# A novel method for integrating genomic and Tn-Seq data to identify common *in vivo* fitness mechanisms across multiple bacterial species

**DOI:** 10.1101/2024.07.24.604993

**Authors:** Derrick E. Fouts, Thomas H. Clarke, Geoffrey B. Severin, Aric N. Brown, Elizabeth N. Ottosen, Caitlyn L. Holmes, Bridget S. Moricz, Sophia Mason, Ritam Sinha, Mark T. Anderson, Victor DiRita, Michael A. Bachman, Harry L. T. Mobley

## Abstract

Sepsis is life-threatening organ dysfunction due to an unregulated immune response to infection. Bacteremia is a leading cause of sepsis, and members of the *Enterobacterales* cause nearly half of bacteremia cases annually. While previous Tn-Seq studies to identify novel bacteremia-fitness genes have provided valuable insight into virulence mechanisms, evidence for common pathways across species is lacking. To identify common fitness pathways in five bacteremia- caused *Enterobacterales* species, we utilized the JCVI pan-genome pipeline to integrate Tn-Seq fitness data with multiple available functional data types. Core genes from species pan-genomes were used to construct a multi-species core pan-genome, producing 2,850 core gene clusters found in four out of the five species. Integration of Tn-Seq fitness data enabled identification of 373 protein clusters that were conserved in all five species. A scoring rubric and filter was applied to these clusters, which incorporated Tn-Seq fitness defects, operon localization, and antibiotic susceptibility data, which reduced the number of bacteremia-fitness genes and identified seven common fitness mechanisms. Independent mutational validation of one prioritized fitness gene, *tatC,* showed reduced fitness *in vivo* and increased susceptibility to beta-lactams that were restored following *tatC* complementation *in trans*. By integrating known operon structures and antibiotic susceptibility with Tn-Seq fitness data, common genes within the core pan-genome emerged and revealed mechanisms that are essential for colonization of, or survival in, the mammalian bloodstream. Our prediction and validation of *tatC* as a common bacteremia fitness factor and contributor of antibiotic resistance supports the utility of this bioinformatic approach. This study represents a major step forward to prioritize potentially novel targets for therapy against these deadly widespread sepsis infections.

**Author Summary:** Bacteremia is a leading cause of sepsis, a life-threatening condition where an unregulated immune response to infection causes systemic organ failure. Nearly half of bacteremia cases are caused by members of the Gram-negative bacterial taxonomic order *Enterobacterales*. Given the public health impact of bacteremia and the reduction of existing antibiotic treatment options, novel strategies are needed to combat these infections. Pan-genome software was used to predict seven shared fitness pathways in these bacteria that may serve as novel targets for treatment of bacteremia. Briefly, a scoring rubric was applied to shared pan-genome clusters, which incorporated multiple data types, including Tn-Seq fitness defects, operon localization, and antibiotic susceptibility data to rank and prioritize fitness genes. To validate one of our predictions, mutations were constructed in *tatC*, which showed both reduced fitness in mice and increased susceptibility to *beta*-lactam antibiotics; complementation restored fitness and antibiotic susceptibility to wild type levels. This study takes a novel bioinformatics approach to build a core pan-genome across multiple distantly related bacteria to integrate computational and experimental data to predict important shared fitness genes and represents a major step forward toward identifying novel targets of therapy against these deadly widespread life-threatening infections.

## Introduction

Sepsis is the life-threatening organ dysfunction resulting from a dysregulated and overwhelming immune response to infection. There are an estimated 49 million cases of sepsis that occur annually worldwide (1) and in the United States sepsis accounts for one in every 2-3 hospital deaths (2). Bacteremia, or the presence of bacteria in the bloodstream, is the leading cause of sepsis with Gram-negative species belonging to the *Enterobacterales* order among the most common causes of bacteremia (3–5). Drug resistance among this cohort of pathogens is a major limitation to effective treatment (4, 6) and contributes to the large burden of antibiotic resistant bacterial threats recognized by the Centers for Disease Control and Prevention (7). Given the public health impact of *Enterobacterales* bacteremia and the waning efficacy of existing treatments, novel strategies are needed to combat these infections. To develop such approaches, it is critical to first understand the genetic and molecular basis by which these bacteria cause bloodstream infections (BSI).

Bacteremia can be divided into three phases of pathogenesis: initial primary site infection, dissemination to the bloodstream, and growth and survival in blood and filtering organs (8). The primary site can also serve as a reservoir for the pathogen that intermittently re-seeds the bloodstream and prolongs systemic infection. We have previously determined that *Enterobacterales* species replicate rapidly in the liver and spleen during bacteremia (9). For some tissue and species combinations, bacterial burdens decrease over time reflecting an ability of the host immune system to overcome this rapid proliferation, while in other instances bacterial growth outpaces clearance and bacterial numbers rapidly expand. This balance between a bacterium’s ability to proliferate and withstand immune-mediated clearance can be collectively described as infection fitness.

To define the genetic requirements for “bacteremia-fitness,” we previously performed a series of transposon sequencing (Tn-Seq) studies in which bacterial genes disrupted by transposon insertion mutations are assessed for their contribution to bacteremia-fitness (10–13). These studies have resulted in the identification and characterization of fitness genes ranked by impact for four *Enterobacterales* BSI pathogens; *Escherichia coli*, *Klebsiella pneumoniae*, *Citrobacter freundii*, and *Serratia marcescens*. Fitness gene identification has also allowed us to build models of metabolism and characterize bacterial pathways required for replication and survival in the host, enabling opportunities for the development of novel therapeutic strategies against these antibiotic resistant pathogens. While the Tn-Seq studies have provided valuable insight on an organismal level, comprehensive information on shared and unique fitness strategies across Gram-negative species is lacking, due in part to the unique nature and depth of each transposon insertion library. By integrating Tn-Seq fitness datasets with comprehensive pan-genomic information on gene content, operon structure, and gene function across species we can gain a better understanding of infection processes for this important group of pathogens.

Our pan-genome pipeline (14) uses PanOCT (15) to cluster orthologous proteins to identify core, variable and singleton gene or protein clusters and provides the data structure to easily link gene functions such as GO terms and antimicrobial resistance (AMR) genes as well as linkage of genes into genomic islands. The core genome of a bacterial pathogen is the collection of genes shared by all or nearly all strains of a species. This core genome is generally enriched for critical functions including energy production, amino acid metabolism, metabolite transport, nucleotide metabolism, and translational machinery (16). On the other hand, the accessory genome is the set of genes that vary across strains of a species. These genes are often involved in protein secretion and defense against innate host immunity, as well as many niche-specific functions that include traits such as fimbriae, toxins, or iron acquisition systems, which reside in so-called “flexible genomic islands” or fGIs (17).

In this report, we first constructed individual species pan-genomes to identify genes shared within *E. coli, K. pneumoniae, C. freundii, S. marcescens*, and *Enterobacter hormaechei* genomes. Each species pan-genome represents the sum of the core and accessory genes for all strains of a species under study. To add biological context, known operon structures, virulence gene predictions, AMR genes and Tn-Seq bacteremia-fitness data were overlaid onto each pan-genome, visually depicted by using the program *PanACEA* (18). Then we took each species’ core genes to build a multi–species pan-genome, which identified the core genes shared across species (19). By integrating our multi-species pan-genome and genome-wide Tn-Seq fitness data, we were able to infer fitness genes in *E. hormaechei,* which lacked Tn-Seq data. We identified 373 protein clusters that were conserved in all five species and predicted to contribute to bacteremia in at least one of them. Applying a scoring rubric to these bacteremia protein clusters, which incorporated the magnitude of a cluster’s fitness defect as predicted by Tn-Seq, its operon localization, and published antibiotic susceptibility data, seven common bacteremia-fitness pathways were identified. Finally, to validate our findings within the prioritized list of bacteremia-fitness genes, the twin-arginine translocation (Tat) system was selected to test independent mutations of four bacteremia-causing *Enterobacterales* for reduced fitness *in vivo* and for increased susceptibility to beta-lactam antibiotics *in vitro*. The TatC transmembrane protein strongly contributes to *C. freundii* bacteremia fitness (11) and was likewise predicted to be a bacteremia-fitness factor in *K. pneumoniae* (13), but not in the *S. marcescens* (10, 20), UPEC *E. coli* (12) or *E. hormaechei*, which lacked Tn-Seq data. Independent insertional inactivation of *tatC* showed reduced fitness *in vivo*, confirming one of our bioinformatics fitness predictions, even in the three species lacking Tn-Seq evidence. The *tatC* mutants also showed increased susceptibility to beta-lactam and beta-lactam plus inhibitor that was restored following *tatC* complementation *in trans*. This raises the exciting possibility that inhibitors of these gene products could simultaneously decrease *in vivo* fitness and sensitize the bacteria to FDA-approved standard-of-care antibiotics. Combined, this study provides an atlas of gene conservation across the *Enterobacterales* and the fitness contribution of these genes to bacteremia, a powerful resource that can be leveraged to advance our understanding of pathogenesis and explore novel therapeutic targets.

## Results

### Species Pan-genomes

Protein-level pan-genomes of five common sepsis-causing *Enterobacterales* species were constructed for two key reasons. First, to integrate multiple data types and Tn-Seq datasets to decipher common bacteremia-fitness factors across species. Second, to generate core pan-genomes that can be used downstream to build a “Multi-Species Core” (MSC) pan-genome that represents highly conserved genes that may be targets of interest for future therapies that would treat infections from multiple species (**Fig 1**). Proteins from representative genomes of *E. coli*, *K. pneumoniae*, *S. marcescens, C. freundii,* and *E. hormaechei* were used to first build species pan-genomes using PanOCT separately within the JCVI pan-genome pipeline (**S1 Table, S1-5 Datasets**). Centroids (*i.e*., orthologous protein cluster representatives) of core and accessory pan-genome clusters were searched against curated databases of bacteremia-fitness genes as defined by transposon insertion-site sequencing (**S2 Table**), classic *E. coli* “essential” genes (**S3 Table**), virulence genes (21) (**S4 Table**), and AMR (22) genes (**Table 1**).

**Fig 1.**
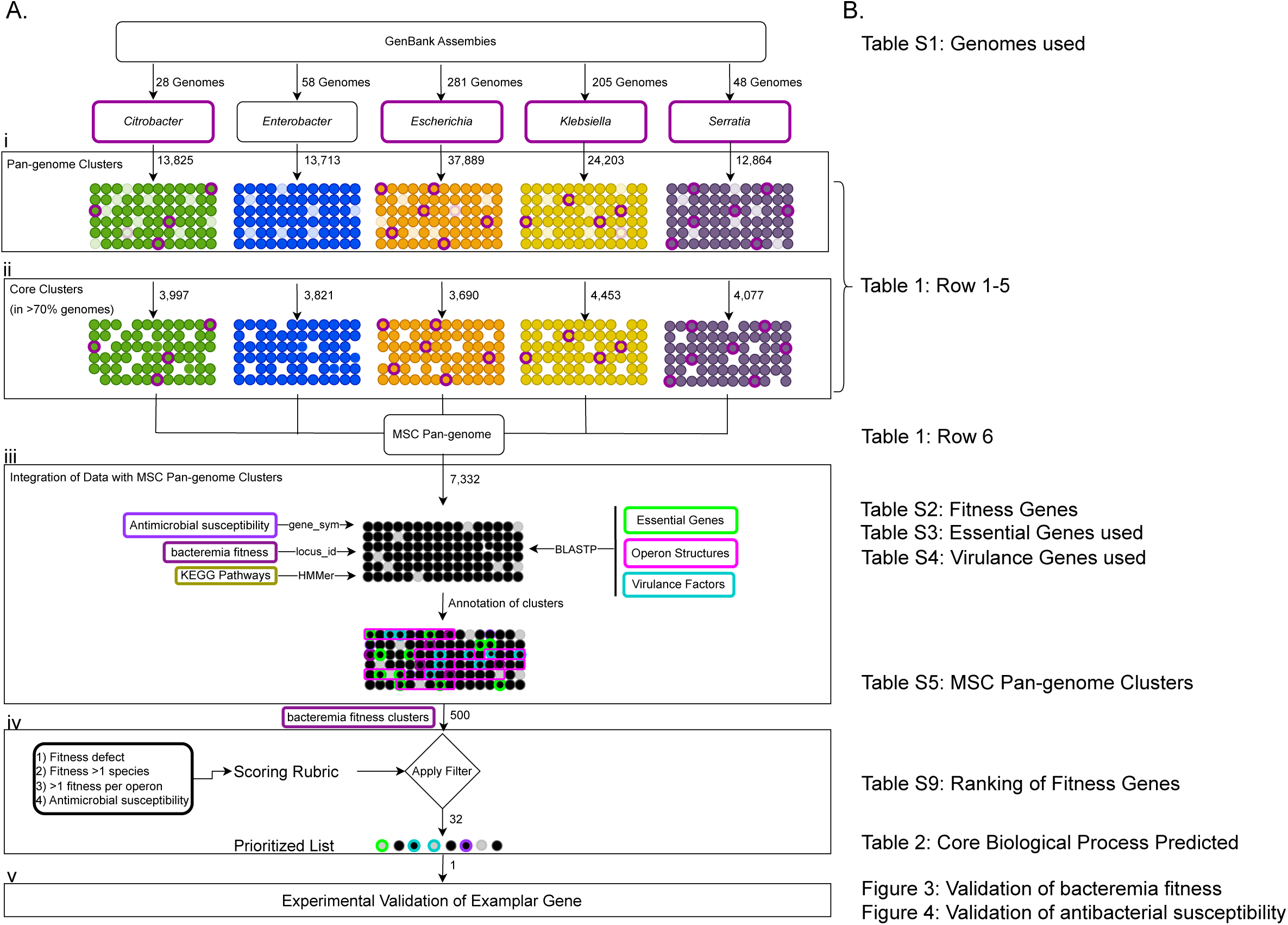
Construction of Genus and Multi-Species Core Pan-Genomes. The workflow of bioinformatic processes used in this study is depicted (**A**). Genome assemblies for the five genera shown were downloaded from NCBI, including those genomes used in Tn-Seq studies to identify fitness genes (purple outline) (10–13). The assemblies were used as input to the JCVI pan-genome pipeline to create pan-genomes for each genus (i). From the resulting pan-genome clusters, core proteins (orthologs present in >70% of the genomes) were selected (ii) as input for the multi-species core pan-genome using the same pipeline. In addition to the TIGRFam and CARD annotations from the pan-genome pipeline, we integrated six annotation features from multiple sources to the resulting MSC pan-genome clusters (iii). *E. coli* essential genes (90–94), operon membership (95, 96), and virulence factor information (21) was assigned through BLASTP, KEGG pathway information was mapped using HMMer, and antibiotic susceptibility (23, 24) and Tn-Seq bacteremia fitness genes were mapped by gene_syms and locus_ids, respectively. To only those MSC pan-genome clusters labeled as Tn-Seq bacteremia fitness, we applied a scoring rubric to filter the list of 500 proteins down to 32 prioritized proteins (iv). Lastly, an exemplar gene, *tatC*, was selected for validation of fitness *in vivo* and antimicrobial susceptibility *in vitro* in all five genera (v). The location of the input and output data tables is also shown (**B**).

**Table 1.**
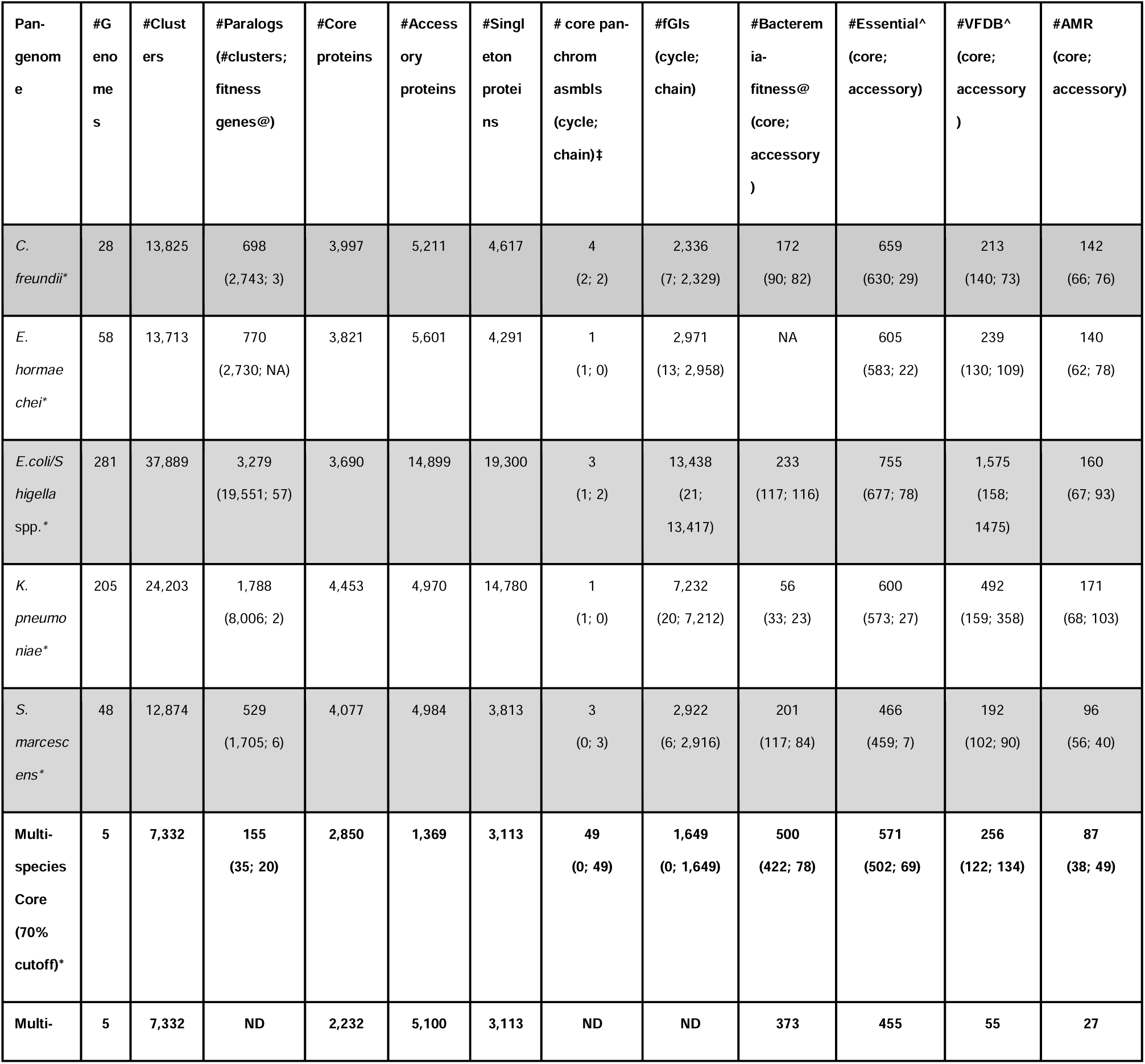
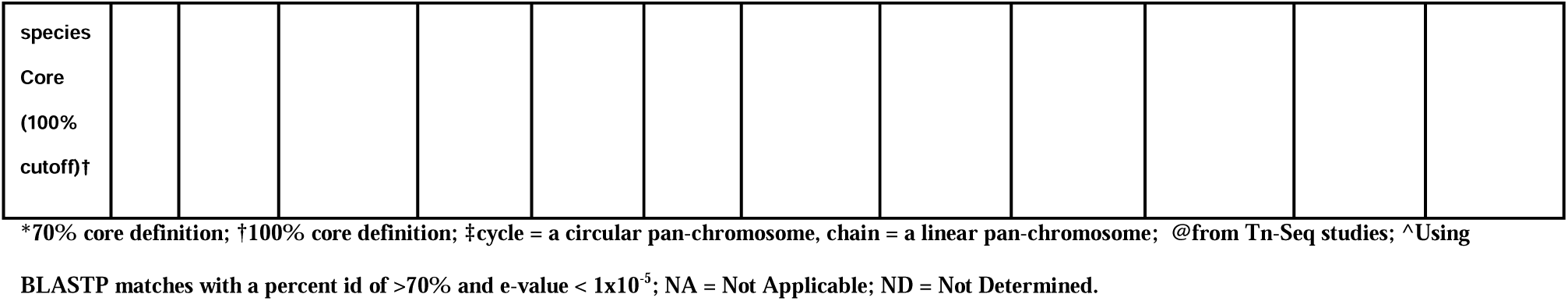
Pan-genome Statistics.

The core genomes of each species varied from 3,690 protein clusters in *E. coli* to 4,453 in *K. pneumoniae*. Based on 281 genomes, *E. coli* had the largest pan-genome, encoding 37,889 protein clusters, which is 8 times the number of proteins encoded by the average *E. coli* genome (*i.e*., 4,661 proteins) (**Table 1, S1 Table**). *K. pneumoniae* had the second largest pan-genome with 24,203 protein clusters across 205 genomes, but the number of proteins of this pan-genome was 4.5 times that of the average *K. pneumoniae* genome (*i.e.*, 5,322 proteins). The remaining species’ pan-genome number of proteins were between 2.7-3.0 times larger than their respective genome averages. The number of strain-specific protein clusters from the species pan-genomes ranged from a minimum of 0 to a maximum of 488 (*Escherichia*: 0 to 488, *Klebsiella*: 1 to 296, *Enterobacter*: 0 to 280, *Citrobacter*: 4 to 449, and *Serratia*: 0 to 449) (**S1 Table**), indicating that each species has a large and variable accessory genome.

Since our pan-genome software, PanOCT, does not collapse or discard paralogs, we were able to determine the extent of paralogy within each species pan-genome. Of the 37,889 total *E. coli* pan-genome protein clusters, 52% were paralogous, reducing the size of the pan-genome to 18,338 protein clusters, which is still considerably larger than the other species pan-genomes (**Table 1**). *Klebsiella* had the second highest number of paralogs (33%) followed by *Enterobacter*/*Citrobacter* (20% each) and *Serratia* (13%).

A total of 569 CARD/RGI-predicted AMR protein clusters were identified across all 4 species with Tn-Seq data (*i.e.*, 160 in *E. coli*, 171 in *K. pneumoniae*, 142 in *C. freundii*, and 96 in *S. marcescens*) (**Table 1**). The distribution of AMR genes across the core and accessory species pan-genomes was roughly even, but with slightly more in the accessory pan-genome (45% core, 55% accessory) except for *S. marcescens* (58% core, 42% accessory). In stark contrast, genes essential for growth under laboratory conditions were skewed towards core (95% core, 5% accessory). Predicted virulence protein clusters varied across species with *E. coli* and *K. pneumoniae* having more clusters in the accessory pan-genome.

### Fitness genes across species

A total of 695 published bacteremia-fitness genes derived from four Tn-Seq studies (10–13) of *E. coli* (247 genes), *K. pneumoniae* (74 genes), *C. freundii* (172 genes), *S. marcescens* (202 genes) (**S2 Table)** were mapped to 662 species pan-genome clusters (**Table 1**). The distribution of species-specific fitness gene clusters across the core and accessory pan-genomes showed slightly more in the core pan-genome (54% core, 46% accessory), with *K. pneumoniae* (59% core, 41% accessory) and *S. marcescens* (58% core, 42% accessory) each having more bias toward core genes. This indicates that many genes important for causing bacteremia are highly conserved within each species and may be conserved across species as well.

### Multi-species Pan-genome

To identify genes conserved across species and determine if they are multi-species bacteremia-fitness factors, core protein sequences from each of the five species-level pan-genomes were combined and analyzed in a second pan-genome run - the MSC pan-genome. For this MSC pan-genome, clusters were labeled as core if they contained proteins from at least four out of five species core pan-genomes. There were 2,232 clusters shared across all five species and 2,850 shared in four of five species (**Table 1**, 100% and 70% cutoff respectively, and **S1A Fig**). *Serratia* had the largest number of unique core protein clusters within its species core genome at 1,124 with *Klebsiella* having the second largest number of unique core clusters at 892 (**S1A Fig**). Despite *Escherichia* having the most accessory genes (**Table 1**) in the species pan-genome, it only has 385 unique core protein clusters within the MSC pan-genome.

### Identification of common bacteremia-fitness and virulence genes

Next, we analyzed the MSC pan-genome for bacteremia-fitness genes (**S5 Table, S6 Dataset**). Using a criterion that at least one member of a MSC pan-genome protein cluster was previously identified by Tn-Seq, 500 clusters were identified as bacteremia-fitness factors with 422 conserved in 4 of 5 species and 373 were conserved across all five species (**Table 1, S1B Fig**). These bacteremia-fitness clusters included proteins from multiple species with published Tn-Seq results implicating them as fitness factors. For example, 11 clusters contained Tn-Seq fitness factors across 3 species: 7 from *C. freundii*, *K. pneumoniae*, and *S. marcescens*, and 4 from *C. freundii*, *E. coli*, and *S. marcescens* (**S6 Table**). Combining these pan-genome clusters with Tn-Seq data can be used to predict potential fitness genes in species and strains that lack experimentally derived data.

MSC pan-genome clusters with homology to known virulence factors were identified by searching against the virulence factor database (VFDB) (21). Out of 256 total clusters assigned to virulence functions (**Table 1**), 55 (21%) were conserved across all five genera (**S1C Fig**). The maximum number of shared virulence factors between four, three and two species was 49 (*Citrobacter*:*Enterobacter*:*Escherichia*:*Serratia*), 12 (*Citrobacter*:*Enterobacter*:*Escherichia*) and 7 (*Citrobacter*:*Escherichia*), respectively. *K. pneumoniae* had the highest number of species-specific virulence factors with 45. We then determined how many MSC pan-genome clusters predicted to be bacteremia-fitness factors were also homologous to known virulence factors (**S7 Table**). Of the 500 bacteremia-fitness clusters and 256 virulence clusters conserved in 4 or more species, 34 were predicted bacteremia-fitness factors associated with known virulence mechanisms (*e.g.*, LPS and capsular production for immune modulation, flagella and fimbriae for motility and adherence, and genes for the acquisition of iron and magnesium).

Using an orthologous approach to identifying multi-species fitness factors, we searched for core operons that contained distinct predicted fitness genes across species. We identified 369 operons that contained at least one bacteremia-fitness factor in the MSC pan-genome (**S8 Table**). Of the 67 operons that contained more than one bacteremia-fitness factor in the MSC pan-genome, 14 operons contained bacteremia-fitness genes from 3 species, 29 from 2 species, and 24 from one species (**S8 Table**). One prominent example was the operon encoding the Enterobacterial common antigen (ECA), a 12-gene locus with 9 bacteremia-fitness genes identified in either *S. marcescens*, *C. freundii*, or *K. pneumoniae* (**Fig 2**). The WzxE ECA flippase within this operon is a predicted fitness factor in all 3 species while WecA (UDP-*N*-acetylglucosamine-UP *N*-acetylglucosaminephosphotransferase) and RffH (glucose-1-phosphate thymidylyltransferase) proteins were fitness factors in two species.

**Fig 2.**
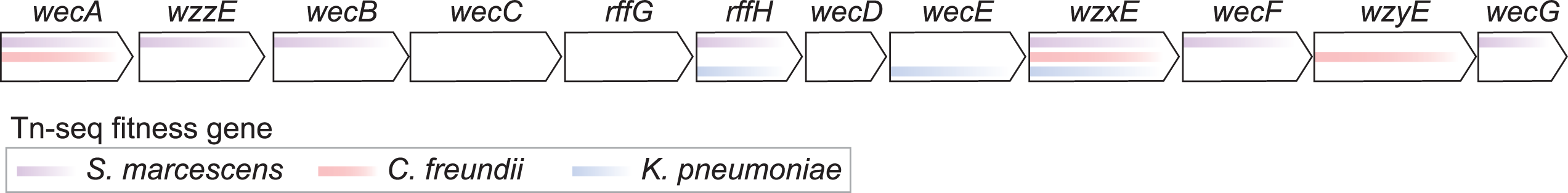
The ECA operon is a shared bacteremia-fitness locus between species. Genes within the conserved ECA operon are represented by arrows and colored lines indicate genes that were identified as significant bacteremia-fitness genes in previously published Tn-Seq studies.

To prioritize genes for future investigation and to identify conserved biological processes contributing to bacteremia we devised a scoring rubric using four genotypic and phenotypic characteristics (see Materials and Methods for details). The MSC pan-genome bacteremia-fitness genes were individually scored based on the magnitude of their published fitness defect predicted in any bacteremia Tn-Seq screen, whether the same gene was predicted to be a bacteremia-fitness factor in multiple Tn-Seq screens (*i.e*., multiple species), if multiple bacteremia-fitness factors were encoded in the same operon, and whether mutation of that gene was previously found to increase antibiotic susceptibility of *E. coli* BW25113 (23, 24). Scores were summed in two ways: first as individual genes to distill the most meaningful bacteremia-fitness factors from the 500 total MSC pan-genome bacteremia-fitness clusters identified, and then as the total score of bacteremia-fitness factors encoded in the same operon to identify biological pathways of interest (**S9 Table**). The rubric produced scores for individual centroids and for operons that most closely mapped to the exponential distribution using the Kolmogorov–Smirnov test with the cutoff capturing the top 10% and 5% of the centroids as shown on the graph (**S2 Fig**).

Application of this scoring rubric filters led to a reduction of potential bacteremia-fitness factors from 500 to 32 (6.4%) and 73 (14.6%) prioritized bacteremia-fitness factors having centroid scores of >=8 and >=6, which correspond to centroids in the top 5% and top 10%, respectively (**S2 Fig**). Additionally, summation of the individual scores for bacteremia-fitness factors encoded within the same operon led to the identification of 11 operons with total scores that were in the top 0.5%. Investigation of the gene functions encoded by these highly-scored individual bacteremia-fitness factors and operons revealed seven biological processes that are predicted to contribute to the full virulence capacity of *Enterobacterales* during bacteremia (**Table 2**). These processes include maintenance of the proton motive force, resistance to antimicrobial peptides and complement, protein and small molecule transport, DNA repair and homologous recombination, shikimate biosynthesis, global gene regulation (pre- and post-transcription), and resistance to oxidative stress. While not an exhaustive accounting of all biological processes that contribute to bacteremia or of the functions encoded within the top scoring genes and operons, these seven biological functions (**Table 2**) include, at a minimum, 19 of the 32 (59%) fitness genes with scores in the top 5% and 40 of the 73 (55%) fitness genes with scores in the top 10% (**S2 Fig**). Additionally, these seven biological functions encompass genetic elements encoded by 10 of the 11 highest scoring bacteremia-fitness factor rich operons identified.

**Table 2.**
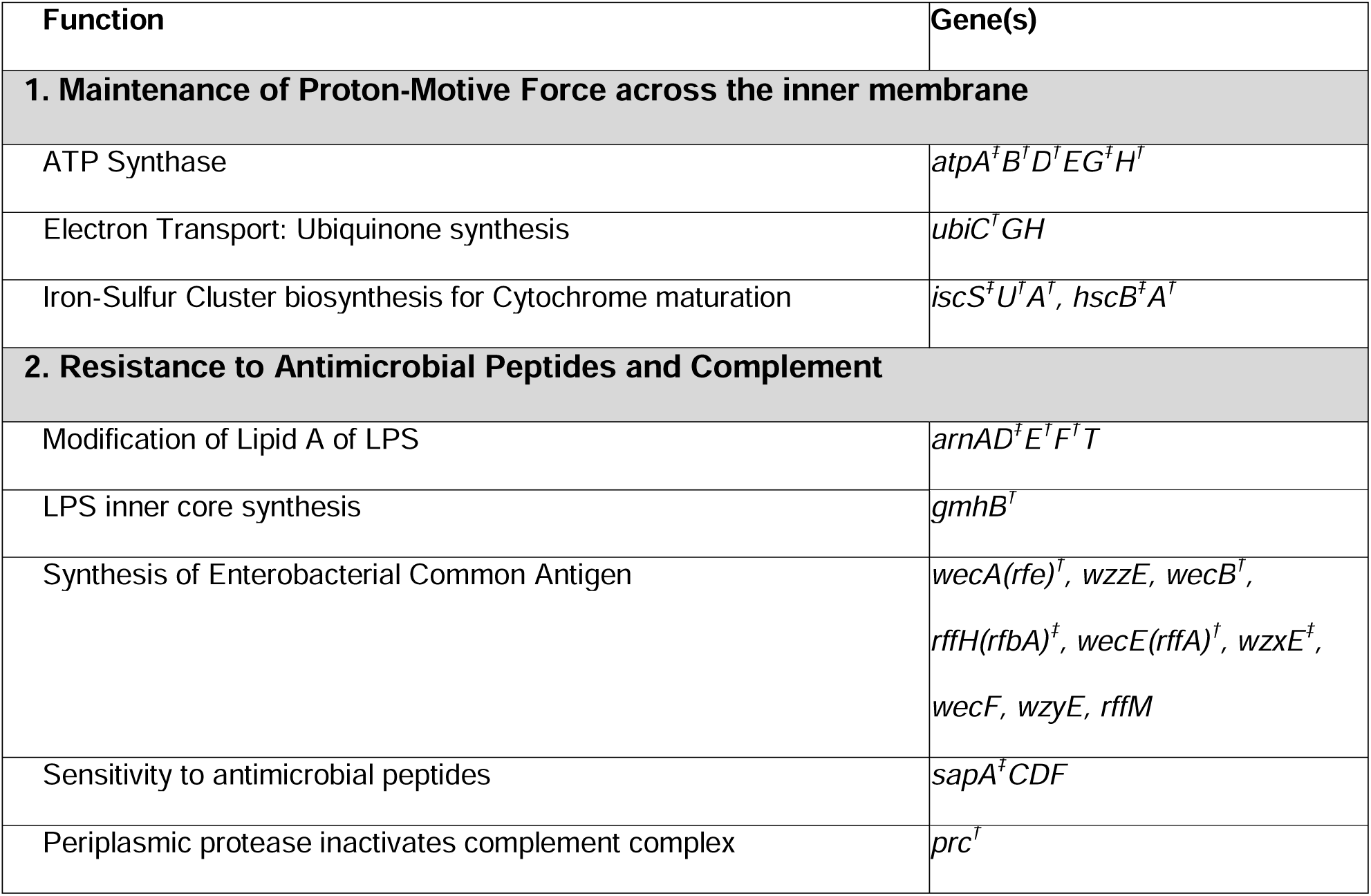
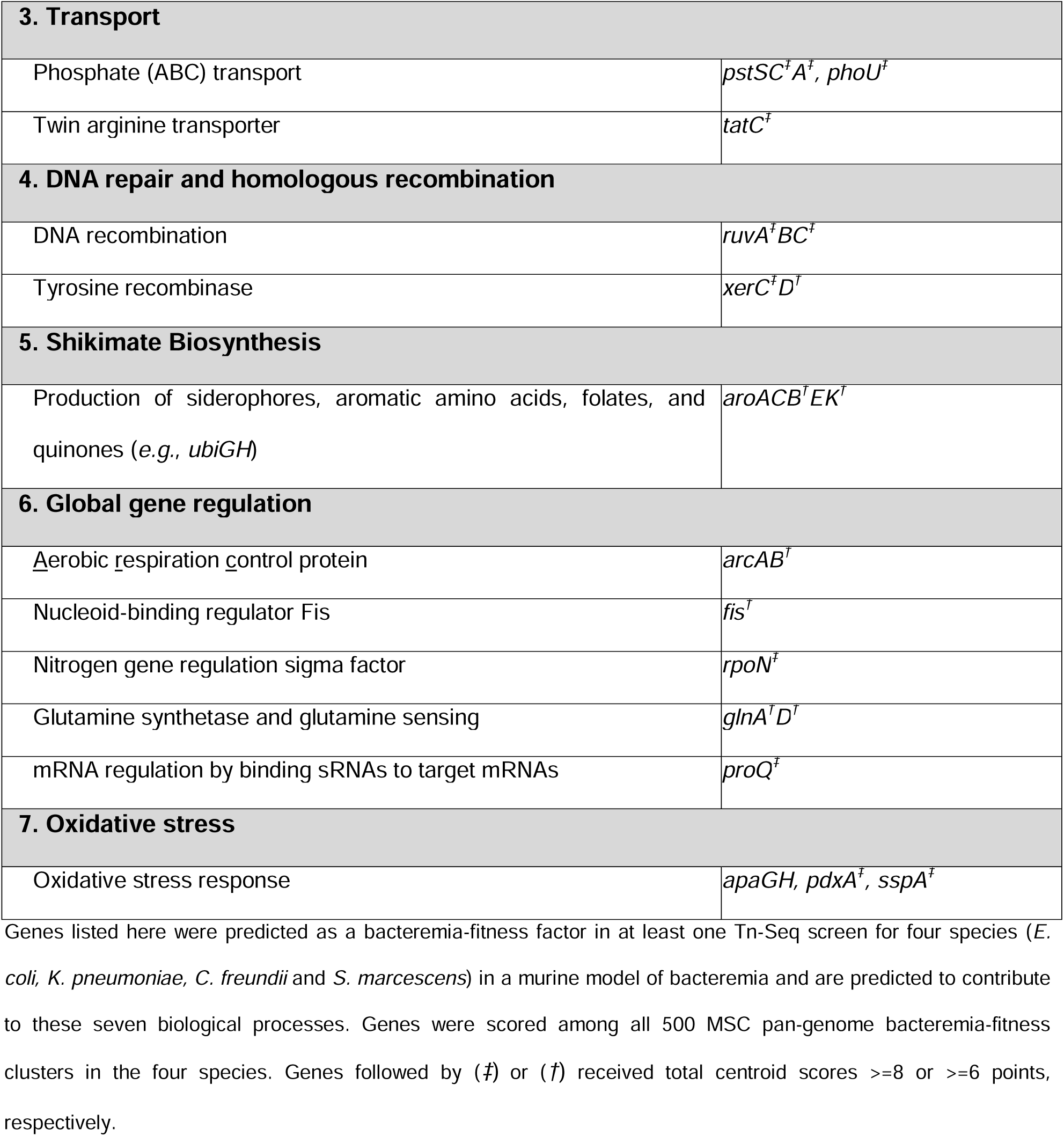
Core Biological Processes Predicted to be Required for Full Virulence Capacity of *Enterobacterales* during Bacteremia using a Multifactorial Prioritization Scoring Rubric*.

#### Validation of a representative core fitness gene function for virulence phenotype in a bacteremia mouse model

In our Tn-Seq screens, numerous genes have been predicted to represent fitness factors during experimental bacteremia. This is a first step in identifying relevant genes that are critical for survival in the bloodstream. To validate our findings within the prioritized MSC genes, the twin-arginine translocation (Tat) system was selected for further study. It was selected because of the variable results (i.e., essentiality/fitness roles) from published Tn-Seq studies (10–13) and its role in virulence and antimicrobial susceptibility in other Gram-negative pathogens (11, 25–29) and certain pathotypes of *E. coli* (*30, 31*). It also tests our operon scoring cut-off since it barely meets the 5% operon score cut-off of 10.28 with a score of 11 (**S9 Table**). The Tat system, under Transport functions (**Table 2**), facilitates translocation of folded proteins across the cytoplasmic membrane in bacteria and while the system itself (TatABC) is widely conserved, the translocated proteins that serve as substrates for the system often vary between species (32) and may therefore alter the requirement for the translocation system in different organisms. The *tatC* allele was one of the highest-scoring predicted bacteremia-fitness factors (**Table 2** and **S9 Table**).

To directly test whether TatC was a conserved bacteremia-fitness factor among the remaining three species of interest, *tatC* mutants were constructed in *S. marcescens*, *K. pneumoniae, E. hormaechei* and *E. coli* followed by competition infections whereby the wild-type strain and the differentially marked *tatC* mutant strain were co-inoculated in a murine infection model. For all four tested species, the *tatC* mutant was significantly outcompeted by the wild-type strain when bacteria were inoculated into the bloodstream via tail vein injection (TVI), as indicated by competitive indices that were significantly below zero (**Fig 3A-D**). Furthermore, *tatC* was required in every tested organ via this model, demonstrating that folded protein translocation was universally important not only across species but within different infection niches.

**Fig 3.**
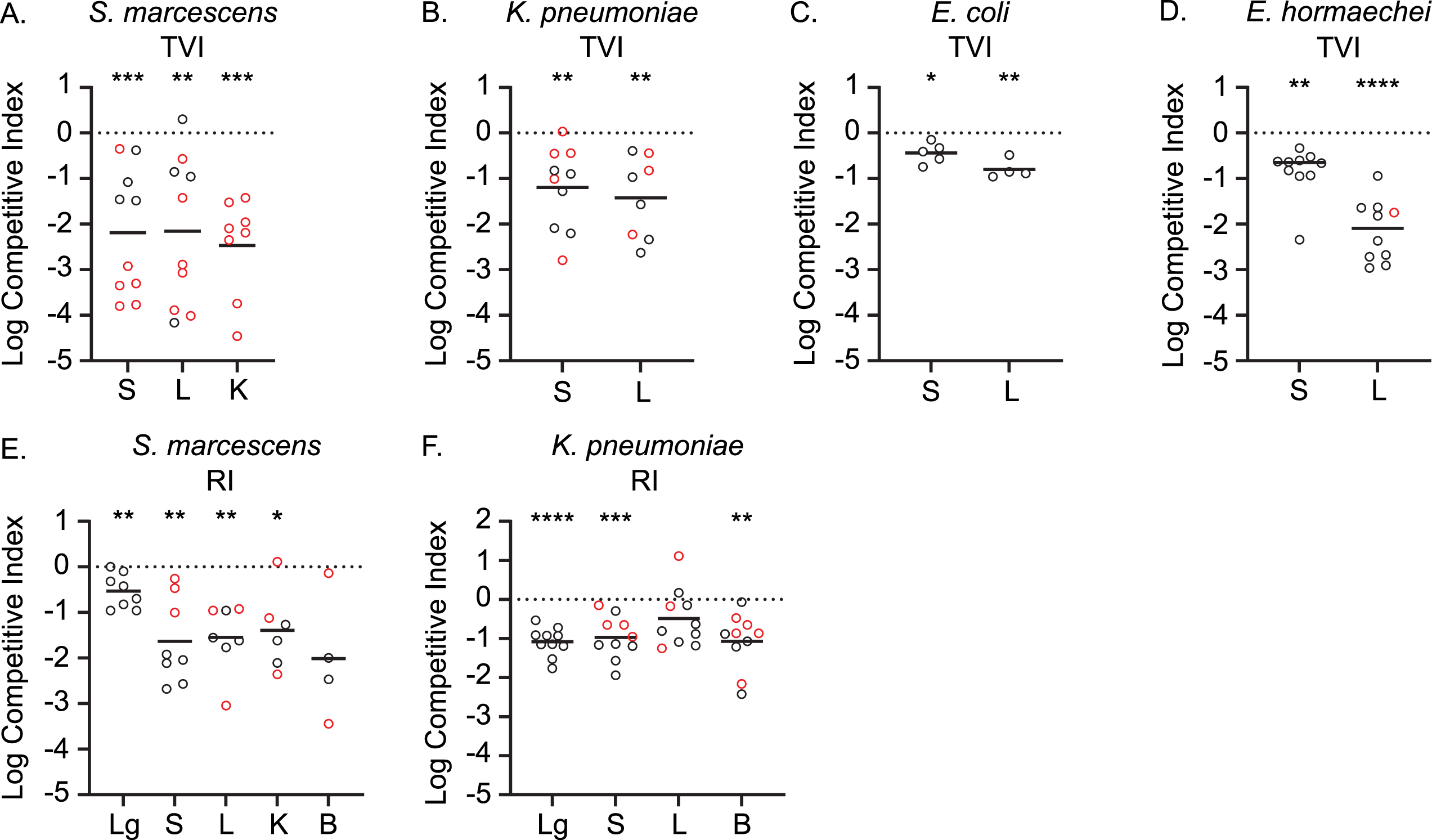
Validation of *tatC* contributions to bacteremia fitness. Mice were inoculated via tail vein injection (TVI) or retropharyngeal inoculation (RI) with a mixture of wild-type and *tatC* mutant derivatives of *S. marcescens* (A and E), *K. pneumoniae* (B and F), *E. coli* (C), or *E. hormaechei* (D). Viable bacteria were enumerated in the spleen (S), liver, (L), kidneys (K), lungs (Lg) or blood (B) at 24 h post-inoculation and the relative fitness of each strain was determined as the log competitive index. Only *S. marcescens* was assayed in the kidneys based on previously determined infection profiles [8]. Mean indices (solid line) that deviated significantly from the hypothetical value of zero (dotted line), which represents neutral fitness, were determined by one-sample t-test: * (P < 0.05), ** (P < 0.01), *** (P < 0.001), **** (P<0.0001). Red symbols designate samples from which *tatC* mutant bacteria were at or below the limit of detection.

Since the TVI route represents direct bacteremia, we additionally sought to determine how TatC contributes to disseminated infection via a bacteremic pneumonia model. Following retropharyngeal inoculation (RI), *S. marcescens* and *K. pneumoniae tatC* mutants were outcompeted in the lungs of infected mice (**Fig 3E-F**). For *S. marcescens*, an even greater competitive disadvantage was observed for bacteria that had escaped the primary infection site and disseminated to the spleen, liver, and kidneys (**Fig 3A and E**). This result was further supported by a trend toward lower recovery of the *tatC* mutant from the bloodstream and suggests that Tat translocation is critically important for escape from the lungs or survival in distal sites for this organism. Similarly, *K. pneumoniae tatC* mutants that disseminated from the primary infection site were also significantly outcompeted by wild-type bacteria (**Fig 3B and F**). Together, these results establish the importance of the Tat translocation system, and very likely its substrate proteins, in Gram-negative bacteremia and demonstrate the value of comparing Tn-Seq fitness hits between species.

#### Contribution of conserved fitness genes to intrinsic antibiotic resistance

Previous work in *E. coli* revealed that many core genes had contributions to antibiotic resistance that were not predictable based on their known functions (23, 24). Genes that contribute to both bacteremia fitness and antibiotic resistance could be attractive therapeutic targets as disruption of their function in combination with antibiotic therapy would be expected to have synergistic efficacy. Therefore, we mapped published data on the antibiotic susceptibility of a comprehensive panel of *E. coli* mutants to multiple classes of antibiotics to our MSC pan-genome. *E. coli* mutants in *tatC* were previously shown to have increased susceptibility to beta-lactam antibiotics. Unlike *E. coli*, the species *K. pneumoniae*, *S. marcescens*, *E. hormaechei* and *C. freundii* have intrinsic resistance to ampicillin due to chromosomally-encoded beta-lactamases. To test whether disruption of *tatC* by either a transposon insertion (*tatC::tn*) or deletion (Δ*tatC::nptII*) could counteract this intrinsic resistance, we used Epsilometer tests to determine the minimum inhibitory concentration (MIC) of ampicillin or piperacillin-tazobactam for the wild-type and *tatC* mutants of these four species. The most striking finding was in *K. pneumoniae*, where disruption of *tatC* by either approach lowered the MIC to ≤ 8, which is a level that would be considered susceptible to ampicillin in clinical care of an infection, and is similar to disruption of the beta-lactamase itself (*bla::tn*) (**Table 3**, **Fig 4**). This increased susceptibility to ampicillin in *K. pneumoniae* was counteracted by provision of either *tatC* (pACYC*tatC*) or the *tatABCD* operon (pACYC*tatABCD*) on a multicopy plasmid, but not the empty vector control (pACYCev). When *tatC* was interrupted in a carbapenem-resistant isolate of *E. hormaechei*, the resistance profile shifted from resistant to sensitive dose dependent (SDD) to piperacillin-tazobactam (**Table 3**). The resistance phenotype of the *tatC* mutant was returned to parental levels when *tatABC* was provided on a plasmid. In the case of *S. marcescens*, which has an extremely high resistance to ampicillin (> 256 µg/ml), mutation of *tatC* caused a substantial reduction in MIC, which could be rescued by provision of the *tatC* in trans from a multicopy plasmid (pBB26) and not the vector control (pBBR1MCS-5) (**Table 3**). In *C. freundii*, we tested piperacillin-tazobactam as a beta-lactam and beta-lactamase inhibitor combination without intrinsic resistance and found that a *tatC* mutation also lowered the MIC and this phenotype was likewise rescued by provision of *tatC* (pBB4 (*tatC*+)) in trans and not the vector control ((pBBR1MCS-5; **Table 3**). Combined with animal models of infection, these data demonstrate that *tatC* represents a conserved bacteremia-fitness factor that also contributes to intrinsic resistance to beta-lactam antibiotics in *Enterobacterales* and has the potential to restore the use of antibiotics to which these species have intrinsic or acquired resistance.

**Fig 4.**
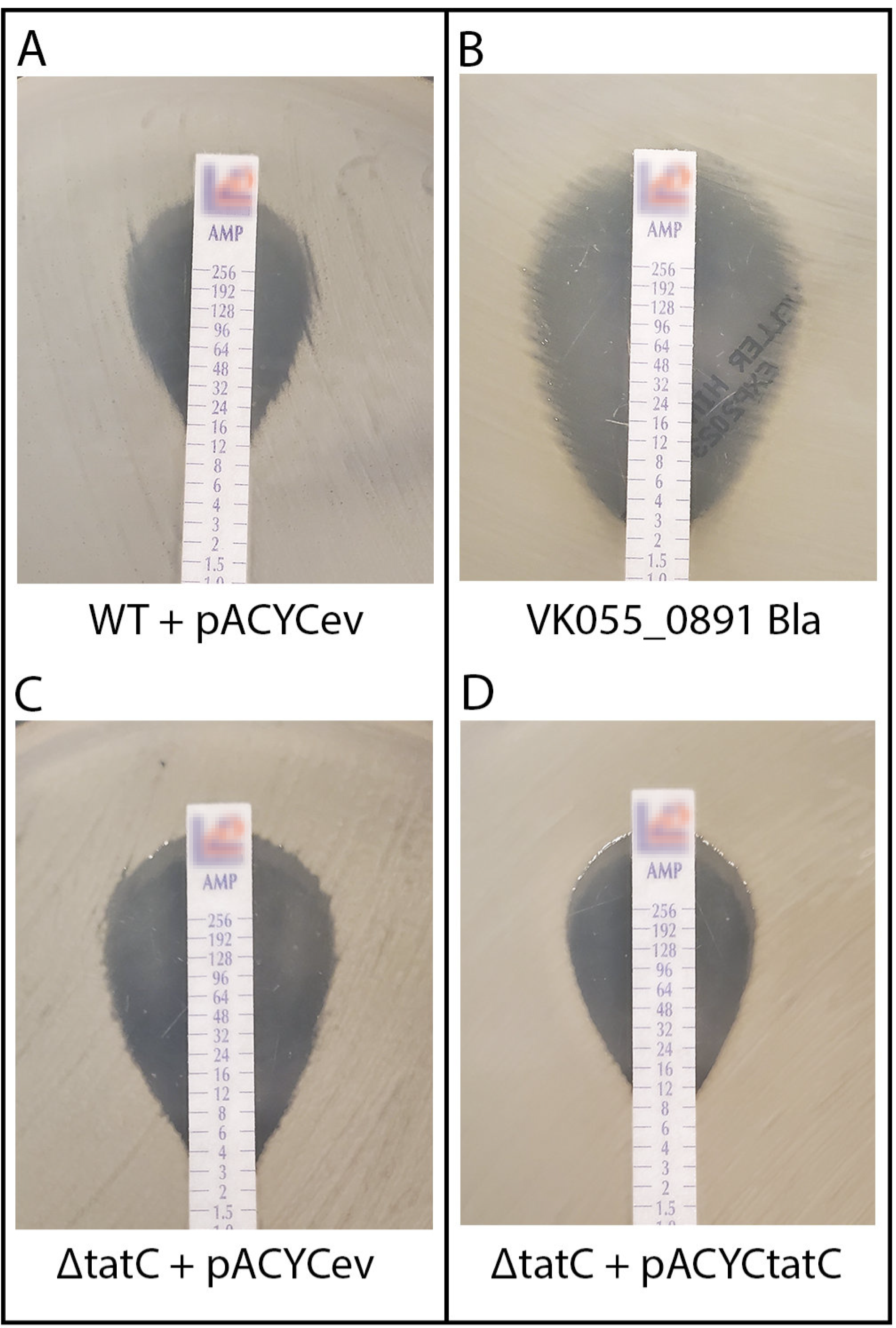
Disruption of *K. pneumoniae tatC* confers susceptibility to ampicillin. Epsilometer (E-test) results for ampicillin are shown with minimum inhibitory concentration (MIC) at the point the zone of clearing touches the test strip for A) WT KPPR1 + pACYCev (pACYC184), B) KPPR1 *bla* transposon mutant VK055_0891_bla, C) KPPR1 Δ*tatC* + pACYCev and D) KPPR1 Δ*tatC* + pACYC*tatC*.

**Table 3.**
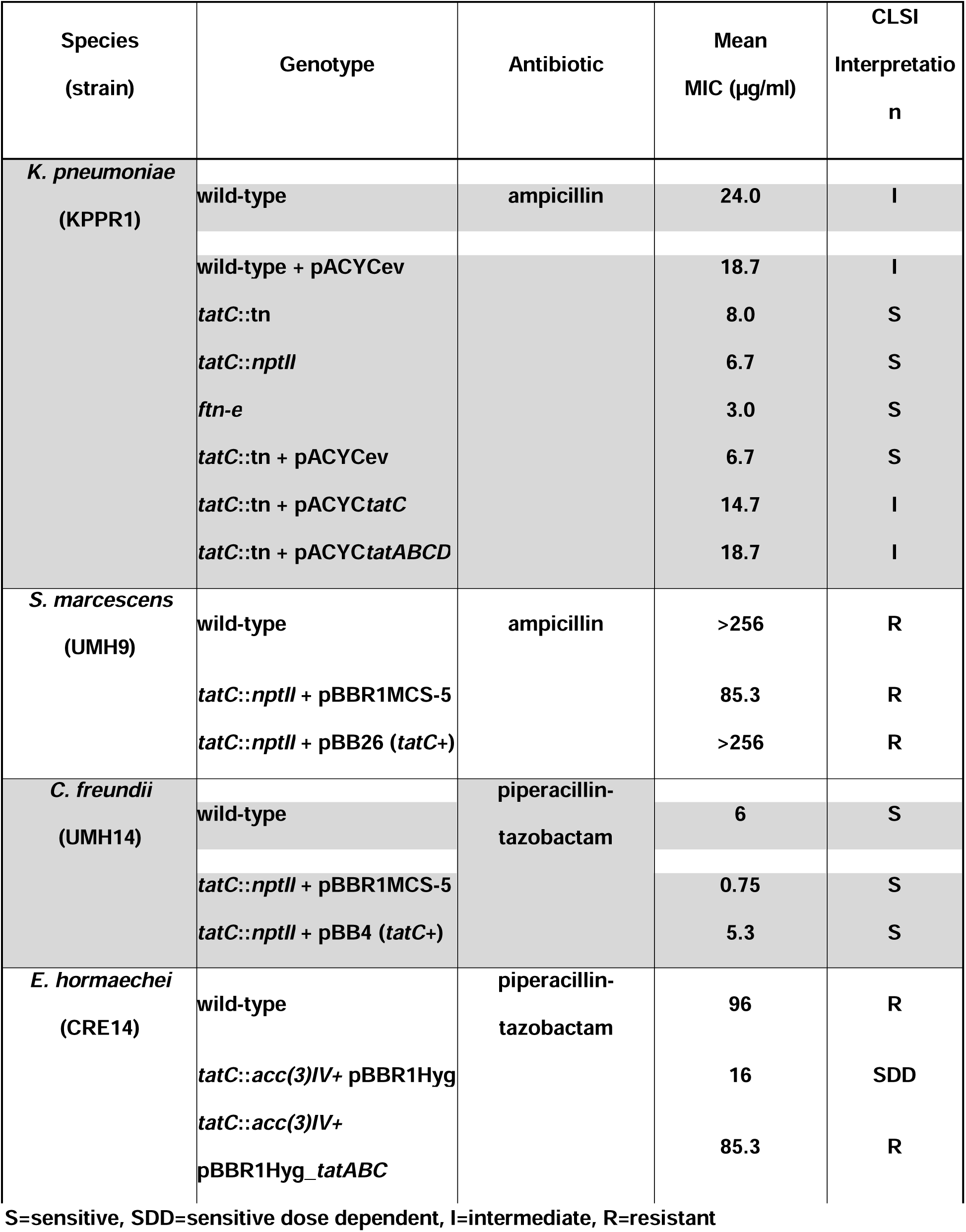
Susceptibility of *tatC* mutants to Beta-lactam antibiotics.

## Discussion

While Tn-Seq has become a powerful tool to simultaneously assess the contribution of all non-essential genes in animal models of infection, mechanistic studies for evaluation of specific genes in animal models of bacteremia have been conducted only sparingly (33). While there have been more than 200 published Tn-Seq studies dealing with bacterial pathogens *in vitro* or *in vivo* (*e.g.*, (12, 34–84)), only a fraction of these have addressed Gram-negative pathogens in animal models of infection; for example: (12, 34, 36, 42, 46, 48, 51, 58, 60, 61, 65, 69, 71–84). In practice, characteristics of Gram-negative bacterial species causing bacteremia have been hard to identify, and only a few studies (58, 85, 86), beyond our work (10, 12, 13, 73, 87–89), have sought fitness and virulence factors in models of bacteremia. Despite the valuable insight Tn-Seq studies have provided at the organismal level, comprehensive information on shared and unique fitness strategies across Gram-negative species is lacking, due in part to the unique nature and depth of each transposon insertion library and a lack of computational tools to compare multiple datasets across pan-genomes..

Because each of our published bacteremia Tn-Seq studies (10–13) seemed to identify different genes (i.e., no single gene was found to be associated with bacteremia-fitness across all studies/organisms), we were unable to identify operons/pathway associations until we took our computational data integration approach presented here. In this report, we leveraged the JCVI pan-genome pipeline as a tool to integrate multiple disparate data types to identify key pathways and mechanisms used by *Enterobacterales* to survive in the bloodstream. Within the core genes (2,850/2,232, 70%/100% core, respectively) shared by *E. coli*/*Shigella spp*.*, K. pneumoniae, S. marcescens*, *C. freundii,* and *E. hormaechei,* statistically significant bacteremia-fitness factors were identified in core clusters (424/373, 70%/100% core, respectively) containing at least one bacteremia-fitness factor identified in any of the four Tn-Seq studies performed in our TVI model of bacteremia. Using a multifactorial scoring rubric, we propose prioritization of 73 of the 500 total conserved bacteremia-fitness factors found in the MSC pan-genome, that have scores in the top 10% for future investigation. Of these 73 prioritized fitness factor genes, 36 were predicted by Tn-Seq to contribute to bacteremia in at least two species and 41 were found to increase antibiotic susceptibility in *E. coli* BW25113 when mutated (23, 24). Summation of individual gene scores encoded in the same operon revealed 27 operons enriched for bacteremia-fitness factors with scores in the top 5%. Investigation of the biological functions encoded by these operons and high scoring bacteremia fitness factors revealed a minimum of seven common biological functions that significantly contribute to *Enterobacterales* fitness during bacteremia. These data support a model in which full virulence of *Enterobacterales* during bacteremia in-part requires the seven biological functions outlined in **Table 2** and discussed below.

To aid in the discovery of potential novel antibiotic targets to clear bacteremia, we conducted species-level and family-level pan-genomic analyses that identified the core pathogenic genome of these *Enterobacterales* species. The identification of unique and common virulence determinants among these Gram-negative bacterial pathogens also provided substantial advances toward determining mechanisms of pathogenesis for these pathogens that disseminate to and survive in the bloodstream. By integrating the pan-*Enterobacterales* genome, genome-wide Tn-Seq fitness data (10–13), and published lists of essential genes (90–94), antibiotic mutational susceptibility (23, 24), virulence (21) and operon data (95, 96), we have identified 373 conserved fitness genes across all five species that were shown to be important in at least one species during experimental bacteremia. Inhibition of these bacteremia-fitness factors *in vivo* could allow the immune system to clear infecting bacteria more rapidly or even kill them directly. Furthermore, 95 mutants in these bacteremia-fitness genes also confer increased susceptibility to common antibiotics in a laboratory strain of *E. coli* (23, 24). This raises the exciting possibility that inhibitors of these gene products can simultaneously decrease *in vivo* fitness and sensitize the bacteria to FDA-approved standard-of-care antibiotics.

The MSC pan-genome produced 500 orthologous protein clusters that contained at least one bacteremia-fitness factor from at least one published Tn-Seq dataset, which is too many to validate. Thus, we devised a subjective scoring rubric that combined four lines of pro-fitness biologically meaningful evidence (e.g., the published fold-change of the fitness defect associated with transposon interruption of a gene in any one species, being identified as a fitness factor in multiple species, whether multiple fitness factors were encoded in the same operon, and if previously found to confer increased antibiotic susceptibility) as a non-statistical way to filter the list of 500 potential bacteremia fitness genes to a reduced and manageable set of 32 genes to independently knock-out in each pathogen and test in the murine model of bacteremia. This resulted in several conserved and shared *Enterobacterales* genes that fall into common bacterial cell functions of which we discuss seven (**Table 2**). These included 1) maintenance of proton-motive force across the inner membrane, 2) resistance to antimicrobial peptides and complement, 3) substrate transport, 4) genome maintenance, 5) shikimate biosynthesis, 6) global gene regulation, and 7) oxidative stress. While it is beyond the scope of this study to discuss all these findings, it is instructive to summarize notable predicted bacteremia-fitness factors below in the context of the seven highlighted common bacterial cell processes that support survival of *Enterobacterales* species in the mammalian bloodstream.

Enterobacterial Common Antigen is a critical outer membrane component in *Enterobacterales* that may affect resistance to antimicrobial peptides and complement (97). Interestingly, while identified by Tn-Seq in *K. pneumoniae*, *C. freundii*, and *S. marcescens* (**Fig 2**), none of the ECA genes were identified by Tn-Seq as fitness factors in *E. coli* suggesting that either the biological role of ECA during infection differs in this species or that ECA genes were not adequately targeted by transposon mutagenesis in *E. coli*. While this discrepancy is currently under investigation, the clear benefit of such a comparison is in forming testable hypotheses regarding the conserved function of ECA or other shared operons across Gram-negative bacteremia-causing genera. Combined, our findings provide strong evidence that production of ECA is an important fitness determinant since mutation of even a single gene in the processive ECA biosynthesis pathway may result in a lack of ECA production or its proper localization in the bacterium. Clearly, surface-exposed macromolecules on the Gram-negative outer membrane, when present or absent or biochemically modified, may modulate surface charge. These changes in charge can confer, in some cases, resistance to binding by complement components and host-derived antimicrobial peptides, thus conferring protection from these host innate immune responses.

Twin-arginine translocation system (98) represents perhaps the most critical transport system and fitness factor overall required for bacteremia. The TatC transmembrane protein was previously shown to contribute to bacteremia fitness in *C. freundii* (11) and *K. pneumoniae* (13) (**S2 Table**). However, neither the *S. marcescens* (10) nor *E. coli* (12) Tn-Seq findings identified TatC as a significant fitness factor, and was even found to be essential in *S. marcescens* for growth on laboratory media (20). Tat mutants of *Salmonella enterica* serovar Typhimurium displayed increased susceptibility to antibiotics that target the cell wall (99) as did a laboratory strain of *E. coli* (23). The tat secretion was also shown to have a role in virulence phenotypes in other Gram-negative pathogens (11, 25–29) and two pathotypes (e.g., EHEC and ExPEC) of *E. coli* (30, 31). It was this discrepancy in essentiality and roles in antibiotic susceptibility and virulence, in addition to the relatively low operon score of 11 (i.e., a score of 10.28 is at the 5% cutoff), that motivated our decision to test independent *tatC* knockouts in *K. pneumoniae*, *S. marcescens*, *E. coli* and *E. hormaechei* for essentiality, fitness and antibiotic susceptibility. Despite this apparent requirement for *tatC*, it is intriguing that we were able to produce viable and stable independent deletions of *tatC* in the four species studied here, suggesting that *tatC* is not essential for growth. Perhaps the discrepancy with previous studies is due to lack of sufficient representation of *tatC* mutants in the inoculum or the initial transposon library, increased susceptibility to the antibiotic used in genetic screens, or strain-specific differences in Tat requirements. Our results showed that the *tatC* allele demonstrated a significant contribution to the fitness of all four species tested in this study in a murine model of bacteremia (**Fig 3**). While it is possible that disruption of the TatC transmembrane component may have unintended consequences, such as altering bacterial envelope integrity, the most likely explanation for this shared fitness phenotype is a requirement for appropriate localization of Tat-secreted proteins for infection fitness. Indeed, we have previously attributed at least some of the fitness defect associated with *C. freundii tatC* mutation to SufI loss of function (11). Whether SufI is required for infection of these additional species remains to be determined. However, it may also be informative in future work to compare the repertoire of Tat-secreted proteins in each species using the conserved N-terminal twin-arginine motif, since it is additionally possible that the requirement for TatC is due to species-specific translocation of different proteins.

### Conclusions

The use of our pan-genome pipeline as a powerful tool for integrating multiple data types and sources to infer biological functions was demonstrated in this study. Comparison of the four *Enterobacterales* Tn-Seq datasets revealed, at first inspection, limited common mutations and functional insight. It wasn’t until those results were mapped to orthologous protein clusters and overlaid upon operon structure that common pathways emerged (**Table 2**). This enabled the prediction of common genes within this core pan-genome and brought to light mechanisms that are essential for colonization of, or survival in, the mammalian bloodstream. This represents a step forward in the quest to identify novel targets of therapy against these deadly widespread infections often accompanied by development of life-threatening sepsis. Our prediction and subsequent validation of *tatC* as a common bacteremia fitness factor (**Fig 3**) and further demonstration of increased antibiotic susceptibility of strains carrying mutations in this allele (**Fig 4**, **Table 3**) supports the utility of our bioinformatic method for integration and prioritizing of other biological datasets to identify genes and pathways of interest. Having predicted the importance of many more genes (**S9 Table**) and pathways, highlighted in **Table 2**, it is now critical to follow up this study by investigating the importance of these bacteremia-fitness factor genes across representative species of the *Enterobacterales* and validate their attenuation in the murine model of bacteremia.

## Materials and Methods

### Species-level pan-genome analysis

The pan-genome for the five *Enterobacteriaceae* species were constructed using all the proteins from a curated list of publicly available genomes for each species using the JCVI pan-genome pipeline (14). Specifically, we used the “user_core_cutoff” branch of the JCVI pan-genome pipeline found on GitHub (100). These included the reference genomes used for Tn-Seq as well as genome quality, UMH and common laboratory strains, and epidemiological importance. The number of genomes in each species-level pan-genome run ranged from a minimum of 28 (*Citrobacter*) to a maximum of 281 (*Escherichia*) and consisted mainly of a single species per genus (*e.g.*, *C. freundii*, *E. hormaechei, K. pneumoniae,* and *Serratia marcescens*) where only the *Escherichia* pan-genome also contained *Shigella* spp. The default parameters for the pan-genome pipeline were used, except for a core cutoff (i.e., run_pangenome.pl --core_cutoff) of 70% of genomes as opposed to the 95% default.

### *Enterobacteriaceae* Multi-Species Core (MSC) Pan-genome analysis

The pan-genome pipeline results were used to create a core “pseudo-genome” for each species from the sequences of the core centroids and their location on the species-level consensus pan-chromosome as shown in the Core.att attribute output file (101). These core pseudo-genomes from the five species were integrated in a MSC pan-genome using the same parameters including the 70% core cutoff. A consensus MSC pan-chromosome was constructed as described for the species-level pan-chromosomes. For all species pan-genomes and the MSC pan-genome, the TIGR roles and AMR were assigned to centroids using the best match for the HMMer3 (102) alignment of the TIGRFAMs (103) and the CARD (22), respectively as part of the pan-genome pipeline (14). Other high-level gene features were integrated into the pan-genomes as described below.

### Integration of Tn-Seq bacteremia-fitness genes with pan-genomes

As the original Tn-Seq fitness experiments were performed on single bacterial strains (10–13) whose genomes were included in the individual species pan-genomes (*e.g.*, *C. freundii* UMH14 biosample SAMN07729546, UPEC *E. coli* CFT073 biosample SAMN02604094, *K. pneumoniae* ATCC 43816 KPPR1 biosample SAMN02982872, and *S. marcescens* UMH9 biosample SAMN06164063), we were able to annotate the fitness gene clusters using their unique gene identifiers (**S2 Table**). If the species pan-genome bacteremia-fitness clusters were core (*i.e.*, orthologous sequences present in greater than 70% of the genomes), they were used in the MSC pan-genome and the MSC pan-genome clusters were labeled a fitness centroid.

### Integration of operon structures, essential and virulence factors, and KEGG ontology

Experimentally-determined operon structures for *E. coli* and *Klebsiella* (95, 96, 104) were obtained and mapped to the MSC pan-genome centroids with BLASTP (reciprocal best matches with e-value < 1×10^-5^). For the *E. coli* operons, additional information, such as the Gene Symbol and the Blattner identifier, were also included (**S5 Table**). To include as many Blattner identifiers as possible for the MSC pan-genome clusters, in instances where the MSC clusters were not in operons, we used the Blattner identifier and the gene symbol of any gene from the genome of *E. coli* str. K-12 substr. MG1655 (BioSample SAMN02604091) with a good BLASTP match (same cutoff as above) to the MSC pan-genome clusters. We also mapped the antibiotic susceptibility of the genes from (23, 24) using the Blattner-derived gene symbols.

Genes that were found to be essential for growth of *E. coli* K-12 and derivatives under laboratory growth conditions were obtained from five different manuscripts/databases (90–94). A unique set of “essential” genes was created (**S3 Table**) from these combined data sets and assigned the unique *E. coli* K-12 composite (ECK*) identifier. BLASTP was used to map the translated essential proteins to the *Escherichia* pan-genome centroids.

Similarly, we used the full set of protein sequences from the Virulence Factor Database (VFDB) (**S4 Table**) (21) downloaded on August 31, 2021, to do a similar BLASTP mapping to species-level pan-genome centroids. If the reciprocal best matches had an e-value < 1×10^-5^ and an additional requirement that the BLASTP match had to have more than 70% identity, then the centroid was annotated as “Essential” or a “Virulence Factor.” The cluster annotation for the species-level pan-genome was passed to the MSC pan-genomes as described for bacteremia-fitness proteins above.

Additionally, KEGG annotations were mapped to the MSC centroids using hmmsearch v 3.3.2 (105) against the KEGG ontology dataset collected in KOfam (106) with an e-value < 1×10^-5^ (**S5 Table**).

### Venn Diagrams

Venn Diagrams were created with R with the Venn Package (107), using data from **S5 Table.** For each of the Venn diagrams comprising **S1 Fig,** a different list of MSC cluster IDs was used as input for each species in the MSC pan-genome to count the numbers of shared orthologs in: A) all MSC clusters; B) all MSC clusters with at least one Tn-Seq bacteremia-fitness protein; and C) all MSC clusters with at least one virulence factor protein.

### Pan-chromosome visualizations

Browsable pan-chromosome visualizations, including the fitness, antimicrobial resistance, and TIGRFam annotations were constructed for each species pan-genome and the MSC pan-genome with PanACEA using default parameters (**S1-5 Dataset, S5 Table**) (18). The species pan-genome comparison was constructed with data from **S5 Table**, using a polar graph from *ggplot2* (108). Orthologs for each MSC cluster were identified in each species as well as the relative position in each species pan-genome relative to *dnaA*. MSC clusters that were not present in a species were noted with a gap. The radial position of orthologs on the figure was the relative location of the *Escherichia* ortholog, while the color was based on the location of ortholog in each species. The fitness clusters with an *Escherichia* ortholog are shown at the *Escherichia* ortholog position.

### Prioritization of fitness genes and operons

We ranked MSC pan-genome bacteremia-fitness clusters and operons identified among the four species Tn-Seq data using a scoring rubric. The rubric was based on four additive criteria: 1) The published fold-change of the fitness defect associated with transposon interruption of a gene in any one species, 2) a single gene being identified as a fitness factor in multiple species, 3) whether multiple fitness factors were encoded in the same operon, and 4) a fitness factor single-gene-knockout was previously found to confer increased antibiotic susceptibility in *E. coli* BW25113 (23, 24).

The top 60 bacteremia-fitness factors in any one species Tn-Seq data set (*i.e.*, the top 60 least fit transposon mutants) were awarded points based on the published magnitude of the fitness defect determined in any one species. The top 1 - 20 least fit bacteremia-fitness factors were awarded 3 points, 21 - 40 were awarded 2 points, and 41 - 60 were awarded 1 point. All scores for each MSC bacteremia-fitness factor were then summed across species. (*e.g.*, *wzxE* received 6 total points in this category by accumulating 3 points each for providing the 6th and 15th greatest fitness defects when mutated in *S. marcescen*s UMH9 and *K. pneumoniae* KPPR1, respectively.) One item of note; *K. pneumoniae* KPPR1 bacteremia Tn-Seq (13) only identified 55 fitness factors, therefore this species was slightly underrepresented in this scoring category.

The MSC pan-genome bacteremia-fitness factor clusters were also awarded points based on their identification in multiple species Tn-Seq datasets, regardless of the magnitude of their defect. If a cluster member was identified as a bacteremia-fitness factor in three species or two species it was awarded 3 points or 2 points, respectively. (*e.g.*, *wzxE* received 3 points for being identified as a bacteremia-fitness factor in Tn-Seq datasets from *C. freundii* UMH14, S. *marcescen*s UMH9, and *K. pneumoniae* KPPR1.)

The MSC pan-genome bacteremia-fitness factor clusters were awarded points based on the identification of at least one other bacteremia-fitness factor encoded in the same operon, regardless of which species any one fitness factor was identified in. Fitness factors found encoded in an operon with 5 to 9 other fitness factors were awarded 3 points, those encoded with 3 to 4 other fitness factors were awarded 2 points, and those encoded in an operon with 1 to 2 other fitness factors were awarded 1 point. An additional point was awarded to fitness factors found in an operon where at least 50% of the genes in that operon were found to be fitness factors. (*e.g.*, *wzxE* is encoded in a 12 gene operon with 9 other fitness factors being identified in at least one species’ Tn-Seq dataset. Therefore, *wzxE* received a total of 4 points; 3 points for being encoded in an operon with 5 to 9 other fitness factors and 1 additional point for greater than 50% of the genes in that operon being identified as fitness factors.) Finally, MSC pan-genome bacteremia-fitness factor clusters were given additional points if mutation of that gene in *E. coli* BW25113 conferred increased susceptibility to an antibiotic (23, 24). Fitness factor clusters received another 2 points if mutation of the orthologous gene in the Keio collection was found to have increased sensitivity to any antibiotics, regardless of the mode of action.

A histogram of the total score of both individual centroids and operons was plotted with ggplot2 (108), and the Kolmogorov–Smirnov test, in the stats library in R (109), was to determine the most representative distribution. The most representative distribution (exponential) was used to find the cutoff values giving the top scoring 5% and 10% of the centroids and operons.

### Generation of *tatC* mutant derivatives and complementation constructs

Mutations in *tatC* were generated by lambda red recombineering using established protocols (10, 11, 73, 110). For *C. freundii*, *E. coli, K. pneumoniae,* and *S. marcescens*, the *nptII* kanamycin resistance cassette was PCR-amplified from pKD4 (110, 111) and targeted to an in-frame deletion of the *tatC* ORF via 5’-end homologous sequences (**S10 Table**). For *E. hormaechei,* the *acc*(*3*)*IV* apramycin resistance cassette was amplified from pUC18-miniTn7T-Apr (112). Recombination was facilitated by Red functions encoded on pKD46, pSIM19, or pSIM18 depending on the species mutated (110, 111). All mutations were confirmed by sequencing of PCR-amplified alleles or by PCR amplicon size and recombineering plasmids were cured prior to phenotypic analysis.

Genetic complementation of the *S. marcescens tatC* mutation was achieved by insertion of the UMH9 *tatC* ORF into plasmid vector pBBR1MCS-5 (113) via isothermal assembly, followed by sequencing verification. The resulting plasmid, pBB26, was transformed into *S. marcescens* via electroporation. Genetic complementation of the *K. pneumoniae tatC* mutation with the pAYCYC_tatC_ plasmid was achieved by using SOE PCR (114) to connect the promoter of the *tat* operon to the *tatC* ORF. The pACYC_tatABCD_ plasmid insert was made by amplifying *tatABCD* including its native promoter. The amplicons were inserted into pACYC184 at the *Hin*dIII and *Sal*I sites and the resulting plasmids were sequenced for verification. Correct plasmids were transformed into *K. pneumoniae* via electroporation. The *E. hormaechei tatC* mutant was complemented by insertion of wild-type *tatABC* into pBBR1Hyg, a hygromycin resistant derivative of pBBR1MCS-5 (this study), by isothermal assembly. The resulting plasmid, pBBR1Hyg_*tatABC* was transformed into *E. hormaechei* by electroporation.

### Murine bacteremia models

All experiments involving the use of laboratory animals were conducted with protocols approved by the University of Michigan or Michigan State University Institutional Animal Care and Use Committee and were performed in accordance with the Office of Laboratory Animal Welfare guidelines. Wild-type and *tatC* mutant derivatives were cultured separately in LB medium, then washed with PBS and normalized to the final density prior to mixing 1:1. Tail vein injections were performed on 7-8 week old C57BL/6J mice as previously described for all tested species (9, 33) and organs were collected for homogenization and CFU determination at 24 h post-injection. For the bacteremic pneumonia model, 1×10^7^ CFU *S. marcescens* or 1x10^6^ CFU *K. pneumoniae* was administered retropharyngeally into lightly anesthetized mice, as previously described (73). Mice were sacrificed at 24 h post-inoculation for CFU determinations. Competitive indices for both infection models were determined as the ratio of mutant/wild-type bacteria after 24 h compared to that same ratio in the inoculum.

### Determination of minimum inhibitory concentration

The minimum inhibitory concentration (MIC) of ampicillin and piperacillin-tazobactam was determined using Epsilometer (E-test) strips. This experiment was carried out following the guidelines of the Clinical Microbiology Procedure Handbook (115). Bacteria were cultured on Lysogeny Broth (LB) agar, then inoculated in 3 mL of LB broth the following day. After reaching OD_600_ of 0.1, cultures were spread onto Mueller-Hinton agar using a sterile cotton-swab and allowed to dry. Then E-test strips were placed on the dried plates and incubated at 37°C overnight. The MIC was determined by observing the point at which the zone of clearing touches the test strip. The Clinical and Laboratory Standards Institute (CLSI) interpretations were obtained from *Performance Standards for Antimicrobial Susceptibility Testing* (116).

## Acknowledgements

We would like to thank all members of the Gram-negative bacilli bacteremia team for their contribution to this manuscript.

## Supporting information

S1 Table. Genomes Used in this Study.

S1 Dataset. *E. coli* PanACEA Output.

S2 Dataset. *K. pneumoniae* PanACEA Output.

S3 Dataset. *E. hormaechei* PanACEA Output.

S4 Dataset. *C. freundii* PanACEA Output.

S5 Dataset. *S. marcescens* PanACEA Output.

**S2 Table. Tn-Seq Fitness Genes used in this Study.**

S3 Table. *E. coli* Essential Genes used in this Study.

**S4 Table. Virulence Genes used in this Study.**

**S5 Table. Multi-species Core Pan-genome Clusters.**

**S6 Dataset. MSC Pan-genome PanACEA Output.**

**S6 Table. Tn-Seq Fitness Factors Shared Across 3 Species.**

**S7 Table. Bacteremia-fitness and Virulence MSC clusters.**

**S8 Table. Fitness Genes Mapped to Known Operons.**

**S9 Table. Ranking of Fitness Genes.**

**S10 Table. Oligonucleotide Primers Used in this Study.**

**S1 Fig. Proteins Shared Across Five bacteremia-causing *Enterobacterales species* in the MSC Pan-genome.** All orthologous protein clusters (**A**), the clusters labeled as “bacteremia-fitness” from Tn-Seq data (**B**), and the clusters labeled as “virulence factor” with matches to the VFDB (**C**) from the MSC pan-genome were binned, counted, and placed into a Venn diagram by whether clusters contained proteins from *C. freundii*, *E. hormaechei, E. coli/Shigella* spp., *K. pneumoniae*, and *S. marcescens*. Singleton clusters, representing species-specific core genes, are noted in the outermost ellipses of the Venn diagram. The Venn diagram is not to scale.

**S2 Fig. Distribution of Bacteremia-fitness factor centroid scores.** Summary of total centroid and total operon scores for all 500 bacteremia-fitness factors (**A**) and 366 bacteremia-fitness operons (**B**) scored in S9 Table. The vertical black dotted lines depict the top 5% cutoff and the gray vertical dotted lines depict the top 10% cutoff using the exponential distribution. Light gray box in (**A**) indicates the 73 bacteremia-fitness factors with a total centroid score in the top 10% of the scores (i.e., genes receiving at least 6 total points, and includes both † and ‡ genes highlighted in Table 2). The dark gray box indicates the 32 bacteremia-fitness factors with a total centroid score in the top 5% (i.e., genes receiving at least 8 total points, including only ‡ genes highlighted in **Table 2**). The colored lines show the different distributions that were compared to the scores, including the exponential (red), poisson (blue) and Gaussian (purple). The Kolmogorov-Smirnov test indicated that the Exponential Distribution was the closest fit and was used.

## References

1. Rudd KE, Johnson SC, Agesa KM, Shackelford KA, Tsoi D, Kievlan DR, Colombara DV, Ikuta KS, Kissoon N, Finfer S, Fleischmann-Struzek C, Machado FR, Reinhart KK, Rowan K, Seymour CW, Watson RS, West TE, Marinho F, Hay SI, Lozano R, Lopez AD, Angus DC, Murray CJL, Naghavi M. 2020. Global, regional, and national sepsis incidence and mortality, 1990-2017: analysis for the Global Burden of Disease Study. Lancet 395:200–211.

2. Liu V, Escobar GJ, Greene JD, Soule J, Whippy A, Angus DC, Iwashyna TJ. 2014. Hospital deaths in patients with sepsis from 2 independent cohorts. JAMA 312:90– 92.

3. Wisplinghoff H, Bischoff T, Tallent SM, Seifert H, Wenzel RP, Edmond MB. 2004. Nosocomial bloodstream infections in US hospitals: analysis of 24,179 cases from a prospective nationwide surveillance study. Clin Infect Dis 39:309–317.

4. Thaden JT, Li Y, Ruffin F, Maskarinec SA, Hill-Rorie JM, Wanda LC, Reed SD, Fowler VG Jr. 2017. Increased Costs Associated with Bloodstream Infections Caused by Multidrug-Resistant Gram-Negative Bacteria Are Due Primarily to Patients with Hospital-Acquired Infections. Antimicrob Agents Chemother 61.

5. Anderson DJ, Moehring RW, Sloane R, Schmader KE, Weber DJ, Fowler VG Jr, Smathers E, Sexton DJ. 2014. Bloodstream infections in community hospitals in the 21st century: a multicenter cohort study. PLoS One 9:e91713.

6. Diekema DJ, Hsueh P-R, Mendes RE, Pfaller MA, Rolston KV, Sader HS, Jones RN. 2019. The Microbiology of Bloodstream Infection: 20-Year Trends from the SENTRY Antimicrobial Surveillance Program. Antimicrob Agents Chemother 63.

7. Centers For Disease Control And Prevention, Centers for Disease Control and Prevention (U.S.). 2019. Antibiotic resistance threats in the United States, 2019 10.15620/cdc:82532.

8. Holmes CL, Anderson MT, Mobley HLT, Bachman MA. 2021. Pathogenesis of Gram-Negative Bacteremia. Clin Microbiol Rev 34.

9. Anderson MT, Brown AN, Pirani A, Smith SN, Photenhauer AL, Sun Y, Snitkin ES, Bachman MA, Mobley HLT. 2021. Replication Dynamics for Six Gram-Negative Bacterial Species during Bloodstream Infection. MBio 12:e0111421.

10. Anderson MT, Mitchell LA, Zhao L, Mobley HLT. 2017. Capsule Production and Glucose Metabolism Dictate Fitness during Bacteremia. MBio 8.

11. Anderson MT, Mitchell LA, Zhao L, Mobley HLT. 2018. *Citrobacter freundii* fitness during bloodstream infection. Sci Rep 8:11792.

12. Subashchandrabose S, Smith SN, Spurbeck RR, Kole MM, Mobley HLT. 2013. Genome-wide detection of fitness genes in uropathogenic *Escherichia coli* during systemic infection. PLoS Pathog 9:e1003788.

13. Holmes CL, Wilcox AE, Forsyth V, Smith SN, Moricz BS, Unverdorben LV, Mason S, Wu W, Zhao L, Mobley HLT, Bachman MA. 2023. *Klebsiella pneumoniae* causes bacteremia using factors that mediate tissue-specific fitness and resistance to oxidative stress. PLoS Pathog 19:e1011233.

14. Inman JM, Sutton GG, Beck E, Brinkac LM, Clarke TH, Fouts DE. 2019. Large-scale comparative analysis of microbial pan-genomes using PanOCT. Bioinformatics 35:1049–1050.

15. Fouts DE, Brinkac L, Beck E, Inman J, Sutton G. 2012. PanOCT: Automated Clustering of Orthologs Using Conserved Gene Neighborhood for Pan-Genomic Analysis of Bacterial Strains and Closely Related Species. Nucleic Acids Res 40:e172.

16. Tettelin H, Masignani V, Cieslewicz MJ, Donati C, Medini D, Ward NL, Angiuoli SV, Crabtree J, Jones AL, Durkin AS, Deboy RT, Davidsen TM, Mora M, Scarselli M, Margarit y Ros I, Peterson JD, Hauser CR, Sundaram JP, Nelson WC, Madupu R, Brinkac LM, Dodson RJ, Rosovitz MJ, Sullivan SA, Daugherty SC, Haft DH, Selengut J, Gwinn ML, Zhou L, Zafar N, Khouri H, Radune D, Dimitrov G, Watkins K, O’Connor KJB, Smith S, Utterback TR, White O, Rubens CE, Grandi G, Madoff LC, Kasper DL, Telford JL, Wessels MR, Rappuoli R, Fraser CM. 2005. Genome analysis of multiple pathogenic isolates of *Streptococcus agalactiae*: implications for the microbial “pan-genome.” Proc Natl Acad Sci U S A 102:13950–13955.

17. Rodriguez-Valera F, Ussery DW. 2012. Is the pan-genome also a pan-selectome? F1000Res 1:16.

18. Clarke TH, Brinkac LM, Inman JM, Sutton G, Fouts DE. 2018. PanACEA: a bioinformatics tool for the exploration and visualization of bacterial pan-chromosomes. BMC Bioinformatics 19:246.

19. Chen Y, Stine OC, Badger JH, Gil AI, Nair GB, Nishibuchi M, Fouts DE. 2011. Comparative genomic analysis of *Vibrio parahaemolyticus*: serotype conversion and virulence. BMC Genomics 12:294.

20. Lazarus JE, Warr AR, Westervelt KA, Hooper DC, Waldor MK. 2021. A Genome-Scale Antibiotic Screen in *Serratia marcescens* Identifies YdgH as a Conserved Modifier of Cephalosporin and Detergent Susceptibility. Antimicrob Agents Chemother 65:e0078621.

21. Chen L, Yang J, Yu J, Yao Z, Sun L, Shen Y, Jin Q. 2005. VFDB: a reference database for bacterial virulence factors. Nucleic Acids Res 33:D325–8.

22. McArthur AG, Waglechner N, Nizam F, Yan A, Azad MA, Baylay AJ, Bhullar K, Canova MJ, De Pascale G, Ejim L, Kalan L, King AM, Koteva K, Morar M, Mulvey MR, O’Brien JS, Pawlowski AC, Piddock LJV, Spanogiannopoulos P, Sutherland AD, Tang I, Taylor PL, Thaker M, Wang W, Yan M, Yu T, Wright GD. 2013. The comprehensive antibiotic resistance database. Antimicrob Agents Chemother 57:3348–3357.

23. Tamae C, Liu A, Kim K, Sitz D, Hong J, Becket E, Bui A, Solaimani P, Tran KP, Yang H, Miller JH. 2008. Determination of antibiotic hypersensitivity among 4,000 single-gene-knockout mutants of *Escherichia coli*. J Bacteriol 190:5981–5988.

24. Liu A, Tran L, Becket E, Lee K, Chinn L, Park E, Tran K, Miller JH. 2010. Antibiotic sensitivity profiles determined with an *Escherichia coli* gene knockout collection: generating an antibiotic bar code. Antimicrob Agents Chemother 54:1393–1403.

25. Ochsner UA, Snyder A, Vasil AI, Vasil ML. 2002. Effects of the twin-arginine translocase on secretion of virulence factors, stress response, and pathogenesis. Proc Natl Acad Sci U S A 99:8312–8317.

26. Lavander M, Ericsson SK, Bröms JE, Forsberg A. 2006. The twin arginine translocation system is essential for virulence of *Yersinia pseudotuberculosis*. Infect Immun 74:1768–1776.

27. Reynolds MM, Bogomolnaya L, Guo J, Aldrich L, Bokhari D, Santiviago CA, McClelland M, Andrews-Polymenis H. 2011. Abrogation of the twin arginine transport system in *Salmonella enterica* serovar Typhimurium leads to colonization defects during infection. PLoS One 6:e15800.

28. Urrutia ÍM, Sabag A, Valenzuela C, Labra B, Álvarez SA, Santiviago CA. 2018. Contribution of the Twin-Arginine Translocation System to the Intracellular Survival of Typhimurium in. Front Microbiol 9:3001.

29. Mickael CS, Lam P-KS, Berberov EM, Allan B, Potter AA, Köster W. 2010. *Salmonella enterica* serovar Enteritidis *tatB* and *tatC* mutants are impaired in Caco-2 cell invasion *in vitro* and show reduced systemic spread in chickens. Infect Immun 78:3493– 3505.

30. Pradel N, Ye C, Livrelli V, Xu J, Joly B, Wu L-F. 2003. Contribution of the twin arginine translocation system to the virulence of enterohemorrhagic *Escherichia coli* O157:H7. Infect Immun 71:4908–4916.

31. Liu J, Yin F, Liu T, Li S, Tan C, Li L, Zhou R, Huang Q. 2020. The Tat system and its dependent cell division proteins are critical for virulence of extra-intestinal pathogenic. Virulence 11:1279–1292.

32. Palmer T, Stansfeld PJ. 2020. Targeting of proteins to the twin-arginine translocation pathway. Mol Microbiol 113:861–871.

33. Smith SN, Hagan EC, Lane MC, Mobley HLT. 2010. Dissemination and systemic colonization of uropathogenic *Escherichia coli* in a murine model of bacteremia. MBio 1.

34. Skurnik D, Roux D, Aschard H, Cattoir V, Yoder-Himes D, Lory S, Pier GB. 2013. A comprehensive analysis of *in vitro* and *in vivo* genetic fitness of *Pseudomonas aeruginosa* using high-throughput sequencing of transposon libraries. PLoS Pathog 9:e1003582.

35. van Opijnen T, Camilli A. 2012. A fine scale phenotype-genotype virulence map of a bacterial pathogen. Genome Res 22:2541–2551.

36. Wang N, Ozer EA, Mandel MJ, Hauser AR. 2014. Genome-wide identification of *Acinetobacter baumannii* genes necessary for persistence in the lung. MBio 5:e01163–14.

37. Baugh L, Gallagher LA, Patrapuvich R, Clifton MC, Gardberg AS, Edwards TE, Armour B, Begley DW, Dieterich SH, Dranow DM, Abendroth J, Fairman JW, Fox D 3rd, Staker BL, Phan I, Gillespie A, Choi R, Nakazawa-Hewitt S, Nguyen MT, Napuli A, Barrett L, Buchko GW, Stacy R, Myler PJ, Stewart LJ, Manoil C, Van Voorhis WC. 2013. Combining functional and structural genomics to sample the essential Burkholderia structome. PLoS One 8:e53851.

38. Brutinel ED, Gralnick JA. 2012. Anomalies of the anaerobic tricarboxylic acid cycle in *Shewanella oneidensis* revealed by Tn-seq. Mol Microbiol 86:273–283.

39. Burghout P, Zomer A, van der Gaast-de Jongh CE, Janssen-Megens EM, Françoijs K-J, Stunnenberg HG, Hermans PWM. 2013. *Streptococcus pneumoniae* folate biosynthesis responds to environmental CO2 levels. J Bacteriol 195:1573–1582.

40. Byrne RT, Chen SH, Wood EA, Cabot EL, Cox MM. 2014. *Escherichia coli* genes and pathways involved in surviving extreme exposure to ionizing radiation. J Bacteriol 196:3534–3545.

41. Carter R, Wolf J, van Opijnen T, Muller M, Obert C, Burnham C, Mann B, Li Y, Hayden RT, Pestina T, Persons D, Camilli A, Flynn PM, Tuomanen EI, Rosch JW. 2014. Genomic analyses of pneumococci from children with sickle cell disease expose host-specific bacterial adaptations and deficits in current interventions. Cell Host Microbe 15:587–599.

42. Chaudhuri RR, Morgan E, Peters SE, Pleasance SJ, Hudson DL, Davies HM, Wang J, van Diemen PM, Buckley AM, Bowen AJ, Pullinger GD, Turner DJ, Langridge GC, Turner AK, Parkhill J, Charles IG, Maskell DJ, Stevens MP. 2013. Comprehensive assignment of roles for *Salmonella typhimurium* genes in intestinal colonization of food-producing animals. PLoS Genet 9:e1003456.

43. de Vries SPW, Eleveld MJ, Hermans PWM, Bootsma HJ. 2013. Characterization of the molecular interplay between *Moraxella catarrhalis* and human respiratory tract epithelial cells. PLoS One 8:e72193.

44. Dembek M, Barquist L, Boinett CJ, Cain AK, Mayho M, Lawley TD, Fairweather NF, Fagan RP. 2015. High-throughput analysis of gene essentiality and sporulation in *Clostridium difficile*. MBio 6:e02383.

45. Dong TG, Ho BT, Yoder-Himes DR, Mekalanos JJ. 2013. Identification of T6SS-dependent effector and immunity proteins by Tn-seq in *Vibrio cholerae*. Proc Natl Acad Sci U S A 110:2623–2628.

46. Fu Y, Waldor MK, Mekalanos JJ. 2013. Tn-Seq analysis of *Vibrio cholerae* intestinal colonization reveals a role for T6SS-mediated antibacterial activity in the host. Cell Host Microbe 14:652–663.

47. Gallagher LA, Shendure J, Manoil C. 2011. Genome-scale identification of resistance functions in *Pseudomonas aeruginosa* using Tn-seq. MBio 2:e00315–10.

48. Gawronski JD, Wong SMS, Giannoukos G, Ward DV, Akerley BJ. 2009. Tracking insertion mutants within libraries by deep sequencing and a genome-wide screen for *Haemophilus* genes required in the lung. Proc Natl Acad Sci U S A 106:16422–16427.

49. Goodman AL, McNulty NP, Zhao Y, Leip D, Mitra RD, Lozupone CA, Knight R, Gordon JI. 2009. Identifying genetic determinants needed to establish a human gut symbiont in its habitat. Cell Host Microbe 6:279–289.

50. Johnson JG, Livny J, Dirita VJ. 2014. High-throughput sequencing of *Campylobacter jejuni* insertion mutant libraries reveals *mapA* as a fitness factor for chicken colonization. J Bacteriol 196:1958–1967.

51. Kamp HD, Patimalla-Dipali B, Lazinski DW, Wallace-Gadsden F, Camilli A. 2013. Gene fitness landscapes of *Vibrio cholerae* at important stages of its life cycle. PLoS Pathog 9:e1003800.

52. Khatiwara A, Jiang T, Sung S-S, Dawoud T, Kim JN, Bhattacharya D, Kim H-B, Ricke SC, Kwon YM. 2012. Genome scanning for conditionally essential genes in *Salmonella enterica* Serotype Typhimurium. Appl Environ Microbiol 78:3098–3107.

53. Langereis JD, Zomer A, Stunnenberg HG, Burghout P, Hermans PWM. 2013. Nontypeable *Haemophilus influenzae* carbonic anhydrase is important for environmental and intracellular survival. J Bacteriol 195:2737–2746.

54. Luan S-L, Chaudhuri RR, Peters SE, Mayho M, Weinert LA, Crowther SA, Wang J, Langford PR, Rycroft A, Wren BW, Tucker AW, Maskell DJ, BRaDP1T Consortium. 2013. Generation of a Tn5 transposon library in Haemophilus parasuis and analysis by transposon-directed insertion-site sequencing (TraDIS). Vet Microbiol 166:558–566.

55. Santa Maria JP Jr, Sadaka A, Moussa SH, Brown S, Zhang YJ, Rubin EJ, Gilmore MS, Walker S. 2014. Compound-gene interaction mapping reveals distinct roles for Staphylococcus aureus teichoic acids. Proc Natl Acad Sci U S A 111:12510–12515.

56. McDonough E, Lazinski DW, Camilli A. 2014. Identification of *in vivo* regulators of the *Vibrio cholerae xds* gene using a high-throughput genetic selection. Mol Microbiol 92:302–315.

57. Moule MG, Hemsley CM, Seet Q, Guerra-Assunção JA, Lim J, Sarkar-Tyson M, Clark TG, Tan PBO, Titball RW, Cuccui J, Wren BW. 2014. Genome-wide saturation mutagenesis of *Burkholderia pseudomallei* K96243 predicts essential genes and novel targets for antimicrobial development. MBio 5:e00926–13.

58. Palace SG, Proulx MK, Lu S, Baker RE, Goguen JD. 2014. Genome-wide mutant fitness profiling identifies nutritional requirements for optimal growth of *Yersinia pestis* in deep tissue. MBio 5.

59. Remmele CW, Xian Y, Albrecht M, Faulstich M, Fraunholz M, Heinrichs E, Dittrich MT, Müller T, Reinhardt R, Rudel T. 2014. Transcriptional landscape and essential genes of *Neisseria gonorrhoeae*. Nucleic Acids Res 42:10579–10595.

60. Troy EB, Lin T, Gao L, Lazinski DW, Camilli A, Norris SJ, Hu LT. 2013. Understanding barriers to *Borrelia burgdorferi* dissemination during infection using massively parallel sequencing. Infect Immun 81:2347–2357.

61. Turner KH, Everett J, Trivedi U, Rumbaugh KP, Whiteley M. 2014. Requirements for *Pseudomonas aeruginosa* acute burn and chronic surgical wound infection. PLoS Genet 10:e1004518.

62. Valentino MD, Foulston L, Sadaka A, Kos VN, Villet RA, Santa Maria J Jr, Lazinski DW, Camilli A, Walker S, Hooper DC, Gilmore MS. 2014. Genes contributing to Staphylococcus aureus fitness in abscess- and infection-related ecologies. MBio 5:e01729– 14.

63. Verhagen LM, de Jonge MI, Burghout P, Schraa K, Spagnuolo L, Mennens S, Eleveld MJ, van der Gaast-de Jongh CE, Zomer A, Hermans PWM, Bootsma HJ. 2014. Genome-wide identification of genes essential for the survival of *Streptococcus pneumoniae* in human saliva. PLoS One 9:e89541.

64. Weerdenburg EM, Abdallah AM, Rangkuti F, Abd El Ghany M, Otto TD, Adroub SA, Molenaar D, Ummels R, Ter Veen K, van Stempvoort G, van der Sar AM, Ali S, Langridge GC, Thomson NR, Pain A, Bitter W. 2015. Genome-wide transposon mutagenesis indicates that Mycobacterium marinum customizes its virulence mechanisms for survival and replication in different hosts. Infect Immun 83:1778–1788.

65. Wiles TJ, Norton JP, Russell CW, Dalley BK, Fischer KF, Mulvey MA. 2013. Combining quantitative genetic footprinting and trait enrichment analysis to identify fitness determinants of a bacterial pathogen. PLoS Genet 9:e1003716.

66. Xiao J, Sun S, Liu Z, Fan C, Zhu B, Zhang D. 2023. Analysis of key genes for the survival of *Pantoea agglomerans* under nutritional stress. Int J Biol Macromol 253:127059.

67. Kado T, Akbary Z, Motooka D, Sparks IL, Melzer ES, Nakamura S, Rojas ER, Morita YS, Siegrist MS. 2023. A cell wall synthase accelerates plasma membrane partitioning in mycobacteria. Elife 12.

68. Ma J, Ahmed MAH, Shao S, Zhang Y, Wang Q, Yin K. 2024. The QseE-QseF two-component system: A key mediator of epinephrine-regulated virulence in the marine pathogen *Edwardsiella piscicida*. Microbiol Res 279:127561.

69. Lo H-Y, Long DR, Holmes EA, Penewit K, Hodgson T, Lewis JD, Waalkes A, Salipante SJ. 2023. Transposon sequencing identifies genes impacting invasion in a human macrophage model. Infect Immun 91:e0022823.

70. Kim G-L, Hooven TA, Norambuena J, Li B, Boyd JM, Yang JH, Parker D. 2021. Growth and Stress Tolerance Comprise Independent Metabolic Strategies Critical for *Staphylococcus aureus* Infection. MBio 12:e0081421.

71. Shull LM, Camilli A. 2018. Transposon Sequencing of *Vibrio cholerae* in the Infant Rabbit Model of Cholera. Methods Mol Biol 1839:103–116.

72. Turner KH, Wessel AK, Palmer GC, Murray JL, Whiteley M. 2015. Essential genome of *Pseudomonas aeruginosa* in cystic fibrosis sputum. Proc Natl Acad Sci U S A 112:4110–4115.

73. Bachman MA, Breen P, Deornellas V, Mu Q, Zhao L, Wu W, Cavalcoli JD, Mobley HLT. 2015. Genome-Wide Identification of *Klebsiella pneumoniae* Fitness Genes during Lung Infection. MBio 6:e00775.

74. Gutierrez MG, Yoder-Himes DR, Warawa JM. 2015. Comprehensive identification of virulence factors required for respiratory melioidosis using Tn-seq mutagenesis. Front Cell Infect Microbiol 5:78.

75. Lourdault K, Matsunaga J, Haake DA. 2016. High-Throughput Parallel Sequencing to Measure Fitness of *Leptospira interrogans* Transposon Insertion Mutants during Acute Infection. PLoS Negl Trop Dis 10:e0005117.

76. Russell CW, Mulvey MA. 2015. The Extraintestinal Pathogenic *Escherichia coli* Factor RqlI Constrains the Genotoxic Effects of the RecQ-Like Helicase RqlH. PLoS Pathog 11:e1005317.

77. Stacy A, Fleming D, Lamont RJ, Rumbaugh KP, Whiteley M. 2016. A Commensal Bacterium Promotes Virulence of an Opportunistic Pathogen via Cross-Respiration. MBio 7.

78. Troy EB, Lin T, Gao L, Lazinski DW, Lundt M, Camilli A, Norris SJ, Hu LT. 2016. Global Tn-seq analysis of carbohydrate utilization and vertebrate infectivity of *Borrelia burgdorferi*. Mol Microbiol 101:1003–1023.

79. Lourdault K, Matsunaga J, Evangelista KV, Haake DA. 2017. High-throughput Parallel Sequencing to Measure Fitness of *Leptospira interrogans* Transposon Insertion Mutants During Golden Syrian Hamster Infection. J Vis Exp 10.3791/56442.

80. Forsyth VS, Mobley HLT, Armbruster CE. 2019. Transposon Insertion Site Sequencing in a Urinary Tract Model. Methods Mol Biol 2021:297–337.

81. Eichelberger KR, Sepúlveda VE, Ford J, Selitsky SR, Mieczkowski PA, Parker JS, Goldman WE. 2020. Tn-Seq Analysis Identifies Genes Important for Yersinia pestis Adherence during Primary Pneumonic Plague. mSphere 5.

82. Perpich JD, Yakoumatos L, Stocke KS, Lewin GR, Ramos A, Yoder-Himes DR, Whiteley M, Lamont RJ. 2022. *Porphyromonas gingivalis* Tyrosine Kinase Is a Fitness Determinant in Polymicrobial Infections. Infect Immun 90:e0017022.

83. Belanger CR, Dostert M, Blimkie TM, Lee AH-Y, Dhillon BK, Wu BC, Akhoundsadegh N, Rahanjam N, Castillo-Arnemann J, Falsafi R, Pletzer D, Haney CH, Hancock REW. 2022. Surviving the host: Microbial metabolic genes required for growth of in physiologically-relevant conditions. Front Microbiol 13:1055512.

84. Price SL, Thibault D, Garrison TM, Brady A, Guo H, Kehl-Fie TE, Garneau-Tsodikova S, Perry RD, van Opijnen T, Lawrenz MB. 2023. Droplet Tn-Seq identifies the primary secretion mechanism for yersiniabactin in *Yersinia pestis*. EMBO Rep 24:e57369.

85. Capel E, Barnier J-P, Zomer AL, Bole-Feysot C, Nussbaumer T, Jamet A, Lécuyer H, Euphrasie D, Virion Z, Frapy E, Pélissier P, Join-Lambert O, Rattei T, Bourdoulous S, Nassif X, Coureuil M. 2017. Peripheral blood vessels are a niche for blood-borne meningococci. Virulence 8:1808–1819.

86. Brauer AL, White AN, Learman BS, Johnson AO, Armbruster CE. 2019. d-Serine Degradation by Proteus mirabilis Contributes to Fitness during Single-Species and Polymicrobial Catheter-Associated Urinary Tract Infection. mSphere 4.

87. Subashchandrabose S, Smith S, DeOrnellas V, Crepin S, Kole M, Zahdeh C, Mobley HLT. 2016. Acinetobacter baumannii Genes Required for Bacterial Survival during Bloodstream Infection. mSphere 1.

88. Anderson MT, Mitchell LA, Mobley HLT. 2017. Cysteine Biosynthesis Controls *Serratia marcescens* Phospholipase Activity. J Bacteriol 199.

89. Armbruster CE, Forsyth VS, Johnson AO, Smith SN, White AN, Brauer AL, Learman BS, Zhao L, Wu W, Anderson MT, Bachman MA, Mobley HLT. 2019. Twin arginine translocation, ammonia incorporation, and polyamine biosynthesis are crucial for *Proteus mirabilis* fitness during bloodstream infection. PLoS Pathog 15:e1007653.

90. Gerdes SY, Scholle MD, Campbell JW, Balázsi G, Ravasz E, Daugherty MD, Somera AL, Kyrpides NC, Anderson I, Gelfand MS, Bhattacharya A, Kapatral V, D’Souza M, Baev MV, Grechkin Y, Mseeh F, Fonstein MY, Overbeek R, Barabási A-L, Oltvai ZN, Osterman AL. 2003. Experimental determination and system level analysis of essential genes in *Escherichia coli* MG1655. J Bacteriol 185:5673–5684.

91. Hashimoto M, Ichimura T, Mizoguchi H, Tanaka K, Fujimitsu K, Keyamura K, Ote T, Yamakawa T, Yamazaki Y, Mori H, Katayama T, Kato J-I. 2005. Cell size and nucleoid organization of engineered *Escherichia coli* cells with a reduced genome. Mol Microbiol 55:137–149.

92. Baba T, Ara T, Hasegawa M, Takai Y, Okumura Y, Baba M, Datsenko KA, Tomita M, Wanner BL, Mori H. 2006. Construction of *Escherichia coli* K-12 in-frame, single-gene knockout mutants: the Keio collection. Mol Syst Biol 2:2006.0008.

93. Goodall ECA, Robinson A, Johnston IG, Jabbari S, Turner KA, Cunningham AF, Lund PA, Cole JA, Henderson IR. 2018. The Essential Genome of K-12. MBio 9.

94. PEC (Profiling of E. coli Chromosome). https://shigen.nig.ac.jp/ecoli/pec/. Retrieved 25 January 2023.

95. Salgado H, Santos-Zavaleta A, Gama-Castro S, Millán-Zárate D, Díaz-Peredo E, Sánchez-Solano F, Pérez-Rueda E, Bonavides-Martínez C, Collado-Vides J. 2001. RegulonDB (version 3.2): transcriptional regulation and operon organization in *Escherichia coli* K-12. Nucleic Acids Res 29:72–74.

96. Seo J-H, Hong JS-J, Kim D, Cho B-K, Huang T-W, Tsai S-F, Palsson BO, Charusanti P. 2012. Multiple-omic data analysis of *Klebsiella pneumoniae* MGH 78578 reveals its transcriptional architecture and regulatory features. BMC Genomics 13:679.

97. Rai AK, Mitchell AM. 2020. Enterobacterial Common Antigen: Synthesis and Function of an Enigmatic Molecule. MBio 11.

98. Berks BC, Sargent F, Palmer T. 2000. The Tat protein export pathway. Mol Microbiol 35:260–274.

99. M Brauer A, R Rogers A, R Ellermeier J. 2021. Twin-arginine translocation (Tat) mutants in serovar Typhimurium have increased susceptibility to cell wall targeting antibiotics. FEMS Microbes 2:xtab004.

100. GitHub - JCVenterInstitute/PanGenomePipeline: PanGenomePipeline. GitHub. https://github.com/JCVenterInstitute/PanGenomePipeline. Retrieved 19 January 2024.

101. Chan AP, Sutton G, DePew J, Krishnakumar R, Choi Y, Huang X-Z, Beck E, Harkins DM, Kim M, Lesho EP, Nikolich MP, Fouts DE. 2015. A novel method of consensus pan-chromosome assembly and large-scale comparative analysis reveal the highly flexible pan-genome of *Acinetobacter baumannii*. Genome Biol 16:143.

102. Mistry J, Finn RD, Eddy SR, Bateman A, Punta M. 2013. Challenges in homology search: HMMER3 and convergent evolution of coiled-coil regions. Nucleic Acids Res 41:e121.

103. Haft DH, Loftus BJ, Richardson DL, Yang F, Eisen JA, Paulsen IT, White O. 2001. TIGRFAMs: a protein family resource for the functional identification of proteins. Nucleic Acids Res 29:41–43.

104. Santos-Zavaleta A, Salgado H, Gama-Castro S, Sánchez-Pérez M, Gómez-Romero L, Ledezma-Tejeida D, García-Sotelo JS, Alquicira-Hernández K, Muñiz-Rascado LJ, Peña-Loredo P, Ishida-Gutiérrez C, Velázquez-Ramírez DA, Del Moral-Chávez V, Bonavides-Martínez C, Méndez-Cruz C-F, Galagan J, Collado-Vides J. 2019. RegulonDB v 10.5: tackling challenges to unify classic and high throughput knowledge of gene regulation in *E. coli* K-12. Nucleic Acids Res 47:D212–D220.

105. Eddy SR. 2011. Accelerated Profile HMM Searches. PLoS Comput Biol 7:e1002195.

106. Aramaki T, Blanc-Mathieu R, Endo H, Ohkubo K, Kanehisa M, Goto S, Ogata H. 2020. KofamKOALA: KEGG Ortholog assignment based on profile HMM and adaptive score threshold. Bioinformatics 36:2251–2252.

107. Dusa A. 2024. Draw Venn Diagrams [R package venn version 1.12].

108. Wickham H. 2009. ggplot2: Elegant Graphics for Data Analysis. Springer Science & Business Media.

109. R Core Team. 2021. R: A Language and Environment for Statistical Computing. R Foundation for Statistical Computing, Vienna, Austria.

110. Datsenko KA, Wanner BL. 2000. One-step inactivation of chromosomal genes in *Escherichia coli* K-12 using PCR products. Proc Natl Acad Sci U S A 97:6640–6645.

111. Datta S, Costantino N, Court DL. 2006. A set of recombineering plasmids for gram-negative bacteria. Gene 379:109–115.

112. Ducas-Mowchun K, De Silva PM, Crisostomo L, Fernando DM, Chao T-C, Pelka P, Schweizer HP, Kumar A. 2019. Next Generation of Tn-Based Single-Copy Insertion Elements for Use in Multi- and Pan-Drug-Resistant Strains of *Acinetobacter baumannii*. Appl Environ Microbiol 85.

113. Kovach ME, Elzer PH, Hill DS, Robertson GT, Farris MA, Roop RM 2nd, Peterson KM. 1995. Four new derivatives of the broad-host-range cloning vector pBBR1MCS, carrying different antibiotic-resistance cassettes. Gene 166:175–176.

114. You C, Zhang Y-HP. 2014. Simple cloning and DNA assembly in *Escherichia coli* by prolonged overlap extension PCR. Methods Mol Biol 1116:183–192.

115. Dwived HP, Brocco F, Matuschek E. 2023. Section 7.5 Gradient Diffusion Tests, p. . In Leber, AL, Burnham, C-AD (eds.), Clinical Microbiology Procedures Handbook. ASM Press.

116. 2023. M100 Performance Standards for Antimicrobial Susceptibility Testing. Clinical and Laboratory Standards Institute.

