## Supplementary material for "A novel method for integrating genomic and Tn-Seq data to identify common *in vivo* fitness mechanisms across multiple bacterial species": S1 Dataset: CL_INS_1.html

Legend

 Mobile +extrachromosomalelementfunctions
 Other
 Hypothetical
 AntibioticResistance
 Transport +binding proteins
 All VFDB Genes

FULL


WINDOWSVGPNG

Trim RowsRemove SingletonsSave Fasta

CL\_13


CL\_13


Break


CL\_13


CL\_13


CL\_13


CL\_13


CL\_1976


CL\_13

HighlightSelectShow Genomes


221

CL\_15


48

CL\_15


1

CL\_15


1

CL\_15


1

CL\_16


1

Break


1

CL\_15


1

CL\_15


1

Break

fGI ID


CL\_INS\_1
CL\_INS\_1
CL\_INS\_1
CL\_INS\_382
CL\_INS\_1
CL\_INS\_1
CL\_INS\_3
CL\_INS\_44
CL\_INS\_44
CL\_INS\_44
CL\_INS\_159
CL\_INS\_1
CL\_INS\_382
CL\_INS\_382
CL\_INS\_1
CL\_INS\_1
CL\_INS\_1
CL\_INS\_1
CL\_INS\_159
CL\_INS\_207
CL\_INS\_207
CL\_INS\_86
CL\_INS\_30
CL\_INS\_352
CL\_INS\_42
CL\_INS\_99
CL\_INS\_272
CL\_INS\_272
CL\_INS\_272
CL\_INS\_272
CL\_INS\_272
CL\_INS\_272
CL\_INS\_272
CL\_INS\_272
CL\_INS\_272
CL\_INS\_272
CL\_INS\_272
CL\_INS\_237
CL\_INS\_382
CL\_INS\_382
CL\_INS\_159
CL\_INS\_159
CL\_INS\_156
CL\_INS\_237
CL\_INS\_382
CL\_INS\_1
CL\_INS\_1
CL\_INS\_159
CL\_INS\_159
CL\_INS\_159
CL\_INS\_159
CL\_INS\_159
CL\_INS\_159
CL\_INS\_1
CL\_INS\_1
CL\_INS\_382
CL\_INS\_382
CL\_INS\_382
CL\_INS\_382
CL\_INS\_382
CL\_INS\_382
CL\_INS\_382
CL\_INS\_382
CL\_INS\_382
CL\_INS\_382
CL\_INS\_382
CL\_INS\_382
CL\_INS\_382
CL\_INS\_382
CL\_INS\_382
CL\_INS\_382
CL\_INS\_382
CL\_INS\_382
CL\_INS\_382
CL\_INS\_159
CL\_INS\_382
CL\_INS\_382
CL\_INS\_382
CL\_INS\_382
CL\_INS\_382
CL\_INS\_382
CL\_INS\_159
CL\_INS\_382
CL\_INS\_1
CL\_INS\_1
CL\_INS\_382
CL\_INS\_382
CL\_INS\_382
CL\_INS\_382
CL\_INS\_382
CL\_INS\_382
CL\_INS\_382
CL\_INS\_382
CL\_INS\_99
CL\_INS\_99
CL\_INS\_382
CL\_INS\_382
CL\_INS\_382
CL\_INS\_382
CL\_INS\_1
CL\_INS\_149
CL\_INS\_1
CL\_INS\_382
CL\_INS\_382
CL\_INS\_382
CL\_INS\_1
CL\_INS\_385
CL\_INS\_385
CL\_INS\_385
CL\_INS\_1
CL\_INS\_123
CL\_INS\_1
CL\_INS\_247
CL\_INS\_1
CL\_INS\_237
CL\_INS\_237
CL\_INS\_237
CL\_INS\_237
CL\_INS\_382
CL\_INS\_382
CL\_INS\_382
CL\_INS\_368
CL\_INS\_1
Cluster ID


CL\_14
CL\_22769
CL\_25942
CL\_11169
CL\_17000
CL\_6564
CL\_6563
CL\_22540
CL\_22541
CL\_22542
CL\_19710
CL\_9919
CL\_19711
CL\_5038
CL\_5037
CL\_5036
CL\_5035
CL\_5034
CL\_5033
CL\_6149
CL\_6150
CL\_1495
CL\_7667
CL\_6833
CL\_4435
CL\_16985
CL\_6334
CL\_6333
CL\_6332
CL\_6331
CL\_6330
CL\_6329
CL\_6328
CL\_6327
CL\_6326
CL\_6325
CL\_6324
CL\_6207
CL\_4973
CL\_4974
CL\_14632
CL\_14631
CL\_14630
CL\_6140
CL\_4240
CL\_20052
CL\_5590
CL\_4258
CL\_4259
CL\_4260
CL\_4261
CL\_5630
CL\_5575
CL\_17002
CL\_17001
CL\_4270
CL\_4271
CL\_5634
CL\_6933
CL\_6934
CL\_5637
CL\_5638
CL\_5639
CL\_4277
CL\_4278
CL\_5567
CL\_5566
CL\_5565
CL\_5564
CL\_5563
CL\_5562
CL\_5561
CL\_5560
CL\_4284
CL\_5651
CL\_5556
CL\_5555
CL\_5554
CL\_5553
CL\_5552
CL\_5551
CL\_5550
CL\_5549
CL\_34933
CL\_34932
CL\_5548
CL\_5662
CL\_4299
CL\_4300
CL\_4301
CL\_4302
CL\_4303
CL\_10406
CL\_14207
CL\_14206
CL\_4276
CL\_4275
CL\_4274
CL\_14183
CL\_16984
CL\_5146
CL\_17003
CL\_10407
CL\_10411
CL\_5600
CL\_26722
CL\_5522
CL\_5521
CL\_5520
CL\_34931
CL\_5297
CL\_25707
CL\_5514
CL\_34930
CL\_5512
CL\_5526
CL\_5527
CL\_5528
CL\_5291
CL\_5290
CL\_5531
CL\_14980
CL\_25663
