## Supplementary material for "A novel method for integrating genomic and Tn-Seq data to identify common *in vivo* fitness mechanisms across multiple bacterial species": S1 Dataset: CL_INS_2.html

Legend

 Mobile +extrachromosomalelementfunctions
 Proteinsynthesis/fate
 Other
 Regulatoryfunctions
 Hypothetical
 All EssentialGenes
 All VFDB Genes

FULL


WINDOWSVGPNG

Trim RowsRemove SingletonsSave Fasta

CL\_15


CL\_15


CL\_15


CL\_15


CL\_15


CL\_15


CL\_13


CL\_15


CL\_43


CL\_15


CL\_15


CL\_15


CL\_15


CL\_15


CL\_15

HighlightSelectShow Genomes


175

CL\_16


76

CL\_16


11

CL\_16


6

CL\_16


1

CL\_16


1

CL\_16


1

CL\_16


1

CL\_16


1

CL\_16


1

CL\_17


1

CL\_16


1

CL\_16


1

CL\_18


1

CL\_18


1

CL\_19

fGI ID


CL\_INS\_2
CL\_INS\_2
CL\_INS\_3
CL\_INS\_3
CL\_INS\_3
CL\_INS\_2
CL\_INS\_2
CL\_INS\_2
CL\_INS\_1
CL\_INS\_3
CL\_INS\_3
CL\_INS\_3
CL\_INS\_3
CL\_INS\_2
CL\_INS\_2
CL\_INS\_2
CL\_INS\_2
CL\_INS\_2
CL\_INS\_2
CL\_INS\_2
CL\_INS\_2
CL\_INS\_2
CL\_INS\_2
Cluster ID


CL\_34946
CL\_30259
CL\_30260
CL\_12919
CL\_6802
CL\_35148
CL\_15110
CL\_34945
CL\_6564
CL\_6563
CL\_6562
CL\_6801
CL\_6800
CL\_23960
CL\_23959
CL\_23958
CL\_23957
CL\_23956
CL\_23955
CL\_23954
CL\_23953
CL\_23952
CL\_23951
