## Supplementary material for "A novel method for integrating genomic and Tn-Seq data to identify common *in vivo* fitness mechanisms across multiple bacterial species": S1 Dataset: CL_INS_3.html

Legend

 Mobile +extrachromosomalelementfunctions
 Proteinsynthesis/fate
 Other
 Regulatoryfunctions
 Hypothetical
 EnergyMetabolism
 All VFDB Genes

FULL


WINDOWSVGPNG

Trim RowsRemove SingletonsSave Fasta

CL\_17


CL\_17


CL\_17


CL\_17


CL\_17


CL\_17


CL\_17


CL\_17


CL\_17


CL\_17


CL\_17


CL\_17


CL\_16


CL\_17


CL\_17


Break


CL\_15


CL\_17


CL\_17


CL\_17


CL\_17


CL\_17


CL\_17


CL\_17


CL\_17


CL\_17


CL\_30

HighlightSelectShow Genomes


86

CL\_18


38

CL\_18


33

CL\_18


30

CL\_18


29

CL\_18


25

CL\_18


7

CL\_18


6

CL\_18


2

CL\_18


2

CL\_18


2

CL\_18


2

CL\_18


2

CL\_18


2

CL\_18


1

CL\_18


1

CL\_18


1

CL\_18


1

CL\_15


1

CL\_18


1

CL\_111


1

CL\_18


1

CL\_18


1

CL\_18


1

CL\_18


1

CL\_18


1

CL\_18


1

CL\_18

fGI ID


CL\_INS\_3
CL\_INS\_3
CL\_INS\_3
CL\_INS\_3
CL\_INS\_3
CL\_INS\_3
CL\_INS\_3
CL\_INS\_3
CL\_INS\_3
CL\_INS\_3
CL\_INS\_3
CL\_INS\_123
CL\_INS\_123
CL\_INS\_123
CL\_INS\_3
CL\_INS\_123
CL\_INS\_3
CL\_INS\_3
CL\_INS\_3
CL\_INS\_3
CL\_INS\_3
Cluster ID


CL\_24151
CL\_11088
CL\_11087
CL\_12230
CL\_6563
CL\_6562
CL\_4320
CL\_6802
CL\_23010
CL\_12919
CL\_30260
CL\_11779
CL\_11778
CL\_10329
CL\_11777
CL\_10328
CL\_11776
CL\_6801
CL\_26628
CL\_6800
CL\_4321
