## Supplementary material for "A novel method for integrating genomic and Tn-Seq data to identify common *in vivo* fitness mechanisms across multiple bacterial species": S1 Dataset: CL_INS_5.html

Legend

 Mobile +extrachromosomalelementfunctions
 Other
 Hypothetical
 All EssentialGenes
 All VFDB Genes

FULL


WINDOWSVGPNG

Trim RowsRemove SingletonsSave Fasta

CL\_43


CL\_43


CL\_43


CL\_43


CL\_43


CL\_28


CL\_43


CL\_43

HighlightSelectShow Genomes


134

CL\_44


87

CL\_44


27

CL\_44


22

CL\_44


3

CL\_44


1

CL\_44


1

CL\_16


1

CL\_44

fGI ID


CL\_INS\_5
CL\_INS\_5
CL\_INS\_5
CL\_INS\_5
Cluster ID


CL\_7948
CL\_7947
CL\_16538
CL\_16539
