## Supplementary material for "A novel method for integrating genomic and Tn-Seq data to identify common *in vivo* fitness mechanisms across multiple bacterial species": S1 Dataset: CL_INS_10.html

Legend

 Mobile +extrachromosomalelementfunctions
 Other
 Hypothetical
 All VFDB Genes
 Transport +binding proteins

FULL


WINDOWSVGPNG

Trim RowsRemove SingletonsSave Fasta

CL\_98


CL\_98


CL\_98

HighlightSelectShow Genomes


270

CL\_99


1

CL\_100


1

CL\_100

fGI ID


CL\_INS\_10
CL\_INS\_10
CL\_INS\_10
CL\_INS\_10
CL\_INS\_10
CL\_INS\_10
CL\_INS\_10
CL\_INS\_10
CL\_INS\_10
CL\_INS\_10
CL\_INS\_10
CL\_INS\_10
CL\_INS\_10
CL\_INS\_10
CL\_INS\_10
CL\_INS\_10
CL\_INS\_10
CL\_INS\_10
CL\_INS\_10
CL\_INS\_10
CL\_INS\_10
CL\_INS\_10
CL\_INS\_10
CL\_INS\_10
CL\_INS\_10
CL\_INS\_10
CL\_INS\_10
CL\_INS\_10
CL\_INS\_10
CL\_INS\_10
CL\_INS\_10
CL\_INS\_10
CL\_INS\_10
CL\_INS\_10
CL\_INS\_10
CL\_INS\_10
CL\_INS\_10
CL\_INS\_10
CL\_INS\_10
CL\_INS\_10
CL\_INS\_10
CL\_INS\_10
CL\_INS\_10
CL\_INS\_10
CL\_INS\_10
CL\_INS\_10
CL\_INS\_10
CL\_INS\_10
CL\_INS\_10
CL\_INS\_10
CL\_INS\_10
CL\_INS\_10
Cluster ID


CL\_9247
CL\_9248
CL\_9249
CL\_4805
CL\_19990
CL\_19989
CL\_27982
CL\_19988
CL\_19987
CL\_32393
CL\_19985
CL\_14419
CL\_19984
CL\_19983
CL\_32394
CL\_32395
CL\_19981
CL\_19979
CL\_19978
CL\_19977
CL\_19976
CL\_9965
CL\_9966
CL\_4788
CL\_4784
CL\_4783
CL\_4782
CL\_4781
CL\_4780
CL\_4779
CL\_19974
CL\_4777
CL\_4776
CL\_4775
CL\_4774
CL\_4773
CL\_4772
CL\_4771
CL\_4770
CL\_4769
CL\_4768
CL\_4767
CL\_4766
CL\_4765
CL\_4764
CL\_12596
CL\_27047
CL\_27046
CL\_19972
CL\_19971
CL\_19970
CL\_19968
