## Supplementary material for "A novel method for integrating genomic and Tn-Seq data to identify common *in vivo* fitness mechanisms across multiple bacterial species": S1 Dataset: CL_INS_11.html

Legend

 Mobile +extrachromosomalelementfunctions
 Other
 Regulatoryfunctions
 Hypothetical
 EnergyMetabolism
 All Fitness Genes
 All VFDB Genes

FULL


WINDOWSVGPNG

Trim RowsRemove SingletonsSave Fasta

CL\_105


CL\_105


CL\_105


CL\_105


CL\_105


CL\_105


CL\_105


CL\_105


CL\_105


CL\_111


CL\_105


CL\_105


CL\_105


CL\_105


CL\_105


CL\_105


CL\_105

HighlightSelectShow Genomes


205

CL\_106


24

CL\_106


21

CL\_106


7

CL\_106


3

CL\_106


3

CL\_106


2

CL\_106


1

CL\_106


1

CL\_106


1

CL\_106


1

CL\_109


1

CL\_106


1

CL\_106


1

CL\_106


1

CL\_106


1

CL\_106


1

CL\_106

fGI ID


CL\_INS\_11
CL\_INS\_11
CL\_INS\_11
CL\_INS\_11
CL\_INS\_11
CL\_INS\_11
CL\_INS\_11
CL\_INS\_11
CL\_INS\_11
CL\_INS\_11
CL\_INS\_11
CL\_INS\_11
CL\_INS\_11
CL\_INS\_11
CL\_INS\_11
CL\_INS\_11
CL\_INS\_11
Cluster ID


CL\_30269
CL\_30268
CL\_30267
CL\_7268
CL\_30264
CL\_30265
CL\_13625
CL\_13626
CL\_7269
CL\_7270
CL\_11086
CL\_7272
CL\_7945
CL\_22211
CL\_11085
CL\_7271
CL\_14353
