## Supplementary material for "A novel method for integrating genomic and Tn-Seq data to identify common *in vivo* fitness mechanisms across multiple bacterial species": S1 Dataset: CL_INS_12.html

Legend

 Mobile +extrachromosomalelementfunctions
 Hypothetical
 All VFDB Genes

FULL


WINDOWSVGPNG

Trim RowsRemove SingletonsSave Fasta

CL\_124


CL\_124


CL\_124


CL\_124


CL\_124


CL\_124

HighlightSelectShow Genomes


204

CL\_125


66

CL\_125


2

CL\_125


1

CL\_125


1

CL\_131


1

CL\_125

fGI ID


CL\_INS\_12
CL\_INS\_12
CL\_INS\_12
CL\_INS\_12
Cluster ID


CL\_28860
CL\_15223
CL\_16540
CL\_11084
