## Supplementary material for "A novel method for integrating genomic and Tn-Seq data to identify common *in vivo* fitness mechanisms across multiple bacterial species": S1 Dataset: CL_INS_15.html

Legend

 Mobile +extrachromosomalelementfunctions
 Regulatoryfunctions
 Hypothetical
 DNA Metabolism
 All EssentialGenes
 Transcription
 All Fitness Genes
 Proteinsynthesis/fate
 Other
 Centralintermediarymetabolism
 Transport +binding proteins
 All VFDB Genes

FULL


WINDOWSVGPNG

Trim RowsRemove SingletonsSave Fasta

CL\_203


CL\_203


CL\_203


CL\_203


CL\_203


CL\_203


CL\_203


CL\_203


CL\_203


CL\_203


CL\_203


CL\_203


CL\_203


CL\_203


CL\_203


CL\_203


CL\_203


CL\_203


CL\_203


CL\_203


CL\_203


CL\_203


CL\_203


CL\_203


CL\_203


CL\_203


CL\_203


CL\_203


CL\_203


CL\_203


CL\_203


CL\_203


CL\_203


CL\_203


CL\_203


CL\_203


CL\_203


CL\_203


CL\_203


CL\_203


CL\_203


CL\_1935


CL\_234


CL\_203


CL\_203


CL\_203


CL\_203


CL\_203


CL\_203


CL\_203


CL\_203


CL\_203


CL\_203


CL\_203


CL\_203


CL\_203


CL\_203


CL\_203


CL\_1935


CL\_3454


CL\_203


CL\_203


CL\_203


CL\_203


CL\_203


CL\_203


CL\_203


CL\_203


CL\_203


CL\_203


CL\_203


CL\_203


CL\_203


CL\_203


CL\_203


CL\_203


CL\_203


CL\_203


CL\_203


CL\_203


CL\_203


CL\_203


CL\_202


CL\_203


CL\_203


CL\_203


CL\_203


CL\_203


CL\_203


CL\_203


CL\_203


CL\_203


CL\_203


CL\_203


CL\_203


CL\_203


CL\_203


CL\_203


CL\_203


CL\_203


CL\_203


CL\_203


CL\_203


CL\_203


CL\_203


CL\_203


CL\_203


CL\_203


CL\_203


CL\_203


CL\_203


CL\_203


CL\_203


CL\_203


CL\_203


CL\_203


CL\_203


CL\_234


CL\_203


CL\_203


CL\_203


CL\_203


CL\_203


CL\_269


CL\_203


CL\_203


CL\_234


CL\_203


CL\_203


CL\_203


CL\_203


CL\_203


CL\_203


CL\_203


CL\_203


CL\_203


CL\_203


CL\_203


CL\_203


CL\_463


CL\_203


CL\_203


CL\_203


CL\_203


CL\_203


CL\_203


CL\_203


CL\_203


CL\_203


CL\_203


CL\_203


CL\_203


CL\_234


CL\_203


CL\_203


CL\_203


CL\_203


CL\_203


CL\_203


CL\_203


CL\_234


CL\_203


CL\_234


CL\_203


CL\_203


CL\_203


CL\_203


CL\_203


CL\_203


CL\_325


CL\_203


CL\_203


CL\_203


CL\_218


CL\_287


CL\_203


CL\_203


CL\_234


CL\_1405


CL\_203

HighlightSelectShow Genomes


54

CL\_206


14

CL\_206


8

CL\_206


8

CL\_206


5

CL\_1935


4

CL\_206


4

CL\_206


4

CL\_206


3

CL\_206


3

CL\_206


3

CL\_206


3

CL\_206


3

CL\_206


3

CL\_206


3

CL\_206


2

CL\_206


2

CL\_206


2

CL\_206


2

CL\_206


2

CL\_206


2

CL\_1935


2

CL\_206


2

CL\_206


2

CL\_206


2

CL\_206


1

CL\_206


1

CL\_206


1

CL\_206


1

CL\_206


1

CL\_206


1

CL\_206


1

CL\_206


1

CL\_206


1

CL\_206


1

CL\_206


1

CL\_206


1

CL\_206


1

CL\_206


1

CL\_206


1

CL\_206


1

CL\_206


1

CL\_206


1

CL\_206


1

CL\_206


1

CL\_206


1

CL\_206


1

CL\_206


1

CL\_206


1

CL\_206


1

CL\_206


1

CL\_206


1

CL\_206


1

CL\_206


1

CL\_206


1

CL\_206


1

CL\_206


1

CL\_206


1

CL\_206


1

CL\_206


1

CL\_206


1

CL\_206


1

CL\_206


1

CL\_206


1

CL\_206


1

CL\_206


1

CL\_206


1

CL\_206


1

CL\_206


1

CL\_206


1

CL\_206


1

CL\_206


1

CL\_206


1

CL\_206


1

CL\_206


1

CL\_1935


1

CL\_206


1

CL\_206


1

CL\_206


1

CL\_206


1

CL\_206


1

CL\_206


1

CL\_206


1

CL\_206


1

CL\_206


1

CL\_206


1

CL\_206


1

CL\_206


1

CL\_206


1

CL\_361


1

CL\_221


1

CL\_206


1

CL\_206


1

CL\_206


1

CL\_206


1

CL\_206


1

CL\_206


1

CL\_206


1

CL\_206


1

CL\_206


1

CL\_206


1

CL\_206


1

CL\_206


1

CL\_230


1

CL\_206


1

Break


1

CL\_206


1

CL\_206


1

CL\_206


1

CL\_206


1

CL\_206


1

CL\_206


1

CL\_206


1

CL\_206


1

CL\_206


1

CL\_206


1

CL\_206


1

CL\_206


1

CL\_206


1

CL\_206


1

CL\_206


1

CL\_206


1

CL\_206


1

CL\_206


1

CL\_206


1

CL\_206


1

CL\_206


1

CL\_206


1

CL\_206


1

CL\_206


1

CL\_206


1

CL\_206


1

CL\_206


1

CL\_206


1

CL\_206


1

CL\_206


1

CL\_206


1

CL\_230


1

CL\_206


1

CL\_206


1

CL\_206


1

CL\_229


1

CL\_206


1

CL\_206


1

CL\_206


1

CL\_206


1

CL\_206


1

CL\_2665


1

CL\_206


1

CL\_206


1

CL\_206


1

CL\_206


1

CL\_206


1

CL\_206


1

CL\_206


1

CL\_206


1

CL\_206


1

CL\_206


1

CL\_296


1

CL\_206


1

CL\_206


1

CL\_206


1

CL\_206


1

CL\_206


1

CL\_206


1

CL\_206


1

CL\_206


1

CL\_208


1

CL\_206


1

CL\_206


1

CL\_206


1

CL\_206


1

CL\_206


1

CL\_206


1

CL\_206


1

CL\_206


1

CL\_206


1

CL\_3455


1

CL\_206


1

CL\_206


1

CL\_2846

fGI ID


CL\_INS\_247
CL\_INS\_70
CL\_INS\_15
CL\_INS\_15
CL\_INS\_15
CL\_INS\_15
CL\_INS\_15
CL\_INS\_15
CL\_INS\_15
CL\_INS\_39
CL\_INS\_38
CL\_INS\_39
CL\_INS\_15
CL\_INS\_237
CL\_INS\_237
CL\_INS\_237
CL\_INS\_382
CL\_INS\_237
CL\_INS\_237
CL\_INS\_247
CL\_INS\_233
CL\_INS\_15
CL\_INS\_15
CL\_INS\_15
CL\_INS\_15
CL\_INS\_15
CL\_INS\_15
CL\_INS\_15
CL\_INS\_15
CL\_INS\_15
CL\_INS\_117
CL\_INS\_117
CL\_INS\_117
CL\_INS\_117
CL\_INS\_117
CL\_INS\_15
CL\_INS\_15
CL\_INS\_247
CL\_INS\_247
CL\_INS\_247
CL\_INS\_70
CL\_INS\_70
CL\_INS\_15
CL\_INS\_70
CL\_INS\_70
CL\_INS\_30
CL\_INS\_30
CL\_INS\_15
CL\_INS\_220
CL\_INS\_15
CL\_INS\_220
CL\_INS\_220
CL\_INS\_220
CL\_INS\_220
CL\_INS\_99
CL\_INS\_15
CL\_INS\_20
CL\_INS\_20
CL\_INS\_20
CL\_INS\_20
CL\_INS\_15
CL\_INS\_15
CL\_INS\_220
CL\_INS\_220
CL\_INS\_220
CL\_INS\_15
CL\_INS\_15
CL\_INS\_15
CL\_INS\_220
CL\_INS\_15
CL\_INS\_220
CL\_INS\_220
CL\_INS\_15
CL\_INS\_15
CL\_INS\_15
CL\_INS\_15
CL\_INS\_15
CL\_INS\_15
CL\_INS\_15
CL\_INS\_15
CL\_INS\_15
CL\_INS\_15
CL\_INS\_15
CL\_INS\_15
CL\_INS\_15
CL\_INS\_15
CL\_INS\_15
CL\_INS\_15
CL\_INS\_15
CL\_INS\_15
CL\_INS\_220
CL\_INS\_15
CL\_INS\_15
CL\_INS\_39
CL\_INS\_15
CL\_INS\_15
CL\_INS\_15
CL\_INS\_15
CL\_INS\_15
CL\_INS\_39
CL\_INS\_39
CL\_INS\_15
CL\_INS\_39
CL\_INS\_15
CL\_INS\_15
CL\_INS\_15
CL\_INS\_15
CL\_INS\_15
CL\_INS\_117
CL\_INS\_15
CL\_INS\_15
CL\_INS\_15
CL\_INS\_15
CL\_INS\_237
CL\_INS\_247
CL\_INS\_30
CL\_INS\_237
CL\_INS\_237
CL\_INS\_70
CL\_INS\_247
CL\_INS\_247
CL\_INS\_247
CL\_INS\_70
CL\_INS\_237
CL\_INS\_20
CL\_INS\_20
CL\_INS\_20
CL\_INS\_20
CL\_INS\_20
CL\_INS\_86
CL\_INS\_204
CL\_INS\_204
CL\_INS\_204
CL\_INS\_204
CL\_INS\_204
CL\_INS\_204
CL\_INS\_204
CL\_INS\_204
CL\_INS\_204
CL\_INS\_204
CL\_INS\_204
CL\_INS\_15
CL\_INS\_15
CL\_INS\_86
CL\_INS\_204
CL\_INS\_204
CL\_INS\_204
CL\_INS\_204
CL\_INS\_204
CL\_INS\_204
CL\_INS\_204
CL\_INS\_204
CL\_INS\_204
CL\_INS\_204
CL\_INS\_204
CL\_INS\_204
CL\_INS\_204
CL\_INS\_204
CL\_INS\_204
CL\_INS\_86
CL\_INS\_382
CL\_INS\_204
CL\_INS\_382
CL\_INS\_382
CL\_INS\_382
CL\_INS\_382
CL\_INS\_382
CL\_INS\_382
CL\_INS\_382
CL\_INS\_382
CL\_INS\_382
CL\_INS\_20
CL\_INS\_237
CL\_INS\_295
CL\_INS\_237
CL\_INS\_70
CL\_INS\_247
CL\_INS\_237
CL\_INS\_237
CL\_INS\_237
CL\_INS\_237
CL\_INS\_70
CL\_INS\_237
CL\_INS\_295
CL\_INS\_70
CL\_INS\_237
CL\_INS\_237
CL\_INS\_20
CL\_INS\_247
CL\_INS\_15
CL\_INS\_20
CL\_INS\_20
CL\_INS\_247
CL\_INS\_15
CL\_INS\_247
CL\_INS\_272
CL\_INS\_272
CL\_INS\_272
CL\_INS\_272
CL\_INS\_272
CL\_INS\_272
CL\_INS\_272
CL\_INS\_272
CL\_INS\_272
CL\_INS\_272
CL\_INS\_272
CL\_INS\_272
CL\_INS\_272
CL\_INS\_273
CL\_INS\_273
CL\_INS\_15
CL\_INS\_272
CL\_INS\_272
CL\_INS\_272
CL\_INS\_272
CL\_INS\_272
CL\_INS\_272
CL\_INS\_272
CL\_INS\_273
CL\_INS\_272
CL\_INS\_15
CL\_INS\_15
CL\_INS\_15
CL\_INS\_15
CL\_INS\_15
CL\_INS\_15
CL\_INS\_15
CL\_INS\_15
CL\_INS\_15
CL\_INS\_237
CL\_INS\_237
CL\_INS\_308
CL\_INS\_15
CL\_INS\_15
CL\_INS\_15
CL\_INS\_15
CL\_INS\_15
CL\_INS\_15
CL\_INS\_15
CL\_INS\_15
CL\_INS\_15
CL\_INS\_15
CL\_INS\_15
CL\_INS\_15
CL\_INS\_15
CL\_INS\_15
CL\_INS\_15
CL\_INS\_15
CL\_INS\_15
CL\_INS\_15
CL\_INS\_15
CL\_INS\_15
CL\_INS\_15
CL\_INS\_15
CL\_INS\_15
CL\_INS\_15
CL\_INS\_15
CL\_INS\_15
CL\_INS\_15
CL\_INS\_123
CL\_INS\_15
CL\_INS\_15
CL\_INS\_15
CL\_INS\_15
CL\_INS\_15
CL\_INS\_15
CL\_INS\_15
CL\_INS\_15
CL\_INS\_15
CL\_INS\_207
CL\_INS\_207
CL\_INS\_15
CL\_INS\_15
CL\_INS\_157
CL\_INS\_15
CL\_INS\_15
CL\_INS\_15
CL\_INS\_15
CL\_INS\_15
CL\_INS\_15
CL\_INS\_237
CL\_INS\_382
CL\_INS\_382
CL\_INS\_352
CL\_INS\_352
CL\_INS\_352
CL\_INS\_352
CL\_INS\_352
CL\_INS\_352
CL\_INS\_352
CL\_INS\_352
CL\_INS\_352
CL\_INS\_352
CL\_INS\_352
CL\_INS\_352
CL\_INS\_352
CL\_INS\_352
CL\_INS\_352
CL\_INS\_352
CL\_INS\_352
CL\_INS\_352
CL\_INS\_352
CL\_INS\_352
CL\_INS\_352
CL\_INS\_352
CL\_INS\_352
CL\_INS\_352
CL\_INS\_352
CL\_INS\_352
CL\_INS\_352
CL\_INS\_352
CL\_INS\_352
CL\_INS\_352
CL\_INS\_352
CL\_INS\_352
CL\_INS\_352
CL\_INS\_352
CL\_INS\_352
CL\_INS\_352
CL\_INS\_352
CL\_INS\_352
CL\_INS\_352
CL\_INS\_237
CL\_INS\_352
CL\_INS\_352
CL\_INS\_15
CL\_INS\_15
CL\_INS\_20
CL\_INS\_382
CL\_INS\_15
CL\_INS\_15
CL\_INS\_20
CL\_INS\_20
CL\_INS\_20
CL\_INS\_20
CL\_INS\_382
CL\_INS\_382
CL\_INS\_382
CL\_INS\_382
CL\_INS\_382
CL\_INS\_382
CL\_INS\_382
CL\_INS\_382
CL\_INS\_382
CL\_INS\_382
CL\_INS\_382
CL\_INS\_382
CL\_INS\_382
CL\_INS\_382
CL\_INS\_382
CL\_INS\_382
CL\_INS\_382
CL\_INS\_382
CL\_INS\_382
CL\_INS\_26
CL\_INS\_20
CL\_INS\_20
CL\_INS\_20
CL\_INS\_20
CL\_INS\_20
CL\_INS\_20
CL\_INS\_20
CL\_INS\_15
CL\_INS\_20
CL\_INS\_20
CL\_INS\_15
CL\_INS\_15
CL\_INS\_15
CL\_INS\_15
CL\_INS\_20
CL\_INS\_15
CL\_INS\_20
CL\_INS\_20
CL\_INS\_20
CL\_INS\_15
CL\_INS\_15
CL\_INS\_15
CL\_INS\_15
CL\_INS\_20
CL\_INS\_15
CL\_INS\_15
CL\_INS\_15
CL\_INS\_15
CL\_INS\_15
CL\_INS\_15
CL\_INS\_15
CL\_INS\_15
CL\_INS\_15
CL\_INS\_15
CL\_INS\_15
CL\_INS\_15
CL\_INS\_15
CL\_INS\_15
CL\_INS\_15
CL\_INS\_15
CL\_INS\_15
CL\_INS\_15
CL\_INS\_15
CL\_INS\_117
CL\_INS\_117
CL\_INS\_117
CL\_INS\_117
CL\_INS\_117
CL\_INS\_15
CL\_INS\_15
CL\_INS\_15
CL\_INS\_15
CL\_INS\_15
CL\_INS\_117
CL\_INS\_117
CL\_INS\_15
CL\_INS\_15
CL\_INS\_15
CL\_INS\_117
CL\_INS\_20
CL\_INS\_15
CL\_INS\_15
CL\_INS\_15
CL\_INS\_237
CL\_INS\_15
CL\_INS\_15
CL\_INS\_15
CL\_INS\_15
CL\_INS\_15
CL\_INS\_15
CL\_INS\_15
CL\_INS\_15
CL\_INS\_117
CL\_INS\_15
CL\_INS\_117
CL\_INS\_117
CL\_INS\_117
CL\_INS\_20
CL\_INS\_117
CL\_INS\_15
CL\_INS\_15
CL\_INS\_15
CL\_INS\_15
CL\_INS\_42
CL\_INS\_117
CL\_INS\_117
CL\_INS\_117
CL\_INS\_15
CL\_INS\_15
CL\_INS\_15
CL\_INS\_15
CL\_INS\_15
CL\_INS\_15
CL\_INS\_15
CL\_INS\_15
CL\_INS\_20
CL\_INS\_15
CL\_INS\_15
CL\_INS\_15
CL\_INS\_15
CL\_INS\_15
CL\_INS\_260
CL\_INS\_15
CL\_INS\_117
CL\_INS\_117
CL\_INS\_117
CL\_INS\_15
CL\_INS\_15
CL\_INS\_15
CL\_INS\_15
CL\_INS\_15
CL\_INS\_247
CL\_INS\_247
CL\_INS\_247
CL\_INS\_247
CL\_INS\_247
CL\_INS\_247
CL\_INS\_247
CL\_INS\_247
CL\_INS\_247
CL\_INS\_382
CL\_INS\_382
CL\_INS\_207
CL\_INS\_207
CL\_INS\_207
CL\_INS\_207
CL\_INS\_207
CL\_INS\_15
CL\_INS\_15
CL\_INS\_15
CL\_INS\_295
CL\_INS\_247
CL\_INS\_15
CL\_INS\_15
CL\_INS\_382
CL\_INS\_106
CL\_INS\_382
CL\_INS\_106
CL\_INS\_132
CL\_INS\_106
CL\_INS\_106
CL\_INS\_106
CL\_INS\_106
CL\_INS\_15
CL\_INS\_382
CL\_INS\_247
CL\_INS\_247
CL\_INS\_247
CL\_INS\_247
CL\_INS\_247
CL\_INS\_382
CL\_INS\_382
CL\_INS\_106
CL\_INS\_295
CL\_INS\_295
CL\_INS\_20
CL\_INS\_207
CL\_INS\_15
CL\_INS\_15
CL\_INS\_15
CL\_INS\_15
CL\_INS\_15
CL\_INS\_15
CL\_INS\_15
CL\_INS\_30
CL\_INS\_30
CL\_INS\_30
CL\_INS\_15
CL\_INS\_237
CL\_INS\_237
CL\_INS\_70
CL\_INS\_70
CL\_INS\_70
CL\_INS\_70
CL\_INS\_70
CL\_INS\_237
CL\_INS\_30
CL\_INS\_30
CL\_INS\_30
CL\_INS\_30
CL\_INS\_30
CL\_INS\_237
CL\_INS\_237
CL\_INS\_237
CL\_INS\_237
CL\_INS\_295
CL\_INS\_30
CL\_INS\_237
CL\_INS\_159
CL\_INS\_159
CL\_INS\_159
CL\_INS\_247
CL\_INS\_247
CL\_INS\_30
CL\_INS\_30
CL\_INS\_30
CL\_INS\_86
CL\_INS\_86
CL\_INS\_30
CL\_INS\_30
CL\_INS\_30
CL\_INS\_15
CL\_INS\_15
CL\_INS\_30
CL\_INS\_30
CL\_INS\_123
CL\_INS\_170
CL\_INS\_30
CL\_INS\_15
CL\_INS\_15
CL\_INS\_15
CL\_INS\_15
CL\_INS\_30
CL\_INS\_15
CL\_INS\_15
CL\_INS\_15
CL\_INS\_15
CL\_INS\_15
CL\_INS\_30
CL\_INS\_30
CL\_INS\_237
CL\_INS\_237
CL\_INS\_237
CL\_INS\_237
CL\_INS\_70
CL\_INS\_237
CL\_INS\_237
CL\_INS\_237
CL\_INS\_237
CL\_INS\_237
CL\_INS\_70
CL\_INS\_237
CL\_INS\_237
CL\_INS\_237
CL\_INS\_237
CL\_INS\_237
CL\_INS\_368
CL\_INS\_368
CL\_INS\_368
CL\_INS\_20
CL\_INS\_20
CL\_INS\_20
CL\_INS\_20
CL\_INS\_20
CL\_INS\_20
CL\_INS\_20
CL\_INS\_20
CL\_INS\_368
CL\_INS\_382
CL\_INS\_15
CL\_INS\_15
CL\_INS\_15
CL\_INS\_15
CL\_INS\_352
CL\_INS\_352
CL\_INS\_15
CL\_INS\_70
CL\_INS\_70
CL\_INS\_70
CL\_INS\_70
CL\_INS\_70
CL\_INS\_70
CL\_INS\_247
CL\_INS\_237
CL\_INS\_237
CL\_INS\_237
CL\_INS\_237
CL\_INS\_70
CL\_INS\_70
CL\_INS\_70
CL\_INS\_70
CL\_INS\_70
CL\_INS\_247
CL\_INS\_237
CL\_INS\_237
CL\_INS\_237
CL\_INS\_237
CL\_INS\_237
CL\_INS\_237
CL\_INS\_237
CL\_INS\_295
CL\_INS\_295
CL\_INS\_70
CL\_INS\_70
CL\_INS\_295
CL\_INS\_295
CL\_INS\_382
CL\_INS\_295
CL\_INS\_295
CL\_INS\_70
CL\_INS\_70
CL\_INS\_70
CL\_INS\_295
CL\_INS\_295
CL\_INS\_295
CL\_INS\_295
CL\_INS\_295
CL\_INS\_30
CL\_INS\_30
CL\_INS\_70
CL\_INS\_295
CL\_INS\_295
CL\_INS\_295
CL\_INS\_295
CL\_INS\_295
CL\_INS\_295
CL\_INS\_295
CL\_INS\_295
CL\_INS\_295
CL\_INS\_295
CL\_INS\_237
CL\_INS\_237
CL\_INS\_295
CL\_INS\_15
CL\_INS\_70
CL\_INS\_15
CL\_INS\_15
CL\_INS\_70
CL\_INS\_70
CL\_INS\_237
CL\_INS\_237
CL\_INS\_237
CL\_INS\_237
CL\_INS\_352
CL\_INS\_15
CL\_INS\_117
CL\_INS\_117
CL\_INS\_117
CL\_INS\_117
CL\_INS\_15
CL\_INS\_295
CL\_INS\_295
CL\_INS\_295
CL\_INS\_237
CL\_INS\_237
CL\_INS\_15
CL\_INS\_295
CL\_INS\_237
CL\_INS\_237
CL\_INS\_237
CL\_INS\_237
CL\_INS\_237
CL\_INS\_237
CL\_INS\_237
CL\_INS\_15
Cluster ID


CL\_9539
CL\_8107
CL\_37072
CL\_37073
CL\_37074
CL\_12245
CL\_12246
CL\_12247
CL\_12248
CL\_30511
CL\_6996
CL\_30510
CL\_26110
CL\_26111
CL\_9376
CL\_9375
CL\_6385
CL\_9374
CL\_26112
CL\_9370
CL\_9369
CL\_28543
CL\_28542
CL\_28541
CL\_28540
CL\_28539
CL\_28538
CL\_28537
CL\_28536
CL\_28535
CL\_10529
CL\_13649
CL\_13650
CL\_13651
CL\_13652
CL\_15225
CL\_34281
CL\_14013
CL\_14014
CL\_14015
CL\_8591
CL\_8592
CL\_34280
CL\_6808
CL\_6809
CL\_1937
CL\_1936
CL\_7943
CL\_7942
CL\_7941
CL\_7940
CL\_7939
CL\_7938
CL\_13041
CL\_13042
CL\_13043
CL\_7056
CL\_7057
CL\_7058
CL\_7059
CL\_28574
CL\_7937
CL\_11609
CL\_11608
CL\_7936
CL\_7935
CL\_16064
CL\_16065
CL\_7934
CL\_7933
CL\_7932
CL\_7931
CL\_11607
CL\_7930
CL\_7929
CL\_11606
CL\_21859
CL\_21860
CL\_11605
CL\_16066
CL\_7928
CL\_7927
CL\_7926
CL\_35146
CL\_35145
CL\_7925
CL\_7924
CL\_11604
CL\_21861
CL\_21862
CL\_11603
CL\_34446
CL\_34445
CL\_7923
CL\_13045
CL\_13046
CL\_13047
CL\_13048
CL\_13049
CL\_30509
CL\_11602
CL\_31451
CL\_11601
CL\_30508
CL\_7922
CL\_7921
CL\_7920
CL\_35144
CL\_7919
CL\_7918
CL\_21863
CL\_21864
CL\_21865
CL\_7970
CL\_7204
CL\_7749
CL\_7206
CL\_7290
CL\_6417
CL\_16835
CL\_20656
CL\_16836
CL\_8642
CL\_4375
CL\_267
CL\_266
CL\_265
CL\_264
CL\_28353
CL\_7344
CL\_6520
CL\_1105
CL\_1104
CL\_1103
CL\_1101
CL\_1100
CL\_5393
CL\_5394
CL\_5395
CL\_5396
CL\_5398
CL\_28534
CL\_28533
CL\_10273
CL\_5401
CL\_5402
CL\_5403
CL\_5404
CL\_5405
CL\_5406
CL\_5407
CL\_5408
CL\_5409
CL\_5410
CL\_1099
CL\_1098
CL\_1097
CL\_5411
CL\_5412
CL\_10976
CL\_5809
CL\_5415
CL\_5416
CL\_7539
CL\_7540
CL\_524
CL\_522
CL\_521
CL\_520
CL\_519
CL\_518
CL\_28354
CL\_7208
CL\_20280
CL\_14318
CL\_10547
CL\_7470
CL\_13846
CL\_6415
CL\_20007
CL\_20009
CL\_8126
CL\_6730
CL\_20279
CL\_6824
CL\_20278
CL\_4374
CL\_4373
CL\_13465
CL\_13464
CL\_4372
CL\_4371
CL\_4370
CL\_9037
CL\_4369
CL\_6323
CL\_6324
CL\_6325
CL\_6326
CL\_6327
CL\_6328
CL\_6329
CL\_6330
CL\_6331
CL\_6332
CL\_6333
CL\_6334
CL\_6335
CL\_21501
CL\_21502
CL\_21503
CL\_6336
CL\_6337
CL\_6338
CL\_6339
CL\_6340
CL\_6341
CL\_6342
CL\_21504
CL\_6344
CL\_26060
CL\_13463
CL\_13462
CL\_9036
CL\_9035
CL\_9034
CL\_9033
CL\_20655
CL\_9032
CL\_9888
CL\_9887
CL\_9886
CL\_9885
CL\_9884
CL\_9883
CL\_9882
CL\_9881
CL\_9880
CL\_9879
CL\_9877
CL\_9876
CL\_9875
CL\_9874
CL\_10261
CL\_10262
CL\_10263
CL\_10264
CL\_10265
CL\_10266
CL\_10267
CL\_10268
CL\_10269
CL\_20654
CL\_20653
CL\_20652
CL\_24495
CL\_24494
CL\_24493
CL\_24492
CL\_8157
CL\_24491
CL\_29807
CL\_9031
CL\_29806
CL\_29805
CL\_29804
CL\_9030
CL\_29803
CL\_9029
CL\_9028
CL\_9027
CL\_9026
CL\_9025
CL\_204
CL\_15451
CL\_15452
CL\_15453
CL\_30946
CL\_22111
CL\_26157
CL\_6413
CL\_7691
CL\_8832
CL\_9439
CL\_9438
CL\_9437
CL\_9436
CL\_9435
CL\_9434
CL\_9433
CL\_9432
CL\_9431
CL\_9430
CL\_9429
CL\_9428
CL\_9427
CL\_9426
CL\_9425
CL\_9424
CL\_9423
CL\_9422
CL\_9421
CL\_9420
CL\_9419
CL\_9418
CL\_9417
CL\_9416
CL\_9415
CL\_9414
CL\_9413
CL\_9412
CL\_9411
CL\_9410
CL\_9409
CL\_9408
CL\_9407
CL\_9406
CL\_9405
CL\_9404
CL\_9403
CL\_9402
CL\_9401
CL\_26156
CL\_9440
CL\_9441
CL\_26155
CL\_26154
CL\_205
CL\_10804
CL\_19176
CL\_13289
CL\_4324
CL\_4325
CL\_4326
CL\_8209
CL\_517
CL\_516
CL\_7366
CL\_7365
CL\_5350
CL\_4410
CL\_4409
CL\_4408
CL\_4407
CL\_6479
CL\_6483
CL\_4402
CL\_10337
CL\_508
CL\_4400
CL\_7821
CL\_5799
CL\_4396
CL\_4395
CL\_4327
CL\_4328
CL\_4329
CL\_4330
CL\_4331
CL\_4332
CL\_4333
CL\_4334
CL\_13290
CL\_4335
CL\_4336
CL\_27634
CL\_29964
CL\_29963
CL\_29962
CL\_4337
CL\_11289
CL\_4338
CL\_4339
CL\_4340
CL\_28177
CL\_25943
CL\_20979
CL\_20980
CL\_4341
CL\_15226
CL\_36653
CL\_36652
CL\_36651
CL\_36650
CL\_36649
CL\_36648
CL\_36647
CL\_36646
CL\_36645
CL\_36644
CL\_36643
CL\_36642
CL\_36641
CL\_36640
CL\_36639
CL\_36638
CL\_36637
CL\_36636
CL\_9176
CL\_10455
CL\_10456
CL\_10457
CL\_10458
CL\_10459
CL\_10460
CL\_27552
CL\_27133
CL\_27132
CL\_13795
CL\_34879
CL\_34216
CL\_34217
CL\_34218
CL\_13437
CL\_4342
CL\_29092
CL\_32577
CL\_32576
CL\_8124
CL\_5790
CL\_5791
CL\_21731
CL\_21732
CL\_21733
CL\_21734
CL\_27131
CL\_27130
CL\_7014
CL\_21844
CL\_7013
CL\_7012
CL\_7011
CL\_5787
CL\_11774
CL\_15454
CL\_9610
CL\_36337
CL\_26059
CL\_4435
CL\_4497
CL\_4498
CL\_4499
CL\_8467
CL\_16543
CL\_16544
CL\_16545
CL\_16546
CL\_6560
CL\_7347
CL\_33371
CL\_5788
CL\_22808
CL\_22807
CL\_29269
CL\_29268
CL\_29267
CL\_5789
CL\_25383
CL\_4343
CL\_1409
CL\_1410
CL\_25382
CL\_7348
CL\_7349
CL\_27774
CL\_27773
CL\_14332
CL\_14333
CL\_14334
CL\_14335
CL\_21989
CL\_21988
CL\_21987
CL\_21986
CL\_21985
CL\_8554
CL\_10807
CL\_8952
CL\_8953
CL\_20646
CL\_20645
CL\_20644
CL\_20643
CL\_27772
CL\_13731
CL\_13732
CL\_7567
CL\_14355
CL\_14356
CL\_8647
CL\_13733
CL\_8648
CL\_8649
CL\_8650
CL\_13734
CL\_13735
CL\_12139
CL\_12138
CL\_13736
CL\_12137
CL\_10496
CL\_8656
CL\_8657
CL\_8658
CL\_8659
CL\_13737
CL\_13738
CL\_13309
CL\_13739
CL\_13740
CL\_8565
CL\_233
CL\_23368
CL\_23367
CL\_23366
CL\_23365
CL\_33969
CL\_33968
CL\_33967
CL\_1934
CL\_235
CL\_7751
CL\_23975
CL\_6813
CL\_13552
CL\_7831
CL\_1933
CL\_1932
CL\_6420
CL\_10546
CL\_11973
CL\_6422
CL\_5241
CL\_6423
CL\_6424
CL\_6425
CL\_20010
CL\_20011
CL\_20012
CL\_15981
CL\_20277
CL\_5246
CL\_11783
CL\_13741
CL\_13742
CL\_13743
CL\_13744
CL\_13745
CL\_5231
CL\_5232
CL\_7752
CL\_5814
CL\_5815
CL\_10984
CL\_7753
CL\_7754
CL\_13746
CL\_13747
CL\_13748
CL\_13749
CL\_10715
CL\_7075
CL\_13750
CL\_35432
CL\_35431
CL\_12249
CL\_12250
CL\_13751
CL\_13752
CL\_13753
CL\_13754
CL\_14357
CL\_14358
CL\_7755
CL\_7756
CL\_13755
CL\_11710
CL\_13756
CL\_11711
CL\_11105
CL\_8127
CL\_11106
CL\_8128
CL\_8556
CL\_8555
CL\_8129
CL\_8130
CL\_8131
CL\_7979
CL\_7980
CL\_7981
CL\_7982
CL\_13757
CL\_13758
CL\_7550
CL\_7551
CL\_7552
CL\_7553
CL\_13759
CL\_13760
CL\_7554
CL\_7555
CL\_7556
CL\_7557
CL\_20651
CL\_20650
CL\_20649
CL\_20648
CL\_10071
CL\_10073
CL\_20647
CL\_6836
CL\_6838
CL\_6839
CL\_6840
CL\_6841
CL\_6842
CL\_6844
CL\_7301
CL\_7988
CL\_7989
CL\_7990
CL\_7672
CL\_7673
CL\_14683
CL\_7234
CL\_7235
CL\_15979
CL\_6845
CL\_6846
CL\_6847
CL\_6848
CL\_6849
CL\_13761
CL\_7872
CL\_13762
CL\_20284
CL\_15973
CL\_18091
CL\_20285
CL\_20286
CL\_8216
CL\_20287
CL\_20288
CL\_11221
CL\_10331
CL\_10330
CL\_20289
CL\_20290
CL\_20291
CL\_20292
CL\_20293
CL\_7667
CL\_11295
CL\_6410
CL\_20294
CL\_20295
CL\_20296
CL\_20297
CL\_20298
CL\_20299
CL\_20300
CL\_20301
CL\_20302
CL\_20303
CL\_20304
CL\_7871
CL\_7870
CL\_13763
CL\_7869
CL\_18302
CL\_18303
CL\_18089
CL\_9371
CL\_8780
CL\_18161
CL\_7236
CL\_18304
CL\_5159
CL\_10315
CL\_4494
CL\_4495
CL\_1411
CL\_10140
CL\_13764
CL\_13765
CL\_13766
CL\_13767
CL\_13768
CL\_13212
CL\_14359
CL\_13769
CL\_13770
CL\_13771
CL\_13772
CL\_13773
CL\_13774
CL\_13775
CL\_13776
CL\_7010
