## Supplementary material for "A novel method for integrating genomic and Tn-Seq data to identify common *in vivo* fitness mechanisms across multiple bacterial species": S1 Dataset: CL_INS_16.html

Legend

 Mobile +extrachromosomalelementfunctions
 Hypothetical
 Other
 All VFDB Genes

FULL


WINDOWSVGPNG

Trim RowsRemove SingletonsSave Fasta

CL\_211


CL\_211


CL\_211


CL\_211


CL\_211


CL\_211


CL\_211


CL\_211


CL\_211


CL\_211


CL\_211

HighlightSelectShow Genomes


138

CL\_214


117

CL\_214


13

CL\_214


2

CL\_216


2

CL\_217


1

CL\_214


1

CL\_217


1

CL\_216


1

CL\_217


1

CL\_334


1

CL\_216

fGI ID


CL\_INS\_16
CL\_INS\_16
CL\_INS\_16
CL\_INS\_16
CL\_INS\_16
CL\_INS\_16
Cluster ID


CL\_212
CL\_215
CL\_213
CL\_24306
CL\_4344
CL\_4345
