## Supplementary material for "A novel method for integrating genomic and Tn-Seq data to identify common *in vivo* fitness mechanisms across multiple bacterial species": S1 Dataset: CL_INS_17.html

Legend

 Mobile +extrachromosomalelementfunctions
 Hypothetical
 Other
 All VFDB Genes

FULL


WINDOWSVGPNG

Trim RowsRemove SingletonsSave Fasta

CL\_214


CL\_214


CL\_214


CL\_214


CL\_214


CL\_214


CL\_211


CL\_214


CL\_214


CL\_214


CL\_214


CL\_214


CL\_214


CL\_211


CL\_214


CL\_214


CL\_214


CL\_214


CL\_214


CL\_214


CL\_211


CL\_211

HighlightSelectShow Genomes


135

CL\_216


76

CL\_216


19

CL\_216


18

CL\_216


3

CL\_216


3

CL\_216


2

CL\_216


2

CL\_217


1

CL\_216


1

CL\_217


1

CL\_221


1

CL\_217


1

CL\_216


1

CL\_216


1

CL\_490


1

CL\_217


1

CL\_217


1

CL\_217


1

CL\_217


1

CL\_216


1

CL\_216


1

CL\_216

fGI ID


CL\_INS\_17
CL\_INS\_17
CL\_INS\_16
CL\_INS\_17
CL\_INS\_382
CL\_INS\_17
CL\_INS\_16
CL\_INS\_17
CL\_INS\_17
CL\_INS\_17
CL\_INS\_16
CL\_INS\_16
CL\_INS\_17
CL\_INS\_17
CL\_INS\_17
CL\_INS\_17
CL\_INS\_17
CL\_INS\_17
CL\_INS\_17
CL\_INS\_17
CL\_INS\_17
CL\_INS\_17
CL\_INS\_17
CL\_INS\_17
CL\_INS\_17
CL\_INS\_17
CL\_INS\_17
CL\_INS\_17
CL\_INS\_17
CL\_INS\_17
CL\_INS\_17
CL\_INS\_17
CL\_INS\_17
CL\_INS\_17
CL\_INS\_17
CL\_INS\_17
CL\_INS\_17
CL\_INS\_17
CL\_INS\_17
CL\_INS\_17
CL\_INS\_17
CL\_INS\_17
CL\_INS\_17
CL\_INS\_17
CL\_INS\_17
CL\_INS\_17
CL\_INS\_17
CL\_INS\_17
CL\_INS\_17
CL\_INS\_17
CL\_INS\_17
Cluster ID


CL\_26058
CL\_11971
CL\_213
CL\_30945
CL\_502
CL\_537
CL\_215
CL\_11970
CL\_7009
CL\_219
CL\_4344
CL\_4345
CL\_5465
CL\_5464
CL\_5463
CL\_5462
CL\_5461
CL\_5460
CL\_5459
CL\_5458
CL\_5457
CL\_5456
CL\_5455
CL\_5454
CL\_5453
CL\_5452
CL\_7626
CL\_5451
CL\_5450
CL\_5449
CL\_5448
CL\_5447
CL\_5446
CL\_5445
CL\_5444
CL\_5443
CL\_5442
CL\_5441
CL\_5440
CL\_5439
CL\_5438
CL\_5437
CL\_5436
CL\_5435
CL\_5434
CL\_5433
CL\_5432
CL\_5431
CL\_7625
CL\_7624
CL\_7623
