## Supplementary material for "A novel method for integrating genomic and Tn-Seq data to identify common *in vivo* fitness mechanisms across multiple bacterial species": S1 Dataset: CL_INS_19.html

Legend

 Hypothetical
 DNA Metabolism
 Other
 All VFDB Genes

FULL


WINDOWSVGPNG

Trim RowsRemove SingletonsSave Fasta

CL\_218


CL\_218


CL\_217


CL\_218


CL\_218


CL\_218


CL\_218


CL\_218


CL\_214

HighlightSelectShow Genomes


147

CL\_221


113

CL\_221


4

CL\_221


3

CL\_221


1

CL\_224


1

CL\_223


1

CL\_223


1

CL\_222


1

CL\_221

fGI ID


CL\_INS\_17
CL\_INS\_17
CL\_INS\_19
Cluster ID


CL\_7009
CL\_219
CL\_220
