## Supplementary material for "A novel method for integrating genomic and Tn-Seq data to identify common *in vivo* fitness mechanisms across multiple bacterial species": S1 Dataset: CL_INS_25.html

Legend

 Mobile +extrachromosomalelementfunctions
 Regulatoryfunctions
 Hypothetical
 DNA Metabolism
 AntibioticResistance
 All EssentialGenes
 All Fitness Genes
 Proteinsynthesis/fate
 Other
 Transport +binding proteins
 All VFDB Genes

FULL


WINDOWSVGPNG

Trim RowsRemove SingletonsSave Fasta

CL\_290


CL\_290


CL\_290


CL\_290


CL\_289


CL\_289


CL\_290


CL\_290


CL\_290


CL\_290


CL\_280


CL\_289


CL\_289


CL\_290


CL\_230


CL\_230


CL\_290


CL\_290


CL\_290


CL\_289


CL\_290


CL\_290


CL\_290


CL\_230


CL\_230


CL\_290


CL\_290


CL\_289


CL\_290

HighlightSelectShow Genomes


138

CL\_291


71

CL\_291


8

CL\_291


4

CL\_291


3

CL\_291


2

CL\_291


2

CL\_291


1

CL\_292


1

CL\_292


1

CL\_291


1

CL\_291


1

CL\_291


1

CL\_291


1

CL\_344


1

CL\_291


1

CL\_291


1

CL\_291


1

Break


1

CL\_291


1

CL\_291


1

CL\_291


1

CL\_291


1

CL\_296


1

CL\_291


1

CL\_291


1

CL\_291


1

CL\_291


1

CL\_291


1

CL\_291

fGI ID


CL\_INS\_25
CL\_INS\_25
CL\_INS\_25
CL\_INS\_247
CL\_INS\_247
CL\_INS\_247
CL\_INS\_247
CL\_INS\_247
CL\_INS\_207
CL\_INS\_20
CL\_INS\_20
CL\_INS\_20
CL\_INS\_20
CL\_INS\_20
CL\_INS\_20
CL\_INS\_20
CL\_INS\_20
CL\_INS\_20
CL\_INS\_20
CL\_INS\_20
CL\_INS\_20
CL\_INS\_20
CL\_INS\_20
CL\_INS\_123
CL\_INS\_20
CL\_INS\_20
CL\_INS\_20
CL\_INS\_20
CL\_INS\_20
CL\_INS\_20
CL\_INS\_20
CL\_INS\_20
CL\_INS\_20
CL\_INS\_20
CL\_INS\_20
CL\_INS\_237
CL\_INS\_237
CL\_INS\_237
CL\_INS\_237
CL\_INS\_237
CL\_INS\_20
CL\_INS\_20
CL\_INS\_20
CL\_INS\_247
CL\_INS\_20
CL\_INS\_20
CL\_INS\_25
CL\_INS\_25
CL\_INS\_382
CL\_INS\_25
CL\_INS\_25
CL\_INS\_20
CL\_INS\_25
CL\_INS\_25
CL\_INS\_25
CL\_INS\_25
CL\_INS\_20
CL\_INS\_207
CL\_INS\_382
CL\_INS\_382
CL\_INS\_20
CL\_INS\_382
CL\_INS\_382
CL\_INS\_207
CL\_INS\_382
CL\_INS\_382
CL\_INS\_382
CL\_INS\_382
CL\_INS\_382
CL\_INS\_20
CL\_INS\_382
CL\_INS\_382
CL\_INS\_382
CL\_INS\_382
CL\_INS\_382
CL\_INS\_20
CL\_INS\_382
CL\_INS\_382
CL\_INS\_382
CL\_INS\_382
CL\_INS\_382
CL\_INS\_106
CL\_INS\_20
CL\_INS\_382
CL\_INS\_204
CL\_INS\_382
CL\_INS\_382
CL\_INS\_207
CL\_INS\_382
CL\_INS\_382
CL\_INS\_204
CL\_INS\_20
CL\_INS\_20
CL\_INS\_204
CL\_INS\_204
CL\_INS\_204
CL\_INS\_204
CL\_INS\_204
CL\_INS\_204
CL\_INS\_204
CL\_INS\_204
CL\_INS\_204
CL\_INS\_204
CL\_INS\_204
CL\_INS\_204
CL\_INS\_204
CL\_INS\_204
CL\_INS\_204
CL\_INS\_204
CL\_INS\_204
CL\_INS\_86
CL\_INS\_204
CL\_INS\_204
CL\_INS\_20
CL\_INS\_204
CL\_INS\_204
CL\_INS\_204
CL\_INS\_204
CL\_INS\_204
CL\_INS\_204
CL\_INS\_207
CL\_INS\_204
CL\_INS\_204
CL\_INS\_204
CL\_INS\_204
CL\_INS\_207
CL\_INS\_20
CL\_INS\_20
CL\_INS\_20
CL\_INS\_20
CL\_INS\_20
CL\_INS\_20
CL\_INS\_20
CL\_INS\_20
CL\_INS\_20
CL\_INS\_20
CL\_INS\_20
CL\_INS\_207
CL\_INS\_207
CL\_INS\_207
CL\_INS\_20
CL\_INS\_30
CL\_INS\_20
CL\_INS\_20
CL\_INS\_20
CL\_INS\_20
CL\_INS\_20
CL\_INS\_247
CL\_INS\_247
CL\_INS\_20
CL\_INS\_20
CL\_INS\_20
CL\_INS\_20
CL\_INS\_20
CL\_INS\_20
CL\_INS\_20
CL\_INS\_20
CL\_INS\_20
CL\_INS\_174
CL\_INS\_382
CL\_INS\_99
CL\_INS\_20
CL\_INS\_20
CL\_INS\_20
CL\_INS\_247
CL\_INS\_20
CL\_INS\_20
CL\_INS\_20
CL\_INS\_20
CL\_INS\_20
CL\_INS\_20
CL\_INS\_20
CL\_INS\_20
CL\_INS\_20
CL\_INS\_20
CL\_INS\_20
CL\_INS\_20
CL\_INS\_20
CL\_INS\_20
CL\_INS\_20
CL\_INS\_20
CL\_INS\_20
CL\_INS\_20
CL\_INS\_20
CL\_INS\_20
CL\_INS\_20
CL\_INS\_20
CL\_INS\_20
CL\_INS\_20
CL\_INS\_20
CL\_INS\_20
CL\_INS\_20
CL\_INS\_20
CL\_INS\_237
CL\_INS\_20
CL\_INS\_20
CL\_INS\_20
CL\_INS\_25
CL\_INS\_25
CL\_INS\_25
CL\_INS\_385
CL\_INS\_247
CL\_INS\_44
CL\_INS\_247
CL\_INS\_247
CL\_INS\_247
CL\_INS\_247
CL\_INS\_26
CL\_INS\_26
CL\_INS\_26
CL\_INS\_26
CL\_INS\_26
CL\_INS\_99
CL\_INS\_99
CL\_INS\_25
CL\_INS\_25
CL\_INS\_25
CL\_INS\_25
CL\_INS\_25
CL\_INS\_25
CL\_INS\_237
CL\_INS\_237
CL\_INS\_237
CL\_INS\_237
CL\_INS\_237
CL\_INS\_237
CL\_INS\_237
CL\_INS\_237
CL\_INS\_382
CL\_INS\_25
CL\_INS\_382
CL\_INS\_25
CL\_INS\_25
CL\_INS\_354
CL\_INS\_354
CL\_INS\_25
CL\_INS\_237
CL\_INS\_25
CL\_INS\_25
CL\_INS\_25
CL\_INS\_237
CL\_INS\_247
CL\_INS\_25
CL\_INS\_237
CL\_INS\_25
CL\_INS\_25
CL\_INS\_382
CL\_INS\_25
CL\_INS\_237
CL\_INS\_237
CL\_INS\_237
CL\_INS\_237
CL\_INS\_25
CL\_INS\_25
CL\_INS\_25
CL\_INS\_237
CL\_INS\_237
CL\_INS\_237
CL\_INS\_237
CL\_INS\_237
CL\_INS\_382
CL\_INS\_382
CL\_INS\_382
CL\_INS\_117
CL\_INS\_247
CL\_INS\_247
CL\_INS\_382
CL\_INS\_382
CL\_INS\_247
CL\_INS\_247
CL\_INS\_247
CL\_INS\_247
CL\_INS\_110
CL\_INS\_382
CL\_INS\_382
CL\_INS\_382
CL\_INS\_368
CL\_INS\_368
CL\_INS\_55
CL\_INS\_55
CL\_INS\_55
CL\_INS\_159
CL\_INS\_159
CL\_INS\_382
CL\_INS\_382
CL\_INS\_382
CL\_INS\_382
CL\_INS\_99
CL\_INS\_159
CL\_INS\_237
CL\_INS\_247
CL\_INS\_382
CL\_INS\_382
CL\_INS\_382
CL\_INS\_382
CL\_INS\_382
CL\_INS\_382
CL\_INS\_382
CL\_INS\_382
CL\_INS\_382
CL\_INS\_382
CL\_INS\_382
CL\_INS\_382
CL\_INS\_247
CL\_INS\_382
CL\_INS\_382
CL\_INS\_382
CL\_INS\_382
CL\_INS\_382
CL\_INS\_25
CL\_INS\_25
CL\_INS\_382
CL\_INS\_25
CL\_INS\_237
CL\_INS\_237
CL\_INS\_25
CL\_INS\_25
CL\_INS\_25
CL\_INS\_237
CL\_INS\_25
CL\_INS\_25
CL\_INS\_237
CL\_INS\_237
CL\_INS\_237
CL\_INS\_237
CL\_INS\_237
CL\_INS\_237
CL\_INS\_237
CL\_INS\_237
CL\_INS\_237
CL\_INS\_25
CL\_INS\_25
CL\_INS\_237
CL\_INS\_25
CL\_INS\_25
CL\_INS\_25
CL\_INS\_382
CL\_INS\_382
CL\_INS\_237
CL\_INS\_237
CL\_INS\_237
CL\_INS\_237
CL\_INS\_237
CL\_INS\_237
CL\_INS\_237
CL\_INS\_237
CL\_INS\_25
CL\_INS\_25
CL\_INS\_123
CL\_INS\_237
CL\_INS\_237
CL\_INS\_237
CL\_INS\_237
CL\_INS\_237
CL\_INS\_237
CL\_INS\_237
Cluster ID


CL\_12927
CL\_20982
CL\_28856
CL\_8955
CL\_8956
CL\_8957
CL\_8958
CL\_8960
CL\_8961
CL\_24490
CL\_24489
CL\_24488
CL\_10039
CL\_24487
CL\_24486
CL\_24485
CL\_24484
CL\_21218
CL\_8090
CL\_8089
CL\_8088
CL\_8087
CL\_21217
CL\_8157
CL\_8084
CL\_21216
CL\_21215
CL\_21214
CL\_7002
CL\_21213
CL\_21212
CL\_21211
CL\_7001
CL\_21210
CL\_21209
CL\_8942
CL\_8941
CL\_8940
CL\_8939
CL\_8938
CL\_10028
CL\_10027
CL\_10026
CL\_2533
CL\_10025
CL\_261
CL\_35124
CL\_35123
CL\_6661
CL\_35122
CL\_35121
CL\_17699
CL\_35120
CL\_35119
CL\_35118
CL\_35117
CL\_263
CL\_5430
CL\_5429
CL\_2285
CL\_21208
CL\_2284
CL\_10270
CL\_19175
CL\_5428
CL\_5427
CL\_5426
CL\_5424
CL\_5423
CL\_10271
CL\_1093
CL\_1094
CL\_1095
CL\_1096
CL\_2280
CL\_21207
CL\_2279
CL\_2278
CL\_5422
CL\_5421
CL\_5420
CL\_17641
CL\_24572
CL\_5416
CL\_5415
CL\_5419
CL\_7339
CL\_7340
CL\_5807
CL\_5808
CL\_5414
CL\_5413
CL\_10272
CL\_5412
CL\_5411
CL\_1097
CL\_1098
CL\_1099
CL\_5410
CL\_5409
CL\_5408
CL\_5407
CL\_5406
CL\_5405
CL\_5404
CL\_5403
CL\_5402
CL\_5401
CL\_5400
CL\_5399
CL\_10273
CL\_5398
CL\_10274
CL\_36492
CL\_5396
CL\_5395
CL\_5394
CL\_5393
CL\_1100
CL\_1101
CL\_5392
CL\_1104
CL\_1103
CL\_2277
CL\_6520
CL\_4717
CL\_24473
CL\_10275
CL\_21206
CL\_10276
CL\_24472
CL\_9022
CL\_9021
CL\_10277
CL\_21205
CL\_21204
CL\_21203
CL\_231
CL\_232
CL\_233
CL\_9020
CL\_235
CL\_236
CL\_237
CL\_238
CL\_239
CL\_24471
CL\_2541
CL\_240
CL\_24483
CL\_24482
CL\_24481
CL\_24480
CL\_24479
CL\_24478
CL\_24477
CL\_24476
CL\_24475
CL\_6192
CL\_6147
CL\_6148
CL\_6201
CL\_6145
CL\_24474
CL\_241
CL\_9019
CL\_6282
CL\_6283
CL\_6284
CL\_6285
CL\_6286
CL\_6287
CL\_6288
CL\_6289
CL\_6290
CL\_6291
CL\_6292
CL\_6293
CL\_6294
CL\_6295
CL\_9018
CL\_9017
CL\_9609
CL\_20642
CL\_9608
CL\_9016
CL\_9015
CL\_9014
CL\_9013
CL\_9012
CL\_9011
CL\_245
CL\_246
CL\_247
CL\_257
CL\_258
CL\_259
CL\_4380
CL\_33358
CL\_32892
CL\_5296
CL\_10421
CL\_10422
CL\_10423
CL\_10382
CL\_10383
CL\_5601
CL\_23690
CL\_23689
CL\_23688
CL\_23686
CL\_34219
CL\_5692
CL\_5693
CL\_4381
CL\_26946
CL\_4382
CL\_4383
CL\_11990
CL\_4384
CL\_20870
CL\_20869
CL\_20868
CL\_20864
CL\_20863
CL\_17620
CL\_17619
CL\_17618
CL\_5013
CL\_20857
CL\_10605
CL\_20851
CL\_20852
CL\_20854
CL\_20853
CL\_20849
CL\_20848
CL\_20844
CL\_20842
CL\_27409
CL\_20821
CL\_7007
CL\_20840
CL\_10590
CL\_20837
CL\_20836
CL\_14193
CL\_27408
CL\_17162
CL\_17161
CL\_17160
CL\_17159
CL\_20835
CL\_20834
CL\_20833
CL\_7684
CL\_7685
CL\_7686
CL\_7687
CL\_10583
CL\_4972
CL\_4974
CL\_5662
CL\_5505
CL\_4233
CL\_4234
CL\_4235
CL\_4236
CL\_5045
CL\_5046
CL\_5047
CL\_5048
CL\_5049
CL\_7692
CL\_7691
CL\_8832
CL\_11816
CL\_11815
CL\_5028
CL\_5029
CL\_14147
CL\_5031
CL\_5032
CL\_19401
CL\_19402
CL\_5050
CL\_5051
CL\_5052
CL\_5053
CL\_11689
CL\_5054
CL\_4254
CL\_11835
CL\_4262
CL\_4263
CL\_5062
CL\_5063
CL\_5064
CL\_5065
CL\_5066
CL\_5067
CL\_5068
CL\_5069
CL\_5070
CL\_5071
CL\_5072
CL\_5073
CL\_5074
CL\_5075
CL\_20826
CL\_20825
CL\_5077
CL\_20824
CL\_17167
CL\_17168
CL\_23536
CL\_20822
CL\_31629
CL\_20820
CL\_20819
CL\_20818
CL\_10591
CL\_20817
CL\_20951
CL\_20815
CL\_10785
CL\_20949
CL\_20814
CL\_20813
CL\_10782
CL\_20812
CL\_20811
CL\_20810
CL\_20809
CL\_20808
CL\_20807
CL\_5093
CL\_5094
CL\_20806
CL\_20804
CL\_14935
CL\_10770
CL\_10769
CL\_14899
CL\_20803
CL\_20802
CL\_20801
CL\_20800
CL\_6957
CL\_20799
CL\_20798
CL\_20797
CL\_20796
CL\_20948
CL\_20795
CL\_20947
