## Supplementary material for "A novel method for integrating genomic and Tn-Seq data to identify common *in vivo* fitness mechanisms across multiple bacterial species": S1 Dataset: CL_INS_28.html

Legend

 Mobile +extrachromosomalelementfunctions
 Hypothetical
 All EssentialGenes
 Other
 All VFDB Genes

FULL


WINDOWSVGPNG

Trim RowsRemove SingletonsSave Fasta

CL\_305


CL\_305


CL\_305


CL\_304


CL\_305


CL\_301


CL\_305


CL\_302


CL\_305


CL\_305


CL\_305


CL\_305


CL\_301


CL\_305


CL\_305

HighlightSelectShow Genomes


131

CL\_306


105

CL\_306


5

CL\_306


3

CL\_306


2

CL\_306


2

CL\_306


2

CL\_306


1

CL\_306


1

CL\_306


1

CL\_308


1

CL\_306


1

CL\_306


1

CL\_306


1

CL\_306


1

CL\_308

fGI ID


CL\_INS\_28
CL\_INS\_28
CL\_INS\_70
CL\_INS\_28
CL\_INS\_28
CL\_INS\_27
CL\_INS\_28
CL\_INS\_27
CL\_INS\_29
CL\_INS\_27
CL\_INS\_29
Cluster ID


CL\_10465
CL\_23646
CL\_7252
CL\_23950
CL\_23949
CL\_4385
CL\_10018
CL\_4386
CL\_4387
CL\_4388
CL\_4389
