## Supplementary material for "A novel method for integrating genomic and Tn-Seq data to identify common *in vivo* fitness mechanisms across multiple bacterial species": S1 Dataset: CL_INS_29.html

Legend

 Mobile +extrachromosomalelementfunctions
 Regulatoryfunctions
 Hypothetical
 All EssentialGenes
 Proteinsynthesis/fate
 Other
 Transport +binding proteins
 All VFDB Genes

FULL


WINDOWSVGPNG

Trim RowsRemove SingletonsSave Fasta

CL\_306


CL\_306


CL\_306


CL\_306


CL\_306


CL\_306


CL\_230


CL\_305


CL\_305

HighlightSelectShow Genomes


177

CL\_308


25

CL\_308


4

CL\_309


2

CL\_308


2

CL\_308


1

CL\_308


1

CL\_308


1

CL\_308


1

CL\_308

fGI ID


CL\_INS\_27
CL\_INS\_29
CL\_INS\_27
CL\_INS\_29
CL\_INS\_29
CL\_INS\_29
CL\_INS\_29
CL\_INS\_20
CL\_INS\_20
CL\_INS\_20
CL\_INS\_20
CL\_INS\_20
CL\_INS\_20
CL\_INS\_20
CL\_INS\_207
CL\_INS\_20
CL\_INS\_20
CL\_INS\_20
CL\_INS\_207
CL\_INS\_207
CL\_INS\_20
CL\_INS\_30
CL\_INS\_20
CL\_INS\_20
CL\_INS\_20
CL\_INS\_20
CL\_INS\_247
CL\_INS\_247
CL\_INS\_247
CL\_INS\_20
CL\_INS\_20
CL\_INS\_20
CL\_INS\_20
CL\_INS\_20
CL\_INS\_20
CL\_INS\_20
CL\_INS\_20
CL\_INS\_20
CL\_INS\_20
CL\_INS\_20
CL\_INS\_20
CL\_INS\_20
CL\_INS\_20
CL\_INS\_20
CL\_INS\_20
CL\_INS\_20
CL\_INS\_149
CL\_INS\_20
CL\_INS\_20
CL\_INS\_20
CL\_INS\_20
CL\_INS\_20
CL\_INS\_20
CL\_INS\_20
CL\_INS\_20
Cluster ID


CL\_4386
CL\_4387
CL\_4388
CL\_4389
CL\_307
CL\_10278
CL\_28576
CL\_9024
CL\_8254
CL\_8252
CL\_8251
CL\_8250
CL\_8249
CL\_8248
CL\_8247
CL\_9023
CL\_9022
CL\_9021
CL\_232
CL\_233
CL\_9020
CL\_235
CL\_236
CL\_237
CL\_238
CL\_239
CL\_2541
CL\_240
CL\_241
CL\_9019
CL\_6282
CL\_6283
CL\_6284
CL\_6285
CL\_6286
CL\_6287
CL\_6288
CL\_6289
CL\_6290
CL\_6291
CL\_6292
CL\_6293
CL\_6294
CL\_6295
CL\_9018
CL\_9017
CL\_4093
CL\_9016
CL\_9015
CL\_9014
CL\_9013
CL\_9012
CL\_9011
CL\_245
CL\_246
