## Supplementary material for "A novel method for integrating genomic and Tn-Seq data to identify common *in vivo* fitness mechanisms across multiple bacterial species": S1 Dataset: CL_INS_33.html

Legend

 Mobile +extrachromosomalelementfunctions
 Hypothetical
 All EssentialGenes
 Other
 All VFDB Genes

FULL


WINDOWSVGPNG

Trim RowsRemove SingletonsSave Fasta

CL\_357


CL\_357


CL\_357


CL\_357


CL\_356


CL\_357


CL\_357


CL\_357

HighlightSelectShow Genomes


188

CL\_359


82

CL\_359


1

CL\_359


1

CL\_359


1

CL\_359


1

CL\_359


1

CL\_359


1

CL\_431

fGI ID


CL\_INS\_33
CL\_INS\_33
CL\_INS\_33
CL\_INS\_33
CL\_INS\_33
Cluster ID


CL\_7350
CL\_30934
CL\_21202
CL\_36063
CL\_358
