## Supplementary material for "A novel method for integrating genomic and Tn-Seq data to identify common *in vivo* fitness mechanisms across multiple bacterial species": S1 Dataset: CL_INS_34.html

Legend

 Hypothetical
 Other
 Fatty acid +phospholipidmetabolism
 All VFDB Genes

FULL


WINDOWSVGPNG

Trim RowsRemove SingletonsSave Fasta

CL\_379


CL\_379


CL\_378


CL\_379


CL\_378

HighlightSelectShow Genomes


220

CL\_380


46

CL\_380


1

CL\_380


1

CL\_380


1

CL\_380

fGI ID


CL\_INS\_34
CL\_INS\_34
Cluster ID


CL\_10467
CL\_30487
