## Supplementary material for "A novel method for integrating genomic and Tn-Seq data to identify common *in vivo* fitness mechanisms across multiple bacterial species": S1 Dataset: CL_INS_35.html

Legend

 Mobile +extrachromosomalelementfunctions
 Hypothetical
 All EssentialGenes
 Other
 All VFDB Genes

FULL


WINDOWSVGPNG

Trim RowsRemove SingletonsSave Fasta

CL\_403


CL\_403


CL\_403


CL\_403


CL\_403


CL\_403


CL\_402


CL\_403


CL\_403


CL\_403

HighlightSelectShow Genomes


197

CL\_404


66

CL\_404


8

CL\_404


1

CL\_404


1

CL\_404


1

CL\_404


1

CL\_404


1

CL\_405


1

CL\_409


1

CL\_404

fGI ID


CL\_INS\_35
CL\_INS\_35
CL\_INS\_35
CL\_INS\_35
CL\_INS\_35
CL\_INS\_35
CL\_INS\_35
CL\_INS\_35
CL\_INS\_35
Cluster ID


CL\_13059
CL\_17795
CL\_28855
CL\_28854
CL\_28853
CL\_4392
CL\_4393
CL\_22998
CL\_4394
