## Supplementary material for "A novel method for integrating genomic and Tn-Seq data to identify common *in vivo* fitness mechanisms across multiple bacterial species": S1 Dataset: CL_INS_36.html

Legend

 Hypothetical
 All EssentialGenes
 Other
 All VFDB Genes

FULL


WINDOWSVGPNG

Trim RowsRemove SingletonsSave Fasta

CL\_455


CL\_455


CL\_453


CL\_455


CL\_455

HighlightSelectShow Genomes


220

CL\_456


47

CL\_456


2

CL\_456


1

CL\_463


1

CL\_457

fGI ID


CL\_INS\_36
CL\_INS\_36
CL\_INS\_38
CL\_INS\_38
CL\_INS\_36
CL\_INS\_38
Cluster ID


CL\_7000
CL\_6999
CL\_10471
CL\_10473
CL\_7915
CL\_6996
