## Supplementary material for "A novel method for integrating genomic and Tn-Seq data to identify common *in vivo* fitness mechanisms across multiple bacterial species": S1 Dataset: CL_INS_38.html

Legend

 Mobile +extrachromosomalelementfunctions
 Regulatoryfunctions
 Hypothetical
 Other
 All VFDB Genes

FULL


WINDOWSVGPNG

Trim RowsRemove SingletonsSave Fasta

CL\_461


CL\_461


CL\_461


CL\_461


CL\_461


CL\_461


CL\_461


CL\_460


CL\_461

HighlightSelectShow Genomes


234

CL\_462


24

CL\_462


8

CL\_462


1

CL\_462


1

CL\_462


1

CL\_462


1

CL\_462


1

CL\_462


1

CL\_463

fGI ID


CL\_INS\_38
CL\_INS\_38
CL\_INS\_38
CL\_INS\_38
CL\_INS\_38
CL\_INS\_38
CL\_INS\_38
CL\_INS\_38
Cluster ID


CL\_6997
CL\_6996
CL\_30485
CL\_30930
CL\_10470
CL\_10471
CL\_10472
CL\_10473
