## Supplementary material for "A novel method for integrating genomic and Tn-Seq data to identify common *in vivo* fitness mechanisms across multiple bacterial species": S1 Dataset: CL_INS_39.html

Legend

 Mobile +extrachromosomalelementfunctions
 Regulatoryfunctions
 Hypothetical
 Other
 Centralintermediarymetabolism
 All VFDB Genes

FULL


WINDOWSVGPNG

Trim RowsRemove SingletonsSave Fasta

CL\_462


CL\_462


CL\_462


CL\_462


CL\_462


CL\_462


CL\_462


CL\_462


CL\_206


CL\_462


CL\_462


CL\_462


CL\_461


CL\_462


CL\_462


CL\_462


CL\_462


CL\_455

HighlightSelectShow Genomes


190

CL\_463


40

CL\_463


16

CL\_463


14

CL\_463


4

CL\_463


2

CL\_463


1

CL\_463


1

CL\_463


1

CL\_463


1

CL\_463


1

CL\_463


1

CL\_463


1

CL\_463


1

CL\_463


1

CL\_339


1

CL\_464


1

CL\_463


1

CL\_463

fGI ID


CL\_INS\_39
CL\_INS\_38
CL\_INS\_39
CL\_INS\_39
CL\_INS\_36
CL\_INS\_38
CL\_INS\_39
CL\_INS\_39
CL\_INS\_39
CL\_INS\_39
CL\_INS\_39
CL\_INS\_38
CL\_INS\_39
CL\_INS\_39
CL\_INS\_39
Cluster ID


CL\_30931
CL\_10473
CL\_13472
CL\_13777
CL\_7915
CL\_6997
CL\_11601
CL\_11602
CL\_30509
CL\_7923
CL\_30510
CL\_6996
CL\_30511
CL\_6995
CL\_21866
