## Supplementary material for "A novel method for integrating genomic and Tn-Seq data to identify common *in vivo* fitness mechanisms across multiple bacterial species": S1 Dataset: CL_INS_42.html

Legend

 Mobile +extrachromosomalelementfunctions
 Regulatoryfunctions
 Hypothetical
 DNA Metabolism
 AntibioticResistance
 All EssentialGenes
 All Fitness Genes
 Proteinsynthesis/fate
 Other
 Transport +binding proteins
 All VFDB Genes

FULL


WINDOWSVGPNG

Trim RowsRemove SingletonsSave Fasta

CL\_500


CL\_500


CL\_500


CL\_500


CL\_500


CL\_500


CL\_500


CL\_4516


CL\_500


CL\_500


CL\_500


CL\_500


CL\_4486


CL\_4516


CL\_500


CL\_500


CL\_500


CL\_500


CL\_500


CL\_500


CL\_4516


CL\_500


CL\_4516


CL\_4427


CL\_500


CL\_500


CL\_500


CL\_500


CL\_500


CL\_500


CL\_4519


CL\_500


CL\_500


CL\_500


CL\_4519


CL\_500


CL\_4519


CL\_500


CL\_500


CL\_4516


CL\_500


CL\_500


CL\_500


CL\_4516


CL\_500


CL\_4516


CL\_500


CL\_4516


CL\_4487


CL\_4519


CL\_500


CL\_500


CL\_500


CL\_500


CL\_4486


CL\_500


CL\_500


CL\_500


CL\_348


CL\_500


CL\_4519


CL\_500


CL\_500


CL\_500


CL\_500


CL\_4516


CL\_500


CL\_500


CL\_4519


CL\_500


CL\_500


CL\_4516


CL\_500


CL\_500


CL\_500


CL\_4487


CL\_500


CL\_500


CL\_500


CL\_500


CL\_4516


CL\_500


CL\_500


CL\_352


CL\_500


CL\_500


CL\_500


CL\_500


CL\_500


CL\_4516


CL\_4519


CL\_500


CL\_4516


CL\_500


CL\_500


Break


CL\_500


CL\_500


CL\_500


CL\_4519


CL\_500


CL\_500


CL\_500


CL\_4516


CL\_500


CL\_500


CL\_500


CL\_500


CL\_4516


CL\_500


CL\_500


CL\_4519


CL\_500


CL\_500


CL\_500


CL\_500


CL\_500


CL\_4519


CL\_500


CL\_500


CL\_4427


CL\_500


CL\_4519


CL\_500


CL\_500


CL\_500


CL\_500


CL\_4516


CL\_4516


CL\_4427


CL\_4516


CL\_500


CL\_500


CL\_500


CL\_500


CL\_500


CL\_500


CL\_500


CL\_500


CL\_500


CL\_500


CL\_500


CL\_500


CL\_500


CL\_4427


CL\_4516


CL\_4516


CL\_500


CL\_4486


CL\_4516


CL\_500


CL\_4516


CL\_4516


CL\_500


CL\_500


CL\_500


CL\_500


CL\_500


CL\_1084


CL\_500


CL\_500


CL\_500


CL\_500


CL\_4516


CL\_500


CL\_500


CL\_4487


CL\_500


CL\_500


Break


CL\_500


CL\_500


CL\_500


CL\_500


CL\_500


CL\_500


CL\_500


CL\_500


CL\_500


CL\_500


CL\_500


CL\_500


CL\_500


CL\_500


CL\_500


CL\_500


CL\_500


CL\_4516

HighlightSelectShow Genomes


63

CL\_538


20

CL\_538


20

CL\_538


16

CL\_538


5

CL\_538


5

CL\_538


4

CL\_538


4

CL\_538


3

CL\_538


3

CL\_541


3

CL\_538


3

CL\_538


2

CL\_538


2

CL\_538


2

CL\_538


2

CL\_538


2

CL\_4427


2

CL\_538


2

CL\_538


2

CL\_538


2

CL\_538


2

CL\_538


2

CL\_538


2

CL\_538


1

CL\_4427


1

CL\_4427


1

CL\_541


1

CL\_4427


1

CL\_538


1

CL\_4427


1

CL\_538


1

CL\_4427


1

CL\_538


1

CL\_4427


1

CL\_538


1

CL\_544


1

CL\_538


1

CL\_538


1

CL\_4427


1

CL\_538


1

CL\_4427


1

CL\_538


1

CL\_538


1

CL\_538


1

CL\_538


1

CL\_538


1

CL\_538


1

CL\_538


1

CL\_538


1

CL\_538


1

CL\_538


1

CL\_538


1

CL\_4427


1

CL\_4427


1

CL\_538


1

CL\_348


1

CL\_4427


1

CL\_541


1

CL\_538


1

CL\_541


1

CL\_538


1

CL\_4427


1

CL\_4427


1

CL\_4427


1

CL\_538


1

CL\_538


1

CL\_221


1

CL\_4427


1

CL\_538


1

CL\_538


1

CL\_4427


1

CL\_538


1

CL\_4427


1

CL\_4427


1

CL\_4519


1

CL\_538


1

CL\_4427


1

CL\_541


1

CL\_4427


1

CL\_4427


1

CL\_538


1

CL\_541


1

CL\_538


1

CL\_538


1

CL\_4427


1

CL\_4427


1

CL\_538


1

CL\_538


1

CL\_538


1

CL\_538


1

CL\_538


1

CL\_541


1

CL\_538


1

CL\_4519


1

CL\_538


1

CL\_538


1

CL\_538


1

CL\_4427


1

CL\_4427


1

CL\_538


1

CL\_4427


1

Break


1

CL\_538


1

CL\_538


1

CL\_538


1

CL\_4427


1

CL\_538


1

CL\_538


1

CL\_538


1

CL\_4427


1

CL\_4427


1

CL\_538


1

CL\_538


1

CL\_541


1

CL\_538


1

CL\_4427


1

CL\_538


1

CL\_538


1

CL\_4427


1

CL\_4427


1

CL\_538


1

CL\_4427


1

CL\_538


1

CL\_420


1

CL\_548


1

CL\_538


1

CL\_538


1

CL\_538


1

CL\_538


1

CL\_538


1

CL\_538


1

CL\_4427


1

CL\_4427


1

CL\_538


1

CL\_538


1

CL\_538


1

CL\_538


1

CL\_4487


1

CL\_618


1

CL\_538


1

CL\_544


1

CL\_538


1

CL\_4427


1

CL\_4427


1

CL\_538


1

CL\_538


1

CL\_538


1

CL\_4427


1

CL\_538


1

CL\_538


1

CL\_4427


1

CL\_538


1

CL\_538


1

CL\_4427


1

CL\_538


1

CL\_538


1

CL\_538


1

CL\_538


1

CL\_538


1

CL\_4427


1

CL\_4427


1

CL\_538


1

CL\_538


1

CL\_538


1

CL\_541


1

CL\_4427


1

CL\_538


1

CL\_4427


1

CL\_538


1

CL\_538


1

CL\_538


1

CL\_4427


1

CL\_538


1

CL\_538


1

CL\_4427


1

CL\_541


1

CL\_538


1

CL\_538


1

CL\_4427


1

CL\_538


1

CL\_538


1

CL\_538


1

CL\_1073


1

CL\_4486


1

CL\_4427


1

CL\_540


1

CL\_538


1

CL\_538

fGI ID


CL\_INS\_42
CL\_INS\_42
CL\_INS\_42
CL\_INS\_42
CL\_INS\_42
CL\_INS\_42
CL\_INS\_42
CL\_INS\_86
CL\_INS\_382
CL\_INS\_99
CL\_INS\_86
CL\_INS\_382
CL\_INS\_382
CL\_INS\_42
CL\_INS\_382
CL\_INS\_382
CL\_INS\_382
CL\_INS\_42
CL\_INS\_42
CL\_INS\_382
CL\_INS\_42
CL\_INS\_382
CL\_INS\_382
CL\_INS\_42
CL\_INS\_42
CL\_INS\_42
CL\_INS\_382
CL\_INS\_42
CL\_INS\_42
CL\_INS\_42
CL\_INS\_42
CL\_INS\_42
CL\_INS\_42
CL\_INS\_382
CL\_INS\_42
CL\_INS\_42
CL\_INS\_42
CL\_INS\_42
CL\_INS\_42
CL\_INS\_42
CL\_INS\_42
CL\_INS\_42
CL\_INS\_42
CL\_INS\_382
CL\_INS\_382
CL\_INS\_42
CL\_INS\_382
CL\_INS\_382
CL\_INS\_382
CL\_INS\_382
CL\_INS\_382
CL\_INS\_382
CL\_INS\_382
CL\_INS\_382
CL\_INS\_382
CL\_INS\_382
CL\_INS\_382
CL\_INS\_382
CL\_INS\_382
CL\_INS\_382
CL\_INS\_382
CL\_INS\_382
CL\_INS\_382
CL\_INS\_382
CL\_INS\_382
CL\_INS\_382
CL\_INS\_382
CL\_INS\_382
CL\_INS\_382
CL\_INS\_382
CL\_INS\_382
CL\_INS\_382
CL\_INS\_382
CL\_INS\_382
CL\_INS\_382
CL\_INS\_382
CL\_INS\_382
CL\_INS\_382
CL\_INS\_382
CL\_INS\_382
CL\_INS\_382
CL\_INS\_382
CL\_INS\_382
CL\_INS\_382
CL\_INS\_382
CL\_INS\_382
CL\_INS\_382
CL\_INS\_382
CL\_INS\_382
CL\_INS\_382
CL\_INS\_382
CL\_INS\_382
CL\_INS\_382
CL\_INS\_382
CL\_INS\_382
CL\_INS\_382
CL\_INS\_42
CL\_INS\_382
CL\_INS\_382
CL\_INS\_382
CL\_INS\_382
CL\_INS\_382
CL\_INS\_382
CL\_INS\_382
CL\_INS\_42
CL\_INS\_42
CL\_INS\_382
CL\_INS\_382
CL\_INS\_382
CL\_INS\_382
CL\_INS\_382
CL\_INS\_382
CL\_INS\_382
CL\_INS\_382
CL\_INS\_382
CL\_INS\_382
CL\_INS\_382
CL\_INS\_382
CL\_INS\_382
CL\_INS\_382
CL\_INS\_382
CL\_INS\_382
CL\_INS\_382
CL\_INS\_382
CL\_INS\_382
CL\_INS\_382
CL\_INS\_382
CL\_INS\_382
CL\_INS\_382
CL\_INS\_382
CL\_INS\_382
CL\_INS\_382
CL\_INS\_382
CL\_INS\_382
CL\_INS\_382
CL\_INS\_382
CL\_INS\_382
CL\_INS\_382
CL\_INS\_382
CL\_INS\_382
CL\_INS\_42
CL\_INS\_382
CL\_INS\_99
CL\_INS\_382
CL\_INS\_382
CL\_INS\_382
CL\_INS\_382
CL\_INS\_382
CL\_INS\_382
CL\_INS\_382
CL\_INS\_382
CL\_INS\_382
CL\_INS\_382
CL\_INS\_42
CL\_INS\_382
CL\_INS\_146
CL\_INS\_146
CL\_INS\_382
CL\_INS\_382
CL\_INS\_382
CL\_INS\_382
CL\_INS\_382
CL\_INS\_382
CL\_INS\_382
CL\_INS\_382
CL\_INS\_382
CL\_INS\_382
CL\_INS\_382
CL\_INS\_382
CL\_INS\_382
CL\_INS\_382
CL\_INS\_382
CL\_INS\_382
CL\_INS\_42
CL\_INS\_382
CL\_INS\_382
CL\_INS\_382
CL\_INS\_382
CL\_INS\_382
CL\_INS\_382
CL\_INS\_382
CL\_INS\_42
CL\_INS\_382
CL\_INS\_382
CL\_INS\_382
CL\_INS\_382
CL\_INS\_382
CL\_INS\_382
CL\_INS\_382
CL\_INS\_382
CL\_INS\_382
CL\_INS\_382
CL\_INS\_382
CL\_INS\_42
CL\_INS\_42
CL\_INS\_382
CL\_INS\_382
CL\_INS\_382
CL\_INS\_382
CL\_INS\_382
CL\_INS\_382
CL\_INS\_382
CL\_INS\_382
CL\_INS\_382
CL\_INS\_382
CL\_INS\_382
CL\_INS\_382
CL\_INS\_382
CL\_INS\_382
CL\_INS\_382
CL\_INS\_382
CL\_INS\_382
CL\_INS\_382
CL\_INS\_382
CL\_INS\_382
CL\_INS\_42
CL\_INS\_42
CL\_INS\_42
CL\_INS\_42
CL\_INS\_42
CL\_INS\_42
CL\_INS\_382
CL\_INS\_382
CL\_INS\_382
CL\_INS\_42
CL\_INS\_42
CL\_INS\_42
CL\_INS\_382
CL\_INS\_382
CL\_INS\_382
CL\_INS\_382
CL\_INS\_382
CL\_INS\_382
CL\_INS\_382
CL\_INS\_382
CL\_INS\_382
CL\_INS\_382
CL\_INS\_382
CL\_INS\_382
CL\_INS\_382
CL\_INS\_382
CL\_INS\_382
CL\_INS\_382
CL\_INS\_382
CL\_INS\_382
CL\_INS\_382
CL\_INS\_382
CL\_INS\_382
CL\_INS\_382
CL\_INS\_42
CL\_INS\_382
CL\_INS\_382
CL\_INS\_382
CL\_INS\_382
CL\_INS\_382
CL\_INS\_382
CL\_INS\_42
CL\_INS\_42
CL\_INS\_42
CL\_INS\_60
CL\_INS\_20
CL\_INS\_20
CL\_INS\_42
CL\_INS\_42
CL\_INS\_42
CL\_INS\_42
CL\_INS\_42
CL\_INS\_42
CL\_INS\_42
CL\_INS\_42
CL\_INS\_42
CL\_INS\_42
CL\_INS\_42
CL\_INS\_382
CL\_INS\_382
CL\_INS\_382
CL\_INS\_382
CL\_INS\_237
CL\_INS\_237
CL\_INS\_237
CL\_INS\_237
CL\_INS\_382
CL\_INS\_42
CL\_INS\_42
CL\_INS\_42
CL\_INS\_382
CL\_INS\_382
CL\_INS\_382
CL\_INS\_382
CL\_INS\_382
CL\_INS\_382
CL\_INS\_42
CL\_INS\_42
CL\_INS\_382
CL\_INS\_42
CL\_INS\_42
CL\_INS\_382
CL\_INS\_382
CL\_INS\_382
CL\_INS\_382
CL\_INS\_382
CL\_INS\_382
CL\_INS\_382
CL\_INS\_382
CL\_INS\_382
CL\_INS\_382
CL\_INS\_382
CL\_INS\_382
CL\_INS\_382
CL\_INS\_42
CL\_INS\_382
CL\_INS\_382
CL\_INS\_42
CL\_INS\_382
CL\_INS\_382
CL\_INS\_382
CL\_INS\_382
CL\_INS\_382
CL\_INS\_382
CL\_INS\_382
CL\_INS\_382
CL\_INS\_382
CL\_INS\_382
CL\_INS\_382
CL\_INS\_42
CL\_INS\_382
CL\_INS\_382
CL\_INS\_382
CL\_INS\_42
CL\_INS\_20
CL\_INS\_207
CL\_INS\_207
CL\_INS\_207
CL\_INS\_385
CL\_INS\_385
CL\_INS\_382
CL\_INS\_382
CL\_INS\_382
CL\_INS\_382
CL\_INS\_382
CL\_INS\_382
CL\_INS\_382
CL\_INS\_382
CL\_INS\_382
CL\_INS\_382
CL\_INS\_382
CL\_INS\_382
CL\_INS\_382
CL\_INS\_382
CL\_INS\_382
CL\_INS\_382
CL\_INS\_204
CL\_INS\_382
CL\_INS\_382
CL\_INS\_382
CL\_INS\_42
CL\_INS\_42
CL\_INS\_382
CL\_INS\_382
CL\_INS\_382
CL\_INS\_382
CL\_INS\_382
CL\_INS\_382
CL\_INS\_382
CL\_INS\_382
CL\_INS\_382
CL\_INS\_382
CL\_INS\_382
CL\_INS\_382
CL\_INS\_382
CL\_INS\_382
CL\_INS\_382
CL\_INS\_382
CL\_INS\_382
CL\_INS\_382
CL\_INS\_382
CL\_INS\_382
CL\_INS\_382
CL\_INS\_382
CL\_INS\_382
CL\_INS\_382
CL\_INS\_382
CL\_INS\_382
CL\_INS\_382
CL\_INS\_382
CL\_INS\_382
CL\_INS\_382
CL\_INS\_382
CL\_INS\_42
CL\_INS\_42
CL\_INS\_382
CL\_INS\_382
CL\_INS\_382
CL\_INS\_382
CL\_INS\_382
CL\_INS\_382
CL\_INS\_382
CL\_INS\_382
CL\_INS\_382
CL\_INS\_382
CL\_INS\_382
CL\_INS\_382
CL\_INS\_382
CL\_INS\_382
CL\_INS\_382
CL\_INS\_382
CL\_INS\_382
CL\_INS\_382
CL\_INS\_382
CL\_INS\_382
CL\_INS\_382
CL\_INS\_382
CL\_INS\_382
CL\_INS\_382
CL\_INS\_382
CL\_INS\_382
CL\_INS\_382
CL\_INS\_382
CL\_INS\_382
CL\_INS\_382
CL\_INS\_382
CL\_INS\_382
CL\_INS\_382
CL\_INS\_42
CL\_INS\_382
CL\_INS\_382
CL\_INS\_382
CL\_INS\_382
CL\_INS\_382
CL\_INS\_382
CL\_INS\_382
CL\_INS\_382
CL\_INS\_382
CL\_INS\_382
CL\_INS\_382
CL\_INS\_382
CL\_INS\_382
CL\_INS\_382
CL\_INS\_382
CL\_INS\_382
CL\_INS\_382
CL\_INS\_382
CL\_INS\_382
CL\_INS\_382
CL\_INS\_382
CL\_INS\_382
CL\_INS\_382
CL\_INS\_382
CL\_INS\_382
CL\_INS\_382
CL\_INS\_382
CL\_INS\_382
CL\_INS\_382
CL\_INS\_382
CL\_INS\_382
CL\_INS\_382
CL\_INS\_42
CL\_INS\_382
CL\_INS\_99
CL\_INS\_382
CL\_INS\_42
CL\_INS\_42
CL\_INS\_42
CL\_INS\_207
CL\_INS\_99
CL\_INS\_99
CL\_INS\_99
CL\_INS\_385
CL\_INS\_385
CL\_INS\_99
CL\_INS\_99
CL\_INS\_99
CL\_INS\_382
CL\_INS\_99
CL\_INS\_99
CL\_INS\_382
CL\_INS\_382
CL\_INS\_382
CL\_INS\_382
CL\_INS\_382
CL\_INS\_99
CL\_INS\_382
CL\_INS\_382
CL\_INS\_136
CL\_INS\_99
CL\_INS\_382
CL\_INS\_99
CL\_INS\_99
CL\_INS\_382
CL\_INS\_382
CL\_INS\_382
CL\_INS\_382
CL\_INS\_42
CL\_INS\_42
CL\_INS\_42
CL\_INS\_42
CL\_INS\_382
CL\_INS\_42
CL\_INS\_42
CL\_INS\_42
CL\_INS\_42
CL\_INS\_86
CL\_INS\_86
CL\_INS\_60
CL\_INS\_99
CL\_INS\_42
CL\_INS\_86
CL\_INS\_20
CL\_INS\_42
CL\_INS\_86
CL\_INS\_146
CL\_INS\_99
CL\_INS\_99
CL\_INS\_86
CL\_INS\_86
CL\_INS\_20
CL\_INS\_20
CL\_INS\_382
CL\_INS\_86
CL\_INS\_86
CL\_INS\_99
CL\_INS\_86
CL\_INS\_385
CL\_INS\_86
CL\_INS\_385
CL\_INS\_385
CL\_INS\_385
CL\_INS\_385
CL\_INS\_385
CL\_INS\_99
CL\_INS\_149
CL\_INS\_87
CL\_INS\_385
CL\_INS\_42
CL\_INS\_385
CL\_INS\_99
CL\_INS\_99
CL\_INS\_385
CL\_INS\_382
CL\_INS\_382
CL\_INS\_382
CL\_INS\_42
CL\_INS\_382
CL\_INS\_382
CL\_INS\_204
CL\_INS\_382
CL\_INS\_382
CL\_INS\_42
CL\_INS\_86
CL\_INS\_382
CL\_INS\_382
CL\_INS\_382
CL\_INS\_382
CL\_INS\_382
CL\_INS\_382
CL\_INS\_382
CL\_INS\_382
CL\_INS\_382
CL\_INS\_382
CL\_INS\_382
CL\_INS\_382
CL\_INS\_382
CL\_INS\_382
CL\_INS\_382
CL\_INS\_42
CL\_INS\_204
CL\_INS\_204
CL\_INS\_204
CL\_INS\_204
CL\_INS\_204
CL\_INS\_204
CL\_INS\_204
CL\_INS\_204
CL\_INS\_204
CL\_INS\_204
CL\_INS\_204
CL\_INS\_204
CL\_INS\_204
CL\_INS\_204
CL\_INS\_204
CL\_INS\_86
CL\_INS\_207
CL\_INS\_204
CL\_INS\_204
CL\_INS\_204
CL\_INS\_204
CL\_INS\_204
CL\_INS\_204
CL\_INS\_204
CL\_INS\_204
CL\_INS\_204
CL\_INS\_207
CL\_INS\_99
CL\_INS\_42
CL\_INS\_20
CL\_INS\_20
CL\_INS\_42
CL\_INS\_385
CL\_INS\_99
CL\_INS\_99
CL\_INS\_385
CL\_INS\_86
CL\_INS\_237
CL\_INS\_385
CL\_INS\_86
CL\_INS\_86
CL\_INS\_382
CL\_INS\_17
CL\_INS\_42
CL\_INS\_42
CL\_INS\_42
Cluster ID


CL\_21200
CL\_16949
CL\_6355
CL\_13915
CL\_7914
CL\_27129
CL\_13482
CL\_6782
CL\_4518
CL\_6747
CL\_4488
CL\_1089
CL\_5798
CL\_501
CL\_502
CL\_9956
CL\_503
CL\_26041
CL\_37545
CL\_504
CL\_14424
CL\_505
CL\_506
CL\_27128
CL\_29676
CL\_27636
CL\_507
CL\_19169
CL\_4435
CL\_13447
CL\_13446
CL\_6552
CL\_31732
CL\_13473
CL\_31731
CL\_31730
CL\_31729
CL\_31728
CL\_31727
CL\_31726
CL\_31725
CL\_31724
CL\_31723
CL\_5429
CL\_2285
CL\_35396
CL\_2284
CL\_10270
CL\_2283
CL\_2282
CL\_17633
CL\_17634
CL\_30576
CL\_30577
CL\_30578
CL\_16807
CL\_4658
CL\_30579
CL\_30580
CL\_30581
CL\_30582
CL\_30583
CL\_30584
CL\_2281
CL\_5427
CL\_5426
CL\_5425
CL\_8576
CL\_8577
CL\_6035
CL\_7543
CL\_7542
CL\_10175
CL\_5424
CL\_5423
CL\_17635
CL\_33135
CL\_8578
CL\_8579
CL\_8580
CL\_1093
CL\_1094
CL\_1095
CL\_1096
CL\_2280
CL\_37260
CL\_37261
CL\_33913
CL\_12066
CL\_12064
CL\_17640
CL\_12766
CL\_19355
CL\_7339
CL\_7541
CL\_7540
CL\_26558
CL\_26559
CL\_7119
CL\_7118
CL\_13636
CL\_7117
CL\_13638
CL\_12119
CL\_22645
CL\_22644
CL\_6036
CL\_6037
CL\_4656
CL\_13564
CL\_4655
CL\_2279
CL\_2278
CL\_5422
CL\_19351
CL\_19352
CL\_5421
CL\_33912
CL\_5420
CL\_5419
CL\_16010
CL\_10168
CL\_16967
CL\_7539
CL\_28352
CL\_13562
CL\_12762
CL\_12760
CL\_12996
CL\_8185
CL\_8184
CL\_12797
CL\_12383
CL\_13524
CL\_9098
CL\_4534
CL\_27281
CL\_30585
CL\_30586
CL\_27280
CL\_22643
CL\_7537
CL\_8183
CL\_7534
CL\_7533
CL\_9123
CL\_15776
CL\_15777
CL\_15778
CL\_15779
CL\_15780
CL\_15781
CL\_4395
CL\_26555
CL\_20109
CL\_26556
CL\_26557
CL\_17794
CL\_4396
CL\_4397
CL\_4398
CL\_4399
CL\_7033
CL\_9953
CL\_13792
CL\_13793
CL\_13794
CL\_5364
CL\_5363
CL\_7032
CL\_5799
CL\_8277
CL\_7821
CL\_20956
CL\_13513
CL\_9955
CL\_8276
CL\_17254
CL\_11963
CL\_10337
CL\_5362
CL\_35839
CL\_5361
CL\_4400
CL\_508
CL\_4401
CL\_5360
CL\_5359
CL\_5358
CL\_5357
CL\_6793
CL\_5356
CL\_8326
CL\_28276
CL\_28277
CL\_6648
CL\_9226
CL\_12059
CL\_6792
CL\_6791
CL\_6790
CL\_6789
CL\_34823
CL\_34822
CL\_6788
CL\_6787
CL\_6786
CL\_5351
CL\_4402
CL\_16844
CL\_16845
CL\_16846
CL\_5800
CL\_6483
CL\_23319
CL\_17545
CL\_31722
CL\_31721
CL\_31720
CL\_31719
CL\_31718
CL\_16570
CL\_8207
CL\_6479
CL\_21867
CL\_21868
CL\_21869
CL\_4403
CL\_4404
CL\_4405
CL\_5801
CL\_26153
CL\_5802
CL\_7362
CL\_21697
CL\_13634
CL\_13635
CL\_4406
CL\_4407
CL\_4408
CL\_4409
CL\_10918
CL\_19805
CL\_29105
CL\_29106
CL\_21886
CL\_21887
CL\_7819
CL\_4410
CL\_31717
CL\_32626
CL\_32627
CL\_32628
CL\_32629
CL\_5350
CL\_5349
CL\_31716
CL\_27921
CL\_27922
CL\_7044
CL\_7045
CL\_7046
CL\_31715
CL\_31714
CL\_16351
CL\_31713
CL\_31712
CL\_31711
CL\_31710
CL\_31709
CL\_31708
CL\_31707
CL\_31706
CL\_5348
CL\_5347
CL\_6658
CL\_6659
CL\_6660
CL\_8319
CL\_9119
CL\_9120
CL\_9095
CL\_32136
CL\_32135
CL\_32134
CL\_5346
CL\_5345
CL\_5344
CL\_15461
CL\_19935
CL\_19934
CL\_21870
CL\_21871
CL\_509
CL\_510
CL\_26919
CL\_18305
CL\_18306
CL\_18307
CL\_7617
CL\_7616
CL\_8206
CL\_11962
CL\_511
CL\_32568
CL\_512
CL\_513
CL\_514
CL\_515
CL\_8469
CL\_4411
CL\_4412
CL\_22887
CL\_8205
CL\_8470
CL\_4086
CL\_11961
CL\_14227
CL\_516
CL\_517
CL\_518
CL\_519
CL\_520
CL\_521
CL\_523
CL\_522
CL\_6785
CL\_524
CL\_19806
CL\_1494
CL\_1493
CL\_1492
CL\_1491
CL\_34825
CL\_1328
CL\_526
CL\_10323
CL\_527
CL\_525
CL\_37120
CL\_4469
CL\_34444
CL\_6784
CL\_7048
CL\_9009
CL\_6783
CL\_15241
CL\_4651
CL\_20785
CL\_11937
CL\_11936
CL\_5924
CL\_6770
CL\_5809
CL\_528
CL\_29677
CL\_29678
CL\_7030
CL\_7818
CL\_7029
CL\_7028
CL\_530
CL\_4413
CL\_13474
CL\_13475
CL\_13476
CL\_13477
CL\_4414
CL\_8471
CL\_531
CL\_532
CL\_5343
CL\_5342
CL\_5341
CL\_6452
CL\_5340
CL\_12006
CL\_12009
CL\_8906
CL\_9944
CL\_13890
CL\_529
CL\_1497
CL\_15242
CL\_8581
CL\_8905
CL\_11960
CL\_8904
CL\_34130
CL\_21872
CL\_4415
CL\_32630
CL\_32631
CL\_32632
CL\_26152
CL\_8647
CL\_8648
CL\_37118
CL\_13510
CL\_8112
CL\_8113
CL\_8655
CL\_13639
CL\_16131
CL\_23647
CL\_12541
CL\_23764
CL\_13310
CL\_23765
CL\_8653
CL\_35423
CL\_16429
CL\_8114
CL\_12137
CL\_13640
CL\_13641
CL\_13642
CL\_13643
CL\_13737
CL\_13738
CL\_17067
CL\_8123
CL\_37117
CL\_35840
CL\_6461
CL\_30587
CL\_21006
CL\_4528
CL\_4527
CL\_4416
CL\_4530
CL\_4417
CL\_4529
CL\_4695
CL\_4418
CL\_13478
CL\_13479
CL\_11914
CL\_13480
CL\_26151
CL\_4419
CL\_8472
CL\_32567
CL\_32566
CL\_32565
CL\_4420
CL\_4526
CL\_4421
CL\_6741
CL\_4629
CL\_4628
CL\_30588
CL\_4523
CL\_11958
CL\_6744
CL\_11913
CL\_27637
CL\_11957
CL\_7112
CL\_4517
CL\_27638
CL\_27639
CL\_27640
CL\_11956
CL\_8586
CL\_8587
CL\_8588
CL\_25199
CL\_35841
CL\_4520
CL\_13789
CL\_13790
CL\_8712
CL\_4620
CL\_4485
CL\_4525
CL\_8582
CL\_8583
CL\_8584
CL\_4423
CL\_17290
CL\_4522
CL\_4521
CL\_4484
CL\_8155
CL\_4424
CL\_23650
CL\_23651
CL\_4425
CL\_4426
CL\_4422
CL\_8473
CL\_8474
CL\_32397
CL\_32396
CL\_21873
CL\_6743
CL\_21874
CL\_21875
CL\_21876
CL\_21877
CL\_4429
CL\_4430
CL\_9516
CL\_4431
CL\_21878
CL\_7025
CL\_10474
CL\_13483
CL\_4515
CL\_8166
CL\_10518
CL\_4489
CL\_1324
CL\_1495
CL\_10475
CL\_10476
CL\_4432
CL\_8585
CL\_10526
CL\_13526
CL\_7022
CL\_13791
CL\_4490
CL\_5810
CL\_5811
CL\_5812
CL\_5813
CL\_22642
CL\_4618
CL\_12560
CL\_4513
CL\_16571
CL\_17793
CL\_17792
CL\_7109
CL\_534
CL\_11286
CL\_5805
CL\_5806
CL\_20418
CL\_31430
CL\_5807
CL\_5808
CL\_5414
CL\_17747
CL\_1496
CL\_28578
CL\_7024
CL\_533
CL\_16567
CL\_17075
CL\_8216
CL\_4533
CL\_37119
CL\_4532
CL\_8903
CL\_8902
CL\_11959
CL\_8901
CL\_8900
CL\_8899
CL\_36064
CL\_4531
CL\_31429
CL\_5412
CL\_5411
CL\_1097
CL\_1098
CL\_1099
CL\_5410
CL\_5409
CL\_5408
CL\_5407
CL\_5406
CL\_5405
CL\_5404
CL\_5403
CL\_5402
CL\_5401
CL\_10273
CL\_11934
CL\_5396
CL\_5395
CL\_5394
CL\_5393
CL\_1100
CL\_1101
CL\_1102
CL\_1103
CL\_1104
CL\_5392
CL\_4433
CL\_4440
CL\_4434
CL\_16568
CL\_9053
CL\_22641
CL\_10342
CL\_8589
CL\_6993
CL\_535
CL\_536
CL\_15244
CL\_5814
CL\_5815
CL\_7557
CL\_537
CL\_15245
CL\_31428
CL\_31427
