## Supplementary material for "A novel method for integrating genomic and Tn-Seq data to identify common *in vivo* fitness mechanisms across multiple bacterial species": S1 Dataset: CL_INS_43.html

Legend

 Mobile +extrachromosomalelementfunctions
 Hypothetical
 DNA Metabolism
 All EssentialGenes
 AntibioticResistance
 Other
 Transport +binding proteins
 All VFDB Genes

FULL


WINDOWSVGPNG

Trim RowsRemove SingletonsSave Fasta

CL\_540


CL\_540


CL\_540


CL\_540


CL\_539


CL\_540


CL\_540


CL\_540


CL\_500


CL\_540


CL\_539


CL\_540


CL\_540


CL\_540


CL\_540


CL\_540


CL\_540


CL\_540


CL\_540


CL\_540


CL\_540


CL\_540


CL\_539


CL\_540


CL\_539


CL\_540


CL\_538


CL\_540


CL\_540


CL\_500


CL\_539


CL\_540


CL\_539


CL\_540


CL\_540


CL\_540


CL\_540


CL\_540


CL\_540


CL\_539


CL\_539


CL\_540


CL\_540


CL\_540


CL\_538


CL\_540


CL\_539


CL\_540


CL\_540


CL\_540


CL\_500


CL\_539


CL\_540


CL\_500


CL\_222


CL\_540


CL\_540


CL\_540


CL\_540


CL\_500


CL\_540


CL\_540


CL\_540


CL\_540


CL\_539


CL\_540


CL\_539


CL\_540


CL\_500


Break


CL\_539


CL\_540


CL\_540


CL\_539


CL\_539


CL\_540


CL\_540


CL\_539


CL\_540


CL\_500


CL\_540


CL\_500


CL\_540


CL\_540


CL\_538


CL\_540


CL\_540


CL\_500


CL\_500


CL\_540


CL\_540


CL\_540

HighlightSelectShow Genomes


102

CL\_541


15

CL\_541


8

CL\_541


6

CL\_541


5

CL\_541


4

CL\_541


4

CL\_541


3

CL\_541


3

CL\_541


3

CL\_541


3

CL\_541


3

CL\_541


3

CL\_541


2

CL\_541


2

CL\_541


2

CL\_541


2

CL\_541


2

CL\_541


2

CL\_541


2

CL\_541


2

CL\_541


2

CL\_541


2

CL\_541


1

CL\_541


1

CL\_541


1

CL\_541


1

CL\_541


1

CL\_541


1

CL\_541


1

CL\_541


1

CL\_541


1

CL\_541


1

CL\_541


1

CL\_541


1

CL\_541


1

CL\_541


1

CL\_541


1

CL\_541


1

CL\_541


1

CL\_541


1

CL\_541


1

CL\_541


1

CL\_541


1

CL\_541


1

CL\_541


1

CL\_541


1

CL\_541


1

CL\_541


1

CL\_541


1

CL\_541


1

CL\_541


1

CL\_541


1

CL\_541


1

CL\_541


1

CL\_541


1

Break


1

CL\_541


1

CL\_541


1

CL\_541


1

CL\_541


1

CL\_541


1

CL\_541


1

CL\_541


1

CL\_541


1

CL\_541


1

CL\_217


1

CL\_541


1

CL\_541


1

CL\_541


1

CL\_541


1

CL\_541


1

CL\_541


1

CL\_541


1

CL\_541


1

CL\_541


1

CL\_541


1

CL\_541


1

CL\_541


1

CL\_541


1

CL\_541


1

CL\_541


1

CL\_541


1

CL\_541


1

CL\_541


1

CL\_541


1

CL\_541


1

CL\_541


1

CL\_541


1

CL\_541


1

CL\_541


1

CL\_541


1

CL\_541

fGI ID


CL\_INS\_382
CL\_INS\_43
CL\_INS\_117
CL\_INS\_117
CL\_INS\_43
CL\_INS\_43
CL\_INS\_43
CL\_INS\_43
CL\_INS\_43
CL\_INS\_43
CL\_INS\_43
CL\_INS\_43
CL\_INS\_43
CL\_INS\_43
CL\_INS\_43
CL\_INS\_43
CL\_INS\_247
CL\_INS\_247
CL\_INS\_382
CL\_INS\_233
CL\_INS\_43
CL\_INS\_43
CL\_INS\_42
CL\_INS\_43
CL\_INS\_43
CL\_INS\_43
CL\_INS\_43
CL\_INS\_43
CL\_INS\_43
CL\_INS\_43
CL\_INS\_146
CL\_INS\_43
CL\_INS\_42
CL\_INS\_42
CL\_INS\_43
CL\_INS\_43
CL\_INS\_43
CL\_INS\_43
CL\_INS\_43
CL\_INS\_42
CL\_INS\_43
CL\_INS\_43
CL\_INS\_43
CL\_INS\_43
CL\_INS\_43
CL\_INS\_43
CL\_INS\_43
CL\_INS\_43
CL\_INS\_43
CL\_INS\_43
CL\_INS\_43
CL\_INS\_43
CL\_INS\_43
CL\_INS\_43
CL\_INS\_43
CL\_INS\_43
CL\_INS\_43
CL\_INS\_43
CL\_INS\_43
CL\_INS\_43
CL\_INS\_43
CL\_INS\_43
CL\_INS\_43
CL\_INS\_43
CL\_INS\_43
CL\_INS\_43
CL\_INS\_43
CL\_INS\_43
CL\_INS\_43
CL\_INS\_43
CL\_INS\_43
CL\_INS\_43
CL\_INS\_43
CL\_INS\_43
CL\_INS\_382
CL\_INS\_382
CL\_INS\_382
CL\_INS\_382
CL\_INS\_382
CL\_INS\_17
CL\_INS\_42
CL\_INS\_382
CL\_INS\_382
CL\_INS\_382
CL\_INS\_382
CL\_INS\_382
CL\_INS\_382
CL\_INS\_382
CL\_INS\_382
CL\_INS\_382
CL\_INS\_382
CL\_INS\_382
CL\_INS\_382
CL\_INS\_382
CL\_INS\_382
CL\_INS\_382
CL\_INS\_382
CL\_INS\_42
CL\_INS\_382
CL\_INS\_382
CL\_INS\_382
CL\_INS\_382
CL\_INS\_382
CL\_INS\_382
CL\_INS\_382
CL\_INS\_382
CL\_INS\_382
CL\_INS\_382
CL\_INS\_382
CL\_INS\_382
CL\_INS\_382
CL\_INS\_382
CL\_INS\_382
CL\_INS\_382
CL\_INS\_382
CL\_INS\_382
CL\_INS\_382
CL\_INS\_382
CL\_INS\_382
CL\_INS\_382
CL\_INS\_382
CL\_INS\_382
CL\_INS\_382
CL\_INS\_382
CL\_INS\_382
CL\_INS\_99
CL\_INS\_382
CL\_INS\_382
CL\_INS\_382
CL\_INS\_382
CL\_INS\_87
CL\_INS\_99
CL\_INS\_99
CL\_INS\_42
CL\_INS\_43
Cluster ID


CL\_9956
CL\_34821
CL\_4498
CL\_4497
CL\_27712
CL\_27711
CL\_27710
CL\_27709
CL\_15246
CL\_15247
CL\_20127
CL\_16572
CL\_15248
CL\_27345
CL\_27344
CL\_27343
CL\_5047
CL\_5046
CL\_13034
CL\_12657
CL\_14093
CL\_14094
CL\_4435
CL\_31626
CL\_17255
CL\_17256
CL\_4436
CL\_4437
CL\_12252
CL\_12251
CL\_5268
CL\_14425
CL\_13447
CL\_13446
CL\_8477
CL\_23456
CL\_8079
CL\_8078
CL\_23455
CL\_6552
CL\_6551
CL\_24112
CL\_8475
CL\_8476
CL\_16573
CL\_16574
CL\_27641
CL\_8478
CL\_4438
CL\_23454
CL\_15249
CL\_15250
CL\_15251
CL\_4439
CL\_32133
CL\_7352
CL\_7353
CL\_34131
CL\_11955
CL\_11954
CL\_11953
CL\_11952
CL\_11951
CL\_11950
CL\_11949
CL\_11948
CL\_34132
CL\_11947
CL\_11946
CL\_11945
CL\_11944
CL\_11943
CL\_11942
CL\_11941
CL\_502
CL\_503
CL\_504
CL\_505
CL\_506
CL\_537
CL\_19169
CL\_6479
CL\_4407
CL\_4408
CL\_4409
CL\_508
CL\_4401
CL\_5360
CL\_5359
CL\_5358
CL\_5357
CL\_507
CL\_4395
CL\_4396
CL\_5799
CL\_7821
CL\_9955
CL\_20956
CL\_511
CL\_512
CL\_513
CL\_514
CL\_515
CL\_516
CL\_517
CL\_518
CL\_519
CL\_520
CL\_521
CL\_5429
CL\_2285
CL\_2284
CL\_2283
CL\_2282
CL\_17634
CL\_6035
CL\_522
CL\_524
CL\_525
CL\_526
CL\_527
CL\_528
CL\_529
CL\_530
CL\_531
CL\_4431
CL\_1496
CL\_533
CL\_4414
CL\_4532
CL\_4513
CL\_534
CL\_4433
CL\_4440
CL\_4441
