## Supplementary material for "A novel method for integrating genomic and Tn-Seq data to identify common *in vivo* fitness mechanisms across multiple bacterial species": S1 Dataset: CL_INS_44.html

Legend

 Mobile +extrachromosomalelementfunctions
 Hypothetical
 AntibioticResistance
 Other
 All VFDB Genes

FULL


WINDOWSVGPNG

Trim RowsRemove SingletonsSave Fasta

CL\_552


CL\_552


CL\_552


CL\_553


CL\_552


CL\_552


CL\_553


CL\_564


CL\_552


CL\_552

HighlightSelectShow Genomes


264

CL\_551


4

CL\_1976


3

CL\_1976


2

CL\_551


1

CL\_549


1

CL\_549


1

CL\_551


1

CL\_551


1

CL\_1976


1

CL\_1976

fGI ID


CL\_INS\_44
CL\_INS\_44
CL\_INS\_247
CL\_INS\_247
CL\_INS\_247
CL\_INS\_247
CL\_INS\_44
CL\_INS\_247
CL\_INS\_44
CL\_INS\_247
CL\_INS\_247
CL\_INS\_247
CL\_INS\_60
CL\_INS\_247
CL\_INS\_123
CL\_INS\_44
CL\_INS\_44
CL\_INS\_44
CL\_INS\_44
CL\_INS\_44
CL\_INS\_44
CL\_INS\_44
CL\_INS\_44
CL\_INS\_44
CL\_INS\_44
CL\_INS\_44
CL\_INS\_247
CL\_INS\_44
CL\_INS\_247
CL\_INS\_247
CL\_INS\_123
CL\_INS\_123
CL\_INS\_44
CL\_INS\_247
CL\_INS\_44
CL\_INS\_44
CL\_INS\_44
CL\_INS\_44
CL\_INS\_44
CL\_INS\_44
CL\_INS\_44
CL\_INS\_44
CL\_INS\_44
CL\_INS\_44
CL\_INS\_44
CL\_INS\_44
Cluster ID


CL\_12281
CL\_22543
CL\_5601
CL\_10383
CL\_10382
CL\_10423
CL\_10422
CL\_10421
CL\_11134
CL\_10642
CL\_10641
CL\_10395
CL\_10394
CL\_10393
CL\_10392
CL\_22786
CL\_22787
CL\_22788
CL\_22789
CL\_22790
CL\_22791
CL\_22792
CL\_22793
CL\_22794
CL\_22795
CL\_22796
CL\_15522
CL\_22797
CL\_15523
CL\_10412
CL\_10715
CL\_5300
CL\_5518
CL\_10388
CL\_22798
CL\_22542
CL\_22541
CL\_22540
CL\_22539
CL\_22538
CL\_22537
CL\_22536
CL\_22535
CL\_22534
CL\_22533
CL\_22532
