## Supplementary material for "A novel method for integrating genomic and Tn-Seq data to identify common *in vivo* fitness mechanisms across multiple bacterial species": S1 Dataset: CL_INS_45.html

Legend

 Mobile +extrachromosomalelementfunctions
 Hypothetical
 Other
 All VFDB Genes

FULL


WINDOWSVGPNG

Trim RowsRemove SingletonsSave Fasta

CL\_552


CL\_552


CL\_552


CL\_552


CL\_552


CL\_551


CL\_552


CL\_552


CL\_552


CL\_552


CL\_551


CL\_344


CL\_552


CL\_552


CL\_552


CL\_552


CL\_552

HighlightSelectShow Genomes


167

CL\_553


39

CL\_553


25

CL\_553


20

CL\_553


16

CL\_553


2

CL\_553


2

CL\_555


1

CL\_553


1

CL\_553


1

CL\_553


1

CL\_553


1

CL\_553


1

CL\_553


1

CL\_555


1

CL\_348


1

CL\_555


1

CL\_553

fGI ID


CL\_INS\_45
CL\_INS\_45
CL\_INS\_45
CL\_INS\_45
CL\_INS\_45
CL\_INS\_45
CL\_INS\_45
CL\_INS\_45
CL\_INS\_159
CL\_INS\_159
CL\_INS\_159
CL\_INS\_159
CL\_INS\_110
Cluster ID


CL\_30928
CL\_26206
CL\_5374
CL\_19743
CL\_8204
CL\_6992
CL\_7354
CL\_27202
CL\_14113
CL\_23395
CL\_23394
CL\_23393
CL\_14120
