## Supplementary material for "A novel method for integrating genomic and Tn-Seq data to identify common *in vivo* fitness mechanisms across multiple bacterial species": S1 Dataset: CL_INS_47.html

Legend

 Mobile +extrachromosomalelementfunctions
 Hypothetical
 All EssentialGenes
 Other
 All VFDB Genes

FULL


WINDOWSVGPNG

Trim RowsRemove SingletonsSave Fasta

CL\_581


CL\_581


CL\_581


CL\_581


CL\_580


CL\_581


CL\_581

HighlightSelectShow Genomes


161

CL\_583


111

CL\_583


1

CL\_818


1

CL\_584


1

CL\_583


1

CL\_583


1

CL\_583

fGI ID


CL\_INS\_47
CL\_INS\_47
CL\_INS\_47
CL\_INS\_47
CL\_INS\_68
Cluster ID


CL\_582
CL\_12948
CL\_16577
CL\_30431
CL\_9938
