## Supplementary material for "A novel method for integrating genomic and Tn-Seq data to identify common *in vivo* fitness mechanisms across multiple bacterial species": S1 Dataset: CL_INS_50.html

Legend

 Mobile +extrachromosomalelementfunctions
 Hypothetical
 All EssentialGenes
 Other
 All VFDB Genes

FULL


WINDOWSVGPNG

Trim RowsRemove SingletonsSave Fasta

CL\_626


CL\_626


CL\_626


CL\_626


CL\_625


CL\_626


CL\_626

HighlightSelectShow Genomes


139

CL\_628


96

CL\_628


34

CL\_628


1

CL\_628


1

CL\_628


1

CL\_628


1

CL\_628

fGI ID


CL\_INS\_50
CL\_INS\_50
CL\_INS\_50
Cluster ID


CL\_627
CL\_4445
CL\_5373
