## Supplementary material for "A novel method for integrating genomic and Tn-Seq data to identify common *in vivo* fitness mechanisms across multiple bacterial species": S1 Dataset: CL_INS_52.html

Legend

 Mobile +extrachromosomalelementfunctions
 Regulatoryfunctions
 Hypothetical
 Proteinsynthesis/fate
 Other
 Transport +binding proteins
 All VFDB Genes

FULL


WINDOWSVGPNG

Trim RowsRemove SingletonsSave Fasta

CL\_639


CL\_639


CL\_639


CL\_639


CL\_639


CL\_639


CL\_639


CL\_639


CL\_639


CL\_639


CL\_639

HighlightSelectShow Genomes


207

CL\_640


49

CL\_640


10

CL\_640


3

CL\_640


1

Break


1

CL\_640


1

CL\_640


1

CL\_640


1

CL\_640


1

CL\_640


1

CL\_640

fGI ID


CL\_INS\_52
CL\_INS\_52
CL\_INS\_52
CL\_INS\_52
CL\_INS\_52
CL\_INS\_52
CL\_INS\_52
CL\_INS\_52
CL\_INS\_52
CL\_INS\_52
CL\_INS\_52
Cluster ID


CL\_14360
CL\_9054
CL\_6991
CL\_30480
CL\_6990
CL\_7913
CL\_7912
CL\_6989
CL\_9055
CL\_7911
CL\_7910
