## Supplementary material for "A novel method for integrating genomic and Tn-Seq data to identify common *in vivo* fitness mechanisms across multiple bacterial species": S1 Dataset: CL_INS_53.html

Legend

 Mobile +extrachromosomalelementfunctions
 Regulatoryfunctions
 Hypothetical
 All EssentialGenes
 All Fitness Genes
 Transport +binding proteins
 All VFDB Genes

FULL


WINDOWSVGPNG

Trim RowsRemove SingletonsSave Fasta

CL\_649


CL\_649


CL\_649


CL\_649


CL\_649


CL\_649


CL\_649


CL\_649


CL\_648


CL\_649


CL\_649


CL\_649


CL\_649


CL\_649


CL\_649

HighlightSelectShow Genomes


168

CL\_651


69

CL\_651


12

CL\_654


3

CL\_651


2

CL\_651


2

CL\_654


1

CL\_656


1

CL\_651


1

CL\_651


1

CL\_652


1

CL\_651


1

CL\_651


1

CL\_652


1

CL\_654


1

CL\_652

fGI ID


CL\_INS\_53
CL\_INS\_53
CL\_INS\_53
CL\_INS\_53
CL\_INS\_53
Cluster ID


CL\_5372
CL\_26843
CL\_650
CL\_6548
CL\_20983
