## Supplementary material for "A novel method for integrating genomic and Tn-Seq data to identify common *in vivo* fitness mechanisms across multiple bacterial species": S1 Dataset: CL_INS_55.html

Legend

 Mobile +extrachromosomalelementfunctions
 Regulatoryfunctions
 Hypothetical
 DNA Metabolism
 All EssentialGenes
 AntibioticResistance
 Transcription
 All Fitness Genes
 Biosynthesis ofcofactors,prostheticgroups, +carriers
 Cell Envelope
 Proteinsynthesis/fate
 Other
 Cellularprocesses
 Transport +binding proteins
 All VFDB Genes

FULL


WINDOWSVGPNG

Trim RowsRemove SingletonsSave Fasta

CL\_660


CL\_660


CL\_660


CL\_660


CL\_660


CL\_660


CL\_660


CL\_660


CL\_660


CL\_660


CL\_660


CL\_660


CL\_660


CL\_660


CL\_660


CL\_660


CL\_660


CL\_660


CL\_660


CL\_660


CL\_660


CL\_660


CL\_660


CL\_660


CL\_660


CL\_660


CL\_660


CL\_660


CL\_660


CL\_660


CL\_660


CL\_660


CL\_660


CL\_660


CL\_660


CL\_660


CL\_660


CL\_660


CL\_660


CL\_660


CL\_660


CL\_660


CL\_660


CL\_660


CL\_660


CL\_660


CL\_660


CL\_660


CL\_660


CL\_660


CL\_660


CL\_660


CL\_660


CL\_660


CL\_660


CL\_660


CL\_787


CL\_660


CL\_660


Break


CL\_660


CL\_660


CL\_660


CL\_660


CL\_660


CL\_660


CL\_660


CL\_660


CL\_660


CL\_660


CL\_660

HighlightSelectShow Genomes


93

CL\_664


64

CL\_664


12

CL\_664


11

CL\_664


8

CL\_664


6

CL\_664


5

CL\_664


5

CL\_664


4

CL\_664


4

CL\_664


3

CL\_664


2

CL\_664


2

Break


2

CL\_664


2

CL\_664


2

CL\_664


2

CL\_664


2

CL\_664


2

CL\_664


1

CL\_664


1

CL\_664


1

CL\_664


1

CL\_664


1

CL\_664


1

CL\_664


1

Break


1

CL\_664


1

CL\_664


1

CL\_664


1

CL\_664


1

CL\_664


1

CL\_664


1

CL\_664


1

CL\_664


1

CL\_664


1

CL\_664


1

CL\_664


1

CL\_664


1

CL\_664


1

CL\_664


1

CL\_664


1

CL\_664


1

CL\_664


1

CL\_665


1

CL\_664


1

CL\_664


1

CL\_664


1

CL\_664


1

CL\_664


1

CL\_664


1

CL\_664


1

CL\_664


1

CL\_664


1

CL\_664


1

CL\_664


1

CL\_752


1

CL\_664


1

CL\_664


1

CL\_664


1

CL\_664


1

CL\_664


1

CL\_664


1

CL\_664


1

CL\_664


1

CL\_664


1

CL\_664


1

CL\_664


1

CL\_664


1

CL\_664


1

CL\_664


1

CL\_664

fGI ID


CL\_INS\_55
CL\_INS\_55
CL\_INS\_55
CL\_INS\_55
CL\_INS\_55
CL\_INS\_55
CL\_INS\_55
CL\_INS\_55
CL\_INS\_55
CL\_INS\_55
CL\_INS\_55
CL\_INS\_55
CL\_INS\_55
CL\_INS\_55
CL\_INS\_55
CL\_INS\_55
CL\_INS\_271
CL\_INS\_55
CL\_INS\_117
CL\_INS\_55
CL\_INS\_55
CL\_INS\_55
CL\_INS\_55
CL\_INS\_55
CL\_INS\_55
CL\_INS\_55
CL\_INS\_55
CL\_INS\_55
CL\_INS\_55
CL\_INS\_117
CL\_INS\_55
CL\_INS\_55
CL\_INS\_237
CL\_INS\_55
CL\_INS\_55
CL\_INS\_55
CL\_INS\_55
CL\_INS\_55
CL\_INS\_55
CL\_INS\_55
CL\_INS\_55
CL\_INS\_55
CL\_INS\_55
CL\_INS\_55
CL\_INS\_55
CL\_INS\_55
CL\_INS\_55
CL\_INS\_55
CL\_INS\_55
CL\_INS\_117
CL\_INS\_55
CL\_INS\_117
CL\_INS\_55
CL\_INS\_55
CL\_INS\_55
CL\_INS\_117
CL\_INS\_117
CL\_INS\_55
CL\_INS\_55
CL\_INS\_55
CL\_INS\_55
CL\_INS\_55
CL\_INS\_55
CL\_INS\_65
CL\_INS\_65
CL\_INS\_65
CL\_INS\_65
CL\_INS\_65
CL\_INS\_65
CL\_INS\_65
CL\_INS\_65
CL\_INS\_65
CL\_INS\_65
CL\_INS\_65
CL\_INS\_65
CL\_INS\_65
CL\_INS\_382
CL\_INS\_65
CL\_INS\_65
CL\_INS\_382
CL\_INS\_65
CL\_INS\_65
CL\_INS\_65
CL\_INS\_65
CL\_INS\_55
CL\_INS\_55
CL\_INS\_237
CL\_INS\_237
CL\_INS\_237
CL\_INS\_237
CL\_INS\_237
CL\_INS\_237
CL\_INS\_237
CL\_INS\_237
CL\_INS\_117
CL\_INS\_237
CL\_INS\_237
CL\_INS\_237
CL\_INS\_55
CL\_INS\_237
CL\_INS\_237
CL\_INS\_271
CL\_INS\_55
CL\_INS\_55
CL\_INS\_55
CL\_INS\_55
CL\_INS\_55
CL\_INS\_271
CL\_INS\_271
CL\_INS\_55
CL\_INS\_55
CL\_INS\_55
CL\_INS\_55
CL\_INS\_368
CL\_INS\_55
CL\_INS\_237
CL\_INS\_237
CL\_INS\_237
CL\_INS\_286
CL\_INS\_247
CL\_INS\_247
CL\_INS\_247
CL\_INS\_247
CL\_INS\_247
CL\_INS\_247
CL\_INS\_382
CL\_INS\_55
CL\_INS\_55
CL\_INS\_55
CL\_INS\_382
CL\_INS\_382
CL\_INS\_382
CL\_INS\_159
CL\_INS\_247
CL\_INS\_55
CL\_INS\_55
CL\_INS\_273
CL\_INS\_286
CL\_INS\_55
CL\_INS\_55
CL\_INS\_55
CL\_INS\_55
CL\_INS\_55
CL\_INS\_55
CL\_INS\_55
CL\_INS\_55
CL\_INS\_55
CL\_INS\_83
CL\_INS\_55
CL\_INS\_55
CL\_INS\_55
CL\_INS\_237
CL\_INS\_55
CL\_INS\_55
CL\_INS\_55
CL\_INS\_247
CL\_INS\_159
CL\_INS\_159
CL\_INS\_55
CL\_INS\_55
CL\_INS\_55
CL\_INS\_247
CL\_INS\_247
CL\_INS\_159
CL\_INS\_382
CL\_INS\_382
CL\_INS\_382
CL\_INS\_55
CL\_INS\_382
CL\_INS\_382
CL\_INS\_382
CL\_INS\_382
CL\_INS\_382
CL\_INS\_55
CL\_INS\_382
CL\_INS\_382
CL\_INS\_382
CL\_INS\_382
CL\_INS\_382
CL\_INS\_382
CL\_INS\_382
CL\_INS\_382
CL\_INS\_382
CL\_INS\_382
CL\_INS\_382
CL\_INS\_382
CL\_INS\_247
CL\_INS\_123
CL\_INS\_247
CL\_INS\_247
CL\_INS\_382
CL\_INS\_382
CL\_INS\_247
CL\_INS\_382
CL\_INS\_382
CL\_INS\_382
CL\_INS\_382
CL\_INS\_382
CL\_INS\_382
CL\_INS\_382
CL\_INS\_382
CL\_INS\_382
CL\_INS\_382
CL\_INS\_382
CL\_INS\_382
CL\_INS\_382
CL\_INS\_382
CL\_INS\_382
CL\_INS\_382
CL\_INS\_382
CL\_INS\_382
CL\_INS\_382
CL\_INS\_382
CL\_INS\_382
CL\_INS\_382
CL\_INS\_382
CL\_INS\_382
CL\_INS\_382
CL\_INS\_382
CL\_INS\_382
CL\_INS\_382
CL\_INS\_382
CL\_INS\_382
CL\_INS\_382
CL\_INS\_382
CL\_INS\_382
CL\_INS\_382
CL\_INS\_382
CL\_INS\_382
CL\_INS\_382
CL\_INS\_382
CL\_INS\_382
CL\_INS\_382
CL\_INS\_382
CL\_INS\_382
CL\_INS\_237
CL\_INS\_382
CL\_INS\_159
CL\_INS\_302
CL\_INS\_382
CL\_INS\_382
CL\_INS\_60
CL\_INS\_159
CL\_INS\_99
CL\_INS\_382
CL\_INS\_382
CL\_INS\_110
CL\_INS\_247
CL\_INS\_247
CL\_INS\_247
CL\_INS\_247
CL\_INS\_247
CL\_INS\_382
CL\_INS\_385
CL\_INS\_385
CL\_INS\_385
CL\_INS\_385
CL\_INS\_123
CL\_INS\_247
CL\_INS\_247
CL\_INS\_247
CL\_INS\_247
CL\_INS\_123
CL\_INS\_207
CL\_INS\_247
CL\_INS\_247
CL\_INS\_247
CL\_INS\_247
CL\_INS\_247
CL\_INS\_247
CL\_INS\_247
CL\_INS\_247
CL\_INS\_247
CL\_INS\_247
CL\_INS\_247
CL\_INS\_272
CL\_INS\_207
CL\_INS\_207
CL\_INS\_207
CL\_INS\_159
CL\_INS\_382
CL\_INS\_233
CL\_INS\_233
CL\_INS\_233
CL\_INS\_382
CL\_INS\_382
CL\_INS\_382
CL\_INS\_382
CL\_INS\_382
CL\_INS\_382
CL\_INS\_382
CL\_INS\_382
CL\_INS\_159
CL\_INS\_382
CL\_INS\_382
CL\_INS\_382
CL\_INS\_382
CL\_INS\_382
CL\_INS\_382
CL\_INS\_382
CL\_INS\_382
CL\_INS\_382
CL\_INS\_382
CL\_INS\_382
CL\_INS\_382
CL\_INS\_382
CL\_INS\_382
CL\_INS\_382
CL\_INS\_382
CL\_INS\_382
CL\_INS\_382
CL\_INS\_382
CL\_INS\_382
CL\_INS\_382
CL\_INS\_382
CL\_INS\_382
CL\_INS\_159
CL\_INS\_385
CL\_INS\_382
CL\_INS\_382
CL\_INS\_382
CL\_INS\_159
CL\_INS\_159
CL\_INS\_382
CL\_INS\_385
CL\_INS\_385
CL\_INS\_382
CL\_INS\_382
CL\_INS\_382
CL\_INS\_382
CL\_INS\_385
CL\_INS\_385
CL\_INS\_159
CL\_INS\_159
CL\_INS\_159
CL\_INS\_159
CL\_INS\_382
CL\_INS\_1
CL\_INS\_55
CL\_INS\_117
CL\_INS\_117
CL\_INS\_55
CL\_INS\_55
CL\_INS\_159
CL\_INS\_237
CL\_INS\_237
CL\_INS\_237
CL\_INS\_237
CL\_INS\_237
CL\_INS\_382
CL\_INS\_382
CL\_INS\_382
CL\_INS\_156
CL\_INS\_156
CL\_INS\_156
CL\_INS\_286
CL\_INS\_237
CL\_INS\_156
CL\_INS\_156
CL\_INS\_156
CL\_INS\_286
CL\_INS\_156
CL\_INS\_156
CL\_INS\_20
CL\_INS\_20
CL\_INS\_70
CL\_INS\_20
CL\_INS\_20
CL\_INS\_117
CL\_INS\_55
CL\_INS\_55
CL\_INS\_55
CL\_INS\_55
CL\_INS\_302
CL\_INS\_156
CL\_INS\_55
CL\_INS\_55
CL\_INS\_55
CL\_INS\_117
CL\_INS\_55
CL\_INS\_55
CL\_INS\_55
CL\_INS\_207
CL\_INS\_207
CL\_INS\_207
CL\_INS\_237
CL\_INS\_237
CL\_INS\_55
CL\_INS\_55
CL\_INS\_55
CL\_INS\_55
CL\_INS\_55
CL\_INS\_237
CL\_INS\_237
CL\_INS\_237
CL\_INS\_237
CL\_INS\_237
CL\_INS\_237
CL\_INS\_237
CL\_INS\_237
CL\_INS\_237
CL\_INS\_237
CL\_INS\_237
CL\_INS\_237
CL\_INS\_237
CL\_INS\_237
CL\_INS\_55
CL\_INS\_55
CL\_INS\_55
CL\_INS\_55
CL\_INS\_55
CL\_INS\_55
CL\_INS\_382
CL\_INS\_237
CL\_INS\_99
CL\_INS\_382
CL\_INS\_99
CL\_INS\_117
CL\_INS\_272
CL\_INS\_382
CL\_INS\_117
CL\_INS\_237
CL\_INS\_237
CL\_INS\_237
CL\_INS\_237
CL\_INS\_237
CL\_INS\_237
CL\_INS\_237
CL\_INS\_237
CL\_INS\_237
CL\_INS\_237
CL\_INS\_237
CL\_INS\_237
CL\_INS\_237
CL\_INS\_237
CL\_INS\_237
CL\_INS\_237
CL\_INS\_237
CL\_INS\_55
CL\_INS\_237
CL\_INS\_237
CL\_INS\_237
CL\_INS\_237
CL\_INS\_267
CL\_INS\_286
CL\_INS\_247
CL\_INS\_286
CL\_INS\_55
CL\_INS\_55
CL\_INS\_55
CL\_INS\_286
CL\_INS\_55
CL\_INS\_70
CL\_INS\_237
CL\_INS\_247
CL\_INS\_247
CL\_INS\_131
CL\_INS\_55
CL\_INS\_237
CL\_INS\_237
CL\_INS\_237
CL\_INS\_237
CL\_INS\_237
CL\_INS\_237
CL\_INS\_237
CL\_INS\_237
CL\_INS\_237
CL\_INS\_237
CL\_INS\_237
CL\_INS\_237
CL\_INS\_237
CL\_INS\_237
CL\_INS\_55
CL\_INS\_55
CL\_INS\_55
CL\_INS\_55
CL\_INS\_55
CL\_INS\_55
CL\_INS\_55
CL\_INS\_55
CL\_INS\_55
CL\_INS\_55
CL\_INS\_55
CL\_INS\_55
CL\_INS\_271
CL\_INS\_55
CL\_INS\_55
CL\_INS\_271
CL\_INS\_55
CL\_INS\_55
CL\_INS\_55
CL\_INS\_237
CL\_INS\_237
CL\_INS\_237
CL\_INS\_237
CL\_INS\_237
CL\_INS\_237
CL\_INS\_237
CL\_INS\_237
CL\_INS\_382
CL\_INS\_237
CL\_INS\_55
CL\_INS\_233
CL\_INS\_174
CL\_INS\_55
CL\_INS\_368
CL\_INS\_352
CL\_INS\_237
CL\_INS\_237
CL\_INS\_237
CL\_INS\_368
CL\_INS\_368
CL\_INS\_237
CL\_INS\_237
CL\_INS\_55
CL\_INS\_55
CL\_INS\_55
CL\_INS\_55
CL\_INS\_286
CL\_INS\_286
CL\_INS\_55
CL\_INS\_237
CL\_INS\_237
CL\_INS\_55
CL\_INS\_159
CL\_INS\_159
CL\_INS\_286
CL\_INS\_237
CL\_INS\_237
CL\_INS\_237
CL\_INS\_55
CL\_INS\_286
CL\_INS\_237
CL\_INS\_237
CL\_INS\_55
CL\_INS\_123
CL\_INS\_55
Cluster ID


CL\_19807
CL\_30479
CL\_30478
CL\_30477
CL\_10966
CL\_10477
CL\_10478
CL\_10479
CL\_17257
CL\_24311
CL\_24312
CL\_12287
CL\_12288
CL\_12286
CL\_11767
CL\_12289
CL\_3631
CL\_15474
CL\_11556
CL\_35424
CL\_7909
CL\_28579
CL\_7908
CL\_21879
CL\_21880
CL\_21881
CL\_21882
CL\_7907
CL\_31426
CL\_11555
CL\_13065
CL\_7906
CL\_7905
CL\_16093
CL\_19658
CL\_11587
CL\_11586
CL\_11585
CL\_11584
CL\_19657
CL\_11583
CL\_11582
CL\_11581
CL\_11580
CL\_19656
CL\_11579
CL\_11578
CL\_11577
CL\_11576
CL\_11575
CL\_16094
CL\_9176
CL\_16095
CL\_16096
CL\_16097
CL\_16098
CL\_7919
CL\_16099
CL\_16100
CL\_16101
CL\_35427
CL\_35426
CL\_35425
CL\_30467
CL\_30466
CL\_30465
CL\_30464
CL\_30463
CL\_30462
CL\_30461
CL\_30460
CL\_30459
CL\_30458
CL\_30457
CL\_30456
CL\_30455
CL\_16006
CL\_30454
CL\_30453
CL\_4547
CL\_30452
CL\_30451
CL\_30450
CL\_30449
CL\_30448
CL\_23935
CL\_11366
CL\_6138
CL\_6137
CL\_6136
CL\_11367
CL\_11368
CL\_11369
CL\_11370
CL\_16259
CL\_11371
CL\_11380
CL\_11381
CL\_34261
CL\_11383
CL\_11384
CL\_9538
CL\_9598
CL\_24294
CL\_6796
CL\_22991
CL\_9537
CL\_9536
CL\_9535
CL\_20461
CL\_20472
CL\_20473
CL\_20474
CL\_20475
CL\_20476
CL\_6761
CL\_6760
CL\_10419
CL\_5295
CL\_5691
CL\_5690
CL\_5689
CL\_5688
CL\_12106
CL\_5019
CL\_9917
CL\_10668
CL\_20477
CL\_10671
CL\_5001
CL\_5000
CL\_10401
CL\_5033
CL\_4233
CL\_13716
CL\_13717
CL\_13718
CL\_13719
CL\_20478
CL\_20479
CL\_20480
CL\_20481
CL\_20482
CL\_20483
CL\_20484
CL\_20485
CL\_20486
CL\_19485
CL\_20487
CL\_20488
CL\_20489
CL\_5697
CL\_20490
CL\_5040
CL\_5039
CL\_4234
CL\_5032
CL\_5031
CL\_14147
CL\_5029
CL\_5028
CL\_5018
CL\_5017
CL\_5506
CL\_5507
CL\_5508
CL\_4310
CL\_14159
CL\_5013
CL\_5123
CL\_5122
CL\_5121
CL\_5120
CL\_6953
CL\_5118
CL\_5117
CL\_5116
CL\_5115
CL\_5114
CL\_5113
CL\_5112
CL\_5111
CL\_5110
CL\_5109
CL\_5108
CL\_5107
CL\_5106
CL\_6957
CL\_6936
CL\_6937
CL\_5105
CL\_5104
CL\_5103
CL\_5102
CL\_5101
CL\_5100
CL\_5099
CL\_5098
CL\_5097
CL\_5096
CL\_5095
CL\_5094
CL\_5093
CL\_5092
CL\_5091
CL\_5090
CL\_5089
CL\_5088
CL\_5087
CL\_5086
CL\_5085
CL\_5084
CL\_5083
CL\_5082
CL\_5080
CL\_5079
CL\_19403
CL\_5078
CL\_5077
CL\_5076
CL\_5075
CL\_5074
CL\_5073
CL\_5072
CL\_5071
CL\_5069
CL\_5068
CL\_5066
CL\_5065
CL\_5064
CL\_5063
CL\_6942
CL\_5061
CL\_5627
CL\_5059
CL\_5058
CL\_4256
CL\_5620
CL\_5056
CL\_4254
CL\_4253
CL\_4252
CL\_5053
CL\_5052
CL\_5051
CL\_5050
CL\_5049
CL\_5048
CL\_5047
CL\_5046
CL\_5045
CL\_6946
CL\_9918
CL\_13406
CL\_13407
CL\_13408
CL\_5296
CL\_5300
CL\_5299
CL\_15544
CL\_15067
CL\_15545
CL\_10392
CL\_10314
CL\_10421
CL\_10423
CL\_10382
CL\_10383
CL\_5601
CL\_10385
CL\_10386
CL\_10387
CL\_10388
CL\_10389
CL\_10390
CL\_5615
CL\_5614
CL\_5613
CL\_5536
CL\_5539
CL\_10411
CL\_5542
CL\_5543
CL\_5544
CL\_4302
CL\_4301
CL\_4300
CL\_4299
CL\_5662
CL\_5548
CL\_4294
CL\_5549
CL\_5550
CL\_5551
CL\_5552
CL\_5553
CL\_5554
CL\_5555
CL\_5556
CL\_5557
CL\_5558
CL\_5559
CL\_4284
CL\_5560
CL\_5561
CL\_5562
CL\_5563
CL\_5564
CL\_5565
CL\_5566
CL\_5567
CL\_4278
CL\_4277
CL\_5639
CL\_5638
CL\_5637
CL\_5572
CL\_5573
CL\_5574
CL\_4271
CL\_4270
CL\_5575
CL\_5577
CL\_5579
CL\_13400
CL\_4266
CL\_4265
CL\_5062
CL\_4263
CL\_4262
CL\_5625
CL\_5584
CL\_4261
CL\_4260
CL\_4259
CL\_4258
CL\_4257
CL\_5590
CL\_5591
CL\_5592
CL\_5593
CL\_20462
CL\_6987
CL\_20463
CL\_5594
CL\_5595
CL\_5596
CL\_5597
CL\_4462
CL\_4974
CL\_4973
CL\_4972
CL\_5474
CL\_5475
CL\_5477
CL\_5480
CL\_6976
CL\_5481
CL\_5482
CL\_5483
CL\_5484
CL\_5486
CL\_5488
CL\_5491
CL\_5492
CL\_5493
CL\_5494
CL\_5495
CL\_5496
CL\_5497
CL\_5498
CL\_5499
CL\_5500
CL\_5501
CL\_5502
CL\_5503
CL\_5043
CL\_5504
CL\_5505
CL\_20464
CL\_20465
CL\_20466
CL\_5509
CL\_5510
CL\_5511
CL\_5512
CL\_5513
CL\_20467
CL\_20468
CL\_20469
CL\_20470
CL\_20471
CL\_9672
CL\_6195
CL\_6194
CL\_6193
CL\_6190
CL\_6189
CL\_6188
CL\_6187
CL\_6186
CL\_6183
CL\_6182
CL\_6181
CL\_11393
CL\_11394
CL\_26646
CL\_26647
CL\_11397
CL\_34260
CL\_34259
CL\_34258
CL\_5364
CL\_6639
CL\_33407
CL\_22329
CL\_11404
CL\_11407
CL\_6180
CL\_6177
CL\_6176
CL\_6175
CL\_6174
CL\_6173
CL\_6172
CL\_6171
CL\_11410
CL\_11411
CL\_11412
CL\_11413
CL\_11414
CL\_11415
CL\_11416
CL\_11417
CL\_11418
CL\_11419
CL\_11421
CL\_11422
CL\_15294
CL\_6163
CL\_11426
CL\_6144
CL\_6143
CL\_20491
CL\_11664
CL\_6844
CL\_11663
CL\_20492
CL\_20493
CL\_6168
CL\_20494
CL\_6305
CL\_6829
CL\_6830
CL\_8282
CL\_15541
CL\_11660
CL\_15295
CL\_11350
CL\_11351
CL\_11352
CL\_15298
CL\_23936
CL\_23937
CL\_11356
CL\_11357
CL\_11358
CL\_11359
CL\_11360
CL\_11361
CL\_11362
CL\_11363
CL\_18418
CL\_18419
CL\_18420
CL\_18421
CL\_18422
CL\_14210
CL\_14211
CL\_18416
CL\_14209
CL\_34262
CL\_661
CL\_12290
CL\_662
CL\_29799
CL\_29798
CL\_663
CL\_10316
CL\_28351
CL\_11661
CL\_11347
CL\_11346
CL\_11345
CL\_11344
CL\_11343
CL\_6139
CL\_6140
CL\_11342
CL\_6141
CL\_6142
CL\_20495
CL\_12653
CL\_9501
CL\_20496
CL\_6018
CL\_6028
CL\_6027
CL\_6026
CL\_6025
CL\_20497
CL\_20498
CL\_6024
CL\_6023
CL\_20499
CL\_20500
CL\_20501
CL\_20502
CL\_20503
CL\_20504
CL\_20505
CL\_6414
CL\_6415
CL\_20506
CL\_11665
CL\_11666
CL\_20507
CL\_11385
CL\_6135
CL\_6134
CL\_6133
CL\_11668
CL\_11388
CL\_6196
CL\_15292
CL\_4315
CL\_15293
