## Supplementary material for "A novel method for integrating genomic and Tn-Seq data to identify common *in vivo* fitness mechanisms across multiple bacterial species": S1 Dataset: CL_INS_58.html

Legend

 Mobile +extrachromosomalelementfunctions
 Regulatoryfunctions
 Hypothetical
 Biosynthesis ofcofactors,prostheticgroups, +carriers
 Other
 EnergyMetabolism
 All VFDB Genes
 Transport +binding proteins

FULL


WINDOWSVGPNG

Trim RowsRemove SingletonsSave Fasta

CL\_686


CL\_686


CL\_686


CL\_686


CL\_686


CL\_686


CL\_686


CL\_686


CL\_686


CL\_686


CL\_686


CL\_686


CL\_686


CL\_686


CL\_686


CL\_686


CL\_682


CL\_685


CL\_686


CL\_686


CL\_686


CL\_686


CL\_686


CL\_686


CL\_686


CL\_686


CL\_686


CL\_686


CL\_686


CL\_686


CL\_686


CL\_686


CL\_686


CL\_686


CL\_686


CL\_686


CL\_686


CL\_686


CL\_686


CL\_685


CL\_686


CL\_686


CL\_686


CL\_685


CL\_686


CL\_686

HighlightSelectShow Genomes


97

CL\_691


43

CL\_691


27

CL\_691


27

CL\_691


13

CL\_691


9

CL\_691


7

CL\_691


6

CL\_691


5

CL\_691


4

CL\_691


3

CL\_691


2

CL\_691


2

CL\_691


2

CL\_691


1

CL\_691


1

CL\_691


1

CL\_691


1

CL\_691


1

CL\_691


1

CL\_691


1

CL\_691


1

CL\_691


1

CL\_691


1

CL\_691


1

CL\_691


1

CL\_691


1

CL\_691


1

CL\_691


1

CL\_691


1

CL\_691


1

CL\_691


1

CL\_692


1

CL\_691


1

CL\_691


1

CL\_691


1

CL\_691


1

CL\_691


1

Break


1

Break


1

CL\_691


1

CL\_691


1

CL\_691


1

CL\_691


1

CL\_691


1

CL\_691


1

CL\_691

fGI ID


CL\_INS\_58
CL\_INS\_58
CL\_INS\_58
CL\_INS\_58
CL\_INS\_58
CL\_INS\_58
CL\_INS\_58
CL\_INS\_58
CL\_INS\_58
CL\_INS\_58
CL\_INS\_58
CL\_INS\_58
CL\_INS\_58
CL\_INS\_58
CL\_INS\_58
CL\_INS\_58
CL\_INS\_58
CL\_INS\_58
CL\_INS\_58
CL\_INS\_58
CL\_INS\_58
CL\_INS\_58
CL\_INS\_58
CL\_INS\_58
CL\_INS\_58
CL\_INS\_58
CL\_INS\_58
CL\_INS\_58
CL\_INS\_58
CL\_INS\_58
CL\_INS\_58
CL\_INS\_58
CL\_INS\_58
CL\_INS\_58
CL\_INS\_58
Cluster ID


CL\_30446
CL\_29940
CL\_29939
CL\_29938
CL\_11766
CL\_687
CL\_9056
CL\_688
CL\_37189
CL\_23171
CL\_689
CL\_7038
CL\_7037
CL\_690
CL\_11285
CL\_11284
CL\_11283
CL\_11282
CL\_11281
CL\_11280
CL\_11279
CL\_10480
CL\_13293
CL\_10481
CL\_10482
CL\_10483
CL\_10484
CL\_10485
CL\_10486
CL\_10487
CL\_10488
CL\_9057
CL\_15257
CL\_9058
CL\_9059
