## Supplementary material for "A novel method for integrating genomic and Tn-Seq data to identify common *in vivo* fitness mechanisms across multiple bacterial species": S1 Dataset: CL_INS_59.html

Legend

 Mobile +extrachromosomalelementfunctions
 Regulatoryfunctions
 Hypothetical
 DNA Metabolism
 All EssentialGenes
 Other
 Transport +binding proteins
 All VFDB Genes

FULL


WINDOWSVGPNG

Trim RowsRemove SingletonsSave Fasta

CL\_719


CL\_719


CL\_719


CL\_719


CL\_800


CL\_2151


CL\_591


CL\_719


CL\_788


CL\_738


CL\_719


CL\_719


CL\_719

HighlightSelectShow Genomes


266

CL\_718


1

CL\_850


1

CL\_718


1

CL\_704


1

CL\_718


1

CL\_718


1

CL\_718


1

CL\_740


1

CL\_718


1

CL\_718


1

CL\_753


1

CL\_2073


1

CL\_1788

fGI ID


CL\_INS\_59
CL\_INS\_61
CL\_INS\_59
CL\_INS\_59
CL\_INS\_59
CL\_INS\_59
CL\_INS\_144
CL\_INS\_146
CL\_INS\_59
CL\_INS\_59
CL\_INS\_128
CL\_INS\_108
CL\_INS\_65
CL\_INS\_65
CL\_INS\_65
CL\_INS\_382
CL\_INS\_61
CL\_INS\_61
CL\_INS\_61
CL\_INS\_61
CL\_INS\_72
CL\_INS\_72
CL\_INS\_59
CL\_INS\_59
CL\_INS\_59
CL\_INS\_72
CL\_INS\_72
CL\_INS\_59
CL\_INS\_59
CL\_INS\_59
CL\_INS\_59
CL\_INS\_382
CL\_INS\_108
CL\_INS\_108
CL\_INS\_108
CL\_INS\_108
CL\_INS\_108
CL\_INS\_59
CL\_INS\_59
CL\_INS\_59
CL\_INS\_59
CL\_INS\_59
CL\_INS\_59
CL\_INS\_59
CL\_INS\_59
CL\_INS\_59
CL\_INS\_60
CL\_INS\_108
CL\_INS\_60
CL\_INS\_60
CL\_INS\_60
CL\_INS\_59
CL\_INS\_204
CL\_INS\_61
CL\_INS\_204
CL\_INS\_204
CL\_INS\_204
CL\_INS\_61
CL\_INS\_61
CL\_INS\_61
CL\_INS\_204
CL\_INS\_61
CL\_INS\_204
CL\_INS\_204
CL\_INS\_204
CL\_INS\_204
CL\_INS\_204
CL\_INS\_204
CL\_INS\_204
CL\_INS\_61
CL\_INS\_61
CL\_INS\_61
CL\_INS\_108
CL\_INS\_59
CL\_INS\_59
CL\_INS\_59
CL\_INS\_204
CL\_INS\_204
CL\_INS\_204
CL\_INS\_204
CL\_INS\_204
CL\_INS\_65
CL\_INS\_65
CL\_INS\_382
CL\_INS\_65
CL\_INS\_106
CL\_INS\_65
CL\_INS\_204
CL\_INS\_204
CL\_INS\_61
CL\_INS\_382
CL\_INS\_136
CL\_INS\_382
CL\_INS\_382
CL\_INS\_136
CL\_INS\_382
CL\_INS\_382
CL\_INS\_72
CL\_INS\_72
CL\_INS\_72
CL\_INS\_382
CL\_INS\_72
CL\_INS\_382
CL\_INS\_382
CL\_INS\_382
CL\_INS\_382
CL\_INS\_59
CL\_INS\_59
CL\_INS\_59
CL\_INS\_59
CL\_INS\_59
CL\_INS\_207
CL\_INS\_170
CL\_INS\_207
CL\_INS\_207
CL\_INS\_207
CL\_INS\_207
CL\_INS\_204
CL\_INS\_204
CL\_INS\_207
CL\_INS\_207
CL\_INS\_207
CL\_INS\_207
CL\_INS\_207
CL\_INS\_207
CL\_INS\_207
CL\_INS\_170
CL\_INS\_170
CL\_INS\_207
CL\_INS\_207
CL\_INS\_66
CL\_INS\_66
CL\_INS\_207
CL\_INS\_207
CL\_INS\_66
CL\_INS\_66
CL\_INS\_66
CL\_INS\_66
CL\_INS\_66
CL\_INS\_66
CL\_INS\_66
CL\_INS\_66
CL\_INS\_382
CL\_INS\_382
CL\_INS\_382
CL\_INS\_382
CL\_INS\_382
CL\_INS\_382
CL\_INS\_65
CL\_INS\_59
CL\_INS\_382
CL\_INS\_59
CL\_INS\_382
CL\_INS\_65
CL\_INS\_382
CL\_INS\_382
CL\_INS\_65
CL\_INS\_382
CL\_INS\_382
CL\_INS\_382
CL\_INS\_65
CL\_INS\_86
CL\_INS\_61
CL\_INS\_61
CL\_INS\_61
CL\_INS\_59
Cluster ID


CL\_22977
CL\_4447
CL\_30824
CL\_30825
CL\_30826
CL\_30827
CL\_12417
CL\_12386
CL\_30174
CL\_30175
CL\_4511
CL\_28820
CL\_10490
CL\_30439
CL\_30440
CL\_12006
CL\_12304
CL\_22980
CL\_22981
CL\_22982
CL\_12969
CL\_13084
CL\_30473
CL\_30472
CL\_30471
CL\_13083
CL\_13082
CL\_12968
CL\_12967
CL\_12966
CL\_12965
CL\_525
CL\_12468
CL\_12467
CL\_12466
CL\_12465
CL\_12464
CL\_12964
CL\_12963
CL\_12962
CL\_12961
CL\_12960
CL\_12959
CL\_12958
CL\_12957
CL\_12956
CL\_12955
CL\_12461
CL\_12460
CL\_12459
CL\_12458
CL\_12954
CL\_12457
CL\_12527
CL\_5897
CL\_5898
CL\_5899
CL\_22983
CL\_22984
CL\_22985
CL\_5906
CL\_22986
CL\_5908
CL\_5909
CL\_5910
CL\_5911
CL\_5912
CL\_5913
CL\_5914
CL\_9221
CL\_22987
CL\_12455
CL\_12454
CL\_12953
CL\_12952
CL\_12303
CL\_5917
CL\_5918
CL\_5919
CL\_5920
CL\_5921
CL\_12471
CL\_30441
CL\_1317
CL\_30442
CL\_9942
CL\_12302
CL\_9222
CL\_5922
CL\_22988
CL\_5340
CL\_1318
CL\_526
CL\_10323
CL\_10525
CL\_4465
CL\_4464
CL\_12719
CL\_13080
CL\_13079
CL\_4463
CL\_12301
CL\_8162
CL\_10939
CL\_10514
CL\_7123
CL\_12300
CL\_12299
CL\_12298
CL\_12297
CL\_12296
CL\_6514
CL\_6515
CL\_6517
CL\_6518
CL\_6519
CL\_5202
CL\_6520
CL\_1105
CL\_6522
CL\_6523
CL\_6524
CL\_6525
CL\_6526
CL\_6527
CL\_6528
CL\_6530
CL\_6531
CL\_11569
CL\_8482
CL\_12987
CL\_12986
CL\_6534
CL\_6535
CL\_12985
CL\_12984
CL\_12983
CL\_12982
CL\_12981
CL\_12980
CL\_12979
CL\_12978
CL\_1311
CL\_12295
CL\_1523
CL\_11926
CL\_11255
CL\_11925
CL\_13081
CL\_12294
CL\_8671
CL\_30176
CL\_1308
CL\_12395
CL\_1306
CL\_5360
CL\_12293
CL\_1303
CL\_1522
CL\_1314
CL\_16182
CL\_8672
CL\_12977
CL\_22989
CL\_22990
CL\_12292
