## Supplementary material for "A novel method for integrating genomic and Tn-Seq data to identify common *in vivo* fitness mechanisms across multiple bacterial species": S1 Dataset: CL_INS_60.html

Legend

 Mobile +extrachromosomalelementfunctions
 Regulatoryfunctions
 Hypothetical
 DNA Metabolism
 AntibioticResistance
 All EssentialGenes
 All Fitness Genes
 Proteinsynthesis/fate
 Other
 Cellularprocesses
 Transport +binding proteins
 All VFDB Genes

FULL


WINDOWSVGPNG

Trim RowsRemove SingletonsSave Fasta

CL\_724


CL\_4516


CL\_4516


CL\_724


CL\_724


CL\_724


CL\_724


CL\_4486


CL\_724


CL\_4486


CL\_724


CL\_724


CL\_4519


CL\_724


CL\_724


CL\_724


CL\_724


CL\_724


CL\_724


CL\_724


Break


CL\_724


CL\_724


CL\_724


CL\_724


CL\_4516


Break


CL\_724


CL\_4519


CL\_724


CL\_724


CL\_724


CL\_724


CL\_724


CL\_724


CL\_4516


CL\_724


CL\_2008


CL\_724


CL\_724


CL\_724


CL\_724


CL\_724


CL\_4486


CL\_720


CL\_724


CL\_724


CL\_724


CL\_724


CL\_724


CL\_724


CL\_724


CL\_4516


CL\_724


CL\_724


CL\_4427


CL\_724


CL\_724


CL\_724


CL\_724


CL\_724


CL\_724


CL\_724


CL\_724


CL\_724


CL\_724


CL\_4486


CL\_724


CL\_724


CL\_724


CL\_724


CL\_724


CL\_724


CL\_724


CL\_724


CL\_4519


CL\_4487


CL\_4516


CL\_724


CL\_724


CL\_4519


CL\_724


CL\_724


Break


CL\_724


CL\_724


CL\_4516


CL\_724


CL\_724


CL\_790

HighlightSelectShow Genomes


204

CL\_725


5

CL\_725


3

CL\_725


2

CL\_725


2

CL\_725


2

CL\_725


1

CL\_4519


1

CL\_725


1

CL\_725


1

CL\_725


1

CL\_4427


1

CL\_725


1

CL\_725


1

CL\_725


1

CL\_725


1

CL\_725


1

CL\_725


1

CL\_4427


1

CL\_725


1

CL\_725


1

CL\_725


1

Break


1

CL\_2007


1

CL\_725


1

CL\_725


1

CL\_725


1

CL\_725


1

CL\_4427


1

CL\_725


1

CL\_4427


1

CL\_725


1

CL\_4427


1

CL\_725


1

CL\_725


1

CL\_725


1

CL\_725


1

CL\_725


1

CL\_725


1

CL\_4427


1

CL\_725


1

CL\_4427


1

Break


1

CL\_4427


1

CL\_725


1

CL\_725


1

CL\_725


1

CL\_4427


1

CL\_725


1

CL\_725


1

CL\_4427


1

CL\_4427


1

CL\_725


1

CL\_725


1

CL\_4427


1

CL\_725


1

CL\_725


1

CL\_4516


1

CL\_4427


1

CL\_725


1

CL\_725


1

CL\_725


1

CL\_725


1

CL\_725


1

CL\_4486


1

CL\_725


1

CL\_4427


1

CL\_725


1

CL\_725


1

CL\_725


1

CL\_4427


1

CL\_725


1

CL\_725


1

CL\_725


1

CL\_725


1

CL\_725


1

CL\_725


1

CL\_725


1

CL\_725


1

CL\_725


1

CL\_725


1

CL\_725


1

CL\_4486


1

CL\_725


1

CL\_725


1

CL\_854


1

CL\_4427


1

CL\_725


1

CL\_725


1

CL\_4427


1

CL\_725

fGI ID


CL\_INS\_60
CL\_INS\_60
CL\_INS\_146
CL\_INS\_86
CL\_INS\_60
CL\_INS\_60
CL\_INS\_60
CL\_INS\_60
CL\_INS\_60
CL\_INS\_60
CL\_INS\_60
CL\_INS\_60
CL\_INS\_60
CL\_INS\_60
CL\_INS\_60
CL\_INS\_60
CL\_INS\_60
CL\_INS\_60
CL\_INS\_60
CL\_INS\_60
CL\_INS\_60
CL\_INS\_60
CL\_INS\_60
CL\_INS\_60
CL\_INS\_60
CL\_INS\_60
CL\_INS\_60
CL\_INS\_60
CL\_INS\_60
CL\_INS\_60
CL\_INS\_60
CL\_INS\_60
CL\_INS\_60
CL\_INS\_60
CL\_INS\_60
CL\_INS\_60
CL\_INS\_60
CL\_INS\_60
CL\_INS\_60
CL\_INS\_60
CL\_INS\_60
CL\_INS\_60
CL\_INS\_60
CL\_INS\_60
CL\_INS\_382
CL\_INS\_382
CL\_INS\_382
CL\_INS\_60
CL\_INS\_60
CL\_INS\_60
CL\_INS\_60
CL\_INS\_382
CL\_INS\_60
CL\_INS\_99
CL\_INS\_99
CL\_INS\_60
CL\_INS\_382
CL\_INS\_382
CL\_INS\_382
CL\_INS\_382
CL\_INS\_382
CL\_INS\_382
CL\_INS\_382
CL\_INS\_382
CL\_INS\_382
CL\_INS\_382
CL\_INS\_382
CL\_INS\_382
CL\_INS\_382
CL\_INS\_382
CL\_INS\_382
CL\_INS\_382
CL\_INS\_382
CL\_INS\_382
CL\_INS\_382
CL\_INS\_382
CL\_INS\_382
CL\_INS\_382
CL\_INS\_382
CL\_INS\_382
CL\_INS\_382
CL\_INS\_382
CL\_INS\_382
CL\_INS\_382
CL\_INS\_382
CL\_INS\_382
CL\_INS\_204
CL\_INS\_60
CL\_INS\_60
CL\_INS\_60
CL\_INS\_60
CL\_INS\_60
CL\_INS\_60
CL\_INS\_60
CL\_INS\_60
CL\_INS\_382
CL\_INS\_382
CL\_INS\_382
CL\_INS\_382
CL\_INS\_382
CL\_INS\_60
CL\_INS\_382
CL\_INS\_382
CL\_INS\_60
CL\_INS\_382
CL\_INS\_382
CL\_INS\_382
CL\_INS\_382
CL\_INS\_382
CL\_INS\_382
CL\_INS\_60
CL\_INS\_382
CL\_INS\_60
CL\_INS\_382
CL\_INS\_237
CL\_INS\_204
CL\_INS\_204
CL\_INS\_204
CL\_INS\_382
CL\_INS\_382
CL\_INS\_60
CL\_INS\_382
CL\_INS\_382
CL\_INS\_382
CL\_INS\_60
CL\_INS\_382
CL\_INS\_146
CL\_INS\_60
CL\_INS\_382
CL\_INS\_60
CL\_INS\_60
CL\_INS\_60
CL\_INS\_382
CL\_INS\_382
CL\_INS\_60
CL\_INS\_382
CL\_INS\_382
CL\_INS\_60
CL\_INS\_382
CL\_INS\_60
CL\_INS\_60
CL\_INS\_382
CL\_INS\_60
CL\_INS\_382
CL\_INS\_60
CL\_INS\_60
CL\_INS\_382
CL\_INS\_382
CL\_INS\_382
CL\_INS\_60
CL\_INS\_382
CL\_INS\_60
CL\_INS\_60
CL\_INS\_60
CL\_INS\_382
CL\_INS\_382
CL\_INS\_382
CL\_INS\_382
CL\_INS\_382
CL\_INS\_60
CL\_INS\_60
CL\_INS\_382
CL\_INS\_382
CL\_INS\_382
CL\_INS\_382
CL\_INS\_382
CL\_INS\_60
CL\_INS\_60
CL\_INS\_60
CL\_INS\_382
CL\_INS\_382
CL\_INS\_382
CL\_INS\_60
CL\_INS\_382
CL\_INS\_382
CL\_INS\_382
CL\_INS\_382
CL\_INS\_382
CL\_INS\_382
CL\_INS\_382
CL\_INS\_60
CL\_INS\_382
CL\_INS\_204
CL\_INS\_382
CL\_INS\_382
CL\_INS\_60
CL\_INS\_60
CL\_INS\_382
CL\_INS\_382
CL\_INS\_382
CL\_INS\_382
CL\_INS\_60
CL\_INS\_382
CL\_INS\_60
CL\_INS\_60
CL\_INS\_60
CL\_INS\_60
CL\_INS\_60
CL\_INS\_60
CL\_INS\_204
CL\_INS\_60
CL\_INS\_382
CL\_INS\_60
CL\_INS\_382
CL\_INS\_382
CL\_INS\_382
CL\_INS\_60
CL\_INS\_382
CL\_INS\_382
CL\_INS\_20
CL\_INS\_20
CL\_INS\_60
CL\_INS\_382
CL\_INS\_382
CL\_INS\_382
CL\_INS\_382
CL\_INS\_382
CL\_INS\_382
CL\_INS\_382
CL\_INS\_382
CL\_INS\_382
CL\_INS\_382
CL\_INS\_60
CL\_INS\_60
CL\_INS\_146
CL\_INS\_382
CL\_INS\_382
CL\_INS\_382
CL\_INS\_20
CL\_INS\_20
CL\_INS\_60
CL\_INS\_60
CL\_INS\_382
CL\_INS\_382
CL\_INS\_382
CL\_INS\_382
CL\_INS\_382
CL\_INS\_382
CL\_INS\_382
CL\_INS\_382
CL\_INS\_237
CL\_INS\_382
CL\_INS\_382
CL\_INS\_382
CL\_INS\_382
CL\_INS\_382
CL\_INS\_382
CL\_INS\_382
CL\_INS\_382
CL\_INS\_382
CL\_INS\_60
CL\_INS\_60
CL\_INS\_60
CL\_INS\_60
CL\_INS\_60
CL\_INS\_207
CL\_INS\_207
CL\_INS\_207
CL\_INS\_237
CL\_INS\_382
CL\_INS\_382
CL\_INS\_20
CL\_INS\_20
CL\_INS\_382
CL\_INS\_382
CL\_INS\_60
CL\_INS\_382
CL\_INS\_60
CL\_INS\_60
CL\_INS\_60
CL\_INS\_60
CL\_INS\_60
CL\_INS\_60
CL\_INS\_60
CL\_INS\_60
CL\_INS\_60
CL\_INS\_60
CL\_INS\_60
CL\_INS\_60
CL\_INS\_60
CL\_INS\_60
CL\_INS\_60
CL\_INS\_60
CL\_INS\_60
CL\_INS\_60
CL\_INS\_60
CL\_INS\_60
CL\_INS\_60
CL\_INS\_60
CL\_INS\_60
CL\_INS\_60
CL\_INS\_60
CL\_INS\_60
CL\_INS\_204
CL\_INS\_204
CL\_INS\_204
CL\_INS\_60
CL\_INS\_20
CL\_INS\_20
CL\_INS\_204
CL\_INS\_204
CL\_INS\_204
CL\_INS\_204
CL\_INS\_204
CL\_INS\_204
CL\_INS\_204
CL\_INS\_204
CL\_INS\_204
CL\_INS\_204
CL\_INS\_204
CL\_INS\_204
CL\_INS\_204
CL\_INS\_204
CL\_INS\_204
CL\_INS\_86
CL\_INS\_207
CL\_INS\_60
CL\_INS\_60
CL\_INS\_207
CL\_INS\_204
CL\_INS\_204
CL\_INS\_204
CL\_INS\_204
CL\_INS\_204
CL\_INS\_204
CL\_INS\_204
CL\_INS\_204
CL\_INS\_204
CL\_INS\_20
CL\_INS\_60
CL\_INS\_60
CL\_INS\_204
CL\_INS\_60
CL\_INS\_60
CL\_INS\_20
CL\_INS\_20
CL\_INS\_60
CL\_INS\_382
CL\_INS\_60
CL\_INS\_106
CL\_INS\_204
CL\_INS\_204
CL\_INS\_204
CL\_INS\_204
CL\_INS\_204
CL\_INS\_204
CL\_INS\_204
CL\_INS\_204
CL\_INS\_204
CL\_INS\_204
CL\_INS\_204
CL\_INS\_204
CL\_INS\_204
CL\_INS\_204
CL\_INS\_204
CL\_INS\_204
CL\_INS\_60
CL\_INS\_60
CL\_INS\_204
CL\_INS\_204
CL\_INS\_204
CL\_INS\_204
CL\_INS\_204
CL\_INS\_204
CL\_INS\_60
CL\_INS\_149
CL\_INS\_60
CL\_INS\_237
CL\_INS\_60
CL\_INS\_237
CL\_INS\_237
CL\_INS\_60
CL\_INS\_60
CL\_INS\_60
CL\_INS\_60
CL\_INS\_237
CL\_INS\_382
CL\_INS\_60
CL\_INS\_60
CL\_INS\_382
CL\_INS\_237
CL\_INS\_20
CL\_INS\_20
CL\_INS\_20
CL\_INS\_60
CL\_INS\_60
CL\_INS\_20
CL\_INS\_20
CL\_INS\_20
CL\_INS\_20
CL\_INS\_20
CL\_INS\_20
CL\_INS\_20
CL\_INS\_20
CL\_INS\_60
CL\_INS\_60
CL\_INS\_60
CL\_INS\_60
CL\_INS\_60
CL\_INS\_20
CL\_INS\_20
CL\_INS\_60
CL\_INS\_60
CL\_INS\_60
CL\_INS\_60
CL\_INS\_20
CL\_INS\_20
CL\_INS\_20
CL\_INS\_60
CL\_INS\_60
CL\_INS\_60
CL\_INS\_60
CL\_INS\_237
CL\_INS\_60
CL\_INS\_60
CL\_INS\_60
CL\_INS\_60
CL\_INS\_60
CL\_INS\_60
CL\_INS\_60
CL\_INS\_60
CL\_INS\_60
CL\_INS\_382
CL\_INS\_382
CL\_INS\_382
CL\_INS\_382
CL\_INS\_382
CL\_INS\_382
CL\_INS\_136
CL\_INS\_99
CL\_INS\_60
CL\_INS\_60
CL\_INS\_86
CL\_INS\_86
CL\_INS\_99
CL\_INS\_382
CL\_INS\_382
CL\_INS\_382
CL\_INS\_382
CL\_INS\_382
CL\_INS\_106
CL\_INS\_20
CL\_INS\_20
CL\_INS\_60
CL\_INS\_60
CL\_INS\_146
CL\_INS\_146
CL\_INS\_382
CL\_INS\_382
CL\_INS\_382
CL\_INS\_382
CL\_INS\_382
CL\_INS\_382
CL\_INS\_382
CL\_INS\_382
CL\_INS\_382
CL\_INS\_382
CL\_INS\_382
CL\_INS\_382
CL\_INS\_382
CL\_INS\_60
CL\_INS\_382
CL\_INS\_382
CL\_INS\_382
CL\_INS\_60
CL\_INS\_382
CL\_INS\_382
CL\_INS\_382
CL\_INS\_382
CL\_INS\_382
CL\_INS\_382
CL\_INS\_382
CL\_INS\_60
CL\_INS\_60
CL\_INS\_60
CL\_INS\_382
CL\_INS\_382
CL\_INS\_382
CL\_INS\_382
CL\_INS\_382
CL\_INS\_382
CL\_INS\_382
CL\_INS\_382
CL\_INS\_382
CL\_INS\_382
CL\_INS\_382
CL\_INS\_382
CL\_INS\_382
CL\_INS\_382
CL\_INS\_382
CL\_INS\_382
CL\_INS\_382
CL\_INS\_382
CL\_INS\_382
CL\_INS\_382
CL\_INS\_382
CL\_INS\_60
CL\_INS\_382
CL\_INS\_60
CL\_INS\_60
CL\_INS\_60
CL\_INS\_382
CL\_INS\_60
CL\_INS\_382
CL\_INS\_382
CL\_INS\_20
CL\_INS\_382
CL\_INS\_382
CL\_INS\_382
CL\_INS\_382
CL\_INS\_382
CL\_INS\_382
CL\_INS\_382
CL\_INS\_60
CL\_INS\_60
CL\_INS\_382
CL\_INS\_60
CL\_INS\_382
CL\_INS\_382
CL\_INS\_382
CL\_INS\_382
CL\_INS\_382
CL\_INS\_382
CL\_INS\_382
CL\_INS\_382
CL\_INS\_382
CL\_INS\_237
CL\_INS\_149
CL\_INS\_170
CL\_INS\_86
CL\_INS\_382
CL\_INS\_382
CL\_INS\_60
CL\_INS\_60
CL\_INS\_60
CL\_INS\_382
CL\_INS\_382
CL\_INS\_60
CL\_INS\_60
CL\_INS\_60
CL\_INS\_60
CL\_INS\_382
CL\_INS\_60
CL\_INS\_60
CL\_INS\_60
CL\_INS\_60
CL\_INS\_60
CL\_INS\_60
CL\_INS\_60
CL\_INS\_60
CL\_INS\_60
CL\_INS\_382
CL\_INS\_382
CL\_INS\_60
CL\_INS\_60
CL\_INS\_60
CL\_INS\_382
CL\_INS\_99
CL\_INS\_86
CL\_INS\_60
CL\_INS\_385
CL\_INS\_60
CL\_INS\_385
CL\_INS\_385
CL\_INS\_385
CL\_INS\_60
CL\_INS\_99
CL\_INS\_382
CL\_INS\_86
CL\_INS\_382
CL\_INS\_99
CL\_INS\_86
CL\_INS\_99
CL\_INS\_87
CL\_INS\_20
CL\_INS\_20
CL\_INS\_20
CL\_INS\_20
CL\_INS\_60
CL\_INS\_86
CL\_INS\_86
CL\_INS\_99
CL\_INS\_60
CL\_INS\_60
CL\_INS\_99
CL\_INS\_99
CL\_INS\_99
CL\_INS\_86
CL\_INS\_20
CL\_INS\_60
CL\_INS\_60
CL\_INS\_99
CL\_INS\_60
CL\_INS\_60
CL\_INS\_382
CL\_INS\_60
CL\_INS\_60
CL\_INS\_60
CL\_INS\_60
CL\_INS\_60
CL\_INS\_60
CL\_INS\_60
CL\_INS\_382
CL\_INS\_20
CL\_INS\_382
CL\_INS\_20
CL\_INS\_20
CL\_INS\_60
CL\_INS\_60
CL\_INS\_20
CL\_INS\_382
CL\_INS\_382
CL\_INS\_382
CL\_INS\_382
CL\_INS\_382
CL\_INS\_382
CL\_INS\_382
CL\_INS\_382
CL\_INS\_382
CL\_INS\_382
CL\_INS\_382
CL\_INS\_382
CL\_INS\_382
CL\_INS\_382
CL\_INS\_382
CL\_INS\_382
CL\_INS\_382
CL\_INS\_382
CL\_INS\_382
CL\_INS\_382
CL\_INS\_382
CL\_INS\_60
CL\_INS\_207
CL\_INS\_382
CL\_INS\_207
CL\_INS\_207
CL\_INS\_207
CL\_INS\_207
CL\_INS\_207
CL\_INS\_382
CL\_INS\_382
CL\_INS\_207
CL\_INS\_207
CL\_INS\_207
CL\_INS\_207
CL\_INS\_99
CL\_INS\_382
CL\_INS\_382
CL\_INS\_382
CL\_INS\_382
CL\_INS\_382
CL\_INS\_382
CL\_INS\_382
CL\_INS\_136
CL\_INS\_136
CL\_INS\_136
CL\_INS\_10
CL\_INS\_99
CL\_INS\_136
CL\_INS\_136
CL\_INS\_136
CL\_INS\_136
CL\_INS\_136
CL\_INS\_87
CL\_INS\_87
CL\_INS\_87
CL\_INS\_136
CL\_INS\_136
CL\_INS\_136
CL\_INS\_136
CL\_INS\_136
CL\_INS\_136
CL\_INS\_136
CL\_INS\_60
CL\_INS\_382
CL\_INS\_382
CL\_INS\_20
CL\_INS\_382
CL\_INS\_382
CL\_INS\_382
CL\_INS\_382
CL\_INS\_382
CL\_INS\_382
CL\_INS\_382
CL\_INS\_382
CL\_INS\_382
CL\_INS\_382
CL\_INS\_382
CL\_INS\_99
CL\_INS\_99
CL\_INS\_60
CL\_INS\_86
CL\_INS\_385
CL\_INS\_382
CL\_INS\_86
CL\_INS\_382
CL\_INS\_60
CL\_INS\_382
CL\_INS\_382
CL\_INS\_382
CL\_INS\_382
CL\_INS\_382
CL\_INS\_382
CL\_INS\_382
CL\_INS\_382
CL\_INS\_382
CL\_INS\_382
CL\_INS\_382
CL\_INS\_382
CL\_INS\_382
CL\_INS\_60
CL\_INS\_60
CL\_INS\_60
CL\_INS\_382
CL\_INS\_382
CL\_INS\_60
CL\_INS\_60
CL\_INS\_60
CL\_INS\_60
CL\_INS\_382
CL\_INS\_382
CL\_INS\_382
CL\_INS\_382
CL\_INS\_382
CL\_INS\_382
CL\_INS\_382
CL\_INS\_382
CL\_INS\_382
CL\_INS\_382
CL\_INS\_382
CL\_INS\_382
CL\_INS\_382
CL\_INS\_382
CL\_INS\_382
CL\_INS\_382
CL\_INS\_382
CL\_INS\_382
CL\_INS\_382
CL\_INS\_382
CL\_INS\_382
CL\_INS\_382
CL\_INS\_382
CL\_INS\_382
CL\_INS\_382
CL\_INS\_382
CL\_INS\_382
CL\_INS\_382
CL\_INS\_382
CL\_INS\_382
CL\_INS\_382
CL\_INS\_382
CL\_INS\_382
CL\_INS\_382
CL\_INS\_382
CL\_INS\_382
CL\_INS\_382
CL\_INS\_382
CL\_INS\_382
CL\_INS\_382
CL\_INS\_382
CL\_INS\_99
CL\_INS\_99
CL\_INS\_382
CL\_INS\_382
CL\_INS\_99
CL\_INS\_86
CL\_INS\_382
CL\_INS\_382
CL\_INS\_382
CL\_INS\_60
CL\_INS\_385
CL\_INS\_385
CL\_INS\_60
CL\_INS\_60
CL\_INS\_60
CL\_INS\_60
CL\_INS\_385
CL\_INS\_60
CL\_INS\_99
CL\_INS\_385
CL\_INS\_385
CL\_INS\_60
CL\_INS\_382
CL\_INS\_382
CL\_INS\_382
CL\_INS\_60
CL\_INS\_382
CL\_INS\_382
CL\_INS\_382
CL\_INS\_382
CL\_INS\_382
CL\_INS\_382
CL\_INS\_382
CL\_INS\_382
CL\_INS\_382
CL\_INS\_382
CL\_INS\_382
CL\_INS\_382
CL\_INS\_382
CL\_INS\_382
CL\_INS\_382
CL\_INS\_382
CL\_INS\_60
CL\_INS\_60
CL\_INS\_237
CL\_INS\_237
CL\_INS\_60
CL\_INS\_60
CL\_INS\_60
CL\_INS\_60
CL\_INS\_60
CL\_INS\_60
CL\_INS\_60
CL\_INS\_60
CL\_INS\_237
CL\_INS\_237
CL\_INS\_237
CL\_INS\_20
CL\_INS\_60
CL\_INS\_60
CL\_INS\_20
CL\_INS\_60
CL\_INS\_60
CL\_INS\_60
CL\_INS\_20
CL\_INS\_20
CL\_INS\_20
CL\_INS\_20
CL\_INS\_20
CL\_INS\_60
CL\_INS\_382
CL\_INS\_382
CL\_INS\_382
CL\_INS\_382
CL\_INS\_60
CL\_INS\_60
CL\_INS\_60
CL\_INS\_60
CL\_INS\_60
CL\_INS\_20
CL\_INS\_20
CL\_INS\_20
CL\_INS\_60
CL\_INS\_237
CL\_INS\_60
CL\_INS\_237
CL\_INS\_60
CL\_INS\_60
CL\_INS\_60
CL\_INS\_60
CL\_INS\_60
CL\_INS\_60
CL\_INS\_60
CL\_INS\_60
CL\_INS\_60
CL\_INS\_60
CL\_INS\_60
CL\_INS\_60
CL\_INS\_60
CL\_INS\_60
CL\_INS\_60
CL\_INS\_60
CL\_INS\_60
CL\_INS\_60
CL\_INS\_60
CL\_INS\_60
CL\_INS\_60
CL\_INS\_60
CL\_INS\_60
CL\_INS\_60
CL\_INS\_60
CL\_INS\_60
CL\_INS\_60
CL\_INS\_60
CL\_INS\_60
CL\_INS\_60
CL\_INS\_60
CL\_INS\_60
CL\_INS\_60
CL\_INS\_60
CL\_INS\_60
CL\_INS\_60
CL\_INS\_60
CL\_INS\_60
CL\_INS\_60
CL\_INS\_60
CL\_INS\_60
CL\_INS\_60
CL\_INS\_60
CL\_INS\_60
CL\_INS\_60
CL\_INS\_60
CL\_INS\_60
CL\_INS\_60
CL\_INS\_60
CL\_INS\_60
CL\_INS\_60
CL\_INS\_60
CL\_INS\_60
CL\_INS\_60
CL\_INS\_60
CL\_INS\_60
CL\_INS\_60
CL\_INS\_60
CL\_INS\_60
CL\_INS\_60
CL\_INS\_60
CL\_INS\_60
CL\_INS\_60
CL\_INS\_60
CL\_INS\_60
CL\_INS\_60
CL\_INS\_60
CL\_INS\_60
CL\_INS\_60
CL\_INS\_60
CL\_INS\_60
CL\_INS\_60
CL\_INS\_60
CL\_INS\_60
CL\_INS\_60
CL\_INS\_60
CL\_INS\_60
CL\_INS\_60
CL\_INS\_60
CL\_INS\_60
CL\_INS\_60
CL\_INS\_60
CL\_INS\_60
CL\_INS\_60
CL\_INS\_60
CL\_INS\_60
CL\_INS\_60
CL\_INS\_60
CL\_INS\_60
CL\_INS\_60
CL\_INS\_60
CL\_INS\_60
CL\_INS\_60
CL\_INS\_60
CL\_INS\_60
CL\_INS\_60
CL\_INS\_20
CL\_INS\_20
CL\_INS\_60
CL\_INS\_60
CL\_INS\_60
CL\_INS\_60
CL\_INS\_60
CL\_INS\_60
CL\_INS\_60
CL\_INS\_60
CL\_INS\_60
CL\_INS\_60
CL\_INS\_60
CL\_INS\_60
CL\_INS\_60
CL\_INS\_60
CL\_INS\_60
CL\_INS\_60
CL\_INS\_60
CL\_INS\_60
CL\_INS\_60
CL\_INS\_60
CL\_INS\_60
CL\_INS\_60
CL\_INS\_60
CL\_INS\_60
CL\_INS\_60
CL\_INS\_60
CL\_INS\_60
CL\_INS\_60
CL\_INS\_60
CL\_INS\_60
CL\_INS\_60
CL\_INS\_60
CL\_INS\_60
CL\_INS\_60
CL\_INS\_60
CL\_INS\_60
CL\_INS\_60
CL\_INS\_60
CL\_INS\_60
CL\_INS\_60
CL\_INS\_60
CL\_INS\_60
CL\_INS\_60
CL\_INS\_247
CL\_INS\_60
CL\_INS\_247
CL\_INS\_123
CL\_INS\_123
CL\_INS\_247
CL\_INS\_123
CL\_INS\_207
CL\_INS\_207
CL\_INS\_207
CL\_INS\_60
CL\_INS\_159
CL\_INS\_159
CL\_INS\_60
CL\_INS\_382
CL\_INS\_382
CL\_INS\_60
CL\_INS\_60
CL\_INS\_60
CL\_INS\_60
CL\_INS\_382
CL\_INS\_382
CL\_INS\_382
CL\_INS\_382
CL\_INS\_382
CL\_INS\_382
CL\_INS\_382
CL\_INS\_382
CL\_INS\_382
CL\_INS\_382
CL\_INS\_382
CL\_INS\_382
CL\_INS\_60
CL\_INS\_60
CL\_INS\_60
CL\_INS\_382
CL\_INS\_382
CL\_INS\_385
CL\_INS\_385
CL\_INS\_385
CL\_INS\_159
CL\_INS\_385
CL\_INS\_385
CL\_INS\_382
CL\_INS\_382
CL\_INS\_382
CL\_INS\_382
CL\_INS\_60
CL\_INS\_60
CL\_INS\_159
CL\_INS\_382
CL\_INS\_382
CL\_INS\_60
CL\_INS\_382
CL\_INS\_382
CL\_INS\_382
CL\_INS\_382
CL\_INS\_382
CL\_INS\_382
CL\_INS\_382
CL\_INS\_382
CL\_INS\_60
CL\_INS\_382
CL\_INS\_382
CL\_INS\_60
CL\_INS\_382
CL\_INS\_382
CL\_INS\_382
CL\_INS\_382
CL\_INS\_382
CL\_INS\_382
CL\_INS\_60
CL\_INS\_159
CL\_INS\_60
CL\_INS\_159
CL\_INS\_159
CL\_INS\_159
CL\_INS\_385
CL\_INS\_159
CL\_INS\_382
CL\_INS\_382
CL\_INS\_99
CL\_INS\_99
CL\_INS\_382
CL\_INS\_237
CL\_INS\_382
CL\_INS\_60
CL\_INS\_60
CL\_INS\_382
CL\_INS\_237
CL\_INS\_60
CL\_INS\_159
CL\_INS\_159
CL\_INS\_60
CL\_INS\_382
CL\_INS\_382
CL\_INS\_382
CL\_INS\_60
CL\_INS\_60
CL\_INS\_60
CL\_INS\_149
CL\_INS\_159
CL\_INS\_159
CL\_INS\_159
CL\_INS\_60
CL\_INS\_60
CL\_INS\_60
CL\_INS\_60
CL\_INS\_60
CL\_INS\_60
CL\_INS\_60
CL\_INS\_60
CL\_INS\_60
CL\_INS\_237
CL\_INS\_237
CL\_INS\_60
CL\_INS\_60
CL\_INS\_60
CL\_INS\_60
CL\_INS\_60
CL\_INS\_60
CL\_INS\_60
CL\_INS\_60
CL\_INS\_60
CL\_INS\_60
CL\_INS\_60
CL\_INS\_60
CL\_INS\_60
CL\_INS\_247
CL\_INS\_60
CL\_INS\_247
CL\_INS\_60
CL\_INS\_60
CL\_INS\_60
CL\_INS\_60
CL\_INS\_60
CL\_INS\_60
CL\_INS\_60
CL\_INS\_60
CL\_INS\_60
CL\_INS\_60
CL\_INS\_60
CL\_INS\_60
CL\_INS\_60
CL\_INS\_60
CL\_INS\_60
CL\_INS\_60
CL\_INS\_60
CL\_INS\_60
CL\_INS\_60
CL\_INS\_60
CL\_INS\_60
CL\_INS\_60
CL\_INS\_60
CL\_INS\_60
CL\_INS\_60
CL\_INS\_60
CL\_INS\_60
CL\_INS\_60
CL\_INS\_60
CL\_INS\_60
CL\_INS\_60
CL\_INS\_60
CL\_INS\_60
CL\_INS\_60
CL\_INS\_60
CL\_INS\_60
CL\_INS\_60
CL\_INS\_60
CL\_INS\_60
CL\_INS\_60
CL\_INS\_60
CL\_INS\_60
CL\_INS\_60
CL\_INS\_60
CL\_INS\_60
CL\_INS\_60
CL\_INS\_60
CL\_INS\_60
CL\_INS\_60
CL\_INS\_60
Cluster ID


CL\_15259
CL\_9516
CL\_8166
CL\_7024
CL\_20748
CL\_20747
CL\_20746
CL\_20745
CL\_20744
CL\_20743
CL\_20742
CL\_20741
CL\_20740
CL\_20739
CL\_20738
CL\_20737
CL\_20736
CL\_20735
CL\_20734
CL\_20733
CL\_20732
CL\_9841
CL\_9842
CL\_9843
CL\_20731
CL\_20730
CL\_20729
CL\_20728
CL\_20727
CL\_20726
CL\_20725
CL\_20724
CL\_20723
CL\_33860
CL\_20985
CL\_27913
CL\_5369
CL\_13085
CL\_13086
CL\_13087
CL\_13088
CL\_852
CL\_13089
CL\_13090
CL\_37190
CL\_9060
CL\_9061
CL\_13485
CL\_13486
CL\_13487
CL\_13488
CL\_5368
CL\_19739
CL\_4489
CL\_13075
CL\_12328
CL\_13874
CL\_8335
CL\_7356
CL\_7357
CL\_7358
CL\_7359
CL\_7360
CL\_7361
CL\_11939
CL\_11938
CL\_2283
CL\_10270
CL\_2282
CL\_2281
CL\_5428
CL\_5427
CL\_5426
CL\_7543
CL\_7542
CL\_1095
CL\_1096
CL\_2280
CL\_2279
CL\_5422
CL\_5421
CL\_5420
CL\_5419
CL\_4651
CL\_11937
CL\_11936
CL\_5924
CL\_17755
CL\_17754
CL\_17753
CL\_17752
CL\_17751
CL\_17750
CL\_17749
CL\_17748
CL\_12766
CL\_1512
CL\_5343
CL\_5342
CL\_5341
CL\_10322
CL\_6452
CL\_5798
CL\_29797
CL\_5799
CL\_4397
CL\_4398
CL\_4399
CL\_13484
CL\_5367
CL\_25047
CL\_6794
CL\_7510
CL\_10317
CL\_11206
CL\_9223
CL\_9222
CL\_5922
CL\_10318
CL\_5366
CL\_16980
CL\_5365
CL\_7618
CL\_5364
CL\_23166
CL\_6638
CL\_12804
CL\_15107
CL\_18912
CL\_28167
CL\_28166
CL\_28165
CL\_29937
CL\_5363
CL\_16866
CL\_11916
CL\_8276
CL\_28164
CL\_10337
CL\_28163
CL\_28350
CL\_23065
CL\_28349
CL\_23064
CL\_33356
CL\_33355
CL\_5362
CL\_5361
CL\_4400
CL\_8278
CL\_508
CL\_34764
CL\_34763
CL\_16366
CL\_4401
CL\_4402
CL\_4403
CL\_13634
CL\_13635
CL\_10319
CL\_10320
CL\_7362
CL\_7363
CL\_7364
CL\_4406
CL\_6483
CL\_7509
CL\_7508
CL\_7507
CL\_16570
CL\_6479
CL\_4407
CL\_34847
CL\_23066
CL\_23067
CL\_4408
CL\_4409
CL\_5360
CL\_7031
CL\_5359
CL\_28737
CL\_5358
CL\_12788
CL\_5357
CL\_12059
CL\_28531
CL\_19885
CL\_11732
CL\_21331
CL\_13888
CL\_23179
CL\_11733
CL\_6793
CL\_29796
CL\_12955
CL\_27101
CL\_12460
CL\_12459
CL\_12458
CL\_12457
CL\_29795
CL\_20417
CL\_35115
CL\_5356
CL\_5355
CL\_5354
CL\_11734
CL\_5353
CL\_5352
CL\_9948
CL\_9947
CL\_9946
CL\_9082
CL\_6655
CL\_9083
CL\_9084
CL\_9085
CL\_9086
CL\_9087
CL\_9088
CL\_9089
CL\_6792
CL\_16585
CL\_16586
CL\_16587
CL\_9076
CL\_9077
CL\_37191
CL\_15105
CL\_15104
CL\_35114
CL\_35113
CL\_6791
CL\_6790
CL\_6788
CL\_6787
CL\_37192
CL\_37193
CL\_29936
CL\_8326
CL\_19187
CL\_6789
CL\_9078
CL\_9079
CL\_9080
CL\_9081
CL\_25380
CL\_11915
CL\_25379
CL\_25378
CL\_26140
CL\_6653
CL\_9116
CL\_27932
CL\_26139
CL\_26138
CL\_26137
CL\_26136
CL\_6660
CL\_6661
CL\_7119
CL\_6662
CL\_6663
CL\_6664
CL\_13890
CL\_36919
CL\_10323
CL\_6666
CL\_6667
CL\_6668
CL\_6669
CL\_6670
CL\_6671
CL\_6672
CL\_6673
CL\_6674
CL\_6675
CL\_6676
CL\_6677
CL\_6678
CL\_6679
CL\_6680
CL\_6681
CL\_6682
CL\_6683
CL\_6684
CL\_6685
CL\_6686
CL\_6687
CL\_15102
CL\_15101
CL\_15100
CL\_15099
CL\_5415
CL\_5414
CL\_10975
CL\_11935
CL\_14636
CL\_14637
CL\_5412
CL\_5411
CL\_1097
CL\_1098
CL\_1099
CL\_5410
CL\_5409
CL\_5408
CL\_5407
CL\_5406
CL\_5405
CL\_5404
CL\_5403
CL\_5402
CL\_5401
CL\_10273
CL\_11934
CL\_17745
CL\_17744
CL\_17743
CL\_5398
CL\_5396
CL\_5395
CL\_5394
CL\_5393
CL\_1100
CL\_1101
CL\_1102
CL\_6594
CL\_20958
CL\_20959
CL\_20960
CL\_1104
CL\_11933
CL\_17742
CL\_2273
CL\_2272
CL\_35112
CL\_9944
CL\_9943
CL\_9942
CL\_5921
CL\_5920
CL\_5919
CL\_5917
CL\_5916
CL\_5915
CL\_5914
CL\_5912
CL\_5911
CL\_5910
CL\_5909
CL\_5908
CL\_5907
CL\_5906
CL\_5905
CL\_5904
CL\_9941
CL\_9940
CL\_5900
CL\_5899
CL\_5898
CL\_5897
CL\_5896
CL\_1103
CL\_9939
CL\_5200
CL\_6454
CL\_8319
CL\_8966
CL\_9119
CL\_9120
CL\_28162
CL\_31705
CL\_31704
CL\_28161
CL\_9121
CL\_5340
CL\_22131
CL\_26135
CL\_5809
CL\_7049
CL\_7050
CL\_7051
CL\_8968
CL\_8317
CL\_8316
CL\_8969
CL\_8970
CL\_7052
CL\_7053
CL\_7054
CL\_7055
CL\_7056
CL\_7057
CL\_36498
CL\_13496
CL\_13497
CL\_7064
CL\_36499
CL\_7058
CL\_26262
CL\_36918
CL\_36917
CL\_13630
CL\_11738
CL\_7065
CL\_7066
CL\_7067
CL\_13498
CL\_14819
CL\_36916
CL\_26133
CL\_19186
CL\_26132
CL\_26131
CL\_32432
CL\_26130
CL\_26129
CL\_32433
CL\_26128
CL\_32434
CL\_32435
CL\_4628
CL\_6743
CL\_11958
CL\_6744
CL\_11913
CL\_8169
CL\_4484
CL\_8168
CL\_28345
CL\_28344
CL\_4429
CL\_4430
CL\_4431
CL\_17391
CL\_6648
CL\_34385
CL\_34384
CL\_9226
CL\_6650
CL\_8324
CL\_8323
CL\_36496
CL\_36497
CL\_11735
CL\_11736
CL\_6786
CL\_6651
CL\_29935
CL\_29934
CL\_29933
CL\_29932
CL\_29931
CL\_29930
CL\_29929
CL\_8206
CL\_26705
CL\_8552
CL\_5351
CL\_26150
CL\_4410
CL\_19805
CL\_509
CL\_10321
CL\_27708
CL\_7617
CL\_7616
CL\_23761
CL\_23762
CL\_511
CL\_512
CL\_14224
CL\_14225
CL\_14226
CL\_513
CL\_514
CL\_515
CL\_23763
CL\_4411
CL\_11961
CL\_14227
CL\_516
CL\_23068
CL\_23069
CL\_517
CL\_518
CL\_519
CL\_520
CL\_5350
CL\_7365
CL\_7366
CL\_15461
CL\_8205
CL\_25377
CL\_25376
CL\_6654
CL\_5349
CL\_9945
CL\_6656
CL\_6657
CL\_6658
CL\_28736
CL\_8965
CL\_6659
CL\_14825
CL\_17392
CL\_9090
CL\_9091
CL\_9092
CL\_9093
CL\_9094
CL\_9095
CL\_32895
CL\_32896
CL\_9096
CL\_32897
CL\_37194
CL\_4469
CL\_12009
CL\_9097
CL\_9009
CL\_529
CL\_530
CL\_9008
CL\_1497
CL\_4514
CL\_4601
CL\_4602
CL\_1495
CL\_8581
CL\_9098
CL\_33911
CL\_33910
CL\_33909
CL\_28347
CL\_25129
CL\_32643
CL\_32644
CL\_32645
CL\_32646
CL\_28529
CL\_33134
CL\_28735
CL\_33908
CL\_33907
CL\_33906
CL\_33905
CL\_33904
CL\_33903
CL\_33902
CL\_22576
CL\_4426
CL\_28346
CL\_33901
CL\_33900
CL\_11957
CL\_6747
CL\_4488
CL\_28732
CL\_9125
CL\_31703
CL\_9127
CL\_9128
CL\_28526
CL\_9129
CL\_7112
CL\_4517
CL\_7025
CL\_4432
CL\_4618
CL\_6782
CL\_10520
CL\_4513
CL\_6781
CL\_6780
CL\_6779
CL\_6778
CL\_6777
CL\_8585
CL\_10526
CL\_9159
CL\_19004
CL\_25891
CL\_8586
CL\_8587
CL\_8588
CL\_7022
CL\_7368
CL\_33899
CL\_17342
CL\_9160
CL\_17343
CL\_33898
CL\_4534
CL\_32898
CL\_32899
CL\_32900
CL\_32901
CL\_32902
CL\_32903
CL\_28530
CL\_5348
CL\_15103
CL\_5347
CL\_5380
CL\_7043
CL\_7044
CL\_16352
CL\_7045
CL\_5346
CL\_5345
CL\_5344
CL\_521
CL\_522
CL\_6785
CL\_25375
CL\_25374
CL\_25373
CL\_7539
CL\_7538
CL\_524
CL\_525
CL\_13636
CL\_27707
CL\_13638
CL\_7118
CL\_7117
CL\_7537
CL\_6784
CL\_11311
CL\_33354
CL\_9122
CL\_7533
CL\_7532
CL\_7531
CL\_7530
CL\_7529
CL\_7528
CL\_7527
CL\_6741
CL\_7526
CL\_7525
CL\_7524
CL\_7523
CL\_7522
CL\_12383
CL\_16565
CL\_27706
CL\_16567
CL\_6783
CL\_8906
CL\_526
CL\_1318
CL\_1319
CL\_1320
CL\_4764
CL\_1326
CL\_4470
CL\_4471
CL\_4472
CL\_4473
CL\_4474
CL\_4475
CL\_4476
CL\_4477
CL\_4478
CL\_4479
CL\_4480
CL\_4481
CL\_4482
CL\_1321
CL\_1322
CL\_4483
CL\_527
CL\_17747
CL\_17746
CL\_7048
CL\_29241
CL\_29240
CL\_528
CL\_531
CL\_532
CL\_4413
CL\_4414
CL\_8905
CL\_8904
CL\_1496
CL\_8589
CL\_534
CL\_30572
CL\_4490
CL\_4491
CL\_533
CL\_1324
CL\_29239
CL\_28348
CL\_4533
CL\_4532
CL\_8903
CL\_8902
CL\_8901
CL\_8900
CL\_8899
CL\_4629
CL\_4531
CL\_4530
CL\_11959
CL\_4529
CL\_7113
CL\_32819
CL\_33133
CL\_33132
CL\_4528
CL\_4527
CL\_28734
CL\_28733
CL\_28528
CL\_28527
CL\_4526
CL\_4415
CL\_4416
CL\_4417
CL\_4695
CL\_8647
CL\_8648
CL\_13639
CL\_23647
CL\_12541
CL\_23764
CL\_13310
CL\_23765
CL\_8653
CL\_8114
CL\_12137
CL\_35423
CL\_16429
CL\_13640
CL\_13641
CL\_13642
CL\_13643
CL\_17067
CL\_8123
CL\_13644
CL\_13645
CL\_13646
CL\_35422
CL\_4418
CL\_4419
CL\_7367
CL\_4420
CL\_29238
CL\_29237
CL\_29236
CL\_4421
CL\_4525
CL\_4524
CL\_4523
CL\_4522
CL\_4521
CL\_4520
CL\_4485
CL\_4518
CL\_8712
CL\_4620
CL\_4515
CL\_9158
CL\_9099
CL\_9100
CL\_33353
CL\_19205
CL\_11304
CL\_28343
CL\_32647
CL\_32648
CL\_32649
CL\_19206
CL\_33131
CL\_33272
CL\_9101
CL\_9102
CL\_9103
CL\_4423
CL\_4424
CL\_4425
CL\_35861
CL\_6636
CL\_9062
CL\_9063
CL\_9064
CL\_9065
CL\_9066
CL\_9067
CL\_9068
CL\_9069
CL\_9070
CL\_9071
CL\_9072
CL\_9073
CL\_9074
CL\_9075
CL\_9953
CL\_9952
CL\_9951
CL\_5389
CL\_5388
CL\_13489
CL\_13490
CL\_13491
CL\_13492
CL\_13493
CL\_13494
CL\_13495
CL\_12057
CL\_5386
CL\_5385
CL\_5384
CL\_6641
CL\_6642
CL\_6643
CL\_14020
CL\_6644
CL\_17676
CL\_15106
CL\_8332
CL\_8330
CL\_5383
CL\_5382
CL\_9950
CL\_25363
CL\_7032
CL\_7821
CL\_9955
CL\_13513
CL\_26145
CL\_26144
CL\_26143
CL\_26142
CL\_26141
CL\_9949
CL\_8962
CL\_8963
CL\_17549
CL\_8328
CL\_8327
CL\_6637
CL\_32633
CL\_32634
CL\_13502
CL\_32635
CL\_32636
CL\_32637
CL\_32638
CL\_32639
CL\_32640
CL\_32641
CL\_32642
CL\_35862
CL\_35863
CL\_35864
CL\_35865
CL\_35866
CL\_35867
CL\_35868
CL\_35869
CL\_35870
CL\_35871
CL\_35872
CL\_35873
CL\_35874
CL\_35875
CL\_35876
CL\_35877
CL\_35878
CL\_35879
CL\_35880
CL\_35881
CL\_35882
CL\_35883
CL\_35884
CL\_35885
CL\_35886
CL\_35887
CL\_35888
CL\_35889
CL\_35890
CL\_35891
CL\_35892
CL\_35893
CL\_35894
CL\_35895
CL\_35896
CL\_35897
CL\_35898
CL\_35899
CL\_35900
CL\_35901
CL\_35902
CL\_35903
CL\_35904
CL\_35905
CL\_35906
CL\_35907
CL\_35908
CL\_35909
CL\_35910
CL\_35911
CL\_35912
CL\_35913
CL\_35914
CL\_35915
CL\_35916
CL\_35917
CL\_35918
CL\_35919
CL\_35920
CL\_35921
CL\_35922
CL\_35923
CL\_35924
CL\_35925
CL\_35926
CL\_35927
CL\_35928
CL\_35929
CL\_35930
CL\_35931
CL\_35932
CL\_35933
CL\_35934
CL\_35935
CL\_35936
CL\_35937
CL\_35938
CL\_35939
CL\_35940
CL\_35941
CL\_35942
CL\_35943
CL\_35944
CL\_35945
CL\_35946
CL\_7046
CL\_7047
CL\_35947
CL\_35948
CL\_35949
CL\_35950
CL\_35951
CL\_35952
CL\_35953
CL\_35954
CL\_35955
CL\_35956
CL\_35957
CL\_35958
CL\_35959
CL\_35960
CL\_35961
CL\_35962
CL\_35963
CL\_35964
CL\_35807
CL\_35806
CL\_35805
CL\_35804
CL\_35803
CL\_35802
CL\_35801
CL\_35800
CL\_35799
CL\_35798
CL\_35797
CL\_35796
CL\_35795
CL\_35794
CL\_35793
CL\_35792
CL\_35791
CL\_35790
CL\_35789
CL\_35788
CL\_35787
CL\_35786
CL\_35785
CL\_10396
CL\_13720
CL\_10395
CL\_10394
CL\_10393
CL\_10392
CL\_5297
CL\_5299
CL\_5300
CL\_5614
CL\_5613
CL\_5536
CL\_5537
CL\_5539
CL\_4307
CL\_5633
CL\_10407
CL\_10406
CL\_15604
CL\_16528
CL\_21766
CL\_21767
CL\_4303
CL\_4302
CL\_4301
CL\_4300
CL\_4299
CL\_4297
CL\_5662
CL\_5548
CL\_4293
CL\_11338
CL\_5659
CL\_5551
CL\_5657
CL\_11829
CL\_11830
CL\_5554
CL\_5555
CL\_5654
CL\_5653
CL\_5652
CL\_5651
CL\_5650
CL\_5649
CL\_4284
CL\_5560
CL\_5561
CL\_5562
CL\_11441
CL\_11440
CL\_5643
CL\_5563
CL\_4279
CL\_5641
CL\_5567
CL\_4278
CL\_4277
CL\_5639
CL\_5638
CL\_5637
CL\_6934
CL\_6933
CL\_6966
CL\_4270
CL\_4269
CL\_5576
CL\_5579
CL\_5580
CL\_4265
CL\_5062
CL\_4263
CL\_4262
CL\_6930
CL\_5630
CL\_6963
CL\_4261
CL\_4260
CL\_4259
CL\_6929
CL\_4258
CL\_4257
CL\_4256
CL\_11832
CL\_11834
CL\_11835
CL\_5055
CL\_4253
CL\_4252
CL\_26440
CL\_4240
CL\_6140
CL\_19342
CL\_14631
CL\_14632
CL\_10399
CL\_4974
CL\_4973
CL\_4972
CL\_10397
CL\_11814
CL\_24017
CL\_11826
CL\_5033
CL\_5032
CL\_5031
CL\_5030
CL\_11825
CL\_11824
CL\_11823
CL\_11822
CL\_11821
CL\_11820
CL\_11819
CL\_11818
CL\_8505
CL\_8506
CL\_11817
CL\_21852
CL\_13615
CL\_10653
CL\_10652
CL\_10651
CL\_10650
CL\_10649
CL\_10648
CL\_10647
CL\_10646
CL\_11135
CL\_10645
CL\_10664
CL\_35784
CL\_10390
CL\_35783
CL\_35782
CL\_35781
CL\_35780
CL\_35779
CL\_35778
CL\_35777
CL\_35776
CL\_35775
CL\_35774
CL\_35773
CL\_35772
CL\_35771
CL\_35770
CL\_35769
CL\_35768
CL\_35767
CL\_35766
CL\_35765
CL\_35764
CL\_35763
CL\_35762
CL\_35761
CL\_35760
CL\_35759
CL\_35758
CL\_35757
CL\_35756
CL\_35755
CL\_35754
CL\_35753
CL\_35752
CL\_35751
CL\_35750
CL\_35749
CL\_35748
CL\_35747
CL\_35746
CL\_35745
CL\_35744
CL\_35743
CL\_35742
CL\_35741
CL\_35740
CL\_35739
CL\_35738
CL\_35737
CL\_35736
CL\_35735
CL\_35734
