## Supplementary material for "A novel method for integrating genomic and Tn-Seq data to identify common *in vivo* fitness mechanisms across multiple bacterial species": S1 Dataset: CL_INS_63.html

Legend

 Mobile +extrachromosomalelementfunctions
 Regulatoryfunctions
 Hypothetical
 Other
 All VFDB Genes

FULL


WINDOWSVGPNG

Trim RowsRemove SingletonsSave Fasta

CL\_769


CL\_769


CL\_769


CL\_769


CL\_769

HighlightSelectShow Genomes


268

CL\_770


1

CL\_770


1

CL\_770


1

CL\_770


1

CL\_770

fGI ID


CL\_INS\_63
CL\_INS\_63
CL\_INS\_63
CL\_INS\_63
CL\_INS\_63
CL\_INS\_63
CL\_INS\_63
CL\_INS\_63
CL\_INS\_63
CL\_INS\_63
CL\_INS\_63
CL\_INS\_63
CL\_INS\_63
CL\_INS\_63
CL\_INS\_63
CL\_INS\_63
CL\_INS\_63
CL\_INS\_63
CL\_INS\_63
CL\_INS\_63
CL\_INS\_63
CL\_INS\_71
CL\_INS\_71
CL\_INS\_71
CL\_INS\_63
CL\_INS\_71
CL\_INS\_71
CL\_INS\_71
CL\_INS\_71
CL\_INS\_71
CL\_INS\_71
CL\_INS\_71
CL\_INS\_71
CL\_INS\_71
CL\_INS\_71
CL\_INS\_71
CL\_INS\_71
CL\_INS\_71
CL\_INS\_71
CL\_INS\_71
CL\_INS\_63
CL\_INS\_71
CL\_INS\_71
CL\_INS\_71
CL\_INS\_71
CL\_INS\_63
CL\_INS\_63
Cluster ID


CL\_23361
CL\_23360
CL\_23359
CL\_23358
CL\_23357
CL\_25216
CL\_25215
CL\_25214
CL\_25213
CL\_23356
CL\_23355
CL\_23354
CL\_23353
CL\_23352
CL\_34898
CL\_34897
CL\_34896
CL\_23351
CL\_23350
CL\_37515
CL\_37514
CL\_31040
CL\_31039
CL\_31038
CL\_37441
CL\_31036
CL\_31035
CL\_31034
CL\_31033
CL\_31032
CL\_31031
CL\_31030
CL\_31029
CL\_31027
CL\_31026
CL\_31025
CL\_31023
CL\_31022
CL\_31021
CL\_31020
CL\_37444
CL\_31018
CL\_31017
CL\_31016
CL\_31015
CL\_37513
CL\_37512
