## Supplementary material for "A novel method for integrating genomic and Tn-Seq data to identify common *in vivo* fitness mechanisms across multiple bacterial species": S1 Dataset: CL_INS_64.html

Legend

 Hypothetical
 Other
 All VFDB Genes

FULL


WINDOWSVGPNG

Trim RowsRemove SingletonsSave Fasta

CL\_773


CL\_773


CL\_773


CL\_773


CL\_773


CL\_773


CL\_773


CL\_754


CL\_773


CL\_773


CL\_773


CL\_772


CL\_773


CL\_773


CL\_773


CL\_773

HighlightSelectShow Genomes


109

CL\_774


38

CL\_774


30

CL\_774


14

CL\_774


3

CL\_775


2

CL\_775


2

CL\_775


1

CL\_774


1

CL\_775


1

CL\_774


1

CL\_774


1

CL\_774


1

CL\_753


1

CL\_774


1

CL\_774


1

CL\_774

fGI ID


CL\_INS\_64
CL\_INS\_64
CL\_INS\_64
CL\_INS\_64
CL\_INS\_64
CL\_INS\_64
CL\_INS\_64
CL\_INS\_64
CL\_INS\_64
CL\_INS\_64
Cluster ID


CL\_29452
CL\_13445
CL\_13444
CL\_7370
CL\_4448
CL\_14083
CL\_10489
CL\_8203
CL\_15261
CL\_8202
