## Supplementary material for "A novel method for integrating genomic and Tn-Seq data to identify common *in vivo* fitness mechanisms across multiple bacterial species": S1 Dataset: CL_INS_65.html

Legend

 Mobile +extrachromosomalelementfunctions
 Regulatoryfunctions
 Hypothetical
 All EssentialGenes
 All Fitness Genes
 Other
 All VFDB Genes

FULL


WINDOWSVGPNG

Trim RowsRemove SingletonsSave Fasta

CL\_787


CL\_787


CL\_786


CL\_786


CL\_787


CL\_718


CL\_786


CL\_785


CL\_787


CL\_787


CL\_787


CL\_787


CL\_784

HighlightSelectShow Genomes


215

CL\_788


42

CL\_788


5

CL\_788


2

CL\_788


1

CL\_788


1

CL\_788


1

CL\_788


1

CL\_788


1

CL\_788


1

CL\_664


1

CL\_789


1

CL\_788


1

CL\_788

fGI ID


CL\_INS\_65
CL\_INS\_65
CL\_INS\_65
CL\_INS\_65
CL\_INS\_65
CL\_INS\_65
CL\_INS\_65
CL\_INS\_65
CL\_INS\_65
CL\_INS\_65
CL\_INS\_65
CL\_INS\_65
CL\_INS\_65
CL\_INS\_65
CL\_INS\_65
CL\_INS\_382
CL\_INS\_65
CL\_INS\_65
CL\_INS\_382
CL\_INS\_65
CL\_INS\_65
CL\_INS\_65
CL\_INS\_65
CL\_INS\_382
CL\_INS\_65
CL\_INS\_382
CL\_INS\_382
CL\_INS\_65
CL\_INS\_382
CL\_INS\_382
CL\_INS\_65
CL\_INS\_382
CL\_INS\_382
CL\_INS\_86
CL\_INS\_65
CL\_INS\_382
CL\_INS\_65
CL\_INS\_382
CL\_INS\_65
CL\_INS\_65
CL\_INS\_204
CL\_INS\_204
CL\_INS\_204
CL\_INS\_65
CL\_INS\_136
CL\_INS\_382
CL\_INS\_65
CL\_INS\_65
CL\_INS\_65
CL\_INS\_65
CL\_INS\_65
CL\_INS\_204
CL\_INS\_204
CL\_INS\_204
CL\_INS\_65
CL\_INS\_65
CL\_INS\_86
CL\_INS\_155
CL\_INS\_155
CL\_INS\_155
CL\_INS\_155
CL\_INS\_65
CL\_INS\_65
CL\_INS\_65
CL\_INS\_65
CL\_INS\_65
CL\_INS\_65
CL\_INS\_65
CL\_INS\_65
CL\_INS\_65
CL\_INS\_65
CL\_INS\_65
CL\_INS\_65
CL\_INS\_65
CL\_INS\_65
CL\_INS\_65
CL\_INS\_65
CL\_INS\_65
CL\_INS\_65
CL\_INS\_65
CL\_INS\_65
CL\_INS\_65
CL\_INS\_65
CL\_INS\_65
CL\_INS\_109
CL\_INS\_65
CL\_INS\_65
CL\_INS\_65
CL\_INS\_65
CL\_INS\_155
CL\_INS\_155
CL\_INS\_155
CL\_INS\_65
CL\_INS\_65
CL\_INS\_65
CL\_INS\_65
CL\_INS\_65
CL\_INS\_182
CL\_INS\_65
CL\_INS\_182
CL\_INS\_65
CL\_INS\_65
Cluster ID


CL\_32563
CL\_30468
CL\_30467
CL\_30466
CL\_30465
CL\_30464
CL\_30463
CL\_30462
CL\_30461
CL\_30460
CL\_30459
CL\_30458
CL\_30457
CL\_30456
CL\_30455
CL\_16006
CL\_30454
CL\_30453
CL\_4547
CL\_30452
CL\_30451
CL\_30450
CL\_30449
CL\_1303
CL\_12293
CL\_5360
CL\_1306
CL\_12395
CL\_1308
CL\_8671
CL\_13081
CL\_12295
CL\_1311
CL\_8672
CL\_16182
CL\_1314
CL\_30442
CL\_1317
CL\_30441
CL\_12471
CL\_5921
CL\_5922
CL\_9222
CL\_12302
CL\_1318
CL\_12006
CL\_30440
CL\_30439
CL\_10490
CL\_10491
CL\_35539
CL\_1105
CL\_1104
CL\_1103
CL\_35540
CL\_35541
CL\_8232
CL\_8231
CL\_8230
CL\_8229
CL\_8228
CL\_35542
CL\_35543
CL\_35544
CL\_35545
CL\_35546
CL\_35547
CL\_35548
CL\_35549
CL\_35550
CL\_35551
CL\_35552
CL\_35553
CL\_35554
CL\_35555
CL\_35556
CL\_35557
CL\_35558
CL\_35559
CL\_35560
CL\_35561
CL\_35562
CL\_35563
CL\_35564
CL\_22580
CL\_24085
CL\_24084
CL\_24083
CL\_24082
CL\_11006
CL\_11005
CL\_11004
CL\_24081
CL\_24080
CL\_24079
CL\_24078
CL\_24077
CL\_10996
CL\_24076
CL\_10994
CL\_24075
CL\_24074
