## Supplementary material for "A novel method for integrating genomic and Tn-Seq data to identify common *in vivo* fitness mechanisms across multiple bacterial species": S1 Dataset: CL_INS_69.html

Legend

 Mobile +extrachromosomalelementfunctions
 Regulatoryfunctions
 Hypothetical
 DNA Metabolism
 All Fitness Genes
 Other
 All VFDB Genes

FULL


WINDOWSVGPNG

Trim RowsRemove SingletonsSave Fasta

CL\_833


CL\_833


CL\_833


CL\_833


CL\_833


CL\_833


CL\_833


CL\_833


CL\_833


CL\_833


CL\_833


CL\_833


CL\_833


CL\_833


CL\_833


CL\_833


CL\_833


CL\_833


CL\_833


CL\_833


CL\_833

HighlightSelectShow Genomes


254

CL\_834


2

CL\_834


2

CL\_834


2

CL\_834


1

CL\_834


1

CL\_834


1

CL\_834


1

CL\_834


1

CL\_834


1

CL\_834


1

CL\_834


1

CL\_834


1

CL\_834


1

CL\_834


1

CL\_834


1

CL\_834


1

CL\_834


1

CL\_836


1

CL\_834


1

CL\_834


1

CL\_834

fGI ID


CL\_INS\_69
CL\_INS\_174
CL\_INS\_69
CL\_INS\_382
CL\_INS\_247
CL\_INS\_106
CL\_INS\_106
CL\_INS\_106
CL\_INS\_382
CL\_INS\_382
CL\_INS\_132
CL\_INS\_106
CL\_INS\_132
CL\_INS\_106
CL\_INS\_106
CL\_INS\_132
CL\_INS\_382
CL\_INS\_382
CL\_INS\_69
CL\_INS\_382
CL\_INS\_382
CL\_INS\_382
CL\_INS\_247
CL\_INS\_382
CL\_INS\_69
CL\_INS\_69
CL\_INS\_69
CL\_INS\_69
CL\_INS\_69
CL\_INS\_69
CL\_INS\_382
CL\_INS\_247
CL\_INS\_69
CL\_INS\_247
CL\_INS\_69
CL\_INS\_247
CL\_INS\_247
CL\_INS\_247
CL\_INS\_382
CL\_INS\_69
CL\_INS\_382
CL\_INS\_382
CL\_INS\_382
CL\_INS\_69
CL\_INS\_69
CL\_INS\_69
CL\_INS\_69
CL\_INS\_69
CL\_INS\_69
CL\_INS\_69
CL\_INS\_69
CL\_INS\_382
CL\_INS\_382
CL\_INS\_247
CL\_INS\_247
CL\_INS\_247
CL\_INS\_247
CL\_INS\_247
CL\_INS\_382
CL\_INS\_382
CL\_INS\_382
CL\_INS\_106
CL\_INS\_69
CL\_INS\_132
CL\_INS\_132
CL\_INS\_174
CL\_INS\_132
CL\_INS\_132
CL\_INS\_132
CL\_INS\_132
CL\_INS\_132
CL\_INS\_132
CL\_INS\_132
CL\_INS\_132
CL\_INS\_382
CL\_INS\_69
CL\_INS\_69
CL\_INS\_69
CL\_INS\_69
CL\_INS\_132
CL\_INS\_174
CL\_INS\_69
CL\_INS\_69
CL\_INS\_69
CL\_INS\_69
CL\_INS\_69
CL\_INS\_69
CL\_INS\_69
CL\_INS\_69
CL\_INS\_69
CL\_INS\_69
CL\_INS\_69
CL\_INS\_69
CL\_INS\_69
CL\_INS\_69
CL\_INS\_69
CL\_INS\_69
CL\_INS\_69
CL\_INS\_69
CL\_INS\_69
CL\_INS\_69
CL\_INS\_69
CL\_INS\_69
CL\_INS\_69
CL\_INS\_69
CL\_INS\_237
CL\_INS\_69
CL\_INS\_382
CL\_INS\_382
CL\_INS\_382
Cluster ID


CL\_34060
CL\_16581
CL\_32131
CL\_8648
CL\_8255
CL\_8649
CL\_16469
CL\_16582
CL\_23647
CL\_12541
CL\_8650
CL\_8651
CL\_8652
CL\_12139
CL\_12138
CL\_13311
CL\_13510
CL\_13310
CL\_32130
CL\_8112
CL\_13512
CL\_8113
CL\_20757
CL\_8114
CL\_32129
CL\_32128
CL\_32127
CL\_32126
CL\_33130
CL\_32125
CL\_12137
CL\_17495
CL\_30648
CL\_8115
CL\_15922
CL\_15923
CL\_15924
CL\_8122
CL\_13640
CL\_34061
CL\_13641
CL\_13642
CL\_13643
CL\_30984
CL\_30985
CL\_30986
CL\_30987
CL\_30988
CL\_30989
CL\_30990
CL\_30991
CL\_8654
CL\_8655
CL\_10496
CL\_8656
CL\_8657
CL\_8658
CL\_8659
CL\_17067
CL\_13737
CL\_13738
CL\_13309
CL\_35808
CL\_10497
CL\_13102
CL\_23120
CL\_10498
CL\_10499
CL\_10500
CL\_10501
CL\_10502
CL\_10503
CL\_10504
CL\_10505
CL\_8123
CL\_26782
CL\_23648
CL\_23649
CL\_34062
CL\_10506
CL\_10507
CL\_23121
CL\_30992
CL\_27865
CL\_30993
CL\_30994
CL\_30995
CL\_30996
CL\_30997
CL\_30998
CL\_30999
CL\_31000
CL\_31001
CL\_31002
CL\_31003
CL\_31004
CL\_31005
CL\_31006
CL\_31007
CL\_6567
CL\_6568
CL\_6569
CL\_27193
CL\_31008
CL\_31009
CL\_31010
CL\_31011
CL\_8216
CL\_4975
CL\_8841
