## Supplementary material for "A novel method for integrating genomic and Tn-Seq data to identify common *in vivo* fitness mechanisms across multiple bacterial species": S1 Dataset: CL_INS_70.html

Legend

 Mobile +extrachromosomalelementfunctions
 Regulatoryfunctions
 Hypothetical
 DNA Metabolism
 All EssentialGenes
 All Fitness Genes
 Cell Envelope
 Proteinsynthesis/fate
 Other
 Centralintermediarymetabolism
 EnergyMetabolism
 Transport +binding proteins
 All VFDB Genes

FULL


WINDOWSVGPNG

Trim RowsRemove SingletonsSave Fasta

CL\_835


CL\_835


CL\_835


CL\_835


CL\_1935


CL\_835


CL\_835


CL\_835


CL\_835


CL\_835


CL\_835


CL\_835


CL\_1935


CL\_833


CL\_835


CL\_835


CL\_835


CL\_835


CL\_835


CL\_835


CL\_835


CL\_835


CL\_835


CL\_835


CL\_835


CL\_234


CL\_834

HighlightSelectShow Genomes


234

CL\_836


11

CL\_836


5

CL\_836


4

CL\_836


3

CL\_836


2

CL\_836


2

CL\_836


1

CL\_1935


1

Break


1

CL\_836


1

CL\_836


1

CL\_836


1

CL\_836


1

CL\_836


1

CL\_1935


1

Break


1

CL\_234


1

CL\_836


1

CL\_836


1

CL\_836


1

CL\_836


1

CL\_836


1

CL\_836


1

CL\_836


1

CL\_234


1

CL\_836


1

CL\_836

fGI ID


CL\_INS\_70
CL\_INS\_70
CL\_INS\_70
CL\_INS\_70
CL\_INS\_70
CL\_INS\_70
CL\_INS\_70
CL\_INS\_70
CL\_INS\_70
CL\_INS\_70
CL\_INS\_70
CL\_INS\_70
CL\_INS\_149
CL\_INS\_70
CL\_INS\_70
CL\_INS\_70
CL\_INS\_70
CL\_INS\_70
CL\_INS\_70
CL\_INS\_70
CL\_INS\_70
CL\_INS\_70
CL\_INS\_70
CL\_INS\_70
CL\_INS\_70
CL\_INS\_70
CL\_INS\_70
CL\_INS\_70
CL\_INS\_70
CL\_INS\_70
CL\_INS\_70
CL\_INS\_70
CL\_INS\_70
CL\_INS\_70
CL\_INS\_70
CL\_INS\_70
CL\_INS\_70
CL\_INS\_70
CL\_INS\_70
CL\_INS\_70
CL\_INS\_70
CL\_INS\_70
CL\_INS\_70
CL\_INS\_70
CL\_INS\_70
CL\_INS\_70
CL\_INS\_70
CL\_INS\_70
CL\_INS\_70
CL\_INS\_70
CL\_INS\_382
CL\_INS\_70
CL\_INS\_70
CL\_INS\_70
CL\_INS\_70
CL\_INS\_70
CL\_INS\_70
CL\_INS\_70
CL\_INS\_70
CL\_INS\_70
CL\_INS\_70
CL\_INS\_70
CL\_INS\_30
CL\_INS\_70
CL\_INS\_70
CL\_INS\_70
CL\_INS\_70
CL\_INS\_70
CL\_INS\_70
CL\_INS\_70
CL\_INS\_70
CL\_INS\_70
CL\_INS\_70
CL\_INS\_70
CL\_INS\_70
CL\_INS\_70
CL\_INS\_70
CL\_INS\_70
CL\_INS\_70
CL\_INS\_70
CL\_INS\_70
CL\_INS\_70
CL\_INS\_70
CL\_INS\_70
CL\_INS\_70
CL\_INS\_70
CL\_INS\_70
CL\_INS\_70
CL\_INS\_70
CL\_INS\_70
CL\_INS\_70
CL\_INS\_70
CL\_INS\_70
CL\_INS\_70
CL\_INS\_70
CL\_INS\_70
CL\_INS\_70
CL\_INS\_70
CL\_INS\_70
CL\_INS\_70
CL\_INS\_70
CL\_INS\_70
CL\_INS\_70
CL\_INS\_70
CL\_INS\_70
CL\_INS\_70
CL\_INS\_70
CL\_INS\_70
CL\_INS\_70
CL\_INS\_70
CL\_INS\_70
CL\_INS\_70
CL\_INS\_70
CL\_INS\_70
CL\_INS\_70
CL\_INS\_70
CL\_INS\_70
CL\_INS\_70
CL\_INS\_30
CL\_INS\_70
CL\_INS\_70
CL\_INS\_70
CL\_INS\_70
CL\_INS\_70
CL\_INS\_70
CL\_INS\_70
CL\_INS\_70
CL\_INS\_149
CL\_INS\_70
CL\_INS\_70
CL\_INS\_70
CL\_INS\_70
CL\_INS\_70
CL\_INS\_70
CL\_INS\_70
CL\_INS\_70
CL\_INS\_70
CL\_INS\_70
CL\_INS\_70
CL\_INS\_70
CL\_INS\_70
CL\_INS\_70
CL\_INS\_70
CL\_INS\_70
CL\_INS\_70
CL\_INS\_70
CL\_INS\_70
CL\_INS\_70
CL\_INS\_70
CL\_INS\_70
CL\_INS\_70
CL\_INS\_70
CL\_INS\_70
CL\_INS\_382
CL\_INS\_382
CL\_INS\_382
CL\_INS\_70
CL\_INS\_70
CL\_INS\_70
CL\_INS\_70
CL\_INS\_70
CL\_INS\_70
CL\_INS\_70
CL\_INS\_70
CL\_INS\_70
CL\_INS\_70
CL\_INS\_70
CL\_INS\_70
CL\_INS\_70
CL\_INS\_70
CL\_INS\_70
CL\_INS\_70
CL\_INS\_70
CL\_INS\_70
CL\_INS\_70
CL\_INS\_70
CL\_INS\_70
CL\_INS\_70
CL\_INS\_70
CL\_INS\_70
CL\_INS\_70
CL\_INS\_70
CL\_INS\_70
CL\_INS\_70
CL\_INS\_70
CL\_INS\_70
CL\_INS\_70
CL\_INS\_70
CL\_INS\_70
CL\_INS\_70
CL\_INS\_70
CL\_INS\_70
CL\_INS\_70
CL\_INS\_70
CL\_INS\_70
CL\_INS\_70
CL\_INS\_70
CL\_INS\_70
CL\_INS\_70
CL\_INS\_70
CL\_INS\_70
CL\_INS\_70
CL\_INS\_70
CL\_INS\_70
CL\_INS\_70
CL\_INS\_70
CL\_INS\_70
CL\_INS\_70
CL\_INS\_70
CL\_INS\_70
CL\_INS\_70
CL\_INS\_70
CL\_INS\_70
CL\_INS\_70
CL\_INS\_70
CL\_INS\_70
CL\_INS\_70
CL\_INS\_70
CL\_INS\_70
CL\_INS\_70
CL\_INS\_70
CL\_INS\_70
CL\_INS\_70
CL\_INS\_70
CL\_INS\_70
CL\_INS\_70
CL\_INS\_70
CL\_INS\_70
CL\_INS\_70
CL\_INS\_70
CL\_INS\_70
CL\_INS\_70
CL\_INS\_70
CL\_INS\_70
CL\_INS\_70
CL\_INS\_70
CL\_INS\_70
CL\_INS\_70
CL\_INS\_70
CL\_INS\_70
CL\_INS\_70
CL\_INS\_70
CL\_INS\_70
CL\_INS\_70
CL\_INS\_70
CL\_INS\_70
CL\_INS\_70
CL\_INS\_70
CL\_INS\_70
CL\_INS\_70
CL\_INS\_70
CL\_INS\_70
CL\_INS\_70
CL\_INS\_70
CL\_INS\_70
CL\_INS\_70
CL\_INS\_382
CL\_INS\_70
CL\_INS\_70
CL\_INS\_70
CL\_INS\_382
CL\_INS\_149
CL\_INS\_70
CL\_INS\_70
CL\_INS\_70
CL\_INS\_70
CL\_INS\_70
CL\_INS\_70
CL\_INS\_70
CL\_INS\_70
CL\_INS\_70
CL\_INS\_70
CL\_INS\_70
CL\_INS\_70
CL\_INS\_70
CL\_INS\_70
CL\_INS\_70
CL\_INS\_70
CL\_INS\_382
CL\_INS\_382
CL\_INS\_70
CL\_INS\_70
CL\_INS\_70
CL\_INS\_70
CL\_INS\_70
CL\_INS\_70
CL\_INS\_70
CL\_INS\_70
CL\_INS\_70
CL\_INS\_70
CL\_INS\_70
CL\_INS\_70
CL\_INS\_70
CL\_INS\_70
CL\_INS\_70
CL\_INS\_70
CL\_INS\_70
CL\_INS\_70
CL\_INS\_30
CL\_INS\_30
CL\_INS\_70
CL\_INS\_70
CL\_INS\_70
CL\_INS\_70
CL\_INS\_30
CL\_INS\_70
CL\_INS\_70
CL\_INS\_30
CL\_INS\_30
CL\_INS\_30
CL\_INS\_30
CL\_INS\_70
CL\_INS\_70
CL\_INS\_30
CL\_INS\_70
CL\_INS\_70
CL\_INS\_70
CL\_INS\_70
CL\_INS\_70
CL\_INS\_30
CL\_INS\_70
CL\_INS\_70
CL\_INS\_70
CL\_INS\_70
CL\_INS\_70
CL\_INS\_70
CL\_INS\_70
CL\_INS\_30
CL\_INS\_70
CL\_INS\_70
CL\_INS\_70
CL\_INS\_70
CL\_INS\_70
CL\_INS\_30
CL\_INS\_30
CL\_INS\_70
CL\_INS\_30
CL\_INS\_30
CL\_INS\_70
CL\_INS\_70
CL\_INS\_30
CL\_INS\_70
CL\_INS\_70
Cluster ID


CL\_16104
CL\_26456
CL\_10492
CL\_17258
CL\_17259
CL\_17260
CL\_9621
CL\_9620
CL\_9619
CL\_9618
CL\_23122
CL\_9617
CL\_5146
CL\_27642
CL\_11991
CL\_26418
CL\_27127
CL\_11992
CL\_37251
CL\_7036
CL\_13627
CL\_7035
CL\_7034
CL\_27126
CL\_29453
CL\_28159
CL\_15745
CL\_7899
CL\_26037
CL\_26036
CL\_26035
CL\_26034
CL\_8037
CL\_30649
CL\_30650
CL\_7895
CL\_17790
CL\_17789
CL\_17788
CL\_17787
CL\_17786
CL\_8591
CL\_8592
CL\_7894
CL\_7893
CL\_7892
CL\_8593
CL\_8594
CL\_8595
CL\_6836
CL\_7557
CL\_6837
CL\_10326
CL\_7672
CL\_6838
CL\_6839
CL\_8596
CL\_8597
CL\_8598
CL\_6840
CL\_6841
CL\_6842
CL\_6856
CL\_5682
CL\_8601
CL\_8602
CL\_5493
CL\_8603
CL\_8605
CL\_8604
CL\_8606
CL\_8607
CL\_8608
CL\_8609
CL\_8610
CL\_8611
CL\_8612
CL\_8613
CL\_8614
CL\_8615
CL\_8616
CL\_8617
CL\_8618
CL\_5391
CL\_30651
CL\_30652
CL\_11221
CL\_7242
CL\_11191
CL\_7241
CL\_6426
CL\_5320
CL\_5321
CL\_10435
CL\_8750
CL\_8749
CL\_8748
CL\_8747
CL\_14344
CL\_4094
CL\_4095
CL\_4096
CL\_4097
CL\_4098
CL\_4099
CL\_4100
CL\_4101
CL\_6831
CL\_6829
CL\_6411
CL\_7898
CL\_7897
CL\_7896
CL\_8590
CL\_26250
CL\_8628
CL\_14649
CL\_17597
CL\_11295
CL\_10581
CL\_15528
CL\_10330
CL\_10331
CL\_10332
CL\_8599
CL\_8600
CL\_6410
CL\_4093
CL\_6828
CL\_7252
CL\_14651
CL\_14652
CL\_14653
CL\_28158
CL\_15746
CL\_15747
CL\_14654
CL\_14655
CL\_14656
CL\_14657
CL\_14658
CL\_14659
CL\_14660
CL\_14661
CL\_14662
CL\_14663
CL\_14664
CL\_15748
CL\_14665
CL\_15749
CL\_15750
CL\_15751
CL\_14666
CL\_8832
CL\_7691
CL\_7692
CL\_7435
CL\_7434
CL\_14667
CL\_14668
CL\_14669
CL\_14670
CL\_14671
CL\_14672
CL\_14673
CL\_14674
CL\_14675
CL\_14676
CL\_14677
CL\_14678
CL\_14679
CL\_14680
CL\_8632
CL\_8631
CL\_8630
CL\_12888
CL\_14681
CL\_13233
CL\_13232
CL\_13231
CL\_7214
CL\_7671
CL\_14682
CL\_7857
CL\_13783
CL\_7856
CL\_7673
CL\_7233
CL\_8781
CL\_15980
CL\_18096
CL\_18095
CL\_18094
CL\_18093
CL\_18092
CL\_15973
CL\_18091
CL\_17096
CL\_18090
CL\_7869
CL\_18089
CL\_18088
CL\_9371
CL\_18087
CL\_18086
CL\_18085
CL\_18084
CL\_18083
CL\_18082
CL\_18081
CL\_18080
CL\_18079
CL\_18078
CL\_18077
CL\_18076
CL\_18075
CL\_18074
CL\_18073
CL\_18072
CL\_18071
CL\_18070
CL\_18069
CL\_18068
CL\_18067
CL\_18066
CL\_18065
CL\_18064
CL\_18063
CL\_18062
CL\_18061
CL\_18060
CL\_18059
CL\_18058
CL\_18057
CL\_18056
CL\_18055
CL\_18054
CL\_18053
CL\_18052
CL\_18051
CL\_18050
CL\_8619
CL\_22734
CL\_30653
CL\_13787
CL\_6418
CL\_1932
CL\_1933
CL\_7831
CL\_8620
CL\_14683
CL\_7234
CL\_7235
CL\_10408
CL\_10409
CL\_14684
CL\_8376
CL\_8779
CL\_7298
CL\_7299
CL\_10807
CL\_11826
CL\_26249
CL\_26248
CL\_26247
CL\_10592
CL\_14685
CL\_6979
CL\_26246
CL\_26245
CL\_26244
CL\_26243
CL\_26242
CL\_26241
CL\_26240
CL\_26239
CL\_25227
CL\_26238
CL\_6177
CL\_6178
CL\_6179
CL\_6980
CL\_6981
CL\_6982
CL\_6983
CL\_22967
CL\_6984
CL\_10832
CL\_15752
CL\_15753
CL\_15754
CL\_7690
CL\_8641
CL\_1931
CL\_15755
CL\_10547
CL\_8508
CL\_9139
CL\_5231
CL\_5230
CL\_9140
CL\_13848
CL\_8642
CL\_8621
CL\_7751
CL\_7829
CL\_26033
CL\_7827
CL\_7750
CL\_7749
CL\_2551
CL\_7824
CL\_26031
CL\_1934
CL\_10610
CL\_18049
CL\_18048
CL\_26032
CL\_8746
CL\_1937
CL\_6809
CL\_6808
CL\_8107
CL\_6417
CL\_6416
CL\_6419
CL\_18098
CL\_18097
CL\_6420
CL\_10546
CL\_8126
CL\_7255
CL\_6824
CL\_7254
CL\_5245
CL\_8129
CL\_6423
CL\_5241
CL\_11105
CL\_6421
CL\_5246
CL\_7253
CL\_6826
