## Supplementary material for "A novel method for integrating genomic and Tn-Seq data to identify common *in vivo* fitness mechanisms across multiple bacterial species": S1 Dataset: CL_INS_71.html

Legend

 Mobile +extrachromosomalelementfunctions
 Regulatoryfunctions
 Hypothetical
 DNA Metabolism
 AntibioticResistance
 Proteinsynthesis/fate
 Other
 Transport +binding proteins
 All VFDB Genes

FULL


WINDOWSVGPNG

Trim RowsRemove SingletonsSave Fasta

CL\_845


CL\_845


CL\_845


CL\_845


CL\_845


CL\_845


CL\_845


CL\_845


CL\_845


CL\_845


CL\_845


CL\_845


CL\_845


CL\_845


CL\_845


CL\_845


CL\_845


CL\_845


CL\_845


CL\_845


CL\_845


CL\_845


CL\_845


CL\_861


CL\_845

HighlightSelectShow Genomes


236

CL\_846


3

CL\_846


3

CL\_846


2

CL\_846


1

CL\_846


1

CL\_846


1

CL\_846


1

CL\_846


1

CL\_846


1

CL\_846


1

Break


1

CL\_860


1

CL\_846


1

CL\_846


1

CL\_846


1

CL\_846


1

CL\_846


1

CL\_846


1

CL\_846


1

CL\_846


1

CL\_846


1

CL\_846


1

CL\_846


1

CL\_846


1

CL\_846

fGI ID


CL\_INS\_71
CL\_INS\_71
CL\_INS\_155
CL\_INS\_155
CL\_INS\_155
CL\_INS\_155
CL\_INS\_155
CL\_INS\_155
CL\_INS\_155
CL\_INS\_155
CL\_INS\_155
CL\_INS\_155
CL\_INS\_74
CL\_INS\_74
CL\_INS\_74
CL\_INS\_74
CL\_INS\_74
CL\_INS\_74
CL\_INS\_74
CL\_INS\_74
CL\_INS\_74
CL\_INS\_74
CL\_INS\_155
CL\_INS\_155
CL\_INS\_155
CL\_INS\_155
CL\_INS\_155
CL\_INS\_155
CL\_INS\_86
CL\_INS\_155
CL\_INS\_86
CL\_INS\_155
CL\_INS\_86
CL\_INS\_155
CL\_INS\_155
CL\_INS\_155
CL\_INS\_155
CL\_INS\_155
CL\_INS\_155
CL\_INS\_155
CL\_INS\_86
CL\_INS\_86
CL\_INS\_86
CL\_INS\_86
CL\_INS\_155
CL\_INS\_155
CL\_INS\_155
CL\_INS\_155
CL\_INS\_86
CL\_INS\_155
CL\_INS\_155
CL\_INS\_155
CL\_INS\_155
CL\_INS\_86
CL\_INS\_71
CL\_INS\_155
CL\_INS\_149
CL\_INS\_149
CL\_INS\_149
CL\_INS\_170
CL\_INS\_149
CL\_INS\_149
CL\_INS\_149
CL\_INS\_149
CL\_INS\_149
CL\_INS\_74
CL\_INS\_149
CL\_INS\_149
CL\_INS\_149
CL\_INS\_149
CL\_INS\_149
CL\_INS\_149
CL\_INS\_170
CL\_INS\_149
CL\_INS\_170
CL\_INS\_149
CL\_INS\_149
CL\_INS\_149
CL\_INS\_149
CL\_INS\_149
CL\_INS\_71
CL\_INS\_71
CL\_INS\_71
CL\_INS\_71
CL\_INS\_382
CL\_INS\_146
CL\_INS\_71
CL\_INS\_71
CL\_INS\_149
CL\_INS\_149
CL\_INS\_149
CL\_INS\_149
CL\_INS\_149
CL\_INS\_149
CL\_INS\_71
CL\_INS\_207
CL\_INS\_71
CL\_INS\_71
CL\_INS\_149
CL\_INS\_149
CL\_INS\_149
CL\_INS\_149
CL\_INS\_71
CL\_INS\_149
CL\_INS\_149
CL\_INS\_71
CL\_INS\_71
CL\_INS\_149
CL\_INS\_149
CL\_INS\_149
CL\_INS\_74
CL\_INS\_71
CL\_INS\_74
CL\_INS\_74
CL\_INS\_74
CL\_INS\_74
CL\_INS\_71
CL\_INS\_71
CL\_INS\_71
CL\_INS\_71
CL\_INS\_71
CL\_INS\_71
CL\_INS\_71
CL\_INS\_71
CL\_INS\_71
CL\_INS\_71
CL\_INS\_71
CL\_INS\_71
CL\_INS\_71
CL\_INS\_71
CL\_INS\_71
CL\_INS\_71
CL\_INS\_71
CL\_INS\_71
CL\_INS\_71
CL\_INS\_71
CL\_INS\_71
CL\_INS\_71
CL\_INS\_71
CL\_INS\_71
CL\_INS\_71
CL\_INS\_71
CL\_INS\_71
CL\_INS\_71
CL\_INS\_71
CL\_INS\_71
CL\_INS\_71
CL\_INS\_71
CL\_INS\_71
CL\_INS\_71
CL\_INS\_71
CL\_INS\_71
CL\_INS\_71
CL\_INS\_71
CL\_INS\_71
CL\_INS\_71
CL\_INS\_71
CL\_INS\_71
CL\_INS\_71
CL\_INS\_71
CL\_INS\_71
CL\_INS\_71
CL\_INS\_71
CL\_INS\_71
CL\_INS\_71
CL\_INS\_149
CL\_INS\_149
CL\_INS\_149
CL\_INS\_149
CL\_INS\_149
CL\_INS\_170
CL\_INS\_149
CL\_INS\_149
CL\_INS\_71
CL\_INS\_71
CL\_INS\_71
CL\_INS\_71
CL\_INS\_207
CL\_INS\_71
CL\_INS\_71
CL\_INS\_149
CL\_INS\_204
CL\_INS\_204
CL\_INS\_71
CL\_INS\_204
CL\_INS\_237
CL\_INS\_149
CL\_INS\_170
CL\_INS\_204
CL\_INS\_74
CL\_INS\_74
CL\_INS\_204
CL\_INS\_71
CL\_INS\_170
CL\_INS\_170
CL\_INS\_149
CL\_INS\_74
CL\_INS\_149
CL\_INS\_170
CL\_INS\_149
CL\_INS\_71
CL\_INS\_149
CL\_INS\_170
CL\_INS\_71
CL\_INS\_71
CL\_INS\_71
CL\_INS\_71
CL\_INS\_71
CL\_INS\_71
CL\_INS\_71
CL\_INS\_71
CL\_INS\_71
CL\_INS\_71
CL\_INS\_149
CL\_INS\_71
CL\_INS\_71
CL\_INS\_71
CL\_INS\_71
CL\_INS\_71
CL\_INS\_71
CL\_INS\_71
CL\_INS\_71
CL\_INS\_149
CL\_INS\_170
CL\_INS\_247
CL\_INS\_247
CL\_INS\_247
CL\_INS\_247
CL\_INS\_247
CL\_INS\_247
CL\_INS\_247
CL\_INS\_60
CL\_INS\_60
CL\_INS\_60
CL\_INS\_60
CL\_INS\_60
CL\_INS\_60
CL\_INS\_60
CL\_INS\_60
CL\_INS\_60
CL\_INS\_86
CL\_INS\_71
CL\_INS\_71
CL\_INS\_71
CL\_INS\_71
CL\_INS\_71
CL\_INS\_71
CL\_INS\_71
CL\_INS\_71
CL\_INS\_71
CL\_INS\_71
CL\_INS\_382
CL\_INS\_382
CL\_INS\_382
CL\_INS\_71
CL\_INS\_71
CL\_INS\_382
CL\_INS\_382
CL\_INS\_71
CL\_INS\_71
CL\_INS\_71
CL\_INS\_71
CL\_INS\_382
CL\_INS\_71
CL\_INS\_71
CL\_INS\_71
CL\_INS\_71
CL\_INS\_71
CL\_INS\_382
CL\_INS\_382
CL\_INS\_382
CL\_INS\_159
CL\_INS\_117
CL\_INS\_117
CL\_INS\_159
CL\_INS\_382
CL\_INS\_382
CL\_INS\_382
CL\_INS\_382
CL\_INS\_382
CL\_INS\_159
CL\_INS\_159
CL\_INS\_159
CL\_INS\_159
CL\_INS\_385
CL\_INS\_71
CL\_INS\_382
CL\_INS\_382
CL\_INS\_382
CL\_INS\_382
CL\_INS\_385
CL\_INS\_385
CL\_INS\_382
CL\_INS\_159
CL\_INS\_159
CL\_INS\_382
CL\_INS\_382
CL\_INS\_382
CL\_INS\_385
CL\_INS\_159
CL\_INS\_382
CL\_INS\_382
CL\_INS\_382
CL\_INS\_382
CL\_INS\_382
CL\_INS\_382
CL\_INS\_382
CL\_INS\_382
CL\_INS\_382
CL\_INS\_382
CL\_INS\_382
CL\_INS\_247
CL\_INS\_275
CL\_INS\_71
CL\_INS\_86
CL\_INS\_215
CL\_INS\_71
CL\_INS\_382
CL\_INS\_382
CL\_INS\_382
CL\_INS\_382
CL\_INS\_382
CL\_INS\_382
CL\_INS\_382
CL\_INS\_382
CL\_INS\_382
CL\_INS\_382
CL\_INS\_382
CL\_INS\_382
CL\_INS\_382
CL\_INS\_382
CL\_INS\_382
CL\_INS\_382
CL\_INS\_382
CL\_INS\_117
CL\_INS\_117
CL\_INS\_233
CL\_INS\_233
CL\_INS\_233
CL\_INS\_233
CL\_INS\_233
CL\_INS\_382
CL\_INS\_159
CL\_INS\_207
CL\_INS\_207
CL\_INS\_207
CL\_INS\_207
CL\_INS\_207
CL\_INS\_207
CL\_INS\_207
CL\_INS\_207
CL\_INS\_237
CL\_INS\_237
CL\_INS\_247
CL\_INS\_247
CL\_INS\_247
CL\_INS\_71
CL\_INS\_99
CL\_INS\_99
CL\_INS\_71
CL\_INS\_71
CL\_INS\_123
CL\_INS\_247
CL\_INS\_123
CL\_INS\_123
CL\_INS\_123
CL\_INS\_247
CL\_INS\_247
Cluster ID


CL\_14644
CL\_14645
CL\_10915
CL\_10914
CL\_10913
CL\_10912
CL\_10911
CL\_10910
CL\_10909
CL\_10907
CL\_10906
CL\_10905
CL\_10904
CL\_10903
CL\_10902
CL\_10901
CL\_10900
CL\_10899
CL\_10898
CL\_10897
CL\_10896
CL\_10895
CL\_10894
CL\_10893
CL\_10892
CL\_10891
CL\_10890
CL\_10889
CL\_10888
CL\_10887
CL\_10886
CL\_10885
CL\_10884
CL\_10883
CL\_10882
CL\_11024
CL\_11025
CL\_10879
CL\_10878
CL\_10877
CL\_11029
CL\_11030
CL\_8226
CL\_8227
CL\_8228
CL\_8229
CL\_8230
CL\_8231
CL\_8232
CL\_10870
CL\_10869
CL\_11035
CL\_11036
CL\_5201
CL\_30659
CL\_8483
CL\_5200
CL\_5199
CL\_4598
CL\_4597
CL\_4596
CL\_14647
CL\_4595
CL\_4594
CL\_11993
CL\_11049
CL\_4593
CL\_4592
CL\_4591
CL\_4590
CL\_4589
CL\_4588
CL\_4587
CL\_6599
CL\_4585
CL\_9190
CL\_9189
CL\_9188
CL\_29792
CL\_11048
CL\_27584
CL\_27583
CL\_11047
CL\_11046
CL\_8683
CL\_11995
CL\_11996
CL\_11997
CL\_11994
CL\_9187
CL\_9186
CL\_4580
CL\_4579
CL\_11045
CL\_11044
CL\_4666
CL\_11998
CL\_11999
CL\_12557
CL\_11043
CL\_4578
CL\_4577
CL\_27259
CL\_8986
CL\_4576
CL\_30678
CL\_30679
CL\_17515
CL\_4575
CL\_4574
CL\_4573
CL\_12555
CL\_11042
CL\_11041
CL\_11040
CL\_11039
CL\_11038
CL\_13297
CL\_11050
CL\_31012
CL\_31013
CL\_31014
CL\_31015
CL\_31016
CL\_31017
CL\_31018
CL\_31019
CL\_31020
CL\_31021
CL\_31022
CL\_31023
CL\_31024
CL\_31025
CL\_31026
CL\_31027
CL\_31028
CL\_31029
CL\_31030
CL\_31031
CL\_31032
CL\_31033
CL\_31034
CL\_31035
CL\_31036
CL\_31037
CL\_31038
CL\_31039
CL\_31040
CL\_31041
CL\_31042
CL\_31043
CL\_31044
CL\_31045
CL\_31046
CL\_31047
CL\_31048
CL\_31049
CL\_31050
CL\_31051
CL\_31052
CL\_25999
CL\_25998
CL\_25997
CL\_25996
CL\_30654
CL\_4613
CL\_9192
CL\_4611
CL\_4610
CL\_4609
CL\_4608
CL\_4607
CL\_4606
CL\_30655
CL\_33129
CL\_33128
CL\_30656
CL\_5392
CL\_30657
CL\_30658
CL\_4605
CL\_4604
CL\_6594
CL\_29794
CL\_1103
CL\_4514
CL\_2276
CL\_7073
CL\_1105
CL\_11365
CL\_27795
CL\_10867
CL\_14646
CL\_5170
CL\_5198
CL\_14492
CL\_13296
CL\_4603
CL\_4602
CL\_4601
CL\_29793
CL\_4600
CL\_4599
CL\_30660
CL\_30661
CL\_30662
CL\_30663
CL\_30664
CL\_30665
CL\_30666
CL\_30667
CL\_30668
CL\_30669
CL\_6529
CL\_30670
CL\_30671
CL\_30672
CL\_30673
CL\_30674
CL\_30675
CL\_30676
CL\_30677
CL\_14648
CL\_5883
CL\_10641
CL\_10642
CL\_10421
CL\_10423
CL\_10382
CL\_10383
CL\_5601
CL\_10645
CL\_10646
CL\_10647
CL\_10648
CL\_10649
CL\_10650
CL\_10651
CL\_10652
CL\_10653
CL\_10420
CL\_10654
CL\_10655
CL\_10656
CL\_10657
CL\_10658
CL\_11136
CL\_10659
CL\_10660
CL\_10662
CL\_10663
CL\_5014
CL\_5015
CL\_5016
CL\_11137
CL\_11138
CL\_5000
CL\_5001
CL\_11139
CL\_11140
CL\_11141
CL\_11142
CL\_11143
CL\_11144
CL\_11145
CL\_11146
CL\_11147
CL\_10661
CL\_4235
CL\_4236
CL\_4237
CL\_6239
CL\_5593
CL\_5592
CL\_5053
CL\_4253
CL\_4254
CL\_4255
CL\_4256
CL\_4257
CL\_4258
CL\_4259
CL\_4260
CL\_4261
CL\_5625
CL\_11131
CL\_5627
CL\_4263
CL\_5062
CL\_4265
CL\_4266
CL\_13400
CL\_5579
CL\_5577
CL\_5575
CL\_4270
CL\_4271
CL\_5574
CL\_5573
CL\_5572
CL\_5637
CL\_5638
CL\_5639
CL\_4277
CL\_4278
CL\_5567
CL\_5566
CL\_5565
CL\_5564
CL\_5563
CL\_5562
CL\_10389
CL\_11196
CL\_5515
CL\_5516
CL\_10427
CL\_11442
CL\_4287
CL\_5556
CL\_5555
CL\_5554
CL\_5553
CL\_5552
CL\_5551
CL\_5659
CL\_11338
CL\_4293
CL\_4294
CL\_5548
CL\_5662
CL\_4297
CL\_4299
CL\_4300
CL\_5664
CL\_6072
CL\_5665
CL\_5544
CL\_5543
CL\_5542
CL\_6935
CL\_5541
CL\_10411
CL\_5539
CL\_5536
CL\_5613
CL\_5614
CL\_22269
CL\_11132
CL\_5509
CL\_5510
CL\_5511
CL\_5512
CL\_5513
CL\_5019
CL\_10412
CL\_10413
CL\_6297
CL\_5692
CL\_5693
CL\_5533
CL\_11133
CL\_5300
CL\_5299
CL\_5298
CL\_5297
CL\_10392
CL\_10393
CL\_10395
