## Supplementary material for "A novel method for integrating genomic and Tn-Seq data to identify common *in vivo* fitness mechanisms across multiple bacterial species": S1 Dataset: CL_INS_76.html

Legend

 Mobile +extrachromosomalelementfunctions
 Regulatoryfunctions
 Hypothetical
 DNA Metabolism
 All EssentialGenes
 All Fitness Genes
 Proteinsynthesis/fate
 Other
 Transport +binding proteins
 All VFDB Genes

FULL


WINDOWSVGPNG

Trim RowsRemove SingletonsSave Fasta

CL\_922


CL\_4516


CL\_922


CL\_922


CL\_1072


CL\_922


CL\_922


CL\_922


CL\_4487


CL\_4519


CL\_922


CL\_922


CL\_922


CL\_1072


CL\_922


CL\_922


CL\_922


CL\_922


CL\_922


CL\_922


CL\_1208


CL\_4487


CL\_922


CL\_922


CL\_922


CL\_922

HighlightSelectShow Genomes


257

CL\_923


1

CL\_923


1

CL\_923


1

CL\_1207


1

CL\_923


1

CL\_4487


1

CL\_923


1

CL\_923


1

CL\_923


1

CL\_923


1

CL\_923


1

CL\_923


1

CL\_4427


1

CL\_923


1

CL\_923


1

CL\_923


1

CL\_4427


1

CL\_958


1

CL\_923


1

CL\_947


1

CL\_923


1

CL\_923


1

CL\_923


1

CL\_4487


1

CL\_923


1

CL\_923

fGI ID


CL\_INS\_76
CL\_INS\_99
CL\_INS\_76
CL\_INS\_76
CL\_INS\_76
CL\_INS\_76
CL\_INS\_76
CL\_INS\_76
CL\_INS\_76
CL\_INS\_76
CL\_INS\_76
CL\_INS\_76
CL\_INS\_76
CL\_INS\_76
CL\_INS\_76
CL\_INS\_76
CL\_INS\_76
CL\_INS\_76
CL\_INS\_76
CL\_INS\_76
CL\_INS\_155
CL\_INS\_237
CL\_INS\_207
CL\_INS\_385
CL\_INS\_155
CL\_INS\_155
CL\_INS\_155
CL\_INS\_76
CL\_INS\_385
CL\_INS\_207
CL\_INS\_207
CL\_INS\_207
CL\_INS\_207
CL\_INS\_76
CL\_INS\_207
CL\_INS\_76
CL\_INS\_76
CL\_INS\_382
CL\_INS\_382
CL\_INS\_382
CL\_INS\_382
CL\_INS\_382
CL\_INS\_83
CL\_INS\_382
CL\_INS\_86
CL\_INS\_99
CL\_INS\_76
CL\_INS\_76
CL\_INS\_382
CL\_INS\_76
CL\_INS\_382
CL\_INS\_382
CL\_INS\_382
CL\_INS\_382
CL\_INS\_99
CL\_INS\_382
CL\_INS\_99
CL\_INS\_99
CL\_INS\_384
CL\_INS\_382
CL\_INS\_99
CL\_INS\_76
CL\_INS\_76
CL\_INS\_76
CL\_INS\_76
CL\_INS\_76
CL\_INS\_76
CL\_INS\_76
CL\_INS\_382
CL\_INS\_382
CL\_INS\_382
CL\_INS\_382
CL\_INS\_86
CL\_INS\_382
CL\_INS\_99
CL\_INS\_99
CL\_INS\_99
CL\_INS\_382
CL\_INS\_382
CL\_INS\_382
CL\_INS\_382
CL\_INS\_382
CL\_INS\_382
CL\_INS\_382
CL\_INS\_382
CL\_INS\_382
CL\_INS\_99
CL\_INS\_83
CL\_INS\_83
CL\_INS\_83
CL\_INS\_384
CL\_INS\_384
CL\_INS\_384
CL\_INS\_384
CL\_INS\_384
CL\_INS\_384
CL\_INS\_99
CL\_INS\_99
CL\_INS\_99
CL\_INS\_79
CL\_INS\_382
CL\_INS\_146
CL\_INS\_146
CL\_INS\_247
CL\_INS\_76
CL\_INS\_76
CL\_INS\_76
CL\_INS\_76
CL\_INS\_79
CL\_INS\_382
CL\_INS\_382
CL\_INS\_382
CL\_INS\_382
CL\_INS\_382
CL\_INS\_382
CL\_INS\_76
CL\_INS\_76
CL\_INS\_76
CL\_INS\_99
CL\_INS\_382
CL\_INS\_382
CL\_INS\_382
CL\_INS\_382
CL\_INS\_146
CL\_INS\_146
CL\_INS\_382
CL\_INS\_382
CL\_INS\_99
CL\_INS\_382
CL\_INS\_76
CL\_INS\_99
CL\_INS\_382
CL\_INS\_76
CL\_INS\_382
CL\_INS\_99
CL\_INS\_99
CL\_INS\_382
CL\_INS\_382
CL\_INS\_382
CL\_INS\_382
CL\_INS\_382
CL\_INS\_382
CL\_INS\_382
CL\_INS\_79
CL\_INS\_382
CL\_INS\_83
CL\_INS\_83
CL\_INS\_83
CL\_INS\_83
CL\_INS\_83
CL\_INS\_83
CL\_INS\_83
CL\_INS\_83
CL\_INS\_83
CL\_INS\_83
CL\_INS\_83
CL\_INS\_83
CL\_INS\_382
CL\_INS\_382
CL\_INS\_83
CL\_INS\_83
CL\_INS\_83
CL\_INS\_83
CL\_INS\_83
CL\_INS\_83
CL\_INS\_382
CL\_INS\_76
CL\_INS\_76
CL\_INS\_382
CL\_INS\_79
CL\_INS\_382
CL\_INS\_382
CL\_INS\_382
CL\_INS\_153
CL\_INS\_382
CL\_INS\_382
CL\_INS\_382
CL\_INS\_382
CL\_INS\_382
CL\_INS\_382
CL\_INS\_99
CL\_INS\_99
CL\_INS\_99
CL\_INS\_99
CL\_INS\_99
CL\_INS\_99
CL\_INS\_99
CL\_INS\_99
CL\_INS\_99
CL\_INS\_99
CL\_INS\_382
CL\_INS\_99
CL\_INS\_382
CL\_INS\_382
CL\_INS\_382
CL\_INS\_382
CL\_INS\_382
CL\_INS\_382
CL\_INS\_382
CL\_INS\_382
CL\_INS\_99
CL\_INS\_382
CL\_INS\_382
CL\_INS\_382
CL\_INS\_382
CL\_INS\_382
CL\_INS\_382
CL\_INS\_382
CL\_INS\_382
CL\_INS\_382
CL\_INS\_382
CL\_INS\_382
CL\_INS\_382
CL\_INS\_382
CL\_INS\_382
CL\_INS\_382
CL\_INS\_382
CL\_INS\_99
CL\_INS\_99
CL\_INS\_382
CL\_INS\_382
CL\_INS\_83
CL\_INS\_83
CL\_INS\_99
CL\_INS\_382
CL\_INS\_382
CL\_INS\_382
CL\_INS\_382
CL\_INS\_382
CL\_INS\_382
CL\_INS\_382
CL\_INS\_382
CL\_INS\_382
CL\_INS\_382
CL\_INS\_382
CL\_INS\_382
CL\_INS\_382
CL\_INS\_382
CL\_INS\_99
CL\_INS\_99
CL\_INS\_382
CL\_INS\_382
CL\_INS\_382
CL\_INS\_382
CL\_INS\_382
CL\_INS\_382
CL\_INS\_10
CL\_INS\_10
CL\_INS\_10
CL\_INS\_99
CL\_INS\_382
CL\_INS\_382
CL\_INS\_382
CL\_INS\_382
CL\_INS\_382
CL\_INS\_382
CL\_INS\_382
CL\_INS\_382
CL\_INS\_382
CL\_INS\_384
CL\_INS\_146
CL\_INS\_207
CL\_INS\_382
CL\_INS\_207
CL\_INS\_207
CL\_INS\_207
CL\_INS\_207
CL\_INS\_207
CL\_INS\_382
CL\_INS\_382
CL\_INS\_207
CL\_INS\_207
CL\_INS\_207
CL\_INS\_207
CL\_INS\_99
CL\_INS\_382
CL\_INS\_382
CL\_INS\_382
CL\_INS\_382
CL\_INS\_382
CL\_INS\_382
CL\_INS\_382
CL\_INS\_76
CL\_INS\_76
CL\_INS\_382
CL\_INS\_128
CL\_INS\_76
CL\_INS\_76
CL\_INS\_76
CL\_INS\_76
CL\_INS\_76
CL\_INS\_76
CL\_INS\_76
CL\_INS\_76
CL\_INS\_76
CL\_INS\_76
CL\_INS\_76
CL\_INS\_76
CL\_INS\_76
CL\_INS\_76
CL\_INS\_99
CL\_INS\_76
CL\_INS\_83
CL\_INS\_76
CL\_INS\_83
CL\_INS\_153
CL\_INS\_76
CL\_INS\_83
CL\_INS\_83
CL\_INS\_76
CL\_INS\_76
CL\_INS\_60
CL\_INS\_76
CL\_INS\_83
CL\_INS\_83
CL\_INS\_76
CL\_INS\_99
CL\_INS\_99
CL\_INS\_382
CL\_INS\_382
CL\_INS\_382
CL\_INS\_382
CL\_INS\_382
CL\_INS\_382
CL\_INS\_382
CL\_INS\_382
CL\_INS\_382
CL\_INS\_382
CL\_INS\_382
CL\_INS\_382
CL\_INS\_207
CL\_INS\_382
CL\_INS\_382
CL\_INS\_382
CL\_INS\_99
CL\_INS\_207
CL\_INS\_382
CL\_INS\_207
CL\_INS\_207
CL\_INS\_207
CL\_INS\_207
CL\_INS\_382
CL\_INS\_99
CL\_INS\_207
CL\_INS\_207
CL\_INS\_382
CL\_INS\_99
CL\_INS\_86
CL\_INS\_384
CL\_INS\_87
CL\_INS\_76
CL\_INS\_76
CL\_INS\_76
CL\_INS\_99
CL\_INS\_99
CL\_INS\_99
CL\_INS\_149
CL\_INS\_149
CL\_INS\_149
CL\_INS\_99
CL\_INS\_99
CL\_INS\_149
CL\_INS\_149
CL\_INS\_99
CL\_INS\_149
CL\_INS\_149
CL\_INS\_149
CL\_INS\_149
CL\_INS\_99
CL\_INS\_99
CL\_INS\_99
CL\_INS\_99
CL\_INS\_99
CL\_INS\_99
CL\_INS\_99
CL\_INS\_99
CL\_INS\_99
CL\_INS\_99
CL\_INS\_99
CL\_INS\_99
CL\_INS\_99
CL\_INS\_99
CL\_INS\_99
CL\_INS\_99
CL\_INS\_99
CL\_INS\_99
CL\_INS\_99
CL\_INS\_99
CL\_INS\_99
CL\_INS\_99
CL\_INS\_99
CL\_INS\_99
CL\_INS\_99
CL\_INS\_99
CL\_INS\_99
CL\_INS\_99
CL\_INS\_99
CL\_INS\_99
CL\_INS\_99
CL\_INS\_99
CL\_INS\_99
CL\_INS\_382
CL\_INS\_382
CL\_INS\_99
CL\_INS\_99
CL\_INS\_382
CL\_INS\_382
CL\_INS\_382
CL\_INS\_382
CL\_INS\_382
CL\_INS\_382
CL\_INS\_382
CL\_INS\_382
CL\_INS\_76
CL\_INS\_382
CL\_INS\_382
CL\_INS\_146
CL\_INS\_76
CL\_INS\_382
CL\_INS\_99
CL\_INS\_76
CL\_INS\_99
CL\_INS\_99
CL\_INS\_76
CL\_INS\_76
CL\_INS\_76
CL\_INS\_76
CL\_INS\_99
CL\_INS\_76
CL\_INS\_76
CL\_INS\_76
CL\_INS\_76
CL\_INS\_76
CL\_INS\_76
CL\_INS\_76
CL\_INS\_76
CL\_INS\_76
CL\_INS\_76
CL\_INS\_76
CL\_INS\_76
CL\_INS\_76
CL\_INS\_76
CL\_INS\_76
CL\_INS\_76
CL\_INS\_76
CL\_INS\_76
CL\_INS\_76
CL\_INS\_76
CL\_INS\_76
CL\_INS\_76
CL\_INS\_76
CL\_INS\_76
CL\_INS\_76
CL\_INS\_76
CL\_INS\_76
CL\_INS\_76
CL\_INS\_76
CL\_INS\_76
CL\_INS\_76
CL\_INS\_76
CL\_INS\_76
CL\_INS\_76
CL\_INS\_76
CL\_INS\_83
CL\_INS\_79
CL\_INS\_76
CL\_INS\_76
CL\_INS\_76
CL\_INS\_153
CL\_INS\_153
CL\_INS\_153
CL\_INS\_153
CL\_INS\_153
CL\_INS\_99
CL\_INS\_153
CL\_INS\_382
CL\_INS\_76
CL\_INS\_76
CL\_INS\_76
CL\_INS\_76
CL\_INS\_76
CL\_INS\_76
CL\_INS\_76
CL\_INS\_76
CL\_INS\_76
CL\_INS\_76
CL\_INS\_76
CL\_INS\_76
CL\_INS\_76
CL\_INS\_76
CL\_INS\_76
CL\_INS\_76
CL\_INS\_76
CL\_INS\_83
CL\_INS\_76
CL\_INS\_76
CL\_INS\_76
CL\_INS\_76
CL\_INS\_153
CL\_INS\_382
CL\_INS\_382
CL\_INS\_76
CL\_INS\_99
CL\_INS\_76
CL\_INS\_76
CL\_INS\_76
CL\_INS\_76
CL\_INS\_76
CL\_INS\_76
CL\_INS\_76
CL\_INS\_76
CL\_INS\_76
CL\_INS\_153
CL\_INS\_76
CL\_INS\_76
CL\_INS\_76
CL\_INS\_76
CL\_INS\_76
CL\_INS\_76
CL\_INS\_76
CL\_INS\_76
CL\_INS\_76
CL\_INS\_76
CL\_INS\_76
CL\_INS\_76
CL\_INS\_76
CL\_INS\_76
CL\_INS\_76
CL\_INS\_76
CL\_INS\_99
CL\_INS\_99
CL\_INS\_382
CL\_INS\_76
CL\_INS\_382
CL\_INS\_60
CL\_INS\_99
CL\_INS\_60
CL\_INS\_76
CL\_INS\_385
CL\_INS\_385
CL\_INS\_79
CL\_INS\_76
CL\_INS\_76
CL\_INS\_79
CL\_INS\_382
CL\_INS\_79
CL\_INS\_79
CL\_INS\_382
CL\_INS\_382
CL\_INS\_76
CL\_INS\_76
CL\_INS\_76
CL\_INS\_76
CL\_INS\_76
CL\_INS\_76
CL\_INS\_76
CL\_INS\_76
CL\_INS\_76
CL\_INS\_76
CL\_INS\_76
CL\_INS\_76
CL\_INS\_382
CL\_INS\_76
CL\_INS\_382
CL\_INS\_382
CL\_INS\_76
CL\_INS\_76
CL\_INS\_382
CL\_INS\_382
CL\_INS\_83
CL\_INS\_146
CL\_INS\_382
CL\_INS\_382
CL\_INS\_83
CL\_INS\_99
CL\_INS\_146
CL\_INS\_83
CL\_INS\_99
CL\_INS\_83
CL\_INS\_83
CL\_INS\_83
CL\_INS\_83
CL\_INS\_83
CL\_INS\_83
CL\_INS\_83
CL\_INS\_83
CL\_INS\_83
CL\_INS\_83
CL\_INS\_83
CL\_INS\_83
CL\_INS\_83
CL\_INS\_83
CL\_INS\_83
CL\_INS\_83
CL\_INS\_83
CL\_INS\_207
CL\_INS\_83
CL\_INS\_83
CL\_INS\_83
CL\_INS\_83
CL\_INS\_83
CL\_INS\_83
CL\_INS\_99
CL\_INS\_83
CL\_INS\_83
CL\_INS\_83
CL\_INS\_83
CL\_INS\_83
CL\_INS\_83
CL\_INS\_83
CL\_INS\_83
CL\_INS\_83
CL\_INS\_76
CL\_INS\_382
CL\_INS\_76
CL\_INS\_76
CL\_INS\_76
CL\_INS\_76
CL\_INS\_76
CL\_INS\_79
CL\_INS\_79
CL\_INS\_79
CL\_INS\_76
CL\_INS\_76
CL\_INS\_76
CL\_INS\_76
CL\_INS\_76
CL\_INS\_76
CL\_INS\_76
CL\_INS\_76
CL\_INS\_76
CL\_INS\_76
CL\_INS\_76
CL\_INS\_76
CL\_INS\_76
CL\_INS\_79
CL\_INS\_79
CL\_INS\_79
CL\_INS\_79
CL\_INS\_79
CL\_INS\_79
CL\_INS\_79
CL\_INS\_79
CL\_INS\_79
CL\_INS\_79
CL\_INS\_79
CL\_INS\_79
CL\_INS\_79
CL\_INS\_79
CL\_INS\_79
CL\_INS\_79
CL\_INS\_79
CL\_INS\_79
CL\_INS\_79
CL\_INS\_79
CL\_INS\_79
CL\_INS\_79
CL\_INS\_79
CL\_INS\_79
CL\_INS\_79
CL\_INS\_79
CL\_INS\_79
CL\_INS\_79
CL\_INS\_382
CL\_INS\_128
CL\_INS\_128
CL\_INS\_128
CL\_INS\_79
CL\_INS\_79
CL\_INS\_382
CL\_INS\_382
CL\_INS\_76
CL\_INS\_382
CL\_INS\_382
CL\_INS\_382
CL\_INS\_382
CL\_INS\_382
CL\_INS\_382
CL\_INS\_382
CL\_INS\_382
CL\_INS\_76
CL\_INS\_76
CL\_INS\_76
CL\_INS\_207
CL\_INS\_76
CL\_INS\_382
CL\_INS\_382
CL\_INS\_382
CL\_INS\_382
CL\_INS\_99
CL\_INS\_382
CL\_INS\_382
CL\_INS\_382
CL\_INS\_384
CL\_INS\_20
CL\_INS\_99
CL\_INS\_382
CL\_INS\_382
CL\_INS\_99
CL\_INS\_86
CL\_INS\_99
CL\_INS\_86
CL\_INS\_382
CL\_INS\_154
CL\_INS\_382
CL\_INS\_382
CL\_INS\_382
CL\_INS\_132
CL\_INS\_132
CL\_INS\_106
CL\_INS\_106
CL\_INS\_382
CL\_INS\_132
CL\_INS\_132
CL\_INS\_132
CL\_INS\_154
CL\_INS\_132
CL\_INS\_132
CL\_INS\_132
CL\_INS\_132
CL\_INS\_132
CL\_INS\_132
CL\_INS\_132
CL\_INS\_132
CL\_INS\_384
CL\_INS\_204
CL\_INS\_204
CL\_INS\_384
CL\_INS\_384
CL\_INS\_99
CL\_INS\_384
CL\_INS\_384
CL\_INS\_146
CL\_INS\_99
CL\_INS\_99
CL\_INS\_99
CL\_INS\_99
CL\_INS\_382
CL\_INS\_382
CL\_INS\_382
CL\_INS\_99
CL\_INS\_99
CL\_INS\_99
CL\_INS\_76
CL\_INS\_76
CL\_INS\_76
CL\_INS\_76
CL\_INS\_76
CL\_INS\_76
CL\_INS\_76
CL\_INS\_76
CL\_INS\_76
CL\_INS\_76
CL\_INS\_76
CL\_INS\_76
CL\_INS\_382
CL\_INS\_76
CL\_INS\_76
CL\_INS\_76
CL\_INS\_76
CL\_INS\_76
CL\_INS\_76
CL\_INS\_76
CL\_INS\_76
CL\_INS\_99
CL\_INS\_99
CL\_INS\_99
CL\_INS\_99
CL\_INS\_99
CL\_INS\_99
CL\_INS\_99
CL\_INS\_86
CL\_INS\_86
CL\_INS\_86
CL\_INS\_76
CL\_INS\_76
CL\_INS\_76
CL\_INS\_76
CL\_INS\_76
CL\_INS\_76
CL\_INS\_76
CL\_INS\_83
CL\_INS\_76
CL\_INS\_76
CL\_INS\_76
CL\_INS\_99
CL\_INS\_382
CL\_INS\_382
CL\_INS\_382
CL\_INS\_79
CL\_INS\_79
CL\_INS\_382
CL\_INS\_382
CL\_INS\_382
CL\_INS\_79
CL\_INS\_99
CL\_INS\_79
CL\_INS\_79
CL\_INS\_79
CL\_INS\_79
CL\_INS\_79
CL\_INS\_79
CL\_INS\_79
CL\_INS\_79
CL\_INS\_79
CL\_INS\_79
CL\_INS\_79
CL\_INS\_79
CL\_INS\_79
CL\_INS\_79
CL\_INS\_79
CL\_INS\_79
CL\_INS\_79
CL\_INS\_79
CL\_INS\_79
CL\_INS\_79
CL\_INS\_79
CL\_INS\_79
CL\_INS\_79
CL\_INS\_79
CL\_INS\_79
CL\_INS\_79
CL\_INS\_79
CL\_INS\_79
CL\_INS\_99
CL\_INS\_382
CL\_INS\_99
CL\_INS\_99
CL\_INS\_99
CL\_INS\_83
CL\_INS\_99
CL\_INS\_131
CL\_INS\_128
CL\_INS\_384
CL\_INS\_384
CL\_INS\_384
CL\_INS\_384
CL\_INS\_99
CL\_INS\_99
CL\_INS\_99
CL\_INS\_382
CL\_INS\_382
CL\_INS\_382
CL\_INS\_99
CL\_INS\_382
CL\_INS\_382
CL\_INS\_99
CL\_INS\_382
CL\_INS\_382
CL\_INS\_382
CL\_INS\_385
CL\_INS\_385
CL\_INS\_385
CL\_INS\_385
CL\_INS\_99
CL\_INS\_385
CL\_INS\_382
CL\_INS\_382
CL\_INS\_384
CL\_INS\_382
CL\_INS\_382
CL\_INS\_384
CL\_INS\_384
CL\_INS\_382
CL\_INS\_83
CL\_INS\_99
CL\_INS\_99
CL\_INS\_99
CL\_INS\_99
CL\_INS\_99
CL\_INS\_382
CL\_INS\_382
CL\_INS\_382
CL\_INS\_382
CL\_INS\_382
CL\_INS\_382
CL\_INS\_382
CL\_INS\_76
CL\_INS\_382
CL\_INS\_382
CL\_INS\_382
CL\_INS\_128
CL\_INS\_382
CL\_INS\_99
CL\_INS\_99
CL\_INS\_382
CL\_INS\_382
CL\_INS\_382
CL\_INS\_382
CL\_INS\_382
CL\_INS\_99
CL\_INS\_382
CL\_INS\_76
CL\_INS\_146
CL\_INS\_382
CL\_INS\_382
CL\_INS\_99
CL\_INS\_99
CL\_INS\_99
CL\_INS\_128
CL\_INS\_382
CL\_INS\_382
CL\_INS\_382
CL\_INS\_382
CL\_INS\_382
CL\_INS\_83
CL\_INS\_382
Cluster ID


CL\_26941
CL\_10521
CL\_20249
CL\_16137
CL\_16138
CL\_16139
CL\_21195
CL\_21194
CL\_21193
CL\_21192
CL\_21191
CL\_21190
CL\_21189
CL\_21188
CL\_21187
CL\_21186
CL\_21185
CL\_21184
CL\_21183
CL\_21182
CL\_6000
CL\_6001
CL\_6002
CL\_9723
CL\_9199
CL\_7780
CL\_17024
CL\_21181
CL\_7778
CL\_6012
CL\_6013
CL\_6014
CL\_6015
CL\_21180
CL\_6017
CL\_21179
CL\_31700
CL\_31699
CL\_31698
CL\_31697
CL\_31696
CL\_31695
CL\_14234
CL\_8201
CL\_8585
CL\_9159
CL\_31690
CL\_16110
CL\_16111
CL\_17263
CL\_16112
CL\_1527
CL\_1526
CL\_8197
CL\_14692
CL\_8196
CL\_14693
CL\_16113
CL\_16114
CL\_15910
CL\_17303
CL\_32904
CL\_31689
CL\_33351
CL\_37262
CL\_37263
CL\_37264
CL\_17304
CL\_17305
CL\_23879
CL\_23880
CL\_4560
CL\_8674
CL\_14237
CL\_18046
CL\_18045
CL\_18044
CL\_5276
CL\_8995
CL\_8996
CL\_8997
CL\_7124
CL\_10937
CL\_9150
CL\_9151
CL\_8675
CL\_15790
CL\_28521
CL\_28520
CL\_28519
CL\_16115
CL\_16116
CL\_16117
CL\_16118
CL\_16119
CL\_16120
CL\_17306
CL\_4558
CL\_4557
CL\_17307
CL\_10938
CL\_17308
CL\_17309
CL\_6844
CL\_28157
CL\_28156
CL\_28155
CL\_33349
CL\_32650
CL\_5282
CL\_6773
CL\_11922
CL\_4546
CL\_17066
CL\_16967
CL\_31414
CL\_31413
CL\_31412
CL\_16661
CL\_17485
CL\_15862
CL\_16419
CL\_16418
CL\_17310
CL\_17311
CL\_5343
CL\_16417
CL\_18043
CL\_8678
CL\_31416
CL\_31415
CL\_4548
CL\_37265
CL\_8679
CL\_18042
CL\_18041
CL\_4552
CL\_4551
CL\_15766
CL\_31694
CL\_31693
CL\_31692
CL\_31691
CL\_17349
CL\_13108
CL\_17350
CL\_17351
CL\_17352
CL\_17353
CL\_17354
CL\_17355
CL\_17356
CL\_17357
CL\_17358
CL\_17359
CL\_17360
CL\_17361
CL\_8690
CL\_27035
CL\_17362
CL\_17363
CL\_17364
CL\_17365
CL\_17366
CL\_17367
CL\_8680
CL\_37266
CL\_12410
CL\_15904
CL\_32681
CL\_4550
CL\_8683
CL\_8684
CL\_17266
CL\_1515
CL\_8688
CL\_32682
CL\_32683
CL\_16126
CL\_16127
CL\_18040
CL\_18039
CL\_18038
CL\_18037
CL\_18036
CL\_18035
CL\_4542
CL\_16660
CL\_18034
CL\_18033
CL\_15997
CL\_18032
CL\_15999
CL\_16000
CL\_9291
CL\_9290
CL\_9289
CL\_9288
CL\_9287
CL\_9286
CL\_9285
CL\_9284
CL\_9283
CL\_9282
CL\_9281
CL\_9280
CL\_9279
CL\_9278
CL\_9277
CL\_9276
CL\_9275
CL\_9274
CL\_9273
CL\_9272
CL\_9271
CL\_9270
CL\_9269
CL\_18031
CL\_9267
CL\_9266
CL\_4644
CL\_36220
CL\_36221
CL\_6043
CL\_4413
CL\_4414
CL\_4532
CL\_4416
CL\_9265
CL\_9264
CL\_9262
CL\_9261
CL\_16006
CL\_16007
CL\_9258
CL\_9257
CL\_16008
CL\_9256
CL\_18030
CL\_18029
CL\_9254
CL\_9253
CL\_9252
CL\_16009
CL\_9251
CL\_9250
CL\_9249
CL\_9248
CL\_9247
CL\_18028
CL\_11920
CL\_14246
CL\_14247
CL\_36211
CL\_36212
CL\_18027
CL\_18026
CL\_9761
CL\_4535
CL\_20250
CL\_16651
CL\_9122
CL\_7533
CL\_7532
CL\_7531
CL\_7530
CL\_7529
CL\_7528
CL\_7527
CL\_6741
CL\_7526
CL\_7525
CL\_7524
CL\_7523
CL\_7522
CL\_4534
CL\_533
CL\_4533
CL\_4415
CL\_6045
CL\_14371
CL\_7116
CL\_31682
CL\_31681
CL\_6461
CL\_26458
CL\_31680
CL\_31679
CL\_33338
CL\_33337
CL\_33336
CL\_31678
CL\_31677
CL\_32111
CL\_32917
CL\_31676
CL\_31675
CL\_32110
CL\_32109
CL\_32108
CL\_22575
CL\_31674
CL\_17384
CL\_31673
CL\_17385
CL\_33335
CL\_32918
CL\_17386
CL\_17387
CL\_32107
CL\_32106
CL\_19004
CL\_32105
CL\_17388
CL\_17389
CL\_32104
CL\_18025
CL\_18024
CL\_4531
CL\_4417
CL\_4418
CL\_4419
CL\_4420
CL\_4421
CL\_4525
CL\_4423
CL\_4424
CL\_4425
CL\_4530
CL\_4529
CL\_17912
CL\_4528
CL\_4527
CL\_4526
CL\_6046
CL\_17913
CL\_4629
CL\_17914
CL\_17915
CL\_17916
CL\_13074
CL\_8169
CL\_4485
CL\_17917
CL\_17918
CL\_4517
CL\_4431
CL\_4515
CL\_28083
CL\_4513
CL\_37267
CL\_37268
CL\_37269
CL\_6047
CL\_6048
CL\_6049
CL\_4689
CL\_4688
CL\_4687
CL\_6050
CL\_6051
CL\_4685
CL\_4684
CL\_6052
CL\_4682
CL\_4681
CL\_4679
CL\_4678
CL\_18023
CL\_18022
CL\_18021
CL\_18020
CL\_18019
CL\_18018
CL\_18017
CL\_18016
CL\_18015
CL\_18014
CL\_18013
CL\_18012
CL\_18011
CL\_18010
CL\_18009
CL\_18008
CL\_18007
CL\_18006
CL\_18005
CL\_18004
CL\_18003
CL\_18002
CL\_18001
CL\_18000
CL\_17999
CL\_17998
CL\_17997
CL\_17996
CL\_17995
CL\_17994
CL\_17993
CL\_17992
CL\_17991
CL\_6042
CL\_14245
CL\_17990
CL\_15900
CL\_16414
CL\_16415
CL\_15800
CL\_4647
CL\_16416
CL\_7539
CL\_16445
CL\_16444
CL\_31411
CL\_12994
CL\_16615
CL\_20782
CL\_20883
CL\_15773
CL\_20884
CL\_31410
CL\_23513
CL\_23514
CL\_20779
CL\_20885
CL\_20886
CL\_20887
CL\_20888
CL\_20889
CL\_20890
CL\_20891
CL\_20892
CL\_20893
CL\_20894
CL\_20895
CL\_20896
CL\_20897
CL\_20898
CL\_20899
CL\_20900
CL\_20901
CL\_20902
CL\_20903
CL\_20904
CL\_20905
CL\_20906
CL\_20907
CL\_20908
CL\_20909
CL\_20910
CL\_20911
CL\_20912
CL\_20913
CL\_20914
CL\_20915
CL\_20916
CL\_20917
CL\_20918
CL\_20919
CL\_20920
CL\_20921
CL\_20922
CL\_16140
CL\_19950
CL\_17312
CL\_33127
CL\_33348
CL\_17313
CL\_17314
CL\_31688
CL\_31687
CL\_31686
CL\_31685
CL\_12382
CL\_31684
CL\_11919
CL\_31409
CL\_31408
CL\_31407
CL\_31406
CL\_31405
CL\_31404
CL\_31403
CL\_31402
CL\_31401
CL\_31400
CL\_31399
CL\_31398
CL\_31397
CL\_31396
CL\_31395
CL\_31394
CL\_31393
CL\_28331
CL\_31392
CL\_31391
CL\_31390
CL\_31389
CL\_31683
CL\_19014
CL\_19013
CL\_28153
CL\_13515
CL\_17315
CL\_17316
CL\_17317
CL\_17318
CL\_17319
CL\_17320
CL\_17321
CL\_17322
CL\_17323
CL\_17324
CL\_17325
CL\_17326
CL\_17327
CL\_17328
CL\_17329
CL\_17330
CL\_17331
CL\_17332
CL\_17333
CL\_17334
CL\_17335
CL\_17336
CL\_17337
CL\_17338
CL\_17339
CL\_17340
CL\_13509
CL\_12791
CL\_8167
CL\_17341
CL\_9124
CL\_17342
CL\_9160
CL\_17343
CL\_17344
CL\_9101
CL\_9102
CL\_31631
CL\_33347
CL\_33346
CL\_32124
CL\_7118
CL\_32651
CL\_19217
CL\_13524
CL\_9098
CL\_32123
CL\_32122
CL\_32121
CL\_32120
CL\_32119
CL\_32118
CL\_32117
CL\_32116
CL\_32115
CL\_32114
CL\_32113
CL\_32112
CL\_8696
CL\_28154
CL\_12999
CL\_531
CL\_33345
CL\_33344
CL\_17368
CL\_7537
CL\_17369
CL\_17370
CL\_8698
CL\_13000
CL\_17371
CL\_17372
CL\_17373
CL\_28097
CL\_28096
CL\_28095
CL\_28094
CL\_28093
CL\_28092
CL\_28091
CL\_28090
CL\_28089
CL\_32697
CL\_28088
CL\_28087
CL\_28086
CL\_28085
CL\_17411
CL\_17412
CL\_17413
CL\_17414
CL\_32698
CL\_32699
CL\_32700
CL\_32701
CL\_32702
CL\_32703
CL\_16628
CL\_17374
CL\_9157
CL\_17375
CL\_17376
CL\_17377
CL\_17378
CL\_17379
CL\_17380
CL\_17381
CL\_17382
CL\_17383
CL\_33343
CL\_28098
CL\_34478
CL\_33342
CL\_33341
CL\_33340
CL\_33339
CL\_32652
CL\_32653
CL\_32654
CL\_32905
CL\_32906
CL\_32907
CL\_32908
CL\_32909
CL\_32910
CL\_32911
CL\_32912
CL\_32913
CL\_33126
CL\_32914
CL\_32915
CL\_32916
CL\_32655
CL\_32656
CL\_32657
CL\_32658
CL\_32659
CL\_32660
CL\_32661
CL\_32662
CL\_32663
CL\_32664
CL\_32665
CL\_32666
CL\_32667
CL\_32668
CL\_32669
CL\_32670
CL\_32671
CL\_32672
CL\_32673
CL\_32674
CL\_32675
CL\_17390
CL\_32677
CL\_32676
CL\_32678
CL\_32679
CL\_32680
CL\_15757
CL\_1525
CL\_17347
CL\_17348
CL\_14240
CL\_23907
CL\_14694
CL\_4543
CL\_4544
CL\_37271
CL\_16011
CL\_27089
CL\_17037
CL\_36206
CL\_36207
CL\_36208
CL\_36209
CL\_36210
CL\_37272
CL\_37273
CL\_37274
CL\_17906
CL\_37275
CL\_4545
CL\_8187
CL\_8186
CL\_8689
CL\_15902
CL\_7540
CL\_10980
CL\_4648
CL\_16128
CL\_16129
CL\_15801
CL\_5340
CL\_526
CL\_8183
CL\_8180
CL\_8179
CL\_8178
CL\_8647
CL\_16130
CL\_8648
CL\_16131
CL\_13512
CL\_8650
CL\_8652
CL\_12139
CL\_12138
CL\_12137
CL\_16132
CL\_10497
CL\_13102
CL\_16133
CL\_10498
CL\_10499
CL\_10500
CL\_10502
CL\_10503
CL\_10504
CL\_10505
CL\_10506
CL\_16134
CL\_1098
CL\_1099
CL\_14704
CL\_14705
CL\_10971
CL\_16135
CL\_16136
CL\_8177
CL\_8176
CL\_8175
CL\_8174
CL\_8173
CL\_8172
CL\_8171
CL\_8170
CL\_32684
CL\_17267
CL\_17268
CL\_17269
CL\_17270
CL\_17271
CL\_17272
CL\_17273
CL\_17274
CL\_17275
CL\_17276
CL\_17277
CL\_17278
CL\_17279
CL\_17280
CL\_7534
CL\_17281
CL\_17282
CL\_17283
CL\_17284
CL\_17285
CL\_17286
CL\_17287
CL\_17288
CL\_17289
CL\_17290
CL\_17291
CL\_17292
CL\_15782
CL\_8155
CL\_7112
CL\_4430
CL\_4488
CL\_1495
CL\_36213
CL\_17293
CL\_17294
CL\_17295
CL\_17296
CL\_17297
CL\_17298
CL\_17299
CL\_17300
CL\_17301
CL\_17302
CL\_32685
CL\_6452
CL\_4646
CL\_8694
CL\_32686
CL\_32687
CL\_12762
CL\_12995
CL\_12996
CL\_16616
CL\_13561
CL\_16350
CL\_27155
CL\_32688
CL\_17242
CL\_32689
CL\_31784
CL\_28309
CL\_31783
CL\_31782
CL\_31781
CL\_31780
CL\_31779
CL\_31778
CL\_31777
CL\_31776
CL\_31775
CL\_31774
CL\_31773
CL\_31772
CL\_32690
CL\_26212
CL\_26213
CL\_21941
CL\_32691
CL\_32692
CL\_32693
CL\_32694
CL\_32695
CL\_17989
CL\_5283
CL\_17988
CL\_17987
CL\_16121
CL\_28518
CL\_15767
CL\_11660
CL\_28304
CL\_16122
CL\_16123
CL\_16124
CL\_16125
CL\_17986
CL\_17985
CL\_17984
CL\_5278
CL\_4559
CL\_10936
CL\_31168
CL\_8676
CL\_19017
CL\_20768
CL\_8677
CL\_16422
CL\_16421
CL\_17491
CL\_17490
CL\_17489
CL\_17488
CL\_17487
CL\_17486
CL\_4556
CL\_10514
CL\_20253
CL\_8681
CL\_8682
CL\_20252
CL\_20251
CL\_4555
CL\_20882
CL\_31167
CL\_17983
CL\_17982
CL\_17981
CL\_17980
CL\_1523
CL\_11926
CL\_11925
CL\_11924
CL\_12697
CL\_13300
CL\_14238
CL\_33350
CL\_14239
CL\_15788
CL\_15789
CL\_14241
CL\_8673
CL\_17979
CL\_17040
CL\_8670
CL\_30168
CL\_7126
CL\_14236
CL\_36204
CL\_4563
CL\_16423
CL\_37270
CL\_19961
CL\_17264
CL\_4562
CL\_1312
CL\_1313
CL\_15759
CL\_17265
CL\_36205
CL\_8199
CL\_14235
CL\_8200
CL\_4565
CL\_17346
CL\_8198
