## Supplementary material for "A novel method for integrating genomic and Tn-Seq data to identify common *in vivo* fitness mechanisms across multiple bacterial species": S1 Dataset: CL_INS_78.html

Legend

 Mobile +extrachromosomalelementfunctions
 Hypothetical
 All EssentialGenes
 Other
 All VFDB Genes

FULL


WINDOWSVGPNG

Trim RowsRemove SingletonsSave Fasta

CL\_953


CL\_953


CL\_951


CL\_953


CL\_953


CL\_953


CL\_953


CL\_953


CL\_953


CL\_940

HighlightSelectShow Genomes


182

CL\_955


84

CL\_955


2

CL\_955


1

CL\_966


1

CL\_946


1

CL\_966


1

CL\_956


1

CL\_955


1

CL\_956


1

CL\_955

fGI ID


CL\_INS\_78
CL\_INS\_78
Cluster ID


CL\_11928
CL\_954
