## Supplementary material for "A novel method for integrating genomic and Tn-Seq data to identify common *in vivo* fitness mechanisms across multiple bacterial species": S1 Dataset: CL_INS_80.html

Legend

 Mobile +extrachromosomalelementfunctions
 Regulatoryfunctions
 Hypothetical
 DNA Metabolism
 All EssentialGenes
 Biosynthesis ofcofactors,prostheticgroups, +carriers
 All Fitness Genes
 Cell Envelope
 Proteinsynthesis/fate
 Other
 Centralintermediarymetabolism
 EnergyMetabolism
 Transport +binding proteins
 All VFDB Genes

FULL


WINDOWSVGPNG

Trim RowsRemove SingletonsSave Fasta

CL\_973


CL\_973


CL\_973


CL\_973


CL\_973


CL\_973


CL\_973


CL\_973


CL\_973


CL\_973


CL\_973


CL\_973


CL\_973


CL\_973


CL\_973


CL\_973


CL\_973


CL\_973


CL\_973

HighlightSelectShow Genomes


214

CL\_974


33

CL\_982


5

CL\_982


2

CL\_982


1

CL\_234


1

CL\_982


1

CL\_1935


1

CL\_983


1

CL\_1935


1

CL\_1935


1

CL\_1935


1

CL\_1935


1

CL\_1935


1

CL\_1935


1

CL\_982


1

CL\_234


1

CL\_1935


1

CL\_1935


1

CL\_1935

fGI ID


CL\_INS\_237
CL\_INS\_80
CL\_INS\_80
CL\_INS\_80
CL\_INS\_70
CL\_INS\_70
CL\_INS\_80
CL\_INS\_80
CL\_INS\_80
CL\_INS\_70
CL\_INS\_80
CL\_INS\_70
CL\_INS\_70
CL\_INS\_70
CL\_INS\_70
CL\_INS\_70
CL\_INS\_70
CL\_INS\_247
CL\_INS\_247
CL\_INS\_247
CL\_INS\_247
CL\_INS\_247
CL\_INS\_247
CL\_INS\_247
CL\_INS\_247
CL\_INS\_247
CL\_INS\_30
CL\_INS\_80
CL\_INS\_237
CL\_INS\_20
CL\_INS\_80
CL\_INS\_80
CL\_INS\_149
CL\_INS\_80
CL\_INS\_80
CL\_INS\_80
CL\_INS\_80
CL\_INS\_174
CL\_INS\_80
CL\_INS\_80
CL\_INS\_70
CL\_INS\_70
CL\_INS\_70
CL\_INS\_30
CL\_INS\_70
CL\_INS\_70
CL\_INS\_207
CL\_INS\_123
CL\_INS\_123
CL\_INS\_347
CL\_INS\_70
CL\_INS\_70
CL\_INS\_70
CL\_INS\_70
CL\_INS\_70
CL\_INS\_70
CL\_INS\_70
CL\_INS\_20
CL\_INS\_70
CL\_INS\_70
CL\_INS\_80
CL\_INS\_80
CL\_INS\_128
CL\_INS\_128
CL\_INS\_128
CL\_INS\_70
CL\_INS\_237
CL\_INS\_70
CL\_INS\_70
CL\_INS\_70
CL\_INS\_70
CL\_INS\_237
CL\_INS\_123
CL\_INS\_149
CL\_INS\_70
CL\_INS\_70
CL\_INS\_70
CL\_INS\_70
CL\_INS\_70
CL\_INS\_70
CL\_INS\_70
CL\_INS\_70
CL\_INS\_70
CL\_INS\_70
CL\_INS\_70
CL\_INS\_70
CL\_INS\_70
CL\_INS\_70
CL\_INS\_70
CL\_INS\_70
CL\_INS\_70
CL\_INS\_237
CL\_INS\_70
CL\_INS\_70
CL\_INS\_70
CL\_INS\_30
CL\_INS\_70
CL\_INS\_70
CL\_INS\_30
CL\_INS\_30
CL\_INS\_30
CL\_INS\_30
CL\_INS\_30
CL\_INS\_30
CL\_INS\_237
CL\_INS\_80
CL\_INS\_70
CL\_INS\_70
CL\_INS\_70
CL\_INS\_70
CL\_INS\_70
CL\_INS\_70
CL\_INS\_70
CL\_INS\_70
CL\_INS\_70
CL\_INS\_70
CL\_INS\_70
CL\_INS\_70
CL\_INS\_70
CL\_INS\_70
CL\_INS\_70
CL\_INS\_70
CL\_INS\_382
CL\_INS\_382
CL\_INS\_382
CL\_INS\_70
CL\_INS\_70
CL\_INS\_70
CL\_INS\_70
CL\_INS\_70
CL\_INS\_70
CL\_INS\_70
CL\_INS\_70
CL\_INS\_70
CL\_INS\_70
CL\_INS\_70
CL\_INS\_70
CL\_INS\_70
CL\_INS\_70
CL\_INS\_70
CL\_INS\_70
CL\_INS\_70
CL\_INS\_70
CL\_INS\_70
CL\_INS\_70
CL\_INS\_70
CL\_INS\_70
CL\_INS\_70
CL\_INS\_70
CL\_INS\_70
CL\_INS\_70
CL\_INS\_70
CL\_INS\_70
CL\_INS\_70
CL\_INS\_70
CL\_INS\_70
CL\_INS\_70
CL\_INS\_70
CL\_INS\_70
CL\_INS\_70
CL\_INS\_70
CL\_INS\_382
CL\_INS\_70
CL\_INS\_70
CL\_INS\_70
CL\_INS\_70
CL\_INS\_70
CL\_INS\_70
CL\_INS\_80
CL\_INS\_270
CL\_INS\_270
CL\_INS\_270
CL\_INS\_270
CL\_INS\_270
CL\_INS\_270
CL\_INS\_270
CL\_INS\_270
CL\_INS\_270
CL\_INS\_270
CL\_INS\_270
CL\_INS\_270
CL\_INS\_270
CL\_INS\_270
CL\_INS\_270
CL\_INS\_270
CL\_INS\_270
CL\_INS\_270
CL\_INS\_237
CL\_INS\_237
CL\_INS\_295
CL\_INS\_70
CL\_INS\_80
CL\_INS\_80
CL\_INS\_110
CL\_INS\_20
CL\_INS\_20
CL\_INS\_20
CL\_INS\_80
CL\_INS\_80
CL\_INS\_80
CL\_INS\_80
CL\_INS\_80
CL\_INS\_80
CL\_INS\_238
CL\_INS\_237
CL\_INS\_117
CL\_INS\_237
CL\_INS\_237
CL\_INS\_70
CL\_INS\_70
CL\_INS\_70
CL\_INS\_237
CL\_INS\_80
CL\_INS\_80
CL\_INS\_159
CL\_INS\_30
CL\_INS\_30
CL\_INS\_70
CL\_INS\_70
CL\_INS\_70
CL\_INS\_70
CL\_INS\_382
CL\_INS\_70
CL\_INS\_70
CL\_INS\_70
CL\_INS\_70
CL\_INS\_70
CL\_INS\_70
CL\_INS\_70
CL\_INS\_70
CL\_INS\_70
CL\_INS\_70
CL\_INS\_70
CL\_INS\_70
CL\_INS\_80
CL\_INS\_80
CL\_INS\_80
CL\_INS\_70
CL\_INS\_70
CL\_INS\_70
CL\_INS\_70
CL\_INS\_70
CL\_INS\_70
CL\_INS\_70
CL\_INS\_70
CL\_INS\_80
CL\_INS\_80
CL\_INS\_247
CL\_INS\_80
CL\_INS\_30
CL\_INS\_30
CL\_INS\_70
CL\_INS\_30
CL\_INS\_30
CL\_INS\_70
CL\_INS\_237
CL\_INS\_237
CL\_INS\_20
CL\_INS\_20
CL\_INS\_80
CL\_INS\_20
CL\_INS\_20
CL\_INS\_20
CL\_INS\_20
CL\_INS\_20
CL\_INS\_20
CL\_INS\_20
CL\_INS\_20
CL\_INS\_20
CL\_INS\_368
CL\_INS\_20
CL\_INS\_20
CL\_INS\_70
CL\_INS\_20
CL\_INS\_20
CL\_INS\_20
CL\_INS\_80
CL\_INS\_30
CL\_INS\_80
CL\_INS\_160
CL\_INS\_80
CL\_INS\_20
CL\_INS\_20
CL\_INS\_80
CL\_INS\_20
CL\_INS\_80
CL\_INS\_80
CL\_INS\_80
CL\_INS\_70
CL\_INS\_237
CL\_INS\_70
CL\_INS\_70
CL\_INS\_70
CL\_INS\_70
CL\_INS\_70
CL\_INS\_80
CL\_INS\_70
CL\_INS\_247
CL\_INS\_30
CL\_INS\_247
CL\_INS\_20
CL\_INS\_70
CL\_INS\_237
Cluster ID


CL\_979
CL\_5339
CL\_5338
CL\_12349
CL\_15745
CL\_7899
CL\_8624
CL\_8625
CL\_8626
CL\_7898
CL\_29454
CL\_7897
CL\_7896
CL\_26035
CL\_26034
CL\_8037
CL\_8628
CL\_22116
CL\_22118
CL\_22119
CL\_22120
CL\_22121
CL\_22122
CL\_22123
CL\_22124
CL\_22125
CL\_10984
CL\_22736
CL\_11715
CL\_26050
CL\_22737
CL\_22738
CL\_5146
CL\_22742
CL\_22741
CL\_22740
CL\_26321
CL\_6192
CL\_22744
CL\_26322
CL\_14649
CL\_8599
CL\_6842
CL\_6856
CL\_5682
CL\_8601
CL\_10217
CL\_10328
CL\_10329
CL\_7265
CL\_10330
CL\_10331
CL\_10332
CL\_8612
CL\_8613
CL\_8614
CL\_8615
CL\_9135
CL\_8616
CL\_8617
CL\_9136
CL\_9137
CL\_10333
CL\_10334
CL\_10335
CL\_11221
CL\_5149
CL\_7242
CL\_11191
CL\_7241
CL\_6426
CL\_6413
CL\_4995
CL\_9138
CL\_5320
CL\_5321
CL\_10435
CL\_8750
CL\_8749
CL\_8748
CL\_8747
CL\_14344
CL\_4094
CL\_4095
CL\_4096
CL\_4097
CL\_4098
CL\_4099
CL\_4100
CL\_4101
CL\_6831
CL\_6830
CL\_6829
CL\_6411
CL\_6828
CL\_11295
CL\_7252
CL\_7253
CL\_5246
CL\_6425
CL\_6424
CL\_6423
CL\_5241
CL\_6422
CL\_6729
CL\_14650
CL\_14651
CL\_14652
CL\_14653
CL\_14654
CL\_14655
CL\_14656
CL\_14657
CL\_14658
CL\_14659
CL\_14660
CL\_14661
CL\_14662
CL\_14663
CL\_14664
CL\_14665
CL\_14666
CL\_8832
CL\_7691
CL\_7692
CL\_7435
CL\_7434
CL\_14667
CL\_14668
CL\_14669
CL\_14670
CL\_14671
CL\_14672
CL\_14673
CL\_14674
CL\_14675
CL\_14676
CL\_14677
CL\_14678
CL\_14679
CL\_14680
CL\_8632
CL\_12888
CL\_14681
CL\_13233
CL\_13232
CL\_13231
CL\_7214
CL\_7671
CL\_14682
CL\_7895
CL\_8591
CL\_8592
CL\_7894
CL\_7893
CL\_7892
CL\_8593
CL\_8594
CL\_8595
CL\_10326
CL\_7672
CL\_7557
CL\_6838
CL\_6839
CL\_6840
CL\_6841
CL\_6410
CL\_8600
CL\_7891
CL\_7890
CL\_7889
CL\_7888
CL\_7887
CL\_7886
CL\_7885
CL\_7884
CL\_7883
CL\_7882
CL\_7881
CL\_7880
CL\_7879
CL\_7878
CL\_7877
CL\_7876
CL\_7875
CL\_7874
CL\_7873
CL\_7872
CL\_7871
CL\_7870
CL\_7869
CL\_7868
CL\_7867
CL\_7866
CL\_7865
CL\_6973
CL\_9476
CL\_27301
CL\_27299
CL\_7864
CL\_7863
CL\_7862
CL\_7861
CL\_7860
CL\_6971
CL\_6970
CL\_7859
CL\_7858
CL\_7857
CL\_7856
CL\_7673
CL\_9131
CL\_9132
CL\_9133
CL\_9134
CL\_5237
CL\_5238
CL\_14683
CL\_7234
CL\_7235
CL\_14684
CL\_8376
CL\_8779
CL\_7298
CL\_14685
CL\_6979
CL\_6980
CL\_6981
CL\_6982
CL\_6983
CL\_6984
CL\_22967
CL\_7690
CL\_8641
CL\_14686
CL\_14687
CL\_14688
CL\_6421
CL\_6420
CL\_1932
CL\_1931
CL\_6419
CL\_10547
CL\_8508
CL\_9139
CL\_14689
CL\_14690
CL\_13465
CL\_14691
CL\_5231
CL\_5230
CL\_9140
CL\_7751
CL\_235
CL\_6416
CL\_7855
CL\_4462
CL\_7854
CL\_7853
CL\_7852
CL\_7851
CL\_12002
CL\_7850
CL\_7849
CL\_7848
CL\_7847
CL\_7846
CL\_7845
CL\_7844
CL\_7843
CL\_5495
CL\_5494
CL\_5493
CL\_5492
CL\_5491
CL\_7842
CL\_7841
CL\_7840
CL\_31974
CL\_6406
CL\_7839
CL\_7838
CL\_7837
CL\_7836
CL\_7835
CL\_7834
CL\_13785
CL\_7833
CL\_6179
CL\_7832
CL\_1933
CL\_13787
CL\_7831
CL\_8620
CL\_6417
CL\_7830
CL\_7829
CL\_7828
CL\_7827
CL\_7826
CL\_7825
CL\_7824
CL\_7823
