## Supplementary material for "A novel method for integrating genomic and Tn-Seq data to identify common *in vivo* fitness mechanisms across multiple bacterial species": S1 Dataset: CL_INS_81.html

Legend

 Mobile +extrachromosomalelementfunctions
 Regulatoryfunctions
 Hypothetical
 DNA Metabolism
 All EssentialGenes
 Biosynthesis ofcofactors,prostheticgroups, +carriers
 All Fitness Genes
 Cell Envelope
 Proteinsynthesis/fate
 Other
 Centralintermediarymetabolism
 EnergyMetabolism
 Transport +binding proteins
 All VFDB Genes

FULL


WINDOWSVGPNG

Trim RowsRemove SingletonsSave Fasta

CL\_976


CL\_976


CL\_973


CL\_1935


CL\_976


CL\_1935


CL\_973


CL\_976


CL\_976


CL\_976


CL\_1935


CL\_976


CL\_976


CL\_973


CL\_975


CL\_976


CL\_976


CL\_976


CL\_1935


CL\_975


CL\_976


CL\_1935


CL\_976


CL\_976


CL\_976


CL\_976


CL\_976


CL\_976


CL\_976


CL\_976


CL\_976


CL\_1935


Break


CL\_976


CL\_976


CL\_976


CL\_976


CL\_976


CL\_976


CL\_976


CL\_976


CL\_976


CL\_975


CL\_976


CL\_974


CL\_976


CL\_969


CL\_976


CL\_976


CL\_976


CL\_976


CL\_976


CL\_976


CL\_976


CL\_976


CL\_976


CL\_976


CL\_976


CL\_976


CL\_976


CL\_976


CL\_976


CL\_974


CL\_974


CL\_976


CL\_973


CL\_971


CL\_976


CL\_976


CL\_1940


CL\_976


CL\_976


CL\_976


CL\_234


CL\_976


CL\_976


CL\_976


CL\_1935


CL\_976


CL\_976


CL\_976


CL\_976


CL\_976


CL\_976


CL\_976


CL\_976


CL\_976


CL\_973


CL\_976


CL\_975

HighlightSelectShow Genomes


63

CL\_982


56

CL\_982


36

CL\_982


9

CL\_982


6

CL\_982


5

CL\_982


5

CL\_982


5

CL\_982


5

CL\_982


4

CL\_982


4

CL\_982


3

CL\_982


3

CL\_982


2

CL\_982


2

CL\_982


2

CL\_982


2

CL\_982


2

CL\_982


2

CL\_982


2

CL\_982


2

CL\_983


2

CL\_982


1

CL\_982


1

CL\_982


1

CL\_234


1

CL\_982


1

CL\_234


1

CL\_982


1

CL\_234


1

CL\_2942


1

CL\_982


1

CL\_982


1

CL\_982


1

CL\_982


1

CL\_234


1

CL\_982


1

CL\_982


1

CL\_982


1

CL\_1935


1

CL\_1935


1

CL\_234


1

CL\_982


1

CL\_982


1

CL\_982


1

CL\_982


1

CL\_982


1

CL\_982


1

CL\_982


1

CL\_982


1

CL\_982


1

CL\_234


1

CL\_982


1

CL\_234


1

CL\_982


1

CL\_982


1

CL\_982


1

CL\_983


1

CL\_2528


1

CL\_1935


1

Break


1

CL\_982


1

CL\_982


1

CL\_982


1

CL\_982


1

CL\_982


1

CL\_982


1

CL\_982


1

CL\_982


1

CL\_982


1

CL\_982


1

CL\_982


1

CL\_982


1

CL\_982


1

CL\_982


1

CL\_982


1

CL\_982


1

CL\_984


1

CL\_982


1

CL\_982


1

CL\_234


1

CL\_982


1

CL\_1928


1

CL\_983


1

CL\_982


1

CL\_982


1

CL\_982


1

CL\_982


1

CL\_982


1

CL\_982


1

CL\_982

fGI ID


CL\_INS\_80
CL\_INS\_81
CL\_INS\_80
CL\_INS\_81
CL\_INS\_81
CL\_INS\_81
CL\_INS\_81
CL\_INS\_81
CL\_INS\_81
CL\_INS\_30
CL\_INS\_70
CL\_INS\_81
CL\_INS\_81
CL\_INS\_247
CL\_INS\_207
CL\_INS\_81
CL\_INS\_81
CL\_INS\_207
CL\_INS\_207
CL\_INS\_382
CL\_INS\_247
CL\_INS\_247
CL\_INS\_247
CL\_INS\_247
CL\_INS\_247
CL\_INS\_247
CL\_INS\_247
CL\_INS\_247
CL\_INS\_247
CL\_INS\_207
CL\_INS\_207
CL\_INS\_207
CL\_INS\_207
CL\_INS\_207
CL\_INS\_81
CL\_INS\_81
CL\_INS\_81
CL\_INS\_81
CL\_INS\_81
CL\_INS\_247
CL\_INS\_30
CL\_INS\_30
CL\_INS\_30
CL\_INS\_30
CL\_INS\_30
CL\_INS\_30
CL\_INS\_159
CL\_INS\_81
CL\_INS\_81
CL\_INS\_159
CL\_INS\_159
CL\_INS\_159
CL\_INS\_159
CL\_INS\_237
CL\_INS\_81
CL\_INS\_81
CL\_INS\_81
CL\_INS\_81
CL\_INS\_81
CL\_INS\_81
CL\_INS\_81
CL\_INS\_81
CL\_INS\_81
CL\_INS\_81
CL\_INS\_81
CL\_INS\_81
CL\_INS\_81
CL\_INS\_81
CL\_INS\_81
CL\_INS\_81
CL\_INS\_81
CL\_INS\_81
CL\_INS\_81
CL\_INS\_81
CL\_INS\_81
CL\_INS\_81
CL\_INS\_81
CL\_INS\_81
CL\_INS\_81
CL\_INS\_81
CL\_INS\_81
CL\_INS\_159
CL\_INS\_387
CL\_INS\_81
CL\_INS\_81
CL\_INS\_81
CL\_INS\_81
CL\_INS\_81
CL\_INS\_81
CL\_INS\_247
CL\_INS\_80
CL\_INS\_80
CL\_INS\_80
CL\_INS\_81
CL\_INS\_70
CL\_INS\_81
CL\_INS\_70
CL\_INS\_81
CL\_INS\_70
CL\_INS\_247
CL\_INS\_247
CL\_INS\_247
CL\_INS\_247
CL\_INS\_207
CL\_INS\_207
CL\_INS\_207
CL\_INS\_207
CL\_INS\_81
CL\_INS\_81
CL\_INS\_81
CL\_INS\_70
CL\_INS\_70
CL\_INS\_70
CL\_INS\_70
CL\_INS\_70
CL\_INS\_70
CL\_INS\_70
CL\_INS\_70
CL\_INS\_70
CL\_INS\_70
CL\_INS\_70
CL\_INS\_70
CL\_INS\_81
CL\_INS\_70
CL\_INS\_382
CL\_INS\_387
CL\_INS\_387
CL\_INS\_387
CL\_INS\_387
CL\_INS\_387
CL\_INS\_70
CL\_INS\_70
CL\_INS\_70
CL\_INS\_70
CL\_INS\_70
CL\_INS\_70
CL\_INS\_70
CL\_INS\_70
CL\_INS\_30
CL\_INS\_70
CL\_INS\_80
CL\_INS\_270
CL\_INS\_270
CL\_INS\_270
CL\_INS\_270
CL\_INS\_270
CL\_INS\_270
CL\_INS\_270
CL\_INS\_270
CL\_INS\_270
CL\_INS\_270
CL\_INS\_270
CL\_INS\_270
CL\_INS\_270
CL\_INS\_270
CL\_INS\_270
CL\_INS\_270
CL\_INS\_270
CL\_INS\_270
CL\_INS\_81
CL\_INS\_237
CL\_INS\_237
CL\_INS\_237
CL\_INS\_237
CL\_INS\_295
CL\_INS\_70
CL\_INS\_81
CL\_INS\_80
CL\_INS\_81
CL\_INS\_81
CL\_INS\_80
CL\_INS\_110
CL\_INS\_20
CL\_INS\_20
CL\_INS\_20
CL\_INS\_20
CL\_INS\_80
CL\_INS\_20
CL\_INS\_20
CL\_INS\_80
CL\_INS\_80
CL\_INS\_80
CL\_INS\_238
CL\_INS\_237
CL\_INS\_117
CL\_INS\_81
CL\_INS\_20
CL\_INS\_237
CL\_INS\_237
CL\_INS\_70
CL\_INS\_80
CL\_INS\_237
CL\_INS\_207
CL\_INS\_80
CL\_INS\_80
CL\_INS\_81
CL\_INS\_80
CL\_INS\_80
CL\_INS\_80
CL\_INS\_237
CL\_INS\_237
CL\_INS\_237
CL\_INS\_237
CL\_INS\_237
CL\_INS\_149
CL\_INS\_247
CL\_INS\_247
CL\_INS\_247
CL\_INS\_123
CL\_INS\_81
CL\_INS\_174
CL\_INS\_80
CL\_INS\_81
CL\_INS\_70
CL\_INS\_81
CL\_INS\_70
CL\_INS\_30
CL\_INS\_70
CL\_INS\_70
CL\_INS\_70
CL\_INS\_70
CL\_INS\_70
CL\_INS\_70
CL\_INS\_70
CL\_INS\_30
CL\_INS\_30
CL\_INS\_30
CL\_INS\_30
CL\_INS\_159
CL\_INS\_30
CL\_INS\_159
CL\_INS\_30
CL\_INS\_30
CL\_INS\_30
CL\_INS\_30
CL\_INS\_30
CL\_INS\_30
CL\_INS\_30
CL\_INS\_70
CL\_INS\_70
CL\_INS\_70
CL\_INS\_70
CL\_INS\_70
CL\_INS\_70
CL\_INS\_70
CL\_INS\_70
CL\_INS\_70
CL\_INS\_70
CL\_INS\_81
CL\_INS\_70
CL\_INS\_70
CL\_INS\_70
CL\_INS\_70
CL\_INS\_70
CL\_INS\_70
CL\_INS\_70
CL\_INS\_83
CL\_INS\_81
CL\_INS\_70
CL\_INS\_70
CL\_INS\_70
CL\_INS\_70
CL\_INS\_70
CL\_INS\_70
CL\_INS\_70
CL\_INS\_81
CL\_INS\_382
CL\_INS\_382
CL\_INS\_382
CL\_INS\_70
CL\_INS\_70
CL\_INS\_70
CL\_INS\_70
CL\_INS\_70
CL\_INS\_70
CL\_INS\_70
CL\_INS\_70
CL\_INS\_70
CL\_INS\_70
CL\_INS\_70
CL\_INS\_70
CL\_INS\_70
CL\_INS\_70
CL\_INS\_70
CL\_INS\_70
CL\_INS\_382
CL\_INS\_237
CL\_INS\_20
CL\_INS\_70
CL\_INS\_295
CL\_INS\_295
CL\_INS\_237
CL\_INS\_237
CL\_INS\_237
CL\_INS\_237
CL\_INS\_237
CL\_INS\_81
CL\_INS\_81
CL\_INS\_81
CL\_INS\_70
CL\_INS\_70
CL\_INS\_70
CL\_INS\_70
CL\_INS\_70
CL\_INS\_81
CL\_INS\_70
CL\_INS\_387
CL\_INS\_70
CL\_INS\_70
CL\_INS\_70
CL\_INS\_70
CL\_INS\_354
CL\_INS\_80
CL\_INS\_81
CL\_INS\_80
CL\_INS\_70
CL\_INS\_70
CL\_INS\_70
CL\_INS\_237
CL\_INS\_81
CL\_INS\_20
CL\_INS\_20
CL\_INS\_80
CL\_INS\_20
CL\_INS\_155
CL\_INS\_204
CL\_INS\_155
CL\_INS\_155
CL\_INS\_155
CL\_INS\_155
CL\_INS\_155
CL\_INS\_86
CL\_INS\_155
CL\_INS\_155
CL\_INS\_155
CL\_INS\_155
CL\_INS\_86
CL\_INS\_86
CL\_INS\_86
CL\_INS\_86
CL\_INS\_155
CL\_INS\_155
CL\_INS\_155
CL\_INS\_155
CL\_INS\_155
CL\_INS\_155
CL\_INS\_155
CL\_INS\_86
CL\_INS\_155
CL\_INS\_86
CL\_INS\_155
CL\_INS\_86
CL\_INS\_155
CL\_INS\_155
CL\_INS\_155
CL\_INS\_155
CL\_INS\_155
CL\_INS\_155
CL\_INS\_74
CL\_INS\_74
CL\_INS\_74
CL\_INS\_74
CL\_INS\_74
CL\_INS\_74
CL\_INS\_74
CL\_INS\_155
CL\_INS\_155
CL\_INS\_155
CL\_INS\_155
CL\_INS\_155
CL\_INS\_155
CL\_INS\_155
CL\_INS\_182
CL\_INS\_155
CL\_INS\_155
CL\_INS\_155
CL\_INS\_155
CL\_INS\_66
CL\_INS\_20
CL\_INS\_20
CL\_INS\_20
CL\_INS\_20
CL\_INS\_20
CL\_INS\_20
CL\_INS\_20
CL\_INS\_20
CL\_INS\_20
CL\_INS\_368
CL\_INS\_81
CL\_INS\_20
CL\_INS\_20
CL\_INS\_70
CL\_INS\_20
CL\_INS\_20
CL\_INS\_20
CL\_INS\_80
CL\_INS\_30
CL\_INS\_160
CL\_INS\_80
CL\_INS\_81
CL\_INS\_20
CL\_INS\_20
CL\_INS\_20
CL\_INS\_80
CL\_INS\_20
CL\_INS\_80
CL\_INS\_80
CL\_INS\_81
CL\_INS\_81
CL\_INS\_80
CL\_INS\_237
CL\_INS\_70
CL\_INS\_81
CL\_INS\_81
CL\_INS\_81
CL\_INS\_70
CL\_INS\_70
CL\_INS\_237
CL\_INS\_237
CL\_INS\_295
CL\_INS\_295
CL\_INS\_224
CL\_INS\_224
CL\_INS\_224
CL\_INS\_224
CL\_INS\_81
CL\_INS\_81
CL\_INS\_224
CL\_INS\_81
CL\_INS\_295
CL\_INS\_224
CL\_INS\_237
CL\_INS\_237
CL\_INS\_224
CL\_INS\_81
CL\_INS\_81
CL\_INS\_224
CL\_INS\_237
CL\_INS\_81
CL\_INS\_81
CL\_INS\_81
CL\_INS\_387
CL\_INS\_81
CL\_INS\_70
CL\_INS\_70
CL\_INS\_70
CL\_INS\_247
CL\_INS\_81
CL\_INS\_70
CL\_INS\_387
CL\_INS\_387
CL\_INS\_387
CL\_INS\_382
CL\_INS\_70
CL\_INS\_237
CL\_INS\_70
CL\_INS\_70
CL\_INS\_70
CL\_INS\_70
CL\_INS\_70
CL\_INS\_70
CL\_INS\_70
CL\_INS\_70
CL\_INS\_70
CL\_INS\_70
CL\_INS\_70
CL\_INS\_70
CL\_INS\_70
CL\_INS\_70
CL\_INS\_70
CL\_INS\_70
CL\_INS\_70
CL\_INS\_70
CL\_INS\_70
CL\_INS\_70
CL\_INS\_70
CL\_INS\_70
CL\_INS\_70
CL\_INS\_70
CL\_INS\_70
CL\_INS\_70
CL\_INS\_70
CL\_INS\_70
CL\_INS\_70
CL\_INS\_70
CL\_INS\_20
CL\_INS\_70
CL\_INS\_70
CL\_INS\_70
CL\_INS\_128
CL\_INS\_128
CL\_INS\_128
CL\_INS\_70
CL\_INS\_70
CL\_INS\_81
CL\_INS\_207
CL\_INS\_123
CL\_INS\_123
CL\_INS\_347
CL\_INS\_70
CL\_INS\_81
CL\_INS\_81
CL\_INS\_20
CL\_INS\_20
CL\_INS\_20
CL\_INS\_20
CL\_INS\_20
CL\_INS\_20
CL\_INS\_70
CL\_INS\_70
CL\_INS\_70
CL\_INS\_81
CL\_INS\_81
CL\_INS\_81
CL\_INS\_81
CL\_INS\_81
CL\_INS\_387
CL\_INS\_387
CL\_INS\_70
CL\_INS\_70
CL\_INS\_70
CL\_INS\_70
CL\_INS\_70
CL\_INS\_70
CL\_INS\_70
CL\_INS\_70
CL\_INS\_70
CL\_INS\_70
CL\_INS\_382
CL\_INS\_382
CL\_INS\_70
CL\_INS\_70
CL\_INS\_70
CL\_INS\_70
CL\_INS\_70
CL\_INS\_70
CL\_INS\_81
CL\_INS\_70
CL\_INS\_70
CL\_INS\_70
CL\_INS\_70
CL\_INS\_70
CL\_INS\_387
CL\_INS\_149
CL\_INS\_382
CL\_INS\_81
CL\_INS\_159
CL\_INS\_156
CL\_INS\_156
CL\_INS\_70
CL\_INS\_81
CL\_INS\_81
CL\_INS\_81
CL\_INS\_81
CL\_INS\_81
CL\_INS\_81
CL\_INS\_81
CL\_INS\_81
CL\_INS\_70
CL\_INS\_81
CL\_INS\_81
CL\_INS\_81
CL\_INS\_81
CL\_INS\_81
CL\_INS\_81
CL\_INS\_81
CL\_INS\_70
CL\_INS\_81
CL\_INS\_70
CL\_INS\_70
CL\_INS\_70
CL\_INS\_70
CL\_INS\_237
CL\_INS\_70
CL\_INS\_70
CL\_INS\_159
CL\_INS\_70
CL\_INS\_30
CL\_INS\_30
CL\_INS\_247
CL\_INS\_81
CL\_INS\_81
CL\_INS\_30
CL\_INS\_30
CL\_INS\_30
CL\_INS\_81
CL\_INS\_30
CL\_INS\_247
CL\_INS\_20
CL\_INS\_70
CL\_INS\_237
CL\_INS\_81
CL\_INS\_30
CL\_INS\_247
CL\_INS\_247
CL\_INS\_30
CL\_INS\_70
CL\_INS\_70
CL\_INS\_237
CL\_INS\_70
CL\_INS\_70
CL\_INS\_70
CL\_INS\_80
CL\_INS\_30
CL\_INS\_30
CL\_INS\_30
CL\_INS\_30
CL\_INS\_30
CL\_INS\_387
CL\_INS\_70
CL\_INS\_70
CL\_INS\_30
CL\_INS\_70
CL\_INS\_70
CL\_INS\_70
CL\_INS\_80
CL\_INS\_70
CL\_INS\_30
CL\_INS\_30
CL\_INS\_159
CL\_INS\_81
CL\_INS\_81
CL\_INS\_81
CL\_INS\_81
Cluster ID


CL\_5338
CL\_31948
CL\_12349
CL\_12350
CL\_20989
CL\_27705
CL\_27704
CL\_28581
CL\_28582
CL\_1938
CL\_7899
CL\_977
CL\_24430
CL\_12000
CL\_7548
CL\_23771
CL\_23772
CL\_23773
CL\_23774
CL\_4086
CL\_14332
CL\_14333
CL\_14334
CL\_14335
CL\_21989
CL\_21988
CL\_21987
CL\_21986
CL\_21985
CL\_8952
CL\_8953
CL\_20646
CL\_20645
CL\_20644
CL\_23775
CL\_23776
CL\_23777
CL\_23778
CL\_23779
CL\_7470
CL\_5232
CL\_5235
CL\_5237
CL\_5238
CL\_5239
CL\_5240
CL\_978
CL\_37479
CL\_37478
CL\_13862
CL\_13863
CL\_13864
CL\_13865
CL\_979
CL\_5797
CL\_9528
CL\_9527
CL\_9526
CL\_15477
CL\_15478
CL\_15479
CL\_15480
CL\_15481
CL\_15482
CL\_15483
CL\_16142
CL\_15484
CL\_15485
CL\_15486
CL\_15487
CL\_15488
CL\_15489
CL\_15490
CL\_15491
CL\_15492
CL\_15493
CL\_15494
CL\_15495
CL\_15496
CL\_15497
CL\_15498
CL\_980
CL\_981
CL\_21178
CL\_21177
CL\_9002
CL\_29235
CL\_29234
CL\_29233
CL\_8623
CL\_8624
CL\_8625
CL\_8626
CL\_8627
CL\_7898
CL\_13778
CL\_7897
CL\_26028
CL\_7896
CL\_15549
CL\_10020
CL\_10021
CL\_9460
CL\_26178
CL\_8139
CL\_2554
CL\_2553
CL\_34479
CL\_35809
CL\_35810
CL\_7895
CL\_17789
CL\_8591
CL\_8592
CL\_7894
CL\_7893
CL\_7892
CL\_8593
CL\_8594
CL\_8595
CL\_10326
CL\_7672
CL\_33398
CL\_6836
CL\_7557
CL\_37737
CL\_37738
CL\_37739
CL\_37740
CL\_37741
CL\_6837
CL\_6838
CL\_6839
CL\_6840
CL\_6841
CL\_6842
CL\_6410
CL\_8600
CL\_6856
CL\_5682
CL\_7891
CL\_7890
CL\_7889
CL\_7888
CL\_7887
CL\_7886
CL\_7885
CL\_7884
CL\_7883
CL\_7882
CL\_7881
CL\_7880
CL\_7879
CL\_7878
CL\_7877
CL\_7876
CL\_7875
CL\_7874
CL\_7873
CL\_23973
CL\_7872
CL\_7871
CL\_14365
CL\_13779
CL\_7870
CL\_7869
CL\_35733
CL\_7868
CL\_35732
CL\_13780
CL\_7867
CL\_7866
CL\_7865
CL\_6972
CL\_6973
CL\_9476
CL\_7864
CL\_6974
CL\_13781
CL\_7863
CL\_7862
CL\_7861
CL\_7860
CL\_6971
CL\_6970
CL\_27298
CL\_13782
CL\_7859
CL\_7858
CL\_8628
CL\_22736
CL\_11715
CL\_9482
CL\_22737
CL\_22738
CL\_22739
CL\_22740
CL\_22741
CL\_22742
CL\_8629
CL\_8131
CL\_7979
CL\_6806
CL\_6805
CL\_5146
CL\_26693
CL\_9539
CL\_11712
CL\_4995
CL\_22743
CL\_6192
CL\_22744
CL\_22745
CL\_14649
CL\_33124
CL\_17597
CL\_11295
CL\_10581
CL\_15528
CL\_8599
CL\_6828
CL\_7252
CL\_7253
CL\_6826
CL\_6425
CL\_6424
CL\_6423
CL\_5241
CL\_5242
CL\_5243
CL\_5244
CL\_5245
CL\_5246
CL\_5247
CL\_5248
CL\_5249
CL\_5250
CL\_6422
CL\_6421
CL\_8619
CL\_14651
CL\_14652
CL\_14653
CL\_15746
CL\_15747
CL\_14654
CL\_14655
CL\_14656
CL\_16627
CL\_14657
CL\_14658
CL\_14659
CL\_14660
CL\_14661
CL\_14662
CL\_14663
CL\_16628
CL\_16629
CL\_14664
CL\_15748
CL\_14665
CL\_15749
CL\_15750
CL\_15751
CL\_14666
CL\_16630
CL\_8832
CL\_7691
CL\_7692
CL\_7435
CL\_7434
CL\_14667
CL\_14668
CL\_14669
CL\_14670
CL\_14671
CL\_14672
CL\_14673
CL\_14674
CL\_14675
CL\_14676
CL\_14677
CL\_14678
CL\_14679
CL\_14680
CL\_13034
CL\_16631
CL\_20259
CL\_8632
CL\_8633
CL\_8634
CL\_8635
CL\_7987
CL\_7986
CL\_8636
CL\_8637
CL\_8638
CL\_8639
CL\_8640
CL\_8631
CL\_8630
CL\_12888
CL\_14681
CL\_13233
CL\_16632
CL\_13232
CL\_22971
CL\_13231
CL\_7214
CL\_7671
CL\_14682
CL\_16633
CL\_27301
CL\_27300
CL\_27299
CL\_7857
CL\_13783
CL\_7856
CL\_7855
CL\_12001
CL\_7854
CL\_7853
CL\_7852
CL\_7851
CL\_11037
CL\_1105
CL\_11036
CL\_8483
CL\_11035
CL\_10869
CL\_10870
CL\_8232
CL\_8231
CL\_8230
CL\_8229
CL\_8228
CL\_8227
CL\_8226
CL\_11030
CL\_11029
CL\_10877
CL\_10878
CL\_10879
CL\_11025
CL\_11024
CL\_10882
CL\_10883
CL\_10884
CL\_10885
CL\_10886
CL\_10887
CL\_10888
CL\_10889
CL\_10890
CL\_10891
CL\_10892
CL\_10893
CL\_10894
CL\_10895
CL\_10896
CL\_10897
CL\_10898
CL\_10899
CL\_10900
CL\_10901
CL\_11005
CL\_11004
CL\_10905
CL\_10906
CL\_10907
CL\_10909
CL\_10910
CL\_10996
CL\_10913
CL\_10914
CL\_10915
CL\_10916
CL\_22135
CL\_12002
CL\_7850
CL\_7849
CL\_7848
CL\_7847
CL\_7846
CL\_7845
CL\_12003
CL\_7844
CL\_7843
CL\_13784
CL\_5495
CL\_5494
CL\_5493
CL\_5492
CL\_5491
CL\_7842
CL\_7841
CL\_7840
CL\_6406
CL\_7839
CL\_13619
CL\_12004
CL\_7838
CL\_7837
CL\_7836
CL\_7835
CL\_7834
CL\_13785
CL\_13786
CL\_12005
CL\_7833
CL\_7832
CL\_7673
CL\_27297
CL\_27296
CL\_27295
CL\_14683
CL\_7234
CL\_17158
CL\_18150
CL\_18151
CL\_18152
CL\_7323
CL\_7322
CL\_7321
CL\_7722
CL\_26027
CL\_26026
CL\_13676
CL\_26025
CL\_18158
CL\_7718
CL\_13820
CL\_7717
CL\_7716
CL\_26024
CL\_26023
CL\_7314
CL\_7236
CL\_23420
CL\_26022
CL\_26021
CL\_22969
CL\_33326
CL\_7235
CL\_10408
CL\_10409
CL\_8932
CL\_33325
CL\_14684
CL\_37742
CL\_37743
CL\_37744
CL\_8376
CL\_8779
CL\_5149
CL\_11221
CL\_7242
CL\_11191
CL\_7241
CL\_6426
CL\_5320
CL\_5321
CL\_10435
CL\_8750
CL\_8749
CL\_8748
CL\_8747
CL\_14344
CL\_4094
CL\_4095
CL\_4096
CL\_4097
CL\_4098
CL\_4099
CL\_4100
CL\_4101
CL\_6831
CL\_6829
CL\_6411
CL\_10331
CL\_10332
CL\_8612
CL\_8613
CL\_8614
CL\_8615
CL\_9135
CL\_8616
CL\_8617
CL\_8618
CL\_10333
CL\_10334
CL\_10335
CL\_6420
CL\_10330
CL\_10327
CL\_10217
CL\_10328
CL\_10329
CL\_7265
CL\_8601
CL\_23348
CL\_27294
CL\_20265
CL\_20264
CL\_20263
CL\_20262
CL\_20261
CL\_20260
CL\_7298
CL\_7299
CL\_14685
CL\_35569
CL\_35570
CL\_35571
CL\_35572
CL\_35573
CL\_30684
CL\_30685
CL\_6979
CL\_26245
CL\_26244
CL\_26243
CL\_26242
CL\_26241
CL\_26240
CL\_26239
CL\_25227
CL\_26238
CL\_6177
CL\_6178
CL\_6179
CL\_6980
CL\_6981
CL\_6982
CL\_6983
CL\_22967
CL\_35574
CL\_7690
CL\_10592
CL\_26247
CL\_26248
CL\_26249
CL\_30686
CL\_11826
CL\_10807
CL\_35575
CL\_11249
CL\_12025
CL\_12024
CL\_8641
CL\_35576
CL\_35577
CL\_35578
CL\_35579
CL\_35580
CL\_35581
CL\_35582
CL\_35583
CL\_1931
CL\_27293
CL\_27292
CL\_27291
CL\_27290
CL\_27289
CL\_27288
CL\_27287
CL\_1932
CL\_14366
CL\_1933
CL\_13787
CL\_6418
CL\_7831
CL\_8564
CL\_8620
CL\_6419
CL\_6727
CL\_8508
CL\_7751
CL\_235
CL\_7828
CL\_14367
CL\_14368
CL\_7827
CL\_1936
CL\_1937
CL\_23972
CL\_7750
CL\_7826
CL\_7825
CL\_7824
CL\_7823
CL\_9525
CL\_7749
CL\_7204
CL\_7203
CL\_2551
CL\_6809
CL\_6808
CL\_7822
CL\_15755
CL\_10547
CL\_9139
CL\_14689
CL\_5231
CL\_5230
CL\_5229
CL\_5228
CL\_5227
CL\_16634
CL\_13848
CL\_9140
CL\_5226
CL\_6416
CL\_8642
CL\_6417
CL\_7830
CL\_7829
CL\_1934
CL\_5225
CL\_5224
CL\_8643
CL\_8644
CL\_8645
CL\_8646
