## Supplementary material for "A novel method for integrating genomic and Tn-Seq data to identify common *in vivo* fitness mechanisms across multiple bacterial species": S1 Dataset: CL_INS_83.html

Legend

 Mobile +extrachromosomalelementfunctions
 Regulatoryfunctions
 Hypothetical
 DNA Metabolism
 AntibioticResistance
 Purines,pyrimidines,nucleosides, +nucleotides
 Transcription
 All EssentialGenes
 Biosynthesis ofcofactors,prostheticgroups, +carriers
 All Fitness Genes
 Proteinsynthesis/fate
 Other
 EnergyMetabolism
 Transport +binding proteins
 All VFDB Genes

FULL


WINDOWSVGPNG

Trim RowsRemove SingletonsSave Fasta

CL\_1072


CL\_4516


CL\_4516


CL\_1072


CL\_1072


CL\_1072


CL\_1820


CL\_1072


CL\_1072


CL\_4427


CL\_1072


CL\_946


CL\_1072


CL\_1072


CL\_1072


CL\_1072


CL\_4516


CL\_957


CL\_1072


CL\_1072


CL\_4516


CL\_1072


CL\_4516


CL\_1072


CL\_1072


CL\_1072


CL\_1072


CL\_1072


CL\_1072


CL\_1072


CL\_1072


CL\_1072


CL\_1072


CL\_4516


CL\_4516


CL\_4516


CL\_1072


CL\_1072


CL\_1072


CL\_1072


CL\_1072


CL\_1072


CL\_1072


CL\_1072


CL\_1917


CL\_1072


CL\_1072


CL\_1072


CL\_1072


CL\_1072


CL\_1072


CL\_1072


CL\_1072


CL\_1072


CL\_4516


CL\_1072


CL\_1072


CL\_4516


CL\_1072


CL\_1072


CL\_1072


CL\_4516


CL\_1072


CL\_1072


CL\_1072


CL\_4516


CL\_1072


CL\_1072


CL\_500


CL\_1072


CL\_4487


CL\_4516


CL\_1072


CL\_4516


CL\_1072


CL\_1072


CL\_1072


CL\_1072


CL\_1072


CL\_4516


CL\_1072


CL\_1072


CL\_1072


CL\_1072


CL\_1072


CL\_4427


CL\_1072


CL\_4516


CL\_1072


CL\_1072


CL\_1072


CL\_1072


CL\_4516

HighlightSelectShow Genomes


211

CL\_1073


2

CL\_1073


2

CL\_1073


2

CL\_1073


2

CL\_4487


1

CL\_923


1

CL\_1073


1

CL\_1073


1

Break


1

CL\_1073


1

CL\_4427


1

CL\_1073


1

CL\_1073


1

CL\_1073


1

CL\_1073


1

CL\_4487


1

CL\_1073


1

CL\_1073


1

CL\_1073


1

CL\_1073


1

CL\_1073


1

CL\_1073


1

CL\_1073


1

CL\_4487


1

CL\_4427


1

CL\_4486


1

CL\_1073


1

CL\_1073


1

CL\_4486


1

CL\_4427


1

CL\_4519


1

CL\_1073


1

CL\_1073


1

CL\_1073


1

CL\_1073


1

CL\_1073


1

CL\_4516


1

CL\_4487


1

CL\_1073


1

CL\_4427


1

CL\_4487


1

CL\_1073


1

CL\_1073


1

CL\_4487


1

CL\_1073


1

CL\_1073


1

CL\_1073


1

CL\_1915


1

CL\_1073


1

CL\_1073


1

CL\_1073


1

CL\_1073


1

CL\_1073


1

CL\_1073


1

CL\_1073


1

CL\_1073


1

CL\_1073


1

CL\_1073


1

CL\_1073


1

CL\_4487


1

CL\_4427


1

CL\_1073


1

CL\_1073


1

CL\_4486


1

CL\_1073


1

CL\_1073


1

CL\_1073


1

CL\_1073


1

CL\_1073


1

CL\_1073


1

CL\_1073


1

CL\_1073


1

CL\_1915


1

CL\_1073


1

CL\_1073


1

CL\_4487


1

CL\_1073


1

CL\_1915


1

CL\_4487


1

CL\_1073


1

CL\_1171


1

CL\_1073


1

CL\_1073


1

Break


1

CL\_923


1

CL\_1073


1

CL\_1915


1

CL\_1073


1

CL\_1073


1

CL\_1073


1

CL\_1073


1

CL\_4516


1

CL\_1073

fGI ID


CL\_INS\_385
CL\_INS\_385
CL\_INS\_382
CL\_INS\_382
CL\_INS\_385
CL\_INS\_385
CL\_INS\_385
CL\_INS\_83
CL\_INS\_382
CL\_INS\_83
CL\_INS\_384
CL\_INS\_384
CL\_INS\_382
CL\_INS\_382
CL\_INS\_382
CL\_INS\_382
CL\_INS\_382
CL\_INS\_382
CL\_INS\_382
CL\_INS\_382
CL\_INS\_382
CL\_INS\_382
CL\_INS\_382
CL\_INS\_382
CL\_INS\_382
CL\_INS\_382
CL\_INS\_382
CL\_INS\_382
CL\_INS\_382
CL\_INS\_382
CL\_INS\_207
CL\_INS\_382
CL\_INS\_382
CL\_INS\_382
CL\_INS\_382
CL\_INS\_382
CL\_INS\_382
CL\_INS\_382
CL\_INS\_382
CL\_INS\_382
CL\_INS\_382
CL\_INS\_382
CL\_INS\_382
CL\_INS\_382
CL\_INS\_382
CL\_INS\_382
CL\_INS\_382
CL\_INS\_382
CL\_INS\_382
CL\_INS\_382
CL\_INS\_382
CL\_INS\_382
CL\_INS\_382
CL\_INS\_382
CL\_INS\_382
CL\_INS\_382
CL\_INS\_382
CL\_INS\_382
CL\_INS\_382
CL\_INS\_60
CL\_INS\_382
CL\_INS\_382
CL\_INS\_382
CL\_INS\_382
CL\_INS\_382
CL\_INS\_382
CL\_INS\_382
CL\_INS\_382
CL\_INS\_83
CL\_INS\_83
CL\_INS\_83
CL\_INS\_382
CL\_INS\_382
CL\_INS\_382
CL\_INS\_382
CL\_INS\_382
CL\_INS\_79
CL\_INS\_79
CL\_INS\_79
CL\_INS\_79
CL\_INS\_79
CL\_INS\_79
CL\_INS\_382
CL\_INS\_79
CL\_INS\_382
CL\_INS\_382
CL\_INS\_237
CL\_INS\_237
CL\_INS\_20
CL\_INS\_79
CL\_INS\_79
CL\_INS\_20
CL\_INS\_20
CL\_INS\_20
CL\_INS\_79
CL\_INS\_79
CL\_INS\_79
CL\_INS\_79
CL\_INS\_79
CL\_INS\_382
CL\_INS\_79
CL\_INS\_382
CL\_INS\_79
CL\_INS\_382
CL\_INS\_79
CL\_INS\_79
CL\_INS\_79
CL\_INS\_382
CL\_INS\_79
CL\_INS\_79
CL\_INS\_146
CL\_INS\_146
CL\_INS\_146
CL\_INS\_79
CL\_INS\_79
CL\_INS\_20
CL\_INS\_20
CL\_INS\_382
CL\_INS\_382
CL\_INS\_382
CL\_INS\_382
CL\_INS\_382
CL\_INS\_382
CL\_INS\_382
CL\_INS\_83
CL\_INS\_99
CL\_INS\_128
CL\_INS\_128
CL\_INS\_83
CL\_INS\_128
CL\_INS\_83
CL\_INS\_79
CL\_INS\_99
CL\_INS\_99
CL\_INS\_79
CL\_INS\_79
CL\_INS\_79
CL\_INS\_382
CL\_INS\_382
CL\_INS\_382
CL\_INS\_83
CL\_INS\_83
CL\_INS\_83
CL\_INS\_83
CL\_INS\_83
CL\_INS\_382
CL\_INS\_382
CL\_INS\_382
CL\_INS\_382
CL\_INS\_382
CL\_INS\_382
CL\_INS\_382
CL\_INS\_382
CL\_INS\_99
CL\_INS\_83
CL\_INS\_382
CL\_INS\_382
CL\_INS\_384
CL\_INS\_160
CL\_INS\_382
CL\_INS\_83
CL\_INS\_382
CL\_INS\_207
CL\_INS\_207
CL\_INS\_99
CL\_INS\_79
CL\_INS\_79
CL\_INS\_79
CL\_INS\_83
CL\_INS\_83
CL\_INS\_382
CL\_INS\_83
CL\_INS\_382
CL\_INS\_382
CL\_INS\_382
CL\_INS\_382
CL\_INS\_83
CL\_INS\_382
CL\_INS\_382
CL\_INS\_99
CL\_INS\_146
CL\_INS\_382
CL\_INS\_83
CL\_INS\_382
CL\_INS\_382
CL\_INS\_128
CL\_INS\_382
CL\_INS\_128
CL\_INS\_382
CL\_INS\_99
CL\_INS\_382
CL\_INS\_83
CL\_INS\_382
CL\_INS\_83
CL\_INS\_382
CL\_INS\_382
CL\_INS\_382
CL\_INS\_382
CL\_INS\_382
CL\_INS\_382
CL\_INS\_382
CL\_INS\_99
CL\_INS\_83
CL\_INS\_83
CL\_INS\_83
CL\_INS\_382
CL\_INS\_382
CL\_INS\_382
CL\_INS\_382
CL\_INS\_382
CL\_INS\_382
CL\_INS\_382
CL\_INS\_83
CL\_INS\_382
CL\_INS\_382
CL\_INS\_83
CL\_INS\_99
CL\_INS\_79
CL\_INS\_382
CL\_INS\_382
CL\_INS\_382
CL\_INS\_99
CL\_INS\_382
CL\_INS\_99
CL\_INS\_99
CL\_INS\_382
CL\_INS\_99
CL\_INS\_99
CL\_INS\_83
CL\_INS\_83
CL\_INS\_83
CL\_INS\_83
CL\_INS\_83
CL\_INS\_99
CL\_INS\_83
CL\_INS\_128
CL\_INS\_99
CL\_INS\_128
CL\_INS\_83
CL\_INS\_83
CL\_INS\_83
CL\_INS\_83
CL\_INS\_83
CL\_INS\_385
CL\_INS\_83
CL\_INS\_83
CL\_INS\_83
CL\_INS\_83
CL\_INS\_83
CL\_INS\_83
CL\_INS\_83
CL\_INS\_83
CL\_INS\_83
CL\_INS\_83
CL\_INS\_83
CL\_INS\_83
CL\_INS\_83
CL\_INS\_83
CL\_INS\_83
CL\_INS\_83
CL\_INS\_83
CL\_INS\_83
CL\_INS\_83
CL\_INS\_83
CL\_INS\_83
CL\_INS\_83
CL\_INS\_83
CL\_INS\_83
CL\_INS\_83
CL\_INS\_83
CL\_INS\_83
CL\_INS\_83
CL\_INS\_83
CL\_INS\_83
CL\_INS\_83
CL\_INS\_83
CL\_INS\_83
CL\_INS\_207
CL\_INS\_207
CL\_INS\_207
CL\_INS\_207
CL\_INS\_207
CL\_INS\_83
CL\_INS\_83
CL\_INS\_382
CL\_INS\_146
CL\_INS\_146
CL\_INS\_207
CL\_INS\_83
CL\_INS\_83
CL\_INS\_83
CL\_INS\_83
CL\_INS\_83
CL\_INS\_83
CL\_INS\_83
CL\_INS\_83
CL\_INS\_382
CL\_INS\_99
CL\_INS\_131
CL\_INS\_382
CL\_INS\_99
CL\_INS\_83
CL\_INS\_83
CL\_INS\_99
CL\_INS\_83
CL\_INS\_83
CL\_INS\_99
CL\_INS\_99
CL\_INS\_382
CL\_INS\_99
CL\_INS\_131
CL\_INS\_83
CL\_INS\_83
CL\_INS\_83
CL\_INS\_83
CL\_INS\_128
CL\_INS\_382
CL\_INS\_382
CL\_INS\_382
CL\_INS\_382
CL\_INS\_382
CL\_INS\_382
CL\_INS\_83
CL\_INS\_382
CL\_INS\_79
CL\_INS\_83
CL\_INS\_384
CL\_INS\_83
CL\_INS\_83
CL\_INS\_83
CL\_INS\_382
CL\_INS\_99
CL\_INS\_79
CL\_INS\_382
CL\_INS\_382
CL\_INS\_382
CL\_INS\_382
CL\_INS\_99
CL\_INS\_99
CL\_INS\_99
CL\_INS\_99
CL\_INS\_99
CL\_INS\_382
CL\_INS\_83
CL\_INS\_83
CL\_INS\_99
CL\_INS\_99
CL\_INS\_83
CL\_INS\_382
CL\_INS\_382
CL\_INS\_382
CL\_INS\_382
CL\_INS\_382
CL\_INS\_382
CL\_INS\_382
CL\_INS\_99
CL\_INS\_382
CL\_INS\_382
CL\_INS\_382
CL\_INS\_86
CL\_INS\_83
CL\_INS\_382
CL\_INS\_99
CL\_INS\_83
CL\_INS\_83
CL\_INS\_83
CL\_INS\_128
CL\_INS\_128
CL\_INS\_128
CL\_INS\_382
CL\_INS\_382
CL\_INS\_382
CL\_INS\_83
CL\_INS\_83
CL\_INS\_382
CL\_INS\_83
CL\_INS\_382
CL\_INS\_171
CL\_INS\_171
CL\_INS\_207
CL\_INS\_83
CL\_INS\_83
CL\_INS\_382
CL\_INS\_149
CL\_INS\_382
CL\_INS\_382
CL\_INS\_382
CL\_INS\_382
CL\_INS\_382
CL\_INS\_382
CL\_INS\_382
CL\_INS\_79
CL\_INS\_382
CL\_INS\_83
CL\_INS\_83
CL\_INS\_83
CL\_INS\_83
CL\_INS\_83
CL\_INS\_83
CL\_INS\_83
CL\_INS\_83
CL\_INS\_83
CL\_INS\_83
CL\_INS\_83
CL\_INS\_83
CL\_INS\_382
CL\_INS\_382
CL\_INS\_204
CL\_INS\_382
CL\_INS\_99
CL\_INS\_83
CL\_INS\_83
CL\_INS\_83
CL\_INS\_83
CL\_INS\_382
CL\_INS\_382
CL\_INS\_83
CL\_INS\_83
CL\_INS\_83
CL\_INS\_83
CL\_INS\_83
CL\_INS\_382
CL\_INS\_83
CL\_INS\_382
CL\_INS\_382
CL\_INS\_382
CL\_INS\_384
CL\_INS\_384
CL\_INS\_382
CL\_INS\_83
CL\_INS\_382
CL\_INS\_382
CL\_INS\_382
CL\_INS\_146
CL\_INS\_83
CL\_INS\_382
CL\_INS\_382
CL\_INS\_83
CL\_INS\_83
CL\_INS\_83
CL\_INS\_83
CL\_INS\_83
CL\_INS\_99
CL\_INS\_83
CL\_INS\_83
CL\_INS\_83
CL\_INS\_83
CL\_INS\_83
CL\_INS\_83
CL\_INS\_83
CL\_INS\_83
CL\_INS\_83
CL\_INS\_83
CL\_INS\_83
CL\_INS\_83
CL\_INS\_83
CL\_INS\_83
CL\_INS\_83
CL\_INS\_83
CL\_INS\_83
CL\_INS\_83
CL\_INS\_83
CL\_INS\_83
CL\_INS\_83
CL\_INS\_83
CL\_INS\_83
CL\_INS\_83
CL\_INS\_83
CL\_INS\_83
CL\_INS\_83
CL\_INS\_83
CL\_INS\_83
CL\_INS\_83
CL\_INS\_83
CL\_INS\_83
CL\_INS\_83
CL\_INS\_83
CL\_INS\_83
CL\_INS\_83
CL\_INS\_83
CL\_INS\_83
CL\_INS\_83
CL\_INS\_83
CL\_INS\_83
CL\_INS\_83
CL\_INS\_83
CL\_INS\_83
CL\_INS\_83
CL\_INS\_83
CL\_INS\_83
CL\_INS\_83
CL\_INS\_83
CL\_INS\_384
CL\_INS\_384
CL\_INS\_382
CL\_INS\_382
CL\_INS\_382
CL\_INS\_382
CL\_INS\_83
CL\_INS\_146
CL\_INS\_382
CL\_INS\_384
CL\_INS\_83
CL\_INS\_83
CL\_INS\_83
CL\_INS\_83
CL\_INS\_83
CL\_INS\_382
CL\_INS\_83
CL\_INS\_99
CL\_INS\_384
CL\_INS\_382
CL\_INS\_382
CL\_INS\_237
CL\_INS\_83
CL\_INS\_382
CL\_INS\_382
CL\_INS\_382
CL\_INS\_382
CL\_INS\_99
CL\_INS\_382
CL\_INS\_382
CL\_INS\_382
CL\_INS\_382
CL\_INS\_382
CL\_INS\_382
CL\_INS\_83
CL\_INS\_83
CL\_INS\_99
CL\_INS\_382
CL\_INS\_83
CL\_INS\_99
CL\_INS\_99
CL\_INS\_83
CL\_INS\_382
CL\_INS\_382
CL\_INS\_382
CL\_INS\_83
CL\_INS\_382
CL\_INS\_83
CL\_INS\_83
CL\_INS\_83
CL\_INS\_79
CL\_INS\_382
CL\_INS\_86
CL\_INS\_204
CL\_INS\_86
CL\_INS\_154
CL\_INS\_99
CL\_INS\_86
CL\_INS\_99
CL\_INS\_83
CL\_INS\_384
CL\_INS\_384
CL\_INS\_83
CL\_INS\_204
CL\_INS\_204
CL\_INS\_384
CL\_INS\_384
CL\_INS\_204
CL\_INS\_204
CL\_INS\_204
CL\_INS\_146
CL\_INS\_204
CL\_INS\_83
CL\_INS\_83
CL\_INS\_83
CL\_INS\_382
CL\_INS\_382
CL\_INS\_106
CL\_INS\_154
CL\_INS\_106
CL\_INS\_106
CL\_INS\_132
CL\_INS\_382
CL\_INS\_247
CL\_INS\_247
CL\_INS\_132
CL\_INS\_132
CL\_INS\_174
CL\_INS\_132
CL\_INS\_132
CL\_INS\_132
CL\_INS\_132
CL\_INS\_132
CL\_INS\_132
CL\_INS\_132
CL\_INS\_99
CL\_INS\_99
CL\_INS\_99
CL\_INS\_99
CL\_INS\_99
CL\_INS\_83
CL\_INS\_83
CL\_INS\_83
CL\_INS\_83
CL\_INS\_83
CL\_INS\_99
CL\_INS\_99
CL\_INS\_99
CL\_INS\_99
CL\_INS\_382
CL\_INS\_20
CL\_INS\_146
CL\_INS\_382
CL\_INS\_83
CL\_INS\_83
CL\_INS\_382
CL\_INS\_207
CL\_INS\_207
CL\_INS\_207
CL\_INS\_207
CL\_INS\_207
CL\_INS\_382
CL\_INS\_382
CL\_INS\_382
CL\_INS\_83
CL\_INS\_83
CL\_INS\_83
CL\_INS\_368
CL\_INS\_83
CL\_INS\_83
CL\_INS\_83
CL\_INS\_83
CL\_INS\_382
CL\_INS\_382
CL\_INS\_382
CL\_INS\_382
CL\_INS\_382
CL\_INS\_99
CL\_INS\_99
CL\_INS\_99
CL\_INS\_99
CL\_INS\_99
CL\_INS\_99
CL\_INS\_146
CL\_INS\_86
CL\_INS\_86
CL\_INS\_385
CL\_INS\_237
CL\_INS\_149
CL\_INS\_170
CL\_INS\_385
CL\_INS\_385
CL\_INS\_10
CL\_INS\_83
CL\_INS\_99
CL\_INS\_146
CL\_INS\_146
CL\_INS\_146
CL\_INS\_99
CL\_INS\_79
CL\_INS\_382
CL\_INS\_382
CL\_INS\_83
CL\_INS\_83
CL\_INS\_83
CL\_INS\_86
CL\_INS\_382
CL\_INS\_83
CL\_INS\_382
CL\_INS\_382
CL\_INS\_382
CL\_INS\_382
CL\_INS\_83
CL\_INS\_382
CL\_INS\_99
CL\_INS\_99
CL\_INS\_83
CL\_INS\_83
CL\_INS\_83
CL\_INS\_83
CL\_INS\_83
CL\_INS\_83
CL\_INS\_83
CL\_INS\_83
CL\_INS\_83
CL\_INS\_83
CL\_INS\_83
CL\_INS\_83
CL\_INS\_83
CL\_INS\_83
CL\_INS\_83
CL\_INS\_83
CL\_INS\_83
CL\_INS\_83
CL\_INS\_83
CL\_INS\_83
CL\_INS\_83
CL\_INS\_83
CL\_INS\_83
CL\_INS\_83
CL\_INS\_83
CL\_INS\_83
CL\_INS\_83
CL\_INS\_83
CL\_INS\_83
CL\_INS\_83
CL\_INS\_83
CL\_INS\_83
CL\_INS\_83
CL\_INS\_83
CL\_INS\_83
CL\_INS\_83
CL\_INS\_83
CL\_INS\_83
CL\_INS\_83
CL\_INS\_128
CL\_INS\_83
CL\_INS\_382
CL\_INS\_99
CL\_INS\_83
CL\_INS\_99
CL\_INS\_382
CL\_INS\_83
CL\_INS\_99
CL\_INS\_83
CL\_INS\_83
CL\_INS\_382
CL\_INS\_382
CL\_INS\_128
CL\_INS\_128
CL\_INS\_128
CL\_INS\_83
CL\_INS\_83
CL\_INS\_83
CL\_INS\_83
CL\_INS\_83
CL\_INS\_83
CL\_INS\_83
CL\_INS\_99
CL\_INS\_382
CL\_INS\_83
CL\_INS\_83
CL\_INS\_382
CL\_INS\_83
CL\_INS\_382
CL\_INS\_382
CL\_INS\_83
CL\_INS\_83
CL\_INS\_83
CL\_INS\_382
CL\_INS\_83
CL\_INS\_146
CL\_INS\_83
CL\_INS\_83
CL\_INS\_83
CL\_INS\_83
CL\_INS\_83
CL\_INS\_83
CL\_INS\_83
CL\_INS\_83
CL\_INS\_83
CL\_INS\_83
CL\_INS\_83
CL\_INS\_83
CL\_INS\_83
CL\_INS\_99
CL\_INS\_99
CL\_INS\_83
CL\_INS\_83
CL\_INS\_83
CL\_INS\_83
CL\_INS\_83
CL\_INS\_83
CL\_INS\_83
CL\_INS\_83
CL\_INS\_83
CL\_INS\_83
CL\_INS\_83
CL\_INS\_83
CL\_INS\_83
CL\_INS\_99
CL\_INS\_83
CL\_INS\_83
CL\_INS\_99
CL\_INS\_207
CL\_INS\_83
CL\_INS\_83
CL\_INS\_83
CL\_INS\_83
CL\_INS\_83
CL\_INS\_83
CL\_INS\_99
CL\_INS\_382
CL\_INS\_99
CL\_INS\_382
CL\_INS\_382
CL\_INS\_382
CL\_INS\_83
CL\_INS\_83
CL\_INS\_382
CL\_INS\_379
CL\_INS\_382
CL\_INS\_83
CL\_INS\_382
CL\_INS\_382
CL\_INS\_83
CL\_INS\_382
CL\_INS\_384
CL\_INS\_382
CL\_INS\_382
CL\_INS\_382
CL\_INS\_382
CL\_INS\_382
CL\_INS\_382
CL\_INS\_382
CL\_INS\_382
CL\_INS\_99
CL\_INS\_99
CL\_INS\_382
CL\_INS\_83
CL\_INS\_382
CL\_INS\_83
CL\_INS\_83
CL\_INS\_83
CL\_INS\_83
CL\_INS\_83
CL\_INS\_83
CL\_INS\_83
CL\_INS\_83
CL\_INS\_83
CL\_INS\_83
CL\_INS\_83
CL\_INS\_83
CL\_INS\_83
CL\_INS\_83
CL\_INS\_83
CL\_INS\_83
CL\_INS\_83
CL\_INS\_83
CL\_INS\_382
CL\_INS\_382
CL\_INS\_382
CL\_INS\_382
CL\_INS\_382
CL\_INS\_382
CL\_INS\_382
CL\_INS\_382
CL\_INS\_382
CL\_INS\_83
CL\_INS\_83
CL\_INS\_382
CL\_INS\_382
CL\_INS\_382
CL\_INS\_382
CL\_INS\_83
CL\_INS\_83
CL\_INS\_83
CL\_INS\_83
CL\_INS\_83
CL\_INS\_83
CL\_INS\_83
CL\_INS\_382
CL\_INS\_382
CL\_INS\_382
CL\_INS\_382
CL\_INS\_382
CL\_INS\_385
CL\_INS\_382
CL\_INS\_382
CL\_INS\_382
CL\_INS\_382
CL\_INS\_382
CL\_INS\_382
CL\_INS\_382
CL\_INS\_382
CL\_INS\_382
CL\_INS\_382
CL\_INS\_382
CL\_INS\_382
CL\_INS\_382
CL\_INS\_382
CL\_INS\_382
CL\_INS\_382
CL\_INS\_384
CL\_INS\_382
CL\_INS\_83
CL\_INS\_83
CL\_INS\_83
CL\_INS\_83
CL\_INS\_83
CL\_INS\_83
CL\_INS\_83
CL\_INS\_83
CL\_INS\_83
CL\_INS\_83
CL\_INS\_83
CL\_INS\_83
CL\_INS\_83
CL\_INS\_83
CL\_INS\_83
CL\_INS\_83
CL\_INS\_83
CL\_INS\_83
CL\_INS\_83
CL\_INS\_382
CL\_INS\_382
CL\_INS\_382
CL\_INS\_382
CL\_INS\_382
CL\_INS\_382
CL\_INS\_382
CL\_INS\_382
CL\_INS\_382
CL\_INS\_382
CL\_INS\_382
CL\_INS\_382
CL\_INS\_382
CL\_INS\_382
CL\_INS\_146
CL\_INS\_146
CL\_INS\_146
CL\_INS\_146
CL\_INS\_382
CL\_INS\_146
CL\_INS\_128
CL\_INS\_382
CL\_INS\_207
CL\_INS\_99
CL\_INS\_207
CL\_INS\_382
CL\_INS\_382
CL\_INS\_382
CL\_INS\_384
CL\_INS\_384
CL\_INS\_382
CL\_INS\_382
CL\_INS\_382
CL\_INS\_207
CL\_INS\_207
CL\_INS\_99
CL\_INS\_207
CL\_INS\_207
CL\_INS\_382
CL\_INS\_382
CL\_INS\_382
CL\_INS\_382
CL\_INS\_207
CL\_INS\_207
CL\_INS\_207
CL\_INS\_207
CL\_INS\_382
CL\_INS\_382
CL\_INS\_382
CL\_INS\_128
CL\_INS\_382
CL\_INS\_99
CL\_INS\_83
CL\_INS\_83
CL\_INS\_83
CL\_INS\_83
CL\_INS\_382
CL\_INS\_83
CL\_INS\_99
CL\_INS\_382
CL\_INS\_99
CL\_INS\_86
CL\_INS\_384
CL\_INS\_99
CL\_INS\_382
CL\_INS\_99
CL\_INS\_83
CL\_INS\_99
CL\_INS\_382
CL\_INS\_382
CL\_INS\_99
CL\_INS\_83
CL\_INS\_99
CL\_INS\_99
CL\_INS\_99
CL\_INS\_83
CL\_INS\_83
CL\_INS\_83
CL\_INS\_83
CL\_INS\_83
CL\_INS\_83
CL\_INS\_382
CL\_INS\_99
CL\_INS\_86
CL\_INS\_385
CL\_INS\_385
CL\_INS\_146
CL\_INS\_83
CL\_INS\_86
CL\_INS\_99
CL\_INS\_87
CL\_INS\_99
CL\_INS\_382
CL\_INS\_99
CL\_INS\_83
CL\_INS\_83
CL\_INS\_237
CL\_INS\_99
CL\_INS\_99
CL\_INS\_99
CL\_INS\_99
CL\_INS\_379
CL\_INS\_379
CL\_INS\_379
CL\_INS\_99
CL\_INS\_382
CL\_INS\_86
CL\_INS\_83
CL\_INS\_99
CL\_INS\_83
CL\_INS\_83
CL\_INS\_83
CL\_INS\_382
CL\_INS\_83
CL\_INS\_382
CL\_INS\_83
CL\_INS\_382
CL\_INS\_382
CL\_INS\_20
CL\_INS\_382
CL\_INS\_86
CL\_INS\_99
CL\_INS\_385
CL\_INS\_86
CL\_INS\_99
CL\_INS\_99
CL\_INS\_99
CL\_INS\_99
CL\_INS\_99
CL\_INS\_99
CL\_INS\_99
CL\_INS\_60
CL\_INS\_83
CL\_INS\_83
CL\_INS\_83
CL\_INS\_83
CL\_INS\_83
CL\_INS\_83
CL\_INS\_83
CL\_INS\_83
CL\_INS\_83
CL\_INS\_83
CL\_INS\_83
CL\_INS\_83
CL\_INS\_83
CL\_INS\_83
CL\_INS\_83
CL\_INS\_83
CL\_INS\_83
CL\_INS\_83
CL\_INS\_83
CL\_INS\_83
CL\_INS\_83
CL\_INS\_83
CL\_INS\_83
CL\_INS\_83
CL\_INS\_83
CL\_INS\_83
CL\_INS\_83
CL\_INS\_83
CL\_INS\_83
CL\_INS\_83
CL\_INS\_83
CL\_INS\_83
CL\_INS\_83
CL\_INS\_83
CL\_INS\_83
CL\_INS\_83
CL\_INS\_83
CL\_INS\_83
CL\_INS\_83
CL\_INS\_83
CL\_INS\_83
CL\_INS\_83
CL\_INS\_83
CL\_INS\_83
CL\_INS\_83
CL\_INS\_83
CL\_INS\_83
CL\_INS\_83
CL\_INS\_83
CL\_INS\_83
CL\_INS\_83
CL\_INS\_83
CL\_INS\_83
CL\_INS\_83
CL\_INS\_83
CL\_INS\_83
CL\_INS\_83
CL\_INS\_83
CL\_INS\_83
CL\_INS\_83
CL\_INS\_83
CL\_INS\_83
CL\_INS\_83
CL\_INS\_83
CL\_INS\_83
CL\_INS\_83
CL\_INS\_83
CL\_INS\_83
CL\_INS\_83
CL\_INS\_83
CL\_INS\_83
CL\_INS\_83
CL\_INS\_83
CL\_INS\_83
CL\_INS\_382
CL\_INS\_382
CL\_INS\_382
CL\_INS\_382
CL\_INS\_382
CL\_INS\_382
CL\_INS\_99
CL\_INS\_71
CL\_INS\_149
CL\_INS\_382
CL\_INS\_382
CL\_INS\_83
CL\_INS\_83
CL\_INS\_83
CL\_INS\_382
CL\_INS\_382
CL\_INS\_382
CL\_INS\_382
CL\_INS\_83
CL\_INS\_382
CL\_INS\_83
CL\_INS\_83
CL\_INS\_83
CL\_INS\_382
CL\_INS\_382
CL\_INS\_99
CL\_INS\_99
CL\_INS\_99
CL\_INS\_99
CL\_INS\_382
CL\_INS\_382
CL\_INS\_382
CL\_INS\_382
CL\_INS\_382
CL\_INS\_382
CL\_INS\_382
CL\_INS\_382
CL\_INS\_382
CL\_INS\_382
CL\_INS\_382
CL\_INS\_382
CL\_INS\_382
CL\_INS\_382
CL\_INS\_83
CL\_INS\_83
CL\_INS\_83
CL\_INS\_83
CL\_INS\_83
CL\_INS\_382
CL\_INS\_146
CL\_INS\_146
CL\_INS\_382
CL\_INS\_382
CL\_INS\_382
CL\_INS\_99
CL\_INS\_382
CL\_INS\_83
CL\_INS\_83
CL\_INS\_83
CL\_INS\_83
CL\_INS\_83
CL\_INS\_83
CL\_INS\_83
CL\_INS\_83
CL\_INS\_83
CL\_INS\_83
CL\_INS\_83
CL\_INS\_83
CL\_INS\_99
CL\_INS\_83
CL\_INS\_83
CL\_INS\_83
CL\_INS\_83
CL\_INS\_83
CL\_INS\_99
CL\_INS\_99
CL\_INS\_382
CL\_INS\_146
CL\_INS\_146
CL\_INS\_382
CL\_INS\_382
CL\_INS\_382
CL\_INS\_382
CL\_INS\_382
CL\_INS\_382
CL\_INS\_146
CL\_INS\_146
CL\_INS\_146
CL\_INS\_146
CL\_INS\_146
CL\_INS\_146
CL\_INS\_146
CL\_INS\_146
CL\_INS\_146
CL\_INS\_146
CL\_INS\_146
CL\_INS\_146
CL\_INS\_146
CL\_INS\_382
CL\_INS\_382
CL\_INS\_382
CL\_INS\_83
CL\_INS\_382
CL\_INS\_83
CL\_INS\_83
CL\_INS\_83
CL\_INS\_83
CL\_INS\_382
CL\_INS\_83
CL\_INS\_382
CL\_INS\_382
CL\_INS\_382
CL\_INS\_382
CL\_INS\_128
CL\_INS\_83
CL\_INS\_83
CL\_INS\_83
CL\_INS\_83
CL\_INS\_83
CL\_INS\_99
CL\_INS\_146
CL\_INS\_382
CL\_INS\_83
CL\_INS\_99
CL\_INS\_146
CL\_INS\_83
CL\_INS\_83
CL\_INS\_99
CL\_INS\_83
CL\_INS\_83
CL\_INS\_83
CL\_INS\_83
CL\_INS\_83
CL\_INS\_83
CL\_INS\_83
CL\_INS\_83
CL\_INS\_83
CL\_INS\_83
CL\_INS\_83
CL\_INS\_83
CL\_INS\_83
CL\_INS\_83
CL\_INS\_83
CL\_INS\_83
CL\_INS\_99
CL\_INS\_83
CL\_INS\_83
CL\_INS\_83
CL\_INS\_83
CL\_INS\_83
CL\_INS\_83
CL\_INS\_83
CL\_INS\_83
CL\_INS\_83
CL\_INS\_83
CL\_INS\_83
CL\_INS\_83
CL\_INS\_83
CL\_INS\_83
CL\_INS\_20
CL\_INS\_20
CL\_INS\_20
CL\_INS\_83
CL\_INS\_83
CL\_INS\_83
CL\_INS\_83
CL\_INS\_207
CL\_INS\_83
CL\_INS\_83
CL\_INS\_83
CL\_INS\_83
CL\_INS\_83
CL\_INS\_79
CL\_INS\_79
CL\_INS\_79
CL\_INS\_79
CL\_INS\_385
CL\_INS\_207
CL\_INS\_207
CL\_INS\_207
CL\_INS\_207
CL\_INS\_382
CL\_INS\_382
CL\_INS\_382
CL\_INS\_382
CL\_INS\_83
CL\_INS\_207
CL\_INS\_237
CL\_INS\_237
CL\_INS\_83
CL\_INS\_83
CL\_INS\_382
CL\_INS\_382
CL\_INS\_382
CL\_INS\_247
CL\_INS\_382
CL\_INS\_382
CL\_INS\_382
CL\_INS\_382
CL\_INS\_247
CL\_INS\_247
CL\_INS\_237
CL\_INS\_247
CL\_INS\_247
CL\_INS\_247
CL\_INS\_247
CL\_INS\_44
CL\_INS\_247
CL\_INS\_44
CL\_INS\_247
CL\_INS\_247
CL\_INS\_247
CL\_INS\_60
CL\_INS\_247
CL\_INS\_123
CL\_INS\_247
CL\_INS\_247
CL\_INS\_247
CL\_INS\_247
CL\_INS\_123
CL\_INS\_83
CL\_INS\_385
CL\_INS\_83
CL\_INS\_83
CL\_INS\_83
CL\_INS\_83
CL\_INS\_99
CL\_INS\_382
CL\_INS\_83
CL\_INS\_343
CL\_INS\_385
CL\_INS\_99
CL\_INS\_83
CL\_INS\_159
CL\_INS\_83
CL\_INS\_83
CL\_INS\_247
CL\_INS\_382
CL\_INS\_382
CL\_INS\_382
CL\_INS\_382
CL\_INS\_382
CL\_INS\_83
CL\_INS\_382
CL\_INS\_385
CL\_INS\_159
CL\_INS\_159
CL\_INS\_159
CL\_INS\_385
CL\_INS\_60
CL\_INS\_382
CL\_INS\_382
CL\_INS\_382
CL\_INS\_382
CL\_INS\_385
CL\_INS\_99
CL\_INS\_382
CL\_INS\_60
CL\_INS\_382
CL\_INS\_382
CL\_INS\_382
CL\_INS\_382
CL\_INS\_382
CL\_INS\_382
CL\_INS\_382
CL\_INS\_382
CL\_INS\_382
CL\_INS\_382
CL\_INS\_382
CL\_INS\_60
CL\_INS\_382
CL\_INS\_382
CL\_INS\_382
CL\_INS\_382
CL\_INS\_382
CL\_INS\_382
CL\_INS\_159
CL\_INS\_385
CL\_INS\_385
CL\_INS\_382
CL\_INS\_382
CL\_INS\_60
CL\_INS\_60
CL\_INS\_60
CL\_INS\_382
CL\_INS\_382
CL\_INS\_382
CL\_INS\_382
CL\_INS\_83
CL\_INS\_382
CL\_INS\_382
CL\_INS\_382
CL\_INS\_382
CL\_INS\_382
CL\_INS\_382
CL\_INS\_382
CL\_INS\_382
CL\_INS\_382
CL\_INS\_382
CL\_INS\_83
Cluster ID


CL\_23895
CL\_7144
CL\_13034
CL\_9124
CL\_9125
CL\_22544
CL\_22545
CL\_29791
CL\_29790
CL\_29000
CL\_29001
CL\_29002
CL\_4670
CL\_28513
CL\_28512
CL\_4667
CL\_21014
CL\_4664
CL\_9745
CL\_4663
CL\_4662
CL\_15867
CL\_15866
CL\_4660
CL\_4659
CL\_4658
CL\_12800
CL\_12379
CL\_12380
CL\_4654
CL\_4653
CL\_4652
CL\_10983
CL\_5368
CL\_5367
CL\_6794
CL\_9953
CL\_7361
CL\_5364
CL\_5363
CL\_5362
CL\_5361
CL\_4400
CL\_508
CL\_4401
CL\_5360
CL\_5359
CL\_5358
CL\_5357
CL\_6793
CL\_8326
CL\_17391
CL\_6648
CL\_9226
CL\_6786
CL\_5351
CL\_4410
CL\_5350
CL\_5349
CL\_6656
CL\_6658
CL\_6659
CL\_17392
CL\_9090
CL\_9091
CL\_9092
CL\_9093
CL\_9094
CL\_17393
CL\_17394
CL\_17395
CL\_5340
CL\_526
CL\_12009
CL\_9098
CL\_27282
CL\_16594
CL\_16595
CL\_16742
CL\_16596
CL\_16597
CL\_16739
CL\_16006
CL\_17550
CL\_8334
CL\_6638
CL\_5389
CL\_5388
CL\_6641
CL\_13503
CL\_13504
CL\_9950
CL\_9949
CL\_8962
CL\_8964
CL\_9229
CL\_31762
CL\_31761
CL\_31760
CL\_9078
CL\_24460
CL\_9080
CL\_28278
CL\_509
CL\_27894
CL\_13507
CL\_21777
CL\_6655
CL\_9117
CL\_31759
CL\_17540
CL\_13972
CL\_17539
CL\_32704
CL\_32705
CL\_13071
CL\_15696
CL\_16444
CL\_12994
CL\_16615
CL\_12995
CL\_12759
CL\_12760
CL\_12996
CL\_12997
CL\_20884
CL\_33112
CL\_33114
CL\_33312
CL\_33116
CL\_12798
CL\_32021
CL\_23513
CL\_23514
CL\_31754
CL\_31753
CL\_16617
CL\_8700
CL\_8701
CL\_8702
CL\_28337
CL\_28336
CL\_28335
CL\_28334
CL\_28333
CL\_8703
CL\_8704
CL\_8705
CL\_8706
CL\_8707
CL\_8708
CL\_8709
CL\_8710
CL\_8711
CL\_28332
CL\_8172
CL\_8171
CL\_18335
CL\_6406
CL\_8170
CL\_28331
CL\_8169
CL\_17917
CL\_17918
CL\_8168
CL\_32706
CL\_32707
CL\_32708
CL\_14233
CL\_14234
CL\_8201
CL\_12993
CL\_8669
CL\_8200
CL\_1527
CL\_4565
CL\_17346
CL\_8198
CL\_15756
CL\_17040
CL\_19961
CL\_8199
CL\_32919
CL\_7126
CL\_1526
CL\_14814
CL\_1525
CL\_17347
CL\_1524
CL\_16113
CL\_5274
CL\_27413
CL\_14238
CL\_34192
CL\_14239
CL\_15788
CL\_15789
CL\_8673
CL\_11924
CL\_12697
CL\_8676
CL\_15908
CL\_27414
CL\_27415
CL\_27416
CL\_5276
CL\_8670
CL\_21792
CL\_17305
CL\_30168
CL\_23879
CL\_8195
CL\_36214
CL\_8671
CL\_8196
CL\_19227
CL\_14693
CL\_14694
CL\_11255
CL\_11926
CL\_11925
CL\_13299
CL\_13300
CL\_4564
CL\_4563
CL\_1311
CL\_1312
CL\_1313
CL\_33323
CL\_33322
CL\_33321
CL\_33320
CL\_33319
CL\_15759
CL\_19960
CL\_19959
CL\_4557
CL\_17265
CL\_28150
CL\_28149
CL\_28148
CL\_28147
CL\_28146
CL\_17490
CL\_28145
CL\_28144
CL\_28143
CL\_28142
CL\_28141
CL\_28140
CL\_28139
CL\_28138
CL\_28137
CL\_28136
CL\_28135
CL\_28134
CL\_28133
CL\_28132
CL\_28131
CL\_28130
CL\_28129
CL\_28128
CL\_28127
CL\_28126
CL\_28125
CL\_28124
CL\_28123
CL\_28122
CL\_28121
CL\_28120
CL\_28119
CL\_28118
CL\_28117
CL\_28116
CL\_28115
CL\_28114
CL\_28113
CL\_17558
CL\_17557
CL\_17556
CL\_17555
CL\_17554
CL\_31836
CL\_28112
CL\_7504
CL\_28111
CL\_28110
CL\_17553
CL\_33310
CL\_28109
CL\_28108
CL\_28107
CL\_28106
CL\_28105
CL\_28104
CL\_28103
CL\_10936
CL\_15760
CL\_13110
CL\_16191
CL\_15761
CL\_15762
CL\_15763
CL\_13303
CL\_15764
CL\_15765
CL\_16150
CL\_16151
CL\_15766
CL\_15767
CL\_11660
CL\_31668
CL\_31667
CL\_31666
CL\_30066
CL\_28304
CL\_16419
CL\_16418
CL\_5343
CL\_16417
CL\_16416
CL\_27034
CL\_16152
CL\_14235
CL\_15757
CL\_15758
CL\_14813
CL\_36620
CL\_16149
CL\_31064
CL\_8197
CL\_14692
CL\_23907
CL\_1523
CL\_1522
CL\_1521
CL\_5277
CL\_17980
CL\_17981
CL\_17982
CL\_17983
CL\_20767
CL\_5278
CL\_20026
CL\_31065
CL\_17984
CL\_17985
CL\_31066
CL\_4562
CL\_4457
CL\_4458
CL\_8995
CL\_4561
CL\_7125
CL\_4559
CL\_31168
CL\_23880
CL\_4560
CL\_30167
CL\_8674
CL\_23906
CL\_8675
CL\_15790
CL\_28521
CL\_28520
CL\_28519
CL\_17348
CL\_14240
CL\_14241
CL\_8996
CL\_8997
CL\_14695
CL\_14696
CL\_14697
CL\_9150
CL\_36215
CL\_6447
CL\_19678
CL\_19679
CL\_7250
CL\_36216
CL\_36217
CL\_9151
CL\_19226
CL\_19225
CL\_19224
CL\_6448
CL\_10510
CL\_10511
CL\_27286
CL\_10512
CL\_17349
CL\_13108
CL\_17350
CL\_17351
CL\_17352
CL\_17353
CL\_17354
CL\_17355
CL\_17356
CL\_17357
CL\_17358
CL\_17359
CL\_17360
CL\_17361
CL\_7124
CL\_7123
CL\_16420
CL\_34870
CL\_8194
CL\_37252
CL\_37253
CL\_37254
CL\_19958
CL\_5281
CL\_5282
CL\_8193
CL\_8192
CL\_8191
CL\_8190
CL\_8189
CL\_5283
CL\_23904
CL\_10981
CL\_8188
CL\_10514
CL\_30412
CL\_30411
CL\_10937
CL\_31669
CL\_1317
CL\_8677
CL\_4556
CL\_17309
CL\_27426
CL\_4555
CL\_4554
CL\_30069
CL\_30068
CL\_30067
CL\_30687
CL\_20882
CL\_16121
CL\_28518
CL\_31067
CL\_31068
CL\_31069
CL\_31070
CL\_31071
CL\_31072
CL\_31073
CL\_31074
CL\_31075
CL\_31076
CL\_31077
CL\_31078
CL\_31079
CL\_31080
CL\_31081
CL\_31082
CL\_31083
CL\_31084
CL\_31085
CL\_31086
CL\_31087
CL\_31088
CL\_31089
CL\_31090
CL\_31091
CL\_31092
CL\_31093
CL\_31094
CL\_31095
CL\_31096
CL\_31097
CL\_31098
CL\_31099
CL\_31100
CL\_31101
CL\_31102
CL\_31103
CL\_31104
CL\_31105
CL\_31106
CL\_31107
CL\_31108
CL\_31109
CL\_31110
CL\_31111
CL\_31112
CL\_31113
CL\_31114
CL\_16124
CL\_16125
CL\_16126
CL\_16127
CL\_16422
CL\_8678
CL\_33318
CL\_12804
CL\_8679
CL\_16123
CL\_33317
CL\_33316
CL\_33315
CL\_33314
CL\_33313
CL\_8680
CL\_23905
CL\_13304
CL\_14698
CL\_10515
CL\_8681
CL\_8434
CL\_11923
CL\_6449
CL\_6773
CL\_11922
CL\_6451
CL\_9000
CL\_4546
CL\_17066
CL\_16967
CL\_7540
CL\_7539
CL\_4644
CL\_36220
CL\_36221
CL\_6043
CL\_16445
CL\_27417
CL\_27418
CL\_27419
CL\_27420
CL\_15773
CL\_15774
CL\_15775
CL\_27421
CL\_8184
CL\_28341
CL\_28340
CL\_28339
CL\_26457
CL\_527
CL\_10976
CL\_10975
CL\_8180
CL\_23903
CL\_8179
CL\_8178
CL\_27796
CL\_23902
CL\_14807
CL\_14805
CL\_23901
CL\_1098
CL\_1099
CL\_14704
CL\_14705
CL\_5410
CL\_5409
CL\_5408
CL\_8177
CL\_1097
CL\_28846
CL\_28845
CL\_28844
CL\_8648
CL\_16131
CL\_16469
CL\_28843
CL\_8649
CL\_8651
CL\_8652
CL\_8113
CL\_20757
CL\_10496
CL\_10497
CL\_13102
CL\_23120
CL\_10498
CL\_10499
CL\_10500
CL\_10502
CL\_10503
CL\_10505
CL\_10506
CL\_10974
CL\_10973
CL\_10972
CL\_10971
CL\_8176
CL\_23900
CL\_23899
CL\_23898
CL\_23897
CL\_23896
CL\_8175
CL\_8174
CL\_8173
CL\_8183
CL\_9944
CL\_8182
CL\_8181
CL\_7534
CL\_30065
CL\_29229
CL\_7533
CL\_7532
CL\_7531
CL\_7530
CL\_7529
CL\_7528
CL\_7527
CL\_9123
CL\_15776
CL\_29228
CL\_29227
CL\_29226
CL\_29225
CL\_29224
CL\_29223
CL\_29222
CL\_29221
CL\_15777
CL\_15778
CL\_15779
CL\_15780
CL\_15781
CL\_17289
CL\_17290
CL\_17291
CL\_17292
CL\_15782
CL\_8155
CL\_28489
CL\_4430
CL\_7025
CL\_21530
CL\_4514
CL\_4601
CL\_4602
CL\_21531
CL\_21532
CL\_4764
CL\_27590
CL\_1326
CL\_28488
CL\_28487
CL\_28486
CL\_13790
CL\_25184
CL\_4548
CL\_4547
CL\_29455
CL\_36218
CL\_36219
CL\_8686
CL\_8687
CL\_28342
CL\_8688
CL\_17433
CL\_17434
CL\_15904
CL\_28102
CL\_19017
CL\_20768
CL\_15907
CL\_31215
CL\_31214
CL\_31213
CL\_31212
CL\_31211
CL\_31210
CL\_31209
CL\_31208
CL\_31207
CL\_31206
CL\_31205
CL\_31204
CL\_31203
CL\_31202
CL\_31201
CL\_31200
CL\_31199
CL\_31198
CL\_31197
CL\_31196
CL\_31195
CL\_31194
CL\_31193
CL\_31192
CL\_31191
CL\_31190
CL\_31189
CL\_31188
CL\_31187
CL\_31186
CL\_31185
CL\_31184
CL\_31183
CL\_31182
CL\_31181
CL\_31180
CL\_31179
CL\_31178
CL\_31177
CL\_28320
CL\_31176
CL\_12360
CL\_13509
CL\_31175
CL\_15906
CL\_15905
CL\_19016
CL\_14812
CL\_19213
CL\_19015
CL\_8186
CL\_8689
CL\_33884
CL\_33883
CL\_33882
CL\_36619
CL\_36618
CL\_36617
CL\_36616
CL\_36615
CL\_36614
CL\_36613
CL\_15902
CL\_8690
CL\_17362
CL\_23512
CL\_4648
CL\_22683
CL\_4647
CL\_14244
CL\_28516
CL\_28515
CL\_28514
CL\_12698
CL\_32187
CL\_20784
CL\_31595
CL\_31596
CL\_31597
CL\_34194
CL\_27427
CL\_27428
CL\_27429
CL\_27430
CL\_27431
CL\_27432
CL\_27433
CL\_27434
CL\_27435
CL\_13560
CL\_20888
CL\_27436
CL\_27437
CL\_27438
CL\_27439
CL\_27440
CL\_27441
CL\_27442
CL\_27443
CL\_27444
CL\_27445
CL\_27446
CL\_27447
CL\_27448
CL\_13789
CL\_27449
CL\_27450
CL\_17053
CL\_17559
CL\_28329
CL\_27451
CL\_27452
CL\_27453
CL\_27454
CL\_19950
CL\_15903
CL\_10980
CL\_8691
CL\_8692
CL\_27285
CL\_27284
CL\_28517
CL\_14370
CL\_10940
CL\_17237
CL\_10941
CL\_30688
CL\_13562
CL\_12762
CL\_19957
CL\_8693
CL\_24313
CL\_7120
CL\_7538
CL\_12006
CL\_7119
CL\_7118
CL\_27283
CL\_12797
CL\_13524
CL\_13561
CL\_28731
CL\_7117
CL\_19956
CL\_8185
CL\_16157
CL\_16158
CL\_16159
CL\_16160
CL\_16161
CL\_16162
CL\_16163
CL\_16164
CL\_16165
CL\_16166
CL\_16167
CL\_16168
CL\_16169
CL\_16170
CL\_16171
CL\_16172
CL\_16173
CL\_16174
CL\_7537
CL\_8696
CL\_6784
CL\_7048
CL\_10323
CL\_6783
CL\_9009
CL\_1497
CL\_1496
CL\_13788
CL\_11921
CL\_11920
CL\_11919
CL\_8697
CL\_13000
CL\_13001
CL\_13002
CL\_13003
CL\_13004
CL\_13005
CL\_13006
CL\_17371
CL\_11918
CL\_7030
CL\_7029
CL\_7028
CL\_532
CL\_14248
CL\_6045
CL\_14371
CL\_12999
CL\_19014
CL\_19013
CL\_4534
CL\_533
CL\_27281
CL\_21008
CL\_27280
CL\_4533
CL\_7116
CL\_4413
CL\_4414
CL\_4532
CL\_4416
CL\_15899
CL\_15898
CL\_21525
CL\_21526
CL\_21527
CL\_17396
CL\_17397
CL\_17398
CL\_17399
CL\_17400
CL\_17401
CL\_17402
CL\_17403
CL\_17404
CL\_17405
CL\_17406
CL\_17407
CL\_17408
CL\_17409
CL\_17410
CL\_19012
CL\_4415
CL\_4531
CL\_4417
CL\_4530
CL\_4418
CL\_4419
CL\_4420
CL\_6461
CL\_8901
CL\_28496
CL\_21006
CL\_21005
CL\_6741
CL\_28495
CL\_28494
CL\_28493
CL\_28492
CL\_28491
CL\_6744
CL\_28490
CL\_26458
CL\_4529
CL\_17912
CL\_6046
CL\_17913
CL\_4528
CL\_7367
CL\_4527
CL\_27588
CL\_27589
CL\_4421
CL\_4526
CL\_4525
CL\_7526
CL\_7525
CL\_21528
CL\_7524
CL\_7523
CL\_4423
CL\_4424
CL\_4425
CL\_4629
CL\_17914
CL\_17915
CL\_17916
CL\_13074
CL\_4524
CL\_4523
CL\_4522
CL\_19951
CL\_11913
CL\_17057
CL\_19011
CL\_19010
CL\_19009
CL\_19008
CL\_4628
CL\_19007
CL\_19006
CL\_4521
CL\_7522
CL\_7027
CL\_21529
CL\_4520
CL\_11957
CL\_7112
CL\_28330
CL\_4485
CL\_4518
CL\_4517
CL\_4431
CL\_31665
CL\_28003
CL\_6747
CL\_12701
CL\_31174
CL\_31173
CL\_31172
CL\_31171
CL\_31170
CL\_31169
CL\_8167
CL\_4489
CL\_1495
CL\_24314
CL\_16571
CL\_8166
CL\_8165
CL\_1324
CL\_10342
CL\_4513
CL\_8589
CL\_8712
CL\_7111
CL\_11917
CL\_19005
CL\_15588
CL\_4620
CL\_17052
CL\_17051
CL\_13075
CL\_15783
CL\_15784
CL\_15785
CL\_12385
CL\_4432
CL\_4515
CL\_33397
CL\_17033
CL\_27422
CL\_27423
CL\_27424
CL\_10982
CL\_27425
CL\_14236
CL\_36621
CL\_14237
CL\_16423
CL\_10474
CL\_9158
CL\_8585
CL\_13526
CL\_13791
CL\_4490
CL\_8586
CL\_8587
CL\_8588
CL\_7109
CL\_534
CL\_4433
CL\_9159
CL\_19004
CL\_19003
CL\_19002
CL\_19001
CL\_19000
CL\_18999
CL\_18998
CL\_18997
CL\_18996
CL\_18995
CL\_18994
CL\_18993
CL\_18992
CL\_18991
CL\_18990
CL\_18989
CL\_18988
CL\_18987
CL\_18986
CL\_18985
CL\_18984
CL\_18983
CL\_18982
CL\_18981
CL\_18980
CL\_18979
CL\_18978
CL\_18977
CL\_18976
CL\_18975
CL\_18974
CL\_18973
CL\_18972
CL\_18971
CL\_18970
CL\_18969
CL\_18968
CL\_18967
CL\_18966
CL\_18965
CL\_18964
CL\_18963
CL\_18962
CL\_18961
CL\_18960
CL\_18959
CL\_18958
CL\_18957
CL\_18956
CL\_18955
CL\_18954
CL\_18953
CL\_18952
CL\_18951
CL\_18950
CL\_18949
CL\_18948
CL\_18947
CL\_18946
CL\_18945
CL\_18944
CL\_18943
CL\_18942
CL\_18941
CL\_18940
CL\_18939
CL\_18938
CL\_18937
CL\_18936
CL\_18935
CL\_18934
CL\_18933
CL\_18932
CL\_18931
CL\_18930
CL\_10938
CL\_4459
CL\_10939
CL\_30166
CL\_30165
CL\_4552
CL\_30692
CL\_11996
CL\_11994
CL\_8683
CL\_8684
CL\_37196
CL\_37197
CL\_37198
CL\_31694
CL\_31693
CL\_31692
CL\_31691
CL\_34193
CL\_8187
CL\_28849
CL\_28848
CL\_28847
CL\_4545
CL\_4544
CL\_16968
CL\_20027
CL\_16969
CL\_20028
CL\_7019
CL\_34871
CL\_33569
CL\_33568
CL\_8909
CL\_10918
CL\_10919
CL\_7820
CL\_7819
CL\_517
CL\_518
CL\_519
CL\_520
CL\_5805
CL\_15768
CL\_15769
CL\_15770
CL\_15771
CL\_15772
CL\_17485
CL\_28511
CL\_28510
CL\_15862
CL\_6452
CL\_4543
CL\_14242
CL\_14243
CL\_14699
CL\_14700
CL\_14701
CL\_14702
CL\_14703
CL\_16153
CL\_16154
CL\_16155
CL\_16156
CL\_28101
CL\_28100
CL\_28099
CL\_17064
CL\_17363
CL\_17364
CL\_17365
CL\_17366
CL\_17367
CL\_17062
CL\_17061
CL\_4646
CL\_4645
CL\_6038
CL\_14245
CL\_6042
CL\_29789
CL\_18027
CL\_18026
CL\_9761
CL\_28509
CL\_28508
CL\_28507
CL\_28506
CL\_28505
CL\_28504
CL\_28503
CL\_28502
CL\_28501
CL\_28500
CL\_28499
CL\_28498
CL\_28497
CL\_14376
CL\_8694
CL\_28098
CL\_31598
CL\_17368
CL\_19955
CL\_19954
CL\_19953
CL\_19952
CL\_531
CL\_12998
CL\_21523
CL\_14246
CL\_14247
CL\_4535
CL\_8050
CL\_29232
CL\_29231
CL\_29230
CL\_21524
CL\_17369
CL\_8695
CL\_17370
CL\_8698
CL\_28338
CL\_17372
CL\_17373
CL\_16628
CL\_17374
CL\_9157
CL\_17375
CL\_17376
CL\_17377
CL\_17378
CL\_17379
CL\_17380
CL\_17381
CL\_17382
CL\_17383
CL\_17384
CL\_17385
CL\_17386
CL\_17387
CL\_17388
CL\_17389
CL\_28097
CL\_28096
CL\_28095
CL\_28094
CL\_28093
CL\_33311
CL\_28092
CL\_28091
CL\_28090
CL\_28089
CL\_32697
CL\_28088
CL\_28087
CL\_28086
CL\_28085
CL\_17411
CL\_7065
CL\_7066
CL\_7067
CL\_17412
CL\_17413
CL\_17414
CL\_32698
CL\_32699
CL\_32700
CL\_32701
CL\_32702
CL\_32703
CL\_17299
CL\_17415
CL\_17416
CL\_17417
CL\_17418
CL\_17419
CL\_17420
CL\_17421
CL\_17422
CL\_17423
CL\_5599
CL\_4310
CL\_5508
CL\_5507
CL\_37203
CL\_5511
CL\_5512
CL\_5513
CL\_35321
CL\_35320
CL\_9687
CL\_9688
CL\_9689
CL\_9690
CL\_9691
CL\_10667
CL\_10666
CL\_10665
CL\_11981
CL\_5601
CL\_5697
CL\_5691
CL\_10383
CL\_10382
CL\_10423
CL\_10422
CL\_10421
CL\_11134
CL\_10642
CL\_10641
CL\_10395
CL\_10394
CL\_10393
CL\_10392
CL\_15545
CL\_15067
CL\_15544
CL\_5299
CL\_5300
CL\_37204
CL\_5296
CL\_19485
CL\_37205
CL\_37206
CL\_13255
CL\_6148
CL\_6147
CL\_14090
CL\_5618
CL\_5619
CL\_14089
CL\_37207
CL\_5053
CL\_20829
CL\_37208
CL\_5054
CL\_4254
CL\_4255
CL\_4256
CL\_4257
CL\_5059
CL\_20953
CL\_14125
CL\_6929
CL\_4259
CL\_4260
CL\_4261
CL\_5584
CL\_6930
CL\_4262
CL\_4263
CL\_5062
CL\_4265
CL\_4266
CL\_6931
CL\_5579
CL\_5576
CL\_4269
CL\_4271
CL\_5634
CL\_6933
CL\_6934
CL\_5637
CL\_5638
CL\_5639
CL\_4277
CL\_4278
CL\_5567
CL\_5641
CL\_4279
CL\_5563
CL\_5562
CL\_5561
CL\_5560
CL\_4284
CL\_5651
CL\_5652
CL\_5654
CL\_5555
CL\_5554
CL\_11830
CL\_11829
CL\_5657
CL\_5551
CL\_5659
CL\_11338
CL\_4293
CL\_18390
CL\_5662
CL\_4297
CL\_4299
CL\_4300
CL\_4301
CL\_4302
CL\_4303
CL\_4304
CL\_4305
CL\_4306
CL\_5540
