## Supplementary material for "A novel method for integrating genomic and Tn-Seq data to identify common *in vivo* fitness mechanisms across multiple bacterial species": S1 Dataset: CL_INS_85.html

Legend

 Mobile +extrachromosomalelementfunctions
 Regulatoryfunctions
 Hypothetical
 DNA Metabolism
 AntibioticResistance
 All EssentialGenes
 Biosynthesis ofcofactors,prostheticgroups, +carriers
 All Fitness Genes
 Proteinsynthesis/fate
 Other
 EnergyMetabolism
 Cellularprocesses
 Transport +binding proteins
 All VFDB Genes

FULL


WINDOWSVGPNG

Trim RowsRemove SingletonsSave Fasta

CL\_1084


CL\_1084


CL\_1084


CL\_1084


CL\_1084


CL\_1084


CL\_4519


CL\_1084


CL\_1084


CL\_1084


CL\_1084


CL\_1084


CL\_1084


CL\_1084


CL\_1084


CL\_1084


CL\_1084


CL\_1084


CL\_1084


CL\_1084


CL\_1084


CL\_1084


CL\_1084


CL\_1084


CL\_1084


CL\_1084


CL\_4516


CL\_1084


CL\_1084


CL\_1084


CL\_1084


CL\_1084


CL\_4516


CL\_1084


CL\_1084


CL\_1084


CL\_1084


CL\_1084


CL\_1084


CL\_1084


CL\_1084


CL\_1084


CL\_1084


CL\_1084


CL\_1084


CL\_1084


CL\_1084


CL\_1084


CL\_1084


Break


CL\_1084


CL\_1084


CL\_1084


CL\_1084


CL\_1084


CL\_1084


CL\_1084


CL\_1084


CL\_1084


CL\_4519


CL\_1084


CL\_1084


CL\_1084


CL\_1084


CL\_1084


CL\_4486


CL\_1084


CL\_1084


CL\_1084


CL\_1084


CL\_1084


CL\_1084


CL\_4516


CL\_1084


CL\_1084


CL\_4516


CL\_1084


CL\_1084


CL\_1084


CL\_1084


CL\_1084


CL\_1084


CL\_1084


CL\_1084


CL\_1084


CL\_1084


CL\_1084


CL\_1084


CL\_4516


CL\_1084


CL\_1084


CL\_1084


CL\_1084


CL\_1084


CL\_1084


CL\_1084


CL\_1084


CL\_1084

HighlightSelectShow Genomes


158

CL\_1109


5

CL\_4427


4

CL\_1109


4

CL\_4427


4

CL\_1109


4

CL\_4427


3

CL\_1109


3

CL\_1110


2

CL\_4427


1

CL\_4427


1

CL\_4427


1

CL\_1109


1

CL\_1110


1

CL\_4427


1

CL\_4427


1

CL\_4427


1

CL\_538


1

CL\_4427


1

CL\_4427


1

CL\_4427


1

CL\_1110


1

CL\_4519


1

CL\_4427


1

CL\_1123


1

CL\_1109


1

CL\_1109


1

CL\_1109


1

CL\_4427


1

CL\_4427


1

CL\_1109


1

CL\_4427


1

CL\_4427


1

CL\_1109


1

CL\_4427


1

CL\_1110


1

CL\_1121


1

CL\_4427


1

CL\_1109


1

CL\_4427


1

CL\_1109


1

CL\_4427


1

CL\_1110


1

CL\_1109


1

CL\_1109


1

CL\_4427


1

CL\_4427


1

CL\_1109


1

CL\_1109


1

CL\_4427


1

CL\_1109


1

CL\_1110


1

CL\_4427


1

CL\_4516


1

CL\_4486


1

CL\_1110


1

CL\_4427


1

Break


1

CL\_4487


1

CL\_4427


1

CL\_1109


1

CL\_1109


1

CL\_1109


1

CL\_1109


1

CL\_1110


1

CL\_4427


1

CL\_1109


1

CL\_4427


1

CL\_4427


1

CL\_4519


1

CL\_1109


1

CL\_1109


1

CL\_1110


1

CL\_1109


1

CL\_1109


1

CL\_4427


1

CL\_1109


1

CL\_1819


1

CL\_4427


1

CL\_4427


1

CL\_4427


1

CL\_1109


1

CL\_1110


1

CL\_4486


1

CL\_1110


1

CL\_1109


1

CL\_1109


1

CL\_1110


1

CL\_4427


1

CL\_1109


1

CL\_1109


1

CL\_4427


1

CL\_4427


1

CL\_4519


1

CL\_4427


1

CL\_4486


1

CL\_1109


1

CL\_4519


1

CL\_4427

fGI ID


CL\_INS\_86
CL\_INS\_85
CL\_INS\_85
CL\_INS\_382
CL\_INS\_86
CL\_INS\_86
CL\_INS\_86
CL\_INS\_86
CL\_INS\_20
CL\_INS\_385
CL\_INS\_99
CL\_INS\_20
CL\_INS\_85
CL\_INS\_99
CL\_INS\_85
CL\_INS\_86
CL\_INS\_382
CL\_INS\_382
CL\_INS\_382
CL\_INS\_149
CL\_INS\_382
CL\_INS\_85
CL\_INS\_85
CL\_INS\_85
CL\_INS\_85
CL\_INS\_382
CL\_INS\_85
CL\_INS\_382
CL\_INS\_85
CL\_INS\_85
CL\_INS\_85
CL\_INS\_85
CL\_INS\_85
CL\_INS\_85
CL\_INS\_85
CL\_INS\_85
CL\_INS\_86
CL\_INS\_85
CL\_INS\_86
CL\_INS\_85
CL\_INS\_382
CL\_INS\_85
CL\_INS\_85
CL\_INS\_382
CL\_INS\_382
CL\_INS\_106
CL\_INS\_153
CL\_INS\_153
CL\_INS\_153
CL\_INS\_153
CL\_INS\_382
CL\_INS\_20
CL\_INS\_207
CL\_INS\_382
CL\_INS\_382
CL\_INS\_382
CL\_INS\_382
CL\_INS\_382
CL\_INS\_382
CL\_INS\_382
CL\_INS\_86
CL\_INS\_86
CL\_INS\_86
CL\_INS\_86
CL\_INS\_86
CL\_INS\_382
CL\_INS\_382
CL\_INS\_85
CL\_INS\_85
CL\_INS\_85
CL\_INS\_85
CL\_INS\_85
CL\_INS\_85
CL\_INS\_85
CL\_INS\_20
CL\_INS\_20
CL\_INS\_382
CL\_INS\_382
CL\_INS\_85
CL\_INS\_20
CL\_INS\_85
CL\_INS\_85
CL\_INS\_85
CL\_INS\_207
CL\_INS\_207
CL\_INS\_153
CL\_INS\_207
CL\_INS\_207
CL\_INS\_85
CL\_INS\_207
CL\_INS\_207
CL\_INS\_153
CL\_INS\_153
CL\_INS\_153
CL\_INS\_153
CL\_INS\_153
CL\_INS\_153
CL\_INS\_153
CL\_INS\_153
CL\_INS\_85
CL\_INS\_85
CL\_INS\_85
CL\_INS\_97
CL\_INS\_97
CL\_INS\_97
CL\_INS\_97
CL\_INS\_85
CL\_INS\_85
CL\_INS\_189
CL\_INS\_85
CL\_INS\_85
CL\_INS\_85
CL\_INS\_85
CL\_INS\_382
CL\_INS\_382
CL\_INS\_382
CL\_INS\_85
CL\_INS\_382
CL\_INS\_382
CL\_INS\_382
CL\_INS\_382
CL\_INS\_382
CL\_INS\_207
CL\_INS\_85
CL\_INS\_204
CL\_INS\_382
CL\_INS\_86
CL\_INS\_86
CL\_INS\_382
CL\_INS\_382
CL\_INS\_86
CL\_INS\_86
CL\_INS\_20
CL\_INS\_382
CL\_INS\_382
CL\_INS\_382
CL\_INS\_86
CL\_INS\_382
CL\_INS\_382
CL\_INS\_153
CL\_INS\_153
CL\_INS\_153
CL\_INS\_153
CL\_INS\_85
CL\_INS\_85
CL\_INS\_86
CL\_INS\_86
CL\_INS\_86
CL\_INS\_86
CL\_INS\_382
CL\_INS\_382
CL\_INS\_382
CL\_INS\_382
CL\_INS\_382
CL\_INS\_382
CL\_INS\_382
CL\_INS\_85
CL\_INS\_85
CL\_INS\_85
CL\_INS\_382
CL\_INS\_106
CL\_INS\_85
CL\_INS\_207
CL\_INS\_207
CL\_INS\_20
CL\_INS\_85
CL\_INS\_86
CL\_INS\_382
CL\_INS\_204
CL\_INS\_85
CL\_INS\_382
CL\_INS\_382
CL\_INS\_382
CL\_INS\_382
CL\_INS\_382
CL\_INS\_382
CL\_INS\_382
CL\_INS\_382
CL\_INS\_382
CL\_INS\_382
CL\_INS\_85
CL\_INS\_85
CL\_INS\_382
CL\_INS\_382
CL\_INS\_382
CL\_INS\_382
CL\_INS\_382
CL\_INS\_382
CL\_INS\_85
CL\_INS\_85
CL\_INS\_382
CL\_INS\_382
CL\_INS\_382
CL\_INS\_382
CL\_INS\_382
CL\_INS\_382
CL\_INS\_382
CL\_INS\_382
CL\_INS\_85
CL\_INS\_85
CL\_INS\_382
CL\_INS\_382
CL\_INS\_146
CL\_INS\_382
CL\_INS\_382
CL\_INS\_85
CL\_INS\_382
CL\_INS\_382
CL\_INS\_382
CL\_INS\_382
CL\_INS\_382
CL\_INS\_382
CL\_INS\_382
CL\_INS\_382
CL\_INS\_85
CL\_INS\_85
CL\_INS\_382
CL\_INS\_382
CL\_INS\_382
CL\_INS\_382
CL\_INS\_382
CL\_INS\_382
CL\_INS\_382
CL\_INS\_382
CL\_INS\_382
CL\_INS\_382
CL\_INS\_382
CL\_INS\_382
CL\_INS\_382
CL\_INS\_382
CL\_INS\_382
CL\_INS\_382
CL\_INS\_382
CL\_INS\_382
CL\_INS\_382
CL\_INS\_382
CL\_INS\_382
CL\_INS\_382
CL\_INS\_382
CL\_INS\_382
CL\_INS\_382
CL\_INS\_382
CL\_INS\_382
CL\_INS\_382
CL\_INS\_382
CL\_INS\_382
CL\_INS\_382
CL\_INS\_382
CL\_INS\_384
CL\_INS\_384
CL\_INS\_382
CL\_INS\_85
CL\_INS\_85
CL\_INS\_382
CL\_INS\_382
CL\_INS\_382
CL\_INS\_60
CL\_INS\_382
CL\_INS\_60
CL\_INS\_382
CL\_INS\_382
CL\_INS\_382
CL\_INS\_42
CL\_INS\_85
CL\_INS\_85
CL\_INS\_85
CL\_INS\_86
CL\_INS\_385
CL\_INS\_385
CL\_INS\_385
CL\_INS\_385
CL\_INS\_385
CL\_INS\_385
CL\_INS\_385
CL\_INS\_237
CL\_INS\_99
CL\_INS\_99
CL\_INS\_20
CL\_INS\_237
CL\_INS\_85
CL\_INS\_85
CL\_INS\_20
CL\_INS\_86
CL\_INS\_86
CL\_INS\_382
CL\_INS\_382
CL\_INS\_382
CL\_INS\_382
CL\_INS\_382
CL\_INS\_382
CL\_INS\_382
CL\_INS\_382
CL\_INS\_382
CL\_INS\_382
CL\_INS\_382
CL\_INS\_382
CL\_INS\_382
CL\_INS\_382
CL\_INS\_382
CL\_INS\_42
CL\_INS\_382
CL\_INS\_382
CL\_INS\_382
CL\_INS\_146
CL\_INS\_146
CL\_INS\_382
CL\_INS\_207
CL\_INS\_382
CL\_INS\_146
CL\_INS\_20
CL\_INS\_146
CL\_INS\_146
CL\_INS\_146
CL\_INS\_382
CL\_INS\_146
CL\_INS\_382
CL\_INS\_382
CL\_INS\_382
CL\_INS\_146
CL\_INS\_146
CL\_INS\_146
CL\_INS\_146
CL\_INS\_382
CL\_INS\_382
CL\_INS\_382
CL\_INS\_382
CL\_INS\_382
CL\_INS\_382
CL\_INS\_85
CL\_INS\_85
CL\_INS\_85
CL\_INS\_382
CL\_INS\_382
CL\_INS\_382
CL\_INS\_382
CL\_INS\_382
CL\_INS\_382
CL\_INS\_382
CL\_INS\_382
CL\_INS\_382
CL\_INS\_382
CL\_INS\_382
CL\_INS\_382
CL\_INS\_382
CL\_INS\_382
CL\_INS\_382
CL\_INS\_382
CL\_INS\_382
CL\_INS\_382
CL\_INS\_382
CL\_INS\_382
CL\_INS\_382
CL\_INS\_53
CL\_INS\_382
CL\_INS\_382
CL\_INS\_382
CL\_INS\_382
CL\_INS\_382
CL\_INS\_382
CL\_INS\_382
CL\_INS\_382
CL\_INS\_85
CL\_INS\_85
CL\_INS\_382
CL\_INS\_382
CL\_INS\_85
CL\_INS\_382
CL\_INS\_382
CL\_INS\_382
CL\_INS\_382
CL\_INS\_382
CL\_INS\_382
CL\_INS\_382
CL\_INS\_382
CL\_INS\_382
CL\_INS\_204
CL\_INS\_382
CL\_INS\_382
CL\_INS\_382
CL\_INS\_20
CL\_INS\_382
CL\_INS\_382
CL\_INS\_382
CL\_INS\_382
CL\_INS\_20
CL\_INS\_86
CL\_INS\_86
CL\_INS\_136
CL\_INS\_136
CL\_INS\_136
CL\_INS\_136
CL\_INS\_136
CL\_INS\_136
CL\_INS\_136
CL\_INS\_85
CL\_INS\_136
CL\_INS\_87
CL\_INS\_87
CL\_INS\_87
CL\_INS\_136
CL\_INS\_136
CL\_INS\_136
CL\_INS\_136
CL\_INS\_136
CL\_INS\_136
CL\_INS\_136
CL\_INS\_60
CL\_INS\_99
CL\_INS\_136
CL\_INS\_382
CL\_INS\_382
CL\_INS\_382
CL\_INS\_99
CL\_INS\_99
CL\_INS\_382
CL\_INS\_382
CL\_INS\_382
CL\_INS\_382
CL\_INS\_382
CL\_INS\_382
CL\_INS\_85
CL\_INS\_85
CL\_INS\_85
CL\_INS\_85
CL\_INS\_382
CL\_INS\_86
CL\_INS\_204
CL\_INS\_86
CL\_INS\_85
CL\_INS\_86
CL\_INS\_86
CL\_INS\_86
CL\_INS\_86
CL\_INS\_86
CL\_INS\_86
CL\_INS\_86
CL\_INS\_86
CL\_INS\_86
CL\_INS\_86
CL\_INS\_86
CL\_INS\_86
CL\_INS\_204
CL\_INS\_204
CL\_INS\_204
CL\_INS\_204
CL\_INS\_204
CL\_INS\_204
CL\_INS\_204
CL\_INS\_204
CL\_INS\_204
CL\_INS\_204
CL\_INS\_85
CL\_INS\_85
CL\_INS\_85
CL\_INS\_204
CL\_INS\_204
CL\_INS\_204
CL\_INS\_204
CL\_INS\_86
CL\_INS\_204
CL\_INS\_86
CL\_INS\_204
CL\_INS\_204
CL\_INS\_204
CL\_INS\_160
CL\_INS\_204
CL\_INS\_204
CL\_INS\_204
CL\_INS\_85
CL\_INS\_85
CL\_INS\_85
CL\_INS\_149
CL\_INS\_204
CL\_INS\_204
CL\_INS\_204
CL\_INS\_86
CL\_INS\_204
CL\_INS\_207
CL\_INS\_204
CL\_INS\_86
CL\_INS\_86
CL\_INS\_85
CL\_INS\_204
CL\_INS\_204
CL\_INS\_204
CL\_INS\_204
CL\_INS\_86
CL\_INS\_86
CL\_INS\_86
CL\_INS\_382
CL\_INS\_382
CL\_INS\_382
CL\_INS\_382
CL\_INS\_382
CL\_INS\_382
CL\_INS\_382
CL\_INS\_382
CL\_INS\_85
CL\_INS\_384
CL\_INS\_382
CL\_INS\_382
CL\_INS\_382
CL\_INS\_382
CL\_INS\_382
CL\_INS\_382
CL\_INS\_85
CL\_INS\_382
CL\_INS\_382
CL\_INS\_382
CL\_INS\_382
CL\_INS\_382
CL\_INS\_382
CL\_INS\_382
CL\_INS\_382
CL\_INS\_382
CL\_INS\_99
CL\_INS\_382
CL\_INS\_382
CL\_INS\_85
CL\_INS\_85
CL\_INS\_382
CL\_INS\_382
CL\_INS\_382
CL\_INS\_382
CL\_INS\_382
CL\_INS\_85
CL\_INS\_382
CL\_INS\_382
CL\_INS\_382
CL\_INS\_382
CL\_INS\_382
CL\_INS\_382
CL\_INS\_382
CL\_INS\_382
CL\_INS\_382
CL\_INS\_382
CL\_INS\_382
CL\_INS\_382
CL\_INS\_382
CL\_INS\_382
CL\_INS\_382
CL\_INS\_382
CL\_INS\_382
CL\_INS\_382
CL\_INS\_382
CL\_INS\_382
CL\_INS\_382
CL\_INS\_382
CL\_INS\_382
CL\_INS\_382
CL\_INS\_382
CL\_INS\_382
CL\_INS\_382
CL\_INS\_382
CL\_INS\_382
CL\_INS\_382
CL\_INS\_382
CL\_INS\_382
CL\_INS\_382
CL\_INS\_382
CL\_INS\_382
CL\_INS\_382
CL\_INS\_382
CL\_INS\_382
CL\_INS\_382
CL\_INS\_382
CL\_INS\_85
CL\_INS\_99
CL\_INS\_85
CL\_INS\_382
CL\_INS\_382
CL\_INS\_382
CL\_INS\_382
CL\_INS\_85
CL\_INS\_85
CL\_INS\_85
CL\_INS\_382
CL\_INS\_382
CL\_INS\_384
CL\_INS\_382
CL\_INS\_382
CL\_INS\_382
CL\_INS\_382
CL\_INS\_382
CL\_INS\_382
CL\_INS\_86
CL\_INS\_86
CL\_INS\_86
CL\_INS\_86
CL\_INS\_136
CL\_INS\_385
CL\_INS\_87
CL\_INS\_385
CL\_INS\_87
CL\_INS\_87
CL\_INS\_87
CL\_INS\_86
CL\_INS\_155
CL\_INS\_85
CL\_INS\_85
CL\_INS\_20
CL\_INS\_155
CL\_INS\_155
CL\_INS\_155
CL\_INS\_155
CL\_INS\_85
CL\_INS\_155
CL\_INS\_155
CL\_INS\_155
CL\_INS\_155
CL\_INS\_155
CL\_INS\_155
CL\_INS\_155
CL\_INS\_155
CL\_INS\_155
CL\_INS\_155
CL\_INS\_155
CL\_INS\_155
CL\_INS\_155
CL\_INS\_155
CL\_INS\_155
CL\_INS\_155
CL\_INS\_155
CL\_INS\_155
CL\_INS\_155
CL\_INS\_155
CL\_INS\_86
CL\_INS\_155
CL\_INS\_86
CL\_INS\_155
CL\_INS\_86
CL\_INS\_155
CL\_INS\_155
CL\_INS\_155
CL\_INS\_155
CL\_INS\_155
CL\_INS\_155
CL\_INS\_155
CL\_INS\_86
CL\_INS\_86
CL\_INS\_86
CL\_INS\_86
CL\_INS\_155
CL\_INS\_155
CL\_INS\_155
CL\_INS\_155
CL\_INS\_86
CL\_INS\_155
CL\_INS\_155
CL\_INS\_155
CL\_INS\_155
CL\_INS\_204
CL\_INS\_204
CL\_INS\_204
CL\_INS\_86
CL\_INS\_86
CL\_INS\_86
CL\_INS\_86
CL\_INS\_86
CL\_INS\_155
CL\_INS\_86
CL\_INS\_86
CL\_INS\_86
CL\_INS\_237
CL\_INS\_86
CL\_INS\_86
CL\_INS\_237
CL\_INS\_237
CL\_INS\_237
CL\_INS\_384
CL\_INS\_86
CL\_INS\_382
CL\_INS\_85
CL\_INS\_85
CL\_INS\_55
CL\_INS\_55
CL\_INS\_55
CL\_INS\_55
CL\_INS\_55
CL\_INS\_55
CL\_INS\_55
CL\_INS\_55
CL\_INS\_55
CL\_INS\_85
CL\_INS\_85
CL\_INS\_85
CL\_INS\_85
CL\_INS\_85
CL\_INS\_85
CL\_INS\_85
CL\_INS\_85
CL\_INS\_85
CL\_INS\_85
CL\_INS\_85
CL\_INS\_85
CL\_INS\_85
CL\_INS\_85
CL\_INS\_85
CL\_INS\_85
CL\_INS\_85
CL\_INS\_85
CL\_INS\_85
CL\_INS\_85
CL\_INS\_85
CL\_INS\_85
CL\_INS\_85
CL\_INS\_85
CL\_INS\_85
CL\_INS\_85
CL\_INS\_85
CL\_INS\_85
CL\_INS\_85
CL\_INS\_85
CL\_INS\_85
CL\_INS\_85
CL\_INS\_85
CL\_INS\_85
CL\_INS\_85
CL\_INS\_85
CL\_INS\_85
CL\_INS\_85
CL\_INS\_85
CL\_INS\_85
CL\_INS\_85
CL\_INS\_85
CL\_INS\_85
CL\_INS\_85
CL\_INS\_85
CL\_INS\_85
CL\_INS\_85
CL\_INS\_85
CL\_INS\_85
CL\_INS\_85
CL\_INS\_85
CL\_INS\_85
CL\_INS\_85
CL\_INS\_85
CL\_INS\_85
CL\_INS\_85
CL\_INS\_85
CL\_INS\_85
CL\_INS\_85
CL\_INS\_85
CL\_INS\_85
CL\_INS\_85
CL\_INS\_85
CL\_INS\_85
CL\_INS\_85
CL\_INS\_247
CL\_INS\_247
CL\_INS\_247
CL\_INS\_247
CL\_INS\_382
CL\_INS\_85
CL\_INS\_85
CL\_INS\_85
CL\_INS\_85
CL\_INS\_85
CL\_INS\_85
CL\_INS\_85
CL\_INS\_85
CL\_INS\_85
CL\_INS\_85
CL\_INS\_85
CL\_INS\_85
CL\_INS\_85
CL\_INS\_85
CL\_INS\_85
CL\_INS\_85
CL\_INS\_85
CL\_INS\_85
CL\_INS\_85
CL\_INS\_85
CL\_INS\_85
CL\_INS\_85
CL\_INS\_85
CL\_INS\_85
CL\_INS\_85
CL\_INS\_85
CL\_INS\_85
CL\_INS\_382
CL\_INS\_85
CL\_INS\_85
CL\_INS\_85
CL\_INS\_85
CL\_INS\_85
CL\_INS\_85
CL\_INS\_85
CL\_INS\_85
CL\_INS\_85
CL\_INS\_85
CL\_INS\_85
CL\_INS\_85
CL\_INS\_85
CL\_INS\_85
CL\_INS\_85
CL\_INS\_85
CL\_INS\_85
CL\_INS\_85
CL\_INS\_85
CL\_INS\_382
CL\_INS\_382
CL\_INS\_382
CL\_INS\_382
CL\_INS\_382
CL\_INS\_382
CL\_INS\_382
CL\_INS\_247
CL\_INS\_382
CL\_INS\_382
CL\_INS\_382
CL\_INS\_382
CL\_INS\_382
CL\_INS\_25
CL\_INS\_25
CL\_INS\_382
CL\_INS\_25
CL\_INS\_237
CL\_INS\_237
CL\_INS\_25
CL\_INS\_25
CL\_INS\_25
CL\_INS\_237
CL\_INS\_25
CL\_INS\_25
CL\_INS\_237
CL\_INS\_237
CL\_INS\_85
CL\_INS\_237
CL\_INS\_237
CL\_INS\_237
CL\_INS\_237
CL\_INS\_237
CL\_INS\_237
CL\_INS\_25
CL\_INS\_25
CL\_INS\_237
CL\_INS\_25
CL\_INS\_25
CL\_INS\_25
CL\_INS\_382
CL\_INS\_382
CL\_INS\_237
CL\_INS\_237
CL\_INS\_237
CL\_INS\_237
CL\_INS\_237
CL\_INS\_237
CL\_INS\_237
CL\_INS\_237
CL\_INS\_237
CL\_INS\_25
CL\_INS\_25
CL\_INS\_123
CL\_INS\_237
CL\_INS\_237
CL\_INS\_237
CL\_INS\_237
CL\_INS\_237
CL\_INS\_237
CL\_INS\_237
CL\_INS\_237
CL\_INS\_237
CL\_INS\_237
CL\_INS\_237
CL\_INS\_237
CL\_INS\_237
CL\_INS\_237
CL\_INS\_237
CL\_INS\_382
CL\_INS\_233
CL\_INS\_25
CL\_INS\_237
CL\_INS\_237
CL\_INS\_237
CL\_INS\_237
CL\_INS\_70
CL\_INS\_237
CL\_INS\_85
CL\_INS\_85
CL\_INS\_117
CL\_INS\_237
CL\_INS\_382
CL\_INS\_354
CL\_INS\_354
CL\_INS\_25
CL\_INS\_237
CL\_INS\_25
CL\_INS\_25
CL\_INS\_237
CL\_INS\_247
CL\_INS\_25
CL\_INS\_237
CL\_INS\_25
CL\_INS\_25
CL\_INS\_382
CL\_INS\_237
CL\_INS\_237
CL\_INS\_237
CL\_INS\_237
CL\_INS\_25
CL\_INS\_25
CL\_INS\_25
CL\_INS\_237
CL\_INS\_237
CL\_INS\_237
CL\_INS\_237
CL\_INS\_382
CL\_INS\_382
CL\_INS\_382
CL\_INS\_117
CL\_INS\_247
CL\_INS\_247
CL\_INS\_382
CL\_INS\_382
CL\_INS\_247
CL\_INS\_247
CL\_INS\_247
CL\_INS\_247
CL\_INS\_110
CL\_INS\_382
CL\_INS\_382
CL\_INS\_382
CL\_INS\_368
CL\_INS\_368
CL\_INS\_55
CL\_INS\_55
CL\_INS\_55
CL\_INS\_159
CL\_INS\_159
CL\_INS\_382
CL\_INS\_382
CL\_INS\_382
CL\_INS\_382
CL\_INS\_99
CL\_INS\_382
CL\_INS\_87
Cluster ID


CL\_9523
CL\_30689
CL\_30823
CL\_12009
CL\_8585
CL\_10526
CL\_6782
CL\_1495
CL\_10475
CL\_11286
CL\_10520
CL\_8279
CL\_21898
CL\_4431
CL\_21899
CL\_1324
CL\_4432
CL\_33569
CL\_33568
CL\_4093
CL\_8909
CL\_1085
CL\_1086
CL\_1087
CL\_14522
CL\_14523
CL\_24429
CL\_8843
CL\_33863
CL\_34697
CL\_34698
CL\_32189
CL\_27581
CL\_27580
CL\_27579
CL\_23515
CL\_23516
CL\_17631
CL\_1088
CL\_14372
CL\_1089
CL\_1090
CL\_31664
CL\_33308
CL\_11406
CL\_21161
CL\_24315
CL\_24316
CL\_24317
CL\_24318
CL\_10270
CL\_7544
CL\_14633
CL\_2283
CL\_2282
CL\_2281
CL\_5427
CL\_5426
CL\_1093
CL\_1094
CL\_1091
CL\_1092
CL\_23517
CL\_23518
CL\_23519
CL\_1095
CL\_1096
CL\_33864
CL\_33865
CL\_33866
CL\_33867
CL\_33868
CL\_33869
CL\_33870
CL\_23543
CL\_23544
CL\_4644
CL\_9266
CL\_33871
CL\_11477
CL\_24325
CL\_24326
CL\_24327
CL\_14846
CL\_14845
CL\_24328
CL\_14843
CL\_14842
CL\_26237
CL\_14841
CL\_14840
CL\_24329
CL\_24330
CL\_24331
CL\_24332
CL\_24333
CL\_24334
CL\_24335
CL\_24336
CL\_24337
CL\_24338
CL\_24339
CL\_8850
CL\_8849
CL\_8848
CL\_8847
CL\_24340
CL\_24341
CL\_24342
CL\_24343
CL\_24344
CL\_24345
CL\_33872
CL\_2280
CL\_2279
CL\_2278
CL\_32192
CL\_5422
CL\_5421
CL\_5420
CL\_5419
CL\_4651
CL\_20411
CL\_27578
CL\_5924
CL\_7339
CL\_23520
CL\_17632
CL\_17633
CL\_17634
CL\_11313
CL\_7621
CL\_17690
CL\_7543
CL\_7542
CL\_5425
CL\_10174
CL\_5423
CL\_17635
CL\_24319
CL\_24320
CL\_24321
CL\_24322
CL\_32190
CL\_32191
CL\_17636
CL\_17637
CL\_17638
CL\_17639
CL\_17640
CL\_12766
CL\_16967
CL\_17234
CL\_17235
CL\_7540
CL\_7539
CL\_32193
CL\_32194
CL\_32195
CL\_1512
CL\_17641
CL\_24323
CL\_10165
CL\_10164
CL\_10163
CL\_24324
CL\_17642
CL\_5416
CL\_5415
CL\_24428
CL\_5798
CL\_21883
CL\_5799
CL\_8277
CL\_5362
CL\_11916
CL\_7821
CL\_9955
CL\_13513
CL\_8276
CL\_10336
CL\_34848
CL\_11963
CL\_7033
CL\_9953
CL\_13792
CL\_13793
CL\_13794
CL\_14373
CL\_14374
CL\_5364
CL\_5363
CL\_7032
CL\_22729
CL\_5361
CL\_4400
CL\_508
CL\_10337
CL\_10338
CL\_34849
CL\_4401
CL\_4402
CL\_30410
CL\_5800
CL\_6483
CL\_29104
CL\_7364
CL\_35354
CL\_27931
CL\_23319
CL\_8207
CL\_23123
CL\_21697
CL\_6479
CL\_26920
CL\_23070
CL\_4403
CL\_7362
CL\_21884
CL\_21885
CL\_4404
CL\_4405
CL\_5801
CL\_5802
CL\_13634
CL\_13635
CL\_4406
CL\_4407
CL\_4408
CL\_4409
CL\_10918
CL\_29105
CL\_29106
CL\_21886
CL\_21887
CL\_21888
CL\_25892
CL\_25893
CL\_14524
CL\_14525
CL\_10919
CL\_37595
CL\_37594
CL\_37593
CL\_5360
CL\_7031
CL\_5359
CL\_5358
CL\_25202
CL\_25203
CL\_5357
CL\_31663
CL\_13514
CL\_6793
CL\_20417
CL\_11732
CL\_11733
CL\_5354
CL\_11734
CL\_5352
CL\_5356
CL\_6792
CL\_28277
CL\_31662
CL\_31661
CL\_31660
CL\_4490
CL\_18913
CL\_21900
CL\_21901
CL\_5810
CL\_5811
CL\_5812
CL\_5813
CL\_5594
CL\_534
CL\_4433
CL\_4434
CL\_536
CL\_28082
CL\_28081
CL\_16568
CL\_5814
CL\_5815
CL\_9076
CL\_9077
CL\_9078
CL\_9079
CL\_9080
CL\_6791
CL\_6790
CL\_6789
CL\_6788
CL\_6787
CL\_6786
CL\_5351
CL\_11915
CL\_4410
CL\_509
CL\_26919
CL\_34480
CL\_34481
CL\_23762
CL\_30409
CL\_30408
CL\_4654
CL\_4653
CL\_4652
CL\_13070
CL\_13071
CL\_30407
CL\_30406
CL\_30405
CL\_7538
CL\_13072
CL\_8184
CL\_6784
CL\_25588
CL\_30404
CL\_30403
CL\_30402
CL\_12386
CL\_17733
CL\_17732
CL\_17731
CL\_31975
CL\_4411
CL\_4412
CL\_26918
CL\_26917
CL\_26916
CL\_31976
CL\_31977
CL\_516
CL\_511
CL\_512
CL\_513
CL\_33307
CL\_514
CL\_515
CL\_7820
CL\_7819
CL\_21889
CL\_21890
CL\_517
CL\_518
CL\_21891
CL\_519
CL\_520
CL\_5350
CL\_5803
CL\_5804
CL\_5372
CL\_8206
CL\_5349
CL\_5348
CL\_5347
CL\_5346
CL\_5345
CL\_34135
CL\_5344
CL\_26915
CL\_35811
CL\_5805
CL\_5343
CL\_36065
CL\_5342
CL\_34136
CL\_5341
CL\_6452
CL\_5340
CL\_5806
CL\_20418
CL\_5807
CL\_5808
CL\_5414
CL\_17730
CL\_25894
CL\_25895
CL\_5413
CL\_17747
CL\_15242
CL\_15241
CL\_21698
CL\_17746
CL\_23521
CL\_23522
CL\_1318
CL\_1319
CL\_1320
CL\_4470
CL\_4471
CL\_4472
CL\_4473
CL\_29003
CL\_4474
CL\_4475
CL\_4476
CL\_4477
CL\_4478
CL\_4479
CL\_4480
CL\_4481
CL\_4482
CL\_1321
CL\_1322
CL\_4483
CL\_4485
CL\_6746
CL\_521
CL\_522
CL\_524
CL\_8586
CL\_8588
CL\_13636
CL\_13637
CL\_13638
CL\_525
CL\_12006
CL\_526
CL\_37115
CL\_37114
CL\_37113
CL\_37112
CL\_5809
CL\_10976
CL\_10975
CL\_8180
CL\_33873
CL\_17643
CL\_8178
CL\_14638
CL\_14639
CL\_17644
CL\_17645
CL\_15467
CL\_17646
CL\_11307
CL\_17647
CL\_17648
CL\_17649
CL\_5412
CL\_5411
CL\_1097
CL\_1098
CL\_1099
CL\_5410
CL\_5409
CL\_5408
CL\_5407
CL\_5406
CL\_33874
CL\_33875
CL\_33876
CL\_5405
CL\_5404
CL\_5403
CL\_5402
CL\_17650
CL\_5401
CL\_10273
CL\_5399
CL\_5398
CL\_10274
CL\_6406
CL\_5396
CL\_5395
CL\_5394
CL\_33877
CL\_33878
CL\_33879
CL\_2276
CL\_4604
CL\_5393
CL\_1100
CL\_17651
CL\_1101
CL\_5392
CL\_1102
CL\_17652
CL\_5201
CL\_27577
CL\_6594
CL\_1103
CL\_1104
CL\_2277
CL\_1106
CL\_1107
CL\_7344
CL\_527
CL\_528
CL\_529
CL\_530
CL\_531
CL\_1496
CL\_533
CL\_4533
CL\_28084
CL\_28083
CL\_21892
CL\_21893
CL\_15898
CL\_21894
CL\_21895
CL\_21896
CL\_36066
CL\_16567
CL\_17075
CL\_8216
CL\_10804
CL\_7030
CL\_22579
CL\_7029
CL\_7028
CL\_7818
CL\_13515
CL\_532
CL\_4413
CL\_36067
CL\_36068
CL\_4414
CL\_4415
CL\_13476
CL\_13477
CL\_4532
CL\_14375
CL\_14376
CL\_4529
CL\_4531
CL\_4530
CL\_6461
CL\_34137
CL\_8901
CL\_4416
CL\_4417
CL\_4695
CL\_8647
CL\_8648
CL\_13639
CL\_23647
CL\_13510
CL\_13512
CL\_8113
CL\_8654
CL\_12137
CL\_13640
CL\_13641
CL\_13642
CL\_13643
CL\_17067
CL\_31978
CL\_13644
CL\_13645
CL\_13646
CL\_4418
CL\_4528
CL\_4527
CL\_4526
CL\_4419
CL\_4420
CL\_4629
CL\_4524
CL\_12007
CL\_7367
CL\_11914
CL\_4421
CL\_37111
CL\_15782
CL\_37110
CL\_4525
CL\_4422
CL\_33306
CL\_33305
CL\_14377
CL\_14378
CL\_14379
CL\_8473
CL\_4423
CL\_25204
CL\_4424
CL\_4425
CL\_4426
CL\_4522
CL\_4521
CL\_11913
CL\_7027
CL\_4429
CL\_4430
CL\_4488
CL\_6769
CL\_6768
CL\_4513
CL\_6993
CL\_20419
CL\_8275
CL\_8274
CL\_7025
CL\_10916
CL\_13517
CL\_13518
CL\_10476
CL\_10915
CL\_10914
CL\_10913
CL\_10912
CL\_13516
CL\_10911
CL\_10910
CL\_10909
CL\_10907
CL\_10906
CL\_10905
CL\_11004
CL\_11005
CL\_11006
CL\_11007
CL\_11008
CL\_11009
CL\_11010
CL\_11011
CL\_10894
CL\_10893
CL\_10892
CL\_10891
CL\_10890
CL\_10889
CL\_10888
CL\_10887
CL\_10886
CL\_10885
CL\_10884
CL\_10883
CL\_10882
CL\_11024
CL\_11025
CL\_10879
CL\_10878
CL\_10877
CL\_11029
CL\_11030
CL\_8226
CL\_8227
CL\_8228
CL\_8229
CL\_8230
CL\_8231
CL\_8232
CL\_10870
CL\_10869
CL\_11035
CL\_8483
CL\_10867
CL\_1105
CL\_6520
CL\_17653
CL\_17654
CL\_17655
CL\_17656
CL\_17657
CL\_11037
CL\_7024
CL\_7023
CL\_7022
CL\_6765
CL\_13647
CL\_7021
CL\_5595
CL\_5596
CL\_6764
CL\_30348
CL\_7020
CL\_7019
CL\_23781
CL\_11511
CL\_14209
CL\_18416
CL\_14211
CL\_14210
CL\_18422
CL\_18421
CL\_18420
CL\_18419
CL\_18418
CL\_23865
CL\_23866
CL\_23847
CL\_19573
CL\_19572
CL\_19571
CL\_19570
CL\_19569
CL\_19568
CL\_19567
CL\_19566
CL\_19565
CL\_19564
CL\_19563
CL\_19562
CL\_19561
CL\_19560
CL\_19559
CL\_19558
CL\_19557
CL\_19556
CL\_19555
CL\_19554
CL\_19553
CL\_19552
CL\_19551
CL\_19550
CL\_19549
CL\_19548
CL\_19547
CL\_23848
CL\_23849
CL\_23154
CL\_23153
CL\_19654
CL\_19653
CL\_19652
CL\_23151
CL\_23850
CL\_19649
CL\_19648
CL\_19647
CL\_19646
CL\_19645
CL\_19644
CL\_19639
CL\_23149
CL\_19638
CL\_19637
CL\_19636
CL\_23851
CL\_19634
CL\_23852
CL\_19633
CL\_19632
CL\_19631
CL\_19630
CL\_19629
CL\_19627
CL\_19623
CL\_19622
CL\_19621
CL\_19620
CL\_19619
CL\_23146
CL\_23853
CL\_21983
CL\_21982
CL\_21981
CL\_10807
CL\_19617
CL\_19616
CL\_19615
CL\_19614
CL\_19613
CL\_23854
CL\_23855
CL\_19612
CL\_19611
CL\_19609
CL\_19608
CL\_19607
CL\_19606
CL\_19605
CL\_19604
CL\_19603
CL\_19602
CL\_19601
CL\_19600
CL\_19599
CL\_19598
CL\_19597
CL\_19596
CL\_19595
CL\_19594
CL\_19593
CL\_19592
CL\_16415
CL\_19591
CL\_19590
CL\_19587
CL\_23163
CL\_23164
CL\_19586
CL\_19585
CL\_19584
CL\_19583
CL\_19582
CL\_19581
CL\_19580
CL\_19579
CL\_19578
CL\_19577
CL\_23159
CL\_19576
CL\_19575
CL\_23856
CL\_4253
CL\_4254
CL\_4255
CL\_5066
CL\_5067
CL\_5068
CL\_5069
CL\_5070
CL\_5071
CL\_5072
CL\_5073
CL\_5074
CL\_4467
CL\_20826
CL\_20825
CL\_5077
CL\_20824
CL\_17167
CL\_17168
CL\_23536
CL\_20822
CL\_31629
CL\_20820
CL\_20819
CL\_20818
CL\_10591
CL\_20817
CL\_20816
CL\_20815
CL\_10785
CL\_20949
CL\_20814
CL\_20813
CL\_10782
CL\_20812
CL\_20811
CL\_20810
CL\_20809
CL\_20808
CL\_20807
CL\_5093
CL\_5094
CL\_20806
CL\_20805
CL\_20804
CL\_10770
CL\_10769
CL\_14899
CL\_20803
CL\_13211
CL\_20802
CL\_20801
CL\_20800
CL\_6957
CL\_20799
CL\_20798
CL\_20797
CL\_20796
CL\_20948
CL\_20795
CL\_20947
CL\_20870
CL\_20869
CL\_20868
CL\_20864
CL\_20863
CL\_17620
CL\_17619
CL\_17618
CL\_5013
CL\_4309
CL\_20857
CL\_7674
CL\_7871
CL\_18866
CL\_18867
CL\_7869
CL\_18424
CL\_31628
CL\_31968
CL\_6969
CL\_17764
CL\_10605
CL\_20854
CL\_20853
CL\_20849
CL\_20848
CL\_20842
CL\_27409
CL\_20821
CL\_7007
CL\_20840
CL\_10590
CL\_20837
CL\_20836
CL\_14193
CL\_17162
CL\_17161
CL\_17160
CL\_17159
CL\_20835
CL\_20834
CL\_20833
CL\_7684
CL\_7685
CL\_7686
CL\_10583
CL\_4972
CL\_4974
CL\_5662
CL\_5505
CL\_4233
CL\_4234
CL\_4235
CL\_4236
CL\_5045
CL\_5046
CL\_5047
CL\_5048
CL\_5049
CL\_7692
CL\_7691
CL\_8832
CL\_11816
CL\_11815
CL\_5028
CL\_5029
CL\_14147
CL\_5031
CL\_5032
CL\_19401
CL\_19402
CL\_5050
CL\_5051
CL\_5052
CL\_4251
CL\_1108
