## Supplementary material for "A novel method for integrating genomic and Tn-Seq data to identify common *in vivo* fitness mechanisms across multiple bacterial species": S1 Dataset: CL_INS_87.html

Legend

 Mobile +extrachromosomalelementfunctions
 Hypothetical
 All Fitness Genes
 Other
 Transport +binding proteins
 All VFDB Genes

FULL


WINDOWSVGPNG

Trim RowsRemove SingletonsSave Fasta

CL\_1119


CL\_1119


CL\_1119


CL\_1119


CL\_1119


CL\_1119


CL\_1119


CL\_1119


CL\_1084


CL\_1119


CL\_1115


CL\_1118


CL\_1119


CL\_1119


CL\_1119


CL\_1119


CL\_1119

HighlightSelectShow Genomes


89

CL\_1121


67

CL\_1121


65

CL\_1121


17

CL\_1121


8

CL\_1122


2

CL\_1121


1

CL\_1121


1

CL\_1121


1

CL\_1121


1

CL\_1121


1

CL\_1121


1

CL\_1121


1

CL\_1121


1

CL\_1121


1

CL\_1121


1

CL\_1121


1

CL\_1121

fGI ID


CL\_INS\_87
CL\_INS\_87
CL\_INS\_87
CL\_INS\_87
CL\_INS\_87
CL\_INS\_87
CL\_INS\_87
CL\_INS\_87
CL\_INS\_87
CL\_INS\_87
CL\_INS\_87
CL\_INS\_87
CL\_INS\_87
CL\_INS\_87
CL\_INS\_87
CL\_INS\_87
Cluster ID


CL\_22254
CL\_22965
CL\_15273
CL\_19737
CL\_4450
CL\_1120
CL\_8778
CL\_29788
CL\_4475
CL\_4476
CL\_4477
CL\_4513
CL\_20419
CL\_8275
CL\_8274
CL\_1108
