## Supplementary material for "A novel method for integrating genomic and Tn-Seq data to identify common *in vivo* fitness mechanisms across multiple bacterial species": S1 Dataset: CL_INS_89.html

Legend

 Mobile +extrachromosomalelementfunctions
 Regulatoryfunctions
 Hypothetical
 DNA Metabolism
 Proteinsynthesis/fate
 Other
 All VFDB Genes

FULL


WINDOWSVGPNG

Trim RowsRemove SingletonsSave Fasta

CL\_1135


CL\_1135


Break


CL\_1135


CL\_1135


CL\_1135


CL\_1135

HighlightSelectShow Genomes


259

CL\_1136


2

CL\_1136


1

CL\_1136


1

CL\_1137


1

CL\_1136


1

Break


1

Break

fGI ID


CL\_INS\_89
CL\_INS\_89
CL\_INS\_89
CL\_INS\_89
CL\_INS\_89
CL\_INS\_89
CL\_INS\_89
CL\_INS\_89
CL\_INS\_382
CL\_INS\_382
CL\_INS\_89
CL\_INS\_89
CL\_INS\_89
CL\_INS\_89
CL\_INS\_89
CL\_INS\_89
CL\_INS\_89
CL\_INS\_89
CL\_INS\_89
CL\_INS\_89
CL\_INS\_89
CL\_INS\_89
CL\_INS\_89
CL\_INS\_233
CL\_INS\_233
CL\_INS\_237
CL\_INS\_89
CL\_INS\_89
CL\_INS\_89
CL\_INS\_382
CL\_INS\_295
CL\_INS\_343
CL\_INS\_343
CL\_INS\_343
CL\_INS\_343
CL\_INS\_385
CL\_INS\_343
CL\_INS\_382
CL\_INS\_343
CL\_INS\_343
CL\_INS\_343
CL\_INS\_343
CL\_INS\_89
CL\_INS\_89
CL\_INS\_89
CL\_INS\_233
CL\_INS\_233
CL\_INS\_89
CL\_INS\_233
CL\_INS\_343
CL\_INS\_233
CL\_INS\_233
CL\_INS\_233
CL\_INS\_233
CL\_INS\_233
CL\_INS\_233
CL\_INS\_89
CL\_INS\_89
CL\_INS\_382
CL\_INS\_159
CL\_INS\_382
CL\_INS\_233
CL\_INS\_233
CL\_INS\_233
CL\_INS\_233
CL\_INS\_233
CL\_INS\_382
CL\_INS\_382
CL\_INS\_233
CL\_INS\_233
CL\_INS\_233
CL\_INS\_382
CL\_INS\_70
CL\_INS\_70
CL\_INS\_70
CL\_INS\_233
CL\_INS\_233
CL\_INS\_233
CL\_INS\_233
CL\_INS\_382
CL\_INS\_382
CL\_INS\_89
CL\_INS\_89
CL\_INS\_89
CL\_INS\_89
CL\_INS\_382
CL\_INS\_382
CL\_INS\_233
CL\_INS\_233
CL\_INS\_89
Cluster ID


CL\_20003
CL\_20039
CL\_20038
CL\_4248
CL\_4247
CL\_4246
CL\_4245
CL\_4244
CL\_4243
CL\_4241
CL\_8662
CL\_8663
CL\_8664
CL\_8665
CL\_8666
CL\_8667
CL\_8668
CL\_13007
CL\_13030
CL\_13008
CL\_13009
CL\_13010
CL\_13011
CL\_12675
CL\_12674
CL\_13012
CL\_13013
CL\_13014
CL\_13015
CL\_6262
CL\_13016
CL\_13017
CL\_13018
CL\_13019
CL\_13020
CL\_5522
CL\_13021
CL\_4975
CL\_13022
CL\_13023
CL\_13024
CL\_13025
CL\_13026
CL\_13027
CL\_13028
CL\_12647
CL\_12648
CL\_13029
CL\_12649
CL\_13031
CL\_12650
CL\_12655
CL\_12656
CL\_12657
CL\_12653
CL\_12659
CL\_13032
CL\_13033
CL\_13034
CL\_5539
CL\_5599
CL\_5541
CL\_6935
CL\_5542
CL\_5543
CL\_5544
CL\_4302
CL\_4301
CL\_5545
CL\_12660
CL\_12661
CL\_4299
CL\_8749
CL\_8748
CL\_8747
CL\_12664
CL\_12666
CL\_12667
CL\_12668
CL\_4236
CL\_4235
CL\_13035
CL\_13036
CL\_13037
CL\_13038
CL\_5015
CL\_5014
CL\_12670
CL\_12671
CL\_13039
