## Supplementary material for "A novel method for integrating genomic and Tn-Seq data to identify common *in vivo* fitness mechanisms across multiple bacterial species": S1 Dataset: CL_INS_90.html

Legend

 Hypothetical
 All Fitness Genes
 Transport +binding proteins
 All VFDB Genes

FULL


WINDOWSVGPNG

Trim RowsRemove SingletonsSave Fasta

CL\_1142


CL\_1142


CL\_1142


CL\_1142


CL\_1142


CL\_1142


CL\_1142


CL\_1141


CL\_1140


CL\_1142

HighlightSelectShow Genomes


223

CL\_1143


17

CL\_1143


14

CL\_1143


14

CL\_1143


2

CL\_1143


1

CL\_1143


1

CL\_1143


1

CL\_1143


1

CL\_1143


1

CL\_1143

fGI ID


CL\_INS\_382
CL\_INS\_90
CL\_INS\_90
CL\_INS\_90
CL\_INS\_90
CL\_INS\_90
CL\_INS\_90
CL\_INS\_90
CL\_INS\_90
CL\_INS\_90
CL\_INS\_90
CL\_INS\_90
CL\_INS\_90
CL\_INS\_90
CL\_INS\_90
CL\_INS\_90
CL\_INS\_90
CL\_INS\_90
CL\_INS\_90
CL\_INS\_90
CL\_INS\_90
CL\_INS\_90
CL\_INS\_90
Cluster ID


CL\_14193
CL\_20248
CL\_13523
CL\_5817
CL\_5816
CL\_10509
CL\_36913
CL\_36912
CL\_36911
CL\_36910
CL\_36909
CL\_36908
CL\_36907
CL\_36906
CL\_36905
CL\_36904
CL\_36903
CL\_36902
CL\_36901
CL\_36900
CL\_36899
CL\_36898
CL\_36897
