## Supplementary material for "A novel method for integrating genomic and Tn-Seq data to identify common *in vivo* fitness mechanisms across multiple bacterial species": S1 Dataset: CL_INS_94.html

Legend

 Mobile +extrachromosomalelementfunctions
 Hypothetical
 All Fitness Genes
 Other
 All VFDB Genes

FULL


WINDOWSVGPNG

Trim RowsRemove SingletonsSave Fasta

CL\_1182


CL\_1182


CL\_1182


CL\_1182


CL\_1182


CL\_1182


CL\_1182


CL\_1182

HighlightSelectShow Genomes


205

CL\_1183


32

CL\_1183


25

CL\_1183


4

CL\_1183


3

CL\_1183


1

Break


1

CL\_1183


1

CL\_1183

fGI ID


CL\_INS\_94
CL\_INS\_94
CL\_INS\_141
CL\_INS\_94
CL\_INS\_94
CL\_INS\_94
CL\_INS\_94
CL\_INS\_94
CL\_INS\_94
CL\_INS\_94
CL\_INS\_94
CL\_INS\_94
Cluster ID


CL\_22462
CL\_7812
CL\_4615
CL\_18926
CL\_18925
CL\_9143
CL\_9144
CL\_9145
CL\_9146
CL\_9147
CL\_9148
CL\_9149
