## Supplementary material for "A novel method for integrating genomic and Tn-Seq data to identify common *in vivo* fitness mechanisms across multiple bacterial species": S1 Dataset: CL_INS_95.html

Legend

 Mobile +extrachromosomalelementfunctions
 Hypothetical
 All Fitness Genes
 All VFDB Genes

FULL


WINDOWSVGPNG

Trim RowsRemove SingletonsSave Fasta

CL\_1185


CL\_1185


CL\_1185


CL\_1185

HighlightSelectShow Genomes


268

CL\_1184


1

CL\_1184


1

CL\_1184


1

CL\_1184

fGI ID


CL\_INS\_95
CL\_INS\_95
CL\_INS\_95
CL\_INS\_194
CL\_INS\_194
CL\_INS\_194
CL\_INS\_194
CL\_INS\_95
CL\_INS\_194
CL\_INS\_382
CL\_INS\_95
CL\_INS\_95
CL\_INS\_95
CL\_INS\_95
CL\_INS\_95
CL\_INS\_95
CL\_INS\_95
CL\_INS\_95
CL\_INS\_95
CL\_INS\_95
CL\_INS\_95
CL\_INS\_95
CL\_INS\_95
Cluster ID


CL\_7499
CL\_7500
CL\_7501
CL\_7502
CL\_6392
CL\_6391
CL\_6389
CL\_7503
CL\_28328
CL\_7504
CL\_37286
CL\_37285
CL\_37284
CL\_37283
CL\_37282
CL\_37281
CL\_37280
CL\_37279
CL\_37278
CL\_37277
CL\_37276
CL\_7505
CL\_7506
