## Supplementary material for "A novel method for integrating genomic and Tn-Seq data to identify common *in vivo* fitness mechanisms across multiple bacterial species": S1 Dataset: CL_INS_97.html

Legend

 Mobile +extrachromosomalelementfunctions
 Regulatoryfunctions
 Hypothetical
 DNA Metabolism
 All Fitness Genes
 Proteinsynthesis/fate
 Other
 Transport +binding proteins
 All VFDB Genes

FULL


WINDOWSVGPNG

Trim RowsRemove SingletonsSave Fasta

CL\_1201


CL\_1200


CL\_1201


CL\_1201


CL\_1201


CL\_1200


CL\_1201


CL\_1201


CL\_1201


CL\_1200


CL\_1201


CL\_1201


CL\_1201


CL\_1201

HighlightSelectShow Genomes


262

CL\_1202


2

CL\_1202


1

CL\_1202


1

CL\_1202


1

CL\_1202


1

CL\_1202


1

CL\_1202


1

CL\_1202


1

CL\_1202


1

CL\_1202


1

CL\_1202


1

CL\_1202


1

CL\_1202


1

CL\_1202

fGI ID


CL\_INS\_97
CL\_INS\_97
CL\_INS\_97
CL\_INS\_97
CL\_INS\_97
CL\_INS\_97
CL\_INS\_97
CL\_INS\_86
CL\_INS\_97
CL\_INS\_97
CL\_INS\_97
CL\_INS\_97
CL\_INS\_97
CL\_INS\_97
CL\_INS\_20
CL\_INS\_20
CL\_INS\_20
CL\_INS\_20
CL\_INS\_97
CL\_INS\_97
CL\_INS\_97
CL\_INS\_97
CL\_INS\_97
CL\_INS\_20
CL\_INS\_20
CL\_INS\_20
CL\_INS\_20
CL\_INS\_97
CL\_INS\_97
CL\_INS\_97
CL\_INS\_20
CL\_INS\_20
CL\_INS\_20
CL\_INS\_20
CL\_INS\_97
CL\_INS\_97
CL\_INS\_97
CL\_INS\_20
CL\_INS\_20
CL\_INS\_97
CL\_INS\_204
CL\_INS\_204
CL\_INS\_204
CL\_INS\_204
CL\_INS\_204
CL\_INS\_204
CL\_INS\_204
CL\_INS\_204
CL\_INS\_204
CL\_INS\_204
CL\_INS\_204
CL\_INS\_204
CL\_INS\_204
CL\_INS\_204
CL\_INS\_204
CL\_INS\_204
CL\_INS\_204
CL\_INS\_204
CL\_INS\_204
CL\_INS\_204
CL\_INS\_20
CL\_INS\_204
CL\_INS\_204
CL\_INS\_97
CL\_INS\_97
CL\_INS\_97
CL\_INS\_97
CL\_INS\_204
CL\_INS\_204
CL\_INS\_204
CL\_INS\_204
CL\_INS\_204
CL\_INS\_204
CL\_INS\_97
CL\_INS\_97
CL\_INS\_146
CL\_INS\_97
CL\_INS\_97
CL\_INS\_97
CL\_INS\_97
CL\_INS\_97
CL\_INS\_97
CL\_INS\_382
CL\_INS\_97
CL\_INS\_97
CL\_INS\_97
CL\_INS\_97
CL\_INS\_97
CL\_INS\_97
CL\_INS\_97
CL\_INS\_97
CL\_INS\_97
CL\_INS\_97
CL\_INS\_97
CL\_INS\_382
CL\_INS\_382
CL\_INS\_97
CL\_INS\_382
CL\_INS\_97
CL\_INS\_97
CL\_INS\_97
CL\_INS\_97
CL\_INS\_382
CL\_INS\_97
CL\_INS\_97
CL\_INS\_97
CL\_INS\_97
CL\_INS\_97
CL\_INS\_97
CL\_INS\_97
CL\_INS\_97
CL\_INS\_97
CL\_INS\_97
CL\_INS\_97
CL\_INS\_97
CL\_INS\_97
CL\_INS\_97
CL\_INS\_97
CL\_INS\_97
CL\_INS\_20
CL\_INS\_20
CL\_INS\_97
CL\_INS\_97
CL\_INS\_97
CL\_INS\_97
CL\_INS\_97
CL\_INS\_97
CL\_INS\_97
CL\_INS\_382
CL\_INS\_97
CL\_INS\_97
CL\_INS\_97
CL\_INS\_97
CL\_INS\_97
CL\_INS\_97
CL\_INS\_97
CL\_INS\_97
CL\_INS\_97
CL\_INS\_97
CL\_INS\_97
CL\_INS\_97
CL\_INS\_97
CL\_INS\_97
CL\_INS\_97
CL\_INS\_97
CL\_INS\_97
CL\_INS\_97
CL\_INS\_97
CL\_INS\_97
CL\_INS\_97
CL\_INS\_97
CL\_INS\_97
CL\_INS\_97
CL\_INS\_97
CL\_INS\_97
CL\_INS\_97
CL\_INS\_97
CL\_INS\_97
CL\_INS\_97
CL\_INS\_97
CL\_INS\_97
CL\_INS\_97
CL\_INS\_97
CL\_INS\_146
CL\_INS\_97
CL\_INS\_146
CL\_INS\_146
CL\_INS\_146
CL\_INS\_146
CL\_INS\_146
CL\_INS\_146
CL\_INS\_146
CL\_INS\_146
CL\_INS\_146
CL\_INS\_146
CL\_INS\_146
CL\_INS\_146
CL\_INS\_146
CL\_INS\_146
CL\_INS\_146
CL\_INS\_146
CL\_INS\_146
CL\_INS\_146
CL\_INS\_146
CL\_INS\_146
CL\_INS\_146
CL\_INS\_146
CL\_INS\_204
CL\_INS\_155
CL\_INS\_86
CL\_INS\_149
CL\_INS\_382
CL\_INS\_382
CL\_INS\_97
CL\_INS\_97
CL\_INS\_97
CL\_INS\_97
CL\_INS\_204
CL\_INS\_97
CL\_INS\_97
CL\_INS\_97
CL\_INS\_97
CL\_INS\_97
CL\_INS\_97
CL\_INS\_97
CL\_INS\_97
CL\_INS\_382
CL\_INS\_382
CL\_INS\_382
CL\_INS\_382
CL\_INS\_382
CL\_INS\_382
CL\_INS\_382
CL\_INS\_382
CL\_INS\_382
CL\_INS\_382
CL\_INS\_382
CL\_INS\_382
CL\_INS\_382
CL\_INS\_382
CL\_INS\_382
CL\_INS\_99
CL\_INS\_97
CL\_INS\_97
CL\_INS\_97
CL\_INS\_97
CL\_INS\_97
CL\_INS\_97
CL\_INS\_382
CL\_INS\_237
CL\_INS\_97
CL\_INS\_97
CL\_INS\_382
CL\_INS\_97
CL\_INS\_97
CL\_INS\_97
CL\_INS\_97
CL\_INS\_97
CL\_INS\_97
CL\_INS\_97
CL\_INS\_97
CL\_INS\_97
CL\_INS\_97
CL\_INS\_97
CL\_INS\_97
CL\_INS\_97
CL\_INS\_97
CL\_INS\_97
CL\_INS\_97
CL\_INS\_97
CL\_INS\_97
CL\_INS\_97
CL\_INS\_97
CL\_INS\_97
CL\_INS\_97
CL\_INS\_97
CL\_INS\_97
CL\_INS\_97
CL\_INS\_97
CL\_INS\_97
CL\_INS\_97
CL\_INS\_97
CL\_INS\_97
CL\_INS\_97
CL\_INS\_97
CL\_INS\_97
CL\_INS\_97
CL\_INS\_97
CL\_INS\_136
CL\_INS\_87
CL\_INS\_97
CL\_INS\_97
CL\_INS\_87
CL\_INS\_87
CL\_INS\_136
CL\_INS\_97
CL\_INS\_97
CL\_INS\_97
CL\_INS\_97
CL\_INS\_97
CL\_INS\_97
CL\_INS\_97
CL\_INS\_97
CL\_INS\_97
CL\_INS\_97
CL\_INS\_97
CL\_INS\_97
CL\_INS\_97
CL\_INS\_97
CL\_INS\_97
CL\_INS\_97
CL\_INS\_97
CL\_INS\_97
CL\_INS\_99
CL\_INS\_207
CL\_INS\_97
CL\_INS\_97
CL\_INS\_97
CL\_INS\_207
CL\_INS\_207
CL\_INS\_97
CL\_INS\_97
CL\_INS\_106
CL\_INS\_106
CL\_INS\_106
CL\_INS\_146
CL\_INS\_97
CL\_INS\_146
CL\_INS\_60
Cluster ID


CL\_23704
CL\_22960
CL\_12487
CL\_12486
CL\_12485
CL\_26324
CL\_26325
CL\_17653
CL\_26326
CL\_6487
CL\_8897
CL\_6486
CL\_13953
CL\_13952
CL\_13951
CL\_13950
CL\_13949
CL\_13948
CL\_13947
CL\_13946
CL\_13945
CL\_13944
CL\_13943
CL\_13942
CL\_13941
CL\_13940
CL\_13939
CL\_13938
CL\_13937
CL\_13936
CL\_13935
CL\_13934
CL\_13933
CL\_13932
CL\_13931
CL\_13930
CL\_13929
CL\_13928
CL\_13927
CL\_36515
CL\_5921
CL\_5920
CL\_5919
CL\_5918
CL\_5917
CL\_5916
CL\_5915
CL\_5914
CL\_5913
CL\_5912
CL\_5911
CL\_5910
CL\_5909
CL\_5908
CL\_7386
CL\_5907
CL\_5906
CL\_5905
CL\_5904
CL\_5903
CL\_13926
CL\_5900
CL\_5899
CL\_36516
CL\_36517
CL\_36518
CL\_36519
CL\_5898
CL\_5897
CL\_9292
CL\_1104
CL\_1103
CL\_2277
CL\_8896
CL\_8895
CL\_8894
CL\_8893
CL\_8892
CL\_6485
CL\_6484
CL\_10090
CL\_10091
CL\_6483
CL\_36507
CL\_36508
CL\_6482
CL\_6481
CL\_6480
CL\_34101
CL\_34100
CL\_23733
CL\_23732
CL\_23731
CL\_19874
CL\_6479
CL\_4407
CL\_19873
CL\_4408
CL\_6478
CL\_6477
CL\_6476
CL\_19872
CL\_10918
CL\_36509
CL\_36510
CL\_19871
CL\_19870
CL\_10092
CL\_8891
CL\_8890
CL\_8889
CL\_10093
CL\_10094
CL\_10095
CL\_10096
CL\_10097
CL\_10098
CL\_8888
CL\_8887
CL\_8324
CL\_8323
CL\_8886
CL\_8885
CL\_8884
CL\_8883
CL\_8882
CL\_8881
CL\_8880
CL\_517
CL\_10099
CL\_10100
CL\_10101
CL\_10102
CL\_10103
CL\_34099
CL\_34098
CL\_6475
CL\_6474
CL\_6473
CL\_34097
CL\_34096
CL\_10104
CL\_10105
CL\_19869
CL\_19868
CL\_19867
CL\_19866
CL\_19865
CL\_19864
CL\_6472
CL\_6471
CL\_6470
CL\_34095
CL\_34094
CL\_34093
CL\_34092
CL\_21297
CL\_21298
CL\_21299
CL\_21300
CL\_21301
CL\_21302
CL\_21303
CL\_14058
CL\_34091
CL\_14057
CL\_14056
CL\_14055
CL\_14054
CL\_14053
CL\_14052
CL\_14051
CL\_21304
CL\_21305
CL\_21306
CL\_21307
CL\_21308
CL\_21309
CL\_21310
CL\_21311
CL\_21312
CL\_21313
CL\_14050
CL\_14049
CL\_14048
CL\_14047
CL\_14046
CL\_10867
CL\_8483
CL\_5201
CL\_5200
CL\_519
CL\_520
CL\_36511
CL\_36512
CL\_36513
CL\_36514
CL\_9223
CL\_6469
CL\_6468
CL\_6467
CL\_6466
CL\_6465
CL\_6464
CL\_6463
CL\_6462
CL\_532
CL\_4413
CL\_4414
CL\_4532
CL\_4531
CL\_4530
CL\_6461
CL\_4528
CL\_4527
CL\_4526
CL\_4629
CL\_4423
CL\_4523
CL\_4522
CL\_4521
CL\_4520
CL\_19863
CL\_10106
CL\_8879
CL\_8878
CL\_8877
CL\_8876
CL\_6661
CL\_8319
CL\_8875
CL\_8874
CL\_6664
CL\_8873
CL\_8872
CL\_8871
CL\_8870
CL\_8869
CL\_8868
CL\_8867
CL\_8866
CL\_8865
CL\_8864
CL\_8863
CL\_8862
CL\_8861
CL\_8860
CL\_8859
CL\_8858
CL\_8857
CL\_8856
CL\_8855
CL\_8854
CL\_8853
CL\_8852
CL\_8851
CL\_8850
CL\_8849
CL\_8848
CL\_8847
CL\_8846
CL\_8845
CL\_8844
CL\_10107
CL\_10108
CL\_10109
CL\_10110
CL\_10111
CL\_4474
CL\_4475
CL\_10112
CL\_10113
CL\_4476
CL\_4477
CL\_7498
CL\_10114
CL\_10115
CL\_10116
CL\_10117
CL\_10118
CL\_10119
CL\_10120
CL\_10121
CL\_10122
CL\_10123
CL\_19862
CL\_19861
CL\_10124
CL\_10125
CL\_10126
CL\_10127
CL\_10128
CL\_10129
CL\_4485
CL\_6460
CL\_10130
CL\_10131
CL\_10132
CL\_6459
CL\_6458
CL\_10133
CL\_10134
CL\_10135
CL\_6457
CL\_6456
CL\_6455
CL\_10136
CL\_9768
CL\_6454
