## Supplementary material for "A novel method for integrating genomic and Tn-Seq data to identify common *in vivo* fitness mechanisms across multiple bacterial species": S1 Dataset: CL_INS_98.html

Legend

 Mobile +extrachromosomalelementfunctions
 Hypothetical
 DNA Metabolism
 All EssentialGenes
 Other
 All VFDB Genes

FULL


WINDOWSVGPNG

Trim RowsRemove SingletonsSave Fasta

CL\_1202


CL\_1202


CL\_1202


CL\_1202


CL\_1202


CL\_1202


CL\_1202


CL\_1202


CL\_1202


CL\_1202


CL\_1202

HighlightSelectShow Genomes


263

CL\_1203


1

CL\_1203


1

CL\_1203


1

CL\_1203


1

CL\_1203


1

CL\_1203


1

CL\_1203


1

CL\_1203


1

CL\_1203


1

CL\_1203


1

CL\_1203

fGI ID


CL\_INS\_98
CL\_INS\_98
CL\_INS\_146
CL\_INS\_98
CL\_INS\_98
CL\_INS\_98
CL\_INS\_79
CL\_INS\_146
CL\_INS\_146
CL\_INS\_146
CL\_INS\_146
CL\_INS\_79
CL\_INS\_79
CL\_INS\_79
CL\_INS\_98
CL\_INS\_79
CL\_INS\_98
CL\_INS\_98
CL\_INS\_146
CL\_INS\_146
CL\_INS\_98
CL\_INS\_98
CL\_INS\_98
CL\_INS\_98
CL\_INS\_146
CL\_INS\_146
CL\_INS\_146
CL\_INS\_146
CL\_INS\_146
CL\_INS\_146
CL\_INS\_146
CL\_INS\_98
CL\_INS\_98
CL\_INS\_98
CL\_INS\_98
Cluster ID


CL\_17729
CL\_30874
CL\_5258
CL\_31658
CL\_30873
CL\_8163
CL\_5337
CL\_7608
CL\_7609
CL\_10349
CL\_10348
CL\_5336
CL\_5335
CL\_5334
CL\_5333
CL\_5332
CL\_26914
CL\_26913
CL\_5331
CL\_5330
CL\_5329
CL\_5328
CL\_5327
CL\_5326
CL\_10347
CL\_5266
CL\_5267
CL\_5268
CL\_5269
CL\_10345
CL\_10346
CL\_14249
CL\_14250
CL\_14251
CL\_5325
