## Supplementary material for "A novel method for integrating genomic and Tn-Seq data to identify common *in vivo* fitness mechanisms across multiple bacterial species": S1 Dataset: CL_INS_100.html

Legend

 Mobile +extrachromosomalelementfunctions
 Regulatoryfunctions
 Hypothetical
 DNA Metabolism
 AntibioticResistance
 Other
 All VFDB Genes

FULL


WINDOWSVGPNG

Trim RowsRemove SingletonsSave Fasta

CL\_1211


CL\_1211


CL\_1211


CL\_1211


CL\_1211


CL\_1211


CL\_1211


CL\_1211


CL\_1211


CL\_1211


CL\_1211


CL\_1211


CL\_1211


CL\_1211


CL\_1211


CL\_1211


CL\_1211


CL\_1211


CL\_1211


CL\_1211


CL\_1211


CL\_1211


CL\_1211


CL\_1211


CL\_1211

HighlightSelectShow Genomes


230

CL\_1210


3

CL\_1207


1

CL\_1207


1

CL\_1207


1

CL\_1207


1

CL\_1207


1

CL\_1207


1

CL\_1207


1

CL\_1207


1

CL\_1207


1

CL\_1207


1

CL\_1207


1

CL\_1207


1

CL\_4486


1

CL\_1207


1

CL\_1207


1

CL\_1207


1

CL\_1207


1

CL\_1207


1

CL\_1207


1

CL\_1207


1

CL\_1208


1

CL\_1207


1

CL\_1207


1

CL\_1207

fGI ID


CL\_INS\_99
CL\_INS\_100
CL\_INS\_99
CL\_INS\_100
CL\_INS\_99
CL\_INS\_99
CL\_INS\_99
CL\_INS\_99
CL\_INS\_99
CL\_INS\_87
CL\_INS\_99
CL\_INS\_99
CL\_INS\_99
CL\_INS\_10
CL\_INS\_99
CL\_INS\_99
CL\_INS\_99
CL\_INS\_99
CL\_INS\_99
CL\_INS\_99
CL\_INS\_99
CL\_INS\_99
CL\_INS\_382
CL\_INS\_99
CL\_INS\_99
CL\_INS\_99
CL\_INS\_99
CL\_INS\_99
CL\_INS\_99
CL\_INS\_382
CL\_INS\_99
CL\_INS\_99
CL\_INS\_99
CL\_INS\_99
CL\_INS\_86
CL\_INS\_99
CL\_INS\_86
CL\_INS\_382
CL\_INS\_99
CL\_INS\_99
CL\_INS\_99
CL\_INS\_99
CL\_INS\_382
CL\_INS\_382
CL\_INS\_382
CL\_INS\_99
CL\_INS\_99
CL\_INS\_99
CL\_INS\_99
CL\_INS\_275
CL\_INS\_99
CL\_INS\_382
CL\_INS\_382
CL\_INS\_382
CL\_INS\_99
CL\_INS\_382
CL\_INS\_382
CL\_INS\_382
CL\_INS\_382
CL\_INS\_382
CL\_INS\_99
CL\_INS\_382
CL\_INS\_382
CL\_INS\_382
CL\_INS\_382
CL\_INS\_382
CL\_INS\_382
CL\_INS\_382
CL\_INS\_382
CL\_INS\_99
CL\_INS\_99
CL\_INS\_382
CL\_INS\_382
CL\_INS\_99
CL\_INS\_382
CL\_INS\_99
CL\_INS\_382
CL\_INS\_382
CL\_INS\_382
CL\_INS\_382
CL\_INS\_382
CL\_INS\_99
CL\_INS\_382
CL\_INS\_382
CL\_INS\_382
CL\_INS\_382
CL\_INS\_382
CL\_INS\_382
CL\_INS\_382
CL\_INS\_382
CL\_INS\_382
CL\_INS\_382
CL\_INS\_382
CL\_INS\_382
CL\_INS\_382
CL\_INS\_382
CL\_INS\_382
CL\_INS\_382
CL\_INS\_382
CL\_INS\_382
CL\_INS\_382
CL\_INS\_382
CL\_INS\_99
CL\_INS\_382
CL\_INS\_382
CL\_INS\_382
CL\_INS\_86
CL\_INS\_382
CL\_INS\_99
CL\_INS\_99
CL\_INS\_99
CL\_INS\_382
CL\_INS\_99
CL\_INS\_382
CL\_INS\_382
CL\_INS\_382
CL\_INS\_382
CL\_INS\_382
CL\_INS\_382
CL\_INS\_382
CL\_INS\_382
CL\_INS\_382
CL\_INS\_382
CL\_INS\_382
CL\_INS\_382
CL\_INS\_382
CL\_INS\_382
CL\_INS\_382
CL\_INS\_382
CL\_INS\_382
CL\_INS\_382
CL\_INS\_382
CL\_INS\_382
CL\_INS\_382
CL\_INS\_382
CL\_INS\_99
CL\_INS\_382
CL\_INS\_382
CL\_INS\_382
CL\_INS\_382
CL\_INS\_382
CL\_INS\_382
CL\_INS\_382
CL\_INS\_382
CL\_INS\_86
CL\_INS\_382
CL\_INS\_86
CL\_INS\_382
CL\_INS\_99
CL\_INS\_99
CL\_INS\_86
CL\_INS\_382
CL\_INS\_382
CL\_INS\_382
CL\_INS\_382
CL\_INS\_382
CL\_INS\_382
Cluster ID


CL\_30871
CL\_35643
CL\_35644
CL\_9169
CL\_9168
CL\_10947
CL\_10521
CL\_6767
CL\_534
CL\_4513
CL\_4618
CL\_7109
CL\_1326
CL\_4764
CL\_35645
CL\_10944
CL\_10945
CL\_10946
CL\_10943
CL\_10942
CL\_9167
CL\_9166
CL\_9165
CL\_9164
CL\_9163
CL\_9162
CL\_9161
CL\_9160
CL\_9159
CL\_9158
CL\_13529
CL\_13528
CL\_13527
CL\_8586
CL\_4515
CL\_6747
CL\_8585
CL\_4432
CL\_4431
CL\_12701
CL\_4620
CL\_35633
CL\_8712
CL\_4518
CL\_12360
CL\_4485
CL\_4520
CL\_5692
CL\_5693
CL\_11196
CL\_13559
CL\_8169
CL\_8170
CL\_8171
CL\_8711
CL\_8710
CL\_8709
CL\_8708
CL\_8707
CL\_8706
CL\_9157
CL\_8705
CL\_8704
CL\_8703
CL\_8702
CL\_8701
CL\_8700
CL\_8699
CL\_8698
CL\_13560
CL\_13561
CL\_8697
CL\_8696
CL\_8695
CL\_7537
CL\_16851
CL\_7117
CL\_7118
CL\_7119
CL\_7120
CL\_8694
CL\_9156
CL\_4543
CL\_4544
CL\_8693
CL\_10941
CL\_10940
CL\_8185
CL\_12996
CL\_12760
CL\_12762
CL\_13562
CL\_7121
CL\_12698
CL\_4646
CL\_4647
CL\_4648
CL\_8690
CL\_8689
CL\_4545
CL\_4546
CL\_8187
CL\_9000
CL\_8186
CL\_8688
CL\_8687
CL\_8686
CL\_4547
CL\_9155
CL\_9154
CL\_9153
CL\_4651
CL\_9152
CL\_9151
CL\_9150
CL\_10939
CL\_4459
CL\_10938
CL\_10937
CL\_10936
CL\_8997
CL\_8996
CL\_8995
CL\_4560
CL\_8195
CL\_1523
CL\_1524
CL\_1525
CL\_5274
CL\_4548
CL\_8684
CL\_8683
CL\_4550
CL\_8682
CL\_8681
CL\_33567
CL\_8680
CL\_4552
CL\_8679
CL\_8678
CL\_4556
CL\_8677
CL\_8676
CL\_8675
CL\_8674
CL\_8673
CL\_8672
CL\_4562
CL\_4563
CL\_4564
CL\_10935
CL\_8196
CL\_8197
CL\_1526
CL\_7126
CL\_8199
CL\_8669
