## Supplementary material for "A novel method for integrating genomic and Tn-Seq data to identify common *in vivo* fitness mechanisms across multiple bacterial species": S1 Dataset: CL_INS_107.html

Legend

 Hypothetical
 All Fitness Genes
 Other
 Transport +binding proteins

FULL


WINDOWSVGPNG

Trim RowsRemove SingletonsSave Fasta

CL\_1298


CL\_1298


CL\_1298


CL\_1296

HighlightSelectShow Genomes


209

CL\_1299


35

CL\_1299


26

CL\_1299


1

CL\_1299

fGI ID


CL\_INS\_107
CL\_INS\_107
Cluster ID


CL\_13307
CL\_7104
