## Supplementary material for "A novel method for integrating genomic and Tn-Seq data to identify common *in vivo* fitness mechanisms across multiple bacterial species": S1 Dataset: CL_INS_109.html

Legend

 Mobile +extrachromosomalelementfunctions
 Regulatoryfunctions
 Hypothetical
 DNA Metabolism
 All EssentialGenes
 All Fitness Genes
 Cell Envelope
 Other
 EnergyMetabolism
 Transport +binding proteins
 All VFDB Genes

FULL


WINDOWSVGPNG

Trim RowsRemove SingletonsSave Fasta

CL\_1338


CL\_1338


CL\_1338


CL\_1338


CL\_1338


CL\_1338


CL\_1338


CL\_1338


CL\_1338


CL\_1338


CL\_1338


CL\_1338


CL\_1338


CL\_1338


CL\_1338


CL\_1338


CL\_1338


CL\_1338


CL\_1338


CL\_1338


CL\_1338


CL\_1587


CL\_1338


CL\_1338


CL\_1338


CL\_1338


CL\_1338


CL\_1338


CL\_1338


CL\_1338


CL\_1338


CL\_1338


CL\_1338


CL\_1338


CL\_1338


CL\_1338


CL\_1338


CL\_1338


CL\_1337


CL\_1338


CL\_1338


CL\_1338


CL\_1338


CL\_1338


CL\_1338


CL\_1338


CL\_1338


CL\_1338


CL\_1338


CL\_1338


CL\_1338


CL\_1338


CL\_1338


CL\_1338


CL\_1338


CL\_1338


CL\_1338


CL\_1338


CL\_1338


CL\_1338


CL\_1338


CL\_1337


CL\_1338


CL\_1338


CL\_1338


CL\_1338


CL\_1338


CL\_1338


CL\_1338

HighlightSelectShow Genomes


80

CL\_1360


57

CL\_1360


17

CL\_1360


16

CL\_1360


12

CL\_1360


11

CL\_1360


7

CL\_1360


4

CL\_1360


3

CL\_1360


2

CL\_1362


2

CL\_1360


2

CL\_1360


2

CL\_1360


2

CL\_1362


1

CL\_1360


1

CL\_1360


1

CL\_1360


1

CL\_1362


1

CL\_1362


1

CL\_1360


1

CL\_1364


1

CL\_1360


1

CL\_1366


1

CL\_1360


1

CL\_1360


1

CL\_1360


1

CL\_1362


1

CL\_1360


1

CL\_1362


1

Break


1

CL\_1365


1

CL\_1360


1

CL\_1362


1

CL\_1360


1

CL\_1360


1

CL\_1362


1

CL\_1374


1

CL\_1360


1

CL\_1360


1

CL\_1360


1

CL\_1360


1

CL\_1366


1

CL\_1360


1

CL\_1364


1

CL\_1360


1

CL\_1366


1

CL\_1360


1

CL\_1360


1

CL\_1362


1

CL\_1360


1

CL\_1360


1

CL\_1360


1

CL\_1360


1

CL\_1360


1

CL\_1586


1

CL\_1360


1

CL\_1360


1

CL\_1362


1

CL\_1360


1

CL\_1360


1

CL\_1360


1

CL\_1360


1

CL\_1360


1

CL\_1362


1

CL\_1360


1

CL\_1367


1

CL\_1360


1

CL\_1360


1

CL\_1360

fGI ID


CL\_INS\_109
CL\_INS\_109
CL\_INS\_109
CL\_INS\_109
CL\_INS\_382
CL\_INS\_109
CL\_INS\_109
CL\_INS\_109
CL\_INS\_109
CL\_INS\_109
CL\_INS\_109
CL\_INS\_109
CL\_INS\_109
CL\_INS\_109
CL\_INS\_155
CL\_INS\_204
CL\_INS\_155
CL\_INS\_109
CL\_INS\_109
CL\_INS\_109
CL\_INS\_109
CL\_INS\_86
CL\_INS\_155
CL\_INS\_155
CL\_INS\_155
CL\_INS\_155
CL\_INS\_86
CL\_INS\_86
CL\_INS\_86
CL\_INS\_86
CL\_INS\_155
CL\_INS\_155
CL\_INS\_155
CL\_INS\_155
CL\_INS\_155
CL\_INS\_155
CL\_INS\_155
CL\_INS\_86
CL\_INS\_155
CL\_INS\_86
CL\_INS\_155
CL\_INS\_86
CL\_INS\_155
CL\_INS\_155
CL\_INS\_155
CL\_INS\_155
CL\_INS\_155
CL\_INS\_155
CL\_INS\_109
CL\_INS\_109
CL\_INS\_155
CL\_INS\_155
CL\_INS\_155
CL\_INS\_155
CL\_INS\_155
CL\_INS\_155
CL\_INS\_155
CL\_INS\_155
CL\_INS\_155
CL\_INS\_155
CL\_INS\_155
CL\_INS\_182
CL\_INS\_155
CL\_INS\_155
CL\_INS\_155
CL\_INS\_155
CL\_INS\_109
CL\_INS\_109
CL\_INS\_109
CL\_INS\_109
CL\_INS\_109
CL\_INS\_109
CL\_INS\_109
CL\_INS\_109
CL\_INS\_109
CL\_INS\_109
CL\_INS\_109
CL\_INS\_109
CL\_INS\_109
CL\_INS\_109
CL\_INS\_109
CL\_INS\_109
CL\_INS\_109
CL\_INS\_109
CL\_INS\_109
CL\_INS\_109
CL\_INS\_109
CL\_INS\_109
CL\_INS\_109
CL\_INS\_109
CL\_INS\_382
CL\_INS\_382
CL\_INS\_382
CL\_INS\_382
CL\_INS\_382
CL\_INS\_382
CL\_INS\_382
CL\_INS\_109
CL\_INS\_109
CL\_INS\_70
CL\_INS\_109
Cluster ID


CL\_13531
CL\_37109
CL\_16898
CL\_16991
CL\_14091
CL\_25828
CL\_35859
CL\_35860
CL\_1339
CL\_1340
CL\_20991
CL\_1341
CL\_1342
CL\_1343
CL\_11037
CL\_10867
CL\_8483
CL\_10868
CL\_29697
CL\_29449
CL\_29450
CL\_8232
CL\_8231
CL\_8230
CL\_8229
CL\_8228
CL\_8227
CL\_8226
CL\_11030
CL\_11029
CL\_10877
CL\_10878
CL\_10879
CL\_11025
CL\_11024
CL\_10882
CL\_10883
CL\_10884
CL\_10885
CL\_10886
CL\_10887
CL\_10888
CL\_10889
CL\_10890
CL\_10891
CL\_10892
CL\_10893
CL\_10894
CL\_22580
CL\_22581
CL\_11009
CL\_11008
CL\_11007
CL\_11006
CL\_11005
CL\_11004
CL\_10905
CL\_10906
CL\_10907
CL\_10909
CL\_10910
CL\_10996
CL\_10913
CL\_10914
CL\_10915
CL\_10916
CL\_1344
CL\_12452
CL\_1345
CL\_1346
CL\_29698
CL\_1347
CL\_1348
CL\_1349
CL\_1350
CL\_1351
CL\_1352
CL\_1353
CL\_24100
CL\_1354
CL\_12432
CL\_1355
CL\_19433
CL\_1356
CL\_13925
CL\_13924
CL\_13923
CL\_6763
CL\_6762
CL\_1357
CL\_8689
CL\_4545
CL\_6451
CL\_1514
CL\_5283
CL\_5282
CL\_10154
CL\_1358
CL\_1359
CL\_6426
CL\_21534
