## Supplementary material for "A novel method for integrating genomic and Tn-Seq data to identify common *in vivo* fitness mechanisms across multiple bacterial species": S1 Dataset: CL_INS_112.html

Legend

 Mobile +extrachromosomalelementfunctions
 Hypothetical
 All EssentialGenes
 All Fitness Genes
 Cell Envelope
 Other
 EnergyMetabolism
 All VFDB Genes

FULL


WINDOWSVGPNG

Trim RowsRemove SingletonsSave Fasta

CL\_1367


CL\_1367


CL\_1367


CL\_1367


CL\_1367


CL\_1367


CL\_1367


CL\_1362


CL\_1367


CL\_1367


CL\_1366


CL\_1367


CL\_1367

HighlightSelectShow Genomes


241

CL\_1368


11

CL\_1368


5

CL\_1368


5

CL\_1368


2

CL\_1368


1

CL\_1368


1

CL\_1369


1

CL\_1368


1

CL\_1370


1

CL\_1368


1

CL\_1368


1

CL\_1373


1

CL\_1368

fGI ID


CL\_INS\_112
CL\_INS\_112
CL\_INS\_112
CL\_INS\_112
CL\_INS\_112
CL\_INS\_112
CL\_INS\_112
CL\_INS\_112
CL\_INS\_112
CL\_INS\_111
CL\_INS\_111
CL\_INS\_112
CL\_INS\_112
Cluster ID


CL\_13308
CL\_10527
CL\_10528
CL\_12714
CL\_12713
CL\_30064
CL\_30063
CL\_30062
CL\_26017
CL\_11756
CL\_11755
CL\_30841
CL\_11754
