## Supplementary material for "A novel method for integrating genomic and Tn-Seq data to identify common *in vivo* fitness mechanisms across multiple bacterial species": S1 Dataset: CL_INS_117.html

Legend

 Mobile +extrachromosomalelementfunctions
 Regulatoryfunctions
 Hypothetical
 DNA Metabolism
 Transcription
 AntibioticResistance
 All EssentialGenes
 All Fitness Genes
 Other
 Centralintermediarymetabolism
 Cellularprocesses
 Transport +binding proteins
 All VFDB Genes

FULL


WINDOWSVGPNG

Trim RowsRemove SingletonsSave Fasta

CL\_1405


CL\_1405


CL\_1405


CL\_1405


CL\_1405


CL\_1405


CL\_1405


CL\_1405


CL\_1405


CL\_1405


CL\_1405


CL\_1405


CL\_1405


CL\_1405


CL\_1405


CL\_1405


CL\_1405


CL\_1405


CL\_1405


CL\_1405


CL\_1405


CL\_1405


CL\_1405


CL\_1405


CL\_1405


CL\_1405


CL\_1405


CL\_1404


CL\_1405


CL\_1405


CL\_1405


CL\_1405


CL\_1405


CL\_1405


CL\_1405


CL\_1405


CL\_1405


CL\_1405


CL\_1404


CL\_1405


CL\_1405


CL\_1405


CL\_1405


CL\_1405


CL\_1405


CL\_1405


CL\_1405


CL\_1405


CL\_1405


CL\_1405


CL\_1405


CL\_1405


CL\_1405


CL\_1405


CL\_471


CL\_1405


CL\_1405


CL\_1405


CL\_1404


CL\_1405


Break


CL\_1405


CL\_1404


CL\_1405


CL\_1404


CL\_1405


CL\_1405


CL\_1405


CL\_1405


CL\_1405


CL\_1405


CL\_1405


CL\_1404


CL\_1405


CL\_1405


CL\_1405


CL\_1405


CL\_1404


CL\_1405


CL\_1291


CL\_1405


CL\_1405


CL\_1405


CL\_1405


CL\_294


CL\_1405


CL\_1405


CL\_1405


CL\_1405


CL\_1405


CL\_1405


CL\_1404


CL\_1405


CL\_1405


CL\_1405


CL\_1405


CL\_1405


CL\_1404


CL\_1405


CL\_1405


CL\_1405


CL\_1405


CL\_1405


CL\_1405


CL\_1405


CL\_1405


CL\_1405


CL\_1405


CL\_1405


CL\_1405


CL\_1405


CL\_1405


CL\_1405


CL\_1405


CL\_1405


CL\_1405


CL\_1405


CL\_3599


CL\_3600


CL\_1405


CL\_1405


CL\_1405


CL\_1405


CL\_1405


CL\_1405


CL\_539


CL\_1405


CL\_1405


CL\_1405


Break


CL\_1405


CL\_1405


CL\_1405


CL\_1405


CL\_1405


CL\_1405


CL\_1405


CL\_1405


CL\_1404


CL\_1405


CL\_1405


CL\_1405

HighlightSelectShow Genomes


21

CL\_1413


17

CL\_1413


16

CL\_1413


11

CL\_1413


9

CL\_1413


9

CL\_1413


7

CL\_1413


6

CL\_1413


5

CL\_1413


5

CL\_1413


5

CL\_1413


4

CL\_1413


4

CL\_1413


4

CL\_1413


4

CL\_1413


3

CL\_1413


3

CL\_1413


3

CL\_1413


3

CL\_1413


3

CL\_1413


3

CL\_1413


3

CL\_1413


3

CL\_1413


2

CL\_1413


2

CL\_1413


2

CL\_1413


2

CL\_1413


2

CL\_1413


2

CL\_1413


2

CL\_1413


2

CL\_1413


2

CL\_1413


2

CL\_1413


1

CL\_1413


1

CL\_1413


1

CL\_1413


1

CL\_1413


1

CL\_1413


1

CL\_1413


1

CL\_1413


1

CL\_1413


1

CL\_1413


1

CL\_1413


1

CL\_1413


1

CL\_1413


1

CL\_1413


1

CL\_1413


1

CL\_1413


1

CL\_1442


1

CL\_1413


1

CL\_1413


1

CL\_206


1

CL\_1413


1

CL\_1413


1

CL\_1413


1

CL\_1413


1

CL\_1413


1

CL\_1413


1

CL\_1413


1

CL\_1413


1

CL\_1413


1

CL\_1413


1

CL\_1413


1

CL\_1421


1

CL\_1413


1

CL\_1413


1

CL\_1413


1

CL\_1413


1

CL\_1413


1

CL\_1413


1

CL\_1413


1

CL\_1413


1

CL\_1413


1

CL\_1413


1

CL\_1489


1

CL\_1413


1

CL\_1413


1

CL\_1413


1

CL\_1413


1

CL\_1413


1

CL\_1252


1

CL\_1413


1

CL\_1413


1

CL\_1413


1

CL\_1413


1

CL\_1413


1

CL\_1413


1

CL\_1413


1

CL\_1413


1

CL\_1413


1

CL\_1413


1

CL\_1413


1

CL\_1413


1

CL\_1413


1

CL\_1413


1

CL\_1413


1

CL\_1413


1

CL\_1413


1

CL\_1413


1

CL\_1413


1

CL\_1413


1

CL\_1413


1

CL\_1413


1

CL\_1413


1

CL\_1413


1

CL\_1413


1

CL\_1421


1

CL\_1414


1

CL\_1413


1

CL\_1413


1

CL\_1413


1

CL\_1413


1

CL\_994


1

CL\_1413


1

CL\_1413


1

CL\_1455


1

Break


1

CL\_1413


1

CL\_1413


1

CL\_1413


1

CL\_475


1

CL\_1413


1

CL\_1413


1

CL\_1413


1

CL\_1413


1

CL\_1413


1

CL\_1413


1

CL\_1413


1

CL\_1413


1

CL\_1413


1

CL\_1413


1

CL\_3772


1

CL\_1413


1

CL\_1413


1

CL\_1413


1

CL\_1413


1

CL\_1413


1

CL\_1413


1

CL\_1413


1

CL\_1413


1

CL\_1413


1

CL\_1413

fGI ID


CL\_INS\_117
CL\_INS\_117
CL\_INS\_106
CL\_INS\_106
CL\_INS\_237
CL\_INS\_237
CL\_INS\_237
CL\_INS\_237
CL\_INS\_117
CL\_INS\_117
CL\_INS\_117
CL\_INS\_117
CL\_INS\_117
CL\_INS\_117
CL\_INS\_117
CL\_INS\_117
CL\_INS\_117
CL\_INS\_117
CL\_INS\_117
CL\_INS\_117
CL\_INS\_117
CL\_INS\_117
CL\_INS\_117
CL\_INS\_117
CL\_INS\_117
CL\_INS\_117
CL\_INS\_117
CL\_INS\_117
CL\_INS\_117
CL\_INS\_117
CL\_INS\_117
CL\_INS\_117
CL\_INS\_117
CL\_INS\_117
CL\_INS\_117
CL\_INS\_117
CL\_INS\_117
CL\_INS\_117
CL\_INS\_117
CL\_INS\_117
CL\_INS\_117
CL\_INS\_117
CL\_INS\_382
CL\_INS\_117
CL\_INS\_117
CL\_INS\_117
CL\_INS\_117
CL\_INS\_117
CL\_INS\_117
CL\_INS\_117
CL\_INS\_117
CL\_INS\_117
CL\_INS\_117
CL\_INS\_117
CL\_INS\_117
CL\_INS\_117
CL\_INS\_117
CL\_INS\_117
CL\_INS\_117
CL\_INS\_207
CL\_INS\_207
CL\_INS\_207
CL\_INS\_382
CL\_INS\_117
CL\_INS\_159
CL\_INS\_382
CL\_INS\_382
CL\_INS\_382
CL\_INS\_117
CL\_INS\_117
CL\_INS\_382
CL\_INS\_233
CL\_INS\_233
CL\_INS\_233
CL\_INS\_382
CL\_INS\_382
CL\_INS\_382
CL\_INS\_382
CL\_INS\_382
CL\_INS\_382
CL\_INS\_382
CL\_INS\_382
CL\_INS\_60
CL\_INS\_60
CL\_INS\_60
CL\_INS\_382
CL\_INS\_382
CL\_INS\_382
CL\_INS\_159
CL\_INS\_382
CL\_INS\_382
CL\_INS\_382
CL\_INS\_382
CL\_INS\_382
CL\_INS\_382
CL\_INS\_382
CL\_INS\_382
CL\_INS\_382
CL\_INS\_382
CL\_INS\_382
CL\_INS\_382
CL\_INS\_382
CL\_INS\_382
CL\_INS\_382
CL\_INS\_382
CL\_INS\_382
CL\_INS\_382
CL\_INS\_382
CL\_INS\_159
CL\_INS\_382
CL\_INS\_385
CL\_INS\_382
CL\_INS\_382
CL\_INS\_382
CL\_INS\_117
CL\_INS\_117
CL\_INS\_117
CL\_INS\_382
CL\_INS\_382
CL\_INS\_382
CL\_INS\_156
CL\_INS\_156
CL\_INS\_156
CL\_INS\_247
CL\_INS\_86
CL\_INS\_149
CL\_INS\_149
CL\_INS\_149
CL\_INS\_149
CL\_INS\_149
CL\_INS\_149
CL\_INS\_57
CL\_INS\_247
CL\_INS\_247
CL\_INS\_247
CL\_INS\_123
CL\_INS\_247
CL\_INS\_123
CL\_INS\_123
CL\_INS\_123
CL\_INS\_247
CL\_INS\_247
CL\_INS\_247
CL\_INS\_247
CL\_INS\_247
CL\_INS\_237
CL\_INS\_237
CL\_INS\_237
CL\_INS\_86
CL\_INS\_117
CL\_INS\_117
CL\_INS\_30
CL\_INS\_70
CL\_INS\_117
CL\_INS\_117
CL\_INS\_117
CL\_INS\_117
CL\_INS\_117
CL\_INS\_237
CL\_INS\_233
CL\_INS\_382
CL\_INS\_117
CL\_INS\_20
CL\_INS\_20
CL\_INS\_20
CL\_INS\_70
CL\_INS\_20
CL\_INS\_117
CL\_INS\_117
CL\_INS\_117
CL\_INS\_121
CL\_INS\_121
CL\_INS\_121
CL\_INS\_10
CL\_INS\_121
CL\_INS\_121
CL\_INS\_121
CL\_INS\_121
CL\_INS\_121
CL\_INS\_121
CL\_INS\_117
CL\_INS\_117
CL\_INS\_117
CL\_INS\_117
CL\_INS\_117
CL\_INS\_117
CL\_INS\_117
CL\_INS\_117
CL\_INS\_117
CL\_INS\_117
CL\_INS\_117
CL\_INS\_117
CL\_INS\_117
CL\_INS\_117
CL\_INS\_117
CL\_INS\_117
CL\_INS\_117
CL\_INS\_117
CL\_INS\_117
CL\_INS\_224
CL\_INS\_224
CL\_INS\_117
CL\_INS\_117
CL\_INS\_117
CL\_INS\_117
CL\_INS\_117
CL\_INS\_117
CL\_INS\_117
CL\_INS\_117
CL\_INS\_117
CL\_INS\_117
CL\_INS\_117
CL\_INS\_117
CL\_INS\_117
CL\_INS\_10
CL\_INS\_117
CL\_INS\_117
CL\_INS\_117
CL\_INS\_117
CL\_INS\_117
CL\_INS\_117
CL\_INS\_117
CL\_INS\_117
CL\_INS\_117
CL\_INS\_117
CL\_INS\_237
CL\_INS\_237
CL\_INS\_237
CL\_INS\_237
CL\_INS\_117
CL\_INS\_247
CL\_INS\_237
CL\_INS\_237
CL\_INS\_237
CL\_INS\_237
CL\_INS\_237
CL\_INS\_237
CL\_INS\_237
CL\_INS\_382
CL\_INS\_237
CL\_INS\_237
CL\_INS\_117
CL\_INS\_237
CL\_INS\_237
CL\_INS\_117
CL\_INS\_117
CL\_INS\_237
CL\_INS\_117
CL\_INS\_237
CL\_INS\_237
CL\_INS\_237
CL\_INS\_237
CL\_INS\_237
CL\_INS\_237
CL\_INS\_237
CL\_INS\_237
CL\_INS\_237
CL\_INS\_117
CL\_INS\_237
CL\_INS\_237
CL\_INS\_117
CL\_INS\_237
CL\_INS\_237
CL\_INS\_237
CL\_INS\_117
CL\_INS\_237
CL\_INS\_117
CL\_INS\_117
CL\_INS\_117
CL\_INS\_117
CL\_INS\_117
CL\_INS\_237
CL\_INS\_237
CL\_INS\_237
CL\_INS\_237
CL\_INS\_237
CL\_INS\_237
CL\_INS\_237
CL\_INS\_237
CL\_INS\_117
CL\_INS\_117
CL\_INS\_237
CL\_INS\_237
CL\_INS\_117
CL\_INS\_117
CL\_INS\_237
CL\_INS\_117
CL\_INS\_237
CL\_INS\_237
CL\_INS\_237
CL\_INS\_237
CL\_INS\_117
CL\_INS\_117
CL\_INS\_237
CL\_INS\_237
CL\_INS\_237
CL\_INS\_237
CL\_INS\_237
CL\_INS\_117
CL\_INS\_237
CL\_INS\_117
CL\_INS\_237
CL\_INS\_117
CL\_INS\_237
CL\_INS\_117
CL\_INS\_117
CL\_INS\_237
CL\_INS\_237
CL\_INS\_237
CL\_INS\_237
CL\_INS\_237
CL\_INS\_117
CL\_INS\_204
CL\_INS\_149
CL\_INS\_170
CL\_INS\_149
CL\_INS\_237
CL\_INS\_237
CL\_INS\_237
CL\_INS\_237
CL\_INS\_237
CL\_INS\_237
CL\_INS\_237
CL\_INS\_237
CL\_INS\_237
CL\_INS\_10
CL\_INS\_10
CL\_INS\_10
CL\_INS\_117
CL\_INS\_117
CL\_INS\_117
CL\_INS\_117
CL\_INS\_117
CL\_INS\_117
CL\_INS\_117
CL\_INS\_117
CL\_INS\_117
CL\_INS\_117
CL\_INS\_117
CL\_INS\_117
CL\_INS\_117
CL\_INS\_117
CL\_INS\_117
CL\_INS\_117
CL\_INS\_117
CL\_INS\_117
CL\_INS\_117
CL\_INS\_117
CL\_INS\_117
CL\_INS\_117
CL\_INS\_117
CL\_INS\_117
CL\_INS\_117
CL\_INS\_117
CL\_INS\_117
CL\_INS\_117
CL\_INS\_117
CL\_INS\_117
CL\_INS\_117
CL\_INS\_117
CL\_INS\_117
CL\_INS\_224
CL\_INS\_117
CL\_INS\_117
CL\_INS\_117
CL\_INS\_117
CL\_INS\_117
CL\_INS\_117
CL\_INS\_117
CL\_INS\_117
CL\_INS\_117
CL\_INS\_117
CL\_INS\_117
CL\_INS\_117
CL\_INS\_117
CL\_INS\_117
CL\_INS\_117
CL\_INS\_117
CL\_INS\_117
CL\_INS\_117
CL\_INS\_117
CL\_INS\_117
CL\_INS\_117
CL\_INS\_117
CL\_INS\_117
CL\_INS\_117
CL\_INS\_117
CL\_INS\_117
CL\_INS\_117
CL\_INS\_117
CL\_INS\_117
CL\_INS\_117
CL\_INS\_117
CL\_INS\_117
CL\_INS\_117
CL\_INS\_117
CL\_INS\_117
CL\_INS\_117
CL\_INS\_117
CL\_INS\_117
CL\_INS\_117
CL\_INS\_117
CL\_INS\_117
CL\_INS\_117
CL\_INS\_117
CL\_INS\_117
CL\_INS\_117
CL\_INS\_117
CL\_INS\_117
CL\_INS\_117
CL\_INS\_117
CL\_INS\_117
CL\_INS\_117
CL\_INS\_117
CL\_INS\_117
CL\_INS\_117
CL\_INS\_117
CL\_INS\_117
CL\_INS\_117
CL\_INS\_117
CL\_INS\_117
CL\_INS\_20
CL\_INS\_117
CL\_INS\_117
CL\_INS\_117
CL\_INS\_117
CL\_INS\_117
CL\_INS\_117
CL\_INS\_343
CL\_INS\_343
CL\_INS\_343
CL\_INS\_343
CL\_INS\_117
CL\_INS\_20
CL\_INS\_117
CL\_INS\_117
CL\_INS\_117
CL\_INS\_117
CL\_INS\_224
CL\_INS\_117
CL\_INS\_117
CL\_INS\_117
CL\_INS\_117
CL\_INS\_117
CL\_INS\_117
CL\_INS\_117
CL\_INS\_117
CL\_INS\_117
CL\_INS\_117
CL\_INS\_117
CL\_INS\_117
CL\_INS\_117
CL\_INS\_117
CL\_INS\_117
CL\_INS\_117
CL\_INS\_117
CL\_INS\_117
Cluster ID


CL\_12436
CL\_11257
CL\_1293
CL\_1294
CL\_6765
CL\_5595
CL\_5596
CL\_6764
CL\_14110
CL\_14109
CL\_14108
CL\_14107
CL\_14106
CL\_14105
CL\_5324
CL\_1406
CL\_9925
CL\_35107
CL\_35106
CL\_10529
CL\_13648
CL\_13649
CL\_13650
CL\_13651
CL\_13652
CL\_10455
CL\_10456
CL\_10457
CL\_10458
CL\_13795
CL\_34879
CL\_7014
CL\_7013
CL\_7012
CL\_7011
CL\_1407
CL\_32054
CL\_28025
CL\_4495
CL\_17032
CL\_17031
CL\_17030
CL\_6385
CL\_4496
CL\_4343
CL\_1409
CL\_1410
CL\_22785
CL\_23567
CL\_23568
CL\_23569
CL\_10141
CL\_13437
CL\_23566
CL\_31848
CL\_31849
CL\_31850
CL\_31851
CL\_31852
CL\_5614
CL\_5613
CL\_5536
CL\_5600
CL\_13399
CL\_4307
CL\_10407
CL\_10406
CL\_4303
CL\_5665
CL\_6072
CL\_5664
CL\_5545
CL\_12660
CL\_12661
CL\_4299
CL\_5662
CL\_5548
CL\_4294
CL\_4293
CL\_11338
CL\_5659
CL\_5551
CL\_5657
CL\_11829
CL\_11830
CL\_5554
CL\_5555
CL\_5556
CL\_5651
CL\_4284
CL\_5560
CL\_5561
CL\_5562
CL\_5563
CL\_5564
CL\_5565
CL\_5566
CL\_5567
CL\_4278
CL\_4277
CL\_5639
CL\_5638
CL\_5637
CL\_6934
CL\_6933
CL\_5634
CL\_4271
CL\_4270
CL\_5575
CL\_5579
CL\_13400
CL\_4265
CL\_5061
CL\_4262
CL\_5582
CL\_5592
CL\_5593
CL\_4974
CL\_4973
CL\_4972
CL\_5474
CL\_5475
CL\_5477
CL\_6749
CL\_5516
CL\_6752
CL\_6753
CL\_6754
CL\_6755
CL\_6756
CL\_6757
CL\_6758
CL\_13409
CL\_11980
CL\_5302
CL\_5300
CL\_5299
CL\_5298
CL\_5297
CL\_10392
CL\_10393
CL\_10395
CL\_10641
CL\_10642
CL\_10421
CL\_6761
CL\_6760
CL\_10419
CL\_10418
CL\_10417
CL\_13402
CL\_11295
CL\_6410
CL\_13401
CL\_6968
CL\_6969
CL\_5496
CL\_6970
CL\_6971
CL\_12667
CL\_6923
CL\_5505
CL\_6973
CL\_5495
CL\_5494
CL\_5493
CL\_5492
CL\_10949
CL\_18923
CL\_16216
CL\_14232
CL\_4807
CL\_4806
CL\_4805
CL\_9958
CL\_4804
CL\_4803
CL\_4802
CL\_4801
CL\_14420
CL\_16217
CL\_16218
CL\_16219
CL\_16220
CL\_16221
CL\_16222
CL\_16223
CL\_16224
CL\_16225
CL\_16226
CL\_16227
CL\_16228
CL\_16229
CL\_16230
CL\_16231
CL\_9969
CL\_9970
CL\_9971
CL\_9972
CL\_12597
CL\_9974
CL\_16232
CL\_16233
CL\_16234
CL\_16235
CL\_16236
CL\_16237
CL\_16238
CL\_16239
CL\_16240
CL\_16241
CL\_16242
CL\_16243
CL\_16244
CL\_4764
CL\_16245
CL\_16246
CL\_16247
CL\_16248
CL\_16249
CL\_11355
CL\_11354
CL\_11353
CL\_16250
CL\_16251
CL\_11352
CL\_11351
CL\_11350
CL\_11349
CL\_11348
CL\_10021
CL\_11347
CL\_11346
CL\_11345
CL\_11344
CL\_11343
CL\_6139
CL\_6140
CL\_6141
CL\_6142
CL\_6143
CL\_11427
CL\_11426
CL\_6163
CL\_11424
CL\_11423
CL\_11422
CL\_11420
CL\_11419
CL\_11418
CL\_11417
CL\_11416
CL\_11415
CL\_11414
CL\_11413
CL\_11412
CL\_11411
CL\_16252
CL\_11410
CL\_6171
CL\_11409
CL\_6172
CL\_6173
CL\_6174
CL\_11408
CL\_6175
CL\_6176
CL\_11407
CL\_16253
CL\_16254
CL\_16255
CL\_11394
CL\_11393
CL\_6181
CL\_6182
CL\_6183
CL\_6186
CL\_6187
CL\_6188
CL\_11392
CL\_11391
CL\_6189
CL\_6190
CL\_11390
CL\_11389
CL\_6193
CL\_16256
CL\_6194
CL\_6195
CL\_6196
CL\_11388
CL\_16257
CL\_16258
CL\_6134
CL\_6135
CL\_11385
CL\_11384
CL\_11383
CL\_11382
CL\_11381
CL\_11372
CL\_11371
CL\_16259
CL\_11370
CL\_16260
CL\_16261
CL\_11368
CL\_11367
CL\_6136
CL\_6137
CL\_6138
CL\_16262
CL\_6520
CL\_4603
CL\_4602
CL\_4601
CL\_4514
CL\_11363
CL\_11362
CL\_11361
CL\_11360
CL\_11359
CL\_11358
CL\_11357
CL\_11356
CL\_9247
CL\_9248
CL\_9249
CL\_16263
CL\_16264
CL\_16265
CL\_16266
CL\_16267
CL\_16268
CL\_16269
CL\_16270
CL\_16271
CL\_16272
CL\_16273
CL\_16274
CL\_16275
CL\_16276
CL\_16277
CL\_16278
CL\_16279
CL\_16280
CL\_16281
CL\_16282
CL\_16283
CL\_16284
CL\_16285
CL\_16286
CL\_16287
CL\_16288
CL\_16289
CL\_16290
CL\_16291
CL\_16292
CL\_16293
CL\_16294
CL\_16295
CL\_9978
CL\_16296
CL\_16297
CL\_16298
CL\_16299
CL\_16300
CL\_16301
CL\_16302
CL\_16303
CL\_16304
CL\_16305
CL\_16306
CL\_16307
CL\_7805
CL\_7804
CL\_30367
CL\_31966
CL\_16098
CL\_1408
CL\_472
CL\_473
CL\_9176
CL\_7102
CL\_5818
CL\_13796
CL\_13797
CL\_21699
CL\_13798
CL\_7101
CL\_10530
CL\_10531
CL\_22136
CL\_11774
CL\_4494
CL\_17659
CL\_10140
CL\_15832
CL\_7803
CL\_7802
CL\_21902
CL\_7919
CL\_7801
CL\_9175
CL\_11568
CL\_11567
CL\_16308
CL\_16309
CL\_16310
CL\_16311
CL\_16312
CL\_9177
CL\_11566
CL\_11565
CL\_11564
CL\_11563
CL\_19724
CL\_11562
CL\_11561
CL\_11560
CL\_11559
CL\_11558
CL\_11557
CL\_19723
CL\_11556
CL\_1411
CL\_11258
CL\_4497
CL\_23909
CL\_23806
CL\_23807
CL\_34282
CL\_4498
CL\_5788
CL\_28024
CL\_4499
CL\_4500
CL\_23705
CL\_4750
CL\_7100
CL\_20158
CL\_20157
CL\_20159
CL\_20160
CL\_11514
CL\_11515
CL\_11517
CL\_11518
CL\_20151
CL\_20152
CL\_20153
CL\_11555
CL\_11575
CL\_30366
CL\_11752
CL\_11751
CL\_1412
