## Supplementary material for "A novel method for integrating genomic and Tn-Seq data to identify common *in vivo* fitness mechanisms across multiple bacterial species": S1 Dataset: CL_INS_118.html

Legend

 Mobile +extrachromosomalelementfunctions
 All EssentialGenes
 All Fitness Genes
 Other
 EnergyMetabolism
 All VFDB Genes

FULL


WINDOWSVGPNG

Trim RowsRemove SingletonsSave Fasta

CL\_1420


CL\_1420


CL\_1420


CL\_1420


CL\_1420


CL\_1420


CL\_1405


CL\_1420


CL\_1300


CL\_1420


CL\_1420


CL\_1463


CL\_1405


CL\_1420

HighlightSelectShow Genomes


84

CL\_1421


58

CL\_1421


44

CL\_1421


2

CL\_1421


1

CL\_1421


1

CL\_1421


1

CL\_1421


1

CL\_1421


1

CL\_1421


1

CL\_1421


1

CL\_1424


1

CL\_1421


1

CL\_1421


1

CL\_1421

fGI ID


CL\_INS\_118
CL\_INS\_118
CL\_INS\_118
CL\_INS\_118
CL\_INS\_118
CL\_INS\_118
CL\_INS\_118
CL\_INS\_118
CL\_INS\_117
CL\_INS\_118
Cluster ID


CL\_9596
CL\_22137
CL\_22685
CL\_30159
CL\_9924
CL\_29787
CL\_34668
CL\_4501
CL\_11257
CL\_12437
