## Supplementary material for "A novel method for integrating genomic and Tn-Seq data to identify common *in vivo* fitness mechanisms across multiple bacterial species": S1 Dataset: CL_INS_120.html

Legend

 Mobile +extrachromosomalelementfunctions
 Other
 All VFDB Genes

FULL


WINDOWSVGPNG

Trim RowsRemove SingletonsSave Fasta

CL\_1430


CL\_1430


CL\_1430


CL\_1430

HighlightSelectShow Genomes


191

CL\_1431


15

CL\_1431


1

CL\_1431


1

CL\_1431

fGI ID


CL\_INS\_120
CL\_INS\_120
CL\_INS\_120
CL\_INS\_120
CL\_INS\_120
CL\_INS\_120
CL\_INS\_120
CL\_INS\_120
CL\_INS\_120
Cluster ID


CL\_16313
CL\_16314
CL\_5819
CL\_5820
CL\_5821
CL\_5822
CL\_5823
CL\_5824
CL\_5825
