## Supplementary material for "A novel method for integrating genomic and Tn-Seq data to identify common *in vivo* fitness mechanisms across multiple bacterial species": S1 Dataset: CL_INS_122.html

Legend

 Mobile +extrachromosomalelementfunctions
 Regulatoryfunctions
 Hypothetical
 DNA Metabolism
 AntibioticResistance
 All Fitness Genes
 Other
 Transport +binding proteins
 All VFDB Genes

FULL


WINDOWSVGPNG

Trim RowsRemove SingletonsSave Fasta

CL\_1453


CL\_1453


CL\_1453


CL\_1453


CL\_1453


CL\_1453


CL\_1453


CL\_1453


CL\_1453


CL\_1453


CL\_1453


CL\_1453


CL\_1453


CL\_1453


CL\_1453


CL\_1453


CL\_1453


CL\_1453


CL\_1452


CL\_1453


CL\_1453


CL\_1453


CL\_4486


CL\_1453


CL\_1453

HighlightSelectShow Genomes


196

CL\_4502


3

CL\_4502


2

CL\_4503


1

CL\_1454


1

CL\_1454


1

Break


1

CL\_4502


1

CL\_1454


1

CL\_1454


1

CL\_1454


1

CL\_1454


1

CL\_1454


1

CL\_1454


1

CL\_1454


1

CL\_1454


1

CL\_1454


1

CL\_1454


1

CL\_1454


1

CL\_4502


1

CL\_1454


1

CL\_1462


1

CL\_1462


1

CL\_4502


1

CL\_1454


1

CL\_1454

fGI ID


CL\_INS\_122
CL\_INS\_123
CL\_INS\_123
CL\_INS\_123
CL\_INS\_122
CL\_INS\_123
CL\_INS\_122
CL\_INS\_123
CL\_INS\_123
CL\_INS\_123
CL\_INS\_123
CL\_INS\_123
CL\_INS\_123
CL\_INS\_86
CL\_INS\_207
CL\_INS\_86
CL\_INS\_86
CL\_INS\_99
CL\_INS\_207
CL\_INS\_207
CL\_INS\_207
CL\_INS\_122
CL\_INS\_123
CL\_INS\_123
CL\_INS\_123
CL\_INS\_123
CL\_INS\_123
CL\_INS\_123
CL\_INS\_123
CL\_INS\_247
CL\_INS\_123
CL\_INS\_123
CL\_INS\_123
CL\_INS\_123
CL\_INS\_123
CL\_INS\_123
CL\_INS\_123
CL\_INS\_123
CL\_INS\_123
CL\_INS\_123
CL\_INS\_123
CL\_INS\_123
CL\_INS\_123
CL\_INS\_123
CL\_INS\_123
CL\_INS\_123
CL\_INS\_149
CL\_INS\_149
CL\_INS\_149
CL\_INS\_149
CL\_INS\_123
CL\_INS\_237
CL\_INS\_237
CL\_INS\_57
CL\_INS\_57
CL\_INS\_149
CL\_INS\_149
CL\_INS\_149
CL\_INS\_149
CL\_INS\_149
CL\_INS\_149
CL\_INS\_57
CL\_INS\_237
CL\_INS\_247
CL\_INS\_382
CL\_INS\_123
CL\_INS\_123
CL\_INS\_70
CL\_INS\_70
CL\_INS\_123
CL\_INS\_123
CL\_INS\_123
CL\_INS\_123
CL\_INS\_123
CL\_INS\_123
CL\_INS\_123
CL\_INS\_123
CL\_INS\_123
CL\_INS\_123
CL\_INS\_123
CL\_INS\_123
CL\_INS\_123
CL\_INS\_123
CL\_INS\_123
CL\_INS\_123
CL\_INS\_123
CL\_INS\_123
CL\_INS\_122
CL\_INS\_122
CL\_INS\_122
CL\_INS\_122
CL\_INS\_122
CL\_INS\_123
CL\_INS\_123
CL\_INS\_123
CL\_INS\_122
CL\_INS\_122
CL\_INS\_122
CL\_INS\_123
CL\_INS\_123
CL\_INS\_123
CL\_INS\_123
CL\_INS\_123
CL\_INS\_123
CL\_INS\_123
CL\_INS\_123
CL\_INS\_123
CL\_INS\_123
CL\_INS\_123
CL\_INS\_123
CL\_INS\_123
CL\_INS\_123
CL\_INS\_123
CL\_INS\_123
CL\_INS\_123
CL\_INS\_122
CL\_INS\_382
CL\_INS\_123
CL\_INS\_123
CL\_INS\_382
CL\_INS\_123
CL\_INS\_382
CL\_INS\_123
CL\_INS\_123
CL\_INS\_123
CL\_INS\_123
CL\_INS\_123
CL\_INS\_123
CL\_INS\_382
CL\_INS\_123
CL\_INS\_123
CL\_INS\_123
CL\_INS\_123
CL\_INS\_123
CL\_INS\_123
CL\_INS\_123
CL\_INS\_123
CL\_INS\_123
CL\_INS\_123
CL\_INS\_123
CL\_INS\_123
CL\_INS\_123
CL\_INS\_123
CL\_INS\_123
CL\_INS\_123
CL\_INS\_123
CL\_INS\_123
CL\_INS\_123
Cluster ID


CL\_9595
CL\_9594
CL\_9593
CL\_9592
CL\_28975
CL\_28976
CL\_35105
CL\_4504
CL\_35104
CL\_35103
CL\_35102
CL\_35101
CL\_35100
CL\_4488
CL\_11956
CL\_10526
CL\_6782
CL\_10520
CL\_1493
CL\_1492
CL\_1491
CL\_14269
CL\_5826
CL\_5827
CL\_5828
CL\_5829
CL\_14270
CL\_14271
CL\_14272
CL\_5514
CL\_15177
CL\_15176
CL\_5300
CL\_10715
CL\_5298
CL\_5297
CL\_10392
CL\_33526
CL\_10142
CL\_14795
CL\_14796
CL\_9893
CL\_9892
CL\_9891
CL\_9890
CL\_35189
CL\_7485
CL\_7486
CL\_7604
CL\_7487
CL\_35188
CL\_6761
CL\_6760
CL\_6759
CL\_6758
CL\_6757
CL\_6756
CL\_6755
CL\_6754
CL\_6753
CL\_6752
CL\_6751
CL\_6750
CL\_6749
CL\_6748
CL\_5323
CL\_5322
CL\_5321
CL\_5320
CL\_5319
CL\_5318
CL\_5317
CL\_5316
CL\_5315
CL\_5314
CL\_5313
CL\_5312
CL\_5311
CL\_12011
CL\_12012
CL\_12013
CL\_12014
CL\_12015
CL\_12016
CL\_12017
CL\_12018
CL\_12019
CL\_20994
CL\_20995
CL\_20996
CL\_20997
CL\_20998
CL\_12020
CL\_12021
CL\_12022
CL\_20246
CL\_20245
CL\_20999
CL\_1456
CL\_1457
CL\_1458
CL\_1459
CL\_1460
CL\_10147
CL\_10148
CL\_10149
CL\_33592
CL\_33591
CL\_33590
CL\_5305
CL\_5304
CL\_8159
CL\_8158
CL\_16971
CL\_16972
CL\_14118
CL\_4235
CL\_8157
CL\_8156
CL\_8843
CL\_8842
CL\_8841
CL\_13922
CL\_13921
CL\_13920
CL\_8840
CL\_8839
CL\_8838
CL\_8909
CL\_9786
CL\_9785
CL\_9782
CL\_8837
CL\_8836
CL\_5310
CL\_14797
CL\_5309
CL\_5308
CL\_5307
CL\_5306
CL\_8835
CL\_8834
CL\_34090
CL\_8833
CL\_13919
CL\_13918
CL\_10152
CL\_10153
