## Supplementary material for "A novel method for integrating genomic and Tn-Seq data to identify common *in vivo* fitness mechanisms across multiple bacterial species": S1 Dataset: CL_INS_123.html

Legend

 Mobile +extrachromosomalelementfunctions
 Regulatoryfunctions
 Hypothetical
 DNA Metabolism
 AntibioticResistance
 All Fitness Genes
 Other
 Transport +binding proteins
 All VFDB Genes

FULL


WINDOWSVGPNG

Trim RowsRemove SingletonsSave Fasta

CL\_4503


CL\_4503


CL\_4503


CL\_4503


CL\_4503


CL\_4503


CL\_4503


CL\_4503


CL\_1453


CL\_4503


CL\_4503


CL\_4503


CL\_4503


CL\_4502


CL\_4503


CL\_1453


CL\_4503


CL\_1453


CL\_4503


CL\_1453


CL\_1537


CL\_1453


CL\_4503


CL\_1453


CL\_1453


CL\_4502


CL\_4503


CL\_1450


CL\_1453


CL\_4503


CL\_4503


CL\_4503


CL\_4503


CL\_4503


CL\_4503


CL\_4503


CL\_1453


CL\_4503


CL\_4502


CL\_4503


CL\_4503


CL\_1452


CL\_4503


CL\_4503


CL\_4503


CL\_4503


CL\_4503


CL\_4503


CL\_4503


CL\_4503


CL\_4503


CL\_4503


CL\_1453


CL\_4503


CL\_4503


CL\_4503


CL\_4503


CL\_4503


CL\_1453


CL\_1453


CL\_4503


CL\_4503


CL\_4503


CL\_4503


CL\_1453


CL\_1453


CL\_4503


CL\_4502


CL\_4503


CL\_4503


CL\_4503


CL\_4503


CL\_1453


CL\_4502


CL\_4503


CL\_1453


CL\_4503


CL\_4503


CL\_4503


CL\_1453


CL\_4502


CL\_4503


CL\_4503


CL\_4503


CL\_4503


CL\_4503


CL\_1452


CL\_4503


CL\_4503

HighlightSelectShow Genomes


102

CL\_1454


6

CL\_1454


6

CL\_1454


5

CL\_1454


5

CL\_1462


5

CL\_1454


5

CL\_1455


3

CL\_1462


3

CL\_1454


3

CL\_1462


2

CL\_1454


2

CL\_1454


2

CL\_1462


2

CL\_1454


2

CL\_1454


1

CL\_1454


1

CL\_1454


1

CL\_1454


1

CL\_1454


1

CL\_1454


1

CL\_1454


1

CL\_1454


1

CL\_1454


1

CL\_1454


1

CL\_1454


1

CL\_1454


1

Break


1

CL\_1454


1

CL\_1454


1

CL\_1462


1

CL\_1454


1

CL\_1454


1

CL\_1454


1

CL\_1454


1

CL\_1454


1

CL\_1454


1

CL\_1454


1

CL\_1454


1

CL\_1454


1

CL\_1454


1

CL\_1454


1

CL\_1454


1

CL\_1454


1

CL\_1455


1

CL\_1454


1

CL\_1454


1

CL\_1535


1

CL\_1454


1

CL\_1454


1

CL\_1454


1

CL\_1454


1

CL\_1454


1

CL\_1454


1

CL\_1454


1

CL\_1455


1

CL\_1454


1

CL\_1454


1

CL\_1454


1

CL\_1454


1

CL\_1454


1

CL\_1454


1

CL\_1454


1

CL\_1462


1

CL\_1454


1

CL\_1454


1

CL\_1454


1

CL\_1455


1

CL\_1454


1

CL\_1454


1

CL\_1455


1

CL\_1454


1

CL\_1454


1

CL\_1454


1

CL\_1454


1

CL\_1462


1

CL\_1454


1

CL\_1454


1

CL\_1454


1

CL\_1465


1

CL\_1454


1

CL\_1454


1

CL\_1465


1

CL\_1454


1

CL\_1454


1

CL\_1454


1

CL\_1454


1

CL\_1454


1

CL\_1454


1

CL\_1454

fGI ID


CL\_INS\_123
CL\_INS\_382
CL\_INS\_123
CL\_INS\_123
CL\_INS\_123
CL\_INS\_123
CL\_INS\_123
CL\_INS\_123
CL\_INS\_123
CL\_INS\_123
CL\_INS\_123
CL\_INS\_123
CL\_INS\_123
CL\_INS\_123
CL\_INS\_123
CL\_INS\_123
CL\_INS\_123
CL\_INS\_123
CL\_INS\_123
CL\_INS\_123
CL\_INS\_123
CL\_INS\_123
CL\_INS\_123
CL\_INS\_123
CL\_INS\_123
CL\_INS\_123
CL\_INS\_123
CL\_INS\_123
CL\_INS\_123
CL\_INS\_123
CL\_INS\_123
CL\_INS\_123
CL\_INS\_123
CL\_INS\_123
CL\_INS\_149
CL\_INS\_149
CL\_INS\_149
CL\_INS\_149
CL\_INS\_123
CL\_INS\_123
CL\_INS\_123
CL\_INS\_123
CL\_INS\_123
CL\_INS\_123
CL\_INS\_123
CL\_INS\_123
CL\_INS\_123
CL\_INS\_123
CL\_INS\_123
CL\_INS\_123
CL\_INS\_123
CL\_INS\_123
CL\_INS\_30
CL\_INS\_70
CL\_INS\_123
CL\_INS\_123
CL\_INS\_123
CL\_INS\_123
CL\_INS\_123
CL\_INS\_123
CL\_INS\_123
CL\_INS\_123
CL\_INS\_123
CL\_INS\_123
CL\_INS\_123
CL\_INS\_123
CL\_INS\_123
CL\_INS\_123
CL\_INS\_123
CL\_INS\_123
CL\_INS\_123
CL\_INS\_123
CL\_INS\_123
CL\_INS\_123
CL\_INS\_123
CL\_INS\_123
CL\_INS\_123
CL\_INS\_123
CL\_INS\_123
CL\_INS\_123
CL\_INS\_123
CL\_INS\_123
CL\_INS\_123
CL\_INS\_123
CL\_INS\_123
CL\_INS\_123
CL\_INS\_123
CL\_INS\_123
CL\_INS\_123
CL\_INS\_123
CL\_INS\_123
CL\_INS\_123
CL\_INS\_123
CL\_INS\_123
CL\_INS\_123
CL\_INS\_123
CL\_INS\_123
CL\_INS\_123
CL\_INS\_123
CL\_INS\_123
CL\_INS\_123
CL\_INS\_123
CL\_INS\_123
CL\_INS\_123
CL\_INS\_123
CL\_INS\_123
CL\_INS\_123
CL\_INS\_123
CL\_INS\_123
CL\_INS\_123
CL\_INS\_123
CL\_INS\_123
CL\_INS\_123
CL\_INS\_123
CL\_INS\_123
CL\_INS\_123
CL\_INS\_123
CL\_INS\_123
CL\_INS\_123
CL\_INS\_123
CL\_INS\_123
CL\_INS\_123
CL\_INS\_123
CL\_INS\_123
CL\_INS\_123
CL\_INS\_123
CL\_INS\_123
CL\_INS\_123
CL\_INS\_123
CL\_INS\_123
CL\_INS\_123
CL\_INS\_123
CL\_INS\_123
CL\_INS\_123
CL\_INS\_123
CL\_INS\_123
CL\_INS\_123
CL\_INS\_123
CL\_INS\_123
CL\_INS\_123
CL\_INS\_123
CL\_INS\_123
CL\_INS\_123
CL\_INS\_123
CL\_INS\_123
CL\_INS\_123
CL\_INS\_123
CL\_INS\_123
CL\_INS\_123
CL\_INS\_123
CL\_INS\_123
CL\_INS\_123
CL\_INS\_123
CL\_INS\_123
CL\_INS\_123
CL\_INS\_123
CL\_INS\_70
CL\_INS\_70
CL\_INS\_123
CL\_INS\_123
CL\_INS\_123
CL\_INS\_123
CL\_INS\_123
CL\_INS\_382
CL\_INS\_123
CL\_INS\_123
CL\_INS\_123
CL\_INS\_123
CL\_INS\_123
CL\_INS\_382
CL\_INS\_123
CL\_INS\_123
CL\_INS\_123
CL\_INS\_382
CL\_INS\_123
CL\_INS\_123
CL\_INS\_123
CL\_INS\_123
CL\_INS\_123
CL\_INS\_123
CL\_INS\_123
CL\_INS\_123
CL\_INS\_123
CL\_INS\_123
CL\_INS\_123
CL\_INS\_123
CL\_INS\_123
CL\_INS\_123
CL\_INS\_123
CL\_INS\_123
CL\_INS\_123
CL\_INS\_123
CL\_INS\_123
CL\_INS\_123
CL\_INS\_123
CL\_INS\_123
CL\_INS\_123
CL\_INS\_123
CL\_INS\_123
CL\_INS\_123
CL\_INS\_123
CL\_INS\_123
CL\_INS\_123
CL\_INS\_123
CL\_INS\_123
CL\_INS\_123
CL\_INS\_123
CL\_INS\_123
CL\_INS\_123
CL\_INS\_123
CL\_INS\_123
CL\_INS\_123
CL\_INS\_123
CL\_INS\_123
CL\_INS\_123
CL\_INS\_123
CL\_INS\_123
CL\_INS\_123
CL\_INS\_123
CL\_INS\_123
CL\_INS\_123
CL\_INS\_123
CL\_INS\_123
CL\_INS\_123
CL\_INS\_123
CL\_INS\_123
CL\_INS\_123
CL\_INS\_123
CL\_INS\_123
CL\_INS\_123
CL\_INS\_123
CL\_INS\_123
CL\_INS\_123
CL\_INS\_123
CL\_INS\_123
CL\_INS\_123
CL\_INS\_123
CL\_INS\_123
CL\_INS\_123
CL\_INS\_123
CL\_INS\_123
CL\_INS\_123
CL\_INS\_123
CL\_INS\_123
CL\_INS\_123
CL\_INS\_123
CL\_INS\_123
CL\_INS\_123
CL\_INS\_123
CL\_INS\_123
CL\_INS\_123
CL\_INS\_123
CL\_INS\_123
CL\_INS\_123
CL\_INS\_123
CL\_INS\_123
CL\_INS\_123
CL\_INS\_123
CL\_INS\_123
CL\_INS\_123
CL\_INS\_123
CL\_INS\_123
CL\_INS\_123
CL\_INS\_123
CL\_INS\_123
CL\_INS\_123
CL\_INS\_123
CL\_INS\_123
CL\_INS\_123
CL\_INS\_123
CL\_INS\_123
CL\_INS\_123
CL\_INS\_123
CL\_INS\_123
CL\_INS\_123
CL\_INS\_123
CL\_INS\_123
CL\_INS\_123
CL\_INS\_123
CL\_INS\_123
CL\_INS\_123
CL\_INS\_123
CL\_INS\_123
CL\_INS\_123
CL\_INS\_123
CL\_INS\_123
CL\_INS\_123
CL\_INS\_123
CL\_INS\_123
CL\_INS\_123
CL\_INS\_123
CL\_INS\_123
CL\_INS\_123
CL\_INS\_123
CL\_INS\_123
CL\_INS\_123
CL\_INS\_123
CL\_INS\_123
CL\_INS\_123
CL\_INS\_123
CL\_INS\_123
CL\_INS\_123
CL\_INS\_123
CL\_INS\_123
CL\_INS\_123
CL\_INS\_123
CL\_INS\_123
CL\_INS\_123
CL\_INS\_123
CL\_INS\_123
CL\_INS\_123
CL\_INS\_123
CL\_INS\_123
CL\_INS\_123
CL\_INS\_123
CL\_INS\_123
CL\_INS\_123
CL\_INS\_123
CL\_INS\_123
CL\_INS\_123
CL\_INS\_123
CL\_INS\_123
CL\_INS\_123
CL\_INS\_123
CL\_INS\_123
CL\_INS\_123
CL\_INS\_123
CL\_INS\_123
CL\_INS\_123
CL\_INS\_123
CL\_INS\_123
CL\_INS\_123
CL\_INS\_123
CL\_INS\_123
CL\_INS\_123
CL\_INS\_123
CL\_INS\_123
CL\_INS\_123
CL\_INS\_123
CL\_INS\_123
CL\_INS\_123
CL\_INS\_123
CL\_INS\_123
CL\_INS\_123
CL\_INS\_123
CL\_INS\_123
CL\_INS\_123
CL\_INS\_123
CL\_INS\_123
CL\_INS\_123
CL\_INS\_123
CL\_INS\_123
CL\_INS\_123
CL\_INS\_123
CL\_INS\_123
CL\_INS\_123
CL\_INS\_123
CL\_INS\_123
CL\_INS\_123
CL\_INS\_123
CL\_INS\_123
CL\_INS\_123
CL\_INS\_123
CL\_INS\_123
CL\_INS\_123
CL\_INS\_123
CL\_INS\_123
CL\_INS\_123
CL\_INS\_123
CL\_INS\_123
CL\_INS\_123
CL\_INS\_123
CL\_INS\_123
CL\_INS\_123
CL\_INS\_123
CL\_INS\_123
CL\_INS\_123
CL\_INS\_123
CL\_INS\_123
CL\_INS\_123
CL\_INS\_123
CL\_INS\_123
CL\_INS\_123
CL\_INS\_123
CL\_INS\_123
CL\_INS\_123
CL\_INS\_123
CL\_INS\_123
CL\_INS\_123
CL\_INS\_123
CL\_INS\_123
CL\_INS\_123
CL\_INS\_123
CL\_INS\_123
CL\_INS\_123
CL\_INS\_123
CL\_INS\_123
CL\_INS\_123
CL\_INS\_123
CL\_INS\_123
CL\_INS\_123
CL\_INS\_123
CL\_INS\_123
CL\_INS\_123
CL\_INS\_123
CL\_INS\_123
CL\_INS\_123
CL\_INS\_123
CL\_INS\_123
CL\_INS\_123
CL\_INS\_123
CL\_INS\_123
CL\_INS\_123
CL\_INS\_123
CL\_INS\_123
CL\_INS\_123
CL\_INS\_123
CL\_INS\_123
CL\_INS\_123
CL\_INS\_123
CL\_INS\_123
CL\_INS\_123
CL\_INS\_123
CL\_INS\_123
CL\_INS\_123
CL\_INS\_123
CL\_INS\_123
CL\_INS\_123
CL\_INS\_123
CL\_INS\_123
CL\_INS\_123
CL\_INS\_123
CL\_INS\_123
CL\_INS\_123
CL\_INS\_123
CL\_INS\_123
CL\_INS\_123
CL\_INS\_123
CL\_INS\_123
CL\_INS\_123
CL\_INS\_123
CL\_INS\_382
CL\_INS\_149
CL\_INS\_123
CL\_INS\_123
CL\_INS\_123
CL\_INS\_123
CL\_INS\_123
CL\_INS\_123
CL\_INS\_123
CL\_INS\_123
CL\_INS\_123
CL\_INS\_123
CL\_INS\_123
CL\_INS\_123
CL\_INS\_123
CL\_INS\_123
CL\_INS\_123
CL\_INS\_123
CL\_INS\_123
CL\_INS\_123
CL\_INS\_123
CL\_INS\_123
CL\_INS\_123
CL\_INS\_123
CL\_INS\_123
CL\_INS\_123
CL\_INS\_123
CL\_INS\_123
CL\_INS\_123
CL\_INS\_123
CL\_INS\_123
CL\_INS\_123
CL\_INS\_123
CL\_INS\_123
CL\_INS\_123
CL\_INS\_123
CL\_INS\_123
CL\_INS\_123
CL\_INS\_123
CL\_INS\_123
CL\_INS\_123
CL\_INS\_123
CL\_INS\_123
CL\_INS\_123
CL\_INS\_123
CL\_INS\_123
CL\_INS\_123
CL\_INS\_123
CL\_INS\_123
CL\_INS\_123
CL\_INS\_123
CL\_INS\_123
CL\_INS\_123
CL\_INS\_123
CL\_INS\_123
CL\_INS\_123
CL\_INS\_123
CL\_INS\_123
CL\_INS\_123
CL\_INS\_123
CL\_INS\_123
CL\_INS\_123
CL\_INS\_123
CL\_INS\_123
CL\_INS\_123
Cluster ID


CL\_36607
CL\_10804
CL\_32052
CL\_32051
CL\_28976
CL\_37077
CL\_37078
CL\_1456
CL\_1457
CL\_31160
CL\_31159
CL\_1458
CL\_1459
CL\_1460
CL\_16999
CL\_22138
CL\_22139
CL\_22140
CL\_24105
CL\_24104
CL\_24103
CL\_24102
CL\_15177
CL\_15176
CL\_5300
CL\_10715
CL\_5298
CL\_5297
CL\_10392
CL\_33526
CL\_33525
CL\_14795
CL\_14796
CL\_35189
CL\_7485
CL\_7486
CL\_7604
CL\_7487
CL\_35188
CL\_9594
CL\_9593
CL\_9592
CL\_24143
CL\_24144
CL\_24145
CL\_24146
CL\_22272
CL\_26601
CL\_26602
CL\_26603
CL\_26604
CL\_26605
CL\_5238
CL\_5682
CL\_5314
CL\_5313
CL\_5312
CL\_5311
CL\_10148
CL\_10149
CL\_33592
CL\_33591
CL\_33590
CL\_10150
CL\_4316
CL\_4315
CL\_4314
CL\_9898
CL\_10151
CL\_5305
CL\_5304
CL\_5303
CL\_4504
CL\_16648
CL\_20420
CL\_16315
CL\_16316
CL\_16317
CL\_16318
CL\_17955
CL\_10142
CL\_11534
CL\_35104
CL\_35103
CL\_35102
CL\_35101
CL\_35100
CL\_35099
CL\_9178
CL\_9179
CL\_9180
CL\_9181
CL\_26938
CL\_26937
CL\_26936
CL\_10532
CL\_10533
CL\_10534
CL\_10535
CL\_10536
CL\_10537
CL\_10538
CL\_10539
CL\_10540
CL\_10541
CL\_10542
CL\_34300
CL\_36891
CL\_17660
CL\_19808
CL\_19809
CL\_19810
CL\_19811
CL\_19812
CL\_19813
CL\_30061
CL\_30060
CL\_30059
CL\_30058
CL\_30057
CL\_30056
CL\_30055
CL\_30054
CL\_9893
CL\_9892
CL\_9891
CL\_9890
CL\_30053
CL\_30052
CL\_30051
CL\_7800
CL\_23334
CL\_23333
CL\_26785
CL\_26786
CL\_26787
CL\_26788
CL\_26789
CL\_25130
CL\_25131
CL\_25132
CL\_25133
CL\_25134
CL\_25135
CL\_10143
CL\_9182
CL\_9183
CL\_10144
CL\_10145
CL\_24418
CL\_6590
CL\_8157
CL\_26935
CL\_8156
CL\_5323
CL\_5322
CL\_5321
CL\_5320
CL\_5319
CL\_5318
CL\_5317
CL\_5316
CL\_5315
CL\_8843
CL\_20718
CL\_8842
CL\_13922
CL\_13921
CL\_13920
CL\_8841
CL\_8840
CL\_8839
CL\_8838
CL\_8909
CL\_9787
CL\_9786
CL\_9785
CL\_9782
CL\_8837
CL\_8836
CL\_5310
CL\_5309
CL\_5308
CL\_30050
CL\_9784
CL\_27591
CL\_27592
CL\_9783
CL\_29456
CL\_5826
CL\_5827
CL\_5828
CL\_5829
CL\_14270
CL\_14271
CL\_14272
CL\_14273
CL\_5830
CL\_5831
CL\_5832
CL\_29457
CL\_29458
CL\_29459
CL\_29460
CL\_29461
CL\_29462
CL\_29463
CL\_29464
CL\_29465
CL\_29466
CL\_29467
CL\_29468
CL\_29469
CL\_29470
CL\_29471
CL\_29472
CL\_29473
CL\_29474
CL\_29475
CL\_29476
CL\_29477
CL\_29478
CL\_29479
CL\_29480
CL\_29481
CL\_29482
CL\_29483
CL\_29484
CL\_29485
CL\_29486
CL\_29487
CL\_29488
CL\_29489
CL\_29490
CL\_29491
CL\_29492
CL\_29493
CL\_29494
CL\_29495
CL\_8159
CL\_21145
CL\_8158
CL\_16971
CL\_16972
CL\_21144
CL\_21143
CL\_5307
CL\_5306
CL\_8835
CL\_4995
CL\_32050
CL\_8834
CL\_34090
CL\_8833
CL\_24417
CL\_24416
CL\_24415
CL\_24414
CL\_24413
CL\_24412
CL\_14797
CL\_20421
CL\_10146
CL\_10147
CL\_13919
CL\_13918
CL\_10152
CL\_24101
CL\_10153
CL\_9591
CL\_27593
CL\_21142
CL\_21141
CL\_21140
CL\_21139
CL\_21138
CL\_21137
CL\_21136
CL\_21135
CL\_21134
CL\_21133
CL\_21132
CL\_21131
CL\_21130
CL\_21129
CL\_21128
CL\_21127
CL\_21126
CL\_21125
CL\_21124
CL\_21123
CL\_21122
CL\_21121
CL\_21120
CL\_21119
CL\_21118
CL\_21117
CL\_21116
CL\_21115
CL\_21114
CL\_21113
CL\_21112
CL\_21111
CL\_21110
CL\_21109
CL\_21108
CL\_21107
CL\_21106
CL\_21105
CL\_21104
CL\_21103
CL\_21102
CL\_21101
CL\_21100
CL\_21099
CL\_21098
CL\_21097
CL\_21096
CL\_21095
CL\_21094
CL\_21093
CL\_21092
CL\_21091
CL\_21090
CL\_21089
CL\_21088
CL\_21087
CL\_12010
CL\_7799
CL\_13653
CL\_27545
CL\_13654
CL\_13655
CL\_7798
CL\_37255
CL\_7096
CL\_7095
CL\_12011
CL\_12012
CL\_12013
CL\_12014
CL\_12015
CL\_12016
CL\_12017
CL\_12018
CL\_12019
CL\_12020
CL\_25136
CL\_22612
CL\_12021
CL\_27594
CL\_27595
CL\_27596
CL\_27597
CL\_27598
CL\_27599
CL\_27600
CL\_27601
CL\_22611
CL\_25917
CL\_25918
CL\_25919
CL\_25920
CL\_25921
CL\_25922
CL\_25923
CL\_25924
CL\_25925
CL\_25926
CL\_25927
CL\_25928
CL\_22610
CL\_22609
CL\_22608
CL\_22607
CL\_22606
CL\_22605
CL\_22604
CL\_22603
CL\_35812
CL\_35813
CL\_35814
CL\_35815
CL\_35816
CL\_35817
CL\_35818
CL\_35819
CL\_35820
CL\_35821
CL\_35822
CL\_35823
CL\_35824
CL\_35825
CL\_35826
CL\_35827
CL\_35828
CL\_35829
CL\_35830
CL\_35831
CL\_35832
CL\_22602
CL\_22601
CL\_22600
CL\_22599
CL\_35833
CL\_12022
CL\_12023
CL\_22598
CL\_22597
CL\_25137
CL\_25138
CL\_25139
CL\_25140
CL\_25141
CL\_25142
CL\_25143
CL\_25144
CL\_25145
CL\_25146
CL\_26790
CL\_25147
CL\_25148
CL\_23792
CL\_24382
CL\_24383
CL\_5629
CL\_23795
CL\_23796
CL\_5727
CL\_24384
CL\_23797
CL\_10677
CL\_10678
CL\_10679
CL\_10680
CL\_10682
CL\_23798
CL\_22379
CL\_23799
CL\_24395
CL\_27273
CL\_16006
CL\_11994
CL\_23800
CL\_25454
CL\_25453
CL\_23803
CL\_24396
CL\_36195
CL\_10704
CL\_36194
CL\_30646
CL\_10703
CL\_23809
CL\_23810
CL\_23811
CL\_23812
CL\_23813
CL\_37209
CL\_37210
CL\_10328
CL\_10329
CL\_11778
CL\_11779
CL\_15016
CL\_36192
CL\_6957
CL\_21850
CL\_23816
CL\_23817
CL\_23818
CL\_23819
CL\_23820
CL\_23821
CL\_23822
CL\_23823
CL\_23824
CL\_23825
CL\_23826
CL\_23827
CL\_23828
CL\_23829
CL\_24400
CL\_23830
CL\_23831
CL\_23832
CL\_23833
CL\_23834
CL\_23835
CL\_23836
CL\_6911
CL\_5721
CL\_5722
CL\_23837
CL\_23782
CL\_23783
CL\_23784
CL\_23785
CL\_23786
CL\_23787
CL\_24403
CL\_23788
CL\_24404
CL\_23789
CL\_23790
CL\_23791
