## Supplementary material for "A novel method for integrating genomic and Tn-Seq data to identify common *in vivo* fitness mechanisms across multiple bacterial species": S1 Dataset: CL_INS_124.html

Legend

 Mobile +extrachromosomalelementfunctions
 Regulatoryfunctions
 Hypothetical
 DNA Metabolism
 AntibioticResistance
 All Fitness Genes
 Proteinsynthesis/fate
 Other
 EnergyMetabolism
 Transport +binding proteins
 All VFDB Genes

FULL


WINDOWSVGPNG

Trim RowsRemove SingletonsSave Fasta

CL\_1455


CL\_1455


CL\_4503


CL\_1455


CL\_1455


CL\_1455


CL\_4503


CL\_4503


CL\_1455


CL\_1455


CL\_1455


CL\_1455


CL\_1455


CL\_4503


CL\_1455


CL\_1455


CL\_1455


CL\_1455


CL\_4503


CL\_1455


CL\_1455


CL\_1454


CL\_1455


CL\_1455


CL\_4503


CL\_1455


CL\_1455


CL\_4502


CL\_1455


CL\_1455


CL\_1455


CL\_1455


CL\_1455


CL\_1455


CL\_1455


CL\_4503


CL\_1455


CL\_1373


CL\_1455


CL\_1455


CL\_1455


CL\_1480


CL\_1440


CL\_1455


CL\_1455


CL\_1455


CL\_1453


CL\_1455


CL\_1455


CL\_1453

HighlightSelectShow Genomes


113

CL\_1462


24

CL\_1462


5

CL\_1462


4

CL\_1462


3

CL\_1462


3

CL\_1462


3

CL\_1462


3

CL\_1462


2

CL\_1462


2

CL\_1462


2

CL\_1462


2

CL\_1463


2

CL\_1462


2

CL\_1462


1

CL\_1462


1

CL\_1462


1

CL\_1462


1

CL\_1462


1

CL\_1462


1

CL\_1462


1

CL\_1462


1

CL\_1462


1

CL\_1462


1

CL\_1479


1

CL\_1462


1

CL\_1462


1

CL\_1462


1

CL\_1462


1

CL\_1462


1

CL\_1462


1

CL\_1462


1

CL\_1462


1

CL\_1462


1

CL\_1462


1

CL\_1462


1

CL\_1462


1

CL\_1462


1

CL\_1462


1

CL\_1462


1

CL\_1472


1

CL\_1462


1

CL\_1462


1

CL\_1462


1

CL\_1462


1

CL\_1465


1

CL\_1405


1

CL\_1462


1

CL\_1462


1

CL\_1462


1

CL\_1462

fGI ID


CL\_INS\_110
CL\_INS\_124
CL\_INS\_124
CL\_INS\_124
CL\_INS\_124
CL\_INS\_123
CL\_INS\_123
CL\_INS\_123
CL\_INS\_124
CL\_INS\_123
CL\_INS\_247
CL\_INS\_382
CL\_INS\_123
CL\_INS\_247
CL\_INS\_123
CL\_INS\_123
CL\_INS\_385
CL\_INS\_286
CL\_INS\_382
CL\_INS\_382
CL\_INS\_382
CL\_INS\_382
CL\_INS\_382
CL\_INS\_124
CL\_INS\_124
CL\_INS\_124
CL\_INS\_385
CL\_INS\_123
CL\_INS\_123
CL\_INS\_123
CL\_INS\_123
CL\_INS\_123
CL\_INS\_123
CL\_INS\_123
CL\_INS\_123
CL\_INS\_122
CL\_INS\_122
CL\_INS\_122
CL\_INS\_122
CL\_INS\_122
CL\_INS\_123
CL\_INS\_123
CL\_INS\_123
CL\_INS\_122
CL\_INS\_122
CL\_INS\_122
CL\_INS\_123
CL\_INS\_124
CL\_INS\_124
CL\_INS\_126
CL\_INS\_126
CL\_INS\_126
CL\_INS\_126
CL\_INS\_126
CL\_INS\_207
CL\_INS\_207
CL\_INS\_207
CL\_INS\_237
CL\_INS\_237
CL\_INS\_247
CL\_INS\_123
CL\_INS\_385
CL\_INS\_123
CL\_INS\_359
CL\_INS\_247
CL\_INS\_359
CL\_INS\_247
CL\_INS\_247
CL\_INS\_369
CL\_INS\_369
CL\_INS\_71
CL\_INS\_247
CL\_INS\_382
CL\_INS\_382
CL\_INS\_382
CL\_INS\_382
CL\_INS\_247
CL\_INS\_382
CL\_INS\_382
CL\_INS\_382
CL\_INS\_124
CL\_INS\_124
CL\_INS\_1
CL\_INS\_382
CL\_INS\_382
CL\_INS\_124
CL\_INS\_124
CL\_INS\_124
CL\_INS\_124
CL\_INS\_124
CL\_INS\_124
CL\_INS\_124
CL\_INS\_124
CL\_INS\_124
CL\_INS\_286
CL\_INS\_286
CL\_INS\_286
CL\_INS\_286
CL\_INS\_286
CL\_INS\_124
CL\_INS\_124
CL\_INS\_124
CL\_INS\_124
CL\_INS\_123
CL\_INS\_123
CL\_INS\_123
CL\_INS\_123
CL\_INS\_124
CL\_INS\_124
CL\_INS\_124
CL\_INS\_92
CL\_INS\_123
CL\_INS\_123
CL\_INS\_123
CL\_INS\_123
CL\_INS\_237
CL\_INS\_237
CL\_INS\_237
CL\_INS\_308
CL\_INS\_15
CL\_INS\_15
CL\_INS\_15
CL\_INS\_15
CL\_INS\_15
CL\_INS\_15
CL\_INS\_15
CL\_INS\_124
CL\_INS\_15
CL\_INS\_15
CL\_INS\_15
CL\_INS\_15
CL\_INS\_124
CL\_INS\_124
CL\_INS\_124
CL\_INS\_237
CL\_INS\_237
CL\_INS\_159
CL\_INS\_124
CL\_INS\_124
CL\_INS\_124
CL\_INS\_124
CL\_INS\_124
CL\_INS\_124
CL\_INS\_124
CL\_INS\_124
CL\_INS\_124
CL\_INS\_124
CL\_INS\_124
CL\_INS\_124
CL\_INS\_124
CL\_INS\_124
CL\_INS\_124
CL\_INS\_124
CL\_INS\_124
CL\_INS\_124
CL\_INS\_124
CL\_INS\_124
CL\_INS\_237
CL\_INS\_237
CL\_INS\_55
CL\_INS\_124
CL\_INS\_382
CL\_INS\_382
CL\_INS\_124
CL\_INS\_359
CL\_INS\_124
CL\_INS\_124
CL\_INS\_124
CL\_INS\_124
CL\_INS\_10
CL\_INS\_149
CL\_INS\_136
CL\_INS\_136
CL\_INS\_136
CL\_INS\_382
CL\_INS\_382
CL\_INS\_382
CL\_INS\_382
CL\_INS\_382
CL\_INS\_382
CL\_INS\_382
CL\_INS\_382
CL\_INS\_382
CL\_INS\_382
CL\_INS\_60
CL\_INS\_60
CL\_INS\_60
CL\_INS\_295
CL\_INS\_124
CL\_INS\_124
CL\_INS\_124
CL\_INS\_124
CL\_INS\_124
CL\_INS\_124
CL\_INS\_124
CL\_INS\_124
CL\_INS\_124
CL\_INS\_124
CL\_INS\_124
CL\_INS\_124
CL\_INS\_124
CL\_INS\_124
CL\_INS\_124
CL\_INS\_124
CL\_INS\_124
CL\_INS\_189
CL\_INS\_189
CL\_INS\_124
CL\_INS\_247
CL\_INS\_273
CL\_INS\_124
CL\_INS\_273
CL\_INS\_273
CL\_INS\_189
CL\_INS\_189
CL\_INS\_273
CL\_INS\_273
CL\_INS\_189
CL\_INS\_189
CL\_INS\_273
CL\_INS\_273
CL\_INS\_286
CL\_INS\_273
CL\_INS\_286
CL\_INS\_286
CL\_INS\_189
CL\_INS\_273
CL\_INS\_189
CL\_INS\_189
CL\_INS\_189
CL\_INS\_273
CL\_INS\_189
CL\_INS\_189
CL\_INS\_189
CL\_INS\_273
CL\_INS\_273
CL\_INS\_382
CL\_INS\_124
CL\_INS\_124
CL\_INS\_124
CL\_INS\_124
CL\_INS\_124
CL\_INS\_124
CL\_INS\_124
CL\_INS\_20
CL\_INS\_124
CL\_INS\_124
CL\_INS\_123
CL\_INS\_123
CL\_INS\_123
CL\_INS\_382
CL\_INS\_123
CL\_INS\_382
CL\_INS\_123
CL\_INS\_123
CL\_INS\_124
CL\_INS\_123
CL\_INS\_247
CL\_INS\_123
CL\_INS\_123
CL\_INS\_123
CL\_INS\_123
CL\_INS\_123
CL\_INS\_123
CL\_INS\_123
CL\_INS\_123
CL\_INS\_123
CL\_INS\_123
CL\_INS\_123
CL\_INS\_123
CL\_INS\_123
CL\_INS\_123
CL\_INS\_123
CL\_INS\_123
CL\_INS\_123
CL\_INS\_123
CL\_INS\_123
CL\_INS\_123
CL\_INS\_124
Cluster ID


CL\_22932
CL\_28023
CL\_28022
CL\_36375
CL\_20924
CL\_20421
CL\_9591
CL\_9592
CL\_16849
CL\_10142
CL\_5302
CL\_5301
CL\_5300
CL\_5299
CL\_5298
CL\_5297
CL\_5296
CL\_5295
CL\_5294
CL\_5293
CL\_5292
CL\_5291
CL\_5290
CL\_5289
CL\_5288
CL\_5287
CL\_5286
CL\_12012
CL\_12013
CL\_12014
CL\_12015
CL\_12016
CL\_12017
CL\_12018
CL\_12019
CL\_20994
CL\_20995
CL\_20996
CL\_20997
CL\_20998
CL\_12020
CL\_12021
CL\_12022
CL\_20246
CL\_20245
CL\_20999
CL\_1456
CL\_36523
CL\_10340
CL\_14444
CL\_4506
CL\_4507
CL\_4508
CL\_4509
CL\_5509
CL\_5510
CL\_5511
CL\_5512
CL\_5513
CL\_5514
CL\_15177
CL\_20144
CL\_15176
CL\_15069
CL\_11649
CL\_11650
CL\_11652
CL\_11653
CL\_11654
CL\_11655
CL\_5533
CL\_11981
CL\_10665
CL\_10666
CL\_10667
CL\_9691
CL\_9690
CL\_9689
CL\_9688
CL\_9687
CL\_9921
CL\_9920
CL\_9919
CL\_9918
CL\_9917
CL\_9916
CL\_9915
CL\_9914
CL\_9913
CL\_9912
CL\_9911
CL\_9910
CL\_9909
CL\_9908
CL\_9907
CL\_9906
CL\_9905
CL\_9904
CL\_9903
CL\_9902
CL\_9901
CL\_9900
CL\_9899
CL\_9898
CL\_4314
CL\_4315
CL\_4316
CL\_9897
CL\_9896
CL\_9895
CL\_9894
CL\_9893
CL\_9892
CL\_9891
CL\_9890
CL\_9889
CL\_9888
CL\_9887
CL\_9886
CL\_9885
CL\_9884
CL\_9883
CL\_9882
CL\_9881
CL\_9880
CL\_9879
CL\_9878
CL\_9877
CL\_9876
CL\_9875
CL\_9874
CL\_9873
CL\_9872
CL\_6275
CL\_6415
CL\_6414
CL\_9871
CL\_9870
CL\_9869
CL\_9868
CL\_9867
CL\_9866
CL\_9865
CL\_9864
CL\_9863
CL\_9862
CL\_9861
CL\_9860
CL\_9859
CL\_9858
CL\_9857
CL\_9856
CL\_9855
CL\_9854
CL\_9853
CL\_9852
CL\_9851
CL\_7986
CL\_7987
CL\_5043
CL\_9850
CL\_5001
CL\_5000
CL\_9849
CL\_9848
CL\_9847
CL\_9846
CL\_9845
CL\_9844
CL\_4764
CL\_4601
CL\_1320
CL\_1319
CL\_1318
CL\_4400
CL\_5361
CL\_5362
CL\_5363
CL\_5364
CL\_7618
CL\_5365
CL\_5366
CL\_5367
CL\_5368
CL\_9843
CL\_9842
CL\_9841
CL\_9840
CL\_6280
CL\_9839
CL\_9838
CL\_9837
CL\_9836
CL\_9835
CL\_9834
CL\_9833
CL\_9832
CL\_9831
CL\_9830
CL\_9829
CL\_9828
CL\_9827
CL\_9826
CL\_9825
CL\_9824
CL\_9823
CL\_9822
CL\_9821
CL\_5160
CL\_9820
CL\_9819
CL\_9818
CL\_9817
CL\_9816
CL\_9815
CL\_9814
CL\_9813
CL\_9812
CL\_9811
CL\_6165
CL\_6306
CL\_6307
CL\_6169
CL\_9810
CL\_9809
CL\_9808
CL\_9807
CL\_9806
CL\_9805
CL\_9804
CL\_9803
CL\_9802
CL\_9801
CL\_9800
CL\_9799
CL\_9798
CL\_8909
CL\_9797
CL\_9796
CL\_9795
CL\_9794
CL\_9793
CL\_9792
CL\_9791
CL\_6201
CL\_9790
CL\_9789
CL\_5304
CL\_8157
CL\_8156
CL\_8843
CL\_8842
CL\_8841
CL\_8840
CL\_8839
CL\_9788
CL\_9787
CL\_5236
CL\_9786
CL\_9785
CL\_9784
CL\_9783
CL\_9782
CL\_8837
CL\_8836
CL\_5310
CL\_5309
CL\_5308
CL\_5307
CL\_8835
CL\_8834
CL\_8833
CL\_1457
CL\_31160
CL\_31159
CL\_1458
CL\_1459
CL\_1460
CL\_1461
