## Supplementary material for "A novel method for integrating genomic and Tn-Seq data to identify common *in vivo* fitness mechanisms across multiple bacterial species": S1 Dataset: CL_INS_126.html

Legend

 Mobile +extrachromosomalelementfunctions
 Regulatoryfunctions
 Hypothetical
 All EssentialGenes
 All Fitness Genes
 Other
 Transport +binding proteins
 All VFDB Genes

FULL


WINDOWSVGPNG

Trim RowsRemove SingletonsSave Fasta

CL\_1478


CL\_1478


CL\_1478


CL\_1478


CL\_1478


CL\_1478


CL\_1478


CL\_1478


CL\_1478


CL\_1478


CL\_1478


CL\_1478


CL\_1478


CL\_1478


CL\_1478


CL\_1455


CL\_1478


CL\_1478


CL\_1478


CL\_1478


CL\_1478


CL\_1478


CL\_1483


CL\_1478

HighlightSelectShow Genomes


122

CL\_1479


116

CL\_1479


7

CL\_1479


5

CL\_1479


4

CL\_1479


4

CL\_1479


1

CL\_1480


1

CL\_1479


1

CL\_1281


1

CL\_1452


1

CL\_1479


1

CL\_1481


1

CL\_1479


1

CL\_1479


1

CL\_1281


1

CL\_1479


1

CL\_1481


1

CL\_1479


1

CL\_1479


1

CL\_1479


1

CL\_1479


1

CL\_1480


1

CL\_1479


1

CL\_1479

fGI ID


CL\_INS\_126
CL\_INS\_126
CL\_INS\_126
CL\_INS\_126
CL\_INS\_126
CL\_INS\_126
CL\_INS\_126
CL\_INS\_126
CL\_INS\_126
CL\_INS\_126
CL\_INS\_126
CL\_INS\_126
Cluster ID


CL\_14444
CL\_17466
CL\_4505
CL\_4506
CL\_4507
CL\_4508
CL\_30049
CL\_4509
CL\_18922
CL\_18921
CL\_18920
CL\_4510
