## Supplementary material for "A novel method for integrating genomic and Tn-Seq data to identify common *in vivo* fitness mechanisms across multiple bacterial species": S1 Dataset: CL_INS_132.html

Legend

 Mobile +extrachromosomalelementfunctions
 Hypothetical
 All EssentialGenes
 All Fitness Genes
 Other
 All VFDB Genes

FULL


WINDOWSVGPNG

Trim RowsRemove SingletonsSave Fasta

CL\_1550


CL\_1550


CL\_1550


CL\_1549


CL\_1550


CL\_1550


CL\_1550


CL\_1550


CL\_1550


CL\_1550


CL\_1550


CL\_1550


CL\_1550


CL\_1550


CL\_1550

HighlightSelectShow Genomes


141

CL\_1551


111

CL\_1551


8

CL\_1552


2

CL\_1551


2

CL\_1552


1

CL\_1552


1

CL\_1552


1

CL\_1552


1

CL\_1552


1

CL\_1551


1

CL\_1552


1

CL\_1552


1

CL\_1552


1

CL\_1552


1

CL\_1552

fGI ID


CL\_INS\_132
CL\_INS\_132
CL\_INS\_132
CL\_INS\_132
CL\_INS\_106
CL\_INS\_132
CL\_INS\_382
CL\_INS\_382
CL\_INS\_132
CL\_INS\_382
CL\_INS\_382
CL\_INS\_132
CL\_INS\_132
CL\_INS\_132
CL\_INS\_132
CL\_INS\_132
CL\_INS\_132
CL\_INS\_132
CL\_INS\_132
CL\_INS\_132
CL\_INS\_132
CL\_INS\_132
CL\_INS\_132
CL\_INS\_132
CL\_INS\_132
CL\_INS\_132
CL\_INS\_132
CL\_INS\_132
CL\_INS\_132
CL\_INS\_132
CL\_INS\_132
CL\_INS\_132
CL\_INS\_132
CL\_INS\_247
CL\_INS\_132
CL\_INS\_132
CL\_INS\_382
CL\_INS\_382
CL\_INS\_382
CL\_INS\_382
CL\_INS\_382
CL\_INS\_132
CL\_INS\_132
CL\_INS\_106
CL\_INS\_132
CL\_INS\_382
CL\_INS\_106
CL\_INS\_382
CL\_INS\_382
CL\_INS\_132
CL\_INS\_132
CL\_INS\_132
CL\_INS\_132
CL\_INS\_132
CL\_INS\_132
CL\_INS\_132
CL\_INS\_106
CL\_INS\_132
CL\_INS\_382
CL\_INS\_382
CL\_INS\_382
CL\_INS\_106
CL\_INS\_382
CL\_INS\_132
CL\_INS\_132
CL\_INS\_132
CL\_INS\_132
CL\_INS\_106
CL\_INS\_132
CL\_INS\_132
CL\_INS\_106
CL\_INS\_132
CL\_INS\_382
Cluster ID


CL\_34381
CL\_5273
CL\_27096
CL\_27063
CL\_13309
CL\_27062
CL\_13738
CL\_13737
CL\_31388
CL\_13642
CL\_13641
CL\_31387
CL\_27057
CL\_35632
CL\_20756
CL\_16132
CL\_10506
CL\_10505
CL\_32197
CL\_32198
CL\_27061
CL\_27060
CL\_27059
CL\_27058
CL\_10504
CL\_10503
CL\_10502
CL\_10501
CL\_10500
CL\_10499
CL\_10498
CL\_13102
CL\_10497
CL\_10496
CL\_32199
CL\_13100
CL\_8655
CL\_8114
CL\_8113
CL\_8112
CL\_13510
CL\_27056
CL\_32200
CL\_12139
CL\_32201
CL\_12137
CL\_12138
CL\_8654
CL\_8653
CL\_31386
CL\_31385
CL\_31384
CL\_8652
CL\_27055
CL\_27054
CL\_10495
CL\_8651
CL\_8650
CL\_13310
CL\_13512
CL\_12541
CL\_8649
CL\_23647
CL\_32202
CL\_31383
CL\_17068
CL\_17069
CL\_16469
CL\_13311
CL\_13312
CL\_16582
CL\_13099
CL\_8648
