## Supplementary material for "A novel method for integrating genomic and Tn-Seq data to identify common *in vivo* fitness mechanisms across multiple bacterial species": S1 Dataset: CL_INS_133.html

Legend

 Mobile +extrachromosomalelementfunctions
 Hypothetical
 All EssentialGenes
 All Fitness Genes
 Other
 All VFDB Genes

FULL


WINDOWSVGPNG

Trim RowsRemove SingletonsSave Fasta

CL\_1651


CL\_1651


CL\_1650


CL\_1651


CL\_1651


CL\_1651

HighlightSelectShow Genomes


247

CL\_1652


10

CL\_1652


1

CL\_1652


1

CL\_1652


1

CL\_1652


1

CL\_1652

fGI ID


CL\_INS\_133
CL\_INS\_133
CL\_INS\_133
CL\_INS\_133
CL\_INS\_133
CL\_INS\_133
Cluster ID


CL\_35642
CL\_13545
CL\_13546
CL\_12710
CL\_13547
CL\_13548
