## Supplementary material for "A novel method for integrating genomic and Tn-Seq data to identify common *in vivo* fitness mechanisms across multiple bacterial species": S1 Dataset: CL_INS_136.html

Legend

 Mobile +extrachromosomalelementfunctions
 Hypothetical
 DNA Metabolism
 AntibioticResistance
 All EssentialGenes
 All Fitness Genes
 Proteinsynthesis/fate
 Other
 Transport +binding proteins
 All VFDB Genes

FULL


WINDOWSVGPNG

Trim RowsRemove SingletonsSave Fasta

CL\_1668


CL\_1668


CL\_4427


CL\_1668


CL\_1668


CL\_1668


CL\_1668


CL\_1668


CL\_1668


CL\_1667


CL\_1668


CL\_4427


CL\_4487


CL\_1668

HighlightSelectShow Genomes


178

CL\_1669


66

CL\_1669


2

CL\_1669


1

CL\_1669


1

CL\_1669


1

CL\_1669


1

CL\_1300


1

CL\_4486


1

CL\_4487


1

CL\_1669


1

CL\_1671


1

CL\_1669


1

CL\_1669


1

CL\_4487

fGI ID


CL\_INS\_136
CL\_INS\_136
CL\_INS\_136
CL\_INS\_136
CL\_INS\_99
CL\_INS\_86
CL\_INS\_86
CL\_INS\_86
CL\_INS\_382
CL\_INS\_20
CL\_INS\_86
CL\_INS\_136
CL\_INS\_99
CL\_INS\_86
CL\_INS\_136
CL\_INS\_99
CL\_INS\_86
CL\_INS\_136
CL\_INS\_86
CL\_INS\_136
CL\_INS\_136
CL\_INS\_136
CL\_INS\_136
CL\_INS\_136
CL\_INS\_136
CL\_INS\_136
CL\_INS\_136
CL\_INS\_136
CL\_INS\_87
CL\_INS\_87
CL\_INS\_87
CL\_INS\_136
CL\_INS\_136
CL\_INS\_136
CL\_INS\_136
CL\_INS\_136
CL\_INS\_136
CL\_INS\_136
CL\_INS\_136
CL\_INS\_382
CL\_INS\_136
CL\_INS\_382
CL\_INS\_382
CL\_INS\_382
CL\_INS\_136
CL\_INS\_382
CL\_INS\_382
CL\_INS\_382
CL\_INS\_382
CL\_INS\_382
CL\_INS\_99
CL\_INS\_99
CL\_INS\_382
CL\_INS\_382
CL\_INS\_382
CL\_INS\_382
CL\_INS\_382
CL\_INS\_382
CL\_INS\_382
CL\_INS\_382
CL\_INS\_136
CL\_INS\_382
CL\_INS\_382
CL\_INS\_382
CL\_INS\_382
CL\_INS\_382
CL\_INS\_382
CL\_INS\_382
CL\_INS\_382
CL\_INS\_382
CL\_INS\_382
CL\_INS\_382
CL\_INS\_382
CL\_INS\_382
CL\_INS\_382
CL\_INS\_382
CL\_INS\_382
CL\_INS\_382
CL\_INS\_382
CL\_INS\_136
CL\_INS\_382
CL\_INS\_382
CL\_INS\_382
CL\_INS\_382
CL\_INS\_382
CL\_INS\_382
CL\_INS\_382
CL\_INS\_382
CL\_INS\_382
CL\_INS\_382
CL\_INS\_382
CL\_INS\_382
CL\_INS\_382
CL\_INS\_382
CL\_INS\_382
CL\_INS\_382
CL\_INS\_382
CL\_INS\_382
CL\_INS\_382
CL\_INS\_382
CL\_INS\_382
CL\_INS\_382
CL\_INS\_382
CL\_INS\_382
CL\_INS\_382
CL\_INS\_382
CL\_INS\_382
CL\_INS\_382
CL\_INS\_382
CL\_INS\_382
CL\_INS\_382
CL\_INS\_382
CL\_INS\_382
CL\_INS\_382
CL\_INS\_382
CL\_INS\_382
CL\_INS\_382
CL\_INS\_136
Cluster ID


CL\_14448
CL\_27993
CL\_33524
CL\_30621
CL\_10520
CL\_6782
CL\_10526
CL\_8585
CL\_4432
CL\_10474
CL\_7025
CL\_33392
CL\_8586
CL\_7024
CL\_6769
CL\_4431
CL\_4430
CL\_6746
CL\_7027
CL\_4484
CL\_1322
CL\_1321
CL\_4482
CL\_4481
CL\_4480
CL\_4479
CL\_4478
CL\_7498
CL\_4477
CL\_4476
CL\_4475
CL\_4474
CL\_4473
CL\_4472
CL\_4471
CL\_4470
CL\_1320
CL\_1319
CL\_1318
CL\_4469
CL\_10525
CL\_4465
CL\_4464
CL\_4463
CL\_20036
CL\_4459
CL\_1317
CL\_1316
CL\_1315
CL\_1314
CL\_1313
CL\_1312
CL\_1311
CL\_1309
CL\_1308
CL\_1307
CL\_1305
CL\_1304
CL\_1303
CL\_1302
CL\_13481
CL\_11913
CL\_4426
CL\_4425
CL\_4424
CL\_4423
CL\_4525
CL\_4421
CL\_4420
CL\_4419
CL\_4418
CL\_4417
CL\_4416
CL\_4415
CL\_4533
CL\_533
CL\_532
CL\_13474
CL\_4413
CL\_34063
CL\_528
CL\_527
CL\_526
CL\_12006
CL\_5340
CL\_6452
CL\_5341
CL\_5342
CL\_5343
CL\_521
CL\_520
CL\_519
CL\_518
CL\_517
CL\_516
CL\_515
CL\_514
CL\_513
CL\_512
CL\_511
CL\_509
CL\_4410
CL\_4409
CL\_4408
CL\_4407
CL\_6479
CL\_6483
CL\_4402
CL\_4401
CL\_508
CL\_4400
CL\_5361
CL\_5362
CL\_5799
CL\_4396
CL\_30622
CL\_30623
CL\_8048
