## Supplementary material for "A novel method for integrating genomic and Tn-Seq data to identify common *in vivo* fitness mechanisms across multiple bacterial species": S1 Dataset: CL_INS_137.html

Legend

 Mobile +extrachromosomalelementfunctions
 Hypothetical
 All Fitness Genes
 Other
 Transport +binding proteins
 All VFDB Genes

FULL


WINDOWSVGPNG

Trim RowsRemove SingletonsSave Fasta

CL\_1677


CL\_1677


CL\_1676


CL\_1677


Break


CL\_1677

HighlightSelectShow Genomes


178

CL\_1679


88

CL\_1679


3

CL\_1679


1

CL\_1679


1

CL\_1679


1

CL\_1680

fGI ID


CL\_INS\_137
CL\_INS\_137
CL\_INS\_137
CL\_INS\_237
Cluster ID


CL\_30379
CL\_1678
CL\_16995
CL\_7986
