## Supplementary material for "A novel method for integrating genomic and Tn-Seq data to identify common *in vivo* fitness mechanisms across multiple bacterial species": S1 Dataset: CL_INS_144.html

Legend

 Mobile +extrachromosomalelementfunctions
 Regulatoryfunctions
 Hypothetical
 DNA Metabolism
 All EssentialGenes
 All Fitness Genes
 Proteinsynthesis/fate
 Other
 Transport +binding proteins
 All VFDB Genes

FULL


WINDOWSVGPNG

Trim RowsRemove SingletonsSave Fasta

CL\_1789


CL\_1285


CL\_4427


CL\_1789


CL\_1789


CL\_1789


CL\_1788


CL\_1789


CL\_1789


CL\_1788


CL\_1789


CL\_1789


CL\_1788


CL\_1124

HighlightSelectShow Genomes


263

CL\_1790


1

CL\_1790


1

CL\_1790


1

CL\_4486


1

CL\_1790


1

CL\_1790


1

CL\_1790


1

CL\_1790


1

CL\_1790


1

CL\_1790


1

CL\_1790


1

CL\_1793


1

CL\_1790


1

CL\_1790

fGI ID


CL\_INS\_144
CL\_INS\_144
CL\_INS\_384
CL\_INS\_99
CL\_INS\_99
CL\_INS\_382
CL\_INS\_144
CL\_INS\_144
CL\_INS\_144
CL\_INS\_61
CL\_INS\_144
CL\_INS\_144
CL\_INS\_144
CL\_INS\_144
CL\_INS\_144
CL\_INS\_144
CL\_INS\_144
CL\_INS\_144
CL\_INS\_144
CL\_INS\_144
CL\_INS\_144
CL\_INS\_60
CL\_INS\_60
CL\_INS\_136
CL\_INS\_136
CL\_INS\_144
CL\_INS\_144
CL\_INS\_382
CL\_INS\_136
CL\_INS\_144
CL\_INS\_144
CL\_INS\_144
CL\_INS\_144
CL\_INS\_144
CL\_INS\_144
CL\_INS\_144
CL\_INS\_144
CL\_INS\_144
CL\_INS\_144
CL\_INS\_144
CL\_INS\_144
CL\_INS\_144
CL\_INS\_382
CL\_INS\_144
CL\_INS\_144
CL\_INS\_144
CL\_INS\_144
CL\_INS\_144
CL\_INS\_144
CL\_INS\_144
CL\_INS\_144
CL\_INS\_144
CL\_INS\_144
CL\_INS\_144
CL\_INS\_131
CL\_INS\_144
CL\_INS\_144
CL\_INS\_382
CL\_INS\_382
CL\_INS\_382
CL\_INS\_382
CL\_INS\_382
CL\_INS\_382
CL\_INS\_382
CL\_INS\_382
CL\_INS\_382
CL\_INS\_382
CL\_INS\_382
CL\_INS\_382
CL\_INS\_382
CL\_INS\_382
CL\_INS\_382
CL\_INS\_382
CL\_INS\_382
CL\_INS\_382
CL\_INS\_382
CL\_INS\_382
CL\_INS\_382
CL\_INS\_382
CL\_INS\_207
CL\_INS\_207
CL\_INS\_144
CL\_INS\_144
CL\_INS\_144
CL\_INS\_144
CL\_INS\_144
CL\_INS\_144
CL\_INS\_99
CL\_INS\_144
CL\_INS\_144
CL\_INS\_144
CL\_INS\_99
CL\_INS\_144
CL\_INS\_144
CL\_INS\_144
CL\_INS\_144
CL\_INS\_144
CL\_INS\_144
CL\_INS\_144
CL\_INS\_144
CL\_INS\_144
CL\_INS\_144
CL\_INS\_144
CL\_INS\_144
CL\_INS\_144
CL\_INS\_144
CL\_INS\_144
CL\_INS\_144
CL\_INS\_144
CL\_INS\_144
CL\_INS\_144
CL\_INS\_144
CL\_INS\_144
CL\_INS\_99
CL\_INS\_382
CL\_INS\_385
CL\_INS\_99
CL\_INS\_385
CL\_INS\_385
CL\_INS\_385
CL\_INS\_385
CL\_INS\_382
CL\_INS\_382
CL\_INS\_382
CL\_INS\_382
CL\_INS\_382
CL\_INS\_128
CL\_INS\_128
CL\_INS\_128
CL\_INS\_382
CL\_INS\_382
CL\_INS\_144
CL\_INS\_144
CL\_INS\_144
CL\_INS\_144
CL\_INS\_144
CL\_INS\_144
CL\_INS\_144
CL\_INS\_144
CL\_INS\_144
CL\_INS\_144
CL\_INS\_61
CL\_INS\_61
CL\_INS\_61
CL\_INS\_144
CL\_INS\_144
CL\_INS\_144
CL\_INS\_144
CL\_INS\_144
CL\_INS\_144
CL\_INS\_144
CL\_INS\_144
CL\_INS\_144
CL\_INS\_144
CL\_INS\_144
CL\_INS\_144
CL\_INS\_144
CL\_INS\_144
CL\_INS\_144
CL\_INS\_144
CL\_INS\_144
CL\_INS\_144
CL\_INS\_144
CL\_INS\_204
CL\_INS\_144
CL\_INS\_144
CL\_INS\_144
CL\_INS\_144
CL\_INS\_144
CL\_INS\_144
CL\_INS\_144
CL\_INS\_144
CL\_INS\_144
CL\_INS\_76
CL\_INS\_144
CL\_INS\_144
CL\_INS\_382
CL\_INS\_144
CL\_INS\_382
CL\_INS\_144
CL\_INS\_144
CL\_INS\_144
CL\_INS\_144
CL\_INS\_144
CL\_INS\_144
CL\_INS\_144
CL\_INS\_144
CL\_INS\_144
CL\_INS\_144
CL\_INS\_144
CL\_INS\_65
CL\_INS\_144
CL\_INS\_382
CL\_INS\_382
CL\_INS\_382
CL\_INS\_382
CL\_INS\_144
CL\_INS\_144
CL\_INS\_144
CL\_INS\_382
CL\_INS\_144
CL\_INS\_382
CL\_INS\_131
CL\_INS\_382
CL\_INS\_382
CL\_INS\_382
CL\_INS\_382
CL\_INS\_382
CL\_INS\_382
CL\_INS\_382
CL\_INS\_382
CL\_INS\_382
CL\_INS\_382
CL\_INS\_382
CL\_INS\_237
CL\_INS\_382
CL\_INS\_382
CL\_INS\_382
CL\_INS\_382
CL\_INS\_144
CL\_INS\_237
CL\_INS\_382
CL\_INS\_382
CL\_INS\_382
CL\_INS\_382
CL\_INS\_382
CL\_INS\_382
CL\_INS\_382
CL\_INS\_382
CL\_INS\_382
CL\_INS\_65
CL\_INS\_86
CL\_INS\_382
CL\_INS\_382
CL\_INS\_382
CL\_INS\_382
CL\_INS\_382
CL\_INS\_382
CL\_INS\_382
CL\_INS\_382
CL\_INS\_382
CL\_INS\_144
CL\_INS\_144
CL\_INS\_144
CL\_INS\_144
CL\_INS\_144
CL\_INS\_144
CL\_INS\_144
CL\_INS\_144
CL\_INS\_382
CL\_INS\_384
CL\_INS\_382
CL\_INS\_382
CL\_INS\_382
CL\_INS\_144
CL\_INS\_144
Cluster ID


CL\_32554
CL\_20244
CL\_20243
CL\_13075
CL\_17051
CL\_4432
CL\_12417
CL\_12418
CL\_30387
CL\_12455
CL\_30388
CL\_30389
CL\_30390
CL\_30391
CL\_30392
CL\_12416
CL\_12741
CL\_12742
CL\_12743
CL\_12744
CL\_12415
CL\_6668
CL\_6667
CL\_4472
CL\_1319
CL\_12414
CL\_30394
CL\_4469
CL\_10525
CL\_12413
CL\_17462
CL\_17461
CL\_17460
CL\_17458
CL\_17457
CL\_17456
CL\_17455
CL\_17454
CL\_32049
CL\_17452
CL\_17451
CL\_17450
CL\_15780
CL\_17449
CL\_17448
CL\_17446
CL\_17445
CL\_17444
CL\_17443
CL\_17442
CL\_17441
CL\_17440
CL\_17439
CL\_17438
CL\_17437
CL\_17436
CL\_17435
CL\_4425
CL\_4424
CL\_4423
CL\_4525
CL\_4421
CL\_4420
CL\_4419
CL\_4418
CL\_4417
CL\_4416
CL\_4532
CL\_4533
CL\_1496
CL\_20242
CL\_11311
CL\_6784
CL\_7537
CL\_7494
CL\_1506
CL\_12996
CL\_12760
CL\_4466
CL\_17225
CL\_17224
CL\_32773
CL\_32774
CL\_32775
CL\_32776
CL\_32777
CL\_32778
CL\_32779
CL\_32780
CL\_32781
CL\_32782
CL\_17292
CL\_32783
CL\_32784
CL\_32785
CL\_32786
CL\_32787
CL\_32788
CL\_32789
CL\_32790
CL\_32791
CL\_32792
CL\_32793
CL\_32794
CL\_32795
CL\_32796
CL\_32797
CL\_32798
CL\_32799
CL\_32800
CL\_32801
CL\_32802
CL\_32803
CL\_16661
CL\_17066
CL\_17486
CL\_17487
CL\_17488
CL\_17489
CL\_17490
CL\_17491
CL\_16421
CL\_16422
CL\_9150
CL\_8997
CL\_8996
CL\_14241
CL\_14240
CL\_17348
CL\_4562
CL\_8198
CL\_32804
CL\_32805
CL\_32806
CL\_32807
CL\_32808
CL\_36222
CL\_36223
CL\_36224
CL\_36225
CL\_36226
CL\_22983
CL\_22984
CL\_22985
CL\_36227
CL\_36228
CL\_36229
CL\_36230
CL\_36231
CL\_36232
CL\_36233
CL\_36234
CL\_36235
CL\_36236
CL\_36237
CL\_36238
CL\_36239
CL\_36240
CL\_36241
CL\_36242
CL\_36243
CL\_36244
CL\_36245
CL\_9222
CL\_36246
CL\_36247
CL\_36248
CL\_36249
CL\_36250
CL\_36251
CL\_36252
CL\_12412
CL\_12411
CL\_12410
CL\_12409
CL\_12408
CL\_8680
CL\_12407
CL\_4552
CL\_12406
CL\_12405
CL\_12404
CL\_12403
CL\_12402
CL\_12401
CL\_12400
CL\_12399
CL\_12398
CL\_12397
CL\_12396
CL\_12395
CL\_36253
CL\_4465
CL\_4464
CL\_4463
CL\_8162
CL\_30395
CL\_30396
CL\_30397
CL\_8676
CL\_12751
CL\_16191
CL\_13110
CL\_13112
CL\_20241
CL\_17434
CL\_17433
CL\_5283
CL\_6773
CL\_5282
CL\_20240
CL\_20239
CL\_20238
CL\_4548
CL\_8434
CL\_13108
CL\_16006
CL\_20237
CL\_7123
CL\_13107
CL\_11403
CL\_8678
CL\_4459
CL\_1317
CL\_1316
CL\_12477
CL\_1315
CL\_1314
CL\_20236
CL\_20235
CL\_16182
CL\_8672
CL\_1311
CL\_8671
CL\_20234
CL\_20233
CL\_1309
CL\_1308
CL\_1307
CL\_1306
CL\_5360
CL\_12745
CL\_12746
CL\_12747
CL\_12748
CL\_12749
CL\_12394
CL\_12393
CL\_12392
CL\_1305
CL\_15285
CL\_1304
CL\_12391
CL\_12390
CL\_12389
CL\_12750
