## Supplementary material for "A novel method for integrating genomic and Tn-Seq data to identify common *in vivo* fitness mechanisms across multiple bacterial species": S1 Dataset: CL_INS_146.html

Legend

 Mobile +extrachromosomalelementfunctions
 Regulatoryfunctions
 Hypothetical
 DNA Metabolism
 All EssentialGenes
 All Fitness Genes
 Proteinsynthesis/fate
 Other
 EnergyMetabolism
 All VFDB Genes
 Transport +binding proteins

FULL


WINDOWSVGPNG

Trim RowsRemove SingletonsSave Fasta

CL\_1819


CL\_1819


CL\_1819


CL\_1819


CL\_1819


CL\_1819


CL\_1819


CL\_1819


CL\_1819


CL\_1819


CL\_1819


CL\_1819


CL\_1819


CL\_1819


CL\_1819


CL\_1819


CL\_1819


CL\_1819


CL\_1819


CL\_4486


CL\_1819


CL\_1819


CL\_4427


CL\_4486


CL\_1819


CL\_1819


CL\_1819


CL\_4427


CL\_1819


CL\_1819


CL\_1819


CL\_4487


CL\_1073


CL\_1819


CL\_1819


CL\_1819


CL\_1819


CL\_1819


CL\_4427


CL\_1819


CL\_1819


CL\_1819


CL\_4487


CL\_1819


CL\_4486


CL\_1819


CL\_1819


CL\_1819

HighlightSelectShow Genomes


232

CL\_1820


2

CL\_1820


2

CL\_1820


1

CL\_1820


1

CL\_1821


1

CL\_4427


1

CL\_1820


1

CL\_4516


1

CL\_1820


1

CL\_1820


1

CL\_1917


1

CL\_1821


1

CL\_1820


1

CL\_1084


1

CL\_1820


1

CL\_1820


1

CL\_1820


1

CL\_1820


1

CL\_4486


1

CL\_1820


1

CL\_1820


1

CL\_1820


1

CL\_1820


1

CL\_1820


1

CL\_4516


1

CL\_1820


1

CL\_1820


1

CL\_1820


1

CL\_4487


1

CL\_1820


1

CL\_1820


1

CL\_1820


1

CL\_1820


1

CL\_1820


1

CL\_1820


1

CL\_1820


1

CL\_4487


1

CL\_1820


1

CL\_1820


1

CL\_1820


1

CL\_1820


1

CL\_1820


1

CL\_1820


1

CL\_1820


1

CL\_1820


1

CL\_1820


1

CL\_4516


1

CL\_1820

fGI ID


CL\_INS\_146
CL\_INS\_146
CL\_INS\_146
CL\_INS\_204
CL\_INS\_146
CL\_INS\_146
CL\_INS\_146
CL\_INS\_146
CL\_INS\_146
CL\_INS\_146
CL\_INS\_146
CL\_INS\_146
CL\_INS\_146
CL\_INS\_146
CL\_INS\_146
CL\_INS\_146
CL\_INS\_146
CL\_INS\_146
CL\_INS\_146
CL\_INS\_146
CL\_INS\_146
CL\_INS\_146
CL\_INS\_146
CL\_INS\_146
CL\_INS\_146
CL\_INS\_146
CL\_INS\_146
CL\_INS\_146
CL\_INS\_146
CL\_INS\_146
CL\_INS\_146
CL\_INS\_146
CL\_INS\_146
CL\_INS\_146
CL\_INS\_146
CL\_INS\_146
CL\_INS\_146
CL\_INS\_146
CL\_INS\_146
CL\_INS\_146
CL\_INS\_146
CL\_INS\_146
CL\_INS\_146
CL\_INS\_146
CL\_INS\_146
CL\_INS\_146
CL\_INS\_146
CL\_INS\_146
CL\_INS\_146
CL\_INS\_382
CL\_INS\_99
CL\_INS\_146
CL\_INS\_207
CL\_INS\_146
CL\_INS\_146
CL\_INS\_146
CL\_INS\_207
CL\_INS\_207
CL\_INS\_207
CL\_INS\_382
CL\_INS\_382
CL\_INS\_146
CL\_INS\_146
CL\_INS\_146
CL\_INS\_146
CL\_INS\_146
CL\_INS\_146
CL\_INS\_146
CL\_INS\_207
CL\_INS\_207
CL\_INS\_207
CL\_INS\_207
CL\_INS\_207
CL\_INS\_382
CL\_INS\_382
CL\_INS\_382
CL\_INS\_146
CL\_INS\_146
CL\_INS\_146
CL\_INS\_146
CL\_INS\_146
CL\_INS\_146
CL\_INS\_146
CL\_INS\_146
CL\_INS\_146
CL\_INS\_146
CL\_INS\_146
CL\_INS\_146
CL\_INS\_87
CL\_INS\_149
CL\_INS\_146
CL\_INS\_86
CL\_INS\_99
CL\_INS\_382
CL\_INS\_99
CL\_INS\_99
CL\_INS\_99
CL\_INS\_99
CL\_INS\_99
CL\_INS\_99
CL\_INS\_99
CL\_INS\_382
CL\_INS\_382
CL\_INS\_382
CL\_INS\_382
CL\_INS\_382
CL\_INS\_382
CL\_INS\_382
CL\_INS\_382
CL\_INS\_207
CL\_INS\_382
CL\_INS\_382
CL\_INS\_99
CL\_INS\_382
CL\_INS\_99
CL\_INS\_146
CL\_INS\_146
CL\_INS\_146
CL\_INS\_146
CL\_INS\_146
CL\_INS\_146
CL\_INS\_146
CL\_INS\_146
CL\_INS\_146
CL\_INS\_146
CL\_INS\_146
CL\_INS\_146
CL\_INS\_146
CL\_INS\_146
CL\_INS\_146
CL\_INS\_146
CL\_INS\_146
CL\_INS\_146
CL\_INS\_146
CL\_INS\_146
CL\_INS\_146
CL\_INS\_146
CL\_INS\_146
CL\_INS\_146
CL\_INS\_146
CL\_INS\_146
CL\_INS\_146
CL\_INS\_146
CL\_INS\_146
CL\_INS\_106
CL\_INS\_382
CL\_INS\_146
CL\_INS\_382
CL\_INS\_382
CL\_INS\_382
CL\_INS\_382
CL\_INS\_146
CL\_INS\_146
CL\_INS\_146
CL\_INS\_146
CL\_INS\_146
CL\_INS\_146
CL\_INS\_146
CL\_INS\_146
CL\_INS\_146
CL\_INS\_146
CL\_INS\_99
CL\_INS\_146
CL\_INS\_146
CL\_INS\_99
CL\_INS\_146
CL\_INS\_146
CL\_INS\_146
CL\_INS\_146
CL\_INS\_146
CL\_INS\_146
CL\_INS\_146
CL\_INS\_146
CL\_INS\_146
CL\_INS\_382
CL\_INS\_146
CL\_INS\_99
CL\_INS\_99
CL\_INS\_99
CL\_INS\_382
CL\_INS\_146
CL\_INS\_146
CL\_INS\_146
CL\_INS\_146
CL\_INS\_146
CL\_INS\_146
CL\_INS\_146
CL\_INS\_146
CL\_INS\_146
CL\_INS\_146
CL\_INS\_146
CL\_INS\_146
CL\_INS\_146
CL\_INS\_146
CL\_INS\_146
CL\_INS\_146
CL\_INS\_146
CL\_INS\_146
CL\_INS\_146
CL\_INS\_146
CL\_INS\_146
CL\_INS\_146
CL\_INS\_146
CL\_INS\_146
CL\_INS\_146
CL\_INS\_382
CL\_INS\_70
CL\_INS\_70
CL\_INS\_146
CL\_INS\_237
CL\_INS\_382
CL\_INS\_382
CL\_INS\_382
CL\_INS\_382
CL\_INS\_382
CL\_INS\_382
CL\_INS\_382
CL\_INS\_382
CL\_INS\_382
CL\_INS\_382
CL\_INS\_382
CL\_INS\_99
CL\_INS\_382
CL\_INS\_99
CL\_INS\_99
CL\_INS\_99
CL\_INS\_99
CL\_INS\_99
CL\_INS\_99
CL\_INS\_99
CL\_INS\_146
CL\_INS\_99
CL\_INS\_146
CL\_INS\_146
CL\_INS\_146
CL\_INS\_146
CL\_INS\_146
CL\_INS\_146
CL\_INS\_382
CL\_INS\_146
CL\_INS\_146
CL\_INS\_146
CL\_INS\_382
CL\_INS\_382
CL\_INS\_146
CL\_INS\_382
CL\_INS\_382
CL\_INS\_146
CL\_INS\_204
CL\_INS\_204
CL\_INS\_204
CL\_INS\_204
CL\_INS\_204
CL\_INS\_86
CL\_INS\_99
CL\_INS\_146
CL\_INS\_86
CL\_INS\_146
CL\_INS\_20
CL\_INS\_146
CL\_INS\_146
CL\_INS\_146
CL\_INS\_146
CL\_INS\_146
CL\_INS\_146
CL\_INS\_382
CL\_INS\_146
CL\_INS\_146
CL\_INS\_146
CL\_INS\_99
CL\_INS\_382
CL\_INS\_99
CL\_INS\_146
CL\_INS\_146
CL\_INS\_146
CL\_INS\_207
CL\_INS\_382
CL\_INS\_146
CL\_INS\_146
CL\_INS\_382
CL\_INS\_382
CL\_INS\_382
CL\_INS\_382
CL\_INS\_146
CL\_INS\_207
CL\_INS\_146
CL\_INS\_146
CL\_INS\_146
CL\_INS\_146
CL\_INS\_106
CL\_INS\_106
CL\_INS\_146
CL\_INS\_146
CL\_INS\_146
CL\_INS\_146
CL\_INS\_146
CL\_INS\_207
CL\_INS\_146
CL\_INS\_146
CL\_INS\_146
CL\_INS\_146
CL\_INS\_146
CL\_INS\_146
CL\_INS\_207
CL\_INS\_146
CL\_INS\_146
CL\_INS\_146
CL\_INS\_146
CL\_INS\_146
CL\_INS\_99
CL\_INS\_99
CL\_INS\_146
CL\_INS\_86
CL\_INS\_99
CL\_INS\_99
CL\_INS\_207
CL\_INS\_207
CL\_INS\_207
CL\_INS\_207
CL\_INS\_207
CL\_INS\_207
CL\_INS\_382
CL\_INS\_99
CL\_INS\_146
CL\_INS\_146
CL\_INS\_146
CL\_INS\_86
CL\_INS\_86
CL\_INS\_20
CL\_INS\_20
CL\_INS\_99
CL\_INS\_99
CL\_INS\_86
CL\_INS\_146
CL\_INS\_146
CL\_INS\_146
CL\_INS\_146
CL\_INS\_146
CL\_INS\_146
CL\_INS\_146
CL\_INS\_146
CL\_INS\_146
CL\_INS\_146
CL\_INS\_146
CL\_INS\_146
CL\_INS\_146
CL\_INS\_146
CL\_INS\_146
CL\_INS\_146
CL\_INS\_146
CL\_INS\_146
CL\_INS\_146
CL\_INS\_146
CL\_INS\_146
CL\_INS\_382
CL\_INS\_382
CL\_INS\_146
CL\_INS\_99
CL\_INS\_382
CL\_INS\_146
CL\_INS\_382
CL\_INS\_382
CL\_INS\_146
CL\_INS\_146
CL\_INS\_146
CL\_INS\_146
CL\_INS\_146
CL\_INS\_99
CL\_INS\_146
CL\_INS\_146
CL\_INS\_146
CL\_INS\_86
CL\_INS\_146
CL\_INS\_99
CL\_INS\_146
CL\_INS\_382
CL\_INS\_146
CL\_INS\_146
CL\_INS\_146
CL\_INS\_146
CL\_INS\_382
CL\_INS\_382
CL\_INS\_382
CL\_INS\_86
CL\_INS\_146
CL\_INS\_146
CL\_INS\_146
CL\_INS\_146
CL\_INS\_382
CL\_INS\_382
CL\_INS\_146
CL\_INS\_146
CL\_INS\_146
CL\_INS\_146
CL\_INS\_146
CL\_INS\_146
CL\_INS\_146
CL\_INS\_146
CL\_INS\_146
CL\_INS\_146
CL\_INS\_146
CL\_INS\_146
CL\_INS\_382
CL\_INS\_146
CL\_INS\_99
CL\_INS\_382
CL\_INS\_382
CL\_INS\_146
CL\_INS\_146
CL\_INS\_146
CL\_INS\_146
CL\_INS\_382
CL\_INS\_382
CL\_INS\_99
CL\_INS\_382
CL\_INS\_146
CL\_INS\_146
CL\_INS\_146
CL\_INS\_86
CL\_INS\_99
CL\_INS\_99
CL\_INS\_99
CL\_INS\_146
CL\_INS\_146
CL\_INS\_146
CL\_INS\_146
CL\_INS\_146
CL\_INS\_146
CL\_INS\_382
CL\_INS\_146
CL\_INS\_146
CL\_INS\_382
CL\_INS\_99
CL\_INS\_99
CL\_INS\_382
CL\_INS\_146
CL\_INS\_146
CL\_INS\_146
CL\_INS\_146
CL\_INS\_146
CL\_INS\_146
CL\_INS\_382
CL\_INS\_146
CL\_INS\_382
CL\_INS\_382
CL\_INS\_382
CL\_INS\_382
CL\_INS\_382
CL\_INS\_382
CL\_INS\_382
CL\_INS\_382
CL\_INS\_382
CL\_INS\_382
CL\_INS\_382
CL\_INS\_382
CL\_INS\_382
CL\_INS\_146
CL\_INS\_146
CL\_INS\_146
CL\_INS\_146
CL\_INS\_146
CL\_INS\_146
CL\_INS\_146
CL\_INS\_146
CL\_INS\_146
CL\_INS\_146
CL\_INS\_146
CL\_INS\_146
CL\_INS\_146
CL\_INS\_146
CL\_INS\_382
CL\_INS\_146
CL\_INS\_382
CL\_INS\_382
CL\_INS\_382
CL\_INS\_382
CL\_INS\_382
CL\_INS\_382
CL\_INS\_382
CL\_INS\_382
CL\_INS\_146
CL\_INS\_146
CL\_INS\_146
CL\_INS\_146
CL\_INS\_146
CL\_INS\_146
CL\_INS\_146
CL\_INS\_146
CL\_INS\_146
CL\_INS\_146
CL\_INS\_146
CL\_INS\_146
CL\_INS\_146
CL\_INS\_382
CL\_INS\_382
CL\_INS\_382
CL\_INS\_237
CL\_INS\_146
CL\_INS\_146
CL\_INS\_382
CL\_INS\_146
CL\_INS\_146
CL\_INS\_146
CL\_INS\_146
CL\_INS\_382
CL\_INS\_382
CL\_INS\_382
CL\_INS\_382
CL\_INS\_382
CL\_INS\_382
CL\_INS\_382
CL\_INS\_382
CL\_INS\_146
CL\_INS\_382
CL\_INS\_146
CL\_INS\_146
CL\_INS\_146
CL\_INS\_146
CL\_INS\_146
CL\_INS\_146
CL\_INS\_146
CL\_INS\_146
CL\_INS\_146
CL\_INS\_146
CL\_INS\_146
CL\_INS\_146
CL\_INS\_146
CL\_INS\_146
CL\_INS\_20
CL\_INS\_207
CL\_INS\_207
CL\_INS\_146
CL\_INS\_146
CL\_INS\_382
CL\_INS\_382
CL\_INS\_146
CL\_INS\_146
CL\_INS\_382
CL\_INS\_382
CL\_INS\_20
CL\_INS\_146
CL\_INS\_382
CL\_INS\_146
CL\_INS\_146
CL\_INS\_146
CL\_INS\_382
CL\_INS\_86
CL\_INS\_382
CL\_INS\_146
CL\_INS\_146
CL\_INS\_146
CL\_INS\_146
CL\_INS\_146
CL\_INS\_146
CL\_INS\_146
CL\_INS\_146
CL\_INS\_382
CL\_INS\_382
CL\_INS\_382
CL\_INS\_382
CL\_INS\_382
CL\_INS\_382
CL\_INS\_382
CL\_INS\_382
CL\_INS\_382
CL\_INS\_382
CL\_INS\_382
CL\_INS\_382
CL\_INS\_382
CL\_INS\_382
CL\_INS\_382
CL\_INS\_382
CL\_INS\_382
CL\_INS\_382
CL\_INS\_382
CL\_INS\_382
CL\_INS\_382
CL\_INS\_99
CL\_INS\_86
CL\_INS\_382
CL\_INS\_382
CL\_INS\_382
CL\_INS\_382
CL\_INS\_382
CL\_INS\_382
CL\_INS\_382
CL\_INS\_382
CL\_INS\_146
CL\_INS\_382
CL\_INS\_382
CL\_INS\_146
CL\_INS\_146
CL\_INS\_146
CL\_INS\_99
CL\_INS\_146
CL\_INS\_146
CL\_INS\_382
CL\_INS\_382
CL\_INS\_382
CL\_INS\_237
CL\_INS\_382
CL\_INS\_146
CL\_INS\_146
CL\_INS\_382
CL\_INS\_382
CL\_INS\_382
CL\_INS\_99
CL\_INS\_99
CL\_INS\_99
CL\_INS\_146
CL\_INS\_382
CL\_INS\_382
CL\_INS\_382
CL\_INS\_382
CL\_INS\_382
CL\_INS\_382
CL\_INS\_382
CL\_INS\_382
CL\_INS\_146
CL\_INS\_146
CL\_INS\_146
CL\_INS\_146
CL\_INS\_382
CL\_INS\_146
CL\_INS\_382
CL\_INS\_146
CL\_INS\_146
CL\_INS\_146
CL\_INS\_382
CL\_INS\_146
CL\_INS\_382
CL\_INS\_382
CL\_INS\_382
CL\_INS\_146
CL\_INS\_146
CL\_INS\_146
CL\_INS\_382
CL\_INS\_382
CL\_INS\_382
CL\_INS\_382
CL\_INS\_99
CL\_INS\_382
CL\_INS\_382
CL\_INS\_382
CL\_INS\_382
CL\_INS\_207
CL\_INS\_207
CL\_INS\_207
CL\_INS\_146
CL\_INS\_382
CL\_INS\_382
CL\_INS\_382
CL\_INS\_382
CL\_INS\_382
CL\_INS\_382
CL\_INS\_382
CL\_INS\_382
CL\_INS\_382
CL\_INS\_382
CL\_INS\_382
CL\_INS\_382
CL\_INS\_146
CL\_INS\_146
CL\_INS\_207
CL\_INS\_207
CL\_INS\_382
CL\_INS\_146
CL\_INS\_146
CL\_INS\_207
CL\_INS\_382
CL\_INS\_382
CL\_INS\_382
CL\_INS\_382
CL\_INS\_146
CL\_INS\_146
CL\_INS\_207
CL\_INS\_382
CL\_INS\_207
CL\_INS\_207
CL\_INS\_146
CL\_INS\_146
CL\_INS\_146
CL\_INS\_382
CL\_INS\_146
CL\_INS\_146
CL\_INS\_382
CL\_INS\_382
CL\_INS\_382
CL\_INS\_146
CL\_INS\_146
CL\_INS\_146
CL\_INS\_146
CL\_INS\_146
CL\_INS\_146
CL\_INS\_146
CL\_INS\_382
CL\_INS\_207
CL\_INS\_146
CL\_INS\_146
CL\_INS\_146
CL\_INS\_382
CL\_INS\_382
CL\_INS\_149
CL\_INS\_382
CL\_INS\_382
CL\_INS\_382
CL\_INS\_207
CL\_INS\_207
CL\_INS\_146
CL\_INS\_146
CL\_INS\_146
CL\_INS\_382
CL\_INS\_382
CL\_INS\_382
CL\_INS\_382
CL\_INS\_382
CL\_INS\_382
CL\_INS\_382
CL\_INS\_207
CL\_INS\_382
CL\_INS\_146
CL\_INS\_146
CL\_INS\_382
CL\_INS\_382
CL\_INS\_382
CL\_INS\_382
CL\_INS\_382
CL\_INS\_382
CL\_INS\_382
CL\_INS\_382
CL\_INS\_382
CL\_INS\_382
CL\_INS\_382
CL\_INS\_382
CL\_INS\_382
CL\_INS\_382
CL\_INS\_382
CL\_INS\_146
CL\_INS\_20
CL\_INS\_20
CL\_INS\_20
CL\_INS\_20
CL\_INS\_146
CL\_INS\_20
CL\_INS\_146
CL\_INS\_146
CL\_INS\_146
CL\_INS\_382
CL\_INS\_382
CL\_INS\_382
CL\_INS\_146
CL\_INS\_146
CL\_INS\_382
CL\_INS\_146
CL\_INS\_382
CL\_INS\_146
CL\_INS\_146
CL\_INS\_382
CL\_INS\_146
CL\_INS\_146
CL\_INS\_146
CL\_INS\_106
CL\_INS\_146
CL\_INS\_146
CL\_INS\_146
CL\_INS\_146
CL\_INS\_382
CL\_INS\_382
CL\_INS\_382
CL\_INS\_382
CL\_INS\_382
CL\_INS\_382
CL\_INS\_146
CL\_INS\_146
CL\_INS\_146
CL\_INS\_146
CL\_INS\_146
CL\_INS\_382
CL\_INS\_382
CL\_INS\_382
CL\_INS\_382
CL\_INS\_146
CL\_INS\_382
CL\_INS\_382
CL\_INS\_382
CL\_INS\_382
CL\_INS\_382
CL\_INS\_382
CL\_INS\_382
CL\_INS\_382
CL\_INS\_382
CL\_INS\_382
CL\_INS\_382
CL\_INS\_146
CL\_INS\_237
CL\_INS\_237
CL\_INS\_237
CL\_INS\_237
CL\_INS\_146
CL\_INS\_146
CL\_INS\_237
CL\_INS\_237
CL\_INS\_146
CL\_INS\_382
CL\_INS\_146
CL\_INS\_146
CL\_INS\_382
CL\_INS\_382
CL\_INS\_382
CL\_INS\_382
CL\_INS\_146
CL\_INS\_146
Cluster ID


CL\_15175
CL\_15174
CL\_15173
CL\_5896
CL\_15172
CL\_5271
CL\_5270
CL\_29496
CL\_29497
CL\_29498
CL\_10345
CL\_10346
CL\_5269
CL\_10347
CL\_5268
CL\_26934
CL\_5267
CL\_7610
CL\_5266
CL\_5330
CL\_5265
CL\_5264
CL\_5263
CL\_5262
CL\_5261
CL\_5260
CL\_5259
CL\_5258
CL\_10348
CL\_10349
CL\_5331
CL\_7609
CL\_7608
CL\_34246
CL\_34762
CL\_34761
CL\_10350
CL\_21070
CL\_21069
CL\_21068
CL\_21067
CL\_21066
CL\_7607
CL\_13141
CL\_30401
CL\_12386
CL\_30402
CL\_30403
CL\_30404
CL\_25588
CL\_7522
CL\_22921
CL\_7523
CL\_13142
CL\_13143
CL\_13144
CL\_7524
CL\_7525
CL\_7526
CL\_6741
CL\_7527
CL\_13145
CL\_13146
CL\_13147
CL\_13148
CL\_13149
CL\_13150
CL\_13151
CL\_7528
CL\_7529
CL\_7530
CL\_7531
CL\_7532
CL\_28720
CL\_4425
CL\_4424
CL\_12387
CL\_30879
CL\_30880
CL\_30881
CL\_30882
CL\_28110
CL\_28111
CL\_28486
CL\_28487
CL\_28488
CL\_13614
CL\_13613
CL\_4513
CL\_12560
CL\_37592
CL\_4515
CL\_17053
CL\_4517
CL\_13790
CL\_13789
CL\_15782
CL\_17292
CL\_17291
CL\_17290
CL\_17289
CL\_15781
CL\_15780
CL\_15779
CL\_15778
CL\_15777
CL\_15776
CL\_9123
CL\_7533
CL\_9122
CL\_12383
CL\_6784
CL\_12382
CL\_7534
CL\_8183
CL\_28970
CL\_15136
CL\_28969
CL\_14044
CL\_14045
CL\_14046
CL\_14047
CL\_14048
CL\_14049
CL\_14050
CL\_21313
CL\_21312
CL\_21311
CL\_21310
CL\_21309
CL\_21308
CL\_21307
CL\_21306
CL\_21305
CL\_21304
CL\_14051
CL\_14052
CL\_14053
CL\_14054
CL\_14055
CL\_14056
CL\_14057
CL\_14058
CL\_28968
CL\_15139
CL\_526
CL\_28967
CL\_5806
CL\_5805
CL\_520
CL\_519
CL\_15171
CL\_15170
CL\_15169
CL\_15168
CL\_15167
CL\_15166
CL\_28966
CL\_28965
CL\_28964
CL\_28963
CL\_10521
CL\_15835
CL\_8166
CL\_4431
CL\_36070
CL\_36071
CL\_36072
CL\_32267
CL\_10969
CL\_10970
CL\_36073
CL\_36074
CL\_36075
CL\_8167
CL\_15836
CL\_6747
CL\_13509
CL\_8168
CL\_8169
CL\_15837
CL\_15838
CL\_15839
CL\_15840
CL\_15841
CL\_15842
CL\_15843
CL\_15844
CL\_15845
CL\_15846
CL\_15847
CL\_15848
CL\_15849
CL\_15850
CL\_15851
CL\_15852
CL\_15853
CL\_15854
CL\_15855
CL\_15856
CL\_15857
CL\_15858
CL\_15859
CL\_15860
CL\_15861
CL\_8170
CL\_7434
CL\_7435
CL\_12757
CL\_7436
CL\_8171
CL\_8172
CL\_8710
CL\_8709
CL\_8708
CL\_8707
CL\_8706
CL\_8705
CL\_8702
CL\_8701
CL\_8700
CL\_12758
CL\_8697
CL\_8173
CL\_8174
CL\_8175
CL\_8176
CL\_10971
CL\_10972
CL\_10973
CL\_36076
CL\_10974
CL\_36077
CL\_36078
CL\_36079
CL\_36080
CL\_36081
CL\_36082
CL\_7538
CL\_30405
CL\_30406
CL\_30407
CL\_8694
CL\_8693
CL\_12381
CL\_7539
CL\_16967
CL\_8177
CL\_5408
CL\_5409
CL\_5410
CL\_1099
CL\_1098
CL\_8178
CL\_8179
CL\_16650
CL\_8180
CL\_8181
CL\_8182
CL\_16651
CL\_16652
CL\_16653
CL\_16654
CL\_16655
CL\_16656
CL\_7120
CL\_16657
CL\_16658
CL\_16659
CL\_16660
CL\_4465
CL\_16661
CL\_16662
CL\_16663
CL\_16664
CL\_4653
CL\_4654
CL\_30408
CL\_30409
CL\_23762
CL\_509
CL\_12380
CL\_12379
CL\_4657
CL\_9771
CL\_9770
CL\_9769
CL\_9768
CL\_6455
CL\_6456
CL\_6457
CL\_10156
CL\_10157
CL\_10158
CL\_10159
CL\_10160
CL\_6458
CL\_9767
CL\_9766
CL\_9765
CL\_9764
CL\_9763
CL\_10161
CL\_6460
CL\_23202
CL\_11305
CL\_12753
CL\_33012
CL\_33013
CL\_28004
CL\_17033
CL\_10967
CL\_7023
CL\_13526
CL\_10518
CL\_28471
CL\_17423
CL\_17422
CL\_17421
CL\_17420
CL\_28470
CL\_9124
CL\_7521
CL\_28469
CL\_28468
CL\_28467
CL\_5815
CL\_5814
CL\_16568
CL\_4434
CL\_4433
CL\_534
CL\_4490
CL\_25149
CL\_25150
CL\_25151
CL\_25152
CL\_25153
CL\_25154
CL\_25155
CL\_25156
CL\_25157
CL\_25158
CL\_25159
CL\_25160
CL\_25161
CL\_25162
CL\_25163
CL\_25164
CL\_25165
CL\_25166
CL\_25167
CL\_25168
CL\_25169
CL\_533
CL\_1496
CL\_17373
CL\_17372
CL\_8698
CL\_17370
CL\_6042
CL\_14245
CL\_6038
CL\_16194
CL\_16193
CL\_23203
CL\_4617
CL\_4618
CL\_4619
CL\_27256
CL\_27255
CL\_4430
CL\_28489
CL\_8155
CL\_28490
CL\_6744
CL\_28491
CL\_28492
CL\_28493
CL\_28494
CL\_28495
CL\_21006
CL\_28496
CL\_4429
CL\_27254
CL\_27253
CL\_27252
CL\_4627
CL\_4628
CL\_8473
CL\_27251
CL\_27250
CL\_27249
CL\_27248
CL\_27247
CL\_27246
CL\_27245
CL\_27244
CL\_27243
CL\_27242
CL\_27241
CL\_27240
CL\_8184
CL\_13072
CL\_20884
CL\_15774
CL\_15773
CL\_20782
CL\_20783
CL\_20784
CL\_23523
CL\_8185
CL\_7119
CL\_12385
CL\_4432
CL\_22922
CL\_12384
CL\_12754
CL\_1495
CL\_17051
CL\_17052
CL\_4620
CL\_4621
CL\_4622
CL\_4623
CL\_4624
CL\_4625
CL\_4626
CL\_8712
CL\_12755
CL\_12756
CL\_4518
CL\_4485
CL\_4520
CL\_4521
CL\_28466
CL\_28465
CL\_28464
CL\_28463
CL\_28462
CL\_28461
CL\_12007
CL\_28460
CL\_4522
CL\_4523
CL\_4423
CL\_4629
CL\_4421
CL\_4420
CL\_4419
CL\_4418
CL\_13640
CL\_7612
CL\_4417
CL\_4416
CL\_4415
CL\_4630
CL\_4631
CL\_4632
CL\_4633
CL\_4634
CL\_4635
CL\_4636
CL\_4637
CL\_4638
CL\_4639
CL\_4640
CL\_4641
CL\_4642
CL\_4643
CL\_4644
CL\_4645
CL\_4524
CL\_4525
CL\_4526
CL\_4527
CL\_4528
CL\_4529
CL\_6461
CL\_14376
CL\_28497
CL\_28498
CL\_28499
CL\_28500
CL\_28501
CL\_28502
CL\_28503
CL\_28504
CL\_28505
CL\_28506
CL\_28507
CL\_28508
CL\_28509
CL\_4646
CL\_4647
CL\_4648
CL\_244
CL\_4649
CL\_4650
CL\_4530
CL\_28459
CL\_28458
CL\_28457
CL\_28456
CL\_4531
CL\_4532
CL\_4533
CL\_4414
CL\_4413
CL\_532
CL\_19013
CL\_19014
CL\_9762
CL\_9761
CL\_9760
CL\_9759
CL\_9758
CL\_9757
CL\_9756
CL\_9755
CL\_9754
CL\_9753
CL\_9752
CL\_9751
CL\_9750
CL\_9749
CL\_9748
CL\_10162
CL\_10163
CL\_10164
CL\_10165
CL\_10166
CL\_10167
CL\_10168
CL\_23312
CL\_36083
CL\_36084
CL\_19352
CL\_12768
CL\_12769
CL\_10169
CL\_10170
CL\_10171
CL\_10172
CL\_10173
CL\_5423
CL\_10174
CL\_10175
CL\_10176
CL\_10177
CL\_10178
CL\_10179
CL\_10180
CL\_10181
CL\_10182
CL\_10183
CL\_4565
CL\_4566
CL\_7115
CL\_21008
CL\_28721
CL\_28722
CL\_28723
CL\_28724
CL\_28725
CL\_28726
CL\_28727
CL\_28728
CL\_28729
CL\_28730
CL\_7116
CL\_14371
CL\_6045
CL\_4534
CL\_9098
CL\_13524
CL\_12797
CL\_28731
CL\_10976
CL\_12999
CL\_11919
CL\_11920
CL\_7537
CL\_7117
CL\_7118
CL\_12759
CL\_8696
CL\_28455
CL\_17368
CL\_28098
CL\_28454
CL\_28453
CL\_28452
CL\_15902
CL\_17311
CL\_17310
CL\_4546
CL\_10981
CL\_6773
CL\_6076
CL\_5282
CL\_17309
CL\_17308
CL\_10938
CL\_10937
CL\_10936
CL\_15759
CL\_1313
CL\_1312
CL\_19961
CL\_14236
CL\_7126
CL\_10982
CL\_10983
CL\_4469
CL\_12996
CL\_12995
CL\_16615
CL\_37591
CL\_37590
CL\_37589
CL\_37588
CL\_12760
CL\_12761
CL\_12762
CL\_12763
CL\_12764
CL\_12765
CL\_12766
CL\_12767
CL\_13562
CL\_6452
CL\_15862
CL\_15863
CL\_28510
CL\_28511
CL\_17485
CL\_4544
CL\_7540
CL\_10980
CL\_15903
CL\_11936
CL\_11937
CL\_4651
CL\_4652
CL\_17563
CL\_17564
CL\_17565
CL\_25170
CL\_5419
CL\_5420
CL\_5421
CL\_5422
CL\_2278
CL\_23311
CL\_2279
CL\_4655
CL\_13564
CL\_4656
CL\_12800
CL\_4658
CL\_12378
CL\_12377
CL\_15864
CL\_15865
CL\_6037
CL\_20787
CL\_16665
CL\_12376
CL\_4659
CL\_4660
CL\_15866
CL\_15867
CL\_16666
CL\_12801
CL\_4661
CL\_4662
CL\_16451
CL\_16452
CL\_21013
CL\_9747
CL\_9746
CL\_9745
CL\_12375
CL\_12374
CL\_4663
CL\_4664
CL\_9744
CL\_9743
CL\_9742
CL\_9741
CL\_9740
CL\_9739
CL\_9738
CL\_9737
CL\_21014
CL\_4665
CL\_12803
CL\_12804
CL\_22574
CL\_10510
CL\_10511
CL\_11994
CL\_8683
CL\_8684
CL\_4548
CL\_15868
CL\_4666
CL\_13152
CL\_13153
CL\_21015
CL\_4667
CL\_28512
CL\_28513
CL\_6036
CL\_8580
CL\_8579
CL\_8578
CL\_14633
CL\_12119
CL\_26557
CL\_26556
CL\_7542
CL\_12121
CL\_7543
CL\_12120
CL\_8576
CL\_2281
CL\_2282
CL\_2283
CL\_20109
CL\_11406
CL\_11405
CL\_17074
CL\_12994
CL\_16444
CL\_16445
CL\_37316
CL\_15697
CL\_15696
CL\_15695
CL\_13071
CL\_13070
CL\_14825
CL\_17539
CL\_13972
CL\_17540
CL\_5347
CL\_5348
CL\_5349
CL\_11736
CL\_11735
CL\_5350
CL\_15150
CL\_6651
CL\_15151
CL\_16587
CL\_4406
CL\_28962
CL\_28961
CL\_28960
CL\_15156
CL\_15158
CL\_15159
CL\_8894
CL\_20104
CL\_508
CL\_4400
CL\_5361
CL\_5362
CL\_5799
CL\_5798
CL\_28959
CL\_28958
CL\_28957
CL\_28956
CL\_15164
CL\_4410
CL\_4408
CL\_4407
CL\_6479
CL\_30410
CL\_4402
CL\_4401
CL\_5351
CL\_6786
CL\_6787
CL\_6788
CL\_6789
CL\_6790
CL\_6791
CL\_6792
CL\_12059
CL\_32838
CL\_5384
CL\_5385
CL\_5386
CL\_5387
CL\_37317
CL\_37318
CL\_5388
CL\_5389
CL\_19855
CL\_6638
CL\_11995
CL\_11729
CL\_11402
CL\_4668
CL\_4669
CL\_4670
CL\_12770
CL\_4671
