## Supplementary material for "A novel method for integrating genomic and Tn-Seq data to identify common *in vivo* fitness mechanisms across multiple bacterial species": S1 Dataset: CL_INS_148.html

Legend

 Mobile +extrachromosomalelementfunctions
 Hypothetical
 All EssentialGenes
 All VFDB Genes

FULL


WINDOWSVGPNG

Trim RowsRemove SingletonsSave Fasta

CL\_1828


CL\_1828


CL\_1826


CL\_1828

HighlightSelectShow Genomes


164

CL\_1830


112

CL\_1830


2

CL\_1830


1

CL\_1830

fGI ID


CL\_INS\_148
CL\_INS\_148
CL\_INS\_148
CL\_INS\_148
CL\_INS\_148
CL\_INS\_148
CL\_INS\_148
Cluster ID


CL\_17504
CL\_17505
CL\_17506
CL\_17507
CL\_17508
CL\_17509
CL\_1829
