## Supplementary material for "A novel method for integrating genomic and Tn-Seq data to identify common *in vivo* fitness mechanisms across multiple bacterial species": S1 Dataset: CL_INS_149.html

Legend

 Mobile +extrachromosomalelementfunctions
 Regulatoryfunctions
 Hypothetical
 DNA Metabolism
 All EssentialGenes
 Cell Envelope
 Proteinsynthesis/fate
 Other
 EnergyMetabolism
 All VFDB Genes
 Transport +binding proteins

FULL


WINDOWSVGPNG

Trim RowsRemove SingletonsSave Fasta

CL\_1859


CL\_1859


CL\_1859


CL\_1859


CL\_1859


CL\_1859


CL\_1859


CL\_1859


CL\_1859


CL\_1859


CL\_1859


CL\_1859


CL\_1859


CL\_1859


CL\_1859


CL\_1859


CL\_1859


CL\_1859


CL\_1859


CL\_1859


CL\_1859


CL\_1859


CL\_1859


CL\_1859


CL\_1859


CL\_1859


CL\_1859


CL\_1859


CL\_1859


CL\_1859


CL\_1859


CL\_1859


CL\_1859


CL\_1858


CL\_1859


CL\_1859


CL\_1859


Break


CL\_1859


CL\_1859


CL\_1859


CL\_1859


CL\_1859


CL\_1858


CL\_1859


CL\_1859


CL\_1859


CL\_1859


CL\_1859


CL\_1859


CL\_1859


CL\_1859


CL\_1859


CL\_1859


CL\_1859


CL\_1859


CL\_1859


CL\_1859


CL\_1859


CL\_1859


CL\_1859


CL\_1859

HighlightSelectShow Genomes


198

CL\_1860


11

CL\_1860


6

CL\_1860


4

CL\_1860


2

CL\_1860


2

CL\_1860


1

CL\_1860


1

CL\_1860


1

CL\_1860


1

CL\_1860


1

CL\_1860


1

CL\_1860


1

CL\_1862


1

CL\_1860


1

CL\_1860


1

CL\_1860


1

CL\_1860


1

CL\_1860


1

CL\_1860


1

CL\_1860


1

CL\_1860


1

CL\_1860


1

CL\_1860


1

CL\_1860


1

CL\_1860


1

CL\_1860


1

CL\_1860


1

CL\_1860


1

Break


1

CL\_1860


1

CL\_1860


1

CL\_1860


1

CL\_1860


1

CL\_1860


1

CL\_1860


1

CL\_1860


1

CL\_1860


1

CL\_1860


1

CL\_1860


1

CL\_1860


1

CL\_1860


1

CL\_1860


1

CL\_1860


1

CL\_1860


1

CL\_1860


1

CL\_1860


1

CL\_1860


1

CL\_1860


1

CL\_1860


1

CL\_1860


1

CL\_1860


1

CL\_1860


1

CL\_1860


1

CL\_1860


1

CL\_1860


1

CL\_1860


1

CL\_1860


1

CL\_1860


1

CL\_1860


1

CL\_1860


1

CL\_1860


1

CL\_1860

fGI ID


CL\_INS\_149
CL\_INS\_149
CL\_INS\_149
CL\_INS\_149
CL\_INS\_149
CL\_INS\_149
CL\_INS\_382
CL\_INS\_149
CL\_INS\_149
CL\_INS\_149
CL\_INS\_149
CL\_INS\_149
CL\_INS\_149
CL\_INS\_149
CL\_INS\_149
CL\_INS\_149
CL\_INS\_149
CL\_INS\_149
CL\_INS\_149
CL\_INS\_149
CL\_INS\_149
CL\_INS\_149
CL\_INS\_149
CL\_INS\_149
CL\_INS\_149
CL\_INS\_149
CL\_INS\_149
CL\_INS\_149
CL\_INS\_149
CL\_INS\_149
CL\_INS\_149
CL\_INS\_149
CL\_INS\_149
CL\_INS\_149
CL\_INS\_149
CL\_INS\_149
CL\_INS\_149
CL\_INS\_149
CL\_INS\_149
CL\_INS\_149
CL\_INS\_149
CL\_INS\_149
CL\_INS\_149
CL\_INS\_149
CL\_INS\_149
CL\_INS\_149
CL\_INS\_149
CL\_INS\_149
CL\_INS\_149
CL\_INS\_149
CL\_INS\_149
CL\_INS\_149
CL\_INS\_149
CL\_INS\_149
CL\_INS\_149
CL\_INS\_382
CL\_INS\_149
CL\_INS\_149
CL\_INS\_149
CL\_INS\_149
CL\_INS\_149
CL\_INS\_149
CL\_INS\_149
CL\_INS\_149
CL\_INS\_149
CL\_INS\_149
CL\_INS\_149
CL\_INS\_149
CL\_INS\_149
CL\_INS\_149
CL\_INS\_149
CL\_INS\_149
CL\_INS\_149
CL\_INS\_149
CL\_INS\_149
CL\_INS\_149
CL\_INS\_149
CL\_INS\_149
CL\_INS\_149
CL\_INS\_149
CL\_INS\_382
CL\_INS\_382
CL\_INS\_149
CL\_INS\_149
CL\_INS\_149
CL\_INS\_149
CL\_INS\_149
CL\_INS\_149
CL\_INS\_149
CL\_INS\_149
CL\_INS\_149
CL\_INS\_149
CL\_INS\_149
CL\_INS\_149
CL\_INS\_149
CL\_INS\_149
CL\_INS\_149
CL\_INS\_149
CL\_INS\_149
CL\_INS\_149
CL\_INS\_149
CL\_INS\_149
CL\_INS\_149
CL\_INS\_149
CL\_INS\_149
CL\_INS\_149
CL\_INS\_149
CL\_INS\_149
CL\_INS\_149
CL\_INS\_149
CL\_INS\_149
CL\_INS\_149
CL\_INS\_149
CL\_INS\_149
CL\_INS\_149
CL\_INS\_149
CL\_INS\_149
CL\_INS\_149
CL\_INS\_149
CL\_INS\_149
CL\_INS\_149
CL\_INS\_149
CL\_INS\_149
CL\_INS\_149
CL\_INS\_149
CL\_INS\_149
CL\_INS\_149
CL\_INS\_149
CL\_INS\_149
CL\_INS\_149
CL\_INS\_149
CL\_INS\_149
CL\_INS\_149
CL\_INS\_149
CL\_INS\_149
CL\_INS\_149
CL\_INS\_149
CL\_INS\_149
CL\_INS\_149
CL\_INS\_149
CL\_INS\_149
CL\_INS\_149
CL\_INS\_149
CL\_INS\_149
CL\_INS\_149
CL\_INS\_149
CL\_INS\_149
CL\_INS\_149
CL\_INS\_149
CL\_INS\_149
CL\_INS\_149
CL\_INS\_149
CL\_INS\_149
CL\_INS\_149
CL\_INS\_149
CL\_INS\_149
CL\_INS\_149
CL\_INS\_149
CL\_INS\_149
CL\_INS\_149
CL\_INS\_149
CL\_INS\_149
CL\_INS\_149
CL\_INS\_149
CL\_INS\_149
CL\_INS\_149
CL\_INS\_149
CL\_INS\_149
CL\_INS\_149
CL\_INS\_149
CL\_INS\_149
CL\_INS\_149
CL\_INS\_149
CL\_INS\_149
CL\_INS\_149
CL\_INS\_149
CL\_INS\_149
CL\_INS\_149
CL\_INS\_149
CL\_INS\_149
CL\_INS\_149
CL\_INS\_149
CL\_INS\_149
CL\_INS\_149
CL\_INS\_149
CL\_INS\_149
CL\_INS\_149
CL\_INS\_149
CL\_INS\_149
CL\_INS\_149
CL\_INS\_149
CL\_INS\_149
CL\_INS\_149
CL\_INS\_149
CL\_INS\_149
CL\_INS\_149
CL\_INS\_149
CL\_INS\_149
CL\_INS\_149
CL\_INS\_149
CL\_INS\_149
CL\_INS\_149
CL\_INS\_149
CL\_INS\_149
CL\_INS\_149
CL\_INS\_149
CL\_INS\_149
CL\_INS\_149
CL\_INS\_149
CL\_INS\_149
CL\_INS\_149
CL\_INS\_149
CL\_INS\_149
CL\_INS\_149
CL\_INS\_149
CL\_INS\_149
CL\_INS\_149
CL\_INS\_149
CL\_INS\_149
CL\_INS\_149
CL\_INS\_149
CL\_INS\_149
CL\_INS\_149
CL\_INS\_149
CL\_INS\_382
CL\_INS\_149
CL\_INS\_149
CL\_INS\_149
CL\_INS\_149
CL\_INS\_149
CL\_INS\_149
CL\_INS\_149
CL\_INS\_149
CL\_INS\_149
CL\_INS\_170
CL\_INS\_149
CL\_INS\_149
CL\_INS\_170
CL\_INS\_149
CL\_INS\_149
CL\_INS\_149
CL\_INS\_149
CL\_INS\_149
CL\_INS\_149
CL\_INS\_149
CL\_INS\_149
CL\_INS\_149
CL\_INS\_149
CL\_INS\_149
CL\_INS\_149
CL\_INS\_149
CL\_INS\_149
CL\_INS\_149
CL\_INS\_149
CL\_INS\_170
CL\_INS\_149
CL\_INS\_149
CL\_INS\_149
CL\_INS\_149
CL\_INS\_149
CL\_INS\_149
CL\_INS\_149
CL\_INS\_149
CL\_INS\_149
CL\_INS\_149
CL\_INS\_170
CL\_INS\_149
CL\_INS\_170
CL\_INS\_170
CL\_INS\_149
CL\_INS\_149
CL\_INS\_204
CL\_INS\_149
CL\_INS\_149
CL\_INS\_149
CL\_INS\_170
CL\_INS\_149
CL\_INS\_204
CL\_INS\_149
CL\_INS\_170
CL\_INS\_149
CL\_INS\_149
CL\_INS\_149
CL\_INS\_149
CL\_INS\_149
CL\_INS\_149
CL\_INS\_149
CL\_INS\_149
CL\_INS\_149
CL\_INS\_149
CL\_INS\_170
CL\_INS\_149
CL\_INS\_149
CL\_INS\_149
CL\_INS\_149
CL\_INS\_149
CL\_INS\_149
CL\_INS\_149
CL\_INS\_149
CL\_INS\_149
CL\_INS\_149
CL\_INS\_149
CL\_INS\_149
CL\_INS\_149
CL\_INS\_149
CL\_INS\_149
CL\_INS\_149
Cluster ID


CL\_11745
CL\_11744
CL\_16954
CL\_24346
CL\_19769
CL\_11826
CL\_10807
CL\_29785
CL\_29784
CL\_13912
CL\_27695
CL\_27694
CL\_30044
CL\_30043
CL\_30042
CL\_19770
CL\_5146
CL\_14063
CL\_14062
CL\_14061
CL\_14060
CL\_19771
CL\_19772
CL\_19773
CL\_19774
CL\_19775
CL\_37476
CL\_37475
CL\_37474
CL\_37473
CL\_37472
CL\_37471
CL\_37470
CL\_37469
CL\_9138
CL\_4093
CL\_26172
CL\_10951
CL\_4678
CL\_4679
CL\_4681
CL\_4682
CL\_4683
CL\_4684
CL\_4685
CL\_10952
CL\_10953
CL\_4687
CL\_4688
CL\_4689
CL\_4690
CL\_4691
CL\_4692
CL\_4693
CL\_4694
CL\_4695
CL\_4696
CL\_4697
CL\_4698
CL\_4699
CL\_4700
CL\_4701
CL\_4702
CL\_4705
CL\_4706
CL\_4707
CL\_4708
CL\_4709
CL\_4710
CL\_4711
CL\_4712
CL\_35071
CL\_35072
CL\_4713
CL\_20927
CL\_10954
CL\_37790
CL\_37789
CL\_37788
CL\_19226
CL\_19225
CL\_19224
CL\_10955
CL\_10956
CL\_32446
CL\_32447
CL\_35631
CL\_10957
CL\_4720
CL\_4721
CL\_4722
CL\_4723
CL\_10958
CL\_10959
CL\_4724
CL\_4725
CL\_4726
CL\_10960
CL\_10961
CL\_4727
CL\_4728
CL\_4729
CL\_14283
CL\_10962
CL\_10963
CL\_4731
CL\_4732
CL\_4733
CL\_4734
CL\_10964
CL\_4735
CL\_10965
CL\_21535
CL\_21536
CL\_21537
CL\_15869
CL\_15870
CL\_15871
CL\_15872
CL\_20232
CL\_15873
CL\_15874
CL\_15875
CL\_15876
CL\_6757
CL\_6756
CL\_6755
CL\_6754
CL\_6753
CL\_6752
CL\_7487
CL\_24347
CL\_7604
CL\_11251
CL\_7603
CL\_7486
CL\_33015
CL\_33016
CL\_7485
CL\_37549
CL\_37550
CL\_25512
CL\_25516
CL\_25517
CL\_7484
CL\_7483
CL\_20231
CL\_7482
CL\_7481
CL\_7480
CL\_34380
CL\_34433
CL\_7479
CL\_7478
CL\_37461
CL\_7477
CL\_7476
CL\_7475
CL\_27693
CL\_27692
CL\_5140
CL\_7474
CL\_36526
CL\_15877
CL\_36524
CL\_36525
CL\_7473
CL\_13908
CL\_13907
CL\_13906
CL\_11250
CL\_7490
CL\_13911
CL\_13910
CL\_13909
CL\_7489
CL\_7488
CL\_37468
CL\_37467
CL\_37466
CL\_37465
CL\_37464
CL\_37463
CL\_37462
CL\_15878
CL\_7606
CL\_7605
CL\_33014
CL\_17510
CL\_17511
CL\_19821
CL\_17512
CL\_17513
CL\_17514
CL\_17515
CL\_33109
CL\_33108
CL\_33107
CL\_33106
CL\_33105
CL\_33104
CL\_33103
CL\_4574
CL\_4575
CL\_4576
CL\_8986
CL\_4577
CL\_4578
CL\_11043
CL\_17516
CL\_16372
CL\_33102
CL\_33101
CL\_14648
CL\_9188
CL\_9189
CL\_11045
CL\_12557
CL\_4579
CL\_4580
CL\_9186
CL\_9187
CL\_29792
CL\_11994
CL\_8683
CL\_16373
CL\_16374
CL\_17517
CL\_33100
CL\_33099
CL\_19822
CL\_11048
CL\_21538
CL\_9190
CL\_4585
CL\_31840
CL\_6599
CL\_4586
CL\_33098
CL\_33097
CL\_6529
CL\_33096
CL\_33095
CL\_33094
CL\_33093
CL\_33092
CL\_33091
CL\_33090
CL\_33089
CL\_14492
CL\_5199
CL\_5200
CL\_14647
CL\_33088
CL\_4587
CL\_4588
CL\_4589
CL\_4590
CL\_4591
CL\_4592
CL\_4593
CL\_11993
CL\_4594
CL\_4595
CL\_4596
CL\_4597
CL\_4598
CL\_4599
CL\_7073
CL\_12560
CL\_2276
CL\_11444
CL\_4600
CL\_8067
CL\_4601
CL\_4602
CL\_4603
CL\_4604
CL\_4605
CL\_5883
CL\_31839
CL\_33087
CL\_33086
CL\_33085
CL\_33084
CL\_33083
CL\_27602
CL\_27603
CL\_4606
CL\_4607
CL\_4608
CL\_4609
CL\_29594
CL\_29595
CL\_4610
CL\_4611
CL\_27930
CL\_28303
CL\_9192
CL\_19823
CL\_19824
CL\_6732
CL\_33220
CL\_4613
CL\_31838
CL\_4614
CL\_17518
