## Supplementary material for "A novel method for integrating genomic and Tn-Seq data to identify common *in vivo* fitness mechanisms across multiple bacterial species": S1 Dataset: CL_INS_154.html

Legend

 Mobile +extrachromosomalelementfunctions
 Regulatoryfunctions
 Hypothetical
 DNA Metabolism
 All EssentialGenes
 Biosynthesis ofcofactors,prostheticgroups, +carriers
 All Fitness Genes
 Proteinsynthesis/fate
 Other
 Transport +binding proteins
 All VFDB Genes

FULL


WINDOWSVGPNG

Trim RowsRemove SingletonsSave Fasta

CL\_1917


CL\_1917


CL\_1917


CL\_1917


CL\_1917


CL\_1917


CL\_1917


CL\_1917


CL\_1917


CL\_1917


CL\_1917


CL\_1917


CL\_1917


CL\_1917


CL\_1917


CL\_1917


CL\_1917


CL\_1917


CL\_1917


CL\_1917


CL\_1917


CL\_1917


CL\_1917


CL\_1917


CL\_1917


CL\_1917


CL\_1917


CL\_1917


CL\_1917


CL\_1917


CL\_1917


CL\_1917


CL\_1917


CL\_1917


CL\_1917


CL\_1917


CL\_1912


CL\_1917


CL\_1917


CL\_1917


CL\_1917


CL\_1915


CL\_1917


CL\_1917


CL\_1917


CL\_1917


CL\_1917


CL\_1917


CL\_1917


CL\_1917


CL\_1917


CL\_1917


CL\_1917


CL\_1917


CL\_1917


CL\_1917


CL\_1917


CL\_1917


CL\_1917


CL\_1917


CL\_1917


CL\_1917


CL\_1917


CL\_1917


CL\_1917


CL\_1917


CL\_1917


CL\_1917


CL\_1917


CL\_1917


CL\_1917


CL\_1917


CL\_1917


CL\_1924


CL\_1917


CL\_1917


CL\_1917


CL\_1917

HighlightSelectShow Genomes


67

CL\_4737


40

CL\_4737


35

CL\_4737


15

CL\_1919


9

CL\_4737


7

CL\_4737


5

CL\_1920


5

CL\_4737


4

CL\_4737


4

CL\_4737


2

CL\_4737


2

CL\_4737


2

CL\_1919


2

CL\_4737


2

CL\_4737


1

CL\_4737


1

CL\_4737


1

CL\_4737


1

CL\_4737


1

CL\_1919


1

CL\_1920


1

CL\_1919


1

CL\_4737


1

CL\_4737


1

CL\_1919


1

CL\_4737


1

CL\_1919


1

CL\_4737


1

CL\_4737


1

CL\_1919


1

CL\_1919


1

CL\_4737


1

CL\_1920


1

CL\_4737


1

CL\_1919


1

Break


1

CL\_4737


1

CL\_4737


1

CL\_4737


1

CL\_1919


1

CL\_1919


1

CL\_4737


1

CL\_1919


1

CL\_4737


1

CL\_4737


1

CL\_4737


1

CL\_4737


1

CL\_4737


1

CL\_4737


1

CL\_4737


1

CL\_4737


1

CL\_1879


1

CL\_4737


1

CL\_1919


1

CL\_4737


1

CL\_1923


1

CL\_1919


1

CL\_4737


1

CL\_1919


1

CL\_1923


1

CL\_4737


1

CL\_1919


1

CL\_4737


1

CL\_4737


1

CL\_4737


1

CL\_1919


1

CL\_4737


1

CL\_1919


1

CL\_1921


1

CL\_1923


1

CL\_1919


1

CL\_1919


1

CL\_4737


1

CL\_4737


1

CL\_1919


1

CL\_4737


1

CL\_4737


1

CL\_4737

fGI ID


CL\_INS\_154
CL\_INS\_154
CL\_INS\_154
CL\_INS\_154
CL\_INS\_154
CL\_INS\_154
CL\_INS\_154
CL\_INS\_154
CL\_INS\_154
CL\_INS\_156
CL\_INS\_156
CL\_INS\_156
CL\_INS\_156
CL\_INS\_156
CL\_INS\_156
CL\_INS\_155
CL\_INS\_155
CL\_INS\_155
CL\_INS\_154
CL\_INS\_154
CL\_INS\_157
CL\_INS\_156
CL\_INS\_207
CL\_INS\_154
CL\_INS\_207
CL\_INS\_207
CL\_INS\_155
CL\_INS\_155
CL\_INS\_154
CL\_INS\_155
CL\_INS\_155
CL\_INS\_155
CL\_INS\_155
CL\_INS\_157
CL\_INS\_157
CL\_INS\_154
CL\_INS\_157
CL\_INS\_154
CL\_INS\_154
CL\_INS\_155
CL\_INS\_155
CL\_INS\_155
CL\_INS\_155
CL\_INS\_155
CL\_INS\_155
CL\_INS\_237
CL\_INS\_207
CL\_INS\_154
CL\_INS\_154
CL\_INS\_207
CL\_INS\_155
CL\_INS\_385
CL\_INS\_385
CL\_INS\_385
CL\_INS\_155
CL\_INS\_155
CL\_INS\_154
CL\_INS\_154
CL\_INS\_237
CL\_INS\_237
CL\_INS\_237
CL\_INS\_237
CL\_INS\_237
CL\_INS\_237
CL\_INS\_154
CL\_INS\_155
CL\_INS\_155
CL\_INS\_385
CL\_INS\_154
CL\_INS\_155
CL\_INS\_155
CL\_INS\_155
CL\_INS\_155
CL\_INS\_155
CL\_INS\_155
CL\_INS\_155
CL\_INS\_155
CL\_INS\_154
CL\_INS\_154
CL\_INS\_154
CL\_INS\_382
CL\_INS\_382
CL\_INS\_154
CL\_INS\_154
CL\_INS\_154
CL\_INS\_146
CL\_INS\_154
CL\_INS\_154
CL\_INS\_132
CL\_INS\_154
CL\_INS\_132
CL\_INS\_132
CL\_INS\_132
CL\_INS\_132
CL\_INS\_132
CL\_INS\_132
CL\_INS\_154
CL\_INS\_132
CL\_INS\_247
CL\_INS\_106
CL\_INS\_106
CL\_INS\_106
CL\_INS\_106
CL\_INS\_106
CL\_INS\_154
CL\_INS\_106
CL\_INS\_382
CL\_INS\_382
CL\_INS\_154
CL\_INS\_154
CL\_INS\_154
CL\_INS\_86
CL\_INS\_146
CL\_INS\_154
CL\_INS\_99
CL\_INS\_382
CL\_INS\_382
CL\_INS\_382
CL\_INS\_382
CL\_INS\_382
CL\_INS\_99
CL\_INS\_382
CL\_INS\_382
CL\_INS\_382
CL\_INS\_154
CL\_INS\_154
CL\_INS\_382
CL\_INS\_79
CL\_INS\_382
CL\_INS\_382
CL\_INS\_382
CL\_INS\_382
CL\_INS\_382
CL\_INS\_382
CL\_INS\_155
CL\_INS\_154
CL\_INS\_154
CL\_INS\_154
CL\_INS\_154
CL\_INS\_154
CL\_INS\_154
CL\_INS\_154
CL\_INS\_154
CL\_INS\_155
CL\_INS\_154
CL\_INS\_156
CL\_INS\_154
CL\_INS\_154
CL\_INS\_154
CL\_INS\_156
CL\_INS\_237
CL\_INS\_154
CL\_INS\_154
CL\_INS\_154
CL\_INS\_154
CL\_INS\_154
CL\_INS\_154
CL\_INS\_154
CL\_INS\_154
CL\_INS\_157
CL\_INS\_157
CL\_INS\_157
CL\_INS\_157
CL\_INS\_20
CL\_INS\_20
CL\_INS\_26
CL\_INS\_20
CL\_INS\_20
CL\_INS\_20
CL\_INS\_20
CL\_INS\_20
CL\_INS\_20
CL\_INS\_20
CL\_INS\_20
CL\_INS\_20
CL\_INS\_20
CL\_INS\_20
CL\_INS\_20
CL\_INS\_20
CL\_INS\_20
CL\_INS\_157
CL\_INS\_157
CL\_INS\_157
CL\_INS\_157
CL\_INS\_157
CL\_INS\_157
CL\_INS\_157
CL\_INS\_157
CL\_INS\_42
CL\_INS\_154
CL\_INS\_154
CL\_INS\_154
CL\_INS\_154
CL\_INS\_154
CL\_INS\_154
CL\_INS\_154
CL\_INS\_154
CL\_INS\_154
CL\_INS\_154
CL\_INS\_154
CL\_INS\_154
CL\_INS\_154
CL\_INS\_154
CL\_INS\_154
CL\_INS\_154
CL\_INS\_154
CL\_INS\_154
CL\_INS\_154
CL\_INS\_154
CL\_INS\_154
CL\_INS\_154
CL\_INS\_154
CL\_INS\_154
CL\_INS\_154
CL\_INS\_154
CL\_INS\_154
CL\_INS\_154
CL\_INS\_154
CL\_INS\_155
CL\_INS\_155
CL\_INS\_155
CL\_INS\_155
CL\_INS\_155
CL\_INS\_154
CL\_INS\_154
CL\_INS\_155
CL\_INS\_155
CL\_INS\_155
CL\_INS\_155
CL\_INS\_155
CL\_INS\_155
CL\_INS\_155
CL\_INS\_155
CL\_INS\_155
CL\_INS\_155
CL\_INS\_155
CL\_INS\_155
CL\_INS\_155
CL\_INS\_155
CL\_INS\_155
CL\_INS\_155
CL\_INS\_155
CL\_INS\_155
CL\_INS\_155
CL\_INS\_155
CL\_INS\_155
CL\_INS\_155
CL\_INS\_155
CL\_INS\_155
CL\_INS\_155
CL\_INS\_171
CL\_INS\_154
CL\_INS\_368
CL\_INS\_368
CL\_INS\_154
CL\_INS\_155
CL\_INS\_155
CL\_INS\_155
CL\_INS\_155
CL\_INS\_155
CL\_INS\_155
CL\_INS\_155
CL\_INS\_155
CL\_INS\_155
CL\_INS\_155
CL\_INS\_155
CL\_INS\_155
CL\_INS\_155
CL\_INS\_155
CL\_INS\_155
CL\_INS\_155
CL\_INS\_155
CL\_INS\_155
CL\_INS\_155
CL\_INS\_155
CL\_INS\_155
CL\_INS\_155
CL\_INS\_155
CL\_INS\_155
CL\_INS\_155
CL\_INS\_155
CL\_INS\_155
CL\_INS\_155
CL\_INS\_155
CL\_INS\_155
CL\_INS\_155
CL\_INS\_155
CL\_INS\_155
CL\_INS\_155
CL\_INS\_155
CL\_INS\_155
CL\_INS\_155
CL\_INS\_155
CL\_INS\_155
CL\_INS\_155
CL\_INS\_155
CL\_INS\_155
CL\_INS\_155
CL\_INS\_155
CL\_INS\_155
CL\_INS\_155
CL\_INS\_155
CL\_INS\_155
CL\_INS\_155
CL\_INS\_155
CL\_INS\_155
CL\_INS\_204
CL\_INS\_155
CL\_INS\_155
CL\_INS\_155
CL\_INS\_155
CL\_INS\_155
CL\_INS\_155
CL\_INS\_155
CL\_INS\_155
CL\_INS\_155
CL\_INS\_155
CL\_INS\_155
CL\_INS\_155
CL\_INS\_155
CL\_INS\_155
CL\_INS\_155
CL\_INS\_155
CL\_INS\_155
CL\_INS\_155
CL\_INS\_155
CL\_INS\_154
CL\_INS\_157
CL\_INS\_156
CL\_INS\_237
CL\_INS\_155
CL\_INS\_154
CL\_INS\_156
CL\_INS\_154
CL\_INS\_154
CL\_INS\_154
CL\_INS\_154
CL\_INS\_154
CL\_INS\_154
CL\_INS\_154
CL\_INS\_154
CL\_INS\_154
CL\_INS\_154
CL\_INS\_154
CL\_INS\_154
CL\_INS\_154
CL\_INS\_154
CL\_INS\_154
CL\_INS\_154
Cluster ID


CL\_30893
CL\_30892
CL\_25258
CL\_9512
CL\_13870
CL\_13869
CL\_13868
CL\_13867
CL\_13866
CL\_24176
CL\_24175
CL\_24174
CL\_24173
CL\_24172
CL\_8299
CL\_9209
CL\_9208
CL\_17029
CL\_9734
CL\_9733
CL\_9732
CL\_9207
CL\_9206
CL\_22746
CL\_9205
CL\_9204
CL\_9203
CL\_7792
CL\_9730
CL\_9202
CL\_7791
CL\_7789
CL\_17028
CL\_9729
CL\_9728
CL\_9727
CL\_9726
CL\_9725
CL\_9724
CL\_6000
CL\_7786
CL\_6003
CL\_17027
CL\_17026
CL\_17025
CL\_6001
CL\_6002
CL\_22747
CL\_9731
CL\_7784
CL\_17024
CL\_7782
CL\_9723
CL\_7781
CL\_9199
CL\_9198
CL\_9722
CL\_9721
CL\_9720
CL\_9719
CL\_9718
CL\_9717
CL\_9716
CL\_9715
CL\_9714
CL\_7780
CL\_17023
CL\_7779
CL\_22748
CL\_9196
CL\_9195
CL\_7777
CL\_9194
CL\_7776
CL\_7775
CL\_17022
CL\_17021
CL\_22749
CL\_22750
CL\_22751
CL\_4517
CL\_4518
CL\_36604
CL\_36603
CL\_14803
CL\_8177
CL\_36602
CL\_36601
CL\_10505
CL\_36600
CL\_10504
CL\_10503
CL\_10502
CL\_10500
CL\_10499
CL\_10498
CL\_16133
CL\_10497
CL\_10496
CL\_12139
CL\_13735
CL\_13734
CL\_8651
CL\_8649
CL\_28843
CL\_16469
CL\_16131
CL\_8648
CL\_16130
CL\_36599
CL\_23903
CL\_8180
CL\_16651
CL\_18336
CL\_8183
CL\_8184
CL\_7119
CL\_4543
CL\_4544
CL\_8186
CL\_14812
CL\_15904
CL\_6773
CL\_6450
CL\_36598
CL\_36597
CL\_8676
CL\_14694
CL\_1524
CL\_1525
CL\_1526
CL\_1527
CL\_10982
CL\_10983
CL\_6439
CL\_12501
CL\_11554
CL\_11553
CL\_11552
CL\_11551
CL\_17954
CL\_17953
CL\_17952
CL\_7374
CL\_33887
CL\_24171
CL\_34003
CL\_34004
CL\_34005
CL\_24169
CL\_8272
CL\_25257
CL\_33720
CL\_33888
CL\_33889
CL\_8271
CL\_24584
CL\_24583
CL\_24582
CL\_24188
CL\_24187
CL\_204
CL\_24186
CL\_4325
CL\_4326
CL\_4327
CL\_4328
CL\_4329
CL\_4330
CL\_4331
CL\_4332
CL\_4333
CL\_4334
CL\_4335
CL\_4336
CL\_4337
CL\_4338
CL\_4340
CL\_4341
CL\_4342
CL\_24185
CL\_24184
CL\_24183
CL\_24182
CL\_24181
CL\_24180
CL\_34002
CL\_24179
CL\_4435
CL\_14576
CL\_14577
CL\_14578
CL\_35395
CL\_35394
CL\_35393
CL\_35392
CL\_35391
CL\_35390
CL\_35389
CL\_35388
CL\_35387
CL\_35386
CL\_35385
CL\_35384
CL\_35383
CL\_35382
CL\_35381
CL\_35380
CL\_35379
CL\_35378
CL\_35377
CL\_35376
CL\_35375
CL\_35374
CL\_35373
CL\_35372
CL\_35371
CL\_35370
CL\_11900
CL\_11899
CL\_10692
CL\_10693
CL\_10694
CL\_34625
CL\_5760
CL\_5758
CL\_5756
CL\_5754
CL\_5753
CL\_11897
CL\_11896
CL\_11895
CL\_11894
CL\_10308
CL\_22457
CL\_22456
CL\_10307
CL\_11893
CL\_11892
CL\_11891
CL\_17524
CL\_22455
CL\_22454
CL\_22453
CL\_22452
CL\_22451
CL\_22450
CL\_22449
CL\_22448
CL\_22447
CL\_21675
CL\_26236
CL\_26235
CL\_26234
CL\_26233
CL\_6438
CL\_6437
CL\_6436
CL\_6435
CL\_6434
CL\_14382
CL\_6433
CL\_6432
CL\_10916
CL\_10915
CL\_10914
CL\_10913
CL\_10912
CL\_10911
CL\_10910
CL\_10909
CL\_10907
CL\_10906
CL\_10905
CL\_11004
CL\_11005
CL\_11006
CL\_11007
CL\_11008
CL\_11009
CL\_11010
CL\_11011
CL\_10894
CL\_10893
CL\_10892
CL\_10891
CL\_10890
CL\_10889
CL\_10887
CL\_10885
CL\_10883
CL\_10882
CL\_11024
CL\_11025
CL\_10879
CL\_10878
CL\_10877
CL\_8228
CL\_8229
CL\_8230
CL\_8231
CL\_10870
CL\_10869
CL\_11035
CL\_11036
CL\_8483
CL\_10867
CL\_11037
CL\_6431
CL\_6430
CL\_6429
CL\_6428
CL\_11910
CL\_11909
CL\_11908
CL\_11907
CL\_11906
CL\_11905
CL\_11904
CL\_11903
CL\_11902
CL\_11901
CL\_6427
CL\_12775
CL\_1918
CL\_17020
CL\_19370
CL\_10351
CL\_11610
CL\_8270
CL\_8269
CL\_31887
CL\_8829
CL\_13436
CL\_31888
CL\_31889
CL\_8268
CL\_8267
CL\_8266
CL\_8265
CL\_8264
CL\_5253
CL\_33020
CL\_33021
CL\_33022
CL\_33023
CL\_36596
CL\_33024
CL\_33025
