## Supplementary material for "A novel method for integrating genomic and Tn-Seq data to identify common *in vivo* fitness mechanisms across multiple bacterial species": S1 Dataset: CL_INS_156.html

Legend

 Mobile +extrachromosomalelementfunctions
 Hypothetical
 All Fitness Genes
 Other
 Transport +binding proteins
 All VFDB Genes

FULL


WINDOWSVGPNG

Trim RowsRemove SingletonsSave Fasta

CL\_1922


CL\_1922


CL\_1922


CL\_1922


CL\_1922


CL\_1922


CL\_1922


CL\_1922


CL\_1922


CL\_1922


CL\_1922


CL\_1922


CL\_1922


CL\_1923


CL\_1922


CL\_234


CL\_1925


CL\_1964


CL\_1923


Break


CL\_1922

HighlightSelectShow Genomes


219

CL\_1921


5

CL\_1925


4

CL\_1925


4

CL\_1925


2

CL\_1921


1

CL\_1924


1

CL\_1921


1

CL\_1920


1

CL\_1924


1

CL\_1921


1

CL\_1921


1

CL\_1925


1

CL\_1921


1

CL\_1921


1

CL\_1921


1

CL\_1921


1

CL\_1921


1

CL\_1921


1

CL\_1921


1

CL\_1921


1

CL\_1925

fGI ID


CL\_INS\_156
CL\_INS\_156
CL\_INS\_156
CL\_INS\_156
CL\_INS\_156
CL\_INS\_156
CL\_INS\_156
CL\_INS\_156
CL\_INS\_156
CL\_INS\_156
CL\_INS\_156
CL\_INS\_237
CL\_INS\_156
CL\_INS\_156
CL\_INS\_156
CL\_INS\_237
CL\_INS\_155
CL\_INS\_156
CL\_INS\_156
CL\_INS\_156
CL\_INS\_156
CL\_INS\_156
CL\_INS\_156
CL\_INS\_156
CL\_INS\_156
CL\_INS\_156
CL\_INS\_156
CL\_INS\_156
CL\_INS\_156
CL\_INS\_156
CL\_INS\_156
CL\_INS\_156
CL\_INS\_156
CL\_INS\_156
CL\_INS\_156
CL\_INS\_30
CL\_INS\_30
CL\_INS\_30
CL\_INS\_30
CL\_INS\_30
CL\_INS\_237
CL\_INS\_237
CL\_INS\_237
CL\_INS\_156
CL\_INS\_30
CL\_INS\_237
CL\_INS\_237
CL\_INS\_70
CL\_INS\_237
CL\_INS\_237
CL\_INS\_30
CL\_INS\_30
CL\_INS\_237
CL\_INS\_237
CL\_INS\_159
CL\_INS\_156
CL\_INS\_70
CL\_INS\_237
CL\_INS\_30
CL\_INS\_30
CL\_INS\_30
CL\_INS\_30
CL\_INS\_156
CL\_INS\_156
CL\_INS\_156
CL\_INS\_159
CL\_INS\_156
CL\_INS\_156
CL\_INS\_156
CL\_INS\_156
CL\_INS\_156
CL\_INS\_156
CL\_INS\_156
CL\_INS\_156
CL\_INS\_156
CL\_INS\_156
CL\_INS\_156
CL\_INS\_156
CL\_INS\_156
CL\_INS\_156
CL\_INS\_156
CL\_INS\_156
CL\_INS\_156
CL\_INS\_156
CL\_INS\_156
CL\_INS\_155
CL\_INS\_155
CL\_INS\_156
CL\_INS\_156
CL\_INS\_207
CL\_INS\_207
CL\_INS\_207
CL\_INS\_155
CL\_INS\_155
CL\_INS\_155
CL\_INS\_155
CL\_INS\_156
CL\_INS\_156
CL\_INS\_155
CL\_INS\_385
CL\_INS\_156
CL\_INS\_156
CL\_INS\_155
CL\_INS\_156
CL\_INS\_207
CL\_INS\_156
CL\_INS\_385
CL\_INS\_156
CL\_INS\_156
CL\_INS\_155
CL\_INS\_155
CL\_INS\_156
CL\_INS\_155
CL\_INS\_155
CL\_INS\_155
CL\_INS\_156
CL\_INS\_155
CL\_INS\_155
CL\_INS\_155
CL\_INS\_382
CL\_INS\_382
CL\_INS\_382
CL\_INS\_382
CL\_INS\_382
CL\_INS\_382
CL\_INS\_382
CL\_INS\_159
CL\_INS\_382
CL\_INS\_382
CL\_INS\_382
CL\_INS\_385
CL\_INS\_159
CL\_INS\_382
CL\_INS\_382
CL\_INS\_382
CL\_INS\_382
CL\_INS\_382
CL\_INS\_382
CL\_INS\_382
CL\_INS\_382
CL\_INS\_382
CL\_INS\_382
CL\_INS\_382
CL\_INS\_382
CL\_INS\_382
CL\_INS\_382
CL\_INS\_159
CL\_INS\_382
CL\_INS\_382
CL\_INS\_382
CL\_INS\_382
CL\_INS\_382
CL\_INS\_382
CL\_INS\_382
CL\_INS\_382
CL\_INS\_382
CL\_INS\_382
CL\_INS\_382
CL\_INS\_382
CL\_INS\_382
CL\_INS\_233
CL\_INS\_233
CL\_INS\_233
CL\_INS\_382
CL\_INS\_382
CL\_INS\_382
CL\_INS\_382
CL\_INS\_382
CL\_INS\_382
CL\_INS\_382
CL\_INS\_156
CL\_INS\_156
CL\_INS\_156
CL\_INS\_156
CL\_INS\_156
CL\_INS\_156
CL\_INS\_156
CL\_INS\_156
CL\_INS\_156
CL\_INS\_156
CL\_INS\_156
CL\_INS\_156
CL\_INS\_156
CL\_INS\_156
CL\_INS\_382
CL\_INS\_156
CL\_INS\_156
CL\_INS\_156
CL\_INS\_156
CL\_INS\_156
CL\_INS\_156
CL\_INS\_156
CL\_INS\_156
CL\_INS\_247
CL\_INS\_247
CL\_INS\_382
CL\_INS\_382
CL\_INS\_156
CL\_INS\_156
CL\_INS\_156
CL\_INS\_156
CL\_INS\_156
CL\_INS\_156
CL\_INS\_156
CL\_INS\_156
CL\_INS\_20
CL\_INS\_156
CL\_INS\_20
CL\_INS\_20
CL\_INS\_70
CL\_INS\_20
CL\_INS\_20
CL\_INS\_156
CL\_INS\_156
CL\_INS\_30
CL\_INS\_30
CL\_INS\_156
CL\_INS\_156
CL\_INS\_156
CL\_INS\_156
CL\_INS\_156
CL\_INS\_156
CL\_INS\_156
CL\_INS\_237
CL\_INS\_237
CL\_INS\_237
CL\_INS\_237
CL\_INS\_156
CL\_INS\_156
CL\_INS\_156
CL\_INS\_237
CL\_INS\_237
CL\_INS\_237
CL\_INS\_237
CL\_INS\_237
CL\_INS\_237
CL\_INS\_237
CL\_INS\_237
CL\_INS\_382
CL\_INS\_382
CL\_INS\_382
CL\_INS\_159
CL\_INS\_159
CL\_INS\_156
CL\_INS\_237
CL\_INS\_382
CL\_INS\_382
CL\_INS\_385
Cluster ID


CL\_28955
CL\_13435
CL\_36153
CL\_36154
CL\_36155
CL\_36156
CL\_36157
CL\_36158
CL\_36159
CL\_36160
CL\_36161
CL\_8272
CL\_13432
CL\_24168
CL\_11610
CL\_8270
CL\_8269
CL\_8829
CL\_13328
CL\_24169
CL\_24170
CL\_24171
CL\_24172
CL\_24173
CL\_24174
CL\_24175
CL\_24176
CL\_34013
CL\_34012
CL\_34011
CL\_34010
CL\_34009
CL\_34008
CL\_34007
CL\_34006
CL\_5228
CL\_5229
CL\_5230
CL\_5231
CL\_5232
CL\_8559
CL\_8558
CL\_15555
CL\_34942
CL\_5240
CL\_8127
CL\_8128
CL\_8129
CL\_8131
CL\_7979
CL\_5233
CL\_8511
CL\_6415
CL\_6414
CL\_9871
CL\_34943
CL\_11105
CL\_6730
CL\_6425
CL\_5246
CL\_5247
CL\_5248
CL\_8299
CL\_12024
CL\_12025
CL\_11249
CL\_12026
CL\_12027
CL\_12028
CL\_12029
CL\_12030
CL\_12031
CL\_12032
CL\_12033
CL\_12034
CL\_12035
CL\_12036
CL\_12037
CL\_12038
CL\_12039
CL\_12040
CL\_12041
CL\_12042
CL\_12043
CL\_12044
CL\_9209
CL\_9208
CL\_7794
CL\_9207
CL\_9206
CL\_9205
CL\_9204
CL\_9203
CL\_17029
CL\_7792
CL\_9202
CL\_9201
CL\_9200
CL\_7786
CL\_11546
CL\_35012
CL\_35013
CL\_7789
CL\_7785
CL\_7784
CL\_7783
CL\_7782
CL\_35010
CL\_35011
CL\_9199
CL\_9198
CL\_9197
CL\_9196
CL\_9195
CL\_7777
CL\_35009
CL\_9194
CL\_7776
CL\_7775
CL\_4262
CL\_4263
CL\_5062
CL\_4265
CL\_5580
CL\_5579
CL\_4269
CL\_5575
CL\_4270
CL\_4271
CL\_5574
CL\_5573
CL\_5572
CL\_5637
CL\_5638
CL\_5639
CL\_4277
CL\_4278
CL\_5567
CL\_5566
CL\_5565
CL\_5564
CL\_5563
CL\_5562
CL\_5561
CL\_5560
CL\_4284
CL\_5651
CL\_5556
CL\_5555
CL\_5554
CL\_5553
CL\_5552
CL\_5551
CL\_5659
CL\_11338
CL\_4293
CL\_4294
CL\_5548
CL\_5662
CL\_4299
CL\_12661
CL\_12660
CL\_5545
CL\_4301
CL\_4302
CL\_4303
CL\_10406
CL\_10407
CL\_10411
CL\_5600
CL\_4987
CL\_4988
CL\_4989
CL\_4990
CL\_25916
CL\_25915
CL\_4991
CL\_16462
CL\_4992
CL\_4993
CL\_4994
CL\_15643
CL\_25914
CL\_25913
CL\_5507
CL\_18395
CL\_14567
CL\_14568
CL\_14569
CL\_14570
CL\_25912
CL\_25911
CL\_25910
CL\_4233
CL\_4234
CL\_4235
CL\_4236
CL\_5502
CL\_25909
CL\_25908
CL\_25907
CL\_25906
CL\_25905
CL\_25904
CL\_25903
CL\_4434
CL\_25902
CL\_5495
CL\_5494
CL\_5493
CL\_5492
CL\_5491
CL\_5488
CL\_5486
CL\_5237
CL\_5238
CL\_5483
CL\_5482
CL\_5481
CL\_5477
CL\_5475
CL\_5474
CL\_25901
CL\_7687
CL\_7686
CL\_7685
CL\_7684
CL\_25900
CL\_25899
CL\_25898
CL\_7710
CL\_7709
CL\_7708
CL\_7707
CL\_7706
CL\_8385
CL\_7305
CL\_5151
CL\_6056
CL\_4973
CL\_4974
CL\_14632
CL\_14631
CL\_14630
CL\_6140
CL\_4240
CL\_5059
CL\_5625
