## Supplementary material for "A novel method for integrating genomic and Tn-Seq data to identify common *in vivo* fitness mechanisms across multiple bacterial species": S1 Dataset: CL_INS_157.html

Legend

 Mobile +extrachromosomalelementfunctions
 Regulatoryfunctions
 Hypothetical
 All EssentialGenes
 All Fitness Genes
 Other
 Transport +binding proteins
 All VFDB Genes

FULL


WINDOWSVGPNG

Trim RowsRemove SingletonsSave Fasta

CL\_1923


CL\_1923


CL\_1923


CL\_1923


CL\_1923


CL\_1923


CL\_1923


CL\_1923


CL\_1923


CL\_1923


CL\_1923


CL\_1924


CL\_1925


CL\_1923


CL\_1923


CL\_1923


CL\_1923


CL\_1923

HighlightSelectShow Genomes


227

CL\_1922


2

CL\_1922


1

CL\_1922


1

CL\_1921


1

CL\_1917


1

CL\_1922


1

CL\_1922


1

CL\_1920


1

CL\_1925


1

CL\_1922


1

CL\_1921


1

CL\_1922


1

CL\_1922


1

CL\_1917


1

CL\_1917


1

CL\_1915


1

CL\_3857


1

CL\_1922

fGI ID


CL\_INS\_155
CL\_INS\_157
CL\_INS\_382
CL\_INS\_157
CL\_INS\_157
CL\_INS\_157
CL\_INS\_157
CL\_INS\_157
CL\_INS\_156
CL\_INS\_237
CL\_INS\_155
CL\_INS\_157
CL\_INS\_157
CL\_INS\_156
CL\_INS\_157
CL\_INS\_157
CL\_INS\_155
CL\_INS\_155
CL\_INS\_155
CL\_INS\_155
CL\_INS\_156
CL\_INS\_157
CL\_INS\_157
CL\_INS\_237
CL\_INS\_237
CL\_INS\_237
CL\_INS\_237
CL\_INS\_157
CL\_INS\_155
CL\_INS\_155
CL\_INS\_385
CL\_INS\_156
CL\_INS\_156
CL\_INS\_207
CL\_INS\_156
CL\_INS\_155
CL\_INS\_385
CL\_INS\_156
CL\_INS\_156
CL\_INS\_155
CL\_INS\_385
CL\_INS\_157
CL\_INS\_207
CL\_INS\_237
CL\_INS\_155
CL\_INS\_157
CL\_INS\_157
CL\_INS\_157
CL\_INS\_157
CL\_INS\_157
CL\_INS\_157
CL\_INS\_155
CL\_INS\_155
CL\_INS\_155
CL\_INS\_155
CL\_INS\_155
CL\_INS\_155
CL\_INS\_157
CL\_INS\_237
CL\_INS\_157
CL\_INS\_42
CL\_INS\_157
CL\_INS\_157
CL\_INS\_157
CL\_INS\_157
CL\_INS\_157
CL\_INS\_157
CL\_INS\_157
CL\_INS\_157
CL\_INS\_20
CL\_INS\_20
CL\_INS\_20
CL\_INS\_20
CL\_INS\_20
CL\_INS\_20
CL\_INS\_20
CL\_INS\_20
CL\_INS\_20
CL\_INS\_20
CL\_INS\_20
CL\_INS\_20
CL\_INS\_20
CL\_INS\_20
CL\_INS\_26
CL\_INS\_20
CL\_INS\_20
CL\_INS\_157
CL\_INS\_157
CL\_INS\_157
CL\_INS\_157
CL\_INS\_157
CL\_INS\_156
CL\_INS\_156
CL\_INS\_156
CL\_INS\_156
CL\_INS\_156
CL\_INS\_156
CL\_INS\_156
CL\_INS\_156
CL\_INS\_156
CL\_INS\_156
CL\_INS\_156
CL\_INS\_156
CL\_INS\_156
CL\_INS\_156
CL\_INS\_156
CL\_INS\_156
CL\_INS\_156
CL\_INS\_156
CL\_INS\_156
CL\_INS\_159
CL\_INS\_156
CL\_INS\_155
CL\_INS\_155
CL\_INS\_155
CL\_INS\_155
CL\_INS\_155
CL\_INS\_155
CL\_INS\_155
CL\_INS\_155
CL\_INS\_155
CL\_INS\_155
CL\_INS\_155
CL\_INS\_155
CL\_INS\_155
CL\_INS\_155
CL\_INS\_155
CL\_INS\_155
CL\_INS\_155
CL\_INS\_155
CL\_INS\_155
CL\_INS\_157
CL\_INS\_157
CL\_INS\_157
CL\_INS\_157
CL\_INS\_157
CL\_INS\_157
CL\_INS\_157
CL\_INS\_157
CL\_INS\_156
CL\_INS\_156
Cluster ID


CL\_1918
CL\_16424
CL\_10804
CL\_8304
CL\_8303
CL\_8302
CL\_8301
CL\_8300
CL\_13328
CL\_8272
CL\_8269
CL\_19436
CL\_10351
CL\_13435
CL\_19435
CL\_22566
CL\_7775
CL\_7776
CL\_9194
CL\_7777
CL\_35009
CL\_22567
CL\_22568
CL\_9719
CL\_9720
CL\_16930
CL\_16931
CL\_22569
CL\_9198
CL\_9199
CL\_7782
CL\_35010
CL\_35011
CL\_7784
CL\_7785
CL\_7786
CL\_11546
CL\_35012
CL\_35013
CL\_7789
CL\_7781
CL\_22570
CL\_6002
CL\_6001
CL\_6000
CL\_22571
CL\_22572
CL\_22573
CL\_9726
CL\_9728
CL\_9729
CL\_9202
CL\_7792
CL\_9203
CL\_17029
CL\_9208
CL\_9209
CL\_9732
CL\_10186
CL\_24178
CL\_4435
CL\_24179
CL\_34002
CL\_24180
CL\_24181
CL\_24182
CL\_24183
CL\_24184
CL\_24185
CL\_4342
CL\_4341
CL\_4340
CL\_4338
CL\_4337
CL\_4336
CL\_4335
CL\_4334
CL\_4333
CL\_4332
CL\_4331
CL\_4330
CL\_4329
CL\_4328
CL\_4327
CL\_4326
CL\_4325
CL\_24186
CL\_204
CL\_24187
CL\_24188
CL\_14383
CL\_12044
CL\_12043
CL\_12042
CL\_12041
CL\_12040
CL\_12039
CL\_12038
CL\_12037
CL\_12036
CL\_12035
CL\_12034
CL\_12033
CL\_12032
CL\_12031
CL\_12030
CL\_12029
CL\_12028
CL\_12027
CL\_12026
CL\_11249
CL\_12025
CL\_10308
CL\_11894
CL\_11897
CL\_5753
CL\_5754
CL\_5756
CL\_5758
CL\_10694
CL\_10693
CL\_10692
CL\_11899
CL\_11900
CL\_11901
CL\_11902
CL\_11903
CL\_11906
CL\_11907
CL\_11909
CL\_11910
CL\_36270
CL\_36269
CL\_36268
CL\_36267
CL\_36266
CL\_36265
CL\_36264
CL\_36263
CL\_12024
CL\_8299
