## Supplementary material for "A novel method for integrating genomic and Tn-Seq data to identify common *in vivo* fitness mechanisms across multiple bacterial species": S1 Dataset: CL_INS_158.html

Legend

 Mobile +extrachromosomalelementfunctions
 Regulatoryfunctions
 Hypothetical
 DNA Metabolism
 All EssentialGenes
 Biosynthesis ofcofactors,prostheticgroups, +carriers
 All Fitness Genes
 Proteinsynthesis/fate
 Other
 Transport +binding proteins
 All VFDB Genes

FULL


WINDOWSVGPNG

Trim RowsRemove SingletonsSave Fasta

CL\_1924


CL\_1924


CL\_1922


CL\_1922


CL\_1922


CL\_1924


CL\_1924


CL\_1924


CL\_1922


CL\_1924


CL\_1924


CL\_1924


CL\_1921


CL\_1924


CL\_1924


CL\_1924


CL\_1924


CL\_1924


CL\_1924


CL\_1924


CL\_1923


CL\_1922


CL\_1924


CL\_1924


CL\_1924


Break


CL\_1924


CL\_1924


CL\_1924


CL\_1924


CL\_1924


CL\_1922


CL\_1924


CL\_1924


CL\_1924

HighlightSelectShow Genomes


224

CL\_1925


8

CL\_1925


5

CL\_1925


4

CL\_1925


4

CL\_1925


2

CL\_1925


1

CL\_1925


1

CL\_1928


1

CL\_1925


1

CL\_1925


1

CL\_1925


1

CL\_1925


1

CL\_1925


1

CL\_2296


1

CL\_1925


1

CL\_1925


1

CL\_1925


1

CL\_1922


1

CL\_1925


1

CL\_1925


1

CL\_1925


1

CL\_1925


1

CL\_1922


1

CL\_1925


1

CL\_1926


1

CL\_1925


1

CL\_1925


1

CL\_1925


1

CL\_4737


1

CL\_1925


1

CL\_234


1

CL\_1925


1

CL\_1925


1

CL\_1925


1

CL\_1925

fGI ID


CL\_INS\_158
CL\_INS\_156
CL\_INS\_156
CL\_INS\_156
CL\_INS\_156
CL\_INS\_156
CL\_INS\_156
CL\_INS\_154
CL\_INS\_154
CL\_INS\_154
CL\_INS\_156
CL\_INS\_156
CL\_INS\_155
CL\_INS\_237
CL\_INS\_156
CL\_INS\_156
CL\_INS\_154
CL\_INS\_158
CL\_INS\_207
CL\_INS\_158
CL\_INS\_158
CL\_INS\_158
CL\_INS\_158
CL\_INS\_158
CL\_INS\_156
CL\_INS\_158
CL\_INS\_158
CL\_INS\_156
CL\_INS\_156
CL\_INS\_158
CL\_INS\_158
CL\_INS\_158
CL\_INS\_158
CL\_INS\_158
CL\_INS\_237
CL\_INS\_158
CL\_INS\_158
CL\_INS\_158
CL\_INS\_158
CL\_INS\_158
CL\_INS\_224
CL\_INS\_158
CL\_INS\_158
CL\_INS\_158
CL\_INS\_158
CL\_INS\_158
CL\_INS\_158
CL\_INS\_158
CL\_INS\_158
CL\_INS\_158
CL\_INS\_158
CL\_INS\_158
CL\_INS\_158
CL\_INS\_158
CL\_INS\_158
CL\_INS\_158
CL\_INS\_158
CL\_INS\_158
CL\_INS\_158
CL\_INS\_156
CL\_INS\_156
CL\_INS\_156
CL\_INS\_159
CL\_INS\_156
CL\_INS\_156
CL\_INS\_156
CL\_INS\_156
CL\_INS\_156
CL\_INS\_156
CL\_INS\_156
CL\_INS\_156
CL\_INS\_156
CL\_INS\_156
CL\_INS\_156
CL\_INS\_156
CL\_INS\_156
CL\_INS\_156
CL\_INS\_156
CL\_INS\_156
CL\_INS\_156
CL\_INS\_156
CL\_INS\_156
CL\_INS\_158
CL\_INS\_158
CL\_INS\_154
CL\_INS\_237
CL\_INS\_157
CL\_INS\_155
CL\_INS\_155
CL\_INS\_155
CL\_INS\_157
CL\_INS\_157
CL\_INS\_157
CL\_INS\_157
CL\_INS\_157
CL\_INS\_157
CL\_INS\_158
CL\_INS\_158
CL\_INS\_155
CL\_INS\_158
CL\_INS\_158
CL\_INS\_385
CL\_INS\_155
CL\_INS\_156
CL\_INS\_207
CL\_INS\_158
CL\_INS\_385
CL\_INS\_153
CL\_INS\_207
CL\_INS\_207
CL\_INS\_155
CL\_INS\_237
CL\_INS\_207
CL\_INS\_157
CL\_INS\_385
CL\_INS\_155
CL\_INS\_385
CL\_INS\_158
CL\_INS\_158
CL\_INS\_155
CL\_INS\_155
CL\_INS\_155
CL\_INS\_155
CL\_INS\_385
CL\_INS\_155
CL\_INS\_155
CL\_INS\_157
CL\_INS\_237
CL\_INS\_237
CL\_INS\_158
CL\_INS\_154
CL\_INS\_154
CL\_INS\_237
CL\_INS\_237
CL\_INS\_157
CL\_INS\_157
CL\_INS\_155
CL\_INS\_237
CL\_INS\_237
CL\_INS\_237
CL\_INS\_237
CL\_INS\_155
CL\_INS\_155
CL\_INS\_158
CL\_INS\_158
CL\_INS\_158
CL\_INS\_158
CL\_INS\_158
CL\_INS\_158
CL\_INS\_158
CL\_INS\_158
CL\_INS\_158
CL\_INS\_158
CL\_INS\_158
CL\_INS\_158
CL\_INS\_158
CL\_INS\_158
CL\_INS\_155
CL\_INS\_155
CL\_INS\_155
CL\_INS\_155
CL\_INS\_155
CL\_INS\_155
CL\_INS\_155
CL\_INS\_155
CL\_INS\_155
CL\_INS\_155
CL\_INS\_155
CL\_INS\_155
CL\_INS\_155
CL\_INS\_155
CL\_INS\_155
CL\_INS\_155
CL\_INS\_155
CL\_INS\_155
CL\_INS\_155
CL\_INS\_155
CL\_INS\_155
CL\_INS\_155
CL\_INS\_155
CL\_INS\_162
CL\_INS\_154
CL\_INS\_154
CL\_INS\_201
CL\_INS\_155
CL\_INS\_155
CL\_INS\_155
CL\_INS\_155
CL\_INS\_158
CL\_INS\_155
CL\_INS\_155
CL\_INS\_155
CL\_INS\_155
CL\_INS\_155
CL\_INS\_155
CL\_INS\_155
CL\_INS\_155
CL\_INS\_155
CL\_INS\_158
CL\_INS\_158
CL\_INS\_158
CL\_INS\_158
CL\_INS\_158
CL\_INS\_158
CL\_INS\_158
CL\_INS\_158
CL\_INS\_158
CL\_INS\_158
CL\_INS\_158
CL\_INS\_155
CL\_INS\_155
CL\_INS\_155
CL\_INS\_158
CL\_INS\_155
CL\_INS\_171
CL\_INS\_171
CL\_INS\_171
CL\_INS\_154
CL\_INS\_368
CL\_INS\_368
CL\_INS\_154
CL\_INS\_158
CL\_INS\_158
CL\_INS\_158
CL\_INS\_158
CL\_INS\_158
CL\_INS\_158
CL\_INS\_123
CL\_INS\_123
CL\_INS\_158
CL\_INS\_158
CL\_INS\_158
CL\_INS\_158
CL\_INS\_158
CL\_INS\_158
CL\_INS\_158
CL\_INS\_386
CL\_INS\_386
CL\_INS\_123
CL\_INS\_123
CL\_INS\_123
CL\_INS\_386
CL\_INS\_386
CL\_INS\_386
CL\_INS\_386
CL\_INS\_386
CL\_INS\_158
CL\_INS\_158
CL\_INS\_157
CL\_INS\_158
CL\_INS\_70
CL\_INS\_70
CL\_INS\_237
CL\_INS\_70
CL\_INS\_70
CL\_INS\_159
CL\_INS\_237
Cluster ID


CL\_24177
CL\_24176
CL\_24175
CL\_24174
CL\_24173
CL\_24172
CL\_24171
CL\_34003
CL\_34004
CL\_34005
CL\_24170
CL\_24169
CL\_8269
CL\_8270
CL\_11610
CL\_24168
CL\_25257
CL\_8830
CL\_10314
CL\_10313
CL\_10312
CL\_10311
CL\_10310
CL\_10309
CL\_13435
CL\_13434
CL\_13433
CL\_13432
CL\_8829
CL\_8828
CL\_13431
CL\_8827
CL\_13430
CL\_13657
CL\_8272
CL\_13658
CL\_13659
CL\_13660
CL\_13661
CL\_13662
CL\_4750
CL\_13663
CL\_13664
CL\_37783
CL\_37782
CL\_13665
CL\_13666
CL\_7774
CL\_7773
CL\_7772
CL\_7771
CL\_7770
CL\_7769
CL\_7768
CL\_7767
CL\_7766
CL\_7765
CL\_7764
CL\_7763
CL\_8299
CL\_12024
CL\_12025
CL\_11249
CL\_12026
CL\_12027
CL\_12028
CL\_12029
CL\_12030
CL\_12031
CL\_12032
CL\_12033
CL\_12034
CL\_12035
CL\_12036
CL\_12037
CL\_12038
CL\_12039
CL\_12040
CL\_12041
CL\_12042
CL\_12043
CL\_12044
CL\_11550
CL\_11549
CL\_9734
CL\_10186
CL\_9732
CL\_9203
CL\_7792
CL\_9202
CL\_9729
CL\_9728
CL\_9726
CL\_22573
CL\_22572
CL\_22571
CL\_10187
CL\_10188
CL\_7789
CL\_11548
CL\_11547
CL\_11546
CL\_7786
CL\_7785
CL\_7784
CL\_11545
CL\_7782
CL\_10189
CL\_10190
CL\_10191
CL\_6000
CL\_6001
CL\_6002
CL\_22570
CL\_7781
CL\_7780
CL\_7779
CL\_11544
CL\_11543
CL\_9196
CL\_9195
CL\_7777
CL\_9194
CL\_9723
CL\_9199
CL\_9198
CL\_22569
CL\_16931
CL\_16930
CL\_10192
CL\_9722
CL\_9721
CL\_9720
CL\_9719
CL\_22568
CL\_22567
CL\_7776
CL\_9718
CL\_9717
CL\_9716
CL\_9715
CL\_7775
CL\_6439
CL\_10193
CL\_10194
CL\_10195
CL\_10196
CL\_10197
CL\_10198
CL\_10199
CL\_10200
CL\_10201
CL\_10202
CL\_10203
CL\_10204
CL\_10205
CL\_10206
CL\_6438
CL\_6437
CL\_6436
CL\_6435
CL\_6434
CL\_6433
CL\_6432
CL\_6431
CL\_6430
CL\_6429
CL\_6428
CL\_11910
CL\_11909
CL\_11908
CL\_11907
CL\_11906
CL\_11905
CL\_11904
CL\_11903
CL\_11902
CL\_11901
CL\_11900
CL\_11899
CL\_5761
CL\_34625
CL\_5760
CL\_5759
CL\_10692
CL\_10693
CL\_10694
CL\_5758
CL\_11898
CL\_5756
CL\_5754
CL\_5753
CL\_11897
CL\_11896
CL\_11895
CL\_11894
CL\_10308
CL\_10307
CL\_10306
CL\_10305
CL\_10304
CL\_10303
CL\_10302
CL\_10301
CL\_10300
CL\_10299
CL\_10298
CL\_10297
CL\_10296
CL\_11893
CL\_11892
CL\_11891
CL\_11890
CL\_11889
CL\_21676
CL\_21675
CL\_21674
CL\_26236
CL\_26235
CL\_26234
CL\_26233
CL\_18337
CL\_18338
CL\_18339
CL\_18340
CL\_18341
CL\_18342
CL\_10148
CL\_10147
CL\_18343
CL\_18344
CL\_18345
CL\_18346
CL\_18347
CL\_18348
CL\_18349
CL\_34626
CL\_34627
CL\_9178
CL\_9179
CL\_9180
CL\_34628
CL\_35229
CL\_34629
CL\_34630
CL\_34631
CL\_34632
CL\_34633
CL\_14383
CL\_26232
CL\_6421
CL\_6420
CL\_28451
CL\_6419
CL\_6418
CL\_6727
CL\_6813
