## Supplementary material for "A novel method for integrating genomic and Tn-Seq data to identify common *in vivo* fitness mechanisms across multiple bacterial species": S1 Dataset: CL_INS_159.html

Legend

 Mobile +extrachromosomalelementfunctions
 Regulatoryfunctions
 Hypothetical
 DNA Metabolism
 All EssentialGenes
 All Fitness Genes
 Biosynthesis ofcofactors,prostheticgroups, +carriers
 Other
 EnergyMetabolism
 Transport +binding proteins
 All VFDB Genes

FULL


WINDOWSVGPNG

Trim RowsRemove SingletonsSave Fasta

CL\_1928


CL\_1935


CL\_1935


CL\_1928


CL\_1928


CL\_1928


CL\_1928


CL\_1928


CL\_1948


CL\_1928


CL\_1928


CL\_1928


CL\_1928


CL\_1928


CL\_1928


CL\_1928


CL\_1935


CL\_1928


CL\_1935


CL\_1928


CL\_1926


CL\_1928


CL\_1928


CL\_1928


CL\_1928


CL\_1928


CL\_1928


CL\_1928


CL\_1928


CL\_1928


CL\_1928


CL\_1928


CL\_1928


CL\_1928


CL\_1928


CL\_339


CL\_1928


CL\_1928


CL\_1928


CL\_1928


CL\_1928


CL\_1928


CL\_1928


CL\_1928


CL\_1928


CL\_1928


CL\_1928


CL\_1928


CL\_1928


CL\_1928


CL\_1928


CL\_1928


CL\_1928


CL\_1928


CL\_1928


CL\_1928


CL\_1928


CL\_1928


CL\_1928


CL\_339


CL\_1928


CL\_1935


CL\_982


CL\_1935


CL\_1928


CL\_1928


CL\_1928


CL\_1928


CL\_1928


CL\_1928


CL\_1928


CL\_1928


CL\_1928


CL\_1935


CL\_1928


CL\_1928


CL\_1928


CL\_1928


CL\_1928


CL\_1928


CL\_1928


CL\_1928


CL\_234


CL\_1928


CL\_1928


CL\_1928


CL\_1928


CL\_1928


CL\_339


CL\_1928


CL\_1928


CL\_1928


CL\_1928


CL\_1928


CL\_3417


CL\_1927


CL\_1928


CL\_1928


CL\_1928


CL\_1928


CL\_1928


CL\_1928


CL\_1928


CL\_1928


CL\_1935


CL\_1928


CL\_1928


CL\_1928


CL\_1928


CL\_1928


CL\_339


CL\_1928


CL\_1928


CL\_1928


CL\_1928


CL\_1928


CL\_234


CL\_339


CL\_1928


CL\_1928


CL\_1928


CL\_1928


Break


CL\_1928


CL\_1928


CL\_1928


CL\_1928


CL\_1928


CL\_1935


CL\_1928


CL\_1928


CL\_1928


CL\_1928


CL\_1928


CL\_1928


CL\_1935


CL\_3417


CL\_1928


CL\_1928


CL\_1928


CL\_1928


CL\_1928


CL\_1928


CL\_234


CL\_1928


CL\_1935


CL\_1928


CL\_1928


CL\_1935


CL\_1928


CL\_1928


CL\_1928


CL\_1928


CL\_1928


CL\_1928


CL\_1928


CL\_1928


CL\_234


CL\_1935


CL\_1928


CL\_1928


CL\_1928


CL\_1928


CL\_1928


CL\_1928


CL\_1925


CL\_1928


CL\_1928


CL\_1928


CL\_1928


CL\_1928


CL\_1928


CL\_234


CL\_1928


CL\_1928


CL\_234


CL\_1928


CL\_1928


CL\_1928


CL\_1928


CL\_1928


CL\_1928


CL\_1928


CL\_1928


CL\_234


CL\_1928


CL\_1928


CL\_1928


CL\_1928


CL\_1928


CL\_1928


CL\_1928


CL\_1928


CL\_1928


CL\_1928


CL\_1928


CL\_1928


CL\_1928


CL\_1928

HighlightSelectShow Genomes


37

CL\_1940


27

CL\_1940


23

CL\_1940


19

CL\_1940


16

CL\_341


12

CL\_341


10

CL\_1941


4

CL\_234


4

CL\_1940


4

CL\_341


3

CL\_1940


3

CL\_1940


2

CL\_1935


2

CL\_1940


2

CL\_1940


2

CL\_3420


2

CL\_1940


2

CL\_1940


1

CL\_1940


1

CL\_1940


1

CL\_1940


1

CL\_1940


1

CL\_1940


1

CL\_1935


1

CL\_1940


1

CL\_1940


1

CL\_1940


1

CL\_1940


1

CL\_1940


1

CL\_1941


1

CL\_1940


1

CL\_1940


1

CL\_1940


1

CL\_1941


1

CL\_1935


1

CL\_1940


1

CL\_234


1

CL\_1935


1

CL\_976


1

CL\_1940


1

CL\_1940


1

CL\_1940


1

CL\_234


1

CL\_1935


1

CL\_234


1

CL\_234


1

CL\_1940


1

CL\_1940


1

CL\_1940


1

CL\_1940


1

CL\_1940


1

CL\_1940


1

CL\_1940


1

CL\_234


1

CL\_1940


1

CL\_1940


1

CL\_234


1

CL\_1940


1

CL\_1940


1

CL\_1940


1

CL\_1940


1

CL\_1940


1

CL\_1940


1

CL\_1940


1

CL\_1940


1

CL\_1940


1

CL\_234


1

CL\_1940


1

CL\_1940


1

CL\_341


1

CL\_234


1

CL\_1940


1

CL\_1940


1

CL\_1940


1

CL\_1940


1

CL\_1940


1

CL\_234


1

CL\_1940


1

CL\_1940


1

CL\_1940


1

CL\_1940


1

CL\_1940


1

CL\_1940


1

CL\_1940


1

CL\_1940


1

CL\_1940


1

CL\_234


1

CL\_1940


1

CL\_1940


1

CL\_234


1

CL\_1935


1

CL\_1940


1

CL\_1940


1

CL\_234


1

CL\_1940


1

CL\_1940


1

CL\_1941


1

CL\_1940


1

CL\_1940


1

CL\_234


1

CL\_1940


1

CL\_1940


1

CL\_1940


1

CL\_1940


1

CL\_1940


1

CL\_1941


1

CL\_1940


1

CL\_1941


1

CL\_1940


1

CL\_1941


1

CL\_1940


1

CL\_341


1

CL\_1940


1

CL\_1940


1

CL\_234


1

CL\_1940


1

CL\_1940


1

CL\_1940


1

CL\_1941


1

CL\_1940


1

CL\_1940


1

CL\_1940


1

CL\_1940


1

CL\_1940


1

CL\_1940


1

CL\_1935


1

CL\_234


1

CL\_1935


1

CL\_1940


1

CL\_234


1

CL\_1940


1

CL\_234


1

CL\_1940


1

CL\_1940


1

CL\_234


1

CL\_1940


1

CL\_1940


1

CL\_1940


1

CL\_1940


1

CL\_1940


1

CL\_1940


1

CL\_1940


1

CL\_1940


1

CL\_1940


1

CL\_1940


1

CL\_1940


1

CL\_1940


1

CL\_1935


1

CL\_1940


1

CL\_1940


1

CL\_1940


1

CL\_1935


1

CL\_1940


1

CL\_1941


1

CL\_1940


1

CL\_1940


1

CL\_1940


1

CL\_1940


1

CL\_1940


1

CL\_1941


1

CL\_1940


1

CL\_234


1

CL\_1935


1

CL\_1940


1

CL\_1940


1

CL\_1940


1

CL\_1940


1

CL\_234


1

CL\_1940


1

CL\_1940


1

CL\_1940


1

CL\_1940


1

CL\_1940


1

CL\_339


1

CL\_1935


1

CL\_1940


1

CL\_234


1

CL\_1940


1

CL\_1940


1

CL\_1935


1

CL\_1940


1

CL\_1940


1

CL\_234


1

CL\_1940


1

CL\_1940


1

CL\_1940


1

CL\_1940


1

CL\_1940


1

CL\_1940


1

CL\_1940


1

CL\_1940


1

CL\_1940


1

CL\_1940


1

CL\_1940


1

CL\_1940


1

CL\_1940


1

CL\_1940


1

CL\_1940


1

CL\_1940

fGI ID


CL\_INS\_159
CL\_INS\_159
CL\_INS\_159
CL\_INS\_159
CL\_INS\_159
CL\_INS\_159
CL\_INS\_159
CL\_INS\_159
CL\_INS\_159
CL\_INS\_159
CL\_INS\_159
CL\_INS\_159
CL\_INS\_159
CL\_INS\_159
CL\_INS\_159
CL\_INS\_159
CL\_INS\_159
CL\_INS\_159
CL\_INS\_159
CL\_INS\_159
CL\_INS\_159
CL\_INS\_159
CL\_INS\_159
CL\_INS\_159
CL\_INS\_159
CL\_INS\_159
CL\_INS\_159
CL\_INS\_159
CL\_INS\_159
CL\_INS\_237
CL\_INS\_159
CL\_INS\_237
CL\_INS\_159
CL\_INS\_237
CL\_INS\_159
CL\_INS\_159
CL\_INS\_159
CL\_INS\_159
CL\_INS\_159
CL\_INS\_159
CL\_INS\_159
CL\_INS\_149
CL\_INS\_70
CL\_INS\_159
CL\_INS\_159
CL\_INS\_159
CL\_INS\_159
CL\_INS\_159
CL\_INS\_159
CL\_INS\_159
CL\_INS\_159
CL\_INS\_247
CL\_INS\_149
CL\_INS\_159
CL\_INS\_31
CL\_INS\_31
CL\_INS\_31
CL\_INS\_31
CL\_INS\_382
CL\_INS\_31
CL\_INS\_237
CL\_INS\_31
CL\_INS\_159
CL\_INS\_159
CL\_INS\_30
CL\_INS\_159
CL\_INS\_159
CL\_INS\_159
CL\_INS\_382
CL\_INS\_159
CL\_INS\_159
CL\_INS\_159
CL\_INS\_159
CL\_INS\_159
CL\_INS\_159
CL\_INS\_159
CL\_INS\_159
CL\_INS\_159
CL\_INS\_159
CL\_INS\_159
CL\_INS\_159
CL\_INS\_159
CL\_INS\_159
CL\_INS\_159
CL\_INS\_159
CL\_INS\_159
CL\_INS\_159
CL\_INS\_159
CL\_INS\_159
CL\_INS\_159
CL\_INS\_159
CL\_INS\_159
CL\_INS\_159
CL\_INS\_159
CL\_INS\_159
CL\_INS\_159
CL\_INS\_159
CL\_INS\_159
CL\_INS\_159
CL\_INS\_159
CL\_INS\_70
CL\_INS\_70
CL\_INS\_70
CL\_INS\_159
CL\_INS\_159
CL\_INS\_30
CL\_INS\_30
CL\_INS\_159
CL\_INS\_30
CL\_INS\_159
CL\_INS\_159
CL\_INS\_30
CL\_INS\_159
CL\_INS\_159
CL\_INS\_70
CL\_INS\_159
CL\_INS\_30
CL\_INS\_30
CL\_INS\_159
CL\_INS\_30
CL\_INS\_30
CL\_INS\_159
CL\_INS\_237
CL\_INS\_70
CL\_INS\_159
CL\_INS\_30
CL\_INS\_159
CL\_INS\_237
CL\_INS\_237
CL\_INS\_159
CL\_INS\_237
CL\_INS\_237
CL\_INS\_237
CL\_INS\_70
CL\_INS\_237
CL\_INS\_237
CL\_INS\_237
CL\_INS\_70
CL\_INS\_70
CL\_INS\_70
CL\_INS\_70
CL\_INS\_70
CL\_INS\_237
CL\_INS\_237
CL\_INS\_70
CL\_INS\_70
CL\_INS\_159
CL\_INS\_159
CL\_INS\_159
CL\_INS\_159
CL\_INS\_159
CL\_INS\_70
CL\_INS\_159
CL\_INS\_70
CL\_INS\_237
CL\_INS\_70
CL\_INS\_70
CL\_INS\_159
CL\_INS\_159
CL\_INS\_237
CL\_INS\_237
CL\_INS\_70
CL\_INS\_70
CL\_INS\_237
CL\_INS\_159
CL\_INS\_159
CL\_INS\_159
CL\_INS\_159
CL\_INS\_70
CL\_INS\_159
CL\_INS\_159
CL\_INS\_237
CL\_INS\_237
CL\_INS\_30
CL\_INS\_30
CL\_INS\_247
CL\_INS\_159
CL\_INS\_159
CL\_INS\_237
CL\_INS\_237
CL\_INS\_30
CL\_INS\_30
CL\_INS\_159
CL\_INS\_159
CL\_INS\_159
CL\_INS\_159
CL\_INS\_237
CL\_INS\_237
CL\_INS\_70
CL\_INS\_70
CL\_INS\_70
CL\_INS\_159
CL\_INS\_70
CL\_INS\_159
CL\_INS\_70
CL\_INS\_237
CL\_INS\_30
CL\_INS\_30
CL\_INS\_237
CL\_INS\_247
CL\_INS\_159
CL\_INS\_159
CL\_INS\_159
CL\_INS\_30
CL\_INS\_30
CL\_INS\_159
CL\_INS\_30
CL\_INS\_159
CL\_INS\_30
CL\_INS\_159
CL\_INS\_30
CL\_INS\_30
CL\_INS\_237
CL\_INS\_30
CL\_INS\_159
CL\_INS\_159
CL\_INS\_159
CL\_INS\_159
CL\_INS\_159
CL\_INS\_159
CL\_INS\_159
CL\_INS\_159
CL\_INS\_159
CL\_INS\_30
CL\_INS\_159
CL\_INS\_159
CL\_INS\_159
CL\_INS\_30
CL\_INS\_237
CL\_INS\_30
CL\_INS\_159
CL\_INS\_159
CL\_INS\_159
CL\_INS\_159
CL\_INS\_30
CL\_INS\_70
CL\_INS\_70
CL\_INS\_247
CL\_INS\_149
CL\_INS\_30
CL\_INS\_30
CL\_INS\_30
CL\_INS\_159
CL\_INS\_159
CL\_INS\_159
CL\_INS\_237
CL\_INS\_237
CL\_INS\_30
CL\_INS\_159
CL\_INS\_159
CL\_INS\_159
CL\_INS\_30
CL\_INS\_86
CL\_INS\_86
CL\_INS\_159
CL\_INS\_30
CL\_INS\_30
CL\_INS\_30
CL\_INS\_30
CL\_INS\_30
CL\_INS\_30
CL\_INS\_30
CL\_INS\_247
CL\_INS\_159
CL\_INS\_159
CL\_INS\_247
CL\_INS\_159
CL\_INS\_159
CL\_INS\_70
CL\_INS\_159
CL\_INS\_159
CL\_INS\_159
CL\_INS\_30
CL\_INS\_159
CL\_INS\_30
CL\_INS\_247
CL\_INS\_247
CL\_INS\_159
CL\_INS\_159
CL\_INS\_159
CL\_INS\_159
CL\_INS\_30
CL\_INS\_159
CL\_INS\_237
CL\_INS\_30
CL\_INS\_159
CL\_INS\_30
CL\_INS\_30
CL\_INS\_70
CL\_INS\_30
CL\_INS\_30
CL\_INS\_30
CL\_INS\_237
CL\_INS\_30
CL\_INS\_30
CL\_INS\_30
CL\_INS\_30
CL\_INS\_237
CL\_INS\_237
CL\_INS\_30
CL\_INS\_159
CL\_INS\_30
CL\_INS\_30
CL\_INS\_30
CL\_INS\_159
CL\_INS\_159
CL\_INS\_159
CL\_INS\_30
CL\_INS\_70
CL\_INS\_30
CL\_INS\_159
CL\_INS\_123
CL\_INS\_382
CL\_INS\_382
CL\_INS\_237
CL\_INS\_159
CL\_INS\_159
CL\_INS\_159
CL\_INS\_159
CL\_INS\_159
CL\_INS\_159
CL\_INS\_70
CL\_INS\_159
CL\_INS\_237
CL\_INS\_159
CL\_INS\_237
CL\_INS\_159
CL\_INS\_159
CL\_INS\_159
CL\_INS\_159
CL\_INS\_237
CL\_INS\_237
CL\_INS\_159
CL\_INS\_159
CL\_INS\_159
CL\_INS\_159
CL\_INS\_70
CL\_INS\_382
CL\_INS\_159
CL\_INS\_159
CL\_INS\_159
CL\_INS\_159
CL\_INS\_159
CL\_INS\_159
CL\_INS\_159
CL\_INS\_159
CL\_INS\_159
CL\_INS\_159
CL\_INS\_159
CL\_INS\_159
CL\_INS\_382
CL\_INS\_382
CL\_INS\_382
CL\_INS\_159
CL\_INS\_159
CL\_INS\_159
CL\_INS\_159
CL\_INS\_159
CL\_INS\_159
CL\_INS\_159
CL\_INS\_382
CL\_INS\_159
CL\_INS\_382
CL\_INS\_159
CL\_INS\_159
CL\_INS\_159
CL\_INS\_159
CL\_INS\_159
CL\_INS\_382
CL\_INS\_382
CL\_INS\_382
CL\_INS\_159
CL\_INS\_159
CL\_INS\_159
CL\_INS\_159
CL\_INS\_159
CL\_INS\_159
CL\_INS\_159
CL\_INS\_159
CL\_INS\_382
CL\_INS\_382
CL\_INS\_159
CL\_INS\_159
CL\_INS\_159
CL\_INS\_159
CL\_INS\_159
CL\_INS\_159
CL\_INS\_159
CL\_INS\_159
CL\_INS\_237
CL\_INS\_159
CL\_INS\_159
CL\_INS\_159
CL\_INS\_159
CL\_INS\_159
CL\_INS\_159
CL\_INS\_159
CL\_INS\_159
CL\_INS\_159
CL\_INS\_159
CL\_INS\_159
CL\_INS\_159
CL\_INS\_159
CL\_INS\_159
CL\_INS\_159
CL\_INS\_159
CL\_INS\_159
CL\_INS\_159
CL\_INS\_159
CL\_INS\_159
CL\_INS\_159
CL\_INS\_159
CL\_INS\_159
CL\_INS\_159
CL\_INS\_123
CL\_INS\_159
CL\_INS\_159
CL\_INS\_247
CL\_INS\_159
CL\_INS\_159
CL\_INS\_159
CL\_INS\_159
CL\_INS\_159
CL\_INS\_159
CL\_INS\_159
CL\_INS\_159
CL\_INS\_159
CL\_INS\_159
CL\_INS\_159
CL\_INS\_382
CL\_INS\_382
CL\_INS\_382
CL\_INS\_233
CL\_INS\_233
CL\_INS\_233
CL\_INS\_233
CL\_INS\_233
CL\_INS\_382
CL\_INS\_382
CL\_INS\_382
CL\_INS\_382
CL\_INS\_382
CL\_INS\_159
CL\_INS\_159
CL\_INS\_382
CL\_INS\_159
CL\_INS\_382
CL\_INS\_382
CL\_INS\_382
CL\_INS\_382
CL\_INS\_382
CL\_INS\_382
CL\_INS\_159
CL\_INS\_382
CL\_INS\_382
CL\_INS\_382
CL\_INS\_382
CL\_INS\_159
CL\_INS\_159
CL\_INS\_382
CL\_INS\_382
CL\_INS\_382
CL\_INS\_382
CL\_INS\_382
CL\_INS\_382
CL\_INS\_382
CL\_INS\_382
CL\_INS\_382
CL\_INS\_382
CL\_INS\_382
CL\_INS\_382
CL\_INS\_382
CL\_INS\_382
CL\_INS\_382
CL\_INS\_159
CL\_INS\_159
CL\_INS\_159
CL\_INS\_382
CL\_INS\_382
CL\_INS\_382
CL\_INS\_382
CL\_INS\_382
CL\_INS\_382
CL\_INS\_159
CL\_INS\_159
CL\_INS\_159
CL\_INS\_159
CL\_INS\_382
CL\_INS\_159
CL\_INS\_159
CL\_INS\_382
CL\_INS\_159
CL\_INS\_159
CL\_INS\_159
CL\_INS\_159
CL\_INS\_159
CL\_INS\_159
CL\_INS\_159
CL\_INS\_159
CL\_INS\_159
CL\_INS\_159
CL\_INS\_159
CL\_INS\_159
CL\_INS\_159
CL\_INS\_159
CL\_INS\_159
CL\_INS\_237
CL\_INS\_159
CL\_INS\_159
CL\_INS\_159
CL\_INS\_159
CL\_INS\_159
CL\_INS\_159
CL\_INS\_159
CL\_INS\_159
CL\_INS\_159
CL\_INS\_159
CL\_INS\_159
CL\_INS\_159
CL\_INS\_159
CL\_INS\_159
CL\_INS\_159
CL\_INS\_159
Cluster ID


CL\_30333
CL\_22914
CL\_15165
CL\_21932
CL\_24107
CL\_1929
CL\_36527
CL\_29337
CL\_29336
CL\_29335
CL\_37578
CL\_4738
CL\_5848
CL\_5849
CL\_5850
CL\_5851
CL\_5852
CL\_5853
CL\_5854
CL\_5855
CL\_5856
CL\_5857
CL\_5858
CL\_5859
CL\_5860
CL\_5861
CL\_5862
CL\_5863
CL\_21065
CL\_8721
CL\_980
CL\_979
CL\_5252
CL\_1930
CL\_36455
CL\_36454
CL\_36453
CL\_36452
CL\_36451
CL\_29596
CL\_9134
CL\_9138
CL\_6426
CL\_5251
CL\_13865
CL\_13864
CL\_13863
CL\_13862
CL\_978
CL\_19211
CL\_19210
CL\_6844
CL\_6732
CL\_10295
CL\_26231
CL\_26230
CL\_26229
CL\_7762
CL\_7761
CL\_7760
CL\_26431
CL\_7759
CL\_7758
CL\_12045
CL\_7757
CL\_12046
CL\_37159
CL\_12047
CL\_7692
CL\_11249
CL\_11248
CL\_33588
CL\_11247
CL\_11246
CL\_11245
CL\_11244
CL\_11243
CL\_11242
CL\_11241
CL\_11240
CL\_11239
CL\_11238
CL\_11237
CL\_11236
CL\_11235
CL\_11234
CL\_11233
CL\_11232
CL\_11231
CL\_11230
CL\_11229
CL\_34188
CL\_11228
CL\_26253
CL\_11227
CL\_11226
CL\_11225
CL\_11224
CL\_11223
CL\_11222
CL\_10330
CL\_10331
CL\_11221
CL\_11220
CL\_33587
CL\_7090
CL\_5250
CL\_24581
CL\_5249
CL\_28954
CL\_13550
CL\_5248
CL\_9511
CL\_9584
CL\_6826
CL\_13551
CL\_7089
CL\_5247
CL\_22752
CL\_5246
CL\_6425
CL\_28863
CL\_6730
CL\_11105
CL\_9871
CL\_6422
CL\_34141
CL\_10822
CL\_7832
CL\_34657
CL\_10548
CL\_11974
CL\_11973
CL\_10546
CL\_7256
CL\_6729
CL\_6728
CL\_6421
CL\_6420
CL\_8508
CL\_15755
CL\_10547
CL\_8505
CL\_8506
CL\_1932
CL\_1933
CL\_37106
CL\_32809
CL\_32810
CL\_32811
CL\_10549
CL\_7831
CL\_25946
CL\_8620
CL\_13552
CL\_8642
CL\_6417
CL\_17662
CL\_17663
CL\_17157
CL\_7207
CL\_6419
CL\_6418
CL\_4375
CL\_6727
CL\_8733
CL\_8734
CL\_8735
CL\_7829
CL\_8736
CL\_8737
CL\_7288
CL\_6813
CL\_7751
CL\_235
CL\_6726
CL\_22753
CL\_6725
CL\_6724
CL\_7205
CL\_2551
CL\_5245
CL\_10545
CL\_8722
CL\_8723
CL\_23878
CL\_6825
CL\_10375
CL\_8037
CL\_6824
CL\_7255
CL\_37107
CL\_8126
CL\_24297
CL\_1931
CL\_12176
CL\_7254
CL\_6424
CL\_16977
CL\_10372
CL\_24296
CL\_8724
CL\_8725
CL\_23183
CL\_23184
CL\_28864
CL\_23185
CL\_5244
CL\_5243
CL\_5242
CL\_6423
CL\_4983
CL\_6731
CL\_5241
CL\_8726
CL\_8727
CL\_8728
CL\_8729
CL\_8730
CL\_8731
CL\_8732
CL\_14384
CL\_14385
CL\_5240
CL\_22565
CL\_22564
CL\_7471
CL\_5239
CL\_11715
CL\_5238
CL\_33419
CL\_29597
CL\_29598
CL\_29599
CL\_5237
CL\_7252
CL\_6828
CL\_5236
CL\_4093
CL\_8511
CL\_12048
CL\_5235
CL\_24348
CL\_24349
CL\_6453
CL\_7088
CL\_7087
CL\_5234
CL\_20025
CL\_13807
CL\_13808
CL\_5233
CL\_5815
CL\_5814
CL\_22546
CL\_8738
CL\_7756
CL\_7755
CL\_7754
CL\_7753
CL\_10984
CL\_7752
CL\_7470
CL\_36595
CL\_36594
CL\_17589
CL\_29783
CL\_15911
CL\_10581
CL\_34064
CL\_34065
CL\_15912
CL\_5232
CL\_15913
CL\_5231
CL\_13745
CL\_13744
CL\_13743
CL\_13742
CL\_13741
CL\_29782
CL\_5230
CL\_7975
CL\_28451
CL\_5229
CL\_35834
CL\_5228
CL\_5227
CL\_6416
CL\_5226
CL\_13809
CL\_1934
CL\_7289
CL\_7827
CL\_7750
CL\_7749
CL\_5225
CL\_6723
CL\_6722
CL\_1936
CL\_5224
CL\_7971
CL\_1937
CL\_1938
CL\_28953
CL\_28952
CL\_28448
CL\_1939
CL\_14344
CL\_18097
CL\_6721
CL\_4995
CL\_7691
CL\_8832
CL\_11110
CL\_20717
CL\_16476
CL\_16477
CL\_16478
CL\_16479
CL\_16881
CL\_6411
CL\_20024
CL\_6414
CL\_28951
CL\_6415
CL\_21064
CL\_21063
CL\_28450
CL\_28449
CL\_10609
CL\_6413
CL\_10294
CL\_10293
CL\_10292
CL\_14059
CL\_7253
CL\_4086
CL\_21539
CL\_21540
CL\_21541
CL\_23409
CL\_23408
CL\_31558
CL\_31557
CL\_31556
CL\_31555
CL\_31554
CL\_31553
CL\_31552
CL\_4984
CL\_4235
CL\_4236
CL\_6239
CL\_14143
CL\_14142
CL\_11837
CL\_11836
CL\_5053
CL\_5620
CL\_4256
CL\_31551
CL\_5059
CL\_5577
CL\_31550
CL\_31549
CL\_31548
CL\_5572
CL\_5571
CL\_5570
CL\_4276
CL\_31547
CL\_31546
CL\_31545
CL\_31544
CL\_31543
CL\_31541
CL\_31540
CL\_31539
CL\_10406
CL\_10407
CL\_4307
CL\_5539
CL\_11666
CL\_11665
CL\_31538
CL\_5031
CL\_5032
CL\_5033
CL\_6967
CL\_31537
CL\_31536
CL\_23393
CL\_23394
CL\_23395
CL\_14113
CL\_31535
CL\_7639
CL\_31534
CL\_31533
CL\_31532
CL\_28652
CL\_28653
CL\_28654
CL\_23412
CL\_28655
CL\_23411
CL\_23410
CL\_31542
CL\_31594
CL\_31593
CL\_31592
CL\_31591
CL\_31590
CL\_13653
CL\_31589
CL\_31588
CL\_7007
CL\_31587
CL\_31586
CL\_31585
CL\_31584
CL\_31583
CL\_31582
CL\_31581
CL\_31580
CL\_31579
CL\_31578
CL\_5506
CL\_4310
CL\_5600
CL\_5599
CL\_5541
CL\_6935
CL\_5542
CL\_5543
CL\_5544
CL\_4302
CL\_4301
CL\_4300
CL\_4299
CL\_5662
CL\_5661
CL\_5660
CL\_5549
CL\_5550
CL\_5551
CL\_5552
CL\_5553
CL\_5554
CL\_5555
CL\_5556
CL\_5651
CL\_4284
CL\_5560
CL\_5561
CL\_5562
CL\_5645
CL\_5643
CL\_5563
CL\_5564
CL\_5565
CL\_5566
CL\_5567
CL\_4278
CL\_4277
CL\_5639
CL\_5638
CL\_5637
CL\_6934
CL\_6933
CL\_5634
CL\_4271
CL\_4270
CL\_5575
CL\_6965
CL\_5630
CL\_5579
CL\_5580
CL\_4265
CL\_5062
CL\_4263
CL\_4262
CL\_4261
CL\_4260
CL\_4259
CL\_4258
CL\_4257
CL\_31576
CL\_23619
CL\_5050
CL\_31575
CL\_23621
CL\_23622
CL\_23623
CL\_31574
CL\_14631
CL\_14632
CL\_31573
CL\_20463
CL\_31572
CL\_26904
CL\_23861
CL\_31571
CL\_19710
CL\_31570
CL\_4462
CL\_31569
CL\_31568
CL\_31567
CL\_31566
CL\_31565
CL\_31564
CL\_31563
CL\_31562
CL\_31561
CL\_31560
CL\_31559
CL\_31577
CL\_5864
CL\_5865
CL\_5866
CL\_5867
