## Supplementary material for "A novel method for integrating genomic and Tn-Seq data to identify common *in vivo* fitness mechanisms across multiple bacterial species": S1 Dataset: CL_INS_164.html

Legend

 Hypothetical
 All Fitness Genes
 All VFDB Genes

FULL


WINDOWSVGPNG

Trim RowsRemove SingletonsSave Fasta

CL\_1993


CL\_1993


CL\_1993


CL\_1994


CL\_1993

HighlightSelectShow Genomes


249

CL\_1992


2

CL\_1964


1

CL\_1964


1

CL\_1992


1

Break

fGI ID


CL\_INS\_164
CL\_INS\_164
CL\_INS\_164
CL\_INS\_164
CL\_INS\_164
CL\_INS\_164
CL\_INS\_164
CL\_INS\_164
CL\_INS\_164
CL\_INS\_164
CL\_INS\_164
CL\_INS\_164
Cluster ID


CL\_35223
CL\_35224
CL\_35225
CL\_35226
CL\_35227
CL\_22554
CL\_22553
CL\_22552
CL\_22551
CL\_22550
CL\_22549
CL\_22548
