## Supplementary material for "A novel method for integrating genomic and Tn-Seq data to identify common *in vivo* fitness mechanisms across multiple bacterial species": S1 Dataset: CL_INS_165.html

Legend

 Hypothetical
 All Fitness Genes
 Other
 All VFDB Genes

FULL


WINDOWSVGPNG

Trim RowsRemove SingletonsSave Fasta

CL\_1994


CL\_1994


CL\_1994


CL\_1994


CL\_1994


CL\_1994


CL\_1994


CL\_1994


CL\_1994


CL\_1994


CL\_1994


CL\_1994


CL\_1994

HighlightSelectShow Genomes


248

CL\_1993


3

CL\_1993


2

CL\_1964


2

CL\_1964


1

CL\_1993


1

CL\_1964


1

CL\_1964


1

CL\_1964


1

CL\_1964


1

CL\_1964


1

CL\_1992


1

CL\_1964


1

CL\_1964

fGI ID


CL\_INS\_165
CL\_INS\_165
CL\_INS\_165
CL\_INS\_165
CL\_INS\_165
CL\_INS\_165
CL\_INS\_165
CL\_INS\_165
CL\_INS\_165
CL\_INS\_165
CL\_INS\_165
CL\_INS\_165
CL\_INS\_165
CL\_INS\_165
CL\_INS\_165
CL\_INS\_165
CL\_INS\_165
CL\_INS\_165
CL\_INS\_165
CL\_INS\_165
CL\_INS\_165
CL\_INS\_165
CL\_INS\_165
CL\_INS\_165
CL\_INS\_165
CL\_INS\_165
CL\_INS\_165
CL\_INS\_165
CL\_INS\_165
CL\_INS\_165
CL\_INS\_165
CL\_INS\_165
CL\_INS\_165
CL\_INS\_165
CL\_INS\_165
CL\_INS\_165
CL\_INS\_165
CL\_INS\_165
CL\_INS\_165
CL\_INS\_165
CL\_INS\_165
CL\_INS\_165
CL\_INS\_165
CL\_INS\_165
CL\_INS\_165
CL\_INS\_165
CL\_INS\_165
Cluster ID


CL\_6706
CL\_6707
CL\_6708
CL\_6709
CL\_6710
CL\_6711
CL\_6712
CL\_14069
CL\_14068
CL\_14067
CL\_14066
CL\_24302
CL\_24301
CL\_24300
CL\_24299
CL\_28943
CL\_28944
CL\_28945
CL\_28946
CL\_28947
CL\_28948
CL\_14070
CL\_34813
CL\_34814
CL\_34815
CL\_34816
CL\_34817
CL\_22777
CL\_22778
CL\_22779
CL\_22780
CL\_22781
CL\_22782
CL\_23940
CL\_23941
CL\_23942
CL\_23943
CL\_23944
CL\_23945
CL\_14065
CL\_37653
CL\_37652
CL\_37651
CL\_37650
CL\_37649
CL\_37648
CL\_37647
