## Supplementary material for "A novel method for integrating genomic and Tn-Seq data to identify common *in vivo* fitness mechanisms across multiple bacterial species": S1 Dataset: CL_INS_166.html

Legend

 Mobile +extrachromosomalelementfunctions
 Hypothetical
 All EssentialGenes
 All Fitness Genes
 Other
 All VFDB Genes

FULL


WINDOWSVGPNG

Trim RowsRemove SingletonsSave Fasta

CL\_1996


CL\_1996


CL\_1996


CL\_1996


CL\_1996


CL\_1964


CL\_1996


CL\_1996


CL\_1996


CL\_1961


CL\_1976


CL\_1996


CL\_1964


CL\_1996


CL\_1994


CL\_1996


CL\_1996


CL\_1996

HighlightSelectShow Genomes


201

CL\_1997


47

CL\_1997


7

CL\_1997


3

CL\_1997


2

CL\_1997


2

CL\_1997


1

CL\_1998


1

CL\_1997


1

CL\_1997


1

CL\_1997


1

CL\_1997


1

CL\_1997


1

CL\_1997


1

CL\_1997


1

CL\_1997


1

CL\_1998


1

CL\_1997


1

CL\_1999

fGI ID


CL\_INS\_166
CL\_INS\_166
CL\_INS\_160
CL\_INS\_160
CL\_INS\_166
CL\_INS\_161
CL\_INS\_161
CL\_INS\_166
CL\_INS\_166
CL\_INS\_166
CL\_INS\_166
CL\_INS\_166
CL\_INS\_166
CL\_INS\_166
CL\_INS\_166
CL\_INS\_166
CL\_INS\_166
CL\_INS\_166
CL\_INS\_166
CL\_INS\_166
Cluster ID


CL\_7740
CL\_7084
CL\_5869
CL\_27006
CL\_8740
CL\_8741
CL\_8742
CL\_30807
CL\_6705
CL\_36150
CL\_37654
CL\_15134
CL\_26226
CL\_26225
CL\_26224
CL\_26223
CL\_26222
CL\_6704
CL\_6703
CL\_24350
