## Supplementary material for "A novel method for integrating genomic and Tn-Seq data to identify common *in vivo* fitness mechanisms across multiple bacterial species": S1 Dataset: CL_INS_167.html

Legend

 Hypothetical
 All VFDB Genes

FULL


WINDOWSVGPNG

Trim RowsRemove SingletonsSave Fasta

CL\_1997


CL\_1997


CL\_1997


CL\_1997


CL\_1997


CL\_1996


CL\_1997


CL\_1996

HighlightSelectShow Genomes


224

CL\_1998


35

CL\_1998


5

CL\_1998


2

CL\_1998


1

CL\_1998


1

CL\_1998


1

CL\_1998


1

CL\_1998

fGI ID


CL\_INS\_167
CL\_INS\_167
CL\_INS\_167
CL\_INS\_166
CL\_INS\_167
Cluster ID


CL\_17942
CL\_17941
CL\_11199
CL\_7084
CL\_7083
