## Supplementary material for "A novel method for integrating genomic and Tn-Seq data to identify common *in vivo* fitness mechanisms across multiple bacterial species": S1 Dataset: CL_INS_168.html

Legend

 Hypothetical
 All EssentialGenes
 All VFDB Genes

FULL


WINDOWSVGPNG

Trim RowsRemove SingletonsSave Fasta

CL\_2004


CL\_2004


CL\_2004


CL\_2003


CL\_2003

HighlightSelectShow Genomes


188

CL\_2006


70

CL\_2006


3

CL\_2007


3

CL\_2006


1

CL\_2006

fGI ID

CL\_INS\_168
Cluster ID

CL\_2005
