## Supplementary material for "A novel method for integrating genomic and Tn-Seq data to identify common *in vivo* fitness mechanisms across multiple bacterial species": S1 Dataset: CL_INS_169.html

Legend

 Mobile +extrachromosomalelementfunctions
 Regulatoryfunctions
 Hypothetical
 DNA Metabolism
 All EssentialGenes
 All Fitness Genes
 Proteinsynthesis/fate
 Other
 Transport +binding proteins
 All VFDB Genes

FULL


WINDOWSVGPNG

Trim RowsRemove SingletonsSave Fasta

CL\_2007


CL\_2007


CL\_2007


CL\_2007


CL\_2007


CL\_2007


CL\_2847


CL\_2007


CL\_2007


CL\_2007


CL\_2007


CL\_2006


CL\_2006


CL\_2006


CL\_725


CL\_2007


CL\_2006


CL\_2007


CL\_2007


CL\_2007

HighlightSelectShow Genomes


244

CL\_2008


3

CL\_2008


2

CL\_2012


2

CL\_2008


1

CL\_2008


1

CL\_4516


1

CL\_2008


1

CL\_2008


1

CL\_2008


1

CL\_2008


1

CL\_2008


1

CL\_2008


1

CL\_2008


1

CL\_2008


1

CL\_2008


1

CL\_2008


1

CL\_2008


1

CL\_2008


1

CL\_2008


1

CL\_724

fGI ID


CL\_INS\_169
CL\_INS\_169
CL\_INS\_169
CL\_INS\_169
CL\_INS\_169
CL\_INS\_169
CL\_INS\_169
CL\_INS\_169
CL\_INS\_169
CL\_INS\_169
CL\_INS\_169
CL\_INS\_169
CL\_INS\_169
CL\_INS\_169
CL\_INS\_169
CL\_INS\_385
CL\_INS\_86
CL\_INS\_99
CL\_INS\_382
CL\_INS\_382
CL\_INS\_382
CL\_INS\_382
CL\_INS\_169
CL\_INS\_169
CL\_INS\_169
CL\_INS\_169
CL\_INS\_99
CL\_INS\_60
CL\_INS\_60
CL\_INS\_99
CL\_INS\_60
CL\_INS\_60
CL\_INS\_382
CL\_INS\_86
CL\_INS\_382
CL\_INS\_99
CL\_INS\_382
CL\_INS\_60
CL\_INS\_60
CL\_INS\_382
CL\_INS\_60
CL\_INS\_60
CL\_INS\_60
CL\_INS\_60
CL\_INS\_60
CL\_INS\_60
CL\_INS\_60
CL\_INS\_382
CL\_INS\_382
CL\_INS\_382
CL\_INS\_60
CL\_INS\_60
CL\_INS\_60
CL\_INS\_382
CL\_INS\_382
CL\_INS\_382
CL\_INS\_382
CL\_INS\_382
CL\_INS\_382
CL\_INS\_382
CL\_INS\_382
CL\_INS\_382
CL\_INS\_382
CL\_INS\_382
CL\_INS\_20
CL\_INS\_20
CL\_INS\_20
CL\_INS\_60
CL\_INS\_20
CL\_INS\_20
CL\_INS\_382
CL\_INS\_237
CL\_INS\_237
CL\_INS\_382
CL\_INS\_382
CL\_INS\_382
CL\_INS\_382
CL\_INS\_382
CL\_INS\_382
CL\_INS\_382
CL\_INS\_382
CL\_INS\_382
CL\_INS\_382
CL\_INS\_382
CL\_INS\_382
CL\_INS\_382
CL\_INS\_382
CL\_INS\_382
CL\_INS\_382
CL\_INS\_382
CL\_INS\_382
CL\_INS\_382
CL\_INS\_382
CL\_INS\_382
CL\_INS\_382
CL\_INS\_382
CL\_INS\_382
CL\_INS\_382
CL\_INS\_382
CL\_INS\_382
CL\_INS\_385
CL\_INS\_385
CL\_INS\_385
CL\_INS\_385
CL\_INS\_169
CL\_INS\_169
CL\_INS\_169
CL\_INS\_382
CL\_INS\_382
CL\_INS\_382
CL\_INS\_382
CL\_INS\_169
CL\_INS\_385
CL\_INS\_382
CL\_INS\_382
CL\_INS\_382
CL\_INS\_86
CL\_INS\_99
CL\_INS\_86
CL\_INS\_86
CL\_INS\_99
CL\_INS\_136
CL\_INS\_382
CL\_INS\_382
CL\_INS\_382
CL\_INS\_382
CL\_INS\_60
CL\_INS\_60
CL\_INS\_60
CL\_INS\_60
CL\_INS\_60
CL\_INS\_60
CL\_INS\_60
CL\_INS\_60
CL\_INS\_60
CL\_INS\_237
CL\_INS\_60
CL\_INS\_237
CL\_INS\_382
CL\_INS\_382
CL\_INS\_382
CL\_INS\_60
CL\_INS\_382
CL\_INS\_237
CL\_INS\_237
CL\_INS\_237
CL\_INS\_237
CL\_INS\_237
CL\_INS\_382
CL\_INS\_382
CL\_INS\_207
CL\_INS\_207
CL\_INS\_207
CL\_INS\_60
CL\_INS\_60
CL\_INS\_60
CL\_INS\_60
CL\_INS\_60
CL\_INS\_382
CL\_INS\_382
CL\_INS\_382
CL\_INS\_382
CL\_INS\_382
CL\_INS\_382
CL\_INS\_237
CL\_INS\_382
CL\_INS\_60
CL\_INS\_237
CL\_INS\_237
CL\_INS\_237
CL\_INS\_237
CL\_INS\_237
CL\_INS\_237
CL\_INS\_237
CL\_INS\_237
CL\_INS\_237
CL\_INS\_382
CL\_INS\_237
CL\_INS\_237
CL\_INS\_237
CL\_INS\_237
CL\_INS\_382
CL\_INS\_60
CL\_INS\_20
CL\_INS\_20
CL\_INS\_60
CL\_INS\_60
CL\_INS\_60
CL\_INS\_60
CL\_INS\_60
CL\_INS\_382
CL\_INS\_382
CL\_INS\_382
CL\_INS\_60
CL\_INS\_60
CL\_INS\_60
CL\_INS\_79
Cluster ID


CL\_12511
CL\_32454
CL\_12786
CL\_22907
CL\_6397
CL\_23662
CL\_9215
CL\_21706
CL\_27691
CL\_27690
CL\_27689
CL\_27688
CL\_17345
CL\_18915
CL\_18914
CL\_18913
CL\_4490
CL\_4618
CL\_532
CL\_15241
CL\_17747
CL\_6664
CL\_27964
CL\_27963
CL\_27962
CL\_27961
CL\_9162
CL\_33898
CL\_17343
CL\_9160
CL\_17342
CL\_33899
CL\_4432
CL\_7025
CL\_4517
CL\_7112
CL\_11957
CL\_33900
CL\_33901
CL\_22576
CL\_33902
CL\_33903
CL\_33904
CL\_33905
CL\_33906
CL\_33907
CL\_33908
CL\_28529
CL\_25129
CL\_28347
CL\_33909
CL\_33910
CL\_33911
CL\_9098
CL\_9009
CL\_9097
CL\_4469
CL\_9096
CL\_9095
CL\_9094
CL\_9093
CL\_9092
CL\_9091
CL\_9090
CL\_7047
CL\_7046
CL\_7045
CL\_16352
CL\_7043
CL\_5380
CL\_5347
CL\_8320
CL\_8321
CL\_5348
CL\_5349
CL\_5350
CL\_4410
CL\_5351
CL\_5352
CL\_5353
CL\_5354
CL\_5355
CL\_5356
CL\_5357
CL\_5358
CL\_5359
CL\_5360
CL\_4401
CL\_508
CL\_4400
CL\_5361
CL\_5362
CL\_5363
CL\_5364
CL\_7618
CL\_5365
CL\_5366
CL\_5367
CL\_13484
CL\_5368
CL\_9102
CL\_9101
CL\_19206
CL\_11304
CL\_27960
CL\_27959
CL\_27958
CL\_12999
CL\_11919
CL\_11920
CL\_7537
CL\_27957
CL\_19205
CL\_9100
CL\_9099
CL\_9158
CL\_4515
CL\_4431
CL\_4430
CL\_4429
CL\_8168
CL\_4484
CL\_6744
CL\_11958
CL\_6743
CL\_4628
CL\_32435
CL\_32434
CL\_26128
CL\_32433
CL\_26129
CL\_26130
CL\_32432
CL\_26131
CL\_26132
CL\_19186
CL\_26133
CL\_7049
CL\_6784
CL\_526
CL\_5809
CL\_26135
CL\_5340
CL\_9121
CL\_9120
CL\_9119
CL\_8319
CL\_6660
CL\_6659
CL\_6658
CL\_26136
CL\_26137
CL\_26138
CL\_26139
CL\_27932
CL\_9116
CL\_6653
CL\_26140
CL\_11915
CL\_25380
CL\_9080
CL\_9079
CL\_9078
CL\_6789
CL\_19187
CL\_8326
CL\_8327
CL\_8328
CL\_11731
CL\_17725
CL\_5384
CL\_5385
CL\_5386
CL\_5387
CL\_8333
CL\_5388
CL\_18912
CL\_5389
CL\_6639
CL\_6637
CL\_11206
CL\_10317
CL\_17549
CL\_8962
CL\_9949
CL\_26141
CL\_26142
CL\_26143
CL\_26144
CL\_26145
CL\_7821
CL\_7032
CL\_6636
CL\_27913
CL\_20985
CL\_33860
CL\_9110
