## Supplementary material for "A novel method for integrating genomic and Tn-Seq data to identify common *in vivo* fitness mechanisms across multiple bacterial species": S1 Dataset: CL_INS_171.html

Legend

 Mobile +extrachromosomalelementfunctions
 Regulatoryfunctions
 Hypothetical
 DNA Metabolism
 All EssentialGenes
 AntibioticResistance
 All Fitness Genes
 Proteinsynthesis/fate
 Other
 Transport +binding proteins
 All VFDB Genes

FULL


WINDOWSVGPNG

Trim RowsRemove SingletonsSave Fasta

CL\_2015


CL\_2015


CL\_2015


CL\_2014


CL\_2015


CL\_2015


CL\_2015


CL\_2015


CL\_2015


CL\_2015


CL\_2015


CL\_2015


CL\_2015


CL\_2015


CL\_2015


CL\_2015


CL\_2015


CL\_2015


CL\_2015


CL\_2015


CL\_2015


CL\_2015


CL\_2015


CL\_2015


CL\_2015


CL\_2015


CL\_2015


Break


CL\_855


CL\_2015


CL\_2015


CL\_2015


CL\_855


Break


CL\_2015


CL\_2015


CL\_2015


CL\_2015


CL\_2015


CL\_2015


CL\_2015


CL\_2015


CL\_2015


CL\_2015


CL\_2014


CL\_2015


CL\_855


CL\_2015


CL\_2015


CL\_2015


CL\_2015


CL\_2015


CL\_2015


CL\_2015


CL\_2015


CL\_2015


CL\_2015


CL\_2015


CL\_2015


CL\_2015

HighlightSelectShow Genomes


197

CL\_2017


26

CL\_2017


2

CL\_2017


2

CL\_2017


1

CL\_2017


1

CL\_2017


1

CL\_2017


1

CL\_2017


1

CL\_2017


1

CL\_2017


1

CL\_2017


1

CL\_2017


1

CL\_2017


1

CL\_2017


1

CL\_2017


1

CL\_2017


1

CL\_2017


1

CL\_2017


1

CL\_2017


1

CL\_2017


1

CL\_2017


1

CL\_2017


1

CL\_2017


1

CL\_2017


1

CL\_2017


1

CL\_2017


1

CL\_2017


1

CL\_2017


1

CL\_2017


1

Break


1

CL\_2017


1

Break


1

CL\_2017


1

CL\_2017


1

CL\_854


1

CL\_854


1

CL\_2017


1

CL\_2017


1

CL\_854


1

CL\_2017


1

CL\_2017


1

CL\_2017


1

CL\_2017


1

CL\_2017


1

CL\_2017


1

CL\_2017


1

CL\_2017


1

CL\_2017


1

CL\_2017


1

CL\_2017


1

CL\_2017


1

CL\_2017


1

CL\_2017


1

CL\_2017


1

CL\_2017


1

CL\_2017


1

CL\_2017


1

CL\_2017


1

CL\_2017


1

CL\_2017

fGI ID


CL\_INS\_171
CL\_INS\_171
CL\_INS\_171
CL\_INS\_171
CL\_INS\_171
CL\_INS\_171
CL\_INS\_171
CL\_INS\_171
CL\_INS\_171
CL\_INS\_170
CL\_INS\_171
CL\_INS\_171
CL\_INS\_171
CL\_INS\_171
CL\_INS\_171
CL\_INS\_171
CL\_INS\_171
CL\_INS\_170
CL\_INS\_170
CL\_INS\_170
CL\_INS\_207
CL\_INS\_171
CL\_INS\_171
CL\_INS\_171
CL\_INS\_170
CL\_INS\_170
CL\_INS\_170
CL\_INS\_171
CL\_INS\_171
CL\_INS\_170
CL\_INS\_171
CL\_INS\_207
CL\_INS\_170
CL\_INS\_171
CL\_INS\_171
CL\_INS\_171
CL\_INS\_171
CL\_INS\_171
CL\_INS\_171
CL\_INS\_171
CL\_INS\_170
CL\_INS\_170
CL\_INS\_170
CL\_INS\_170
CL\_INS\_170
CL\_INS\_207
CL\_INS\_171
CL\_INS\_171
CL\_INS\_171
CL\_INS\_171
CL\_INS\_171
CL\_INS\_171
CL\_INS\_237
CL\_INS\_237
CL\_INS\_204
CL\_INS\_171
CL\_INS\_204
CL\_INS\_155
CL\_INS\_204
CL\_INS\_86
CL\_INS\_237
CL\_INS\_204
CL\_INS\_204
CL\_INS\_207
CL\_INS\_171
CL\_INS\_171
CL\_INS\_171
CL\_INS\_171
CL\_INS\_171
CL\_INS\_171
CL\_INS\_171
CL\_INS\_171
CL\_INS\_171
CL\_INS\_171
CL\_INS\_171
CL\_INS\_171
CL\_INS\_171
CL\_INS\_171
CL\_INS\_382
CL\_INS\_382
CL\_INS\_171
CL\_INS\_99
CL\_INS\_70
CL\_INS\_70
CL\_INS\_149
CL\_INS\_170
CL\_INS\_149
CL\_INS\_382
CL\_INS\_171
CL\_INS\_171
CL\_INS\_171
CL\_INS\_207
CL\_INS\_171
CL\_INS\_171
CL\_INS\_149
CL\_INS\_70
CL\_INS\_70
CL\_INS\_171
CL\_INS\_171
CL\_INS\_171
CL\_INS\_149
CL\_INS\_170
CL\_INS\_149
CL\_INS\_171
CL\_INS\_247
CL\_INS\_247
CL\_INS\_247
CL\_INS\_247
CL\_INS\_247
CL\_INS\_247
CL\_INS\_247
CL\_INS\_247
CL\_INS\_247
CL\_INS\_123
CL\_INS\_123
CL\_INS\_123
CL\_INS\_149
CL\_INS\_149
CL\_INS\_149
CL\_INS\_149
CL\_INS\_149
CL\_INS\_57
CL\_INS\_57
CL\_INS\_171
CL\_INS\_171
CL\_INS\_171
CL\_INS\_171
CL\_INS\_171
CL\_INS\_171
CL\_INS\_171
CL\_INS\_171
CL\_INS\_171
CL\_INS\_171
CL\_INS\_385
CL\_INS\_385
CL\_INS\_385
CL\_INS\_385
CL\_INS\_171
CL\_INS\_247
CL\_INS\_207
CL\_INS\_149
CL\_INS\_170
CL\_INS\_86
CL\_INS\_149
CL\_INS\_149
CL\_INS\_170
CL\_INS\_170
CL\_INS\_170
CL\_INS\_171
CL\_INS\_207
CL\_INS\_171
CL\_INS\_171
CL\_INS\_171
CL\_INS\_171
CL\_INS\_171
CL\_INS\_171
CL\_INS\_207
CL\_INS\_149
CL\_INS\_171
CL\_INS\_171
CL\_INS\_171
CL\_INS\_207
CL\_INS\_170
CL\_INS\_171
CL\_INS\_170
CL\_INS\_171
CL\_INS\_171
CL\_INS\_171
CL\_INS\_171
CL\_INS\_171
CL\_INS\_171
CL\_INS\_171
CL\_INS\_207
CL\_INS\_170
CL\_INS\_171
CL\_INS\_171
CL\_INS\_171
CL\_INS\_207
CL\_INS\_171
CL\_INS\_207
CL\_INS\_207
CL\_INS\_207
CL\_INS\_149
CL\_INS\_171
CL\_INS\_171
CL\_INS\_171
CL\_INS\_171
CL\_INS\_171
CL\_INS\_171
CL\_INS\_171
CL\_INS\_170
CL\_INS\_171
CL\_INS\_170
CL\_INS\_171
CL\_INS\_171
CL\_INS\_170
CL\_INS\_171
CL\_INS\_171
CL\_INS\_171
CL\_INS\_171
CL\_INS\_171
CL\_INS\_171
CL\_INS\_171
CL\_INS\_171
CL\_INS\_171
CL\_INS\_170
CL\_INS\_171
CL\_INS\_171
CL\_INS\_171
CL\_INS\_171
CL\_INS\_171
CL\_INS\_171
CL\_INS\_171
CL\_INS\_171
CL\_INS\_171
CL\_INS\_171
CL\_INS\_171
CL\_INS\_171
CL\_INS\_171
CL\_INS\_171
CL\_INS\_171
CL\_INS\_171
CL\_INS\_171
CL\_INS\_170
CL\_INS\_171
CL\_INS\_149
CL\_INS\_149
CL\_INS\_171
CL\_INS\_170
CL\_INS\_170
CL\_INS\_171
CL\_INS\_171
CL\_INS\_171
CL\_INS\_171
CL\_INS\_171
CL\_INS\_171
CL\_INS\_171
CL\_INS\_170
CL\_INS\_171
CL\_INS\_170
CL\_INS\_170
CL\_INS\_171
CL\_INS\_171
CL\_INS\_171
CL\_INS\_171
CL\_INS\_171
CL\_INS\_149
CL\_INS\_171
CL\_INS\_171
CL\_INS\_171
CL\_INS\_171
CL\_INS\_171
CL\_INS\_171
CL\_INS\_170
CL\_INS\_207
CL\_INS\_170
CL\_INS\_149
CL\_INS\_171
CL\_INS\_171
CL\_INS\_171
CL\_INS\_171
CL\_INS\_171
CL\_INS\_170
CL\_INS\_207
CL\_INS\_171
CL\_INS\_207
CL\_INS\_171
CL\_INS\_171
CL\_INS\_171
CL\_INS\_171
CL\_INS\_171
CL\_INS\_171
CL\_INS\_171
CL\_INS\_171
CL\_INS\_171
CL\_INS\_171
CL\_INS\_171
CL\_INS\_171
CL\_INS\_171
CL\_INS\_207
CL\_INS\_171
CL\_INS\_171
CL\_INS\_171
CL\_INS\_170
CL\_INS\_171
CL\_INS\_171
CL\_INS\_171
CL\_INS\_171
CL\_INS\_171
CL\_INS\_171
CL\_INS\_170
CL\_INS\_171
CL\_INS\_171
CL\_INS\_171
CL\_INS\_171
CL\_INS\_171
CL\_INS\_171
CL\_INS\_171
CL\_INS\_171
CL\_INS\_171
CL\_INS\_171
CL\_INS\_171
CL\_INS\_171
CL\_INS\_171
CL\_INS\_171
CL\_INS\_171
CL\_INS\_171
CL\_INS\_171
CL\_INS\_171
CL\_INS\_171
CL\_INS\_171
CL\_INS\_171
CL\_INS\_171
CL\_INS\_171
CL\_INS\_171
CL\_INS\_171
CL\_INS\_171
CL\_INS\_171
CL\_INS\_170
CL\_INS\_170
CL\_INS\_171
CL\_INS\_171
CL\_INS\_171
CL\_INS\_171
CL\_INS\_171
CL\_INS\_171
CL\_INS\_171
CL\_INS\_170
CL\_INS\_152
CL\_INS\_170
CL\_INS\_171
CL\_INS\_171
CL\_INS\_171
CL\_INS\_170
CL\_INS\_171
CL\_INS\_170
CL\_INS\_171
CL\_INS\_170
CL\_INS\_171
CL\_INS\_171
CL\_INS\_170
CL\_INS\_171
CL\_INS\_171
CL\_INS\_171
CL\_INS\_170
CL\_INS\_171
CL\_INS\_171
CL\_INS\_171
CL\_INS\_171
CL\_INS\_171
CL\_INS\_170
CL\_INS\_172
CL\_INS\_170
CL\_INS\_171
CL\_INS\_171
CL\_INS\_171
CL\_INS\_171
CL\_INS\_171
CL\_INS\_149
CL\_INS\_171
CL\_INS\_171
CL\_INS\_171
CL\_INS\_171
CL\_INS\_171
CL\_INS\_171
CL\_INS\_171
CL\_INS\_171
CL\_INS\_171
Cluster ID


CL\_10289
CL\_29837
CL\_29836
CL\_29835
CL\_29834
CL\_22682
CL\_22681
CL\_14493
CL\_26221
CL\_2016
CL\_33581
CL\_33580
CL\_14451
CL\_24108
CL\_24109
CL\_14452
CL\_32203
CL\_5208
CL\_5207
CL\_5883
CL\_6518
CL\_20139
CL\_23877
CL\_23876
CL\_4608
CL\_7078
CL\_5206
CL\_22692
CL\_5205
CL\_5204
CL\_29334
CL\_11275
CL\_7077
CL\_13365
CL\_29333
CL\_29332
CL\_29331
CL\_20398
CL\_22693
CL\_22694
CL\_7076
CL\_7075
CL\_7074
CL\_5203
CL\_36593
CL\_5202
CL\_12512
CL\_12513
CL\_12514
CL\_12515
CL\_12516
CL\_12517
CL\_24099
CL\_24098
CL\_6520
CL\_15330
CL\_1105
CL\_8483
CL\_1104
CL\_1106
CL\_4514
CL\_2277
CL\_1103
CL\_5884
CL\_35002
CL\_24110
CL\_34704
CL\_23258
CL\_32204
CL\_32205
CL\_16679
CL\_23875
CL\_5885
CL\_5886
CL\_16857
CL\_8066
CL\_8146
CL\_8145
CL\_21780
CL\_21781
CL\_12787
CL\_8194
CL\_5320
CL\_5321
CL\_10957
CL\_4602
CL\_4603
CL\_1519
CL\_19926
CL\_27797
CL\_23231
CL\_7250
CL\_19679
CL\_19678
CL\_8067
CL\_10408
CL\_10409
CL\_30320
CL\_30319
CL\_30318
CL\_4600
CL\_4599
CL\_4598
CL\_25262
CL\_5601
CL\_10383
CL\_10382
CL\_10423
CL\_10421
CL\_10642
CL\_10641
CL\_10395
CL\_10393
CL\_10392
CL\_5297
CL\_5298
CL\_6753
CL\_6754
CL\_6755
CL\_6756
CL\_6757
CL\_6758
CL\_6759
CL\_25532
CL\_25531
CL\_25530
CL\_20559
CL\_20560
CL\_20561
CL\_20562
CL\_20563
CL\_20564
CL\_13405
CL\_13406
CL\_13407
CL\_13408
CL\_5296
CL\_25529
CL\_10388
CL\_10314
CL\_2276
CL\_7073
CL\_5201
CL\_5200
CL\_5199
CL\_5198
CL\_4597
CL\_7072
CL\_29833
CL\_6521
CL\_35001
CL\_16858
CL\_15331
CL\_15332
CL\_8068
CL\_20137
CL\_5197
CL\_4596
CL\_26220
CL\_26219
CL\_26218
CL\_6522
CL\_5196
CL\_29329
CL\_5195
CL\_32206
CL\_32207
CL\_32208
CL\_32209
CL\_32210
CL\_32211
CL\_32212
CL\_6523
CL\_8069
CL\_20136
CL\_22680
CL\_20135
CL\_10567
CL\_29832
CL\_6526
CL\_10566
CL\_6528
CL\_6529
CL\_25250
CL\_25249
CL\_25248
CL\_25247
CL\_25246
CL\_25245
CL\_25244
CL\_5194
CL\_24096
CL\_5193
CL\_24111
CL\_27342
CL\_5192
CL\_24095
CL\_24094
CL\_24093
CL\_24092
CL\_24091
CL\_24090
CL\_24089
CL\_23232
CL\_23233
CL\_5191
CL\_29761
CL\_29760
CL\_29759
CL\_29758
CL\_29757
CL\_29756
CL\_29755
CL\_29754
CL\_29753
CL\_29752
CL\_29751
CL\_29750
CL\_22700
CL\_22701
CL\_22702
CL\_22703
CL\_22704
CL\_5190
CL\_20133
CL\_4590
CL\_4589
CL\_22679
CL\_4587
CL\_5189
CL\_30606
CL\_30607
CL\_30608
CL\_30609
CL\_30610
CL\_23234
CL\_23235
CL\_5188
CL\_5187
CL\_6530
CL\_6531
CL\_12518
CL\_5186
CL\_32213
CL\_32214
CL\_25243
CL\_6599
CL\_29328
CL\_23236
CL\_23237
CL\_5185
CL\_20132
CL\_22705
CL\_4586
CL\_8482
CL\_5887
CL\_9190
CL\_27341
CL\_22678
CL\_22677
CL\_22676
CL\_14455
CL\_4585
CL\_5184
CL\_29831
CL\_9005
CL\_29830
CL\_35000
CL\_34999
CL\_25059
CL\_25060
CL\_9109
CL\_24088
CL\_26217
CL\_26216
CL\_23238
CL\_19927
CL\_23239
CL\_26461
CL\_10564
CL\_10377
CL\_23240
CL\_23241
CL\_10376
CL\_5183
CL\_5182
CL\_5181
CL\_5180
CL\_5179
CL\_20130
CL\_4584
CL\_12519
CL\_19677
CL\_7594
CL\_7593
CL\_26254
CL\_20118
CL\_16859
CL\_16860
CL\_5888
CL\_14453
CL\_14454
CL\_19942
CL\_23124
CL\_23125
CL\_23126
CL\_22706
CL\_22707
CL\_22708
CL\_22709
CL\_22209
CL\_27610
CL\_27611
CL\_27612
CL\_27613
CL\_21674
CL\_21675
CL\_21676
CL\_7070
CL\_7069
CL\_22521
CL\_8144
CL\_8143
CL\_8142
CL\_25528
CL\_8070
CL\_8071
CL\_5889
CL\_5890
CL\_5178
CL\_14456
CL\_14457
CL\_14458
CL\_5177
CL\_8072
CL\_5176
CL\_8074
CL\_5175
CL\_14459
CL\_29327
CL\_5174
CL\_20128
CL\_7592
CL\_6610
CL\_5173
CL\_8075
CL\_6612
CL\_30317
CL\_30316
CL\_30315
CL\_5172
CL\_5171
CL\_5170
CL\_30601
CL\_30602
CL\_30603
CL\_30604
CL\_30605
CL\_14492
CL\_24097
CL\_29330
CL\_22695
CL\_22696
CL\_19943
CL\_22697
CL\_22698
CL\_22699
CL\_2020
