## Supplementary material for "A novel method for integrating genomic and Tn-Seq data to identify common *in vivo* fitness mechanisms across multiple bacterial species": S1 Dataset: CL_INS_172.html

Legend

 Mobile +extrachromosomalelementfunctions
 Regulatoryfunctions
 Hypothetical
 All EssentialGenes
 Other
 All VFDB Genes

FULL


WINDOWSVGPNG

Trim RowsRemove SingletonsSave Fasta

CL\_2018


CL\_2018


CL\_2018


CL\_2018


CL\_2018


CL\_2018


CL\_2018


CL\_2014


CL\_2018


CL\_2018


CL\_2014


CL\_2018

HighlightSelectShow Genomes


261

CL\_2019


9

CL\_2027


1

CL\_2027


1

CL\_2027


1

CL\_2027


1

CL\_2019


1

CL\_2027


1

CL\_2019


1

CL\_2027


1

CL\_2019


1

CL\_2019


1

CL\_2019

fGI ID


CL\_INS\_172
CL\_INS\_172
CL\_INS\_172
CL\_INS\_172
CL\_INS\_172
CL\_INS\_172
CL\_INS\_172
CL\_INS\_172
CL\_INS\_172
CL\_INS\_172
CL\_INS\_170
CL\_INS\_170
CL\_INS\_170
CL\_INS\_170
CL\_INS\_170
CL\_INS\_170
CL\_INS\_170
CL\_INS\_170
CL\_INS\_170
CL\_INS\_170
CL\_INS\_170
CL\_INS\_170
CL\_INS\_170
CL\_INS\_170
CL\_INS\_170
CL\_INS\_170
CL\_INS\_170
CL\_INS\_170
CL\_INS\_170
CL\_INS\_170
CL\_INS\_170
CL\_INS\_170
CL\_INS\_170
CL\_INS\_170
CL\_INS\_170
CL\_INS\_170
CL\_INS\_170
CL\_INS\_170
CL\_INS\_170
CL\_INS\_170
CL\_INS\_170
CL\_INS\_170
CL\_INS\_170
CL\_INS\_170
CL\_INS\_170
CL\_INS\_170
CL\_INS\_170
CL\_INS\_170
CL\_INS\_170
CL\_INS\_170
CL\_INS\_170
CL\_INS\_170
CL\_INS\_170
CL\_INS\_170
CL\_INS\_170
CL\_INS\_170
CL\_INS\_170
CL\_INS\_170
CL\_INS\_172
Cluster ID


CL\_25869
CL\_33514
CL\_6702
CL\_6701
CL\_6700
CL\_6699
CL\_6698
CL\_6697
CL\_6696
CL\_6695
CL\_7080
CL\_7079
CL\_2016
CL\_5208
CL\_7078
CL\_7077
CL\_7076
CL\_7075
CL\_7074
CL\_7073
CL\_6515
CL\_5883
CL\_4608
CL\_5203
CL\_4602
CL\_5170
CL\_7072
CL\_5198
CL\_4597
CL\_5196
CL\_5195
CL\_5194
CL\_5193
CL\_7071
CL\_5192
CL\_5191
CL\_5190
CL\_5189
CL\_5188
CL\_6530
CL\_4587
CL\_4586
CL\_7070
CL\_7069
CL\_6531
CL\_4585
CL\_4584
CL\_36865
CL\_36864
CL\_10376
CL\_5889
CL\_5178
CL\_5177
CL\_5176
CL\_5175
CL\_5174
CL\_5173
CL\_5172
CL\_5171
