## Supplementary material for "A novel method for integrating genomic and Tn-Seq data to identify common *in vivo* fitness mechanisms across multiple bacterial species": S1 Dataset: CL_INS_173.html

Legend

 Mobile +extrachromosomalelementfunctions
 Hypothetical
 Other
 All VFDB Genes

FULL


WINDOWSVGPNG

Trim RowsRemove SingletonsSave Fasta

CL\_2027


CL\_2027


CL\_2027


CL\_2027


CL\_2027


CL\_2027


CL\_2027


CL\_2027


CL\_2027


CL\_2027

HighlightSelectShow Genomes


238

CL\_2026


17

CL\_2026


9

CL\_2018


2

CL\_2026


1

CL\_2018


1

CL\_2026


1

CL\_2018


1

CL\_2018


1

CL\_2018


1

CL\_2018

fGI ID


CL\_INS\_173
CL\_INS\_173
CL\_INS\_173
CL\_INS\_173
CL\_INS\_173
CL\_INS\_172
CL\_INS\_172
CL\_INS\_172
CL\_INS\_172
CL\_INS\_172
CL\_INS\_172
CL\_INS\_172
Cluster ID


CL\_14386
CL\_20224
CL\_7382
CL\_6694
CL\_35185
CL\_6695
CL\_6696
CL\_6697
CL\_6698
CL\_6699
CL\_6700
CL\_6701
