## Supplementary material for "A novel method for integrating genomic and Tn-Seq data to identify common *in vivo* fitness mechanisms across multiple bacterial species": S1 Dataset: CL_INS_176.html

Legend

 Mobile +extrachromosomalelementfunctions
 Regulatoryfunctions
 Hypothetical
 DNA Metabolism
 All EssentialGenes
 All Fitness Genes
 Proteinsynthesis/fate
 Other
 All VFDB Genes
 Transport +binding proteins

FULL


WINDOWSVGPNG

Trim RowsRemove SingletonsSave Fasta

CL\_2053


CL\_2044


CL\_2053


CL\_2053


CL\_2053


CL\_2053


CL\_2053


CL\_2053


CL\_2052


CL\_2053


CL\_2053


CL\_2050


CL\_2053


CL\_2053


CL\_2053


CL\_2053


CL\_2053


CL\_2052


CL\_2053


CL\_2053


CL\_2053


CL\_2053


CL\_2053


CL\_2053


CL\_2053


CL\_2053


CL\_2053


CL\_2053

HighlightSelectShow Genomes


237

CL\_2054


1

CL\_2054


1

CL\_2055


1

CL\_2054


1

CL\_2054


1

CL\_2054


1

CL\_2054


1

CL\_2054


1

CL\_2054


1

CL\_2054


1

CL\_2054


1

CL\_2054


1

CL\_2054


1

CL\_2054


1

CL\_2049


1

CL\_2054


1

CL\_2054


1

CL\_2054


1

CL\_2043


1

CL\_2054


1

CL\_2060


1

CL\_2054


1

CL\_2054


1

CL\_2054


1

CL\_2054


1

CL\_2054


1

CL\_2054


1

CL\_2054

fGI ID


CL\_INS\_176
CL\_INS\_176
CL\_INS\_237
CL\_INS\_382
CL\_INS\_382
CL\_INS\_382
CL\_INS\_382
CL\_INS\_382
CL\_INS\_382
CL\_INS\_207
CL\_INS\_207
CL\_INS\_382
CL\_INS\_382
CL\_INS\_382
CL\_INS\_382
CL\_INS\_176
CL\_INS\_99
CL\_INS\_204
CL\_INS\_99
CL\_INS\_99
CL\_INS\_176
CL\_INS\_176
CL\_INS\_176
CL\_INS\_153
CL\_INS\_176
CL\_INS\_176
CL\_INS\_176
CL\_INS\_176
CL\_INS\_176
CL\_INS\_176
CL\_INS\_176
CL\_INS\_61
CL\_INS\_176
CL\_INS\_176
CL\_INS\_176
CL\_INS\_176
CL\_INS\_176
CL\_INS\_176
CL\_INS\_176
CL\_INS\_146
CL\_INS\_146
CL\_INS\_70
CL\_INS\_176
CL\_INS\_176
CL\_INS\_237
CL\_INS\_99
CL\_INS\_176
CL\_INS\_176
CL\_INS\_76
CL\_INS\_76
CL\_INS\_76
CL\_INS\_76
CL\_INS\_76
CL\_INS\_76
CL\_INS\_83
CL\_INS\_76
CL\_INS\_76
CL\_INS\_76
CL\_INS\_76
CL\_INS\_76
CL\_INS\_76
CL\_INS\_76
CL\_INS\_76
CL\_INS\_176
CL\_INS\_76
CL\_INS\_76
CL\_INS\_76
CL\_INS\_76
CL\_INS\_76
CL\_INS\_76
CL\_INS\_76
CL\_INS\_76
CL\_INS\_76
CL\_INS\_128
CL\_INS\_99
CL\_INS\_176
CL\_INS\_60
CL\_INS\_382
CL\_INS\_76
CL\_INS\_176
CL\_INS\_176
CL\_INS\_176
CL\_INS\_176
CL\_INS\_176
CL\_INS\_237
CL\_INS\_176
CL\_INS\_176
CL\_INS\_176
CL\_INS\_176
CL\_INS\_176
CL\_INS\_176
CL\_INS\_176
CL\_INS\_382
CL\_INS\_382
CL\_INS\_382
CL\_INS\_99
CL\_INS\_83
CL\_INS\_384
CL\_INS\_382
CL\_INS\_382
CL\_INS\_382
CL\_INS\_382
CL\_INS\_176
CL\_INS\_76
CL\_INS\_99
CL\_INS\_382
CL\_INS\_382
CL\_INS\_382
CL\_INS\_153
CL\_INS\_382
CL\_INS\_382
CL\_INS\_382
CL\_INS\_382
CL\_INS\_382
CL\_INS\_382
CL\_INS\_382
CL\_INS\_382
CL\_INS\_128
CL\_INS\_99
CL\_INS\_382
CL\_INS\_382
CL\_INS\_382
CL\_INS\_176
CL\_INS\_176
CL\_INS\_382
CL\_INS\_382
CL\_INS\_176
CL\_INS\_76
CL\_INS\_176
CL\_INS\_176
CL\_INS\_176
CL\_INS\_86
CL\_INS\_382
CL\_INS\_99
CL\_INS\_83
CL\_INS\_382
CL\_INS\_99
CL\_INS\_99
CL\_INS\_382
CL\_INS\_382
CL\_INS\_382
CL\_INS\_99
CL\_INS\_99
CL\_INS\_382
CL\_INS\_382
CL\_INS\_382
CL\_INS\_382
CL\_INS\_382
CL\_INS\_382
CL\_INS\_382
CL\_INS\_176
CL\_INS\_382
CL\_INS\_382
CL\_INS\_382
CL\_INS\_382
CL\_INS\_382
CL\_INS\_382
CL\_INS\_382
CL\_INS\_382
CL\_INS\_382
CL\_INS\_176
CL\_INS\_382
CL\_INS\_382
CL\_INS\_176
CL\_INS\_176
CL\_INS\_176
CL\_INS\_176
CL\_INS\_382
CL\_INS\_382
CL\_INS\_382
CL\_INS\_382
CL\_INS\_382
CL\_INS\_382
CL\_INS\_382
CL\_INS\_382
CL\_INS\_382
CL\_INS\_382
CL\_INS\_382
CL\_INS\_382
CL\_INS\_176
CL\_INS\_129
CL\_INS\_382
CL\_INS\_382
CL\_INS\_99
CL\_INS\_382
CL\_INS\_382
CL\_INS\_382
CL\_INS\_382
CL\_INS\_382
CL\_INS\_382
CL\_INS\_382
CL\_INS\_382
CL\_INS\_382
CL\_INS\_382
CL\_INS\_382
CL\_INS\_99
CL\_INS\_176
CL\_INS\_176
CL\_INS\_176
CL\_INS\_176
CL\_INS\_176
CL\_INS\_176
CL\_INS\_176
CL\_INS\_382
CL\_INS\_176
CL\_INS\_176
CL\_INS\_176
CL\_INS\_176
CL\_INS\_176
CL\_INS\_176
CL\_INS\_176
CL\_INS\_176
CL\_INS\_382
CL\_INS\_382
CL\_INS\_382
CL\_INS\_382
CL\_INS\_382
CL\_INS\_382
CL\_INS\_382
CL\_INS\_382
CL\_INS\_176
CL\_INS\_176
CL\_INS\_176
CL\_INS\_99
CL\_INS\_99
CL\_INS\_382
CL\_INS\_99
CL\_INS\_382
CL\_INS\_129
CL\_INS\_79
CL\_INS\_99
CL\_INS\_382
CL\_INS\_382
CL\_INS\_76
CL\_INS\_176
CL\_INS\_382
CL\_INS\_176
CL\_INS\_146
CL\_INS\_146
CL\_INS\_146
CL\_INS\_382
CL\_INS\_99
CL\_INS\_382
CL\_INS\_176
CL\_INS\_176
CL\_INS\_176
CL\_INS\_176
CL\_INS\_176
CL\_INS\_176
CL\_INS\_176
CL\_INS\_176
CL\_INS\_176
CL\_INS\_176
CL\_INS\_176
CL\_INS\_176
CL\_INS\_176
CL\_INS\_176
CL\_INS\_176
CL\_INS\_176
CL\_INS\_176
CL\_INS\_176
CL\_INS\_176
CL\_INS\_176
CL\_INS\_176
CL\_INS\_176
CL\_INS\_382
CL\_INS\_382
CL\_INS\_79
CL\_INS\_79
CL\_INS\_79
CL\_INS\_99
CL\_INS\_99
CL\_INS\_176
CL\_INS\_79
CL\_INS\_176
CL\_INS\_153
CL\_INS\_176
CL\_INS\_176
CL\_INS\_176
CL\_INS\_176
CL\_INS\_176
CL\_INS\_176
CL\_INS\_176
CL\_INS\_60
CL\_INS\_382
CL\_INS\_176
CL\_INS\_207
CL\_INS\_207
CL\_INS\_207
CL\_INS\_60
CL\_INS\_79
CL\_INS\_382
CL\_INS\_176
CL\_INS\_176
CL\_INS\_176
CL\_INS\_176
CL\_INS\_99
CL\_INS\_382
CL\_INS\_176
CL\_INS\_382
CL\_INS\_382
CL\_INS\_382
CL\_INS\_176
CL\_INS\_382
CL\_INS\_382
CL\_INS\_176
CL\_INS\_57
CL\_INS\_382
CL\_INS\_382
CL\_INS\_207
CL\_INS\_207
CL\_INS\_207
CL\_INS\_176
CL\_INS\_176
CL\_INS\_176
CL\_INS\_176
CL\_INS\_146
CL\_INS\_146
CL\_INS\_382
CL\_INS\_382
CL\_INS\_382
CL\_INS\_146
CL\_INS\_207
CL\_INS\_382
CL\_INS\_382
CL\_INS\_382
CL\_INS\_382
CL\_INS\_382
CL\_INS\_382
CL\_INS\_382
CL\_INS\_382
CL\_INS\_382
CL\_INS\_382
CL\_INS\_382
CL\_INS\_176
CL\_INS\_382
CL\_INS\_382
CL\_INS\_146
CL\_INS\_382
CL\_INS\_382
CL\_INS\_382
CL\_INS\_146
CL\_INS\_382
CL\_INS\_176
CL\_INS\_176
CL\_INS\_382
CL\_INS\_176
CL\_INS\_382
CL\_INS\_368
CL\_INS\_382
CL\_INS\_382
CL\_INS\_382
CL\_INS\_146
CL\_INS\_382
CL\_INS\_382
CL\_INS\_382
CL\_INS\_382
CL\_INS\_382
CL\_INS\_382
CL\_INS\_382
CL\_INS\_382
CL\_INS\_382
CL\_INS\_382
CL\_INS\_176
CL\_INS\_20
CL\_INS\_382
CL\_INS\_20
CL\_INS\_20
CL\_INS\_20
CL\_INS\_20
CL\_INS\_382
CL\_INS\_176
CL\_INS\_20
CL\_INS\_176
CL\_INS\_20
CL\_INS\_176
CL\_INS\_20
CL\_INS\_20
CL\_INS\_176
CL\_INS\_176
CL\_INS\_382
CL\_INS\_146
CL\_INS\_146
CL\_INS\_207
CL\_INS\_176
CL\_INS\_382
CL\_INS\_146
CL\_INS\_382
CL\_INS\_382
CL\_INS\_207
CL\_INS\_176
CL\_INS\_382
CL\_INS\_382
CL\_INS\_382
CL\_INS\_382
CL\_INS\_382
CL\_INS\_207
CL\_INS\_207
CL\_INS\_207
CL\_INS\_146
CL\_INS\_146
CL\_INS\_146
CL\_INS\_382
CL\_INS\_382
CL\_INS\_146
CL\_INS\_146
CL\_INS\_382
CL\_INS\_382
CL\_INS\_382
CL\_INS\_382
CL\_INS\_176
CL\_INS\_176
CL\_INS\_340
CL\_INS\_176
CL\_INS\_176
CL\_INS\_382
CL\_INS\_382
CL\_INS\_60
CL\_INS\_60
CL\_INS\_382
CL\_INS\_60
CL\_INS\_176
CL\_INS\_176
CL\_INS\_176
CL\_INS\_176
CL\_INS\_176
CL\_INS\_382
CL\_INS\_382
CL\_INS\_382
CL\_INS\_382
CL\_INS\_106
CL\_INS\_189
CL\_INS\_176
CL\_INS\_176
CL\_INS\_176
CL\_INS\_382
CL\_INS\_382
CL\_INS\_176
CL\_INS\_176
CL\_INS\_176
CL\_INS\_382
CL\_INS\_20
CL\_INS\_146
CL\_INS\_146
CL\_INS\_146
CL\_INS\_79
CL\_INS\_79
CL\_INS\_382
CL\_INS\_79
CL\_INS\_79
CL\_INS\_79
CL\_INS\_382
CL\_INS\_79
CL\_INS\_176
CL\_INS\_382
CL\_INS\_382
CL\_INS\_79
CL\_INS\_382
CL\_INS\_42
CL\_INS\_382
CL\_INS\_382
CL\_INS\_42
CL\_INS\_42
CL\_INS\_79
CL\_INS\_79
CL\_INS\_79
CL\_INS\_79
CL\_INS\_237
CL\_INS\_131
CL\_INS\_85
CL\_INS\_60
CL\_INS\_382
CL\_INS\_382
CL\_INS\_382
CL\_INS\_382
CL\_INS\_79
CL\_INS\_60
CL\_INS\_20
CL\_INS\_176
CL\_INS\_20
CL\_INS\_379
CL\_INS\_60
CL\_INS\_379
CL\_INS\_176
CL\_INS\_176
CL\_INS\_176
CL\_INS\_176
CL\_INS\_99
CL\_INS\_99
CL\_INS\_83
CL\_INS\_83
CL\_INS\_176
CL\_INS\_176
CL\_INS\_176
CL\_INS\_83
CL\_INS\_99
CL\_INS\_83
CL\_INS\_99
CL\_INS\_176
CL\_INS\_146
CL\_INS\_128
CL\_INS\_176
CL\_INS\_99
CL\_INS\_385
CL\_INS\_99
CL\_INS\_99
CL\_INS\_146
CL\_INS\_99
CL\_INS\_99
CL\_INS\_86
CL\_INS\_86
CL\_INS\_99
CL\_INS\_99
CL\_INS\_382
CL\_INS\_86
CL\_INS\_99
CL\_INS\_176
CL\_INS\_176
CL\_INS\_99
CL\_INS\_176
CL\_INS\_382
CL\_INS\_176
CL\_INS\_99
CL\_INS\_176
CL\_INS\_99
CL\_INS\_99
CL\_INS\_99
CL\_INS\_99
CL\_INS\_99
CL\_INS\_99
CL\_INS\_99
CL\_INS\_382
CL\_INS\_382
CL\_INS\_382
CL\_INS\_382
CL\_INS\_382
CL\_INS\_382
CL\_INS\_382
CL\_INS\_176
CL\_INS\_176
CL\_INS\_176
CL\_INS\_128
CL\_INS\_176
CL\_INS\_176
CL\_INS\_176
CL\_INS\_176
CL\_INS\_176
CL\_INS\_176
CL\_INS\_176
CL\_INS\_176
CL\_INS\_176
CL\_INS\_176
CL\_INS\_382
CL\_INS\_176
CL\_INS\_144
CL\_INS\_176
CL\_INS\_176
CL\_INS\_176
CL\_INS\_176
CL\_INS\_176
CL\_INS\_382
CL\_INS\_176
CL\_INS\_176
CL\_INS\_176
CL\_INS\_176
CL\_INS\_176
CL\_INS\_176
CL\_INS\_176
CL\_INS\_176
CL\_INS\_99
CL\_INS\_176
CL\_INS\_176
CL\_INS\_176
CL\_INS\_176
CL\_INS\_176
CL\_INS\_176
CL\_INS\_176
CL\_INS\_176
CL\_INS\_176
CL\_INS\_176
CL\_INS\_176
CL\_INS\_176
CL\_INS\_176
CL\_INS\_176
CL\_INS\_176
CL\_INS\_176
CL\_INS\_176
CL\_INS\_176
CL\_INS\_176
CL\_INS\_176
CL\_INS\_176
CL\_INS\_176
CL\_INS\_176
CL\_INS\_176
CL\_INS\_176
CL\_INS\_176
CL\_INS\_176
CL\_INS\_176
CL\_INS\_176
CL\_INS\_176
CL\_INS\_176
CL\_INS\_176
CL\_INS\_176
CL\_INS\_176
CL\_INS\_176
CL\_INS\_176
CL\_INS\_176
CL\_INS\_176
CL\_INS\_176
CL\_INS\_176
CL\_INS\_176
CL\_INS\_176
CL\_INS\_176
CL\_INS\_176
CL\_INS\_176
CL\_INS\_176
CL\_INS\_176
CL\_INS\_176
CL\_INS\_176
CL\_INS\_176
CL\_INS\_42
CL\_INS\_382
CL\_INS\_382
CL\_INS\_176
CL\_INS\_382
CL\_INS\_382
CL\_INS\_382
CL\_INS\_382
CL\_INS\_382
CL\_INS\_382
CL\_INS\_382
CL\_INS\_382
CL\_INS\_176
CL\_INS\_382
CL\_INS\_382
CL\_INS\_382
CL\_INS\_382
CL\_INS\_382
CL\_INS\_382
CL\_INS\_382
CL\_INS\_382
CL\_INS\_382
CL\_INS\_176
CL\_INS\_176
CL\_INS\_176
CL\_INS\_382
CL\_INS\_382
CL\_INS\_176
CL\_INS\_382
CL\_INS\_382
CL\_INS\_382
CL\_INS\_382
CL\_INS\_382
CL\_INS\_382
CL\_INS\_176
CL\_INS\_60
CL\_INS\_382
CL\_INS\_382
CL\_INS\_382
CL\_INS\_176
CL\_INS\_176
CL\_INS\_382
CL\_INS\_382
CL\_INS\_382
CL\_INS\_176
CL\_INS\_123
CL\_INS\_382
CL\_INS\_382
CL\_INS\_382
CL\_INS\_176
CL\_INS\_176
Cluster ID


CL\_13176
CL\_30308
CL\_9121
CL\_9094
CL\_9093
CL\_9092
CL\_9091
CL\_9090
CL\_17392
CL\_7531
CL\_7532
CL\_7533
CL\_15241
CL\_13890
CL\_9944
CL\_22903
CL\_13559
CL\_12788
CL\_6767
CL\_534
CL\_27233
CL\_30307
CL\_30306
CL\_12525
CL\_13177
CL\_13178
CL\_13179
CL\_30305
CL\_30304
CL\_30303
CL\_12526
CL\_12527
CL\_12528
CL\_22905
CL\_12529
CL\_12530
CL\_12531
CL\_12532
CL\_12533
CL\_9740
CL\_9738
CL\_7252
CL\_12789
CL\_12790
CL\_4514
CL\_4489
CL\_27954
CL\_32215
CL\_20922
CL\_20921
CL\_20920
CL\_31389
CL\_31391
CL\_31392
CL\_28331
CL\_31393
CL\_31394
CL\_31395
CL\_31396
CL\_31397
CL\_31398
CL\_31399
CL\_31400
CL\_32216
CL\_31401
CL\_31402
CL\_31403
CL\_31404
CL\_31405
CL\_31406
CL\_31407
CL\_31408
CL\_31409
CL\_27953
CL\_9162
CL\_27952
CL\_17342
CL\_9124
CL\_17341
CL\_32040
CL\_32039
CL\_32038
CL\_32037
CL\_32036
CL\_15588
CL\_32035
CL\_32034
CL\_32033
CL\_32032
CL\_32031
CL\_32030
CL\_32029
CL\_12999
CL\_11919
CL\_11920
CL\_12382
CL\_12798
CL\_23978
CL\_8696
CL\_7537
CL\_17368
CL\_28098
CL\_32028
CL\_17269
CL\_17267
CL\_4545
CL\_8688
CL\_1515
CL\_17266
CL\_8684
CL\_8683
CL\_4550
CL\_4551
CL\_4552
CL\_8679
CL\_8678
CL\_4556
CL\_17265
CL\_15759
CL\_4560
CL\_8195
CL\_23879
CL\_32027
CL\_32026
CL\_7126
CL\_10982
CL\_32025
CL\_16110
CL\_32024
CL\_32023
CL\_32022
CL\_8585
CL\_4432
CL\_4620
CL\_11917
CL\_8167
CL\_12791
CL\_13509
CL\_12360
CL\_8712
CL\_4518
CL\_4485
CL\_4520
CL\_4521
CL\_4425
CL\_4522
CL\_4523
CL\_4524
CL\_4525
CL\_4421
CL\_12792
CL\_4526
CL\_4527
CL\_4528
CL\_4418
CL\_4629
CL\_21005
CL\_21006
CL\_7113
CL\_6461
CL\_21007
CL\_4529
CL\_4530
CL\_12793
CL\_12794
CL\_12795
CL\_12796
CL\_4531
CL\_4532
CL\_4414
CL\_4417
CL\_21008
CL\_7116
CL\_14371
CL\_4413
CL\_1496
CL\_4533
CL\_533
CL\_4534
CL\_21009
CL\_16618
CL\_12009
CL\_9098
CL\_17372
CL\_13524
CL\_12383
CL\_12797
CL\_8184
CL\_8185
CL\_7538
CL\_7120
CL\_7539
CL\_8169
CL\_8170
CL\_8171
CL\_8711
CL\_27951
CL\_28286
CL\_27950
CL\_27949
CL\_27948
CL\_27947
CL\_27946
CL\_8172
CL\_32269
CL\_32270
CL\_32271
CL\_32272
CL\_32273
CL\_32274
CL\_32275
CL\_32276
CL\_8710
CL\_8709
CL\_8708
CL\_8707
CL\_8706
CL\_8705
CL\_8704
CL\_8703
CL\_28285
CL\_28284
CL\_28283
CL\_12758
CL\_20774
CL\_8699
CL\_13560
CL\_8697
CL\_17073
CL\_16350
CL\_13561
CL\_15775
CL\_15774
CL\_20779
CL\_20780
CL\_15773
CL\_20781
CL\_20782
CL\_20783
CL\_20784
CL\_16445
CL\_23514
CL\_8702
CL\_27945
CL\_27944
CL\_27943
CL\_27942
CL\_27941
CL\_27940
CL\_27939
CL\_27938
CL\_32277
CL\_32278
CL\_32279
CL\_32280
CL\_32281
CL\_32282
CL\_32283
CL\_32284
CL\_32285
CL\_32286
CL\_32287
CL\_32288
CL\_32289
CL\_32290
CL\_8701
CL\_8700
CL\_16617
CL\_31753
CL\_31754
CL\_23513
CL\_20884
CL\_28282
CL\_16616
CL\_28281
CL\_17314
CL\_28280
CL\_33029
CL\_27937
CL\_27936
CL\_27935
CL\_27934
CL\_27933
CL\_27932
CL\_5349
CL\_28279
CL\_26136
CL\_26137
CL\_26138
CL\_26139
CL\_32021
CL\_12996
CL\_35470
CL\_35469
CL\_35468
CL\_35467
CL\_13563
CL\_524
CL\_35466
CL\_12760
CL\_12762
CL\_13562
CL\_21010
CL\_6452
CL\_7540
CL\_12799
CL\_6751
CL\_20785
CL\_4651
CL\_17563
CL\_17564
CL\_17565
CL\_35465
CL\_35464
CL\_21011
CL\_21012
CL\_21013
CL\_9747
CL\_9745
CL\_10168
CL\_4652
CL\_13070
CL\_4653
CL\_4654
CL\_5419
CL\_5420
CL\_5421
CL\_19352
CL\_19351
CL\_12768
CL\_5422
CL\_2278
CL\_2279
CL\_4655
CL\_20786
CL\_13564
CL\_12379
CL\_4657
CL\_12800
CL\_4656
CL\_6037
CL\_20787
CL\_6036
CL\_27471
CL\_27472
CL\_12119
CL\_27473
CL\_10270
CL\_17186
CL\_7542
CL\_12121
CL\_7543
CL\_10177
CL\_12120
CL\_8576
CL\_2281
CL\_2282
CL\_2283
CL\_20109
CL\_11406
CL\_11405
CL\_16615
CL\_12994
CL\_33193
CL\_15697
CL\_16444
CL\_15696
CL\_15695
CL\_13071
CL\_15694
CL\_16010
CL\_27470
CL\_15692
CL\_32291
CL\_15691
CL\_32292
CL\_15690
CL\_12769
CL\_32293
CL\_32294
CL\_4658
CL\_12378
CL\_12377
CL\_12376
CL\_22902
CL\_4659
CL\_12801
CL\_15866
CL\_15867
CL\_4661
CL\_12802
CL\_4662
CL\_4663
CL\_4664
CL\_9744
CL\_21014
CL\_4665
CL\_15868
CL\_4666
CL\_13152
CL\_13153
CL\_21015
CL\_4667
CL\_4670
CL\_12803
CL\_12804
CL\_10510
CL\_5423
CL\_5424
CL\_10175
CL\_32295
CL\_32296
CL\_21606
CL\_33192
CL\_33191
CL\_6658
CL\_5347
CL\_6657
CL\_6656
CL\_5348
CL\_28349
CL\_31803
CL\_33190
CL\_33189
CL\_33188
CL\_33187
CL\_5350
CL\_4410
CL\_11915
CL\_5351
CL\_6650
CL\_6649
CL\_33186
CL\_33185
CL\_33184
CL\_5355
CL\_5356
CL\_33183
CL\_33182
CL\_33181
CL\_7031
CL\_14825
CL\_17539
CL\_13972
CL\_17540
CL\_31759
CL\_9117
CL\_6655
CL\_21777
CL\_13507
CL\_27894
CL\_509
CL\_28278
CL\_32020
CL\_9080
CL\_9079
CL\_24460
CL\_9078
CL\_17545
CL\_23319
CL\_27931
CL\_28277
CL\_28276
CL\_31760
CL\_31761
CL\_31762
CL\_9229
CL\_11731
CL\_23314
CL\_13514
CL\_6644
CL\_5357
CL\_5358
CL\_5359
CL\_5360
CL\_8964
CL\_17549
CL\_8962
CL\_28440
CL\_9949
CL\_31763
CL\_26142
CL\_31764
CL\_12524
CL\_31805
CL\_31806
CL\_31807
CL\_12385
CL\_13075
CL\_31114
CL\_31113
CL\_32263
CL\_32264
CL\_32265
CL\_31174
CL\_6747
CL\_31175
CL\_29360
CL\_32266
CL\_32267
CL\_28320
CL\_32268
CL\_13527
CL\_11286
CL\_13526
CL\_10518
CL\_28486
CL\_17051
CL\_17052
CL\_7025
CL\_4430
CL\_7521
CL\_17053
CL\_4517
CL\_4429
CL\_13789
CL\_28817
CL\_30624
CL\_19006
CL\_30625
CL\_4628
CL\_21004
CL\_4431
CL\_33197
CL\_13790
CL\_8155
CL\_15782
CL\_17292
CL\_17291
CL\_17290
CL\_17289
CL\_15781
CL\_15780
CL\_15779
CL\_15778
CL\_15777
CL\_15776
CL\_9123
CL\_34537
CL\_34538
CL\_34539
CL\_32745
CL\_34540
CL\_34541
CL\_34542
CL\_34543
CL\_34544
CL\_34545
CL\_34546
CL\_34547
CL\_34548
CL\_28068
CL\_7534
CL\_28069
CL\_17443
CL\_28070
CL\_28071
CL\_33196
CL\_33195
CL\_33194
CL\_12995
CL\_34549
CL\_34550
CL\_34551
CL\_34552
CL\_34553
CL\_34554
CL\_34555
CL\_34556
CL\_32085
CL\_34557
CL\_34558
CL\_34559
CL\_34560
CL\_34561
CL\_34562
CL\_34563
CL\_34564
CL\_34565
CL\_34566
CL\_34567
CL\_34568
CL\_34569
CL\_34570
CL\_34571
CL\_34572
CL\_34573
CL\_34574
CL\_34575
CL\_34576
CL\_34577
CL\_34578
CL\_34579
CL\_34580
CL\_34581
CL\_34582
CL\_34583
CL\_34584
CL\_34585
CL\_34586
CL\_34587
CL\_34588
CL\_34589
CL\_34590
CL\_34591
CL\_34592
CL\_34593
CL\_34594
CL\_34595
CL\_34596
CL\_34597
CL\_34598
CL\_34599
CL\_34600
CL\_34601
CL\_34602
CL\_34603
CL\_34604
CL\_34605
CL\_34606
CL\_16351
CL\_522
CL\_521
CL\_34607
CL\_519
CL\_518
CL\_517
CL\_516
CL\_515
CL\_514
CL\_513
CL\_512
CL\_34608
CL\_8206
CL\_4409
CL\_4408
CL\_4407
CL\_4406
CL\_13635
CL\_13634
CL\_4403
CL\_4402
CL\_33180
CL\_33179
CL\_33178
CL\_4401
CL\_23064
CL\_28275
CL\_11916
CL\_508
CL\_4400
CL\_5361
CL\_5362
CL\_17074
CL\_35463
CL\_13486
CL\_5363
CL\_6638
CL\_7361
CL\_33177
CL\_33176
CL\_5799
CL\_10511
CL\_10512
CL\_33175
CL\_27273
CL\_11402
CL\_4668
CL\_4669
CL\_12534
CL\_12535
