## Supplementary material for "A novel method for integrating genomic and Tn-Seq data to identify common *in vivo* fitness mechanisms across multiple bacterial species": S1 Dataset: CL_INS_178.html

Legend

 Mobile +extrachromosomalelementfunctions
 Hypothetical
 Proteinsynthesis/fate
 Other
 EnergyMetabolism
 All VFDB Genes

FULL


WINDOWSVGPNG

Trim RowsRemove SingletonsSave Fasta

CL\_2081


CL\_2081


CL\_2081


CL\_2081


CL\_2080


CL\_2081

HighlightSelectShow Genomes


237

CL\_2082


7

CL\_2082


1

CL\_2083


1

CL\_2082


1

CL\_2082


1

CL\_4754

fGI ID


CL\_INS\_178
CL\_INS\_178
CL\_INS\_178
CL\_INS\_178
CL\_INS\_178
Cluster ID


CL\_30300
CL\_20397
CL\_20396
CL\_20395
CL\_20394
