## Supplementary material for "A novel method for integrating genomic and Tn-Seq data to identify common *in vivo* fitness mechanisms across multiple bacterial species": S1 Dataset: CL_INS_179.html

Legend

 Mobile +extrachromosomalelementfunctions
 Regulatoryfunctions
 Hypothetical
 DNA Metabolism
 All EssentialGenes
 All Fitness Genes
 Proteinsynthesis/fate
 Other
 Transport +binding proteins
 All VFDB Genes

FULL


WINDOWSVGPNG

Trim RowsRemove SingletonsSave Fasta

CL\_2116


CL\_2116


CL\_2116


CL\_2116


CL\_2116


CL\_2116


CL\_2116


CL\_2116


CL\_2116


CL\_2116


CL\_2115


CL\_2116


CL\_2116


CL\_4519


CL\_2116


CL\_2116


CL\_2116


CL\_2116


CL\_2116


CL\_2116


CL\_2116


CL\_2116


CL\_2116


CL\_2116


CL\_2116


CL\_2116


CL\_2116


CL\_2116


CL\_2116


CL\_2116


CL\_2116

HighlightSelectShow Genomes


137

CL\_2118


110

CL\_2118


3

CL\_2118


1

CL\_2119


1

CL\_2118


1

CL\_2118


1

CL\_2118


1

CL\_2118


1

CL\_2118


1

CL\_2118


1

CL\_2118


1

CL\_2118


1

CL\_2118


1

CL\_2118


1

CL\_2118


1

CL\_2118


1

CL\_4516


1

CL\_2118


1

CL\_2118


1

CL\_2118


1

CL\_2118


1

CL\_2118


1

CL\_2118


1

CL\_2118


1

CL\_2118


1

CL\_2118


1

CL\_2119


1

CL\_2118


1

CL\_2118


1

CL\_2118


1

CL\_2118

fGI ID


CL\_INS\_179
CL\_INS\_179
CL\_INS\_179
CL\_INS\_385
CL\_INS\_20
CL\_INS\_86
CL\_INS\_179
CL\_INS\_179
CL\_INS\_179
CL\_INS\_179
CL\_INS\_179
CL\_INS\_179
CL\_INS\_106
CL\_INS\_179
CL\_INS\_382
CL\_INS\_106
CL\_INS\_247
CL\_INS\_106
CL\_INS\_382
CL\_INS\_132
CL\_INS\_179
CL\_INS\_207
CL\_INS\_179
CL\_INS\_179
CL\_INS\_179
CL\_INS\_247
CL\_INS\_247
CL\_INS\_247
CL\_INS\_179
CL\_INS\_179
CL\_INS\_179
CL\_INS\_179
CL\_INS\_106
CL\_INS\_132
CL\_INS\_382
CL\_INS\_382
CL\_INS\_382
CL\_INS\_106
CL\_INS\_382
CL\_INS\_179
CL\_INS\_69
CL\_INS\_69
CL\_INS\_69
CL\_INS\_69
CL\_INS\_69
CL\_INS\_382
CL\_INS\_382
CL\_INS\_247
CL\_INS\_69
CL\_INS\_382
CL\_INS\_382
CL\_INS\_382
CL\_INS\_382
CL\_INS\_382
CL\_INS\_382
CL\_INS\_382
CL\_INS\_132
CL\_INS\_179
CL\_INS\_179
CL\_INS\_106
CL\_INS\_106
CL\_INS\_382
CL\_INS\_179
CL\_INS\_179
CL\_INS\_132
CL\_INS\_69
CL\_INS\_247
CL\_INS\_247
CL\_INS\_247
CL\_INS\_179
CL\_INS\_179
CL\_INS\_179
CL\_INS\_179
CL\_INS\_20
CL\_INS\_247
CL\_INS\_247
CL\_INS\_247
CL\_INS\_247
CL\_INS\_247
CL\_INS\_247
CL\_INS\_382
CL\_INS\_207
CL\_INS\_179
CL\_INS\_179
CL\_INS\_179
CL\_INS\_179
CL\_INS\_179
CL\_INS\_179
CL\_INS\_179
CL\_INS\_179
CL\_INS\_179
CL\_INS\_382
CL\_INS\_179
CL\_INS\_106
CL\_INS\_106
CL\_INS\_179
CL\_INS\_179
CL\_INS\_179
CL\_INS\_179
CL\_INS\_179
CL\_INS\_179
CL\_INS\_179
CL\_INS\_179
CL\_INS\_382
CL\_INS\_382
CL\_INS\_382
CL\_INS\_382
CL\_INS\_382
CL\_INS\_382
CL\_INS\_382
CL\_INS\_382
CL\_INS\_179
CL\_INS\_382
CL\_INS\_179
CL\_INS\_382
CL\_INS\_382
CL\_INS\_382
CL\_INS\_99
CL\_INS\_382
CL\_INS\_382
CL\_INS\_99
CL\_INS\_99
CL\_INS\_382
CL\_INS\_382
CL\_INS\_382
CL\_INS\_382
CL\_INS\_382
CL\_INS\_382
CL\_INS\_382
CL\_INS\_382
CL\_INS\_382
CL\_INS\_382
CL\_INS\_382
CL\_INS\_382
CL\_INS\_382
CL\_INS\_382
CL\_INS\_382
CL\_INS\_86
CL\_INS\_382
CL\_INS\_382
CL\_INS\_382
CL\_INS\_382
CL\_INS\_207
CL\_INS\_179
CL\_INS\_179
CL\_INS\_179
CL\_INS\_179
CL\_INS\_179
CL\_INS\_207
CL\_INS\_207
CL\_INS\_179
CL\_INS\_179
CL\_INS\_155
CL\_INS\_204
CL\_INS\_146
CL\_INS\_146
CL\_INS\_146
CL\_INS\_146
CL\_INS\_146
CL\_INS\_146
CL\_INS\_146
CL\_INS\_146
CL\_INS\_146
CL\_INS\_146
CL\_INS\_146
CL\_INS\_146
CL\_INS\_146
CL\_INS\_146
CL\_INS\_146
CL\_INS\_146
CL\_INS\_146
CL\_INS\_146
CL\_INS\_146
CL\_INS\_146
CL\_INS\_146
CL\_INS\_146
CL\_INS\_146
CL\_INS\_97
CL\_INS\_97
CL\_INS\_97
CL\_INS\_97
CL\_INS\_97
CL\_INS\_97
CL\_INS\_97
CL\_INS\_179
CL\_INS\_179
CL\_INS\_179
CL\_INS\_179
CL\_INS\_179
CL\_INS\_179
CL\_INS\_179
CL\_INS\_179
CL\_INS\_179
CL\_INS\_179
CL\_INS\_179
CL\_INS\_179
CL\_INS\_146
CL\_INS\_179
CL\_INS\_179
CL\_INS\_179
CL\_INS\_179
CL\_INS\_179
CL\_INS\_382
CL\_INS\_179
CL\_INS\_179
CL\_INS\_385
CL\_INS\_179
Cluster ID


CL\_2117
CL\_30910
CL\_34142
CL\_18913
CL\_1494
CL\_1324
CL\_12539
CL\_33724
CL\_15921
CL\_12540
CL\_33725
CL\_20934
CL\_8649
CL\_31499
CL\_13310
CL\_16469
CL\_8255
CL\_16582
CL\_12541
CL\_10495
CL\_12542
CL\_12543
CL\_12544
CL\_31498
CL\_30031
CL\_14015
CL\_14014
CL\_14013
CL\_30030
CL\_30029
CL\_12545
CL\_31381
CL\_8651
CL\_8652
CL\_13512
CL\_8113
CL\_8114
CL\_13735
CL\_8653
CL\_32815
CL\_32129
CL\_32128
CL\_32127
CL\_32126
CL\_32125
CL\_8654
CL\_12137
CL\_17495
CL\_30648
CL\_13640
CL\_16429
CL\_13641
CL\_13642
CL\_13643
CL\_13737
CL\_13738
CL\_8650
CL\_28908
CL\_28907
CL\_12139
CL\_12138
CL\_8655
CL\_31497
CL\_31496
CL\_10497
CL\_15922
CL\_15923
CL\_15924
CL\_8122
CL\_20935
CL\_28906
CL\_28905
CL\_28904
CL\_11159
CL\_10496
CL\_8656
CL\_8657
CL\_8658
CL\_12136
CL\_8659
CL\_17067
CL\_15935
CL\_28903
CL\_28902
CL\_17928
CL\_17927
CL\_17926
CL\_17925
CL\_17924
CL\_28901
CL\_28900
CL\_8123
CL\_17923
CL\_15925
CL\_12546
CL\_15926
CL\_30028
CL\_30027
CL\_30026
CL\_30025
CL\_16430
CL\_5169
CL\_15339
CL\_4521
CL\_4425
CL\_6743
CL\_4524
CL\_4525
CL\_7527
CL\_4420
CL\_4419
CL\_36124
CL\_4530
CL\_36123
CL\_4532
CL\_4533
CL\_1496
CL\_27276
CL\_11311
CL\_6784
CL\_12382
CL\_23470
CL\_7538
CL\_7539
CL\_7540
CL\_7339
CL\_5419
CL\_5420
CL\_5421
CL\_5422
CL\_2278
CL\_2279
CL\_2280
CL\_1096
CL\_5423
CL\_5424
CL\_10175
CL\_11313
CL\_2281
CL\_2282
CL\_2283
CL\_20109
CL\_20110
CL\_36122
CL\_36121
CL\_36120
CL\_36119
CL\_36118
CL\_21316
CL\_9771
CL\_21315
CL\_21314
CL\_8483
CL\_10867
CL\_14046
CL\_14047
CL\_14048
CL\_14049
CL\_14050
CL\_21313
CL\_21312
CL\_21311
CL\_21310
CL\_21309
CL\_21308
CL\_21307
CL\_21306
CL\_21305
CL\_21304
CL\_14051
CL\_14052
CL\_14053
CL\_14054
CL\_14055
CL\_14056
CL\_14057
CL\_14058
CL\_21303
CL\_21302
CL\_21301
CL\_21300
CL\_21299
CL\_21298
CL\_21297
CL\_21296
CL\_21295
CL\_21294
CL\_21293
CL\_21292
CL\_21291
CL\_21290
CL\_21289
CL\_21288
CL\_21287
CL\_21286
CL\_21285
CL\_10178
CL\_21284
CL\_21283
CL\_21282
CL\_21281
CL\_21280
CL\_11314
CL\_21279
CL\_21278
CL\_21277
CL\_21276
