## Supplementary material for "A novel method for integrating genomic and Tn-Seq data to identify common *in vivo* fitness mechanisms across multiple bacterial species": S1 Dataset: CL_INS_181.html

Legend

 Regulatoryfunctions
 Hypothetical
 All EssentialGenes
 All VFDB Genes

FULL


WINDOWSVGPNG

Trim RowsRemove SingletonsSave Fasta

CL\_2155


CL\_2155

HighlightSelectShow Genomes


267

CL\_2156


11

CL\_2156

fGI ID


CL\_INS\_181
CL\_INS\_181
CL\_INS\_181
Cluster ID


CL\_13565
CL\_13566
CL\_13567
