## Supplementary material for "A novel method for integrating genomic and Tn-Seq data to identify common *in vivo* fitness mechanisms across multiple bacterial species": S1 Dataset: CL_INS_182.html

Legend

 Mobile +extrachromosomalelementfunctions
 Regulatoryfunctions
 Hypothetical
 All EssentialGenes
 All Fitness Genes
 Other
 All VFDB Genes

FULL


WINDOWSVGPNG

Trim RowsRemove SingletonsSave Fasta

CL\_2156


CL\_2156


CL\_2156


CL\_2156


CL\_2156

HighlightSelectShow Genomes


246

CL\_2157


4

CL\_2157


2

CL\_2158


1

CL\_2157


1

CL\_860

fGI ID


CL\_INS\_182
CL\_INS\_182
CL\_INS\_182
CL\_INS\_182
CL\_INS\_182
CL\_INS\_182
CL\_INS\_182
CL\_INS\_182
CL\_INS\_182
CL\_INS\_182
CL\_INS\_182
CL\_INS\_182
CL\_INS\_182
CL\_INS\_182
CL\_INS\_155
CL\_INS\_155
CL\_INS\_155
CL\_INS\_155
CL\_INS\_155
CL\_INS\_155
CL\_INS\_155
CL\_INS\_155
CL\_INS\_182
CL\_INS\_182
CL\_INS\_182
CL\_INS\_182
CL\_INS\_182
CL\_INS\_182
CL\_INS\_182
CL\_INS\_182
CL\_INS\_182
CL\_INS\_182
CL\_INS\_182
CL\_INS\_182
CL\_INS\_155
CL\_INS\_155
CL\_INS\_182
CL\_INS\_182
CL\_INS\_182
CL\_INS\_86
CL\_INS\_86
CL\_INS\_86
CL\_INS\_86
CL\_INS\_155
CL\_INS\_155
CL\_INS\_155
CL\_INS\_182
CL\_INS\_182
CL\_INS\_182
CL\_INS\_182
CL\_INS\_155
CL\_INS\_155
CL\_INS\_155
CL\_INS\_204
Cluster ID


CL\_17922
CL\_15342
CL\_10992
CL\_10993
CL\_10994
CL\_10995
CL\_10996
CL\_10997
CL\_10998
CL\_10999
CL\_11000
CL\_11001
CL\_11002
CL\_11003
CL\_11004
CL\_11005
CL\_11006
CL\_11007
CL\_11008
CL\_11009
CL\_11010
CL\_11011
CL\_11012
CL\_11013
CL\_11014
CL\_11015
CL\_11016
CL\_11017
CL\_11018
CL\_11019
CL\_11020
CL\_11021
CL\_11022
CL\_11023
CL\_11024
CL\_11025
CL\_11026
CL\_11027
CL\_11028
CL\_11029
CL\_11030
CL\_8226
CL\_8227
CL\_8228
CL\_8229
CL\_8230
CL\_11031
CL\_11032
CL\_11033
CL\_11034
CL\_11035
CL\_8483
CL\_11036
CL\_1105
