## Supplementary material for "A novel method for integrating genomic and Tn-Seq data to identify common *in vivo* fitness mechanisms across multiple bacterial species": S1 Dataset: CL_INS_184.html

Legend

 All VFDB Genes

FULL


WINDOWSVGPNG

Trim RowsRemove SingletonsSave Fasta

CL\_2167


CL\_2167


CL\_2167


CL\_2163


CL\_2167

HighlightSelectShow Genomes


138

CL\_2169


132

CL\_2169


2

CL\_2170


1

CL\_2169


1

CL\_2174

fGI ID

CL\_INS\_184
Cluster ID

CL\_2168
