## Supplementary material for "A novel method for integrating genomic and Tn-Seq data to identify common *in vivo* fitness mechanisms across multiple bacterial species": S1 Dataset: CL_INS_189.html

Legend

 Mobile +extrachromosomalelementfunctions
 Regulatoryfunctions
 Hypothetical
 DNA Metabolism
 AntibioticResistance
 All EssentialGenes
 All Fitness Genes
 Proteinsynthesis/fate
 Other
 EnergyMetabolism
 Cellularprocesses
 Transport +binding proteins
 All VFDB Genes

FULL


WINDOWSVGPNG

Trim RowsRemove SingletonsSave Fasta

CL\_2270


CL\_2270


CL\_2270


CL\_2270


CL\_2270


CL\_2270


CL\_2270


CL\_2270


CL\_2270


CL\_2270


CL\_2270


CL\_2270


CL\_2270


CL\_2270


CL\_2270


CL\_2270


CL\_2270


CL\_2270


CL\_2270


CL\_2270


CL\_2270


CL\_2270


CL\_2270


CL\_2270


CL\_2270


CL\_2270


CL\_2270


CL\_2270


CL\_2270


CL\_2270


CL\_2270


CL\_2270


CL\_2270


CL\_2270


CL\_2270


CL\_2270


CL\_2270


CL\_2270


CL\_2270


CL\_2270


CL\_2270


CL\_2270


CL\_2270


CL\_2270


CL\_2270


CL\_2270


CL\_2270


CL\_2270


CL\_2270


CL\_2270


CL\_2270


CL\_2270


CL\_2270


Break


CL\_2270


CL\_2270


CL\_2270


CL\_2269


CL\_2269


CL\_2270


CL\_2269


CL\_2270


CL\_2270


CL\_2270


CL\_2270


CL\_2270


CL\_2270


CL\_2270


CL\_2270


CL\_2270


CL\_2270


CL\_2270


CL\_2270


CL\_2270


CL\_2270


CL\_2270


CL\_2270


CL\_2270


CL\_2270


CL\_2270


CL\_2270


CL\_2270


CL\_2270


CL\_2270


CL\_2270


CL\_2270


CL\_2270


CL\_2270


CL\_2270


CL\_2270


CL\_2270


CL\_2270


CL\_2270


CL\_4487


CL\_2270


CL\_1976


CL\_2270


CL\_2270


CL\_2270


CL\_2270


CL\_2270


CL\_2270


CL\_2270


CL\_2270


CL\_2270


CL\_2270


CL\_2270


CL\_2270


CL\_2270


CL\_2270


CL\_2270


CL\_2270


CL\_2270


CL\_2270


CL\_2270


CL\_2270


CL\_2270


CL\_2270


CL\_2270


CL\_2270


CL\_2270


CL\_2270


CL\_2270


CL\_2270


CL\_2312


CL\_2270


CL\_2270


CL\_2270


CL\_2270


CL\_2270

HighlightSelectShow Genomes


45

CL\_2289


39

CL\_2289


20

CL\_2289


17

CL\_2289


10

CL\_2289


6

CL\_2289


5

CL\_2289


5

CL\_2289


3

CL\_2289


3

CL\_2289


2

CL\_2289


2

CL\_2290


2

CL\_2289


2

CL\_2290


2

CL\_2289


2

CL\_2289


2

CL\_2289


1

CL\_2289


1

CL\_2289


1

CL\_2289


1

CL\_2289


1

CL\_2289


1

CL\_2289


1

CL\_549


1

CL\_2289


1

CL\_2289


1

CL\_2289


1

CL\_2289


1

CL\_2289


1

CL\_2289


1

CL\_2289


1

CL\_2289


1

CL\_2289


1

CL\_2289


1

CL\_2289


1

CL\_2289


1

CL\_2289


1

CL\_2289


1

CL\_2289


1

CL\_2289


1

CL\_2310


1

CL\_2289


1

CL\_2289


1

CL\_2289


1

CL\_2289


1

CL\_2289


1

CL\_2289


1

CL\_2289


1

CL\_2289


1

CL\_2289


1

CL\_2289


1

CL\_2289


1

CL\_2289


1

CL\_2289


1

CL\_2289


1

CL\_2289


1

CL\_2289


1

CL\_2289


1

CL\_2289


1

CL\_2289


1

CL\_2289


1

CL\_2289


1

CL\_2289


1

Break


1

CL\_2289


1

CL\_2289


1

CL\_2289


1

CL\_2290


1

CL\_2289


1

CL\_2289


1

CL\_2289


1

CL\_2289


1

CL\_2289


1

CL\_2289


1

CL\_2289


1

CL\_2289


1

CL\_2289


1

CL\_2732


1

CL\_2290


1

CL\_2289


1

CL\_2295


1

CL\_2289


1

CL\_2289


1

CL\_2289


1

CL\_2289


1

CL\_2289


1

CL\_4516


1

CL\_2290


1

CL\_2289


1

CL\_2289


1

CL\_2289


1

CL\_2289


1

CL\_2289


1

CL\_2289


1

CL\_2289


1

CL\_2289


1

CL\_2289


1

CL\_2289


1

CL\_2289


1

CL\_2289


1

CL\_2289


1

CL\_2290


1

CL\_2289


1

CL\_2289


1

Break


1

CL\_2289


1

CL\_2289


1

CL\_2289


1

CL\_2289


1

CL\_2290


1

CL\_2289


1

CL\_2289


1

CL\_2289


1

CL\_2289


1

CL\_2289


1

CL\_2289


1

CL\_2289


1

CL\_2289


1

CL\_2289


1

CL\_2289


1

CL\_2289


1

CL\_2292


1

CL\_2289


1

CL\_2289


1

CL\_2289


1

CL\_2289


1

CL\_2257


1

CL\_2289


1

CL\_2289


1

CL\_2289

fGI ID


CL\_INS\_189
CL\_INS\_60
CL\_INS\_189
CL\_INS\_189
CL\_INS\_189
CL\_INS\_189
CL\_INS\_189
CL\_INS\_189
CL\_INS\_189
CL\_INS\_189
CL\_INS\_189
CL\_INS\_189
CL\_INS\_189
CL\_INS\_189
CL\_INS\_189
CL\_INS\_382
CL\_INS\_382
CL\_INS\_382
CL\_INS\_382
CL\_INS\_382
CL\_INS\_382
CL\_INS\_382
CL\_INS\_189
CL\_INS\_382
CL\_INS\_382
CL\_INS\_382
CL\_INS\_60
CL\_INS\_60
CL\_INS\_60
CL\_INS\_382
CL\_INS\_382
CL\_INS\_189
CL\_INS\_189
CL\_INS\_97
CL\_INS\_97
CL\_INS\_97
CL\_INS\_189
CL\_INS\_189
CL\_INS\_189
CL\_INS\_189
CL\_INS\_189
CL\_INS\_189
CL\_INS\_97
CL\_INS\_99
CL\_INS\_382
CL\_INS\_382
CL\_INS\_384
CL\_INS\_189
CL\_INS\_382
CL\_INS\_382
CL\_INS\_99
CL\_INS\_99
CL\_INS\_99
CL\_INS\_99
CL\_INS\_99
CL\_INS\_189
CL\_INS\_189
CL\_INS\_189
CL\_INS\_189
CL\_INS\_189
CL\_INS\_189
CL\_INS\_189
CL\_INS\_189
CL\_INS\_189
CL\_INS\_189
CL\_INS\_153
CL\_INS\_153
CL\_INS\_153
CL\_INS\_382
CL\_INS\_189
CL\_INS\_189
CL\_INS\_189
CL\_INS\_146
CL\_INS\_204
CL\_INS\_204
CL\_INS\_204
CL\_INS\_204
CL\_INS\_86
CL\_INS\_99
CL\_INS\_86
CL\_INS\_123
CL\_INS\_20
CL\_INS\_99
CL\_INS\_382
CL\_INS\_382
CL\_INS\_204
CL\_INS\_204
CL\_INS\_204
CL\_INS\_204
CL\_INS\_189
CL\_INS\_129
CL\_INS\_129
CL\_INS\_189
CL\_INS\_129
CL\_INS\_129
CL\_INS\_129
CL\_INS\_129
CL\_INS\_189
CL\_INS\_189
CL\_INS\_189
CL\_INS\_189
CL\_INS\_189
CL\_INS\_189
CL\_INS\_189
CL\_INS\_189
CL\_INS\_189
CL\_INS\_189
CL\_INS\_189
CL\_INS\_189
CL\_INS\_189
CL\_INS\_189
CL\_INS\_189
CL\_INS\_189
CL\_INS\_189
CL\_INS\_189
CL\_INS\_189
CL\_INS\_189
CL\_INS\_189
CL\_INS\_189
CL\_INS\_189
CL\_INS\_189
CL\_INS\_189
CL\_INS\_189
CL\_INS\_189
CL\_INS\_382
CL\_INS\_382
CL\_INS\_189
CL\_INS\_129
CL\_INS\_129
CL\_INS\_129
CL\_INS\_129
CL\_INS\_129
CL\_INS\_129
CL\_INS\_79
CL\_INS\_79
CL\_INS\_189
CL\_INS\_189
CL\_INS\_189
CL\_INS\_189
CL\_INS\_189
CL\_INS\_131
CL\_INS\_131
CL\_INS\_79
CL\_INS\_129
CL\_INS\_129
CL\_INS\_129
CL\_INS\_129
CL\_INS\_129
CL\_INS\_129
CL\_INS\_189
CL\_INS\_129
CL\_INS\_129
CL\_INS\_129
CL\_INS\_129
CL\_INS\_79
CL\_INS\_86
CL\_INS\_382
CL\_INS\_189
CL\_INS\_189
CL\_INS\_189
CL\_INS\_20
CL\_INS\_20
CL\_INS\_189
CL\_INS\_60
CL\_INS\_189
CL\_INS\_189
CL\_INS\_189
CL\_INS\_189
CL\_INS\_189
CL\_INS\_20
CL\_INS\_20
CL\_INS\_20
CL\_INS\_189
CL\_INS\_382
CL\_INS\_189
CL\_INS\_189
CL\_INS\_189
CL\_INS\_189
CL\_INS\_189
CL\_INS\_189
CL\_INS\_189
CL\_INS\_189
CL\_INS\_189
CL\_INS\_20
CL\_INS\_20
CL\_INS\_189
CL\_INS\_79
CL\_INS\_189
CL\_INS\_382
CL\_INS\_237
CL\_INS\_237
CL\_INS\_382
CL\_INS\_382
CL\_INS\_189
CL\_INS\_189
CL\_INS\_97
CL\_INS\_79
CL\_INS\_129
CL\_INS\_79
CL\_INS\_99
CL\_INS\_79
CL\_INS\_382
CL\_INS\_382
CL\_INS\_382
CL\_INS\_382
CL\_INS\_382
CL\_INS\_42
CL\_INS\_20
CL\_INS\_20
CL\_INS\_382
CL\_INS\_382
CL\_INS\_189
CL\_INS\_79
CL\_INS\_189
CL\_INS\_189
CL\_INS\_189
CL\_INS\_189
CL\_INS\_189
CL\_INS\_189
CL\_INS\_189
CL\_INS\_20
CL\_INS\_60
CL\_INS\_60
CL\_INS\_20
CL\_INS\_189
CL\_INS\_20
CL\_INS\_60
CL\_INS\_189
CL\_INS\_60
CL\_INS\_384
CL\_INS\_384
CL\_INS\_384
CL\_INS\_189
CL\_INS\_189
CL\_INS\_382
CL\_INS\_79
CL\_INS\_189
CL\_INS\_189
CL\_INS\_60
CL\_INS\_382
CL\_INS\_60
CL\_INS\_189
CL\_INS\_189
CL\_INS\_382
CL\_INS\_189
CL\_INS\_189
CL\_INS\_382
CL\_INS\_20
CL\_INS\_382
CL\_INS\_146
CL\_INS\_146
CL\_INS\_189
CL\_INS\_189
CL\_INS\_189
CL\_INS\_189
CL\_INS\_79
CL\_INS\_60
CL\_INS\_79
CL\_INS\_189
CL\_INS\_42
CL\_INS\_42
CL\_INS\_189
CL\_INS\_189
CL\_INS\_189
CL\_INS\_189
CL\_INS\_189
CL\_INS\_20
CL\_INS\_382
CL\_INS\_384
CL\_INS\_146
CL\_INS\_189
CL\_INS\_189
CL\_INS\_189
CL\_INS\_189
CL\_INS\_106
CL\_INS\_189
CL\_INS\_146
CL\_INS\_189
CL\_INS\_106
CL\_INS\_189
CL\_INS\_189
CL\_INS\_382
CL\_INS\_384
CL\_INS\_189
CL\_INS\_20
CL\_INS\_189
CL\_INS\_382
CL\_INS\_382
CL\_INS\_189
CL\_INS\_60
CL\_INS\_189
CL\_INS\_189
CL\_INS\_382
CL\_INS\_60
CL\_INS\_382
CL\_INS\_20
CL\_INS\_189
CL\_INS\_189
CL\_INS\_20
CL\_INS\_146
CL\_INS\_60
CL\_INS\_60
CL\_INS\_20
CL\_INS\_382
CL\_INS\_382
CL\_INS\_382
CL\_INS\_382
CL\_INS\_382
CL\_INS\_189
CL\_INS\_20
CL\_INS\_20
CL\_INS\_189
CL\_INS\_20
CL\_INS\_189
CL\_INS\_189
CL\_INS\_189
CL\_INS\_382
CL\_INS\_60
CL\_INS\_237
CL\_INS\_382
CL\_INS\_189
CL\_INS\_189
CL\_INS\_189
CL\_INS\_237
CL\_INS\_189
CL\_INS\_189
CL\_INS\_382
CL\_INS\_382
CL\_INS\_79
CL\_INS\_382
CL\_INS\_382
CL\_INS\_189
CL\_INS\_382
CL\_INS\_189
CL\_INS\_189
CL\_INS\_382
CL\_INS\_20
CL\_INS\_20
CL\_INS\_189
CL\_INS\_20
CL\_INS\_60
CL\_INS\_382
CL\_INS\_189
CL\_INS\_189
CL\_INS\_189
CL\_INS\_146
CL\_INS\_20
CL\_INS\_189
CL\_INS\_189
CL\_INS\_189
CL\_INS\_189
CL\_INS\_189
CL\_INS\_189
CL\_INS\_189
CL\_INS\_189
CL\_INS\_189
CL\_INS\_189
CL\_INS\_189
CL\_INS\_189
CL\_INS\_189
CL\_INS\_79
CL\_INS\_20
CL\_INS\_189
CL\_INS\_20
CL\_INS\_20
CL\_INS\_20
CL\_INS\_60
CL\_INS\_60
CL\_INS\_237
CL\_INS\_237
CL\_INS\_237
CL\_INS\_237
CL\_INS\_189
CL\_INS\_189
CL\_INS\_189
CL\_INS\_237
CL\_INS\_189
CL\_INS\_189
CL\_INS\_189
CL\_INS\_189
CL\_INS\_189
CL\_INS\_189
CL\_INS\_189
CL\_INS\_189
CL\_INS\_189
CL\_INS\_382
CL\_INS\_382
CL\_INS\_382
CL\_INS\_382
CL\_INS\_382
CL\_INS\_189
CL\_INS\_189
CL\_INS\_189
CL\_INS\_20
CL\_INS\_60
CL\_INS\_60
CL\_INS\_60
CL\_INS\_60
CL\_INS\_60
CL\_INS\_382
CL\_INS\_382
CL\_INS\_189
CL\_INS\_128
CL\_INS\_382
CL\_INS\_382
CL\_INS\_79
CL\_INS\_131
CL\_INS\_237
CL\_INS\_382
CL\_INS\_60
CL\_INS\_384
CL\_INS\_60
CL\_INS\_189
CL\_INS\_20
CL\_INS\_79
CL\_INS\_79
CL\_INS\_20
CL\_INS\_60
CL\_INS\_189
CL\_INS\_237
CL\_INS\_189
CL\_INS\_189
CL\_INS\_189
CL\_INS\_237
CL\_INS\_382
CL\_INS\_204
CL\_INS\_237
CL\_INS\_146
CL\_INS\_382
CL\_INS\_146
CL\_INS\_189
CL\_INS\_201
CL\_INS\_382
CL\_INS\_189
CL\_INS\_189
CL\_INS\_382
CL\_INS\_189
CL\_INS\_146
CL\_INS\_382
CL\_INS\_237
CL\_INS\_189
CL\_INS\_97
CL\_INS\_382
CL\_INS\_79
CL\_INS\_237
CL\_INS\_189
CL\_INS\_382
CL\_INS\_237
CL\_INS\_60
CL\_INS\_382
CL\_INS\_201
CL\_INS\_201
CL\_INS\_128
CL\_INS\_131
CL\_INS\_128
CL\_INS\_128
CL\_INS\_189
CL\_INS\_201
CL\_INS\_201
CL\_INS\_99
CL\_INS\_189
CL\_INS\_189
CL\_INS\_60
CL\_INS\_79
CL\_INS\_79
CL\_INS\_79
CL\_INS\_79
CL\_INS\_79
CL\_INS\_131
CL\_INS\_131
CL\_INS\_189
CL\_INS\_189
CL\_INS\_131
CL\_INS\_131
CL\_INS\_189
CL\_INS\_189
CL\_INS\_189
CL\_INS\_189
CL\_INS\_189
CL\_INS\_189
CL\_INS\_189
CL\_INS\_189
CL\_INS\_189
CL\_INS\_189
CL\_INS\_189
CL\_INS\_189
CL\_INS\_189
CL\_INS\_189
CL\_INS\_189
CL\_INS\_131
CL\_INS\_382
CL\_INS\_189
CL\_INS\_189
CL\_INS\_189
CL\_INS\_189
CL\_INS\_189
CL\_INS\_382
CL\_INS\_382
CL\_INS\_189
CL\_INS\_189
CL\_INS\_189
CL\_INS\_189
CL\_INS\_189
CL\_INS\_20
CL\_INS\_382
CL\_INS\_382
CL\_INS\_382
CL\_INS\_189
CL\_INS\_189
CL\_INS\_189
CL\_INS\_189
CL\_INS\_189
CL\_INS\_189
CL\_INS\_189
CL\_INS\_189
CL\_INS\_189
CL\_INS\_189
CL\_INS\_189
CL\_INS\_382
CL\_INS\_382
CL\_INS\_189
CL\_INS\_189
CL\_INS\_20
CL\_INS\_86
CL\_INS\_86
CL\_INS\_382
CL\_INS\_189
CL\_INS\_382
CL\_INS\_189
CL\_INS\_189
CL\_INS\_189
CL\_INS\_189
CL\_INS\_189
CL\_INS\_189
CL\_INS\_189
CL\_INS\_189
CL\_INS\_189
CL\_INS\_189
CL\_INS\_189
CL\_INS\_189
CL\_INS\_189
CL\_INS\_189
CL\_INS\_189
CL\_INS\_189
CL\_INS\_189
CL\_INS\_189
CL\_INS\_189
CL\_INS\_189
CL\_INS\_189
CL\_INS\_189
CL\_INS\_189
CL\_INS\_189
CL\_INS\_189
CL\_INS\_189
CL\_INS\_189
CL\_INS\_189
CL\_INS\_189
CL\_INS\_189
CL\_INS\_189
CL\_INS\_79
CL\_INS\_189
CL\_INS\_189
CL\_INS\_79
CL\_INS\_79
CL\_INS\_189
CL\_INS\_189
CL\_INS\_189
CL\_INS\_189
CL\_INS\_384
CL\_INS\_189
CL\_INS\_20
CL\_INS\_189
CL\_INS\_189
CL\_INS\_86
CL\_INS\_385
CL\_INS\_385
CL\_INS\_385
CL\_INS\_86
CL\_INS\_189
CL\_INS\_189
CL\_INS\_189
CL\_INS\_189
CL\_INS\_189
CL\_INS\_189
CL\_INS\_189
CL\_INS\_189
CL\_INS\_189
CL\_INS\_189
CL\_INS\_189
CL\_INS\_189
CL\_INS\_189
CL\_INS\_207
CL\_INS\_189
CL\_INS\_207
CL\_INS\_207
CL\_INS\_207
CL\_INS\_207
CL\_INS\_207
CL\_INS\_207
CL\_INS\_189
CL\_INS\_189
CL\_INS\_189
CL\_INS\_123
CL\_INS\_189
CL\_INS\_189
CL\_INS\_189
CL\_INS\_189
CL\_INS\_189
CL\_INS\_189
CL\_INS\_189
CL\_INS\_189
CL\_INS\_189
CL\_INS\_189
CL\_INS\_189
CL\_INS\_189
CL\_INS\_189
CL\_INS\_189
CL\_INS\_189
CL\_INS\_20
CL\_INS\_382
CL\_INS\_86
CL\_INS\_382
CL\_INS\_99
CL\_INS\_99
CL\_INS\_99
CL\_INS\_207
CL\_INS\_207
CL\_INS\_207
CL\_INS\_207
CL\_INS\_382
CL\_INS\_382
CL\_INS\_207
CL\_INS\_207
CL\_INS\_207
CL\_INS\_207
CL\_INS\_207
CL\_INS\_382
CL\_INS\_207
CL\_INS\_207
CL\_INS\_382
CL\_INS\_189
CL\_INS\_189
CL\_INS\_189
CL\_INS\_189
CL\_INS\_189
CL\_INS\_189
CL\_INS\_189
CL\_INS\_189
CL\_INS\_382
CL\_INS\_189
CL\_INS\_189
CL\_INS\_189
CL\_INS\_189
CL\_INS\_189
CL\_INS\_189
CL\_INS\_189
CL\_INS\_189
CL\_INS\_237
CL\_INS\_237
CL\_INS\_237
CL\_INS\_237
CL\_INS\_237
CL\_INS\_237
CL\_INS\_237
CL\_INS\_237
CL\_INS\_237
CL\_INS\_237
CL\_INS\_237
CL\_INS\_237
CL\_INS\_247
CL\_INS\_155
CL\_INS\_155
CL\_INS\_155
CL\_INS\_155
CL\_INS\_155
CL\_INS\_155
CL\_INS\_155
CL\_INS\_155
CL\_INS\_155
CL\_INS\_155
CL\_INS\_155
CL\_INS\_20
CL\_INS\_20
CL\_INS\_189
CL\_INS\_189
CL\_INS\_189
CL\_INS\_382
CL\_INS\_382
CL\_INS\_189
CL\_INS\_189
CL\_INS\_189
CL\_INS\_189
CL\_INS\_189
CL\_INS\_247
CL\_INS\_247
CL\_INS\_247
CL\_INS\_247
CL\_INS\_189
CL\_INS\_247
CL\_INS\_247
CL\_INS\_247
CL\_INS\_247
CL\_INS\_247
CL\_INS\_247
CL\_INS\_247
CL\_INS\_247
CL\_INS\_247
CL\_INS\_247
CL\_INS\_247
CL\_INS\_247
CL\_INS\_247
CL\_INS\_247
CL\_INS\_247
CL\_INS\_247
CL\_INS\_20
CL\_INS\_20
CL\_INS\_189
CL\_INS\_189
CL\_INS\_123
CL\_INS\_189
CL\_INS\_189
CL\_INS\_189
CL\_INS\_149
CL\_INS\_204
CL\_INS\_382
CL\_INS\_382
CL\_INS\_189
CL\_INS\_189
CL\_INS\_189
CL\_INS\_189
CL\_INS\_189
CL\_INS\_189
CL\_INS\_189
CL\_INS\_189
CL\_INS\_189
CL\_INS\_189
CL\_INS\_20
CL\_INS\_189
CL\_INS\_189
CL\_INS\_189
CL\_INS\_204
CL\_INS\_189
CL\_INS\_20
CL\_INS\_189
CL\_INS\_189
CL\_INS\_60
CL\_INS\_189
CL\_INS\_20
CL\_INS\_189
CL\_INS\_189
CL\_INS\_204
CL\_INS\_204
CL\_INS\_204
CL\_INS\_204
CL\_INS\_204
CL\_INS\_204
CL\_INS\_204
CL\_INS\_204
CL\_INS\_189
CL\_INS\_189
CL\_INS\_189
CL\_INS\_204
CL\_INS\_189
CL\_INS\_204
CL\_INS\_86
CL\_INS\_189
CL\_INS\_204
CL\_INS\_204
CL\_INS\_189
CL\_INS\_20
CL\_INS\_189
CL\_INS\_189
CL\_INS\_189
CL\_INS\_204
CL\_INS\_204
CL\_INS\_204
CL\_INS\_86
CL\_INS\_189
CL\_INS\_189
CL\_INS\_189
CL\_INS\_189
CL\_INS\_189
CL\_INS\_189
CL\_INS\_189
CL\_INS\_189
CL\_INS\_189
CL\_INS\_189
CL\_INS\_189
CL\_INS\_20
CL\_INS\_204
CL\_INS\_189
CL\_INS\_189
CL\_INS\_189
CL\_INS\_207
CL\_INS\_20
CL\_INS\_20
CL\_INS\_204
CL\_INS\_189
CL\_INS\_189
CL\_INS\_106
CL\_INS\_189
CL\_INS\_189
CL\_INS\_106
CL\_INS\_189
CL\_INS\_189
CL\_INS\_189
CL\_INS\_60
CL\_INS\_60
CL\_INS\_60
CL\_INS\_60
CL\_INS\_60
CL\_INS\_60
CL\_INS\_60
CL\_INS\_60
CL\_INS\_60
CL\_INS\_60
CL\_INS\_60
CL\_INS\_60
CL\_INS\_60
CL\_INS\_60
CL\_INS\_60
CL\_INS\_60
CL\_INS\_60
CL\_INS\_60
CL\_INS\_60
CL\_INS\_60
CL\_INS\_60
CL\_INS\_60
CL\_INS\_189
CL\_INS\_146
CL\_INS\_189
CL\_INS\_189
CL\_INS\_189
CL\_INS\_189
CL\_INS\_189
CL\_INS\_60
CL\_INS\_204
CL\_INS\_189
CL\_INS\_204
CL\_INS\_204
CL\_INS\_204
CL\_INS\_204
CL\_INS\_20
CL\_INS\_204
CL\_INS\_204
CL\_INS\_79
CL\_INS\_79
CL\_INS\_79
CL\_INS\_189
CL\_INS\_189
CL\_INS\_204
CL\_INS\_204
CL\_INS\_204
CL\_INS\_204
CL\_INS\_204
CL\_INS\_204
CL\_INS\_204
CL\_INS\_204
CL\_INS\_204
CL\_INS\_204
CL\_INS\_204
CL\_INS\_204
CL\_INS\_204
CL\_INS\_204
CL\_INS\_61
CL\_INS\_204
CL\_INS\_204
CL\_INS\_204
CL\_INS\_204
CL\_INS\_204
CL\_INS\_204
CL\_INS\_106
CL\_INS\_106
CL\_INS\_382
CL\_INS\_189
CL\_INS\_204
CL\_INS\_189
CL\_INS\_189
CL\_INS\_204
CL\_INS\_204
CL\_INS\_204
CL\_INS\_204
CL\_INS\_204
CL\_INS\_189
CL\_INS\_189
CL\_INS\_382
CL\_INS\_189
CL\_INS\_189
CL\_INS\_189
CL\_INS\_189
CL\_INS\_189
CL\_INS\_189
CL\_INS\_189
CL\_INS\_189
CL\_INS\_189
CL\_INS\_189
CL\_INS\_189
CL\_INS\_189
CL\_INS\_189
CL\_INS\_382
CL\_INS\_382
CL\_INS\_382
CL\_INS\_382
CL\_INS\_382
CL\_INS\_382
CL\_INS\_382
CL\_INS\_20
CL\_INS\_189
CL\_INS\_189
CL\_INS\_189
CL\_INS\_189
CL\_INS\_189
CL\_INS\_20
CL\_INS\_20
CL\_INS\_20
CL\_INS\_20
CL\_INS\_20
CL\_INS\_189
CL\_INS\_20
CL\_INS\_189
CL\_INS\_189
CL\_INS\_189
CL\_INS\_189
CL\_INS\_189
CL\_INS\_189
CL\_INS\_189
CL\_INS\_189
CL\_INS\_189
CL\_INS\_189
CL\_INS\_189
CL\_INS\_189
CL\_INS\_189
CL\_INS\_189
CL\_INS\_189
CL\_INS\_189
CL\_INS\_189
CL\_INS\_189
CL\_INS\_60
CL\_INS\_60
CL\_INS\_189
CL\_INS\_20
CL\_INS\_20
CL\_INS\_189
CL\_INS\_189
CL\_INS\_20
CL\_INS\_20
CL\_INS\_20
CL\_INS\_20
CL\_INS\_20
CL\_INS\_20
CL\_INS\_20
CL\_INS\_20
CL\_INS\_20
CL\_INS\_189
CL\_INS\_20
CL\_INS\_189
CL\_INS\_382
CL\_INS\_189
CL\_INS\_340
CL\_INS\_79
CL\_INS\_189
CL\_INS\_382
CL\_INS\_382
CL\_INS\_189
CL\_INS\_237
CL\_INS\_189
CL\_INS\_189
CL\_INS\_189
CL\_INS\_189
CL\_INS\_189
CL\_INS\_189
CL\_INS\_189
CL\_INS\_189
CL\_INS\_189
CL\_INS\_189
CL\_INS\_189
CL\_INS\_237
CL\_INS\_20
CL\_INS\_86
CL\_INS\_60
CL\_INS\_189
CL\_INS\_189
CL\_INS\_189
CL\_INS\_189
CL\_INS\_189
CL\_INS\_189
CL\_INS\_189
CL\_INS\_189
CL\_INS\_189
CL\_INS\_189
CL\_INS\_189
CL\_INS\_79
CL\_INS\_79
CL\_INS\_79
CL\_INS\_79
CL\_INS\_79
CL\_INS\_79
CL\_INS\_79
CL\_INS\_79
CL\_INS\_79
CL\_INS\_79
CL\_INS\_79
CL\_INS\_79
CL\_INS\_79
CL\_INS\_79
CL\_INS\_79
CL\_INS\_79
CL\_INS\_79
CL\_INS\_79
CL\_INS\_79
CL\_INS\_79
CL\_INS\_79
CL\_INS\_382
CL\_INS\_189
CL\_INS\_189
CL\_INS\_189
CL\_INS\_189
CL\_INS\_146
CL\_INS\_382
CL\_INS\_382
CL\_INS\_382
CL\_INS\_146
CL\_INS\_189
CL\_INS\_146
CL\_INS\_144
CL\_INS\_382
CL\_INS\_237
CL\_INS\_237
CL\_INS\_20
CL\_INS\_189
CL\_INS\_189
CL\_INS\_189
CL\_INS\_189
CL\_INS\_189
CL\_INS\_189
CL\_INS\_189
CL\_INS\_189
CL\_INS\_189
CL\_INS\_79
CL\_INS\_382
CL\_INS\_382
CL\_INS\_146
CL\_INS\_382
CL\_INS\_382
CL\_INS\_382
CL\_INS\_382
CL\_INS\_79
CL\_INS\_382
CL\_INS\_382
CL\_INS\_189
CL\_INS\_42
CL\_INS\_42
CL\_INS\_42
CL\_INS\_189
CL\_INS\_79
CL\_INS\_189
CL\_INS\_237
CL\_INS\_189
CL\_INS\_382
CL\_INS\_382
CL\_INS\_79
CL\_INS\_60
CL\_INS\_189
CL\_INS\_189
CL\_INS\_189
CL\_INS\_189
CL\_INS\_189
CL\_INS\_189
CL\_INS\_189
CL\_INS\_189
CL\_INS\_189
CL\_INS\_189
CL\_INS\_189
CL\_INS\_382
CL\_INS\_382
CL\_INS\_382
CL\_INS\_382
CL\_INS\_189
CL\_INS\_189
CL\_INS\_189
CL\_INS\_382
CL\_INS\_60
CL\_INS\_382
CL\_INS\_382
CL\_INS\_382
CL\_INS\_382
CL\_INS\_382
CL\_INS\_382
CL\_INS\_189
CL\_INS\_189
CL\_INS\_60
CL\_INS\_189
CL\_INS\_189
CL\_INS\_189
CL\_INS\_189
CL\_INS\_189
CL\_INS\_189
CL\_INS\_189
CL\_INS\_189
CL\_INS\_189
CL\_INS\_189
CL\_INS\_189
CL\_INS\_189
CL\_INS\_189
CL\_INS\_189
CL\_INS\_189
CL\_INS\_189
CL\_INS\_189
CL\_INS\_189
CL\_INS\_189
CL\_INS\_189
CL\_INS\_189
CL\_INS\_189
CL\_INS\_189
CL\_INS\_189
CL\_INS\_189
CL\_INS\_189
CL\_INS\_189
CL\_INS\_189
CL\_INS\_189
CL\_INS\_189
CL\_INS\_97
CL\_INS\_97
CL\_INS\_97
CL\_INS\_97
CL\_INS\_97
CL\_INS\_97
CL\_INS\_97
CL\_INS\_189
CL\_INS\_106
CL\_INS\_106
CL\_INS\_106
CL\_INS\_106
CL\_INS\_106
CL\_INS\_106
CL\_INS\_189
CL\_INS\_97
CL\_INS\_97
CL\_INS\_97
CL\_INS\_97
CL\_INS\_189
CL\_INS\_189
CL\_INS\_97
CL\_INS\_97
CL\_INS\_189
CL\_INS\_189
CL\_INS\_97
CL\_INS\_189
CL\_INS\_189
CL\_INS\_97
CL\_INS\_97
CL\_INS\_189
CL\_INS\_189
CL\_INS\_189
CL\_INS\_189
CL\_INS\_189
CL\_INS\_189
CL\_INS\_189
CL\_INS\_189
CL\_INS\_189
CL\_INS\_189
CL\_INS\_189
CL\_INS\_189
CL\_INS\_189
CL\_INS\_97
CL\_INS\_97
CL\_INS\_189
CL\_INS\_189
CL\_INS\_189
CL\_INS\_189
CL\_INS\_189
CL\_INS\_189
CL\_INS\_97
CL\_INS\_189
CL\_INS\_189
CL\_INS\_189
CL\_INS\_20
CL\_INS\_189
CL\_INS\_189
CL\_INS\_97
CL\_INS\_20
CL\_INS\_20
CL\_INS\_20
CL\_INS\_189
CL\_INS\_189
CL\_INS\_97
CL\_INS\_97
CL\_INS\_60
CL\_INS\_97
CL\_INS\_189
CL\_INS\_189
CL\_INS\_146
CL\_INS\_189
CL\_INS\_97
CL\_INS\_97
CL\_INS\_189
CL\_INS\_189
CL\_INS\_189
CL\_INS\_146
CL\_INS\_189
CL\_INS\_189
CL\_INS\_97
CL\_INS\_189
CL\_INS\_189
CL\_INS\_189
CL\_INS\_189
CL\_INS\_97
CL\_INS\_189
CL\_INS\_189
CL\_INS\_97
CL\_INS\_97
CL\_INS\_146
CL\_INS\_146
CL\_INS\_382
CL\_INS\_189
CL\_INS\_189
CL\_INS\_189
CL\_INS\_189
CL\_INS\_189
CL\_INS\_189
CL\_INS\_189
CL\_INS\_189
CL\_INS\_189
CL\_INS\_189
CL\_INS\_189
CL\_INS\_189
CL\_INS\_189
CL\_INS\_189
CL\_INS\_189
CL\_INS\_189
CL\_INS\_189
CL\_INS\_189
CL\_INS\_384
CL\_INS\_237
CL\_INS\_189
CL\_INS\_189
CL\_INS\_189
CL\_INS\_189
CL\_INS\_189
CL\_INS\_189
CL\_INS\_189
CL\_INS\_368
CL\_INS\_368
CL\_INS\_368
CL\_INS\_189
CL\_INS\_189
CL\_INS\_189
CL\_INS\_189
CL\_INS\_189
CL\_INS\_189
CL\_INS\_189
CL\_INS\_189
CL\_INS\_189
CL\_INS\_189
CL\_INS\_155
CL\_INS\_189
CL\_INS\_207
CL\_INS\_207
CL\_INS\_237
CL\_INS\_207
CL\_INS\_385
CL\_INS\_155
CL\_INS\_155
CL\_INS\_385
CL\_INS\_385
CL\_INS\_207
CL\_INS\_207
CL\_INS\_207
CL\_INS\_207
CL\_INS\_76
CL\_INS\_207
CL\_INS\_76
CL\_INS\_189
CL\_INS\_189
CL\_INS\_189
CL\_INS\_189
CL\_INS\_189
CL\_INS\_189
CL\_INS\_189
CL\_INS\_207
CL\_INS\_207
CL\_INS\_207
CL\_INS\_207
CL\_INS\_207
CL\_INS\_207
CL\_INS\_207
CL\_INS\_207
CL\_INS\_189
CL\_INS\_189
CL\_INS\_189
CL\_INS\_189
CL\_INS\_189
CL\_INS\_189
CL\_INS\_207
CL\_INS\_207
CL\_INS\_207
CL\_INS\_207
CL\_INS\_207
CL\_INS\_207
CL\_INS\_207
CL\_INS\_207
CL\_INS\_207
CL\_INS\_189
CL\_INS\_207
CL\_INS\_207
CL\_INS\_189
CL\_INS\_189
CL\_INS\_189
CL\_INS\_189
CL\_INS\_189
CL\_INS\_189
CL\_INS\_189
CL\_INS\_189
CL\_INS\_189
CL\_INS\_189
CL\_INS\_189
CL\_INS\_189
CL\_INS\_189
CL\_INS\_189
CL\_INS\_189
CL\_INS\_207
CL\_INS\_189
CL\_INS\_189
CL\_INS\_247
CL\_INS\_20
CL\_INS\_20
CL\_INS\_20
CL\_INS\_20
CL\_INS\_20
CL\_INS\_20
CL\_INS\_20
CL\_INS\_207
CL\_INS\_189
CL\_INS\_189
CL\_INS\_189
CL\_INS\_189
CL\_INS\_189
CL\_INS\_189
CL\_INS\_189
CL\_INS\_189
CL\_INS\_382
CL\_INS\_382
CL\_INS\_189
CL\_INS\_382
CL\_INS\_159
CL\_INS\_189
CL\_INS\_189
CL\_INS\_189
CL\_INS\_189
CL\_INS\_382
CL\_INS\_382
CL\_INS\_189
CL\_INS\_189
CL\_INS\_189
CL\_INS\_189
CL\_INS\_189
CL\_INS\_189
CL\_INS\_237
CL\_INS\_189
CL\_INS\_189
CL\_INS\_237
CL\_INS\_237
CL\_INS\_237
CL\_INS\_189
CL\_INS\_149
CL\_INS\_149
CL\_INS\_189
CL\_INS\_159
CL\_INS\_189
CL\_INS\_149
CL\_INS\_70
CL\_INS\_286
CL\_INS\_286
CL\_INS\_237
CL\_INS\_247
CL\_INS\_189
CL\_INS\_189
CL\_INS\_70
CL\_INS\_70
CL\_INS\_30
CL\_INS\_30
CL\_INS\_30
CL\_INS\_237
CL\_INS\_70
CL\_INS\_189
CL\_INS\_189
CL\_INS\_149
CL\_INS\_382
CL\_INS\_382
CL\_INS\_189
CL\_INS\_189
CL\_INS\_189
CL\_INS\_189
CL\_INS\_189
CL\_INS\_189
CL\_INS\_189
CL\_INS\_189
CL\_INS\_189
CL\_INS\_189
CL\_INS\_189
CL\_INS\_189
CL\_INS\_368
CL\_INS\_237
CL\_INS\_237
CL\_INS\_237
CL\_INS\_237
CL\_INS\_70
CL\_INS\_237
CL\_INS\_237
CL\_INS\_237
CL\_INS\_237
CL\_INS\_237
CL\_INS\_70
CL\_INS\_237
CL\_INS\_189
CL\_INS\_30
CL\_INS\_237
CL\_INS\_237
CL\_INS\_237
CL\_INS\_368
CL\_INS\_189
CL\_INS\_30
CL\_INS\_189
CL\_INS\_189
CL\_INS\_170
CL\_INS\_123
CL\_INS\_30
CL\_INS\_237
CL\_INS\_237
CL\_INS\_237
CL\_INS\_237
CL\_INS\_70
CL\_INS\_189
CL\_INS\_189
CL\_INS\_189
CL\_INS\_189
CL\_INS\_189
CL\_INS\_189
CL\_INS\_189
CL\_INS\_189
CL\_INS\_70
CL\_INS\_70
CL\_INS\_30
CL\_INS\_30
CL\_INS\_70
CL\_INS\_70
CL\_INS\_237
CL\_INS\_247
CL\_INS\_237
CL\_INS\_237
CL\_INS\_247
CL\_INS\_30
CL\_INS\_30
CL\_INS\_159
CL\_INS\_189
CL\_INS\_189
CL\_INS\_189
CL\_INS\_189
CL\_INS\_189
CL\_INS\_189
CL\_INS\_189
CL\_INS\_20
CL\_INS\_189
CL\_INS\_189
CL\_INS\_189
CL\_INS\_189
CL\_INS\_60
CL\_INS\_60
CL\_INS\_189
CL\_INS\_189
CL\_INS\_20
CL\_INS\_20
CL\_INS\_60
CL\_INS\_189
CL\_INS\_189
CL\_INS\_189
CL\_INS\_189
CL\_INS\_189
CL\_INS\_20
CL\_INS\_189
CL\_INS\_189
CL\_INS\_382
CL\_INS\_189
CL\_INS\_20
CL\_INS\_189
CL\_INS\_189
CL\_INS\_189
CL\_INS\_189
CL\_INS\_189
CL\_INS\_60
CL\_INS\_189
CL\_INS\_20
CL\_INS\_189
CL\_INS\_189
CL\_INS\_189
CL\_INS\_60
CL\_INS\_189
CL\_INS\_189
CL\_INS\_60
CL\_INS\_189
CL\_INS\_20
CL\_INS\_20
CL\_INS\_189
CL\_INS\_189
CL\_INS\_189
CL\_INS\_189
CL\_INS\_189
CL\_INS\_189
CL\_INS\_189
CL\_INS\_189
CL\_INS\_189
CL\_INS\_189
CL\_INS\_189
CL\_INS\_189
CL\_INS\_189
CL\_INS\_189
CL\_INS\_189
CL\_INS\_189
CL\_INS\_189
CL\_INS\_189
CL\_INS\_273
CL\_INS\_273
CL\_INS\_273
CL\_INS\_273
CL\_INS\_273
CL\_INS\_273
CL\_INS\_273
CL\_INS\_189
CL\_INS\_273
CL\_INS\_189
CL\_INS\_189
CL\_INS\_273
CL\_INS\_273
CL\_INS\_189
CL\_INS\_189
CL\_INS\_273
CL\_INS\_273
CL\_INS\_189
CL\_INS\_189
CL\_INS\_189
CL\_INS\_189
CL\_INS\_189
CL\_INS\_273
CL\_INS\_189
CL\_INS\_273
CL\_INS\_273
CL\_INS\_189
CL\_INS\_273
CL\_INS\_273
CL\_INS\_189
CL\_INS\_189
CL\_INS\_273
CL\_INS\_273
CL\_INS\_273
CL\_INS\_273
CL\_INS\_189
CL\_INS\_273
CL\_INS\_273
CL\_INS\_273
CL\_INS\_189
CL\_INS\_189
CL\_INS\_189
CL\_INS\_189
CL\_INS\_273
CL\_INS\_273
CL\_INS\_273
CL\_INS\_189
CL\_INS\_273
CL\_INS\_273
CL\_INS\_273
CL\_INS\_189
CL\_INS\_189
CL\_INS\_189
CL\_INS\_273
CL\_INS\_189
CL\_INS\_273
CL\_INS\_189
CL\_INS\_273
CL\_INS\_189
CL\_INS\_273
CL\_INS\_273
CL\_INS\_273
CL\_INS\_189
CL\_INS\_189
CL\_INS\_189
CL\_INS\_189
CL\_INS\_189
CL\_INS\_273
CL\_INS\_273
CL\_INS\_189
CL\_INS\_273
CL\_INS\_273
CL\_INS\_273
CL\_INS\_273
CL\_INS\_189
CL\_INS\_189
CL\_INS\_189
CL\_INS\_189
CL\_INS\_189
CL\_INS\_273
CL\_INS\_273
CL\_INS\_273
CL\_INS\_273
CL\_INS\_273
CL\_INS\_273
CL\_INS\_189
CL\_INS\_189
CL\_INS\_189
CL\_INS\_273
CL\_INS\_273
CL\_INS\_273
CL\_INS\_189
CL\_INS\_273
CL\_INS\_273
CL\_INS\_273
CL\_INS\_273
CL\_INS\_273
CL\_INS\_273
CL\_INS\_273
CL\_INS\_99
CL\_INS\_189
CL\_INS\_273
CL\_INS\_189
CL\_INS\_273
CL\_INS\_189
CL\_INS\_273
CL\_INS\_273
CL\_INS\_189
CL\_INS\_189
CL\_INS\_189
CL\_INS\_273
CL\_INS\_189
CL\_INS\_189
CL\_INS\_57
CL\_INS\_57
CL\_INS\_149
CL\_INS\_149
CL\_INS\_149
CL\_INS\_149
CL\_INS\_149
CL\_INS\_149
CL\_INS\_57
CL\_INS\_273
CL\_INS\_273
CL\_INS\_189
CL\_INS\_189
CL\_INS\_273
CL\_INS\_273
CL\_INS\_189
CL\_INS\_189
CL\_INS\_273
CL\_INS\_273
CL\_INS\_286
CL\_INS\_273
CL\_INS\_286
CL\_INS\_286
CL\_INS\_189
CL\_INS\_189
CL\_INS\_189
CL\_INS\_273
CL\_INS\_189
CL\_INS\_189
CL\_INS\_189
CL\_INS\_273
CL\_INS\_189
CL\_INS\_189
CL\_INS\_189
CL\_INS\_273
CL\_INS\_273
CL\_INS\_273
CL\_INS\_189
CL\_INS\_273
CL\_INS\_273
CL\_INS\_189
CL\_INS\_189
CL\_INS\_189
CL\_INS\_189
CL\_INS\_273
CL\_INS\_189
CL\_INS\_171
CL\_INS\_171
CL\_INS\_171
CL\_INS\_171
CL\_INS\_171
CL\_INS\_189
CL\_INS\_189
CL\_INS\_272
CL\_INS\_272
CL\_INS\_272
CL\_INS\_272
CL\_INS\_272
CL\_INS\_272
CL\_INS\_272
CL\_INS\_272
CL\_INS\_272
CL\_INS\_272
CL\_INS\_272
CL\_INS\_272
CL\_INS\_272
CL\_INS\_272
CL\_INS\_272
CL\_INS\_272
CL\_INS\_272
CL\_INS\_272
CL\_INS\_272
CL\_INS\_272
CL\_INS\_272
CL\_INS\_272
CL\_INS\_272
CL\_INS\_272
CL\_INS\_368
CL\_INS\_368
CL\_INS\_273
CL\_INS\_189
CL\_INS\_207
CL\_INS\_189
CL\_INS\_189
CL\_INS\_189
CL\_INS\_189
CL\_INS\_189
CL\_INS\_382
CL\_INS\_382
CL\_INS\_382
CL\_INS\_247
CL\_INS\_382
CL\_INS\_207
CL\_INS\_207
CL\_INS\_207
CL\_INS\_207
CL\_INS\_189
CL\_INS\_247
CL\_INS\_189
CL\_INS\_247
CL\_INS\_247
CL\_INS\_123
CL\_INS\_247
CL\_INS\_123
CL\_INS\_247
CL\_INS\_189
CL\_INS\_189
CL\_INS\_189
CL\_INS\_189
CL\_INS\_189
CL\_INS\_273
CL\_INS\_273
CL\_INS\_189
CL\_INS\_273
CL\_INS\_189
CL\_INS\_189
CL\_INS\_189
CL\_INS\_189
CL\_INS\_189
CL\_INS\_189
CL\_INS\_189
CL\_INS\_273
CL\_INS\_273
CL\_INS\_207
CL\_INS\_70
CL\_INS\_70
CL\_INS\_70
CL\_INS\_70
CL\_INS\_70
CL\_INS\_70
CL\_INS\_70
CL\_INS\_70
CL\_INS\_70
CL\_INS\_70
CL\_INS\_70
CL\_INS\_70
CL\_INS\_70
CL\_INS\_70
CL\_INS\_20
CL\_INS\_20
CL\_INS\_20
CL\_INS\_20
CL\_INS\_20
CL\_INS\_20
CL\_INS\_189
CL\_INS\_273
CL\_INS\_273
CL\_INS\_273
CL\_INS\_189
CL\_INS\_189
CL\_INS\_189
CL\_INS\_189
CL\_INS\_189
CL\_INS\_189
CL\_INS\_189
Cluster ID


CL\_22891
CL\_13498
CL\_33970
CL\_12815
CL\_12814
CL\_12813
CL\_10543
CL\_10544
CL\_29828
CL\_29827
CL\_29826
CL\_29825
CL\_29824
CL\_29823
CL\_29822
CL\_518
CL\_519
CL\_520
CL\_521
CL\_522
CL\_524
CL\_525
CL\_22888
CL\_4411
CL\_4407
CL\_6479
CL\_7507
CL\_7508
CL\_7509
CL\_6483
CL\_4402
CL\_34143
CL\_34144
CL\_8847
CL\_8848
CL\_8849
CL\_37628
CL\_37627
CL\_37626
CL\_37625
CL\_37624
CL\_37623
CL\_8850
CL\_4485
CL\_8169
CL\_8170
CL\_18335
CL\_37622
CL\_8171
CL\_8172
CL\_8173
CL\_8174
CL\_8175
CL\_8176
CL\_10971
CL\_37621
CL\_37620
CL\_37619
CL\_37618
CL\_37617
CL\_37616
CL\_37615
CL\_37614
CL\_37613
CL\_37612
CL\_34677
CL\_34678
CL\_34679
CL\_5807
CL\_37611
CL\_37610
CL\_37609
CL\_8177
CL\_5408
CL\_5409
CL\_5410
CL\_1099
CL\_8178
CL\_8179
CL\_8180
CL\_21105
CL\_11477
CL\_9267
CL\_9266
CL\_4644
CL\_1098
CL\_1097
CL\_5411
CL\_5412
CL\_32323
CL\_16324
CL\_16325
CL\_17536
CL\_16326
CL\_16327
CL\_16328
CL\_16621
CL\_16684
CL\_16685
CL\_16686
CL\_16687
CL\_32324
CL\_32325
CL\_32326
CL\_32327
CL\_32328
CL\_32329
CL\_32330
CL\_32331
CL\_32332
CL\_32333
CL\_32334
CL\_32335
CL\_32336
CL\_32337
CL\_32338
CL\_32339
CL\_32340
CL\_32341
CL\_32342
CL\_32343
CL\_32344
CL\_32345
CL\_32346
CL\_7539
CL\_7540
CL\_32347
CL\_16688
CL\_16333
CL\_16334
CL\_16335
CL\_16336
CL\_16337
CL\_17537
CL\_17538
CL\_16689
CL\_16690
CL\_16691
CL\_16692
CL\_16693
CL\_16338
CL\_16339
CL\_16340
CL\_16341
CL\_16342
CL\_16343
CL\_16344
CL\_16345
CL\_16346
CL\_34610
CL\_16347
CL\_16348
CL\_16349
CL\_16618
CL\_27306
CL\_10976
CL\_6664
CL\_34424
CL\_34423
CL\_34422
CL\_6663
CL\_6662
CL\_33450
CL\_8966
CL\_13968
CL\_13969
CL\_13970
CL\_13971
CL\_36446
CL\_15697
CL\_15696
CL\_15695
CL\_9224
CL\_6661
CL\_15141
CL\_15142
CL\_15143
CL\_15144
CL\_15145
CL\_15146
CL\_15147
CL\_15148
CL\_15149
CL\_7047
CL\_7046
CL\_27120
CL\_9117
CL\_9225
CL\_8965
CL\_8319
CL\_6660
CL\_13890
CL\_9944
CL\_24458
CL\_24459
CL\_13930
CL\_16617
CL\_17073
CL\_16350
CL\_13561
CL\_16616
CL\_12995
CL\_16615
CL\_12994
CL\_16444
CL\_16445
CL\_16351
CL\_13071
CL\_14825
CL\_6659
CL\_6658
CL\_32832
CL\_32093
CL\_27929
CL\_27928
CL\_27927
CL\_27926
CL\_27925
CL\_27924
CL\_27923
CL\_7045
CL\_16352
CL\_7044
CL\_7043
CL\_25174
CL\_5380
CL\_6657
CL\_34611
CL\_6656
CL\_34375
CL\_11210
CL\_20987
CL\_25871
CL\_34842
CL\_6655
CL\_21777
CL\_13974
CL\_7042
CL\_6654
CL\_5348
CL\_9945
CL\_17541
CL\_17542
CL\_5349
CL\_8322
CL\_9115
CL\_9082
CL\_9947
CL\_9081
CL\_11736
CL\_11735
CL\_26712
CL\_26713
CL\_26714
CL\_26715
CL\_13507
CL\_6653
CL\_17543
CL\_17544
CL\_27922
CL\_27921
CL\_27920
CL\_27919
CL\_27918
CL\_11209
CL\_17673
CL\_6652
CL\_6651
CL\_8325
CL\_15151
CL\_15152
CL\_15153
CL\_15154
CL\_15155
CL\_15156
CL\_15157
CL\_15158
CL\_22672
CL\_6650
CL\_13975
CL\_13976
CL\_9075
CL\_34374
CL\_25177
CL\_5381
CL\_6649
CL\_5353
CL\_5354
CL\_12058
CL\_11733
CL\_17674
CL\_17675
CL\_11732
CL\_17676
CL\_5355
CL\_6645
CL\_23871
CL\_23870
CL\_11207
CL\_16587
CL\_16586
CL\_16585
CL\_7041
CL\_5352
CL\_13888
CL\_21331
CL\_9226
CL\_6648
CL\_30680
CL\_15104
CL\_15105
CL\_25365
CL\_14826
CL\_6647
CL\_6646
CL\_23872
CL\_8326
CL\_8327
CL\_8328
CL\_12059
CL\_37752
CL\_37751
CL\_25364
CL\_17725
CL\_9227
CL\_9228
CL\_6787
CL\_6788
CL\_27917
CL\_6791
CL\_6792
CL\_16584
CL\_5356
CL\_14827
CL\_33578
CL\_6793
CL\_14020
CL\_8330
CL\_8331
CL\_8332
CL\_15106
CL\_9074
CL\_25872
CL\_32836
CL\_32837
CL\_32838
CL\_5382
CL\_33509
CL\_33508
CL\_33507
CL\_32839
CL\_14828
CL\_32840
CL\_32841
CL\_32842
CL\_32843
CL\_32844
CL\_32845
CL\_32846
CL\_32847
CL\_8964
CL\_8329
CL\_9230
CL\_8963
CL\_8962
CL\_5383
CL\_6643
CL\_6642
CL\_5384
CL\_5385
CL\_5386
CL\_5387
CL\_37448
CL\_37447
CL\_37446
CL\_8333
CL\_9231
CL\_9232
CL\_9233
CL\_9234
CL\_9235
CL\_9236
CL\_9237
CL\_9238
CL\_9239
CL\_8335
CL\_5798
CL\_5367
CL\_6794
CL\_13874
CL\_11529
CL\_9240
CL\_9241
CL\_9949
CL\_26141
CL\_26142
CL\_26143
CL\_26144
CL\_26145
CL\_7821
CL\_7032
CL\_34612
CL\_31641
CL\_8682
CL\_13108
CL\_25184
CL\_31642
CL\_11206
CL\_7361
CL\_25047
CL\_6635
CL\_20985
CL\_34613
CL\_9950
CL\_13504
CL\_13503
CL\_6641
CL\_13502
CL\_14829
CL\_5388
CL\_27237
CL\_27620
CL\_27621
CL\_6639
CL\_5364
CL\_5935
CL\_5389
CL\_12804
CL\_6448
CL\_19855
CL\_19854
CL\_6640
CL\_6638
CL\_13988
CL\_13989
CL\_10317
CL\_25178
CL\_11729
CL\_11402
CL\_11728
CL\_11727
CL\_8881
CL\_8334
CL\_17550
CL\_5149
CL\_17551
CL\_16006
CL\_6637
CL\_25363
CL\_6636
CL\_13628
CL\_13337
CL\_31643
CL\_31644
CL\_31645
CL\_31646
CL\_17552
CL\_21018
CL\_21017
CL\_27916
CL\_27915
CL\_27914
CL\_27913
CL\_16600
CL\_16599
CL\_16598
CL\_16597
CL\_16596
CL\_16694
CL\_16695
CL\_16696
CL\_16697
CL\_16698
CL\_16699
CL\_16700
CL\_16701
CL\_16702
CL\_16703
CL\_16704
CL\_16705
CL\_16706
CL\_16707
CL\_16708
CL\_16709
CL\_16710
CL\_16711
CL\_16712
CL\_16713
CL\_16714
CL\_11660
CL\_4651
CL\_34145
CL\_34146
CL\_34147
CL\_34148
CL\_34149
CL\_10168
CL\_12122
CL\_35221
CL\_35220
CL\_35219
CL\_35218
CL\_35217
CL\_26547
CL\_5423
CL\_5424
CL\_5425
CL\_34150
CL\_34151
CL\_34152
CL\_34153
CL\_34154
CL\_34155
CL\_34156
CL\_34157
CL\_34158
CL\_37601
CL\_37600
CL\_11406
CL\_11405
CL\_37599
CL\_37598
CL\_17690
CL\_7621
CL\_11313
CL\_17634
CL\_17689
CL\_5426
CL\_35216
CL\_35215
CL\_35214
CL\_35213
CL\_35212
CL\_35211
CL\_35210
CL\_16715
CL\_16716
CL\_16717
CL\_16718
CL\_16719
CL\_16720
CL\_16721
CL\_16722
CL\_16723
CL\_16724
CL\_16725
CL\_16726
CL\_16727
CL\_16728
CL\_16729
CL\_16730
CL\_16731
CL\_16732
CL\_16733
CL\_16734
CL\_16735
CL\_16736
CL\_16737
CL\_16738
CL\_16739
CL\_16740
CL\_16741
CL\_16742
CL\_16595
CL\_16743
CL\_16744
CL\_34159
CL\_16745
CL\_2287
CL\_26933
CL\_2271
CL\_20687
CL\_20686
CL\_17653
CL\_34378
CL\_34377
CL\_34376
CL\_8585
CL\_34691
CL\_34692
CL\_15349
CL\_11535
CL\_23447
CL\_11218
CL\_21677
CL\_10865
CL\_10864
CL\_23462
CL\_23463
CL\_23464
CL\_23465
CL\_10863
CL\_9496
CL\_10862
CL\_10861
CL\_10860
CL\_10859
CL\_10858
CL\_10857
CL\_10856
CL\_10855
CL\_19676
CL\_11534
CL\_11533
CL\_11532
CL\_11531
CL\_34079
CL\_33451
CL\_16861
CL\_27687
CL\_27686
CL\_27685
CL\_27684
CL\_13573
CL\_36044
CL\_27239
CL\_27238
CL\_27576
CL\_35222
CL\_4432
CL\_4488
CL\_4517
CL\_7112
CL\_8155
CL\_7522
CL\_7523
CL\_7524
CL\_7525
CL\_7526
CL\_6741
CL\_7527
CL\_7528
CL\_7529
CL\_7530
CL\_7531
CL\_7532
CL\_7533
CL\_9122
CL\_12124
CL\_5809
CL\_27616
CL\_33512
CL\_33511
CL\_33510
CL\_27617
CL\_13954
CL\_25630
CL\_23169
CL\_5366
CL\_13572
CL\_36043
CL\_37334
CL\_35369
CL\_37335
CL\_37336
CL\_37337
CL\_37338
CL\_18879
CL\_18878
CL\_18877
CL\_18876
CL\_18875
CL\_18874
CL\_18873
CL\_18872
CL\_18871
CL\_18870
CL\_18869
CL\_18868
CL\_8932
CL\_6438
CL\_6437
CL\_6436
CL\_6435
CL\_6434
CL\_6433
CL\_6432
CL\_6431
CL\_6430
CL\_6429
CL\_6428
CL\_7838
CL\_7837
CL\_37339
CL\_37340
CL\_37341
CL\_8554
CL\_10807
CL\_37342
CL\_37343
CL\_37344
CL\_37345
CL\_6357
CL\_4842
CL\_4841
CL\_5947
CL\_4840
CL\_4839
CL\_4838
CL\_4837
CL\_4836
CL\_4835
CL\_4834
CL\_4833
CL\_4832
CL\_2743
CL\_2742
CL\_2741
CL\_2740
CL\_2739
CL\_2737
CL\_2736
CL\_2735
CL\_2734
CL\_2272
CL\_2273
CL\_37508
CL\_37507
CL\_4995
CL\_23297
CL\_2274
CL\_2275
CL\_2276
CL\_2277
CL\_2278
CL\_2279
CL\_34705
CL\_25870
CL\_17671
CL\_17672
CL\_8975
CL\_8974
CL\_8309
CL\_12055
CL\_34395
CL\_34394
CL\_6693
CL\_17721
CL\_17720
CL\_17719
CL\_21566
CL\_11739
CL\_7067
CL\_34393
CL\_28435
CL\_14819
CL\_12056
CL\_7066
CL\_14820
CL\_14821
CL\_6594
CL\_1102
CL\_1101
CL\_1100
CL\_5393
CL\_5394
CL\_5395
CL\_5396
CL\_17718
CL\_17717
CL\_17716
CL\_10274
CL\_36450
CL\_5398
CL\_10273
CL\_16862
CL\_5399
CL\_5400
CL\_17715
CL\_17714
CL\_17713
CL\_17712
CL\_17711
CL\_5403
CL\_5404
CL\_5405
CL\_11307
CL\_17710
CL\_17709
CL\_17708
CL\_17707
CL\_17706
CL\_17705
CL\_17704
CL\_17703
CL\_17702
CL\_17701
CL\_17700
CL\_17699
CL\_5407
CL\_36449
CL\_36448
CL\_36447
CL\_11310
CL\_21509
CL\_21508
CL\_9292
CL\_26215
CL\_26214
CL\_6692
CL\_37655
CL\_24342
CL\_6691
CL\_6690
CL\_6689
CL\_6688
CL\_6687
CL\_6686
CL\_6685
CL\_6684
CL\_6683
CL\_6682
CL\_6681
CL\_6680
CL\_6679
CL\_6678
CL\_6677
CL\_6676
CL\_6675
CL\_6674
CL\_6673
CL\_6672
CL\_6671
CL\_6670
CL\_6669
CL\_6668
CL\_6667
CL\_6666
CL\_6665
CL\_15136
CL\_15137
CL\_15138
CL\_22675
CL\_22674
CL\_13633
CL\_6454
CL\_9217
CL\_9218
CL\_5897
CL\_5898
CL\_5899
CL\_5900
CL\_13926
CL\_23874
CL\_21709
CL\_21941
CL\_26213
CL\_26212
CL\_9219
CL\_9220
CL\_5903
CL\_5904
CL\_5905
CL\_5906
CL\_5907
CL\_7386
CL\_5908
CL\_5909
CL\_5910
CL\_5911
CL\_5912
CL\_5913
CL\_5914
CL\_5915
CL\_9221
CL\_5916
CL\_5917
CL\_5918
CL\_5919
CL\_5920
CL\_5921
CL\_9942
CL\_15139
CL\_526
CL\_15140
CL\_5922
CL\_23873
CL\_33579
CL\_9222
CL\_9223
CL\_5414
CL\_5415
CL\_5924
CL\_17698
CL\_17697
CL\_16967
CL\_37608
CL\_37607
CL\_37606
CL\_37605
CL\_37604
CL\_37603
CL\_37602
CL\_17696
CL\_17695
CL\_17694
CL\_17693
CL\_17692
CL\_17691
CL\_2280
CL\_2281
CL\_2282
CL\_2283
CL\_2284
CL\_2285
CL\_1096
CL\_34767
CL\_17723
CL\_17722
CL\_13632
CL\_32830
CL\_33173
CL\_7058
CL\_7057
CL\_11157
CL\_7056
CL\_7055
CL\_26711
CL\_7054
CL\_17727
CL\_13955
CL\_13956
CL\_13957
CL\_13958
CL\_13959
CL\_13960
CL\_13961
CL\_13962
CL\_13963
CL\_13964
CL\_13965
CL\_13966
CL\_13967
CL\_8312
CL\_8313
CL\_8314
CL\_8315
CL\_8316
CL\_8317
CL\_19856
CL\_7053
CL\_7052
CL\_30611
CL\_30612
CL\_8970
CL\_16592
CL\_16591
CL\_16590
CL\_8969
CL\_8968
CL\_7051
CL\_16589
CL\_21778
CL\_22673
CL\_7050
CL\_8967
CL\_6783
CL\_8318
CL\_22766
CL\_13629
CL\_14824
CL\_12383
CL\_7048
CL\_32831
CL\_7049
CL\_7459
CL\_9497
CL\_7458
CL\_21678
CL\_21679
CL\_21680
CL\_35209
CL\_22671
CL\_22670
CL\_33172
CL\_7457
CL\_5673
CL\_7368
CL\_4490
CL\_9129
CL\_27912
CL\_27911
CL\_27910
CL\_27909
CL\_27908
CL\_27907
CL\_27906
CL\_27905
CL\_27904
CL\_27903
CL\_27902
CL\_27901
CL\_16614
CL\_16613
CL\_16612
CL\_12345
CL\_12344
CL\_12343
CL\_16611
CL\_12341
CL\_16610
CL\_16609
CL\_12340
CL\_12339
CL\_12338
CL\_12337
CL\_12336
CL\_16606
CL\_16605
CL\_16604
CL\_16603
CL\_16602
CL\_11938
CL\_32348
CL\_32349
CL\_32350
CL\_32351
CL\_17539
CL\_9086
CL\_9085
CL\_9084
CL\_13972
CL\_13973
CL\_17540
CL\_12412
CL\_5347
CL\_8320
CL\_8321
CL\_15103
CL\_32833
CL\_32834
CL\_32835
CL\_27900
CL\_27899
CL\_27898
CL\_27897
CL\_27896
CL\_27895
CL\_27894
CL\_5350
CL\_4410
CL\_15150
CL\_11915
CL\_5351
CL\_6786
CL\_9080
CL\_24460
CL\_9079
CL\_9078
CL\_26211
CL\_17545
CL\_28277
CL\_28276
CL\_17546
CL\_17547
CL\_17548
CL\_11731
CL\_11730
CL\_23319
CL\_27931
CL\_9229
CL\_6644
CL\_13977
CL\_13978
CL\_13979
CL\_13980
CL\_13981
CL\_13982
CL\_13983
CL\_13984
CL\_13985
CL\_13986
CL\_13987
CL\_5357
CL\_5358
CL\_5359
CL\_5360
CL\_16863
CL\_16864
CL\_16865
CL\_11916
CL\_16866
CL\_4401
CL\_4400
CL\_5361
CL\_5362
CL\_5363
CL\_10318
CL\_11530
CL\_22889
CL\_17549
CL\_27893
CL\_27892
CL\_25510
CL\_16746
CL\_37455
CL\_37454
CL\_37453
CL\_37452
CL\_37451
CL\_37450
CL\_37449
CL\_28437
CL\_28436
CL\_34432
CL\_34431
CL\_34430
CL\_16747
CL\_16748
CL\_16749
CL\_16750
CL\_16751
CL\_34429
CL\_16752
CL\_16753
CL\_34428
CL\_20093
CL\_20094
CL\_20095
CL\_16754
CL\_16755
CL\_8856
CL\_8857
CL\_8858
CL\_8859
CL\_8860
CL\_8861
CL\_8862
CL\_34427
CL\_14772
CL\_14771
CL\_14770
CL\_14769
CL\_14768
CL\_14767
CL\_34426
CL\_8863
CL\_8864
CL\_8865
CL\_8866
CL\_20096
CL\_16756
CL\_8868
CL\_8869
CL\_20097
CL\_20098
CL\_10109
CL\_20099
CL\_16757
CL\_8871
CL\_8872
CL\_34425
CL\_20100
CL\_16758
CL\_16759
CL\_16760
CL\_16761
CL\_16762
CL\_16763
CL\_16764
CL\_20101
CL\_34421
CL\_34420
CL\_16765
CL\_8877
CL\_8879
CL\_34419
CL\_34418
CL\_34417
CL\_34416
CL\_34415
CL\_34414
CL\_19870
CL\_16766
CL\_16767
CL\_16768
CL\_15462
CL\_16769
CL\_16770
CL\_8886
CL\_8323
CL\_8324
CL\_9948
CL\_25175
CL\_25176
CL\_8888
CL\_8889
CL\_11734
CL\_8890
CL\_20102
CL\_20103
CL\_15159
CL\_15160
CL\_8895
CL\_8896
CL\_15161
CL\_15162
CL\_15163
CL\_15164
CL\_16771
CL\_16772
CL\_8887
CL\_16773
CL\_16774
CL\_16775
CL\_16776
CL\_8891
CL\_34413
CL\_16777
CL\_8892
CL\_8893
CL\_8894
CL\_20104
CL\_508
CL\_16778
CL\_16779
CL\_16780
CL\_16781
CL\_20105
CL\_20106
CL\_20107
CL\_16782
CL\_16783
CL\_16784
CL\_16785
CL\_34412
CL\_34411
CL\_34410
CL\_34409
CL\_16786
CL\_16787
CL\_20108
CL\_2286
CL\_1930
CL\_10555
CL\_10556
CL\_10557
CL\_10558
CL\_10559
CL\_7465
CL\_8257
CL\_13711
CL\_11386
CL\_11387
CL\_25242
CL\_25241
CL\_25240
CL\_25239
CL\_25238
CL\_25237
CL\_25236
CL\_25235
CL\_25234
CL\_25233
CL\_7789
CL\_25232
CL\_10190
CL\_10191
CL\_6001
CL\_6002
CL\_9723
CL\_9199
CL\_7780
CL\_7779
CL\_7778
CL\_6012
CL\_6013
CL\_6014
CL\_6015
CL\_21180
CL\_6017
CL\_21179
CL\_22440
CL\_22439
CL\_22438
CL\_22437
CL\_22436
CL\_22435
CL\_22434
CL\_7590
CL\_7589
CL\_7588
CL\_19851
CL\_10037
CL\_10036
CL\_10035
CL\_10034
CL\_22433
CL\_22432
CL\_22431
CL\_22430
CL\_22429
CL\_7726
CL\_6370
CL\_6369
CL\_7416
CL\_6367
CL\_6366
CL\_7415
CL\_7414
CL\_7413
CL\_7412
CL\_22428
CL\_7411
CL\_7410
CL\_22427
CL\_22426
CL\_22425
CL\_22424
CL\_22423
CL\_22422
CL\_22421
CL\_22420
CL\_22419
CL\_22418
CL\_22417
CL\_22416
CL\_22415
CL\_22414
CL\_22413
CL\_9701
CL\_26716
CL\_8256
CL\_8255
CL\_8254
CL\_8253
CL\_8252
CL\_8251
CL\_8250
CL\_8249
CL\_8248
CL\_8247
CL\_8246
CL\_8245
CL\_24303
CL\_26717
CL\_26718
CL\_7464
CL\_7463
CL\_7462
CL\_8832
CL\_7691
CL\_24351
CL\_7692
CL\_6721
CL\_7461
CL\_7460
CL\_7737
CL\_28898
CL\_1497
CL\_529
CL\_29054
CL\_30287
CL\_25522
CL\_34206
CL\_34207
CL\_34208
CL\_14117
CL\_25521
CL\_25520
CL\_6244
CL\_6245
CL\_6246
CL\_25518
CL\_25517
CL\_25516
CL\_25519
CL\_1929
CL\_34209
CL\_6732
CL\_9140
CL\_11664
CL\_11663
CL\_4462
CL\_6844
CL\_25515
CL\_25835
CL\_7253
CL\_6826
CL\_5245
CL\_6424
CL\_6422
CL\_6729
CL\_6421
CL\_25514
CL\_25513
CL\_25512
CL\_4975
CL\_8216
CL\_18447
CL\_18448
CL\_18449
CL\_18450
CL\_18451
CL\_18452
CL\_18453
CL\_18454
CL\_18455
CL\_35626
CL\_35625
CL\_34210
CL\_13757
CL\_32150
CL\_7979
CL\_8131
CL\_8130
CL\_8129
CL\_8555
CL\_8556
CL\_8128
CL\_11106
CL\_8127
CL\_11105
CL\_11711
CL\_35624
CL\_8511
CL\_11710
CL\_11709
CL\_11708
CL\_33404
CL\_35623
CL\_5233
CL\_34211
CL\_34212
CL\_7075
CL\_10715
CL\_13749
CL\_8558
CL\_8559
CL\_8560
CL\_7832
CL\_15755
CL\_32848
CL\_32849
CL\_32850
CL\_32851
CL\_32852
CL\_32853
CL\_32854
CL\_32855
CL\_6420
CL\_6419
CL\_8907
CL\_8738
CL\_8508
CL\_6416
CL\_7291
CL\_7828
CL\_6724
CL\_7205
CL\_6726
CL\_1934
CL\_5225
CL\_5224
CL\_25511
CL\_4811
CL\_4812
CL\_30286
CL\_4813
CL\_4814
CL\_2288
CL\_7065
CL\_23296
CL\_23295
CL\_23294
CL\_23293
CL\_13497
CL\_13496
CL\_27618
CL\_27619
CL\_25496
CL\_11212
CL\_7064
CL\_7063
CL\_7062
CL\_35462
CL\_35461
CL\_35460
CL\_7061
CL\_22562
CL\_36042
CL\_7060
CL\_8973
CL\_26262
CL\_8972
CL\_11216
CL\_11215
CL\_11214
CL\_13631
CL\_13630
CL\_14822
CL\_14823
CL\_11213
CL\_8310
CL\_8311
CL\_12057
CL\_25172
CL\_25173
CL\_11738
CL\_11737
CL\_8971
CL\_7059
CL\_25062
CL\_25063
CL\_25064
CL\_25065
CL\_25066
CL\_25067
CL\_25068
CL\_25069
CL\_25070
CL\_25071
CL\_25072
CL\_25073
CL\_25074
CL\_25075
CL\_25076
CL\_25077
CL\_25078
CL\_20599
CL\_20600
CL\_20601
CL\_20602
CL\_20603
CL\_20604
CL\_20605
CL\_20606
CL\_20607
CL\_20608
CL\_20609
CL\_20610
CL\_20611
CL\_20612
CL\_20613
CL\_20615
CL\_20616
CL\_20617
CL\_20618
CL\_20619
CL\_20620
CL\_20622
CL\_20623
CL\_20624
CL\_20625
CL\_20626
CL\_20627
CL\_20628
CL\_20629
CL\_20630
CL\_20631
CL\_20632
CL\_20633
CL\_20634
CL\_20635
CL\_20636
CL\_20520
CL\_20521
CL\_20522
CL\_20523
CL\_20524
CL\_20525
CL\_20526
CL\_20527
CL\_20528
CL\_20530
CL\_20533
CL\_24876
CL\_20535
CL\_20536
CL\_20537
CL\_20538
CL\_20539
CL\_24875
CL\_25079
CL\_25080
CL\_25081
CL\_20541
CL\_20544
CL\_20545
CL\_25082
CL\_25083
CL\_24820
CL\_25084
CL\_25085
CL\_25086
CL\_25087
CL\_25088
CL\_24818
CL\_24817
CL\_24816
CL\_24815
CL\_24814
CL\_24813
CL\_24812
CL\_24810
CL\_24809
CL\_24806
CL\_25089
CL\_25090
CL\_24803
CL\_24801
CL\_24799
CL\_25091
CL\_24798
CL\_24797
CL\_25092
CL\_25093
CL\_25094
CL\_24795
CL\_25095
CL\_24793
CL\_24792
CL\_25096
CL\_24791
CL\_24790
CL\_24789
CL\_24788
CL\_25097
CL\_24787
CL\_14149
CL\_25098
CL\_24786
CL\_24785
CL\_25099
CL\_24783
CL\_24778
CL\_24777
CL\_25100
CL\_24776
CL\_24775
CL\_9820
CL\_25101
CL\_25102
CL\_6759
CL\_6758
CL\_6757
CL\_6756
CL\_6755
CL\_6754
CL\_6753
CL\_6752
CL\_6751
CL\_9818
CL\_9817
CL\_9816
CL\_9815
CL\_9814
CL\_9813
CL\_9812
CL\_9811
CL\_6165
CL\_6306
CL\_6307
CL\_6169
CL\_9810
CL\_9809
CL\_9808
CL\_25103
CL\_25104
CL\_9807
CL\_9806
CL\_9805
CL\_9804
CL\_9803
CL\_9802
CL\_9801
CL\_9800
CL\_9799
CL\_9798
CL\_25105
CL\_25106
CL\_24245
CL\_24244
CL\_25107
CL\_25108
CL\_25109
CL\_25110
CL\_24242
CL\_25111
CL\_20559
CL\_20560
CL\_20561
CL\_20563
CL\_20564
CL\_9822
CL\_9823
CL\_6342
CL\_6341
CL\_6340
CL\_6339
CL\_6338
CL\_6337
CL\_6336
CL\_6335
CL\_6334
CL\_6333
CL\_6332
CL\_6331
CL\_6330
CL\_6329
CL\_6328
CL\_6327
CL\_6326
CL\_6325
CL\_6324
CL\_6323
CL\_6322
CL\_6321
CL\_6320
CL\_6319
CL\_6318
CL\_8488
CL\_6317
CL\_24240
CL\_9677
CL\_25112
CL\_25113
CL\_25114
CL\_25115
CL\_25116
CL\_9687
CL\_9688
CL\_9689
CL\_9690
CL\_9691
CL\_9692
CL\_9693
CL\_9694
CL\_9695
CL\_25117
CL\_10421
CL\_22504
CL\_10395
CL\_10393
CL\_10392
CL\_15067
CL\_5297
CL\_5299
CL\_15066
CL\_15065
CL\_24235
CL\_24234
CL\_24233
CL\_24232
CL\_24231
CL\_24230
CL\_24229
CL\_25118
CL\_25119
CL\_25120
CL\_20579
CL\_25121
CL\_24932
CL\_25122
CL\_24228
CL\_24227
CL\_25123
CL\_14680
CL\_14679
CL\_14678
CL\_14677
CL\_14676
CL\_14675
CL\_14674
CL\_14673
CL\_14672
CL\_14671
CL\_14670
CL\_14669
CL\_14668
CL\_14667
CL\_20260
CL\_20261
CL\_20262
CL\_20263
CL\_20264
CL\_20265
CL\_20583
CL\_20585
CL\_20586
CL\_20594
CL\_20595
CL\_20596
CL\_25124
CL\_25125
CL\_25126
CL\_25127
CL\_25128
