## Supplementary material for "A novel method for integrating genomic and Tn-Seq data to identify common *in vivo* fitness mechanisms across multiple bacterial species": S1 Dataset: CL_INS_192.html

Legend

 Hypothetical
 Other
 All VFDB Genes

FULL


WINDOWSVGPNG

Trim RowsRemove SingletonsSave Fasta

CL\_2338


CL\_2338


CL\_2338


CL\_2338


CL\_2337

HighlightSelectShow Genomes


219

CL\_2340


47

CL\_2340


2

CL\_2340


1

Break


1

CL\_2340

fGI ID


CL\_INS\_192
CL\_INS\_192
CL\_INS\_192
CL\_INS\_382
Cluster ID


CL\_2339
CL\_20971
CL\_16988
CL\_6664
