## Supplementary material for "A novel method for integrating genomic and Tn-Seq data to identify common *in vivo* fitness mechanisms across multiple bacterial species": S1 Dataset: CL_INS_196.html

Legend

 Mobile +extrachromosomalelementfunctions
 Hypothetical
 Other

FULL


WINDOWSVGPNG

Trim RowsRemove SingletonsSave Fasta

CL\_2393


CL\_2393


CL\_2393


CL\_2393


CL\_2393


CL\_2393


CL\_2393


CL\_2393


CL\_2393


CL\_2392


CL\_2392


CL\_2392


CL\_2393


CL\_2393


CL\_2393


CL\_2392


CL\_2393

HighlightSelectShow Genomes


86

CL\_2397


70

CL\_2397


15

CL\_2397


13

CL\_2397


11

CL\_2397


4

CL\_2397


2

CL\_2397


1

CL\_2404


1

CL\_2399


1

CL\_2397


1

CL\_2397


1

CL\_2397


1

CL\_2399


1

CL\_2397


1

CL\_2400


1

CL\_2397


1

CL\_2397

fGI ID


CL\_INS\_196
CL\_INS\_196
CL\_INS\_196
CL\_INS\_197
CL\_INS\_196
Cluster ID


CL\_2394
CL\_15927
CL\_2395
CL\_2396
CL\_22881
