## Supplementary material for "A novel method for integrating genomic and Tn-Seq data to identify common *in vivo* fitness mechanisms across multiple bacterial species": S1 Dataset: CL_INS_199.html

Legend

 Mobile +extrachromosomalelementfunctions
 Other
 All VFDB Genes

FULL


WINDOWSVGPNG

Trim RowsRemove SingletonsSave Fasta

CL\_2402


CL\_2402


CL\_2402


CL\_2400


CL\_2402


CL\_2402


CL\_2402


CL\_2402


CL\_2393

HighlightSelectShow Genomes


185

CL\_2404


9

CL\_2404


4

CL\_2405


3

CL\_2404


1

CL\_2404


1

CL\_2404


1

CL\_2404


1

Break


1

CL\_2404

fGI ID


CL\_INS\_199
CL\_INS\_199
CL\_INS\_199
CL\_INS\_199
Cluster ID


CL\_35208
CL\_2403
CL\_12573
CL\_33506
