## Supplementary material for "A novel method for integrating genomic and Tn-Seq data to identify common *in vivo* fitness mechanisms across multiple bacterial species": S1 Dataset: CL_INS_200.html

Legend

 Mobile +extrachromosomalelementfunctions
 Hypothetical
 All VFDB Genes

FULL


WINDOWSVGPNG

Trim RowsRemove SingletonsSave Fasta

CL\_2415


CL\_2415


CL\_2414


CL\_2415


CL\_2414


CL\_2415


CL\_2414


CL\_2415


CL\_2414


CL\_2414


CL\_2415


CL\_2414


CL\_2414

HighlightSelectShow Genomes


142

CL\_2417


112

CL\_2417


5

CL\_2417


1

CL\_2417


1

CL\_2417


1

CL\_2417


1

CL\_2417


1

CL\_2417


1

CL\_2417


1

CL\_2417


1

CL\_2419


1

CL\_2417


1

CL\_2417

fGI ID


CL\_INS\_200
CL\_INS\_200
CL\_INS\_200
CL\_INS\_200
CL\_INS\_200
CL\_INS\_200
CL\_INS\_200
Cluster ID


CL\_6634
CL\_21259
CL\_21258
CL\_9709
CL\_9708
CL\_24304
CL\_2416
