## Supplementary material for "A novel method for integrating genomic and Tn-Seq data to identify common *in vivo* fitness mechanisms across multiple bacterial species": S1 Dataset: CL_INS_204.html

Legend

 Mobile +extrachromosomalelementfunctions
 Hypothetical
 Other
 All VFDB Genes

FULL


WINDOWSVGPNG

Trim RowsRemove SingletonsSave Fasta

CL\_2475


CL\_2507


CL\_2475


CL\_2475


CL\_2475


CL\_2475


CL\_2475


CL\_2475


CL\_2475


CL\_2475


CL\_2475


CL\_2475


CL\_2475


CL\_2475


CL\_2475


CL\_2475


CL\_2475

HighlightSelectShow Genomes


260

CL\_2474


1

CL\_2474


1

CL\_2474


1

CL\_2474


1

CL\_2528


1

CL\_2474


1

CL\_2474


1

Break


1

CL\_2474


1

CL\_2474


1

CL\_2474


1

CL\_2474


1

CL\_2474


1

CL\_2474


1

CL\_2474


1

CL\_2474


1

CL\_2474

fGI ID


CL\_INS\_204
CL\_INS\_204
CL\_INS\_204
CL\_INS\_204
CL\_INS\_204
CL\_INS\_204
CL\_INS\_204
CL\_INS\_204
CL\_INS\_204
CL\_INS\_204
CL\_INS\_204
CL\_INS\_204
CL\_INS\_204
CL\_INS\_204
CL\_INS\_204
CL\_INS\_204
CL\_INS\_204
CL\_INS\_204
CL\_INS\_204
CL\_INS\_204
CL\_INS\_204
CL\_INS\_204
CL\_INS\_204
CL\_INS\_204
CL\_INS\_204
CL\_INS\_204
CL\_INS\_204
CL\_INS\_204
CL\_INS\_204
CL\_INS\_204
CL\_INS\_204
CL\_INS\_204
CL\_INS\_204
CL\_INS\_204
CL\_INS\_204
CL\_INS\_204
CL\_INS\_204
CL\_INS\_204
CL\_INS\_204
CL\_INS\_204
CL\_INS\_204
CL\_INS\_204
CL\_INS\_204
CL\_INS\_204
CL\_INS\_204
CL\_INS\_204
CL\_INS\_204
CL\_INS\_204
CL\_INS\_204
CL\_INS\_204
CL\_INS\_204
CL\_INS\_204
CL\_INS\_204
CL\_INS\_204
CL\_INS\_204
CL\_INS\_204
CL\_INS\_204
CL\_INS\_204
CL\_INS\_204
CL\_INS\_204
CL\_INS\_204
CL\_INS\_204
CL\_INS\_204
CL\_INS\_204
CL\_INS\_204
CL\_INS\_204
CL\_INS\_204
CL\_INS\_204
CL\_INS\_204
CL\_INS\_204
CL\_INS\_204
CL\_INS\_204
CL\_INS\_204
CL\_INS\_204
CL\_INS\_204
CL\_INS\_204
CL\_INS\_204
CL\_INS\_204
CL\_INS\_204
CL\_INS\_204
CL\_INS\_204
CL\_INS\_204
CL\_INS\_204
CL\_INS\_204
CL\_INS\_204
CL\_INS\_204
CL\_INS\_204
CL\_INS\_204
CL\_INS\_204
CL\_INS\_204
CL\_INS\_204
CL\_INS\_204
CL\_INS\_204
CL\_INS\_204
CL\_INS\_204
CL\_INS\_204
CL\_INS\_204
CL\_INS\_204
CL\_INS\_204
CL\_INS\_204
CL\_INS\_204
CL\_INS\_204
CL\_INS\_204
CL\_INS\_204
CL\_INS\_204
CL\_INS\_204
CL\_INS\_204
CL\_INS\_204
CL\_INS\_204
CL\_INS\_204
CL\_INS\_204
CL\_INS\_204
CL\_INS\_204
CL\_INS\_204
CL\_INS\_204
CL\_INS\_204
CL\_INS\_204
CL\_INS\_204
CL\_INS\_204
CL\_INS\_204
CL\_INS\_204
CL\_INS\_204
CL\_INS\_204
CL\_INS\_204
CL\_INS\_204
CL\_INS\_204
CL\_INS\_204
CL\_INS\_204
CL\_INS\_204
CL\_INS\_204
CL\_INS\_204
CL\_INS\_204
CL\_INS\_204
CL\_INS\_204
CL\_INS\_204
CL\_INS\_204
CL\_INS\_204
CL\_INS\_204
CL\_INS\_204
CL\_INS\_204
CL\_INS\_204
CL\_INS\_204
CL\_INS\_204
CL\_INS\_204
CL\_INS\_204
CL\_INS\_204
CL\_INS\_204
CL\_INS\_204
CL\_INS\_204
CL\_INS\_204
CL\_INS\_204
CL\_INS\_204
CL\_INS\_204
CL\_INS\_204
CL\_INS\_204
CL\_INS\_204
CL\_INS\_204
CL\_INS\_204
CL\_INS\_204
CL\_INS\_204
CL\_INS\_204
CL\_INS\_204
CL\_INS\_204
Cluster ID


CL\_13201
CL\_12457
CL\_13200
CL\_13199
CL\_13198
CL\_13197
CL\_12574
CL\_5940
CL\_30139
CL\_5939
CL\_12577
CL\_12576
CL\_12575
CL\_5938
CL\_32015
CL\_32016
CL\_32017
CL\_5937
CL\_21711
CL\_26794
CL\_21569
CL\_34067
CL\_7393
CL\_36276
CL\_36275
CL\_21570
CL\_5936
CL\_21568
CL\_16420
CL\_7392
CL\_7391
CL\_5935
CL\_5934
CL\_26793
CL\_5933
CL\_5932
CL\_21710
CL\_7390
CL\_18904
CL\_18905
CL\_5931
CL\_5930
CL\_5929
CL\_26792
CL\_5928
CL\_5927
CL\_18906
CL\_18907
CL\_7389
CL\_7388
CL\_7387
CL\_18908
CL\_18909
CL\_36274
CL\_5926
CL\_5925
CL\_18910
CL\_5924
CL\_5415
CL\_5923
CL\_5414
CL\_10975
CL\_5412
CL\_5411
CL\_1097
CL\_1098
CL\_1099
CL\_5410
CL\_5409
CL\_5408
CL\_5407
CL\_5406
CL\_5405
CL\_5404
CL\_5403
CL\_5402
CL\_5401
CL\_5400
CL\_5399
CL\_5398
CL\_10274
CL\_5396
CL\_5395
CL\_5394
CL\_5393
CL\_1100
CL\_1101
CL\_1102
CL\_9223
CL\_9222
CL\_5922
CL\_5921
CL\_5920
CL\_5919
CL\_5918
CL\_5917
CL\_5916
CL\_5915
CL\_5914
CL\_5913
CL\_5912
CL\_5911
CL\_5910
CL\_5909
CL\_5908
CL\_7386
CL\_5907
CL\_5906
CL\_5905
CL\_5904
CL\_5903
CL\_5902
CL\_5901
CL\_21567
CL\_21709
CL\_26791
CL\_34066
CL\_23874
CL\_5900
CL\_5899
CL\_5898
CL\_5897
CL\_30140
CL\_12788
CL\_13334
CL\_22636
CL\_22637
CL\_32018
CL\_11444
CL\_5896
CL\_9292
CL\_21566
CL\_9217
CL\_21565
CL\_1103
CL\_5895
CL\_6594
CL\_2277
CL\_1104
CL\_1105
CL\_6520
CL\_5894
CL\_4604
CL\_18911
CL\_7385
CL\_23996
CL\_23997
CL\_23998
CL\_23999
CL\_24000
CL\_24001
CL\_24002
CL\_24003
CL\_10867
CL\_24004
CL\_24005
CL\_24006
CL\_24007
CL\_24008
CL\_24009
CL\_24010
CL\_24011
CL\_24012
