## Supplementary material for "A novel method for integrating genomic and Tn-Seq data to identify common *in vivo* fitness mechanisms across multiple bacterial species": S1 Dataset: CL_INS_207.html

Legend

 Mobile +extrachromosomalelementfunctions
 Regulatoryfunctions
 Hypothetical
 DNA Metabolism
 AntibioticResistance
 All EssentialGenes
 All Fitness Genes
 Biosynthesis ofcofactors,prostheticgroups, +carriers
 Proteinsynthesis/fate
 Other
 EnergyMetabolism
 Transport +binding proteins
 All VFDB Genes

FULL


WINDOWSVGPNG

Trim RowsRemove SingletonsSave Fasta

CL\_2528


CL\_2528


CL\_2528


CL\_2528


CL\_2528


CL\_2528


CL\_2528


CL\_2528


CL\_2528


CL\_2528


CL\_2528


CL\_2528


CL\_2528


CL\_2528


CL\_4487


CL\_2528


CL\_2528


CL\_2528


CL\_2528


CL\_2528


CL\_2528


CL\_2528


CL\_2528


CL\_2528


CL\_2528


CL\_2528


CL\_287


CL\_2528


CL\_2528


CL\_2528


CL\_2528


CL\_2528


CL\_2528


CL\_2528


CL\_2528


CL\_2528


CL\_2528


CL\_2528


CL\_2528


CL\_2528


CL\_2528


CL\_2528


CL\_1935


CL\_2528


CL\_2528


CL\_2528


CL\_2528


CL\_2528


CL\_2528


CL\_2528


CL\_2528


CL\_2528


CL\_4085


CL\_2528


CL\_2528


CL\_2528


CL\_2528


CL\_2528


CL\_2528


CL\_2528


CL\_2528


CL\_2528


CL\_2528


CL\_2528


CL\_2528


CL\_2528


CL\_2528


CL\_2528


CL\_2528


CL\_2528


CL\_2528


CL\_2528


CL\_2528


CL\_2528


CL\_2528


CL\_2528


CL\_2528


CL\_2528


CL\_2528


CL\_2528


CL\_2528


CL\_1124


CL\_2528


CL\_2528


CL\_2528


CL\_2528


CL\_2528


CL\_2528


CL\_2528


CL\_2528


CL\_2528


CL\_2528


CL\_2528


CL\_2528


CL\_2528


CL\_2528


CL\_2528


CL\_2528


CL\_2528


CL\_2528


CL\_2528


CL\_2528


CL\_2528


CL\_2528


CL\_2528


CL\_2528


CL\_2528


CL\_2528


CL\_2528


CL\_2528


CL\_2528


CL\_2528


CL\_2528


CL\_2528


CL\_2528


CL\_2528


CL\_2528


CL\_4486


CL\_2528


CL\_4486


CL\_2528


CL\_2528


CL\_2528


CL\_2528


CL\_2528


CL\_2528


CL\_4427


CL\_2528


CL\_2528


CL\_2528


CL\_2528


CL\_2528


CL\_2528


CL\_2528


CL\_2528


CL\_2528


CL\_2528


CL\_4486


CL\_2528


CL\_2528


CL\_269


CL\_2528


CL\_2528


CL\_2528


CL\_2528


CL\_2528


CL\_2528


CL\_2528


CL\_2528


CL\_2528


CL\_2528


CL\_2528


CL\_2528


CL\_2528


CL\_2528


CL\_2528


CL\_2528


CL\_2528


CL\_2528


CL\_2528


CL\_2528


CL\_2528


CL\_2528


CL\_4487


CL\_2528


CL\_2528


CL\_2528


CL\_2528


CL\_2528


CL\_2528

HighlightSelectShow Genomes


33

CL\_2555


21

CL\_2555


12

CL\_2555


10

CL\_2555


10

CL\_2555


8

CL\_2555


7

CL\_2555


6

CL\_2555


5

CL\_2555


4

CL\_4516


4

CL\_2555


2

CL\_2555


2

CL\_2555


2

CL\_2555


2

CL\_2555


2

CL\_2555


2

CL\_2555


2

CL\_2555


2

CL\_2555


2

CL\_2555


2

CL\_2555


2

CL\_2555


1

CL\_2555


1

CL\_4516


1

CL\_2555


1

CL\_2555


1

CL\_2555


1

CL\_2555


1

CL\_2555


1

CL\_2560


1

CL\_2555


1

CL\_2555


1

CL\_2555


1

CL\_2555


1

CL\_3275


1

CL\_2555


1

CL\_2555


1

CL\_2555


1

CL\_2555


1

CL\_2555


1

Break


1

CL\_234


1

CL\_2555


1

CL\_2555


1

CL\_2555


1

CL\_2555


1

CL\_2555


1

CL\_2555


1

CL\_2555


1

CL\_2555


1

CL\_2555


1

CL\_2555


1

CL\_2555


1

CL\_2555


1

CL\_2555


1

CL\_2555


1

CL\_1125


1

CL\_2555


1

CL\_2555


1

CL\_2555


1

CL\_2555


1

CL\_2555


1

CL\_2555


1

CL\_2555


1

CL\_2555


1

CL\_2555


1

CL\_2555


1

CL\_4516


1

CL\_2555


1

CL\_2555


1

CL\_2555


1

CL\_2555


1

CL\_2555


1

CL\_2555


1

CL\_2555


1

CL\_2475


1

CL\_2555


1

CL\_2555


1

CL\_2731


1

CL\_2555


1

CL\_2555


1

CL\_2555


1

CL\_2555


1

CL\_2555


1

CL\_2555


1

CL\_2555


1

CL\_2556


1

CL\_2555


1

CL\_2555


1

CL\_2555


1

CL\_2555


1

CL\_2555


1

CL\_2555


1

CL\_2555


1

CL\_2555


1

CL\_976


1

CL\_2555


1

CL\_2555


1

CL\_2555


1

CL\_2555


1

CL\_2555


1

CL\_2555


1

CL\_2555


1

CL\_2555


1

CL\_2555


1

CL\_2555


1

CL\_2555


1

CL\_2555


1

CL\_2555


1

CL\_2555


1

CL\_2555


1

CL\_2555


1

CL\_2555


1

CL\_2555


1

CL\_4516


1

CL\_2555


1

CL\_2555


1

CL\_2555


1

CL\_2555


1

CL\_2555


1

CL\_2556


1

CL\_2555


1

CL\_2555


1

CL\_230


1

CL\_2555


1

CL\_2555


1

CL\_2555


1

CL\_2555


1

CL\_2555


1

CL\_2555


1

CL\_2555


1

CL\_2555


1

CL\_2555


1

CL\_2555


1

CL\_2555


1

CL\_2555


1

CL\_2556


1

CL\_2555


1

CL\_2555


1

CL\_2555


1

CL\_2555


1

CL\_2555


1

CL\_2555


1

CL\_2555


1

CL\_2555


1

CL\_2555


1

CL\_2556


1

CL\_2555


1

CL\_2555


1

CL\_2555


1

CL\_2555


1

CL\_2555


1

CL\_2555


1

CL\_2555


1

CL\_2555


1

CL\_2555


1

CL\_2555


1

CL\_289


1

CL\_2555


1

CL\_2555


1

CL\_2555


1

CL\_2555


1

CL\_2555


1

CL\_2555


1

CL\_2555


1

CL\_2555


1

CL\_2555


1

CL\_2555


1

CL\_2555


1

CL\_2555

fGI ID


CL\_INS\_207
CL\_INS\_207
CL\_INS\_204
CL\_INS\_207
CL\_INS\_207
CL\_INS\_207
CL\_INS\_207
CL\_INS\_207
CL\_INS\_207
CL\_INS\_207
CL\_INS\_207
CL\_INS\_207
CL\_INS\_207
CL\_INS\_207
CL\_INS\_207
CL\_INS\_207
CL\_INS\_207
CL\_INS\_207
CL\_INS\_207
CL\_INS\_247
CL\_INS\_70
CL\_INS\_70
CL\_INS\_70
CL\_INS\_70
CL\_INS\_207
CL\_INS\_207
CL\_INS\_207
CL\_INS\_247
CL\_INS\_207
CL\_INS\_207
CL\_INS\_207
CL\_INS\_207
CL\_INS\_207
CL\_INS\_207
CL\_INS\_207
CL\_INS\_207
CL\_INS\_207
CL\_INS\_207
CL\_INS\_207
CL\_INS\_207
CL\_INS\_207
CL\_INS\_207
CL\_INS\_207
CL\_INS\_207
CL\_INS\_207
CL\_INS\_207
CL\_INS\_207
CL\_INS\_207
CL\_INS\_207
CL\_INS\_207
CL\_INS\_207
CL\_INS\_207
CL\_INS\_207
CL\_INS\_207
CL\_INS\_207
CL\_INS\_207
CL\_INS\_207
CL\_INS\_207
CL\_INS\_207
CL\_INS\_207
CL\_INS\_207
CL\_INS\_207
CL\_INS\_207
CL\_INS\_207
CL\_INS\_207
CL\_INS\_207
CL\_INS\_207
CL\_INS\_207
CL\_INS\_207
CL\_INS\_207
CL\_INS\_207
CL\_INS\_170
CL\_INS\_207
CL\_INS\_207
CL\_INS\_207
CL\_INS\_207
CL\_INS\_207
CL\_INS\_207
CL\_INS\_207
CL\_INS\_207
CL\_INS\_207
CL\_INS\_207
CL\_INS\_207
CL\_INS\_207
CL\_INS\_207
CL\_INS\_207
CL\_INS\_207
CL\_INS\_207
CL\_INS\_207
CL\_INS\_207
CL\_INS\_207
CL\_INS\_247
CL\_INS\_207
CL\_INS\_207
CL\_INS\_207
CL\_INS\_247
CL\_INS\_247
CL\_INS\_247
CL\_INS\_247
CL\_INS\_247
CL\_INS\_247
CL\_INS\_247
CL\_INS\_247
CL\_INS\_207
CL\_INS\_247
CL\_INS\_247
CL\_INS\_237
CL\_INS\_237
CL\_INS\_237
CL\_INS\_237
CL\_INS\_207
CL\_INS\_207
CL\_INS\_207
CL\_INS\_207
CL\_INS\_247
CL\_INS\_155
CL\_INS\_155
CL\_INS\_207
CL\_INS\_155
CL\_INS\_155
CL\_INS\_207
CL\_INS\_207
CL\_INS\_207
CL\_INS\_207
CL\_INS\_99
CL\_INS\_86
CL\_INS\_86
CL\_INS\_207
CL\_INS\_207
CL\_INS\_207
CL\_INS\_207
CL\_INS\_207
CL\_INS\_207
CL\_INS\_207
CL\_INS\_207
CL\_INS\_207
CL\_INS\_87
CL\_INS\_86
CL\_INS\_99
CL\_INS\_86
CL\_INS\_99
CL\_INS\_99
CL\_INS\_99
CL\_INS\_99
CL\_INS\_99
CL\_INS\_99
CL\_INS\_207
CL\_INS\_207
CL\_INS\_207
CL\_INS\_207
CL\_INS\_207
CL\_INS\_382
CL\_INS\_382
CL\_INS\_207
CL\_INS\_207
CL\_INS\_207
CL\_INS\_207
CL\_INS\_207
CL\_INS\_382
CL\_INS\_207
CL\_INS\_207
CL\_INS\_382
CL\_INS\_382
CL\_INS\_382
CL\_INS\_99
CL\_INS\_99
CL\_INS\_207
CL\_INS\_207
CL\_INS\_207
CL\_INS\_207
CL\_INS\_207
CL\_INS\_207
CL\_INS\_207
CL\_INS\_99
CL\_INS\_237
CL\_INS\_382
CL\_INS\_382
CL\_INS\_207
CL\_INS\_382
CL\_INS\_207
CL\_INS\_207
CL\_INS\_207
CL\_INS\_207
CL\_INS\_207
CL\_INS\_207
CL\_INS\_207
CL\_INS\_207
CL\_INS\_207
CL\_INS\_207
CL\_INS\_207
CL\_INS\_207
CL\_INS\_207
CL\_INS\_207
CL\_INS\_207
CL\_INS\_207
CL\_INS\_207
CL\_INS\_207
CL\_INS\_207
CL\_INS\_207
CL\_INS\_204
CL\_INS\_207
CL\_INS\_207
CL\_INS\_204
CL\_INS\_204
CL\_INS\_149
CL\_INS\_86
CL\_INS\_207
CL\_INS\_247
CL\_INS\_207
CL\_INS\_170
CL\_INS\_207
CL\_INS\_207
CL\_INS\_207
CL\_INS\_207
CL\_INS\_170
CL\_INS\_170
CL\_INS\_170
CL\_INS\_170
CL\_INS\_170
CL\_INS\_170
CL\_INS\_207
CL\_INS\_207
CL\_INS\_207
CL\_INS\_207
CL\_INS\_86
CL\_INS\_207
CL\_INS\_207
CL\_INS\_207
CL\_INS\_207
CL\_INS\_207
CL\_INS\_207
CL\_INS\_155
CL\_INS\_149
CL\_INS\_207
CL\_INS\_70
CL\_INS\_70
CL\_INS\_149
CL\_INS\_207
CL\_INS\_207
CL\_INS\_207
CL\_INS\_207
CL\_INS\_207
CL\_INS\_170
CL\_INS\_207
CL\_INS\_207
CL\_INS\_207
CL\_INS\_99
CL\_INS\_237
CL\_INS\_99
CL\_INS\_86
CL\_INS\_99
CL\_INS\_382
CL\_INS\_86
CL\_INS\_382
CL\_INS\_382
CL\_INS\_382
CL\_INS\_382
CL\_INS\_382
CL\_INS\_382
CL\_INS\_382
CL\_INS\_382
CL\_INS\_382
CL\_INS\_382
CL\_INS\_382
CL\_INS\_382
CL\_INS\_382
CL\_INS\_382
CL\_INS\_382
CL\_INS\_382
CL\_INS\_207
CL\_INS\_207
CL\_INS\_382
CL\_INS\_382
CL\_INS\_207
CL\_INS\_382
CL\_INS\_207
CL\_INS\_207
CL\_INS\_99
CL\_INS\_382
CL\_INS\_382
CL\_INS\_382
CL\_INS\_382
CL\_INS\_207
CL\_INS\_207
CL\_INS\_207
CL\_INS\_207
CL\_INS\_207
CL\_INS\_207
CL\_INS\_207
CL\_INS\_207
CL\_INS\_207
CL\_INS\_207
CL\_INS\_207
CL\_INS\_207
CL\_INS\_207
CL\_INS\_207
CL\_INS\_207
CL\_INS\_207
CL\_INS\_207
CL\_INS\_207
CL\_INS\_207
CL\_INS\_207
CL\_INS\_207
CL\_INS\_207
CL\_INS\_207
CL\_INS\_207
CL\_INS\_207
CL\_INS\_207
CL\_INS\_207
CL\_INS\_207
CL\_INS\_207
CL\_INS\_382
CL\_INS\_382
CL\_INS\_382
CL\_INS\_382
CL\_INS\_382
CL\_INS\_382
CL\_INS\_207
CL\_INS\_207
CL\_INS\_207
CL\_INS\_207
CL\_INS\_207
CL\_INS\_207
CL\_INS\_207
CL\_INS\_207
CL\_INS\_207
CL\_INS\_207
CL\_INS\_207
CL\_INS\_207
CL\_INS\_99
CL\_INS\_382
CL\_INS\_382
CL\_INS\_382
CL\_INS\_207
CL\_INS\_207
CL\_INS\_207
CL\_INS\_207
CL\_INS\_382
CL\_INS\_207
CL\_INS\_382
CL\_INS\_207
CL\_INS\_207
CL\_INS\_207
CL\_INS\_207
CL\_INS\_207
CL\_INS\_207
CL\_INS\_207
CL\_INS\_207
CL\_INS\_207
CL\_INS\_149
CL\_INS\_149
CL\_INS\_149
CL\_INS\_149
CL\_INS\_207
CL\_INS\_99
CL\_INS\_149
CL\_INS\_149
CL\_INS\_99
CL\_INS\_99
CL\_INS\_149
CL\_INS\_149
CL\_INS\_170
CL\_INS\_149
CL\_INS\_149
CL\_INS\_207
CL\_INS\_207
CL\_INS\_207
CL\_INS\_207
CL\_INS\_207
CL\_INS\_207
CL\_INS\_207
CL\_INS\_207
CL\_INS\_170
CL\_INS\_207
CL\_INS\_207
CL\_INS\_207
CL\_INS\_207
CL\_INS\_207
CL\_INS\_207
CL\_INS\_207
CL\_INS\_207
CL\_INS\_207
CL\_INS\_170
CL\_INS\_170
CL\_INS\_170
CL\_INS\_170
CL\_INS\_207
CL\_INS\_207
CL\_INS\_207
CL\_INS\_207
CL\_INS\_207
CL\_INS\_207
CL\_INS\_207
CL\_INS\_207
CL\_INS\_207
CL\_INS\_207
CL\_INS\_207
CL\_INS\_207
CL\_INS\_207
CL\_INS\_207
CL\_INS\_207
CL\_INS\_207
CL\_INS\_207
CL\_INS\_207
CL\_INS\_207
CL\_INS\_207
CL\_INS\_207
CL\_INS\_149
CL\_INS\_170
CL\_INS\_170
CL\_INS\_149
CL\_INS\_170
CL\_INS\_170
CL\_INS\_170
CL\_INS\_207
CL\_INS\_207
CL\_INS\_207
CL\_INS\_170
CL\_INS\_207
CL\_INS\_207
CL\_INS\_207
CL\_INS\_207
CL\_INS\_207
CL\_INS\_207
CL\_INS\_207
CL\_INS\_207
CL\_INS\_207
CL\_INS\_207
CL\_INS\_207
CL\_INS\_207
CL\_INS\_207
CL\_INS\_207
CL\_INS\_207
CL\_INS\_207
CL\_INS\_207
CL\_INS\_207
CL\_INS\_207
CL\_INS\_207
CL\_INS\_207
CL\_INS\_207
CL\_INS\_207
CL\_INS\_207
CL\_INS\_207
CL\_INS\_247
CL\_INS\_247
CL\_INS\_207
CL\_INS\_207
CL\_INS\_247
CL\_INS\_207
CL\_INS\_247
CL\_INS\_207
CL\_INS\_207
CL\_INS\_247
CL\_INS\_207
CL\_INS\_207
CL\_INS\_207
CL\_INS\_207
CL\_INS\_207
CL\_INS\_207
CL\_INS\_207
CL\_INS\_207
CL\_INS\_207
CL\_INS\_207
CL\_INS\_207
CL\_INS\_207
CL\_INS\_207
CL\_INS\_207
CL\_INS\_207
CL\_INS\_207
CL\_INS\_207
CL\_INS\_207
CL\_INS\_207
CL\_INS\_207
CL\_INS\_207
CL\_INS\_207
CL\_INS\_207
CL\_INS\_207
CL\_INS\_207
CL\_INS\_207
CL\_INS\_207
CL\_INS\_207
CL\_INS\_207
CL\_INS\_207
CL\_INS\_207
CL\_INS\_207
CL\_INS\_207
CL\_INS\_207
CL\_INS\_207
CL\_INS\_207
CL\_INS\_207
CL\_INS\_207
CL\_INS\_207
CL\_INS\_207
CL\_INS\_207
CL\_INS\_207
CL\_INS\_207
CL\_INS\_207
CL\_INS\_207
CL\_INS\_207
CL\_INS\_207
CL\_INS\_207
CL\_INS\_86
CL\_INS\_86
CL\_INS\_207
CL\_INS\_207
CL\_INS\_204
CL\_INS\_204
CL\_INS\_86
CL\_INS\_207
CL\_INS\_204
CL\_INS\_204
CL\_INS\_170
CL\_INS\_207
CL\_INS\_204
CL\_INS\_207
CL\_INS\_207
CL\_INS\_207
CL\_INS\_207
CL\_INS\_207
CL\_INS\_207
CL\_INS\_204
CL\_INS\_204
CL\_INS\_204
CL\_INS\_204
CL\_INS\_204
CL\_INS\_207
CL\_INS\_207
CL\_INS\_207
CL\_INS\_204
CL\_INS\_204
CL\_INS\_207
CL\_INS\_207
CL\_INS\_207
CL\_INS\_207
CL\_INS\_207
CL\_INS\_207
CL\_INS\_207
CL\_INS\_207
CL\_INS\_207
CL\_INS\_207
CL\_INS\_207
CL\_INS\_207
CL\_INS\_207
CL\_INS\_207
CL\_INS\_207
CL\_INS\_207
CL\_INS\_207
CL\_INS\_207
CL\_INS\_207
CL\_INS\_207
CL\_INS\_207
CL\_INS\_207
CL\_INS\_207
CL\_INS\_207
CL\_INS\_207
CL\_INS\_207
CL\_INS\_155
CL\_INS\_207
CL\_INS\_207
CL\_INS\_204
CL\_INS\_204
CL\_INS\_86
CL\_INS\_204
CL\_INS\_207
CL\_INS\_204
CL\_INS\_204
CL\_INS\_204
CL\_INS\_204
CL\_INS\_86
CL\_INS\_207
CL\_INS\_207
CL\_INS\_99
CL\_INS\_99
CL\_INS\_207
CL\_INS\_204
CL\_INS\_204
CL\_INS\_204
CL\_INS\_204
CL\_INS\_86
CL\_INS\_86
CL\_INS\_86
CL\_INS\_204
CL\_INS\_204
CL\_INS\_204
CL\_INS\_204
CL\_INS\_204
CL\_INS\_204
CL\_INS\_207
CL\_INS\_207
CL\_INS\_207
CL\_INS\_204
CL\_INS\_207
CL\_INS\_207
CL\_INS\_207
CL\_INS\_207
CL\_INS\_204
CL\_INS\_204
CL\_INS\_204
CL\_INS\_382
CL\_INS\_382
CL\_INS\_207
CL\_INS\_207
CL\_INS\_382
CL\_INS\_207
CL\_INS\_207
CL\_INS\_207
CL\_INS\_207
CL\_INS\_382
CL\_INS\_382
CL\_INS\_207
CL\_INS\_382
CL\_INS\_382
CL\_INS\_382
CL\_INS\_382
CL\_INS\_86
CL\_INS\_207
CL\_INS\_207
CL\_INS\_207
CL\_INS\_207
CL\_INS\_207
CL\_INS\_86
CL\_INS\_382
CL\_INS\_382
CL\_INS\_207
CL\_INS\_382
CL\_INS\_382
CL\_INS\_86
CL\_INS\_86
CL\_INS\_382
CL\_INS\_382
CL\_INS\_382
CL\_INS\_382
CL\_INS\_382
CL\_INS\_382
CL\_INS\_382
CL\_INS\_382
CL\_INS\_382
CL\_INS\_86
CL\_INS\_86
CL\_INS\_382
CL\_INS\_382
CL\_INS\_382
CL\_INS\_382
CL\_INS\_382
CL\_INS\_382
CL\_INS\_382
CL\_INS\_382
CL\_INS\_382
CL\_INS\_382
CL\_INS\_382
CL\_INS\_382
CL\_INS\_382
CL\_INS\_207
CL\_INS\_207
CL\_INS\_207
CL\_INS\_207
CL\_INS\_207
CL\_INS\_207
CL\_INS\_207
CL\_INS\_207
CL\_INS\_207
CL\_INS\_207
CL\_INS\_207
CL\_INS\_382
CL\_INS\_247
CL\_INS\_207
CL\_INS\_207
CL\_INS\_207
CL\_INS\_382
CL\_INS\_207
CL\_INS\_382
CL\_INS\_382
CL\_INS\_382
CL\_INS\_382
CL\_INS\_382
CL\_INS\_207
CL\_INS\_207
CL\_INS\_382
CL\_INS\_382
CL\_INS\_382
CL\_INS\_382
CL\_INS\_382
CL\_INS\_382
CL\_INS\_99
CL\_INS\_99
CL\_INS\_99
CL\_INS\_382
CL\_INS\_382
CL\_INS\_382
CL\_INS\_382
CL\_INS\_382
CL\_INS\_382
CL\_INS\_382
CL\_INS\_382
CL\_INS\_382
CL\_INS\_382
CL\_INS\_382
CL\_INS\_382
CL\_INS\_382
CL\_INS\_382
CL\_INS\_382
CL\_INS\_382
CL\_INS\_382
CL\_INS\_382
CL\_INS\_382
CL\_INS\_382
CL\_INS\_382
CL\_INS\_382
CL\_INS\_382
CL\_INS\_382
CL\_INS\_382
CL\_INS\_382
CL\_INS\_99
CL\_INS\_382
CL\_INS\_382
CL\_INS\_382
CL\_INS\_382
CL\_INS\_382
CL\_INS\_382
CL\_INS\_207
CL\_INS\_207
CL\_INS\_207
CL\_INS\_207
CL\_INS\_207
CL\_INS\_207
CL\_INS\_207
CL\_INS\_207
CL\_INS\_207
CL\_INS\_207
CL\_INS\_207
CL\_INS\_207
CL\_INS\_207
CL\_INS\_207
CL\_INS\_207
CL\_INS\_207
CL\_INS\_207
CL\_INS\_207
CL\_INS\_207
CL\_INS\_207
CL\_INS\_207
CL\_INS\_207
CL\_INS\_207
CL\_INS\_207
CL\_INS\_207
CL\_INS\_207
CL\_INS\_207
CL\_INS\_207
CL\_INS\_207
CL\_INS\_382
CL\_INS\_382
CL\_INS\_382
CL\_INS\_207
CL\_INS\_382
CL\_INS\_99
CL\_INS\_207
CL\_INS\_207
CL\_INS\_382
CL\_INS\_382
CL\_INS\_382
CL\_INS\_382
CL\_INS\_382
CL\_INS\_382
CL\_INS\_207
CL\_INS\_382
CL\_INS\_382
CL\_INS\_207
CL\_INS\_382
CL\_INS\_207
CL\_INS\_207
CL\_INS\_207
CL\_INS\_382
CL\_INS\_207
CL\_INS\_207
CL\_INS\_207
CL\_INS\_207
CL\_INS\_207
CL\_INS\_207
CL\_INS\_207
CL\_INS\_382
CL\_INS\_382
CL\_INS\_382
CL\_INS\_382
CL\_INS\_207
CL\_INS\_382
CL\_INS\_382
CL\_INS\_382
CL\_INS\_382
CL\_INS\_207
CL\_INS\_207
CL\_INS\_207
CL\_INS\_207
CL\_INS\_207
CL\_INS\_382
CL\_INS\_382
CL\_INS\_382
CL\_INS\_382
CL\_INS\_382
CL\_INS\_382
CL\_INS\_207
CL\_INS\_207
CL\_INS\_382
CL\_INS\_207
CL\_INS\_207
CL\_INS\_382
CL\_INS\_382
CL\_INS\_207
CL\_INS\_207
CL\_INS\_207
CL\_INS\_207
CL\_INS\_207
CL\_INS\_382
CL\_INS\_382
CL\_INS\_382
CL\_INS\_382
CL\_INS\_207
CL\_INS\_207
CL\_INS\_207
CL\_INS\_207
CL\_INS\_207
CL\_INS\_207
CL\_INS\_207
CL\_INS\_207
CL\_INS\_207
CL\_INS\_207
CL\_INS\_207
CL\_INS\_207
CL\_INS\_207
CL\_INS\_207
CL\_INS\_382
CL\_INS\_207
CL\_INS\_207
CL\_INS\_207
CL\_INS\_207
CL\_INS\_207
CL\_INS\_207
CL\_INS\_207
CL\_INS\_207
CL\_INS\_207
CL\_INS\_207
CL\_INS\_207
CL\_INS\_207
CL\_INS\_207
CL\_INS\_207
CL\_INS\_207
CL\_INS\_207
CL\_INS\_207
CL\_INS\_207
CL\_INS\_207
CL\_INS\_207
CL\_INS\_207
CL\_INS\_207
CL\_INS\_207
CL\_INS\_207
CL\_INS\_207
CL\_INS\_207
CL\_INS\_207
CL\_INS\_207
CL\_INS\_207
CL\_INS\_207
CL\_INS\_207
CL\_INS\_207
CL\_INS\_207
CL\_INS\_207
CL\_INS\_207
CL\_INS\_207
CL\_INS\_207
CL\_INS\_207
CL\_INS\_207
CL\_INS\_207
CL\_INS\_207
CL\_INS\_207
CL\_INS\_207
CL\_INS\_207
CL\_INS\_207
CL\_INS\_207
CL\_INS\_207
CL\_INS\_207
CL\_INS\_207
CL\_INS\_207
CL\_INS\_207
CL\_INS\_207
CL\_INS\_207
CL\_INS\_207
CL\_INS\_207
CL\_INS\_207
CL\_INS\_207
CL\_INS\_207
CL\_INS\_207
CL\_INS\_207
CL\_INS\_207
CL\_INS\_207
CL\_INS\_207
CL\_INS\_207
CL\_INS\_207
CL\_INS\_207
CL\_INS\_207
CL\_INS\_86
CL\_INS\_99
CL\_INS\_207
CL\_INS\_207
CL\_INS\_207
CL\_INS\_207
CL\_INS\_207
CL\_INS\_207
CL\_INS\_207
CL\_INS\_207
CL\_INS\_237
CL\_INS\_237
CL\_INS\_247
CL\_INS\_247
CL\_INS\_247
CL\_INS\_207
CL\_INS\_207
CL\_INS\_207
CL\_INS\_207
CL\_INS\_207
CL\_INS\_207
CL\_INS\_207
CL\_INS\_207
CL\_INS\_207
CL\_INS\_207
CL\_INS\_207
CL\_INS\_207
CL\_INS\_207
CL\_INS\_207
CL\_INS\_207
CL\_INS\_207
CL\_INS\_207
CL\_INS\_207
CL\_INS\_207
CL\_INS\_207
CL\_INS\_207
CL\_INS\_207
CL\_INS\_207
CL\_INS\_207
CL\_INS\_207
CL\_INS\_207
CL\_INS\_207
CL\_INS\_207
CL\_INS\_207
CL\_INS\_207
CL\_INS\_207
CL\_INS\_207
CL\_INS\_207
CL\_INS\_207
CL\_INS\_207
CL\_INS\_207
CL\_INS\_207
CL\_INS\_207
CL\_INS\_207
CL\_INS\_207
CL\_INS\_207
CL\_INS\_207
CL\_INS\_207
CL\_INS\_207
CL\_INS\_207
CL\_INS\_207
CL\_INS\_207
CL\_INS\_207
CL\_INS\_207
CL\_INS\_207
CL\_INS\_207
CL\_INS\_207
CL\_INS\_207
CL\_INS\_207
CL\_INS\_207
CL\_INS\_207
CL\_INS\_207
CL\_INS\_207
CL\_INS\_207
CL\_INS\_207
CL\_INS\_207
CL\_INS\_207
CL\_INS\_207
CL\_INS\_207
CL\_INS\_207
CL\_INS\_207
CL\_INS\_207
CL\_INS\_207
CL\_INS\_207
CL\_INS\_207
CL\_INS\_207
CL\_INS\_207
CL\_INS\_207
CL\_INS\_70
CL\_INS\_207
CL\_INS\_207
CL\_INS\_207
CL\_INS\_207
CL\_INS\_207
CL\_INS\_207
CL\_INS\_207
CL\_INS\_207
CL\_INS\_237
CL\_INS\_207
CL\_INS\_155
CL\_INS\_237
CL\_INS\_207
CL\_INS\_155
CL\_INS\_207
CL\_INS\_207
CL\_INS\_207
CL\_INS\_207
CL\_INS\_207
CL\_INS\_207
CL\_INS\_207
CL\_INS\_207
CL\_INS\_207
CL\_INS\_207
CL\_INS\_207
CL\_INS\_207
CL\_INS\_207
CL\_INS\_207
CL\_INS\_207
CL\_INS\_207
CL\_INS\_207
CL\_INS\_207
CL\_INS\_207
CL\_INS\_207
CL\_INS\_207
CL\_INS\_207
CL\_INS\_207
CL\_INS\_247
CL\_INS\_207
CL\_INS\_247
CL\_INS\_247
CL\_INS\_247
CL\_INS\_247
CL\_INS\_247
CL\_INS\_247
CL\_INS\_247
CL\_INS\_207
CL\_INS\_207
CL\_INS\_207
CL\_INS\_207
CL\_INS\_207
CL\_INS\_207
CL\_INS\_207
CL\_INS\_247
CL\_INS\_207
CL\_INS\_207
CL\_INS\_207
CL\_INS\_207
CL\_INS\_207
CL\_INS\_207
CL\_INS\_207
CL\_INS\_207
CL\_INS\_207
CL\_INS\_207
CL\_INS\_207
CL\_INS\_207
CL\_INS\_207
CL\_INS\_207
CL\_INS\_207
CL\_INS\_207
CL\_INS\_207
CL\_INS\_207
CL\_INS\_207
CL\_INS\_207
CL\_INS\_207
CL\_INS\_207
CL\_INS\_207
CL\_INS\_207
CL\_INS\_247
CL\_INS\_247
CL\_INS\_247
CL\_INS\_247
CL\_INS\_247
CL\_INS\_207
CL\_INS\_207
CL\_INS\_247
CL\_INS\_207
CL\_INS\_207
CL\_INS\_207
CL\_INS\_207
CL\_INS\_247
CL\_INS\_247
CL\_INS\_30
CL\_INS\_207
CL\_INS\_207
CL\_INS\_207
CL\_INS\_207
CL\_INS\_207
CL\_INS\_207
CL\_INS\_207
CL\_INS\_207
CL\_INS\_207
CL\_INS\_207
CL\_INS\_207
CL\_INS\_207
CL\_INS\_207
CL\_INS\_207
CL\_INS\_207
CL\_INS\_207
CL\_INS\_207
CL\_INS\_207
CL\_INS\_207
CL\_INS\_207
CL\_INS\_207
CL\_INS\_207
CL\_INS\_207
CL\_INS\_207
CL\_INS\_207
CL\_INS\_207
CL\_INS\_207
CL\_INS\_207
CL\_INS\_207
CL\_INS\_207
CL\_INS\_207
CL\_INS\_207
CL\_INS\_207
CL\_INS\_207
CL\_INS\_207
CL\_INS\_207
CL\_INS\_207
CL\_INS\_207
CL\_INS\_207
CL\_INS\_207
CL\_INS\_207
CL\_INS\_207
CL\_INS\_207
CL\_INS\_207
CL\_INS\_207
CL\_INS\_207
CL\_INS\_207
CL\_INS\_207
CL\_INS\_207
CL\_INS\_207
CL\_INS\_207
CL\_INS\_207
CL\_INS\_207
CL\_INS\_207
CL\_INS\_207
CL\_INS\_247
CL\_INS\_247
CL\_INS\_247
CL\_INS\_247
CL\_INS\_382
CL\_INS\_382
CL\_INS\_382
CL\_INS\_247
CL\_INS\_207
CL\_INS\_207
CL\_INS\_382
CL\_INS\_207
CL\_INS\_207
CL\_INS\_207
CL\_INS\_207
CL\_INS\_207
CL\_INS\_207
CL\_INS\_207
CL\_INS\_207
CL\_INS\_207
CL\_INS\_207
CL\_INS\_207
CL\_INS\_207
CL\_INS\_207
CL\_INS\_207
CL\_INS\_207
CL\_INS\_207
CL\_INS\_237
CL\_INS\_237
CL\_INS\_237
CL\_INS\_237
CL\_INS\_237
CL\_INS\_207
CL\_INS\_207
CL\_INS\_207
CL\_INS\_207
CL\_INS\_207
CL\_INS\_382
CL\_INS\_247
CL\_INS\_382
CL\_INS\_382
CL\_INS\_382
CL\_INS\_207
CL\_INS\_207
CL\_INS\_207
CL\_INS\_207
CL\_INS\_207
CL\_INS\_207
CL\_INS\_207
CL\_INS\_207
CL\_INS\_207
CL\_INS\_207
CL\_INS\_207
CL\_INS\_207
CL\_INS\_207
CL\_INS\_207
CL\_INS\_207
CL\_INS\_207
CL\_INS\_247
CL\_INS\_247
CL\_INS\_247
CL\_INS\_247
CL\_INS\_247
CL\_INS\_247
CL\_INS\_247
CL\_INS\_247
CL\_INS\_149
CL\_INS\_247
CL\_INS\_247
CL\_INS\_207
CL\_INS\_207
CL\_INS\_207
CL\_INS\_237
CL\_INS\_247
CL\_INS\_207
CL\_INS\_70
CL\_INS\_70
CL\_INS\_207
CL\_INS\_207
CL\_INS\_237
CL\_INS\_70
CL\_INS\_70
CL\_INS\_70
CL\_INS\_207
CL\_INS\_70
CL\_INS\_70
CL\_INS\_70
CL\_INS\_70
CL\_INS\_70
CL\_INS\_70
CL\_INS\_70
CL\_INS\_70
CL\_INS\_70
CL\_INS\_70
CL\_INS\_70
CL\_INS\_70
CL\_INS\_70
CL\_INS\_70
CL\_INS\_207
CL\_INS\_70
CL\_INS\_207
CL\_INS\_70
CL\_INS\_70
CL\_INS\_70
CL\_INS\_70
CL\_INS\_207
CL\_INS\_207
CL\_INS\_207
CL\_INS\_70
CL\_INS\_207
CL\_INS\_207
CL\_INS\_207
CL\_INS\_207
CL\_INS\_207
CL\_INS\_207
CL\_INS\_207
CL\_INS\_207
CL\_INS\_207
CL\_INS\_207
CL\_INS\_207
CL\_INS\_207
CL\_INS\_207
CL\_INS\_207
CL\_INS\_207
CL\_INS\_247
CL\_INS\_247
CL\_INS\_247
CL\_INS\_247
CL\_INS\_247
CL\_INS\_247
CL\_INS\_247
CL\_INS\_247
CL\_INS\_247
CL\_INS\_247
CL\_INS\_382
CL\_INS\_382
CL\_INS\_207
CL\_INS\_207
CL\_INS\_70
CL\_INS\_247
CL\_INS\_247
CL\_INS\_247
CL\_INS\_247
CL\_INS\_247
CL\_INS\_247
CL\_INS\_207
CL\_INS\_247
CL\_INS\_247
CL\_INS\_247
CL\_INS\_247
CL\_INS\_247
CL\_INS\_247
CL\_INS\_207
CL\_INS\_207
CL\_INS\_247
CL\_INS\_247
CL\_INS\_247
CL\_INS\_382
CL\_INS\_382
CL\_INS\_207
CL\_INS\_207
CL\_INS\_207
CL\_INS\_207
CL\_INS\_207
CL\_INS\_207
CL\_INS\_207
CL\_INS\_70
CL\_INS\_70
CL\_INS\_70
CL\_INS\_70
CL\_INS\_237
CL\_INS\_149
CL\_INS\_149
CL\_INS\_149
CL\_INS\_207
CL\_INS\_207
CL\_INS\_247
CL\_INS\_382
CL\_INS\_382
CL\_INS\_70
CL\_INS\_70
CL\_INS\_237
CL\_INS\_207
CL\_INS\_207
CL\_INS\_207
CL\_INS\_207
CL\_INS\_207
CL\_INS\_207
CL\_INS\_207
CL\_INS\_237
CL\_INS\_237
CL\_INS\_237
CL\_INS\_207
CL\_INS\_207
CL\_INS\_149
CL\_INS\_207
CL\_INS\_70
CL\_INS\_207
CL\_INS\_207
CL\_INS\_70
CL\_INS\_247
CL\_INS\_207
CL\_INS\_30
CL\_INS\_247
CL\_INS\_70
CL\_INS\_237
CL\_INS\_30
CL\_INS\_237
CL\_INS\_30
CL\_INS\_70
CL\_INS\_237
CL\_INS\_352
CL\_INS\_70
CL\_INS\_247
CL\_INS\_70
CL\_INS\_70
CL\_INS\_70
CL\_INS\_247
CL\_INS\_207
Cluster ID


CL\_35410
CL\_30782
CL\_5939
CL\_25506
CL\_15510
CL\_15511
CL\_15512
CL\_32455
CL\_30009
CL\_30008
CL\_30007
CL\_36982
CL\_36981
CL\_36980
CL\_36979
CL\_36978
CL\_36977
CL\_7395
CL\_7250
CL\_5236
CL\_4094
CL\_4095
CL\_4096
CL\_4097
CL\_15133
CL\_15132
CL\_11989
CL\_7007
CL\_7008
CL\_21845
CL\_36203
CL\_36202
CL\_36201
CL\_10863
CL\_10862
CL\_10861
CL\_10860
CL\_10859
CL\_10858
CL\_10857
CL\_36200
CL\_36199
CL\_36198
CL\_36197
CL\_11881
CL\_11880
CL\_11879
CL\_11878
CL\_11877
CL\_37135
CL\_11876
CL\_11875
CL\_11874
CL\_11873
CL\_11872
CL\_11871
CL\_11870
CL\_11869
CL\_11868
CL\_11867
CL\_11866
CL\_11865
CL\_11864
CL\_11863
CL\_11862
CL\_11861
CL\_11860
CL\_11859
CL\_34163
CL\_11858
CL\_11857
CL\_7075
CL\_11856
CL\_11855
CL\_11854
CL\_33032
CL\_17225
CL\_17224
CL\_28027
CL\_32699
CL\_33033
CL\_33034
CL\_33035
CL\_28471
CL\_17423
CL\_17422
CL\_17421
CL\_17420
CL\_28470
CL\_29056
CL\_29057
CL\_10372
CL\_29058
CL\_29059
CL\_29060
CL\_28878
CL\_28879
CL\_28880
CL\_28881
CL\_28882
CL\_28883
CL\_28884
CL\_28885
CL\_29061
CL\_8427
CL\_8423
CL\_8422
CL\_8421
CL\_8420
CL\_8419
CL\_29062
CL\_29063
CL\_29064
CL\_29065
CL\_28886
CL\_22453
CL\_22452
CL\_29066
CL\_22451
CL\_22450
CL\_14388
CL\_1491
CL\_1492
CL\_1493
CL\_10520
CL\_6782
CL\_10526
CL\_11956
CL\_10968
CL\_17772
CL\_17773
CL\_27004
CL\_23466
CL\_23467
CL\_23468
CL\_23469
CL\_4513
CL\_1324
CL\_4489
CL\_4488
CL\_6747
CL\_7112
CL\_4485
CL\_4520
CL\_8155
CL\_7522
CL\_7523
CL\_27003
CL\_7524
CL\_7525
CL\_7526
CL\_6741
CL\_7527
CL\_7528
CL\_7529
CL\_7530
CL\_7531
CL\_7532
CL\_7533
CL\_9122
CL\_12124
CL\_11311
CL\_6784
CL\_526
CL\_12382
CL\_23470
CL\_17553
CL\_17554
CL\_17555
CL\_17556
CL\_17557
CL\_17558
CL\_17559
CL\_17053
CL\_15588
CL\_8712
CL\_4518
CL\_32856
CL\_12360
CL\_12062
CL\_14077
CL\_14078
CL\_14079
CL\_7346
CL\_35018
CL\_21316
CL\_35017
CL\_9771
CL\_14833
CL\_14834
CL\_14835
CL\_14836
CL\_6458
CL\_6459
CL\_14837
CL\_6460
CL\_26664
CL\_26665
CL\_26666
CL\_6520
CL\_8296
CL\_9530
CL\_1105
CL\_2277
CL\_2276
CL\_5201
CL\_13990
CL\_13991
CL\_6514
CL\_6515
CL\_6516
CL\_6517
CL\_33505
CL\_6518
CL\_5203
CL\_5204
CL\_5883
CL\_5206
CL\_5207
CL\_5208
CL\_6591
CL\_6519
CL\_5202
CL\_23287
CL\_1106
CL\_13992
CL\_11275
CL\_5884
CL\_33504
CL\_33503
CL\_33502
CL\_8483
CL\_5200
CL\_24067
CL\_10408
CL\_10409
CL\_5199
CL\_33501
CL\_33500
CL\_33499
CL\_33498
CL\_33497
CL\_5198
CL\_5197
CL\_16438
CL\_17919
CL\_4433
CL\_4514
CL\_7521
CL\_4515
CL\_4431
CL\_4517
CL\_4429
CL\_11957
CL\_11913
CL\_4521
CL\_4425
CL\_4424
CL\_4423
CL\_4525
CL\_4421
CL\_4420
CL\_4419
CL\_4418
CL\_4417
CL\_4416
CL\_4415
CL\_4533
CL\_533
CL\_17918
CL\_17917
CL\_8169
CL\_8170
CL\_14838
CL\_8171
CL\_32857
CL\_32858
CL\_8711
CL\_8710
CL\_8709
CL\_8708
CL\_8707
CL\_14839
CL\_14840
CL\_14841
CL\_14842
CL\_14843
CL\_14844
CL\_14845
CL\_14846
CL\_14847
CL\_14848
CL\_14849
CL\_14850
CL\_14851
CL\_14852
CL\_10164
CL\_10165
CL\_14853
CL\_14854
CL\_14855
CL\_14856
CL\_14857
CL\_14858
CL\_14859
CL\_14860
CL\_14861
CL\_14862
CL\_14863
CL\_14864
CL\_14865
CL\_8706
CL\_8705
CL\_8704
CL\_8703
CL\_8702
CL\_8700
CL\_32859
CL\_32860
CL\_32861
CL\_32862
CL\_32863
CL\_32864
CL\_32865
CL\_32866
CL\_32867
CL\_32868
CL\_32869
CL\_32870
CL\_8183
CL\_12006
CL\_7538
CL\_13562
CL\_13074
CL\_17916
CL\_17915
CL\_17914
CL\_4629
CL\_17913
CL\_4527
CL\_17912
CL\_17911
CL\_17910
CL\_17909
CL\_6053
CL\_16439
CL\_16440
CL\_16441
CL\_16442
CL\_4678
CL\_4679
CL\_4681
CL\_4682
CL\_33726
CL\_6052
CL\_4684
CL\_4685
CL\_6051
CL\_6050
CL\_4687
CL\_4688
CL\_7072
CL\_4689
CL\_4598
CL\_6521
CL\_26318
CL\_26317
CL\_29323
CL\_19381
CL\_30613
CL\_6522
CL\_24356
CL\_5196
CL\_6523
CL\_8290
CL\_6524
CL\_26316
CL\_6525
CL\_6526
CL\_24354
CL\_24355
CL\_10567
CL\_5193
CL\_5194
CL\_5195
CL\_8069
CL\_33496
CL\_33495
CL\_33494
CL\_33493
CL\_33492
CL\_33491
CL\_33490
CL\_33489
CL\_33488
CL\_33487
CL\_33486
CL\_33485
CL\_33484
CL\_33483
CL\_33482
CL\_33481
CL\_33480
CL\_33479
CL\_6527
CL\_6528
CL\_10566
CL\_6529
CL\_6530
CL\_6531
CL\_6599
CL\_4586
CL\_4585
CL\_4584
CL\_29926
CL\_29927
CL\_29928
CL\_5887
CL\_11569
CL\_8482
CL\_5184
CL\_26315
CL\_31846
CL\_17738
CL\_29322
CL\_29321
CL\_29320
CL\_29319
CL\_9004
CL\_9005
CL\_9006
CL\_15744
CL\_19884
CL\_13993
CL\_6534
CL\_10564
CL\_10565
CL\_24353
CL\_6535
CL\_6536
CL\_13443
CL\_6537
CL\_6538
CL\_6539
CL\_6540
CL\_10562
CL\_10563
CL\_6541
CL\_17739
CL\_6542
CL\_13343
CL\_13344
CL\_6543
CL\_13994
CL\_13995
CL\_13996
CL\_21793
CL\_33577
CL\_29318
CL\_13997
CL\_8349
CL\_13998
CL\_13999
CL\_14000
CL\_26836
CL\_26835
CL\_26834
CL\_16868
CL\_16869
CL\_10037
CL\_10036
CL\_10035
CL\_10034
CL\_23578
CL\_19850
CL\_16870
CL\_16871
CL\_16872
CL\_16873
CL\_16874
CL\_21725
CL\_21724
CL\_14001
CL\_14002
CL\_14003
CL\_27890
CL\_27889
CL\_27888
CL\_27887
CL\_27886
CL\_27885
CL\_27884
CL\_27883
CL\_26138
CL\_27882
CL\_19382
CL\_19383
CL\_12063
CL\_7345
CL\_4717
CL\_36041
CL\_4490
CL\_1495
CL\_15108
CL\_26932
CL\_6594
CL\_1102
CL\_7344
CL\_7343
CL\_1103
CL\_1104
CL\_4599
CL\_5392
CL\_1101
CL\_17677
CL\_17678
CL\_17679
CL\_17680
CL\_17681
CL\_17682
CL\_1100
CL\_5393
CL\_5394
CL\_5395
CL\_5396
CL\_11934
CL\_17743
CL\_26931
CL\_10274
CL\_5398
CL\_13585
CL\_7588
CL\_7587
CL\_7586
CL\_7585
CL\_7584
CL\_10361
CL\_10362
CL\_14292
CL\_9574
CL\_33449
CL\_33448
CL\_33447
CL\_14293
CL\_14294
CL\_9573
CL\_9572
CL\_9571
CL\_7583
CL\_19852
CL\_19851
CL\_8040
CL\_23869
CL\_7589
CL\_7590
CL\_27881
CL\_7789
CL\_10190
CL\_10191
CL\_5399
CL\_5400
CL\_10273
CL\_5401
CL\_17683
CL\_5402
CL\_5403
CL\_5404
CL\_5405
CL\_11307
CL\_11308
CL\_11309
CL\_10973
CL\_10974
CL\_11310
CL\_5406
CL\_5407
CL\_5408
CL\_5409
CL\_17644
CL\_14639
CL\_14638
CL\_5410
CL\_1099
CL\_1098
CL\_1097
CL\_5411
CL\_5412
CL\_34923
CL\_33079
CL\_19359
CL\_10975
CL\_36277
CL\_26667
CL\_7342
CL\_7341
CL\_5414
CL\_5415
CL\_5924
CL\_11936
CL\_11937
CL\_21681
CL\_21682
CL\_5416
CL\_20411
CL\_11312
CL\_26668
CL\_26669
CL\_5808
CL\_5807
CL\_7340
CL\_7339
CL\_19355
CL\_12766
CL\_17640
CL\_17639
CL\_36278
CL\_36279
CL\_36280
CL\_36281
CL\_36282
CL\_10174
CL\_5421
CL\_12064
CL\_12065
CL\_12066
CL\_5422
CL\_17638
CL\_17637
CL\_2278
CL\_2279
CL\_2280
CL\_17635
CL\_1096
CL\_5423
CL\_5424
CL\_10175
CL\_5425
CL\_7621
CL\_11313
CL\_17634
CL\_5426
CL\_5427
CL\_5428
CL\_2283
CL\_11938
CL\_11406
CL\_11405
CL\_10511
CL\_11402
CL\_2284
CL\_2285
CL\_5429
CL\_5430
CL\_36283
CL\_36284
CL\_23582
CL\_23583
CL\_15934
CL\_15935
CL\_36285
CL\_12543
CL\_15937
CL\_15938
CL\_12541
CL\_8255
CL\_15940
CL\_12279
CL\_23547
CL\_20109
CL\_20110
CL\_1095
CL\_7542
CL\_7543
CL\_1094
CL\_1093
CL\_14633
CL\_19175
CL\_7338
CL\_2281
CL\_2282
CL\_10270
CL\_12067
CL\_7337
CL\_6049
CL\_6048
CL\_6047
CL\_4528
CL\_4529
CL\_4530
CL\_4532
CL\_4414
CL\_4413
CL\_1496
CL\_531
CL\_7495
CL\_1499
CL\_1500
CL\_1502
CL\_1503
CL\_1504
CL\_1505
CL\_7494
CL\_1506
CL\_1507
CL\_1508
CL\_1509
CL\_1510
CL\_1511
CL\_1497
CL\_8699
CL\_8698
CL\_8697
CL\_13560
CL\_11920
CL\_7537
CL\_7118
CL\_12996
CL\_8694
CL\_12698
CL\_17560
CL\_17561
CL\_16443
CL\_15699
CL\_15698
CL\_17908
CL\_17907
CL\_17906
CL\_17905
CL\_17904
CL\_17903
CL\_17902
CL\_17901
CL\_17900
CL\_17899
CL\_17898
CL\_17897
CL\_17896
CL\_17895
CL\_17894
CL\_17893
CL\_17892
CL\_17891
CL\_17890
CL\_17889
CL\_17888
CL\_17887
CL\_17886
CL\_17885
CL\_15866
CL\_15867
CL\_532
CL\_33727
CL\_4535
CL\_6043
CL\_33728
CL\_33729
CL\_4644
CL\_16444
CL\_16445
CL\_7539
CL\_7540
CL\_17235
CL\_27355
CL\_17234
CL\_16967
CL\_12123
CL\_17774
CL\_16446
CL\_16447
CL\_16448
CL\_4651
CL\_17562
CL\_17563
CL\_17564
CL\_17565
CL\_17566
CL\_17567
CL\_17568
CL\_1512
CL\_5419
CL\_5420
CL\_4652
CL\_4653
CL\_4654
CL\_12380
CL\_12379
CL\_4658
CL\_12376
CL\_15864
CL\_15865
CL\_16449
CL\_16450
CL\_4655
CL\_13564
CL\_4656
CL\_6037
CL\_6036
CL\_4659
CL\_33730
CL\_4661
CL\_4662
CL\_16451
CL\_16452
CL\_4663
CL\_4664
CL\_4665
CL\_16453
CL\_15868
CL\_4666
CL\_15623
CL\_4667
CL\_4668
CL\_4669
CL\_12068
CL\_27356
CL\_27357
CL\_27002
CL\_20111
CL\_20112
CL\_32871
CL\_33078
CL\_33077
CL\_33076
CL\_21683
CL\_21251
CL\_21250
CL\_5125
CL\_26930
CL\_14389
CL\_14390
CL\_26670
CL\_26671
CL\_7336
CL\_7335
CL\_7334
CL\_7333
CL\_12069
CL\_2529
CL\_32456
CL\_32457
CL\_9356
CL\_9357
CL\_20685
CL\_33554
CL\_13419
CL\_8141
CL\_31890
CL\_31891
CL\_31892
CL\_31893
CL\_31894
CL\_31895
CL\_31896
CL\_31897
CL\_31898
CL\_7784
CL\_31899
CL\_22510
CL\_22511
CL\_28271
CL\_28270
CL\_37436
CL\_19941
CL\_19940
CL\_19939
CL\_19938
CL\_8247
CL\_8338
CL\_8339
CL\_8340
CL\_8341
CL\_8342
CL\_8343
CL\_8344
CL\_8345
CL\_8346
CL\_8347
CL\_8348
CL\_20683
CL\_20682
CL\_20681
CL\_21257
CL\_21256
CL\_21255
CL\_21254
CL\_21253
CL\_21252
CL\_23128
CL\_23129
CL\_23130
CL\_16867
CL\_30631
CL\_30632
CL\_30633
CL\_30634
CL\_30635
CL\_5516
CL\_5692
CL\_5536
CL\_5613
CL\_5614
CL\_22269
CL\_11132
CL\_5509
CL\_5510
CL\_5511
CL\_5512
CL\_5513
CL\_5019
CL\_10412
CL\_10413
CL\_10314
CL\_9706
CL\_9705
CL\_21571
CL\_21572
CL\_21573
CL\_9704
CL\_6370
CL\_6369
CL\_6368
CL\_7416
CL\_14295
CL\_14296
CL\_24560
CL\_6367
CL\_9703
CL\_9702
CL\_6366
CL\_6365
CL\_14297
CL\_7582
CL\_7415
CL\_17569
CL\_7414
CL\_24559
CL\_21723
CL\_7413
CL\_17570
CL\_7412
CL\_7411
CL\_7410
CL\_9701
CL\_9570
CL\_9569
CL\_20684
CL\_14298
CL\_14299
CL\_9700
CL\_9699
CL\_24558
CL\_24557
CL\_24556
CL\_19384
CL\_19385
CL\_19181
CL\_19180
CL\_19179
CL\_19937
CL\_18125
CL\_18126
CL\_16974
CL\_13414
CL\_4817
CL\_33075
CL\_8350
CL\_4818
CL\_7456
CL\_8039
CL\_7455
CL\_8038
CL\_7454
CL\_7453
CL\_7452
CL\_7451
CL\_7450
CL\_7449
CL\_7448
CL\_7447
CL\_7446
CL\_7445
CL\_7444
CL\_7443
CL\_7442
CL\_8037
CL\_15357
CL\_9206
CL\_9205
CL\_9204
CL\_18235
CL\_18234
CL\_18233
CL\_5997
CL\_5998
CL\_5999
CL\_6000
CL\_6001
CL\_6002
CL\_6003
CL\_6004
CL\_6005
CL\_6006
CL\_6007
CL\_6008
CL\_6009
CL\_6010
CL\_23131
CL\_6011
CL\_23132
CL\_6012
CL\_6013
CL\_6014
CL\_6015
CL\_6016
CL\_6017
CL\_23133
CL\_2530
CL\_2531
CL\_28890
CL\_2532
CL\_10212
CL\_10213
CL\_2533
CL\_8961
CL\_8960
CL\_8959
CL\_8958
CL\_8957
CL\_8956
CL\_8955
CL\_8954
CL\_2534
CL\_2535
CL\_2536
CL\_2537
CL\_2538
CL\_2539
CL\_2540
CL\_2541
CL\_27532
CL\_27531
CL\_2542
CL\_2543
CL\_2544
CL\_2545
CL\_2546
CL\_2547
CL\_9028
CL\_9027
CL\_27539
CL\_27538
CL\_27537
CL\_27536
CL\_27535
CL\_27534
CL\_27533
CL\_26672
CL\_26673
CL\_26674
CL\_26675
CL\_26676
CL\_26677
CL\_26678
CL\_241
CL\_240
CL\_9471
CL\_9472
CL\_9473
CL\_26679
CL\_26680
CL\_2548
CL\_233
CL\_232
CL\_231
CL\_2549
CL\_10219
CL\_2550
CL\_2551
CL\_26681
CL\_26682
CL\_26683
CL\_26684
CL\_2552
CL\_25978
CL\_25977
CL\_25976
CL\_25975
CL\_5972
CL\_15358
CL\_13418
CL\_13417
CL\_37346
CL\_13416
CL\_15359
CL\_15360
CL\_15361
CL\_13415
CL\_5973
CL\_5974
CL\_22512
CL\_22513
CL\_5975
CL\_5976
CL\_5977
CL\_28269
CL\_20644
CL\_20645
CL\_20646
CL\_8953
CL\_8952
CL\_8951
CL\_8950
CL\_8949
CL\_8948
CL\_8947
CL\_8946
CL\_8945
CL\_26929
CL\_26928
CL\_26927
CL\_26926
CL\_26925
CL\_6916
CL\_6915
CL\_5978
CL\_5979
CL\_5980
CL\_8140
CL\_15362
CL\_2553
CL\_12578
CL\_12579
CL\_2554
CL\_8108
CL\_8109
CL\_8110
CL\_8111
CL\_8112
CL\_8113
CL\_8114
CL\_15924
CL\_34408
CL\_34407
CL\_8123
CL\_34164
CL\_4819
CL\_24561
CL\_8139
CL\_32014
CL\_11853
CL\_10560
CL\_24352
CL\_6547
CL\_6545
CL\_10561
CL\_5943
CL\_9698
CL\_6897
CL\_6896
CL\_6889
CL\_6888
CL\_6887
CL\_6886
CL\_6885
CL\_9697
CL\_9696
CL\_9695
CL\_9694
CL\_9693
CL\_9692
CL\_9691
CL\_9690
CL\_9689
CL\_9688
CL\_9687
CL\_9686
CL\_9685
CL\_9684
CL\_9683
CL\_9682
CL\_9681
CL\_9680
CL\_9679
CL\_9678
CL\_9677
CL\_6270
CL\_9676
CL\_6150
CL\_6149
CL\_9675
CL\_26178
CL\_9460
CL\_10021
CL\_10020
CL\_15549
CL\_15548
CL\_15547
CL\_15546
CL\_11712
CL\_5146
CL\_8623
CL\_12000
CL\_7548
CL\_23773
CL\_23774
CL\_11713
CL\_11714
CL\_26177
CL\_8781
CL\_15980
CL\_27474
CL\_27475
CL\_7985
CL\_7252
CL\_6828
CL\_6411
CL\_27476
CL\_7435
CL\_14667
CL\_14668
CL\_14669
CL\_14670
CL\_14671
CL\_14672
CL\_14673
CL\_14674
CL\_14675
CL\_14676
CL\_14677
CL\_14678
CL\_14679
CL\_10217
CL\_14680
CL\_25123
CL\_7690
CL\_8641
CL\_7896
CL\_7895
CL\_33731
CL\_33732
CL\_33733
CL\_8628
CL\_9492
CL\_9491
CL\_9490
CL\_9489
CL\_9488
CL\_9487
CL\_9486
CL\_9485
CL\_9484
CL\_9483
CL\_9482
CL\_9481
CL\_9480
CL\_9479
CL\_9478
CL\_22116
CL\_22117
CL\_22118
CL\_22119
CL\_22120
CL\_22121
CL\_22122
CL\_22123
CL\_22124
CL\_22125
CL\_4086
CL\_7557
CL\_32013
CL\_28434
CL\_7214
CL\_7215
CL\_18854
CL\_7567
CL\_28433
CL\_7202
CL\_28432
CL\_28431
CL\_14332
CL\_14333
CL\_14334
CL\_14335
CL\_21989
CL\_21988
CL\_37083
CL\_37084
CL\_21987
CL\_21986
CL\_21985
CL\_8554
CL\_10807
CL\_37167
CL\_37168
CL\_37169
CL\_37170
CL\_37171
CL\_37172
CL\_37173
CL\_6420
CL\_1932
CL\_1933
CL\_8620
CL\_4375
CL\_7490
CL\_7489
CL\_7488
CL\_33734
CL\_33735
CL\_6844
CL\_6177
CL\_6178
CL\_6179
CL\_1931
CL\_16977
CL\_33171
CL\_33170
CL\_33169
CL\_33168
CL\_33167
CL\_33166
CL\_33165
CL\_9119
CL\_8319
CL\_6660
CL\_26136
CL\_26137
CL\_4093
CL\_33164
CL\_8508
CL\_37174
CL\_29067
CL\_9140
CL\_6726
CL\_37175
CL\_1934
CL\_7828
CL\_10610
CL\_10819
CL\_7749
CL\_10818
CL\_1937
CL\_6809
CL\_7970
CL\_7969
CL\_6808
CL\_7204
CL\_8746
CL\_26031
CL\_8107
CL\_7203
CL\_28430
