## Supplementary material for "A novel method for integrating genomic and Tn-Seq data to identify common *in vivo* fitness mechanisms across multiple bacterial species": S1 Dataset: CL_INS_208.html

Legend

 Hypothetical
 Regulatoryfunctions
 All EssentialGenes
 Other

FULL


WINDOWSVGPNG

Trim RowsRemove SingletonsSave Fasta

CL\_2568


CL\_2568


CL\_2568


CL\_2568


CL\_2568


CL\_2567

HighlightSelectShow Genomes


202

CL\_2569


52

CL\_2569


22

CL\_2569


1

CL\_2569


1

CL\_2569


1

CL\_2569

fGI ID


CL\_INS\_208
CL\_INS\_208
CL\_INS\_208
Cluster ID


CL\_7332
CL\_7331
CL\_7330
