## Supplementary material for "A novel method for integrating genomic and Tn-Seq data to identify common *in vivo* fitness mechanisms across multiple bacterial species": S1 Dataset: CL_INS_209.html

Legend

 Hypothetical
 All VFDB Genes

FULL


WINDOWSVGPNG

Trim RowsRemove SingletonsSave Fasta

CL\_2577


CL\_2577


CL\_2577

HighlightSelectShow Genomes


168

CL\_2579


110

CL\_2579


1

CL\_2580

fGI ID

CL\_INS\_209
Cluster ID

CL\_2578
