## Supplementary material for "A novel method for integrating genomic and Tn-Seq data to identify common *in vivo* fitness mechanisms across multiple bacterial species": S1 Dataset: CL_INS_212.html

Legend

 Mobile +extrachromosomalelementfunctions
 Hypothetical
 All EssentialGenes
 Other
 All VFDB Genes

FULL


WINDOWSVGPNG

Trim RowsRemove SingletonsSave Fasta

CL\_2603


CL\_2603


CL\_2603


CL\_2603


CL\_2603


CL\_2602


CL\_2603


CL\_2603

HighlightSelectShow Genomes


234

CL\_2604


28

CL\_2607


5

CL\_2607


4

CL\_2607


2

CL\_2607


1

CL\_2604


1

CL\_2607


1

CL\_2607

fGI ID


CL\_INS\_212
CL\_INS\_212
CL\_INS\_212
CL\_INS\_212
CL\_INS\_212
CL\_INS\_212
Cluster ID


CL\_9363
CL\_9364
CL\_13819
CL\_7732
CL\_5944
CL\_5945
