## Supplementary material for "A novel method for integrating genomic and Tn-Seq data to identify common *in vivo* fitness mechanisms across multiple bacterial species": S1 Dataset: CL_INS_213.html

Legend

 Mobile +extrachromosomalelementfunctions
 Regulatoryfunctions
 Hypothetical
 All EssentialGenes
 Other
 Transport +binding proteins
 All VFDB Genes

FULL


WINDOWSVGPNG

Trim RowsRemove SingletonsSave Fasta

CL\_2606


CL\_2606


CL\_2603


CL\_2606


CL\_2606


CL\_2606


CL\_2606


CL\_2603


CL\_2606


CL\_2606


CL\_2603


CL\_2604


CL\_2603


CL\_2606


CL\_2606


CL\_2606


CL\_2606


CL\_2603


CL\_2605


CL\_2604


CL\_2603


CL\_2606

HighlightSelectShow Genomes


100

CL\_2607


54

CL\_2607


28

CL\_2607


23

CL\_2607


20

CL\_2607


10

CL\_2607


9

CL\_2607


5

CL\_2607


4

CL\_2607


4

CL\_2607


4

CL\_2607


2

CL\_2607


2

CL\_2607


2

CL\_2607


1

CL\_2607


1

CL\_2607


1

CL\_2607


1

CL\_2607


1

CL\_2607


1

CL\_2607


1

CL\_2607


1

CL\_2607

fGI ID


CL\_INS\_213
CL\_INS\_213
CL\_INS\_213
CL\_INS\_212
CL\_INS\_212
CL\_INS\_212
CL\_INS\_212
CL\_INS\_213
CL\_INS\_213
CL\_INS\_213
CL\_INS\_182
CL\_INS\_182
CL\_INS\_381
CL\_INS\_213
CL\_INS\_213
CL\_INS\_182
CL\_INS\_182
CL\_INS\_182
CL\_INS\_155
CL\_INS\_155
CL\_INS\_155
CL\_INS\_155
CL\_INS\_155
CL\_INS\_155
CL\_INS\_155
CL\_INS\_74
CL\_INS\_74
CL\_INS\_74
CL\_INS\_155
CL\_INS\_155
CL\_INS\_155
CL\_INS\_155
CL\_INS\_155
CL\_INS\_155
CL\_INS\_213
CL\_INS\_155
CL\_INS\_86
CL\_INS\_155
CL\_INS\_86
CL\_INS\_155
CL\_INS\_74
CL\_INS\_155
CL\_INS\_155
CL\_INS\_155
CL\_INS\_155
CL\_INS\_74
CL\_INS\_86
CL\_INS\_74
CL\_INS\_74
CL\_INS\_297
CL\_INS\_74
CL\_INS\_155
CL\_INS\_155
CL\_INS\_155
CL\_INS\_86
CL\_INS\_155
CL\_INS\_155
CL\_INS\_155
CL\_INS\_155
CL\_INS\_204
CL\_INS\_74
Cluster ID


CL\_13587
CL\_8036
CL\_20113
CL\_9364
CL\_7732
CL\_5944
CL\_5945
CL\_30275
CL\_7329
CL\_31900
CL\_10992
CL\_10993
CL\_31901
CL\_31902
CL\_31903
CL\_10998
CL\_10999
CL\_11001
CL\_10906
CL\_10905
CL\_11004
CL\_11005
CL\_11006
CL\_11007
CL\_11009
CL\_10897
CL\_10896
CL\_10895
CL\_10894
CL\_10893
CL\_10892
CL\_10891
CL\_10890
CL\_10889
CL\_31904
CL\_10887
CL\_10886
CL\_10885
CL\_10884
CL\_10883
CL\_31905
CL\_11024
CL\_11025
CL\_10879
CL\_10878
CL\_31906
CL\_11029
CL\_10875
CL\_10874
CL\_31907
CL\_10873
CL\_8229
CL\_8230
CL\_8231
CL\_8232
CL\_10870
CL\_10869
CL\_11035
CL\_8483
CL\_10867
CL\_10866
