## Supplementary material for "A novel method for integrating genomic and Tn-Seq data to identify common *in vivo* fitness mechanisms across multiple bacterial species": S1 Dataset: CL_INS_218.html

Legend

 Mobile +extrachromosomalelementfunctions
 Regulatoryfunctions
 Hypothetical
 All EssentialGenes
 AntibioticResistance
 Other
 Transport +binding proteins
 All VFDB Genes

FULL


WINDOWSVGPNG

Trim RowsRemove SingletonsSave Fasta

CL\_2643


CL\_2643


CL\_2643


CL\_2643


CL\_2643


CL\_2643


CL\_2643


CL\_2643


CL\_2643


CL\_2643


CL\_2643


CL\_2643


CL\_2643


CL\_2643


CL\_2643


CL\_2643


CL\_2643


CL\_2641


CL\_2643


CL\_2643


CL\_2643


CL\_2643


CL\_2643


CL\_2643


CL\_2622


CL\_2643


CL\_2643


CL\_2643


CL\_2643


CL\_2643


CL\_2643


CL\_2643


CL\_2643


CL\_2643


CL\_2643


CL\_2643


CL\_2643


CL\_2643


CL\_2643


CL\_2643


CL\_2643


CL\_2643


CL\_2643


CL\_2643


CL\_2643


CL\_2643


CL\_2643


CL\_2643


CL\_2643


CL\_2643


CL\_2948


CL\_2643


CL\_2643


CL\_2643


CL\_2643


CL\_2643


CL\_2643


CL\_2643

HighlightSelectShow Genomes


82

CL\_4831


79

CL\_4831


29

CL\_2652


14

CL\_4831


6

CL\_4831


6

CL\_4831


5

CL\_4831


4

CL\_4831


3

CL\_4831


2

CL\_2652


2

CL\_2652


2

CL\_4831


2

CL\_4831


2

CL\_4831


1

CL\_2652


1

CL\_4831


1

CL\_2652


1

CL\_4831


1

CL\_4831


1

CL\_4831


1

CL\_2652


1

CL\_4831


1

CL\_2652


1

CL\_2652


1

CL\_4831


1

CL\_4831


1

CL\_2652


1

CL\_2652


1

CL\_4831


1

CL\_4831


1

CL\_4831


1

CL\_4831


1

CL\_2652


1

CL\_4831


1

CL\_2652


1

CL\_4831


1

CL\_2652


1

CL\_4831


1

CL\_2652


1

CL\_2652


1

CL\_2652


1

CL\_4831


1

CL\_2652


1

CL\_2652


1

CL\_4831


1

CL\_4831


1

CL\_2652


1

CL\_2652


1

CL\_2652


1

CL\_4831


1

CL\_4831


1

CL\_4831


1

CL\_2631


1

CL\_4831


1

CL\_2704


1

CL\_4831


1

Break


1

CL\_2652

fGI ID


CL\_INS\_218
CL\_INS\_215
CL\_INS\_215
CL\_INS\_247
CL\_INS\_215
CL\_INS\_218
CL\_INS\_224
CL\_INS\_224
CL\_INS\_218
CL\_INS\_219
CL\_INS\_219
CL\_INS\_219
CL\_INS\_219
CL\_INS\_219
CL\_INS\_219
CL\_INS\_218
CL\_INS\_219
CL\_INS\_218
CL\_INS\_218
CL\_INS\_218
CL\_INS\_219
CL\_INS\_219
CL\_INS\_219
CL\_INS\_219
CL\_INS\_218
CL\_INS\_219
CL\_INS\_219
CL\_INS\_219
CL\_INS\_219
CL\_INS\_219
CL\_INS\_219
CL\_INS\_219
CL\_INS\_219
CL\_INS\_247
CL\_INS\_57
CL\_INS\_247
CL\_INS\_247
CL\_INS\_247
CL\_INS\_247
CL\_INS\_247
CL\_INS\_247
CL\_INS\_247
CL\_INS\_247
CL\_INS\_247
CL\_INS\_44
CL\_INS\_247
CL\_INS\_44
CL\_INS\_247
CL\_INS\_247
CL\_INS\_217
CL\_INS\_247
CL\_INS\_60
CL\_INS\_247
CL\_INS\_123
CL\_INS\_123
CL\_INS\_123
CL\_INS\_247
CL\_INS\_247
CL\_INS\_217
CL\_INS\_57
Cluster ID


CL\_8244
CL\_34069
CL\_34070
CL\_10390
CL\_10427
CL\_2644
CL\_12837
CL\_7311
CL\_12605
CL\_2645
CL\_4826
CL\_2646
CL\_2647
CL\_4827
CL\_4828
CL\_12839
CL\_4829
CL\_23579
CL\_34373
CL\_8243
CL\_4830
CL\_2648
CL\_37176
CL\_17571
CL\_25505
CL\_2649
CL\_2650
CL\_15131
CL\_2651
CL\_6361
CL\_6360
CL\_29069
CL\_29068
CL\_10389
CL\_6751
CL\_10388
CL\_10387
CL\_10386
CL\_10385
CL\_10384
CL\_5601
CL\_10383
CL\_10382
CL\_10423
CL\_10422
CL\_10421
CL\_11134
CL\_10642
CL\_10641
CL\_23459
CL\_10395
CL\_10394
CL\_10393
CL\_10392
CL\_5297
CL\_5298
CL\_5299
CL\_5019
CL\_4823
CL\_6759
