## Supplementary material for "A novel method for integrating genomic and Tn-Seq data to identify common *in vivo* fitness mechanisms across multiple bacterial species": S1 Dataset: CL_INS_220.html

Legend

 Mobile +extrachromosomalelementfunctions
 Regulatoryfunctions
 Hypothetical
 All EssentialGenes
 Other
 Transport +binding proteins
 All VFDB Genes

FULL


WINDOWSVGPNG

Trim RowsRemove SingletonsSave Fasta

CL\_2665


CL\_2665


CL\_2665


CL\_2655


CL\_2665


CL\_2665


CL\_2664


CL\_2665


CL\_2665


CL\_230


CL\_2664


CL\_2665


CL\_2665


CL\_2663


CL\_2664


CL\_2665


CL\_2665


CL\_2665


CL\_2665


CL\_2663


CL\_2658


CL\_2665

HighlightSelectShow Genomes


184

CL\_2666


43

CL\_2666


3

CL\_2666


2

CL\_2666


2

CL\_2666


2

CL\_2666


1

CL\_2666


1

Break


1

CL\_2666


1

CL\_2666


1

CL\_2666


1

CL\_2682


1

CL\_2666


1

CL\_2666


1

CL\_2666


1

CL\_203


1

CL\_2669


1

CL\_2672


1

CL\_2666


1

CL\_2666


1

CL\_2666


1

CL\_2666

fGI ID


CL\_INS\_220
CL\_INS\_220
CL\_INS\_220
CL\_INS\_220
CL\_INS\_220
CL\_INS\_220
CL\_INS\_220
CL\_INS\_220
CL\_INS\_20
CL\_INS\_20
CL\_INS\_220
CL\_INS\_220
CL\_INS\_237
CL\_INS\_237
CL\_INS\_237
CL\_INS\_237
CL\_INS\_237
CL\_INS\_220
CL\_INS\_220
CL\_INS\_220
CL\_INS\_220
CL\_INS\_220
CL\_INS\_220
CL\_INS\_220
CL\_INS\_220
CL\_INS\_220
CL\_INS\_220
CL\_INS\_220
CL\_INS\_220
CL\_INS\_220
Cluster ID


CL\_30776
CL\_27880
CL\_10215
CL\_8242
CL\_12604
CL\_7396
CL\_37571
CL\_6633
CL\_13291
CL\_30497
CL\_30496
CL\_27804
CL\_8938
CL\_8939
CL\_8940
CL\_8941
CL\_8942
CL\_30273
CL\_11603
CL\_7931
CL\_7932
CL\_7934
CL\_7936
CL\_11608
CL\_11609
CL\_13041
CL\_7938
CL\_7939
CL\_7940
CL\_7942
