## Supplementary material for "A novel method for integrating genomic and Tn-Seq data to identify common *in vivo* fitness mechanisms across multiple bacterial species": S1 Dataset: CL_INS_221.html

Legend

 Mobile +extrachromosomalelementfunctions
 Regulatoryfunctions
 Hypothetical
 All EssentialGenes
 Other
 Transport +binding proteins
 All VFDB Genes

FULL


WINDOWSVGPNG

Trim RowsRemove SingletonsSave Fasta

CL\_2666


CL\_2666


CL\_2666


CL\_2666


CL\_2666


CL\_2666


CL\_2666


CL\_2666


CL\_2666


CL\_2666


CL\_2666


CL\_2666


CL\_2663


CL\_2666


CL\_2663


CL\_2666

HighlightSelectShow Genomes


87

CL\_2668


77

CL\_2668


47

CL\_2668


27

CL\_2669


16

CL\_2669


4

CL\_2668


1

CL\_2669


1

CL\_2669


1

CL\_2669


1

CL\_2669


1

CL\_2668


1

CL\_2669


1

CL\_2668


1

CL\_2669


1

CL\_2668


1

CL\_2668

fGI ID


CL\_INS\_221
CL\_INS\_221
CL\_INS\_221
CL\_INS\_220
CL\_INS\_221
CL\_INS\_221
CL\_INS\_221
CL\_INS\_220
CL\_INS\_221
CL\_INS\_237
CL\_INS\_221
CL\_INS\_237
CL\_INS\_221
CL\_INS\_221
CL\_INS\_221
CL\_INS\_221
CL\_INS\_221
Cluster ID


CL\_22878
CL\_2667
CL\_12603
CL\_27880
CL\_30775
CL\_30187
CL\_7327
CL\_7396
CL\_10569
CL\_8758
CL\_8759
CL\_8760
CL\_35089
CL\_35088
CL\_35087
CL\_35086
CL\_35085
