## Supplementary material for "A novel method for integrating genomic and Tn-Seq data to identify common *in vivo* fitness mechanisms across multiple bacterial species": S1 Dataset: CL_INS_222.html

Legend

 Other
 All VFDB Genes

FULL


WINDOWSVGPNG

Trim RowsRemove SingletonsSave Fasta

CL\_2690


CL\_2690


CL\_2690


CL\_2690

HighlightSelectShow Genomes


249

CL\_2691


16

CL\_2692


2

CL\_2692


1

CL\_2692

fGI ID


CL\_INS\_222
CL\_INS\_222
CL\_INS\_222
CL\_INS\_222
Cluster ID


CL\_5166
CL\_5165
CL\_5164
CL\_5163
