## Supplementary material for "A novel method for integrating genomic and Tn-Seq data to identify common *in vivo* fitness mechanisms across multiple bacterial species": S1 Dataset: CL_INS_223.html

Legend

 Other
 All VFDB Genes

FULL


WINDOWSVGPNG

Trim RowsRemove SingletonsSave Fasta

CL\_2692


CL\_2692


CL\_2692


CL\_2692


CL\_2692

HighlightSelectShow Genomes


246

CL\_2691


17

CL\_2690


2

CL\_2690


1

CL\_2690


1

Break

fGI ID


CL\_INS\_223
CL\_INS\_223
CL\_INS\_222
CL\_INS\_222
CL\_INS\_222
Cluster ID


CL\_16952
CL\_8133
CL\_5163
CL\_5164
CL\_5165
