## Supplementary material for "A novel method for integrating genomic and Tn-Seq data to identify common *in vivo* fitness mechanisms across multiple bacterial species": S1 Dataset: CL_INS_228.html

Legend

 Mobile +extrachromosomalelementfunctions
 Hypothetical
 Other
 EnergyMetabolism
 All VFDB Genes

FULL


WINDOWSVGPNG

Trim RowsRemove SingletonsSave Fasta

CL\_2797


CL\_2797


CL\_2797


CL\_2797


CL\_2796


CL\_2797


CL\_2797

HighlightSelectShow Genomes


228

CL\_2798


8

CL\_2798


3

CL\_2798


2

CL\_2802


2

CL\_2798


1

CL\_2798


1

CL\_2801

fGI ID


CL\_INS\_228
CL\_INS\_228
CL\_INS\_228
CL\_INS\_228
Cluster ID


CL\_30768
CL\_11193
CL\_11192
CL\_11851
