## Supplementary material for "A novel method for integrating genomic and Tn-Seq data to identify common *in vivo* fitness mechanisms across multiple bacterial species": S1 Dataset: CL_INS_229.html

Legend

 Hypothetical
 Other
 All VFDB Genes

FULL


WINDOWSVGPNG

Trim RowsRemove SingletonsSave Fasta

CL\_2799


CL\_2798


CL\_2799


CL\_2799


CL\_2797

HighlightSelectShow Genomes


179

CL\_2801


9

CL\_2801


5

CL\_2801


1

CL\_2802


1

CL\_2801

fGI ID

CL\_INS\_229
Cluster ID

CL\_2800
