## Supplementary material for "A novel method for integrating genomic and Tn-Seq data to identify common *in vivo* fitness mechanisms across multiple bacterial species": S1 Dataset: CL_INS_230.html

Legend

 Hypothetical
 Other
 All VFDB Genes

FULL


WINDOWSVGPNG

Trim RowsRemove SingletonsSave Fasta

CL\_2809


CL\_2809


CL\_2809


CL\_2809


CL\_2809


Break


CL\_2809


CL\_2809


CL\_2809


CL\_2809

HighlightSelectShow Genomes


175

CL\_2810


90

CL\_2810


3

CL\_2810


2

CL\_2810


2

CL\_2810


1

CL\_2810


1

CL\_2810


1

CL\_2810


1

CL\_2811


1

CL\_2810

fGI ID


CL\_INS\_230
CL\_INS\_230
CL\_INS\_230
CL\_INS\_230
CL\_INS\_230
CL\_INS\_230
Cluster ID


CL\_15370
CL\_4845
CL\_4846
CL\_4847
CL\_4848
CL\_4849
