## Supplementary material for "A novel method for integrating genomic and Tn-Seq data to identify common *in vivo* fitness mechanisms across multiple bacterial species": S1 Dataset: CL_INS_231.html

Legend

 Mobile +extrachromosomalelementfunctions
 Hypothetical
 All EssentialGenes
 Proteinsynthesis/fate
 Other
 Transport +binding proteins
 All VFDB Genes

FULL


WINDOWSVGPNG

Trim RowsRemove SingletonsSave Fasta

CL\_2815


CL\_2815


CL\_2815


CL\_2815


CL\_2815


CL\_2815


CL\_2815


CL\_2815


CL\_2815


CL\_2813


CL\_2815


CL\_2815


Break


CL\_2814


CL\_2815

HighlightSelectShow Genomes


155

CL\_2816


87

CL\_2816


20

CL\_2816


2

CL\_2816


1

CL\_2816


1

Break


1

CL\_2816


1

CL\_2816


1

CL\_2816


1

CL\_2816


1

CL\_2816


1

CL\_2817


1

CL\_2816


1

CL\_2816


1

CL\_2816

fGI ID


CL\_INS\_231
CL\_INS\_231
CL\_INS\_231
CL\_INS\_231
CL\_INS\_231
CL\_INS\_237
CL\_INS\_70
CL\_INS\_237
CL\_INS\_237
CL\_INS\_237
CL\_INS\_70
CL\_INS\_70
CL\_INS\_70
CL\_INS\_237
CL\_INS\_237
CL\_INS\_237
CL\_INS\_231
CL\_INS\_382
CL\_INS\_382
CL\_INS\_231
CL\_INS\_231
CL\_INS\_231
CL\_INS\_231
CL\_INS\_232
Cluster ID


CL\_30770
CL\_8536
CL\_17009
CL\_16906
CL\_16905
CL\_7213
CL\_7214
CL\_8517
CL\_8518
CL\_13378
CL\_7673
CL\_14683
CL\_7234
CL\_16459
CL\_16458
CL\_8780
CL\_16904
CL\_5341
CL\_6452
CL\_7308
CL\_34000
CL\_30503
CL\_21684
CL\_15371
