## Supplementary material for "A novel method for integrating genomic and Tn-Seq data to identify common *in vivo* fitness mechanisms across multiple bacterial species": S1 Dataset: CL_INS_232.html

Legend

 Mobile +extrachromosomalelementfunctions
 Hypothetical
 Proteinsynthesis/fate
 Other
 EnergyMetabolism
 Transport +binding proteins
 All VFDB Genes

FULL


WINDOWSVGPNG

Trim RowsRemove SingletonsSave Fasta

CL\_2817


CL\_2817


CL\_2817


CL\_2817


CL\_2817


CL\_2817


CL\_2817


CL\_2817


CL\_2817


CL\_2817

HighlightSelectShow Genomes


227

CL\_2818


30

CL\_2818


13

CL\_2818


2

CL\_2818


2

CL\_2818


1

CL\_2818


1

CL\_2818


1

CL\_2818


1

CL\_2819


1

CL\_2818

fGI ID


CL\_INS\_232
CL\_INS\_232
CL\_INS\_232
CL\_INS\_232
CL\_INS\_232
CL\_INS\_232
CL\_INS\_232
CL\_INS\_232
CL\_INS\_232
CL\_INS\_232
CL\_INS\_247
Cluster ID


CL\_8819
CL\_8535
CL\_15371
CL\_7398
CL\_7399
CL\_7400
CL\_7401
CL\_7402
CL\_7403
CL\_7404
CL\_6844
