## Supplementary material for "A novel method for integrating genomic and Tn-Seq data to identify common *in vivo* fitness mechanisms across multiple bacterial species": S1 Dataset: CL_INS_233.html

Legend

 Mobile +extrachromosomalelementfunctions
 Regulatoryfunctions
 Hypothetical
 All Fitness Genes
 Proteinsynthesis/fate
 Other
 Transport +binding proteins
 All VFDB Genes

FULL


WINDOWSVGPNG

Trim RowsRemove SingletonsSave Fasta

CL\_2818


CL\_2818


CL\_2818


CL\_2852


CL\_2818


CL\_2818


CL\_2818


CL\_2817

HighlightSelectShow Genomes


184

CL\_2819


87

CL\_2819


3

CL\_2820


1

CL\_2819


1

CL\_2846


1

CL\_2819


1

Break


1

CL\_2819

fGI ID


CL\_INS\_233
CL\_INS\_233
CL\_INS\_233
CL\_INS\_233
CL\_INS\_233
CL\_INS\_233
CL\_INS\_233
CL\_INS\_247
CL\_INS\_247
CL\_INS\_247
CL\_INS\_233
CL\_INS\_233
CL\_INS\_233
CL\_INS\_233
CL\_INS\_233
CL\_INS\_233
CL\_INS\_233
CL\_INS\_233
CL\_INS\_233
CL\_INS\_233
CL\_INS\_233
CL\_INS\_233
CL\_INS\_233
CL\_INS\_233
CL\_INS\_233
CL\_INS\_233
CL\_INS\_233
CL\_INS\_233
CL\_INS\_233
CL\_INS\_233
CL\_INS\_233
CL\_INS\_233
CL\_INS\_233
CL\_INS\_233
CL\_INS\_233
CL\_INS\_233
CL\_INS\_233
CL\_INS\_233
CL\_INS\_233
CL\_INS\_233
CL\_INS\_233
CL\_INS\_30
CL\_INS\_233
CL\_INS\_233
CL\_INS\_233
CL\_INS\_233
CL\_INS\_247
CL\_INS\_233
CL\_INS\_233
CL\_INS\_233
CL\_INS\_233
CL\_INS\_233
CL\_INS\_233
CL\_INS\_233
CL\_INS\_233
CL\_INS\_233
CL\_INS\_233
CL\_INS\_233
CL\_INS\_233
CL\_INS\_233
CL\_INS\_233
CL\_INS\_233
CL\_INS\_233
CL\_INS\_233
CL\_INS\_233
CL\_INS\_382
CL\_INS\_382
CL\_INS\_233
CL\_INS\_233
CL\_INS\_233
CL\_INS\_233
CL\_INS\_233
CL\_INS\_382
CL\_INS\_382
CL\_INS\_233
CL\_INS\_233
CL\_INS\_233
CL\_INS\_233
CL\_INS\_382
CL\_INS\_382
CL\_INS\_233
CL\_INS\_70
CL\_INS\_70
CL\_INS\_70
CL\_INS\_233
CL\_INS\_233
CL\_INS\_233
CL\_INS\_233
CL\_INS\_233
CL\_INS\_233
CL\_INS\_233
CL\_INS\_382
CL\_INS\_382
CL\_INS\_382
CL\_INS\_382
CL\_INS\_233
CL\_INS\_233
CL\_INS\_233
CL\_INS\_233
CL\_INS\_233
CL\_INS\_233
CL\_INS\_233
CL\_INS\_233
CL\_INS\_382
CL\_INS\_382
CL\_INS\_382
CL\_INS\_382
CL\_INS\_233
CL\_INS\_233
CL\_INS\_233
CL\_INS\_233
CL\_INS\_233
CL\_INS\_233
CL\_INS\_233
CL\_INS\_382
CL\_INS\_233
CL\_INS\_233
CL\_INS\_233
CL\_INS\_233
CL\_INS\_247
Cluster ID


CL\_30771
CL\_4850
CL\_28603
CL\_28604
CL\_28605
CL\_28606
CL\_7325
CL\_4840
CL\_5947
CL\_4841
CL\_12610
CL\_12611
CL\_12612
CL\_12613
CL\_12614
CL\_10578
CL\_12615
CL\_12616
CL\_12617
CL\_12618
CL\_12619
CL\_12620
CL\_12621
CL\_12622
CL\_12623
CL\_12624
CL\_12625
CL\_12626
CL\_12627
CL\_12628
CL\_12629
CL\_12630
CL\_12631
CL\_12632
CL\_12633
CL\_12634
CL\_12635
CL\_12636
CL\_12637
CL\_12638
CL\_12639
CL\_5237
CL\_12640
CL\_12641
CL\_12642
CL\_9369
CL\_9370
CL\_12643
CL\_12644
CL\_12645
CL\_12646
CL\_12647
CL\_12648
CL\_12649
CL\_12650
CL\_12651
CL\_12652
CL\_12653
CL\_12654
CL\_12655
CL\_12656
CL\_12657
CL\_12658
CL\_12659
CL\_4309
CL\_5600
CL\_5599
CL\_5541
CL\_6935
CL\_5542
CL\_5543
CL\_5544
CL\_4302
CL\_4301
CL\_5545
CL\_12660
CL\_12661
CL\_12662
CL\_5662
CL\_5548
CL\_12663
CL\_8749
CL\_8748
CL\_8747
CL\_12664
CL\_10801
CL\_12665
CL\_12666
CL\_12667
CL\_12668
CL\_12669
CL\_4236
CL\_4235
CL\_5015
CL\_5014
CL\_12670
CL\_12671
CL\_12672
CL\_12673
CL\_12674
CL\_12675
CL\_12676
CL\_12677
CL\_4253
CL\_4251
CL\_6927
CL\_6926
CL\_12678
CL\_12679
CL\_12680
CL\_12681
CL\_12682
CL\_12683
CL\_12684
CL\_10807
CL\_12685
CL\_12686
CL\_12687
CL\_12688
CL\_4838
