## Supplementary material for "A novel method for integrating genomic and Tn-Seq data to identify common *in vivo* fitness mechanisms across multiple bacterial species": S1 Dataset: CL_INS_236.html

Legend

 Mobile +extrachromosomalelementfunctions
 Hypothetical
 All EssentialGenes
 Cell Envelope
 Other
 All VFDB Genes

FULL


WINDOWSVGPNG

Trim RowsRemove SingletonsSave Fasta

CL\_2835


CL\_2835


CL\_2835


CL\_2835


CL\_2835


CL\_2835


CL\_2835


CL\_2835


CL\_2835


CL\_2835


CL\_2835


CL\_2835


CL\_2835


CL\_2835


CL\_2835


CL\_234

HighlightSelectShow Genomes


177

CL\_2836


81

CL\_2836


2

CL\_2836


1

CL\_2836


1

CL\_2836


1

CL\_2836


1

CL\_2836


1

CL\_2837


1

CL\_2846


1

CL\_2837


1

CL\_2836


1

CL\_2837


1

CL\_2836


1

CL\_2836


1

CL\_2836


1

CL\_2836

fGI ID


CL\_INS\_236
CL\_INS\_236
CL\_INS\_236
CL\_INS\_236
CL\_INS\_236
CL\_INS\_237
CL\_INS\_236
CL\_INS\_236
CL\_INS\_382
CL\_INS\_382
CL\_INS\_382
CL\_INS\_123
CL\_INS\_237
CL\_INS\_382
CL\_INS\_70
CL\_INS\_70
CL\_INS\_70
CL\_INS\_237
CL\_INS\_237
CL\_INS\_237
CL\_INS\_237
CL\_INS\_237
CL\_INS\_237
CL\_INS\_70
CL\_INS\_70
CL\_INS\_237
CL\_INS\_70
CL\_INS\_70
CL\_INS\_70
CL\_INS\_70
CL\_INS\_30
CL\_INS\_30
CL\_INS\_237
CL\_INS\_236
CL\_INS\_30
CL\_INS\_30
CL\_INS\_159
CL\_INS\_30
CL\_INS\_159
CL\_INS\_30
CL\_INS\_30
CL\_INS\_30
CL\_INS\_236
CL\_INS\_236
CL\_INS\_237
CL\_INS\_236
CL\_INS\_236
CL\_INS\_236
CL\_INS\_156
CL\_INS\_156
CL\_INS\_156
CL\_INS\_156
CL\_INS\_156
CL\_INS\_156
CL\_INS\_156
CL\_INS\_156
CL\_INS\_156
CL\_INS\_236
CL\_INS\_236
CL\_INS\_237
CL\_INS\_237
CL\_INS\_237
CL\_INS\_236
CL\_INS\_236
CL\_INS\_236
CL\_INS\_236
CL\_INS\_236
Cluster ID


CL\_22872
CL\_27878
CL\_12843
CL\_12842
CL\_8360
CL\_8361
CL\_13590
CL\_30502
CL\_8832
CL\_7691
CL\_7692
CL\_4995
CL\_7297
CL\_8376
CL\_8779
CL\_7298
CL\_7299
CL\_7300
CL\_6244
CL\_6245
CL\_6246
CL\_5147
CL\_5148
CL\_6417
CL\_8642
CL\_13552
CL\_7831
CL\_1933
CL\_1932
CL\_1931
CL\_5233
CL\_5234
CL\_7087
CL\_16466
CL\_5240
CL\_5241
CL\_5242
CL\_5243
CL\_5244
CL\_5245
CL\_5246
CL\_5247
CL\_16465
CL\_16464
CL\_11715
CL\_4985
CL\_16463
CL\_4986
CL\_4987
CL\_4988
CL\_4989
CL\_4990
CL\_16462
CL\_4992
CL\_4993
CL\_4994
CL\_15643
CL\_16461
CL\_16460
CL\_16459
CL\_16458
CL\_8780
CL\_16457
CL\_16456
CL\_12841
CL\_8362
CL\_12128
