## Supplementary material for "A novel method for integrating genomic and Tn-Seq data to identify common *in vivo* fitness mechanisms across multiple bacterial species": S1 Dataset: CL_INS_237.html

Legend

 Mobile +extrachromosomalelementfunctions
 Regulatoryfunctions
 Hypothetical
 DNA Metabolism
 AntibioticResistance
 All EssentialGenes
 Biosynthesis ofcofactors,prostheticgroups, +carriers
 All Fitness Genes
 Cell Envelope
 Proteinsynthesis/fate
 Other
 Centralintermediarymetabolism
 EnergyMetabolism
 Transport +binding proteins
 All VFDB Genes

FULL


WINDOWSVGPNG

Trim RowsRemove SingletonsSave Fasta

CL\_2846


CL\_1935


CL\_2846


CL\_2846


CL\_2846


CL\_1935


CL\_2846


CL\_2846


CL\_2846


CL\_2846


CL\_1935


CL\_1935


CL\_234


CL\_2846


CL\_2846


CL\_234


CL\_2846


CL\_2846


CL\_2846


CL\_2846


CL\_234


CL\_1935


CL\_2846


CL\_2846


CL\_2846


CL\_2846


CL\_2846


CL\_2846


CL\_2846


CL\_2846


CL\_2846


CL\_2846


CL\_2846


CL\_2846


CL\_234


CL\_2846


CL\_1935


CL\_2846


CL\_2846


CL\_1935


CL\_2846


CL\_2845


CL\_2846


CL\_2846


CL\_2846


CL\_2846


CL\_2846


CL\_1935


CL\_2846


CL\_2846


CL\_2846


CL\_2846


CL\_2846


CL\_2846


CL\_2846


CL\_2846


CL\_2846


CL\_1935


CL\_2846


CL\_2846


CL\_1935


CL\_2846


CL\_1935


CL\_2846


CL\_2846


CL\_4118


CL\_2846


CL\_2846


CL\_2846


CL\_2846


CL\_2846


CL\_1935


CL\_2846


CL\_2846


CL\_2846


CL\_2846


CL\_2846


CL\_2846


CL\_2846


CL\_2846


CL\_234


CL\_234


CL\_2846


CL\_234


CL\_1935


CL\_2845


CL\_2846


CL\_2846


CL\_234


CL\_2846


CL\_2846


CL\_2846


CL\_2846


CL\_2846


CL\_2846


CL\_1935


CL\_2846


CL\_1935


CL\_2846


CL\_2846


CL\_2846


CL\_2846


CL\_2846


CL\_2846


CL\_2846


CL\_2846


CL\_2846


CL\_2846


CL\_2846


CL\_2846


CL\_2846


CL\_2846


CL\_1935


CL\_2846


CL\_2846


CL\_2846


CL\_2846


CL\_2846


CL\_2846


CL\_2846


CL\_2846


CL\_234


Break


CL\_2846


CL\_2846


CL\_2846


CL\_2846


CL\_2846


CL\_2846


CL\_2846


CL\_2846


CL\_2846


CL\_2846


CL\_2846


CL\_2846


CL\_2846


CL\_2846


CL\_234


CL\_2846


CL\_2846


CL\_2846


CL\_2846


CL\_2846


CL\_2846


CL\_2846


CL\_1935


CL\_2846


CL\_1935


CL\_2846


CL\_1935


CL\_1935


CL\_2846


CL\_234


CL\_1935


CL\_2846


CL\_2846


CL\_2846


CL\_1935


CL\_2846


CL\_2846


CL\_2846


CL\_1935


CL\_2846


CL\_2846


CL\_1942


CL\_2846


CL\_1935


CL\_2846


CL\_2008


CL\_2846


CL\_2846


CL\_2846


CL\_2846


CL\_234


CL\_2846


CL\_2846


CL\_234


CL\_2846


CL\_2846


CL\_1935


CL\_2846


CL\_1935


CL\_2846


CL\_234


CL\_2846


CL\_2846


CL\_2846


CL\_2846


CL\_2846


CL\_2846


CL\_2846


CL\_2846


CL\_2846


CL\_2846


CL\_2846


CL\_2846


CL\_1935


CL\_2846


CL\_2846


CL\_2846


CL\_2846


CL\_2846


CL\_2846


CL\_2846


CL\_234


CL\_2846


CL\_323


CL\_2846


CL\_2846


CL\_1935


CL\_2846


CL\_2846


CL\_2846


CL\_2846


CL\_1935


CL\_1935


CL\_2846


CL\_2846


CL\_2846


CL\_2846


CL\_2846


CL\_2846


CL\_2846


CL\_2846


CL\_234


CL\_2846


CL\_2846


CL\_2846


CL\_2846


CL\_234


CL\_4119


CL\_2846


CL\_2846


CL\_2846


CL\_2846


CL\_2846


CL\_2846


CL\_2846

HighlightSelectShow Genomes


72

CL\_2847


9

CL\_2847


6

CL\_2847


4

CL\_2866


4

CL\_2145


3

CL\_2847


2

CL\_2853


2

CL\_2847


2

CL\_1935


2

CL\_2866


2

CL\_2847


2

CL\_2847


2

CL\_2847


1

CL\_234


1

CL\_2847


1

CL\_2847


1

CL\_2866


1

CL\_4861


1

CL\_2852


1

CL\_2847


1

CL\_2847


1

CL\_2847


1

CL\_234


1

CL\_2847


1

CL\_2866


1

CL\_2847


1

CL\_1935


1

CL\_2847


1

CL\_234


1

CL\_234


1

CL\_2847


1

CL\_2847


1

CL\_1442


1

CL\_2847


1

CL\_2847


1

CL\_203


1

CL\_2847


1

CL\_2847


1

CL\_2847


1

CL\_2847


1

CL\_2847


1

CL\_2847


1

CL\_2853


1

CL\_2847


1

CL\_1935


1

CL\_2847


1

CL\_2847


1

CL\_2847


1

CL\_2853


1

CL\_234


1

CL\_4853


1

CL\_2847


1

CL\_234


1

CL\_2921


1

CL\_2847


1

CL\_2866


1

CL\_1935


1

CL\_2847


1

CL\_2847


1

CL\_2847


1

CL\_2847


1

CL\_323


1

CL\_2847


1

CL\_234


1

CL\_234


1

CL\_2847


1

CL\_2847


1

CL\_1935


1

CL\_234


1

CL\_2847


1

CL\_234


1

CL\_2847


1

CL\_2847


1

CL\_2145


1

CL\_2865


1

CL\_2853


1

CL\_1935


1

CL\_2866


1

CL\_2847


1

CL\_2847


1

CL\_2847


1

CL\_2847


1

CL\_2847


1

CL\_2847


1

CL\_2847


1

CL\_2847


1

CL\_2835


1

CL\_2847


1

CL\_2847


1

CL\_234


1

CL\_2847


1

CL\_234


1

CL\_2866


1

CL\_2887


1

CL\_2847


1

CL\_2847


1

CL\_234


1

CL\_2847


1

CL\_1935


1

CL\_1935


1

CL\_2847


1

CL\_2847


1

CL\_2847


1

CL\_2847


1

CL\_2847


1

CL\_2847


1

CL\_2847


1

CL\_234


1

CL\_2813


1

CL\_2847


1

CL\_234


1

CL\_1935


1

CL\_2847


1

CL\_234


1

CL\_2847


1

CL\_2847


1

CL\_234


1

CL\_1935


1

CL\_2847


1

CL\_234


1

CL\_234


1

CL\_2847


1

CL\_2847


1

CL\_2847


1

CL\_234


1

CL\_2847


1

CL\_1935


1

CL\_2847


1

CL\_2847


1

CL\_4863


1

CL\_2847


1

CL\_2866


1

CL\_2818


1

CL\_234


1

CL\_2847


1

CL\_2847


1

CL\_2862


1

CL\_2847


1

CL\_234


1

CL\_2847


1

CL\_2847


1

CL\_2847


1

CL\_2847


1

CL\_2847


1

CL\_234


1

CL\_2847


1

CL\_2847


1

CL\_2847


1

CL\_2847


1

CL\_2847


1

CL\_2847


1

CL\_2847


1

CL\_2847


1

CL\_2847


1

CL\_234


1

CL\_234


1

CL\_2847


1

CL\_2847


1

CL\_2852


1

CL\_1935


1

CL\_2847


1

CL\_2847


1

CL\_234


1

CL\_2852


1

CL\_2847


1

CL\_234


1

CL\_2847


1

CL\_2852


1

CL\_2847


1

CL\_234


1

CL\_2852


1

CL\_234


1

CL\_2852


1

CL\_2847


1

CL\_2847


1

CL\_234


1

CL\_2847


1

CL\_2847


1

CL\_2847


1

CL\_2847


1

CL\_2847


1

CL\_2847


1

CL\_2852


1

CL\_2847


1

CL\_2847


1

CL\_2847


1

CL\_2847


1

CL\_2847


1

CL\_2847


1

CL\_1935


1

CL\_2847


1

CL\_234


1

CL\_234


1

CL\_2847


1

CL\_4856


1

CL\_2847


1

CL\_2847


1

CL\_1935


1

Break


1

CL\_2847


1

CL\_2847


1

CL\_2852


1

CL\_2866


1

CL\_2847


1

CL\_2847


1

CL\_2847


1

CL\_2847


1

CL\_1935


1

CL\_2862


1

CL\_2847


1

CL\_1935


1

CL\_1935


1

CL\_2847


1

CL\_2847


1

CL\_2847


1

CL\_2847


1

CL\_2847


1

CL\_234


1

CL\_2879


1

CL\_1935


1

CL\_3958


1

CL\_2847


1

CL\_2853


1

CL\_2847


1

CL\_2847


1

CL\_234


1

CL\_234


1

CL\_2847


1

CL\_2847


1

CL\_2847


1

CL\_2847


1

CL\_2847


1

CL\_234


1

CL\_2866


1

CL\_4119


1

CL\_2866


1

CL\_2853


1

CL\_2852

fGI ID


CL\_INS\_237
CL\_INS\_237
CL\_INS\_233
CL\_INS\_233
CL\_INS\_233
CL\_INS\_233
CL\_INS\_155
CL\_INS\_237
CL\_INS\_237
CL\_INS\_155
CL\_INS\_382
CL\_INS\_382
CL\_INS\_382
CL\_INS\_382
CL\_INS\_237
CL\_INS\_237
CL\_INS\_237
CL\_INS\_237
CL\_INS\_237
CL\_INS\_237
CL\_INS\_237
CL\_INS\_237
CL\_INS\_237
CL\_INS\_237
CL\_INS\_237
CL\_INS\_237
CL\_INS\_237
CL\_INS\_382
CL\_INS\_237
CL\_INS\_382
CL\_INS\_382
CL\_INS\_382
CL\_INS\_382
CL\_INS\_382
CL\_INS\_382
CL\_INS\_382
CL\_INS\_382
CL\_INS\_237
CL\_INS\_237
CL\_INS\_382
CL\_INS\_382
CL\_INS\_382
CL\_INS\_237
CL\_INS\_237
CL\_INS\_237
CL\_INS\_237
CL\_INS\_237
CL\_INS\_382
CL\_INS\_382
CL\_INS\_382
CL\_INS\_237
CL\_INS\_237
CL\_INS\_237
CL\_INS\_237
CL\_INS\_237
CL\_INS\_237
CL\_INS\_237
CL\_INS\_237
CL\_INS\_237
CL\_INS\_237
CL\_INS\_237
CL\_INS\_237
CL\_INS\_237
CL\_INS\_237
CL\_INS\_237
CL\_INS\_237
CL\_INS\_237
CL\_INS\_237
CL\_INS\_237
CL\_INS\_237
CL\_INS\_237
CL\_INS\_237
CL\_INS\_237
CL\_INS\_237
CL\_INS\_237
CL\_INS\_237
CL\_INS\_237
CL\_INS\_237
CL\_INS\_237
CL\_INS\_237
CL\_INS\_237
CL\_INS\_237
CL\_INS\_237
CL\_INS\_237
CL\_INS\_237
CL\_INS\_237
CL\_INS\_237
CL\_INS\_237
CL\_INS\_237
CL\_INS\_237
CL\_INS\_237
CL\_INS\_237
CL\_INS\_237
CL\_INS\_237
CL\_INS\_237
CL\_INS\_237
CL\_INS\_237
CL\_INS\_237
CL\_INS\_237
CL\_INS\_237
CL\_INS\_237
CL\_INS\_237
CL\_INS\_237
CL\_INS\_237
CL\_INS\_237
CL\_INS\_237
CL\_INS\_237
CL\_INS\_237
CL\_INS\_237
CL\_INS\_237
CL\_INS\_237
CL\_INS\_237
CL\_INS\_237
CL\_INS\_237
CL\_INS\_237
CL\_INS\_237
CL\_INS\_237
CL\_INS\_237
CL\_INS\_237
CL\_INS\_237
CL\_INS\_237
CL\_INS\_237
CL\_INS\_237
CL\_INS\_237
CL\_INS\_237
CL\_INS\_237
CL\_INS\_237
CL\_INS\_237
CL\_INS\_237
CL\_INS\_237
CL\_INS\_237
CL\_INS\_237
CL\_INS\_237
CL\_INS\_237
CL\_INS\_237
CL\_INS\_237
CL\_INS\_237
CL\_INS\_237
CL\_INS\_237
CL\_INS\_237
CL\_INS\_237
CL\_INS\_237
CL\_INS\_237
CL\_INS\_237
CL\_INS\_237
CL\_INS\_237
CL\_INS\_382
CL\_INS\_382
CL\_INS\_382
CL\_INS\_382
CL\_INS\_237
CL\_INS\_247
CL\_INS\_382
CL\_INS\_382
CL\_INS\_382
CL\_INS\_237
CL\_INS\_237
CL\_INS\_237
CL\_INS\_237
CL\_INS\_237
CL\_INS\_247
CL\_INS\_237
CL\_INS\_382
CL\_INS\_382
CL\_INS\_237
CL\_INS\_382
CL\_INS\_237
CL\_INS\_382
CL\_INS\_237
CL\_INS\_382
CL\_INS\_382
CL\_INS\_382
CL\_INS\_382
CL\_INS\_382
CL\_INS\_382
CL\_INS\_382
CL\_INS\_237
CL\_INS\_382
CL\_INS\_382
CL\_INS\_237
CL\_INS\_382
CL\_INS\_247
CL\_INS\_382
CL\_INS\_382
CL\_INS\_382
CL\_INS\_382
CL\_INS\_382
CL\_INS\_382
CL\_INS\_382
CL\_INS\_237
CL\_INS\_237
CL\_INS\_237
CL\_INS\_237
CL\_INS\_237
CL\_INS\_237
CL\_INS\_237
CL\_INS\_237
CL\_INS\_237
CL\_INS\_237
CL\_INS\_237
CL\_INS\_237
CL\_INS\_237
CL\_INS\_237
CL\_INS\_237
CL\_INS\_237
CL\_INS\_237
CL\_INS\_237
CL\_INS\_237
CL\_INS\_237
CL\_INS\_237
CL\_INS\_237
CL\_INS\_237
CL\_INS\_237
CL\_INS\_237
CL\_INS\_237
CL\_INS\_237
CL\_INS\_237
CL\_INS\_237
CL\_INS\_237
CL\_INS\_237
CL\_INS\_237
CL\_INS\_237
CL\_INS\_237
CL\_INS\_237
CL\_INS\_237
CL\_INS\_237
CL\_INS\_237
CL\_INS\_237
CL\_INS\_237
CL\_INS\_237
CL\_INS\_237
CL\_INS\_237
CL\_INS\_237
CL\_INS\_237
CL\_INS\_237
CL\_INS\_237
CL\_INS\_237
CL\_INS\_237
CL\_INS\_237
CL\_INS\_237
CL\_INS\_237
CL\_INS\_237
CL\_INS\_237
CL\_INS\_237
CL\_INS\_237
CL\_INS\_237
CL\_INS\_237
CL\_INS\_237
CL\_INS\_237
CL\_INS\_237
CL\_INS\_237
CL\_INS\_237
CL\_INS\_237
CL\_INS\_237
CL\_INS\_237
CL\_INS\_237
CL\_INS\_237
CL\_INS\_237
CL\_INS\_123
CL\_INS\_237
CL\_INS\_237
CL\_INS\_237
CL\_INS\_237
CL\_INS\_237
CL\_INS\_237
CL\_INS\_237
CL\_INS\_237
CL\_INS\_237
CL\_INS\_237
CL\_INS\_123
CL\_INS\_237
CL\_INS\_237
CL\_INS\_237
CL\_INS\_237
CL\_INS\_237
CL\_INS\_237
CL\_INS\_237
CL\_INS\_123
CL\_INS\_237
CL\_INS\_247
CL\_INS\_237
CL\_INS\_237
CL\_INS\_237
CL\_INS\_237
CL\_INS\_237
CL\_INS\_237
CL\_INS\_237
CL\_INS\_237
CL\_INS\_237
CL\_INS\_237
CL\_INS\_237
CL\_INS\_237
CL\_INS\_237
CL\_INS\_237
CL\_INS\_237
CL\_INS\_237
CL\_INS\_237
CL\_INS\_237
CL\_INS\_237
CL\_INS\_237
CL\_INS\_237
CL\_INS\_237
CL\_INS\_237
CL\_INS\_237
CL\_INS\_237
CL\_INS\_237
CL\_INS\_237
CL\_INS\_237
CL\_INS\_237
CL\_INS\_237
CL\_INS\_237
CL\_INS\_237
CL\_INS\_237
CL\_INS\_237
CL\_INS\_237
CL\_INS\_237
CL\_INS\_237
CL\_INS\_237
CL\_INS\_237
CL\_INS\_237
CL\_INS\_237
CL\_INS\_237
CL\_INS\_237
CL\_INS\_237
CL\_INS\_237
CL\_INS\_237
CL\_INS\_237
CL\_INS\_237
CL\_INS\_237
CL\_INS\_237
CL\_INS\_237
CL\_INS\_237
CL\_INS\_237
CL\_INS\_237
CL\_INS\_237
CL\_INS\_237
CL\_INS\_237
CL\_INS\_237
CL\_INS\_237
CL\_INS\_237
CL\_INS\_237
CL\_INS\_237
CL\_INS\_237
CL\_INS\_237
CL\_INS\_237
CL\_INS\_237
CL\_INS\_237
CL\_INS\_237
CL\_INS\_237
CL\_INS\_237
CL\_INS\_237
CL\_INS\_237
CL\_INS\_237
CL\_INS\_237
CL\_INS\_237
CL\_INS\_237
CL\_INS\_237
CL\_INS\_237
CL\_INS\_237
CL\_INS\_237
CL\_INS\_237
CL\_INS\_237
CL\_INS\_237
CL\_INS\_237
CL\_INS\_237
CL\_INS\_237
CL\_INS\_237
CL\_INS\_237
CL\_INS\_237
CL\_INS\_237
CL\_INS\_237
CL\_INS\_237
CL\_INS\_237
CL\_INS\_237
CL\_INS\_237
CL\_INS\_237
CL\_INS\_237
CL\_INS\_237
CL\_INS\_237
CL\_INS\_237
CL\_INS\_237
CL\_INS\_237
CL\_INS\_237
CL\_INS\_237
CL\_INS\_237
CL\_INS\_237
CL\_INS\_237
CL\_INS\_237
CL\_INS\_237
CL\_INS\_237
CL\_INS\_237
CL\_INS\_237
CL\_INS\_237
CL\_INS\_237
CL\_INS\_237
CL\_INS\_237
CL\_INS\_237
CL\_INS\_237
CL\_INS\_237
CL\_INS\_237
CL\_INS\_237
CL\_INS\_237
CL\_INS\_237
CL\_INS\_237
CL\_INS\_237
CL\_INS\_237
CL\_INS\_237
CL\_INS\_237
CL\_INS\_237
CL\_INS\_237
CL\_INS\_237
CL\_INS\_237
CL\_INS\_237
CL\_INS\_237
CL\_INS\_237
CL\_INS\_237
CL\_INS\_237
CL\_INS\_237
CL\_INS\_237
CL\_INS\_237
CL\_INS\_382
CL\_INS\_382
CL\_INS\_237
CL\_INS\_237
CL\_INS\_237
CL\_INS\_237
CL\_INS\_237
CL\_INS\_237
CL\_INS\_237
CL\_INS\_237
CL\_INS\_237
CL\_INS\_237
CL\_INS\_237
CL\_INS\_237
CL\_INS\_237
CL\_INS\_237
CL\_INS\_237
CL\_INS\_237
CL\_INS\_237
CL\_INS\_237
CL\_INS\_237
CL\_INS\_237
CL\_INS\_237
CL\_INS\_237
CL\_INS\_237
CL\_INS\_237
CL\_INS\_237
CL\_INS\_237
CL\_INS\_237
CL\_INS\_237
CL\_INS\_237
CL\_INS\_237
CL\_INS\_237
CL\_INS\_237
CL\_INS\_237
CL\_INS\_237
CL\_INS\_237
CL\_INS\_237
CL\_INS\_237
CL\_INS\_237
CL\_INS\_237
CL\_INS\_237
CL\_INS\_247
CL\_INS\_237
CL\_INS\_237
CL\_INS\_237
CL\_INS\_237
CL\_INS\_237
CL\_INS\_237
CL\_INS\_237
CL\_INS\_247
CL\_INS\_237
CL\_INS\_237
CL\_INS\_237
CL\_INS\_237
CL\_INS\_237
CL\_INS\_237
CL\_INS\_237
CL\_INS\_237
CL\_INS\_237
CL\_INS\_237
CL\_INS\_382
CL\_INS\_237
CL\_INS\_237
CL\_INS\_237
CL\_INS\_237
CL\_INS\_237
CL\_INS\_237
CL\_INS\_237
CL\_INS\_237
CL\_INS\_237
CL\_INS\_237
CL\_INS\_237
CL\_INS\_382
CL\_INS\_237
CL\_INS\_237
CL\_INS\_237
CL\_INS\_237
CL\_INS\_237
CL\_INS\_237
CL\_INS\_237
CL\_INS\_237
CL\_INS\_237
CL\_INS\_237
CL\_INS\_237
CL\_INS\_237
CL\_INS\_237
CL\_INS\_237
CL\_INS\_237
CL\_INS\_237
CL\_INS\_237
CL\_INS\_237
CL\_INS\_237
CL\_INS\_237
CL\_INS\_237
CL\_INS\_237
CL\_INS\_237
CL\_INS\_237
CL\_INS\_149
CL\_INS\_237
CL\_INS\_237
CL\_INS\_237
CL\_INS\_237
CL\_INS\_237
CL\_INS\_237
CL\_INS\_237
CL\_INS\_237
CL\_INS\_237
CL\_INS\_237
CL\_INS\_237
CL\_INS\_237
CL\_INS\_237
CL\_INS\_237
CL\_INS\_237
CL\_INS\_237
CL\_INS\_237
CL\_INS\_237
CL\_INS\_237
CL\_INS\_237
CL\_INS\_237
CL\_INS\_237
CL\_INS\_237
CL\_INS\_237
CL\_INS\_237
CL\_INS\_247
CL\_INS\_247
CL\_INS\_237
CL\_INS\_237
CL\_INS\_237
CL\_INS\_237
CL\_INS\_237
CL\_INS\_237
CL\_INS\_237
CL\_INS\_237
CL\_INS\_237
CL\_INS\_237
CL\_INS\_237
CL\_INS\_237
CL\_INS\_237
CL\_INS\_237
CL\_INS\_382
CL\_INS\_237
CL\_INS\_237
CL\_INS\_237
CL\_INS\_237
CL\_INS\_237
CL\_INS\_237
CL\_INS\_237
CL\_INS\_237
CL\_INS\_237
CL\_INS\_237
CL\_INS\_237
CL\_INS\_237
CL\_INS\_237
CL\_INS\_237
CL\_INS\_237
CL\_INS\_237
CL\_INS\_237
CL\_INS\_237
CL\_INS\_237
CL\_INS\_237
CL\_INS\_237
CL\_INS\_237
CL\_INS\_237
CL\_INS\_30
CL\_INS\_247
CL\_INS\_237
CL\_INS\_237
CL\_INS\_237
CL\_INS\_237
CL\_INS\_237
CL\_INS\_237
CL\_INS\_237
CL\_INS\_70
CL\_INS\_237
CL\_INS\_237
CL\_INS\_237
CL\_INS\_237
CL\_INS\_237
CL\_INS\_237
CL\_INS\_237
CL\_INS\_237
CL\_INS\_237
CL\_INS\_237
CL\_INS\_237
CL\_INS\_237
CL\_INS\_237
CL\_INS\_237
CL\_INS\_237
CL\_INS\_237
CL\_INS\_70
CL\_INS\_237
CL\_INS\_237
CL\_INS\_237
CL\_INS\_237
CL\_INS\_237
CL\_INS\_270
CL\_INS\_270
CL\_INS\_270
CL\_INS\_270
CL\_INS\_270
CL\_INS\_270
CL\_INS\_270
CL\_INS\_270
CL\_INS\_270
CL\_INS\_270
CL\_INS\_270
CL\_INS\_270
CL\_INS\_270
CL\_INS\_270
CL\_INS\_270
CL\_INS\_270
CL\_INS\_270
CL\_INS\_270
CL\_INS\_30
CL\_INS\_30
CL\_INS\_237
CL\_INS\_237
CL\_INS\_237
CL\_INS\_237
CL\_INS\_237
CL\_INS\_237
CL\_INS\_237
CL\_INS\_30
CL\_INS\_237
CL\_INS\_237
CL\_INS\_237
CL\_INS\_237
CL\_INS\_237
CL\_INS\_237
CL\_INS\_149
CL\_INS\_237
CL\_INS\_237
CL\_INS\_237
CL\_INS\_247
CL\_INS\_237
CL\_INS\_237
CL\_INS\_237
CL\_INS\_237
CL\_INS\_237
CL\_INS\_237
CL\_INS\_237
CL\_INS\_237
CL\_INS\_237
CL\_INS\_237
CL\_INS\_237
CL\_INS\_237
CL\_INS\_237
CL\_INS\_237
CL\_INS\_237
CL\_INS\_237
CL\_INS\_237
CL\_INS\_237
CL\_INS\_237
CL\_INS\_237
CL\_INS\_237
CL\_INS\_237
CL\_INS\_237
CL\_INS\_237
CL\_INS\_237
CL\_INS\_237
CL\_INS\_237
CL\_INS\_237
CL\_INS\_237
CL\_INS\_237
CL\_INS\_237
CL\_INS\_237
CL\_INS\_237
CL\_INS\_237
CL\_INS\_237
CL\_INS\_237
CL\_INS\_237
CL\_INS\_237
CL\_INS\_237
CL\_INS\_237
CL\_INS\_247
CL\_INS\_237
CL\_INS\_237
CL\_INS\_237
CL\_INS\_237
CL\_INS\_237
CL\_INS\_237
CL\_INS\_237
CL\_INS\_237
CL\_INS\_237
CL\_INS\_237
CL\_INS\_70
CL\_INS\_237
CL\_INS\_237
CL\_INS\_237
CL\_INS\_237
CL\_INS\_237
CL\_INS\_237
CL\_INS\_237
CL\_INS\_237
CL\_INS\_237
CL\_INS\_237
CL\_INS\_237
CL\_INS\_237
CL\_INS\_237
CL\_INS\_237
CL\_INS\_237
CL\_INS\_237
CL\_INS\_237
CL\_INS\_237
CL\_INS\_237
CL\_INS\_237
CL\_INS\_237
CL\_INS\_237
CL\_INS\_237
CL\_INS\_237
CL\_INS\_237
CL\_INS\_237
CL\_INS\_237
CL\_INS\_237
CL\_INS\_237
CL\_INS\_237
CL\_INS\_237
CL\_INS\_237
CL\_INS\_237
CL\_INS\_237
CL\_INS\_237
CL\_INS\_237
CL\_INS\_237
CL\_INS\_237
CL\_INS\_237
CL\_INS\_237
CL\_INS\_237
CL\_INS\_237
CL\_INS\_237
CL\_INS\_237
CL\_INS\_237
CL\_INS\_237
CL\_INS\_237
CL\_INS\_237
CL\_INS\_237
CL\_INS\_237
CL\_INS\_237
CL\_INS\_237
CL\_INS\_237
CL\_INS\_237
CL\_INS\_237
CL\_INS\_70
CL\_INS\_70
CL\_INS\_70
CL\_INS\_237
CL\_INS\_237
CL\_INS\_30
CL\_INS\_237
CL\_INS\_237
CL\_INS\_237
CL\_INS\_70
CL\_INS\_70
CL\_INS\_70
CL\_INS\_237
CL\_INS\_237
CL\_INS\_237
CL\_INS\_237
CL\_INS\_237
CL\_INS\_237
CL\_INS\_237
CL\_INS\_237
CL\_INS\_237
CL\_INS\_237
CL\_INS\_237
CL\_INS\_237
CL\_INS\_237
CL\_INS\_237
CL\_INS\_237
CL\_INS\_237
CL\_INS\_237
CL\_INS\_237
CL\_INS\_237
CL\_INS\_70
CL\_INS\_70
CL\_INS\_247
CL\_INS\_70
CL\_INS\_70
CL\_INS\_237
CL\_INS\_237
CL\_INS\_237
CL\_INS\_237
CL\_INS\_237
CL\_INS\_70
CL\_INS\_237
CL\_INS\_247
CL\_INS\_204
CL\_INS\_237
CL\_INS\_237
CL\_INS\_247
CL\_INS\_247
CL\_INS\_247
CL\_INS\_237
CL\_INS\_237
CL\_INS\_237
CL\_INS\_237
CL\_INS\_237
CL\_INS\_237
CL\_INS\_237
CL\_INS\_237
CL\_INS\_237
CL\_INS\_237
CL\_INS\_237
CL\_INS\_237
CL\_INS\_237
CL\_INS\_237
CL\_INS\_237
CL\_INS\_237
CL\_INS\_237
CL\_INS\_237
CL\_INS\_237
CL\_INS\_237
CL\_INS\_237
CL\_INS\_237
CL\_INS\_30
CL\_INS\_237
CL\_INS\_237
CL\_INS\_237
CL\_INS\_237
CL\_INS\_237
CL\_INS\_237
CL\_INS\_237
CL\_INS\_237
CL\_INS\_237
CL\_INS\_382
CL\_INS\_237
CL\_INS\_382
CL\_INS\_237
CL\_INS\_382
CL\_INS\_382
CL\_INS\_237
CL\_INS\_237
CL\_INS\_237
CL\_INS\_237
CL\_INS\_237
CL\_INS\_237
CL\_INS\_237
CL\_INS\_237
CL\_INS\_237
CL\_INS\_237
CL\_INS\_237
CL\_INS\_237
CL\_INS\_70
CL\_INS\_237
CL\_INS\_237
CL\_INS\_70
CL\_INS\_237
CL\_INS\_237
CL\_INS\_237
CL\_INS\_237
CL\_INS\_237
CL\_INS\_237
CL\_INS\_237
CL\_INS\_237
CL\_INS\_237
CL\_INS\_70
CL\_INS\_70
CL\_INS\_247
CL\_INS\_247
CL\_INS\_247
CL\_INS\_247
CL\_INS\_247
CL\_INS\_247
CL\_INS\_247
CL\_INS\_247
CL\_INS\_247
CL\_INS\_247
CL\_INS\_247
CL\_INS\_247
CL\_INS\_247
CL\_INS\_247
CL\_INS\_247
CL\_INS\_237
CL\_INS\_237
CL\_INS\_237
CL\_INS\_247
CL\_INS\_247
CL\_INS\_247
CL\_INS\_247
CL\_INS\_237
CL\_INS\_237
CL\_INS\_237
CL\_INS\_237
CL\_INS\_237
CL\_INS\_237
CL\_INS\_237
CL\_INS\_237
CL\_INS\_237
CL\_INS\_237
CL\_INS\_237
CL\_INS\_237
CL\_INS\_237
CL\_INS\_237
CL\_INS\_237
CL\_INS\_237
CL\_INS\_237
CL\_INS\_237
CL\_INS\_237
CL\_INS\_237
CL\_INS\_237
CL\_INS\_237
CL\_INS\_237
CL\_INS\_237
CL\_INS\_237
CL\_INS\_237
CL\_INS\_237
CL\_INS\_237
CL\_INS\_237
CL\_INS\_237
CL\_INS\_237
CL\_INS\_237
CL\_INS\_237
CL\_INS\_237
CL\_INS\_237
CL\_INS\_237
CL\_INS\_237
CL\_INS\_237
CL\_INS\_237
CL\_INS\_237
CL\_INS\_237
CL\_INS\_237
CL\_INS\_237
CL\_INS\_237
CL\_INS\_237
CL\_INS\_237
CL\_INS\_237
CL\_INS\_237
CL\_INS\_237
CL\_INS\_237
CL\_INS\_237
CL\_INS\_237
CL\_INS\_237
CL\_INS\_237
CL\_INS\_237
CL\_INS\_247
CL\_INS\_247
CL\_INS\_247
CL\_INS\_247
CL\_INS\_237
CL\_INS\_237
CL\_INS\_237
CL\_INS\_237
CL\_INS\_237
CL\_INS\_237
CL\_INS\_237
CL\_INS\_237
CL\_INS\_237
CL\_INS\_237
CL\_INS\_237
CL\_INS\_237
CL\_INS\_237
CL\_INS\_237
CL\_INS\_237
CL\_INS\_237
CL\_INS\_237
CL\_INS\_237
CL\_INS\_237
CL\_INS\_237
CL\_INS\_237
CL\_INS\_237
CL\_INS\_237
CL\_INS\_237
CL\_INS\_237
CL\_INS\_237
CL\_INS\_237
CL\_INS\_237
CL\_INS\_237
CL\_INS\_237
CL\_INS\_382
CL\_INS\_247
CL\_INS\_237
CL\_INS\_57
CL\_INS\_149
CL\_INS\_149
CL\_INS\_149
CL\_INS\_149
CL\_INS\_149
CL\_INS\_149
CL\_INS\_57
CL\_INS\_57
CL\_INS\_382
CL\_INS\_237
CL\_INS\_237
CL\_INS\_237
CL\_INS\_237
CL\_INS\_237
CL\_INS\_237
CL\_INS\_237
CL\_INS\_237
CL\_INS\_237
CL\_INS\_237
CL\_INS\_237
CL\_INS\_237
CL\_INS\_237
CL\_INS\_237
CL\_INS\_237
CL\_INS\_237
CL\_INS\_237
CL\_INS\_237
CL\_INS\_237
CL\_INS\_247
CL\_INS\_237
CL\_INS\_237
CL\_INS\_123
CL\_INS\_237
CL\_INS\_237
CL\_INS\_237
CL\_INS\_237
CL\_INS\_237
CL\_INS\_237
CL\_INS\_237
CL\_INS\_237
CL\_INS\_237
CL\_INS\_237
CL\_INS\_237
CL\_INS\_237
CL\_INS\_237
CL\_INS\_237
CL\_INS\_237
CL\_INS\_237
CL\_INS\_237
CL\_INS\_382
CL\_INS\_237
CL\_INS\_237
CL\_INS\_237
CL\_INS\_237
CL\_INS\_237
CL\_INS\_70
CL\_INS\_237
CL\_INS\_237
CL\_INS\_237
CL\_INS\_70
CL\_INS\_237
CL\_INS\_237
CL\_INS\_70
CL\_INS\_237
CL\_INS\_237
CL\_INS\_237
CL\_INS\_237
CL\_INS\_237
CL\_INS\_237
CL\_INS\_237
CL\_INS\_352
CL\_INS\_237
CL\_INS\_149
CL\_INS\_237
CL\_INS\_237
CL\_INS\_237
CL\_INS\_70
CL\_INS\_237
CL\_INS\_70
CL\_INS\_247
CL\_INS\_70
CL\_INS\_237
CL\_INS\_70
CL\_INS\_70
CL\_INS\_70
CL\_INS\_237
CL\_INS\_237
CL\_INS\_237
CL\_INS\_237
CL\_INS\_237
CL\_INS\_237
CL\_INS\_237
CL\_INS\_382
CL\_INS\_382
CL\_INS\_382
CL\_INS\_382
CL\_INS\_382
CL\_INS\_237
CL\_INS\_237
CL\_INS\_237
CL\_INS\_237
CL\_INS\_237
CL\_INS\_237
CL\_INS\_237
CL\_INS\_237
CL\_INS\_237
CL\_INS\_237
CL\_INS\_237
CL\_INS\_237
CL\_INS\_237
CL\_INS\_237
CL\_INS\_237
CL\_INS\_237
CL\_INS\_237
CL\_INS\_237
CL\_INS\_237
CL\_INS\_237
CL\_INS\_237
CL\_INS\_382
CL\_INS\_382
CL\_INS\_70
CL\_INS\_237
CL\_INS\_70
CL\_INS\_70
CL\_INS\_237
CL\_INS\_237
CL\_INS\_237
CL\_INS\_70
CL\_INS\_237
CL\_INS\_237
CL\_INS\_237
CL\_INS\_237
CL\_INS\_237
CL\_INS\_237
CL\_INS\_237
CL\_INS\_237
CL\_INS\_237
CL\_INS\_247
CL\_INS\_237
CL\_INS\_237
CL\_INS\_70
CL\_INS\_382
CL\_INS\_382
CL\_INS\_382
CL\_INS\_70
CL\_INS\_237
CL\_INS\_237
CL\_INS\_237
CL\_INS\_237
CL\_INS\_237
CL\_INS\_237
CL\_INS\_237
CL\_INS\_70
CL\_INS\_237
CL\_INS\_237
CL\_INS\_237
CL\_INS\_237
CL\_INS\_237
CL\_INS\_237
CL\_INS\_237
CL\_INS\_237
CL\_INS\_237
CL\_INS\_237
CL\_INS\_237
CL\_INS\_237
CL\_INS\_237
CL\_INS\_237
CL\_INS\_382
CL\_INS\_237
CL\_INS\_237
CL\_INS\_237
CL\_INS\_237
CL\_INS\_237
CL\_INS\_237
CL\_INS\_237
CL\_INS\_237
CL\_INS\_237
CL\_INS\_237
CL\_INS\_237
CL\_INS\_237
CL\_INS\_237
CL\_INS\_237
CL\_INS\_237
CL\_INS\_237
CL\_INS\_237
CL\_INS\_237
CL\_INS\_237
CL\_INS\_237
CL\_INS\_237
CL\_INS\_237
CL\_INS\_237
CL\_INS\_237
CL\_INS\_149
CL\_INS\_237
CL\_INS\_237
CL\_INS\_237
CL\_INS\_237
CL\_INS\_237
CL\_INS\_237
CL\_INS\_237
CL\_INS\_237
CL\_INS\_237
CL\_INS\_237
CL\_INS\_237
CL\_INS\_237
CL\_INS\_237
CL\_INS\_237
CL\_INS\_237
CL\_INS\_237
CL\_INS\_237
CL\_INS\_237
CL\_INS\_237
CL\_INS\_237
CL\_INS\_237
CL\_INS\_237
CL\_INS\_237
CL\_INS\_237
CL\_INS\_237
CL\_INS\_237
CL\_INS\_237
CL\_INS\_237
CL\_INS\_237
CL\_INS\_237
CL\_INS\_237
CL\_INS\_237
CL\_INS\_237
CL\_INS\_237
CL\_INS\_237
CL\_INS\_237
CL\_INS\_237
CL\_INS\_237
CL\_INS\_237
CL\_INS\_237
CL\_INS\_237
CL\_INS\_237
CL\_INS\_237
CL\_INS\_237
CL\_INS\_237
CL\_INS\_237
CL\_INS\_237
CL\_INS\_237
CL\_INS\_237
CL\_INS\_237
CL\_INS\_237
CL\_INS\_237
CL\_INS\_237
CL\_INS\_70
CL\_INS\_70
CL\_INS\_70
CL\_INS\_70
CL\_INS\_70
CL\_INS\_70
CL\_INS\_237
CL\_INS\_237
CL\_INS\_237
CL\_INS\_237
CL\_INS\_237
CL\_INS\_237
CL\_INS\_237
CL\_INS\_237
CL\_INS\_237
CL\_INS\_237
CL\_INS\_237
CL\_INS\_237
CL\_INS\_382
CL\_INS\_237
CL\_INS\_237
CL\_INS\_237
CL\_INS\_237
CL\_INS\_237
CL\_INS\_237
CL\_INS\_237
CL\_INS\_247
CL\_INS\_70
CL\_INS\_70
CL\_INS\_70
CL\_INS\_70
CL\_INS\_70
CL\_INS\_70
CL\_INS\_237
CL\_INS\_237
CL\_INS\_237
CL\_INS\_237
CL\_INS\_237
CL\_INS\_237
CL\_INS\_237
CL\_INS\_237
CL\_INS\_237
CL\_INS\_237
CL\_INS\_237
CL\_INS\_237
CL\_INS\_237
CL\_INS\_237
CL\_INS\_237
CL\_INS\_237
CL\_INS\_237
CL\_INS\_237
CL\_INS\_237
CL\_INS\_30
CL\_INS\_30
CL\_INS\_237
CL\_INS\_237
CL\_INS\_70
CL\_INS\_237
CL\_INS\_237
CL\_INS\_237
CL\_INS\_237
CL\_INS\_237
CL\_INS\_237
CL\_INS\_237
CL\_INS\_237
CL\_INS\_237
CL\_INS\_30
CL\_INS\_237
CL\_INS\_30
CL\_INS\_237
CL\_INS\_237
CL\_INS\_237
CL\_INS\_70
CL\_INS\_237
CL\_INS\_30
CL\_INS\_30
CL\_INS\_237
CL\_INS\_237
CL\_INS\_237
CL\_INS\_237
CL\_INS\_237
CL\_INS\_237
CL\_INS\_70
CL\_INS\_70
CL\_INS\_382
CL\_INS\_237
CL\_INS\_237
CL\_INS\_237
CL\_INS\_237
CL\_INS\_247
CL\_INS\_30
CL\_INS\_30
CL\_INS\_30
CL\_INS\_30
CL\_INS\_30
CL\_INS\_382
CL\_INS\_237
CL\_INS\_237
CL\_INS\_237
CL\_INS\_237
CL\_INS\_237
CL\_INS\_237
CL\_INS\_237
CL\_INS\_237
CL\_INS\_70
CL\_INS\_237
CL\_INS\_237
CL\_INS\_70
CL\_INS\_237
CL\_INS\_237
CL\_INS\_237
CL\_INS\_237
CL\_INS\_247
CL\_INS\_382
CL\_INS\_382
CL\_INS\_237
CL\_INS\_237
CL\_INS\_382
CL\_INS\_237
CL\_INS\_237
CL\_INS\_237
CL\_INS\_237
CL\_INS\_237
CL\_INS\_237
CL\_INS\_237
CL\_INS\_237
CL\_INS\_237
CL\_INS\_237
CL\_INS\_237
CL\_INS\_237
CL\_INS\_237
CL\_INS\_237
CL\_INS\_237
CL\_INS\_70
CL\_INS\_237
CL\_INS\_70
CL\_INS\_70
CL\_INS\_237
CL\_INS\_237
CL\_INS\_237
CL\_INS\_237
CL\_INS\_237
CL\_INS\_237
CL\_INS\_237
CL\_INS\_237
CL\_INS\_149
CL\_INS\_237
CL\_INS\_237
CL\_INS\_237
CL\_INS\_237
CL\_INS\_237
CL\_INS\_237
CL\_INS\_237
CL\_INS\_237
CL\_INS\_237
CL\_INS\_237
CL\_INS\_237
CL\_INS\_237
CL\_INS\_237
CL\_INS\_237
CL\_INS\_237
CL\_INS\_237
CL\_INS\_237
CL\_INS\_237
CL\_INS\_237
CL\_INS\_87
CL\_INS\_237
CL\_INS\_237
CL\_INS\_237
CL\_INS\_237
CL\_INS\_237
CL\_INS\_237
CL\_INS\_237
CL\_INS\_237
CL\_INS\_237
CL\_INS\_30
CL\_INS\_237
CL\_INS\_237
CL\_INS\_237
CL\_INS\_237
CL\_INS\_237
CL\_INS\_237
CL\_INS\_237
CL\_INS\_237
CL\_INS\_30
CL\_INS\_30
CL\_INS\_237
CL\_INS\_237
CL\_INS\_237
CL\_INS\_352
CL\_INS\_237
CL\_INS\_237
CL\_INS\_237
CL\_INS\_237
CL\_INS\_237
CL\_INS\_237
CL\_INS\_237
CL\_INS\_237
CL\_INS\_352
CL\_INS\_352
CL\_INS\_237
CL\_INS\_352
CL\_INS\_237
CL\_INS\_352
CL\_INS\_237
CL\_INS\_237
CL\_INS\_237
CL\_INS\_237
CL\_INS\_237
CL\_INS\_237
CL\_INS\_237
CL\_INS\_237
CL\_INS\_237
CL\_INS\_237
CL\_INS\_237
CL\_INS\_237
CL\_INS\_237
CL\_INS\_237
CL\_INS\_237
CL\_INS\_237
CL\_INS\_237
CL\_INS\_237
CL\_INS\_237
CL\_INS\_237
CL\_INS\_237
CL\_INS\_237
CL\_INS\_237
CL\_INS\_237
CL\_INS\_237
CL\_INS\_237
CL\_INS\_155
CL\_INS\_237
CL\_INS\_237
CL\_INS\_237
CL\_INS\_237
CL\_INS\_237
CL\_INS\_237
CL\_INS\_237
CL\_INS\_237
CL\_INS\_237
CL\_INS\_237
CL\_INS\_237
CL\_INS\_237
CL\_INS\_237
CL\_INS\_237
CL\_INS\_237
CL\_INS\_237
CL\_INS\_237
CL\_INS\_237
CL\_INS\_237
CL\_INS\_237
CL\_INS\_237
CL\_INS\_237
CL\_INS\_70
CL\_INS\_70
CL\_INS\_352
CL\_INS\_352
CL\_INS\_352
CL\_INS\_237
CL\_INS\_237
CL\_INS\_237
CL\_INS\_237
CL\_INS\_237
CL\_INS\_70
CL\_INS\_237
CL\_INS\_237
CL\_INS\_237
CL\_INS\_237
CL\_INS\_237
CL\_INS\_247
CL\_INS\_247
CL\_INS\_237
CL\_INS\_237
CL\_INS\_237
CL\_INS\_237
CL\_INS\_237
CL\_INS\_237
CL\_INS\_237
CL\_INS\_237
CL\_INS\_237
CL\_INS\_237
CL\_INS\_237
CL\_INS\_237
CL\_INS\_237
CL\_INS\_237
CL\_INS\_237
CL\_INS\_237
CL\_INS\_237
CL\_INS\_237
CL\_INS\_237
CL\_INS\_237
CL\_INS\_237
CL\_INS\_237
CL\_INS\_237
CL\_INS\_237
CL\_INS\_237
CL\_INS\_237
CL\_INS\_237
CL\_INS\_237
CL\_INS\_247
CL\_INS\_247
CL\_INS\_237
CL\_INS\_237
CL\_INS\_233
CL\_INS\_237
CL\_INS\_237
CL\_INS\_237
CL\_INS\_237
CL\_INS\_237
CL\_INS\_237
CL\_INS\_247
CL\_INS\_70
CL\_INS\_237
CL\_INS\_237
CL\_INS\_237
CL\_INS\_237
CL\_INS\_237
CL\_INS\_237
CL\_INS\_237
CL\_INS\_70
CL\_INS\_237
CL\_INS\_237
CL\_INS\_237
CL\_INS\_237
CL\_INS\_237
CL\_INS\_237
CL\_INS\_237
CL\_INS\_237
CL\_INS\_237
CL\_INS\_237
CL\_INS\_237
CL\_INS\_237
CL\_INS\_237
CL\_INS\_237
CL\_INS\_237
CL\_INS\_237
CL\_INS\_237
CL\_INS\_237
CL\_INS\_237
CL\_INS\_237
CL\_INS\_237
CL\_INS\_237
CL\_INS\_237
CL\_INS\_237
CL\_INS\_247
CL\_INS\_237
CL\_INS\_237
CL\_INS\_237
CL\_INS\_237
CL\_INS\_237
CL\_INS\_237
CL\_INS\_237
CL\_INS\_237
CL\_INS\_237
CL\_INS\_237
CL\_INS\_237
CL\_INS\_237
CL\_INS\_237
CL\_INS\_237
CL\_INS\_237
CL\_INS\_237
CL\_INS\_237
CL\_INS\_237
CL\_INS\_237
CL\_INS\_237
CL\_INS\_237
CL\_INS\_237
CL\_INS\_70
CL\_INS\_233
CL\_INS\_237
CL\_INS\_237
CL\_INS\_237
CL\_INS\_247
CL\_INS\_237
CL\_INS\_247
CL\_INS\_237
CL\_INS\_237
CL\_INS\_237
CL\_INS\_237
CL\_INS\_237
CL\_INS\_237
CL\_INS\_237
CL\_INS\_237
CL\_INS\_237
CL\_INS\_237
CL\_INS\_70
CL\_INS\_70
CL\_INS\_70
CL\_INS\_70
CL\_INS\_70
CL\_INS\_70
CL\_INS\_70
CL\_INS\_70
CL\_INS\_70
CL\_INS\_70
CL\_INS\_70
CL\_INS\_70
CL\_INS\_155
CL\_INS\_237
CL\_INS\_237
CL\_INS\_237
CL\_INS\_237
CL\_INS\_237
CL\_INS\_237
CL\_INS\_237
CL\_INS\_237
CL\_INS\_237
CL\_INS\_237
CL\_INS\_237
CL\_INS\_237
CL\_INS\_237
CL\_INS\_237
CL\_INS\_237
CL\_INS\_237
CL\_INS\_237
CL\_INS\_237
CL\_INS\_237
CL\_INS\_237
CL\_INS\_237
CL\_INS\_237
CL\_INS\_237
CL\_INS\_237
CL\_INS\_237
CL\_INS\_237
CL\_INS\_237
CL\_INS\_237
CL\_INS\_237
CL\_INS\_237
CL\_INS\_237
CL\_INS\_237
CL\_INS\_237
CL\_INS\_237
CL\_INS\_237
CL\_INS\_123
CL\_INS\_237
CL\_INS\_237
CL\_INS\_237
CL\_INS\_237
CL\_INS\_237
CL\_INS\_237
CL\_INS\_237
CL\_INS\_237
CL\_INS\_237
CL\_INS\_70
CL\_INS\_70
CL\_INS\_247
CL\_INS\_237
CL\_INS\_237
CL\_INS\_237
CL\_INS\_237
CL\_INS\_237
CL\_INS\_237
CL\_INS\_237
CL\_INS\_237
CL\_INS\_382
CL\_INS\_237
CL\_INS\_237
CL\_INS\_237
CL\_INS\_237
CL\_INS\_237
CL\_INS\_237
CL\_INS\_237
CL\_INS\_70
CL\_INS\_70
CL\_INS\_237
CL\_INS\_237
CL\_INS\_237
CL\_INS\_237
CL\_INS\_237
CL\_INS\_237
CL\_INS\_237
CL\_INS\_237
CL\_INS\_237
CL\_INS\_237
CL\_INS\_237
CL\_INS\_237
CL\_INS\_237
CL\_INS\_237
CL\_INS\_237
CL\_INS\_237
CL\_INS\_237
CL\_INS\_237
CL\_INS\_237
CL\_INS\_237
CL\_INS\_237
CL\_INS\_237
CL\_INS\_237
CL\_INS\_237
CL\_INS\_237
CL\_INS\_237
CL\_INS\_237
CL\_INS\_237
CL\_INS\_237
CL\_INS\_237
CL\_INS\_237
CL\_INS\_237
CL\_INS\_237
CL\_INS\_237
CL\_INS\_237
CL\_INS\_237
CL\_INS\_237
CL\_INS\_237
CL\_INS\_237
CL\_INS\_237
CL\_INS\_237
CL\_INS\_237
CL\_INS\_237
CL\_INS\_237
CL\_INS\_237
CL\_INS\_237
CL\_INS\_237
CL\_INS\_237
CL\_INS\_237
CL\_INS\_237
CL\_INS\_237
CL\_INS\_237
CL\_INS\_237
CL\_INS\_237
CL\_INS\_237
CL\_INS\_237
CL\_INS\_237
CL\_INS\_237
CL\_INS\_237
CL\_INS\_237
CL\_INS\_237
CL\_INS\_237
CL\_INS\_237
CL\_INS\_382
CL\_INS\_237
CL\_INS\_70
CL\_INS\_237
CL\_INS\_237
CL\_INS\_237
CL\_INS\_237
CL\_INS\_237
CL\_INS\_237
CL\_INS\_237
CL\_INS\_237
CL\_INS\_237
CL\_INS\_237
CL\_INS\_237
CL\_INS\_237
CL\_INS\_247
CL\_INS\_237
CL\_INS\_237
CL\_INS\_247
CL\_INS\_237
CL\_INS\_237
CL\_INS\_237
CL\_INS\_237
CL\_INS\_237
CL\_INS\_70
CL\_INS\_70
CL\_INS\_237
CL\_INS\_237
CL\_INS\_237
CL\_INS\_237
CL\_INS\_237
CL\_INS\_70
CL\_INS\_70
CL\_INS\_70
CL\_INS\_70
CL\_INS\_70
CL\_INS\_237
CL\_INS\_70
CL\_INS\_237
CL\_INS\_237
CL\_INS\_237
CL\_INS\_70
CL\_INS\_30
CL\_INS\_237
CL\_INS\_237
CL\_INS\_237
CL\_INS\_237
CL\_INS\_237
CL\_INS\_237
CL\_INS\_237
CL\_INS\_237
CL\_INS\_237
CL\_INS\_237
CL\_INS\_30
CL\_INS\_30
CL\_INS\_237
CL\_INS\_237
CL\_INS\_30
CL\_INS\_30
CL\_INS\_237
CL\_INS\_237
CL\_INS\_237
CL\_INS\_237
CL\_INS\_237
CL\_INS\_237
CL\_INS\_237
CL\_INS\_237
CL\_INS\_237
CL\_INS\_70
CL\_INS\_70
CL\_INS\_237
CL\_INS\_70
CL\_INS\_237
CL\_INS\_237
CL\_INS\_237
CL\_INS\_30
CL\_INS\_237
CL\_INS\_237
CL\_INS\_237
CL\_INS\_70
CL\_INS\_237
CL\_INS\_237
CL\_INS\_237
CL\_INS\_237
CL\_INS\_70
CL\_INS\_237
CL\_INS\_237
CL\_INS\_237
CL\_INS\_237
CL\_INS\_237
CL\_INS\_70
CL\_INS\_237
CL\_INS\_70
CL\_INS\_237
CL\_INS\_70
CL\_INS\_237
CL\_INS\_70
CL\_INS\_237
CL\_INS\_237
CL\_INS\_70
CL\_INS\_237
CL\_INS\_70
CL\_INS\_237
CL\_INS\_237
CL\_INS\_30
CL\_INS\_237
CL\_INS\_70
CL\_INS\_237
CL\_INS\_70
CL\_INS\_237
CL\_INS\_237
CL\_INS\_70
CL\_INS\_70
CL\_INS\_237
CL\_INS\_237
CL\_INS\_237
CL\_INS\_237
CL\_INS\_237
CL\_INS\_237
CL\_INS\_237
CL\_INS\_237
CL\_INS\_237
CL\_INS\_237
CL\_INS\_247
CL\_INS\_247
CL\_INS\_237
CL\_INS\_237
CL\_INS\_237
CL\_INS\_237
CL\_INS\_123
CL\_INS\_237
CL\_INS\_237
CL\_INS\_237
CL\_INS\_237
CL\_INS\_237
CL\_INS\_237
CL\_INS\_237
CL\_INS\_237
CL\_INS\_237
CL\_INS\_237
CL\_INS\_237
CL\_INS\_237
CL\_INS\_237
CL\_INS\_237
CL\_INS\_237
CL\_INS\_237
CL\_INS\_237
CL\_INS\_237
CL\_INS\_237
CL\_INS\_237
CL\_INS\_237
CL\_INS\_237
CL\_INS\_237
CL\_INS\_237
CL\_INS\_237
CL\_INS\_237
CL\_INS\_237
CL\_INS\_237
CL\_INS\_237
CL\_INS\_30
CL\_INS\_30
CL\_INS\_237
CL\_INS\_237
CL\_INS\_30
CL\_INS\_237
CL\_INS\_237
CL\_INS\_70
CL\_INS\_237
CL\_INS\_237
CL\_INS\_237
CL\_INS\_237
CL\_INS\_30
CL\_INS\_70
CL\_INS\_70
CL\_INS\_237
CL\_INS\_237
CL\_INS\_237
CL\_INS\_237
CL\_INS\_247
CL\_INS\_237
CL\_INS\_237
CL\_INS\_30
CL\_INS\_237
CL\_INS\_237
CL\_INS\_237
CL\_INS\_237
CL\_INS\_237
CL\_INS\_237
CL\_INS\_30
CL\_INS\_237
CL\_INS\_237
CL\_INS\_30
CL\_INS\_247
CL\_INS\_237
CL\_INS\_237
CL\_INS\_237
CL\_INS\_237
CL\_INS\_352
CL\_INS\_352
CL\_INS\_352
CL\_INS\_352
CL\_INS\_352
CL\_INS\_352
CL\_INS\_352
CL\_INS\_352
CL\_INS\_352
CL\_INS\_352
CL\_INS\_352
CL\_INS\_352
CL\_INS\_352
CL\_INS\_352
CL\_INS\_352
CL\_INS\_352
CL\_INS\_352
CL\_INS\_352
CL\_INS\_352
CL\_INS\_352
CL\_INS\_352
CL\_INS\_352
CL\_INS\_352
CL\_INS\_352
CL\_INS\_352
CL\_INS\_352
CL\_INS\_352
CL\_INS\_352
CL\_INS\_352
CL\_INS\_352
CL\_INS\_352
CL\_INS\_352
CL\_INS\_352
CL\_INS\_352
CL\_INS\_352
CL\_INS\_352
CL\_INS\_352
CL\_INS\_352
CL\_INS\_352
CL\_INS\_352
CL\_INS\_352
CL\_INS\_237
CL\_INS\_237
CL\_INS\_352
CL\_INS\_237
CL\_INS\_352
CL\_INS\_352
CL\_INS\_237
CL\_INS\_247
CL\_INS\_237
CL\_INS\_237
CL\_INS\_237
CL\_INS\_237
CL\_INS\_237
CL\_INS\_237
CL\_INS\_237
CL\_INS\_237
CL\_INS\_237
CL\_INS\_237
CL\_INS\_237
CL\_INS\_237
CL\_INS\_237
CL\_INS\_237
CL\_INS\_237
CL\_INS\_237
CL\_INS\_382
CL\_INS\_237
CL\_INS\_237
CL\_INS\_382
CL\_INS\_237
CL\_INS\_237
CL\_INS\_237
CL\_INS\_237
CL\_INS\_237
CL\_INS\_237
CL\_INS\_237
CL\_INS\_237
CL\_INS\_237
CL\_INS\_237
CL\_INS\_237
CL\_INS\_149
CL\_INS\_237
CL\_INS\_237
CL\_INS\_237
CL\_INS\_237
CL\_INS\_237
CL\_INS\_237
CL\_INS\_30
CL\_INS\_30
CL\_INS\_237
CL\_INS\_237
CL\_INS\_247
CL\_INS\_237
CL\_INS\_237
CL\_INS\_237
CL\_INS\_237
CL\_INS\_247
CL\_INS\_237
CL\_INS\_70
CL\_INS\_237
CL\_INS\_237
CL\_INS\_237
CL\_INS\_237
CL\_INS\_237
CL\_INS\_237
CL\_INS\_237
CL\_INS\_237
CL\_INS\_237
CL\_INS\_237
CL\_INS\_237
CL\_INS\_237
CL\_INS\_237
CL\_INS\_237
CL\_INS\_237
CL\_INS\_237
CL\_INS\_247
CL\_INS\_237
CL\_INS\_247
CL\_INS\_247
CL\_INS\_247
CL\_INS\_237
CL\_INS\_247
CL\_INS\_237
CL\_INS\_237
CL\_INS\_247
CL\_INS\_237
CL\_INS\_237
CL\_INS\_237
CL\_INS\_237
CL\_INS\_237
CL\_INS\_237
CL\_INS\_237
CL\_INS\_237
CL\_INS\_237
CL\_INS\_237
CL\_INS\_237
CL\_INS\_237
CL\_INS\_237
CL\_INS\_237
CL\_INS\_237
CL\_INS\_237
CL\_INS\_237
CL\_INS\_237
CL\_INS\_237
CL\_INS\_237
CL\_INS\_237
CL\_INS\_237
CL\_INS\_237
CL\_INS\_237
CL\_INS\_237
CL\_INS\_237
CL\_INS\_237
CL\_INS\_237
CL\_INS\_237
CL\_INS\_237
CL\_INS\_237
CL\_INS\_237
CL\_INS\_237
CL\_INS\_237
CL\_INS\_237
CL\_INS\_237
CL\_INS\_237
CL\_INS\_237
CL\_INS\_237
CL\_INS\_237
CL\_INS\_237
CL\_INS\_237
CL\_INS\_237
CL\_INS\_237
CL\_INS\_237
CL\_INS\_237
CL\_INS\_237
CL\_INS\_247
CL\_INS\_237
CL\_INS\_237
CL\_INS\_237
CL\_INS\_237
CL\_INS\_237
CL\_INS\_237
CL\_INS\_237
CL\_INS\_237
CL\_INS\_247
CL\_INS\_237
CL\_INS\_237
CL\_INS\_237
CL\_INS\_237
CL\_INS\_237
CL\_INS\_237
CL\_INS\_237
CL\_INS\_237
CL\_INS\_237
CL\_INS\_237
CL\_INS\_237
CL\_INS\_237
CL\_INS\_237
CL\_INS\_237
CL\_INS\_237
CL\_INS\_237
CL\_INS\_237
CL\_INS\_237
CL\_INS\_237
CL\_INS\_237
CL\_INS\_237
CL\_INS\_237
CL\_INS\_237
CL\_INS\_237
CL\_INS\_237
CL\_INS\_237
CL\_INS\_237
CL\_INS\_237
CL\_INS\_237
CL\_INS\_247
CL\_INS\_237
CL\_INS\_237
CL\_INS\_237
CL\_INS\_237
CL\_INS\_237
CL\_INS\_237
CL\_INS\_237
CL\_INS\_237
CL\_INS\_237
CL\_INS\_237
CL\_INS\_237
CL\_INS\_237
CL\_INS\_237
CL\_INS\_237
CL\_INS\_237
CL\_INS\_237
CL\_INS\_237
CL\_INS\_247
CL\_INS\_237
CL\_INS\_237
CL\_INS\_237
CL\_INS\_237
CL\_INS\_237
CL\_INS\_237
CL\_INS\_237
CL\_INS\_237
CL\_INS\_237
CL\_INS\_237
CL\_INS\_237
CL\_INS\_237
CL\_INS\_237
CL\_INS\_237
CL\_INS\_237
CL\_INS\_237
CL\_INS\_70
CL\_INS\_237
CL\_INS\_237
CL\_INS\_237
CL\_INS\_70
CL\_INS\_237
CL\_INS\_237
CL\_INS\_237
CL\_INS\_237
CL\_INS\_237
CL\_INS\_237
CL\_INS\_30
CL\_INS\_237
CL\_INS\_237
CL\_INS\_237
CL\_INS\_30
CL\_INS\_237
CL\_INS\_70
CL\_INS\_382
CL\_INS\_30
CL\_INS\_237
CL\_INS\_30
CL\_INS\_237
CL\_INS\_237
CL\_INS\_237
CL\_INS\_237
CL\_INS\_237
CL\_INS\_237
CL\_INS\_237
CL\_INS\_237
CL\_INS\_237
CL\_INS\_237
CL\_INS\_70
CL\_INS\_237
CL\_INS\_237
CL\_INS\_237
CL\_INS\_70
CL\_INS\_247
CL\_INS\_237
Cluster ID


CL\_20307
CL\_28607
CL\_7325
CL\_28606
CL\_28605
CL\_28604
CL\_6439
CL\_8272
CL\_8270
CL\_8269
CL\_5368
CL\_13484
CL\_5367
CL\_10317
CL\_11206
CL\_6637
CL\_6639
CL\_5389
CL\_5388
CL\_8333
CL\_5387
CL\_5386
CL\_5385
CL\_5384
CL\_17725
CL\_11731
CL\_8328
CL\_8326
CL\_19187
CL\_6789
CL\_9078
CL\_9079
CL\_9080
CL\_5351
CL\_4410
CL\_5350
CL\_5349
CL\_8321
CL\_8320
CL\_5347
CL\_6658
CL\_6659
CL\_6660
CL\_8319
CL\_9119
CL\_9120
CL\_9121
CL\_5340
CL\_526
CL\_6784
CL\_7049
CL\_19186
CL\_4346
CL\_4347
CL\_4348
CL\_36530
CL\_36531
CL\_36532
CL\_36533
CL\_36534
CL\_36535
CL\_31794
CL\_4116
CL\_4115
CL\_4114
CL\_4113
CL\_4112
CL\_4111
CL\_4110
CL\_4109
CL\_4108
CL\_4107
CL\_4106
CL\_4105
CL\_4104
CL\_4103
CL\_4102
CL\_9541
CL\_7259
CL\_7258
CL\_9542
CL\_6723
CL\_6724
CL\_24053
CL\_24052
CL\_9672
CL\_5673
CL\_24051
CL\_24050
CL\_24049
CL\_24048
CL\_24047
CL\_6888
CL\_6887
CL\_6886
CL\_6885
CL\_9697
CL\_24046
CL\_33074
CL\_8363
CL\_17879
CL\_17878
CL\_26431
CL\_17124
CL\_17125
CL\_17126
CL\_17127
CL\_17128
CL\_31275
CL\_17877
CL\_17876
CL\_17875
CL\_17874
CL\_8364
CL\_8365
CL\_24066
CL\_24065
CL\_24064
CL\_24063
CL\_24062
CL\_24061
CL\_24060
CL\_24059
CL\_24058
CL\_24057
CL\_24056
CL\_14320
CL\_14321
CL\_14322
CL\_21946
CL\_21947
CL\_21948
CL\_21949
CL\_20947
CL\_20870
CL\_20869
CL\_20868
CL\_20865
CL\_20864
CL\_20863
CL\_17620
CL\_20862
CL\_17619
CL\_17618
CL\_20861
CL\_20860
CL\_5013
CL\_5014
CL\_5015
CL\_5016
CL\_35487
CL\_5018
CL\_19401
CL\_19402
CL\_5050
CL\_25427
CL\_35488
CL\_35489
CL\_14141
CL\_11689
CL\_5054
CL\_5055
CL\_4255
CL\_5057
CL\_5058
CL\_5059
CL\_35490
CL\_5060
CL\_23701
CL\_5627
CL\_5061
CL\_6942
CL\_5063
CL\_5064
CL\_5065
CL\_5066
CL\_6939
CL\_5067
CL\_5068
CL\_14165
CL\_5069
CL\_5070
CL\_5071
CL\_5072
CL\_5073
CL\_5074
CL\_5075
CL\_5076
CL\_5077
CL\_35491
CL\_35492
CL\_35316
CL\_19404
CL\_20821
CL\_20820
CL\_35315
CL\_17168
CL\_17167
CL\_17166
CL\_17165
CL\_17164
CL\_35314
CL\_35313
CL\_35312
CL\_20817
CL\_20951
CL\_20815
CL\_10785
CL\_20949
CL\_20814
CL\_20813
CL\_10782
CL\_35311
CL\_20810
CL\_35493
CL\_35494
CL\_35495
CL\_22350
CL\_35496
CL\_35497
CL\_35498
CL\_35499
CL\_35500
CL\_35501
CL\_35502
CL\_35503
CL\_35504
CL\_35505
CL\_35506
CL\_35507
CL\_35508
CL\_35509
CL\_35510
CL\_35511
CL\_35512
CL\_35513
CL\_35514
CL\_35515
CL\_35516
CL\_35517
CL\_35518
CL\_35519
CL\_35520
CL\_35521
CL\_35522
CL\_35523
CL\_31010
CL\_35524
CL\_35525
CL\_35526
CL\_35527
CL\_35528
CL\_35529
CL\_35530
CL\_20795
CL\_21950
CL\_21951
CL\_21952
CL\_21850
CL\_21849
CL\_21848
CL\_21953
CL\_26312
CL\_22582
CL\_20948
CL\_20796
CL\_20797
CL\_20799
CL\_20798
CL\_6957
CL\_22147
CL\_20802
CL\_21954
CL\_21955
CL\_17873
CL\_21956
CL\_21957
CL\_5298
CL\_21958
CL\_11649
CL\_21959
CL\_21960
CL\_21961
CL\_21962
CL\_21963
CL\_21964
CL\_21965
CL\_21966
CL\_9887
CL\_9888
CL\_9889
CL\_17872
CL\_17871
CL\_14888
CL\_17870
CL\_17869
CL\_17868
CL\_17867
CL\_17866
CL\_17865
CL\_17864
CL\_8396
CL\_8397
CL\_22247
CL\_21967
CL\_14323
CL\_14324
CL\_14325
CL\_21968
CL\_21969
CL\_21970
CL\_8391
CL\_21971
CL\_21972
CL\_21973
CL\_21974
CL\_21975
CL\_21976
CL\_21977
CL\_21978
CL\_21979
CL\_22246
CL\_22245
CL\_22244
CL\_22243
CL\_22242
CL\_22241
CL\_22240
CL\_22239
CL\_22238
CL\_8398
CL\_24045
CL\_24044
CL\_17863
CL\_17862
CL\_22237
CL\_8399
CL\_8400
CL\_17861
CL\_17860
CL\_17859
CL\_17858
CL\_17857
CL\_17856
CL\_17855
CL\_8401
CL\_8402
CL\_8403
CL\_24043
CL\_24042
CL\_24041
CL\_24040
CL\_8404
CL\_20848
CL\_8405
CL\_8406
CL\_8407
CL\_24039
CL\_17854
CL\_17853
CL\_17852
CL\_17851
CL\_17850
CL\_17849
CL\_17848
CL\_17847
CL\_17846
CL\_17845
CL\_17844
CL\_17843
CL\_17842
CL\_17841
CL\_17840
CL\_17839
CL\_17838
CL\_17837
CL\_8408
CL\_8409
CL\_8410
CL\_8411
CL\_17829
CL\_17828
CL\_17827
CL\_17826
CL\_17825
CL\_17824
CL\_17823
CL\_17822
CL\_17821
CL\_17820
CL\_17819
CL\_17818
CL\_17817
CL\_17816
CL\_17815
CL\_17814
CL\_17813
CL\_17812
CL\_8394
CL\_8395
CL\_8393
CL\_8392
CL\_17811
CL\_17810
CL\_17809
CL\_17808
CL\_17807
CL\_17806
CL\_17805
CL\_17804
CL\_8366
CL\_8367
CL\_8368
CL\_8369
CL\_8370
CL\_8371
CL\_8372
CL\_8373
CL\_8374
CL\_8375
CL\_7761
CL\_8376
CL\_7297
CL\_14412
CL\_8377
CL\_8378
CL\_8379
CL\_8380
CL\_8381
CL\_8382
CL\_8383
CL\_8384
CL\_17803
CL\_17802
CL\_17801
CL\_17800
CL\_17799
CL\_8412
CL\_8413
CL\_8414
CL\_24038
CL\_536
CL\_8415
CL\_8416
CL\_22236
CL\_8417
CL\_17832
CL\_17831
CL\_17830
CL\_8418
CL\_24037
CL\_14880
CL\_22235
CL\_14879
CL\_8419
CL\_17836
CL\_17835
CL\_17834
CL\_17833
CL\_8420
CL\_8421
CL\_8422
CL\_8423
CL\_24036
CL\_24035
CL\_24034
CL\_22234
CL\_8424
CL\_8425
CL\_8426
CL\_8427
CL\_22233
CL\_8428
CL\_24033
CL\_8429
CL\_24032
CL\_8430
CL\_8431
CL\_8432
CL\_22232
CL\_22231
CL\_8682
CL\_8433
CL\_8434
CL\_26209
CL\_8435
CL\_26311
CL\_11728
CL\_22230
CL\_22229
CL\_11403
CL\_22228
CL\_22227
CL\_10515
CL\_8436
CL\_8437
CL\_8438
CL\_8439
CL\_8440
CL\_14331
CL\_26208
CL\_8441
CL\_18180
CL\_18179
CL\_22226
CL\_8442
CL\_26310
CL\_5145
CL\_5144
CL\_5143
CL\_21249
CL\_21248
CL\_21247
CL\_21246
CL\_21245
CL\_21244
CL\_5142
CL\_5141
CL\_5140
CL\_5139
CL\_5138
CL\_5137
CL\_5136
CL\_5135
CL\_26309
CL\_37530
CL\_26308
CL\_37435
CL\_19788
CL\_34289
CL\_34290
CL\_34291
CL\_34292
CL\_26307
CL\_26306
CL\_37570
CL\_26305
CL\_26304
CL\_26303
CL\_26302
CL\_26301
CL\_26300
CL\_26299
CL\_26298
CL\_14334
CL\_14335
CL\_14336
CL\_7579
CL\_14337
CL\_14338
CL\_14339
CL\_15538
CL\_26297
CL\_26296
CL\_26295
CL\_26294
CL\_26293
CL\_26292
CL\_26291
CL\_23913
CL\_8554
CL\_4951
CL\_9557
CL\_8518
CL\_18148
CL\_4950
CL\_26290
CL\_4948
CL\_4949
CL\_26289
CL\_23915
CL\_23916
CL\_4945
CL\_9560
CL\_9559
CL\_9558
CL\_4944
CL\_4943
CL\_15588
CL\_23917
CL\_6764
CL\_15589
CL\_26288
CL\_26287
CL\_7667
CL\_9370
CL\_26112
CL\_20008
CL\_20007
CL\_6415
CL\_18883
CL\_18882
CL\_18881
CL\_6426
CL\_18880
CL\_18879
CL\_18878
CL\_18877
CL\_18876
CL\_18875
CL\_18874
CL\_18873
CL\_18872
CL\_18871
CL\_18870
CL\_18869
CL\_18868
CL\_18867
CL\_18866
CL\_18865
CL\_7869
CL\_18864
CL\_14365
CL\_7872
CL\_13761
CL\_13779
CL\_7873
CL\_7874
CL\_7875
CL\_7876
CL\_7877
CL\_7878
CL\_7879
CL\_7880
CL\_7881
CL\_7882
CL\_7883
CL\_7884
CL\_7885
CL\_7886
CL\_7887
CL\_7888
CL\_7889
CL\_7890
CL\_11296
CL\_11295
CL\_36441
CL\_18863
CL\_6849
CL\_17170
CL\_21575
CL\_6848
CL\_6847
CL\_6856
CL\_8818
CL\_6846
CL\_7986
CL\_7985
CL\_7987
CL\_16063
CL\_11826
CL\_15590
CL\_26286
CL\_12241
CL\_5160
CL\_26285
CL\_26284
CL\_26283
CL\_22038
CL\_22039
CL\_22040
CL\_26282
CL\_21998
CL\_21999
CL\_15584
CL\_15583
CL\_15582
CL\_15581
CL\_15580
CL\_15579
CL\_15578
CL\_10446
CL\_10445
CL\_10444
CL\_10443
CL\_10442
CL\_10441
CL\_10440
CL\_10439
CL\_10438
CL\_10437
CL\_8760
CL\_8758
CL\_22000
CL\_22001
CL\_34724
CL\_22002
CL\_6845
CL\_7245
CL\_12078
CL\_22412
CL\_5148
CL\_25949
CL\_29308
CL\_29307
CL\_7998
CL\_17687
CL\_7997
CL\_7996
CL\_7995
CL\_7994
CL\_7993
CL\_7992
CL\_17175
CL\_17174
CL\_17173
CL\_10581
CL\_13776
CL\_13775
CL\_13774
CL\_13773
CL\_36445
CL\_36444
CL\_36443
CL\_36442
CL\_13772
CL\_13771
CL\_17131
CL\_17132
CL\_13770
CL\_37215
CL\_37216
CL\_37217
CL\_37218
CL\_13768
CL\_31992
CL\_31993
CL\_31994
CL\_31995
CL\_31996
CL\_10582
CL\_27877
CL\_7678
CL\_7677
CL\_7676
CL\_12088
CL\_14595
CL\_7675
CL\_7984
CL\_7674
CL\_7679
CL\_7682
CL\_7681
CL\_7680
CL\_7683
CL\_7684
CL\_14594
CL\_12086
CL\_7685
CL\_7686
CL\_7687
CL\_35343
CL\_35342
CL\_35341
CL\_35340
CL\_35339
CL\_35338
CL\_35337
CL\_35336
CL\_35335
CL\_35334
CL\_979
CL\_6410
CL\_8600
CL\_8599
CL\_8817
CL\_8816
CL\_5247
CL\_7243
CL\_26521
CL\_26522
CL\_7242
CL\_11191
CL\_7241
CL\_12129
CL\_11190
CL\_11189
CL\_7240
CL\_7239
CL\_7238
CL\_7441
CL\_29996
CL\_29995
CL\_29994
CL\_29993
CL\_29992
CL\_7440
CL\_7237
CL\_7439
CL\_18304
CL\_7236
CL\_7581
CL\_7438
CL\_7235
CL\_7234
CL\_15979
CL\_15980
CL\_14683
CL\_15981
CL\_20012
CL\_20011
CL\_20010
CL\_15982
CL\_7253
CL\_27324
CL\_5236
CL\_5398
CL\_25950
CL\_19725
CL\_12106
CL\_5688
CL\_5689
CL\_12104
CL\_12103
CL\_12102
CL\_12101
CL\_12100
CL\_12099
CL\_12098
CL\_12097
CL\_12096
CL\_12095
CL\_12094
CL\_30006
CL\_30005
CL\_30004
CL\_30003
CL\_30002
CL\_30001
CL\_30000
CL\_29999
CL\_29998
CL\_29997
CL\_27334
CL\_8553
CL\_27333
CL\_27332
CL\_27331
CL\_27330
CL\_27329
CL\_27328
CL\_15627
CL\_27327
CL\_27326
CL\_6097
CL\_15625
CL\_6099
CL\_20055
CL\_6100
CL\_6101
CL\_6102
CL\_6103
CL\_6104
CL\_6105
CL\_6106
CL\_6107
CL\_6108
CL\_6109
CL\_6110
CL\_6111
CL\_27325
CL\_7300
CL\_7299
CL\_11188
CL\_8780
CL\_9371
CL\_9372
CL\_9373
CL\_33162
CL\_18424
CL\_16458
CL\_16459
CL\_18150
CL\_17158
CL\_23663
CL\_8781
CL\_7233
CL\_8782
CL\_7232
CL\_8783
CL\_8784
CL\_8785
CL\_7231
CL\_8786
CL\_7230
CL\_7229
CL\_8787
CL\_7228
CL\_7227
CL\_8788
CL\_7226
CL\_7225
CL\_7224
CL\_7223
CL\_7222
CL\_7221
CL\_8789
CL\_7220
CL\_7219
CL\_7218
CL\_7217
CL\_7216
CL\_35729
CL\_23995
CL\_23994
CL\_33991
CL\_33990
CL\_23993
CL\_23992
CL\_23991
CL\_23990
CL\_23989
CL\_23988
CL\_23987
CL\_35333
CL\_35332
CL\_35331
CL\_35330
CL\_35329
CL\_14393
CL\_35328
CL\_35327
CL\_35326
CL\_7565
CL\_7306
CL\_19831
CL\_21794
CL\_21795
CL\_21796
CL\_244
CL\_23274
CL\_34371
CL\_34370
CL\_34369
CL\_34368
CL\_34367
CL\_34366
CL\_34365
CL\_34364
CL\_34363
CL\_34362
CL\_34361
CL\_22666
CL\_8942
CL\_8941
CL\_8940
CL\_8939
CL\_8938
CL\_8937
CL\_8936
CL\_8935
CL\_8934
CL\_8933
CL\_13588
CL\_22665
CL\_22664
CL\_22663
CL\_22662
CL\_13368
CL\_13369
CL\_13370
CL\_12907
CL\_12908
CL\_13371
CL\_23012
CL\_33740
CL\_13372
CL\_13373
CL\_13374
CL\_27775
CL\_13610
CL\_7710
CL\_7709
CL\_7708
CL\_36721
CL\_36720
CL\_7707
CL\_7706
CL\_22225
CL\_22224
CL\_22223
CL\_8385
CL\_7305
CL\_7304
CL\_6760
CL\_10419
CL\_6761
CL\_5697
CL\_6748
CL\_6749
CL\_6750
CL\_6751
CL\_6752
CL\_6753
CL\_6754
CL\_6755
CL\_6756
CL\_6757
CL\_6758
CL\_6759
CL\_6056
CL\_27358
CL\_7303
CL\_8386
CL\_14326
CL\_14327
CL\_14328
CL\_14329
CL\_14330
CL\_8387
CL\_8388
CL\_8389
CL\_8390
CL\_35322
CL\_26529
CL\_10584
CL\_36127
CL\_36126
CL\_36125
CL\_10835
CL\_19733
CL\_36095
CL\_10834
CL\_4995
CL\_14401
CL\_17133
CL\_17134
CL\_7296
CL\_10585
CL\_10833
CL\_10586
CL\_10587
CL\_10588
CL\_30636
CL\_12176
CL\_12175
CL\_6967
CL\_35325
CL\_10831
CL\_10589
CL\_17159
CL\_13034
CL\_10590
CL\_10830
CL\_10591
CL\_14394
CL\_14395
CL\_8632
CL\_13593
CL\_17169
CL\_10829
CL\_10592
CL\_22730
CL\_10828
CL\_7690
CL\_14117
CL\_7689
CL\_7688
CL\_12085
CL\_10593
CL\_13212
CL\_13213
CL\_10594
CL\_12894
CL\_4093
CL\_17172
CL\_17171
CL\_10595
CL\_8631
CL\_16631
CL\_8630
CL\_6844
CL\_10832
CL\_37134
CL\_12888
CL\_13233
CL\_13231
CL\_247
CL\_15115
CL\_15470
CL\_15471
CL\_15472
CL\_15473
CL\_21826
CL\_5090
CL\_5091
CL\_5092
CL\_5093
CL\_5094
CL\_20806
CL\_20805
CL\_20804
CL\_14935
CL\_10770
CL\_10769
CL\_14899
CL\_20803
CL\_13211
CL\_12904
CL\_7705
CL\_5528
CL\_5527
CL\_5526
CL\_5525
CL\_5512
CL\_5513
CL\_7704
CL\_7703
CL\_14593
CL\_7702
CL\_8909
CL\_10807
CL\_10435
CL\_22005
CL\_7856
CL\_7857
CL\_7858
CL\_7859
CL\_6971
CL\_7671
CL\_14411
CL\_22559
CL\_34807
CL\_23986
CL\_23985
CL\_23984
CL\_23983
CL\_23982
CL\_23981
CL\_19734
CL\_24099
CL\_24098
CL\_5682
CL\_5292
CL\_5293
CL\_5294
CL\_15528
CL\_14340
CL\_8517
CL\_8132
CL\_8516
CL\_21021
CL\_8515
CL\_8514
CL\_7214
CL\_15526
CL\_7213
CL\_25361
CL\_14341
CL\_14342
CL\_14400
CL\_22558
CL\_8210
CL\_7294
CL\_6163
CL\_11426
CL\_6144
CL\_6143
CL\_6142
CL\_6141
CL\_11342
CL\_6140
CL\_6139
CL\_11343
CL\_11344
CL\_11345
CL\_11346
CL\_11347
CL\_11349
CL\_11350
CL\_11351
CL\_11352
CL\_15298
CL\_23936
CL\_23937
CL\_11356
CL\_11357
CL\_11358
CL\_11359
CL\_11360
CL\_11361
CL\_11362
CL\_11363
CL\_4514
CL\_2276
CL\_11366
CL\_6138
CL\_6137
CL\_6136
CL\_11367
CL\_11368
CL\_11369
CL\_11370
CL\_34812
CL\_11371
CL\_11380
CL\_11381
CL\_11383
CL\_11384
CL\_11385
CL\_6135
CL\_6134
CL\_34811
CL\_34810
CL\_11388
CL\_6196
CL\_6195
CL\_6194
CL\_6193
CL\_6190
CL\_6189
CL\_6188
CL\_6187
CL\_6186
CL\_6183
CL\_6182
CL\_6181
CL\_11393
CL\_11394
CL\_34809
CL\_6175
CL\_6174
CL\_6173
CL\_6172
CL\_6171
CL\_11410
CL\_11411
CL\_11412
CL\_11413
CL\_11414
CL\_11415
CL\_11416
CL\_11417
CL\_11418
CL\_11419
CL\_11421
CL\_11422
CL\_34808
CL\_6984
CL\_6983
CL\_6982
CL\_6981
CL\_6980
CL\_6979
CL\_16977
CL\_10596
CL\_10597
CL\_10598
CL\_10599
CL\_10600
CL\_10601
CL\_10602
CL\_10603
CL\_10604
CL\_10827
CL\_10826
CL\_10605
CL\_14867
CL\_14868
CL\_14869
CL\_14870
CL\_14871
CL\_14872
CL\_6805
CL\_10372
CL\_7893
CL\_7892
CL\_8593
CL\_8594
CL\_8595
CL\_10326
CL\_35457
CL\_26166
CL\_26165
CL\_27805
CL\_27806
CL\_26164
CL\_26163
CL\_35456
CL\_6413
CL\_15963
CL\_15964
CL\_15965
CL\_15966
CL\_37667
CL\_10373
CL\_10374
CL\_19192
CL\_19193
CL\_19194
CL\_7254
CL\_6423
CL\_6731
CL\_26167
CL\_6824
CL\_19195
CL\_1930
CL\_10825
CL\_10824
CL\_8281
CL\_23134
CL\_8513
CL\_10823
CL\_8512
CL\_8511
CL\_21022
CL\_5233
CL\_14541
CL\_14542
CL\_21582
CL\_6828
CL\_21583
CL\_13751
CL\_13750
CL\_21584
CL\_21585
CL\_21586
CL\_21587
CL\_8510
CL\_8509
CL\_10547
CL\_8126
CL\_6385
CL\_13214
CL\_37393
CL\_20310
CL\_8125
CL\_7470
CL\_5232
CL\_5231
CL\_5230
CL\_5229
CL\_5228
CL\_8832
CL\_8361
CL\_10822
CL\_10820
CL\_10609
CL\_37133
CL\_37132
CL\_23071
CL\_13821
CL\_7434
CL\_7433
CL\_7432
CL\_7435
CL\_29305
CL\_7717
CL\_13820
CL\_22659
CL\_14396
CL\_8216
CL\_4975
CL\_27622
CL\_23072
CL\_8841
CL\_36435
CL\_36434
CL\_36433
CL\_13234
CL\_27359
CL\_36719
CL\_27360
CL\_7436
CL\_10548
CL\_31999
CL\_35324
CL\_13552
CL\_33065
CL\_33064
CL\_33063
CL\_8642
CL\_28451
CL\_9140
CL\_7829
CL\_6978
CL\_5594
CL\_5595
CL\_6765
CL\_30531
CL\_5596
CL\_5597
CL\_5147
CL\_5146
CL\_18171
CL\_6246
CL\_6245
CL\_6244
CL\_7713
CL\_12079
CL\_7712
CL\_32012
CL\_6075
CL\_26539
CL\_26540
CL\_10583
CL\_34165
CL\_11850
CL\_21797
CL\_21798
CL\_21799
CL\_21800
CL\_21801
CL\_8778
CL\_26832
CL\_26831
CL\_26830
CL\_7711
CL\_25189
CL\_7292
CL\_4374
CL\_11707
CL\_14318
CL\_5226
CL\_4375
CL\_20308
CL\_7208
CL\_20257
CL\_7291
CL\_26685
CL\_7290
CL\_15525
CL\_7750
CL\_7749
CL\_7568
CL\_10789
CL\_25972
CL\_8504
CL\_8239
CL\_8124
CL\_26176
CL\_10611
CL\_10817
CL\_10816
CL\_10612
CL\_8505
CL\_7969
CL\_8503
CL\_15563
CL\_8502
CL\_15562
CL\_5159
CL\_32875
CL\_32876
CL\_21996
CL\_21997
CL\_24555
CL\_24554
CL\_24553
CL\_24552
CL\_24551
CL\_24550
CL\_24549
CL\_24548
CL\_24547
CL\_24546
CL\_24545
CL\_24544
CL\_24543
CL\_24542
CL\_24541
CL\_24540
CL\_29070
CL\_29071
CL\_29072
CL\_29073
CL\_5998
CL\_29074
CL\_6000
CL\_6001
CL\_29075
CL\_16932
CL\_16931
CL\_16930
CL\_9720
CL\_9719
CL\_9718
CL\_9717
CL\_9716
CL\_9715
CL\_21644
CL\_21643
CL\_21642
CL\_29633
CL\_29634
CL\_29635
CL\_29636
CL\_26801
CL\_6207
CL\_16929
CL\_5158
CL\_5321
CL\_5320
CL\_6833
CL\_6029
CL\_6028
CL\_6027
CL\_6026
CL\_6025
CL\_6024
CL\_6023
CL\_6411
CL\_20017
CL\_20016
CL\_20015
CL\_8534
CL\_26520
CL\_14332
CL\_14333
CL\_27337
CL\_27336
CL\_27335
CL\_14306
CL\_14307
CL\_14308
CL\_14309
CL\_14310
CL\_14311
CL\_20304
CL\_37214
CL\_35041
CL\_10836
CL\_34745
CL\_34744
CL\_34743
CL\_34742
CL\_34741
CL\_15537
CL\_15536
CL\_13467
CL\_15535
CL\_15534
CL\_15533
CL\_15532
CL\_15531
CL\_15530
CL\_15529
CL\_16926
CL\_8533
CL\_14349
CL\_7307
CL\_9369
CL\_15967
CL\_15968
CL\_15969
CL\_15970
CL\_15971
CL\_15972
CL\_14911
CL\_15973
CL\_15974
CL\_15975
CL\_15976
CL\_15977
CL\_15978
CL\_17572
CL\_17573
CL\_7252
CL\_6827
CL\_6414
CL\_26467
CL\_26468
CL\_8721
CL\_11262
CL\_36536
CL\_36537
CL\_36538
CL\_36539
CL\_36540
CL\_36541
CL\_36542
CL\_36543
CL\_36544
CL\_36545
CL\_36546
CL\_36547
CL\_36548
CL\_36549
CL\_36550
CL\_36551
CL\_36552
CL\_17574
CL\_8532
CL\_8531
CL\_8530
CL\_8529
CL\_8528
CL\_8527
CL\_8526
CL\_8525
CL\_8524
CL\_8523
CL\_10805
CL\_8637
CL\_8636
CL\_8635
CL\_8522
CL\_6976
CL\_32362
CL\_32363
CL\_7855
CL\_8521
CL\_21020
CL\_8520
CL\_8519
CL\_7673
CL\_10801
CL\_11715
CL\_37391
CL\_37392
CL\_11714
CL\_11713
CL\_11712
CL\_9374
CL\_9375
CL\_9131
CL\_9376
CL\_26111
CL\_9377
CL\_33161
CL\_31766
CL\_33160
CL\_33159
CL\_4094
CL\_8747
CL\_8748
CL\_8749
CL\_8750
CL\_4095
CL\_4096
CL\_4097
CL\_4098
CL\_4099
CL\_4100
CL\_4101
CL\_7374
CL\_13375
CL\_13376
CL\_13378
CL\_13377
CL\_26541
CL\_26542
CL\_15527
CL\_6806
CL\_33041
CL\_33042
CL\_33043
CL\_33044
CL\_33045
CL\_33046
CL\_33047
CL\_13217
CL\_13218
CL\_32509
CL\_32508
CL\_18808
CL\_33073
CL\_33072
CL\_33071
CL\_33070
CL\_18856
CL\_34740
CL\_34739
CL\_34738
CL\_34737
CL\_34736
CL\_34735
CL\_34734
CL\_34733
CL\_21592
CL\_20448
CL\_6590
CL\_11800
CL\_4089
CL\_4090
CL\_4092
CL\_18430
CL\_7871
CL\_4091
CL\_22719
CL\_34732
CL\_10408
CL\_10409
CL\_8932
CL\_34731
CL\_34730
CL\_34729
CL\_34728
CL\_34727
CL\_34726
CL\_34725
CL\_7244
CL\_10804
CL\_26530
CL\_12077
CL\_18903
CL\_18902
CL\_18901
CL\_18900
CL\_18899
CL\_7298
CL\_8779
CL\_18161
CL\_20014
CL\_18898
CL\_18897
CL\_18896
CL\_18895
CL\_10186
CL\_18894
CL\_19189
CL\_19190
CL\_19191
CL\_8241
CL\_8240
CL\_7426
CL\_7425
CL\_7427
CL\_7428
CL\_7429
CL\_7430
CL\_7431
CL\_18893
CL\_18892
CL\_18891
CL\_18890
CL\_18889
CL\_18888
CL\_18887
CL\_18886
CL\_18885
CL\_18884
CL\_13012
CL\_7991
CL\_5157
CL\_28990
CL\_5156
CL\_13591
CL\_26523
CL\_26524
CL\_26525
CL\_26526
CL\_26527
CL\_26528
CL\_25190
CL\_25191
CL\_13592
CL\_5155
CL\_5154
CL\_5153
CL\_5152
CL\_21802
CL\_5151
CL\_5150
CL\_25192
CL\_5149
CL\_22661
CL\_17163
CL\_17162
CL\_17161
CL\_17160
CL\_22660
CL\_34950
CL\_8555
CL\_8556
CL\_4086
CL\_13846
CL\_1931
CL\_23976
CL\_35459
CL\_35458
CL\_7990
CL\_36037
CL\_7989
CL\_7988
CL\_7301
CL\_7302
CL\_16843
CL\_7437
CL\_20270
CL\_17759
CL\_17760
CL\_17761
CL\_17762
CL\_17763
CL\_17764
CL\_17087
CL\_17088
CL\_20013
CL\_18095
CL\_18096
CL\_18428
CL\_25193
CL\_31997
CL\_31998
CL\_18429
CL\_6842
CL\_6841
CL\_6840
CL\_6839
CL\_6838
CL\_14866
CL\_6836
CL\_14537
CL\_36440
CL\_11783
CL\_6826
CL\_6425
CL\_36553
CL\_36554
CL\_36555
CL\_36556
CL\_36557
CL\_36558
CL\_36559
CL\_36560
CL\_33069
CL\_33068
CL\_6424
CL\_5241
CL\_22733
CL\_33067
CL\_5246
CL\_5245
CL\_21576
CL\_21577
CL\_21578
CL\_21579
CL\_21580
CL\_21581
CL\_6730
CL\_6825
CL\_10375
CL\_7255
CL\_6829
CL\_6830
CL\_6831
CL\_10606
CL\_10607
CL\_10608
CL\_6422
CL\_7256
CL\_6729
CL\_6728
CL\_6421
CL\_27681
CL\_27680
CL\_11974
CL\_11973
CL\_10546
CL\_36439
CL\_36438
CL\_36437
CL\_36436
CL\_11972
CL\_6420
CL\_10821
CL\_1932
CL\_29306
CL\_6179
CL\_27776
CL\_1933
CL\_8910
CL\_33066
CL\_7831
CL\_33391
CL\_8620
CL\_34071
CL\_37131
CL\_5227
CL\_37130
CL\_6417
CL\_36561
CL\_6418
CL\_21632
CL\_8564
CL\_6416
CL\_7672
CL\_11725
CL\_11724
CL\_11723
CL\_11722
CL\_11721
CL\_11720
CL\_11719
CL\_11718
CL\_11717
CL\_11716
CL\_18854
CL\_7215
CL\_7670
CL\_7669
CL\_25973
CL\_6076
CL\_9785
CL\_7668
CL\_8790
CL\_8791
CL\_8792
CL\_8793
CL\_8212
CL\_11187
CL\_7295
CL\_8794
CL\_21803
CL\_21804
CL\_22658
CL\_7666
CL\_35455
CL\_12110
CL\_7665
CL\_7664
CL\_7212
CL\_23926
CL\_23927
CL\_7211
CL\_7210
CL\_7293
CL\_33741
CL\_33742
CL\_33743
CL\_14399
CL\_7209
CL\_6813
CL\_7751
CL\_235
CL\_8507
CL\_8506
CL\_1934
CL\_7206
CL\_7207
CL\_10610
CL\_35073
CL\_10819
CL\_10818
CL\_21980
CL\_2551
CL\_8746
CL\_6809
CL\_7970
CL\_17577
CL\_17578
CL\_17579
CL\_7204
CL\_17157
CL\_8815
CL\_7827
CL\_37129
CL\_7289
CL\_7288
CL\_7205
CL\_6722
CL\_18099
CL\_1939
CL\_18100
CL\_18101
CL\_7971
CL\_7567
CL\_33062
CL\_17580
CL\_17581
CL\_17582
CL\_17583
CL\_17584
CL\_9401
CL\_9402
CL\_9403
CL\_9404
CL\_9405
CL\_9406
CL\_9407
CL\_9408
CL\_9409
CL\_9410
CL\_9411
CL\_9412
CL\_9413
CL\_9414
CL\_9415
CL\_9416
CL\_9417
CL\_9418
CL\_9419
CL\_9420
CL\_9421
CL\_9422
CL\_9423
CL\_9424
CL\_9425
CL\_9426
CL\_9427
CL\_9428
CL\_9429
CL\_9430
CL\_9431
CL\_9432
CL\_9433
CL\_9434
CL\_9435
CL\_9436
CL\_9437
CL\_9438
CL\_9439
CL\_32150
CL\_32149
CL\_9440
CL\_26156
CL\_9441
CL\_17585
CL\_31793
CL\_7007
CL\_31792
CL\_33061
CL\_4462
CL\_7701
CL\_12081
CL\_17129
CL\_17130
CL\_12082
CL\_7700
CL\_7699
CL\_7698
CL\_7697
CL\_7696
CL\_7695
CL\_7694
CL\_7693
CL\_7692
CL\_12083
CL\_8629
CL\_7691
CL\_32354
CL\_32355
CL\_32356
CL\_32357
CL\_32358
CL\_32359
CL\_32360
CL\_32361
CL\_9477
CL\_11110
CL\_15962
CL\_6732
CL\_17575
CL\_17576
CL\_30637
CL\_30638
CL\_30639
CL\_30640
CL\_1936
CL\_1937
CL\_13822
CL\_7823
CL\_7203
CL\_7822
CL\_17156
CL\_17155
CL\_17154
CL\_7202
CL\_13709
CL\_6808
CL\_10461
CL\_25971
CL\_25970
CL\_25969
CL\_25968
CL\_32458
CL\_25967
CL\_25966
CL\_25965
CL\_32459
CL\_32460
CL\_32461
CL\_32462
CL\_22214
CL\_22213
CL\_23443
CL\_7287
CL\_14619
CL\_8501
CL\_7286
CL\_8500
CL\_8499
CL\_8498
CL\_26531
CL\_26532
CL\_8497
CL\_27575
CL\_27574
CL\_17684
CL\_7905
CL\_27530
CL\_27529
CL\_27528
CL\_14543
CL\_14544
CL\_14545
CL\_14546
CL\_14547
CL\_14548
CL\_14549
CL\_29637
CL\_24197
CL\_13679
CL\_8814
CL\_24198
CL\_8813
CL\_8812
CL\_26469
CL\_26470
CL\_8811
CL\_24199
CL\_8810
CL\_8809
CL\_24200
CL\_24201
CL\_24202
CL\_27679
CL\_27678
CL\_26471
CL\_8808
CL\_8807
CL\_8806
CL\_8805
CL\_8804
CL\_8803
CL\_13680
CL\_13681
CL\_13682
CL\_11186
CL\_10613
CL\_10614
CL\_10615
CL\_13355
CL\_13356
CL\_10616
CL\_10815
CL\_20974
CL\_10814
CL\_10813
CL\_11185
CL\_11184
CL\_11183
CL\_14009
CL\_26795
CL\_26796
CL\_29014
CL\_11182
CL\_13823
CL\_13824
CL\_13825
CL\_12070
CL\_12071
CL\_12072
CL\_12073
CL\_12074
CL\_7285
CL\_8496
CL\_8495
CL\_14397
CL\_8494
CL\_13683
CL\_17686
CL\_32507
CL\_32506
CL\_32505
CL\_32504
CL\_23474
CL\_25194
CL\_11181
CL\_33390
CL\_33389
CL\_26281
CL\_7284
CL\_11706
CL\_11849
CL\_11848
CL\_11705
CL\_23445
CL\_23444
CL\_35043
CL\_23664
CL\_23665
CL\_23666
CL\_23667
CL\_23668
CL\_27350
CL\_10617
CL\_23242
CL\_23243
CL\_23244
CL\_7283
CL\_17685
CL\_8802
CL\_8801
CL\_27677
CL\_8800
CL\_13826
CL\_7981
CL\_14538
CL\_7980
CL\_7979
CL\_8131
CL\_14539
CL\_14540
CL\_27323
CL\_20278
CL\_8130
CL\_8129
CL\_8128
CL\_11106
CL\_8127
CL\_11105
CL\_11711
CL\_13756
CL\_11710
CL\_11709
CL\_11708
CL\_13755
CL\_5240
CL\_7088
CL\_20009
CL\_7087
CL\_5234
CL\_15555
CL\_5391
CL\_7019
CL\_5238
CL\_27682
CL\_5237
CL\_14312
CL\_14313
CL\_14314
CL\_14315
CL\_14316
CL\_14317
CL\_8558
CL\_8559
CL\_8560
CL\_7832
CL\_6419
CL\_20006
CL\_20005
CL\_20004
CL\_8508
CL\_6726
CL\_15983
