## Supplementary material for "A novel method for integrating genomic and Tn-Seq data to identify common *in vivo* fitness mechanisms across multiple bacterial species": S1 Dataset: CL_INS_239.html

Legend

 Mobile +extrachromosomalelementfunctions
 Regulatoryfunctions
 Hypothetical
 DNA Metabolism
 Proteinsynthesis/fate
 Other
 Transport +binding proteins
 All VFDB Genes

FULL


WINDOWSVGPNG

Trim RowsRemove SingletonsSave Fasta

CL\_2866


CL\_2866


CL\_2866


CL\_2866


CL\_2866


CL\_2866


CL\_2866


CL\_2866


CL\_2866


CL\_2866


CL\_2866


CL\_2866


CL\_2866


CL\_2866


CL\_2866

HighlightSelectShow Genomes


239

CL\_2865


2

CL\_2846


2

CL\_2846


1

CL\_2846


1

CL\_2846


1

CL\_2846


1

CL\_2846


1

CL\_234


1

CL\_2864


1

CL\_2846


1

CL\_234


1

CL\_2846


1

CL\_2846


1

CL\_2846


1

CL\_2846

fGI ID


CL\_INS\_352
CL\_INS\_237
CL\_INS\_237
CL\_INS\_239
CL\_INS\_237
CL\_INS\_237
CL\_INS\_237
CL\_INS\_237
CL\_INS\_237
CL\_INS\_237
CL\_INS\_237
CL\_INS\_237
CL\_INS\_237
CL\_INS\_70
CL\_INS\_247
CL\_INS\_233
CL\_INS\_239
CL\_INS\_237
CL\_INS\_237
CL\_INS\_237
CL\_INS\_237
CL\_INS\_237
CL\_INS\_237
CL\_INS\_237
CL\_INS\_237
CL\_INS\_237
CL\_INS\_237
CL\_INS\_237
CL\_INS\_239
CL\_INS\_247
CL\_INS\_237
CL\_INS\_237
CL\_INS\_237
CL\_INS\_237
CL\_INS\_237
CL\_INS\_237
CL\_INS\_237
CL\_INS\_237
CL\_INS\_237
CL\_INS\_237
CL\_INS\_237
CL\_INS\_237
CL\_INS\_237
CL\_INS\_237
CL\_INS\_237
CL\_INS\_237
CL\_INS\_237
CL\_INS\_237
CL\_INS\_237
CL\_INS\_237
CL\_INS\_237
CL\_INS\_237
CL\_INS\_237
CL\_INS\_237
CL\_INS\_237
CL\_INS\_239
CL\_INS\_237
CL\_INS\_237
CL\_INS\_247
CL\_INS\_247
CL\_INS\_247
CL\_INS\_247
CL\_INS\_237
CL\_INS\_247
CL\_INS\_247
CL\_INS\_247
CL\_INS\_239
CL\_INS\_247
CL\_INS\_247
CL\_INS\_247
CL\_INS\_30
CL\_INS\_239
CL\_INS\_237
CL\_INS\_237
CL\_INS\_237
CL\_INS\_237
CL\_INS\_237
CL\_INS\_237
CL\_INS\_237
CL\_INS\_237
CL\_INS\_237
CL\_INS\_237
CL\_INS\_237
CL\_INS\_247
CL\_INS\_247
CL\_INS\_247
CL\_INS\_237
CL\_INS\_247
CL\_INS\_237
CL\_INS\_70
CL\_INS\_70
CL\_INS\_70
CL\_INS\_237
CL\_INS\_237
CL\_INS\_237
CL\_INS\_237
CL\_INS\_237
CL\_INS\_237
CL\_INS\_237
CL\_INS\_237
CL\_INS\_237
CL\_INS\_70
CL\_INS\_70
CL\_INS\_237
CL\_INS\_237
CL\_INS\_237
CL\_INS\_237
CL\_INS\_237
CL\_INS\_237
CL\_INS\_237
CL\_INS\_237
CL\_INS\_237
CL\_INS\_237
CL\_INS\_237
CL\_INS\_237
CL\_INS\_237
CL\_INS\_237
CL\_INS\_237
CL\_INS\_149
CL\_INS\_237
CL\_INS\_237
CL\_INS\_237
CL\_INS\_237
CL\_INS\_237
CL\_INS\_149
CL\_INS\_237
CL\_INS\_237
CL\_INS\_237
CL\_INS\_237
CL\_INS\_237
CL\_INS\_237
CL\_INS\_237
CL\_INS\_237
CL\_INS\_237
CL\_INS\_237
CL\_INS\_237
CL\_INS\_237
Cluster ID


CL\_6833
CL\_23274
CL\_13467
CL\_33158
CL\_33159
CL\_33160
CL\_31766
CL\_33161
CL\_9377
CL\_9376
CL\_9375
CL\_9374
CL\_33162
CL\_9371
CL\_9370
CL\_9369
CL\_34360
CL\_34361
CL\_34362
CL\_34363
CL\_34364
CL\_34365
CL\_34366
CL\_34367
CL\_34368
CL\_34369
CL\_34370
CL\_34371
CL\_8931
CL\_8932
CL\_8933
CL\_8934
CL\_8935
CL\_8936
CL\_8937
CL\_8938
CL\_8939
CL\_8940
CL\_8941
CL\_8942
CL\_244
CL\_10437
CL\_10438
CL\_10439
CL\_10440
CL\_10441
CL\_10442
CL\_10443
CL\_10444
CL\_10445
CL\_10446
CL\_21999
CL\_21998
CL\_21997
CL\_21996
CL\_11847
CL\_11848
CL\_11849
CL\_7284
CL\_7283
CL\_8497
CL\_8498
CL\_8499
CL\_8500
CL\_7286
CL\_8501
CL\_36096
CL\_7287
CL\_7203
CL\_7204
CL\_1936
CL\_12093
CL\_12094
CL\_12095
CL\_12096
CL\_12097
CL\_12098
CL\_12099
CL\_12100
CL\_12101
CL\_12102
CL\_12103
CL\_12104
CL\_5689
CL\_5688
CL\_12106
CL\_19725
CL\_5236
CL\_11189
CL\_7241
CL\_11191
CL\_7242
CL\_7425
CL\_7426
CL\_7427
CL\_7428
CL\_7429
CL\_7430
CL\_7431
CL\_7432
CL\_7433
CL\_7434
CL\_7435
CL\_7436
CL\_7437
CL\_7438
CL\_7236
CL\_7439
CL\_7237
CL\_7440
CL\_7441
CL\_7239
CL\_7240
CL\_5135
CL\_5136
CL\_5137
CL\_5138
CL\_5139
CL\_5140
CL\_5141
CL\_5142
CL\_5143
CL\_5144
CL\_5145
CL\_5146
CL\_5147
CL\_5148
CL\_5149
CL\_5150
CL\_5151
CL\_5152
CL\_5153
CL\_5154
CL\_5155
CL\_5156
CL\_5157
CL\_5158
