## Supplementary material for "A novel method for integrating genomic and Tn-Seq data to identify common *in vivo* fitness mechanisms across multiple bacterial species": S1 Dataset: CL_INS_241.html

Legend

 Mobile +extrachromosomalelementfunctions
 Hypothetical
 Regulatoryfunctions
 Other
 All VFDB Genes

FULL


WINDOWSVGPNG

Trim RowsRemove SingletonsSave Fasta

CL\_2890


CL\_2890


CL\_2890


CL\_2890


CL\_2890


CL\_2890


CL\_2890


CL\_2890

HighlightSelectShow Genomes


189

CL\_2891


68

CL\_2891


6

CL\_2891


4

CL\_2891


1

CL\_2897


1

CL\_2891


1

CL\_2891


1

CL\_2891

fGI ID


CL\_INS\_241
CL\_INS\_241
CL\_INS\_241
CL\_INS\_242
CL\_INS\_242
CL\_INS\_242
CL\_INS\_241
CL\_INS\_241
CL\_INS\_241
CL\_INS\_241
CL\_INS\_70
CL\_INS\_70
CL\_INS\_70
CL\_INS\_70
CL\_INS\_70
CL\_INS\_70
CL\_INS\_70
CL\_INS\_241
CL\_INS\_241
CL\_INS\_241
CL\_INS\_241
CL\_INS\_241
CL\_INS\_241
CL\_INS\_241
CL\_INS\_241
CL\_INS\_241
Cluster ID


CL\_7282
CL\_7281
CL\_7663
CL\_7662
CL\_7661
CL\_7280
CL\_7660
CL\_29991
CL\_29990
CL\_29989
CL\_4095
CL\_4096
CL\_4097
CL\_4098
CL\_4099
CL\_4100
CL\_4101
CL\_6247
CL\_29988
CL\_29987
CL\_29986
CL\_29985
CL\_29984
CL\_29983
CL\_29982
CL\_29981
