## Supplementary material for "A novel method for integrating genomic and Tn-Seq data to identify common *in vivo* fitness mechanisms across multiple bacterial species": S1 Dataset: CL_INS_242.html

Legend

 Hypothetical
 All EssentialGenes
 Other
 All VFDB Genes

FULL


WINDOWSVGPNG

Trim RowsRemove SingletonsSave Fasta

CL\_2893


CL\_2893


CL\_2893


CL\_2893


CL\_2893


CL\_2893


CL\_2890


CL\_2893


CL\_2892


CL\_2892

HighlightSelectShow Genomes


96

CL\_2897


86

CL\_2897


83

CL\_2897


2

CL\_2897


2

CL\_2897


2

CL\_2897


1

CL\_2897


1

CL\_2897


1

CL\_2897


1

CL\_2897

fGI ID


CL\_INS\_242
CL\_INS\_242
CL\_INS\_242
CL\_INS\_242
CL\_INS\_242
CL\_INS\_242
CL\_INS\_242
Cluster ID


CL\_15377
CL\_2894
CL\_7662
CL\_7661
CL\_7280
CL\_2895
CL\_2896
