## Supplementary material for "A novel method for integrating genomic and Tn-Seq data to identify common *in vivo* fitness mechanisms across multiple bacterial species": S1 Dataset: CL_INS_245.html

Legend

 Mobile +extrachromosomalelementfunctions
 Hypothetical
 All EssentialGenes
 Other
 Transport +binding proteins
 All VFDB Genes

FULL


WINDOWSVGPNG

Trim RowsRemove SingletonsSave Fasta

CL\_2913


CL\_2913


CL\_2913


CL\_2912


CL\_2917


CL\_2913

HighlightSelectShow Genomes


148

CL\_2915


66

CL\_2915


37

CL\_2917


1

CL\_2915


1

CL\_2915


1

CL\_2915

fGI ID


CL\_INS\_245
CL\_INS\_246
Cluster ID


CL\_2914
CL\_2918
