## Supplementary material for "A novel method for integrating genomic and Tn-Seq data to identify common *in vivo* fitness mechanisms across multiple bacterial species": S1 Dataset: CL_INS_246.html

Legend

 Mobile +extrachromosomalelementfunctions
 All EssentialGenes
 Cell Envelope
 Other
 All VFDB Genes

FULL


WINDOWSVGPNG

Trim RowsRemove SingletonsSave Fasta

CL\_2917


CL\_2917


CL\_2917


CL\_2917


CL\_2917


CL\_2917


CL\_2917


CL\_2916


CL\_2917


CL\_2917


CL\_2916


CL\_2917


CL\_2917


CL\_3860


CL\_2917


CL\_2917


CL\_2917


CL\_2917


CL\_2917


CL\_2917

HighlightSelectShow Genomes


172

CL\_2920


18

CL\_2920


11

CL\_2921


2

CL\_2920


1

CL\_2915


1

CL\_2921


1

CL\_2921


1

CL\_2920


1

CL\_2920


1

CL\_2920


1

CL\_2920


1

CL\_2920


1

CL\_2920


1

CL\_2920


1

CL\_2920


1

CL\_2921


1

CL\_2920


1

CL\_2920


1

Break


1

CL\_2920

fGI ID


CL\_INS\_246
CL\_INS\_246
CL\_INS\_246
CL\_INS\_246
CL\_INS\_246
CL\_INS\_246
CL\_INS\_246
CL\_INS\_246
Cluster ID


CL\_16903
CL\_22557
CL\_24539
CL\_22516
CL\_6629
CL\_8799
CL\_2918
CL\_2919
