## Supplementary material for "A novel method for integrating genomic and Tn-Seq data to identify common *in vivo* fitness mechanisms across multiple bacterial species": S1 Dataset: CL_INS_248.html

Legend

 Mobile +extrachromosomalelementfunctions
 Regulatoryfunctions
 Hypothetical
 DNA Metabolism
 Other
 Transport +binding proteins
 All VFDB Genes

FULL


WINDOWSVGPNG

Trim RowsRemove SingletonsSave Fasta

CL\_2942


CL\_2942


CL\_2941


CL\_2940

HighlightSelectShow Genomes


246

CL\_2943


31

CL\_2943


1

CL\_2943


1

CL\_2943

fGI ID


CL\_INS\_248
CL\_INS\_247
CL\_INS\_70
CL\_INS\_247
CL\_INS\_30
CL\_INS\_70
CL\_INS\_247
CL\_INS\_247
CL\_INS\_70
CL\_INS\_70
CL\_INS\_70
CL\_INS\_70
CL\_INS\_70
CL\_INS\_247
CL\_INS\_247
CL\_INS\_247
CL\_INS\_247
CL\_INS\_247
CL\_INS\_247
CL\_INS\_247
CL\_INS\_247
CL\_INS\_247
CL\_INS\_247
CL\_INS\_247
CL\_INS\_247
CL\_INS\_247
CL\_INS\_247
CL\_INS\_382
CL\_INS\_247
CL\_INS\_247
CL\_INS\_247
CL\_INS\_247
CL\_INS\_247
CL\_INS\_247
CL\_INS\_247
Cluster ID


CL\_7275
CL\_7203
CL\_7824
CL\_7204
CL\_7749
CL\_10610
CL\_7828
CL\_6726
CL\_8508
CL\_10547
CL\_1931
CL\_8641
CL\_7690
CL\_5236
CL\_21981
CL\_21982
CL\_21983
CL\_23853
CL\_21985
CL\_21986
CL\_21987
CL\_21988
CL\_21989
CL\_14335
CL\_14334
CL\_14333
CL\_14332
CL\_4086
CL\_11712
CL\_15546
CL\_15547
CL\_15548
CL\_15549
CL\_10021
CL\_9460
