## Supplementary material for "A novel method for integrating genomic and Tn-Seq data to identify common *in vivo* fitness mechanisms across multiple bacterial species": S1 Dataset: CL_INS_249.html

Legend

 Hypothetical
 All EssentialGenes
 Other

FULL


WINDOWSVGPNG

Trim RowsRemove SingletonsSave Fasta

CL\_2954


CL\_2954


CL\_2954


CL\_2954


CL\_2954

HighlightSelectShow Genomes


112

CL\_2956


97

CL\_2956


2

CL\_2957


1

CL\_2956


1

CL\_2956

fGI ID


CL\_INS\_249
CL\_INS\_249
CL\_INS\_249
Cluster ID


CL\_16797
CL\_2955
CL\_15386
