## Supplementary material for "A novel method for integrating genomic and Tn-Seq data to identify common *in vivo* fitness mechanisms across multiple bacterial species": S1 Dataset: CL_INS_253.html

Legend

 Mobile +extrachromosomalelementfunctions
 Regulatoryfunctions
 Hypothetical
 All EssentialGenes
 Other
 Transport +binding proteins
 All VFDB Genes

FULL


WINDOWSVGPNG

Trim RowsRemove SingletonsSave Fasta

CL\_3007


CL\_3007


CL\_3007


CL\_3007


CL\_3007


CL\_3004


CL\_2995


CL\_3007


CL\_3007


CL\_3007


CL\_3007


CL\_3004


CL\_3007


CL\_3119


CL\_3120


CL\_2994


CL\_3003


CL\_3007


CL\_2998


CL\_3007


CL\_3007


CL\_3007


CL\_3007


CL\_3007


CL\_3007


CL\_3003


CL\_3007


CL\_3007


CL\_3003


CL\_3006


CL\_3007


CL\_3007


CL\_3007


CL\_3003

HighlightSelectShow Genomes


118

CL\_3012


108

CL\_3012


10

CL\_3012


3

CL\_3012


3

CL\_3012


2

CL\_3012


2

CL\_3012


2

CL\_3012


2

CL\_3012


2

CL\_3012


2

CL\_3012


1

CL\_3012


1

CL\_3012


1

CL\_3012


1

CL\_3012


1

CL\_3012


1

CL\_3012


1

CL\_3012


1

CL\_3012


1

CL\_3012


1

CL\_3012


1

CL\_3012


1

CL\_3012


1

CL\_3012


1

CL\_3014


1

CL\_3012


1

CL\_3012


1

CL\_3012


1

CL\_3012


1

CL\_3012


1

CL\_3012


1

CL\_3012


1

CL\_3012


1

CL\_3012

fGI ID


CL\_INS\_253
CL\_INS\_253
CL\_INS\_253
CL\_INS\_253
CL\_INS\_253
CL\_INS\_253
CL\_INS\_253
CL\_INS\_253
CL\_INS\_253
CL\_INS\_253
CL\_INS\_253
CL\_INS\_253
CL\_INS\_253
CL\_INS\_253
CL\_INS\_253
CL\_INS\_253
CL\_INS\_253
CL\_INS\_253
CL\_INS\_253
CL\_INS\_253
CL\_INS\_253
CL\_INS\_253
CL\_INS\_253
CL\_INS\_253
CL\_INS\_253
CL\_INS\_253
Cluster ID


CL\_30758
CL\_28608
CL\_28609
CL\_31467
CL\_33048
CL\_23595
CL\_12145
CL\_12146
CL\_5957
CL\_8444
CL\_3008
CL\_3009
CL\_34857
CL\_25504
CL\_3010
CL\_29820
CL\_5956
CL\_12850
CL\_5955
CL\_30130
CL\_22215
CL\_30131
CL\_29819
CL\_29818
CL\_30132
CL\_3011
