## Supplementary material for "A novel method for integrating genomic and Tn-Seq data to identify common *in vivo* fitness mechanisms across multiple bacterial species": S1 Dataset: CL_INS_257.html

Legend

 Hypothetical
 All EssentialGenes
 All VFDB Genes

FULL


WINDOWSVGPNG

Trim RowsRemove SingletonsSave Fasta

CL\_3116


CL\_3116


CL\_3116


CL\_3116

HighlightSelectShow Genomes


230

CL\_3117


43

CL\_3117


2

CL\_3117


1

CL\_3118

fGI ID


CL\_INS\_257
CL\_INS\_257
Cluster ID


CL\_15394
CL\_5950
