## Supplementary material for "A novel method for integrating genomic and Tn-Seq data to identify common *in vivo* fitness mechanisms across multiple bacterial species": S1 Dataset: CL_INS_258.html

Legend

 Mobile +extrachromosomalelementfunctions
 Regulatoryfunctions
 Hypothetical
 All EssentialGenes
 Other
 All VFDB Genes

FULL


WINDOWSVGPNG

Trim RowsRemove SingletonsSave Fasta

CL\_3118


CL\_3118


CL\_3118


CL\_3118


CL\_3118


CL\_3118


CL\_3118


CL\_3118


CL\_3012


CL\_3118


CL\_3118

HighlightSelectShow Genomes


187

CL\_3119


79

CL\_3119


3

CL\_3119


1

CL\_3119


1

CL\_3119


1

CL\_4119


1

CL\_3119


1

CL\_3119


1

CL\_3119


1

CL\_2746


1

CL\_3119

fGI ID


CL\_INS\_253
CL\_INS\_258
CL\_INS\_258
CL\_INS\_226
CL\_INS\_226
CL\_INS\_258
CL\_INS\_258
CL\_INS\_258
CL\_INS\_368
CL\_INS\_253
CL\_INS\_253
CL\_INS\_253
CL\_INS\_258
Cluster ID


CL\_12850
CL\_5951
CL\_5952
CL\_12845
CL\_10216
CL\_5953
CL\_30519
CL\_5954
CL\_10436
CL\_5955
CL\_5956
CL\_5957
CL\_5958
