## Supplementary material for "A novel method for integrating genomic and Tn-Seq data to identify common *in vivo* fitness mechanisms across multiple bacterial species": S1 Dataset: CL_INS_261.html

Legend

 Mobile +extrachromosomalelementfunctions
 Regulatoryfunctions
 Hypothetical
 All EssentialGenes
 Other
 All VFDB Genes

FULL


WINDOWSVGPNG

Trim RowsRemove SingletonsSave Fasta

CL\_3742


CL\_3742


CL\_3742


CL\_3742


CL\_3742


CL\_3742


CL\_3742


CL\_3742


CL\_3742


Break


CL\_3742


CL\_3742


CL\_3556


CL\_3742


CL\_3742

HighlightSelectShow Genomes


129

CL\_3736


123

CL\_3736


8

CL\_3735


4

CL\_3736


2

CL\_3736


2

CL\_3736


1

CL\_3736


1

CL\_3736


1

CL\_3728


1

CL\_3736


1

CL\_3736


1

CL\_3734


1

CL\_3736


1

CL\_3558


1

CL\_3736

fGI ID


CL\_INS\_261
CL\_INS\_261
CL\_INS\_261
CL\_INS\_282
CL\_INS\_261
CL\_INS\_282
CL\_INS\_261
Cluster ID


CL\_30524
CL\_24203
CL\_3741
CL\_3740
CL\_3739
CL\_3738
CL\_3737
