## Supplementary material for "A novel method for integrating genomic and Tn-Seq data to identify common *in vivo* fitness mechanisms across multiple bacterial species": S1 Dataset: CL_INS_263.html

Legend

 Hypothetical
 Other
 EnergyMetabolism
 All VFDB Genes

FULL


WINDOWSVGPNG

Trim RowsRemove SingletonsSave Fasta

CL\_3709


CL\_3709


CL\_3709


CL\_3709

HighlightSelectShow Genomes


192

CL\_3708


77

CL\_3708


6

CL\_3708


1

CL\_3705

fGI ID


CL\_INS\_263
CL\_INS\_263
Cluster ID


CL\_4879
CL\_4880
