## Supplementary material for "A novel method for integrating genomic and Tn-Seq data to identify common *in vivo* fitness mechanisms across multiple bacterial species": S1 Dataset: CL_INS_264.html

Legend

 Mobile +extrachromosomalelementfunctions
 Hypothetical
 Other
 All VFDB Genes

FULL


WINDOWSVGPNG

Trim RowsRemove SingletonsSave Fasta

CL\_4881


CL\_4881


CL\_4881


CL\_4881


CL\_3687


CL\_4881


CL\_4881


CL\_4881


CL\_3687


CL\_4881


CL\_3687


CL\_4881


CL\_3687


CL\_3687


CL\_3688


CL\_4881


CL\_3687


CL\_4881


CL\_4881


CL\_4881


CL\_3688


CL\_3687


CL\_4881


CL\_3521


CL\_4881


CL\_4881


CL\_4881


CL\_4881


CL\_3687


CL\_3687


CL\_4881


CL\_4881


CL\_4881


CL\_3687


CL\_4881


CL\_4881


CL\_4881

HighlightSelectShow Genomes


86

CL\_3686


30

CL\_3686


28

CL\_3686


22

CL\_3686


10

CL\_3686


7

CL\_3686


5

CL\_3686


5

CL\_3686


3

CL\_3686


3

CL\_3686


2

CL\_3686


2

CL\_3686


2

CL\_3686


2

CL\_3686


2

CL\_3686


2

CL\_3686


2

CL\_3686


1

CL\_3686


1

CL\_3686


1

CL\_3686


1

CL\_3686


1

CL\_3686


1

CL\_3686


1

CL\_3686


1

CL\_3686


1

CL\_3686


1

CL\_3686


1

CL\_3686


1

CL\_3686


1

CL\_3686


1

CL\_3686


1

CL\_3686


1

CL\_3686


1

CL\_3686


1

CL\_3686


1

CL\_3686


1

CL\_3686

fGI ID


CL\_INS\_264
CL\_INS\_264
CL\_INS\_264
CL\_INS\_264
CL\_INS\_264
CL\_INS\_264
CL\_INS\_264
CL\_INS\_264
CL\_INS\_264
CL\_INS\_264
CL\_INS\_264
CL\_INS\_264
CL\_INS\_264
CL\_INS\_264
CL\_INS\_264
CL\_INS\_264
CL\_INS\_264
CL\_INS\_264
CL\_INS\_264
CL\_INS\_264
CL\_INS\_264
CL\_INS\_264
CL\_INS\_264
CL\_INS\_264
CL\_INS\_264
CL\_INS\_264
Cluster ID


CL\_30536
CL\_12218
CL\_14466
CL\_6626
CL\_4882
CL\_4883
CL\_27873
CL\_4884
CL\_24527
CL\_24526
CL\_12857
CL\_12856
CL\_12855
CL\_23029
CL\_23030
CL\_23031
CL\_23032
CL\_12153
CL\_7176
CL\_7654
CL\_7653
CL\_7175
CL\_36097
CL\_7174
CL\_7173
CL\_7652
