## Supplementary material for "A novel method for integrating genomic and Tn-Seq data to identify common *in vivo* fitness mechanisms across multiple bacterial species": S1 Dataset: CL_INS_265.html

Legend

 Mobile +extrachromosomalelementfunctions
 Other
 Transport +binding proteins

FULL


WINDOWSVGPNG

Trim RowsRemove SingletonsSave Fasta

CL\_3680


CL\_3680


CL\_3680


CL\_3680

HighlightSelectShow Genomes


183

CL\_3679


87

CL\_3679


3

CL\_3679


1

CL\_3679

fGI ID


CL\_INS\_265
CL\_INS\_265
CL\_INS\_265
Cluster ID


CL\_5959
CL\_5960
CL\_30753
