## Supplementary material for "A novel method for integrating genomic and Tn-Seq data to identify common *in vivo* fitness mechanisms across multiple bacterial species": S1 Dataset: CL_INS_267.html

Legend

 Mobile +extrachromosomalelementfunctions
 Hypothetical
 Other
 All VFDB Genes

FULL


WINDOWSVGPNG

Trim RowsRemove SingletonsSave Fasta

CL\_3674


CL\_3674


CL\_3674


CL\_3674


CL\_3674


CL\_3674


CL\_3674


CL\_3675


CL\_3674


CL\_3674


CL\_3674


CL\_3674


CL\_3674


CL\_3674


CL\_3674


CL\_3674


CL\_3674


CL\_3674


CL\_3674


CL\_3675


CL\_3674


CL\_3674


CL\_3674

HighlightSelectShow Genomes


124

CL\_3670


19

CL\_3670


18

CL\_3670


12

CL\_3670


8

CL\_3670


5

CL\_3669


4

CL\_3669


3

CL\_3670


3

CL\_3670


3

CL\_3670


2

CL\_3670


1

CL\_3670


1

CL\_3669


1

CL\_3670


1

CL\_3670


1

CL\_3669


1

CL\_3669


1

CL\_3670


1

CL\_3670


1

CL\_3670


1

CL\_3670


1

CL\_3670


1

CL\_3669

fGI ID


CL\_INS\_267
CL\_INS\_267
CL\_INS\_267
CL\_INS\_267
CL\_INS\_267
CL\_INS\_267
CL\_INS\_267
CL\_INS\_267
CL\_INS\_247
CL\_INS\_267
CL\_INS\_267
CL\_INS\_267
Cluster ID


CL\_15550
CL\_24525
CL\_12154
CL\_12155
CL\_3673
CL\_12858
CL\_3672
CL\_20491
CL\_6844
CL\_3671
CL\_6625
CL\_20305
