## Supplementary material for "A novel method for integrating genomic and Tn-Seq data to identify common *in vivo* fitness mechanisms across multiple bacterial species": S1 Dataset: CL_INS_268.html

Legend

 Hypothetical
 Other
 All VFDB Genes

FULL


WINDOWSVGPNG

Trim RowsRemove SingletonsSave Fasta

CL\_3657


CL\_3657


CL\_3657


CL\_3657


CL\_3657


CL\_3657


CL\_3658


CL\_3658


CL\_3657

HighlightSelectShow Genomes


208

CL\_3656


62

CL\_3656


3

CL\_3656


1

CL\_3655


1

CL\_3656


1

CL\_3655


1

CL\_3656


1

CL\_3656


1

CL\_3656

fGI ID


CL\_INS\_268
CL\_INS\_268
CL\_INS\_268
CL\_INS\_269
Cluster ID


CL\_27478
CL\_10620
CL\_5962
CL\_5963
