## Supplementary material for "A novel method for integrating genomic and Tn-Seq data to identify common *in vivo* fitness mechanisms across multiple bacterial species": S1 Dataset: CL_INS_269.html

Legend

 Hypothetical
 Other
 All VFDB Genes
 Transport +binding proteins

FULL


WINDOWSVGPNG

Trim RowsRemove SingletonsSave Fasta

CL\_3656


CL\_3656


CL\_3656


CL\_3657


CL\_3657

HighlightSelectShow Genomes


259

CL\_3655


16

CL\_3655


2

CL\_3655


1

CL\_3655


1

CL\_3655

fGI ID


CL\_INS\_269
CL\_INS\_269
CL\_INS\_269
CL\_INS\_269
CL\_INS\_269
CL\_INS\_269
CL\_INS\_269
CL\_INS\_269
Cluster ID


CL\_15403
CL\_5963
CL\_7172
CL\_7171
CL\_7170
CL\_7169
CL\_7168
CL\_7167
