## Supplementary material for "A novel method for integrating genomic and Tn-Seq data to identify common *in vivo* fitness mechanisms across multiple bacterial species": S1 Dataset: CL_INS_271.html

Legend

 Mobile +extrachromosomalelementfunctions
 Regulatoryfunctions
 Hypothetical
 All EssentialGenes
 All Fitness Genes
 Other
 Transport +binding proteins
 All VFDB Genes

FULL


WINDOWSVGPNG

Trim RowsRemove SingletonsSave Fasta

CL\_3635


CL\_3635


CL\_3635


CL\_3635


CL\_3635


CL\_3635


CL\_3635


CL\_3635


CL\_3635


CL\_3635


CL\_3635


CL\_3635


CL\_3635


CL\_3635


CL\_3635


CL\_3635


CL\_3637


CL\_3635


CL\_1440


CL\_3635


CL\_3635


CL\_3635


CL\_3635


CL\_3635


CL\_3635


CL\_3635


CL\_3456


CL\_3635


CL\_3635


CL\_3635


CL\_3635


CL\_3635


CL\_3635


CL\_3635


CL\_3635


CL\_3635


CL\_3635


CL\_3635

HighlightSelectShow Genomes


55

CL\_3630


50

CL\_3630


47

CL\_3630


34

CL\_3630


19

CL\_3630


13

CL\_3630


11

CL\_3630


8

CL\_3630


5

CL\_3630


5

CL\_3630


4

CL\_3630


2

CL\_3630


2

CL\_3630


2

CL\_3630


1

CL\_3630


1

CL\_3630


1

CL\_3630


1

CL\_3630


1

CL\_3630


1

CL\_3630


1

CL\_3630


1

CL\_3630


1

CL\_3630


1

CL\_3630


1

CL\_3623


1

CL\_3630


1

CL\_3630


1

CL\_3630


1

CL\_3630


1

CL\_3630


1

Break


1

CL\_3512


1

CL\_3629


1

CL\_3687


1

CL\_3630


1

CL\_3566


1

CL\_3630


1

CL\_3630

fGI ID


CL\_INS\_271
CL\_INS\_271
CL\_INS\_271
CL\_INS\_271
CL\_INS\_271
CL\_INS\_271
CL\_INS\_271
CL\_INS\_271
CL\_INS\_271
CL\_INS\_271
CL\_INS\_271
CL\_INS\_271
CL\_INS\_271
CL\_INS\_271
CL\_INS\_271
CL\_INS\_271
CL\_INS\_271
CL\_INS\_271
CL\_INS\_271
CL\_INS\_271
CL\_INS\_271
CL\_INS\_271
CL\_INS\_271
CL\_INS\_271
CL\_INS\_271
CL\_INS\_271
CL\_INS\_271
CL\_INS\_271
Cluster ID


CL\_662
CL\_3520
CL\_3518
CL\_32496
CL\_3634
CL\_3633
CL\_3632
CL\_3631
CL\_663
CL\_9538
CL\_9536
CL\_9535
CL\_11697
CL\_3519
CL\_5130
CL\_11696
CL\_7651
CL\_30751
CL\_30750
CL\_7166
CL\_7165
CL\_7164
CL\_7163
CL\_7162
CL\_7161
CL\_4885
CL\_4886
CL\_9459
