## Supplementary material for "A novel method for integrating genomic and Tn-Seq data to identify common *in vivo* fitness mechanisms across multiple bacterial species": S1 Dataset: CL_INS_272.html

Legend

 Mobile +extrachromosomalelementfunctions
 Regulatoryfunctions
 Hypothetical
 DNA Metabolism
 AntibioticResistance
 Other
 Transport +binding proteins
 All VFDB Genes

FULL


WINDOWSVGPNG

Trim RowsRemove SingletonsSave Fasta

CL\_3628


CL\_3628


CL\_3628


CL\_3628


CL\_3629


CL\_3628


Break


CL\_3628


CL\_3628


CL\_3628


CL\_3628


CL\_3628


CL\_3628


CL\_3636


CL\_3628


CL\_3628


CL\_3628


CL\_3628


CL\_3628


CL\_3628


CL\_3628


CL\_3628


CL\_3628


Break


CL\_3628

HighlightSelectShow Genomes


120

CL\_3626


61

CL\_3626


56

CL\_3626


12

CL\_3626


3

CL\_3626


3

CL\_3626


2

CL\_3626


2

CL\_3626


2

CL\_3625


1

CL\_3625


1

CL\_4129


1

CL\_3457


1

CL\_3625


1

CL\_3626


1

CL\_3626


1

CL\_3626


1

Break


1

CL\_3626


1

CL\_3626


1

CL\_3625


1

CL\_3626


1

CL\_3626


1

CL\_3625


1

CL\_3626


1

CL\_3625

fGI ID


CL\_INS\_272
CL\_INS\_272
CL\_INS\_271
CL\_INS\_271
CL\_INS\_271
CL\_INS\_272
CL\_INS\_272
CL\_INS\_272
CL\_INS\_272
CL\_INS\_272
CL\_INS\_382
CL\_INS\_382
CL\_INS\_272
CL\_INS\_272
CL\_INS\_272
CL\_INS\_272
CL\_INS\_294
CL\_INS\_272
CL\_INS\_272
CL\_INS\_294
CL\_INS\_294
CL\_INS\_294
CL\_INS\_294
CL\_INS\_272
CL\_INS\_272
CL\_INS\_272
CL\_INS\_272
CL\_INS\_237
CL\_INS\_272
CL\_INS\_272
CL\_INS\_272
CL\_INS\_272
CL\_INS\_272
CL\_INS\_272
CL\_INS\_272
CL\_INS\_272
CL\_INS\_272
CL\_INS\_272
CL\_INS\_272
CL\_INS\_272
CL\_INS\_272
CL\_INS\_272
CL\_INS\_272
CL\_INS\_272
CL\_INS\_272
CL\_INS\_272
CL\_INS\_272
CL\_INS\_272
CL\_INS\_272
CL\_INS\_272
CL\_INS\_272
CL\_INS\_272
CL\_INS\_272
CL\_INS\_272
CL\_INS\_272
CL\_INS\_272
CL\_INS\_272
CL\_INS\_272
CL\_INS\_272
CL\_INS\_272
CL\_INS\_272
CL\_INS\_368
CL\_INS\_368
CL\_INS\_272
CL\_INS\_272
CL\_INS\_272
CL\_INS\_272
CL\_INS\_272
CL\_INS\_272
CL\_INS\_272
CL\_INS\_272
CL\_INS\_272
CL\_INS\_272
CL\_INS\_272
CL\_INS\_247
CL\_INS\_247
CL\_INS\_86
CL\_INS\_237
CL\_INS\_86
CL\_INS\_237
CL\_INS\_237
CL\_INS\_237
CL\_INS\_86
CL\_INS\_247
CL\_INS\_247
CL\_INS\_382
CL\_INS\_382
CL\_INS\_382
CL\_INS\_382
CL\_INS\_247
CL\_INS\_382
CL\_INS\_382
CL\_INS\_382
CL\_INS\_272
CL\_INS\_207
CL\_INS\_207
CL\_INS\_207
CL\_INS\_159
CL\_INS\_382
CL\_INS\_382
CL\_INS\_382
CL\_INS\_382
CL\_INS\_382
CL\_INS\_382
CL\_INS\_382
CL\_INS\_382
CL\_INS\_382
CL\_INS\_382
CL\_INS\_382
CL\_INS\_382
CL\_INS\_159
CL\_INS\_382
CL\_INS\_382
CL\_INS\_382
CL\_INS\_382
CL\_INS\_382
CL\_INS\_382
CL\_INS\_159
CL\_INS\_382
CL\_INS\_382
CL\_INS\_382
CL\_INS\_382
CL\_INS\_382
CL\_INS\_382
CL\_INS\_382
CL\_INS\_382
CL\_INS\_382
CL\_INS\_382
CL\_INS\_382
CL\_INS\_382
CL\_INS\_382
CL\_INS\_382
CL\_INS\_159
CL\_INS\_385
CL\_INS\_382
CL\_INS\_382
CL\_INS\_382
CL\_INS\_159
CL\_INS\_159
CL\_INS\_382
CL\_INS\_385
CL\_INS\_385
CL\_INS\_382
CL\_INS\_382
CL\_INS\_382
CL\_INS\_382
CL\_INS\_385
CL\_INS\_368
CL\_INS\_159
CL\_INS\_159
CL\_INS\_159
CL\_INS\_382
CL\_INS\_159
CL\_INS\_382
CL\_INS\_99
CL\_INS\_99
CL\_INS\_382
CL\_INS\_382
CL\_INS\_382
CL\_INS\_159
CL\_INS\_99
CL\_INS\_382
CL\_INS\_382
CL\_INS\_159
CL\_INS\_159
CL\_INS\_272
CL\_INS\_272
CL\_INS\_156
CL\_INS\_272
CL\_INS\_207
CL\_INS\_272
CL\_INS\_272
CL\_INS\_382
CL\_INS\_382
CL\_INS\_382
CL\_INS\_247
CL\_INS\_382
CL\_INS\_382
CL\_INS\_382
CL\_INS\_382
CL\_INS\_382
CL\_INS\_382
CL\_INS\_382
CL\_INS\_70
CL\_INS\_272
CL\_INS\_237
CL\_INS\_237
CL\_INS\_237
CL\_INS\_20
CL\_INS\_20
CL\_INS\_237
CL\_INS\_237
CL\_INS\_382
CL\_INS\_382
CL\_INS\_237
CL\_INS\_247
CL\_INS\_382
CL\_INS\_272
CL\_INS\_272
CL\_INS\_272
CL\_INS\_272
CL\_INS\_70
CL\_INS\_70
CL\_INS\_70
CL\_INS\_70
Cluster ID


CL\_12075
CL\_27872
CL\_7163
CL\_7162
CL\_7161
CL\_14156
CL\_14154
CL\_14153
CL\_14152
CL\_17006
CL\_8554
CL\_10807
CL\_17011
CL\_4887
CL\_26273
CL\_13595
CL\_3627
CL\_15404
CL\_25962
CL\_23751
CL\_7637
CL\_30544
CL\_7153
CL\_13226
CL\_13227
CL\_13228
CL\_13229
CL\_1930
CL\_9630
CL\_6343
CL\_6344
CL\_6345
CL\_6342
CL\_6341
CL\_6340
CL\_6339
CL\_6338
CL\_6337
CL\_6336
CL\_6335
CL\_6334
CL\_6333
CL\_6332
CL\_10238
CL\_10237
CL\_6331
CL\_6330
CL\_6329
CL\_6328
CL\_6327
CL\_6326
CL\_6325
CL\_6324
CL\_6323
CL\_9629
CL\_6322
CL\_6321
CL\_6320
CL\_6319
CL\_21243
CL\_21242
CL\_6318
CL\_8488
CL\_15129
CL\_15128
CL\_15127
CL\_15126
CL\_15125
CL\_15124
CL\_15123
CL\_15122
CL\_15121
CL\_15120
CL\_15119
CL\_5601
CL\_5678
CL\_10414
CL\_5697
CL\_10420
CL\_6761
CL\_6760
CL\_10419
CL\_10418
CL\_10664
CL\_11981
CL\_10665
CL\_10666
CL\_10667
CL\_9691
CL\_9690
CL\_9689
CL\_9688
CL\_9687
CL\_5615
CL\_5614
CL\_5613
CL\_5536
CL\_5539
CL\_10411
CL\_10407
CL\_10406
CL\_4303
CL\_4302
CL\_4301
CL\_4300
CL\_4299
CL\_5662
CL\_5548
CL\_4294
CL\_5549
CL\_5550
CL\_5551
CL\_5552
CL\_5553
CL\_5554
CL\_5555
CL\_5556
CL\_5651
CL\_4284
CL\_5560
CL\_5561
CL\_5562
CL\_5563
CL\_5564
CL\_5565
CL\_5566
CL\_5567
CL\_4278
CL\_4277
CL\_5639
CL\_5638
CL\_5637
CL\_5572
CL\_5573
CL\_5574
CL\_4271
CL\_4270
CL\_5575
CL\_5577
CL\_5579
CL\_13400
CL\_4266
CL\_4265
CL\_5062
CL\_4263
CL\_4262
CL\_5625
CL\_5585
CL\_4261
CL\_4260
CL\_4259
CL\_14125
CL\_4258
CL\_4256
CL\_14149
CL\_11834
CL\_11835
CL\_4254
CL\_4253
CL\_5053
CL\_5052
CL\_5051
CL\_5050
CL\_11836
CL\_11837
CL\_11838
CL\_11839
CL\_5502
CL\_15118
CL\_10314
CL\_15117
CL\_15116
CL\_10404
CL\_5038
CL\_10403
CL\_6844
CL\_5001
CL\_5000
CL\_10401
CL\_9918
CL\_9917
CL\_6177
CL\_6178
CL\_6179
CL\_6180
CL\_6181
CL\_6182
CL\_244
CL\_245
CL\_246
CL\_247
CL\_15115
CL\_4974
CL\_4973
CL\_6413
CL\_10372
CL\_10605
CL\_15114
CL\_15113
CL\_15112
CL\_15111
CL\_4098
CL\_4099
CL\_4100
CL\_4101
