## Supplementary material for "A novel method for integrating genomic and Tn-Seq data to identify common *in vivo* fitness mechanisms across multiple bacterial species": S1 Dataset: CL_INS_274.html

Legend

 Mobile +extrachromosomalelementfunctions
 Hypothetical
 Regulatoryfunctions
 All EssentialGenes
 Biosynthesis ofcofactors,prostheticgroups, +carriers
 Other
 Transport +binding proteins
 All VFDB Genes

FULL


WINDOWSVGPNG

Trim RowsRemove SingletonsSave Fasta

CL\_3613


CL\_3613


CL\_3613


CL\_3613


CL\_3613


CL\_3613


CL\_3613


CL\_3613


CL\_3613


CL\_3613


CL\_3613


CL\_3614


CL\_3547


CL\_3613


CL\_3613


CL\_3613


CL\_3613


CL\_3613


CL\_3615


CL\_3613


CL\_3613


CL\_3613

HighlightSelectShow Genomes


85

CL\_3610


65

CL\_3610


58

CL\_3610


49

CL\_3610


2

CL\_3609


2

CL\_3610


1

CL\_3610


1

CL\_3610


1

CL\_3610


1

CL\_3616


1

CL\_3610


1

CL\_3610


1

CL\_3610


1

CL\_3610


1

CL\_3610


1

CL\_3610


1

CL\_3609


1

CL\_3610


1

CL\_3610


1

CL\_3610


1

CL\_3608


1

CL\_3610

fGI ID


CL\_INS\_237
CL\_INS\_274
CL\_INS\_274
CL\_INS\_274
CL\_INS\_274
CL\_INS\_274
CL\_INS\_274
CL\_INS\_274
CL\_INS\_274
CL\_INS\_274
CL\_INS\_274
CL\_INS\_275
CL\_INS\_275
CL\_INS\_275
CL\_INS\_275
CL\_INS\_275
CL\_INS\_275
CL\_INS\_275
CL\_INS\_55
CL\_INS\_55
CL\_INS\_55
CL\_INS\_55
CL\_INS\_55
CL\_INS\_55
CL\_INS\_55
CL\_INS\_286
CL\_INS\_273
CL\_INS\_274
CL\_INS\_274
CL\_INS\_274
CL\_INS\_274
CL\_INS\_274
Cluster ID


CL\_6207
CL\_7394
CL\_4889
CL\_30543
CL\_4890
CL\_23752
CL\_15406
CL\_12863
CL\_3612
CL\_3611
CL\_11843
CL\_7160
CL\_7650
CL\_7158
CL\_7649
CL\_7648
CL\_7647
CL\_7646
CL\_20484
CL\_20483
CL\_20482
CL\_20481
CL\_20480
CL\_20479
CL\_20478
CL\_13719
CL\_13718
CL\_28616
CL\_28617
CL\_28618
CL\_28619
CL\_28620
