## Supplementary material for "A novel method for integrating genomic and Tn-Seq data to identify common *in vivo* fitness mechanisms across multiple bacterial species": S1 Dataset: CL_INS_275.html

Legend

 Mobile +extrachromosomalelementfunctions
 Regulatoryfunctions
 Hypothetical
 All EssentialGenes
 All VFDB Genes

FULL


WINDOWSVGPNG

Trim RowsRemove SingletonsSave Fasta

CL\_3609


CL\_3609


CL\_3609


CL\_3609


CL\_3609


CL\_3609


CL\_3613


CL\_3609


CL\_3609


CL\_3610


CL\_3609

HighlightSelectShow Genomes


156

CL\_3608


82

CL\_3608


27

CL\_3608


4

CL\_3608


1

CL\_3607


1

CL\_3608


1

CL\_3608


1

CL\_3608


1

CL\_3608


1

CL\_3608


1

CL\_3607

fGI ID


CL\_INS\_275
CL\_INS\_275
CL\_INS\_275
CL\_INS\_275
CL\_INS\_275
CL\_INS\_275
CL\_INS\_275
CL\_INS\_275
CL\_INS\_275
CL\_INS\_275
CL\_INS\_275
CL\_INS\_275
Cluster ID


CL\_7160
CL\_7650
CL\_16491
CL\_7159
CL\_7158
CL\_7649
CL\_7648
CL\_7647
CL\_7646
CL\_7645
CL\_11196
CL\_19699
