## Supplementary material for "A novel method for integrating genomic and Tn-Seq data to identify common *in vivo* fitness mechanisms across multiple bacterial species": S1 Dataset: CL_INS_280.html

Legend

 Hypothetical
 Other
 All VFDB Genes

FULL


WINDOWSVGPNG

Trim RowsRemove SingletonsSave Fasta

CL\_3572


CL\_3572


CL\_3572


CL\_3572


CL\_3572


CL\_3572

HighlightSelectShow Genomes


185

CL\_3570


53

CL\_3570


34

CL\_3570


3

CL\_3569


1

CL\_3566


1

CL\_3569

fGI ID


CL\_INS\_280
CL\_INS\_280
CL\_INS\_280
CL\_INS\_280
CL\_INS\_280
Cluster ID


CL\_7157
CL\_3571
CL\_5964
CL\_5965
CL\_5967
