## Supplementary material for "A novel method for integrating genomic and Tn-Seq data to identify common *in vivo* fitness mechanisms across multiple bacterial species": S1 Dataset: CL_INS_281.html

Legend

 Other
 All VFDB Genes

FULL


WINDOWSVGPNG

Trim RowsRemove SingletonsSave Fasta

CL\_3569


CL\_3569


CL\_3569


CL\_3569


CL\_3569


CL\_3569


CL\_3573


CL\_3569


CL\_3570


CL\_3569


CL\_3572

HighlightSelectShow Genomes


217

CL\_3568


49

CL\_3568


2

CL\_3568


2

CL\_3568


1

CL\_3567


1

CL\_3568


1

CL\_3568


1

CL\_3568


1

CL\_3568


1

CL\_3568


1

CL\_3568

fGI ID


CL\_INS\_280
CL\_INS\_280
CL\_INS\_281
CL\_INS\_280
CL\_INS\_281
Cluster ID


CL\_5964
CL\_5965
CL\_5966
CL\_5967
CL\_5968
