## Supplementary material for "A novel method for integrating genomic and Tn-Seq data to identify common *in vivo* fitness mechanisms across multiple bacterial species": S1 Dataset: CL_INS_282.html

Legend

 Mobile +extrachromosomalelementfunctions
 Hypothetical
 All EssentialGenes
 Other
 All VFDB Genes

FULL


WINDOWSVGPNG

Trim RowsRemove SingletonsSave Fasta

CL\_3558


CL\_3558


CL\_3559


CL\_3558


CL\_3558


CL\_3558


CL\_3558


CL\_3558


CL\_3558


CL\_3736


CL\_3559


CL\_3539

HighlightSelectShow Genomes


154

CL\_3556


98

CL\_3556


7

CL\_3556


2

CL\_3554


1

CL\_3556


1

CL\_3556


1

CL\_3556


1

CL\_3742


1

CL\_3596


1

CL\_3556


1

CL\_3556


1

CL\_3556

fGI ID


CL\_INS\_282
CL\_INS\_282
CL\_INS\_282
CL\_INS\_282
CL\_INS\_282
CL\_INS\_282
CL\_INS\_282
CL\_INS\_282
CL\_INS\_282
Cluster ID


CL\_12865
CL\_32463
CL\_26161
CL\_3738
CL\_3557
CL\_29640
CL\_3740
CL\_13412
CL\_13411
