## Supplementary material for "A novel method for integrating genomic and Tn-Seq data to identify common *in vivo* fitness mechanisms across multiple bacterial species": S1 Dataset: CL_INS_286.html

Legend

 Mobile +extrachromosomalelementfunctions
 Regulatoryfunctions
 Hypothetical
 DNA Metabolism
 AntibioticResistance
 All EssentialGenes
 Cell Envelope
 Other
 Transport +binding proteins
 All VFDB Genes

FULL


WINDOWSVGPNG

Trim RowsRemove SingletonsSave Fasta

CL\_3521


CL\_3521


CL\_3521


CL\_3521


CL\_3521


CL\_3521


CL\_3521


CL\_3521


CL\_3521


CL\_3521


CL\_3521


CL\_3521


CL\_3521


CL\_3521


CL\_3521


CL\_3521


CL\_3521


CL\_3521


CL\_3521


CL\_3521


CL\_3521


CL\_3521


CL\_3521


CL\_3521


CL\_3521


CL\_3521


CL\_3521


CL\_3522


CL\_3521


CL\_3521


CL\_3521


CL\_3521


CL\_3521


CL\_3521


CL\_3521


CL\_3521


CL\_3521


CL\_3521


CL\_3521


CL\_3521


CL\_3521


CL\_3521


CL\_3521


CL\_3521


CL\_3521


CL\_3521


CL\_3521


CL\_3521


CL\_3521


CL\_3521


CL\_3521


CL\_3521


CL\_3521


CL\_3521


CL\_3521


CL\_3521


CL\_3521


CL\_3521


CL\_3194


CL\_3521


CL\_3521


CL\_3521


CL\_3521


CL\_3521


CL\_3521


CL\_3521

HighlightSelectShow Genomes


92

CL\_3516


32

CL\_3516


19

CL\_3516


12

CL\_3516


11

CL\_3516


7

CL\_3516


5

CL\_3516


5

CL\_3516


5

CL\_3516


5

CL\_3516


5

CL\_3516


4

CL\_3516


4

CL\_3516


4

CL\_3516


4

CL\_3516


3

CL\_3516


3

CL\_3516


2

CL\_3516


2

CL\_3516


2

CL\_3516


2

CL\_3516


2

CL\_3516


2

CL\_3516


1

CL\_3516


1

CL\_3516


1

CL\_3552


1

CL\_3516


1

CL\_3516


1

CL\_3516


1

CL\_3516


1

CL\_3516


1

CL\_3515


1

CL\_3508


1

CL\_3515


1

CL\_3516


1

CL\_3516


1

CL\_3516


1

CL\_3516


1

CL\_3516


1

CL\_3516


1

CL\_3516


1

CL\_3516


1

CL\_3516


1

CL\_3516


1

CL\_3686


1

CL\_3516


1

Break


1

CL\_3516


1

CL\_3516


1

Break


1

CL\_3516


1

CL\_3193


1

CL\_3516


1

CL\_3516


1

CL\_3516


1

CL\_3516


1

CL\_3516


1

CL\_4900


1

CL\_3516


1

CL\_3516


1

CL\_3516


1

CL\_3516


1

CL\_3516


1

CL\_3516


1

CL\_3516


1

CL\_3516

fGI ID


CL\_INS\_286
CL\_INS\_264
CL\_INS\_264
CL\_INS\_286
CL\_INS\_286
CL\_INS\_286
CL\_INS\_286
CL\_INS\_271
CL\_INS\_271
CL\_INS\_271
CL\_INS\_271
CL\_INS\_271
CL\_INS\_351
CL\_INS\_351
CL\_INS\_351
CL\_INS\_351
CL\_INS\_351
CL\_INS\_351
CL\_INS\_351
CL\_INS\_351
CL\_INS\_351
CL\_INS\_351
CL\_INS\_286
CL\_INS\_159
CL\_INS\_286
CL\_INS\_286
CL\_INS\_156
CL\_INS\_156
CL\_INS\_156
CL\_INS\_286
CL\_INS\_156
CL\_INS\_20
CL\_INS\_271
CL\_INS\_271
CL\_INS\_271
CL\_INS\_271
CL\_INS\_286
CL\_INS\_286
CL\_INS\_286
CL\_INS\_286
CL\_INS\_271
CL\_INS\_286
CL\_INS\_286
CL\_INS\_286
CL\_INS\_271
CL\_INS\_286
CL\_INS\_286
CL\_INS\_271
CL\_INS\_271
CL\_INS\_286
CL\_INS\_286
CL\_INS\_286
CL\_INS\_286
CL\_INS\_286
CL\_INS\_286
CL\_INS\_159
CL\_INS\_159
CL\_INS\_286
CL\_INS\_237
CL\_INS\_237
CL\_INS\_237
CL\_INS\_286
CL\_INS\_237
CL\_INS\_286
CL\_INS\_286
CL\_INS\_237
CL\_INS\_237
CL\_INS\_237
CL\_INS\_237
CL\_INS\_237
CL\_INS\_174
CL\_INS\_286
CL\_INS\_286
CL\_INS\_286
CL\_INS\_286
CL\_INS\_286
CL\_INS\_237
CL\_INS\_237
CL\_INS\_237
CL\_INS\_237
CL\_INS\_272
CL\_INS\_70
CL\_INS\_382
CL\_INS\_382
CL\_INS\_117
CL\_INS\_237
CL\_INS\_237
CL\_INS\_237
CL\_INS\_237
CL\_INS\_286
CL\_INS\_286
CL\_INS\_382
CL\_INS\_207
CL\_INS\_207
CL\_INS\_207
CL\_INS\_237
CL\_INS\_237
CL\_INS\_385
CL\_INS\_385
CL\_INS\_385
CL\_INS\_385
CL\_INS\_286
CL\_INS\_247
CL\_INS\_57
CL\_INS\_286
CL\_INS\_237
CL\_INS\_237
CL\_INS\_385
CL\_INS\_247
CL\_INS\_71
CL\_INS\_86
CL\_INS\_237
CL\_INS\_237
CL\_INS\_237
CL\_INS\_286
CL\_INS\_217
CL\_INS\_247
CL\_INS\_117
CL\_INS\_123
CL\_INS\_247
CL\_INS\_286
CL\_INS\_286
CL\_INS\_286
CL\_INS\_237
CL\_INS\_70
CL\_INS\_286
CL\_INS\_286
CL\_INS\_286
CL\_INS\_286
CL\_INS\_131
CL\_INS\_237
CL\_INS\_237
CL\_INS\_237
CL\_INS\_237
CL\_INS\_237
CL\_INS\_237
CL\_INS\_237
CL\_INS\_237
CL\_INS\_382
CL\_INS\_237
CL\_INS\_237
CL\_INS\_286
CL\_INS\_237
CL\_INS\_286
Cluster ID


CL\_30535
CL\_30536
CL\_12218
CL\_11695
CL\_11694
CL\_11693
CL\_20002
CL\_3634
CL\_663
CL\_9538
CL\_9535
CL\_11697
CL\_11119
CL\_11122
CL\_11121
CL\_13366
CL\_7643
CL\_7642
CL\_7641
CL\_7561
CL\_7560
CL\_7559
CL\_7640
CL\_7639
CL\_34805
CL\_5480
CL\_5481
CL\_5482
CL\_5483
CL\_5484
CL\_5486
CL\_9476
CL\_3633
CL\_3632
CL\_3631
CL\_3519
CL\_24519
CL\_24518
CL\_24517
CL\_4894
CL\_3518
CL\_29077
CL\_12868
CL\_5131
CL\_5130
CL\_3517
CL\_28424
CL\_3520
CL\_662
CL\_29814
CL\_9458
CL\_9810
CL\_9809
CL\_11663
CL\_11664
CL\_11665
CL\_11666
CL\_20507
CL\_11385
CL\_6135
CL\_6134
CL\_11668
CL\_9672
CL\_19786
CL\_13719
CL\_11388
CL\_6196
CL\_6195
CL\_6194
CL\_6193
CL\_6192
CL\_9903
CL\_9904
CL\_9905
CL\_9906
CL\_9907
CL\_6190
CL\_6189
CL\_6182
CL\_6181
CL\_6180
CL\_6179
CL\_6178
CL\_6177
CL\_6176
CL\_6175
CL\_6174
CL\_6173
CL\_6172
CL\_34693
CL\_34694
CL\_4972
CL\_5509
CL\_5510
CL\_5511
CL\_5512
CL\_5513
CL\_5522
CL\_5521
CL\_5520
CL\_5519
CL\_34695
CL\_10388
CL\_6751
CL\_19701
CL\_5528
CL\_5527
CL\_5296
CL\_5514
CL\_5515
CL\_5516
CL\_6761
CL\_6760
CL\_10419
CL\_5295
CL\_23459
CL\_5302
CL\_10417
CL\_5298
CL\_5299
CL\_11646
CL\_6307
CL\_20494
CL\_23274
CL\_14344
CL\_18446
CL\_18445
CL\_18444
CL\_18443
CL\_11660
CL\_11347
CL\_11346
CL\_11345
CL\_11344
CL\_11343
CL\_6139
CL\_6140
CL\_11342
CL\_6141
CL\_6142
CL\_6143
CL\_20503
CL\_7223
CL\_20504
