## Supplementary material for "A novel method for integrating genomic and Tn-Seq data to identify common *in vivo* fitness mechanisms across multiple bacterial species": S1 Dataset: CL_INS_287.html

Legend

 Mobile +extrachromosomalelementfunctions
 Hypothetical
 Other

FULL


WINDOWSVGPNG

Trim RowsRemove SingletonsSave Fasta

CL\_3510


CL\_3510


CL\_3384


CL\_3511


CL\_3510


CL\_3510

HighlightSelectShow Genomes


179

CL\_4895


32

CL\_4895


1

CL\_4895


1

CL\_4895


1

CL\_4895


1

CL\_3507

fGI ID


CL\_INS\_287
CL\_INS\_287
CL\_INS\_307
Cluster ID


CL\_6622
CL\_24029
CL\_10431
