## Supplementary material for "A novel method for integrating genomic and Tn-Seq data to identify common *in vivo* fitness mechanisms across multiple bacterial species": S1 Dataset: CL_INS_288.html

Legend

 Mobile +extrachromosomalelementfunctions
 Hypothetical
 All VFDB Genes

FULL


WINDOWSVGPNG

Trim RowsRemove SingletonsSave Fasta

CL\_3495


CL\_3495


CL\_3495


CL\_3495


CL\_3495

HighlightSelectShow Genomes


204

CL\_3493


69

CL\_3493


2

CL\_3493


1

CL\_3493


1

CL\_3486

fGI ID


CL\_INS\_288
CL\_INS\_288
CL\_INS\_288
CL\_INS\_288
Cluster ID


CL\_35184
CL\_3494
CL\_9627
CL\_4897
