## Supplementary material for "A novel method for integrating genomic and Tn-Seq data to identify common *in vivo* fitness mechanisms across multiple bacterial species": S1 Dataset: CL_INS_289.html

Legend

 Hypothetical
 All EssentialGenes
 Other
 All VFDB Genes

FULL


WINDOWSVGPNG

Trim RowsRemove SingletonsSave Fasta

CL\_3491


CL\_3491


CL\_3491


CL\_3491


CL\_3491


CL\_3498


CL\_3491


CL\_3491


CL\_3496


CL\_3491


CL\_3492


CL\_3491


CL\_3491


CL\_3496


CL\_3491


CL\_3491


CL\_3491


CL\_3491


CL\_3491


CL\_3491

HighlightSelectShow Genomes


111

CL\_3487


55

CL\_3486


42

CL\_3487


33

CL\_3487


21

CL\_3487


2

CL\_3487


2

CL\_3487


2

CL\_3486


1

CL\_3487


1

CL\_3487


1

CL\_3487


1

CL\_3486


1

CL\_3487


1

CL\_3487


1

CL\_3487


1

CL\_3487


1

CL\_2502


1

Break


1

CL\_3486


1

CL\_3487

fGI ID


CL\_INS\_289
CL\_INS\_289
CL\_INS\_289
CL\_INS\_289
CL\_INS\_289
CL\_INS\_289
CL\_INS\_289
CL\_INS\_290
CL\_INS\_289
CL\_INS\_290
CL\_INS\_288
CL\_INS\_290
CL\_INS\_289
Cluster ID


CL\_3490
CL\_22222
CL\_3489
CL\_15180
CL\_15181
CL\_15182
CL\_5129
CL\_5128
CL\_7424
CL\_5127
CL\_4897
CL\_4898
CL\_3488
