## Supplementary material for "A novel method for integrating genomic and Tn-Seq data to identify common *in vivo* fitness mechanisms across multiple bacterial species": S1 Dataset: CL_INS_291.html

Legend

 Hypothetical
 All VFDB Genes

FULL


WINDOWSVGPNG

Trim RowsRemove SingletonsSave Fasta

CL\_3485


CL\_3485


CL\_3485


CL\_3485

HighlightSelectShow Genomes


220

CL\_3482


36

CL\_3482


21

CL\_3482


1

CL\_3482

fGI ID


CL\_INS\_291
CL\_INS\_291
CL\_INS\_291
Cluster ID


CL\_3484
CL\_3483
CL\_7580
