## Supplementary material for "A novel method for integrating genomic and Tn-Seq data to identify common *in vivo* fitness mechanisms across multiple bacterial species": S1 Dataset: CL_INS_294.html

Legend

 Mobile +extrachromosomalelementfunctions
 Regulatoryfunctions
 Hypothetical
 All EssentialGenes
 Other
 EnergyMetabolism
 Transport +binding proteins
 All VFDB Genes

FULL


WINDOWSVGPNG

Trim RowsRemove SingletonsSave Fasta

CL\_3457


CL\_3457


CL\_3457


CL\_3457


CL\_3457


CL\_3457


CL\_3457


CL\_3457


CL\_3457


CL\_3457


CL\_3457


CL\_3457


CL\_3457


CL\_3625


CL\_3457


CL\_3457


CL\_3457


CL\_3457


CL\_3457

HighlightSelectShow Genomes


162

CL\_3456


86

CL\_3456


6

CL\_3456


3

CL\_3455


1

CL\_3456


1

CL\_3628


1

CL\_3456


1

CL\_3455


1

CL\_3455


1

CL\_3456


1

CL\_3451


1

CL\_3456


1

CL\_3454


1

CL\_3456


1

CL\_3942


1

CL\_3456


1

CL\_3455


1

CL\_3434


1

CL\_3456

fGI ID


CL\_INS\_294
CL\_INS\_294
CL\_INS\_294
CL\_INS\_294
CL\_INS\_237
CL\_INS\_123
CL\_INS\_368
CL\_INS\_368
CL\_INS\_368
CL\_INS\_294
CL\_INS\_294
CL\_INS\_207
CL\_INS\_382
CL\_INS\_382
CL\_INS\_70
CL\_INS\_70
CL\_INS\_304
CL\_INS\_294
CL\_INS\_294
CL\_INS\_294
CL\_INS\_294
CL\_INS\_294
CL\_INS\_294
CL\_INS\_294
CL\_INS\_294
CL\_INS\_294
CL\_INS\_294
CL\_INS\_294
CL\_INS\_294
CL\_INS\_294
Cluster ID


CL\_10240
CL\_29641
CL\_17150
CL\_28870
CL\_1930
CL\_4995
CL\_10371
CL\_10370
CL\_10369
CL\_10368
CL\_36488
CL\_5125
CL\_8832
CL\_7691
CL\_6828
CL\_7252
CL\_3428
CL\_37060
CL\_7154
CL\_7638
CL\_7153
CL\_30544
CL\_7637
CL\_23751
CL\_3627
CL\_7636
CL\_7152
CL\_7151
CL\_7150
CL\_11692
