## Supplementary material for "A novel method for integrating genomic and Tn-Seq data to identify common *in vivo* fitness mechanisms across multiple bacterial species": S1 Dataset: CL_INS_295.html

Legend

 Mobile +extrachromosomalelementfunctions
 Regulatoryfunctions
 Hypothetical
 DNA Metabolism
 All EssentialGenes
 AntibioticResistance
 All Fitness Genes
 Proteinsynthesis/fate
 Other
 Centralintermediarymetabolism
 EnergyMetabolism
 Transport +binding proteins
 All VFDB Genes

FULL


WINDOWSVGPNG

Trim RowsRemove SingletonsSave Fasta

CL\_3455


CL\_3455


CL\_3455


CL\_1935


CL\_234


CL\_3455


CL\_3455


CL\_3455


CL\_3455


CL\_234


CL\_234


CL\_3455


CL\_234


CL\_3455


CL\_1935


CL\_3455


CL\_234


CL\_3455


CL\_3455


CL\_3455


CL\_3455


CL\_3455


CL\_3455


CL\_3455


CL\_3455


CL\_3455


CL\_3455


CL\_3457


CL\_3455


CL\_3455


CL\_3455


CL\_3455


CL\_3455


CL\_3455


CL\_3455


CL\_3455


CL\_3455


CL\_3455


CL\_3455


CL\_3455


CL\_206


CL\_3455


CL\_3455


CL\_3455


CL\_3455


CL\_3455


CL\_3455


CL\_3455


CL\_3455


CL\_3455


CL\_3455


CL\_1935


CL\_3455


CL\_3455


CL\_3455


CL\_3455


CL\_3455


CL\_3455


CL\_3455


CL\_3455


CL\_3455


CL\_3455


CL\_3455


CL\_3455


CL\_3455


CL\_3455


CL\_3455


CL\_3455


CL\_3455


CL\_3455


CL\_3455


CL\_3455


CL\_3456


CL\_3455


CL\_3455


CL\_3455


CL\_3455


CL\_3455


CL\_3455


CL\_3455


CL\_3455


CL\_3455


CL\_3455


CL\_3455


CL\_3455


CL\_3455


CL\_3455


CL\_3455


CL\_3456


CL\_3455


CL\_3455


CL\_3455


CL\_3455


CL\_3455


CL\_3455


CL\_3455


Break


CL\_3455


CL\_3455


CL\_3455


CL\_3455


CL\_3455


CL\_3455


CL\_3455


CL\_3455


CL\_234

HighlightSelectShow Genomes


162

CL\_3454


6

CL\_3452


3

CL\_3454


3

CL\_3454


3

CL\_3454


2

CL\_3452


2

CL\_3452


2

CL\_3454


2

CL\_234


2

CL\_3454


2

CL\_3454


2

CL\_3452


2

CL\_3454


2

CL\_3452


1

CL\_3454


1

CL\_234


1

CL\_3454


1

CL\_3454


1

CL\_3454


1

CL\_3454


1

CL\_3454


1

CL\_3452


1

CL\_3454


1

CL\_1935


1

CL\_3454


1

CL\_3452


1

CL\_3454


1

CL\_3454


1

CL\_3454


1

CL\_3454


1

CL\_3454


1

CL\_3454


1

CL\_3454


1

CL\_3452


1

CL\_3452


1

CL\_3452


1

CL\_234


1

CL\_234


1

CL\_3454


1

CL\_3454


1

CL\_3454


1

CL\_3454


1

CL\_3452


1

CL\_3452


1

CL\_3452


1

CL\_3454


1

CL\_3454


1

CL\_3454


1

CL\_234


1

CL\_3452


1

CL\_234


1

CL\_3454


1

CL\_3454


1

CL\_1935


1

CL\_3452


1

CL\_3452


1

CL\_234


1

CL\_3454


1

CL\_3454


1

CL\_3454


1

CL\_234


1

CL\_3454


1

CL\_3452


1

CL\_3452


1

CL\_3454


1

CL\_3454


1

CL\_3452


1

Break


1

CL\_3452


1

CL\_3454


1

CL\_3454


1

CL\_3454


1

CL\_3454


1

CL\_3454


1

CL\_3450


1

CL\_3454


1

CL\_3452


1

CL\_234


1

CL\_234


1

CL\_3454


1

CL\_3454


1

CL\_3452


1

CL\_3454


1

CL\_3454


1

CL\_3454


1

CL\_3454


1

CL\_3452


1

CL\_3452


1

CL\_3454


1

CL\_3452


1

CL\_234


1

CL\_3454


1

CL\_3452


1

CL\_3452


1

CL\_203


1

CL\_3452


1

CL\_3454


1

CL\_3454


1

CL\_3454


1

CL\_3454


1

CL\_3454


1

CL\_3454


1

CL\_3454


1

CL\_3454


1

CL\_234


1

CL\_3454

fGI ID


CL\_INS\_294
CL\_INS\_295
CL\_INS\_295
CL\_INS\_295
CL\_INS\_295
CL\_INS\_295
CL\_INS\_295
CL\_INS\_295
CL\_INS\_294
CL\_INS\_294
CL\_INS\_294
CL\_INS\_294
CL\_INS\_294
CL\_INS\_294
CL\_INS\_295
CL\_INS\_295
CL\_INS\_295
CL\_INS\_295
CL\_INS\_295
CL\_INS\_295
CL\_INS\_295
CL\_INS\_295
CL\_INS\_207
CL\_INS\_207
CL\_INS\_295
CL\_INS\_237
CL\_INS\_237
CL\_INS\_295
CL\_INS\_295
CL\_INS\_295
CL\_INS\_207
CL\_INS\_295
CL\_INS\_295
CL\_INS\_207
CL\_INS\_207
CL\_INS\_20
CL\_INS\_207
CL\_INS\_295
CL\_INS\_207
CL\_INS\_295
CL\_INS\_295
CL\_INS\_295
CL\_INS\_207
CL\_INS\_207
CL\_INS\_207
CL\_INS\_368
CL\_INS\_271
CL\_INS\_70
CL\_INS\_295
CL\_INS\_207
CL\_INS\_207
CL\_INS\_295
CL\_INS\_295
CL\_INS\_295
CL\_INS\_352
CL\_INS\_352
CL\_INS\_352
CL\_INS\_352
CL\_INS\_352
CL\_INS\_352
CL\_INS\_352
CL\_INS\_352
CL\_INS\_352
CL\_INS\_352
CL\_INS\_352
CL\_INS\_352
CL\_INS\_352
CL\_INS\_352
CL\_INS\_352
CL\_INS\_352
CL\_INS\_352
CL\_INS\_352
CL\_INS\_352
CL\_INS\_352
CL\_INS\_352
CL\_INS\_352
CL\_INS\_352
CL\_INS\_352
CL\_INS\_352
CL\_INS\_352
CL\_INS\_352
CL\_INS\_352
CL\_INS\_352
CL\_INS\_352
CL\_INS\_352
CL\_INS\_352
CL\_INS\_352
CL\_INS\_352
CL\_INS\_352
CL\_INS\_352
CL\_INS\_352
CL\_INS\_352
CL\_INS\_352
CL\_INS\_352
CL\_INS\_352
CL\_INS\_207
CL\_INS\_295
CL\_INS\_295
CL\_INS\_295
CL\_INS\_295
CL\_INS\_20
CL\_INS\_295
CL\_INS\_295
CL\_INS\_295
CL\_INS\_295
CL\_INS\_295
CL\_INS\_295
CL\_INS\_295
CL\_INS\_295
CL\_INS\_295
CL\_INS\_295
CL\_INS\_295
CL\_INS\_295
CL\_INS\_295
CL\_INS\_295
CL\_INS\_295
CL\_INS\_295
CL\_INS\_295
CL\_INS\_295
CL\_INS\_20
CL\_INS\_20
CL\_INS\_295
CL\_INS\_207
CL\_INS\_20
CL\_INS\_207
CL\_INS\_295
CL\_INS\_295
CL\_INS\_295
CL\_INS\_295
CL\_INS\_295
CL\_INS\_295
CL\_INS\_295
CL\_INS\_295
CL\_INS\_295
CL\_INS\_295
CL\_INS\_295
CL\_INS\_123
CL\_INS\_123
CL\_INS\_123
CL\_INS\_295
CL\_INS\_295
CL\_INS\_295
CL\_INS\_295
CL\_INS\_368
CL\_INS\_368
CL\_INS\_295
CL\_INS\_295
CL\_INS\_368
CL\_INS\_368
CL\_INS\_368
CL\_INS\_368
CL\_INS\_207
CL\_INS\_295
CL\_INS\_295
CL\_INS\_295
CL\_INS\_294
CL\_INS\_368
CL\_INS\_368
CL\_INS\_368
CL\_INS\_368
CL\_INS\_368
CL\_INS\_295
CL\_INS\_237
CL\_INS\_237
CL\_INS\_237
CL\_INS\_237
CL\_INS\_237
CL\_INS\_237
CL\_INS\_295
CL\_INS\_295
CL\_INS\_295
CL\_INS\_295
CL\_INS\_295
CL\_INS\_295
CL\_INS\_20
CL\_INS\_295
CL\_INS\_295
CL\_INS\_295
CL\_INS\_295
CL\_INS\_295
CL\_INS\_295
CL\_INS\_295
CL\_INS\_295
CL\_INS\_295
CL\_INS\_295
CL\_INS\_70
CL\_INS\_295
CL\_INS\_295
CL\_INS\_295
CL\_INS\_295
CL\_INS\_295
CL\_INS\_295
CL\_INS\_295
CL\_INS\_295
CL\_INS\_295
CL\_INS\_247
CL\_INS\_295
CL\_INS\_60
CL\_INS\_294
CL\_INS\_295
CL\_INS\_295
CL\_INS\_295
CL\_INS\_295
CL\_INS\_295
CL\_INS\_295
CL\_INS\_295
CL\_INS\_295
CL\_INS\_295
CL\_INS\_295
CL\_INS\_295
CL\_INS\_237
CL\_INS\_295
CL\_INS\_295
CL\_INS\_20
CL\_INS\_247
CL\_INS\_368
CL\_INS\_295
CL\_INS\_295
CL\_INS\_295
CL\_INS\_382
CL\_INS\_295
CL\_INS\_237
CL\_INS\_237
CL\_INS\_295
CL\_INS\_237
CL\_INS\_237
CL\_INS\_237
CL\_INS\_237
CL\_INS\_270
CL\_INS\_270
CL\_INS\_270
CL\_INS\_270
CL\_INS\_270
CL\_INS\_270
CL\_INS\_270
CL\_INS\_270
CL\_INS\_270
CL\_INS\_270
CL\_INS\_270
CL\_INS\_270
CL\_INS\_270
CL\_INS\_270
CL\_INS\_270
CL\_INS\_270
CL\_INS\_270
CL\_INS\_270
CL\_INS\_295
CL\_INS\_237
CL\_INS\_237
CL\_INS\_237
CL\_INS\_237
CL\_INS\_247
CL\_INS\_247
CL\_INS\_237
CL\_INS\_70
CL\_INS\_237
CL\_INS\_295
CL\_INS\_382
CL\_INS\_382
CL\_INS\_295
CL\_INS\_295
CL\_INS\_295
CL\_INS\_295
CL\_INS\_382
CL\_INS\_295
CL\_INS\_295
CL\_INS\_295
CL\_INS\_295
CL\_INS\_295
CL\_INS\_237
CL\_INS\_237
CL\_INS\_237
CL\_INS\_237
CL\_INS\_237
CL\_INS\_237
CL\_INS\_237
CL\_INS\_237
CL\_INS\_237
CL\_INS\_237
CL\_INS\_237
CL\_INS\_237
CL\_INS\_368
CL\_INS\_295
CL\_INS\_295
CL\_INS\_237
CL\_INS\_237
CL\_INS\_237
CL\_INS\_295
CL\_INS\_295
CL\_INS\_295
CL\_INS\_156
CL\_INS\_156
CL\_INS\_156
CL\_INS\_156
CL\_INS\_295
CL\_INS\_295
CL\_INS\_295
CL\_INS\_295
CL\_INS\_295
CL\_INS\_154
CL\_INS\_154
CL\_INS\_154
CL\_INS\_343
CL\_INS\_295
CL\_INS\_295
CL\_INS\_295
CL\_INS\_295
CL\_INS\_237
CL\_INS\_295
CL\_INS\_295
CL\_INS\_159
CL\_INS\_247
CL\_INS\_237
CL\_INS\_295
CL\_INS\_247
CL\_INS\_247
CL\_INS\_237
CL\_INS\_237
CL\_INS\_295
CL\_INS\_237
CL\_INS\_295
CL\_INS\_295
CL\_INS\_295
CL\_INS\_237
CL\_INS\_295
CL\_INS\_295
CL\_INS\_295
CL\_INS\_295
CL\_INS\_295
CL\_INS\_295
CL\_INS\_70
CL\_INS\_295
CL\_INS\_295
CL\_INS\_295
CL\_INS\_295
CL\_INS\_247
CL\_INS\_295
CL\_INS\_123
CL\_INS\_123
CL\_INS\_247
CL\_INS\_123
CL\_INS\_385
CL\_INS\_247
CL\_INS\_86
CL\_INS\_117
CL\_INS\_86
CL\_INS\_237
CL\_INS\_237
CL\_INS\_237
CL\_INS\_247
CL\_INS\_247
CL\_INS\_237
CL\_INS\_159
CL\_INS\_70
CL\_INS\_70
CL\_INS\_237
CL\_INS\_382
CL\_INS\_44
CL\_INS\_247
CL\_INS\_247
CL\_INS\_149
CL\_INS\_149
CL\_INS\_149
CL\_INS\_149
CL\_INS\_149
CL\_INS\_149
CL\_INS\_57
CL\_INS\_295
CL\_INS\_237
CL\_INS\_237
CL\_INS\_237
CL\_INS\_295
CL\_INS\_237
CL\_INS\_368
CL\_INS\_368
CL\_INS\_30
CL\_INS\_247
CL\_INS\_247
CL\_INS\_247
CL\_INS\_237
CL\_INS\_237
CL\_INS\_237
CL\_INS\_237
CL\_INS\_237
CL\_INS\_237
CL\_INS\_237
CL\_INS\_237
CL\_INS\_237
CL\_INS\_237
CL\_INS\_237
CL\_INS\_239
CL\_INS\_368
CL\_INS\_368
CL\_INS\_368
CL\_INS\_237
CL\_INS\_237
CL\_INS\_295
CL\_INS\_237
CL\_INS\_237
CL\_INS\_295
CL\_INS\_295
CL\_INS\_295
CL\_INS\_295
CL\_INS\_295
CL\_INS\_295
CL\_INS\_237
CL\_INS\_237
CL\_INS\_295
CL\_INS\_295
CL\_INS\_295
CL\_INS\_70
CL\_INS\_70
CL\_INS\_70
CL\_INS\_237
CL\_INS\_247
CL\_INS\_295
CL\_INS\_295
CL\_INS\_295
CL\_INS\_295
CL\_INS\_237
CL\_INS\_295
CL\_INS\_295
CL\_INS\_110
CL\_INS\_286
CL\_INS\_286
CL\_INS\_159
CL\_INS\_149
CL\_INS\_295
CL\_INS\_295
CL\_INS\_295
CL\_INS\_295
CL\_INS\_295
CL\_INS\_368
CL\_INS\_70
CL\_INS\_237
CL\_INS\_295
CL\_INS\_295
CL\_INS\_295
CL\_INS\_247
CL\_INS\_247
CL\_INS\_247
CL\_INS\_247
CL\_INS\_295
CL\_INS\_295
CL\_INS\_295
CL\_INS\_295
CL\_INS\_295
CL\_INS\_295
CL\_INS\_295
CL\_INS\_295
CL\_INS\_295
CL\_INS\_295
CL\_INS\_295
CL\_INS\_20
CL\_INS\_20
CL\_INS\_20
CL\_INS\_20
CL\_INS\_20
CL\_INS\_20
CL\_INS\_20
CL\_INS\_20
CL\_INS\_20
CL\_INS\_20
CL\_INS\_20
CL\_INS\_20
CL\_INS\_20
CL\_INS\_20
CL\_INS\_20
CL\_INS\_20
CL\_INS\_295
CL\_INS\_20
CL\_INS\_20
CL\_INS\_23
CL\_INS\_247
CL\_INS\_247
CL\_INS\_247
CL\_INS\_20
CL\_INS\_20
CL\_INS\_20
CL\_INS\_352
CL\_INS\_352
CL\_INS\_237
CL\_INS\_237
CL\_INS\_237
CL\_INS\_237
CL\_INS\_237
CL\_INS\_237
CL\_INS\_237
CL\_INS\_237
CL\_INS\_237
CL\_INS\_237
CL\_INS\_237
CL\_INS\_295
CL\_INS\_295
CL\_INS\_295
CL\_INS\_382
CL\_INS\_382
CL\_INS\_295
CL\_INS\_70
CL\_INS\_70
CL\_INS\_237
CL\_INS\_237
CL\_INS\_295
CL\_INS\_237
CL\_INS\_237
CL\_INS\_237
CL\_INS\_237
CL\_INS\_237
CL\_INS\_382
CL\_INS\_237
CL\_INS\_237
CL\_INS\_295
CL\_INS\_295
CL\_INS\_295
CL\_INS\_237
CL\_INS\_237
CL\_INS\_237
CL\_INS\_237
CL\_INS\_237
CL\_INS\_237
CL\_INS\_237
CL\_INS\_237
CL\_INS\_368
CL\_INS\_368
CL\_INS\_368
CL\_INS\_368
CL\_INS\_368
CL\_INS\_368
CL\_INS\_368
CL\_INS\_368
CL\_INS\_368
CL\_INS\_295
CL\_INS\_295
CL\_INS\_295
CL\_INS\_295
CL\_INS\_295
CL\_INS\_295
CL\_INS\_295
CL\_INS\_295
CL\_INS\_295
CL\_INS\_295
CL\_INS\_295
CL\_INS\_343
CL\_INS\_295
CL\_INS\_295
CL\_INS\_295
CL\_INS\_295
CL\_INS\_295
CL\_INS\_295
CL\_INS\_295
CL\_INS\_295
CL\_INS\_295
CL\_INS\_295
CL\_INS\_295
CL\_INS\_295
CL\_INS\_295
CL\_INS\_295
CL\_INS\_295
CL\_INS\_295
CL\_INS\_295
CL\_INS\_295
CL\_INS\_295
CL\_INS\_295
CL\_INS\_295
CL\_INS\_295
CL\_INS\_295
CL\_INS\_295
CL\_INS\_237
CL\_INS\_237
CL\_INS\_295
CL\_INS\_295
CL\_INS\_295
CL\_INS\_295
CL\_INS\_295
CL\_INS\_295
CL\_INS\_295
CL\_INS\_295
CL\_INS\_295
CL\_INS\_237
CL\_INS\_237
CL\_INS\_237
CL\_INS\_237
CL\_INS\_237
CL\_INS\_295
CL\_INS\_295
CL\_INS\_295
CL\_INS\_295
CL\_INS\_295
CL\_INS\_295
CL\_INS\_237
CL\_INS\_295
CL\_INS\_70
CL\_INS\_70
CL\_INS\_247
CL\_INS\_247
CL\_INS\_247
CL\_INS\_247
CL\_INS\_247
CL\_INS\_247
CL\_INS\_247
CL\_INS\_247
CL\_INS\_247
CL\_INS\_247
CL\_INS\_247
CL\_INS\_247
CL\_INS\_247
CL\_INS\_247
CL\_INS\_70
CL\_INS\_70
CL\_INS\_295
CL\_INS\_295
CL\_INS\_70
CL\_INS\_70
CL\_INS\_237
CL\_INS\_295
CL\_INS\_70
CL\_INS\_70
CL\_INS\_30
CL\_INS\_159
CL\_INS\_30
CL\_INS\_295
CL\_INS\_237
CL\_INS\_237
CL\_INS\_237
CL\_INS\_70
CL\_INS\_247
CL\_INS\_237
CL\_INS\_295
CL\_INS\_295
CL\_INS\_70
CL\_INS\_237
CL\_INS\_237
CL\_INS\_237
CL\_INS\_295
CL\_INS\_237
CL\_INS\_295
CL\_INS\_237
CL\_INS\_237
CL\_INS\_237
CL\_INS\_237
CL\_INS\_237
CL\_INS\_237
CL\_INS\_70
CL\_INS\_368
CL\_INS\_368
CL\_INS\_237
CL\_INS\_237
CL\_INS\_237
CL\_INS\_237
CL\_INS\_295
CL\_INS\_20
CL\_INS\_30
CL\_INS\_237
CL\_INS\_237
CL\_INS\_237
CL\_INS\_295
CL\_INS\_295
CL\_INS\_70
CL\_INS\_295
CL\_INS\_237
CL\_INS\_295
CL\_INS\_70
CL\_INS\_30
CL\_INS\_295
CL\_INS\_247
CL\_INS\_237
CL\_INS\_237
CL\_INS\_237
CL\_INS\_237
CL\_INS\_237
CL\_INS\_70
CL\_INS\_295
CL\_INS\_295
CL\_INS\_295
CL\_INS\_295
CL\_INS\_295
CL\_INS\_70
CL\_INS\_237
CL\_INS\_237
CL\_INS\_237
CL\_INS\_237
CL\_INS\_237
CL\_INS\_70
CL\_INS\_295
CL\_INS\_70
CL\_INS\_70
CL\_INS\_295
CL\_INS\_237
CL\_INS\_159
CL\_INS\_70
CL\_INS\_70
CL\_INS\_70
CL\_INS\_159
CL\_INS\_237
CL\_INS\_70
CL\_INS\_70
CL\_INS\_70
CL\_INS\_70
CL\_INS\_30
CL\_INS\_30
CL\_INS\_295
CL\_INS\_70
CL\_INS\_70
CL\_INS\_70
CL\_INS\_295
CL\_INS\_237
CL\_INS\_295
CL\_INS\_295
CL\_INS\_295
CL\_INS\_295
CL\_INS\_70
CL\_INS\_237
CL\_INS\_237
CL\_INS\_237
CL\_INS\_237
CL\_INS\_237
CL\_INS\_237
CL\_INS\_237
CL\_INS\_237
CL\_INS\_295
CL\_INS\_237
CL\_INS\_237
CL\_INS\_237
CL\_INS\_237
CL\_INS\_237
CL\_INS\_70
CL\_INS\_237
CL\_INS\_237
CL\_INS\_237
CL\_INS\_237
CL\_INS\_295
CL\_INS\_295
CL\_INS\_295
CL\_INS\_295
CL\_INS\_295
CL\_INS\_295
CL\_INS\_295
CL\_INS\_382
CL\_INS\_70
CL\_INS\_295
CL\_INS\_70
CL\_INS\_295
CL\_INS\_295
CL\_INS\_295
CL\_INS\_368
CL\_INS\_20
CL\_INS\_237
CL\_INS\_237
CL\_INS\_237
CL\_INS\_295
CL\_INS\_237
CL\_INS\_237
CL\_INS\_221
CL\_INS\_237
CL\_INS\_237
CL\_INS\_237
CL\_INS\_237
CL\_INS\_295
CL\_INS\_237
CL\_INS\_237
CL\_INS\_237
CL\_INS\_237
CL\_INS\_237
CL\_INS\_295
CL\_INS\_237
CL\_INS\_237
CL\_INS\_295
CL\_INS\_295
CL\_INS\_295
CL\_INS\_70
CL\_INS\_295
CL\_INS\_237
CL\_INS\_295
CL\_INS\_295
CL\_INS\_295
CL\_INS\_237
CL\_INS\_237
CL\_INS\_237
CL\_INS\_70
CL\_INS\_237
CL\_INS\_295
CL\_INS\_295
CL\_INS\_295
CL\_INS\_295
CL\_INS\_295
CL\_INS\_237
CL\_INS\_382
CL\_INS\_295
CL\_INS\_295
CL\_INS\_295
CL\_INS\_295
CL\_INS\_295
CL\_INS\_295
CL\_INS\_295
CL\_INS\_295
CL\_INS\_295
CL\_INS\_295
CL\_INS\_295
CL\_INS\_237
CL\_INS\_247
CL\_INS\_247
CL\_INS\_247
CL\_INS\_247
CL\_INS\_237
CL\_INS\_247
CL\_INS\_295
CL\_INS\_237
CL\_INS\_295
CL\_INS\_237
CL\_INS\_295
CL\_INS\_295
CL\_INS\_237
CL\_INS\_295
CL\_INS\_295
CL\_INS\_295
CL\_INS\_295
CL\_INS\_295
CL\_INS\_295
CL\_INS\_295
CL\_INS\_295
CL\_INS\_295
CL\_INS\_295
CL\_INS\_295
CL\_INS\_237
CL\_INS\_295
CL\_INS\_295
CL\_INS\_295
CL\_INS\_295
CL\_INS\_295
CL\_INS\_295
CL\_INS\_295
CL\_INS\_295
CL\_INS\_295
CL\_INS\_295
CL\_INS\_70
CL\_INS\_70
CL\_INS\_70
CL\_INS\_295
CL\_INS\_295
CL\_INS\_70
CL\_INS\_30
CL\_INS\_295
CL\_INS\_30
CL\_INS\_295
CL\_INS\_295
CL\_INS\_295
CL\_INS\_295
CL\_INS\_295
CL\_INS\_70
CL\_INS\_159
CL\_INS\_159
CL\_INS\_159
CL\_INS\_159
CL\_INS\_159
CL\_INS\_159
CL\_INS\_159
CL\_INS\_159
CL\_INS\_159
CL\_INS\_159
CL\_INS\_159
CL\_INS\_159
CL\_INS\_159
CL\_INS\_159
CL\_INS\_159
CL\_INS\_159
CL\_INS\_70
CL\_INS\_70
CL\_INS\_295
CL\_INS\_295
CL\_INS\_382
CL\_INS\_295
CL\_INS\_295
CL\_INS\_295
CL\_INS\_295
CL\_INS\_295
CL\_INS\_295
CL\_INS\_295
CL\_INS\_123
CL\_INS\_237
CL\_INS\_237
CL\_INS\_123
CL\_INS\_382
CL\_INS\_295
CL\_INS\_295
CL\_INS\_295
CL\_INS\_382
CL\_INS\_237
CL\_INS\_237
CL\_INS\_295
CL\_INS\_295
CL\_INS\_237
CL\_INS\_295
CL\_INS\_295
CL\_INS\_237
CL\_INS\_237
CL\_INS\_237
CL\_INS\_233
CL\_INS\_295
CL\_INS\_149
CL\_INS\_295
CL\_INS\_295
CL\_INS\_237
CL\_INS\_237
CL\_INS\_237
CL\_INS\_237
CL\_INS\_237
CL\_INS\_237
CL\_INS\_70
CL\_INS\_70
CL\_INS\_70
CL\_INS\_70
CL\_INS\_70
CL\_INS\_295
CL\_INS\_295
CL\_INS\_70
CL\_INS\_70
CL\_INS\_295
CL\_INS\_295
CL\_INS\_237
CL\_INS\_237
CL\_INS\_295
CL\_INS\_237
CL\_INS\_237
CL\_INS\_237
CL\_INS\_295
CL\_INS\_237
CL\_INS\_70
CL\_INS\_70
CL\_INS\_382
CL\_INS\_237
CL\_INS\_237
CL\_INS\_123
CL\_INS\_237
CL\_INS\_237
CL\_INS\_237
CL\_INS\_237
CL\_INS\_295
CL\_INS\_295
CL\_INS\_295
CL\_INS\_295
CL\_INS\_295
CL\_INS\_295
CL\_INS\_295
CL\_INS\_237
CL\_INS\_70
CL\_INS\_237
CL\_INS\_237
CL\_INS\_237
CL\_INS\_237
CL\_INS\_295
CL\_INS\_224
CL\_INS\_237
CL\_INS\_237
CL\_INS\_224
CL\_INS\_295
CL\_INS\_295
CL\_INS\_224
CL\_INS\_295
CL\_INS\_295
CL\_INS\_295
CL\_INS\_224
CL\_INS\_295
CL\_INS\_295
CL\_INS\_295
CL\_INS\_295
CL\_INS\_295
CL\_INS\_295
CL\_INS\_224
CL\_INS\_233
CL\_INS\_295
CL\_INS\_295
CL\_INS\_237
CL\_INS\_237
CL\_INS\_237
CL\_INS\_237
CL\_INS\_295
CL\_INS\_295
CL\_INS\_70
CL\_INS\_70
CL\_INS\_70
CL\_INS\_70
CL\_INS\_295
CL\_INS\_295
CL\_INS\_295
CL\_INS\_295
CL\_INS\_295
CL\_INS\_295
CL\_INS\_295
CL\_INS\_295
CL\_INS\_295
CL\_INS\_237
CL\_INS\_295
CL\_INS\_237
CL\_INS\_237
CL\_INS\_237
CL\_INS\_30
CL\_INS\_295
CL\_INS\_237
CL\_INS\_237
CL\_INS\_295
CL\_INS\_295
CL\_INS\_237
CL\_INS\_237
CL\_INS\_237
CL\_INS\_295
CL\_INS\_295
CL\_INS\_295
CL\_INS\_30
CL\_INS\_70
CL\_INS\_368
CL\_INS\_20
CL\_INS\_20
CL\_INS\_20
CL\_INS\_20
CL\_INS\_20
CL\_INS\_20
CL\_INS\_20
CL\_INS\_20
CL\_INS\_368
CL\_INS\_368
CL\_INS\_237
CL\_INS\_237
CL\_INS\_237
CL\_INS\_237
CL\_INS\_295
CL\_INS\_295
CL\_INS\_295
CL\_INS\_237
CL\_INS\_295
CL\_INS\_295
CL\_INS\_295
CL\_INS\_70
CL\_INS\_237
CL\_INS\_237
CL\_INS\_237
CL\_INS\_237
CL\_INS\_237
CL\_INS\_70
CL\_INS\_237
CL\_INS\_237
CL\_INS\_237
CL\_INS\_237
CL\_INS\_30
CL\_INS\_30
CL\_INS\_30
CL\_INS\_30
CL\_INS\_30
CL\_INS\_86
CL\_INS\_86
CL\_INS\_30
CL\_INS\_30
CL\_INS\_30
CL\_INS\_247
CL\_INS\_247
CL\_INS\_159
CL\_INS\_159
CL\_INS\_159
CL\_INS\_237
CL\_INS\_70
CL\_INS\_237
CL\_INS\_295
CL\_INS\_295
CL\_INS\_247
CL\_INS\_30
CL\_INS\_30
CL\_INS\_30
CL\_INS\_30
CL\_INS\_295
CL\_INS\_295
CL\_INS\_237
CL\_INS\_30
CL\_INS\_247
CL\_INS\_368
CL\_INS\_295
CL\_INS\_237
CL\_INS\_237
CL\_INS\_70
CL\_INS\_295
CL\_INS\_237
CL\_INS\_247
CL\_INS\_70
CL\_INS\_237
CL\_INS\_70
CL\_INS\_207
CL\_INS\_20
CL\_INS\_295
CL\_INS\_295
CL\_INS\_295
Cluster ID


CL\_10240
CL\_12870
CL\_27860
CL\_27859
CL\_27858
CL\_27857
CL\_27856
CL\_27855
CL\_7638
CL\_7153
CL\_7152
CL\_7151
CL\_7150
CL\_11692
CL\_18117
CL\_18118
CL\_18119
CL\_18121
CL\_18122
CL\_18120
CL\_18123
CL\_18124
CL\_18125
CL\_18126
CL\_24028
CL\_13776
CL\_13775
CL\_21993
CL\_33049
CL\_33050
CL\_5972
CL\_33051
CL\_33052
CL\_5975
CL\_5976
CL\_8451
CL\_5977
CL\_8452
CL\_5978
CL\_32881
CL\_32880
CL\_32879
CL\_16974
CL\_13414
CL\_8140
CL\_6018
CL\_3520
CL\_14685
CL\_33053
CL\_5979
CL\_5980
CL\_15419
CL\_20116
CL\_20117
CL\_9401
CL\_9402
CL\_9403
CL\_9404
CL\_9405
CL\_9406
CL\_9407
CL\_9408
CL\_9409
CL\_9410
CL\_9411
CL\_9412
CL\_9413
CL\_9414
CL\_9415
CL\_9416
CL\_9417
CL\_9418
CL\_9419
CL\_9420
CL\_9421
CL\_9422
CL\_9423
CL\_9424
CL\_9425
CL\_9426
CL\_9427
CL\_9428
CL\_9429
CL\_9430
CL\_9431
CL\_9432
CL\_9433
CL\_9434
CL\_9435
CL\_9436
CL\_9437
CL\_9438
CL\_9439
CL\_9440
CL\_9441
CL\_5125
CL\_9625
CL\_31789
CL\_33576
CL\_13410
CL\_26270
CL\_15418
CL\_19929
CL\_19930
CL\_27009
CL\_27008
CL\_21024
CL\_21025
CL\_8449
CL\_8450
CL\_25195
CL\_25196
CL\_5969
CL\_5970
CL\_5971
CL\_20114
CL\_20115
CL\_24359
CL\_24360
CL\_7628
CL\_8457
CL\_21026
CL\_5973
CL\_15986
CL\_5974
CL\_21027
CL\_14470
CL\_14471
CL\_14472
CL\_14473
CL\_14474
CL\_14475
CL\_14476
CL\_34859
CL\_13829
CL\_13830
CL\_9180
CL\_9179
CL\_9178
CL\_29514
CL\_29515
CL\_29516
CL\_29517
CL\_12911
CL\_13831
CL\_29518
CL\_29519
CL\_13832
CL\_13833
CL\_13834
CL\_13835
CL\_7442
CL\_10365
CL\_10366
CL\_10367
CL\_10368
CL\_10369
CL\_10370
CL\_10371
CL\_14500
CL\_20669
CL\_20668
CL\_15473
CL\_15472
CL\_15471
CL\_15470
CL\_15115
CL\_247
CL\_18861
CL\_18860
CL\_18859
CL\_18858
CL\_18857
CL\_24205
CL\_21214
CL\_36141
CL\_24206
CL\_24207
CL\_24208
CL\_24209
CL\_24210
CL\_24211
CL\_24212
CL\_24213
CL\_24214
CL\_10581
CL\_19847
CL\_19846
CL\_10251
CL\_10250
CL\_10249
CL\_10248
CL\_10247
CL\_10246
CL\_16998
CL\_11980
CL\_9840
CL\_9841
CL\_17150
CL\_17149
CL\_17148
CL\_33156
CL\_33155
CL\_33154
CL\_33153
CL\_33152
CL\_33151
CL\_33150
CL\_33149
CL\_28263
CL\_11110
CL\_28262
CL\_28261
CL\_4377
CL\_2550
CL\_13836
CL\_13837
CL\_13838
CL\_13839
CL\_7557
CL\_17147
CL\_14365
CL\_7872
CL\_20277
CL\_15981
CL\_20012
CL\_20011
CL\_20010
CL\_7873
CL\_7874
CL\_7875
CL\_7876
CL\_7877
CL\_7878
CL\_7879
CL\_7880
CL\_7881
CL\_7882
CL\_7883
CL\_7884
CL\_7885
CL\_7886
CL\_7887
CL\_7888
CL\_7889
CL\_7890
CL\_17146
CL\_17167
CL\_17168
CL\_10590
CL\_10830
CL\_6844
CL\_7007
CL\_17169
CL\_8632
CL\_17170
CL\_5124
CL\_8832
CL\_7691
CL\_14553
CL\_14554
CL\_14555
CL\_14556
CL\_14193
CL\_14557
CL\_14558
CL\_14559
CL\_14560
CL\_14561
CL\_7701
CL\_12081
CL\_12082
CL\_7700
CL\_7699
CL\_7698
CL\_7697
CL\_7696
CL\_7695
CL\_7694
CL\_7693
CL\_12083
CL\_12084
CL\_14562
CL\_14563
CL\_14411
CL\_22559
CL\_7302
CL\_14564
CL\_14565
CL\_14566
CL\_14567
CL\_14568
CL\_14569
CL\_14570
CL\_14571
CL\_14572
CL\_14573
CL\_14574
CL\_14575
CL\_14576
CL\_14577
CL\_14578
CL\_14579
CL\_14580
CL\_14581
CL\_14582
CL\_14583
CL\_7688
CL\_14584
CL\_14585
CL\_7639
CL\_10372
CL\_6413
CL\_13016
CL\_16926
CL\_14396
CL\_18856
CL\_7984
CL\_18855
CL\_7674
CL\_14586
CL\_14587
CL\_14588
CL\_7579
CL\_33746
CL\_33747
CL\_33748
CL\_33749
CL\_33750
CL\_33751
CL\_8781
CL\_33752
CL\_33753
CL\_33754
CL\_33755
CL\_10395
CL\_22259
CL\_10392
CL\_5298
CL\_5299
CL\_5300
CL\_5296
CL\_5678
CL\_5516
CL\_10417
CL\_10418
CL\_10419
CL\_6760
CL\_6761
CL\_10664
CL\_11981
CL\_10373
CL\_1929
CL\_6829
CL\_6411
CL\_10374
CL\_5301
CL\_5518
CL\_10393
CL\_5302
CL\_6757
CL\_6756
CL\_6755
CL\_6754
CL\_6753
CL\_6752
CL\_6751
CL\_36149
CL\_13368
CL\_13369
CL\_13370
CL\_13596
CL\_6976
CL\_12108
CL\_12107
CL\_8553
CL\_12106
CL\_5688
CL\_5689
CL\_12104
CL\_12103
CL\_12102
CL\_12100
CL\_12101
CL\_12099
CL\_12098
CL\_12097
CL\_12096
CL\_12095
CL\_12094
CL\_12093
CL\_12092
CL\_12091
CL\_12090
CL\_7306
CL\_13610
CL\_14403
CL\_7709
CL\_7708
CL\_35323
CL\_14406
CL\_14407
CL\_14408
CL\_14409
CL\_14410
CL\_35322
CL\_26529
CL\_10811
CL\_7578
CL\_7577
CL\_7242
CL\_11191
CL\_7241
CL\_11189
CL\_5236
CL\_7576
CL\_20975
CL\_21685
CL\_34267
CL\_4462
CL\_15612
CL\_20976
CL\_29376
CL\_11663
CL\_11664
CL\_11665
CL\_6732
CL\_26747
CL\_29642
CL\_29643
CL\_29644
CL\_29645
CL\_6832
CL\_5321
CL\_244
CL\_37555
CL\_37556
CL\_34760
CL\_8960
CL\_8959
CL\_8958
CL\_8957
CL\_24027
CL\_24026
CL\_24025
CL\_24024
CL\_24023
CL\_24022
CL\_24021
CL\_24020
CL\_14477
CL\_14478
CL\_14479
CL\_9019
CL\_6283
CL\_6284
CL\_6285
CL\_6286
CL\_6287
CL\_6288
CL\_6289
CL\_6290
CL\_6291
CL\_6292
CL\_6293
CL\_6294
CL\_6295
CL\_9018
CL\_9017
CL\_9624
CL\_9623
CL\_19845
CL\_19844
CL\_241
CL\_240
CL\_2541
CL\_239
CL\_238
CL\_237
CL\_6029
CL\_6028
CL\_6027
CL\_6026
CL\_6025
CL\_6024
CL\_6023
CL\_8132
CL\_8516
CL\_21021
CL\_8515
CL\_8513
CL\_10823
CL\_10810
CL\_10809
CL\_10808
CL\_10807
CL\_8554
CL\_14011
CL\_7435
CL\_7434
CL\_5149
CL\_8530
CL\_10806
CL\_8529
CL\_8528
CL\_10805
CL\_8637
CL\_8636
CL\_10804
CL\_15536
CL\_13467
CL\_21589
CL\_21590
CL\_21591
CL\_21592
CL\_20448
CL\_11800
CL\_4089
CL\_4090
CL\_4091
CL\_22719
CL\_34732
CL\_20449
CL\_17781
CL\_17782
CL\_17783
CL\_20450
CL\_20451
CL\_20452
CL\_20453
CL\_20454
CL\_16927
CL\_21593
CL\_33388
CL\_10754
CL\_10753
CL\_10752
CL\_10751
CL\_10750
CL\_10749
CL\_10748
CL\_10747
CL\_5618
CL\_10746
CL\_10745
CL\_10744
CL\_10743
CL\_10742
CL\_10741
CL\_10740
CL\_10739
CL\_10738
CL\_10737
CL\_10736
CL\_10735
CL\_10734
CL\_10733
CL\_10732
CL\_10731
CL\_10730
CL\_10729
CL\_10728
CL\_10727
CL\_10726
CL\_10424
CL\_21594
CL\_21595
CL\_12907
CL\_12908
CL\_20667
CL\_20666
CL\_20665
CL\_20664
CL\_20663
CL\_20662
CL\_20661
CL\_20660
CL\_20659
CL\_13371
CL\_13372
CL\_13373
CL\_13374
CL\_13376
CL\_21596
CL\_21597
CL\_21598
CL\_21599
CL\_21600
CL\_21601
CL\_17158
CL\_21602
CL\_7234
CL\_7233
CL\_8782
CL\_7232
CL\_8783
CL\_8784
CL\_8785
CL\_7231
CL\_8786
CL\_7229
CL\_8787
CL\_7228
CL\_7227
CL\_8788
CL\_7226
CL\_7225
CL\_14683
CL\_7673
CL\_33756
CL\_33757
CL\_8748
CL\_8747
CL\_14340
CL\_37569
CL\_7253
CL\_6826
CL\_5245
CL\_10545
CL\_7254
CL\_17145
CL\_13378
CL\_9557
CL\_8518
CL\_7672
CL\_18854
CL\_8517
CL\_34072
CL\_34073
CL\_7214
CL\_21803
CL\_6076
CL\_7213
CL\_14398
CL\_14342
CL\_35403
CL\_7664
CL\_7212
CL\_7211
CL\_7293
CL\_7292
CL\_4374
CL\_6417
CL\_21603
CL\_21604
CL\_7294
CL\_7210
CL\_14399
CL\_7209
CL\_33387
CL\_236
CL\_6425
CL\_20278
CL\_6825
CL\_10375
CL\_27319
CL\_27318
CL\_6824
CL\_20279
CL\_6730
CL\_10796
CL\_7255
CL\_6422
CL\_17144
CL\_7470
CL\_11974
CL\_11973
CL\_7256
CL\_6729
CL\_6728
CL\_6421
CL\_10795
CL\_10794
CL\_10793
CL\_10792
CL\_20309
CL\_8126
CL\_20009
CL\_20007
CL\_13214
CL\_20310
CL\_8125
CL\_10546
CL\_35622
CL\_6420
CL\_10547
CL\_34636
CL\_10548
CL\_10549
CL\_6419
CL\_8508
CL\_6418
CL\_6727
CL\_4375
CL\_1932
CL\_1933
CL\_7831
CL\_8620
CL\_5246
CL\_5247
CL\_27527
CL\_4099
CL\_4100
CL\_4101
CL\_35406
CL\_13377
CL\_33758
CL\_33759
CL\_33760
CL\_35405
CL\_7671
CL\_7670
CL\_7669
CL\_8790
CL\_8791
CL\_8792
CL\_8793
CL\_8212
CL\_11187
CL\_35404
CL\_14401
CL\_7296
CL\_25361
CL\_7295
CL\_8794
CL\_6426
CL\_7665
CL\_12110
CL\_14400
CL\_8210
CL\_34720
CL\_34719
CL\_34718
CL\_27526
CL\_27525
CL\_27524
CL\_27523
CL\_7692
CL\_6828
CL\_32376
CL\_7252
CL\_21994
CL\_21995
CL\_17122
CL\_11816
CL\_7838
CL\_21996
CL\_21997
CL\_21998
CL\_34722
CL\_21999
CL\_8760
CL\_8759
CL\_8758
CL\_22000
CL\_22001
CL\_22002
CL\_27522
CL\_13774
CL\_13773
CL\_13772
CL\_13771
CL\_13770
CL\_13769
CL\_13212
CL\_13768
CL\_13767
CL\_13766
CL\_13765
CL\_7869
CL\_7870
CL\_7871
CL\_27521
CL\_27520
CL\_27519
CL\_6414
CL\_6415
CL\_13846
CL\_1931
CL\_16977
CL\_27518
CL\_27517
CL\_27516
CL\_27515
CL\_27514
CL\_12241
CL\_7761
CL\_27513
CL\_27512
CL\_27511
CL\_27510
CL\_27509
CL\_27508
CL\_27507
CL\_27506
CL\_27505
CL\_27504
CL\_27503
CL\_7223
CL\_7221
CL\_8789
CL\_7220
CL\_7219
CL\_7216
CL\_7215
CL\_22006
CL\_15526
CL\_33761
CL\_23926
CL\_33762
CL\_26095
CL\_6830
CL\_27502
CL\_27501
CL\_27500
CL\_27499
CL\_27498
CL\_27497
CL\_27496
CL\_27495
CL\_27494
CL\_27493
CL\_27492
CL\_20304
CL\_20303
CL\_20302
CL\_20301
CL\_20300
CL\_20299
CL\_20298
CL\_20297
CL\_20296
CL\_20295
CL\_20294
CL\_6410
CL\_8600
CL\_8599
CL\_27491
CL\_27490
CL\_6842
CL\_11295
CL\_14589
CL\_7667
CL\_20293
CL\_20292
CL\_20291
CL\_20290
CL\_20289
CL\_10330
CL\_11223
CL\_11224
CL\_11225
CL\_11226
CL\_11227
CL\_11228
CL\_11229
CL\_11230
CL\_11231
CL\_11232
CL\_11233
CL\_11234
CL\_11235
CL\_11236
CL\_11238
CL\_11239
CL\_10331
CL\_11221
CL\_20288
CL\_20287
CL\_8216
CL\_7575
CL\_7574
CL\_7573
CL\_7572
CL\_7571
CL\_7570
CL\_7569
CL\_6590
CL\_6805
CL\_6806
CL\_9785
CL\_4975
CL\_24215
CL\_24216
CL\_24217
CL\_8841
CL\_8635
CL\_1930
CL\_8634
CL\_8633
CL\_7987
CL\_10803
CL\_10802
CL\_7986
CL\_8505
CL\_8506
CL\_10801
CL\_10800
CL\_4093
CL\_32374
CL\_32375
CL\_7240
CL\_7239
CL\_7441
CL\_7440
CL\_7237
CL\_7439
CL\_4094
CL\_4095
CL\_4096
CL\_4097
CL\_4098
CL\_20286
CL\_20285
CL\_18091
CL\_15973
CL\_20284
CL\_13762
CL\_13761
CL\_6849
CL\_34721
CL\_22005
CL\_18863
CL\_7990
CL\_9052
CL\_6244
CL\_7299
CL\_7298
CL\_8376
CL\_7297
CL\_14412
CL\_4995
CL\_7989
CL\_7988
CL\_7301
CL\_6848
CL\_31309
CL\_31310
CL\_31311
CL\_31312
CL\_31313
CL\_31314
CL\_31315
CL\_7236
CL\_7235
CL\_18161
CL\_8780
CL\_16459
CL\_8239
CL\_18159
CL\_7716
CL\_7717
CL\_13820
CL\_7718
CL\_18158
CL\_18157
CL\_13676
CL\_27567
CL\_27568
CL\_27569
CL\_10853
CL\_31316
CL\_27570
CL\_27571
CL\_27572
CL\_27573
CL\_18152
CL\_7723
CL\_7325
CL\_30532
CL\_18151
CL\_18150
CL\_6847
CL\_6846
CL\_6845
CL\_22003
CL\_22004
CL\_6841
CL\_6840
CL\_6839
CL\_6838
CL\_27489
CL\_27488
CL\_27487
CL\_27486
CL\_27485
CL\_27484
CL\_27483
CL\_27482
CL\_27481
CL\_14318
CL\_20280
CL\_7208
CL\_7207
CL\_7206
CL\_7749
CL\_27779
CL\_7291
CL\_7290
CL\_24019
CL\_24018
CL\_6724
CL\_7289
CL\_7205
CL\_10791
CL\_10790
CL\_16953
CL\_5226
CL\_6836
CL\_7556
CL\_7555
CL\_7554
CL\_13760
CL\_13759
CL\_7553
CL\_7552
CL\_7551
CL\_7550
CL\_13758
CL\_7982
CL\_7981
CL\_7980
CL\_7979
CL\_8131
CL\_10799
CL\_22731
CL\_22732
CL\_22733
CL\_23204
CL\_10798
CL\_10797
CL\_8129
CL\_8555
CL\_8556
CL\_8128
CL\_11106
CL\_8127
CL\_11105
CL\_11711
CL\_13756
CL\_11710
CL\_13755
CL\_7756
CL\_7755
CL\_7754
CL\_7753
CL\_10984
CL\_5815
CL\_5814
CL\_7752
CL\_5232
CL\_5231
CL\_13745
CL\_13744
CL\_13743
CL\_13742
CL\_13741
CL\_13552
CL\_8642
CL\_6813
CL\_13597
CL\_27317
CL\_6726
CL\_7751
CL\_235
CL\_1934
CL\_5225
CL\_34074
CL\_27480
CL\_7288
CL\_1937
CL\_7204
CL\_6807
CL\_7149
CL\_7970
CL\_6722
CL\_6809
CL\_14590
CL\_7568
CL\_7567
CL\_6808
CL\_10789
CL\_8107
CL\_233
CL\_8565
CL\_13740
CL\_13739
CL\_13732
