## Supplementary material for "A novel method for integrating genomic and Tn-Seq data to identify common *in vivo* fitness mechanisms across multiple bacterial species": S1 Dataset: CL_INS_299.html

Legend

 Mobile +extrachromosomalelementfunctions
 Hypothetical
 All EssentialGenes
 Other
 Transport +binding proteins
 All VFDB Genes

FULL


WINDOWSVGPNG

Trim RowsRemove SingletonsSave Fasta

CL\_3446


CL\_3446


CL\_4900


CL\_3446


CL\_3447


CL\_3446


CL\_3446

HighlightSelectShow Genomes


262

CL\_3445


5

CL\_3445


1

CL\_3445


1

CL\_3445


1

CL\_3445


1

CL\_3442


1

CL\_3445

fGI ID


CL\_INS\_299
CL\_INS\_299
CL\_INS\_299
CL\_INS\_207
CL\_INS\_207
CL\_INS\_207
CL\_INS\_207
CL\_INS\_299
CL\_INS\_299
Cluster ID


CL\_11691
CL\_16493
CL\_16494
CL\_9489
CL\_9490
CL\_9491
CL\_9492
CL\_16495
CL\_16496
