## Supplementary material for "A novel method for integrating genomic and Tn-Seq data to identify common *in vivo* fitness mechanisms across multiple bacterial species": S1 Dataset: CL_INS_300.html

Legend

 Mobile +extrachromosomalelementfunctions
 Hypothetical
 Other
 All VFDB Genes

FULL


WINDOWSVGPNG

Trim RowsRemove SingletonsSave Fasta

CL\_3442


CL\_3442


CL\_3442


CL\_3442


CL\_3442


CL\_3442


CL\_3442


CL\_3447


CL\_3442


CL\_3442


CL\_3442

HighlightSelectShow Genomes


205

CL\_3441


40

CL\_3441


20

CL\_3441


4

CL\_3441


4

CL\_3441


2

CL\_3441


1

CL\_3440


1

CL\_3441


1

CL\_3441


1

CL\_3441


1

CL\_3441

fGI ID


CL\_INS\_300
CL\_INS\_300
CL\_INS\_300
CL\_INS\_300
CL\_INS\_300
CL\_INS\_300
CL\_INS\_300
Cluster ID


CL\_30738
CL\_30521
CL\_13840
CL\_7632
CL\_10635
CL\_10636
CL\_33148
