## Supplementary material for "A novel method for integrating genomic and Tn-Seq data to identify common *in vivo* fitness mechanisms across multiple bacterial species": S1 Dataset: CL_INS_301.html

Legend

 Mobile +extrachromosomalelementfunctions
 Hypothetical
 Other
 Cellularprocesses
 All VFDB Genes
 Transport +binding proteins

FULL


WINDOWSVGPNG

Trim RowsRemove SingletonsSave Fasta

CL\_3439


CL\_3439


CL\_3439


CL\_3439


CL\_3439


CL\_3439


CL\_3439


CL\_3439

HighlightSelectShow Genomes


198

CL\_3438


6

CL\_3438


3

CL\_3438


2

CL\_3438


1

CL\_3438


1

CL\_3437


1

CL\_3438


1

CL\_3438

fGI ID


CL\_INS\_299
CL\_INS\_301
CL\_INS\_301
CL\_INS\_301
CL\_INS\_301
CL\_INS\_301
CL\_INS\_121
CL\_INS\_121
CL\_INS\_121
CL\_INS\_10
CL\_INS\_121
CL\_INS\_121
CL\_INS\_121
CL\_INS\_121
CL\_INS\_121
CL\_INS\_121
CL\_INS\_10
CL\_INS\_301
CL\_INS\_121
CL\_INS\_121
CL\_INS\_121
CL\_INS\_301
CL\_INS\_121
CL\_INS\_121
CL\_INS\_121
CL\_INS\_121
CL\_INS\_121
CL\_INS\_10
CL\_INS\_121
CL\_INS\_301
CL\_INS\_10
CL\_INS\_10
CL\_INS\_10
CL\_INS\_10
CL\_INS\_10
CL\_INS\_10
CL\_INS\_121
CL\_INS\_10
CL\_INS\_10
CL\_INS\_10
CL\_INS\_10
CL\_INS\_10
CL\_INS\_10
CL\_INS\_10
CL\_INS\_10
CL\_INS\_10
CL\_INS\_10
CL\_INS\_10
CL\_INS\_10
CL\_INS\_10
CL\_INS\_170
CL\_INS\_121
Cluster ID


CL\_11691
CL\_22520
CL\_34939
CL\_15422
CL\_15423
CL\_15424
CL\_14232
CL\_4807
CL\_4806
CL\_4805
CL\_9958
CL\_4804
CL\_4803
CL\_4802
CL\_4801
CL\_14420
CL\_14419
CL\_14418
CL\_4796
CL\_4795
CL\_4793
CL\_4792
CL\_4791
CL\_4790
CL\_14415
CL\_14414
CL\_4789
CL\_4788
CL\_4787
CL\_34868
CL\_4784
CL\_4783
CL\_4782
CL\_4781
CL\_4780
CL\_4779
CL\_4778
CL\_4777
CL\_4776
CL\_4775
CL\_4774
CL\_4773
CL\_4772
CL\_4771
CL\_4770
CL\_4769
CL\_4768
CL\_4767
CL\_4766
CL\_4765
CL\_4602
CL\_4760
