## Supplementary material for "A novel method for integrating genomic and Tn-Seq data to identify common *in vivo* fitness mechanisms across multiple bacterial species": S1 Dataset: CL_INS_303.html

Legend

 Mobile +extrachromosomalelementfunctions
 Hypothetical
 Other
 All VFDB Genes
 Transport +binding proteins

FULL


WINDOWSVGPNG

Trim RowsRemove SingletonsSave Fasta

CL\_3430


CL\_3430


CL\_3434


CL\_3431


CL\_3434


CL\_3430


CL\_3434


CL\_3434


CL\_3687


CL\_3430


Break


CL\_3434


CL\_3430


CL\_3434

HighlightSelectShow Genomes


86

CL\_3429


76

CL\_3429


16

CL\_3429


5

CL\_3429


4

CL\_3429


4

CL\_3426


1

CL\_3429


1

CL\_3429


1

CL\_3429


1

CL\_3429


1

CL\_3429


1

CL\_3429


1

CL\_3427


1

CL\_3429

fGI ID


CL\_INS\_303
CL\_INS\_264
CL\_INS\_303
CL\_INS\_141
CL\_INS\_141
CL\_INS\_303
CL\_INS\_382
CL\_INS\_247
CL\_INS\_304
Cluster ID


CL\_30737
CL\_4883
CL\_30546
CL\_4615
CL\_4616
CL\_16990
CL\_7557
CL\_5045
CL\_6615
