## Supplementary material for "A novel method for integrating genomic and Tn-Seq data to identify common *in vivo* fitness mechanisms across multiple bacterial species": S1 Dataset: CL_INS_311.html

Legend

 Mobile +extrachromosomalelementfunctions
 All EssentialGenes
 Other
 All VFDB Genes

FULL


WINDOWSVGPNG

Trim RowsRemove SingletonsSave Fasta

CL\_3342


CL\_3342


CL\_3342


CL\_3342


CL\_3342


CL\_3342

HighlightSelectShow Genomes


143

CL\_3341


128

CL\_3341


2

CL\_3341


1

CL\_3340


1

CL\_3338


1

CL\_3341

fGI ID


CL\_INS\_311
CL\_INS\_311
Cluster ID


CL\_4906
CL\_12877
