## Supplementary material for "A novel method for integrating genomic and Tn-Seq data to identify common *in vivo* fitness mechanisms across multiple bacterial species": S1 Dataset: CL_INS_312.html

Legend

 Mobile +extrachromosomalelementfunctions
 Hypothetical
 Other
 All VFDB Genes

FULL


WINDOWSVGPNG

Trim RowsRemove SingletonsSave Fasta

CL\_3320


CL\_3320


CL\_3321


CL\_3321


CL\_3320


CL\_3321

HighlightSelectShow Genomes


160

CL\_3319


37

CL\_3319


1

CL\_3319


1

CL\_3319


1

CL\_3319


1

CL\_3319

fGI ID


CL\_INS\_312
CL\_INS\_312
CL\_INS\_312
CL\_INS\_312
CL\_INS\_312
CL\_INS\_312
Cluster ID


CL\_11130
CL\_26446
CL\_13240
CL\_13239
CL\_30552
CL\_30553
