## Supplementary material for "A novel method for integrating genomic and Tn-Seq data to identify common *in vivo* fitness mechanisms across multiple bacterial species": S1 Dataset: CL_INS_313.html

Legend

 Mobile +extrachromosomalelementfunctions
 Hypothetical
 Other
 All VFDB Genes

FULL


WINDOWSVGPNG

Trim RowsRemove SingletonsSave Fasta

CL\_3314


CL\_3314


CL\_3314


CL\_3314


CL\_3314


CL\_3314


CL\_3314


CL\_3314


CL\_3314


CL\_3314


CL\_3316


CL\_3314


CL\_3316

HighlightSelectShow Genomes


103

CL\_3313


74

CL\_3313


34

CL\_3313


23

CL\_3313


23

CL\_3313


10

CL\_3313


5

CL\_3313


2

CL\_3313


1

CL\_3313


1

CL\_3313


1

CL\_3313


1

CL\_3313


1

CL\_3313

fGI ID


CL\_INS\_313
CL\_INS\_313
CL\_INS\_313
CL\_INS\_313
CL\_INS\_313
CL\_INS\_313
CL\_INS\_313
CL\_INS\_313
CL\_INS\_313
CL\_INS\_313
Cluster ID


CL\_23932
CL\_4907
CL\_13599
CL\_12205
CL\_5473
CL\_4908
CL\_4909
CL\_4910
CL\_4911
CL\_23931
