## Supplementary material for "A novel method for integrating genomic and Tn-Seq data to identify common *in vivo* fitness mechanisms across multiple bacterial species": S1 Dataset: CL_INS_314.html

Legend

 Mobile +extrachromosomalelementfunctions
 Hypothetical
 All EssentialGenes
 Other
 Transport +binding proteins

FULL


WINDOWSVGPNG

Trim RowsRemove SingletonsSave Fasta

CL\_3308


CL\_3308


CL\_3308


CL\_3308


CL\_3308


CL\_3309

HighlightSelectShow Genomes


145

CL\_3307


107

CL\_3307


13

CL\_3306


8

CL\_3303


1

CL\_3307


1

CL\_3307

fGI ID


CL\_INS\_314
CL\_INS\_314
CL\_INS\_316
CL\_INS\_316
Cluster ID


CL\_23038
CL\_7147
CL\_4912
CL\_4913
