## Supplementary material for "A novel method for integrating genomic and Tn-Seq data to identify common *in vivo* fitness mechanisms across multiple bacterial species": S1 Dataset: CL_INS_317.html

Legend

 Transport +binding proteins
 All VFDB Genes

FULL


WINDOWSVGPNG

Trim RowsRemove SingletonsSave Fasta

CL\_3299


CL\_3299

HighlightSelectShow Genomes


238

CL\_3298


36

CL\_3298

fGI ID

CL\_INS\_317
Cluster ID

CL\_9444
