## Supplementary material for "A novel method for integrating genomic and Tn-Seq data to identify common *in vivo* fitness mechanisms across multiple bacterial species": S1 Dataset: CL_INS_318.html

Legend

 Mobile +extrachromosomalelementfunctions
 Regulatoryfunctions
 Hypothetical
 AntibioticResistance
 Proteinsynthesis/fate
 Other
 Transport +binding proteins
 All VFDB Genes

FULL


WINDOWSVGPNG

Trim RowsRemove SingletonsSave Fasta

CL\_3293


CL\_3293


CL\_3293


CL\_3293


CL\_3293


CL\_3293


CL\_3293


CL\_3293


CL\_3293


CL\_3293


CL\_3293


CL\_3293


CL\_3293


CL\_3293


CL\_3293


CL\_3293


CL\_3293


CL\_3294


CL\_3293


CL\_3293


CL\_3293


CL\_3294


CL\_3293

HighlightSelectShow Genomes


156

CL\_3292


29

CL\_3292


22

CL\_3292


19

CL\_3292


11

CL\_3292


10

CL\_3292


7

CL\_3292


6

CL\_3292


2

CL\_3292


1

CL\_3292


1

CL\_3292


1

CL\_3292


1

CL\_3292


1

CL\_3292


1

CL\_3292


1

CL\_3291


1

CL\_3292


1

CL\_3292


1

CL\_3292


1

CL\_3291


1

CL\_3292


1

CL\_3292


1

CL\_3292

fGI ID


CL\_INS\_318
CL\_INS\_318
CL\_INS\_318
CL\_INS\_237
CL\_INS\_318
CL\_INS\_318
CL\_INS\_385
CL\_INS\_318
CL\_INS\_318
CL\_INS\_318
CL\_INS\_318
CL\_INS\_318
CL\_INS\_318
CL\_INS\_318
CL\_INS\_318
CL\_INS\_318
CL\_INS\_318
CL\_INS\_318
CL\_INS\_318
CL\_INS\_318
CL\_INS\_318
CL\_INS\_318
CL\_INS\_318
CL\_INS\_318
Cluster ID


CL\_30122
CL\_26269
CL\_7146
CL\_5149
CL\_13600
CL\_7145
CL\_7144
CL\_36709
CL\_7143
CL\_7142
CL\_7141
CL\_7140
CL\_7139
CL\_7138
CL\_8026
CL\_8025
CL\_9445
CL\_7137
CL\_7136
CL\_7135
CL\_26534
CL\_26535
CL\_26536
CL\_7134
