## Supplementary material for "A novel method for integrating genomic and Tn-Seq data to identify common *in vivo* fitness mechanisms across multiple bacterial species": S1 Dataset: CL_INS_319.html

Legend

 Hypothetical
 Proteinsynthesis/fate
 Other
 All VFDB Genes

FULL


WINDOWSVGPNG

Trim RowsRemove SingletonsSave Fasta

CL\_3291


CL\_3291


CL\_3291


CL\_3291


CL\_3291


CL\_3291


CL\_3292


CL\_3292

HighlightSelectShow Genomes


185

CL\_3290


81

CL\_3290


2

CL\_3290


2

CL\_3290


2

CL\_3290


2

CL\_3290


1

CL\_3290


1

CL\_3290

fGI ID


CL\_INS\_319
CL\_INS\_319
CL\_INS\_319
CL\_INS\_319
CL\_INS\_319
CL\_INS\_319
CL\_INS\_319
Cluster ID


CL\_15432
CL\_7133
CL\_7132
CL\_7131
CL\_7130
CL\_7129
CL\_8024
