## Supplementary material for "A novel method for integrating genomic and Tn-Seq data to identify common *in vivo* fitness mechanisms across multiple bacterial species": S1 Dataset: CL_INS_322.html

Legend

 Mobile +extrachromosomalelementfunctions
 Regulatoryfunctions
 Hypothetical
 Other
 All VFDB Genes

FULL


WINDOWSVGPNG

Trim RowsRemove SingletonsSave Fasta

CL\_3243


CL\_3243


CL\_3256


CL\_3243


CL\_3243


CL\_3243


CL\_3257


CL\_3243


CL\_3243


CL\_3243


CL\_3243


CL\_3243


CL\_3243


CL\_3243


CL\_3256


CL\_3243

HighlightSelectShow Genomes


174

CL\_3244


85

CL\_3247


41

CL\_3244


4

CL\_3247


2

CL\_3245


2

CL\_3247


1

CL\_3244


1

CL\_3256


1

CL\_3247


1

CL\_3259


1

CL\_3256


1

CL\_3272


1

CL\_3247


1

CL\_3257


1

CL\_3244


1

CL\_3246

fGI ID


CL\_INS\_322
CL\_INS\_322
CL\_INS\_322
CL\_INS\_321
CL\_INS\_321
CL\_INS\_321
CL\_INS\_321
CL\_INS\_321
CL\_INS\_321
CL\_INS\_321
CL\_INS\_321
CL\_INS\_321
CL\_INS\_321
Cluster ID


CL\_30743
CL\_30742
CL\_7178
CL\_9039
CL\_3254
CL\_12878
CL\_9040
CL\_9041
CL\_11327
CL\_11326
CL\_11325
CL\_11324
CL\_11323
