## Supplementary material for "A novel method for integrating genomic and Tn-Seq data to identify common *in vivo* fitness mechanisms across multiple bacterial species": S1 Dataset: CL_INS_323.html

Legend

 Mobile +extrachromosomalelementfunctions
 Regulatoryfunctions
 Hypothetical
 Other
 All VFDB Genes

FULL


WINDOWSVGPNG

Trim RowsRemove SingletonsSave Fasta

CL\_3240


CL\_3240


CL\_3240


CL\_3240


CL\_3240


CL\_3240


CL\_3240


CL\_3240


CL\_3240


CL\_3240


CL\_3240


CL\_3240


CL\_3240


CL\_3240


CL\_3240


CL\_3240


CL\_3240


CL\_3240


CL\_3240


CL\_3240


CL\_3240


CL\_3240


CL\_3240

HighlightSelectShow Genomes


90

CL\_3239


75

CL\_3239


31

CL\_3239


24

CL\_3239


16

CL\_3239


9

CL\_3239


7

CL\_3239


4

CL\_3239


2

CL\_3239


2

CL\_3239


2

CL\_3239


1

CL\_3239


1

CL\_3239


1

CL\_3239


1

CL\_3239


1

CL\_3239


1

CL\_3239


1

CL\_3239


1

CL\_3238


1

CL\_3239


1

CL\_3239


1

CL\_3239


1

CL\_3239

fGI ID


CL\_INS\_323
CL\_INS\_323
CL\_INS\_323
CL\_INS\_323
CL\_INS\_323
CL\_INS\_323
CL\_INS\_323
CL\_INS\_323
CL\_INS\_323
CL\_INS\_323
CL\_INS\_323
CL\_INS\_323
CL\_INS\_323
CL\_INS\_323
CL\_INS\_323
CL\_INS\_323
CL\_INS\_323
Cluster ID


CL\_12879
CL\_12195
CL\_25882
CL\_25883
CL\_35364
CL\_36707
CL\_36706
CL\_26447
CL\_9562
CL\_15185
CL\_4916
CL\_4917
CL\_4918
CL\_23930
CL\_7179
CL\_8022
CL\_7180
