## Supplementary material for "A novel method for integrating genomic and Tn-Seq data to identify common *in vivo* fitness mechanisms across multiple bacterial species": S1 Dataset: CL_INS_326.html

Legend

 Mobile +extrachromosomalelementfunctions
 Regulatoryfunctions
 Hypothetical
 DNA Metabolism
 AntibioticResistance
 All Fitness Genes
 Proteinsynthesis/fate
 Other
 Transport +binding proteins
 All VFDB Genes

FULL


WINDOWSVGPNG

Trim RowsRemove SingletonsSave Fasta

CL\_3216


CL\_3216


CL\_3216


CL\_3216


CL\_3216


CL\_3216


CL\_3216


Break


CL\_3216


CL\_3216


CL\_3216


CL\_3216


CL\_3216


CL\_3216


CL\_3216


CL\_3216


CL\_3216


CL\_3216


CL\_3216


CL\_3216


CL\_3216


CL\_3216


CL\_3216


CL\_3216


CL\_3216


CL\_3216


CL\_3216


CL\_3216


CL\_3216


CL\_3216


CL\_3216


CL\_3216


CL\_3216


CL\_3216


CL\_3216


CL\_3216


CL\_3216

HighlightSelectShow Genomes


243

CL\_3215


2

CL\_3215


1

Break


1

CL\_3215


1

CL\_3215


1

CL\_3215


1

CL\_3215


1

CL\_3215


1

CL\_3215


1

CL\_3215


1

CL\_3215


1

CL\_3215


1

CL\_3215


1

CL\_3215


1

CL\_3215


1

CL\_3215


1

CL\_3215


1

CL\_3215


1

CL\_3215


1

CL\_3215


1

CL\_3215


1

CL\_3215


1

CL\_3215


1

CL\_3215


1

CL\_3215


1

CL\_3215


1

CL\_3215


1

CL\_3215


1

CL\_3215


1

CL\_3215


1

CL\_3215


1

CL\_3215


1

CL\_3215


1

CL\_3215


1

CL\_3215


1

CL\_3215


1

CL\_3215

fGI ID


CL\_INS\_326
CL\_INS\_171
CL\_INS\_326
CL\_INS\_326
CL\_INS\_170
CL\_INS\_326
CL\_INS\_326
CL\_INS\_171
CL\_INS\_326
CL\_INS\_171
CL\_INS\_326
CL\_INS\_171
CL\_INS\_326
CL\_INS\_170
CL\_INS\_170
CL\_INS\_170
CL\_INS\_170
CL\_INS\_326
CL\_INS\_326
CL\_INS\_326
CL\_INS\_326
CL\_INS\_170
CL\_INS\_152
CL\_INS\_171
CL\_INS\_326
CL\_INS\_326
CL\_INS\_326
CL\_INS\_379
CL\_INS\_379
CL\_INS\_379
CL\_INS\_171
CL\_INS\_171
CL\_INS\_73
CL\_INS\_73
CL\_INS\_326
CL\_INS\_326
CL\_INS\_326
CL\_INS\_326
CL\_INS\_326
CL\_INS\_326
CL\_INS\_326
CL\_INS\_74
CL\_INS\_326
CL\_INS\_326
CL\_INS\_326
CL\_INS\_326
CL\_INS\_326
CL\_INS\_326
CL\_INS\_326
CL\_INS\_326
CL\_INS\_326
CL\_INS\_326
CL\_INS\_149
CL\_INS\_149
CL\_INS\_326
CL\_INS\_326
CL\_INS\_326
CL\_INS\_326
CL\_INS\_326
CL\_INS\_326
CL\_INS\_326
CL\_INS\_326
CL\_INS\_149
CL\_INS\_326
CL\_INS\_326
CL\_INS\_326
CL\_INS\_326
CL\_INS\_326
CL\_INS\_170
CL\_INS\_326
CL\_INS\_171
CL\_INS\_66
CL\_INS\_66
CL\_INS\_326
CL\_INS\_326
CL\_INS\_326
CL\_INS\_207
CL\_INS\_326
CL\_INS\_326
CL\_INS\_207
CL\_INS\_326
CL\_INS\_326
CL\_INS\_170
CL\_INS\_171
CL\_INS\_171
CL\_INS\_170
CL\_INS\_171
CL\_INS\_326
CL\_INS\_326
CL\_INS\_326
CL\_INS\_326
CL\_INS\_326
CL\_INS\_326
CL\_INS\_326
CL\_INS\_326
CL\_INS\_326
CL\_INS\_326
CL\_INS\_170
CL\_INS\_326
CL\_INS\_326
CL\_INS\_170
CL\_INS\_326
CL\_INS\_170
CL\_INS\_170
CL\_INS\_326
CL\_INS\_326
CL\_INS\_326
CL\_INS\_170
CL\_INS\_326
CL\_INS\_149
CL\_INS\_171
CL\_INS\_326
CL\_INS\_326
CL\_INS\_326
CL\_INS\_326
CL\_INS\_326
CL\_INS\_171
CL\_INS\_326
CL\_INS\_171
CL\_INS\_170
CL\_INS\_326
CL\_INS\_326
CL\_INS\_326
CL\_INS\_170
CL\_INS\_170
CL\_INS\_326
CL\_INS\_326
CL\_INS\_326
CL\_INS\_170
CL\_INS\_326
CL\_INS\_326
CL\_INS\_170
CL\_INS\_170
CL\_INS\_326
CL\_INS\_170
CL\_INS\_170
CL\_INS\_326
CL\_INS\_326
CL\_INS\_326
CL\_INS\_326
CL\_INS\_170
CL\_INS\_171
CL\_INS\_171
CL\_INS\_326
CL\_INS\_207
CL\_INS\_170
CL\_INS\_170
CL\_INS\_326
CL\_INS\_149
CL\_INS\_207
CL\_INS\_170
CL\_INS\_149
CL\_INS\_326
CL\_INS\_149
CL\_INS\_155
CL\_INS\_204
CL\_INS\_204
CL\_INS\_204
CL\_INS\_204
CL\_INS\_326
CL\_INS\_326
CL\_INS\_326
CL\_INS\_86
CL\_INS\_326
CL\_INS\_149
CL\_INS\_237
CL\_INS\_326
CL\_INS\_247
CL\_INS\_247
CL\_INS\_123
CL\_INS\_123
CL\_INS\_123
CL\_INS\_247
CL\_INS\_247
CL\_INS\_326
CL\_INS\_326
CL\_INS\_247
CL\_INS\_86
CL\_INS\_207
CL\_INS\_207
CL\_INS\_207
CL\_INS\_170
CL\_INS\_170
CL\_INS\_170
CL\_INS\_170
CL\_INS\_170
CL\_INS\_171
CL\_INS\_171
CL\_INS\_326
CL\_INS\_326
CL\_INS\_170
CL\_INS\_170
CL\_INS\_170
CL\_INS\_170
CL\_INS\_149
CL\_INS\_149
CL\_INS\_149
CL\_INS\_149
CL\_INS\_149
CL\_INS\_149
CL\_INS\_149
CL\_INS\_326
CL\_INS\_326
CL\_INS\_326
CL\_INS\_326
CL\_INS\_149
CL\_INS\_149
CL\_INS\_149
CL\_INS\_170
CL\_INS\_326
CL\_INS\_149
CL\_INS\_170
CL\_INS\_326
CL\_INS\_149
CL\_INS\_326
CL\_INS\_326
CL\_INS\_170
CL\_INS\_171
CL\_INS\_171
CL\_INS\_170
CL\_INS\_20
CL\_INS\_326
CL\_INS\_326
CL\_INS\_149
CL\_INS\_204
CL\_INS\_204
CL\_INS\_149
CL\_INS\_149
CL\_INS\_149
CL\_INS\_171
CL\_INS\_326
CL\_INS\_73
CL\_INS\_326
CL\_INS\_326
CL\_INS\_170
CL\_INS\_170
CL\_INS\_170
CL\_INS\_171
CL\_INS\_171
CL\_INS\_326
CL\_INS\_326
CL\_INS\_326
CL\_INS\_171
CL\_INS\_171
CL\_INS\_207
CL\_INS\_326
CL\_INS\_326
CL\_INS\_326
CL\_INS\_326
CL\_INS\_326
CL\_INS\_326
CL\_INS\_326
CL\_INS\_326
CL\_INS\_326
CL\_INS\_326
CL\_INS\_326
CL\_INS\_326
CL\_INS\_326
CL\_INS\_326
CL\_INS\_326
CL\_INS\_326
CL\_INS\_326
CL\_INS\_326
CL\_INS\_326
CL\_INS\_326
CL\_INS\_326
CL\_INS\_326
CL\_INS\_326
CL\_INS\_326
CL\_INS\_326
CL\_INS\_326
CL\_INS\_326
CL\_INS\_326
CL\_INS\_326
CL\_INS\_326
CL\_INS\_326
CL\_INS\_326
CL\_INS\_71
CL\_INS\_326
CL\_INS\_326
CL\_INS\_326
CL\_INS\_207
CL\_INS\_207
CL\_INS\_326
CL\_INS\_326
CL\_INS\_326
CL\_INS\_149
CL\_INS\_170
CL\_INS\_149
CL\_INS\_149
CL\_INS\_149
CL\_INS\_326
CL\_INS\_326
CL\_INS\_326
CL\_INS\_326
CL\_INS\_326
CL\_INS\_159
CL\_INS\_159
CL\_INS\_326
CL\_INS\_30
CL\_INS\_326
CL\_INS\_326
CL\_INS\_326
CL\_INS\_326
CL\_INS\_326
CL\_INS\_326
CL\_INS\_326
CL\_INS\_326
CL\_INS\_326
CL\_INS\_326
CL\_INS\_326
CL\_INS\_247
CL\_INS\_382
CL\_INS\_382
CL\_INS\_326
CL\_INS\_382
CL\_INS\_326
CL\_INS\_326
CL\_INS\_326
CL\_INS\_382
CL\_INS\_326
CL\_INS\_326
CL\_INS\_326
CL\_INS\_326
CL\_INS\_326
CL\_INS\_123
CL\_INS\_326
CL\_INS\_343
CL\_INS\_295
CL\_INS\_326
CL\_INS\_326
CL\_INS\_326
CL\_INS\_326
CL\_INS\_326
CL\_INS\_326
CL\_INS\_326
CL\_INS\_326
CL\_INS\_382
CL\_INS\_382
CL\_INS\_382
CL\_INS\_326
CL\_INS\_326
CL\_INS\_326
CL\_INS\_326
CL\_INS\_326
CL\_INS\_326
CL\_INS\_326
CL\_INS\_326
CL\_INS\_326
CL\_INS\_237
CL\_INS\_237
CL\_INS\_237
CL\_INS\_237
CL\_INS\_326
CL\_INS\_326
CL\_INS\_382
CL\_INS\_326
CL\_INS\_382
CL\_INS\_382
CL\_INS\_382
CL\_INS\_382
CL\_INS\_382
CL\_INS\_237
CL\_INS\_237
CL\_INS\_382
CL\_INS\_247
CL\_INS\_326
CL\_INS\_326
CL\_INS\_326
CL\_INS\_326
CL\_INS\_326
CL\_INS\_326
CL\_INS\_326
CL\_INS\_326
CL\_INS\_237
CL\_INS\_326
CL\_INS\_326
CL\_INS\_326
CL\_INS\_207
CL\_INS\_326
CL\_INS\_326
CL\_INS\_326
CL\_INS\_326
CL\_INS\_326
CL\_INS\_326
CL\_INS\_326
CL\_INS\_326
CL\_INS\_326
CL\_INS\_326
CL\_INS\_326
CL\_INS\_326
CL\_INS\_237
CL\_INS\_237
CL\_INS\_237
CL\_INS\_326
CL\_INS\_326
CL\_INS\_326
CL\_INS\_201
CL\_INS\_382
CL\_INS\_382
CL\_INS\_382
CL\_INS\_99
CL\_INS\_117
CL\_INS\_326
CL\_INS\_326
CL\_INS\_99
CL\_INS\_384
CL\_INS\_382
CL\_INS\_326
CL\_INS\_326
CL\_INS\_326
CL\_INS\_326
CL\_INS\_326
CL\_INS\_326
CL\_INS\_326
CL\_INS\_326
CL\_INS\_326
CL\_INS\_326
CL\_INS\_326
CL\_INS\_326
CL\_INS\_326
CL\_INS\_326
CL\_INS\_326
CL\_INS\_37
CL\_INS\_326
CL\_INS\_326
CL\_INS\_326
CL\_INS\_237
CL\_INS\_326
CL\_INS\_326
CL\_INS\_326
CL\_INS\_326
CL\_INS\_326
CL\_INS\_326
CL\_INS\_326
CL\_INS\_326
CL\_INS\_326
CL\_INS\_326
CL\_INS\_326
CL\_INS\_326
CL\_INS\_326
CL\_INS\_326
CL\_INS\_237
CL\_INS\_326
CL\_INS\_237
CL\_INS\_237
CL\_INS\_237
CL\_INS\_237
CL\_INS\_237
CL\_INS\_237
CL\_INS\_326
CL\_INS\_237
CL\_INS\_237
CL\_INS\_237
CL\_INS\_237
CL\_INS\_237
CL\_INS\_326
CL\_INS\_326
CL\_INS\_326
CL\_INS\_326
CL\_INS\_237
CL\_INS\_326
CL\_INS\_326
CL\_INS\_326
CL\_INS\_326
CL\_INS\_237
CL\_INS\_326
CL\_INS\_326
CL\_INS\_326
CL\_INS\_326
CL\_INS\_326
CL\_INS\_326
CL\_INS\_326
CL\_INS\_326
CL\_INS\_326
CL\_INS\_326
CL\_INS\_326
CL\_INS\_326
CL\_INS\_162
CL\_INS\_162
CL\_INS\_162
CL\_INS\_162
CL\_INS\_326
CL\_INS\_326
CL\_INS\_326
CL\_INS\_123
CL\_INS\_326
CL\_INS\_326
CL\_INS\_207
CL\_INS\_207
CL\_INS\_326
CL\_INS\_237
CL\_INS\_237
CL\_INS\_237
CL\_INS\_237
CL\_INS\_237
CL\_INS\_326
CL\_INS\_326
CL\_INS\_326
CL\_INS\_326
CL\_INS\_149
CL\_INS\_326
CL\_INS\_326
CL\_INS\_326
CL\_INS\_326
CL\_INS\_71
CL\_INS\_326
CL\_INS\_326
CL\_INS\_326
CL\_INS\_326
CL\_INS\_326
CL\_INS\_326
CL\_INS\_326
CL\_INS\_326
CL\_INS\_326
CL\_INS\_326
CL\_INS\_326
CL\_INS\_326
CL\_INS\_326
CL\_INS\_326
CL\_INS\_326
CL\_INS\_326
CL\_INS\_326
CL\_INS\_326
CL\_INS\_326
CL\_INS\_326
CL\_INS\_326
CL\_INS\_326
CL\_INS\_326
CL\_INS\_326
CL\_INS\_326
CL\_INS\_326
CL\_INS\_326
CL\_INS\_326
CL\_INS\_326
CL\_INS\_326
CL\_INS\_326
CL\_INS\_326
CL\_INS\_326
CL\_INS\_326
CL\_INS\_326
CL\_INS\_326
CL\_INS\_326
CL\_INS\_326
CL\_INS\_326
CL\_INS\_326
CL\_INS\_326
CL\_INS\_326
CL\_INS\_326
CL\_INS\_326
CL\_INS\_326
CL\_INS\_123
CL\_INS\_123
CL\_INS\_123
CL\_INS\_123
CL\_INS\_123
CL\_INS\_326
CL\_INS\_326
CL\_INS\_326
CL\_INS\_326
CL\_INS\_326
CL\_INS\_326
CL\_INS\_326
CL\_INS\_326
CL\_INS\_326
CL\_INS\_326
CL\_INS\_326
CL\_INS\_326
CL\_INS\_326
CL\_INS\_326
CL\_INS\_326
CL\_INS\_326
CL\_INS\_326
CL\_INS\_326
CL\_INS\_326
CL\_INS\_326
CL\_INS\_326
CL\_INS\_326
CL\_INS\_326
CL\_INS\_326
CL\_INS\_326
CL\_INS\_382
CL\_INS\_189
CL\_INS\_189
CL\_INS\_123
CL\_INS\_247
CL\_INS\_247
CL\_INS\_295
CL\_INS\_326
CL\_INS\_326
CL\_INS\_326
CL\_INS\_326
CL\_INS\_247
CL\_INS\_247
CL\_INS\_247
CL\_INS\_359
CL\_INS\_117
CL\_INS\_382
CL\_INS\_326
CL\_INS\_326
CL\_INS\_326
CL\_INS\_326
CL\_INS\_326
CL\_INS\_326
CL\_INS\_326
CL\_INS\_326
CL\_INS\_326
CL\_INS\_326
CL\_INS\_326
CL\_INS\_326
CL\_INS\_326
CL\_INS\_326
CL\_INS\_326
CL\_INS\_326
CL\_INS\_326
CL\_INS\_326
CL\_INS\_326
CL\_INS\_326
CL\_INS\_326
CL\_INS\_326
CL\_INS\_326
CL\_INS\_326
CL\_INS\_326
CL\_INS\_326
CL\_INS\_326
CL\_INS\_326
CL\_INS\_326
CL\_INS\_326
CL\_INS\_326
CL\_INS\_326
CL\_INS\_326
CL\_INS\_326
CL\_INS\_326
CL\_INS\_326
CL\_INS\_326
CL\_INS\_326
CL\_INS\_326
CL\_INS\_326
CL\_INS\_326
CL\_INS\_326
CL\_INS\_326
CL\_INS\_326
CL\_INS\_326
CL\_INS\_326
CL\_INS\_326
CL\_INS\_326
CL\_INS\_326
CL\_INS\_326
CL\_INS\_326
CL\_INS\_326
CL\_INS\_326
CL\_INS\_326
CL\_INS\_326
CL\_INS\_326
CL\_INS\_326
CL\_INS\_207
CL\_INS\_326
CL\_INS\_326
CL\_INS\_326
CL\_INS\_326
CL\_INS\_326
CL\_INS\_326
CL\_INS\_326
Cluster ID


CL\_6613
CL\_6612
CL\_26160
CL\_6611
CL\_5173
CL\_23253
CL\_23254
CL\_14457
CL\_29297
CL\_6610
CL\_35074
CL\_7592
CL\_11813
CL\_5174
CL\_5175
CL\_5176
CL\_5177
CL\_23255
CL\_35075
CL\_35076
CL\_35077
CL\_5178
CL\_5890
CL\_5179
CL\_29296
CL\_29295
CL\_29294
CL\_23526
CL\_12363
CL\_23525
CL\_22708
CL\_22707
CL\_19377
CL\_19378
CL\_19720
CL\_19719
CL\_19718
CL\_19717
CL\_19716
CL\_6609
CL\_35621
CL\_4573
CL\_6608
CL\_24138
CL\_6607
CL\_6606
CL\_21238
CL\_24137
CL\_6605
CL\_6604
CL\_24136
CL\_24135
CL\_9188
CL\_9189
CL\_24134
CL\_21237
CL\_21236
CL\_6603
CL\_24511
CL\_26159
CL\_21235
CL\_21234
CL\_9190
CL\_6600
CL\_24133
CL\_24132
CL\_24131
CL\_22410
CL\_5889
CL\_22409
CL\_5888
CL\_22616
CL\_22615
CL\_13362
CL\_26537
CL\_13363
CL\_8482
CL\_13364
CL\_35078
CL\_5184
CL\_6602
CL\_6601
CL\_10376
CL\_10377
CL\_22521
CL\_7069
CL\_20118
CL\_11812
CL\_11811
CL\_8453
CL\_8454
CL\_22585
CL\_10378
CL\_10379
CL\_10380
CL\_35620
CL\_35619
CL\_4584
CL\_20119
CL\_23256
CL\_5887
CL\_33618
CL\_4586
CL\_6531
CL\_20120
CL\_20121
CL\_20122
CL\_4585
CL\_29293
CL\_6599
CL\_5185
CL\_32217
CL\_32218
CL\_14483
CL\_14484
CL\_14485
CL\_5186
CL\_35079
CL\_5187
CL\_5188
CL\_14486
CL\_14487
CL\_14488
CL\_5189
CL\_5190
CL\_33617
CL\_33616
CL\_33615
CL\_5191
CL\_14489
CL\_14490
CL\_5192
CL\_8069
CL\_14491
CL\_5193
CL\_5194
CL\_32219
CL\_32220
CL\_32221
CL\_32222
CL\_5195
CL\_16858
CL\_8068
CL\_29292
CL\_6521
CL\_5196
CL\_7072
CL\_29291
CL\_14492
CL\_5197
CL\_5198
CL\_5199
CL\_23257
CL\_5200
CL\_8483
CL\_1103
CL\_1104
CL\_1105
CL\_6520
CL\_22522
CL\_22523
CL\_22524
CL\_5201
CL\_33613
CL\_2276
CL\_4514
CL\_26106
CL\_5019
CL\_5299
CL\_5298
CL\_5297
CL\_10392
CL\_10393
CL\_15520
CL\_19698
CL\_26105
CL\_5601
CL\_1106
CL\_5202
CL\_5884
CL\_11275
CL\_7077
CL\_7078
CL\_5203
CL\_5204
CL\_5883
CL\_5205
CL\_14493
CL\_9352
CL\_9353
CL\_5206
CL\_5207
CL\_5208
CL\_4587
CL\_4588
CL\_4589
CL\_4590
CL\_4591
CL\_4592
CL\_4593
CL\_11993
CL\_6598
CL\_22408
CL\_6597
CL\_6596
CL\_4594
CL\_4595
CL\_4596
CL\_4597
CL\_33614
CL\_4598
CL\_5170
CL\_29290
CL\_4601
CL\_6595
CL\_24510
CL\_7073
CL\_23231
CL\_27797
CL\_4599
CL\_22407
CL\_32223
CL\_32224
CL\_4600
CL\_6594
CL\_4604
CL\_4605
CL\_4606
CL\_8067
CL\_20398
CL\_29289
CL\_20138
CL\_34867
CL\_34866
CL\_7074
CL\_7075
CL\_7076
CL\_13365
CL\_23258
CL\_22584
CL\_33054
CL\_10381
CL\_19678
CL\_19679
CL\_7250
CL\_35618
CL\_35617
CL\_35616
CL\_35615
CL\_35614
CL\_35613
CL\_35612
CL\_35611
CL\_35610
CL\_35609
CL\_35608
CL\_35607
CL\_35606
CL\_35605
CL\_35604
CL\_35603
CL\_35602
CL\_35601
CL\_35600
CL\_35599
CL\_35598
CL\_35597
CL\_35596
CL\_24130
CL\_35595
CL\_24129
CL\_24128
CL\_24127
CL\_24126
CL\_24125
CL\_24124
CL\_24123
CL\_14646
CL\_6593
CL\_24122
CL\_24121
CL\_6518
CL\_6519
CL\_22458
CL\_24120
CL\_6592
CL\_4607
CL\_4608
CL\_4609
CL\_4610
CL\_4611
CL\_22401
CL\_22400
CL\_11844
CL\_11845
CL\_20568
CL\_5032
CL\_5033
CL\_22399
CL\_5237
CL\_22398
CL\_22397
CL\_22396
CL\_22395
CL\_22394
CL\_22393
CL\_22392
CL\_22391
CL\_22390
CL\_22389
CL\_22388
CL\_5045
CL\_19401
CL\_19402
CL\_22387
CL\_5050
CL\_19705
CL\_22386
CL\_22385
CL\_4236
CL\_22384
CL\_22383
CL\_22382
CL\_22381
CL\_22380
CL\_22379
CL\_22378
CL\_5618
CL\_10748
CL\_22377
CL\_22376
CL\_22375
CL\_22374
CL\_10757
CL\_22373
CL\_22372
CL\_22371
CL\_5073
CL\_5074
CL\_5075
CL\_22370
CL\_22369
CL\_22368
CL\_22367
CL\_22366
CL\_22365
CL\_22364
CL\_22363
CL\_22362
CL\_10785
CL\_20949
CL\_20814
CL\_10782
CL\_22361
CL\_22360
CL\_5092
CL\_22359
CL\_5094
CL\_5095
CL\_5096
CL\_5097
CL\_5098
CL\_10770
CL\_10769
CL\_5102
CL\_5103
CL\_22358
CL\_22357
CL\_22356
CL\_22355
CL\_22354
CL\_22353
CL\_22352
CL\_22351
CL\_22350
CL\_22349
CL\_22348
CL\_22347
CL\_4666
CL\_22346
CL\_22345
CL\_22344
CL\_22343
CL\_22342
CL\_22341
CL\_11673
CL\_22340
CL\_22339
CL\_22338
CL\_22337
CL\_22336
CL\_6181
CL\_11393
CL\_11394
CL\_11395
CL\_22335
CL\_22334
CL\_22333
CL\_8682
CL\_16006
CL\_17074
CL\_11404
CL\_11407
CL\_22332
CL\_22331
CL\_16121
CL\_13501
CL\_22329
CL\_22328
CL\_22327
CL\_22326
CL\_22325
CL\_22324
CL\_22323
CL\_22322
CL\_22321
CL\_22320
CL\_22319
CL\_22318
CL\_22317
CL\_22316
CL\_22315
CL\_22314
CL\_6998
CL\_22313
CL\_22312
CL\_22311
CL\_11417
CL\_22310
CL\_22309
CL\_22308
CL\_22307
CL\_22306
CL\_22305
CL\_22304
CL\_22303
CL\_22302
CL\_22301
CL\_22300
CL\_22299
CL\_22298
CL\_22297
CL\_11343
CL\_22296
CL\_11344
CL\_11345
CL\_11346
CL\_11347
CL\_11356
CL\_11357
CL\_22295
CL\_11358
CL\_11360
CL\_11361
CL\_11362
CL\_11363
CL\_22294
CL\_22293
CL\_22292
CL\_22291
CL\_11367
CL\_22290
CL\_11770
CL\_22289
CL\_22288
CL\_11383
CL\_22287
CL\_22286
CL\_22285
CL\_22284
CL\_22283
CL\_22282
CL\_22281
CL\_22280
CL\_22279
CL\_22278
CL\_22277
CL\_22276
CL\_5736
CL\_5735
CL\_5734
CL\_5733
CL\_22275
CL\_22274
CL\_22273
CL\_22272
CL\_22271
CL\_22270
CL\_22269
CL\_11132
CL\_22268
CL\_6888
CL\_6887
CL\_6886
CL\_6885
CL\_9697
CL\_22267
CL\_22266
CL\_22265
CL\_22264
CL\_5146
CL\_22263
CL\_22262
CL\_22261
CL\_22260
CL\_5515
CL\_22330
CL\_14979
CL\_21494
CL\_21493
CL\_21491
CL\_21490
CL\_21489
CL\_21480
CL\_21479
CL\_21478
CL\_21477
CL\_21476
CL\_21475
CL\_21474
CL\_21473
CL\_21472
CL\_21469
CL\_21468
CL\_14981
CL\_14983
CL\_14984
CL\_14985
CL\_14986
CL\_14987
CL\_14988
CL\_21467
CL\_14990
CL\_14991
CL\_14992
CL\_14993
CL\_14994
CL\_14995
CL\_14996
CL\_14997
CL\_14998
CL\_14999
CL\_15000
CL\_15001
CL\_15009
CL\_15010
CL\_15011
CL\_15012
CL\_15013
CL\_15014
CL\_15015
CL\_10328
CL\_10329
CL\_11778
CL\_11779
CL\_15016
CL\_15018
CL\_15031
CL\_15032
CL\_15033
CL\_15034
CL\_15039
CL\_15040
CL\_15041
CL\_15042
CL\_15043
CL\_15044
CL\_15045
CL\_15046
CL\_15047
CL\_15048
CL\_15054
CL\_15055
CL\_15056
CL\_15057
CL\_15058
CL\_15059
CL\_15061
CL\_15062
CL\_15063
CL\_15064
CL\_10208
CL\_15065
CL\_15066
CL\_10715
CL\_10412
CL\_15522
CL\_22259
CL\_22258
CL\_22257
CL\_22256
CL\_22255
CL\_10386
CL\_10385
CL\_10384
CL\_13730
CL\_10417
CL\_9687
CL\_15071
CL\_15073
CL\_15074
CL\_15076
CL\_15077
CL\_21461
CL\_15078
CL\_15079
CL\_15080
CL\_15081
CL\_15082
CL\_15083
CL\_15085
CL\_15086
CL\_15087
CL\_15088
CL\_15089
CL\_15090
CL\_15091
CL\_15092
CL\_15093
CL\_15094
CL\_15095
CL\_15096
CL\_15097
CL\_15098
CL\_14945
CL\_14946
CL\_14947
CL\_14949
CL\_14950
CL\_14952
CL\_14953
CL\_14954
CL\_14955
CL\_14956
CL\_14957
CL\_14958
CL\_14960
CL\_14961
CL\_14962
CL\_14963
CL\_14964
CL\_14965
CL\_14966
CL\_14967
CL\_14968
CL\_14969
CL\_14970
CL\_14971
CL\_14972
CL\_14974
CL\_14975
CL\_14976
CL\_14977
CL\_14978
CL\_22509
CL\_6591
CL\_12561
CL\_12562
CL\_21233
CL\_22525
CL\_20123
CL\_35594
CL\_22508
