## Supplementary material for "A novel method for integrating genomic and Tn-Seq data to identify common *in vivo* fitness mechanisms across multiple bacterial species": S1 Dataset: CL_INS_328.html

Legend

 Mobile +extrachromosomalelementfunctions
 Hypothetical
 Other
 EnergyMetabolism
 Transport +binding proteins
 All VFDB Genes

FULL


WINDOWSVGPNG

Trim RowsRemove SingletonsSave Fasta

CL\_3190


CL\_3190


CL\_3190


CL\_3190


CL\_3190


CL\_3190


CL\_3190


CL\_3190


CL\_3190


CL\_3190


CL\_3190


CL\_3190


CL\_3190

HighlightSelectShow Genomes


147

CL\_3189


61

CL\_3189


39

CL\_3189


11

CL\_3189


4

CL\_3189


4

CL\_3189


3

CL\_3189


2

CL\_3189


2

CL\_3189


1

CL\_3189


1

CL\_3186


1

CL\_3187


1

CL\_3189

fGI ID


CL\_INS\_328
CL\_INS\_328
CL\_INS\_328
CL\_INS\_328
CL\_INS\_328
CL\_INS\_328
CL\_INS\_328
CL\_INS\_328
CL\_INS\_328
CL\_INS\_328
Cluster ID


CL\_12191
CL\_12881
CL\_30560
CL\_7181
CL\_7182
CL\_7183
CL\_7184
CL\_8020
CL\_23382
CL\_23381
