## Supplementary material for "A novel method for integrating genomic and Tn-Seq data to identify common *in vivo* fitness mechanisms across multiple bacterial species": S1 Dataset: CL_INS_329.html

Legend

 Mobile +extrachromosomalelementfunctions
 Regulatoryfunctions
 Hypothetical
 Other
 All VFDB Genes

FULL


WINDOWSVGPNG

Trim RowsRemove SingletonsSave Fasta

CL\_3171


CL\_3171


CL\_3171


CL\_3171


CL\_3171


CL\_3171


CL\_3171


CL\_3171


CL\_3171


CL\_3171

HighlightSelectShow Genomes


163

CL\_3170


83

CL\_3170


18

CL\_3170


7

CL\_3170


2

CL\_3170


1

CL\_3170


1

CL\_3170


1

CL\_3170


1

CL\_3170


1

CL\_3170

fGI ID


CL\_INS\_329
CL\_INS\_329
CL\_INS\_329
CL\_INS\_329
CL\_INS\_329
CL\_INS\_329
CL\_INS\_329
CL\_INS\_329
CL\_INS\_329
CL\_INS\_329
CL\_INS\_329
CL\_INS\_329
CL\_INS\_329
Cluster ID


CL\_13242
CL\_7185
CL\_11809
CL\_11808
CL\_11807
CL\_11806
CL\_11805
CL\_11804
CL\_11803
CL\_11802
CL\_4922
CL\_26984
CL\_4923
