## Supplementary material for "A novel method for integrating genomic and Tn-Seq data to identify common *in vivo* fitness mechanisms across multiple bacterial species": S1 Dataset: CL_INS_330.html

Legend

 Hypothetical
 All EssentialGenes
 Other

FULL


WINDOWSVGPNG

Trim RowsRemove SingletonsSave Fasta

CL\_3169


CL\_3169


CL\_3169


CL\_3169


CL\_3169


CL\_3169


CL\_3169


CL\_3169


CL\_3169

HighlightSelectShow Genomes


193

CL\_3168


77

CL\_3168


2

CL\_3168


1

CL\_3168


1

CL\_3194


1

CL\_3168


1

Break


1

CL\_3167


1

CL\_3168

fGI ID


CL\_INS\_330
CL\_INS\_330
CL\_INS\_330
CL\_INS\_330
CL\_INS\_330
Cluster ID


CL\_16804
CL\_28260
CL\_36676
CL\_8018
CL\_7186
