## Supplementary material for "A novel method for integrating genomic and Tn-Seq data to identify common *in vivo* fitness mechanisms across multiple bacterial species": S1 Dataset: CL_INS_331.html

Legend

 Mobile +extrachromosomalelementfunctions
 Hypothetical
 All EssentialGenes
 Other
 All VFDB Genes

FULL


WINDOWSVGPNG

Trim RowsRemove SingletonsSave Fasta

CL\_3152


CL\_3152


CL\_3152


CL\_3154


CL\_3152


CL\_3152


CL\_3152


CL\_3152


CL\_3152


CL\_3152


CL\_3152

HighlightSelectShow Genomes


154

CL\_3150


107

CL\_3150


4

CL\_3150


3

CL\_3150


2

CL\_3149


1

CL\_3150


1

Break


1

CL\_3150


1

CL\_3150


1

CL\_3150


1

CL\_3150

fGI ID


CL\_INS\_331
CL\_INS\_331
CL\_INS\_331
CL\_INS\_331
CL\_INS\_331
CL\_INS\_331
CL\_INS\_331
CL\_INS\_331
CL\_INS\_331
CL\_INS\_331
CL\_INS\_331
Cluster ID


CL\_30748
CL\_13601
CL\_24218
CL\_13602
CL\_13603
CL\_12190
CL\_12189
CL\_12188
CL\_3151
CL\_28259
CL\_30121
