## Supplementary material for "A novel method for integrating genomic and Tn-Seq data to identify common *in vivo* fitness mechanisms across multiple bacterial species": S1 Dataset: CL_INS_332.html

Legend

 Mobile +extrachromosomalelementfunctions
 Regulatoryfunctions
 Hypothetical
 DNA Metabolism
 Purines,pyrimidines,nucleosides, +nucleotides
 All EssentialGenes
 AntibioticResistance
 Biosynthesis ofcofactors,prostheticgroups, +carriers
 All Fitness Genes
 Cell Envelope
 Proteinsynthesis/fate
 Other
 Cellularprocesses
 Transport +binding proteins
 All VFDB Genes

FULL


WINDOWSVGPNG

Trim RowsRemove SingletonsSave Fasta

CL\_3133


CL\_3133


CL\_3164


CL\_3832


CL\_3133


CL\_3133


CL\_3553


CL\_3133


CL\_3133


CL\_4487


CL\_3133


CL\_3133


CL\_3133

HighlightSelectShow Genomes


148

CL\_3835


105

CL\_3835


4

CL\_3835


2

CL\_3835


2

CL\_3836


2

CL\_3775


1

CL\_3835


1

Break


1

Break


1

CL\_3835


1

CL\_3835


1

CL\_3775


1

Break

fGI ID


CL\_INS\_237
CL\_INS\_92
CL\_INS\_332
CL\_INS\_332
CL\_INS\_332
CL\_INS\_332
CL\_INS\_382
CL\_INS\_382
CL\_INS\_382
CL\_INS\_382
CL\_INS\_382
CL\_INS\_382
CL\_INS\_382
CL\_INS\_382
CL\_INS\_382
CL\_INS\_382
CL\_INS\_382
CL\_INS\_382
CL\_INS\_382
CL\_INS\_99
CL\_INS\_99
CL\_INS\_382
CL\_INS\_382
CL\_INS\_332
CL\_INS\_382
CL\_INS\_382
CL\_INS\_99
CL\_INS\_99
CL\_INS\_99
CL\_INS\_382
CL\_INS\_382
CL\_INS\_382
CL\_INS\_382
CL\_INS\_382
CL\_INS\_382
CL\_INS\_382
CL\_INS\_382
CL\_INS\_382
CL\_INS\_382
CL\_INS\_382
CL\_INS\_382
CL\_INS\_382
CL\_INS\_382
CL\_INS\_382
CL\_INS\_382
CL\_INS\_99
CL\_INS\_99
CL\_INS\_382
CL\_INS\_382
CL\_INS\_99
CL\_INS\_382
CL\_INS\_237
CL\_INS\_237
CL\_INS\_237
CL\_INS\_237
CL\_INS\_70
CL\_INS\_237
CL\_INS\_70
CL\_INS\_70
CL\_INS\_70
CL\_INS\_70
CL\_INS\_70
CL\_INS\_70
CL\_INS\_237
CL\_INS\_237
CL\_INS\_237
CL\_INS\_237
CL\_INS\_237
CL\_INS\_382
CL\_INS\_237
CL\_INS\_55
CL\_INS\_55
CL\_INS\_55
CL\_INS\_332
CL\_INS\_55
CL\_INS\_55
CL\_INS\_55
CL\_INS\_55
CL\_INS\_55
CL\_INS\_55
CL\_INS\_332
CL\_INS\_332
CL\_INS\_332
CL\_INS\_332
CL\_INS\_247
CL\_INS\_332
CL\_INS\_70
CL\_INS\_70
CL\_INS\_70
CL\_INS\_332
CL\_INS\_332
CL\_INS\_70
CL\_INS\_332
CL\_INS\_332
CL\_INS\_332
CL\_INS\_385
CL\_INS\_332
CL\_INS\_286
CL\_INS\_332
CL\_INS\_332
CL\_INS\_55
CL\_INS\_332
CL\_INS\_55
CL\_INS\_302
CL\_INS\_156
CL\_INS\_382
CL\_INS\_382
CL\_INS\_159
CL\_INS\_382
CL\_INS\_332
CL\_INS\_382
CL\_INS\_71
CL\_INS\_71
CL\_INS\_71
CL\_INS\_71
CL\_INS\_382
CL\_INS\_71
CL\_INS\_71
CL\_INS\_382
CL\_INS\_332
CL\_INS\_332
CL\_INS\_237
CL\_INS\_295
CL\_INS\_332
CL\_INS\_332
CL\_INS\_332
CL\_INS\_332
CL\_INS\_332
CL\_INS\_332
CL\_INS\_332
CL\_INS\_332
CL\_INS\_332
CL\_INS\_332
CL\_INS\_332
CL\_INS\_332
CL\_INS\_382
CL\_INS\_382
CL\_INS\_60
CL\_INS\_60
CL\_INS\_368
CL\_INS\_368
CL\_INS\_30
CL\_INS\_30
CL\_INS\_30
CL\_INS\_30
CL\_INS\_30
CL\_INS\_60
CL\_INS\_60
CL\_INS\_60
CL\_INS\_60
CL\_INS\_368
CL\_INS\_382
CL\_INS\_237
CL\_INS\_237
CL\_INS\_237
CL\_INS\_237
CL\_INS\_237
CL\_INS\_237
CL\_INS\_237
CL\_INS\_332
CL\_INS\_332
CL\_INS\_332
CL\_INS\_332
CL\_INS\_332
CL\_INS\_332
CL\_INS\_55
CL\_INS\_60
CL\_INS\_159
CL\_INS\_159
CL\_INS\_159
CL\_INS\_332
CL\_INS\_60
CL\_INS\_60
CL\_INS\_60
CL\_INS\_60
CL\_INS\_60
CL\_INS\_60
CL\_INS\_60
CL\_INS\_60
CL\_INS\_60
CL\_INS\_368
CL\_INS\_368
CL\_INS\_247
CL\_INS\_382
CL\_INS\_382
CL\_INS\_382
CL\_INS\_382
CL\_INS\_247
CL\_INS\_382
CL\_INS\_382
CL\_INS\_382
CL\_INS\_71
CL\_INS\_71
CL\_INS\_117
CL\_INS\_117
CL\_INS\_159
CL\_INS\_382
CL\_INS\_382
CL\_INS\_382
CL\_INS\_382
CL\_INS\_1
CL\_INS\_382
CL\_INS\_159
CL\_INS\_159
CL\_INS\_159
CL\_INS\_368
CL\_INS\_385
CL\_INS\_382
CL\_INS\_159
CL\_INS\_382
CL\_INS\_233
CL\_INS\_207
CL\_INS\_207
CL\_INS\_207
CL\_INS\_332
CL\_INS\_332
CL\_INS\_382
CL\_INS\_382
CL\_INS\_382
CL\_INS\_382
CL\_INS\_382
CL\_INS\_332
CL\_INS\_382
CL\_INS\_382
CL\_INS\_382
CL\_INS\_382
CL\_INS\_382
CL\_INS\_382
CL\_INS\_382
CL\_INS\_382
CL\_INS\_382
CL\_INS\_382
CL\_INS\_382
CL\_INS\_60
CL\_INS\_382
CL\_INS\_382
CL\_INS\_382
CL\_INS\_382
CL\_INS\_382
CL\_INS\_382
CL\_INS\_385
CL\_INS\_385
CL\_INS\_159
CL\_INS\_385
CL\_INS\_385
CL\_INS\_385
CL\_INS\_382
CL\_INS\_382
CL\_INS\_60
CL\_INS\_60
CL\_INS\_60
CL\_INS\_382
CL\_INS\_332
CL\_INS\_382
CL\_INS\_382
CL\_INS\_83
CL\_INS\_382
CL\_INS\_382
CL\_INS\_382
CL\_INS\_233
CL\_INS\_233
CL\_INS\_233
CL\_INS\_382
CL\_INS\_382
CL\_INS\_233
CL\_INS\_233
CL\_INS\_233
CL\_INS\_233
CL\_INS\_60
CL\_INS\_60
CL\_INS\_382
CL\_INS\_382
CL\_INS\_382
CL\_INS\_382
CL\_INS\_386
CL\_INS\_233
CL\_INS\_233
CL\_INS\_332
CL\_INS\_332
CL\_INS\_117
CL\_INS\_332
CL\_INS\_332
CL\_INS\_156
CL\_INS\_332
CL\_INS\_332
CL\_INS\_332
CL\_INS\_332
CL\_INS\_332
CL\_INS\_70
CL\_INS\_359
CL\_INS\_332
CL\_INS\_247
CL\_INS\_332
CL\_INS\_332
CL\_INS\_332
CL\_INS\_332
CL\_INS\_237
CL\_INS\_332
CL\_INS\_332
CL\_INS\_332
CL\_INS\_332
CL\_INS\_332
CL\_INS\_332
CL\_INS\_332
CL\_INS\_332
CL\_INS\_332
CL\_INS\_123
CL\_INS\_123
CL\_INS\_332
CL\_INS\_332
CL\_INS\_332
CL\_INS\_332
CL\_INS\_332
CL\_INS\_162
CL\_INS\_162
CL\_INS\_162
CL\_INS\_332
CL\_INS\_99
CL\_INS\_162
CL\_INS\_162
CL\_INS\_162
CL\_INS\_162
CL\_INS\_162
CL\_INS\_154
CL\_INS\_201
CL\_INS\_99
CL\_INS\_99
CL\_INS\_99
CL\_INS\_99
CL\_INS\_332
CL\_INS\_237
CL\_INS\_332
CL\_INS\_332
CL\_INS\_332
CL\_INS\_332
CL\_INS\_332
CL\_INS\_368
CL\_INS\_332
CL\_INS\_359
CL\_INS\_332
CL\_INS\_332
CL\_INS\_326
CL\_INS\_332
CL\_INS\_332
CL\_INS\_332
CL\_INS\_272
CL\_INS\_272
CL\_INS\_272
CL\_INS\_272
CL\_INS\_272
CL\_INS\_272
CL\_INS\_272
CL\_INS\_272
CL\_INS\_272
CL\_INS\_272
CL\_INS\_272
CL\_INS\_369
CL\_INS\_247
CL\_INS\_247
CL\_INS\_359
CL\_INS\_359
CL\_INS\_247
CL\_INS\_359
CL\_INS\_247
CL\_INS\_247
CL\_INS\_247
CL\_INS\_44
CL\_INS\_247
CL\_INS\_332
CL\_INS\_332
CL\_INS\_247
CL\_INS\_247
CL\_INS\_247
CL\_INS\_247
CL\_INS\_237
CL\_INS\_237
CL\_INS\_237
CL\_INS\_237
CL\_INS\_237
CL\_INS\_237
CL\_INS\_237
CL\_INS\_382
CL\_INS\_149
CL\_INS\_332
CL\_INS\_332
CL\_INS\_332
CL\_INS\_332
CL\_INS\_332
CL\_INS\_332
CL\_INS\_332
CL\_INS\_332
CL\_INS\_332
CL\_INS\_332
CL\_INS\_332
CL\_INS\_332
CL\_INS\_332
CL\_INS\_332
CL\_INS\_332
CL\_INS\_332
CL\_INS\_332
CL\_INS\_332
CL\_INS\_332
CL\_INS\_332
CL\_INS\_86
CL\_INS\_71
CL\_INS\_385
CL\_INS\_332
CL\_INS\_83
CL\_INS\_332
CL\_INS\_332
CL\_INS\_332
CL\_INS\_332
CL\_INS\_332
CL\_INS\_332
CL\_INS\_332
CL\_INS\_332
CL\_INS\_332
CL\_INS\_332
CL\_INS\_332
CL\_INS\_332
CL\_INS\_332
CL\_INS\_332
CL\_INS\_332
CL\_INS\_332
CL\_INS\_332
CL\_INS\_332
CL\_INS\_332
CL\_INS\_332
CL\_INS\_332
CL\_INS\_332
CL\_INS\_332
CL\_INS\_332
CL\_INS\_332
CL\_INS\_332
CL\_INS\_332
CL\_INS\_332
CL\_INS\_332
CL\_INS\_332
CL\_INS\_332
CL\_INS\_332
CL\_INS\_332
CL\_INS\_332
CL\_INS\_332
CL\_INS\_332
CL\_INS\_332
CL\_INS\_332
CL\_INS\_332
CL\_INS\_332
CL\_INS\_332
CL\_INS\_332
CL\_INS\_332
CL\_INS\_332
CL\_INS\_332
CL\_INS\_332
CL\_INS\_332
CL\_INS\_332
CL\_INS\_332
CL\_INS\_332
CL\_INS\_332
CL\_INS\_332
CL\_INS\_332
CL\_INS\_332
CL\_INS\_332
CL\_INS\_332
CL\_INS\_332
CL\_INS\_332
CL\_INS\_332
CL\_INS\_332
CL\_INS\_332
CL\_INS\_332
CL\_INS\_85
CL\_INS\_85
CL\_INS\_85
CL\_INS\_85
CL\_INS\_85
CL\_INS\_85
CL\_INS\_85
CL\_INS\_85
CL\_INS\_85
CL\_INS\_85
CL\_INS\_85
CL\_INS\_85
CL\_INS\_85
CL\_INS\_85
CL\_INS\_85
CL\_INS\_85
CL\_INS\_85
CL\_INS\_85
CL\_INS\_85
CL\_INS\_85
CL\_INS\_85
CL\_INS\_85
CL\_INS\_85
CL\_INS\_85
CL\_INS\_85
CL\_INS\_85
CL\_INS\_85
CL\_INS\_332
CL\_INS\_85
CL\_INS\_85
CL\_INS\_85
CL\_INS\_85
CL\_INS\_85
CL\_INS\_85
CL\_INS\_85
CL\_INS\_85
CL\_INS\_85
CL\_INS\_85
CL\_INS\_85
CL\_INS\_85
CL\_INS\_237
CL\_INS\_326
CL\_INS\_70
CL\_INS\_382
CL\_INS\_382
CL\_INS\_85
CL\_INS\_332
CL\_INS\_332
CL\_INS\_85
CL\_INS\_85
CL\_INS\_382
CL\_INS\_85
CL\_INS\_85
CL\_INS\_85
CL\_INS\_85
CL\_INS\_85
CL\_INS\_85
CL\_INS\_85
CL\_INS\_85
CL\_INS\_85
CL\_INS\_85
CL\_INS\_85
CL\_INS\_85
CL\_INS\_85
CL\_INS\_85
CL\_INS\_85
CL\_INS\_85
CL\_INS\_85
CL\_INS\_85
CL\_INS\_332
CL\_INS\_85
CL\_INS\_85
CL\_INS\_85
CL\_INS\_85
CL\_INS\_85
CL\_INS\_85
CL\_INS\_85
CL\_INS\_332
CL\_INS\_85
CL\_INS\_85
CL\_INS\_85
CL\_INS\_85
CL\_INS\_85
CL\_INS\_332
CL\_INS\_332
CL\_INS\_332
CL\_INS\_85
CL\_INS\_332
CL\_INS\_85
CL\_INS\_85
CL\_INS\_85
CL\_INS\_85
CL\_INS\_85
CL\_INS\_85
CL\_INS\_332
CL\_INS\_85
CL\_INS\_85
CL\_INS\_85
CL\_INS\_85
CL\_INS\_354
CL\_INS\_354
CL\_INS\_354
CL\_INS\_354
CL\_INS\_237
CL\_INS\_237
CL\_INS\_85
CL\_INS\_85
CL\_INS\_85
CL\_INS\_85
CL\_INS\_85
CL\_INS\_85
CL\_INS\_332
CL\_INS\_332
CL\_INS\_332
CL\_INS\_85
CL\_INS\_85
CL\_INS\_85
CL\_INS\_332
Cluster ID


CL\_16977
CL\_34869
CL\_30724
CL\_14494
CL\_14495
CL\_14496
CL\_7818
CL\_1496
CL\_4533
CL\_7116
CL\_7115
CL\_4529
CL\_7113
CL\_4527
CL\_4526
CL\_4629
CL\_4524
CL\_4523
CL\_4522
CL\_4520
CL\_13789
CL\_12762
CL\_12760
CL\_34873
CL\_12996
CL\_7118
CL\_13561
CL\_28731
CL\_13560
CL\_8698
CL\_8700
CL\_8701
CL\_8702
CL\_8703
CL\_8704
CL\_8705
CL\_8706
CL\_8707
CL\_8708
CL\_8709
CL\_8710
CL\_8172
CL\_8171
CL\_8170
CL\_8169
CL\_4485
CL\_29360
CL\_4518
CL\_8712
CL\_4620
CL\_13034
CL\_10589
CL\_10590
CL\_10591
CL\_4462
CL\_8632
CL\_10595
CL\_6984
CL\_6983
CL\_6982
CL\_6981
CL\_6980
CL\_6979
CL\_10588
CL\_10587
CL\_10586
CL\_10585
CL\_10584
CL\_10804
CL\_10583
CL\_14211
CL\_18416
CL\_14209
CL\_18417
CL\_18418
CL\_18419
CL\_18420
CL\_18421
CL\_18422
CL\_14210
CL\_18388
CL\_18357
CL\_18358
CL\_18359
CL\_7007
CL\_18360
CL\_7252
CL\_6828
CL\_6411
CL\_18361
CL\_18362
CL\_7435
CL\_18363
CL\_18364
CL\_18365
CL\_5520
CL\_18366
CL\_11663
CL\_6944
CL\_6945
CL\_5499
CL\_18367
CL\_5500
CL\_5501
CL\_5502
CL\_4236
CL\_4237
CL\_6239
CL\_4235
CL\_10402
CL\_5001
CL\_11139
CL\_11140
CL\_11141
CL\_11142
CL\_11143
CL\_11147
CL\_10661
CL\_5000
CL\_18368
CL\_18369
CL\_9477
CL\_16998
CL\_18370
CL\_18371
CL\_18372
CL\_18373
CL\_18374
CL\_18375
CL\_18376
CL\_18377
CL\_18378
CL\_18379
CL\_18380
CL\_18381
CL\_4973
CL\_4972
CL\_10397
CL\_11814
CL\_11815
CL\_11816
CL\_7667
CL\_7840
CL\_11295
CL\_11296
CL\_8553
CL\_11818
CL\_11820
CL\_11821
CL\_11822
CL\_13617
CL\_4974
CL\_5151
CL\_7305
CL\_5153
CL\_5154
CL\_5155
CL\_5156
CL\_5157
CL\_18382
CL\_18383
CL\_18384
CL\_18385
CL\_18386
CL\_18387
CL\_5029
CL\_5030
CL\_5031
CL\_5032
CL\_5033
CL\_11827
CL\_10653
CL\_10652
CL\_10651
CL\_10650
CL\_10649
CL\_10648
CL\_10647
CL\_10646
CL\_10645
CL\_10644
CL\_10643
CL\_10664
CL\_10665
CL\_10666
CL\_10667
CL\_9691
CL\_9690
CL\_9689
CL\_9688
CL\_9687
CL\_11137
CL\_11138
CL\_5593
CL\_5592
CL\_5053
CL\_4253
CL\_4254
CL\_4255
CL\_4256
CL\_5590
CL\_4257
CL\_4258
CL\_4259
CL\_4261
CL\_5585
CL\_5625
CL\_5059
CL\_5630
CL\_4262
CL\_4309
CL\_5536
CL\_5613
CL\_5614
CL\_34874
CL\_3834
CL\_4263
CL\_5062
CL\_4265
CL\_5580
CL\_5579
CL\_6932
CL\_4270
CL\_4271
CL\_5574
CL\_15681
CL\_15682
CL\_5637
CL\_5638
CL\_5639
CL\_4277
CL\_4278
CL\_5567
CL\_5641
CL\_4279
CL\_5563
CL\_5562
CL\_5561
CL\_5560
CL\_4284
CL\_5649
CL\_5650
CL\_5651
CL\_5652
CL\_5653
CL\_5654
CL\_5555
CL\_5554
CL\_11830
CL\_11829
CL\_5657
CL\_5551
CL\_18389
CL\_5549
CL\_4294
CL\_18390
CL\_5662
CL\_4297
CL\_4299
CL\_12661
CL\_12660
CL\_5545
CL\_4301
CL\_4302
CL\_5544
CL\_5543
CL\_5542
CL\_6935
CL\_16528
CL\_15604
CL\_10406
CL\_10407
CL\_10411
CL\_5600
CL\_11428
CL\_12643
CL\_12644
CL\_18391
CL\_18392
CL\_6072
CL\_18393
CL\_18394
CL\_18395
CL\_18396
CL\_18397
CL\_18398
CL\_18399
CL\_18400
CL\_5682
CL\_5681
CL\_5680
CL\_5678
CL\_5677
CL\_5676
CL\_5675
CL\_5674
CL\_5673
CL\_5672
CL\_5671
CL\_5670
CL\_5669
CL\_5668
CL\_5667
CL\_5666
CL\_19451
CL\_19452
CL\_5722
CL\_5721
CL\_5720
CL\_5719
CL\_5718
CL\_5717
CL\_5716
CL\_5715
CL\_5714
CL\_5713
CL\_5712
CL\_5711
CL\_5765
CL\_5764
CL\_5763
CL\_5762
CL\_5761
CL\_5760
CL\_5759
CL\_5702
CL\_5701
CL\_5700
CL\_5699
CL\_5698
CL\_5697
CL\_5696
CL\_5695
CL\_5686
CL\_5685
CL\_5684
CL\_5683
CL\_19453
CL\_19454
CL\_19455
CL\_19456
CL\_11673
CL\_19457
CL\_19458
CL\_19459
CL\_6333
CL\_6332
CL\_6331
CL\_6330
CL\_6329
CL\_6328
CL\_6327
CL\_6326
CL\_6325
CL\_6324
CL\_6323
CL\_11654
CL\_11653
CL\_11652
CL\_11651
CL\_11650
CL\_11649
CL\_15069
CL\_15545
CL\_15067
CL\_15544
CL\_5518
CL\_5691
CL\_19460
CL\_19461
CL\_19462
CL\_19463
CL\_6879
CL\_6880
CL\_11347
CL\_11346
CL\_11345
CL\_11344
CL\_11343
CL\_6139
CL\_6140
CL\_6141
CL\_4093
CL\_19464
CL\_19465
CL\_19466
CL\_19467
CL\_19468
CL\_19469
CL\_19470
CL\_19471
CL\_19472
CL\_19473
CL\_19474
CL\_19475
CL\_19476
CL\_19477
CL\_19478
CL\_19479
CL\_19480
CL\_19481
CL\_19482
CL\_19483
CL\_5516
CL\_5515
CL\_5296
CL\_19484
CL\_19485
CL\_19486
CL\_19487
CL\_19488
CL\_19489
CL\_19490
CL\_19491
CL\_19492
CL\_19493
CL\_19494
CL\_19495
CL\_19496
CL\_19497
CL\_19498
CL\_19499
CL\_19500
CL\_19501
CL\_19502
CL\_19503
CL\_19504
CL\_19505
CL\_19506
CL\_19507
CL\_19508
CL\_19509
CL\_19510
CL\_19511
CL\_19512
CL\_19513
CL\_19514
CL\_19515
CL\_19516
CL\_19517
CL\_19518
CL\_19519
CL\_19520
CL\_19521
CL\_19522
CL\_19523
CL\_19524
CL\_19525
CL\_19526
CL\_19527
CL\_19528
CL\_19529
CL\_19530
CL\_19531
CL\_19532
CL\_19533
CL\_19534
CL\_19535
CL\_19536
CL\_19537
CL\_19538
CL\_19539
CL\_19540
CL\_19541
CL\_19542
CL\_19543
CL\_19544
CL\_19545
CL\_3833
CL\_19546
CL\_19547
CL\_19548
CL\_19549
CL\_19550
CL\_19551
CL\_19552
CL\_19553
CL\_19554
CL\_19555
CL\_19556
CL\_19557
CL\_19558
CL\_19559
CL\_19560
CL\_19561
CL\_19562
CL\_19563
CL\_19564
CL\_19565
CL\_19566
CL\_19567
CL\_19568
CL\_19569
CL\_19570
CL\_19571
CL\_19572
CL\_19573
CL\_19574
CL\_19575
CL\_19576
CL\_19577
CL\_19578
CL\_19579
CL\_19580
CL\_19581
CL\_19582
CL\_19583
CL\_19584
CL\_19585
CL\_19586
CL\_11394
CL\_11395
CL\_6179
CL\_6178
CL\_6177
CL\_19587
CL\_19588
CL\_19589
CL\_19590
CL\_19591
CL\_16415
CL\_19592
CL\_19593
CL\_19594
CL\_19595
CL\_19596
CL\_19597
CL\_19598
CL\_19599
CL\_19600
CL\_19601
CL\_19602
CL\_19603
CL\_19604
CL\_19605
CL\_19606
CL\_19607
CL\_19608
CL\_19609
CL\_19610
CL\_19611
CL\_19612
CL\_19613
CL\_19614
CL\_19615
CL\_19616
CL\_19617
CL\_19618
CL\_19619
CL\_19620
CL\_19621
CL\_19622
CL\_19623
CL\_19624
CL\_19625
CL\_19626
CL\_19627
CL\_19628
CL\_19629
CL\_19630
CL\_19631
CL\_19632
CL\_19633
CL\_19634
CL\_19635
CL\_19636
CL\_19637
CL\_19638
CL\_19639
CL\_19640
CL\_19641
CL\_19642
CL\_19643
CL\_11393
CL\_6181
CL\_19644
CL\_19645
CL\_19646
CL\_19647
CL\_19648
CL\_19649
CL\_10640
CL\_19650
CL\_19651
CL\_19652
CL\_19653
CL\_19654
CL\_19655
