## Supplementary material for "A novel method for integrating genomic and Tn-Seq data to identify common *in vivo* fitness mechanisms across multiple bacterial species": S1 Dataset: CL_INS_334.html

Legend

 Hypothetical
 Transcription
 Proteinsynthesis/fate
 Other
 EnergyMetabolism
 All VFDB Genes
 Transport +binding proteins

FULL


WINDOWSVGPNG

Trim RowsRemove SingletonsSave Fasta

CL\_3842


CL\_3842


CL\_3842


CL\_3841


CL\_3842


CL\_3839


CL\_3842


CL\_3842


CL\_3842

HighlightSelectShow Genomes


195

CL\_3843


41

CL\_3843


29

CL\_3843


1

CL\_3843


1

CL\_3845


1

CL\_3843


1

CL\_3843


1

CL\_3844


1

CL\_3844

fGI ID


CL\_INS\_334
CL\_INS\_334
CL\_INS\_334
CL\_INS\_334
CL\_INS\_334
CL\_INS\_334
CL\_INS\_334
Cluster ID


CL\_36957
CL\_7187
CL\_7188
CL\_5984
CL\_5985
CL\_5988
CL\_5989
