## Supplementary material for "A novel method for integrating genomic and Tn-Seq data to identify common *in vivo* fitness mechanisms across multiple bacterial species": S1 Dataset: CL_INS_335.html

Legend

 Mobile +extrachromosomalelementfunctions
 Hypothetical
 Regulatoryfunctions
 Transcription
 Proteinsynthesis/fate
 Other
 Transport +binding proteins
 All VFDB Genes

FULL


WINDOWSVGPNG

Trim RowsRemove SingletonsSave Fasta

CL\_3844


CL\_3844


CL\_3844


CL\_3844


CL\_3844


CL\_3844


CL\_3844


CL\_3844


CL\_3844


CL\_3844


CL\_3844


CL\_3843


CL\_3844


CL\_3844


CL\_3844


CL\_3844


CL\_3842


CL\_3843


CL\_3844

HighlightSelectShow Genomes


124

CL\_3845


115

CL\_3845


13

CL\_3845


5

CL\_3845


4

CL\_3845


2

CL\_3845


1

CL\_3845


1

CL\_3845


1

CL\_3846


1

CL\_3846


1

CL\_3845


1

CL\_3845


1

CL\_3845


1

CL\_3845


1

CL\_3845


1

CL\_3845


1

CL\_3845


1

CL\_3845


1

CL\_3845

fGI ID


CL\_INS\_335
CL\_INS\_335
CL\_INS\_334
CL\_INS\_335
CL\_INS\_334
CL\_INS\_335
CL\_INS\_335
CL\_INS\_334
CL\_INS\_334
CL\_INS\_335
CL\_INS\_335
Cluster ID


CL\_11322
CL\_33921
CL\_5984
CL\_30247
CL\_5985
CL\_5986
CL\_5987
CL\_5988
CL\_5989
CL\_5990
CL\_21720
