## Supplementary material for "A novel method for integrating genomic and Tn-Seq data to identify common *in vivo* fitness mechanisms across multiple bacterial species": S1 Dataset: CL_INS_337.html

Legend

 Hypothetical
 All EssentialGenes
 All VFDB Genes

FULL


WINDOWSVGPNG

Trim RowsRemove SingletonsSave Fasta

CL\_3853


CL\_3853


CL\_3852


CL\_3853

HighlightSelectShow Genomes


210

CL\_3855


67

CL\_3855


2

CL\_3855


1

CL\_3855

fGI ID


CL\_INS\_337
CL\_INS\_337
Cluster ID


CL\_3854
CL\_16505
