## Supplementary material for "A novel method for integrating genomic and Tn-Seq data to identify common *in vivo* fitness mechanisms across multiple bacterial species": S1 Dataset: CL_INS_338.html

Legend

 All EssentialGenes
 Other
 All VFDB Genes

FULL


WINDOWSVGPNG

Trim RowsRemove SingletonsSave Fasta

CL\_3855


CL\_3855


CL\_3855


CL\_3855


CL\_1923


CL\_3855

HighlightSelectShow Genomes


164

CL\_3857


109

CL\_3857


1

CL\_3860


1

CL\_3857


1

CL\_3857


1

CL\_3861

fGI ID


CL\_INS\_338
CL\_INS\_338
Cluster ID


CL\_3856
CL\_24509
