## Supplementary material for "A novel method for integrating genomic and Tn-Seq data to identify common *in vivo* fitness mechanisms across multiple bacterial species": S1 Dataset: CL_INS_339.html

Legend

 Mobile +extrachromosomalelementfunctions
 Hypothetical
 Biosynthesis ofcofactors,prostheticgroups, +carriers
 Transport +binding proteins
 All VFDB Genes

FULL


WINDOWSVGPNG

Trim RowsRemove SingletonsSave Fasta

CL\_3862


CL\_3862


CL\_3862


CL\_3862


CL\_3862


CL\_3593


CL\_3862


CL\_3862


CL\_3862


CL\_3862


CL\_3879


CL\_3862


CL\_3860

HighlightSelectShow Genomes


204

CL\_3863


57

CL\_3863


3

CL\_3863


2

CL\_3863


1

CL\_3863


1

CL\_3863


1

CL\_3863


1

CL\_3863


1

CL\_2844


1

CL\_3863


1

CL\_3863


1

CL\_3863


1

CL\_3863

fGI ID


CL\_INS\_339
CL\_INS\_339
CL\_INS\_339
CL\_INS\_284
CL\_INS\_339
CL\_INS\_339
CL\_INS\_339
CL\_INS\_339
CL\_INS\_339
Cluster ID


CL\_12882
CL\_33545
CL\_25051
CL\_3528
CL\_21232
CL\_10074
CL\_7566
CL\_15194
CL\_12184
