## Supplementary material for "A novel method for integrating genomic and Tn-Seq data to identify common *in vivo* fitness mechanisms across multiple bacterial species": S1 Dataset: CL_INS_341.html

Legend

 All EssentialGenes
 Other
 Transport +binding proteins
 All VFDB Genes

FULL


WINDOWSVGPNG

Trim RowsRemove SingletonsSave Fasta

CL\_3875


CL\_3875


CL\_3875


CL\_3875


CL\_3875

HighlightSelectShow Genomes


222

CL\_3876


28

CL\_3876


20

CL\_3876


1

CL\_3876


1

CL\_3876

fGI ID


CL\_INS\_341
CL\_INS\_341
CL\_INS\_341
CL\_INS\_341
CL\_INS\_341
CL\_INS\_341
CL\_INS\_341
Cluster ID


CL\_11126
CL\_7189
CL\_11125
CL\_11124
CL\_7190
CL\_7191
CL\_7192
