## Supplementary material for "A novel method for integrating genomic and Tn-Seq data to identify common *in vivo* fitness mechanisms across multiple bacterial species": S1 Dataset: CL_INS_342.html

Legend

 Hypothetical
 All EssentialGenes
 Other
 All VFDB Genes

FULL


WINDOWSVGPNG

Trim RowsRemove SingletonsSave Fasta

CL\_3877


CL\_3877


CL\_3877


CL\_3876


CL\_3877


CL\_3877

HighlightSelectShow Genomes


213

CL\_3878


56

CL\_3878


1

CL\_3880


1

CL\_3878


1

CL\_3879


1

CL\_3879

fGI ID


CL\_INS\_342
CL\_INS\_342
CL\_INS\_343
CL\_INS\_343
CL\_INS\_343
CL\_INS\_343
CL\_INS\_343
CL\_INS\_343
CL\_INS\_343
CL\_INS\_343
Cluster ID


CL\_7193
CL\_9450
CL\_7194
CL\_7195
CL\_7196
CL\_8017
CL\_8016
CL\_8015
CL\_8014
CL\_7197
