## Supplementary material for "A novel method for integrating genomic and Tn-Seq data to identify common *in vivo* fitness mechanisms across multiple bacterial species": S1 Dataset: CL_INS_343.html

Legend

 Mobile +extrachromosomalelementfunctions
 Regulatoryfunctions
 Hypothetical
 All EssentialGenes
 Biosynthesis ofcofactors,prostheticgroups, +carriers
 All Fitness Genes
 Other
 Transport +binding proteins
 All VFDB Genes

FULL


WINDOWSVGPNG

Trim RowsRemove SingletonsSave Fasta

CL\_3878


CL\_3878


CL\_3878


CL\_3878


CL\_3878


CL\_3878


CL\_3877


CL\_3878


CL\_3878


CL\_3878


CL\_3878


CL\_3878


CL\_3878


CL\_3878


CL\_3878


CL\_3878


CL\_3878


CL\_3878


CL\_3877


CL\_3878


CL\_3958


CL\_3878


CL\_3863


CL\_3878

HighlightSelectShow Genomes


133

CL\_3879


59

CL\_3879


58

CL\_3879


4

CL\_3880


1

CL\_3879


1

CL\_3879


1

CL\_3879


1

CL\_3879


1

Break


1

CL\_234


1

CL\_3879


1

CL\_3879


1

CL\_3879


1

CL\_3879


1

CL\_3879


1

CL\_3879


1

CL\_3879


1

CL\_3880


1

CL\_3879


1

CL\_3881


1

CL\_3879


1

CL\_3880


1

CL\_3879


1

CL\_3879

fGI ID


CL\_INS\_343
CL\_INS\_343
CL\_INS\_343
CL\_INS\_343
CL\_INS\_343
CL\_INS\_343
CL\_INS\_343
CL\_INS\_343
CL\_INS\_343
CL\_INS\_343
CL\_INS\_343
CL\_INS\_343
CL\_INS\_343
CL\_INS\_70
CL\_INS\_237
CL\_INS\_343
CL\_INS\_343
CL\_INS\_343
CL\_INS\_343
CL\_INS\_343
CL\_INS\_343
CL\_INS\_343
CL\_INS\_343
CL\_INS\_343
CL\_INS\_343
CL\_INS\_237
CL\_INS\_237
CL\_INS\_237
CL\_INS\_237
CL\_INS\_237
CL\_INS\_237
CL\_INS\_237
CL\_INS\_237
CL\_INS\_237
CL\_INS\_237
CL\_INS\_237
CL\_INS\_237
CL\_INS\_237
CL\_INS\_237
CL\_INS\_237
CL\_INS\_237
CL\_INS\_237
CL\_INS\_237
CL\_INS\_382
CL\_INS\_237
CL\_INS\_237
CL\_INS\_237
CL\_INS\_382
CL\_INS\_237
CL\_INS\_237
CL\_INS\_237
CL\_INS\_237
CL\_INS\_343
CL\_INS\_343
CL\_INS\_343
CL\_INS\_343
CL\_INS\_343
CL\_INS\_343
CL\_INS\_343
CL\_INS\_343
CL\_INS\_343
CL\_INS\_343
CL\_INS\_343
CL\_INS\_343
CL\_INS\_343
CL\_INS\_343
CL\_INS\_343
CL\_INS\_343
CL\_INS\_233
CL\_INS\_30
CL\_INS\_233
CL\_INS\_343
CL\_INS\_233
CL\_INS\_343
CL\_INS\_343
CL\_INS\_343
CL\_INS\_233
CL\_INS\_233
CL\_INS\_233
CL\_INS\_233
CL\_INS\_159
CL\_INS\_382
CL\_INS\_233
CL\_INS\_233
CL\_INS\_343
CL\_INS\_233
CL\_INS\_233
CL\_INS\_382
CL\_INS\_382
CL\_INS\_233
CL\_INS\_233
CL\_INS\_233
CL\_INS\_70
CL\_INS\_343
CL\_INS\_70
CL\_INS\_70
CL\_INS\_70
CL\_INS\_70
CL\_INS\_233
CL\_INS\_233
CL\_INS\_233
CL\_INS\_233
CL\_INS\_233
CL\_INS\_343
CL\_INS\_382
CL\_INS\_382
CL\_INS\_343
CL\_INS\_382
CL\_INS\_382
CL\_INS\_233
CL\_INS\_233
CL\_INS\_382
CL\_INS\_382
CL\_INS\_382
CL\_INS\_233
CL\_INS\_233
CL\_INS\_233
CL\_INS\_233
CL\_INS\_233
CL\_INS\_343
CL\_INS\_343
CL\_INS\_343
CL\_INS\_343
CL\_INS\_343
CL\_INS\_343
CL\_INS\_343
CL\_INS\_343
CL\_INS\_343
CL\_INS\_343
CL\_INS\_343
CL\_INS\_343
CL\_INS\_30
CL\_INS\_70
CL\_INS\_343
CL\_INS\_343
CL\_INS\_343
CL\_INS\_382
CL\_INS\_343
CL\_INS\_343
CL\_INS\_343
CL\_INS\_343
CL\_INS\_343
CL\_INS\_343
CL\_INS\_233
CL\_INS\_233
CL\_INS\_233
CL\_INS\_233
CL\_INS\_343
CL\_INS\_233
CL\_INS\_343
CL\_INS\_343
CL\_INS\_343
CL\_INS\_382
CL\_INS\_343
CL\_INS\_343
CL\_INS\_343
CL\_INS\_233
CL\_INS\_233
CL\_INS\_233
CL\_INS\_247
CL\_INS\_247
CL\_INS\_247
CL\_INS\_247
CL\_INS\_233
CL\_INS\_233
CL\_INS\_233
CL\_INS\_233
CL\_INS\_233
CL\_INS\_233
CL\_INS\_233
CL\_INS\_233
CL\_INS\_233
CL\_INS\_233
CL\_INS\_233
CL\_INS\_233
CL\_INS\_233
CL\_INS\_233
CL\_INS\_233
CL\_INS\_233
CL\_INS\_233
CL\_INS\_233
CL\_INS\_233
CL\_INS\_233
CL\_INS\_233
CL\_INS\_233
CL\_INS\_233
CL\_INS\_233
CL\_INS\_233
CL\_INS\_233
CL\_INS\_233
CL\_INS\_343
CL\_INS\_233
CL\_INS\_233
CL\_INS\_237
Cluster ID


CL\_12183
CL\_37548
CL\_34967
CL\_23756
CL\_34282
CL\_23807
CL\_23806
CL\_23909
CL\_4927
CL\_10241
CL\_7194
CL\_7195
CL\_32001
CL\_8037
CL\_7980
CL\_32002
CL\_32003
CL\_32004
CL\_32005
CL\_32006
CL\_32007
CL\_32008
CL\_32009
CL\_32010
CL\_32011
CL\_4374
CL\_4375
CL\_5148
CL\_5147
CL\_7713
CL\_26539
CL\_26540
CL\_32012
CL\_6075
CL\_7711
CL\_7710
CL\_7709
CL\_7708
CL\_7707
CL\_7706
CL\_8385
CL\_7305
CL\_7304
CL\_6056
CL\_7705
CL\_7704
CL\_7703
CL\_8909
CL\_7687
CL\_7686
CL\_7685
CL\_7684
CL\_14579
CL\_7196
CL\_8017
CL\_8016
CL\_8015
CL\_8014
CL\_7197
CL\_8013
CL\_30961
CL\_30962
CL\_30963
CL\_30964
CL\_30965
CL\_30966
CL\_30967
CL\_30968
CL\_12639
CL\_5237
CL\_12640
CL\_22859
CL\_12642
CL\_30969
CL\_30970
CL\_30971
CL\_12656
CL\_12655
CL\_12657
CL\_12653
CL\_5539
CL\_5599
CL\_5541
CL\_6935
CL\_30972
CL\_5543
CL\_5544
CL\_4302
CL\_4301
CL\_5545
CL\_12660
CL\_12661
CL\_10435
CL\_22836
CL\_8750
CL\_8749
CL\_8748
CL\_8747
CL\_12664
CL\_10801
CL\_12666
CL\_12667
CL\_12668
CL\_30973
CL\_4236
CL\_4235
CL\_5618
CL\_5015
CL\_5014
CL\_12671
CL\_12674
CL\_4251
CL\_6927
CL\_6926
CL\_12678
CL\_12679
CL\_12659
CL\_12680
CL\_12682
CL\_30974
CL\_13265
CL\_13266
CL\_30975
CL\_13256
CL\_13285
CL\_30976
CL\_22855
CL\_22856
CL\_13017
CL\_13018
CL\_13019
CL\_7667
CL\_6426
CL\_13020
CL\_13253
CL\_13021
CL\_4975
CL\_13022
CL\_13023
CL\_13024
CL\_30977
CL\_13025
CL\_30978
CL\_12646
CL\_12647
CL\_12648
CL\_12649
CL\_13031
CL\_12650
CL\_30979
CL\_30980
CL\_30981
CL\_4306
CL\_30982
CL\_30221
CL\_30983
CL\_12685
CL\_12686
CL\_12688
CL\_4838
CL\_4840
CL\_5947
CL\_4841
CL\_12610
CL\_12611
CL\_12612
CL\_12613
CL\_12614
CL\_10578
CL\_12615
CL\_12616
CL\_12617
CL\_12618
CL\_12619
CL\_12620
CL\_12622
CL\_12623
CL\_12624
CL\_12625
CL\_12626
CL\_12627
CL\_12628
CL\_12629
CL\_12630
CL\_12631
CL\_12632
CL\_12633
CL\_12634
CL\_12635
CL\_12636
CL\_13274
CL\_12637
CL\_12638
CL\_12241
