## Supplementary material for "A novel method for integrating genomic and Tn-Seq data to identify common *in vivo* fitness mechanisms across multiple bacterial species": S1 Dataset: CL_INS_345.html

Legend

 Regulatoryfunctions
 Hypothetical
 Other

FULL


WINDOWSVGPNG

Trim RowsRemove SingletonsSave Fasta

CL\_3887


CL\_3887


CL\_3887

HighlightSelectShow Genomes


274

CL\_3888


3

CL\_3888


1

CL\_3888

fGI ID


CL\_INS\_345
CL\_INS\_345
CL\_INS\_345
Cluster ID


CL\_15196
CL\_15197
CL\_15198
