## Supplementary material for "A novel method for integrating genomic and Tn-Seq data to identify common *in vivo* fitness mechanisms across multiple bacterial species": S1 Dataset: CL_INS_346.html

Legend

 Mobile +extrachromosomalelementfunctions
 Hypothetical
 Other
 Transport +binding proteins
 All VFDB Genes

FULL


WINDOWSVGPNG

Trim RowsRemove SingletonsSave Fasta

CL\_3889


CL\_3889


CL\_3889


CL\_3889


CL\_3889


CL\_3889


CL\_3888


CL\_3889


CL\_3889


CL\_3889

HighlightSelectShow Genomes


160

CL\_3891


60

CL\_3891


44

CL\_3891


4

CL\_3891


2

CL\_3892


1

CL\_3891


1

CL\_3891


1

CL\_3907


1

CL\_3892


1

CL\_3452

fGI ID


CL\_INS\_346
CL\_INS\_346
CL\_INS\_346
CL\_INS\_237
CL\_INS\_237
CL\_INS\_346
CL\_INS\_346
CL\_INS\_346
CL\_INS\_346
CL\_INS\_346
CL\_INS\_346
Cluster ID


CL\_3890
CL\_10255
CL\_30709
CL\_7245
CL\_8124
CL\_8012
CL\_8011
CL\_7198
CL\_23041
CL\_3910
CL\_3909
