## Supplementary material for "A novel method for integrating genomic and Tn-Seq data to identify common *in vivo* fitness mechanisms across multiple bacterial species": S1 Dataset: CL_INS_348.html

Legend

 Mobile +extrachromosomalelementfunctions
 Regulatoryfunctions
 Hypothetical
 All EssentialGenes
 Other
 Transport +binding proteins
 All VFDB Genes

FULL


WINDOWSVGPNG

Trim RowsRemove SingletonsSave Fasta

CL\_3907


CL\_3907


CL\_3907


CL\_3907


CL\_3907


CL\_3907


CL\_3907


CL\_3907


CL\_3907


CL\_3906


CL\_4929


CL\_3907


CL\_3906


CL\_3907


CL\_3907


CL\_3907


CL\_3906


CL\_3906


CL\_3907


CL\_3907


CL\_3907


CL\_3907


CL\_3907


CL\_3907


CL\_3907


CL\_3907


CL\_3906


CL\_3907


CL\_3907


CL\_3907


CL\_3907

HighlightSelectShow Genomes


133

CL\_3914


82

CL\_3914


15

CL\_3914


3

CL\_3914


3

CL\_3915


2

CL\_3914


2

CL\_3914


2

CL\_3915


2

CL\_3915


1

CL\_3914


1

CL\_3914


1

CL\_3914


1

CL\_3914


1

CL\_4001


1

CL\_3915


1

CL\_3914


1

CL\_3914


1

CL\_3914


1

CL\_3889


1

CL\_3914


1

CL\_3914


1

CL\_3923


1

CL\_3914


1

CL\_3914


1

CL\_3914


1

CL\_3914


1

CL\_3914


1

CL\_3916


1

CL\_3914


1

CL\_3915


1

CL\_3914

fGI ID


CL\_INS\_348
CL\_INS\_348
CL\_INS\_348
CL\_INS\_348
CL\_INS\_348
CL\_INS\_346
CL\_INS\_297
CL\_INS\_297
CL\_INS\_297
CL\_INS\_297
CL\_INS\_297
CL\_INS\_346
CL\_INS\_348
CL\_INS\_297
CL\_INS\_297
CL\_INS\_348
CL\_INS\_348
CL\_INS\_348
CL\_INS\_348
CL\_INS\_348
CL\_INS\_356
CL\_INS\_356
CL\_INS\_356
CL\_INS\_356
CL\_INS\_70
CL\_INS\_356
CL\_INS\_70
CL\_INS\_356
CL\_INS\_356
CL\_INS\_356
Cluster ID


CL\_12902
CL\_12901
CL\_32407
CL\_14497
CL\_4928
CL\_3909
CL\_12181
CL\_12180
CL\_12179
CL\_12178
CL\_15553
CL\_3910
CL\_37661
CL\_15554
CL\_3911
CL\_3908
CL\_3912
CL\_3913
CL\_23928
CL\_12883
CL\_12884
CL\_12885
CL\_12886
CL\_12887
CL\_12888
CL\_12889
CL\_8632
CL\_12890
CL\_12891
CL\_12892
