## Supplementary material for "A novel method for integrating genomic and Tn-Seq data to identify common *in vivo* fitness mechanisms across multiple bacterial species": S1 Dataset: CL_INS_351.html

Legend

 Mobile +extrachromosomalelementfunctions
 Hypothetical
 Regulatoryfunctions
 Transcription
 Cell Envelope
 Other
 All VFDB Genes
 Transport +binding proteins

FULL


WINDOWSVGPNG

Trim RowsRemove SingletonsSave Fasta

CL\_3949


CL\_3949


CL\_3949


CL\_3948


CL\_3949


CL\_3949


CL\_3949


CL\_3949


CL\_3949


CL\_3949


CL\_3949


CL\_3949


CL\_3948


CL\_3949


CL\_4028

HighlightSelectShow Genomes


237

CL\_3950


14

CL\_3950


9

CL\_3950


2

CL\_3950


2

CL\_3950


1

CL\_4027


1

CL\_3950


1

CL\_3950


1

CL\_3950


1

CL\_3950


1

CL\_3950


1

CL\_3957


1

CL\_3950


1

CL\_3845


1

CL\_3950

fGI ID


CL\_INS\_351
CL\_INS\_351
CL\_INS\_351
CL\_INS\_351
CL\_INS\_351
CL\_INS\_351
CL\_INS\_351
CL\_INS\_351
CL\_INS\_351
CL\_INS\_351
CL\_INS\_351
CL\_INS\_351
CL\_INS\_351
CL\_INS\_351
CL\_INS\_351
CL\_INS\_351
CL\_INS\_351
CL\_INS\_351
CL\_INS\_159
CL\_INS\_351
CL\_INS\_351
Cluster ID


CL\_29288
CL\_29287
CL\_11123
CL\_11122
CL\_11121
CL\_11120
CL\_13366
CL\_19661
CL\_19660
CL\_19659
CL\_20001
CL\_7643
CL\_7642
CL\_7641
CL\_7561
CL\_7560
CL\_7559
CL\_11119
CL\_7639
CL\_11118
CL\_11117
