## Supplementary material for "A novel method for integrating genomic and Tn-Seq data to identify common *in vivo* fitness mechanisms across multiple bacterial species": S1 Dataset: CL_INS_352.html

Legend

 Mobile +extrachromosomalelementfunctions
 Hypothetical
 Other
 All VFDB Genes

FULL


WINDOWSVGPNG

Trim RowsRemove SingletonsSave Fasta

CL\_3950


CL\_3950


CL\_3950


CL\_3950


CL\_3950


CL\_3950


CL\_3950


CL\_3950


CL\_3950


CL\_3950

HighlightSelectShow Genomes


259

CL\_3951


3

CL\_3951


1

CL\_3951


1

CL\_1935


1

CL\_3958


1

CL\_3951


1

CL\_3951


1

CL\_3958


1

CL\_3958


1

CL\_3958

fGI ID


CL\_INS\_352
CL\_INS\_352
CL\_INS\_352
CL\_INS\_352
CL\_INS\_352
CL\_INS\_382
CL\_INS\_382
CL\_INS\_382
CL\_INS\_70
CL\_INS\_70
CL\_INS\_352
CL\_INS\_352
CL\_INS\_352
CL\_INS\_352
CL\_INS\_352
CL\_INS\_352
CL\_INS\_352
CL\_INS\_352
CL\_INS\_352
CL\_INS\_352
CL\_INS\_352
CL\_INS\_352
CL\_INS\_352
CL\_INS\_352
CL\_INS\_70
CL\_INS\_149
CL\_INS\_70
CL\_INS\_352
CL\_INS\_352
CL\_INS\_352
CL\_INS\_352
CL\_INS\_352
CL\_INS\_352
CL\_INS\_352
CL\_INS\_352
CL\_INS\_352
CL\_INS\_352
CL\_INS\_352
CL\_INS\_352
CL\_INS\_352
CL\_INS\_352
CL\_INS\_352
CL\_INS\_352
CL\_INS\_352
CL\_INS\_352
CL\_INS\_352
CL\_INS\_352
CL\_INS\_352
CL\_INS\_352
CL\_INS\_352
CL\_INS\_352
CL\_INS\_352
CL\_INS\_352
CL\_INS\_352
CL\_INS\_352
CL\_INS\_352
CL\_INS\_352
CL\_INS\_352
CL\_INS\_352
CL\_INS\_352
CL\_INS\_352
CL\_INS\_352
CL\_INS\_352
CL\_INS\_352
CL\_INS\_352
CL\_INS\_352
CL\_INS\_352
CL\_INS\_352
CL\_INS\_352
CL\_INS\_352
CL\_INS\_352
CL\_INS\_352
CL\_INS\_352
CL\_INS\_352
CL\_INS\_352
CL\_INS\_352
CL\_INS\_352
CL\_INS\_352
CL\_INS\_352
CL\_INS\_352
CL\_INS\_352
CL\_INS\_352
CL\_INS\_352
CL\_INS\_352
CL\_INS\_352
CL\_INS\_352
CL\_INS\_70
CL\_INS\_30
Cluster ID


CL\_30729
CL\_13243
CL\_15202
CL\_27363
CL\_10594
CL\_8216
CL\_4975
CL\_8841
CL\_10592
CL\_7690
CL\_10073
CL\_10072
CL\_10071
CL\_10070
CL\_10069
CL\_10068
CL\_10067
CL\_10066
CL\_10065
CL\_10064
CL\_10063
CL\_10062
CL\_6028
CL\_6029
CL\_5321
CL\_5146
CL\_5320
CL\_6833
CL\_5159
CL\_10061
CL\_10060
CL\_10059
CL\_10058
CL\_10057
CL\_10056
CL\_10055
CL\_10054
CL\_10053
CL\_17585
CL\_9441
CL\_9440
CL\_9439
CL\_9438
CL\_9437
CL\_9436
CL\_9435
CL\_9434
CL\_9433
CL\_9432
CL\_9431
CL\_9430
CL\_9429
CL\_9428
CL\_9427
CL\_9426
CL\_9425
CL\_9424
CL\_9423
CL\_9422
CL\_9421
CL\_9420
CL\_9419
CL\_9418
CL\_9417
CL\_9416
CL\_9415
CL\_9414
CL\_9413
CL\_9412
CL\_9411
CL\_9410
CL\_9409
CL\_9408
CL\_9407
CL\_9406
CL\_9405
CL\_9404
CL\_9403
CL\_9402
CL\_9401
CL\_17584
CL\_17583
CL\_8502
CL\_8503
CL\_7969
CL\_8504
CL\_6809
CL\_1937
