## Supplementary material for "A novel method for integrating genomic and Tn-Seq data to identify common *in vivo* fitness mechanisms across multiple bacterial species": S1 Dataset: CL_INS_353.html

Legend

 Mobile +extrachromosomalelementfunctions
 Regulatoryfunctions
 Hypothetical
 Other
 Centralintermediarymetabolism
 Transport +binding proteins
 All VFDB Genes

FULL


WINDOWSVGPNG

Trim RowsRemove SingletonsSave Fasta

CL\_3956


CL\_3956


CL\_3955


CL\_3956


CL\_3956


CL\_3956


CL\_3956


CL\_3956


CL\_3949


CL\_3956


CL\_3956


CL\_3956


CL\_3944

HighlightSelectShow Genomes


190

CL\_3957


57

CL\_3957


2

CL\_3957


1

CL\_3957


1

CL\_3958


1

CL\_4175


1

CL\_3958


1

CL\_3958


1

CL\_3957


1

CL\_3958


1

CL\_3958


1

CL\_1935


1

CL\_3957

fGI ID


CL\_INS\_353
CL\_INS\_353
CL\_INS\_353
CL\_INS\_353
CL\_INS\_353
CL\_INS\_353
CL\_INS\_353
CL\_INS\_353
CL\_INS\_353
CL\_INS\_353
CL\_INS\_353
CL\_INS\_353
CL\_INS\_353
CL\_INS\_353
CL\_INS\_353
CL\_INS\_237
CL\_INS\_237
CL\_INS\_237
CL\_INS\_237
CL\_INS\_237
CL\_INS\_237
CL\_INS\_354
CL\_INS\_354
CL\_INS\_237
CL\_INS\_237
CL\_INS\_237
CL\_INS\_237
CL\_INS\_237
CL\_INS\_155
CL\_INS\_237
CL\_INS\_237
CL\_INS\_237
CL\_INS\_237
CL\_INS\_237
CL\_INS\_237
CL\_INS\_354
CL\_INS\_354
CL\_INS\_247
CL\_INS\_237
CL\_INS\_247
CL\_INS\_237
CL\_INS\_247
CL\_INS\_247
CL\_INS\_237
CL\_INS\_237
CL\_INS\_237
CL\_INS\_237
CL\_INS\_237
CL\_INS\_237
CL\_INS\_237
CL\_INS\_237
CL\_INS\_237
CL\_INS\_237
CL\_INS\_237
CL\_INS\_237
CL\_INS\_237
CL\_INS\_70
CL\_INS\_237
CL\_INS\_237
CL\_INS\_237
CL\_INS\_237
CL\_INS\_237
CL\_INS\_237
CL\_INS\_237
CL\_INS\_237
CL\_INS\_237
CL\_INS\_237
CL\_INS\_30
CL\_INS\_70
CL\_INS\_368
CL\_INS\_368
CL\_INS\_368
CL\_INS\_368
CL\_INS\_368
CL\_INS\_368
CL\_INS\_368
CL\_INS\_368
CL\_INS\_368
CL\_INS\_368
CL\_INS\_155
CL\_INS\_237
CL\_INS\_237
CL\_INS\_237
CL\_INS\_247
CL\_INS\_237
CL\_INS\_237
CL\_INS\_237
CL\_INS\_247
CL\_INS\_247
CL\_INS\_247
CL\_INS\_247
CL\_INS\_237
CL\_INS\_237
CL\_INS\_237
CL\_INS\_247
CL\_INS\_247
CL\_INS\_247
CL\_INS\_247
CL\_INS\_247
CL\_INS\_247
CL\_INS\_247
CL\_INS\_247
CL\_INS\_247
CL\_INS\_247
CL\_INS\_247
CL\_INS\_247
CL\_INS\_247
CL\_INS\_247
CL\_INS\_247
CL\_INS\_70
CL\_INS\_70
CL\_INS\_70
CL\_INS\_70
CL\_INS\_237
CL\_INS\_237
CL\_INS\_237
CL\_INS\_237
CL\_INS\_237
CL\_INS\_237
CL\_INS\_70
CL\_INS\_70
CL\_INS\_237
CL\_INS\_237
CL\_INS\_237
CL\_INS\_237
CL\_INS\_382
CL\_INS\_70
CL\_INS\_70
CL\_INS\_70
CL\_INS\_237
CL\_INS\_237
CL\_INS\_237
CL\_INS\_237
CL\_INS\_237
CL\_INS\_237
CL\_INS\_237
CL\_INS\_87
Cluster ID


CL\_28993
CL\_5471
CL\_23051
CL\_23050
CL\_25930
CL\_25931
CL\_25932
CL\_25933
CL\_25934
CL\_25935
CL\_25936
CL\_25937
CL\_25938
CL\_25939
CL\_25940
CL\_9715
CL\_9716
CL\_9717
CL\_9718
CL\_9719
CL\_9720
CL\_29649
CL\_29650
CL\_16930
CL\_16931
CL\_16932
CL\_29075
CL\_6001
CL\_6000
CL\_29074
CL\_5998
CL\_29073
CL\_29072
CL\_29071
CL\_29070
CL\_29651
CL\_29652
CL\_7202
CL\_10789
CL\_7567
CL\_7568
CL\_7203
CL\_7204
CL\_7205
CL\_7206
CL\_7207
CL\_7208
CL\_4375
CL\_7209
CL\_7210
CL\_4374
CL\_7292
CL\_7293
CL\_7211
CL\_7212
CL\_7213
CL\_7214
CL\_7664
CL\_7665
CL\_8794
CL\_7295
CL\_8212
CL\_8793
CL\_8792
CL\_8791
CL\_8790
CL\_7666
CL\_7667
CL\_5682
CL\_9546
CL\_9547
CL\_9548
CL\_9549
CL\_9550
CL\_9551
CL\_9553
CL\_9554
CL\_9555
CL\_9556
CL\_7374
CL\_6076
CL\_7669
CL\_7670
CL\_7215
CL\_7216
CL\_7217
CL\_7218
CL\_7219
CL\_7220
CL\_8789
CL\_7221
CL\_7222
CL\_7223
CL\_7224
CL\_7225
CL\_7226
CL\_8788
CL\_7227
CL\_7228
CL\_8787
CL\_7229
CL\_7230
CL\_8786
CL\_7231
CL\_8785
CL\_8784
CL\_8783
CL\_7232
CL\_8782
CL\_7233
CL\_7234
CL\_7235
CL\_8781
CL\_8780
CL\_7236
CL\_7237
CL\_7238
CL\_7239
CL\_7240
CL\_7241
CL\_7242
CL\_7243
CL\_7244
CL\_7245
CL\_7297
CL\_8376
CL\_8779
CL\_7298
CL\_7299
CL\_7300
CL\_6244
CL\_6245
CL\_6246
CL\_5147
CL\_5148
CL\_5149
CL\_8778
