## Supplementary material for "A novel method for integrating genomic and Tn-Seq data to identify common *in vivo* fitness mechanisms across multiple bacterial species": S1 Dataset: CL_INS_356.html

Legend

 Mobile +extrachromosomalelementfunctions
 Hypothetical
 Other
 All VFDB Genes

FULL


WINDOWSVGPNG

Trim RowsRemove SingletonsSave Fasta

CL\_4001


CL\_4001


CL\_4001


CL\_4001


CL\_4001


CL\_4001


CL\_4001


CL\_4001


CL\_4001


CL\_4001

HighlightSelectShow Genomes


241

CL\_4002


12

CL\_4004


9

CL\_4002


2

CL\_4002


1

CL\_4008


1

CL\_4003


1

CL\_4002


1

CL\_4003


1

CL\_4002


1

CL\_3907

fGI ID


CL\_INS\_356
CL\_INS\_356
CL\_INS\_357
CL\_INS\_356
CL\_INS\_357
CL\_INS\_357
CL\_INS\_356
CL\_INS\_356
CL\_INS\_356
CL\_INS\_356
CL\_INS\_70
CL\_INS\_356
CL\_INS\_70
CL\_INS\_356
CL\_INS\_356
CL\_INS\_356
CL\_INS\_356
Cluster ID


CL\_8777
CL\_10256
CL\_8776
CL\_24503
CL\_8775
CL\_8774
CL\_12893
CL\_12892
CL\_12891
CL\_12890
CL\_8632
CL\_12889
CL\_12888
CL\_12887
CL\_12886
CL\_12885
CL\_12884
