## Supplementary material for "A novel method for integrating genomic and Tn-Seq data to identify common *in vivo* fitness mechanisms across multiple bacterial species": S1 Dataset: CL_INS_357.html

Legend

 Other

FULL


WINDOWSVGPNG

Trim RowsRemove SingletonsSave Fasta

CL\_4004


CL\_4004


CL\_4005

HighlightSelectShow Genomes


253

CL\_4003


11

CL\_4001


1

CL\_4003

fGI ID


CL\_INS\_357
CL\_INS\_357
CL\_INS\_357
Cluster ID


CL\_8774
CL\_8775
CL\_8776
