## Supplementary material for "A novel method for integrating genomic and Tn-Seq data to identify common *in vivo* fitness mechanisms across multiple bacterial species": S1 Dataset: CL_INS_363.html

Legend

 Mobile +extrachromosomalelementfunctions
 Regulatoryfunctions
 Hypothetical
 All EssentialGenes
 Other
 Transport +binding proteins
 All VFDB Genes

FULL


WINDOWSVGPNG

Trim RowsRemove SingletonsSave Fasta

CL\_4042


CL\_4042


CL\_4042


CL\_4042


CL\_4042


CL\_4042


CL\_4042


CL\_4040


CL\_4042


CL\_4042


CL\_4042


CL\_4042

HighlightSelectShow Genomes


139

CL\_4045


115

CL\_4045


11

CL\_4045


6

CL\_4045


1

CL\_4045


1

CL\_4045


1

CL\_4045


1

CL\_4045


1

CL\_4045


1

CL\_4045


1

CL\_4045


1

CL\_4045

fGI ID


CL\_INS\_363
CL\_INS\_363
CL\_INS\_363
CL\_INS\_363
CL\_INS\_363
CL\_INS\_363
CL\_INS\_363
CL\_INS\_363
CL\_INS\_363
CL\_INS\_149
CL\_INS\_149
CL\_INS\_149
CL\_INS\_149
CL\_INS\_149
CL\_INS\_149
CL\_INS\_363
CL\_INS\_170
CL\_INS\_363
CL\_INS\_363
CL\_INS\_363
CL\_INS\_363
CL\_INS\_86
CL\_INS\_155
CL\_INS\_149
CL\_INS\_363
CL\_INS\_363
CL\_INS\_363
CL\_INS\_363
CL\_INS\_363
CL\_INS\_363
CL\_INS\_363
CL\_INS\_363
CL\_INS\_363
CL\_INS\_363
CL\_INS\_363
CL\_INS\_71
CL\_INS\_71
CL\_INS\_363
CL\_INS\_363
CL\_INS\_363
CL\_INS\_363
CL\_INS\_207
CL\_INS\_363
CL\_INS\_170
CL\_INS\_149
CL\_INS\_149
CL\_INS\_149
CL\_INS\_149
CL\_INS\_204
CL\_INS\_237
CL\_INS\_149
CL\_INS\_170
CL\_INS\_149
CL\_INS\_149
CL\_INS\_170
CL\_INS\_149
CL\_INS\_363
CL\_INS\_363
CL\_INS\_363
CL\_INS\_363
CL\_INS\_363
CL\_INS\_363
CL\_INS\_363
CL\_INS\_363
CL\_INS\_363
CL\_INS\_363
CL\_INS\_363
CL\_INS\_363
CL\_INS\_170
CL\_INS\_149
CL\_INS\_149
CL\_INS\_149
CL\_INS\_149
CL\_INS\_149
CL\_INS\_149
CL\_INS\_149
CL\_INS\_149
CL\_INS\_149
CL\_INS\_149
CL\_INS\_149
CL\_INS\_170
CL\_INS\_149
CL\_INS\_170
CL\_INS\_149
CL\_INS\_149
CL\_INS\_149
CL\_INS\_149
CL\_INS\_363
CL\_INS\_363
CL\_INS\_363
CL\_INS\_363
CL\_INS\_149
CL\_INS\_71
CL\_INS\_71
CL\_INS\_363
CL\_INS\_149
CL\_INS\_149
CL\_INS\_149
CL\_INS\_149
CL\_INS\_363
CL\_INS\_207
CL\_INS\_71
CL\_INS\_71
CL\_INS\_149
CL\_INS\_363
CL\_INS\_363
CL\_INS\_363
CL\_INS\_363
CL\_INS\_149
CL\_INS\_149
CL\_INS\_149
CL\_INS\_363
CL\_INS\_363
CL\_INS\_149
CL\_INS\_149
CL\_INS\_149
CL\_INS\_363
CL\_INS\_363
CL\_INS\_363
CL\_INS\_363
CL\_INS\_363
CL\_INS\_363
CL\_INS\_363
CL\_INS\_363
CL\_INS\_363
Cluster ID


CL\_12076
CL\_13710
CL\_12228
CL\_24119
CL\_24118
CL\_36674
CL\_36673
CL\_36672
CL\_26088
CL\_17518
CL\_4614
CL\_4613
CL\_4611
CL\_4610
CL\_4609
CL\_26087
CL\_5883
CL\_26086
CL\_26085
CL\_26084
CL\_9191
CL\_5201
CL\_11036
CL\_5199
CL\_36671
CL\_36670
CL\_36669
CL\_36668
CL\_36667
CL\_36666
CL\_36665
CL\_36664
CL\_36663
CL\_36662
CL\_36661
CL\_11996
CL\_11997
CL\_36660
CL\_36659
CL\_26083
CL\_26082
CL\_13992
CL\_26081
CL\_4608
CL\_4607
CL\_4606
CL\_4605
CL\_8067
CL\_4604
CL\_4514
CL\_4603
CL\_4602
CL\_4601
CL\_4600
CL\_4599
CL\_4598
CL\_26080
CL\_26079
CL\_26078
CL\_26077
CL\_26076
CL\_26075
CL\_26074
CL\_26073
CL\_26072
CL\_26071
CL\_26070
CL\_26069
CL\_4597
CL\_4596
CL\_4595
CL\_4594
CL\_11993
CL\_4593
CL\_4592
CL\_4591
CL\_4590
CL\_6529
CL\_4589
CL\_4588
CL\_4587
CL\_6599
CL\_4585
CL\_9190
CL\_16374
CL\_9187
CL\_9186
CL\_17181
CL\_17180
CL\_26068
CL\_26067
CL\_11048
CL\_27584
CL\_27583
CL\_35362
CL\_4580
CL\_4579
CL\_11045
CL\_12557
CL\_33634
CL\_4666
CL\_11998
CL\_11999
CL\_4578
CL\_26066
CL\_36658
CL\_36657
CL\_26065
CL\_4577
CL\_8986
CL\_4576
CL\_26064
CL\_26063
CL\_4575
CL\_4574
CL\_17515
CL\_24117
CL\_4571
CL\_36656
CL\_4570
CL\_4043
CL\_4044
CL\_35361
CL\_35360
CL\_35359
