## Supplementary material for "A novel method for integrating genomic and Tn-Seq data to identify common *in vivo* fitness mechanisms across multiple bacterial species": S1 Dataset: CL_INS_367.html

Legend

 Mobile +extrachromosomalelementfunctions
 Regulatoryfunctions
 Hypothetical
 All EssentialGenes
 AntibioticResistance
 Other
 Transport +binding proteins
 All VFDB Genes

FULL


WINDOWSVGPNG

Trim RowsRemove SingletonsSave Fasta

CL\_4079


CL\_4079


CL\_4078


CL\_4079


CL\_4079


CL\_4079


CL\_4079


CL\_4079


CL\_4079


CL\_4079


CL\_4079


CL\_4078


CL\_3958


CL\_4079


CL\_4079

HighlightSelectShow Genomes


166

CL\_4085


81

CL\_4085


4

CL\_4085


2

CL\_4085


2

CL\_4085


1

CL\_4057


1

CL\_4128


1

CL\_4119


1

CL\_4085


1

CL\_4118


1

CL\_234


1

CL\_4085


1

CL\_4085


1

CL\_4118


1

CL\_4123

fGI ID


CL\_INS\_367
CL\_INS\_368
CL\_INS\_20
CL\_INS\_368
CL\_INS\_247
CL\_INS\_368
CL\_INS\_87
CL\_INS\_237
CL\_INS\_237
CL\_INS\_354
CL\_INS\_354
CL\_INS\_354
CL\_INS\_354
CL\_INS\_367
CL\_INS\_237
CL\_INS\_237
CL\_INS\_237
CL\_INS\_367
CL\_INS\_367
CL\_INS\_237
CL\_INS\_237
CL\_INS\_237
CL\_INS\_368
CL\_INS\_368
CL\_INS\_368
CL\_INS\_368
CL\_INS\_368
CL\_INS\_368
CL\_INS\_368
CL\_INS\_20
CL\_INS\_237
CL\_INS\_237
CL\_INS\_237
CL\_INS\_237
CL\_INS\_237
CL\_INS\_237
CL\_INS\_237
CL\_INS\_237
CL\_INS\_237
CL\_INS\_237
CL\_INS\_237
CL\_INS\_237
CL\_INS\_382
CL\_INS\_382
CL\_INS\_149
CL\_INS\_70
CL\_INS\_70
CL\_INS\_70
CL\_INS\_70
CL\_INS\_70
CL\_INS\_70
CL\_INS\_70
CL\_INS\_155
CL\_INS\_237
CL\_INS\_237
CL\_INS\_237
CL\_INS\_237
CL\_INS\_237
CL\_INS\_70
CL\_INS\_237
CL\_INS\_237
CL\_INS\_237
CL\_INS\_247
CL\_INS\_368
CL\_INS\_380
CL\_INS\_237
CL\_INS\_367
CL\_INS\_237
CL\_INS\_237
CL\_INS\_237
CL\_INS\_367
CL\_INS\_367
CL\_INS\_367
CL\_INS\_367
CL\_INS\_367
CL\_INS\_367
CL\_INS\_367
CL\_INS\_367
CL\_INS\_367
CL\_INS\_367
CL\_INS\_367
CL\_INS\_367
CL\_INS\_367
CL\_INS\_367
CL\_INS\_367
CL\_INS\_367
CL\_INS\_367
CL\_INS\_367
CL\_INS\_367
CL\_INS\_367
CL\_INS\_367
CL\_INS\_367
CL\_INS\_367
CL\_INS\_367
CL\_INS\_367
CL\_INS\_237
CL\_INS\_367
CL\_INS\_367
CL\_INS\_237
Cluster ID


CL\_4080
CL\_8287
CL\_8286
CL\_8285
CL\_6749
CL\_8284
CL\_8778
CL\_12078
CL\_5148
CL\_22105
CL\_4081
CL\_4082
CL\_4083
CL\_4084
CL\_4943
CL\_4944
CL\_4945
CL\_4946
CL\_4947
CL\_4948
CL\_4949
CL\_4950
CL\_5994
CL\_31464
CL\_31463
CL\_31462
CL\_31461
CL\_31460
CL\_32580
CL\_4350
CL\_7991
CL\_5157
CL\_5156
CL\_5155
CL\_5153
CL\_5152
CL\_5151
CL\_5150
CL\_7990
CL\_7989
CL\_7988
CL\_7301
CL\_8216
CL\_4975
CL\_4093
CL\_4095
CL\_4096
CL\_4097
CL\_4098
CL\_4099
CL\_4100
CL\_4101
CL\_7374
CL\_13375
CL\_13376
CL\_13377
CL\_13378
CL\_14340
CL\_7671
CL\_7670
CL\_7669
CL\_6076
CL\_5236
CL\_16880
CL\_22042
CL\_5147
CL\_22041
CL\_22040
CL\_22039
CL\_22038
CL\_22037
CL\_22036
CL\_22035
CL\_22034
CL\_22033
CL\_22032
CL\_22031
CL\_22030
CL\_22029
CL\_22028
CL\_22027
CL\_22026
CL\_22025
CL\_22024
CL\_22023
CL\_22022
CL\_22021
CL\_22020
CL\_22019
CL\_22018
CL\_22017
CL\_22016
CL\_22015
CL\_22014
CL\_22013
CL\_7666
CL\_22012
CL\_22011
CL\_4375
