## Supplementary material for "A novel method for integrating genomic and Tn-Seq data to identify common *in vivo* fitness mechanisms across multiple bacterial species": S1 Dataset: CL_INS_368.html

Legend

 Mobile +extrachromosomalelementfunctions
 Regulatoryfunctions
 Hypothetical
 DNA Metabolism
 Transcription
 All EssentialGenes
 AntibioticResistance
 All Fitness Genes
 Proteinsynthesis/fate
 Other
 Centralintermediarymetabolism
 EnergyMetabolism
 Transport +binding proteins
 All VFDB Genes

FULL


WINDOWSVGPNG

Trim RowsRemove SingletonsSave Fasta

CL\_1935


CL\_234


CL\_4085


CL\_4085


CL\_4085


CL\_1935


CL\_4085


CL\_234


CL\_4085


CL\_4085


CL\_4085


CL\_4085


CL\_4085


CL\_4085


CL\_4085


CL\_4085


CL\_4085


CL\_4085


CL\_234


CL\_1935


CL\_4085


CL\_4085


CL\_4085


CL\_4085


CL\_4085


CL\_4085


CL\_4085


CL\_4085


CL\_4085


CL\_4085


CL\_4085


CL\_4085


CL\_4085


CL\_4085


CL\_4085


CL\_4085


CL\_4085


CL\_4085


CL\_4085


CL\_4085


CL\_4085


CL\_4085


CL\_4085


CL\_4085


CL\_4085


CL\_4085


CL\_2847


CL\_4085


CL\_4085


CL\_4085


CL\_4085


CL\_4085


CL\_4085


CL\_4085


CL\_4085


CL\_1935


CL\_4085


CL\_4085


CL\_4085


CL\_4085


CL\_4085


CL\_4085


CL\_4085


CL\_4085


CL\_4085


CL\_4085


CL\_4085


CL\_4085


CL\_4085


CL\_4085


CL\_4085


CL\_4085


CL\_4085


CL\_4085


CL\_4085


CL\_4085


CL\_4085


CL\_4085


CL\_4085


CL\_4085


CL\_4085


CL\_4085


CL\_4085


CL\_4085


CL\_4085


CL\_1935


CL\_4085


CL\_4085


CL\_4085


CL\_4085


CL\_4085


CL\_4085


CL\_4085


CL\_4085


CL\_4085


CL\_4085


CL\_4085


CL\_234


CL\_4085


CL\_4085


CL\_4085


CL\_4085


CL\_234


CL\_4085


CL\_4085


Break


CL\_234


CL\_4160


CL\_4085


CL\_4085


CL\_4085


CL\_4085


CL\_4085


CL\_4085


CL\_4085


CL\_4085


CL\_4085


CL\_4085


CL\_4085


CL\_4085


CL\_234


CL\_4085


CL\_4085


CL\_1935


CL\_4085


CL\_4085


CL\_4085


CL\_4085


CL\_4085


CL\_4085


CL\_4085


CL\_4085


CL\_4085


CL\_4085


CL\_4085


CL\_4085


CL\_4085


CL\_4085


CL\_4085


CL\_4074


CL\_4085


CL\_4085


CL\_4085


CL\_4085


CL\_4085


CL\_4085


CL\_4085


CL\_4085


CL\_4085


CL\_4085


CL\_4085


CL\_4085


CL\_4085


CL\_4085


CL\_4085


CL\_4085


CL\_4085


CL\_4085


CL\_4060


CL\_4085


CL\_4085


CL\_4085


CL\_4085


CL\_3873


CL\_4085


CL\_4085


CL\_1935


CL\_1935


CL\_4085


CL\_4085


CL\_4085


CL\_4085


CL\_4085


CL\_4085


CL\_4085


CL\_4085


CL\_4085


CL\_4085


CL\_1935


CL\_4085


CL\_1935


CL\_4085


CL\_4085


CL\_4085


CL\_4085


CL\_4085


CL\_1935


CL\_4085


CL\_4085


CL\_4085


CL\_1935


CL\_4085


CL\_4085


CL\_4085


CL\_4085


CL\_4085


CL\_4085


CL\_4085


CL\_4085


CL\_4085


CL\_4085


CL\_4085


CL\_4085


CL\_4085


CL\_4085


CL\_1935


CL\_3958


CL\_4085


CL\_4085


CL\_4085


CL\_4085


CL\_1935


CL\_4085


CL\_4085


CL\_1935


CL\_4085


CL\_4085


CL\_4085


CL\_4085


CL\_4085


CL\_4085


CL\_4079


CL\_4085


CL\_4085


CL\_4085


CL\_4085


CL\_4085


CL\_4085


CL\_4085


CL\_4085


CL\_4085


CL\_4085


CL\_4085


CL\_4085


CL\_4085


CL\_4079


CL\_4085


CL\_4085


CL\_4085


CL\_4085


CL\_4085


CL\_4085


CL\_4085


CL\_4085


CL\_4085


CL\_4085


CL\_4085


CL\_4085


CL\_4085


CL\_4085


CL\_4085


CL\_4085


CL\_4085


CL\_4085


CL\_4085


CL\_4085


CL\_4085


CL\_4085


CL\_4085


CL\_4085


CL\_4085


CL\_4085


CL\_4085


CL\_4085


CL\_1935


CL\_4085


CL\_4085


CL\_4085


CL\_1935


CL\_4085


CL\_234


CL\_4085


CL\_4085


CL\_4085


CL\_4085


CL\_4085


CL\_4085


CL\_3958


CL\_1935


CL\_4085


CL\_4085


CL\_4085


CL\_4085


CL\_4085


CL\_2192


CL\_4085

HighlightSelectShow Genomes


8

CL\_4118


5

CL\_4118


3

CL\_4128


3

CL\_4118


3

CL\_4118


3

CL\_4118


2

CL\_4118


2

CL\_4118


2

CL\_4118


2

CL\_4118


2

CL\_4118


2

CL\_4118


2

CL\_4118


2

CL\_4118


2

CL\_234


2

CL\_4118


2

CL\_234


2

CL\_4118


2

CL\_4118


2

CL\_4118


2

CL\_4118


2

CL\_4118


2

CL\_4118


2

CL\_4118


1

CL\_234


1

CL\_4118


1

CL\_4118


1

CL\_234


1

CL\_4118


1

CL\_4118


1

CL\_4118


1

CL\_4118


1

CL\_4118


1

CL\_4118


1

CL\_4118


1

Break


1

CL\_4118


1

CL\_4118


1

CL\_4118


1

CL\_234


1

CL\_4118


1

CL\_4121


1

CL\_4118


1

CL\_4118


1

CL\_4118


1

CL\_4118


1

CL\_4118


1

CL\_4118


1

CL\_4118


1

CL\_4118


1

CL\_4118


1

CL\_4118


1

CL\_234


1

CL\_4118


1

CL\_4127


1

CL\_4118


1

Break


1

CL\_4118


1

CL\_4118


1

CL\_4118


1

CL\_4118


1

CL\_4118


1

CL\_4118


1

CL\_4130


1

CL\_4118


1

CL\_1935


1

CL\_234


1

CL\_4118


1

CL\_4118


1

CL\_4118


1

CL\_4118


1

CL\_4127


1

CL\_4118


1

CL\_234


1

CL\_4118


1

CL\_4150


1

CL\_4130


1

CL\_4118


1

CL\_234


1

CL\_4118


1

CL\_4118


1

CL\_4119


1

CL\_4118


1

CL\_4126


1

CL\_4118


1

CL\_4118


1

CL\_4128


1

CL\_1153


1

CL\_4118


1

CL\_4118


1

CL\_4118


1

CL\_4118


1

CL\_4126


1

CL\_4118


1

CL\_4129


1

CL\_4118


1

CL\_4118


1

CL\_4118


1

CL\_234


1

CL\_4118


1

CL\_4118


1

CL\_4118


1

CL\_4118


1

CL\_4118


1

CL\_4119


1

CL\_4118


1

CL\_4118


1

CL\_4118


1

CL\_4118


1

CL\_4118


1

CL\_4118


1

CL\_4118


1

CL\_4127


1

CL\_4118


1

CL\_4118


1

CL\_4118


1

CL\_234


1

CL\_4118


1

CL\_4118


1

CL\_4118


1

CL\_4118


1

CL\_4118


1

CL\_4118


1

CL\_4118


1

CL\_4118


1

CL\_1935


1

CL\_4126


1

CL\_4120


1

CL\_4118


1

CL\_4119


1

CL\_4127


1

CL\_3957


1

CL\_234


1

CL\_4118


1

CL\_4118


1

CL\_4123


1

CL\_234


1

CL\_4118


1

CL\_4118


1

CL\_4118


1

CL\_4150


1

CL\_4118


1

CL\_4118


1

CL\_4118


1

CL\_234


1

CL\_4118


1

CL\_4118


1

CL\_4118


1

CL\_4129


1

CL\_4129


1

CL\_4118


1

CL\_3870


1

CL\_4118


1

CL\_4127


1

CL\_4118


1

CL\_234


1

CL\_4118


1

CL\_1935


1

CL\_4118


1

CL\_4118


1

CL\_4118


1

CL\_4118


1

CL\_4118


1

CL\_4118


1

CL\_4118


1

CL\_4118


1

CL\_4118


1

CL\_4118


1

CL\_4121


1

CL\_4118


1

CL\_4118


1

CL\_4118


1

CL\_4146


1

CL\_4119


1

CL\_234


1

CL\_234


1

CL\_4118


1

CL\_4118


1

CL\_4118


1

CL\_4118


1

CL\_4118


1

CL\_234


1

CL\_4130


1

CL\_4118


1

CL\_234


1

CL\_4118


1

CL\_4118


1

CL\_1935


1

CL\_4118


1

CL\_234


1

CL\_4118


1

CL\_234


1

CL\_4118


1

CL\_234


1

CL\_234


1

CL\_4118


1

CL\_4118


1

CL\_1935


1

CL\_4118


1

CL\_4118


1

CL\_4118


1

CL\_4119


1

CL\_4118


1

CL\_4119


1

CL\_4130


1

CL\_4118


1

CL\_4118


1

CL\_234


1

CL\_234


1

CL\_4118


1

Break


1

CL\_4118


1

CL\_4118


1

CL\_234


1

CL\_4118


1

CL\_2555


1

CL\_4118


1

CL\_4118


1

CL\_4118


1

CL\_1935


1

CL\_4118


1

CL\_4118


1

CL\_4119


1

CL\_4118


1

CL\_4118


1

CL\_4118


1

CL\_4118


1

CL\_4118


1

CL\_4118


1

CL\_234


1

CL\_4118


1

CL\_234


1

CL\_4118


1

CL\_4118


1

CL\_4118


1

CL\_4118


1

CL\_4118


1

CL\_4118


1

CL\_4118


1

CL\_4119


1

CL\_4118


1

CL\_4118


1

CL\_4118


1

CL\_4118


1

CL\_234


1

CL\_4118


1

CL\_4118


1

CL\_4118


1

CL\_4118


1

CL\_4118


1

CL\_4118


1

Break


1

CL\_3873


1

CL\_4118


1

CL\_4118


1

CL\_4118


1

CL\_4118


1

CL\_234


1

CL\_4118


1

CL\_4118


1

CL\_4118


1

CL\_4118


1

CL\_4118


1

CL\_4118


1

CL\_4118


1

CL\_4118


1

CL\_1935


1

CL\_4118


1

CL\_4118


1

CL\_1935


1

CL\_4118


1

CL\_234


1

CL\_4118


1

CL\_4118


1

CL\_4118


1

CL\_234


1

CL\_3846


1

CL\_4118


1

CL\_4118


1

CL\_4118


1

CL\_4118


1

CL\_4118


1

CL\_1935


1

CL\_4118


1

CL\_4118


1

CL\_4118

fGI ID


CL\_INS\_368
CL\_INS\_368
CL\_INS\_368
CL\_INS\_368
CL\_INS\_368
CL\_INS\_368
CL\_INS\_368
CL\_INS\_368
CL\_INS\_368
CL\_INS\_368
CL\_INS\_368
CL\_INS\_368
CL\_INS\_368
CL\_INS\_368
CL\_INS\_368
CL\_INS\_368
CL\_INS\_368
CL\_INS\_368
CL\_INS\_123
CL\_INS\_368
CL\_INS\_368
CL\_INS\_20
CL\_INS\_20
CL\_INS\_20
CL\_INS\_368
CL\_INS\_237
CL\_INS\_368
CL\_INS\_368
CL\_INS\_368
CL\_INS\_368
CL\_INS\_368
CL\_INS\_368
CL\_INS\_368
CL\_INS\_237
CL\_INS\_237
CL\_INS\_237
CL\_INS\_237
CL\_INS\_70
CL\_INS\_70
CL\_INS\_70
CL\_INS\_382
CL\_INS\_237
CL\_INS\_237
CL\_INS\_146
CL\_INS\_99
CL\_INS\_20
CL\_INS\_99
CL\_INS\_99
CL\_INS\_368
CL\_INS\_99
CL\_INS\_368
CL\_INS\_382
CL\_INS\_99
CL\_INS\_368
CL\_INS\_368
CL\_INS\_207
CL\_INS\_207
CL\_INS\_382
CL\_INS\_368
CL\_INS\_368
CL\_INS\_368
CL\_INS\_368
CL\_INS\_368
CL\_INS\_368
CL\_INS\_368
CL\_INS\_368
CL\_INS\_368
CL\_INS\_368
CL\_INS\_368
CL\_INS\_368
CL\_INS\_368
CL\_INS\_20
CL\_INS\_20
CL\_INS\_20
CL\_INS\_20
CL\_INS\_20
CL\_INS\_368
CL\_INS\_20
CL\_INS\_20
CL\_INS\_207
CL\_INS\_20
CL\_INS\_20
CL\_INS\_237
CL\_INS\_368
CL\_INS\_237
CL\_INS\_368
CL\_INS\_237
CL\_INS\_368
CL\_INS\_368
CL\_INS\_368
CL\_INS\_368
CL\_INS\_368
CL\_INS\_207
CL\_INS\_207
CL\_INS\_207
CL\_INS\_207
CL\_INS\_207
CL\_INS\_368
CL\_INS\_207
CL\_INS\_207
CL\_INS\_368
CL\_INS\_368
CL\_INS\_368
CL\_INS\_368
CL\_INS\_368
CL\_INS\_368
CL\_INS\_207
CL\_INS\_20
CL\_INS\_368
CL\_INS\_207
CL\_INS\_368
CL\_INS\_368
CL\_INS\_368
CL\_INS\_368
CL\_INS\_247
CL\_INS\_237
CL\_INS\_237
CL\_INS\_237
CL\_INS\_237
CL\_INS\_237
CL\_INS\_237
CL\_INS\_247
CL\_INS\_247
CL\_INS\_237
CL\_INS\_247
CL\_INS\_247
CL\_INS\_247
CL\_INS\_237
CL\_INS\_368
CL\_INS\_368
CL\_INS\_368
CL\_INS\_368
CL\_INS\_368
CL\_INS\_368
CL\_INS\_368
CL\_INS\_368
CL\_INS\_368
CL\_INS\_368
CL\_INS\_368
CL\_INS\_368
CL\_INS\_368
CL\_INS\_368
CL\_INS\_368
CL\_INS\_368
CL\_INS\_368
CL\_INS\_368
CL\_INS\_368
CL\_INS\_368
CL\_INS\_368
CL\_INS\_368
CL\_INS\_368
CL\_INS\_368
CL\_INS\_368
CL\_INS\_368
CL\_INS\_368
CL\_INS\_368
CL\_INS\_368
CL\_INS\_368
CL\_INS\_368
CL\_INS\_368
CL\_INS\_368
CL\_INS\_368
CL\_INS\_368
CL\_INS\_368
CL\_INS\_368
CL\_INS\_368
CL\_INS\_368
CL\_INS\_368
CL\_INS\_368
CL\_INS\_368
CL\_INS\_368
CL\_INS\_368
CL\_INS\_368
CL\_INS\_207
CL\_INS\_207
CL\_INS\_207
CL\_INS\_368
CL\_INS\_368
CL\_INS\_368
CL\_INS\_368
CL\_INS\_368
CL\_INS\_368
CL\_INS\_368
CL\_INS\_368
CL\_INS\_237
CL\_INS\_368
CL\_INS\_237
CL\_INS\_237
CL\_INS\_237
CL\_INS\_237
CL\_INS\_237
CL\_INS\_237
CL\_INS\_368
CL\_INS\_237
CL\_INS\_368
CL\_INS\_368
CL\_INS\_368
CL\_INS\_368
CL\_INS\_368
CL\_INS\_368
CL\_INS\_368
CL\_INS\_368
CL\_INS\_368
CL\_INS\_368
CL\_INS\_368
CL\_INS\_237
CL\_INS\_237
CL\_INS\_247
CL\_INS\_368
CL\_INS\_247
CL\_INS\_368
CL\_INS\_368
CL\_INS\_368
CL\_INS\_368
CL\_INS\_368
CL\_INS\_368
CL\_INS\_368
CL\_INS\_368
CL\_INS\_368
CL\_INS\_368
CL\_INS\_368
CL\_INS\_368
CL\_INS\_368
CL\_INS\_368
CL\_INS\_368
CL\_INS\_368
CL\_INS\_368
CL\_INS\_368
CL\_INS\_368
CL\_INS\_368
CL\_INS\_368
CL\_INS\_368
CL\_INS\_368
CL\_INS\_368
CL\_INS\_368
CL\_INS\_368
CL\_INS\_368
CL\_INS\_368
CL\_INS\_368
CL\_INS\_368
CL\_INS\_368
CL\_INS\_20
CL\_INS\_368
CL\_INS\_368
CL\_INS\_20
CL\_INS\_20
CL\_INS\_20
CL\_INS\_368
CL\_INS\_368
CL\_INS\_368
CL\_INS\_368
CL\_INS\_368
CL\_INS\_368
CL\_INS\_368
CL\_INS\_368
CL\_INS\_368
CL\_INS\_368
CL\_INS\_352
CL\_INS\_352
CL\_INS\_237
CL\_INS\_237
CL\_INS\_237
CL\_INS\_368
CL\_INS\_368
CL\_INS\_237
CL\_INS\_237
CL\_INS\_237
CL\_INS\_368
CL\_INS\_368
CL\_INS\_368
CL\_INS\_368
CL\_INS\_368
CL\_INS\_368
CL\_INS\_368
CL\_INS\_368
CL\_INS\_368
CL\_INS\_368
CL\_INS\_237
CL\_INS\_221
CL\_INS\_237
CL\_INS\_368
CL\_INS\_368
CL\_INS\_368
CL\_INS\_237
CL\_INS\_237
CL\_INS\_237
CL\_INS\_237
CL\_INS\_368
CL\_INS\_368
CL\_INS\_368
CL\_INS\_368
CL\_INS\_368
CL\_INS\_368
CL\_INS\_368
CL\_INS\_368
CL\_INS\_237
CL\_INS\_247
CL\_INS\_368
CL\_INS\_237
CL\_INS\_368
CL\_INS\_368
CL\_INS\_70
CL\_INS\_70
CL\_INS\_368
CL\_INS\_30
CL\_INS\_30
CL\_INS\_30
CL\_INS\_70
CL\_INS\_70
CL\_INS\_70
CL\_INS\_70
CL\_INS\_368
CL\_INS\_368
CL\_INS\_368
CL\_INS\_368
CL\_INS\_70
CL\_INS\_70
CL\_INS\_368
CL\_INS\_368
CL\_INS\_368
CL\_INS\_368
CL\_INS\_368
CL\_INS\_368
CL\_INS\_368
CL\_INS\_368
CL\_INS\_368
CL\_INS\_368
CL\_INS\_368
CL\_INS\_368
CL\_INS\_70
CL\_INS\_70
CL\_INS\_237
CL\_INS\_368
CL\_INS\_368
CL\_INS\_368
CL\_INS\_368
CL\_INS\_368
CL\_INS\_352
CL\_INS\_70
CL\_INS\_70
CL\_INS\_368
CL\_INS\_368
CL\_INS\_70
CL\_INS\_368
CL\_INS\_70
CL\_INS\_70
CL\_INS\_70
CL\_INS\_70
CL\_INS\_368
CL\_INS\_149
CL\_INS\_368
CL\_INS\_368
CL\_INS\_368
CL\_INS\_70
CL\_INS\_368
CL\_INS\_368
CL\_INS\_70
CL\_INS\_368
CL\_INS\_368
CL\_INS\_368
CL\_INS\_368
CL\_INS\_368
CL\_INS\_368
CL\_INS\_368
CL\_INS\_368
CL\_INS\_368
CL\_INS\_369
CL\_INS\_368
CL\_INS\_368
CL\_INS\_368
CL\_INS\_368
CL\_INS\_368
CL\_INS\_368
CL\_INS\_368
CL\_INS\_368
CL\_INS\_368
CL\_INS\_368
CL\_INS\_368
CL\_INS\_368
CL\_INS\_368
CL\_INS\_368
CL\_INS\_368
CL\_INS\_368
CL\_INS\_368
CL\_INS\_368
CL\_INS\_368
CL\_INS\_368
CL\_INS\_368
CL\_INS\_20
CL\_INS\_368
CL\_INS\_368
CL\_INS\_207
CL\_INS\_237
CL\_INS\_237
CL\_INS\_237
CL\_INS\_237
CL\_INS\_368
CL\_INS\_368
CL\_INS\_368
CL\_INS\_368
CL\_INS\_368
CL\_INS\_368
CL\_INS\_368
CL\_INS\_368
CL\_INS\_368
CL\_INS\_368
CL\_INS\_368
CL\_INS\_368
CL\_INS\_368
CL\_INS\_368
CL\_INS\_70
CL\_INS\_368
CL\_INS\_368
CL\_INS\_368
CL\_INS\_368
CL\_INS\_368
CL\_INS\_368
CL\_INS\_368
CL\_INS\_368
CL\_INS\_207
CL\_INS\_368
CL\_INS\_368
CL\_INS\_207
CL\_INS\_207
CL\_INS\_207
CL\_INS\_207
CL\_INS\_207
CL\_INS\_368
CL\_INS\_368
CL\_INS\_368
CL\_INS\_207
CL\_INS\_207
CL\_INS\_207
CL\_INS\_207
CL\_INS\_207
CL\_INS\_368
CL\_INS\_368
CL\_INS\_368
CL\_INS\_368
CL\_INS\_368
CL\_INS\_368
CL\_INS\_368
CL\_INS\_368
CL\_INS\_368
CL\_INS\_207
CL\_INS\_207
CL\_INS\_207
CL\_INS\_207
CL\_INS\_207
CL\_INS\_207
CL\_INS\_368
CL\_INS\_368
CL\_INS\_368
CL\_INS\_368
CL\_INS\_368
CL\_INS\_368
CL\_INS\_368
CL\_INS\_368
CL\_INS\_368
CL\_INS\_20
CL\_INS\_237
CL\_INS\_237
CL\_INS\_237
CL\_INS\_237
CL\_INS\_237
CL\_INS\_368
CL\_INS\_368
CL\_INS\_368
CL\_INS\_247
CL\_INS\_247
CL\_INS\_247
CL\_INS\_247
CL\_INS\_368
CL\_INS\_368
CL\_INS\_368
CL\_INS\_368
CL\_INS\_368
CL\_INS\_368
CL\_INS\_368
CL\_INS\_368
CL\_INS\_368
CL\_INS\_368
CL\_INS\_368
CL\_INS\_368
CL\_INS\_368
CL\_INS\_207
CL\_INS\_207
CL\_INS\_207
CL\_INS\_207
CL\_INS\_207
CL\_INS\_207
CL\_INS\_207
CL\_INS\_207
CL\_INS\_368
CL\_INS\_368
CL\_INS\_368
CL\_INS\_368
CL\_INS\_368
CL\_INS\_368
CL\_INS\_368
CL\_INS\_368
CL\_INS\_368
CL\_INS\_368
CL\_INS\_368
CL\_INS\_368
CL\_INS\_368
CL\_INS\_207
CL\_INS\_237
CL\_INS\_207
CL\_INS\_368
CL\_INS\_155
CL\_INS\_237
CL\_INS\_207
CL\_INS\_155
CL\_INS\_207
CL\_INS\_207
CL\_INS\_207
CL\_INS\_368
CL\_INS\_368
CL\_INS\_207
CL\_INS\_207
CL\_INS\_207
CL\_INS\_207
CL\_INS\_207
CL\_INS\_207
CL\_INS\_207
CL\_INS\_207
CL\_INS\_207
CL\_INS\_207
CL\_INS\_207
CL\_INS\_247
CL\_INS\_237
CL\_INS\_237
CL\_INS\_237
CL\_INS\_237
CL\_INS\_237
CL\_INS\_237
CL\_INS\_237
CL\_INS\_237
CL\_INS\_237
CL\_INS\_237
CL\_INS\_237
CL\_INS\_237
CL\_INS\_368
CL\_INS\_368
CL\_INS\_368
CL\_INS\_368
CL\_INS\_368
CL\_INS\_368
CL\_INS\_368
CL\_INS\_368
CL\_INS\_30
CL\_INS\_368
CL\_INS\_368
CL\_INS\_368
CL\_INS\_368
CL\_INS\_368
CL\_INS\_368
CL\_INS\_368
CL\_INS\_70
CL\_INS\_368
CL\_INS\_368
CL\_INS\_368
CL\_INS\_368
CL\_INS\_368
CL\_INS\_382
CL\_INS\_382
CL\_INS\_382
CL\_INS\_382
CL\_INS\_382
CL\_INS\_382
CL\_INS\_368
CL\_INS\_71
CL\_INS\_368
CL\_INS\_368
CL\_INS\_368
CL\_INS\_237
CL\_INS\_368
CL\_INS\_368
CL\_INS\_368
CL\_INS\_237
CL\_INS\_237
CL\_INS\_237
CL\_INS\_20
CL\_INS\_20
CL\_INS\_20
CL\_INS\_237
CL\_INS\_237
CL\_INS\_368
CL\_INS\_237
CL\_INS\_237
CL\_INS\_237
CL\_INS\_237
CL\_INS\_368
CL\_INS\_237
CL\_INS\_368
CL\_INS\_368
CL\_INS\_368
CL\_INS\_368
CL\_INS\_368
CL\_INS\_368
CL\_INS\_368
CL\_INS\_368
CL\_INS\_368
CL\_INS\_368
CL\_INS\_368
CL\_INS\_368
CL\_INS\_368
CL\_INS\_368
CL\_INS\_368
CL\_INS\_368
CL\_INS\_368
CL\_INS\_368
CL\_INS\_368
CL\_INS\_368
CL\_INS\_368
CL\_INS\_368
CL\_INS\_368
CL\_INS\_368
CL\_INS\_368
CL\_INS\_368
CL\_INS\_368
CL\_INS\_368
CL\_INS\_368
CL\_INS\_368
CL\_INS\_368
CL\_INS\_368
CL\_INS\_368
CL\_INS\_368
CL\_INS\_368
CL\_INS\_368
CL\_INS\_368
CL\_INS\_368
CL\_INS\_368
CL\_INS\_237
CL\_INS\_382
CL\_INS\_382
CL\_INS\_382
CL\_INS\_368
CL\_INS\_368
CL\_INS\_237
CL\_INS\_368
CL\_INS\_368
CL\_INS\_368
CL\_INS\_368
CL\_INS\_368
CL\_INS\_368
CL\_INS\_368
CL\_INS\_368
CL\_INS\_368
CL\_INS\_237
CL\_INS\_237
CL\_INS\_237
CL\_INS\_237
CL\_INS\_368
CL\_INS\_237
CL\_INS\_237
CL\_INS\_237
CL\_INS\_237
CL\_INS\_237
CL\_INS\_237
CL\_INS\_247
CL\_INS\_247
CL\_INS\_247
CL\_INS\_247
CL\_INS\_237
CL\_INS\_237
CL\_INS\_368
CL\_INS\_368
CL\_INS\_368
CL\_INS\_351
CL\_INS\_368
CL\_INS\_368
CL\_INS\_351
CL\_INS\_351
CL\_INS\_368
CL\_INS\_368
CL\_INS\_382
CL\_INS\_368
CL\_INS\_237
CL\_INS\_237
CL\_INS\_237
CL\_INS\_237
CL\_INS\_237
CL\_INS\_237
CL\_INS\_237
CL\_INS\_368
CL\_INS\_368
CL\_INS\_237
CL\_INS\_237
CL\_INS\_237
CL\_INS\_237
CL\_INS\_237
CL\_INS\_368
CL\_INS\_237
CL\_INS\_237
CL\_INS\_237
CL\_INS\_237
CL\_INS\_237
CL\_INS\_237
CL\_INS\_237
CL\_INS\_368
CL\_INS\_368
CL\_INS\_368
CL\_INS\_368
CL\_INS\_368
CL\_INS\_237
CL\_INS\_237
CL\_INS\_237
CL\_INS\_237
CL\_INS\_237
CL\_INS\_237
CL\_INS\_237
CL\_INS\_237
CL\_INS\_237
CL\_INS\_368
CL\_INS\_87
CL\_INS\_237
CL\_INS\_237
CL\_INS\_237
CL\_INS\_237
CL\_INS\_237
CL\_INS\_237
CL\_INS\_237
CL\_INS\_237
CL\_INS\_237
CL\_INS\_368
CL\_INS\_237
CL\_INS\_382
CL\_INS\_237
CL\_INS\_368
CL\_INS\_237
CL\_INS\_237
CL\_INS\_237
CL\_INS\_382
CL\_INS\_237
CL\_INS\_237
CL\_INS\_368
CL\_INS\_368
CL\_INS\_368
CL\_INS\_237
CL\_INS\_237
CL\_INS\_237
CL\_INS\_237
CL\_INS\_237
CL\_INS\_237
CL\_INS\_237
CL\_INS\_237
CL\_INS\_237
CL\_INS\_237
CL\_INS\_237
CL\_INS\_237
CL\_INS\_368
CL\_INS\_237
CL\_INS\_368
CL\_INS\_368
CL\_INS\_368
CL\_INS\_368
CL\_INS\_368
CL\_INS\_20
CL\_INS\_368
CL\_INS\_247
CL\_INS\_369
CL\_INS\_368
CL\_INS\_368
CL\_INS\_174
CL\_INS\_20
CL\_INS\_20
CL\_INS\_20
CL\_INS\_20
CL\_INS\_20
CL\_INS\_20
CL\_INS\_20
CL\_INS\_20
CL\_INS\_20
CL\_INS\_20
CL\_INS\_368
CL\_INS\_368
CL\_INS\_368
CL\_INS\_369
CL\_INS\_247
CL\_INS\_368
CL\_INS\_368
CL\_INS\_207
CL\_INS\_368
CL\_INS\_123
CL\_INS\_123
CL\_INS\_123
CL\_INS\_123
CL\_INS\_99
CL\_INS\_368
CL\_INS\_368
CL\_INS\_99
CL\_INS\_99
CL\_INS\_368
CL\_INS\_99
CL\_INS\_99
CL\_INS\_123
CL\_INS\_99
CL\_INS\_99
CL\_INS\_99
CL\_INS\_99
CL\_INS\_368
CL\_INS\_368
CL\_INS\_99
CL\_INS\_99
CL\_INS\_99
CL\_INS\_99
CL\_INS\_99
CL\_INS\_99
CL\_INS\_99
CL\_INS\_99
CL\_INS\_99
CL\_INS\_155
CL\_INS\_155
CL\_INS\_155
CL\_INS\_99
CL\_INS\_99
CL\_INS\_99
CL\_INS\_99
CL\_INS\_99
CL\_INS\_99
CL\_INS\_99
CL\_INS\_99
CL\_INS\_99
CL\_INS\_368
CL\_INS\_368
CL\_INS\_123
CL\_INS\_382
CL\_INS\_71
CL\_INS\_71
CL\_INS\_382
CL\_INS\_382
CL\_INS\_382
CL\_INS\_382
CL\_INS\_60
CL\_INS\_60
CL\_INS\_368
CL\_INS\_368
CL\_INS\_368
CL\_INS\_60
CL\_INS\_60
CL\_INS\_60
CL\_INS\_60
CL\_INS\_368
CL\_INS\_117
CL\_INS\_368
CL\_INS\_368
CL\_INS\_368
CL\_INS\_368
CL\_INS\_368
CL\_INS\_368
CL\_INS\_247
CL\_INS\_247
CL\_INS\_247
CL\_INS\_247
CL\_INS\_247
CL\_INS\_247
CL\_INS\_247
CL\_INS\_247
CL\_INS\_123
CL\_INS\_123
CL\_INS\_123
CL\_INS\_247
CL\_INS\_123
CL\_INS\_159
CL\_INS\_159
CL\_INS\_159
CL\_INS\_368
CL\_INS\_117
CL\_INS\_368
CL\_INS\_117
CL\_INS\_117
CL\_INS\_368
CL\_INS\_368
CL\_INS\_368
CL\_INS\_368
CL\_INS\_368
CL\_INS\_368
CL\_INS\_368
CL\_INS\_382
CL\_INS\_385
CL\_INS\_159
CL\_INS\_382
CL\_INS\_382
CL\_INS\_382
CL\_INS\_382
CL\_INS\_382
CL\_INS\_382
CL\_INS\_60
CL\_INS\_382
CL\_INS\_382
CL\_INS\_382
CL\_INS\_382
CL\_INS\_382
CL\_INS\_382
CL\_INS\_71
CL\_INS\_382
CL\_INS\_382
CL\_INS\_382
CL\_INS\_382
CL\_INS\_368
CL\_INS\_60
CL\_INS\_60
CL\_INS\_60
CL\_INS\_60
CL\_INS\_60
CL\_INS\_60
CL\_INS\_60
CL\_INS\_60
CL\_INS\_60
CL\_INS\_368
CL\_INS\_368
CL\_INS\_247
CL\_INS\_247
CL\_INS\_247
CL\_INS\_207
CL\_INS\_207
CL\_INS\_207
CL\_INS\_207
CL\_INS\_207
CL\_INS\_207
CL\_INS\_237
CL\_INS\_237
CL\_INS\_247
CL\_INS\_30
CL\_INS\_368
CL\_INS\_368
CL\_INS\_368
CL\_INS\_368
CL\_INS\_368
CL\_INS\_368
CL\_INS\_70
CL\_INS\_207
CL\_INS\_207
CL\_INS\_159
CL\_INS\_382
CL\_INS\_382
CL\_INS\_368
CL\_INS\_365
CL\_INS\_365
CL\_INS\_365
CL\_INS\_365
CL\_INS\_368
CL\_INS\_368
CL\_INS\_20
CL\_INS\_368
CL\_INS\_382
CL\_INS\_368
CL\_INS\_382
CL\_INS\_382
CL\_INS\_368
CL\_INS\_368
CL\_INS\_368
CL\_INS\_368
CL\_INS\_368
CL\_INS\_368
CL\_INS\_368
CL\_INS\_368
CL\_INS\_368
CL\_INS\_237
CL\_INS\_237
CL\_INS\_237
CL\_INS\_368
CL\_INS\_237
CL\_INS\_237
CL\_INS\_237
CL\_INS\_368
CL\_INS\_237
CL\_INS\_70
CL\_INS\_247
CL\_INS\_247
CL\_INS\_247
CL\_INS\_247
CL\_INS\_247
CL\_INS\_247
CL\_INS\_247
CL\_INS\_247
CL\_INS\_247
CL\_INS\_247
CL\_INS\_247
CL\_INS\_247
CL\_INS\_247
CL\_INS\_247
CL\_INS\_247
CL\_INS\_237
CL\_INS\_70
CL\_INS\_123
CL\_INS\_368
CL\_INS\_368
CL\_INS\_70
CL\_INS\_237
CL\_INS\_368
CL\_INS\_368
CL\_INS\_368
CL\_INS\_247
CL\_INS\_247
CL\_INS\_368
CL\_INS\_368
CL\_INS\_20
CL\_INS\_20
CL\_INS\_20
CL\_INS\_247
CL\_INS\_247
CL\_INS\_70
CL\_INS\_70
CL\_INS\_237
CL\_INS\_57
CL\_INS\_149
CL\_INS\_149
CL\_INS\_149
CL\_INS\_149
CL\_INS\_149
CL\_INS\_149
CL\_INS\_57
CL\_INS\_57
CL\_INS\_368
CL\_INS\_368
CL\_INS\_237
CL\_INS\_237
CL\_INS\_237
CL\_INS\_237
CL\_INS\_237
CL\_INS\_237
CL\_INS\_237
CL\_INS\_368
CL\_INS\_237
CL\_INS\_237
CL\_INS\_237
CL\_INS\_368
CL\_INS\_237
CL\_INS\_237
CL\_INS\_237
CL\_INS\_368
CL\_INS\_368
CL\_INS\_237
CL\_INS\_368
CL\_INS\_368
CL\_INS\_368
CL\_INS\_239
CL\_INS\_237
CL\_INS\_237
CL\_INS\_237
CL\_INS\_237
CL\_INS\_237
CL\_INS\_237
CL\_INS\_237
CL\_INS\_237
CL\_INS\_237
CL\_INS\_237
CL\_INS\_237
CL\_INS\_368
CL\_INS\_247
CL\_INS\_247
CL\_INS\_247
CL\_INS\_30
CL\_INS\_382
CL\_INS\_237
CL\_INS\_368
CL\_INS\_368
CL\_INS\_30
CL\_INS\_70
CL\_INS\_368
CL\_INS\_368
CL\_INS\_368
CL\_INS\_368
CL\_INS\_368
CL\_INS\_368
CL\_INS\_368
CL\_INS\_237
CL\_INS\_123
CL\_INS\_368
CL\_INS\_237
CL\_INS\_237
CL\_INS\_237
CL\_INS\_237
CL\_INS\_237
CL\_INS\_237
CL\_INS\_237
CL\_INS\_237
CL\_INS\_368
CL\_INS\_368
CL\_INS\_237
CL\_INS\_368
CL\_INS\_368
CL\_INS\_237
CL\_INS\_20
CL\_INS\_237
CL\_INS\_368
CL\_INS\_368
CL\_INS\_368
CL\_INS\_368
CL\_INS\_70
CL\_INS\_20
CL\_INS\_207
CL\_INS\_247
CL\_INS\_368
CL\_INS\_207
CL\_INS\_368
CL\_INS\_237
CL\_INS\_237
CL\_INS\_237
CL\_INS\_237
CL\_INS\_368
CL\_INS\_368
CL\_INS\_368
CL\_INS\_368
CL\_INS\_368
CL\_INS\_368
CL\_INS\_368
CL\_INS\_368
CL\_INS\_368
CL\_INS\_368
CL\_INS\_368
CL\_INS\_368
CL\_INS\_368
CL\_INS\_368
CL\_INS\_368
CL\_INS\_237
CL\_INS\_237
CL\_INS\_237
CL\_INS\_237
CL\_INS\_237
CL\_INS\_237
CL\_INS\_20
CL\_INS\_70
CL\_INS\_247
CL\_INS\_237
CL\_INS\_352
CL\_INS\_368
CL\_INS\_237
CL\_INS\_237
CL\_INS\_247
CL\_INS\_237
CL\_INS\_368
CL\_INS\_368
CL\_INS\_368
CL\_INS\_368
CL\_INS\_368
CL\_INS\_20
CL\_INS\_20
CL\_INS\_20
CL\_INS\_20
CL\_INS\_20
CL\_INS\_20
CL\_INS\_20
CL\_INS\_20
CL\_INS\_368
CL\_INS\_207
CL\_INS\_368
CL\_INS\_368
CL\_INS\_368
CL\_INS\_368
CL\_INS\_368
CL\_INS\_368
CL\_INS\_237
CL\_INS\_237
CL\_INS\_237
CL\_INS\_237
CL\_INS\_237
CL\_INS\_368
CL\_INS\_237
CL\_INS\_237
CL\_INS\_237
CL\_INS\_368
CL\_INS\_368
CL\_INS\_368
CL\_INS\_368
CL\_INS\_368
CL\_INS\_368
CL\_INS\_368
CL\_INS\_237
CL\_INS\_237
CL\_INS\_368
CL\_INS\_368
CL\_INS\_368
CL\_INS\_237
CL\_INS\_237
CL\_INS\_237
CL\_INS\_368
CL\_INS\_368
CL\_INS\_237
CL\_INS\_237
CL\_INS\_237
CL\_INS\_237
CL\_INS\_237
CL\_INS\_237
CL\_INS\_237
CL\_INS\_237
CL\_INS\_237
CL\_INS\_237
CL\_INS\_237
CL\_INS\_237
CL\_INS\_237
CL\_INS\_20
CL\_INS\_237
CL\_INS\_237
CL\_INS\_237
CL\_INS\_368
CL\_INS\_368
CL\_INS\_20
CL\_INS\_20
CL\_INS\_20
CL\_INS\_20
CL\_INS\_20
CL\_INS\_20
CL\_INS\_20
CL\_INS\_20
CL\_INS\_20
CL\_INS\_368
CL\_INS\_368
CL\_INS\_368
CL\_INS\_368
CL\_INS\_368
CL\_INS\_368
CL\_INS\_368
CL\_INS\_237
CL\_INS\_247
CL\_INS\_247
CL\_INS\_247
CL\_INS\_368
CL\_INS\_368
CL\_INS\_368
CL\_INS\_368
CL\_INS\_368
CL\_INS\_237
CL\_INS\_237
CL\_INS\_237
CL\_INS\_368
CL\_INS\_369
CL\_INS\_237
CL\_INS\_237
CL\_INS\_237
CL\_INS\_237
CL\_INS\_237
CL\_INS\_237
CL\_INS\_237
CL\_INS\_368
CL\_INS\_368
CL\_INS\_382
CL\_INS\_382
CL\_INS\_237
CL\_INS\_237
CL\_INS\_237
CL\_INS\_382
CL\_INS\_368
CL\_INS\_60
CL\_INS\_382
CL\_INS\_382
CL\_INS\_382
CL\_INS\_382
CL\_INS\_382
CL\_INS\_382
CL\_INS\_382
CL\_INS\_382
CL\_INS\_382
CL\_INS\_382
CL\_INS\_382
CL\_INS\_382
CL\_INS\_382
CL\_INS\_382
CL\_INS\_382
CL\_INS\_382
CL\_INS\_159
CL\_INS\_159
CL\_INS\_368
CL\_INS\_368
CL\_INS\_385
CL\_INS\_382
CL\_INS\_382
CL\_INS\_382
CL\_INS\_382
CL\_INS\_385
CL\_INS\_368
CL\_INS\_159
CL\_INS\_159
CL\_INS\_159
CL\_INS\_159
CL\_INS\_382
CL\_INS\_382
CL\_INS\_382
CL\_INS\_382
CL\_INS\_382
CL\_INS\_99
CL\_INS\_382
CL\_INS\_159
CL\_INS\_117
CL\_INS\_117
CL\_INS\_159
CL\_INS\_382
CL\_INS\_382
CL\_INS\_382
CL\_INS\_382
CL\_INS\_382
CL\_INS\_247
CL\_INS\_368
CL\_INS\_368
CL\_INS\_149
CL\_INS\_70
CL\_INS\_70
CL\_INS\_70
CL\_INS\_70
CL\_INS\_207
CL\_INS\_207
CL\_INS\_207
CL\_INS\_207
CL\_INS\_70
CL\_INS\_368
CL\_INS\_70
CL\_INS\_70
CL\_INS\_70
CL\_INS\_368
CL\_INS\_368
CL\_INS\_368
CL\_INS\_155
CL\_INS\_368
CL\_INS\_368
CL\_INS\_368
CL\_INS\_368
CL\_INS\_368
CL\_INS\_368
CL\_INS\_368
CL\_INS\_368
CL\_INS\_368
CL\_INS\_368
CL\_INS\_368
CL\_INS\_368
CL\_INS\_368
CL\_INS\_368
CL\_INS\_237
CL\_INS\_237
CL\_INS\_237
CL\_INS\_237
CL\_INS\_237
CL\_INS\_237
CL\_INS\_237
CL\_INS\_237
CL\_INS\_368
CL\_INS\_70
CL\_INS\_368
CL\_INS\_237
CL\_INS\_368
CL\_INS\_368
CL\_INS\_368
CL\_INS\_368
CL\_INS\_368
CL\_INS\_368
CL\_INS\_70
CL\_INS\_70
CL\_INS\_237
CL\_INS\_70
CL\_INS\_70
CL\_INS\_368
CL\_INS\_237
CL\_INS\_368
CL\_INS\_70
CL\_INS\_368
CL\_INS\_368
CL\_INS\_368
CL\_INS\_368
CL\_INS\_368
CL\_INS\_368
CL\_INS\_70
CL\_INS\_368
CL\_INS\_368
CL\_INS\_368
CL\_INS\_368
CL\_INS\_382
CL\_INS\_368
CL\_INS\_382
CL\_INS\_368
CL\_INS\_368
CL\_INS\_368
CL\_INS\_368
CL\_INS\_368
CL\_INS\_368
CL\_INS\_368
CL\_INS\_368
CL\_INS\_368
CL\_INS\_368
CL\_INS\_237
CL\_INS\_368
CL\_INS\_368
CL\_INS\_368
CL\_INS\_368
CL\_INS\_368
CL\_INS\_368
CL\_INS\_207
CL\_INS\_368
CL\_INS\_237
CL\_INS\_237
CL\_INS\_368
CL\_INS\_368
CL\_INS\_253
CL\_INS\_70
CL\_INS\_70
CL\_INS\_368
CL\_INS\_247
CL\_INS\_237
CL\_INS\_70
CL\_INS\_237
CL\_INS\_368
CL\_INS\_237
CL\_INS\_70
CL\_INS\_368
CL\_INS\_368
CL\_INS\_368
CL\_INS\_70
CL\_INS\_30
CL\_INS\_368
CL\_INS\_368
CL\_INS\_30
CL\_INS\_368
CL\_INS\_237
CL\_INS\_159
CL\_INS\_30
CL\_INS\_159
CL\_INS\_237
CL\_INS\_42
CL\_INS\_368
CL\_INS\_368
CL\_INS\_30
CL\_INS\_159
CL\_INS\_30
CL\_INS\_30
CL\_INS\_368
CL\_INS\_237
CL\_INS\_237
CL\_INS\_368
CL\_INS\_70
CL\_INS\_368
CL\_INS\_368
CL\_INS\_368
CL\_INS\_368
CL\_INS\_368
CL\_INS\_70
CL\_INS\_237
CL\_INS\_237
CL\_INS\_237
CL\_INS\_237
CL\_INS\_237
CL\_INS\_70
CL\_INS\_237
CL\_INS\_237
CL\_INS\_237
CL\_INS\_237
CL\_INS\_237
CL\_INS\_237
CL\_INS\_368
CL\_INS\_237
CL\_INS\_368
CL\_INS\_237
CL\_INS\_368
CL\_INS\_30
CL\_INS\_368
CL\_INS\_368
CL\_INS\_30
CL\_INS\_237
CL\_INS\_237
CL\_INS\_70
CL\_INS\_237
CL\_INS\_237
CL\_INS\_368
CL\_INS\_368
CL\_INS\_237
CL\_INS\_368
CL\_INS\_70
CL\_INS\_237
CL\_INS\_30
CL\_INS\_368
CL\_INS\_368
CL\_INS\_368
CL\_INS\_368
CL\_INS\_368
CL\_INS\_368
CL\_INS\_368
CL\_INS\_368
CL\_INS\_368
CL\_INS\_368
CL\_INS\_30
CL\_INS\_30
CL\_INS\_368
CL\_INS\_368
CL\_INS\_159
CL\_INS\_30
CL\_INS\_368
CL\_INS\_368
CL\_INS\_237
CL\_INS\_237
CL\_INS\_237
CL\_INS\_247
CL\_INS\_70
CL\_INS\_70
CL\_INS\_368
CL\_INS\_70
CL\_INS\_70
CL\_INS\_368
CL\_INS\_368
CL\_INS\_368
CL\_INS\_237
CL\_INS\_368
CL\_INS\_30
CL\_INS\_30
CL\_INS\_237
CL\_INS\_70
CL\_INS\_70
CL\_INS\_70
CL\_INS\_368
CL\_INS\_368
CL\_INS\_368
CL\_INS\_368
CL\_INS\_368
CL\_INS\_368
CL\_INS\_159
CL\_INS\_70
CL\_INS\_237
CL\_INS\_368
CL\_INS\_368
CL\_INS\_368
CL\_INS\_30
CL\_INS\_30
CL\_INS\_237
CL\_INS\_247
CL\_INS\_247
CL\_INS\_159
CL\_INS\_237
CL\_INS\_237
CL\_INS\_368
CL\_INS\_368
CL\_INS\_30
CL\_INS\_30
CL\_INS\_70
CL\_INS\_30
CL\_INS\_237
CL\_INS\_368
CL\_INS\_368
CL\_INS\_70
CL\_INS\_70
CL\_INS\_368
CL\_INS\_382
CL\_INS\_368
CL\_INS\_70
CL\_INS\_70
CL\_INS\_368
CL\_INS\_368
CL\_INS\_237
CL\_INS\_70
CL\_INS\_368
CL\_INS\_368
CL\_INS\_352
CL\_INS\_352
CL\_INS\_352
CL\_INS\_352
CL\_INS\_352
CL\_INS\_352
CL\_INS\_352
CL\_INS\_352
CL\_INS\_352
CL\_INS\_352
CL\_INS\_352
CL\_INS\_352
CL\_INS\_352
CL\_INS\_352
CL\_INS\_352
CL\_INS\_352
CL\_INS\_352
CL\_INS\_352
CL\_INS\_352
CL\_INS\_352
CL\_INS\_352
CL\_INS\_352
CL\_INS\_352
CL\_INS\_352
CL\_INS\_352
CL\_INS\_352
CL\_INS\_352
CL\_INS\_352
CL\_INS\_352
CL\_INS\_352
CL\_INS\_352
CL\_INS\_352
CL\_INS\_352
CL\_INS\_352
CL\_INS\_352
CL\_INS\_352
CL\_INS\_352
CL\_INS\_352
CL\_INS\_352
CL\_INS\_352
CL\_INS\_352
CL\_INS\_352
CL\_INS\_368
CL\_INS\_368
CL\_INS\_368
CL\_INS\_70
CL\_INS\_368
CL\_INS\_368
CL\_INS\_368
CL\_INS\_70
CL\_INS\_368
CL\_INS\_368
CL\_INS\_368
CL\_INS\_382
CL\_INS\_368
CL\_INS\_368
CL\_INS\_368
CL\_INS\_368
CL\_INS\_368
CL\_INS\_368
CL\_INS\_368
CL\_INS\_368
CL\_INS\_368
CL\_INS\_368
CL\_INS\_368
CL\_INS\_382
CL\_INS\_368
CL\_INS\_70
CL\_INS\_237
CL\_INS\_237
CL\_INS\_237
CL\_INS\_237
CL\_INS\_237
CL\_INS\_237
CL\_INS\_237
CL\_INS\_237
CL\_INS\_237
CL\_INS\_237
CL\_INS\_237
CL\_INS\_237
CL\_INS\_237
CL\_INS\_237
CL\_INS\_237
CL\_INS\_382
CL\_INS\_237
CL\_INS\_70
CL\_INS\_368
CL\_INS\_237
CL\_INS\_237
CL\_INS\_237
CL\_INS\_237
CL\_INS\_237
CL\_INS\_70
CL\_INS\_70
CL\_INS\_70
CL\_INS\_70
CL\_INS\_70
CL\_INS\_70
CL\_INS\_30
CL\_INS\_70
CL\_INS\_368
CL\_INS\_159
Cluster ID


CL\_29971
CL\_29970
CL\_29969
CL\_13711
CL\_11386
CL\_11387
CL\_36956
CL\_36955
CL\_17111
CL\_17110
CL\_17109
CL\_17108
CL\_17107
CL\_17106
CL\_17105
CL\_32145
CL\_32144
CL\_32143
CL\_6590
CL\_12906
CL\_36143
CL\_24148
CL\_24149
CL\_24150
CL\_9452
CL\_8910
CL\_37179
CL\_33375
CL\_32383
CL\_32384
CL\_32385
CL\_26266
CL\_26265
CL\_6246
CL\_6245
CL\_6244
CL\_7300
CL\_7299
CL\_7298
CL\_8779
CL\_8376
CL\_7297
CL\_7236
CL\_11305
CL\_13527
CL\_10476
CL\_13526
CL\_10518
CL\_32581
CL\_12791
CL\_23673
CL\_12360
CL\_4520
CL\_29273
CL\_37242
CL\_7525
CL\_7526
CL\_4525
CL\_29225
CL\_37581
CL\_37240
CL\_37239
CL\_37238
CL\_37237
CL\_37580
CL\_22106
CL\_4087
CL\_22811
CL\_22526
CL\_22527
CL\_34937
CL\_8254
CL\_8253
CL\_8252
CL\_8251
CL\_8250
CL\_22810
CL\_8249
CL\_8248
CL\_8247
CL\_22528
CL\_22529
CL\_5149
CL\_23015
CL\_10593
CL\_23016
CL\_12904
CL\_22809
CL\_20416
CL\_20415
CL\_20414
CL\_20413
CL\_5972
CL\_5973
CL\_5974
CL\_5975
CL\_5976
CL\_8460
CL\_5977
CL\_5978
CL\_8461
CL\_8462
CL\_20873
CL\_20874
CL\_20875
CL\_37419
CL\_5979
CL\_17797
CL\_26109
CL\_5980
CL\_7627
CL\_8103
CL\_36654
CL\_19911
CL\_7284
CL\_7285
CL\_12074
CL\_12073
CL\_12072
CL\_12071
CL\_12070
CL\_8497
CL\_8498
CL\_8499
CL\_8500
CL\_7286
CL\_8501
CL\_10612
CL\_37127
CL\_37126
CL\_37125
CL\_6588
CL\_6587
CL\_6586
CL\_6585
CL\_6584
CL\_9454
CL\_6583
CL\_6582
CL\_6581
CL\_9455
CL\_6580
CL\_6579
CL\_6578
CL\_6577
CL\_6576
CL\_6575
CL\_6574
CL\_9456
CL\_37124
CL\_6573
CL\_11642
CL\_11641
CL\_6589
CL\_11300
CL\_11299
CL\_16512
CL\_8773
CL\_8772
CL\_11638
CL\_8771
CL\_8770
CL\_11637
CL\_8769
CL\_11636
CL\_11635
CL\_11634
CL\_11633
CL\_11632
CL\_11631
CL\_18846
CL\_18845
CL\_18844
CL\_7590
CL\_7589
CL\_7588
CL\_31618
CL\_18843
CL\_18842
CL\_11630
CL\_11629
CL\_11628
CL\_11627
CL\_11626
CL\_8406
CL\_18841
CL\_8407
CL\_8408
CL\_8409
CL\_8410
CL\_8411
CL\_8412
CL\_11625
CL\_8414
CL\_11624
CL\_11623
CL\_34838
CL\_34837
CL\_31619
CL\_31620
CL\_31621
CL\_18840
CL\_18839
CL\_11622
CL\_11621
CL\_8420
CL\_8421
CL\_8423
CL\_11620
CL\_8427
CL\_11619
CL\_31622
CL\_11618
CL\_11617
CL\_11616
CL\_11615
CL\_11614
CL\_11613
CL\_18838
CL\_18837
CL\_18836
CL\_7422
CL\_7421
CL\_7420
CL\_31623
CL\_31624
CL\_31625
CL\_16880
CL\_18835
CL\_18834
CL\_18833
CL\_18832
CL\_18831
CL\_18830
CL\_6018
CL\_25885
CL\_25886
CL\_6019
CL\_6020
CL\_6021
CL\_6022
CL\_13854
CL\_13855
CL\_13856
CL\_13857
CL\_13858
CL\_13859
CL\_13860
CL\_13861
CL\_18829
CL\_18828
CL\_18827
CL\_26846
CL\_26845
CL\_26844
CL\_18826
CL\_11612
CL\_6029
CL\_6028
CL\_6027
CL\_6026
CL\_6025
CL\_20497
CL\_20498
CL\_6024
CL\_6023
CL\_8132
CL\_13842
CL\_8768
CL\_8767
CL\_8766
CL\_8765
CL\_7843
CL\_8764
CL\_8763
CL\_8762
CL\_8761
CL\_8760
CL\_8759
CL\_8758
CL\_8757
CL\_8756
CL\_8755
CL\_6849
CL\_6848
CL\_6847
CL\_6846
CL\_11116
CL\_11115
CL\_11114
CL\_11113
CL\_11112
CL\_11298
CL\_11297
CL\_11111
CL\_6845
CL\_6844
CL\_6843
CL\_11110
CL\_11109
CL\_11108
CL\_6842
CL\_1931
CL\_25941
CL\_7750
CL\_7749
CL\_2551
CL\_6410
CL\_8600
CL\_6841
CL\_6840
CL\_8754
CL\_8753
CL\_8752
CL\_8751
CL\_6839
CL\_6838
CL\_17136
CL\_19784
CL\_19783
CL\_21829
CL\_21830
CL\_21831
CL\_21832
CL\_21833
CL\_21834
CL\_21835
CL\_21836
CL\_21837
CL\_6837
CL\_6836
CL\_11783
CL\_6835
CL\_27723
CL\_27722
CL\_29342
CL\_6834
CL\_6833
CL\_6426
CL\_5320
CL\_29090
CL\_6832
CL\_5321
CL\_13611
CL\_8750
CL\_8749
CL\_8748
CL\_8747
CL\_14345
CL\_11826
CL\_14346
CL\_14347
CL\_14348
CL\_10435
CL\_21030
CL\_34936
CL\_14344
CL\_5994
CL\_31464
CL\_31463
CL\_31462
CL\_31461
CL\_31460
CL\_32580
CL\_10050
CL\_36655
CL\_24116
CL\_28243
CL\_28242
CL\_28241
CL\_28240
CL\_28239
CL\_28238
CL\_17186
CL\_28237
CL\_28236
CL\_28235
CL\_28234
CL\_28233
CL\_28232
CL\_28231
CL\_28230
CL\_26107
CL\_26108
CL\_32408
CL\_33920
CL\_8455
CL\_8456
CL\_8457
CL\_8458
CL\_8459
CL\_9701
CL\_15963
CL\_15964
CL\_15965
CL\_15966
CL\_20669
CL\_27554
CL\_20872
CL\_8104
CL\_20282
CL\_25229
CL\_13385
CL\_13386
CL\_13387
CL\_21920
CL\_25228
CL\_13388
CL\_13389
CL\_13390
CL\_25227
CL\_25226
CL\_25225
CL\_23528
CL\_23529
CL\_23530
CL\_7407
CL\_7408
CL\_7409
CL\_7410
CL\_31915
CL\_31916
CL\_21723
CL\_7411
CL\_7412
CL\_7413
CL\_7414
CL\_29275
CL\_29276
CL\_29277
CL\_7415
CL\_17568
CL\_21724
CL\_21725
CL\_16874
CL\_31917
CL\_31918
CL\_31919
CL\_31920
CL\_21726
CL\_21727
CL\_21728
CL\_21729
CL\_21730
CL\_6366
CL\_6367
CL\_7416
CL\_6368
CL\_6369
CL\_6370
CL\_7417
CL\_7418
CL\_29278
CL\_29279
CL\_19183
CL\_19182
CL\_7631
CL\_7630
CL\_7629
CL\_7628
CL\_14310
CL\_14309
CL\_14308
CL\_14307
CL\_14306
CL\_14499
CL\_23181
CL\_22219
CL\_8960
CL\_8959
CL\_8958
CL\_8957
CL\_22218
CL\_14500
CL\_35016
CL\_35015
CL\_35014
CL\_14501
CL\_14502
CL\_14503
CL\_32885
CL\_32886
CL\_32887
CL\_32888
CL\_32889
CL\_19181
CL\_19180
CL\_19179
CL\_18125
CL\_18126
CL\_16974
CL\_13414
CL\_8140
CL\_4158
CL\_4157
CL\_4156
CL\_4155
CL\_4154
CL\_4153
CL\_21031
CL\_28570
CL\_28571
CL\_28572
CL\_28573
CL\_5995
CL\_5996
CL\_5997
CL\_5998
CL\_5999
CL\_29520
CL\_6000
CL\_6001
CL\_6002
CL\_6003
CL\_6004
CL\_6005
CL\_6006
CL\_29521
CL\_29522
CL\_6007
CL\_6008
CL\_6009
CL\_6010
CL\_6011
CL\_6012
CL\_6013
CL\_6014
CL\_6015
CL\_6016
CL\_6017
CL\_7998
CL\_7997
CL\_7996
CL\_7995
CL\_7994
CL\_7993
CL\_7992
CL\_7991
CL\_5157
CL\_5156
CL\_5155
CL\_5154
CL\_5153
CL\_37733
CL\_37734
CL\_6861
CL\_6860
CL\_6859
CL\_6858
CL\_6857
CL\_27721
CL\_6856
CL\_18851
CL\_18850
CL\_18849
CL\_18848
CL\_11640
CL\_11639
CL\_18847
CL\_8599
CL\_30641
CL\_30642
CL\_5683
CL\_30643
CL\_24249
CL\_5294
CL\_5293
CL\_5292
CL\_5291
CL\_5290
CL\_5531
CL\_14980
CL\_5533
CL\_20475
CL\_23210
CL\_30644
CL\_12077
CL\_37182
CL\_37181
CL\_37180
CL\_17160
CL\_17161
CL\_17162
CL\_7260
CL\_7261
CL\_11090
CL\_12078
CL\_11800
CL\_4088
CL\_4089
CL\_4090
CL\_4091
CL\_4092
CL\_22712
CL\_9477
CL\_11799
CL\_11798
CL\_11797
CL\_31373
CL\_10046
CL\_10045
CL\_20755
CL\_28547
CL\_28546
CL\_28545
CL\_28544
CL\_11796
CL\_11795
CL\_11794
CL\_11793
CL\_20751
CL\_20752
CL\_20753
CL\_20754
CL\_11792
CL\_11791
CL\_11790
CL\_11789
CL\_11788
CL\_33138
CL\_31379
CL\_31378
CL\_33137
CL\_31377
CL\_31376
CL\_31375
CL\_31374
CL\_20749
CL\_20750
CL\_28548
CL\_11787
CL\_11786
CL\_11785
CL\_11784
CL\_7565
CL\_4550
CL\_11314
CL\_11315
CL\_22764
CL\_11318
CL\_20448
CL\_20449
CL\_17781
CL\_17782
CL\_17783
CL\_20450
CL\_20451
CL\_20452
CL\_20453
CL\_20454
CL\_4943
CL\_4944
CL\_4945
CL\_9560
CL\_26956
CL\_9559
CL\_9558
CL\_4948
CL\_4949
CL\_4950
CL\_4951
CL\_14220
CL\_14219
CL\_14218
CL\_14217
CL\_9557
CL\_8518
CL\_7564
CL\_7563
CL\_7562
CL\_7561
CL\_29809
CL\_29808
CL\_7560
CL\_7559
CL\_34075
CL\_7558
CL\_7557
CL\_33406
CL\_5148
CL\_7306
CL\_13368
CL\_13369
CL\_13370
CL\_12907
CL\_12908
CL\_12909
CL\_12910
CL\_13371
CL\_13372
CL\_13373
CL\_13374
CL\_5147
CL\_25197
CL\_7243
CL\_26539
CL\_26540
CL\_7713
CL\_7712
CL\_12079
CL\_10583
CL\_26805
CL\_26806
CL\_26807
CL\_26808
CL\_26809
CL\_5152
CL\_5151
CL\_5150
CL\_7990
CL\_7989
CL\_7988
CL\_7301
CL\_7872
CL\_7302
CL\_25198
CL\_8778
CL\_11850
CL\_7711
CL\_7710
CL\_7709
CL\_7708
CL\_7707
CL\_7706
CL\_8385
CL\_7305
CL\_26810
CL\_7304
CL\_6056
CL\_7705
CL\_12080
CL\_7704
CL\_7703
CL\_14593
CL\_8909
CL\_7702
CL\_4462
CL\_32409
CL\_32410
CL\_32411
CL\_7701
CL\_12081
CL\_12082
CL\_7700
CL\_7699
CL\_7698
CL\_7697
CL\_7696
CL\_7695
CL\_7694
CL\_7693
CL\_12083
CL\_12084
CL\_8629
CL\_8287
CL\_24502
CL\_24501
CL\_24500
CL\_14086
CL\_8286
CL\_8285
CL\_6749
CL\_21220
CL\_33374
CL\_33373
CL\_11596
CL\_11594
CL\_11593
CL\_11592
CL\_11591
CL\_11590
CL\_11589
CL\_7515
CL\_7514
CL\_7513
CL\_7512
CL\_13382
CL\_13383
CL\_8284
CL\_8283
CL\_8282
CL\_33544
CL\_7249
CL\_7250
CL\_9453
CL\_10680
CL\_10679
CL\_10678
CL\_10677
CL\_11433
CL\_11432
CL\_5779
CL\_10676
CL\_10675
CL\_10718
CL\_10717
CL\_10716
CL\_10715
CL\_10714
CL\_10713
CL\_10712
CL\_10711
CL\_26879
CL\_10699
CL\_26878
CL\_10698
CL\_10697
CL\_10696
CL\_10695
CL\_5699
CL\_5700
CL\_5701
CL\_5702
CL\_10694
CL\_10693
CL\_10692
CL\_10691
CL\_10690
CL\_10689
CL\_5711
CL\_10688
CL\_10687
CL\_10686
CL\_10685
CL\_10684
CL\_11436
CL\_25758
CL\_10682
CL\_4974
CL\_11137
CL\_11138
CL\_5000
CL\_5001
CL\_4973
CL\_4972
CL\_10397
CL\_11814
CL\_34959
CL\_11815
CL\_11816
CL\_11818
CL\_11820
CL\_11821
CL\_11822
CL\_13617
CL\_4343
CL\_10363
CL\_8488
CL\_6318
CL\_10369
CL\_10370
CL\_10371
CL\_10383
CL\_10382
CL\_10423
CL\_10421
CL\_10642
CL\_10641
CL\_10395
CL\_10393
CL\_10392
CL\_5297
CL\_5298
CL\_5299
CL\_5300
CL\_5031
CL\_5032
CL\_5033
CL\_34958
CL\_11514
CL\_11519
CL\_11518
CL\_11517
CL\_11516
CL\_34957
CL\_34956
CL\_34955
CL\_22592
CL\_22590
CL\_22591
CL\_5574
CL\_5573
CL\_5572
CL\_5637
CL\_5638
CL\_5639
CL\_4277
CL\_4278
CL\_5567
CL\_5641
CL\_4279
CL\_5563
CL\_5562
CL\_5561
CL\_5560
CL\_4284
CL\_11442
CL\_4287
CL\_5556
CL\_5555
CL\_5554
CL\_23075
CL\_10653
CL\_10652
CL\_10651
CL\_10650
CL\_10649
CL\_10648
CL\_10647
CL\_10646
CL\_10645
CL\_10644
CL\_10643
CL\_10664
CL\_13409
CL\_10390
CL\_5614
CL\_22269
CL\_11132
CL\_5509
CL\_5510
CL\_5511
CL\_5512
CL\_5513
CL\_5019
CL\_11296
CL\_35446
CL\_35447
CL\_35448
CL\_35449
CL\_35450
CL\_35451
CL\_7829
CL\_5613
CL\_5536
CL\_5539
CL\_10411
CL\_10407
CL\_24256
CL\_8000
CL\_8001
CL\_8002
CL\_8003
CL\_24257
CL\_24258
CL\_6201
CL\_24259
CL\_10406
CL\_23073
CL\_4303
CL\_10804
CL\_29280
CL\_29281
CL\_29282
CL\_29283
CL\_29284
CL\_29285
CL\_29286
CL\_7419
CL\_12087
CL\_7677
CL\_7676
CL\_12088
CL\_12089
CL\_14595
CL\_7675
CL\_7984
CL\_7983
CL\_7674
CL\_7233
CL\_8782
CL\_7232
CL\_8783
CL\_8784
CL\_8785
CL\_7231
CL\_8786
CL\_7230
CL\_7229
CL\_8787
CL\_7228
CL\_7227
CL\_8788
CL\_7226
CL\_7225
CL\_7224
CL\_7673
CL\_4995
CL\_20455
CL\_20456
CL\_7672
CL\_8517
CL\_33971
CL\_33136
CL\_31372
CL\_4369
CL\_4370
CL\_25218
CL\_25217
CL\_4371
CL\_4372
CL\_4373
CL\_18854
CL\_7215
CL\_7671
CL\_7214
CL\_15526
CL\_6751
CL\_6752
CL\_6753
CL\_6754
CL\_6755
CL\_6756
CL\_6757
CL\_6758
CL\_6759
CL\_21603
CL\_21604
CL\_7294
CL\_7670
CL\_7669
CL\_8790
CL\_8791
CL\_8792
CL\_8793
CL\_36287
CL\_6076
CL\_8212
CL\_7295
CL\_36286
CL\_7666
CL\_8794
CL\_7665
CL\_9545
CL\_8211
CL\_8210
CL\_12090
CL\_12091
CL\_12092
CL\_12093
CL\_12094
CL\_12095
CL\_12096
CL\_12097
CL\_12098
CL\_12099
CL\_12100
CL\_12101
CL\_12102
CL\_12103
CL\_12104
CL\_12105
CL\_5689
CL\_5688
CL\_12106
CL\_8553
CL\_8554
CL\_6976
CL\_12107
CL\_12108
CL\_7667
CL\_5682
CL\_6855
CL\_6854
CL\_6853
CL\_6852
CL\_6851
CL\_21827
CL\_6850
CL\_7223
CL\_9785
CL\_12109
CL\_7213
CL\_14342
CL\_12110
CL\_7664
CL\_7212
CL\_7211
CL\_7293
CL\_7292
CL\_13379
CL\_13380
CL\_4374
CL\_17628
CL\_17629
CL\_4375
CL\_4376
CL\_7208
CL\_13850
CL\_13851
CL\_13852
CL\_13853
CL\_6416
CL\_4377
CL\_233
CL\_2550
CL\_13836
CL\_231
CL\_13384
CL\_14868
CL\_14869
CL\_14870
CL\_14871
CL\_21919
CL\_8102
CL\_8101
CL\_8100
CL\_8099
CL\_8098
CL\_8097
CL\_8096
CL\_17233
CL\_20283
CL\_14504
CL\_8095
CL\_8094
CL\_8093
CL\_14596
CL\_7207
CL\_7289
CL\_7288
CL\_7206
CL\_7205
CL\_7568
CL\_9607
CL\_6809
CL\_7567
CL\_7970
CL\_7969
CL\_6572
CL\_8124
CL\_7822
CL\_8932
CL\_22214
CL\_23205
CL\_23206
CL\_22212
CL\_33405
CL\_7556
CL\_7555
CL\_7554
CL\_13760
CL\_13759
CL\_7553
CL\_7552
CL\_7551
CL\_7550
CL\_7549
CL\_7548
CL\_7547
CL\_7546
CL\_7545
CL\_13758
CL\_7982
CL\_13757
CL\_7981
CL\_7980
CL\_14539
CL\_14540
CL\_7979
CL\_23014
CL\_8131
CL\_8130
CL\_22213
CL\_8463
CL\_8464
CL\_8465
CL\_8466
CL\_11097
CL\_11096
CL\_11095
CL\_6805
CL\_6806
CL\_11094
CL\_11093
CL\_9543
CL\_9542
CL\_7258
CL\_7259
CL\_35398
CL\_35397
CL\_9541
CL\_4102
CL\_4103
CL\_4104
CL\_4105
CL\_4106
CL\_4107
CL\_4108
CL\_4109
CL\_4110
CL\_4111
CL\_4112
CL\_4113
CL\_4350
CL\_4114
CL\_4115
CL\_4116
CL\_23674
CL\_11782
CL\_4357
CL\_4358
CL\_4359
CL\_4360
CL\_4361
CL\_4362
CL\_4363
CL\_4364
CL\_4365
CL\_23176
CL\_26235
CL\_26234
CL\_23190
CL\_23191
CL\_23192
CL\_35042
CL\_7222
CL\_7221
CL\_8789
CL\_7220
CL\_14597
CL\_12111
CL\_10436
CL\_6030
CL\_5468
CL\_10437
CL\_10438
CL\_10439
CL\_26062
CL\_10259
CL\_10440
CL\_10441
CL\_10442
CL\_10443
CL\_10444
CL\_10445
CL\_10446
CL\_10447
CL\_17785
CL\_8216
CL\_4975
CL\_7987
CL\_7986
CL\_7985
CL\_8841
CL\_34864
CL\_16528
CL\_4302
CL\_4301
CL\_4300
CL\_4299
CL\_4297
CL\_5662
CL\_5548
CL\_4294
CL\_4293
CL\_11338
CL\_5659
CL\_5551
CL\_5552
CL\_5553
CL\_4271
CL\_4270
CL\_5575
CL\_5577
CL\_14175
CL\_11341
CL\_4266
CL\_4265
CL\_5062
CL\_4263
CL\_4262
CL\_5625
CL\_5585
CL\_4261
CL\_4260
CL\_4259
CL\_4258
CL\_4257
CL\_4256
CL\_4255
CL\_4254
CL\_11835
CL\_11834
CL\_4253
CL\_5053
CL\_5592
CL\_5593
CL\_6239
CL\_4237
CL\_4236
CL\_4235
CL\_9918
CL\_9917
CL\_5236
CL\_9613
CL\_13249
CL\_4093
CL\_4094
CL\_4095
CL\_4096
CL\_4097
CL\_15133
CL\_15132
CL\_10361
CL\_14292
CL\_4098
CL\_37123
CL\_4099
CL\_4100
CL\_4101
CL\_30726
CL\_34935
CL\_22107
CL\_7374
CL\_9556
CL\_9555
CL\_9554
CL\_9553
CL\_9552
CL\_9551
CL\_9550
CL\_9549
CL\_9548
CL\_9547
CL\_9546
CL\_8215
CL\_8214
CL\_8213
CL\_13375
CL\_13376
CL\_13377
CL\_13378
CL\_26541
CL\_26542
CL\_15527
CL\_14340
CL\_7251
CL\_6831
CL\_23910
CL\_6830
CL\_21838
CL\_21839
CL\_21840
CL\_21841
CL\_21842
CL\_21843
CL\_1933
CL\_7831
CL\_13552
CL\_8620
CL\_6829
CL\_14893
CL\_8281
CL\_8280
CL\_7255
CL\_25224
CL\_25223
CL\_25222
CL\_25221
CL\_25220
CL\_25219
CL\_6179
CL\_10048
CL\_10047
CL\_35193
CL\_25887
CL\_6177
CL\_10049
CL\_6178
CL\_9038
CL\_25888
CL\_25889
CL\_31951
CL\_25890
CL\_12911
CL\_12912
CL\_12913
CL\_6248
CL\_12914
CL\_23012
CL\_23013
CL\_13831
CL\_13832
CL\_13833
CL\_13834
CL\_13835
CL\_7442
CL\_6807
CL\_10611
CL\_10817
CL\_22217
CL\_22216
CL\_22215
CL\_6411
CL\_6828
CL\_32890
CL\_8533
CL\_14349
CL\_7252
CL\_10820
CL\_37183
CL\_6827
CL\_7253
CL\_32891
CL\_11294
CL\_11293
CL\_6826
CL\_5245
CL\_33635
CL\_33636
CL\_7254
CL\_7978
CL\_12176
CL\_28864
CL\_23184
CL\_28863
CL\_10374
CL\_13915
CL\_34862
CL\_4920
CL\_5247
CL\_22752
CL\_5246
CL\_6425
CL\_34170
CL\_10825
CL\_6825
CL\_11975
CL\_6824
CL\_13843
CL\_13844
CL\_6823
CL\_6822
CL\_11107
CL\_8129
CL\_8555
CL\_8556
CL\_8128
CL\_11106
CL\_8127
CL\_11105
CL\_11711
CL\_13756
CL\_11710
CL\_13755
CL\_11709
CL\_11708
CL\_33404
CL\_7832
CL\_33403
CL\_7256
CL\_13845
CL\_6424
CL\_11292
CL\_11291
CL\_6423
CL\_6731
CL\_6730
CL\_8126
CL\_13214
CL\_8125
CL\_36030
CL\_36029
CL\_10822
CL\_28862
CL\_10546
CL\_11972
CL\_5241
CL\_34171
CL\_34172
CL\_6821
CL\_6820
CL\_6819
CL\_6818
CL\_6817
CL\_6816
CL\_6815
CL\_6814
CL\_6422
CL\_5232
CL\_7977
CL\_7976
CL\_7975
CL\_5229
CL\_7974
CL\_7973
CL\_11974
CL\_11973
CL\_13846
CL\_7470
CL\_10547
CL\_9139
CL\_13847
CL\_13848
CL\_8642
CL\_13849
CL\_23188
CL\_23189
CL\_6729
CL\_8908
CL\_8738
CL\_8907
CL\_6728
CL\_6421
CL\_8619
CL\_22734
CL\_11104
CL\_11103
CL\_11102
CL\_11101
CL\_11100
CL\_11099
CL\_6727
CL\_8508
CL\_6813
CL\_6812
CL\_6811
CL\_6810
CL\_7751
CL\_235
CL\_8815
CL\_7828
CL\_6726
CL\_6725
CL\_6724
CL\_6723
CL\_7257
CL\_11098
CL\_1936
CL\_7971
CL\_7824
CL\_1937
CL\_6722
CL\_35047
CL\_35048
CL\_8746
CL\_6808
CL\_9544
CL\_4086
CL\_13381
CL\_6420
CL\_1932
CL\_19349
CL\_19350
CL\_6415
CL\_6419
CL\_37184
CL\_37185
CL\_9401
CL\_9402
CL\_9403
CL\_9404
CL\_9405
CL\_9406
CL\_9407
CL\_9408
CL\_9409
CL\_9410
CL\_9411
CL\_9412
CL\_9413
CL\_9414
CL\_9415
CL\_9416
CL\_9417
CL\_9418
CL\_9419
CL\_9420
CL\_9421
CL\_9422
CL\_9423
CL\_9424
CL\_9425
CL\_9426
CL\_9427
CL\_9428
CL\_9429
CL\_9430
CL\_9431
CL\_9432
CL\_9433
CL\_9434
CL\_9435
CL\_9436
CL\_9437
CL\_9438
CL\_9439
CL\_9440
CL\_9441
CL\_17585
CL\_37186
CL\_37187
CL\_37188
CL\_6418
CL\_34076
CL\_33402
CL\_33401
CL\_6417
CL\_7972
CL\_35045
CL\_35046
CL\_7692
CL\_6505
CL\_6506
CL\_6507
CL\_6508
CL\_6509
CL\_6510
CL\_10927
CL\_10926
CL\_10925
CL\_10924
CL\_23140
CL\_7691
CL\_21828
CL\_7690
CL\_14117
CL\_7689
CL\_7688
CL\_12085
CL\_7687
CL\_7686
CL\_7685
CL\_7684
CL\_14594
CL\_12086
CL\_7683
CL\_7682
CL\_7680
CL\_7679
CL\_7678
CL\_8832
CL\_8780
CL\_8781
CL\_34863
CL\_5597
CL\_5596
CL\_5595
CL\_5594
CL\_6978
CL\_6979
CL\_6980
CL\_6981
CL\_6982
CL\_6983
CL\_6984
CL\_11295
CL\_10581
CL\_4117
CL\_1929
