## Supplementary material for "A novel method for integrating genomic and Tn-Seq data to identify common *in vivo* fitness mechanisms across multiple bacterial species": S1 Dataset: CL_INS_369.html

Legend

 Mobile +extrachromosomalelementfunctions
 Hypothetical
 Other
 Transport +binding proteins
 All VFDB Genes

FULL


WINDOWSVGPNG

Trim RowsRemove SingletonsSave Fasta

CL\_4120


CL\_4120


CL\_4120


CL\_4120


CL\_4120


CL\_4120


CL\_4120


CL\_4120


CL\_4085


CL\_4120


CL\_4120


CL\_4120


CL\_4120


CL\_4085


CL\_4120


CL\_4120


CL\_4120

HighlightSelectShow Genomes


184

CL\_4121


19

CL\_4122


5

CL\_4130


4

CL\_4121


2

CL\_4121


2

CL\_4122


2

CL\_4121


2

CL\_4121


1

CL\_4121


1

CL\_4121


1

CL\_4122


1

CL\_4953


1

Break


1

CL\_4121


1

CL\_4130


1

CL\_4121


1

CL\_4122

fGI ID


CL\_INS\_369
CL\_INS\_369
CL\_INS\_369
CL\_INS\_369
CL\_INS\_369
CL\_INS\_369
CL\_INS\_247
CL\_INS\_369
CL\_INS\_369
CL\_INS\_247
CL\_INS\_369
CL\_INS\_99
CL\_INS\_99
CL\_INS\_247
CL\_INS\_369
CL\_INS\_369
CL\_INS\_369
CL\_INS\_207
CL\_INS\_207
CL\_INS\_207
CL\_INS\_207
CL\_INS\_247
CL\_INS\_70
CL\_INS\_70
CL\_INS\_70
CL\_INS\_70
CL\_INS\_70
CL\_INS\_70
CL\_INS\_70
CL\_INS\_70
Cluster ID


CL\_11290
CL\_10259
CL\_26186
CL\_15209
CL\_13248
CL\_11092
CL\_6749
CL\_21220
CL\_8283
CL\_8282
CL\_34927
CL\_6747
CL\_4489
CL\_11653
CL\_11654
CL\_11655
CL\_24116
CL\_5975
CL\_5976
CL\_5977
CL\_5978
CL\_5236
CL\_4094
CL\_4095
CL\_4096
CL\_4097
CL\_4098
CL\_4099
CL\_4100
CL\_4101
