## Supplementary material for "A novel method for integrating genomic and Tn-Seq data to identify common *in vivo* fitness mechanisms across multiple bacterial species": S1 Dataset: CL_INS_370.html

Legend

 Hypothetical
 All VFDB Genes

FULL


WINDOWSVGPNG

Trim RowsRemove SingletonsSave Fasta

CL\_4122


CL\_4122


CL\_4122


CL\_4122


CL\_4122

HighlightSelectShow Genomes


207

CL\_4121


19

CL\_4120


1

CL\_4120


1

CL\_4120


1

CL\_4120

fGI ID


CL\_INS\_292
CL\_INS\_369
Cluster ID


CL\_11091
CL\_11092
