## Supplementary material for "A novel method for integrating genomic and Tn-Seq data to identify common *in vivo* fitness mechanisms across multiple bacterial species": S1 Dataset: CL_INS_372.html

Legend

 Regulatoryfunctions
 Hypothetical
 Other
 EnergyMetabolism
 All VFDB Genes
 Transport +binding proteins

FULL


WINDOWSVGPNG

Trim RowsRemove SingletonsSave Fasta

CL\_4136


CL\_4136


CL\_4136


CL\_4136


CL\_4136


CL\_4136


CL\_4136


CL\_4136


CL\_4136


CL\_4136


CL\_4136


CL\_4136


CL\_4136


CL\_4136


CL\_4136

HighlightSelectShow Genomes


246

CL\_4137


5

CL\_4137


4

CL\_4137


3

CL\_4138


3

CL\_4137


1

CL\_4137


1

CL\_4137


1

CL\_4137


1

CL\_4137


1

CL\_4138


1

CL\_4138


1

CL\_4137


1

CL\_4137


1

CL\_4137


1

CL\_4137

fGI ID


CL\_INS\_372
CL\_INS\_372
CL\_INS\_372
CL\_INS\_372
CL\_INS\_372
CL\_INS\_372
CL\_INS\_372
CL\_INS\_372
CL\_INS\_372
CL\_INS\_372
CL\_INS\_372
CL\_INS\_372
CL\_INS\_372
CL\_INS\_372
CL\_INS\_372
CL\_INS\_372
CL\_INS\_372
Cluster ID


CL\_7968
CL\_7967
CL\_7966
CL\_7965
CL\_7964
CL\_7963
CL\_12112
CL\_7962
CL\_7961
CL\_7960
CL\_12113
CL\_7959
CL\_34753
CL\_7958
CL\_12114
CL\_7957
CL\_7956
