## Supplementary material for "A novel method for integrating genomic and Tn-Seq data to identify common *in vivo* fitness mechanisms across multiple bacterial species": S1 Dataset: CL_INS_374.html

Legend

 Mobile +extrachromosomalelementfunctions
 Hypothetical
 Regulatoryfunctions
 DNA Metabolism
 AntibioticResistance
 Transcription
 Other
 All VFDB Genes

FULL


WINDOWSVGPNG

Trim RowsRemove SingletonsSave Fasta

CL\_4953


CL\_4953


CL\_4148


CL\_4147


CL\_4953


CL\_4953


CL\_4953


CL\_4953


CL\_4085


CL\_4953


CL\_4953


CL\_4148


CL\_4953


CL\_4953

HighlightSelectShow Genomes


165

CL\_4150


33

CL\_4150


6

CL\_4150


3

CL\_4150


3

CL\_4160


2

CL\_4160


1

CL\_4160


1

CL\_4150


1

CL\_4150


1

CL\_4160


1

CL\_4120


1

CL\_4150


1

CL\_4160


1

CL\_4160

fGI ID


CL\_INS\_374
CL\_INS\_374
CL\_INS\_374
CL\_INS\_374
CL\_INS\_368
CL\_INS\_368
CL\_INS\_368
CL\_INS\_374
CL\_INS\_359
CL\_INS\_374
CL\_INS\_368
CL\_INS\_69
CL\_INS\_20
CL\_INS\_20
CL\_INS\_20
CL\_INS\_374
CL\_INS\_374
CL\_INS\_86
CL\_INS\_247
CL\_INS\_247
CL\_INS\_374
CL\_INS\_374
CL\_INS\_374
CL\_INS\_237
CL\_INS\_237
CL\_INS\_237
CL\_INS\_247
CL\_INS\_189
CL\_INS\_247
CL\_INS\_247
CL\_INS\_123
CL\_INS\_374
CL\_INS\_374
CL\_INS\_247
CL\_INS\_247
CL\_INS\_123
CL\_INS\_382
CL\_INS\_247
CL\_INS\_374
CL\_INS\_247
CL\_INS\_247
CL\_INS\_69
CL\_INS\_69
CL\_INS\_374
Cluster ID


CL\_33637
CL\_4149
CL\_23044
CL\_17784
CL\_17783
CL\_17782
CL\_17781
CL\_23045
CL\_6570
CL\_17780
CL\_4155
CL\_6569
CL\_7260
CL\_7261
CL\_11090
CL\_11089
CL\_30645
CL\_5516
CL\_10388
CL\_10387
CL\_22507
CL\_22506
CL\_22505
CL\_6750
CL\_6760
CL\_6761
CL\_10421
CL\_22504
CL\_10395
CL\_10393
CL\_10392
CL\_22503
CL\_22502
CL\_15544
CL\_5299
CL\_5300
CL\_5301
CL\_5302
CL\_22501
CL\_11980
CL\_13409
CL\_6568
CL\_6567
CL\_15214
