## Supplementary material for "A novel method for integrating genomic and Tn-Seq data to identify common *in vivo* fitness mechanisms across multiple bacterial species": S1 Dataset: CL_INS_375.html

Legend

 Mobile +extrachromosomalelementfunctions
 Regulatoryfunctions
 Hypothetical
 DNA Metabolism
 AntibioticResistance
 Transcription
 All EssentialGenes
 Other
 Transport +binding proteins
 All VFDB Genes

FULL


WINDOWSVGPNG

Trim RowsRemove SingletonsSave Fasta

CL\_4148


CL\_4150


CL\_4150


CL\_4150


CL\_4150


CL\_4150


CL\_4150


CL\_4150


CL\_4150


CL\_4150


CL\_4150


CL\_4953


CL\_4953


CL\_4150


CL\_4150


CL\_4150


CL\_4150


CL\_4150


CL\_4039


CL\_4150


CL\_4150


CL\_4148


CL\_4024


CL\_4150


CL\_4150


CL\_4150


CL\_4150


CL\_4150


CL\_4150


CL\_4150


CL\_4150


CL\_4150


CL\_4150


CL\_4150


CL\_4146


CL\_4150


CL\_4953


CL\_4148


CL\_4148


CL\_4150


CL\_4148


CL\_4150


CL\_4135


CL\_4150


CL\_4150


CL\_4150


CL\_4150


CL\_4150


CL\_4150


CL\_4150


CL\_4150


CL\_4150


CL\_4150


CL\_4150


CL\_4150


CL\_4953


CL\_4150


CL\_4150


CL\_4953


CL\_4150


CL\_4118


CL\_4148


CL\_4150


CL\_4150


CL\_4953


CL\_4150


CL\_4150


CL\_4150


CL\_4133


CL\_4150


CL\_4150


CL\_4148


CL\_4150


CL\_4150

HighlightSelectShow Genomes


48

CL\_4160


48

CL\_4160


31

CL\_4160


18

CL\_4160


17

CL\_4160


11

CL\_4160


10

CL\_4160


9

CL\_4160


8

CL\_4160


5

CL\_4160


3

CL\_4160


3

CL\_4160


2

CL\_4160


2

CL\_4160


2

CL\_4160


2

CL\_4160


2

CL\_4160


2

CL\_4160


1

CL\_4160


1

CL\_4160


1

CL\_4160


1

CL\_4160


1

CL\_4160


1

CL\_4160


1

CL\_4160


1

CL\_4160


1

CL\_4160


1

CL\_4161


1

CL\_4160


1

CL\_4085


1

CL\_4160


1

CL\_4160


1

CL\_4160


1

CL\_4160


1

CL\_4160


1

CL\_4160


1

CL\_4160


1

CL\_4160


1

CL\_4160


1

CL\_4161


1

CL\_4160


1

CL\_4160


1

CL\_4160


1

CL\_4160


1

CL\_4160


1

CL\_4160


1

CL\_4160


1

CL\_4160


1

CL\_4038


1

CL\_4160


1

CL\_4161


1

CL\_4160


1

CL\_4160


1

CL\_4160


1

CL\_4160


1

CL\_4160


1

CL\_4160


1

CL\_4160


1

CL\_4160


1

CL\_4160


1

CL\_4160


1

CL\_4160


1

CL\_4160


1

CL\_4160


1

CL\_4160


1

CL\_4160


1

CL\_4160


1

CL\_4160


1

CL\_4160


1

CL\_4160


1

CL\_4160


1

CL\_4160


1

CL\_4160


1

CL\_4160

fGI ID


CL\_INS\_375
CL\_INS\_375
CL\_INS\_375
CL\_INS\_374
CL\_INS\_359
CL\_INS\_374
CL\_INS\_368
CL\_INS\_368
CL\_INS\_368
CL\_INS\_374
CL\_INS\_375
CL\_INS\_375
CL\_INS\_375
CL\_INS\_359
CL\_INS\_374
CL\_INS\_374
CL\_INS\_375
CL\_INS\_368
CL\_INS\_70
CL\_INS\_70
CL\_INS\_70
CL\_INS\_237
CL\_INS\_237
CL\_INS\_237
CL\_INS\_237
CL\_INS\_237
CL\_INS\_375
CL\_INS\_375
CL\_INS\_375
CL\_INS\_247
CL\_INS\_247
CL\_INS\_207
CL\_INS\_237
CL\_INS\_237
CL\_INS\_237
CL\_INS\_237
CL\_INS\_237
CL\_INS\_237
CL\_INS\_237
CL\_INS\_237
CL\_INS\_237
CL\_INS\_237
CL\_INS\_237
CL\_INS\_237
CL\_INS\_237
CL\_INS\_237
CL\_INS\_237
CL\_INS\_237
CL\_INS\_237
CL\_INS\_237
CL\_INS\_368
CL\_INS\_368
CL\_INS\_368
CL\_INS\_368
CL\_INS\_375
CL\_INS\_20
CL\_INS\_375
CL\_INS\_375
CL\_INS\_20
CL\_INS\_69
CL\_INS\_375
CL\_INS\_368
CL\_INS\_368
CL\_INS\_368
CL\_INS\_375
CL\_INS\_247
CL\_INS\_247
CL\_INS\_247
CL\_INS\_247
CL\_INS\_368
CL\_INS\_368
CL\_INS\_368
CL\_INS\_71
CL\_INS\_368
CL\_INS\_382
CL\_INS\_382
CL\_INS\_382
CL\_INS\_382
CL\_INS\_382
CL\_INS\_382
CL\_INS\_217
CL\_INS\_374
CL\_INS\_374
CL\_INS\_374
CL\_INS\_237
CL\_INS\_237
CL\_INS\_20
CL\_INS\_374
CL\_INS\_373
CL\_INS\_374
CL\_INS\_237
CL\_INS\_247
CL\_INS\_189
CL\_INS\_247
CL\_INS\_247
CL\_INS\_123
CL\_INS\_374
CL\_INS\_374
CL\_INS\_247
CL\_INS\_247
CL\_INS\_123
CL\_INS\_382
CL\_INS\_247
CL\_INS\_374
CL\_INS\_247
CL\_INS\_247
CL\_INS\_69
CL\_INS\_69
CL\_INS\_375
CL\_INS\_375
CL\_INS\_374
CL\_INS\_373
Cluster ID


CL\_12917
CL\_4954
CL\_30955
CL\_23044
CL\_13250
CL\_17784
CL\_17783
CL\_17782
CL\_17781
CL\_23045
CL\_12061
CL\_15989
CL\_22108
CL\_6570
CL\_17780
CL\_33637
CL\_4151
CL\_21030
CL\_10435
CL\_5321
CL\_5320
CL\_9477
CL\_4092
CL\_4091
CL\_4090
CL\_4089
CL\_7955
CL\_7954
CL\_4152
CL\_10412
CL\_10413
CL\_10314
CL\_4116
CL\_4115
CL\_4114
CL\_4113
CL\_4112
CL\_4111
CL\_4110
CL\_4109
CL\_4108
CL\_4107
CL\_4106
CL\_4105
CL\_4104
CL\_4103
CL\_4102
CL\_9541
CL\_7259
CL\_7258
CL\_21031
CL\_4153
CL\_4154
CL\_4155
CL\_28189
CL\_7260
CL\_9457
CL\_13466
CL\_7261
CL\_6569
CL\_29091
CL\_4156
CL\_4157
CL\_4158
CL\_4159
CL\_10388
CL\_10387
CL\_10664
CL\_10390
CL\_30644
CL\_23210
CL\_20475
CL\_5533
CL\_14980
CL\_5531
CL\_5290
CL\_5291
CL\_5292
CL\_5293
CL\_5294
CL\_23459
CL\_22507
CL\_22506
CL\_22505
CL\_6750
CL\_6760
CL\_11090
CL\_11089
CL\_6804
CL\_30645
CL\_6761
CL\_10421
CL\_22504
CL\_10395
CL\_10393
CL\_10392
CL\_22503
CL\_22502
CL\_15544
CL\_5299
CL\_5300
CL\_5301
CL\_5302
CL\_22501
CL\_11980
CL\_13409
CL\_6568
CL\_6567
CL\_6566
CL\_23758
CL\_15214
CL\_6803
