## Supplementary material for "A novel method for integrating genomic and Tn-Seq data to identify common *in vivo* fitness mechanisms across multiple bacterial species": S1 Dataset: CL_INS_378.html

Legend

 Mobile +extrachromosomalelementfunctions
 Regulatoryfunctions
 Hypothetical
 Other
 Transport +binding proteins
 All VFDB Genes

FULL


WINDOWSVGPNG

Trim RowsRemove SingletonsSave Fasta

CL\_4165


CL\_4165


CL\_4166


CL\_4165


CL\_4165


CL\_4165


CL\_4165


CL\_4165


CL\_4167


CL\_4165


CL\_4165


CL\_4165

HighlightSelectShow Genomes


192

CL\_4164


49

CL\_4163


4

CL\_4164


1

CL\_4163


1

CL\_4163


1

CL\_4163


1

CL\_4164


1

CL\_4163


1

CL\_4164


1

CL\_4163


1

CL\_4164


1

CL\_4163

fGI ID


CL\_INS\_237
CL\_INS\_378
CL\_INS\_377
CL\_INS\_377
CL\_INS\_376
CL\_INS\_376
Cluster ID


CL\_244
CL\_34752
CL\_7950
CL\_26543
CL\_7951
CL\_7952
