## Supplementary material for "A novel method for integrating genomic and Tn-Seq data to identify common *in vivo* fitness mechanisms across multiple bacterial species": S1 Dataset: CL_INS_379.html

Legend

 Mobile +extrachromosomalelementfunctions
 Regulatoryfunctions
 Hypothetical
 DNA Metabolism
 All EssentialGenes
 All Fitness Genes
 Biosynthesis ofcofactors,prostheticgroups, +carriers
 Proteinsynthesis/fate
 Other
 EnergyMetabolism
 Transport +binding proteins
 All VFDB Genes

FULL


WINDOWSVGPNG

Trim RowsRemove SingletonsSave Fasta

CL\_4180


CL\_4516


CL\_4180


CL\_4516


CL\_4180


CL\_4180


CL\_4180


CL\_4180


CL\_4180


CL\_4180


CL\_4180


CL\_4180


CL\_4516


CL\_4180


CL\_4487


CL\_4180


CL\_4180


CL\_4427


CL\_4516


CL\_4180


CL\_4180


CL\_4180


CL\_4519


CL\_4486


CL\_4180


CL\_4180


CL\_4486


CL\_4516


CL\_4180


CL\_4180


CL\_4180


CL\_4516


CL\_4519


CL\_4180


CL\_4516


CL\_4180


CL\_4180


CL\_4180


CL\_4180


CL\_4180


CL\_4180


CL\_4487


CL\_4516


CL\_4126


CL\_4180


CL\_4516


CL\_4180


CL\_4516


CL\_4180


CL\_4180


CL\_4486


CL\_4516


CL\_4180


CL\_4180


CL\_4180


CL\_4180


CL\_4486


CL\_4516


CL\_4180


CL\_4180


CL\_4180


CL\_4180


CL\_4180


CL\_4180


CL\_4180


CL\_4180


CL\_4180


CL\_4180


CL\_4487


CL\_4180

HighlightSelectShow Genomes


223

CL\_4181


1

CL\_4181


1

CL\_4516


1

CL\_4181


1

CL\_3873


1

CL\_4427


1

CL\_4486


1

CL\_4427


1

CL\_4181


1

CL\_4073


1

CL\_4181


1

CL\_4181


1

CL\_4181


1

CL\_4181


1

CL\_4181


1

CL\_4181


1

CL\_4181


1

CL\_4181


1

CL\_4181


1

CL\_4181


1

CL\_4181


1

CL\_4486


1

CL\_4181


1

CL\_4181


1

CL\_4487


1

CL\_4427


1

CL\_4181


1

CL\_4181


1

CL\_4486


1

CL\_4181


1

CL\_4486


1

CL\_4181


1

CL\_4181


1

CL\_1915


1

CL\_4181


1

CL\_4181


1

CL\_4181


1

CL\_4181


1

CL\_4516


1

CL\_4427


1

CL\_4181


1

CL\_4181


1

CL\_4181


1

CL\_4181


1

CL\_4181


1

CL\_4181


1

CL\_4486


1

CL\_4181


1

CL\_4427


1

CL\_4487


1

CL\_4181


1

CL\_4181


1

CL\_4181


1

CL\_4486


1

CL\_4181


1

CL\_4427


1

CL\_4181


1

CL\_4181


1

CL\_4486


1

CL\_4181


1

CL\_4427


1

CL\_4181


1

CL\_4181


1

CL\_4427


1

CL\_4181


1

CL\_4181


1

CL\_4181


1

CL\_4427


1

CL\_4181


1

CL\_4181

fGI ID


CL\_INS\_385
CL\_INS\_385
CL\_INS\_385
CL\_INS\_385
CL\_INS\_99
CL\_INS\_379
CL\_INS\_379
CL\_INS\_379
CL\_INS\_379
CL\_INS\_379
CL\_INS\_379
CL\_INS\_379
CL\_INS\_379
CL\_INS\_379
CL\_INS\_379
CL\_INS\_382
CL\_INS\_379
CL\_INS\_379
CL\_INS\_382
CL\_INS\_379
CL\_INS\_379
CL\_INS\_379
CL\_INS\_382
CL\_INS\_379
CL\_INS\_379
CL\_INS\_382
CL\_INS\_379
CL\_INS\_382
CL\_INS\_379
CL\_INS\_379
CL\_INS\_379
CL\_INS\_379
CL\_INS\_382
CL\_INS\_382
CL\_INS\_379
CL\_INS\_379
CL\_INS\_379
CL\_INS\_379
CL\_INS\_379
CL\_INS\_382
CL\_INS\_60
CL\_INS\_382
CL\_INS\_382
CL\_INS\_382
CL\_INS\_382
CL\_INS\_382
CL\_INS\_379
CL\_INS\_60
CL\_INS\_379
CL\_INS\_20
CL\_INS\_20
CL\_INS\_60
CL\_INS\_237
CL\_INS\_79
CL\_INS\_79
CL\_INS\_79
CL\_INS\_79
CL\_INS\_382
CL\_INS\_79
CL\_INS\_382
CL\_INS\_382
CL\_INS\_79
CL\_INS\_382
CL\_INS\_79
CL\_INS\_79
CL\_INS\_79
CL\_INS\_382
CL\_INS\_79
CL\_INS\_79
CL\_INS\_146
CL\_INS\_146
CL\_INS\_146
CL\_INS\_20
CL\_INS\_20
CL\_INS\_20
CL\_INS\_379
CL\_INS\_382
CL\_INS\_379
CL\_INS\_382
CL\_INS\_237
CL\_INS\_382
CL\_INS\_382
CL\_INS\_382
CL\_INS\_382
CL\_INS\_379
CL\_INS\_379
CL\_INS\_379
CL\_INS\_379
CL\_INS\_379
CL\_INS\_79
CL\_INS\_79
CL\_INS\_79
CL\_INS\_79
CL\_INS\_79
CL\_INS\_79
CL\_INS\_79
CL\_INS\_79
CL\_INS\_79
CL\_INS\_79
CL\_INS\_79
CL\_INS\_79
CL\_INS\_79
CL\_INS\_79
CL\_INS\_79
CL\_INS\_79
CL\_INS\_20
CL\_INS\_20
CL\_INS\_20
CL\_INS\_20
CL\_INS\_379
CL\_INS\_379
CL\_INS\_379
CL\_INS\_379
CL\_INS\_79
CL\_INS\_79
CL\_INS\_79
CL\_INS\_382
CL\_INS\_382
CL\_INS\_379
CL\_INS\_379
CL\_INS\_379
CL\_INS\_382
CL\_INS\_382
CL\_INS\_382
CL\_INS\_382
CL\_INS\_382
CL\_INS\_382
CL\_INS\_382
CL\_INS\_382
CL\_INS\_382
CL\_INS\_382
CL\_INS\_379
CL\_INS\_379
CL\_INS\_379
CL\_INS\_379
CL\_INS\_379
CL\_INS\_79
CL\_INS\_379
CL\_INS\_379
CL\_INS\_379
CL\_INS\_379
CL\_INS\_379
CL\_INS\_379
CL\_INS\_379
CL\_INS\_379
CL\_INS\_379
CL\_INS\_379
CL\_INS\_379
CL\_INS\_379
CL\_INS\_379
CL\_INS\_379
CL\_INS\_42
CL\_INS\_382
CL\_INS\_382
CL\_INS\_382
CL\_INS\_382
CL\_INS\_382
CL\_INS\_382
CL\_INS\_382
CL\_INS\_382
CL\_INS\_382
CL\_INS\_379
CL\_INS\_79
CL\_INS\_79
CL\_INS\_79
CL\_INS\_79
CL\_INS\_99
CL\_INS\_382
CL\_INS\_99
CL\_INS\_382
CL\_INS\_99
CL\_INS\_79
CL\_INS\_79
CL\_INS\_382
CL\_INS\_382
CL\_INS\_382
CL\_INS\_382
CL\_INS\_382
CL\_INS\_382
CL\_INS\_379
CL\_INS\_379
CL\_INS\_379
CL\_INS\_379
CL\_INS\_379
CL\_INS\_379
CL\_INS\_379
CL\_INS\_379
CL\_INS\_379
CL\_INS\_382
CL\_INS\_379
CL\_INS\_379
CL\_INS\_379
CL\_INS\_379
CL\_INS\_379
CL\_INS\_379
CL\_INS\_379
CL\_INS\_379
CL\_INS\_379
CL\_INS\_379
CL\_INS\_379
CL\_INS\_379
CL\_INS\_379
CL\_INS\_146
CL\_INS\_146
CL\_INS\_379
CL\_INS\_146
CL\_INS\_379
CL\_INS\_379
CL\_INS\_379
CL\_INS\_379
CL\_INS\_379
CL\_INS\_379
CL\_INS\_379
CL\_INS\_379
CL\_INS\_379
CL\_INS\_379
CL\_INS\_379
CL\_INS\_379
CL\_INS\_379
CL\_INS\_379
CL\_INS\_382
CL\_INS\_382
CL\_INS\_382
CL\_INS\_382
CL\_INS\_382
CL\_INS\_382
CL\_INS\_382
CL\_INS\_97
CL\_INS\_97
CL\_INS\_97
CL\_INS\_97
CL\_INS\_379
CL\_INS\_379
CL\_INS\_379
CL\_INS\_379
CL\_INS\_379
CL\_INS\_379
CL\_INS\_379
CL\_INS\_379
CL\_INS\_379
CL\_INS\_379
CL\_INS\_128
CL\_INS\_385
CL\_INS\_385
CL\_INS\_379
CL\_INS\_379
CL\_INS\_379
CL\_INS\_379
CL\_INS\_379
CL\_INS\_379
CL\_INS\_379
CL\_INS\_379
CL\_INS\_379
CL\_INS\_379
CL\_INS\_379
CL\_INS\_379
CL\_INS\_379
CL\_INS\_379
CL\_INS\_379
CL\_INS\_379
CL\_INS\_379
CL\_INS\_352
CL\_INS\_237
CL\_INS\_237
CL\_INS\_379
CL\_INS\_237
CL\_INS\_237
CL\_INS\_237
CL\_INS\_379
CL\_INS\_379
CL\_INS\_379
CL\_INS\_379
CL\_INS\_379
CL\_INS\_379
CL\_INS\_379
CL\_INS\_379
CL\_INS\_379
CL\_INS\_379
CL\_INS\_379
CL\_INS\_379
CL\_INS\_379
CL\_INS\_379
CL\_INS\_379
CL\_INS\_379
CL\_INS\_379
CL\_INS\_379
CL\_INS\_379
CL\_INS\_379
CL\_INS\_379
CL\_INS\_379
CL\_INS\_379
CL\_INS\_379
CL\_INS\_379
CL\_INS\_379
CL\_INS\_379
CL\_INS\_379
CL\_INS\_379
CL\_INS\_379
CL\_INS\_379
CL\_INS\_379
CL\_INS\_379
CL\_INS\_379
CL\_INS\_379
CL\_INS\_379
CL\_INS\_379
CL\_INS\_379
CL\_INS\_379
CL\_INS\_379
CL\_INS\_379
CL\_INS\_379
CL\_INS\_379
CL\_INS\_379
CL\_INS\_379
CL\_INS\_379
CL\_INS\_379
CL\_INS\_379
CL\_INS\_379
CL\_INS\_379
CL\_INS\_379
CL\_INS\_379
CL\_INS\_379
CL\_INS\_379
CL\_INS\_379
CL\_INS\_379
CL\_INS\_379
CL\_INS\_382
CL\_INS\_382
CL\_INS\_382
CL\_INS\_382
CL\_INS\_379
CL\_INS\_368
CL\_INS\_382
CL\_INS\_382
CL\_INS\_382
CL\_INS\_207
CL\_INS\_382
CL\_INS\_382
CL\_INS\_379
CL\_INS\_382
CL\_INS\_379
CL\_INS\_379
CL\_INS\_379
CL\_INS\_379
CL\_INS\_379
CL\_INS\_382
CL\_INS\_86
CL\_INS\_20
CL\_INS\_382
CL\_INS\_86
CL\_INS\_382
CL\_INS\_382
CL\_INS\_382
CL\_INS\_86
CL\_INS\_86
CL\_INS\_382
CL\_INS\_382
CL\_INS\_382
CL\_INS\_382
CL\_INS\_382
CL\_INS\_382
CL\_INS\_129
CL\_INS\_382
CL\_INS\_382
CL\_INS\_382
CL\_INS\_379
CL\_INS\_379
CL\_INS\_379
CL\_INS\_382
CL\_INS\_382
CL\_INS\_382
CL\_INS\_382
CL\_INS\_382
CL\_INS\_382
CL\_INS\_382
CL\_INS\_382
CL\_INS\_382
CL\_INS\_382
CL\_INS\_382
CL\_INS\_382
CL\_INS\_379
CL\_INS\_379
CL\_INS\_382
CL\_INS\_382
CL\_INS\_382
CL\_INS\_382
CL\_INS\_382
CL\_INS\_382
CL\_INS\_382
CL\_INS\_382
CL\_INS\_382
CL\_INS\_382
CL\_INS\_382
CL\_INS\_382
CL\_INS\_382
CL\_INS\_382
CL\_INS\_382
CL\_INS\_382
CL\_INS\_382
CL\_INS\_382
CL\_INS\_382
CL\_INS\_382
CL\_INS\_382
CL\_INS\_382
CL\_INS\_382
CL\_INS\_382
CL\_INS\_382
CL\_INS\_382
CL\_INS\_382
CL\_INS\_382
CL\_INS\_382
CL\_INS\_382
CL\_INS\_382
CL\_INS\_382
CL\_INS\_382
CL\_INS\_382
CL\_INS\_382
CL\_INS\_382
CL\_INS\_382
CL\_INS\_382
CL\_INS\_382
CL\_INS\_382
CL\_INS\_382
CL\_INS\_382
CL\_INS\_382
CL\_INS\_382
CL\_INS\_382
CL\_INS\_382
CL\_INS\_382
CL\_INS\_382
CL\_INS\_382
CL\_INS\_382
CL\_INS\_382
CL\_INS\_382
CL\_INS\_382
CL\_INS\_382
CL\_INS\_382
CL\_INS\_382
CL\_INS\_10
CL\_INS\_10
CL\_INS\_10
CL\_INS\_382
CL\_INS\_382
CL\_INS\_379
CL\_INS\_382
CL\_INS\_382
CL\_INS\_379
CL\_INS\_382
CL\_INS\_382
CL\_INS\_382
CL\_INS\_382
CL\_INS\_382
CL\_INS\_382
CL\_INS\_86
CL\_INS\_382
CL\_INS\_382
CL\_INS\_379
CL\_INS\_379
CL\_INS\_379
CL\_INS\_379
CL\_INS\_379
CL\_INS\_379
CL\_INS\_379
CL\_INS\_379
CL\_INS\_379
CL\_INS\_379
CL\_INS\_379
CL\_INS\_379
CL\_INS\_379
CL\_INS\_379
CL\_INS\_382
CL\_INS\_379
CL\_INS\_382
CL\_INS\_382
CL\_INS\_382
CL\_INS\_382
CL\_INS\_382
CL\_INS\_382
CL\_INS\_382
CL\_INS\_382
CL\_INS\_382
CL\_INS\_382
CL\_INS\_382
CL\_INS\_382
CL\_INS\_382
CL\_INS\_382
CL\_INS\_382
CL\_INS\_382
CL\_INS\_382
CL\_INS\_382
CL\_INS\_379
CL\_INS\_379
CL\_INS\_379
CL\_INS\_379
CL\_INS\_128
CL\_INS\_128
CL\_INS\_382
CL\_INS\_382
CL\_INS\_207
CL\_INS\_382
CL\_INS\_382
CL\_INS\_379
CL\_INS\_379
CL\_INS\_379
CL\_INS\_379
CL\_INS\_379
CL\_INS\_382
CL\_INS\_382
CL\_INS\_382
CL\_INS\_382
CL\_INS\_99
CL\_INS\_379
CL\_INS\_379
CL\_INS\_86
CL\_INS\_99
CL\_INS\_86
CL\_INS\_204
CL\_INS\_204
CL\_INS\_204
CL\_INS\_204
CL\_INS\_204
CL\_INS\_204
CL\_INS\_204
CL\_INS\_146
CL\_INS\_99
CL\_INS\_99
CL\_INS\_99
CL\_INS\_99
CL\_INS\_146
CL\_INS\_379
CL\_INS\_379
CL\_INS\_379
CL\_INS\_382
CL\_INS\_99
CL\_INS\_379
CL\_INS\_382
CL\_INS\_207
CL\_INS\_382
CL\_INS\_382
CL\_INS\_382
CL\_INS\_382
CL\_INS\_382
CL\_INS\_382
CL\_INS\_382
CL\_INS\_20
CL\_INS\_207
CL\_INS\_207
CL\_INS\_20
CL\_INS\_382
CL\_INS\_379
CL\_INS\_379
CL\_INS\_379
CL\_INS\_379
CL\_INS\_379
CL\_INS\_382
CL\_INS\_379
CL\_INS\_382
CL\_INS\_382
CL\_INS\_382
CL\_INS\_379
CL\_INS\_379
CL\_INS\_379
CL\_INS\_379
CL\_INS\_379
CL\_INS\_146
CL\_INS\_382
CL\_INS\_382
CL\_INS\_382
CL\_INS\_382
CL\_INS\_382
CL\_INS\_382
CL\_INS\_382
CL\_INS\_382
CL\_INS\_382
CL\_INS\_382
CL\_INS\_382
CL\_INS\_382
CL\_INS\_379
CL\_INS\_79
CL\_INS\_99
CL\_INS\_128
CL\_INS\_379
CL\_INS\_379
CL\_INS\_99
CL\_INS\_20
CL\_INS\_379
CL\_INS\_379
CL\_INS\_379
CL\_INS\_379
CL\_INS\_385
CL\_INS\_99
CL\_INS\_99
CL\_INS\_382
CL\_INS\_379
CL\_INS\_382
CL\_INS\_382
CL\_INS\_382
CL\_INS\_382
CL\_INS\_207
CL\_INS\_382
CL\_INS\_204
CL\_INS\_382
CL\_INS\_382
CL\_INS\_382
CL\_INS\_382
CL\_INS\_382
CL\_INS\_382
CL\_INS\_204
CL\_INS\_207
CL\_INS\_204
CL\_INS\_379
CL\_INS\_204
CL\_INS\_207
CL\_INS\_99
CL\_INS\_99
CL\_INS\_207
CL\_INS\_207
CL\_INS\_86
CL\_INS\_204
CL\_INS\_204
CL\_INS\_204
CL\_INS\_379
CL\_INS\_379
CL\_INS\_204
CL\_INS\_20
CL\_INS\_379
CL\_INS\_379
CL\_INS\_379
CL\_INS\_204
CL\_INS\_204
CL\_INS\_204
CL\_INS\_207
CL\_INS\_204
CL\_INS\_20
CL\_INS\_20
CL\_INS\_379
CL\_INS\_204
CL\_INS\_204
CL\_INS\_20
CL\_INS\_20
CL\_INS\_204
CL\_INS\_20
CL\_INS\_379
CL\_INS\_382
CL\_INS\_20
CL\_INS\_382
CL\_INS\_379
CL\_INS\_207
CL\_INS\_379
CL\_INS\_382
CL\_INS\_382
CL\_INS\_382
CL\_INS\_382
CL\_INS\_382
CL\_INS\_382
CL\_INS\_382
CL\_INS\_382
CL\_INS\_379
CL\_INS\_379
CL\_INS\_379
CL\_INS\_379
CL\_INS\_379
CL\_INS\_207
CL\_INS\_207
CL\_INS\_207
CL\_INS\_207
CL\_INS\_207
CL\_INS\_379
CL\_INS\_382
CL\_INS\_382
CL\_INS\_207
CL\_INS\_207
CL\_INS\_379
CL\_INS\_379
CL\_INS\_379
CL\_INS\_207
CL\_INS\_368
CL\_INS\_136
CL\_INS\_207
CL\_INS\_99
CL\_INS\_99
CL\_INS\_382
CL\_INS\_382
CL\_INS\_382
CL\_INS\_382
CL\_INS\_382
CL\_INS\_382
CL\_INS\_379
CL\_INS\_382
CL\_INS\_382
CL\_INS\_382
CL\_INS\_382
CL\_INS\_382
CL\_INS\_382
CL\_INS\_382
CL\_INS\_382
CL\_INS\_379
CL\_INS\_379
CL\_INS\_379
CL\_INS\_379
CL\_INS\_379
CL\_INS\_382
CL\_INS\_382
CL\_INS\_382
CL\_INS\_382
CL\_INS\_99
CL\_INS\_99
CL\_INS\_99
CL\_INS\_99
CL\_INS\_149
CL\_INS\_149
CL\_INS\_149
CL\_INS\_99
CL\_INS\_99
CL\_INS\_149
CL\_INS\_149
CL\_INS\_99
CL\_INS\_149
CL\_INS\_149
CL\_INS\_149
CL\_INS\_149
CL\_INS\_382
CL\_INS\_382
CL\_INS\_379
CL\_INS\_379
CL\_INS\_382
CL\_INS\_382
CL\_INS\_382
CL\_INS\_379
CL\_INS\_379
CL\_INS\_382
CL\_INS\_382
CL\_INS\_382
CL\_INS\_382
CL\_INS\_382
CL\_INS\_382
CL\_INS\_382
CL\_INS\_379
CL\_INS\_382
CL\_INS\_86
CL\_INS\_99
CL\_INS\_86
CL\_INS\_382
CL\_INS\_99
CL\_INS\_99
CL\_INS\_99
CL\_INS\_99
CL\_INS\_379
CL\_INS\_99
CL\_INS\_86
CL\_INS\_99
CL\_INS\_86
CL\_INS\_86
CL\_INS\_149
CL\_INS\_86
CL\_INS\_20
CL\_INS\_385
CL\_INS\_99
CL\_INS\_385
CL\_INS\_99
CL\_INS\_20
CL\_INS\_385
CL\_INS\_20
CL\_INS\_207
CL\_INS\_207
CL\_INS\_207
CL\_INS\_379
CL\_INS\_379
CL\_INS\_379
CL\_INS\_379
CL\_INS\_207
CL\_INS\_99
CL\_INS\_86
CL\_INS\_86
CL\_INS\_99
CL\_INS\_385
CL\_INS\_385
CL\_INS\_379
CL\_INS\_379
CL\_INS\_379
CL\_INS\_379
CL\_INS\_146
CL\_INS\_385
CL\_INS\_385
CL\_INS\_146
CL\_INS\_385
CL\_INS\_385
CL\_INS\_385
CL\_INS\_379
CL\_INS\_385
CL\_INS\_385
CL\_INS\_379
CL\_INS\_382
CL\_INS\_382
CL\_INS\_86
CL\_INS\_382
CL\_INS\_99
CL\_INS\_379
CL\_INS\_99
CL\_INS\_99
CL\_INS\_99
CL\_INS\_379
CL\_INS\_86
CL\_INS\_237
CL\_INS\_87
CL\_INS\_128
CL\_INS\_379
CL\_INS\_379
CL\_INS\_379
CL\_INS\_382
CL\_INS\_382
CL\_INS\_379
CL\_INS\_382
CL\_INS\_382
CL\_INS\_382
CL\_INS\_382
CL\_INS\_382
CL\_INS\_382
CL\_INS\_382
CL\_INS\_382
CL\_INS\_382
CL\_INS\_382
CL\_INS\_379
CL\_INS\_382
CL\_INS\_382
CL\_INS\_382
CL\_INS\_379
CL\_INS\_379
CL\_INS\_379
CL\_INS\_379
CL\_INS\_379
CL\_INS\_379
CL\_INS\_379
CL\_INS\_379
CL\_INS\_99
CL\_INS\_382
CL\_INS\_86
CL\_INS\_86
CL\_INS\_99
CL\_INS\_99
CL\_INS\_207
CL\_INS\_207
CL\_INS\_146
CL\_INS\_207
CL\_INS\_86
CL\_INS\_385
CL\_INS\_382
CL\_INS\_382
CL\_INS\_382
CL\_INS\_382
Cluster ID


CL\_31736
CL\_31735
CL\_31734
CL\_31733
CL\_12701
CL\_28187
CL\_28186
CL\_28185
CL\_28184
CL\_28183
CL\_28182
CL\_28181
CL\_28180
CL\_28179
CL\_12115
CL\_13612
CL\_33638
CL\_33639
CL\_6031
CL\_33446
CL\_33056
CL\_33057
CL\_23375
CL\_17779
CL\_17778
CL\_1519
CL\_23759
CL\_34077
CL\_29274
CL\_29968
CL\_29967
CL\_29966
CL\_16516
CL\_20412
CL\_30135
CL\_30134
CL\_6032
CL\_6033
CL\_12116
CL\_11402
CL\_13486
CL\_17074
CL\_5363
CL\_5362
CL\_5361
CL\_4400
CL\_31764
CL\_26142
CL\_31763
CL\_9949
CL\_8962
CL\_17549
CL\_11731
CL\_9229
CL\_31762
CL\_31761
CL\_31760
CL\_9078
CL\_24460
CL\_9080
CL\_5351
CL\_28278
CL\_509
CL\_27894
CL\_13507
CL\_21777
CL\_6655
CL\_9117
CL\_31759
CL\_17540
CL\_13972
CL\_17539
CL\_9599
CL\_9600
CL\_9601
CL\_28191
CL\_6034
CL\_33058
CL\_20109
CL\_8434
CL\_8335
CL\_2284
CL\_21645
CL\_12118
CL\_28192
CL\_28193
CL\_28194
CL\_28195
CL\_28196
CL\_16606
CL\_12336
CL\_12337
CL\_12338
CL\_12339
CL\_12340
CL\_16609
CL\_16610
CL\_12341
CL\_16611
CL\_12343
CL\_12344
CL\_12345
CL\_16612
CL\_16613
CL\_16614
CL\_14825
CL\_13071
CL\_15696
CL\_15697
CL\_31758
CL\_31757
CL\_31756
CL\_31755
CL\_31754
CL\_31753
CL\_16617
CL\_8700
CL\_1496
CL\_23049
CL\_23048
CL\_23047
CL\_4533
CL\_4415
CL\_4416
CL\_4417
CL\_27463
CL\_34078
CL\_6740
CL\_7372
CL\_27462
CL\_8701
CL\_31455
CL\_31454
CL\_31453
CL\_31452
CL\_23526
CL\_28309
CL\_31752
CL\_31751
CL\_31750
CL\_31749
CL\_31748
CL\_31747
CL\_31746
CL\_31745
CL\_31744
CL\_31743
CL\_31742
CL\_31741
CL\_31740
CL\_31739
CL\_16351
CL\_16445
CL\_32386
CL\_32387
CL\_32388
CL\_32389
CL\_15773
CL\_15774
CL\_15775
CL\_16444
CL\_27159
CL\_27158
CL\_27157
CL\_27156
CL\_16616
CL\_13561
CL\_11920
CL\_13560
CL\_8698
CL\_12758
CL\_27155
CL\_26457
CL\_20242
CL\_4534
CL\_533
CL\_6045
CL\_4414
CL\_4532
CL\_27154
CL\_27153
CL\_27152
CL\_27151
CL\_27150
CL\_27149
CL\_27148
CL\_27147
CL\_27146
CL\_4628
CL\_27145
CL\_27144
CL\_27143
CL\_27142
CL\_27141
CL\_27140
CL\_27139
CL\_27138
CL\_27137
CL\_27136
CL\_27135
CL\_27134
CL\_6055
CL\_20784
CL\_20783
CL\_28197
CL\_20782
CL\_28198
CL\_28199
CL\_28200
CL\_28201
CL\_28202
CL\_28203
CL\_28204
CL\_28205
CL\_28206
CL\_28207
CL\_28208
CL\_28209
CL\_28210
CL\_28211
CL\_8707
CL\_8708
CL\_8709
CL\_8710
CL\_8172
CL\_8171
CL\_8170
CL\_8850
CL\_8849
CL\_8848
CL\_8847
CL\_37422
CL\_37423
CL\_32392
CL\_28212
CL\_28213
CL\_28214
CL\_28215
CL\_28216
CL\_28217
CL\_28218
CL\_28219
CL\_17468
CL\_9128
CL\_28220
CL\_28221
CL\_28222
CL\_28223
CL\_28224
CL\_28225
CL\_28226
CL\_28227
CL\_33640
CL\_33641
CL\_33642
CL\_33643
CL\_33644
CL\_33645
CL\_33646
CL\_33647
CL\_33648
CL\_6028
CL\_6027
CL\_6026
CL\_33649
CL\_6025
CL\_6024
CL\_6023
CL\_33650
CL\_33651
CL\_33652
CL\_33653
CL\_33654
CL\_33655
CL\_33656
CL\_33657
CL\_33658
CL\_33659
CL\_33660
CL\_33661
CL\_33662
CL\_33663
CL\_33664
CL\_33665
CL\_33666
CL\_33667
CL\_33668
CL\_33669
CL\_33670
CL\_33671
CL\_33672
CL\_33673
CL\_33674
CL\_33675
CL\_33676
CL\_33677
CL\_33678
CL\_33679
CL\_33680
CL\_33681
CL\_33682
CL\_33683
CL\_33684
CL\_33685
CL\_33686
CL\_33687
CL\_33688
CL\_33689
CL\_33690
CL\_33691
CL\_33692
CL\_33693
CL\_33694
CL\_33695
CL\_33696
CL\_33697
CL\_33698
CL\_33699
CL\_33700
CL\_33701
CL\_33702
CL\_33703
CL\_33704
CL\_33705
CL\_33706
CL\_15990
CL\_15991
CL\_12117
CL\_11939
CL\_25502
CL\_17186
CL\_10270
CL\_11938
CL\_2283
CL\_14633
CL\_5427
CL\_5426
CL\_17777
CL\_2282
CL\_29660
CL\_29659
CL\_29658
CL\_29657
CL\_29656
CL\_2281
CL\_7621
CL\_17690
CL\_5425
CL\_10174
CL\_6035
CL\_12120
CL\_7338
CL\_23518
CL\_23519
CL\_7543
CL\_12121
CL\_7542
CL\_1095
CL\_1096
CL\_2280
CL\_27162
CL\_4658
CL\_12800
CL\_12119
CL\_17776
CL\_17775
CL\_16806
CL\_15992
CL\_15993
CL\_8576
CL\_8577
CL\_10175
CL\_5424
CL\_5423
CL\_15994
CL\_1093
CL\_1094
CL\_6036
CL\_16807
CL\_16808
CL\_16809
CL\_6037
CL\_4656
CL\_13564
CL\_15995
CL\_15996
CL\_15997
CL\_15998
CL\_15999
CL\_16000
CL\_9291
CL\_9290
CL\_9289
CL\_9288
CL\_9287
CL\_9286
CL\_10804
CL\_16001
CL\_16002
CL\_16003
CL\_9284
CL\_9283
CL\_9282
CL\_9281
CL\_9280
CL\_9279
CL\_9278
CL\_9277
CL\_9276
CL\_9275
CL\_9274
CL\_9273
CL\_9272
CL\_9271
CL\_9270
CL\_9269
CL\_16004
CL\_16005
CL\_9266
CL\_4644
CL\_9265
CL\_9264
CL\_9263
CL\_9262
CL\_9261
CL\_16006
CL\_16007
CL\_9258
CL\_9257
CL\_16008
CL\_9256
CL\_9254
CL\_9253
CL\_9252
CL\_16009
CL\_9251
CL\_9250
CL\_9249
CL\_9248
CL\_9247
CL\_4655
CL\_12768
CL\_30614
CL\_19351
CL\_2279
CL\_16810
CL\_2278
CL\_19352
CL\_21646
CL\_21647
CL\_5422
CL\_5421
CL\_7024
CL\_12122
CL\_21648
CL\_16811
CL\_16812
CL\_16813
CL\_16814
CL\_16815
CL\_16816
CL\_16817
CL\_16818
CL\_16819
CL\_16820
CL\_16821
CL\_16822
CL\_16823
CL\_16824
CL\_10170
CL\_30615
CL\_5420
CL\_5419
CL\_1511
CL\_16517
CL\_1510
CL\_1509
CL\_1508
CL\_1507
CL\_1506
CL\_26544
CL\_7494
CL\_1505
CL\_1504
CL\_1503
CL\_1502
CL\_1500
CL\_1499
CL\_7495
CL\_23374
CL\_23373
CL\_23372
CL\_23371
CL\_8050
CL\_8051
CL\_531
CL\_4651
CL\_20411
CL\_19353
CL\_19354
CL\_35442
CL\_35441
CL\_35440
CL\_36289
CL\_36288
CL\_7119
CL\_8185
CL\_8184
CL\_8696
CL\_8183
CL\_37420
CL\_37421
CL\_8180
CL\_8179
CL\_8178
CL\_1098
CL\_5395
CL\_5394
CL\_1099
CL\_5410
CL\_5409
CL\_5408
CL\_8177
CL\_8176
CL\_8175
CL\_8174
CL\_8173
CL\_6038
CL\_6039
CL\_6040
CL\_6041
CL\_6042
CL\_6043
CL\_6044
CL\_4535
CL\_12123
CL\_16967
CL\_17234
CL\_17235
CL\_16010
CL\_10168
CL\_8689
CL\_7339
CL\_21611
CL\_10165
CL\_10164
CL\_10163
CL\_17774
CL\_27161
CL\_27160
CL\_31459
CL\_31458
CL\_31457
CL\_9096
CL\_31456
CL\_16011
CL\_7540
CL\_7539
CL\_17236
CL\_17237
CL\_17238
CL\_17239
CL\_17240
CL\_12761
CL\_13562
CL\_12762
CL\_12760
CL\_12996
CL\_7538
CL\_526
CL\_7118
CL\_7117
CL\_7537
CL\_10323
CL\_12006
CL\_4469
CL\_17241
CL\_17242
CL\_17243
CL\_17244
CL\_17245
CL\_17246
CL\_17247
CL\_10474
CL\_17248
CL\_17249
CL\_17250
CL\_17251
CL\_17252
CL\_23470
CL\_12382
CL\_6784
CL\_29655
CL\_13000
CL\_7534
CL\_17730
CL\_11311
CL\_12124
CL\_5416
CL\_5415
CL\_6770
CL\_6664
CL\_9009
CL\_6783
CL\_529
CL\_1497
CL\_5414
CL\_26667
CL\_5412
CL\_20410
CL\_5411
CL\_11310
CL\_10974
CL\_10973
CL\_11309
CL\_11308
CL\_11307
CL\_5405
CL\_5404
CL\_5403
CL\_33919
CL\_33918
CL\_5400
CL\_26629
CL\_33917
CL\_33916
CL\_33915
CL\_5393
CL\_1100
CL\_1101
CL\_5392
CL\_1104
CL\_19416
CL\_19417
CL\_20409
CL\_1103
CL\_2277
CL\_26630
CL\_26631
CL\_4604
CL\_21665
CL\_33914
CL\_17747
CL\_17746
CL\_15241
CL\_37243
CL\_9122
CL\_23370
CL\_7533
CL\_9123
CL\_15776
CL\_15777
CL\_15778
CL\_15779
CL\_15780
CL\_15781
CL\_35439
CL\_35438
CL\_35437
CL\_35436
CL\_37244
CL\_7532
CL\_7531
CL\_7530
CL\_7529
CL\_7528
CL\_37245
CL\_7527
CL\_6741
CL\_7526
CL\_7525
CL\_29965
CL\_37246
CL\_37247
CL\_7524
CL\_29273
CL\_4484
CL\_7523
CL\_7522
CL\_8155
CL\_16012
CL\_12698
CL\_16013
CL\_16014
CL\_16015
CL\_16016
CL\_36395
CL\_16017
CL\_16018
CL\_16019
CL\_16020
CL\_16021
CL\_16022
CL\_16023
CL\_16024
CL\_16825
CL\_36394
CL\_36393
CL\_36392
CL\_36391
CL\_4531
CL\_4530
CL\_4529
CL\_4528
CL\_6046
CL\_6047
CL\_6048
CL\_6049
CL\_4689
CL\_4688
CL\_4687
CL\_6050
CL\_6051
CL\_4685
CL\_4684
CL\_6052
CL\_4682
CL\_4681
CL\_4679
CL\_4678
CL\_7115
CL\_6461
CL\_16671
CL\_16670
CL\_4526
CL\_4422
CL\_6742
CL\_35435
CL\_35434
CL\_4629
CL\_4525
CL\_4524
CL\_6743
CL\_7371
CL\_4523
CL\_4522
CL\_35433
CL\_11913
CL\_7027
CL\_13790
CL\_4430
CL\_4521
CL\_4520
CL\_13789
CL\_4485
CL\_7112
CL\_29272
CL\_6747
CL\_4488
CL\_4489
CL\_1324
CL\_4490
CL\_12560
CL\_1495
CL\_10475
CL\_11286
CL\_7521
CL\_16850
CL\_4618
CL\_1494
CL\_20457
CL\_6766
CL\_1493
CL\_1492
CL\_1491
CL\_29271
CL\_29270
CL\_34214
CL\_34215
CL\_11956
CL\_8586
CL\_10526
CL\_6782
CL\_10520
CL\_31340
CL\_37805
CL\_12362
CL\_12363
CL\_23525
CL\_12364
CL\_13613
CL\_21277
CL\_29671
CL\_13614
CL\_12125
CL\_12126
CL\_12127
CL\_37248
CL\_13791
CL\_25199
CL\_25200
CL\_4518
CL\_4517
CL\_7025
CL\_8712
CL\_4620
CL\_32390
CL\_17033
CL\_13075
CL\_12385
CL\_32391
CL\_4515
CL\_4514
CL\_4513
CL\_10341
CL\_15783
CL\_15784
CL\_15785
CL\_16025
CL\_16026
CL\_16826
CL\_4418
CL\_4419
CL\_4420
CL\_4421
CL\_21649
CL\_21650
CL\_21651
CL\_21652
CL\_21653
CL\_16027
CL\_16827
CL\_4527
CL\_16028
CL\_16029
CL\_16828
CL\_16829
CL\_16830
CL\_16831
CL\_16832
CL\_16833
CL\_16030
CL\_16834
CL\_4431
CL\_4432
CL\_8585
CL\_7023
CL\_8587
CL\_8588
CL\_17773
CL\_17772
CL\_10967
CL\_6053
CL\_6054
CL\_16031
CL\_4423
CL\_4424
CL\_4425
CL\_4426
