## Supplementary material for "A novel method for integrating genomic and Tn-Seq data to identify common *in vivo* fitness mechanisms across multiple bacterial species": S1 Dataset: CL_INS_380.html

Legend

 Mobile +extrachromosomalelementfunctions
 Regulatoryfunctions
 Hypothetical
 All EssentialGenes
 Other
 All VFDB Genes

FULL


WINDOWSVGPNG

Trim RowsRemove SingletonsSave Fasta

CL\_4191


CL\_4191


CL\_4191


CL\_4191


CL\_4191


CL\_4191


CL\_4191


CL\_4191


CL\_4078


CL\_4191


CL\_4191

HighlightSelectShow Genomes


121

CL\_4193


71

CL\_4193


59

CL\_4193


7

Break


4

CL\_4194


3

Break


2

CL\_4193


1

CL\_4193


1

CL\_4193


1

CL\_4193


1

CL\_4165

fGI ID


CL\_INS\_380
CL\_INS\_380
CL\_INS\_380
CL\_INS\_380
CL\_INS\_380
CL\_INS\_380
Cluster ID


CL\_30953
CL\_22042
CL\_14413
CL\_4192
CL\_27587
CL\_5467
