## Supplementary material for "A novel method for integrating genomic and Tn-Seq data to identify common *in vivo* fitness mechanisms across multiple bacterial species": S1 Dataset: CL_INS_381.html

Legend

 Mobile +extrachromosomalelementfunctions
 Regulatoryfunctions
 Hypothetical
 Other
 All VFDB Genes

FULL


WINDOWSVGPNG

Trim RowsRemove SingletonsSave Fasta

CL\_4197


CL\_4197


CL\_4197


CL\_4197


CL\_4197


CL\_4197


CL\_4197


CL\_4197


CL\_4197


CL\_4197


CL\_4194


CL\_4197


CL\_4197


CL\_4197


CL\_4197


CL\_4197

HighlightSelectShow Genomes


171

CL\_4198


30

CL\_4198


18

CL\_4198


12

CL\_4198


8

Break


5

CL\_4198


3

CL\_4198


2

CL\_4198


2

Break


1

CL\_4198


1

CL\_4198


1

CL\_4198


1

CL\_4199


1

CL\_4198


1

CL\_4198


1

CL\_4198

fGI ID


CL\_INS\_381
CL\_INS\_381
CL\_INS\_381
CL\_INS\_381
CL\_INS\_381
CL\_INS\_381
CL\_INS\_381
CL\_INS\_381
CL\_INS\_381
CL\_INS\_381
CL\_INS\_381
CL\_INS\_381
CL\_INS\_155
CL\_INS\_204
CL\_INS\_204
CL\_INS\_381
CL\_INS\_155
CL\_INS\_182
CL\_INS\_182
CL\_INS\_182
CL\_INS\_182
CL\_INS\_381
CL\_INS\_381
CL\_INS\_381
CL\_INS\_86
CL\_INS\_381
CL\_INS\_381
CL\_INS\_381
CL\_INS\_182
CL\_INS\_182
CL\_INS\_182
CL\_INS\_155
CL\_INS\_155
CL\_INS\_182
CL\_INS\_182
CL\_INS\_182
CL\_INS\_182
CL\_INS\_182
CL\_INS\_381
CL\_INS\_182
CL\_INS\_182
CL\_INS\_182
CL\_INS\_182
CL\_INS\_182
CL\_INS\_182
CL\_INS\_182
CL\_INS\_155
CL\_INS\_155
CL\_INS\_155
CL\_INS\_155
CL\_INS\_155
CL\_INS\_155
CL\_INS\_155
CL\_INS\_155
CL\_INS\_182
CL\_INS\_182
CL\_INS\_182
CL\_INS\_182
CL\_INS\_182
CL\_INS\_182
CL\_INS\_155
CL\_INS\_155
CL\_INS\_155
CL\_INS\_381
CL\_INS\_182
Cluster ID


CL\_7262
CL\_7263
CL\_23141
CL\_13393
CL\_13394
CL\_10448
CL\_10449
CL\_10450
CL\_10451
CL\_10452
CL\_10453
CL\_11781
CL\_11037
CL\_1105
CL\_10867
CL\_36028
CL\_11035
CL\_11034
CL\_11033
CL\_11032
CL\_11031
CL\_36027
CL\_36026
CL\_36025
CL\_8227
CL\_36024
CL\_36023
CL\_36022
CL\_11028
CL\_11027
CL\_11026
CL\_11025
CL\_11024
CL\_11023
CL\_11022
CL\_11021
CL\_11020
CL\_11019
CL\_36004
CL\_11018
CL\_11017
CL\_11016
CL\_11015
CL\_11014
CL\_11013
CL\_11012
CL\_11011
CL\_11010
CL\_11009
CL\_11008
CL\_11007
CL\_11006
CL\_11005
CL\_11004
CL\_11003
CL\_11002
CL\_11001
CL\_11000
CL\_10999
CL\_10998
CL\_10911
CL\_10912
CL\_10913
CL\_31901
CL\_10993
