## Supplementary material for "A novel method for integrating genomic and Tn-Seq data to identify common *in vivo* fitness mechanisms across multiple bacterial species": S1 Dataset: CL_INS_383.html

Legend

 Mobile +extrachromosomalelementfunctions
 Regulatoryfunctions
 Hypothetical
 DNA Metabolism
 All EssentialGenes
 AntibioticResistance
 All Fitness Genes
 Proteinsynthesis/fate
 Other
 EnergyMetabolism
 Transport +binding proteins
 All VFDB Genes

FULL


WINDOWSVGPNG

Trim RowsRemove SingletonsSave Fasta

CL\_4519


CL\_4519


CL\_4519


CL\_4519


CL\_4519


CL\_4427


CL\_4519


CL\_1529


CL\_1300


CL\_4519


CL\_4519


CL\_1820


CL\_4519


CL\_4180


CL\_1300


CL\_4180


CL\_1084


CL\_3870


CL\_4519


CL\_1820


CL\_1207


CL\_4519


CL\_1072


CL\_1072


CL\_4180


CL\_1529


CL\_1529


CL\_230


CL\_4519


CL\_4519


CL\_4519


CL\_4427


CL\_1300


CL\_230


CL\_4519


CL\_4519


CL\_1823


CL\_4519


CL\_4519


CL\_1529


CL\_1207


CL\_1529


CL\_1529


CL\_230


CL\_1529


CL\_1300


CL\_500


CL\_2555


CL\_4519


CL\_1300


CL\_4519


CL\_1207


CL\_4427


CL\_2555


CL\_4519


CL\_4180


CL\_4519


CL\_1300


CL\_4519


CL\_4519


CL\_724


CL\_1300


CL\_4181


CL\_1529


CL\_1300


CL\_4519


CL\_4519


CL\_1300


CL\_1529


CL\_3870


CL\_1072


CL\_1531


CL\_1300


CL\_3870


CL\_4519


CL\_4519


CL\_1820


CL\_4519


CL\_1207


CL\_4519


CL\_1207


CL\_230


CL\_1300


CL\_1529


CL\_4519


CL\_1300


CL\_1529


CL\_1917


CL\_4519


CL\_1529


CL\_1529


CL\_4519


CL\_1300


CL\_4519


CL\_4519


CL\_4519


CL\_4519


CL\_1300


CL\_1300


CL\_1300


CL\_1300


CL\_4074


CL\_4519


CL\_1084


CL\_230


CL\_1529


CL\_1529


CL\_1529


CL\_1300


CL\_1529


CL\_4519


CL\_4519


CL\_4519


CL\_1300


CL\_1084


CL\_1300


CL\_1529


CL\_1300


CL\_1529


CL\_1452


CL\_230


CL\_4519


CL\_4519


CL\_4519


CL\_4519


CL\_4180


CL\_4519


CL\_4519


CL\_724


CL\_230


CL\_1917


CL\_230


CL\_4519


CL\_1300


CL\_4519


CL\_1529


CL\_1207


CL\_4180


CL\_1529


CL\_4519


CL\_4180


CL\_1529


CL\_4519


CL\_2555


CL\_1529


CL\_4519

HighlightSelectShow Genomes


125

CL\_4486


9

CL\_4487


7

CL\_1109


2

CL\_4487


2

CL\_4516


2

CL\_4486


2

CL\_1208


2

CL\_4486


2

CL\_4486


1

CL\_725


1

CL\_923


1

CL\_4486


1

CL\_270


1

CL\_4486


1

CL\_4486


1

CL\_4486


1

CL\_4486


1

CL\_4486


1

CL\_1330


1

CL\_4486


1

CL\_4486


1

CL\_1110


1

CL\_4486


1

CL\_4486


1

CL\_4486


1

CL\_4486


1

CL\_4486


1

CL\_4486


1

CL\_1490


1

CL\_725


1

CL\_1490


1

CL\_4486


1

CL\_4486


1

CL\_4486


1

CL\_4516


1

CL\_538


1

CL\_4486


1

Break


1

CL\_538


1

CL\_4486


1

CL\_4486


1

CL\_4486


1

CL\_4486


1

CL\_4486


1

CL\_4486


1

CL\_4486


1

CL\_4486


1

CL\_4486


1

CL\_1109


1

CL\_4486


1

CL\_1208


1

CL\_4486


1

CL\_4486


1

CL\_4486


1

CL\_538


1

CL\_4486


1

CL\_1110


1

CL\_4486


1

CL\_1825


1

CL\_538


1

CL\_4486


1

CL\_4486


1

CL\_4486


1

CL\_4486


1

CL\_4486


1

CL\_1330


1

CL\_4487


1

CL\_4486


1

CL\_4486


1

CL\_4486


1

CL\_4486


1

CL\_4486


1

CL\_4486


1

CL\_4486


1

CL\_538


1

CL\_4516


1

CL\_4486


1

CL\_1490


1

CL\_4486


1

Break


1

CL\_4486


1

CL\_4486


1

CL\_4486


1

CL\_4486


1

CL\_4516


1

CL\_4486


1

CL\_4486


1

CL\_4486


1

CL\_4181


1

CL\_4486


1

CL\_4486


1

CL\_1208


1

CL\_4486


1

CL\_538


1

CL\_538


1

CL\_1208


1

CL\_1110


1

CL\_4486


1

CL\_4486


1

CL\_4486


1

CL\_4486


1

CL\_4486


1

CL\_1490


1

CL\_4486


1

CL\_4486


1

CL\_4486


1

CL\_4486


1

CL\_4486


1

CL\_4486


1

CL\_4486


1

CL\_1208


1

CL\_538


1

CL\_1490


1

CL\_4486


1

CL\_4486


1

CL\_4486


1

CL\_4486


1

CL\_4486


1

CL\_4486


1

CL\_4486


1

CL\_4486


1

CL\_4487


1

CL\_725


1

CL\_1915


1

CL\_538


1

CL\_4486


1

CL\_1110


1

CL\_538


1

CL\_4486


1

CL\_4486


1

CL\_4486


1

CL\_4486


1

CL\_538


1

CL\_4486


1

CL\_1209


1

CL\_4486


1

CL\_4486


1

CL\_4486


1

CL\_4486


1

CL\_725


1

CL\_4486


1

CL\_4486


1

CL\_1915


1

CL\_4486


1

CL\_4486


1

CL\_4181

fGI ID


CL\_INS\_383
CL\_INS\_147
CL\_INS\_383
CL\_INS\_383
CL\_INS\_383
CL\_INS\_136
CL\_INS\_99
CL\_INS\_383
CL\_INS\_85
CL\_INS\_99
CL\_INS\_382
CL\_INS\_99
CL\_INS\_99
CL\_INS\_99
CL\_INS\_99
CL\_INS\_99
CL\_INS\_382
CL\_INS\_382
CL\_INS\_382
CL\_INS\_382
CL\_INS\_146
CL\_INS\_146
CL\_INS\_146
CL\_INS\_207
CL\_INS\_146
CL\_INS\_146
CL\_INS\_146
CL\_INS\_146
CL\_INS\_146
CL\_INS\_146
CL\_INS\_146
CL\_INS\_383
CL\_INS\_20
CL\_INS\_20
CL\_INS\_20
CL\_INS\_247
CL\_INS\_247
CL\_INS\_247
CL\_INS\_247
CL\_INS\_247
CL\_INS\_247
CL\_INS\_247
CL\_INS\_20
CL\_INS\_20
CL\_INS\_20
CL\_INS\_247
CL\_INS\_247
CL\_INS\_247
CL\_INS\_247
CL\_INS\_70
CL\_INS\_70
CL\_INS\_247
CL\_INS\_247
CL\_INS\_247
CL\_INS\_247
CL\_INS\_247
CL\_INS\_247
CL\_INS\_247
CL\_INS\_247
CL\_INS\_247
CL\_INS\_247
CL\_INS\_30
CL\_INS\_70
CL\_INS\_146
CL\_INS\_382
CL\_INS\_382
CL\_INS\_382
CL\_INS\_207
CL\_INS\_382
CL\_INS\_382
CL\_INS\_207
CL\_INS\_146
CL\_INS\_146
CL\_INS\_382
CL\_INS\_382
CL\_INS\_382
CL\_INS\_382
CL\_INS\_382
CL\_INS\_382
CL\_INS\_382
CL\_INS\_382
CL\_INS\_382
CL\_INS\_382
CL\_INS\_20
CL\_INS\_379
CL\_INS\_379
CL\_INS\_379
CL\_INS\_379
CL\_INS\_382
CL\_INS\_207
CL\_INS\_368
CL\_INS\_340
CL\_INS\_207
CL\_INS\_207
CL\_INS\_207
CL\_INS\_207
CL\_INS\_207
CL\_INS\_382
CL\_INS\_379
CL\_INS\_379
CL\_INS\_379
CL\_INS\_382
CL\_INS\_382
CL\_INS\_340
CL\_INS\_237
CL\_INS\_207
CL\_INS\_207
CL\_INS\_207
CL\_INS\_207
CL\_INS\_207
CL\_INS\_207
CL\_INS\_207
CL\_INS\_382
CL\_INS\_382
CL\_INS\_207
CL\_INS\_382
CL\_INS\_384
CL\_INS\_382
CL\_INS\_382
CL\_INS\_382
CL\_INS\_382
CL\_INS\_382
CL\_INS\_382
CL\_INS\_379
CL\_INS\_382
CL\_INS\_384
CL\_INS\_382
CL\_INS\_382
CL\_INS\_128
CL\_INS\_128
CL\_INS\_128
CL\_INS\_128
CL\_INS\_382
CL\_INS\_382
CL\_INS\_382
CL\_INS\_382
CL\_INS\_382
CL\_INS\_382
CL\_INS\_382
CL\_INS\_382
CL\_INS\_382
CL\_INS\_382
CL\_INS\_382
CL\_INS\_382
CL\_INS\_382
CL\_INS\_382
CL\_INS\_382
CL\_INS\_382
CL\_INS\_382
CL\_INS\_382
CL\_INS\_382
CL\_INS\_382
CL\_INS\_382
CL\_INS\_382
CL\_INS\_382
CL\_INS\_382
CL\_INS\_382
CL\_INS\_382
CL\_INS\_382
CL\_INS\_382
CL\_INS\_382
CL\_INS\_382
CL\_INS\_382
CL\_INS\_382
CL\_INS\_382
CL\_INS\_382
CL\_INS\_382
CL\_INS\_382
CL\_INS\_382
CL\_INS\_382
CL\_INS\_382
CL\_INS\_382
CL\_INS\_382
CL\_INS\_382
CL\_INS\_382
CL\_INS\_382
CL\_INS\_382
CL\_INS\_382
CL\_INS\_382
CL\_INS\_382
CL\_INS\_382
CL\_INS\_382
CL\_INS\_85
CL\_INS\_382
CL\_INS\_382
CL\_INS\_382
CL\_INS\_382
CL\_INS\_382
CL\_INS\_382
CL\_INS\_382
CL\_INS\_382
CL\_INS\_382
CL\_INS\_382
CL\_INS\_382
CL\_INS\_382
CL\_INS\_382
CL\_INS\_382
CL\_INS\_382
CL\_INS\_382
CL\_INS\_382
CL\_INS\_382
CL\_INS\_382
CL\_INS\_382
CL\_INS\_382
CL\_INS\_85
CL\_INS\_382
CL\_INS\_382
CL\_INS\_382
CL\_INS\_382
CL\_INS\_382
CL\_INS\_382
CL\_INS\_382
CL\_INS\_382
CL\_INS\_42
CL\_INS\_382
CL\_INS\_382
CL\_INS\_382
CL\_INS\_382
CL\_INS\_382
CL\_INS\_382
CL\_INS\_382
CL\_INS\_382
CL\_INS\_382
CL\_INS\_382
CL\_INS\_382
CL\_INS\_382
CL\_INS\_382
CL\_INS\_382
CL\_INS\_382
CL\_INS\_382
CL\_INS\_382
CL\_INS\_382
CL\_INS\_108
CL\_INS\_382
CL\_INS\_382
CL\_INS\_382
CL\_INS\_86
CL\_INS\_204
CL\_INS\_86
CL\_INS\_99
CL\_INS\_86
CL\_INS\_99
CL\_INS\_99
CL\_INS\_99
CL\_INS\_99
CL\_INS\_99
CL\_INS\_99
CL\_INS\_99
CL\_INS\_99
CL\_INS\_382
CL\_INS\_146
CL\_INS\_146
CL\_INS\_382
CL\_INS\_382
CL\_INS\_382
CL\_INS\_382
CL\_INS\_382
CL\_INS\_382
CL\_INS\_382
CL\_INS\_382
CL\_INS\_382
CL\_INS\_382
CL\_INS\_384
CL\_INS\_382
CL\_INS\_382
CL\_INS\_382
CL\_INS\_382
CL\_INS\_382
CL\_INS\_382
CL\_INS\_382
CL\_INS\_382
CL\_INS\_382
CL\_INS\_382
CL\_INS\_382
CL\_INS\_382
CL\_INS\_128
CL\_INS\_382
CL\_INS\_384
CL\_INS\_382
CL\_INS\_382
CL\_INS\_384
CL\_INS\_99
CL\_INS\_99
CL\_INS\_99
CL\_INS\_382
CL\_INS\_382
CL\_INS\_382
CL\_INS\_382
CL\_INS\_382
CL\_INS\_382
CL\_INS\_382
CL\_INS\_382
CL\_INS\_384
CL\_INS\_382
CL\_INS\_108
CL\_INS\_108
CL\_INS\_108
CL\_INS\_237
CL\_INS\_108
CL\_INS\_136
CL\_INS\_384
CL\_INS\_108
CL\_INS\_108
CL\_INS\_382
CL\_INS\_108
CL\_INS\_382
CL\_INS\_382
CL\_INS\_382
CL\_INS\_99
CL\_INS\_382
CL\_INS\_382
CL\_INS\_382
CL\_INS\_382
CL\_INS\_382
CL\_INS\_128
CL\_INS\_128
CL\_INS\_128
CL\_INS\_128
CL\_INS\_20
CL\_INS\_20
CL\_INS\_20
CL\_INS\_20
CL\_INS\_108
CL\_INS\_108
CL\_INS\_108
CL\_INS\_382
CL\_INS\_382
CL\_INS\_382
CL\_INS\_382
CL\_INS\_382
CL\_INS\_382
CL\_INS\_382
CL\_INS\_382
CL\_INS\_382
CL\_INS\_382
CL\_INS\_382
CL\_INS\_382
CL\_INS\_382
CL\_INS\_382
CL\_INS\_382
CL\_INS\_382
CL\_INS\_382
CL\_INS\_382
CL\_INS\_99
CL\_INS\_382
CL\_INS\_99
CL\_INS\_99
CL\_INS\_99
CL\_INS\_99
CL\_INS\_382
CL\_INS\_382
CL\_INS\_382
CL\_INS\_379
CL\_INS\_99
CL\_INS\_384
CL\_INS\_384
CL\_INS\_382
CL\_INS\_382
CL\_INS\_382
CL\_INS\_382
CL\_INS\_382
CL\_INS\_382
CL\_INS\_382
CL\_INS\_382
CL\_INS\_382
CL\_INS\_382
CL\_INS\_382
CL\_INS\_382
CL\_INS\_382
CL\_INS\_382
CL\_INS\_382
CL\_INS\_20
CL\_INS\_20
CL\_INS\_382
CL\_INS\_129
CL\_INS\_382
CL\_INS\_383
CL\_INS\_207
CL\_INS\_60
CL\_INS\_207
CL\_INS\_382
CL\_INS\_382
CL\_INS\_383
CL\_INS\_383
CL\_INS\_383
CL\_INS\_382
CL\_INS\_382
CL\_INS\_382
CL\_INS\_99
CL\_INS\_99
CL\_INS\_382
CL\_INS\_382
CL\_INS\_382
CL\_INS\_382
CL\_INS\_382
CL\_INS\_382
CL\_INS\_382
CL\_INS\_382
CL\_INS\_382
CL\_INS\_382
CL\_INS\_136
CL\_INS\_384
CL\_INS\_108
CL\_INS\_108
CL\_INS\_204
CL\_INS\_382
CL\_INS\_204
CL\_INS\_382
CL\_INS\_382
CL\_INS\_382
CL\_INS\_108
CL\_INS\_108
CL\_INS\_108
CL\_INS\_382
CL\_INS\_382
CL\_INS\_382
CL\_INS\_108
CL\_INS\_382
CL\_INS\_99
CL\_INS\_154
CL\_INS\_146
CL\_INS\_382
CL\_INS\_382
CL\_INS\_382
CL\_INS\_136
CL\_INS\_136
CL\_INS\_108
CL\_INS\_384
CL\_INS\_136
CL\_INS\_108
CL\_INS\_136
CL\_INS\_136
CL\_INS\_108
CL\_INS\_108
CL\_INS\_136
CL\_INS\_136
CL\_INS\_136
CL\_INS\_87
CL\_INS\_87
CL\_INS\_87
CL\_INS\_136
CL\_INS\_136
CL\_INS\_136
CL\_INS\_136
CL\_INS\_136
CL\_INS\_108
CL\_INS\_136
CL\_INS\_136
CL\_INS\_136
CL\_INS\_60
CL\_INS\_108
CL\_INS\_108
CL\_INS\_108
CL\_INS\_382
CL\_INS\_382
CL\_INS\_382
CL\_INS\_382
CL\_INS\_382
CL\_INS\_382
CL\_INS\_382
CL\_INS\_382
CL\_INS\_382
CL\_INS\_85
CL\_INS\_382
CL\_INS\_382
CL\_INS\_382
CL\_INS\_382
CL\_INS\_382
CL\_INS\_382
CL\_INS\_382
CL\_INS\_382
CL\_INS\_382
CL\_INS\_382
CL\_INS\_382
CL\_INS\_382
CL\_INS\_382
CL\_INS\_99
CL\_INS\_382
CL\_INS\_382
CL\_INS\_382
CL\_INS\_382
CL\_INS\_382
CL\_INS\_382
CL\_INS\_382
CL\_INS\_382
CL\_INS\_382
CL\_INS\_382
CL\_INS\_382
CL\_INS\_382
CL\_INS\_382
CL\_INS\_382
CL\_INS\_382
CL\_INS\_382
CL\_INS\_382
CL\_INS\_42
CL\_INS\_382
CL\_INS\_382
CL\_INS\_99
CL\_INS\_99
CL\_INS\_382
CL\_INS\_382
CL\_INS\_382
CL\_INS\_86
CL\_INS\_382
CL\_INS\_108
CL\_INS\_382
CL\_INS\_382
CL\_INS\_382
CL\_INS\_106
CL\_INS\_106
CL\_INS\_382
CL\_INS\_382
CL\_INS\_382
CL\_INS\_382
CL\_INS\_382
CL\_INS\_382
CL\_INS\_382
CL\_INS\_382
CL\_INS\_382
CL\_INS\_382
CL\_INS\_382
CL\_INS\_247
CL\_INS\_247
CL\_INS\_382
CL\_INS\_382
CL\_INS\_382
CL\_INS\_382
CL\_INS\_382
CL\_INS\_382
CL\_INS\_108
CL\_INS\_108
CL\_INS\_108
CL\_INS\_108
CL\_INS\_108
CL\_INS\_108
CL\_INS\_382
CL\_INS\_382
CL\_INS\_382
CL\_INS\_382
CL\_INS\_382
CL\_INS\_128
CL\_INS\_85
CL\_INS\_108
CL\_INS\_108
CL\_INS\_108
CL\_INS\_108
CL\_INS\_136
CL\_INS\_99
CL\_INS\_99
CL\_INS\_382
CL\_INS\_382
CL\_INS\_128
CL\_INS\_123
CL\_INS\_128
CL\_INS\_382
CL\_INS\_128
CL\_INS\_382
CL\_INS\_382
CL\_INS\_382
CL\_INS\_382
CL\_INS\_382
CL\_INS\_382
CL\_INS\_128
CL\_INS\_384
CL\_INS\_382
CL\_INS\_128
CL\_INS\_128
CL\_INS\_382
CL\_INS\_382
CL\_INS\_382
CL\_INS\_382
CL\_INS\_382
CL\_INS\_99
CL\_INS\_128
CL\_INS\_128
CL\_INS\_99
CL\_INS\_382
CL\_INS\_382
CL\_INS\_382
CL\_INS\_382
CL\_INS\_382
CL\_INS\_382
CL\_INS\_99
CL\_INS\_99
CL\_INS\_382
CL\_INS\_382
CL\_INS\_99
CL\_INS\_99
CL\_INS\_99
CL\_INS\_382
CL\_INS\_83
CL\_INS\_382
CL\_INS\_99
CL\_INS\_382
CL\_INS\_382
CL\_INS\_382
CL\_INS\_146
CL\_INS\_382
CL\_INS\_99
CL\_INS\_382
CL\_INS\_99
CL\_INS\_99
CL\_INS\_99
CL\_INS\_382
CL\_INS\_128
CL\_INS\_99
CL\_INS\_99
CL\_INS\_99
CL\_INS\_128
CL\_INS\_128
CL\_INS\_128
CL\_INS\_128
CL\_INS\_128
CL\_INS\_128
CL\_INS\_128
CL\_INS\_128
CL\_INS\_128
CL\_INS\_128
CL\_INS\_128
CL\_INS\_128
CL\_INS\_128
CL\_INS\_128
CL\_INS\_128
CL\_INS\_128
CL\_INS\_128
CL\_INS\_382
CL\_INS\_99
CL\_INS\_382
CL\_INS\_99
CL\_INS\_382
CL\_INS\_382
CL\_INS\_128
CL\_INS\_128
CL\_INS\_128
CL\_INS\_128
CL\_INS\_128
CL\_INS\_128
CL\_INS\_128
CL\_INS\_128
CL\_INS\_128
CL\_INS\_128
CL\_INS\_128
CL\_INS\_128
CL\_INS\_385
CL\_INS\_385
CL\_INS\_128
CL\_INS\_128
CL\_INS\_128
CL\_INS\_128
CL\_INS\_128
CL\_INS\_128
CL\_INS\_382
CL\_INS\_382
CL\_INS\_382
CL\_INS\_382
CL\_INS\_382
CL\_INS\_99
CL\_INS\_382
CL\_INS\_382
CL\_INS\_128
CL\_INS\_382
CL\_INS\_382
CL\_INS\_83
CL\_INS\_382
CL\_INS\_382
CL\_INS\_382
CL\_INS\_382
CL\_INS\_382
CL\_INS\_382
CL\_INS\_382
CL\_INS\_382
CL\_INS\_382
CL\_INS\_149
CL\_INS\_382
CL\_INS\_382
CL\_INS\_382
CL\_INS\_128
CL\_INS\_128
CL\_INS\_382
CL\_INS\_368
CL\_INS\_382
CL\_INS\_382
CL\_INS\_382
CL\_INS\_20
CL\_INS\_207
CL\_INS\_207
CL\_INS\_382
CL\_INS\_382
CL\_INS\_382
CL\_INS\_382
CL\_INS\_379
CL\_INS\_379
CL\_INS\_382
CL\_INS\_382
CL\_INS\_382
CL\_INS\_382
CL\_INS\_382
CL\_INS\_382
CL\_INS\_382
CL\_INS\_382
CL\_INS\_382
CL\_INS\_379
CL\_INS\_382
CL\_INS\_383
CL\_INS\_383
CL\_INS\_383
CL\_INS\_383
CL\_INS\_383
CL\_INS\_382
CL\_INS\_20
CL\_INS\_382
CL\_INS\_382
CL\_INS\_382
CL\_INS\_382
CL\_INS\_86
CL\_INS\_382
CL\_INS\_86
CL\_INS\_382
CL\_INS\_20
CL\_INS\_20
CL\_INS\_20
CL\_INS\_20
CL\_INS\_207
CL\_INS\_20
CL\_INS\_382
CL\_INS\_340
CL\_INS\_340
CL\_INS\_86
CL\_INS\_86
CL\_INS\_382
CL\_INS\_382
CL\_INS\_382
CL\_INS\_86
CL\_INS\_86
CL\_INS\_86
CL\_INS\_382
CL\_INS\_382
CL\_INS\_382
CL\_INS\_382
CL\_INS\_382
CL\_INS\_382
CL\_INS\_379
CL\_INS\_382
CL\_INS\_382
CL\_INS\_382
CL\_INS\_382
CL\_INS\_382
CL\_INS\_382
CL\_INS\_382
CL\_INS\_382
CL\_INS\_382
CL\_INS\_20
CL\_INS\_20
CL\_INS\_382
CL\_INS\_382
CL\_INS\_128
CL\_INS\_128
CL\_INS\_128
CL\_INS\_207
CL\_INS\_382
CL\_INS\_382
CL\_INS\_382
CL\_INS\_382
CL\_INS\_99
CL\_INS\_99
CL\_INS\_379
CL\_INS\_379
CL\_INS\_128
CL\_INS\_128
CL\_INS\_128
CL\_INS\_170
CL\_INS\_20
CL\_INS\_20
CL\_INS\_382
CL\_INS\_382
CL\_INS\_382
CL\_INS\_128
CL\_INS\_128
CL\_INS\_382
CL\_INS\_382
CL\_INS\_382
CL\_INS\_382
CL\_INS\_382
CL\_INS\_128
CL\_INS\_128
CL\_INS\_382
CL\_INS\_382
CL\_INS\_382
CL\_INS\_382
CL\_INS\_382
CL\_INS\_382
CL\_INS\_382
CL\_INS\_382
CL\_INS\_382
CL\_INS\_382
CL\_INS\_382
CL\_INS\_207
CL\_INS\_382
CL\_INS\_382
CL\_INS\_382
CL\_INS\_382
CL\_INS\_382
CL\_INS\_382
CL\_INS\_382
CL\_INS\_382
CL\_INS\_382
CL\_INS\_382
CL\_INS\_382
CL\_INS\_382
CL\_INS\_382
CL\_INS\_382
CL\_INS\_382
CL\_INS\_382
CL\_INS\_382
CL\_INS\_382
CL\_INS\_382
CL\_INS\_382
CL\_INS\_383
CL\_INS\_128
CL\_INS\_128
CL\_INS\_128
CL\_INS\_128
CL\_INS\_128
CL\_INS\_128
CL\_INS\_128
CL\_INS\_207
CL\_INS\_207
CL\_INS\_60
CL\_INS\_207
CL\_INS\_207
CL\_INS\_207
CL\_INS\_382
CL\_INS\_382
CL\_INS\_146
CL\_INS\_146
CL\_INS\_146
CL\_INS\_207
CL\_INS\_207
CL\_INS\_379
CL\_INS\_368
CL\_INS\_207
CL\_INS\_207
CL\_INS\_146
CL\_INS\_99
CL\_INS\_99
CL\_INS\_99
CL\_INS\_86
CL\_INS\_382
CL\_INS\_99
CL\_INS\_379
CL\_INS\_99
CL\_INS\_99
CL\_INS\_99
CL\_INS\_379
CL\_INS\_379
CL\_INS\_153
CL\_INS\_153
CL\_INS\_153
CL\_INS\_153
CL\_INS\_153
CL\_INS\_153
CL\_INS\_153
CL\_INS\_382
CL\_INS\_86
CL\_INS\_108
CL\_INS\_136
CL\_INS\_86
CL\_INS\_86
CL\_INS\_86
CL\_INS\_86
CL\_INS\_20
CL\_INS\_146
CL\_INS\_86
CL\_INS\_86
CL\_INS\_128
CL\_INS\_86
CL\_INS\_86
CL\_INS\_237
CL\_INS\_237
CL\_INS\_237
CL\_INS\_237
CL\_INS\_86
CL\_INS\_382
CL\_INS\_207
CL\_INS\_383
CL\_INS\_383
CL\_INS\_383
CL\_INS\_383
CL\_INS\_99
CL\_INS\_99
CL\_INS\_383
CL\_INS\_383
CL\_INS\_99
CL\_INS\_99
CL\_INS\_86
CL\_INS\_237
CL\_INS\_385
CL\_INS\_17
CL\_INS\_99
CL\_INS\_20
CL\_INS\_20
CL\_INS\_42
CL\_INS\_385
CL\_INS\_99
CL\_INS\_99
CL\_INS\_385
CL\_INS\_99
CL\_INS\_99
CL\_INS\_383
CL\_INS\_383
CL\_INS\_99
CL\_INS\_99
CL\_INS\_86
CL\_INS\_87
CL\_INS\_204
CL\_INS\_379
CL\_INS\_385
CL\_INS\_42
CL\_INS\_385
CL\_INS\_385
CL\_INS\_99
CL\_INS\_128
CL\_INS\_121
CL\_INS\_121
CL\_INS\_10
CL\_INS\_121
CL\_INS\_121
CL\_INS\_128
CL\_INS\_128
CL\_INS\_121
CL\_INS\_128
CL\_INS\_121
CL\_INS\_121
CL\_INS\_128
CL\_INS\_128
CL\_INS\_121
CL\_INS\_128
CL\_INS\_128
CL\_INS\_10
CL\_INS\_10
CL\_INS\_10
CL\_INS\_128
CL\_INS\_128
CL\_INS\_121
CL\_INS\_10
CL\_INS\_117
CL\_INS\_117
CL\_INS\_117
CL\_INS\_117
CL\_INS\_128
CL\_INS\_224
CL\_INS\_224
CL\_INS\_224
CL\_INS\_224
CL\_INS\_10
CL\_INS\_10
CL\_INS\_10
CL\_INS\_224
CL\_INS\_224
CL\_INS\_10
CL\_INS\_10
CL\_INS\_10
CL\_INS\_10
CL\_INS\_128
CL\_INS\_128
CL\_INS\_128
CL\_INS\_128
CL\_INS\_128
CL\_INS\_128
CL\_INS\_128
CL\_INS\_128
CL\_INS\_99
CL\_INS\_86
CL\_INS\_99
CL\_INS\_207
CL\_INS\_99
CL\_INS\_99
CL\_INS\_146
CL\_INS\_128
CL\_INS\_99
CL\_INS\_99
CL\_INS\_99
CL\_INS\_85
CL\_INS\_382
CL\_INS\_86
CL\_INS\_86
CL\_INS\_99
CL\_INS\_60
CL\_INS\_60
CL\_INS\_382
CL\_INS\_385
CL\_INS\_385
CL\_INS\_99
CL\_INS\_149
CL\_INS\_86
CL\_INS\_20
CL\_INS\_20
CL\_INS\_99
CL\_INS\_20
CL\_INS\_237
CL\_INS\_385
CL\_INS\_383
CL\_INS\_207
CL\_INS\_207
CL\_INS\_207
CL\_INS\_207
CL\_INS\_207
CL\_INS\_207
CL\_INS\_207
CL\_INS\_207
CL\_INS\_385
CL\_INS\_20
CL\_INS\_20
CL\_INS\_20
CL\_INS\_207
CL\_INS\_207
CL\_INS\_207
CL\_INS\_86
CL\_INS\_86
CL\_INS\_86
CL\_INS\_385
CL\_INS\_385
CL\_INS\_385
CL\_INS\_385
CL\_INS\_385
CL\_INS\_385
CL\_INS\_385
CL\_INS\_237
CL\_INS\_237
CL\_INS\_237
CL\_INS\_237
CL\_INS\_237
CL\_INS\_382
CL\_INS\_237
CL\_INS\_237
CL\_INS\_237
CL\_INS\_237
CL\_INS\_237
CL\_INS\_237
CL\_INS\_237
CL\_INS\_237
CL\_INS\_237
CL\_INS\_237
CL\_INS\_237
CL\_INS\_237
CL\_INS\_237
CL\_INS\_237
CL\_INS\_237
CL\_INS\_237
CL\_INS\_237
CL\_INS\_237
CL\_INS\_237
CL\_INS\_237
CL\_INS\_237
CL\_INS\_237
CL\_INS\_237
CL\_INS\_237
CL\_INS\_149
CL\_INS\_237
CL\_INS\_237
CL\_INS\_237
CL\_INS\_237
CL\_INS\_237
CL\_INS\_237
CL\_INS\_237
CL\_INS\_237
CL\_INS\_237
CL\_INS\_237
CL\_INS\_237
CL\_INS\_237
CL\_INS\_237
CL\_INS\_237
CL\_INS\_237
CL\_INS\_237
CL\_INS\_237
CL\_INS\_237
CL\_INS\_237
CL\_INS\_237
CL\_INS\_237
CL\_INS\_237
CL\_INS\_237
CL\_INS\_237
CL\_INS\_237
CL\_INS\_237
CL\_INS\_237
CL\_INS\_237
CL\_INS\_237
CL\_INS\_237
CL\_INS\_237
CL\_INS\_237
CL\_INS\_237
CL\_INS\_237
CL\_INS\_237
CL\_INS\_237
CL\_INS\_237
CL\_INS\_237
CL\_INS\_237
CL\_INS\_237
CL\_INS\_237
CL\_INS\_237
CL\_INS\_237
CL\_INS\_237
CL\_INS\_237
CL\_INS\_237
CL\_INS\_237
CL\_INS\_237
CL\_INS\_237
CL\_INS\_237
CL\_INS\_237
CL\_INS\_237
CL\_INS\_86
CL\_INS\_237
CL\_INS\_237
CL\_INS\_237
CL\_INS\_247
CL\_INS\_247
CL\_INS\_247
CL\_INS\_247
CL\_INS\_247
CL\_INS\_123
CL\_INS\_123
CL\_INS\_247
CL\_INS\_123
CL\_INS\_71
CL\_INS\_71
CL\_INS\_247
CL\_INS\_247
CL\_INS\_247
CL\_INS\_382
CL\_INS\_382
CL\_INS\_382
CL\_INS\_382
CL\_INS\_247
CL\_INS\_382
CL\_INS\_382
CL\_INS\_382
CL\_INS\_383
CL\_INS\_271
Cluster ID


CL\_30343
CL\_30342
CL\_30345
CL\_30346
CL\_30347
CL\_6746
CL\_13790
CL\_21897
CL\_21898
CL\_7112
CL\_4517
CL\_12701
CL\_23652
CL\_23653
CL\_7111
CL\_7110
CL\_4518
CL\_4670
CL\_4669
CL\_4668
CL\_13153
CL\_13152
CL\_22921
CL\_4666
CL\_13151
CL\_13150
CL\_13149
CL\_13148
CL\_13147
CL\_13146
CL\_13145
CL\_29665
CL\_14214
CL\_14215
CL\_14216
CL\_14218
CL\_14219
CL\_14220
CL\_8957
CL\_8958
CL\_8959
CL\_8960
CL\_14221
CL\_14222
CL\_14223
CL\_17101
CL\_9539
CL\_10020
CL\_15549
CL\_7896
CL\_8628
CL\_22116
CL\_22117
CL\_22118
CL\_22119
CL\_22120
CL\_22121
CL\_22122
CL\_22123
CL\_22124
CL\_22125
CL\_10984
CL\_1931
CL\_12375
CL\_9745
CL\_4663
CL\_4662
CL\_4661
CL\_4660
CL\_4659
CL\_12376
CL\_12377
CL\_12378
CL\_4658
CL\_12379
CL\_12380
CL\_4654
CL\_4652
CL\_13612
CL\_11939
CL\_11938
CL\_5429
CL\_2285
CL\_26545
CL\_29274
CL\_29968
CL\_29967
CL\_29966
CL\_10983
CL\_27356
CL\_11318
CL\_11317
CL\_5943
CL\_2554
CL\_2553
CL\_21250
CL\_21251
CL\_6031
CL\_17779
CL\_17778
CL\_12116
CL\_12117
CL\_6034
CL\_22008
CL\_8434
CL\_4819
CL\_4818
CL\_8350
CL\_4817
CL\_2529
CL\_20112
CL\_20111
CL\_12068
CL\_7337
CL\_20110
CL\_20109
CL\_11316
CL\_11315
CL\_11314
CL\_8335
CL\_2284
CL\_21645
CL\_2283
CL\_17777
CL\_2282
CL\_16123
CL\_6638
CL\_8200
CL\_17502
CL\_17501
CL\_17500
CL\_17499
CL\_8669
CL\_8199
CL\_7126
CL\_5368
CL\_5367
CL\_10317
CL\_10318
CL\_5363
CL\_5362
CL\_1089
CL\_5798
CL\_7033
CL\_7032
CL\_5361
CL\_502
CL\_503
CL\_504
CL\_505
CL\_506
CL\_507
CL\_4395
CL\_4396
CL\_5799
CL\_7821
CL\_4400
CL\_508
CL\_4401
CL\_5360
CL\_7031
CL\_5359
CL\_5358
CL\_5357
CL\_12059
CL\_6793
CL\_5356
CL\_5355
CL\_5354
CL\_5353
CL\_5352
CL\_6792
CL\_6791
CL\_6790
CL\_6789
CL\_6788
CL\_6787
CL\_6786
CL\_5351
CL\_4402
CL\_6483
CL\_6479
CL\_23070
CL\_4403
CL\_4404
CL\_4405
CL\_4406
CL\_4407
CL\_4408
CL\_4409
CL\_10918
CL\_10919
CL\_7820
CL\_7819
CL\_4410
CL\_5350
CL\_5349
CL\_5348
CL\_5347
CL\_5346
CL\_5345
CL\_5344
CL\_521
CL\_522
CL\_35811
CL\_524
CL\_525
CL\_509
CL\_511
CL\_512
CL\_513
CL\_4411
CL\_11961
CL\_8469
CL\_8470
CL\_516
CL\_517
CL\_518
CL\_21891
CL\_519
CL\_520
CL\_5805
CL\_5806
CL\_5807
CL\_5808
CL\_5809
CL\_5343
CL\_5342
CL\_5341
CL\_6452
CL\_5340
CL\_526
CL\_34439
CL\_527
CL\_12383
CL\_529
CL\_10976
CL\_10975
CL\_8180
CL\_8179
CL\_8178
CL\_10974
CL\_10973
CL\_10972
CL\_10971
CL\_8176
CL\_8175
CL\_8174
CL\_8173
CL\_8172
CL\_10970
CL\_10969
CL\_7030
CL\_7818
CL\_7029
CL\_7028
CL\_532
CL\_528
CL\_531
CL\_1302
CL\_1303
CL\_1304
CL\_15285
CL\_1305
CL\_1306
CL\_1307
CL\_1308
CL\_1309
CL\_4562
CL\_8195
CL\_23880
CL\_4560
CL\_4561
CL\_7125
CL\_4559
CL\_17265
CL\_9150
CL\_1310
CL\_1311
CL\_8995
CL\_4456
CL\_1312
CL\_1313
CL\_15759
CL\_11761
CL\_4457
CL\_8996
CL\_4458
CL\_1314
CL\_1315
CL\_1316
CL\_1317
CL\_19912
CL\_4459
CL\_37510
CL\_4460
CL\_4461
CL\_4462
CL\_20037
CL\_20036
CL\_20035
CL\_6775
CL\_6774
CL\_4548
CL\_22684
CL\_14254
CL\_14255
CL\_8162
CL\_14149
CL\_8997
CL\_7124
CL\_7123
CL\_12008
CL\_7122
CL\_33962
CL\_33963
CL\_37540
CL\_23658
CL\_6781
CL\_6780
CL\_6779
CL\_6778
CL\_10522
CL\_10523
CL\_10524
CL\_10514
CL\_8680
CL\_6449
CL\_6450
CL\_6773
CL\_4546
CL\_4545
CL\_8689
CL\_8690
CL\_4648
CL\_7121
CL\_4647
CL\_14244
CL\_4644
CL\_14245
CL\_6042
CL\_14246
CL\_14247
CL\_8691
CL\_8692
CL\_28485
CL\_28484
CL\_28483
CL\_28482
CL\_7120
CL\_10980
CL\_4646
CL\_17237
CL\_10979
CL\_10978
CL\_10977
CL\_10940
CL\_10941
CL\_8693
CL\_8694
CL\_7119
CL\_8185
CL\_7118
CL\_7117
CL\_12009
CL\_8184
CL\_12797
CL\_13524
CL\_9098
CL\_7537
CL\_8696
CL\_7536
CL\_7535
CL\_6784
CL\_16618
CL\_11311
CL\_36954
CL\_12124
CL\_33354
CL\_9122
CL\_7533
CL\_9123
CL\_36953
CL\_36952
CL\_36951
CL\_17035
CL\_24295
CL\_4534
CL\_28481
CL\_13515
CL\_19820
CL\_19819
CL\_19946
CL\_19945
CL\_19944
CL\_5281
CL\_11758
CL\_4463
CL\_4464
CL\_4465
CL\_10525
CL\_19913
CL\_6772
CL\_6771
CL\_5924
CL\_6770
CL\_5414
CL\_17747
CL\_6664
CL\_6783
CL\_34442
CL\_34441
CL\_34440
CL\_4466
CL\_4467
CL\_4468
CL\_20140
CL\_12006
CL\_8183
CL\_18336
CL\_16651
CL\_9944
CL\_7534
CL\_4469
CL\_1318
CL\_1319
CL\_9520
CL\_9519
CL\_1320
CL\_25372
CL\_4470
CL\_4471
CL\_34438
CL\_34437
CL\_4472
CL\_4473
CL\_4474
CL\_4475
CL\_4476
CL\_4477
CL\_7498
CL\_4478
CL\_4479
CL\_4480
CL\_4481
CL\_8161
CL\_4482
CL\_1321
CL\_1322
CL\_4483
CL\_37804
CL\_37803
CL\_37802
CL\_1496
CL\_4533
CL\_533
CL\_4413
CL\_14371
CL\_7116
CL\_21008
CL\_4414
CL\_4415
CL\_14375
CL\_14376
CL\_4418
CL\_7372
CL\_27462
CL\_12007
CL\_7367
CL\_11914
CL\_8471
CL\_4532
CL\_4531
CL\_7115
CL\_7114
CL\_7113
CL\_22578
CL\_4416
CL\_4417
CL\_4530
CL\_4529
CL\_4528
CL\_4527
CL\_4526
CL\_4629
CL\_4419
CL\_4420
CL\_8472
CL\_4421
CL\_4525
CL\_4628
CL\_6743
CL\_11958
CL\_8473
CL\_8474
CL\_4522
CL\_4426
CL\_4520
CL\_22575
CL\_4521
CL\_11913
CL\_11957
CL\_7027
CL\_6045
CL\_37801
CL\_8647
CL\_8648
CL\_16131
CL\_16469
CL\_16582
CL\_13510
CL\_8112
CL\_8113
CL\_23647
CL\_12541
CL\_23764
CL\_13310
CL\_23765
CL\_8653
CL\_8114
CL\_12137
CL\_17495
CL\_15924
CL\_13640
CL\_13641
CL\_13642
CL\_13643
CL\_17067
CL\_8123
CL\_37800
CL\_37799
CL\_37798
CL\_37797
CL\_37796
CL\_37795
CL\_4422
CL\_4423
CL\_4424
CL\_4425
CL\_6744
CL\_6745
CL\_14377
CL\_37794
CL\_37793
CL\_37792
CL\_37791
CL\_4484
CL\_8168
CL\_8155
CL\_4567
CL\_4566
CL\_17498
CL\_4995
CL\_37155
CL\_4565
CL\_28971
CL\_14798
CL\_1526
CL\_1525
CL\_1524
CL\_1523
CL\_14236
CL\_14814
CL\_14813
CL\_14237
CL\_32769
CL\_32768
CL\_11255
CL\_11926
CL\_11925
CL\_11924
CL\_8676
CL\_15908
CL\_32767
CL\_32766
CL\_20767
CL\_10936
CL\_10937
CL\_4556
CL\_4555
CL\_10938
CL\_19017
CL\_15907
CL\_15906
CL\_15905
CL\_15904
CL\_14812
CL\_20768
CL\_20769
CL\_4547
CL\_29455
CL\_8188
CL\_9000
CL\_8187
CL\_8186
CL\_4544
CL\_12381
CL\_4543
CL\_14242
CL\_14243
CL\_20770
CL\_27998
CL\_20771
CL\_8688
CL\_32765
CL\_27999
CL\_17062
CL\_17061
CL\_32764
CL\_32763
CL\_32762
CL\_28001
CL\_32761
CL\_32760
CL\_32759
CL\_32758
CL\_32757
CL\_32756
CL\_32755
CL\_32754
CL\_32753
CL\_32752
CL\_32751
CL\_32750
CL\_32749
CL\_21005
CL\_28480
CL\_4524
CL\_22577
CL\_22576
CL\_4523
CL\_32748
CL\_32747
CL\_32746
CL\_32745
CL\_32744
CL\_32743
CL\_32742
CL\_32741
CL\_32740
CL\_32739
CL\_32738
CL\_28219
CL\_17468
CL\_9128
CL\_17467
CL\_32737
CL\_32736
CL\_32735
CL\_32734
CL\_32733
CL\_10982
CL\_15910
CL\_1527
CL\_8670
CL\_8671
CL\_13299
CL\_13300
CL\_5274
CL\_31425
CL\_8197
CL\_8196
CL\_19227
CL\_11253
CL\_11254
CL\_5275
CL\_5276
CL\_1522
CL\_1521
CL\_5277
CL\_5278
CL\_9151
CL\_19226
CL\_19225
CL\_19224
CL\_6447
CL\_11912
CL\_19916
CL\_6448
CL\_17186
CL\_10270
CL\_17633
CL\_17634
CL\_26546
CL\_19175
CL\_14633
CL\_1093
CL\_1094
CL\_12118
CL\_12119
CL\_17776
CL\_17775
CL\_12120
CL\_7543
CL\_12121
CL\_7542
CL\_6036
CL\_6037
CL\_4656
CL\_4655
CL\_12768
CL\_30614
CL\_2281
CL\_29666
CL\_29667
CL\_29668
CL\_29669
CL\_29670
CL\_6035
CL\_7544
CL\_7338
CL\_8577
CL\_10175
CL\_5424
CL\_7621
CL\_5425
CL\_10174
CL\_5423
CL\_26547
CL\_26548
CL\_26549
CL\_26550
CL\_12065
CL\_26551
CL\_17635
CL\_36291
CL\_36290
CL\_23518
CL\_23519
CL\_1095
CL\_1096
CL\_2280
CL\_17637
CL\_17638
CL\_17639
CL\_17640
CL\_12766
CL\_2279
CL\_2278
CL\_5422
CL\_10170
CL\_30615
CL\_5421
CL\_5420
CL\_5419
CL\_7339
CL\_7541
CL\_17774
CL\_12122
CL\_10168
CL\_4651
CL\_26552
CL\_26553
CL\_19353
CL\_19354
CL\_33884
CL\_33883
CL\_33882
CL\_12123
CL\_7540
CL\_7539
CL\_5416
CL\_7538
CL\_23470
CL\_12382
CL\_36289
CL\_36288
CL\_11256
CL\_32406
CL\_32405
CL\_7075
CL\_19963
CL\_19962
CL\_5282
CL\_5283
CL\_10981
CL\_33964
CL\_33965
CL\_11922
CL\_10939
CL\_11748
CL\_11749
CL\_1520
CL\_37154
CL\_13543
CL\_1519
CL\_1518
CL\_1517
CL\_1516
CL\_1515
CL\_1514
CL\_6451
CL\_1513
CL\_1512
CL\_16967
CL\_17234
CL\_27355
CL\_17235
CL\_1511
CL\_16517
CL\_1510
CL\_1509
CL\_1508
CL\_1507
CL\_1506
CL\_7494
CL\_1505
CL\_1504
CL\_1503
CL\_1502
CL\_1501
CL\_1500
CL\_1499
CL\_1498
CL\_1497
CL\_8581
CL\_7495
CL\_21700
CL\_15834
CL\_15833
CL\_14274
CL\_8050
CL\_8051
CL\_17497
CL\_17496
CL\_7532
CL\_7531
CL\_9516
CL\_7530
CL\_7529
CL\_7528
CL\_7527
CL\_6741
CL\_13144
CL\_13143
CL\_13142
CL\_7526
CL\_7525
CL\_29965
CL\_29273
CL\_7524
CL\_7523
CL\_13141
CL\_7522
CL\_13789
CL\_4485
CL\_4429
CL\_8712
CL\_4620
CL\_32390
CL\_17033
CL\_13075
CL\_12385
CL\_32391
CL\_32392
CL\_21927
CL\_21926
CL\_21925
CL\_21924
CL\_21923
CL\_21922
CL\_16668
CL\_19201
CL\_4430
CL\_28972
CL\_6769
CL\_10920
CL\_10921
CL\_10922
CL\_7025
CL\_10474
CL\_8166
CL\_7024
CL\_10526
CL\_13538
CL\_7023
CL\_7022
CL\_6765
CL\_5595
CL\_5596
CL\_6764
CL\_7020
CL\_7019
CL\_10968
CL\_29688
CL\_29689
CL\_29690
CL\_29691
CL\_10518
CL\_13526
CL\_29692
CL\_29693
CL\_10520
CL\_534
CL\_535
CL\_536
CL\_15244
CL\_537
CL\_4433
CL\_4434
CL\_16568
CL\_9053
CL\_22641
CL\_10342
CL\_8589
CL\_6993
CL\_6767
CL\_10521
CL\_29694
CL\_29695
CL\_10947
CL\_4489
CL\_1324
CL\_4513
CL\_12788
CL\_6055
CL\_16571
CL\_17793
CL\_17792
CL\_1328
CL\_7109
CL\_21076
CL\_4807
CL\_4806
CL\_4805
CL\_9958
CL\_4802
CL\_21077
CL\_9959
CL\_4801
CL\_21078
CL\_4796
CL\_4795
CL\_21079
CL\_4794
CL\_4793
CL\_9963
CL\_9964
CL\_9965
CL\_9966
CL\_4788
CL\_9967
CL\_9968
CL\_4787
CL\_4784
CL\_9969
CL\_9970
CL\_9971
CL\_9972
CL\_9973
CL\_9974
CL\_9975
CL\_9976
CL\_9977
CL\_4774
CL\_4773
CL\_4772
CL\_9978
CL\_9979
CL\_4769
CL\_4768
CL\_4767
CL\_4766
CL\_9980
CL\_21080
CL\_21081
CL\_21082
CL\_21083
CL\_21084
CL\_21085
CL\_21086
CL\_6747
CL\_4488
CL\_7521
CL\_11956
CL\_8586
CL\_8587
CL\_10967
CL\_13537
CL\_4431
CL\_27275
CL\_27274
CL\_21899
CL\_4432
CL\_6782
CL\_8585
CL\_9159
CL\_19004
CL\_25891
CL\_9100
CL\_19205
CL\_19206
CL\_4618
CL\_12560
CL\_1495
CL\_10475
CL\_10476
CL\_8588
CL\_1494
CL\_4346
CL\_34640
CL\_34641
CL\_19181
CL\_19180
CL\_19179
CL\_18125
CL\_18126
CL\_16974
CL\_13414
CL\_8140
CL\_34642
CL\_264
CL\_265
CL\_266
CL\_1493
CL\_1492
CL\_1491
CL\_4515
CL\_4490
CL\_6054
CL\_18913
CL\_21900
CL\_21901
CL\_5810
CL\_5811
CL\_5812
CL\_5813
CL\_6163
CL\_11426
CL\_6144
CL\_6143
CL\_6142
CL\_6141
CL\_11342
CL\_6140
CL\_6139
CL\_11343
CL\_11344
CL\_11345
CL\_11346
CL\_11347
CL\_11349
CL\_11350
CL\_11351
CL\_11352
CL\_15298
CL\_23936
CL\_23937
CL\_11356
CL\_11357
CL\_11358
CL\_11359
CL\_11360
CL\_11361
CL\_11362
CL\_11363
CL\_4514
CL\_2276
CL\_11366
CL\_6138
CL\_6137
CL\_6136
CL\_11367
CL\_11368
CL\_11369
CL\_11370
CL\_34812
CL\_11371
CL\_11380
CL\_11381
CL\_11383
CL\_11384
CL\_11385
CL\_6135
CL\_6134
CL\_34811
CL\_34810
CL\_11388
CL\_6196
CL\_6195
CL\_6194
CL\_6193
CL\_6190
CL\_6189
CL\_6188
CL\_6187
CL\_6186
CL\_6183
CL\_6182
CL\_6181
CL\_11393
CL\_11394
CL\_34809
CL\_6175
CL\_6174
CL\_6173
CL\_6172
CL\_6171
CL\_11410
CL\_11411
CL\_11412
CL\_11413
CL\_11414
CL\_11415
CL\_11416
CL\_11417
CL\_11418
CL\_11419
CL\_11421
CL\_11422
CL\_10418
CL\_10419
CL\_6760
CL\_6761
CL\_10421
CL\_10642
CL\_10641
CL\_10395
CL\_10393
CL\_10392
CL\_5297
CL\_5299
CL\_5300
CL\_11133
CL\_5533
CL\_5302
CL\_11980
CL\_13409
CL\_10665
CL\_10666
CL\_10667
CL\_9691
CL\_9690
CL\_9689
CL\_9688
CL\_9687
CL\_34824
CL\_3634
