## Supplementary material for "A novel method for integrating genomic and Tn-Seq data to identify common *in vivo* fitness mechanisms across multiple bacterial species": S1 Dataset: CL_INS_384.html

Legend

 Mobile +extrachromosomalelementfunctions
 Regulatoryfunctions
 Hypothetical
 DNA Metabolism
 AntibioticResistance
 All EssentialGenes
 All Fitness Genes
 Proteinsynthesis/fate
 Other
 All VFDB Genes
 Transport +binding proteins

FULL


WINDOWSVGPNG

Trim RowsRemove SingletonsSave Fasta

CL\_4486


CL\_4486


CL\_4519


CL\_4486


CL\_4486


CL\_4486


CL\_4486


CL\_4486


CL\_2555


CL\_4519


CL\_4486


CL\_1072


CL\_4486


CL\_4427


CL\_4486


CL\_4486


CL\_4519


CL\_1917


CL\_4486


CL\_1072


CL\_1450


CL\_1917


CL\_1529


CL\_4486


CL\_500


CL\_4486


CL\_1669


CL\_4486


CL\_4486


CL\_4486


CL\_4486


CL\_2289


CL\_4486


CL\_4180


CL\_1529


CL\_4486


CL\_4486


CL\_4486


CL\_1207


CL\_4486


CL\_4486


CL\_1917


CL\_1529


CL\_4427


CL\_4486


CL\_1917


CL\_1820


CL\_1917


CL\_4486


CL\_1820


CL\_1529


CL\_1529


CL\_4486


CL\_4486


CL\_1917


CL\_1072


CL\_946


CL\_4486


CL\_4486


CL\_1072


CL\_4486


CL\_4519


CL\_3870


CL\_1207


CL\_1917


CL\_1917


CL\_4486


CL\_4486


CL\_4486


CL\_4486


CL\_4180


CL\_4486


CL\_1300


CL\_1300


CL\_1207


CL\_1300


CL\_4486


CL\_4486


CL\_4486


CL\_922


CL\_1917


CL\_4486


CL\_922


CL\_4486


CL\_1084


CL\_3870


CL\_4486


CL\_1917


CL\_4486


CL\_1207


CL\_1529


CL\_4486


CL\_4486


CL\_1529


CL\_4179


CL\_4486


CL\_4486


CL\_1072


CL\_1072


CL\_4486


CL\_1072


CL\_1917


CL\_3870


CL\_4486


CL\_2555


CL\_1072


CL\_1300


CL\_4486


CL\_4486


CL\_1529


CL\_1072


CL\_1529


CL\_1529

HighlightSelectShow Genomes


138

CL\_4487


14

CL\_1110


9

CL\_4487


8

CL\_1110


7

CL\_1208


4

CL\_4516


3

CL\_1110


3

CL\_1110


2

CL\_4487


2

CL\_4487


2

CL\_538


2

CL\_4487


1

CL\_4516


1

CL\_4487


1

CL\_538


1

CL\_725


1

CL\_4487


1

CL\_4487


1

CL\_1490


1

CL\_4487


1

CL\_4487


1

CL\_4487


1

CL\_4487


1

CL\_1110


1

CL\_4487


1

CL\_1490


1

CL\_4487


1

CL\_4181


1

CL\_3873


1

CL\_4181


1

CL\_269


1

CL\_4487


1

CL\_1119


1

CL\_4487


1

CL\_4487


1

CL\_1211


1

CL\_1490


1

CL\_1330


1

CL\_4487


1

CL\_725


1

CL\_1208


1

CL\_4487


1

CL\_4487


1

CL\_4487


1

CL\_4181


1

CL\_4487


1

CL\_4487


1

CL\_4487


1

CL\_1330


1

CL\_4487


1

CL\_4487


1

CL\_4487


1

CL\_1819


1

CL\_1110


1

CL\_4487


1

CL\_4487


1

CL\_4487


1

CL\_4181


1

CL\_538


1

CL\_4487


1

CL\_725


1

CL\_4487


1

CL\_4487


1

CL\_4487


1

CL\_4487


1

CL\_4487


1

CL\_1110


1

CL\_1208


1

CL\_1110


1

CL\_269


1

CL\_4487


1

CL\_269


1

CL\_4487


1

CL\_4487


1

CL\_4487


1

CL\_4487


1

CL\_1208


1

CL\_725


1

CL\_3873


1

CL\_4487


1

CL\_4487


1

CL\_1490


1

CL\_4487


1

CL\_269


1

CL\_4487


1

CL\_4487


1

CL\_1668


1

CL\_4487


1

CL\_4516


1

CL\_4487


1

CL\_4487


1

CL\_1109


1

CL\_4502


1

CL\_4487


1

CL\_4487


1

CL\_269


CL\_10920
CL\_10921
CL\_10922
CL\_29200
CL\_29201
CL\_29202
CL\_29203
CL\_29204
CL\_29205
CL\_29206
CL\_20111
CL\_12116
CL\_12068
CL\_7337
CL\_20110
CL\_29001
CL\_29002
CL\_8201
CL\_8200
CL\_4430
CL\_16518
CL\_9516
CL\_7026
CL\_7025
CL\_10474
CL\_7024
CL\_7023
CL\_7022
CL\_7021
CL\_6765
CL\_30135
CL\_30134
CL\_6034
CL\_8434
CL\_10983
CL\_1302
CL\_1303
CL\_1304
CL\_1305
CL\_15285
CL\_1306
CL\_1307
CL\_1308
CL\_1309
CL\_15286
CL\_1310
CL\_1311
CL\_4456
CL\_11761
CL\_4457
CL\_4458
CL\_23661
CL\_23660
CL\_6747
CL\_22922
CL\_7521
CL\_4488
CL\_9124
CL\_9125
CL\_17468
CL\_33881
CL\_33880
CL\_9128
CL\_11956
CL\_13791
CL\_25199
CL\_25200
CL\_4489
CL\_1324
CL\_7109
CL\_4432
CL\_12385
CL\_17051
CL\_8585
CL\_10526
CL\_8586
CL\_8587
CL\_8588
CL\_17773
CL\_10967
CL\_4618
CL\_10342
CL\_8589
CL\_1495
CL\_10475
CL\_11286
CL\_535
CL\_10520
CL\_11305
CL\_13613
CL\_13614
CL\_1494
CL\_20457
CL\_1493
CL\_1492
CL\_17792
CL\_1328
CL\_534
CL\_6767
CL\_4433
CL\_4434
CL\_536
CL\_28083
CL\_4490
CL\_5810
CL\_5811
CL\_5812
CL\_5813
CL\_5594
CL\_5595
CL\_5596
CL\_6764
CL\_7020
CL\_7019
CL\_7817
CL\_5814
CL\_5815
CL\_30348
CL\_35222
CL\_22407
CL\_2276
CL\_4514
CL\_23468
CL\_23467
CL\_23466
CL\_26207
CL\_6782
CL\_4513
CL\_7520
CL\_7519
CL\_7518
CL\_7517
CL\_7516
CL\_7515
CL\_7514
CL\_7513
CL\_7512
CL\_6781
CL\_6780
CL\_6779
CL\_6778
CL\_4346
CL\_8539
CL\_8540
CL\_8541
CL\_4116
CL\_8542
CL\_8543
CL\_8544
CL\_8545
CL\_8546
CL\_8547
CL\_8548
CL\_8549
CL\_8550
CL\_8551
CL\_8553
CL\_8554
CL\_7979
CL\_8131
CL\_8130
CL\_8129
CL\_8128
CL\_8127
CL\_5240
CL\_15555
CL\_8558
CL\_8559
CL\_8560
CL\_7832
CL\_1933
CL\_7831
CL\_6417
CL\_7291
CL\_7828
CL\_6416
CL\_1934
CL\_233
CL\_8565
CL\_232
CL\_231
CL\_264
CL\_265
CL\_266
CL\_267
CL\_11316
CL\_30623
CL\_30622
CL\_4396
CL\_23979
CL\_11315
CL\_34186
CL\_11314
CL\_8335
CL\_25184
CL\_2284
CL\_2283
CL\_12118
CL\_12119
CL\_7543
CL\_7542
CL\_37219
CL\_37220
CL\_12120
CL\_37221
CL\_37222
CL\_37223
CL\_37224
CL\_37225
CL\_37226
CL\_37227
CL\_37228
CL\_37229
CL\_37230
CL\_37231
CL\_37232
CL\_37233
CL\_2282
CL\_2281
CL\_7621
CL\_5427
CL\_5426
CL\_5425
CL\_10174
CL\_5423
CL\_5424
CL\_1096
CL\_2280
CL\_11313
CL\_17690
CL\_1093
CL\_1094
CL\_6036
CL\_6037
CL\_4656
CL\_4670
CL\_4669
CL\_4668
CL\_4667
CL\_4666
CL\_15868
CL\_21014
CL\_9744
CL\_4548
CL\_15287
CL\_15288
CL\_15289
CL\_4463
CL\_4464
CL\_4465
CL\_10525
CL\_19913
CL\_5368
CL\_5367
CL\_6636
CL\_2287
CL\_2286
CL\_6635
CL\_1089
CL\_5798
CL\_7033
CL\_9953
CL\_7361
CL\_6638
CL\_13501
CL\_13502
CL\_6641
CL\_13503
CL\_13504
CL\_9950
CL\_9949
CL\_8962
CL\_11731
CL\_13792
CL\_13793
CL\_13794
CL\_5364
CL\_5388
CL\_5387
CL\_5386
CL\_5385
CL\_5384
CL\_5383
CL\_5382
CL\_34374
CL\_9075
CL\_9229
CL\_28276
CL\_28277
CL\_9078
CL\_24460
CL\_6650
CL\_8325
CL\_8324
CL\_8323
CL\_20987
CL\_11210
CL\_34375
CL\_6656
CL\_6658
CL\_7043
CL\_7044
CL\_7045
CL\_7046
CL\_13508
CL\_525
CL\_8185
CL\_23978
CL\_12382
CL\_8184
CL\_7047
CL\_8319
CL\_6662
CL\_6663
CL\_6664
CL\_7049
CL\_5412
CL\_5411
CL\_1097
CL\_5363
CL\_8669
CL\_8199
CL\_5799
CL\_5362
CL\_5361
CL\_4400
CL\_508
CL\_4401
CL\_5360
CL\_5359
CL\_5358
CL\_25202
CL\_25203
CL\_6793
CL\_5356
CL\_6792
CL\_6791
CL\_13505
CL\_13506
CL\_6648
CL\_9226
CL\_5352
CL\_9081
CL\_9082
CL\_13507
CL\_6655
CL\_6790
CL\_6789
CL\_6788
CL\_6787
CL\_6786
CL\_5351
CL\_4410
CL\_5350
CL\_5372
CL\_5349
CL\_5348
CL\_5347
CL\_5346
CL\_5345
CL\_5344
CL\_35839
CL\_8276
CL\_11963
CL\_10337
CL\_4402
CL\_6483
CL\_5800
CL\_4403
CL\_7362
CL\_21697
CL\_6479
CL\_4407
CL\_4408
CL\_4409
CL\_10918
CL\_509
CL\_511
CL\_512
CL\_513
CL\_514
CL\_515
CL\_4411
CL\_8470
CL\_516
CL\_517
CL\_518
CL\_519
CL\_520
CL\_521
CL\_5343
CL\_5342
CL\_6452
CL\_5341
CL\_524
CL\_7030
CL\_7818
CL\_4413
CL\_13474
CL\_4414
CL\_4651
CL\_12123
CL\_37234
CL\_37235
CL\_20785
CL\_11937
CL\_11936
CL\_5924
CL\_6770
CL\_5809
CL\_528
CL\_34063
CL\_531
CL\_1496
CL\_533
CL\_16111
CL\_16112
CL\_4567
CL\_4566
CL\_31988
CL\_31987
CL\_13075
CL\_20243
CL\_31986
CL\_31985
CL\_31984
CL\_10982
CL\_15910
CL\_7126
CL\_8670
CL\_8671
CL\_14236
CL\_14814
CL\_14813
CL\_23879
CL\_23880
CL\_13300
CL\_19961
CL\_1312
CL\_1313
CL\_1314
CL\_1315
CL\_1316
CL\_1317
CL\_19912
CL\_4459
CL\_23659
CL\_23658
CL\_20036
CL\_20035
CL\_15759
CL\_19959
CL\_4557
CL\_7123
CL\_35190
CL\_8194
CL\_19916
CL\_4555
CL\_35629
CL\_31983
CL\_4551
CL\_31475
CL\_16126
CL\_16127
CL\_8187
CL\_8186
CL\_8689
CL\_23883
CL\_23884
CL\_23885
CL\_23886
CL\_23887
CL\_15902
CL\_8690
CL\_10980
CL\_10979
CL\_10978
CL\_10977
CL\_10940
CL\_23657
CL\_10941
CL\_4466
CL\_4467
CL\_4468
CL\_4469
CL\_7540
CL\_7539
CL\_4544
CL\_4543
CL\_4644
CL\_14245
CL\_6042
CL\_21523
CL\_4535
CL\_20250
CL\_16651
CL\_19222
CL\_16011
CL\_4648
CL\_16128
CL\_16129
CL\_15801
CL\_5340
CL\_12006
CL\_527
CL\_10976
CL\_10975
CL\_526
CL\_1318
CL\_1319
CL\_9519
CL\_1320
CL\_4470
CL\_4471
CL\_4472
CL\_4473
CL\_4474
CL\_4475
CL\_4476
CL\_4477
CL\_7498
CL\_4478
CL\_4479
CL\_4480
CL\_4481
CL\_4482
CL\_1321
CL\_1322
CL\_4483
CL\_4484
CL\_8183
CL\_18336
CL\_8182
CL\_8181
CL\_8180
CL\_8179
CL\_23888
CL\_23889
CL\_23890
CL\_23891
CL\_23892
CL\_23893
CL\_8178
CL\_27796
CL\_10974
CL\_10973
CL\_10972
CL\_10971
CL\_23894
CL\_8647
CL\_16130
CL\_8648
CL\_13639
CL\_23647
CL\_12541
CL\_23764
CL\_13310
CL\_23765
CL\_8653
CL\_35423
CL\_16429
CL\_13640
CL\_13641
CL\_13642
CL\_13643
CL\_17067
CL\_8123
CL\_35840
CL\_6461
CL\_8901
CL\_4419
CL\_4420
CL\_4421
CL\_16131
CL\_13512
CL\_8650
CL\_8652
CL\_12139
CL\_12138
CL\_12137
CL\_16132
CL\_10497
CL\_13102
CL\_16133
CL\_10498
CL\_10499
CL\_10500
CL\_10502
CL\_10503
CL\_10504
CL\_10505
CL\_10506
CL\_16134
CL\_1098
CL\_14807
CL\_1099
CL\_5410
CL\_14805
CL\_14804
CL\_5409
CL\_5408
CL\_14704
CL\_14705
CL\_16135
CL\_16136
CL\_8177
CL\_8176
CL\_8175
CL\_8174
CL\_8173
CL\_8172
CL\_10970
CL\_10969
CL\_8171
CL\_18335
CL\_21673
CL\_8170
CL\_8169
CL\_8168
CL\_13509
CL\_4647
CL\_4646
CL\_4645
CL\_6038
CL\_7120
CL\_8691
CL\_8692
CL\_8693
CL\_24313
CL\_8694
CL\_7538
CL\_6784
CL\_11311
CL\_9122
CL\_7119
CL\_7118
CL\_7117
CL\_13562
CL\_12762
CL\_12760
CL\_12996
CL\_7537
CL\_7536
CL\_7535
CL\_11920
CL\_11919
CL\_12999
CL\_17372
CL\_9098
CL\_8696
CL\_8697
CL\_8698
CL\_8700
CL\_8701
CL\_8702
CL\_8703
CL\_8704
CL\_8705
CL\_9157
CL\_10516
CL\_10517
CL\_8706
CL\_8707
CL\_8708
CL\_8709
CL\_8710
CL\_8711
CL\_26457
CL\_20242
CL\_4534
CL\_6045
CL\_4533
CL\_7116
CL\_21008
CL\_4415
CL\_4416
CL\_4417
CL\_4695
CL\_7372
CL\_4532
CL\_15899
CL\_15898
CL\_4531
CL\_4530
CL\_4418
CL\_4529
CL\_4528
CL\_4527
CL\_27588
CL\_27589
CL\_4526
CL\_4422
CL\_4524
CL\_4525
CL\_4423
CL\_4424
CL\_4425
CL\_4426
CL\_25204
CL\_4522
CL\_4521
CL\_4629
CL\_4628
CL\_4523
CL\_11958
CL\_6744
CL\_6745
CL\_4520
CL\_12360
CL\_23673
CL\_31982
CL\_6743
CL\_7371
CL\_31981
CL\_19951
CL\_11913
CL\_13481
CL\_13482
CL\_25205
CL\_17057
CL\_11957
CL\_4429
CL\_4518
CL\_4517
CL\_7111
CL\_12701
CL\_4431
CL\_6769
CL\_17053
CL\_31980
CL\_31979
CL\_31425
CL\_4565
CL\_1527
CL\_1526
CL\_4564
CL\_4563
CL\_1525
CL\_13299
CL\_8197
CL\_14692
CL\_8196
CL\_1524
CL\_1523
CL\_8195
CL\_4560
CL\_4562
CL\_8995
CL\_14693
CL\_14694
CL\_11926
CL\_11925
CL\_11924
CL\_12697
CL\_8676
CL\_15908
CL\_15907
CL\_15906
CL\_15905
CL\_8677
CL\_4552
CL\_4556
CL\_16113
CL\_4561
CL\_7125
CL\_29207
CL\_29208
CL\_29209
CL\_29210
CL\_29211
CL\_29212
CL\_29213
CL\_29214
CL\_29215
CL\_29216
CL\_29217
CL\_29218
CL\_4617
CL\_29219
CL\_4515
CL\_4620
CL\_16114
CL\_14239
CL\_15788
CL\_15789
CL\_14241
CL\_8996
CL\_4559
CL\_7124
CL\_8997
CL\_10936
CL\_10937
CL\_9150
CL\_8599
CL\_13301
CL\_13302
CL\_13303
CL\_9152
CL\_9151
CL\_19226
CL\_19225
CL\_19224
CL\_16115
CL\_16116
CL\_16117
CL\_16118
CL\_16119
CL\_16120
CL\_16121
CL\_16122
CL\_16123
CL\_16124
CL\_16125
CL\_5274
CL\_11253
CL\_11254
CL\_11255
CL\_5275
CL\_5276
CL\_1522
CL\_1521
CL\_5277
CL\_5278
CL\_6447
CL\_21781
CL\_21780
CL\_22955
CL\_10344
CL\_35417
CL\_8685
CL\_36623
CL\_11256
CL\_5281
CL\_5282
CL\_22954
CL\_6448
CL\_10510
CL\_22574
CL\_12804
CL\_12803
CL\_4665
CL\_4664
CL\_9745
CL\_4663
CL\_16452
CL\_16451
CL\_4662
CL\_4661
CL\_12801
CL\_4659
CL\_4658
CL\_12800
CL\_12379
CL\_13564
CL\_4655
CL\_12768
CL\_19351
CL\_19352
CL\_10170
CL\_12122
CL\_2279
CL\_2278
CL\_5422
CL\_5421
CL\_5420
CL\_5419
CL\_16967
CL\_17234
CL\_17235
CL\_7339
CL\_17774
CL\_35993
CL\_4654
CL\_4653
CL\_4652
CL\_10168
CL\_15903
CL\_10511
CL\_11405
CL\_11406
CL\_20109
CL\_10512
CL\_10513
CL\_10514
CL\_30412
CL\_30411
CL\_23881
CL\_23882
CL\_4547
CL\_20253
CL\_8681
CL\_8682
CL\_20252
CL\_20251
CL\_35630
CL\_8680
CL\_13304
CL\_14698
CL\_13305
CL\_13306
CL\_8684
CL\_10515
CL\_6449
CL\_6450
CL\_1519
CL\_1518
CL\_1517
CL\_1516
CL\_1515
CL\_6773
CL\_15904
CL\_14812
CL\_11922
CL\_4546
CL\_4545
CL\_10939
CL\_5283
CL\_8188
CL\_10981
CL\_9000
CL\_1514
CL\_6451
CL\_1513
CL\_1512
CL\_19197
CL\_1511
CL\_16517
CL\_1510
CL\_1509
CL\_1508
CL\_1507
CL\_1506
CL\_34669
CL\_7494
CL\_1505
CL\_12009
CL\_37236
CL\_10323
CL\_6783
CL\_12124
CL\_1504
CL\_1503
CL\_1502
CL\_1501
CL\_1500
CL\_1499
CL\_9001
CL\_1498
CL\_1497
CL\_7495
CL\_8050
CL\_23656
CL\_23655
CL\_23654
CL\_7533
CL\_37237
CL\_37238
CL\_37239
CL\_37240
CL\_8051
CL\_7532
CL\_7531
CL\_7530
CL\_7529
CL\_7528
CL\_7527
CL\_29225
CL\_37241
CL\_6741
CL\_7526
CL\_21528
CL\_7525
CL\_36888
CL\_36889
CL\_37242
CL\_29273
CL\_7524
CL\_7523
CL\_7522
CL\_8155
CL\_4485
CL\_21529
CL\_6746
CL\_7112
