## Supplementary material for "A novel method for integrating genomic and Tn-Seq data to identify common *in vivo* fitness mechanisms across multiple bacterial species": S1 Dataset: CL_INS_385.html

Legend

 Mobile +extrachromosomalelementfunctions
 Regulatoryfunctions
 Hypothetical
 DNA Metabolism
 All EssentialGenes
 AntibioticResistance
 All Fitness Genes
 Proteinsynthesis/fate
 Other
 Transport +binding proteins
 All VFDB Genes

FULL


WINDOWSVGPNG

Trim RowsRemove SingletonsSave Fasta

CL\_4516


CL\_4516


CL\_4516


CL\_4516


CL\_4516


CL\_4516


CL\_4516


CL\_4516


CL\_4516


CL\_4516


CL\_4516


CL\_4516


CL\_4516


CL\_4516


CL\_4516


CL\_4516


CL\_4516


CL\_4516


CL\_4516


CL\_4516


CL\_4516


CL\_4516


CL\_4516


CL\_4516


CL\_4516


CL\_4516


CL\_4516


CL\_4516


CL\_4516


CL\_4516


CL\_4516


CL\_4516


CL\_4516


CL\_4516


CL\_4516


CL\_4516


CL\_4516


CL\_4516


CL\_4516


CL\_4516


CL\_4516


CL\_4516


CL\_4516


CL\_4516


CL\_4516


CL\_4516


CL\_4516


CL\_4516


CL\_4516


CL\_4516


CL\_4516


CL\_4516


CL\_4516


CL\_4516


CL\_4516


CL\_4516


CL\_4516


CL\_4516


CL\_4516


CL\_4516


CL\_4516


CL\_4516


CL\_4516


CL\_4516


CL\_4516


CL\_4516


CL\_4516


CL\_4516


CL\_4516


CL\_4516


CL\_4516


CL\_4516


CL\_4516


CL\_4516


CL\_4516


CL\_4516


CL\_4516


CL\_4516


CL\_4516


CL\_4516


CL\_4516


CL\_4516


CL\_4516


CL\_4516


CL\_4516


CL\_4516


CL\_4516


CL\_4516


CL\_4516


CL\_4516


CL\_4516


CL\_4516


CL\_4516


CL\_4516


CL\_4516


CL\_4516


CL\_4516


CL\_4516


CL\_4516


CL\_4516


CL\_4516


CL\_4516


CL\_4516


CL\_4516


CL\_4516


CL\_4516


CL\_4516


CL\_4516


CL\_4516


CL\_4516


CL\_4516


CL\_4516


CL\_4516


CL\_4516


CL\_4516


CL\_4516


CL\_4516


CL\_4516


CL\_4516


CL\_4516


CL\_4516


CL\_4516


CL\_4516


CL\_4516


CL\_4516


CL\_4516


CL\_4516


CL\_4516


CL\_4516


CL\_4516


CL\_4516


CL\_4516


CL\_4516


CL\_4516


CL\_4516


CL\_4516


CL\_4516


CL\_4516


CL\_4516


CL\_4516


CL\_4516


CL\_4516

HighlightSelectShow Genomes


13

CL\_1490


12

CL\_1490


6

CL\_1915


5

CL\_725


4

CL\_2528


4

CL\_538


4

CL\_1490


3

CL\_725


3

CL\_1490


2

CL\_1330


2

CL\_1073


2

CL\_538


2

CL\_1490


2

CL\_1073


2

CL\_1110


2

CL\_3873


2

CL\_1330


2

CL\_1330


2

CL\_1330


2

CL\_538


2

CL\_1490


2

CL\_538


2

CL\_1330


1

CL\_4181


1

CL\_1073


1

CL\_538


1

CL\_538


1

CL\_269


1

CL\_1915


1

CL\_1330


1

CL\_1109


1

CL\_4181


1

CL\_1489


1

CL\_1109


1

CL\_1490


1

CL\_270


1

CL\_1330


1

CL\_1915


1

CL\_1330


1

CL\_2007


1

CL\_2528


1

CL\_1073


1

CL\_538


1

CL\_1073


1

CL\_538


1

CL\_538


1

CL\_4181


1

CL\_538


1

CL\_1915


1

CL\_538


1

CL\_1490


1

CL\_947


1

CL\_1490


1

CL\_1126


1

CL\_1330


1

CL\_1330


1

CL\_1490


1

CL\_1490


1

CL\_2528


1

CL\_1330


1

CL\_538


1

CL\_725


1

CL\_1330


1

CL\_1488


1

CL\_1208


1

CL\_538


1

CL\_1110


1

CL\_538


1

Break


1

Break


1

CL\_1915


1

CL\_1490


1

CL\_725


1

CL\_923


1

CL\_1330


1

CL\_1915


1

CL\_538


1

CL\_538


1

CL\_3873


1

CL\_4181


1

CL\_2270


1

CL\_1915


1

CL\_1109


1

CL\_1208


1

CL\_1490


1

CL\_3873


1

CL\_4181


1

CL\_1073


1

CL\_1073


1

CL\_4181


1

CL\_1208


1

CL\_2528


1

CL\_1490


1

CL\_4181


1

CL\_538


1

CL\_4181


1

CL\_1330


1

CL\_1073


1

CL\_538


1

CL\_1126


1

CL\_4181


1

CL\_1073


1

CL\_1073


1

CL\_1819


1

CL\_1915


1

CL\_4181


1

CL\_725


1

CL\_538


1

CL\_1073


1

CL\_725


1

CL\_1208


1

CL\_1490


1

CL\_725


1

CL\_1073


1

CL\_538


1

CL\_1109


1

CL\_4181


1

CL\_1109


1

CL\_1073


1

CL\_1915


1

CL\_2116


1

CL\_1489


1

CL\_1490


1

CL\_1330


1

CL\_1490


1

CL\_4181


1

CL\_1110


1

CL\_1073


1

CL\_1915


1

CL\_1490


1

CL\_538


1

CL\_1208


1

CL\_1073


1

CL\_1073


1

CL\_538


1

CL\_1819


1

CL\_538


1

CL\_3873


1

CL\_538


1

CL\_538


1

CL\_1073


1

CL\_1819

fGI ID


CL\_INS\_385
CL\_INS\_87
CL\_INS\_87
CL\_INS\_87
CL\_INS\_86
CL\_INS\_385
CL\_INS\_385
CL\_INS\_385
CL\_INS\_385
CL\_INS\_385
CL\_INS\_385
CL\_INS\_385
CL\_INS\_146
CL\_INS\_99
CL\_INS\_99
CL\_INS\_385
CL\_INS\_136
CL\_INS\_86
CL\_INS\_99
CL\_INS\_382
CL\_INS\_146
CL\_INS\_146
CL\_INS\_146
CL\_INS\_207
CL\_INS\_382
CL\_INS\_207
CL\_INS\_385
CL\_INS\_385
CL\_INS\_146
CL\_INS\_99
CL\_INS\_146
CL\_INS\_385
CL\_INS\_123
CL\_INS\_385
CL\_INS\_86
CL\_INS\_382
CL\_INS\_385
CL\_INS\_385
CL\_INS\_385
CL\_INS\_86
CL\_INS\_86
CL\_INS\_385
CL\_INS\_99
CL\_INS\_99
CL\_INS\_146
CL\_INS\_146
CL\_INS\_20
CL\_INS\_86
CL\_INS\_385
CL\_INS\_385
CL\_INS\_20
CL\_INS\_385
CL\_INS\_20
CL\_INS\_99
CL\_INS\_146
CL\_INS\_146
CL\_INS\_146
CL\_INS\_385
CL\_INS\_385
CL\_INS\_385
CL\_INS\_385
CL\_INS\_385
CL\_INS\_385
CL\_INS\_385
CL\_INS\_385
CL\_INS\_385
CL\_INS\_385
CL\_INS\_385
CL\_INS\_385
CL\_INS\_385
CL\_INS\_385
CL\_INS\_385
CL\_INS\_385
CL\_INS\_385
CL\_INS\_99
CL\_INS\_99
CL\_INS\_275
CL\_INS\_237
CL\_INS\_385
CL\_INS\_385
CL\_INS\_86
CL\_INS\_86
CL\_INS\_237
CL\_INS\_237
CL\_INS\_237
CL\_INS\_382
CL\_INS\_20
CL\_INS\_86
CL\_INS\_86
CL\_INS\_20
CL\_INS\_20
CL\_INS\_237
CL\_INS\_237
CL\_INS\_237
CL\_INS\_20
CL\_INS\_20
CL\_INS\_86
CL\_INS\_86
CL\_INS\_99
CL\_INS\_20
CL\_INS\_385
CL\_INS\_385
CL\_INS\_385
CL\_INS\_385
CL\_INS\_385
CL\_INS\_86
CL\_INS\_385
CL\_INS\_385
CL\_INS\_385
CL\_INS\_385
CL\_INS\_385
CL\_INS\_385
CL\_INS\_99
CL\_INS\_20
CL\_INS\_99
CL\_INS\_86
CL\_INS\_99
CL\_INS\_385
CL\_INS\_385
CL\_INS\_385
CL\_INS\_20
CL\_INS\_20
CL\_INS\_385
CL\_INS\_99
CL\_INS\_99
CL\_INS\_99
CL\_INS\_385
CL\_INS\_237
CL\_INS\_385
CL\_INS\_17
CL\_INS\_385
CL\_INS\_155
CL\_INS\_155
CL\_INS\_155
CL\_INS\_155
CL\_INS\_155
CL\_INS\_385
CL\_INS\_385
CL\_INS\_155
CL\_INS\_385
CL\_INS\_385
CL\_INS\_385
CL\_INS\_385
CL\_INS\_385
CL\_INS\_385
CL\_INS\_385
CL\_INS\_385
CL\_INS\_385
CL\_INS\_385
CL\_INS\_385
CL\_INS\_385
CL\_INS\_385
CL\_INS\_385
CL\_INS\_385
CL\_INS\_385
CL\_INS\_155
CL\_INS\_385
CL\_INS\_155
CL\_INS\_385
CL\_INS\_385
CL\_INS\_155
CL\_INS\_155
CL\_INS\_155
CL\_INS\_385
CL\_INS\_385
CL\_INS\_385
CL\_INS\_385
CL\_INS\_155
CL\_INS\_155
CL\_INS\_155
CL\_INS\_155
CL\_INS\_385
CL\_INS\_385
CL\_INS\_385
CL\_INS\_385
CL\_INS\_99
CL\_INS\_99
CL\_INS\_385
CL\_INS\_385
CL\_INS\_385
CL\_INS\_207
CL\_INS\_207
CL\_INS\_382
CL\_INS\_385
CL\_INS\_385
CL\_INS\_385
CL\_INS\_237
CL\_INS\_385
CL\_INS\_385
CL\_INS\_149
CL\_INS\_170
CL\_INS\_10
CL\_INS\_99
CL\_INS\_385
CL\_INS\_99
CL\_INS\_87
CL\_INS\_385
CL\_INS\_385
CL\_INS\_385
CL\_INS\_207
CL\_INS\_385
CL\_INS\_385
CL\_INS\_385
CL\_INS\_385
CL\_INS\_86
CL\_INS\_149
CL\_INS\_86
CL\_INS\_385
CL\_INS\_385
CL\_INS\_385
CL\_INS\_385
CL\_INS\_385
CL\_INS\_385
CL\_INS\_385
CL\_INS\_385
CL\_INS\_385
CL\_INS\_385
CL\_INS\_385
CL\_INS\_385
CL\_INS\_385
CL\_INS\_207
CL\_INS\_207
CL\_INS\_207
CL\_INS\_207
CL\_INS\_207
CL\_INS\_207
CL\_INS\_207
CL\_INS\_207
CL\_INS\_207
CL\_INS\_207
CL\_INS\_207
CL\_INS\_207
CL\_INS\_207
CL\_INS\_207
CL\_INS\_207
CL\_INS\_207
CL\_INS\_207
CL\_INS\_207
CL\_INS\_207
CL\_INS\_207
CL\_INS\_207
CL\_INS\_20
CL\_INS\_237
CL\_INS\_385
CL\_INS\_207
CL\_INS\_207
CL\_INS\_207
CL\_INS\_207
CL\_INS\_207
CL\_INS\_207
CL\_INS\_207
CL\_INS\_207
CL\_INS\_385
CL\_INS\_20
CL\_INS\_20
CL\_INS\_20
CL\_INS\_20
CL\_INS\_207
CL\_INS\_385
CL\_INS\_385
CL\_INS\_207
CL\_INS\_385
CL\_INS\_207
CL\_INS\_385
CL\_INS\_385
CL\_INS\_382
CL\_INS\_385
CL\_INS\_382
CL\_INS\_385
CL\_INS\_382
CL\_INS\_382
CL\_INS\_382
CL\_INS\_382
CL\_INS\_382
CL\_INS\_382
CL\_INS\_382
CL\_INS\_99
CL\_INS\_382
CL\_INS\_382
CL\_INS\_382
CL\_INS\_385
CL\_INS\_385
CL\_INS\_385
CL\_INS\_385
CL\_INS\_99
CL\_INS\_385
CL\_INS\_382
CL\_INS\_99
CL\_INS\_382
CL\_INS\_382
CL\_INS\_382
CL\_INS\_385
CL\_INS\_382
CL\_INS\_385
CL\_INS\_382
CL\_INS\_382
CL\_INS\_99
CL\_INS\_385
CL\_INS\_99
CL\_INS\_385
CL\_INS\_385
CL\_INS\_99
CL\_INS\_382
CL\_INS\_382
CL\_INS\_382
CL\_INS\_385
CL\_INS\_382
CL\_INS\_382
CL\_INS\_382
CL\_INS\_382
CL\_INS\_99
CL\_INS\_382
CL\_INS\_385
CL\_INS\_385
CL\_INS\_385
CL\_INS\_382
CL\_INS\_382
CL\_INS\_382
CL\_INS\_99
CL\_INS\_99
CL\_INS\_385
CL\_INS\_385
CL\_INS\_99
CL\_INS\_385
CL\_INS\_385
CL\_INS\_385
CL\_INS\_207
CL\_INS\_207
CL\_INS\_385
CL\_INS\_385
CL\_INS\_385
CL\_INS\_385
CL\_INS\_382
CL\_INS\_233
CL\_INS\_382
CL\_INS\_382
CL\_INS\_382
CL\_INS\_382
CL\_INS\_385
CL\_INS\_385
CL\_INS\_385
CL\_INS\_385
CL\_INS\_385
CL\_INS\_385
CL\_INS\_385
CL\_INS\_343
CL\_INS\_385
CL\_INS\_237
CL\_INS\_247
CL\_INS\_237
CL\_INS\_382
CL\_INS\_99
CL\_INS\_99
CL\_INS\_382
CL\_INS\_382
CL\_INS\_159
CL\_INS\_385
CL\_INS\_159
CL\_INS\_159
CL\_INS\_159
CL\_INS\_385
CL\_INS\_385
CL\_INS\_382
CL\_INS\_382
CL\_INS\_382
CL\_INS\_382
CL\_INS\_385
CL\_INS\_385
CL\_INS\_382
CL\_INS\_99
CL\_INS\_99
CL\_INS\_382
CL\_INS\_382
CL\_INS\_382
CL\_INS\_382
CL\_INS\_385
CL\_INS\_159
CL\_INS\_382
CL\_INS\_382
CL\_INS\_382
CL\_INS\_382
CL\_INS\_382
CL\_INS\_382
CL\_INS\_382
CL\_INS\_382
CL\_INS\_382
CL\_INS\_382
CL\_INS\_159
CL\_INS\_382
CL\_INS\_385
CL\_INS\_159
CL\_INS\_382
CL\_INS\_382
CL\_INS\_382
CL\_INS\_382
CL\_INS\_385
CL\_INS\_385
CL\_INS\_159
CL\_INS\_385
CL\_INS\_385
CL\_INS\_385
CL\_INS\_382
CL\_INS\_382
CL\_INS\_382
CL\_INS\_382
CL\_INS\_159
CL\_INS\_382
CL\_INS\_382
CL\_INS\_382
CL\_INS\_382
CL\_INS\_382
CL\_INS\_382
CL\_INS\_382
CL\_INS\_233
CL\_INS\_233
CL\_INS\_233
Cluster ID


CL\_13791
CL\_20419
CL\_8275
CL\_8274
CL\_4515
CL\_23895
CL\_21530
CL\_31737
CL\_31736
CL\_31735
CL\_31734
CL\_31733
CL\_8166
CL\_10518
CL\_13526
CL\_22953
CL\_6769
CL\_7024
CL\_7521
CL\_9124
CL\_28486
CL\_28111
CL\_28110
CL\_10968
CL\_4432
CL\_6053
CL\_16031
CL\_14248
CL\_12384
CL\_12385
CL\_12386
CL\_20144
CL\_15177
CL\_5286
CL\_8585
CL\_9099
CL\_34376
CL\_34377
CL\_34378
CL\_17653
CL\_7023
CL\_6768
CL\_8586
CL\_8587
CL\_10967
CL\_23203
CL\_10474
CL\_1495
CL\_24314
CL\_16571
CL\_10475
CL\_11286
CL\_10476
CL\_8588
CL\_13613
CL\_13614
CL\_11305
CL\_21277
CL\_29671
CL\_25199
CL\_35841
CL\_22642
CL\_25586
CL\_25585
CL\_25584
CL\_25583
CL\_25582
CL\_25581
CL\_25580
CL\_25579
CL\_5522
CL\_5521
CL\_5520
CL\_5519
CL\_5692
CL\_5693
CL\_11196
CL\_6765
CL\_19367
CL\_19368
CL\_13647
CL\_7021
CL\_5595
CL\_5596
CL\_6764
CL\_7019
CL\_22649
CL\_5814
CL\_5815
CL\_11079
CL\_11078
CL\_8935
CL\_8934
CL\_8933
CL\_9606
CL\_9605
CL\_10526
CL\_6782
CL\_10520
CL\_8279
CL\_12125
CL\_12126
CL\_12127
CL\_31340
CL\_37805
CL\_7022
CL\_11757
CL\_22686
CL\_5810
CL\_5811
CL\_5812
CL\_5813
CL\_6767
CL\_6766
CL\_534
CL\_535
CL\_4433
CL\_33304
CL\_33303
CL\_33302
CL\_4434
CL\_16568
CL\_22641
CL\_10342
CL\_8713
CL\_8589
CL\_6993
CL\_536
CL\_15244
CL\_537
CL\_34204
CL\_7775
CL\_7776
CL\_9194
CL\_7777
CL\_9195
CL\_7778
CL\_7779
CL\_7780
CL\_7781
CL\_34203
CL\_9723
CL\_7782
CL\_13805
CL\_34202
CL\_27038
CL\_27039
CL\_27040
CL\_27041
CL\_34201
CL\_34200
CL\_27042
CL\_27043
CL\_24191
CL\_24192
CL\_7791
CL\_33561
CL\_17025
CL\_34199
CL\_34198
CL\_17027
CL\_6003
CL\_7786
CL\_11546
CL\_13803
CL\_13802
CL\_13801
CL\_7789
CL\_9202
CL\_7792
CL\_9203
CL\_27044
CL\_34197
CL\_34196
CL\_34195
CL\_4489
CL\_10521
CL\_7144
CL\_16850
CL\_28028
CL\_28027
CL\_17224
CL\_13034
CL\_22544
CL\_22545
CL\_34140
CL\_4514
CL\_21531
CL\_21532
CL\_4601
CL\_4602
CL\_4764
CL\_1326
CL\_1327
CL\_4618
CL\_4513
CL\_4512
CL\_34852
CL\_25587
CL\_11956
CL\_27036
CL\_17419
CL\_27037
CL\_17252
CL\_1324
CL\_12560
CL\_4490
CL\_20457
CL\_18913
CL\_21900
CL\_21901
CL\_19205
CL\_11304
CL\_19206
CL\_9101
CL\_9102
CL\_4491
CL\_19914
CL\_26016
CL\_7497
CL\_4717
CL\_6915
CL\_6916
CL\_9701
CL\_7410
CL\_7411
CL\_7412
CL\_7413
CL\_7414
CL\_7415
CL\_6366
CL\_6367
CL\_7416
CL\_6369
CL\_6370
CL\_21252
CL\_21253
CL\_21254
CL\_21255
CL\_21256
CL\_21257
CL\_1494
CL\_4346
CL\_34640
CL\_19181
CL\_19180
CL\_19179
CL\_18125
CL\_18126
CL\_16974
CL\_13414
CL\_8140
CL\_34642
CL\_264
CL\_265
CL\_266
CL\_267
CL\_1493
CL\_14445
CL\_17792
CL\_1492
CL\_1328
CL\_1491
CL\_34825
CL\_17494
CL\_10982
CL\_17493
CL\_14798
CL\_17492
CL\_14237
CL\_13300
CL\_11255
CL\_4561
CL\_7125
CL\_4559
CL\_8675
CL\_15790
CL\_8677
CL\_16422
CL\_16421
CL\_17491
CL\_17490
CL\_17489
CL\_17488
CL\_17487
CL\_17486
CL\_17066
CL\_16661
CL\_17485
CL\_15862
CL\_6452
CL\_17484
CL\_4646
CL\_17483
CL\_7119
CL\_8185
CL\_12382
CL\_17482
CL\_17481
CL\_17480
CL\_17479
CL\_17478
CL\_1497
CL\_532
CL\_4413
CL\_17477
CL\_4532
CL\_4531
CL\_4530
CL\_4529
CL\_17476
CL\_4527
CL\_17475
CL\_17474
CL\_17473
CL\_4523
CL\_4522
CL\_4521
CL\_4520
CL\_13789
CL\_17472
CL\_17471
CL\_4620
CL\_17470
CL\_17469
CL\_9125
CL\_17422
CL\_17423
CL\_9127
CL\_17468
CL\_9128
CL\_28526
CL\_9100
CL\_5541
CL\_5600
CL\_4310
CL\_5508
CL\_5014
CL\_25206
CL\_25207
CL\_25208
CL\_5296
CL\_13408
CL\_13407
CL\_13406
CL\_5618
CL\_5619
CL\_11689
CL\_5054
CL\_5055
CL\_11835
CL\_11834
CL\_14149
CL\_4256
CL\_4257
CL\_4258
CL\_6929
CL\_4259
CL\_4260
CL\_4261
CL\_5584
CL\_5625
CL\_4262
CL\_4263
CL\_5062
CL\_4265
CL\_4266
CL\_13400
CL\_5579
CL\_21032
CL\_14903
CL\_4269
CL\_4270
CL\_4271
CL\_5574
CL\_5573
CL\_5572
CL\_5637
CL\_5638
CL\_5639
CL\_4277
CL\_4278
CL\_5567
CL\_5566
CL\_5565
CL\_5564
CL\_5563
CL\_5643
CL\_4086
CL\_5644
CL\_5645
CL\_5562
CL\_5561
CL\_5560
CL\_4284
CL\_5649
CL\_5650
CL\_5651
CL\_5652
CL\_5653
CL\_5654
CL\_5555
CL\_5554
CL\_5552
CL\_5551
CL\_5550
CL\_5549
CL\_4294
CL\_5548
CL\_5662
CL\_4297
CL\_4299
CL\_4300
CL\_5544
CL\_5542
CL\_6935
