## Supplementary material for "A novel method for integrating genomic and Tn-Seq data to identify common *in vivo* fitness mechanisms across multiple bacterial species": S1 Dataset: CL_INS_388.html

Legend

 Mobile +extrachromosomalelementfunctions
 Regulatoryfunctions
 Hypothetical
 DNA Metabolism
 All EssentialGenes
 All Fitness Genes
 Proteinsynthesis/fate
 Other
 Transport +binding proteins
 All VFDB Genes

FULL


WINDOWSVGPNG

Trim RowsRemove SingletonsSave Fasta

CL\_1935


CL\_1935


CL\_1935


CL\_1935


CL\_1935


CL\_1935


CL\_1935


CL\_1935


CL\_1935


CL\_1935


CL\_1935


CL\_1935


CL\_1935


CL\_1935


CL\_1935


CL\_1935


CL\_1935


CL\_1935


CL\_1935


CL\_1935


CL\_1935


CL\_1935


CL\_1935


CL\_1935


CL\_1935


CL\_1935


CL\_1935


CL\_1935


CL\_1935


CL\_1935


CL\_1935


CL\_1935


CL\_1935


CL\_1935


CL\_1935


CL\_1935


CL\_1935


CL\_1935


CL\_1935


CL\_1935


CL\_1935


CL\_1935


CL\_1935


CL\_1935


CL\_1935


CL\_1935


CL\_1935


CL\_1935


CL\_1935


CL\_1935


CL\_1935


CL\_1935


CL\_1935


CL\_1935


CL\_1935


CL\_1935


CL\_1935


CL\_1935


CL\_1935


CL\_1935


CL\_1935


CL\_1935


CL\_1935


CL\_1935


CL\_1935


CL\_1935


CL\_1935


CL\_1935


CL\_1935


CL\_1935


CL\_1935


CL\_1935


CL\_1935


CL\_1935


CL\_1935


CL\_1935


CL\_1935


CL\_1935


CL\_1935


CL\_1935


CL\_1935


CL\_1935


CL\_1935


CL\_1935


CL\_1935


CL\_1935


CL\_1935


CL\_1935


CL\_1935


CL\_1935


CL\_1935


CL\_1935


CL\_1935


CL\_1935


CL\_1935


CL\_1935


CL\_1935


CL\_1935


CL\_1935


CL\_1935


CL\_1935


CL\_1935


CL\_1935

HighlightSelectShow Genomes


27

CL\_1940


23

CL\_1940


11

CL\_3454


9

CL\_982


9

CL\_2847


8

CL\_4118


5

CL\_982


4

CL\_982


4

CL\_230


3

CL\_2847


3

CL\_4118


2

CL\_3957


2

CL\_982


2

CL\_4017


2

CL\_4118


2

CL\_3957


2

CL\_982


2

CL\_1940


2

CL\_2847


2

CL\_2847


1

CL\_2941


1

CL\_4118


1

CL\_4118


1

CL\_2847


1

CL\_2847


1

CL\_1940


1

CL\_2847


1

CL\_4118


1

CL\_2847


1

CL\_2847


1

CL\_982


1

CL\_2847


1

CL\_4118


1

CL\_836


1

CL\_1940


1

CL\_2847


1

CL\_2941


1

CL\_2847


1

CL\_3454


1

CL\_4118


1

CL\_203


1

CL\_1940


1

CL\_2847


1

CL\_982


1

CL\_2847


1

CL\_3957


1

CL\_269


1

CL\_4118


1

CL\_2847


1

CL\_2847


1

CL\_4118


1

CL\_1940


1

CL\_4118


1

CL\_4118


1

CL\_2852


1

CL\_2847


1

CL\_2847


1

CL\_2847


1

CL\_2941


1

CL\_3957


1

CL\_1940


1

CL\_3957


1

CL\_1940


1

CL\_2847


1

CL\_4118


1

CL\_2847


1

CL\_4118


1

CL\_3957


1

Break


1

CL\_269


1

CL\_4017


1

CL\_2847


1

CL\_3454


1

CL\_2867


1

CL\_3957


1

CL\_3957


1

CL\_3957


1

CL\_2847


1

CL\_2847


1

CL\_4851


1

CL\_4118


1

CL\_1940


1

CL\_3957


1

CL\_1940


1

CL\_234


1

CL\_269


1

CL\_4118


1

CL\_2847


1

CL\_2847


1

CL\_1940


1

CL\_2847


1

CL\_3957


1

CL\_983


1

CL\_2941


1

CL\_4118


1

CL\_3956


1

CL\_2555


1

CL\_2847


1

CL\_3950


1

CL\_4118


1

CL\_2847


1

CL\_2847


1

CL\_1940

fGI ID


CL\_INS\_159
CL\_INS\_237
CL\_INS\_237
CL\_INS\_30
CL\_INS\_247
CL\_INS\_20
CL\_INS\_20
CL\_INS\_30
CL\_INS\_368
CL\_INS\_368
CL\_INS\_368
CL\_INS\_368
CL\_INS\_368
CL\_INS\_30
CL\_INS\_388
CL\_INS\_237
CL\_INS\_70
CL\_INS\_70
CL\_INS\_70
CL\_INS\_388
CL\_INS\_388
CL\_INS\_237
CL\_INS\_237
CL\_INS\_30
CL\_INS\_30
CL\_INS\_237
CL\_INS\_237
CL\_INS\_30
CL\_INS\_159
CL\_INS\_159
CL\_INS\_30
CL\_INS\_30
CL\_INS\_237
CL\_INS\_237
CL\_INS\_237
CL\_INS\_30
CL\_INS\_70
CL\_INS\_237
CL\_INS\_237
CL\_INS\_368
CL\_INS\_368
CL\_INS\_368
CL\_INS\_368
CL\_INS\_368
CL\_INS\_237
CL\_INS\_237
CL\_INS\_237
CL\_INS\_237
CL\_INS\_237
CL\_INS\_237
CL\_INS\_237
CL\_INS\_237
CL\_INS\_237
CL\_INS\_237
CL\_INS\_237
CL\_INS\_237
CL\_INS\_368
CL\_INS\_368
CL\_INS\_368
CL\_INS\_382
CL\_INS\_382
CL\_INS\_70
CL\_INS\_368
CL\_INS\_237
CL\_INS\_237
CL\_INS\_237
CL\_INS\_20
CL\_INS\_20
CL\_INS\_20
CL\_INS\_20
CL\_INS\_20
CL\_INS\_20
CL\_INS\_20
CL\_INS\_20
CL\_INS\_20
CL\_INS\_20
CL\_INS\_20
CL\_INS\_20
CL\_INS\_20
CL\_INS\_20
CL\_INS\_20
CL\_INS\_26
CL\_INS\_20
CL\_INS\_20
CL\_INS\_20
CL\_INS\_20
CL\_INS\_70
CL\_INS\_382
CL\_INS\_237
CL\_INS\_237
CL\_INS\_368
CL\_INS\_70
CL\_INS\_70
CL\_INS\_70
CL\_INS\_70
CL\_INS\_70
CL\_INS\_70
CL\_INS\_70
CL\_INS\_70
CL\_INS\_237
CL\_INS\_70
CL\_INS\_70
CL\_INS\_70
CL\_INS\_70
CL\_INS\_368
CL\_INS\_237
CL\_INS\_237
CL\_INS\_237
CL\_INS\_237
CL\_INS\_237
CL\_INS\_237
CL\_INS\_237
CL\_INS\_237
CL\_INS\_237
CL\_INS\_237
CL\_INS\_237
CL\_INS\_237
CL\_INS\_237
CL\_INS\_237
CL\_INS\_237
CL\_INS\_237
CL\_INS\_237
CL\_INS\_237
CL\_INS\_237
CL\_INS\_237
CL\_INS\_368
CL\_INS\_368
CL\_INS\_237
CL\_INS\_247
CL\_INS\_247
CL\_INS\_237
CL\_INS\_237
CL\_INS\_237
CL\_INS\_247
CL\_INS\_237
CL\_INS\_237
CL\_INS\_237
CL\_INS\_237
CL\_INS\_237
CL\_INS\_237
CL\_INS\_237
CL\_INS\_237
CL\_INS\_237
CL\_INS\_237
CL\_INS\_237
CL\_INS\_237
CL\_INS\_237
CL\_INS\_237
CL\_INS\_237
CL\_INS\_237
CL\_INS\_247
CL\_INS\_237
CL\_INS\_237
CL\_INS\_237
CL\_INS\_237
CL\_INS\_237
CL\_INS\_237
CL\_INS\_237
CL\_INS\_237
CL\_INS\_237
CL\_INS\_237
CL\_INS\_237
CL\_INS\_237
CL\_INS\_237
CL\_INS\_237
CL\_INS\_237
CL\_INS\_237
CL\_INS\_237
CL\_INS\_237
CL\_INS\_237
CL\_INS\_237
CL\_INS\_237
CL\_INS\_237
CL\_INS\_237
CL\_INS\_237
CL\_INS\_237
CL\_INS\_237
CL\_INS\_237
CL\_INS\_237
CL\_INS\_237
CL\_INS\_237
CL\_INS\_237
CL\_INS\_237
CL\_INS\_237
CL\_INS\_237
CL\_INS\_237
CL\_INS\_247
CL\_INS\_237
CL\_INS\_237
CL\_INS\_237
CL\_INS\_237
CL\_INS\_237
CL\_INS\_247
CL\_INS\_237
CL\_INS\_247
CL\_INS\_237
CL\_INS\_237
CL\_INS\_237
CL\_INS\_237
CL\_INS\_237
CL\_INS\_237
CL\_INS\_237
CL\_INS\_237
CL\_INS\_237
CL\_INS\_237
CL\_INS\_237
CL\_INS\_237
CL\_INS\_237
CL\_INS\_237
CL\_INS\_237
CL\_INS\_237
CL\_INS\_237
CL\_INS\_237
CL\_INS\_237
CL\_INS\_237
CL\_INS\_247
CL\_INS\_247
CL\_INS\_247
CL\_INS\_247
CL\_INS\_237
CL\_INS\_237
CL\_INS\_237
CL\_INS\_237
CL\_INS\_237
CL\_INS\_237
CL\_INS\_237
CL\_INS\_70
CL\_INS\_70
CL\_INS\_237
CL\_INS\_237
CL\_INS\_237
CL\_INS\_237
CL\_INS\_237
CL\_INS\_237
CL\_INS\_237
CL\_INS\_237
CL\_INS\_237
CL\_INS\_30
CL\_INS\_237
CL\_INS\_237
CL\_INS\_237
CL\_INS\_30
CL\_INS\_70
CL\_INS\_247
CL\_INS\_382
CL\_INS\_368
CL\_INS\_368
CL\_INS\_368
CL\_INS\_237
CL\_INS\_368
CL\_INS\_368
CL\_INS\_368
CL\_INS\_368
CL\_INS\_368
CL\_INS\_368
CL\_INS\_352
CL\_INS\_352
CL\_INS\_247
CL\_INS\_207
CL\_INS\_368
CL\_INS\_368
CL\_INS\_368
CL\_INS\_368
CL\_INS\_368
CL\_INS\_368
CL\_INS\_368
CL\_INS\_368
CL\_INS\_368
CL\_INS\_368
CL\_INS\_368
CL\_INS\_237
CL\_INS\_237
CL\_INS\_247
CL\_INS\_352
CL\_INS\_352
CL\_INS\_352
CL\_INS\_352
CL\_INS\_352
CL\_INS\_352
CL\_INS\_352
CL\_INS\_352
CL\_INS\_352
CL\_INS\_352
CL\_INS\_352
CL\_INS\_352
CL\_INS\_352
CL\_INS\_352
CL\_INS\_352
CL\_INS\_352
CL\_INS\_352
CL\_INS\_352
CL\_INS\_352
CL\_INS\_352
CL\_INS\_352
CL\_INS\_352
CL\_INS\_352
CL\_INS\_352
CL\_INS\_352
CL\_INS\_352
CL\_INS\_352
CL\_INS\_352
CL\_INS\_352
CL\_INS\_352
CL\_INS\_352
CL\_INS\_352
CL\_INS\_352
CL\_INS\_352
CL\_INS\_352
CL\_INS\_352
CL\_INS\_352
CL\_INS\_352
CL\_INS\_352
CL\_INS\_352
CL\_INS\_352
CL\_INS\_352
CL\_INS\_352
CL\_INS\_352
CL\_INS\_352
CL\_INS\_247
CL\_INS\_237
CL\_INS\_20
CL\_INS\_207
CL\_INS\_207
CL\_INS\_20
CL\_INS\_20
CL\_INS\_20
CL\_INS\_237
CL\_INS\_20
CL\_INS\_20
CL\_INS\_20
CL\_INS\_20
CL\_INS\_20
CL\_INS\_20
CL\_INS\_20
CL\_INS\_20
CL\_INS\_20
CL\_INS\_20
CL\_INS\_20
CL\_INS\_20
CL\_INS\_237
CL\_INS\_247
CL\_INS\_247
CL\_INS\_247
CL\_INS\_237
CL\_INS\_247
Cluster ID


CL\_5224
CL\_10819
CL\_10818
CL\_7750
CL\_7826
CL\_20256
CL\_7825
CL\_7749
CL\_11098
CL\_11097
CL\_11096
CL\_11095
CL\_7257
CL\_1936
CL\_36563
CL\_6722
CL\_8746
CL\_26031
CL\_8107
CL\_13215
CL\_13216
CL\_13217
CL\_13218
CL\_7971
CL\_1937
CL\_13822
CL\_7823
CL\_1938
CL\_28953
CL\_28952
CL\_1939
CL\_8738
CL\_6415
CL\_6414
CL\_6413
CL\_5237
CL\_6809
CL\_10789
CL\_7970
CL\_8463
CL\_12111
CL\_10436
CL\_6030
CL\_5468
CL\_10437
CL\_10438
CL\_10439
CL\_10440
CL\_10441
CL\_10442
CL\_10443
CL\_10444
CL\_10445
CL\_10446
CL\_8124
CL\_26176
CL\_8464
CL\_8465
CL\_8466
CL\_6177
CL\_6178
CL\_6179
CL\_6807
CL\_10817
CL\_10612
CL\_10816
CL\_4342
CL\_4341
CL\_4340
CL\_4339
CL\_4338
CL\_4337
CL\_4336
CL\_4335
CL\_4334
CL\_4333
CL\_4332
CL\_4331
CL\_4330
CL\_4329
CL\_4328
CL\_4327
CL\_4326
CL\_4325
CL\_4324
CL\_205
CL\_6808
CL\_8841
CL\_36435
CL\_36434
CL\_9544
CL\_4095
CL\_4096
CL\_4097
CL\_4098
CL\_4099
CL\_4100
CL\_4101
CL\_6831
CL\_6830
CL\_6829
CL\_6411
CL\_6828
CL\_7252
CL\_9543
CL\_6723
CL\_9542
CL\_7258
CL\_7259
CL\_9541
CL\_4102
CL\_4103
CL\_4104
CL\_4105
CL\_4106
CL\_4107
CL\_4108
CL\_4109
CL\_4110
CL\_4111
CL\_4112
CL\_4113
CL\_4114
CL\_4115
CL\_4116
CL\_23674
CL\_4117
CL\_7205
CL\_7204
CL\_7203
CL\_7822
CL\_17156
CL\_17155
CL\_8497
CL\_23474
CL\_25194
CL\_27530
CL\_27529
CL\_27528
CL\_32507
CL\_32506
CL\_32505
CL\_32504
CL\_10815
CL\_10814
CL\_10813
CL\_11186
CL\_11185
CL\_11184
CL\_11183
CL\_14009
CL\_26795
CL\_26796
CL\_29014
CL\_11182
CL\_11181
CL\_12070
CL\_12071
CL\_12072
CL\_12073
CL\_12074
CL\_7285
CL\_14543
CL\_14544
CL\_14545
CL\_14546
CL\_14547
CL\_14548
CL\_14549
CL\_10613
CL\_10614
CL\_10615
CL\_10616
CL\_24197
CL\_8814
CL\_24198
CL\_8813
CL\_8812
CL\_8811
CL\_24199
CL\_8806
CL\_8809
CL\_24200
CL\_24201
CL\_24202
CL\_8803
CL\_7284
CL\_11706
CL\_11849
CL\_11848
CL\_27350
CL\_10617
CL\_7283
CL\_8802
CL\_7202
CL\_13709
CL\_8364
CL\_8365
CL\_14320
CL\_14321
CL\_14322
CL\_14323
CL\_14324
CL\_14325
CL\_8386
CL\_14326
CL\_14327
CL\_14328
CL\_14329
CL\_14330
CL\_8440
CL\_14331
CL\_8441
CL\_8442
CL\_7245
CL\_14332
CL\_14333
CL\_14334
CL\_14335
CL\_14336
CL\_14337
CL\_14338
CL\_14339
CL\_13377
CL\_13378
CL\_14340
CL\_7671
CL\_7214
CL\_7213
CL\_14341
CL\_14342
CL\_7211
CL\_7293
CL\_7292
CL\_4374
CL\_4375
CL\_7208
CL\_1934
CL\_7206
CL\_7288
CL\_7568
CL\_2551
CL\_7824
CL\_7567
CL\_4086
CL\_13381
CL\_13382
CL\_13383
CL\_12078
CL\_13384
CL\_13385
CL\_13386
CL\_13387
CL\_13388
CL\_13389
CL\_8504
CL\_7969
CL\_28432
CL\_28431
CL\_6018
CL\_8102
CL\_8101
CL\_8100
CL\_8099
CL\_8098
CL\_8097
CL\_8096
CL\_8095
CL\_8094
CL\_8093
CL\_6805
CL\_6806
CL\_5236
CL\_8503
CL\_8502
CL\_17583
CL\_17584
CL\_9401
CL\_9402
CL\_9403
CL\_9404
CL\_9405
CL\_9406
CL\_9407
CL\_9408
CL\_9409
CL\_9410
CL\_9411
CL\_9412
CL\_9413
CL\_9414
CL\_9415
CL\_9416
CL\_9417
CL\_9418
CL\_9419
CL\_9420
CL\_9421
CL\_9422
CL\_9423
CL\_9424
CL\_9425
CL\_9426
CL\_9427
CL\_9428
CL\_9429
CL\_9430
CL\_9431
CL\_9432
CL\_9433
CL\_9434
CL\_9435
CL\_9436
CL\_9437
CL\_9438
CL\_9439
CL\_9440
CL\_9441
CL\_7287
CL\_14619
CL\_14223
CL\_231
CL\_232
CL\_9601
CL\_9600
CL\_9599
CL\_4346
CL\_22805
CL\_9024
CL\_22804
CL\_22803
CL\_22802
CL\_22801
CL\_22800
CL\_22799
CL\_264
CL\_265
CL\_266
CL\_267
CL\_10611
CL\_8501
CL\_7286
CL\_8500
CL\_8499
CL\_8498
