## Supplementary figures and images for "A novel method for integrating genomic and Tn-Seq data to identify common *in vivo* fitness mechanisms across multiple bacterial species"

### S1 Fig

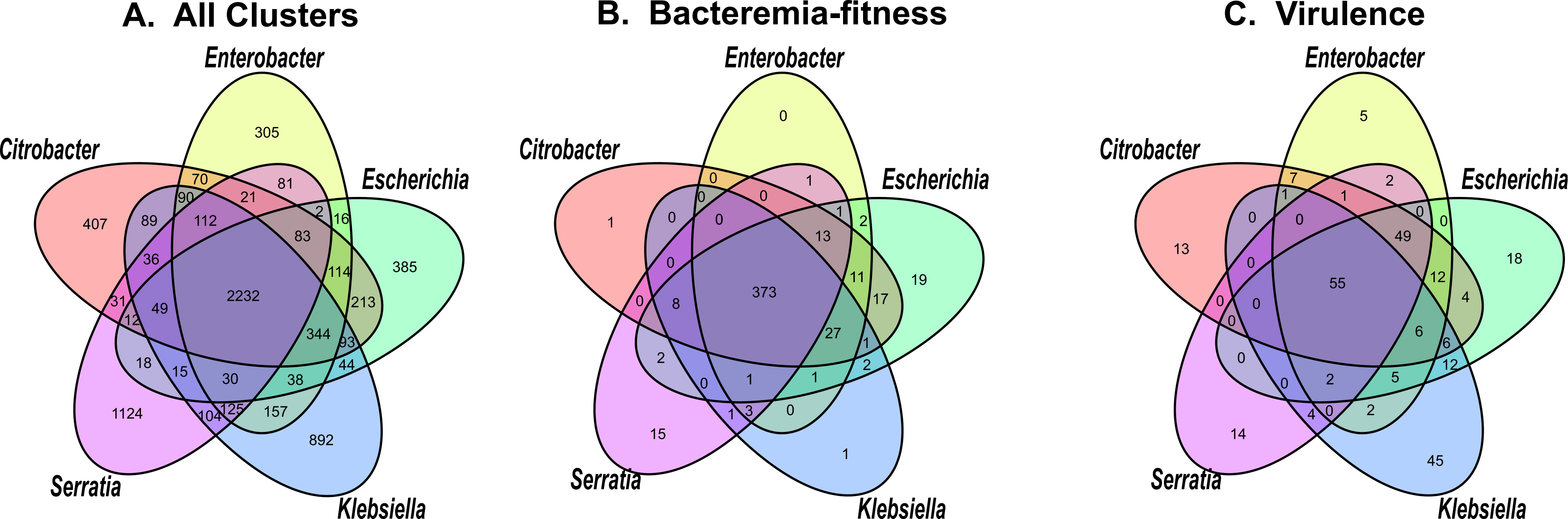

### S2 Fig

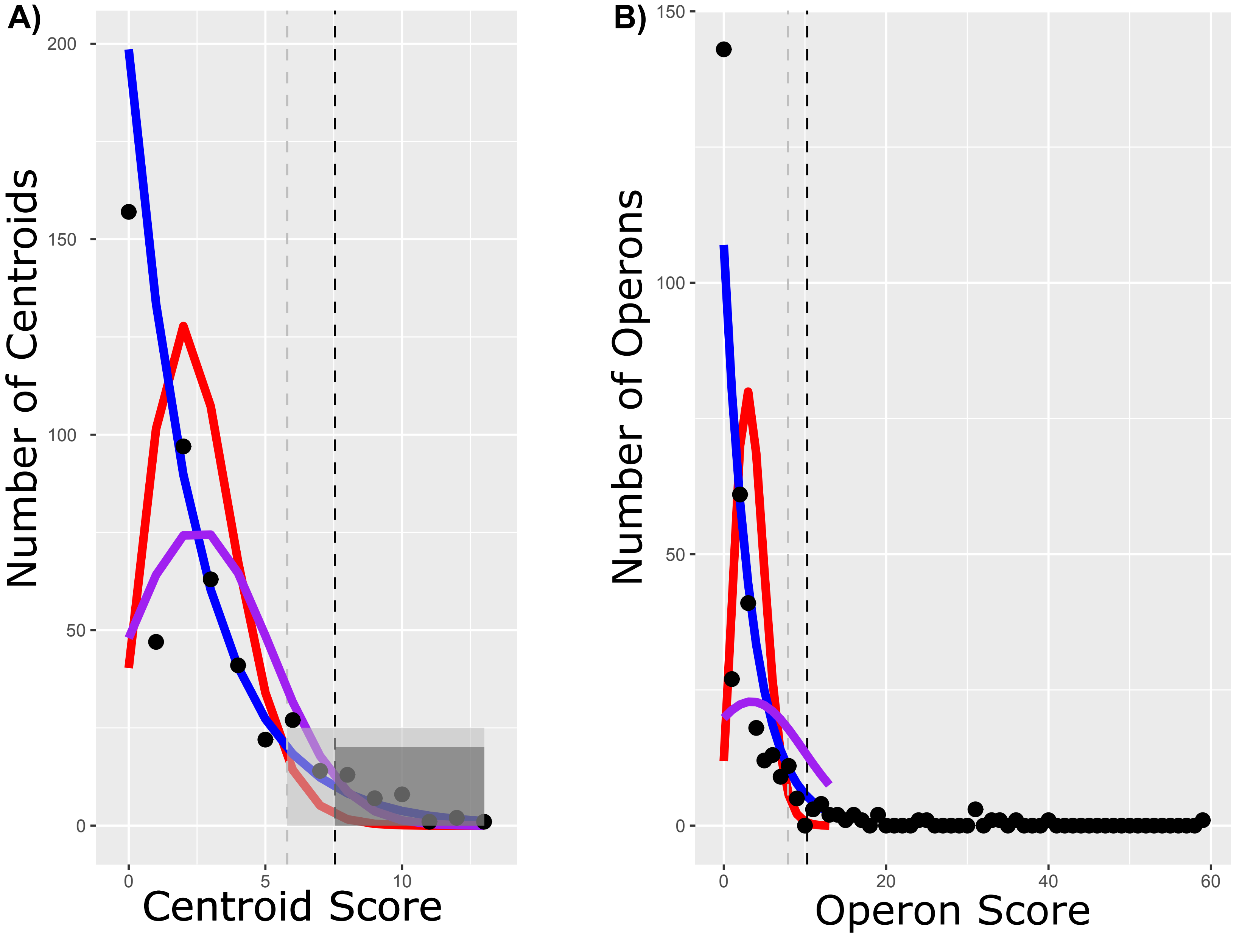
