## Supplementary material for "A novel method for integrating genomic and Tn-Seq data to identify common *in vivo* fitness mechanisms across multiple bacterial species": S2 Dataset: CL_INS_1.html

Legend

 Mobile +extrachromosomalelementfunctions
 Other
 Regulatoryfunctions
 Hypothetical
 Biosynthesis ofcofactors,prostheticgroups, +carriers
 All VFDB Genes

FULL


WINDOWSVGPNG

Trim RowsRemove SingletonsSave Fasta

CL\_204


CL\_204


Break


CL\_204


CL\_204


CL\_204


CL\_204


Break


Break


Break


Break


CL\_204


CL\_204

HighlightSelectShow Genomes


128

CL\_205


43

CL\_205


9

CL\_205


6

CL\_205


4

Break


1

Break


1

CL\_205


1

CL\_205


1

CL\_205


1

CL\_205


1

CL\_205


1

Break


1

CL\_205

fGI ID


CL\_INS\_1
CL\_INS\_1
CL\_INS\_1
CL\_INS\_1
CL\_INS\_1
CL\_INS\_1
CL\_INS\_1
CL\_INS\_278
CL\_INS\_58
CL\_INS\_278
CL\_INS\_318
CL\_INS\_318
CL\_INS\_318
CL\_INS\_318
CL\_INS\_318
CL\_INS\_318
CL\_INS\_1
CL\_INS\_212
CL\_INS\_212
CL\_INS\_278
CL\_INS\_212
CL\_INS\_212
CL\_INS\_278
CL\_INS\_212
CL\_INS\_212
CL\_INS\_212
CL\_INS\_212
CL\_INS\_212
CL\_INS\_212
CL\_INS\_212
CL\_INS\_212
CL\_INS\_212
CL\_INS\_212
CL\_INS\_271
CL\_INS\_212
CL\_INS\_212
CL\_INS\_212
CL\_INS\_212
CL\_INS\_212
CL\_INS\_1
CL\_INS\_318
CL\_INS\_385
CL\_INS\_278
CL\_INS\_278
CL\_INS\_278
CL\_INS\_278
CL\_INS\_278
CL\_INS\_278
CL\_INS\_1
CL\_INS\_1
CL\_INS\_1
CL\_INS\_278
CL\_INS\_278
CL\_INS\_278
CL\_INS\_278
CL\_INS\_278
CL\_INS\_278
CL\_INS\_278
CL\_INS\_278
CL\_INS\_278
CL\_INS\_278
CL\_INS\_278
CL\_INS\_278
CL\_INS\_278
CL\_INS\_278
CL\_INS\_278
CL\_INS\_278
Cluster ID


CL\_17797
CL\_6524
CL\_23546
CL\_17852
CL\_17851
CL\_17850
CL\_12293
CL\_5987
CL\_5986
CL\_6829
CL\_6102
CL\_6101
CL\_6100
CL\_6099
CL\_6098
CL\_6097
CL\_5971
CL\_12418
CL\_12417
CL\_12338
CL\_12416
CL\_12415
CL\_12336
CL\_12414
CL\_12413
CL\_12412
CL\_12411
CL\_12410
CL\_12409
CL\_12408
CL\_12407
CL\_12406
CL\_12405
CL\_5229
CL\_12404
CL\_12403
CL\_12402
CL\_12401
CL\_12400
CL\_12379
CL\_7112
CL\_12378
CL\_6745
CL\_6744
CL\_6743
CL\_6742
CL\_6741
CL\_6740
CL\_6739
CL\_6738
CL\_6737
CL\_6736
CL\_6735
CL\_6734
CL\_6733
CL\_6732
CL\_6763
CL\_6762
CL\_6761
CL\_6760
CL\_6759
CL\_6758
CL\_6757
CL\_6756
CL\_6755
CL\_6754
CL\_6753
