## Supplementary material for "A novel method for integrating genomic and Tn-Seq data to identify common *in vivo* fitness mechanisms across multiple bacterial species": S2 Dataset: CL_INS_2.html

Legend

 Other
 Hypothetical
 All EssentialGenes
 All VFDB Genes

FULL


WINDOWSVGPNG

Trim RowsRemove SingletonsSave Fasta

CL\_213


CL\_213


CL\_213


CL\_212

HighlightSelectShow Genomes


126

CL\_214


71

CL\_214


2

CL\_215


1

CL\_214

fGI ID

CL\_INS\_2
Cluster ID

CL\_5970
