## Supplementary material for "A novel method for integrating genomic and Tn-Seq data to identify common *in vivo* fitness mechanisms across multiple bacterial species": S2 Dataset: CL_INS_3.html

Legend

 Other
 Hypothetical
 All VFDB Genes

FULL


WINDOWSVGPNG

Trim RowsRemove SingletonsSave Fasta

CL\_229


CL\_229


CL\_229


CL\_229

HighlightSelectShow Genomes


190

CL\_230


6

CL\_230


1

CL\_230


1

CL\_230

fGI ID


CL\_INS\_3
CL\_INS\_3
CL\_INS\_3
Cluster ID


CL\_6352
CL\_6351
CL\_6350
