## Supplementary material for "A novel method for integrating genomic and Tn-Seq data to identify common *in vivo* fitness mechanisms across multiple bacterial species": S2 Dataset: CL_INS_5.html

Legend

 Other
 Hypothetical
 All EssentialGenes
 Transport +binding proteins
 All VFDB Genes

FULL


WINDOWSVGPNG

Trim RowsRemove SingletonsSave Fasta

CL\_244


CL\_244

HighlightSelectShow Genomes


192

CL\_245


12

CL\_245

fGI ID


CL\_INS\_5
CL\_INS\_5
CL\_INS\_5
Cluster ID


CL\_7933
CL\_7932
CL\_7931
