## Supplementary material for "A novel method for integrating genomic and Tn-Seq data to identify common *in vivo* fitness mechanisms across multiple bacterial species": S2 Dataset: CL_INS_11.html

Legend

 Mobile +extrachromosomalelementfunctions
 Regulatoryfunctions
 Hypothetical
 All EssentialGenes

FULL


WINDOWSVGPNG

Trim RowsRemove SingletonsSave Fasta

CL\_367


CL\_367


CL\_367


CL\_367

HighlightSelectShow Genomes


114

CL\_369


87

CL\_369


1

CL\_369


1

Break

fGI ID


CL\_INS\_11
CL\_INS\_11
Cluster ID


CL\_11527
CL\_368
