## Supplementary material for "A novel method for integrating genomic and Tn-Seq data to identify common *in vivo* fitness mechanisms across multiple bacterial species": S2 Dataset: CL_INS_12.html

Legend

 Regulatoryfunctions
 Hypothetical
 All Fitness Genes
 Other
 Transport +binding proteins
 All VFDB Genes

FULL


WINDOWSVGPNG

Trim RowsRemove SingletonsSave Fasta

CL\_410


CL\_410


CL\_410


CL\_410


CL\_410


Break


Break


Break


CL\_410


CL\_410


Break


CL\_410

HighlightSelectShow Genomes


147

CL\_411


37

CL\_411


3

Break


3

CL\_411


3

CL\_411


1

CL\_411


1

CL\_411


1

CL\_411


1

Break


1

CL\_411


1

CL\_411


1

CL\_411

fGI ID


CL\_INS\_12
CL\_INS\_12
CL\_INS\_12
CL\_INS\_12
CL\_INS\_307
CL\_INS\_278
CL\_INS\_12
CL\_INS\_12
CL\_INS\_12
CL\_INS\_12
CL\_INS\_278
CL\_INS\_12
CL\_INS\_263
CL\_INS\_263
CL\_INS\_12
CL\_INS\_12
CL\_INS\_263
CL\_INS\_132
CL\_INS\_132
CL\_INS\_30
CL\_INS\_30
Cluster ID


CL\_14437
CL\_18361
CL\_6103
CL\_6542
CL\_8177
CL\_5935
CL\_6110
CL\_6109
CL\_6108
CL\_6107
CL\_6137
CL\_5871
CL\_5467
CL\_5468
CL\_6785
CL\_6784
CL\_5470
CL\_6872
CL\_6871
CL\_6870
CL\_6869
