## Supplementary material for "A novel method for integrating genomic and Tn-Seq data to identify common *in vivo* fitness mechanisms across multiple bacterial species": S2 Dataset: CL_INS_15.html

Legend

 Hypothetical
 All EssentialGenes
 All VFDB Genes

FULL


WINDOWSVGPNG

Trim RowsRemove SingletonsSave Fasta

CL\_458


CL\_458


CL\_458


CL\_457


CL\_458


CL\_458


CL\_458


CL\_458


CL\_457


CL\_458


CL\_458

HighlightSelectShow Genomes


96

CL\_460


85

CL\_460


7

CL\_460


1

CL\_460


1

CL\_460


1

CL\_5289


1

CL\_464


1

CL\_460


1

CL\_460


1

CL\_487


1

CL\_5289

fGI ID


CL\_INS\_15
CL\_INS\_15
CL\_INS\_15
CL\_INS\_15
CL\_INS\_15
CL\_INS\_318
CL\_INS\_318
CL\_INS\_15
Cluster ID


CL\_23146
CL\_7043
CL\_20528
CL\_459
CL\_15239
CL\_8142
CL\_8143
CL\_18536
