## Supplementary material for "A novel method for integrating genomic and Tn-Seq data to identify common *in vivo* fitness mechanisms across multiple bacterial species": S2 Dataset: CL_INS_16.html

Legend

 Mobile +extrachromosomalelementfunctions
 Hypothetical
 All EssentialGenes
 All VFDB Genes

FULL


WINDOWSVGPNG

Trim RowsRemove SingletonsSave Fasta

CL\_461


CL\_461


CL\_5289


CL\_461


CL\_461


CL\_461

HighlightSelectShow Genomes


110

CL\_463


84

CL\_463


1

CL\_463


1

CL\_463


1

CL\_465


1

CL\_464

fGI ID


CL\_INS\_16
CL\_INS\_16
Cluster ID


CL\_9364
CL\_462
