## Supplementary material for "A novel method for integrating genomic and Tn-Seq data to identify common *in vivo* fitness mechanisms across multiple bacterial species": S2 Dataset: CL_INS_17.html

Legend

 Hypothetical
 All EssentialGenes
 Other
 All VFDB Genes

FULL


WINDOWSVGPNG

Trim RowsRemove SingletonsSave Fasta

CL\_464


CL\_464


CL\_463


CL\_461


CL\_429


CL\_464


CL\_463

HighlightSelectShow Genomes


124

CL\_465


68

CL\_465


3

CL\_465


1

CL\_465


1

CL\_465


1

CL\_466


1

CL\_465

fGI ID


CL\_INS\_17
CL\_INS\_17
Cluster ID


CL\_14581
CL\_5862
