## Supplementary material for "A novel method for integrating genomic and Tn-Seq data to identify common *in vivo* fitness mechanisms across multiple bacterial species": S2 Dataset: CL_INS_19.html

Legend

 Mobile +extrachromosomalelementfunctions
 Hypothetical
 All VFDB Genes

FULL


WINDOWSVGPNG

Trim RowsRemove SingletonsSave Fasta

CL\_482


CL\_482


CL\_482


CL\_482


CL\_482


Break

HighlightSelectShow Genomes


114

CL\_484


86

CL\_484


1

CL\_486


1

CL\_486


1

Break


1

CL\_484

fGI ID


CL\_INS\_204
CL\_INS\_19
Cluster ID


CL\_5953
CL\_483
