## Supplementary material for "A novel method for integrating genomic and Tn-Seq data to identify common *in vivo* fitness mechanisms across multiple bacterial species": S2 Dataset: CL_INS_25.html

Legend

 Mobile +extrachromosomalelementfunctions
 Regulatoryfunctions
 Hypothetical
 All VFDB Genes

FULL


WINDOWSVGPNG

Trim RowsRemove SingletonsSave Fasta

CL\_531


CL\_531


CL\_531


CL\_531


CL\_530


CL\_531

HighlightSelectShow Genomes


118

CL\_532


78

CL\_532


2

CL\_533


1

CL\_532


1

CL\_532


1

CL\_532

fGI ID


CL\_INS\_25
CL\_INS\_25
CL\_INS\_25
CL\_INS\_25
CL\_INS\_25
CL\_INS\_26
CL\_INS\_26
CL\_INS\_26
Cluster ID


CL\_14223
CL\_14224
CL\_14225
CL\_5286
CL\_16432
CL\_11117
CL\_5858
CL\_5857
