## Supplementary material for "A novel method for integrating genomic and Tn-Seq data to identify common *in vivo* fitness mechanisms across multiple bacterial species": S2 Dataset: CL_INS_28.html

Legend

 Hypothetical
 Other
 All VFDB Genes

FULL


WINDOWSVGPNG

Trim RowsRemove SingletonsSave Fasta

CL\_546


CL\_546


CL\_546


CL\_544


CL\_545


CL\_546

HighlightSelectShow Genomes


114

CL\_547


44

CL\_547


1

CL\_547


1

CL\_547


1

CL\_547


1

CL\_553

fGI ID


CL\_INS\_28
CL\_INS\_27
Cluster ID


CL\_5853
CL\_13783
