## Supplementary material for "A novel method for integrating genomic and Tn-Seq data to identify common *in vivo* fitness mechanisms across multiple bacterial species": S2 Dataset: CL_INS_29.html

Legend

 Mobile +extrachromosomalelementfunctions
 Hypothetical
 All VFDB Genes

FULL


WINDOWSVGPNG

Trim RowsRemove SingletonsSave Fasta

CL\_549


CL\_549


CL\_549


CL\_549


CL\_552


CL\_549


CL\_549


CL\_549

HighlightSelectShow Genomes


157

CL\_548


22

CL\_542


8

CL\_542


5

CL\_542


1

CL\_548


1

CL\_542


1

CL\_548


1

CL\_548

fGI ID


CL\_INS\_29
CL\_INS\_27
CL\_INS\_27
CL\_INS\_27
CL\_INS\_27
Cluster ID


CL\_18599
CL\_13783
CL\_5284
CL\_7400
CL\_5285
