## Supplementary material for "A novel method for integrating genomic and Tn-Seq data to identify common *in vivo* fitness mechanisms across multiple bacterial species": S2 Dataset: CL_INS_34.html

Legend

 Hypothetical
 All Fitness Genes
 Other
 All VFDB Genes

FULL


WINDOWSVGPNG

Trim RowsRemove SingletonsSave Fasta

CL\_603


CL\_603


CL\_603

HighlightSelectShow Genomes


199

CL\_604


2

CL\_604


1

CL\_604

fGI ID


CL\_INS\_34
CL\_INS\_34
CL\_INS\_34
CL\_INS\_34
CL\_INS\_34
CL\_INS\_34
CL\_INS\_34
Cluster ID


CL\_8378
CL\_8377
CL\_8376
CL\_11512
CL\_8375
CL\_8374
CL\_8373
