## Supplementary material for "A novel method for integrating genomic and Tn-Seq data to identify common *in vivo* fitness mechanisms across multiple bacterial species": S2 Dataset: CL_INS_35.html

Legend

 Mobile +extrachromosomalelementfunctions
 Other
 Transport +binding proteins
 All VFDB Genes

FULL


WINDOWSVGPNG

Trim RowsRemove SingletonsSave Fasta

CL\_615


CL\_615


CL\_615


CL\_615


CL\_615


CL\_615

HighlightSelectShow Genomes


102

CL\_616


95

CL\_616


4

CL\_616


2

CL\_616


1

CL\_616


1

CL\_616

fGI ID


CL\_INS\_35
CL\_INS\_35
CL\_INS\_35
CL\_INS\_35
Cluster ID


CL\_5282
CL\_5281
CL\_21296
CL\_5280
