## Supplementary material for "A novel method for integrating genomic and Tn-Seq data to identify common *in vivo* fitness mechanisms across multiple bacterial species": S2 Dataset: CL_INS_36.html

Legend

 Hypothetical
 All EssentialGenes
 Other
 Transport +binding proteins
 All VFDB Genes

FULL


WINDOWSVGPNG

Trim RowsRemove SingletonsSave Fasta

CL\_616


CL\_616


CL\_616


CL\_616


CL\_521


CL\_616


CL\_616


CL\_616

HighlightSelectShow Genomes


188

CL\_617


9

CL\_617


3

CL\_617


1

CL\_617


1

CL\_617


1

CL\_617


1

CL\_617


1

CL\_520

fGI ID


CL\_INS\_36
CL\_INS\_36
CL\_INS\_36
CL\_INS\_36
CL\_INS\_36
CL\_INS\_10
CL\_INS\_36
CL\_INS\_36
CL\_INS\_10
CL\_INS\_36
Cluster ID


CL\_11116
CL\_11115
CL\_11114
CL\_11113
CL\_7713
CL\_4594
CL\_12010
CL\_23545
CL\_4593
CL\_7712
