## Supplementary material for "A novel method for integrating genomic and Tn-Seq data to identify common *in vivo* fitness mechanisms across multiple bacterial species": S2 Dataset: CL_INS_39.html

Legend

 Regulatoryfunctions
 Hypothetical
 EnergyMetabolism

FULL


WINDOWSVGPNG

Trim RowsRemove SingletonsSave Fasta

CL\_633


CL\_633

HighlightSelectShow Genomes


145

CL\_635


60

CL\_635

fGI ID

CL\_INS\_39
Cluster ID

CL\_634
