## Supplementary material for "A novel method for integrating genomic and Tn-Seq data to identify common *in vivo* fitness mechanisms across multiple bacterial species": S2 Dataset: CL_INS_41.html

Legend

 Mobile +extrachromosomalelementfunctions
 Regulatoryfunctions
 Hypothetical
 All EssentialGenes
 Other
 All VFDB Genes

FULL


WINDOWSVGPNG

Trim RowsRemove SingletonsSave Fasta

CL\_663


CL\_663


CL\_663


CL\_663


CL\_663

HighlightSelectShow Genomes


96

CL\_664


61

CL\_664


45

CL\_664


2

CL\_664


1

CL\_664

fGI ID


CL\_INS\_41
CL\_INS\_41
CL\_INS\_41
CL\_INS\_41
CL\_INS\_41
CL\_INS\_41
CL\_INS\_41
Cluster ID


CL\_7763
CL\_18552
CL\_6333
CL\_6332
CL\_6331
CL\_6330
CL\_6329
