## Supplementary material for "A novel method for integrating genomic and Tn-Seq data to identify common *in vivo* fitness mechanisms across multiple bacterial species": S2 Dataset: CL_INS_42.html

Legend

 Mobile +extrachromosomalelementfunctions
 Regulatoryfunctions
 Hypothetical
 Other
 All VFDB Genes

FULL


WINDOWSVGPNG

Trim RowsRemove SingletonsSave Fasta

CL\_676


CL\_676


CL\_676


CL\_676


CL\_676


CL\_676


Break


CL\_676


CL\_676


CL\_676


CL\_676


CL\_676

HighlightSelectShow Genomes


186

CL\_677


5

CL\_677


3

CL\_677


2

Break


2

CL\_677


2

CL\_677


1

CL\_677


1

CL\_677


1

Break


1

CL\_677


1

Break


1

Break

fGI ID


CL\_INS\_42
CL\_INS\_42
CL\_INS\_278
CL\_INS\_278
CL\_INS\_278
CL\_INS\_278
CL\_INS\_278
CL\_INS\_42
CL\_INS\_42
CL\_INS\_42
CL\_INS\_42
CL\_INS\_42
CL\_INS\_278
CL\_INS\_42
CL\_INS\_42
CL\_INS\_42
CL\_INS\_42
CL\_INS\_42
CL\_INS\_42
CL\_INS\_42
Cluster ID


CL\_5584
CL\_23195
CL\_6810
CL\_7063
CL\_7064
CL\_5990
CL\_5989
CL\_7795
CL\_7796
CL\_7797
CL\_13643
CL\_13644
CL\_8141
CL\_6327
CL\_6326
CL\_20445
CL\_6804
CL\_6805
CL\_6685
CL\_20444
