## Supplementary material for "A novel method for integrating genomic and Tn-Seq data to identify common *in vivo* fitness mechanisms across multiple bacterial species": S2 Dataset: CL_INS_43.html

Legend

 Hypothetical
 Other
 All VFDB Genes

FULL


WINDOWSVGPNG

Trim RowsRemove SingletonsSave Fasta

CL\_678


CL\_678


CL\_678

HighlightSelectShow Genomes


111

CL\_680


91

CL\_680


1

CL\_681

fGI ID

CL\_INS\_43
Cluster ID

CL\_679
