## Supplementary material for "A novel method for integrating genomic and Tn-Seq data to identify common *in vivo* fitness mechanisms across multiple bacterial species": S2 Dataset: CL_INS_44.html

Legend

 Mobile +extrachromosomalelementfunctions
 Regulatoryfunctions
 Hypothetical
 DNA Metabolism
 All EssentialGenes
 Proteinsynthesis/fate
 Other
 EnergyMetabolism
 Transport +binding proteins
 All VFDB Genes

FULL


WINDOWSVGPNG

Trim RowsRemove SingletonsSave Fasta

CL\_691


CL\_691


CL\_691


CL\_691


CL\_691


CL\_691


CL\_691


CL\_691


CL\_691


CL\_691


CL\_691


CL\_691


CL\_691


CL\_691


CL\_691


CL\_691


CL\_691


Break


CL\_691


CL\_691


CL\_691


CL\_691


CL\_691


CL\_691


CL\_691


CL\_691


CL\_691


CL\_691


CL\_691


CL\_691


CL\_691


CL\_691


CL\_691


CL\_691


CL\_691


CL\_691


CL\_691


CL\_691


CL\_691

HighlightSelectShow Genomes


155

CL\_692


6

CL\_758


5

CL\_692


2

CL\_758


2

CL\_759


1

CL\_759


1

CL\_759


1

CL\_758


1

CL\_758


1

CL\_758


1

CL\_758


1

CL\_759


1

CL\_758


1

CL\_758


1

CL\_758


1

CL\_758


1

Break


1

CL\_692


1

CL\_758


1

CL\_759


1

CL\_758


1

CL\_758


1

CL\_758


1

CL\_758


1

CL\_759


1

CL\_758


1

CL\_758


1

Break


1

CL\_758


1

CL\_758


1

CL\_759


1

CL\_758


1

CL\_759


1

CL\_758


1

CL\_758


1

CL\_758


1

CL\_758


1

CL\_692


1

CL\_758

fGI ID


CL\_INS\_44
CL\_INS\_44
CL\_INS\_44
CL\_INS\_44
CL\_INS\_44
CL\_INS\_44
CL\_INS\_44
CL\_INS\_44
CL\_INS\_44
CL\_INS\_44
CL\_INS\_44
CL\_INS\_44
CL\_INS\_44
CL\_INS\_278
CL\_INS\_278
CL\_INS\_278
CL\_INS\_278
CL\_INS\_278
CL\_INS\_278
CL\_INS\_44
CL\_INS\_278
CL\_INS\_44
CL\_INS\_44
CL\_INS\_44
CL\_INS\_44
CL\_INS\_385
CL\_INS\_132
CL\_INS\_44
CL\_INS\_44
CL\_INS\_44
CL\_INS\_44
CL\_INS\_44
CL\_INS\_44
CL\_INS\_44
CL\_INS\_44
CL\_INS\_44
CL\_INS\_44
CL\_INS\_44
CL\_INS\_44
CL\_INS\_44
CL\_INS\_44
CL\_INS\_44
CL\_INS\_44
CL\_INS\_44
CL\_INS\_321
CL\_INS\_44
CL\_INS\_44
CL\_INS\_44
CL\_INS\_44
CL\_INS\_44
CL\_INS\_44
CL\_INS\_44
CL\_INS\_44
CL\_INS\_44
CL\_INS\_44
CL\_INS\_44
CL\_INS\_44
CL\_INS\_44
CL\_INS\_44
CL\_INS\_44
CL\_INS\_44
CL\_INS\_44
CL\_INS\_44
CL\_INS\_44
CL\_INS\_44
CL\_INS\_44
CL\_INS\_44
CL\_INS\_44
CL\_INS\_44
CL\_INS\_44
CL\_INS\_44
CL\_INS\_321
CL\_INS\_44
CL\_INS\_321
CL\_INS\_321
CL\_INS\_44
CL\_INS\_321
CL\_INS\_57
CL\_INS\_44
CL\_INS\_44
CL\_INS\_44
CL\_INS\_44
CL\_INS\_44
CL\_INS\_44
CL\_INS\_44
CL\_INS\_44
CL\_INS\_44
CL\_INS\_44
CL\_INS\_44
CL\_INS\_44
CL\_INS\_44
CL\_INS\_44
CL\_INS\_44
CL\_INS\_44
CL\_INS\_44
CL\_INS\_44
CL\_INS\_44
CL\_INS\_44
CL\_INS\_44
CL\_INS\_44
CL\_INS\_44
CL\_INS\_351
CL\_INS\_44
CL\_INS\_44
CL\_INS\_44
CL\_INS\_44
CL\_INS\_44
CL\_INS\_44
CL\_INS\_44
CL\_INS\_44
CL\_INS\_44
CL\_INS\_44
CL\_INS\_44
CL\_INS\_44
CL\_INS\_44
CL\_INS\_44
CL\_INS\_44
CL\_INS\_44
CL\_INS\_307
CL\_INS\_74
CL\_INS\_74
CL\_INS\_44
CL\_INS\_74
CL\_INS\_74
CL\_INS\_74
CL\_INS\_44
CL\_INS\_74
CL\_INS\_44
CL\_INS\_44
CL\_INS\_44
CL\_INS\_74
CL\_INS\_74
CL\_INS\_44
CL\_INS\_351
CL\_INS\_351
CL\_INS\_74
CL\_INS\_351
CL\_INS\_74
CL\_INS\_74
CL\_INS\_74
CL\_INS\_74
CL\_INS\_74
CL\_INS\_74
CL\_INS\_44
CL\_INS\_44
CL\_INS\_74
CL\_INS\_44
CL\_INS\_44
CL\_INS\_44
CL\_INS\_44
CL\_INS\_44
CL\_INS\_44
CL\_INS\_44
CL\_INS\_44
CL\_INS\_44
CL\_INS\_44
CL\_INS\_44
CL\_INS\_44
CL\_INS\_44
CL\_INS\_44
CL\_INS\_44
CL\_INS\_44
CL\_INS\_44
CL\_INS\_44
CL\_INS\_351
CL\_INS\_44
CL\_INS\_351
CL\_INS\_351
CL\_INS\_44
CL\_INS\_44
CL\_INS\_44
CL\_INS\_44
CL\_INS\_44
CL\_INS\_351
CL\_INS\_351
CL\_INS\_351
CL\_INS\_351
CL\_INS\_351
CL\_INS\_44
CL\_INS\_44
CL\_INS\_44
CL\_INS\_44
CL\_INS\_44
CL\_INS\_351
CL\_INS\_351
CL\_INS\_351
CL\_INS\_351
CL\_INS\_351
CL\_INS\_351
CL\_INS\_351
CL\_INS\_351
CL\_INS\_44
CL\_INS\_44
CL\_INS\_44
CL\_INS\_44
CL\_INS\_132
CL\_INS\_44
CL\_INS\_44
CL\_INS\_44
CL\_INS\_44
CL\_INS\_44
CL\_INS\_44
CL\_INS\_44
CL\_INS\_44
CL\_INS\_44
CL\_INS\_44
CL\_INS\_351
CL\_INS\_351
CL\_INS\_44
CL\_INS\_44
CL\_INS\_44
CL\_INS\_44
CL\_INS\_44
CL\_INS\_44
CL\_INS\_44
CL\_INS\_44
CL\_INS\_44
CL\_INS\_44
CL\_INS\_44
CL\_INS\_44
CL\_INS\_44
CL\_INS\_44
CL\_INS\_44
CL\_INS\_44
CL\_INS\_44
CL\_INS\_44
CL\_INS\_44
CL\_INS\_44
CL\_INS\_44
CL\_INS\_44
CL\_INS\_44
CL\_INS\_44
CL\_INS\_204
CL\_INS\_204
CL\_INS\_204
CL\_INS\_44
CL\_INS\_44
CL\_INS\_44
CL\_INS\_44
CL\_INS\_321
CL\_INS\_44
CL\_INS\_44
CL\_INS\_321
CL\_INS\_321
CL\_INS\_44
CL\_INS\_321
CL\_INS\_321
CL\_INS\_321
CL\_INS\_321
CL\_INS\_44
CL\_INS\_44
CL\_INS\_321
CL\_INS\_321
CL\_INS\_44
CL\_INS\_44
CL\_INS\_44
CL\_INS\_44
CL\_INS\_44
CL\_INS\_91
CL\_INS\_44
CL\_INS\_44
CL\_INS\_44
CL\_INS\_44
CL\_INS\_44
CL\_INS\_44
CL\_INS\_44
CL\_INS\_44
CL\_INS\_44
CL\_INS\_44
CL\_INS\_44
CL\_INS\_44
CL\_INS\_44
CL\_INS\_44
CL\_INS\_44
CL\_INS\_44
CL\_INS\_44
CL\_INS\_44
CL\_INS\_44
CL\_INS\_44
CL\_INS\_44
CL\_INS\_44
CL\_INS\_44
CL\_INS\_44
CL\_INS\_44
CL\_INS\_44
CL\_INS\_207
CL\_INS\_278
CL\_INS\_44
CL\_INS\_44
CL\_INS\_44
CL\_INS\_44
CL\_INS\_44
CL\_INS\_44
CL\_INS\_44
CL\_INS\_57
CL\_INS\_57
CL\_INS\_57
CL\_INS\_57
CL\_INS\_57
CL\_INS\_57
CL\_INS\_57
CL\_INS\_57
CL\_INS\_44
CL\_INS\_44
CL\_INS\_246
CL\_INS\_44
CL\_INS\_246
CL\_INS\_246
CL\_INS\_57
CL\_INS\_57
CL\_INS\_44
CL\_INS\_57
CL\_INS\_246
CL\_INS\_44
CL\_INS\_44
CL\_INS\_246
CL\_INS\_44
CL\_INS\_44
CL\_INS\_44
CL\_INS\_44
CL\_INS\_44
CL\_INS\_44
CL\_INS\_57
CL\_INS\_57
CL\_INS\_57
CL\_INS\_246
CL\_INS\_246
CL\_INS\_246
CL\_INS\_334
CL\_INS\_44
CL\_INS\_44
CL\_INS\_44
CL\_INS\_44
CL\_INS\_44
CL\_INS\_44
CL\_INS\_57
CL\_INS\_44
CL\_INS\_57
Cluster ID


CL\_17870
CL\_5277
CL\_5276
CL\_5275
CL\_5274
CL\_5273
CL\_13075
CL\_13074
CL\_13073
CL\_13072
CL\_13071
CL\_13070
CL\_13069
CL\_6731
CL\_6730
CL\_6729
CL\_6728
CL\_6877
CL\_6876
CL\_6875
CL\_6874
CL\_6873
CL\_11835
CL\_5846
CL\_5845
CL\_7810
CL\_6822
CL\_5844
CL\_10538
CL\_10537
CL\_5843
CL\_5842
CL\_5841
CL\_5840
CL\_23544
CL\_23543
CL\_23542
CL\_23541
CL\_23540
CL\_23539
CL\_23538
CL\_14280
CL\_14281
CL\_14282
CL\_6339
CL\_10287
CL\_10286
CL\_10285
CL\_10284
CL\_19137
CL\_11511
CL\_11510
CL\_19136
CL\_19135
CL\_19134
CL\_11509
CL\_11508
CL\_20194
CL\_10281
CL\_10280
CL\_10279
CL\_10278
CL\_22589
CL\_10277
CL\_10282
CL\_10283
CL\_10276
CL\_20031
CL\_12172
CL\_6538
CL\_6537
CL\_6536
CL\_6535
CL\_6534
CL\_6533
CL\_6532
CL\_6531
CL\_746
CL\_20784
CL\_20783
CL\_8350
CL\_24200
CL\_24199
CL\_24198
CL\_24197
CL\_24196
CL\_24195
CL\_24194
CL\_14061
CL\_8372
CL\_8371
CL\_8368
CL\_13807
CL\_8365
CL\_8364
CL\_14060
CL\_23696
CL\_12924
CL\_12925
CL\_12927
CL\_22274
CL\_5561
CL\_17104
CL\_20619
CL\_20618
CL\_20617
CL\_20616
CL\_20615
CL\_7917
CL\_7916
CL\_7915
CL\_7918
CL\_17684
CL\_20614
CL\_17687
CL\_18730
CL\_18731
CL\_18732
CL\_6821
CL\_18733
CL\_18734
CL\_18735
CL\_18736
CL\_5568
CL\_5569
CL\_18737
CL\_13265
CL\_18738
CL\_21238
CL\_21237
CL\_7140
CL\_9061
CL\_16473
CL\_7142
CL\_18649
CL\_7143
CL\_7466
CL\_9064
CL\_7144
CL\_7146
CL\_7147
CL\_7467
CL\_7148
CL\_7468
CL\_7469
CL\_7151
CL\_21236
CL\_18739
CL\_12011
CL\_5839
CL\_17972
CL\_17971
CL\_17970
CL\_17969
CL\_17968
CL\_17967
CL\_17966
CL\_17965
CL\_17964
CL\_17963
CL\_17962
CL\_17961
CL\_17960
CL\_17959
CL\_11608
CL\_20997
CL\_11607
CL\_15323
CL\_11606
CL\_11605
CL\_20996
CL\_20995
CL\_20994
CL\_11604
CL\_11603
CL\_11602
CL\_15459
CL\_18220
CL\_20993
CL\_20992
CL\_20991
CL\_20990
CL\_20989
CL\_11601
CL\_12045
CL\_11600
CL\_11599
CL\_11598
CL\_12046
CL\_12047
CL\_7152
CL\_17958
CL\_14283
CL\_14284
CL\_14285
CL\_6840
CL\_14286
CL\_14287
CL\_17957
CL\_7531
CL\_6530
CL\_6529
CL\_6528
CL\_6527
CL\_6526
CL\_6525
CL\_11597
CL\_11596
CL\_11595
CL\_11594
CL\_11593
CL\_11592
CL\_7041
CL\_11591
CL\_8842
CL\_11590
CL\_11589
CL\_11588
CL\_11587
CL\_11586
CL\_11585
CL\_11584
CL\_11112
CL\_11111
CL\_11110
CL\_11109
CL\_11108
CL\_11107
CL\_11106
CL\_11105
CL\_11104
CL\_11103
CL\_5475
CL\_6159
CL\_5524
CL\_11102
CL\_11101
CL\_11100
CL\_11099
CL\_7428
CL\_20613
CL\_20612
CL\_7421
CL\_7419
CL\_9166
CL\_8355
CL\_8354
CL\_8353
CL\_8352
CL\_8351
CL\_20611
CL\_7427
CL\_7426
CL\_10242
CL\_11098
CL\_11097
CL\_11096
CL\_11095
CL\_6027
CL\_11094
CL\_11093
CL\_11092
CL\_6394
CL\_23487
CL\_18653
CL\_12855
CL\_23486
CL\_23485
CL\_23484
CL\_23483
CL\_23482
CL\_18654
CL\_18655
CL\_18656
CL\_18657
CL\_18658
CL\_18659
CL\_18660
CL\_18661
CL\_18662
CL\_18663
CL\_18664
CL\_18665
CL\_18666
CL\_18667
CL\_9529
CL\_6081
CL\_18668
CL\_18669
CL\_18670
CL\_18671
CL\_18672
CL\_18673
CL\_18674
CL\_721
CL\_722
CL\_723
CL\_724
CL\_725
CL\_726
CL\_727
CL\_728
CL\_8989
CL\_18675
CL\_6427
CL\_18676
CL\_6429
CL\_6430
CL\_733
CL\_734
CL\_18677
CL\_735
CL\_737
CL\_18678
CL\_18679
CL\_6432
CL\_18680
CL\_18681
CL\_18682
CL\_18683
CL\_18684
CL\_18685
CL\_741
CL\_742
CL\_743
CL\_6438
CL\_6439
CL\_6440
CL\_1165
CL\_8830
CL\_15458
CL\_696
CL\_5838
CL\_753
CL\_754
CL\_755
CL\_7399
CL\_756
