## Supplementary material for "A novel method for integrating genomic and Tn-Seq data to identify common *in vivo* fitness mechanisms across multiple bacterial species": S2 Dataset: CL_INS_45.html

Legend

 Mobile +extrachromosomalelementfunctions
 Regulatoryfunctions
 Hypothetical
 DNA Metabolism
 All EssentialGenes
 Proteinsynthesis/fate
 Other
 EnergyMetabolism
 Transport +binding proteins
 All VFDB Genes

FULL


WINDOWSVGPNG

Trim RowsRemove SingletonsSave Fasta

CL\_692


CL\_691


CL\_692


CL\_692


CL\_692


CL\_692


CL\_692


Break


CL\_692


CL\_692


CL\_692


CL\_692


CL\_692


CL\_692


CL\_692


CL\_692


CL\_692


CL\_692


CL\_691


CL\_692


CL\_691


CL\_691


CL\_692


CL\_692


CL\_692


CL\_691


CL\_691


CL\_692


CL\_692


CL\_692


CL\_692


CL\_691


CL\_692


CL\_692


CL\_692


CL\_692


CL\_692


CL\_692


CL\_692


CL\_692


CL\_690


CL\_692


CL\_692


CL\_692


CL\_692


CL\_692


CL\_691


CL\_691


CL\_691


CL\_692


CL\_692


CL\_691


CL\_692


CL\_692


CL\_692


CL\_692


CL\_692


CL\_692


CL\_692


CL\_692


CL\_692


CL\_692


CL\_692


CL\_692


CL\_692


CL\_692


CL\_692


CL\_865


CL\_692


CL\_692


CL\_692


CL\_692


CL\_691


CL\_691


CL\_692


CL\_691


CL\_692


CL\_692


CL\_692


CL\_692


CL\_692


CL\_692


CL\_692


CL\_692


CL\_691


CL\_692


CL\_692


CL\_692


CL\_692


CL\_692


CL\_692


CL\_692


CL\_692


CL\_691


CL\_692


CL\_692


CL\_692


CL\_692


CL\_692


CL\_691


CL\_692


CL\_692


CL\_692


CL\_692


CL\_692


CL\_692


CL\_692


CL\_692


CL\_692


CL\_691


CL\_692


CL\_692


CL\_692


CL\_692


CL\_692


CL\_692


CL\_692


CL\_692


CL\_691


CL\_691


CL\_692


CL\_691


CL\_692


CL\_692


CL\_692


CL\_692


CL\_691


CL\_692


CL\_692


CL\_692


CL\_692


CL\_692


CL\_692


CL\_692


CL\_691


CL\_692


CL\_692


CL\_692


CL\_692


CL\_692


CL\_692


CL\_692


CL\_692


CL\_692


CL\_692


CL\_692


CL\_692


CL\_692


CL\_692


CL\_692


CL\_692


CL\_692


CL\_692


CL\_692


CL\_692


CL\_692


CL\_691


CL\_692


CL\_692


CL\_691


CL\_692


CL\_692

HighlightSelectShow Genomes


7

CL\_758


6

CL\_758


4

CL\_758


4

CL\_758


3

CL\_758


3

CL\_758


3

CL\_758


3

CL\_758


3

CL\_758


2

CL\_758


2

CL\_758


2

CL\_758


2

CL\_758


2

CL\_759


2

CL\_758


2

CL\_761


2

CL\_758


2

CL\_758


2

CL\_758


1

CL\_758


1

CL\_758


1

CL\_758


1

CL\_758


1

CL\_758


1

CL\_758


1

CL\_758


1

CL\_758


1

Break


1

CL\_758


1

CL\_758


1

CL\_758


1

CL\_758


1

CL\_758


1

CL\_758


1

CL\_758


1

CL\_758


1

CL\_758


1

CL\_758


1

CL\_758


1

Break


1

CL\_758


1

Break


1

CL\_758


1

CL\_758


1

CL\_758


1

CL\_758


1

CL\_758


1

CL\_758


1

CL\_758


1

CL\_758


1

CL\_758


1

CL\_758


1

CL\_758


1

CL\_758


1

CL\_758


1

CL\_758


1

CL\_758


1

CL\_758


1

CL\_758


1

CL\_758


1

CL\_758


1

CL\_758


1

CL\_758


1

CL\_758


1

CL\_758


1

CL\_758


1

CL\_758


1

CL\_758


1

CL\_758


1

CL\_862


1

CL\_758


1

CL\_758


1

CL\_758


1

CL\_758


1

CL\_758


1

CL\_758


1

CL\_758


1

CL\_758


1

CL\_758


1

CL\_758


1

CL\_759


1

CL\_758


1

CL\_758


1

CL\_758


1

CL\_758


1

CL\_758


1

CL\_758


1

CL\_759


1

CL\_758


1

CL\_758


1

CL\_758


1

CL\_758


1

CL\_758


1

CL\_758


1

CL\_758


1

CL\_758


1

CL\_758


1

CL\_758


1

CL\_758


1

CL\_758


1

CL\_758


1

CL\_758


1

CL\_759


1

CL\_758


1

CL\_759


1

CL\_758


1

CL\_758


1

CL\_758


1

CL\_758


1

CL\_758


1

CL\_758


1

CL\_758


1

CL\_758


1

CL\_758


1

CL\_758


1

CL\_758


1

CL\_758


1

CL\_758


1

CL\_758


1

CL\_758


1

CL\_758


1

CL\_758


1

CL\_758


1

CL\_758


1

CL\_758


1

CL\_758


1

CL\_758


1

CL\_758


1

CL\_758


1

CL\_758


1

CL\_758


1

CL\_758


1

CL\_758


1

Break


1

CL\_758


1

CL\_758


1

CL\_758


1

CL\_758


1

CL\_758


1

CL\_758


1

CL\_758


1

CL\_758


1

CL\_758


1

CL\_758


1

CL\_759


1

CL\_758


1

CL\_758


1

CL\_758


1

CL\_758


1

CL\_758


1

CL\_758


1

CL\_759


1

CL\_758


1

CL\_758


1

CL\_758


1

CL\_758


1

CL\_758


1

CL\_758


1

CL\_758


1

CL\_758


1

CL\_758


1

CL\_758

fGI ID


CL\_INS\_385
CL\_INS\_44
CL\_INS\_44
CL\_INS\_44
CL\_INS\_44
CL\_INS\_44
CL\_INS\_45
CL\_INS\_45
CL\_INS\_74
CL\_INS\_45
CL\_INS\_74
CL\_INS\_45
CL\_INS\_45
CL\_INS\_45
CL\_INS\_45
CL\_INS\_45
CL\_INS\_45
CL\_INS\_45
CL\_INS\_44
CL\_INS\_44
CL\_INS\_44
CL\_INS\_44
CL\_INS\_44
CL\_INS\_44
CL\_INS\_45
CL\_INS\_45
CL\_INS\_44
CL\_INS\_44
CL\_INS\_44
CL\_INS\_44
CL\_INS\_44
CL\_INS\_44
CL\_INS\_44
CL\_INS\_44
CL\_INS\_44
CL\_INS\_44
CL\_INS\_44
CL\_INS\_44
CL\_INS\_44
CL\_INS\_44
CL\_INS\_44
CL\_INS\_44
CL\_INS\_44
CL\_INS\_45
CL\_INS\_45
CL\_INS\_45
CL\_INS\_45
CL\_INS\_45
CL\_INS\_351
CL\_INS\_74
CL\_INS\_321
CL\_INS\_278
CL\_INS\_132
CL\_INS\_278
CL\_INS\_278
CL\_INS\_278
CL\_INS\_278
CL\_INS\_278
CL\_INS\_132
CL\_INS\_45
CL\_INS\_401
CL\_INS\_278
CL\_INS\_278
CL\_INS\_278
CL\_INS\_45
CL\_INS\_45
CL\_INS\_132
CL\_INS\_307
CL\_INS\_307
CL\_INS\_45
CL\_INS\_45
CL\_INS\_45
CL\_INS\_45
CL\_INS\_45
CL\_INS\_45
CL\_INS\_45
CL\_INS\_45
CL\_INS\_45
CL\_INS\_45
CL\_INS\_45
CL\_INS\_45
CL\_INS\_45
CL\_INS\_45
CL\_INS\_45
CL\_INS\_45
CL\_INS\_45
CL\_INS\_155
CL\_INS\_155
CL\_INS\_155
CL\_INS\_155
CL\_INS\_155
CL\_INS\_44
CL\_INS\_44
CL\_INS\_307
CL\_INS\_74
CL\_INS\_74
CL\_INS\_44
CL\_INS\_74
CL\_INS\_74
CL\_INS\_74
CL\_INS\_44
CL\_INS\_74
CL\_INS\_44
CL\_INS\_44
CL\_INS\_45
CL\_INS\_44
CL\_INS\_44
CL\_INS\_44
CL\_INS\_44
CL\_INS\_44
CL\_INS\_74
CL\_INS\_74
CL\_INS\_44
CL\_INS\_351
CL\_INS\_351
CL\_INS\_74
CL\_INS\_351
CL\_INS\_74
CL\_INS\_74
CL\_INS\_74
CL\_INS\_74
CL\_INS\_74
CL\_INS\_74
CL\_INS\_44
CL\_INS\_44
CL\_INS\_74
CL\_INS\_44
CL\_INS\_44
CL\_INS\_44
CL\_INS\_45
CL\_INS\_44
CL\_INS\_44
CL\_INS\_44
CL\_INS\_44
CL\_INS\_44
CL\_INS\_44
CL\_INS\_44
CL\_INS\_44
CL\_INS\_44
CL\_INS\_44
CL\_INS\_44
CL\_INS\_44
CL\_INS\_44
CL\_INS\_351
CL\_INS\_351
CL\_INS\_44
CL\_INS\_44
CL\_INS\_44
CL\_INS\_44
CL\_INS\_44
CL\_INS\_351
CL\_INS\_351
CL\_INS\_351
CL\_INS\_351
CL\_INS\_351
CL\_INS\_44
CL\_INS\_44
CL\_INS\_44
CL\_INS\_44
CL\_INS\_44
CL\_INS\_351
CL\_INS\_351
CL\_INS\_351
CL\_INS\_351
CL\_INS\_351
CL\_INS\_351
CL\_INS\_351
CL\_INS\_351
CL\_INS\_44
CL\_INS\_44
CL\_INS\_44
CL\_INS\_44
CL\_INS\_132
CL\_INS\_44
CL\_INS\_44
CL\_INS\_44
CL\_INS\_351
CL\_INS\_351
CL\_INS\_44
CL\_INS\_45
CL\_INS\_321
CL\_INS\_155
CL\_INS\_45
CL\_INS\_45
CL\_INS\_45
CL\_INS\_45
CL\_INS\_45
CL\_INS\_45
CL\_INS\_45
CL\_INS\_45
CL\_INS\_45
CL\_INS\_45
CL\_INS\_45
CL\_INS\_45
CL\_INS\_45
CL\_INS\_45
CL\_INS\_44
CL\_INS\_44
CL\_INS\_44
CL\_INS\_44
CL\_INS\_44
CL\_INS\_44
CL\_INS\_44
CL\_INS\_44
CL\_INS\_44
CL\_INS\_26
CL\_INS\_26
CL\_INS\_26
CL\_INS\_278
CL\_INS\_278
CL\_INS\_45
CL\_INS\_45
CL\_INS\_45
CL\_INS\_45
CL\_INS\_44
CL\_INS\_132
CL\_INS\_44
CL\_INS\_45
CL\_INS\_44
CL\_INS\_45
CL\_INS\_45
CL\_INS\_45
CL\_INS\_45
CL\_INS\_45
CL\_INS\_45
CL\_INS\_321
CL\_INS\_321
CL\_INS\_45
CL\_INS\_45
CL\_INS\_45
CL\_INS\_318
CL\_INS\_318
CL\_INS\_318
CL\_INS\_318
CL\_INS\_45
CL\_INS\_45
CL\_INS\_318
CL\_INS\_318
CL\_INS\_57
CL\_INS\_321
CL\_INS\_45
CL\_INS\_45
CL\_INS\_45
CL\_INS\_45
CL\_INS\_45
CL\_INS\_321
CL\_INS\_45
CL\_INS\_334
CL\_INS\_334
CL\_INS\_74
CL\_INS\_74
CL\_INS\_74
CL\_INS\_74
CL\_INS\_74
CL\_INS\_74
CL\_INS\_321
CL\_INS\_132
CL\_INS\_132
CL\_INS\_132
CL\_INS\_45
CL\_INS\_10
CL\_INS\_45
CL\_INS\_45
CL\_INS\_45
CL\_INS\_44
CL\_INS\_45
CL\_INS\_44
CL\_INS\_44
CL\_INS\_44
CL\_INS\_44
CL\_INS\_44
CL\_INS\_44
CL\_INS\_44
CL\_INS\_44
CL\_INS\_44
CL\_INS\_45
CL\_INS\_132
CL\_INS\_321
CL\_INS\_321
CL\_INS\_321
CL\_INS\_321
CL\_INS\_334
CL\_INS\_74
CL\_INS\_45
CL\_INS\_351
CL\_INS\_45
CL\_INS\_45
CL\_INS\_45
CL\_INS\_45
CL\_INS\_45
CL\_INS\_91
CL\_INS\_91
CL\_INS\_45
CL\_INS\_45
CL\_INS\_45
CL\_INS\_45
CL\_INS\_351
CL\_INS\_351
CL\_INS\_351
CL\_INS\_45
CL\_INS\_45
CL\_INS\_45
CL\_INS\_45
CL\_INS\_45
CL\_INS\_45
CL\_INS\_45
CL\_INS\_45
CL\_INS\_45
CL\_INS\_45
CL\_INS\_45
CL\_INS\_278
CL\_INS\_278
CL\_INS\_278
CL\_INS\_278
CL\_INS\_45
CL\_INS\_385
CL\_INS\_278
CL\_INS\_278
CL\_INS\_321
CL\_INS\_321
CL\_INS\_321
CL\_INS\_321
CL\_INS\_45
CL\_INS\_45
CL\_INS\_321
CL\_INS\_45
CL\_INS\_321
CL\_INS\_321
CL\_INS\_278
CL\_INS\_45
CL\_INS\_45
CL\_INS\_45
CL\_INS\_321
CL\_INS\_321
CL\_INS\_321
CL\_INS\_321
CL\_INS\_321
CL\_INS\_321
CL\_INS\_321
CL\_INS\_321
CL\_INS\_45
CL\_INS\_45
CL\_INS\_45
CL\_INS\_45
CL\_INS\_45
CL\_INS\_278
CL\_INS\_45
CL\_INS\_45
CL\_INS\_45
CL\_INS\_45
CL\_INS\_45
CL\_INS\_45
CL\_INS\_74
CL\_INS\_45
CL\_INS\_359
CL\_INS\_359
CL\_INS\_26
CL\_INS\_26
CL\_INS\_45
CL\_INS\_45
CL\_INS\_45
CL\_INS\_45
CL\_INS\_45
CL\_INS\_321
CL\_INS\_351
CL\_INS\_45
CL\_INS\_45
CL\_INS\_45
CL\_INS\_45
CL\_INS\_45
CL\_INS\_45
CL\_INS\_45
CL\_INS\_45
CL\_INS\_45
CL\_INS\_400
CL\_INS\_45
CL\_INS\_26
CL\_INS\_26
CL\_INS\_26
CL\_INS\_26
CL\_INS\_26
CL\_INS\_351
CL\_INS\_351
CL\_INS\_351
CL\_INS\_45
CL\_INS\_45
CL\_INS\_45
CL\_INS\_45
CL\_INS\_45
CL\_INS\_351
CL\_INS\_351
CL\_INS\_44
CL\_INS\_26
CL\_INS\_26
CL\_INS\_45
CL\_INS\_44
CL\_INS\_45
CL\_INS\_45
CL\_INS\_44
CL\_INS\_45
CL\_INS\_45
CL\_INS\_45
CL\_INS\_45
CL\_INS\_44
CL\_INS\_44
CL\_INS\_44
CL\_INS\_44
CL\_INS\_44
CL\_INS\_45
CL\_INS\_91
CL\_INS\_45
CL\_INS\_45
CL\_INS\_45
CL\_INS\_45
CL\_INS\_45
CL\_INS\_45
CL\_INS\_26
CL\_INS\_45
CL\_INS\_45
CL\_INS\_45
CL\_INS\_45
CL\_INS\_278
CL\_INS\_278
CL\_INS\_278
CL\_INS\_278
CL\_INS\_391
CL\_INS\_45
CL\_INS\_45
CL\_INS\_45
CL\_INS\_45
CL\_INS\_45
CL\_INS\_45
CL\_INS\_45
CL\_INS\_351
CL\_INS\_351
CL\_INS\_45
CL\_INS\_351
CL\_INS\_351
CL\_INS\_45
CL\_INS\_45
CL\_INS\_45
CL\_INS\_45
CL\_INS\_45
CL\_INS\_45
CL\_INS\_45
CL\_INS\_45
CL\_INS\_45
CL\_INS\_45
CL\_INS\_45
CL\_INS\_307
CL\_INS\_45
CL\_INS\_321
CL\_INS\_321
CL\_INS\_307
CL\_INS\_307
CL\_INS\_321
CL\_INS\_307
CL\_INS\_45
CL\_INS\_45
CL\_INS\_334
CL\_INS\_334
CL\_INS\_334
CL\_INS\_334
CL\_INS\_334
CL\_INS\_334
CL\_INS\_334
CL\_INS\_45
CL\_INS\_45
CL\_INS\_45
CL\_INS\_45
CL\_INS\_45
CL\_INS\_45
CL\_INS\_45
CL\_INS\_45
CL\_INS\_45
CL\_INS\_45
CL\_INS\_45
CL\_INS\_45
CL\_INS\_45
CL\_INS\_45
CL\_INS\_45
CL\_INS\_334
CL\_INS\_334
CL\_INS\_334
CL\_INS\_334
CL\_INS\_45
CL\_INS\_45
CL\_INS\_45
CL\_INS\_45
CL\_INS\_45
CL\_INS\_45
CL\_INS\_45
CL\_INS\_45
CL\_INS\_321
CL\_INS\_334
CL\_INS\_334
CL\_INS\_307
CL\_INS\_45
CL\_INS\_45
CL\_INS\_45
CL\_INS\_45
CL\_INS\_45
CL\_INS\_45
CL\_INS\_45
CL\_INS\_45
CL\_INS\_45
CL\_INS\_45
CL\_INS\_45
CL\_INS\_45
CL\_INS\_334
CL\_INS\_334
CL\_INS\_321
CL\_INS\_334
CL\_INS\_334
CL\_INS\_334
CL\_INS\_334
CL\_INS\_45
CL\_INS\_45
CL\_INS\_45
CL\_INS\_45
CL\_INS\_45
CL\_INS\_45
CL\_INS\_45
CL\_INS\_45
CL\_INS\_45
CL\_INS\_45
CL\_INS\_44
CL\_INS\_45
CL\_INS\_45
CL\_INS\_45
CL\_INS\_45
CL\_INS\_155
CL\_INS\_139
CL\_INS\_278
CL\_INS\_278
CL\_INS\_278
CL\_INS\_401
CL\_INS\_401
CL\_INS\_401
CL\_INS\_278
CL\_INS\_278
CL\_INS\_278
CL\_INS\_45
CL\_INS\_44
CL\_INS\_278
CL\_INS\_45
CL\_INS\_44
CL\_INS\_44
CL\_INS\_321
CL\_INS\_321
CL\_INS\_321
CL\_INS\_74
CL\_INS\_45
CL\_INS\_45
CL\_INS\_45
CL\_INS\_44
CL\_INS\_321
CL\_INS\_321
CL\_INS\_321
CL\_INS\_45
CL\_INS\_321
CL\_INS\_44
CL\_INS\_321
CL\_INS\_321
CL\_INS\_321
CL\_INS\_321
CL\_INS\_321
CL\_INS\_45
CL\_INS\_45
CL\_INS\_45
CL\_INS\_45
CL\_INS\_45
CL\_INS\_45
CL\_INS\_45
CL\_INS\_45
CL\_INS\_45
CL\_INS\_45
CL\_INS\_51
CL\_INS\_51
CL\_INS\_51
CL\_INS\_45
CL\_INS\_45
CL\_INS\_44
CL\_INS\_44
CL\_INS\_44
CL\_INS\_45
CL\_INS\_45
CL\_INS\_44
CL\_INS\_45
CL\_INS\_45
CL\_INS\_45
CL\_INS\_45
CL\_INS\_45
CL\_INS\_45
CL\_INS\_45
CL\_INS\_45
CL\_INS\_26
CL\_INS\_45
CL\_INS\_26
CL\_INS\_26
CL\_INS\_44
CL\_INS\_44
CL\_INS\_44
CL\_INS\_45
CL\_INS\_321
CL\_INS\_321
CL\_INS\_321
CL\_INS\_401
CL\_INS\_45
CL\_INS\_321
CL\_INS\_321
CL\_INS\_321
CL\_INS\_321
CL\_INS\_321
CL\_INS\_45
CL\_INS\_44
CL\_INS\_321
CL\_INS\_351
CL\_INS\_321
CL\_INS\_321
CL\_INS\_45
CL\_INS\_45
CL\_INS\_45
CL\_INS\_45
CL\_INS\_45
CL\_INS\_45
CL\_INS\_45
CL\_INS\_45
CL\_INS\_45
CL\_INS\_45
CL\_INS\_45
CL\_INS\_45
CL\_INS\_45
CL\_INS\_45
CL\_INS\_45
CL\_INS\_45
CL\_INS\_45
CL\_INS\_45
CL\_INS\_45
CL\_INS\_45
CL\_INS\_45
CL\_INS\_45
CL\_INS\_45
CL\_INS\_45
CL\_INS\_45
CL\_INS\_45
CL\_INS\_45
CL\_INS\_45
CL\_INS\_45
CL\_INS\_74
CL\_INS\_74
CL\_INS\_74
CL\_INS\_74
CL\_INS\_74
CL\_INS\_45
CL\_INS\_45
CL\_INS\_45
CL\_INS\_74
CL\_INS\_45
CL\_INS\_45
CL\_INS\_45
CL\_INS\_278
CL\_INS\_278
CL\_INS\_45
CL\_INS\_45
CL\_INS\_45
CL\_INS\_45
CL\_INS\_45
CL\_INS\_155
CL\_INS\_45
CL\_INS\_45
CL\_INS\_45
CL\_INS\_45
CL\_INS\_321
CL\_INS\_45
CL\_INS\_74
CL\_INS\_45
CL\_INS\_74
CL\_INS\_45
CL\_INS\_44
CL\_INS\_44
CL\_INS\_44
CL\_INS\_44
CL\_INS\_44
CL\_INS\_44
CL\_INS\_45
CL\_INS\_45
CL\_INS\_45
CL\_INS\_45
CL\_INS\_45
CL\_INS\_45
CL\_INS\_45
CL\_INS\_45
CL\_INS\_45
CL\_INS\_45
CL\_INS\_45
CL\_INS\_26
CL\_INS\_45
CL\_INS\_45
CL\_INS\_26
CL\_INS\_45
CL\_INS\_91
CL\_INS\_91
CL\_INS\_91
CL\_INS\_91
CL\_INS\_91
CL\_INS\_26
CL\_INS\_45
CL\_INS\_26
CL\_INS\_74
CL\_INS\_26
CL\_INS\_321
CL\_INS\_321
CL\_INS\_321
CL\_INS\_45
CL\_INS\_45
CL\_INS\_321
CL\_INS\_321
CL\_INS\_45
CL\_INS\_45
CL\_INS\_45
CL\_INS\_74
CL\_INS\_351
CL\_INS\_45
CL\_INS\_45
CL\_INS\_45
CL\_INS\_45
CL\_INS\_45
CL\_INS\_45
CL\_INS\_45
CL\_INS\_45
CL\_INS\_45
CL\_INS\_321
CL\_INS\_321
CL\_INS\_321
CL\_INS\_321
CL\_INS\_45
CL\_INS\_45
CL\_INS\_45
CL\_INS\_45
CL\_INS\_45
CL\_INS\_45
CL\_INS\_45
CL\_INS\_44
CL\_INS\_26
CL\_INS\_321
CL\_INS\_45
CL\_INS\_26
CL\_INS\_26
CL\_INS\_45
CL\_INS\_45
CL\_INS\_45
CL\_INS\_45
CL\_INS\_45
CL\_INS\_45
CL\_INS\_204
CL\_INS\_45
CL\_INS\_287
CL\_INS\_321
CL\_INS\_45
CL\_INS\_321
CL\_INS\_321
CL\_INS\_45
CL\_INS\_44
CL\_INS\_44
CL\_INS\_45
CL\_INS\_45
CL\_INS\_239
CL\_INS\_44
CL\_INS\_45
CL\_INS\_44
CL\_INS\_44
CL\_INS\_44
CL\_INS\_45
CL\_INS\_45
CL\_INS\_45
CL\_INS\_44
CL\_INS\_44
CL\_INS\_44
CL\_INS\_45
CL\_INS\_45
CL\_INS\_45
CL\_INS\_45
CL\_INS\_45
CL\_INS\_45
CL\_INS\_45
CL\_INS\_45
CL\_INS\_44
CL\_INS\_45
CL\_INS\_45
CL\_INS\_44
CL\_INS\_45
CL\_INS\_45
CL\_INS\_45
CL\_INS\_45
CL\_INS\_45
CL\_INS\_45
CL\_INS\_44
CL\_INS\_44
CL\_INS\_45
CL\_INS\_45
CL\_INS\_45
CL\_INS\_45
CL\_INS\_45
CL\_INS\_45
CL\_INS\_44
CL\_INS\_44
CL\_INS\_45
CL\_INS\_45
CL\_INS\_44
CL\_INS\_44
CL\_INS\_44
CL\_INS\_44
CL\_INS\_45
CL\_INS\_45
CL\_INS\_45
CL\_INS\_45
CL\_INS\_44
CL\_INS\_44
CL\_INS\_44
CL\_INS\_44
CL\_INS\_44
CL\_INS\_44
CL\_INS\_44
CL\_INS\_44
CL\_INS\_44
CL\_INS\_44
CL\_INS\_44
CL\_INS\_44
CL\_INS\_44
CL\_INS\_44
CL\_INS\_44
CL\_INS\_207
CL\_INS\_278
CL\_INS\_44
CL\_INS\_44
CL\_INS\_44
CL\_INS\_44
CL\_INS\_44
CL\_INS\_44
CL\_INS\_44
CL\_INS\_57
CL\_INS\_57
CL\_INS\_57
CL\_INS\_57
CL\_INS\_57
CL\_INS\_57
CL\_INS\_57
CL\_INS\_57
CL\_INS\_57
CL\_INS\_57
CL\_INS\_57
CL\_INS\_57
CL\_INS\_44
CL\_INS\_44
CL\_INS\_246
CL\_INS\_44
CL\_INS\_246
CL\_INS\_246
CL\_INS\_57
CL\_INS\_57
CL\_INS\_44
CL\_INS\_57
CL\_INS\_57
CL\_INS\_57
CL\_INS\_246
CL\_INS\_57
CL\_INS\_57
CL\_INS\_57
CL\_INS\_57
CL\_INS\_44
CL\_INS\_44
CL\_INS\_246
CL\_INS\_44
CL\_INS\_44
CL\_INS\_44
CL\_INS\_44
CL\_INS\_44
CL\_INS\_44
CL\_INS\_57
CL\_INS\_57
CL\_INS\_57
CL\_INS\_57
CL\_INS\_57
CL\_INS\_57
CL\_INS\_57
CL\_INS\_45
CL\_INS\_57
CL\_INS\_57
CL\_INS\_57
CL\_INS\_57
CL\_INS\_45
CL\_INS\_45
CL\_INS\_45
CL\_INS\_45
CL\_INS\_45
CL\_INS\_44
CL\_INS\_44
CL\_INS\_45
CL\_INS\_44
CL\_INS\_246
CL\_INS\_246
CL\_INS\_246
CL\_INS\_334
CL\_INS\_45
CL\_INS\_44
CL\_INS\_45
CL\_INS\_45
CL\_INS\_45
CL\_INS\_45
CL\_INS\_45
CL\_INS\_45
CL\_INS\_45
CL\_INS\_45
CL\_INS\_44
CL\_INS\_44
CL\_INS\_44
CL\_INS\_45
CL\_INS\_45
CL\_INS\_45
CL\_INS\_45
CL\_INS\_45
CL\_INS\_45
CL\_INS\_45
CL\_INS\_45
CL\_INS\_45
CL\_INS\_45
CL\_INS\_45
CL\_INS\_45
CL\_INS\_45
CL\_INS\_45
CL\_INS\_45
CL\_INS\_44
CL\_INS\_44
CL\_INS\_44
CL\_INS\_44
CL\_INS\_44
CL\_INS\_44
CL\_INS\_45
CL\_INS\_45
CL\_INS\_45
CL\_INS\_45
CL\_INS\_45
CL\_INS\_45
CL\_INS\_45
CL\_INS\_45
CL\_INS\_45
CL\_INS\_45
CL\_INS\_321
CL\_INS\_321
CL\_INS\_321
CL\_INS\_321
CL\_INS\_321
CL\_INS\_45
CL\_INS\_45
CL\_INS\_57
CL\_INS\_44
CL\_INS\_44
CL\_INS\_44
CL\_INS\_204
CL\_INS\_45
CL\_INS\_45
CL\_INS\_45
CL\_INS\_45
CL\_INS\_45
CL\_INS\_45
CL\_INS\_45
CL\_INS\_204
CL\_INS\_204
CL\_INS\_44
CL\_INS\_44
CL\_INS\_44
CL\_INS\_44
CL\_INS\_321
CL\_INS\_321
CL\_INS\_321
CL\_INS\_44
CL\_INS\_44
CL\_INS\_44
CL\_INS\_44
CL\_INS\_44
CL\_INS\_91
CL\_INS\_44
CL\_INS\_44
CL\_INS\_44
CL\_INS\_385
CL\_INS\_385
CL\_INS\_401
CL\_INS\_45
CL\_INS\_45
CL\_INS\_385
CL\_INS\_307
CL\_INS\_307
CL\_INS\_307
CL\_INS\_307
CL\_INS\_307
CL\_INS\_307
CL\_INS\_307
CL\_INS\_307
CL\_INS\_307
CL\_INS\_307
CL\_INS\_307
CL\_INS\_307
CL\_INS\_307
CL\_INS\_74
CL\_INS\_307
CL\_INS\_307
CL\_INS\_307
CL\_INS\_45
CL\_INS\_318
CL\_INS\_307
CL\_INS\_307
CL\_INS\_307
CL\_INS\_307
CL\_INS\_45
CL\_INS\_57
CL\_INS\_44
CL\_INS\_45
CL\_INS\_57
CL\_INS\_45
Cluster ID


CL\_7810
CL\_5277
CL\_5276
CL\_5275
CL\_5274
CL\_5273
CL\_23889
CL\_23890
CL\_22226
CL\_23891
CL\_22225
CL\_14579
CL\_14578
CL\_14577
CL\_14576
CL\_14575
CL\_14574
CL\_7042
CL\_23543
CL\_23542
CL\_23541
CL\_23540
CL\_23539
CL\_23538
CL\_21394
CL\_21395
CL\_23487
CL\_10286
CL\_10285
CL\_10284
CL\_11511
CL\_11510
CL\_11509
CL\_11508
CL\_20194
CL\_10281
CL\_10282
CL\_10283
CL\_10276
CL\_14281
CL\_14282
CL\_20784
CL\_20783
CL\_21607
CL\_13162
CL\_13161
CL\_7534
CL\_7533
CL\_5561
CL\_13169
CL\_11630
CL\_6047
CL\_6048
CL\_7592
CL\_6050
CL\_6051
CL\_6052
CL\_6053
CL\_6054
CL\_5933
CL\_6858
CL\_5935
CL\_5936
CL\_5937
CL\_12432
CL\_12431
CL\_8246
CL\_8245
CL\_8244
CL\_22033
CL\_22032
CL\_12849
CL\_12850
CL\_19214
CL\_19215
CL\_23268
CL\_23267
CL\_19216
CL\_19217
CL\_19218
CL\_19219
CL\_19220
CL\_19221
CL\_19222
CL\_19223
CL\_14815
CL\_12009
CL\_12008
CL\_12007
CL\_12006
CL\_12005
CL\_18731
CL\_18732
CL\_6821
CL\_18733
CL\_18734
CL\_18735
CL\_18736
CL\_5568
CL\_5569
CL\_18737
CL\_13265
CL\_18738
CL\_20997
CL\_20443
CL\_20619
CL\_20618
CL\_20617
CL\_20615
CL\_21237
CL\_7140
CL\_9061
CL\_16473
CL\_7142
CL\_18649
CL\_7143
CL\_7466
CL\_9064
CL\_7144
CL\_7146
CL\_7147
CL\_7467
CL\_7148
CL\_7468
CL\_7469
CL\_7151
CL\_21236
CL\_18739
CL\_12011
CL\_23595
CL\_17971
CL\_17970
CL\_17969
CL\_17968
CL\_17967
CL\_17966
CL\_17965
CL\_17964
CL\_17963
CL\_17962
CL\_17961
CL\_17960
CL\_17959
CL\_11607
CL\_15323
CL\_11606
CL\_11605
CL\_20996
CL\_20995
CL\_20994
CL\_11604
CL\_11603
CL\_11602
CL\_15459
CL\_18220
CL\_20993
CL\_20992
CL\_20991
CL\_20990
CL\_20989
CL\_11601
CL\_12045
CL\_11600
CL\_11599
CL\_11598
CL\_12046
CL\_12047
CL\_7152
CL\_17958
CL\_14283
CL\_14284
CL\_14285
CL\_6840
CL\_14286
CL\_14287
CL\_17957
CL\_11597
CL\_11596
CL\_11595
CL\_20654
CL\_12174
CL\_12913
CL\_12914
CL\_12915
CL\_12916
CL\_12917
CL\_16958
CL\_16956
CL\_16957
CL\_16955
CL\_16954
CL\_16953
CL\_16952
CL\_16951
CL\_16950
CL\_16949
CL\_11111
CL\_11110
CL\_11109
CL\_11108
CL\_11107
CL\_11106
CL\_11105
CL\_11104
CL\_11103
CL\_12012
CL\_12013
CL\_12014
CL\_5633
CL\_6666
CL\_23846
CL\_23845
CL\_23844
CL\_23843
CL\_11586
CL\_8211
CL\_11585
CL\_24070
CL\_11584
CL\_23842
CL\_23841
CL\_23840
CL\_23839
CL\_23838
CL\_23837
CL\_17338
CL\_17336
CL\_9181
CL\_9180
CL\_9179
CL\_9178
CL\_9177
CL\_10455
CL\_9176
CL\_9175
CL\_21629
CL\_10454
CL\_9174
CL\_21716
CL\_12144
CL\_23831
CL\_23830
CL\_14070
CL\_23829
CL\_23828
CL\_7524
CL\_9930
CL\_8657
CL\_6135
CL\_9929
CL\_9928
CL\_9927
CL\_9926
CL\_9925
CL\_9924
CL\_5298
CL\_6320
CL\_6321
CL\_6322
CL\_18812
CL\_14693
CL\_18811
CL\_22222
CL\_22223
CL\_17104
CL\_9923
CL\_5846
CL\_5845
CL\_5844
CL\_5843
CL\_5842
CL\_5841
CL\_5840
CL\_10538
CL\_10537
CL\_14816
CL\_5765
CL\_8270
CL\_12143
CL\_8272
CL\_8273
CL\_8274
CL\_14069
CL\_10434
CL\_10433
CL\_22554
CL\_22553
CL\_19701
CL\_19702
CL\_17566
CL\_14311
CL\_14312
CL\_17994
CL\_22552
CL\_19703
CL\_19704
CL\_10432
CL\_10431
CL\_10430
CL\_10429
CL\_10428
CL\_10427
CL\_10426
CL\_10425
CL\_10424
CL\_10423
CL\_10782
CL\_10781
CL\_10780
CL\_10779
CL\_9786
CL\_9785
CL\_9784
CL\_9783
CL\_10778
CL\_10777
CL\_6672
CL\_6673
CL\_10776
CL\_10775
CL\_10774
CL\_10773
CL\_15019
CL\_15020
CL\_10772
CL\_15021
CL\_10771
CL\_10770
CL\_9776
CL\_15022
CL\_15023
CL\_15024
CL\_10769
CL\_10768
CL\_10767
CL\_10766
CL\_10765
CL\_10764
CL\_10763
CL\_10762
CL\_16431
CL\_16430
CL\_16429
CL\_16428
CL\_16427
CL\_10902
CL\_17105
CL\_16426
CL\_16425
CL\_16424
CL\_16423
CL\_8363
CL\_8362
CL\_18200
CL\_19015
CL\_19014
CL\_18199
CL\_18198
CL\_18197
CL\_18196
CL\_18195
CL\_22139
CL\_22140
CL\_17335
CL\_14217
CL\_9361
CL\_9360
CL\_9359
CL\_9358
CL\_20442
CL\_20441
CL\_20440
CL\_22104
CL\_20439
CL\_17721
CL\_22103
CL\_21493
CL\_21494
CL\_21495
CL\_21496
CL\_21497
CL\_19694
CL\_19695
CL\_19696
CL\_19697
CL\_19698
CL\_19699
CL\_19700
CL\_21295
CL\_14216
CL\_14215
CL\_8372
CL\_18205
CL\_18204
CL\_18203
CL\_8371
CL\_13809
CL\_13808
CL\_8368
CL\_8370
CL\_8369
CL\_8367
CL\_8366
CL\_13807
CL\_8365
CL\_8364
CL\_14060
CL\_23696
CL\_9173
CL\_14310
CL\_9172
CL\_9171
CL\_9170
CL\_21294
CL\_21293
CL\_21292
CL\_12365
CL\_23596
CL\_23597
CL\_23598
CL\_23599
CL\_110
CL\_109
CL\_108
CL\_107
CL\_6936
CL\_22102
CL\_22101
CL\_22100
CL\_22099
CL\_21291
CL\_21290
CL\_21289
CL\_14199
CL\_14198
CL\_21288
CL\_14196
CL\_14195
CL\_18126
CL\_23885
CL\_18127
CL\_18128
CL\_18129
CL\_18130
CL\_18131
CL\_18132
CL\_15769
CL\_15768
CL\_7762
CL\_7761
CL\_17796
CL\_9282
CL\_9283
CL\_7760
CL\_7759
CL\_7758
CL\_7757
CL\_15767
CL\_15766
CL\_7756
CL\_7755
CL\_7754
CL\_9267
CL\_8658
CL\_8276
CL\_8277
CL\_18895
CL\_18894
CL\_18893
CL\_18892
CL\_18266
CL\_18265
CL\_12522
CL\_12142
CL\_12141
CL\_12140
CL\_12139
CL\_14068
CL\_14067
CL\_23827
CL\_14066
CL\_7294
CL\_7295
CL\_7296
CL\_7297
CL\_18264
CL\_18263
CL\_18262
CL\_18261
CL\_14065
CL\_14064
CL\_14063
CL\_14062
CL\_9285
CL\_7753
CL\_7752
CL\_7751
CL\_15183
CL\_7750
CL\_17795
CL\_7749
CL\_7748
CL\_7747
CL\_7746
CL\_7745
CL\_7744
CL\_7743
CL\_7742
CL\_15184
CL\_9268
CL\_9269
CL\_9289
CL\_9270
CL\_9273
CL\_9274
CL\_9275
CL\_15765
CL\_15764
CL\_15763
CL\_15762
CL\_15761
CL\_15760
CL\_15759
CL\_15758
CL\_15757
CL\_6541
CL\_20031
CL\_12629
CL\_6540
CL\_22350
CL\_6539
CL\_5387
CL\_12280
CL\_5973
CL\_5974
CL\_5975
CL\_8206
CL\_8205
CL\_8204
CL\_5979
CL\_5980
CL\_5991
CL\_12630
CL\_12172
CL\_6079
CL\_12491
CL\_6538
CL\_6537
CL\_6339
CL\_6340
CL\_6536
CL\_14573
CL\_14572
CL\_14571
CL\_14570
CL\_6535
CL\_6534
CL\_6533
CL\_6341
CL\_20653
CL\_13616
CL\_6532
CL\_6342
CL\_6343
CL\_6344
CL\_6345
CL\_6531
CL\_18360
CL\_7532
CL\_12631
CL\_20652
CL\_7929
CL\_7928
CL\_7927
CL\_7926
CL\_7925
CL\_7924
CL\_7923
CL\_7922
CL\_7921
CL\_7920
CL\_7919
CL\_7918
CL\_17684
CL\_20614
CL\_17685
CL\_17686
CL\_17687
CL\_21694
CL\_21693
CL\_21692
CL\_21691
CL\_17688
CL\_17689
CL\_17690
CL\_17691
CL\_18202
CL\_22138
CL\_19016
CL\_18201
CL\_7917
CL\_7916
CL\_7915
CL\_7914
CL\_7423
CL\_7422
CL\_7421
CL\_10422
CL\_10421
CL\_8358
CL\_8357
CL\_8356
CL\_7420
CL\_7419
CL\_23883
CL\_9166
CL\_8355
CL\_14193
CL\_7418
CL\_7417
CL\_12525
CL\_12524
CL\_12523
CL\_15326
CL\_15327
CL\_15328
CL\_15329
CL\_15330
CL\_15331
CL\_15332
CL\_15333
CL\_15334
CL\_15335
CL\_15336
CL\_15337
CL\_15338
CL\_15339
CL\_15340
CL\_15341
CL\_15342
CL\_15343
CL\_15344
CL\_15345
CL\_15346
CL\_15347
CL\_15348
CL\_15349
CL\_15350
CL\_15351
CL\_11493
CL\_11495
CL\_9355
CL\_8361
CL\_11496
CL\_8839
CL\_15565
CL\_8360
CL\_9169
CL\_9168
CL\_20438
CL\_23600
CL\_8053
CL\_8052
CL\_9167
CL\_21630
CL\_18133
CL\_18134
CL\_18135
CL\_18136
CL\_20437
CL\_18137
CL\_23355
CL\_23884
CL\_8359
CL\_9354
CL\_8838
CL\_8837
CL\_8836
CL\_8835
CL\_13074
CL\_13073
CL\_13072
CL\_13071
CL\_13070
CL\_13069
CL\_13785
CL\_13786
CL\_13787
CL\_13788
CL\_13789
CL\_13790
CL\_13791
CL\_13792
CL\_13793
CL\_13794
CL\_13795
CL\_13796
CL\_20938
CL\_13797
CL\_13798
CL\_13799
CL\_14313
CL\_14314
CL\_14315
CL\_14316
CL\_14317
CL\_17624
CL\_17622
CL\_17623
CL\_10420
CL\_17621
CL\_7416
CL\_8353
CL\_8352
CL\_9165
CL\_9164
CL\_10419
CL\_7413
CL\_23836
CL\_23835
CL\_23834
CL\_7913
CL\_8834
CL\_21952
CL\_21951
CL\_21950
CL\_21949
CL\_21948
CL\_21947
CL\_12003
CL\_12002
CL\_12001
CL\_9353
CL\_9352
CL\_9351
CL\_9350
CL\_8833
CL\_8832
CL\_13160
CL\_13159
CL\_8831
CL\_17993
CL\_7912
CL\_5839
CL\_20937
CL\_7412
CL\_21976
CL\_17618
CL\_17617
CL\_13800
CL\_13801
CL\_13802
CL\_13803
CL\_13804
CL\_13805
CL\_8086
CL\_13806
CL\_7011
CL\_8354
CL\_23601
CL\_7415
CL\_7414
CL\_22098
CL\_8351
CL\_8350
CL\_12856
CL\_12857
CL\_9257
CL\_14061
CL\_11754
CL\_6530
CL\_6529
CL\_6528
CL\_20936
CL\_20935
CL\_20934
CL\_6527
CL\_6526
CL\_6525
CL\_7037
CL\_19224
CL\_19225
CL\_5272
CL\_21491
CL\_8847
CL\_8846
CL\_8845
CL\_11594
CL\_21490
CL\_21489
CL\_11593
CL\_13737
CL\_13738
CL\_22046
CL\_12004
CL\_9357
CL\_9356
CL\_11592
CL\_7041
CL\_7040
CL\_7039
CL\_8844
CL\_8843
CL\_20651
CL\_20650
CL\_11591
CL\_8842
CL\_8841
CL\_8840
CL\_11590
CL\_11589
CL\_11588
CL\_11587
CL\_18206
CL\_693
CL\_694
CL\_695
CL\_18653
CL\_18654
CL\_18655
CL\_18656
CL\_18657
CL\_18658
CL\_18659
CL\_18660
CL\_18661
CL\_18662
CL\_18663
CL\_18664
CL\_18665
CL\_18666
CL\_18667
CL\_9529
CL\_6081
CL\_18668
CL\_18669
CL\_18670
CL\_18671
CL\_18672
CL\_18673
CL\_18674
CL\_721
CL\_722
CL\_723
CL\_724
CL\_725
CL\_726
CL\_727
CL\_728
CL\_729
CL\_730
CL\_731
CL\_732
CL\_8989
CL\_18675
CL\_6427
CL\_18676
CL\_6429
CL\_6430
CL\_733
CL\_734
CL\_18677
CL\_735
CL\_736
CL\_6173
CL\_737
CL\_738
CL\_18101
CL\_739
CL\_740
CL\_18678
CL\_18679
CL\_6432
CL\_18680
CL\_18681
CL\_18682
CL\_18683
CL\_18684
CL\_18685
CL\_741
CL\_742
CL\_743
CL\_744
CL\_745
CL\_746
CL\_747
CL\_748
CL\_749
CL\_8061
CL\_18102
CL\_21717
CL\_750
CL\_751
CL\_752
CL\_12922
CL\_12923
CL\_12924
CL\_12925
CL\_12926
CL\_12927
CL\_6438
CL\_6439
CL\_6440
CL\_1165
CL\_7038
CL\_7531
CL\_22245
CL\_15572
CL\_15571
CL\_15570
CL\_15569
CL\_15568
CL\_15567
CL\_15566
CL\_8830
CL\_15458
CL\_696
CL\_12851
CL\_12918
CL\_697
CL\_12919
CL\_12920
CL\_12921
CL\_698
CL\_699
CL\_700
CL\_701
CL\_702
CL\_12852
CL\_12853
CL\_12854
CL\_22031
CL\_12855
CL\_23486
CL\_23485
CL\_23484
CL\_23483
CL\_23482
CL\_703
CL\_704
CL\_705
CL\_706
CL\_707
CL\_708
CL\_709
CL\_710
CL\_711
CL\_712
CL\_713
CL\_714
CL\_715
CL\_716
CL\_717
CL\_718
CL\_719
CL\_720
CL\_5838
CL\_753
CL\_754
CL\_5475
CL\_7393
CL\_7392
CL\_7391
CL\_7390
CL\_7389
CL\_22106
CL\_22105
CL\_6159
CL\_5524
CL\_11102
CL\_11101
CL\_11100
CL\_11099
CL\_7428
CL\_7427
CL\_7426
CL\_10242
CL\_11098
CL\_11097
CL\_11096
CL\_11095
CL\_6027
CL\_11094
CL\_11093
CL\_11092
CL\_7128
CL\_7126
CL\_7124
CL\_8099
CL\_91
CL\_90
CL\_89
CL\_87
CL\_86
CL\_85
CL\_84
CL\_83
CL\_82
CL\_81
CL\_80
CL\_79
CL\_77
CL\_76
CL\_75
CL\_74
CL\_73
CL\_72
CL\_5951
CL\_15178
CL\_9416
CL\_5941
CL\_5940
CL\_5939
CL\_5938
CL\_16488
CL\_755
CL\_7399
CL\_22436
CL\_756
CL\_757
