## Supplementary material for "A novel method for integrating genomic and Tn-Seq data to identify common *in vivo* fitness mechanisms across multiple bacterial species": S2 Dataset: CL_INS_47.html

Legend

 Hypothetical
 Other
 All VFDB Genes

FULL


WINDOWSVGPNG

Trim RowsRemove SingletonsSave Fasta

CL\_761


CL\_761


CL\_761


CL\_760


CL\_761


Break


CL\_761


CL\_761


CL\_761

HighlightSelectShow Genomes


148

CL\_762


41

CL\_762


3

Break


1

CL\_762


1

Break


1

CL\_762


1

Break


1

CL\_762


1

CL\_776

fGI ID


CL\_INS\_204
CL\_INS\_47
CL\_INS\_47
Cluster ID


CL\_5475
CL\_7741
CL\_11737
