## Supplementary material for "A novel method for integrating genomic and Tn-Seq data to identify common *in vivo* fitness mechanisms across multiple bacterial species": S2 Dataset: CL_INS_50.html

Legend

 Hypothetical
 Regulatoryfunctions
 Proteinsynthesis/fate
 Other
 All VFDB Genes

FULL


WINDOWSVGPNG

Trim RowsRemove SingletonsSave Fasta

CL\_793


CL\_793


CL\_793


CL\_793


CL\_793


CL\_793

HighlightSelectShow Genomes


176

CL\_792


20

CL\_788


2

CL\_788


2

CL\_788


1

CL\_787


1

CL\_788

fGI ID


CL\_INS\_50
CL\_INS\_50
CL\_INS\_50
CL\_INS\_50
CL\_INS\_50
CL\_INS\_50
CL\_INS\_50
CL\_INS\_50
CL\_INS\_50
CL\_INS\_50
CL\_INS\_50
CL\_INS\_50
CL\_INS\_50
CL\_INS\_50
CL\_INS\_50
CL\_INS\_50
CL\_INS\_50
Cluster ID


CL\_7725
CL\_7726
CL\_7727
CL\_7728
CL\_7729
CL\_7730
CL\_7731
CL\_7732
CL\_7733
CL\_22826
CL\_7734
CL\_7735
CL\_7736
CL\_7737
CL\_7738
CL\_7739
CL\_7740
