## Supplementary material for "A novel method for integrating genomic and Tn-Seq data to identify common *in vivo* fitness mechanisms across multiple bacterial species": S2 Dataset: CL_INS_52.html

Legend

 Mobile +extrachromosomalelementfunctions
 Hypothetical
 Other
 All VFDB Genes

FULL


WINDOWSVGPNG

Trim RowsRemove SingletonsSave Fasta

CL\_814


CL\_814


CL\_814


CL\_814


CL\_814


CL\_814


CL\_814


CL\_814


CL\_814


CL\_814

HighlightSelectShow Genomes


148

CL\_815


5

CL\_817


4

CL\_817


2

CL\_815


2

CL\_817


2

CL\_817


1

CL\_817


1

CL\_817


1

CL\_817


1

CL\_817

fGI ID


CL\_INS\_52
CL\_INS\_52
CL\_INS\_53
CL\_INS\_53
CL\_INS\_53
CL\_INS\_53
CL\_INS\_53
CL\_INS\_53
CL\_INS\_53
CL\_INS\_53
CL\_INS\_53
CL\_INS\_53
Cluster ID


CL\_9345
CL\_10275
CL\_10274
CL\_10273
CL\_21681
CL\_21680
CL\_21679
CL\_10272
CL\_11834
CL\_11833
CL\_11832
CL\_11831
