## Supplementary material for "A novel method for integrating genomic and Tn-Seq data to identify common *in vivo* fitness mechanisms across multiple bacterial species": S2 Dataset: CL_INS_53.html

Legend

 Mobile +extrachromosomalelementfunctions
 Hypothetical
 Other
 Proteinsynthesis/fate
 All VFDB Genes

FULL


WINDOWSVGPNG

Trim RowsRemove SingletonsSave Fasta

CL\_817


CL\_817


CL\_817


CL\_817


CL\_817


CL\_817


CL\_817


CL\_817


CL\_817


CL\_817

HighlightSelectShow Genomes


160

CL\_816


5

CL\_814


4

CL\_814


3

CL\_814


2

CL\_814


1

CL\_814


1

CL\_788


1

CL\_814


1

CL\_814


1

CL\_814

fGI ID


CL\_INS\_53
CL\_INS\_53
CL\_INS\_53
CL\_INS\_53
CL\_INS\_53
CL\_INS\_53
CL\_INS\_53
CL\_INS\_53
CL\_INS\_53
CL\_INS\_53
CL\_INS\_53
CL\_INS\_53
CL\_INS\_53
CL\_INS\_53
CL\_INS\_53
CL\_INS\_53
CL\_INS\_53
CL\_INS\_53
CL\_INS\_53
CL\_INS\_53
CL\_INS\_53
CL\_INS\_53
CL\_INS\_53
CL\_INS\_53
CL\_INS\_53
CL\_INS\_53
CL\_INS\_53
CL\_INS\_53
Cluster ID


CL\_19171
CL\_11831
CL\_21679
CL\_21680
CL\_21681
CL\_11832
CL\_11833
CL\_11834
CL\_10272
CL\_10273
CL\_10274
CL\_8187
CL\_8186
CL\_8185
CL\_8184
CL\_10809
CL\_10808
CL\_10807
CL\_10806
CL\_10805
CL\_10804
CL\_10803
CL\_10802
CL\_10801
CL\_10800
CL\_10799
CL\_10798
CL\_13476
