## Supplementary material for "A novel method for integrating genomic and Tn-Seq data to identify common *in vivo* fitness mechanisms across multiple bacterial species": S2 Dataset: CL_INS_55.html

Legend

 Regulatoryfunctions
 Hypothetical
 Other
 All VFDB Genes

FULL


WINDOWSVGPNG

Trim RowsRemove SingletonsSave Fasta

CL\_828


CL\_828


CL\_828


CL\_828


CL\_828


CL\_828


CL\_828


CL\_828


CL\_828


CL\_828


CL\_828


CL\_828

HighlightSelectShow Genomes


57

CL\_831


32

CL\_831


30

CL\_831


25

CL\_831


18

CL\_831


14

CL\_831


14

CL\_831


11

CL\_831


1

Break


1

CL\_831


1

CL\_832


1

CL\_831

fGI ID


CL\_INS\_55
CL\_INS\_55
CL\_INS\_55
CL\_INS\_55
CL\_INS\_55
CL\_INS\_55
CL\_INS\_55
CL\_INS\_55
CL\_INS\_55
CL\_INS\_55
CL\_INS\_55
Cluster ID


CL\_829
CL\_830
CL\_18891
CL\_5835
CL\_8829
CL\_8828
CL\_8827
CL\_8826
CL\_8825
CL\_8824
CL\_6150
