## Supplementary material for "A novel method for integrating genomic and Tn-Seq data to identify common *in vivo* fitness mechanisms across multiple bacterial species": S2 Dataset: CL_INS_58.html

Legend

 Mobile +extrachromosomalelementfunctions
 Regulatoryfunctions
 Hypothetical
 DNA Metabolism
 AntibioticResistance
 Proteinsynthesis/fate
 Other
 EnergyMetabolism
 Transport +binding proteins
 All VFDB Genes

FULL


WINDOWSVGPNG

Trim RowsRemove SingletonsSave Fasta

CL\_897


CL\_897


CL\_897


CL\_897


CL\_897

HighlightSelectShow Genomes


121

CL\_898


80

CL\_898


1

CL\_899


1

Break


1

CL\_899

fGI ID


CL\_INS\_58
CL\_INS\_58
CL\_INS\_58
CL\_INS\_58
CL\_INS\_58
CL\_INS\_58
CL\_INS\_58
CL\_INS\_58
CL\_INS\_58
CL\_INS\_58
CL\_INS\_58
CL\_INS\_58
CL\_INS\_58
CL\_INS\_58
CL\_INS\_58
CL\_INS\_58
CL\_INS\_58
CL\_INS\_58
CL\_INS\_132
CL\_INS\_278
CL\_INS\_132
CL\_INS\_132
CL\_INS\_132
CL\_INS\_160
CL\_INS\_132
CL\_INS\_132
CL\_INS\_132
CL\_INS\_132
CL\_INS\_132
CL\_INS\_132
CL\_INS\_132
CL\_INS\_307
CL\_INS\_307
CL\_INS\_307
CL\_INS\_278
CL\_INS\_278
CL\_INS\_278
CL\_INS\_278
CL\_INS\_278
CL\_INS\_278
CL\_INS\_278
CL\_INS\_278
CL\_INS\_278
CL\_INS\_132
CL\_INS\_278
CL\_INS\_132
CL\_INS\_132
CL\_INS\_278
CL\_INS\_278
CL\_INS\_278
CL\_INS\_278
CL\_INS\_58
CL\_INS\_58
CL\_INS\_278
CL\_INS\_278
CL\_INS\_278
CL\_INS\_278
CL\_INS\_278
CL\_INS\_278
CL\_INS\_278
CL\_INS\_278
CL\_INS\_278
CL\_INS\_278
CL\_INS\_278
CL\_INS\_278
CL\_INS\_278
CL\_INS\_58
CL\_INS\_58
CL\_INS\_58
CL\_INS\_58
CL\_INS\_58
CL\_INS\_160
CL\_INS\_160
CL\_INS\_160
CL\_INS\_160
CL\_INS\_58
CL\_INS\_58
CL\_INS\_58
CL\_INS\_58
CL\_INS\_58
CL\_INS\_58
CL\_INS\_30
CL\_INS\_30
CL\_INS\_132
CL\_INS\_132
CL\_INS\_58
CL\_INS\_58
CL\_INS\_30
CL\_INS\_30
CL\_INS\_30
CL\_INS\_30
CL\_INS\_30
CL\_INS\_30
CL\_INS\_278
CL\_INS\_58
CL\_INS\_58
CL\_INS\_278
CL\_INS\_58
CL\_INS\_278
CL\_INS\_278
CL\_INS\_178
CL\_INS\_58
CL\_INS\_58
CL\_INS\_278
CL\_INS\_278
CL\_INS\_278
CL\_INS\_212
CL\_INS\_278
CL\_INS\_278
CL\_INS\_278
CL\_INS\_160
CL\_INS\_58
CL\_INS\_58
CL\_INS\_278
CL\_INS\_58
CL\_INS\_58
CL\_INS\_278
CL\_INS\_278
CL\_INS\_278
CL\_INS\_58
CL\_INS\_58
CL\_INS\_278
CL\_INS\_160
CL\_INS\_160
CL\_INS\_278
Cluster ID


CL\_5265
CL\_12930
CL\_12931
CL\_12932
CL\_12933
CL\_12934
CL\_12935
CL\_12936
CL\_12937
CL\_12938
CL\_12939
CL\_12940
CL\_12941
CL\_12942
CL\_12943
CL\_12944
CL\_12945
CL\_12946
CL\_8171
CL\_7577
CL\_6822
CL\_6788
CL\_6881
CL\_6174
CL\_6175
CL\_6176
CL\_6177
CL\_6178
CL\_6179
CL\_6180
CL\_6181
CL\_3387
CL\_3386
CL\_6182
CL\_6183
CL\_6184
CL\_6185
CL\_6186
CL\_6187
CL\_6874
CL\_12259
CL\_6876
CL\_6877
CL\_6840
CL\_6839
CL\_6838
CL\_6837
CL\_6836
CL\_6834
CL\_6833
CL\_6832
CL\_12947
CL\_12948
CL\_7947
CL\_6885
CL\_12260
CL\_7948
CL\_6883
CL\_6882
CL\_7984
CL\_12261
CL\_6842
CL\_6843
CL\_6844
CL\_6845
CL\_6846
CL\_8653
CL\_12949
CL\_12262
CL\_12371
CL\_12263
CL\_6848
CL\_6849
CL\_6850
CL\_6851
CL\_6852
CL\_7988
CL\_6854
CL\_12264
CL\_12265
CL\_12266
CL\_6869
CL\_6870
CL\_6871
CL\_6872
CL\_12372
CL\_12373
CL\_7955
CL\_6860
CL\_6859
CL\_7997
CL\_7959
CL\_7998
CL\_5981
CL\_8000
CL\_8001
CL\_6829
CL\_5986
CL\_5987
CL\_5988
CL\_6828
CL\_6827
CL\_6826
CL\_5991
CL\_5992
CL\_5993
CL\_6825
CL\_5995
CL\_5996
CL\_5997
CL\_7963
CL\_12950
CL\_7585
CL\_6001
CL\_10833
CL\_10832
CL\_6004
CL\_6005
CL\_6006
CL\_6880
CL\_12267
CL\_6009
CL\_8019
CL\_8020
CL\_6013
