## Supplementary material for "A novel method for integrating genomic and Tn-Seq data to identify common *in vivo* fitness mechanisms across multiple bacterial species": S2 Dataset: CL_INS_59.html

Legend

 Regulatoryfunctions
 Hypothetical
 All EssentialGenes

FULL


WINDOWSVGPNG

Trim RowsRemove SingletonsSave Fasta

CL\_914


CL\_914

HighlightSelectShow Genomes


118

CL\_916


87

CL\_916

fGI ID

CL\_INS\_59
Cluster ID

CL\_915
