## Supplementary material for "A novel method for integrating genomic and Tn-Seq data to identify common *in vivo* fitness mechanisms across multiple bacterial species": S2 Dataset: CL_INS_60.html

Legend

 Mobile +extrachromosomalelementfunctions
 Hypothetical
 All EssentialGenes
 Other
 Transport +binding proteins
 All VFDB Genes

FULL


WINDOWSVGPNG

Trim RowsRemove SingletonsSave Fasta

CL\_925


CL\_925


CL\_925


CL\_922


CL\_925


CL\_925


CL\_925


CL\_925


CL\_925


CL\_925

HighlightSelectShow Genomes


110

CL\_926


85

CL\_926


3

CL\_926


1

CL\_926


1

CL\_926


1

CL\_926


1

CL\_926


1

CL\_926


1

CL\_926


1

CL\_926

fGI ID


CL\_INS\_60
CL\_INS\_60
CL\_INS\_60
CL\_INS\_60
CL\_INS\_60
CL\_INS\_60
CL\_INS\_60
CL\_INS\_60
CL\_INS\_60
CL\_INS\_60
CL\_INS\_60
CL\_INS\_60
CL\_INS\_60
CL\_INS\_60
Cluster ID


CL\_21098
CL\_21099
CL\_21100
CL\_5264
CL\_5263
CL\_5262
CL\_5261
CL\_5260
CL\_5259
CL\_5258
CL\_5257
CL\_5256
CL\_5255
CL\_5254
