## Supplementary material for "A novel method for integrating genomic and Tn-Seq data to identify common *in vivo* fitness mechanisms across multiple bacterial species": S2 Dataset: CL_INS_63.html

Legend

 Hypothetical
 Other
 Transport +binding proteins

FULL


WINDOWSVGPNG

Trim RowsRemove SingletonsSave Fasta

CL\_958


CL\_958


CL\_958

HighlightSelectShow Genomes


113

CL\_960


87

CL\_960


1

CL\_963

fGI ID

CL\_INS\_63
Cluster ID

CL\_959
