## Supplementary material for "A novel method for integrating genomic and Tn-Seq data to identify common *in vivo* fitness mechanisms across multiple bacterial species": S2 Dataset: CL_INS_65.html

Legend

 Regulatoryfunctions
 Hypothetical
 Other

FULL


WINDOWSVGPNG

Trim RowsRemove SingletonsSave Fasta

CL\_973


CL\_973


CL\_973


CL\_973

HighlightSelectShow Genomes


136

CL\_974


67

CL\_974


1

CL\_975


1

CL\_975

fGI ID

CL\_INS\_65
Cluster ID

CL\_5836
