## Supplementary material for "A novel method for integrating genomic and Tn-Seq data to identify common *in vivo* fitness mechanisms across multiple bacterial species": S2 Dataset: CL_INS_69.html

Legend

 Hypothetical
 Other
 All VFDB Genes

FULL


WINDOWSVGPNG

Trim RowsRemove SingletonsSave Fasta

CL\_1060


CL\_1060


CL\_1060


CL\_1060


CL\_1060

HighlightSelectShow Genomes


134

CL\_1062


62

CL\_1062


7

CL\_1062


1

CL\_1063


1

CL\_1062

fGI ID


CL\_INS\_69
CL\_INS\_69
CL\_INS\_69
Cluster ID


CL\_6151
CL\_1061
CL\_12639
