## Supplementary material for "A novel method for integrating genomic and Tn-Seq data to identify common *in vivo* fitness mechanisms across multiple bacterial species": S2 Dataset: CL_INS_70.html

Legend

 Hypothetical
 All EssentialGenes
 All VFDB Genes

FULL


WINDOWSVGPNG

Trim RowsRemove SingletonsSave Fasta

CL\_1084


CL\_1084

HighlightSelectShow Genomes


130

CL\_1085


75

CL\_1085

fGI ID

CL\_INS\_70
Cluster ID

CL\_5837
