## Supplementary material for "A novel method for integrating genomic and Tn-Seq data to identify common *in vivo* fitness mechanisms across multiple bacterial species": S2 Dataset: CL_INS_71.html

Legend

 Regulatoryfunctions
 Hypothetical
 Other
 All VFDB Genes

FULL


WINDOWSVGPNG

Trim RowsRemove SingletonsSave Fasta

CL\_1143


CL\_1143


CL\_1143


CL\_1143


CL\_1143

HighlightSelectShow Genomes


186

CL\_1144


8

CL\_1144


7

CL\_1144


3

CL\_1144


1

CL\_1144

fGI ID


CL\_INS\_71
CL\_INS\_71
CL\_INS\_71
CL\_INS\_71
CL\_INS\_71
CL\_INS\_71
CL\_INS\_71
CL\_INS\_71
CL\_INS\_71
CL\_INS\_71
CL\_INS\_71
CL\_INS\_71
CL\_INS\_71
CL\_INS\_71
CL\_INS\_71
CL\_INS\_71
CL\_INS\_71
CL\_INS\_71
CL\_INS\_71
Cluster ID


CL\_9917
CL\_9916
CL\_9915
CL\_9914
CL\_9913
CL\_15025
CL\_9912
CL\_11506
CL\_17956
CL\_11505
CL\_11504
CL\_11503
CL\_17955
CL\_11502
CL\_11501
CL\_11500
CL\_11499
CL\_11498
CL\_11497
