## Supplementary material for "A novel method for integrating genomic and Tn-Seq data to identify common *in vivo* fitness mechanisms across multiple bacterial species": S2 Dataset: CL_INS_76.html

Legend

 Mobile +extrachromosomalelementfunctions
 Hypothetical
 DNA Metabolism
 Transcription
 Other
 Cellularprocesses
 All VFDB Genes

FULL


WINDOWSVGPNG

Trim RowsRemove SingletonsSave Fasta

CL\_1174


CL\_1174


CL\_1174


CL\_1174


CL\_1174


CL\_1174


CL\_1174


Break


Break


CL\_1174


CL\_1174


CL\_1174


CL\_1174


CL\_1173


Break


Break

HighlightSelectShow Genomes


84

CL\_1176


83

CL\_1176


13

CL\_1176


7

CL\_1176


5

CL\_1176


2

CL\_1176


2

CL\_1176


1

CL\_1176


1

CL\_1176


1

CL\_1176


1

Break


1

Break


1

Break


1

CL\_1176


1

CL\_1176


1

CL\_1176

fGI ID


CL\_INS\_76
CL\_INS\_105
CL\_INS\_278
CL\_INS\_278
CL\_INS\_278
CL\_INS\_263
CL\_INS\_263
CL\_INS\_76
CL\_INS\_278
CL\_INS\_76
CL\_INS\_132
CL\_INS\_132
CL\_INS\_278
CL\_INS\_76
CL\_INS\_278
CL\_INS\_278
CL\_INS\_278
CL\_INS\_30
CL\_INS\_30
CL\_INS\_132
CL\_INS\_132
CL\_INS\_401
CL\_INS\_278
CL\_INS\_132
CL\_INS\_278
CL\_INS\_278
CL\_INS\_132
CL\_INS\_132
CL\_INS\_132
CL\_INS\_132
CL\_INS\_76
CL\_INS\_132
CL\_INS\_76
CL\_INS\_105
CL\_INS\_298
CL\_INS\_1
CL\_INS\_318
CL\_INS\_385
CL\_INS\_76
CL\_INS\_197
CL\_INS\_278
CL\_INS\_278
CL\_INS\_278
CL\_INS\_278
CL\_INS\_278
CL\_INS\_278
CL\_INS\_278
CL\_INS\_197
CL\_INS\_197
CL\_INS\_197
CL\_INS\_197
CL\_INS\_197
CL\_INS\_197
CL\_INS\_197
CL\_INS\_197
CL\_INS\_278
CL\_INS\_57
CL\_INS\_57
CL\_INS\_246
CL\_INS\_246
CL\_INS\_246
CL\_INS\_246
CL\_INS\_246
CL\_INS\_246
CL\_INS\_246
CL\_INS\_246
CL\_INS\_57
CL\_INS\_246
CL\_INS\_57
CL\_INS\_57
CL\_INS\_246
CL\_INS\_246
CL\_INS\_246
CL\_INS\_246
CL\_INS\_246
CL\_INS\_246
CL\_INS\_246
CL\_INS\_246
CL\_INS\_57
CL\_INS\_57
CL\_INS\_57
CL\_INS\_57
CL\_INS\_57
CL\_INS\_57
CL\_INS\_57
CL\_INS\_246
CL\_INS\_246
CL\_INS\_246
CL\_INS\_246
CL\_INS\_246
CL\_INS\_246
CL\_INS\_359
CL\_INS\_359
CL\_INS\_359
CL\_INS\_359
CL\_INS\_359
CL\_INS\_359
CL\_INS\_359
CL\_INS\_359
CL\_INS\_359
CL\_INS\_359
CL\_INS\_359
CL\_INS\_359
CL\_INS\_359
CL\_INS\_359
CL\_INS\_359
CL\_INS\_359
CL\_INS\_359
CL\_INS\_359
CL\_INS\_359
CL\_INS\_359
CL\_INS\_359
CL\_INS\_359
CL\_INS\_359
CL\_INS\_44
Cluster ID


CL\_20428
CL\_6044
CL\_6043
CL\_6042
CL\_6041
CL\_5463
CL\_5462
CL\_6692
CL\_6005
CL\_13717
CL\_6806
CL\_6823
CL\_6800
CL\_6861
CL\_5973
CL\_5974
CL\_5975
CL\_6860
CL\_6859
CL\_7058
CL\_6008
CL\_6010
CL\_6011
CL\_6012
CL\_6013
CL\_6014
CL\_6016
CL\_6017
CL\_6018
CL\_6019
CL\_7062
CL\_6022
CL\_1175
CL\_6198
CL\_7910
CL\_12379
CL\_7112
CL\_12378
CL\_12490
CL\_6820
CL\_6819
CL\_6818
CL\_6817
CL\_6137
CL\_6816
CL\_6815
CL\_6814
CL\_6812
CL\_6811
CL\_10120
CL\_8193
CL\_8192
CL\_8191
CL\_8190
CL\_8189
CL\_12342
CL\_742
CL\_741
CL\_6437
CL\_6436
CL\_6435
CL\_6434
CL\_6433
CL\_6432
CL\_12211
CL\_737
CL\_735
CL\_6431
CL\_734
CL\_733
CL\_6430
CL\_6429
CL\_6428
CL\_6427
CL\_6426
CL\_6425
CL\_6424
CL\_6423
CL\_727
CL\_726
CL\_725
CL\_724
CL\_723
CL\_722
CL\_721
CL\_6422
CL\_12210
CL\_6421
CL\_12209
CL\_6420
CL\_6419
CL\_6418
CL\_6417
CL\_6416
CL\_6415
CL\_6414
CL\_6413
CL\_6412
CL\_6411
CL\_6410
CL\_6409
CL\_6408
CL\_6407
CL\_6406
CL\_6405
CL\_6404
CL\_6402
CL\_6401
CL\_6400
CL\_6399
CL\_6398
CL\_6397
CL\_6396
CL\_6395
CL\_6394
