## Supplementary material for "A novel method for integrating genomic and Tn-Seq data to identify common *in vivo* fitness mechanisms across multiple bacterial species": S2 Dataset: CL_INS_78.html

Legend

 AntibioticResistance
 EnergyMetabolism

FULL


WINDOWSVGPNG

Trim RowsRemove SingletonsSave Fasta

CL\_1194


CL\_1194


CL\_1194

HighlightSelectShow Genomes


115

CL\_1196


86

CL\_1196


1

CL\_1197

fGI ID

CL\_INS\_78
Cluster ID

CL\_1195
