## Supplementary material for "A novel method for integrating genomic and Tn-Seq data to identify common *in vivo* fitness mechanisms across multiple bacterial species": S2 Dataset: CL_INS_80.html

Legend

 Mobile +extrachromosomalelementfunctions
 Hypothetical
 Other
 All VFDB Genes

FULL


WINDOWSVGPNG

Trim RowsRemove SingletonsSave Fasta

CL\_1206


CL\_1206


CL\_1206


CL\_1205


CL\_1206

HighlightSelectShow Genomes


106

CL\_1207


90

CL\_1207


3

CL\_1207


2

CL\_1207


1

Break

fGI ID


CL\_INS\_80
CL\_INS\_80
CL\_INS\_321
CL\_INS\_321
Cluster ID


CL\_18890
CL\_5253
CL\_6344
CL\_6343
