## Supplementary material for "A novel method for integrating genomic and Tn-Seq data to identify common *in vivo* fitness mechanisms across multiple bacterial species": S2 Dataset: CL_INS_81.html

Legend

 Mobile +extrachromosomalelementfunctions
 Hypothetical
 All VFDB Genes

FULL


WINDOWSVGPNG

Trim RowsRemove SingletonsSave Fasta

CL\_1244


CL\_1244


CL\_1244


CL\_1244

HighlightSelectShow Genomes


141

CL\_1246


61

CL\_1246


1

CL\_1246


1

CL\_1246

fGI ID


CL\_INS\_81
CL\_INS\_81
CL\_INS\_81
CL\_INS\_81
Cluster ID


CL\_14694
CL\_1245
CL\_16292
CL\_16291
