## Supplementary material for "A novel method for integrating genomic and Tn-Seq data to identify common *in vivo* fitness mechanisms across multiple bacterial species": S2 Dataset: CL_INS_83.html

Legend

 Mobile +extrachromosomalelementfunctions
 Regulatoryfunctions
 Hypothetical
 Other
 All VFDB Genes

FULL


WINDOWSVGPNG

Trim RowsRemove SingletonsSave Fasta

CL\_1311


CL\_1311


CL\_1311


CL\_1311


CL\_1311


CL\_1311


CL\_1311


CL\_1311

HighlightSelectShow Genomes


183

CL\_1312


15

CL\_1312


1

CL\_1312


1

CL\_1312


1

CL\_1312


1

CL\_1312


1

CL\_1312


1

CL\_1312

fGI ID


CL\_INS\_83
CL\_INS\_83
CL\_INS\_74
CL\_INS\_83
CL\_INS\_83
CL\_INS\_83
CL\_INS\_83
CL\_INS\_83
CL\_INS\_83
CL\_INS\_83
CL\_INS\_83
CL\_INS\_83
CL\_INS\_83
CL\_INS\_83
CL\_INS\_83
CL\_INS\_83
CL\_INS\_83
CL\_INS\_83
CL\_INS\_83
CL\_INS\_83
CL\_INS\_83
CL\_INS\_83
CL\_INS\_83
CL\_INS\_83
CL\_INS\_83
CL\_INS\_83
CL\_INS\_83
CL\_INS\_83
CL\_INS\_83
CL\_INS\_83
CL\_INS\_83
CL\_INS\_83
CL\_INS\_83
CL\_INS\_83
Cluster ID


CL\_17303
CL\_16491
CL\_6857
CL\_23481
CL\_7909
CL\_7908
CL\_7907
CL\_7906
CL\_7905
CL\_20988
CL\_20987
CL\_20986
CL\_20985
CL\_17302
CL\_17301
CL\_17300
CL\_17299
CL\_17298
CL\_17297
CL\_17296
CL\_17295
CL\_17294
CL\_17293
CL\_19376
CL\_19375
CL\_19374
CL\_19373
CL\_19372
CL\_19371
CL\_19370
CL\_19369
CL\_19368
CL\_19367
CL\_19366
