## Supplementary material for "A novel method for integrating genomic and Tn-Seq data to identify common *in vivo* fitness mechanisms across multiple bacterial species": S2 Dataset: CL_INS_87.html

Legend

 Hypothetical
 Other
 All VFDB Genes

FULL


WINDOWSVGPNG

Trim RowsRemove SingletonsSave Fasta

CL\_1354


CL\_1354


CL\_1354


CL\_1354

HighlightSelectShow Genomes


113

CL\_1357


85

CL\_1357


2

CL\_1357


1

CL\_1358

fGI ID


CL\_INS\_87
CL\_INS\_87
Cluster ID


CL\_1355
CL\_1356
