## Supplementary material for "A novel method for integrating genomic and Tn-Seq data to identify common *in vivo* fitness mechanisms across multiple bacterial species": S2 Dataset: CL_INS_89.html

Legend

 Regulatoryfunctions
 Hypothetical
 All EssentialGenes
 Other
 All VFDB Genes

FULL


WINDOWSVGPNG

Trim RowsRemove SingletonsSave Fasta

CL\_1374


CL\_1374


CL\_1374


CL\_1374

HighlightSelectShow Genomes


194

CL\_1393


8

CL\_1393


1

CL\_1393


1

CL\_1393

fGI ID


CL\_INS\_89
CL\_INS\_89
CL\_INS\_89
CL\_INS\_89
CL\_INS\_89
CL\_INS\_89
CL\_INS\_89
CL\_INS\_89
CL\_INS\_89
CL\_INS\_89
CL\_INS\_89
CL\_INS\_89
CL\_INS\_89
CL\_INS\_89
CL\_INS\_89
CL\_INS\_89
CL\_INS\_89
CL\_INS\_89
CL\_INS\_89
Cluster ID


CL\_1375
CL\_1376
CL\_1377
CL\_1378
CL\_1379
CL\_1380
CL\_1381
CL\_1382
CL\_1383
CL\_1384
CL\_1385
CL\_22092
CL\_1386
CL\_1387
CL\_1388
CL\_1389
CL\_1390
CL\_1391
CL\_1392
