## Supplementary material for "A novel method for integrating genomic and Tn-Seq data to identify common *in vivo* fitness mechanisms across multiple bacterial species": S2 Dataset: CL_INS_90.html

Legend

 Hypothetical
 All EssentialGenes

FULL


WINDOWSVGPNG

Trim RowsRemove SingletonsSave Fasta

CL\_1398


CL\_1398

HighlightSelectShow Genomes


194

CL\_1402


9

CL\_1402

fGI ID


CL\_INS\_90
CL\_INS\_90
CL\_INS\_90
Cluster ID


CL\_1399
CL\_1400
CL\_1401
