## Supplementary material for "A novel method for integrating genomic and Tn-Seq data to identify common *in vivo* fitness mechanisms across multiple bacterial species": S2 Dataset: CL_INS_94.html

Legend

 Mobile +extrachromosomalelementfunctions
 Hypothetical
 All VFDB Genes

FULL


WINDOWSVGPNG

Trim RowsRemove SingletonsSave Fasta

CL\_1417


CL\_1417


CL\_1417


CL\_1417

HighlightSelectShow Genomes


174

CL\_1418


25

CL\_1418


2

CL\_1419


1

CL\_1418

fGI ID


CL\_INS\_94
CL\_INS\_94
Cluster ID


CL\_19354
CL\_5712
