## Supplementary material for "A novel method for integrating genomic and Tn-Seq data to identify common *in vivo* fitness mechanisms across multiple bacterial species": S2 Dataset: CL_INS_98.html

Legend

 Hypothetical
 All VFDB Genes

FULL


WINDOWSVGPNG

Trim RowsRemove SingletonsSave Fasta

CL\_1445


CL\_1445


CL\_1445


CL\_1444

HighlightSelectShow Genomes


115

CL\_1447


87

CL\_1447


2

CL\_1447


1

CL\_1447

fGI ID


CL\_INS\_98
CL\_INS\_98
Cluster ID


CL\_1446
CL\_6603
