## Supplementary material for "A novel method for integrating genomic and Tn-Seq data to identify common *in vivo* fitness mechanisms across multiple bacterial species": S2 Dataset: CL_INS_100.html

Legend

 Regulatoryfunctions
 Hypothetical
 Other
 All VFDB Genes
 Transport +binding proteins

FULL


WINDOWSVGPNG

Trim RowsRemove SingletonsSave Fasta

CL\_1460


CL\_1460


CL\_1460


CL\_1460


CL\_1459


CL\_1460

HighlightSelectShow Genomes


102

CL\_1461


86

CL\_1461


6

CL\_1461


2

CL\_1461


1

CL\_1461


1

CL\_1461

fGI ID


CL\_INS\_100
CL\_INS\_100
CL\_INS\_100
CL\_INS\_100
CL\_INS\_100
CL\_INS\_100
CL\_INS\_100
CL\_INS\_100
CL\_INS\_100
CL\_INS\_100
CL\_INS\_100
CL\_INS\_100
CL\_INS\_100
Cluster ID


CL\_5179
CL\_7903
CL\_7902
CL\_7901
CL\_7900
CL\_7899
CL\_7898
CL\_7897
CL\_7896
CL\_7895
CL\_7894
CL\_17170
CL\_5178
