## Supplementary material for "A novel method for integrating genomic and Tn-Seq data to identify common *in vivo* fitness mechanisms across multiple bacterial species": S2 Dataset: CL_INS_109.html

Legend

 Hypothetical
 Other
 All VFDB Genes

FULL


WINDOWSVGPNG

Trim RowsRemove SingletonsSave Fasta

CL\_1530


CL\_1530


CL\_1530


CL\_1529


Break


CL\_1530

HighlightSelectShow Genomes


134

CL\_1532


55

CL\_1537


11

CL\_1532


1

CL\_1532


1

CL\_1532


1

CL\_1533

fGI ID


CL\_INS\_109
CL\_INS\_109
Cluster ID


CL\_1531
CL\_7518
