## Supplementary material for "A novel method for integrating genomic and Tn-Seq data to identify common *in vivo* fitness mechanisms across multiple bacterial species": S2 Dataset: CL_INS_112.html

Legend

 Regulatoryfunctions
 Hypothetical
 Other
 All VFDB Genes

FULL


WINDOWSVGPNG

Trim RowsRemove SingletonsSave Fasta

CL\_1553


CL\_1553


CL\_1553


CL\_1553

HighlightSelectShow Genomes


178

CL\_1554


9

CL\_1554


5

CL\_1554


1

CL\_1554

fGI ID


CL\_INS\_112
CL\_INS\_112
CL\_INS\_112
CL\_INS\_112
CL\_INS\_112
CL\_INS\_112
CL\_INS\_112
CL\_INS\_112
CL\_INS\_112
CL\_INS\_112
CL\_INS\_112
Cluster ID


CL\_5177
CL\_5176
CL\_5175
CL\_5174
CL\_5173
CL\_5172
CL\_5171
CL\_5170
CL\_16290
CL\_5169
CL\_5168
