## Supplementary material for "A novel method for integrating genomic and Tn-Seq data to identify common *in vivo* fitness mechanisms across multiple bacterial species": S2 Dataset: CL_INS_117.html

Legend

 Regulatoryfunctions
 Other
 Transport +binding proteins
 All VFDB Genes

FULL


WINDOWSVGPNG

Trim RowsRemove SingletonsSave Fasta

CL\_1686


CL\_1686


CL\_1686


CL\_1686

HighlightSelectShow Genomes


191

CL\_1687


12

CL\_1687


1

CL\_1687


1

CL\_1687

fGI ID


CL\_INS\_117
CL\_INS\_117
CL\_INS\_117
CL\_INS\_117
CL\_INS\_117
CL\_INS\_36
CL\_INS\_117
CL\_INS\_10
CL\_INS\_10
CL\_INS\_36
Cluster ID


CL\_7718
CL\_7717
CL\_7716
CL\_7715
CL\_7714
CL\_7713
CL\_12859
CL\_4594
CL\_4593
CL\_7712
