## Supplementary material for "A novel method for integrating genomic and Tn-Seq data to identify common *in vivo* fitness mechanisms across multiple bacterial species": S2 Dataset: CL_INS_120.html

Legend

 Hypothetical
 Other
 All VFDB Genes

FULL


WINDOWSVGPNG

Trim RowsRemove SingletonsSave Fasta

CL\_1706


CL\_1706


CL\_1706


CL\_1706


CL\_1706


CL\_1706


CL\_1706


CL\_1706

HighlightSelectShow Genomes


101

CL\_1708


85

CL\_1708


8

CL\_1708


3

CL\_1708


2

CL\_1708


2

CL\_1708


1

CL\_1721


1

CL\_1708

fGI ID


CL\_INS\_120
CL\_INS\_120
CL\_INS\_120
CL\_INS\_120
Cluster ID


CL\_7893
CL\_19174
CL\_1707
CL\_14453
