## Supplementary material for "A novel method for integrating genomic and Tn-Seq data to identify common *in vivo* fitness mechanisms across multiple bacterial species": S2 Dataset: CL_INS_122.html

Legend

 Hypothetical
 Other
 All VFDB Genes

FULL


WINDOWSVGPNG

Trim RowsRemove SingletonsSave Fasta

CL\_1728


CL\_1728


CL\_1728


CL\_1728


CL\_1728


CL\_1728

HighlightSelectShow Genomes


153

CL\_1729


41

CL\_1729


5

CL\_1729


2

CL\_1729


2

CL\_1729


1

CL\_1729

fGI ID


CL\_INS\_122
CL\_INS\_122
CL\_INS\_122
CL\_INS\_122
Cluster ID


CL\_6190
CL\_5708
CL\_9821
CL\_18356
