## Supplementary material for "A novel method for integrating genomic and Tn-Seq data to identify common *in vivo* fitness mechanisms across multiple bacterial species": S2 Dataset: CL_INS_123.html

Legend

 Hypothetical
 Other
 All VFDB Genes

FULL


WINDOWSVGPNG

Trim RowsRemove SingletonsSave Fasta

CL\_1729


CL\_1729


CL\_1729


CL\_1729

HighlightSelectShow Genomes


113

CL\_1730


90

CL\_1730


1

CL\_1731


1

CL\_1730

fGI ID


CL\_INS\_123
CL\_INS\_123
Cluster ID


CL\_12823
CL\_5707
