## Supplementary material for "A novel method for integrating genomic and Tn-Seq data to identify common *in vivo* fitness mechanisms across multiple bacterial species": S2 Dataset: CL_INS_124.html

Legend

 Hypothetical
 Biosynthesis ofcofactors,prostheticgroups, +carriers
 Other

FULL


WINDOWSVGPNG

Trim RowsRemove SingletonsSave Fasta

CL\_1733


CL\_1733

HighlightSelectShow Genomes


130

CL\_1734


75

CL\_1734

fGI ID

CL\_INS\_124
Cluster ID

CL\_5706
