## Supplementary material for "A novel method for integrating genomic and Tn-Seq data to identify common *in vivo* fitness mechanisms across multiple bacterial species": S2 Dataset: CL_INS_126.html

Legend

 Hypothetical
 Other
 All VFDB Genes

FULL


WINDOWSVGPNG

Trim RowsRemove SingletonsSave Fasta

CL\_1749


CL\_1749


CL\_1749


CL\_1749


CL\_1748


CL\_1749

HighlightSelectShow Genomes


114

CL\_1751


84

CL\_1751


2

CL\_1752


1

CL\_1751


1

CL\_1751


1

CL\_1752

fGI ID


CL\_INS\_126
CL\_INS\_126
Cluster ID


CL\_1750
CL\_17282
