## Supplementary material for "A novel method for integrating genomic and Tn-Seq data to identify common *in vivo* fitness mechanisms across multiple bacterial species": S2 Dataset: CL_INS_133.html

Legend

 Hypothetical
 Other

FULL


WINDOWSVGPNG

Trim RowsRemove SingletonsSave Fasta

CL\_1849


CL\_1849


CL\_1849

HighlightSelectShow Genomes


116

CL\_1851


87

CL\_1851


1

CL\_1851

fGI ID


CL\_INS\_133
CL\_INS\_133
Cluster ID


CL\_1850
CL\_14044
