## Supplementary material for "A novel method for integrating genomic and Tn-Seq data to identify common *in vivo* fitness mechanisms across multiple bacterial species": S2 Dataset: CL_INS_136.html

Legend

 Hypothetical
 Cellularprocesses
 All VFDB Genes

FULL


WINDOWSVGPNG

Trim RowsRemove SingletonsSave Fasta

CL\_1913


CL\_1913


CL\_1913


CL\_1913


CL\_1913

HighlightSelectShow Genomes


115

CL\_1915


86

CL\_1915


1

CL\_1915


1

Break


1

CL\_1915

fGI ID


CL\_INS\_136
CL\_INS\_307
CL\_INS\_136
CL\_INS\_136
Cluster ID


CL\_21284
CL\_81
CL\_9817
CL\_1914
