## Supplementary material for "A novel method for integrating genomic and Tn-Seq data to identify common *in vivo* fitness mechanisms across multiple bacterial species": S2 Dataset: CL_INS_137.html

Legend

 Mobile +extrachromosomalelementfunctions
 Hypothetical
 All EssentialGenes
 Other
 All VFDB Genes

FULL


WINDOWSVGPNG

Trim RowsRemove SingletonsSave Fasta

CL\_1920


CL\_1920


CL\_1920


CL\_1920


CL\_1920


CL\_1920


CL\_1920


CL\_1920


CL\_1920


CL\_1920


CL\_1920


CL\_1920


CL\_1920

HighlightSelectShow Genomes


161

CL\_1921


32

CL\_1921


1

CL\_1921


1

CL\_1921


1

CL\_1921


1

CL\_1921


1

CL\_1921


1

Break


1

CL\_1921


1

CL\_1921


1

CL\_1921


1

CL\_1921


1

CL\_1921

fGI ID


CL\_INS\_287
CL\_INS\_137
CL\_INS\_137
CL\_INS\_137
CL\_INS\_137
CL\_INS\_137
CL\_INS\_137
CL\_INS\_137
CL\_INS\_137
CL\_INS\_137
CL\_INS\_137
CL\_INS\_137
CL\_INS\_137
CL\_INS\_137
CL\_INS\_137
Cluster ID


CL\_7011
CL\_9127
CL\_16523
CL\_23607
CL\_5623
CL\_5624
CL\_22027
CL\_19343
CL\_19342
CL\_19341
CL\_5625
CL\_5626
CL\_5627
CL\_5628
CL\_5629
