## Supplementary material for "A novel method for integrating genomic and Tn-Seq data to identify common *in vivo* fitness mechanisms across multiple bacterial species": S2 Dataset: CL_INS_140.html

Legend

 Hypothetical
 All EssentialGenes
 Other
 All VFDB Genes
 Transport +binding proteins

FULL


WINDOWSVGPNG

Trim RowsRemove SingletonsSave Fasta

CL\_1943


CL\_1943


CL\_1943


CL\_1943


CL\_1943


CL\_1943


CL\_1943


CL\_1943


CL\_1943

HighlightSelectShow Genomes


145

CL\_6355


34

CL\_1931


2

CL\_1931


2

CL\_6355


1

CL\_1928


1

CL\_1931


1

CL\_1928


1

CL\_1931


1

CL\_1931

fGI ID


CL\_INS\_140
CL\_INS\_139
CL\_INS\_139
CL\_INS\_139
CL\_INS\_139
CL\_INS\_139
CL\_INS\_139
CL\_INS\_139
CL\_INS\_139
CL\_INS\_139
CL\_INS\_139
CL\_INS\_139
CL\_INS\_138
Cluster ID


CL\_1942
CL\_1941
CL\_23406
CL\_1940
CL\_1939
CL\_1938
CL\_1937
CL\_1936
CL\_1935
CL\_1934
CL\_1933
CL\_1932
CL\_1930
