## Supplementary material for "A novel method for integrating genomic and Tn-Seq data to identify common *in vivo* fitness mechanisms across multiple bacterial species": S2 Dataset: CL_INS_144.html

Legend

 Mobile +extrachromosomalelementfunctions
 Hypothetical
 All EssentialGenes
 Other
 All VFDB Genes

FULL


WINDOWSVGPNG

Trim RowsRemove SingletonsSave Fasta

CL\_1982


CL\_1982


CL\_1982


CL\_1982


CL\_1982


CL\_1982


CL\_1982


CL\_1982


CL\_1981


CL\_1982


CL\_1982


Break


CL\_1982


CL\_1982


CL\_1982


Break


CL\_1982


CL\_1982


CL\_1982


CL\_1982

HighlightSelectShow Genomes


102

CL\_1984


74

CL\_1984


6

CL\_1984


3

CL\_1984


3

CL\_1984


3

CL\_1984


2

Break


1

CL\_1984


1

CL\_1984


1

CL\_1984


1

CL\_1984


1

CL\_1984


1

CL\_1984


1

CL\_1984


1

CL\_1984


1

CL\_1984


1

CL\_1984


1

CL\_1984


1

Break


1

CL\_1984

fGI ID


CL\_INS\_318
CL\_INS\_144
CL\_INS\_144
CL\_INS\_144
CL\_INS\_144
CL\_INS\_160
CL\_INS\_144
CL\_INS\_144
CL\_INS\_144
CL\_INS\_26
CL\_INS\_144
CL\_INS\_144
CL\_INS\_144
CL\_INS\_144
CL\_INS\_144
CL\_INS\_144
CL\_INS\_144
CL\_INS\_278
CL\_INS\_278
CL\_INS\_132
CL\_INS\_132
CL\_INS\_132
CL\_INS\_359
CL\_INS\_359
CL\_INS\_359
CL\_INS\_278
CL\_INS\_278
CL\_INS\_160
CL\_INS\_385
CL\_INS\_385
CL\_INS\_385
CL\_INS\_401
Cluster ID


CL\_10121
CL\_15627
CL\_12426
CL\_1983
CL\_22355
CL\_7082
CL\_7010
CL\_8798
CL\_9126
CL\_8797
CL\_8796
CL\_8795
CL\_8794
CL\_8793
CL\_8792
CL\_8791
CL\_8790
CL\_6013
CL\_6014
CL\_6015
CL\_6016
CL\_6017
CL\_4356
CL\_4355
CL\_4354
CL\_11802
CL\_12216
CL\_68
CL\_69
CL\_12217
CL\_6693
CL\_6040
