## Supplementary material for "A novel method for integrating genomic and Tn-Seq data to identify common *in vivo* fitness mechanisms across multiple bacterial species": S2 Dataset: CL_INS_146.html

Legend

 Hypothetical
 All EssentialGenes
 All VFDB Genes

FULL


WINDOWSVGPNG

Trim RowsRemove SingletonsSave Fasta

CL\_1999


CL\_1999


CL\_1997

HighlightSelectShow Genomes


116

CL\_2001


87

CL\_2001


1

CL\_2001

fGI ID

CL\_INS\_146
Cluster ID

CL\_2000
