## Supplementary material for "A novel method for integrating genomic and Tn-Seq data to identify common *in vivo* fitness mechanisms across multiple bacterial species": S2 Dataset: CL_INS_148.html

Legend

 Mobile +extrachromosomalelementfunctions
 Regulatoryfunctions
 Hypothetical
 DNA Metabolism
 Other
 All VFDB Genes
 Transport +binding proteins

FULL


WINDOWSVGPNG

Trim RowsRemove SingletonsSave Fasta

CL\_2038


CL\_2038


CL\_2037


CL\_2038


CL\_2037


CL\_2038


CL\_2038

HighlightSelectShow Genomes


109

CL\_2040


83

CL\_2040


3

CL\_2040


1

CL\_2042


1

CL\_2040


1

CL\_2040


1

CL\_2040

fGI ID


CL\_INS\_148
CL\_INS\_148
CL\_INS\_148
CL\_INS\_148
CL\_INS\_149
CL\_INS\_149
CL\_INS\_148
CL\_INS\_148
CL\_INS\_148
CL\_INS\_148
CL\_INS\_148
Cluster ID


CL\_12661
CL\_12662
CL\_2039
CL\_23523
CL\_23522
CL\_23521
CL\_15272
CL\_15273
CL\_15274
CL\_15275
CL\_15276
