## Supplementary material for "A novel method for integrating genomic and Tn-Seq data to identify common *in vivo* fitness mechanisms across multiple bacterial species": S2 Dataset: CL_INS_149.html

Legend

 Hypothetical
 DNA Metabolism
 All VFDB Genes

FULL


WINDOWSVGPNG

Trim RowsRemove SingletonsSave Fasta

CL\_2040


CL\_2040


CL\_2038

HighlightSelectShow Genomes


114

CL\_2042


88

CL\_2042


1

CL\_2042

fGI ID


CL\_INS\_149
CL\_INS\_149
CL\_INS\_149
CL\_INS\_149
Cluster ID


CL\_2041
CL\_23522
CL\_23521
CL\_23520
