## Supplementary material for "A novel method for integrating genomic and Tn-Seq data to identify common *in vivo* fitness mechanisms across multiple bacterial species": S2 Dataset: CL_INS_154.html

Legend

 Hypothetical
 All EssentialGenes
 Other
 All VFDB Genes

FULL


WINDOWSVGPNG

Trim RowsRemove SingletonsSave Fasta

CL\_2067


CL\_2067


CL\_2067


CL\_2067


CL\_2067


CL\_2067


CL\_2067


CL\_2067


CL\_2067

HighlightSelectShow Genomes


154

CL\_2070


34

CL\_2070


3

CL\_2070


3

CL\_2070


2

CL\_2070


2

CL\_2070


1

CL\_2070


1

CL\_2070


1

CL\_2070

fGI ID


CL\_INS\_154
CL\_INS\_154
CL\_INS\_154
CL\_INS\_154
CL\_INS\_154
Cluster ID


CL\_2068
CL\_7211
CL\_2069
CL\_8789
CL\_20415
