## Supplementary material for "A novel method for integrating genomic and Tn-Seq data to identify common *in vivo* fitness mechanisms across multiple bacterial species": S2 Dataset: CL_INS_156.html

Legend

 Hypothetical
 Other
 All VFDB Genes

FULL


WINDOWSVGPNG

Trim RowsRemove SingletonsSave Fasta

CL\_5158


CL\_5158


CL\_5158


CL\_5158


CL\_5158

HighlightSelectShow Genomes


69

CL\_5155


55

CL\_5155


47

CL\_5155


27

CL\_5155


3

CL\_5155

fGI ID


CL\_INS\_156
CL\_INS\_156
CL\_INS\_156
CL\_INS\_156
CL\_INS\_156
CL\_INS\_156
Cluster ID


CL\_5157
CL\_5156
CL\_10375
CL\_10374
CL\_10373
CL\_10372
