## Supplementary material for "A novel method for integrating genomic and Tn-Seq data to identify common *in vivo* fitness mechanisms across multiple bacterial species": S2 Dataset: CL_INS_157.html

Legend

 Mobile +extrachromosomalelementfunctions
 Hypothetical
 DNA Metabolism
 Other
 All VFDB Genes

FULL


WINDOWSVGPNG

Trim RowsRemove SingletonsSave Fasta

CL\_5154


CL\_5154


CL\_5154


CL\_5155


CL\_2080

HighlightSelectShow Genomes


110

CL\_5152


86

CL\_5152


1

CL\_5152


1

CL\_5152


1

CL\_5152

fGI ID


CL\_INS\_157
CL\_INS\_157
CL\_INS\_155
CL\_INS\_155
CL\_INS\_263
CL\_INS\_263
CL\_INS\_263
CL\_INS\_155
CL\_INS\_155
CL\_INS\_155
CL\_INS\_155
Cluster ID


CL\_21420
CL\_5153
CL\_2082
CL\_5638
CL\_5639
CL\_5641
CL\_8721
CL\_19226
CL\_7473
CL\_8783
CL\_5670
