## Supplementary material for "A novel method for integrating genomic and Tn-Seq data to identify common *in vivo* fitness mechanisms across multiple bacterial species": S2 Dataset: CL_INS_159.html

Legend

 Mobile +extrachromosomalelementfunctions
 Regulatoryfunctions
 Hypothetical
 DNA Metabolism
 All EssentialGenes
 All Fitness Genes
 Other
 Proteinsynthesis/fate
 All VFDB Genes

FULL


WINDOWSVGPNG

Trim RowsRemove SingletonsSave Fasta

CL\_5143


CL\_5143


CL\_5143


CL\_5143


CL\_5143


CL\_5143


CL\_5141

HighlightSelectShow Genomes


148

CL\_5144


42

CL\_5145


1

CL\_5144


1

CL\_2080


1

CL\_5144


1

CL\_5144


1

CL\_5144

fGI ID


CL\_INS\_159
CL\_INS\_159
CL\_INS\_159
CL\_INS\_159
CL\_INS\_159
CL\_INS\_155
CL\_INS\_155
CL\_INS\_155
CL\_INS\_155
CL\_INS\_155
CL\_INS\_155
CL\_INS\_155
CL\_INS\_155
CL\_INS\_155
CL\_INS\_155
CL\_INS\_155
CL\_INS\_155
CL\_INS\_155
CL\_INS\_155
CL\_INS\_155
CL\_INS\_155
CL\_INS\_155
CL\_INS\_155
CL\_INS\_155
CL\_INS\_155
CL\_INS\_155
CL\_INS\_155
CL\_INS\_155
CL\_INS\_155
CL\_INS\_155
CL\_INS\_155
CL\_INS\_155
CL\_INS\_155
CL\_INS\_263
CL\_INS\_155
CL\_INS\_155
CL\_INS\_155
CL\_INS\_155
CL\_INS\_155
CL\_INS\_155
CL\_INS\_155
CL\_INS\_155
CL\_INS\_155
CL\_INS\_155
CL\_INS\_155
CL\_INS\_155
CL\_INS\_155
CL\_INS\_155
CL\_INS\_155
CL\_INS\_155
CL\_INS\_155
CL\_INS\_155
CL\_INS\_155
CL\_INS\_155
CL\_INS\_155
Cluster ID


CL\_23685
CL\_21359
CL\_21358
CL\_21357
CL\_17004
CL\_7257
CL\_7256
CL\_7255
CL\_7475
CL\_7254
CL\_7253
CL\_7252
CL\_7476
CL\_7477
CL\_7478
CL\_7479
CL\_7606
CL\_7607
CL\_7608
CL\_7609
CL\_7610
CL\_7483
CL\_7611
CL\_7612
CL\_7485
CL\_7887
CL\_7614
CL\_7238
CL\_7237
CL\_7236
CL\_7501
CL\_7502
CL\_7503
CL\_7504
CL\_7505
CL\_7233
CL\_7232
CL\_7231
CL\_7230
CL\_7229
CL\_7228
CL\_7227
CL\_7226
CL\_18103
CL\_8788
CL\_7222
CL\_7221
CL\_7220
CL\_7219
CL\_7218
CL\_7217
CL\_7215
CL\_7214
CL\_7515
CL\_7212
