## Supplementary material for "A novel method for integrating genomic and Tn-Seq data to identify common *in vivo* fitness mechanisms across multiple bacterial species": S2 Dataset: CL_INS_164.html

Legend

 Hypothetical
 Other
 Cellularprocesses
 All VFDB Genes

FULL


WINDOWSVGPNG

Trim RowsRemove SingletonsSave Fasta

CL\_2135


CL\_2135


CL\_2134


CL\_2135

HighlightSelectShow Genomes


116

CL\_2137


83

CL\_2137


2

CL\_2137


1

CL\_2138

fGI ID

CL\_INS\_164
Cluster ID

CL\_2136
