## Supplementary material for "A novel method for integrating genomic and Tn-Seq data to identify common *in vivo* fitness mechanisms across multiple bacterial species": S2 Dataset: CL_INS_165.html

Legend

 Hypothetical
 Cellularprocesses
 All VFDB Genes

FULL


WINDOWSVGPNG

Trim RowsRemove SingletonsSave Fasta

CL\_2137


CL\_2137


CL\_2137


CL\_2135

HighlightSelectShow Genomes


188

CL\_2138


11

CL\_2140


3

CL\_2140


1

CL\_2138

fGI ID

CL\_INS\_129
Cluster ID

CL\_8782
