## Supplementary material for "A novel method for integrating genomic and Tn-Seq data to identify common *in vivo* fitness mechanisms across multiple bacterial species": S2 Dataset: CL_INS_166.html

Legend

 Mobile +extrachromosomalelementfunctions
 Regulatoryfunctions
 Hypothetical
 Other
 All VFDB Genes

FULL


WINDOWSVGPNG

Trim RowsRemove SingletonsSave Fasta

CL\_2149


CL\_2149


CL\_2149


CL\_2149


CL\_2149


CL\_2149


CL\_2149


CL\_2149

HighlightSelectShow Genomes


137

CL\_5127


53

CL\_5127


1

CL\_5127


1

CL\_5127


1

CL\_5127


1

CL\_5127


1

CL\_5127


1

CL\_5127

fGI ID


CL\_INS\_166
CL\_INS\_166
CL\_INS\_166
CL\_INS\_166
CL\_INS\_166
CL\_INS\_166
CL\_INS\_166
CL\_INS\_166
CL\_INS\_166
CL\_INS\_166
Cluster ID


CL\_5135
CL\_5134
CL\_18595
CL\_5133
CL\_5132
CL\_14538
CL\_5131
CL\_5130
CL\_5129
CL\_5128
