## Supplementary material for "A novel method for integrating genomic and Tn-Seq data to identify common *in vivo* fitness mechanisms across multiple bacterial species": S2 Dataset: CL_INS_167.html

Legend

 Regulatoryfunctions
 Hypothetical
 All EssentialGenes
 Other
 Transport +binding proteins
 All VFDB Genes

FULL


WINDOWSVGPNG

Trim RowsRemove SingletonsSave Fasta

CL\_2152


CL\_2152


CL\_2152


CL\_2152


CL\_2152


CL\_2152


CL\_2152


CL\_2152


Break


CL\_2152


Break


CL\_2152

HighlightSelectShow Genomes


139

CL\_2153


47

CL\_2153


2

CL\_2153


2

CL\_2154


2

Break


1

CL\_2153


1

CL\_2153


1

CL\_4858


1

CL\_2153


1

Break


1

CL\_2153


1

Break

fGI ID


CL\_INS\_167
CL\_INS\_329
CL\_INS\_167
CL\_INS\_167
CL\_INS\_263
CL\_INS\_263
CL\_INS\_263
CL\_INS\_263
CL\_INS\_58
CL\_INS\_132
CL\_INS\_385
CL\_INS\_167
CL\_INS\_167
CL\_INS\_167
CL\_INS\_167
CL\_INS\_167
CL\_INS\_167
CL\_INS\_26
CL\_INS\_204
CL\_INS\_204
CL\_INS\_278
CL\_INS\_278
Cluster ID


CL\_13351
CL\_12028
CL\_17855
CL\_6170
CL\_5457
CL\_5456
CL\_5455
CL\_5454
CL\_6880
CL\_6823
CL\_6693
CL\_6571
CL\_17798
CL\_17226
CL\_17227
CL\_17228
CL\_17229
CL\_14253
CL\_6062
CL\_6063
CL\_6710
CL\_7577
