## Supplementary material for "A novel method for integrating genomic and Tn-Seq data to identify common *in vivo* fitness mechanisms across multiple bacterial species": S2 Dataset: CL_INS_168.html

Legend

 Hypothetical
 All EssentialGenes
 Transport +binding proteins

FULL


WINDOWSVGPNG

Trim RowsRemove SingletonsSave Fasta

CL\_2177


CL\_2177


CL\_2177

HighlightSelectShow Genomes


115

CL\_2179


86

CL\_2179


1

CL\_2180

fGI ID

CL\_INS\_168
Cluster ID

CL\_2178
