## Supplementary material for "A novel method for integrating genomic and Tn-Seq data to identify common *in vivo* fitness mechanisms across multiple bacterial species": S2 Dataset: CL_INS_171.html

Legend

 Mobile +extrachromosomalelementfunctions
 Regulatoryfunctions
 Hypothetical
 DNA Metabolism
 All EssentialGenes
 Other
 All VFDB Genes

FULL


WINDOWSVGPNG

Trim RowsRemove SingletonsSave Fasta

CL\_2213


CL\_2213


CL\_2213


CL\_2213


CL\_2213


CL\_2213


CL\_2213


CL\_2213


CL\_2213


CL\_2213


CL\_2213


CL\_2213


CL\_2213


CL\_2213


CL\_2213


CL\_2213


CL\_2213


CL\_2213


CL\_2213


CL\_2213

HighlightSelectShow Genomes


132

CL\_2214


21

CL\_2214


13

CL\_2214


12

CL\_2214


6

CL\_2214


4

CL\_2214


3

CL\_2214


1

CL\_2214


1

CL\_2214


1

CL\_2214


1

CL\_2214


1

CL\_2214


1

CL\_2214


1

Break


1

CL\_2214


1

CL\_2214


1

CL\_2214


1

CL\_2214


1

CL\_2214


1

CL\_2214

fGI ID


CL\_INS\_174
CL\_INS\_171
CL\_INS\_171
CL\_INS\_171
CL\_INS\_171
CL\_INS\_171
CL\_INS\_171
CL\_INS\_171
CL\_INS\_171
CL\_INS\_287
CL\_INS\_171
CL\_INS\_171
CL\_INS\_171
CL\_INS\_171
CL\_INS\_351
CL\_INS\_171
CL\_INS\_171
CL\_INS\_171
CL\_INS\_171
CL\_INS\_171
CL\_INS\_44
CL\_INS\_74
CL\_INS\_74
CL\_INS\_74
CL\_INS\_74
CL\_INS\_74
CL\_INS\_74
CL\_INS\_74
CL\_INS\_74
CL\_INS\_74
CL\_INS\_74
CL\_INS\_74
CL\_INS\_171
CL\_INS\_171
CL\_INS\_351
CL\_INS\_171
CL\_INS\_307
CL\_INS\_171
CL\_INS\_171
CL\_INS\_171
CL\_INS\_171
CL\_INS\_351
CL\_INS\_44
CL\_INS\_351
CL\_INS\_351
CL\_INS\_351
CL\_INS\_351
CL\_INS\_351
CL\_INS\_351
CL\_INS\_171
CL\_INS\_351
CL\_INS\_351
CL\_INS\_351
CL\_INS\_351
CL\_INS\_351
CL\_INS\_351
CL\_INS\_351
CL\_INS\_293
CL\_INS\_351
CL\_INS\_351
CL\_INS\_171
CL\_INS\_171
CL\_INS\_171
CL\_INS\_171
CL\_INS\_171
CL\_INS\_178
CL\_INS\_278
CL\_INS\_58
CL\_INS\_278
CL\_INS\_58
CL\_INS\_171
CL\_INS\_171
CL\_INS\_30
CL\_INS\_30
CL\_INS\_278
CL\_INS\_278
CL\_INS\_278
CL\_INS\_160
CL\_INS\_171
CL\_INS\_171
CL\_INS\_171
CL\_INS\_204
CL\_INS\_171
CL\_INS\_171
CL\_INS\_171
CL\_INS\_385
CL\_INS\_132
CL\_INS\_278
CL\_INS\_278
CL\_INS\_278
CL\_INS\_160
CL\_INS\_278
CL\_INS\_278
CL\_INS\_278
CL\_INS\_105
CL\_INS\_278
CL\_INS\_278
CL\_INS\_278
CL\_INS\_401
CL\_INS\_278
CL\_INS\_278
CL\_INS\_278
CL\_INS\_278
CL\_INS\_160
CL\_INS\_278
CL\_INS\_278
CL\_INS\_278
CL\_INS\_278
CL\_INS\_278
CL\_INS\_278
CL\_INS\_278
CL\_INS\_278
CL\_INS\_132
CL\_INS\_132
CL\_INS\_132
CL\_INS\_278
CL\_INS\_171
CL\_INS\_171
CL\_INS\_278
CL\_INS\_160
CL\_INS\_160
CL\_INS\_307
CL\_INS\_171
CL\_INS\_171
CL\_INS\_385
CL\_INS\_171
CL\_INS\_278
CL\_INS\_132
CL\_INS\_171
CL\_INS\_171
CL\_INS\_132
CL\_INS\_132
CL\_INS\_132
CL\_INS\_132
CL\_INS\_171
CL\_INS\_171
CL\_INS\_171
CL\_INS\_171
CL\_INS\_171
CL\_INS\_171
CL\_INS\_171
CL\_INS\_132
CL\_INS\_278
CL\_INS\_278
CL\_INS\_160
CL\_INS\_278
CL\_INS\_401
CL\_INS\_278
CL\_INS\_132
CL\_INS\_278
CL\_INS\_171
CL\_INS\_278
CL\_INS\_278
CL\_INS\_278
CL\_INS\_278
CL\_INS\_278
CL\_INS\_278
CL\_INS\_278
CL\_INS\_278
CL\_INS\_278
CL\_INS\_278
CL\_INS\_212
CL\_INS\_278
CL\_INS\_278
CL\_INS\_171
CL\_INS\_58
CL\_INS\_178
CL\_INS\_171
CL\_INS\_171
CL\_INS\_171
CL\_INS\_171
CL\_INS\_171
CL\_INS\_171
CL\_INS\_171
Cluster ID


CL\_6937
CL\_14832
CL\_11780
CL\_14701
CL\_7711
CL\_7710
CL\_19569
CL\_7709
CL\_15555
CL\_7011
CL\_15999
CL\_15998
CL\_15997
CL\_15996
CL\_11608
CL\_22288
CL\_20649
CL\_16476
CL\_16475
CL\_16474
CL\_16473
CL\_7143
CL\_9064
CL\_7144
CL\_7145
CL\_7146
CL\_7147
CL\_15441
CL\_12846
CL\_12845
CL\_9065
CL\_7151
CL\_18222
CL\_18221
CL\_11607
CL\_15554
CL\_6821
CL\_15553
CL\_15552
CL\_15551
CL\_15550
CL\_12042
CL\_11605
CL\_12043
CL\_12044
CL\_11604
CL\_18220
CL\_15459
CL\_11603
CL\_22289
CL\_11602
CL\_12045
CL\_11601
CL\_11600
CL\_11599
CL\_11598
CL\_11597
CL\_7942
CL\_11596
CL\_16472
CL\_20297
CL\_20298
CL\_20321
CL\_20315
CL\_20316
CL\_7960
CL\_5987
CL\_5986
CL\_6829
CL\_8001
CL\_20317
CL\_20318
CL\_7998
CL\_7959
CL\_5978
CL\_5977
CL\_5976
CL\_10071
CL\_18623
CL\_20319
CL\_20320
CL\_6066
CL\_15667
CL\_15666
CL\_8651
CL\_6156
CL\_6054
CL\_6053
CL\_6052
CL\_6051
CL\_6851
CL\_7592
CL\_7593
CL\_6047
CL\_6044
CL\_6043
CL\_6042
CL\_6041
CL\_6040
CL\_6845
CL\_6844
CL\_6843
CL\_6842
CL\_6841
CL\_6882
CL\_6883
CL\_7948
CL\_6885
CL\_7947
CL\_6879
CL\_6830
CL\_6831
CL\_8207
CL\_8208
CL\_8209
CL\_6874
CL\_7940
CL\_10906
CL\_6186
CL\_9219
CL\_9218
CL\_3386
CL\_7821
CL\_11713
CL\_8212
CL\_20322
CL\_6106
CL\_6022
CL\_6021
CL\_6020
CL\_6019
CL\_6018
CL\_6017
CL\_6016
CL\_20323
CL\_20324
CL\_20325
CL\_20326
CL\_20327
CL\_20328
CL\_20329
CL\_6015
CL\_6014
CL\_6013
CL\_8020
CL\_6011
CL\_6010
CL\_6009
CL\_6008
CL\_6006
CL\_10925
CL\_6004
CL\_6003
CL\_6002
CL\_6001
CL\_6000
CL\_5999
CL\_5998
CL\_5997
CL\_5996
CL\_5995
CL\_6825
CL\_5993
CL\_5992
CL\_10837
CL\_6826
CL\_6828
CL\_20330
CL\_20331
CL\_20332
CL\_20333
CL\_20334
CL\_20335
CL\_20336
