## Supplementary material for "A novel method for integrating genomic and Tn-Seq data to identify common *in vivo* fitness mechanisms across multiple bacterial species": S2 Dataset: CL_INS_176.html

Legend

 Hypothetical
 Other
 All VFDB Genes

FULL


WINDOWSVGPNG

Trim RowsRemove SingletonsSave Fasta

CL\_2273


CL\_2273


CL\_2273


CL\_2273


CL\_2273


CL\_2273


CL\_2273


CL\_2273


CL\_2273


CL\_2273


CL\_2272


CL\_2272


CL\_2273


CL\_2273

HighlightSelectShow Genomes


96

CL\_2285


57

CL\_2285


34

CL\_2285


2

CL\_2285


2

CL\_2285


2

CL\_2285


1

CL\_2285


1

CL\_2285


1

CL\_2285


1

CL\_2285


1

CL\_2285


1

CL\_2285


1

CL\_2285


1

CL\_2285

fGI ID


CL\_INS\_176
CL\_INS\_176
CL\_INS\_176
CL\_INS\_176
CL\_INS\_176
CL\_INS\_176
CL\_INS\_176
CL\_INS\_176
CL\_INS\_176
CL\_INS\_176
CL\_INS\_176
CL\_INS\_176
Cluster ID


CL\_2274
CL\_2275
CL\_2276
CL\_2277
CL\_2278
CL\_2279
CL\_2280
CL\_15549
CL\_2281
CL\_2282
CL\_2283
CL\_2284
