## Supplementary material for "A novel method for integrating genomic and Tn-Seq data to identify common *in vivo* fitness mechanisms across multiple bacterial species": S2 Dataset: CL_INS_178.html

Legend

 Mobile +extrachromosomalelementfunctions
 Hypothetical
 DNA Metabolism
 All EssentialGenes
 Other
 All VFDB Genes

FULL


WINDOWSVGPNG

Trim RowsRemove SingletonsSave Fasta

CL\_2299


CL\_2299


CL\_2299


CL\_2299


CL\_2299


CL\_2299


CL\_2299


CL\_2299


CL\_2299


CL\_2299


CL\_2299


CL\_2299


CL\_2299


CL\_2299


CL\_2299


CL\_2299


CL\_2299


CL\_2299


CL\_2299


CL\_2299


CL\_2299


CL\_2299


CL\_2299


CL\_2299


CL\_2299


CL\_2299


CL\_2299


CL\_2299


CL\_2299


CL\_2299


CL\_2299


CL\_2299


CL\_2299


CL\_2299


CL\_2262


CL\_2299


CL\_2299


CL\_2299


CL\_2299


CL\_2299


CL\_2299


CL\_2298


CL\_2299


CL\_2299


CL\_2298


CL\_2299


CL\_2299


CL\_2299


CL\_2299


CL\_2292


CL\_2299


CL\_2299


CL\_2299


CL\_2299


Break


CL\_2299


CL\_2299


CL\_2299


CL\_2299


CL\_2299


CL\_2290


CL\_2299


CL\_2298


CL\_2299


CL\_2299


CL\_2299


CL\_2299


CL\_2299


CL\_2299


CL\_2299


CL\_2299


CL\_2299


CL\_2299


Break


CL\_2299


Break


CL\_2299


CL\_2299


CL\_2298


CL\_2299


CL\_2299


CL\_2298


CL\_2299


CL\_2299


CL\_2299


CL\_2299


CL\_2299


CL\_2299


CL\_2299


CL\_2299


CL\_2299

HighlightSelectShow Genomes


22

CL\_2304


19

CL\_2304


13

CL\_2304


13

CL\_2304


12

CL\_2304


10

CL\_2304


7

CL\_2304


4

CL\_2304


3

CL\_2304


3

CL\_2304


3

Break


3

CL\_2304


3

CL\_2304


3

CL\_2304


2

CL\_2304


2

CL\_2304


2

CL\_2304


2

CL\_2304


2

CL\_2304


2

CL\_2304


2

CL\_2304


2

CL\_2304


2

CL\_2304


1

CL\_2304


1

CL\_2304


1

CL\_2304


1

CL\_2304


1

CL\_2326


1

CL\_2304


1

CL\_2304


1

CL\_2304


1

CL\_2304


1

Break


1

CL\_2304


1

CL\_2304


1

CL\_2304


1

CL\_2304


1

CL\_2305


1

CL\_2304


1

CL\_2304


1

CL\_2304


1

CL\_2304


1

CL\_2304


1

CL\_2304


1

CL\_2304


1

CL\_2304


1

CL\_2304


1

CL\_2304


1

CL\_2304


1

CL\_2304


1

CL\_2304


1

CL\_2304


1

CL\_2304


1

CL\_2304


1

CL\_2304


1

CL\_2304


1

CL\_2304


1

CL\_2304


1

CL\_2304


1

Break


1

CL\_2304


1

CL\_2304


1

CL\_2304


1

CL\_2304


1

CL\_2305


1

CL\_2304


1

CL\_2304


1

CL\_2304


1

CL\_2305


1

CL\_2305


1

CL\_2304


1

CL\_2305


1

CL\_2304


1

CL\_2304


1

CL\_2304


1

CL\_2304


1

CL\_2304


1

CL\_2304


1

CL\_2304


1

CL\_2304


1

CL\_2333


1

CL\_2304


1

CL\_2304


1

CL\_2304


1

CL\_2304


1

CL\_2304


1

CL\_2304


1

CL\_2304


1

CL\_2304


1

CL\_2304


1

CL\_2304

fGI ID


CL\_INS\_178
CL\_INS\_178
CL\_INS\_204
CL\_INS\_178
CL\_INS\_178
CL\_INS\_178
CL\_INS\_178
CL\_INS\_204
CL\_INS\_178
CL\_INS\_178
CL\_INS\_178
CL\_INS\_178
CL\_INS\_178
CL\_INS\_178
CL\_INS\_178
CL\_INS\_278
CL\_INS\_278
CL\_INS\_278
CL\_INS\_178
CL\_INS\_178
CL\_INS\_178
CL\_INS\_178
CL\_INS\_385
CL\_INS\_178
CL\_INS\_178
CL\_INS\_178
CL\_INS\_178
CL\_INS\_178
CL\_INS\_178
CL\_INS\_132
CL\_INS\_132
CL\_INS\_132
CL\_INS\_132
CL\_INS\_132
CL\_INS\_132
CL\_INS\_132
CL\_INS\_178
CL\_INS\_178
CL\_INS\_178
CL\_INS\_178
CL\_INS\_178
CL\_INS\_178
CL\_INS\_178
CL\_INS\_178
CL\_INS\_178
CL\_INS\_178
CL\_INS\_178
CL\_INS\_178
CL\_INS\_178
CL\_INS\_178
CL\_INS\_178
CL\_INS\_178
CL\_INS\_178
CL\_INS\_178
CL\_INS\_178
CL\_INS\_178
CL\_INS\_178
CL\_INS\_178
CL\_INS\_178
CL\_INS\_178
CL\_INS\_178
CL\_INS\_178
CL\_INS\_178
CL\_INS\_178
CL\_INS\_178
CL\_INS\_178
CL\_INS\_178
CL\_INS\_178
CL\_INS\_178
CL\_INS\_178
CL\_INS\_278
CL\_INS\_307
CL\_INS\_91
CL\_INS\_178
CL\_INS\_263
CL\_INS\_178
CL\_INS\_178
CL\_INS\_178
CL\_INS\_178
CL\_INS\_178
CL\_INS\_155
CL\_INS\_307
CL\_INS\_307
CL\_INS\_178
CL\_INS\_178
CL\_INS\_178
CL\_INS\_91
CL\_INS\_91
CL\_INS\_91
CL\_INS\_91
CL\_INS\_91
CL\_INS\_91
CL\_INS\_91
CL\_INS\_91
CL\_INS\_91
CL\_INS\_91
CL\_INS\_91
CL\_INS\_91
CL\_INS\_91
CL\_INS\_91
CL\_INS\_91
CL\_INS\_91
CL\_INS\_307
CL\_INS\_155
CL\_INS\_263
CL\_INS\_307
CL\_INS\_307
CL\_INS\_307
CL\_INS\_155
CL\_INS\_178
CL\_INS\_178
CL\_INS\_178
CL\_INS\_178
CL\_INS\_178
CL\_INS\_178
CL\_INS\_174
CL\_INS\_174
CL\_INS\_174
CL\_INS\_178
CL\_INS\_178
CL\_INS\_178
CL\_INS\_174
CL\_INS\_178
Cluster ID


CL\_7260
CL\_7261
CL\_8086
CL\_12380
CL\_10001
CL\_2300
CL\_2301
CL\_5953
CL\_6828
CL\_7960
CL\_7053
CL\_7052
CL\_7051
CL\_5985
CL\_5984
CL\_5983
CL\_5982
CL\_5981
CL\_7704
CL\_7703
CL\_11779
CL\_7702
CL\_7810
CL\_7708
CL\_7707
CL\_7706
CL\_7705
CL\_7701
CL\_5125
CL\_9985
CL\_8242
CL\_12269
CL\_7944
CL\_7945
CL\_7946
CL\_6806
CL\_20184
CL\_5124
CL\_9329
CL\_9328
CL\_6523
CL\_5675
CL\_10240
CL\_7472
CL\_11467
CL\_11466
CL\_11465
CL\_11464
CL\_11463
CL\_11462
CL\_11461
CL\_11460
CL\_11459
CL\_11458
CL\_11457
CL\_5673
CL\_11456
CL\_11455
CL\_11454
CL\_11453
CL\_5674
CL\_6136
CL\_5123
CL\_5122
CL\_5121
CL\_13630
CL\_13631
CL\_13632
CL\_8390
CL\_8389
CL\_8388
CL\_8387
CL\_10368
CL\_9123
CL\_8335
CL\_9122
CL\_10367
CL\_9120
CL\_9119
CL\_9118
CL\_6997
CL\_6996
CL\_6995
CL\_10366
CL\_10365
CL\_10364
CL\_6611
CL\_6612
CL\_8327
CL\_6613
CL\_6614
CL\_6615
CL\_6616
CL\_6617
CL\_6618
CL\_6619
CL\_6620
CL\_6621
CL\_6622
CL\_6985
CL\_6623
CL\_6646
CL\_6645
CL\_6984
CL\_5385
CL\_12093
CL\_10954
CL\_10955
CL\_8325
CL\_7700
CL\_10356
CL\_7699
CL\_13838
CL\_13839
CL\_7698
CL\_7697
CL\_7696
CL\_6937
CL\_2302
CL\_14711
CL\_7883
CL\_9812
CL\_2303
