## Supplementary material for "A novel method for integrating genomic and Tn-Seq data to identify common *in vivo* fitness mechanisms across multiple bacterial species": S2 Dataset: CL_INS_179.html

Legend

 Hypothetical
 All VFDB Genes

FULL


WINDOWSVGPNG

Trim RowsRemove SingletonsSave Fasta

CL\_2305


CL\_2305


CL\_2305


CL\_2305


CL\_2305


CL\_2305

HighlightSelectShow Genomes


94

CL\_2308


43

CL\_2308


27

CL\_2308


3

CL\_2308


1

CL\_2308


1

CL\_2309

fGI ID


CL\_INS\_179
CL\_INS\_179
CL\_INS\_385
Cluster ID


CL\_2306
CL\_2307
CL\_7810
