## Supplementary material for "A novel method for integrating genomic and Tn-Seq data to identify common *in vivo* fitness mechanisms across multiple bacterial species": S2 Dataset: CL_INS_181.html

Legend

 Hypothetical
 Other
 All VFDB Genes

FULL


WINDOWSVGPNG

Trim RowsRemove SingletonsSave Fasta

CL\_2342


CL\_2342


CL\_2341


CL\_2341

HighlightSelectShow Genomes


130

CL\_2343


73

CL\_2343


1

CL\_2343


1

CL\_2343

fGI ID

CL\_INS\_181
Cluster ID

CL\_5677
