## Supplementary material for "A novel method for integrating genomic and Tn-Seq data to identify common *in vivo* fitness mechanisms across multiple bacterial species": S2 Dataset: CL_INS_182.html

Legend

 Mobile +extrachromosomalelementfunctions
 Hypothetical
 Other
 All VFDB Genes

FULL


WINDOWSVGPNG

Trim RowsRemove SingletonsSave Fasta

CL\_2350


CL\_2350


CL\_2350


CL\_2350


CL\_2349

HighlightSelectShow Genomes


193

CL\_2351


1

CL\_2351


1

CL\_2351


1

CL\_2352


1

CL\_2351

fGI ID


CL\_INS\_182
CL\_INS\_182
CL\_INS\_182
CL\_INS\_182
CL\_INS\_182
CL\_INS\_182
CL\_INS\_182
CL\_INS\_182
CL\_INS\_182
CL\_INS\_182
CL\_INS\_182
CL\_INS\_182
CL\_INS\_334
CL\_INS\_182
Cluster ID


CL\_17277
CL\_13214
CL\_21462
CL\_13215
CL\_13216
CL\_13217
CL\_13218
CL\_13219
CL\_13220
CL\_13221
CL\_13222
CL\_24190
CL\_1165
CL\_13223
