## Supplementary material for "A novel method for integrating genomic and Tn-Seq data to identify common *in vivo* fitness mechanisms across multiple bacterial species": S2 Dataset: CL_INS_184.html

Legend

 Hypothetical
 EnergyMetabolism
 All VFDB Genes

FULL


WINDOWSVGPNG

Trim RowsRemove SingletonsSave Fasta

CL\_2379


CL\_2379


CL\_2379


CL\_2379

HighlightSelectShow Genomes


118

CL\_2381


82

CL\_2381


3

CL\_2382


1

CL\_2382

fGI ID

CL\_INS\_184
Cluster ID

CL\_2380
