## Supplementary material for "A novel method for integrating genomic and Tn-Seq data to identify common *in vivo* fitness mechanisms across multiple bacterial species": S2 Dataset: CL_INS_189.html

Legend

 Regulatoryfunctions
 Hypothetical
 Other
 EnergyMetabolism
 All VFDB Genes

FULL


WINDOWSVGPNG

Trim RowsRemove SingletonsSave Fasta

CL\_2445


CL\_2445


CL\_2445


CL\_2445


CL\_2445


CL\_2445


CL\_2444

HighlightSelectShow Genomes


144

CL\_2446


18

CL\_2446


18

CL\_2446


14

CL\_2446


6

CL\_2446


1

CL\_2447


1

CL\_2446

fGI ID


CL\_INS\_189
CL\_INS\_189
CL\_INS\_189
CL\_INS\_189
Cluster ID


CL\_7293
CL\_6979
CL\_6978
CL\_6977
