## Supplementary material for "A novel method for integrating genomic and Tn-Seq data to identify common *in vivo* fitness mechanisms across multiple bacterial species": S2 Dataset: CL_INS_190.html

Legend

 Hypothetical
 Proteinsynthesis/fate
 Other

FULL


WINDOWSVGPNG

Trim RowsRemove SingletonsSave Fasta

CL\_2459


CL\_2459


CL\_2459

HighlightSelectShow Genomes


115

CL\_2462


79

CL\_2462


7

CL\_2462

fGI ID


CL\_INS\_190
CL\_INS\_190
Cluster ID


CL\_2460
CL\_2461
