## Supplementary material for "A novel method for integrating genomic and Tn-Seq data to identify common *in vivo* fitness mechanisms across multiple bacterial species": S2 Dataset: CL_INS_192.html

Legend

 Mobile +extrachromosomalelementfunctions
 Hypothetical
 All VFDB Genes

FULL


WINDOWSVGPNG

Trim RowsRemove SingletonsSave Fasta

CL\_2489


CL\_2489


CL\_2489


CL\_2489


CL\_2489


CL\_2488


CL\_2488


CL\_2488


CL\_2488


CL\_2489


CL\_2489


CL\_2489


CL\_2488


CL\_2489


CL\_2488


CL\_2488


CL\_2488


CL\_2489

HighlightSelectShow Genomes


44

CL\_2492


39

CL\_2492


39

CL\_2492


35

CL\_2492


21

CL\_2492


6

CL\_2492


3

CL\_2492


2

CL\_2492


1

CL\_2492


1

CL\_2497


1

CL\_2493


1

CL\_2492


1

CL\_2492


1

CL\_2492


1

CL\_2492


1

CL\_2492


1

CL\_2492


1

CL\_2492

fGI ID


CL\_INS\_191
CL\_INS\_192
CL\_INS\_192
CL\_INS\_192
CL\_INS\_192
Cluster ID


CL\_9080
CL\_12185
CL\_2490
CL\_2491
CL\_5680
