## Supplementary material for "A novel method for integrating genomic and Tn-Seq data to identify common *in vivo* fitness mechanisms across multiple bacterial species": S2 Dataset: CL_INS_196.html

Legend

 Hypothetical
 Other
 All VFDB Genes

FULL


WINDOWSVGPNG

Trim RowsRemove SingletonsSave Fasta

CL\_2530


CL\_2530


CL\_2529


CL\_2530


CL\_2529


CL\_2530


CL\_2530


CL\_2530

HighlightSelectShow Genomes


103

CL\_2533


80

CL\_2533


1

CL\_2533


1

CL\_2533


1

CL\_2533


1

CL\_2534


1

CL\_2533


1

Break

fGI ID


CL\_INS\_196
CL\_INS\_196
CL\_INS\_196
CL\_INS\_196
CL\_INS\_196
CL\_INS\_196
Cluster ID


CL\_2531
CL\_2532
CL\_6165
CL\_6164
CL\_6163
CL\_6162
