## Supplementary material for "A novel method for integrating genomic and Tn-Seq data to identify common *in vivo* fitness mechanisms across multiple bacterial species": S2 Dataset: CL_INS_199.html

Legend

 Hypothetical
 Other
 All VFDB Genes

FULL


WINDOWSVGPNG

Trim RowsRemove SingletonsSave Fasta

CL\_2577


CL\_2577


CL\_2577

HighlightSelectShow Genomes


143

CL\_2578


55

CL\_2578


1

CL\_2579

fGI ID

CL\_INS\_199
Cluster ID

CL\_6160
