## Supplementary material for "A novel method for integrating genomic and Tn-Seq data to identify common *in vivo* fitness mechanisms across multiple bacterial species": S2 Dataset: CL_INS_200.html

Legend

 Hypothetical
 All VFDB Genes

FULL


WINDOWSVGPNG

Trim RowsRemove SingletonsSave Fasta

CL\_2587


CL\_2587


CL\_2587

HighlightSelectShow Genomes


118

CL\_2589


84

CL\_2589


1

CL\_2589

fGI ID


CL\_INS\_200
CL\_INS\_200
Cluster ID


CL\_20519
CL\_2588
