## Supplementary material for "A novel method for integrating genomic and Tn-Seq data to identify common *in vivo* fitness mechanisms across multiple bacterial species": S2 Dataset: CL_INS_204.html

Legend

 Mobile +extrachromosomalelementfunctions
 Regulatoryfunctions
 Hypothetical
 DNA Metabolism
 AntibioticResistance
 Proteinsynthesis/fate
 Other
 EnergyMetabolism
 All VFDB Genes
 Transport +binding proteins

FULL


WINDOWSVGPNG

Trim RowsRemove SingletonsSave Fasta

CL\_2616


CL\_2616


CL\_2616


CL\_2616


CL\_2608


CL\_2616


CL\_2616


CL\_2616


CL\_2616


CL\_2616


CL\_2608


CL\_2608


CL\_2616


CL\_2616


CL\_2616


CL\_2608


CL\_2616


CL\_2616


CL\_2616


CL\_2616


CL\_2616


CL\_2616


CL\_2616


CL\_2616


CL\_2616


CL\_2616


CL\_2615


CL\_2616


CL\_2616


CL\_2616


CL\_2616


CL\_2616


CL\_2616


CL\_2612


CL\_2616


CL\_2616


CL\_2616


CL\_2616


CL\_2608


CL\_2616


CL\_2616


CL\_2608


CL\_2616


CL\_2616


CL\_2616


CL\_2616


CL\_2616


CL\_2964


CL\_2608


CL\_2616


CL\_2616


CL\_2616


CL\_2616


CL\_2608


CL\_2616


CL\_2612


CL\_2616


CL\_2616


CL\_2608


CL\_2599


CL\_2615


CL\_2599


CL\_2616


CL\_2616


CL\_2616


CL\_2616


CL\_2608


CL\_2616


CL\_2616


CL\_2616


CL\_2608


CL\_2616


CL\_2608


CL\_2616


CL\_2616


CL\_2608


CL\_2616


CL\_2616


CL\_2616


CL\_2616


CL\_2616


CL\_2616


CL\_2616


CL\_2599


Break


CL\_2616


CL\_2616


CL\_2612


CL\_2616


CL\_2616


CL\_2616


CL\_2616


CL\_2608


CL\_2616


CL\_2616


CL\_2616


CL\_2616


CL\_2599


CL\_2608


CL\_2616


CL\_2608


CL\_2608


CL\_2616


CL\_2603


CL\_2616


CL\_2608


CL\_2616


CL\_2616


CL\_2616

HighlightSelectShow Genomes


14

CL\_5520


12

CL\_5520


11

CL\_5520


10

CL\_5520


7

CL\_5520


6

CL\_5518


4

CL\_5520


4

CL\_5520


3

CL\_5520


3

CL\_5520


3

CL\_5520


3

CL\_5520


3

CL\_5520


3

CL\_5520


2

CL\_5520


2

CL\_5520


2

CL\_5520


2

CL\_5520


2

CL\_5519


2

CL\_5520


2

CL\_5520


2

CL\_2617


2

CL\_5520


1

CL\_5516


1

CL\_5520


1

CL\_5520


1

CL\_5520


1

CL\_2641


1

CL\_5519


1

CL\_5519


1

CL\_5520


1

CL\_5520


1

CL\_5520


1

CL\_5520


1

CL\_5520


1

CL\_5520


1

CL\_5520


1

CL\_5520


1

CL\_5520


1

CL\_5520


1

CL\_5520


1

CL\_5520


1

CL\_5520


1

CL\_5520


1

CL\_5519


1

CL\_5520


1

CL\_5520


1

CL\_5520


1

CL\_5520


1

CL\_5520


1

CL\_5520


1

CL\_5520


1

CL\_5518


1

CL\_5520


1

CL\_2641


1

CL\_5520


1

CL\_5520


1

CL\_5520


1

CL\_5520


1

CL\_5520


1

CL\_5520


1

CL\_5520


1

CL\_5520


1

CL\_5515


1

CL\_5520


1

CL\_5518


1

CL\_5520


1

CL\_5520


1

CL\_5520


1

CL\_2618


1

CL\_5520


1

CL\_5520


1

CL\_5520


1

CL\_5520


1

CL\_5520


1

CL\_5520


1

CL\_5520


1

CL\_5519


1

CL\_5520


1

CL\_5520


1

CL\_2617


1

CL\_5515


1

CL\_5519


1

CL\_5520


1

CL\_5520


1

CL\_5520


1

CL\_5520


1

CL\_5520


1

CL\_5520


1

CL\_2647


1

CL\_5520


1

CL\_5520


1

CL\_5520


1

CL\_5520


1

CL\_5517


1

CL\_5520


1

CL\_5520


1

CL\_5520


1

CL\_5520


1

CL\_5520


1

CL\_5520


1

CL\_5520


1

CL\_5520


1

CL\_5520


1

CL\_5520


1

CL\_5520


1

CL\_5519


1

CL\_2618


1

CL\_5520

fGI ID


CL\_INS\_204
CL\_INS\_204
CL\_INS\_204
CL\_INS\_204
CL\_INS\_204
CL\_INS\_204
CL\_INS\_204
CL\_INS\_204
CL\_INS\_204
CL\_INS\_204
CL\_INS\_204
CL\_INS\_204
CL\_INS\_204
CL\_INS\_204
CL\_INS\_204
CL\_INS\_204
CL\_INS\_204
CL\_INS\_204
CL\_INS\_204
CL\_INS\_204
CL\_INS\_204
CL\_INS\_204
CL\_INS\_204
CL\_INS\_204
CL\_INS\_204
CL\_INS\_204
CL\_INS\_204
CL\_INS\_204
CL\_INS\_204
CL\_INS\_204
CL\_INS\_204
CL\_INS\_204
CL\_INS\_204
CL\_INS\_204
CL\_INS\_204
CL\_INS\_204
CL\_INS\_204
CL\_INS\_204
CL\_INS\_204
CL\_INS\_204
CL\_INS\_204
CL\_INS\_204
CL\_INS\_204
CL\_INS\_204
CL\_INS\_204
CL\_INS\_204
CL\_INS\_204
CL\_INS\_204
CL\_INS\_204
CL\_INS\_204
CL\_INS\_204
CL\_INS\_204
CL\_INS\_204
CL\_INS\_204
CL\_INS\_204
CL\_INS\_204
CL\_INS\_204
CL\_INS\_204
CL\_INS\_204
CL\_INS\_204
CL\_INS\_204
CL\_INS\_204
CL\_INS\_204
CL\_INS\_204
CL\_INS\_204
CL\_INS\_204
CL\_INS\_204
CL\_INS\_204
CL\_INS\_204
CL\_INS\_204
CL\_INS\_204
CL\_INS\_204
CL\_INS\_204
CL\_INS\_204
CL\_INS\_204
CL\_INS\_204
CL\_INS\_204
CL\_INS\_204
CL\_INS\_204
CL\_INS\_204
CL\_INS\_204
CL\_INS\_204
CL\_INS\_204
CL\_INS\_204
CL\_INS\_204
CL\_INS\_204
CL\_INS\_204
CL\_INS\_204
CL\_INS\_204
CL\_INS\_204
CL\_INS\_204
CL\_INS\_204
CL\_INS\_204
CL\_INS\_204
CL\_INS\_204
CL\_INS\_204
CL\_INS\_204
CL\_INS\_204
CL\_INS\_204
CL\_INS\_204
CL\_INS\_204
CL\_INS\_204
CL\_INS\_204
CL\_INS\_204
CL\_INS\_204
CL\_INS\_204
CL\_INS\_204
CL\_INS\_204
CL\_INS\_204
CL\_INS\_204
CL\_INS\_204
CL\_INS\_204
CL\_INS\_204
CL\_INS\_204
CL\_INS\_204
CL\_INS\_204
CL\_INS\_204
CL\_INS\_204
CL\_INS\_204
CL\_INS\_204
CL\_INS\_204
CL\_INS\_204
CL\_INS\_204
CL\_INS\_204
CL\_INS\_204
CL\_INS\_204
CL\_INS\_204
CL\_INS\_204
CL\_INS\_204
CL\_INS\_204
CL\_INS\_204
CL\_INS\_204
CL\_INS\_204
CL\_INS\_204
CL\_INS\_204
CL\_INS\_204
CL\_INS\_204
CL\_INS\_204
CL\_INS\_204
CL\_INS\_204
CL\_INS\_204
CL\_INS\_204
CL\_INS\_204
CL\_INS\_204
CL\_INS\_204
CL\_INS\_204
CL\_INS\_204
CL\_INS\_204
CL\_INS\_204
CL\_INS\_204
CL\_INS\_204
CL\_INS\_204
CL\_INS\_204
CL\_INS\_204
CL\_INS\_204
CL\_INS\_204
CL\_INS\_204
CL\_INS\_204
CL\_INS\_204
CL\_INS\_204
CL\_INS\_204
CL\_INS\_204
CL\_INS\_204
CL\_INS\_204
CL\_INS\_204
CL\_INS\_204
CL\_INS\_204
CL\_INS\_204
CL\_INS\_204
CL\_INS\_204
CL\_INS\_204
CL\_INS\_204
CL\_INS\_204
CL\_INS\_204
CL\_INS\_204
CL\_INS\_204
CL\_INS\_204
CL\_INS\_204
CL\_INS\_204
CL\_INS\_204
CL\_INS\_204
CL\_INS\_204
CL\_INS\_204
CL\_INS\_204
CL\_INS\_204
CL\_INS\_204
CL\_INS\_204
CL\_INS\_204
CL\_INS\_204
CL\_INS\_204
CL\_INS\_204
CL\_INS\_204
CL\_INS\_204
CL\_INS\_204
CL\_INS\_204
CL\_INS\_204
CL\_INS\_204
CL\_INS\_204
CL\_INS\_204
CL\_INS\_204
CL\_INS\_204
CL\_INS\_204
CL\_INS\_204
CL\_INS\_204
CL\_INS\_204
CL\_INS\_204
CL\_INS\_204
CL\_INS\_204
CL\_INS\_204
CL\_INS\_204
CL\_INS\_204
CL\_INS\_204
CL\_INS\_204
CL\_INS\_204
CL\_INS\_204
CL\_INS\_204
CL\_INS\_204
CL\_INS\_204
CL\_INS\_204
CL\_INS\_204
CL\_INS\_204
CL\_INS\_204
CL\_INS\_204
CL\_INS\_204
CL\_INS\_204
CL\_INS\_204
CL\_INS\_204
CL\_INS\_204
CL\_INS\_204
CL\_INS\_204
CL\_INS\_204
CL\_INS\_204
CL\_INS\_204
CL\_INS\_204
CL\_INS\_204
CL\_INS\_204
CL\_INS\_204
Cluster ID


CL\_8762
CL\_11441
CL\_11440
CL\_11439
CL\_11438
CL\_11451
CL\_11040
CL\_11039
CL\_11038
CL\_11037
CL\_11036
CL\_21454
CL\_21196
CL\_21195
CL\_21194
CL\_21193
CL\_21192
CL\_21191
CL\_21190
CL\_21189
CL\_21188
CL\_6067
CL\_6066
CL\_6065
CL\_6064
CL\_6063
CL\_6062
CL\_6061
CL\_6060
CL\_6059
CL\_6058
CL\_5528
CL\_5527
CL\_5475
CL\_9313
CL\_9312
CL\_5526
CL\_19275
CL\_19274
CL\_19273
CL\_5525
CL\_5685
CL\_5686
CL\_5687
CL\_5688
CL\_5689
CL\_5690
CL\_5691
CL\_5692
CL\_5693
CL\_5694
CL\_11046
CL\_5695
CL\_5696
CL\_5697
CL\_5698
CL\_5699
CL\_5700
CL\_24051
CL\_24050
CL\_11778
CL\_5701
CL\_15442
CL\_5702
CL\_5953
CL\_18191
CL\_18190
CL\_18189
CL\_18188
CL\_18187
CL\_18186
CL\_18185
CL\_18184
CL\_18183
CL\_18182
CL\_18181
CL\_18180
CL\_18179
CL\_18178
CL\_18177
CL\_18176
CL\_18175
CL\_18174
CL\_18173
CL\_18172
CL\_18171
CL\_18170
CL\_18169
CL\_18168
CL\_18167
CL\_5522
CL\_10239
CL\_9079
CL\_9078
CL\_9077
CL\_9076
CL\_9075
CL\_7471
CL\_16061
CL\_12886
CL\_17272
CL\_7690
CL\_17152
CL\_17151
CL\_17150
CL\_17149
CL\_17148
CL\_17147
CL\_17146
CL\_17145
CL\_17144
CL\_17143
CL\_17142
CL\_17141
CL\_17140
CL\_17139
CL\_17138
CL\_17137
CL\_17136
CL\_17135
CL\_19272
CL\_8605
CL\_19271
CL\_10354
CL\_19270
CL\_19269
CL\_19268
CL\_19267
CL\_19266
CL\_19265
CL\_19264
CL\_10353
CL\_10352
CL\_10351
CL\_9305
CL\_5521
CL\_20648
CL\_6159
CL\_5524
CL\_21946
CL\_17453
CL\_18884
CL\_7882
CL\_7881
CL\_7880
CL\_7879
CL\_7878
CL\_7877
CL\_7876
CL\_7875
CL\_13848
CL\_12832
CL\_14428
CL\_8086
CL\_8188
CL\_16934
CL\_16933
CL\_16932
CL\_16931
CL\_16930
CL\_16929
CL\_8210
CL\_9311
CL\_9310
CL\_9309
CL\_9308
CL\_9307
CL\_9306
CL\_5523
CL\_15990
CL\_6123
CL\_6122
CL\_21453
CL\_21452
CL\_21451
CL\_7695
CL\_7694
CL\_7693
CL\_7692
CL\_7691
CL\_16546
CL\_16547
CL\_16548
CL\_16549
CL\_16550
CL\_16551
CL\_16552
CL\_20559
CL\_16553
CL\_16554
CL\_16555
CL\_16556
CL\_16557
CL\_16558
CL\_16559
CL\_20560
CL\_16560
CL\_16561
CL\_16562
CL\_16563
CL\_16564
CL\_16565
CL\_16566
CL\_20561
CL\_16567
CL\_20562
CL\_20563
CL\_20564
CL\_20565
CL\_20566
CL\_16568
CL\_16569
CL\_16570
CL\_20567
CL\_16571
CL\_16572
CL\_16573
CL\_16574
CL\_16575
CL\_16576
CL\_16577
CL\_16578
CL\_16579
CL\_16580
CL\_16581
CL\_16582
CL\_16583
CL\_16584
CL\_16585
CL\_16586
CL\_16587
CL\_16588
CL\_16589
CL\_16590
CL\_16591
CL\_16592
CL\_16593
