## Supplementary material for "A novel method for integrating genomic and Tn-Seq data to identify common *in vivo* fitness mechanisms across multiple bacterial species": S2 Dataset: CL_INS_207.html

Legend

 Mobile +extrachromosomalelementfunctions
 Regulatoryfunctions
 Hypothetical
 DNA Metabolism
 All EssentialGenes
 Other
 Transport +binding proteins
 All VFDB Genes

FULL


WINDOWSVGPNG

Trim RowsRemove SingletonsSave Fasta

CL\_2622


CL\_2622


CL\_2622


CL\_2622


CL\_2622


CL\_2622

HighlightSelectShow Genomes


160

CL\_2621


22

CL\_2620


2

CL\_2620


1

Break


1

CL\_2620


1

CL\_2620

fGI ID


CL\_INS\_207
CL\_INS\_270
CL\_INS\_207
CL\_INS\_270
CL\_INS\_207
CL\_INS\_155
CL\_INS\_197
CL\_INS\_197
CL\_INS\_197
CL\_INS\_197
CL\_INS\_197
CL\_INS\_278
CL\_INS\_207
CL\_INS\_207
CL\_INS\_207
CL\_INS\_207
CL\_INS\_207
CL\_INS\_207
CL\_INS\_207
CL\_INS\_207
CL\_INS\_207
CL\_INS\_207
CL\_INS\_207
CL\_INS\_207
CL\_INS\_207
CL\_INS\_207
CL\_INS\_207
CL\_INS\_207
CL\_INS\_207
CL\_INS\_207
CL\_INS\_207
CL\_INS\_207
CL\_INS\_207
CL\_INS\_207
CL\_INS\_132
CL\_INS\_278
CL\_INS\_278
CL\_INS\_278
CL\_INS\_278
CL\_INS\_278
CL\_INS\_278
Cluster ID


CL\_18850
CL\_8063
CL\_18851
CL\_8064
CL\_8065
CL\_5388
CL\_8189
CL\_8190
CL\_8191
CL\_8192
CL\_8193
CL\_6001
CL\_7111
CL\_9408
CL\_9407
CL\_9529
CL\_9530
CL\_10820
CL\_9531
CL\_9532
CL\_9533
CL\_9536
CL\_10319
CL\_10318
CL\_9537
CL\_9538
CL\_9541
CL\_9542
CL\_9543
CL\_9544
CL\_9545
CL\_9547
CL\_9548
CL\_10819
CL\_7987
CL\_7593
CL\_7592
CL\_6050
CL\_6051
CL\_6052
CL\_6053
