## Supplementary material for "A novel method for integrating genomic and Tn-Seq data to identify common *in vivo* fitness mechanisms across multiple bacterial species": S2 Dataset: CL_INS_209.html

Legend

 Hypothetical
 All EssentialGenes
 AntibioticResistance
 All VFDB Genes

FULL


WINDOWSVGPNG

Trim RowsRemove SingletonsSave Fasta

CL\_2624


CL\_2624


CL\_2624


CL\_2622

HighlightSelectShow Genomes


142

CL\_2626


1

CL\_2627


1

CL\_2626


1

CL\_2626

fGI ID

CL\_INS\_209
Cluster ID

CL\_2625
