## Supplementary material for "A novel method for integrating genomic and Tn-Seq data to identify common *in vivo* fitness mechanisms across multiple bacterial species": S2 Dataset: CL_INS_212.html

Legend

 Mobile +extrachromosomalelementfunctions
 Regulatoryfunctions
 Hypothetical
 DNA Metabolism
 Other
 Transport +binding proteins
 All VFDB Genes

FULL


WINDOWSVGPNG

Trim RowsRemove SingletonsSave Fasta

CL\_2642


CL\_2642


CL\_2642


CL\_2642


Break


CL\_3365


CL\_2642


CL\_2642


CL\_2642


CL\_2642


CL\_2643


CL\_2642

HighlightSelectShow Genomes


173

CL\_2641


8

CL\_2641


2

CL\_2641


1

CL\_2636


1

CL\_2641


1

CL\_2641


1

CL\_2641


1

Break


1

Break


1

Break


1

CL\_2641


1

Break

fGI ID


CL\_INS\_212
CL\_INS\_212
CL\_INS\_132
CL\_INS\_278
CL\_INS\_212
CL\_INS\_278
CL\_INS\_385
CL\_INS\_212
CL\_INS\_212
CL\_INS\_212
CL\_INS\_212
CL\_INS\_212
CL\_INS\_212
CL\_INS\_38
CL\_INS\_212
CL\_INS\_212
CL\_INS\_211
CL\_INS\_278
CL\_INS\_278
CL\_INS\_278
CL\_INS\_278
CL\_INS\_278
CL\_INS\_132
CL\_INS\_278
CL\_INS\_278
CL\_INS\_278
CL\_INS\_278
CL\_INS\_212
CL\_INS\_160
CL\_INS\_160
CL\_INS\_212
CL\_INS\_160
CL\_INS\_160
CL\_INS\_160
CL\_INS\_212
CL\_INS\_212
CL\_INS\_212
CL\_INS\_212
CL\_INS\_212
CL\_INS\_212
CL\_INS\_212
CL\_INS\_212
CL\_INS\_212
CL\_INS\_212
CL\_INS\_212
CL\_INS\_212
CL\_INS\_212
CL\_INS\_212
CL\_INS\_212
CL\_INS\_212
CL\_INS\_212
CL\_INS\_212
CL\_INS\_271
CL\_INS\_212
CL\_INS\_212
CL\_INS\_212
CL\_INS\_212
CL\_INS\_212
CL\_INS\_212
CL\_INS\_212
CL\_INS\_212
CL\_INS\_212
CL\_INS\_212
CL\_INS\_278
CL\_INS\_212
CL\_INS\_212
CL\_INS\_278
CL\_INS\_212
CL\_INS\_212
Cluster ID


CL\_19182
CL\_13849
CL\_6823
CL\_6005
CL\_6825
CL\_6155
CL\_6156
CL\_3171
CL\_3174
CL\_5607
CL\_3176
CL\_3177
CL\_5606
CL\_5605
CL\_5604
CL\_5603
CL\_6157
CL\_12439
CL\_12438
CL\_12437
CL\_8167
CL\_8141
CL\_8219
CL\_6844
CL\_6843
CL\_6842
CL\_7984
CL\_7983
CL\_7982
CL\_7981
CL\_7980
CL\_12487
CL\_7979
CL\_7978
CL\_12486
CL\_12485
CL\_7975
CL\_7974
CL\_7973
CL\_7972
CL\_7970
CL\_9397
CL\_10540
CL\_12482
CL\_12483
CL\_12484
CL\_12436
CL\_12400
CL\_12401
CL\_12402
CL\_12403
CL\_12404
CL\_5229
CL\_12405
CL\_12406
CL\_12407
CL\_12408
CL\_12409
CL\_12410
CL\_12411
CL\_12412
CL\_12413
CL\_12414
CL\_12336
CL\_12415
CL\_12416
CL\_12338
CL\_12417
CL\_12418
