## Supplementary material for "A novel method for integrating genomic and Tn-Seq data to identify common *in vivo* fitness mechanisms across multiple bacterial species": S2 Dataset: CL_INS_213.html

Legend

 Hypothetical
 Other
 All VFDB Genes

FULL


WINDOWSVGPNG

Trim RowsRemove SingletonsSave Fasta

CL\_2649


CL\_2649


CL\_2649


CL\_2649


CL\_2649


CL\_2617

HighlightSelectShow Genomes


123

CL\_2650


29

CL\_2650


13

CL\_2650


12

CL\_2650


1

CL\_2650


1

CL\_2650

fGI ID


CL\_INS\_213
CL\_INS\_213
CL\_INS\_213
Cluster ID


CL\_5514
CL\_5621
CL\_23872
