## Supplementary material for "A novel method for integrating genomic and Tn-Seq data to identify common *in vivo* fitness mechanisms across multiple bacterial species": S2 Dataset: CL_INS_218.html

Legend

 AntibioticResistance
 Other
 All VFDB Genes

FULL


WINDOWSVGPNG

Trim RowsRemove SingletonsSave Fasta

CL\_2674


CL\_2674


CL\_2672


CL\_2674

HighlightSelectShow Genomes


81

CL\_2675


69

CL\_2675


3

CL\_2675


1

CL\_2681

fGI ID

CL\_INS\_218
Cluster ID

CL\_6572
