## Supplementary material for "A novel method for integrating genomic and Tn-Seq data to identify common *in vivo* fitness mechanisms across multiple bacterial species": S2 Dataset: CL_INS_220.html

Legend

 Mobile +extrachromosomalelementfunctions
 Hypothetical
 AntibioticResistance
 Other
 All VFDB Genes

FULL


WINDOWSVGPNG

Trim RowsRemove SingletonsSave Fasta

CL\_2681


CL\_2681


CL\_2681


CL\_2681


CL\_2681


CL\_2681


CL\_2681


CL\_2681

HighlightSelectShow Genomes


184

CL\_2680


2

CL\_2680


2

CL\_2679


2

CL\_2680


1

CL\_2674


1

CL\_2677


1

CL\_2680


1

CL\_2636

fGI ID


CL\_INS\_211
CL\_INS\_385
CL\_INS\_220
CL\_INS\_220
CL\_INS\_220
CL\_INS\_220
CL\_INS\_219
Cluster ID


CL\_2638
CL\_12091
CL\_14985
CL\_20645
CL\_20646
CL\_13226
CL\_13225
