## Supplementary material for "A novel method for integrating genomic and Tn-Seq data to identify common *in vivo* fitness mechanisms across multiple bacterial species": S2 Dataset: CL_INS_221.html

Legend

 Mobile +extrachromosomalelementfunctions
 Hypothetical
 DNA Metabolism
 Other
 All VFDB Genes
 Transport +binding proteins

FULL


WINDOWSVGPNG

Trim RowsRemove SingletonsSave Fasta

CL\_2682


CL\_2682


CL\_2682


CL\_2682


CL\_2682


CL\_2682


CL\_2682


CL\_2682


CL\_2682


CL\_2682


CL\_2682


CL\_2636


CL\_2670


CL\_2682


CL\_2682


CL\_2682


CL\_2676


CL\_2682


CL\_2682


CL\_2682


CL\_2682


CL\_2682


CL\_2682


CL\_2682


CL\_2682

HighlightSelectShow Genomes


48

CL\_2684


44

CL\_2684


27

CL\_2684


20

CL\_2684


16

CL\_2684


12

CL\_2684


9

CL\_2684


8

CL\_2684


2

CL\_2684


2

CL\_2684


2

CL\_2684


1

CL\_2684


1

CL\_2684


1

CL\_2684


1

CL\_2684


1

CL\_2684


1

CL\_2684


1

CL\_2684


1

CL\_2684


1

CL\_2684


1

CL\_2684


1

CL\_2684


1

CL\_2684


1

CL\_2684


1

CL\_2684

fGI ID


CL\_INS\_221
CL\_INS\_221
CL\_INS\_385
CL\_INS\_221
CL\_INS\_221
CL\_INS\_221
CL\_INS\_221
CL\_INS\_221
CL\_INS\_221
CL\_INS\_221
CL\_INS\_221
CL\_INS\_221
CL\_INS\_221
CL\_INS\_334
CL\_INS\_334
CL\_INS\_334
CL\_INS\_334
CL\_INS\_221
CL\_INS\_221
CL\_INS\_221
CL\_INS\_221
CL\_INS\_221
CL\_INS\_221
CL\_INS\_221
CL\_INS\_221
Cluster ID


CL\_13850
CL\_22303
CL\_12091
CL\_2683
CL\_22581
CL\_22580
CL\_22579
CL\_5512
CL\_8761
CL\_19072
CL\_15982
CL\_5620
CL\_22545
CL\_7294
CL\_7295
CL\_7296
CL\_7297
CL\_7298
CL\_6575
CL\_6576
CL\_6577
CL\_6578
CL\_5511
CL\_5510
CL\_6579
