## Supplementary material for "A novel method for integrating genomic and Tn-Seq data to identify common *in vivo* fitness mechanisms across multiple bacterial species": S2 Dataset: CL_INS_222.html

Legend

 Regulatoryfunctions
 AntibioticResistance
 Centralintermediarymetabolism

FULL


WINDOWSVGPNG

Trim RowsRemove SingletonsSave Fasta

CL\_2687


CL\_2687

HighlightSelectShow Genomes


124

CL\_2688


79

CL\_2688

fGI ID

CL\_INS\_222
Cluster ID

CL\_5619
