## Supplementary material for "A novel method for integrating genomic and Tn-Seq data to identify common *in vivo* fitness mechanisms across multiple bacterial species": S2 Dataset: CL_INS_230.html

Legend

 Mobile +extrachromosomalelementfunctions
 Regulatoryfunctions
 Hypothetical
 Other
 All VFDB Genes

FULL


WINDOWSVGPNG

Trim RowsRemove SingletonsSave Fasta

CL\_2747


CL\_2747


CL\_2747


CL\_2747


CL\_2747


CL\_2746


CL\_2747

HighlightSelectShow Genomes


124

CL\_2748


40

CL\_2748


21

CL\_2748


6

CL\_2748


2

CL\_2748


1

CL\_2748


1

CL\_2748

fGI ID


CL\_INS\_230
CL\_INS\_230
CL\_INS\_230
CL\_INS\_230
CL\_INS\_230
CL\_INS\_230
CL\_INS\_230
CL\_INS\_230
CL\_INS\_230
CL\_INS\_230
CL\_INS\_230
CL\_INS\_230
Cluster ID


CL\_5617
CL\_13851
CL\_5616
CL\_11450
CL\_11449
CL\_11448
CL\_11447
CL\_11446
CL\_11445
CL\_11444
CL\_11443
CL\_11442
