## Supplementary material for "A novel method for integrating genomic and Tn-Seq data to identify common *in vivo* fitness mechanisms across multiple bacterial species": S2 Dataset: CL_INS_231.html

Legend

 Hypothetical
 Other
 Proteinsynthesis/fate

FULL


WINDOWSVGPNG

Trim RowsRemove SingletonsSave Fasta

CL\_2753


CL\_2753


CL\_2753

HighlightSelectShow Genomes


113

CL\_2755


87

CL\_2755


1

CL\_2756

fGI ID

CL\_INS\_231
Cluster ID

CL\_2754
