## Supplementary material for "A novel method for integrating genomic and Tn-Seq data to identify common *in vivo* fitness mechanisms across multiple bacterial species": S2 Dataset: CL_INS_232.html

Legend

 Mobile +extrachromosomalelementfunctions
 Hypothetical
 Other
 All VFDB Genes

FULL


WINDOWSVGPNG

Trim RowsRemove SingletonsSave Fasta

CL\_2756


CL\_2756


CL\_2755


CL\_2756


CL\_2756


CL\_2755

HighlightSelectShow Genomes


107

CL\_2757


88

CL\_2757


2

CL\_2757


1

CL\_2757


1

CL\_2757


1

CL\_2757

fGI ID


CL\_INS\_232
CL\_INS\_232
CL\_INS\_278
CL\_INS\_278
CL\_INS\_278
CL\_INS\_160
CL\_INS\_278
CL\_INS\_278
CL\_INS\_278
CL\_INS\_278
CL\_INS\_232
CL\_INS\_278
CL\_INS\_132
CL\_INS\_232
CL\_INS\_91
CL\_INS\_278
CL\_INS\_278
CL\_INS\_278
Cluster ID


CL\_7300
CL\_5509
CL\_15652
CL\_7577
CL\_13334
CL\_68
CL\_15654
CL\_11243
CL\_15655
CL\_6789
CL\_15656
CL\_15658
CL\_8078
CL\_8077
CL\_6886
CL\_15675
CL\_15676
CL\_6079
