## Supplementary material for "A novel method for integrating genomic and Tn-Seq data to identify common *in vivo* fitness mechanisms across multiple bacterial species": S2 Dataset: CL_INS_233.html

Legend

 Mobile +extrachromosomalelementfunctions
 Other
 EnergyMetabolism

FULL


WINDOWSVGPNG

Trim RowsRemove SingletonsSave Fasta

CL\_2770


CL\_2770


CL\_2770

HighlightSelectShow Genomes


109

CL\_2772


92

CL\_2772


1

CL\_2794

fGI ID

CL\_INS\_233
Cluster ID

CL\_2771
