## Supplementary material for "A novel method for integrating genomic and Tn-Seq data to identify common *in vivo* fitness mechanisms across multiple bacterial species": S2 Dataset: CL_INS_237.html

Legend

 Hypothetical
 Other
 All VFDB Genes

FULL


WINDOWSVGPNG

Trim RowsRemove SingletonsSave Fasta

CL\_2819


CL\_2819


CL\_2819


CL\_2819

HighlightSelectShow Genomes


115

CL\_2821


85

CL\_2821


1

CL\_2823


1

CL\_2822

fGI ID

CL\_INS\_237
Cluster ID

CL\_2820
