## Supplementary material for "A novel method for integrating genomic and Tn-Seq data to identify common *in vivo* fitness mechanisms across multiple bacterial species": S2 Dataset: CL_INS_239.html

Legend

 Mobile +extrachromosomalelementfunctions
 Hypothetical
 Other
 EnergyMetabolism
 All VFDB Genes

FULL


WINDOWSVGPNG

Trim RowsRemove SingletonsSave Fasta

CL\_5614


CL\_5614


CL\_2912


CL\_5614


CL\_5614


CL\_2912


CL\_2912


CL\_5614


CL\_2912


CL\_5614


CL\_5614


CL\_5614


CL\_5614


CL\_2912


CL\_2912


CL\_5614


CL\_5614


CL\_2912


CL\_5614


CL\_2912


CL\_5614


CL\_5614


CL\_5614

HighlightSelectShow Genomes


49

CL\_2916


41

CL\_2916


25

CL\_2916


25

CL\_2916


17

CL\_2916


12

CL\_2916


6

CL\_2916


4

CL\_2916


4

CL\_2916


2

CL\_2916


2

CL\_2916


1

CL\_2917


1

CL\_2916


1

CL\_2916


1

CL\_2916


1

CL\_2916


1

CL\_2917


1

CL\_2916


1

CL\_2916


1

CL\_2916


1

CL\_2916


1

CL\_2916


1

CL\_2916

fGI ID


CL\_INS\_239
CL\_INS\_239
CL\_INS\_239
CL\_INS\_239
CL\_INS\_239
CL\_INS\_239
CL\_INS\_239
CL\_INS\_307
CL\_INS\_239
CL\_INS\_307
CL\_INS\_239
CL\_INS\_239
CL\_INS\_239
CL\_INS\_239
CL\_INS\_239
CL\_INS\_239
CL\_INS\_239
CL\_INS\_239
CL\_INS\_239
Cluster ID


CL\_14650
CL\_2913
CL\_17426
CL\_14986
CL\_9257
CL\_22637
CL\_12090
CL\_11725
CL\_11722
CL\_11723
CL\_22578
CL\_22577
CL\_18152
CL\_18153
CL\_22576
CL\_22636
CL\_11731
CL\_2914
CL\_2915
