## Supplementary material for "A novel method for integrating genomic and Tn-Seq data to identify common *in vivo* fitness mechanisms across multiple bacterial species": S2 Dataset: CL_INS_242.html

Legend

 Hypothetical
 Other
 All VFDB Genes

FULL


WINDOWSVGPNG

Trim RowsRemove SingletonsSave Fasta

CL\_2935


CL\_2935


CL\_2935


CL\_2935

HighlightSelectShow Genomes


130

CL\_2937


50

CL\_2937


10

CL\_2938


1

CL\_2939

fGI ID


CL\_INS\_242
CL\_INS\_243
CL\_INS\_243
Cluster ID


CL\_2936
CL\_6971
CL\_6970
