## Supplementary material for "A novel method for integrating genomic and Tn-Seq data to identify common *in vivo* fitness mechanisms across multiple bacterial species": S2 Dataset: CL_INS_246.html

Legend

 Mobile +extrachromosomalelementfunctions
 Regulatoryfunctions
 Hypothetical
 DNA Metabolism
 Other
 Transport +binding proteins
 All VFDB Genes

FULL


WINDOWSVGPNG

Trim RowsRemove SingletonsSave Fasta

CL\_2964


CL\_2964


CL\_2964


CL\_2964


CL\_2964


CL\_2964


CL\_2964


CL\_2964


CL\_2964


CL\_2964


CL\_2964


CL\_2964


CL\_2964


CL\_2964


CL\_2964


CL\_2964


CL\_2964


CL\_2964


CL\_2964


CL\_2964


CL\_2964


CL\_2964


CL\_2964


CL\_2964


CL\_2964


CL\_2964


CL\_2964


CL\_2964


CL\_2964


CL\_2964


CL\_2938


CL\_2964


CL\_2964


CL\_2964


CL\_2964


CL\_2964


CL\_2964


CL\_2964


CL\_2964


CL\_2964


CL\_2964


CL\_2964


CL\_2964


CL\_2964


CL\_2964


CL\_2964


CL\_2964


CL\_2964


CL\_2964


CL\_2964


CL\_2964


CL\_2964


CL\_2964


CL\_2964


CL\_2964


CL\_2964


CL\_2964


CL\_2964


CL\_2964


CL\_2964


CL\_2964


CL\_2964


CL\_2964


CL\_2964


CL\_2964


CL\_2964


CL\_2964


CL\_2964


CL\_2964


CL\_2964


CL\_2964


CL\_2964


CL\_2964


CL\_2920


Break


CL\_2964


CL\_2964


CL\_5516


CL\_2964


CL\_2964


CL\_2964


CL\_2964


CL\_2964


CL\_2964


CL\_2964


CL\_2964


CL\_2964


CL\_2964


CL\_2964


CL\_2964


CL\_2938


CL\_2964


CL\_2964

HighlightSelectShow Genomes


23

CL\_2968


15

CL\_2968


12

CL\_2968


11

CL\_2968


8

CL\_2968


8

CL\_2968


8

CL\_2968


7

CL\_2968


5

CL\_2968


4

CL\_2968


3

CL\_2968


3

CL\_2968


3

CL\_2969


3

CL\_2968


3

CL\_2968


3

CL\_2968


2

Break


2

CL\_2968


2

CL\_2968


2

CL\_2968


2

CL\_2968


2

CL\_2968


2

CL\_2968


2

CL\_2968


1

CL\_2968


1

CL\_2968


1

CL\_2968


1

CL\_2968


1

CL\_2969


1

CL\_2968


1

CL\_2968


1

CL\_2968


1

CL\_2968


1

CL\_2968


1

CL\_2968


1

CL\_2968


1

CL\_2968


1

Break


1

CL\_2968


1

Break


1

CL\_2969


1

CL\_2968


1

CL\_2968


1

CL\_2968


1

CL\_2968


1

CL\_2995


1

CL\_2968


1

CL\_2968


1

CL\_2968


1

CL\_2968


1

CL\_2968


1

CL\_2968


1

CL\_2968


1

CL\_2968


1

CL\_2968


1

CL\_2968


1

CL\_2968


1

CL\_2968


1

CL\_2968


1

CL\_2968


1

CL\_2968


1

CL\_2968


1

CL\_2968


1

CL\_2968


1

CL\_2968


1

CL\_5520


1

CL\_2968


1

CL\_2968


1

CL\_2968


1

CL\_2968


1

CL\_2968


1

CL\_2968


1

CL\_2968


1

CL\_2968


1

CL\_2968


1

CL\_2968


1

CL\_2968


1

CL\_2968


1

CL\_2968


1

CL\_2968


1

CL\_2968


1

CL\_2968


1

CL\_2969


1

CL\_2968


1

CL\_1720


1

CL\_2968


1

CL\_2968


1

CL\_2968


1

CL\_2968


1

CL\_2968


1

CL\_2968


1

CL\_2968


1

CL\_2968

fGI ID


CL\_INS\_246
CL\_INS\_246
CL\_INS\_246
CL\_INS\_246
CL\_INS\_246
CL\_INS\_246
CL\_INS\_246
CL\_INS\_204
CL\_INS\_204
CL\_INS\_204
CL\_INS\_246
CL\_INS\_246
CL\_INS\_246
CL\_INS\_246
CL\_INS\_246
CL\_INS\_246
CL\_INS\_246
CL\_INS\_246
CL\_INS\_246
CL\_INS\_246
CL\_INS\_246
CL\_INS\_246
CL\_INS\_246
CL\_INS\_246
CL\_INS\_246
CL\_INS\_246
CL\_INS\_246
CL\_INS\_246
CL\_INS\_246
CL\_INS\_246
CL\_INS\_246
CL\_INS\_246
CL\_INS\_246
CL\_INS\_246
CL\_INS\_246
CL\_INS\_243
CL\_INS\_243
CL\_INS\_246
CL\_INS\_246
CL\_INS\_246
CL\_INS\_246
CL\_INS\_243
CL\_INS\_243
CL\_INS\_243
CL\_INS\_243
CL\_INS\_246
CL\_INS\_243
CL\_INS\_246
CL\_INS\_246
CL\_INS\_246
CL\_INS\_246
CL\_INS\_243
CL\_INS\_246
CL\_INS\_246
CL\_INS\_246
CL\_INS\_246
CL\_INS\_246
CL\_INS\_246
CL\_INS\_246
CL\_INS\_246
CL\_INS\_246
CL\_INS\_246
CL\_INS\_246
CL\_INS\_246
CL\_INS\_246
CL\_INS\_246
CL\_INS\_246
CL\_INS\_246
CL\_INS\_246
CL\_INS\_246
CL\_INS\_246
CL\_INS\_246
CL\_INS\_246
CL\_INS\_246
CL\_INS\_246
CL\_INS\_246
CL\_INS\_246
CL\_INS\_246
CL\_INS\_246
CL\_INS\_243
CL\_INS\_246
CL\_INS\_246
CL\_INS\_243
CL\_INS\_246
CL\_INS\_246
CL\_INS\_243
CL\_INS\_263
CL\_INS\_246
CL\_INS\_263
CL\_INS\_263
CL\_INS\_263
CL\_INS\_263
CL\_INS\_263
CL\_INS\_263
CL\_INS\_263
CL\_INS\_263
CL\_INS\_263
CL\_INS\_263
CL\_INS\_263
CL\_INS\_263
CL\_INS\_263
CL\_INS\_263
CL\_INS\_246
CL\_INS\_246
CL\_INS\_246
CL\_INS\_57
CL\_INS\_57
CL\_INS\_57
CL\_INS\_246
CL\_INS\_246
CL\_INS\_246
CL\_INS\_246
CL\_INS\_246
CL\_INS\_246
CL\_INS\_246
CL\_INS\_246
CL\_INS\_57
CL\_INS\_246
CL\_INS\_57
CL\_INS\_57
CL\_INS\_246
CL\_INS\_246
CL\_INS\_246
CL\_INS\_246
CL\_INS\_246
CL\_INS\_246
CL\_INS\_246
CL\_INS\_246
CL\_INS\_57
CL\_INS\_57
CL\_INS\_57
CL\_INS\_57
CL\_INS\_57
CL\_INS\_57
CL\_INS\_57
CL\_INS\_246
CL\_INS\_246
CL\_INS\_246
CL\_INS\_246
CL\_INS\_246
CL\_INS\_246
Cluster ID


CL\_23817
CL\_14987
CL\_17425
CL\_2965
CL\_2966
CL\_2967
CL\_10231
CL\_11441
CL\_11440
CL\_11439
CL\_11767
CL\_10230
CL\_10229
CL\_10228
CL\_23251
CL\_10227
CL\_11436
CL\_11435
CL\_11434
CL\_11433
CL\_11432
CL\_11431
CL\_11430
CL\_11429
CL\_11428
CL\_11427
CL\_11426
CL\_11425
CL\_11229
CL\_11230
CL\_11231
CL\_11232
CL\_11424
CL\_11423
CL\_11422
CL\_5613
CL\_5612
CL\_10544
CL\_22091
CL\_5611
CL\_8760
CL\_8759
CL\_8758
CL\_8757
CL\_8756
CL\_5610
CL\_6638
CL\_14178
CL\_14179
CL\_14180
CL\_14181
CL\_6639
CL\_22249
CL\_8062
CL\_22304
CL\_6968
CL\_6967
CL\_6966
CL\_6965
CL\_6964
CL\_6963
CL\_6962
CL\_6961
CL\_6960
CL\_5492
CL\_23464
CL\_23190
CL\_23191
CL\_23192
CL\_8898
CL\_8899
CL\_14159
CL\_17454
CL\_5491
CL\_15980
CL\_5490
CL\_21795
CL\_13051
CL\_5489
CL\_12366
CL\_6637
CL\_5488
CL\_14630
CL\_5487
CL\_5486
CL\_5485
CL\_6726
CL\_6802
CL\_6801
CL\_5448
CL\_5449
CL\_5450
CL\_5452
CL\_5453
CL\_5454
CL\_5455
CL\_5456
CL\_5457
CL\_5458
CL\_5459
CL\_5460
CL\_5461
CL\_6440
CL\_6439
CL\_6438
CL\_743
CL\_742
CL\_741
CL\_6437
CL\_6436
CL\_6435
CL\_6434
CL\_6433
CL\_6432
CL\_12211
CL\_737
CL\_735
CL\_6431
CL\_734
CL\_733
CL\_6430
CL\_6429
CL\_6428
CL\_6427
CL\_6426
CL\_6425
CL\_6424
CL\_6423
CL\_727
CL\_726
CL\_725
CL\_724
CL\_723
CL\_722
CL\_721
CL\_6422
CL\_12210
CL\_6421
CL\_12209
CL\_6420
CL\_6419
