## Supplementary material for "A novel method for integrating genomic and Tn-Seq data to identify common *in vivo* fitness mechanisms across multiple bacterial species": S2 Dataset: CL_INS_248.html

Legend

 Regulatoryfunctions
 Hypothetical
 Other
 All VFDB Genes

FULL


WINDOWSVGPNG

Trim RowsRemove SingletonsSave Fasta

CL\_3002


CL\_3002


CL\_3002


CL\_3002


CL\_3001

HighlightSelectShow Genomes


195

CL\_3003


6

CL\_3006


1

CL\_3005


1

CL\_3005


1

CL\_3003

fGI ID


CL\_INS\_248
CL\_INS\_248
CL\_INS\_249
CL\_INS\_249
CL\_INS\_249
CL\_INS\_249
CL\_INS\_249
CL\_INS\_248
CL\_INS\_248
CL\_INS\_248
CL\_INS\_248
CL\_INS\_248
CL\_INS\_248
CL\_INS\_248
CL\_INS\_248
CL\_INS\_248
CL\_INS\_248
CL\_INS\_248
CL\_INS\_248
Cluster ID


CL\_20182
CL\_9805
CL\_9804
CL\_9803
CL\_9802
CL\_9801
CL\_9800
CL\_20768
CL\_20767
CL\_20766
CL\_20765
CL\_23612
CL\_20181
CL\_20180
CL\_20179
CL\_20764
CL\_20763
CL\_20762
CL\_20761
