## Supplementary material for "A novel method for integrating genomic and Tn-Seq data to identify common *in vivo* fitness mechanisms across multiple bacterial species": S2 Dataset: CL_INS_249.html

Legend

 Regulatoryfunctions
 Hypothetical
 All VFDB Genes

FULL


WINDOWSVGPNG

Trim RowsRemove SingletonsSave Fasta

CL\_3006


CL\_3006


CL\_3006

HighlightSelectShow Genomes


189

CL\_3005


6

CL\_3002


1

CL\_3005

fGI ID


CL\_INS\_249
CL\_INS\_249
CL\_INS\_249
CL\_INS\_249
CL\_INS\_249
CL\_INS\_249
CL\_INS\_249
Cluster ID


CL\_12683
CL\_9799
CL\_9800
CL\_9801
CL\_9802
CL\_9803
CL\_9804
