## Supplementary material for "A novel method for integrating genomic and Tn-Seq data to identify common *in vivo* fitness mechanisms across multiple bacterial species": S2 Dataset: CL_INS_253.html

Legend

 Mobile +extrachromosomalelementfunctions
 Regulatoryfunctions
 Hypothetical
 DNA Metabolism
 AntibioticResistance
 Other
 Cellularprocesses
 Transport +binding proteins
 All VFDB Genes

FULL


WINDOWSVGPNG

Trim RowsRemove SingletonsSave Fasta

CL\_3070


CL\_3070


CL\_3070

HighlightSelectShow Genomes


116

CL\_3072


87

CL\_3072


1

Break

fGI ID


CL\_INS\_253
CL\_INS\_253
CL\_INS\_253
CL\_INS\_253
CL\_INS\_253
CL\_INS\_253
CL\_INS\_253
CL\_INS\_253
CL\_INS\_253
CL\_INS\_253
CL\_INS\_253
CL\_INS\_253
CL\_INS\_253
CL\_INS\_253
CL\_INS\_253
CL\_INS\_253
CL\_INS\_253
CL\_INS\_253
CL\_INS\_253
CL\_INS\_253
CL\_INS\_253
CL\_INS\_253
CL\_INS\_253
CL\_INS\_253
CL\_INS\_253
CL\_INS\_253
CL\_INS\_253
CL\_INS\_253
CL\_INS\_253
CL\_INS\_253
CL\_INS\_253
CL\_INS\_253
CL\_INS\_253
CL\_INS\_253
CL\_INS\_253
CL\_INS\_253
CL\_INS\_253
CL\_INS\_253
CL\_INS\_253
CL\_INS\_253
CL\_INS\_253
CL\_INS\_253
CL\_INS\_253
CL\_INS\_253
CL\_INS\_253
CL\_INS\_253
CL\_INS\_366
CL\_INS\_253
CL\_INS\_253
CL\_INS\_253
CL\_INS\_253
CL\_INS\_253
CL\_INS\_253
CL\_INS\_253
CL\_INS\_253
CL\_INS\_253
CL\_INS\_253
CL\_INS\_253
CL\_INS\_253
CL\_INS\_253
CL\_INS\_253
CL\_INS\_253
CL\_INS\_253
CL\_INS\_253
CL\_INS\_253
CL\_INS\_253
CL\_INS\_253
CL\_INS\_253
CL\_INS\_253
CL\_INS\_253
CL\_INS\_253
CL\_INS\_253
CL\_INS\_253
CL\_INS\_253
CL\_INS\_253
CL\_INS\_253
CL\_INS\_253
CL\_INS\_253
CL\_INS\_278
CL\_INS\_278
CL\_INS\_253
CL\_INS\_253
CL\_INS\_253
CL\_INS\_253
CL\_INS\_253
CL\_INS\_278
CL\_INS\_278
CL\_INS\_253
CL\_INS\_253
CL\_INS\_253
CL\_INS\_253
CL\_INS\_253
CL\_INS\_253
CL\_INS\_253
CL\_INS\_253
CL\_INS\_253
CL\_INS\_253
CL\_INS\_105
CL\_INS\_253
CL\_INS\_253
CL\_INS\_155
CL\_INS\_57
CL\_INS\_253
CL\_INS\_253
CL\_INS\_253
CL\_INS\_253
CL\_INS\_253
CL\_INS\_253
CL\_INS\_160
CL\_INS\_278
CL\_INS\_278
CL\_INS\_253
CL\_INS\_253
CL\_INS\_132
CL\_INS\_307
CL\_INS\_307
CL\_INS\_307
CL\_INS\_253
CL\_INS\_253
CL\_INS\_253
CL\_INS\_253
CL\_INS\_253
CL\_INS\_253
CL\_INS\_253
CL\_INS\_160
CL\_INS\_132
CL\_INS\_132
CL\_INS\_132
CL\_INS\_132
CL\_INS\_132
CL\_INS\_132
CL\_INS\_132
CL\_INS\_132
CL\_INS\_253
CL\_INS\_278
CL\_INS\_278
CL\_INS\_278
CL\_INS\_278
CL\_INS\_278
CL\_INS\_278
CL\_INS\_278
CL\_INS\_278
CL\_INS\_278
CL\_INS\_278
CL\_INS\_278
CL\_INS\_278
CL\_INS\_278
CL\_INS\_278
CL\_INS\_278
CL\_INS\_278
CL\_INS\_278
CL\_INS\_278
CL\_INS\_105
CL\_INS\_278
CL\_INS\_278
CL\_INS\_278
CL\_INS\_278
CL\_INS\_278
CL\_INS\_278
CL\_INS\_278
CL\_INS\_204
CL\_INS\_278
CL\_INS\_253
CL\_INS\_204
CL\_INS\_204
CL\_INS\_278
CL\_INS\_278
CL\_INS\_278
CL\_INS\_204
CL\_INS\_204
CL\_INS\_204
CL\_INS\_204
CL\_INS\_204
CL\_INS\_253
CL\_INS\_74
CL\_INS\_253
CL\_INS\_253
CL\_INS\_385
CL\_INS\_253
CL\_INS\_212
CL\_INS\_212
CL\_INS\_253
CL\_INS\_212
CL\_INS\_278
CL\_INS\_253
CL\_INS\_253
CL\_INS\_253
CL\_INS\_253
CL\_INS\_253
CL\_INS\_253
CL\_INS\_253
CL\_INS\_253
CL\_INS\_253
CL\_INS\_253
CL\_INS\_253
CL\_INS\_253
CL\_INS\_160
CL\_INS\_58
CL\_INS\_58
CL\_INS\_253
CL\_INS\_30
CL\_INS\_160
CL\_INS\_160
CL\_INS\_160
CL\_INS\_160
CL\_INS\_160
CL\_INS\_160
CL\_INS\_253
CL\_INS\_253
CL\_INS\_253
CL\_INS\_160
CL\_INS\_160
CL\_INS\_160
CL\_INS\_278
CL\_INS\_278
CL\_INS\_253
CL\_INS\_197
CL\_INS\_197
CL\_INS\_253
CL\_INS\_253
CL\_INS\_253
CL\_INS\_253
CL\_INS\_253
CL\_INS\_278
CL\_INS\_278
CL\_INS\_278
CL\_INS\_278
CL\_INS\_197
Cluster ID


CL\_3071
CL\_11342
CL\_11343
CL\_11344
CL\_11345
CL\_11346
CL\_11347
CL\_11348
CL\_11349
CL\_11350
CL\_11351
CL\_11352
CL\_11353
CL\_11354
CL\_11355
CL\_11356
CL\_11357
CL\_11358
CL\_11359
CL\_11360
CL\_11361
CL\_11363
CL\_11364
CL\_11365
CL\_11366
CL\_11367
CL\_11368
CL\_11369
CL\_11370
CL\_11371
CL\_11372
CL\_11373
CL\_11374
CL\_11375
CL\_11376
CL\_11377
CL\_11378
CL\_11341
CL\_11301
CL\_11302
CL\_11303
CL\_11304
CL\_11305
CL\_11306
CL\_11307
CL\_11308
CL\_11309
CL\_11310
CL\_11311
CL\_11312
CL\_11313
CL\_11314
CL\_11315
CL\_11316
CL\_11317
CL\_11318
CL\_11319
CL\_11320
CL\_11321
CL\_11322
CL\_11323
CL\_11324
CL\_11325
CL\_11326
CL\_11327
CL\_11328
CL\_11329
CL\_11330
CL\_11331
CL\_11332
CL\_11333
CL\_11334
CL\_11335
CL\_11336
CL\_11337
CL\_11338
CL\_11339
CL\_11340
CL\_9246
CL\_9249
CL\_20487
CL\_10116
CL\_20488
CL\_20486
CL\_21589
CL\_8650
CL\_7588
CL\_15807
CL\_22256
CL\_22257
CL\_22258
CL\_18468
CL\_12750
CL\_12749
CL\_12748
CL\_12747
CL\_12746
CL\_6865
CL\_12745
CL\_12744
CL\_5387
CL\_6173
CL\_22259
CL\_22260
CL\_22261
CL\_22262
CL\_22263
CL\_22264
CL\_6711
CL\_6710
CL\_8159
CL\_12741
CL\_7939
CL\_6181
CL\_3387
CL\_3386
CL\_6182
CL\_9983
CL\_7078
CL\_9984
CL\_7079
CL\_12740
CL\_12739
CL\_12738
CL\_7080
CL\_8243
CL\_9985
CL\_8242
CL\_7944
CL\_7945
CL\_7946
CL\_6806
CL\_8219
CL\_12766
CL\_6879
CL\_7947
CL\_6885
CL\_12260
CL\_7948
CL\_6883
CL\_6882
CL\_7984
CL\_12261
CL\_6842
CL\_6843
CL\_6844
CL\_6845
CL\_6846
CL\_7595
CL\_6041
CL\_6042
CL\_6043
CL\_6044
CL\_6047
CL\_7593
CL\_7592
CL\_6050
CL\_6051
CL\_6052
CL\_6053
CL\_6058
CL\_12765
CL\_12764
CL\_6059
CL\_6060
CL\_12763
CL\_12762
CL\_12761
CL\_6062
CL\_6063
CL\_6064
CL\_6065
CL\_6066
CL\_22255
CL\_6857
CL\_8101
CL\_18433
CL\_6156
CL\_19430
CL\_7972
CL\_7973
CL\_20041
CL\_7975
CL\_6833
CL\_18388
CL\_14484
CL\_18387
CL\_18386
CL\_18385
CL\_18384
CL\_18383
CL\_18382
CL\_18381
CL\_18380
CL\_18379
CL\_18378
CL\_6851
CL\_6852
CL\_7988
CL\_18377
CL\_6869
CL\_11883
CL\_11882
CL\_11881
CL\_15101
CL\_11290
CL\_10042
CL\_10098
CL\_20043
CL\_20042
CL\_11297
CL\_7059
CL\_8020
CL\_8167
CL\_8141
CL\_13994
CL\_6811
CL\_10120
CL\_10117
CL\_10114
CL\_10112
CL\_10111
CL\_10110
CL\_6137
CL\_6816
CL\_6815
CL\_6814
CL\_6812
