## Supplementary material for "A novel method for integrating genomic and Tn-Seq data to identify common *in vivo* fitness mechanisms across multiple bacterial species": S2 Dataset: CL_INS_257.html

Legend

 Hypothetical
 All EssentialGenes
 All VFDB Genes

FULL


WINDOWSVGPNG

Trim RowsRemove SingletonsSave Fasta

CL\_3114


CL\_3114


CL\_3113


CL\_3113

HighlightSelectShow Genomes


115

CL\_3116


87

CL\_3116


1

CL\_3116


1

CL\_3116

fGI ID

CL\_INS\_257
Cluster ID

CL\_3115
