## Supplementary material for "A novel method for integrating genomic and Tn-Seq data to identify common *in vivo* fitness mechanisms across multiple bacterial species": S2 Dataset: CL_INS_258.html

Legend

 Hypothetical
 All VFDB Genes

FULL


WINDOWSVGPNG

Trim RowsRemove SingletonsSave Fasta

CL\_3127


CL\_3127


CL\_3127

HighlightSelectShow Genomes


116

CL\_3129


86

CL\_3129


1

Break

fGI ID

CL\_INS\_258
Cluster ID

CL\_3128
