## Supplementary material for "A novel method for integrating genomic and Tn-Seq data to identify common *in vivo* fitness mechanisms across multiple bacterial species": S2 Dataset: CL_INS_261.html

Legend

 Mobile +extrachromosomalelementfunctions
 Regulatoryfunctions
 Hypothetical
 DNA Metabolism
 All EssentialGenes
 AntibioticResistance
 Proteinsynthesis/fate
 Other
 EnergyMetabolism
 Cellularprocesses
 Transport +binding proteins
 All VFDB Genes

FULL


WINDOWSVGPNG

Trim RowsRemove SingletonsSave Fasta

CL\_3168


CL\_3168


CL\_3168


CL\_3168


CL\_3168


CL\_3168


CL\_3168


CL\_3168


CL\_3168


CL\_3168


CL\_3168


CL\_3168


CL\_3168


CL\_3168


CL\_3168


CL\_3168


CL\_3168


CL\_3168


CL\_3168


CL\_3168


CL\_3168


CL\_3168


CL\_3168


CL\_3168


CL\_3168


CL\_3168


CL\_3168


CL\_3168


CL\_3168


CL\_3168


CL\_3168


CL\_3168


CL\_3168


CL\_3168


CL\_3168


CL\_3168


CL\_3168


CL\_3168


CL\_3168


CL\_3168


CL\_3168


CL\_3168


CL\_3168


CL\_3168


CL\_3168


CL\_3168


CL\_3168


CL\_3168


CL\_3168


CL\_3168


CL\_3168


CL\_3168


CL\_3168


CL\_3168


CL\_3168


CL\_3168


CL\_3168


CL\_3168


CL\_3168


CL\_3168


CL\_3166


CL\_3168


CL\_3168


CL\_3168


CL\_3168


CL\_3168


CL\_3168


CL\_3168


CL\_3168


CL\_3168


CL\_3168


CL\_3168


Break


CL\_3168


CL\_3168


CL\_3168


CL\_3168


CL\_3168


CL\_3168


CL\_3168


CL\_3168


CL\_3168


CL\_3168


CL\_3168


CL\_3168


CL\_3168


CL\_1409


CL\_3168


CL\_3168


CL\_3168


CL\_3168


CL\_3168


CL\_3168


CL\_3168


CL\_3168


CL\_3168


CL\_3168


CL\_3168


CL\_3168


CL\_3168


CL\_3168


CL\_3168


CL\_3168


CL\_3168


CL\_3168


CL\_3168


CL\_3168


CL\_3168


CL\_3168


CL\_3168


CL\_3168


CL\_3168

HighlightSelectShow Genomes


23

CL\_3178


11

CL\_3178


7

CL\_3179


7

CL\_3178


5

CL\_3178


4

CL\_3178


4

CL\_3178


4

CL\_3178


3

CL\_3178


3

CL\_3178


3

CL\_3178


3

CL\_3178


3

CL\_3178


3

CL\_3178


3

CL\_3178


3

CL\_3179


3

CL\_3178


3

CL\_3178


2

CL\_3178


2

CL\_3179


2

CL\_3178


2

CL\_3178


2

CL\_3178


2

CL\_3178


2

CL\_3178


2

Break


2

CL\_3178


2

CL\_3178


2

CL\_3178


1

CL\_3178


1

CL\_3178


1

CL\_3178


1

CL\_3210


1

CL\_3180


1

CL\_3178


1

CL\_3179


1

CL\_3178


1

CL\_3179


1

CL\_3178


1

CL\_3178


1

CL\_3179


1

CL\_3179


1

CL\_3178


1

Break


1

CL\_3178


1

CL\_3178


1

CL\_3179


1

CL\_3178


1

CL\_3178


1

CL\_3178


1

CL\_3204


1

CL\_3474


1

CL\_3178


1

CL\_3178


1

CL\_3178


1

CL\_3178


1

CL\_3178


1

CL\_3178


1

CL\_3183


1

CL\_3178


1

CL\_3178


1

CL\_3178


1

CL\_3178


1

CL\_3178


1

CL\_3178


1

CL\_3178


1

CL\_3178


1

CL\_3178


1

CL\_3178


1

Break


1

CL\_3178


1

CL\_3178


1

CL\_3178


1

CL\_3178


1

CL\_3178


1

CL\_3178


1

CL\_3209


1

CL\_3183


1

CL\_3178


1

CL\_3183


1

CL\_3179


1

CL\_3178


1

CL\_3179


1

CL\_3178


1

CL\_3178


1

CL\_3178


1

CL\_3178


1

CL\_3178


1

CL\_3178


1

Break


1

CL\_3183


1

CL\_3178


1

CL\_3179


1

CL\_3178


1

CL\_3179


1

CL\_3178


1

CL\_3178


1

CL\_3178


1

CL\_3179


1

CL\_3178


1

CL\_3178


1

CL\_3178


1

CL\_3178


1

CL\_3183


1

CL\_3178


1

CL\_3179


1

CL\_3178


1

CL\_3183


1

CL\_3178


1

CL\_3178


1

CL\_3178


1

CL\_3178

fGI ID


CL\_INS\_261
CL\_INS\_261
CL\_INS\_261
CL\_INS\_261
CL\_INS\_267
CL\_INS\_261
CL\_INS\_261
CL\_INS\_267
CL\_INS\_261
CL\_INS\_261
CL\_INS\_261
CL\_INS\_261
CL\_INS\_261
CL\_INS\_261
CL\_INS\_261
CL\_INS\_261
CL\_INS\_26
CL\_INS\_261
CL\_INS\_261
CL\_INS\_321
CL\_INS\_321
CL\_INS\_321
CL\_INS\_321
CL\_INS\_261
CL\_INS\_278
CL\_INS\_278
CL\_INS\_278
CL\_INS\_278
CL\_INS\_261
CL\_INS\_261
CL\_INS\_261
CL\_INS\_212
CL\_INS\_212
CL\_INS\_261
CL\_INS\_212
CL\_INS\_261
CL\_INS\_261
CL\_INS\_212
CL\_INS\_212
CL\_INS\_261
CL\_INS\_282
CL\_INS\_212
CL\_INS\_38
CL\_INS\_212
CL\_INS\_212
CL\_INS\_261
CL\_INS\_261
CL\_INS\_261
CL\_INS\_261
CL\_INS\_38
CL\_INS\_38
CL\_INS\_38
CL\_INS\_58
CL\_INS\_58
CL\_INS\_278
CL\_INS\_278
CL\_INS\_132
CL\_INS\_278
CL\_INS\_278
CL\_INS\_132
CL\_INS\_132
CL\_INS\_132
CL\_INS\_132
CL\_INS\_132
CL\_INS\_278
CL\_INS\_278
CL\_INS\_278
CL\_INS\_278
CL\_INS\_278
CL\_INS\_278
CL\_INS\_278
CL\_INS\_132
CL\_INS\_318
CL\_INS\_318
CL\_INS\_400
CL\_INS\_278
CL\_INS\_400
CL\_INS\_400
CL\_INS\_400
CL\_INS\_400
CL\_INS\_400
CL\_INS\_400
CL\_INS\_261
CL\_INS\_261
CL\_INS\_261
CL\_INS\_132
CL\_INS\_278
CL\_INS\_278
CL\_INS\_278
CL\_INS\_132
CL\_INS\_132
CL\_INS\_307
CL\_INS\_307
CL\_INS\_160
CL\_INS\_160
CL\_INS\_278
CL\_INS\_171
CL\_INS\_171
CL\_INS\_278
CL\_INS\_132
CL\_INS\_132
CL\_INS\_132
CL\_INS\_132
CL\_INS\_132
CL\_INS\_261
CL\_INS\_132
CL\_INS\_253
CL\_INS\_278
CL\_INS\_278
CL\_INS\_278
CL\_INS\_278
CL\_INS\_278
CL\_INS\_278
CL\_INS\_278
CL\_INS\_278
CL\_INS\_278
CL\_INS\_278
CL\_INS\_278
CL\_INS\_278
CL\_INS\_278
CL\_INS\_278
CL\_INS\_278
CL\_INS\_58
CL\_INS\_278
CL\_INS\_278
CL\_INS\_278
CL\_INS\_105
CL\_INS\_160
CL\_INS\_160
CL\_INS\_160
CL\_INS\_160
CL\_INS\_58
CL\_INS\_58
CL\_INS\_58
CL\_INS\_58
CL\_INS\_204
CL\_INS\_261
CL\_INS\_261
CL\_INS\_261
CL\_INS\_58
CL\_INS\_30
CL\_INS\_30
CL\_INS\_132
CL\_INS\_132
CL\_INS\_278
CL\_INS\_278
CL\_INS\_278
CL\_INS\_30
CL\_INS\_30
CL\_INS\_30
CL\_INS\_278
CL\_INS\_278
CL\_INS\_278
CL\_INS\_278
CL\_INS\_278
CL\_INS\_278
CL\_INS\_58
CL\_INS\_278
CL\_INS\_278
CL\_INS\_178
CL\_INS\_58
CL\_INS\_58
CL\_INS\_278
CL\_INS\_278
CL\_INS\_278
CL\_INS\_212
CL\_INS\_278
CL\_INS\_278
CL\_INS\_278
CL\_INS\_278
CL\_INS\_278
CL\_INS\_278
CL\_INS\_278
CL\_INS\_278
CL\_INS\_278
CL\_INS\_261
CL\_INS\_278
CL\_INS\_278
CL\_INS\_132
CL\_INS\_278
CL\_INS\_278
CL\_INS\_261
CL\_INS\_385
CL\_INS\_318
CL\_INS\_307
CL\_INS\_385
CL\_INS\_278
CL\_INS\_261
CL\_INS\_261
CL\_INS\_307
CL\_INS\_385
CL\_INS\_261
CL\_INS\_261
CL\_INS\_261
CL\_INS\_261
CL\_INS\_261
CL\_INS\_261
CL\_INS\_261
CL\_INS\_261
CL\_INS\_261
CL\_INS\_261
CL\_INS\_261
CL\_INS\_261
CL\_INS\_318
CL\_INS\_318
CL\_INS\_334
CL\_INS\_318
CL\_INS\_391
CL\_INS\_261
CL\_INS\_132
CL\_INS\_261
CL\_INS\_261
CL\_INS\_261
CL\_INS\_385
CL\_INS\_318
CL\_INS\_318
CL\_INS\_204
CL\_INS\_246
CL\_INS\_278
CL\_INS\_278
CL\_INS\_385
CL\_INS\_278
CL\_INS\_278
CL\_INS\_385
CL\_INS\_261
CL\_INS\_261
CL\_INS\_91
CL\_INS\_385
CL\_INS\_278
CL\_INS\_278
CL\_INS\_278
CL\_INS\_385
CL\_INS\_261
CL\_INS\_261
CL\_INS\_278
CL\_INS\_385
CL\_INS\_318
CL\_INS\_204
CL\_INS\_278
CL\_INS\_261
CL\_INS\_261
CL\_INS\_261
CL\_INS\_261
CL\_INS\_261
CL\_INS\_261
CL\_INS\_261
CL\_INS\_261
CL\_INS\_261
CL\_INS\_261
CL\_INS\_261
CL\_INS\_261
CL\_INS\_261
CL\_INS\_278
CL\_INS\_278
CL\_INS\_278
CL\_INS\_30
CL\_INS\_278
CL\_INS\_385
CL\_INS\_278
CL\_INS\_385
CL\_INS\_261
CL\_INS\_261
CL\_INS\_261
CL\_INS\_278
CL\_INS\_318
CL\_INS\_318
CL\_INS\_318
CL\_INS\_318
CL\_INS\_318
CL\_INS\_318
CL\_INS\_318
CL\_INS\_318
CL\_INS\_318
CL\_INS\_318
CL\_INS\_318
CL\_INS\_318
CL\_INS\_385
CL\_INS\_385
CL\_INS\_57
CL\_INS\_38
CL\_INS\_261
CL\_INS\_261
CL\_INS\_261
CL\_INS\_261
CL\_INS\_261
CL\_INS\_261
CL\_INS\_261
CL\_INS\_261
CL\_INS\_261
CL\_INS\_261
CL\_INS\_261
CL\_INS\_261
CL\_INS\_261
CL\_INS\_261
CL\_INS\_261
CL\_INS\_261
CL\_INS\_261
CL\_INS\_261
CL\_INS\_318
CL\_INS\_385
CL\_INS\_307
CL\_INS\_307
CL\_INS\_307
CL\_INS\_385
CL\_INS\_385
CL\_INS\_261
CL\_INS\_261
CL\_INS\_261
CL\_INS\_385
CL\_INS\_385
CL\_INS\_385
CL\_INS\_385
CL\_INS\_385
CL\_INS\_385
CL\_INS\_318
CL\_INS\_318
CL\_INS\_129
CL\_INS\_385
CL\_INS\_385
CL\_INS\_261
CL\_INS\_401
CL\_INS\_401
CL\_INS\_261
CL\_INS\_53
CL\_INS\_53
CL\_INS\_53
CL\_INS\_53
CL\_INS\_53
CL\_INS\_53
CL\_INS\_261
CL\_INS\_155
CL\_INS\_155
CL\_INS\_155
CL\_INS\_155
CL\_INS\_263
CL\_INS\_155
CL\_INS\_155
CL\_INS\_261
CL\_INS\_261
CL\_INS\_261
CL\_INS\_261
CL\_INS\_261
CL\_INS\_351
CL\_INS\_351
CL\_INS\_351
CL\_INS\_351
CL\_INS\_351
CL\_INS\_351
CL\_INS\_351
CL\_INS\_211
CL\_INS\_318
CL\_INS\_385
CL\_INS\_385
CL\_INS\_278
CL\_INS\_278
CL\_INS\_261
CL\_INS\_278
CL\_INS\_261
CL\_INS\_261
CL\_INS\_261
CL\_INS\_261
CL\_INS\_287
CL\_INS\_401
CL\_INS\_261
CL\_INS\_385
CL\_INS\_385
CL\_INS\_261
CL\_INS\_261
CL\_INS\_261
CL\_INS\_261
CL\_INS\_278
CL\_INS\_204
CL\_INS\_261
CL\_INS\_261
CL\_INS\_261
CL\_INS\_261
CL\_INS\_261
Cluster ID


CL\_5482
CL\_7308
CL\_7309
CL\_7310
CL\_7311
CL\_7312
CL\_17424
CL\_7313
CL\_11566
CL\_7314
CL\_10226
CL\_18046
CL\_18045
CL\_18044
CL\_18043
CL\_3169
CL\_12365
CL\_18751
CL\_3170
CL\_6345
CL\_6344
CL\_6343
CL\_6342
CL\_6641
CL\_6764
CL\_6765
CL\_6766
CL\_6767
CL\_18166
CL\_16922
CL\_12686
CL\_3171
CL\_3174
CL\_3175
CL\_5607
CL\_3172
CL\_3173
CL\_3176
CL\_3177
CL\_23409
CL\_10905
CL\_5606
CL\_5605
CL\_5604
CL\_5603
CL\_5602
CL\_12187
CL\_13853
CL\_18589
CL\_5601
CL\_5600
CL\_5599
CL\_6880
CL\_12267
CL\_6009
CL\_6011
CL\_6012
CL\_6013
CL\_6014
CL\_6015
CL\_6016
CL\_6017
CL\_6018
CL\_6019
CL\_6810
CL\_7063
CL\_7064
CL\_9944
CL\_6106
CL\_6105
CL\_6104
CL\_9205
CL\_8142
CL\_8143
CL\_9203
CL\_8213
CL\_9202
CL\_9201
CL\_8158
CL\_8157
CL\_8156
CL\_6080
CL\_16362
CL\_16361
CL\_16360
CL\_13749
CL\_13750
CL\_13751
CL\_6089
CL\_13772
CL\_6181
CL\_3387
CL\_3386
CL\_9218
CL\_9219
CL\_6186
CL\_10906
CL\_7940
CL\_6874
CL\_8209
CL\_8208
CL\_8207
CL\_7946
CL\_6806
CL\_12767
CL\_8219
CL\_12766
CL\_6879
CL\_7947
CL\_6885
CL\_12260
CL\_7948
CL\_6883
CL\_6882
CL\_7984
CL\_12261
CL\_6842
CL\_6843
CL\_6844
CL\_6845
CL\_6846
CL\_7595
CL\_8653
CL\_6041
CL\_6042
CL\_6043
CL\_6044
CL\_6848
CL\_6849
CL\_6850
CL\_6851
CL\_6852
CL\_7988
CL\_6854
CL\_12264
CL\_6060
CL\_16359
CL\_16358
CL\_16357
CL\_12266
CL\_6869
CL\_6870
CL\_6871
CL\_6872
CL\_5973
CL\_5974
CL\_5975
CL\_6860
CL\_6859
CL\_7997
CL\_5979
CL\_5980
CL\_5981
CL\_5982
CL\_5983
CL\_6829
CL\_5986
CL\_5987
CL\_5988
CL\_6828
CL\_6827
CL\_6826
CL\_5991
CL\_5992
CL\_5993
CL\_6825
CL\_5995
CL\_5996
CL\_5997
CL\_5998
CL\_5999
CL\_6000
CL\_6001
CL\_6002
CL\_6003
CL\_16356
CL\_6004
CL\_6005
CL\_6823
CL\_6868
CL\_101
CL\_14947
CL\_65
CL\_13967
CL\_8858
CL\_64
CL\_8050
CL\_14857
CL\_18092
CL\_8857
CL\_6156
CL\_8928
CL\_8927
CL\_8926
CL\_8925
CL\_8924
CL\_8923
CL\_8922
CL\_8921
CL\_8920
CL\_8918
CL\_8917
CL\_8916
CL\_8915
CL\_12168
CL\_7068
CL\_9405
CL\_6936
CL\_9401
CL\_9400
CL\_9399
CL\_9398
CL\_12169
CL\_13445
CL\_13446
CL\_121
CL\_6123
CL\_8898
CL\_10666
CL\_11879
CL\_16355
CL\_11877
CL\_10452
CL\_14943
CL\_14743
CL\_16354
CL\_8954
CL\_117
CL\_116
CL\_115
CL\_114
CL\_7770
CL\_13930
CL\_8955
CL\_113
CL\_112
CL\_111
CL\_5953
CL\_110
CL\_14952
CL\_14953
CL\_14954
CL\_14955
CL\_14956
CL\_14957
CL\_14958
CL\_14959
CL\_14960
CL\_14961
CL\_14962
CL\_14963
CL\_13929
CL\_109
CL\_108
CL\_107
CL\_105
CL\_104
CL\_13928
CL\_103
CL\_7536
CL\_16353
CL\_16352
CL\_16351
CL\_102
CL\_10436
CL\_10437
CL\_10438
CL\_10439
CL\_10440
CL\_10441
CL\_10442
CL\_10443
CL\_10444
CL\_10445
CL\_10446
CL\_10447
CL\_7092
CL\_16350
CL\_8061
CL\_5598
CL\_7689
CL\_23394
CL\_23393
CL\_23392
CL\_7688
CL\_9301
CL\_22820
CL\_10196
CL\_19616
CL\_19615
CL\_19614
CL\_7680
CL\_19613
CL\_19612
CL\_7683
CL\_22821
CL\_22822
CL\_13926
CL\_9517
CL\_90
CL\_78
CL\_76
CL\_5951
CL\_8874
CL\_10453
CL\_16349
CL\_16348
CL\_16347
CL\_7126
CL\_12156
CL\_12157
CL\_16346
CL\_16345
CL\_7096
CL\_7095
CL\_9191
CL\_8888
CL\_12158
CL\_8855
CL\_18084
CL\_8180
CL\_8181
CL\_18085
CL\_10807
CL\_10808
CL\_10809
CL\_8184
CL\_8185
CL\_8186
CL\_18086
CL\_7237
CL\_7236
CL\_10260
CL\_12559
CL\_7504
CL\_7505
CL\_7233
CL\_18087
CL\_18088
CL\_18089
CL\_18090
CL\_18091
CL\_9390
CL\_9391
CL\_9392
CL\_9393
CL\_9394
CL\_9395
CL\_9396
CL\_6157
CL\_120
CL\_119
CL\_118
CL\_131
CL\_132
CL\_7822
CL\_7785
CL\_18093
CL\_7971
CL\_18094
CL\_18095
CL\_6023
CL\_6024
CL\_18096
CL\_14746
CL\_14747
CL\_18097
CL\_18098
CL\_18099
CL\_18100
CL\_133
CL\_8086
CL\_16344
CL\_14948
CL\_5481
CL\_5480
CL\_19585
