## Supplementary material for "A novel method for integrating genomic and Tn-Seq data to identify common *in vivo* fitness mechanisms across multiple bacterial species": S2 Dataset: CL_INS_263.html

Legend

 Mobile +extrachromosomalelementfunctions
 Regulatoryfunctions
 Hypothetical
 DNA Metabolism
 Transcription
 All EssentialGenes
 AntibioticResistance
 Proteinsynthesis/fate
 Other
 Transport +binding proteins
 All VFDB Genes

FULL


WINDOWSVGPNG

Trim RowsRemove SingletonsSave Fasta

CL\_3186


CL\_3185


CL\_3186


CL\_3186


CL\_3186


CL\_3186


CL\_3186


CL\_3186


CL\_3186


CL\_3186


CL\_3186


CL\_3186


CL\_3186


CL\_3186


CL\_3186


CL\_3186


CL\_3186


CL\_3186


CL\_3186


CL\_3185


CL\_3186


CL\_3186

HighlightSelectShow Genomes


177

CL\_3187


2

CL\_3187


1

CL\_3188


1

CL\_3188


1

CL\_3188


1

CL\_3187


1

CL\_3188


1

CL\_3187


1

CL\_1403


1

CL\_3187


1

CL\_3189


1

CL\_3188


1

CL\_3188


1

CL\_3188


1

CL\_3188


1

CL\_3188


1

CL\_3187


1

CL\_3188


1

CL\_3188


1

CL\_3187


1

CL\_3188


1

CL\_3188

fGI ID


CL\_INS\_263
CL\_INS\_263
CL\_INS\_263
CL\_INS\_263
CL\_INS\_263
CL\_INS\_263
CL\_INS\_263
CL\_INS\_263
CL\_INS\_263
CL\_INS\_263
CL\_INS\_263
CL\_INS\_263
CL\_INS\_263
CL\_INS\_263
CL\_INS\_263
CL\_INS\_263
CL\_INS\_263
CL\_INS\_263
CL\_INS\_263
CL\_INS\_263
CL\_INS\_263
CL\_INS\_263
CL\_INS\_263
CL\_INS\_263
CL\_INS\_263
CL\_INS\_263
CL\_INS\_263
CL\_INS\_263
CL\_INS\_263
CL\_INS\_263
CL\_INS\_263
CL\_INS\_263
CL\_INS\_263
CL\_INS\_263
CL\_INS\_263
CL\_INS\_263
CL\_INS\_263
CL\_INS\_263
CL\_INS\_263
CL\_INS\_263
CL\_INS\_263
CL\_INS\_263
CL\_INS\_263
CL\_INS\_263
CL\_INS\_263
CL\_INS\_263
CL\_INS\_263
CL\_INS\_263
CL\_INS\_263
CL\_INS\_263
CL\_INS\_263
CL\_INS\_263
CL\_INS\_263
CL\_INS\_263
CL\_INS\_263
CL\_INS\_263
CL\_INS\_263
CL\_INS\_263
CL\_INS\_263
CL\_INS\_263
CL\_INS\_263
CL\_INS\_263
CL\_INS\_263
CL\_INS\_263
CL\_INS\_263
CL\_INS\_263
CL\_INS\_263
CL\_INS\_263
CL\_INS\_263
CL\_INS\_263
CL\_INS\_263
CL\_INS\_263
CL\_INS\_263
CL\_INS\_263
CL\_INS\_263
CL\_INS\_263
CL\_INS\_263
CL\_INS\_263
CL\_INS\_263
CL\_INS\_263
CL\_INS\_263
CL\_INS\_263
CL\_INS\_263
CL\_INS\_263
CL\_INS\_263
CL\_INS\_263
CL\_INS\_263
CL\_INS\_263
CL\_INS\_263
CL\_INS\_263
CL\_INS\_263
CL\_INS\_263
CL\_INS\_263
CL\_INS\_263
CL\_INS\_263
CL\_INS\_263
CL\_INS\_263
CL\_INS\_263
CL\_INS\_263
CL\_INS\_263
CL\_INS\_263
CL\_INS\_263
CL\_INS\_263
CL\_INS\_263
CL\_INS\_263
CL\_INS\_263
CL\_INS\_263
CL\_INS\_263
CL\_INS\_263
CL\_INS\_263
CL\_INS\_263
CL\_INS\_263
CL\_INS\_263
CL\_INS\_263
CL\_INS\_263
CL\_INS\_263
CL\_INS\_263
CL\_INS\_263
CL\_INS\_263
CL\_INS\_263
CL\_INS\_263
CL\_INS\_263
CL\_INS\_263
CL\_INS\_263
CL\_INS\_263
CL\_INS\_263
CL\_INS\_263
CL\_INS\_263
CL\_INS\_263
CL\_INS\_263
CL\_INS\_263
CL\_INS\_263
CL\_INS\_263
CL\_INS\_263
CL\_INS\_263
CL\_INS\_263
CL\_INS\_263
CL\_INS\_263
CL\_INS\_263
CL\_INS\_263
CL\_INS\_263
CL\_INS\_263
CL\_INS\_263
CL\_INS\_263
CL\_INS\_263
CL\_INS\_263
CL\_INS\_263
CL\_INS\_263
CL\_INS\_263
CL\_INS\_263
CL\_INS\_263
CL\_INS\_263
CL\_INS\_263
CL\_INS\_263
CL\_INS\_263
CL\_INS\_263
CL\_INS\_263
CL\_INS\_263
CL\_INS\_263
CL\_INS\_263
CL\_INS\_263
CL\_INS\_263
CL\_INS\_263
CL\_INS\_263
CL\_INS\_263
CL\_INS\_263
CL\_INS\_263
CL\_INS\_263
CL\_INS\_263
CL\_INS\_263
CL\_INS\_263
CL\_INS\_263
CL\_INS\_263
CL\_INS\_263
CL\_INS\_263
CL\_INS\_263
CL\_INS\_263
CL\_INS\_263
CL\_INS\_263
CL\_INS\_263
CL\_INS\_263
CL\_INS\_263
CL\_INS\_263
CL\_INS\_263
CL\_INS\_263
CL\_INS\_263
CL\_INS\_263
CL\_INS\_263
CL\_INS\_263
CL\_INS\_263
CL\_INS\_263
CL\_INS\_263
CL\_INS\_204
CL\_INS\_263
CL\_INS\_263
CL\_INS\_263
CL\_INS\_263
CL\_INS\_263
CL\_INS\_263
CL\_INS\_263
CL\_INS\_263
CL\_INS\_263
CL\_INS\_263
CL\_INS\_263
CL\_INS\_263
CL\_INS\_263
CL\_INS\_263
CL\_INS\_263
CL\_INS\_263
CL\_INS\_263
CL\_INS\_263
CL\_INS\_263
CL\_INS\_263
CL\_INS\_263
CL\_INS\_263
CL\_INS\_263
CL\_INS\_263
CL\_INS\_263
CL\_INS\_263
CL\_INS\_263
CL\_INS\_263
CL\_INS\_263
CL\_INS\_263
CL\_INS\_263
CL\_INS\_263
CL\_INS\_263
CL\_INS\_263
CL\_INS\_263
CL\_INS\_263
CL\_INS\_263
CL\_INS\_263
CL\_INS\_263
CL\_INS\_263
CL\_INS\_263
CL\_INS\_263
CL\_INS\_263
CL\_INS\_263
CL\_INS\_263
CL\_INS\_263
CL\_INS\_263
CL\_INS\_263
CL\_INS\_263
CL\_INS\_263
CL\_INS\_263
CL\_INS\_263
CL\_INS\_263
CL\_INS\_263
CL\_INS\_263
CL\_INS\_263
CL\_INS\_263
CL\_INS\_263
CL\_INS\_263
CL\_INS\_263
CL\_INS\_263
CL\_INS\_263
CL\_INS\_263
CL\_INS\_263
CL\_INS\_263
CL\_INS\_263
CL\_INS\_263
CL\_INS\_263
CL\_INS\_263
CL\_INS\_263
CL\_INS\_263
CL\_INS\_263
CL\_INS\_263
CL\_INS\_263
CL\_INS\_263
CL\_INS\_263
CL\_INS\_263
CL\_INS\_263
CL\_INS\_263
CL\_INS\_263
CL\_INS\_263
CL\_INS\_263
CL\_INS\_263
CL\_INS\_263
CL\_INS\_263
CL\_INS\_263
CL\_INS\_263
CL\_INS\_263
CL\_INS\_263
CL\_INS\_263
CL\_INS\_263
CL\_INS\_263
CL\_INS\_263
CL\_INS\_263
CL\_INS\_263
CL\_INS\_263
CL\_INS\_263
CL\_INS\_263
CL\_INS\_263
CL\_INS\_263
CL\_INS\_263
CL\_INS\_263
CL\_INS\_263
CL\_INS\_263
CL\_INS\_263
CL\_INS\_263
CL\_INS\_263
CL\_INS\_263
CL\_INS\_263
CL\_INS\_263
CL\_INS\_263
CL\_INS\_263
CL\_INS\_263
CL\_INS\_263
CL\_INS\_263
CL\_INS\_263
CL\_INS\_263
CL\_INS\_263
CL\_INS\_263
CL\_INS\_263
CL\_INS\_263
CL\_INS\_263
CL\_INS\_263
CL\_INS\_263
CL\_INS\_263
CL\_INS\_263
CL\_INS\_263
CL\_INS\_263
CL\_INS\_263
CL\_INS\_263
CL\_INS\_263
CL\_INS\_263
CL\_INS\_263
CL\_INS\_263
CL\_INS\_263
CL\_INS\_263
CL\_INS\_263
CL\_INS\_263
CL\_INS\_263
CL\_INS\_263
CL\_INS\_263
CL\_INS\_263
CL\_INS\_263
CL\_INS\_263
CL\_INS\_263
CL\_INS\_263
CL\_INS\_263
CL\_INS\_263
CL\_INS\_263
CL\_INS\_263
CL\_INS\_263
CL\_INS\_263
CL\_INS\_263
CL\_INS\_263
CL\_INS\_263
CL\_INS\_263
CL\_INS\_263
CL\_INS\_263
CL\_INS\_263
CL\_INS\_263
CL\_INS\_263
CL\_INS\_263
CL\_INS\_263
CL\_INS\_263
CL\_INS\_263
CL\_INS\_263
CL\_INS\_263
CL\_INS\_263
CL\_INS\_263
CL\_INS\_263
CL\_INS\_263
CL\_INS\_263
CL\_INS\_263
CL\_INS\_263
CL\_INS\_263
CL\_INS\_263
CL\_INS\_263
CL\_INS\_263
CL\_INS\_263
CL\_INS\_263
CL\_INS\_263
CL\_INS\_263
CL\_INS\_263
CL\_INS\_263
CL\_INS\_263
CL\_INS\_263
CL\_INS\_263
CL\_INS\_263
CL\_INS\_263
CL\_INS\_263
CL\_INS\_263
CL\_INS\_263
CL\_INS\_263
CL\_INS\_263
CL\_INS\_263
CL\_INS\_263
CL\_INS\_263
CL\_INS\_263
CL\_INS\_263
CL\_INS\_263
CL\_INS\_263
CL\_INS\_263
CL\_INS\_263
CL\_INS\_263
CL\_INS\_263
CL\_INS\_263
CL\_INS\_263
CL\_INS\_263
CL\_INS\_263
CL\_INS\_263
CL\_INS\_263
CL\_INS\_263
CL\_INS\_263
CL\_INS\_263
CL\_INS\_263
CL\_INS\_263
CL\_INS\_263
CL\_INS\_263
CL\_INS\_263
CL\_INS\_263
CL\_INS\_263
CL\_INS\_263
CL\_INS\_263
CL\_INS\_263
CL\_INS\_263
CL\_INS\_263
CL\_INS\_263
CL\_INS\_263
CL\_INS\_263
CL\_INS\_263
CL\_INS\_263
CL\_INS\_263
CL\_INS\_263
CL\_INS\_263
CL\_INS\_263
CL\_INS\_263
CL\_INS\_263
CL\_INS\_263
CL\_INS\_263
CL\_INS\_263
CL\_INS\_263
CL\_INS\_263
CL\_INS\_263
CL\_INS\_263
CL\_INS\_263
CL\_INS\_263
CL\_INS\_263
CL\_INS\_263
CL\_INS\_263
CL\_INS\_263
CL\_INS\_263
CL\_INS\_263
CL\_INS\_263
CL\_INS\_263
CL\_INS\_263
CL\_INS\_263
CL\_INS\_263
CL\_INS\_263
CL\_INS\_263
CL\_INS\_263
CL\_INS\_263
CL\_INS\_263
CL\_INS\_263
CL\_INS\_263
CL\_INS\_263
CL\_INS\_263
CL\_INS\_263
CL\_INS\_263
CL\_INS\_263
CL\_INS\_263
CL\_INS\_263
CL\_INS\_263
CL\_INS\_263
CL\_INS\_263
CL\_INS\_263
CL\_INS\_263
CL\_INS\_263
CL\_INS\_263
CL\_INS\_263
CL\_INS\_263
CL\_INS\_263
CL\_INS\_263
CL\_INS\_263
CL\_INS\_263
CL\_INS\_263
CL\_INS\_263
CL\_INS\_263
CL\_INS\_263
CL\_INS\_263
CL\_INS\_263
CL\_INS\_263
CL\_INS\_263
CL\_INS\_263
CL\_INS\_263
CL\_INS\_263
CL\_INS\_263
CL\_INS\_263
CL\_INS\_263
CL\_INS\_263
CL\_INS\_263
CL\_INS\_263
CL\_INS\_263
CL\_INS\_263
CL\_INS\_263
CL\_INS\_263
CL\_INS\_263
CL\_INS\_263
CL\_INS\_263
CL\_INS\_263
CL\_INS\_263
CL\_INS\_263
CL\_INS\_263
CL\_INS\_263
CL\_INS\_263
CL\_INS\_263
CL\_INS\_263
CL\_INS\_263
CL\_INS\_263
CL\_INS\_263
CL\_INS\_263
CL\_INS\_263
CL\_INS\_263
CL\_INS\_263
CL\_INS\_263
CL\_INS\_263
CL\_INS\_263
CL\_INS\_263
CL\_INS\_263
CL\_INS\_263
CL\_INS\_263
CL\_INS\_263
CL\_INS\_263
CL\_INS\_263
CL\_INS\_263
CL\_INS\_263
CL\_INS\_263
CL\_INS\_263
CL\_INS\_263
CL\_INS\_263
CL\_INS\_263
CL\_INS\_263
CL\_INS\_263
CL\_INS\_263
CL\_INS\_263
CL\_INS\_263
CL\_INS\_263
CL\_INS\_263
CL\_INS\_263
CL\_INS\_263
CL\_INS\_263
CL\_INS\_263
CL\_INS\_263
CL\_INS\_263
CL\_INS\_263
CL\_INS\_263
CL\_INS\_263
CL\_INS\_263
CL\_INS\_263
CL\_INS\_263
CL\_INS\_263
CL\_INS\_263
CL\_INS\_263
CL\_INS\_263
CL\_INS\_263
CL\_INS\_263
CL\_INS\_263
CL\_INS\_263
CL\_INS\_263
CL\_INS\_263
CL\_INS\_263
CL\_INS\_263
CL\_INS\_263
CL\_INS\_263
CL\_INS\_263
CL\_INS\_263
CL\_INS\_263
CL\_INS\_263
CL\_INS\_263
CL\_INS\_263
CL\_INS\_263
CL\_INS\_263
CL\_INS\_263
CL\_INS\_263
CL\_INS\_263
CL\_INS\_263
CL\_INS\_263
CL\_INS\_263
CL\_INS\_263
CL\_INS\_263
CL\_INS\_263
CL\_INS\_263
CL\_INS\_263
CL\_INS\_263
CL\_INS\_263
CL\_INS\_263
CL\_INS\_263
CL\_INS\_263
CL\_INS\_263
CL\_INS\_263
CL\_INS\_263
CL\_INS\_263
CL\_INS\_263
CL\_INS\_263
CL\_INS\_263
CL\_INS\_263
CL\_INS\_263
CL\_INS\_263
CL\_INS\_263
CL\_INS\_263
CL\_INS\_263
CL\_INS\_263
CL\_INS\_263
CL\_INS\_263
CL\_INS\_263
CL\_INS\_263
CL\_INS\_263
CL\_INS\_263
CL\_INS\_263
CL\_INS\_263
CL\_INS\_263
CL\_INS\_263
CL\_INS\_263
CL\_INS\_263
CL\_INS\_263
CL\_INS\_263
CL\_INS\_263
CL\_INS\_263
CL\_INS\_263
CL\_INS\_263
Cluster ID


CL\_6077
CL\_6076
CL\_13228
CL\_13229
CL\_6689
CL\_6688
CL\_14628
CL\_19586
CL\_19587
CL\_14625
CL\_17134
CL\_17133
CL\_17132
CL\_5479
CL\_14627
CL\_14626
CL\_17455
CL\_5181
CL\_7842
CL\_21572
CL\_21573
CL\_21574
CL\_12652
CL\_21447
CL\_21446
CL\_21445
CL\_21444
CL\_22239
CL\_8753
CL\_8752
CL\_8751
CL\_8750
CL\_8749
CL\_5182
CL\_12881
CL\_5183
CL\_22186
CL\_22185
CL\_22184
CL\_20420
CL\_7196
CL\_21443
CL\_7195
CL\_7841
CL\_7840
CL\_18806
CL\_8741
CL\_9719
CL\_9718
CL\_9717
CL\_8740
CL\_8739
CL\_8738
CL\_8737
CL\_8736
CL\_8735
CL\_8734
CL\_8733
CL\_8732
CL\_8731
CL\_8730
CL\_8729
CL\_8728
CL\_19358
CL\_12876
CL\_5213
CL\_5214
CL\_20975
CL\_21442
CL\_21441
CL\_8727
CL\_10407
CL\_19077
CL\_19589
CL\_6988
CL\_19590
CL\_19591
CL\_6556
CL\_23413
CL\_5184
CL\_5185
CL\_5186
CL\_5187
CL\_23463
CL\_7198
CL\_5188
CL\_5189
CL\_6228
CL\_6229
CL\_16342
CL\_16341
CL\_15258
CL\_15257
CL\_6230
CL\_17456
CL\_17457
CL\_5190
CL\_5191
CL\_23410
CL\_23411
CL\_23412
CL\_22043
CL\_8748
CL\_8747
CL\_8746
CL\_8745
CL\_8744
CL\_8743
CL\_8742
CL\_11623
CL\_11622
CL\_5192
CL\_5193
CL\_21202
CL\_6233
CL\_6234
CL\_6235
CL\_15628
CL\_22183
CL\_22182
CL\_22181
CL\_22180
CL\_5199
CL\_5200
CL\_5201
CL\_17458
CL\_5202
CL\_5203
CL\_5204
CL\_19712
CL\_19711
CL\_19710
CL\_5205
CL\_5206
CL\_5207
CL\_5208
CL\_23462
CL\_7189
CL\_7188
CL\_7187
CL\_7186
CL\_15629
CL\_15630
CL\_17459
CL\_5210
CL\_5211
CL\_5212
CL\_6237
CL\_11663
CL\_6239
CL\_5217
CL\_5218
CL\_5219
CL\_18458
CL\_8726
CL\_22179
CL\_6727
CL\_6726
CL\_8725
CL\_8724
CL\_12540
CL\_6238
CL\_7837
CL\_14559
CL\_14560
CL\_11021
CL\_5478
CL\_22305
CL\_22306
CL\_22307
CL\_8766
CL\_8767
CL\_8768
CL\_8769
CL\_8770
CL\_22308
CL\_22309
CL\_22310
CL\_22311
CL\_22312
CL\_22313
CL\_22314
CL\_22315
CL\_22316
CL\_22317
CL\_22318
CL\_22319
CL\_22320
CL\_22321
CL\_16343
CL\_5477
CL\_5476
CL\_5475
CL\_5474
CL\_5473
CL\_5472
CL\_17131
CL\_16071
CL\_17130
CL\_17129
CL\_17128
CL\_17127
CL\_17126
CL\_5471
CL\_14624
CL\_5470
CL\_20977
CL\_14623
CL\_5469
CL\_15611
CL\_5468
CL\_5467
CL\_5466
CL\_5465
CL\_5464
CL\_5463
CL\_5462
CL\_5461
CL\_17125
CL\_5460
CL\_5459
CL\_17124
CL\_5458
CL\_5457
CL\_5456
CL\_17123
CL\_17122
CL\_5455
CL\_5454
CL\_15610
CL\_15609
CL\_15608
CL\_15607
CL\_15606
CL\_15605
CL\_15604
CL\_14821
CL\_10179
CL\_15603
CL\_15602
CL\_15601
CL\_15600
CL\_15599
CL\_5453
CL\_17121
CL\_5452
CL\_5451
CL\_5450
CL\_17120
CL\_17119
CL\_6801
CL\_17118
CL\_17117
CL\_17116
CL\_17115
CL\_17114
CL\_5449
CL\_19588
CL\_20976
CL\_5448
CL\_5447
CL\_14622
CL\_5446
CL\_5445
CL\_17423
CL\_17422
CL\_8696
CL\_17421
CL\_17420
CL\_17419
CL\_17418
CL\_17417
CL\_17416
CL\_17415
CL\_17414
CL\_17413
CL\_17412
CL\_17411
CL\_17410
CL\_17409
CL\_17408
CL\_17407
CL\_17406
CL\_17405
CL\_17404
CL\_17403
CL\_17402
CL\_17401
CL\_17400
CL\_17399
CL\_17398
CL\_17397
CL\_17396
CL\_17395
CL\_17394
CL\_17393
CL\_17392
CL\_17391
CL\_17390
CL\_17389
CL\_17388
CL\_17387
CL\_17386
CL\_17385
CL\_17384
CL\_17383
CL\_17382
CL\_17381
CL\_17380
CL\_9920
CL\_9921
CL\_5385
CL\_5664
CL\_7287
CL\_7286
CL\_7285
CL\_7284
CL\_7283
CL\_7282
CL\_7281
CL\_13230
CL\_13231
CL\_13232
CL\_5384
CL\_5383
CL\_5382
CL\_5662
CL\_9640
CL\_5660
CL\_5378
CL\_5377
CL\_5376
CL\_5375
CL\_5374
CL\_5373
CL\_7277
CL\_8772
CL\_13233
CL\_5659
CL\_13234
CL\_13235
CL\_13236
CL\_13237
CL\_13238
CL\_13239
CL\_13240
CL\_13241
CL\_13242
CL\_13243
CL\_9111
CL\_22322
CL\_22323
CL\_22324
CL\_22325
CL\_22326
CL\_22327
CL\_13244
CL\_9091
CL\_13245
CL\_13246
CL\_13247
CL\_13248
CL\_13249
CL\_13250
CL\_13251
CL\_13252
CL\_13253
CL\_9096
CL\_13254
CL\_13255
CL\_13256
CL\_13257
CL\_13258
CL\_13259
CL\_13260
CL\_9621
CL\_13261
CL\_13262
CL\_8779
CL\_6591
CL\_6590
CL\_6589
CL\_6588
CL\_13846
CL\_13845
CL\_13844
CL\_13843
CL\_7273
CL\_13842
CL\_17087
CL\_17088
CL\_7504
CL\_6243
CL\_5444
CL\_5651
CL\_17951
CL\_22178
CL\_22177
CL\_22176
CL\_6241
CL\_18526
CL\_18525
CL\_5221
CL\_11620
CL\_5222
CL\_5223
CL\_17113
CL\_21440
CL\_5225
CL\_21439
CL\_5643
CL\_13188
CL\_5224
CL\_5438
CL\_5437
CL\_5436
CL\_12645
CL\_5435
CL\_20098
CL\_23414
CL\_23415
CL\_23416
CL\_23417
CL\_23418
CL\_5237
CL\_21999
CL\_7834
CL\_19592
CL\_19593
CL\_19594
CL\_17112
CL\_17111
CL\_17110
CL\_17109
CL\_17108
CL\_5226
CL\_17460
CL\_17461
CL\_17462
CL\_12538
CL\_5639
CL\_5640
CL\_5641
CL\_5341
CL\_5434
CL\_5433
CL\_5432
CL\_5431
CL\_5236
CL\_5430
CL\_6246
CL\_6247
CL\_6248
CL\_9708
CL\_5443
CL\_5442
CL\_15598
CL\_5441
CL\_11577
CL\_11578
CL\_22175
CL\_22174
CL\_6992
CL\_22173
CL\_19229
CL\_15872
CL\_21834
CL\_22172
CL\_21417
CL\_8721
CL\_17463
CL\_8315
CL\_5644
CL\_11614
CL\_15597
CL\_5440
CL\_15596
CL\_5439
CL\_15595
CL\_15594
CL\_15593
CL\_15592
CL\_15591
CL\_5232
CL\_5233
CL\_11020
CL\_11019
CL\_5234
CL\_5235
CL\_15590
CL\_6559
CL\_6560
CL\_6561
CL\_8314
CL\_6250
CL\_23419
CL\_23420
CL\_5426
CL\_5244
CL\_22171
CL\_11611
CL\_11610
CL\_15589
CL\_11612
CL\_23357
CL\_5425
CL\_8717
CL\_5245
CL\_12643
CL\_5247
CL\_7179
CL\_8716
CL\_8715
CL\_8714
CL\_8713
CL\_23421
CL\_16493
CL\_5424
CL\_11580
CL\_11581
CL\_8712
CL\_5423
CL\_23422
CL\_5422
CL\_5421
CL\_6564
CL\_15970
CL\_6254
CL\_9703
CL\_9702
CL\_6255
CL\_6256
CL\_8711
CL\_21436
CL\_5420
CL\_5248
CL\_16492
CL\_5250
CL\_21437
CL\_23423
CL\_23424
CL\_5419
CL\_17379
CL\_14970
CL\_2097
CL\_12539
CL\_5418
CL\_14621
CL\_22328
CL\_8332
CL\_13818
CL\_15169
CL\_20240
CL\_6285
CL\_22329
CL\_8317
CL\_7184
CL\_8316
CL\_22330
CL\_22331
CL\_17378
CL\_17377
CL\_5231
CL\_12535
CL\_12534
CL\_12533
CL\_22332
CL\_22333
CL\_13816
CL\_12532
CL\_12531
CL\_14620
CL\_8335
CL\_13815
CL\_22334
CL\_22335
CL\_8336
CL\_14619
CL\_12866
CL\_13814
CL\_13813
CL\_5429
CL\_5428
CL\_19363
CL\_19364
CL\_5427
CL\_21438
CL\_23358
CL\_5238
CL\_5239
CL\_19595
CL\_6251
CL\_19596
CL\_19597
CL\_19598
CL\_19599
CL\_19600
CL\_19601
CL\_19602
CL\_19603
CL\_19604
CL\_14616
CL\_7831
CL\_21631
CL\_7830
CL\_5242
CL\_17376
CL\_17375
CL\_12530
CL\_14618
CL\_12529
CL\_12528
CL\_7265
CL\_14617
CL\_5417
CL\_5416
CL\_5415
CL\_17464
CL\_5414
CL\_5413
CL\_5412
CL\_5411
