## Supplementary material for "A novel method for integrating genomic and Tn-Seq data to identify common *in vivo* fitness mechanisms across multiple bacterial species": S2 Dataset: CL_INS_264.html

Legend

 Mobile +extrachromosomalelementfunctions
 Regulatoryfunctions
 Hypothetical
 DNA Metabolism
 AntibioticResistance
 All EssentialGenes
 Proteinsynthesis/fate
 Other
 Transport +binding proteins
 All VFDB Genes

FULL


WINDOWSVGPNG

Trim RowsRemove SingletonsSave Fasta

CL\_3188


CL\_3188


CL\_3188


CL\_3188


CL\_3188


CL\_3188


CL\_3188


CL\_3188


CL\_3188


CL\_3188


CL\_3188


CL\_3188


CL\_3188


CL\_3188


CL\_3188

HighlightSelectShow Genomes


183

CL\_3187


1

CL\_3186


1

CL\_1404


1

CL\_3186


1

CL\_3186


1

CL\_3186


1

CL\_3186


1

CL\_3186


1

CL\_3186


1

CL\_3186


1

CL\_3186


1

CL\_3186


1

CL\_3186


1

CL\_3186


1

CL\_3186

fGI ID


CL\_INS\_263
CL\_INS\_263
CL\_INS\_263
CL\_INS\_263
CL\_INS\_263
CL\_INS\_263
CL\_INS\_263
CL\_INS\_263
CL\_INS\_263
CL\_INS\_263
CL\_INS\_263
CL\_INS\_263
CL\_INS\_263
CL\_INS\_263
CL\_INS\_263
CL\_INS\_263
CL\_INS\_263
CL\_INS\_263
CL\_INS\_263
CL\_INS\_263
CL\_INS\_263
CL\_INS\_263
CL\_INS\_263
CL\_INS\_263
CL\_INS\_263
CL\_INS\_263
CL\_INS\_263
CL\_INS\_263
CL\_INS\_263
CL\_INS\_263
CL\_INS\_263
CL\_INS\_263
CL\_INS\_263
CL\_INS\_263
CL\_INS\_263
CL\_INS\_263
CL\_INS\_263
CL\_INS\_263
CL\_INS\_263
CL\_INS\_263
CL\_INS\_263
CL\_INS\_263
CL\_INS\_263
CL\_INS\_263
CL\_INS\_263
CL\_INS\_263
CL\_INS\_263
CL\_INS\_263
CL\_INS\_263
CL\_INS\_263
CL\_INS\_91
CL\_INS\_91
CL\_INS\_263
CL\_INS\_263
CL\_INS\_263
CL\_INS\_263
CL\_INS\_263
CL\_INS\_263
CL\_INS\_263
CL\_INS\_263
CL\_INS\_263
CL\_INS\_263
CL\_INS\_263
CL\_INS\_263
CL\_INS\_263
CL\_INS\_263
CL\_INS\_263
CL\_INS\_263
CL\_INS\_263
CL\_INS\_263
CL\_INS\_263
CL\_INS\_263
CL\_INS\_263
CL\_INS\_263
CL\_INS\_263
CL\_INS\_263
CL\_INS\_263
CL\_INS\_263
CL\_INS\_263
CL\_INS\_263
CL\_INS\_263
CL\_INS\_263
CL\_INS\_263
CL\_INS\_263
CL\_INS\_263
CL\_INS\_263
CL\_INS\_263
CL\_INS\_263
CL\_INS\_263
CL\_INS\_263
CL\_INS\_263
CL\_INS\_263
CL\_INS\_263
CL\_INS\_263
CL\_INS\_263
CL\_INS\_91
CL\_INS\_91
CL\_INS\_263
CL\_INS\_263
CL\_INS\_263
CL\_INS\_263
CL\_INS\_263
CL\_INS\_263
CL\_INS\_263
CL\_INS\_263
CL\_INS\_263
CL\_INS\_263
CL\_INS\_263
CL\_INS\_263
CL\_INS\_263
CL\_INS\_263
CL\_INS\_263
CL\_INS\_263
CL\_INS\_263
CL\_INS\_263
CL\_INS\_263
CL\_INS\_263
CL\_INS\_263
CL\_INS\_263
CL\_INS\_263
CL\_INS\_263
CL\_INS\_263
CL\_INS\_263
CL\_INS\_263
CL\_INS\_263
CL\_INS\_271
CL\_INS\_271
CL\_INS\_91
CL\_INS\_91
CL\_INS\_91
CL\_INS\_91
CL\_INS\_91
CL\_INS\_263
CL\_INS\_263
CL\_INS\_263
CL\_INS\_263
CL\_INS\_263
CL\_INS\_263
CL\_INS\_263
CL\_INS\_263
CL\_INS\_263
CL\_INS\_263
CL\_INS\_263
CL\_INS\_263
CL\_INS\_263
CL\_INS\_263
CL\_INS\_263
CL\_INS\_263
CL\_INS\_263
CL\_INS\_91
CL\_INS\_307
CL\_INS\_91
CL\_INS\_91
CL\_INS\_91
CL\_INS\_91
CL\_INS\_91
CL\_INS\_91
CL\_INS\_204
CL\_INS\_263
CL\_INS\_263
CL\_INS\_263
CL\_INS\_263
CL\_INS\_263
CL\_INS\_263
CL\_INS\_263
CL\_INS\_263
CL\_INS\_263
CL\_INS\_263
CL\_INS\_263
CL\_INS\_263
CL\_INS\_263
CL\_INS\_263
CL\_INS\_263
CL\_INS\_263
CL\_INS\_263
CL\_INS\_263
CL\_INS\_263
CL\_INS\_263
CL\_INS\_263
CL\_INS\_263
CL\_INS\_263
CL\_INS\_263
CL\_INS\_263
CL\_INS\_263
CL\_INS\_263
CL\_INS\_263
CL\_INS\_263
CL\_INS\_263
CL\_INS\_263
CL\_INS\_263
CL\_INS\_263
CL\_INS\_263
CL\_INS\_263
CL\_INS\_263
CL\_INS\_263
CL\_INS\_263
CL\_INS\_263
CL\_INS\_263
CL\_INS\_263
CL\_INS\_263
CL\_INS\_263
CL\_INS\_263
CL\_INS\_263
CL\_INS\_263
CL\_INS\_263
CL\_INS\_263
CL\_INS\_263
CL\_INS\_263
CL\_INS\_263
CL\_INS\_263
CL\_INS\_263
CL\_INS\_263
CL\_INS\_263
CL\_INS\_263
CL\_INS\_263
CL\_INS\_263
CL\_INS\_263
CL\_INS\_263
CL\_INS\_263
CL\_INS\_263
CL\_INS\_263
CL\_INS\_263
CL\_INS\_263
CL\_INS\_263
CL\_INS\_263
CL\_INS\_263
CL\_INS\_263
CL\_INS\_263
CL\_INS\_263
CL\_INS\_263
CL\_INS\_263
CL\_INS\_263
CL\_INS\_263
CL\_INS\_263
CL\_INS\_263
CL\_INS\_263
CL\_INS\_263
CL\_INS\_263
CL\_INS\_263
CL\_INS\_263
CL\_INS\_263
CL\_INS\_263
CL\_INS\_263
CL\_INS\_263
CL\_INS\_263
CL\_INS\_263
CL\_INS\_263
CL\_INS\_263
CL\_INS\_263
CL\_INS\_263
CL\_INS\_263
CL\_INS\_263
CL\_INS\_263
CL\_INS\_263
CL\_INS\_263
CL\_INS\_263
CL\_INS\_263
CL\_INS\_263
CL\_INS\_263
CL\_INS\_263
CL\_INS\_263
CL\_INS\_263
CL\_INS\_263
CL\_INS\_263
CL\_INS\_263
CL\_INS\_263
CL\_INS\_263
CL\_INS\_263
CL\_INS\_263
CL\_INS\_263
CL\_INS\_263
CL\_INS\_263
CL\_INS\_263
CL\_INS\_263
CL\_INS\_263
CL\_INS\_263
CL\_INS\_263
CL\_INS\_263
CL\_INS\_263
CL\_INS\_263
CL\_INS\_263
CL\_INS\_263
CL\_INS\_263
CL\_INS\_263
CL\_INS\_263
CL\_INS\_263
CL\_INS\_263
CL\_INS\_263
CL\_INS\_263
CL\_INS\_263
CL\_INS\_263
CL\_INS\_263
CL\_INS\_263
CL\_INS\_263
CL\_INS\_263
CL\_INS\_263
CL\_INS\_263
CL\_INS\_263
CL\_INS\_263
CL\_INS\_263
CL\_INS\_263
CL\_INS\_263
CL\_INS\_263
CL\_INS\_263
CL\_INS\_263
CL\_INS\_263
CL\_INS\_263
CL\_INS\_263
CL\_INS\_263
CL\_INS\_263
CL\_INS\_263
CL\_INS\_263
CL\_INS\_263
CL\_INS\_263
CL\_INS\_263
CL\_INS\_263
CL\_INS\_263
CL\_INS\_263
CL\_INS\_263
CL\_INS\_263
CL\_INS\_263
CL\_INS\_263
CL\_INS\_263
CL\_INS\_263
CL\_INS\_263
CL\_INS\_263
CL\_INS\_263
CL\_INS\_91
CL\_INS\_263
CL\_INS\_263
CL\_INS\_263
CL\_INS\_91
CL\_INS\_263
CL\_INS\_263
CL\_INS\_263
CL\_INS\_263
CL\_INS\_263
CL\_INS\_263
CL\_INS\_263
CL\_INS\_263
CL\_INS\_263
CL\_INS\_263
CL\_INS\_263
CL\_INS\_263
CL\_INS\_263
CL\_INS\_263
CL\_INS\_263
CL\_INS\_263
CL\_INS\_263
CL\_INS\_91
CL\_INS\_91
CL\_INS\_263
CL\_INS\_263
CL\_INS\_263
CL\_INS\_91
CL\_INS\_263
CL\_INS\_263
CL\_INS\_91
CL\_INS\_91
CL\_INS\_91
CL\_INS\_91
CL\_INS\_263
CL\_INS\_263
CL\_INS\_263
CL\_INS\_263
CL\_INS\_263
CL\_INS\_263
CL\_INS\_263
CL\_INS\_263
CL\_INS\_263
CL\_INS\_263
CL\_INS\_263
CL\_INS\_263
CL\_INS\_263
CL\_INS\_307
CL\_INS\_263
CL\_INS\_263
CL\_INS\_263
CL\_INS\_263
CL\_INS\_263
CL\_INS\_263
CL\_INS\_91
CL\_INS\_91
CL\_INS\_91
CL\_INS\_263
CL\_INS\_263
CL\_INS\_263
CL\_INS\_91
CL\_INS\_263
CL\_INS\_263
CL\_INS\_263
CL\_INS\_263
CL\_INS\_263
CL\_INS\_204
CL\_INS\_263
CL\_INS\_263
CL\_INS\_263
CL\_INS\_263
CL\_INS\_263
CL\_INS\_263
CL\_INS\_263
CL\_INS\_263
CL\_INS\_263
CL\_INS\_263
CL\_INS\_263
CL\_INS\_263
CL\_INS\_263
CL\_INS\_263
CL\_INS\_263
CL\_INS\_263
CL\_INS\_263
CL\_INS\_263
CL\_INS\_263
CL\_INS\_263
CL\_INS\_263
CL\_INS\_263
CL\_INS\_263
CL\_INS\_263
CL\_INS\_263
CL\_INS\_263
CL\_INS\_263
CL\_INS\_263
CL\_INS\_263
CL\_INS\_263
CL\_INS\_263
CL\_INS\_263
CL\_INS\_263
CL\_INS\_263
CL\_INS\_263
CL\_INS\_263
CL\_INS\_263
CL\_INS\_263
CL\_INS\_263
CL\_INS\_263
CL\_INS\_263
CL\_INS\_263
CL\_INS\_263
CL\_INS\_263
CL\_INS\_263
CL\_INS\_263
CL\_INS\_263
CL\_INS\_263
CL\_INS\_263
CL\_INS\_263
CL\_INS\_263
CL\_INS\_263
CL\_INS\_263
CL\_INS\_263
CL\_INS\_263
CL\_INS\_263
CL\_INS\_263
CL\_INS\_263
CL\_INS\_263
CL\_INS\_263
CL\_INS\_263
CL\_INS\_263
CL\_INS\_263
CL\_INS\_263
CL\_INS\_263
CL\_INS\_263
CL\_INS\_263
CL\_INS\_263
CL\_INS\_263
CL\_INS\_263
CL\_INS\_263
CL\_INS\_263
CL\_INS\_263
CL\_INS\_263
Cluster ID


CL\_5411
CL\_5412
CL\_14616
CL\_14617
CL\_7265
CL\_5413
CL\_17464
CL\_21436
CL\_5414
CL\_5415
CL\_5416
CL\_5417
CL\_8711
CL\_6256
CL\_6255
CL\_5248
CL\_6254
CL\_21437
CL\_8712
CL\_8713
CL\_8714
CL\_8715
CL\_8716
CL\_8717
CL\_11581
CL\_11580
CL\_12528
CL\_12529
CL\_14618
CL\_12530
CL\_22335
CL\_22334
CL\_13815
CL\_13813
CL\_13814
CL\_12866
CL\_14619
CL\_8336
CL\_8335
CL\_13816
CL\_22333
CL\_22332
CL\_14620
CL\_17375
CL\_17376
CL\_5238
CL\_5237
CL\_11019
CL\_11020
CL\_5429
CL\_17198
CL\_17197
CL\_5236
CL\_5235
CL\_5234
CL\_5430
CL\_5431
CL\_5432
CL\_5433
CL\_5434
CL\_12645
CL\_5435
CL\_5436
CL\_5437
CL\_5438
CL\_12531
CL\_12532
CL\_12533
CL\_12534
CL\_12535
CL\_20240
CL\_15169
CL\_22331
CL\_22330
CL\_8316
CL\_7184
CL\_8317
CL\_22329
CL\_6285
CL\_13188
CL\_5231
CL\_5643
CL\_13818
CL\_12539
CL\_8332
CL\_22328
CL\_21439
CL\_5225
CL\_21440
CL\_17377
CL\_17378
CL\_14621
CL\_5418
CL\_15970
CL\_6564
CL\_17200
CL\_17199
CL\_5419
CL\_5420
CL\_5421
CL\_5422
CL\_5423
CL\_5245
CL\_16493
CL\_5424
CL\_5244
CL\_5425
CL\_5426
CL\_21438
CL\_5427
CL\_5428
CL\_7179
CL\_19364
CL\_19363
CL\_5242
CL\_21631
CL\_7831
CL\_5239
CL\_11612
CL\_15589
CL\_6561
CL\_6560
CL\_6559
CL\_6246
CL\_5233
CL\_15434
CL\_5343
CL\_17196
CL\_17195
CL\_17194
CL\_17193
CL\_17192
CL\_21834
CL\_15872
CL\_19229
CL\_22173
CL\_6992
CL\_15590
CL\_5232
CL\_8315
CL\_17463
CL\_8721
CL\_17462
CL\_17461
CL\_17460
CL\_5226
CL\_7834
CL\_5224
CL\_6556
CL\_6555
CL\_9714
CL\_17191
CL\_17190
CL\_17189
CL\_17188
CL\_17187
CL\_17186
CL\_8086
CL\_11578
CL\_11577
CL\_5223
CL\_5222
CL\_5221
CL\_5651
CL\_17951
CL\_8724
CL\_8725
CL\_6726
CL\_14622
CL\_21441
CL\_21442
CL\_5214
CL\_5213
CL\_20976
CL\_19588
CL\_5212
CL\_5211
CL\_8726
CL\_6239
CL\_11663
CL\_6238
CL\_6237
CL\_15630
CL\_15629
CL\_5210
CL\_23462
CL\_7186
CL\_7187
CL\_7188
CL\_7189
CL\_17459
CL\_5208
CL\_19710
CL\_19711
CL\_19712
CL\_5207
CL\_5206
CL\_5205
CL\_5204
CL\_5203
CL\_5202
CL\_17458
CL\_5201
CL\_15628
CL\_5200
CL\_5199
CL\_6235
CL\_8741
CL\_18806
CL\_7840
CL\_7841
CL\_7195
CL\_9717
CL\_9718
CL\_9719
CL\_21443
CL\_7196
CL\_5188
CL\_5187
CL\_23463
CL\_6234
CL\_6233
CL\_17457
CL\_17456
CL\_6230
CL\_6229
CL\_6228
CL\_5189
CL\_5186
CL\_5185
CL\_5184
CL\_5183
CL\_5182
CL\_21444
CL\_21445
CL\_21446
CL\_21447
CL\_5181
CL\_17455
CL\_15591
CL\_15592
CL\_15593
CL\_15594
CL\_15595
CL\_5439
CL\_15596
CL\_5440
CL\_15597
CL\_5441
CL\_6988
CL\_5442
CL\_15598
CL\_5443
CL\_2097
CL\_12540
CL\_18458
CL\_5219
CL\_5218
CL\_14970
CL\_22327
CL\_22326
CL\_22325
CL\_22324
CL\_22323
CL\_22322
CL\_9111
CL\_22321
CL\_22320
CL\_22319
CL\_22318
CL\_22317
CL\_22316
CL\_22315
CL\_22314
CL\_22313
CL\_22312
CL\_22311
CL\_22310
CL\_22309
CL\_22308
CL\_8770
CL\_8769
CL\_8768
CL\_8767
CL\_8766
CL\_22307
CL\_22306
CL\_22305
CL\_17379
CL\_5444
CL\_5217
CL\_20975
CL\_5445
CL\_12876
CL\_19358
CL\_8728
CL\_8729
CL\_8730
CL\_8731
CL\_8732
CL\_8733
CL\_8734
CL\_8735
CL\_8736
CL\_8737
CL\_8738
CL\_8739
CL\_8740
CL\_11622
CL\_11623
CL\_8742
CL\_8744
CL\_8745
CL\_8746
CL\_8747
CL\_8748
CL\_8750
CL\_8751
CL\_8752
CL\_12652
CL\_21574
CL\_21573
CL\_21572
CL\_5446
CL\_5447
CL\_5448
CL\_5449
CL\_17185
CL\_5450
CL\_5451
CL\_5452
CL\_17184
CL\_5453
CL\_5454
CL\_15599
CL\_15600
CL\_15601
CL\_15602
CL\_15603
CL\_10179
CL\_14821
CL\_15604
CL\_15605
CL\_15606
CL\_15607
CL\_15608
CL\_15609
CL\_15610
CL\_5455
CL\_17183
CL\_17182
CL\_5456
CL\_5457
CL\_5458
CL\_17181
CL\_5459
CL\_5460
CL\_17180
CL\_17179
CL\_17178
CL\_17177
CL\_5461
CL\_5462
CL\_5463
CL\_5464
CL\_5465
CL\_5466
CL\_15257
CL\_15258
CL\_16341
CL\_16342
CL\_5467
CL\_5468
CL\_20977
CL\_17176
CL\_14623
CL\_5469
CL\_15611
CL\_5470
CL\_14624
CL\_5471
CL\_17175
CL\_17174
CL\_17173
CL\_5472
CL\_5473
CL\_14625
CL\_17172
CL\_14626
CL\_14627
CL\_5479
CL\_5474
CL\_7842
CL\_5475
CL\_5476
CL\_5477
CL\_16343
CL\_5478
CL\_6243
CL\_7504
CL\_17088
CL\_17087
CL\_13842
CL\_7273
CL\_13843
CL\_13844
CL\_13845
CL\_13846
CL\_7277
CL\_5373
CL\_5374
CL\_5375
CL\_5376
CL\_5377
CL\_5378
CL\_5660
CL\_9640
CL\_5662
CL\_5382
CL\_5383
CL\_5384
CL\_5385
CL\_9921
CL\_9920
CL\_17380
CL\_17381
CL\_17382
CL\_17383
CL\_17384
CL\_17385
CL\_17386
CL\_17387
CL\_17388
CL\_17389
CL\_17390
CL\_17391
CL\_17392
CL\_17393
CL\_17394
CL\_17395
CL\_17396
CL\_17397
CL\_17398
CL\_17399
CL\_17400
CL\_17401
CL\_17402
CL\_17403
CL\_17404
CL\_17405
CL\_17406
CL\_17407
CL\_17408
CL\_17409
CL\_17410
CL\_17411
CL\_17412
CL\_17413
CL\_17414
CL\_17415
CL\_17416
CL\_17417
CL\_17418
CL\_17419
CL\_17420
CL\_17421
CL\_8696
CL\_17422
