## Supplementary material for "A novel method for integrating genomic and Tn-Seq data to identify common *in vivo* fitness mechanisms across multiple bacterial species": S2 Dataset: CL_INS_265.html

Legend

 Hypothetical
 Other

FULL


WINDOWSVGPNG

Trim RowsRemove SingletonsSave Fasta

CL\_3198


CL\_3198


CL\_3197


CL\_3198

HighlightSelectShow Genomes


107

CL\_3200


82

CL\_3200


2

CL\_3200


1

CL\_3201

fGI ID

CL\_INS\_265
Cluster ID

CL\_3199
