## Supplementary material for "A novel method for integrating genomic and Tn-Seq data to identify common *in vivo* fitness mechanisms across multiple bacterial species": S2 Dataset: CL_INS_267.html

Legend

 Hypothetical
 Other
 All VFDB Genes

FULL


WINDOWSVGPNG

Trim RowsRemove SingletonsSave Fasta

CL\_3207


CL\_3207


CL\_3207


CL\_3207


CL\_3207


CL\_3166


CL\_3207


CL\_3168

HighlightSelectShow Genomes


141

CL\_3209


51

CL\_3209


3

CL\_3209


2

CL\_3209


1

CL\_3209


1

CL\_3209


1

CL\_3209


1

CL\_3209

fGI ID


CL\_INS\_267
CL\_INS\_267
CL\_INS\_267
CL\_INS\_267
CL\_INS\_267
CL\_INS\_267
CL\_INS\_267
CL\_INS\_267
Cluster ID


CL\_16468
CL\_10506
CL\_21711
CL\_23250
CL\_7311
CL\_7313
CL\_3208
CL\_20412
