## Supplementary material for "A novel method for integrating genomic and Tn-Seq data to identify common *in vivo* fitness mechanisms across multiple bacterial species": S2 Dataset: CL_INS_268.html

Legend

 Regulatoryfunctions
 Hypothetical
 All VFDB Genes

FULL


WINDOWSVGPNG

Trim RowsRemove SingletonsSave Fasta

CL\_3220


CL\_3220


CL\_3220

HighlightSelectShow Genomes


123

CL\_3222


80

CL\_3222


1

CL\_3223

fGI ID

CL\_INS\_268
Cluster ID

CL\_3221
