## Supplementary material for "A novel method for integrating genomic and Tn-Seq data to identify common *in vivo* fitness mechanisms across multiple bacterial species": S2 Dataset: CL_INS_269.html

Legend

 Mobile +extrachromosomalelementfunctions
 Hypothetical
 EnergyMetabolism
 All VFDB Genes

FULL


WINDOWSVGPNG

Trim RowsRemove SingletonsSave Fasta

CL\_3242


CL\_3242


CL\_3242

HighlightSelectShow Genomes


112

CL\_3244


87

CL\_3244


1

CL\_3244

fGI ID


CL\_INS\_269
CL\_INS\_132
CL\_INS\_269
Cluster ID


CL\_3243
CL\_6359
CL\_21367
