## Supplementary material for "A novel method for integrating genomic and Tn-Seq data to identify common *in vivo* fitness mechanisms across multiple bacterial species": S2 Dataset: CL_INS_271.html

Legend

 Mobile +extrachromosomalelementfunctions
 Regulatoryfunctions
 Hypothetical
 DNA Metabolism
 All EssentialGenes
 Cell Envelope
 Other
 All VFDB Genes

FULL


WINDOWSVGPNG

Trim RowsRemove SingletonsSave Fasta

CL\_3272


CL\_3272


CL\_3271


CL\_3272


CL\_3272


CL\_3272


CL\_3254


CL\_3272


CL\_3272


CL\_3272


CL\_3272


CL\_3272


CL\_3271

HighlightSelectShow Genomes


145

CL\_3273


35

CL\_3273


11

CL\_3273


2

CL\_3273


1

CL\_3273


1

CL\_3273


1

CL\_3273


1

CL\_3273


1

CL\_3273


1

CL\_3273


1

CL\_3273


1

CL\_3273


1

CL\_3273

fGI ID


CL\_INS\_271
CL\_INS\_271
CL\_INS\_271
CL\_INS\_271
CL\_INS\_271
CL\_INS\_270
CL\_INS\_271
CL\_INS\_270
CL\_INS\_263
CL\_INS\_263
CL\_INS\_271
CL\_INS\_263
CL\_INS\_263
CL\_INS\_263
CL\_INS\_263
CL\_INS\_263
CL\_INS\_271
CL\_INS\_271
CL\_INS\_271
CL\_INS\_271
CL\_INS\_263
CL\_INS\_263
CL\_INS\_263
CL\_INS\_263
CL\_INS\_263
CL\_INS\_271
CL\_INS\_263
CL\_INS\_271
CL\_INS\_263
CL\_INS\_271
CL\_INS\_271
CL\_INS\_271
CL\_INS\_271
CL\_INS\_271
CL\_INS\_263
CL\_INS\_271
CL\_INS\_271
CL\_INS\_271
CL\_INS\_271
CL\_INS\_271
CL\_INS\_271
CL\_INS\_271
CL\_INS\_271
CL\_INS\_271
CL\_INS\_271
CL\_INS\_271
CL\_INS\_271
CL\_INS\_271
CL\_INS\_271
CL\_INS\_263
CL\_INS\_271
CL\_INS\_271
CL\_INS\_271
CL\_INS\_271
CL\_INS\_271
CL\_INS\_271
CL\_INS\_271
CL\_INS\_263
CL\_INS\_263
CL\_INS\_263
CL\_INS\_263
CL\_INS\_263
CL\_INS\_271
CL\_INS\_271
CL\_INS\_271
CL\_INS\_271
CL\_INS\_271
CL\_INS\_271
CL\_INS\_271
CL\_INS\_271
CL\_INS\_263
CL\_INS\_271
CL\_INS\_263
CL\_INS\_271
CL\_INS\_271
Cluster ID


CL\_16920
CL\_14712
CL\_5595
CL\_12819
CL\_12818
CL\_12817
CL\_23904
CL\_12816
CL\_5479
CL\_5181
CL\_23576
CL\_5183
CL\_5184
CL\_5185
CL\_5186
CL\_7196
CL\_18250
CL\_5194
CL\_21685
CL\_21684
CL\_6235
CL\_15628
CL\_5201
CL\_5202
CL\_5203
CL\_23575
CL\_5204
CL\_11028
CL\_7188
CL\_11027
CL\_11026
CL\_11025
CL\_11024
CL\_23574
CL\_5212
CL\_23573
CL\_23572
CL\_23571
CL\_23570
CL\_23569
CL\_23568
CL\_23567
CL\_23566
CL\_23565
CL\_23564
CL\_23563
CL\_23562
CL\_23561
CL\_8723
CL\_5640
CL\_23560
CL\_5229
CL\_11979
CL\_23559
CL\_5343
CL\_15434
CL\_23558
CL\_6247
CL\_6248
CL\_5237
CL\_15589
CL\_11612
CL\_23318
CL\_23557
CL\_23321
CL\_23556
CL\_23322
CL\_23323
CL\_23324
CL\_23325
CL\_8711
CL\_23555
CL\_5226
CL\_23554
CL\_12815
