## Supplementary material for "A novel method for integrating genomic and Tn-Seq data to identify common *in vivo* fitness mechanisms across multiple bacterial species": S2 Dataset: CL_INS_274.html

Legend

 Mobile +extrachromosomalelementfunctions
 Regulatoryfunctions
 Hypothetical
 DNA Metabolism
 Other
 All VFDB Genes

FULL


WINDOWSVGPNG

Trim RowsRemove SingletonsSave Fasta

CL\_3301


CL\_3301


CL\_3301


CL\_3301

HighlightSelectShow Genomes


200

CL\_3300


1

CL\_3300


1

CL\_3299


1

CL\_3300

fGI ID


CL\_INS\_274
CL\_INS\_273
CL\_INS\_274
CL\_INS\_274
CL\_INS\_273
CL\_INS\_74
CL\_INS\_274
CL\_INS\_273
CL\_INS\_273
CL\_INS\_273
CL\_INS\_273
CL\_INS\_273
CL\_INS\_274
CL\_INS\_273
CL\_INS\_273
CL\_INS\_273
CL\_INS\_273
CL\_INS\_274
CL\_INS\_263
CL\_INS\_263
CL\_INS\_263
CL\_INS\_263
CL\_INS\_307
CL\_INS\_263
CL\_INS\_263
CL\_INS\_263
CL\_INS\_307
CL\_INS\_263
CL\_INS\_263
CL\_INS\_307
CL\_INS\_263
CL\_INS\_91
CL\_INS\_91
CL\_INS\_263
CL\_INS\_263
CL\_INS\_263
CL\_INS\_263
CL\_INS\_263
CL\_INS\_263
CL\_INS\_263
CL\_INS\_263
CL\_INS\_307
CL\_INS\_307
CL\_INS\_263
CL\_INS\_263
CL\_INS\_263
CL\_INS\_263
CL\_INS\_263
CL\_INS\_307
CL\_INS\_263
CL\_INS\_263
CL\_INS\_263
CL\_INS\_91
CL\_INS\_263
CL\_INS\_263
CL\_INS\_91
CL\_INS\_273
CL\_INS\_271
CL\_INS\_271
CL\_INS\_271
CL\_INS\_271
CL\_INS\_263
CL\_INS\_271
CL\_INS\_263
CL\_INS\_263
CL\_INS\_91
CL\_INS\_263
CL\_INS\_263
CL\_INS\_263
CL\_INS\_263
CL\_INS\_263
CL\_INS\_91
CL\_INS\_273
CL\_INS\_263
CL\_INS\_263
CL\_INS\_307
CL\_INS\_91
CL\_INS\_263
CL\_INS\_263
CL\_INS\_263
CL\_INS\_263
CL\_INS\_263
CL\_INS\_263
CL\_INS\_263
CL\_INS\_263
CL\_INS\_263
Cluster ID


CL\_17371
CL\_10493
CL\_17372
CL\_23850
CL\_10496
CL\_7141
CL\_23851
CL\_10500
CL\_10501
CL\_10497
CL\_10498
CL\_10499
CL\_17373
CL\_10502
CL\_10503
CL\_10504
CL\_10505
CL\_17374
CL\_5411
CL\_5412
CL\_5413
CL\_5416
CL\_11016
CL\_6256
CL\_5248
CL\_5247
CL\_5246
CL\_5245
CL\_7179
CL\_5243
CL\_5242
CL\_11017
CL\_11018
CL\_5239
CL\_5238
CL\_5237
CL\_11019
CL\_11020
CL\_5233
CL\_5232
CL\_5231
CL\_9711
CL\_6245
CL\_8332
CL\_5225
CL\_5224
CL\_5223
CL\_5222
CL\_9714
CL\_5443
CL\_5444
CL\_11021
CL\_11022
CL\_5214
CL\_5213
CL\_7185
CL\_11023
CL\_11024
CL\_11025
CL\_11026
CL\_11027
CL\_7188
CL\_11028
CL\_5204
CL\_5203
CL\_11029
CL\_5202
CL\_5201
CL\_5200
CL\_5199
CL\_6235
CL\_7192
CL\_11030
CL\_6229
CL\_6228
CL\_11031
CL\_8323
CL\_5189
CL\_5188
CL\_7198
CL\_5186
CL\_5185
CL\_5184
CL\_5183
CL\_5182
CL\_5181
