## Supplementary material for "A novel method for integrating genomic and Tn-Seq data to identify common *in vivo* fitness mechanisms across multiple bacterial species": S2 Dataset: CL_INS_275.html

Legend

 Hypothetical
 All EssentialGenes
 Other

FULL


WINDOWSVGPNG

Trim RowsRemove SingletonsSave Fasta

CL\_3314


CL\_3314


CL\_3314


CL\_3314


CL\_3314

HighlightSelectShow Genomes


83

CL\_3316


70

CL\_3316


49

CL\_3316


1

CL\_3317


1

CL\_3328

fGI ID


CL\_INS\_275
CL\_INS\_275
CL\_INS\_275
Cluster ID


CL\_7856
CL\_5593
CL\_3315
