## Supplementary material for "A novel method for integrating genomic and Tn-Seq data to identify common *in vivo* fitness mechanisms across multiple bacterial species": S2 Dataset: CL_INS_280.html

Legend

 Mobile +extrachromosomalelementfunctions
 Regulatoryfunctions
 Hypothetical
 DNA Metabolism
 AntibioticResistance
 All Fitness Genes
 Biosynthesis ofcofactors,prostheticgroups, +carriers
 Proteinsynthesis/fate
 Other
 Centralintermediarymetabolism
 EnergyMetabolism
 Transport +binding proteins
 All VFDB Genes

FULL


WINDOWSVGPNG

Trim RowsRemove SingletonsSave Fasta

CL\_3357


CL\_3357


CL\_3357


CL\_3357


CL\_3357


CL\_3357


CL\_3357


CL\_3357


CL\_3357


CL\_3357


Break


CL\_3357


CL\_3357


CL\_3357


CL\_3357


CL\_3357


CL\_3357


CL\_3357


CL\_3357


CL\_3357


CL\_3357


CL\_3357


CL\_3357


CL\_3357


CL\_3357


CL\_3357


CL\_3357


CL\_3357


CL\_3357


CL\_3357


CL\_3357

HighlightSelectShow Genomes


165

CL\_3364


2

Break


2

CL\_3363


2

CL\_3364


1

CL\_3363


1

CL\_3364


1

CL\_3364


1

CL\_3364


1

CL\_3364


1

CL\_3364


1

CL\_3364


1

CL\_3364


1

CL\_3364


1

CL\_3364


1

CL\_3363


1

CL\_3364


1

CL\_3364


1

CL\_3364


1

CL\_3364


1

CL\_3422


1

CL\_3364


1

CL\_3364


1

CL\_3364


1

CL\_3364


1

CL\_3364


1

CL\_3364


1

CL\_3364


1

CL\_3364


1

CL\_3363


1

CL\_3364


1

CL\_3364

fGI ID


CL\_INS\_280
CL\_INS\_280
CL\_INS\_278
CL\_INS\_278
CL\_INS\_278
CL\_INS\_278
CL\_INS\_278
CL\_INS\_278
CL\_INS\_278
CL\_INS\_278
CL\_INS\_278
CL\_INS\_278
CL\_INS\_278
CL\_INS\_278
CL\_INS\_278
CL\_INS\_278
CL\_INS\_278
CL\_INS\_278
CL\_INS\_278
CL\_INS\_278
CL\_INS\_278
CL\_INS\_278
CL\_INS\_278
CL\_INS\_278
CL\_INS\_278
CL\_INS\_278
CL\_INS\_278
CL\_INS\_278
CL\_INS\_278
CL\_INS\_132
CL\_INS\_280
CL\_INS\_278
CL\_INS\_278
CL\_INS\_278
CL\_INS\_278
CL\_INS\_278
CL\_INS\_280
CL\_INS\_278
CL\_INS\_278
CL\_INS\_278
CL\_INS\_278
CL\_INS\_278
CL\_INS\_280
CL\_INS\_278
CL\_INS\_278
CL\_INS\_278
CL\_INS\_278
CL\_INS\_278
CL\_INS\_278
CL\_INS\_278
CL\_INS\_287
CL\_INS\_280
CL\_INS\_278
CL\_INS\_280
CL\_INS\_280
CL\_INS\_280
CL\_INS\_280
CL\_INS\_280
CL\_INS\_280
CL\_INS\_280
CL\_INS\_280
CL\_INS\_280
CL\_INS\_280
CL\_INS\_278
CL\_INS\_278
CL\_INS\_278
CL\_INS\_278
CL\_INS\_278
CL\_INS\_278
CL\_INS\_278
CL\_INS\_278
CL\_INS\_278
CL\_INS\_278
CL\_INS\_278
CL\_INS\_278
CL\_INS\_278
CL\_INS\_278
CL\_INS\_278
CL\_INS\_278
CL\_INS\_278
CL\_INS\_280
CL\_INS\_307
CL\_INS\_307
CL\_INS\_280
CL\_INS\_280
CL\_INS\_280
CL\_INS\_278
CL\_INS\_278
CL\_INS\_280
CL\_INS\_280
CL\_INS\_280
CL\_INS\_280
CL\_INS\_280
CL\_INS\_280
CL\_INS\_278
CL\_INS\_278
CL\_INS\_278
CL\_INS\_278
CL\_INS\_278
CL\_INS\_278
CL\_INS\_278
CL\_INS\_278
CL\_INS\_278
CL\_INS\_280
CL\_INS\_278
CL\_INS\_278
CL\_INS\_278
CL\_INS\_278
CL\_INS\_278
CL\_INS\_280
CL\_INS\_278
CL\_INS\_278
CL\_INS\_278
CL\_INS\_278
CL\_INS\_278
CL\_INS\_278
CL\_INS\_278
CL\_INS\_278
CL\_INS\_278
CL\_INS\_280
CL\_INS\_280
CL\_INS\_280
CL\_INS\_278
CL\_INS\_278
CL\_INS\_278
CL\_INS\_278
CL\_INS\_278
CL\_INS\_278
CL\_INS\_278
CL\_INS\_278
CL\_INS\_278
CL\_INS\_278
CL\_INS\_280
CL\_INS\_278
CL\_INS\_278
CL\_INS\_278
CL\_INS\_278
CL\_INS\_278
CL\_INS\_278
CL\_INS\_278
CL\_INS\_278
CL\_INS\_278
CL\_INS\_278
CL\_INS\_278
CL\_INS\_278
CL\_INS\_278
CL\_INS\_280
CL\_INS\_280
CL\_INS\_280
CL\_INS\_280
CL\_INS\_280
CL\_INS\_280
CL\_INS\_280
CL\_INS\_280
CL\_INS\_280
CL\_INS\_280
CL\_INS\_280
CL\_INS\_278
CL\_INS\_278
CL\_INS\_278
CL\_INS\_278
CL\_INS\_278
CL\_INS\_278
CL\_INS\_278
CL\_INS\_278
CL\_INS\_278
CL\_INS\_280
CL\_INS\_278
CL\_INS\_278
CL\_INS\_278
CL\_INS\_278
CL\_INS\_278
CL\_INS\_280
CL\_INS\_278
CL\_INS\_280
CL\_INS\_280
CL\_INS\_204
CL\_INS\_293
CL\_INS\_278
CL\_INS\_280
CL\_INS\_280
CL\_INS\_280
CL\_INS\_278
CL\_INS\_280
CL\_INS\_278
CL\_INS\_278
CL\_INS\_278
CL\_INS\_278
CL\_INS\_278
CL\_INS\_278
CL\_INS\_278
CL\_INS\_278
CL\_INS\_278
CL\_INS\_280
CL\_INS\_278
CL\_INS\_278
CL\_INS\_278
CL\_INS\_278
CL\_INS\_278
CL\_INS\_278
CL\_INS\_278
CL\_INS\_278
CL\_INS\_280
CL\_INS\_278
CL\_INS\_278
CL\_INS\_278
CL\_INS\_278
CL\_INS\_278
CL\_INS\_278
CL\_INS\_278
CL\_INS\_278
CL\_INS\_278
CL\_INS\_278
CL\_INS\_278
CL\_INS\_278
CL\_INS\_280
CL\_INS\_280
CL\_INS\_280
CL\_INS\_280
CL\_INS\_278
CL\_INS\_278
CL\_INS\_280
CL\_INS\_278
CL\_INS\_280
CL\_INS\_280
CL\_INS\_278
CL\_INS\_278
CL\_INS\_278
CL\_INS\_278
CL\_INS\_278
CL\_INS\_278
CL\_INS\_278
CL\_INS\_278
CL\_INS\_278
CL\_INS\_278
CL\_INS\_280
CL\_INS\_278
CL\_INS\_280
CL\_INS\_280
CL\_INS\_280
CL\_INS\_278
CL\_INS\_278
CL\_INS\_278
CL\_INS\_278
CL\_INS\_278
CL\_INS\_278
CL\_INS\_280
CL\_INS\_278
CL\_INS\_278
Cluster ID


CL\_8007
CL\_7318
CL\_8060
CL\_8059
CL\_8058
CL\_7319
CL\_7320
CL\_7321
CL\_7322
CL\_7323
CL\_7324
CL\_7325
CL\_53
CL\_7326
CL\_7327
CL\_7328
CL\_7329
CL\_7330
CL\_7331
CL\_7332
CL\_49
CL\_48
CL\_47
CL\_46
CL\_45
CL\_44
CL\_7333
CL\_43
CL\_42
CL\_8211
CL\_7334
CL\_41
CL\_40
CL\_7335
CL\_7336
CL\_7337
CL\_7338
CL\_7339
CL\_11976
CL\_11975
CL\_11974
CL\_11973
CL\_11972
CL\_11971
CL\_11970
CL\_11969
CL\_11968
CL\_11967
CL\_11966
CL\_11965
CL\_11964
CL\_11963
CL\_8310
CL\_15968
CL\_15967
CL\_15966
CL\_15965
CL\_15964
CL\_15963
CL\_15962
CL\_15961
CL\_15960
CL\_15959
CL\_3368
CL\_3369
CL\_3370
CL\_3371
CL\_3372
CL\_3373
CL\_3374
CL\_3375
CL\_3376
CL\_3377
CL\_3378
CL\_3379
CL\_3380
CL\_3381
CL\_3382
CL\_3383
CL\_3384
CL\_3385
CL\_3386
CL\_3387
CL\_3388
CL\_3389
CL\_3390
CL\_3391
CL\_3392
CL\_3393
CL\_3394
CL\_3395
CL\_3396
CL\_3397
CL\_3398
CL\_3399
CL\_3400
CL\_3401
CL\_12351
CL\_12350
CL\_3402
CL\_3403
CL\_3404
CL\_3405
CL\_15958
CL\_3406
CL\_3407
CL\_3408
CL\_5592
CL\_3409
CL\_16917
CL\_3410
CL\_7463
CL\_3411
CL\_3412
CL\_3413
CL\_5591
CL\_11609
CL\_3414
CL\_3415
CL\_15867
CL\_15866
CL\_23020
CL\_3416
CL\_15957
CL\_15956
CL\_15955
CL\_15954
CL\_15953
CL\_15952
CL\_15951
CL\_15950
CL\_15949
CL\_15948
CL\_15947
CL\_7462
CL\_7461
CL\_7460
CL\_7459
CL\_7458
CL\_7457
CL\_15588
CL\_7456
CL\_7455
CL\_7454
CL\_7453
CL\_7452
CL\_3417
CL\_3418
CL\_3419
CL\_3420
CL\_3421
CL\_15541
CL\_7687
CL\_7686
CL\_7685
CL\_7684
CL\_11766
CL\_8057
CL\_8056
CL\_8055
CL\_8054
CL\_8053
CL\_8052
CL\_8051
CL\_13048
CL\_13047
CL\_22913
CL\_13046
CL\_13045
CL\_13044
CL\_13043
CL\_13042
CL\_15865
CL\_13041
CL\_22912
CL\_22911
CL\_5475
CL\_7942
CL\_11962
CL\_7340
CL\_7341
CL\_7342
CL\_7343
CL\_18588
CL\_7344
CL\_7345
CL\_7346
CL\_7347
CL\_7348
CL\_7349
CL\_7350
CL\_16916
CL\_7351
CL\_16915
CL\_7352
CL\_7353
CL\_7354
CL\_7355
CL\_7356
CL\_7357
CL\_7358
CL\_7359
CL\_7360
CL\_7361
CL\_7362
CL\_7363
CL\_18108
CL\_3367
CL\_6155
CL\_6661
CL\_6662
CL\_9794
CL\_5635
CL\_5634
CL\_5633
CL\_15547
CL\_15546
CL\_15545
CL\_15544
CL\_6663
CL\_6664
CL\_6665
CL\_6666
CL\_16062
CL\_16767
CL\_6667
CL\_6668
CL\_6669
CL\_6670
CL\_6671
CL\_6672
CL\_6673
CL\_9665
CL\_9775
CL\_9774
CL\_16063
CL\_9773
CL\_15543
CL\_6674
CL\_6675
CL\_9772
CL\_6676
CL\_6677
CL\_6678
CL\_6679
CL\_6680
CL\_15542
CL\_6681
CL\_6682
