## Supplementary material for "A novel method for integrating genomic and Tn-Seq data to identify common *in vivo* fitness mechanisms across multiple bacterial species": S2 Dataset: CL_INS_281.html

Legend

 Mobile +extrachromosomalelementfunctions
 Regulatoryfunctions
 Hypothetical
 DNA Metabolism
 Biosynthesis ofcofactors,prostheticgroups, +carriers
 Cell Envelope
 Proteinsynthesis/fate
 Other
 All VFDB Genes

FULL


WINDOWSVGPNG

Trim RowsRemove SingletonsSave Fasta

CL\_3358


CL\_3358


CL\_3358


CL\_3358


CL\_3422


Break


CL\_3358


CL\_3358


CL\_3358


CL\_3358


CL\_3358


CL\_3430


CL\_3358


CL\_3358


CL\_3422

HighlightSelectShow Genomes


180

CL\_3365


6

CL\_3365


3

CL\_3422


2

Break


1

CL\_3365


1

CL\_3365


1

CL\_3365


1

Break


1

CL\_3365


1

CL\_3365


1

CL\_3365


1

CL\_3365


1

CL\_3365


1

CL\_3365


1

CL\_3365

fGI ID


CL\_INS\_321
CL\_INS\_212
CL\_INS\_321
CL\_INS\_321
CL\_INS\_178
CL\_INS\_178
CL\_INS\_278
CL\_INS\_282
CL\_INS\_282
CL\_INS\_282
CL\_INS\_281
CL\_INS\_212
CL\_INS\_212
CL\_INS\_212
CL\_INS\_212
CL\_INS\_212
CL\_INS\_212
CL\_INS\_212
CL\_INS\_160
CL\_INS\_160
CL\_INS\_160
CL\_INS\_212
CL\_INS\_160
CL\_INS\_160
CL\_INS\_212
CL\_INS\_278
CL\_INS\_278
CL\_INS\_278
CL\_INS\_278
CL\_INS\_278
CL\_INS\_278
CL\_INS\_278
CL\_INS\_278
CL\_INS\_278
CL\_INS\_278
CL\_INS\_278
CL\_INS\_278
CL\_INS\_280
CL\_INS\_280
CL\_INS\_278
CL\_INS\_278
CL\_INS\_278
CL\_INS\_278
CL\_INS\_278
CL\_INS\_278
CL\_INS\_278
CL\_INS\_278
CL\_INS\_278
CL\_INS\_278
CL\_INS\_278
CL\_INS\_278
CL\_INS\_138
CL\_INS\_278
CL\_INS\_278
CL\_INS\_278
CL\_INS\_278
CL\_INS\_278
CL\_INS\_278
CL\_INS\_278
CL\_INS\_278
CL\_INS\_278
CL\_INS\_278
CL\_INS\_278
CL\_INS\_278
CL\_INS\_280
CL\_INS\_282
CL\_INS\_278
CL\_INS\_278
CL\_INS\_282
CL\_INS\_282
CL\_INS\_282
CL\_INS\_278
CL\_INS\_282
CL\_INS\_282
CL\_INS\_278
CL\_INS\_278
CL\_INS\_278
CL\_INS\_278
CL\_INS\_278
CL\_INS\_278
CL\_INS\_278
CL\_INS\_278
CL\_INS\_282
CL\_INS\_282
CL\_INS\_282
CL\_INS\_282
CL\_INS\_282
CL\_INS\_282
CL\_INS\_282
CL\_INS\_282
CL\_INS\_282
CL\_INS\_282
CL\_INS\_282
CL\_INS\_282
CL\_INS\_282
CL\_INS\_278
CL\_INS\_278
CL\_INS\_278
CL\_INS\_278
CL\_INS\_278
CL\_INS\_278
CL\_INS\_278
CL\_INS\_278
CL\_INS\_278
CL\_INS\_278
CL\_INS\_280
CL\_INS\_278
CL\_INS\_278
CL\_INS\_278
CL\_INS\_278
CL\_INS\_278
CL\_INS\_278
CL\_INS\_278
CL\_INS\_282
CL\_INS\_278
CL\_INS\_278
CL\_INS\_278
CL\_INS\_278
CL\_INS\_278
CL\_INS\_278
CL\_INS\_278
CL\_INS\_278
CL\_INS\_278
CL\_INS\_278
CL\_INS\_278
CL\_INS\_280
CL\_INS\_280
CL\_INS\_280
CL\_INS\_282
CL\_INS\_278
CL\_INS\_278
CL\_INS\_278
CL\_INS\_278
CL\_INS\_278
CL\_INS\_278
CL\_INS\_278
CL\_INS\_278
CL\_INS\_278
CL\_INS\_278
CL\_INS\_278
CL\_INS\_278
CL\_INS\_278
CL\_INS\_278
CL\_INS\_278
CL\_INS\_278
CL\_INS\_278
CL\_INS\_278
CL\_INS\_278
CL\_INS\_278
CL\_INS\_278
CL\_INS\_278
CL\_INS\_278
CL\_INS\_278
CL\_INS\_282
CL\_INS\_278
CL\_INS\_278
CL\_INS\_278
CL\_INS\_278
CL\_INS\_278
CL\_INS\_278
CL\_INS\_278
CL\_INS\_278
CL\_INS\_278
CL\_INS\_278
CL\_INS\_278
CL\_INS\_278
CL\_INS\_278
CL\_INS\_278
CL\_INS\_278
CL\_INS\_278
CL\_INS\_278
CL\_INS\_278
CL\_INS\_278
CL\_INS\_278
CL\_INS\_278
CL\_INS\_282
CL\_INS\_282
CL\_INS\_282
CL\_INS\_281
CL\_INS\_282
CL\_INS\_281
CL\_INS\_282
CL\_INS\_282
CL\_INS\_282
CL\_INS\_282
CL\_INS\_282
CL\_INS\_282
CL\_INS\_281
CL\_INS\_282
CL\_INS\_282
CL\_INS\_282
CL\_INS\_282
CL\_INS\_281
CL\_INS\_282
CL\_INS\_281
CL\_INS\_281
Cluster ID


CL\_6341
CL\_9397
CL\_12218
CL\_12219
CL\_12380
CL\_10001
CL\_8310
CL\_8309
CL\_8308
CL\_21710
CL\_8307
CL\_7970
CL\_7972
CL\_7973
CL\_7974
CL\_7975
CL\_12485
CL\_12486
CL\_7978
CL\_7979
CL\_12487
CL\_7980
CL\_7981
CL\_7982
CL\_7983
CL\_7984
CL\_6842
CL\_6843
CL\_6844
CL\_6682
CL\_6681
CL\_6680
CL\_6679
CL\_6678
CL\_6677
CL\_6676
CL\_9772
CL\_6675
CL\_6674
CL\_9665
CL\_6673
CL\_6672
CL\_6671
CL\_6670
CL\_6669
CL\_6668
CL\_6667
CL\_6666
CL\_6664
CL\_6663
CL\_5633
CL\_15786
CL\_5634
CL\_5635
CL\_9794
CL\_6662
CL\_6661
CL\_6155
CL\_3415
CL\_3414
CL\_3413
CL\_3412
CL\_3411
CL\_3410
CL\_16917
CL\_20389
CL\_3409
CL\_3408
CL\_20390
CL\_20391
CL\_20392
CL\_12350
CL\_20393
CL\_20394
CL\_3407
CL\_3406
CL\_3405
CL\_3404
CL\_3403
CL\_3402
CL\_3401
CL\_3400
CL\_20395
CL\_20396
CL\_20397
CL\_20398
CL\_20399
CL\_20400
CL\_20401
CL\_20402
CL\_20403
CL\_20404
CL\_20405
CL\_20406
CL\_20407
CL\_3367
CL\_3399
CL\_3384
CL\_3383
CL\_3382
CL\_3381
CL\_3380
CL\_7363
CL\_7362
CL\_7361
CL\_7360
CL\_7359
CL\_7358
CL\_7357
CL\_7356
CL\_7355
CL\_7354
CL\_7353
CL\_21703
CL\_7352
CL\_7351
CL\_16916
CL\_7350
CL\_7349
CL\_7348
CL\_7347
CL\_7346
CL\_7345
CL\_7344
CL\_7343
CL\_7342
CL\_7341
CL\_7340
CL\_21704
CL\_3379
CL\_3378
CL\_3377
CL\_3376
CL\_3375
CL\_3374
CL\_3373
CL\_3372
CL\_3371
CL\_3370
CL\_3369
CL\_3368
CL\_7339
CL\_7337
CL\_7336
CL\_7335
CL\_40
CL\_41
CL\_42
CL\_43
CL\_7333
CL\_44
CL\_45
CL\_46
CL\_21705
CL\_47
CL\_48
CL\_49
CL\_7332
CL\_7331
CL\_7330
CL\_7329
CL\_7328
CL\_7327
CL\_7326
CL\_53
CL\_7325
CL\_7324
CL\_7323
CL\_7322
CL\_7321
CL\_7320
CL\_7319
CL\_8058
CL\_8059
CL\_8060
CL\_21706
CL\_21707
CL\_8306
CL\_11421
CL\_8305
CL\_11420
CL\_8304
CL\_8303
CL\_8302
CL\_8301
CL\_21708
CL\_8300
CL\_11419
CL\_8299
CL\_21709
CL\_8298
CL\_8297
CL\_11418
CL\_8296
CL\_21655
CL\_8295
