## Supplementary material for "A novel method for integrating genomic and Tn-Seq data to identify common *in vivo* fitness mechanisms across multiple bacterial species": S2 Dataset: CL_INS_282.html

Legend

 Mobile +extrachromosomalelementfunctions
 Regulatoryfunctions
 Hypothetical
 DNA Metabolism
 All EssentialGenes
 AntibioticResistance
 All Fitness Genes
 Biosynthesis ofcofactors,prostheticgroups, +carriers
 Proteinsynthesis/fate
 Other
 Transport +binding proteins
 All VFDB Genes

FULL


WINDOWSVGPNG

Trim RowsRemove SingletonsSave Fasta

CL\_3365


CL\_3365


CL\_3365


CL\_3365


CL\_3365


CL\_3365


CL\_3365


CL\_3358


CL\_3365


CL\_3365


CL\_3365


CL\_3365


CL\_3365


CL\_3365


CL\_3365


CL\_3365


CL\_3365


CL\_3365


CL\_3365


CL\_3365


CL\_3365


CL\_3365


CL\_3365


CL\_3365


CL\_3365


CL\_3365


CL\_3365


CL\_3365


CL\_3365


CL\_3365


CL\_3365


CL\_3365


CL\_3365


CL\_3365


CL\_3365


CL\_3365


CL\_3365


CL\_3365


CL\_3365


CL\_3365


CL\_3365


CL\_3365


CL\_3365


CL\_3365


CL\_3365


CL\_3365


CL\_3365


CL\_3365


CL\_3365


CL\_3365


CL\_3365


CL\_3365


CL\_3365


CL\_3365


CL\_3365


CL\_3365


CL\_3365


CL\_3365


CL\_3365


CL\_3365


CL\_3365


CL\_3365


CL\_3365


CL\_3365


CL\_3365


CL\_3365


CL\_3357


CL\_3365


CL\_3365


CL\_3365


CL\_3365


CL\_3365


CL\_3365


CL\_3365


CL\_3365


CL\_3365


CL\_3365


CL\_3365


CL\_3365

HighlightSelectShow Genomes


59

CL\_3422


48

CL\_3422


6

CL\_3422


4

CL\_3422


3

CL\_3422


3

CL\_3422


2

CL\_3422


2

CL\_3422


2

CL\_3422


2

CL\_3422


2

CL\_3422


2

CL\_3422


2

CL\_3422


2

CL\_3422


1

CL\_3422


1

CL\_3422


1

CL\_3422


1

CL\_3422


1

CL\_3422


1

CL\_3422


1

CL\_3422


1

CL\_3422


1

CL\_3422


1

CL\_3422


1

CL\_3422


1

CL\_3422


1

Break


1

CL\_3422


1

CL\_3422


1

CL\_3422


1

CL\_3422


1

CL\_3422


1

CL\_3422


1

CL\_3422


1

CL\_3422


1

CL\_3358


1

CL\_3422


1

CL\_3422


1

CL\_3358


1

CL\_3422


1

CL\_3422


1

CL\_3422


1

CL\_3422


1

CL\_3422


1

CL\_3422


1

CL\_3422


1

CL\_3422


1

CL\_3422


1

CL\_3422


1

CL\_3422


1

CL\_3422


1

CL\_3423


1

CL\_3422


1

CL\_3422


1

CL\_3422


1

CL\_3422


1

CL\_3422


1

CL\_3422


1

CL\_3422


1

CL\_3422


1

CL\_3422


1

CL\_3422


1

CL\_3422


1

CL\_3422


1

CL\_3422


1

CL\_3423


1

CL\_3422


1

CL\_3422


1

CL\_3422


1

CL\_3422


1

CL\_3422


1

CL\_3422


1

CL\_2641


1

CL\_3422


1

CL\_3422


1

CL\_3422


1

CL\_3422


1

CL\_3422


1

CL\_3422

fGI ID


CL\_INS\_282
CL\_INS\_282
CL\_INS\_282
CL\_INS\_282
CL\_INS\_282
CL\_INS\_282
CL\_INS\_282
CL\_INS\_282
CL\_INS\_282
CL\_INS\_282
CL\_INS\_282
CL\_INS\_282
CL\_INS\_282
CL\_INS\_282
CL\_INS\_282
CL\_INS\_282
CL\_INS\_282
CL\_INS\_282
CL\_INS\_282
CL\_INS\_282
CL\_INS\_282
CL\_INS\_282
CL\_INS\_282
CL\_INS\_282
CL\_INS\_282
CL\_INS\_282
CL\_INS\_282
CL\_INS\_282
CL\_INS\_282
CL\_INS\_282
CL\_INS\_278
CL\_INS\_282
CL\_INS\_282
CL\_INS\_282
CL\_INS\_282
CL\_INS\_282
CL\_INS\_282
CL\_INS\_282
CL\_INS\_282
CL\_INS\_282
CL\_INS\_282
CL\_INS\_282
CL\_INS\_282
CL\_INS\_282
CL\_INS\_278
CL\_INS\_278
CL\_INS\_278
CL\_INS\_278
CL\_INS\_278
CL\_INS\_282
CL\_INS\_282
CL\_INS\_282
CL\_INS\_293
CL\_INS\_278
CL\_INS\_278
CL\_INS\_278
CL\_INS\_278
CL\_INS\_278
CL\_INS\_278
CL\_INS\_278
CL\_INS\_278
CL\_INS\_278
CL\_INS\_278
CL\_INS\_278
CL\_INS\_278
CL\_INS\_278
CL\_INS\_282
CL\_INS\_282
CL\_INS\_282
CL\_INS\_278
CL\_INS\_278
CL\_INS\_282
CL\_INS\_278
CL\_INS\_278
CL\_INS\_282
CL\_INS\_282
CL\_INS\_278
CL\_INS\_307
CL\_INS\_278
CL\_INS\_307
CL\_INS\_278
CL\_INS\_278
CL\_INS\_282
CL\_INS\_282
CL\_INS\_282
CL\_INS\_278
CL\_INS\_280
CL\_INS\_132
CL\_INS\_278
CL\_INS\_278
CL\_INS\_278
CL\_INS\_278
CL\_INS\_278
CL\_INS\_280
CL\_INS\_278
CL\_INS\_278
CL\_INS\_278
CL\_INS\_278
CL\_INS\_278
CL\_INS\_278
CL\_INS\_278
CL\_INS\_278
CL\_INS\_278
CL\_INS\_278
CL\_INS\_278
CL\_INS\_278
CL\_INS\_282
CL\_INS\_282
CL\_INS\_278
CL\_INS\_282
CL\_INS\_278
CL\_INS\_282
CL\_INS\_278
CL\_INS\_278
CL\_INS\_278
CL\_INS\_278
CL\_INS\_278
CL\_INS\_280
CL\_INS\_307
CL\_INS\_307
CL\_INS\_280
CL\_INS\_280
CL\_INS\_280
CL\_INS\_278
CL\_INS\_278
CL\_INS\_280
CL\_INS\_280
CL\_INS\_280
CL\_INS\_280
CL\_INS\_280
CL\_INS\_280
CL\_INS\_278
CL\_INS\_278
CL\_INS\_282
CL\_INS\_282
CL\_INS\_278
CL\_INS\_282
CL\_INS\_282
CL\_INS\_282
CL\_INS\_278
CL\_INS\_282
CL\_INS\_278
CL\_INS\_278
CL\_INS\_282
CL\_INS\_278
CL\_INS\_278
CL\_INS\_278
CL\_INS\_278
CL\_INS\_278
CL\_INS\_282
CL\_INS\_278
CL\_INS\_278
CL\_INS\_278
CL\_INS\_280
CL\_INS\_278
CL\_INS\_278
CL\_INS\_278
CL\_INS\_278
CL\_INS\_278
CL\_INS\_278
CL\_INS\_278
CL\_INS\_278
CL\_INS\_280
CL\_INS\_280
CL\_INS\_278
CL\_INS\_321
CL\_INS\_321
CL\_INS\_321
CL\_INS\_282
CL\_INS\_280
CL\_INS\_280
CL\_INS\_280
CL\_INS\_280
CL\_INS\_280
CL\_INS\_282
CL\_INS\_282
CL\_INS\_282
CL\_INS\_282
CL\_INS\_282
CL\_INS\_282
CL\_INS\_282
CL\_INS\_278
CL\_INS\_278
CL\_INS\_278
CL\_INS\_278
CL\_INS\_278
CL\_INS\_278
CL\_INS\_278
CL\_INS\_280
CL\_INS\_278
CL\_INS\_278
CL\_INS\_278
CL\_INS\_278
CL\_INS\_278
CL\_INS\_278
CL\_INS\_278
CL\_INS\_278
CL\_INS\_278
CL\_INS\_278
CL\_INS\_282
CL\_INS\_282
CL\_INS\_278
CL\_INS\_282
CL\_INS\_278
CL\_INS\_278
CL\_INS\_278
CL\_INS\_278
CL\_INS\_278
CL\_INS\_278
CL\_INS\_278
CL\_INS\_278
CL\_INS\_278
CL\_INS\_278
CL\_INS\_278
CL\_INS\_278
CL\_INS\_278
CL\_INS\_280
CL\_INS\_278
CL\_INS\_280
CL\_INS\_280
CL\_INS\_280
CL\_INS\_280
CL\_INS\_280
CL\_INS\_282
CL\_INS\_282
CL\_INS\_282
CL\_INS\_282
CL\_INS\_282
CL\_INS\_282
CL\_INS\_282
CL\_INS\_282
CL\_INS\_282
CL\_INS\_359
CL\_INS\_359
CL\_INS\_282
CL\_INS\_359
CL\_INS\_282
CL\_INS\_282
CL\_INS\_282
CL\_INS\_359
CL\_INS\_282
CL\_INS\_282
CL\_INS\_282
CL\_INS\_282
CL\_INS\_278
CL\_INS\_278
CL\_INS\_278
CL\_INS\_278
CL\_INS\_278
CL\_INS\_278
CL\_INS\_278
CL\_INS\_278
CL\_INS\_280
CL\_INS\_280
CL\_INS\_280
CL\_INS\_278
CL\_INS\_211
CL\_INS\_280
CL\_INS\_278
CL\_INS\_278
CL\_INS\_278
CL\_INS\_278
CL\_INS\_278
CL\_INS\_278
CL\_INS\_278
CL\_INS\_278
CL\_INS\_278
CL\_INS\_280
CL\_INS\_278
CL\_INS\_282
CL\_INS\_278
CL\_INS\_278
CL\_INS\_278
CL\_INS\_278
CL\_INS\_278
CL\_INS\_278
CL\_INS\_278
CL\_INS\_280
CL\_INS\_278
CL\_INS\_278
CL\_INS\_278
CL\_INS\_278
CL\_INS\_278
CL\_INS\_282
CL\_INS\_282
CL\_INS\_282
CL\_INS\_282
CL\_INS\_282
CL\_INS\_282
CL\_INS\_282
CL\_INS\_282
CL\_INS\_282
CL\_INS\_282
CL\_INS\_282
CL\_INS\_282
CL\_INS\_282
CL\_INS\_282
CL\_INS\_282
CL\_INS\_282
CL\_INS\_282
CL\_INS\_282
CL\_INS\_282
CL\_INS\_132
CL\_INS\_278
CL\_INS\_385
CL\_INS\_212
CL\_INS\_212
CL\_INS\_212
CL\_INS\_212
CL\_INS\_212
CL\_INS\_212
CL\_INS\_38
CL\_INS\_212
CL\_INS\_212
CL\_INS\_278
CL\_INS\_278
CL\_INS\_278
CL\_INS\_278
CL\_INS\_278
CL\_INS\_278
CL\_INS\_278
CL\_INS\_280
CL\_INS\_278
CL\_INS\_278
CL\_INS\_278
CL\_INS\_278
CL\_INS\_278
CL\_INS\_278
CL\_INS\_278
CL\_INS\_278
CL\_INS\_278
CL\_INS\_280
CL\_INS\_280
CL\_INS\_278
CL\_INS\_278
CL\_INS\_278
CL\_INS\_278
CL\_INS\_278
CL\_INS\_278
CL\_INS\_278
Cluster ID


CL\_23985
CL\_20407
CL\_20406
CL\_20405
CL\_20404
CL\_20403
CL\_20402
CL\_20401
CL\_20400
CL\_20399
CL\_20398
CL\_20397
CL\_20396
CL\_20395
CL\_8296
CL\_8297
CL\_8298
CL\_21709
CL\_8299
CL\_8300
CL\_21708
CL\_8301
CL\_8302
CL\_8303
CL\_8304
CL\_8305
CL\_8306
CL\_21707
CL\_21706
CL\_3366
CL\_8060
CL\_8411
CL\_8410
CL\_8409
CL\_8408
CL\_8407
CL\_8406
CL\_8405
CL\_8404
CL\_8403
CL\_21636
CL\_8402
CL\_8401
CL\_21637
CL\_8059
CL\_8058
CL\_7319
CL\_7320
CL\_7321
CL\_21710
CL\_8308
CL\_8309
CL\_7942
CL\_7322
CL\_7323
CL\_7324
CL\_7325
CL\_53
CL\_7326
CL\_7327
CL\_7328
CL\_7329
CL\_7330
CL\_7331
CL\_7332
CL\_49
CL\_21688
CL\_21687
CL\_21686
CL\_48
CL\_47
CL\_21705
CL\_46
CL\_45
CL\_18105
CL\_23814
CL\_44
CL\_8858
CL\_8050
CL\_8857
CL\_7333
CL\_43
CL\_18106
CL\_23813
CL\_18107
CL\_42
CL\_7334
CL\_8211
CL\_40
CL\_41
CL\_7335
CL\_7336
CL\_7337
CL\_7338
CL\_7339
CL\_3368
CL\_3369
CL\_3370
CL\_3371
CL\_3372
CL\_3373
CL\_3374
CL\_3375
CL\_3376
CL\_3377
CL\_3378
CL\_13972
CL\_13973
CL\_108
CL\_13974
CL\_3379
CL\_21704
CL\_3380
CL\_3381
CL\_3382
CL\_3383
CL\_3384
CL\_3385
CL\_3386
CL\_3387
CL\_3388
CL\_3389
CL\_3390
CL\_3391
CL\_3392
CL\_3393
CL\_3394
CL\_3395
CL\_3396
CL\_3397
CL\_3398
CL\_3399
CL\_3400
CL\_20394
CL\_20393
CL\_12350
CL\_20392
CL\_20391
CL\_20390
CL\_3401
CL\_20520
CL\_3402
CL\_3403
CL\_16918
CL\_3404
CL\_3405
CL\_3406
CL\_3407
CL\_3408
CL\_20389
CL\_5592
CL\_3409
CL\_7463
CL\_16917
CL\_3410
CL\_3411
CL\_3412
CL\_3413
CL\_5591
CL\_11609
CL\_3414
CL\_3415
CL\_15867
CL\_15866
CL\_3416
CL\_5590
CL\_5589
CL\_5588
CL\_5587
CL\_15541
CL\_7687
CL\_7686
CL\_7685
CL\_7684
CL\_17208
CL\_17209
CL\_17210
CL\_17211
CL\_17212
CL\_17213
CL\_17214
CL\_13048
CL\_13047
CL\_13046
CL\_13045
CL\_13044
CL\_13043
CL\_13042
CL\_15865
CL\_13041
CL\_7462
CL\_20409
CL\_7461
CL\_7460
CL\_7459
CL\_7458
CL\_7457
CL\_20408
CL\_15588
CL\_20521
CL\_10905
CL\_7456
CL\_23743
CL\_7455
CL\_7454
CL\_7453
CL\_7452
CL\_15957
CL\_15956
CL\_15955
CL\_15954
CL\_15953
CL\_15952
CL\_15951
CL\_15950
CL\_15949
CL\_15948
CL\_15947
CL\_3417
CL\_3418
CL\_3419
CL\_3420
CL\_3421
CL\_8294
CL\_8293
CL\_8292
CL\_8291
CL\_8290
CL\_8289
CL\_8288
CL\_7451
CL\_7450
CL\_7449
CL\_10350
CL\_23984
CL\_10349
CL\_10348
CL\_10347
CL\_10345
CL\_10346
CL\_23983
CL\_23982
CL\_7448
CL\_7447
CL\_8057
CL\_8056
CL\_8055
CL\_8054
CL\_8053
CL\_8052
CL\_8051
CL\_6945
CL\_7340
CL\_7341
CL\_7342
CL\_7343
CL\_6157
CL\_18588
CL\_7344
CL\_7345
CL\_7346
CL\_7347
CL\_7348
CL\_7349
CL\_7350
CL\_16916
CL\_7351
CL\_16915
CL\_7352
CL\_21703
CL\_7353
CL\_7354
CL\_7355
CL\_7356
CL\_7357
CL\_7358
CL\_7359
CL\_7360
CL\_7361
CL\_7362
CL\_7363
CL\_18108
CL\_3367
CL\_6959
CL\_6958
CL\_6957
CL\_6956
CL\_6955
CL\_13854
CL\_13855
CL\_6954
CL\_6953
CL\_13856
CL\_23249
CL\_22407
CL\_6952
CL\_6951
CL\_6950
CL\_13857
CL\_6949
CL\_6948
CL\_6947
CL\_6946
CL\_6155
CL\_6156
CL\_3171
CL\_3174
CL\_5607
CL\_3176
CL\_3177
CL\_5606
CL\_5605
CL\_5604
CL\_5603
CL\_6661
CL\_6662
CL\_5635
CL\_5634
CL\_5633
CL\_6663
CL\_6664
CL\_6665
CL\_6666
CL\_6667
CL\_6668
CL\_6669
CL\_6670
CL\_6671
CL\_6672
CL\_6673
CL\_9665
CL\_6674
CL\_6675
CL\_6676
CL\_6677
CL\_6678
CL\_6679
CL\_6680
CL\_6681
CL\_6682
