## Supplementary material for "A novel method for integrating genomic and Tn-Seq data to identify common *in vivo* fitness mechanisms across multiple bacterial species": S2 Dataset: CL_INS_286.html

Legend

 Regulatoryfunctions
 Hypothetical
 Other
 All VFDB Genes

FULL


WINDOWSVGPNG

Trim RowsRemove SingletonsSave Fasta

CL\_3445


CL\_3445


CL\_3445


CL\_3445

HighlightSelectShow Genomes


198

CL\_3446


4

CL\_3446


2

CL\_3446


1

CL\_3446

fGI ID


CL\_INS\_42
CL\_INS\_286
CL\_INS\_286
CL\_INS\_286
CL\_INS\_286
CL\_INS\_286
CL\_INS\_286
CL\_INS\_286
CL\_INS\_286
CL\_INS\_286
CL\_INS\_286
CL\_INS\_286
CL\_INS\_286
CL\_INS\_286
CL\_INS\_286
CL\_INS\_286
CL\_INS\_286
CL\_INS\_286
CL\_INS\_286
CL\_INS\_286
CL\_INS\_286
Cluster ID


CL\_5584
CL\_13858
CL\_13859
CL\_13860
CL\_13861
CL\_13862
CL\_13863
CL\_13864
CL\_13865
CL\_13866
CL\_13867
CL\_13868
CL\_13869
CL\_13870
CL\_13871
CL\_13872
CL\_13873
CL\_13874
CL\_13875
CL\_13876
CL\_13877
