## Supplementary material for "A novel method for integrating genomic and Tn-Seq data to identify common *in vivo* fitness mechanisms across multiple bacterial species": S2 Dataset: CL_INS_288.html

Legend

 Mobile +extrachromosomalelementfunctions
 Hypothetical
 Other
 All VFDB Genes

FULL


WINDOWSVGPNG

Trim RowsRemove SingletonsSave Fasta

CL\_3466


CL\_3466


CL\_3466


CL\_3466


CL\_3466


CL\_3457


CL\_3466


CL\_3457


CL\_3466


CL\_3466


CL\_3457


CL\_3466


CL\_3457


CL\_3457


CL\_3457


CL\_3457


CL\_3466


CL\_3466


CL\_3491


CL\_3457


CL\_3466


CL\_3457


CL\_3466


CL\_3457


CL\_3466


CL\_3466


CL\_3466


CL\_3466


CL\_3466


CL\_3466


CL\_3466


CL\_3466


CL\_3466

HighlightSelectShow Genomes


134

CL\_3468


17

CL\_3468


11

CL\_3468


10

CL\_3468


8

CL\_3457


4

CL\_3468


4

CL\_3468


3

CL\_3468


2

CL\_3468


1

CL\_3468


1

CL\_3468


1

CL\_3468


1

CL\_3468


1

CL\_3468


1

CL\_3468


1

CL\_3468


1

CL\_3457


1

CL\_3468


1

CL\_3468


1

CL\_3468


1

CL\_3468


1

CL\_3468


1

CL\_3468


1

CL\_3468


1

CL\_3470


1

CL\_3468


1

CL\_3474


1

CL\_3468


1

CL\_3457


1

CL\_3468


1

CL\_3470


1

CL\_3468


1

CL\_3457

fGI ID


CL\_INS\_287
CL\_INS\_288
CL\_INS\_288
CL\_INS\_287
CL\_INS\_288
CL\_INS\_288
CL\_INS\_288
CL\_INS\_288
CL\_INS\_288
CL\_INS\_287
CL\_INS\_288
CL\_INS\_288
CL\_INS\_288
CL\_INS\_288
CL\_INS\_288
CL\_INS\_288
CL\_INS\_288
CL\_INS\_288
CL\_INS\_288
CL\_INS\_288
CL\_INS\_288
CL\_INS\_288
CL\_INS\_287
CL\_INS\_287
CL\_INS\_287
CL\_INS\_287
CL\_INS\_287
CL\_INS\_287
CL\_INS\_287
CL\_INS\_288
CL\_INS\_287
CL\_INS\_287
CL\_INS\_287
CL\_INS\_287
CL\_INS\_287
CL\_INS\_287
CL\_INS\_287
CL\_INS\_287
CL\_INS\_288
CL\_INS\_278
CL\_INS\_287
CL\_INS\_278
CL\_INS\_278
CL\_INS\_288
Cluster ID


CL\_8047
CL\_8046
CL\_3467
CL\_12188
CL\_18585
CL\_13878
CL\_14861
CL\_15780
CL\_8049
CL\_9298
CL\_5405
CL\_5404
CL\_5403
CL\_15587
CL\_18752
CL\_18753
CL\_18754
CL\_14845
CL\_14846
CL\_14847
CL\_14848
CL\_14849
CL\_14990
CL\_14991
CL\_10344
CL\_10343
CL\_6943
CL\_6942
CL\_13879
CL\_17367
CL\_16064
CL\_6941
CL\_6940
CL\_10342
CL\_10341
CL\_6264
CL\_6265
CL\_6939
CL\_14992
CL\_109
CL\_14182
CL\_8050
CL\_108
CL\_6938
