## Supplementary material for "A novel method for integrating genomic and Tn-Seq data to identify common *in vivo* fitness mechanisms across multiple bacterial species": S2 Dataset: CL_INS_289.html

Legend

 Mobile +extrachromosomalelementfunctions
 Hypothetical
 Other
 All VFDB Genes

FULL


WINDOWSVGPNG

Trim RowsRemove SingletonsSave Fasta

CL\_3468


CL\_3468


CL\_3468


CL\_3468


CL\_3468


CL\_3468


CL\_3468


CL\_3468


CL\_3466


CL\_3468


CL\_3468


CL\_3468


CL\_3468


CL\_3466

HighlightSelectShow Genomes


93

CL\_3470


90

CL\_3470


2

CL\_3471


2

CL\_3471


2

CL\_3471


1

CL\_3471


1

CL\_3470


1

CL\_3471


1

CL\_3470


1

CL\_3471


1

CL\_3470


1

CL\_3470


1

CL\_3471


1

CL\_3470

fGI ID


CL\_INS\_391
CL\_INS\_289
CL\_INS\_290
CL\_INS\_290
CL\_INS\_287
CL\_INS\_290
CL\_INS\_290
CL\_INS\_290
CL\_INS\_290
CL\_INS\_66
CL\_INS\_290
CL\_INS\_290
CL\_INS\_290
CL\_INS\_290
CL\_INS\_290
CL\_INS\_287
CL\_INS\_288
CL\_INS\_288
CL\_INS\_290
CL\_INS\_307
CL\_INS\_278
CL\_INS\_278
CL\_INS\_318
CL\_INS\_290
CL\_INS\_288
Cluster ID


CL\_6936
CL\_3469
CL\_10977
CL\_20522
CL\_16064
CL\_6258
CL\_6259
CL\_6260
CL\_6261
CL\_8045
CL\_14183
CL\_8044
CL\_8043
CL\_8042
CL\_8041
CL\_13879
CL\_17367
CL\_13878
CL\_22336
CL\_8857
CL\_8050
CL\_107
CL\_9517
CL\_22337
CL\_14849
