## Supplementary material for "A novel method for integrating genomic and Tn-Seq data to identify common *in vivo* fitness mechanisms across multiple bacterial species": S2 Dataset: CL_INS_291.html

Legend

 Mobile +extrachromosomalelementfunctions
 Hypothetical
 All VFDB Genes

FULL


WINDOWSVGPNG

Trim RowsRemove SingletonsSave Fasta

CL\_3471


CL\_3471


CL\_3471


CL\_3471


CL\_3471


CL\_3471


CL\_3471


CL\_3466


CL\_3471


CL\_3470


CL\_3471


CL\_3471


CL\_3471


CL\_3471

HighlightSelectShow Genomes


86

CL\_3474


42

CL\_3474


40

CL\_3474


21

CL\_3474


7

CL\_3474


1

CL\_3474


1

CL\_3474


1

CL\_3474


1

CL\_3474


1

CL\_3474


1

CL\_3490


1

CL\_3474


1

CL\_3474


1

CL\_3474

fGI ID


CL\_INS\_291
CL\_INS\_291
CL\_INS\_288
CL\_INS\_290
CL\_INS\_290
CL\_INS\_287
CL\_INS\_287
CL\_INS\_290
CL\_INS\_291
CL\_INS\_290
CL\_INS\_291
CL\_INS\_291
CL\_INS\_291
CL\_INS\_291
CL\_INS\_291
CL\_INS\_291
CL\_INS\_291
CL\_INS\_291
CL\_INS\_291
Cluster ID


CL\_6935
CL\_6934
CL\_5404
CL\_12286
CL\_6263
CL\_6264
CL\_6265
CL\_6266
CL\_7446
CL\_3472
CL\_15781
CL\_15782
CL\_15783
CL\_15784
CL\_14859
CL\_14858
CL\_3473
CL\_17366
CL\_22237
