## Supplementary material for "A novel method for integrating genomic and Tn-Seq data to identify common *in vivo* fitness mechanisms across multiple bacterial species": S2 Dataset: CL_INS_294.html

Legend

 Hypothetical
 Other
 Proteinsynthesis/fate
 Transport +binding proteins
 All VFDB Genes

FULL


WINDOWSVGPNG

Trim RowsRemove SingletonsSave Fasta

CL\_3501


CL\_3501

HighlightSelectShow Genomes


199

CL\_3502


3

CL\_3502

fGI ID


CL\_INS\_294
CL\_INS\_294
CL\_INS\_294
CL\_INS\_294
CL\_INS\_294
CL\_INS\_294
Cluster ID


CL\_14862
CL\_14863
CL\_14864
CL\_14865
CL\_14866
CL\_14867
