## Supplementary material for "A novel method for integrating genomic and Tn-Seq data to identify common *in vivo* fitness mechanisms across multiple bacterial species": S2 Dataset: CL_INS_299.html

Legend

 Hypothetical
 Other
 All VFDB Genes

FULL


WINDOWSVGPNG

Trim RowsRemove SingletonsSave Fasta

CL\_3581


CL\_3581


CL\_3581


CL\_3581


CL\_3580


CL\_3581


CL\_3580

HighlightSelectShow Genomes


101

CL\_3582


49

CL\_3582


47

CL\_3582


3

CL\_3582


2

CL\_3582


1

CL\_3583


1

CL\_3582

fGI ID


CL\_INS\_299
CL\_INS\_299
CL\_INS\_299
CL\_INS\_299
CL\_INS\_299
Cluster ID


CL\_17353
CL\_17352
CL\_14184
CL\_5393
CL\_5392
