## Supplementary material for "A novel method for integrating genomic and Tn-Seq data to identify common *in vivo* fitness mechanisms across multiple bacterial species": S2 Dataset: CL_INS_301.html

Legend

 Mobile +extrachromosomalelementfunctions
 Regulatoryfunctions
 Hypothetical
 DNA Metabolism
 All EssentialGenes
 Proteinsynthesis/fate
 Other
 EnergyMetabolism
 Transport +binding proteins
 All VFDB Genes

FULL


WINDOWSVGPNG

Trim RowsRemove SingletonsSave Fasta

CL\_3605


CL\_3605


CL\_3605


CL\_3605


CL\_3605


CL\_3605


CL\_3605


CL\_3605


CL\_3605


CL\_3605


CL\_3605

HighlightSelectShow Genomes


178

CL\_3606


14

CL\_3606


1

CL\_3606


1

CL\_3606


1

CL\_3606


1

CL\_3606


1

CL\_3606


1

CL\_3606


1

CL\_3606


1

CL\_3606


1

CL\_3606

fGI ID


CL\_INS\_74
CL\_INS\_74
CL\_INS\_74
CL\_INS\_301
CL\_INS\_301
CL\_INS\_74
CL\_INS\_301
CL\_INS\_301
CL\_INS\_301
CL\_INS\_301
CL\_INS\_301
CL\_INS\_301
CL\_INS\_301
CL\_INS\_301
CL\_INS\_301
CL\_INS\_301
CL\_INS\_301
CL\_INS\_313
CL\_INS\_313
CL\_INS\_313
CL\_INS\_313
CL\_INS\_313
CL\_INS\_313
CL\_INS\_313
CL\_INS\_313
CL\_INS\_313
CL\_INS\_313
CL\_INS\_313
CL\_INS\_313
CL\_INS\_301
CL\_INS\_301
CL\_INS\_301
CL\_INS\_301
CL\_INS\_301
CL\_INS\_301
CL\_INS\_301
CL\_INS\_301
CL\_INS\_301
CL\_INS\_301
CL\_INS\_301
CL\_INS\_301
CL\_INS\_301
CL\_INS\_301
CL\_INS\_301
CL\_INS\_301
CL\_INS\_301
CL\_INS\_301
CL\_INS\_301
CL\_INS\_385
CL\_INS\_301
CL\_INS\_301
CL\_INS\_301
CL\_INS\_301
CL\_INS\_301
CL\_INS\_301
CL\_INS\_301
CL\_INS\_155
CL\_INS\_155
CL\_INS\_155
CL\_INS\_155
CL\_INS\_155
CL\_INS\_155
CL\_INS\_155
CL\_INS\_155
CL\_INS\_277
CL\_INS\_301
CL\_INS\_277
CL\_INS\_277
CL\_INS\_277
CL\_INS\_277
CL\_INS\_277
CL\_INS\_277
CL\_INS\_277
CL\_INS\_301
CL\_INS\_301
CL\_INS\_301
CL\_INS\_301
CL\_INS\_155
CL\_INS\_301
CL\_INS\_301
CL\_INS\_301
CL\_INS\_301
CL\_INS\_301
CL\_INS\_321
CL\_INS\_307
CL\_INS\_301
CL\_INS\_155
CL\_INS\_155
CL\_INS\_155
CL\_INS\_155
CL\_INS\_301
CL\_INS\_301
CL\_INS\_301
CL\_INS\_301
CL\_INS\_301
CL\_INS\_301
CL\_INS\_301
CL\_INS\_301
CL\_INS\_155
CL\_INS\_155
CL\_INS\_155
CL\_INS\_301
CL\_INS\_155
CL\_INS\_301
CL\_INS\_301
CL\_INS\_301
CL\_INS\_301
CL\_INS\_263
CL\_INS\_301
CL\_INS\_301
CL\_INS\_155
CL\_INS\_301
CL\_INS\_301
CL\_INS\_155
CL\_INS\_155
CL\_INS\_263
CL\_INS\_263
CL\_INS\_263
CL\_INS\_263
CL\_INS\_263
CL\_INS\_263
CL\_INS\_155
CL\_INS\_155
CL\_INS\_263
CL\_INS\_263
CL\_INS\_155
CL\_INS\_301
CL\_INS\_301
CL\_INS\_263
CL\_INS\_155
CL\_INS\_155
CL\_INS\_155
CL\_INS\_155
CL\_INS\_155
CL\_INS\_155
CL\_INS\_155
CL\_INS\_155
CL\_INS\_155
CL\_INS\_155
CL\_INS\_155
CL\_INS\_155
CL\_INS\_155
CL\_INS\_155
CL\_INS\_263
CL\_INS\_155
CL\_INS\_155
CL\_INS\_263
CL\_INS\_301
CL\_INS\_155
CL\_INS\_155
CL\_INS\_155
CL\_INS\_155
CL\_INS\_155
CL\_INS\_301
CL\_INS\_301
CL\_INS\_155
CL\_INS\_301
CL\_INS\_155
CL\_INS\_155
CL\_INS\_155
CL\_INS\_155
CL\_INS\_155
CL\_INS\_155
CL\_INS\_155
CL\_INS\_301
CL\_INS\_155
CL\_INS\_278
CL\_INS\_155
CL\_INS\_301
CL\_INS\_155
CL\_INS\_321
CL\_INS\_301
CL\_INS\_301
CL\_INS\_301
CL\_INS\_155
CL\_INS\_155
CL\_INS\_321
CL\_INS\_321
CL\_INS\_321
CL\_INS\_321
CL\_INS\_321
CL\_INS\_321
CL\_INS\_301
CL\_INS\_155
CL\_INS\_155
CL\_INS\_155
CL\_INS\_155
CL\_INS\_263
CL\_INS\_155
CL\_INS\_155
CL\_INS\_155
CL\_INS\_155
CL\_INS\_155
CL\_INS\_155
CL\_INS\_155
CL\_INS\_155
CL\_INS\_155
CL\_INS\_155
CL\_INS\_155
CL\_INS\_155
CL\_INS\_155
CL\_INS\_301
CL\_INS\_155
CL\_INS\_155
CL\_INS\_155
CL\_INS\_155
CL\_INS\_155
CL\_INS\_263
CL\_INS\_271
CL\_INS\_277
CL\_INS\_263
CL\_INS\_155
CL\_INS\_155
CL\_INS\_301
CL\_INS\_301
CL\_INS\_263
CL\_INS\_263
CL\_INS\_301
CL\_INS\_301
CL\_INS\_301
CL\_INS\_301
CL\_INS\_301
CL\_INS\_301
CL\_INS\_301
Cluster ID


CL\_10974
CL\_11765
CL\_11764
CL\_20633
CL\_20632
CL\_11763
CL\_11762
CL\_11761
CL\_11760
CL\_23431
CL\_11759
CL\_11758
CL\_11757
CL\_10973
CL\_19617
CL\_10972
CL\_10971
CL\_10970
CL\_10969
CL\_10968
CL\_10967
CL\_10966
CL\_10965
CL\_10964
CL\_10963
CL\_10962
CL\_10961
CL\_10960
CL\_10959
CL\_10958
CL\_10957
CL\_19618
CL\_19619
CL\_19620
CL\_19621
CL\_19622
CL\_19623
CL\_19624
CL\_19625
CL\_19626
CL\_19627
CL\_19628
CL\_19629
CL\_19630
CL\_19631
CL\_19632
CL\_19633
CL\_19634
CL\_6156
CL\_16618
CL\_16619
CL\_16620
CL\_16621
CL\_16622
CL\_16623
CL\_16624
CL\_7253
CL\_7252
CL\_7251
CL\_7250
CL\_7249
CL\_7248
CL\_7247
CL\_7246
CL\_15306
CL\_16625
CL\_15308
CL\_15309
CL\_15310
CL\_15311
CL\_15312
CL\_15313
CL\_15314
CL\_16626
CL\_16627
CL\_16628
CL\_16629
CL\_7234
CL\_16630
CL\_16631
CL\_16632
CL\_7677
CL\_8703
CL\_6584
CL\_7676
CL\_17016
CL\_6582
CL\_6581
CL\_5388
CL\_5387
CL\_8702
CL\_8701
CL\_23744
CL\_23745
CL\_23746
CL\_23747
CL\_23748
CL\_23749
CL\_23048
CL\_23049
CL\_23050
CL\_23750
CL\_8325
CL\_8700
CL\_8699
CL\_8698
CL\_8697
CL\_8696
CL\_23751
CL\_23752
CL\_7675
CL\_7674
CL\_17017
CL\_7288
CL\_6644
CL\_5384
CL\_7287
CL\_5383
CL\_7286
CL\_7285
CL\_7284
CL\_9106
CL\_13193
CL\_7283
CL\_7282
CL\_8695
CL\_8694
CL\_8693
CL\_7281
CL\_7280
CL\_7279
CL\_7673
CL\_7672
CL\_7671
CL\_7670
CL\_7669
CL\_5368
CL\_7668
CL\_7272
CL\_5364
CL\_6605
CL\_5359
CL\_5358
CL\_9111
CL\_5357
CL\_5356
CL\_5651
CL\_8692
CL\_8691
CL\_8690
CL\_5656
CL\_5655
CL\_7667
CL\_23753
CL\_23754
CL\_5653
CL\_23755
CL\_9090
CL\_6986
CL\_2109
CL\_2108
CL\_2107
CL\_2106
CL\_2105
CL\_8689
CL\_7666
CL\_7665
CL\_7664
CL\_23756
CL\_7663
CL\_16633
CL\_17018
CL\_17019
CL\_17020
CL\_2103
CL\_2102
CL\_16634
CL\_16635
CL\_16636
CL\_16637
CL\_16638
CL\_16639
CL\_23757
CL\_2101
CL\_2100
CL\_2099
CL\_2098
CL\_2097
CL\_2096
CL\_2095
CL\_2094
CL\_2093
CL\_17021
CL\_17022
CL\_2092
CL\_7662
CL\_7661
CL\_2089
CL\_2091
CL\_2090
CL\_19333
CL\_23758
CL\_2088
CL\_2087
CL\_2086
CL\_2085
CL\_2084
CL\_7265
CL\_5343
CL\_8688
CL\_13188
CL\_14893
CL\_7660
CL\_23759
CL\_23760
CL\_5415
CL\_6285
CL\_16640
CL\_7659
CL\_7658
CL\_17023
CL\_17024
CL\_7657
CL\_23761
