## Supplementary material for "A novel method for integrating genomic and Tn-Seq data to identify common *in vivo* fitness mechanisms across multiple bacterial species": S2 Dataset: CL_INS_303.html

Legend

 Hypothetical
 All VFDB Genes

FULL


WINDOWSVGPNG

Trim RowsRemove SingletonsSave Fasta

CL\_3634


CL\_3634


CL\_3634

HighlightSelectShow Genomes


115

CL\_3636


86

CL\_3636


1

CL\_3636

fGI ID


CL\_INS\_303
CL\_INS\_303
Cluster ID


CL\_3635
CL\_21370
