## Supplementary material for "A novel method for integrating genomic and Tn-Seq data to identify common *in vivo* fitness mechanisms across multiple bacterial species": S2 Dataset: CL_INS_311.html

Legend

 Hypothetical
 Proteinsynthesis/fate
 All VFDB Genes

FULL


WINDOWSVGPNG

Trim RowsRemove SingletonsSave Fasta

CL\_3794


CL\_3794

HighlightSelectShow Genomes


117

CL\_3796


87

CL\_3796

fGI ID

CL\_INS\_311
Cluster ID

CL\_3795
