## Supplementary material for "A novel method for integrating genomic and Tn-Seq data to identify common *in vivo* fitness mechanisms across multiple bacterial species": S2 Dataset: CL_INS_312.html

Legend

 Mobile +extrachromosomalelementfunctions
 Regulatoryfunctions
 Hypothetical
 DNA Metabolism
 All EssentialGenes
 Biosynthesis ofcofactors,prostheticgroups, +carriers
 Other
 EnergyMetabolism
 Cellularprocesses
 All VFDB Genes
 Transport +binding proteins

FULL


WINDOWSVGPNG

Trim RowsRemove SingletonsSave Fasta

CL\_3813


CL\_3813


CL\_3813


CL\_3813


CL\_3813


CL\_3813


CL\_3813


CL\_3813


CL\_3813


CL\_3813


CL\_3813


CL\_3813


CL\_3811


CL\_3813


CL\_3813


CL\_3813


CL\_3813


CL\_3813


CL\_3813


CL\_3813


CL\_3813


CL\_3812


CL\_3813


CL\_4246


CL\_3813


CL\_3811


CL\_3813


CL\_3813


CL\_3813


CL\_3813


Break


CL\_3813


CL\_3813


CL\_3813


CL\_3813


CL\_3813


CL\_3813


CL\_3807


CL\_3813


CL\_3813


CL\_3813


CL\_3813


CL\_3813


CL\_3813


CL\_3813


CL\_3813


CL\_3813


CL\_3813


CL\_3813


CL\_3813


CL\_3813


CL\_3813


CL\_3813


CL\_3813


CL\_3811

HighlightSelectShow Genomes


139

CL\_3814


3

CL\_3814


3

CL\_3814


2

CL\_3814


1

CL\_3814


1

CL\_3814


1

CL\_3814


1

CL\_3814


1

CL\_3814


1

CL\_3814


1

Break


1

CL\_3814


1

CL\_3814


1

CL\_3814


1

CL\_3814


1

CL\_3814


1

CL\_3814


1

CL\_3814


1

CL\_3814


1

CL\_3814


1

CL\_3814


1

CL\_3814


1

CL\_3814


1

CL\_3814


1

Break


1

CL\_3814


1

CL\_3814


1

CL\_3814


1

CL\_3814


1

CL\_3814


1

CL\_3814


1

CL\_3814


1

CL\_3814


1

CL\_3814


1

CL\_3814


1

CL\_3814


1

CL\_3814


1

CL\_3814


1

CL\_3814


1

CL\_3814


1

CL\_3814


1

CL\_3814


1

CL\_3814


1

Break


1

CL\_3814


1

CL\_3814


1

CL\_3814


1

CL\_3814


1

CL\_3814


1

CL\_3814


1

CL\_3814


1

CL\_3814


1

Break


1

CL\_3814


1

CL\_3814

fGI ID


CL\_INS\_312
CL\_INS\_321
CL\_INS\_312
CL\_INS\_312
CL\_INS\_312
CL\_INS\_312
CL\_INS\_312
CL\_INS\_312
CL\_INS\_312
CL\_INS\_312
CL\_INS\_312
CL\_INS\_312
CL\_INS\_312
CL\_INS\_312
CL\_INS\_312
CL\_INS\_74
CL\_INS\_74
CL\_INS\_74
CL\_INS\_74
CL\_INS\_312
CL\_INS\_312
CL\_INS\_312
CL\_INS\_312
CL\_INS\_312
CL\_INS\_312
CL\_INS\_312
CL\_INS\_312
CL\_INS\_312
CL\_INS\_312
CL\_INS\_312
CL\_INS\_312
CL\_INS\_312
CL\_INS\_312
CL\_INS\_312
CL\_INS\_312
CL\_INS\_312
CL\_INS\_312
CL\_INS\_312
CL\_INS\_312
CL\_INS\_312
CL\_INS\_312
CL\_INS\_312
CL\_INS\_312
CL\_INS\_312
CL\_INS\_312
CL\_INS\_312
CL\_INS\_312
CL\_INS\_321
CL\_INS\_312
CL\_INS\_312
CL\_INS\_321
CL\_INS\_321
CL\_INS\_321
CL\_INS\_321
CL\_INS\_321
CL\_INS\_321
CL\_INS\_321
CL\_INS\_321
CL\_INS\_312
CL\_INS\_312
CL\_INS\_312
CL\_INS\_312
CL\_INS\_334
CL\_INS\_312
CL\_INS\_312
CL\_INS\_312
CL\_INS\_312
CL\_INS\_312
CL\_INS\_312
CL\_INS\_312
CL\_INS\_312
CL\_INS\_312
CL\_INS\_312
CL\_INS\_312
CL\_INS\_312
CL\_INS\_312
CL\_INS\_312
CL\_INS\_312
CL\_INS\_312
CL\_INS\_312
CL\_INS\_312
CL\_INS\_312
CL\_INS\_312
CL\_INS\_312
CL\_INS\_312
CL\_INS\_312
CL\_INS\_312
CL\_INS\_312
CL\_INS\_312
CL\_INS\_312
CL\_INS\_312
CL\_INS\_312
CL\_INS\_312
CL\_INS\_276
CL\_INS\_312
CL\_INS\_312
CL\_INS\_312
CL\_INS\_312
CL\_INS\_312
CL\_INS\_312
CL\_INS\_312
CL\_INS\_312
CL\_INS\_312
CL\_INS\_312
CL\_INS\_91
CL\_INS\_312
CL\_INS\_312
CL\_INS\_312
CL\_INS\_91
CL\_INS\_91
CL\_INS\_139
CL\_INS\_91
CL\_INS\_91
CL\_INS\_312
CL\_INS\_312
CL\_INS\_312
CL\_INS\_312
CL\_INS\_312
CL\_INS\_312
CL\_INS\_312
CL\_INS\_312
CL\_INS\_312
CL\_INS\_312
CL\_INS\_312
CL\_INS\_10
CL\_INS\_312
CL\_INS\_312
CL\_INS\_312
CL\_INS\_312
CL\_INS\_312
CL\_INS\_312
CL\_INS\_312
CL\_INS\_313
CL\_INS\_312
CL\_INS\_312
CL\_INS\_91
CL\_INS\_312
CL\_INS\_312
CL\_INS\_313
CL\_INS\_313
CL\_INS\_312
CL\_INS\_312
CL\_INS\_312
CL\_INS\_312
CL\_INS\_312
CL\_INS\_312
CL\_INS\_312
CL\_INS\_312
CL\_INS\_312
CL\_INS\_312
CL\_INS\_312
CL\_INS\_312
CL\_INS\_312
CL\_INS\_312
CL\_INS\_312
CL\_INS\_312
CL\_INS\_312
CL\_INS\_312
CL\_INS\_312
CL\_INS\_312
CL\_INS\_312
CL\_INS\_312
CL\_INS\_312
CL\_INS\_312
CL\_INS\_312
CL\_INS\_312
CL\_INS\_312
CL\_INS\_312
CL\_INS\_312
CL\_INS\_312
CL\_INS\_312
CL\_INS\_312
CL\_INS\_312
CL\_INS\_312
CL\_INS\_312
CL\_INS\_277
CL\_INS\_312
CL\_INS\_312
CL\_INS\_312
CL\_INS\_312
CL\_INS\_312
CL\_INS\_139
CL\_INS\_312
CL\_INS\_347
CL\_INS\_312
CL\_INS\_347
CL\_INS\_312
CL\_INS\_347
CL\_INS\_312
CL\_INS\_312
CL\_INS\_347
CL\_INS\_347
CL\_INS\_347
CL\_INS\_91
CL\_INS\_307
CL\_INS\_271
CL\_INS\_312
CL\_INS\_307
CL\_INS\_312
CL\_INS\_312
CL\_INS\_278
CL\_INS\_278
CL\_INS\_278
CL\_INS\_278
CL\_INS\_278
CL\_INS\_132
CL\_INS\_132
CL\_INS\_312
CL\_INS\_312
CL\_INS\_312
CL\_INS\_312
CL\_INS\_312
CL\_INS\_312
CL\_INS\_312
CL\_INS\_312
CL\_INS\_273
CL\_INS\_312
CL\_INS\_347
CL\_INS\_312
CL\_INS\_312
CL\_INS\_312
CL\_INS\_307
CL\_INS\_312
CL\_INS\_312
CL\_INS\_91
CL\_INS\_312
CL\_INS\_312
CL\_INS\_263
CL\_INS\_263
CL\_INS\_263
CL\_INS\_263
CL\_INS\_312
CL\_INS\_271
CL\_INS\_312
CL\_INS\_312
CL\_INS\_312
CL\_INS\_263
CL\_INS\_273
CL\_INS\_312
CL\_INS\_91
CL\_INS\_91
CL\_INS\_312
CL\_INS\_263
CL\_INS\_312
CL\_INS\_312
CL\_INS\_155
CL\_INS\_312
CL\_INS\_307
CL\_INS\_312
CL\_INS\_91
CL\_INS\_312
CL\_INS\_312
CL\_INS\_263
CL\_INS\_312
CL\_INS\_312
CL\_INS\_307
CL\_INS\_263
CL\_INS\_312
CL\_INS\_312
CL\_INS\_91
CL\_INS\_312
CL\_INS\_263
CL\_INS\_321
CL\_INS\_307
CL\_INS\_307
CL\_INS\_307
CL\_INS\_312
CL\_INS\_347
CL\_INS\_312
CL\_INS\_312
CL\_INS\_312
CL\_INS\_312
CL\_INS\_312
CL\_INS\_312
CL\_INS\_347
CL\_INS\_312
CL\_INS\_312
CL\_INS\_312
CL\_INS\_347
CL\_INS\_347
CL\_INS\_347
CL\_INS\_347
CL\_INS\_347
CL\_INS\_312
CL\_INS\_312
CL\_INS\_347
CL\_INS\_312
CL\_INS\_312
CL\_INS\_312
CL\_INS\_312
CL\_INS\_347
CL\_INS\_347
CL\_INS\_347
CL\_INS\_347
CL\_INS\_347
CL\_INS\_312
CL\_INS\_91
CL\_INS\_347
CL\_INS\_312
CL\_INS\_312
CL\_INS\_347
CL\_INS\_312
CL\_INS\_312
CL\_INS\_347
CL\_INS\_312
CL\_INS\_278
CL\_INS\_312
CL\_INS\_385
CL\_INS\_312
CL\_INS\_278
CL\_INS\_312
CL\_INS\_278
CL\_INS\_312
CL\_INS\_278
CL\_INS\_278
CL\_INS\_278
CL\_INS\_278
CL\_INS\_278
CL\_INS\_278
CL\_INS\_278
CL\_INS\_132
CL\_INS\_307
CL\_INS\_278
CL\_INS\_197
CL\_INS\_278
CL\_INS\_278
CL\_INS\_278
CL\_INS\_278
CL\_INS\_278
CL\_INS\_278
CL\_INS\_278
CL\_INS\_401
CL\_INS\_197
CL\_INS\_197
CL\_INS\_278
CL\_INS\_312
CL\_INS\_312
CL\_INS\_278
CL\_INS\_278
CL\_INS\_278
CL\_INS\_278
CL\_INS\_278
CL\_INS\_278
CL\_INS\_278
CL\_INS\_278
CL\_INS\_278
CL\_INS\_278
CL\_INS\_312
CL\_INS\_278
CL\_INS\_278
CL\_INS\_278
CL\_INS\_278
CL\_INS\_278
CL\_INS\_278
CL\_INS\_312
CL\_INS\_312
CL\_INS\_278
CL\_INS\_312
CL\_INS\_278
CL\_INS\_278
CL\_INS\_278
CL\_INS\_278
CL\_INS\_278
CL\_INS\_278
CL\_INS\_278
CL\_INS\_278
CL\_INS\_278
CL\_INS\_278
CL\_INS\_278
CL\_INS\_278
CL\_INS\_278
CL\_INS\_278
CL\_INS\_278
CL\_INS\_278
CL\_INS\_278
CL\_INS\_278
CL\_INS\_278
CL\_INS\_278
CL\_INS\_278
CL\_INS\_278
CL\_INS\_278
CL\_INS\_278
CL\_INS\_278
CL\_INS\_278
CL\_INS\_278
CL\_INS\_278
CL\_INS\_278
CL\_INS\_278
CL\_INS\_278
CL\_INS\_312
CL\_INS\_312
CL\_INS\_385
CL\_INS\_307
CL\_INS\_312
CL\_INS\_263
CL\_INS\_263
CL\_INS\_312
Cluster ID


CL\_20974
CL\_6923
CL\_20604
CL\_20603
CL\_20744
CL\_20743
CL\_20742
CL\_20741
CL\_20740
CL\_17977
CL\_17976
CL\_17975
CL\_14082
CL\_14081
CL\_14080
CL\_14079
CL\_14078
CL\_14077
CL\_14076
CL\_17974
CL\_14075
CL\_14074
CL\_15540
CL\_13486
CL\_23077
CL\_23459
CL\_23458
CL\_15539
CL\_15538
CL\_15537
CL\_15536
CL\_15535
CL\_15534
CL\_15533
CL\_15532
CL\_23076
CL\_15531
CL\_15530
CL\_15529
CL\_23075
CL\_15528
CL\_15527
CL\_14073
CL\_14072
CL\_14071
CL\_16905
CL\_16904
CL\_12060
CL\_16903
CL\_16902
CL\_12058
CL\_12057
CL\_12056
CL\_717
CL\_716
CL\_715
CL\_714
CL\_713
CL\_16901
CL\_16900
CL\_16899
CL\_16898
CL\_12050
CL\_16897
CL\_15285
CL\_20973
CL\_5557
CL\_13030
CL\_5556
CL\_5555
CL\_12189
CL\_12190
CL\_6276
CL\_6277
CL\_6278
CL\_12191
CL\_6279
CL\_6280
CL\_6281
CL\_5554
CL\_5553
CL\_12423
CL\_16439
CL\_16172
CL\_23613
CL\_18109
CL\_14609
CL\_23157
CL\_15779
CL\_15778
CL\_18759
CL\_22504
CL\_20629
CL\_11015
CL\_20972
CL\_20971
CL\_20970
CL\_20851
CL\_20850
CL\_15196
CL\_15197
CL\_15198
CL\_19642
CL\_19643
CL\_5550
CL\_5549
CL\_5548
CL\_22088
CL\_11675
CL\_11676
CL\_12319
CL\_11672
CL\_11673
CL\_15286
CL\_12249
CL\_18758
CL\_6275
CL\_19941
CL\_14608
CL\_14607
CL\_20523
CL\_19942
CL\_19943
CL\_20524
CL\_14693
CL\_19944
CL\_22244
CL\_5552
CL\_14606
CL\_14605
CL\_16438
CL\_5551
CL\_17103
CL\_19644
CL\_19645
CL\_5547
CL\_14604
CL\_15777
CL\_10962
CL\_10961
CL\_5546
CL\_5545
CL\_19646
CL\_19647
CL\_19648
CL\_5544
CL\_5543
CL\_5542
CL\_5541
CL\_5540
CL\_5539
CL\_5538
CL\_5537
CL\_5536
CL\_16173
CL\_5535
CL\_5534
CL\_5533
CL\_12193
CL\_19945
CL\_12248
CL\_12247
CL\_12246
CL\_12245
CL\_12244
CL\_12243
CL\_12242
CL\_12241
CL\_12240
CL\_12239
CL\_12238
CL\_12237
CL\_12192
CL\_12236
CL\_5532
CL\_5531
CL\_5530
CL\_16174
CL\_18110
CL\_18549
CL\_6282
CL\_6283
CL\_12194
CL\_5796
CL\_13029
CL\_5797
CL\_18548
CL\_5798
CL\_18583
CL\_12195
CL\_5799
CL\_5800
CL\_5802
CL\_5803
CL\_12063
CL\_5229
CL\_14603
CL\_7832
CL\_19649
CL\_6284
CL\_8141
CL\_8167
CL\_12437
CL\_12438
CL\_12439
CL\_6320
CL\_6319
CL\_18582
CL\_18581
CL\_18580
CL\_18579
CL\_18578
CL\_12196
CL\_18577
CL\_18576
CL\_6288
CL\_15776
CL\_5801
CL\_16437
CL\_16641
CL\_16642
CL\_14561
CL\_23614
CL\_22087
CL\_5227
CL\_15287
CL\_15288
CL\_17461
CL\_17462
CL\_5643
CL\_5231
CL\_17784
CL\_8723
CL\_18111
CL\_15199
CL\_18547
CL\_5341
CL\_15870
CL\_18112
CL\_5228
CL\_19650
CL\_19651
CL\_6285
CL\_15775
CL\_15774
CL\_5642
CL\_18757
CL\_9709
CL\_20628
CL\_12294
CL\_20627
CL\_6286
CL\_5436
CL\_20969
CL\_20968
CL\_18728
CL\_8315
CL\_6287
CL\_12197
CL\_8318
CL\_13028
CL\_5644
CL\_13027
CL\_9711
CL\_5342
CL\_19946
CL\_6289
CL\_5804
CL\_12198
CL\_12199
CL\_12200
CL\_12201
CL\_12202
CL\_12203
CL\_5805
CL\_13026
CL\_18113
CL\_13025
CL\_5806
CL\_5807
CL\_5808
CL\_5809
CL\_5810
CL\_19652
CL\_12420
CL\_5811
CL\_18114
CL\_6290
CL\_12204
CL\_6291
CL\_5812
CL\_5813
CL\_5814
CL\_5815
CL\_5816
CL\_15773
CL\_5817
CL\_5818
CL\_15200
CL\_19653
CL\_5819
CL\_18756
CL\_6292
CL\_5820
CL\_5821
CL\_6885
CL\_6783
CL\_6782
CL\_6781
CL\_6635
CL\_6634
CL\_6633
CL\_6632
CL\_6631
CL\_6630
CL\_6629
CL\_6628
CL\_6627
CL\_6626
CL\_6625
CL\_6822
CL\_6821
CL\_6079
CL\_6820
CL\_6819
CL\_6818
CL\_6817
CL\_6137
CL\_6816
CL\_6815
CL\_6814
CL\_6813
CL\_6812
CL\_6811
CL\_6500
CL\_6499
CL\_6498
CL\_6497
CL\_6496
CL\_6495
CL\_6494
CL\_6493
CL\_6492
CL\_6491
CL\_6490
CL\_6489
CL\_6488
CL\_6487
CL\_6486
CL\_6485
CL\_6484
CL\_6483
CL\_6482
CL\_6481
CL\_6480
CL\_6479
CL\_6478
CL\_6477
CL\_6476
CL\_6475
CL\_6474
CL\_6473
CL\_6472
CL\_6471
CL\_6470
CL\_6469
CL\_6468
CL\_6467
CL\_6466
CL\_6465
CL\_6464
CL\_6463
CL\_6462
CL\_6461
CL\_6460
CL\_6459
CL\_6458
CL\_6457
CL\_6658
CL\_6657
CL\_6656
CL\_6655
CL\_6654
CL\_6653
CL\_6652
CL\_6651
CL\_6650
CL\_6649
CL\_6648
CL\_6695
CL\_6694
CL\_6693
CL\_6691
CL\_6690
CL\_6689
CL\_6688
CL\_6687
