## Supplementary material for "A novel method for integrating genomic and Tn-Seq data to identify common *in vivo* fitness mechanisms across multiple bacterial species": S2 Dataset: CL_INS_313.html

Legend

 Mobile +extrachromosomalelementfunctions
 Hypothetical
 AntibioticResistance
 Other
 All VFDB Genes

FULL


WINDOWSVGPNG

Trim RowsRemove SingletonsSave Fasta

CL\_3818


CL\_3818


CL\_3818


CL\_3818


CL\_3818


Break


CL\_3818


CL\_3818


CL\_3818


CL\_3817


CL\_3818


CL\_3818


CL\_3818


CL\_3818


CL\_3818


CL\_3818


CL\_3818


CL\_3818


CL\_3818


CL\_3818


CL\_3818

HighlightSelectShow Genomes


153

CL\_3819


18

CL\_3819


12

CL\_3819


2

CL\_3819


2

CL\_3819


1

CL\_3819


1

CL\_3820


1

CL\_3819


1

CL\_3819


1

CL\_3819


1

CL\_3819


1

CL\_3819


1

CL\_3819


1

CL\_3819


1

CL\_3819


1

CL\_3819


1

CL\_3819


1

CL\_3819


1

CL\_3819


1

CL\_3819


1

CL\_3819

fGI ID


CL\_INS\_91
CL\_INS\_91
CL\_INS\_313
CL\_INS\_91
CL\_INS\_91
CL\_INS\_91
CL\_INS\_91
CL\_INS\_91
CL\_INS\_91
CL\_INS\_91
CL\_INS\_91
CL\_INS\_91
CL\_INS\_91
CL\_INS\_91
CL\_INS\_74
CL\_INS\_313
CL\_INS\_313
CL\_INS\_313
CL\_INS\_313
CL\_INS\_313
CL\_INS\_313
CL\_INS\_313
CL\_INS\_91
CL\_INS\_313
CL\_INS\_313
CL\_INS\_313
CL\_INS\_313
CL\_INS\_313
CL\_INS\_313
CL\_INS\_313
CL\_INS\_313
CL\_INS\_313
CL\_INS\_91
CL\_INS\_91
CL\_INS\_91
CL\_INS\_313
CL\_INS\_91
CL\_INS\_313
CL\_INS\_313
CL\_INS\_313
CL\_INS\_313
CL\_INS\_91
CL\_INS\_91
CL\_INS\_91
CL\_INS\_91
CL\_INS\_91
CL\_INS\_91
CL\_INS\_313
CL\_INS\_313
CL\_INS\_313
CL\_INS\_313
CL\_INS\_313
CL\_INS\_313
CL\_INS\_313
CL\_INS\_313
CL\_INS\_91
CL\_INS\_313
CL\_INS\_313
CL\_INS\_313
CL\_INS\_313
CL\_INS\_313
CL\_INS\_313
CL\_INS\_313
CL\_INS\_313
CL\_INS\_313
CL\_INS\_313
CL\_INS\_313
CL\_INS\_313
CL\_INS\_313
CL\_INS\_313
CL\_INS\_313
CL\_INS\_313
CL\_INS\_313
CL\_INS\_313
CL\_INS\_313
CL\_INS\_263
CL\_INS\_263
CL\_INS\_263
CL\_INS\_263
CL\_INS\_263
CL\_INS\_313
CL\_INS\_313
CL\_INS\_74
CL\_INS\_74
CL\_INS\_74
CL\_INS\_313
CL\_INS\_91
CL\_INS\_91
CL\_INS\_91
CL\_INS\_91
CL\_INS\_91
CL\_INS\_91
CL\_INS\_91
CL\_INS\_91
CL\_INS\_313
CL\_INS\_313
CL\_INS\_91
CL\_INS\_91
CL\_INS\_313
CL\_INS\_91
CL\_INS\_313
CL\_INS\_313
CL\_INS\_313
CL\_INS\_91
CL\_INS\_91
CL\_INS\_91
CL\_INS\_91
CL\_INS\_91
CL\_INS\_313
CL\_INS\_313
CL\_INS\_313
CL\_INS\_91
CL\_INS\_91
CL\_INS\_263
CL\_INS\_263
CL\_INS\_263
CL\_INS\_313
CL\_INS\_91
CL\_INS\_313
CL\_INS\_91
CL\_INS\_263
CL\_INS\_313
CL\_INS\_313
CL\_INS\_313
CL\_INS\_321
CL\_INS\_91
CL\_INS\_91
CL\_INS\_313
CL\_INS\_91
CL\_INS\_91
CL\_INS\_91
CL\_INS\_313
CL\_INS\_313
CL\_INS\_313
CL\_INS\_313
CL\_INS\_313
CL\_INS\_313
CL\_INS\_313
CL\_INS\_313
Cluster ID


CL\_11671
CL\_5550
CL\_17103
CL\_11672
CL\_11673
CL\_5547
CL\_11674
CL\_11675
CL\_11676
CL\_11677
CL\_11678
CL\_11679
CL\_11680
CL\_11681
CL\_11814
CL\_12833
CL\_12834
CL\_12835
CL\_11941
CL\_24185
CL\_24184
CL\_24183
CL\_5714
CL\_24182
CL\_24181
CL\_24180
CL\_24179
CL\_24178
CL\_24177
CL\_24176
CL\_24175
CL\_24174
CL\_24068
CL\_11927
CL\_11926
CL\_22502
CL\_11925
CL\_11940
CL\_11939
CL\_11938
CL\_11937
CL\_11936
CL\_11935
CL\_11934
CL\_11933
CL\_11932
CL\_11931
CL\_11930
CL\_11929
CL\_11928
CL\_21822
CL\_7656
CL\_12498
CL\_7655
CL\_12499
CL\_10194
CL\_10959
CL\_10960
CL\_10961
CL\_10962
CL\_19654
CL\_19655
CL\_19656
CL\_10963
CL\_10964
CL\_10965
CL\_10966
CL\_10967
CL\_10968
CL\_10969
CL\_10970
CL\_21821
CL\_21820
CL\_21819
CL\_17781
CL\_5468
CL\_5467
CL\_5466
CL\_5465
CL\_5464
CL\_21818
CL\_21817
CL\_9342
CL\_9341
CL\_9340
CL\_22899
CL\_14828
CL\_10193
CL\_12836
CL\_10192
CL\_12837
CL\_12838
CL\_10191
CL\_10190
CL\_22023
CL\_10189
CL\_12839
CL\_14825
CL\_22503
CL\_10188
CL\_10187
CL\_10186
CL\_10185
CL\_12840
CL\_10184
CL\_11682
CL\_11683
CL\_10183
CL\_15410
CL\_18755
CL\_15411
CL\_12841
CL\_10182
CL\_15605
CL\_15604
CL\_14821
CL\_24173
CL\_10181
CL\_12842
CL\_10180
CL\_10179
CL\_22926
CL\_21816
CL\_22898
CL\_6534
CL\_10178
CL\_10177
CL\_19657
CL\_12843
CL\_10176
CL\_10175
CL\_10174
CL\_11684
CL\_11924
CL\_12844
CL\_24172
CL\_19658
CL\_10173
CL\_10172
