## Supplementary material for "A novel method for integrating genomic and Tn-Seq data to identify common *in vivo* fitness mechanisms across multiple bacterial species": S2 Dataset: CL_INS_314.html

Legend

 Hypothetical
 Other
 All VFDB Genes

FULL


WINDOWSVGPNG

Trim RowsRemove SingletonsSave Fasta

CL\_3825


CL\_3825


CL\_3825

HighlightSelectShow Genomes


134

CL\_3826


70

CL\_3826


1

CL\_3828

fGI ID

CL\_INS\_314
Cluster ID

CL\_5822
