## Supplementary material for "A novel method for integrating genomic and Tn-Seq data to identify common *in vivo* fitness mechanisms across multiple bacterial species": S2 Dataset: CL_INS_317.html

Legend

 Mobile +extrachromosomalelementfunctions
 Hypothetical
 All Fitness Genes
 All VFDB Genes

FULL


WINDOWSVGPNG

Trim RowsRemove SingletonsSave Fasta

CL\_3857


CL\_3857


CL\_3856


CL\_3857

HighlightSelectShow Genomes


164

CL\_3858


36

CL\_3858


2

CL\_3858


1

CL\_3858

fGI ID


CL\_INS\_317
CL\_INS\_317
CL\_INS\_317
Cluster ID


CL\_20626
CL\_20625
CL\_5823
