## Supplementary material for "A novel method for integrating genomic and Tn-Seq data to identify common *in vivo* fitness mechanisms across multiple bacterial species": S2 Dataset: CL_INS_318.html

Legend

 Mobile +extrachromosomalelementfunctions
 Regulatoryfunctions
 Hypothetical
 DNA Metabolism
 All EssentialGenes
 AntibioticResistance
 All Fitness Genes
 Biosynthesis ofcofactors,prostheticgroups, +carriers
 Cell Envelope
 Proteinsynthesis/fate
 Other
 EnergyMetabolism
 Transport +binding proteins
 All VFDB Genes

FULL


WINDOWSVGPNG

Trim RowsRemove SingletonsSave Fasta

CL\_3887


CL\_3887


CL\_3887


Break


CL\_3887


CL\_3887


CL\_3887


CL\_3887


CL\_3887


CL\_3887


Break


CL\_3887


CL\_3887


CL\_3887

HighlightSelectShow Genomes


160

CL\_3888


29

CL\_3888


4

Break


1

CL\_3888


1

Break


1

Break


1

Break


1

CL\_1112


1

Break


1

CL\_3888


1

CL\_3888


1

CL\_3888


1

Break


1

Break

fGI ID


CL\_INS\_318
CL\_INS\_318
CL\_INS\_318
CL\_INS\_318
CL\_INS\_160
CL\_INS\_318
CL\_INS\_160
CL\_INS\_318
CL\_INS\_318
CL\_INS\_318
CL\_INS\_318
CL\_INS\_318
CL\_INS\_318
CL\_INS\_318
CL\_INS\_318
CL\_INS\_318
CL\_INS\_318
CL\_INS\_318
CL\_INS\_318
CL\_INS\_318
CL\_INS\_318
CL\_INS\_318
CL\_INS\_91
CL\_INS\_342
CL\_INS\_400
CL\_INS\_243
CL\_INS\_243
CL\_INS\_318
CL\_INS\_318
CL\_INS\_400
CL\_INS\_401
CL\_INS\_401
CL\_INS\_401
CL\_INS\_401
CL\_INS\_401
CL\_INS\_318
CL\_INS\_318
CL\_INS\_318
CL\_INS\_278
CL\_INS\_385
CL\_INS\_385
CL\_INS\_368
CL\_INS\_368
CL\_INS\_368
CL\_INS\_368
CL\_INS\_368
CL\_INS\_318
CL\_INS\_385
CL\_INS\_385
CL\_INS\_385
CL\_INS\_385
CL\_INS\_385
CL\_INS\_318
CL\_INS\_385
CL\_INS\_385
CL\_INS\_385
CL\_INS\_132
CL\_INS\_321
CL\_INS\_321
CL\_INS\_385
CL\_INS\_385
CL\_INS\_318
CL\_INS\_318
CL\_INS\_318
CL\_INS\_318
CL\_INS\_318
CL\_INS\_318
CL\_INS\_318
CL\_INS\_318
CL\_INS\_318
CL\_INS\_318
CL\_INS\_318
CL\_INS\_318
CL\_INS\_400
CL\_INS\_400
CL\_INS\_400
CL\_INS\_400
CL\_INS\_400
CL\_INS\_400
CL\_INS\_278
CL\_INS\_400
CL\_INS\_132
CL\_INS\_132
CL\_INS\_132
CL\_INS\_278
CL\_INS\_278
CL\_INS\_278
CL\_INS\_278
CL\_INS\_278
CL\_INS\_132
CL\_INS\_132
CL\_INS\_400
CL\_INS\_132
CL\_INS\_385
CL\_INS\_318
CL\_INS\_385
CL\_INS\_385
CL\_INS\_318
CL\_INS\_318
CL\_INS\_318
CL\_INS\_132
CL\_INS\_132
CL\_INS\_132
CL\_INS\_132
CL\_INS\_132
CL\_INS\_132
CL\_INS\_132
CL\_INS\_132
CL\_INS\_132
CL\_INS\_132
CL\_INS\_132
CL\_INS\_132
CL\_INS\_132
CL\_INS\_132
CL\_INS\_160
CL\_INS\_278
CL\_INS\_278
CL\_INS\_278
CL\_INS\_278
CL\_INS\_278
CL\_INS\_132
CL\_INS\_132
CL\_INS\_132
CL\_INS\_132
CL\_INS\_132
CL\_INS\_132
CL\_INS\_307
CL\_INS\_307
CL\_INS\_132
CL\_INS\_132
CL\_INS\_318
CL\_INS\_318
CL\_INS\_318
CL\_INS\_401
CL\_INS\_278
CL\_INS\_278
CL\_INS\_278
CL\_INS\_132
CL\_INS\_132
CL\_INS\_132
CL\_INS\_132
CL\_INS\_318
CL\_INS\_318
CL\_INS\_318
CL\_INS\_318
CL\_INS\_318
CL\_INS\_318
CL\_INS\_318
CL\_INS\_318
CL\_INS\_318
CL\_INS\_318
CL\_INS\_278
CL\_INS\_318
CL\_INS\_132
CL\_INS\_318
CL\_INS\_318
CL\_INS\_318
CL\_INS\_318
CL\_INS\_318
CL\_INS\_318
CL\_INS\_132
CL\_INS\_278
CL\_INS\_278
CL\_INS\_132
CL\_INS\_401
CL\_INS\_401
CL\_INS\_318
CL\_INS\_401
CL\_INS\_278
CL\_INS\_401
CL\_INS\_278
CL\_INS\_278
CL\_INS\_278
CL\_INS\_401
CL\_INS\_278
CL\_INS\_132
CL\_INS\_132
CL\_INS\_132
CL\_INS\_278
CL\_INS\_278
CL\_INS\_278
CL\_INS\_401
CL\_INS\_278
CL\_INS\_318
CL\_INS\_318
CL\_INS\_278
CL\_INS\_278
CL\_INS\_278
CL\_INS\_278
CL\_INS\_278
CL\_INS\_401
CL\_INS\_307
CL\_INS\_401
CL\_INS\_278
CL\_INS\_278
CL\_INS\_278
CL\_INS\_278
CL\_INS\_278
CL\_INS\_278
CL\_INS\_401
CL\_INS\_58
CL\_INS\_318
CL\_INS\_178
CL\_INS\_58
CL\_INS\_278
CL\_INS\_278
CL\_INS\_58
CL\_INS\_278
CL\_INS\_278
CL\_INS\_318
CL\_INS\_178
CL\_INS\_178
CL\_INS\_178
CL\_INS\_178
CL\_INS\_178
CL\_INS\_278
CL\_INS\_278
CL\_INS\_278
CL\_INS\_278
CL\_INS\_132
CL\_INS\_132
CL\_INS\_132
CL\_INS\_278
CL\_INS\_278
CL\_INS\_278
CL\_INS\_401
CL\_INS\_278
CL\_INS\_204
CL\_INS\_204
CL\_INS\_318
CL\_INS\_204
CL\_INS\_204
CL\_INS\_204
CL\_INS\_278
CL\_INS\_204
CL\_INS\_204
CL\_INS\_204
CL\_INS\_253
CL\_INS\_204
CL\_INS\_132
CL\_INS\_132
CL\_INS\_318
CL\_INS\_132
CL\_INS\_318
CL\_INS\_30
CL\_INS\_401
CL\_INS\_401
CL\_INS\_401
CL\_INS\_278
CL\_INS\_278
CL\_INS\_278
CL\_INS\_278
CL\_INS\_278
CL\_INS\_278
CL\_INS\_278
CL\_INS\_401
CL\_INS\_105
CL\_INS\_278
CL\_INS\_278
CL\_INS\_278
CL\_INS\_278
CL\_INS\_318
CL\_INS\_318
CL\_INS\_318
CL\_INS\_318
CL\_INS\_204
CL\_INS\_318
CL\_INS\_318
CL\_INS\_318
CL\_INS\_318
CL\_INS\_318
CL\_INS\_58
CL\_INS\_318
CL\_INS\_318
CL\_INS\_318
CL\_INS\_58
CL\_INS\_58
CL\_INS\_160
CL\_INS\_132
CL\_INS\_132
CL\_INS\_132
CL\_INS\_278
CL\_INS\_278
CL\_INS\_278
CL\_INS\_278
CL\_INS\_278
CL\_INS\_160
CL\_INS\_278
CL\_INS\_278
CL\_INS\_278
CL\_INS\_278
CL\_INS\_278
CL\_INS\_278
CL\_INS\_278
CL\_INS\_278
CL\_INS\_132
CL\_INS\_318
CL\_INS\_318
CL\_INS\_318
CL\_INS\_318
CL\_INS\_318
CL\_INS\_318
CL\_INS\_132
CL\_INS\_132
CL\_INS\_278
CL\_INS\_171
CL\_INS\_171
CL\_INS\_278
CL\_INS\_160
CL\_INS\_160
CL\_INS\_318
CL\_INS\_278
CL\_INS\_278
CL\_INS\_318
CL\_INS\_318
CL\_INS\_318
CL\_INS\_318
CL\_INS\_318
CL\_INS\_318
CL\_INS\_318
CL\_INS\_318
CL\_INS\_318
CL\_INS\_318
CL\_INS\_318
CL\_INS\_318
CL\_INS\_318
CL\_INS\_318
CL\_INS\_318
CL\_INS\_132
CL\_INS\_278
CL\_INS\_318
CL\_INS\_318
CL\_INS\_318
CL\_INS\_318
CL\_INS\_318
CL\_INS\_318
CL\_INS\_318
CL\_INS\_318
CL\_INS\_318
CL\_INS\_318
CL\_INS\_318
CL\_INS\_318
CL\_INS\_318
CL\_INS\_318
CL\_INS\_318
CL\_INS\_318
CL\_INS\_318
CL\_INS\_318
CL\_INS\_318
CL\_INS\_318
CL\_INS\_318
CL\_INS\_318
CL\_INS\_318
CL\_INS\_318
CL\_INS\_318
CL\_INS\_318
CL\_INS\_318
CL\_INS\_318
CL\_INS\_318
CL\_INS\_318
CL\_INS\_318
CL\_INS\_318
CL\_INS\_318
CL\_INS\_318
CL\_INS\_318
CL\_INS\_253
CL\_INS\_253
CL\_INS\_253
CL\_INS\_253
CL\_INS\_253
CL\_INS\_253
CL\_INS\_253
CL\_INS\_253
CL\_INS\_253
CL\_INS\_253
CL\_INS\_253
CL\_INS\_253
CL\_INS\_253
CL\_INS\_253
CL\_INS\_253
CL\_INS\_253
CL\_INS\_253
CL\_INS\_253
CL\_INS\_253
CL\_INS\_253
CL\_INS\_253
CL\_INS\_253
CL\_INS\_253
CL\_INS\_253
CL\_INS\_253
CL\_INS\_253
CL\_INS\_253
CL\_INS\_253
CL\_INS\_253
CL\_INS\_253
CL\_INS\_253
CL\_INS\_253
CL\_INS\_253
CL\_INS\_253
CL\_INS\_253
CL\_INS\_253
CL\_INS\_253
CL\_INS\_366
CL\_INS\_253
CL\_INS\_253
CL\_INS\_253
CL\_INS\_253
CL\_INS\_253
CL\_INS\_253
CL\_INS\_253
CL\_INS\_253
CL\_INS\_253
CL\_INS\_253
CL\_INS\_253
CL\_INS\_253
CL\_INS\_253
CL\_INS\_253
CL\_INS\_253
CL\_INS\_253
CL\_INS\_253
CL\_INS\_253
CL\_INS\_253
CL\_INS\_253
CL\_INS\_253
CL\_INS\_253
CL\_INS\_253
CL\_INS\_253
CL\_INS\_318
CL\_INS\_253
CL\_INS\_253
CL\_INS\_253
CL\_INS\_253
CL\_INS\_253
CL\_INS\_253
CL\_INS\_253
CL\_INS\_253
CL\_INS\_253
CL\_INS\_253
CL\_INS\_253
CL\_INS\_253
CL\_INS\_318
CL\_INS\_318
CL\_INS\_318
CL\_INS\_318
CL\_INS\_318
CL\_INS\_318
CL\_INS\_318
CL\_INS\_318
CL\_INS\_318
CL\_INS\_318
CL\_INS\_318
CL\_INS\_318
CL\_INS\_318
CL\_INS\_318
CL\_INS\_318
CL\_INS\_318
CL\_INS\_318
CL\_INS\_318
CL\_INS\_318
CL\_INS\_318
CL\_INS\_334
CL\_INS\_318
CL\_INS\_318
CL\_INS\_318
CL\_INS\_318
CL\_INS\_318
CL\_INS\_318
CL\_INS\_318
CL\_INS\_278
CL\_INS\_278
CL\_INS\_278
CL\_INS\_385
CL\_INS\_318
CL\_INS\_318
CL\_INS\_385
CL\_INS\_287
CL\_INS\_385
CL\_INS\_318
CL\_INS\_318
CL\_INS\_318
CL\_INS\_307
CL\_INS\_307
CL\_INS\_278
CL\_INS\_278
CL\_INS\_385
CL\_INS\_318
CL\_INS\_318
CL\_INS\_318
CL\_INS\_318
CL\_INS\_318
CL\_INS\_318
CL\_INS\_318
CL\_INS\_318
CL\_INS\_318
CL\_INS\_318
CL\_INS\_318
CL\_INS\_385
CL\_INS\_385
CL\_INS\_385
CL\_INS\_278
CL\_INS\_278
CL\_INS\_278
CL\_INS\_385
CL\_INS\_278
CL\_INS\_385
CL\_INS\_318
CL\_INS\_204
CL\_INS\_278
CL\_INS\_278
CL\_INS\_278
CL\_INS\_30
CL\_INS\_278
CL\_INS\_278
CL\_INS\_263
CL\_INS\_263
CL\_INS\_278
CL\_INS\_132
CL\_INS\_132
CL\_INS\_132
CL\_INS\_318
CL\_INS\_278
CL\_INS\_385
CL\_INS\_385
CL\_INS\_278
CL\_INS\_278
CL\_INS\_278
CL\_INS\_318
CL\_INS\_318
CL\_INS\_318
CL\_INS\_318
CL\_INS\_318
CL\_INS\_318
CL\_INS\_318
CL\_INS\_318
CL\_INS\_318
CL\_INS\_318
CL\_INS\_318
CL\_INS\_318
CL\_INS\_278
CL\_INS\_385
CL\_INS\_318
CL\_INS\_318
CL\_INS\_385
CL\_INS\_385
CL\_INS\_385
CL\_INS\_385
CL\_INS\_385
CL\_INS\_385
CL\_INS\_385
CL\_INS\_204
CL\_INS\_321
CL\_INS\_321
CL\_INS\_321
CL\_INS\_307
CL\_INS\_307
CL\_INS\_385
CL\_INS\_318
CL\_INS\_171
CL\_INS\_318
CL\_INS\_385
CL\_INS\_278
CL\_INS\_278
CL\_INS\_385
CL\_INS\_91
CL\_INS\_318
CL\_INS\_318
CL\_INS\_318
CL\_INS\_318
CL\_INS\_318
CL\_INS\_318
CL\_INS\_278
CL\_INS\_293
CL\_INS\_318
CL\_INS\_318
CL\_INS\_318
CL\_INS\_318
CL\_INS\_385
CL\_INS\_318
CL\_INS\_318
CL\_INS\_334
CL\_INS\_318
CL\_INS\_318
CL\_INS\_318
CL\_INS\_132
CL\_INS\_132
CL\_INS\_318
CL\_INS\_318
CL\_INS\_401
CL\_INS\_307
CL\_INS\_307
CL\_INS\_307
CL\_INS\_307
CL\_INS\_307
CL\_INS\_307
CL\_INS\_307
CL\_INS\_307
CL\_INS\_278
CL\_INS\_385
CL\_INS\_57
CL\_INS\_385
CL\_INS\_318
CL\_INS\_401
CL\_INS\_401
CL\_INS\_318
CL\_INS\_318
CL\_INS\_30
CL\_INS\_30
CL\_INS\_30
CL\_INS\_204
CL\_INS\_307
CL\_INS\_307
CL\_INS\_307
CL\_INS\_307
CL\_INS\_307
CL\_INS\_307
CL\_INS\_307
CL\_INS\_307
CL\_INS\_307
CL\_INS\_307
CL\_INS\_307
CL\_INS\_307
CL\_INS\_307
CL\_INS\_307
CL\_INS\_307
CL\_INS\_74
CL\_INS\_307
CL\_INS\_307
CL\_INS\_307
CL\_INS\_307
CL\_INS\_318
CL\_INS\_318
CL\_INS\_129
CL\_INS\_307
CL\_INS\_385
CL\_INS\_385
CL\_INS\_318
CL\_INS\_318
CL\_INS\_318
CL\_INS\_318
CL\_INS\_318
CL\_INS\_318
CL\_INS\_204
CL\_INS\_246
CL\_INS\_246
CL\_INS\_318
CL\_INS\_318
CL\_INS\_318
CL\_INS\_318
CL\_INS\_204
Cluster ID


CL\_7370
CL\_6925
CL\_9700
CL\_9699
CL\_7963
CL\_8013
CL\_8012
CL\_13662
CL\_13663
CL\_13664
CL\_13665
CL\_13666
CL\_13667
CL\_13668
CL\_13669
CL\_13670
CL\_13671
CL\_13672
CL\_13673
CL\_13674
CL\_13675
CL\_13676
CL\_6027
CL\_12215
CL\_8215
CL\_6120
CL\_6121
CL\_13427
CL\_12566
CL\_6033
CL\_7091
CL\_6034
CL\_6037
CL\_6038
CL\_6039
CL\_23335
CL\_20685
CL\_23336
CL\_6089
CL\_15224
CL\_8228
CL\_8125
CL\_8126
CL\_8127
CL\_8128
CL\_8129
CL\_23337
CL\_8212
CL\_12399
CL\_12398
CL\_12397
CL\_12396
CL\_14666
CL\_12394
CL\_14667
CL\_11247
CL\_6793
CL\_12219
CL\_12218
CL\_6693
CL\_12217
CL\_20916
CL\_20917
CL\_22706
CL\_11260
CL\_11261
CL\_11262
CL\_11259
CL\_12972
CL\_12971
CL\_12970
CL\_22705
CL\_22704
CL\_6080
CL\_8156
CL\_8157
CL\_8158
CL\_9201
CL\_9202
CL\_8213
CL\_9203
CL\_8230
CL\_8231
CL\_8232
CL\_6081
CL\_6082
CL\_6084
CL\_6085
CL\_6086
CL\_6087
CL\_13353
CL\_8233
CL\_8229
CL\_106
CL\_23338
CL\_9941
CL\_7810
CL\_14670
CL\_13507
CL\_13505
CL\_6806
CL\_8219
CL\_13760
CL\_8220
CL\_8221
CL\_8222
CL\_8223
CL\_6712
CL\_6713
CL\_8224
CL\_8225
CL\_8226
CL\_8227
CL\_8166
CL\_68
CL\_8141
CL\_6070
CL\_6071
CL\_6072
CL\_6073
CL\_7946
CL\_7945
CL\_7944
CL\_12269
CL\_8242
CL\_8243
CL\_8244
CL\_8245
CL\_8246
CL\_11233
CL\_10856
CL\_17932
CL\_7112
CL\_7065
CL\_7064
CL\_7063
CL\_6810
CL\_6019
CL\_6018
CL\_6017
CL\_6016
CL\_11708
CL\_11709
CL\_11710
CL\_9416
CL\_9417
CL\_9418
CL\_9016
CL\_10865
CL\_11711
CL\_10866
CL\_10868
CL\_10867
CL\_9205
CL\_6102
CL\_6101
CL\_6100
CL\_6099
CL\_6098
CL\_6097
CL\_6015
CL\_6014
CL\_6013
CL\_6012
CL\_7965
CL\_7966
CL\_7967
CL\_8202
CL\_8200
CL\_8201
CL\_8199
CL\_8198
CL\_6011
CL\_6010
CL\_6009
CL\_6008
CL\_7058
CL\_6823
CL\_6005
CL\_6004
CL\_6003
CL\_8203
CL\_6002
CL\_10834
CL\_12489
CL\_6001
CL\_6000
CL\_5999
CL\_5998
CL\_5997
CL\_7055
CL\_7056
CL\_7057
CL\_5996
CL\_5995
CL\_5994
CL\_5993
CL\_5992
CL\_5991
CL\_7054
CL\_6826
CL\_12759
CL\_6828
CL\_6827
CL\_5988
CL\_5987
CL\_5986
CL\_6829
CL\_5983
CL\_7586
CL\_5985
CL\_7051
CL\_7052
CL\_7053
CL\_7960
CL\_5982
CL\_5981
CL\_5980
CL\_5979
CL\_7958
CL\_7957
CL\_7956
CL\_5975
CL\_5974
CL\_5973
CL\_7050
CL\_8650
CL\_6067
CL\_6066
CL\_23339
CL\_6064
CL\_6063
CL\_6062
CL\_12763
CL\_6060
CL\_6061
CL\_6059
CL\_12764
CL\_6058
CL\_7951
CL\_7952
CL\_7953
CL\_6872
CL\_7954
CL\_7955
CL\_8206
CL\_8205
CL\_8204
CL\_6053
CL\_6052
CL\_6051
CL\_6050
CL\_7592
CL\_7593
CL\_6047
CL\_6045
CL\_6044
CL\_6043
CL\_6042
CL\_6041
CL\_7595
CL\_6824
CL\_23340
CL\_23341
CL\_23342
CL\_6065
CL\_23343
CL\_23344
CL\_23345
CL\_23346
CL\_23347
CL\_12264
CL\_23348
CL\_10121
CL\_20578
CL\_7988
CL\_6852
CL\_6851
CL\_6049
CL\_6048
CL\_7987
CL\_6846
CL\_6845
CL\_6844
CL\_6843
CL\_6842
CL\_6841
CL\_6882
CL\_6883
CL\_7948
CL\_6885
CL\_7947
CL\_6879
CL\_6830
CL\_6831
CL\_8207
CL\_20859
CL\_20858
CL\_20857
CL\_20856
CL\_20855
CL\_8218
CL\_8208
CL\_8209
CL\_6874
CL\_7940
CL\_10906
CL\_6186
CL\_9219
CL\_9218
CL\_11712
CL\_8383
CL\_8384
CL\_8385
CL\_8386
CL\_23349
CL\_20872
CL\_20873
CL\_20874
CL\_20875
CL\_20876
CL\_20877
CL\_20878
CL\_20879
CL\_20880
CL\_20881
CL\_20882
CL\_20883
CL\_6881
CL\_7577
CL\_18990
CL\_14381
CL\_14379
CL\_14378
CL\_14377
CL\_14376
CL\_14375
CL\_14374
CL\_14373
CL\_14372
CL\_14371
CL\_14370
CL\_14369
CL\_14368
CL\_14367
CL\_14398
CL\_14417
CL\_14416
CL\_14415
CL\_14414
CL\_14413
CL\_14412
CL\_14411
CL\_14410
CL\_14408
CL\_13606
CL\_13546
CL\_13547
CL\_13548
CL\_14382
CL\_14383
CL\_14384
CL\_14385
CL\_14386
CL\_18975
CL\_11349
CL\_11348
CL\_11347
CL\_11346
CL\_11345
CL\_11344
CL\_11343
CL\_11342
CL\_11340
CL\_11339
CL\_11338
CL\_11337
CL\_11336
CL\_11335
CL\_11334
CL\_11333
CL\_11332
CL\_11331
CL\_11330
CL\_11329
CL\_11328
CL\_11327
CL\_11326
CL\_11325
CL\_11324
CL\_11323
CL\_11321
CL\_11320
CL\_11319
CL\_11318
CL\_11317
CL\_11316
CL\_11315
CL\_11314
CL\_11312
CL\_11311
CL\_11310
CL\_11309
CL\_11308
CL\_11307
CL\_11306
CL\_11305
CL\_11304
CL\_11303
CL\_11302
CL\_11301
CL\_11341
CL\_11378
CL\_11377
CL\_11376
CL\_11375
CL\_11374
CL\_11372
CL\_11371
CL\_11370
CL\_11369
CL\_11368
CL\_11367
CL\_11366
CL\_11365
CL\_11364
CL\_11363
CL\_11362
CL\_11361
CL\_11360
CL\_11359
CL\_11358
CL\_11357
CL\_11356
CL\_11355
CL\_11354
CL\_11353
CL\_11352
CL\_11351
CL\_11350
CL\_14388
CL\_14389
CL\_14390
CL\_14391
CL\_14392
CL\_18980
CL\_18981
CL\_18982
CL\_18983
CL\_17897
CL\_18984
CL\_18878
CL\_8142
CL\_8143
CL\_18985
CL\_23350
CL\_23351
CL\_20684
CL\_20683
CL\_20682
CL\_17783
CL\_9517
CL\_9174
CL\_10454
CL\_9176
CL\_10455
CL\_9177
CL\_9178
CL\_133
CL\_132
CL\_131
CL\_13445
CL\_13446
CL\_121
CL\_10456
CL\_12392
CL\_15221
CL\_8613
CL\_8612
CL\_23367
CL\_5932
CL\_8387
CL\_5936
CL\_5937
CL\_10457
CL\_10458
CL\_10459
CL\_10460
CL\_10461
CL\_10462
CL\_10463
CL\_10464
CL\_10465
CL\_120
CL\_13447
CL\_10466
CL\_119
CL\_118
CL\_117
CL\_116
CL\_115
CL\_114
CL\_7770
CL\_113
CL\_112
CL\_111
CL\_5953
CL\_109
CL\_108
CL\_107
CL\_105
CL\_6074
CL\_6075
CL\_6076
CL\_6077
CL\_6078
CL\_8237
CL\_8171
CL\_13749
CL\_23368
CL\_13751
CL\_13928
CL\_13927
CL\_103
CL\_104
CL\_102
CL\_10436
CL\_10437
CL\_10438
CL\_10439
CL\_10440
CL\_10441
CL\_10442
CL\_10443
CL\_10444
CL\_10445
CL\_10446
CL\_10447
CL\_24
CL\_7096
CL\_7095
CL\_9191
CL\_8887
CL\_8886
CL\_8885
CL\_8884
CL\_8883
CL\_8882
CL\_8881
CL\_5475
CL\_7426
CL\_7427
CL\_7428
CL\_3386
CL\_3387
CL\_13428
CL\_12797
CL\_7821
CL\_23329
CL\_16350
CL\_6673
CL\_6672
CL\_10777
CL\_8954
CL\_8953
CL\_8952
CL\_14938
CL\_14939
CL\_7104
CL\_14940
CL\_10452
CL\_7942
CL\_10451
CL\_10450
CL\_10449
CL\_10448
CL\_8880
CL\_23330
CL\_14756
CL\_7068
CL\_9405
CL\_9521
CL\_9520
CL\_8211
CL\_6054
CL\_6055
CL\_6056
CL\_6057
CL\_5938
CL\_5939
CL\_5940
CL\_5941
CL\_5942
CL\_17694
CL\_5944
CL\_8857
CL\_8050
CL\_5943
CL\_8061
CL\_90
CL\_22220
CL\_6025
CL\_6026
CL\_22219
CL\_22217
CL\_6028
CL\_6029
CL\_6030
CL\_8086
CL\_89
CL\_87
CL\_86
CL\_85
CL\_84
CL\_88
CL\_83
CL\_82
CL\_81
CL\_80
CL\_79
CL\_78
CL\_77
CL\_75
CL\_76
CL\_74
CL\_73
CL\_72
CL\_5951
CL\_7938
CL\_14769
CL\_8938
CL\_8888
CL\_8858
CL\_64
CL\_65
CL\_13967
CL\_66
CL\_23331
CL\_8915
CL\_12168
CL\_14776
CL\_6123
CL\_8898
CL\_8899
CL\_23332
CL\_23333
CL\_23334
CL\_9420
CL\_21188
