## Supplementary material for "A novel method for integrating genomic and Tn-Seq data to identify common *in vivo* fitness mechanisms across multiple bacterial species": S2 Dataset: CL_INS_319.html

Legend

 Mobile +extrachromosomalelementfunctions
 Regulatoryfunctions
 Hypothetical
 All EssentialGenes
 Other
 All VFDB Genes

FULL


WINDOWSVGPNG

Trim RowsRemove SingletonsSave Fasta

CL\_3901


CL\_3901


CL\_3901


CL\_3901


CL\_3901


CL\_3901


CL\_3901


CL\_3901

HighlightSelectShow Genomes


123

CL\_3902


75

CL\_3902


1

CL\_3902


1

CL\_3902


1

CL\_3902


1

CL\_3902


1

CL\_3902


1

CL\_3902

fGI ID


CL\_INS\_319
CL\_INS\_319
CL\_INS\_319
CL\_INS\_319
CL\_INS\_319
CL\_INS\_319
CL\_INS\_319
CL\_INS\_319
CL\_INS\_319
CL\_INS\_319
CL\_INS\_319
CL\_INS\_319
CL\_INS\_319
CL\_INS\_319
CL\_INS\_319
CL\_INS\_319
CL\_INS\_319
CL\_INS\_319
CL\_INS\_319
CL\_INS\_319
CL\_INS\_319
CL\_INS\_319
CL\_INS\_319
CL\_INS\_319
CL\_INS\_319
CL\_INS\_319
CL\_INS\_319
CL\_INS\_319
CL\_INS\_319
CL\_INS\_319
CL\_INS\_319
CL\_INS\_319
CL\_INS\_319
CL\_INS\_319
CL\_INS\_319
CL\_INS\_319
CL\_INS\_319
CL\_INS\_319
CL\_INS\_319
CL\_INS\_319
CL\_INS\_319
CL\_INS\_319
CL\_INS\_319
CL\_INS\_319
CL\_INS\_319
CL\_INS\_319
CL\_INS\_319
CL\_INS\_319
CL\_INS\_319
CL\_INS\_319
CL\_INS\_319
CL\_INS\_319
CL\_INS\_319
CL\_INS\_319
CL\_INS\_319
CL\_INS\_319
CL\_INS\_319
CL\_INS\_319
Cluster ID


CL\_5334
CL\_12705
CL\_16175
CL\_16176
CL\_12706
CL\_19068
CL\_12707
CL\_12708
CL\_5927
CL\_5926
CL\_5925
CL\_5924
CL\_13740
CL\_12709
CL\_5923
CL\_5921
CL\_12710
CL\_12711
CL\_5920
CL\_11078
CL\_12712
CL\_12713
CL\_5915
CL\_7210
CL\_5913
CL\_5912
CL\_5911
CL\_5910
CL\_5909
CL\_9647
CL\_9648
CL\_5906
CL\_11068
CL\_11067
CL\_5751
CL\_5750
CL\_5905
CL\_5904
CL\_5903
CL\_5902
CL\_5901
CL\_5900
CL\_5899
CL\_8684
CL\_11128
CL\_11174
CL\_5748
CL\_6384
CL\_12714
CL\_8687
CL\_5897
CL\_5896
CL\_5895
CL\_5894
CL\_5893
CL\_5892
CL\_5891
CL\_5890
