## Supplementary material for "A novel method for integrating genomic and Tn-Seq data to identify common *in vivo* fitness mechanisms across multiple bacterial species": S2 Dataset: CL_INS_322.html

Legend

 Mobile +extrachromosomalelementfunctions
 Regulatoryfunctions
 Hypothetical
 Other
 Transport +binding proteins
 All VFDB Genes

FULL


WINDOWSVGPNG

Trim RowsRemove SingletonsSave Fasta

CL\_3918


CL\_3918


CL\_3918


CL\_3918


CL\_3918


CL\_3918


CL\_3918


CL\_3918


CL\_3918


CL\_3918


CL\_3918


CL\_3918

HighlightSelectShow Genomes


167

CL\_3917


2

CL\_3916


1

CL\_3916


1

CL\_3916


1

CL\_3916


1

CL\_3916


1

CL\_3916


1

CL\_3916


1

CL\_3916


1

CL\_3916


1

CL\_3916


1

CL\_3916

fGI ID


CL\_INS\_321
CL\_INS\_321
CL\_INS\_204
CL\_INS\_321
CL\_INS\_321
CL\_INS\_321
CL\_INS\_321
CL\_INS\_321
CL\_INS\_321
CL\_INS\_321
CL\_INS\_321
CL\_INS\_321
CL\_INS\_321
CL\_INS\_321
CL\_INS\_321
CL\_INS\_321
CL\_INS\_321
CL\_INS\_321
CL\_INS\_321
CL\_INS\_321
CL\_INS\_321
CL\_INS\_321
CL\_INS\_321
CL\_INS\_321
CL\_INS\_321
CL\_INS\_321
CL\_INS\_322
CL\_INS\_321
CL\_INS\_321
CL\_INS\_321
CL\_INS\_321
CL\_INS\_321
CL\_INS\_321
CL\_INS\_321
CL\_INS\_321
CL\_INS\_321
CL\_INS\_321
CL\_INS\_334
CL\_INS\_334
CL\_INS\_334
CL\_INS\_334
CL\_INS\_321
CL\_INS\_321
CL\_INS\_321
CL\_INS\_321
CL\_INS\_321
CL\_INS\_321
CL\_INS\_322
CL\_INS\_321
CL\_INS\_321
CL\_INS\_321
CL\_INS\_321
CL\_INS\_321
CL\_INS\_321
CL\_INS\_321
CL\_INS\_321
CL\_INS\_321
CL\_INS\_321
CL\_INS\_321
CL\_INS\_321
CL\_INS\_321
CL\_INS\_132
CL\_INS\_132
CL\_INS\_132
CL\_INS\_132
CL\_INS\_132
CL\_INS\_132
CL\_INS\_132
CL\_INS\_321
CL\_INS\_132
CL\_INS\_132
CL\_INS\_132
CL\_INS\_132
CL\_INS\_132
CL\_INS\_132
CL\_INS\_132
CL\_INS\_132
CL\_INS\_132
CL\_INS\_132
CL\_INS\_132
CL\_INS\_132
CL\_INS\_132
CL\_INS\_132
CL\_INS\_319
CL\_INS\_319
CL\_INS\_132
CL\_INS\_319
CL\_INS\_132
CL\_INS\_132
CL\_INS\_132
CL\_INS\_132
CL\_INS\_132
CL\_INS\_132
CL\_INS\_132
CL\_INS\_132
CL\_INS\_132
CL\_INS\_132
CL\_INS\_132
CL\_INS\_321
Cluster ID


CL\_6922
CL\_6923
CL\_8086
CL\_5327
CL\_5328
CL\_5329
CL\_23677
CL\_5330
CL\_5331
CL\_5332
CL\_14191
CL\_14190
CL\_14189
CL\_14188
CL\_14187
CL\_14186
CL\_21427
CL\_6346
CL\_6345
CL\_6344
CL\_6343
CL\_6342
CL\_6341
CL\_6340
CL\_11630
CL\_11629
CL\_10168
CL\_10169
CL\_10170
CL\_10171
CL\_5720
CL\_12144
CL\_7524
CL\_8270
CL\_12143
CL\_8272
CL\_8273
CL\_8274
CL\_8658
CL\_8276
CL\_8277
CL\_21165
CL\_21166
CL\_21167
CL\_10470
CL\_10471
CL\_10472
CL\_5786
CL\_5785
CL\_5784
CL\_5783
CL\_5782
CL\_5781
CL\_5333
CL\_5780
CL\_5779
CL\_5778
CL\_5777
CL\_5776
CL\_5775
CL\_5774
CL\_5773
CL\_5772
CL\_5771
CL\_5770
CL\_5769
CL\_5768
CL\_5767
CL\_5766
CL\_5765
CL\_5764
CL\_5763
CL\_5762
CL\_5761
CL\_5760
CL\_5759
CL\_5758
CL\_5757
CL\_5756
CL\_5755
CL\_5754
CL\_5753
CL\_5752
CL\_5751
CL\_5750
CL\_5749
CL\_5748
CL\_5747
CL\_5746
CL\_5745
CL\_5744
CL\_5743
CL\_5742
CL\_5741
CL\_5740
CL\_5739
CL\_5738
CL\_5737
CL\_5736
