## Supplementary material for "A novel method for integrating genomic and Tn-Seq data to identify common *in vivo* fitness mechanisms across multiple bacterial species": S2 Dataset: CL_INS_323.html

Legend

 Regulatoryfunctions
 Hypothetical
 DNA Metabolism
 All EssentialGenes
 Other
 Transport +binding proteins
 All VFDB Genes

FULL


WINDOWSVGPNG

Trim RowsRemove SingletonsSave Fasta

CL\_3920


CL\_3920


CL\_3920


CL\_3920


CL\_3920


CL\_3920


CL\_3920


CL\_3920


CL\_3920

HighlightSelectShow Genomes


180

CL\_3919


1

CL\_3916


1

CL\_3916


1

CL\_3916


1

CL\_3916


1

CL\_3916


1

CL\_3916


1

CL\_3916


1

CL\_3916

fGI ID


CL\_INS\_323
CL\_INS\_321
CL\_INS\_323
CL\_INS\_321
CL\_INS\_321
CL\_INS\_321
CL\_INS\_321
CL\_INS\_321
CL\_INS\_321
CL\_INS\_321
CL\_INS\_321
CL\_INS\_321
CL\_INS\_321
CL\_INS\_321
CL\_INS\_321
CL\_INS\_321
CL\_INS\_321
CL\_INS\_321
CL\_INS\_323
CL\_INS\_321
CL\_INS\_321
CL\_INS\_321
CL\_INS\_307
CL\_INS\_321
CL\_INS\_321
CL\_INS\_334
CL\_INS\_334
CL\_INS\_321
CL\_INS\_321
CL\_INS\_321
CL\_INS\_321
CL\_INS\_321
CL\_INS\_321
CL\_INS\_321
CL\_INS\_321
CL\_INS\_321
CL\_INS\_321
CL\_INS\_334
CL\_INS\_334
CL\_INS\_334
CL\_INS\_321
CL\_INS\_321
CL\_INS\_321
CL\_INS\_334
CL\_INS\_321
CL\_INS\_321
CL\_INS\_321
CL\_INS\_321
CL\_INS\_321
CL\_INS\_321
CL\_INS\_321
CL\_INS\_321
CL\_INS\_321
CL\_INS\_321
CL\_INS\_321
CL\_INS\_321
CL\_INS\_321
CL\_INS\_321
CL\_INS\_321
CL\_INS\_321
CL\_INS\_321
CL\_INS\_321
CL\_INS\_321
CL\_INS\_321
CL\_INS\_321
CL\_INS\_321
CL\_INS\_321
CL\_INS\_321
CL\_INS\_321
CL\_INS\_321
CL\_INS\_321
CL\_INS\_321
CL\_INS\_321
CL\_INS\_321
CL\_INS\_321
CL\_INS\_321
CL\_INS\_321
CL\_INS\_321
CL\_INS\_321
CL\_INS\_321
CL\_INS\_321
CL\_INS\_321
CL\_INS\_321
CL\_INS\_321
CL\_INS\_321
CL\_INS\_321
CL\_INS\_321
CL\_INS\_321
Cluster ID


CL\_23008
CL\_23009
CL\_14601
CL\_14435
CL\_6346
CL\_6345
CL\_6344
CL\_6343
CL\_6342
CL\_6341
CL\_6340
CL\_11630
CL\_11629
CL\_14602
CL\_17523
CL\_5588
CL\_5589
CL\_5590
CL\_9278
CL\_9279
CL\_9280
CL\_9281
CL\_7761
CL\_9282
CL\_9283
CL\_7755
CL\_7754
CL\_9284
CL\_9285
CL\_9286
CL\_9287
CL\_9288
CL\_9289
CL\_9290
CL\_9291
CL\_9292
CL\_9293
CL\_9294
CL\_9295
CL\_9296
CL\_13125
CL\_13126
CL\_13127
CL\_1165
CL\_13128
CL\_13129
CL\_13130
CL\_12060
CL\_13131
CL\_7412
CL\_8352
CL\_7413
CL\_7414
CL\_8353
CL\_10479
CL\_8354
CL\_8355
CL\_10480
CL\_10481
CL\_10482
CL\_17522
CL\_17521
CL\_10483
CL\_7415
CL\_7416
CL\_7417
CL\_7418
CL\_7419
CL\_7420
CL\_7421
CL\_7422
CL\_7423
CL\_11752
CL\_11751
CL\_13132
CL\_11750
CL\_10705
CL\_10704
CL\_13133
CL\_9351
CL\_9352
CL\_9353
CL\_13134
CL\_7433
CL\_7432
CL\_7431
CL\_17520
CL\_13135
