## Supplementary material for "A novel method for integrating genomic and Tn-Seq data to identify common *in vivo* fitness mechanisms across multiple bacterial species": S2 Dataset: CL_INS_326.html

Legend

 Mobile +extrachromosomalelementfunctions
 Regulatoryfunctions
 Hypothetical
 All EssentialGenes
 Other
 Transport +binding proteins
 All VFDB Genes

FULL


WINDOWSVGPNG

Trim RowsRemove SingletonsSave Fasta

CL\_3957


CL\_3957


CL\_3957


CL\_3956


CL\_3957


CL\_3957


CL\_3916


CL\_3916


CL\_3923

HighlightSelectShow Genomes


128

CL\_3960


58

CL\_3960


3

CL\_3960


1

CL\_3960


1

CL\_3960


1

CL\_3960


1

CL\_3960


1

CL\_3960


1

CL\_3960

fGI ID


CL\_INS\_326
CL\_INS\_326
CL\_INS\_321
CL\_INS\_321
CL\_INS\_321
CL\_INS\_321
CL\_INS\_321
CL\_INS\_321
CL\_INS\_321
CL\_INS\_321
CL\_INS\_263
CL\_INS\_263
CL\_INS\_307
CL\_INS\_155
CL\_INS\_155
CL\_INS\_155
CL\_INS\_155
CL\_INS\_155
CL\_INS\_155
CL\_INS\_155
CL\_INS\_155
CL\_INS\_155
CL\_INS\_155
CL\_INS\_155
CL\_INS\_155
CL\_INS\_155
CL\_INS\_155
CL\_INS\_155
CL\_INS\_155
CL\_INS\_155
CL\_INS\_321
CL\_INS\_321
CL\_INS\_321
CL\_INS\_321
CL\_INS\_155
CL\_INS\_155
CL\_INS\_321
CL\_INS\_155
CL\_INS\_321
CL\_INS\_321
CL\_INS\_321
CL\_INS\_321
CL\_INS\_321
CL\_INS\_321
CL\_INS\_321
CL\_INS\_155
CL\_INS\_155
CL\_INS\_155
CL\_INS\_155
CL\_INS\_155
CL\_INS\_155
CL\_INS\_155
CL\_INS\_155
CL\_INS\_321
CL\_INS\_321
CL\_INS\_263
CL\_INS\_321
CL\_INS\_321
CL\_INS\_321
CL\_INS\_321
CL\_INS\_321
CL\_INS\_321
CL\_INS\_321
CL\_INS\_321
CL\_INS\_321
CL\_INS\_321
CL\_INS\_321
CL\_INS\_321
CL\_INS\_321
CL\_INS\_321
CL\_INS\_321
CL\_INS\_326
Cluster ID


CL\_3958
CL\_18575
CL\_19391
CL\_19390
CL\_6339
CL\_19660
CL\_19661
CL\_19662
CL\_19663
CL\_19664
CL\_5385
CL\_5664
CL\_6645
CL\_11051
CL\_11052
CL\_7250
CL\_11053
CL\_7248
CL\_7247
CL\_7246
CL\_7245
CL\_7244
CL\_7243
CL\_7242
CL\_7241
CL\_7240
CL\_7239
CL\_11054
CL\_11055
CL\_2110
CL\_19665
CL\_19666
CL\_19667
CL\_18825
CL\_7664
CL\_7663
CL\_16633
CL\_2102
CL\_16634
CL\_16635
CL\_19668
CL\_16636
CL\_16637
CL\_16638
CL\_16639
CL\_2093
CL\_2091
CL\_7661
CL\_2088
CL\_2087
CL\_2086
CL\_2085
CL\_2084
CL\_13027
CL\_19669
CL\_8721
CL\_19670
CL\_5333
CL\_11903
CL\_11902
CL\_11630
CL\_5298
CL\_6536
CL\_6343
CL\_6344
CL\_6345
CL\_6346
CL\_22825
CL\_11901
CL\_14435
CL\_19671
CL\_3959
