## Supplementary material for "A novel method for integrating genomic and Tn-Seq data to identify common *in vivo* fitness mechanisms across multiple bacterial species": S2 Dataset: CL_INS_328.html

Legend

 Hypothetical
 All VFDB Genes

FULL


WINDOWSVGPNG

Trim RowsRemove SingletonsSave Fasta

CL\_3989


CL\_3989


CL\_3989

HighlightSelectShow Genomes


116

CL\_3991


87

CL\_3991


1

CL\_3992

fGI ID

CL\_INS\_328
Cluster ID

CL\_3990
