## Supplementary material for "A novel method for integrating genomic and Tn-Seq data to identify common *in vivo* fitness mechanisms across multiple bacterial species": S2 Dataset: CL_INS_329.html

Legend

 Mobile +extrachromosomalelementfunctions
 Regulatoryfunctions
 Hypothetical
 Other
 All VFDB Genes

FULL


WINDOWSVGPNG

Trim RowsRemove SingletonsSave Fasta

CL\_3999


CL\_3999


CL\_3998


CL\_3999


CL\_3999

HighlightSelectShow Genomes


127

CL\_4001


70

CL\_4001


2

CL\_4001


1

Break


1

Break

fGI ID


CL\_INS\_329
CL\_INS\_329
CL\_INS\_263
CL\_INS\_263
CL\_INS\_263
CL\_INS\_263
CL\_INS\_263
CL\_INS\_263
CL\_INS\_263
CL\_INS\_263
CL\_INS\_263
CL\_INS\_263
CL\_INS\_263
CL\_INS\_263
CL\_INS\_263
CL\_INS\_263
CL\_INS\_246
CL\_INS\_263
CL\_INS\_263
CL\_INS\_263
CL\_INS\_329
CL\_INS\_329
CL\_INS\_321
CL\_INS\_321
CL\_INS\_321
CL\_INS\_321
CL\_INS\_321
CL\_INS\_321
CL\_INS\_321
CL\_INS\_321
CL\_INS\_321
CL\_INS\_321
CL\_INS\_321
CL\_INS\_321
CL\_INS\_321
CL\_INS\_321
CL\_INS\_263
CL\_INS\_263
CL\_INS\_263
CL\_INS\_263
CL\_INS\_329
CL\_INS\_329
CL\_INS\_329
CL\_INS\_329
Cluster ID


CL\_4000
CL\_12028
CL\_5461
CL\_5460
CL\_5459
CL\_5458
CL\_5457
CL\_5456
CL\_5455
CL\_5454
CL\_5453
CL\_5452
CL\_5450
CL\_5449
CL\_5448
CL\_6801
CL\_6802
CL\_6726
CL\_5218
CL\_5219
CL\_12182
CL\_12181
CL\_6725
CL\_6724
CL\_6723
CL\_6602
CL\_6600
CL\_6599
CL\_6598
CL\_6597
CL\_6596
CL\_6595
CL\_8073
CL\_6594
CL\_6593
CL\_6592
CL\_6591
CL\_6590
CL\_6589
CL\_6588
CL\_6587
CL\_8780
CL\_6586
CL\_6585
