## Supplementary material for "A novel method for integrating genomic and Tn-Seq data to identify common *in vivo* fitness mechanisms across multiple bacterial species": S2 Dataset: CL_INS_330.html

Legend

 Hypothetical
 All VFDB Genes

FULL


WINDOWSVGPNG

Trim RowsRemove SingletonsSave Fasta

CL\_4001


CL\_4001


CL\_4001

HighlightSelectShow Genomes


131

CL\_4002


73

CL\_4002


1

CL\_4002

fGI ID


CL\_INS\_330
CL\_INS\_330
Cluster ID


CL\_5787
CL\_12512
