## Supplementary material for "A novel method for integrating genomic and Tn-Seq data to identify common *in vivo* fitness mechanisms across multiple bacterial species": S2 Dataset: CL_INS_331.html

Legend

 Hypothetical
 All VFDB Genes

FULL


WINDOWSVGPNG

Trim RowsRemove SingletonsSave Fasta

CL\_4008


CL\_4008


CL\_4008


CL\_4007

HighlightSelectShow Genomes


103

CL\_4009


90

CL\_4009


9

CL\_4009


3

CL\_4009

fGI ID


CL\_INS\_331
CL\_INS\_331
Cluster ID


CL\_5788
CL\_11410
