## Supplementary material for "A novel method for integrating genomic and Tn-Seq data to identify common *in vivo* fitness mechanisms across multiple bacterial species": S2 Dataset: CL_INS_332.html

Legend

 Hypothetical
 All VFDB Genes
 Transport +binding proteins

FULL


WINDOWSVGPNG

Trim RowsRemove SingletonsSave Fasta

CL\_4024


CL\_4024

HighlightSelectShow Genomes


118

CL\_4026


85

CL\_4026

fGI ID

CL\_INS\_332
Cluster ID

CL\_4025
