## Supplementary material for "A novel method for integrating genomic and Tn-Seq data to identify common *in vivo* fitness mechanisms across multiple bacterial species": S2 Dataset: CL_INS_334.html

Legend

 Mobile +extrachromosomalelementfunctions
 Regulatoryfunctions
 Hypothetical
 DNA Metabolism
 All EssentialGenes
 Proteinsynthesis/fate
 Other
 All VFDB Genes
 Transport +binding proteins

FULL


WINDOWSVGPNG

Trim RowsRemove SingletonsSave Fasta

CL\_4066


CL\_4066


CL\_4066


CL\_4066


CL\_4066


CL\_4066


CL\_4066


CL\_4066


CL\_4066


CL\_4066


CL\_4066


CL\_4066


CL\_4066


CL\_3709


CL\_4066

HighlightSelectShow Genomes


149

CL\_5326


1

CL\_4067


1

CL\_4067


1

CL\_4067


1

CL\_5325


1

CL\_4067


1

CL\_4067


1

CL\_4067


1

CL\_4067


1

CL\_4067


1

CL\_4067


1

CL\_4067


1

CL\_4067


1

CL\_5326


1

CL\_4067

fGI ID


CL\_INS\_334
CL\_INS\_334
CL\_INS\_334
CL\_INS\_334
CL\_INS\_334
CL\_INS\_334
CL\_INS\_334
CL\_INS\_334
CL\_INS\_334
CL\_INS\_334
CL\_INS\_334
CL\_INS\_334
CL\_INS\_334
CL\_INS\_334
CL\_INS\_334
CL\_INS\_334
CL\_INS\_334
CL\_INS\_334
CL\_INS\_334
CL\_INS\_334
CL\_INS\_334
CL\_INS\_334
CL\_INS\_334
CL\_INS\_334
CL\_INS\_334
CL\_INS\_334
CL\_INS\_334
CL\_INS\_334
CL\_INS\_334
CL\_INS\_334
CL\_INS\_334
CL\_INS\_334
CL\_INS\_334
CL\_INS\_334
CL\_INS\_334
CL\_INS\_334
CL\_INS\_334
CL\_INS\_334
CL\_INS\_334
CL\_INS\_334
CL\_INS\_334
CL\_INS\_334
CL\_INS\_334
CL\_INS\_334
CL\_INS\_334
CL\_INS\_334
CL\_INS\_334
CL\_INS\_334
CL\_INS\_334
CL\_INS\_334
CL\_INS\_334
CL\_INS\_334
CL\_INS\_334
CL\_INS\_334
CL\_INS\_334
CL\_INS\_334
CL\_INS\_334
CL\_INS\_334
CL\_INS\_334
CL\_INS\_334
CL\_INS\_334
CL\_INS\_334
CL\_INS\_334
CL\_INS\_334
CL\_INS\_334
CL\_INS\_334
CL\_INS\_334
CL\_INS\_334
CL\_INS\_334
CL\_INS\_334
CL\_INS\_334
CL\_INS\_334
CL\_INS\_334
CL\_INS\_334
CL\_INS\_334
CL\_INS\_334
CL\_INS\_334
CL\_INS\_334
CL\_INS\_334
CL\_INS\_334
CL\_INS\_334
CL\_INS\_334
CL\_INS\_334
CL\_INS\_334
CL\_INS\_334
CL\_INS\_334
CL\_INS\_334
CL\_INS\_334
CL\_INS\_334
CL\_INS\_334
Cluster ID


CL\_21187
CL\_21186
CL\_21185
CL\_21184
CL\_21183
CL\_21182
CL\_9296
CL\_9295
CL\_9294
CL\_21181
CL\_7297
CL\_7296
CL\_7295
CL\_7294
CL\_23007
CL\_13024
CL\_13023
CL\_13022
CL\_13021
CL\_7068
CL\_13020
CL\_1165
CL\_21164
CL\_21163
CL\_21162
CL\_21161
CL\_21160
CL\_21159
CL\_21158
CL\_21157
CL\_21156
CL\_21155
CL\_21154
CL\_21153
CL\_12050
CL\_9276
CL\_8680
CL\_8679
CL\_8678
CL\_8677
CL\_8676
CL\_8675
CL\_8674
CL\_8673
CL\_8672
CL\_8671
CL\_8670
CL\_8669
CL\_8668
CL\_8667
CL\_8666
CL\_8665
CL\_8664
CL\_8663
CL\_8662
CL\_7752
CL\_9275
CL\_9274
CL\_9273
CL\_9272
CL\_9271
CL\_9270
CL\_9269
CL\_9268
CL\_7753
CL\_7754
CL\_7755
CL\_7756
CL\_17327
CL\_17326
CL\_17325
CL\_17324
CL\_9267
CL\_9266
CL\_9265
CL\_9264
CL\_9263
CL\_20597
CL\_8661
CL\_8660
CL\_8659
CL\_8277
CL\_8276
CL\_8658
CL\_8274
CL\_8657
CL\_6135
CL\_17783
CL\_8266
CL\_8263
