## Supplementary material for "A novel method for integrating genomic and Tn-Seq data to identify common *in vivo* fitness mechanisms across multiple bacterial species": S2 Dataset: CL_INS_335.html

Legend

 Mobile +extrachromosomalelementfunctions
 Regulatoryfunctions
 Hypothetical
 DNA Metabolism
 All EssentialGenes
 Proteinsynthesis/fate
 Other
 All VFDB Genes

FULL


WINDOWSVGPNG

Trim RowsRemove SingletonsSave Fasta

CL\_5324


CL\_5324


CL\_5324


CL\_4066


CL\_4066


CL\_5324


CL\_4066


CL\_5324


CL\_5324


CL\_4066


CL\_4066


CL\_5324


CL\_4066


CL\_5324


CL\_4066


CL\_5324


CL\_4066


CL\_4066


CL\_5324


CL\_4066


CL\_4066


CL\_5324


CL\_4066

HighlightSelectShow Genomes


137

CL\_4067


9

CL\_4067


2

CL\_4067


1

CL\_4067


1

CL\_4067


1

CL\_4067


1

CL\_4067


1

CL\_4067


1

CL\_4068


1

CL\_4067


1

CL\_4067


1

CL\_4067


1

CL\_4067


1

CL\_4067


1

CL\_4067


1

CL\_4067


1

CL\_4067


1

CL\_4067


1

CL\_4067


1

CL\_4067


1

CL\_4067


1

CL\_4067


1

CL\_4067

fGI ID


CL\_INS\_335
CL\_INS\_335
CL\_INS\_335
CL\_INS\_335
CL\_INS\_334
CL\_INS\_334
CL\_INS\_334
CL\_INS\_334
CL\_INS\_334
CL\_INS\_334
CL\_INS\_335
CL\_INS\_335
CL\_INS\_335
CL\_INS\_335
CL\_INS\_334
CL\_INS\_334
CL\_INS\_334
CL\_INS\_334
CL\_INS\_334
CL\_INS\_334
CL\_INS\_334
CL\_INS\_334
CL\_INS\_334
CL\_INS\_334
CL\_INS\_334
CL\_INS\_334
CL\_INS\_334
CL\_INS\_335
CL\_INS\_335
CL\_INS\_334
CL\_INS\_132
CL\_INS\_335
CL\_INS\_335
CL\_INS\_335
CL\_INS\_335
CL\_INS\_335
CL\_INS\_335
CL\_INS\_335
CL\_INS\_335
CL\_INS\_335
CL\_INS\_335
CL\_INS\_335
CL\_INS\_335
CL\_INS\_335
CL\_INS\_335
CL\_INS\_335
CL\_INS\_335
CL\_INS\_335
CL\_INS\_335
CL\_INS\_335
CL\_INS\_334
CL\_INS\_334
CL\_INS\_334
CL\_INS\_334
CL\_INS\_334
CL\_INS\_334
CL\_INS\_334
CL\_INS\_334
CL\_INS\_334
CL\_INS\_334
CL\_INS\_334
CL\_INS\_334
CL\_INS\_334
CL\_INS\_334
CL\_INS\_334
CL\_INS\_334
CL\_INS\_334
CL\_INS\_334
CL\_INS\_334
CL\_INS\_334
CL\_INS\_334
CL\_INS\_132
CL\_INS\_334
CL\_INS\_334
CL\_INS\_334
CL\_INS\_334
CL\_INS\_334
CL\_INS\_334
CL\_INS\_334
CL\_INS\_334
CL\_INS\_334
CL\_INS\_321
CL\_INS\_335
CL\_INS\_334
CL\_INS\_334
CL\_INS\_334
CL\_INS\_334
CL\_INS\_334
CL\_INS\_335
CL\_INS\_334
CL\_INS\_334
CL\_INS\_334
CL\_INS\_334
CL\_INS\_334
CL\_INS\_334
CL\_INS\_334
CL\_INS\_334
CL\_INS\_334
CL\_INS\_334
CL\_INS\_334
CL\_INS\_334
CL\_INS\_334
CL\_INS\_334
CL\_INS\_334
CL\_INS\_321
CL\_INS\_321
CL\_INS\_335
CL\_INS\_321
CL\_INS\_335
CL\_INS\_335
CL\_INS\_335
CL\_INS\_307
CL\_INS\_307
CL\_INS\_307
CL\_INS\_307
CL\_INS\_307
CL\_INS\_45
CL\_INS\_335
CL\_INS\_335
CL\_INS\_335
CL\_INS\_335
CL\_INS\_334
CL\_INS\_334
CL\_INS\_334
CL\_INS\_334
CL\_INS\_335
CL\_INS\_335
CL\_INS\_334
CL\_INS\_204
Cluster ID


CL\_11756
CL\_13612
CL\_16334
CL\_13611
CL\_13023
CL\_13022
CL\_13021
CL\_7068
CL\_13020
CL\_1165
CL\_20601
CL\_20600
CL\_20599
CL\_20598
CL\_21164
CL\_21163
CL\_21162
CL\_21161
CL\_21160
CL\_21159
CL\_21158
CL\_21157
CL\_21156
CL\_21155
CL\_21154
CL\_21153
CL\_12050
CL\_8279
CL\_8278
CL\_13024
CL\_6840
CL\_18028
CL\_18027
CL\_18026
CL\_18025
CL\_18024
CL\_18023
CL\_18022
CL\_18021
CL\_18020
CL\_18019
CL\_18018
CL\_18017
CL\_18016
CL\_18015
CL\_18014
CL\_18013
CL\_18012
CL\_18011
CL\_13019
CL\_9276
CL\_8680
CL\_8679
CL\_8678
CL\_8677
CL\_8676
CL\_8675
CL\_8674
CL\_8673
CL\_8672
CL\_8671
CL\_8670
CL\_8669
CL\_8668
CL\_8667
CL\_8666
CL\_8665
CL\_8664
CL\_8663
CL\_8662
CL\_7752
CL\_8229
CL\_9275
CL\_9274
CL\_9273
CL\_9272
CL\_9271
CL\_9270
CL\_9269
CL\_9268
CL\_7753
CL\_9285
CL\_18224
CL\_8661
CL\_8660
CL\_8659
CL\_8277
CL\_8276
CL\_8275
CL\_8658
CL\_9267
CL\_9266
CL\_9265
CL\_9264
CL\_7754
CL\_7755
CL\_7756
CL\_17327
CL\_17326
CL\_17325
CL\_17324
CL\_9263
CL\_20597
CL\_8274
CL\_8273
CL\_8272
CL\_8271
CL\_8270
CL\_8269
CL\_8268
CL\_8267
CL\_7757
CL\_11525
CL\_7759
CL\_7760
CL\_7761
CL\_14070
CL\_13610
CL\_13609
CL\_13608
CL\_13607
CL\_8657
CL\_6135
CL\_17783
CL\_8266
CL\_8265
CL\_8264
CL\_8263
CL\_8188
