## Supplementary material for "A novel method for integrating genomic and Tn-Seq data to identify common *in vivo* fitness mechanisms across multiple bacterial species": S2 Dataset: CL_INS_337.html

Legend

 Mobile +extrachromosomalelementfunctions
 Hypothetical
 Other
 All VFDB Genes

FULL


WINDOWSVGPNG

Trim RowsRemove SingletonsSave Fasta

CL\_4094


CL\_4094


CL\_4094

HighlightSelectShow Genomes


131

CL\_4095


72

CL\_4095


1

CL\_4095

fGI ID


CL\_INS\_337
CL\_INS\_337
Cluster ID


CL\_18544
CL\_5323
