## Supplementary material for "A novel method for integrating genomic and Tn-Seq data to identify common *in vivo* fitness mechanisms across multiple bacterial species": S2 Dataset: CL_INS_338.html

Legend

 Mobile +extrachromosomalelementfunctions
 Hypothetical
 Proteinsynthesis/fate
 Other
 All VFDB Genes

FULL


WINDOWSVGPNG

Trim RowsRemove SingletonsSave Fasta

CL\_4115


CL\_4115


CL\_4115


CL\_4115


CL\_4115


CL\_4115

HighlightSelectShow Genomes


196

CL\_4116


2

CL\_4117


1

CL\_4117


1

CL\_4117


1

CL\_4117


1

CL\_4117

fGI ID


CL\_INS\_338
CL\_INS\_339
CL\_INS\_339
CL\_INS\_263
CL\_INS\_263
CL\_INS\_339
CL\_INS\_313
CL\_INS\_263
CL\_INS\_263
CL\_INS\_339
CL\_INS\_339
CL\_INS\_263
CL\_INS\_263
CL\_INS\_263
CL\_INS\_91
CL\_INS\_263
CL\_INS\_263
CL\_INS\_263
CL\_INS\_263
CL\_INS\_339
CL\_INS\_263
CL\_INS\_263
CL\_INS\_263
CL\_INS\_338
CL\_INS\_339
CL\_INS\_339
CL\_INS\_132
CL\_INS\_132
CL\_INS\_132
CL\_INS\_385
CL\_INS\_321
CL\_INS\_321
CL\_INS\_385
CL\_INS\_385
CL\_INS\_368
CL\_INS\_368
CL\_INS\_368
CL\_INS\_368
CL\_INS\_368
CL\_INS\_339
CL\_INS\_339
CL\_INS\_339
CL\_INS\_339
Cluster ID


CL\_16436
CL\_16435
CL\_16434
CL\_15608
CL\_15607
CL\_17782
CL\_17781
CL\_5468
CL\_5467
CL\_17780
CL\_17779
CL\_15606
CL\_15605
CL\_15604
CL\_11925
CL\_14821
CL\_10179
CL\_15603
CL\_15602
CL\_16433
CL\_15601
CL\_15600
CL\_15599
CL\_14030
CL\_14029
CL\_14028
CL\_12981
CL\_8079
CL\_6714
CL\_8080
CL\_12219
CL\_12218
CL\_6693
CL\_8228
CL\_8125
CL\_8126
CL\_8127
CL\_8128
CL\_8129
CL\_12599
CL\_8130
CL\_17323
CL\_12601
