## Supplementary material for "A novel method for integrating genomic and Tn-Seq data to identify common *in vivo* fitness mechanisms across multiple bacterial species": S2 Dataset: CL_INS_341.html

Legend

 Hypothetical
 Other
 All VFDB Genes

FULL


WINDOWSVGPNG

Trim RowsRemove SingletonsSave Fasta

CL\_4127


CL\_4127


CL\_4127


CL\_4127

HighlightSelectShow Genomes


121

CL\_4129


81

CL\_4129


2

CL\_4130


1

CL\_4130

fGI ID

CL\_INS\_341
Cluster ID

CL\_4128
