## Supplementary material for "A novel method for integrating genomic and Tn-Seq data to identify common *in vivo* fitness mechanisms across multiple bacterial species": S2 Dataset: CL_INS_342.html

Legend

 Hypothetical
 Other
 All VFDB Genes

FULL


WINDOWSVGPNG

Trim RowsRemove SingletonsSave Fasta

CL\_4141


CL\_4141


CL\_4141


CL\_4141


CL\_4141


CL\_4141


CL\_4141


CL\_4141


CL\_4141


CL\_4141


CL\_4141


CL\_4141


CL\_4141

HighlightSelectShow Genomes


164

CL\_4142


13

CL\_4142


5

CL\_4142


4

CL\_4142


4

CL\_4142


2

CL\_4142


2

CL\_4142


2

CL\_4142


1

CL\_4142


1

CL\_4142


1

CL\_4142


1

Break


1

CL\_4142

fGI ID


CL\_INS\_342
CL\_INS\_132
CL\_INS\_278
CL\_INS\_278
CL\_INS\_278
CL\_INS\_342
CL\_INS\_342
CL\_INS\_342
CL\_INS\_342
CL\_INS\_342
CL\_INS\_342
CL\_INS\_342
CL\_INS\_342
CL\_INS\_342
CL\_INS\_342
CL\_INS\_342
CL\_INS\_342
CL\_INS\_342
CL\_INS\_342
CL\_INS\_342
CL\_INS\_342
CL\_INS\_342
CL\_INS\_342
CL\_INS\_342
CL\_INS\_342
CL\_INS\_342
CL\_INS\_342
CL\_INS\_342
CL\_INS\_342
CL\_INS\_342
Cluster ID


CL\_12215
CL\_11233
CL\_5973
CL\_5974
CL\_5975
CL\_7169
CL\_7168
CL\_7167
CL\_7166
CL\_7165
CL\_11755
CL\_11408
CL\_11407
CL\_11406
CL\_11405
CL\_11404
CL\_7164
CL\_7163
CL\_7162
CL\_7161
CL\_7160
CL\_7159
CL\_7158
CL\_7157
CL\_7156
CL\_7155
CL\_7154
CL\_7653
CL\_7153
CL\_7652
