## Supplementary material for "A novel method for integrating genomic and Tn-Seq data to identify common *in vivo* fitness mechanisms across multiple bacterial species": S2 Dataset: CL_INS_346.html

Legend

 Hypothetical
 Other
 All VFDB Genes

FULL


WINDOWSVGPNG

Trim RowsRemove SingletonsSave Fasta

CL\_4226


CL\_4226


CL\_4226


CL\_4225

HighlightSelectShow Genomes


107

CL\_4228


80

CL\_4228


3

CL\_4229


1

CL\_4228

fGI ID

CL\_INS\_346
Cluster ID

CL\_4227
