## Supplementary material for "A novel method for integrating genomic and Tn-Seq data to identify common *in vivo* fitness mechanisms across multiple bacterial species": S2 Dataset: CL_INS_351.html

Legend

 Mobile +extrachromosomalelementfunctions
 Regulatoryfunctions
 Hypothetical
 DNA Metabolism
 All EssentialGenes
 Cell Envelope
 Other
 Transport +binding proteins
 All VFDB Genes

FULL


WINDOWSVGPNG

Trim RowsRemove SingletonsSave Fasta

CL\_4278


CL\_4278


CL\_4278


CL\_4278


CL\_4278


CL\_4281


CL\_4278


CL\_4278


CL\_4278


CL\_4278


CL\_4278


CL\_4278


CL\_4278


CL\_4278


CL\_4278


CL\_4278


CL\_4278


CL\_4278


CL\_4278


CL\_4278


CL\_4278


CL\_4278


CL\_4276


CL\_4276


CL\_4278

HighlightSelectShow Genomes


177

CL\_4279


4

CL\_4279


2

CL\_4279


1

CL\_4279


1

CL\_4279


1

CL\_4279


1

CL\_4279


1

CL\_4279


1

CL\_4279


1

CL\_4279


1

CL\_4279


1

CL\_4279


1

CL\_4279


1

CL\_4279


1

CL\_4279


1

CL\_4279


1

CL\_4279


1

CL\_4279


1

CL\_4279


1

CL\_4279


1

CL\_4279


1

CL\_4279


1

CL\_4279


1

CL\_4279


1

CL\_4279

fGI ID


CL\_INS\_352
CL\_INS\_352
CL\_INS\_352
CL\_INS\_352
CL\_INS\_351
CL\_INS\_351
CL\_INS\_351
CL\_INS\_351
CL\_INS\_351
CL\_INS\_351
CL\_INS\_351
CL\_INS\_351
CL\_INS\_351
CL\_INS\_351
CL\_INS\_351
CL\_INS\_351
CL\_INS\_74
CL\_INS\_351
CL\_INS\_74
CL\_INS\_74
CL\_INS\_74
CL\_INS\_74
CL\_INS\_74
CL\_INS\_74
CL\_INS\_74
CL\_INS\_321
CL\_INS\_74
CL\_INS\_351
CL\_INS\_351
CL\_INS\_351
CL\_INS\_351
CL\_INS\_351
CL\_INS\_351
CL\_INS\_351
CL\_INS\_351
CL\_INS\_351
CL\_INS\_351
CL\_INS\_351
CL\_INS\_351
CL\_INS\_351
CL\_INS\_351
CL\_INS\_351
CL\_INS\_351
CL\_INS\_351
CL\_INS\_351
CL\_INS\_351
CL\_INS\_351
CL\_INS\_351
CL\_INS\_351
CL\_INS\_351
CL\_INS\_307
CL\_INS\_307
CL\_INS\_351
CL\_INS\_351
CL\_INS\_352
CL\_INS\_352
CL\_INS\_352
CL\_INS\_352
CL\_INS\_352
CL\_INS\_352
CL\_INS\_352
CL\_INS\_352
CL\_INS\_352
CL\_INS\_352
CL\_INS\_352
CL\_INS\_352
CL\_INS\_352
CL\_INS\_352
CL\_INS\_352
CL\_INS\_352
CL\_INS\_352
CL\_INS\_352
CL\_INS\_352
CL\_INS\_351
CL\_INS\_351
CL\_INS\_351
CL\_INS\_351
CL\_INS\_351
CL\_INS\_351
CL\_INS\_351
CL\_INS\_351
CL\_INS\_351
CL\_INS\_351
CL\_INS\_351
CL\_INS\_351
CL\_INS\_351
CL\_INS\_351
CL\_INS\_351
CL\_INS\_351
CL\_INS\_351
CL\_INS\_351
CL\_INS\_351
CL\_INS\_351
CL\_INS\_351
CL\_INS\_351
CL\_INS\_351
CL\_INS\_351
CL\_INS\_351
CL\_INS\_351
CL\_INS\_351
CL\_INS\_351
CL\_INS\_351
CL\_INS\_351
CL\_INS\_351
CL\_INS\_351
CL\_INS\_351
CL\_INS\_351
CL\_INS\_74
CL\_INS\_74
CL\_INS\_74
CL\_INS\_74
CL\_INS\_74
CL\_INS\_74
CL\_INS\_74
CL\_INS\_74
CL\_INS\_74
CL\_INS\_74
CL\_INS\_74
CL\_INS\_351
CL\_INS\_351
CL\_INS\_74
CL\_INS\_351
CL\_INS\_351
CL\_INS\_351
CL\_INS\_351
CL\_INS\_351
CL\_INS\_74
CL\_INS\_74
CL\_INS\_74
CL\_INS\_351
CL\_INS\_351
CL\_INS\_74
CL\_INS\_351
CL\_INS\_74
CL\_INS\_74
CL\_INS\_351
CL\_INS\_74
CL\_INS\_74
CL\_INS\_74
CL\_INS\_351
CL\_INS\_351
CL\_INS\_351
CL\_INS\_74
CL\_INS\_74
CL\_INS\_351
CL\_INS\_351
CL\_INS\_351
CL\_INS\_351
CL\_INS\_74
CL\_INS\_351
CL\_INS\_351
CL\_INS\_351
CL\_INS\_351
CL\_INS\_351
CL\_INS\_351
CL\_INS\_351
CL\_INS\_351
CL\_INS\_351
CL\_INS\_351
CL\_INS\_351
CL\_INS\_351
CL\_INS\_351
CL\_INS\_351
CL\_INS\_351
CL\_INS\_351
CL\_INS\_351
CL\_INS\_351
CL\_INS\_351
CL\_INS\_351
CL\_INS\_351
CL\_INS\_351
CL\_INS\_351
CL\_INS\_321
CL\_INS\_321
CL\_INS\_351
CL\_INS\_321
CL\_INS\_321
CL\_INS\_321
CL\_INS\_351
CL\_INS\_321
CL\_INS\_351
CL\_INS\_91
CL\_INS\_91
CL\_INS\_351
CL\_INS\_351
CL\_INS\_351
CL\_INS\_351
CL\_INS\_351
CL\_INS\_351
CL\_INS\_351
CL\_INS\_351
CL\_INS\_351
CL\_INS\_351
CL\_INS\_351
CL\_INS\_351
CL\_INS\_351
CL\_INS\_351
CL\_INS\_351
CL\_INS\_351
CL\_INS\_351
CL\_INS\_351
CL\_INS\_351
CL\_INS\_351
CL\_INS\_351
CL\_INS\_351
CL\_INS\_351
CL\_INS\_351
CL\_INS\_351
CL\_INS\_351
CL\_INS\_351
CL\_INS\_351
CL\_INS\_351
CL\_INS\_351
CL\_INS\_351
CL\_INS\_351
CL\_INS\_351
CL\_INS\_351
CL\_INS\_351
CL\_INS\_351
CL\_INS\_351
CL\_INS\_351
CL\_INS\_385
CL\_INS\_351
CL\_INS\_351
CL\_INS\_351
CL\_INS\_351
Cluster ID


CL\_6902
CL\_6903
CL\_22081
CL\_22082
CL\_20729
CL\_20728
CL\_20727
CL\_20726
CL\_20725
CL\_20724
CL\_20723
CL\_20722
CL\_20721
CL\_20720
CL\_13124
CL\_13123
CL\_5958
CL\_13122
CL\_5959
CL\_5960
CL\_5961
CL\_5962
CL\_7525
CL\_5964
CL\_5965
CL\_7524
CL\_7523
CL\_10166
CL\_10165
CL\_15322
CL\_10164
CL\_10163
CL\_10162
CL\_10161
CL\_10160
CL\_10159
CL\_10158
CL\_10157
CL\_19115
CL\_19114
CL\_19113
CL\_19112
CL\_19111
CL\_19110
CL\_19109
CL\_19108
CL\_19107
CL\_19106
CL\_19105
CL\_19104
CL\_5939
CL\_5938
CL\_19103
CL\_19102
CL\_6919
CL\_6918
CL\_6917
CL\_6916
CL\_6915
CL\_6914
CL\_6913
CL\_6912
CL\_6911
CL\_6910
CL\_6909
CL\_6908
CL\_6907
CL\_5726
CL\_5727
CL\_5728
CL\_5729
CL\_5730
CL\_6906
CL\_6905
CL\_11179
CL\_11178
CL\_15518
CL\_11177
CL\_15517
CL\_15516
CL\_11176
CL\_11175
CL\_7152
CL\_9697
CL\_16472
CL\_11596
CL\_12047
CL\_12046
CL\_11597
CL\_21983
CL\_21982
CL\_11598
CL\_11599
CL\_11600
CL\_11601
CL\_12045
CL\_11602
CL\_24171
CL\_11603
CL\_11604
CL\_12044
CL\_12043
CL\_12042
CL\_11607
CL\_15324
CL\_11608
CL\_15325
CL\_7151
CL\_12845
CL\_12846
CL\_9065
CL\_7150
CL\_7149
CL\_7148
CL\_7467
CL\_7147
CL\_7146
CL\_7144
CL\_9696
CL\_7466
CL\_7145
CL\_15459
CL\_15323
CL\_24170
CL\_24169
CL\_18220
CL\_21152
CL\_9064
CL\_7143
CL\_18649
CL\_7142
CL\_9063
CL\_9695
CL\_9062
CL\_9061
CL\_5561
CL\_18534
CL\_7141
CL\_7140
CL\_12847
CL\_12848
CL\_18650
CL\_9060
CL\_9059
CL\_7139
CL\_7138
CL\_7137
CL\_8834
CL\_7913
CL\_17819
CL\_17820
CL\_17821
CL\_20955
CL\_20954
CL\_20953
CL\_17823
CL\_17824
CL\_20952
CL\_20951
CL\_17825
CL\_17826
CL\_17827
CL\_17828
CL\_20950
CL\_17829
CL\_17830
CL\_17831
CL\_17832
CL\_20949
CL\_20948
CL\_20947
CL\_20946
CL\_8352
CL\_7414
CL\_21509
CL\_10419
CL\_8353
CL\_8354
CL\_14193
CL\_8355
CL\_21508
CL\_14312
CL\_14311
CL\_14194
CL\_14195
CL\_14196
CL\_10433
CL\_10432
CL\_10431
CL\_10430
CL\_21507
CL\_21506
CL\_19696
CL\_19695
CL\_19694
CL\_21505
CL\_14197
CL\_14198
CL\_14199
CL\_14200
CL\_14201
CL\_14202
CL\_14203
CL\_14204
CL\_14205
CL\_14206
CL\_14207
CL\_14208
CL\_14209
CL\_14210
CL\_14211
CL\_14212
CL\_14213
CL\_9396
CL\_9395
CL\_9394
CL\_9393
CL\_9392
CL\_9391
CL\_9390
CL\_14214
CL\_8855
CL\_14215
CL\_14216
CL\_14217
CL\_13121
