## Supplementary material for "A novel method for integrating genomic and Tn-Seq data to identify common *in vivo* fitness mechanisms across multiple bacterial species": S2 Dataset: CL_INS_352.html

Legend

 Hypothetical
 Cell Envelope
 Other
 All VFDB Genes

FULL


WINDOWSVGPNG

Trim RowsRemove SingletonsSave Fasta

CL\_4279


CL\_4279


CL\_4279


CL\_4279


CL\_4279

HighlightSelectShow Genomes


98

CL\_4281


78

CL\_4281


28

CL\_4281


1

CL\_4281


1

CL\_4276

fGI ID


CL\_INS\_352
CL\_INS\_352
CL\_INS\_352
CL\_INS\_352
CL\_INS\_352
CL\_INS\_352
CL\_INS\_352
CL\_INS\_352
CL\_INS\_352
CL\_INS\_352
CL\_INS\_352
CL\_INS\_352
CL\_INS\_352
CL\_INS\_352
CL\_INS\_352
CL\_INS\_352
CL\_INS\_352
CL\_INS\_352
CL\_INS\_352
CL\_INS\_352
CL\_INS\_352
CL\_INS\_352
CL\_INS\_352
CL\_INS\_352
CL\_INS\_352
CL\_INS\_352
CL\_INS\_352
Cluster ID


CL\_4280
CL\_6904
CL\_6906
CL\_5730
CL\_5729
CL\_22082
CL\_22081
CL\_6903
CL\_6902
CL\_6901
CL\_13066
CL\_5728
CL\_5727
CL\_5726
CL\_6907
CL\_6908
CL\_6909
CL\_6910
CL\_6911
CL\_6912
CL\_6913
CL\_6914
CL\_6915
CL\_6916
CL\_6917
CL\_6918
CL\_6919
