## Supplementary material for "A novel method for integrating genomic and Tn-Seq data to identify common *in vivo* fitness mechanisms across multiple bacterial species": S2 Dataset: CL_INS_356.html

Legend

 Mobile +extrachromosomalelementfunctions
 Hypothetical
 All VFDB Genes

FULL


WINDOWSVGPNG

Trim RowsRemove SingletonsSave Fasta

CL\_4334


CL\_4334


CL\_4334

HighlightSelectShow Genomes


109

CL\_4336


92

CL\_4336


2

CL\_4336

fGI ID


CL\_INS\_356
CL\_INS\_356
Cluster ID


CL\_14722
CL\_4335
