## Supplementary material for "A novel method for integrating genomic and Tn-Seq data to identify common *in vivo* fitness mechanisms across multiple bacterial species": S2 Dataset: CL_INS_357.html

Legend

 Hypothetical
 AntibioticResistance
 Other
 All VFDB Genes

FULL


WINDOWSVGPNG

Trim RowsRemove SingletonsSave Fasta

CL\_4341


CL\_4341


CL\_4341


CL\_4341


CL\_4341


CL\_4341


CL\_4341

HighlightSelectShow Genomes


127

CL\_4345


61

CL\_4345


12

CL\_4345


2

CL\_4345


1

CL\_4345


1

CL\_4345


1

CL\_4345

fGI ID


CL\_INS\_357
CL\_INS\_357
CL\_INS\_357
CL\_INS\_357
CL\_INS\_357
CL\_INS\_357
CL\_INS\_357
Cluster ID


CL\_4342
CL\_4343
CL\_4344
CL\_6900
CL\_9058
CL\_6899
CL\_17317
