## Supplementary material for "A novel method for integrating genomic and Tn-Seq data to identify common *in vivo* fitness mechanisms across multiple bacterial species": S2 Dataset: CL_INS_363.html

Legend

 Mobile +extrachromosomalelementfunctions
 Regulatoryfunctions
 Hypothetical
 AntibioticResistance
 Other
 All VFDB Genes

FULL


WINDOWSVGPNG

Trim RowsRemove SingletonsSave Fasta

CL\_4406


CL\_4406

HighlightSelectShow Genomes


201

CL\_4405


1

CL\_4405

fGI ID


CL\_INS\_363
CL\_INS\_363
CL\_INS\_363
CL\_INS\_363
CL\_INS\_363
CL\_INS\_363
CL\_INS\_363
CL\_INS\_363
CL\_INS\_363
CL\_INS\_363
CL\_INS\_363
CL\_INS\_363
CL\_INS\_363
CL\_INS\_363
CL\_INS\_363
CL\_INS\_363
CL\_INS\_363
CL\_INS\_363
CL\_INS\_363
CL\_INS\_363
CL\_INS\_363
CL\_INS\_363
CL\_INS\_363
CL\_INS\_363
CL\_INS\_363
CL\_INS\_363
CL\_INS\_363
CL\_INS\_363
CL\_INS\_363
CL\_INS\_363
CL\_INS\_363
CL\_INS\_363
CL\_INS\_363
CL\_INS\_363
CL\_INS\_363
CL\_INS\_363
CL\_INS\_363
CL\_INS\_363
CL\_INS\_363
CL\_INS\_363
CL\_INS\_363
CL\_INS\_363
CL\_INS\_363
CL\_INS\_363
CL\_INS\_363
CL\_INS\_363
CL\_INS\_363
CL\_INS\_363
CL\_INS\_363
CL\_INS\_363
CL\_INS\_363
Cluster ID


CL\_16690
CL\_16689
CL\_16688
CL\_16687
CL\_16686
CL\_16685
CL\_16684
CL\_16683
CL\_9874
CL\_9875
CL\_16682
CL\_9878
CL\_9879
CL\_9880
CL\_16681
CL\_9881
CL\_16680
CL\_9883
CL\_9884
CL\_16679
CL\_16678
CL\_16677
CL\_16676
CL\_16675
CL\_16674
CL\_16673
CL\_16672
CL\_16671
CL\_16670
CL\_16669
CL\_16668
CL\_16667
CL\_16666
CL\_16665
CL\_16664
CL\_16663
CL\_16662
CL\_16661
CL\_16660
CL\_16659
CL\_16658
CL\_16657
CL\_16656
CL\_16655
CL\_16654
CL\_16653
CL\_16652
CL\_16651
CL\_16650
CL\_16649
CL\_16648
