## Supplementary material for "A novel method for integrating genomic and Tn-Seq data to identify common *in vivo* fitness mechanisms across multiple bacterial species": S2 Dataset: CL_INS_367.html

Legend

 Mobile +extrachromosomalelementfunctions
 Regulatoryfunctions
 Hypothetical
 DNA Metabolism
 All EssentialGenes
 Proteinsynthesis/fate
 Other
 Transport +binding proteins
 All VFDB Genes

FULL


WINDOWSVGPNG

Trim RowsRemove SingletonsSave Fasta

CL\_4426


CL\_4426


CL\_4426


CL\_4426


CL\_4426


CL\_4426


CL\_4426


CL\_4426


CL\_4426


CL\_4426


CL\_4426


CL\_4426


CL\_4426


CL\_4426


CL\_4426


CL\_4426


CL\_4426


CL\_4426


CL\_4426


CL\_4426


CL\_4426


CL\_4426


CL\_4426


CL\_4426


CL\_4426


CL\_4426


CL\_4426


CL\_4426


CL\_4426


CL\_4426


CL\_4426


CL\_4426


CL\_4426


CL\_4426


CL\_4426


CL\_4426


CL\_4426


CL\_4426


CL\_4426


CL\_4426


CL\_4426


CL\_4426


CL\_4426


CL\_4426


CL\_4426


CL\_4426


CL\_4426


CL\_4426


CL\_4426

HighlightSelectShow Genomes


131

CL\_4427


5

CL\_4428


4

CL\_4427


4

CL\_4427


3

CL\_4427


2

CL\_4427


2

CL\_4427


1

CL\_4428


1

CL\_4427


1

Break


1

CL\_4428


1

CL\_4428


1

CL\_4428


1

CL\_4427


1

CL\_4427


1

CL\_4428


1

CL\_4427


1

CL\_4428


1

CL\_4427


1

CL\_4428


1

CL\_4428


1

CL\_4428


1

CL\_4427


1

CL\_4428


1

CL\_4427


1

CL\_4428


1

CL\_456


1

CL\_4427


1

CL\_4427


1

CL\_4427


1

CL\_4428


1

CL\_4430


1

CL\_4427


1

CL\_4427


1

CL\_4430


1

CL\_4427


1

CL\_4427


1

CL\_4427


1

CL\_4428


1

CL\_4427


1

CL\_4427


1

CL\_4430


1

CL\_4427


1

CL\_4427


1

CL\_4430


1

CL\_4428


1

CL\_4428


1

CL\_4427


1

CL\_4432

fGI ID


CL\_INS\_44
CL\_INS\_381
CL\_INS\_367
CL\_INS\_367
CL\_INS\_367
CL\_INS\_367
CL\_INS\_367
CL\_INS\_155
CL\_INS\_155
CL\_INS\_155
CL\_INS\_155
CL\_INS\_155
CL\_INS\_155
CL\_INS\_155
CL\_INS\_155
CL\_INS\_367
CL\_INS\_367
CL\_INS\_367
CL\_INS\_368
CL\_INS\_155
CL\_INS\_155
CL\_INS\_381
CL\_INS\_381
CL\_INS\_368
CL\_INS\_367
CL\_INS\_367
CL\_INS\_367
CL\_INS\_367
CL\_INS\_367
CL\_INS\_367
CL\_INS\_367
CL\_INS\_367
CL\_INS\_367
CL\_INS\_367
CL\_INS\_367
CL\_INS\_385
CL\_INS\_367
CL\_INS\_367
CL\_INS\_155
CL\_INS\_367
CL\_INS\_367
CL\_INS\_381
CL\_INS\_367
CL\_INS\_367
CL\_INS\_367
CL\_INS\_368
CL\_INS\_368
CL\_INS\_368
CL\_INS\_368
CL\_INS\_368
CL\_INS\_368
CL\_INS\_368
CL\_INS\_368
CL\_INS\_368
CL\_INS\_368
CL\_INS\_368
CL\_INS\_367
CL\_INS\_368
CL\_INS\_367
CL\_INS\_368
CL\_INS\_368
CL\_INS\_368
CL\_INS\_45
CL\_INS\_26
CL\_INS\_26
CL\_INS\_74
CL\_INS\_321
CL\_INS\_321
CL\_INS\_321
CL\_INS\_321
CL\_INS\_321
CL\_INS\_385
CL\_INS\_368
CL\_INS\_368
CL\_INS\_368
CL\_INS\_368
CL\_INS\_368
CL\_INS\_385
CL\_INS\_204
CL\_INS\_368
CL\_INS\_368
CL\_INS\_368
CL\_INS\_368
CL\_INS\_368
CL\_INS\_368
CL\_INS\_368
CL\_INS\_321
CL\_INS\_321
CL\_INS\_367
CL\_INS\_367
CL\_INS\_367
CL\_INS\_239
CL\_INS\_367
CL\_INS\_321
CL\_INS\_321
CL\_INS\_321
CL\_INS\_321
CL\_INS\_368
CL\_INS\_321
CL\_INS\_321
CL\_INS\_321
CL\_INS\_321
CL\_INS\_321
CL\_INS\_321
CL\_INS\_368
CL\_INS\_368
CL\_INS\_351
CL\_INS\_351
CL\_INS\_351
CL\_INS\_351
CL\_INS\_367
CL\_INS\_368
CL\_INS\_368
CL\_INS\_368
CL\_INS\_368
CL\_INS\_368
CL\_INS\_368
CL\_INS\_368
CL\_INS\_368
CL\_INS\_368
CL\_INS\_368
CL\_INS\_368
CL\_INS\_321
CL\_INS\_321
CL\_INS\_132
CL\_INS\_132
CL\_INS\_132
CL\_INS\_132
CL\_INS\_132
CL\_INS\_132
CL\_INS\_132
CL\_INS\_321
CL\_INS\_367
CL\_INS\_367
CL\_INS\_367
CL\_INS\_367
CL\_INS\_367
CL\_INS\_367
CL\_INS\_367
CL\_INS\_321
CL\_INS\_319
CL\_INS\_319
CL\_INS\_367
CL\_INS\_367
CL\_INS\_319
CL\_INS\_319
CL\_INS\_132
CL\_INS\_132
CL\_INS\_368
CL\_INS\_319
CL\_INS\_319
CL\_INS\_319
CL\_INS\_319
CL\_INS\_319
CL\_INS\_319
CL\_INS\_319
CL\_INS\_367
CL\_INS\_319
CL\_INS\_367
CL\_INS\_367
CL\_INS\_367
CL\_INS\_367
CL\_INS\_367
CL\_INS\_367
CL\_INS\_367
CL\_INS\_367
CL\_INS\_367
CL\_INS\_321
CL\_INS\_321
CL\_INS\_367
CL\_INS\_319
CL\_INS\_319
CL\_INS\_368
CL\_INS\_368
CL\_INS\_368
CL\_INS\_367
CL\_INS\_367
CL\_INS\_319
CL\_INS\_132
CL\_INS\_319
CL\_INS\_319
CL\_INS\_319
CL\_INS\_132
CL\_INS\_132
CL\_INS\_132
CL\_INS\_132
CL\_INS\_132
CL\_INS\_132
CL\_INS\_132
CL\_INS\_132
CL\_INS\_132
CL\_INS\_132
CL\_INS\_132
CL\_INS\_132
CL\_INS\_132
CL\_INS\_132
CL\_INS\_132
CL\_INS\_132
CL\_INS\_132
CL\_INS\_132
CL\_INS\_367
CL\_INS\_132
CL\_INS\_367
CL\_INS\_367
CL\_INS\_367
CL\_INS\_132
CL\_INS\_132
CL\_INS\_367
CL\_INS\_367
CL\_INS\_132
CL\_INS\_368
CL\_INS\_368
CL\_INS\_368
CL\_INS\_321
CL\_INS\_132
CL\_INS\_132
CL\_INS\_132
CL\_INS\_132
CL\_INS\_132
CL\_INS\_132
CL\_INS\_321
CL\_INS\_321
CL\_INS\_321
CL\_INS\_368
CL\_INS\_321
CL\_INS\_368
CL\_INS\_321
CL\_INS\_319
CL\_INS\_319
CL\_INS\_132
CL\_INS\_132
CL\_INS\_368
CL\_INS\_368
CL\_INS\_319
CL\_INS\_367
CL\_INS\_321
CL\_INS\_321
CL\_INS\_367
CL\_INS\_367
CL\_INS\_132
CL\_INS\_132
CL\_INS\_367
CL\_INS\_367
CL\_INS\_319
CL\_INS\_319
CL\_INS\_319
CL\_INS\_319
CL\_INS\_319
CL\_INS\_367
CL\_INS\_367
CL\_INS\_319
CL\_INS\_319
CL\_INS\_319
CL\_INS\_319
CL\_INS\_319
CL\_INS\_319
CL\_INS\_319
CL\_INS\_132
CL\_INS\_319
CL\_INS\_319
CL\_INS\_367
CL\_INS\_367
CL\_INS\_367
CL\_INS\_367
CL\_INS\_367
CL\_INS\_367
CL\_INS\_367
CL\_INS\_367
CL\_INS\_367
CL\_INS\_367
CL\_INS\_367
CL\_INS\_367
CL\_INS\_368
CL\_INS\_134
CL\_INS\_319
CL\_INS\_134
CL\_INS\_134
CL\_INS\_134
CL\_INS\_368
CL\_INS\_367
CL\_INS\_367
CL\_INS\_368
CL\_INS\_368
CL\_INS\_319
CL\_INS\_319
CL\_INS\_368
CL\_INS\_368
CL\_INS\_319
CL\_INS\_132
CL\_INS\_132
CL\_INS\_319
CL\_INS\_367
CL\_INS\_319
CL\_INS\_367
CL\_INS\_319
CL\_INS\_319
CL\_INS\_132
CL\_INS\_368
CL\_INS\_132
CL\_INS\_319
CL\_INS\_319
CL\_INS\_319
CL\_INS\_132
CL\_INS\_368
CL\_INS\_132
CL\_INS\_368
CL\_INS\_367
CL\_INS\_367
CL\_INS\_367
CL\_INS\_367
CL\_INS\_367
CL\_INS\_385
CL\_INS\_105
CL\_INS\_367
CL\_INS\_105
CL\_INS\_278
CL\_INS\_385
CL\_INS\_105
CL\_INS\_368
CL\_INS\_368
CL\_INS\_368
CL\_INS\_368
CL\_INS\_368
CL\_INS\_368
CL\_INS\_368
CL\_INS\_368
CL\_INS\_368
CL\_INS\_368
CL\_INS\_368
CL\_INS\_368
CL\_INS\_368
CL\_INS\_368
CL\_INS\_368
CL\_INS\_368
CL\_INS\_367
CL\_INS\_400
CL\_INS\_367
CL\_INS\_367
CL\_INS\_385
CL\_INS\_367
CL\_INS\_367
CL\_INS\_367
CL\_INS\_367
CL\_INS\_367
CL\_INS\_91
CL\_INS\_232
CL\_INS\_132
CL\_INS\_132
CL\_INS\_367
CL\_INS\_278
CL\_INS\_367
CL\_INS\_243
CL\_INS\_278
CL\_INS\_66
CL\_INS\_253
CL\_INS\_253
CL\_INS\_253
CL\_INS\_253
CL\_INS\_253
CL\_INS\_253
CL\_INS\_253
CL\_INS\_253
CL\_INS\_253
CL\_INS\_253
CL\_INS\_253
CL\_INS\_253
CL\_INS\_253
CL\_INS\_253
CL\_INS\_253
CL\_INS\_253
CL\_INS\_253
CL\_INS\_318
CL\_INS\_253
CL\_INS\_253
CL\_INS\_253
CL\_INS\_253
CL\_INS\_253
CL\_INS\_253
CL\_INS\_253
CL\_INS\_253
CL\_INS\_253
CL\_INS\_253
CL\_INS\_253
CL\_INS\_253
CL\_INS\_253
CL\_INS\_253
CL\_INS\_253
CL\_INS\_253
CL\_INS\_253
CL\_INS\_253
CL\_INS\_253
CL\_INS\_253
CL\_INS\_253
CL\_INS\_253
CL\_INS\_253
CL\_INS\_253
CL\_INS\_253
CL\_INS\_253
CL\_INS\_253
CL\_INS\_253
CL\_INS\_253
CL\_INS\_253
CL\_INS\_253
CL\_INS\_253
CL\_INS\_253
CL\_INS\_367
CL\_INS\_253
CL\_INS\_253
CL\_INS\_253
CL\_INS\_253
CL\_INS\_253
CL\_INS\_253
CL\_INS\_253
CL\_INS\_253
CL\_INS\_253
CL\_INS\_160
CL\_INS\_253
CL\_INS\_253
CL\_INS\_253
CL\_INS\_253
CL\_INS\_253
CL\_INS\_253
CL\_INS\_278
Cluster ID


CL\_8351
CL\_6447
CL\_20944
CL\_15895
CL\_15894
CL\_15893
CL\_15420
CL\_6448
CL\_7488
CL\_6201
CL\_6202
CL\_6203
CL\_6204
CL\_6205
CL\_6206
CL\_20943
CL\_16691
CL\_16692
CL\_15419
CL\_7490
CL\_7492
CL\_17877
CL\_17876
CL\_15418
CL\_20596
CL\_20595
CL\_20594
CL\_20593
CL\_20592
CL\_20591
CL\_20590
CL\_20589
CL\_20588
CL\_20587
CL\_20586
CL\_6156
CL\_20585
CL\_20584
CL\_7493
CL\_20583
CL\_20582
CL\_17875
CL\_20942
CL\_20581
CL\_21599
CL\_21598
CL\_21597
CL\_21596
CL\_21595
CL\_21594
CL\_21593
CL\_15417
CL\_15416
CL\_21592
CL\_15415
CL\_15414
CL\_20941
CL\_15413
CL\_20580
CL\_15412
CL\_21591
CL\_21590
CL\_11754
CL\_17617
CL\_17618
CL\_7913
CL\_8352
CL\_10419
CL\_8353
CL\_8354
CL\_8355
CL\_8228
CL\_8125
CL\_8126
CL\_8127
CL\_8128
CL\_8129
CL\_8212
CL\_5953
CL\_23074
CL\_23073
CL\_23072
CL\_23071
CL\_23070
CL\_23069
CL\_23068
CL\_7412
CL\_7413
CL\_9260
CL\_9259
CL\_9258
CL\_9257
CL\_9256
CL\_7414
CL\_7415
CL\_7416
CL\_7417
CL\_11753
CL\_7421
CL\_7422
CL\_7423
CL\_11752
CL\_11751
CL\_11750
CL\_11749
CL\_11748
CL\_10433
CL\_10432
CL\_10431
CL\_10430
CL\_23723
CL\_11747
CL\_11746
CL\_11745
CL\_11744
CL\_11743
CL\_11742
CL\_11741
CL\_11740
CL\_11739
CL\_9255
CL\_22150
CL\_6390
CL\_5736
CL\_6389
CL\_6388
CL\_6322
CL\_6321
CL\_6320
CL\_6319
CL\_6318
CL\_6386
CL\_17883
CL\_17882
CL\_18339
CL\_18338
CL\_18337
CL\_14596
CL\_23847
CL\_6385
CL\_6384
CL\_5748
CL\_20381
CL\_20380
CL\_5890
CL\_5899
CL\_13741
CL\_10411
CL\_9651
CL\_5891
CL\_5892
CL\_5893
CL\_5894
CL\_5895
CL\_5896
CL\_5897
CL\_5898
CL\_8687
CL\_9650
CL\_9649
CL\_23676
CL\_23675
CL\_12905
CL\_23809
CL\_23808
CL\_23807
CL\_23806
CL\_8686
CL\_8685
CL\_15848
CL\_11174
CL\_11128
CL\_11173
CL\_23863
CL\_23862
CL\_17025
CL\_17026
CL\_8684
CL\_5749
CL\_5750
CL\_5902
CL\_5903
CL\_5754
CL\_5755
CL\_5756
CL\_6376
CL\_5757
CL\_6375
CL\_6308
CL\_6307
CL\_6306
CL\_6305
CL\_5758
CL\_6374
CL\_5760
CL\_5761
CL\_5762
CL\_5763
CL\_5764
CL\_5765
CL\_21642
CL\_12368
CL\_15846
CL\_15845
CL\_15844
CL\_10413
CL\_7209
CL\_20379
CL\_8334
CL\_7207
CL\_23861
CL\_23860
CL\_23859
CL\_8682
CL\_6366
CL\_5769
CL\_5770
CL\_5771
CL\_5772
CL\_5773
CL\_5774
CL\_5775
CL\_5776
CL\_11171
CL\_5778
CL\_11170
CL\_5779
CL\_5900
CL\_5901
CL\_6312
CL\_6311
CL\_12906
CL\_23858
CL\_5751
CL\_22242
CL\_12052
CL\_12053
CL\_21979
CL\_21641
CL\_15847
CL\_6309
CL\_14930
CL\_17027
CL\_11067
CL\_5904
CL\_5905
CL\_11068
CL\_5906
CL\_5907
CL\_5908
CL\_9648
CL\_9647
CL\_5909
CL\_5910
CL\_5911
CL\_5912
CL\_5913
CL\_5914
CL\_7210
CL\_5915
CL\_22243
CL\_12907
CL\_17881
CL\_21978
CL\_21977
CL\_5916
CL\_5917
CL\_5918
CL\_9646
CL\_5919
CL\_23805
CL\_23804
CL\_17723
CL\_11077
CL\_11078
CL\_11079
CL\_11080
CL\_11081
CL\_14595
CL\_14594
CL\_14593
CL\_18336
CL\_15011
CL\_12712
CL\_12713
CL\_17028
CL\_17029
CL\_5920
CL\_9818
CL\_14592
CL\_5921
CL\_5922
CL\_13740
CL\_17030
CL\_5923
CL\_5924
CL\_6361
CL\_11172
CL\_6300
CL\_5925
CL\_5926
CL\_5927
CL\_5928
CL\_5929
CL\_5930
CL\_5931
CL\_12908
CL\_23803
CL\_21643
CL\_20378
CL\_14931
CL\_6864
CL\_6865
CL\_6866
CL\_6867
CL\_6868
CL\_8643
CL\_6855
CL\_11169
CL\_11168
CL\_11167
CL\_11166
CL\_11738
CL\_11165
CL\_11164
CL\_11163
CL\_11162
CL\_11161
CL\_11160
CL\_11159
CL\_19185
CL\_22151
CL\_9933
CL\_9934
CL\_17031
CL\_9203
CL\_14136
CL\_14137
CL\_7810
CL\_17055
CL\_17056
CL\_17057
CL\_17058
CL\_8075
CL\_6886
CL\_8077
CL\_8078
CL\_8079
CL\_16972
CL\_11243
CL\_8115
CL\_6120
CL\_6789
CL\_14668
CL\_11341
CL\_11378
CL\_11377
CL\_11376
CL\_11375
CL\_11374
CL\_11373
CL\_11372
CL\_11371
CL\_11370
CL\_11369
CL\_11368
CL\_11367
CL\_11366
CL\_11365
CL\_11364
CL\_11363
CL\_11362
CL\_11361
CL\_11360
CL\_11359
CL\_11358
CL\_11357
CL\_11356
CL\_11352
CL\_11351
CL\_11350
CL\_11349
CL\_11348
CL\_11347
CL\_11345
CL\_11344
CL\_11343
CL\_11342
CL\_11340
CL\_11339
CL\_11338
CL\_11337
CL\_11336
CL\_11335
CL\_11334
CL\_11333
CL\_11332
CL\_11331
CL\_11330
CL\_11329
CL\_11328
CL\_11327
CL\_11326
CL\_11325
CL\_11324
CL\_13100
CL\_11323
CL\_11321
CL\_11320
CL\_11319
CL\_11318
CL\_11317
CL\_11316
CL\_11314
CL\_11312
CL\_11882
CL\_11306
CL\_11305
CL\_11304
CL\_11303
CL\_11302
CL\_11301
CL\_8213
