## Supplementary material for "A novel method for integrating genomic and Tn-Seq data to identify common *in vivo* fitness mechanisms across multiple bacterial species": S2 Dataset: CL_INS_368.html

Legend

 Mobile +extrachromosomalelementfunctions
 Regulatoryfunctions
 Hypothetical
 DNA Metabolism
 All EssentialGenes
 Other
 Transport +binding proteins
 All VFDB Genes

FULL


WINDOWSVGPNG

Trim RowsRemove SingletonsSave Fasta

CL\_4428


CL\_4428


CL\_4428


CL\_4428


CL\_4428


CL\_4428


CL\_4428


CL\_4428


CL\_4428


CL\_4428


CL\_4428


CL\_4428


CL\_4428


CL\_4428


CL\_4428


CL\_4428


CL\_4428


CL\_4428


CL\_4428

HighlightSelectShow Genomes


166

CL\_4427


5

CL\_4426


1

CL\_4426


1

CL\_4426


1

CL\_4426


1

CL\_4426


1

CL\_4426


1

CL\_457


1

CL\_4426


1

CL\_4426


1

CL\_4426


1

CL\_4426


1

CL\_4426


1

CL\_4426


1

CL\_4426


1

CL\_4426


1

CL\_4426


1

CL\_4426


1

Break

fGI ID


CL\_INS\_368
CL\_INS\_368
CL\_INS\_368
CL\_INS\_368
CL\_INS\_368
CL\_INS\_368
CL\_INS\_368
CL\_INS\_368
CL\_INS\_368
CL\_INS\_368
CL\_INS\_368
CL\_INS\_368
CL\_INS\_368
CL\_INS\_368
CL\_INS\_368
CL\_INS\_368
CL\_INS\_368
CL\_INS\_368
CL\_INS\_368
CL\_INS\_368
CL\_INS\_368
CL\_INS\_368
CL\_INS\_368
CL\_INS\_368
CL\_INS\_368
CL\_INS\_368
CL\_INS\_368
CL\_INS\_155
CL\_INS\_155
CL\_INS\_368
CL\_INS\_155
CL\_INS\_155
CL\_INS\_155
CL\_INS\_155
CL\_INS\_155
CL\_INS\_155
CL\_INS\_155
CL\_INS\_155
CL\_INS\_368
CL\_INS\_368
CL\_INS\_368
CL\_INS\_368
CL\_INS\_368
CL\_INS\_368
CL\_INS\_368
CL\_INS\_368
CL\_INS\_368
CL\_INS\_368
CL\_INS\_368
CL\_INS\_368
CL\_INS\_368
CL\_INS\_368
CL\_INS\_368
CL\_INS\_204
CL\_INS\_385
CL\_INS\_368
CL\_INS\_368
CL\_INS\_368
CL\_INS\_368
CL\_INS\_368
CL\_INS\_385
CL\_INS\_321
CL\_INS\_321
CL\_INS\_321
CL\_INS\_321
CL\_INS\_321
CL\_INS\_74
CL\_INS\_26
CL\_INS\_26
CL\_INS\_368
CL\_INS\_368
CL\_INS\_368
CL\_INS\_368
CL\_INS\_368
CL\_INS\_368
CL\_INS\_368
CL\_INS\_368
CL\_INS\_368
CL\_INS\_351
CL\_INS\_351
CL\_INS\_351
CL\_INS\_351
CL\_INS\_368
CL\_INS\_368
CL\_INS\_321
CL\_INS\_321
CL\_INS\_321
CL\_INS\_321
CL\_INS\_321
CL\_INS\_321
CL\_INS\_368
CL\_INS\_321
CL\_INS\_321
CL\_INS\_321
CL\_INS\_321
CL\_INS\_321
CL\_INS\_321
CL\_INS\_368
CL\_INS\_368
CL\_INS\_368
CL\_INS\_132
CL\_INS\_368
CL\_INS\_132
CL\_INS\_132
CL\_INS\_368
CL\_INS\_132
CL\_INS\_319
CL\_INS\_319
CL\_INS\_319
CL\_INS\_368
CL\_INS\_368
CL\_INS\_368
CL\_INS\_368
CL\_INS\_319
CL\_INS\_319
CL\_INS\_368
CL\_INS\_319
CL\_INS\_368
CL\_INS\_319
CL\_INS\_319
CL\_INS\_319
CL\_INS\_319
CL\_INS\_319
CL\_INS\_319
CL\_INS\_319
CL\_INS\_319
CL\_INS\_319
CL\_INS\_319
CL\_INS\_319
CL\_INS\_319
CL\_INS\_319
CL\_INS\_319
CL\_INS\_368
CL\_INS\_319
CL\_INS\_368
CL\_INS\_132
CL\_INS\_132
CL\_INS\_132
CL\_INS\_319
CL\_INS\_319
CL\_INS\_321
CL\_INS\_321
CL\_INS\_368
CL\_INS\_321
CL\_INS\_321
CL\_INS\_321
CL\_INS\_132
CL\_INS\_132
CL\_INS\_132
CL\_INS\_132
CL\_INS\_132
CL\_INS\_132
CL\_INS\_321
CL\_INS\_132
CL\_INS\_132
CL\_INS\_368
CL\_INS\_368
CL\_INS\_368
CL\_INS\_132
CL\_INS\_132
CL\_INS\_132
CL\_INS\_132
CL\_INS\_132
CL\_INS\_132
CL\_INS\_132
CL\_INS\_132
CL\_INS\_132
CL\_INS\_132
CL\_INS\_132
CL\_INS\_132
CL\_INS\_132
CL\_INS\_132
CL\_INS\_132
CL\_INS\_132
CL\_INS\_132
CL\_INS\_132
CL\_INS\_319
CL\_INS\_319
CL\_INS\_319
CL\_INS\_319
CL\_INS\_132
CL\_INS\_319
CL\_INS\_319
CL\_INS\_368
CL\_INS\_319
CL\_INS\_368
CL\_INS\_368
CL\_INS\_319
CL\_INS\_319
CL\_INS\_319
CL\_INS\_319
CL\_INS\_319
CL\_INS\_319
CL\_INS\_319
CL\_INS\_319
CL\_INS\_368
CL\_INS\_132
CL\_INS\_132
CL\_INS\_132
CL\_INS\_132
CL\_INS\_132
CL\_INS\_132
CL\_INS\_132
CL\_INS\_321
Cluster ID


CL\_9935
CL\_9934
CL\_9933
CL\_15412
CL\_15413
CL\_19185
CL\_22151
CL\_11159
CL\_11160
CL\_11161
CL\_11162
CL\_11163
CL\_11164
CL\_11738
CL\_11165
CL\_11166
CL\_11167
CL\_11168
CL\_11169
CL\_21590
CL\_21591
CL\_15414
CL\_15415
CL\_15416
CL\_21592
CL\_15417
CL\_15418
CL\_7492
CL\_7490
CL\_15419
CL\_6206
CL\_6205
CL\_6204
CL\_6203
CL\_6202
CL\_6201
CL\_7488
CL\_6448
CL\_21593
CL\_21594
CL\_21595
CL\_21596
CL\_21597
CL\_21598
CL\_22150
CL\_9255
CL\_23068
CL\_23069
CL\_23070
CL\_23071
CL\_23072
CL\_23073
CL\_23074
CL\_5953
CL\_8212
CL\_8129
CL\_8128
CL\_8127
CL\_8126
CL\_8125
CL\_8228
CL\_8355
CL\_8354
CL\_8353
CL\_10419
CL\_8352
CL\_7913
CL\_17618
CL\_17617
CL\_11739
CL\_11740
CL\_11741
CL\_11742
CL\_11743
CL\_11744
CL\_11745
CL\_11746
CL\_11747
CL\_10430
CL\_10431
CL\_10432
CL\_10433
CL\_11748
CL\_11749
CL\_11750
CL\_11751
CL\_11752
CL\_7423
CL\_7422
CL\_7421
CL\_11753
CL\_7417
CL\_7416
CL\_7415
CL\_7414
CL\_7413
CL\_7412
CL\_23857
CL\_5931
CL\_11170
CL\_5930
CL\_5929
CL\_5928
CL\_6300
CL\_11172
CL\_6361
CL\_5924
CL\_5923
CL\_5921
CL\_17029
CL\_17028
CL\_17723
CL\_18336
CL\_5920
CL\_12712
CL\_15011
CL\_12713
CL\_14595
CL\_5915
CL\_7210
CL\_5913
CL\_5912
CL\_5911
CL\_5910
CL\_5909
CL\_9647
CL\_9648
CL\_5906
CL\_11068
CL\_5905
CL\_5904
CL\_11067
CL\_12906
CL\_5751
CL\_23858
CL\_6311
CL\_6312
CL\_10411
CL\_5899
CL\_5890
CL\_5779
CL\_5778
CL\_11171
CL\_5776
CL\_5775
CL\_5774
CL\_5773
CL\_5772
CL\_5771
CL\_5770
CL\_5769
CL\_6366
CL\_8682
CL\_5765
CL\_5764
CL\_23859
CL\_23860
CL\_23861
CL\_7209
CL\_10413
CL\_5763
CL\_5762
CL\_5761
CL\_5760
CL\_6374
CL\_6375
CL\_5757
CL\_6376
CL\_5756
CL\_5755
CL\_5754
CL\_5758
CL\_6305
CL\_6306
CL\_6307
CL\_6308
CL\_5903
CL\_5902
CL\_5750
CL\_5900
CL\_5749
CL\_8684
CL\_11128
CL\_11173
CL\_11174
CL\_23862
CL\_23863
CL\_8687
CL\_5897
CL\_5896
CL\_5895
CL\_5894
CL\_5893
CL\_5892
CL\_5891
CL\_9651
CL\_6318
CL\_6319
CL\_6320
CL\_6321
CL\_6322
CL\_6388
CL\_6389
CL\_5736
