## Supplementary material for "A novel method for integrating genomic and Tn-Seq data to identify common *in vivo* fitness mechanisms across multiple bacterial species": S2 Dataset: CL_INS_369.html

Legend

 Hypothetical
 EnergyMetabolism
 All VFDB Genes

FULL


WINDOWSVGPNG

Trim RowsRemove SingletonsSave Fasta

CL\_4450


CL\_4450


CL\_4449

HighlightSelectShow Genomes


129

CL\_4451


72

CL\_4451


1

CL\_4451

fGI ID

CL\_INS\_369
Cluster ID

CL\_7411
