## Supplementary material for "A novel method for integrating genomic and Tn-Seq data to identify common *in vivo* fitness mechanisms across multiple bacterial species": S2 Dataset: CL_INS_370.html

Legend

 Hypothetical
 Other
 All VFDB Genes

FULL


WINDOWSVGPNG

Trim RowsRemove SingletonsSave Fasta

CL\_4452


CL\_4452


CL\_4452


CL\_4451

HighlightSelectShow Genomes


195

CL\_4453


8

CL\_4453


1

CL\_4453


1

CL\_4453

fGI ID


CL\_INS\_370
CL\_INS\_370
CL\_INS\_370
CL\_INS\_370
CL\_INS\_370
CL\_INS\_370
CL\_INS\_370
CL\_INS\_370
CL\_INS\_370
Cluster ID


CL\_12806
CL\_7650
CL\_7649
CL\_7648
CL\_7647
CL\_7646
CL\_7645
CL\_7644
CL\_7643
