## Supplementary material for "A novel method for integrating genomic and Tn-Seq data to identify common *in vivo* fitness mechanisms across multiple bacterial species": S2 Dataset: CL_INS_372.html

Legend

 Hypothetical
 Other
 All VFDB Genes

FULL


WINDOWSVGPNG

Trim RowsRemove SingletonsSave Fasta

CL\_4486


CL\_4486


CL\_4486


CL\_4486

HighlightSelectShow Genomes


115

CL\_4488


86

CL\_4488


1

CL\_4489


1

CL\_4488

fGI ID


CL\_INS\_372
CL\_INS\_373
Cluster ID


CL\_4487
CL\_13018
