## Supplementary material for "A novel method for integrating genomic and Tn-Seq data to identify common *in vivo* fitness mechanisms across multiple bacterial species": S2 Dataset: CL_INS_374.html

Legend

 Regulatoryfunctions
 Hypothetical
 All EssentialGenes
 Other
 All VFDB Genes
 Transport +binding proteins

FULL


WINDOWSVGPNG

Trim RowsRemove SingletonsSave Fasta

CL\_4494


CL\_4494


CL\_4494


CL\_4494


CL\_4494


CL\_4494

HighlightSelectShow Genomes


106

CL\_4509


52

CL\_4509


40

CL\_4509


1

CL\_4509


1

CL\_4509


1

CL\_4509

fGI ID


CL\_INS\_374
CL\_INS\_374
CL\_INS\_374
CL\_INS\_374
CL\_INS\_374
CL\_INS\_374
CL\_INS\_374
CL\_INS\_374
CL\_INS\_374
CL\_INS\_374
CL\_INS\_374
CL\_INS\_374
CL\_INS\_374
CL\_INS\_374
CL\_INS\_374
Cluster ID


CL\_4495
CL\_4496
CL\_4497
CL\_20377
CL\_4498
CL\_4499
CL\_4500
CL\_4501
CL\_4502
CL\_4503
CL\_4504
CL\_4505
CL\_4506
CL\_4507
CL\_4508
