## Supplementary material for "A novel method for integrating genomic and Tn-Seq data to identify common *in vivo* fitness mechanisms across multiple bacterial species": S2 Dataset: CL_INS_375.html

Legend

 Regulatoryfunctions
 Hypothetical
 All EssentialGenes
 Other

FULL


WINDOWSVGPNG

Trim RowsRemove SingletonsSave Fasta

CL\_4523


CL\_4523


CL\_4523


CL\_4523


CL\_4523


CL\_4523


CL\_4523

HighlightSelectShow Genomes


155

CL\_4524


43

CL\_4524


2

CL\_4524


1

CL\_4524


1

CL\_4524


1

CL\_4525


1

CL\_4524

fGI ID


CL\_INS\_375
CL\_INS\_375
CL\_INS\_375
CL\_INS\_375
CL\_INS\_375
CL\_INS\_375
CL\_INS\_375
CL\_INS\_375
CL\_INS\_375
CL\_INS\_375
CL\_INS\_375
CL\_INS\_375
CL\_INS\_375
Cluster ID


CL\_5318
CL\_5317
CL\_5316
CL\_5315
CL\_5314
CL\_5313
CL\_5312
CL\_5311
CL\_5310
CL\_5309
CL\_5308
CL\_5307
CL\_5306
