## Supplementary material for "A novel method for integrating genomic and Tn-Seq data to identify common *in vivo* fitness mechanisms across multiple bacterial species": S2 Dataset: CL_INS_378.html

Legend

 Hypothetical
 All Fitness Genes
 Other
 All VFDB Genes

FULL


WINDOWSVGPNG

Trim RowsRemove SingletonsSave Fasta

CL\_4575


CL\_4575


CL\_1022


CL\_4575


CL\_4574


CL\_4574

HighlightSelectShow Genomes


117

CL\_4577


83

CL\_4577


1

CL\_4577


1

CL\_1021


1

CL\_4577


1

CL\_4577

fGI ID


CL\_INS\_378
CL\_INS\_378
Cluster ID


CL\_4576
CL\_8087
