## Supplementary material for "A novel method for integrating genomic and Tn-Seq data to identify common *in vivo* fitness mechanisms across multiple bacterial species": S2 Dataset: CL_INS_379.html

Legend

 Mobile +extrachromosomalelementfunctions
 Hypothetical
 Regulatoryfunctions
 Other
 All VFDB Genes
 Transport +binding proteins

FULL


WINDOWSVGPNG

Trim RowsRemove SingletonsSave Fasta

CL\_4591


CL\_4591


CL\_4591


CL\_4591


CL\_4591


CL\_4591


CL\_4591


CL\_4591


CL\_4591


CL\_4591


CL\_4591


CL\_4591

HighlightSelectShow Genomes


137

CL\_4601


53

CL\_4601


3

CL\_4601


3

CL\_4601


1

CL\_4601


1

CL\_4601


1

CL\_4601


1

CL\_4601


1

CL\_4601


1

CL\_4601


1

CL\_4601


1

CL\_4601

fGI ID


CL\_INS\_379
CL\_INS\_379
CL\_INS\_379
CL\_INS\_10
CL\_INS\_10
CL\_INS\_10
CL\_INS\_379
CL\_INS\_379
CL\_INS\_379
CL\_INS\_379
CL\_INS\_379
CL\_INS\_379
CL\_INS\_379
CL\_INS\_379
Cluster ID


CL\_4592
CL\_12909
CL\_13017
CL\_4593
CL\_4594
CL\_4595
CL\_4596
CL\_4597
CL\_4598
CL\_4599
CL\_4600
CL\_18520
CL\_18519
CL\_18518
