## Supplementary material for "A novel method for integrating genomic and Tn-Seq data to identify common *in vivo* fitness mechanisms across multiple bacterial species": S2 Dataset: CL_INS_381.html

Legend

 Mobile +extrachromosomalelementfunctions
 Hypothetical
 All EssentialGenes
 Other
 All VFDB Genes

FULL


WINDOWSVGPNG

Trim RowsRemove SingletonsSave Fasta

CL\_4628


CL\_4628


CL\_4628


CL\_4628


CL\_4628


CL\_4628


CL\_4628


CL\_4628


CL\_4628


CL\_4628


CL\_4628


CL\_4628


CL\_4628


CL\_4628


CL\_4628


CL\_4628


CL\_4628


CL\_4628


CL\_4628


CL\_4628

HighlightSelectShow Genomes


154

CL\_4629


15

CL\_4629


10

CL\_4629


8

CL\_4629


1

CL\_4629


1

CL\_4629


1

CL\_4629


1

CL\_4629


1

CL\_4629


1

CL\_4629


1

Break


1

CL\_4629


1

CL\_4629


1

CL\_4629


1

CL\_4629


1

CL\_4629


1

CL\_4629


1

CL\_4629


1

CL\_4629


1

CL\_4629

fGI ID


CL\_INS\_381
CL\_INS\_381
CL\_INS\_381
CL\_INS\_381
CL\_INS\_381
CL\_INS\_381
CL\_INS\_381
CL\_INS\_381
CL\_INS\_381
CL\_INS\_381
CL\_INS\_381
CL\_INS\_381
CL\_INS\_381
CL\_INS\_381
CL\_INS\_381
CL\_INS\_381
CL\_INS\_381
CL\_INS\_381
CL\_INS\_381
CL\_INS\_381
CL\_INS\_381
CL\_INS\_381
CL\_INS\_381
CL\_INS\_381
CL\_INS\_381
CL\_INS\_381
CL\_INS\_381
CL\_INS\_381
CL\_INS\_381
CL\_INS\_381
CL\_INS\_381
CL\_INS\_381
CL\_INS\_381
CL\_INS\_381
CL\_INS\_381
CL\_INS\_381
CL\_INS\_381
CL\_INS\_381
CL\_INS\_155
CL\_INS\_381
CL\_INS\_381
CL\_INS\_381
CL\_INS\_381
CL\_INS\_155
CL\_INS\_381
CL\_INS\_155
CL\_INS\_381
CL\_INS\_381
CL\_INS\_381
CL\_INS\_381
CL\_INS\_381
CL\_INS\_381
CL\_INS\_381
CL\_INS\_155
CL\_INS\_155
CL\_INS\_381
CL\_INS\_381
CL\_INS\_155
CL\_INS\_155
CL\_INS\_155
CL\_INS\_155
CL\_INS\_155
CL\_INS\_381
CL\_INS\_155
CL\_INS\_155
CL\_INS\_381
CL\_INS\_381
CL\_INS\_381
CL\_INS\_381
CL\_INS\_381
CL\_INS\_381
CL\_INS\_381
CL\_INS\_381
CL\_INS\_381
CL\_INS\_381
CL\_INS\_381
CL\_INS\_381
CL\_INS\_381
CL\_INS\_381
CL\_INS\_381
CL\_INS\_381
CL\_INS\_381
CL\_INS\_155
CL\_INS\_381
CL\_INS\_381
CL\_INS\_155
CL\_INS\_381
CL\_INS\_381
CL\_INS\_381
CL\_INS\_381
CL\_INS\_381
CL\_INS\_381
CL\_INS\_278
CL\_INS\_278
CL\_INS\_212
CL\_INS\_278
CL\_INS\_278
CL\_INS\_278
CL\_INS\_58
CL\_INS\_58
CL\_INS\_178
CL\_INS\_278
CL\_INS\_278
CL\_INS\_58
CL\_INS\_278
CL\_INS\_278
CL\_INS\_401
CL\_INS\_278
CL\_INS\_278
CL\_INS\_278
CL\_INS\_278
CL\_INS\_278
CL\_INS\_278
CL\_INS\_278
CL\_INS\_278
CL\_INS\_278
CL\_INS\_278
CL\_INS\_278
CL\_INS\_278
Cluster ID


CL\_22080
CL\_22079
CL\_22078
CL\_22077
CL\_22076
CL\_22075
CL\_22074
CL\_22073
CL\_22072
CL\_22071
CL\_24136
CL\_24135
CL\_24134
CL\_24133
CL\_24132
CL\_24131
CL\_24130
CL\_24129
CL\_24128
CL\_24127
CL\_24126
CL\_24125
CL\_24124
CL\_24123
CL\_24122
CL\_24121
CL\_24120
CL\_17879
CL\_18010
CL\_18009
CL\_18008
CL\_18007
CL\_18006
CL\_18005
CL\_18004
CL\_18003
CL\_16998
CL\_18002
CL\_7492
CL\_17877
CL\_17876
CL\_17875
CL\_23593
CL\_7493
CL\_18001
CL\_7495
CL\_14220
CL\_14221
CL\_18000
CL\_6447
CL\_14218
CL\_14219
CL\_19006
CL\_6448
CL\_7488
CL\_12212
CL\_6200
CL\_6201
CL\_6202
CL\_6203
CL\_6204
CL\_6205
CL\_14647
CL\_6206
CL\_7490
CL\_17878
CL\_6207
CL\_19005
CL\_19004
CL\_19003
CL\_19002
CL\_19001
CL\_6208
CL\_20569
CL\_6209
CL\_6210
CL\_6211
CL\_12213
CL\_6212
CL\_6213
CL\_6214
CL\_17999
CL\_6215
CL\_6216
CL\_8030
CL\_6217
CL\_17874
CL\_19000
CL\_14222
CL\_21059
CL\_6218
CL\_6219
CL\_6882
CL\_6883
CL\_6825
CL\_5993
CL\_5992
CL\_5991
CL\_6826
CL\_6827
CL\_6828
CL\_5988
CL\_5987
CL\_5986
CL\_6829
CL\_5983
CL\_6010
CL\_6768
CL\_6769
CL\_6770
CL\_6771
CL\_6772
CL\_6773
CL\_6774
CL\_6775
CL\_6776
CL\_6777
CL\_6778
CL\_6779
