## Supplementary material for "A novel method for integrating genomic and Tn-Seq data to identify common *in vivo* fitness mechanisms across multiple bacterial species": S2 Dataset: CL_INS_383.html

Legend

 Hypothetical
 All Fitness Genes
 Other
 All VFDB Genes

FULL


WINDOWSVGPNG

Trim RowsRemove SingletonsSave Fasta

CL\_4749


CL\_4749


CL\_4749


CL\_4749


CL\_4749


CL\_4749

HighlightSelectShow Genomes


127

CL\_4750


63

CL\_4750


8

CL\_4750


5

CL\_4750


1

CL\_4750


1

CL\_4751

fGI ID


CL\_INS\_383
CL\_INS\_383
CL\_INS\_383
Cluster ID


CL\_5305
CL\_5304
CL\_17524
