## Supplementary material for "A novel method for integrating genomic and Tn-Seq data to identify common *in vivo* fitness mechanisms across multiple bacterial species": S2 Dataset: CL_INS_384.html

Legend

 Hypothetical
 Other

FULL


WINDOWSVGPNG

Trim RowsRemove SingletonsSave Fasta

CL\_4777


CL\_4777


CL\_4777

HighlightSelectShow Genomes


113

CL\_4778


91

CL\_4778


1

CL\_4778

fGI ID


CL\_INS\_384
CL\_INS\_384
Cluster ID


CL\_5831
CL\_12730
