## Supplementary material for "A novel method for integrating genomic and Tn-Seq data to identify common *in vivo* fitness mechanisms across multiple bacterial species": S2 Dataset: CL_INS_385.html

Legend

 Mobile +extrachromosomalelementfunctions
 Regulatoryfunctions
 Hypothetical
 DNA Metabolism
 AntibioticResistance
 All EssentialGenes
 Biosynthesis ofcofactors,prostheticgroups, +carriers
 Proteinsynthesis/fate
 Other
 Cellularprocesses
 Transport +binding proteins
 All VFDB Genes

FULL


WINDOWSVGPNG

Trim RowsRemove SingletonsSave Fasta

CL\_4780


CL\_4780


CL\_4780


CL\_4780

HighlightSelectShow Genomes


174

CL\_4781


29

CL\_4781


1

Break


1

Break

fGI ID


CL\_INS\_385
CL\_INS\_385
CL\_INS\_278
CL\_INS\_385
CL\_INS\_278
CL\_INS\_278
CL\_INS\_385
CL\_INS\_307
CL\_INS\_385
CL\_INS\_385
CL\_INS\_385
CL\_INS\_385
CL\_INS\_385
CL\_INS\_385
CL\_INS\_385
CL\_INS\_385
CL\_INS\_385
CL\_INS\_385
CL\_INS\_385
CL\_INS\_385
CL\_INS\_385
CL\_INS\_385
CL\_INS\_385
CL\_INS\_385
CL\_INS\_385
CL\_INS\_385
CL\_INS\_385
CL\_INS\_385
CL\_INS\_385
CL\_INS\_385
CL\_INS\_385
CL\_INS\_385
CL\_INS\_385
CL\_INS\_385
CL\_INS\_385
CL\_INS\_385
CL\_INS\_385
CL\_INS\_385
CL\_INS\_385
CL\_INS\_385
CL\_INS\_385
CL\_INS\_385
CL\_INS\_385
CL\_INS\_385
CL\_INS\_385
CL\_INS\_385
CL\_INS\_385
CL\_INS\_385
CL\_INS\_385
CL\_INS\_385
CL\_INS\_385
CL\_INS\_385
CL\_INS\_385
CL\_INS\_385
CL\_INS\_385
CL\_INS\_385
CL\_INS\_385
CL\_INS\_385
CL\_INS\_385
CL\_INS\_385
CL\_INS\_385
CL\_INS\_385
CL\_INS\_385
CL\_INS\_385
CL\_INS\_385
CL\_INS\_385
CL\_INS\_385
CL\_INS\_385
CL\_INS\_385
CL\_INS\_385
CL\_INS\_385
CL\_INS\_385
CL\_INS\_385
CL\_INS\_385
CL\_INS\_278
CL\_INS\_385
CL\_INS\_278
CL\_INS\_385
CL\_INS\_385
CL\_INS\_385
CL\_INS\_278
CL\_INS\_278
CL\_INS\_385
CL\_INS\_57
CL\_INS\_385
CL\_INS\_307
CL\_INS\_307
CL\_INS\_307
CL\_INS\_307
CL\_INS\_307
CL\_INS\_307
CL\_INS\_307
CL\_INS\_307
CL\_INS\_385
CL\_INS\_385
CL\_INS\_385
CL\_INS\_385
CL\_INS\_385
CL\_INS\_385
CL\_INS\_385
CL\_INS\_385
CL\_INS\_385
CL\_INS\_385
CL\_INS\_385
CL\_INS\_385
CL\_INS\_385
CL\_INS\_385
CL\_INS\_385
CL\_INS\_321
CL\_INS\_321
CL\_INS\_385
CL\_INS\_385
CL\_INS\_385
CL\_INS\_385
CL\_INS\_385
CL\_INS\_204
CL\_INS\_385
CL\_INS\_385
CL\_INS\_385
CL\_INS\_385
CL\_INS\_385
CL\_INS\_385
CL\_INS\_385
CL\_INS\_278
CL\_INS\_278
CL\_INS\_385
CL\_INS\_278
CL\_INS\_278
CL\_INS\_385
CL\_INS\_278
CL\_INS\_385
CL\_INS\_278
CL\_INS\_278
CL\_INS\_278
CL\_INS\_278
CL\_INS\_385
CL\_INS\_278
CL\_INS\_278
CL\_INS\_278
CL\_INS\_278
CL\_INS\_278
CL\_INS\_278
CL\_INS\_278
CL\_INS\_385
CL\_INS\_385
CL\_INS\_385
CL\_INS\_385
CL\_INS\_385
CL\_INS\_307
CL\_INS\_278
CL\_INS\_385
CL\_INS\_278
CL\_INS\_385
CL\_INS\_385
CL\_INS\_278
CL\_INS\_278
CL\_INS\_307
CL\_INS\_307
CL\_INS\_307
CL\_INS\_307
CL\_INS\_307
CL\_INS\_385
CL\_INS\_307
CL\_INS\_385
CL\_INS\_385
CL\_INS\_385
CL\_INS\_385
CL\_INS\_385
CL\_INS\_385
CL\_INS\_385
CL\_INS\_385
CL\_INS\_385
CL\_INS\_385
CL\_INS\_385
CL\_INS\_385
CL\_INS\_385
CL\_INS\_385
CL\_INS\_385
CL\_INS\_385
CL\_INS\_385
CL\_INS\_385
CL\_INS\_385
CL\_INS\_385
CL\_INS\_385
CL\_INS\_385
CL\_INS\_385
CL\_INS\_385
CL\_INS\_307
CL\_INS\_278
CL\_INS\_307
CL\_INS\_385
CL\_INS\_385
CL\_INS\_385
CL\_INS\_385
CL\_INS\_385
CL\_INS\_385
CL\_INS\_385
CL\_INS\_385
CL\_INS\_385
CL\_INS\_385
CL\_INS\_385
CL\_INS\_385
CL\_INS\_53
CL\_INS\_53
CL\_INS\_53
CL\_INS\_53
CL\_INS\_385
CL\_INS\_385
CL\_INS\_385
CL\_INS\_385
CL\_INS\_385
CL\_INS\_385
CL\_INS\_385
CL\_INS\_385
CL\_INS\_385
CL\_INS\_385
CL\_INS\_53
CL\_INS\_385
CL\_INS\_385
CL\_INS\_385
CL\_INS\_385
CL\_INS\_385
CL\_INS\_385
CL\_INS\_385
CL\_INS\_385
CL\_INS\_385
CL\_INS\_385
CL\_INS\_385
CL\_INS\_278
CL\_INS\_278
CL\_INS\_385
CL\_INS\_385
CL\_INS\_385
CL\_INS\_385
CL\_INS\_385
CL\_INS\_385
CL\_INS\_385
CL\_INS\_278
CL\_INS\_278
CL\_INS\_278
CL\_INS\_385
CL\_INS\_278
CL\_INS\_385
CL\_INS\_385
CL\_INS\_278
CL\_INS\_278
CL\_INS\_385
CL\_INS\_30
CL\_INS\_278
CL\_INS\_385
CL\_INS\_385
CL\_INS\_385
CL\_INS\_385
CL\_INS\_385
CL\_INS\_385
CL\_INS\_385
CL\_INS\_385
CL\_INS\_385
CL\_INS\_385
CL\_INS\_385
CL\_INS\_385
CL\_INS\_385
CL\_INS\_132
CL\_INS\_132
CL\_INS\_385
CL\_INS\_385
CL\_INS\_385
CL\_INS\_385
CL\_INS\_91
CL\_INS\_385
CL\_INS\_385
CL\_INS\_385
CL\_INS\_385
CL\_INS\_132
CL\_INS\_278
CL\_INS\_278
CL\_INS\_132
CL\_INS\_278
CL\_INS\_385
CL\_INS\_307
CL\_INS\_307
CL\_INS\_132
CL\_INS\_307
CL\_INS\_307
CL\_INS\_132
CL\_INS\_132
CL\_INS\_132
CL\_INS\_132
CL\_INS\_132
CL\_INS\_132
CL\_INS\_132
CL\_INS\_132
CL\_INS\_385
CL\_INS\_385
CL\_INS\_385
CL\_INS\_385
CL\_INS\_385
CL\_INS\_385
CL\_INS\_385
CL\_INS\_385
CL\_INS\_132
CL\_INS\_385
CL\_INS\_385
CL\_INS\_385
CL\_INS\_278
CL\_INS\_278
CL\_INS\_278
CL\_INS\_278
CL\_INS\_278
CL\_INS\_197
CL\_INS\_197
CL\_INS\_197
CL\_INS\_197
CL\_INS\_278
CL\_INS\_278
CL\_INS\_385
CL\_INS\_385
CL\_INS\_385
CL\_INS\_385
CL\_INS\_385
CL\_INS\_385
CL\_INS\_385
CL\_INS\_385
CL\_INS\_385
CL\_INS\_385
CL\_INS\_243
CL\_INS\_243
CL\_INS\_385
CL\_INS\_385
Cluster ID


CL\_8027
CL\_15232
CL\_6089
CL\_15224
CL\_13751
CL\_13750
CL\_15225
CL\_6821
CL\_15226
CL\_15227
CL\_15228
CL\_15229
CL\_15230
CL\_15231
CL\_14107
CL\_14109
CL\_14110
CL\_14111
CL\_14112
CL\_14113
CL\_14114
CL\_14115
CL\_14116
CL\_14118
CL\_14119
CL\_14120
CL\_14121
CL\_14122
CL\_14123
CL\_14124
CL\_14125
CL\_14126
CL\_14127
CL\_14128
CL\_14129
CL\_14130
CL\_14131
CL\_14132
CL\_14133
CL\_14134
CL\_14108
CL\_14100
CL\_14101
CL\_14102
CL\_14103
CL\_15233
CL\_14104
CL\_14105
CL\_14106
CL\_13335
CL\_14667
CL\_12394
CL\_12396
CL\_12397
CL\_12398
CL\_12399
CL\_8212
CL\_10853
CL\_10854
CL\_10855
CL\_10457
CL\_10456
CL\_15221
CL\_13493
CL\_13494
CL\_7794
CL\_9192
CL\_8873
CL\_23956
CL\_23957
CL\_23958
CL\_14464
CL\_6127
CL\_6091
CL\_103
CL\_7536
CL\_102
CL\_7092
CL\_16350
CL\_23907
CL\_6673
CL\_6672
CL\_10777
CL\_8061
CL\_90
CL\_87
CL\_86
CL\_83
CL\_82
CL\_81
CL\_73
CL\_72
CL\_5951
CL\_8874
CL\_8875
CL\_8876
CL\_8877
CL\_8878
CL\_8880
CL\_8881
CL\_8882
CL\_8883
CL\_8884
CL\_8885
CL\_8886
CL\_8887
CL\_8228
CL\_15222
CL\_12218
CL\_12219
CL\_10453
CL\_9385
CL\_7128
CL\_7127
CL\_7126
CL\_5475
CL\_12156
CL\_12157
CL\_16346
CL\_16345
CL\_7096
CL\_12158
CL\_8855
CL\_135
CL\_136
CL\_137
CL\_138
CL\_139
CL\_23908
CL\_141
CL\_142
CL\_143
CL\_144
CL\_145
CL\_146
CL\_147
CL\_148
CL\_149
CL\_150
CL\_151
CL\_152
CL\_153
CL\_154
CL\_7789
CL\_10313
CL\_10312
CL\_7576
CL\_7575
CL\_11688
CL\_10452
CL\_14943
CL\_11877
CL\_11878
CL\_16355
CL\_11879
CL\_10666
CL\_5938
CL\_5939
CL\_5940
CL\_5941
CL\_5942
CL\_5943
CL\_17694
CL\_8929
CL\_22865
CL\_22864
CL\_8930
CL\_8931
CL\_8932
CL\_8933
CL\_8934
CL\_8935
CL\_8936
CL\_8937
CL\_9510
CL\_9511
CL\_9512
CL\_9513
CL\_9514
CL\_9515
CL\_9516
CL\_23909
CL\_7776
CL\_23910
CL\_23911
CL\_65
CL\_64
CL\_8858
CL\_8050
CL\_8857
CL\_6156
CL\_23912
CL\_23913
CL\_10541
CL\_13384
CL\_18320
CL\_13432
CL\_13938
CL\_9452
CL\_14746
CL\_14747
CL\_12091
CL\_13476
CL\_10798
CL\_10799
CL\_10800
CL\_18324
CL\_18325
CL\_18326
CL\_18327
CL\_18328
CL\_18329
CL\_18330
CL\_18331
CL\_18332
CL\_18333
CL\_10809
CL\_10810
CL\_10811
CL\_10812
CL\_10813
CL\_10814
CL\_10815
CL\_9369
CL\_9368
CL\_7129
CL\_9367
CL\_9366
CL\_132
CL\_131
CL\_13445
CL\_119
CL\_12170
CL\_118
CL\_117
CL\_18962
CL\_9592
CL\_116
CL\_115
CL\_114
CL\_7770
CL\_113
CL\_112
CL\_23914
CL\_108
CL\_107
CL\_106
CL\_105
CL\_104
CL\_13928
CL\_13927
CL\_23946
CL\_23947
CL\_23948
CL\_23949
CL\_23950
CL\_23951
CL\_23952
CL\_23953
CL\_23954
CL\_23955
CL\_8080
CL\_6714
CL\_8079
CL\_6782
CL\_8118
CL\_8119
CL\_8120
CL\_6886
CL\_8121
CL\_8122
CL\_8123
CL\_8124
CL\_9205
CL\_6104
CL\_6105
CL\_8211
CL\_6106
CL\_13428
CL\_3387
CL\_3386
CL\_8246
CL\_8245
CL\_8244
CL\_8243
CL\_8242
CL\_12269
CL\_7944
CL\_7945
CL\_7946
CL\_6806
CL\_8219
CL\_7810
CL\_9941
CL\_12378
CL\_69
CL\_12217
CL\_6693
CL\_12421
CL\_12422
CL\_6793
CL\_15223
CL\_11247
CL\_6864
CL\_6817
CL\_6137
CL\_6816
CL\_6815
CL\_6814
CL\_6812
CL\_6811
CL\_10120
CL\_6820
CL\_6819
CL\_6818
CL\_15234
CL\_15235
CL\_15236
CL\_15237
CL\_15238
CL\_11241
CL\_23940
CL\_8643
CL\_23941
CL\_23942
CL\_6121
CL\_6120
CL\_23943
CL\_11245
