## Supplementary material for "A novel method for integrating genomic and Tn-Seq data to identify common *in vivo* fitness mechanisms across multiple bacterial species": S2 Dataset: CL_INS_388.html

Legend

 Mobile +extrachromosomalelementfunctions
 Hypothetical
 All EssentialGenes
 Proteinsynthesis/fate
 All VFDB Genes

FULL


WINDOWSVGPNG

Trim RowsRemove SingletonsSave Fasta

CL\_4858


CL\_4858


CL\_4858


CL\_4858


CL\_4858


CL\_4858


Break


Break


CL\_4858

HighlightSelectShow Genomes


132

CL\_4860


56

CL\_4860


5

CL\_4860


4

Break


1

CL\_4860


1

CL\_2152


1

CL\_4860


1

CL\_4860


1

Break

fGI ID


CL\_INS\_132
CL\_INS\_132
CL\_INS\_388
CL\_INS\_388
CL\_INS\_388
CL\_INS\_388
CL\_INS\_388
Cluster ID


CL\_6881
CL\_8219
CL\_6169
CL\_4859
CL\_13655
CL\_13654
CL\_13712
