## Supplementary material for "A novel method for integrating genomic and Tn-Seq data to identify common *in vivo* fitness mechanisms across multiple bacterial species": S2 Dataset: CL_INS_392.html

Legend

 Hypothetical
 Other
 All VFDB Genes

FULL


WINDOWSVGPNG

Trim RowsRemove SingletonsSave Fasta

CL\_4896


CL\_4896

HighlightSelectShow Genomes


190

CL\_4897


13

CL\_4897

fGI ID


CL\_INS\_392
CL\_INS\_392
CL\_INS\_392
CL\_INS\_392
CL\_INS\_392
CL\_INS\_392
CL\_INS\_392
CL\_INS\_392
Cluster ID


CL\_6897
CL\_6896
CL\_6895
CL\_6894
CL\_6893
CL\_6892
CL\_6891
CL\_6890
