## Supplementary material for "A novel method for integrating genomic and Tn-Seq data to identify common *in vivo* fitness mechanisms across multiple bacterial species": S2 Dataset: CL_INS_393.html

Legend

 Hypothetical
 Other
 Transport +binding proteins
 All VFDB Genes

FULL


WINDOWSVGPNG

Trim RowsRemove SingletonsSave Fasta

CL\_4899


CL\_4899


CL\_4899


CL\_4899

HighlightSelectShow Genomes


118

CL\_4900


80

CL\_4900


6

CL\_4900


1

CL\_4900

fGI ID


CL\_INS\_393
CL\_INS\_393
CL\_INS\_393
Cluster ID


CL\_18275
CL\_5302
CL\_5301
