## Supplementary material for "A novel method for integrating genomic and Tn-Seq data to identify common *in vivo* fitness mechanisms across multiple bacterial species": S2 Dataset: CL_INS_394.html

Legend

 Hypothetical
 All VFDB Genes

FULL


WINDOWSVGPNG

Trim RowsRemove SingletonsSave Fasta

CL\_4922


CL\_4922


CL\_4922

HighlightSelectShow Genomes


181

CL\_4923


19

CL\_4926


1

CL\_4928

fGI ID


CL\_INS\_394
CL\_INS\_394
Cluster ID


CL\_7132
CL\_7131
