## Supplementary material for "A novel method for integrating genomic and Tn-Seq data to identify common *in vivo* fitness mechanisms across multiple bacterial species": S2 Dataset: CL_INS_395.html

Legend

 Hypothetical
 All VFDB Genes

FULL


WINDOWSVGPNG

Trim RowsRemove SingletonsSave Fasta

CL\_4926


CL\_4926


CL\_4926


CL\_4926

HighlightSelectShow Genomes


181

CL\_4925


19

CL\_4922


1

CL\_4925


1

CL\_4925

fGI ID


CL\_INS\_395
CL\_INS\_394
CL\_INS\_395
CL\_INS\_395
Cluster ID


CL\_7130
CL\_7131
CL\_12733
CL\_12732
