## Supplementary material for "A novel method for integrating genomic and Tn-Seq data to identify common *in vivo* fitness mechanisms across multiple bacterial species": S2 Dataset: CL_INS_396.html

Legend

 Hypothetical
 All EssentialGenes
 Other

FULL


WINDOWSVGPNG

Trim RowsRemove SingletonsSave Fasta

CL\_4934


CL\_4934


CL\_4934

HighlightSelectShow Genomes


201

CL\_4935


3

CL\_4935


1

CL\_4935

fGI ID


CL\_INS\_396
CL\_INS\_396
CL\_INS\_396
CL\_INS\_396
Cluster ID


CL\_5829
CL\_5828
CL\_5827
CL\_5826
