## Supplementary material for "A novel method for integrating genomic and Tn-Seq data to identify common *in vivo* fitness mechanisms across multiple bacterial species": S2 Dataset: CL_INS_399.html

Legend

 Mobile +extrachromosomalelementfunctions
 Hypothetical
 AntibioticResistance
 Other
 All VFDB Genes

FULL


WINDOWSVGPNG

Trim RowsRemove SingletonsSave Fasta

CL\_5300


CL\_5300


CL\_4946


CL\_4944


CL\_4946


CL\_5300


CL\_4946


CL\_5300


CL\_5300


CL\_4943


CL\_4946

HighlightSelectShow Genomes


97

CL\_4949


79

CL\_4949


4

CL\_4949


4

CL\_4949


4

CL\_4949


4

CL\_4949


1

CL\_4949


1

Break


1

CL\_4949


1

CL\_4949


1

CL\_4949

fGI ID


CL\_INS\_399
CL\_INS\_399
CL\_INS\_398
CL\_INS\_398
CL\_INS\_399
Cluster ID


CL\_12514
CL\_6456
CL\_9253
CL\_14590
CL\_4948
