## Supplementary material for "A novel method for integrating genomic and Tn-Seq data to identify common *in vivo* fitness mechanisms across multiple bacterial species": S2 Dataset: CL_INS_400.html

Legend

 Mobile +extrachromosomalelementfunctions
 Regulatoryfunctions
 Hypothetical
 DNA Metabolism
 Other
 Transport +binding proteins
 All VFDB Genes

FULL


WINDOWSVGPNG

Trim RowsRemove SingletonsSave Fasta

CL\_4949


CL\_4949


CL\_4949


CL\_4949


CL\_4949


CL\_4949


CL\_4949


CL\_4949


CL\_4949


CL\_4949


CL\_4949


CL\_4949


CL\_4949


CL\_4949


CL\_4949


CL\_4949

HighlightSelectShow Genomes


107

CL\_4951


81

CL\_4951


1

CL\_4951


1

CL\_4952


1

CL\_4951


1

CL\_4951


1

CL\_4951


1

CL\_4952


1

CL\_4951


1

CL\_4951


1

CL\_4951


1

CL\_4951


1

CL\_4951


1

CL\_4951


1

CL\_4952


1

CL\_4951

fGI ID


CL\_INS\_400
CL\_INS\_400
CL\_INS\_400
CL\_INS\_400
CL\_INS\_400
CL\_INS\_400
CL\_INS\_400
CL\_INS\_351
CL\_INS\_400
CL\_INS\_400
CL\_INS\_400
CL\_INS\_400
CL\_INS\_400
CL\_INS\_400
CL\_INS\_400
CL\_INS\_400
CL\_INS\_400
CL\_INS\_400
CL\_INS\_400
CL\_INS\_400
CL\_INS\_400
CL\_INS\_400
CL\_INS\_400
CL\_INS\_400
CL\_INS\_400
CL\_INS\_400
CL\_INS\_278
CL\_INS\_385
CL\_INS\_385
CL\_INS\_278
CL\_INS\_400
CL\_INS\_400
CL\_INS\_400
CL\_INS\_400
CL\_INS\_400
CL\_INS\_400
CL\_INS\_400
CL\_INS\_313
CL\_INS\_400
CL\_INS\_313
CL\_INS\_91
CL\_INS\_91
CL\_INS\_91
CL\_INS\_321
CL\_INS\_263
CL\_INS\_91
CL\_INS\_91
CL\_INS\_263
CL\_INS\_91
CL\_INS\_313
CL\_INS\_313
CL\_INS\_400
CL\_INS\_400
CL\_INS\_400
CL\_INS\_400
CL\_INS\_400
CL\_INS\_400
CL\_INS\_400
CL\_INS\_91
CL\_INS\_313
CL\_INS\_313
CL\_INS\_91
CL\_INS\_400
CL\_INS\_91
CL\_INS\_400
CL\_INS\_400
CL\_INS\_400
CL\_INS\_400
CL\_INS\_91
CL\_INS\_91
CL\_INS\_91
CL\_INS\_400
CL\_INS\_91
CL\_INS\_400
CL\_INS\_400
CL\_INS\_400
CL\_INS\_400
CL\_INS\_400
CL\_INS\_400
CL\_INS\_400
CL\_INS\_400
CL\_INS\_400
CL\_INS\_400
CL\_INS\_400
CL\_INS\_400
CL\_INS\_400
CL\_INS\_400
CL\_INS\_400
CL\_INS\_278
CL\_INS\_400
CL\_INS\_278
CL\_INS\_400
CL\_INS\_400
CL\_INS\_400
CL\_INS\_400
CL\_INS\_400
CL\_INS\_400
CL\_INS\_243
CL\_INS\_243
CL\_INS\_400
CL\_INS\_368
CL\_INS\_368
CL\_INS\_278
CL\_INS\_132
CL\_INS\_278
CL\_INS\_278
CL\_INS\_278
CL\_INS\_400
CL\_INS\_278
CL\_INS\_132
CL\_INS\_278
CL\_INS\_263
Cluster ID


CL\_13316
CL\_13317
CL\_17313
CL\_17312
CL\_13318
CL\_13319
CL\_13320
CL\_7138
CL\_17311
CL\_13321
CL\_13322
CL\_13323
CL\_13324
CL\_22501
CL\_22500
CL\_22499
CL\_22498
CL\_22497
CL\_23067
CL\_23066
CL\_23065
CL\_23064
CL\_23063
CL\_23062
CL\_23061
CL\_17721
CL\_11877
CL\_11878
CL\_16355
CL\_11879
CL\_23060
CL\_23059
CL\_23058
CL\_23057
CL\_23056
CL\_23055
CL\_4950
CL\_11924
CL\_23969
CL\_10174
CL\_10175
CL\_10176
CL\_10178
CL\_6534
CL\_10179
CL\_10180
CL\_11925
CL\_14821
CL\_10181
CL\_11929
CL\_11930
CL\_22496
CL\_22495
CL\_22494
CL\_22493
CL\_22492
CL\_22491
CL\_22490
CL\_10182
CL\_15411
CL\_15410
CL\_10183
CL\_14723
CL\_10184
CL\_14724
CL\_19678
CL\_19679
CL\_19680
CL\_12840
CL\_14725
CL\_19681
CL\_16960
CL\_16959
CL\_14726
CL\_20579
CL\_6033
CL\_21412
CL\_21411
CL\_21410
CL\_21409
CL\_21408
CL\_21407
CL\_21406
CL\_21405
CL\_21404
CL\_21403
CL\_21402
CL\_8233
CL\_6089
CL\_9203
CL\_8213
CL\_9202
CL\_9201
CL\_8158
CL\_8157
CL\_8156
CL\_6080
CL\_6120
CL\_6121
CL\_8215
CL\_8125
CL\_8126
CL\_8238
CL\_6087
CL\_6086
CL\_6085
CL\_6084
CL\_21401
CL\_6081
CL\_8237
CL\_6078
CL\_6077
