## Supplementary material for "A novel method for integrating genomic and Tn-Seq data to identify common *in vivo* fitness mechanisms across multiple bacterial species": S2 Dataset: CL_INS_402.html

Legend

 Hypothetical
 All VFDB Genes

FULL


WINDOWSVGPNG

Trim RowsRemove SingletonsSave Fasta

CL\_5000


CL\_5000


CL\_5000


CL\_5000


CL\_5000

HighlightSelectShow Genomes


78

CL\_5003


75

CL\_5003


42

CL\_5003


8

CL\_5003


2

CL\_5004

fGI ID


CL\_INS\_402
CL\_INS\_402
CL\_INS\_402
Cluster ID


CL\_5825
CL\_5001
CL\_5002
