## Supplementary material for "A novel method for integrating genomic and Tn-Seq data to identify common *in vivo* fitness mechanisms across multiple bacterial species": S2 Dataset: CL_INS_403.html

Legend

 Regulatoryfunctions
 Hypothetical
 All VFDB Genes

FULL


WINDOWSVGPNG

Trim RowsRemove SingletonsSave Fasta

CL\_5026


CL\_5026


CL\_5026


CL\_5025

HighlightSelectShow Genomes


131

CL\_5027


71

CL\_5027


1

Break


1

CL\_5027

fGI ID

CL\_INS\_403
Cluster ID

CL\_5824
