## Supplementary material for "A novel method for integrating genomic and Tn-Seq data to identify common *in vivo* fitness mechanisms across multiple bacterial species": S2 Dataset: CL_INS_404.html

Legend

 Hypothetical
 Other
 Cellularprocesses
 All VFDB Genes

FULL


WINDOWSVGPNG

Trim RowsRemove SingletonsSave Fasta

CL\_5039


CL\_5039


CL\_5039


CL\_5038


CL\_5039


CL\_5039


CL\_5038

HighlightSelectShow Genomes


112

CL\_5041


86

CL\_5041


1

CL\_5041


1

CL\_5041


1

CL\_5041


1

CL\_5041


1

CL\_5041

fGI ID


CL\_INS\_404
CL\_INS\_404
CL\_INS\_404
CL\_INS\_404
Cluster ID


CL\_19682
CL\_11528
CL\_5040
CL\_9614
