## Supplementary material for "A novel method for integrating genomic and Tn-Seq data to identify common *in vivo* fitness mechanisms across multiple bacterial species": S2 Dataset: CL_INS_407.html

Legend

 Mobile +extrachromosomalelementfunctions
 Hypothetical
 All Fitness Genes
 Other
 Transport +binding proteins

FULL


WINDOWSVGPNG

Trim RowsRemove SingletonsSave Fasta

CL\_5077


CL\_5077


CL\_5077


CL\_5077


CL\_5077


CL\_5077


CL\_5077


CL\_5077


CL\_5077

HighlightSelectShow Genomes


156

CL\_5078


9

CL\_5078


3

CL\_5078


1

CL\_5078


1

CL\_5078


1

CL\_5078


1

CL\_5078


1

CL\_5078


1

CL\_5078

fGI ID


CL\_INS\_407
CL\_INS\_407
CL\_INS\_407
CL\_INS\_45
CL\_INS\_45
CL\_INS\_45
CL\_INS\_407
CL\_INS\_407
CL\_INS\_407
CL\_INS\_407
CL\_INS\_407
CL\_INS\_407
CL\_INS\_407
CL\_INS\_407
CL\_INS\_407
CL\_INS\_407
CL\_INS\_407
CL\_INS\_407
CL\_INS\_407
CL\_INS\_407
CL\_INS\_407
CL\_INS\_407
CL\_INS\_407
CL\_INS\_407
CL\_INS\_407
CL\_INS\_407
CL\_INS\_407
CL\_INS\_407
Cluster ID


CL\_12910
CL\_12911
CL\_7410
CL\_7744
CL\_7743
CL\_7742
CL\_23550
CL\_23549
CL\_7409
CL\_7408
CL\_7407
CL\_12025
CL\_14933
CL\_12024
CL\_12023
CL\_7406
CL\_7405
CL\_12022
CL\_12021
CL\_12020
CL\_7404
CL\_12912
CL\_7403
CL\_23548
CL\_23547
CL\_7402
CL\_7401
CL\_12019
