## Supplementary material for "A novel method for integrating genomic and Tn-Seq data to identify common *in vivo* fitness mechanisms across multiple bacterial species": S2 Dataset: CL_INS_408.html

Legend

 Hypothetical
 All EssentialGenes
 Other

FULL


WINDOWSVGPNG

Trim RowsRemove SingletonsSave Fasta

CL\_5088


CL\_5088


CL\_5087


CL\_5088

HighlightSelectShow Genomes


93

CL\_5090


72

CL\_5090


16

CL\_5090


1

CL\_5090

fGI ID


CL\_INS\_408
CL\_INS\_408
Cluster ID


CL\_21241
CL\_5089
