## Supplementary material for "A novel method for integrating genomic and Tn-Seq data to identify common *in vivo* fitness mechanisms across multiple bacterial species": S2 Dataset: CL_INS_409.html

Legend

 Hypothetical
 All EssentialGenes

FULL


WINDOWSVGPNG

Trim RowsRemove SingletonsSave Fasta

CL\_5097


CL\_5097


CL\_5097


CL\_5097

HighlightSelectShow Genomes


84

CL\_5100


56

CL\_5100


3

CL\_5100


1

Break

fGI ID


CL\_INS\_409
CL\_INS\_409
Cluster ID


CL\_5098
CL\_5099
