## Supplementary material for "A novel method for integrating genomic and Tn-Seq data to identify common *in vivo* fitness mechanisms across multiple bacterial species": S3 Dataset: CL_INS_1.html

Legend

 Mobile +extrachromosomalelementfunctions
 Regulatoryfunctions
 Hypothetical
 DNA Metabolism
 AntibioticResistance
 All EssentialGenes
 Biosynthesis ofcofactors,prostheticgroups, +carriers
 Other
 Proteinsynthesis/fate
 EnergyMetabolism
 All VFDB Genes
 Transport +binding proteins

FULL


WINDOWSVGPNG

Trim RowsRemove SingletonsSave Fasta

CL\_28


CL\_28


CL\_28


CL\_28


CL\_28


CL\_28


CL\_28


CL\_28


CL\_28


CL\_28


CL\_28


CL\_28


CL\_28


CL\_28


CL\_28


CL\_28


CL\_28


CL\_28


CL\_28


CL\_30


CL\_28


CL\_4308


CL\_28


CL\_28


CL\_28


CL\_28


CL\_28


CL\_28


CL\_28


CL\_28


CL\_28


CL\_28


CL\_28


Break

HighlightSelectShow Genomes


3

CL\_4333


2

CL\_4333


2

CL\_4333


2

CL\_4333


2

CL\_4333


2

CL\_4333


2

CL\_4333


1

CL\_4333


1

Break


1

CL\_4333


1

CL\_4333


1

CL\_4333


1

CL\_4309


1

CL\_4333


1

CL\_4333


1

CL\_4333


1

CL\_4333


1

CL\_4333


1

CL\_4333


1

CL\_4333


1

CL\_4333


1

CL\_4333


1

CL\_4333


1

CL\_4333


1

Break


1

CL\_4333


1

CL\_4333


1

CL\_4333


1

CL\_4333


1

CL\_4333


1

CL\_4333


1

Break


1

CL\_4333


1

CL\_4333

fGI ID


CL\_INS\_1
CL\_INS\_1
CL\_INS\_1
CL\_INS\_1
CL\_INS\_1
CL\_INS\_1
CL\_INS\_1
CL\_INS\_1
CL\_INS\_1
CL\_INS\_1
CL\_INS\_1
CL\_INS\_1
CL\_INS\_1
CL\_INS\_1
CL\_INS\_1
CL\_INS\_1
CL\_INS\_1
CL\_INS\_1
CL\_INS\_1
CL\_INS\_1
CL\_INS\_1
CL\_INS\_1
CL\_INS\_1
CL\_INS\_1
CL\_INS\_1
CL\_INS\_1
CL\_INS\_1
CL\_INS\_1
CL\_INS\_1
CL\_INS\_1
CL\_INS\_1
CL\_INS\_1
CL\_INS\_1
CL\_INS\_1
CL\_INS\_1
CL\_INS\_1
CL\_INS\_1
CL\_INS\_1
CL\_INS\_1
CL\_INS\_1
CL\_INS\_1
CL\_INS\_1
CL\_INS\_1
CL\_INS\_1
CL\_INS\_1
CL\_INS\_1
CL\_INS\_1
CL\_INS\_1
CL\_INS\_1
CL\_INS\_1
CL\_INS\_1
CL\_INS\_1
CL\_INS\_1
CL\_INS\_1
CL\_INS\_1
CL\_INS\_1
CL\_INS\_1
CL\_INS\_1
CL\_INS\_1
CL\_INS\_1
CL\_INS\_1
CL\_INS\_1
CL\_INS\_1
CL\_INS\_1
CL\_INS\_1
CL\_INS\_1
CL\_INS\_1
CL\_INS\_1
CL\_INS\_1
CL\_INS\_1
CL\_INS\_1
CL\_INS\_1
CL\_INS\_1
CL\_INS\_1
CL\_INS\_1
CL\_INS\_1
CL\_INS\_1
CL\_INS\_1
CL\_INS\_1
CL\_INS\_1
CL\_INS\_1
CL\_INS\_1
CL\_INS\_1
CL\_INS\_1
CL\_INS\_1
CL\_INS\_1
CL\_INS\_1
CL\_INS\_1
CL\_INS\_1
CL\_INS\_1
CL\_INS\_1
CL\_INS\_1
CL\_INS\_1
CL\_INS\_1
CL\_INS\_1
CL\_INS\_1
CL\_INS\_1
CL\_INS\_1
CL\_INS\_1
CL\_INS\_1
CL\_INS\_1
CL\_INS\_1
CL\_INS\_1
CL\_INS\_1
CL\_INS\_1
CL\_INS\_1
CL\_INS\_1
CL\_INS\_1
CL\_INS\_1
CL\_INS\_1
CL\_INS\_1
CL\_INS\_1
CL\_INS\_1
CL\_INS\_1
CL\_INS\_1
CL\_INS\_1
CL\_INS\_1
CL\_INS\_1
CL\_INS\_1
CL\_INS\_1
CL\_INS\_1
CL\_INS\_1
CL\_INS\_1
CL\_INS\_1
CL\_INS\_1
CL\_INS\_1
CL\_INS\_1
CL\_INS\_1
CL\_INS\_1
CL\_INS\_1
CL\_INS\_1
CL\_INS\_1
CL\_INS\_1
CL\_INS\_1
CL\_INS\_1
CL\_INS\_1
CL\_INS\_1
CL\_INS\_1
CL\_INS\_1
CL\_INS\_83
CL\_INS\_83
CL\_INS\_83
CL\_INS\_1
CL\_INS\_1
CL\_INS\_83
CL\_INS\_83
CL\_INS\_83
CL\_INS\_83
CL\_INS\_83
CL\_INS\_83
CL\_INS\_83
CL\_INS\_1
CL\_INS\_1
CL\_INS\_1
CL\_INS\_1
CL\_INS\_1
CL\_INS\_1
CL\_INS\_1
CL\_INS\_1
CL\_INS\_1
CL\_INS\_1
CL\_INS\_1
CL\_INS\_1
CL\_INS\_1
CL\_INS\_1
CL\_INS\_1
CL\_INS\_1
CL\_INS\_1
CL\_INS\_1
CL\_INS\_1
CL\_INS\_1
CL\_INS\_1
CL\_INS\_1
CL\_INS\_1
CL\_INS\_1
CL\_INS\_1
CL\_INS\_1
CL\_INS\_1
CL\_INS\_1
CL\_INS\_1
CL\_INS\_1
CL\_INS\_1
CL\_INS\_1
CL\_INS\_1
CL\_INS\_1
CL\_INS\_1
CL\_INS\_1
CL\_INS\_1
CL\_INS\_1
CL\_INS\_1
CL\_INS\_1
CL\_INS\_1
CL\_INS\_1
CL\_INS\_1
CL\_INS\_1
CL\_INS\_1
CL\_INS\_1
CL\_INS\_1
CL\_INS\_1
CL\_INS\_1
CL\_INS\_1
CL\_INS\_1
CL\_INS\_1
CL\_INS\_1
CL\_INS\_1
CL\_INS\_1
CL\_INS\_1
CL\_INS\_1
CL\_INS\_1
CL\_INS\_1
CL\_INS\_1
CL\_INS\_83
CL\_INS\_1
CL\_INS\_1
CL\_INS\_1
CL\_INS\_1
CL\_INS\_1
CL\_INS\_1
CL\_INS\_1
CL\_INS\_1
CL\_INS\_1
CL\_INS\_1
CL\_INS\_1
CL\_INS\_1
CL\_INS\_1
CL\_INS\_1
CL\_INS\_1
CL\_INS\_1
CL\_INS\_1
CL\_INS\_1
CL\_INS\_1
CL\_INS\_1
CL\_INS\_1
CL\_INS\_1
CL\_INS\_1
CL\_INS\_1
CL\_INS\_1
CL\_INS\_1
CL\_INS\_1
CL\_INS\_1
CL\_INS\_1
CL\_INS\_1
CL\_INS\_1
CL\_INS\_1
CL\_INS\_1
CL\_INS\_1
CL\_INS\_1
CL\_INS\_1
CL\_INS\_1
CL\_INS\_1
CL\_INS\_1
CL\_INS\_1
CL\_INS\_1
CL\_INS\_1
CL\_INS\_1
CL\_INS\_1
CL\_INS\_1
CL\_INS\_1
CL\_INS\_1
CL\_INS\_1
CL\_INS\_1
CL\_INS\_1
CL\_INS\_1
CL\_INS\_1
CL\_INS\_1
CL\_INS\_1
CL\_INS\_1
CL\_INS\_1
CL\_INS\_1
CL\_INS\_1
CL\_INS\_1
CL\_INS\_1
CL\_INS\_1
CL\_INS\_1
CL\_INS\_1
CL\_INS\_1
CL\_INS\_83
CL\_INS\_83
CL\_INS\_1
CL\_INS\_1
CL\_INS\_1
CL\_INS\_1
CL\_INS\_1
CL\_INS\_1
CL\_INS\_1
CL\_INS\_1
CL\_INS\_1
CL\_INS\_1
CL\_INS\_1
CL\_INS\_1
CL\_INS\_1
CL\_INS\_1
CL\_INS\_1
CL\_INS\_1
CL\_INS\_1
CL\_INS\_1
CL\_INS\_1
CL\_INS\_1
CL\_INS\_1
CL\_INS\_1
CL\_INS\_1
CL\_INS\_1
CL\_INS\_1
CL\_INS\_1
CL\_INS\_1
CL\_INS\_1
CL\_INS\_1
CL\_INS\_1
CL\_INS\_1
CL\_INS\_1
CL\_INS\_1
CL\_INS\_1
CL\_INS\_1
CL\_INS\_1
CL\_INS\_1
CL\_INS\_1
CL\_INS\_1
CL\_INS\_1
CL\_INS\_1
CL\_INS\_1
CL\_INS\_1
CL\_INS\_1
CL\_INS\_1
CL\_INS\_1
CL\_INS\_1
CL\_INS\_1
CL\_INS\_1
CL\_INS\_1
CL\_INS\_1
CL\_INS\_1
CL\_INS\_1
CL\_INS\_1
CL\_INS\_1
CL\_INS\_1
CL\_INS\_1
CL\_INS\_1
CL\_INS\_1
CL\_INS\_1
CL\_INS\_1
CL\_INS\_1
CL\_INS\_1
CL\_INS\_1
CL\_INS\_1
CL\_INS\_1
CL\_INS\_1
CL\_INS\_1
CL\_INS\_1
CL\_INS\_1
CL\_INS\_1
CL\_INS\_1
CL\_INS\_1
CL\_INS\_1
CL\_INS\_1
CL\_INS\_1
CL\_INS\_1
CL\_INS\_1
CL\_INS\_1
CL\_INS\_1
CL\_INS\_1
CL\_INS\_1
CL\_INS\_1
CL\_INS\_1
CL\_INS\_1
CL\_INS\_1
CL\_INS\_1
CL\_INS\_1
CL\_INS\_1
CL\_INS\_1
CL\_INS\_1
CL\_INS\_1
CL\_INS\_1
CL\_INS\_1
CL\_INS\_1
CL\_INS\_1
CL\_INS\_1
CL\_INS\_1
CL\_INS\_1
CL\_INS\_1
CL\_INS\_1
CL\_INS\_1
CL\_INS\_1
CL\_INS\_1
CL\_INS\_1
CL\_INS\_1
CL\_INS\_1
CL\_INS\_1
CL\_INS\_1
CL\_INS\_1
CL\_INS\_1
CL\_INS\_1
CL\_INS\_1
CL\_INS\_1
CL\_INS\_1
CL\_INS\_1
CL\_INS\_1
CL\_INS\_1
CL\_INS\_1
CL\_INS\_1
CL\_INS\_1
CL\_INS\_1
CL\_INS\_1
CL\_INS\_1
CL\_INS\_1
CL\_INS\_1
CL\_INS\_1
CL\_INS\_1
CL\_INS\_1
CL\_INS\_1
CL\_INS\_1
CL\_INS\_1
CL\_INS\_1
CL\_INS\_1
CL\_INS\_1
CL\_INS\_1
CL\_INS\_1
CL\_INS\_1
CL\_INS\_1
CL\_INS\_1
CL\_INS\_1
CL\_INS\_1
CL\_INS\_1
CL\_INS\_1
CL\_INS\_1
CL\_INS\_1
CL\_INS\_1
CL\_INS\_1
CL\_INS\_1
CL\_INS\_1
CL\_INS\_1
CL\_INS\_1
CL\_INS\_1
CL\_INS\_1
CL\_INS\_1
CL\_INS\_1
CL\_INS\_1
CL\_INS\_1
CL\_INS\_1
CL\_INS\_1
CL\_INS\_1
CL\_INS\_1
CL\_INS\_1
CL\_INS\_1
CL\_INS\_1
CL\_INS\_1
CL\_INS\_1
CL\_INS\_1
CL\_INS\_1
CL\_INS\_1
CL\_INS\_1
CL\_INS\_1
CL\_INS\_1
CL\_INS\_1
CL\_INS\_1
CL\_INS\_1
CL\_INS\_1
CL\_INS\_1
CL\_INS\_1
CL\_INS\_1
CL\_INS\_1
CL\_INS\_1
CL\_INS\_1
CL\_INS\_1
CL\_INS\_1
CL\_INS\_1
CL\_INS\_1
CL\_INS\_1
CL\_INS\_1
CL\_INS\_1
CL\_INS\_1
CL\_INS\_1
CL\_INS\_1
CL\_INS\_1
CL\_INS\_1
CL\_INS\_1
CL\_INS\_1
CL\_INS\_1
CL\_INS\_1
CL\_INS\_1
CL\_INS\_1
CL\_INS\_1
CL\_INS\_1
CL\_INS\_1
CL\_INS\_1
CL\_INS\_1
CL\_INS\_1
CL\_INS\_1
CL\_INS\_1
CL\_INS\_153
CL\_INS\_153
CL\_INS\_1
CL\_INS\_1
CL\_INS\_1
CL\_INS\_133
CL\_INS\_133
CL\_INS\_1
CL\_INS\_1
CL\_INS\_1
CL\_INS\_1
CL\_INS\_1
CL\_INS\_1
CL\_INS\_1
CL\_INS\_1
CL\_INS\_1
CL\_INS\_1
CL\_INS\_1
CL\_INS\_1
CL\_INS\_1
CL\_INS\_1
CL\_INS\_1
CL\_INS\_1
CL\_INS\_1
CL\_INS\_1
CL\_INS\_1
CL\_INS\_1
CL\_INS\_1
CL\_INS\_1
CL\_INS\_1
CL\_INS\_1
CL\_INS\_1
CL\_INS\_1
CL\_INS\_1
CL\_INS\_1
CL\_INS\_153
CL\_INS\_153
CL\_INS\_153
CL\_INS\_153
CL\_INS\_153
CL\_INS\_153
CL\_INS\_153
CL\_INS\_153
CL\_INS\_153
CL\_INS\_153
CL\_INS\_153
CL\_INS\_153
CL\_INS\_153
CL\_INS\_153
CL\_INS\_153
CL\_INS\_153
CL\_INS\_153
CL\_INS\_153
CL\_INS\_153
CL\_INS\_153
CL\_INS\_153
CL\_INS\_153
CL\_INS\_1
CL\_INS\_153
CL\_INS\_133
CL\_INS\_1
CL\_INS\_1
CL\_INS\_1
CL\_INS\_1
CL\_INS\_1
CL\_INS\_1
CL\_INS\_1
CL\_INS\_1
CL\_INS\_1
CL\_INS\_1
CL\_INS\_1
CL\_INS\_1
CL\_INS\_1
CL\_INS\_1
CL\_INS\_1
CL\_INS\_1
CL\_INS\_1
CL\_INS\_1
CL\_INS\_1
CL\_INS\_1
CL\_INS\_1
CL\_INS\_133
CL\_INS\_1
CL\_INS\_1
CL\_INS\_1
CL\_INS\_1
CL\_INS\_1
CL\_INS\_1
CL\_INS\_1
CL\_INS\_1
CL\_INS\_1
CL\_INS\_1
CL\_INS\_1
CL\_INS\_1
CL\_INS\_1
CL\_INS\_1
CL\_INS\_1
CL\_INS\_1
CL\_INS\_1
CL\_INS\_1
CL\_INS\_1
CL\_INS\_133
CL\_INS\_1
CL\_INS\_1
CL\_INS\_1
CL\_INS\_1
CL\_INS\_1
CL\_INS\_1
CL\_INS\_1
CL\_INS\_1
CL\_INS\_1
CL\_INS\_1
CL\_INS\_1
CL\_INS\_1
CL\_INS\_1
CL\_INS\_1
CL\_INS\_1
CL\_INS\_1
CL\_INS\_1
CL\_INS\_1
CL\_INS\_1
CL\_INS\_1
CL\_INS\_1
CL\_INS\_1
CL\_INS\_1
CL\_INS\_1
CL\_INS\_1
CL\_INS\_1
CL\_INS\_1
CL\_INS\_1
CL\_INS\_1
CL\_INS\_1
CL\_INS\_1
CL\_INS\_1
CL\_INS\_1
CL\_INS\_1
CL\_INS\_1
CL\_INS\_1
CL\_INS\_1
CL\_INS\_1
CL\_INS\_1
CL\_INS\_1
CL\_INS\_1
CL\_INS\_1
CL\_INS\_1
CL\_INS\_1
CL\_INS\_1
CL\_INS\_1
CL\_INS\_1
CL\_INS\_1
CL\_INS\_1
CL\_INS\_1
CL\_INS\_1
CL\_INS\_1
CL\_INS\_1
CL\_INS\_1
CL\_INS\_1
CL\_INS\_1
CL\_INS\_1
CL\_INS\_1
CL\_INS\_1
CL\_INS\_1
CL\_INS\_1
CL\_INS\_1
CL\_INS\_1
CL\_INS\_1
CL\_INS\_1
CL\_INS\_1
CL\_INS\_1
CL\_INS\_1
CL\_INS\_1
CL\_INS\_1
CL\_INS\_1
CL\_INS\_1
CL\_INS\_1
CL\_INS\_1
CL\_INS\_1
CL\_INS\_204
CL\_INS\_1
CL\_INS\_1
CL\_INS\_1
CL\_INS\_1
CL\_INS\_1
CL\_INS\_1
CL\_INS\_1
CL\_INS\_1
CL\_INS\_1
CL\_INS\_1
CL\_INS\_1
CL\_INS\_1
CL\_INS\_1
CL\_INS\_1
CL\_INS\_1
CL\_INS\_1
CL\_INS\_1
CL\_INS\_1
CL\_INS\_1
CL\_INS\_1
CL\_INS\_1
CL\_INS\_1
CL\_INS\_1
CL\_INS\_1
CL\_INS\_1
CL\_INS\_1
CL\_INS\_1
CL\_INS\_1
CL\_INS\_1
CL\_INS\_1
CL\_INS\_1
CL\_INS\_1
CL\_INS\_1
CL\_INS\_1
CL\_INS\_1
CL\_INS\_1
CL\_INS\_1
CL\_INS\_1
CL\_INS\_1
CL\_INS\_1
CL\_INS\_1
CL\_INS\_1
CL\_INS\_1
CL\_INS\_1
CL\_INS\_1
CL\_INS\_1
CL\_INS\_1
CL\_INS\_1
CL\_INS\_1
CL\_INS\_1
CL\_INS\_1
CL\_INS\_1
CL\_INS\_1
CL\_INS\_1
CL\_INS\_1
CL\_INS\_1
CL\_INS\_1
CL\_INS\_1
CL\_INS\_1
CL\_INS\_1
CL\_INS\_1
CL\_INS\_1
CL\_INS\_1
CL\_INS\_1
CL\_INS\_1
CL\_INS\_1
CL\_INS\_1
CL\_INS\_1
CL\_INS\_1
CL\_INS\_1
CL\_INS\_1
CL\_INS\_1
CL\_INS\_1
CL\_INS\_1
CL\_INS\_1
CL\_INS\_1
CL\_INS\_1
CL\_INS\_1
CL\_INS\_1
CL\_INS\_1
CL\_INS\_1
CL\_INS\_1
CL\_INS\_1
CL\_INS\_1
CL\_INS\_1
CL\_INS\_1
CL\_INS\_1
CL\_INS\_1
CL\_INS\_1
CL\_INS\_1
CL\_INS\_1
CL\_INS\_1
CL\_INS\_1
CL\_INS\_1
CL\_INS\_1
Cluster ID


CL\_10026
CL\_10027
CL\_10028
CL\_10029
CL\_10030
CL\_10031
CL\_10032
CL\_10033
CL\_10034
CL\_5748
CL\_5747
CL\_5746
CL\_5745
CL\_5744
CL\_5743
CL\_5742
CL\_5741
CL\_5740
CL\_5739
CL\_5738
CL\_10035
CL\_10036
CL\_10037
CL\_13338
CL\_12524
CL\_26
CL\_25
CL\_12687
CL\_12686
CL\_11684
CL\_11685
CL\_11686
CL\_24
CL\_8135
CL\_13339
CL\_5115
CL\_5114
CL\_5113
CL\_5112
CL\_5111
CL\_5110
CL\_5109
CL\_6564
CL\_9806
CL\_13698
CL\_27
CL\_6920
CL\_5894
CL\_5895
CL\_5896
CL\_22
CL\_21
CL\_20
CL\_19
CL\_10541
CL\_6919
CL\_5196
CL\_10170
CL\_5160
CL\_4784
CL\_4783
CL\_4782
CL\_4781
CL\_4780
CL\_4779
CL\_4778
CL\_4777
CL\_4776
CL\_4775
CL\_4774
CL\_4773
CL\_4772
CL\_4771
CL\_4770
CL\_4769
CL\_4768
CL\_4767
CL\_4766
CL\_4765
CL\_4764
CL\_4763
CL\_4762
CL\_4761
CL\_4760
CL\_4759
CL\_4758
CL\_4757
CL\_4756
CL\_4755
CL\_4754
CL\_4753
CL\_4752
CL\_4751
CL\_4750
CL\_4749
CL\_4748
CL\_4747
CL\_4746
CL\_4745
CL\_4744
CL\_4743
CL\_4742
CL\_4741
CL\_4740
CL\_4739
CL\_4738
CL\_4737
CL\_4736
CL\_4735
CL\_4734
CL\_4733
CL\_4732
CL\_4731
CL\_4730
CL\_4729
CL\_4728
CL\_4727
CL\_4726
CL\_4725
CL\_4724
CL\_4723
CL\_4722
CL\_4721
CL\_4720
CL\_4719
CL\_4718
CL\_4717
CL\_4716
CL\_4715
CL\_4714
CL\_4713
CL\_4712
CL\_4711
CL\_4710
CL\_4709
CL\_4708
CL\_4707
CL\_4706
CL\_4705
CL\_4704
CL\_4703
CL\_4702
CL\_4701
CL\_4700
CL\_4699
CL\_4698
CL\_4697
CL\_4696
CL\_4695
CL\_4694
CL\_4693
CL\_4692
CL\_4691
CL\_4690
CL\_4689
CL\_4688
CL\_4687
CL\_4686
CL\_4685
CL\_4684
CL\_4683
CL\_4682
CL\_4681
CL\_4680
CL\_4679
CL\_4678
CL\_4677
CL\_4676
CL\_4675
CL\_4674
CL\_4673
CL\_4672
CL\_4671
CL\_4670
CL\_4669
CL\_4668
CL\_4667
CL\_4666
CL\_4665
CL\_4664
CL\_4663
CL\_4662
CL\_4661
CL\_4660
CL\_4659
CL\_4658
CL\_4657
CL\_4656
CL\_4655
CL\_4654
CL\_4653
CL\_4652
CL\_4651
CL\_4650
CL\_4649
CL\_4648
CL\_4647
CL\_4646
CL\_4645
CL\_4644
CL\_4643
CL\_4642
CL\_4641
CL\_4640
CL\_4639
CL\_4638
CL\_4637
CL\_4636
CL\_4635
CL\_4634
CL\_4633
CL\_4632
CL\_23
CL\_13697
CL\_5108
CL\_6563
CL\_9089
CL\_7892
CL\_6910
CL\_6909
CL\_7
CL\_6
CL\_5
CL\_3
CL\_2
CL\_10161
CL\_10162
CL\_10163
CL\_10164
CL\_10165
CL\_10166
CL\_10167
CL\_10168
CL\_10169
CL\_13388
CL\_5107
CL\_5106
CL\_11210
CL\_13695
CL\_9805
CL\_9804
CL\_9803
CL\_9802
CL\_13387
CL\_5105
CL\_10064
CL\_10063
CL\_10062
CL\_10061
CL\_10060
CL\_10059
CL\_10058
CL\_10057
CL\_10056
CL\_10055
CL\_10054
CL\_10053
CL\_10052
CL\_10051
CL\_10050
CL\_10049
CL\_10048
CL\_10047
CL\_10046
CL\_10045
CL\_10044
CL\_10043
CL\_10042
CL\_10041
CL\_10040
CL\_3936
CL\_3937
CL\_3938
CL\_3939
CL\_6155
CL\_3940
CL\_3941
CL\_3942
CL\_89
CL\_10039
CL\_4631
CL\_4630
CL\_4629
CL\_4628
CL\_4627
CL\_4626
CL\_4625
CL\_4624
CL\_4623
CL\_4622
CL\_4621
CL\_4620
CL\_4619
CL\_4618
CL\_4617
CL\_4616
CL\_4615
CL\_4614
CL\_4613
CL\_4612
CL\_4611
CL\_4610
CL\_4609
CL\_4608
CL\_4607
CL\_4606
CL\_8112
CL\_8111
CL\_8110
CL\_4605
CL\_5722
CL\_4604
CL\_10249
CL\_4603
CL\_4602
CL\_4601
CL\_4600
CL\_4599
CL\_4598
CL\_4597
CL\_4596
CL\_4595
CL\_4594
CL\_4593
CL\_4592
CL\_4591
CL\_4590
CL\_4589
CL\_4588
CL\_4587
CL\_4586
CL\_4585
CL\_4584
CL\_4583
CL\_4582
CL\_4581
CL\_4580
CL\_4579
CL\_4578
CL\_4577
CL\_5469
CL\_5471
CL\_5472
CL\_11195
CL\_11196
CL\_11197
CL\_11198
CL\_4576
CL\_4575
CL\_4574
CL\_4573
CL\_4572
CL\_4571
CL\_4570
CL\_4569
CL\_4568
CL\_4567
CL\_4566
CL\_4565
CL\_4564
CL\_4563
CL\_4562
CL\_4561
CL\_4560
CL\_4559
CL\_4558
CL\_4557
CL\_4556
CL\_4555
CL\_4554
CL\_4553
CL\_4552
CL\_4551
CL\_4550
CL\_4549
CL\_4548
CL\_4547
CL\_4546
CL\_4545
CL\_4544
CL\_4543
CL\_4542
CL\_4541
CL\_4540
CL\_4539
CL\_10246
CL\_4538
CL\_4537
CL\_4536
CL\_4535
CL\_4534
CL\_4533
CL\_4532
CL\_4531
CL\_4530
CL\_4529
CL\_4528
CL\_8997
CL\_4527
CL\_4526
CL\_4525
CL\_4524
CL\_4523
CL\_4522
CL\_4521
CL\_4520
CL\_4519
CL\_4518
CL\_4517
CL\_4516
CL\_4515
CL\_4514
CL\_4513
CL\_4512
CL\_4511
CL\_4510
CL\_10767
CL\_4509
CL\_4508
CL\_4507
CL\_4506
CL\_4505
CL\_4504
CL\_4503
CL\_4502
CL\_4501
CL\_4500
CL\_4499
CL\_4498
CL\_4497
CL\_10766
CL\_4496
CL\_4495
CL\_4494
CL\_4493
CL\_4492
CL\_4491
CL\_4490
CL\_4489
CL\_4488
CL\_4487
CL\_4486
CL\_4485
CL\_4484
CL\_4483
CL\_5222
CL\_5223
CL\_5468
CL\_5467
CL\_5466
CL\_5465
CL\_4482
CL\_4481
CL\_4480
CL\_10245
CL\_4479
CL\_4478
CL\_4477
CL\_4476
CL\_4475
CL\_4474
CL\_4473
CL\_4472
CL\_4471
CL\_4470
CL\_4469
CL\_4468
CL\_4467
CL\_4466
CL\_4465
CL\_4464
CL\_4463
CL\_4462
CL\_4461
CL\_10244
CL\_10243
CL\_10242
CL\_4460
CL\_4459
CL\_4458
CL\_4457
CL\_4456
CL\_4455
CL\_5321
CL\_4454
CL\_4453
CL\_4452
CL\_5322
CL\_5323
CL\_5324
CL\_5325
CL\_5326
CL\_5327
CL\_5328
CL\_11173
CL\_11174
CL\_11175
CL\_11176
CL\_11177
CL\_11178
CL\_11179
CL\_11180
CL\_11181
CL\_11182
CL\_8852
CL\_10765
CL\_10764
CL\_10763
CL\_10762
CL\_10761
CL\_10760
CL\_10759
CL\_10758
CL\_10757
CL\_10756
CL\_10755
CL\_10754
CL\_11202
CL\_10241
CL\_10240
CL\_10239
CL\_10238
CL\_10237
CL\_10753
CL\_10236
CL\_10235
CL\_10234
CL\_10233
CL\_10752
CL\_10232
CL\_10231
CL\_10230
CL\_10229
CL\_10228
CL\_10227
CL\_10226
CL\_10225
CL\_10224
CL\_10223
CL\_10222
CL\_10221
CL\_10220
CL\_10219
CL\_10218
CL\_10217
CL\_10216
CL\_4451
CL\_5838
CL\_4450
CL\_4449
CL\_8853
CL\_4448
CL\_4447
CL\_4446
CL\_4445
CL\_10215
CL\_5329
CL\_5330
CL\_5331
CL\_4444
CL\_4443
CL\_4442
CL\_4441
CL\_4440
CL\_4439
CL\_4438
CL\_4437
CL\_4436
CL\_4435
CL\_4434
CL\_10741
CL\_5837
CL\_10742
CL\_4433
CL\_10441
CL\_4432
CL\_4431
CL\_4430
CL\_4429
CL\_4428
CL\_4427
CL\_4426
CL\_4425
CL\_4424
CL\_6077
CL\_6076
CL\_6075
CL\_6074
CL\_5202
CL\_11183
CL\_11184
CL\_11185
CL\_10277
CL\_10276
CL\_11186
CL\_8946
CL\_8945
CL\_8944
CL\_8943
CL\_8942
CL\_8941
CL\_8940
CL\_6918
CL\_6917
CL\_6916
CL\_6915
CL\_6914
CL\_8939
CL\_8938
CL\_10285
CL\_10284
CL\_10283
CL\_10282
CL\_11187
CL\_10281
CL\_10280
CL\_11188
CL\_11189
CL\_11190
CL\_11191
CL\_11192
CL\_11193
CL\_11194
CL\_5813
CL\_5812
CL\_8109
CL\_8395
CL\_10740
CL\_7567
CL\_4423
CL\_4422
CL\_4421
CL\_4420
CL\_4419
CL\_4418
CL\_4417
CL\_4416
CL\_10751
CL\_10750
CL\_10749
CL\_10748
CL\_10747
CL\_10746
CL\_10745
CL\_10744
CL\_10743
CL\_4415
CL\_10265
CL\_4414
CL\_4413
CL\_6913
CL\_18
CL\_17
CL\_16
CL\_15
CL\_14
CL\_13
CL\_12
CL\_11
CL\_10
CL\_9
CL\_5163
CL\_13696
CL\_12671
CL\_6912
CL\_6911
CL\_1
CL\_5104
CL\_5261
CL\_5260
CL\_5259
CL\_4412
CL\_4411
CL\_4410
CL\_4409
CL\_10739
CL\_10738
CL\_4408
CL\_4407
CL\_4406
CL\_4405
CL\_4404
CL\_4403
CL\_4402
CL\_4401
CL\_4400
CL\_4399
CL\_4398
CL\_4397
CL\_4396
CL\_4395
CL\_4394
CL\_4393
CL\_4392
CL\_4391
CL\_4390
CL\_4389
CL\_4388
CL\_4387
CL\_4386
CL\_4385
CL\_7175
CL\_4384
CL\_4383
CL\_4382
CL\_11201
CL\_4381
CL\_4380
CL\_10737
CL\_4379
CL\_4378
CL\_4377
CL\_4376
CL\_4375
CL\_4374
CL\_4373
CL\_4372
CL\_4371
CL\_4370
CL\_4369
CL\_4368
CL\_4367
CL\_4366
CL\_4365
CL\_4364
CL\_4363
CL\_4362
CL\_4361
CL\_4360
CL\_4359
CL\_4358
CL\_10736
CL\_4357
CL\_4356
CL\_10735
CL\_4355
CL\_4354
CL\_4353
CL\_4352
CL\_4351
CL\_4350
CL\_4349
CL\_4348
CL\_4347
CL\_4346
CL\_4345
CL\_4344
CL\_4343
CL\_4342
CL\_4341
CL\_4340
CL\_4339
CL\_10734
CL\_4338
CL\_4337
CL\_4336
CL\_4335
CL\_10264
CL\_6067
CL\_4334
