## Supplementary material for "A novel method for integrating genomic and Tn-Seq data to identify common *in vivo* fitness mechanisms across multiple bacterial species": S3 Dataset: CL_INS_2.html

Legend

 Mobile +extrachromosomalelementfunctions
 Hypothetical

FULL


WINDOWSVGPNG

Trim RowsRemove SingletonsSave Fasta

CL\_28


CL\_28

HighlightSelectShow Genomes


35

CL\_29


6

CL\_29

fGI ID

CL\_INS\_2
Cluster ID

CL\_5116
