## Supplementary material for "A novel method for integrating genomic and Tn-Seq data to identify common *in vivo* fitness mechanisms across multiple bacterial species": S3 Dataset: CL_INS_4.html

Legend

 Regulatoryfunctions
 All EssentialGenes
 All VFDB Genes

FULL


WINDOWSVGPNG

Trim RowsRemove SingletonsSave Fasta

CL\_43


CL\_43


CL\_43


CL\_43

HighlightSelectShow Genomes


35

CL\_50


19

CL\_50


1

CL\_50


1

CL\_50

fGI ID


CL\_INS\_4
CL\_INS\_4
CL\_INS\_4
CL\_INS\_4
CL\_INS\_4
CL\_INS\_4
Cluster ID


CL\_44
CL\_45
CL\_46
CL\_47
CL\_48
CL\_49
