## Supplementary material for "A novel method for integrating genomic and Tn-Seq data to identify common *in vivo* fitness mechanisms across multiple bacterial species": S3 Dataset: CL_INS_5.html

Legend

 Mobile +extrachromosomalelementfunctions
 All EssentialGenes
 Proteinsynthesis/fate
 All VFDB Genes

FULL


WINDOWSVGPNG

Trim RowsRemove SingletonsSave Fasta

CL\_82


CL\_82


CL\_82


CL\_82

HighlightSelectShow Genomes


45

CL\_83


9

CL\_83


2

CL\_83


1

CL\_83

fGI ID


CL\_INS\_5
CL\_INS\_5
CL\_INS\_5
Cluster ID


CL\_9505
CL\_6922
CL\_6923
