## Supplementary material for "A novel method for integrating genomic and Tn-Seq data to identify common *in vivo* fitness mechanisms across multiple bacterial species": S3 Dataset: CL_INS_6.html

Legend

 Mobile +extrachromosomalelementfunctions
 Regulatoryfunctions
 Hypothetical
 DNA Metabolism
 All EssentialGenes
 Transcription
 Proteinsynthesis/fate
 Other
 EnergyMetabolism
 Transport +binding proteins
 All VFDB Genes

FULL


WINDOWSVGPNG

Trim RowsRemove SingletonsSave Fasta

CL\_87


CL\_87


CL\_87


CL\_87


CL\_87


CL\_87


CL\_87


CL\_87


CL\_87


CL\_87


CL\_87


CL\_87


CL\_87


CL\_87


CL\_87


CL\_87


CL\_87


CL\_87


CL\_87


CL\_87


CL\_87


CL\_87


CL\_87


CL\_87


CL\_87


CL\_87


CL\_87


CL\_87


CL\_87


CL\_87


CL\_87


CL\_87

HighlightSelectShow Genomes


16

CL\_107


5

CL\_107


3

CL\_107


2

CL\_107


2

CL\_108


2

CL\_107


1

CL\_107


1

CL\_107


1

CL\_107


1

CL\_107


1

CL\_107


1

CL\_107


1

CL\_107


1

CL\_107


1

CL\_107


1

CL\_107


1

CL\_107


1

CL\_107


1

CL\_107


1

CL\_107


1

CL\_107


1

CL\_107


1

CL\_107


1

CL\_107


1

CL\_107


1

CL\_107


1

CL\_107


1

CL\_107


1

CL\_107


1

CL\_107


1

CL\_107


1

CL\_107

fGI ID


CL\_INS\_6
CL\_INS\_6
CL\_INS\_6
CL\_INS\_6
CL\_INS\_6
CL\_INS\_6
CL\_INS\_6
CL\_INS\_6
CL\_INS\_220
CL\_INS\_6
CL\_INS\_6
CL\_INS\_65
CL\_INS\_237
CL\_INS\_1
CL\_INS\_1
CL\_INS\_83
CL\_INS\_83
CL\_INS\_83
CL\_INS\_6
CL\_INS\_6
CL\_INS\_220
CL\_INS\_220
CL\_INS\_219
CL\_INS\_219
CL\_INS\_219
CL\_INS\_219
CL\_INS\_219
CL\_INS\_220
CL\_INS\_220
CL\_INS\_220
CL\_INS\_220
CL\_INS\_220
CL\_INS\_219
CL\_INS\_219
CL\_INS\_6
CL\_INS\_219
CL\_INS\_219
CL\_INS\_219
CL\_INS\_219
CL\_INS\_6
CL\_INS\_219
CL\_INS\_6
CL\_INS\_219
CL\_INS\_219
CL\_INS\_219
CL\_INS\_219
CL\_INS\_219
CL\_INS\_219
CL\_INS\_219
CL\_INS\_219
CL\_INS\_219
CL\_INS\_220
CL\_INS\_96
CL\_INS\_6
CL\_INS\_220
CL\_INS\_6
CL\_INS\_6
CL\_INS\_220
CL\_INS\_220
CL\_INS\_6
CL\_INS\_220
CL\_INS\_198
CL\_INS\_219
CL\_INS\_219
CL\_INS\_219
CL\_INS\_220
CL\_INS\_220
CL\_INS\_220
CL\_INS\_220
CL\_INS\_220
CL\_INS\_220
CL\_INS\_220
CL\_INS\_220
CL\_INS\_220
CL\_INS\_220
CL\_INS\_220
CL\_INS\_219
CL\_INS\_220
CL\_INS\_6
CL\_INS\_218
CL\_INS\_6
CL\_INS\_6
CL\_INS\_6
CL\_INS\_6
CL\_INS\_6
CL\_INS\_220
CL\_INS\_6
CL\_INS\_6
CL\_INS\_6
CL\_INS\_220
CL\_INS\_6
CL\_INS\_220
CL\_INS\_220
CL\_INS\_220
CL\_INS\_220
CL\_INS\_219
CL\_INS\_133
CL\_INS\_133
CL\_INS\_6
CL\_INS\_6
CL\_INS\_220
CL\_INS\_6
CL\_INS\_6
CL\_INS\_6
CL\_INS\_6
CL\_INS\_6
CL\_INS\_6
CL\_INS\_6
CL\_INS\_6
CL\_INS\_6
CL\_INS\_6
CL\_INS\_6
CL\_INS\_6
CL\_INS\_6
CL\_INS\_6
CL\_INS\_6
CL\_INS\_6
CL\_INS\_6
CL\_INS\_6
CL\_INS\_6
CL\_INS\_6
CL\_INS\_219
CL\_INS\_218
CL\_INS\_219
CL\_INS\_219
CL\_INS\_219
CL\_INS\_219
CL\_INS\_219
CL\_INS\_219
CL\_INS\_219
CL\_INS\_219
CL\_INS\_219
CL\_INS\_219
CL\_INS\_219
CL\_INS\_6
CL\_INS\_6
CL\_INS\_218
CL\_INS\_6
CL\_INS\_6
CL\_INS\_6
CL\_INS\_6
CL\_INS\_6
CL\_INS\_6
CL\_INS\_6
CL\_INS\_6
CL\_INS\_6
CL\_INS\_6
CL\_INS\_219
CL\_INS\_219
CL\_INS\_219
CL\_INS\_219
CL\_INS\_219
CL\_INS\_219
CL\_INS\_219
CL\_INS\_6
CL\_INS\_219
CL\_INS\_219
CL\_INS\_219
CL\_INS\_6
CL\_INS\_6
CL\_INS\_6
CL\_INS\_6
CL\_INS\_6
CL\_INS\_6
CL\_INS\_6
CL\_INS\_6
CL\_INS\_6
CL\_INS\_6
CL\_INS\_6
CL\_INS\_220
CL\_INS\_6
CL\_INS\_6
CL\_INS\_6
CL\_INS\_6
CL\_INS\_6
CL\_INS\_6
CL\_INS\_13
CL\_INS\_6
CL\_INS\_13
CL\_INS\_13
CL\_INS\_6
CL\_INS\_6
CL\_INS\_6
CL\_INS\_6
CL\_INS\_6
CL\_INS\_13
CL\_INS\_13
CL\_INS\_13
CL\_INS\_13
CL\_INS\_13
CL\_INS\_6
CL\_INS\_13
CL\_INS\_6
CL\_INS\_13
CL\_INS\_13
CL\_INS\_13
CL\_INS\_13
CL\_INS\_13
CL\_INS\_13
CL\_INS\_13
CL\_INS\_6
CL\_INS\_6
CL\_INS\_6
CL\_INS\_6
CL\_INS\_6
CL\_INS\_6
CL\_INS\_6
CL\_INS\_6
CL\_INS\_6
CL\_INS\_6
CL\_INS\_6
CL\_INS\_6
CL\_INS\_6
CL\_INS\_83
CL\_INS\_83
CL\_INS\_83
CL\_INS\_1
CL\_INS\_1
CL\_INS\_83
CL\_INS\_6
CL\_INS\_6
CL\_INS\_6
CL\_INS\_1
CL\_INS\_6
CL\_INS\_6
CL\_INS\_6
CL\_INS\_83
CL\_INS\_6
CL\_INS\_83
CL\_INS\_6
CL\_INS\_6
CL\_INS\_6
CL\_INS\_6
CL\_INS\_6
CL\_INS\_6
CL\_INS\_6
CL\_INS\_220
CL\_INS\_220
CL\_INS\_6
CL\_INS\_220
CL\_INS\_83
CL\_INS\_83
CL\_INS\_83
CL\_INS\_83
CL\_INS\_83
CL\_INS\_83
CL\_INS\_6
CL\_INS\_6
CL\_INS\_6
CL\_INS\_6
CL\_INS\_6
CL\_INS\_6
CL\_INS\_6
CL\_INS\_83
CL\_INS\_6
CL\_INS\_220
CL\_INS\_220
CL\_INS\_220
CL\_INS\_6
CL\_INS\_6
CL\_INS\_6
CL\_INS\_6
CL\_INS\_6
CL\_INS\_6
CL\_INS\_6
CL\_INS\_6
CL\_INS\_6
CL\_INS\_6
CL\_INS\_6
CL\_INS\_6
CL\_INS\_6
CL\_INS\_6
CL\_INS\_6
CL\_INS\_6
CL\_INS\_6
CL\_INS\_6
CL\_INS\_6
CL\_INS\_6
CL\_INS\_6
CL\_INS\_6
CL\_INS\_6
CL\_INS\_178
CL\_INS\_178
CL\_INS\_178
CL\_INS\_178
CL\_INS\_6
CL\_INS\_6
CL\_INS\_6
CL\_INS\_6
CL\_INS\_6
CL\_INS\_6
CL\_INS\_6
CL\_INS\_220
CL\_INS\_6
CL\_INS\_220
CL\_INS\_220
CL\_INS\_220
CL\_INS\_220
CL\_INS\_220
CL\_INS\_220
CL\_INS\_220
CL\_INS\_220
CL\_INS\_220
CL\_INS\_220
CL\_INS\_220
CL\_INS\_220
CL\_INS\_220
CL\_INS\_6
CL\_INS\_83
CL\_INS\_6
CL\_INS\_220
CL\_INS\_6
CL\_INS\_6
CL\_INS\_6
CL\_INS\_6
CL\_INS\_6
CL\_INS\_6
CL\_INS\_6
CL\_INS\_6
CL\_INS\_6
CL\_INS\_6
CL\_INS\_6
CL\_INS\_6
CL\_INS\_220
CL\_INS\_220
CL\_INS\_220
CL\_INS\_220
CL\_INS\_220
CL\_INS\_220
CL\_INS\_220
CL\_INS\_220
CL\_INS\_220
CL\_INS\_220
CL\_INS\_6
CL\_INS\_220
CL\_INS\_220
CL\_INS\_220
CL\_INS\_220
CL\_INS\_220
CL\_INS\_220
CL\_INS\_220
CL\_INS\_220
CL\_INS\_220
CL\_INS\_220
CL\_INS\_220
CL\_INS\_220
CL\_INS\_220
CL\_INS\_220
CL\_INS\_220
CL\_INS\_220
CL\_INS\_220
CL\_INS\_220
CL\_INS\_220
CL\_INS\_220
CL\_INS\_220
CL\_INS\_220
CL\_INS\_6
CL\_INS\_6
CL\_INS\_6
CL\_INS\_6
CL\_INS\_6
Cluster ID


CL\_11841
CL\_11840
CL\_11839
CL\_11024
CL\_11025
CL\_11026
CL\_11027
CL\_6924
CL\_7732
CL\_13699
CL\_8134
CL\_88
CL\_10038
CL\_10039
CL\_89
CL\_90
CL\_91
CL\_92
CL\_9088
CL\_12260
CL\_6565
CL\_6566
CL\_6411
CL\_6412
CL\_6413
CL\_6414
CL\_6415
CL\_6567
CL\_6568
CL\_6569
CL\_6570
CL\_6571
CL\_6416
CL\_6417
CL\_10971
CL\_6418
CL\_6419
CL\_6420
CL\_6421
CL\_7452
CL\_6422
CL\_11611
CL\_6426
CL\_10655
CL\_10656
CL\_10657
CL\_10658
CL\_10659
CL\_10660
CL\_10661
CL\_10662
CL\_6572
CL\_6573
CL\_8607
CL\_8606
CL\_8605
CL\_8783
CL\_8782
CL\_6574
CL\_8603
CL\_8604
CL\_6575
CL\_6576
CL\_6577
CL\_6578
CL\_6579
CL\_6580
CL\_6581
CL\_6582
CL\_6583
CL\_6584
CL\_6585
CL\_6586
CL\_6587
CL\_6588
CL\_6589
CL\_6590
CL\_6591
CL\_8602
CL\_7841
CL\_10649
CL\_6592
CL\_10648
CL\_10647
CL\_6593
CL\_6594
CL\_10394
CL\_10395
CL\_10396
CL\_6595
CL\_10646
CL\_6596
CL\_6597
CL\_6598
CL\_6599
CL\_6600
CL\_8867
CL\_8868
CL\_10397
CL\_10398
CL\_6601
CL\_6602
CL\_6603
CL\_8778
CL\_6604
CL\_10645
CL\_10644
CL\_10643
CL\_10642
CL\_10641
CL\_10640
CL\_10639
CL\_10638
CL\_10637
CL\_10636
CL\_10635
CL\_10634
CL\_10633
CL\_10632
CL\_10631
CL\_10630
CL\_6427
CL\_6428
CL\_6429
CL\_6430
CL\_6431
CL\_6432
CL\_6433
CL\_6434
CL\_6435
CL\_6436
CL\_6437
CL\_6438
CL\_6439
CL\_8133
CL\_8132
CL\_7840
CL\_10972
CL\_8781
CL\_8780
CL\_8779
CL\_11610
CL\_11609
CL\_10629
CL\_10628
CL\_10627
CL\_10626
CL\_9480
CL\_8131
CL\_10668
CL\_10669
CL\_10670
CL\_10671
CL\_8130
CL\_11608
CL\_8129
CL\_8128
CL\_8127
CL\_8126
CL\_8125
CL\_8124
CL\_8123
CL\_11607
CL\_12395
CL\_5118
CL\_5119
CL\_11606
CL\_11605
CL\_11604
CL\_6605
CL\_9922
CL\_9084
CL\_10973
CL\_6925
CL\_6926
CL\_9087
CL\_6927
CL\_6928
CL\_6709
CL\_10410
CL\_6929
CL\_6930
CL\_6931
CL\_6932
CL\_6933
CL\_6708
CL\_6707
CL\_6706
CL\_6705
CL\_6704
CL\_6934
CL\_6703
CL\_6935
CL\_10411
CL\_6936
CL\_6937
CL\_6938
CL\_6939
CL\_6940
CL\_6941
CL\_6942
CL\_12185
CL\_6943
CL\_6944
CL\_6945
CL\_6946
CL\_6947
CL\_6948
CL\_6949
CL\_6950
CL\_6951
CL\_6952
CL\_9086
CL\_3943
CL\_3942
CL\_3941
CL\_3940
CL\_6155
CL\_6154
CL\_13556
CL\_13555
CL\_6152
CL\_3939
CL\_7686
CL\_93
CL\_10542
CL\_94
CL\_10543
CL\_95
CL\_96
CL\_97
CL\_98
CL\_99
CL\_100
CL\_101
CL\_13134
CL\_10399
CL\_10400
CL\_10401
CL\_10402
CL\_10403
CL\_10404
CL\_10969
CL\_6151
CL\_8777
CL\_6150
CL\_13554
CL\_13553
CL\_13552
CL\_13551
CL\_13550
CL\_13549
CL\_13548
CL\_13547
CL\_13135
CL\_10024
CL\_10023
CL\_10022
CL\_10544
CL\_10545
CL\_10546
CL\_10547
CL\_10548
CL\_10549
CL\_10550
CL\_10551
CL\_10552
CL\_10553
CL\_10554
CL\_10555
CL\_10556
CL\_10557
CL\_13136
CL\_10558
CL\_10559
CL\_10560
CL\_10561
CL\_10562
CL\_10563
CL\_10564
CL\_10565
CL\_6061
CL\_6062
CL\_6063
CL\_6064
CL\_10566
CL\_8776
CL\_8775
CL\_8774
CL\_8773
CL\_8772
CL\_8771
CL\_5166
CL\_5167
CL\_5168
CL\_5170
CL\_5171
CL\_5172
CL\_5173
CL\_5174
CL\_5175
CL\_5176
CL\_10405
CL\_5177
CL\_5178
CL\_5179
CL\_5180
CL\_5181
CL\_102
CL\_103
CL\_8160
CL\_104
CL\_13546
CL\_10567
CL\_8770
CL\_10406
CL\_8769
CL\_8768
CL\_8767
CL\_105
CL\_106
CL\_10407
CL\_10408
CL\_7730
CL\_7729
CL\_7728
CL\_7727
CL\_7726
CL\_7725
CL\_7724
CL\_7723
CL\_7722
CL\_7721
CL\_13336
CL\_7720
CL\_7719
CL\_7718
CL\_7717
CL\_7716
CL\_7715
CL\_7714
CL\_7713
CL\_7712
CL\_7711
CL\_7710
CL\_7709
CL\_7708
CL\_7707
CL\_7706
CL\_9085
CL\_7705
CL\_7704
CL\_7703
CL\_7702
CL\_7701
CL\_7700
CL\_7453
CL\_7454
CL\_8766
CL\_10624
CL\_10625
