## Supplementary material for "A novel method for integrating genomic and Tn-Seq data to identify common *in vivo* fitness mechanisms across multiple bacterial species": S3 Dataset: CL_INS_8.html

Legend

 Hypothetical
 Other
 All VFDB Genes

FULL


WINDOWSVGPNG

Trim RowsRemove SingletonsSave Fasta

CL\_153


CL\_153

HighlightSelectShow Genomes


53

CL\_154


3

CL\_154

fGI ID


CL\_INS\_8
CL\_INS\_8
CL\_INS\_8
CL\_INS\_8
Cluster ID


CL\_7893
CL\_7894
CL\_7895
CL\_7896
