## Supplementary material for "A novel method for integrating genomic and Tn-Seq data to identify common *in vivo* fitness mechanisms across multiple bacterial species": S3 Dataset: CL_INS_10.html

Legend

 Hypothetical
 All EssentialGenes
 Other

FULL


WINDOWSVGPNG

Trim RowsRemove SingletonsSave Fasta

CL\_259


CL\_259


CL\_259

HighlightSelectShow Genomes


38

CL\_260


18

CL\_260


1

CL\_260

fGI ID


CL\_INS\_10
CL\_INS\_10
Cluster ID


CL\_10160
CL\_6611
