## Supplementary material for "A novel method for integrating genomic and Tn-Seq data to identify common *in vivo* fitness mechanisms across multiple bacterial species": S3 Dataset: CL_INS_11.html

Legend

 Hypothetical
 Proteinsynthesis/fate
 All VFDB Genes

FULL


WINDOWSVGPNG

Trim RowsRemove SingletonsSave Fasta

CL\_274


CL\_274

HighlightSelectShow Genomes


38

CL\_275


18

CL\_275

fGI ID

CL\_INS\_11
Cluster ID

CL\_8122
