## Supplementary material for "A novel method for integrating genomic and Tn-Seq data to identify common *in vivo* fitness mechanisms across multiple bacterial species": S3 Dataset: CL_INS_15.html

Legend

 Hypothetical
 Proteinsynthesis/fate
 EnergyMetabolism

FULL


WINDOWSVGPNG

Trim RowsRemove SingletonsSave Fasta

CL\_403


CL\_403

HighlightSelectShow Genomes


30

CL\_404


27

CL\_404

fGI ID

CL\_INS\_15
Cluster ID

CL\_5141
