## Supplementary material for "A novel method for integrating genomic and Tn-Seq data to identify common *in vivo* fitness mechanisms across multiple bacterial species": S3 Dataset: CL_INS_16.html

Legend

 Mobile +extrachromosomalelementfunctions
 Hypothetical
 Other
 Transport +binding proteins

FULL


WINDOWSVGPNG

Trim RowsRemove SingletonsSave Fasta

CL\_429


CL\_429


CL\_429


CL\_429

HighlightSelectShow Genomes


39

CL\_432


10

CL\_432


7

CL\_432


1

CL\_432

fGI ID


CL\_INS\_16
CL\_INS\_16
CL\_INS\_16
Cluster ID


CL\_9507
CL\_430
CL\_431
