## Supplementary material for "A novel method for integrating genomic and Tn-Seq data to identify common *in vivo* fitness mechanisms across multiple bacterial species": S3 Dataset: CL_INS_18.html

Legend

 Hypothetical
 Other
 All VFDB Genes

FULL


WINDOWSVGPNG

Trim RowsRemove SingletonsSave Fasta

CL\_490


CL\_490


CL\_490


CL\_490


CL\_512


CL\_490

HighlightSelectShow Genomes


30

CL\_498


24

CL\_498


1

CL\_498


1

CL\_502


1

CL\_498


1

CL\_498

fGI ID


CL\_INS\_133
CL\_INS\_18
CL\_INS\_18
CL\_INS\_18
CL\_INS\_18
CL\_INS\_18
CL\_INS\_18
CL\_INS\_18
CL\_INS\_18
Cluster ID


CL\_4451
CL\_491
CL\_492
CL\_493
CL\_494
CL\_495
CL\_496
CL\_497
CL\_8372
