## Supplementary material for "A novel method for integrating genomic and Tn-Seq data to identify common *in vivo* fitness mechanisms across multiple bacterial species": S3 Dataset: CL_INS_19.html

Legend

 Mobile +extrachromosomalelementfunctions
 Hypothetical
 Regulatoryfunctions
 DNA Metabolism
 AntibioticResistance
 All EssentialGenes
 Biosynthesis ofcofactors,prostheticgroups, +carriers
 Proteinsynthesis/fate
 Other
 EnergyMetabolism
 Transport +binding proteins
 All VFDB Genes

FULL


WINDOWSVGPNG

Trim RowsRemove SingletonsSave Fasta

CL\_502


CL\_502


CL\_502


CL\_502


CL\_502


CL\_1138


CL\_502


CL\_4308


CL\_502


CL\_502


CL\_1629


CL\_502


Break


CL\_502


CL\_502


CL\_498


CL\_502


CL\_502


CL\_2669


CL\_2800


CL\_502

HighlightSelectShow Genomes


11

CL\_512


5

CL\_512


5

CL\_530


4

CL\_512


3

CL\_512


2

CL\_512


1

CL\_512


1

CL\_512


1

Break


1

CL\_512


1

CL\_512


1

CL\_519


1

CL\_512


1

CL\_490


1

CL\_512


1

CL\_512


1

CL\_512


1

CL\_512


1

CL\_512


1

CL\_512


1

CL\_512

fGI ID


CL\_INS\_19
CL\_INS\_153
CL\_INS\_18
CL\_INS\_18
CL\_INS\_18
CL\_INS\_18
CL\_INS\_18
CL\_INS\_18
CL\_INS\_18
CL\_INS\_19
CL\_INS\_20
CL\_INS\_20
CL\_INS\_20
CL\_INS\_19
CL\_INS\_237
CL\_INS\_237
CL\_INS\_237
CL\_INS\_237
CL\_INS\_237
CL\_INS\_237
CL\_INS\_237
CL\_INS\_237
CL\_INS\_237
CL\_INS\_237
CL\_INS\_237
CL\_INS\_237
CL\_INS\_237
CL\_INS\_237
CL\_INS\_237
CL\_INS\_237
CL\_INS\_237
CL\_INS\_56
CL\_INS\_237
CL\_INS\_237
CL\_INS\_237
CL\_INS\_237
CL\_INS\_237
CL\_INS\_237
CL\_INS\_237
CL\_INS\_237
CL\_INS\_83
CL\_INS\_83
CL\_INS\_83
CL\_INS\_83
CL\_INS\_83
CL\_INS\_83
CL\_INS\_83
CL\_INS\_83
CL\_INS\_83
CL\_INS\_83
CL\_INS\_83
CL\_INS\_83
CL\_INS\_83
CL\_INS\_83
CL\_INS\_83
CL\_INS\_83
CL\_INS\_83
CL\_INS\_83
CL\_INS\_83
CL\_INS\_83
CL\_INS\_83
CL\_INS\_83
CL\_INS\_83
CL\_INS\_83
CL\_INS\_83
CL\_INS\_83
CL\_INS\_83
CL\_INS\_83
CL\_INS\_83
CL\_INS\_83
CL\_INS\_83
CL\_INS\_83
CL\_INS\_83
CL\_INS\_83
CL\_INS\_83
CL\_INS\_83
CL\_INS\_83
CL\_INS\_83
CL\_INS\_83
CL\_INS\_83
CL\_INS\_83
CL\_INS\_83
CL\_INS\_83
CL\_INS\_19
CL\_INS\_19
CL\_INS\_156
CL\_INS\_156
CL\_INS\_156
CL\_INS\_1
CL\_INS\_153
CL\_INS\_1
CL\_INS\_1
CL\_INS\_1
CL\_INS\_1
CL\_INS\_1
CL\_INS\_1
CL\_INS\_1
CL\_INS\_19
CL\_INS\_1
CL\_INS\_1
CL\_INS\_1
CL\_INS\_1
CL\_INS\_1
CL\_INS\_1
CL\_INS\_1
CL\_INS\_1
CL\_INS\_1
CL\_INS\_1
CL\_INS\_1
CL\_INS\_1
CL\_INS\_1
CL\_INS\_1
CL\_INS\_1
CL\_INS\_1
CL\_INS\_1
CL\_INS\_1
CL\_INS\_1
CL\_INS\_1
CL\_INS\_1
CL\_INS\_1
CL\_INS\_1
CL\_INS\_1
CL\_INS\_1
CL\_INS\_1
CL\_INS\_1
CL\_INS\_1
CL\_INS\_1
CL\_INS\_1
CL\_INS\_1
CL\_INS\_1
CL\_INS\_1
CL\_INS\_20
CL\_INS\_20
CL\_INS\_133
CL\_INS\_20
CL\_INS\_20
CL\_INS\_20
CL\_INS\_20
CL\_INS\_20
CL\_INS\_20
CL\_INS\_20
CL\_INS\_19
CL\_INS\_19
CL\_INS\_19
CL\_INS\_19
CL\_INS\_19
CL\_INS\_19
CL\_INS\_19
CL\_INS\_19
CL\_INS\_133
CL\_INS\_19
CL\_INS\_19
CL\_INS\_19
CL\_INS\_19
CL\_INS\_19
CL\_INS\_19
CL\_INS\_19
CL\_INS\_19
CL\_INS\_19
CL\_INS\_1
CL\_INS\_1
CL\_INS\_20
CL\_INS\_20
CL\_INS\_20
CL\_INS\_133
CL\_INS\_20
CL\_INS\_20
CL\_INS\_20
CL\_INS\_133
CL\_INS\_19
CL\_INS\_20
CL\_INS\_20
CL\_INS\_20
CL\_INS\_20
CL\_INS\_20
CL\_INS\_20
CL\_INS\_20
CL\_INS\_133
CL\_INS\_1
CL\_INS\_1
CL\_INS\_1
CL\_INS\_1
CL\_INS\_1
CL\_INS\_1
CL\_INS\_1
CL\_INS\_1
CL\_INS\_19
CL\_INS\_1
CL\_INS\_1
CL\_INS\_1
CL\_INS\_1
CL\_INS\_1
CL\_INS\_1
CL\_INS\_1
CL\_INS\_1
CL\_INS\_1
CL\_INS\_1
CL\_INS\_1
CL\_INS\_1
CL\_INS\_1
CL\_INS\_1
CL\_INS\_19
CL\_INS\_1
CL\_INS\_1
CL\_INS\_1
CL\_INS\_1
CL\_INS\_1
CL\_INS\_1
CL\_INS\_1
CL\_INS\_1
CL\_INS\_1
CL\_INS\_1
CL\_INS\_1
CL\_INS\_1
CL\_INS\_1
CL\_INS\_1
CL\_INS\_1
CL\_INS\_1
CL\_INS\_1
CL\_INS\_1
CL\_INS\_1
CL\_INS\_1
CL\_INS\_1
CL\_INS\_1
CL\_INS\_19
CL\_INS\_1
CL\_INS\_1
CL\_INS\_1
CL\_INS\_1
CL\_INS\_1
CL\_INS\_1
CL\_INS\_1
CL\_INS\_1
CL\_INS\_1
CL\_INS\_1
CL\_INS\_1
CL\_INS\_20
CL\_INS\_55
CL\_INS\_19
CL\_INS\_1
CL\_INS\_133
CL\_INS\_133
CL\_INS\_133
CL\_INS\_133
CL\_INS\_56
CL\_INS\_19
CL\_INS\_19
CL\_INS\_19
CL\_INS\_19
CL\_INS\_19
CL\_INS\_133
CL\_INS\_1
CL\_INS\_156
CL\_INS\_1
CL\_INS\_1
CL\_INS\_1
CL\_INS\_1
CL\_INS\_1
CL\_INS\_1
CL\_INS\_1
CL\_INS\_1
CL\_INS\_1
CL\_INS\_1
CL\_INS\_1
CL\_INS\_1
CL\_INS\_1
CL\_INS\_1
CL\_INS\_1
CL\_INS\_1
CL\_INS\_1
CL\_INS\_1
CL\_INS\_1
CL\_INS\_1
CL\_INS\_1
CL\_INS\_1
CL\_INS\_1
CL\_INS\_1
CL\_INS\_1
CL\_INS\_1
CL\_INS\_34
CL\_INS\_34
CL\_INS\_156
CL\_INS\_34
CL\_INS\_156
CL\_INS\_156
CL\_INS\_156
CL\_INS\_156
CL\_INS\_83
CL\_INS\_83
CL\_INS\_83
CL\_INS\_83
CL\_INS\_83
CL\_INS\_83
CL\_INS\_19
CL\_INS\_237
CL\_INS\_237
CL\_INS\_19
CL\_INS\_19
CL\_INS\_19
CL\_INS\_19
CL\_INS\_19
CL\_INS\_19
CL\_INS\_19
CL\_INS\_19
CL\_INS\_19
CL\_INS\_19
CL\_INS\_19
CL\_INS\_19
CL\_INS\_19
CL\_INS\_19
CL\_INS\_19
CL\_INS\_19
CL\_INS\_19
CL\_INS\_19
CL\_INS\_19
CL\_INS\_19
CL\_INS\_19
CL\_INS\_19
CL\_INS\_19
CL\_INS\_168
CL\_INS\_19
CL\_INS\_19
CL\_INS\_19
CL\_INS\_19
CL\_INS\_19
CL\_INS\_19
CL\_INS\_168
CL\_INS\_19
CL\_INS\_20
CL\_INS\_20
CL\_INS\_20
CL\_INS\_20
CL\_INS\_20
CL\_INS\_1
CL\_INS\_19
CL\_INS\_133
CL\_INS\_20
CL\_INS\_19
CL\_INS\_1
CL\_INS\_1
CL\_INS\_1
CL\_INS\_19
CL\_INS\_1
CL\_INS\_1
CL\_INS\_1
CL\_INS\_1
CL\_INS\_1
CL\_INS\_1
CL\_INS\_1
CL\_INS\_1
CL\_INS\_168
CL\_INS\_1
CL\_INS\_20
CL\_INS\_1
CL\_INS\_83
CL\_INS\_83
CL\_INS\_19
CL\_INS\_19
CL\_INS\_19
CL\_INS\_19
CL\_INS\_19
CL\_INS\_83
CL\_INS\_19
CL\_INS\_1
CL\_INS\_1
CL\_INS\_1
CL\_INS\_1
CL\_INS\_1
CL\_INS\_1
CL\_INS\_19
CL\_INS\_1
CL\_INS\_1
CL\_INS\_1
CL\_INS\_1
CL\_INS\_153
CL\_INS\_153
CL\_INS\_19
CL\_INS\_19
CL\_INS\_19
CL\_INS\_19
CL\_INS\_19
CL\_INS\_19
CL\_INS\_19
CL\_INS\_1
CL\_INS\_1
CL\_INS\_1
CL\_INS\_133
CL\_INS\_156
CL\_INS\_156
CL\_INS\_1
CL\_INS\_133
CL\_INS\_20
CL\_INS\_20
CL\_INS\_20
CL\_INS\_20
CL\_INS\_45
CL\_INS\_45
CL\_INS\_45
CL\_INS\_34
CL\_INS\_34
CL\_INS\_34
CL\_INS\_34
CL\_INS\_1
CL\_INS\_1
CL\_INS\_1
CL\_INS\_1
CL\_INS\_1
CL\_INS\_1
CL\_INS\_1
CL\_INS\_1
CL\_INS\_1
CL\_INS\_156
Cluster ID


CL\_12258
CL\_8121
CL\_8372
CL\_497
CL\_496
CL\_495
CL\_494
CL\_493
CL\_492
CL\_5143
CL\_5144
CL\_5145
CL\_5146
CL\_10415
CL\_5709
CL\_5710
CL\_11135
CL\_11136
CL\_11137
CL\_11138
CL\_11139
CL\_11140
CL\_11141
CL\_8936
CL\_6100
CL\_6099
CL\_6098
CL\_6097
CL\_6096
CL\_6095
CL\_6094
CL\_6092
CL\_11142
CL\_11143
CL\_11144
CL\_11145
CL\_11146
CL\_11147
CL\_11148
CL\_11149
CL\_1627
CL\_1626
CL\_1625
CL\_1624
CL\_9617
CL\_90
CL\_9616
CL\_91
CL\_6154
CL\_12424
CL\_12425
CL\_4894
CL\_12426
CL\_12427
CL\_12428
CL\_12429
CL\_12430
CL\_12431
CL\_10403
CL\_10404
CL\_10969
CL\_6151
CL\_8777
CL\_6150
CL\_12432
CL\_12433
CL\_6148
CL\_9435
CL\_9434
CL\_9433
CL\_9432
CL\_12434
CL\_9431
CL\_9430
CL\_9429
CL\_9428
CL\_9427
CL\_4632
CL\_9425
CL\_9424
CL\_9423
CL\_9422
CL\_9421
CL\_11928
CL\_11927
CL\_2667
CL\_2665
CL\_2664
CL\_4452
CL\_5322
CL\_4587
CL\_4586
CL\_4585
CL\_4584
CL\_4583
CL\_4582
CL\_4581
CL\_10868
CL\_4580
CL\_4579
CL\_4578
CL\_4577
CL\_4576
CL\_4575
CL\_4574
CL\_4573
CL\_4572
CL\_4571
CL\_4570
CL\_4569
CL\_4568
CL\_4567
CL\_4566
CL\_4565
CL\_4564
CL\_4563
CL\_4562
CL\_4561
CL\_4560
CL\_4559
CL\_4558
CL\_4557
CL\_4556
CL\_4555
CL\_4554
CL\_4553
CL\_4552
CL\_4551
CL\_4550
CL\_4549
CL\_4548
CL\_5353
CL\_5828
CL\_5198
CL\_5829
CL\_5830
CL\_5831
CL\_5832
CL\_5833
CL\_5834
CL\_5835
CL\_10867
CL\_10866
CL\_10865
CL\_10864
CL\_10863
CL\_10862
CL\_10861
CL\_9263
CL\_6967
CL\_10860
CL\_10859
CL\_10858
CL\_10857
CL\_10856
CL\_10855
CL\_10854
CL\_10853
CL\_10250
CL\_4433
CL\_4434
CL\_5335
CL\_9148
CL\_9149
CL\_5884
CL\_5826
CL\_9150
CL\_5336
CL\_5337
CL\_8996
CL\_5343
CL\_5344
CL\_5345
CL\_7634
CL\_7635
CL\_7636
CL\_7637
CL\_5157
CL\_4542
CL\_4541
CL\_4540
CL\_4539
CL\_4538
CL\_4537
CL\_4536
CL\_4535
CL\_10852
CL\_4534
CL\_4533
CL\_4531
CL\_4530
CL\_4529
CL\_4528
CL\_4527
CL\_4526
CL\_4525
CL\_4524
CL\_4523
CL\_4522
CL\_4521
CL\_4520
CL\_10851
CL\_4519
CL\_4518
CL\_4516
CL\_4515
CL\_4514
CL\_4513
CL\_4512
CL\_4511
CL\_4510
CL\_4509
CL\_4508
CL\_4507
CL\_4506
CL\_4505
CL\_4504
CL\_4503
CL\_4502
CL\_4501
CL\_4500
CL\_4499
CL\_4498
CL\_4497
CL\_10850
CL\_4494
CL\_4493
CL\_4492
CL\_4491
CL\_4490
CL\_4489
CL\_4488
CL\_4487
CL\_4486
CL\_4485
CL\_4484
CL\_5224
CL\_5806
CL\_10849
CL\_5812
CL\_5201
CL\_5202
CL\_5203
CL\_5204
CL\_5205
CL\_5206
CL\_5207
CL\_5208
CL\_5209
CL\_5210
CL\_5328
CL\_5813
CL\_8855
CL\_6077
CL\_4424
CL\_5331
CL\_4422
CL\_4421
CL\_4420
CL\_4419
CL\_4418
CL\_4417
CL\_4416
CL\_4415
CL\_4414
CL\_4413
CL\_6913
CL\_5261
CL\_5260
CL\_5259
CL\_4601
CL\_4602
CL\_4603
CL\_4604
CL\_4606
CL\_4444
CL\_4443
CL\_4442
CL\_4440
CL\_6090
CL\_6089
CL\_8856
CL\_6088
CL\_6087
CL\_8857
CL\_8858
CL\_8859
CL\_506
CL\_507
CL\_508
CL\_10981
CL\_10982
CL\_10983
CL\_8120
CL\_509
CL\_510
CL\_511
CL\_6979
CL\_10848
CL\_10847
CL\_10846
CL\_5221
CL\_10845
CL\_10844
CL\_10843
CL\_10842
CL\_10841
CL\_10840
CL\_10839
CL\_10838
CL\_10837
CL\_10836
CL\_10835
CL\_10834
CL\_10833
CL\_10832
CL\_10831
CL\_10830
CL\_10829
CL\_10828
CL\_10827
CL\_10826
CL\_10825
CL\_10824
CL\_10823
CL\_9270
CL\_8081
CL\_10822
CL\_5342
CL\_5341
CL\_5340
CL\_5339
CL\_5338
CL\_5837
CL\_8925
CL\_6969
CL\_6968
CL\_10821
CL\_4481
CL\_4479
CL\_4478
CL\_10820
CL\_4461
CL\_4462
CL\_4463
CL\_4464
CL\_4465
CL\_4466
CL\_4467
CL\_4468
CL\_5320
CL\_5160
CL\_5159
CL\_4469
CL\_503
CL\_504
CL\_12688
CL\_8764
CL\_8763
CL\_8762
CL\_12689
CL\_505
CL\_10570
CL\_10242
CL\_4458
CL\_4459
CL\_4460
CL\_4457
CL\_4456
CL\_10819
CL\_4455
CL\_4454
CL\_5321
CL\_4453
CL\_5323
CL\_8601
CL\_8600
CL\_8599
CL\_8598
CL\_8597
CL\_8596
CL\_8595
CL\_8594
CL\_5324
CL\_5325
CL\_5326
CL\_5327
CL\_8850
CL\_8851
CL\_8852
CL\_4451
CL\_5287
CL\_5286
CL\_5285
CL\_5283
CL\_5282
CL\_5281
CL\_5280
CL\_5279
CL\_5278
CL\_5277
CL\_6086
CL\_4450
CL\_4449
CL\_8853
CL\_4448
CL\_4447
CL\_4446
CL\_4445
CL\_5329
CL\_5330
CL\_8854
