## Supplementary material for "A novel method for integrating genomic and Tn-Seq data to identify common *in vivo* fitness mechanisms across multiple bacterial species": S3 Dataset: CL_INS_20.html

Legend

 Mobile +extrachromosomalelementfunctions
 Hypothetical
 Regulatoryfunctions
 DNA Metabolism
 AntibioticResistance
 All EssentialGenes
 Biosynthesis ofcofactors,prostheticgroups, +carriers
 Proteinsynthesis/fate
 Other
 EnergyMetabolism
 All VFDB Genes
 Transport +binding proteins

FULL


WINDOWSVGPNG

Trim RowsRemove SingletonsSave Fasta

CL\_522


CL\_522


CL\_502


CL\_522


CL\_522


CL\_522


CL\_522


CL\_522


CL\_522


Break


CL\_522


CL\_522


CL\_522


CL\_522


CL\_522


CL\_522


CL\_522


CL\_522


CL\_522


CL\_522


CL\_522


CL\_522


CL\_522

HighlightSelectShow Genomes


17

CL\_530


7

CL\_530


5

CL\_530


2

CL\_530


2

CL\_1137


1

CL\_530


1

Break


1

CL\_530


1

CL\_2663


1

CL\_530


1

Break


1

CL\_530


1

Break


1

CL\_1623


1

Break


1

Break


1

Break


1

CL\_530


1

CL\_2807


1

Break


1

CL\_4309


1

Break


1

CL\_531

fGI ID


CL\_INS\_20
CL\_INS\_20
CL\_INS\_20
CL\_INS\_20
CL\_INS\_20
CL\_INS\_20
CL\_INS\_20
CL\_INS\_20
CL\_INS\_20
CL\_INS\_20
CL\_INS\_20
CL\_INS\_20
CL\_INS\_20
CL\_INS\_20
CL\_INS\_20
CL\_INS\_20
CL\_INS\_20
CL\_INS\_20
CL\_INS\_237
CL\_INS\_20
CL\_INS\_20
CL\_INS\_237
CL\_INS\_20
CL\_INS\_237
CL\_INS\_20
CL\_INS\_20
CL\_INS\_20
CL\_INS\_20
CL\_INS\_20
CL\_INS\_20
CL\_INS\_20
CL\_INS\_20
CL\_INS\_20
CL\_INS\_20
CL\_INS\_20
CL\_INS\_20
CL\_INS\_20
CL\_INS\_20
CL\_INS\_20
CL\_INS\_20
CL\_INS\_20
CL\_INS\_20
CL\_INS\_20
CL\_INS\_20
CL\_INS\_1
CL\_INS\_1
CL\_INS\_1
CL\_INS\_1
CL\_INS\_20
CL\_INS\_20
CL\_INS\_20
CL\_INS\_20
CL\_INS\_20
CL\_INS\_20
CL\_INS\_20
CL\_INS\_20
CL\_INS\_20
CL\_INS\_20
CL\_INS\_20
CL\_INS\_20
CL\_INS\_20
CL\_INS\_20
CL\_INS\_1
CL\_INS\_20
CL\_INS\_20
CL\_INS\_20
CL\_INS\_20
CL\_INS\_20
CL\_INS\_20
CL\_INS\_20
CL\_INS\_20
CL\_INS\_20
CL\_INS\_20
CL\_INS\_20
CL\_INS\_20
CL\_INS\_1
CL\_INS\_1
CL\_INS\_1
CL\_INS\_1
CL\_INS\_1
CL\_INS\_1
CL\_INS\_1
CL\_INS\_20
CL\_INS\_20
CL\_INS\_20
CL\_INS\_20
CL\_INS\_20
CL\_INS\_20
CL\_INS\_20
CL\_INS\_20
CL\_INS\_20
CL\_INS\_133
CL\_INS\_20
CL\_INS\_20
CL\_INS\_133
CL\_INS\_20
CL\_INS\_133
CL\_INS\_133
CL\_INS\_133
CL\_INS\_133
CL\_INS\_20
CL\_INS\_133
CL\_INS\_133
CL\_INS\_133
CL\_INS\_133
CL\_INS\_1
CL\_INS\_1
CL\_INS\_1
CL\_INS\_1
CL\_INS\_1
CL\_INS\_1
CL\_INS\_1
CL\_INS\_1
CL\_INS\_1
CL\_INS\_1
CL\_INS\_1
CL\_INS\_1
CL\_INS\_1
CL\_INS\_1
CL\_INS\_1
CL\_INS\_1
CL\_INS\_1
CL\_INS\_133
CL\_INS\_20
CL\_INS\_1
CL\_INS\_20
CL\_INS\_20
CL\_INS\_20
CL\_INS\_20
CL\_INS\_20
CL\_INS\_20
CL\_INS\_20
CL\_INS\_20
CL\_INS\_20
CL\_INS\_1
CL\_INS\_20
CL\_INS\_20
CL\_INS\_20
CL\_INS\_20
CL\_INS\_20
CL\_INS\_20
CL\_INS\_20
CL\_INS\_1
CL\_INS\_20
CL\_INS\_20
CL\_INS\_1
CL\_INS\_20
CL\_INS\_45
CL\_INS\_45
CL\_INS\_20
CL\_INS\_20
CL\_INS\_20
CL\_INS\_20
CL\_INS\_20
CL\_INS\_20
CL\_INS\_237
CL\_INS\_237
CL\_INS\_237
CL\_INS\_237
CL\_INS\_237
CL\_INS\_237
CL\_INS\_237
CL\_INS\_237
CL\_INS\_237
CL\_INS\_237
CL\_INS\_237
CL\_INS\_20
CL\_INS\_237
CL\_INS\_237
CL\_INS\_237
CL\_INS\_237
CL\_INS\_237
CL\_INS\_237
CL\_INS\_237
CL\_INS\_237
CL\_INS\_237
CL\_INS\_237
CL\_INS\_237
CL\_INS\_237
CL\_INS\_237
CL\_INS\_237
CL\_INS\_237
CL\_INS\_237
CL\_INS\_237
CL\_INS\_237
CL\_INS\_237
CL\_INS\_237
CL\_INS\_237
CL\_INS\_237
CL\_INS\_237
CL\_INS\_237
CL\_INS\_237
CL\_INS\_237
CL\_INS\_237
CL\_INS\_237
CL\_INS\_237
CL\_INS\_237
CL\_INS\_237
CL\_INS\_237
CL\_INS\_237
CL\_INS\_237
CL\_INS\_237
CL\_INS\_237
CL\_INS\_237
CL\_INS\_20
CL\_INS\_237
CL\_INS\_237
CL\_INS\_237
CL\_INS\_237
CL\_INS\_237
CL\_INS\_237
CL\_INS\_237
CL\_INS\_237
CL\_INS\_237
CL\_INS\_237
CL\_INS\_237
CL\_INS\_237
CL\_INS\_237
CL\_INS\_237
CL\_INS\_20
CL\_INS\_20
CL\_INS\_20
CL\_INS\_133
CL\_INS\_1
CL\_INS\_1
CL\_INS\_1
CL\_INS\_1
CL\_INS\_1
CL\_INS\_1
CL\_INS\_1
CL\_INS\_20
CL\_INS\_1
CL\_INS\_1
CL\_INS\_1
CL\_INS\_1
CL\_INS\_1
CL\_INS\_1
CL\_INS\_1
CL\_INS\_1
CL\_INS\_1
CL\_INS\_1
CL\_INS\_1
CL\_INS\_1
CL\_INS\_1
CL\_INS\_1
CL\_INS\_1
CL\_INS\_1
CL\_INS\_1
CL\_INS\_1
CL\_INS\_1
CL\_INS\_1
CL\_INS\_1
CL\_INS\_1
CL\_INS\_1
CL\_INS\_1
CL\_INS\_1
CL\_INS\_1
CL\_INS\_1
CL\_INS\_1
CL\_INS\_83
CL\_INS\_83
CL\_INS\_83
CL\_INS\_83
CL\_INS\_83
CL\_INS\_83
CL\_INS\_83
CL\_INS\_1
CL\_INS\_1
CL\_INS\_83
CL\_INS\_83
CL\_INS\_83
CL\_INS\_20
CL\_INS\_20
CL\_INS\_20
CL\_INS\_1
CL\_INS\_1
CL\_INS\_1
CL\_INS\_1
CL\_INS\_1
CL\_INS\_1
CL\_INS\_1
CL\_INS\_1
CL\_INS\_1
CL\_INS\_1
CL\_INS\_1
CL\_INS\_1
CL\_INS\_1
CL\_INS\_1
CL\_INS\_1
CL\_INS\_1
CL\_INS\_1
CL\_INS\_1
CL\_INS\_1
CL\_INS\_1
CL\_INS\_1
CL\_INS\_1
CL\_INS\_1
CL\_INS\_1
CL\_INS\_1
CL\_INS\_1
CL\_INS\_1
CL\_INS\_1
CL\_INS\_1
CL\_INS\_1
CL\_INS\_1
CL\_INS\_1
CL\_INS\_1
CL\_INS\_1
CL\_INS\_1
CL\_INS\_1
CL\_INS\_1
CL\_INS\_1
CL\_INS\_1
CL\_INS\_1
CL\_INS\_1
CL\_INS\_1
CL\_INS\_1
CL\_INS\_1
CL\_INS\_1
CL\_INS\_1
CL\_INS\_1
CL\_INS\_1
CL\_INS\_20
CL\_INS\_1
CL\_INS\_20
CL\_INS\_1
CL\_INS\_1
CL\_INS\_1
CL\_INS\_1
CL\_INS\_1
CL\_INS\_1
CL\_INS\_1
CL\_INS\_153
CL\_INS\_153
CL\_INS\_1
CL\_INS\_1
CL\_INS\_133
CL\_INS\_20
CL\_INS\_20
CL\_INS\_20
CL\_INS\_1
CL\_INS\_1
CL\_INS\_1
CL\_INS\_1
CL\_INS\_1
CL\_INS\_1
CL\_INS\_1
CL\_INS\_1
CL\_INS\_1
CL\_INS\_1
CL\_INS\_1
CL\_INS\_1
CL\_INS\_1
CL\_INS\_1
CL\_INS\_1
CL\_INS\_1
CL\_INS\_20
CL\_INS\_20
CL\_INS\_20
CL\_INS\_1
CL\_INS\_1
CL\_INS\_1
CL\_INS\_1
CL\_INS\_1
CL\_INS\_1
CL\_INS\_20
CL\_INS\_1
CL\_INS\_1
CL\_INS\_1
CL\_INS\_1
CL\_INS\_1
CL\_INS\_1
CL\_INS\_1
CL\_INS\_1
CL\_INS\_1
CL\_INS\_1
CL\_INS\_1
CL\_INS\_1
CL\_INS\_1
CL\_INS\_1
CL\_INS\_1
CL\_INS\_1
CL\_INS\_1
CL\_INS\_1
CL\_INS\_1
CL\_INS\_1
CL\_INS\_1
CL\_INS\_1
CL\_INS\_1
CL\_INS\_1
CL\_INS\_1
CL\_INS\_1
CL\_INS\_1
CL\_INS\_1
CL\_INS\_1
CL\_INS\_1
CL\_INS\_1
CL\_INS\_20
CL\_INS\_1
CL\_INS\_1
CL\_INS\_1
CL\_INS\_1
CL\_INS\_1
CL\_INS\_1
CL\_INS\_1
CL\_INS\_1
CL\_INS\_1
CL\_INS\_1
CL\_INS\_1
CL\_INS\_1
CL\_INS\_1
CL\_INS\_1
CL\_INS\_1
CL\_INS\_1
CL\_INS\_1
CL\_INS\_1
CL\_INS\_1
CL\_INS\_1
CL\_INS\_20
CL\_INS\_1
CL\_INS\_1
CL\_INS\_1
CL\_INS\_1
CL\_INS\_1
CL\_INS\_1
CL\_INS\_1
CL\_INS\_1
CL\_INS\_1
CL\_INS\_1
CL\_INS\_1
CL\_INS\_1
CL\_INS\_1
CL\_INS\_1
CL\_INS\_1
CL\_INS\_1
CL\_INS\_1
CL\_INS\_1
CL\_INS\_1
CL\_INS\_1
CL\_INS\_1
CL\_INS\_1
CL\_INS\_1
CL\_INS\_20
CL\_INS\_1
CL\_INS\_1
CL\_INS\_1
CL\_INS\_1
CL\_INS\_1
CL\_INS\_1
CL\_INS\_1
CL\_INS\_1
CL\_INS\_1
CL\_INS\_1
CL\_INS\_1
CL\_INS\_20
CL\_INS\_1
CL\_INS\_1
CL\_INS\_1
CL\_INS\_1
CL\_INS\_1
CL\_INS\_1
CL\_INS\_1
CL\_INS\_1
CL\_INS\_1
CL\_INS\_1
CL\_INS\_1
CL\_INS\_1
CL\_INS\_1
CL\_INS\_1
CL\_INS\_1
CL\_INS\_1
CL\_INS\_1
CL\_INS\_1
CL\_INS\_1
CL\_INS\_1
CL\_INS\_20
CL\_INS\_1
CL\_INS\_1
CL\_INS\_204
CL\_INS\_1
CL\_INS\_1
CL\_INS\_1
CL\_INS\_20
CL\_INS\_20
CL\_INS\_20
CL\_INS\_20
CL\_INS\_20
CL\_INS\_1
CL\_INS\_1
CL\_INS\_1
CL\_INS\_1
CL\_INS\_1
CL\_INS\_1
CL\_INS\_1
CL\_INS\_1
CL\_INS\_1
CL\_INS\_1
CL\_INS\_1
CL\_INS\_1
CL\_INS\_1
CL\_INS\_1
CL\_INS\_1
CL\_INS\_1
CL\_INS\_1
CL\_INS\_20
CL\_INS\_20
CL\_INS\_20
CL\_INS\_20
CL\_INS\_20
CL\_INS\_20
CL\_INS\_1
CL\_INS\_20
CL\_INS\_20
CL\_INS\_20
CL\_INS\_20
CL\_INS\_20
CL\_INS\_20
CL\_INS\_20
CL\_INS\_20
CL\_INS\_20
CL\_INS\_20
CL\_INS\_20
CL\_INS\_20
CL\_INS\_20
CL\_INS\_20
CL\_INS\_20
CL\_INS\_20
CL\_INS\_20
CL\_INS\_20
CL\_INS\_20
CL\_INS\_20
CL\_INS\_20
CL\_INS\_20
CL\_INS\_20
CL\_INS\_20
CL\_INS\_20
CL\_INS\_20
CL\_INS\_20
CL\_INS\_20
CL\_INS\_20
CL\_INS\_20
CL\_INS\_20
CL\_INS\_20
CL\_INS\_20
CL\_INS\_20
CL\_INS\_20
CL\_INS\_20
CL\_INS\_20
CL\_INS\_20
CL\_INS\_20
CL\_INS\_20
CL\_INS\_20
CL\_INS\_20
CL\_INS\_20
CL\_INS\_20
CL\_INS\_20
CL\_INS\_20
CL\_INS\_20
CL\_INS\_20
CL\_INS\_20
CL\_INS\_20
CL\_INS\_20
CL\_INS\_20
CL\_INS\_20
CL\_INS\_20
CL\_INS\_20
CL\_INS\_20
CL\_INS\_20
CL\_INS\_20
CL\_INS\_20
CL\_INS\_20
CL\_INS\_20
CL\_INS\_20
CL\_INS\_20
CL\_INS\_20
CL\_INS\_20
CL\_INS\_20
CL\_INS\_20
CL\_INS\_20
CL\_INS\_20
CL\_INS\_20
CL\_INS\_20
CL\_INS\_20
CL\_INS\_20
CL\_INS\_20
CL\_INS\_20
CL\_INS\_20
CL\_INS\_20
CL\_INS\_20
CL\_INS\_20
CL\_INS\_20
CL\_INS\_20
CL\_INS\_20
CL\_INS\_20
CL\_INS\_20
CL\_INS\_20
CL\_INS\_20
CL\_INS\_20
CL\_INS\_20
CL\_INS\_20
CL\_INS\_20
CL\_INS\_20
CL\_INS\_20
CL\_INS\_20
CL\_INS\_20
CL\_INS\_20
CL\_INS\_20
CL\_INS\_20
CL\_INS\_20
CL\_INS\_20
CL\_INS\_20
CL\_INS\_20
CL\_INS\_20
CL\_INS\_20
CL\_INS\_20
CL\_INS\_20
CL\_INS\_20
CL\_INS\_20
CL\_INS\_83
CL\_INS\_83
CL\_INS\_20
CL\_INS\_20
CL\_INS\_20
CL\_INS\_20
CL\_INS\_20
CL\_INS\_20
CL\_INS\_20
CL\_INS\_20
CL\_INS\_20
CL\_INS\_20
CL\_INS\_20
CL\_INS\_20
CL\_INS\_20
CL\_INS\_20
CL\_INS\_20
CL\_INS\_20
CL\_INS\_20
CL\_INS\_20
CL\_INS\_20
CL\_INS\_20
CL\_INS\_20
CL\_INS\_20
CL\_INS\_20
CL\_INS\_20
CL\_INS\_20
CL\_INS\_20
CL\_INS\_20
CL\_INS\_20
CL\_INS\_20
CL\_INS\_20
CL\_INS\_20
CL\_INS\_20
CL\_INS\_20
CL\_INS\_20
CL\_INS\_20
CL\_INS\_20
CL\_INS\_20
CL\_INS\_20
CL\_INS\_20
CL\_INS\_20
CL\_INS\_20
CL\_INS\_20
CL\_INS\_20
CL\_INS\_20
CL\_INS\_20
CL\_INS\_20
CL\_INS\_20
CL\_INS\_1
CL\_INS\_20
CL\_INS\_20
CL\_INS\_20
CL\_INS\_20
CL\_INS\_20
CL\_INS\_20
CL\_INS\_20
CL\_INS\_20
CL\_INS\_20
CL\_INS\_20
CL\_INS\_20
CL\_INS\_20
CL\_INS\_20
CL\_INS\_20
CL\_INS\_20
CL\_INS\_20
CL\_INS\_20
CL\_INS\_20
CL\_INS\_20
CL\_INS\_20
CL\_INS\_20
CL\_INS\_20
CL\_INS\_20
CL\_INS\_20
CL\_INS\_20
CL\_INS\_20
CL\_INS\_20
CL\_INS\_20
CL\_INS\_20
CL\_INS\_20
CL\_INS\_20
CL\_INS\_20
CL\_INS\_20
CL\_INS\_20
CL\_INS\_20
CL\_INS\_20
CL\_INS\_20
CL\_INS\_20
CL\_INS\_20
CL\_INS\_20
CL\_INS\_20
CL\_INS\_20
CL\_INS\_20
CL\_INS\_20
CL\_INS\_20
CL\_INS\_20
CL\_INS\_20
CL\_INS\_20
CL\_INS\_20
CL\_INS\_20
CL\_INS\_1
CL\_INS\_20
CL\_INS\_20
CL\_INS\_20
CL\_INS\_20
CL\_INS\_1
CL\_INS\_1
CL\_INS\_20
CL\_INS\_20
CL\_INS\_20
CL\_INS\_20
CL\_INS\_20
CL\_INS\_20
CL\_INS\_20
CL\_INS\_20
CL\_INS\_20
CL\_INS\_20
CL\_INS\_133
CL\_INS\_20
CL\_INS\_133
CL\_INS\_20
CL\_INS\_20
CL\_INS\_1
CL\_INS\_1
CL\_INS\_20
CL\_INS\_1
CL\_INS\_20
CL\_INS\_20
CL\_INS\_20
CL\_INS\_20
CL\_INS\_20
CL\_INS\_20
CL\_INS\_133
CL\_INS\_133
CL\_INS\_133
CL\_INS\_20
CL\_INS\_20
CL\_INS\_20
CL\_INS\_20
CL\_INS\_20
CL\_INS\_20
CL\_INS\_20
CL\_INS\_133
CL\_INS\_20
CL\_INS\_20
CL\_INS\_20
CL\_INS\_20
CL\_INS\_20
CL\_INS\_20
CL\_INS\_20
CL\_INS\_20
CL\_INS\_20
CL\_INS\_20
CL\_INS\_20
CL\_INS\_20
CL\_INS\_20
CL\_INS\_20
CL\_INS\_20
CL\_INS\_20
CL\_INS\_20
CL\_INS\_20
CL\_INS\_20
CL\_INS\_20
CL\_INS\_20
CL\_INS\_20
CL\_INS\_20
CL\_INS\_20
CL\_INS\_20
CL\_INS\_20
CL\_INS\_20
CL\_INS\_20
CL\_INS\_20
CL\_INS\_20
CL\_INS\_20
CL\_INS\_20
CL\_INS\_20
CL\_INS\_20
CL\_INS\_20
CL\_INS\_20
CL\_INS\_20
CL\_INS\_20
CL\_INS\_20
CL\_INS\_20
CL\_INS\_20
CL\_INS\_20
CL\_INS\_20
CL\_INS\_20
CL\_INS\_20
CL\_INS\_20
CL\_INS\_20
CL\_INS\_20
CL\_INS\_20
CL\_INS\_20
CL\_INS\_20
CL\_INS\_20
CL\_INS\_20
CL\_INS\_20
CL\_INS\_20
CL\_INS\_20
CL\_INS\_20
CL\_INS\_20
CL\_INS\_20
CL\_INS\_20
CL\_INS\_20
CL\_INS\_20
CL\_INS\_20
CL\_INS\_20
CL\_INS\_20
CL\_INS\_20
CL\_INS\_20
CL\_INS\_20
CL\_INS\_20
CL\_INS\_20
CL\_INS\_20
CL\_INS\_20
CL\_INS\_20
CL\_INS\_20
CL\_INS\_20
CL\_INS\_20
CL\_INS\_20
CL\_INS\_20
CL\_INS\_20
CL\_INS\_20
CL\_INS\_20
CL\_INS\_20
CL\_INS\_20
CL\_INS\_20
CL\_INS\_20
CL\_INS\_20
CL\_INS\_20
CL\_INS\_20
CL\_INS\_20
CL\_INS\_20
CL\_INS\_20
CL\_INS\_20
CL\_INS\_20
CL\_INS\_20
CL\_INS\_20
CL\_INS\_20
CL\_INS\_20
CL\_INS\_20
CL\_INS\_20
CL\_INS\_20
CL\_INS\_20
CL\_INS\_20
CL\_INS\_20
CL\_INS\_20
CL\_INS\_20
CL\_INS\_20
CL\_INS\_20
CL\_INS\_20
CL\_INS\_20
CL\_INS\_20
CL\_INS\_20
CL\_INS\_20
CL\_INS\_1
CL\_INS\_1
CL\_INS\_20
CL\_INS\_20
CL\_INS\_1
CL\_INS\_20
CL\_INS\_20
CL\_INS\_20
CL\_INS\_20
CL\_INS\_20
CL\_INS\_20
CL\_INS\_20
CL\_INS\_20
CL\_INS\_20
CL\_INS\_20
CL\_INS\_20
CL\_INS\_20
CL\_INS\_20
CL\_INS\_20
CL\_INS\_20
CL\_INS\_20
CL\_INS\_20
CL\_INS\_20
CL\_INS\_20
CL\_INS\_20
CL\_INS\_20
CL\_INS\_20
CL\_INS\_20
CL\_INS\_20
CL\_INS\_20
CL\_INS\_20
CL\_INS\_20
CL\_INS\_20
CL\_INS\_20
CL\_INS\_20
CL\_INS\_20
CL\_INS\_20
CL\_INS\_20
CL\_INS\_1
CL\_INS\_20
CL\_INS\_20
CL\_INS\_20
CL\_INS\_20
CL\_INS\_20
CL\_INS\_20
CL\_INS\_20
CL\_INS\_20
CL\_INS\_20
CL\_INS\_20
CL\_INS\_20
CL\_INS\_20
CL\_INS\_20
CL\_INS\_20
CL\_INS\_20
CL\_INS\_20
CL\_INS\_20
CL\_INS\_20
CL\_INS\_20
CL\_INS\_20
CL\_INS\_20
CL\_INS\_20
CL\_INS\_20
CL\_INS\_20
CL\_INS\_20
CL\_INS\_20
CL\_INS\_20
CL\_INS\_20
CL\_INS\_20
CL\_INS\_20
CL\_INS\_20
CL\_INS\_20
CL\_INS\_20
CL\_INS\_20
CL\_INS\_20
CL\_INS\_20
CL\_INS\_20
CL\_INS\_20
CL\_INS\_20
CL\_INS\_20
CL\_INS\_20
CL\_INS\_20
CL\_INS\_20
CL\_INS\_20
CL\_INS\_20
CL\_INS\_20
CL\_INS\_20
CL\_INS\_20
CL\_INS\_20
Cluster ID


CL\_5144
CL\_5145
CL\_5146
CL\_8901
CL\_8900
CL\_8899
CL\_8898
CL\_523
CL\_8115
CL\_8116
CL\_11926
CL\_11925
CL\_11924
CL\_11923
CL\_11922
CL\_11921
CL\_11920
CL\_11919
CL\_11150
CL\_8118
CL\_6073
CL\_5726
CL\_11293
CL\_6071
CL\_6069
CL\_11294
CL\_11295
CL\_11296
CL\_11297
CL\_11298
CL\_5287
CL\_524
CL\_525
CL\_9811
CL\_526
CL\_8860
CL\_8861
CL\_527
CL\_528
CL\_529
CL\_6959
CL\_10818
CL\_10817
CL\_10816
CL\_10215
CL\_5329
CL\_5330
CL\_5331
CL\_5186
CL\_9170
CL\_9171
CL\_9172
CL\_9173
CL\_9174
CL\_9175
CL\_9176
CL\_9177
CL\_9178
CL\_9142
CL\_5187
CL\_5188
CL\_5189
CL\_4633
CL\_5190
CL\_11918
CL\_11917
CL\_11916
CL\_11915
CL\_11914
CL\_5191
CL\_5192
CL\_5193
CL\_5194
CL\_5195
CL\_10984
CL\_4638
CL\_4639
CL\_4640
CL\_4641
CL\_4642
CL\_4643
CL\_4644
CL\_7634
CL\_10985
CL\_10986
CL\_10987
CL\_10988
CL\_10989
CL\_7635
CL\_7636
CL\_7637
CL\_5328
CL\_10247
CL\_10248
CL\_5198
CL\_5828
CL\_5199
CL\_5200
CL\_4435
CL\_5201
CL\_5889
CL\_5202
CL\_5203
CL\_5204
CL\_5157
CL\_4543
CL\_4542
CL\_4541
CL\_4540
CL\_4539
CL\_4538
CL\_4537
CL\_4536
CL\_4535
CL\_4534
CL\_4533
CL\_4532
CL\_4531
CL\_4530
CL\_4529
CL\_4528
CL\_8997
CL\_5337
CL\_5336
CL\_5837
CL\_5836
CL\_5835
CL\_5834
CL\_5833
CL\_5832
CL\_5831
CL\_5830
CL\_5338
CL\_5339
CL\_5813
CL\_5814
CL\_5815
CL\_5816
CL\_5817
CL\_5818
CL\_5819
CL\_5820
CL\_5812
CL\_5887
CL\_5464
CL\_4595
CL\_7638
CL\_5281
CL\_5282
CL\_5283
CL\_11299
CL\_11300
CL\_11301
CL\_11302
CL\_11303
CL\_5752
CL\_5756
CL\_11151
CL\_5757
CL\_5758
CL\_5759
CL\_5760
CL\_11152
CL\_11153
CL\_11154
CL\_11155
CL\_11304
CL\_5762
CL\_5763
CL\_5764
CL\_11156
CL\_5765
CL\_5766
CL\_5767
CL\_5768
CL\_5769
CL\_5770
CL\_11157
CL\_5771
CL\_5772
CL\_5773
CL\_5774
CL\_5775
CL\_5776
CL\_5777
CL\_5778
CL\_5779
CL\_5780
CL\_11158
CL\_11159
CL\_11160
CL\_11161
CL\_11162
CL\_11163
CL\_11164
CL\_11165
CL\_11166
CL\_11167
CL\_11168
CL\_11169
CL\_5781
CL\_5782
CL\_5783
CL\_5784
CL\_11305
CL\_5785
CL\_11170
CL\_5786
CL\_5787
CL\_5788
CL\_5789
CL\_5790
CL\_11171
CL\_5791
CL\_5792
CL\_5793
CL\_11172
CL\_5794
CL\_5795
CL\_5284
CL\_5285
CL\_5286
CL\_4451
CL\_4659
CL\_4660
CL\_4661
CL\_4662
CL\_4663
CL\_4664
CL\_4665
CL\_7639
CL\_4667
CL\_4668
CL\_4669
CL\_4670
CL\_4671
CL\_4672
CL\_4673
CL\_4674
CL\_4675
CL\_4676
CL\_4677
CL\_4678
CL\_4679
CL\_4680
CL\_4681
CL\_4682
CL\_4683
CL\_5894
CL\_5895
CL\_5896
CL\_4685
CL\_4686
CL\_4687
CL\_4688
CL\_4689
CL\_4690
CL\_4691
CL\_4692
CL\_4693
CL\_4694
CL\_4695
CL\_4696
CL\_4697
CL\_4698
CL\_4699
CL\_4700
CL\_4701
CL\_4702
CL\_4703
CL\_4704
CL\_12523
CL\_7640
CL\_7641
CL\_4527
CL\_4526
CL\_4525
CL\_4524
CL\_4523
CL\_4522
CL\_4521
CL\_4520
CL\_4519
CL\_4518
CL\_4517
CL\_4516
CL\_4515
CL\_4514
CL\_4513
CL\_4512
CL\_4511
CL\_4510
CL\_4509
CL\_4508
CL\_4507
CL\_4506
CL\_4505
CL\_4504
CL\_4503
CL\_4502
CL\_4501
CL\_4500
CL\_4499
CL\_4498
CL\_4497
CL\_4496
CL\_4495
CL\_4494
CL\_4493
CL\_4492
CL\_4491
CL\_4490
CL\_4489
CL\_4488
CL\_4487
CL\_4486
CL\_4480
CL\_4479
CL\_4478
CL\_4477
CL\_4476
CL\_4469
CL\_5159
CL\_5160
CL\_5161
CL\_4457
CL\_4456
CL\_4455
CL\_5321
CL\_4454
CL\_4453
CL\_4452
CL\_5322
CL\_5323
CL\_5324
CL\_5326
CL\_5327
CL\_5224
CL\_5810
CL\_5811
CL\_4707
CL\_4708
CL\_4709
CL\_4710
CL\_4711
CL\_4712
CL\_4713
CL\_4714
CL\_4715
CL\_4716
CL\_4717
CL\_4718
CL\_4719
CL\_4720
CL\_4721
CL\_4722
CL\_7642
CL\_7643
CL\_7644
CL\_4725
CL\_4726
CL\_4727
CL\_4728
CL\_4729
CL\_4730
CL\_7645
CL\_4732
CL\_4733
CL\_4734
CL\_4735
CL\_4736
CL\_4737
CL\_4738
CL\_4739
CL\_4740
CL\_4741
CL\_4742
CL\_4743
CL\_4744
CL\_4745
CL\_4746
CL\_4747
CL\_4748
CL\_4749
CL\_22
CL\_21
CL\_20
CL\_19
CL\_5196
CL\_18
CL\_17
CL\_16
CL\_4750
CL\_4751
CL\_4752
CL\_4753
CL\_12524
CL\_7646
CL\_4386
CL\_4385
CL\_7175
CL\_4384
CL\_4383
CL\_4382
CL\_4381
CL\_4380
CL\_4379
CL\_4378
CL\_4377
CL\_4376
CL\_4375
CL\_4374
CL\_4373
CL\_4372
CL\_4371
CL\_4370
CL\_4369
CL\_4368
CL\_6992
CL\_4367
CL\_4366
CL\_4365
CL\_4364
CL\_4363
CL\_4362
CL\_4361
CL\_4360
CL\_4359
CL\_4358
CL\_4357
CL\_4356
CL\_4355
CL\_4354
CL\_4353
CL\_4352
CL\_4351
CL\_4350
CL\_4349
CL\_4348
CL\_4347
CL\_4346
CL\_4345
CL\_10815
CL\_4344
CL\_4343
CL\_4342
CL\_4341
CL\_4340
CL\_4339
CL\_4338
CL\_4337
CL\_4336
CL\_4335
CL\_4334
CL\_7647
CL\_4388
CL\_4389
CL\_4390
CL\_4391
CL\_4392
CL\_4760
CL\_4394
CL\_4395
CL\_4396
CL\_4397
CL\_4398
CL\_4399
CL\_4400
CL\_4768
CL\_4402
CL\_4403
CL\_4404
CL\_4405
CL\_4406
CL\_4407
CL\_7648
CL\_4409
CL\_4410
CL\_6912
CL\_4414
CL\_10265
CL\_4606
CL\_6082
CL\_6081
CL\_6080
CL\_6079
CL\_6078
CL\_4439
CL\_4438
CL\_6077
CL\_6076
CL\_6075
CL\_6074
CL\_5259
CL\_5260
CL\_5261
CL\_5722
CL\_4422
CL\_4421
CL\_4420
CL\_4419
CL\_4418
CL\_4601
CL\_4602
CL\_12525
CL\_7649
CL\_7650
CL\_7651
CL\_7652
CL\_7653
CL\_4783
CL\_5183
CL\_7945
CL\_7946
CL\_7947
CL\_10095
CL\_7948
CL\_7949
CL\_7950
CL\_7951
CL\_7952
CL\_7953
CL\_7954
CL\_7955
CL\_7956
CL\_7957
CL\_10093
CL\_7959
CL\_7961
CL\_7962
CL\_7963
CL\_7964
CL\_7965
CL\_7966
CL\_7967
CL\_7968
CL\_7969
CL\_7971
CL\_7972
CL\_7973
CL\_10724
CL\_7974
CL\_7976
CL\_7977
CL\_7978
CL\_7979
CL\_13465
CL\_13466
CL\_7912
CL\_10100
CL\_10976
CL\_7910
CL\_13467
CL\_7908
CL\_7907
CL\_7906
CL\_7905
CL\_7904
CL\_7903
CL\_7902
CL\_7901
CL\_7900
CL\_7899
CL\_7970
CL\_8016
CL\_8015
CL\_8014
CL\_8013
CL\_8012
CL\_8011
CL\_8010
CL\_8009
CL\_8008
CL\_8007
CL\_8006
CL\_8005
CL\_8004
CL\_8003
CL\_8002
CL\_8001
CL\_8000
CL\_7999
CL\_7998
CL\_7997
CL\_7996
CL\_7995
CL\_7994
CL\_7993
CL\_7992
CL\_10975
CL\_7991
CL\_7990
CL\_7989
CL\_7988
CL\_7987
CL\_7986
CL\_5557
CL\_7985
CL\_7984
CL\_7983
CL\_7982
CL\_7981
CL\_7980
CL\_10727
CL\_13468
CL\_7915
CL\_7916
CL\_7917
CL\_7918
CL\_7919
CL\_7920
CL\_7921
CL\_7922
CL\_10099
CL\_10097
CL\_7924
CL\_10978
CL\_10979
CL\_6715
CL\_6714
CL\_7925
CL\_7926
CL\_7927
CL\_7929
CL\_7930
CL\_7931
CL\_7932
CL\_7933
CL\_7934
CL\_7935
CL\_7936
CL\_7937
CL\_7938
CL\_7939
CL\_7940
CL\_7941
CL\_7942
CL\_7943
CL\_7944
CL\_10778
CL\_10779
CL\_10780
CL\_10781
CL\_10782
CL\_10783
CL\_10784
CL\_10785
CL\_10786
CL\_10787
CL\_10788
CL\_10789
CL\_10790
CL\_10791
CL\_10792
CL\_10793
CL\_10794
CL\_10795
CL\_10796
CL\_7101
CL\_7102
CL\_10797
CL\_10799
CL\_10800
CL\_10801
CL\_10802
CL\_10803
CL\_10804
CL\_8853
CL\_8390
CL\_8391
CL\_8392
CL\_10294
CL\_12265
CL\_7569
CL\_7570
CL\_7571
CL\_13469
CL\_5350
CL\_5349
CL\_5347
CL\_5346
CL\_5345
CL\_5344
CL\_5343
CL\_5342
CL\_5341
CL\_5340
CL\_5348
CL\_10806
CL\_10807
CL\_10808
CL\_10809
CL\_10810
CL\_10811
CL\_10812
CL\_10813
CL\_10814
CL\_13470
CL\_10768
CL\_10769
CL\_5366
CL\_10770
CL\_10771
CL\_10772
CL\_10773
CL\_10774
CL\_10776
CL\_10777
CL\_10445
CL\_11827
CL\_11828
CL\_5948
CL\_5949
CL\_5950
CL\_5951
CL\_6028
CL\_12526
CL\_10999
CL\_4426
CL\_10254
CL\_10253
CL\_5829
CL\_7590
CL\_4432
CL\_4433
CL\_8388
CL\_5821
CL\_5822
CL\_5823
CL\_5824
CL\_5825
CL\_5826
CL\_5335
CL\_9148
CL\_9149
CL\_5884
CL\_9150
CL\_6967
CL\_8927
CL\_10260
CL\_4434
CL\_6919
CL\_7573
CL\_6916
CL\_10259
CL\_7574
CL\_10258
CL\_7575
CL\_7576
CL\_7577
CL\_7578
CL\_8396
CL\_8397
CL\_8398
CL\_8399
CL\_8400
CL\_8402
CL\_8404
CL\_12527
CL\_6027
CL\_6969
CL\_6968
CL\_9143
CL\_9144
CL\_9145
CL\_8926
CL\_9146
CL\_6026
CL\_6025
CL\_6024
CL\_6023
CL\_6022
CL\_9141
CL\_6021
CL\_6020
CL\_6019
CL\_12528
CL\_12529
CL\_12530
CL\_6018
CL\_6017
CL\_6016
CL\_9140
CL\_9139
CL\_5999
CL\_5998
CL\_12531
CL\_5996
CL\_5997
CL\_5995
CL\_5994
CL\_5993
CL\_5992
CL\_5991
CL\_5990
CL\_5989
CL\_5988
CL\_5987
CL\_5986
CL\_5985
CL\_5984
CL\_5983
CL\_5982
CL\_5981
CL\_5980
CL\_12532
CL\_5979
CL\_5978
CL\_5977
CL\_5976
CL\_5975
CL\_5974
CL\_5973
CL\_5972
CL\_5971
CL\_5970
CL\_5969
CL\_5968
CL\_5967
CL\_5966
CL\_5965
CL\_5964
CL\_5963
CL\_7239
CL\_7191
CL\_7192
CL\_7193
CL\_7194
CL\_7195
CL\_7196
CL\_7197
CL\_7198
CL\_7199
CL\_7200
CL\_7201
CL\_7202
CL\_7203
CL\_7204
CL\_7205
CL\_7206
CL\_5962
CL\_5961
CL\_5960
CL\_5959
CL\_5958
CL\_9147
CL\_5353
CL\_5352
CL\_5351
CL\_5957
CL\_5956
CL\_5955
CL\_5954
CL\_5953
CL\_6001
CL\_6037
CL\_6036
CL\_6035
CL\_7207
CL\_7208
CL\_7209
CL\_7210
CL\_7211
CL\_7212
CL\_7213
CL\_7214
CL\_6034
CL\_6033
CL\_6032
CL\_12533
CL\_6030
CL\_6029
CL\_6031
CL\_5471
CL\_5472
CL\_8018
CL\_6625
CL\_5469
CL\_7215
CL\_7216
CL\_7217
CL\_7218
CL\_7219
CL\_7220
CL\_7221
CL\_7222
CL\_7223
CL\_7224
CL\_7225
CL\_7226
CL\_7227
CL\_7228
CL\_7229
CL\_7230
CL\_7231
CL\_7232
CL\_7233
CL\_7234
CL\_7235
CL\_7236
CL\_7237
CL\_7238
CL\_5877
CL\_5876
CL\_5873
CL\_7240
CL\_5872
CL\_5871
CL\_7241
CL\_7242
CL\_7243
CL\_4666
CL\_7244
CL\_7245
CL\_7246
CL\_7247
CL\_7248
CL\_7249
CL\_7250
CL\_7251
CL\_7252
CL\_7253
CL\_7254
CL\_5890
CL\_7255
CL\_7256
CL\_7257
CL\_7258
CL\_7259
CL\_7260
CL\_7261
CL\_7262
CL\_7263
CL\_7264
CL\_7265
CL\_7266
CL\_7267
CL\_7268
CL\_7269
CL\_7270
CL\_7271
CL\_7272
CL\_7273
CL\_7274
CL\_7275
CL\_7276
CL\_7277
CL\_7278
CL\_7279
CL\_7280
CL\_7281
CL\_7282
CL\_7283
CL\_7284
CL\_7285
CL\_7286
CL\_7287
CL\_7288
CL\_7289
CL\_7290
CL\_7291
