## Supplementary material for "A novel method for integrating genomic and Tn-Seq data to identify common *in vivo* fitness mechanisms across multiple bacterial species": S3 Dataset: CL_INS_21.html

Legend

 Hypothetical
 DNA Metabolism
 Other
 All VFDB Genes

FULL


WINDOWSVGPNG

Trim RowsRemove SingletonsSave Fasta

CL\_584


CL\_584


CL\_584

HighlightSelectShow Genomes


46

CL\_585


5

CL\_596


1

CL\_596

fGI ID


CL\_INS\_21
CL\_INS\_21
Cluster ID


CL\_6615
CL\_594
