## Supplementary material for "A novel method for integrating genomic and Tn-Seq data to identify common *in vivo* fitness mechanisms across multiple bacterial species": S3 Dataset: CL_INS_24.html

Legend

 Other
 EnergyMetabolism
 Transport +binding proteins
 All VFDB Genes

FULL


WINDOWSVGPNG

Trim RowsRemove SingletonsSave Fasta

CL\_636


CL\_636


CL\_636


CL\_636


CL\_635


CL\_636

HighlightSelectShow Genomes


23

CL\_637


15

CL\_637


14

CL\_637


3

CL\_637


1

CL\_637


1

CL\_637

fGI ID


CL\_INS\_24
CL\_INS\_24
CL\_INS\_24
CL\_INS\_24
CL\_INS\_24
CL\_INS\_24
CL\_INS\_24
Cluster ID


CL\_5150
CL\_9397
CL\_9398
CL\_9399
CL\_9400
CL\_9401
CL\_5151
