## Supplementary material for "A novel method for integrating genomic and Tn-Seq data to identify common *in vivo* fitness mechanisms across multiple bacterial species": S3 Dataset: CL_INS_25.html

Legend

 Regulatoryfunctions
 Hypothetical
 All EssentialGenes
 All VFDB Genes

FULL


WINDOWSVGPNG

Trim RowsRemove SingletonsSave Fasta

CL\_642


CL\_642


CL\_642

HighlightSelectShow Genomes


37

CL\_643


19

CL\_643


1

CL\_643

fGI ID


CL\_INS\_25
CL\_INS\_25
Cluster ID


CL\_6617
CL\_9812
