## Supplementary material for "A novel method for integrating genomic and Tn-Seq data to identify common *in vivo* fitness mechanisms across multiple bacterial species": S3 Dataset: CL_INS_27.html

Legend

 Hypothetical
 Other
 All VFDB Genes

FULL


WINDOWSVGPNG

Trim RowsRemove SingletonsSave Fasta

CL\_671


CL\_671


CL\_671

HighlightSelectShow Genomes


38

CL\_672


18

CL\_672


1

CL\_673

fGI ID

CL\_INS\_27
Cluster ID

CL\_6618
