## Supplementary material for "A novel method for integrating genomic and Tn-Seq data to identify common *in vivo* fitness mechanisms across multiple bacterial species": S3 Dataset: CL_INS_28.html

Legend

 Hypothetical
 Other
 Transport +binding proteins
 All VFDB Genes

FULL


WINDOWSVGPNG

Trim RowsRemove SingletonsSave Fasta

CL\_680


CL\_680


CL\_680


CL\_680

HighlightSelectShow Genomes


51

CL\_681


3

CL\_684


1

CL\_683


1

CL\_681

fGI ID


CL\_INS\_28
CL\_INS\_28
CL\_INS\_29
CL\_INS\_29
CL\_INS\_29
Cluster ID


CL\_11781
CL\_5152
CL\_5153
CL\_5154
CL\_5155
