## Supplementary material for "A novel method for integrating genomic and Tn-Seq data to identify common *in vivo* fitness mechanisms across multiple bacterial species": S3 Dataset: CL_INS_29.html

Legend

 Other
 All VFDB Genes

FULL


WINDOWSVGPNG

Trim RowsRemove SingletonsSave Fasta

CL\_684


CL\_684

HighlightSelectShow Genomes


53

CL\_683


3

CL\_680

fGI ID


CL\_INS\_29
CL\_INS\_29
CL\_INS\_29
CL\_INS\_29
Cluster ID


CL\_5156
CL\_5155
CL\_5154
CL\_5153
