## Supplementary material for "A novel method for integrating genomic and Tn-Seq data to identify common *in vivo* fitness mechanisms across multiple bacterial species": S3 Dataset: CL_INS_31.html

Legend

 Regulatoryfunctions
 Hypothetical
 Other
 All VFDB Genes

FULL


WINDOWSVGPNG

Trim RowsRemove SingletonsSave Fasta

CL\_699


CL\_699


CL\_699


CL\_699


CL\_699


CL\_699

HighlightSelectShow Genomes


35

CL\_702


17

CL\_702


2

CL\_703


1

CL\_703


1

Break


1

Break

fGI ID


CL\_INS\_31
CL\_INS\_31
CL\_INS\_31
CL\_INS\_31
CL\_INS\_31
CL\_INS\_31
CL\_INS\_31
Cluster ID


CL\_11208
CL\_11207
CL\_11206
CL\_700
CL\_701
CL\_6965
CL\_6966
