## Supplementary material for "A novel method for integrating genomic and Tn-Seq data to identify common *in vivo* fitness mechanisms across multiple bacterial species": S3 Dataset: CL_INS_33.html

Legend

 Hypothetical
 Other
 All VFDB Genes

FULL


WINDOWSVGPNG

Trim RowsRemove SingletonsSave Fasta

CL\_722


CL\_722


CL\_722

HighlightSelectShow Genomes


34

CL\_723


19

CL\_723


1

Break

fGI ID

CL\_INS\_33
Cluster ID

CL\_10157
