## Supplementary material for "A novel method for integrating genomic and Tn-Seq data to identify common *in vivo* fitness mechanisms across multiple bacterial species": S3 Dataset: CL_INS_34.html

Legend

 Mobile +extrachromosomalelementfunctions
 Regulatoryfunctions
 Hypothetical
 Other
 Transport +binding proteins
 All VFDB Genes

FULL


WINDOWSVGPNG

Trim RowsRemove SingletonsSave Fasta

CL\_758


CL\_758


CL\_758


CL\_758


CL\_758


CL\_758


CL\_758


CL\_758

HighlightSelectShow Genomes


23

CL\_759


18

CL\_759


7

Break


1

CL\_759


1

CL\_759


1

CL\_759


1

CL\_759


1

CL\_759

fGI ID


CL\_INS\_34
CL\_INS\_34
CL\_INS\_34
CL\_INS\_34
CL\_INS\_34
CL\_INS\_34
CL\_INS\_34
CL\_INS\_20
CL\_INS\_20
CL\_INS\_20
CL\_INS\_20
CL\_INS\_20
CL\_INS\_45
CL\_INS\_45
CL\_INS\_45
CL\_INS\_34
CL\_INS\_34
CL\_INS\_34
CL\_INS\_34
CL\_INS\_1
CL\_INS\_34
CL\_INS\_34
CL\_INS\_34
CL\_INS\_1
CL\_INS\_1
CL\_INS\_1
CL\_INS\_1
CL\_INS\_34
Cluster ID


CL\_5802
CL\_5803
CL\_11204
CL\_5804
CL\_6621
CL\_6622
CL\_13658
CL\_5287
CL\_5286
CL\_5285
CL\_5284
CL\_5283
CL\_5282
CL\_5281
CL\_5280
CL\_5279
CL\_5278
CL\_5277
CL\_6086
CL\_4594
CL\_6088
CL\_6089
CL\_6090
CL\_4440
CL\_4442
CL\_4443
CL\_4444
CL\_5805
