## Supplementary material for "A novel method for integrating genomic and Tn-Seq data to identify common *in vivo* fitness mechanisms across multiple bacterial species": S3 Dataset: CL_INS_35.html

Legend

 Regulatoryfunctions
 Hypothetical
 All EssentialGenes
 Other
 All VFDB Genes

FULL


WINDOWSVGPNG

Trim RowsRemove SingletonsSave Fasta

CL\_778


CL\_778


CL\_778

HighlightSelectShow Genomes


42

CL\_779


8

CL\_781


6

CL\_779

fGI ID


CL\_INS\_35
CL\_INS\_35
CL\_INS\_35
CL\_INS\_36
CL\_INS\_36
CL\_INS\_36
CL\_INS\_36
Cluster ID


CL\_4785
CL\_4786
CL\_6626
CL\_6627
CL\_6628
CL\_6629
CL\_6630
