## Supplementary material for "A novel method for integrating genomic and Tn-Seq data to identify common *in vivo* fitness mechanisms across multiple bacterial species": S3 Dataset: CL_INS_36.html

Legend

 Hypothetical
 All EssentialGenes
 Other
 All VFDB Genes

FULL


WINDOWSVGPNG

Trim RowsRemove SingletonsSave Fasta

CL\_781


CL\_781

HighlightSelectShow Genomes


46

CL\_780


8

CL\_778

fGI ID


CL\_INS\_36
CL\_INS\_36
CL\_INS\_36
CL\_INS\_36
CL\_INS\_36
Cluster ID


CL\_6631
CL\_6630
CL\_6629
CL\_6628
CL\_6627
