## Supplementary material for "A novel method for integrating genomic and Tn-Seq data to identify common *in vivo* fitness mechanisms across multiple bacterial species": S3 Dataset: CL_INS_37.html

Legend

 All EssentialGenes
 Other
 Transport +binding proteins

FULL


WINDOWSVGPNG

Trim RowsRemove SingletonsSave Fasta

CL\_791


CL\_791

HighlightSelectShow Genomes


54

CL\_792


3

CL\_792

fGI ID


CL\_INS\_37
CL\_INS\_37
CL\_INS\_37
CL\_INS\_37
Cluster ID


CL\_6632
CL\_6633
CL\_6634
CL\_6635
