## Supplementary material for "A novel method for integrating genomic and Tn-Seq data to identify common *in vivo* fitness mechanisms across multiple bacterial species": S3 Dataset: CL_INS_38.html

Legend

 Other
 All VFDB Genes

FULL


WINDOWSVGPNG

Trim RowsRemove SingletonsSave Fasta

CL\_794


CL\_794


CL\_794

HighlightSelectShow Genomes


51

CL\_795


5

CL\_795


1

CL\_795

fGI ID


CL\_INS\_38
CL\_INS\_38
CL\_INS\_38
Cluster ID


CL\_7292
CL\_7293
CL\_7294
