## Supplementary material for "A novel method for integrating genomic and Tn-Seq data to identify common *in vivo* fitness mechanisms across multiple bacterial species": S3 Dataset: CL_INS_41.html

Legend

 Hypothetical
 All EssentialGenes
 Other
 All VFDB Genes

FULL


WINDOWSVGPNG

Trim RowsRemove SingletonsSave Fasta

CL\_803


CL\_803


CL\_803


CL\_803


CL\_803


CL\_803


CL\_803


CL\_803

HighlightSelectShow Genomes


21

CL\_807


19

CL\_807


7

CL\_807


5

CL\_807


3

CL\_807


1

CL\_809


1

CL\_807


1

CL\_808

fGI ID


CL\_INS\_41
CL\_INS\_41
CL\_INS\_41
CL\_INS\_41
CL\_INS\_41
Cluster ID


CL\_5475
CL\_5476
CL\_804
CL\_805
CL\_806
