## Supplementary material for "A novel method for integrating genomic and Tn-Seq data to identify common *in vivo* fitness mechanisms across multiple bacterial species": S3 Dataset: CL_INS_42.html

Legend

 Hypothetical
 Transcription
 All EssentialGenes
 All VFDB Genes
 Transport +binding proteins

FULL


WINDOWSVGPNG

Trim RowsRemove SingletonsSave Fasta

CL\_854


CL\_854


CL\_854


CL\_854

HighlightSelectShow Genomes


49

CL\_855


5

CL\_857


1

CL\_857


1

CL\_857

fGI ID


CL\_INS\_42
CL\_INS\_42
CL\_INS\_42
CL\_INS\_42
Cluster ID


CL\_9065
CL\_9581
CL\_9580
CL\_9064
