## Supplementary material for "A novel method for integrating genomic and Tn-Seq data to identify common *in vivo* fitness mechanisms across multiple bacterial species": S3 Dataset: CL_INS_43.html

Legend

 Hypothetical
 Transcription
 Transport +binding proteins
 All VFDB Genes

FULL


WINDOWSVGPNG

Trim RowsRemove SingletonsSave Fasta

CL\_856


CL\_856


CL\_854


CL\_854


CL\_854

HighlightSelectShow Genomes


23

CL\_857


21

CL\_857


5

CL\_857


1

CL\_857


1

CL\_857

fGI ID


CL\_INS\_43
CL\_INS\_43
CL\_INS\_42
CL\_INS\_42
CL\_INS\_43
Cluster ID


CL\_6052
CL\_6053
CL\_9580
CL\_9064
CL\_9063
