## Supplementary material for "A novel method for integrating genomic and Tn-Seq data to identify common *in vivo* fitness mechanisms across multiple bacterial species": S3 Dataset: CL_INS_44.html

Legend

 Mobile +extrachromosomalelementfunctions
 Regulatoryfunctions
 Hypothetical
 DNA Metabolism
 Transcription
 Biosynthesis ofcofactors,prostheticgroups, +carriers
 Other
 Transport +binding proteins
 All VFDB Genes

FULL


WINDOWSVGPNG

Trim RowsRemove SingletonsSave Fasta

CL\_870


CL\_870


CL\_870


CL\_870


CL\_870


CL\_870


CL\_870


CL\_870

HighlightSelectShow Genomes


23

CL\_876


11

CL\_876


8

CL\_876


7

CL\_876


3

CL\_876


1

CL\_876


1

CL\_879


1

CL\_876

fGI ID


CL\_INS\_44
CL\_INS\_44
CL\_INS\_44
CL\_INS\_44
CL\_INS\_46
CL\_INS\_44
CL\_INS\_44
CL\_INS\_44
CL\_INS\_45
CL\_INS\_45
CL\_INS\_45
CL\_INS\_1
CL\_INS\_1
CL\_INS\_1
CL\_INS\_1
CL\_INS\_1
CL\_INS\_1
CL\_INS\_1
CL\_INS\_1
CL\_INS\_1
CL\_INS\_1
CL\_INS\_1
CL\_INS\_45
Cluster ID


CL\_4787
CL\_4788
CL\_11314
CL\_871
CL\_872
CL\_873
CL\_874
CL\_875
CL\_5282
CL\_5281
CL\_5280
CL\_4689
CL\_4688
CL\_4687
CL\_4686
CL\_4685
CL\_4684
CL\_4683
CL\_4682
CL\_4681
CL\_4680
CL\_4679
CL\_5487
