## Supplementary material for "A novel method for integrating genomic and Tn-Seq data to identify common *in vivo* fitness mechanisms across multiple bacterial species": S3 Dataset: CL_INS_45.html

Legend

 Mobile +extrachromosomalelementfunctions
 Regulatoryfunctions
 Hypothetical
 DNA Metabolism
 Transcription
 Biosynthesis ofcofactors,prostheticgroups, +carriers
 Proteinsynthesis/fate
 Other
 All VFDB Genes
 Transport +binding proteins

FULL


WINDOWSVGPNG

Trim RowsRemove SingletonsSave Fasta

CL\_876


CL\_876


CL\_876


CL\_876


CL\_876


CL\_876


CL\_876


CL\_851


CL\_876


CL\_876

HighlightSelectShow Genomes


42

CL\_877


5

CL\_879


1

CL\_879


1

CL\_879


1

CL\_879


1

CL\_879


1

CL\_879


1

CL\_877


1

CL\_879


1

CL\_879

fGI ID


CL\_INS\_45
CL\_INS\_45
CL\_INS\_45
CL\_INS\_45
CL\_INS\_45
CL\_INS\_45
CL\_INS\_83
CL\_INS\_83
CL\_INS\_45
CL\_INS\_45
CL\_INS\_83
CL\_INS\_83
CL\_INS\_83
CL\_INS\_83
CL\_INS\_83
CL\_INS\_83
CL\_INS\_83
CL\_INS\_45
CL\_INS\_83
CL\_INS\_45
CL\_INS\_83
CL\_INS\_45
CL\_INS\_45
CL\_INS\_45
CL\_INS\_45
CL\_INS\_1
CL\_INS\_1
CL\_INS\_1
CL\_INS\_1
CL\_INS\_1
CL\_INS\_1
CL\_INS\_1
CL\_INS\_1
CL\_INS\_1
CL\_INS\_1
CL\_INS\_1
CL\_INS\_45
CL\_INS\_45
CL\_INS\_45
CL\_INS\_45
CL\_INS\_83
CL\_INS\_83
CL\_INS\_83
CL\_INS\_45
CL\_INS\_83
CL\_INS\_45
CL\_INS\_45
CL\_INS\_83
CL\_INS\_45
CL\_INS\_45
CL\_INS\_45
CL\_INS\_45
CL\_INS\_45
CL\_INS\_45
CL\_INS\_45
CL\_INS\_45
CL\_INS\_45
CL\_INS\_45
CL\_INS\_45
CL\_INS\_45
CL\_INS\_1
CL\_INS\_1
CL\_INS\_1
CL\_INS\_45
CL\_INS\_45
CL\_INS\_45
CL\_INS\_1
CL\_INS\_45
CL\_INS\_45
CL\_INS\_45
CL\_INS\_45
CL\_INS\_45
CL\_INS\_45
CL\_INS\_45
CL\_INS\_45
CL\_INS\_45
Cluster ID


CL\_5477
CL\_10571
CL\_7654
CL\_5478
CL\_5479
CL\_5480
CL\_3942
CL\_3941
CL\_10416
CL\_10869
CL\_90
CL\_9616
CL\_91
CL\_92
CL\_5481
CL\_5482
CL\_5483
CL\_5484
CL\_5485
CL\_12534
CL\_6649
CL\_5486
CL\_5282
CL\_5281
CL\_5280
CL\_4689
CL\_4688
CL\_4687
CL\_4686
CL\_4685
CL\_4684
CL\_4683
CL\_4682
CL\_4681
CL\_4680
CL\_4679
CL\_11834
CL\_11835
CL\_5487
CL\_5488
CL\_6154
CL\_6153
CL\_9615
CL\_10870
CL\_6150
CL\_10871
CL\_10872
CL\_6148
CL\_10608
CL\_10873
CL\_7665
CL\_7664
CL\_7663
CL\_7662
CL\_7661
CL\_7660
CL\_7659
CL\_7658
CL\_7657
CL\_7656
CL\_5896
CL\_5895
CL\_5894
CL\_10874
CL\_10875
CL\_10876
CL\_5838
CL\_10877
CL\_10878
CL\_10879
CL\_10880
CL\_10881
CL\_10882
CL\_10883
CL\_10884
CL\_10885
