## Supplementary material for "A novel method for integrating genomic and Tn-Seq data to identify common *in vivo* fitness mechanisms across multiple bacterial species": S3 Dataset: CL_INS_46.html

Legend

 Mobile +extrachromosomalelementfunctions
 Regulatoryfunctions
 Hypothetical
 DNA Metabolism
 Transcription
 Biosynthesis ofcofactors,prostheticgroups, +carriers
 Proteinsynthesis/fate
 Other
 Transport +binding proteins
 All VFDB Genes

FULL


WINDOWSVGPNG

Trim RowsRemove SingletonsSave Fasta

CL\_877


CL\_877


CL\_876


CL\_877


CL\_877


CL\_877


CL\_877


CL\_877


CL\_877


CL\_877


CL\_866


CL\_877


CL\_876


CL\_877


CL\_823


CL\_876


CL\_877


CL\_877


CL\_877


CL\_866


CL\_876


CL\_876


CL\_876


CL\_877


CL\_876


CL\_876


CL\_870


CL\_877

HighlightSelectShow Genomes


12

CL\_879


6

CL\_879


5

CL\_879


4

CL\_879


3

CL\_879


3

CL\_879


2

CL\_879


2

CL\_879


2

CL\_879


2

CL\_879


1

CL\_879


1

CL\_821


1

CL\_879


1

CL\_879


1

CL\_879


1

CL\_879


1

CL\_879


1

CL\_886


1

CL\_879


1

CL\_879


1

CL\_879


1

CL\_879


1

CL\_879


1

CL\_879


1

CL\_879


1

CL\_879


1

CL\_879


1

CL\_879

fGI ID


CL\_INS\_46
CL\_INS\_46
CL\_INS\_46
CL\_INS\_46
CL\_INS\_46
CL\_INS\_46
CL\_INS\_46
CL\_INS\_45
CL\_INS\_46
CL\_INS\_45
CL\_INS\_46
CL\_INS\_46
CL\_INS\_46
CL\_INS\_46
CL\_INS\_46
CL\_INS\_46
CL\_INS\_46
CL\_INS\_46
CL\_INS\_46
CL\_INS\_65
CL\_INS\_65
CL\_INS\_65
CL\_INS\_65
CL\_INS\_46
CL\_INS\_46
CL\_INS\_46
CL\_INS\_46
CL\_INS\_46
CL\_INS\_46
CL\_INS\_46
CL\_INS\_46
CL\_INS\_46
CL\_INS\_46
CL\_INS\_46
CL\_INS\_46
CL\_INS\_46
CL\_INS\_83
CL\_INS\_46
CL\_INS\_83
CL\_INS\_83
CL\_INS\_46
CL\_INS\_46
CL\_INS\_220
CL\_INS\_1
CL\_INS\_46
CL\_INS\_46
CL\_INS\_45
CL\_INS\_45
CL\_INS\_45
CL\_INS\_83
CL\_INS\_83
CL\_INS\_45
CL\_INS\_45
CL\_INS\_83
CL\_INS\_83
CL\_INS\_83
CL\_INS\_83
CL\_INS\_83
CL\_INS\_83
CL\_INS\_83
CL\_INS\_46
CL\_INS\_45
CL\_INS\_83
CL\_INS\_45
CL\_INS\_45
CL\_INS\_83
CL\_INS\_45
CL\_INS\_45
CL\_INS\_45
CL\_INS\_1
CL\_INS\_1
CL\_INS\_1
CL\_INS\_1
CL\_INS\_1
CL\_INS\_1
CL\_INS\_1
CL\_INS\_1
CL\_INS\_1
CL\_INS\_1
CL\_INS\_1
CL\_INS\_45
CL\_INS\_45
CL\_INS\_20
CL\_INS\_46
CL\_INS\_45
CL\_INS\_45
CL\_INS\_83
CL\_INS\_83
CL\_INS\_83
CL\_INS\_45
CL\_INS\_83
CL\_INS\_45
CL\_INS\_45
CL\_INS\_83
CL\_INS\_45
CL\_INS\_45
CL\_INS\_45
CL\_INS\_45
CL\_INS\_45
CL\_INS\_45
CL\_INS\_45
CL\_INS\_45
CL\_INS\_45
CL\_INS\_45
CL\_INS\_45
CL\_INS\_45
CL\_INS\_1
CL\_INS\_1
CL\_INS\_1
CL\_INS\_45
CL\_INS\_45
CL\_INS\_45
CL\_INS\_1
CL\_INS\_45
CL\_INS\_45
CL\_INS\_45
CL\_INS\_45
CL\_INS\_45
CL\_INS\_45
CL\_INS\_45
CL\_INS\_45
CL\_INS\_45
CL\_INS\_46
Cluster ID


CL\_13333
CL\_9275
CL\_7295
CL\_9402
CL\_12321
CL\_9925
CL\_9926
CL\_10571
CL\_10572
CL\_7654
CL\_7655
CL\_9179
CL\_9180
CL\_11030
CL\_11031
CL\_11032
CL\_9579
CL\_9584
CL\_9585
CL\_8264
CL\_8265
CL\_8266
CL\_8267
CL\_9586
CL\_9587
CL\_9588
CL\_8583
CL\_6054
CL\_9589
CL\_9590
CL\_9591
CL\_872
CL\_11415
CL\_878
CL\_8592
CL\_8591
CL\_8590
CL\_8589
CL\_8588
CL\_8587
CL\_8586
CL\_8585
CL\_8584
CL\_89
CL\_9578
CL\_9577
CL\_5478
CL\_5479
CL\_5480
CL\_3942
CL\_3941
CL\_10416
CL\_10869
CL\_90
CL\_9616
CL\_91
CL\_92
CL\_5481
CL\_5482
CL\_5483
CL\_11414
CL\_5484
CL\_5485
CL\_12534
CL\_5486
CL\_6649
CL\_5282
CL\_5281
CL\_5280
CL\_4689
CL\_4688
CL\_4687
CL\_4686
CL\_4685
CL\_4684
CL\_4683
CL\_4682
CL\_4681
CL\_4680
CL\_4679
CL\_11834
CL\_11835
CL\_8118
CL\_8117
CL\_5487
CL\_5488
CL\_6154
CL\_6153
CL\_9615
CL\_10870
CL\_6150
CL\_10871
CL\_10872
CL\_6148
CL\_10608
CL\_10873
CL\_7665
CL\_7664
CL\_7663
CL\_7662
CL\_7661
CL\_7660
CL\_7659
CL\_7658
CL\_7657
CL\_7656
CL\_5896
CL\_5895
CL\_5894
CL\_10874
CL\_10875
CL\_10876
CL\_5838
CL\_10877
CL\_10878
CL\_10879
CL\_10880
CL\_10881
CL\_10882
CL\_10883
CL\_10884
CL\_10885
CL\_5489
