## Supplementary material for "A novel method for integrating genomic and Tn-Seq data to identify common *in vivo* fitness mechanisms across multiple bacterial species": S3 Dataset: CL_INS_49.html

Legend

 Mobile +extrachromosomalelementfunctions
 Hypothetical
 Transcription
 Other

FULL


WINDOWSVGPNG

Trim RowsRemove SingletonsSave Fasta

CL\_1043


CL\_1043


CL\_1043

HighlightSelectShow Genomes


31

CL\_1046


25

CL\_1046


1

CL\_1046

fGI ID


CL\_INS\_49
CL\_INS\_49
CL\_INS\_49
CL\_INS\_49
CL\_INS\_49
CL\_INS\_49
Cluster ID


CL\_1044
CL\_1045
CL\_6059
CL\_6060
CL\_6663
CL\_6664
