## Supplementary material for "A novel method for integrating genomic and Tn-Seq data to identify common *in vivo* fitness mechanisms across multiple bacterial species": S3 Dataset: CL_INS_50.html

Legend

 Hypothetical
 Biosynthesis ofcofactors,prostheticgroups, +carriers
 Other
 Transport +binding proteins

FULL


WINDOWSVGPNG

Trim RowsRemove SingletonsSave Fasta

CL\_1061


CL\_1061

HighlightSelectShow Genomes


47

CL\_1062


9

CL\_1062

fGI ID


CL\_INS\_50
CL\_INS\_50
Cluster ID


CL\_8339
CL\_8338
