## Supplementary material for "A novel method for integrating genomic and Tn-Seq data to identify common *in vivo* fitness mechanisms across multiple bacterial species": S3 Dataset: CL_INS_52.html

Legend

 Mobile +extrachromosomalelementfunctions
 Hypothetical
 All EssentialGenes
 Other
 All VFDB Genes

FULL


WINDOWSVGPNG

Trim RowsRemove SingletonsSave Fasta

CL\_1082


CL\_1082


CL\_1082


CL\_1082

HighlightSelectShow Genomes


53

CL\_1083


2

CL\_1083


1

CL\_1083


1

CL\_1083

fGI ID


CL\_INS\_52
CL\_INS\_52
CL\_INS\_52
CL\_INS\_52
CL\_INS\_52
CL\_INS\_52
CL\_INS\_52
CL\_INS\_52
CL\_INS\_52
CL\_INS\_52
CL\_INS\_52
CL\_INS\_52
CL\_INS\_52
CL\_INS\_52
CL\_INS\_52
CL\_INS\_52
CL\_INS\_52
CL\_INS\_52
CL\_INS\_52
CL\_INS\_52
CL\_INS\_52
CL\_INS\_52
CL\_INS\_52
CL\_INS\_52
Cluster ID


CL\_11313
CL\_11312
CL\_11311
CL\_11310
CL\_11836
CL\_11837
CL\_11838
CL\_11309
CL\_11308
CL\_11307
CL\_11306
CL\_13138
CL\_13139
CL\_13140
CL\_13141
CL\_13142
CL\_13143
CL\_13144
CL\_13145
CL\_13146
CL\_13147
CL\_13148
CL\_13149
CL\_13150
