## Supplementary material for "A novel method for integrating genomic and Tn-Seq data to identify common *in vivo* fitness mechanisms across multiple bacterial species": S3 Dataset: CL_INS_53.html

Legend

 Regulatoryfunctions
 Hypothetical
 DNA Metabolism
 Other
 All VFDB Genes

FULL


WINDOWSVGPNG

Trim RowsRemove SingletonsSave Fasta

CL\_1099


CL\_1099


CL\_1098


CL\_1099


CL\_1099


CL\_1099

HighlightSelectShow Genomes


40

CL\_1100


13

CL\_1100


1

CL\_1100


1

CL\_1100


1

CL\_1115


1

CL\_1138

fGI ID


CL\_INS\_53
CL\_INS\_53
CL\_INS\_53
CL\_INS\_53
CL\_INS\_53
CL\_INS\_53
CL\_INS\_53
CL\_INS\_53
CL\_INS\_53
CL\_INS\_53
CL\_INS\_53
CL\_INS\_53
CL\_INS\_53
CL\_INS\_53
CL\_INS\_53
CL\_INS\_53
CL\_INS\_53
CL\_INS\_53
CL\_INS\_53
CL\_INS\_53
CL\_INS\_53
Cluster ID


CL\_12163
CL\_5247
CL\_12162
CL\_10148
CL\_6558
CL\_6557
CL\_6556
CL\_6555
CL\_6554
CL\_6553
CL\_6552
CL\_6551
CL\_6550
CL\_6549
CL\_6548
CL\_6547
CL\_6546
CL\_6545
CL\_6544
CL\_6543
CL\_10147
