## Supplementary material for "A novel method for integrating genomic and Tn-Seq data to identify common *in vivo* fitness mechanisms across multiple bacterial species": S3 Dataset: CL_INS_54.html

Legend

 All EssentialGenes
 Other
 All VFDB Genes

FULL


WINDOWSVGPNG

Trim RowsRemove SingletonsSave Fasta

CL\_1107


CL\_1107


CL\_1107


CL\_1107


CL\_1099


CL\_1107

HighlightSelectShow Genomes


27

CL\_1115


26

CL\_1115


1

Break


1

CL\_1115


1

CL\_1115


1

CL\_1115

fGI ID


CL\_INS\_53
CL\_INS\_54
CL\_INS\_54
CL\_INS\_54
CL\_INS\_54
CL\_INS\_54
CL\_INS\_54
CL\_INS\_54
Cluster ID


CL\_12162
CL\_1108
CL\_1109
CL\_1110
CL\_1111
CL\_1112
CL\_1113
CL\_1114
