## Supplementary material for "A novel method for integrating genomic and Tn-Seq data to identify common *in vivo* fitness mechanisms across multiple bacterial species": S3 Dataset: CL_INS_55.html

Legend

 Mobile +extrachromosomalelementfunctions
 Regulatoryfunctions
 Hypothetical
 AntibioticResistance
 Other
 Transport +binding proteins
 All VFDB Genes

FULL


WINDOWSVGPNG

Trim RowsRemove SingletonsSave Fasta

CL\_1136


CL\_1136


CL\_1136


Break


CL\_1136


CL\_1136


CL\_1136


CL\_1135

HighlightSelectShow Genomes


27

CL\_1137


11

CL\_1137


5

CL\_300


1

CL\_1137


1

CL\_1138


1

CL\_1138


1

CL\_1138


1

CL\_1137

fGI ID


CL\_INS\_55
CL\_INS\_55
CL\_INS\_55
CL\_INS\_55
CL\_INS\_55
CL\_INS\_55
CL\_INS\_55
CL\_INS\_20
CL\_INS\_1
CL\_INS\_55
CL\_INS\_55
CL\_INS\_55
CL\_INS\_55
CL\_INS\_133
CL\_INS\_55
CL\_INS\_55
CL\_INS\_55
CL\_INS\_237
CL\_INS\_237
CL\_INS\_237
CL\_INS\_237
CL\_INS\_237
CL\_INS\_237
CL\_INS\_237
CL\_INS\_237
CL\_INS\_237
CL\_INS\_237
CL\_INS\_55
CL\_INS\_55
CL\_INS\_237
CL\_INS\_237
CL\_INS\_55
CL\_INS\_204
CL\_INS\_1
CL\_INS\_1
CL\_INS\_1
CL\_INS\_20
CL\_INS\_20
CL\_INS\_55
CL\_INS\_55
CL\_INS\_237
CL\_INS\_237
CL\_INS\_237
CL\_INS\_237
CL\_INS\_237
CL\_INS\_237
CL\_INS\_237
CL\_INS\_237
CL\_INS\_237
CL\_INS\_237
CL\_INS\_237
CL\_INS\_55
CL\_INS\_55
CL\_INS\_83
CL\_INS\_55
CL\_INS\_55
CL\_INS\_55
CL\_INS\_55
CL\_INS\_55
CL\_INS\_55
CL\_INS\_55
CL\_INS\_55
CL\_INS\_55
CL\_INS\_55
CL\_INS\_55
CL\_INS\_55
CL\_INS\_55
CL\_INS\_55
CL\_INS\_55
CL\_INS\_55
CL\_INS\_55
CL\_INS\_55
CL\_INS\_55
CL\_INS\_55
CL\_INS\_55
CL\_INS\_55
CL\_INS\_55
CL\_INS\_55
CL\_INS\_55
CL\_INS\_55
Cluster ID


CL\_10622
CL\_9592
CL\_9593
CL\_6624
CL\_6623
CL\_11832
CL\_13713
CL\_8926
CL\_8109
CL\_5806
CL\_5807
CL\_5808
CL\_5251
CL\_7578
CL\_6084
CL\_13712
CL\_13711
CL\_5724
CL\_6093
CL\_6094
CL\_6095
CL\_6096
CL\_6097
CL\_6098
CL\_6099
CL\_6100
CL\_12388
CL\_13710
CL\_6101
CL\_5710
CL\_5709
CL\_13709
CL\_6912
CL\_5469
CL\_5471
CL\_5472
CL\_8018
CL\_6625
CL\_13708
CL\_13707
CL\_5796
CL\_5795
CL\_5794
CL\_5793
CL\_5792
CL\_5791
CL\_5790
CL\_8937
CL\_5789
CL\_5788
CL\_5787
CL\_12535
CL\_12536
CL\_9198
CL\_12537
CL\_12538
CL\_12539
CL\_12540
CL\_12541
CL\_12542
CL\_12543
CL\_12544
CL\_12545
CL\_12546
CL\_12547
CL\_12548
CL\_12549
CL\_12550
CL\_12551
CL\_12552
CL\_12553
CL\_12554
CL\_12555
CL\_12556
CL\_12557
CL\_12558
CL\_12559
CL\_12560
CL\_12561
CL\_12562
