## Supplementary material for "A novel method for integrating genomic and Tn-Seq data to identify common *in vivo* fitness mechanisms across multiple bacterial species": S3 Dataset: CL_INS_56.html

Legend

 Mobile +extrachromosomalelementfunctions
 Regulatoryfunctions
 Hypothetical
 DNA Metabolism
 AntibioticResistance
 All EssentialGenes
 Biosynthesis ofcofactors,prostheticgroups, +carriers
 Proteinsynthesis/fate
 Other
 EnergyMetabolism
 Transport +binding proteins
 All VFDB Genes

FULL


WINDOWSVGPNG

Trim RowsRemove SingletonsSave Fasta

CL\_1137


CL\_512


CL\_1137


CL\_301


CL\_1137


CL\_1136


CL\_1136


CL\_1099


CL\_1136


CL\_1137


CL\_1137


CL\_1137


CL\_1137


CL\_1137

HighlightSelectShow Genomes


38

CL\_1138


2

CL\_1138


2

CL\_522


1

CL\_1138


1

CL\_1138


1

CL\_1138


1

CL\_1138


1

CL\_1138


1

CL\_1138


1

CL\_1138


1

CL\_1138


1

CL\_1138


1

CL\_1138


1

CL\_731

fGI ID


CL\_INS\_55
CL\_INS\_56
CL\_INS\_53
CL\_INS\_53
CL\_INS\_53
CL\_INS\_53
CL\_INS\_53
CL\_INS\_53
CL\_INS\_53
CL\_INS\_53
CL\_INS\_53
CL\_INS\_53
CL\_INS\_53
CL\_INS\_53
CL\_INS\_53
CL\_INS\_53
CL\_INS\_53
CL\_INS\_53
CL\_INS\_53
CL\_INS\_237
CL\_INS\_237
CL\_INS\_237
CL\_INS\_83
CL\_INS\_83
CL\_INS\_83
CL\_INS\_83
CL\_INS\_83
CL\_INS\_83
CL\_INS\_1
CL\_INS\_56
CL\_INS\_56
CL\_INS\_56
CL\_INS\_56
CL\_INS\_56
CL\_INS\_56
CL\_INS\_56
CL\_INS\_56
CL\_INS\_56
CL\_INS\_83
CL\_INS\_56
CL\_INS\_56
CL\_INS\_56
CL\_INS\_83
CL\_INS\_56
CL\_INS\_56
CL\_INS\_56
CL\_INS\_56
CL\_INS\_56
CL\_INS\_56
CL\_INS\_56
CL\_INS\_56
CL\_INS\_56
CL\_INS\_56
CL\_INS\_56
CL\_INS\_56
CL\_INS\_56
CL\_INS\_56
CL\_INS\_56
CL\_INS\_56
CL\_INS\_56
CL\_INS\_237
CL\_INS\_1
CL\_INS\_1
CL\_INS\_83
CL\_INS\_83
CL\_INS\_1
CL\_INS\_1
CL\_INS\_1
CL\_INS\_1
CL\_INS\_1
CL\_INS\_1
CL\_INS\_1
CL\_INS\_56
CL\_INS\_56
CL\_INS\_56
CL\_INS\_56
CL\_INS\_56
CL\_INS\_1
CL\_INS\_1
CL\_INS\_1
CL\_INS\_1
CL\_INS\_1
CL\_INS\_1
CL\_INS\_1
CL\_INS\_1
CL\_INS\_56
CL\_INS\_1
CL\_INS\_1
CL\_INS\_1
CL\_INS\_1
CL\_INS\_1
CL\_INS\_1
CL\_INS\_56
CL\_INS\_1
CL\_INS\_1
CL\_INS\_1
CL\_INS\_1
CL\_INS\_1
CL\_INS\_1
CL\_INS\_1
CL\_INS\_1
CL\_INS\_1
CL\_INS\_1
CL\_INS\_1
CL\_INS\_56
CL\_INS\_56
CL\_INS\_56
CL\_INS\_56
CL\_INS\_1
CL\_INS\_1
CL\_INS\_1
CL\_INS\_237
CL\_INS\_237
CL\_INS\_237
CL\_INS\_237
CL\_INS\_237
CL\_INS\_237
CL\_INS\_237
CL\_INS\_237
CL\_INS\_237
CL\_INS\_237
CL\_INS\_237
CL\_INS\_55
CL\_INS\_55
CL\_INS\_83
CL\_INS\_55
CL\_INS\_55
CL\_INS\_55
CL\_INS\_55
CL\_INS\_55
CL\_INS\_55
CL\_INS\_55
CL\_INS\_55
CL\_INS\_55
CL\_INS\_55
CL\_INS\_55
CL\_INS\_55
CL\_INS\_55
CL\_INS\_55
CL\_INS\_55
CL\_INS\_55
CL\_INS\_55
CL\_INS\_55
CL\_INS\_55
CL\_INS\_55
CL\_INS\_55
CL\_INS\_55
CL\_INS\_55
CL\_INS\_55
CL\_INS\_55
CL\_INS\_55
CL\_INS\_178
CL\_INS\_178
CL\_INS\_178
CL\_INS\_178
CL\_INS\_237
CL\_INS\_237
CL\_INS\_20
CL\_INS\_237
CL\_INS\_237
CL\_INS\_237
CL\_INS\_237
CL\_INS\_237
CL\_INS\_237
CL\_INS\_237
CL\_INS\_237
CL\_INS\_237
CL\_INS\_237
CL\_INS\_237
CL\_INS\_56
CL\_INS\_237
CL\_INS\_237
CL\_INS\_237
CL\_INS\_237
CL\_INS\_237
CL\_INS\_237
CL\_INS\_237
CL\_INS\_237
CL\_INS\_237
CL\_INS\_237
CL\_INS\_237
CL\_INS\_237
CL\_INS\_20
CL\_INS\_56
CL\_INS\_56
CL\_INS\_237
CL\_INS\_56
CL\_INS\_56
CL\_INS\_1
CL\_INS\_56
CL\_INS\_237
CL\_INS\_237
CL\_INS\_237
CL\_INS\_237
CL\_INS\_237
CL\_INS\_237
CL\_INS\_56
CL\_INS\_56
CL\_INS\_56
CL\_INS\_56
CL\_INS\_56
CL\_INS\_56
CL\_INS\_56
CL\_INS\_56
CL\_INS\_56
CL\_INS\_56
CL\_INS\_56
CL\_INS\_237
CL\_INS\_237
CL\_INS\_237
CL\_INS\_237
CL\_INS\_20
CL\_INS\_20
CL\_INS\_20
CL\_INS\_20
CL\_INS\_20
CL\_INS\_237
CL\_INS\_237
CL\_INS\_237
CL\_INS\_1
CL\_INS\_1
CL\_INS\_1
CL\_INS\_1
CL\_INS\_1
CL\_INS\_1
CL\_INS\_1
CL\_INS\_1
CL\_INS\_1
CL\_INS\_1
CL\_INS\_237
CL\_INS\_1
CL\_INS\_1
CL\_INS\_1
CL\_INS\_1
CL\_INS\_1
CL\_INS\_1
CL\_INS\_1
CL\_INS\_1
CL\_INS\_1
CL\_INS\_1
CL\_INS\_237
CL\_INS\_237
CL\_INS\_237
CL\_INS\_237
CL\_INS\_237
CL\_INS\_237
CL\_INS\_237
CL\_INS\_237
CL\_INS\_237
CL\_INS\_237
CL\_INS\_20
CL\_INS\_56
CL\_INS\_56
CL\_INS\_237
CL\_INS\_20
CL\_INS\_56
CL\_INS\_237
CL\_INS\_56
CL\_INS\_56
CL\_INS\_56
CL\_INS\_56
CL\_INS\_56
CL\_INS\_237
CL\_INS\_237
CL\_INS\_20
CL\_INS\_20
CL\_INS\_56
CL\_INS\_56
CL\_INS\_56
CL\_INS\_56
CL\_INS\_56
CL\_INS\_56
CL\_INS\_56
CL\_INS\_56
CL\_INS\_56
CL\_INS\_56
CL\_INS\_56
CL\_INS\_56
CL\_INS\_56
CL\_INS\_56
CL\_INS\_56
CL\_INS\_1
CL\_INS\_1
CL\_INS\_1
CL\_INS\_1
CL\_INS\_1
CL\_INS\_1
CL\_INS\_20
CL\_INS\_20
CL\_INS\_20
CL\_INS\_20
CL\_INS\_20
CL\_INS\_56
CL\_INS\_20
CL\_INS\_1
CL\_INS\_20
CL\_INS\_20
CL\_INS\_1
CL\_INS\_56
CL\_INS\_56
CL\_INS\_56
CL\_INS\_1
CL\_INS\_1
CL\_INS\_1
CL\_INS\_56
CL\_INS\_56
CL\_INS\_1
CL\_INS\_1
CL\_INS\_1
CL\_INS\_1
CL\_INS\_56
CL\_INS\_1
CL\_INS\_56
CL\_INS\_1
CL\_INS\_56
CL\_INS\_56
CL\_INS\_20
CL\_INS\_20
CL\_INS\_20
CL\_INS\_56
CL\_INS\_56
CL\_INS\_20
CL\_INS\_20
CL\_INS\_1
CL\_INS\_1
CL\_INS\_56
CL\_INS\_56
CL\_INS\_56
CL\_INS\_56
CL\_INS\_56
CL\_INS\_1
CL\_INS\_83
CL\_INS\_83
CL\_INS\_237
CL\_INS\_83
CL\_INS\_83
CL\_INS\_83
CL\_INS\_83
CL\_INS\_83
CL\_INS\_56
CL\_INS\_1
CL\_INS\_1
CL\_INS\_1
CL\_INS\_1
CL\_INS\_1
CL\_INS\_1
CL\_INS\_20
CL\_INS\_20
CL\_INS\_20
CL\_INS\_56
CL\_INS\_20
CL\_INS\_20
CL\_INS\_20
CL\_INS\_20
CL\_INS\_20
CL\_INS\_20
CL\_INS\_20
CL\_INS\_20
CL\_INS\_56
CL\_INS\_20
CL\_INS\_20
CL\_INS\_20
CL\_INS\_20
CL\_INS\_20
CL\_INS\_20
CL\_INS\_20
CL\_INS\_20
CL\_INS\_20
CL\_INS\_20
CL\_INS\_20
CL\_INS\_20
CL\_INS\_20
CL\_INS\_20
CL\_INS\_20
CL\_INS\_20
CL\_INS\_20
CL\_INS\_20
CL\_INS\_20
CL\_INS\_20
CL\_INS\_20
CL\_INS\_20
CL\_INS\_20
CL\_INS\_20
CL\_INS\_20
CL\_INS\_20
CL\_INS\_20
CL\_INS\_20
CL\_INS\_20
CL\_INS\_20
CL\_INS\_20
CL\_INS\_20
CL\_INS\_20
CL\_INS\_20
CL\_INS\_20
CL\_INS\_20
CL\_INS\_20
CL\_INS\_20
CL\_INS\_20
CL\_INS\_133
CL\_INS\_20
CL\_INS\_20
CL\_INS\_20
CL\_INS\_20
CL\_INS\_1
CL\_INS\_1
CL\_INS\_1
CL\_INS\_55
CL\_INS\_1
CL\_INS\_1
CL\_INS\_56
CL\_INS\_20
CL\_INS\_20
CL\_INS\_20
CL\_INS\_20
CL\_INS\_20
CL\_INS\_20
CL\_INS\_20
CL\_INS\_20
CL\_INS\_20
CL\_INS\_20
CL\_INS\_45
CL\_INS\_45
CL\_INS\_45
CL\_INS\_34
CL\_INS\_34
CL\_INS\_34
CL\_INS\_34
CL\_INS\_1
CL\_INS\_156
CL\_INS\_34
CL\_INS\_34
CL\_INS\_34
CL\_INS\_1
CL\_INS\_1
CL\_INS\_1
CL\_INS\_1
CL\_INS\_1
CL\_INS\_56
CL\_INS\_56
CL\_INS\_237
CL\_INS\_237
CL\_INS\_237
CL\_INS\_237
CL\_INS\_237
CL\_INS\_237
CL\_INS\_237
CL\_INS\_237
CL\_INS\_237
CL\_INS\_1
CL\_INS\_1
CL\_INS\_1
CL\_INS\_1
CL\_INS\_1
CL\_INS\_1
CL\_INS\_1
CL\_INS\_1
CL\_INS\_1
CL\_INS\_1
CL\_INS\_56
CL\_INS\_1
CL\_INS\_1
CL\_INS\_1
CL\_INS\_1
CL\_INS\_1
CL\_INS\_56
CL\_INS\_237
CL\_INS\_237
CL\_INS\_237
CL\_INS\_237
CL\_INS\_237
CL\_INS\_237
CL\_INS\_237
CL\_INS\_237
CL\_INS\_237
CL\_INS\_237
CL\_INS\_55
CL\_INS\_237
CL\_INS\_237
CL\_INS\_237
CL\_INS\_56
CL\_INS\_56
CL\_INS\_56
CL\_INS\_56
CL\_INS\_56
CL\_INS\_56
CL\_INS\_56
CL\_INS\_56
CL\_INS\_56
CL\_INS\_56
CL\_INS\_56
CL\_INS\_56
CL\_INS\_56
CL\_INS\_56
CL\_INS\_56
CL\_INS\_56
CL\_INS\_56
CL\_INS\_56
CL\_INS\_56
CL\_INS\_56
CL\_INS\_56
CL\_INS\_56
CL\_INS\_56
CL\_INS\_56
CL\_INS\_56
CL\_INS\_56
CL\_INS\_56
CL\_INS\_56
CL\_INS\_133
CL\_INS\_56
CL\_INS\_133
CL\_INS\_237
CL\_INS\_237
CL\_INS\_133
CL\_INS\_133
Cluster ID


CL\_9593
CL\_9594
CL\_6558
CL\_6557
CL\_6556
CL\_6555
CL\_6554
CL\_6553
CL\_6552
CL\_6551
CL\_6550
CL\_6549
CL\_6548
CL\_6547
CL\_6546
CL\_6545
CL\_6544
CL\_6543
CL\_10147
CL\_6542
CL\_510
CL\_509
CL\_508
CL\_507
CL\_506
CL\_505
CL\_504
CL\_503
CL\_8853
CL\_11052
CL\_11053
CL\_11054
CL\_11055
CL\_11056
CL\_11057
CL\_11058
CL\_11059
CL\_11060
CL\_6150
CL\_11061
CL\_11062
CL\_9284
CL\_5485
CL\_9283
CL\_11063
CL\_11064
CL\_11065
CL\_11066
CL\_11067
CL\_11068
CL\_11069
CL\_11070
CL\_11071
CL\_11072
CL\_11073
CL\_11074
CL\_11075
CL\_11076
CL\_11077
CL\_13000
CL\_10038
CL\_10039
CL\_89
CL\_3942
CL\_3941
CL\_3940
CL\_6155
CL\_3939
CL\_3938
CL\_3937
CL\_3936
CL\_10040
CL\_13001
CL\_13002
CL\_13003
CL\_13004
CL\_13005
CL\_10041
CL\_10042
CL\_10043
CL\_10044
CL\_10045
CL\_10046
CL\_10047
CL\_10048
CL\_13006
CL\_10049
CL\_10050
CL\_10051
CL\_10052
CL\_10053
CL\_10054
CL\_13007
CL\_10055
CL\_10056
CL\_10057
CL\_10058
CL\_10059
CL\_10060
CL\_10061
CL\_10062
CL\_10063
CL\_10064
CL\_5105
CL\_13008
CL\_13009
CL\_8949
CL\_13010
CL\_10037
CL\_10036
CL\_10035
CL\_5796
CL\_5795
CL\_5794
CL\_5793
CL\_5792
CL\_5791
CL\_5790
CL\_8937
CL\_5789
CL\_5788
CL\_5787
CL\_12535
CL\_12536
CL\_9198
CL\_12537
CL\_12538
CL\_12539
CL\_12540
CL\_12541
CL\_12542
CL\_12543
CL\_12544
CL\_12545
CL\_12546
CL\_12547
CL\_12548
CL\_12549
CL\_12550
CL\_12551
CL\_12552
CL\_12553
CL\_12554
CL\_12555
CL\_12556
CL\_12557
CL\_12558
CL\_12559
CL\_12560
CL\_12561
CL\_12562
CL\_6061
CL\_6062
CL\_6063
CL\_6064
CL\_5786
CL\_5785
CL\_11305
CL\_5784
CL\_5783
CL\_5782
CL\_5781
CL\_5780
CL\_5779
CL\_5778
CL\_5777
CL\_5776
CL\_5775
CL\_5774
CL\_10585
CL\_5773
CL\_5772
CL\_5771
CL\_5770
CL\_5769
CL\_5768
CL\_5767
CL\_5766
CL\_5765
CL\_5764
CL\_5763
CL\_5762
CL\_11304
CL\_10586
CL\_10587
CL\_5761
CL\_6065
CL\_6066
CL\_6067
CL\_6068
CL\_5760
CL\_5759
CL\_5758
CL\_5757
CL\_5756
CL\_5755
CL\_10588
CL\_10589
CL\_10590
CL\_10591
CL\_10592
CL\_10593
CL\_10594
CL\_10595
CL\_10596
CL\_10597
CL\_10598
CL\_5754
CL\_5162
CL\_5753
CL\_5752
CL\_11303
CL\_11302
CL\_11301
CL\_11300
CL\_11299
CL\_5751
CL\_5750
CL\_5749
CL\_5748
CL\_10034
CL\_10033
CL\_10032
CL\_10031
CL\_10030
CL\_10029
CL\_10028
CL\_10027
CL\_10026
CL\_10025
CL\_5747
CL\_5746
CL\_5745
CL\_5744
CL\_5743
CL\_5742
CL\_5741
CL\_5740
CL\_5739
CL\_5738
CL\_5737
CL\_5736
CL\_5735
CL\_5734
CL\_5733
CL\_5732
CL\_5731
CL\_5730
CL\_5729
CL\_5728
CL\_6069
CL\_12987
CL\_6070
CL\_6071
CL\_11293
CL\_6072
CL\_5726
CL\_10599
CL\_10600
CL\_10601
CL\_10602
CL\_10603
CL\_10604
CL\_5724
CL\_6073
CL\_8118
CL\_11078
CL\_11079
CL\_11080
CL\_11081
CL\_11082
CL\_11083
CL\_11084
CL\_11085
CL\_11086
CL\_11087
CL\_11088
CL\_11089
CL\_11090
CL\_11091
CL\_11092
CL\_6074
CL\_6075
CL\_6076
CL\_6077
CL\_4438
CL\_4439
CL\_6078
CL\_6079
CL\_6080
CL\_6081
CL\_6082
CL\_6083
CL\_5189
CL\_4633
CL\_5190
CL\_5890
CL\_4640
CL\_13706
CL\_13705
CL\_5893
CL\_5894
CL\_5895
CL\_5896
CL\_5314
CL\_5315
CL\_4644
CL\_4659
CL\_4660
CL\_4661
CL\_5898
CL\_4665
CL\_5899
CL\_4667
CL\_5869
CL\_5870
CL\_5871
CL\_5872
CL\_5873
CL\_5874
CL\_5875
CL\_5876
CL\_5877
CL\_4674
CL\_4675
CL\_5878
CL\_5879
CL\_5880
CL\_5881
CL\_5882
CL\_4531
CL\_4693
CL\_4694
CL\_5727
CL\_4695
CL\_4696
CL\_4697
CL\_4698
CL\_4699
CL\_13704
CL\_5222
CL\_5223
CL\_5468
CL\_5467
CL\_5466
CL\_5465
CL\_5964
CL\_5965
CL\_5966
CL\_13703
CL\_5967
CL\_5968
CL\_5969
CL\_5970
CL\_5971
CL\_5972
CL\_5973
CL\_5974
CL\_13702
CL\_5976
CL\_5977
CL\_5978
CL\_5979
CL\_5980
CL\_5981
CL\_5982
CL\_5983
CL\_5984
CL\_5985
CL\_5986
CL\_5987
CL\_5988
CL\_5989
CL\_5990
CL\_5991
CL\_5992
CL\_5993
CL\_5994
CL\_5995
CL\_5996
CL\_5997
CL\_5998
CL\_5999
CL\_9139
CL\_9140
CL\_6016
CL\_6017
CL\_6018
CL\_6019
CL\_6020
CL\_6021
CL\_9141
CL\_6022
CL\_6023
CL\_6024
CL\_6025
CL\_6026
CL\_6027
CL\_6969
CL\_6968
CL\_9143
CL\_9144
CL\_9145
CL\_4422
CL\_4601
CL\_4602
CL\_6084
CL\_4605
CL\_4609
CL\_6085
CL\_5287
CL\_11298
CL\_11297
CL\_11296
CL\_11295
CL\_11294
CL\_5286
CL\_5285
CL\_5284
CL\_5283
CL\_5282
CL\_5281
CL\_5280
CL\_5279
CL\_5278
CL\_5277
CL\_6086
CL\_4594
CL\_6087
CL\_6088
CL\_6089
CL\_6090
CL\_4440
CL\_4442
CL\_4443
CL\_4444
CL\_4606
CL\_6091
CL\_6092
CL\_6093
CL\_6094
CL\_6095
CL\_6096
CL\_6097
CL\_6098
CL\_6099
CL\_10605
CL\_10606
CL\_8938
CL\_8939
CL\_6918
CL\_8940
CL\_8941
CL\_8942
CL\_8943
CL\_8944
CL\_8945
CL\_8946
CL\_8947
CL\_4683
CL\_4682
CL\_4681
CL\_4680
CL\_4679
CL\_8948
CL\_5721
CL\_5720
CL\_5719
CL\_5718
CL\_5717
CL\_5716
CL\_5715
CL\_5714
CL\_5713
CL\_6100
CL\_6101
CL\_5710
CL\_5709
CL\_5708
CL\_6102
CL\_6103
CL\_6104
CL\_6105
CL\_6106
CL\_6107
CL\_6108
CL\_6109
CL\_6110
CL\_6111
CL\_6112
CL\_6113
CL\_6114
CL\_6115
CL\_6116
CL\_6117
CL\_6118
CL\_6119
CL\_6120
CL\_6121
CL\_6122
CL\_6123
CL\_6124
CL\_6125
CL\_6126
CL\_6127
CL\_6128
CL\_5205
CL\_5204
CL\_13701
CL\_5202
CL\_5712
CL\_5711
CL\_5201
CL\_4435
