## Supplementary material for "A novel method for integrating genomic and Tn-Seq data to identify common *in vivo* fitness mechanisms across multiple bacterial species": S3 Dataset: CL_INS_58.html

Legend

 Mobile +extrachromosomalelementfunctions
 Hypothetical
 Other
 All VFDB Genes

FULL


WINDOWSVGPNG

Trim RowsRemove SingletonsSave Fasta

CL\_1212


CL\_1212


CL\_1212

HighlightSelectShow Genomes


54

CL\_1213


2

CL\_1213


1

CL\_1213

fGI ID


CL\_INS\_58
CL\_INS\_58
CL\_INS\_58
CL\_INS\_58
CL\_INS\_58
CL\_INS\_58
Cluster ID


CL\_9403
CL\_9404
CL\_9405
CL\_9406
CL\_12563
CL\_9407
