## Supplementary material for "A novel method for integrating genomic and Tn-Seq data to identify common *in vivo* fitness mechanisms across multiple bacterial species": S3 Dataset: CL_INS_59.html

Legend

 Hypothetical
 All VFDB Genes

FULL


WINDOWSVGPNG

Trim RowsRemove SingletonsSave Fasta

CL\_1219


CL\_1219


CL\_1219

HighlightSelectShow Genomes


34

CL\_1220


16

CL\_1220


7

CL\_1220

fGI ID


CL\_INS\_230
CL\_INS\_230
CL\_INS\_59
Cluster ID


CL\_5490
CL\_5491
CL\_4793
