## Supplementary material for "A novel method for integrating genomic and Tn-Seq data to identify common *in vivo* fitness mechanisms across multiple bacterial species": S3 Dataset: CL_INS_60.html

Legend

 Other
 All VFDB Genes

FULL


WINDOWSVGPNG

Trim RowsRemove SingletonsSave Fasta

CL\_1220


CL\_1220


CL\_1220


CL\_1220


CL\_1220

HighlightSelectShow Genomes


33

CL\_1227


20

CL\_1227


2

CL\_1227


1

CL\_1227


1

CL\_1227

fGI ID


CL\_INS\_60
CL\_INS\_60
CL\_INS\_60
CL\_INS\_60
CL\_INS\_60
CL\_INS\_60
CL\_INS\_60
Cluster ID


CL\_6667
CL\_1221
CL\_1222
CL\_1223
CL\_1224
CL\_1225
CL\_1226
