## Supplementary material for "A novel method for integrating genomic and Tn-Seq data to identify common *in vivo* fitness mechanisms across multiple bacterial species": S3 Dataset: CL_INS_61.html

Legend

 Mobile +extrachromosomalelementfunctions
 Regulatoryfunctions
 Hypothetical
 Other
 All VFDB Genes
 Transport +binding proteins

FULL


WINDOWSVGPNG

Trim RowsRemove SingletonsSave Fasta

CL\_1254


CL\_1254


CL\_1254


CL\_1254


CL\_1254

HighlightSelectShow Genomes


27

CL\_1260


22

CL\_1260


6

CL\_1260


1

CL\_1260


1

CL\_1260

fGI ID


CL\_INS\_61
CL\_INS\_61
CL\_INS\_61
CL\_INS\_61
CL\_INS\_61
CL\_INS\_61
Cluster ID


CL\_1255
CL\_1256
CL\_1257
CL\_1258
CL\_11291
CL\_1259
