## Supplementary material for "A novel method for integrating genomic and Tn-Seq data to identify common *in vivo* fitness mechanisms across multiple bacterial species": S3 Dataset: CL_INS_62.html

Legend

 Hypothetical
 Transport +binding proteins
 All VFDB Genes

FULL


WINDOWSVGPNG

Trim RowsRemove SingletonsSave Fasta

CL\_1261


CL\_1261


CL\_1261


CL\_1261


CL\_1261

HighlightSelectShow Genomes


37

CL\_1262


12

CL\_1262


3

CL\_1262


2

CL\_1262


1

CL\_1262

fGI ID


CL\_INS\_62
CL\_INS\_62
CL\_INS\_62
CL\_INS\_62
Cluster ID


CL\_6130
CL\_6131
CL\_6132
CL\_6133
