## Supplementary material for "A novel method for integrating genomic and Tn-Seq data to identify common *in vivo* fitness mechanisms across multiple bacterial species": S3 Dataset: CL_INS_63.html

Legend

 Hypothetical
 All VFDB Genes

FULL


WINDOWSVGPNG

Trim RowsRemove SingletonsSave Fasta

CL\_1280


CL\_1280


CL\_1280

HighlightSelectShow Genomes


22

CL\_1283


20

CL\_1283


15

CL\_1283

fGI ID


CL\_INS\_63
CL\_INS\_63
Cluster ID


CL\_1281
CL\_1282
