## Supplementary material for "A novel method for integrating genomic and Tn-Seq data to identify common *in vivo* fitness mechanisms across multiple bacterial species": S3 Dataset: CL_INS_64.html

Legend

 Mobile +extrachromosomalelementfunctions
 Hypothetical
 Regulatoryfunctions
 DNA Metabolism
 Transcription
 All EssentialGenes
 Proteinsynthesis/fate
 Other
 Transport +binding proteins
 All VFDB Genes

FULL


WINDOWSVGPNG

Trim RowsRemove SingletonsSave Fasta

CL\_1286


CL\_1286


CL\_1286


CL\_1286


CL\_1286


CL\_1286


CL\_1286


CL\_1286


CL\_1286


CL\_1286


CL\_1286


CL\_1286


CL\_1286


CL\_1286


CL\_1286


CL\_1286


CL\_1286


CL\_1286


CL\_1286


CL\_1286


CL\_1286


CL\_1286


CL\_1286


CL\_1286


CL\_1286


CL\_1286


CL\_1286


CL\_1286


CL\_1286


CL\_1286


CL\_1286


CL\_1286


CL\_1286


CL\_1286


CL\_1286


CL\_1286


CL\_1286


CL\_1286


CL\_1286


CL\_1286


CL\_1286


CL\_1286


CL\_1286

HighlightSelectShow Genomes


4

CL\_1293


4

CL\_1293


3

CL\_1294


3

CL\_1293


2

CL\_1293


2

CL\_1293


2

CL\_1294


2

CL\_1293


1

CL\_1293


1

CL\_1293


1

CL\_1293


1

CL\_1293


1

CL\_1293


1

CL\_1293


1

CL\_1293


1

CL\_1293


1

CL\_1296


1

CL\_1293


1

CL\_1293


1

CL\_1293


1

CL\_1293


1

CL\_1293


1

CL\_1293


1

CL\_1293


1

CL\_1294


1

CL\_1293


1

CL\_1294


1

CL\_1293


1

CL\_1294


1

CL\_1293


1

CL\_1293


1

CL\_1293


1

CL\_1293


1

CL\_1294


1

CL\_1294


1

CL\_1294


1

CL\_1293


1

CL\_1293


1

CL\_1294


1

CL\_1294


1

CL\_1293


1

CL\_1293


1

CL\_1303

fGI ID


CL\_INS\_64
CL\_INS\_64
CL\_INS\_64
CL\_INS\_64
CL\_INS\_64
CL\_INS\_64
CL\_INS\_64
CL\_INS\_64
CL\_INS\_64
CL\_INS\_64
CL\_INS\_64
CL\_INS\_64
CL\_INS\_64
CL\_INS\_64
CL\_INS\_65
CL\_INS\_65
CL\_INS\_64
CL\_INS\_64
CL\_INS\_64
CL\_INS\_64
CL\_INS\_64
CL\_INS\_64
CL\_INS\_64
CL\_INS\_64
CL\_INS\_64
CL\_INS\_64
CL\_INS\_64
CL\_INS\_64
CL\_INS\_64
CL\_INS\_64
CL\_INS\_64
CL\_INS\_64
CL\_INS\_64
CL\_INS\_64
CL\_INS\_64
CL\_INS\_64
CL\_INS\_64
CL\_INS\_64
CL\_INS\_64
CL\_INS\_64
CL\_INS\_64
CL\_INS\_64
CL\_INS\_64
CL\_INS\_64
CL\_INS\_64
CL\_INS\_64
CL\_INS\_83
CL\_INS\_64
CL\_INS\_64
CL\_INS\_64
CL\_INS\_64
CL\_INS\_64
CL\_INS\_64
CL\_INS\_64
CL\_INS\_64
CL\_INS\_64
CL\_INS\_64
CL\_INS\_65
CL\_INS\_65
CL\_INS\_64
CL\_INS\_64
CL\_INS\_64
CL\_INS\_64
CL\_INS\_64
CL\_INS\_83
CL\_INS\_64
CL\_INS\_230
CL\_INS\_230
CL\_INS\_230
CL\_INS\_64
CL\_INS\_230
CL\_INS\_230
CL\_INS\_64
CL\_INS\_65
CL\_INS\_65
CL\_INS\_65
CL\_INS\_65
CL\_INS\_65
CL\_INS\_64
CL\_INS\_64
CL\_INS\_83
CL\_INS\_83
CL\_INS\_83
CL\_INS\_83
CL\_INS\_65
CL\_INS\_230
CL\_INS\_83
CL\_INS\_64
CL\_INS\_64
CL\_INS\_65
CL\_INS\_83
CL\_INS\_64
CL\_INS\_64
CL\_INS\_64
CL\_INS\_64
CL\_INS\_64
CL\_INS\_64
CL\_INS\_230
CL\_INS\_64
CL\_INS\_64
CL\_INS\_64
CL\_INS\_64
CL\_INS\_237
CL\_INS\_64
CL\_INS\_237
CL\_INS\_237
CL\_INS\_64
CL\_INS\_64
CL\_INS\_64
CL\_INS\_230
CL\_INS\_230
CL\_INS\_83
CL\_INS\_230
CL\_INS\_64
CL\_INS\_64
CL\_INS\_64
CL\_INS\_64
CL\_INS\_64
CL\_INS\_64
CL\_INS\_64
CL\_INS\_64
CL\_INS\_64
CL\_INS\_64
CL\_INS\_64
CL\_INS\_64
CL\_INS\_64
CL\_INS\_64
CL\_INS\_220
CL\_INS\_64
CL\_INS\_64
CL\_INS\_64
CL\_INS\_64
CL\_INS\_64
CL\_INS\_64
CL\_INS\_64
CL\_INS\_64
CL\_INS\_64
CL\_INS\_83
CL\_INS\_64
CL\_INS\_64
CL\_INS\_64
CL\_INS\_64
CL\_INS\_64
CL\_INS\_64
CL\_INS\_64
CL\_INS\_64
CL\_INS\_56
CL\_INS\_64
CL\_INS\_64
CL\_INS\_64
CL\_INS\_64
CL\_INS\_64
CL\_INS\_64
CL\_INS\_64
CL\_INS\_64
CL\_INS\_64
CL\_INS\_64
CL\_INS\_64
CL\_INS\_64
CL\_INS\_64
CL\_INS\_64
CL\_INS\_64
CL\_INS\_64
CL\_INS\_64
CL\_INS\_64
CL\_INS\_64
CL\_INS\_64
CL\_INS\_64
CL\_INS\_64
CL\_INS\_64
CL\_INS\_64
CL\_INS\_64
CL\_INS\_64
CL\_INS\_64
CL\_INS\_64
CL\_INS\_64
CL\_INS\_64
CL\_INS\_64
CL\_INS\_64
CL\_INS\_64
CL\_INS\_64
CL\_INS\_64
CL\_INS\_64
CL\_INS\_45
CL\_INS\_45
CL\_INS\_64
CL\_INS\_64
CL\_INS\_64
CL\_INS\_83
CL\_INS\_64
CL\_INS\_6
CL\_INS\_83
CL\_INS\_64
CL\_INS\_64
CL\_INS\_64
CL\_INS\_83
CL\_INS\_6
CL\_INS\_6
CL\_INS\_64
CL\_INS\_83
CL\_INS\_64
CL\_INS\_83
CL\_INS\_64
CL\_INS\_64
CL\_INS\_83
CL\_INS\_83
CL\_INS\_64
CL\_INS\_6
CL\_INS\_83
CL\_INS\_83
CL\_INS\_1
CL\_INS\_1
CL\_INS\_83
CL\_INS\_83
CL\_INS\_64
CL\_INS\_64
CL\_INS\_64
CL\_INS\_64
CL\_INS\_64
CL\_INS\_64
CL\_INS\_64
CL\_INS\_64
CL\_INS\_64
CL\_INS\_64
CL\_INS\_45
CL\_INS\_64
CL\_INS\_64
CL\_INS\_64
CL\_INS\_64
CL\_INS\_64
CL\_INS\_45
CL\_INS\_45
CL\_INS\_45
CL\_INS\_45
CL\_INS\_45
CL\_INS\_64
CL\_INS\_64
CL\_INS\_220
CL\_INS\_45
CL\_INS\_45
CL\_INS\_45
CL\_INS\_45
CL\_INS\_64
CL\_INS\_64
CL\_INS\_64
CL\_INS\_64
CL\_INS\_64
CL\_INS\_64
CL\_INS\_64
CL\_INS\_64
CL\_INS\_64
CL\_INS\_64
CL\_INS\_64
CL\_INS\_64
CL\_INS\_64
CL\_INS\_64
CL\_INS\_64
CL\_INS\_64
CL\_INS\_64
CL\_INS\_64
CL\_INS\_64
CL\_INS\_64
CL\_INS\_64
CL\_INS\_6
CL\_INS\_220
CL\_INS\_220
CL\_INS\_64
CL\_INS\_64
CL\_INS\_64
CL\_INS\_64
CL\_INS\_83
CL\_INS\_83
CL\_INS\_83
CL\_INS\_83
CL\_INS\_64
CL\_INS\_64
CL\_INS\_64
CL\_INS\_83
CL\_INS\_45
CL\_INS\_83
CL\_INS\_56
CL\_INS\_64
CL\_INS\_64
CL\_INS\_64
CL\_INS\_64
CL\_INS\_83
CL\_INS\_83
CL\_INS\_83
CL\_INS\_83
CL\_INS\_83
CL\_INS\_64
CL\_INS\_1
CL\_INS\_65
CL\_INS\_65
CL\_INS\_65
CL\_INS\_65
CL\_INS\_65
CL\_INS\_65
CL\_INS\_65
CL\_INS\_65
CL\_INS\_65
CL\_INS\_65
CL\_INS\_65
CL\_INS\_65
CL\_INS\_65
CL\_INS\_65
CL\_INS\_65
CL\_INS\_64
CL\_INS\_64
CL\_INS\_64
CL\_INS\_64
CL\_INS\_65
CL\_INS\_65
CL\_INS\_65
CL\_INS\_65
CL\_INS\_64
CL\_INS\_64
CL\_INS\_64
CL\_INS\_64
CL\_INS\_64
CL\_INS\_64
CL\_INS\_64
CL\_INS\_65
CL\_INS\_65
CL\_INS\_65
CL\_INS\_65
CL\_INS\_65
CL\_INS\_64
CL\_INS\_65
CL\_INS\_64
CL\_INS\_65
CL\_INS\_65
CL\_INS\_65
CL\_INS\_65
CL\_INS\_83
CL\_INS\_64
CL\_INS\_64
CL\_INS\_83
CL\_INS\_64
CL\_INS\_64
CL\_INS\_65
CL\_INS\_64
CL\_INS\_83
CL\_INS\_83
CL\_INS\_83
CL\_INS\_65
CL\_INS\_65
CL\_INS\_65
CL\_INS\_65
CL\_INS\_83
CL\_INS\_83
CL\_INS\_64
CL\_INS\_64
CL\_INS\_64
CL\_INS\_64
CL\_INS\_65
CL\_INS\_65
CL\_INS\_65
CL\_INS\_65
CL\_INS\_65
CL\_INS\_65
Cluster ID


CL\_12437
CL\_12436
CL\_12036
CL\_12035
CL\_12034
CL\_12033
CL\_12032
CL\_12322
CL\_9062
CL\_12323
CL\_12324
CL\_12325
CL\_12326
CL\_9927
CL\_9928
CL\_9929
CL\_13520
CL\_13519
CL\_13518
CL\_13517
CL\_13516
CL\_13515
CL\_13514
CL\_13513
CL\_8206
CL\_10146
CL\_9814
CL\_9815
CL\_10145
CL\_9816
CL\_9817
CL\_9818
CL\_9819
CL\_9408
CL\_9409
CL\_9410
CL\_9411
CL\_9412
CL\_9413
CL\_9414
CL\_9415
CL\_9416
CL\_9417
CL\_11033
CL\_11034
CL\_11035
CL\_8952
CL\_9602
CL\_9603
CL\_9604
CL\_9605
CL\_9606
CL\_9607
CL\_9608
CL\_9609
CL\_9610
CL\_7296
CL\_7297
CL\_12464
CL\_11413
CL\_9181
CL\_9182
CL\_9183
CL\_9184
CL\_9185
CL\_8207
CL\_5091
CL\_5090
CL\_5089
CL\_8208
CL\_5087
CL\_5086
CL\_4795
CL\_4796
CL\_5248
CL\_4797
CL\_9800
CL\_5492
CL\_8897
CL\_5493
CL\_5494
CL\_5495
CL\_7298
CL\_5652
CL\_5651
CL\_5088
CL\_5656
CL\_11036
CL\_11037
CL\_10421
CL\_5655
CL\_8896
CL\_8895
CL\_8894
CL\_10422
CL\_10423
CL\_10424
CL\_5085
CL\_11038
CL\_9186
CL\_9187
CL\_8893
CL\_1287
CL\_9611
CL\_1288
CL\_1289
CL\_1290
CL\_1291
CL\_9188
CL\_5084
CL\_5083
CL\_5496
CL\_5081
CL\_9061
CL\_5497
CL\_5498
CL\_5499
CL\_5500
CL\_5501
CL\_5502
CL\_5503
CL\_5504
CL\_5505
CL\_6135
CL\_13332
CL\_13331
CL\_13330
CL\_11688
CL\_11290
CL\_11289
CL\_6668
CL\_6669
CL\_6670
CL\_6671
CL\_6672
CL\_6673
CL\_6674
CL\_6675
CL\_13087
CL\_9276
CL\_9277
CL\_9278
CL\_9279
CL\_9280
CL\_9281
CL\_9282
CL\_9283
CL\_13628
CL\_13627
CL\_13626
CL\_13625
CL\_13624
CL\_13623
CL\_13622
CL\_13621
CL\_13620
CL\_13019
CL\_13020
CL\_13021
CL\_13022
CL\_13023
CL\_13024
CL\_13025
CL\_13026
CL\_13027
CL\_13028
CL\_13029
CL\_13030
CL\_13031
CL\_13032
CL\_13033
CL\_13034
CL\_13035
CL\_6136
CL\_6137
CL\_6138
CL\_6139
CL\_6140
CL\_12257
CL\_6141
CL\_6142
CL\_6143
CL\_6144
CL\_10873
CL\_10608
CL\_6145
CL\_6146
CL\_6147
CL\_6148
CL\_10609
CL\_96
CL\_95
CL\_10610
CL\_10611
CL\_10612
CL\_94
CL\_10542
CL\_93
CL\_11412
CL\_12433
CL\_6149
CL\_6150
CL\_9286
CL\_9287
CL\_8777
CL\_6151
CL\_13662
CL\_6152
CL\_6153
CL\_6154
CL\_6155
CL\_3940
CL\_3941
CL\_3942
CL\_9288
CL\_9289
CL\_9290
CL\_10886
CL\_10887
CL\_10888
CL\_10889
CL\_10890
CL\_10891
CL\_10892
CL\_7656
CL\_9191
CL\_9192
CL\_9193
CL\_5506
CL\_5507
CL\_7657
CL\_7658
CL\_7659
CL\_7660
CL\_7661
CL\_9189
CL\_9190
CL\_8159
CL\_7662
CL\_7663
CL\_7664
CL\_7665
CL\_10607
CL\_7666
CL\_7667
CL\_7668
CL\_7669
CL\_7670
CL\_7671
CL\_7672
CL\_7673
CL\_7674
CL\_7675
CL\_7676
CL\_7677
CL\_7678
CL\_7679
CL\_7680
CL\_7681
CL\_7682
CL\_7683
CL\_7684
CL\_7685
CL\_7686
CL\_7447
CL\_7448
CL\_10425
CL\_10426
CL\_10427
CL\_10428
CL\_10429
CL\_10430
CL\_10431
CL\_10432
CL\_10433
CL\_10434
CL\_10435
CL\_6649
CL\_5486
CL\_5485
CL\_9284
CL\_9285
CL\_13619
CL\_13618
CL\_10436
CL\_5483
CL\_5482
CL\_91
CL\_90
CL\_3943
CL\_6156
CL\_89
CL\_8295
CL\_8294
CL\_8293
CL\_8292
CL\_8291
CL\_8290
CL\_8289
CL\_8288
CL\_8287
CL\_8342
CL\_8286
CL\_8285
CL\_8284
CL\_8283
CL\_8282
CL\_10893
CL\_10894
CL\_10895
CL\_10896
CL\_8281
CL\_8280
CL\_8279
CL\_8278
CL\_10897
CL\_10898
CL\_10899
CL\_10900
CL\_10901
CL\_10902
CL\_10903
CL\_8277
CL\_8276
CL\_8275
CL\_8274
CL\_88
CL\_1292
CL\_8273
CL\_10904
CL\_8362
CL\_8272
CL\_8271
CL\_8270
CL\_6676
CL\_6677
CL\_6678
CL\_12564
CL\_12565
CL\_12566
CL\_8269
CL\_13088
CL\_8268
CL\_8587
CL\_8588
CL\_8267
CL\_8266
CL\_8265
CL\_8264
CL\_8263
CL\_8590
CL\_13089
CL\_13090
CL\_13091
CL\_13092
CL\_9060
CL\_9059
CL\_5510
CL\_5511
CL\_5512
CL\_5513
