## Supplementary material for "A novel method for integrating genomic and Tn-Seq data to identify common *in vivo* fitness mechanisms across multiple bacterial species": S3 Dataset: CL_INS_65.html

Legend

 Mobile +extrachromosomalelementfunctions
 Regulatoryfunctions
 Hypothetical
 Transcription
 Proteinsynthesis/fate
 Other
 Transport +binding proteins
 All VFDB Genes

FULL


WINDOWSVGPNG

Trim RowsRemove SingletonsSave Fasta

CL\_1293


CL\_1293


CL\_1293


CL\_1293


CL\_1293


CL\_1286


CL\_1293


CL\_1286


CL\_1293


CL\_1293


CL\_1286


CL\_1293


CL\_1286


CL\_1293


CL\_1293


CL\_1293


CL\_1286


CL\_1286


CL\_1293


CL\_1286


CL\_1286


CL\_1286


CL\_1286

HighlightSelectShow Genomes


8

CL\_1294


6

CL\_1294


6

CL\_1294


6

CL\_1294


5

CL\_1294


3

CL\_1294


3

CL\_1294


2

CL\_1294


2

CL\_1294


1

CL\_1294


1

CL\_1294


1

CL\_1294


1

CL\_1294


1

CL\_1311


1

CL\_1294


1

CL\_1294


1

CL\_1294


1

CL\_1294


1

CL\_1294


1

CL\_1294


1

CL\_1294


1

CL\_1294


1

CL\_1294

fGI ID


CL\_INS\_65
CL\_INS\_65
CL\_INS\_65
CL\_INS\_65
CL\_INS\_65
CL\_INS\_65
CL\_INS\_65
CL\_INS\_65
CL\_INS\_65
CL\_INS\_65
CL\_INS\_65
CL\_INS\_65
CL\_INS\_65
CL\_INS\_65
CL\_INS\_65
CL\_INS\_65
CL\_INS\_65
CL\_INS\_65
CL\_INS\_65
CL\_INS\_65
CL\_INS\_65
CL\_INS\_65
CL\_INS\_83
CL\_INS\_65
CL\_INS\_83
CL\_INS\_83
CL\_INS\_83
CL\_INS\_83
CL\_INS\_65
CL\_INS\_230
CL\_INS\_230
CL\_INS\_230
CL\_INS\_83
CL\_INS\_230
CL\_INS\_65
CL\_INS\_65
CL\_INS\_65
CL\_INS\_65
CL\_INS\_65
CL\_INS\_65
CL\_INS\_65
CL\_INS\_65
CL\_INS\_65
CL\_INS\_65
CL\_INS\_65
CL\_INS\_65
CL\_INS\_65
CL\_INS\_65
CL\_INS\_65
CL\_INS\_65
CL\_INS\_65
CL\_INS\_65
CL\_INS\_65
CL\_INS\_65
CL\_INS\_65
CL\_INS\_65
CL\_INS\_65
CL\_INS\_65
CL\_INS\_65
CL\_INS\_65
CL\_INS\_65
CL\_INS\_65
CL\_INS\_65
CL\_INS\_83
CL\_INS\_65
CL\_INS\_83
CL\_INS\_65
CL\_INS\_65
CL\_INS\_65
CL\_INS\_65
CL\_INS\_83
CL\_INS\_65
CL\_INS\_65
CL\_INS\_65
CL\_INS\_65
CL\_INS\_65
CL\_INS\_65
CL\_INS\_65
CL\_INS\_65
CL\_INS\_65
CL\_INS\_65
Cluster ID


CL\_4796
CL\_4797
CL\_5248
CL\_9291
CL\_9292
CL\_9293
CL\_9058
CL\_9057
CL\_9056
CL\_9928
CL\_9929
CL\_9060
CL\_9059
CL\_4798
CL\_5508
CL\_6157
CL\_6158
CL\_5509
CL\_7297
CL\_12464
CL\_9800
CL\_5492
CL\_5656
CL\_10421
CL\_5494
CL\_5495
CL\_7298
CL\_5652
CL\_5651
CL\_5085
CL\_5084
CL\_5083
CL\_5496
CL\_5081
CL\_8295
CL\_8294
CL\_8293
CL\_8292
CL\_8291
CL\_8290
CL\_8289
CL\_8288
CL\_8287
CL\_8342
CL\_8286
CL\_8285
CL\_8284
CL\_8283
CL\_8282
CL\_8281
CL\_8280
CL\_8279
CL\_8278
CL\_8277
CL\_8276
CL\_8275
CL\_8274
CL\_88
CL\_8273
CL\_8362
CL\_8272
CL\_8271
CL\_8270
CL\_6676
CL\_8269
CL\_8268
CL\_8267
CL\_8266
CL\_8265
CL\_8264
CL\_8263
CL\_5510
CL\_5511
CL\_13036
CL\_5512
CL\_10144
CL\_9821
CL\_9822
CL\_9823
CL\_5513
CL\_5514
