## Supplementary material for "A novel method for integrating genomic and Tn-Seq data to identify common *in vivo* fitness mechanisms across multiple bacterial species": S3 Dataset: CL_INS_69.html

Legend

 Hypothetical
 Transcription
 Other
 All VFDB Genes

FULL


WINDOWSVGPNG

Trim RowsRemove SingletonsSave Fasta

CL\_1310


CL\_1310


CL\_1310


CL\_1293

HighlightSelectShow Genomes


53

CL\_1311


2

CL\_1311


1

CL\_1311


1

CL\_1311

fGI ID


CL\_INS\_69
CL\_INS\_65
CL\_INS\_65
CL\_INS\_65
CL\_INS\_65
CL\_INS\_65
CL\_INS\_69
Cluster ID


CL\_9820
CL\_5512
CL\_10144
CL\_9821
CL\_9822
CL\_9823
CL\_9824
