## Supplementary material for "A novel method for integrating genomic and Tn-Seq data to identify common *in vivo* fitness mechanisms across multiple bacterial species": S3 Dataset: CL_INS_70.html

Legend

 Mobile +extrachromosomalelementfunctions
 Other
 All VFDB Genes

FULL


WINDOWSVGPNG

Trim RowsRemove SingletonsSave Fasta

CL\_1319


CL\_1319


CL\_1319

HighlightSelectShow Genomes


37

CL\_1320


18

CL\_1320


1

CL\_1320

fGI ID


CL\_INS\_70
CL\_INS\_70
CL\_INS\_70
Cluster ID


CL\_11776
CL\_6159
CL\_6160
