## Supplementary material for "A novel method for integrating genomic and Tn-Seq data to identify common *in vivo* fitness mechanisms across multiple bacterial species": S3 Dataset: CL_INS_72.html

Legend

 Hypothetical
 Other
 Transport +binding proteins

FULL


WINDOWSVGPNG

Trim RowsRemove SingletonsSave Fasta

CL\_1325


CL\_1325


CL\_1325

HighlightSelectShow Genomes


32

CL\_1326


15

CL\_1328


9

CL\_1326

fGI ID


CL\_INS\_72
CL\_INS\_72
Cluster ID


CL\_5515
CL\_5516
