## Supplementary material for "A novel method for integrating genomic and Tn-Seq data to identify common *in vivo* fitness mechanisms across multiple bacterial species": S3 Dataset: CL_INS_74.html

Legend

 Hypothetical
 All VFDB Genes

FULL


WINDOWSVGPNG

Trim RowsRemove SingletonsSave Fasta

CL\_1330


CL\_1330


CL\_1330

HighlightSelectShow Genomes


30

CL\_1333


15

CL\_1333


12

CL\_1333

fGI ID


CL\_INS\_74
CL\_INS\_74
CL\_INS\_74
Cluster ID


CL\_4801
CL\_1331
CL\_1332
