## Supplementary material for "A novel method for integrating genomic and Tn-Seq data to identify common *in vivo* fitness mechanisms across multiple bacterial species": S3 Dataset: CL_INS_75.html

Legend

 Mobile +extrachromosomalelementfunctions
 Other
 All VFDB Genes

FULL


WINDOWSVGPNG

Trim RowsRemove SingletonsSave Fasta

CL\_1379


CL\_1379


CL\_1379


CL\_1379


CL\_1379


CL\_1379


CL\_1379


CL\_1379

HighlightSelectShow Genomes


37

CL\_1380


10

CL\_1380


4

CL\_1380


2

CL\_1380


1

CL\_1380


1

CL\_1380


1

CL\_1382


1

CL\_1380

fGI ID


CL\_INS\_75
CL\_INS\_75
CL\_INS\_75
CL\_INS\_9
CL\_INS\_9
CL\_INS\_9
CL\_INS\_75
CL\_INS\_75
CL\_INS\_75
Cluster ID


CL\_6163
CL\_13663
CL\_7689
CL\_6610
CL\_5137
CL\_5138
CL\_6164
CL\_6165
CL\_12256
