## Supplementary material for "A novel method for integrating genomic and Tn-Seq data to identify common *in vivo* fitness mechanisms across multiple bacterial species": S3 Dataset: CL_INS_77.html

Legend

 All EssentialGenes
 EnergyMetabolism
 All VFDB Genes
 Transport +binding proteins

FULL


WINDOWSVGPNG

Trim RowsRemove SingletonsSave Fasta

CL\_1418


CL\_1418


CL\_1418


CL\_1418

HighlightSelectShow Genomes


28

CL\_1427


27

CL\_1427


1

CL\_1427


1

CL\_1427

fGI ID


CL\_INS\_77
CL\_INS\_77
CL\_INS\_77
CL\_INS\_77
CL\_INS\_77
CL\_INS\_77
CL\_INS\_77
CL\_INS\_77
Cluster ID


CL\_1419
CL\_1420
CL\_1421
CL\_1422
CL\_1423
CL\_1424
CL\_1425
CL\_1426
