## Supplementary material for "A novel method for integrating genomic and Tn-Seq data to identify common *in vivo* fitness mechanisms across multiple bacterial species": S3 Dataset: CL_INS_78.html

Legend

 Other
 Transport +binding proteins
 All VFDB Genes

FULL


WINDOWSVGPNG

Trim RowsRemove SingletonsSave Fasta

CL\_1454


CL\_1454


CL\_1454

HighlightSelectShow Genomes


38

CL\_1456


18

CL\_1456


1

CL\_1456

fGI ID


CL\_INS\_78
CL\_INS\_78
Cluster ID


CL\_9055
CL\_1455
