## Supplementary material for "A novel method for integrating genomic and Tn-Seq data to identify common *in vivo* fitness mechanisms across multiple bacterial species": S3 Dataset: CL_INS_80.html

Legend

 All VFDB Genes

FULL


WINDOWSVGPNG

Trim RowsRemove SingletonsSave Fasta

CL\_1515


CL\_1515


CL\_1515


CL\_1515


CL\_1515

HighlightSelectShow Genomes


29

CL\_1517


20

CL\_1517


5

CL\_1517


2

CL\_1518


1

CL\_1523

fGI ID


CL\_INS\_80
CL\_INS\_80
CL\_INS\_80
Cluster ID


CL\_6712
CL\_4804
CL\_1516
