## Supplementary material for "A novel method for integrating genomic and Tn-Seq data to identify common *in vivo* fitness mechanisms across multiple bacterial species": S3 Dataset: CL_INS_81.html

Legend

 Other
 All VFDB Genes

FULL


WINDOWSVGPNG

Trim RowsRemove SingletonsSave Fasta

CL\_1533


CL\_1533


CL\_1533


CL\_1533


CL\_1533


CL\_1532

HighlightSelectShow Genomes


28

CL\_1538


25

CL\_1538


1

CL\_1538


1

CL\_1539


1

CL\_1539


1

CL\_1538

fGI ID


CL\_INS\_81
CL\_INS\_81
CL\_INS\_81
CL\_INS\_81
Cluster ID


CL\_1534
CL\_1535
CL\_1536
CL\_1537
