## Supplementary material for "A novel method for integrating genomic and Tn-Seq data to identify common *in vivo* fitness mechanisms across multiple bacterial species": S3 Dataset: CL_INS_82.html

Legend

 Hypothetical
 Regulatoryfunctions
 All EssentialGenes
 Other
 Centralintermediarymetabolism
 All VFDB Genes

FULL


WINDOWSVGPNG

Trim RowsRemove SingletonsSave Fasta

CL\_1596


CL\_1596


CL\_1596


CL\_1596


CL\_1596


CL\_1596


CL\_1596


CL\_1596


CL\_1596


CL\_1596

HighlightSelectShow Genomes


33

CL\_1604


10

CL\_1604


5

CL\_1604


2

CL\_1604


2

CL\_1604


1

CL\_1604


1

CL\_1604


1

CL\_1604


1

CL\_1604


1

CL\_1604

fGI ID


CL\_INS\_82
CL\_INS\_82
CL\_INS\_82
CL\_INS\_82
CL\_INS\_82
CL\_INS\_82
CL\_INS\_82
CL\_INS\_82
CL\_INS\_82
CL\_INS\_82
CL\_INS\_82
CL\_INS\_82
CL\_INS\_82
CL\_INS\_82
CL\_INS\_82
CL\_INS\_82
CL\_INS\_82
CL\_INS\_82
CL\_INS\_82
CL\_INS\_82
CL\_INS\_82
CL\_INS\_82
CL\_INS\_82
CL\_INS\_82
CL\_INS\_82
CL\_INS\_82
CL\_INS\_82
CL\_INS\_82
CL\_INS\_82
CL\_INS\_82
CL\_INS\_82
CL\_INS\_82
CL\_INS\_82
CL\_INS\_82
CL\_INS\_82
Cluster ID


CL\_1597
CL\_1598
CL\_1599
CL\_1600
CL\_1601
CL\_1602
CL\_1603
CL\_6168
CL\_6169
CL\_6170
CL\_6171
CL\_6172
CL\_6173
CL\_6174
CL\_6175
CL\_6176
CL\_6177
CL\_6178
CL\_6179
CL\_6180
CL\_6181
CL\_6182
CL\_6183
CL\_6184
CL\_6185
CL\_6186
CL\_6187
CL\_6188
CL\_6189
CL\_6190
CL\_6191
CL\_6192
CL\_6193
CL\_6194
CL\_6195
