## Supplementary material for "A novel method for integrating genomic and Tn-Seq data to identify common *in vivo* fitness mechanisms across multiple bacterial species": S3 Dataset: CL_INS_83.html

Legend

 Mobile +extrachromosomalelementfunctions
 Regulatoryfunctions
 Hypothetical
 DNA Metabolism
 All EssentialGenes
 Biosynthesis ofcofactors,prostheticgroups, +carriers
 Other
 Transport +binding proteins
 All VFDB Genes

FULL


WINDOWSVGPNG

Trim RowsRemove SingletonsSave Fasta

CL\_1623


CL\_1623


CL\_1623


CL\_1623


CL\_1623


CL\_1623


CL\_1623


CL\_1623


CL\_1623


CL\_1623


CL\_1623


CL\_1623


CL\_1623


CL\_1623


CL\_1623


CL\_1623


CL\_1623


CL\_1623


CL\_1623


CL\_1623


CL\_1623


CL\_1623


CL\_1623


CL\_1623


CL\_1623


CL\_1623


CL\_1623


CL\_1623


CL\_1623


CL\_1623


CL\_512


CL\_1623


CL\_1623


CL\_1623


CL\_1623


CL\_1623


CL\_1623


CL\_1623


CL\_1623


CL\_1623

HighlightSelectShow Genomes


4

CL\_1629


4

CL\_1631


3

CL\_1629


3

CL\_1629


2

CL\_1629


2

CL\_4828


2

CL\_1629


2

CL\_1629


2

CL\_4828


2

CL\_1629


2

CL\_1629


2

CL\_1629


1

CL\_1629


1

CL\_1629


1

CL\_1629


1

CL\_1629


1

CL\_1633


1

CL\_1629


1

CL\_1629


1

CL\_1629


1

CL\_1629


1

CL\_1629


1

CL\_1629


1

CL\_1629


1

CL\_522


1

CL\_1629


1

CL\_1629


1

CL\_1629


1

CL\_1629


1

CL\_1629


1

CL\_1629


1

CL\_1629


1

CL\_1631


1

CL\_1629


1

CL\_1629


1

CL\_1629


1

CL\_1629


1

CL\_1629


1

CL\_1631


1

CL\_1629

fGI ID


CL\_INS\_83
CL\_INS\_83
CL\_INS\_83
CL\_INS\_83
CL\_INS\_83
CL\_INS\_83
CL\_INS\_83
CL\_INS\_83
CL\_INS\_83
CL\_INS\_83
CL\_INS\_83
CL\_INS\_83
CL\_INS\_83
CL\_INS\_83
CL\_INS\_83
CL\_INS\_83
CL\_INS\_83
CL\_INS\_83
CL\_INS\_83
CL\_INS\_83
CL\_INS\_83
CL\_INS\_83
CL\_INS\_83
CL\_INS\_83
CL\_INS\_83
CL\_INS\_83
CL\_INS\_83
CL\_INS\_83
CL\_INS\_83
CL\_INS\_83
CL\_INS\_83
CL\_INS\_83
CL\_INS\_83
CL\_INS\_83
CL\_INS\_83
CL\_INS\_83
CL\_INS\_83
CL\_INS\_83
CL\_INS\_83
CL\_INS\_83
CL\_INS\_83
CL\_INS\_83
CL\_INS\_83
CL\_INS\_83
CL\_INS\_83
CL\_INS\_83
CL\_INS\_83
CL\_INS\_83
CL\_INS\_83
CL\_INS\_83
CL\_INS\_83
CL\_INS\_83
CL\_INS\_83
CL\_INS\_83
CL\_INS\_83
CL\_INS\_83
CL\_INS\_83
CL\_INS\_83
CL\_INS\_83
CL\_INS\_83
CL\_INS\_83
CL\_INS\_83
CL\_INS\_83
CL\_INS\_83
CL\_INS\_83
CL\_INS\_83
CL\_INS\_83
CL\_INS\_83
CL\_INS\_83
CL\_INS\_83
CL\_INS\_83
CL\_INS\_83
CL\_INS\_83
CL\_INS\_83
CL\_INS\_83
CL\_INS\_83
CL\_INS\_83
CL\_INS\_83
CL\_INS\_83
CL\_INS\_83
CL\_INS\_83
CL\_INS\_83
CL\_INS\_83
CL\_INS\_83
CL\_INS\_83
CL\_INS\_83
CL\_INS\_83
CL\_INS\_83
CL\_INS\_83
CL\_INS\_83
CL\_INS\_83
CL\_INS\_83
CL\_INS\_83
CL\_INS\_83
CL\_INS\_83
CL\_INS\_83
CL\_INS\_83
CL\_INS\_83
CL\_INS\_83
CL\_INS\_83
CL\_INS\_83
CL\_INS\_83
CL\_INS\_83
CL\_INS\_83
CL\_INS\_83
CL\_INS\_83
CL\_INS\_83
CL\_INS\_83
CL\_INS\_83
CL\_INS\_83
CL\_INS\_83
CL\_INS\_83
CL\_INS\_83
CL\_INS\_83
CL\_INS\_83
CL\_INS\_83
CL\_INS\_83
CL\_INS\_83
CL\_INS\_83
CL\_INS\_83
CL\_INS\_83
CL\_INS\_83
CL\_INS\_83
CL\_INS\_83
CL\_INS\_83
CL\_INS\_83
CL\_INS\_83
CL\_INS\_83
CL\_INS\_83
CL\_INS\_83
CL\_INS\_83
CL\_INS\_83
CL\_INS\_83
CL\_INS\_83
CL\_INS\_83
CL\_INS\_83
CL\_INS\_83
CL\_INS\_83
CL\_INS\_83
CL\_INS\_83
CL\_INS\_83
CL\_INS\_83
CL\_INS\_83
CL\_INS\_133
CL\_INS\_83
CL\_INS\_83
CL\_INS\_83
CL\_INS\_83
CL\_INS\_83
CL\_INS\_83
CL\_INS\_83
CL\_INS\_83
CL\_INS\_83
CL\_INS\_83
CL\_INS\_83
CL\_INS\_83
CL\_INS\_83
CL\_INS\_83
CL\_INS\_83
CL\_INS\_83
CL\_INS\_83
CL\_INS\_83
CL\_INS\_83
CL\_INS\_83
CL\_INS\_83
CL\_INS\_83
CL\_INS\_83
CL\_INS\_83
CL\_INS\_83
CL\_INS\_83
CL\_INS\_83
CL\_INS\_83
CL\_INS\_83
CL\_INS\_83
CL\_INS\_83
CL\_INS\_83
CL\_INS\_83
CL\_INS\_83
CL\_INS\_83
CL\_INS\_83
CL\_INS\_83
CL\_INS\_83
CL\_INS\_83
CL\_INS\_83
CL\_INS\_83
CL\_INS\_83
CL\_INS\_83
CL\_INS\_83
CL\_INS\_83
CL\_INS\_83
CL\_INS\_83
CL\_INS\_83
CL\_INS\_83
CL\_INS\_83
CL\_INS\_83
CL\_INS\_83
CL\_INS\_83
CL\_INS\_83
CL\_INS\_83
CL\_INS\_83
CL\_INS\_83
CL\_INS\_83
CL\_INS\_83
CL\_INS\_83
CL\_INS\_83
CL\_INS\_83
CL\_INS\_83
CL\_INS\_83
CL\_INS\_83
CL\_INS\_83
CL\_INS\_83
CL\_INS\_83
CL\_INS\_83
CL\_INS\_83
CL\_INS\_133
CL\_INS\_133
CL\_INS\_83
CL\_INS\_83
CL\_INS\_83
CL\_INS\_83
CL\_INS\_83
CL\_INS\_83
CL\_INS\_83
CL\_INS\_83
CL\_INS\_83
CL\_INS\_83
CL\_INS\_83
CL\_INS\_83
CL\_INS\_83
CL\_INS\_83
CL\_INS\_83
CL\_INS\_83
CL\_INS\_83
CL\_INS\_83
CL\_INS\_83
CL\_INS\_83
CL\_INS\_83
CL\_INS\_83
CL\_INS\_83
CL\_INS\_83
CL\_INS\_83
CL\_INS\_83
CL\_INS\_83
CL\_INS\_83
CL\_INS\_83
CL\_INS\_83
CL\_INS\_83
CL\_INS\_83
CL\_INS\_83
CL\_INS\_83
CL\_INS\_83
CL\_INS\_83
CL\_INS\_83
CL\_INS\_83
CL\_INS\_133
CL\_INS\_83
CL\_INS\_83
CL\_INS\_83
CL\_INS\_83
CL\_INS\_83
CL\_INS\_83
CL\_INS\_83
CL\_INS\_83
CL\_INS\_83
CL\_INS\_83
CL\_INS\_83
CL\_INS\_83
CL\_INS\_83
CL\_INS\_83
CL\_INS\_83
CL\_INS\_83
CL\_INS\_83
CL\_INS\_83
CL\_INS\_83
CL\_INS\_83
CL\_INS\_83
CL\_INS\_83
CL\_INS\_83
CL\_INS\_83
CL\_INS\_83
CL\_INS\_83
CL\_INS\_83
CL\_INS\_83
CL\_INS\_83
CL\_INS\_83
CL\_INS\_83
CL\_INS\_83
CL\_INS\_83
CL\_INS\_83
CL\_INS\_83
CL\_INS\_133
CL\_INS\_83
CL\_INS\_83
CL\_INS\_83
CL\_INS\_83
CL\_INS\_83
CL\_INS\_83
CL\_INS\_83
CL\_INS\_83
CL\_INS\_83
CL\_INS\_83
CL\_INS\_83
CL\_INS\_83
CL\_INS\_83
CL\_INS\_83
CL\_INS\_83
CL\_INS\_83
CL\_INS\_83
CL\_INS\_83
CL\_INS\_83
CL\_INS\_83
CL\_INS\_83
CL\_INS\_83
CL\_INS\_83
CL\_INS\_83
CL\_INS\_83
CL\_INS\_83
CL\_INS\_83
CL\_INS\_133
CL\_INS\_83
CL\_INS\_83
CL\_INS\_83
CL\_INS\_83
CL\_INS\_230
CL\_INS\_230
CL\_INS\_88
CL\_INS\_83
CL\_INS\_83
CL\_INS\_83
CL\_INS\_83
CL\_INS\_83
CL\_INS\_83
CL\_INS\_83
CL\_INS\_83
CL\_INS\_83
CL\_INS\_83
CL\_INS\_83
CL\_INS\_83
CL\_INS\_83
CL\_INS\_83
CL\_INS\_83
CL\_INS\_83
CL\_INS\_83
CL\_INS\_83
CL\_INS\_83
CL\_INS\_83
CL\_INS\_83
CL\_INS\_83
CL\_INS\_83
CL\_INS\_83
CL\_INS\_83
CL\_INS\_83
CL\_INS\_83
CL\_INS\_83
CL\_INS\_83
CL\_INS\_83
CL\_INS\_83
CL\_INS\_83
CL\_INS\_83
CL\_INS\_83
CL\_INS\_83
CL\_INS\_83
CL\_INS\_83
CL\_INS\_83
CL\_INS\_83
CL\_INS\_83
CL\_INS\_83
CL\_INS\_83
CL\_INS\_83
CL\_INS\_83
CL\_INS\_83
CL\_INS\_83
CL\_INS\_83
CL\_INS\_83
CL\_INS\_83
CL\_INS\_83
CL\_INS\_83
CL\_INS\_83
CL\_INS\_83
CL\_INS\_83
CL\_INS\_83
CL\_INS\_83
CL\_INS\_83
CL\_INS\_83
CL\_INS\_83
CL\_INS\_83
CL\_INS\_83
CL\_INS\_83
CL\_INS\_83
CL\_INS\_83
CL\_INS\_83
CL\_INS\_83
CL\_INS\_83
CL\_INS\_83
CL\_INS\_83
CL\_INS\_83
CL\_INS\_83
CL\_INS\_83
CL\_INS\_83
CL\_INS\_83
CL\_INS\_83
CL\_INS\_83
CL\_INS\_83
CL\_INS\_83
CL\_INS\_83
CL\_INS\_83
CL\_INS\_83
CL\_INS\_83
CL\_INS\_83
CL\_INS\_83
CL\_INS\_83
CL\_INS\_83
CL\_INS\_83
CL\_INS\_83
CL\_INS\_83
CL\_INS\_83
CL\_INS\_83
CL\_INS\_83
CL\_INS\_83
CL\_INS\_83
CL\_INS\_83
CL\_INS\_83
CL\_INS\_83
CL\_INS\_83
CL\_INS\_83
CL\_INS\_83
CL\_INS\_83
CL\_INS\_83
CL\_INS\_83
CL\_INS\_83
CL\_INS\_83
CL\_INS\_83
CL\_INS\_83
CL\_INS\_83
CL\_INS\_83
CL\_INS\_83
CL\_INS\_83
CL\_INS\_83
CL\_INS\_83
CL\_INS\_83
CL\_INS\_83
CL\_INS\_83
CL\_INS\_83
CL\_INS\_83
CL\_INS\_83
CL\_INS\_83
CL\_INS\_83
CL\_INS\_83
CL\_INS\_83
CL\_INS\_83
CL\_INS\_83
CL\_INS\_83
CL\_INS\_83
CL\_INS\_83
CL\_INS\_83
CL\_INS\_83
CL\_INS\_83
CL\_INS\_83
CL\_INS\_83
CL\_INS\_83
CL\_INS\_83
CL\_INS\_83
CL\_INS\_83
CL\_INS\_83
CL\_INS\_83
CL\_INS\_83
CL\_INS\_83
CL\_INS\_83
CL\_INS\_83
CL\_INS\_83
CL\_INS\_83
CL\_INS\_83
CL\_INS\_83
CL\_INS\_83
CL\_INS\_83
CL\_INS\_83
CL\_INS\_83
CL\_INS\_83
CL\_INS\_83
CL\_INS\_83
CL\_INS\_83
CL\_INS\_83
CL\_INS\_83
CL\_INS\_83
CL\_INS\_83
CL\_INS\_83
CL\_INS\_83
CL\_INS\_83
CL\_INS\_83
CL\_INS\_83
CL\_INS\_83
CL\_INS\_83
CL\_INS\_83
CL\_INS\_83
CL\_INS\_83
CL\_INS\_83
CL\_INS\_83
CL\_INS\_83
CL\_INS\_83
CL\_INS\_83
CL\_INS\_83
CL\_INS\_83
CL\_INS\_83
CL\_INS\_83
CL\_INS\_83
CL\_INS\_83
CL\_INS\_83
CL\_INS\_83
CL\_INS\_83
CL\_INS\_83
Cluster ID


CL\_9931
CL\_9932
CL\_9933
CL\_9934
CL\_8756
CL\_6713
CL\_3943
CL\_9613
CL\_9614
CL\_8261
CL\_8260
CL\_8259
CL\_8258
CL\_8257
CL\_8341
CL\_8256
CL\_8255
CL\_8254
CL\_8253
CL\_8252
CL\_8251
CL\_8250
CL\_8249
CL\_10983
CL\_10982
CL\_10981
CL\_508
CL\_507
CL\_506
CL\_505
CL\_504
CL\_503
CL\_9421
CL\_9422
CL\_9423
CL\_9424
CL\_9425
CL\_9426
CL\_4632
CL\_9427
CL\_9428
CL\_9429
CL\_9430
CL\_9431
CL\_12434
CL\_9432
CL\_9433
CL\_9434
CL\_9435
CL\_6148
CL\_12433
CL\_12432
CL\_95
CL\_94
CL\_9615
CL\_13617
CL\_13616
CL\_13615
CL\_13614
CL\_13613
CL\_13612
CL\_13611
CL\_13610
CL\_13609
CL\_13608
CL\_13607
CL\_13606
CL\_13605
CL\_13604
CL\_13603
CL\_9617
CL\_1624
CL\_13093
CL\_13094
CL\_13095
CL\_7299
CL\_7300
CL\_7301
CL\_1625
CL\_1626
CL\_1627
CL\_1628
CL\_3942
CL\_3941
CL\_91
CL\_6714
CL\_6715
CL\_6716
CL\_9616
CL\_90
CL\_6154
CL\_6153
CL\_10403
CL\_12431
CL\_12430
CL\_12429
CL\_12428
CL\_12427
CL\_12426
CL\_4894
CL\_12425
CL\_12424
CL\_10404
CL\_10969
CL\_6151
CL\_8777
CL\_6150
CL\_7690
CL\_11411
CL\_11410
CL\_11279
CL\_7354
CL\_8809
CL\_11481
CL\_6725
CL\_5518
CL\_7691
CL\_7692
CL\_7693
CL\_7694
CL\_7695
CL\_7696
CL\_7697
CL\_7698
CL\_7353
CL\_3267
CL\_5521
CL\_3265
CL\_10472
CL\_11409
CL\_11408
CL\_11407
CL\_11406
CL\_11405
CL\_11277
CL\_3257
CL\_11404
CL\_11403
CL\_11402
CL\_11401
CL\_11400
CL\_11399
CL\_11398
CL\_5528
CL\_5529
CL\_5530
CL\_5531
CL\_5532
CL\_11397
CL\_11396
CL\_11395
CL\_11394
CL\_11393
CL\_11392
CL\_11391
CL\_11390
CL\_11389
CL\_11388
CL\_11387
CL\_11386
CL\_11385
CL\_3264
CL\_6726
CL\_3263
CL\_3262
CL\_6727
CL\_10020
CL\_3260
CL\_12572
CL\_12573
CL\_12574
CL\_6728
CL\_6729
CL\_6730
CL\_6732
CL\_6734
CL\_3255
CL\_3254
CL\_10299
CL\_6205
CL\_3251
CL\_10300
CL\_5526
CL\_12575
CL\_12576
CL\_10301
CL\_3249
CL\_3248
CL\_8817
CL\_8731
CL\_8730
CL\_3243
CL\_5534
CL\_12577
CL\_8725
CL\_7822
CL\_3237
CL\_3236
CL\_3235
CL\_9008
CL\_9678
CL\_9679
CL\_9680
CL\_6213
CL\_6214
CL\_6215
CL\_10302
CL\_10303
CL\_3238
CL\_6217
CL\_6218
CL\_10304
CL\_6219
CL\_3234
CL\_5542
CL\_5543
CL\_11384
CL\_8823
CL\_5541
CL\_3232
CL\_3231
CL\_6739
CL\_3230
CL\_8722
CL\_12578
CL\_12579
CL\_12580
CL\_12581
CL\_11344
CL\_11343
CL\_11342
CL\_11341
CL\_11340
CL\_11339
CL\_11338
CL\_11337
CL\_11336
CL\_11335
CL\_11334
CL\_11333
CL\_11332
CL\_11331
CL\_11330
CL\_11329
CL\_11328
CL\_11326
CL\_11325
CL\_11324
CL\_11323
CL\_11322
CL\_12582
CL\_10929
CL\_10930
CL\_11773
CL\_4813
CL\_4814
CL\_12583
CL\_6241
CL\_8950
CL\_8951
CL\_8952
CL\_5656
CL\_5494
CL\_5495
CL\_7298
CL\_5652
CL\_5650
CL\_5649
CL\_9185
CL\_10931
CL\_10932
CL\_10933
CL\_5655
CL\_10934
CL\_5653
CL\_10935
CL\_10936
CL\_10937
CL\_5496
CL\_12584
CL\_12585
CL\_8953
CL\_8751
CL\_9825
CL\_8750
CL\_10616
CL\_8954
CL\_8974
CL\_7384
CL\_6220
CL\_10305
CL\_8723
CL\_6221
CL\_3229
CL\_7415
CL\_3228
CL\_6224
CL\_7414
CL\_6370
CL\_6369
CL\_6368
CL\_6367
CL\_6366
CL\_6365
CL\_6364
CL\_6363
CL\_6362
CL\_6361
CL\_6360
CL\_6359
CL\_6358
CL\_6357
CL\_6356
CL\_6355
CL\_6354
CL\_6353
CL\_6352
CL\_6351
CL\_6350
CL\_11383
CL\_11382
CL\_11381
CL\_11380
CL\_5076
CL\_5075
CL\_6196
CL\_9198
CL\_10432
CL\_10431
CL\_10430
CL\_10429
CL\_13547
CL\_102
CL\_13602
CL\_13601
CL\_13600
CL\_13599
CL\_13598
CL\_13597
CL\_13596
CL\_12564
CL\_6676
CL\_6675
CL\_8268
CL\_8587
CL\_8588
CL\_8263
CL\_8590
CL\_6225
CL\_6226
CL\_6227
CL\_10306
CL\_6229
CL\_6230
CL\_6231
CL\_6232
CL\_6233
CL\_6743
CL\_3214
CL\_9872
CL\_3213
CL\_3212
CL\_3211
CL\_3210
CL\_8720
CL\_3206
CL\_3205
CL\_3204
CL\_3203
CL\_10307
CL\_10308
CL\_10309
CL\_4805
CL\_10613
CL\_10614
CL\_10615
CL\_8755
CL\_8754
CL\_8753
CL\_8752
CL\_9921
CL\_8891
CL\_13545
CL\_9418
CL\_9419
CL\_9420
CL\_6513
CL\_11285
CL\_11284
CL\_11283
CL\_11282
CL\_4693
CL\_4694
CL\_4695
CL\_4696
CL\_4697
CL\_4698
CL\_4699
CL\_4702
CL\_4703
CL\_4704
CL\_6649
CL\_11281
CL\_5485
CL\_11280
CL\_5483
CL\_5482
CL\_5481
CL\_92
CL\_4806
CL\_9920
CL\_9919
CL\_9918
CL\_9917
CL\_9916
CL\_9915
CL\_9913
CL\_9912
CL\_13329
CL\_8890
CL\_8889
CL\_8888
CL\_8887
CL\_8886
CL\_8885
CL\_8884
CL\_8883
CL\_8882
CL\_8881
CL\_8880
CL\_8879
CL\_8878
CL\_8877
CL\_6509
CL\_10534
CL\_8164
CL\_6506
CL\_10533
CL\_10532
CL\_10531
CL\_10530
CL\_13512
CL\_10525
CL\_10524
CL\_10523
CL\_6496
CL\_6495
CL\_6494
CL\_6493
CL\_6492
CL\_6491
CL\_6490
CL\_6489
CL\_4807
CL\_4808
CL\_10310
CL\_10311
CL\_10905
CL\_10312
CL\_10313
CL\_10906
CL\_10314
CL\_12327
CL\_10907
CL\_10908
CL\_10909
CL\_11775
CL\_11774
CL\_10910
CL\_10911
CL\_10912
CL\_10913
CL\_10914
CL\_10915
CL\_10916
CL\_10917
CL\_10918
CL\_10919
CL\_10920
CL\_10921
CL\_10922
CL\_10923
CL\_10924
CL\_10925
CL\_10926
CL\_10927
CL\_10928
CL\_4809
CL\_4810
CL\_4811
CL\_4812
CL\_12328
CL\_12329
CL\_10315
CL\_10316
CL\_4815
CL\_4816
CL\_4817
CL\_9197
CL\_4818
CL\_4819
CL\_4821
CL\_4820
CL\_10317
CL\_4822
CL\_4823
CL\_4824
CL\_4825
CL\_4826
CL\_4827
