## Supplementary material for "A novel method for integrating genomic and Tn-Seq data to identify common *in vivo* fitness mechanisms across multiple bacterial species": S3 Dataset: CL_INS_84.html

Legend

 Hypothetical
 Other

FULL


WINDOWSVGPNG

Trim RowsRemove SingletonsSave Fasta

CL\_4828


CL\_4828


CL\_4828

HighlightSelectShow Genomes


44

CL\_1629


2

CL\_1623


2

CL\_1623

fGI ID


CL\_INS\_84
CL\_INS\_83
CL\_INS\_83
CL\_INS\_83
CL\_INS\_83
CL\_INS\_83
CL\_INS\_83
CL\_INS\_83
CL\_INS\_83
CL\_INS\_83
Cluster ID


CL\_11039
CL\_7698
CL\_7697
CL\_7696
CL\_7695
CL\_7694
CL\_7693
CL\_7692
CL\_7691
CL\_5518
