## Supplementary material for "A novel method for integrating genomic and Tn-Seq data to identify common *in vivo* fitness mechanisms across multiple bacterial species": S3 Dataset: CL_INS_85.html

Legend

 Mobile +extrachromosomalelementfunctions
 Hypothetical
 All EssentialGenes
 Other
 All VFDB Genes

FULL


WINDOWSVGPNG

Trim RowsRemove SingletonsSave Fasta

CL\_1631


CL\_1631


CL\_1631


CL\_1631

HighlightSelectShow Genomes


50

CL\_1630


4

CL\_1623


1

CL\_1623


1

CL\_1623

fGI ID


CL\_INS\_85
CL\_INS\_83
CL\_INS\_83
CL\_INS\_83
CL\_INS\_83
CL\_INS\_83
CL\_INS\_83
CL\_INS\_83
CL\_INS\_86
CL\_INS\_83
CL\_INS\_83
CL\_INS\_83
CL\_INS\_83
CL\_INS\_83
CL\_INS\_83
CL\_INS\_83
CL\_INS\_83
CL\_INS\_83
CL\_INS\_83
CL\_INS\_83
CL\_INS\_83
CL\_INS\_83
Cluster ID


CL\_6717
CL\_6716
CL\_6715
CL\_6714
CL\_91
CL\_3941
CL\_3942
CL\_3943
CL\_8248
CL\_8249
CL\_8250
CL\_8251
CL\_8252
CL\_8253
CL\_8254
CL\_8255
CL\_8256
CL\_8341
CL\_8257
CL\_8258
CL\_8259
CL\_8260
