## Supplementary material for "A novel method for integrating genomic and Tn-Seq data to identify common *in vivo* fitness mechanisms across multiple bacterial species": S3 Dataset: CL_INS_87.html

Legend

 Hypothetical
 Other
 All VFDB Genes

FULL


WINDOWSVGPNG

Trim RowsRemove SingletonsSave Fasta

CL\_1635


CL\_1635

HighlightSelectShow Genomes


52

CL\_1636


4

CL\_1636

fGI ID


CL\_INS\_87
CL\_INS\_87
CL\_INS\_87
CL\_INS\_87
CL\_INS\_87
CL\_INS\_87
CL\_INS\_87
CL\_INS\_87
CL\_INS\_87
CL\_INS\_87
CL\_INS\_87
CL\_INS\_87
CL\_INS\_87
CL\_INS\_87
CL\_INS\_87
CL\_INS\_87
CL\_INS\_87
Cluster ID


CL\_9826
CL\_9827
CL\_9828
CL\_9829
CL\_9830
CL\_9831
CL\_9832
CL\_9833
CL\_9834
CL\_9835
CL\_9836
CL\_9837
CL\_9838
CL\_9839
CL\_9840
CL\_9841
CL\_9842
