## Supplementary material for "A novel method for integrating genomic and Tn-Seq data to identify common *in vivo* fitness mechanisms across multiple bacterial species": S3 Dataset: CL_INS_90.html

Legend

 Regulatoryfunctions
 Other
 Transport +binding proteins

FULL


WINDOWSVGPNG

Trim RowsRemove SingletonsSave Fasta

CL\_1692


CL\_1692


CL\_1692

HighlightSelectShow Genomes


31

CL\_1697


23

CL\_1697


1

CL\_1697

fGI ID


CL\_INS\_90
CL\_INS\_90
CL\_INS\_90
CL\_INS\_90
Cluster ID


CL\_1693
CL\_1694
CL\_1695
CL\_1696
