## Supplementary material for "A novel method for integrating genomic and Tn-Seq data to identify common *in vivo* fitness mechanisms across multiple bacterial species": S3 Dataset: CL_INS_94.html

Legend

 Mobile +extrachromosomalelementfunctions
 Regulatoryfunctions
 Hypothetical
 DNA Metabolism
 All VFDB Genes

FULL


WINDOWSVGPNG

Trim RowsRemove SingletonsSave Fasta

CL\_1737


CL\_1737


CL\_1737

HighlightSelectShow Genomes


47

CL\_1738


6

CL\_1738


3

CL\_1738

fGI ID


CL\_INS\_94
CL\_INS\_94
CL\_INS\_94
CL\_INS\_94
CL\_INS\_94
CL\_INS\_94
CL\_INS\_94
CL\_INS\_94
CL\_INS\_94
CL\_INS\_94
CL\_INS\_94
CL\_INS\_94
Cluster ID


CL\_9843
CL\_10142
CL\_10141
CL\_10140
CL\_10139
CL\_10138
CL\_10137
CL\_10136
CL\_10135
CL\_10134
CL\_10133
CL\_10132
