## Supplementary material for "A novel method for integrating genomic and Tn-Seq data to identify common *in vivo* fitness mechanisms across multiple bacterial species": S3 Dataset: CL_INS_95.html

Legend

 Regulatoryfunctions
 Other
 All VFDB Genes
 Transport +binding proteins

FULL


WINDOWSVGPNG

Trim RowsRemove SingletonsSave Fasta

CL\_1743


CL\_1743


CL\_1742

HighlightSelectShow Genomes


30

CL\_1746


23

CL\_1746


2

CL\_1746

fGI ID


CL\_INS\_95
CL\_INS\_95
Cluster ID


CL\_1744
CL\_1745
