## Supplementary material for "A novel method for integrating genomic and Tn-Seq data to identify common *in vivo* fitness mechanisms across multiple bacterial species": S3 Dataset: CL_INS_96.html

Legend

 Mobile +extrachromosomalelementfunctions
 Regulatoryfunctions
 Hypothetical
 Other
 All VFDB Genes

FULL


WINDOWSVGPNG

Trim RowsRemove SingletonsSave Fasta

CL\_1758


CL\_1758


CL\_1758

HighlightSelectShow Genomes


47

CL\_1759


2

CL\_1759


1

CL\_1759

fGI ID


CL\_INS\_96
CL\_INS\_96
CL\_INS\_96
CL\_INS\_96
CL\_INS\_96
CL\_INS\_96
CL\_INS\_96
CL\_INS\_96
CL\_INS\_96
CL\_INS\_96
CL\_INS\_96
CL\_INS\_96
CL\_INS\_96
CL\_INS\_96
CL\_INS\_96
CL\_INS\_96
CL\_INS\_96
CL\_INS\_96
CL\_INS\_96
CL\_INS\_96
CL\_INS\_96
CL\_INS\_96
CL\_INS\_96
CL\_INS\_96
CL\_INS\_96
CL\_INS\_96
CL\_INS\_96
CL\_INS\_96
CL\_INS\_96
CL\_INS\_96
CL\_INS\_96
CL\_INS\_96
CL\_INS\_96
CL\_INS\_96
CL\_INS\_96
CL\_INS\_96
CL\_INS\_96
CL\_INS\_96
CL\_INS\_96
CL\_INS\_96
CL\_INS\_96
CL\_INS\_96
CL\_INS\_96
CL\_INS\_96
CL\_INS\_96
CL\_INS\_96
CL\_INS\_96
CL\_INS\_96
CL\_INS\_96
CL\_INS\_96
CL\_INS\_96
CL\_INS\_96
Cluster ID


CL\_13096
CL\_7302
CL\_7303
CL\_7304
CL\_7305
CL\_7306
CL\_7307
CL\_7308
CL\_7309
CL\_7310
CL\_7311
CL\_7312
CL\_7313
CL\_7314
CL\_7315
CL\_7316
CL\_7317
CL\_7318
CL\_7319
CL\_7320
CL\_7321
CL\_7322
CL\_7323
CL\_7324
CL\_7325
CL\_7326
CL\_7327
CL\_7328
CL\_7329
CL\_7330
CL\_7331
CL\_7332
CL\_7333
CL\_7334
CL\_7335
CL\_7336
CL\_7337
CL\_7338
CL\_7339
CL\_7340
CL\_7341
CL\_7342
CL\_7343
CL\_7344
CL\_7345
CL\_7346
CL\_7347
CL\_7348
CL\_7349
CL\_7350
CL\_6573
CL\_7351
