## Supplementary material for "A novel method for integrating genomic and Tn-Seq data to identify common *in vivo* fitness mechanisms across multiple bacterial species": S3 Dataset: CL_INS_97.html

Legend

 Mobile +extrachromosomalelementfunctions
 Regulatoryfunctions
 Hypothetical
 Other
 All VFDB Genes

FULL


WINDOWSVGPNG

Trim RowsRemove SingletonsSave Fasta

CL\_1775


CL\_1775

HighlightSelectShow Genomes


54

CL\_1776


2

CL\_1776

fGI ID


CL\_INS\_97
CL\_INS\_97
CL\_INS\_97
CL\_INS\_97
CL\_INS\_97
CL\_INS\_97
CL\_INS\_220
CL\_INS\_97
CL\_INS\_97
CL\_INS\_220
CL\_INS\_97
CL\_INS\_219
CL\_INS\_219
CL\_INS\_219
CL\_INS\_219
CL\_INS\_97
CL\_INS\_219
CL\_INS\_97
CL\_INS\_97
CL\_INS\_97
CL\_INS\_97
CL\_INS\_220
CL\_INS\_220
CL\_INS\_220
CL\_INS\_220
CL\_INS\_220
CL\_INS\_219
CL\_INS\_220
CL\_INS\_97
CL\_INS\_97
CL\_INS\_97
CL\_INS\_97
CL\_INS\_97
CL\_INS\_97
CL\_INS\_97
CL\_INS\_97
CL\_INS\_97
CL\_INS\_97
CL\_INS\_97
CL\_INS\_97
CL\_INS\_97
Cluster ID


CL\_13097
CL\_11956
CL\_11957
CL\_11958
CL\_11959
CL\_11960
CL\_6569
CL\_11961
CL\_11962
CL\_8782
CL\_13098
CL\_6419
CL\_6576
CL\_6577
CL\_6578
CL\_11963
CL\_6427
CL\_13099
CL\_13100
CL\_13101
CL\_13102
CL\_6585
CL\_6586
CL\_6587
CL\_6588
CL\_6589
CL\_6438
CL\_6591
CL\_11968
CL\_11969
CL\_11970
CL\_13103
CL\_13104
CL\_13105
CL\_11974
CL\_11975
CL\_11976
CL\_11977
CL\_13106
CL\_11979
CL\_11980
