## Supplementary material for "A novel method for integrating genomic and Tn-Seq data to identify common *in vivo* fitness mechanisms across multiple bacterial species": S3 Dataset: CL_INS_98.html

Legend

 Mobile +extrachromosomalelementfunctions
 Regulatoryfunctions
 Hypothetical
 DNA Metabolism
 All EssentialGenes
 Other
 EnergyMetabolism
 Transport +binding proteins
 All VFDB Genes

FULL


WINDOWSVGPNG

Trim RowsRemove SingletonsSave Fasta

CL\_1807


CL\_1807


CL\_1807


CL\_1807


CL\_1807


CL\_1807


CL\_1807


CL\_1807


CL\_1807


CL\_1807


CL\_1807


CL\_1807


CL\_1807


CL\_1807


CL\_1807


CL\_1807


CL\_1807


CL\_1807


CL\_1807


CL\_1807


CL\_1807


CL\_1807

HighlightSelectShow Genomes


23

CL\_1808


9

CL\_1808


3

CL\_1808


2

CL\_1808


2

CL\_1808


2

CL\_1808


1

CL\_1808


1

CL\_1808


1

CL\_1808


1

CL\_1808


1

CL\_1808


1

CL\_1808


1

CL\_1808


1

CL\_1808


1

CL\_1808


1

CL\_1808


1

CL\_1808


1

CL\_1808


1

CL\_1808


1

CL\_1808


1

CL\_1808


1

CL\_1808

fGI ID


CL\_INS\_98
CL\_INS\_98
CL\_INS\_98
CL\_INS\_98
CL\_INS\_98
CL\_INS\_98
CL\_INS\_98
CL\_INS\_98
CL\_INS\_98
CL\_INS\_98
CL\_INS\_98
CL\_INS\_98
CL\_INS\_98
CL\_INS\_98
CL\_INS\_83
CL\_INS\_98
CL\_INS\_98
CL\_INS\_98
CL\_INS\_98
CL\_INS\_98
CL\_INS\_98
CL\_INS\_98
CL\_INS\_98
CL\_INS\_83
CL\_INS\_83
CL\_INS\_98
CL\_INS\_98
CL\_INS\_83
CL\_INS\_83
CL\_INS\_83
CL\_INS\_98
CL\_INS\_98
CL\_INS\_98
CL\_INS\_98
CL\_INS\_98
CL\_INS\_98
CL\_INS\_98
CL\_INS\_98
CL\_INS\_98
CL\_INS\_98
CL\_INS\_98
CL\_INS\_98
CL\_INS\_98
CL\_INS\_98
CL\_INS\_98
CL\_INS\_83
CL\_INS\_83
CL\_INS\_217
CL\_INS\_83
CL\_INS\_98
CL\_INS\_83
CL\_INS\_83
CL\_INS\_98
CL\_INS\_98
CL\_INS\_98
CL\_INS\_98
CL\_INS\_98
CL\_INS\_98
CL\_INS\_98
CL\_INS\_83
CL\_INS\_98
CL\_INS\_98
CL\_INS\_83
CL\_INS\_83
CL\_INS\_98
CL\_INS\_98
CL\_INS\_98
CL\_INS\_98
CL\_INS\_98
CL\_INS\_98
CL\_INS\_98
CL\_INS\_98
CL\_INS\_98
CL\_INS\_83
CL\_INS\_83
CL\_INS\_83
CL\_INS\_83
CL\_INS\_83
CL\_INS\_98
CL\_INS\_83
CL\_INS\_98
CL\_INS\_83
CL\_INS\_217
CL\_INS\_98
CL\_INS\_98
CL\_INS\_98
CL\_INS\_98
CL\_INS\_98
CL\_INS\_217
CL\_INS\_98
CL\_INS\_98
CL\_INS\_98
CL\_INS\_98
CL\_INS\_98
CL\_INS\_98
CL\_INS\_98
CL\_INS\_98
CL\_INS\_98
CL\_INS\_98
CL\_INS\_98
CL\_INS\_98
CL\_INS\_98
CL\_INS\_98
CL\_INS\_98
CL\_INS\_98
CL\_INS\_98
CL\_INS\_98
CL\_INS\_98
CL\_INS\_98
CL\_INS\_98
CL\_INS\_98
CL\_INS\_98
CL\_INS\_98
CL\_INS\_98
CL\_INS\_83
CL\_INS\_98
CL\_INS\_98
CL\_INS\_98
CL\_INS\_217
CL\_INS\_217
CL\_INS\_98
CL\_INS\_98
CL\_INS\_98
CL\_INS\_98
CL\_INS\_98
CL\_INS\_98
CL\_INS\_98
CL\_INS\_83
CL\_INS\_83
CL\_INS\_98
CL\_INS\_98
CL\_INS\_98
CL\_INS\_98
CL\_INS\_98
CL\_INS\_98
CL\_INS\_83
CL\_INS\_217
CL\_INS\_98
CL\_INS\_98
CL\_INS\_83
CL\_INS\_98
CL\_INS\_98
CL\_INS\_133
CL\_INS\_133
CL\_INS\_98
CL\_INS\_133
CL\_INS\_133
CL\_INS\_98
CL\_INS\_217
CL\_INS\_133
CL\_INS\_83
CL\_INS\_83
CL\_INS\_217
CL\_INS\_98
CL\_INS\_98
CL\_INS\_98
CL\_INS\_98
CL\_INS\_98
CL\_INS\_98
CL\_INS\_98
CL\_INS\_217
CL\_INS\_217
CL\_INS\_83
CL\_INS\_83
CL\_INS\_98
CL\_INS\_98
CL\_INS\_98
CL\_INS\_98
CL\_INS\_98
CL\_INS\_217
CL\_INS\_83
CL\_INS\_98
CL\_INS\_83
CL\_INS\_133
CL\_INS\_98
CL\_INS\_98
CL\_INS\_98
CL\_INS\_98
CL\_INS\_83
CL\_INS\_98
CL\_INS\_98
CL\_INS\_98
CL\_INS\_98
CL\_INS\_98
CL\_INS\_98
CL\_INS\_98
CL\_INS\_98
CL\_INS\_98
CL\_INS\_98
CL\_INS\_98
CL\_INS\_83
CL\_INS\_98
CL\_INS\_98
CL\_INS\_98
CL\_INS\_98
CL\_INS\_98
CL\_INS\_98
CL\_INS\_98
CL\_INS\_98
CL\_INS\_98
CL\_INS\_98
CL\_INS\_98
CL\_INS\_98
CL\_INS\_98
CL\_INS\_98
CL\_INS\_98
CL\_INS\_98
CL\_INS\_98
CL\_INS\_98
CL\_INS\_98
CL\_INS\_98
CL\_INS\_98
CL\_INS\_98
CL\_INS\_98
CL\_INS\_98
CL\_INS\_98
CL\_INS\_98
CL\_INS\_98
CL\_INS\_98
CL\_INS\_98
CL\_INS\_98
CL\_INS\_98
CL\_INS\_98
CL\_INS\_98
CL\_INS\_98
CL\_INS\_98
CL\_INS\_98
CL\_INS\_98
CL\_INS\_98
CL\_INS\_98
CL\_INS\_83
CL\_INS\_98
CL\_INS\_98
CL\_INS\_98
CL\_INS\_98
CL\_INS\_98
CL\_INS\_98
CL\_INS\_98
CL\_INS\_83
CL\_INS\_83
CL\_INS\_98
CL\_INS\_98
CL\_INS\_98
CL\_INS\_83
CL\_INS\_83
CL\_INS\_83
CL\_INS\_83
CL\_INS\_83
CL\_INS\_83
CL\_INS\_98
CL\_INS\_98
CL\_INS\_98
CL\_INS\_98
CL\_INS\_83
CL\_INS\_98
CL\_INS\_98
CL\_INS\_83
CL\_INS\_83
CL\_INS\_98
CL\_INS\_133
CL\_INS\_98
CL\_INS\_98
CL\_INS\_98
CL\_INS\_98
CL\_INS\_98
CL\_INS\_98
CL\_INS\_98
CL\_INS\_98
CL\_INS\_98
CL\_INS\_98
CL\_INS\_98
CL\_INS\_98
CL\_INS\_98
CL\_INS\_98
CL\_INS\_98
CL\_INS\_133
CL\_INS\_98
CL\_INS\_98
CL\_INS\_98
CL\_INS\_83
CL\_INS\_83
CL\_INS\_83
CL\_INS\_83
CL\_INS\_83
CL\_INS\_98
CL\_INS\_83
CL\_INS\_98
CL\_INS\_98
CL\_INS\_98
CL\_INS\_83
CL\_INS\_98
CL\_INS\_98
CL\_INS\_98
CL\_INS\_98
CL\_INS\_98
CL\_INS\_83
CL\_INS\_83
CL\_INS\_83
CL\_INS\_83
CL\_INS\_83
CL\_INS\_83
CL\_INS\_98
CL\_INS\_98
CL\_INS\_83
CL\_INS\_98
CL\_INS\_98
CL\_INS\_98
CL\_INS\_83
CL\_INS\_83
CL\_INS\_83
CL\_INS\_83
CL\_INS\_83
CL\_INS\_83
CL\_INS\_83
CL\_INS\_83
CL\_INS\_83
CL\_INS\_83
CL\_INS\_83
CL\_INS\_83
CL\_INS\_83
CL\_INS\_83
CL\_INS\_83
CL\_INS\_83
CL\_INS\_83
CL\_INS\_98
CL\_INS\_83
CL\_INS\_83
CL\_INS\_83
CL\_INS\_83
CL\_INS\_83
CL\_INS\_98
CL\_INS\_98
CL\_INS\_83
CL\_INS\_98
CL\_INS\_217
CL\_INS\_217
CL\_INS\_98
CL\_INS\_83
CL\_INS\_83
CL\_INS\_83
CL\_INS\_83
CL\_INS\_83
CL\_INS\_98
CL\_INS\_98
CL\_INS\_98
CL\_INS\_83
CL\_INS\_98
CL\_INS\_98
CL\_INS\_98
CL\_INS\_83
CL\_INS\_83
CL\_INS\_83
CL\_INS\_83
CL\_INS\_98
CL\_INS\_83
CL\_INS\_98
CL\_INS\_98
CL\_INS\_98
CL\_INS\_98
CL\_INS\_98
CL\_INS\_98
CL\_INS\_98
CL\_INS\_98
CL\_INS\_98
CL\_INS\_98
CL\_INS\_98
CL\_INS\_98
CL\_INS\_98
CL\_INS\_98
CL\_INS\_98
CL\_INS\_98
CL\_INS\_98
CL\_INS\_98
CL\_INS\_98
CL\_INS\_98
CL\_INS\_98
CL\_INS\_98
CL\_INS\_98
CL\_INS\_98
CL\_INS\_98
CL\_INS\_98
CL\_INS\_98
CL\_INS\_98
CL\_INS\_98
CL\_INS\_98
CL\_INS\_98
CL\_INS\_98
CL\_INS\_83
CL\_INS\_98
CL\_INS\_83
CL\_INS\_83
CL\_INS\_98
CL\_INS\_98
CL\_INS\_83
CL\_INS\_83
CL\_INS\_83
CL\_INS\_98
CL\_INS\_98
CL\_INS\_98
CL\_INS\_98
CL\_INS\_83
CL\_INS\_98
CL\_INS\_98
CL\_INS\_98
CL\_INS\_98
CL\_INS\_98
CL\_INS\_98
CL\_INS\_83
CL\_INS\_83
CL\_INS\_83
CL\_INS\_98
CL\_INS\_83
CL\_INS\_98
CL\_INS\_98
CL\_INS\_98
CL\_INS\_98
CL\_INS\_98
CL\_INS\_98
CL\_INS\_98
CL\_INS\_98
CL\_INS\_98
CL\_INS\_98
CL\_INS\_98
CL\_INS\_98
CL\_INS\_98
CL\_INS\_133
CL\_INS\_83
CL\_INS\_133
CL\_INS\_217
CL\_INS\_98
CL\_INS\_98
CL\_INS\_98
CL\_INS\_98
CL\_INS\_83
CL\_INS\_83
CL\_INS\_133
CL\_INS\_133
CL\_INS\_98
CL\_INS\_83
CL\_INS\_133
CL\_INS\_133
CL\_INS\_133
CL\_INS\_98
CL\_INS\_98
CL\_INS\_98
CL\_INS\_98
CL\_INS\_98
CL\_INS\_98
CL\_INS\_83
CL\_INS\_83
CL\_INS\_83
CL\_INS\_83
CL\_INS\_83
CL\_INS\_83
CL\_INS\_83
CL\_INS\_83
CL\_INS\_83
CL\_INS\_83
CL\_INS\_83
CL\_INS\_83
CL\_INS\_83
CL\_INS\_83
CL\_INS\_83
CL\_INS\_83
CL\_INS\_98
CL\_INS\_83
CL\_INS\_83
CL\_INS\_83
CL\_INS\_83
CL\_INS\_133
CL\_INS\_83
CL\_INS\_83
CL\_INS\_83
CL\_INS\_98
CL\_INS\_98
CL\_INS\_98
CL\_INS\_98
Cluster ID


CL\_9052
CL\_9051
CL\_10319
CL\_10320
CL\_10321
CL\_10322
CL\_10323
CL\_4831
CL\_4832
CL\_4833
CL\_13107
CL\_13108
CL\_5517
CL\_11379
CL\_5518
CL\_6723
CL\_6724
CL\_13152
CL\_9199
CL\_5519
CL\_11378
CL\_11377
CL\_7352
CL\_11279
CL\_7354
CL\_4931
CL\_11278
CL\_7353
CL\_3267
CL\_5521
CL\_9618
CL\_9295
CL\_9200
CL\_6202
CL\_12791
CL\_12792
CL\_12793
CL\_12794
CL\_12353
CL\_12795
CL\_12796
CL\_12797
CL\_12798
CL\_12799
CL\_12800
CL\_11277
CL\_3257
CL\_5520
CL\_3265
CL\_9294
CL\_6725
CL\_6726
CL\_9201
CL\_9202
CL\_5522
CL\_10131
CL\_7830
CL\_7831
CL\_10721
CL\_3264
CL\_10473
CL\_10130
CL\_3263
CL\_3262
CL\_9619
CL\_9620
CL\_9621
CL\_9622
CL\_9623
CL\_9624
CL\_9625
CL\_3261
CL\_9884
CL\_6727
CL\_3260
CL\_6728
CL\_6729
CL\_6730
CL\_6731
CL\_6732
CL\_6733
CL\_6734
CL\_3259
CL\_9296
CL\_9297
CL\_9298
CL\_3258
CL\_5523
CL\_8811
CL\_11372
CL\_11371
CL\_11370
CL\_11369
CL\_11368
CL\_11367
CL\_11366
CL\_11365
CL\_11364
CL\_11363
CL\_11362
CL\_11361
CL\_11360
CL\_11359
CL\_11358
CL\_11357
CL\_11356
CL\_11355
CL\_11354
CL\_11353
CL\_11352
CL\_11351
CL\_11350
CL\_11349
CL\_11348
CL\_8723
CL\_6203
CL\_6204
CL\_13165
CL\_8812
CL\_3256
CL\_5524
CL\_5525
CL\_8981
CL\_8980
CL\_9626
CL\_9627
CL\_9628
CL\_3255
CL\_3254
CL\_6735
CL\_6736
CL\_8736
CL\_12255
CL\_8737
CL\_3252
CL\_6205
CL\_9674
CL\_13153
CL\_9959
CL\_10300
CL\_13664
CL\_11641
CL\_6207
CL\_6208
CL\_6209
CL\_6210
CL\_6211
CL\_6206
CL\_9239
CL\_5528
CL\_5529
CL\_5530
CL\_3247
CL\_13154
CL\_13155
CL\_13156
CL\_13157
CL\_13158
CL\_13159
CL\_13160
CL\_9784
CL\_6212
CL\_6213
CL\_6214
CL\_11376
CL\_11375
CL\_11374
CL\_11373
CL\_12802
CL\_8727
CL\_8725
CL\_9207
CL\_5534
CL\_5535
CL\_5536
CL\_5537
CL\_5538
CL\_5539
CL\_6215
CL\_11276
CL\_11275
CL\_11274
CL\_11273
CL\_11272
CL\_10126
CL\_10125
CL\_10124
CL\_6216
CL\_10720
CL\_10123
CL\_3238
CL\_12803
CL\_12804
CL\_12805
CL\_12806
CL\_12807
CL\_12808
CL\_12809
CL\_12810
CL\_12811
CL\_12812
CL\_12813
CL\_12814
CL\_12815
CL\_12816
CL\_12817
CL\_12818
CL\_12819
CL\_12820
CL\_12821
CL\_12822
CL\_12823
CL\_12824
CL\_12825
CL\_12826
CL\_12827
CL\_12828
CL\_12829
CL\_12830
CL\_12831
CL\_12832
CL\_12833
CL\_12834
CL\_12835
CL\_12836
CL\_12837
CL\_12838
CL\_7827
CL\_9203
CL\_11248
CL\_5526
CL\_6737
CL\_12801
CL\_5527
CL\_12254
CL\_8734
CL\_8733
CL\_12253
CL\_8731
CL\_8730
CL\_10129
CL\_10128
CL\_10006
CL\_3249
CL\_9678
CL\_9679
CL\_9680
CL\_3248
CL\_8817
CL\_8732
CL\_9629
CL\_9630
CL\_10127
CL\_5531
CL\_13328
CL\_13327
CL\_5532
CL\_3243
CL\_6738
CL\_3242
CL\_9631
CL\_9632
CL\_9633
CL\_9634
CL\_9635
CL\_9636
CL\_9204
CL\_9205
CL\_9206
CL\_8729
CL\_12252
CL\_5533
CL\_13326
CL\_13325
CL\_13324
CL\_5561
CL\_11942
CL\_11941
CL\_5540
CL\_3237
CL\_3236
CL\_3235
CL\_9008
CL\_6217
CL\_8975
CL\_6218
CL\_9208
CL\_10122
CL\_9333
CL\_6219
CL\_11271
CL\_11270
CL\_9879
CL\_9878
CL\_9877
CL\_3234
CL\_6739
CL\_7384
CL\_3232
CL\_3231
CL\_3230
CL\_9637
CL\_9209
CL\_6221
CL\_11347
CL\_11346
CL\_11345
CL\_11344
CL\_11343
CL\_11342
CL\_11341
CL\_11340
CL\_11339
CL\_11338
CL\_11337
CL\_11336
CL\_11335
CL\_11334
CL\_11333
CL\_11332
CL\_11331
CL\_11330
CL\_11329
CL\_11328
CL\_11327
CL\_11326
CL\_11325
CL\_11324
CL\_11323
CL\_11322
CL\_11321
CL\_11320
CL\_3229
CL\_5545
CL\_6222
CL\_6223
CL\_9334
CL\_7415
CL\_6224
CL\_7414
CL\_6226
CL\_6227
CL\_9210
CL\_6228
CL\_10121
CL\_6229
CL\_9211
CL\_9212
CL\_9213
CL\_6230
CL\_6231
CL\_6232
CL\_6233
CL\_10719
CL\_3228
CL\_3226
CL\_3227
CL\_5546
CL\_10718
CL\_6740
CL\_6741
CL\_6742
CL\_3225
CL\_3224
CL\_11269
CL\_5547
CL\_5548
CL\_3223
CL\_3222
CL\_3221
CL\_3220
CL\_3219
CL\_3218
CL\_5549
CL\_10717
CL\_3217
CL\_9638
CL\_11268
CL\_9005
CL\_13162
CL\_11267
CL\_9003
CL\_11266
CL\_11265
CL\_11264
CL\_11113
CL\_3216
CL\_6743
CL\_11640
CL\_3214
CL\_3213
CL\_6234
CL\_6235
CL\_3212
CL\_3211
CL\_3210
CL\_9971
CL\_5550
CL\_6236
CL\_3207
CL\_8720
CL\_5551
CL\_11263
CL\_11262
CL\_11261
CL\_11260
CL\_11259
CL\_3206
CL\_3205
CL\_3204
CL\_12251
CL\_3203
CL\_6744
CL\_6745
CL\_9639
CL\_9640
CL\_9214
CL\_9215
CL\_9216
CL\_5552
CL\_10120
CL\_6237
CL\_6238
CL\_6239
CL\_10716
CL\_4813
CL\_4814
CL\_4930
CL\_6345
CL\_3199
CL\_3198
CL\_9641
CL\_6240
CL\_6241
CL\_5541
CL\_5542
CL\_5543
CL\_5544
CL\_6220
CL\_7821
CL\_7820
CL\_5586
CL\_13161
CL\_7819
CL\_7818
CL\_13323
CL\_13322
CL\_7816
CL\_6225
CL\_6370
CL\_6369
CL\_6368
CL\_6367
CL\_6366
CL\_6365
CL\_6364
CL\_6363
CL\_6362
CL\_6361
CL\_6360
CL\_6359
CL\_6358
CL\_6357
CL\_6356
CL\_13321
CL\_6354
CL\_6353
CL\_6352
CL\_6351
CL\_6350
CL\_11383
CL\_11382
CL\_11381
CL\_3197
CL\_3182
CL\_7393
CL\_3180
