## Supplementary material for "A novel method for integrating genomic and Tn-Seq data to identify common *in vivo* fitness mechanisms across multiple bacterial species": S3 Dataset: CL_INS_100.html

Legend

 Hypothetical
 Other
 All VFDB Genes

FULL


WINDOWSVGPNG

Trim RowsRemove SingletonsSave Fasta

CL\_1832


CL\_1832


CL\_1832


CL\_1832


CL\_1832

HighlightSelectShow Genomes


47

CL\_1833


5

CL\_1839


2

CL\_1839


1

CL\_1839


1

CL\_1839

fGI ID


CL\_INS\_100
CL\_INS\_100
CL\_INS\_100
CL\_INS\_100
CL\_INS\_100
CL\_INS\_100
CL\_INS\_100
CL\_INS\_100
CL\_INS\_100
CL\_INS\_100
CL\_INS\_100
CL\_INS\_100
CL\_INS\_100
CL\_INS\_100
CL\_INS\_100
CL\_INS\_100
CL\_INS\_100
CL\_INS\_100
CL\_INS\_100
CL\_INS\_100
CL\_INS\_100
CL\_INS\_100
Cluster ID


CL\_1834
CL\_6747
CL\_13595
CL\_13594
CL\_4835
CL\_4838
CL\_6242
CL\_13666
CL\_6243
CL\_6244
CL\_10324
CL\_9437
CL\_10325
CL\_10326
CL\_10327
CL\_10328
CL\_10329
CL\_4839
CL\_4840
CL\_4841
CL\_9442
CL\_4843
