## Supplementary material for "A novel method for integrating genomic and Tn-Seq data to identify common *in vivo* fitness mechanisms across multiple bacterial species": S3 Dataset: CL_INS_101.html

Legend

 Mobile +extrachromosomalelementfunctions
 Hypothetical
 Other
 Transport +binding proteins
 All VFDB Genes

FULL


WINDOWSVGPNG

Trim RowsRemove SingletonsSave Fasta

CL\_1833


CL\_1832


CL\_1833


CL\_1833


CL\_1833


CL\_1833


CL\_1833


CL\_1833


CL\_1832


CL\_1833


CL\_1832


CL\_1833


CL\_1833


CL\_1833


CL\_1833


CL\_1833


CL\_1833


CL\_1833


CL\_1833


CL\_1833


CL\_1832


CL\_1833


CL\_1833

HighlightSelectShow Genomes


18

CL\_1839


5

CL\_1839


4

CL\_1839


3

CL\_1839


3

CL\_1839


3

CL\_1839


2

CL\_1839


2

CL\_1839


2

CL\_1839


2

CL\_1839


1

CL\_1839


1

CL\_1839


1

CL\_1839


1

CL\_1839


1

CL\_1839


1

CL\_1839


1

CL\_1839


1

CL\_1839


1

CL\_1839


1

CL\_1839


1

CL\_1839


1

CL\_1839


1

CL\_1839

fGI ID


CL\_INS\_101
CL\_INS\_101
CL\_INS\_101
CL\_INS\_100
CL\_INS\_101
CL\_INS\_101
CL\_INS\_101
CL\_INS\_101
CL\_INS\_100
CL\_INS\_101
CL\_INS\_101
CL\_INS\_101
CL\_INS\_100
CL\_INS\_100
CL\_INS\_100
CL\_INS\_100
CL\_INS\_100
CL\_INS\_101
CL\_INS\_101
CL\_INS\_101
CL\_INS\_101
CL\_INS\_101
CL\_INS\_101
CL\_INS\_100
CL\_INS\_100
CL\_INS\_101
CL\_INS\_101
CL\_INS\_101
CL\_INS\_101
CL\_INS\_100
CL\_INS\_101
CL\_INS\_100
CL\_INS\_100
CL\_INS\_101
CL\_INS\_100
CL\_INS\_100
CL\_INS\_100
CL\_INS\_100
CL\_INS\_100
CL\_INS\_100
CL\_INS\_100
CL\_INS\_100
CL\_INS\_101
CL\_INS\_101
CL\_INS\_101
CL\_INS\_101
CL\_INS\_101
CL\_INS\_101
CL\_INS\_101
CL\_INS\_168
CL\_INS\_168
CL\_INS\_101
CL\_INS\_101
CL\_INS\_101
CL\_INS\_101
CL\_INS\_101
CL\_INS\_101
CL\_INS\_100
CL\_INS\_100
CL\_INS\_101
CL\_INS\_101
Cluster ID


CL\_9050
CL\_10619
CL\_9642
CL\_1834
CL\_1835
CL\_6746
CL\_1836
CL\_1837
CL\_4835
CL\_4836
CL\_4837
CL\_9436
CL\_4838
CL\_6242
CL\_13666
CL\_6243
CL\_6244
CL\_12027
CL\_10938
CL\_10939
CL\_10940
CL\_10941
CL\_10942
CL\_10324
CL\_9437
CL\_9438
CL\_9439
CL\_9440
CL\_9441
CL\_6747
CL\_7355
CL\_13595
CL\_13594
CL\_6748
CL\_10325
CL\_10326
CL\_10327
CL\_10328
CL\_10329
CL\_4839
CL\_4840
CL\_4841
CL\_9935
CL\_7699
CL\_13320
CL\_13109
CL\_13110
CL\_9844
CL\_9845
CL\_9846
CL\_9847
CL\_9848
CL\_10119
CL\_9849
CL\_9850
CL\_1838
CL\_4842
CL\_9442
CL\_4843
CL\_4844
CL\_4845
