## Supplementary material for "A novel method for integrating genomic and Tn-Seq data to identify common *in vivo* fitness mechanisms across multiple bacterial species": S3 Dataset: CL_INS_103.html

Legend

 Hypothetical
 Other

FULL


WINDOWSVGPNG

Trim RowsRemove SingletonsSave Fasta

CL\_1876


CL\_1876

HighlightSelectShow Genomes


50

CL\_1877


8

CL\_1877

fGI ID


CL\_INS\_103
CL\_INS\_103
CL\_INS\_103
Cluster ID


CL\_4850
CL\_4851
CL\_4852
