## Supplementary material for "A novel method for integrating genomic and Tn-Seq data to identify common *in vivo* fitness mechanisms across multiple bacterial species": S3 Dataset: CL_INS_104.html

Legend

 Hypothetical
 Other
 All VFDB Genes

FULL


WINDOWSVGPNG

Trim RowsRemove SingletonsSave Fasta

CL\_1880


CL\_1880


CL\_1880

HighlightSelectShow Genomes


42

CL\_1882


6

CL\_1882


1

CL\_1883

fGI ID

CL\_INS\_104
Cluster ID

CL\_1881
