## Supplementary material for "A novel method for integrating genomic and Tn-Seq data to identify common *in vivo* fitness mechanisms across multiple bacterial species": S3 Dataset: CL_INS_105.html

Legend

 Hypothetical
 Other

FULL


WINDOWSVGPNG

Trim RowsRemove SingletonsSave Fasta

CL\_1889


CL\_1889


CL\_1889


CL\_1889


CL\_1889


CL\_1889


CL\_1889

HighlightSelectShow Genomes


18

CL\_1890


14

CL\_1890


8

CL\_1890


8

CL\_1890


6

CL\_1890


2

CL\_1890


1

CL\_1890

fGI ID


CL\_INS\_105
CL\_INS\_105
CL\_INS\_105
CL\_INS\_105
CL\_INS\_105
Cluster ID


CL\_4853
CL\_4854
CL\_4855
CL\_8247
CL\_8955
