## Supplementary material for "A novel method for integrating genomic and Tn-Seq data to identify common *in vivo* fitness mechanisms across multiple bacterial species": S3 Dataset: CL_INS_107.html

Legend

 Hypothetical
 All VFDB Genes

FULL


WINDOWSVGPNG

Trim RowsRemove SingletonsSave Fasta

CL\_1935


CL\_1935


CL\_1935

HighlightSelectShow Genomes


50

CL\_1936


6

CL\_1936


1

CL\_1936

fGI ID


CL\_INS\_107
CL\_INS\_107
Cluster ID


CL\_6749
CL\_6750
