## Supplementary material for "A novel method for integrating genomic and Tn-Seq data to identify common *in vivo* fitness mechanisms across multiple bacterial species": S3 Dataset: CL_INS_108.html

Legend

 Hypothetical
 All EssentialGenes
 All VFDB Genes

FULL


WINDOWSVGPNG

Trim RowsRemove SingletonsSave Fasta

CL\_1940


CL\_1940

HighlightSelectShow Genomes


41

CL\_1941


16

CL\_1941

fGI ID

CL\_INS\_106
Cluster ID

CL\_6245
