## Supplementary material for "A novel method for integrating genomic and Tn-Seq data to identify common *in vivo* fitness mechanisms across multiple bacterial species": S3 Dataset: CL_INS_110.html

Legend

 Hypothetical
 All EssentialGenes
 Other
 All VFDB Genes

FULL


WINDOWSVGPNG

Trim RowsRemove SingletonsSave Fasta

CL\_1953


CL\_1953

HighlightSelectShow Genomes


45

CL\_1954


5

CL\_1954

fGI ID


CL\_INS\_110
CL\_INS\_110
CL\_INS\_110
Cluster ID


CL\_7356
CL\_7357
CL\_7358
