## Supplementary material for "A novel method for integrating genomic and Tn-Seq data to identify common *in vivo* fitness mechanisms across multiple bacterial species": S3 Dataset: CL_INS_116.html

Legend

 Hypothetical
 Other
 All VFDB Genes

FULL


WINDOWSVGPNG

Trim RowsRemove SingletonsSave Fasta

CL\_2075


CL\_2075


CL\_2075


CL\_2075

HighlightSelectShow Genomes


47

CL\_2076


5

CL\_2085


2

CL\_2085


1

CL\_2085

fGI ID


CL\_INS\_116
CL\_INS\_117
CL\_INS\_117
CL\_INS\_117
CL\_INS\_117
CL\_INS\_117
Cluster ID


CL\_9299
CL\_2077
CL\_2078
CL\_2079
CL\_2081
CL\_2082
