## Supplementary material for "A novel method for integrating genomic and Tn-Seq data to identify common *in vivo* fitness mechanisms across multiple bacterial species": S3 Dataset: CL_INS_117.html

Legend

 Mobile +extrachromosomalelementfunctions
 Hypothetical
 Other
 Transport +binding proteins
 All VFDB Genes

FULL


WINDOWSVGPNG

Trim RowsRemove SingletonsSave Fasta

CL\_2076


CL\_2076


CL\_2076


CL\_2075


CL\_2075


CL\_2075


CL\_2076


CL\_2076

HighlightSelectShow Genomes


21

CL\_2085


19

CL\_2085


6

CL\_2085


5

CL\_2085


2

CL\_2085


1

CL\_2085


1

CL\_2085


1

CL\_2085

fGI ID


CL\_INS\_117
CL\_INS\_117
CL\_INS\_117
CL\_INS\_117
CL\_INS\_117
CL\_INS\_117
CL\_INS\_117
CL\_INS\_117
CL\_INS\_117
CL\_INS\_117
CL\_INS\_117
CL\_INS\_117
CL\_INS\_117
CL\_INS\_117
CL\_INS\_117
CL\_INS\_117
CL\_INS\_117
CL\_INS\_117
CL\_INS\_117
CL\_INS\_117
CL\_INS\_117
CL\_INS\_117
CL\_INS\_117
CL\_INS\_117
CL\_INS\_117
CL\_INS\_117
CL\_INS\_117
CL\_INS\_168
CL\_INS\_168
CL\_INS\_117
CL\_INS\_117
CL\_INS\_117
CL\_INS\_117
CL\_INS\_1
CL\_INS\_117
CL\_INS\_117
CL\_INS\_117
CL\_INS\_117
CL\_INS\_117
CL\_INS\_117
CL\_INS\_117
CL\_INS\_117
CL\_INS\_117
CL\_INS\_117
CL\_INS\_117
CL\_INS\_117
CL\_INS\_117
CL\_INS\_117
Cluster ID


CL\_7734
CL\_2077
CL\_2078
CL\_2079
CL\_2080
CL\_2081
CL\_2082
CL\_2083
CL\_2084
CL\_11770
CL\_11769
CL\_6900
CL\_6902
CL\_6903
CL\_6904
CL\_6905
CL\_6906
CL\_11768
CL\_11767
CL\_11766
CL\_11765
CL\_11764
CL\_11763
CL\_11762
CL\_11761
CL\_11760
CL\_11759
CL\_9846
CL\_9847
CL\_11758
CL\_11757
CL\_11756
CL\_11755
CL\_5163
CL\_11754
CL\_11753
CL\_11752
CL\_11751
CL\_11750
CL\_11749
CL\_11748
CL\_11747
CL\_11746
CL\_11745
CL\_11744
CL\_11743
CL\_11742
CL\_11741
