## Supplementary material for "A novel method for integrating genomic and Tn-Seq data to identify common *in vivo* fitness mechanisms across multiple bacterial species": S3 Dataset: CL_INS_118.html

Legend

 Regulatoryfunctions
 Hypothetical
 All EssentialGenes
 Other
 Transport +binding proteins
 All VFDB Genes

FULL


WINDOWSVGPNG

Trim RowsRemove SingletonsSave Fasta

CL\_2118


CL\_2118


CL\_2118


CL\_2118


CL\_2118


CL\_2118


CL\_2115


CL\_2118


CL\_2118


CL\_2118


CL\_2118


CL\_2116


CL\_2118


CL\_2118


CL\_2118


CL\_2116

HighlightSelectShow Genomes


15

CL\_2121


14

CL\_2121


4

CL\_2121


3

CL\_2121


3

CL\_2121


2

CL\_2121


2

CL\_2121


2

CL\_2121


1

CL\_2121


1

CL\_2121


1

CL\_2121


1

CL\_2121


1

CL\_2139


1

CL\_2121


1

CL\_2121


1

CL\_2121

fGI ID


CL\_INS\_118
CL\_INS\_6
CL\_INS\_118
CL\_INS\_118
CL\_INS\_118
CL\_INS\_118
CL\_INS\_118
CL\_INS\_118
CL\_INS\_118
CL\_INS\_118
CL\_INS\_118
CL\_INS\_118
CL\_INS\_118
CL\_INS\_118
CL\_INS\_118
CL\_INS\_118
Cluster ID


CL\_6754
CL\_11025
CL\_12140
CL\_2119
CL\_7735
CL\_2120
CL\_13112
CL\_10330
CL\_4858
CL\_11319
CL\_10331
CL\_10332
CL\_10333
CL\_4859
CL\_4860
CL\_4861
