## Supplementary material for "A novel method for integrating genomic and Tn-Seq data to identify common *in vivo* fitness mechanisms across multiple bacterial species": S3 Dataset: CL_INS_119.html

Legend

 Hypothetical
 All VFDB Genes

FULL


WINDOWSVGPNG

Trim RowsRemove SingletonsSave Fasta

CL\_2126


CL\_2126


CL\_2126

HighlightSelectShow Genomes


36

CL\_2127


20

CL\_2127


1

CL\_2128

fGI ID

CL\_INS\_119
Cluster ID

CL\_6248
