## Supplementary material for "A novel method for integrating genomic and Tn-Seq data to identify common *in vivo* fitness mechanisms across multiple bacterial species": S3 Dataset: CL_INS_120.html

Legend

 Mobile +extrachromosomalelementfunctions
 Hypothetical
 Other
 All VFDB Genes

FULL


WINDOWSVGPNG

Trim RowsRemove SingletonsSave Fasta

CL\_2164


CL\_2164


CL\_2164


CL\_2164


CL\_2164


CL\_2164

HighlightSelectShow Genomes


43

CL\_2166


10

CL\_2166


1

CL\_2166


1

CL\_2166


1

CL\_2166


1

CL\_2166

fGI ID


CL\_INS\_120
CL\_INS\_120
CL\_INS\_120
CL\_INS\_120
Cluster ID


CL\_11739
CL\_2165
CL\_9936
CL\_7359
