## Supplementary material for "A novel method for integrating genomic and Tn-Seq data to identify common *in vivo* fitness mechanisms across multiple bacterial species": S3 Dataset: CL_INS_121.html

Legend

 Hypothetical
 All EssentialGenes
 Other
 All VFDB Genes

FULL


WINDOWSVGPNG

Trim RowsRemove SingletonsSave Fasta

CL\_2176


CL\_2176


CL\_2176


CL\_2176


CL\_2175


CL\_2176


CL\_2176

HighlightSelectShow Genomes


27

CL\_2177


25

CL\_2177


1

Break


1

CL\_2177


1

CL\_2177


1

CL\_2177


1

CL\_2177

fGI ID


CL\_INS\_107
CL\_INS\_121
CL\_INS\_121
CL\_INS\_121
CL\_INS\_121
CL\_INS\_121
CL\_INS\_121
CL\_INS\_121
CL\_INS\_121
CL\_INS\_121
CL\_INS\_121
CL\_INS\_121
Cluster ID


CL\_6749
CL\_12839
CL\_11258
CL\_4862
CL\_9648
CL\_9649
CL\_9650
CL\_9651
CL\_9652
CL\_9653
CL\_9654
CL\_9655
