## Supplementary material for "A novel method for integrating genomic and Tn-Seq data to identify common *in vivo* fitness mechanisms across multiple bacterial species": S3 Dataset: CL_INS_124.html

Legend

 Regulatoryfunctions
 Hypothetical
 Other
 Transport +binding proteins

FULL


WINDOWSVGPNG

Trim RowsRemove SingletonsSave Fasta

CL\_2257


CL\_2257


CL\_2256

HighlightSelectShow Genomes


47

CL\_2258


8

CL\_2258


1

CL\_2258

fGI ID

CL\_INS\_124
Cluster ID

CL\_4864
