## Supplementary material for "A novel method for integrating genomic and Tn-Seq data to identify common *in vivo* fitness mechanisms across multiple bacterial species": S3 Dataset: CL_INS_125.html

Legend

 Hypothetical
 All VFDB Genes

FULL


WINDOWSVGPNG

Trim RowsRemove SingletonsSave Fasta

CL\_2259


CL\_2259


CL\_2259


CL\_2259


CL\_2259

HighlightSelectShow Genomes


26

CL\_2263


24

CL\_2263


3

CL\_2264


3

CL\_2263


1

CL\_2263

fGI ID


CL\_INS\_125
CL\_INS\_125
CL\_INS\_125
Cluster ID


CL\_2260
CL\_2261
CL\_2262
