## Supplementary material for "A novel method for integrating genomic and Tn-Seq data to identify common *in vivo* fitness mechanisms across multiple bacterial species": S3 Dataset: CL_INS_126.html

Legend

 Hypothetical
 Other

FULL


WINDOWSVGPNG

Trim RowsRemove SingletonsSave Fasta

CL\_4865


CL\_4865


CL\_4865

HighlightSelectShow Genomes


42

CL\_2267


12

CL\_2267


1

CL\_2268

fGI ID


CL\_INS\_20
CL\_INS\_126
Cluster ID


CL\_5557
CL\_5558
