## Supplementary material for "A novel method for integrating genomic and Tn-Seq data to identify common *in vivo* fitness mechanisms across multiple bacterial species": S3 Dataset: CL_INS_127.html

Legend

 Hypothetical
 Other
 All VFDB Genes

FULL


WINDOWSVGPNG

Trim RowsRemove SingletonsSave Fasta

CL\_2273


CL\_2273


CL\_2273


CL\_2273

HighlightSelectShow Genomes


47

CL\_2274


5

CL\_2274


2

CL\_2274


1

CL\_2274

fGI ID


CL\_INS\_127
CL\_INS\_127
CL\_INS\_127
Cluster ID


CL\_7364
CL\_12465
CL\_7365
