## Supplementary material for "A novel method for integrating genomic and Tn-Seq data to identify common *in vivo* fitness mechanisms across multiple bacterial species": S3 Dataset: CL_INS_130.html

Legend

 Mobile +extrachromosomalelementfunctions
 All EssentialGenes
 Other

FULL


WINDOWSVGPNG

Trim RowsRemove SingletonsSave Fasta

CL\_2378


CL\_2378

HighlightSelectShow Genomes


30

CL\_2380


25

CL\_2380

fGI ID

CL\_INS\_130
Cluster ID

CL\_2379
