## Supplementary material for "A novel method for integrating genomic and Tn-Seq data to identify common *in vivo* fitness mechanisms across multiple bacterial species": S3 Dataset: CL_INS_131.html

Legend

 Mobile +extrachromosomalelementfunctions
 Regulatoryfunctions
 Hypothetical
 DNA Metabolism
 All EssentialGenes
 Proteinsynthesis/fate
 Other
 EnergyMetabolism
 Transport +binding proteins
 All VFDB Genes

FULL


WINDOWSVGPNG

Trim RowsRemove SingletonsSave Fasta

CL\_2389


CL\_2389


CL\_2389


CL\_2389


CL\_2389


CL\_2389


CL\_2389


CL\_2389


CL\_2389


CL\_2389


CL\_2389


CL\_2389

HighlightSelectShow Genomes


45

CL\_2390


1

CL\_2390


1

CL\_2390


1

CL\_2390


1

CL\_2390


1

Break


1

CL\_2390


1

CL\_2390


1

CL\_2390


1

CL\_2390


1

CL\_2390


1

CL\_2390

fGI ID


CL\_INS\_131
CL\_INS\_131
CL\_INS\_131
CL\_INS\_198
CL\_INS\_131
CL\_INS\_131
CL\_INS\_131
CL\_INS\_131
CL\_INS\_131
CL\_INS\_131
CL\_INS\_131
CL\_INS\_131
CL\_INS\_131
CL\_INS\_131
CL\_INS\_133
CL\_INS\_131
CL\_INS\_131
CL\_INS\_131
CL\_INS\_133
CL\_INS\_133
CL\_INS\_133
CL\_INS\_133
CL\_INS\_133
CL\_INS\_133
CL\_INS\_133
CL\_INS\_133
CL\_INS\_133
CL\_INS\_133
CL\_INS\_133
CL\_INS\_133
CL\_INS\_133
CL\_INS\_133
CL\_INS\_217
CL\_INS\_131
CL\_INS\_133
CL\_INS\_133
CL\_INS\_131
CL\_INS\_131
CL\_INS\_198
CL\_INS\_198
CL\_INS\_198
CL\_INS\_198
CL\_INS\_198
CL\_INS\_198
CL\_INS\_198
CL\_INS\_198
CL\_INS\_198
CL\_INS\_198
CL\_INS\_198
CL\_INS\_198
CL\_INS\_198
CL\_INS\_198
CL\_INS\_198
CL\_INS\_198
CL\_INS\_198
CL\_INS\_198
CL\_INS\_198
CL\_INS\_219
CL\_INS\_198
CL\_INS\_131
CL\_INS\_131
CL\_INS\_198
CL\_INS\_198
CL\_INS\_198
CL\_INS\_217
CL\_INS\_133
CL\_INS\_133
CL\_INS\_196
CL\_INS\_133
CL\_INS\_133
CL\_INS\_133
CL\_INS\_133
CL\_INS\_133
CL\_INS\_133
CL\_INS\_133
CL\_INS\_133
CL\_INS\_133
CL\_INS\_133
CL\_INS\_133
CL\_INS\_133
CL\_INS\_133
CL\_INS\_133
CL\_INS\_133
CL\_INS\_133
CL\_INS\_133
CL\_INS\_133
CL\_INS\_133
CL\_INS\_133
CL\_INS\_131
CL\_INS\_131
CL\_INS\_131
CL\_INS\_131
CL\_INS\_131
CL\_INS\_131
CL\_INS\_131
CL\_INS\_131
CL\_INS\_131
CL\_INS\_131
CL\_INS\_131
CL\_INS\_131
CL\_INS\_131
CL\_INS\_131
CL\_INS\_131
CL\_INS\_131
CL\_INS\_217
CL\_INS\_217
CL\_INS\_217
CL\_INS\_131
CL\_INS\_131
CL\_INS\_131
CL\_INS\_131
CL\_INS\_131
CL\_INS\_217
CL\_INS\_131
CL\_INS\_131
CL\_INS\_131
CL\_INS\_131
CL\_INS\_131
CL\_INS\_217
CL\_INS\_198
CL\_INS\_131
CL\_INS\_131
CL\_INS\_131
CL\_INS\_131
CL\_INS\_131
CL\_INS\_131
CL\_INS\_131
CL\_INS\_131
CL\_INS\_133
CL\_INS\_217
CL\_INS\_131
CL\_INS\_131
CL\_INS\_220
CL\_INS\_220
CL\_INS\_220
CL\_INS\_220
CL\_INS\_220
CL\_INS\_131
CL\_INS\_133
CL\_INS\_133
CL\_INS\_133
CL\_INS\_133
CL\_INS\_133
CL\_INS\_133
CL\_INS\_133
CL\_INS\_133
CL\_INS\_133
CL\_INS\_133
CL\_INS\_133
CL\_INS\_131
CL\_INS\_131
CL\_INS\_131
CL\_INS\_131
CL\_INS\_198
CL\_INS\_198
CL\_INS\_198
CL\_INS\_198
CL\_INS\_131
CL\_INS\_131
CL\_INS\_131
CL\_INS\_131
CL\_INS\_131
CL\_INS\_220
CL\_INS\_133
CL\_INS\_133
CL\_INS\_133
CL\_INS\_133
CL\_INS\_220
CL\_INS\_131
CL\_INS\_196
CL\_INS\_133
CL\_INS\_133
CL\_INS\_133
CL\_INS\_133
CL\_INS\_133
CL\_INS\_133
CL\_INS\_133
CL\_INS\_133
CL\_INS\_131
CL\_INS\_220
CL\_INS\_133
CL\_INS\_133
CL\_INS\_133
CL\_INS\_133
CL\_INS\_98
CL\_INS\_131
CL\_INS\_98
CL\_INS\_133
CL\_INS\_137
CL\_INS\_217
CL\_INS\_131
CL\_INS\_133
CL\_INS\_131
CL\_INS\_83
CL\_INS\_83
CL\_INS\_131
Cluster ID


CL\_13037
CL\_13038
CL\_12346
CL\_9368
CL\_13511
CL\_13544
CL\_13543
CL\_9035
CL\_13593
CL\_11639
CL\_9034
CL\_9033
CL\_13542
CL\_11987
CL\_10715
CL\_11986
CL\_11985
CL\_11984
CL\_5566
CL\_5568
CL\_8712
CL\_5570
CL\_5571
CL\_5572
CL\_5573
CL\_5577
CL\_8640
CL\_5579
CL\_7742
CL\_5582
CL\_6378
CL\_6377
CL\_6376
CL\_12347
CL\_5585
CL\_5586
CL\_9018
CL\_9017
CL\_8203
CL\_7867
CL\_7866
CL\_7865
CL\_7864
CL\_7863
CL\_7862
CL\_7861
CL\_7860
CL\_7859
CL\_7858
CL\_7857
CL\_7856
CL\_7855
CL\_7854
CL\_7853
CL\_7852
CL\_7851
CL\_6575
CL\_6418
CL\_8568
CL\_12348
CL\_12349
CL\_8567
CL\_7847
CL\_7846
CL\_7418
CL\_7417
CL\_7416
CL\_10517
CL\_8319
CL\_9222
CL\_8643
CL\_6468
CL\_6467
CL\_6466
CL\_8238
CL\_8237
CL\_8236
CL\_7753
CL\_8235
CL\_8234
CL\_5600
CL\_5601
CL\_5603
CL\_5604
CL\_5605
CL\_8745
CL\_5607
CL\_5608
CL\_9032
CL\_9031
CL\_9030
CL\_9029
CL\_9300
CL\_9301
CL\_9302
CL\_9028
CL\_9303
CL\_13510
CL\_9027
CL\_11638
CL\_9026
CL\_9025
CL\_13509
CL\_13508
CL\_6392
CL\_6391
CL\_6390
CL\_9024
CL\_11637
CL\_11636
CL\_11635
CL\_11634
CL\_7424
CL\_13507
CL\_9023
CL\_9022
CL\_9021
CL\_9020
CL\_6384
CL\_8195
CL\_9304
CL\_9305
CL\_9306
CL\_9307
CL\_13506
CL\_13505
CL\_13504
CL\_9019
CL\_7795
CL\_6379
CL\_11633
CL\_11632
CL\_7873
CL\_7872
CL\_7871
CL\_7870
CL\_7869
CL\_9308
CL\_10202
CL\_10201
CL\_10200
CL\_10199
CL\_10198
CL\_10197
CL\_10196
CL\_10195
CL\_10194
CL\_8312
CL\_8311
CL\_12132
CL\_13541
CL\_13540
CL\_11631
CL\_11630
CL\_11629
CL\_11628
CL\_11627
CL\_11626
CL\_11625
CL\_11624
CL\_11623
CL\_11622
CL\_10653
CL\_7756
CL\_7757
CL\_7758
CL\_7759
CL\_10654
CL\_11621
CL\_11620
CL\_5609
CL\_6462
CL\_9328
CL\_6460
CL\_5610
CL\_5611
CL\_6350
CL\_8864
CL\_11983
CL\_6454
CL\_6349
CL\_6348
CL\_6347
CL\_5614
CL\_3182
CL\_13503
CL\_3180
CL\_7388
CL\_7389
CL\_7390
CL\_11982
CL\_5616
CL\_11981
CL\_6714
CL\_6715
CL\_9016
