## Supplementary material for "A novel method for integrating genomic and Tn-Seq data to identify common *in vivo* fitness mechanisms across multiple bacterial species": S3 Dataset: CL_INS_133.html

Legend

 Mobile +extrachromosomalelementfunctions
 Regulatoryfunctions
 Hypothetical
 DNA Metabolism
 AntibioticResistance
 Proteinsynthesis/fate
 Other
 EnergyMetabolism
 Transport +binding proteins
 All VFDB Genes

FULL


WINDOWSVGPNG

Trim RowsRemove SingletonsSave Fasta

CL\_2406


CL\_2406


CL\_2406


CL\_2406


CL\_2406


CL\_2406


CL\_2406


CL\_2406


CL\_2406


CL\_2406


CL\_2406


CL\_3297


Break


CL\_2406


CL\_2406


CL\_2406


CL\_2406


CL\_2406


CL\_2406


CL\_2406


CL\_2406


CL\_2406


CL\_2406

HighlightSelectShow Genomes


31

CL\_2407


5

CL\_2407


2

CL\_2407


1

CL\_2407


1

CL\_2407


1

CL\_2407


1

CL\_2407


1

CL\_2407


1

CL\_2407


1

CL\_2407


1

CL\_2407


1

CL\_2407


1

CL\_2407


1

Break


1

CL\_2407


1

CL\_2407


1

CL\_2407


1

CL\_2407


1

CL\_3704


1

CL\_2407


1

CL\_2407


1

CL\_2407


1

CL\_2407

fGI ID


CL\_INS\_133
CL\_INS\_133
CL\_INS\_133
CL\_INS\_133
CL\_INS\_133
CL\_INS\_133
CL\_INS\_133
CL\_INS\_133
CL\_INS\_133
CL\_INS\_133
CL\_INS\_133
CL\_INS\_133
CL\_INS\_133
CL\_INS\_133
CL\_INS\_133
CL\_INS\_133
CL\_INS\_133
CL\_INS\_133
CL\_INS\_133
CL\_INS\_133
CL\_INS\_133
CL\_INS\_133
CL\_INS\_133
CL\_INS\_133
CL\_INS\_133
CL\_INS\_133
CL\_INS\_133
CL\_INS\_133
CL\_INS\_133
CL\_INS\_133
CL\_INS\_133
CL\_INS\_133
CL\_INS\_133
CL\_INS\_133
CL\_INS\_133
CL\_INS\_133
CL\_INS\_133
CL\_INS\_133
CL\_INS\_133
CL\_INS\_133
CL\_INS\_133
CL\_INS\_133
CL\_INS\_133
CL\_INS\_133
CL\_INS\_133
CL\_INS\_133
CL\_INS\_133
CL\_INS\_133
CL\_INS\_133
CL\_INS\_133
CL\_INS\_133
CL\_INS\_133
CL\_INS\_133
CL\_INS\_133
CL\_INS\_133
CL\_INS\_133
CL\_INS\_133
CL\_INS\_133
CL\_INS\_133
CL\_INS\_133
CL\_INS\_133
CL\_INS\_133
CL\_INS\_133
CL\_INS\_133
CL\_INS\_133
CL\_INS\_133
CL\_INS\_133
CL\_INS\_133
CL\_INS\_133
CL\_INS\_133
CL\_INS\_133
CL\_INS\_133
CL\_INS\_133
CL\_INS\_133
CL\_INS\_133
CL\_INS\_133
CL\_INS\_133
CL\_INS\_133
CL\_INS\_133
CL\_INS\_133
CL\_INS\_133
CL\_INS\_133
CL\_INS\_133
CL\_INS\_133
CL\_INS\_133
CL\_INS\_133
CL\_INS\_133
CL\_INS\_133
CL\_INS\_133
CL\_INS\_133
CL\_INS\_133
CL\_INS\_133
CL\_INS\_133
CL\_INS\_133
CL\_INS\_133
CL\_INS\_133
CL\_INS\_133
CL\_INS\_133
CL\_INS\_133
CL\_INS\_133
CL\_INS\_133
CL\_INS\_133
CL\_INS\_133
CL\_INS\_133
CL\_INS\_133
CL\_INS\_133
CL\_INS\_133
CL\_INS\_133
CL\_INS\_133
CL\_INS\_133
CL\_INS\_133
CL\_INS\_133
CL\_INS\_133
CL\_INS\_133
CL\_INS\_133
CL\_INS\_133
CL\_INS\_133
CL\_INS\_133
CL\_INS\_133
CL\_INS\_133
CL\_INS\_133
CL\_INS\_133
CL\_INS\_133
CL\_INS\_133
CL\_INS\_133
CL\_INS\_133
CL\_INS\_133
CL\_INS\_133
CL\_INS\_133
CL\_INS\_133
CL\_INS\_133
CL\_INS\_133
CL\_INS\_133
CL\_INS\_133
CL\_INS\_133
CL\_INS\_133
CL\_INS\_133
CL\_INS\_133
CL\_INS\_133
CL\_INS\_133
CL\_INS\_133
CL\_INS\_133
CL\_INS\_133
CL\_INS\_133
CL\_INS\_133
CL\_INS\_133
CL\_INS\_133
CL\_INS\_133
CL\_INS\_133
CL\_INS\_133
CL\_INS\_133
CL\_INS\_133
CL\_INS\_133
CL\_INS\_133
CL\_INS\_133
CL\_INS\_133
CL\_INS\_133
CL\_INS\_133
CL\_INS\_133
CL\_INS\_133
CL\_INS\_133
CL\_INS\_133
CL\_INS\_133
CL\_INS\_133
CL\_INS\_133
CL\_INS\_133
CL\_INS\_133
CL\_INS\_133
CL\_INS\_133
CL\_INS\_133
CL\_INS\_133
CL\_INS\_133
CL\_INS\_133
CL\_INS\_133
CL\_INS\_133
CL\_INS\_133
CL\_INS\_133
CL\_INS\_133
CL\_INS\_133
CL\_INS\_133
CL\_INS\_133
CL\_INS\_133
CL\_INS\_133
CL\_INS\_133
CL\_INS\_133
CL\_INS\_133
CL\_INS\_133
CL\_INS\_133
CL\_INS\_133
CL\_INS\_133
CL\_INS\_133
CL\_INS\_133
CL\_INS\_133
CL\_INS\_133
CL\_INS\_133
CL\_INS\_133
CL\_INS\_133
CL\_INS\_133
CL\_INS\_133
CL\_INS\_133
CL\_INS\_133
CL\_INS\_133
CL\_INS\_133
CL\_INS\_133
CL\_INS\_133
CL\_INS\_133
CL\_INS\_133
CL\_INS\_133
CL\_INS\_133
CL\_INS\_133
CL\_INS\_133
CL\_INS\_133
CL\_INS\_133
CL\_INS\_133
CL\_INS\_133
CL\_INS\_133
CL\_INS\_133
CL\_INS\_133
CL\_INS\_133
CL\_INS\_133
CL\_INS\_133
CL\_INS\_133
CL\_INS\_133
CL\_INS\_133
CL\_INS\_133
CL\_INS\_133
CL\_INS\_133
CL\_INS\_133
CL\_INS\_133
CL\_INS\_133
CL\_INS\_133
CL\_INS\_133
CL\_INS\_133
CL\_INS\_133
CL\_INS\_133
CL\_INS\_133
CL\_INS\_133
CL\_INS\_133
CL\_INS\_133
CL\_INS\_133
CL\_INS\_133
CL\_INS\_133
CL\_INS\_133
CL\_INS\_133
CL\_INS\_133
CL\_INS\_133
CL\_INS\_133
CL\_INS\_133
CL\_INS\_133
CL\_INS\_133
CL\_INS\_133
CL\_INS\_133
CL\_INS\_133
CL\_INS\_133
CL\_INS\_133
CL\_INS\_133
CL\_INS\_133
CL\_INS\_133
CL\_INS\_133
CL\_INS\_133
CL\_INS\_133
CL\_INS\_133
CL\_INS\_133
CL\_INS\_133
CL\_INS\_133
CL\_INS\_133
CL\_INS\_133
CL\_INS\_133
CL\_INS\_133
CL\_INS\_133
CL\_INS\_133
CL\_INS\_133
CL\_INS\_133
CL\_INS\_133
CL\_INS\_133
CL\_INS\_133
CL\_INS\_133
CL\_INS\_133
CL\_INS\_133
CL\_INS\_133
CL\_INS\_133
CL\_INS\_133
CL\_INS\_133
CL\_INS\_133
CL\_INS\_133
CL\_INS\_133
CL\_INS\_133
CL\_INS\_133
CL\_INS\_133
CL\_INS\_133
CL\_INS\_133
CL\_INS\_133
CL\_INS\_133
CL\_INS\_133
CL\_INS\_133
CL\_INS\_133
CL\_INS\_133
CL\_INS\_133
CL\_INS\_133
CL\_INS\_133
CL\_INS\_133
CL\_INS\_133
CL\_INS\_133
CL\_INS\_133
CL\_INS\_133
CL\_INS\_133
CL\_INS\_133
CL\_INS\_133
CL\_INS\_133
CL\_INS\_133
CL\_INS\_133
CL\_INS\_133
CL\_INS\_133
CL\_INS\_133
CL\_INS\_133
CL\_INS\_133
CL\_INS\_133
CL\_INS\_133
CL\_INS\_133
CL\_INS\_133
CL\_INS\_133
CL\_INS\_133
CL\_INS\_133
CL\_INS\_133
CL\_INS\_133
CL\_INS\_133
CL\_INS\_133
CL\_INS\_133
CL\_INS\_133
CL\_INS\_133
CL\_INS\_133
CL\_INS\_133
CL\_INS\_133
CL\_INS\_133
CL\_INS\_133
CL\_INS\_133
CL\_INS\_133
CL\_INS\_133
CL\_INS\_133
CL\_INS\_133
CL\_INS\_133
CL\_INS\_133
CL\_INS\_133
CL\_INS\_133
CL\_INS\_133
CL\_INS\_133
CL\_INS\_133
CL\_INS\_133
CL\_INS\_133
CL\_INS\_133
CL\_INS\_133
CL\_INS\_133
CL\_INS\_133
CL\_INS\_133
CL\_INS\_133
CL\_INS\_133
CL\_INS\_133
CL\_INS\_133
CL\_INS\_133
CL\_INS\_133
CL\_INS\_133
CL\_INS\_133
CL\_INS\_133
CL\_INS\_133
CL\_INS\_133
CL\_INS\_133
CL\_INS\_133
CL\_INS\_133
CL\_INS\_133
CL\_INS\_133
CL\_INS\_133
CL\_INS\_133
CL\_INS\_133
CL\_INS\_133
CL\_INS\_133
CL\_INS\_133
CL\_INS\_133
CL\_INS\_133
CL\_INS\_133
CL\_INS\_133
CL\_INS\_133
CL\_INS\_133
CL\_INS\_133
CL\_INS\_133
CL\_INS\_133
CL\_INS\_133
CL\_INS\_133
CL\_INS\_133
CL\_INS\_133
CL\_INS\_133
CL\_INS\_133
CL\_INS\_133
CL\_INS\_133
CL\_INS\_133
CL\_INS\_133
CL\_INS\_133
CL\_INS\_133
CL\_INS\_133
CL\_INS\_133
CL\_INS\_133
CL\_INS\_133
CL\_INS\_133
CL\_INS\_133
CL\_INS\_133
CL\_INS\_133
CL\_INS\_133
CL\_INS\_133
CL\_INS\_133
CL\_INS\_133
CL\_INS\_133
CL\_INS\_133
CL\_INS\_133
Cluster ID


CL\_9656
CL\_7361
CL\_8697
CL\_8696
CL\_8695
CL\_8694
CL\_6485
CL\_6484
CL\_8693
CL\_8692
CL\_8691
CL\_8690
CL\_8689
CL\_8688
CL\_8687
CL\_8686
CL\_6477
CL\_6476
CL\_6475
CL\_6474
CL\_8685
CL\_8684
CL\_8683
CL\_8682
CL\_8681
CL\_8680
CL\_8679
CL\_8678
CL\_8677
CL\_8676
CL\_8675
CL\_8674
CL\_8673
CL\_8672
CL\_8671
CL\_8670
CL\_8669
CL\_8668
CL\_8667
CL\_5559
CL\_5560
CL\_9309
CL\_9310
CL\_3242
CL\_5561
CL\_7736
CL\_7737
CL\_8728
CL\_10339
CL\_12841
CL\_12842
CL\_13303
CL\_12843
CL\_5562
CL\_5563
CL\_5564
CL\_10214
CL\_9217
CL\_9218
CL\_10713
CL\_11943
CL\_5535
CL\_8246
CL\_10715
CL\_10714
CL\_8245
CL\_8875
CL\_8874
CL\_8873
CL\_11041
CL\_5565
CL\_5566
CL\_8244
CL\_8631
CL\_8632
CL\_5567
CL\_13302
CL\_13301
CL\_11738
CL\_7738
CL\_8713
CL\_5568
CL\_7739
CL\_5569
CL\_11042
CL\_8871
CL\_7882
CL\_7881
CL\_8712
CL\_5570
CL\_5571
CL\_11043
CL\_8872
CL\_12844
CL\_12845
CL\_12846
CL\_12350
CL\_13453
CL\_13452
CL\_6207
CL\_8243
CL\_8242
CL\_6208
CL\_11044
CL\_11045
CL\_13451
CL\_6210
CL\_6211
CL\_5528
CL\_8241
CL\_7880
CL\_7879
CL\_5572
CL\_5573
CL\_7828
CL\_5574
CL\_13039
CL\_7740
CL\_5575
CL\_7741
CL\_12847
CL\_12848
CL\_12849
CL\_12850
CL\_8708
CL\_5580
CL\_8240
CL\_5542
CL\_5543
CL\_7821
CL\_7820
CL\_8870
CL\_8869
CL\_5581
CL\_9311
CL\_9312
CL\_7795
CL\_9313
CL\_9314
CL\_11046
CL\_8868
CL\_8867
CL\_5582
CL\_5583
CL\_9219
CL\_6472
CL\_6378
CL\_6377
CL\_6471
CL\_5576
CL\_5577
CL\_8640
CL\_5578
CL\_5579
CL\_7742
CL\_13040
CL\_7743
CL\_13450
CL\_7744
CL\_7745
CL\_5584
CL\_10305
CL\_5585
CL\_10213
CL\_10212
CL\_10211
CL\_10210
CL\_10209
CL\_10208
CL\_10207
CL\_10206
CL\_10205
CL\_10204
CL\_10203
CL\_10202
CL\_10201
CL\_10200
CL\_10199
CL\_10198
CL\_10197
CL\_10196
CL\_10195
CL\_10194
CL\_8312
CL\_8311
CL\_8310
CL\_8309
CL\_8308
CL\_8866
CL\_8306
CL\_8305
CL\_8304
CL\_8865
CL\_9220
CL\_9221
CL\_10386
CL\_7417
CL\_7416
CL\_8319
CL\_7868
CL\_8318
CL\_8317
CL\_8316
CL\_8315
CL\_8314
CL\_8313
CL\_10341
CL\_10342
CL\_12851
CL\_12852
CL\_12853
CL\_12854
CL\_12855
CL\_12856
CL\_12857
CL\_12858
CL\_12859
CL\_12860
CL\_12861
CL\_12862
CL\_12863
CL\_12864
CL\_12865
CL\_10954
CL\_12866
CL\_6464
CL\_12867
CL\_9222
CL\_5586
CL\_13300
CL\_8239
CL\_7746
CL\_10340
CL\_5587
CL\_9315
CL\_9316
CL\_11047
CL\_8200
CL\_11048
CL\_11049
CL\_8201
CL\_9317
CL\_7749
CL\_6466
CL\_13298
CL\_13297
CL\_13296
CL\_8238
CL\_7750
CL\_8237
CL\_8236
CL\_11050
CL\_6469
CL\_11051
CL\_8643
CL\_13449
CL\_13448
CL\_5588
CL\_13299
CL\_13041
CL\_8832
CL\_6468
CL\_6467
CL\_7747
CL\_7748
CL\_5589
CL\_5590
CL\_5591
CL\_5592
CL\_5593
CL\_5594
CL\_5595
CL\_12351
CL\_5596
CL\_5597
CL\_5598
CL\_5599
CL\_10992
CL\_10994
CL\_10995
CL\_10996
CL\_5157
CL\_5204
CL\_5203
CL\_5202
CL\_5201
CL\_4435
CL\_5200
CL\_5199
CL\_5198
CL\_5328
CL\_6967
CL\_10805
CL\_13390
CL\_5337
CL\_6969
CL\_4451
CL\_5884
CL\_7591
CL\_7578
CL\_8396
CL\_8397
CL\_8421
CL\_5327
CL\_8422
CL\_8423
CL\_9318
CL\_9319
CL\_10955
CL\_9320
CL\_7751
CL\_7752
CL\_7753
CL\_9321
CL\_8235
CL\_8748
CL\_8747
CL\_8746
CL\_9322
CL\_7754
CL\_7755
CL\_7756
CL\_7757
CL\_7758
CL\_7759
CL\_6465
CL\_8234
CL\_13295
CL\_10712
CL\_10711
CL\_9323
CL\_9324
CL\_9325
CL\_9326
CL\_10516
CL\_9327
CL\_13042
CL\_13043
CL\_13044
CL\_13045
CL\_9657
CL\_9658
CL\_4813
CL\_9659
CL\_4930
CL\_5600
CL\_5601
CL\_5602
CL\_5603
CL\_5604
CL\_5605
CL\_8233
CL\_8666
CL\_8665
CL\_8664
CL\_8663
CL\_8662
CL\_8661
CL\_8660
CL\_8745
CL\_10710
CL\_10709
CL\_10708
CL\_5606
CL\_5607
CL\_5608
CL\_8744
CL\_5609
CL\_8834
CL\_8835
CL\_8836
CL\_8837
CL\_6462
CL\_9328
CL\_6460
CL\_8303
CL\_10707
CL\_10706
CL\_8964
CL\_6459
CL\_8864
CL\_10193
CL\_5610
CL\_7760
CL\_5611
CL\_6350
CL\_6349
CL\_8743
CL\_8742
CL\_6348
CL\_5612
CL\_5613
CL\_9223
CL\_9224
CL\_6346
CL\_6458
CL\_7761
CL\_10705
CL\_10704
CL\_8863
CL\_6347
CL\_7388
CL\_7412
CL\_7362
CL\_5045
CL\_5044
CL\_5043
CL\_5042
CL\_6456
CL\_5614
CL\_5615
CL\_5616
