## Supplementary material for "A novel method for integrating genomic and Tn-Seq data to identify common *in vivo* fitness mechanisms across multiple bacterial species": S3 Dataset: CL_INS_134.html

Legend

 Regulatoryfunctions
 Hypothetical
 Other
 EnergyMetabolism
 Transport +binding proteins
 All VFDB Genes

FULL


WINDOWSVGPNG

Trim RowsRemove SingletonsSave Fasta

CL\_2420


CL\_2420


CL\_2420

HighlightSelectShow Genomes


49

CL\_2421


5

CL\_2421


3

CL\_2421

fGI ID


CL\_INS\_134
CL\_INS\_134
CL\_INS\_134
CL\_INS\_134
CL\_INS\_134
CL\_INS\_134
CL\_INS\_134
CL\_INS\_134
CL\_INS\_134
CL\_INS\_134
CL\_INS\_134
CL\_INS\_134
CL\_INS\_134
Cluster ID


CL\_6758
CL\_7366
CL\_7367
CL\_7368
CL\_7369
CL\_7370
CL\_6759
CL\_6760
CL\_6761
CL\_6762
CL\_6763
CL\_6764
CL\_6765
