## Supplementary material for "A novel method for integrating genomic and Tn-Seq data to identify common *in vivo* fitness mechanisms across multiple bacterial species": S3 Dataset: CL_INS_136.html

Legend

 Hypothetical
 All EssentialGenes

FULL


WINDOWSVGPNG

Trim RowsRemove SingletonsSave Fasta

CL\_2465


CL\_2465


CL\_2465

HighlightSelectShow Genomes


34

CL\_2466


17

CL\_2466


6

CL\_2466

fGI ID


CL\_INS\_136
CL\_INS\_136
Cluster ID


CL\_10192
CL\_4869
