## Supplementary material for "A novel method for integrating genomic and Tn-Seq data to identify common *in vivo* fitness mechanisms across multiple bacterial species": S3 Dataset: CL_INS_137.html

Legend

 Mobile +extrachromosomalelementfunctions
 Regulatoryfunctions
 Hypothetical
 DNA Metabolism
 All EssentialGenes
 Other
 All VFDB Genes

FULL


WINDOWSVGPNG

Trim RowsRemove SingletonsSave Fasta

CL\_2469


CL\_2469


CL\_2469


CL\_2469

HighlightSelectShow Genomes


52

CL\_2470


2

CL\_2470


2

CL\_2470


1

CL\_2470

fGI ID


CL\_INS\_137
CL\_INS\_137
CL\_INS\_137
CL\_INS\_137
CL\_INS\_137
CL\_INS\_137
CL\_INS\_137
CL\_INS\_137
CL\_INS\_137
CL\_INS\_217
CL\_INS\_217
CL\_INS\_217
CL\_INS\_137
CL\_INS\_137
CL\_INS\_217
CL\_INS\_217
CL\_INS\_217
CL\_INS\_137
CL\_INS\_137
CL\_INS\_83
CL\_INS\_83
CL\_INS\_83
CL\_INS\_137
CL\_INS\_137
CL\_INS\_137
CL\_INS\_137
CL\_INS\_217
CL\_INS\_217
CL\_INS\_217
CL\_INS\_217
CL\_INS\_137
CL\_INS\_137
CL\_INS\_137
CL\_INS\_137
CL\_INS\_137
CL\_INS\_217
CL\_INS\_217
CL\_INS\_217
CL\_INS\_83
CL\_INS\_83
CL\_INS\_217
CL\_INS\_217
CL\_INS\_217
CL\_INS\_137
CL\_INS\_137
CL\_INS\_217
CL\_INS\_137
CL\_INS\_217
CL\_INS\_137
CL\_INS\_137
CL\_INS\_137
CL\_INS\_137
CL\_INS\_137
CL\_INS\_83
CL\_INS\_83
CL\_INS\_83
CL\_INS\_83
CL\_INS\_133
CL\_INS\_133
CL\_INS\_133
CL\_INS\_133
CL\_INS\_133
CL\_INS\_137
CL\_INS\_217
CL\_INS\_137
CL\_INS\_137
CL\_INS\_133
CL\_INS\_217
CL\_INS\_137
CL\_INS\_98
CL\_INS\_137
Cluster ID


CL\_7371
CL\_7372
CL\_7373
CL\_7374
CL\_7375
CL\_7376
CL\_7377
CL\_7378
CL\_7379
CL\_11694
CL\_7423
CL\_6384
CL\_7380
CL\_7381
CL\_6383
CL\_6382
CL\_6381
CL\_7382
CL\_7383
CL\_7384
CL\_6220
CL\_3230
CL\_7385
CL\_7386
CL\_12466
CL\_7387
CL\_6405
CL\_6404
CL\_6403
CL\_6402
CL\_10703
CL\_10702
CL\_10701
CL\_10700
CL\_10699
CL\_6398
CL\_6397
CL\_6396
CL\_5531
CL\_5532
CL\_6393
CL\_6392
CL\_7425
CL\_10698
CL\_10697
CL\_6389
CL\_10696
CL\_10695
CL\_10694
CL\_10693
CL\_10692
CL\_10691
CL\_10690
CL\_6354
CL\_6353
CL\_6352
CL\_6351
CL\_6350
CL\_6349
CL\_6348
CL\_6347
CL\_7388
CL\_7389
CL\_7390
CL\_7391
CL\_10689
CL\_4930
CL\_6345
CL\_7392
CL\_7393
CL\_7394
