## Supplementary material for "A novel method for integrating genomic and Tn-Seq data to identify common *in vivo* fitness mechanisms across multiple bacterial species": S3 Dataset: CL_INS_138.html

Legend

 Hypothetical
 Other

FULL


WINDOWSVGPNG

Trim RowsRemove SingletonsSave Fasta

CL\_2485


CL\_2485


CL\_2485


CL\_2485

HighlightSelectShow Genomes


44

CL\_2486


8

CL\_2486


1

CL\_2486


1

CL\_2486

fGI ID


CL\_INS\_138
CL\_INS\_138
CL\_INS\_138
CL\_INS\_138
CL\_INS\_138
CL\_INS\_138
CL\_INS\_138
CL\_INS\_138
Cluster ID


CL\_4870
CL\_9225
CL\_10293
CL\_9226
CL\_9227
CL\_9228
CL\_9229
CL\_9230
