## Supplementary material for "A novel method for integrating genomic and Tn-Seq data to identify common *in vivo* fitness mechanisms across multiple bacterial species": S3 Dataset: CL_INS_139.html

Legend

 Regulatoryfunctions
 Hypothetical
 All EssentialGenes
 AntibioticResistance
 All VFDB Genes

FULL


WINDOWSVGPNG

Trim RowsRemove SingletonsSave Fasta

CL\_2503


CL\_2503


CL\_2503


CL\_2503


CL\_2503


CL\_2503


CL\_2112


CL\_2502

HighlightSelectShow Genomes


16

CL\_2506


12

CL\_2506


9

CL\_2506


7

CL\_2506


7

CL\_2506


4

CL\_2506


1

CL\_2506


1

CL\_2506

fGI ID


CL\_INS\_139
CL\_INS\_133
CL\_INS\_1
CL\_INS\_20
CL\_INS\_139
CL\_INS\_139
CL\_INS\_139
CL\_INS\_139
Cluster ID


CL\_5885
CL\_5884
CL\_4595
CL\_7638
CL\_13669
CL\_2504
CL\_2505
CL\_6257
