## Supplementary material for "A novel method for integrating genomic and Tn-Seq data to identify common *in vivo* fitness mechanisms across multiple bacterial species": S3 Dataset: CL_INS_140.html

Legend

 Hypothetical
 All VFDB Genes

FULL


WINDOWSVGPNG

Trim RowsRemove SingletonsSave Fasta

CL\_2524


CL\_2524


CL\_2520

HighlightSelectShow Genomes


32

CL\_2525


16

CL\_2525


4

CL\_2525

fGI ID

CL\_INS\_140
Cluster ID

CL\_8340
