## Supplementary material for "A novel method for integrating genomic and Tn-Seq data to identify common *in vivo* fitness mechanisms across multiple bacterial species": S3 Dataset: CL_INS_141.html

Legend

 Hypothetical
 All VFDB Genes

FULL


WINDOWSVGPNG

Trim RowsRemove SingletonsSave Fasta

CL\_2525


CL\_2525

HighlightSelectShow Genomes


54

CL\_2526


3

CL\_2526

fGI ID


CL\_INS\_141
CL\_INS\_141
CL\_INS\_141
Cluster ID


CL\_9444
CL\_9445
CL\_9446
