## Supplementary material for "A novel method for integrating genomic and Tn-Seq data to identify common *in vivo* fitness mechanisms across multiple bacterial species": S3 Dataset: CL_INS_143.html

Legend

 Mobile +extrachromosomalelementfunctions
 Regulatoryfunctions
 Hypothetical
 DNA Metabolism
 All EssentialGenes
 Other
 EnergyMetabolism
 All VFDB Genes
 Transport +binding proteins

FULL


WINDOWSVGPNG

Trim RowsRemove SingletonsSave Fasta

CL\_2575


CL\_2575


CL\_2575


CL\_2575


CL\_2575


CL\_2575


CL\_2575


CL\_2575


CL\_2575


CL\_2575

HighlightSelectShow Genomes


47

CL\_2576


1

CL\_3370


1

CL\_2577


1

CL\_2577


1

CL\_3860


1

CL\_2577


1

CL\_2577


1

CL\_2577


1

CL\_3179


1

Break

fGI ID


CL\_INS\_143
CL\_INS\_144
CL\_INS\_83
CL\_INS\_217
CL\_INS\_143
CL\_INS\_143
CL\_INS\_199
CL\_INS\_217
CL\_INS\_193
CL\_INS\_83
CL\_INS\_83
CL\_INS\_144
CL\_INS\_83
CL\_INS\_217
CL\_INS\_144
CL\_INS\_83
CL\_INS\_194
CL\_INS\_217
CL\_INS\_83
CL\_INS\_83
CL\_INS\_144
CL\_INS\_83
CL\_INS\_98
CL\_INS\_144
CL\_INS\_83
CL\_INS\_83
CL\_INS\_193
CL\_INS\_98
CL\_INS\_217
CL\_INS\_217
CL\_INS\_217
CL\_INS\_217
CL\_INS\_217
CL\_INS\_217
CL\_INS\_217
CL\_INS\_217
CL\_INS\_217
CL\_INS\_217
CL\_INS\_217
CL\_INS\_217
CL\_INS\_83
CL\_INS\_143
CL\_INS\_194
CL\_INS\_144
CL\_INS\_144
CL\_INS\_217
CL\_INS\_217
CL\_INS\_217
CL\_INS\_83
CL\_INS\_83
CL\_INS\_83
CL\_INS\_194
CL\_INS\_194
CL\_INS\_83
CL\_INS\_83
CL\_INS\_83
CL\_INS\_133
CL\_INS\_83
CL\_INS\_83
CL\_INS\_83
CL\_INS\_217
CL\_INS\_217
CL\_INS\_217
CL\_INS\_217
CL\_INS\_217
CL\_INS\_217
CL\_INS\_217
CL\_INS\_217
CL\_INS\_217
CL\_INS\_217
CL\_INS\_217
CL\_INS\_217
CL\_INS\_217
CL\_INS\_217
CL\_INS\_217
CL\_INS\_217
CL\_INS\_217
CL\_INS\_217
CL\_INS\_217
CL\_INS\_217
CL\_INS\_217
CL\_INS\_217
CL\_INS\_217
CL\_INS\_217
CL\_INS\_217
CL\_INS\_217
CL\_INS\_217
CL\_INS\_217
CL\_INS\_217
CL\_INS\_217
CL\_INS\_217
CL\_INS\_217
CL\_INS\_217
CL\_INS\_217
CL\_INS\_83
CL\_INS\_83
CL\_INS\_217
CL\_INS\_83
CL\_INS\_217
CL\_INS\_217
CL\_INS\_217
CL\_INS\_83
CL\_INS\_83
CL\_INS\_83
CL\_INS\_83
CL\_INS\_133
CL\_INS\_133
CL\_INS\_133
CL\_INS\_133
CL\_INS\_217
CL\_INS\_133
CL\_INS\_144
CL\_INS\_144
CL\_INS\_144
CL\_INS\_144
CL\_INS\_83
CL\_INS\_193
CL\_INS\_98
CL\_INS\_83
CL\_INS\_194
CL\_INS\_199
CL\_INS\_199
CL\_INS\_199
CL\_INS\_199
CL\_INS\_199
CL\_INS\_199
CL\_INS\_199
CL\_INS\_199
CL\_INS\_199
CL\_INS\_199
CL\_INS\_199
CL\_INS\_98
CL\_INS\_98
CL\_INS\_98
CL\_INS\_144
CL\_INS\_144
CL\_INS\_98
CL\_INS\_98
CL\_INS\_98
CL\_INS\_194
CL\_INS\_98
CL\_INS\_193
CL\_INS\_194
CL\_INS\_144
CL\_INS\_144
CL\_INS\_144
CL\_INS\_144
CL\_INS\_194
CL\_INS\_194
CL\_INS\_83
CL\_INS\_144
CL\_INS\_144
CL\_INS\_98
CL\_INS\_83
CL\_INS\_83
CL\_INS\_217
CL\_INS\_217
CL\_INS\_98
CL\_INS\_83
CL\_INS\_217
CL\_INS\_217
CL\_INS\_144
CL\_INS\_144
CL\_INS\_144
CL\_INS\_144
CL\_INS\_83
CL\_INS\_217
CL\_INS\_83
CL\_INS\_83
CL\_INS\_143
CL\_INS\_98
CL\_INS\_144
CL\_INS\_144
CL\_INS\_144
CL\_INS\_144
CL\_INS\_144
CL\_INS\_144
CL\_INS\_144
CL\_INS\_144
CL\_INS\_144
CL\_INS\_144
CL\_INS\_133
CL\_INS\_83
CL\_INS\_217
CL\_INS\_217
CL\_INS\_83
CL\_INS\_83
CL\_INS\_144
CL\_INS\_144
CL\_INS\_144
CL\_INS\_144
CL\_INS\_144
CL\_INS\_144
CL\_INS\_98
CL\_INS\_217
CL\_INS\_144
CL\_INS\_83
CL\_INS\_144
CL\_INS\_144
CL\_INS\_144
CL\_INS\_199
CL\_INS\_199
CL\_INS\_199
CL\_INS\_199
CL\_INS\_199
CL\_INS\_199
CL\_INS\_199
CL\_INS\_194
CL\_INS\_83
CL\_INS\_193
CL\_INS\_193
CL\_INS\_193
CL\_INS\_193
CL\_INS\_193
CL\_INS\_144
CL\_INS\_98
CL\_INS\_144
CL\_INS\_133
CL\_INS\_133
CL\_INS\_133
CL\_INS\_98
CL\_INS\_98
CL\_INS\_98
CL\_INS\_83
CL\_INS\_98
CL\_INS\_83
CL\_INS\_98
CL\_INS\_144
CL\_INS\_83
CL\_INS\_83
CL\_INS\_83
CL\_INS\_83
CL\_INS\_144
CL\_INS\_144
CL\_INS\_144
CL\_INS\_144
CL\_INS\_144
CL\_INS\_193
CL\_INS\_83
CL\_INS\_83
CL\_INS\_83
CL\_INS\_83
CL\_INS\_133
CL\_INS\_133
CL\_INS\_133
CL\_INS\_133
CL\_INS\_143
CL\_INS\_144
CL\_INS\_193
CL\_INS\_193
CL\_INS\_98
CL\_INS\_98
CL\_INS\_98
CL\_INS\_98
CL\_INS\_144
CL\_INS\_144
CL\_INS\_144
CL\_INS\_144
CL\_INS\_144
CL\_INS\_83
CL\_INS\_83
CL\_INS\_83
CL\_INS\_144
CL\_INS\_83
CL\_INS\_83
CL\_INS\_83
CL\_INS\_83
CL\_INS\_83
CL\_INS\_98
CL\_INS\_83
CL\_INS\_83
CL\_INS\_193
CL\_INS\_193
CL\_INS\_193
CL\_INS\_193
CL\_INS\_193
CL\_INS\_98
CL\_INS\_194
CL\_INS\_83
CL\_INS\_98
CL\_INS\_83
CL\_INS\_98
CL\_INS\_98
CL\_INS\_98
CL\_INS\_144
CL\_INS\_98
CL\_INS\_194
CL\_INS\_194
CL\_INS\_194
CL\_INS\_133
CL\_INS\_83
CL\_INS\_83
CL\_INS\_98
CL\_INS\_144
CL\_INS\_98
CL\_INS\_144
CL\_INS\_144
CL\_INS\_144
CL\_INS\_144
CL\_INS\_144
CL\_INS\_98
CL\_INS\_98
CL\_INS\_98
CL\_INS\_98
CL\_INS\_98
CL\_INS\_98
CL\_INS\_194
CL\_INS\_194
CL\_INS\_194
CL\_INS\_194
CL\_INS\_98
CL\_INS\_98
CL\_INS\_98
CL\_INS\_199
CL\_INS\_83
CL\_INS\_83
CL\_INS\_144
CL\_INS\_144
CL\_INS\_144
CL\_INS\_144
CL\_INS\_144
CL\_INS\_144
CL\_INS\_144
CL\_INS\_144
CL\_INS\_144
CL\_INS\_144
CL\_INS\_83
CL\_INS\_83
CL\_INS\_83
CL\_INS\_83
CL\_INS\_98
CL\_INS\_83
CL\_INS\_83
CL\_INS\_83
CL\_INS\_83
CL\_INS\_83
CL\_INS\_193
CL\_INS\_199
CL\_INS\_199
CL\_INS\_98
CL\_INS\_98
CL\_INS\_98
CL\_INS\_133
CL\_INS\_194
CL\_INS\_98
CL\_INS\_194
CL\_INS\_98
CL\_INS\_194
CL\_INS\_194
CL\_INS\_194
CL\_INS\_144
CL\_INS\_144
CL\_INS\_144
CL\_INS\_144
CL\_INS\_144
CL\_INS\_144
CL\_INS\_144
CL\_INS\_194
CL\_INS\_144
CL\_INS\_144
CL\_INS\_194
CL\_INS\_144
CL\_INS\_194
CL\_INS\_144
CL\_INS\_144
CL\_INS\_144
CL\_INS\_194
CL\_INS\_194
CL\_INS\_194
CL\_INS\_194
CL\_INS\_217
CL\_INS\_133
CL\_INS\_217
CL\_INS\_133
CL\_INS\_133
CL\_INS\_133
CL\_INS\_133
Cluster ID


CL\_9015
CL\_8739
CL\_7353
CL\_9014
CL\_9013
CL\_9012
CL\_9945
CL\_5520
CL\_11482
CL\_11481
CL\_3267
CL\_8738
CL\_5521
CL\_9661
CL\_11735
CL\_7354
CL\_3266
CL\_9662
CL\_3265
CL\_6726
CL\_13163
CL\_3264
CL\_5522
CL\_13164
CL\_6727
CL\_3260
CL\_11479
CL\_10473
CL\_9663
CL\_9664
CL\_9665
CL\_9666
CL\_9667
CL\_9668
CL\_9669
CL\_9670
CL\_9671
CL\_9672
CL\_9673
CL\_9674
CL\_5526
CL\_9009
CL\_11477
CL\_13170
CL\_13046
CL\_9675
CL\_9676
CL\_9677
CL\_3249
CL\_3248
CL\_5530
CL\_3245
CL\_3244
CL\_5531
CL\_5532
CL\_3243
CL\_7737
CL\_9678
CL\_9679
CL\_9680
CL\_9681
CL\_9682
CL\_9683
CL\_9684
CL\_9685
CL\_9686
CL\_9687
CL\_9688
CL\_9689
CL\_9690
CL\_9691
CL\_9692
CL\_9693
CL\_9694
CL\_9695
CL\_9696
CL\_9697
CL\_9698
CL\_9699
CL\_9700
CL\_9701
CL\_9702
CL\_9703
CL\_9704
CL\_9705
CL\_9706
CL\_9707
CL\_9708
CL\_9709
CL\_9710
CL\_9711
CL\_9712
CL\_9713
CL\_9714
CL\_6358
CL\_6357
CL\_9715
CL\_6355
CL\_9716
CL\_9135
CL\_9134
CL\_6354
CL\_6353
CL\_6352
CL\_6351
CL\_6350
CL\_6349
CL\_6348
CL\_6347
CL\_9717
CL\_6346
CL\_11954
CL\_9946
CL\_12352
CL\_9947
CL\_3263
CL\_11480
CL\_7830
CL\_3262
CL\_9011
CL\_9948
CL\_9949
CL\_9950
CL\_9951
CL\_9952
CL\_9953
CL\_9954
CL\_9955
CL\_9956
CL\_9957
CL\_9958
CL\_9959
CL\_12353
CL\_3261
CL\_12354
CL\_12355
CL\_6203
CL\_13165
CL\_6204
CL\_11245
CL\_5524
CL\_11478
CL\_11246
CL\_12356
CL\_12357
CL\_12358
CL\_12359
CL\_8815
CL\_8816
CL\_8817
CL\_13171
CL\_13172
CL\_8732
CL\_8731
CL\_8730
CL\_9784
CL\_3259
CL\_3258
CL\_3257
CL\_8811
CL\_8812
CL\_13166
CL\_13167
CL\_13168
CL\_13169
CL\_10300
CL\_3256
CL\_3255
CL\_3254
CL\_9010
CL\_3252
CL\_11953
CL\_11952
CL\_11951
CL\_11950
CL\_11949
CL\_11948
CL\_11947
CL\_11946
CL\_11945
CL\_11944
CL\_5528
CL\_5529
CL\_3247
CL\_6212
CL\_6213
CL\_6214
CL\_13173
CL\_13174
CL\_13175
CL\_13176
CL\_13177
CL\_12934
CL\_5533
CL\_8727
CL\_8976
CL\_5534
CL\_9329
CL\_9330
CL\_9331
CL\_9960
CL\_9961
CL\_9962
CL\_9963
CL\_9964
CL\_9965
CL\_9966
CL\_9967
CL\_6215
CL\_11476
CL\_11475
CL\_11474
CL\_11473
CL\_11472
CL\_13047
CL\_8729
CL\_9332
CL\_5535
CL\_11943
CL\_5561
CL\_11942
CL\_11941
CL\_5540
CL\_6217
CL\_8975
CL\_6218
CL\_9333
CL\_13048
CL\_3237
CL\_3236
CL\_3235
CL\_9008
CL\_10480
CL\_13178
CL\_13179
CL\_13180
CL\_10481
CL\_11471
CL\_6219
CL\_3234
CL\_3232
CL\_3231
CL\_5542
CL\_5543
CL\_7821
CL\_7820
CL\_9007
CL\_8724
CL\_11470
CL\_11469
CL\_11270
CL\_9879
CL\_9878
CL\_9877
CL\_13181
CL\_13182
CL\_13183
CL\_13184
CL\_13185
CL\_6225
CL\_6226
CL\_6227
CL\_8721
CL\_6229
CL\_6230
CL\_6231
CL\_6232
CL\_6233
CL\_6234
CL\_7384
CL\_6220
CL\_11468
CL\_11467
CL\_11466
CL\_11465
CL\_11464
CL\_7816
CL\_7815
CL\_3230
CL\_9209
CL\_3228
CL\_6740
CL\_6741
CL\_6742
CL\_9336
CL\_3225
CL\_9968
CL\_9969
CL\_9970
CL\_10305
CL\_8723
CL\_3229
CL\_9334
CL\_9335
CL\_10719
CL\_11940
CL\_12360
CL\_11939
CL\_11938
CL\_11937
CL\_3223
CL\_3222
CL\_3221
CL\_3220
CL\_3219
CL\_5549
CL\_11452
CL\_11453
CL\_11454
CL\_11455
CL\_3218
CL\_3217
CL\_3216
CL\_9873
CL\_3214
CL\_9872
CL\_9995
CL\_9994
CL\_9992
CL\_13186
CL\_13187
CL\_13188
CL\_13189
CL\_13190
CL\_13191
CL\_13192
CL\_3213
CL\_3212
CL\_3211
CL\_3210
CL\_9971
CL\_8720
CL\_3206
CL\_3205
CL\_3204
CL\_3203
CL\_11463
CL\_9972
CL\_9973
CL\_3199
CL\_3198
CL\_3197
CL\_4930
CL\_9006
CL\_9005
CL\_9004
CL\_9003
CL\_9002
CL\_9001
CL\_10483
CL\_12361
CL\_12362
CL\_9133
CL\_13049
CL\_11731
CL\_11730
CL\_10487
CL\_11117
CL\_10489
CL\_10490
CL\_10491
CL\_9131
CL\_9130
CL\_12363
CL\_12364
CL\_9129
CL\_9128
CL\_9127
CL\_9126
CL\_9125
CL\_9742
CL\_6458
CL\_9741
CL\_6456
CL\_5613
CL\_5614
CL\_7412
