## Supplementary material for "A novel method for integrating genomic and Tn-Seq data to identify common *in vivo* fitness mechanisms across multiple bacterial species": S3 Dataset: CL_INS_144.html

Legend

 Mobile +extrachromosomalelementfunctions
 Regulatoryfunctions
 Hypothetical
 DNA Metabolism
 All EssentialGenes
 Other
 EnergyMetabolism
 Transport +binding proteins
 All VFDB Genes

FULL


WINDOWSVGPNG

Trim RowsRemove SingletonsSave Fasta

CL\_2577


CL\_2577


CL\_2577


CL\_2577


CL\_2577


CL\_2577


CL\_2581


CL\_2577


CL\_2579


CL\_2577


CL\_2577


CL\_2577


CL\_2577

HighlightSelectShow Genomes


46

CL\_2576


1

CL\_2575


1

CL\_2575


1

CL\_2575


1

CL\_3371


1

CL\_3271


1

CL\_2576


1

CL\_2575


1

CL\_2576


1

CL\_2572


1


CL\_12115
CL\_6350
CL\_12116
CL\_6354
CL\_12117
CL\_12118
CL\_12119
CL\_6356
CL\_12120
CL\_10210
CL\_6363
CL\_11732
CL\_11733
CL\_9337
CL\_9336
CL\_9124
CL\_9984
CL\_9985
CL\_9986
CL\_9987
CL\_9988
CL\_9989
CL\_9990
CL\_9991
CL\_3207
CL\_4930
CL\_13192
CL\_13191
CL\_13190
CL\_13189
CL\_13188
CL\_13187
CL\_13186
CL\_9992
CL\_9993
CL\_9994
CL\_9995
CL\_9996
CL\_9997
CL\_9998
CL\_3214
CL\_3216
CL\_6234
CL\_6233
CL\_6232
CL\_6231
CL\_6230
CL\_6229
CL\_9999
CL\_8721
CL\_6227
CL\_6226
CL\_7414
CL\_6224
CL\_7415
CL\_10000
CL\_6225
CL\_13185
CL\_13184
CL\_13183
CL\_13182
CL\_13181
CL\_9877
CL\_9878
CL\_9879
CL\_11448
CL\_11271
CL\_11270
CL\_5615
CL\_6346
CL\_7412
CL\_5614
CL\_6456
CL\_9741
CL\_6458
CL\_5613
CL\_9742
CL\_9743
CL\_9744
CL\_9745
CL\_9746
CL\_9747
CL\_9748
CL\_9749
CL\_9750
CL\_9751
CL\_9752
CL\_9753
CL\_9754
CL\_9755
CL\_9756
CL\_9757
CL\_9758
CL\_9759
CL\_9760
CL\_9761
CL\_9762
CL\_9763
CL\_9764
CL\_9765
CL\_9766
CL\_9767
CL\_9768
CL\_9769
CL\_9770
CL\_9771
CL\_9772
CL\_9773
CL\_9774
CL\_9775
CL\_9776
CL\_9777
CL\_9778
CL\_9779
CL\_9780
CL\_9781
CL\_9782
CL\_9125
CL\_9126
CL\_9127
CL\_12364
CL\_12363
CL\_9128
CL\_9129
CL\_9130
CL\_11458
CL\_13049
CL\_9133
CL\_9134
CL\_9135
CL\_9136
CL\_9137
CL\_9138
CL\_12362
CL\_12361
CL\_9131
CL\_9132
CL\_10491
CL\_11457
CL\_6347
CL\_10490
CL\_10489
CL\_11117
CL\_10487
CL\_11730
CL\_11731
CL\_10483
CL\_9001
CL\_9002
CL\_9003
CL\_11456
CL\_9004
CL\_9005
CL\_9006
CL\_3218
CL\_3219
CL\_3220
CL\_3221
CL\_3222
CL\_11455
CL\_11454
CL\_11453
CL\_11452
CL\_11451
CL\_6742
CL\_7816
CL\_9335
CL\_9334
CL\_3229
CL\_11450
CL\_11449
CL\_3223
CL\_3225
CL\_6741
CL\_6740
CL\_12360
CL\_3228
CL\_9637
CL\_9209
CL\_3230
CL\_3231
CL\_3232
CL\_8823
CL\_5541
CL\_11937
CL\_11938
CL\_11939
CL\_11940
CL\_10719
CL\_8723
CL\_10305
CL\_6220
CL\_5543
CL\_5542
CL\_3234
CL\_9333
CL\_7384
CL\_8724
CL\_13180
CL\_13179
CL\_13178
CL\_6219
CL\_13048
CL\_6218
CL\_8975
CL\_6217
CL\_3238
CL\_10001
CL\_10002
CL\_11447
CL\_11446
CL\_11445
CL\_9309
CL\_10481
CL\_10480
CL\_12934
CL\_13177
CL\_13176
CL\_13175
CL\_13174
CL\_13173
CL\_3237
CL\_12121
CL\_12122
CL\_12123
CL\_12124
CL\_5540
CL\_8976
CL\_8727
CL\_9783
CL\_5533
CL\_9332
CL\_9331
CL\_9330
CL\_9329
CL\_8729
CL\_13047
CL\_6215
CL\_11941
CL\_11942
CL\_5561
CL\_11943
CL\_5535
CL\_5534
CL\_3243
CL\_3244
CL\_8730
CL\_8731
CL\_5532
CL\_10003
CL\_5531
CL\_5530
CL\_10004
CL\_10005
CL\_10006
CL\_3248
CL\_3249
CL\_13170
CL\_13046
CL\_5526
CL\_10007
CL\_10008
CL\_10009
CL\_10010
CL\_10011
CL\_10012
CL\_10013
CL\_10014
CL\_10015
CL\_10016
CL\_10017
CL\_10018
CL\_10019
CL\_10300
CL\_13169
CL\_13168
CL\_13167
CL\_13166
CL\_9674
CL\_3252
CL\_3254
CL\_11443
CL\_3256
CL\_5523
CL\_3258
CL\_8812
CL\_13165
CL\_8811
CL\_6214
CL\_13172
CL\_13171
CL\_6213
CL\_6212
CL\_9784
CL\_8817
CL\_9678
CL\_8816
CL\_8815
CL\_7827
CL\_11444
CL\_12359
CL\_12358
CL\_12357
CL\_12356
CL\_11246
CL\_11245
CL\_6204
CL\_6203
CL\_12355
CL\_12354
CL\_3247
CL\_5529
CL\_5528
CL\_9239
CL\_9785
CL\_6205
CL\_9786
CL\_11944
CL\_11945
CL\_11946
CL\_11947
CL\_11948
CL\_11949
CL\_11950
CL\_11951
CL\_11952
CL\_11953
CL\_3259
CL\_11109
CL\_3260
CL\_6727
CL\_13164
CL\_5522
CL\_9787
CL\_9788
CL\_9789
CL\_9790
CL\_3261
CL\_12353
CL\_3262
CL\_8984
CL\_3263
CL\_9947
CL\_12352
CL\_10473
CL\_6725
CL\_9618
CL\_11442
CL\_3269
CL\_3264
CL\_13163
CL\_3265
CL\_10020
CL\_8738
CL\_5521
CL\_3267
CL\_7353
CL\_8739
CL\_5520
CL\_6407
CL\_7830
CL\_9946
CL\_3266
CL\_11954
CL\_7354
CL\_9661
CL\_11735
CL\_9014
