## Supplementary material for "A novel method for integrating genomic and Tn-Seq data to identify common *in vivo* fitness mechanisms across multiple bacterial species": S3 Dataset: CL_INS_145.html

Legend

 Mobile +extrachromosomalelementfunctions
 Regulatoryfunctions
 Hypothetical
 DNA Metabolism
 All EssentialGenes
 Other
 EnergyMetabolism
 All VFDB Genes

FULL


WINDOWSVGPNG

Trim RowsRemove SingletonsSave Fasta

CL\_2584


CL\_2584


CL\_2581


CL\_2584


CL\_2584


CL\_2584

HighlightSelectShow Genomes


29

CL\_2587


25

CL\_2587


1

CL\_2587


1

CL\_2587


1

CL\_2587


1

CL\_2587

fGI ID


CL\_INS\_145
CL\_INS\_145
CL\_INS\_137
CL\_INS\_137
CL\_INS\_137
CL\_INS\_217
CL\_INS\_217
CL\_INS\_217
CL\_INS\_145
CL\_INS\_145
CL\_INS\_217
CL\_INS\_217
CL\_INS\_217
CL\_INS\_217
CL\_INS\_217
CL\_INS\_217
CL\_INS\_217
CL\_INS\_217
CL\_INS\_145
CL\_INS\_145
CL\_INS\_145
CL\_INS\_145
CL\_INS\_145
CL\_INS\_217
CL\_INS\_217
CL\_INS\_145
CL\_INS\_217
CL\_INS\_217
CL\_INS\_145
CL\_INS\_217
CL\_INS\_217
CL\_INS\_145
CL\_INS\_145
CL\_INS\_145
CL\_INS\_217
CL\_INS\_217
CL\_INS\_217
CL\_INS\_217
CL\_INS\_217
CL\_INS\_217
CL\_INS\_217
CL\_INS\_217
CL\_INS\_133
CL\_INS\_198
CL\_INS\_217
CL\_INS\_133
CL\_INS\_133
CL\_INS\_198
CL\_INS\_198
CL\_INS\_217
CL\_INS\_145
CL\_INS\_217
CL\_INS\_145
CL\_INS\_217
CL\_INS\_217
CL\_INS\_217
CL\_INS\_217
CL\_INS\_217
CL\_INS\_217
CL\_INS\_217
CL\_INS\_217
CL\_INS\_145
CL\_INS\_217
CL\_INS\_217
CL\_INS\_217
CL\_INS\_217
CL\_INS\_217
CL\_INS\_217
CL\_INS\_217
CL\_INS\_145
CL\_INS\_145
CL\_INS\_145
CL\_INS\_217
CL\_INS\_217
CL\_INS\_133
CL\_INS\_133
CL\_INS\_217
CL\_INS\_217
CL\_INS\_133
CL\_INS\_133
CL\_INS\_133
CL\_INS\_133
CL\_INS\_133
CL\_INS\_133
CL\_INS\_133
CL\_INS\_133
CL\_INS\_217
CL\_INS\_133
CL\_INS\_145
CL\_INS\_217
CL\_INS\_137
CL\_INS\_137
CL\_INS\_145
CL\_INS\_145
CL\_INS\_83
CL\_INS\_83
CL\_INS\_145
CL\_INS\_145
CL\_INS\_145
CL\_INS\_145
CL\_INS\_145
Cluster ID


CL\_6767
CL\_6768
CL\_7371
CL\_7372
CL\_7373
CL\_6405
CL\_6404
CL\_7431
CL\_9851
CL\_9852
CL\_6398
CL\_6402
CL\_7430
CL\_7428
CL\_6397
CL\_6396
CL\_7802
CL\_7801
CL\_9853
CL\_9854
CL\_9855
CL\_9856
CL\_9857
CL\_6387
CL\_6385
CL\_7796
CL\_7427
CL\_7426
CL\_13050
CL\_6392
CL\_8170
CL\_13051
CL\_13052
CL\_13053
CL\_7424
CL\_11694
CL\_7423
CL\_6384
CL\_6383
CL\_6382
CL\_6381
CL\_6380
CL\_7795
CL\_10186
CL\_6379
CL\_6378
CL\_6377
CL\_9368
CL\_10187
CL\_6376
CL\_9858
CL\_6375
CL\_10188
CL\_6372
CL\_6371
CL\_7793
CL\_7792
CL\_7791
CL\_7790
CL\_7789
CL\_7788
CL\_9859
CL\_7787
CL\_9860
CL\_7786
CL\_7785
CL\_7784
CL\_7783
CL\_7782
CL\_9861
CL\_13054
CL\_13055
CL\_7780
CL\_6463
CL\_5609
CL\_5610
CL\_9862
CL\_9863
CL\_5611
CL\_5612
CL\_5613
CL\_6350
CL\_6349
CL\_6348
CL\_6347
CL\_7388
CL\_9387
CL\_5045
CL\_13056
CL\_7390
CL\_7391
CL\_7392
CL\_9864
CL\_9865
CL\_6714
CL\_6715
CL\_10070
CL\_13057
CL\_2585
CL\_6769
CL\_2586
