## Supplementary material for "A novel method for integrating genomic and Tn-Seq data to identify common *in vivo* fitness mechanisms across multiple bacterial species": S3 Dataset: CL_INS_146.html

Legend

 Mobile +extrachromosomalelementfunctions
 Hypothetical
 DNA Metabolism
 Transport +binding proteins
 All VFDB Genes

FULL


WINDOWSVGPNG

Trim RowsRemove SingletonsSave Fasta

CL\_2607


CL\_2607


CL\_2607


CL\_2607


CL\_2607

HighlightSelectShow Genomes


48

CL\_2608


5

CL\_2671


1

CL\_2671


1

CL\_2671


1

CL\_2671

fGI ID


CL\_INS\_146
CL\_INS\_146
CL\_INS\_157
CL\_INS\_157
CL\_INS\_157
CL\_INS\_157
Cluster ID


CL\_5249
CL\_4897
CL\_4898
CL\_4899
CL\_4900
CL\_4901
