## Supplementary material for "A novel method for integrating genomic and Tn-Seq data to identify common *in vivo* fitness mechanisms across multiple bacterial species": S3 Dataset: CL_INS_147.html

Legend

 Hypothetical
 DNA Metabolism
 Other
 All VFDB Genes

FULL


WINDOWSVGPNG

Trim RowsRemove SingletonsSave Fasta

CL\_2616


CL\_2616


CL\_2615

HighlightSelectShow Genomes


25

CL\_2618


24

CL\_2618


8

CL\_2618

fGI ID

CL\_INS\_147
Cluster ID

CL\_2617
