## Supplementary material for "A novel method for integrating genomic and Tn-Seq data to identify common *in vivo* fitness mechanisms across multiple bacterial species": S3 Dataset: CL_INS_149.html

Legend

 Hypothetical
 Other
 Proteinsynthesis/fate
 All VFDB Genes

FULL


WINDOWSVGPNG

Trim RowsRemove SingletonsSave Fasta

CL\_2627


CL\_2627


CL\_2627


CL\_2626

HighlightSelectShow Genomes


45

CL\_2628


9

CL\_2628


2

CL\_2628


1

CL\_2628

fGI ID


CL\_INS\_149
CL\_INS\_149
CL\_INS\_149
CL\_INS\_149
Cluster ID


CL\_9123
CL\_6770
CL\_6771
CL\_6772
