## Supplementary material for "A novel method for integrating genomic and Tn-Seq data to identify common *in vivo* fitness mechanisms across multiple bacterial species": S3 Dataset: CL_INS_150.html

Legend

 Hypothetical
 All EssentialGenes
 Other
 All VFDB Genes

FULL


WINDOWSVGPNG

Trim RowsRemove SingletonsSave Fasta

CL\_2633


CL\_2633


CL\_2633


CL\_2633

HighlightSelectShow Genomes


54

CL\_2634


1

CL\_2634


1

CL\_2647


1

CL\_2634

fGI ID


CL\_INS\_150
CL\_INS\_153
CL\_INS\_153
CL\_INS\_153
CL\_INS\_153
CL\_INS\_153
CL\_INS\_153
CL\_INS\_150
CL\_INS\_150
CL\_INS\_150
CL\_INS\_150
CL\_INS\_150
CL\_INS\_150
CL\_INS\_150
CL\_INS\_150
CL\_INS\_150
CL\_INS\_150
Cluster ID


CL\_12112
CL\_12111
CL\_12110
CL\_12109
CL\_12108
CL\_12107
CL\_12106
CL\_13592
CL\_13654
CL\_13591
CL\_13590
CL\_13589
CL\_13588
CL\_13587
CL\_13586
CL\_13585
CL\_13584
