## Supplementary material for "A novel method for integrating genomic and Tn-Seq data to identify common *in vivo* fitness mechanisms across multiple bacterial species": S3 Dataset: CL_INS_152.html

Legend

 Hypothetical
 AntibioticResistance
 All EssentialGenes

FULL


WINDOWSVGPNG

Trim RowsRemove SingletonsSave Fasta

CL\_2641


CL\_2641

HighlightSelectShow Genomes


32

CL\_2643


24

CL\_2643

fGI ID

CL\_INS\_152
Cluster ID

CL\_2642
