## Supplementary material for "A novel method for integrating genomic and Tn-Seq data to identify common *in vivo* fitness mechanisms across multiple bacterial species": S3 Dataset: CL_INS_153.html

Legend

 Mobile +extrachromosomalelementfunctions
 Regulatoryfunctions
 Hypothetical
 All EssentialGenes
 Proteinsynthesis/fate
 Other
 Transport +binding proteins
 All VFDB Genes

FULL


WINDOWSVGPNG

Trim RowsRemove SingletonsSave Fasta

CL\_2646


CL\_2646


CL\_2646


CL\_2645


CL\_2646


CL\_2646


CL\_2646


CL\_2645


CL\_2633


CL\_2646

HighlightSelectShow Genomes


26

CL\_2647


22

CL\_2647


2

CL\_2647


2

CL\_2647


1

CL\_2647


1

CL\_2647


1

CL\_2647


1

CL\_2647


1

CL\_2647


1

CL\_2647

fGI ID


CL\_INS\_153
CL\_INS\_153
CL\_INS\_153
CL\_INS\_153
CL\_INS\_153
CL\_INS\_153
CL\_INS\_153
CL\_INS\_153
CL\_INS\_153
CL\_INS\_153
CL\_INS\_153
CL\_INS\_153
CL\_INS\_153
CL\_INS\_153
CL\_INS\_153
CL\_INS\_153
CL\_INS\_153
CL\_INS\_153
CL\_INS\_153
CL\_INS\_133
CL\_INS\_153
CL\_INS\_153
CL\_INS\_153
CL\_INS\_153
CL\_INS\_153
CL\_INS\_153
CL\_INS\_153
CL\_INS\_153
CL\_INS\_153
CL\_INS\_153
CL\_INS\_153
CL\_INS\_153
CL\_INS\_153
CL\_INS\_153
CL\_INS\_153
CL\_INS\_153
CL\_INS\_153
CL\_INS\_153
CL\_INS\_153
CL\_INS\_153
CL\_INS\_153
CL\_INS\_153
CL\_INS\_153
CL\_INS\_153
CL\_INS\_153
CL\_INS\_153
CL\_INS\_153
CL\_INS\_153
CL\_INS\_153
CL\_INS\_153
Cluster ID


CL\_12111
CL\_12110
CL\_12109
CL\_12108
CL\_12107
CL\_12106
CL\_12105
CL\_6775
CL\_6776
CL\_6777
CL\_10943
CL\_6778
CL\_6779
CL\_10688
CL\_10687
CL\_10686
CL\_6780
CL\_6781
CL\_5617
CL\_4451
CL\_10216
CL\_10218
CL\_10219
CL\_10220
CL\_10221
CL\_10222
CL\_10223
CL\_10224
CL\_13583
CL\_10225
CL\_10226
CL\_10227
CL\_10228
CL\_10229
CL\_10230
CL\_10231
CL\_10232
CL\_10752
CL\_10233
CL\_10234
CL\_10235
CL\_10236
CL\_10753
CL\_10237
CL\_8601
CL\_13655
CL\_13656
CL\_5323
CL\_5322
CL\_8121
