## Supplementary material for "A novel method for integrating genomic and Tn-Seq data to identify common *in vivo* fitness mechanisms across multiple bacterial species": S3 Dataset: CL_INS_154.html

Legend

 All EssentialGenes
 Other
 All VFDB Genes

FULL


WINDOWSVGPNG

Trim RowsRemove SingletonsSave Fasta

CL\_2647


CL\_2647


CL\_2647

HighlightSelectShow Genomes


40

CL\_2649


14

CL\_2649


4

CL\_2649

fGI ID


CL\_INS\_154
CL\_INS\_154
Cluster ID


CL\_2648
CL\_5618
