## Supplementary material for "A novel method for integrating genomic and Tn-Seq data to identify common *in vivo* fitness mechanisms across multiple bacterial species": S3 Dataset: CL_INS_155.html

Legend

 Other
 All VFDB Genes

FULL


WINDOWSVGPNG

Trim RowsRemove SingletonsSave Fasta

CL\_2655


CL\_2655


CL\_2655


CL\_2655

HighlightSelectShow Genomes


28

CL\_2658


25

CL\_2658


3

CL\_2658


1

CL\_2658

fGI ID


CL\_INS\_155
CL\_INS\_155
Cluster ID


CL\_2656
CL\_2657
