## Supplementary material for "A novel method for integrating genomic and Tn-Seq data to identify common *in vivo* fitness mechanisms across multiple bacterial species": S3 Dataset: CL_INS_156.html

Legend

 Mobile +extrachromosomalelementfunctions
 Regulatoryfunctions
 Hypothetical
 AntibioticResistance
 Proteinsynthesis/fate
 Other
 Transport +binding proteins
 All VFDB Genes

FULL


WINDOWSVGPNG

Trim RowsRemove SingletonsSave Fasta

CL\_2663


CL\_2663


CL\_2663


CL\_2663


CL\_2663


CL\_2663


CL\_2663


CL\_2663


CL\_2663


CL\_2663


CL\_2663


CL\_2663


CL\_512


CL\_3356

HighlightSelectShow Genomes


18

CL\_2669


8

CL\_2669


8

CL\_2669


7

CL\_2669


6

CL\_2669


3

CL\_2669


2

CL\_2669


1

CL\_522


1

CL\_2669


1

CL\_2669


1

CL\_2669


1

CL\_2669


1

CL\_2669


1

CL\_2669

fGI ID


CL\_INS\_83
CL\_INS\_156
CL\_INS\_156
CL\_INS\_156
CL\_INS\_156
CL\_INS\_156
CL\_INS\_156
CL\_INS\_156
CL\_INS\_20
CL\_INS\_20
CL\_INS\_20
CL\_INS\_20
CL\_INS\_20
CL\_INS\_133
CL\_INS\_133
CL\_INS\_133
CL\_INS\_133
CL\_INS\_133
CL\_INS\_156
CL\_INS\_156
CL\_INS\_133
CL\_INS\_133
CL\_INS\_133
CL\_INS\_133
CL\_INS\_133
CL\_INS\_133
CL\_INS\_237
CL\_INS\_237
CL\_INS\_83
CL\_INS\_83
CL\_INS\_83
CL\_INS\_156
CL\_INS\_156
CL\_INS\_156
CL\_INS\_156
CL\_INS\_34
CL\_INS\_156
CL\_INS\_34
CL\_INS\_34
CL\_INS\_1
CL\_INS\_1
CL\_INS\_1
CL\_INS\_1
CL\_INS\_1
CL\_INS\_1
CL\_INS\_1
CL\_INS\_1
CL\_INS\_1
CL\_INS\_1
CL\_INS\_1
CL\_INS\_1
CL\_INS\_1
CL\_INS\_1
CL\_INS\_1
CL\_INS\_1
CL\_INS\_1
CL\_INS\_1
CL\_INS\_1
CL\_INS\_1
CL\_INS\_1
CL\_INS\_1
CL\_INS\_1
CL\_INS\_1
CL\_INS\_1
CL\_INS\_1
CL\_INS\_156
CL\_INS\_1
CL\_INS\_1
CL\_INS\_156
CL\_INS\_1
CL\_INS\_1
CL\_INS\_1
CL\_INS\_1
CL\_INS\_1
CL\_INS\_1
CL\_INS\_1
CL\_INS\_1
CL\_INS\_1
CL\_INS\_34
CL\_INS\_34
CL\_INS\_34
CL\_INS\_34
CL\_INS\_45
CL\_INS\_45
CL\_INS\_45
CL\_INS\_20
CL\_INS\_20
CL\_INS\_20
CL\_INS\_20
CL\_INS\_133
CL\_INS\_1
CL\_INS\_156
CL\_INS\_156
CL\_INS\_133
CL\_INS\_1
CL\_INS\_1
CL\_INS\_1
CL\_INS\_153
CL\_INS\_153
CL\_INS\_1
CL\_INS\_156
CL\_INS\_156
CL\_INS\_156
CL\_INS\_156
CL\_INS\_156
CL\_INS\_156
Cluster ID


CL\_4894
CL\_4895
CL\_6258
CL\_6259
CL\_6260
CL\_9740
CL\_4896
CL\_8862
CL\_8861
CL\_8860
CL\_526
CL\_525
CL\_524
CL\_5567
CL\_11043
CL\_7879
CL\_5575
CL\_5576
CL\_12220
CL\_12221
CL\_7820
CL\_5586
CL\_5591
CL\_7749
CL\_9318
CL\_8666
CL\_510
CL\_509
CL\_508
CL\_507
CL\_506
CL\_8859
CL\_8858
CL\_8857
CL\_6087
CL\_6088
CL\_8856
CL\_6089
CL\_6090
CL\_4440
CL\_4442
CL\_4443
CL\_4444
CL\_4606
CL\_4604
CL\_4603
CL\_4602
CL\_4601
CL\_5259
CL\_5260
CL\_5261
CL\_6913
CL\_4413
CL\_4414
CL\_4415
CL\_4416
CL\_4417
CL\_4418
CL\_4419
CL\_4420
CL\_4421
CL\_4422
CL\_5331
CL\_4424
CL\_6077
CL\_8855
CL\_5813
CL\_5812
CL\_8854
CL\_5330
CL\_5329
CL\_4445
CL\_4446
CL\_4447
CL\_4448
CL\_8853
CL\_4449
CL\_4450
CL\_6086
CL\_5277
CL\_5278
CL\_5279
CL\_5280
CL\_5281
CL\_5282
CL\_5283
CL\_5285
CL\_5286
CL\_5287
CL\_4451
CL\_8852
CL\_8851
CL\_8850
CL\_5327
CL\_5326
CL\_5325
CL\_5324
CL\_5323
CL\_5322
CL\_4452
CL\_2664
CL\_2665
CL\_5619
CL\_2666
CL\_2667
CL\_2668
