## Supplementary material for "A novel method for integrating genomic and Tn-Seq data to identify common *in vivo* fitness mechanisms across multiple bacterial species": S3 Dataset: CL_INS_157.html

Legend

 Hypothetical
 DNA Metabolism
 Other
 Transport +binding proteins

FULL


WINDOWSVGPNG

Trim RowsRemove SingletonsSave Fasta

CL\_2671


CL\_2671


CL\_2671


CL\_2671


CL\_2671

HighlightSelectShow Genomes


49

CL\_2670


5

CL\_2607


1

CL\_2607


1

CL\_2607


1

CL\_2607

fGI ID


CL\_INS\_157
CL\_INS\_157
CL\_INS\_157
CL\_INS\_157
CL\_INS\_157
Cluster ID


CL\_4902
CL\_4901
CL\_4900
CL\_4899
CL\_4898
