## Supplementary material for "A novel method for integrating genomic and Tn-Seq data to identify common *in vivo* fitness mechanisms across multiple bacterial species": S3 Dataset: CL_INS_158.html

Legend

 Regulatoryfunctions
 Hypothetical
 Cell Envelope
 Other
 Proteinsynthesis/fate
 All VFDB Genes

FULL


WINDOWSVGPNG

Trim RowsRemove SingletonsSave Fasta

CL\_2675


CL\_2675


CL\_2675


CL\_2675


CL\_2674


CL\_2675

HighlightSelectShow Genomes


28

CL\_2676


24

CL\_2676


2

CL\_2676


1

CL\_2676


1

CL\_2676


1

CL\_2676

fGI ID


CL\_INS\_158
CL\_INS\_158
CL\_INS\_158
CL\_INS\_158
CL\_INS\_158
CL\_INS\_158
CL\_INS\_158
CL\_INS\_158
CL\_INS\_158
CL\_INS\_158
CL\_INS\_1
CL\_INS\_1
CL\_INS\_1
CL\_INS\_1
CL\_INS\_1
CL\_INS\_1
CL\_INS\_1
CL\_INS\_1
CL\_INS\_1
CL\_INS\_1
CL\_INS\_158
Cluster ID


CL\_4903
CL\_13011
CL\_6782
CL\_6783
CL\_6784
CL\_6785
CL\_6786
CL\_6787
CL\_8959
CL\_8960
CL\_4609
CL\_4606
CL\_4604
CL\_4603
CL\_4602
CL\_4601
CL\_5259
CL\_5260
CL\_5261
CL\_6913
CL\_8961
