## Supplementary material for "A novel method for integrating genomic and Tn-Seq data to identify common *in vivo* fitness mechanisms across multiple bacterial species": S3 Dataset: CL_INS_159.html

Legend

 Hypothetical
 Other
 EnergyMetabolism
 All VFDB Genes

FULL


WINDOWSVGPNG

Trim RowsRemove SingletonsSave Fasta

CL\_2684


CL\_2684


CL\_2684


CL\_2683


CL\_2684


CL\_2684

HighlightSelectShow Genomes


31

CL\_2685


11

CL\_2685


11

CL\_2685


1

CL\_2685


1

CL\_2686


1

CL\_2685

fGI ID


CL\_INS\_159
CL\_INS\_159
CL\_INS\_159
Cluster ID


CL\_7762
CL\_7763
CL\_8231
