## Supplementary material for "A novel method for integrating genomic and Tn-Seq data to identify common *in vivo* fitness mechanisms across multiple bacterial species": S3 Dataset: CL_INS_163.html

Legend

 Regulatoryfunctions
 Hypothetical
 All VFDB Genes

FULL


WINDOWSVGPNG

Trim RowsRemove SingletonsSave Fasta

CL\_2737


CL\_2737


CL\_2737


CL\_2737


CL\_2737


CL\_2947


CL\_2737

HighlightSelectShow Genomes


25

CL\_2738


13

CL\_2738


11

CL\_2738


4

CL\_2738


2

CL\_2738


1

CL\_2738


1

CL\_2738

fGI ID


CL\_INS\_163
CL\_INS\_163
CL\_INS\_163
CL\_INS\_163
CL\_INS\_142
CL\_INS\_142
CL\_INS\_178
CL\_INS\_178
CL\_INS\_171
Cluster ID


CL\_11459
CL\_5620
CL\_6261
CL\_9738
CL\_2945
CL\_2944
CL\_6832
CL\_6831
CL\_4919
