## Supplementary material for "A novel method for integrating genomic and Tn-Seq data to identify common *in vivo* fitness mechanisms across multiple bacterial species": S3 Dataset: CL_INS_166.html

Legend

 Mobile +extrachromosomalelementfunctions
 Regulatoryfunctions
 Hypothetical
 Other
 Transport +binding proteins
 All VFDB Genes

FULL


WINDOWSVGPNG

Trim RowsRemove SingletonsSave Fasta

CL\_2789


CL\_2789


CL\_2789


CL\_2789


CL\_2789


CL\_2789


CL\_2789


CL\_2789


CL\_2789


CL\_2789


CL\_2789


CL\_2789


CL\_2789


CL\_2789

HighlightSelectShow Genomes


18

CL\_2794


9

CL\_2794


8

CL\_2794


7

CL\_2794


3

CL\_2794


3

CL\_2794


2

CL\_2794


1

CL\_2794


1

CL\_2794


1

CL\_2794


1

CL\_2794


1

CL\_2794


1

CL\_2794


1

CL\_2794

fGI ID


CL\_INS\_166
CL\_INS\_166
CL\_INS\_166
CL\_INS\_166
CL\_INS\_166
CL\_INS\_166
CL\_INS\_166
CL\_INS\_166
CL\_INS\_166
CL\_INS\_166
CL\_INS\_166
CL\_INS\_166
CL\_INS\_166
CL\_INS\_166
CL\_INS\_166
CL\_INS\_166
CL\_INS\_166
CL\_INS\_166
CL\_INS\_166
CL\_INS\_166
CL\_INS\_166
CL\_INS\_166
CL\_INS\_166
CL\_INS\_166
CL\_INS\_166
CL\_INS\_166
CL\_INS\_166
CL\_INS\_166
CL\_INS\_166
CL\_INS\_166
CL\_INS\_166
CL\_INS\_166
CL\_INS\_166
CL\_INS\_166
CL\_INS\_166
CL\_INS\_166
CL\_INS\_166
CL\_INS\_166
CL\_INS\_166
CL\_INS\_166
CL\_INS\_166
CL\_INS\_166
CL\_INS\_166
CL\_INS\_166
CL\_INS\_166
CL\_INS\_166
CL\_INS\_166
CL\_INS\_166
CL\_INS\_166
CL\_INS\_166
CL\_INS\_166
CL\_INS\_166
Cluster ID


CL\_9737
CL\_6829
CL\_6262
CL\_6263
CL\_2790
CL\_2791
CL\_2792
CL\_2793
CL\_12588
CL\_12589
CL\_12590
CL\_12591
CL\_12592
CL\_12593
CL\_12594
CL\_12595
CL\_12596
CL\_12597
CL\_12598
CL\_12599
CL\_12600
CL\_12601
CL\_12602
CL\_12603
CL\_12604
CL\_12605
CL\_12606
CL\_12607
CL\_12608
CL\_12609
CL\_12610
CL\_12611
CL\_12612
CL\_12613
CL\_12614
CL\_12615
CL\_12616
CL\_12617
CL\_12618
CL\_12619
CL\_12620
CL\_12621
CL\_12622
CL\_12623
CL\_12624
CL\_12625
CL\_12626
CL\_12627
CL\_12628
CL\_12629
CL\_12630
CL\_12631
