## Supplementary material for "A novel method for integrating genomic and Tn-Seq data to identify common *in vivo* fitness mechanisms across multiple bacterial species": S3 Dataset: CL_INS_167.html

Legend

 Mobile +extrachromosomalelementfunctions
 Regulatoryfunctions
 Hypothetical
 DNA Metabolism
 AntibioticResistance
 Other
 Transport +binding proteins
 All VFDB Genes

FULL


WINDOWSVGPNG

Trim RowsRemove SingletonsSave Fasta

CL\_2800


CL\_2800


CL\_2800


CL\_2800


CL\_2800


CL\_2800


CL\_2800


CL\_2800

HighlightSelectShow Genomes


46

CL\_2801


2

CL\_2807


1

Break


1

CL\_2807


1

CL\_512


1

CL\_2807


1

CL\_2807


1

CL\_2807

fGI ID


CL\_INS\_168
CL\_INS\_168
CL\_INS\_169
CL\_INS\_168
CL\_INS\_169
CL\_INS\_169
CL\_INS\_169
CL\_INS\_169
CL\_INS\_168
CL\_INS\_168
CL\_INS\_168
CL\_INS\_168
CL\_INS\_168
CL\_INS\_168
CL\_INS\_168
CL\_INS\_168
CL\_INS\_168
CL\_INS\_168
CL\_INS\_168
CL\_INS\_168
CL\_INS\_168
CL\_INS\_168
CL\_INS\_168
CL\_INS\_168
CL\_INS\_168
CL\_INS\_168
CL\_INS\_168
CL\_INS\_168
CL\_INS\_168
CL\_INS\_168
CL\_INS\_55
CL\_INS\_55
CL\_INS\_55
CL\_INS\_20
CL\_INS\_167
CL\_INS\_19
CL\_INS\_19
CL\_INS\_1
CL\_INS\_1
CL\_INS\_156
CL\_INS\_1
CL\_INS\_1
CL\_INS\_133
CL\_INS\_133
CL\_INS\_133
CL\_INS\_133
CL\_INS\_56
CL\_INS\_19
CL\_INS\_19
CL\_INS\_19
CL\_INS\_19
CL\_INS\_19
CL\_INS\_133
CL\_INS\_133
CL\_INS\_1
CL\_INS\_1
CL\_INS\_1
CL\_INS\_153
CL\_INS\_153
CL\_INS\_1
CL\_INS\_1
CL\_INS\_1
CL\_INS\_1
CL\_INS\_1
CL\_INS\_1
CL\_INS\_1
CL\_INS\_1
CL\_INS\_1
CL\_INS\_1
CL\_INS\_1
CL\_INS\_1
CL\_INS\_1
CL\_INS\_1
CL\_INS\_1
CL\_INS\_1
CL\_INS\_1
CL\_INS\_1
CL\_INS\_168
CL\_INS\_1
CL\_INS\_20
CL\_INS\_1
CL\_INS\_168
CL\_INS\_168
CL\_INS\_168
CL\_INS\_168
CL\_INS\_168
CL\_INS\_168
CL\_INS\_168
CL\_INS\_168
CL\_INS\_168
CL\_INS\_168
CL\_INS\_168
CL\_INS\_168
CL\_INS\_168
CL\_INS\_168
CL\_INS\_167
CL\_INS\_168
CL\_INS\_168
CL\_INS\_168
CL\_INS\_168
CL\_INS\_168
CL\_INS\_168
CL\_INS\_168
CL\_INS\_168
CL\_INS\_168
CL\_INS\_168
CL\_INS\_168
CL\_INS\_168
CL\_INS\_168
CL\_INS\_168
CL\_INS\_168
CL\_INS\_168
CL\_INS\_168
CL\_INS\_168
CL\_INS\_168
CL\_INS\_168
CL\_INS\_168
CL\_INS\_83
CL\_INS\_83
CL\_INS\_83
CL\_INS\_83
CL\_INS\_83
CL\_INS\_83
CL\_INS\_237
CL\_INS\_237
Cluster ID


CL\_2802
CL\_7766
CL\_6826
CL\_7767
CL\_6824
CL\_6823
CL\_6822
CL\_7768
CL\_10493
CL\_7770
CL\_7769
CL\_10494
CL\_10495
CL\_10496
CL\_10497
CL\_10498
CL\_10499
CL\_10500
CL\_10501
CL\_10502
CL\_10503
CL\_10504
CL\_10505
CL\_10506
CL\_9847
CL\_9846
CL\_2805
CL\_5250
CL\_5253
CL\_5254
CL\_5251
CL\_5808
CL\_5807
CL\_5161
CL\_11929
CL\_11928
CL\_11927
CL\_4424
CL\_6077
CL\_8855
CL\_5813
CL\_5812
CL\_5201
CL\_5202
CL\_5203
CL\_5204
CL\_5205
CL\_5206
CL\_5207
CL\_5208
CL\_5209
CL\_5210
CL\_5328
CL\_5327
CL\_5326
CL\_5325
CL\_5324
CL\_5323
CL\_5322
CL\_4452
CL\_4453
CL\_4454
CL\_5321
CL\_4455
CL\_4456
CL\_4457
CL\_4458
CL\_4459
CL\_4460
CL\_4461
CL\_4462
CL\_4463
CL\_4464
CL\_4465
CL\_4466
CL\_4467
CL\_4468
CL\_5320
CL\_5160
CL\_5159
CL\_4469
CL\_8229
CL\_8228
CL\_8227
CL\_8226
CL\_8225
CL\_8224
CL\_8223
CL\_8222
CL\_8221
CL\_8220
CL\_8219
CL\_8218
CL\_8217
CL\_8216
CL\_8343
CL\_8215
CL\_8214
CL\_8213
CL\_8361
CL\_8360
CL\_8359
CL\_8358
CL\_8357
CL\_8356
CL\_8355
CL\_8354
CL\_8353
CL\_8352
CL\_8351
CL\_8350
CL\_8349
CL\_8348
CL\_8347
CL\_8212
CL\_8211
CL\_8210
CL\_503
CL\_504
CL\_505
CL\_506
CL\_507
CL\_508
CL\_509
CL\_510
