## Supplementary material for "A novel method for integrating genomic and Tn-Seq data to identify common *in vivo* fitness mechanisms across multiple bacterial species": S3 Dataset: CL_INS_168.html

Legend

 Mobile +extrachromosomalelementfunctions
 Regulatoryfunctions
 Hypothetical
 DNA Metabolism
 All EssentialGenes
 AntibioticResistance
 Proteinsynthesis/fate
 Other
 All VFDB Genes
 Transport +binding proteins

FULL


WINDOWSVGPNG

Trim RowsRemove SingletonsSave Fasta

CL\_2801


CL\_2801


CL\_2801


CL\_2801


CL\_2801


CL\_2800


CL\_2801


CL\_2800


CL\_2801


CL\_2801


CL\_2801


CL\_2801


CL\_2800


CL\_2801


CL\_2801


CL\_2801


CL\_2801


CL\_2801


Break


CL\_2801


CL\_2801


CL\_2801


CL\_522


CL\_2801


CL\_2801


CL\_2798


CL\_2801


CL\_2801


CL\_2801


Break


CL\_2801


CL\_2801


CL\_2801


CL\_2800


CL\_2801


CL\_2801


CL\_2801


CL\_2800


CL\_2801

HighlightSelectShow Genomes


11

CL\_2807


4

CL\_2807


3

CL\_2807


2

CL\_2807


2

CL\_2807


2

CL\_2807


2

CL\_2807


1

CL\_2807


1

CL\_2807


1

CL\_2807


1

CL\_2807


1

CL\_2807


1

CL\_2807


1

CL\_2807


1

CL\_2807


1

CL\_2807


1

CL\_2807


1

CL\_2807


1

CL\_2807


1

CL\_2807


1

CL\_2807


1

CL\_2807


1

CL\_2807


1

CL\_2811


1

CL\_2807


1

CL\_2807


1

CL\_2807


1

CL\_2807


1

CL\_2807


1

CL\_2807


1

CL\_2807


1

CL\_2807


1

CL\_2807


1

CL\_2807


1

Break


1

CL\_2807


1

CL\_2807


1

CL\_2807


1

CL\_2807

fGI ID


CL\_INS\_20
CL\_INS\_1
CL\_INS\_1
CL\_INS\_1
CL\_INS\_1
CL\_INS\_1
CL\_INS\_1
CL\_INS\_168
CL\_INS\_168
CL\_INS\_168
CL\_INS\_168
CL\_INS\_168
CL\_INS\_168
CL\_INS\_168
CL\_INS\_168
CL\_INS\_168
CL\_INS\_168
CL\_INS\_168
CL\_INS\_168
CL\_INS\_168
CL\_INS\_168
CL\_INS\_168
CL\_INS\_168
CL\_INS\_168
CL\_INS\_168
CL\_INS\_168
CL\_INS\_168
CL\_INS\_168
CL\_INS\_168
CL\_INS\_168
CL\_INS\_168
CL\_INS\_168
CL\_INS\_168
CL\_INS\_168
CL\_INS\_168
CL\_INS\_168
CL\_INS\_168
CL\_INS\_168
CL\_INS\_168
CL\_INS\_168
CL\_INS\_168
CL\_INS\_168
CL\_INS\_168
CL\_INS\_168
CL\_INS\_168
CL\_INS\_168
CL\_INS\_168
CL\_INS\_168
CL\_INS\_168
CL\_INS\_168
CL\_INS\_168
CL\_INS\_168
CL\_INS\_168
CL\_INS\_168
CL\_INS\_168
CL\_INS\_168
CL\_INS\_55
CL\_INS\_55
CL\_INS\_55
CL\_INS\_55
CL\_INS\_20
CL\_INS\_1
CL\_INS\_20
CL\_INS\_1
CL\_INS\_168
CL\_INS\_168
CL\_INS\_168
CL\_INS\_168
CL\_INS\_168
CL\_INS\_168
CL\_INS\_168
CL\_INS\_168
CL\_INS\_168
CL\_INS\_168
CL\_INS\_168
CL\_INS\_168
CL\_INS\_168
CL\_INS\_168
CL\_INS\_168
CL\_INS\_168
CL\_INS\_168
CL\_INS\_168
CL\_INS\_168
CL\_INS\_168
CL\_INS\_168
CL\_INS\_168
CL\_INS\_168
CL\_INS\_168
CL\_INS\_168
CL\_INS\_168
CL\_INS\_168
CL\_INS\_168
CL\_INS\_168
CL\_INS\_168
CL\_INS\_168
CL\_INS\_168
CL\_INS\_168
CL\_INS\_168
CL\_INS\_168
CL\_INS\_168
CL\_INS\_1
CL\_INS\_168
CL\_INS\_168
CL\_INS\_1
CL\_INS\_1
CL\_INS\_168
CL\_INS\_1
CL\_INS\_168
CL\_INS\_1
CL\_INS\_1
CL\_INS\_1
CL\_INS\_1
CL\_INS\_168
CL\_INS\_1
CL\_INS\_168
CL\_INS\_1
CL\_INS\_1
CL\_INS\_1
CL\_INS\_168
CL\_INS\_168
CL\_INS\_1
CL\_INS\_168
CL\_INS\_168
CL\_INS\_168
CL\_INS\_168
CL\_INS\_168
CL\_INS\_168
CL\_INS\_168
CL\_INS\_168
CL\_INS\_168
CL\_INS\_168
CL\_INS\_168
CL\_INS\_168
CL\_INS\_168
CL\_INS\_168
CL\_INS\_168
CL\_INS\_168
CL\_INS\_1
CL\_INS\_1
CL\_INS\_1
CL\_INS\_1
CL\_INS\_20
CL\_INS\_168
CL\_INS\_168
CL\_INS\_168
CL\_INS\_20
CL\_INS\_1
CL\_INS\_1
CL\_INS\_168
CL\_INS\_168
CL\_INS\_20
CL\_INS\_168
CL\_INS\_168
CL\_INS\_168
CL\_INS\_168
CL\_INS\_168
CL\_INS\_168
CL\_INS\_168
CL\_INS\_168
CL\_INS\_168
CL\_INS\_168
CL\_INS\_168
CL\_INS\_168
CL\_INS\_20
CL\_INS\_168
CL\_INS\_20
CL\_INS\_168
CL\_INS\_168
CL\_INS\_168
CL\_INS\_168
CL\_INS\_168
CL\_INS\_168
CL\_INS\_168
CL\_INS\_168
CL\_INS\_168
CL\_INS\_168
CL\_INS\_168
CL\_INS\_168
CL\_INS\_168
CL\_INS\_168
CL\_INS\_168
CL\_INS\_168
CL\_INS\_168
CL\_INS\_168
CL\_INS\_168
CL\_INS\_168
CL\_INS\_168
CL\_INS\_168
CL\_INS\_168
CL\_INS\_168
CL\_INS\_168
CL\_INS\_168
CL\_INS\_168
CL\_INS\_168
CL\_INS\_168
CL\_INS\_168
CL\_INS\_168
CL\_INS\_168
CL\_INS\_168
CL\_INS\_168
CL\_INS\_168
CL\_INS\_168
CL\_INS\_1
CL\_INS\_168
CL\_INS\_168
CL\_INS\_20
CL\_INS\_20
CL\_INS\_20
CL\_INS\_20
CL\_INS\_20
CL\_INS\_20
CL\_INS\_20
CL\_INS\_20
CL\_INS\_20
CL\_INS\_1
CL\_INS\_20
CL\_INS\_20
CL\_INS\_20
CL\_INS\_20
CL\_INS\_20
CL\_INS\_20
CL\_INS\_168
CL\_INS\_168
CL\_INS\_168
CL\_INS\_20
CL\_INS\_168
CL\_INS\_168
CL\_INS\_1
CL\_INS\_1
CL\_INS\_1
CL\_INS\_1
CL\_INS\_1
CL\_INS\_1
CL\_INS\_1
CL\_INS\_1
CL\_INS\_1
CL\_INS\_1
CL\_INS\_1
CL\_INS\_1
CL\_INS\_1
CL\_INS\_1
CL\_INS\_1
CL\_INS\_168
CL\_INS\_168
CL\_INS\_168
CL\_INS\_168
CL\_INS\_168
CL\_INS\_168
CL\_INS\_168
CL\_INS\_168
CL\_INS\_168
CL\_INS\_168
CL\_INS\_168
CL\_INS\_168
CL\_INS\_168
CL\_INS\_168
CL\_INS\_168
CL\_INS\_168
CL\_INS\_168
CL\_INS\_168
CL\_INS\_168
CL\_INS\_168
CL\_INS\_168
CL\_INS\_168
CL\_INS\_168
CL\_INS\_168
CL\_INS\_168
CL\_INS\_168
CL\_INS\_168
CL\_INS\_168
CL\_INS\_168
CL\_INS\_168
CL\_INS\_1
CL\_INS\_1
CL\_INS\_1
CL\_INS\_1
CL\_INS\_1
CL\_INS\_1
CL\_INS\_1
CL\_INS\_1
CL\_INS\_1
CL\_INS\_1
CL\_INS\_168
CL\_INS\_168
CL\_INS\_168
CL\_INS\_1
CL\_INS\_20
CL\_INS\_168
CL\_INS\_20
CL\_INS\_20
CL\_INS\_20
CL\_INS\_20
CL\_INS\_20
CL\_INS\_20
CL\_INS\_20
CL\_INS\_20
CL\_INS\_168
CL\_INS\_133
CL\_INS\_20
CL\_INS\_168
CL\_INS\_168
CL\_INS\_168
CL\_INS\_168
CL\_INS\_20
CL\_INS\_55
CL\_INS\_55
CL\_INS\_55
CL\_INS\_168
CL\_INS\_169
CL\_INS\_169
CL\_INS\_168
CL\_INS\_169
CL\_INS\_169
CL\_INS\_169
CL\_INS\_168
CL\_INS\_168
CL\_INS\_168
CL\_INS\_168
CL\_INS\_168
CL\_INS\_220
CL\_INS\_220
CL\_INS\_168
CL\_INS\_168
CL\_INS\_168
CL\_INS\_168
CL\_INS\_168
CL\_INS\_168
CL\_INS\_168
CL\_INS\_168
CL\_INS\_168
CL\_INS\_168
CL\_INS\_168
CL\_INS\_168
CL\_INS\_168
CL\_INS\_168
CL\_INS\_168
CL\_INS\_168
CL\_INS\_168
CL\_INS\_168
CL\_INS\_168
CL\_INS\_168
CL\_INS\_168
CL\_INS\_168
CL\_INS\_168
CL\_INS\_168
CL\_INS\_168
CL\_INS\_168
CL\_INS\_168
CL\_INS\_168
CL\_INS\_168
CL\_INS\_168
CL\_INS\_168
CL\_INS\_168
CL\_INS\_168
CL\_INS\_168
CL\_INS\_169
CL\_INS\_169
CL\_INS\_169
CL\_INS\_169
CL\_INS\_169
CL\_INS\_169
CL\_INS\_169
CL\_INS\_169
CL\_INS\_169
CL\_INS\_169
CL\_INS\_169
CL\_INS\_169
CL\_INS\_168
CL\_INS\_168
CL\_INS\_168
CL\_INS\_168
CL\_INS\_168
CL\_INS\_168
CL\_INS\_168
CL\_INS\_168
CL\_INS\_168
CL\_INS\_1
CL\_INS\_168
CL\_INS\_168
CL\_INS\_168
CL\_INS\_168
CL\_INS\_168
CL\_INS\_168
CL\_INS\_168
CL\_INS\_168
CL\_INS\_168
CL\_INS\_168
CL\_INS\_168
CL\_INS\_168
CL\_INS\_168
CL\_INS\_168
CL\_INS\_168
CL\_INS\_168
CL\_INS\_168
CL\_INS\_168
CL\_INS\_168
CL\_INS\_168
CL\_INS\_168
CL\_INS\_168
CL\_INS\_168
CL\_INS\_168
CL\_INS\_168
CL\_INS\_168
CL\_INS\_168
CL\_INS\_168
CL\_INS\_168
CL\_INS\_168
CL\_INS\_168
CL\_INS\_168
CL\_INS\_1
CL\_INS\_1
CL\_INS\_1
CL\_INS\_1
CL\_INS\_1
CL\_INS\_1
CL\_INS\_168
CL\_INS\_168
CL\_INS\_168
CL\_INS\_168
CL\_INS\_168
CL\_INS\_168
CL\_INS\_168
CL\_INS\_168
CL\_INS\_168
CL\_INS\_168
CL\_INS\_168
CL\_INS\_168
CL\_INS\_168
CL\_INS\_1
CL\_INS\_1
CL\_INS\_56
CL\_INS\_56
CL\_INS\_168
CL\_INS\_168
CL\_INS\_168
CL\_INS\_168
CL\_INS\_168
CL\_INS\_168
CL\_INS\_168
Cluster ID


CL\_7265
CL\_4463
CL\_4464
CL\_4465
CL\_4466
CL\_4467
CL\_4468
CL\_5320
CL\_2802
CL\_2803
CL\_2804
CL\_13058
CL\_13059
CL\_13060
CL\_13061
CL\_7764
CL\_7765
CL\_12632
CL\_12633
CL\_5250
CL\_5621
CL\_5622
CL\_5623
CL\_5252
CL\_5253
CL\_5254
CL\_9338
CL\_9339
CL\_9340
CL\_9341
CL\_9342
CL\_9343
CL\_9344
CL\_9345
CL\_9346
CL\_9347
CL\_9348
CL\_9349
CL\_9350
CL\_9351
CL\_9352
CL\_9353
CL\_9354
CL\_9355
CL\_9356
CL\_9357
CL\_9358
CL\_9359
CL\_9360
CL\_9361
CL\_9362
CL\_9363
CL\_9364
CL\_9365
CL\_6828
CL\_6827
CL\_5251
CL\_5808
CL\_5807
CL\_5806
CL\_5161
CL\_5160
CL\_5159
CL\_4469
CL\_8229
CL\_8228
CL\_8227
CL\_8226
CL\_8225
CL\_8224
CL\_8223
CL\_8222
CL\_8221
CL\_8220
CL\_8219
CL\_8218
CL\_8217
CL\_8216
CL\_8215
CL\_8361
CL\_8360
CL\_8359
CL\_8358
CL\_8357
CL\_8356
CL\_8355
CL\_8354
CL\_8353
CL\_8352
CL\_8351
CL\_8350
CL\_8349
CL\_8348
CL\_8347
CL\_5847
CL\_5846
CL\_12299
CL\_12298
CL\_5841
CL\_5839
CL\_4747
CL\_12297
CL\_12296
CL\_4744
CL\_4743
CL\_12295
CL\_4740
CL\_12294
CL\_4737
CL\_4736
CL\_4735
CL\_4734
CL\_12293
CL\_4729
CL\_12292
CL\_4728
CL\_4727
CL\_4725
CL\_12291
CL\_12290
CL\_4722
CL\_5932
CL\_5931
CL\_5930
CL\_12289
CL\_12288
CL\_12287
CL\_12286
CL\_5925
CL\_5924
CL\_8441
CL\_8442
CL\_8443
CL\_8444
CL\_8445
CL\_12285
CL\_8446
CL\_4676
CL\_4675
CL\_4674
CL\_4673
CL\_5876
CL\_12284
CL\_12283
CL\_8448
CL\_7243
CL\_4666
CL\_4665
CL\_12282
CL\_11679
CL\_7248
CL\_12281
CL\_12280
CL\_12279
CL\_12278
CL\_12277
CL\_12276
CL\_12275
CL\_12274
CL\_12273
CL\_12272
CL\_5462
CL\_5463
CL\_5464
CL\_8019
CL\_7638
CL\_5428
CL\_5429
CL\_5430
CL\_5431
CL\_5432
CL\_5433
CL\_5434
CL\_5435
CL\_5436
CL\_5437
CL\_5438
CL\_5439
CL\_5440
CL\_5441
CL\_5442
CL\_5443
CL\_5444
CL\_5445
CL\_5446
CL\_5447
CL\_5448
CL\_5449
CL\_5450
CL\_5451
CL\_5452
CL\_5453
CL\_5454
CL\_5455
CL\_5456
CL\_5457
CL\_5458
CL\_5459
CL\_5460
CL\_5461
CL\_5311
CL\_5310
CL\_4640
CL\_5309
CL\_5308
CL\_8116
CL\_11926
CL\_11925
CL\_11924
CL\_11923
CL\_11922
CL\_11921
CL\_11920
CL\_11919
CL\_4633
CL\_5190
CL\_11918
CL\_11917
CL\_11916
CL\_11915
CL\_11914
CL\_13535
CL\_13534
CL\_11913
CL\_8115
CL\_8114
CL\_8113
CL\_8112
CL\_8111
CL\_8110
CL\_4439
CL\_4438
CL\_4436
CL\_6077
CL\_6076
CL\_6075
CL\_6074
CL\_8109
CL\_7567
CL\_4432
CL\_4433
CL\_4434
CL\_8108
CL\_8107
CL\_8106
CL\_8105
CL\_8104
CL\_8103
CL\_8102
CL\_8101
CL\_8100
CL\_8099
CL\_8098
CL\_8097
CL\_8096
CL\_8095
CL\_8094
CL\_8093
CL\_8092
CL\_8091
CL\_8090
CL\_8089
CL\_8088
CL\_8087
CL\_8086
CL\_8085
CL\_8084
CL\_8083
CL\_8082
CL\_8081
CL\_8080
CL\_8079
CL\_4605
CL\_4606
CL\_4604
CL\_4603
CL\_4602
CL\_4601
CL\_5259
CL\_5260
CL\_5261
CL\_5722
CL\_8078
CL\_5229
CL\_8077
CL\_4595
CL\_5350
CL\_12266
CL\_7571
CL\_7570
CL\_7569
CL\_12265
CL\_10294
CL\_8392
CL\_8391
CL\_8390
CL\_11010
CL\_6969
CL\_6968
CL\_10828
CL\_12264
CL\_12263
CL\_5848
CL\_6625
CL\_11832
CL\_6623
CL\_6624
CL\_7766
CL\_6826
CL\_6825
CL\_7767
CL\_6824
CL\_6823
CL\_6822
CL\_10346
CL\_6821
CL\_6820
CL\_6819
CL\_6818
CL\_6817
CL\_6816
CL\_12634
CL\_12635
CL\_12636
CL\_12637
CL\_12638
CL\_12639
CL\_12640
CL\_12641
CL\_12642
CL\_6815
CL\_6814
CL\_6813
CL\_6812
CL\_6811
CL\_6810
CL\_6809
CL\_6808
CL\_6807
CL\_6806
CL\_6805
CL\_6804
CL\_6803
CL\_6802
CL\_6801
CL\_6800
CL\_6799
CL\_6798
CL\_6797
CL\_6796
CL\_6795
CL\_6794
CL\_6793
CL\_6792
CL\_6791
CL\_7768
CL\_8849
CL\_8848
CL\_8847
CL\_8846
CL\_8845
CL\_8844
CL\_8843
CL\_8842
CL\_8841
CL\_8840
CL\_8839
CL\_10347
CL\_10348
CL\_10349
CL\_10350
CL\_10351
CL\_10352
CL\_10353
CL\_10354
CL\_10355
CL\_24
CL\_10356
CL\_10357
CL\_9447
CL\_10493
CL\_13538
CL\_13537
CL\_13536
CL\_12262
CL\_7769
CL\_10494
CL\_10495
CL\_10496
CL\_10497
CL\_10498
CL\_10499
CL\_10500
CL\_10501
CL\_10502
CL\_10503
CL\_10504
CL\_10505
CL\_10506
CL\_9847
CL\_9846
CL\_2805
CL\_7770
CL\_7771
CL\_11619
CL\_7772
CL\_7773
CL\_7774
CL\_7775
CL\_5465
CL\_5466
CL\_5467
CL\_5468
CL\_5223
CL\_5222
CL\_8076
CL\_8075
CL\_12271
CL\_12270
CL\_12269
CL\_12268
CL\_12267
CL\_8492
CL\_8491
CL\_8074
CL\_8073
CL\_8072
CL\_8071
CL\_4659
CL\_4644
CL\_5315
CL\_5314
CL\_5312
CL\_8214
CL\_8213
CL\_8212
CL\_8211
CL\_8210
CL\_2806
