## Supplementary material for "A novel method for integrating genomic and Tn-Seq data to identify common *in vivo* fitness mechanisms across multiple bacterial species": S3 Dataset: CL_INS_169.html

Legend

 Mobile +extrachromosomalelementfunctions
 Hypothetical
 DNA Metabolism
 Other
 All VFDB Genes

FULL


WINDOWSVGPNG

Trim RowsRemove SingletonsSave Fasta

CL\_2810


CL\_2810


CL\_2801

HighlightSelectShow Genomes


46

CL\_2811


8

CL\_2811


1

CL\_2811

fGI ID


CL\_INS\_169
CL\_INS\_169
CL\_INS\_169
CL\_INS\_169
CL\_INS\_169
CL\_INS\_169
CL\_INS\_169
CL\_INS\_169
CL\_INS\_169
CL\_INS\_169
CL\_INS\_169
CL\_INS\_169
CL\_INS\_169
CL\_INS\_169
CL\_INS\_169
CL\_INS\_169
CL\_INS\_169
CL\_INS\_169
CL\_INS\_169
CL\_INS\_169
CL\_INS\_169
CL\_INS\_169
CL\_INS\_169
CL\_INS\_169
CL\_INS\_169
CL\_INS\_169
CL\_INS\_169
CL\_INS\_169
Cluster ID


CL\_4909
CL\_4910
CL\_4911
CL\_4912
CL\_4913
CL\_4914
CL\_4915
CL\_4916
CL\_4917
CL\_4918
CL\_6826
CL\_6825
CL\_6824
CL\_6823
CL\_6822
CL\_7768
CL\_8849
CL\_8848
CL\_8847
CL\_8846
CL\_8845
CL\_8844
CL\_8843
CL\_8842
CL\_8841
CL\_8840
CL\_8839
CL\_8838
