## Supplementary material for "A novel method for integrating genomic and Tn-Seq data to identify common *in vivo* fitness mechanisms across multiple bacterial species": S3 Dataset: CL_INS_170.html

Legend

 Regulatoryfunctions
 Hypothetical
 All VFDB Genes

FULL


WINDOWSVGPNG

Trim RowsRemove SingletonsSave Fasta

CL\_2820


CL\_2820


CL\_2820


CL\_2820


CL\_2820


CL\_2819

HighlightSelectShow Genomes


27

CL\_2823


23

CL\_2823


3

CL\_2823


1

CL\_2824


1

CL\_2823


1

CL\_2823

fGI ID


CL\_INS\_170
CL\_INS\_170
Cluster ID


CL\_2821
CL\_2822
