## Supplementary material for "A novel method for integrating genomic and Tn-Seq data to identify common *in vivo* fitness mechanisms across multiple bacterial species": S3 Dataset: CL_INS_171.html

Legend

 Mobile +extrachromosomalelementfunctions
 Hypothetical
 Other
 All VFDB Genes

FULL


WINDOWSVGPNG

Trim RowsRemove SingletonsSave Fasta

CL\_2854


CL\_2854


CL\_2854

HighlightSelectShow Genomes


43

CL\_2855


8

CL\_2855


1

CL\_2855

fGI ID


CL\_INS\_171
CL\_INS\_171
CL\_INS\_171
CL\_INS\_171
CL\_INS\_171
CL\_INS\_171
Cluster ID


CL\_4919
CL\_4920
CL\_11722
CL\_11721
CL\_11720
CL\_11719
