## Supplementary material for "A novel method for integrating genomic and Tn-Seq data to identify common *in vivo* fitness mechanisms across multiple bacterial species": S3 Dataset: CL_INS_172.html

Legend

 Mobile +extrachromosomalelementfunctions
 Hypothetical
 EnergyMetabolism
 All VFDB Genes

FULL


WINDOWSVGPNG

Trim RowsRemove SingletonsSave Fasta

CL\_2860


CL\_2860


CL\_2860


CL\_2860


CL\_2860


CL\_2860

HighlightSelectShow Genomes


24

CL\_2863


20

CL\_2863


9

CL\_2863


1

CL\_2863


1

CL\_2863


1

CL\_2863

fGI ID


CL\_INS\_172
CL\_INS\_172
CL\_INS\_172
CL\_INS\_172
CL\_INS\_172
Cluster ID


CL\_7397
CL\_2861
CL\_6790
CL\_8962
CL\_2862
