## Supplementary material for "A novel method for integrating genomic and Tn-Seq data to identify common *in vivo* fitness mechanisms across multiple bacterial species": S3 Dataset: CL_INS_174.html

Legend

 Hypothetical
 Proteinsynthesis/fate
 Other
 All VFDB Genes

FULL


WINDOWSVGPNG

Trim RowsRemove SingletonsSave Fasta

CL\_2875


CL\_2875


CL\_2875


CL\_2853

HighlightSelectShow Genomes


21

CL\_4922


18

CL\_4922


1

CL\_4922


1

CL\_4922

fGI ID


CL\_INS\_174
CL\_INS\_174
CL\_INS\_174
CL\_INS\_174
Cluster ID


CL\_12094
CL\_4921
CL\_13294
CL\_13293
