## Supplementary material for "A novel method for integrating genomic and Tn-Seq data to identify common *in vivo* fitness mechanisms across multiple bacterial species": S3 Dataset: CL_INS_175.html

Legend

 Regulatoryfunctions
 Other

FULL


WINDOWSVGPNG

Trim RowsRemove SingletonsSave Fasta

CL\_2877


CL\_2877

HighlightSelectShow Genomes


47

CL\_2878


8

CL\_2878

fGI ID


CL\_INS\_175
CL\_INS\_175
Cluster ID


CL\_4923
CL\_4924
