## Supplementary material for "A novel method for integrating genomic and Tn-Seq data to identify common *in vivo* fitness mechanisms across multiple bacterial species": S3 Dataset: CL_INS_176.html

Legend

 Mobile +extrachromosomalelementfunctions
 Regulatoryfunctions
 Hypothetical
 DNA Metabolism
 All EssentialGenes
 Other
 EnergyMetabolism
 All VFDB Genes

FULL


WINDOWSVGPNG

Trim RowsRemove SingletonsSave Fasta

CL\_2922


CL\_2922


CL\_2922


CL\_2922


CL\_2922


CL\_2922


CL\_2922


CL\_2922


CL\_2922


CL\_2922


CL\_2922


CL\_2922


CL\_2922


CL\_2922


CL\_2922


CL\_2922


CL\_2922

HighlightSelectShow Genomes


26

CL\_2924


8

CL\_2924


4

CL\_2924


3

CL\_2924


3

CL\_2924


2

CL\_2924


1

CL\_2924


1

CL\_2924


1

CL\_2924


1

CL\_2924


1

CL\_2924


1

CL\_2924


1

CL\_2924


1

CL\_2924


1

CL\_2924


1

CL\_2924


1

CL\_2924

fGI ID


CL\_INS\_176
CL\_INS\_176
CL\_INS\_176
CL\_INS\_176
CL\_INS\_137
CL\_INS\_176
CL\_INS\_176
CL\_INS\_176
CL\_INS\_176
CL\_INS\_98
CL\_INS\_176
CL\_INS\_176
CL\_INS\_217
CL\_INS\_217
CL\_INS\_176
CL\_INS\_176
CL\_INS\_176
CL\_INS\_145
CL\_INS\_176
CL\_INS\_176
CL\_INS\_176
CL\_INS\_176
CL\_INS\_176
CL\_INS\_137
CL\_INS\_137
CL\_INS\_176
CL\_INS\_176
CL\_INS\_217
CL\_INS\_217
CL\_INS\_133
CL\_INS\_133
CL\_INS\_176
CL\_INS\_176
CL\_INS\_133
CL\_INS\_217
CL\_INS\_133
CL\_INS\_133
CL\_INS\_133
CL\_INS\_176
CL\_INS\_133
CL\_INS\_133
CL\_INS\_133
CL\_INS\_217
CL\_INS\_217
CL\_INS\_176
CL\_INS\_217
CL\_INS\_176
CL\_INS\_176
CL\_INS\_217
CL\_INS\_217
CL\_INS\_217
CL\_INS\_217
CL\_INS\_217
CL\_INS\_83
CL\_INS\_217
CL\_INS\_217
CL\_INS\_217
CL\_INS\_217
CL\_INS\_217
CL\_INS\_217
CL\_INS\_217
CL\_INS\_217
CL\_INS\_217
CL\_INS\_217
CL\_INS\_145
CL\_INS\_217
CL\_INS\_176
CL\_INS\_217
CL\_INS\_176
CL\_INS\_198
CL\_INS\_133
CL\_INS\_133
CL\_INS\_198
CL\_INS\_176
CL\_INS\_217
CL\_INS\_198
CL\_INS\_133
CL\_INS\_217
CL\_INS\_145
CL\_INS\_176
CL\_INS\_217
CL\_INS\_217
CL\_INS\_217
CL\_INS\_145
CL\_INS\_145
CL\_INS\_145
CL\_INS\_145
CL\_INS\_176
CL\_INS\_145
CL\_INS\_176
CL\_INS\_176
CL\_INS\_176
CL\_INS\_176
CL\_INS\_1
CL\_INS\_176
CL\_INS\_176
CL\_INS\_176
CL\_INS\_217
CL\_INS\_217
CL\_INS\_83
CL\_INS\_83
CL\_INS\_133
CL\_INS\_217
CL\_INS\_176
CL\_INS\_217
CL\_INS\_217
CL\_INS\_217
CL\_INS\_176
CL\_INS\_176
CL\_INS\_217
CL\_INS\_176
CL\_INS\_176
CL\_INS\_176
CL\_INS\_176
CL\_INS\_176
CL\_INS\_176
CL\_INS\_217
CL\_INS\_217
CL\_INS\_137
CL\_INS\_137
CL\_INS\_176
Cluster ID


CL\_9734
CL\_13672
CL\_6265
CL\_9235
CL\_7394
CL\_6266
CL\_6267
CL\_6268
CL\_6269
CL\_7393
CL\_9236
CL\_9237
CL\_10191
CL\_8790
CL\_10190
CL\_10189
CL\_7776
CL\_2586
CL\_7777
CL\_7778
CL\_11014
CL\_11015
CL\_11016
CL\_7392
CL\_7391
CL\_9366
CL\_13454
CL\_7390
CL\_7363
CL\_7388
CL\_6347
CL\_11017
CL\_11018
CL\_6348
CL\_11697
CL\_6349
CL\_6350
CL\_6460
CL\_7779
CL\_6462
CL\_5611
CL\_5610
CL\_6463
CL\_7780
CL\_9238
CL\_7781
CL\_11019
CL\_12368
CL\_7782
CL\_7783
CL\_7784
CL\_7785
CL\_7786
CL\_6362
CL\_9860
CL\_7787
CL\_7788
CL\_7789
CL\_7790
CL\_7791
CL\_7792
CL\_7793
CL\_6371
CL\_6372
CL\_10188
CL\_6375
CL\_7794
CL\_6376
CL\_9367
CL\_10187
CL\_6377
CL\_6378
CL\_9368
CL\_9369
CL\_6379
CL\_10186
CL\_7795
CL\_6380
CL\_7796
CL\_9370
CL\_6385
CL\_6386
CL\_6387
CL\_9857
CL\_9856
CL\_9855
CL\_9854
CL\_10185
CL\_9853
CL\_13455
CL\_7797
CL\_7798
CL\_11020
CL\_5107
CL\_7799
CL\_7800
CL\_11021
CL\_7801
CL\_7802
CL\_5530
CL\_5529
CL\_5528
CL\_9239
CL\_9240
CL\_6396
CL\_6397
CL\_6398
CL\_12369
CL\_7803
CL\_7431
CL\_10184
CL\_10183
CL\_10182
CL\_10181
CL\_11022
CL\_11023
CL\_6404
CL\_6405
CL\_7373
CL\_7372
CL\_2923
