## Supplementary material for "A novel method for integrating genomic and Tn-Seq data to identify common *in vivo* fitness mechanisms across multiple bacterial species": S3 Dataset: CL_INS_178.html

Legend

 Mobile +extrachromosomalelementfunctions
 Regulatoryfunctions
 Hypothetical
 DNA Metabolism
 AntibioticResistance
 All EssentialGenes
 Other
 Transport +binding proteins
 All VFDB Genes

FULL


WINDOWSVGPNG

Trim RowsRemove SingletonsSave Fasta

CL\_2927


CL\_2927


CL\_2926


CL\_2927


CL\_2927


CL\_2927


CL\_2927


CL\_2927


CL\_2927


CL\_2927


CL\_2927


CL\_2926


CL\_2927


CL\_2927


CL\_2927


CL\_2927


CL\_2927


CL\_2927


CL\_2927


CL\_2927


CL\_2927


CL\_2927


CL\_2927


CL\_2927


CL\_2927


CL\_2927


CL\_2738


CL\_2927


CL\_2927

HighlightSelectShow Genomes


11

CL\_2947


5

CL\_2947


5

CL\_2947


4

CL\_2947


3

CL\_2947


2

CL\_2947


2

CL\_2947


2

CL\_2947


1

CL\_2947


1

CL\_2947


1

CL\_2947


1

CL\_2947


1

CL\_2947


1

CL\_2735


1

CL\_2947


1

CL\_2947


1

CL\_2947


1

CL\_2947


1

CL\_2947


1

CL\_2947


1

CL\_2947


1

CL\_2947


1

CL\_2947


1

CL\_2947


1

CL\_2947


1

CL\_2947


1

CL\_2947


1

CL\_2947


1

CL\_2948

fGI ID


CL\_INS\_178
CL\_INS\_178
CL\_INS\_178
CL\_INS\_178
CL\_INS\_178
CL\_INS\_178
CL\_INS\_178
CL\_INS\_178
CL\_INS\_178
CL\_INS\_178
CL\_INS\_178
CL\_INS\_178
CL\_INS\_178
CL\_INS\_178
CL\_INS\_178
CL\_INS\_178
CL\_INS\_178
CL\_INS\_178
CL\_INS\_178
CL\_INS\_142
CL\_INS\_178
CL\_INS\_178
CL\_INS\_178
CL\_INS\_178
CL\_INS\_178
CL\_INS\_178
CL\_INS\_178
CL\_INS\_178
CL\_INS\_178
CL\_INS\_178
CL\_INS\_178
CL\_INS\_178
CL\_INS\_178
CL\_INS\_178
CL\_INS\_178
CL\_INS\_178
CL\_INS\_178
CL\_INS\_178
CL\_INS\_178
CL\_INS\_178
CL\_INS\_178
CL\_INS\_178
CL\_INS\_178
CL\_INS\_178
CL\_INS\_178
CL\_INS\_178
CL\_INS\_178
CL\_INS\_178
CL\_INS\_178
CL\_INS\_178
CL\_INS\_178
CL\_INS\_178
CL\_INS\_178
CL\_INS\_142
CL\_INS\_142
CL\_INS\_178
CL\_INS\_142
CL\_INS\_178
CL\_INS\_142
CL\_INS\_178
CL\_INS\_178
CL\_INS\_178
CL\_INS\_178
CL\_INS\_178
CL\_INS\_178
CL\_INS\_178
CL\_INS\_178
CL\_INS\_178
CL\_INS\_178
CL\_INS\_178
CL\_INS\_178
CL\_INS\_178
CL\_INS\_178
CL\_INS\_178
CL\_INS\_178
CL\_INS\_142
CL\_INS\_178
CL\_INS\_142
CL\_INS\_142
CL\_INS\_142
CL\_INS\_142
CL\_INS\_142
CL\_INS\_178
CL\_INS\_178
CL\_INS\_178
CL\_INS\_178
CL\_INS\_1
CL\_INS\_1
CL\_INS\_1
CL\_INS\_1
CL\_INS\_1
CL\_INS\_1
CL\_INS\_133
CL\_INS\_133
CL\_INS\_20
CL\_INS\_133
CL\_INS\_1
CL\_INS\_20
CL\_INS\_1
CL\_INS\_20
CL\_INS\_178
CL\_INS\_1
CL\_INS\_178
CL\_INS\_178
CL\_INS\_178
CL\_INS\_178
CL\_INS\_178
CL\_INS\_178
CL\_INS\_178
CL\_INS\_178
CL\_INS\_178
CL\_INS\_178
CL\_INS\_178
CL\_INS\_178
CL\_INS\_204
CL\_INS\_178
CL\_INS\_178
CL\_INS\_20
CL\_INS\_20
CL\_INS\_20
CL\_INS\_20
CL\_INS\_20
CL\_INS\_45
CL\_INS\_45
CL\_INS\_45
CL\_INS\_34
CL\_INS\_34
CL\_INS\_34
CL\_INS\_178
CL\_INS\_178
CL\_INS\_178
CL\_INS\_178
CL\_INS\_178
CL\_INS\_178
CL\_INS\_178
CL\_INS\_178
CL\_INS\_178
CL\_INS\_178
CL\_INS\_178
CL\_INS\_178
CL\_INS\_178
CL\_INS\_178
CL\_INS\_178
CL\_INS\_178
CL\_INS\_178
CL\_INS\_178
CL\_INS\_178
CL\_INS\_178
CL\_INS\_178
CL\_INS\_178
CL\_INS\_178
CL\_INS\_178
CL\_INS\_178
CL\_INS\_178
CL\_INS\_178
CL\_INS\_178
CL\_INS\_178
CL\_INS\_178
CL\_INS\_178
CL\_INS\_178
CL\_INS\_178
CL\_INS\_178
CL\_INS\_178
CL\_INS\_178
Cluster ID


CL\_6831
CL\_6832
CL\_2928
CL\_2929
CL\_10359
CL\_13292
CL\_7804
CL\_7805
CL\_7806
CL\_7807
CL\_5625
CL\_5626
CL\_5627
CL\_5628
CL\_5629
CL\_5630
CL\_5631
CL\_2930
CL\_2931
CL\_4888
CL\_7808
CL\_2932
CL\_9448
CL\_9449
CL\_9450
CL\_11718
CL\_11717
CL\_9451
CL\_2933
CL\_2934
CL\_2935
CL\_2936
CL\_4927
CL\_2937
CL\_2938
CL\_2939
CL\_9241
CL\_9242
CL\_11460
CL\_9243
CL\_9244
CL\_11256
CL\_9245
CL\_9246
CL\_9452
CL\_9453
CL\_9454
CL\_9455
CL\_11716
CL\_9456
CL\_9457
CL\_9458
CL\_9459
CL\_5632
CL\_4886
CL\_13062
CL\_9234
CL\_5633
CL\_4884
CL\_13063
CL\_13064
CL\_13065
CL\_9733
CL\_7809
CL\_7810
CL\_5634
CL\_13291
CL\_13290
CL\_13289
CL\_5635
CL\_2940
CL\_9731
CL\_9730
CL\_2941
CL\_4928
CL\_4873
CL\_9983
CL\_2942
CL\_2943
CL\_2944
CL\_2945
CL\_2946
CL\_12643
CL\_12644
CL\_12645
CL\_12646
CL\_4439
CL\_4440
CL\_4441
CL\_4442
CL\_4443
CL\_4444
CL\_5202
CL\_5201
CL\_5889
CL\_4435
CL\_10741
CL\_5836
CL\_10441
CL\_5835
CL\_12647
CL\_7567
CL\_12648
CL\_12649
CL\_12650
CL\_12651
CL\_8158
CL\_12652
CL\_12653
CL\_12654
CL\_12655
CL\_12656
CL\_12657
CL\_12658
CL\_6912
CL\_12659
CL\_12660
CL\_5287
CL\_5286
CL\_5285
CL\_5284
CL\_5283
CL\_5282
CL\_5281
CL\_5280
CL\_5279
CL\_5278
CL\_5277
CL\_5276
CL\_7511
CL\_7510
CL\_7509
CL\_7508
CL\_12661
CL\_7507
CL\_7506
CL\_7505
CL\_7504
CL\_7503
CL\_7502
CL\_7501
CL\_12662
CL\_12663
CL\_12664
CL\_6064
CL\_6063
CL\_6062
CL\_6061
CL\_9732
CL\_12665
CL\_12666
CL\_12667
CL\_12668
CL\_7498
CL\_7496
CL\_7495
CL\_7494
CL\_7493
CL\_7492
CL\_7491
CL\_7490
CL\_7489
CL\_12669
CL\_12670
