## Supplementary material for "A novel method for integrating genomic and Tn-Seq data to identify common *in vivo* fitness mechanisms across multiple bacterial species": S3 Dataset: CL_INS_179.html

Legend

 All VFDB Genes

FULL


WINDOWSVGPNG

Trim RowsRemove SingletonsSave Fasta

CL\_2978


CL\_2978


CL\_2978

HighlightSelectShow Genomes


38

CL\_2981


18

CL\_2981


1

CL\_2981

fGI ID


CL\_INS\_179
CL\_INS\_179
Cluster ID


CL\_2979
CL\_2980
