## Supplementary material for "A novel method for integrating genomic and Tn-Seq data to identify common *in vivo* fitness mechanisms across multiple bacterial species": S3 Dataset: CL_INS_181.html

Legend

 Mobile +extrachromosomalelementfunctions
 Hypothetical
 Regulatoryfunctions
 All EssentialGenes
 Other
 All VFDB Genes
 Transport +binding proteins

FULL


WINDOWSVGPNG

Trim RowsRemove SingletonsSave Fasta

CL\_3004


CL\_3004


CL\_3004


CL\_3004


CL\_3004


CL\_3004


CL\_3004


CL\_3004

HighlightSelectShow Genomes


47

CL\_3003


4

CL\_3003


1

CL\_3003


1

CL\_3003


1

CL\_3003


1

CL\_3003


1

CL\_3003


1

CL\_3003

fGI ID


CL\_INS\_181
CL\_INS\_181
CL\_INS\_181
CL\_INS\_181
CL\_INS\_181
CL\_INS\_181
CL\_INS\_181
CL\_INS\_181
CL\_INS\_181
CL\_INS\_181
CL\_INS\_181
CL\_INS\_181
CL\_INS\_64
CL\_INS\_230
CL\_INS\_64
CL\_INS\_230
CL\_INS\_230
CL\_INS\_230
CL\_INS\_64
CL\_INS\_230
CL\_INS\_230
CL\_INS\_230
CL\_INS\_181
CL\_INS\_181
CL\_INS\_181
CL\_INS\_64
CL\_INS\_83
CL\_INS\_83
CL\_INS\_83
CL\_INS\_83
CL\_INS\_65
CL\_INS\_83
CL\_INS\_83
CL\_INS\_181
CL\_INS\_181
CL\_INS\_181
Cluster ID


CL\_13277
CL\_13278
CL\_13279
CL\_13280
CL\_13281
CL\_13282
CL\_13283
CL\_13284
CL\_13285
CL\_13286
CL\_13287
CL\_13288
CL\_8206
CL\_5093
CL\_8207
CL\_5091
CL\_5090
CL\_5089
CL\_8208
CL\_5087
CL\_5086
CL\_5085
CL\_10360
CL\_8787
CL\_10952
CL\_5493
CL\_5494
CL\_5495
CL\_7298
CL\_5652
CL\_5651
CL\_5650
CL\_5649
CL\_5648
CL\_8209
CL\_8656
