## Supplementary material for "A novel method for integrating genomic and Tn-Seq data to identify common *in vivo* fitness mechanisms across multiple bacterial species": S3 Dataset: CL_INS_182.html

Legend

 Mobile +extrachromosomalelementfunctions
 Hypothetical
 Other
 Transport +binding proteins
 All VFDB Genes

FULL


WINDOWSVGPNG

Trim RowsRemove SingletonsSave Fasta

CL\_3021


CL\_3021


CL\_3021


CL\_3021


CL\_3021


CL\_3021


CL\_3021


CL\_3021


CL\_3021

HighlightSelectShow Genomes


46

CL\_3022


3

CL\_3022


2

CL\_3022


1

CL\_3022


1

CL\_3022


1

CL\_3022


1

CL\_3023


1

CL\_3022


1

CL\_3022

fGI ID


CL\_INS\_182
CL\_INS\_182
CL\_INS\_182
CL\_INS\_182
CL\_INS\_1
CL\_INS\_1
CL\_INS\_1
CL\_INS\_1
CL\_INS\_1
CL\_INS\_1
CL\_INS\_1
CL\_INS\_1
CL\_INS\_1
CL\_INS\_182
CL\_INS\_182
CL\_INS\_182
CL\_INS\_182
CL\_INS\_182
CL\_INS\_182
CL\_INS\_182
CL\_INS\_182
CL\_INS\_182
CL\_INS\_182
CL\_INS\_182
CL\_INS\_182
CL\_INS\_182
CL\_INS\_182
CL\_INS\_182
CL\_INS\_182
CL\_INS\_182
CL\_INS\_182
CL\_INS\_182
CL\_INS\_182
CL\_INS\_182
CL\_INS\_182
CL\_INS\_182
CL\_INS\_182
CL\_INS\_182
CL\_INS\_182
CL\_INS\_182
CL\_INS\_182
CL\_INS\_182
CL\_INS\_182
CL\_INS\_182
CL\_INS\_182
CL\_INS\_182
CL\_INS\_182
CL\_INS\_182
CL\_INS\_182
CL\_INS\_182
CL\_INS\_182
CL\_INS\_182
CL\_INS\_182
CL\_INS\_182
CL\_INS\_182
CL\_INS\_182
CL\_INS\_182
CL\_INS\_182
CL\_INS\_182
CL\_INS\_182
CL\_INS\_182
CL\_INS\_182
CL\_INS\_182
CL\_INS\_182
CL\_INS\_182
CL\_INS\_182
CL\_INS\_182
CL\_INS\_182
CL\_INS\_182
CL\_INS\_182
CL\_INS\_182
CL\_INS\_182
CL\_INS\_182
Cluster ID


CL\_10361
CL\_6441
CL\_6442
CL\_9866
CL\_4411
CL\_5261
CL\_5260
CL\_5259
CL\_4601
CL\_4602
CL\_4603
CL\_4604
CL\_4605
CL\_9460
CL\_10685
CL\_6271
CL\_6272
CL\_6273
CL\_9867
CL\_9868
CL\_6274
CL\_6275
CL\_6276
CL\_6277
CL\_6278
CL\_6279
CL\_6280
CL\_13673
CL\_6281
CL\_9461
CL\_13674
CL\_6282
CL\_6283
CL\_6284
CL\_6285
CL\_9462
CL\_10684
CL\_6286
CL\_6287
CL\_6288
CL\_6289
CL\_6290
CL\_6291
CL\_6292
CL\_6293
CL\_6294
CL\_6295
CL\_6296
CL\_6297
CL\_6298
CL\_6299
CL\_9463
CL\_9464
CL\_9465
CL\_10683
CL\_10682
CL\_10681
CL\_10680
CL\_10679
CL\_10678
CL\_6300
CL\_6301
CL\_6302
CL\_6303
CL\_6304
CL\_6445
CL\_6305
CL\_6306
CL\_6307
CL\_13675
CL\_6308
CL\_6309
CL\_6310
