## Supplementary material for "A novel method for integrating genomic and Tn-Seq data to identify common *in vivo* fitness mechanisms across multiple bacterial species": S3 Dataset: CL_INS_185.html

Legend

 Mobile +extrachromosomalelementfunctions
 Hypothetical
 All EssentialGenes
 Other
 All VFDB Genes

FULL


WINDOWSVGPNG

Trim RowsRemove SingletonsSave Fasta

CL\_3076


CL\_3076


CL\_3076


CL\_3076


CL\_3076


CL\_3076


CL\_3076


CL\_3076


CL\_3076


CL\_3076


CL\_3076


CL\_3076


CL\_3076


CL\_3076


CL\_3076


CL\_3076


CL\_3076


CL\_3076


CL\_3076


CL\_3076


CL\_3076


CL\_3076


CL\_3076


CL\_3076


CL\_3076


CL\_3076


CL\_3076


CL\_3076


CL\_3076


CL\_3076


CL\_3076


CL\_3076


CL\_3076


CL\_3076


CL\_3076


CL\_3076


CL\_3076

HighlightSelectShow Genomes


8

CL\_3089


3

CL\_3089


2

CL\_3089


2

CL\_3089


2

CL\_3089


2

CL\_3089


2

CL\_3089


2

CL\_3089


2

CL\_3089


2

CL\_3089


2

CL\_3089


2

CL\_3089


2

CL\_3089


1

CL\_3089


1

CL\_3089


1

CL\_3089


1

CL\_3089


1

CL\_3089


1

CL\_3089


1

CL\_3089


1

CL\_3089


1

CL\_3089


1

CL\_3089


1

CL\_3089


1

CL\_3089


1

CL\_3089


1

CL\_3089


1

CL\_3089


1

CL\_3089


1

CL\_3089


1

CL\_3089


1

CL\_3089


1

CL\_3089


1

CL\_3089


1

CL\_3089


1

CL\_3089


1

CL\_3089

fGI ID


CL\_INS\_185
CL\_INS\_185
CL\_INS\_185
CL\_INS\_185
CL\_INS\_185
CL\_INS\_185
CL\_INS\_185
CL\_INS\_185
CL\_INS\_185
CL\_INS\_185
CL\_INS\_185
CL\_INS\_185
CL\_INS\_185
CL\_INS\_185
CL\_INS\_185
CL\_INS\_185
CL\_INS\_185
CL\_INS\_185
CL\_INS\_185
CL\_INS\_185
CL\_INS\_185
CL\_INS\_185
CL\_INS\_185
CL\_INS\_185
CL\_INS\_185
CL\_INS\_185
CL\_INS\_185
CL\_INS\_185
CL\_INS\_185
CL\_INS\_185
CL\_INS\_185
CL\_INS\_185
CL\_INS\_185
CL\_INS\_185
CL\_INS\_185
CL\_INS\_185
CL\_INS\_185
CL\_INS\_185
CL\_INS\_185
CL\_INS\_185
CL\_INS\_185
CL\_INS\_185
CL\_INS\_185
CL\_INS\_185
CL\_INS\_185
CL\_INS\_185
CL\_INS\_185
CL\_INS\_185
CL\_INS\_185
CL\_INS\_185
CL\_INS\_185
CL\_INS\_185
CL\_INS\_185
CL\_INS\_185
CL\_INS\_185
CL\_INS\_185
CL\_INS\_185
CL\_INS\_185
CL\_INS\_185
CL\_INS\_185
CL\_INS\_185
CL\_INS\_185
CL\_INS\_185
CL\_INS\_185
CL\_INS\_185
CL\_INS\_185
CL\_INS\_185
CL\_INS\_185
CL\_INS\_185
CL\_INS\_185
Cluster ID


CL\_4934
CL\_4935
CL\_4936
CL\_4937
CL\_4938
CL\_4939
CL\_4940
CL\_4941
CL\_4942
CL\_4943
CL\_4944
CL\_4945
CL\_4946
CL\_4947
CL\_4948
CL\_4949
CL\_4950
CL\_4951
CL\_11708
CL\_11707
CL\_4952
CL\_4953
CL\_13676
CL\_4954
CL\_11715
CL\_11714
CL\_11461
CL\_4955
CL\_4956
CL\_4957
CL\_4958
CL\_3077
CL\_3078
CL\_13680
CL\_3088
CL\_6312
CL\_3086
CL\_11462
CL\_3087
CL\_13677
CL\_13678
CL\_13679
CL\_6313
CL\_4959
CL\_10677
CL\_13681
CL\_11713
CL\_11712
CL\_11711
CL\_11710
CL\_11709
CL\_3079
CL\_9466
CL\_3080
CL\_3081
CL\_3082
CL\_3083
CL\_5636
CL\_3084
CL\_3085
CL\_11255
CL\_9467
CL\_4960
CL\_6834
CL\_4961
CL\_7398
CL\_4962
CL\_9869
CL\_6835
CL\_4963
