## Supplementary material for "A novel method for integrating genomic and Tn-Seq data to identify common *in vivo* fitness mechanisms across multiple bacterial species": S3 Dataset: CL_INS_186.html

Legend

 Hypothetical
 Other
 All VFDB Genes

FULL


WINDOWSVGPNG

Trim RowsRemove SingletonsSave Fasta

CL\_3089


CL\_3089


CL\_3089


CL\_3089


CL\_3089


CL\_3089

HighlightSelectShow Genomes


24

CL\_3090


21

CL\_3090


8

CL\_3090


2

CL\_3090


1

CL\_3090


1

CL\_3098

fGI ID


CL\_INS\_186
CL\_INS\_186
CL\_INS\_186
CL\_INS\_187
CL\_INS\_187
CL\_INS\_187
CL\_INS\_187
CL\_INS\_187
CL\_INS\_187
Cluster ID


CL\_7399
CL\_9247
CL\_4964
CL\_3091
CL\_3093
CL\_8963
CL\_3094
CL\_3095
CL\_3096
