## Supplementary material for "A novel method for integrating genomic and Tn-Seq data to identify common *in vivo* fitness mechanisms across multiple bacterial species": S3 Dataset: CL_INS_189.html

Legend

 Hypothetical
 All VFDB Genes

FULL


WINDOWSVGPNG

Trim RowsRemove SingletonsSave Fasta

CL\_3109


CL\_3109


CL\_3108


CL\_3109

HighlightSelectShow Genomes


40

CL\_3110


9

CL\_3110


4

CL\_3110


1

CL\_3111

fGI ID


CL\_INS\_190
CL\_INS\_189
Cluster ID


CL\_4966
CL\_7811
