## Supplementary material for "A novel method for integrating genomic and Tn-Seq data to identify common *in vivo* fitness mechanisms across multiple bacterial species": S3 Dataset: CL_INS_190.html

Legend

 Hypothetical
 All VFDB Genes

FULL


WINDOWSVGPNG

Trim RowsRemove SingletonsSave Fasta

CL\_3110


CL\_3110


CL\_3110


CL\_3108

HighlightSelectShow Genomes


45

CL\_3111


5

CL\_3111


4

CL\_3111


3

CL\_3111

fGI ID


CL\_INS\_190
CL\_INS\_190
CL\_INS\_190
Cluster ID


CL\_9468
CL\_4966
CL\_4967
