## Supplementary material for "A novel method for integrating genomic and Tn-Seq data to identify common *in vivo* fitness mechanisms across multiple bacterial species": S3 Dataset: CL_INS_191.html

Legend

 All VFDB Genes

FULL


WINDOWSVGPNG

Trim RowsRemove SingletonsSave Fasta

CL\_3112


CL\_3112

HighlightSelectShow Genomes


33

CL\_3114


21

CL\_3114

fGI ID


CL\_INS\_191
CL\_INS\_191
CL\_INS\_191
Cluster ID


CL\_3113
CL\_4968
CL\_4969
