## Supplementary material for "A novel method for integrating genomic and Tn-Seq data to identify common *in vivo* fitness mechanisms across multiple bacterial species": S3 Dataset: CL_INS_192.html

Legend

 Hypothetical
 Other
 All VFDB Genes

FULL


WINDOWSVGPNG

Trim RowsRemove SingletonsSave Fasta

CL\_3124


CL\_3124


CL\_3124


CL\_3123

HighlightSelectShow Genomes


27

CL\_3126


27

CL\_3126


2

CL\_3126


1

CL\_3126

fGI ID


CL\_INS\_192
CL\_INS\_192
CL\_INS\_192
Cluster ID


CL\_3125
CL\_9729
CL\_9728
