## Supplementary material for "A novel method for integrating genomic and Tn-Seq data to identify common *in vivo* fitness mechanisms across multiple bacterial species": S3 Dataset: CL_INS_194.html


CL\_7411
CL\_7410
CL\_7409
CL\_12467
CL\_7408
CL\_7407
CL\_7406
CL\_7405
CL\_7404
CL\_7403
CL\_7402
CL\_3270
CL\_3269
CL\_11244
CL\_7832
CL\_7831
CL\_13014
CL\_11442
CL\_9618
CL\_6725
CL\_10473
CL\_8984
CL\_11109
CL\_7353
CL\_3268
CL\_3267
CL\_6202
CL\_3266
CL\_3265
CL\_7830
CL\_9201
CL\_9202
CL\_11898
CL\_9011
CL\_6727
CL\_8983
CL\_10020
CL\_3264
CL\_5522
CL\_3263
CL\_3262
CL\_7829
CL\_7828
CL\_13442
CL\_3261
CL\_3260
CL\_11899
CL\_6203
CL\_6204
CL\_11245
CL\_11246
CL\_11247
CL\_11248
CL\_8979
CL\_11900
CL\_11901
CL\_11902
CL\_11903
CL\_11904
CL\_11905
CL\_11906
CL\_9676
CL\_9677
CL\_3259
CL\_3258
CL\_5523
CL\_11110
CL\_3257
CL\_3256
CL\_11443
CL\_3255
CL\_3254
CL\_6735
CL\_11444
CL\_7827
CL\_7826
CL\_5528
CL\_8978
CL\_5529
CL\_8815
CL\_8816
CL\_5526
CL\_11477
CL\_3253
CL\_3252
CL\_3251
CL\_3250
CL\_3249
CL\_9678
CL\_8817
CL\_8731
CL\_8730
CL\_8727
CL\_3248
CL\_5530
CL\_7802
CL\_7801
CL\_11249
CL\_11250
CL\_10478
CL\_5532
CL\_3247
CL\_3246
CL\_3245
CL\_3244
CL\_3243
CL\_8873
CL\_11111
CL\_7825
CL\_7824
CL\_9309
CL\_11445
CL\_11446
CL\_11447
CL\_5534
CL\_6215
CL\_13443
CL\_13444
CL\_13445
CL\_8725
CL\_9207
CL\_11907
CL\_8819
CL\_11908
CL\_11909
CL\_11910
CL\_5538
CL\_7823
CL\_7822
CL\_13013
CL\_5540
CL\_11911
CL\_3242
CL\_3241
CL\_3240
CL\_3239
CL\_3238
CL\_6217
CL\_6218
CL\_9208
CL\_6219
CL\_11271
CL\_11448
CL\_9879
CL\_9878
CL\_9877
CL\_9637
CL\_11912
CL\_9333
CL\_3237
CL\_3236
CL\_3235
CL\_9008
CL\_3234
CL\_11112
CL\_9967
CL\_5541
CL\_5542
CL\_5543
CL\_7821
CL\_7820
CL\_7819
CL\_7818
CL\_3233
CL\_3232
CL\_3231
CL\_3230
CL\_11251
CL\_11252
CL\_11253
CL\_3229
CL\_6222
CL\_6223
CL\_6224
CL\_6225
CL\_3228
CL\_11449
CL\_11450
CL\_6742
CL\_11451
CL\_11452
CL\_11453
CL\_11454
CL\_11455
CL\_7817
CL\_7816
CL\_7815
CL\_7814
CL\_7414
CL\_6226
CL\_6227
CL\_11254
CL\_13012
CL\_9210
CL\_13446
CL\_6229
CL\_9211
CL\_9212
CL\_9213
CL\_6230
CL\_6231
CL\_6232
CL\_6233
CL\_6740
CL\_6741
CL\_3227
CL\_3226
CL\_3225
CL\_9968
CL\_9969
CL\_9970
CL\_3224
CL\_3223
CL\_3222
CL\_3221
CL\_3220
CL\_3219
CL\_5549
CL\_3218
CL\_9006
CL\_9005
CL\_9004
CL\_11456
CL\_9003
CL\_9002
CL\_9001
CL\_10483
CL\_6347
CL\_11457
CL\_10491
CL\_11458
CL\_9130
CL\_9128
CL\_9127
CL\_9126
CL\_9125
CL\_9742
CL\_5613
CL\_5614
CL\_6346
CL\_3217
CL\_6234
CL\_3216
CL\_3215
CL\_3214
CL\_3213
CL\_3212
CL\_3211
CL\_3210
CL\_11113
CL\_9971
CL\_8720
CL\_5551
CL\_3209
CL\_3208
CL\_3207
CL\_3206
CL\_3205
CL\_3204
CL\_3203
CL\_11114
CL\_11115
CL\_11116
CL\_11117
CL\_7813
CL\_7812
CL\_4932
CL\_4931
CL\_3202
CL\_3201
CL\_3200
CL\_3199
CL\_3198
CL\_3197
CL\_3196
CL\_3195
CL\_3194
CL\_3193
CL\_3192
CL\_3191
CL\_3190
CL\_3189
CL\_3188
CL\_3187
CL\_3186
CL\_3185
CL\_3184
CL\_3183
CL\_3182
CL\_7393
CL\_3181
