## Supplementary material for "A novel method for integrating genomic and Tn-Seq data to identify common *in vivo* fitness mechanisms across multiple bacterial species": S3 Dataset: CL_INS_195.html

Legend

 Hypothetical
 Other
 All VFDB Genes

FULL


WINDOWSVGPNG

Trim RowsRemove SingletonsSave Fasta

CL\_3279


CL\_3279


CL\_3279


CL\_3279


CL\_3279


CL\_3279


CL\_3279


CL\_3279


CL\_3279


CL\_3279

HighlightSelectShow Genomes


34

CL\_3280


6

CL\_3280


5

CL\_3280


3

CL\_3280


2

CL\_3280


2

CL\_3280


2

CL\_3280


1

CL\_3280


1

CL\_3280


1

CL\_3280

fGI ID


CL\_INS\_195
CL\_INS\_195
CL\_INS\_195
CL\_INS\_195
CL\_INS\_195
CL\_INS\_195
CL\_INS\_195
CL\_INS\_195
CL\_INS\_195
Cluster ID


CL\_4971
CL\_4972
CL\_6318
CL\_4973
CL\_6319
CL\_6320
CL\_4974
CL\_4975
CL\_4976
