## Supplementary material for "A novel method for integrating genomic and Tn-Seq data to identify common *in vivo* fitness mechanisms across multiple bacterial species": S3 Dataset: CL_INS_196.html

Legend

 Mobile +extrachromosomalelementfunctions
 Regulatoryfunctions
 Hypothetical
 DNA Metabolism
 All EssentialGenes
 Proteinsynthesis/fate
 Other
 EnergyMetabolism
 All VFDB Genes

FULL


WINDOWSVGPNG

Trim RowsRemove SingletonsSave Fasta

CL\_3296


CL\_3296


CL\_3296


CL\_3296


CL\_2407


CL\_3296


CL\_3296


CL\_3296


CL\_3296


CL\_3296


CL\_3296


CL\_3296


CL\_3296


CL\_3296


CL\_3296


CL\_3296


CL\_3296

HighlightSelectShow Genomes


40

CL\_3297


2

CL\_3297


2

CL\_3297


1

CL\_3297


1

CL\_3297


1

CL\_3297


1

CL\_3705


1

CL\_3297


1

CL\_3297


1

CL\_3297


1

CL\_3297


1

CL\_3297


1

CL\_3298


1

CL\_3297


1

CL\_3297


1

CL\_3297


1

CL\_3297

fGI ID


CL\_INS\_196
CL\_INS\_196
CL\_INS\_196
CL\_INS\_196
CL\_INS\_196
CL\_INS\_196
CL\_INS\_196
CL\_INS\_196
CL\_INS\_196
CL\_INS\_196
CL\_INS\_196
CL\_INS\_196
CL\_INS\_133
CL\_INS\_133
CL\_INS\_133
CL\_INS\_133
CL\_INS\_133
CL\_INS\_133
CL\_INS\_133
CL\_INS\_133
CL\_INS\_133
CL\_INS\_217
CL\_INS\_196
CL\_INS\_217
CL\_INS\_196
CL\_INS\_196
CL\_INS\_133
CL\_INS\_133
CL\_INS\_133
CL\_INS\_220
CL\_INS\_220
CL\_INS\_196
CL\_INS\_196
CL\_INS\_196
CL\_INS\_196
CL\_INS\_196
CL\_INS\_133
CL\_INS\_196
CL\_INS\_133
CL\_INS\_133
CL\_INS\_133
CL\_INS\_133
CL\_INS\_133
CL\_INS\_133
CL\_INS\_133
CL\_INS\_196
CL\_INS\_133
CL\_INS\_133
CL\_INS\_133
CL\_INS\_133
CL\_INS\_133
CL\_INS\_133
CL\_INS\_133
CL\_INS\_133
CL\_INS\_133
CL\_INS\_196
CL\_INS\_196
CL\_INS\_196
CL\_INS\_196
CL\_INS\_133
CL\_INS\_133
CL\_INS\_196
CL\_INS\_196
CL\_INS\_196
CL\_INS\_196
CL\_INS\_196
CL\_INS\_133
CL\_INS\_133
CL\_INS\_196
CL\_INS\_133
CL\_INS\_196
CL\_INS\_133
CL\_INS\_133
CL\_INS\_133
CL\_INS\_133
CL\_INS\_133
CL\_INS\_133
CL\_INS\_133
CL\_INS\_133
CL\_INS\_133
CL\_INS\_133
CL\_INS\_133
CL\_INS\_133
CL\_INS\_133
CL\_INS\_133
CL\_INS\_133
CL\_INS\_133
CL\_INS\_133
CL\_INS\_133
CL\_INS\_133
CL\_INS\_133
CL\_INS\_133
CL\_INS\_133
CL\_INS\_133
CL\_INS\_133
CL\_INS\_133
CL\_INS\_133
CL\_INS\_196
CL\_INS\_133
CL\_INS\_133
CL\_INS\_133
CL\_INS\_133
CL\_INS\_133
CL\_INS\_196
CL\_INS\_133
CL\_INS\_133
CL\_INS\_196
CL\_INS\_133
CL\_INS\_196
CL\_INS\_196
CL\_INS\_133
CL\_INS\_133
CL\_INS\_196
CL\_INS\_220
CL\_INS\_217
CL\_INS\_133
CL\_INS\_133
CL\_INS\_133
CL\_INS\_196
CL\_INS\_133
CL\_INS\_133
CL\_INS\_198
CL\_INS\_133
CL\_INS\_133
CL\_INS\_133
CL\_INS\_196
CL\_INS\_196
CL\_INS\_196
CL\_INS\_196
CL\_INS\_196
CL\_INS\_196
CL\_INS\_133
CL\_INS\_196
CL\_INS\_196
CL\_INS\_196
CL\_INS\_196
CL\_INS\_196
CL\_INS\_196
CL\_INS\_196
CL\_INS\_196
CL\_INS\_196
CL\_INS\_133
CL\_INS\_133
CL\_INS\_133
CL\_INS\_133
CL\_INS\_133
CL\_INS\_133
CL\_INS\_133
CL\_INS\_133
CL\_INS\_133
CL\_INS\_133
CL\_INS\_196
CL\_INS\_133
CL\_INS\_196
CL\_INS\_196
CL\_INS\_133
CL\_INS\_133
CL\_INS\_133
CL\_INS\_133
CL\_INS\_133
CL\_INS\_196
CL\_INS\_196
CL\_INS\_196
CL\_INS\_196
CL\_INS\_196
CL\_INS\_133
CL\_INS\_133
CL\_INS\_133
CL\_INS\_133
CL\_INS\_133
CL\_INS\_133
CL\_INS\_133
CL\_INS\_196
CL\_INS\_196
CL\_INS\_196
CL\_INS\_198
CL\_INS\_133
CL\_INS\_133
CL\_INS\_133
CL\_INS\_133
CL\_INS\_196
CL\_INS\_196
CL\_INS\_133
CL\_INS\_133
CL\_INS\_133
CL\_INS\_133
CL\_INS\_133
CL\_INS\_196
CL\_INS\_133
CL\_INS\_133
CL\_INS\_196
CL\_INS\_133
CL\_INS\_196
CL\_INS\_133
CL\_INS\_133
CL\_INS\_133
CL\_INS\_196
CL\_INS\_196
CL\_INS\_196
CL\_INS\_196
CL\_INS\_196
CL\_INS\_196
CL\_INS\_196
CL\_INS\_133
CL\_INS\_133
CL\_INS\_196
CL\_INS\_196
CL\_INS\_196
CL\_INS\_196
CL\_INS\_196
CL\_INS\_196
CL\_INS\_196
CL\_INS\_133
CL\_INS\_133
CL\_INS\_133
CL\_INS\_196
CL\_INS\_196
CL\_INS\_133
CL\_INS\_133
CL\_INS\_133
CL\_INS\_196
CL\_INS\_196
CL\_INS\_196
CL\_INS\_196
CL\_INS\_196
CL\_INS\_196
CL\_INS\_196
CL\_INS\_196
CL\_INS\_196
CL\_INS\_196
CL\_INS\_133
CL\_INS\_196
CL\_INS\_196
CL\_INS\_196
CL\_INS\_196
CL\_INS\_196
CL\_INS\_196
CL\_INS\_196
CL\_INS\_133
CL\_INS\_133
CL\_INS\_196
CL\_INS\_133
CL\_INS\_133
CL\_INS\_133
CL\_INS\_133
CL\_INS\_133
CL\_INS\_133
CL\_INS\_133
CL\_INS\_133
CL\_INS\_133
CL\_INS\_133
CL\_INS\_196
CL\_INS\_196
CL\_INS\_133
CL\_INS\_133
CL\_INS\_133
CL\_INS\_133
CL\_INS\_133
CL\_INS\_133
CL\_INS\_196
CL\_INS\_196
CL\_INS\_196
CL\_INS\_196
CL\_INS\_133
CL\_INS\_196
CL\_INS\_196
CL\_INS\_196
CL\_INS\_196
CL\_INS\_196
CL\_INS\_133
CL\_INS\_196
CL\_INS\_196
CL\_INS\_133
CL\_INS\_133
CL\_INS\_133
CL\_INS\_133
CL\_INS\_133
CL\_INS\_133
CL\_INS\_217
CL\_INS\_133
CL\_INS\_133
CL\_INS\_196
CL\_INS\_196
CL\_INS\_196
CL\_INS\_133
CL\_INS\_133
CL\_INS\_133
CL\_INS\_196
CL\_INS\_196
CL\_INS\_196
CL\_INS\_196
CL\_INS\_196
CL\_INS\_196
CL\_INS\_133
CL\_INS\_196
CL\_INS\_196
CL\_INS\_133
CL\_INS\_133
CL\_INS\_133
CL\_INS\_196
CL\_INS\_196
CL\_INS\_196
CL\_INS\_196
CL\_INS\_217
CL\_INS\_220
CL\_INS\_196
CL\_INS\_220
CL\_INS\_133
CL\_INS\_133
CL\_INS\_133
CL\_INS\_133
CL\_INS\_133
CL\_INS\_133
CL\_INS\_133
CL\_INS\_133
CL\_INS\_133
CL\_INS\_133
CL\_INS\_196
CL\_INS\_196
CL\_INS\_196
CL\_INS\_196
CL\_INS\_196
CL\_INS\_196
CL\_INS\_196
CL\_INS\_220
CL\_INS\_220
CL\_INS\_220
CL\_INS\_220
CL\_INS\_220
CL\_INS\_220
CL\_INS\_220
CL\_INS\_220
CL\_INS\_220
CL\_INS\_220
CL\_INS\_220
CL\_INS\_220
CL\_INS\_220
CL\_INS\_196
CL\_INS\_133
CL\_INS\_196
CL\_INS\_196
CL\_INS\_196
CL\_INS\_133
CL\_INS\_196
CL\_INS\_196
CL\_INS\_196
CL\_INS\_133
CL\_INS\_133
CL\_INS\_133
CL\_INS\_133
CL\_INS\_196
CL\_INS\_133
CL\_INS\_196
Cluster ID


CL\_9371
CL\_9372
CL\_9373
CL\_9374
CL\_9375
CL\_9376
CL\_9377
CL\_9378
CL\_9379
CL\_9380
CL\_9381
CL\_9382
CL\_6346
CL\_5045
CL\_8660
CL\_8661
CL\_8662
CL\_8663
CL\_8664
CL\_8665
CL\_7412
CL\_7390
CL\_11441
CL\_7413
CL\_10508
CL\_10509
CL\_5615
CL\_7388
CL\_8864
CL\_10381
CL\_10382
CL\_10510
CL\_10511
CL\_10512
CL\_10513
CL\_10514
CL\_9327
CL\_10515
CL\_10516
CL\_9326
CL\_9325
CL\_9324
CL\_9323
CL\_6347
CL\_6348
CL\_8655
CL\_6459
CL\_8964
CL\_8303
CL\_6462
CL\_6349
CL\_6350
CL\_5611
CL\_5610
CL\_5609
CL\_8654
CL\_8653
CL\_8652
CL\_8651
CL\_5608
CL\_8666
CL\_10676
CL\_10675
CL\_10674
CL\_10673
CL\_13274
CL\_5607
CL\_5606
CL\_11440
CL\_8745
CL\_10953
CL\_10954
CL\_8233
CL\_8667
CL\_8668
CL\_8669
CL\_8670
CL\_8671
CL\_8672
CL\_8673
CL\_8674
CL\_8675
CL\_8676
CL\_8677
CL\_8678
CL\_8679
CL\_8680
CL\_8681
CL\_5605
CL\_5604
CL\_5603
CL\_5601
CL\_5600
CL\_5599
CL\_5598
CL\_5597
CL\_5596
CL\_12871
CL\_5594
CL\_5593
CL\_5592
CL\_5591
CL\_10342
CL\_11243
CL\_5590
CL\_5589
CL\_12872
CL\_6469
CL\_9982
CL\_12873
CL\_11048
CL\_10213
CL\_13270
CL\_10387
CL\_6376
CL\_7821
CL\_5543
CL\_5542
CL\_10672
CL\_5585
CL\_6377
CL\_8198
CL\_6471
CL\_7745
CL\_7744
CL\_9117
CL\_9116
CL\_9115
CL\_9114
CL\_9113
CL\_9112
CL\_6472
CL\_10958
CL\_10959
CL\_8831
CL\_8830
CL\_8829
CL\_8828
CL\_10960
CL\_10961
CL\_10962
CL\_8692
CL\_8693
CL\_5584
CL\_5583
CL\_9219
CL\_6378
CL\_5582
CL\_8867
CL\_8868
CL\_7742
CL\_8641
CL\_8640
CL\_8639
CL\_8638
CL\_5579
CL\_5578
CL\_7741
CL\_5576
CL\_5575
CL\_8637
CL\_8636
CL\_8635
CL\_8634
CL\_8633
CL\_7881
CL\_5574
CL\_5573
CL\_5572
CL\_5571
CL\_5570
CL\_5577
CL\_12874
CL\_12875
CL\_12876
CL\_8711
CL\_7879
CL\_7880
CL\_8712
CL\_5568
CL\_11439
CL\_11616
CL\_5567
CL\_5566
CL\_11041
CL\_10715
CL\_8246
CL\_11615
CL\_8728
CL\_5561
CL\_11614
CL\_5560
CL\_11613
CL\_7882
CL\_8632
CL\_8631
CL\_8630
CL\_8629
CL\_8628
CL\_12877
CL\_12878
CL\_12879
CL\_12880
CL\_5535
CL\_11943
CL\_12881
CL\_12882
CL\_12883
CL\_12884
CL\_12885
CL\_12886
CL\_12887
CL\_5613
CL\_5612
CL\_8837
CL\_13276
CL\_13275
CL\_8836
CL\_8835
CL\_8834
CL\_11618
CL\_12888
CL\_11620
CL\_12889
CL\_12890
CL\_12891
CL\_12892
CL\_12893
CL\_12894
CL\_12895
CL\_8234
CL\_8650
CL\_8649
CL\_8648
CL\_8647
CL\_8646
CL\_8645
CL\_8644
CL\_7754
CL\_8235
CL\_8833
CL\_7753
CL\_7752
CL\_7751
CL\_9320
CL\_10955
CL\_9318
CL\_7749
CL\_8236
CL\_8237
CL\_7750
CL\_13273
CL\_13272
CL\_8238
CL\_6466
CL\_6467
CL\_6468
CL\_8832
CL\_8643
CL\_8965
CL\_12896
CL\_12897
CL\_12898
CL\_5595
CL\_12899
CL\_12900
CL\_12901
CL\_12902
CL\_12903
CL\_5588
CL\_13115
CL\_8642
CL\_5587
CL\_7820
CL\_8682
CL\_8683
CL\_8684
CL\_8685
CL\_6373
CL\_10210
CL\_10211
CL\_10956
CL\_10957
CL\_11617
CL\_5586
CL\_9222
CL\_8319
CL\_9122
CL\_9121
CL\_9120
CL\_9119
CL\_9118
CL\_10517
CL\_7416
CL\_13271
CL\_8320
CL\_7417
CL\_10386
CL\_9221
CL\_11242
CL\_11241
CL\_11240
CL\_11239
CL\_6379
CL\_10388
CL\_11238
CL\_10389
CL\_6474
CL\_6475
CL\_6476
CL\_6477
CL\_8686
CL\_8687
CL\_8688
CL\_8689
CL\_8690
CL\_8691
CL\_10518
CL\_10519
CL\_9039
CL\_10520
CL\_9041
CL\_9042
CL\_9043
CL\_10390
CL\_10391
CL\_10392
CL\_10393
CL\_6478
CL\_8966
CL\_8967
CL\_6481
CL\_6479
CL\_6480
CL\_6482
CL\_6483
CL\_8968
CL\_10521
CL\_6484
CL\_9111
CL\_9110
CL\_10963
CL\_8697
CL\_9109
CL\_9108
CL\_9107
CL\_6485
CL\_8694
CL\_8695
CL\_8696
CL\_8969
CL\_7361
CL\_8698
