## Supplementary material for "A novel method for integrating genomic and Tn-Seq data to identify common *in vivo* fitness mechanisms across multiple bacterial species": S3 Dataset: CL_INS_199.html

Legend

 Mobile +extrachromosomalelementfunctions
 Regulatoryfunctions
 Hypothetical
 DNA Metabolism
 All EssentialGenes
 Other
 EnergyMetabolism
 Transport +binding proteins
 All VFDB Genes

FULL


WINDOWSVGPNG

Trim RowsRemove SingletonsSave Fasta

CL\_3371


CL\_3371


CL\_3371


CL\_3371


CL\_3371


CL\_3371


CL\_3371


CL\_3371


CL\_3371


CL\_3371


CL\_2575

HighlightSelectShow Genomes


48

CL\_3370


1

CL\_3370


1

CL\_3370


1

CL\_3370


1

CL\_2577


1

CL\_3370


1

CL\_3370


1

CL\_3370


1

CL\_3370


1

CL\_3370


1

CL\_3370

fGI ID


CL\_INS\_217
CL\_INS\_199
CL\_INS\_83
CL\_INS\_194
CL\_INS\_144
CL\_INS\_144
CL\_INS\_83
CL\_INS\_199
CL\_INS\_144
CL\_INS\_199
CL\_INS\_199
CL\_INS\_98
CL\_INS\_199
CL\_INS\_199
CL\_INS\_199
CL\_INS\_98
CL\_INS\_98
CL\_INS\_83
CL\_INS\_83
CL\_INS\_83
CL\_INS\_83
CL\_INS\_144
CL\_INS\_199
CL\_INS\_83
CL\_INS\_199
CL\_INS\_83
CL\_INS\_194
CL\_INS\_98
CL\_INS\_83
CL\_INS\_199
CL\_INS\_199
CL\_INS\_199
CL\_INS\_199
CL\_INS\_199
CL\_INS\_199
CL\_INS\_199
CL\_INS\_199
CL\_INS\_199
CL\_INS\_199
CL\_INS\_199
CL\_INS\_98
CL\_INS\_83
CL\_INS\_194
CL\_INS\_83
CL\_INS\_98
CL\_INS\_83
CL\_INS\_199
CL\_INS\_144
CL\_INS\_199
CL\_INS\_83
CL\_INS\_98
CL\_INS\_144
CL\_INS\_144
CL\_INS\_144
CL\_INS\_144
CL\_INS\_144
CL\_INS\_144
CL\_INS\_144
CL\_INS\_144
CL\_INS\_144
CL\_INS\_144
CL\_INS\_144
CL\_INS\_144
CL\_INS\_199
CL\_INS\_98
CL\_INS\_217
CL\_INS\_217
CL\_INS\_98
CL\_INS\_98
CL\_INS\_98
CL\_INS\_98
CL\_INS\_199
CL\_INS\_217
CL\_INS\_217
CL\_INS\_83
CL\_INS\_83
CL\_INS\_199
CL\_INS\_83
CL\_INS\_144
CL\_INS\_199
CL\_INS\_98
CL\_INS\_199
CL\_INS\_199
CL\_INS\_199
CL\_INS\_199
CL\_INS\_194
CL\_INS\_194
CL\_INS\_98
CL\_INS\_199
CL\_INS\_98
CL\_INS\_98
CL\_INS\_194
CL\_INS\_98
CL\_INS\_199
CL\_INS\_83
CL\_INS\_133
CL\_INS\_194
CL\_INS\_83
CL\_INS\_83
CL\_INS\_144
CL\_INS\_83
CL\_INS\_83
CL\_INS\_83
CL\_INS\_144
CL\_INS\_144
CL\_INS\_133
CL\_INS\_133
CL\_INS\_83
CL\_INS\_217
CL\_INS\_217
CL\_INS\_217
CL\_INS\_98
CL\_INS\_98
CL\_INS\_98
CL\_INS\_98
CL\_INS\_144
CL\_INS\_144
CL\_INS\_83
CL\_INS\_83
CL\_INS\_83
CL\_INS\_217
CL\_INS\_199
CL\_INS\_199
CL\_INS\_199
CL\_INS\_199
CL\_INS\_199
CL\_INS\_199
CL\_INS\_144
CL\_INS\_98
CL\_INS\_83
CL\_INS\_83
CL\_INS\_199
CL\_INS\_194
CL\_INS\_194
CL\_INS\_83
CL\_INS\_199
CL\_INS\_194
CL\_INS\_199
CL\_INS\_98
CL\_INS\_133
CL\_INS\_199
CL\_INS\_199
CL\_INS\_194
CL\_INS\_199
CL\_INS\_199
CL\_INS\_83
CL\_INS\_199
CL\_INS\_199
CL\_INS\_199
CL\_INS\_199
CL\_INS\_199
CL\_INS\_199
CL\_INS\_199
CL\_INS\_194
CL\_INS\_199
CL\_INS\_199
CL\_INS\_199
CL\_INS\_199
CL\_INS\_98
CL\_INS\_133
CL\_INS\_217
CL\_INS\_144
CL\_INS\_199
CL\_INS\_83
CL\_INS\_199
CL\_INS\_98
CL\_INS\_199
CL\_INS\_199
CL\_INS\_98
CL\_INS\_199
CL\_INS\_199
CL\_INS\_83
CL\_INS\_199
CL\_INS\_199
CL\_INS\_98
CL\_INS\_83
CL\_INS\_98
CL\_INS\_83
CL\_INS\_98
CL\_INS\_83
CL\_INS\_144
CL\_INS\_98
CL\_INS\_83
CL\_INS\_83
CL\_INS\_83
CL\_INS\_83
CL\_INS\_83
CL\_INS\_83
CL\_INS\_83
CL\_INS\_144
CL\_INS\_83
CL\_INS\_133
CL\_INS\_133
CL\_INS\_133
CL\_INS\_133
CL\_INS\_199
CL\_INS\_98
CL\_INS\_98
CL\_INS\_98
CL\_INS\_98
CL\_INS\_199
CL\_INS\_98
CL\_INS\_199
CL\_INS\_199
CL\_INS\_199
CL\_INS\_199
CL\_INS\_83
CL\_INS\_144
CL\_INS\_199
CL\_INS\_199
CL\_INS\_199
CL\_INS\_199
CL\_INS\_199
CL\_INS\_199
CL\_INS\_199
CL\_INS\_199
CL\_INS\_199
CL\_INS\_199
CL\_INS\_199
CL\_INS\_199
CL\_INS\_199
CL\_INS\_199
CL\_INS\_199
CL\_INS\_199
CL\_INS\_199
CL\_INS\_199
CL\_INS\_199
CL\_INS\_199
CL\_INS\_199
CL\_INS\_199
CL\_INS\_199
CL\_INS\_199
CL\_INS\_199
CL\_INS\_199
CL\_INS\_199
CL\_INS\_199
CL\_INS\_199
CL\_INS\_98
CL\_INS\_98
CL\_INS\_98
CL\_INS\_98
CL\_INS\_98
CL\_INS\_199
CL\_INS\_144
CL\_INS\_199
CL\_INS\_83
CL\_INS\_83
CL\_INS\_133
CL\_INS\_83
CL\_INS\_83
CL\_INS\_217
CL\_INS\_199
CL\_INS\_83
CL\_INS\_199
CL\_INS\_83
CL\_INS\_98
CL\_INS\_83
CL\_INS\_83
CL\_INS\_98
CL\_INS\_98
CL\_INS\_83
CL\_INS\_199
CL\_INS\_98
CL\_INS\_98
CL\_INS\_194
CL\_INS\_194
CL\_INS\_194
CL\_INS\_199
CL\_INS\_199
CL\_INS\_98
CL\_INS\_98
CL\_INS\_98
CL\_INS\_98
CL\_INS\_98
CL\_INS\_98
CL\_INS\_199
CL\_INS\_199
CL\_INS\_98
CL\_INS\_83
CL\_INS\_83
CL\_INS\_144
CL\_INS\_194
CL\_INS\_144
CL\_INS\_199
CL\_INS\_83
CL\_INS\_199
CL\_INS\_199
CL\_INS\_199
CL\_INS\_83
CL\_INS\_83
CL\_INS\_83
CL\_INS\_83
CL\_INS\_144
CL\_INS\_144
CL\_INS\_144
CL\_INS\_144
CL\_INS\_144
CL\_INS\_144
CL\_INS\_144
CL\_INS\_98
CL\_INS\_144
CL\_INS\_144
CL\_INS\_144
CL\_INS\_144
CL\_INS\_144
CL\_INS\_144
CL\_INS\_144
CL\_INS\_144
CL\_INS\_98
CL\_INS\_98
CL\_INS\_98
CL\_INS\_98
CL\_INS\_98
CL\_INS\_194
CL\_INS\_199
CL\_INS\_83
CL\_INS\_83
CL\_INS\_83
CL\_INS\_83
CL\_INS\_98
CL\_INS\_83
CL\_INS\_83
CL\_INS\_199
CL\_INS\_98
CL\_INS\_83
CL\_INS\_83
CL\_INS\_83
CL\_INS\_83
CL\_INS\_83
CL\_INS\_194
CL\_INS\_199
CL\_INS\_199
CL\_INS\_199
CL\_INS\_83
CL\_INS\_199
CL\_INS\_199
CL\_INS\_199
CL\_INS\_199
CL\_INS\_199
CL\_INS\_199
CL\_INS\_199
CL\_INS\_199
CL\_INS\_199
CL\_INS\_194
CL\_INS\_98
CL\_INS\_98
CL\_INS\_98
CL\_INS\_133
CL\_INS\_98
CL\_INS\_98
CL\_INS\_98
Cluster ID


CL\_9014
CL\_9945
CL\_7354
CL\_3266
CL\_9946
CL\_9947
CL\_3263
CL\_8740
CL\_8739
CL\_13463
CL\_8807
CL\_7352
CL\_8986
CL\_8808
CL\_9885
CL\_6724
CL\_6202
CL\_7353
CL\_3267
CL\_8809
CL\_10020
CL\_8738
CL\_12382
CL\_3265
CL\_8985
CL\_6725
CL\_8984
CL\_9202
CL\_3262
CL\_9948
CL\_9949
CL\_9950
CL\_9951
CL\_9952
CL\_9953
CL\_9954
CL\_9955
CL\_9956
CL\_9957
CL\_9958
CL\_9959
CL\_6727
CL\_8983
CL\_3260
CL\_3261
CL\_3264
CL\_12943
CL\_10019
CL\_8810
CL\_6726
CL\_10130
CL\_10018
CL\_10017
CL\_10016
CL\_10015
CL\_10014
CL\_10013
CL\_10012
CL\_10011
CL\_10010
CL\_10009
CL\_10008
CL\_10007
CL\_12381
CL\_9884
CL\_3259
CL\_8811
CL\_6203
CL\_6204
CL\_3258
CL\_5523
CL\_12380
CL\_8812
CL\_3256
CL\_11403
CL\_3255
CL\_12379
CL\_3254
CL\_12358
CL\_13462
CL\_6735
CL\_12942
CL\_12941
CL\_8813
CL\_8814
CL\_8815
CL\_8816
CL\_8737
CL\_8982
CL\_8981
CL\_8980
CL\_8979
CL\_8736
CL\_8735
CL\_6205
CL\_5528
CL\_8978
CL\_5529
CL\_5530
CL\_10003
CL\_5531
CL\_5532
CL\_6214
CL\_10002
CL\_10001
CL\_8867
CL\_8868
CL\_5526
CL\_9675
CL\_9676
CL\_9677
CL\_5527
CL\_8734
CL\_8733
CL\_10006
CL\_10005
CL\_10004
CL\_3249
CL\_3248
CL\_8817
CL\_3247
CL\_12940
CL\_12939
CL\_12938
CL\_12937
CL\_12936
CL\_12935
CL\_12934
CL\_8732
CL\_8731
CL\_8730
CL\_12378
CL\_3245
CL\_3244
CL\_3243
CL\_13461
CL\_7824
CL\_13460
CL\_5537
CL\_3242
CL\_8977
CL\_8818
CL\_8819
CL\_8820
CL\_8821
CL\_5534
CL\_9960
CL\_9961
CL\_9962
CL\_9963
CL\_9964
CL\_9965
CL\_9966
CL\_9967
CL\_13523
CL\_13524
CL\_13525
CL\_13526
CL\_8729
CL\_8728
CL\_8727
CL\_8976
CL\_8726
CL\_8725
CL\_9883
CL\_6216
CL\_9882
CL\_9881
CL\_5538
CL\_13459
CL\_8822
CL\_3238
CL\_13527
CL\_9880
CL\_5540
CL\_6217
CL\_8975
CL\_6218
CL\_10122
CL\_6219
CL\_8724
CL\_9333
CL\_3234
CL\_8974
CL\_5541
CL\_8823
CL\_3232
CL\_3231
CL\_3230
CL\_10000
CL\_7415
CL\_5542
CL\_5543
CL\_7821
CL\_7820
CL\_13528
CL\_9879
CL\_9878
CL\_9877
CL\_9637
CL\_9876
CL\_11376
CL\_12377
CL\_12376
CL\_12375
CL\_12374
CL\_3237
CL\_10480
CL\_12933
CL\_12932
CL\_12931
CL\_12930
CL\_12929
CL\_12928
CL\_12927
CL\_12926
CL\_12925
CL\_12924
CL\_12923
CL\_12922
CL\_12921
CL\_12920
CL\_12919
CL\_12918
CL\_12917
CL\_12916
CL\_12915
CL\_12914
CL\_12913
CL\_12912
CL\_12911
CL\_12910
CL\_12909
CL\_12908
CL\_12907
CL\_12906
CL\_12905
CL\_11262
CL\_11261
CL\_11260
CL\_11259
CL\_6237
CL\_12904
CL\_10481
CL\_10482
CL\_7384
CL\_6220
CL\_10305
CL\_8723
CL\_3229
CL\_6223
CL\_13458
CL\_8722
CL\_12373
CL\_3228
CL\_3227
CL\_6224
CL\_6225
CL\_6740
CL\_6741
CL\_7414
CL\_9875
CL\_5546
CL\_3225
CL\_9968
CL\_9969
CL\_9970
CL\_8824
CL\_9874
CL\_3223
CL\_3222
CL\_3221
CL\_3220
CL\_3219
CL\_3218
CL\_12372
CL\_12371
CL\_6742
CL\_6226
CL\_6227
CL\_9999
CL\_13446
CL\_8721
CL\_12370
CL\_6229
CL\_8973
CL\_8972
CL\_8971
CL\_6230
CL\_6231
CL\_6232
CL\_6233
CL\_9998
CL\_9997
CL\_9996
CL\_9995
CL\_9994
CL\_9993
CL\_9992
CL\_3207
CL\_9991
CL\_9990
CL\_9989
CL\_9988
CL\_9987
CL\_9986
CL\_9985
CL\_9984
CL\_9211
CL\_9212
CL\_9213
CL\_3217
CL\_3216
CL\_3215
CL\_9873
CL\_3214
CL\_9872
CL\_3213
CL\_3212
CL\_6234
CL\_3211
CL\_3210
CL\_9871
CL\_9971
CL\_8720
CL\_3206
CL\_3205
CL\_3204
CL\_3203
CL\_7813
CL\_13457
CL\_13456
CL\_8970
CL\_6241
CL\_9972
CL\_9973
CL\_8825
CL\_8826
CL\_8827
CL\_8719
CL\_13529
CL\_8718
CL\_9870
CL\_4932
CL\_4931
CL\_3199
CL\_3198
CL\_4930
CL\_3197
CL\_3182
CL\_3180
