## Supplementary material for "A novel method for integrating genomic and Tn-Seq data to identify common *in vivo* fitness mechanisms across multiple bacterial species": S3 Dataset: CL_INS_200.html

Legend

 Regulatoryfunctions
 Hypothetical
 Other
 All VFDB Genes

FULL


WINDOWSVGPNG

Trim RowsRemove SingletonsSave Fasta

CL\_3415


CL\_3415


CL\_3415


CL\_3415


CL\_3415


CL\_3415


CL\_3415


CL\_3415


CL\_3415

HighlightSelectShow Genomes


25

CL\_3423


14

CL\_3423


11

CL\_3423


2

CL\_3423


1

CL\_3423


1

CL\_3423


1

CL\_3423


1

CL\_3423


1

CL\_3423

fGI ID


CL\_INS\_200
CL\_INS\_200
CL\_INS\_200
CL\_INS\_200
CL\_INS\_200
CL\_INS\_200
CL\_INS\_200
CL\_INS\_200
CL\_INS\_200
Cluster ID


CL\_5637
CL\_5638
CL\_3416
CL\_3417
CL\_3418
CL\_3419
CL\_3420
CL\_3421
CL\_3422
