## Supplementary material for "A novel method for integrating genomic and Tn-Seq data to identify common *in vivo* fitness mechanisms across multiple bacterial species": S3 Dataset: CL_INS_201.html

Legend

 AntibioticResistance
 Other

FULL


WINDOWSVGPNG

Trim RowsRemove SingletonsSave Fasta

CL\_3433


CL\_3433


CL\_3433

HighlightSelectShow Genomes


41

CL\_3434


5

CL\_3434


2

CL\_3434

fGI ID

CL\_INS\_201
Cluster ID

CL\_5006
