## Supplementary material for "A novel method for integrating genomic and Tn-Seq data to identify common *in vivo* fitness mechanisms across multiple bacterial species": S3 Dataset: CL_INS_202.html

Legend

 Mobile +extrachromosomalelementfunctions
 Hypothetical
 DNA Metabolism
 Other
 All VFDB Genes

FULL


WINDOWSVGPNG

Trim RowsRemove SingletonsSave Fasta

CL\_3439


CL\_3439

HighlightSelectShow Genomes


52

CL\_3438


2

CL\_3438

fGI ID


CL\_INS\_202
CL\_INS\_202
CL\_INS\_194
CL\_INS\_98
CL\_INS\_83
CL\_INS\_83
CL\_INS\_83
CL\_INS\_83
CL\_INS\_98
CL\_INS\_98
CL\_INS\_98
CL\_INS\_98
CL\_INS\_202
CL\_INS\_202
CL\_INS\_98
CL\_INS\_98
CL\_INS\_98
CL\_INS\_98
Cluster ID


CL\_13632
CL\_13633
CL\_13014
CL\_7831
CL\_3265
CL\_3264
CL\_6726
CL\_3262
CL\_3261
CL\_9884
CL\_3258
CL\_5523
CL\_13634
CL\_13635
CL\_3199
CL\_3198
CL\_3197
CL\_3180
