## Supplementary material for "A novel method for integrating genomic and Tn-Seq data to identify common *in vivo* fitness mechanisms across multiple bacterial species": S3 Dataset: CL_INS_204.html

Legend

 Mobile +extrachromosomalelementfunctions
 Hypothetical
 Cell Envelope
 Other
 All VFDB Genes

FULL


WINDOWSVGPNG

Trim RowsRemove SingletonsSave Fasta

CL\_3467


CL\_3467


CL\_3458


CL\_3458


CL\_3458


CL\_3458


CL\_3458


CL\_3458


CL\_3467


CL\_3458


CL\_3458


CL\_3467


CL\_3458


CL\_3458


CL\_3467


CL\_3467


CL\_3458


CL\_3458


CL\_3458

HighlightSelectShow Genomes


12

CL\_3468


12

CL\_3468


7

CL\_3468


4

CL\_3468


3

CL\_3468


2

CL\_3468


2

CL\_3468


2

CL\_3468


2

CL\_3468


2

CL\_3468


1

CL\_3468


1

CL\_3468


1

CL\_3468


1

CL\_3468


1

CL\_3468


1

CL\_3468


1

CL\_3468


1

CL\_3468


1

CL\_3468

fGI ID


CL\_INS\_204
CL\_INS\_204
CL\_INS\_204
CL\_INS\_204
CL\_INS\_204
CL\_INS\_204
CL\_INS\_204
CL\_INS\_204
CL\_INS\_204
CL\_INS\_204
CL\_INS\_204
CL\_INS\_204
CL\_INS\_204
CL\_INS\_204
CL\_INS\_204
CL\_INS\_204
CL\_INS\_204
CL\_INS\_204
CL\_INS\_204
CL\_INS\_204
CL\_INS\_204
CL\_INS\_204
CL\_INS\_204
CL\_INS\_204
CL\_INS\_204
CL\_INS\_204
CL\_INS\_204
CL\_INS\_204
CL\_INS\_204
CL\_INS\_204
CL\_INS\_204
CL\_INS\_204
CL\_INS\_204
CL\_INS\_204
CL\_INS\_204
CL\_INS\_204
CL\_INS\_204
CL\_INS\_204
CL\_INS\_204
CL\_INS\_204
CL\_INS\_204
CL\_INS\_204
CL\_INS\_204
CL\_INS\_204
CL\_INS\_204
CL\_INS\_204
CL\_INS\_204
CL\_INS\_204
CL\_INS\_204
CL\_INS\_204
CL\_INS\_204
CL\_INS\_204
CL\_INS\_204
CL\_INS\_204
CL\_INS\_204
CL\_INS\_204
CL\_INS\_204
CL\_INS\_204
CL\_INS\_204
CL\_INS\_204
CL\_INS\_204
CL\_INS\_204
CL\_INS\_204
CL\_INS\_204
CL\_INS\_204
CL\_INS\_204
CL\_INS\_204
CL\_INS\_204
CL\_INS\_204
CL\_INS\_204
CL\_INS\_204
CL\_INS\_204
CL\_INS\_204
CL\_INS\_204
CL\_INS\_204
CL\_INS\_204
CL\_INS\_204
CL\_INS\_204
CL\_INS\_204
CL\_INS\_204
CL\_INS\_204
CL\_INS\_204
CL\_INS\_204
CL\_INS\_204
CL\_INS\_204
CL\_INS\_204
CL\_INS\_204
CL\_INS\_204
CL\_INS\_204
CL\_INS\_204
CL\_INS\_204
CL\_INS\_204
CL\_INS\_204
CL\_INS\_204
Cluster ID


CL\_13265
CL\_13264
CL\_13263
CL\_13262
CL\_9470
CL\_9471
CL\_9472
CL\_9473
CL\_9474
CL\_7834
CL\_10021
CL\_7835
CL\_7836
CL\_7837
CL\_7838
CL\_6912
CL\_6838
CL\_11437
CL\_11436
CL\_11435
CL\_11434
CL\_11433
CL\_6839
CL\_11107
CL\_11106
CL\_11105
CL\_11104
CL\_11103
CL\_6840
CL\_6841
CL\_6842
CL\_6843
CL\_6844
CL\_6845
CL\_9725
CL\_9724
CL\_9723
CL\_9722
CL\_9721
CL\_9720
CL\_9719
CL\_8180
CL\_8179
CL\_8178
CL\_8177
CL\_8176
CL\_8175
CL\_8174
CL\_13489
CL\_13488
CL\_13487
CL\_13486
CL\_13485
CL\_13484
CL\_13483
CL\_13482
CL\_13481
CL\_13480
CL\_13479
CL\_12383
CL\_12463
CL\_12462
CL\_12461
CL\_12460
CL\_12459
CL\_12458
CL\_12457
CL\_12456
CL\_12455
CL\_12454
CL\_12453
CL\_10364
CL\_10365
CL\_10366
CL\_10367
CL\_10368
CL\_10369
CL\_10370
CL\_10371
CL\_10372
CL\_10373
CL\_10374
CL\_5007
CL\_5008
CL\_5009
CL\_5010
CL\_5011
CL\_5012
CL\_5013
CL\_11200
CL\_11199
CL\_5014
CL\_5015
CL\_5016
