## Supplementary material for "A novel method for integrating genomic and Tn-Seq data to identify common *in vivo* fitness mechanisms across multiple bacterial species": S3 Dataset: CL_INS_205.html

Legend

 Hypothetical
 Other
 All VFDB Genes

FULL


WINDOWSVGPNG

Trim RowsRemove SingletonsSave Fasta

CL\_3501


CL\_3501


CL\_3501

HighlightSelectShow Genomes


40

CL\_3503


16

CL\_3503


1

CL\_3504

fGI ID


CL\_INS\_205
CL\_INS\_205
Cluster ID


CL\_5017
CL\_3502
