## Supplementary material for "A novel method for integrating genomic and Tn-Seq data to identify common *in vivo* fitness mechanisms across multiple bacterial species": S3 Dataset: CL_INS_207.html

Legend

 Hypothetical
 Other
 All VFDB Genes

FULL


WINDOWSVGPNG

Trim RowsRemove SingletonsSave Fasta

CL\_3504


CL\_3504


CL\_3504


CL\_3504


CL\_3504


CL\_3504


CL\_3504


CL\_3504

HighlightSelectShow Genomes


28

CL\_3519


19

CL\_3519


5

CL\_3519


1

CL\_3519


1

CL\_3519


1

CL\_3519


1

CL\_3519


1

CL\_3519

fGI ID


CL\_INS\_207
CL\_INS\_207
CL\_INS\_207
CL\_INS\_207
CL\_INS\_207
CL\_INS\_207
CL\_INS\_207
CL\_INS\_207
CL\_INS\_207
CL\_INS\_207
CL\_INS\_207
CL\_INS\_207
CL\_INS\_207
CL\_INS\_207
CL\_INS\_207
Cluster ID


CL\_13261
CL\_3505
CL\_3506
CL\_3507
CL\_3508
CL\_3509
CL\_3510
CL\_3511
CL\_3512
CL\_3513
CL\_3514
CL\_3515
CL\_3516
CL\_3517
CL\_3518
