## Supplementary material for "A novel method for integrating genomic and Tn-Seq data to identify common *in vivo* fitness mechanisms across multiple bacterial species": S3 Dataset: CL_INS_208.html

Legend

 Mobile +extrachromosomalelementfunctions
 Hypothetical
 Biosynthesis ofcofactors,prostheticgroups, +carriers
 All VFDB Genes
 Transport +binding proteins

FULL


WINDOWSVGPNG

Trim RowsRemove SingletonsSave Fasta

CL\_3525


CL\_3525


CL\_3525


CL\_3525


CL\_3525


CL\_3525


CL\_3525


CL\_3525


CL\_3497


CL\_3525


CL\_3525

HighlightSelectShow Genomes


35

CL\_3526


5

CL\_3526


2

CL\_3526


1

CL\_3526


1

CL\_3526


1

CL\_3528


1

CL\_3526


1

CL\_3526


1

CL\_3526


1

CL\_3526


1

CL\_3526

fGI ID


CL\_INS\_208
CL\_INS\_208
CL\_INS\_208
CL\_INS\_208
CL\_INS\_208
CL\_INS\_208
CL\_INS\_208
CL\_INS\_208
CL\_INS\_208
CL\_INS\_208
CL\_INS\_208
CL\_INS\_208
CL\_INS\_209
CL\_INS\_209
CL\_INS\_208
CL\_INS\_209
CL\_INS\_209
CL\_INS\_209
CL\_INS\_209
CL\_INS\_209
CL\_INS\_208
CL\_INS\_209
CL\_INS\_209
CL\_INS\_209
CL\_INS\_208
CL\_INS\_181
CL\_INS\_83
CL\_INS\_83
CL\_INS\_65
CL\_INS\_83
CL\_INS\_83
CL\_INS\_208
CL\_INS\_83
CL\_INS\_83
CL\_INS\_208
CL\_INS\_208
CL\_INS\_230
CL\_INS\_231
CL\_INS\_83
CL\_INS\_208
CL\_INS\_83
CL\_INS\_83
CL\_INS\_83
CL\_INS\_208
CL\_INS\_208
CL\_INS\_208
CL\_INS\_83
CL\_INS\_83
CL\_INS\_208
CL\_INS\_208
CL\_INS\_230
CL\_INS\_230
CL\_INS\_230
CL\_INS\_230
CL\_INS\_230
CL\_INS\_230
CL\_INS\_83
CL\_INS\_83
CL\_INS\_65
CL\_INS\_231
CL\_INS\_208
CL\_INS\_208
CL\_INS\_88
CL\_INS\_208
CL\_INS\_64
CL\_INS\_209
CL\_INS\_208
CL\_INS\_208
CL\_INS\_208
CL\_INS\_208
CL\_INS\_208
CL\_INS\_208
CL\_INS\_208
CL\_INS\_208
CL\_INS\_208
CL\_INS\_208
CL\_INS\_208
CL\_INS\_208
CL\_INS\_208
CL\_INS\_208
CL\_INS\_208
CL\_INS\_208
CL\_INS\_208
CL\_INS\_208
CL\_INS\_208
Cluster ID


CL\_11703
CL\_11702
CL\_12212
CL\_12211
CL\_12210
CL\_12209
CL\_12208
CL\_12207
CL\_12206
CL\_12205
CL\_9384
CL\_6846
CL\_6847
CL\_6848
CL\_9385
CL\_6849
CL\_6850
CL\_6851
CL\_6852
CL\_6853
CL\_9386
CL\_6854
CL\_6855
CL\_6856
CL\_5647
CL\_5648
CL\_5649
CL\_5650
CL\_5651
CL\_5652
CL\_5653
CL\_5654
CL\_5655
CL\_5656
CL\_5657
CL\_5658
CL\_5093
CL\_5094
CL\_5496
CL\_13116
CL\_8252
CL\_8253
CL\_8254
CL\_13117
CL\_13118
CL\_13119
CL\_8257
CL\_8258
CL\_13120
CL\_13121
CL\_5083
CL\_5084
CL\_5085
CL\_5086
CL\_5087
CL\_5088
CL\_5495
CL\_5494
CL\_10421
CL\_8788
CL\_12191
CL\_12452
CL\_6196
CL\_12451
CL\_7296
CL\_5659
CL\_5019
CL\_5020
CL\_5021
CL\_5256
CL\_5022
CL\_5023
CL\_5024
CL\_5025
CL\_5026
CL\_5027
CL\_5028
CL\_5029
CL\_5030
CL\_5031
CL\_5032
CL\_5033
CL\_5034
CL\_5035
CL\_5036
