## Supplementary material for "A novel method for integrating genomic and Tn-Seq data to identify common *in vivo* fitness mechanisms across multiple bacterial species": S3 Dataset: CL_INS_209.html

Legend

 Mobile +extrachromosomalelementfunctions
 Hypothetical
 Biosynthesis ofcofactors,prostheticgroups, +carriers
 All VFDB Genes

FULL


WINDOWSVGPNG

Trim RowsRemove SingletonsSave Fasta

CL\_3526


CL\_3526


CL\_3526


CL\_3526


CL\_3525

HighlightSelectShow Genomes


38

CL\_3528


8

CL\_3528


2

CL\_3528


1

CL\_3529


1

CL\_3528

fGI ID


CL\_INS\_209
CL\_INS\_209
CL\_INS\_209
CL\_INS\_209
CL\_INS\_209
CL\_INS\_209
CL\_INS\_209
CL\_INS\_209
CL\_INS\_209
CL\_INS\_209
CL\_INS\_209
CL\_INS\_209
CL\_INS\_209
Cluster ID


CL\_5037
CL\_3527
CL\_6847
CL\_6848
CL\_6849
CL\_6850
CL\_6851
CL\_6852
CL\_6853
CL\_6854
CL\_6855
CL\_6856
CL\_5659
