## Supplementary material for "A novel method for integrating genomic and Tn-Seq data to identify common *in vivo* fitness mechanisms across multiple bacterial species": S3 Dataset: CL_INS_211.html

Legend

 Hypothetical
 All EssentialGenes
 Other

FULL


WINDOWSVGPNG

Trim RowsRemove SingletonsSave Fasta

CL\_3559


CL\_3559

HighlightSelectShow Genomes


37

CL\_3560


20

CL\_3560

fGI ID


CL\_INS\_211
CL\_INS\_211
Cluster ID


CL\_6335
CL\_6336
