## Supplementary material for "A novel method for integrating genomic and Tn-Seq data to identify common *in vivo* fitness mechanisms across multiple bacterial species": S3 Dataset: CL_INS_212.html

Legend

 Hypothetical
 All VFDB Genes

FULL


WINDOWSVGPNG

Trim RowsRemove SingletonsSave Fasta

CL\_3729


CL\_3729


CL\_3728


CL\_3729

HighlightSelectShow Genomes


37

CL\_3730


18

CL\_3730


1

CL\_3730


1

CL\_3730

fGI ID


CL\_INS\_212
CL\_INS\_212
Cluster ID


CL\_6340
CL\_6341
