## Supplementary material for "A novel method for integrating genomic and Tn-Seq data to identify common *in vivo* fitness mechanisms across multiple bacterial species": S3 Dataset: CL_INS_213.html

Legend

 Hypothetical
 Other
 All VFDB Genes

FULL


WINDOWSVGPNG

Trim RowsRemove SingletonsSave Fasta

CL\_3747


CL\_3747


CL\_3747


CL\_3746


CL\_3747

HighlightSelectShow Genomes


32

CL\_3755


15

CL\_3755


8

CL\_3755


1

CL\_3755


1

CL\_3756

fGI ID


CL\_INS\_213
CL\_INS\_213
CL\_INS\_213
CL\_INS\_213
CL\_INS\_213
CL\_INS\_213
CL\_INS\_213
Cluster ID


CL\_3748
CL\_3749
CL\_3750
CL\_3751
CL\_3752
CL\_3753
CL\_3754
