## Supplementary material for "A novel method for integrating genomic and Tn-Seq data to identify common *in vivo* fitness mechanisms across multiple bacterial species": S3 Dataset: CL_INS_214.html

Legend

 Hypothetical
 All EssentialGenes
 Other
 All VFDB Genes

FULL


WINDOWSVGPNG

Trim RowsRemove SingletonsSave Fasta

CL\_3788


CL\_3788


CL\_3788


CL\_3788


CL\_3788

HighlightSelectShow Genomes


33

CL\_3789


17

CL\_3789


3

CL\_3798


1

CL\_3798


1

CL\_3789

fGI ID


CL\_INS\_214
CL\_INS\_214
CL\_INS\_214
CL\_INS\_214
CL\_INS\_215
CL\_INS\_215
CL\_INS\_215
CL\_INS\_215
CL\_INS\_215
CL\_INS\_215
CL\_INS\_215
CL\_INS\_215
CL\_INS\_215
Cluster ID


CL\_6857
CL\_6343
CL\_6344
CL\_9895
CL\_9896
CL\_9897
CL\_9898
CL\_9899
CL\_9900
CL\_9901
CL\_9902
CL\_9903
CL\_9904
