## Supplementary material for "A novel method for integrating genomic and Tn-Seq data to identify common *in vivo* fitness mechanisms across multiple bacterial species": S3 Dataset: CL_INS_217.html

Legend

 Mobile +extrachromosomalelementfunctions
 Regulatoryfunctions
 Hypothetical
 DNA Metabolism
 All EssentialGenes
 Other
 EnergyMetabolism
 All VFDB Genes

FULL


WINDOWSVGPNG

Trim RowsRemove SingletonsSave Fasta

CL\_3860


CL\_3860


CL\_3860


CL\_3860


CL\_3860


CL\_3860


CL\_3860


CL\_3860


CL\_3860


CL\_3860


CL\_3860


CL\_3860


CL\_3860


CL\_3860


CL\_3860


CL\_2577

HighlightSelectShow Genomes


38

CL\_3861


4

CL\_3861


2

CL\_3861


1

CL\_3861


1

CL\_3861


1

CL\_3861


1

CL\_3861


1

CL\_3861


CL\_7412
CL\_7413
CL\_5042
CL\_5043
CL\_10535
CL\_8791
CL\_5044
CL\_5045
CL\_5046
CL\_5047
CL\_11701
CL\_7363
CL\_9387
CL\_9388
CL\_10191
CL\_8790
CL\_5613
CL\_7362
CL\_11700
CL\_11699
CL\_11698
CL\_11697
CL\_5612
CL\_8837
CL\_10536
CL\_10537
CL\_6462
CL\_5611
CL\_9863
CL\_9862
CL\_5610
CL\_6463
CL\_7780
CL\_7781
CL\_7782
CL\_7783
CL\_7784
CL\_7785
CL\_7786
CL\_9860
CL\_7787
CL\_7788
CL\_7789
CL\_7790
CL\_7791
CL\_7792
CL\_7793
CL\_6371
CL\_6372
CL\_12508
CL\_11047
CL\_5586
CL\_6373
CL\_6374
CL\_6375
CL\_6223
CL\_9389
CL\_6222
CL\_11696
CL\_6376
CL\_6377
CL\_13357
CL\_6378
CL\_6379
CL\_7419
CL\_9390
CL\_9391
CL\_9392
CL\_11695
CL\_6380
CL\_7420
CL\_6218
CL\_12509
CL\_12510
CL\_12511
CL\_7421
CL\_7422
CL\_6381
CL\_6382
CL\_6383
CL\_6384
CL\_6385
CL\_6386
CL\_6387
CL\_6388
CL\_6389
CL\_7425
CL\_12512
CL\_9309
CL\_5562
CL\_8170
CL\_6390
CL\_6391
CL\_7423
CL\_11694
CL\_7424
CL\_8172
CL\_8171
CL\_9393
CL\_13356
CL\_13355
CL\_13354
CL\_13353
CL\_13352
CL\_13351
CL\_7801
CL\_7802
CL\_10695
CL\_11693
CL\_11692
CL\_11691
CL\_6392
CL\_6393
CL\_6394
CL\_6395
CL\_5532
CL\_7426
CL\_7427
CL\_5531
CL\_6396
CL\_6397
CL\_6398
CL\_13350
CL\_13349
CL\_13348
CL\_6399
CL\_6400
CL\_7428
CL\_7429
CL\_7430
CL\_13522
CL\_7431
CL\_6401
CL\_6402
CL\_6403
CL\_6404
CL\_6405
CL\_13347
CL\_7432
CL\_6406
CL\_5615
CL\_5614
CL\_6456
CL\_9741
CL\_6458
CL\_9742
CL\_9743
CL\_9744
CL\_9745
CL\_9746
CL\_9747
CL\_9748
CL\_9749
CL\_9750
CL\_9751
CL\_9752
CL\_9753
CL\_9754
CL\_9755
CL\_9756
CL\_9757
CL\_9758
CL\_9759
CL\_9760
CL\_9761
CL\_9762
CL\_9763
CL\_9764
CL\_9765
CL\_9766
CL\_9767
CL\_9768
CL\_9769
CL\_9770
CL\_9771
CL\_9772
CL\_9773
CL\_9774
CL\_9775
CL\_9776
CL\_9777
CL\_9778
CL\_9779
CL\_9780
CL\_9781
CL\_9782
CL\_8727
CL\_9783
CL\_6214
CL\_6213
CL\_6212
CL\_9784
CL\_3247
CL\_5529
CL\_5528
CL\_9239
CL\_9785
CL\_6205
CL\_9786
CL\_3254
CL\_3256
CL\_8812
CL\_8811
CL\_3259
CL\_3260
CL\_6727
CL\_9787
CL\_9788
CL\_9789
CL\_9790
CL\_5520
CL\_6407
CL\_7433
CL\_6408
CL\_6345
CL\_8173
CL\_6346
CL\_7390
CL\_12507
CL\_9717
CL\_6347
CL\_6348
CL\_6349
CL\_6350
CL\_6351
CL\_6352
CL\_6353
CL\_6354
CL\_9134
CL\_9135
CL\_9716
CL\_6355
CL\_6356
CL\_9715
CL\_6357
CL\_6358
CL\_6359
CL\_6360
CL\_6361
CL\_6362
CL\_6363
CL\_6364
CL\_6365
CL\_6366
CL\_6367
CL\_6368
CL\_6369
CL\_6370
CL\_7414
CL\_6224
CL\_7415
CL\_7416
CL\_7417
CL\_7418
CL\_5585
CL\_9714
CL\_9713
CL\_9712
CL\_9711
CL\_9710
CL\_9709
CL\_9708
CL\_9707
CL\_9706
CL\_9705
CL\_9704
CL\_9703
CL\_9702
CL\_9701
CL\_9700
CL\_9699
CL\_9698
CL\_9697
CL\_9696
CL\_9695
CL\_9694
CL\_9693
CL\_9692
CL\_9691
CL\_9690
CL\_9689
CL\_9688
CL\_9687
CL\_9686
CL\_9685
CL\_9684
CL\_9683
CL\_9682
CL\_9681
CL\_9680
CL\_9679
CL\_9678
CL\_3249
CL\_9677
CL\_9676
CL\_9675
CL\_5526
CL\_9674
CL\_9673
CL\_9672
CL\_9671
CL\_9670
CL\_9669
CL\_9668
CL\_9667
CL\_9666
CL\_9665
CL\_9664
CL\_9663
CL\_3265
CL\_9662
CL\_7354
CL\_9661
CL\_9014
