## Supplementary material for "A novel method for integrating genomic and Tn-Seq data to identify common *in vivo* fitness mechanisms across multiple bacterial species": S3 Dataset: CL_INS_218.html

Legend

 Mobile +extrachromosomalelementfunctions
 Hypothetical
 Other
 All VFDB Genes

FULL


WINDOWSVGPNG

Trim RowsRemove SingletonsSave Fasta

CL\_3885


CL\_3885


CL\_3885


CL\_3885


CL\_3885


CL\_3885


CL\_3885


CL\_3885


CL\_3885


CL\_3885

HighlightSelectShow Genomes


44

CL\_3884


4

CL\_3884


1

CL\_3884


1

CL\_3884


1

CL\_3883


1

CL\_3884


1

CL\_3884


1

CL\_3884


1

CL\_3884


1

CL\_3884

fGI ID


CL\_INS\_218
CL\_INS\_218
CL\_INS\_218
CL\_INS\_218
CL\_INS\_218
CL\_INS\_218
CL\_INS\_218
CL\_INS\_218
CL\_INS\_182
CL\_INS\_182
CL\_INS\_218
CL\_INS\_218
CL\_INS\_218
CL\_INS\_218
CL\_INS\_218
CL\_INS\_218
CL\_INS\_218
CL\_INS\_218
CL\_INS\_182
CL\_INS\_182
CL\_INS\_218
CL\_INS\_218
CL\_INS\_219
CL\_INS\_219
CL\_INS\_219
CL\_INS\_219
CL\_INS\_219
CL\_INS\_219
CL\_INS\_219
CL\_INS\_219
CL\_INS\_219
CL\_INS\_219
CL\_INS\_219
CL\_INS\_218
CL\_INS\_219
CL\_INS\_219
CL\_INS\_218
CL\_INS\_219
CL\_INS\_219
CL\_INS\_219
CL\_INS\_218
CL\_INS\_218
CL\_INS\_218
CL\_INS\_219
CL\_INS\_219
CL\_INS\_219
CL\_INS\_219
CL\_INS\_219
CL\_INS\_219
CL\_INS\_218
CL\_INS\_219
CL\_INS\_219
CL\_INS\_219
CL\_INS\_219
CL\_INS\_219
CL\_INS\_219
CL\_INS\_219
CL\_INS\_220
CL\_INS\_219
CL\_INS\_218
CL\_INS\_218
CL\_INS\_218
Cluster ID


CL\_5050
CL\_5049
CL\_6451
CL\_6450
CL\_6449
CL\_6448
CL\_6447
CL\_6446
CL\_6445
CL\_6304
CL\_13425
CL\_13077
CL\_13076
CL\_7842
CL\_6444
CL\_6443
CL\_7841
CL\_7840
CL\_6442
CL\_6441
CL\_6440
CL\_13685
CL\_6439
CL\_6438
CL\_6437
CL\_6436
CL\_6435
CL\_6434
CL\_6433
CL\_6432
CL\_6431
CL\_6430
CL\_6429
CL\_6428
CL\_6427
CL\_6426
CL\_13075
CL\_6578
CL\_6577
CL\_6576
CL\_6425
CL\_6424
CL\_6423
CL\_6422
CL\_6421
CL\_6420
CL\_6419
CL\_6418
CL\_6417
CL\_13684
CL\_6416
CL\_6415
CL\_6414
CL\_6413
CL\_6412
CL\_6411
CL\_6410
CL\_6566
CL\_5048
CL\_12685
CL\_12684
CL\_6409
