## Supplementary material for "A novel method for integrating genomic and Tn-Seq data to identify common *in vivo* fitness mechanisms across multiple bacterial species": S3 Dataset: CL_INS_219.html

Legend

 Mobile +extrachromosomalelementfunctions
 Regulatoryfunctions
 Hypothetical
 All EssentialGenes
 Other

FULL


WINDOWSVGPNG

Trim RowsRemove SingletonsSave Fasta

CL\_3892


CL\_3892


CL\_3892


CL\_3892


CL\_3892

HighlightSelectShow Genomes


50

CL\_3893


2

CL\_3893


2

CL\_3893


2

CL\_3893


1

CL\_3893

fGI ID


CL\_INS\_219
CL\_INS\_219
CL\_INS\_219
CL\_INS\_219
CL\_INS\_219
CL\_INS\_219
CL\_INS\_219
CL\_INS\_219
CL\_INS\_219
CL\_INS\_219
CL\_INS\_219
CL\_INS\_219
CL\_INS\_219
CL\_INS\_219
CL\_INS\_219
CL\_INS\_219
CL\_INS\_219
CL\_INS\_219
CL\_INS\_219
CL\_INS\_219
CL\_INS\_219
CL\_INS\_219
CL\_INS\_219
CL\_INS\_219
CL\_INS\_219
CL\_INS\_219
CL\_INS\_219
CL\_INS\_219
CL\_INS\_219
CL\_INS\_219
CL\_INS\_219
CL\_INS\_219
CL\_INS\_219
CL\_INS\_219
CL\_INS\_219
CL\_INS\_219
CL\_INS\_219
CL\_INS\_219
CL\_INS\_219
CL\_INS\_219
CL\_INS\_219
CL\_INS\_219
CL\_INS\_219
CL\_INS\_219
CL\_INS\_219
CL\_INS\_219
CL\_INS\_219
CL\_INS\_219
CL\_INS\_219
CL\_INS\_219
CL\_INS\_219
CL\_INS\_219
CL\_INS\_219
CL\_INS\_219
CL\_INS\_219
CL\_INS\_219
CL\_INS\_219
CL\_INS\_219
CL\_INS\_219
CL\_INS\_219
CL\_INS\_219
CL\_INS\_219
CL\_INS\_219
CL\_INS\_219
Cluster ID


CL\_11236
CL\_11235
CL\_9259
CL\_9258
CL\_9477
CL\_9256
CL\_6600
CL\_10671
CL\_10670
CL\_10669
CL\_10668
CL\_10667
CL\_8127
CL\_8128
CL\_8129
CL\_9478
CL\_8130
CL\_9479
CL\_10666
CL\_8131
CL\_9480
CL\_6439
CL\_6590
CL\_10665
CL\_10664
CL\_10663
CL\_10662
CL\_10661
CL\_10660
CL\_10659
CL\_10658
CL\_10657
CL\_10656
CL\_10655
CL\_6438
CL\_6437
CL\_6436
CL\_6435
CL\_6434
CL\_6433
CL\_6432
CL\_6431
CL\_6430
CL\_6429
CL\_6427
CL\_6426
CL\_6422
CL\_13122
CL\_6421
CL\_6420
CL\_6578
CL\_6577
CL\_6576
CL\_6419
CL\_6418
CL\_6417
CL\_6416
CL\_6415
CL\_6414
CL\_6413
CL\_6412
CL\_6411
CL\_6410
CL\_5048
