## Supplementary material for "A novel method for integrating genomic and Tn-Seq data to identify common *in vivo* fitness mechanisms across multiple bacterial species": S3 Dataset: CL_INS_220.html

Legend

 Mobile +extrachromosomalelementfunctions
 Regulatoryfunctions
 Hypothetical
 DNA Metabolism
 All EssentialGenes
 Proteinsynthesis/fate
 Other
 EnergyMetabolism
 Transport +binding proteins
 All VFDB Genes

FULL


WINDOWSVGPNG

Trim RowsRemove SingletonsSave Fasta

CL\_3907


CL\_3907


CL\_3907


CL\_3907


CL\_3907


CL\_3907


CL\_3907


CL\_3907


CL\_3907


CL\_3907


CL\_3907


CL\_3907


CL\_3907


CL\_3907


CL\_3907


CL\_3907


CL\_3907


CL\_3907


CL\_3907


CL\_3907


CL\_3907


CL\_3907


CL\_3907


CL\_3907


CL\_3907


CL\_3907


CL\_3907


CL\_3907


CL\_3907


CL\_3907


CL\_3907


CL\_3907


CL\_3907


CL\_3907


CL\_3907


CL\_3907


CL\_3907


CL\_3907


CL\_3907


CL\_3907


CL\_3907


CL\_3907


CL\_3907


CL\_3907


CL\_3907


CL\_3907

HighlightSelectShow Genomes


4

CL\_4012


3

CL\_4011


2

CL\_4019


2

CL\_4011


2

CL\_4011


2

CL\_4024


2

CL\_4011


2

CL\_4011


1

CL\_4011


1

CL\_4011


1

CL\_4025


1

CL\_4012


1

CL\_4011


1

CL\_4011


1

CL\_4011


1

CL\_4011


1

CL\_4011


1

CL\_4011


1

CL\_4011


1

CL\_4011


1

CL\_4011


1

CL\_4011


1

CL\_4011


1

CL\_4021


1

CL\_4011


1

CL\_4011


1

CL\_4011


1

CL\_4011


1

CL\_4011


1

CL\_4011


1

CL\_4011


1

CL\_4011


1

CL\_4011


1

CL\_4012


1

CL\_4012


1

CL\_4011


1

CL\_4011


1

CL\_4011


1

CL\_4014


1

CL\_4011


1

CL\_4021


1

CL\_4011


1

CL\_4011


1

CL\_4025


1

CL\_4011


1

CL\_4011

fGI ID


CL\_INS\_220
CL\_INS\_220
CL\_INS\_220
CL\_INS\_220
CL\_INS\_220
CL\_INS\_220
CL\_INS\_220
CL\_INS\_198
CL\_INS\_198
CL\_INS\_198
CL\_INS\_198
CL\_INS\_220
CL\_INS\_198
CL\_INS\_220
CL\_INS\_220
CL\_INS\_220
CL\_INS\_220
CL\_INS\_220
CL\_INS\_133
CL\_INS\_220
CL\_INS\_133
CL\_INS\_133
CL\_INS\_133
CL\_INS\_220
CL\_INS\_133
CL\_INS\_217
CL\_INS\_133
CL\_INS\_133
CL\_INS\_220
CL\_INS\_133
CL\_INS\_133
CL\_INS\_133
CL\_INS\_133
CL\_INS\_220
CL\_INS\_220
CL\_INS\_220
CL\_INS\_220
CL\_INS\_220
CL\_INS\_220
CL\_INS\_220
CL\_INS\_220
CL\_INS\_220
CL\_INS\_220
CL\_INS\_220
CL\_INS\_220
CL\_INS\_220
CL\_INS\_220
CL\_INS\_220
CL\_INS\_220
CL\_INS\_220
CL\_INS\_220
CL\_INS\_220
CL\_INS\_220
CL\_INS\_220
CL\_INS\_220
CL\_INS\_220
CL\_INS\_220
CL\_INS\_220
CL\_INS\_220
CL\_INS\_220
CL\_INS\_220
CL\_INS\_220
CL\_INS\_220
CL\_INS\_220
CL\_INS\_220
CL\_INS\_220
CL\_INS\_220
CL\_INS\_220
CL\_INS\_220
CL\_INS\_220
CL\_INS\_220
CL\_INS\_220
CL\_INS\_220
CL\_INS\_220
CL\_INS\_220
CL\_INS\_1
CL\_INS\_220
CL\_INS\_220
CL\_INS\_220
CL\_INS\_220
CL\_INS\_220
CL\_INS\_220
CL\_INS\_220
CL\_INS\_220
CL\_INS\_220
CL\_INS\_220
CL\_INS\_220
CL\_INS\_220
CL\_INS\_220
CL\_INS\_220
CL\_INS\_220
CL\_INS\_220
CL\_INS\_198
CL\_INS\_198
CL\_INS\_198
CL\_INS\_198
CL\_INS\_198
CL\_INS\_198
CL\_INS\_198
CL\_INS\_198
CL\_INS\_198
CL\_INS\_198
CL\_INS\_198
CL\_INS\_198
CL\_INS\_198
CL\_INS\_198
CL\_INS\_198
CL\_INS\_198
CL\_INS\_198
CL\_INS\_198
CL\_INS\_198
CL\_INS\_198
CL\_INS\_220
CL\_INS\_220
CL\_INS\_220
CL\_INS\_220
CL\_INS\_83
CL\_INS\_220
CL\_INS\_220
CL\_INS\_133
CL\_INS\_220
CL\_INS\_220
CL\_INS\_220
CL\_INS\_220
CL\_INS\_220
CL\_INS\_220
CL\_INS\_220
CL\_INS\_220
CL\_INS\_220
CL\_INS\_220
CL\_INS\_220
CL\_INS\_133
CL\_INS\_220
CL\_INS\_220
CL\_INS\_220
CL\_INS\_220
CL\_INS\_133
CL\_INS\_133
CL\_INS\_133
CL\_INS\_220
CL\_INS\_220
CL\_INS\_220
CL\_INS\_220
CL\_INS\_220
CL\_INS\_220
CL\_INS\_220
CL\_INS\_220
CL\_INS\_220
CL\_INS\_220
CL\_INS\_219
CL\_INS\_219
CL\_INS\_219
CL\_INS\_220
CL\_INS\_220
CL\_INS\_220
CL\_INS\_220
CL\_INS\_220
CL\_INS\_220
CL\_INS\_220
CL\_INS\_220
CL\_INS\_220
CL\_INS\_220
CL\_INS\_220
CL\_INS\_220
CL\_INS\_220
CL\_INS\_220
CL\_INS\_219
CL\_INS\_220
CL\_INS\_220
CL\_INS\_220
CL\_INS\_220
CL\_INS\_220
CL\_INS\_220
CL\_INS\_220
CL\_INS\_220
CL\_INS\_220
CL\_INS\_220
CL\_INS\_220
CL\_INS\_220
CL\_INS\_219
CL\_INS\_219
CL\_INS\_220
CL\_INS\_219
CL\_INS\_219
CL\_INS\_220
CL\_INS\_220
CL\_INS\_220
CL\_INS\_220
CL\_INS\_220
CL\_INS\_220
CL\_INS\_220
CL\_INS\_220
CL\_INS\_220
CL\_INS\_220
CL\_INS\_220
CL\_INS\_220
CL\_INS\_220
CL\_INS\_220
CL\_INS\_220
CL\_INS\_220
CL\_INS\_220
CL\_INS\_220
CL\_INS\_220
CL\_INS\_220
CL\_INS\_220
CL\_INS\_220
CL\_INS\_220
CL\_INS\_220
CL\_INS\_220
CL\_INS\_220
CL\_INS\_220
CL\_INS\_220
CL\_INS\_220
CL\_INS\_220
CL\_INS\_220
CL\_INS\_220
CL\_INS\_220
CL\_INS\_220
CL\_INS\_220
CL\_INS\_220
CL\_INS\_220
CL\_INS\_220
CL\_INS\_220
CL\_INS\_220
CL\_INS\_220
CL\_INS\_220
CL\_INS\_220
CL\_INS\_220
CL\_INS\_220
CL\_INS\_220
CL\_INS\_220
CL\_INS\_220
CL\_INS\_220
CL\_INS\_220
CL\_INS\_220
CL\_INS\_220
CL\_INS\_220
CL\_INS\_220
CL\_INS\_220
CL\_INS\_220
CL\_INS\_220
CL\_INS\_220
CL\_INS\_220
CL\_INS\_83
CL\_INS\_83
CL\_INS\_83
CL\_INS\_83
CL\_INS\_220
CL\_INS\_220
CL\_INS\_220
CL\_INS\_220
CL\_INS\_65
CL\_INS\_83
CL\_INS\_220
CL\_INS\_65
CL\_INS\_83
CL\_INS\_83
CL\_INS\_83
CL\_INS\_83
CL\_INS\_220
CL\_INS\_220
CL\_INS\_220
CL\_INS\_220
CL\_INS\_220
CL\_INS\_220
CL\_INS\_220
CL\_INS\_220
CL\_INS\_220
CL\_INS\_220
CL\_INS\_220
CL\_INS\_220
CL\_INS\_220
CL\_INS\_220
CL\_INS\_220
CL\_INS\_220
CL\_INS\_220
CL\_INS\_220
CL\_INS\_220
CL\_INS\_220
CL\_INS\_220
CL\_INS\_83
CL\_INS\_222
CL\_INS\_220
CL\_INS\_222
CL\_INS\_222
CL\_INS\_222
CL\_INS\_222
CL\_INS\_222
CL\_INS\_220
CL\_INS\_220
CL\_INS\_220
CL\_INS\_220
CL\_INS\_220
CL\_INS\_220
CL\_INS\_220
CL\_INS\_220
CL\_INS\_220
CL\_INS\_220
CL\_INS\_220
CL\_INS\_1
CL\_INS\_83
CL\_INS\_45
CL\_INS\_220
CL\_INS\_220
CL\_INS\_220
CL\_INS\_45
CL\_INS\_45
CL\_INS\_45
CL\_INS\_45
CL\_INS\_220
CL\_INS\_220
CL\_INS\_220
CL\_INS\_220
CL\_INS\_220
CL\_INS\_220
CL\_INS\_220
CL\_INS\_220
CL\_INS\_220
CL\_INS\_220
CL\_INS\_220
CL\_INS\_220
CL\_INS\_220
CL\_INS\_220
CL\_INS\_220
CL\_INS\_220
CL\_INS\_83
CL\_INS\_83
CL\_INS\_83
CL\_INS\_83
CL\_INS\_83
CL\_INS\_83
CL\_INS\_83
CL\_INS\_83
CL\_INS\_83
CL\_INS\_220
CL\_INS\_220
CL\_INS\_220
CL\_INS\_220
CL\_INS\_83
CL\_INS\_83
CL\_INS\_220
CL\_INS\_83
CL\_INS\_83
CL\_INS\_83
CL\_INS\_83
CL\_INS\_83
CL\_INS\_83
CL\_INS\_220
CL\_INS\_220
CL\_INS\_220
CL\_INS\_220
CL\_INS\_220
CL\_INS\_220
CL\_INS\_220
CL\_INS\_220
CL\_INS\_220
CL\_INS\_220
CL\_INS\_220
CL\_INS\_220
CL\_INS\_220
CL\_INS\_220
CL\_INS\_220
CL\_INS\_220
CL\_INS\_220
CL\_INS\_220
CL\_INS\_83
CL\_INS\_83
CL\_INS\_220
CL\_INS\_220
CL\_INS\_220
CL\_INS\_83
CL\_INS\_220
CL\_INS\_83
CL\_INS\_220
CL\_INS\_220
CL\_INS\_83
CL\_INS\_220
CL\_INS\_83
CL\_INS\_220
CL\_INS\_220
CL\_INS\_220
CL\_INS\_220
CL\_INS\_220
CL\_INS\_220
CL\_INS\_220
CL\_INS\_220
CL\_INS\_220
CL\_INS\_220
CL\_INS\_220
CL\_INS\_83
CL\_INS\_133
CL\_INS\_98
CL\_INS\_220
CL\_INS\_220
CL\_INS\_220
CL\_INS\_220
CL\_INS\_220
CL\_INS\_83
CL\_INS\_220
CL\_INS\_220
CL\_INS\_220
CL\_INS\_220
CL\_INS\_220
CL\_INS\_220
CL\_INS\_220
CL\_INS\_220
CL\_INS\_220
CL\_INS\_220
CL\_INS\_220
CL\_INS\_220
CL\_INS\_220
CL\_INS\_220
CL\_INS\_220
CL\_INS\_220
CL\_INS\_220
CL\_INS\_220
CL\_INS\_220
CL\_INS\_220
CL\_INS\_220
CL\_INS\_220
CL\_INS\_220
CL\_INS\_220
CL\_INS\_220
CL\_INS\_220
CL\_INS\_220
CL\_INS\_220
CL\_INS\_220
CL\_INS\_220
CL\_INS\_220
CL\_INS\_220
CL\_INS\_1
CL\_INS\_1
CL\_INS\_1
CL\_INS\_1
CL\_INS\_1
CL\_INS\_83
CL\_INS\_83
CL\_INS\_83
CL\_INS\_220
CL\_INS\_220
CL\_INS\_220
CL\_INS\_220
CL\_INS\_133
CL\_INS\_217
CL\_INS\_133
CL\_INS\_217
CL\_INS\_220
CL\_INS\_220
CL\_INS\_220
CL\_INS\_133
CL\_INS\_133
CL\_INS\_133
CL\_INS\_133
CL\_INS\_133
CL\_INS\_133
CL\_INS\_133
CL\_INS\_133
CL\_INS\_220
CL\_INS\_220
CL\_INS\_133
CL\_INS\_133
CL\_INS\_133
CL\_INS\_133
CL\_INS\_133
CL\_INS\_220
CL\_INS\_220
CL\_INS\_133
CL\_INS\_133
CL\_INS\_220
CL\_INS\_217
CL\_INS\_133
CL\_INS\_133
CL\_INS\_217
CL\_INS\_220
CL\_INS\_220
CL\_INS\_133
CL\_INS\_133
CL\_INS\_133
CL\_INS\_133
CL\_INS\_220
CL\_INS\_220
CL\_INS\_220
CL\_INS\_220
CL\_INS\_220
CL\_INS\_220
CL\_INS\_220
CL\_INS\_220
CL\_INS\_220
CL\_INS\_220
CL\_INS\_220
CL\_INS\_220
CL\_INS\_133
CL\_INS\_220
CL\_INS\_133
CL\_INS\_220
CL\_INS\_220
CL\_INS\_220
CL\_INS\_220
CL\_INS\_220
CL\_INS\_220
CL\_INS\_220
CL\_INS\_220
CL\_INS\_220
CL\_INS\_220
CL\_INS\_220
CL\_INS\_220
CL\_INS\_220
CL\_INS\_220
CL\_INS\_220
CL\_INS\_220
CL\_INS\_220
CL\_INS\_220
CL\_INS\_220
CL\_INS\_220
CL\_INS\_220
CL\_INS\_220
CL\_INS\_220
CL\_INS\_220
CL\_INS\_220
CL\_INS\_220
CL\_INS\_220
CL\_INS\_220
CL\_INS\_220
CL\_INS\_220
CL\_INS\_220
CL\_INS\_220
CL\_INS\_220
CL\_INS\_220
CL\_INS\_220
CL\_INS\_220
CL\_INS\_220
CL\_INS\_220
CL\_INS\_220
CL\_INS\_220
CL\_INS\_220
CL\_INS\_220
CL\_INS\_220
CL\_INS\_220
CL\_INS\_220
CL\_INS\_220
CL\_INS\_220
CL\_INS\_220
CL\_INS\_220
CL\_INS\_220
CL\_INS\_220
CL\_INS\_220
CL\_INS\_220
CL\_INS\_220
CL\_INS\_220
CL\_INS\_220
CL\_INS\_220
CL\_INS\_220
CL\_INS\_220
CL\_INS\_220
CL\_INS\_220
CL\_INS\_220
CL\_INS\_220
CL\_INS\_220
CL\_INS\_220
CL\_INS\_220
CL\_INS\_220
CL\_INS\_220
CL\_INS\_220
CL\_INS\_220
CL\_INS\_220
CL\_INS\_220
CL\_INS\_220
CL\_INS\_220
CL\_INS\_220
CL\_INS\_220
CL\_INS\_220
CL\_INS\_220
CL\_INS\_220
CL\_INS\_220
CL\_INS\_220
CL\_INS\_220
CL\_INS\_220
CL\_INS\_220
CL\_INS\_220
CL\_INS\_220
CL\_INS\_220
CL\_INS\_220
CL\_INS\_220
CL\_INS\_220
Cluster ID


CL\_6866
CL\_7843
CL\_7844
CL\_9481
CL\_9482
CL\_9483
CL\_7845
CL\_7846
CL\_7847
CL\_7848
CL\_7849
CL\_9484
CL\_7850
CL\_6452
CL\_13686
CL\_6453
CL\_6454
CL\_6455
CL\_6456
CL\_6457
CL\_6458
CL\_6459
CL\_6460
CL\_6461
CL\_6462
CL\_6463
CL\_6464
CL\_6465
CL\_10654
CL\_7759
CL\_7758
CL\_7757
CL\_7756
CL\_10653
CL\_10652
CL\_10651
CL\_11234
CL\_11432
CL\_11233
CL\_11232
CL\_11231
CL\_11230
CL\_11229
CL\_11431
CL\_11430
CL\_11429
CL\_11428
CL\_11427
CL\_11228
CL\_11227
CL\_11226
CL\_11225
CL\_11224
CL\_13630
CL\_11426
CL\_11425
CL\_11424
CL\_11223
CL\_6867
CL\_6868
CL\_6869
CL\_6870
CL\_13629
CL\_6871
CL\_6872
CL\_6873
CL\_6874
CL\_6875
CL\_6876
CL\_11423
CL\_11422
CL\_6877
CL\_11222
CL\_11221
CL\_6878
CL\_24
CL\_11871
CL\_11870
CL\_6565
CL\_6566
CL\_6567
CL\_6568
CL\_6569
CL\_6570
CL\_6571
CL\_6572
CL\_8606
CL\_8604
CL\_10178
CL\_10177
CL\_8782
CL\_6574
CL\_6575
CL\_7851
CL\_7852
CL\_7853
CL\_7854
CL\_7855
CL\_9485
CL\_7856
CL\_7857
CL\_7858
CL\_7859
CL\_7860
CL\_7861
CL\_7862
CL\_7863
CL\_7864
CL\_7865
CL\_7866
CL\_7867
CL\_8203
CL\_9486
CL\_9487
CL\_9488
CL\_9489
CL\_4816
CL\_8321
CL\_9490
CL\_7868
CL\_7869
CL\_7870
CL\_7871
CL\_7872
CL\_7873
CL\_7874
CL\_7875
CL\_7876
CL\_8323
CL\_7877
CL\_7878
CL\_7879
CL\_8326
CL\_8327
CL\_8328
CL\_9491
CL\_7880
CL\_7881
CL\_7882
CL\_7883
CL\_7884
CL\_7885
CL\_7886
CL\_7887
CL\_7888
CL\_7889
CL\_9492
CL\_7890
CL\_7891
CL\_6576
CL\_6577
CL\_6578
CL\_9253
CL\_9254
CL\_10176
CL\_6579
CL\_6580
CL\_6581
CL\_6582
CL\_6583
CL\_6584
CL\_6585
CL\_6586
CL\_6587
CL\_6588
CL\_6589
CL\_6438
CL\_6591
CL\_9255
CL\_10175
CL\_10174
CL\_6594
CL\_10173
CL\_6595
CL\_10172
CL\_6596
CL\_6597
CL\_6598
CL\_6599
CL\_6600
CL\_9256
CL\_9257
CL\_9258
CL\_9259
CL\_9260
CL\_9261
CL\_10171
CL\_7732
CL\_7730
CL\_7729
CL\_7728
CL\_7727
CL\_7726
CL\_7725
CL\_7724
CL\_7723
CL\_7722
CL\_7721
CL\_7720
CL\_7719
CL\_7718
CL\_7717
CL\_7716
CL\_7715
CL\_7714
CL\_7713
CL\_7712
CL\_7711
CL\_7710
CL\_7709
CL\_7708
CL\_7707
CL\_7706
CL\_9085
CL\_7705
CL\_7704
CL\_11134
CL\_7703
CL\_7702
CL\_7701
CL\_7700
CL\_6601
CL\_6605
CL\_3908
CL\_8169
CL\_8168
CL\_8167
CL\_8166
CL\_8165
CL\_9100
CL\_9099
CL\_9098
CL\_9097
CL\_9096
CL\_11690
CL\_11689
CL\_11688
CL\_13346
CL\_13345
CL\_13344
CL\_13343
CL\_13342
CL\_13341
CL\_5483
CL\_5482
CL\_5481
CL\_92
CL\_13340
CL\_10538
CL\_10539
CL\_9791
CL\_8270
CL\_6676
CL\_9093
CL\_8269
CL\_8268
CL\_8587
CL\_8588
CL\_8263
CL\_5051
CL\_5052
CL\_5053
CL\_5054
CL\_5055
CL\_5056
CL\_5057
CL\_5058
CL\_5059
CL\_5060
CL\_5061
CL\_5062
CL\_5063
CL\_5064
CL\_5065
CL\_5066
CL\_6817
CL\_6816
CL\_10022
CL\_10023
CL\_10024
CL\_6150
CL\_7434
CL\_8161
CL\_7435
CL\_7436
CL\_7437
CL\_7438
CL\_13521
CL\_7439
CL\_7440
CL\_7441
CL\_7442
CL\_7443
CL\_7444
CL\_7445
CL\_7446
CL\_7447
CL\_7448
CL\_8160
CL\_5838
CL\_102
CL\_7656
CL\_8159
CL\_8610
CL\_8609
CL\_7661
CL\_7663
CL\_7664
CL\_7665
CL\_5180
CL\_5179
CL\_5178
CL\_5177
CL\_10405
CL\_5176
CL\_5175
CL\_5174
CL\_5173
CL\_5172
CL\_5171
CL\_5170
CL\_5169
CL\_5168
CL\_5166
CL\_10968
CL\_8777
CL\_9615
CL\_6153
CL\_6154
CL\_9616
CL\_6151
CL\_10969
CL\_10404
CL\_10403
CL\_10402
CL\_10970
CL\_10400
CL\_10399
CL\_6489
CL\_6490
CL\_9394
CL\_6491
CL\_6492
CL\_6493
CL\_6494
CL\_6495
CL\_6496
CL\_13123
CL\_9395
CL\_6497
CL\_6498
CL\_6499
CL\_9396
CL\_6500
CL\_11219
CL\_12513
CL\_11417
CL\_11416
CL\_6501
CL\_13688
CL\_6881
CL\_6502
CL\_6503
CL\_6504
CL\_6505
CL\_6506
CL\_8164
CL\_8163
CL\_6507
CL\_6508
CL\_6509
CL\_6510
CL\_4806
CL\_6511
CL\_6512
CL\_6513
CL\_8162
CL\_4805
CL\_9095
CL\_9094
CL\_11213
CL\_12514
CL\_12515
CL\_12516
CL\_12517
CL\_12518
CL\_12519
CL\_12520
CL\_7450
CL\_4814
CL\_4813
CL\_6239
CL\_7451
CL\_6488
CL\_9908
CL\_9909
CL\_8584
CL\_8590
CL\_10540
CL\_9792
CL\_9493
CL\_3909
CL\_3910
CL\_3911
CL\_3912
CL\_3913
CL\_3914
CL\_8994
CL\_8995
CL\_3915
CL\_3916
CL\_3917
CL\_3918
CL\_3919
CL\_3920
CL\_3921
CL\_3922
CL\_3923
CL\_3924
CL\_3925
CL\_3926
CL\_3927
CL\_3928
CL\_3929
CL\_3930
CL\_3931
CL\_3932
CL\_3933
CL\_3934
CL\_3935
CL\_3936
CL\_3937
CL\_3938
CL\_3939
CL\_3940
CL\_3941
CL\_3942
CL\_3943
CL\_5067
CL\_5257
CL\_5068
CL\_7449
CL\_5042
CL\_8791
CL\_5044
CL\_9387
CL\_10381
CL\_10382
CL\_10383
CL\_8237
CL\_8238
CL\_6466
CL\_6467
CL\_6468
CL\_5590
CL\_5589
CL\_6469
CL\_10384
CL\_6470
CL\_5586
CL\_5585
CL\_6471
CL\_6472
CL\_5582
CL\_6473
CL\_10385
CL\_10386
CL\_10213
CL\_10387
CL\_6376
CL\_6377
CL\_6378
CL\_6379
CL\_10388
CL\_10389
CL\_6474
CL\_6475
CL\_6476
CL\_6477
CL\_10390
CL\_10391
CL\_10392
CL\_10393
CL\_6478
CL\_6479
CL\_6480
CL\_8966
CL\_8967
CL\_6481
CL\_6482
CL\_6483
CL\_6484
CL\_8968
CL\_6485
CL\_13687
CL\_6486
CL\_6487
CL\_3944
CL\_3945
CL\_3946
CL\_3947
CL\_3948
CL\_3949
CL\_3950
CL\_3951
CL\_3952
CL\_3953
CL\_3954
CL\_3955
CL\_3956
CL\_3957
CL\_3958
CL\_3959
CL\_3960
CL\_3961
CL\_3962
CL\_3963
CL\_3964
CL\_3965
CL\_3966
CL\_3967
CL\_3968
CL\_3969
CL\_3970
CL\_3971
CL\_3972
CL\_3973
CL\_3974
CL\_3975
CL\_3976
CL\_3977
CL\_3978
CL\_3979
CL\_3980
CL\_3981
CL\_3982
CL\_3983
CL\_3984
CL\_3985
CL\_3986
CL\_3987
CL\_3988
CL\_3989
CL\_3990
CL\_3991
CL\_3992
CL\_3993
CL\_3994
CL\_3995
CL\_3996
CL\_3997
CL\_3998
CL\_3999
CL\_4000
CL\_4001
CL\_4002
CL\_4003
CL\_4004
CL\_5706
CL\_4005
CL\_4006
CL\_4007
CL\_4008
CL\_6514
CL\_6515
CL\_6516
CL\_6517
CL\_6518
CL\_9793
CL\_6519
CL\_6520
CL\_6521
CL\_6522
CL\_6523
CL\_6524
CL\_6525
CL\_9262
CL\_9494
CL\_9495
CL\_9496
CL\_6526
CL\_6527
CL\_4009
CL\_4010
