## Supplementary material for "A novel method for integrating genomic and Tn-Seq data to identify common *in vivo* fitness mechanisms across multiple bacterial species": S3 Dataset: CL_INS_221.html

Legend

 Mobile +extrachromosomalelementfunctions
 Hypothetical
 DNA Metabolism
 Other
 Proteinsynthesis/fate
 Transport +binding proteins
 All VFDB Genes

FULL


WINDOWSVGPNG

Trim RowsRemove SingletonsSave Fasta

CL\_4012


CL\_4012


CL\_4012


CL\_4012


CL\_4012

HighlightSelectShow Genomes


41

CL\_4011


4

CL\_3907


1

CL\_3907


1

CL\_3907


1

CL\_3907

fGI ID


CL\_INS\_178
CL\_INS\_45
CL\_INS\_45
CL\_INS\_45
CL\_INS\_45
CL\_INS\_220
CL\_INS\_220
CL\_INS\_220
CL\_INS\_45
CL\_INS\_1
CL\_INS\_83
CL\_INS\_220
CL\_INS\_222
CL\_INS\_220
CL\_INS\_220
CL\_INS\_83
CL\_INS\_83
CL\_INS\_220
CL\_INS\_83
CL\_INS\_220
CL\_INS\_83
CL\_INS\_83
CL\_INS\_220
CL\_INS\_220
CL\_INS\_220
CL\_INS\_220
CL\_INS\_220
CL\_INS\_220
CL\_INS\_220
CL\_INS\_220
CL\_INS\_220
CL\_INS\_220
CL\_INS\_220
CL\_INS\_220
CL\_INS\_220
CL\_INS\_220
CL\_INS\_220
CL\_INS\_220
CL\_INS\_220
CL\_INS\_220
CL\_INS\_220
CL\_INS\_220
CL\_INS\_220
CL\_INS\_220
CL\_INS\_220
CL\_INS\_220
CL\_INS\_220
CL\_INS\_220
CL\_INS\_220
CL\_INS\_220
CL\_INS\_220
CL\_INS\_220
CL\_INS\_83
CL\_INS\_83
CL\_INS\_83
CL\_INS\_83
CL\_INS\_220
CL\_INS\_220
CL\_INS\_220
CL\_INS\_220
CL\_INS\_220
CL\_INS\_220
Cluster ID


CL\_8158
CL\_7665
CL\_7664
CL\_7663
CL\_7661
CL\_8609
CL\_8610
CL\_8159
CL\_7656
CL\_5838
CL\_102
CL\_8160
CL\_7435
CL\_8161
CL\_8162
CL\_6513
CL\_4806
CL\_6510
CL\_6509
CL\_8163
CL\_8164
CL\_6506
CL\_6505
CL\_6504
CL\_6503
CL\_6881
CL\_6501
CL\_8165
CL\_8166
CL\_8167
CL\_8168
CL\_8169
CL\_4010
CL\_4009
CL\_4008
CL\_4007
CL\_4006
CL\_4005
CL\_5706
CL\_4004
CL\_4003
CL\_4002
CL\_4001
CL\_4000
CL\_3999
CL\_3949
CL\_3948
CL\_3947
CL\_3946
CL\_3945
CL\_3944
CL\_13340
CL\_92
CL\_5481
CL\_5482
CL\_5483
CL\_13341
CL\_13342
CL\_13343
CL\_13344
CL\_13345
CL\_13346
