## Supplementary material for "A novel method for integrating genomic and Tn-Seq data to identify common *in vivo* fitness mechanisms across multiple bacterial species": S3 Dataset: CL_INS_222.html

Legend

 Hypothetical
 Regulatoryfunctions
 Other
 All VFDB Genes

FULL


WINDOWSVGPNG

Trim RowsRemove SingletonsSave Fasta

CL\_4012


CL\_4012


CL\_4012


CL\_3907

HighlightSelectShow Genomes


36

CL\_4014


6

CL\_4015


4

CL\_4014


1

CL\_4014

fGI ID


CL\_INS\_222
CL\_INS\_222
CL\_INS\_222
CL\_INS\_222
CL\_INS\_222
CL\_INS\_222
CL\_INS\_222
CL\_INS\_222
CL\_INS\_222
Cluster ID


CL\_9497
CL\_9498
CL\_7434
CL\_7435
CL\_7436
CL\_7437
CL\_7438
CL\_13521
CL\_4013
