## Supplementary material for "A novel method for integrating genomic and Tn-Seq data to identify common *in vivo* fitness mechanisms across multiple bacterial species": S3 Dataset: CL_INS_223.html

Legend

 Regulatoryfunctions
 Hypothetical
 Other
 All VFDB Genes

FULL


WINDOWSVGPNG

Trim RowsRemove SingletonsSave Fasta

CL\_4015


CL\_4015

HighlightSelectShow Genomes


41

CL\_4014


6

CL\_4012

fGI ID


CL\_INS\_223
CL\_INS\_222
Cluster ID


CL\_9499
CL\_9498
