## Supplementary material for "A novel method for integrating genomic and Tn-Seq data to identify common *in vivo* fitness mechanisms across multiple bacterial species": S3 Dataset: CL_INS_224.html

Legend

 Hypothetical
 Other
 All VFDB Genes

FULL


WINDOWSVGPNG

Trim RowsRemove SingletonsSave Fasta

CL\_4018


CL\_4018


CL\_4017


CL\_3907

HighlightSelectShow Genomes


27

CL\_4019


14

CL\_4019


3

CL\_4019


2

CL\_4019

fGI ID


CL\_INS\_224
CL\_INS\_224
CL\_INS\_224
CL\_INS\_224
CL\_INS\_224
CL\_INS\_220
CL\_INS\_220
CL\_INS\_220
CL\_INS\_133
CL\_INS\_220
CL\_INS\_133
CL\_INS\_133
CL\_INS\_133
CL\_INS\_220
CL\_INS\_133
CL\_INS\_217
CL\_INS\_133
CL\_INS\_133
CL\_INS\_220
CL\_INS\_133
CL\_INS\_133
CL\_INS\_133
CL\_INS\_133
CL\_INS\_220
CL\_INS\_220
CL\_INS\_220
CL\_INS\_224
Cluster ID


CL\_6528
CL\_6529
CL\_6530
CL\_6531
CL\_6532
CL\_6453
CL\_6454
CL\_6455
CL\_6456
CL\_6457
CL\_6458
CL\_6459
CL\_6460
CL\_6461
CL\_6462
CL\_6463
CL\_6464
CL\_6465
CL\_10654
CL\_7759
CL\_7758
CL\_7757
CL\_7756
CL\_10653
CL\_10652
CL\_10651
CL\_10650
