## Supplementary material for "A novel method for integrating genomic and Tn-Seq data to identify common *in vivo* fitness mechanisms across multiple bacterial species": S3 Dataset: CL_INS_226.html

Legend

 All EssentialGenes
 Other
 All VFDB Genes

FULL


WINDOWSVGPNG

Trim RowsRemove SingletonsSave Fasta

CL\_4088


CL\_4088


CL\_4087

HighlightSelectShow Genomes


37

CL\_4089


19

CL\_4089


1

CL\_4089

fGI ID


CL\_INS\_226
CL\_INS\_226
CL\_INS\_226
CL\_INS\_226
Cluster ID


CL\_6533
CL\_6534
CL\_6535
CL\_6536
