## Supplementary material for "A novel method for integrating genomic and Tn-Seq data to identify common *in vivo* fitness mechanisms across multiple bacterial species": S3 Dataset: CL_INS_227.html

Legend

 Mobile +extrachromosomalelementfunctions
 Regulatoryfunctions
 Hypothetical
 DNA Metabolism
 Transcription
 Other
 All VFDB Genes
 Transport +binding proteins

FULL


WINDOWSVGPNG

Trim RowsRemove SingletonsSave Fasta

CL\_4090


CL\_4090


CL\_4090


CL\_4090


CL\_4090


CL\_4090

HighlightSelectShow Genomes


42

CL\_4091


11

CL\_4091


1

CL\_4091


1

CL\_4091


1

CL\_4091


1

CL\_4091

fGI ID


CL\_INS\_220
CL\_INS\_220
CL\_INS\_220
CL\_INS\_220
CL\_INS\_227
CL\_INS\_227
CL\_INS\_227
CL\_INS\_227
CL\_INS\_227
CL\_INS\_227
CL\_INS\_220
CL\_INS\_227
CL\_INS\_227
CL\_INS\_227
CL\_INS\_227
CL\_INS\_227
CL\_INS\_227
CL\_INS\_227
CL\_INS\_227
CL\_INS\_1
CL\_INS\_227
CL\_INS\_1
CL\_INS\_1
CL\_INS\_1
CL\_INS\_1
CL\_INS\_1
CL\_INS\_227
CL\_INS\_227
CL\_INS\_227
CL\_INS\_227
CL\_INS\_227
CL\_INS\_227
CL\_INS\_227
CL\_INS\_227
CL\_INS\_227
CL\_INS\_227
CL\_INS\_220
CL\_INS\_220
CL\_INS\_220
CL\_INS\_227
CL\_INS\_220
CL\_INS\_220
CL\_INS\_220
CL\_INS\_227
CL\_INS\_220
CL\_INS\_220
CL\_INS\_227
CL\_INS\_227
CL\_INS\_227
CL\_INS\_227
CL\_INS\_227
CL\_INS\_227
CL\_INS\_227
CL\_INS\_227
CL\_INS\_227
CL\_INS\_227
CL\_INS\_227
CL\_INS\_227
CL\_INS\_220
CL\_INS\_227
CL\_INS\_227
Cluster ID


CL\_9100
CL\_9099
CL\_9098
CL\_9097
CL\_6880
CL\_6537
CL\_6538
CL\_11421
CL\_11420
CL\_11419
CL\_11219
CL\_11218
CL\_11217
CL\_11216
CL\_11215
CL\_11214
CL\_11418
CL\_13424
CL\_13423
CL\_8941
CL\_13422
CL\_8942
CL\_8946
CL\_8943
CL\_8944
CL\_8945
CL\_13421
CL\_13420
CL\_13419
CL\_13418
CL\_13417
CL\_13416
CL\_13415
CL\_13414
CL\_13413
CL\_13412
CL\_11417
CL\_11416
CL\_6501
CL\_11220
CL\_6881
CL\_6502
CL\_6503
CL\_6882
CL\_6504
CL\_6505
CL\_6883
CL\_6884
CL\_6885
CL\_6886
CL\_6887
CL\_6888
CL\_6889
CL\_6890
CL\_6891
CL\_6892
CL\_6893
CL\_6894
CL\_11213
CL\_11212
CL\_6895
