## Supplementary material for "A novel method for integrating genomic and Tn-Seq data to identify common *in vivo* fitness mechanisms across multiple bacterial species": S3 Dataset: CL_INS_230.html

Legend

 Mobile +extrachromosomalelementfunctions
 Hypothetical
 DNA Metabolism
 All EssentialGenes
 Transport +binding proteins
 All VFDB Genes

FULL


WINDOWSVGPNG

Trim RowsRemove SingletonsSave Fasta

CL\_4212


CL\_4212


CL\_4212


CL\_4212


CL\_4212


CL\_4212


CL\_4212


CL\_4212


CL\_4212

HighlightSelectShow Genomes


42

CL\_4213


3

CL\_4215


2

CL\_4215


2

CL\_4215


1

CL\_4215


1

CL\_4215


1

CL\_4215


1

CL\_4215


1

CL\_4215

fGI ID


CL\_INS\_230
CL\_INS\_230
CL\_INS\_230
CL\_INS\_230
CL\_INS\_230
CL\_INS\_230
CL\_INS\_230
CL\_INS\_230
CL\_INS\_230
CL\_INS\_230
CL\_INS\_230
CL\_INS\_230
CL\_INS\_230
CL\_INS\_230
CL\_INS\_230
CL\_INS\_230
CL\_INS\_230
CL\_INS\_230
CL\_INS\_230
CL\_INS\_230
CL\_INS\_230
CL\_INS\_230
CL\_INS\_230
CL\_INS\_230
CL\_INS\_230
CL\_INS\_230
CL\_INS\_230
CL\_INS\_230
CL\_INS\_230
CL\_INS\_230
CL\_INS\_230
CL\_INS\_230
Cluster ID


CL\_9501
CL\_9502
CL\_9503
CL\_5707
CL\_4214
CL\_8344
CL\_5490
CL\_5491
CL\_5074
CL\_5075
CL\_5076
CL\_5077
CL\_5078
CL\_5079
CL\_5080
CL\_5081
CL\_13125
CL\_13126
CL\_13127
CL\_13128
CL\_5082
CL\_5083
CL\_5084
CL\_5085
CL\_5086
CL\_5087
CL\_5088
CL\_5089
CL\_5090
CL\_5091
CL\_5092
CL\_5093
