## Supplementary material for "A novel method for integrating genomic and Tn-Seq data to identify common *in vivo* fitness mechanisms across multiple bacterial species": S3 Dataset: CL_INS_231.html

Legend

 Mobile +extrachromosomalelementfunctions
 Hypothetical
 DNA Metabolism
 All EssentialGenes
 Other
 Transport +binding proteins
 All VFDB Genes

FULL


WINDOWSVGPNG

Trim RowsRemove SingletonsSave Fasta

CL\_4213


CL\_4213


CL\_4213


CL\_4213


CL\_4213


CL\_4213


CL\_4213


CL\_4212


CL\_4213


CL\_4212


CL\_4212


CL\_4213


CL\_4213


CL\_4212


CL\_4212


CL\_4213


CL\_4212


CL\_4213


CL\_4213


CL\_4213


CL\_4208


CL\_4212


CL\_4213


CL\_4213


CL\_4212

HighlightSelectShow Genomes


6

CL\_4215


6

CL\_4215


5

CL\_4215


5

CL\_4215


5

CL\_4215


4

CL\_4215


3

CL\_4215


3

CL\_4215


2

CL\_4215


2

CL\_4215


2

CL\_4215


1

CL\_4215


1

CL\_4215


1

CL\_4215


1

CL\_4215


1

CL\_4215


1

CL\_4215


1

CL\_4215


1

CL\_4215


1

CL\_4215


1

CL\_4215


1

CL\_4215


1

CL\_4215


1

CL\_4215


1

CL\_4215

fGI ID


CL\_INS\_231
CL\_INS\_230
CL\_INS\_230
CL\_INS\_231
CL\_INS\_230
CL\_INS\_230
CL\_INS\_230
CL\_INS\_230
CL\_INS\_153
CL\_INS\_153
CL\_INS\_153
CL\_INS\_153
CL\_INS\_231
CL\_INS\_231
CL\_INS\_230
CL\_INS\_231
CL\_INS\_230
CL\_INS\_230
CL\_INS\_230
CL\_INS\_231
CL\_INS\_231
CL\_INS\_83
CL\_INS\_83
CL\_INS\_231
CL\_INS\_83
CL\_INS\_83
CL\_INS\_83
CL\_INS\_83
CL\_INS\_83
CL\_INS\_83
CL\_INS\_83
CL\_INS\_83
CL\_INS\_83
CL\_INS\_230
CL\_INS\_230
CL\_INS\_230
CL\_INS\_230
CL\_INS\_231
CL\_INS\_231
CL\_INS\_230
CL\_INS\_230
CL\_INS\_230
CL\_INS\_230
CL\_INS\_230
CL\_INS\_231
CL\_INS\_83
CL\_INS\_83
CL\_INS\_230
CL\_INS\_230
CL\_INS\_230
CL\_INS\_230
CL\_INS\_65
CL\_INS\_83
CL\_INS\_83
CL\_INS\_230
CL\_INS\_230
CL\_INS\_230
CL\_INS\_83
CL\_INS\_83
CL\_INS\_231
CL\_INS\_231
CL\_INS\_64
CL\_INS\_181
CL\_INS\_231
CL\_INS\_231
CL\_INS\_231
CL\_INS\_64
CL\_INS\_231
CL\_INS\_231
CL\_INS\_231
CL\_INS\_231
CL\_INS\_231
CL\_INS\_230
CL\_INS\_230
CL\_INS\_230
CL\_INS\_230
CL\_INS\_230
CL\_INS\_65
CL\_INS\_231
CL\_INS\_231
Cluster ID


CL\_11687
CL\_5490
CL\_5491
CL\_9092
CL\_9502
CL\_9503
CL\_5707
CL\_4214
CL\_6781
CL\_6780
CL\_10686
CL\_10687
CL\_9091
CL\_9090
CL\_8344
CL\_13691
CL\_5074
CL\_5075
CL\_5076
CL\_9910
CL\_9911
CL\_9912
CL\_9913
CL\_9914
CL\_9915
CL\_9916
CL\_9917
CL\_9918
CL\_9919
CL\_9920
CL\_4806
CL\_9921
CL\_4805
CL\_5077
CL\_5078
CL\_5079
CL\_5080
CL\_12384
CL\_9794
CL\_5081
CL\_13125
CL\_13126
CL\_13127
CL\_13128
CL\_13129
CL\_5496
CL\_5649
CL\_5082
CL\_5083
CL\_5084
CL\_5085
CL\_5651
CL\_5652
CL\_7298
CL\_5086
CL\_5087
CL\_5088
CL\_5495
CL\_5494
CL\_8788
CL\_12944
CL\_5493
CL\_8787
CL\_8786
CL\_8785
CL\_8784
CL\_4795
CL\_9795
CL\_9796
CL\_9797
CL\_9798
CL\_9799
CL\_5089
CL\_5090
CL\_5091
CL\_5092
CL\_5093
CL\_9800
CL\_9801
CL\_5094
