## Supplementary material for "A novel method for integrating genomic and Tn-Seq data to identify common *in vivo* fitness mechanisms across multiple bacterial species": S3 Dataset: CL_INS_233.html

Legend

 Hypothetical
 All EssentialGenes
 Other
 EnergyMetabolism
 Transport +binding proteins
 All VFDB Genes

FULL


WINDOWSVGPNG

Trim RowsRemove SingletonsSave Fasta

CL\_4252


CL\_4252


CL\_4252

HighlightSelectShow Genomes


43

CL\_4253


13

CL\_4253


1

CL\_4253

fGI ID


CL\_INS\_233
CL\_INS\_233
CL\_INS\_233
CL\_INS\_233
CL\_INS\_233
CL\_INS\_233
Cluster ID


CL\_5096
CL\_5097
CL\_5098
CL\_5099
CL\_5100
CL\_5101
