## Supplementary material for "A novel method for integrating genomic and Tn-Seq data to identify common *in vivo* fitness mechanisms across multiple bacterial species": S3 Dataset: CL_INS_236.html

Legend

 Mobile +extrachromosomalelementfunctions
 Hypothetical
 All EssentialGenes
 Other
 All VFDB Genes

FULL


WINDOWSVGPNG

Trim RowsRemove SingletonsSave Fasta

CL\_4275


CL\_4275


CL\_4275

HighlightSelectShow Genomes


55

CL\_4276


1

CL\_4276


1

CL\_4276

fGI ID


CL\_INS\_236
CL\_INS\_236
CL\_INS\_236
CL\_INS\_117
CL\_INS\_236
CL\_INS\_117
CL\_INS\_117
CL\_INS\_117
CL\_INS\_117
CL\_INS\_117
Cluster ID


CL\_6897
CL\_6898
CL\_6899
CL\_6900
CL\_6901
CL\_6902
CL\_6903
CL\_6904
CL\_6905
CL\_6906
