## Supplementary material for "A novel method for integrating genomic and Tn-Seq data to identify common *in vivo* fitness mechanisms across multiple bacterial species": S3 Dataset: CL_INS_237.html

Legend

 Mobile +extrachromosomalelementfunctions
 Regulatoryfunctions
 Hypothetical
 DNA Metabolism
 All EssentialGenes
 Other
 Transport +binding proteins
 All VFDB Genes

FULL


WINDOWSVGPNG

Trim RowsRemove SingletonsSave Fasta

CL\_4308


CL\_4308


CL\_4308


CL\_4308


CL\_4308


CL\_4308


CL\_4308


CL\_4308


CL\_4308


CL\_4308


CL\_4308


CL\_4308


CL\_28


CL\_522


CL\_4308


CL\_4308


CL\_4308

HighlightSelectShow Genomes


31

CL\_4309


7

CL\_4309


5

CL\_4309


2

CL\_4309


1

CL\_4310


1

CL\_4309


1

CL\_4309


1

CL\_4309


1

CL\_4333


1

CL\_4309


1

CL\_512


1

CL\_4309


1

CL\_4309


1

CL\_4309


1

CL\_4315


1

CL\_4309


1

CL\_4309

fGI ID


CL\_INS\_237
CL\_INS\_237
CL\_INS\_237
CL\_INS\_237
CL\_INS\_237
CL\_INS\_237
CL\_INS\_237
CL\_INS\_237
CL\_INS\_237
CL\_INS\_237
CL\_INS\_53
CL\_INS\_237
CL\_INS\_237
CL\_INS\_237
CL\_INS\_237
CL\_INS\_237
CL\_INS\_1
CL\_INS\_1
CL\_INS\_1
CL\_INS\_1
CL\_INS\_1
CL\_INS\_1
CL\_INS\_1
CL\_INS\_1
CL\_INS\_1
CL\_INS\_237
CL\_INS\_237
CL\_INS\_237
CL\_INS\_237
CL\_INS\_237
CL\_INS\_237
CL\_INS\_237
CL\_INS\_237
CL\_INS\_237
CL\_INS\_237
CL\_INS\_237
CL\_INS\_237
CL\_INS\_237
CL\_INS\_237
CL\_INS\_237
CL\_INS\_237
CL\_INS\_237
CL\_INS\_237
CL\_INS\_237
CL\_INS\_237
CL\_INS\_237
CL\_INS\_237
CL\_INS\_237
CL\_INS\_237
CL\_INS\_237
CL\_INS\_237
CL\_INS\_237
CL\_INS\_237
CL\_INS\_237
CL\_INS\_237
CL\_INS\_237
CL\_INS\_237
CL\_INS\_237
CL\_INS\_237
CL\_INS\_237
CL\_INS\_237
CL\_INS\_237
CL\_INS\_237
CL\_INS\_237
CL\_INS\_237
CL\_INS\_237
CL\_INS\_53
CL\_INS\_53
CL\_INS\_237
CL\_INS\_53
CL\_INS\_53
CL\_INS\_53
CL\_INS\_53
CL\_INS\_53
CL\_INS\_53
CL\_INS\_53
CL\_INS\_53
CL\_INS\_53
CL\_INS\_53
CL\_INS\_53
CL\_INS\_53
CL\_INS\_53
CL\_INS\_237
CL\_INS\_237
CL\_INS\_237
CL\_INS\_237
CL\_INS\_237
CL\_INS\_1
CL\_INS\_1
CL\_INS\_1
CL\_INS\_1
CL\_INS\_1
CL\_INS\_1
CL\_INS\_1
CL\_INS\_1
CL\_INS\_1
CL\_INS\_83
CL\_INS\_1
CL\_INS\_1
CL\_INS\_1
CL\_INS\_1
CL\_INS\_1
CL\_INS\_1
CL\_INS\_1
CL\_INS\_1
CL\_INS\_1
CL\_INS\_1
CL\_INS\_1
CL\_INS\_1
CL\_INS\_1
CL\_INS\_1
CL\_INS\_1
CL\_INS\_1
CL\_INS\_1
CL\_INS\_1
CL\_INS\_1
CL\_INS\_1
CL\_INS\_1
CL\_INS\_1
CL\_INS\_1
CL\_INS\_1
CL\_INS\_1
CL\_INS\_1
CL\_INS\_1
CL\_INS\_1
CL\_INS\_1
CL\_INS\_1
CL\_INS\_1
CL\_INS\_1
CL\_INS\_1
CL\_INS\_1
CL\_INS\_1
CL\_INS\_1
CL\_INS\_83
CL\_INS\_83
CL\_INS\_1
CL\_INS\_1
CL\_INS\_237
CL\_INS\_237
CL\_INS\_237
CL\_INS\_237
CL\_INS\_237
CL\_INS\_237
CL\_INS\_237
CL\_INS\_237
CL\_INS\_237
CL\_INS\_237
CL\_INS\_237
CL\_INS\_237
CL\_INS\_237
CL\_INS\_237
CL\_INS\_237
CL\_INS\_237
CL\_INS\_237
CL\_INS\_237
CL\_INS\_237
CL\_INS\_237
CL\_INS\_237
CL\_INS\_237
CL\_INS\_237
CL\_INS\_237
CL\_INS\_237
CL\_INS\_237
CL\_INS\_83
CL\_INS\_83
CL\_INS\_83
CL\_INS\_83
CL\_INS\_83
CL\_INS\_83
CL\_INS\_237
CL\_INS\_237
CL\_INS\_237
CL\_INS\_237
CL\_INS\_237
CL\_INS\_237
CL\_INS\_237
CL\_INS\_237
CL\_INS\_237
CL\_INS\_133
CL\_INS\_237
CL\_INS\_237
CL\_INS\_237
CL\_INS\_237
CL\_INS\_237
CL\_INS\_237
CL\_INS\_237
CL\_INS\_237
CL\_INS\_237
CL\_INS\_1
CL\_INS\_1
CL\_INS\_1
CL\_INS\_1
CL\_INS\_1
CL\_INS\_1
CL\_INS\_1
CL\_INS\_1
CL\_INS\_237
CL\_INS\_237
CL\_INS\_237
CL\_INS\_237
CL\_INS\_237
CL\_INS\_237
CL\_INS\_237
CL\_INS\_237
CL\_INS\_237
CL\_INS\_237
CL\_INS\_237
CL\_INS\_83
CL\_INS\_237
CL\_INS\_237
CL\_INS\_237
CL\_INS\_237
CL\_INS\_237
CL\_INS\_237
CL\_INS\_237
CL\_INS\_237
CL\_INS\_237
CL\_INS\_237
CL\_INS\_1
CL\_INS\_1
CL\_INS\_1
CL\_INS\_1
CL\_INS\_1
CL\_INS\_1
CL\_INS\_1
CL\_INS\_1
CL\_INS\_1
CL\_INS\_1
CL\_INS\_1
CL\_INS\_1
CL\_INS\_1
CL\_INS\_1
CL\_INS\_1
CL\_INS\_1
CL\_INS\_237
CL\_INS\_237
CL\_INS\_237
CL\_INS\_237
CL\_INS\_237
CL\_INS\_237
CL\_INS\_237
CL\_INS\_237
CL\_INS\_237
CL\_INS\_237
CL\_INS\_237
CL\_INS\_237
CL\_INS\_237
CL\_INS\_237
CL\_INS\_237
CL\_INS\_237
CL\_INS\_237
CL\_INS\_237
CL\_INS\_237
CL\_INS\_237
CL\_INS\_237
CL\_INS\_237
CL\_INS\_237
CL\_INS\_237
CL\_INS\_237
CL\_INS\_237
CL\_INS\_237
CL\_INS\_237
CL\_INS\_237
CL\_INS\_237
CL\_INS\_237
CL\_INS\_237
CL\_INS\_237
CL\_INS\_237
CL\_INS\_237
CL\_INS\_237
CL\_INS\_237
CL\_INS\_237
CL\_INS\_237
CL\_INS\_237
CL\_INS\_237
CL\_INS\_237
CL\_INS\_237
CL\_INS\_237
CL\_INS\_237
CL\_INS\_237
CL\_INS\_237
CL\_INS\_237
CL\_INS\_237
CL\_INS\_237
CL\_INS\_237
CL\_INS\_237
CL\_INS\_237
CL\_INS\_237
CL\_INS\_237
CL\_INS\_237
CL\_INS\_237
CL\_INS\_237
CL\_INS\_237
CL\_INS\_237
CL\_INS\_237
CL\_INS\_237
CL\_INS\_237
CL\_INS\_237
CL\_INS\_237
CL\_INS\_237
CL\_INS\_237
CL\_INS\_237
CL\_INS\_237
CL\_INS\_237
CL\_INS\_237
CL\_INS\_237
CL\_INS\_237
CL\_INS\_237
CL\_INS\_237
CL\_INS\_237
CL\_INS\_237
CL\_INS\_237
CL\_INS\_237
Cluster ID


CL\_13561
CL\_13560
CL\_13559
CL\_13558
CL\_13557
CL\_1289
CL\_1288
CL\_1287
CL\_13693
CL\_6542
CL\_6543
CL\_8157
CL\_8156
CL\_8155
CL\_8154
CL\_10025
CL\_10026
CL\_10027
CL\_10028
CL\_10029
CL\_10030
CL\_10031
CL\_10032
CL\_10033
CL\_10034
CL\_11861
CL\_11860
CL\_8153
CL\_8152
CL\_8151
CL\_8150
CL\_8149
CL\_11859
CL\_8148
CL\_8147
CL\_8146
CL\_5000
CL\_4999
CL\_4998
CL\_8608
CL\_8145
CL\_8144
CL\_8143
CL\_8142
CL\_8141
CL\_8140
CL\_8139
CL\_8138
CL\_8137
CL\_8136
CL\_11858
CL\_11857
CL\_11856
CL\_11855
CL\_11854
CL\_11853
CL\_11852
CL\_11851
CL\_11850
CL\_11849
CL\_11848
CL\_11847
CL\_11846
CL\_11845
CL\_11844
CL\_11843
CL\_6544
CL\_6545
CL\_13694
CL\_6546
CL\_6547
CL\_6548
CL\_6549
CL\_6550
CL\_6551
CL\_6552
CL\_6553
CL\_6554
CL\_6555
CL\_6556
CL\_6557
CL\_6558
CL\_6559
CL\_6560
CL\_6561
CL\_11842
CL\_9504
CL\_26
CL\_25
CL\_5115
CL\_5114
CL\_5113
CL\_5112
CL\_5111
CL\_5110
CL\_5109
CL\_4632
CL\_23
CL\_5107
CL\_5106
CL\_9805
CL\_5105
CL\_10064
CL\_10063
CL\_10062
CL\_10061
CL\_10060
CL\_10059
CL\_10058
CL\_10057
CL\_10056
CL\_10055
CL\_10054
CL\_10053
CL\_10052
CL\_10051
CL\_10050
CL\_10049
CL\_10048
CL\_10047
CL\_10046
CL\_10045
CL\_10044
CL\_10043
CL\_10042
CL\_10041
CL\_10040
CL\_3936
CL\_3937
CL\_3938
CL\_3939
CL\_6155
CL\_3940
CL\_3941
CL\_3942
CL\_89
CL\_10039
CL\_10038
CL\_11150
CL\_5708
CL\_11135
CL\_11136
CL\_11137
CL\_11138
CL\_11139
CL\_11140
CL\_11141
CL\_8936
CL\_6100
CL\_5709
CL\_5710
CL\_10606
CL\_10605
CL\_6099
CL\_6098
CL\_11142
CL\_11143
CL\_11144
CL\_11145
CL\_11146
CL\_11147
CL\_11148
CL\_11149
CL\_503
CL\_504
CL\_505
CL\_506
CL\_507
CL\_508
CL\_509
CL\_510
CL\_6097
CL\_6096
CL\_6095
CL\_6094
CL\_6093
CL\_5711
CL\_5712
CL\_5202
CL\_5713
CL\_5714
CL\_5715
CL\_5716
CL\_5717
CL\_5718
CL\_5719
CL\_5720
CL\_5721
CL\_5722
CL\_5261
CL\_5260
CL\_5259
CL\_4601
CL\_4602
CL\_4603
CL\_4604
CL\_5723
CL\_5724
CL\_10604
CL\_12385
CL\_12386
CL\_12387
CL\_5725
CL\_5726
CL\_6071
CL\_12388
CL\_5727
CL\_4694
CL\_5728
CL\_5729
CL\_5730
CL\_5731
CL\_5732
CL\_5733
CL\_5734
CL\_5735
CL\_5736
CL\_5737
CL\_5738
CL\_10035
CL\_10036
CL\_10037
CL\_2
CL\_1
CL\_5739
CL\_5740
CL\_5741
CL\_5742
CL\_5743
CL\_5744
CL\_5745
CL\_5746
CL\_5747
CL\_5748
CL\_5749
CL\_5750
CL\_5751
CL\_5752
CL\_5753
CL\_5162
CL\_5754
CL\_5755
CL\_5756
CL\_11151
CL\_5757
CL\_5758
CL\_5759
CL\_5760
CL\_11152
CL\_11153
CL\_11154
CL\_11155
CL\_12389
CL\_12390
CL\_12391
CL\_12392
CL\_12393
CL\_12394
CL\_5761
CL\_5762
CL\_5763
CL\_5764
CL\_11156
CL\_5765
CL\_5766
CL\_5767
CL\_5768
CL\_5769
CL\_5770
CL\_11157
CL\_5771
CL\_5772
CL\_5773
CL\_5774
CL\_5775
CL\_5776
CL\_5777
CL\_5778
CL\_5779
CL\_5780
CL\_11158
CL\_11159
CL\_11160
CL\_11161
CL\_11162
CL\_11163
CL\_11164
CL\_11165
CL\_11166
CL\_11167
CL\_11168
CL\_11169
CL\_5781
CL\_5782
CL\_5783
CL\_5784
CL\_5785
CL\_11170
CL\_5786
CL\_5787
CL\_5788
CL\_5789
CL\_8937
CL\_5790
CL\_11171
CL\_5791
CL\_5792
CL\_5793
CL\_11172
CL\_5794
CL\_5795
CL\_5796
CL\_5102
