## Supplementary material for "A novel method for integrating genomic and Tn-Seq data to identify common *in vivo* fitness mechanisms across multiple bacterial species": S4 Dataset: CL_INS_1.html

Legend

 Other
 Regulatoryfunctions
 All VFDB Genes
 Transport +binding proteins

FULL


WINDOWSVGPNG

Trim RowsRemove SingletonsSave Fasta

CL\_11


CL\_11


CL\_11

HighlightSelectShow Genomes


16

CL\_12


6

CL\_12


6

CL\_12

fGI ID


CL\_INS\_1
CL\_INS\_1
CL\_INS\_1
CL\_INS\_1
CL\_INS\_1
CL\_INS\_1
CL\_INS\_1
CL\_INS\_1
CL\_INS\_1
CL\_INS\_1
Cluster ID


CL\_9199
CL\_9200
CL\_9201
CL\_5150
CL\_5151
CL\_5152
CL\_5153
CL\_5154
CL\_5155
CL\_5156
