## Supplementary material for "A novel method for integrating genomic and Tn-Seq data to identify common *in vivo* fitness mechanisms across multiple bacterial species": S4 Dataset: CL_INS_2.html

Legend

 Hypothetical
 Cellularprocesses

FULL


WINDOWSVGPNG

Trim RowsRemove SingletonsSave Fasta

CL\_21


CL\_21


CL\_21

HighlightSelectShow Genomes


15

CL\_22


7

CL\_22


2

CL\_22

fGI ID


CL\_INS\_2
CL\_INS\_2
Cluster ID


CL\_6504
CL\_12618
