## Supplementary material for "A novel method for integrating genomic and Tn-Seq data to identify common *in vivo* fitness mechanisms across multiple bacterial species": S4 Dataset: CL_INS_3.html

Legend

 Other
 Hypothetical

FULL


WINDOWSVGPNG

Trim RowsRemove SingletonsSave Fasta

CL\_33


CL\_33


CL\_33


CL\_33

HighlightSelectShow Genomes


14

CL\_36


11

CL\_36


2

CL\_36


1

CL\_36

fGI ID


CL\_INS\_3
CL\_INS\_3
CL\_INS\_3
CL\_INS\_3
CL\_INS\_3
Cluster ID


CL\_12684
CL\_34
CL\_35
CL\_6503
CL\_6502
