## Supplementary material for "A novel method for integrating genomic and Tn-Seq data to identify common *in vivo* fitness mechanisms across multiple bacterial species": S4 Dataset: CL_INS_4.html

Legend

 Mobile +extrachromosomalelementfunctions
 Regulatoryfunctions
 Hypothetical
 DNA Metabolism
 Biosynthesis ofcofactors,prostheticgroups, +carriers
 Other
 All VFDB Genes
 Transport +binding proteins

FULL


WINDOWSVGPNG

Trim RowsRemove SingletonsSave Fasta

CL\_52


CL\_52


CL\_52

HighlightSelectShow Genomes


21

CL\_53


6

CL\_53


1

CL\_74

fGI ID


CL\_INS\_4
CL\_INS\_4
CL\_INS\_4
CL\_INS\_4
CL\_INS\_4
CL\_INS\_4
CL\_INS\_4
CL\_INS\_6
CL\_INS\_6
CL\_INS\_365
CL\_INS\_6
CL\_INS\_6
CL\_INS\_365
CL\_INS\_365
CL\_INS\_365
CL\_INS\_365
CL\_INS\_365
CL\_INS\_365
CL\_INS\_365
CL\_INS\_365
CL\_INS\_6
CL\_INS\_6
CL\_INS\_6
CL\_INS\_6
Cluster ID


CL\_9202
CL\_9203
CL\_9204
CL\_9205
CL\_9206
CL\_9207
CL\_10785
CL\_7859
CL\_11022
CL\_10137
CL\_6641
CL\_6640
CL\_8378
CL\_8379
CL\_8380
CL\_7680
CL\_2880
CL\_2881
CL\_2882
CL\_2883
CL\_11023
CL\_11024
CL\_11025
CL\_11026
