## Supplementary material for "A novel method for integrating genomic and Tn-Seq data to identify common *in vivo* fitness mechanisms across multiple bacterial species": S4 Dataset: CL_INS_5.html

Legend

 Mobile +extrachromosomalelementfunctions
 Regulatoryfunctions
 Hypothetical
 Transcription
 Other
 Transport +binding proteins
 All VFDB Genes

FULL


WINDOWSVGPNG

Trim RowsRemove SingletonsSave Fasta

CL\_61


CL\_61


CL\_61


CL\_61


CL\_4566


CL\_61

HighlightSelectShow Genomes


20

CL\_62


3

CL\_74


2

CL\_74


1

CL\_4567


1

CL\_62


1

CL\_74

fGI ID


CL\_INS\_5
CL\_INS\_5
CL\_INS\_5
CL\_INS\_5
CL\_INS\_5
CL\_INS\_5
CL\_INS\_5
CL\_INS\_5
CL\_INS\_5
CL\_INS\_5
CL\_INS\_5
CL\_INS\_5
CL\_INS\_5
CL\_INS\_5
CL\_INS\_5
CL\_INS\_365
CL\_INS\_5
CL\_INS\_5
CL\_INS\_5
CL\_INS\_5
CL\_INS\_5
CL\_INS\_5
CL\_INS\_270
CL\_INS\_270
CL\_INS\_270
CL\_INS\_270
CL\_INS\_5
CL\_INS\_5
CL\_INS\_5
CL\_INS\_270
CL\_INS\_5
CL\_INS\_5
Cluster ID


CL\_5157
CL\_5158
CL\_5159
CL\_5160
CL\_5682
CL\_5681
CL\_5680
CL\_5679
CL\_5678
CL\_5677
CL\_5676
CL\_5675
CL\_5674
CL\_5673
CL\_5672
CL\_5671
CL\_5670
CL\_6501
CL\_6500
CL\_9197
CL\_9196
CL\_9195
CL\_1202
CL\_1203
CL\_66
CL\_67
CL\_13671
CL\_1209
CL\_1210
CL\_1211
CL\_9194
CL\_9193
