## Supplementary material for "A novel method for integrating genomic and Tn-Seq data to identify common *in vivo* fitness mechanisms across multiple bacterial species": S4 Dataset: CL_INS_12.html

Legend

 Hypothetical
 All Fitness Genes
 All VFDB Genes

FULL


WINDOWSVGPNG

Trim RowsRemove SingletonsSave Fasta

CL\_104


CL\_104


CL\_104

HighlightSelectShow Genomes


14

CL\_105


10

CL\_105


4

CL\_105

fGI ID


CL\_INS\_12
CL\_INS\_12
Cluster ID


CL\_5192
CL\_6496
