## Supplementary material for "A novel method for integrating genomic and Tn-Seq data to identify common *in vivo* fitness mechanisms across multiple bacterial species": S4 Dataset: CL_INS_15.html

Legend

 Mobile +extrachromosomalelementfunctions
 Regulatoryfunctions
 Hypothetical
 Other
 Transport +binding proteins
 All VFDB Genes

FULL


WINDOWSVGPNG

Trim RowsRemove SingletonsSave Fasta

CL\_136


CL\_136


CL\_136

HighlightSelectShow Genomes


15

CL\_137


12

CL\_137


1

CL\_137

fGI ID


CL\_INS\_15
CL\_INS\_15
CL\_INS\_15
CL\_INS\_15
CL\_INS\_15
CL\_INS\_15
CL\_INS\_15
CL\_INS\_15
CL\_INS\_181
CL\_INS\_181
CL\_INS\_181
CL\_INS\_181
CL\_INS\_181
CL\_INS\_181
CL\_INS\_181
CL\_INS\_181
CL\_INS\_15
CL\_INS\_365
CL\_INS\_15
CL\_INS\_15
CL\_INS\_15
CL\_INS\_15
CL\_INS\_15
CL\_INS\_15
CL\_INS\_15
CL\_INS\_15
CL\_INS\_15
CL\_INS\_15
CL\_INS\_15
CL\_INS\_15
CL\_INS\_15
CL\_INS\_15
CL\_INS\_15
CL\_INS\_365
CL\_INS\_365
CL\_INS\_365
CL\_INS\_15
CL\_INS\_132
CL\_INS\_132
CL\_INS\_365
CL\_INS\_365
CL\_INS\_365
CL\_INS\_365
CL\_INS\_365
CL\_INS\_365
CL\_INS\_365
CL\_INS\_365
CL\_INS\_365
CL\_INS\_15
CL\_INS\_365
CL\_INS\_15
CL\_INS\_15
CL\_INS\_15
Cluster ID


CL\_6495
CL\_12354
CL\_12353
CL\_12352
CL\_12351
CL\_12350
CL\_12349
CL\_6606
CL\_8202
CL\_8203
CL\_8204
CL\_8205
CL\_7698
CL\_7697
CL\_7696
CL\_7695
CL\_7694
CL\_5020
CL\_6605
CL\_6604
CL\_6603
CL\_12348
CL\_7683
CL\_7684
CL\_7685
CL\_7686
CL\_7687
CL\_7688
CL\_7689
CL\_7690
CL\_7691
CL\_7692
CL\_7693
CL\_875
CL\_874
CL\_873
CL\_12347
CL\_5138
CL\_5137
CL\_5136
CL\_5847
CL\_7882
CL\_7883
CL\_7884
CL\_7885
CL\_8373
CL\_7886
CL\_5022
CL\_5128
CL\_5764
CL\_10774
CL\_10773
CL\_12346
