## Supplementary material for "A novel method for integrating genomic and Tn-Seq data to identify common *in vivo* fitness mechanisms across multiple bacterial species": S4 Dataset: CL_INS_16.html

Legend

 Hypothetical
 Other
 All VFDB Genes

FULL


WINDOWSVGPNG

Trim RowsRemove SingletonsSave Fasta

CL\_138


CL\_138


CL\_138


CL\_138


CL\_138

HighlightSelectShow Genomes


19

CL\_139


4

CL\_139


3

CL\_139


1

CL\_139


1

CL\_139

fGI ID


CL\_INS\_16
CL\_INS\_16
CL\_INS\_16
CL\_INS\_16
CL\_INS\_16
CL\_INS\_16
CL\_INS\_16
CL\_INS\_16
CL\_INS\_16
CL\_INS\_16
Cluster ID


CL\_9208
CL\_9209
CL\_11027
CL\_12617
CL\_12616
CL\_11028
CL\_11029
CL\_11030
CL\_11031
CL\_9210
