## Supplementary material for "A novel method for integrating genomic and Tn-Seq data to identify common *in vivo* fitness mechanisms across multiple bacterial species": S4 Dataset: CL_INS_19.html

Legend

 Hypothetical
 Other
 All VFDB Genes

FULL


WINDOWSVGPNG

Trim RowsRemove SingletonsSave Fasta

CL\_189


CL\_189


CL\_189

HighlightSelectShow Genomes


17

CL\_190


9

CL\_190


2

CL\_190

fGI ID


CL\_INS\_19
CL\_INS\_19
CL\_INS\_19
Cluster ID


CL\_13347
CL\_13348
CL\_6492
