## Supplementary material for "A novel method for integrating genomic and Tn-Seq data to identify common *in vivo* fitness mechanisms across multiple bacterial species": S4 Dataset: CL_INS_25.html

Legend

 Hypothetical

FULL


WINDOWSVGPNG

Trim RowsRemove SingletonsSave Fasta

CL\_287


CL\_287


CL\_287

HighlightSelectShow Genomes


18

CL\_288


8

CL\_288


2

CL\_288

fGI ID


CL\_INS\_25
CL\_INS\_25
Cluster ID


CL\_6489
CL\_11521
