## Supplementary material for "A novel method for integrating genomic and Tn-Seq data to identify common *in vivo* fitness mechanisms across multiple bacterial species": S4 Dataset: CL_INS_28.html

Legend

 Hypothetical
 Other
 All VFDB Genes

FULL


WINDOWSVGPNG

Trim RowsRemove SingletonsSave Fasta

CL\_300


CL\_300


CL\_300

HighlightSelectShow Genomes


23

CL\_301


3

CL\_305


2

CL\_305

fGI ID


CL\_INS\_28
CL\_INS\_29
CL\_INS\_29
CL\_INS\_29
Cluster ID


CL\_9211
CL\_11520
CL\_11519
CL\_11518
