## Supplementary material for "A novel method for integrating genomic and Tn-Seq data to identify common *in vivo* fitness mechanisms across multiple bacterial species": S4 Dataset: CL_INS_29.html

Legend

 Regulatoryfunctions
 Hypothetical
 All VFDB Genes

FULL


WINDOWSVGPNG

Trim RowsRemove SingletonsSave Fasta

CL\_305


CL\_305


CL\_305


CL\_305

HighlightSelectShow Genomes


22

CL\_304


3

CL\_300


2

CL\_300


1

CL\_303

fGI ID


CL\_INS\_29
CL\_INS\_29
CL\_INS\_29
CL\_INS\_29
Cluster ID


CL\_9212
CL\_11518
CL\_11519
CL\_11520
