## Supplementary material for "A novel method for integrating genomic and Tn-Seq data to identify common *in vivo* fitness mechanisms across multiple bacterial species": S4 Dataset: CL_INS_33.html

Legend

 Hypothetical
 EnergyMetabolism
 All VFDB Genes

FULL


WINDOWSVGPNG

Trim RowsRemove SingletonsSave Fasta

CL\_339


CL\_339

HighlightSelectShow Genomes


14

CL\_342


14

CL\_342

fGI ID


CL\_INS\_33
CL\_INS\_33
Cluster ID


CL\_340
CL\_341
