## Supplementary material for "A novel method for integrating genomic and Tn-Seq data to identify common *in vivo* fitness mechanisms across multiple bacterial species": S4 Dataset: CL_INS_34.html

Legend

 Hypothetical
 Other

FULL


WINDOWSVGPNG

Trim RowsRemove SingletonsSave Fasta

CL\_371


CL\_371


CL\_371

HighlightSelectShow Genomes


23

CL\_372


3

CL\_372


2

CL\_372

fGI ID


CL\_INS\_34
CL\_INS\_34
Cluster ID


CL\_6486
CL\_13235
