## Supplementary material for "A novel method for integrating genomic and Tn-Seq data to identify common *in vivo* fitness mechanisms across multiple bacterial species": S4 Dataset: CL_INS_36.html

Legend

 Mobile +extrachromosomalelementfunctions
 Regulatoryfunctions
 Hypothetical
 All Fitness Genes
 Other
 All VFDB Genes

FULL


WINDOWSVGPNG

Trim RowsRemove SingletonsSave Fasta

CL\_459


CL\_459


CL\_459


CL\_459


CL\_459


CL\_459


CL\_459


CL\_459

HighlightSelectShow Genomes


18

CL\_473


3

CL\_473


2

CL\_473


1

CL\_473


1

CL\_473


1

CL\_473


1

CL\_473


1

CL\_473

fGI ID


CL\_INS\_36
CL\_INS\_36
CL\_INS\_36
CL\_INS\_36
CL\_INS\_36
CL\_INS\_36
CL\_INS\_36
CL\_INS\_36
CL\_INS\_36
CL\_INS\_36
CL\_INS\_36
CL\_INS\_36
CL\_INS\_36
CL\_INS\_36
CL\_INS\_36
CL\_INS\_36
CL\_INS\_36
CL\_INS\_36
CL\_INS\_36
CL\_INS\_36
CL\_INS\_36
CL\_INS\_36
CL\_INS\_36
CL\_INS\_36
CL\_INS\_36
CL\_INS\_36
CL\_INS\_36
CL\_INS\_36
CL\_INS\_36
CL\_INS\_36
CL\_INS\_36
CL\_INS\_36
CL\_INS\_36
CL\_INS\_36
CL\_INS\_36
CL\_INS\_36
CL\_INS\_36
CL\_INS\_36
CL\_INS\_36
CL\_INS\_36
CL\_INS\_36
CL\_INS\_36
CL\_INS\_36
CL\_INS\_36
CL\_INS\_36
CL\_INS\_36
CL\_INS\_36
CL\_INS\_36
CL\_INS\_36
CL\_INS\_36
CL\_INS\_36
CL\_INS\_36
CL\_INS\_36
CL\_INS\_36
CL\_INS\_36
CL\_INS\_36
Cluster ID


CL\_460
CL\_10102
CL\_7988
CL\_461
CL\_13236
CL\_9186
CL\_7355
CL\_462
CL\_463
CL\_464
CL\_9185
CL\_7354
CL\_7353
CL\_465
CL\_13237
CL\_466
CL\_467
CL\_10103
CL\_468
CL\_469
CL\_470
CL\_471
CL\_472
CL\_13238
CL\_9184
CL\_9183
CL\_9182
CL\_9181
CL\_5186
CL\_5185
CL\_5184
CL\_5183
CL\_5182
CL\_5181
CL\_7352
CL\_7351
CL\_7350
CL\_7349
CL\_7348
CL\_7347
CL\_5180
CL\_9180
CL\_5179
CL\_5178
CL\_5177
CL\_5176
CL\_5175
CL\_7346
CL\_5173
CL\_5172
CL\_13239
CL\_13240
CL\_5171
CL\_10104
CL\_5169
CL\_7345
