## Supplementary material for "A novel method for integrating genomic and Tn-Seq data to identify common *in vivo* fitness mechanisms across multiple bacterial species": S4 Dataset: CL_INS_38.html

Legend

 Hypothetical
 All Fitness Genes
 Proteinsynthesis/fate

FULL


WINDOWSVGPNG

Trim RowsRemove SingletonsSave Fasta

CL\_505


CL\_505

HighlightSelectShow Genomes


18

CL\_506


10

CL\_506

fGI ID

CL\_INS\_38
Cluster ID

CL\_6482
