## Supplementary material for "A novel method for integrating genomic and Tn-Seq data to identify common *in vivo* fitness mechanisms across multiple bacterial species": S4 Dataset: CL_INS_39.html

Legend

 Hypothetical
 Other
 Centralintermediarymetabolism

FULL


WINDOWSVGPNG

Trim RowsRemove SingletonsSave Fasta

CL\_518


CL\_518

HighlightSelectShow Genomes


23

CL\_519


5

CL\_519

fGI ID

CL\_INS\_39
Cluster ID

CL\_7987
