## Supplementary material for "A novel method for integrating genomic and Tn-Seq data to identify common *in vivo* fitness mechanisms across multiple bacterial species": S4 Dataset: CL_INS_44.html

Legend

 Hypothetical
 Regulatoryfunctions
 Other
 Transport +binding proteins
 All VFDB Genes

FULL


WINDOWSVGPNG

Trim RowsRemove SingletonsSave Fasta

CL\_580


CL\_580


CL\_580


CL\_580


CL\_580

HighlightSelectShow Genomes


16

CL\_581


8

CL\_581


2

CL\_597


1

CL\_597


1

CL\_597

fGI ID


CL\_INS\_44
CL\_INS\_44
CL\_INS\_44
CL\_INS\_44
CL\_INS\_44
CL\_INS\_44
CL\_INS\_44
CL\_INS\_44
CL\_INS\_44
CL\_INS\_44
CL\_INS\_44
CL\_INS\_44
CL\_INS\_44
CL\_INS\_44
CL\_INS\_44
CL\_INS\_44
CL\_INS\_44
CL\_INS\_44
Cluster ID


CL\_9178
CL\_9177
CL\_9176
CL\_9175
CL\_9174
CL\_9216
CL\_9217
CL\_9218
CL\_9219
CL\_9220
CL\_9221
CL\_9222
CL\_9223
CL\_9224
CL\_9225
CL\_9226
CL\_9227
CL\_9228
