## Supplementary material for "A novel method for integrating genomic and Tn-Seq data to identify common *in vivo* fitness mechanisms across multiple bacterial species": S4 Dataset: CL_INS_45.html

Legend

 Regulatoryfunctions
 Hypothetical
 Other
 Transport +binding proteins
 All VFDB Genes

FULL


WINDOWSVGPNG

Trim RowsRemove SingletonsSave Fasta

CL\_597


CL\_597


CL\_597


CL\_597


CL\_597

HighlightSelectShow Genomes


20

CL\_596


2

CL\_580


1

CL\_580


1

CL\_580


1

CL\_596

fGI ID


CL\_INS\_45
CL\_INS\_45
CL\_INS\_44
CL\_INS\_44
CL\_INS\_44
CL\_INS\_44
CL\_INS\_44
CL\_INS\_44
CL\_INS\_44
CL\_INS\_44
CL\_INS\_44
CL\_INS\_44
CL\_INS\_44
CL\_INS\_44
CL\_INS\_44
Cluster ID


CL\_10105
CL\_9229
CL\_9228
CL\_9227
CL\_9226
CL\_9225
CL\_9224
CL\_9223
CL\_9222
CL\_9221
CL\_9220
CL\_9219
CL\_9218
CL\_9217
CL\_9216
