## Supplementary material for "A novel method for integrating genomic and Tn-Seq data to identify common *in vivo* fitness mechanisms across multiple bacterial species": S4 Dataset: CL_INS_52.html

Legend

 Biosynthesis ofcofactors,prostheticgroups, +carriers
 Cell Envelope
 Proteinsynthesis/fate
 All VFDB Genes

FULL


WINDOWSVGPNG

Trim RowsRemove SingletonsSave Fasta

CL\_733


CL\_733

HighlightSelectShow Genomes


19

CL\_734


9

CL\_734

fGI ID


CL\_INS\_52
CL\_INS\_52
CL\_INS\_52
CL\_INS\_52
Cluster ID


CL\_6525
CL\_6524
CL\_6523
CL\_6522
