## Supplementary material for "A novel method for integrating genomic and Tn-Seq data to identify common *in vivo* fitness mechanisms across multiple bacterial species": S4 Dataset: CL_INS_54.html

Legend

 Hypothetical
 All VFDB Genes

FULL


WINDOWSVGPNG

Trim RowsRemove SingletonsSave Fasta

CL\_753


CL\_753


CL\_753

HighlightSelectShow Genomes


19

CL\_754


8

CL\_754


1

CL\_754

fGI ID


CL\_INS\_54
CL\_INS\_54
CL\_INS\_54
Cluster ID


CL\_9231
CL\_9232
CL\_9233
