## Supplementary material for "A novel method for integrating genomic and Tn-Seq data to identify common *in vivo* fitness mechanisms across multiple bacterial species": S4 Dataset: CL_INS_58.html

Legend

 Hypothetical
 All VFDB Genes

FULL


WINDOWSVGPNG

Trim RowsRemove SingletonsSave Fasta

CL\_803


CL\_803

HighlightSelectShow Genomes


25

CL\_804


3

CL\_804

fGI ID


CL\_INS\_58
CL\_INS\_58
CL\_INS\_58
Cluster ID


CL\_13263
CL\_13264
CL\_13265
