## Supplementary material for "A novel method for integrating genomic and Tn-Seq data to identify common *in vivo* fitness mechanisms across multiple bacterial species": S4 Dataset: CL_INS_59.html

Legend

 Hypothetical
 Other
 All VFDB Genes

FULL


WINDOWSVGPNG

Trim RowsRemove SingletonsSave Fasta

CL\_813


CL\_813


CL\_813

HighlightSelectShow Genomes


17

CL\_814


10

CL\_814


1

CL\_814

fGI ID


CL\_INS\_59
CL\_INS\_59
Cluster ID


CL\_13580
CL\_7317
