## Supplementary material for "A novel method for integrating genomic and Tn-Seq data to identify common *in vivo* fitness mechanisms across multiple bacterial species": S4 Dataset: CL_INS_63.html

Legend

 Mobile +extrachromosomalelementfunctions
 Regulatoryfunctions
 Hypothetical
 Other
 All VFDB Genes

FULL


WINDOWSVGPNG

Trim RowsRemove SingletonsSave Fasta

CL\_824


CL\_824


CL\_824

HighlightSelectShow Genomes


24

CL\_876


3

CL\_876


1

CL\_876

fGI ID


CL\_INS\_63
CL\_INS\_63
CL\_INS\_63
CL\_INS\_63
CL\_INS\_63
CL\_INS\_63
CL\_INS\_63
CL\_INS\_63
CL\_INS\_63
CL\_INS\_63
CL\_INS\_63
CL\_INS\_63
CL\_INS\_63
CL\_INS\_63
CL\_INS\_63
CL\_INS\_63
CL\_INS\_63
CL\_INS\_63
CL\_INS\_63
CL\_INS\_63
CL\_INS\_365
CL\_INS\_365
CL\_INS\_63
CL\_INS\_365
CL\_INS\_365
CL\_INS\_365
CL\_INS\_365
CL\_INS\_365
CL\_INS\_365
CL\_INS\_365
CL\_INS\_365
CL\_INS\_365
CL\_INS\_365
CL\_INS\_365
CL\_INS\_365
CL\_INS\_365
CL\_INS\_365
CL\_INS\_365
CL\_INS\_365
CL\_INS\_63
CL\_INS\_63
CL\_INS\_63
CL\_INS\_63
CL\_INS\_365
CL\_INS\_365
CL\_INS\_365
CL\_INS\_365
CL\_INS\_365
CL\_INS\_365
CL\_INS\_365
CL\_INS\_365
CL\_INS\_365
CL\_INS\_365
CL\_INS\_365
CL\_INS\_365
CL\_INS\_365
CL\_INS\_365
Cluster ID


CL\_13266
CL\_13267
CL\_13268
CL\_13269
CL\_13270
CL\_13271
CL\_825
CL\_826
CL\_827
CL\_828
CL\_829
CL\_830
CL\_831
CL\_832
CL\_833
CL\_834
CL\_835
CL\_836
CL\_837
CL\_838
CL\_839
CL\_840
CL\_841
CL\_842
CL\_843
CL\_844
CL\_845
CL\_846
CL\_847
CL\_848
CL\_849
CL\_850
CL\_851
CL\_852
CL\_853
CL\_854
CL\_855
CL\_856
CL\_857
CL\_858
CL\_859
CL\_860
CL\_861
CL\_862
CL\_863
CL\_864
CL\_865
CL\_866
CL\_867
CL\_868
CL\_869
CL\_870
CL\_871
CL\_872
CL\_873
CL\_874
CL\_875
