## Supplementary material for "A novel method for integrating genomic and Tn-Seq data to identify common *in vivo* fitness mechanisms across multiple bacterial species": S4 Dataset: CL_INS_65.html

Legend

 Hypothetical
 Other
 All VFDB Genes

FULL


WINDOWSVGPNG

Trim RowsRemove SingletonsSave Fasta

CL\_897


CL\_897


CL\_897


CL\_897


CL\_897


CL\_897

HighlightSelectShow Genomes


19

CL\_898


2

CL\_898


1

CL\_898


1

CL\_898


1

CL\_898


1

CL\_898

fGI ID


CL\_INS\_65
CL\_INS\_65
CL\_INS\_16
CL\_INS\_65
CL\_INS\_65
CL\_INS\_16
CL\_INS\_65
CL\_INS\_65
CL\_INS\_65
CL\_INS\_65
Cluster ID


CL\_11512
CL\_11511
CL\_11028
CL\_11510
CL\_11509
CL\_11027
CL\_12614
CL\_11508
CL\_11507
CL\_11506
