## Supplementary material for "A novel method for integrating genomic and Tn-Seq data to identify common *in vivo* fitness mechanisms across multiple bacterial species": S4 Dataset: CL_INS_70.html

Legend

 Mobile +extrachromosomalelementfunctions
 Hypothetical
 Cell Envelope
 Other
 All VFDB Genes

FULL


WINDOWSVGPNG

Trim RowsRemove SingletonsSave Fasta

CL\_981


CL\_981


CL\_981


CL\_980

HighlightSelectShow Genomes


15

CL\_993


11

CL\_993


1

CL\_993


1

CL\_993

fGI ID


CL\_INS\_70
CL\_INS\_70
CL\_INS\_70
CL\_INS\_70
CL\_INS\_70
CL\_INS\_70
CL\_INS\_70
CL\_INS\_70
CL\_INS\_70
CL\_INS\_70
CL\_INS\_70
Cluster ID


CL\_982
CL\_983
CL\_984
CL\_985
CL\_986
CL\_987
CL\_988
CL\_989
CL\_990
CL\_991
CL\_992
