## Supplementary material for "A novel method for integrating genomic and Tn-Seq data to identify common *in vivo* fitness mechanisms across multiple bacterial species": S4 Dataset: CL_INS_71.html

Legend

 Mobile +extrachromosomalelementfunctions
 Hypothetical
 Other
 All VFDB Genes

FULL


WINDOWSVGPNG

Trim RowsRemove SingletonsSave Fasta

CL\_1020


CL\_1020


CL\_1020


CL\_1020


CL\_1020


CL\_1020

HighlightSelectShow Genomes


21

CL\_1021


2

CL\_1021


2

CL\_1021


1

CL\_1022


1

CL\_1021


1

CL\_1021

fGI ID


CL\_INS\_71
CL\_INS\_71
CL\_INS\_71
CL\_INS\_71
CL\_INS\_71
CL\_INS\_71
CL\_INS\_71
CL\_INS\_71
CL\_INS\_71
CL\_INS\_71
CL\_INS\_71
CL\_INS\_71
CL\_INS\_71
CL\_INS\_71
CL\_INS\_71
CL\_INS\_71
CL\_INS\_71
CL\_INS\_71
CL\_INS\_71
CL\_INS\_71
CL\_INS\_71
CL\_INS\_71
CL\_INS\_71
CL\_INS\_71
CL\_INS\_71
CL\_INS\_71
CL\_INS\_71
CL\_INS\_71
CL\_INS\_71
CL\_INS\_71
CL\_INS\_71
CL\_INS\_71
CL\_INS\_71
CL\_INS\_71
CL\_INS\_71
CL\_INS\_71
CL\_INS\_71
CL\_INS\_71
CL\_INS\_71
CL\_INS\_71
CL\_INS\_71
CL\_INS\_71
CL\_INS\_71
CL\_INS\_71
CL\_INS\_71
CL\_INS\_71
CL\_INS\_71
CL\_INS\_71
CL\_INS\_71
CL\_INS\_71
CL\_INS\_71
CL\_INS\_71
CL\_INS\_71
CL\_INS\_71
Cluster ID


CL\_5228
CL\_12293
CL\_13274
CL\_12292
CL\_12291
CL\_12290
CL\_6322
CL\_6321
CL\_6320
CL\_6319
CL\_6318
CL\_6317
CL\_6316
CL\_6315
CL\_6314
CL\_6313
CL\_6312
CL\_6311
CL\_6310
CL\_6309
CL\_6308
CL\_6307
CL\_6306
CL\_6305
CL\_6304
CL\_6303
CL\_6302
CL\_6301
CL\_6300
CL\_6299
CL\_6298
CL\_6297
CL\_6296
CL\_6295
CL\_6294
CL\_6293
CL\_6292
CL\_6291
CL\_6290
CL\_6289
CL\_6288
CL\_6287
CL\_6286
CL\_6285
CL\_6284
CL\_6283
CL\_6282
CL\_6281
CL\_6280
CL\_6279
CL\_6278
CL\_6277
CL\_6276
CL\_6275
