## Supplementary material for "A novel method for integrating genomic and Tn-Seq data to identify common *in vivo* fitness mechanisms across multiple bacterial species": S4 Dataset: CL_INS_74.html

Legend

 Mobile +extrachromosomalelementfunctions
 Regulatoryfunctions
 Hypothetical
 DNA Metabolism
 Other
 All VFDB Genes
 Transport +binding proteins

FULL


WINDOWSVGPNG

Trim RowsRemove SingletonsSave Fasta

CL\_1057


CL\_1057


CL\_1057


CL\_1057


CL\_1057


CL\_1057


CL\_1057


CL\_1057


CL\_1057

HighlightSelectShow Genomes


12

CL\_1068


7

CL\_1068


2

CL\_1068


2

CL\_1068


1

CL\_1068


1

CL\_1068


1

CL\_1068


1

CL\_1068


1

CL\_1068

fGI ID


CL\_INS\_74
CL\_INS\_74
CL\_INS\_74
CL\_INS\_74
CL\_INS\_74
CL\_INS\_74
CL\_INS\_74
CL\_INS\_74
CL\_INS\_74
CL\_INS\_74
CL\_INS\_74
CL\_INS\_74
CL\_INS\_74
CL\_INS\_270
CL\_INS\_270
CL\_INS\_270
CL\_INS\_270
CL\_INS\_74
CL\_INS\_74
CL\_INS\_74
CL\_INS\_74
CL\_INS\_74
CL\_INS\_74
CL\_INS\_74
CL\_INS\_74
CL\_INS\_74
CL\_INS\_74
CL\_INS\_74
CL\_INS\_74
CL\_INS\_74
CL\_INS\_74
CL\_INS\_74
CL\_INS\_74
CL\_INS\_74
CL\_INS\_74
CL\_INS\_74
CL\_INS\_74
CL\_INS\_74
CL\_INS\_74
CL\_INS\_74
CL\_INS\_74
CL\_INS\_74
CL\_INS\_74
CL\_INS\_74
CL\_INS\_74
CL\_INS\_74
CL\_INS\_74
CL\_INS\_74
CL\_INS\_74
CL\_INS\_74
CL\_INS\_74
CL\_INS\_74
CL\_INS\_74
CL\_INS\_74
CL\_INS\_74
CL\_INS\_74
CL\_INS\_74
CL\_INS\_74
CL\_INS\_74
CL\_INS\_74
CL\_INS\_74
CL\_INS\_74
CL\_INS\_74
CL\_INS\_74
CL\_INS\_74
CL\_INS\_74
CL\_INS\_74
CL\_INS\_74
CL\_INS\_74
CL\_INS\_74
CL\_INS\_74
CL\_INS\_74
CL\_INS\_74
CL\_INS\_74
Cluster ID


CL\_12715
CL\_12716
CL\_12717
CL\_11497
CL\_12718
CL\_12719
CL\_5250
CL\_5251
CL\_11495
CL\_12720
CL\_12721
CL\_13275
CL\_13276
CL\_8340
CL\_8341
CL\_8342
CL\_8343
CL\_13277
CL\_13278
CL\_13279
CL\_9246
CL\_9247
CL\_9248
CL\_12722
CL\_10402
CL\_10403
CL\_10404
CL\_10405
CL\_10406
CL\_10407
CL\_10408
CL\_10409
CL\_10410
CL\_10411
CL\_10412
CL\_10413
CL\_10414
CL\_10415
CL\_10416
CL\_10417
CL\_10418
CL\_10419
CL\_10420
CL\_7267
CL\_7266
CL\_7265
CL\_7264
CL\_7263
CL\_7262
CL\_7261
CL\_7260
CL\_7259
CL\_7258
CL\_7257
CL\_7256
CL\_7255
CL\_7254
CL\_7253
CL\_7252
CL\_7251
CL\_7250
CL\_7249
CL\_7248
CL\_1058
CL\_1059
CL\_1060
CL\_5229
CL\_1061
CL\_1062
CL\_1063
CL\_1064
CL\_1065
CL\_1066
CL\_1067
