## Supplementary material for "A novel method for integrating genomic and Tn-Seq data to identify common *in vivo* fitness mechanisms across multiple bacterial species": S4 Dataset: CL_INS_80.html

Legend

 Hypothetical
 Other
 All VFDB Genes

FULL


WINDOWSVGPNG

Trim RowsRemove SingletonsSave Fasta

CL\_1150


CL\_1150

HighlightSelectShow Genomes


21

CL\_1149


6

CL\_1143

fGI ID


CL\_INS\_80
CL\_INS\_79
CL\_INS\_79
CL\_INS\_79
Cluster ID


CL\_9253
CL\_9252
CL\_9251
CL\_9250
