## Supplementary material for "A novel method for integrating genomic and Tn-Seq data to identify common *in vivo* fitness mechanisms across multiple bacterial species": S4 Dataset: CL_INS_81.html

Legend

 Mobile +extrachromosomalelementfunctions
 Hypothetical
 All VFDB Genes

FULL


WINDOWSVGPNG

Trim RowsRemove SingletonsSave Fasta

CL\_1158


CL\_1158


CL\_1158


CL\_1158

HighlightSelectShow Genomes


11

CL\_1159


9

CL\_1159


3

CL\_1159


1

CL\_1159

fGI ID


CL\_INS\_81
CL\_INS\_81
Cluster ID


CL\_5237
CL\_9133
