## Supplementary material for "A novel method for integrating genomic and Tn-Seq data to identify common *in vivo* fitness mechanisms across multiple bacterial species": S4 Dataset: CL_INS_83.html

Legend

 Mobile +extrachromosomalelementfunctions
 Regulatoryfunctions
 Hypothetical
 DNA Metabolism
 Other
 Transport +binding proteins
 All VFDB Genes

FULL


WINDOWSVGPNG

Trim RowsRemove SingletonsSave Fasta

CL\_1197


CL\_1197


CL\_1197


CL\_1197


CL\_1197


CL\_1186


CL\_1174


CL\_1173


CL\_1197


CL\_1197


CL\_1197


CL\_1197


CL\_1197


CL\_1197


CL\_1197


Break


CL\_1197


CL\_1197


CL\_1197


CL\_1197

HighlightSelectShow Genomes


3

CL\_1213


2

CL\_1213


2

CL\_1213


2

CL\_1213


2

CL\_1213


1

CL\_1213


1

CL\_1213


1

CL\_1213


1

CL\_1213


1

CL\_1214


1

CL\_1214


1

CL\_1213


1

CL\_1213


1

CL\_5662


1

CL\_1214


1

CL\_1213


1

CL\_1213


1

CL\_1213


1

CL\_1213


1

Break

fGI ID


CL\_INS\_83
CL\_INS\_83
CL\_INS\_83
CL\_INS\_83
CL\_INS\_83
CL\_INS\_83
CL\_INS\_83
CL\_INS\_83
CL\_INS\_83
CL\_INS\_83
CL\_INS\_83
CL\_INS\_83
CL\_INS\_83
CL\_INS\_83
CL\_INS\_83
CL\_INS\_371
CL\_INS\_83
CL\_INS\_83
CL\_INS\_83
CL\_INS\_83
CL\_INS\_83
CL\_INS\_83
CL\_INS\_83
CL\_INS\_83
CL\_INS\_270
CL\_INS\_270
CL\_INS\_83
CL\_INS\_83
CL\_INS\_83
CL\_INS\_83
CL\_INS\_83
CL\_INS\_5
CL\_INS\_5
CL\_INS\_270
CL\_INS\_83
CL\_INS\_83
CL\_INS\_371
CL\_INS\_270
CL\_INS\_83
CL\_INS\_83
CL\_INS\_83
CL\_INS\_83
CL\_INS\_83
CL\_INS\_83
CL\_INS\_83
CL\_INS\_83
CL\_INS\_83
CL\_INS\_83
CL\_INS\_83
CL\_INS\_83
CL\_INS\_83
CL\_INS\_83
CL\_INS\_83
CL\_INS\_83
CL\_INS\_83
CL\_INS\_83
CL\_INS\_83
CL\_INS\_83
CL\_INS\_83
CL\_INS\_74
CL\_INS\_83
CL\_INS\_83
CL\_INS\_83
CL\_INS\_83
CL\_INS\_83
CL\_INS\_83
CL\_INS\_83
CL\_INS\_83
CL\_INS\_83
CL\_INS\_83
CL\_INS\_83
CL\_INS\_74
CL\_INS\_74
CL\_INS\_83
CL\_INS\_83
CL\_INS\_83
CL\_INS\_83
CL\_INS\_83
CL\_INS\_83
CL\_INS\_83
CL\_INS\_83
CL\_INS\_83
CL\_INS\_74
CL\_INS\_83
CL\_INS\_83
CL\_INS\_83
CL\_INS\_83
CL\_INS\_83
CL\_INS\_83
CL\_INS\_270
CL\_INS\_270
CL\_INS\_270
CL\_INS\_83
CL\_INS\_83
CL\_INS\_83
CL\_INS\_83
CL\_INS\_83
CL\_INS\_83
CL\_INS\_83
CL\_INS\_83
CL\_INS\_83
CL\_INS\_83
CL\_INS\_83
CL\_INS\_83
CL\_INS\_83
CL\_INS\_83
CL\_INS\_83
CL\_INS\_83
CL\_INS\_83
CL\_INS\_83
CL\_INS\_83
CL\_INS\_83
CL\_INS\_83
CL\_INS\_83
CL\_INS\_83
CL\_INS\_83
CL\_INS\_83
CL\_INS\_83
CL\_INS\_83
CL\_INS\_83
CL\_INS\_83
CL\_INS\_83
CL\_INS\_83
CL\_INS\_222
CL\_INS\_222
CL\_INS\_83
CL\_INS\_222
CL\_INS\_222
CL\_INS\_222
CL\_INS\_222
CL\_INS\_222
CL\_INS\_83
CL\_INS\_222
CL\_INS\_222
CL\_INS\_83
CL\_INS\_83
CL\_INS\_83
CL\_INS\_83
CL\_INS\_83
CL\_INS\_83
CL\_INS\_221
CL\_INS\_222
CL\_INS\_221
CL\_INS\_221
CL\_INS\_221
CL\_INS\_221
CL\_INS\_221
CL\_INS\_83
CL\_INS\_221
CL\_INS\_83
CL\_INS\_83
CL\_INS\_221
CL\_INS\_221
CL\_INS\_83
CL\_INS\_222
CL\_INS\_222
CL\_INS\_221
CL\_INS\_221
CL\_INS\_83
CL\_INS\_83
CL\_INS\_222
CL\_INS\_222
CL\_INS\_222
CL\_INS\_222
CL\_INS\_221
CL\_INS\_221
Cluster ID


CL\_9124
CL\_10224
CL\_11066
CL\_11067
CL\_11826
CL\_11825
CL\_12287
CL\_12286
CL\_12285
CL\_12284
CL\_12283
CL\_1198
CL\_11824
CL\_11823
CL\_11822
CL\_1199
CL\_13579
CL\_13578
CL\_13577
CL\_13576
CL\_13575
CL\_1200
CL\_12917
CL\_1201
CL\_1202
CL\_1203
CL\_1204
CL\_1205
CL\_1206
CL\_1207
CL\_1208
CL\_1209
CL\_1210
CL\_1211
CL\_13670
CL\_1212
CL\_9132
CL\_3904
CL\_5239
CL\_4313
CL\_5240
CL\_5241
CL\_5242
CL\_5243
CL\_5244
CL\_11505
CL\_12613
CL\_12612
CL\_12611
CL\_12610
CL\_12609
CL\_12608
CL\_11504
CL\_11503
CL\_11502
CL\_11501
CL\_11500
CL\_11499
CL\_11498
CL\_11497
CL\_5245
CL\_5246
CL\_5247
CL\_5248
CL\_5249
CL\_7408
CL\_12723
CL\_12724
CL\_12725
CL\_5255
CL\_5256
CL\_5250
CL\_5251
CL\_5252
CL\_5253
CL\_5254
CL\_10147
CL\_10148
CL\_10149
CL\_10150
CL\_10151
CL\_11496
CL\_11495
CL\_11494
CL\_11493
CL\_11492
CL\_11491
CL\_11490
CL\_11489
CL\_8354
CL\_8355
CL\_8356
CL\_6269
CL\_6268
CL\_6267
CL\_6266
CL\_6265
CL\_6264
CL\_6263
CL\_6262
CL\_6261
CL\_6260
CL\_6259
CL\_6258
CL\_6257
CL\_6256
CL\_6255
CL\_6254
CL\_6253
CL\_6252
CL\_6251
CL\_6250
CL\_6249
CL\_6248
CL\_6247
CL\_6246
CL\_6245
CL\_6244
CL\_6243
CL\_6242
CL\_6241
CL\_6240
CL\_12516
CL\_5506
CL\_5504
CL\_11820
CL\_669
CL\_670
CL\_9173
CL\_672
CL\_673
CL\_11818
CL\_674
CL\_11817
CL\_12515
CL\_12514
CL\_12513
CL\_12512
CL\_12511
CL\_12510
CL\_3243
CL\_3242
CL\_3241
CL\_3240
CL\_3239
CL\_3238
CL\_3237
CL\_12509
CL\_3235
CL\_12508
CL\_12507
CL\_689
CL\_3232
CL\_12506
CL\_3230
CL\_3229
CL\_3228
CL\_3227
CL\_10705
CL\_12505
CL\_3224
CL\_3223
CL\_3222
CL\_7381
CL\_3220
CL\_3219
