## Supplementary material for "A novel method for integrating genomic and Tn-Seq data to identify common *in vivo* fitness mechanisms across multiple bacterial species": S4 Dataset: CL_INS_85.html

Legend

 Other
 All VFDB Genes
 Transport +binding proteins

FULL


WINDOWSVGPNG

Trim RowsRemove SingletonsSave Fasta

CL\_1252


CL\_1252


CL\_1252

HighlightSelectShow Genomes


14

CL\_1255


12

CL\_1255


2

CL\_1255

fGI ID


CL\_INS\_85
CL\_INS\_85
Cluster ID


CL\_1253
CL\_1254
