## Supplementary material for "A novel method for integrating genomic and Tn-Seq data to identify common *in vivo* fitness mechanisms across multiple bacterial species": S4 Dataset: CL_INS_90.html

Legend

 Regulatoryfunctions
 Hypothetical
 All VFDB Genes

FULL


WINDOWSVGPNG

Trim RowsRemove SingletonsSave Fasta

CL\_1352


CL\_1352

HighlightSelectShow Genomes


20

CL\_1354


8

CL\_1354

fGI ID

CL\_INS\_90
Cluster ID

CL\_1353
