## Supplementary material for "A novel method for integrating genomic and Tn-Seq data to identify common *in vivo* fitness mechanisms across multiple bacterial species": S4 Dataset: CL_INS_98.html

Legend

 Hypothetical
 All VFDB Genes

FULL


WINDOWSVGPNG

Trim RowsRemove SingletonsSave Fasta

CL\_1424


CL\_1424


CL\_1424

HighlightSelectShow Genomes


15

CL\_1425


12

CL\_1425


1

CL\_1425

fGI ID


CL\_INS\_98
CL\_INS\_98
Cluster ID


CL\_7239
CL\_6201
