## Supplementary material for "A novel method for integrating genomic and Tn-Seq data to identify common *in vivo* fitness mechanisms across multiple bacterial species": S4 Dataset: CL_INS_100.html

Legend

 Hypothetical
 Other
 EnergyMetabolism
 All VFDB Genes

FULL


WINDOWSVGPNG

Trim RowsRemove SingletonsSave Fasta

CL\_1431


CL\_1431


CL\_1431


CL\_1431


CL\_1431

HighlightSelectShow Genomes


20

CL\_1432


4

CL\_1432


2

CL\_1432


1

CL\_1432


1

CL\_1432

fGI ID


CL\_INS\_100
CL\_INS\_100
CL\_INS\_100
CL\_INS\_100
CL\_INS\_100
CL\_INS\_100
Cluster ID


CL\_9300
CL\_9301
CL\_9302
CL\_11070
CL\_11071
CL\_9303
