## Supplementary material for "A novel method for integrating genomic and Tn-Seq data to identify common *in vivo* fitness mechanisms across multiple bacterial species": S4 Dataset: CL_INS_107.html

Legend

 Mobile +extrachromosomalelementfunctions
 Regulatoryfunctions
 Hypothetical
 DNA Metabolism
 Transcription
 Other
 EnergyMetabolism
 Transport +binding proteins
 All VFDB Genes

FULL


WINDOWSVGPNG

Trim RowsRemove SingletonsSave Fasta

CL\_1508


CL\_1508


CL\_1508


CL\_1508


CL\_1508


CL\_1508


CL\_1508


CL\_1508


CL\_1504


CL\_1508

HighlightSelectShow Genomes


18

CL\_1509


2

CL\_1509


1

CL\_1509


1

CL\_1509


1

CL\_1509


1

CL\_1509


1

CL\_1509


1

CL\_1509


1

CL\_1509


1

CL\_1509

fGI ID


CL\_INS\_105
CL\_INS\_105
CL\_INS\_105
CL\_INS\_105
CL\_INS\_107
CL\_INS\_107
CL\_INS\_150
CL\_INS\_150
CL\_INS\_150
CL\_INS\_150
CL\_INS\_150
CL\_INS\_107
CL\_INS\_150
CL\_INS\_107
CL\_INS\_107
CL\_INS\_107
CL\_INS\_107
CL\_INS\_107
CL\_INS\_107
CL\_INS\_107
CL\_INS\_49
CL\_INS\_49
CL\_INS\_49
CL\_INS\_49
CL\_INS\_49
CL\_INS\_49
CL\_INS\_49
CL\_INS\_107
CL\_INS\_107
CL\_INS\_107
CL\_INS\_107
CL\_INS\_107
CL\_INS\_107
CL\_INS\_107
CL\_INS\_107
CL\_INS\_107
CL\_INS\_107
CL\_INS\_107
CL\_INS\_107
CL\_INS\_107
CL\_INS\_107
CL\_INS\_107
CL\_INS\_107
CL\_INS\_107
CL\_INS\_107
CL\_INS\_107
CL\_INS\_107
CL\_INS\_107
CL\_INS\_107
CL\_INS\_366
CL\_INS\_107
CL\_INS\_107
CL\_INS\_107
CL\_INS\_107
CL\_INS\_107
CL\_INS\_141
CL\_INS\_141
CL\_INS\_141
CL\_INS\_141
CL\_INS\_107
CL\_INS\_368
CL\_INS\_141
CL\_INS\_141
CL\_INS\_141
CL\_INS\_141
CL\_INS\_141
CL\_INS\_141
CL\_INS\_141
CL\_INS\_107
CL\_INS\_107
CL\_INS\_141
CL\_INS\_141
CL\_INS\_141
CL\_INS\_325
CL\_INS\_141
CL\_INS\_107
CL\_INS\_204
CL\_INS\_107
CL\_INS\_365
CL\_INS\_107
CL\_INS\_107
CL\_INS\_107
CL\_INS\_107
CL\_INS\_107
CL\_INS\_107
CL\_INS\_107
CL\_INS\_107
CL\_INS\_107
CL\_INS\_107
CL\_INS\_107
CL\_INS\_141
CL\_INS\_141
CL\_INS\_141
CL\_INS\_141
CL\_INS\_107
CL\_INS\_107
CL\_INS\_107
CL\_INS\_107
CL\_INS\_107
CL\_INS\_107
CL\_INS\_107
CL\_INS\_141
CL\_INS\_325
CL\_INS\_107
CL\_INS\_107
CL\_INS\_107
CL\_INS\_107
CL\_INS\_107
CL\_INS\_107
CL\_INS\_107
CL\_INS\_141
CL\_INS\_204
CL\_INS\_204
CL\_INS\_204
CL\_INS\_204
CL\_INS\_204
CL\_INS\_107
CL\_INS\_141
CL\_INS\_141
CL\_INS\_141
CL\_INS\_141
CL\_INS\_141
CL\_INS\_141
CL\_INS\_141
CL\_INS\_325
CL\_INS\_325
CL\_INS\_325
CL\_INS\_325
CL\_INS\_325
CL\_INS\_325
CL\_INS\_325
CL\_INS\_325
CL\_INS\_107
CL\_INS\_325
CL\_INS\_325
CL\_INS\_325
CL\_INS\_325
CL\_INS\_325
CL\_INS\_325
CL\_INS\_107
CL\_INS\_325
CL\_INS\_204
CL\_INS\_141
CL\_INS\_141
CL\_INS\_141
CL\_INS\_141
CL\_INS\_141
CL\_INS\_107
CL\_INS\_141
CL\_INS\_141
CL\_INS\_141
CL\_INS\_141
CL\_INS\_141
CL\_INS\_141
CL\_INS\_141
CL\_INS\_141
CL\_INS\_141
CL\_INS\_368
CL\_INS\_141
CL\_INS\_115
CL\_INS\_107
CL\_INS\_107
CL\_INS\_107
CL\_INS\_107
CL\_INS\_107
CL\_INS\_107
CL\_INS\_107
CL\_INS\_107
CL\_INS\_107
CL\_INS\_107
CL\_INS\_107
CL\_INS\_107
CL\_INS\_107
CL\_INS\_107
CL\_INS\_107
CL\_INS\_107
CL\_INS\_107
CL\_INS\_107
CL\_INS\_93
CL\_INS\_107
CL\_INS\_107
CL\_INS\_107
CL\_INS\_107
CL\_INS\_107
CL\_INS\_107
CL\_INS\_107
CL\_INS\_107
CL\_INS\_107
CL\_INS\_107
CL\_INS\_107
CL\_INS\_366
CL\_INS\_107
CL\_INS\_107
CL\_INS\_366
CL\_INS\_107
CL\_INS\_107
CL\_INS\_366
CL\_INS\_107
CL\_INS\_107
CL\_INS\_107
CL\_INS\_107
CL\_INS\_107
CL\_INS\_107
CL\_INS\_107
CL\_INS\_107
CL\_INS\_107
CL\_INS\_107
CL\_INS\_107
CL\_INS\_107
CL\_INS\_107
CL\_INS\_107
CL\_INS\_107
CL\_INS\_107
CL\_INS\_107
CL\_INS\_107
CL\_INS\_107
CL\_INS\_107
CL\_INS\_107
CL\_INS\_107
CL\_INS\_107
CL\_INS\_107
CL\_INS\_204
CL\_INS\_107
CL\_INS\_115
CL\_INS\_107
CL\_INS\_270
CL\_INS\_366
CL\_INS\_107
CL\_INS\_366
CL\_INS\_107
CL\_INS\_107
CL\_INS\_107
CL\_INS\_107
CL\_INS\_107
CL\_INS\_107
CL\_INS\_107
CL\_INS\_107
CL\_INS\_368
CL\_INS\_178
CL\_INS\_107
CL\_INS\_107
CL\_INS\_368
CL\_INS\_107
CL\_INS\_178
CL\_INS\_107
CL\_INS\_368
CL\_INS\_178
CL\_INS\_178
CL\_INS\_178
CL\_INS\_178
CL\_INS\_178
CL\_INS\_107
CL\_INS\_107
CL\_INS\_107
CL\_INS\_107
CL\_INS\_107
CL\_INS\_107
CL\_INS\_107
CL\_INS\_107
CL\_INS\_107
CL\_INS\_107
CL\_INS\_107
CL\_INS\_107
CL\_INS\_366
CL\_INS\_270
CL\_INS\_270
CL\_INS\_107
CL\_INS\_368
CL\_INS\_107
CL\_INS\_107
CL\_INS\_107
CL\_INS\_107
CL\_INS\_107
CL\_INS\_107
CL\_INS\_107
CL\_INS\_107
CL\_INS\_107
CL\_INS\_107
CL\_INS\_107
CL\_INS\_107
CL\_INS\_107
CL\_INS\_107
CL\_INS\_107
CL\_INS\_107
CL\_INS\_107
CL\_INS\_107
CL\_INS\_107
CL\_INS\_107
CL\_INS\_107
CL\_INS\_107
CL\_INS\_107
CL\_INS\_107
CL\_INS\_107
CL\_INS\_107
CL\_INS\_107
CL\_INS\_107
CL\_INS\_107
CL\_INS\_107
CL\_INS\_107
CL\_INS\_107
Cluster ID


CL\_7235
CL\_7234
CL\_7233
CL\_7232
CL\_12604
CL\_12603
CL\_11471
CL\_11470
CL\_11469
CL\_11468
CL\_11467
CL\_12602
CL\_11466
CL\_13698
CL\_13699
CL\_12601
CL\_12600
CL\_10153
CL\_10154
CL\_10155
CL\_4306
CL\_4307
CL\_4308
CL\_4309
CL\_4310
CL\_4311
CL\_4312
CL\_11079
CL\_11080
CL\_11081
CL\_11082
CL\_11083
CL\_11084
CL\_11085
CL\_11086
CL\_11087
CL\_11088
CL\_11089
CL\_11090
CL\_11091
CL\_11092
CL\_10156
CL\_10157
CL\_11093
CL\_7231
CL\_5257
CL\_5258
CL\_5259
CL\_5260
CL\_5261
CL\_5262
CL\_5263
CL\_5264
CL\_5265
CL\_5266
CL\_5267
CL\_5268
CL\_5269
CL\_1285
CL\_1286
CL\_1287
CL\_1288
CL\_1289
CL\_5270
CL\_1292
CL\_2023
CL\_2024
CL\_2025
CL\_5271
CL\_5272
CL\_1297
CL\_1298
CL\_2029
CL\_5273
CL\_2863
CL\_5274
CL\_5275
CL\_10424
CL\_8766
CL\_10425
CL\_5416
CL\_10426
CL\_10427
CL\_10428
CL\_10429
CL\_10430
CL\_10431
CL\_10432
CL\_10433
CL\_10434
CL\_7969
CL\_10435
CL\_10436
CL\_5701
CL\_10437
CL\_10438
CL\_10439
CL\_10440
CL\_10441
CL\_10442
CL\_10443
CL\_7173
CL\_5709
CL\_10444
CL\_10445
CL\_10446
CL\_10447
CL\_10448
CL\_10449
CL\_10450
CL\_10451
CL\_5276
CL\_5277
CL\_5278
CL\_5279
CL\_5280
CL\_5281
CL\_2858
CL\_9410
CL\_9411
CL\_9412
CL\_9413
CL\_9414
CL\_5282
CL\_5283
CL\_5284
CL\_5285
CL\_5286
CL\_5287
CL\_5288
CL\_5289
CL\_5290
CL\_5291
CL\_5292
CL\_5293
CL\_5294
CL\_5295
CL\_5296
CL\_5297
CL\_5298
CL\_5299
CL\_3154
CL\_3153
CL\_3152
CL\_3151
CL\_3150
CL\_3149
CL\_10452
CL\_3148
CL\_3147
CL\_3146
CL\_1319
CL\_3143
CL\_3142
CL\_2059
CL\_2060
CL\_6140
CL\_5728
CL\_2062
CL\_1637
CL\_12282
CL\_12281
CL\_10612
CL\_12280
CL\_10176
CL\_10175
CL\_12726
CL\_12727
CL\_5440
CL\_11695
CL\_12728
CL\_11420
CL\_5439
CL\_5438
CL\_10610
CL\_12279
CL\_11422
CL\_10180
CL\_8296
CL\_5437
CL\_5436
CL\_12729
CL\_5434
CL\_5432
CL\_5431
CL\_5430
CL\_10608
CL\_10607
CL\_12278
CL\_12277
CL\_10604
CL\_12276
CL\_12275
CL\_9038
CL\_12274
CL\_12273
CL\_10538
CL\_12272
CL\_12271
CL\_12270
CL\_12269
CL\_12268
CL\_12267
CL\_10594
CL\_12730
CL\_10973
CL\_10186
CL\_12731
CL\_10187
CL\_10188
CL\_5424
CL\_12732
CL\_12733
CL\_12734
CL\_12735
CL\_5420
CL\_12736
CL\_12737
CL\_12738
CL\_12739
CL\_12740
CL\_1278
CL\_12741
CL\_7224
CL\_12742
CL\_3944
CL\_7128
CL\_12266
CL\_7127
CL\_12265
CL\_5405
CL\_12264
CL\_12263
CL\_5401
CL\_12262
CL\_12261
CL\_12260
CL\_5397
CL\_7115
CL\_11706
CL\_11707
CL\_10584
CL\_11708
CL\_5390
CL\_5389
CL\_7108
CL\_5387
CL\_5386
CL\_5385
CL\_5384
CL\_5383
CL\_5382
CL\_12259
CL\_5381
CL\_5380
CL\_12258
CL\_5379
CL\_5378
CL\_12257
CL\_12256
CL\_12255
CL\_12254
CL\_12253
CL\_7126
CL\_6963
CL\_5710
CL\_12743
CL\_10481
CL\_12744
CL\_12745
CL\_12746
CL\_12747
CL\_12748
CL\_12749
CL\_12750
CL\_12751
CL\_12752
CL\_12753
CL\_12754
CL\_12755
CL\_12756
CL\_12757
CL\_12758
CL\_12759
CL\_12760
CL\_12761
CL\_12762
CL\_12763
CL\_12764
CL\_12765
CL\_12766
CL\_12767
CL\_12768
CL\_12769
CL\_12770
CL\_12771
CL\_12772
CL\_12773
CL\_12774
CL\_12252
