## Supplementary material for "A novel method for integrating genomic and Tn-Seq data to identify common *in vivo* fitness mechanisms across multiple bacterial species": S4 Dataset: CL_INS_109.html

Legend

 Hypothetical
 Other
 All VFDB Genes

FULL


WINDOWSVGPNG

Trim RowsRemove SingletonsSave Fasta

CL\_1524


CL\_1524

HighlightSelectShow Genomes


21

CL\_1525


7

CL\_1525

fGI ID


CL\_INS\_109
CL\_INS\_109
CL\_INS\_109
CL\_INS\_109
Cluster ID


CL\_6194
CL\_6193
CL\_6192
CL\_6191
