## Supplementary material for "A novel method for integrating genomic and Tn-Seq data to identify common *in vivo* fitness mechanisms across multiple bacterial species": S4 Dataset: CL_INS_112.html

Legend

 Hypothetical
 Other
 All VFDB Genes

FULL


WINDOWSVGPNG

Trim RowsRemove SingletonsSave Fasta

CL\_1592


CL\_1592


CL\_1592

HighlightSelectShow Genomes


22

CL\_1593


2

CL\_1593


1

Break

fGI ID


CL\_INS\_112
CL\_INS\_112
CL\_INS\_112
CL\_INS\_112
Cluster ID


CL\_13290
CL\_13291
CL\_13292
CL\_13293
