## Supplementary material for "A novel method for integrating genomic and Tn-Seq data to identify common *in vivo* fitness mechanisms across multiple bacterial species": S4 Dataset: CL_INS_117.html

Legend

 Mobile +extrachromosomalelementfunctions
 Hypothetical
 DNA Metabolism
 Other
 EnergyMetabolism
 Transport +binding proteins
 All VFDB Genes

FULL


WINDOWSVGPNG

Trim RowsRemove SingletonsSave Fasta

CL\_1660


CL\_1660


CL\_1660


CL\_1660


CL\_1660


CL\_1660


CL\_1660


CL\_1660

HighlightSelectShow Genomes


19

CL\_5300


2

CL\_5300


2

CL\_5300


1

CL\_5300


1

CL\_5300


1

CL\_5300


1

CL\_1676


1

CL\_5300

fGI ID


CL\_INS\_117
CL\_INS\_117
CL\_INS\_117
CL\_INS\_117
CL\_INS\_117
CL\_INS\_117
CL\_INS\_117
CL\_INS\_117
CL\_INS\_117
CL\_INS\_117
CL\_INS\_117
CL\_INS\_117
CL\_INS\_117
CL\_INS\_117
CL\_INS\_117
CL\_INS\_117
CL\_INS\_117
CL\_INS\_117
CL\_INS\_117
CL\_INS\_117
CL\_INS\_117
CL\_INS\_117
CL\_INS\_117
CL\_INS\_117
CL\_INS\_117
CL\_INS\_117
CL\_INS\_117
CL\_INS\_117
CL\_INS\_117
CL\_INS\_117
CL\_INS\_117
Cluster ID


CL\_1661
CL\_6182
CL\_6181
CL\_6180
CL\_6179
CL\_9535
CL\_9534
CL\_9531
CL\_9097
CL\_9096
CL\_9095
CL\_9094
CL\_1662
CL\_9098
CL\_1663
CL\_1664
CL\_1665
CL\_1666
CL\_12653
CL\_1667
CL\_6178
CL\_6177
CL\_1668
CL\_1669
CL\_1670
CL\_1671
CL\_1672
CL\_1673
CL\_1674
CL\_11716
CL\_1675
