## Supplementary material for "A novel method for integrating genomic and Tn-Seq data to identify common *in vivo* fitness mechanisms across multiple bacterial species": S4 Dataset: CL_INS_120.html

Legend

 Hypothetical
 Other
 Proteinsynthesis/fate
 EnergyMetabolism
 All VFDB Genes

FULL


WINDOWSVGPNG

Trim RowsRemove SingletonsSave Fasta

CL\_1691


CL\_1691


CL\_1691


CL\_1691


CL\_1690


CL\_1691

HighlightSelectShow Genomes


15

CL\_1696


8

CL\_1696


1

CL\_1696


1

CL\_1696


1

CL\_1696


1

CL\_1696

fGI ID


CL\_INS\_120
CL\_INS\_120
CL\_INS\_120
CL\_INS\_120
CL\_INS\_120
Cluster ID


CL\_5302
CL\_1692
CL\_1693
CL\_1694
CL\_1695
