## Supplementary material for "A novel method for integrating genomic and Tn-Seq data to identify common *in vivo* fitness mechanisms across multiple bacterial species": S4 Dataset: CL_INS_121.html

Legend

 Hypothetical
 Proteinsynthesis/fate
 All VFDB Genes

FULL


WINDOWSVGPNG

Trim RowsRemove SingletonsSave Fasta

CL\_1696


CL\_1696

HighlightSelectShow Genomes


16

CL\_1697


12

CL\_1697

fGI ID

CL\_INS\_121
Cluster ID

CL\_6174
