## Supplementary material for "A novel method for integrating genomic and Tn-Seq data to identify common *in vivo* fitness mechanisms across multiple bacterial species": S4 Dataset: CL_INS_122.html

Legend

 Hypothetical
 Other
 All VFDB Genes

FULL


WINDOWSVGPNG

Trim RowsRemove SingletonsSave Fasta

CL\_1716


CL\_1716


CL\_1716

HighlightSelectShow Genomes


16

CL\_1717


11

CL\_1717


1

CL\_1718

fGI ID

CL\_INS\_122
Cluster ID

CL\_10453
