## Supplementary material for "A novel method for integrating genomic and Tn-Seq data to identify common *in vivo* fitness mechanisms across multiple bacterial species": S4 Dataset: CL_INS_123.html

Legend

 All VFDB Genes

FULL


WINDOWSVGPNG

Trim RowsRemove SingletonsSave Fasta

CL\_1736


CL\_1736


CL\_1736

HighlightSelectShow Genomes


13

CL\_1738


12

CL\_1738


3

CL\_1739

fGI ID


CL\_INS\_123
CL\_INS\_123
Cluster ID


CL\_1737
CL\_6173
