## Supplementary material for "A novel method for integrating genomic and Tn-Seq data to identify common *in vivo* fitness mechanisms across multiple bacterial species": S4 Dataset: CL_INS_124.html

Legend

 Cell Envelope
 All VFDB Genes

FULL


WINDOWSVGPNG

Trim RowsRemove SingletonsSave Fasta

CL\_1739


CL\_1739


CL\_1739


CL\_1739


CL\_1738

HighlightSelectShow Genomes


12

CL\_1741


11

CL\_1741


3

CL\_1741


1

CL\_1741


1

CL\_1741

fGI ID


CL\_INS\_124
CL\_INS\_124
CL\_INS\_124
CL\_INS\_124
Cluster ID


CL\_10454
CL\_13084
CL\_6172
CL\_1740
