## Supplementary material for "A novel method for integrating genomic and Tn-Seq data to identify common *in vivo* fitness mechanisms across multiple bacterial species": S4 Dataset: CL_INS_132.html

Legend

 Mobile +extrachromosomalelementfunctions
 Regulatoryfunctions
 Hypothetical
 DNA Metabolism
 AntibioticResistance
 Proteinsynthesis/fate
 Other
 Transport +binding proteins
 All VFDB Genes

FULL


WINDOWSVGPNG

Trim RowsRemove SingletonsSave Fasta

CL\_1911


CL\_1911


CL\_5729


CL\_1911


CL\_1911


CL\_1911

HighlightSelectShow Genomes


15

CL\_6159


9

CL\_6159


1

CL\_6159


1

CL\_1912


1

CL\_6159


1

Break

fGI ID


CL\_INS\_132
CL\_INS\_132
CL\_INS\_132
CL\_INS\_132
CL\_INS\_132
CL\_INS\_132
CL\_INS\_132
CL\_INS\_132
CL\_INS\_132
CL\_INS\_132
CL\_INS\_132
CL\_INS\_132
CL\_INS\_132
CL\_INS\_132
CL\_INS\_132
CL\_INS\_366
CL\_INS\_204
CL\_INS\_366
CL\_INS\_366
CL\_INS\_366
CL\_INS\_366
CL\_INS\_204
CL\_INS\_132
CL\_INS\_204
CL\_INS\_204
CL\_INS\_204
CL\_INS\_204
CL\_INS\_366
CL\_INS\_132
CL\_INS\_204
CL\_INS\_204
CL\_INS\_132
CL\_INS\_132
CL\_INS\_132
CL\_INS\_204
CL\_INS\_366
CL\_INS\_132
CL\_INS\_132
CL\_INS\_204
CL\_INS\_204
CL\_INS\_132
CL\_INS\_366
CL\_INS\_204
CL\_INS\_132
CL\_INS\_132
CL\_INS\_132
CL\_INS\_132
CL\_INS\_366
CL\_INS\_132
CL\_INS\_132
CL\_INS\_141
CL\_INS\_132
CL\_INS\_132
CL\_INS\_204
CL\_INS\_204
CL\_INS\_141
CL\_INS\_132
CL\_INS\_132
CL\_INS\_132
CL\_INS\_132
CL\_INS\_132
CL\_INS\_132
CL\_INS\_132
CL\_INS\_132
CL\_INS\_132
CL\_INS\_132
CL\_INS\_132
CL\_INS\_132
CL\_INS\_132
CL\_INS\_132
CL\_INS\_132
CL\_INS\_132
CL\_INS\_132
CL\_INS\_132
CL\_INS\_132
CL\_INS\_132
CL\_INS\_132
CL\_INS\_132
CL\_INS\_132
CL\_INS\_132
CL\_INS\_132
CL\_INS\_132
CL\_INS\_132
CL\_INS\_132
CL\_INS\_132
CL\_INS\_132
CL\_INS\_132
CL\_INS\_132
CL\_INS\_132
CL\_INS\_132
CL\_INS\_132
CL\_INS\_132
CL\_INS\_132
CL\_INS\_132
CL\_INS\_132
CL\_INS\_132
CL\_INS\_132
CL\_INS\_132
CL\_INS\_132
CL\_INS\_132
CL\_INS\_132
CL\_INS\_132
CL\_INS\_132
CL\_INS\_132
CL\_INS\_132
CL\_INS\_132
CL\_INS\_365
CL\_INS\_132
CL\_INS\_132
CL\_INS\_132
CL\_INS\_132
CL\_INS\_132
CL\_INS\_365
CL\_INS\_132
CL\_INS\_365
CL\_INS\_132
CL\_INS\_132
CL\_INS\_132
CL\_INS\_132
CL\_INS\_132
CL\_INS\_132
CL\_INS\_132
CL\_INS\_132
CL\_INS\_132
CL\_INS\_365
CL\_INS\_132
CL\_INS\_132
CL\_INS\_132
CL\_INS\_132
CL\_INS\_132
CL\_INS\_132
CL\_INS\_132
CL\_INS\_132
CL\_INS\_132
CL\_INS\_132
CL\_INS\_132
CL\_INS\_132
CL\_INS\_132
CL\_INS\_132
CL\_INS\_132
CL\_INS\_132
CL\_INS\_132
CL\_INS\_132
CL\_INS\_132
CL\_INS\_132
CL\_INS\_132
CL\_INS\_132
CL\_INS\_132
CL\_INS\_132
CL\_INS\_132
CL\_INS\_132
CL\_INS\_132
CL\_INS\_132
CL\_INS\_132
CL\_INS\_132
CL\_INS\_132
CL\_INS\_132
CL\_INS\_132
CL\_INS\_132
CL\_INS\_132
CL\_INS\_132
CL\_INS\_132
CL\_INS\_132
CL\_INS\_132
CL\_INS\_132
CL\_INS\_132
CL\_INS\_132
CL\_INS\_132
CL\_INS\_132
CL\_INS\_132
CL\_INS\_132
CL\_INS\_132
CL\_INS\_132
CL\_INS\_132
CL\_INS\_132
CL\_INS\_132
CL\_INS\_132
CL\_INS\_132
CL\_INS\_132
CL\_INS\_132
CL\_INS\_132
CL\_INS\_132
CL\_INS\_132
CL\_INS\_132
CL\_INS\_132
CL\_INS\_132
CL\_INS\_132
CL\_INS\_132
CL\_INS\_132
CL\_INS\_132
CL\_INS\_132
CL\_INS\_132
CL\_INS\_132
CL\_INS\_132
CL\_INS\_132
CL\_INS\_132
CL\_INS\_132
CL\_INS\_132
CL\_INS\_132
CL\_INS\_132
CL\_INS\_132
CL\_INS\_132
CL\_INS\_365
CL\_INS\_132
CL\_INS\_365
CL\_INS\_365
CL\_INS\_365
CL\_INS\_365
CL\_INS\_365
CL\_INS\_365
CL\_INS\_365
CL\_INS\_365
CL\_INS\_365
CL\_INS\_365
CL\_INS\_365
CL\_INS\_365
CL\_INS\_365
CL\_INS\_365
CL\_INS\_365
CL\_INS\_132
CL\_INS\_132
CL\_INS\_132
CL\_INS\_132
CL\_INS\_132
CL\_INS\_132
CL\_INS\_132
CL\_INS\_132
CL\_INS\_132
CL\_INS\_132
CL\_INS\_132
CL\_INS\_132
CL\_INS\_132
CL\_INS\_132
CL\_INS\_132
CL\_INS\_132
CL\_INS\_132
CL\_INS\_132
CL\_INS\_132
CL\_INS\_132
CL\_INS\_132
CL\_INS\_270
CL\_INS\_132
CL\_INS\_132
CL\_INS\_132
CL\_INS\_132
CL\_INS\_132
CL\_INS\_132
CL\_INS\_132
CL\_INS\_132
CL\_INS\_132
CL\_INS\_132
CL\_INS\_132
CL\_INS\_132
CL\_INS\_132
CL\_INS\_132
CL\_INS\_132
CL\_INS\_132
CL\_INS\_132
CL\_INS\_132
CL\_INS\_132
CL\_INS\_132
CL\_INS\_132
CL\_INS\_132
CL\_INS\_132
CL\_INS\_132
CL\_INS\_132
CL\_INS\_132
CL\_INS\_132
CL\_INS\_132
CL\_INS\_132
CL\_INS\_132
CL\_INS\_132
CL\_INS\_132
CL\_INS\_132
Cluster ID


CL\_10788
CL\_10789
CL\_10790
CL\_2823
CL\_10791
CL\_10792
CL\_10793
CL\_10794
CL\_10795
CL\_10796
CL\_10797
CL\_10798
CL\_10799
CL\_9082
CL\_11134
CL\_2891
CL\_2890
CL\_2887
CL\_2886
CL\_2885
CL\_2884
CL\_7940
CL\_11135
CL\_2889
CL\_11136
CL\_6136
CL\_11137
CL\_6135
CL\_11138
CL\_6133
CL\_9388
CL\_9389
CL\_9390
CL\_11139
CL\_2876
CL\_2875
CL\_9391
CL\_9392
CL\_2874
CL\_2873
CL\_11140
CL\_2872
CL\_6127
CL\_9393
CL\_9394
CL\_9395
CL\_9396
CL\_2871
CL\_11141
CL\_11142
CL\_7207
CL\_11143
CL\_11144
CL\_7205
CL\_7204
CL\_7203
CL\_11286
CL\_11287
CL\_11288
CL\_11289
CL\_11290
CL\_11291
CL\_11292
CL\_11293
CL\_11294
CL\_11295
CL\_11296
CL\_11297
CL\_11298
CL\_11299
CL\_11300
CL\_11301
CL\_11302
CL\_11303
CL\_11304
CL\_11305
CL\_11306
CL\_11307
CL\_11308
CL\_11309
CL\_6568
CL\_6569
CL\_6570
CL\_6571
CL\_6572
CL\_6573
CL\_11310
CL\_11311
CL\_11312
CL\_11313
CL\_11314
CL\_6576
CL\_6578
CL\_11315
CL\_6579
CL\_11316
CL\_11317
CL\_11318
CL\_11319
CL\_11320
CL\_11321
CL\_11322
CL\_11323
CL\_11324
CL\_11145
CL\_11146
CL\_7864
CL\_7865
CL\_11147
CL\_11148
CL\_5138
CL\_5137
CL\_5136
CL\_5135
CL\_7882
CL\_11149
CL\_11150
CL\_11151
CL\_11152
CL\_6550
CL\_11153
CL\_11154
CL\_11155
CL\_7958
CL\_6662
CL\_5879
CL\_5880
CL\_5881
CL\_5882
CL\_5884
CL\_5886
CL\_5887
CL\_5888
CL\_5889
CL\_5890
CL\_7480
CL\_11156
CL\_11157
CL\_11158
CL\_11159
CL\_11160
CL\_7486
CL\_7487
CL\_7488
CL\_7491
CL\_7492
CL\_7493
CL\_7508
CL\_7509
CL\_7510
CL\_7512
CL\_7513
CL\_7514
CL\_11161
CL\_5790
CL\_11162
CL\_5797
CL\_5798
CL\_5799
CL\_5801
CL\_5802
CL\_5803
CL\_5804
CL\_5807
CL\_5808
CL\_5809
CL\_5810
CL\_5811
CL\_5812
CL\_5813
CL\_5814
CL\_5815
CL\_5816
CL\_5818
CL\_5819
CL\_5820
CL\_5822
CL\_5823
CL\_5824
CL\_5825
CL\_5826
CL\_5827
CL\_5828
CL\_5829
CL\_5830
CL\_5831
CL\_5832
CL\_5833
CL\_11163
CL\_6596
CL\_6597
CL\_11164
CL\_11165
CL\_11166
CL\_11167
CL\_11168
CL\_11169
CL\_11170
CL\_11171
CL\_11172
CL\_11173
CL\_11174
CL\_6704
CL\_11175
CL\_7061
CL\_6628
CL\_6629
CL\_6630
CL\_6631
CL\_6632
CL\_4880
CL\_4881
CL\_4882
CL\_4883
CL\_4884
CL\_4885
CL\_7071
CL\_4879
CL\_6657
CL\_11176
CL\_11177
CL\_11178
CL\_11179
CL\_11180
CL\_11181
CL\_11182
CL\_11183
CL\_11184
CL\_11185
CL\_11186
CL\_11187
CL\_11188
CL\_11189
CL\_11190
CL\_11191
CL\_11192
CL\_11193
CL\_11194
CL\_11195
CL\_11196
CL\_7678
CL\_7677
CL\_7676
CL\_5835
CL\_11197
CL\_11198
CL\_11199
CL\_11200
CL\_11201
CL\_11202
CL\_11203
CL\_11204
CL\_5838
CL\_5839
CL\_5841
CL\_11205
CL\_11206
CL\_11207
CL\_11208
CL\_11209
CL\_11210
CL\_11211
CL\_11212
CL\_11213
CL\_11214
CL\_11215
CL\_7675
CL\_7674
CL\_7673
CL\_7672
CL\_7671
CL\_7670
CL\_7668
CL\_7667
