## Supplementary material for "A novel method for integrating genomic and Tn-Seq data to identify common *in vivo* fitness mechanisms across multiple bacterial species": S4 Dataset: CL_INS_133.html

Legend

 Mobile +extrachromosomalelementfunctions
 Hypothetical
 AntibioticResistance
 All VFDB Genes

FULL


WINDOWSVGPNG

Trim RowsRemove SingletonsSave Fasta

CL\_6159


CL\_6159


CL\_1911

HighlightSelectShow Genomes


23

CL\_1912


3

CL\_1912


1

CL\_1912

fGI ID


CL\_INS\_133
CL\_INS\_133
Cluster ID


CL\_13068
CL\_5324
