## Supplementary material for "A novel method for integrating genomic and Tn-Seq data to identify common *in vivo* fitness mechanisms across multiple bacterial species": S4 Dataset: CL_INS_137.html

Legend

 Mobile +extrachromosomalelementfunctions
 Regulatoryfunctions
 Hypothetical
 DNA Metabolism
 Cell Envelope
 Proteinsynthesis/fate
 Other
 EnergyMetabolism
 All VFDB Genes

FULL


WINDOWSVGPNG

Trim RowsRemove SingletonsSave Fasta

CL\_1954


CL\_1954


CL\_1954


CL\_1954


CL\_1954


CL\_5729


CL\_1954


CL\_1954


CL\_1954

HighlightSelectShow Genomes


19

CL\_1955


3

CL\_1955


1

CL\_1955


1

CL\_1955


1

CL\_1955


1

CL\_1955


1

CL\_1955


1

CL\_5729


1

CL\_1956

fGI ID


CL\_INS\_141
CL\_INS\_369
CL\_INS\_137
CL\_INS\_137
CL\_INS\_137
CL\_INS\_137
CL\_INS\_137
CL\_INS\_137
CL\_INS\_137
CL\_INS\_137
CL\_INS\_137
CL\_INS\_137
CL\_INS\_137
CL\_INS\_137
CL\_INS\_368
CL\_INS\_368
CL\_INS\_368
CL\_INS\_368
CL\_INS\_366
CL\_INS\_368
CL\_INS\_368
CL\_INS\_137
CL\_INS\_137
CL\_INS\_137
CL\_INS\_137
CL\_INS\_137
CL\_INS\_141
CL\_INS\_368
CL\_INS\_141
CL\_INS\_141
CL\_INS\_141
CL\_INS\_141
CL\_INS\_368
CL\_INS\_107
CL\_INS\_137
CL\_INS\_141
CL\_INS\_141
CL\_INS\_141
CL\_INS\_137
CL\_INS\_141
CL\_INS\_141
CL\_INS\_141
CL\_INS\_141
CL\_INS\_141
CL\_INS\_141
CL\_INS\_141
CL\_INS\_141
CL\_INS\_137
CL\_INS\_137
CL\_INS\_141
CL\_INS\_368
CL\_INS\_137
CL\_INS\_141
CL\_INS\_368
CL\_INS\_141
CL\_INS\_141
CL\_INS\_141
CL\_INS\_141
CL\_INS\_141
CL\_INS\_368
CL\_INS\_368
CL\_INS\_141
CL\_INS\_368
CL\_INS\_368
CL\_INS\_137
CL\_INS\_137
CL\_INS\_204
CL\_INS\_141
CL\_INS\_141
CL\_INS\_141
CL\_INS\_141
CL\_INS\_141
CL\_INS\_141
CL\_INS\_141
CL\_INS\_204
CL\_INS\_204
CL\_INS\_204
CL\_INS\_368
CL\_INS\_368
CL\_INS\_204
CL\_INS\_204
CL\_INS\_141
CL\_INS\_204
CL\_INS\_368
CL\_INS\_368
CL\_INS\_368
CL\_INS\_368
CL\_INS\_141
CL\_INS\_141
CL\_INS\_141
CL\_INS\_141
CL\_INS\_141
CL\_INS\_137
CL\_INS\_141
CL\_INS\_204
CL\_INS\_141
CL\_INS\_204
CL\_INS\_204
CL\_INS\_204
CL\_INS\_204
CL\_INS\_204
CL\_INS\_141
CL\_INS\_141
CL\_INS\_137
CL\_INS\_137
CL\_INS\_137
CL\_INS\_137
CL\_INS\_137
CL\_INS\_137
CL\_INS\_137
CL\_INS\_137
CL\_INS\_137
CL\_INS\_137
CL\_INS\_137
CL\_INS\_137
CL\_INS\_137
CL\_INS\_137
CL\_INS\_137
CL\_INS\_137
CL\_INS\_137
CL\_INS\_137
CL\_INS\_137
CL\_INS\_137
CL\_INS\_137
CL\_INS\_137
CL\_INS\_137
CL\_INS\_137
CL\_INS\_137
CL\_INS\_137
CL\_INS\_137
CL\_INS\_137
CL\_INS\_137
CL\_INS\_137
CL\_INS\_137
CL\_INS\_137
CL\_INS\_137
CL\_INS\_137
CL\_INS\_137
CL\_INS\_137
CL\_INS\_137
CL\_INS\_137
CL\_INS\_137
CL\_INS\_137
CL\_INS\_137
CL\_INS\_137
CL\_INS\_137
CL\_INS\_137
CL\_INS\_137
CL\_INS\_137
CL\_INS\_137
CL\_INS\_137
CL\_INS\_137
CL\_INS\_137
CL\_INS\_175
CL\_INS\_175
CL\_INS\_175
CL\_INS\_137
CL\_INS\_175
CL\_INS\_175
CL\_INS\_137
CL\_INS\_137
CL\_INS\_137
CL\_INS\_137
CL\_INS\_137
CL\_INS\_137
CL\_INS\_137
CL\_INS\_137
CL\_INS\_137
CL\_INS\_137
CL\_INS\_137
Cluster ID


CL\_9005
CL\_6111
CL\_12469
CL\_13067
CL\_9081
CL\_9080
CL\_9397
CL\_13066
CL\_13065
CL\_13064
CL\_13063
CL\_13062
CL\_13061
CL\_13060
CL\_9398
CL\_9399
CL\_12487
CL\_12486
CL\_12485
CL\_12484
CL\_12483
CL\_9400
CL\_9401
CL\_9402
CL\_9403
CL\_9404
CL\_1280
CL\_1281
CL\_9266
CL\_11973
CL\_1285
CL\_7221
CL\_1287
CL\_5266
CL\_9405
CL\_2015
CL\_2016
CL\_2017
CL\_9406
CL\_1288
CL\_1289
CL\_5270
CL\_1290
CL\_1291
CL\_1292
CL\_2023
CL\_2024
CL\_1293
CL\_1294
CL\_2025
CL\_12482
CL\_9407
CL\_1297
CL\_12481
CL\_1298
CL\_2029
CL\_2030
CL\_7960
CL\_12480
CL\_12479
CL\_12478
CL\_10451
CL\_12477
CL\_12476
CL\_9408
CL\_9409
CL\_5276
CL\_2858
CL\_9410
CL\_9411
CL\_9412
CL\_9413
CL\_9414
CL\_5282
CL\_5716
CL\_3154
CL\_5717
CL\_12475
CL\_12474
CL\_5720
CL\_7023
CL\_12045
CL\_5721
CL\_12473
CL\_12472
CL\_12471
CL\_12470
CL\_3153
CL\_3152
CL\_3151
CL\_3150
CL\_3149
CL\_9415
CL\_3148
CL\_7025
CL\_9416
CL\_7026
CL\_7027
CL\_7028
CL\_7029
CL\_7030
CL\_9417
CL\_2062
CL\_9418
CL\_9419
CL\_9420
CL\_10164
CL\_10165
CL\_10166
CL\_10167
CL\_10168
CL\_10169
CL\_9079
CL\_9078
CL\_9077
CL\_9076
CL\_9075
CL\_9074
CL\_9073
CL\_9072
CL\_9071
CL\_9070
CL\_9421
CL\_10800
CL\_9069
CL\_9422
CL\_9068
CL\_9067
CL\_9423
CL\_9066
CL\_10801
CL\_10802
CL\_9065
CL\_9064
CL\_5118
CL\_9063
CL\_9062
CL\_9061
CL\_9060
CL\_9059
CL\_9058
CL\_9057
CL\_9056
CL\_9055
CL\_9054
CL\_9053
CL\_9052
CL\_9424
CL\_10803
CL\_10804
CL\_10805
CL\_10806
CL\_10807
CL\_7053
CL\_7052
CL\_7051
CL\_9425
CL\_7050
CL\_7049
CL\_9051
CL\_9050
CL\_9049
CL\_12059
CL\_12058
CL\_9426
CL\_9427
CL\_9428
CL\_9429
CL\_9430
CL\_10808
