## Supplementary material for "A novel method for integrating genomic and Tn-Seq data to identify common *in vivo* fitness mechanisms across multiple bacterial species": S4 Dataset: CL_INS_141.html

Legend

 Mobile +extrachromosomalelementfunctions
 Regulatoryfunctions
 Hypothetical
 DNA Metabolism
 All Fitness Genes
 Proteinsynthesis/fate
 Other
 EnergyMetabolism
 Cellularprocesses
 Transport +binding proteins
 All VFDB Genes

FULL


WINDOWSVGPNG

Trim RowsRemove SingletonsSave Fasta

CL\_2002


CL\_2866


CL\_2866


CL\_2002


CL\_2002


CL\_2002


CL\_2002


CL\_2002


CL\_2002


CL\_2866


CL\_2002


CL\_2002


CL\_2002


CL\_2002


CL\_2002


CL\_2002


CL\_2866


CL\_2002


CL\_2866


CL\_2002


CL\_2002


CL\_5729


CL\_2002

HighlightSelectShow Genomes


12

CL\_2067


1

CL\_2067


1

CL\_2067


1

CL\_2821


1

CL\_2867


1

CL\_2067


1

CL\_2867


1

CL\_2067


1

CL\_2067


1

CL\_2067


1

CL\_2067


1

CL\_2867


1

CL\_2067


1

CL\_2821


1

CL\_2067


1

CL\_2821


1

CL\_2067


1

CL\_2067


1

CL\_2067


1

CL\_2067


1

CL\_2067


1

CL\_2067


1

CL\_2067

fGI ID


CL\_INS\_141
CL\_INS\_141
CL\_INS\_141
CL\_INS\_141
CL\_INS\_141
CL\_INS\_141
CL\_INS\_141
CL\_INS\_141
CL\_INS\_141
CL\_INS\_141
CL\_INS\_141
CL\_INS\_141
CL\_INS\_141
CL\_INS\_141
CL\_INS\_141
CL\_INS\_141
CL\_INS\_141
CL\_INS\_141
CL\_INS\_141
CL\_INS\_141
CL\_INS\_141
CL\_INS\_141
CL\_INS\_141
CL\_INS\_141
CL\_INS\_141
CL\_INS\_141
CL\_INS\_141
CL\_INS\_141
CL\_INS\_141
CL\_INS\_141
CL\_INS\_141
CL\_INS\_141
CL\_INS\_141
CL\_INS\_141
CL\_INS\_141
CL\_INS\_141
CL\_INS\_141
CL\_INS\_141
CL\_INS\_141
CL\_INS\_141
CL\_INS\_204
CL\_INS\_141
CL\_INS\_141
CL\_INS\_141
CL\_INS\_141
CL\_INS\_141
CL\_INS\_141
CL\_INS\_141
CL\_INS\_141
CL\_INS\_141
CL\_INS\_141
CL\_INS\_141
CL\_INS\_141
CL\_INS\_141
CL\_INS\_141
CL\_INS\_141
CL\_INS\_141
CL\_INS\_141
CL\_INS\_141
CL\_INS\_141
CL\_INS\_141
CL\_INS\_141
CL\_INS\_141
CL\_INS\_141
CL\_INS\_141
CL\_INS\_141
CL\_INS\_141
CL\_INS\_141
CL\_INS\_141
CL\_INS\_141
CL\_INS\_141
CL\_INS\_141
CL\_INS\_141
CL\_INS\_141
CL\_INS\_141
CL\_INS\_141
CL\_INS\_141
CL\_INS\_141
CL\_INS\_141
CL\_INS\_141
CL\_INS\_141
CL\_INS\_141
CL\_INS\_141
CL\_INS\_141
CL\_INS\_141
CL\_INS\_141
CL\_INS\_141
CL\_INS\_141
CL\_INS\_141
CL\_INS\_141
CL\_INS\_141
CL\_INS\_141
CL\_INS\_141
CL\_INS\_141
CL\_INS\_141
CL\_INS\_141
CL\_INS\_141
CL\_INS\_141
CL\_INS\_141
CL\_INS\_141
CL\_INS\_141
CL\_INS\_141
CL\_INS\_141
CL\_INS\_141
CL\_INS\_141
CL\_INS\_141
CL\_INS\_141
CL\_INS\_141
CL\_INS\_141
CL\_INS\_141
CL\_INS\_141
CL\_INS\_141
CL\_INS\_141
CL\_INS\_141
CL\_INS\_141
CL\_INS\_141
CL\_INS\_141
CL\_INS\_141
CL\_INS\_141
CL\_INS\_141
CL\_INS\_141
CL\_INS\_141
CL\_INS\_141
CL\_INS\_141
CL\_INS\_141
CL\_INS\_141
CL\_INS\_141
CL\_INS\_141
CL\_INS\_141
CL\_INS\_141
CL\_INS\_141
CL\_INS\_141
CL\_INS\_141
CL\_INS\_141
CL\_INS\_141
CL\_INS\_141
CL\_INS\_141
CL\_INS\_141
CL\_INS\_141
CL\_INS\_141
CL\_INS\_141
CL\_INS\_141
CL\_INS\_141
CL\_INS\_141
CL\_INS\_141
CL\_INS\_141
CL\_INS\_141
CL\_INS\_141
CL\_INS\_141
CL\_INS\_141
CL\_INS\_141
CL\_INS\_141
CL\_INS\_141
CL\_INS\_141
CL\_INS\_141
CL\_INS\_141
CL\_INS\_141
CL\_INS\_141
CL\_INS\_141
CL\_INS\_141
CL\_INS\_141
CL\_INS\_141
CL\_INS\_141
CL\_INS\_141
CL\_INS\_141
CL\_INS\_141
CL\_INS\_141
CL\_INS\_141
CL\_INS\_141
CL\_INS\_141
CL\_INS\_141
CL\_INS\_141
CL\_INS\_141
CL\_INS\_141
CL\_INS\_141
CL\_INS\_141
CL\_INS\_141
CL\_INS\_141
CL\_INS\_141
CL\_INS\_141
CL\_INS\_141
CL\_INS\_141
CL\_INS\_141
CL\_INS\_141
CL\_INS\_141
CL\_INS\_141
CL\_INS\_141
CL\_INS\_141
CL\_INS\_141
CL\_INS\_141
CL\_INS\_141
CL\_INS\_141
CL\_INS\_141
CL\_INS\_141
CL\_INS\_141
CL\_INS\_141
CL\_INS\_141
CL\_INS\_141
CL\_INS\_141
CL\_INS\_141
CL\_INS\_141
CL\_INS\_141
CL\_INS\_141
CL\_INS\_141
CL\_INS\_141
CL\_INS\_141
CL\_INS\_141
CL\_INS\_141
CL\_INS\_141
CL\_INS\_141
CL\_INS\_141
CL\_INS\_141
CL\_INS\_141
CL\_INS\_141
CL\_INS\_141
CL\_INS\_204
CL\_INS\_204
CL\_INS\_204
CL\_INS\_141
CL\_INS\_204
CL\_INS\_204
CL\_INS\_204
CL\_INS\_204
CL\_INS\_204
CL\_INS\_204
CL\_INS\_204
CL\_INS\_141
CL\_INS\_141
CL\_INS\_141
CL\_INS\_141
CL\_INS\_141
CL\_INS\_141
CL\_INS\_141
CL\_INS\_141
CL\_INS\_141
CL\_INS\_141
CL\_INS\_141
CL\_INS\_141
CL\_INS\_141
CL\_INS\_141
CL\_INS\_141
CL\_INS\_141
CL\_INS\_141
CL\_INS\_141
CL\_INS\_141
CL\_INS\_141
CL\_INS\_141
CL\_INS\_141
CL\_INS\_141
CL\_INS\_141
CL\_INS\_141
CL\_INS\_141
CL\_INS\_141
CL\_INS\_141
CL\_INS\_141
CL\_INS\_141
CL\_INS\_141
CL\_INS\_141
CL\_INS\_141
CL\_INS\_141
CL\_INS\_141
CL\_INS\_141
CL\_INS\_141
CL\_INS\_141
CL\_INS\_141
CL\_INS\_141
CL\_INS\_141
CL\_INS\_141
CL\_INS\_141
CL\_INS\_141
CL\_INS\_141
CL\_INS\_141
CL\_INS\_141
CL\_INS\_141
CL\_INS\_141
CL\_INS\_141
CL\_INS\_141
CL\_INS\_141
CL\_INS\_204
CL\_INS\_204
CL\_INS\_204
CL\_INS\_204
CL\_INS\_141
CL\_INS\_141
CL\_INS\_141
CL\_INS\_141
CL\_INS\_141
CL\_INS\_141
CL\_INS\_141
CL\_INS\_141
CL\_INS\_141
CL\_INS\_141
CL\_INS\_141
CL\_INS\_141
CL\_INS\_141
CL\_INS\_141
CL\_INS\_141
CL\_INS\_141
CL\_INS\_141
CL\_INS\_141
CL\_INS\_141
CL\_INS\_141
CL\_INS\_141
CL\_INS\_141
CL\_INS\_141
CL\_INS\_141
CL\_INS\_141
CL\_INS\_141
CL\_INS\_141
CL\_INS\_141
CL\_INS\_141
CL\_INS\_141
CL\_INS\_141
CL\_INS\_141
CL\_INS\_365
CL\_INS\_141
CL\_INS\_141
CL\_INS\_141
CL\_INS\_141
CL\_INS\_141
CL\_INS\_141
CL\_INS\_141
CL\_INS\_141
CL\_INS\_141
CL\_INS\_141
CL\_INS\_141
CL\_INS\_141
CL\_INS\_204
CL\_INS\_204
CL\_INS\_204
CL\_INS\_204
CL\_INS\_204
CL\_INS\_204
CL\_INS\_141
CL\_INS\_204
CL\_INS\_141
CL\_INS\_141
CL\_INS\_141
CL\_INS\_141
CL\_INS\_141
CL\_INS\_204
CL\_INS\_141
CL\_INS\_204
CL\_INS\_204
CL\_INS\_204
CL\_INS\_204
CL\_INS\_204
CL\_INS\_141
CL\_INS\_204
CL\_INS\_204
CL\_INS\_141
CL\_INS\_141
CL\_INS\_141
CL\_INS\_141
CL\_INS\_141
CL\_INS\_141
CL\_INS\_141
CL\_INS\_141
CL\_INS\_141
CL\_INS\_141
CL\_INS\_141
CL\_INS\_141
CL\_INS\_141
CL\_INS\_141
CL\_INS\_141
CL\_INS\_365
CL\_INS\_365
CL\_INS\_365
CL\_INS\_365
CL\_INS\_365
CL\_INS\_365
CL\_INS\_365
CL\_INS\_365
CL\_INS\_365
CL\_INS\_365
CL\_INS\_365
CL\_INS\_365
CL\_INS\_365
CL\_INS\_365
CL\_INS\_365
CL\_INS\_365
CL\_INS\_365
CL\_INS\_365
CL\_INS\_141
CL\_INS\_141
CL\_INS\_141
CL\_INS\_141
CL\_INS\_141
CL\_INS\_141
CL\_INS\_141
CL\_INS\_141
CL\_INS\_141
CL\_INS\_141
CL\_INS\_141
CL\_INS\_141
CL\_INS\_141
CL\_INS\_141
CL\_INS\_141
CL\_INS\_141
CL\_INS\_204
CL\_INS\_204
CL\_INS\_141
CL\_INS\_204
CL\_INS\_204
CL\_INS\_204
CL\_INS\_141
CL\_INS\_141
CL\_INS\_141
CL\_INS\_141
CL\_INS\_141
CL\_INS\_141
CL\_INS\_141
CL\_INS\_141
CL\_INS\_141
CL\_INS\_141
CL\_INS\_141
CL\_INS\_141
CL\_INS\_141
CL\_INS\_141
CL\_INS\_141
CL\_INS\_141
CL\_INS\_141
CL\_INS\_141
CL\_INS\_141
CL\_INS\_141
CL\_INS\_141
CL\_INS\_141
CL\_INS\_141
CL\_INS\_141
CL\_INS\_141
CL\_INS\_141
CL\_INS\_141
CL\_INS\_141
CL\_INS\_141
CL\_INS\_141
CL\_INS\_141
CL\_INS\_141
Cluster ID


CL\_9005
CL\_2003
CL\_2004
CL\_10810
CL\_7182
CL\_10811
CL\_7181
CL\_7180
CL\_6152
CL\_7179
CL\_7178
CL\_10813
CL\_7177
CL\_7176
CL\_6151
CL\_10812
CL\_3184
CL\_3183
CL\_3182
CL\_6150
CL\_3181
CL\_3180
CL\_6149
CL\_6148
CL\_3177
CL\_7175
CL\_3175
CL\_3174
CL\_3173
CL\_6147
CL\_6146
CL\_6145
CL\_10814
CL\_10815
CL\_10816
CL\_10817
CL\_11724
CL\_6144
CL\_6143
CL\_10818
CL\_6127
CL\_7174
CL\_13709
CL\_13710
CL\_13814
CL\_13711
CL\_12181
CL\_12182
CL\_12183
CL\_12184
CL\_12185
CL\_13353
CL\_13354
CL\_13355
CL\_13356
CL\_3172
CL\_10819
CL\_3170
CL\_3169
CL\_3168
CL\_3167
CL\_3166
CL\_3165
CL\_3164
CL\_13655
CL\_3163
CL\_3162
CL\_3161
CL\_3160
CL\_3159
CL\_3158
CL\_3157
CL\_3156
CL\_3155
CL\_7209
CL\_1298
CL\_7208
CL\_7207
CL\_5689
CL\_12057
CL\_12056
CL\_12055
CL\_5695
CL\_5690
CL\_5691
CL\_7972
CL\_12919
CL\_7971
CL\_12920
CL\_7970
CL\_7969
CL\_10435
CL\_10436
CL\_7968
CL\_7967
CL\_7966
CL\_7965
CL\_7964
CL\_9431
CL\_9432
CL\_9433
CL\_9434
CL\_5697
CL\_5698
CL\_12054
CL\_12053
CL\_12052
CL\_12051
CL\_9268
CL\_12050
CL\_12049
CL\_12048
CL\_12047
CL\_7223
CL\_7222
CL\_12234
CL\_5267
CL\_5268
CL\_5269
CL\_5701
CL\_7221
CL\_7220
CL\_5702
CL\_5703
CL\_9435
CL\_9436
CL\_9437
CL\_5707
CL\_7217
CL\_10085
CL\_10084
CL\_7173
CL\_7172
CL\_7171
CL\_7170
CL\_7169
CL\_7168
CL\_7167
CL\_7166
CL\_7165
CL\_7164
CL\_7163
CL\_7162
CL\_7161
CL\_7160
CL\_7159
CL\_7158
CL\_7157
CL\_7156
CL\_7155
CL\_7154
CL\_7153
CL\_7152
CL\_7151
CL\_9438
CL\_5708
CL\_9439
CL\_9440
CL\_10083
CL\_12046
CL\_9441
CL\_9442
CL\_9443
CL\_2005
CL\_13660
CL\_13659
CL\_13658
CL\_13657
CL\_1280
CL\_9266
CL\_11973
CL\_1285
CL\_7963
CL\_1288
CL\_1289
CL\_1290
CL\_1291
CL\_5270
CL\_1292
CL\_2006
CL\_2007
CL\_2008
CL\_2009
CL\_2010
CL\_2011
CL\_2012
CL\_2013
CL\_2014
CL\_2015
CL\_2016
CL\_2017
CL\_2018
CL\_2019
CL\_2020
CL\_2021
CL\_2022
CL\_7219
CL\_12233
CL\_12232
CL\_12231
CL\_7211
CL\_2023
CL\_2024
CL\_2025
CL\_7962
CL\_7961
CL\_13656
CL\_1297
CL\_2026
CL\_2027
CL\_2028
CL\_2029
CL\_2030
CL\_7960
CL\_12480
CL\_7020
CL\_7021
CL\_5275
CL\_10451
CL\_5276
CL\_5277
CL\_5278
CL\_5279
CL\_5280
CL\_5715
CL\_5716
CL\_2031
CL\_2032
CL\_2033
CL\_7959
CL\_2034
CL\_2035
CL\_2036
CL\_2037
CL\_2038
CL\_2826
CL\_2825
CL\_2824
CL\_2039
CL\_2040
CL\_2041
CL\_2042
CL\_2043
CL\_2044
CL\_2045
CL\_2046
CL\_2047
CL\_2048
CL\_2049
CL\_2050
CL\_2051
CL\_2052
CL\_2053
CL\_2054
CL\_2055
CL\_2056
CL\_2057
CL\_2058
CL\_2864
CL\_2863
CL\_9444
CL\_9445
CL\_1302
CL\_8180
CL\_8178
CL\_8177
CL\_8176
CL\_8175
CL\_8174
CL\_8173
CL\_8172
CL\_8171
CL\_8170
CL\_8169
CL\_8168
CL\_8167
CL\_8166
CL\_10820
CL\_2862
CL\_6124
CL\_6123
CL\_2861
CL\_10821
CL\_10822
CL\_10823
CL\_10824
CL\_10825
CL\_10826
CL\_10827
CL\_10828
CL\_10829
CL\_10830
CL\_10831
CL\_10832
CL\_10833
CL\_10834
CL\_10835
CL\_10836
CL\_10837
CL\_10838
CL\_10839
CL\_11723
CL\_10840
CL\_10841
CL\_10842
CL\_10843
CL\_10844
CL\_10845
CL\_11722
CL\_10846
CL\_10847
CL\_11721
CL\_10848
CL\_2860
CL\_6122
CL\_2859
CL\_2858
CL\_12449
CL\_12448
CL\_12447
CL\_12446
CL\_9410
CL\_9411
CL\_9412
CL\_9413
CL\_9414
CL\_5282
CL\_3154
CL\_5717
CL\_5718
CL\_5719
CL\_5720
CL\_7023
CL\_12045
CL\_5721
CL\_7103
CL\_7102
CL\_7101
CL\_7100
CL\_13654
CL\_7025
CL\_9416
CL\_7026
CL\_7027
CL\_7028
CL\_7029
CL\_7030
CL\_9417
CL\_7031
CL\_7032
CL\_7150
CL\_7149
CL\_3153
CL\_3152
CL\_3151
CL\_3150
CL\_3149
CL\_12655
CL\_13357
CL\_3148
CL\_3147
CL\_3146
CL\_3145
CL\_6142
CL\_6141
CL\_2857
CL\_2856
CL\_2855
CL\_6120
CL\_2854
CL\_2853
CL\_2852
CL\_2851
CL\_2850
CL\_2849
CL\_2848
CL\_6119
CL\_2847
CL\_2846
CL\_2845
CL\_2844
CL\_2843
CL\_2842
CL\_2841
CL\_2840
CL\_2839
CL\_2838
CL\_2837
CL\_2836
CL\_2835
CL\_2834
CL\_2833
CL\_2832
CL\_2831
CL\_2830
CL\_2829
CL\_2828
CL\_7941
CL\_7942
CL\_7205
CL\_7204
CL\_7203
CL\_7202
CL\_7201
CL\_7200
CL\_7199
CL\_7198
CL\_7197
CL\_7196
CL\_1314
CL\_1315
CL\_8151
CL\_1316
CL\_1317
CL\_1318
CL\_1319
CL\_9446
CL\_7943
CL\_9010
CL\_9009
CL\_7944
CL\_7945
CL\_3143
CL\_3142
CL\_2059
CL\_13358
CL\_13359
CL\_2060
CL\_9447
CL\_6140
CL\_2061
CL\_2062
CL\_2063
CL\_2064
CL\_1326
CL\_2065
CL\_2066
