## Supplementary material for "A novel method for integrating genomic and Tn-Seq data to identify common *in vivo* fitness mechanisms across multiple bacterial species": S4 Dataset: CL_INS_144.html

Legend

 Hypothetical
 DNA Metabolism
 Other
 All VFDB Genes

FULL


WINDOWSVGPNG

Trim RowsRemove SingletonsSave Fasta

CL\_2091


CL\_2091


CL\_2091


CL\_2091

HighlightSelectShow Genomes


21

CL\_2092


3

CL\_2092


2

CL\_2092


2

CL\_2092

fGI ID


CL\_INS\_144
CL\_INS\_144
CL\_INS\_144
CL\_INS\_144
CL\_INS\_144
CL\_INS\_144
CL\_INS\_144
CL\_INS\_144
CL\_INS\_144
CL\_INS\_144
CL\_INS\_144
CL\_INS\_144
CL\_INS\_144
Cluster ID


CL\_5325
CL\_5326
CL\_5327
CL\_13712
CL\_13713
CL\_5204
CL\_607
CL\_608
CL\_5328
CL\_13059
CL\_5329
CL\_5330
CL\_5331
