## Supplementary material for "A novel method for integrating genomic and Tn-Seq data to identify common *in vivo* fitness mechanisms across multiple bacterial species": S4 Dataset: CL_INS_146.html

Legend

 Mobile +extrachromosomalelementfunctions
 Hypothetical
 Proteinsynthesis/fate
 Cellularprocesses
 All VFDB Genes

FULL


WINDOWSVGPNG

Trim RowsRemove SingletonsSave Fasta

CL\_2104


CL\_2104


CL\_2104


CL\_2104


CL\_2104


CL\_2104


CL\_2104


CL\_2104

HighlightSelectShow Genomes


21

CL\_2135


1

CL\_2135


1

CL\_2135


1

CL\_2135


1

CL\_2135


1

CL\_2135


1

CL\_2135


1

CL\_2135

fGI ID


CL\_INS\_146
CL\_INS\_146
CL\_INS\_146
CL\_INS\_146
CL\_INS\_146
CL\_INS\_146
CL\_INS\_146
CL\_INS\_146
CL\_INS\_146
CL\_INS\_146
CL\_INS\_146
CL\_INS\_146
CL\_INS\_146
CL\_INS\_146
CL\_INS\_146
CL\_INS\_146
CL\_INS\_146
CL\_INS\_146
CL\_INS\_146
CL\_INS\_146
CL\_INS\_146
CL\_INS\_146
CL\_INS\_146
CL\_INS\_146
CL\_INS\_146
CL\_INS\_146
CL\_INS\_146
CL\_INS\_146
CL\_INS\_146
CL\_INS\_146
CL\_INS\_146
CL\_INS\_146
CL\_INS\_146
CL\_INS\_146
Cluster ID


CL\_2105
CL\_7946
CL\_2106
CL\_2107
CL\_2108
CL\_2109
CL\_2110
CL\_2111
CL\_2112
CL\_10171
CL\_2113
CL\_2114
CL\_2115
CL\_2116
CL\_2117
CL\_2118
CL\_2119
CL\_2120
CL\_2121
CL\_2122
CL\_2123
CL\_2124
CL\_2125
CL\_7947
CL\_2126
CL\_2127
CL\_2128
CL\_2129
CL\_2130
CL\_2131
CL\_10172
CL\_2132
CL\_2133
CL\_2134
