## Supplementary material for "A novel method for integrating genomic and Tn-Seq data to identify common *in vivo* fitness mechanisms across multiple bacterial species": S4 Dataset: CL_INS_148.html

Legend

 Hypothetical
 Transport +binding proteins
 All VFDB Genes

FULL


WINDOWSVGPNG

Trim RowsRemove SingletonsSave Fasta

CL\_2147


CL\_2147

HighlightSelectShow Genomes


24

CL\_2148


3

CL\_2148

fGI ID

CL\_INS\_148
Cluster ID

CL\_8213
