## Supplementary material for "A novel method for integrating genomic and Tn-Seq data to identify common *in vivo* fitness mechanisms across multiple bacterial species": S4 Dataset: CL_INS_149.html

Legend

 Mobile +extrachromosomalelementfunctions
 Hypothetical
 All Fitness Genes
 Transport +binding proteins
 All VFDB Genes

FULL


WINDOWSVGPNG

Trim RowsRemove SingletonsSave Fasta

CL\_2192


CL\_2192


CL\_2192

HighlightSelectShow Genomes


26

CL\_5332


1

CL\_2199


1

CL\_2195

fGI ID


CL\_INS\_149
CL\_INS\_149
CL\_INS\_149
CL\_INS\_149
Cluster ID


CL\_2193
CL\_9477
CL\_9478
CL\_9479
