## Supplementary material for "A novel method for integrating genomic and Tn-Seq data to identify common *in vivo* fitness mechanisms across multiple bacterial species": S4 Dataset: CL_INS_156.html

Legend

 Hypothetical
 Other
 All VFDB Genes

FULL


WINDOWSVGPNG

Trim RowsRemove SingletonsSave Fasta

CL\_2281


CL\_2281


CL\_2281

HighlightSelectShow Genomes


16

CL\_2282


11

CL\_2282


1

CL\_2283

fGI ID

CL\_INS\_156
Cluster ID

CL\_5341
