## Supplementary material for "A novel method for integrating genomic and Tn-Seq data to identify common *in vivo* fitness mechanisms across multiple bacterial species": S4 Dataset: CL_INS_158.html

Legend

 All Fitness Genes
 All VFDB Genes

FULL


WINDOWSVGPNG

Trim RowsRemove SingletonsSave Fasta

CL\_2294


CL\_2294


CL\_2294

HighlightSelectShow Genomes


17

CL\_2292


8

CL\_2291


3

CL\_2292

fGI ID

CL\_INS\_365
Cluster ID

CL\_2293
