## Supplementary material for "A novel method for integrating genomic and Tn-Seq data to identify common *in vivo* fitness mechanisms across multiple bacterial species": S4 Dataset: CL_INS_163.html

Legend

 Mobile +extrachromosomalelementfunctions
 Hypothetical
 All VFDB Genes

FULL


WINDOWSVGPNG

Trim RowsRemove SingletonsSave Fasta

CL\_2366


CL\_2366


CL\_2366


CL\_2366


CL\_2366


CL\_2365

HighlightSelectShow Genomes


15

CL\_2373


5

CL\_2373


3

CL\_2373


3

CL\_2373


1

CL\_2373


1

CL\_2373

fGI ID


CL\_INS\_163
CL\_INS\_163
CL\_INS\_163
CL\_INS\_163
CL\_INS\_163
CL\_INS\_163
CL\_INS\_163
CL\_INS\_163
Cluster ID


CL\_2367
CL\_7077
CL\_7076
CL\_2368
CL\_2369
CL\_2370
CL\_2371
CL\_2372
