## Supplementary material for "A novel method for integrating genomic and Tn-Seq data to identify common *in vivo* fitness mechanisms across multiple bacterial species": S4 Dataset: CL_INS_164.html

Legend

 Mobile +extrachromosomalelementfunctions
 Regulatoryfunctions
 Hypothetical
 DNA Metabolism
 Other
 EnergyMetabolism
 All VFDB Genes

FULL


WINDOWSVGPNG

Trim RowsRemove SingletonsSave Fasta

CL\_2392


CL\_2392


CL\_2392


CL\_2392

HighlightSelectShow Genomes


14

CL\_2393


10

CL\_2393


2

CL\_2393


1

CL\_2393

fGI ID


CL\_INS\_164
CL\_INS\_164
CL\_INS\_164
CL\_INS\_164
CL\_INS\_164
CL\_INS\_164
CL\_INS\_164
CL\_INS\_164
CL\_INS\_164
CL\_INS\_164
CL\_INS\_366
CL\_INS\_164
CL\_INS\_164
CL\_INS\_164
CL\_INS\_164
CL\_INS\_164
CL\_INS\_164
CL\_INS\_164
CL\_INS\_164
CL\_INS\_164
CL\_INS\_164
CL\_INS\_164
CL\_INS\_164
CL\_INS\_164
CL\_INS\_164
CL\_INS\_164
CL\_INS\_164
CL\_INS\_164
CL\_INS\_164
CL\_INS\_164
CL\_INS\_164
CL\_INS\_270
CL\_INS\_270
CL\_INS\_270
CL\_INS\_164
CL\_INS\_164
CL\_INS\_164
CL\_INS\_164
CL\_INS\_164
CL\_INS\_164
CL\_INS\_270
CL\_INS\_270
CL\_INS\_270
CL\_INS\_270
CL\_INS\_270
CL\_INS\_270
CL\_INS\_270
CL\_INS\_164
CL\_INS\_270
CL\_INS\_270
CL\_INS\_270
CL\_INS\_270
CL\_INS\_270
CL\_INS\_164
CL\_INS\_164
CL\_INS\_164
CL\_INS\_164
CL\_INS\_164
CL\_INS\_164
CL\_INS\_164
CL\_INS\_164
CL\_INS\_164
CL\_INS\_164
CL\_INS\_164
CL\_INS\_164
CL\_INS\_164
Cluster ID


CL\_11398
CL\_11397
CL\_11396
CL\_11395
CL\_11394
CL\_11393
CL\_11392
CL\_11391
CL\_11390
CL\_11389
CL\_7146
CL\_11388
CL\_11387
CL\_11386
CL\_11385
CL\_11384
CL\_11383
CL\_11382
CL\_11381
CL\_11380
CL\_11379
CL\_11378
CL\_11377
CL\_11376
CL\_11375
CL\_11374
CL\_11373
CL\_11372
CL\_11371
CL\_11370
CL\_11369
CL\_3940
CL\_3939
CL\_3938
CL\_11368
CL\_11367
CL\_11366
CL\_11365
CL\_11364
CL\_11363
CL\_11362
CL\_11361
CL\_11360
CL\_11359
CL\_11358
CL\_11357
CL\_11356
CL\_11355
CL\_11354
CL\_11353
CL\_11352
CL\_11351
CL\_11350
CL\_11349
CL\_11348
CL\_11347
CL\_11346
CL\_11345
CL\_11344
CL\_11343
CL\_11342
CL\_11341
CL\_11340
CL\_11339
CL\_6098
CL\_5363
