## Supplementary material for "A novel method for integrating genomic and Tn-Seq data to identify common *in vivo* fitness mechanisms across multiple bacterial species": S4 Dataset: CL_INS_165.html

Legend

 Hypothetical
 Other
 All VFDB Genes

FULL


WINDOWSVGPNG

Trim RowsRemove SingletonsSave Fasta

CL\_2398


CL\_2398


CL\_2398


CL\_2398


CL\_2398


CL\_2398

HighlightSelectShow Genomes


9

CL\_6096


6

CL\_6096


3

CL\_2399


3

CL\_6096


2

CL\_6096


1

CL\_6096

fGI ID


CL\_INS\_165
CL\_INS\_165
CL\_INS\_165
Cluster ID


CL\_6097
CL\_5364
CL\_5365
