## Supplementary material for "A novel method for integrating genomic and Tn-Seq data to identify common *in vivo* fitness mechanisms across multiple bacterial species": S4 Dataset: CL_INS_166.html

Legend

 Mobile +extrachromosomalelementfunctions
 Hypothetical
 All VFDB Genes

FULL


WINDOWSVGPNG

Trim RowsRemove SingletonsSave Fasta

CL\_2419


CL\_2419


CL\_2419


CL\_2419

HighlightSelectShow Genomes


22

CL\_2420


2

CL\_2420


2

CL\_2420


1

CL\_2420

fGI ID


CL\_INS\_166
CL\_INS\_166
CL\_INS\_166
CL\_INS\_166
CL\_INS\_166
Cluster ID


CL\_9483
CL\_9484
CL\_9485
CL\_9486
CL\_9487
