## Supplementary material for "A novel method for integrating genomic and Tn-Seq data to identify common *in vivo* fitness mechanisms across multiple bacterial species": S4 Dataset: CL_INS_167.html

Legend

 Hypothetical
 Regulatoryfunctions
 Other
 All VFDB Genes

FULL


WINDOWSVGPNG

Trim RowsRemove SingletonsSave Fasta

CL\_2422


CL\_2422


CL\_2422


CL\_2422

HighlightSelectShow Genomes


11

CL\_2423


6

CL\_2423


6

CL\_2423


5

CL\_2423

fGI ID


CL\_INS\_167
CL\_INS\_167
CL\_INS\_167
CL\_INS\_167
Cluster ID


CL\_5366
CL\_6095
CL\_6094
CL\_6093
