## Supplementary material for "A novel method for integrating genomic and Tn-Seq data to identify common *in vivo* fitness mechanisms across multiple bacterial species": S4 Dataset: CL_INS_169.html

Legend

 Mobile +extrachromosomalelementfunctions
 Hypothetical
 Other
 All VFDB Genes

FULL


WINDOWSVGPNG

Trim RowsRemove SingletonsSave Fasta

CL\_2425


CL\_2425


CL\_2425


CL\_2425


CL\_2425

HighlightSelectShow Genomes


14

CL\_2428


11

CL\_2428


1

CL\_2428


1

CL\_2428


1

CL\_2429

fGI ID


CL\_INS\_169
CL\_INS\_169
CL\_INS\_169
Cluster ID


CL\_2426
CL\_13736
CL\_2427
