## Supplementary material for "A novel method for integrating genomic and Tn-Seq data to identify common *in vivo* fitness mechanisms across multiple bacterial species": S4 Dataset: CL_INS_171.html

Legend

 Hypothetical
 Other
 All VFDB Genes

FULL


WINDOWSVGPNG

Trim RowsRemove SingletonsSave Fasta

CL\_2438


CL\_2438


CL\_2438


CL\_2438

HighlightSelectShow Genomes


14

CL\_2441


10

CL\_2441


2

CL\_2441


2

CL\_2441

fGI ID


CL\_INS\_171
CL\_INS\_171
CL\_INS\_171
Cluster ID


CL\_12220
CL\_2439
CL\_2440
