## Supplementary material for "A novel method for integrating genomic and Tn-Seq data to identify common *in vivo* fitness mechanisms across multiple bacterial species": S4 Dataset: CL_INS_173.html

Legend

 Mobile +extrachromosomalelementfunctions
 Hypothetical
 Other
 All VFDB Genes

FULL


WINDOWSVGPNG

Trim RowsRemove SingletonsSave Fasta

CL\_2471


CL\_2471


CL\_2471


CL\_2471

HighlightSelectShow Genomes


18

CL\_2472


4

CL\_2472


3

CL\_2478


1

CL\_2472

fGI ID


CL\_INS\_173
CL\_INS\_173
CL\_INS\_174
CL\_INS\_174
CL\_INS\_173
Cluster ID


CL\_10859
CL\_6091
CL\_6090
CL\_6089
CL\_6088
