## Supplementary material for "A novel method for integrating genomic and Tn-Seq data to identify common *in vivo* fitness mechanisms across multiple bacterial species": S4 Dataset: CL_INS_176.html

Legend

 Mobile +extrachromosomalelementfunctions
 Hypothetical
 Regulatoryfunctions
 Other
 All VFDB Genes

FULL


WINDOWSVGPNG

Trim RowsRemove SingletonsSave Fasta

CL\_2482


CL\_2482


CL\_2482


Break

HighlightSelectShow Genomes


20

CL\_2483


6

CL\_2483


1

CL\_2483


1

CL\_2483

fGI ID


CL\_INS\_176
CL\_INS\_176
CL\_INS\_176
CL\_INS\_176
CL\_INS\_176
CL\_INS\_176
CL\_INS\_176
CL\_INS\_176
CL\_INS\_176
Cluster ID


CL\_9519
CL\_9520
CL\_9521
CL\_9522
CL\_9523
CL\_9524
CL\_9525
CL\_9526
CL\_9527
