## Supplementary material for "A novel method for integrating genomic and Tn-Seq data to identify common *in vivo* fitness mechanisms across multiple bacterial species": S4 Dataset: CL_INS_179.html

Legend

 Mobile +extrachromosomalelementfunctions
 Regulatoryfunctions
 Hypothetical
 DNA Metabolism
 Proteinsynthesis/fate
 Other
 All VFDB Genes

FULL


WINDOWSVGPNG

Trim RowsRemove SingletonsSave Fasta

CL\_2505


CL\_2505


CL\_2505


CL\_2505


CL\_2505


CL\_2505

HighlightSelectShow Genomes


18

CL\_2504


4

CL\_2503


2

CL\_2503


1

CL\_2504


1

CL\_2503


1

CL\_2503

fGI ID


CL\_INS\_179
CL\_INS\_179
CL\_INS\_178
CL\_INS\_366
CL\_INS\_366
CL\_INS\_366
CL\_INS\_366
CL\_INS\_366
CL\_INS\_178
CL\_INS\_178
CL\_INS\_366
CL\_INS\_178
CL\_INS\_178
CL\_INS\_366
CL\_INS\_366
CL\_INS\_366
CL\_INS\_178
CL\_INS\_178
CL\_INS\_178
CL\_INS\_178
CL\_INS\_270
CL\_INS\_178
CL\_INS\_366
CL\_INS\_366
CL\_INS\_366
CL\_INS\_178
CL\_INS\_366
CL\_INS\_178
CL\_INS\_178
CL\_INS\_178
CL\_INS\_178
CL\_INS\_178
CL\_INS\_178
CL\_INS\_178
CL\_INS\_178
CL\_INS\_178
CL\_INS\_178
CL\_INS\_178
CL\_INS\_178
CL\_INS\_178
CL\_INS\_178
CL\_INS\_178
CL\_INS\_178
CL\_INS\_178
CL\_INS\_178
CL\_INS\_178
CL\_INS\_178
CL\_INS\_178
CL\_INS\_178
CL\_INS\_178
CL\_INS\_178
CL\_INS\_178
CL\_INS\_141
CL\_INS\_141
CL\_INS\_178
CL\_INS\_141
CL\_INS\_141
CL\_INS\_178
CL\_INS\_178
CL\_INS\_178
CL\_INS\_178
Cluster ID


CL\_10250
CL\_10893
CL\_10892
CL\_7147
CL\_10891
CL\_7146
CL\_7145
CL\_10890
CL\_10889
CL\_10888
CL\_9045
CL\_10887
CL\_10886
CL\_9038
CL\_10536
CL\_10537
CL\_10885
CL\_10884
CL\_10883
CL\_10882
CL\_10881
CL\_10880
CL\_7130
CL\_7129
CL\_7128
CL\_10879
CL\_7127
CL\_10878
CL\_7125
CL\_7124
CL\_7123
CL\_7122
CL\_10877
CL\_10876
CL\_7117
CL\_7116
CL\_7115
CL\_10875
CL\_7112
CL\_7111
CL\_7110
CL\_7109
CL\_5390
CL\_5387
CL\_5386
CL\_5385
CL\_5384
CL\_5383
CL\_10874
CL\_10873
CL\_10872
CL\_10871
CL\_7103
CL\_7102
CL\_10870
CL\_7101
CL\_7100
CL\_10869
CL\_10868
CL\_7096
CL\_10867
