## Supplementary material for "A novel method for integrating genomic and Tn-Seq data to identify common *in vivo* fitness mechanisms across multiple bacterial species": S4 Dataset: CL_INS_181.html

Legend

 Mobile +extrachromosomalelementfunctions
 Hypothetical
 Other
 Transport +binding proteins
 All VFDB Genes

FULL


WINDOWSVGPNG

Trim RowsRemove SingletonsSave Fasta

CL\_2516


CL\_2516


CL\_2516


CL\_2516

HighlightSelectShow Genomes


12

CL\_2518


12

CL\_2518


3

CL\_2518


1

CL\_2518

fGI ID


CL\_INS\_181
CL\_INS\_181
CL\_INS\_181
CL\_INS\_181
CL\_INS\_181
CL\_INS\_181
CL\_INS\_181
CL\_INS\_181
CL\_INS\_181
CL\_INS\_181
CL\_INS\_181
CL\_INS\_181
CL\_INS\_181
CL\_INS\_181
CL\_INS\_181
CL\_INS\_181
CL\_INS\_181
CL\_INS\_365
CL\_INS\_365
CL\_INS\_365
CL\_INS\_365
CL\_INS\_365
CL\_INS\_365
CL\_INS\_365
CL\_INS\_181
Cluster ID


CL\_2517
CL\_12962
CL\_12961
CL\_12960
CL\_12959
CL\_12958
CL\_12957
CL\_12956
CL\_12955
CL\_8202
CL\_8203
CL\_8204
CL\_8205
CL\_7698
CL\_7697
CL\_7696
CL\_7695
CL\_5020
CL\_8206
CL\_8207
CL\_4892
CL\_8208
CL\_8209
CL\_8210
CL\_8211
