## Supplementary material for "A novel method for integrating genomic and Tn-Seq data to identify common *in vivo* fitness mechanisms across multiple bacterial species": S4 Dataset: CL_INS_182.html

Legend

 Mobile +extrachromosomalelementfunctions
 Hypothetical
 Proteinsynthesis/fate
 All VFDB Genes

FULL


WINDOWSVGPNG

Trim RowsRemove SingletonsSave Fasta

CL\_2530


CL\_2530


CL\_2530


CL\_2530


CL\_2530

HighlightSelectShow Genomes


23

CL\_2531


2

CL\_2531


1

CL\_2531


1

CL\_2531


1

CL\_2531

fGI ID


CL\_INS\_182
CL\_INS\_182
CL\_INS\_182
CL\_INS\_182
CL\_INS\_182
CL\_INS\_182
CL\_INS\_182
CL\_INS\_182
CL\_INS\_182
CL\_INS\_182
CL\_INS\_182
CL\_INS\_182
CL\_INS\_182
CL\_INS\_182
CL\_INS\_182
CL\_INS\_182
CL\_INS\_182
CL\_INS\_182
CL\_INS\_182
CL\_INS\_182
Cluster ID


CL\_9529
CL\_9530
CL\_7948
CL\_12217
CL\_12216
CL\_12215
CL\_12214
CL\_12213
CL\_12212
CL\_7949
CL\_7082
CL\_7950
CL\_7084
CL\_7951
CL\_7952
CL\_7953
CL\_7954
CL\_7955
CL\_7956
CL\_7957
