## Supplementary material for "A novel method for integrating genomic and Tn-Seq data to identify common *in vivo* fitness mechanisms across multiple bacterial species": S4 Dataset: CL_INS_184.html

Legend

 Mobile +extrachromosomalelementfunctions
 Hypothetical
 DNA Metabolism
 Other
 EnergyMetabolism
 Transport +binding proteins
 All VFDB Genes

FULL


WINDOWSVGPNG

Trim RowsRemove SingletonsSave Fasta

CL\_2558


CL\_2558


CL\_2558

HighlightSelectShow Genomes


26

CL\_2557


1

CL\_2557


1

CL\_2557

fGI ID


CL\_INS\_184
CL\_INS\_117
CL\_INS\_117
CL\_INS\_184
CL\_INS\_184
CL\_INS\_117
CL\_INS\_117
CL\_INS\_117
CL\_INS\_117
CL\_INS\_117
CL\_INS\_184
CL\_INS\_117
CL\_INS\_117
CL\_INS\_117
CL\_INS\_184
CL\_INS\_184
CL\_INS\_184
CL\_INS\_117
CL\_INS\_117
CL\_INS\_117
CL\_INS\_117
Cluster ID


CL\_9536
CL\_9535
CL\_9534
CL\_9533
CL\_9532
CL\_9531
CL\_9097
CL\_1667
CL\_1668
CL\_1669
CL\_11712
CL\_1670
CL\_1671
CL\_1672
CL\_11713
CL\_11714
CL\_11715
CL\_1673
CL\_1674
CL\_11716
CL\_1675
