## Supplementary material for "A novel method for integrating genomic and Tn-Seq data to identify common *in vivo* fitness mechanisms across multiple bacterial species": S4 Dataset: CL_INS_189.html

Legend

 Mobile +extrachromosomalelementfunctions
 Hypothetical
 Other
 EnergyMetabolism
 All VFDB Genes

FULL


WINDOWSVGPNG

Trim RowsRemove SingletonsSave Fasta

CL\_2654


CL\_2654


CL\_2654


CL\_2654


CL\_2653


CL\_2654


CL\_2654

HighlightSelectShow Genomes


11

CL\_2658


10

CL\_2658


3

CL\_2658


1

CL\_2665


1

CL\_2658


1

CL\_2658


1

CL\_2658

fGI ID


CL\_INS\_189
CL\_INS\_189
CL\_INS\_189
CL\_INS\_191
CL\_INS\_189
Cluster ID


CL\_10251
CL\_5376
CL\_2655
CL\_2656
CL\_2657
