## Supplementary material for "A novel method for integrating genomic and Tn-Seq data to identify common *in vivo* fitness mechanisms across multiple bacterial species": S4 Dataset: CL_INS_195.html

Legend

 Mobile +extrachromosomalelementfunctions
 Regulatoryfunctions
 Hypothetical
 DNA Metabolism
 All Fitness Genes
 Proteinsynthesis/fate
 Other
 All VFDB Genes

FULL


WINDOWSVGPNG

Trim RowsRemove SingletonsSave Fasta

CL\_2714


CL\_2714


CL\_2714


CL\_2714


CL\_2714


CL\_2714


CL\_2714


CL\_2714


CL\_2714


CL\_2714


CL\_2168


CL\_2714


CL\_2714

HighlightSelectShow Genomes


14

CL\_2715


1

CL\_2715


1

CL\_2715


1

CL\_2715


1

CL\_2715


1

CL\_2715


1

CL\_2715


1

CL\_2715


1

CL\_2715


1

CL\_2715


1

CL\_2715


1

CL\_2167


1

CL\_2715

fGI ID


CL\_INS\_195
CL\_INS\_195
CL\_INS\_195
CL\_INS\_195
CL\_INS\_195
CL\_INS\_195
CL\_INS\_195
CL\_INS\_195
CL\_INS\_195
CL\_INS\_195
CL\_INS\_195
CL\_INS\_195
CL\_INS\_195
CL\_INS\_195
CL\_INS\_195
CL\_INS\_195
CL\_INS\_195
CL\_INS\_195
CL\_INS\_364
CL\_INS\_195
CL\_INS\_195
CL\_INS\_195
CL\_INS\_195
CL\_INS\_107
CL\_INS\_107
CL\_INS\_195
CL\_INS\_195
CL\_INS\_195
CL\_INS\_195
CL\_INS\_195
CL\_INS\_195
CL\_INS\_195
CL\_INS\_195
CL\_INS\_195
CL\_INS\_195
CL\_INS\_195
CL\_INS\_195
CL\_INS\_195
CL\_INS\_195
CL\_INS\_195
CL\_INS\_195
CL\_INS\_195
CL\_INS\_195
CL\_INS\_195
CL\_INS\_195
CL\_INS\_195
CL\_INS\_195
CL\_INS\_195
CL\_INS\_195
CL\_INS\_195
CL\_INS\_195
CL\_INS\_195
CL\_INS\_195
CL\_INS\_195
CL\_INS\_195
CL\_INS\_195
CL\_INS\_195
CL\_INS\_195
CL\_INS\_195
CL\_INS\_195
CL\_INS\_195
CL\_INS\_195
CL\_INS\_195
CL\_INS\_195
CL\_INS\_195
CL\_INS\_195
CL\_INS\_195
CL\_INS\_195
CL\_INS\_195
CL\_INS\_195
CL\_INS\_195
CL\_INS\_195
CL\_INS\_195
CL\_INS\_195
CL\_INS\_195
CL\_INS\_195
CL\_INS\_195
CL\_INS\_195
CL\_INS\_195
CL\_INS\_195
CL\_INS\_195
CL\_INS\_195
CL\_INS\_195
CL\_INS\_195
CL\_INS\_195
CL\_INS\_195
CL\_INS\_195
CL\_INS\_195
CL\_INS\_195
CL\_INS\_195
CL\_INS\_195
CL\_INS\_195
CL\_INS\_195
CL\_INS\_195
CL\_INS\_195
CL\_INS\_195
CL\_INS\_107
CL\_INS\_107
CL\_INS\_178
CL\_INS\_107
CL\_INS\_195
CL\_INS\_195
CL\_INS\_178
CL\_INS\_178
CL\_INS\_195
CL\_INS\_195
CL\_INS\_178
CL\_INS\_178
CL\_INS\_178
CL\_INS\_195
CL\_INS\_107
CL\_INS\_178
CL\_INS\_107
CL\_INS\_368
CL\_INS\_368
CL\_INS\_107
CL\_INS\_107
CL\_INS\_195
CL\_INS\_368
CL\_INS\_368
CL\_INS\_178
CL\_INS\_195
CL\_INS\_195
CL\_INS\_195
CL\_INS\_195
CL\_INS\_195
CL\_INS\_195
CL\_INS\_368
CL\_INS\_368
CL\_INS\_368
CL\_INS\_368
CL\_INS\_195
CL\_INS\_195
CL\_INS\_195
CL\_INS\_107
CL\_INS\_195
CL\_INS\_368
CL\_INS\_270
CL\_INS\_368
CL\_INS\_366
CL\_INS\_366
CL\_INS\_366
CL\_INS\_195
CL\_INS\_195
CL\_INS\_195
CL\_INS\_270
CL\_INS\_195
CL\_INS\_195
CL\_INS\_107
CL\_INS\_195
CL\_INS\_195
CL\_INS\_195
CL\_INS\_195
CL\_INS\_195
CL\_INS\_195
CL\_INS\_195
CL\_INS\_107
CL\_INS\_195
CL\_INS\_270
CL\_INS\_195
CL\_INS\_204
CL\_INS\_195
CL\_INS\_195
CL\_INS\_195
CL\_INS\_195
CL\_INS\_195
CL\_INS\_195
CL\_INS\_195
CL\_INS\_195
CL\_INS\_366
CL\_INS\_366
CL\_INS\_366
CL\_INS\_366
CL\_INS\_195
CL\_INS\_195
CL\_INS\_195
CL\_INS\_195
CL\_INS\_204
CL\_INS\_195
CL\_INS\_195
CL\_INS\_195
CL\_INS\_195
CL\_INS\_366
CL\_INS\_366
CL\_INS\_195
CL\_INS\_195
CL\_INS\_195
CL\_INS\_195
CL\_INS\_195
CL\_INS\_366
CL\_INS\_195
CL\_INS\_195
CL\_INS\_107
CL\_INS\_107
CL\_INS\_195
CL\_INS\_195
CL\_INS\_93
CL\_INS\_195
CL\_INS\_195
CL\_INS\_107
CL\_INS\_195
CL\_INS\_195
CL\_INS\_195
CL\_INS\_107
CL\_INS\_195
CL\_INS\_195
CL\_INS\_195
CL\_INS\_107
CL\_INS\_195
CL\_INS\_195
CL\_INS\_195
CL\_INS\_195
CL\_INS\_195
CL\_INS\_107
CL\_INS\_107
CL\_INS\_107
CL\_INS\_107
CL\_INS\_195
CL\_INS\_195
CL\_INS\_195
CL\_INS\_107
CL\_INS\_195
CL\_INS\_195
CL\_INS\_107
CL\_INS\_195
CL\_INS\_107
CL\_INS\_195
CL\_INS\_107
CL\_INS\_195
CL\_INS\_195
CL\_INS\_195
CL\_INS\_195
CL\_INS\_195
CL\_INS\_195
CL\_INS\_195
CL\_INS\_195
CL\_INS\_195
CL\_INS\_195
CL\_INS\_195
CL\_INS\_195
CL\_INS\_195
CL\_INS\_195
CL\_INS\_195
CL\_INS\_195
CL\_INS\_195
CL\_INS\_195
CL\_INS\_195
CL\_INS\_195
CL\_INS\_195
CL\_INS\_195
CL\_INS\_195
CL\_INS\_195
CL\_INS\_195
CL\_INS\_195
CL\_INS\_195
CL\_INS\_195
CL\_INS\_195
CL\_INS\_195
CL\_INS\_195
CL\_INS\_195
CL\_INS\_195
CL\_INS\_195
CL\_INS\_195
CL\_INS\_195
CL\_INS\_195
CL\_INS\_195
CL\_INS\_195
CL\_INS\_195
CL\_INS\_195
CL\_INS\_195
CL\_INS\_195
CL\_INS\_195
CL\_INS\_195
CL\_INS\_195
CL\_INS\_195
CL\_INS\_195
CL\_INS\_195
CL\_INS\_195
CL\_INS\_195
CL\_INS\_195
CL\_INS\_195
CL\_INS\_195
CL\_INS\_195
CL\_INS\_195
CL\_INS\_107
CL\_INS\_107
CL\_INS\_195
CL\_INS\_107
CL\_INS\_107
CL\_INS\_107
CL\_INS\_107
CL\_INS\_195
CL\_INS\_195
CL\_INS\_195
CL\_INS\_195
CL\_INS\_107
CL\_INS\_195
CL\_INS\_195
CL\_INS\_195
CL\_INS\_195
CL\_INS\_107
CL\_INS\_195
CL\_INS\_195
CL\_INS\_195
CL\_INS\_195
CL\_INS\_195
Cluster ID


CL\_12205
CL\_12204
CL\_12203
CL\_12202
CL\_12201
CL\_11431
CL\_6077
CL\_13613
CL\_11711
CL\_6076
CL\_6075
CL\_11336
CL\_11335
CL\_6074
CL\_6073
CL\_12016
CL\_11710
CL\_11709
CL\_4842
CL\_10579
CL\_5377
CL\_9537
CL\_9538
CL\_5378
CL\_5379
CL\_9539
CL\_9540
CL\_9541
CL\_9542
CL\_9543
CL\_9544
CL\_9545
CL\_9546
CL\_9547
CL\_9548
CL\_9549
CL\_9550
CL\_9551
CL\_9552
CL\_9553
CL\_9554
CL\_9555
CL\_9556
CL\_9557
CL\_9558
CL\_9559
CL\_9560
CL\_9561
CL\_9562
CL\_9563
CL\_9564
CL\_9565
CL\_9566
CL\_9567
CL\_9568
CL\_9569
CL\_9570
CL\_9571
CL\_9572
CL\_9573
CL\_9574
CL\_9575
CL\_9576
CL\_9577
CL\_9578
CL\_9579
CL\_9580
CL\_9581
CL\_9582
CL\_9583
CL\_9584
CL\_9585
CL\_9586
CL\_9587
CL\_9588
CL\_9589
CL\_9590
CL\_9591
CL\_9592
CL\_9593
CL\_9594
CL\_9595
CL\_9596
CL\_9597
CL\_9598
CL\_9599
CL\_9600
CL\_9601
CL\_9602
CL\_9603
CL\_9604
CL\_9605
CL\_9606
CL\_9607
CL\_9608
CL\_9609
CL\_5380
CL\_5381
CL\_7116
CL\_5382
CL\_11430
CL\_10580
CL\_5383
CL\_5384
CL\_10581
CL\_10582
CL\_5385
CL\_5386
CL\_5387
CL\_5388
CL\_5389
CL\_5390
CL\_11708
CL\_10583
CL\_10584
CL\_11707
CL\_11706
CL\_11705
CL\_10585
CL\_10586
CL\_7115
CL\_5391
CL\_5392
CL\_5393
CL\_5394
CL\_5395
CL\_5396
CL\_5397
CL\_10587
CL\_10588
CL\_10589
CL\_5398
CL\_5399
CL\_5400
CL\_5401
CL\_11704
CL\_10503
CL\_5710
CL\_10502
CL\_7126
CL\_7127
CL\_7128
CL\_11703
CL\_10590
CL\_10591
CL\_5402
CL\_5403
CL\_5404
CL\_5405
CL\_10592
CL\_5406
CL\_10593
CL\_5407
CL\_5408
CL\_5409
CL\_5410
CL\_10594
CL\_11702
CL\_3944
CL\_10595
CL\_7014
CL\_11429
CL\_11428
CL\_11427
CL\_11426
CL\_11701
CL\_11700
CL\_11699
CL\_11698
CL\_10538
CL\_10537
CL\_10536
CL\_9038
CL\_10596
CL\_10597
CL\_10598
CL\_10599
CL\_1278
CL\_10600
CL\_10601
CL\_10602
CL\_10603
CL\_9039
CL\_9040
CL\_11425
CL\_11424
CL\_11423
CL\_11697
CL\_11696
CL\_10604
CL\_10605
CL\_10606
CL\_10607
CL\_10608
CL\_5411
CL\_5412
CL\_5413
CL\_5414
CL\_5415
CL\_5416
CL\_5417
CL\_5418
CL\_5419
CL\_5420
CL\_5421
CL\_5422
CL\_5423
CL\_5424
CL\_5425
CL\_5426
CL\_5427
CL\_5428
CL\_5429
CL\_5430
CL\_5431
CL\_10180
CL\_11422
CL\_10179
CL\_10178
CL\_10609
CL\_10610
CL\_11421
CL\_10611
CL\_5432
CL\_5433
CL\_5434
CL\_5435
CL\_5436
CL\_9610
CL\_9611
CL\_9612
CL\_9613
CL\_9614
CL\_9615
CL\_9616
CL\_9617
CL\_9618
CL\_9619
CL\_9620
CL\_9621
CL\_9622
CL\_9623
CL\_9624
CL\_9625
CL\_9626
CL\_9627
CL\_9628
CL\_9629
CL\_9630
CL\_9631
CL\_9632
CL\_9633
CL\_9634
CL\_9635
CL\_9636
CL\_9637
CL\_9638
CL\_9639
CL\_9640
CL\_9641
CL\_9642
CL\_9643
CL\_9644
CL\_9645
CL\_9646
CL\_9647
CL\_9648
CL\_9649
CL\_9650
CL\_9651
CL\_9652
CL\_9653
CL\_9654
CL\_9655
CL\_9656
CL\_9657
CL\_9658
CL\_9659
CL\_9660
CL\_9661
CL\_9662
CL\_9663
CL\_9664
CL\_9665
CL\_5437
CL\_5438
CL\_13612
CL\_5439
CL\_11420
CL\_11695
CL\_5440
CL\_11419
CL\_11694
CL\_11693
CL\_11692
CL\_10176
CL\_5441
CL\_9666
CL\_5442
CL\_11418
CL\_10612
CL\_5443
CL\_9667
CL\_5444
CL\_5445
CL\_5446
