## Supplementary material for "A novel method for integrating genomic and Tn-Seq data to identify common *in vivo* fitness mechanisms across multiple bacterial species": S4 Dataset: CL_INS_200.html

Legend

 Mobile +extrachromosomalelementfunctions
 Other
 All VFDB Genes

FULL


WINDOWSVGPNG

Trim RowsRemove SingletonsSave Fasta

CL\_2807


CL\_2807


CL\_2807


CL\_2807

HighlightSelectShow Genomes


13

CL\_2808


10

CL\_2808


1

CL\_2808


1

CL\_2808

fGI ID


CL\_INS\_200
CL\_INS\_200
CL\_INS\_207
Cluster ID


CL\_13742
CL\_5450
CL\_6043
