## Supplementary material for "A novel method for integrating genomic and Tn-Seq data to identify common *in vivo* fitness mechanisms across multiple bacterial species": S4 Dataset: CL_INS_202.html

Legend

 Mobile +extrachromosomalelementfunctions
 Regulatoryfunctions
 Hypothetical
 DNA Metabolism
 All Fitness Genes
 Other
 Transport +binding proteins
 All VFDB Genes

FULL


WINDOWSVGPNG

Trim RowsRemove SingletonsSave Fasta

CL\_2814


CL\_2814


CL\_2814


CL\_2814


CL\_2814


CL\_5729


CL\_2814

HighlightSelectShow Genomes


21

CL\_2815


3

CL\_2815


1

CL\_2815


1

CL\_2815


1

CL\_2815


1

CL\_2815


1

CL\_5729

fGI ID


CL\_INS\_202
CL\_INS\_202
CL\_INS\_202
CL\_INS\_202
CL\_INS\_202
CL\_INS\_202
CL\_INS\_369
CL\_INS\_369
CL\_INS\_141
CL\_INS\_270
CL\_INS\_202
CL\_INS\_270
CL\_INS\_202
CL\_INS\_365
CL\_INS\_312
CL\_INS\_312
CL\_INS\_312
CL\_INS\_83
CL\_INS\_83
CL\_INS\_83
CL\_INS\_202
CL\_INS\_202
CL\_INS\_202
CL\_INS\_202
CL\_INS\_202
CL\_INS\_202
CL\_INS\_202
CL\_INS\_115
CL\_INS\_115
CL\_INS\_202
CL\_INS\_202
CL\_INS\_368
CL\_INS\_204
CL\_INS\_204
CL\_INS\_204
CL\_INS\_202
CL\_INS\_204
CL\_INS\_204
CL\_INS\_204
CL\_INS\_204
CL\_INS\_204
CL\_INS\_204
CL\_INS\_204
CL\_INS\_202
CL\_INS\_202
CL\_INS\_204
CL\_INS\_204
CL\_INS\_204
CL\_INS\_204
CL\_INS\_204
CL\_INS\_204
CL\_INS\_204
CL\_INS\_204
CL\_INS\_204
CL\_INS\_204
CL\_INS\_204
CL\_INS\_204
CL\_INS\_368
CL\_INS\_270
CL\_INS\_368
CL\_INS\_368
CL\_INS\_368
CL\_INS\_366
CL\_INS\_368
CL\_INS\_325
CL\_INS\_115
CL\_INS\_141
CL\_INS\_141
CL\_INS\_325
CL\_INS\_141
CL\_INS\_141
CL\_INS\_141
CL\_INS\_141
CL\_INS\_141
CL\_INS\_368
CL\_INS\_202
CL\_INS\_368
CL\_INS\_368
CL\_INS\_368
CL\_INS\_202
CL\_INS\_202
CL\_INS\_202
CL\_INS\_368
CL\_INS\_368
CL\_INS\_202
Cluster ID


CL\_6067
CL\_6066
CL\_6065
CL\_6064
CL\_6063
CL\_10915
CL\_12682
CL\_6111
CL\_9005
CL\_5520
CL\_10916
CL\_9151
CL\_10917
CL\_4900
CL\_4903
CL\_4902
CL\_4901
CL\_6243
CL\_6242
CL\_6241
CL\_10918
CL\_11985
CL\_11984
CL\_11983
CL\_11982
CL\_11981
CL\_11980
CL\_1636
CL\_1635
CL\_11979
CL\_11978
CL\_12680
CL\_12406
CL\_12405
CL\_11977
CL\_11976
CL\_9286
CL\_9285
CL\_9284
CL\_9283
CL\_9282
CL\_9281
CL\_9280
CL\_11975
CL\_11974
CL\_9279
CL\_9278
CL\_9277
CL\_9276
CL\_9275
CL\_9274
CL\_7200
CL\_7201
CL\_7202
CL\_9273
CL\_7204
CL\_7205
CL\_12679
CL\_5402
CL\_12678
CL\_12677
CL\_12676
CL\_7210
CL\_12675
CL\_8182
CL\_7218
CL\_7219
CL\_2022
CL\_8183
CL\_5703
CL\_7220
CL\_7221
CL\_5701
CL\_11973
CL\_11972
CL\_11971
CL\_12674
CL\_12673
CL\_11970
CL\_11969
CL\_11968
CL\_11967
CL\_12672
CL\_12671
CL\_12670
