## Supplementary material for "A novel method for integrating genomic and Tn-Seq data to identify common *in vivo* fitness mechanisms across multiple bacterial species": S4 Dataset: CL_INS_204.html

Legend

 Mobile +extrachromosomalelementfunctions
 Regulatoryfunctions
 Hypothetical
 DNA Metabolism
 Other
 Transport +binding proteins
 All VFDB Genes

FULL


WINDOWSVGPNG

Trim RowsRemove SingletonsSave Fasta

CL\_2899


CL\_2899


CL\_2867


CL\_2899


CL\_2867

HighlightSelectShow Genomes


26

CL\_2900


1

CL\_2866


1

CL\_2900


1

CL\_2866


1

CL\_2900

fGI ID


CL\_INS\_204
CL\_INS\_204
CL\_INS\_204
CL\_INS\_204
CL\_INS\_204
CL\_INS\_204
CL\_INS\_204
CL\_INS\_204
CL\_INS\_204
CL\_INS\_204
CL\_INS\_204
CL\_INS\_204
CL\_INS\_204
CL\_INS\_204
CL\_INS\_204
CL\_INS\_204
CL\_INS\_204
CL\_INS\_204
CL\_INS\_204
CL\_INS\_204
CL\_INS\_204
CL\_INS\_204
CL\_INS\_204
CL\_INS\_204
CL\_INS\_204
CL\_INS\_204
CL\_INS\_204
CL\_INS\_204
CL\_INS\_204
CL\_INS\_204
CL\_INS\_204
CL\_INS\_204
CL\_INS\_204
CL\_INS\_204
CL\_INS\_204
CL\_INS\_204
CL\_INS\_204
CL\_INS\_204
CL\_INS\_204
CL\_INS\_204
CL\_INS\_204
CL\_INS\_204
CL\_INS\_204
CL\_INS\_204
CL\_INS\_204
CL\_INS\_204
CL\_INS\_204
CL\_INS\_204
CL\_INS\_204
CL\_INS\_204
CL\_INS\_204
CL\_INS\_204
CL\_INS\_204
CL\_INS\_204
CL\_INS\_204
CL\_INS\_204
CL\_INS\_204
CL\_INS\_204
CL\_INS\_204
CL\_INS\_204
CL\_INS\_204
CL\_INS\_204
CL\_INS\_204
CL\_INS\_204
CL\_INS\_204
CL\_INS\_204
CL\_INS\_204
CL\_INS\_204
CL\_INS\_204
CL\_INS\_204
CL\_INS\_204
CL\_INS\_204
CL\_INS\_204
CL\_INS\_204
CL\_INS\_204
CL\_INS\_204
CL\_INS\_204
CL\_INS\_204
CL\_INS\_204
CL\_INS\_204
CL\_INS\_204
CL\_INS\_204
CL\_INS\_204
CL\_INS\_204
CL\_INS\_204
CL\_INS\_204
CL\_INS\_204
CL\_INS\_204
CL\_INS\_204
CL\_INS\_204
CL\_INS\_204
CL\_INS\_204
CL\_INS\_204
CL\_INS\_204
CL\_INS\_204
CL\_INS\_204
CL\_INS\_204
CL\_INS\_204
Cluster ID


CL\_6126
CL\_6127
CL\_2873
CL\_2874
CL\_7131
CL\_2876
CL\_12404
CL\_12403
CL\_9388
CL\_6133
CL\_7940
CL\_11137
CL\_6136
CL\_11136
CL\_2889
CL\_2890
CL\_12402
CL\_7019
CL\_7018
CL\_7017
CL\_7016
CL\_7015
CL\_7014
CL\_7013
CL\_7012
CL\_7011
CL\_1278
CL\_7010
CL\_7009
CL\_7008
CL\_7007
CL\_7006
CL\_7005
CL\_7004
CL\_7003
CL\_7002
CL\_7001
CL\_7000
CL\_6999
CL\_6998
CL\_6997
CL\_6996
CL\_7034
CL\_7033
CL\_7032
CL\_12407
CL\_12406
CL\_12405
CL\_11977
CL\_9286
CL\_9285
CL\_9284
CL\_9283
CL\_9282
CL\_9281
CL\_9280
CL\_9279
CL\_9278
CL\_9277
CL\_9276
CL\_9275
CL\_9274
CL\_7200
CL\_7201
CL\_7202
CL\_9273
CL\_7204
CL\_7205
CL\_2861
CL\_6123
CL\_6124
CL\_2862
CL\_7031
CL\_7030
CL\_7029
CL\_7028
CL\_7027
CL\_7026
CL\_7025
CL\_5721
CL\_7024
CL\_7023
CL\_5720
CL\_5719
CL\_5718
CL\_5717
CL\_3154
CL\_5716
CL\_5715
CL\_7022
CL\_5280
CL\_5279
CL\_5278
CL\_5277
CL\_5276
CL\_5275
CL\_7021
CL\_7020
