## Supplementary material for "A novel method for integrating genomic and Tn-Seq data to identify common *in vivo* fitness mechanisms across multiple bacterial species": S4 Dataset: CL_INS_207.html

Legend

 Mobile +extrachromosomalelementfunctions
 Regulatoryfunctions
 Hypothetical
 DNA Metabolism
 All Fitness Genes
 Proteinsynthesis/fate
 Other
 Fatty acid +phospholipidmetabolism
 EnergyMetabolism
 All VFDB Genes
 Transport +binding proteins

FULL


WINDOWSVGPNG

Trim RowsRemove SingletonsSave Fasta

CL\_2930


CL\_2930


CL\_2930


CL\_2930


CL\_2930


CL\_2930


CL\_2930


CL\_2894


CL\_2930


CL\_2930


CL\_2930


CL\_2930


CL\_2930


CL\_2930


CL\_2930

HighlightSelectShow Genomes


11

CL\_2937


3

CL\_2945


2

CL\_2937


2

CL\_2937


1

CL\_2937


1

CL\_2945


1

CL\_2937


1

CL\_2937


1

CL\_2945


1

CL\_2937


1

CL\_2937


1

CL\_2945


1

CL\_2937


1

CL\_2821


1

CL\_2937

fGI ID


CL\_INS\_207
CL\_INS\_207
CL\_INS\_207
CL\_INS\_207
CL\_INS\_207
CL\_INS\_207
CL\_INS\_207
CL\_INS\_207
CL\_INS\_207
CL\_INS\_207
CL\_INS\_207
CL\_INS\_207
CL\_INS\_207
CL\_INS\_207
CL\_INS\_207
CL\_INS\_207
CL\_INS\_207
CL\_INS\_207
CL\_INS\_207
CL\_INS\_207
CL\_INS\_207
CL\_INS\_207
CL\_INS\_365
CL\_INS\_207
CL\_INS\_207
CL\_INS\_207
CL\_INS\_207
CL\_INS\_207
CL\_INS\_207
CL\_INS\_207
CL\_INS\_207
CL\_INS\_207
CL\_INS\_207
CL\_INS\_366
CL\_INS\_207
CL\_INS\_207
CL\_INS\_141
CL\_INS\_141
CL\_INS\_141
CL\_INS\_207
CL\_INS\_132
CL\_INS\_207
CL\_INS\_207
CL\_INS\_207
CL\_INS\_207
CL\_INS\_207
CL\_INS\_207
CL\_INS\_207
CL\_INS\_207
CL\_INS\_207
CL\_INS\_207
CL\_INS\_207
CL\_INS\_207
CL\_INS\_207
CL\_INS\_207
CL\_INS\_207
CL\_INS\_207
CL\_INS\_207
CL\_INS\_207
CL\_INS\_207
CL\_INS\_207
CL\_INS\_207
CL\_INS\_207
CL\_INS\_207
CL\_INS\_207
CL\_INS\_207
CL\_INS\_207
CL\_INS\_207
CL\_INS\_207
CL\_INS\_207
CL\_INS\_207
CL\_INS\_207
CL\_INS\_207
CL\_INS\_207
CL\_INS\_207
CL\_INS\_207
CL\_INS\_207
CL\_INS\_366
CL\_INS\_366
CL\_INS\_204
CL\_INS\_366
CL\_INS\_366
CL\_INS\_366
CL\_INS\_366
CL\_INS\_366
CL\_INS\_204
CL\_INS\_207
CL\_INS\_366
CL\_INS\_366
CL\_INS\_366
CL\_INS\_366
CL\_INS\_366
CL\_INS\_204
CL\_INS\_204
CL\_INS\_366
CL\_INS\_366
CL\_INS\_366
CL\_INS\_207
CL\_INS\_141
CL\_INS\_141
CL\_INS\_207
CL\_INS\_207
CL\_INS\_207
CL\_INS\_207
CL\_INS\_207
CL\_INS\_207
CL\_INS\_207
CL\_INS\_207
CL\_INS\_207
CL\_INS\_207
Cluster ID


CL\_6043
CL\_6042
CL\_6041
CL\_6040
CL\_6039
CL\_7939
CL\_2931
CL\_2932
CL\_2933
CL\_2934
CL\_2935
CL\_2936
CL\_5451
CL\_5452
CL\_5454
CL\_5455
CL\_8947
CL\_5453
CL\_13743
CL\_5456
CL\_5457
CL\_5458
CL\_5459
CL\_5460
CL\_5461
CL\_5462
CL\_6995
CL\_6994
CL\_5463
CL\_5464
CL\_8948
CL\_13744
CL\_8949
CL\_6132
CL\_8950
CL\_8951
CL\_2037
CL\_2038
CL\_2826
CL\_8952
CL\_2823
CL\_9670
CL\_9671
CL\_9672
CL\_9673
CL\_9674
CL\_9675
CL\_9676
CL\_9677
CL\_9678
CL\_9679
CL\_9680
CL\_9681
CL\_9682
CL\_9683
CL\_9684
CL\_9685
CL\_9686
CL\_9687
CL\_9688
CL\_9689
CL\_13611
CL\_13610
CL\_13609
CL\_13608
CL\_13607
CL\_13606
CL\_13605
CL\_13604
CL\_13603
CL\_13602
CL\_13601
CL\_13600
CL\_13599
CL\_13598
CL\_13597
CL\_13596
CL\_2892
CL\_2891
CL\_2890
CL\_2887
CL\_2886
CL\_2885
CL\_6135
CL\_6134
CL\_6133
CL\_8946
CL\_6131
CL\_6130
CL\_6129
CL\_6128
CL\_2875
CL\_2874
CL\_2873
CL\_2872
CL\_2871
CL\_2870
CL\_8945
CL\_3168
CL\_3167
CL\_8944
CL\_8943
CL\_8942
CL\_8941
CL\_8940
CL\_8939
CL\_8938
CL\_8937
CL\_8936
CL\_8935
