## Supplementary material for "A novel method for integrating genomic and Tn-Seq data to identify common *in vivo* fitness mechanisms across multiple bacterial species": S4 Dataset: CL_INS_208.html

Legend

 Mobile +extrachromosomalelementfunctions
 Hypothetical
 Proteinsynthesis/fate
 Other
 Fatty acid +phospholipidmetabolism
 All VFDB Genes
 Transport +binding proteins

FULL


WINDOWSVGPNG

Trim RowsRemove SingletonsSave Fasta

CL\_2943


CL\_2943


CL\_2943


CL\_2930


CL\_2943


CL\_2943


CL\_2943


CL\_2930


CL\_2943


CL\_2930


CL\_2943


CL\_2943


CL\_2930

HighlightSelectShow Genomes


5

CL\_2945


4

CL\_2945


3

CL\_2945


3

CL\_2945


3

CL\_2945


2

CL\_2945


2

CL\_2945


1

CL\_2945


1

CL\_2945


1

CL\_2945


1

CL\_2945


1

CL\_2945


1

CL\_2945

fGI ID


CL\_INS\_207
CL\_INS\_207
CL\_INS\_207
CL\_INS\_207
CL\_INS\_208
CL\_INS\_208
CL\_INS\_208
CL\_INS\_208
CL\_INS\_208
CL\_INS\_208
CL\_INS\_208
CL\_INS\_208
CL\_INS\_208
CL\_INS\_208
CL\_INS\_208
CL\_INS\_208
CL\_INS\_365
CL\_INS\_208
CL\_INS\_208
CL\_INS\_208
CL\_INS\_207
CL\_INS\_207
CL\_INS\_207
CL\_INS\_365
CL\_INS\_207
CL\_INS\_207
CL\_INS\_207
CL\_INS\_207
CL\_INS\_208
CL\_INS\_207
CL\_INS\_207
CL\_INS\_207
CL\_INS\_207
CL\_INS\_207
CL\_INS\_207
CL\_INS\_207
CL\_INS\_207
CL\_INS\_207
CL\_INS\_207
CL\_INS\_207
CL\_INS\_207
CL\_INS\_207
CL\_INS\_207
CL\_INS\_207
CL\_INS\_207
CL\_INS\_207
CL\_INS\_207
CL\_INS\_207
CL\_INS\_208
CL\_INS\_207
CL\_INS\_207
CL\_INS\_207
CL\_INS\_207
CL\_INS\_207
CL\_INS\_207
CL\_INS\_207
CL\_INS\_207
CL\_INS\_207
CL\_INS\_207
CL\_INS\_207
CL\_INS\_207
CL\_INS\_207
CL\_INS\_207
CL\_INS\_207
CL\_INS\_207
CL\_INS\_208
Cluster ID


CL\_6042
CL\_6041
CL\_6040
CL\_6039
CL\_6038
CL\_7938
CL\_2944
CL\_10613
CL\_10614
CL\_10919
CL\_10920
CL\_10921
CL\_12199
CL\_12198
CL\_12197
CL\_12196
CL\_12561
CL\_12195
CL\_12194
CL\_12193
CL\_5452
CL\_5453
CL\_5457
CL\_5459
CL\_5461
CL\_5462
CL\_6995
CL\_6994
CL\_6993
CL\_9671
CL\_9672
CL\_9673
CL\_9674
CL\_9675
CL\_9676
CL\_9677
CL\_9678
CL\_9679
CL\_9680
CL\_9681
CL\_9682
CL\_9683
CL\_9684
CL\_9685
CL\_9686
CL\_9687
CL\_9688
CL\_9689
CL\_9690
CL\_13611
CL\_13610
CL\_13609
CL\_13608
CL\_13607
CL\_13606
CL\_13605
CL\_13604
CL\_13603
CL\_13602
CL\_13601
CL\_13600
CL\_13599
CL\_13598
CL\_13597
CL\_13596
CL\_13595
