## Supplementary material for "A novel method for integrating genomic and Tn-Seq data to identify common *in vivo* fitness mechanisms across multiple bacterial species": S4 Dataset: CL_INS_214.html

Legend

 Hypothetical
 Other
 Transport +binding proteins
 All VFDB Genes

FULL


WINDOWSVGPNG

Trim RowsRemove SingletonsSave Fasta

CL\_3039


CL\_3039

HighlightSelectShow Genomes


19

CL\_3049


9

CL\_3049

fGI ID


CL\_INS\_214
CL\_INS\_214
CL\_INS\_214
CL\_INS\_214
CL\_INS\_214
CL\_INS\_214
CL\_INS\_214
CL\_INS\_214
CL\_INS\_214
Cluster ID


CL\_3040
CL\_3041
CL\_3042
CL\_3043
CL\_3044
CL\_3045
CL\_3046
CL\_3047
CL\_3048
