## Supplementary material for "A novel method for integrating genomic and Tn-Seq data to identify common *in vivo* fitness mechanisms across multiple bacterial species": S4 Dataset: CL_INS_218.html

Legend

 Mobile +extrachromosomalelementfunctions
 Hypothetical
 DNA Metabolism
 AntibioticResistance
 Proteinsynthesis/fate
 Other
 All VFDB Genes

FULL


WINDOWSVGPNG

Trim RowsRemove SingletonsSave Fasta

CL\_3189


CL\_3189

HighlightSelectShow Genomes


18

CL\_3190


2

CL\_3190

fGI ID


CL\_INS\_218
CL\_INS\_337
CL\_INS\_337
CL\_INS\_337
CL\_INS\_337
CL\_INS\_337
CL\_INS\_337
CL\_INS\_337
CL\_INS\_337
CL\_INS\_337
CL\_INS\_337
CL\_INS\_218
Cluster ID


CL\_13127
CL\_13128
CL\_10781
CL\_10782
CL\_5901
CL\_5900
CL\_7806
CL\_10784
CL\_7808
CL\_7809
CL\_13129
CL\_13130
