## Supplementary material for "A novel method for integrating genomic and Tn-Seq data to identify common *in vivo* fitness mechanisms across multiple bacterial species": S4 Dataset: CL_INS_221.html

Legend

 Mobile +extrachromosomalelementfunctions
 Regulatoryfunctions
 Hypothetical
 Other
 All VFDB Genes

FULL


WINDOWSVGPNG

Trim RowsRemove SingletonsSave Fasta

CL\_3217


CL\_3217


CL\_3217


CL\_3217


CL\_3217


CL\_3217

HighlightSelectShow Genomes


20

CL\_3255


3

CL\_3255


2

CL\_3255


1

CL\_3255


1

CL\_3255


1

CL\_3255

fGI ID


CL\_INS\_221
CL\_INS\_222
CL\_INS\_222
CL\_INS\_222
CL\_INS\_221
CL\_INS\_221
CL\_INS\_222
CL\_INS\_222
CL\_INS\_222
CL\_INS\_222
CL\_INS\_222
CL\_INS\_221
CL\_INS\_222
CL\_INS\_221
CL\_INS\_221
CL\_INS\_222
CL\_INS\_222
CL\_INS\_222
CL\_INS\_222
CL\_INS\_222
CL\_INS\_222
CL\_INS\_221
CL\_INS\_221
CL\_INS\_222
CL\_INS\_222
CL\_INS\_222
CL\_INS\_221
CL\_INS\_221
CL\_INS\_221
CL\_INS\_221
CL\_INS\_221
CL\_INS\_221
CL\_INS\_221
CL\_INS\_221
CL\_INS\_221
CL\_INS\_221
CL\_INS\_222
CL\_INS\_222
CL\_INS\_222
CL\_INS\_222
CL\_INS\_221
CL\_INS\_221
CL\_INS\_221
CL\_INS\_221
CL\_INS\_222
CL\_INS\_222
CL\_INS\_221
CL\_INS\_221
CL\_INS\_221
CL\_INS\_221
CL\_INS\_221
CL\_INS\_221
CL\_INS\_221
CL\_INS\_221
CL\_INS\_221
CL\_INS\_221
CL\_INS\_221
CL\_INS\_221
CL\_INS\_221
CL\_INS\_222
CL\_INS\_221
CL\_INS\_221
CL\_INS\_221
CL\_INS\_221
CL\_INS\_221
CL\_INS\_221
CL\_INS\_221
CL\_INS\_221
CL\_INS\_221
CL\_INS\_221
CL\_INS\_221
CL\_INS\_221
CL\_INS\_221
CL\_INS\_221
CL\_INS\_221
CL\_INS\_221
CL\_INS\_221
CL\_INS\_221
CL\_INS\_221
CL\_INS\_221
CL\_INS\_221
CL\_INS\_221
Cluster ID


CL\_13131
CL\_5469
CL\_5470
CL\_5471
CL\_13132
CL\_13133
CL\_5474
CL\_5475
CL\_5476
CL\_5478
CL\_5479
CL\_6465
CL\_5480
CL\_13134
CL\_13135
CL\_5483
CL\_5484
CL\_5485
CL\_5486
CL\_5487
CL\_5488
CL\_13136
CL\_13137
CL\_5491
CL\_5492
CL\_5493
CL\_13138
CL\_13139
CL\_6235
CL\_6236
CL\_6237
CL\_6238
CL\_3218
CL\_3219
CL\_3220
CL\_3221
CL\_7381
CL\_3222
CL\_3223
CL\_3224
CL\_3225
CL\_3226
CL\_3227
CL\_3228
CL\_3229
CL\_3230
CL\_3231
CL\_3232
CL\_3233
CL\_689
CL\_688
CL\_3234
CL\_3235
CL\_3236
CL\_3237
CL\_3238
CL\_3239
CL\_3240
CL\_3241
CL\_3242
CL\_3243
CL\_3244
CL\_3245
CL\_3246
CL\_3247
CL\_3248
CL\_3249
CL\_11417
CL\_11416
CL\_11415
CL\_11414
CL\_11413
CL\_11412
CL\_11411
CL\_11410
CL\_11409
CL\_11408
CL\_3250
CL\_3251
CL\_3252
CL\_3253
CL\_3254
