## Supplementary material for "A novel method for integrating genomic and Tn-Seq data to identify common *in vivo* fitness mechanisms across multiple bacterial species": S4 Dataset: CL_INS_222.html

Legend

 Mobile +extrachromosomalelementfunctions
 Regulatoryfunctions
 Hypothetical
 DNA Metabolism
 All Fitness Genes
 Other
 All VFDB Genes
 Transport +binding proteins

FULL


WINDOWSVGPNG

Trim RowsRemove SingletonsSave Fasta

CL\_3286


CL\_3286


CL\_3286


CL\_3286


CL\_3286


CL\_3286

HighlightSelectShow Genomes


15

CL\_3287


8

CL\_3287


2

CL\_3287


1

CL\_3287


1

CL\_3287


1

CL\_3287

fGI ID


CL\_INS\_222
CL\_INS\_222
CL\_INS\_222
CL\_INS\_222
CL\_INS\_222
CL\_INS\_222
CL\_INS\_222
CL\_INS\_222
CL\_INS\_222
CL\_INS\_222
CL\_INS\_222
CL\_INS\_222
CL\_INS\_222
CL\_INS\_222
CL\_INS\_222
CL\_INS\_222
CL\_INS\_222
CL\_INS\_222
CL\_INS\_222
CL\_INS\_222
CL\_INS\_222
CL\_INS\_222
CL\_INS\_222
CL\_INS\_222
CL\_INS\_222
CL\_INS\_222
CL\_INS\_222
CL\_INS\_222
CL\_INS\_222
CL\_INS\_222
CL\_INS\_222
CL\_INS\_222
CL\_INS\_222
CL\_INS\_222
CL\_INS\_222
CL\_INS\_222
CL\_INS\_222
CL\_INS\_222
CL\_INS\_222
CL\_INS\_222
CL\_INS\_222
CL\_INS\_222
CL\_INS\_222
CL\_INS\_222
CL\_INS\_222
CL\_INS\_222
CL\_INS\_222
CL\_INS\_222
CL\_INS\_222
CL\_INS\_222
CL\_INS\_222
CL\_INS\_222
CL\_INS\_222
CL\_INS\_222
CL\_INS\_222
CL\_INS\_222
CL\_INS\_222
CL\_INS\_222
CL\_INS\_222
CL\_INS\_222
CL\_INS\_222
CL\_INS\_222
CL\_INS\_222
CL\_INS\_222
CL\_INS\_222
CL\_INS\_222
CL\_INS\_222
CL\_INS\_222
CL\_INS\_222
CL\_INS\_222
CL\_INS\_222
CL\_INS\_222
CL\_INS\_222
Cluster ID


CL\_11407
CL\_11406
CL\_7381
CL\_3222
CL\_3223
CL\_3224
CL\_12180
CL\_12179
CL\_12178
CL\_12177
CL\_3229
CL\_3230
CL\_12176
CL\_12175
CL\_3242
CL\_12174
CL\_12173
CL\_12172
CL\_11815
CL\_11817
CL\_674
CL\_9172
CL\_673
CL\_672
CL\_9173
CL\_11819
CL\_670
CL\_5467
CL\_5468
CL\_5469
CL\_5470
CL\_5471
CL\_6463
CL\_5472
CL\_5473
CL\_5474
CL\_5475
CL\_5476
CL\_5477
CL\_5478
CL\_5479
CL\_5480
CL\_5481
CL\_5482
CL\_5483
CL\_5484
CL\_5485
CL\_5486
CL\_5487
CL\_5488
CL\_5489
CL\_5490
CL\_5491
CL\_5492
CL\_5493
CL\_5494
CL\_5495
CL\_5496
CL\_5497
CL\_5498
CL\_5499
CL\_5500
CL\_5501
CL\_5502
CL\_11484
CL\_669
CL\_5503
CL\_5504
CL\_12171
CL\_11962
CL\_5505
CL\_5506
CL\_7997
