## Supplementary material for "A novel method for integrating genomic and Tn-Seq data to identify common *in vivo* fitness mechanisms across multiple bacterial species": S4 Dataset: CL_INS_223.html

Legend

 Mobile +extrachromosomalelementfunctions
 Hypothetical
 Other
 Transport +binding proteins
 All VFDB Genes

FULL


WINDOWSVGPNG

Trim RowsRemove SingletonsSave Fasta

CL\_3301


CL\_3301


CL\_3301


CL\_3301


CL\_3301


CL\_3301


CL\_3301


CL\_3301


CL\_3301


CL\_3301

HighlightSelectShow Genomes


18

CL\_3319


2

CL\_3319


1

CL\_3319


1

CL\_3319


1

CL\_3319


1

CL\_3319


1

CL\_3319


1

CL\_3319


1

CL\_3319


1

CL\_3319

fGI ID


CL\_INS\_223
CL\_INS\_223
CL\_INS\_223
CL\_INS\_223
CL\_INS\_223
CL\_INS\_223
CL\_INS\_223
CL\_INS\_223
CL\_INS\_223
CL\_INS\_223
CL\_INS\_223
CL\_INS\_223
CL\_INS\_223
CL\_INS\_223
CL\_INS\_223
CL\_INS\_223
CL\_INS\_365
CL\_INS\_365
CL\_INS\_223
CL\_INS\_223
CL\_INS\_223
CL\_INS\_223
CL\_INS\_223
CL\_INS\_223
CL\_INS\_223
CL\_INS\_223
CL\_INS\_223
CL\_INS\_223
CL\_INS\_223
Cluster ID


CL\_6990
CL\_7996
CL\_3302
CL\_11961
CL\_3303
CL\_3304
CL\_3305
CL\_3306
CL\_3307
CL\_3308
CL\_3309
CL\_12922
CL\_5507
CL\_3310
CL\_3311
CL\_8152
CL\_8398
CL\_8397
CL\_3312
CL\_3313
CL\_3314
CL\_3315
CL\_3316
CL\_8927
CL\_13746
CL\_13747
CL\_5508
CL\_3317
CL\_3318
