## Supplementary material for "A novel method for integrating genomic and Tn-Seq data to identify common *in vivo* fitness mechanisms across multiple bacterial species": S4 Dataset: CL_INS_229.html

Legend

 Mobile +extrachromosomalelementfunctions
 Hypothetical
 Other
 All VFDB Genes

FULL


WINDOWSVGPNG

Trim RowsRemove SingletonsSave Fasta

CL\_3341


CL\_3341


CL\_3341

HighlightSelectShow Genomes


25

CL\_3343


2

CL\_3343


1

CL\_3343

fGI ID


CL\_INS\_229
CL\_INS\_229
Cluster ID


CL\_10927
CL\_3342
