## Supplementary material for "A novel method for integrating genomic and Tn-Seq data to identify common *in vivo* fitness mechanisms across multiple bacterial species": S4 Dataset: CL_INS_230.html

Legend

 Regulatoryfunctions
 Hypothetical
 Other

FULL


WINDOWSVGPNG

Trim RowsRemove SingletonsSave Fasta

CL\_3356


CL\_3356

HighlightSelectShow Genomes


16

CL\_3360


12

CL\_3360

fGI ID


CL\_INS\_230
CL\_INS\_230
CL\_INS\_230
Cluster ID


CL\_3357
CL\_3358
CL\_3359
