## Supplementary material for "A novel method for integrating genomic and Tn-Seq data to identify common *in vivo* fitness mechanisms across multiple bacterial species": S4 Dataset: CL_INS_231.html

Legend

 Hypothetical
 Biosynthesis ofcofactors,prostheticgroups, +carriers
 Other

FULL


WINDOWSVGPNG

Trim RowsRemove SingletonsSave Fasta

CL\_3375


CL\_3375

HighlightSelectShow Genomes


20

CL\_3376


8

CL\_3376

fGI ID

CL\_INS\_231
Cluster ID

CL\_6031
