## Supplementary material for "A novel method for integrating genomic and Tn-Seq data to identify common *in vivo* fitness mechanisms across multiple bacterial species": S4 Dataset: CL_INS_232.html

Legend

 Hypothetical
 Other
 All VFDB Genes

FULL


WINDOWSVGPNG

Trim RowsRemove SingletonsSave Fasta

CL\_3377


CL\_3377


CL\_3377

HighlightSelectShow Genomes


15

CL\_3378


12

CL\_3378


1

CL\_3378

fGI ID


CL\_INS\_232
CL\_INS\_232
Cluster ID


CL\_13748
CL\_6030
