## Supplementary material for "A novel method for integrating genomic and Tn-Seq data to identify common *in vivo* fitness mechanisms across multiple bacterial species": S4 Dataset: CL_INS_233.html

Legend

 Mobile +extrachromosomalelementfunctions
 Hypothetical
 Other
 All VFDB Genes

FULL


WINDOWSVGPNG

Trim RowsRemove SingletonsSave Fasta

CL\_3390


CL\_3390


CL\_3390


CL\_3390

HighlightSelectShow Genomes


15

CL\_3392


8

CL\_3392


4

CL\_3392


1

CL\_3392

fGI ID


CL\_INS\_233
CL\_INS\_233
CL\_INS\_233
CL\_INS\_233
CL\_INS\_233
CL\_INS\_233
Cluster ID


CL\_5510
CL\_3391
CL\_6029
CL\_6028
CL\_6027
CL\_6026
