## Supplementary material for "A novel method for integrating genomic and Tn-Seq data to identify common *in vivo* fitness mechanisms across multiple bacterial species": S4 Dataset: CL_INS_241.html

Legend

 Hypothetical
 All VFDB Genes

FULL


WINDOWSVGPNG

Trim RowsRemove SingletonsSave Fasta

CL\_3483


CL\_3483


CL\_3483


Break

HighlightSelectShow Genomes


10

CL\_3484


10

CL\_3484


7

CL\_3484


1

CL\_3484

fGI ID


CL\_INS\_241
CL\_INS\_241
Cluster ID


CL\_6021
CL\_12646
