## Supplementary material for "A novel method for integrating genomic and Tn-Seq data to identify common *in vivo* fitness mechanisms across multiple bacterial species": S4 Dataset: CL_INS_242.html

Legend

 Hypothetical
 Centralintermediarymetabolism
 All VFDB Genes

FULL


WINDOWSVGPNG

Trim RowsRemove SingletonsSave Fasta

CL\_3491


CL\_3491

HighlightSelectShow Genomes


25

CL\_3492


3

CL\_3492

fGI ID

CL\_INS\_242
Cluster ID

CL\_8358
