## Supplementary material for "A novel method for integrating genomic and Tn-Seq data to identify common *in vivo* fitness mechanisms across multiple bacterial species": S4 Dataset: CL_INS_253.html

Legend

 Hypothetical
 Other

FULL


WINDOWSVGPNG

Trim RowsRemove SingletonsSave Fasta

CL\_3616


CL\_3616


CL\_3616


CL\_3616


CL\_3616

HighlightSelectShow Genomes


21

CL\_3617


2

CL\_3617


2

CL\_3617


1

CL\_3617


1

CL\_3617

fGI ID


CL\_INS\_253
CL\_INS\_253
CL\_INS\_253
CL\_INS\_253
Cluster ID


CL\_8885
CL\_5514
CL\_5515
CL\_5516
