## Supplementary material for "A novel method for integrating genomic and Tn-Seq data to identify common *in vivo* fitness mechanisms across multiple bacterial species": S4 Dataset: CL_INS_254.html

Legend

 Hypothetical
 Proteinsynthesis/fate
 All VFDB Genes

FULL


WINDOWSVGPNG

Trim RowsRemove SingletonsSave Fasta

CL\_3618


CL\_3618

HighlightSelectShow Genomes


25

CL\_3619


3

CL\_3619

fGI ID


CL\_INS\_254
CL\_INS\_254
Cluster ID


CL\_13143
CL\_13144
