## Supplementary material for "A novel method for integrating genomic and Tn-Seq data to identify common *in vivo* fitness mechanisms across multiple bacterial species": S4 Dataset: CL_INS_258.html

Legend

 Mobile +extrachromosomalelementfunctions
 Regulatoryfunctions
 Hypothetical
 DNA Metabolism
 All VFDB Genes

FULL


WINDOWSVGPNG

Trim RowsRemove SingletonsSave Fasta

CL\_3697


CL\_3697


CL\_3697


CL\_3697

HighlightSelectShow Genomes


24

CL\_3698


2

CL\_3698


1

CL\_3698


1

CL\_3698

fGI ID


CL\_INS\_258
CL\_INS\_258
CL\_INS\_258
CL\_INS\_258
CL\_INS\_258
CL\_INS\_258
CL\_INS\_258
CL\_INS\_258
CL\_INS\_258
CL\_INS\_258
CL\_INS\_258
CL\_INS\_258
CL\_INS\_258
CL\_INS\_258
CL\_INS\_258
CL\_INS\_258
CL\_INS\_258
CL\_INS\_258
CL\_INS\_258
CL\_INS\_258
CL\_INS\_144
CL\_INS\_258
CL\_INS\_258
Cluster ID


CL\_12669
CL\_12668
CL\_12667
CL\_12666
CL\_12665
CL\_12664
CL\_12663
CL\_12662
CL\_13145
CL\_13146
CL\_13147
CL\_13148
CL\_13365
CL\_13366
CL\_13149
CL\_13150
CL\_13151
CL\_13152
CL\_13153
CL\_13154
CL\_608
CL\_13155
CL\_12661
