## Supplementary material for "A novel method for integrating genomic and Tn-Seq data to identify common *in vivo* fitness mechanisms across multiple bacterial species": S4 Dataset: CL_INS_263.html

Legend

 Hypothetical
 All VFDB Genes

FULL


WINDOWSVGPNG

Trim RowsRemove SingletonsSave Fasta

CL\_3752


CL\_3752

HighlightSelectShow Genomes


20

CL\_3753


8

CL\_3753

fGI ID


CL\_INS\_263
CL\_INS\_263
Cluster ID


CL\_9693
CL\_9694
