## Supplementary material for "A novel method for integrating genomic and Tn-Seq data to identify common *in vivo* fitness mechanisms across multiple bacterial species": S4 Dataset: CL_INS_264.html

Legend

 Hypothetical
 All VFDB Genes

FULL


WINDOWSVGPNG

Trim RowsRemove SingletonsSave Fasta

CL\_3785


CL\_3785


CL\_3784


CL\_3784


CL\_3784


CL\_3785

HighlightSelectShow Genomes


22

CL\_3786


2

CL\_3786


1

CL\_3786


1

CL\_3786


1

CL\_3786


1

CL\_3786

fGI ID


CL\_INS\_264
CL\_INS\_264
CL\_INS\_264
Cluster ID


CL\_8154
CL\_9696
CL\_12167
