## Supplementary material for "A novel method for integrating genomic and Tn-Seq data to identify common *in vivo* fitness mechanisms across multiple bacterial species": S4 Dataset: CL_INS_265.html

Legend

 Mobile +extrachromosomalelementfunctions
 Regulatoryfunctions
 Hypothetical
 Other

FULL


WINDOWSVGPNG

Trim RowsRemove SingletonsSave Fasta

CL\_3806


CL\_3806


CL\_3806


CL\_3806


CL\_3806


CL\_3806

HighlightSelectShow Genomes


13

CL\_3810


8

CL\_3810


4

CL\_3810


1

CL\_3810


1

CL\_3810


1

CL\_3810

fGI ID


CL\_INS\_265
CL\_INS\_265
CL\_INS\_265
CL\_INS\_265
CL\_INS\_265
CL\_INS\_265
CL\_INS\_265
CL\_INS\_265
CL\_INS\_265
CL\_INS\_265
Cluster ID


CL\_12166
CL\_6003
CL\_6002
CL\_6001
CL\_6984
CL\_6983
CL\_9699
CL\_3807
CL\_3808
CL\_3809
