## Supplementary material for "A novel method for integrating genomic and Tn-Seq data to identify common *in vivo* fitness mechanisms across multiple bacterial species": S4 Dataset: CL_INS_267.html

Legend

 Regulatoryfunctions
 All VFDB Genes

FULL


WINDOWSVGPNG

Trim RowsRemove SingletonsSave Fasta

CL\_3827


CL\_3827


CL\_3827

HighlightSelectShow Genomes


14

CL\_3834


13

CL\_3834


1

CL\_3834

fGI ID


CL\_INS\_267
CL\_INS\_267
CL\_INS\_267
CL\_INS\_267
CL\_INS\_267
CL\_INS\_267
Cluster ID


CL\_3828
CL\_3829
CL\_3830
CL\_3831
CL\_3832
CL\_3833
