## Supplementary material for "A novel method for integrating genomic and Tn-Seq data to identify common *in vivo* fitness mechanisms across multiple bacterial species": S4 Dataset: CL_INS_268.html

Legend

 Mobile +extrachromosomalelementfunctions
 Regulatoryfunctions
 Hypothetical
 DNA Metabolism
 Other
 All VFDB Genes

FULL


WINDOWSVGPNG

Trim RowsRemove SingletonsSave Fasta

CL\_3847


CL\_3847


CL\_3847


CL\_3847


CL\_3847

HighlightSelectShow Genomes


13

CL\_3850


8

CL\_3850


2

CL\_3850


1

CL\_3866


1

CL\_3969

fGI ID


CL\_INS\_268
CL\_INS\_268
CL\_INS\_268
CL\_INS\_268
CL\_INS\_268
CL\_INS\_270
CL\_INS\_270
CL\_INS\_270
CL\_INS\_270
CL\_INS\_270
CL\_INS\_270
CL\_INS\_270
CL\_INS\_270
CL\_INS\_270
CL\_INS\_270
CL\_INS\_270
CL\_INS\_270
CL\_INS\_270
CL\_INS\_270
CL\_INS\_270
CL\_INS\_270
CL\_INS\_270
CL\_INS\_270
CL\_INS\_270
CL\_INS\_270
CL\_INS\_270
Cluster ID


CL\_9700
CL\_3848
CL\_3849
CL\_13753
CL\_13754
CL\_11960
CL\_8356
CL\_8355
CL\_8354
CL\_8353
CL\_8352
CL\_8351
CL\_8350
CL\_8349
CL\_11959
CL\_11958
CL\_11957
CL\_8343
CL\_8342
CL\_8341
CL\_8340
CL\_8339
CL\_8338
CL\_8337
CL\_6379
CL\_11956
