## Supplementary material for "A novel method for integrating genomic and Tn-Seq data to identify common *in vivo* fitness mechanisms across multiple bacterial species": S4 Dataset: CL_INS_269.html

Legend

 Mobile +extrachromosomalelementfunctions
 Hypothetical
 Other

FULL


WINDOWSVGPNG

Trim RowsRemove SingletonsSave Fasta

CL\_3855


CL\_3855


CL\_3855

HighlightSelectShow Genomes


20

CL\_3856


2

CL\_3856


1

CL\_3856

fGI ID


CL\_INS\_269
CL\_INS\_269
CL\_INS\_269
Cluster ID


CL\_10255
CL\_10256
CL\_10257
