## Supplementary material for "A novel method for integrating genomic and Tn-Seq data to identify common *in vivo* fitness mechanisms across multiple bacterial species": S4 Dataset: CL_INS_271.html

Legend

 Mobile +extrachromosomalelementfunctions
 DNA Metabolism
 All VFDB Genes

FULL


WINDOWSVGPNG

Trim RowsRemove SingletonsSave Fasta

CL\_3988


CL\_3988


CL\_3987

HighlightSelectShow Genomes


25

CL\_3989


2

CL\_3989


1

CL\_3989

fGI ID

CL\_INS\_271
Cluster ID

CL\_7993
