## Supplementary material for "A novel method for integrating genomic and Tn-Seq data to identify common *in vivo* fitness mechanisms across multiple bacterial species": S4 Dataset: CL_INS_272.html

Legend

 Mobile +extrachromosomalelementfunctions
 Hypothetical
 Other
 All VFDB Genes

FULL


WINDOWSVGPNG

Trim RowsRemove SingletonsSave Fasta

CL\_3989


CL\_3989


CL\_3989


CL\_3989


CL\_3989


CL\_3989


CL\_3989


CL\_3989


CL\_3989

HighlightSelectShow Genomes


13

CL\_3990


6

CL\_3990


3

CL\_3990


1

CL\_3990


1

CL\_3991


1

CL\_3990


1

CL\_3990


1

CL\_3990


1

CL\_3990

fGI ID


CL\_INS\_272
CL\_INS\_272
CL\_INS\_272
CL\_INS\_272
CL\_INS\_272
CL\_INS\_272
CL\_INS\_272
CL\_INS\_272
CL\_INS\_272
CL\_INS\_272
CL\_INS\_272
CL\_INS\_272
CL\_INS\_272
CL\_INS\_272
CL\_INS\_272
CL\_INS\_272
CL\_INS\_272
CL\_INS\_272
CL\_INS\_272
CL\_INS\_272
CL\_INS\_272
CL\_INS\_272
CL\_INS\_272
CL\_INS\_272
CL\_INS\_272
CL\_INS\_272
CL\_INS\_272
CL\_INS\_272
CL\_INS\_272
CL\_INS\_272
CL\_INS\_272
CL\_INS\_272
CL\_INS\_272
CL\_INS\_272
CL\_INS\_272
CL\_INS\_272
CL\_INS\_272
CL\_INS\_272
CL\_INS\_272
CL\_INS\_272
CL\_INS\_272
CL\_INS\_272
CL\_INS\_272
CL\_INS\_272
CL\_INS\_272
CL\_INS\_272
CL\_INS\_272
CL\_INS\_272
Cluster ID


CL\_5534
CL\_9702
CL\_5535
CL\_5536
CL\_5537
CL\_5538
CL\_5539
CL\_5540
CL\_5541
CL\_5542
CL\_5543
CL\_5544
CL\_5545
CL\_5546
CL\_5547
CL\_5548
CL\_5549
CL\_5550
CL\_5551
CL\_5552
CL\_5553
CL\_5554
CL\_5555
CL\_5556
CL\_5557
CL\_5558
CL\_5559
CL\_5560
CL\_9703
CL\_11683
CL\_9704
CL\_5561
CL\_5562
CL\_5563
CL\_5564
CL\_5565
CL\_5566
CL\_5567
CL\_5568
CL\_5569
CL\_5570
CL\_5571
CL\_5572
CL\_5573
CL\_5574
CL\_5575
CL\_5576
CL\_5577
