## Supplementary material for "A novel method for integrating genomic and Tn-Seq data to identify common *in vivo* fitness mechanisms across multiple bacterial species": S4 Dataset: CL_INS_274.html

Legend

 Hypothetical
 Other
 All VFDB Genes

FULL


WINDOWSVGPNG

Trim RowsRemove SingletonsSave Fasta

CL\_4071


CL\_4071


CL\_4071


CL\_4068


CL\_4071


CL\_4071

HighlightSelectShow Genomes


12

CL\_4073


10

CL\_4073


2

CL\_4073


1

CL\_4073


1

CL\_4073


1

CL\_4073

fGI ID


CL\_INS\_274
CL\_INS\_274
CL\_INS\_274
CL\_INS\_274
Cluster ID


CL\_8333
CL\_4072
CL\_6956
CL\_6955
