## Supplementary material for "A novel method for integrating genomic and Tn-Seq data to identify common *in vivo* fitness mechanisms across multiple bacterial species": S4 Dataset: CL_INS_281.html

Legend

 Regulatoryfunctions
 Hypothetical
 Other
 All VFDB Genes

FULL


WINDOWSVGPNG

Trim RowsRemove SingletonsSave Fasta

CL\_4101


CL\_4101


CL\_4101

HighlightSelectShow Genomes


21

CL\_4102


6

CL\_4102


1

CL\_4102

fGI ID


CL\_INS\_281
CL\_INS\_281
CL\_INS\_281
Cluster ID


CL\_13187
CL\_5584
CL\_5585
