## Supplementary material for "A novel method for integrating genomic and Tn-Seq data to identify common *in vivo* fitness mechanisms across multiple bacterial species": S4 Dataset: CL_INS_282.html

Legend

 Hypothetical
 Biosynthesis ofcofactors,prostheticgroups, +carriers
 All VFDB Genes

FULL


WINDOWSVGPNG

Trim RowsRemove SingletonsSave Fasta

CL\_4107


CL\_4107

HighlightSelectShow Genomes


19

CL\_4108


9

CL\_4108

fGI ID

CL\_INS\_282
Cluster ID

CL\_5978
