## Supplementary material for "A novel method for integrating genomic and Tn-Seq data to identify common *in vivo* fitness mechanisms across multiple bacterial species": S4 Dataset: CL_INS_287.html

Legend

 Hypothetical
 Other
 Transport +binding proteins

FULL


WINDOWSVGPNG

Trim RowsRemove SingletonsSave Fasta

CL\_4167


CL\_4167

HighlightSelectShow Genomes


13

CL\_4168


12

CL\_4168

fGI ID

CL\_INS\_287
Cluster ID

CL\_5975
