## Supplementary material for "A novel method for integrating genomic and Tn-Seq data to identify common *in vivo* fitness mechanisms across multiple bacterial species": S4 Dataset: CL_INS_289.html

Legend

 Other
 EnergyMetabolism
 Transport +binding proteins
 All VFDB Genes

FULL


WINDOWSVGPNG

Trim RowsRemove SingletonsSave Fasta

CL\_4185


CL\_4185

HighlightSelectShow Genomes


25

CL\_4186


3

CL\_4186

fGI ID


CL\_INS\_289
CL\_INS\_289
CL\_INS\_289
CL\_INS\_289
CL\_INS\_289
CL\_INS\_289
CL\_INS\_289
CL\_INS\_289
Cluster ID


CL\_10674
CL\_10675
CL\_10676
CL\_10677
CL\_10678
CL\_10679
CL\_10680
CL\_10681
