## Supplementary material for "A novel method for integrating genomic and Tn-Seq data to identify common *in vivo* fitness mechanisms across multiple bacterial species": S4 Dataset: CL_INS_291.html

Legend

 Hypothetical
 All VFDB Genes

FULL


WINDOWSVGPNG

Trim RowsRemove SingletonsSave Fasta

CL\_4224


CL\_4224


CL\_4224


Break

HighlightSelectShow Genomes


18

CL\_4225


7

CL\_4225


1

CL\_4225


1

CL\_4225

fGI ID


CL\_INS\_291
CL\_INS\_291
Cluster ID


CL\_8156
CL\_5971
