## Supplementary material for "A novel method for integrating genomic and Tn-Seq data to identify common *in vivo* fitness mechanisms across multiple bacterial species": S4 Dataset: CL_INS_293.html

Legend

 Hypothetical
 Other
 Transport +binding proteins

FULL


WINDOWSVGPNG

Trim RowsRemove SingletonsSave Fasta

CL\_4249


CL\_4249

HighlightSelectShow Genomes


23

CL\_4251


3

CL\_4251

fGI ID

CL\_INS\_293
Cluster ID

CL\_4250
