## Supplementary material for "A novel method for integrating genomic and Tn-Seq data to identify common *in vivo* fitness mechanisms across multiple bacterial species": S4 Dataset: CL_INS_295.html

Legend

 Regulatoryfunctions
 Hypothetical
 DNA Metabolism
 Transcription
 Transport +binding proteins

FULL


WINDOWSVGPNG

Trim RowsRemove SingletonsSave Fasta

CL\_4268


CL\_4268


CL\_4268


CL\_4268


CL\_4268


CL\_4268


CL\_4268

HighlightSelectShow Genomes


18

CL\_4269


3

CL\_4269


3

CL\_4269


1

CL\_4269


1

CL\_4269


1

CL\_4269


1

CL\_4269

fGI ID


CL\_INS\_295
CL\_INS\_295
CL\_INS\_295
CL\_INS\_295
CL\_INS\_295
CL\_INS\_295
CL\_INS\_295
CL\_INS\_295
CL\_INS\_295
CL\_INS\_295
CL\_INS\_295
CL\_INS\_295
CL\_INS\_295
CL\_INS\_295
CL\_INS\_295
CL\_INS\_295
Cluster ID


CL\_10997
CL\_10998
CL\_10999
CL\_12152
CL\_12151
CL\_12150
CL\_11000
CL\_11001
CL\_11002
CL\_12874
CL\_12875
CL\_12876
CL\_12877
CL\_12878
CL\_12879
CL\_11003
