## Supplementary material for "A novel method for integrating genomic and Tn-Seq data to identify common *in vivo* fitness mechanisms across multiple bacterial species": S4 Dataset: CL_INS_300.html

Legend

 Hypothetical
 All VFDB Genes

FULL


WINDOWSVGPNG

Trim RowsRemove SingletonsSave Fasta

CL\_4337


CL\_4337


CL\_4337

HighlightSelectShow Genomes


23

CL\_4338


3

CL\_4341


2

CL\_4338

fGI ID


CL\_INS\_300
CL\_INS\_300
CL\_INS\_300
Cluster ID


CL\_5615
CL\_13197
CL\_13198
