## Supplementary material for "A novel method for integrating genomic and Tn-Seq data to identify common *in vivo* fitness mechanisms across multiple bacterial species": S4 Dataset: CL_INS_301.html

Legend

 Hypothetical
 Other
 All VFDB Genes

FULL


WINDOWSVGPNG

Trim RowsRemove SingletonsSave Fasta

CL\_4340


CL\_4340


CL\_4337


CL\_4340

HighlightSelectShow Genomes


18

CL\_4341


6

CL\_4341


3

CL\_4341


1

CL\_4342

fGI ID


CL\_INS\_301
CL\_INS\_300
CL\_INS\_301
Cluster ID


CL\_8852
CL\_13198
CL\_13199
