## Supplementary material for "A novel method for integrating genomic and Tn-Seq data to identify common *in vivo* fitness mechanisms across multiple bacterial species": S4 Dataset: CL_INS_303.html

Legend

 Mobile +extrachromosomalelementfunctions
 Regulatoryfunctions
 Hypothetical
 Other
 Transport +binding proteins

FULL


WINDOWSVGPNG

Trim RowsRemove SingletonsSave Fasta

CL\_4344


CL\_4344


CL\_4344


CL\_4344


CL\_4344

HighlightSelectShow Genomes


13

CL\_4351


10

CL\_4351


3

CL\_4351


1

CL\_4351


1

CL\_4351

fGI ID


CL\_INS\_303
CL\_INS\_303
CL\_INS\_303
CL\_INS\_303
CL\_INS\_303
CL\_INS\_303
CL\_INS\_303
Cluster ID


CL\_4345
CL\_4346
CL\_12365
CL\_4347
CL\_4348
CL\_4349
CL\_4350
