## Supplementary material for "A novel method for integrating genomic and Tn-Seq data to identify common *in vivo* fitness mechanisms across multiple bacterial species": S4 Dataset: CL_INS_311.html

Legend

 Mobile +extrachromosomalelementfunctions
 Hypothetical
 All VFDB Genes

FULL


WINDOWSVGPNG

Trim RowsRemove SingletonsSave Fasta

CL\_4446


CL\_4446


CL\_4446

HighlightSelectShow Genomes


16

CL\_4447


11

CL\_4447


1

CL\_4447

fGI ID


CL\_INS\_311
CL\_INS\_311
CL\_INS\_311
Cluster ID


CL\_6531
CL\_5626
CL\_5625
