## Supplementary material for "A novel method for integrating genomic and Tn-Seq data to identify common *in vivo* fitness mechanisms across multiple bacterial species": S4 Dataset: CL_INS_312.html

Legend

 Mobile +extrachromosomalelementfunctions
 Regulatoryfunctions
 Hypothetical
 DNA Metabolism
 AntibioticResistance
 Biosynthesis ofcofactors,prostheticgroups, +carriers
 Proteinsynthesis/fate
 Other
 EnergyMetabolism
 Transport +binding proteins
 All VFDB Genes

FULL


WINDOWSVGPNG

Trim RowsRemove SingletonsSave Fasta

CL\_4456


CL\_4456


CL\_5662


CL\_4456


CL\_4456


CL\_4456


CL\_4456


CL\_4456


Break


CL\_5662


CL\_5662


CL\_4456


CL\_4456


CL\_4456


CL\_4456


CL\_4456


CL\_4456


CL\_5661


CL\_4456


CL\_3862

HighlightSelectShow Genomes


12

CL\_4457


3

CL\_4457


1

CL\_4457


1

CL\_4457


1

CL\_4459


1

CL\_4457


1

CL\_5662


1

CL\_5661


1

CL\_4457


1

CL\_4457


1

CL\_4457


1

CL\_76


1

CL\_4457


1

CL\_256


1

CL\_4457


1

CL\_4458


1

CL\_4457


1

CL\_4457


1

CL\_256


1

CL\_4457

fGI ID


CL\_INS\_312
CL\_INS\_270
CL\_INS\_312
CL\_INS\_312
CL\_INS\_8
CL\_INS\_8
CL\_INS\_8
CL\_INS\_312
CL\_INS\_312
CL\_INS\_312
CL\_INS\_312
CL\_INS\_312
CL\_INS\_312
CL\_INS\_312
CL\_INS\_312
CL\_INS\_312
CL\_INS\_312
CL\_INS\_312
CL\_INS\_312
CL\_INS\_312
CL\_INS\_312
CL\_INS\_312
CL\_INS\_312
CL\_INS\_312
CL\_INS\_297
CL\_INS\_297
CL\_INS\_297
CL\_INS\_312
CL\_INS\_312
CL\_INS\_312
CL\_INS\_312
CL\_INS\_312
CL\_INS\_312
CL\_INS\_312
CL\_INS\_312
CL\_INS\_312
CL\_INS\_312
CL\_INS\_312
CL\_INS\_312
CL\_INS\_312
CL\_INS\_312
CL\_INS\_312
CL\_INS\_312
CL\_INS\_312
CL\_INS\_312
CL\_INS\_312
CL\_INS\_312
CL\_INS\_365
CL\_INS\_365
CL\_INS\_365
CL\_INS\_365
CL\_INS\_365
CL\_INS\_365
CL\_INS\_365
CL\_INS\_312
CL\_INS\_312
CL\_INS\_312
CL\_INS\_312
CL\_INS\_312
CL\_INS\_312
CL\_INS\_312
CL\_INS\_312
CL\_INS\_312
CL\_INS\_312
CL\_INS\_312
CL\_INS\_365
CL\_INS\_365
CL\_INS\_365
CL\_INS\_365
CL\_INS\_365
CL\_INS\_365
CL\_INS\_365
CL\_INS\_365
CL\_INS\_365
CL\_INS\_365
CL\_INS\_365
CL\_INS\_365
CL\_INS\_365
CL\_INS\_365
CL\_INS\_365
CL\_INS\_365
CL\_INS\_365
CL\_INS\_365
CL\_INS\_365
CL\_INS\_365
CL\_INS\_365
CL\_INS\_365
CL\_INS\_365
CL\_INS\_365
CL\_INS\_365
CL\_INS\_312
CL\_INS\_131
CL\_INS\_131
CL\_INS\_131
CL\_INS\_131
CL\_INS\_131
CL\_INS\_132
CL\_INS\_131
CL\_INS\_132
CL\_INS\_131
CL\_INS\_132
CL\_INS\_132
CL\_INS\_131
CL\_INS\_131
CL\_INS\_132
CL\_INS\_131
CL\_INS\_132
CL\_INS\_131
CL\_INS\_312
CL\_INS\_131
CL\_INS\_131
CL\_INS\_312
CL\_INS\_312
CL\_INS\_312
CL\_INS\_312
CL\_INS\_365
CL\_INS\_312
CL\_INS\_312
CL\_INS\_312
CL\_INS\_312
CL\_INS\_312
CL\_INS\_312
CL\_INS\_312
CL\_INS\_312
CL\_INS\_312
CL\_INS\_312
CL\_INS\_312
CL\_INS\_312
CL\_INS\_312
CL\_INS\_312
CL\_INS\_312
CL\_INS\_312
CL\_INS\_312
CL\_INS\_312
CL\_INS\_312
CL\_INS\_312
CL\_INS\_312
CL\_INS\_312
CL\_INS\_312
CL\_INS\_312
CL\_INS\_312
CL\_INS\_312
CL\_INS\_312
CL\_INS\_312
CL\_INS\_312
CL\_INS\_312
CL\_INS\_312
CL\_INS\_312
CL\_INS\_312
CL\_INS\_312
CL\_INS\_312
CL\_INS\_312
CL\_INS\_312
CL\_INS\_312
CL\_INS\_312
CL\_INS\_312
CL\_INS\_312
CL\_INS\_312
CL\_INS\_312
CL\_INS\_312
CL\_INS\_312
CL\_INS\_312
CL\_INS\_270
CL\_INS\_270
CL\_INS\_270
CL\_INS\_270
CL\_INS\_270
CL\_INS\_270
CL\_INS\_270
CL\_INS\_270
CL\_INS\_270
CL\_INS\_207
CL\_INS\_312
CL\_INS\_312
CL\_INS\_312
CL\_INS\_297
CL\_INS\_312
CL\_INS\_365
CL\_INS\_312
CL\_INS\_366
CL\_INS\_335
CL\_INS\_335
CL\_INS\_335
CL\_INS\_335
CL\_INS\_335
CL\_INS\_335
CL\_INS\_312
CL\_INS\_312
CL\_INS\_312
CL\_INS\_312
CL\_INS\_312
CL\_INS\_312
CL\_INS\_312
CL\_INS\_312
CL\_INS\_312
CL\_INS\_312
CL\_INS\_312
CL\_INS\_312
CL\_INS\_312
CL\_INS\_312
CL\_INS\_312
CL\_INS\_312
CL\_INS\_312
CL\_INS\_312
CL\_INS\_312
CL\_INS\_312
CL\_INS\_312
CL\_INS\_297
CL\_INS\_297
CL\_INS\_297
CL\_INS\_297
CL\_INS\_270
CL\_INS\_270
CL\_INS\_365
CL\_INS\_270
CL\_INS\_365
CL\_INS\_270
CL\_INS\_270
CL\_INS\_270
CL\_INS\_297
CL\_INS\_297
CL\_INS\_297
CL\_INS\_297
CL\_INS\_297
CL\_INS\_49
CL\_INS\_297
CL\_INS\_312
CL\_INS\_312
CL\_INS\_312
CL\_INS\_312
CL\_INS\_270
CL\_INS\_312
CL\_INS\_312
CL\_INS\_312
CL\_INS\_312
CL\_INS\_312
CL\_INS\_312
CL\_INS\_312
CL\_INS\_312
CL\_INS\_312
CL\_INS\_312
CL\_INS\_312
CL\_INS\_312
CL\_INS\_312
CL\_INS\_312
CL\_INS\_365
CL\_INS\_312
CL\_INS\_312
CL\_INS\_312
CL\_INS\_312
CL\_INS\_365
CL\_INS\_365
CL\_INS\_132
CL\_INS\_312
CL\_INS\_132
CL\_INS\_132
CL\_INS\_132
CL\_INS\_132
CL\_INS\_132
CL\_INS\_132
CL\_INS\_132
CL\_INS\_132
CL\_INS\_132
CL\_INS\_132
CL\_INS\_132
CL\_INS\_132
CL\_INS\_312
CL\_INS\_132
CL\_INS\_132
CL\_INS\_312
CL\_INS\_132
CL\_INS\_132
CL\_INS\_132
CL\_INS\_132
CL\_INS\_132
CL\_INS\_132
CL\_INS\_132
CL\_INS\_132
CL\_INS\_312
CL\_INS\_312
CL\_INS\_312
CL\_INS\_312
CL\_INS\_312
CL\_INS\_312
CL\_INS\_312
CL\_INS\_312
CL\_INS\_312
CL\_INS\_312
CL\_INS\_312
CL\_INS\_312
CL\_INS\_312
CL\_INS\_312
CL\_INS\_312
CL\_INS\_312
CL\_INS\_312
CL\_INS\_312
CL\_INS\_312
CL\_INS\_312
CL\_INS\_312
CL\_INS\_312
CL\_INS\_312
CL\_INS\_312
CL\_INS\_312
CL\_INS\_312
CL\_INS\_312
CL\_INS\_312
CL\_INS\_312
CL\_INS\_312
CL\_INS\_312
CL\_INS\_312
CL\_INS\_312
CL\_INS\_312
CL\_INS\_312
CL\_INS\_312
CL\_INS\_312
CL\_INS\_312
CL\_INS\_312
CL\_INS\_312
CL\_INS\_312
CL\_INS\_312
CL\_INS\_312
CL\_INS\_312
CL\_INS\_312
CL\_INS\_312
CL\_INS\_312
CL\_INS\_312
CL\_INS\_312
CL\_INS\_312
CL\_INS\_312
CL\_INS\_312
CL\_INS\_365
CL\_INS\_312
CL\_INS\_312
CL\_INS\_312
CL\_INS\_312
CL\_INS\_312
CL\_INS\_312
CL\_INS\_312
CL\_INS\_312
CL\_INS\_312
CL\_INS\_312
CL\_INS\_312
CL\_INS\_312
CL\_INS\_312
CL\_INS\_312
CL\_INS\_312
CL\_INS\_312
CL\_INS\_312
CL\_INS\_312
CL\_INS\_312
CL\_INS\_312
CL\_INS\_312
CL\_INS\_312
CL\_INS\_312
CL\_INS\_132
CL\_INS\_132
CL\_INS\_132
CL\_INS\_132
CL\_INS\_132
CL\_INS\_365
CL\_INS\_365
CL\_INS\_365
CL\_INS\_365
CL\_INS\_365
CL\_INS\_365
CL\_INS\_365
CL\_INS\_365
CL\_INS\_365
CL\_INS\_365
CL\_INS\_365
CL\_INS\_365
CL\_INS\_365
CL\_INS\_365
CL\_INS\_365
CL\_INS\_365
CL\_INS\_365
CL\_INS\_365
CL\_INS\_365
CL\_INS\_365
CL\_INS\_365
CL\_INS\_365
CL\_INS\_365
CL\_INS\_365
CL\_INS\_365
CL\_INS\_365
CL\_INS\_365
CL\_INS\_365
CL\_INS\_365
CL\_INS\_312
CL\_INS\_312
CL\_INS\_365
CL\_INS\_365
CL\_INS\_365
CL\_INS\_365
CL\_INS\_365
CL\_INS\_365
CL\_INS\_365
CL\_INS\_365
CL\_INS\_365
CL\_INS\_365
CL\_INS\_365
CL\_INS\_365
CL\_INS\_365
CL\_INS\_365
CL\_INS\_365
CL\_INS\_365
CL\_INS\_365
CL\_INS\_365
CL\_INS\_365
CL\_INS\_365
CL\_INS\_365
CL\_INS\_365
CL\_INS\_365
CL\_INS\_365
CL\_INS\_365
CL\_INS\_365
CL\_INS\_365
CL\_INS\_365
CL\_INS\_365
CL\_INS\_365
CL\_INS\_365
CL\_INS\_365
CL\_INS\_365
CL\_INS\_365
CL\_INS\_365
CL\_INS\_365
CL\_INS\_365
CL\_INS\_365
CL\_INS\_365
CL\_INS\_365
CL\_INS\_132
CL\_INS\_132
CL\_INS\_132
CL\_INS\_132
CL\_INS\_132
CL\_INS\_132
CL\_INS\_132
CL\_INS\_132
CL\_INS\_312
CL\_INS\_132
CL\_INS\_132
CL\_INS\_132
CL\_INS\_132
CL\_INS\_132
CL\_INS\_132
CL\_INS\_132
CL\_INS\_132
CL\_INS\_132
CL\_INS\_132
CL\_INS\_132
CL\_INS\_312
CL\_INS\_312
CL\_INS\_312
CL\_INS\_312
CL\_INS\_312
CL\_INS\_312
CL\_INS\_312
CL\_INS\_365
CL\_INS\_365
CL\_INS\_365
CL\_INS\_365
CL\_INS\_365
CL\_INS\_365
CL\_INS\_365
CL\_INS\_365
CL\_INS\_365
CL\_INS\_365
CL\_INS\_365
CL\_INS\_312
CL\_INS\_312
CL\_INS\_132
CL\_INS\_132
CL\_INS\_132
CL\_INS\_132
CL\_INS\_132
CL\_INS\_132
CL\_INS\_132
CL\_INS\_132
CL\_INS\_132
CL\_INS\_132
CL\_INS\_132
CL\_INS\_132
CL\_INS\_132
CL\_INS\_132
CL\_INS\_312
CL\_INS\_132
CL\_INS\_312
CL\_INS\_312
CL\_INS\_365
CL\_INS\_312
CL\_INS\_312
CL\_INS\_365
CL\_INS\_312
CL\_INS\_312
CL\_INS\_312
CL\_INS\_312
CL\_INS\_312
CL\_INS\_312
CL\_INS\_312
CL\_INS\_312
CL\_INS\_312
CL\_INS\_312
CL\_INS\_312
CL\_INS\_312
CL\_INS\_312
CL\_INS\_312
CL\_INS\_312
CL\_INS\_312
CL\_INS\_312
CL\_INS\_312
CL\_INS\_312
CL\_INS\_312
CL\_INS\_312
CL\_INS\_312
CL\_INS\_312
CL\_INS\_312
CL\_INS\_312
CL\_INS\_312
CL\_INS\_312
CL\_INS\_312
CL\_INS\_312
CL\_INS\_312
CL\_INS\_312
CL\_INS\_312
CL\_INS\_312
CL\_INS\_312
CL\_INS\_312
CL\_INS\_312
CL\_INS\_312
CL\_INS\_312
CL\_INS\_365
CL\_INS\_365
CL\_INS\_365
CL\_INS\_365
CL\_INS\_312
CL\_INS\_312
CL\_INS\_365
CL\_INS\_365
CL\_INS\_312
CL\_INS\_365
CL\_INS\_365
CL\_INS\_365
CL\_INS\_365
CL\_INS\_365
CL\_INS\_365
CL\_INS\_312
CL\_INS\_365
CL\_INS\_365
CL\_INS\_312
CL\_INS\_312
CL\_INS\_312
CL\_INS\_312
CL\_INS\_312
CL\_INS\_312
CL\_INS\_312
CL\_INS\_312
CL\_INS\_312
CL\_INS\_132
CL\_INS\_312
CL\_INS\_312
CL\_INS\_175
CL\_INS\_175
CL\_INS\_175
CL\_INS\_365
CL\_INS\_312
CL\_INS\_312
CL\_INS\_312
CL\_INS\_312
CL\_INS\_312
CL\_INS\_175
CL\_INS\_175
CL\_INS\_175
CL\_INS\_175
CL\_INS\_175
CL\_INS\_175
CL\_INS\_175
CL\_INS\_175
CL\_INS\_312
CL\_INS\_365
CL\_INS\_365
CL\_INS\_312
CL\_INS\_365
CL\_INS\_365
CL\_INS\_365
CL\_INS\_365
CL\_INS\_365
CL\_INS\_365
CL\_INS\_365
CL\_INS\_312
CL\_INS\_312
CL\_INS\_371
CL\_INS\_371
CL\_INS\_371
CL\_INS\_371
CL\_INS\_371
CL\_INS\_371
CL\_INS\_371
CL\_INS\_371
CL\_INS\_365
CL\_INS\_312
CL\_INS\_312
CL\_INS\_312
CL\_INS\_312
CL\_INS\_312
CL\_INS\_312
CL\_INS\_312
CL\_INS\_312
CL\_INS\_312
CL\_INS\_365
CL\_INS\_365
CL\_INS\_181
CL\_INS\_181
CL\_INS\_181
CL\_INS\_181
CL\_INS\_312
CL\_INS\_312
CL\_INS\_312
CL\_INS\_312
CL\_INS\_312
CL\_INS\_312
CL\_INS\_365
CL\_INS\_312
CL\_INS\_312
CL\_INS\_312
CL\_INS\_312
CL\_INS\_312
CL\_INS\_312
CL\_INS\_312
CL\_INS\_312
CL\_INS\_312
CL\_INS\_312
CL\_INS\_365
CL\_INS\_312
CL\_INS\_365
CL\_INS\_365
CL\_INS\_365
CL\_INS\_365
CL\_INS\_365
CL\_INS\_365
CL\_INS\_365
CL\_INS\_365
CL\_INS\_365
CL\_INS\_365
CL\_INS\_15
CL\_INS\_365
CL\_INS\_365
CL\_INS\_365
CL\_INS\_365
CL\_INS\_365
CL\_INS\_365
CL\_INS\_365
CL\_INS\_365
CL\_INS\_365
CL\_INS\_312
CL\_INS\_137
CL\_INS\_312
CL\_INS\_312
CL\_INS\_312
CL\_INS\_312
CL\_INS\_312
CL\_INS\_312
CL\_INS\_312
CL\_INS\_312
CL\_INS\_312
CL\_INS\_132
CL\_INS\_365
CL\_INS\_312
CL\_INS\_365
CL\_INS\_365
CL\_INS\_181
CL\_INS\_181
CL\_INS\_181
CL\_INS\_15
CL\_INS\_312
CL\_INS\_312
CL\_INS\_83
CL\_INS\_312
CL\_INS\_312
CL\_INS\_312
CL\_INS\_312
CL\_INS\_312
CL\_INS\_312
CL\_INS\_312
CL\_INS\_312
CL\_INS\_297
CL\_INS\_297
CL\_INS\_297
CL\_INS\_312
CL\_INS\_312
CL\_INS\_312
CL\_INS\_312
CL\_INS\_312
CL\_INS\_312
CL\_INS\_312
CL\_INS\_312
CL\_INS\_312
CL\_INS\_312
CL\_INS\_312
CL\_INS\_312
CL\_INS\_312
CL\_INS\_312
CL\_INS\_312
CL\_INS\_312
CL\_INS\_312
CL\_INS\_312
CL\_INS\_312
CL\_INS\_312
CL\_INS\_312
CL\_INS\_312
CL\_INS\_312
CL\_INS\_312
CL\_INS\_312
CL\_INS\_312
CL\_INS\_312
CL\_INS\_312
CL\_INS\_312
CL\_INS\_312
CL\_INS\_312
CL\_INS\_312
CL\_INS\_312
CL\_INS\_312
CL\_INS\_312
CL\_INS\_312
CL\_INS\_312
CL\_INS\_312
CL\_INS\_312
CL\_INS\_312
CL\_INS\_312
CL\_INS\_312
CL\_INS\_312
CL\_INS\_312
CL\_INS\_312
CL\_INS\_131
CL\_INS\_312
CL\_INS\_312
CL\_INS\_312
CL\_INS\_312
CL\_INS\_312
CL\_INS\_312
CL\_INS\_312
CL\_INS\_312
CL\_INS\_131
CL\_INS\_131
CL\_INS\_131
CL\_INS\_131
CL\_INS\_131
CL\_INS\_131
CL\_INS\_312
CL\_INS\_312
CL\_INS\_312
CL\_INS\_365
CL\_INS\_312
CL\_INS\_312
CL\_INS\_312
CL\_INS\_312
CL\_INS\_365
CL\_INS\_312
CL\_INS\_312
CL\_INS\_312
CL\_INS\_365
CL\_INS\_365
CL\_INS\_365
CL\_INS\_365
CL\_INS\_312
CL\_INS\_312
CL\_INS\_312
CL\_INS\_312
CL\_INS\_312
CL\_INS\_312
CL\_INS\_312
CL\_INS\_312
CL\_INS\_312
CL\_INS\_312
CL\_INS\_312
CL\_INS\_312
CL\_INS\_365
CL\_INS\_312
CL\_INS\_312
CL\_INS\_365
CL\_INS\_365
CL\_INS\_312
CL\_INS\_312
CL\_INS\_312
CL\_INS\_312
CL\_INS\_312
CL\_INS\_312
CL\_INS\_312
CL\_INS\_312
CL\_INS\_312
CL\_INS\_365
CL\_INS\_312
CL\_INS\_312
CL\_INS\_312
CL\_INS\_312
CL\_INS\_312
CL\_INS\_312
CL\_INS\_270
CL\_INS\_270
CL\_INS\_270
CL\_INS\_270
CL\_INS\_312
CL\_INS\_365
CL\_INS\_312
CL\_INS\_312
CL\_INS\_365
CL\_INS\_365
CL\_INS\_312
CL\_INS\_312
CL\_INS\_365
CL\_INS\_365
Cluster ID


CL\_9788
CL\_9789
CL\_9790
CL\_5627
CL\_5628
CL\_5629
CL\_5630
CL\_7405
CL\_7404
CL\_7403
CL\_7402
CL\_7401
CL\_13201
CL\_13202
CL\_13203
CL\_13204
CL\_10117
CL\_10116
CL\_10115
CL\_10114
CL\_10113
CL\_13205
CL\_13206
CL\_13207
CL\_13208
CL\_13209
CL\_13210
CL\_13211
CL\_13212
CL\_13213
CL\_10055
CL\_10056
CL\_10057
CL\_10058
CL\_10059
CL\_10060
CL\_10061
CL\_10062
CL\_10063
CL\_10064
CL\_10065
CL\_10066
CL\_10067
CL\_10068
CL\_10069
CL\_10070
CL\_7318
CL\_11827
CL\_11828
CL\_11829
CL\_11830
CL\_11831
CL\_11832
CL\_13761
CL\_10071
CL\_10072
CL\_10073
CL\_4994
CL\_4995
CL\_4996
CL\_4997
CL\_10074
CL\_10075
CL\_10076
CL\_10077
CL\_10078
CL\_8851
CL\_10389
CL\_13762
CL\_10391
CL\_13763
CL\_13764
CL\_13765
CL\_13766
CL\_13767
CL\_13768
CL\_13769
CL\_13770
CL\_13771
CL\_9840
CL\_7061
CL\_6627
CL\_6628
CL\_6629
CL\_6630
CL\_6631
CL\_6632
CL\_13821
CL\_13772
CL\_13773
CL\_8850
CL\_8228
CL\_8229
CL\_8230
CL\_8231
CL\_8232
CL\_6568
CL\_8233
CL\_6570
CL\_8234
CL\_6572
CL\_6573
CL\_6574
CL\_6575
CL\_6576
CL\_6577
CL\_6579
CL\_8235
CL\_12148
CL\_8237
CL\_8238
CL\_12147
CL\_12146
CL\_12145
CL\_12144
CL\_9135
CL\_9136
CL\_9138
CL\_9139
CL\_9140
CL\_9731
CL\_9732
CL\_9733
CL\_9734
CL\_9735
CL\_9736
CL\_9737
CL\_9738
CL\_9739
CL\_9740
CL\_9741
CL\_9742
CL\_9743
CL\_9744
CL\_9745
CL\_9746
CL\_9747
CL\_9748
CL\_9749
CL\_9750
CL\_9751
CL\_9752
CL\_9753
CL\_9754
CL\_9755
CL\_9756
CL\_9757
CL\_9758
CL\_9759
CL\_9760
CL\_9761
CL\_9762
CL\_9763
CL\_9764
CL\_9765
CL\_9766
CL\_9767
CL\_9768
CL\_9769
CL\_9148
CL\_9149
CL\_9150
CL\_3888
CL\_9151
CL\_10694
CL\_5520
CL\_3886
CL\_3885
CL\_3884
CL\_3883
CL\_3882
CL\_8948
CL\_10695
CL\_10696
CL\_10697
CL\_10043
CL\_9152
CL\_2883
CL\_9153
CL\_2879
CL\_9154
CL\_9155
CL\_9156
CL\_9157
CL\_9158
CL\_9159
CL\_10698
CL\_9147
CL\_9146
CL\_9145
CL\_9770
CL\_9143
CL\_9144
CL\_9142
CL\_9141
CL\_9771
CL\_9772
CL\_9773
CL\_9774
CL\_9775
CL\_9776
CL\_9777
CL\_9778
CL\_9779
CL\_9780
CL\_9781
CL\_9782
CL\_5606
CL\_5605
CL\_9160
CL\_9161
CL\_3891
CL\_5756
CL\_5755
CL\_5754
CL\_5753
CL\_5752
CL\_5751
CL\_3893
CL\_9162
CL\_9163
CL\_9164
CL\_5750
CL\_5749
CL\_5748
CL\_5604
CL\_9783
CL\_9784
CL\_9785
CL\_9786
CL\_6327
CL\_9787
CL\_11944
CL\_11943
CL\_11942
CL\_11941
CL\_11940
CL\_11939
CL\_11938
CL\_11937
CL\_11936
CL\_11935
CL\_11934
CL\_11933
CL\_11932
CL\_10399
CL\_11931
CL\_11930
CL\_11929
CL\_11928
CL\_8206
CL\_8207
CL\_5820
CL\_5821
CL\_5822
CL\_5823
CL\_5824
CL\_5825
CL\_5826
CL\_5827
CL\_5828
CL\_5829
CL\_5830
CL\_5831
CL\_5832
CL\_5833
CL\_5834
CL\_5835
CL\_11198
CL\_13797
CL\_11199
CL\_11200
CL\_11201
CL\_11202
CL\_11203
CL\_11204
CL\_5838
CL\_5839
CL\_5840
CL\_5842
CL\_7518
CL\_13796
CL\_5845
CL\_7553
CL\_7554
CL\_7555
CL\_7556
CL\_7557
CL\_7560
CL\_7561
CL\_7562
CL\_7563
CL\_7564
CL\_7565
CL\_7566
CL\_7567
CL\_7568
CL\_7569
CL\_7570
CL\_7571
CL\_7572
CL\_7573
CL\_7574
CL\_7575
CL\_7576
CL\_7577
CL\_7578
CL\_7579
CL\_13795
CL\_7580
CL\_7581
CL\_7583
CL\_7584
CL\_7599
CL\_7600
CL\_7601
CL\_7602
CL\_7517
CL\_7468
CL\_7469
CL\_7470
CL\_7471
CL\_7472
CL\_7473
CL\_7474
CL\_7475
CL\_7476
CL\_5850
CL\_5857
CL\_5858
CL\_5859
CL\_5860
CL\_5861
CL\_5862
CL\_5863
CL\_5865
CL\_5866
CL\_13794
CL\_5868
CL\_5869
CL\_5870
CL\_5871
CL\_5872
CL\_7477
CL\_5873
CL\_5874
CL\_13793
CL\_13792
CL\_13791
CL\_13790
CL\_5875
CL\_7478
CL\_5876
CL\_5878
CL\_5879
CL\_5880
CL\_5881
CL\_5882
CL\_5886
CL\_867
CL\_864
CL\_8568
CL\_8569
CL\_8570
CL\_8571
CL\_8572
CL\_8573
CL\_8574
CL\_8575
CL\_8576
CL\_8577
CL\_8578
CL\_8579
CL\_8580
CL\_8581
CL\_8582
CL\_8583
CL\_8584
CL\_8585
CL\_8586
CL\_8587
CL\_8588
CL\_6861
CL\_6860
CL\_6859
CL\_6858
CL\_8589
CL\_8590
CL\_13789
CL\_13788
CL\_8591
CL\_8592
CL\_8593
CL\_8594
CL\_8595
CL\_8596
CL\_8597
CL\_8598
CL\_8599
CL\_8600
CL\_8601
CL\_8602
CL\_8603
CL\_8604
CL\_8605
CL\_8606
CL\_8607
CL\_8608
CL\_8609
CL\_8610
CL\_856
CL\_855
CL\_854
CL\_853
CL\_852
CL\_851
CL\_849
CL\_848
CL\_8469
CL\_847
CL\_846
CL\_844
CL\_843
CL\_8466
CL\_8465
CL\_8464
CL\_8463
CL\_8462
CL\_8461
CL\_8460
CL\_5887
CL\_5888
CL\_5889
CL\_5890
CL\_7480
CL\_11156
CL\_11157
CL\_11158
CL\_13787
CL\_11159
CL\_11160
CL\_7486
CL\_7487
CL\_7488
CL\_7491
CL\_7492
CL\_7493
CL\_7508
CL\_7509
CL\_7510
CL\_10768
CL\_10769
CL\_13786
CL\_13785
CL\_13784
CL\_13783
CL\_13782
CL\_6701
CL\_9996
CL\_9995
CL\_6703
CL\_9993
CL\_6702
CL\_5124
CL\_9992
CL\_9991
CL\_9989
CL\_9988
CL\_13781
CL\_13780
CL\_5801
CL\_5802
CL\_5803
CL\_5804
CL\_5807
CL\_5808
CL\_5809
CL\_5810
CL\_5811
CL\_5812
CL\_5813
CL\_5814
CL\_5815
CL\_5816
CL\_7516
CL\_5819
CL\_8705
CL\_9812
CL\_4968
CL\_9813
CL\_8710
CL\_4966
CL\_8711
CL\_4963
CL\_9814
CL\_9815
CL\_4960
CL\_4959
CL\_9816
CL\_4958
CL\_8717
CL\_8718
CL\_9817
CL\_8722
CL\_8723
CL\_4953
CL\_4952
CL\_4951
CL\_4950
CL\_9818
CL\_9819
CL\_9820
CL\_9821
CL\_4949
CL\_4947
CL\_4946
CL\_4945
CL\_4944
CL\_4943
CL\_4942
CL\_9822
CL\_4940
CL\_4939
CL\_4938
CL\_4937
CL\_9823
CL\_4935
CL\_9824
CL\_9825
CL\_4931
CL\_4930
CL\_4929
CL\_4928
CL\_4927
CL\_4925
CL\_4924
CL\_4923
CL\_4922
CL\_12914
CL\_4921
CL\_4920
CL\_4919
CL\_4918
CL\_4917
CL\_4916
CL\_4912
CL\_4911
CL\_4910
CL\_4909
CL\_4907
CL\_4906
CL\_4904
CL\_4903
CL\_12143
CL\_12142
CL\_12141
CL\_12140
CL\_7670
CL\_12139
CL\_12138
CL\_7068
CL\_7069
CL\_7070
CL\_4881
CL\_4902
CL\_4901
CL\_8849
CL\_8848
CL\_8847
CL\_8846
CL\_8845
CL\_8844
CL\_8843
CL\_8842
CL\_8841
CL\_8840
CL\_8839
CL\_8838
CL\_8837
CL\_8836
CL\_8835
CL\_8834
CL\_8833
CL\_8832
CL\_6453
CL\_6454
CL\_6455
CL\_6326
CL\_8831
CL\_8830
CL\_5747
CL\_5746
CL\_5745
CL\_5744
CL\_8829
CL\_8828
CL\_8827
CL\_5743
CL\_4900
CL\_13811
CL\_13810
CL\_4898
CL\_4897
CL\_4896
CL\_4895
CL\_4894
CL\_12890
CL\_4893
CL\_4892
CL\_8208
CL\_8203
CL\_8204
CL\_8205
CL\_7698
CL\_4891
CL\_4890
CL\_4889
CL\_4887
CL\_6651
CL\_4886
CL\_4885
CL\_12137
CL\_12136
CL\_12135
CL\_12134
CL\_12133
CL\_12132
CL\_12131
CL\_12130
CL\_12129
CL\_12128
CL\_4884
CL\_13825
CL\_4882
CL\_4883
CL\_13822
CL\_13823
CL\_4880
CL\_4879
CL\_6657
CL\_5768
CL\_7860
CL\_6659
CL\_5128
CL\_7886
CL\_8373
CL\_7885
CL\_7884
CL\_7883
CL\_7882
CL\_5847
CL\_7881
CL\_7880
CL\_7879
CL\_5118
CL\_7876
CL\_7874
CL\_7873
CL\_7871
CL\_7870
CL\_7869
CL\_7868
CL\_7867
CL\_7866
CL\_7865
CL\_7864
CL\_12907
CL\_7863
CL\_6660
CL\_7697
CL\_7696
CL\_7695
CL\_7694
CL\_11927
CL\_11926
CL\_6240
CL\_11925
CL\_11924
CL\_11923
CL\_11922
CL\_11921
CL\_11920
CL\_11919
CL\_11918
CL\_11917
CL\_11916
CL\_11915
CL\_11914
CL\_11913
CL\_11912
CL\_11911
CL\_11910
CL\_11909
CL\_11908
CL\_11907
CL\_11906
CL\_11905
CL\_11904
CL\_11903
CL\_11902
CL\_11901
CL\_11900
CL\_11899
CL\_11898
CL\_11897
CL\_11896
CL\_11895
CL\_11894
CL\_11893
CL\_11892
CL\_11891
CL\_11890
CL\_11889
CL\_11888
CL\_11887
CL\_11886
CL\_11885
CL\_11884
CL\_11883
CL\_11882
CL\_11881
CL\_11880
CL\_11879
CL\_11878
CL\_11877
CL\_11876
CL\_11875
CL\_11874
CL\_11873
CL\_11872
CL\_11871
CL\_11870
CL\_3086
CL\_3087
CL\_3088
CL\_3090
CL\_11869
CL\_11868
CL\_11867
CL\_11866
CL\_11865
CL\_11112
CL\_11111
CL\_11110
CL\_11109
CL\_11108
CL\_11107
CL\_11864
CL\_7861
CL\_5014
CL\_5013
CL\_5012
CL\_5011
CL\_5010
CL\_5035
CL\_5009
CL\_12905
CL\_10352
CL\_10351
CL\_10349
CL\_12904
CL\_10348
CL\_7531
CL\_5933
CL\_7532
CL\_7896
CL\_7895
CL\_13807
CL\_13806
CL\_13805
CL\_13804
CL\_13803
CL\_13802
CL\_5935
CL\_13801
CL\_7795
CL\_9847
CL\_9846
CL\_7794
CL\_7535
CL\_4868
CL\_4869
CL\_4870
CL\_4871
CL\_4872
CL\_4873
CL\_4874
CL\_4875
CL\_4876
CL\_4877
CL\_12362
CL\_5763
CL\_5762
CL\_5761
CL\_12361
CL\_12360
CL\_5760
CL\_5759
CL\_5758
CL\_5757
CL\_12359
CL\_4878
CL\_13800
CL\_7643
CL\_7682
CL\_5924
CL\_13799
CL\_13798
CL\_11664
CL\_8369
