## Supplementary material for "A novel method for integrating genomic and Tn-Seq data to identify common *in vivo* fitness mechanisms across multiple bacterial species": S4 Dataset: CL_INS_313.html

Legend

 Hypothetical
 All VFDB Genes

FULL


WINDOWSVGPNG

Trim RowsRemove SingletonsSave Fasta

CL\_4494


CL\_4494


CL\_4494

HighlightSelectShow Genomes


24

CL\_4495


3

CL\_4495


1

CL\_4495

fGI ID


CL\_INS\_313
CL\_INS\_313
CL\_INS\_313
CL\_INS\_313
Cluster ID


CL\_10699
CL\_10700
CL\_5742
CL\_5741
