## Supplementary material for "A novel method for integrating genomic and Tn-Seq data to identify common *in vivo* fitness mechanisms across multiple bacterial species": S4 Dataset: CL_INS_314.html

Legend

 Hypothetical
 All VFDB Genes

FULL


WINDOWSVGPNG

Trim RowsRemove SingletonsSave Fasta

CL\_4499


CL\_4499

HighlightSelectShow Genomes


25

CL\_4500


3

CL\_4500

fGI ID


CL\_INS\_314
CL\_INS\_314
CL\_INS\_314
CL\_INS\_314
CL\_INS\_314
CL\_INS\_314
CL\_INS\_314
Cluster ID


CL\_13214
CL\_13215
CL\_13216
CL\_13217
CL\_13218
CL\_13219
CL\_13220
