## Supplementary material for "A novel method for integrating genomic and Tn-Seq data to identify common *in vivo* fitness mechanisms across multiple bacterial species": S4 Dataset: CL_INS_319.html

Legend

 Mobile +extrachromosomalelementfunctions
 Hypothetical
 Centralintermediarymetabolism
 All VFDB Genes

FULL


WINDOWSVGPNG

Trim RowsRemove SingletonsSave Fasta

CL\_4518


CL\_4518


CL\_4513

HighlightSelectShow Genomes


19

CL\_4519


8

CL\_4519


1

CL\_4519

fGI ID


CL\_INS\_319
CL\_INS\_319
CL\_INS\_319
Cluster ID


CL\_7399
CL\_7398
CL\_6528
