## Supplementary material for "A novel method for integrating genomic and Tn-Seq data to identify common *in vivo* fitness mechanisms across multiple bacterial species": S4 Dataset: CL_INS_323.html

Legend

 Mobile +extrachromosomalelementfunctions
 Regulatoryfunctions
 Hypothetical
 DNA Metabolism
 Other
 Transport +binding proteins
 All VFDB Genes

FULL


WINDOWSVGPNG

Trim RowsRemove SingletonsSave Fasta

CL\_4522


CL\_4522


CL\_4522

HighlightSelectShow Genomes


14

CL\_4523


13

CL\_4523


1

CL\_4523

fGI ID


CL\_INS\_323
CL\_INS\_323
CL\_INS\_323
CL\_INS\_323
CL\_INS\_323
CL\_INS\_323
CL\_INS\_323
CL\_INS\_323
CL\_INS\_323
CL\_INS\_323
CL\_INS\_323
CL\_INS\_323
CL\_INS\_323
CL\_INS\_323
Cluster ID


CL\_5732
CL\_7397
CL\_7396
CL\_7395
CL\_7394
CL\_7393
CL\_7392
CL\_7391
CL\_7390
CL\_7389
CL\_7388
CL\_7387
CL\_7386
CL\_7385
