## Supplementary material for "A novel method for integrating genomic and Tn-Seq data to identify common *in vivo* fitness mechanisms across multiple bacterial species": S4 Dataset: CL_INS_326.html

Legend

 Hypothetical
 All Fitness Genes
 Transport +binding proteins

FULL


WINDOWSVGPNG

Trim RowsRemove SingletonsSave Fasta

CL\_4536


CL\_4536

HighlightSelectShow Genomes


24

CL\_4537


4

CL\_4537

fGI ID

CL\_INS\_326
Cluster ID

CL\_12926
