## Supplementary material for "A novel method for integrating genomic and Tn-Seq data to identify common *in vivo* fitness mechanisms across multiple bacterial species": S4 Dataset: CL_INS_328.html

Legend

 Hypothetical
 Biosynthesis ofcofactors,prostheticgroups, +carriers
 Transport +binding proteins
 All VFDB Genes

FULL


WINDOWSVGPNG

Trim RowsRemove SingletonsSave Fasta

CL\_4541


CL\_4541


CL\_4541


CL\_4541


CL\_4541

HighlightSelectShow Genomes


20

CL\_4542


3

CL\_4542


3

CL\_4542


1

CL\_4542


1

CL\_4542

fGI ID


CL\_INS\_328
CL\_INS\_328
CL\_INS\_328
CL\_INS\_328
Cluster ID


CL\_9797
CL\_11855
CL\_9798
CL\_9799
