## Supplementary material for "A novel method for integrating genomic and Tn-Seq data to identify common *in vivo* fitness mechanisms across multiple bacterial species": S4 Dataset: CL_INS_329.html

Legend

 Other
 All VFDB Genes

FULL


WINDOWSVGPNG

Trim RowsRemove SingletonsSave Fasta

CL\_4547


CL\_4547


CL\_4547

HighlightSelectShow Genomes


17

CL\_4550


10

CL\_4550


1

CL\_4550

fGI ID


CL\_INS\_329
CL\_INS\_329
CL\_INS\_329
Cluster ID


CL\_10702
CL\_4548
CL\_4549
