## Supplementary material for "A novel method for integrating genomic and Tn-Seq data to identify common *in vivo* fitness mechanisms across multiple bacterial species": S4 Dataset: CL_INS_331.html

Legend

 Hypothetical
 Other
 All VFDB Genes

FULL


WINDOWSVGPNG

Trim RowsRemove SingletonsSave Fasta

CL\_4560


CL\_4560

HighlightSelectShow Genomes


23

CL\_5685


4

CL\_4566

fGI ID


CL\_INS\_331
CL\_INS\_331
CL\_INS\_331
CL\_INS\_331
Cluster ID


CL\_4561
CL\_4562
CL\_4563
CL\_4564
