## Supplementary material for "A novel method for integrating genomic and Tn-Seq data to identify common *in vivo* fitness mechanisms across multiple bacterial species": S4 Dataset: CL_INS_332.html

Legend

 Regulatoryfunctions
 Hypothetical
 Other
 All VFDB Genes

FULL


WINDOWSVGPNG

Trim RowsRemove SingletonsSave Fasta

CL\_5685


CL\_5685


CL\_4560

HighlightSelectShow Genomes


14

CL\_4566


10

CL\_4566


4

CL\_4566

fGI ID


CL\_INS\_332
CL\_INS\_331
CL\_INS\_331
CL\_INS\_331
CL\_INS\_332
Cluster ID


CL\_5684
CL\_4562
CL\_4563
CL\_4564
CL\_4565
