## Supplementary material for "A novel method for integrating genomic and Tn-Seq data to identify common *in vivo* fitness mechanisms across multiple bacterial species": S4 Dataset: CL_INS_334.html

Legend

 Mobile +extrachromosomalelementfunctions
 Hypothetical
 All VFDB Genes

FULL


WINDOWSVGPNG

Trim RowsRemove SingletonsSave Fasta

CL\_4576


CL\_4576


CL\_4576

HighlightSelectShow Genomes


18

CL\_4577


2

CL\_4577


1

CL\_4577

fGI ID


CL\_INS\_334
CL\_INS\_334
CL\_INS\_334
Cluster ID


CL\_5164
CL\_5165
CL\_5166
