## Supplementary material for "A novel method for integrating genomic and Tn-Seq data to identify common *in vivo* fitness mechanisms across multiple bacterial species": S4 Dataset: CL_INS_335.html

Legend

 Mobile +extrachromosomalelementfunctions
 Hypothetical
 Proteinsynthesis/fate
 Other
 All VFDB Genes

FULL


WINDOWSVGPNG

Trim RowsRemove SingletonsSave Fasta

CL\_4577


CL\_4577


CL\_4577


CL\_4577


CL\_4577


CL\_4577

HighlightSelectShow Genomes


11

CL\_4578


4

CL\_4581


2

CL\_4578


2

CL\_4578


1

CL\_4578


1

CL\_4578

fGI ID


CL\_INS\_335
CL\_INS\_335
CL\_INS\_335
CL\_INS\_335
CL\_INS\_335
CL\_INS\_335
CL\_INS\_335
CL\_INS\_335
CL\_INS\_335
Cluster ID


CL\_11013
CL\_4579
CL\_12534
CL\_9159
CL\_9158
CL\_9157
CL\_9156
CL\_9155
CL\_9154
