## Supplementary material for "A novel method for integrating genomic and Tn-Seq data to identify common *in vivo* fitness mechanisms across multiple bacterial species": S4 Dataset: CL_INS_337.html

Legend

 Mobile +extrachromosomalelementfunctions
 Hypothetical
 DNA Metabolism
 AntibioticResistance
 Other
 EnergyMetabolism
 All VFDB Genes

FULL


WINDOWSVGPNG

Trim RowsRemove SingletonsSave Fasta

CL\_4584


CL\_4584


CL\_4584


CL\_4584

HighlightSelectShow Genomes


22

CL\_4585


4

CL\_4585


1

CL\_4586


1

CL\_4586

fGI ID


CL\_INS\_337
CL\_INS\_337
CL\_INS\_337
CL\_INS\_337
CL\_INS\_337
CL\_INS\_337
CL\_INS\_337
CL\_INS\_337
CL\_INS\_337
CL\_INS\_337
CL\_INS\_337
CL\_INS\_337
CL\_INS\_337
Cluster ID


CL\_6521
CL\_6520
CL\_13345
CL\_13128
CL\_10781
CL\_10782
CL\_5901
CL\_5900
CL\_7806
CL\_10784
CL\_7808
CL\_7809
CL\_13129
