## Supplementary material for "A novel method for integrating genomic and Tn-Seq data to identify common *in vivo* fitness mechanisms across multiple bacterial species": S4 Dataset: CL_INS_342.html

Legend

 Mobile +extrachromosomalelementfunctions
 Other

FULL


WINDOWSVGPNG

Trim RowsRemove SingletonsSave Fasta

CL\_4648


CL\_4648

HighlightSelectShow Genomes


25

CL\_4649


3

CL\_4649

fGI ID

CL\_INS\_342
Cluster ID

CL\_13221
