## Supplementary material for "A novel method for integrating genomic and Tn-Seq data to identify common *in vivo* fitness mechanisms across multiple bacterial species": S4 Dataset: CL_INS_343.html

Legend

 Hypothetical
 Other
 All VFDB Genes

FULL


WINDOWSVGPNG

Trim RowsRemove SingletonsSave Fasta

CL\_4651


CL\_4651


CL\_4651

HighlightSelectShow Genomes


18

CL\_4652


9

CL\_4652


1

CL\_4653

fGI ID


CL\_INS\_343
CL\_INS\_343
Cluster ID


CL\_6517
CL\_13776
