## Supplementary material for "A novel method for integrating genomic and Tn-Seq data to identify common *in vivo* fitness mechanisms across multiple bacterial species": S4 Dataset: CL_INS_348.html

Legend

 Mobile +extrachromosomalelementfunctions
 Regulatoryfunctions
 Hypothetical
 Other
 Transport +binding proteins
 All VFDB Genes

FULL


WINDOWSVGPNG

Trim RowsRemove SingletonsSave Fasta

CL\_4673


CL\_4673


CL\_4673


CL\_4673


CL\_4673

HighlightSelectShow Genomes


18

CL\_4674


7

CL\_4674


1

CL\_4674


1

CL\_4674


1

CL\_4674

fGI ID


CL\_INS\_348
CL\_INS\_348
CL\_INS\_348
CL\_INS\_348
CL\_INS\_348
CL\_INS\_348
CL\_INS\_348
CL\_INS\_222
CL\_INS\_222
CL\_INS\_222
CL\_INS\_222
CL\_INS\_348
CL\_INS\_221
CL\_INS\_83
CL\_INS\_348
CL\_INS\_348
CL\_INS\_221
CL\_INS\_221
CL\_INS\_222
CL\_INS\_222
CL\_INS\_348
CL\_INS\_348
CL\_INS\_348
CL\_INS\_348
CL\_INS\_221
CL\_INS\_348
CL\_INS\_348
CL\_INS\_348
CL\_INS\_348
CL\_INS\_221
CL\_INS\_221
CL\_INS\_221
CL\_INS\_221
CL\_INS\_221
CL\_INS\_222
CL\_INS\_221
CL\_INS\_348
CL\_INS\_348
CL\_INS\_348
CL\_INS\_221
CL\_INS\_348
CL\_INS\_348
CL\_INS\_348
CL\_INS\_348
CL\_INS\_348
CL\_INS\_348
CL\_INS\_348
CL\_INS\_348
CL\_INS\_348
CL\_INS\_348
CL\_INS\_348
CL\_INS\_348
CL\_INS\_348
CL\_INS\_348
CL\_INS\_348
CL\_INS\_348
CL\_INS\_348
CL\_INS\_348
CL\_INS\_348
CL\_INS\_348
Cluster ID


CL\_6515
CL\_6514
CL\_10703
CL\_10704
CL\_7384
CL\_7383
CL\_7382
CL\_3224
CL\_3223
CL\_3222
CL\_7381
CL\_7380
CL\_3220
CL\_10705
CL\_11014
CL\_7379
CL\_3227
CL\_3228
CL\_3229
CL\_3230
CL\_7378
CL\_7377
CL\_7376
CL\_7375
CL\_3235
CL\_7374
CL\_11015
CL\_11016
CL\_7373
CL\_3237
CL\_3238
CL\_3239
CL\_3240
CL\_3241
CL\_3242
CL\_3243
CL\_7372
CL\_7371
CL\_7370
CL\_3246
CL\_7369
CL\_10706
CL\_10707
CL\_10708
CL\_11017
CL\_11018
CL\_7368
CL\_7367
CL\_7366
CL\_7365
CL\_7364
CL\_7363
CL\_7362
CL\_7361
CL\_11019
CL\_10709
CL\_7360
CL\_10710
CL\_7359
CL\_7358
