## Supplementary material for "A novel method for integrating genomic and Tn-Seq data to identify common *in vivo* fitness mechanisms across multiple bacterial species": S4 Dataset: CL_INS_349.html

Legend

 Regulatoryfunctions
 Hypothetical
 Other
 All VFDB Genes

FULL


WINDOWSVGPNG

Trim RowsRemove SingletonsSave Fasta

CL\_4676


CL\_4676


CL\_4676

HighlightSelectShow Genomes


21

CL\_4677


4

CL\_4677


3

CL\_4677

fGI ID


CL\_INS\_349
CL\_INS\_349
CL\_INS\_349
CL\_INS\_349
CL\_INS\_349
Cluster ID


CL\_6513
CL\_6512
CL\_6511
CL\_13224
CL\_13225
