## Supplementary material for "A novel method for integrating genomic and Tn-Seq data to identify common *in vivo* fitness mechanisms across multiple bacterial species": S4 Dataset: CL_INS_351.html

Legend

 Hypothetical
 Transport +binding proteins
 All VFDB Genes

FULL


WINDOWSVGPNG

Trim RowsRemove SingletonsSave Fasta

CL\_4681


CL\_4681

HighlightSelectShow Genomes


25

CL\_4682


3

CL\_4682

fGI ID


CL\_INS\_351
CL\_INS\_351
Cluster ID


CL\_6510
CL\_6509
