## Supplementary material for "A novel method for integrating genomic and Tn-Seq data to identify common *in vivo* fitness mechanisms across multiple bacterial species": S4 Dataset: CL_INS_352.html

Legend

 Hypothetical
 Transport +binding proteins

FULL


WINDOWSVGPNG

Trim RowsRemove SingletonsSave Fasta

CL\_4691


CL\_4691

HighlightSelectShow Genomes


15

CL\_4696


13

CL\_4696

fGI ID


CL\_INS\_352
CL\_INS\_352
CL\_INS\_352
CL\_INS\_352
Cluster ID


CL\_4692
CL\_4693
CL\_4694
CL\_4695
