## Supplementary material for "A novel method for integrating genomic and Tn-Seq data to identify common *in vivo* fitness mechanisms across multiple bacterial species": S4 Dataset: CL_INS_356.html

Legend

 Other
 All VFDB Genes

FULL


WINDOWSVGPNG

Trim RowsRemove SingletonsSave Fasta

CL\_4715


CL\_4715

HighlightSelectShow Genomes


25

CL\_4716


2

CL\_4711

fGI ID


CL\_INS\_356
CL\_INS\_356
CL\_INS\_356
Cluster ID


CL\_12356
CL\_12357
CL\_12358
