## Supplementary material for "A novel method for integrating genomic and Tn-Seq data to identify common *in vivo* fitness mechanisms across multiple bacterial species": S4 Dataset: CL_INS_357.html

Legend

 Hypothetical
 All Fitness Genes
 All VFDB Genes

FULL


WINDOWSVGPNG

Trim RowsRemove SingletonsSave Fasta

CL\_4723


CL\_4723

HighlightSelectShow Genomes


24

CL\_4724


4

CL\_4724

fGI ID

CL\_INS\_357
Cluster ID

CL\_12355
