## Supplementary material for "A novel method for integrating genomic and Tn-Seq data to identify common *in vivo* fitness mechanisms across multiple bacterial species": S4 Dataset: CL_INS_358.html

Legend

 Hypothetical
 Proteinsynthesis/fate
 Other

FULL


WINDOWSVGPNG

Trim RowsRemove SingletonsSave Fasta

CL\_4757


CL\_4757


CL\_4757


CL\_4757

HighlightSelectShow Genomes


23

CL\_4758


2

CL\_4758


2

CL\_4758


1

CL\_4758

fGI ID


CL\_INS\_358
CL\_INS\_358
Cluster ID


CL\_9801
CL\_9802
