## Supplementary material for "A novel method for integrating genomic and Tn-Seq data to identify common *in vivo* fitness mechanisms across multiple bacterial species": S4 Dataset: CL_INS_364.html

Legend

 Mobile +extrachromosomalelementfunctions
 Hypothetical
 AntibioticResistance
 All Fitness Genes
 Other
 Transport +binding proteins
 All VFDB Genes

FULL


WINDOWSVGPNG

Trim RowsRemove SingletonsSave Fasta

CL\_4837


CL\_4837


CL\_4837


CL\_4837


CL\_4837


CL\_4837


CL\_4837

HighlightSelectShow Genomes


14

CL\_4853


7

CL\_4853


1

Break


1

CL\_4853


1

CL\_4853


1

CL\_4853


1

CL\_4853

fGI ID


CL\_INS\_364
CL\_INS\_364
CL\_INS\_364
CL\_INS\_364
CL\_INS\_364
CL\_INS\_364
CL\_INS\_364
CL\_INS\_364
CL\_INS\_364
CL\_INS\_364
CL\_INS\_364
CL\_INS\_364
CL\_INS\_364
CL\_INS\_364
CL\_INS\_364
CL\_INS\_364
CL\_INS\_364
CL\_INS\_365
CL\_INS\_364
CL\_INS\_365
CL\_INS\_364
CL\_INS\_364
Cluster ID


CL\_9803
CL\_9804
CL\_9805
CL\_9806
CL\_4838
CL\_4839
CL\_4840
CL\_4841
CL\_4842
CL\_13778
CL\_13779
CL\_4843
CL\_4844
CL\_4845
CL\_9198
CL\_4846
CL\_4847
CL\_4848
CL\_4849
CL\_4850
CL\_4851
CL\_4852
