## Supplementary material for "A novel method for integrating genomic and Tn-Seq data to identify common *in vivo* fitness mechanisms across multiple bacterial species": S4 Dataset: CL_INS_367.html

Legend

 Mobile +extrachromosomalelementfunctions
 Hypothetical
 Proteinsynthesis/fate
 Other
 Transport +binding proteins

FULL


WINDOWSVGPNG

Trim RowsRemove SingletonsSave Fasta

CL\_2867


CL\_2867


CL\_2867

HighlightSelectShow Genomes


22

CL\_2866


2

CL\_2246


1

CL\_2246

fGI ID


CL\_INS\_367
CL\_INS\_141
CL\_INS\_152
CL\_INS\_325
CL\_INS\_141
CL\_INS\_141
CL\_INS\_141
CL\_INS\_141
CL\_INS\_141
CL\_INS\_141
CL\_INS\_141
CL\_INS\_141
CL\_INS\_141
CL\_INS\_141
CL\_INS\_141
CL\_INS\_141
CL\_INS\_141
CL\_INS\_141
CL\_INS\_141
CL\_INS\_141
CL\_INS\_325
CL\_INS\_141
CL\_INS\_325
CL\_INS\_152
CL\_INS\_152
CL\_INS\_152
CL\_INS\_152
CL\_INS\_152
CL\_INS\_152
CL\_INS\_152
CL\_INS\_152
Cluster ID


CL\_13026
CL\_7207
CL\_13025
CL\_8179
CL\_8178
CL\_8177
CL\_8176
CL\_8175
CL\_8174
CL\_8173
CL\_8172
CL\_8171
CL\_8170
CL\_8169
CL\_8168
CL\_8167
CL\_8166
CL\_1315
CL\_1316
CL\_1317
CL\_8162
CL\_7945
CL\_8160
CL\_13024
CL\_13023
CL\_13022
CL\_13021
CL\_13020
CL\_13019
CL\_13018
CL\_13017
