## Supplementary material for "A novel method for integrating genomic and Tn-Seq data to identify common *in vivo* fitness mechanisms across multiple bacterial species": S4 Dataset: CL_INS_368.html

Legend

 Mobile +extrachromosomalelementfunctions
 Regulatoryfunctions
 Hypothetical
 DNA Metabolism
 All Fitness Genes
 Cell Envelope
 Proteinsynthesis/fate
 Other
 EnergyMetabolism
 Transport +binding proteins
 All VFDB Genes

FULL


WINDOWSVGPNG

Trim RowsRemove SingletonsSave Fasta

CL\_2094


CL\_2866


CL\_2866


CL\_2866


CL\_2866


CL\_2866


CL\_4533


CL\_4533


CL\_2866


CL\_2866


CL\_2866


CL\_2815


CL\_2866


CL\_2866


CL\_2866


CL\_4533


CL\_1954


CL\_2866


CL\_2866


CL\_2866


CL\_2866


CL\_2866


CL\_2866


CL\_2245


CL\_2866


CL\_2094


CL\_4533


CL\_2094


CL\_2866


CL\_2866

HighlightSelectShow Genomes


2

CL\_5729


2

CL\_5729


2

CL\_5729


1

CL\_2899


1

CL\_2067


1

CL\_2067


1

CL\_5729


1

CL\_5729


1

CL\_5729


1

CL\_2095


1

CL\_2899


1

CL\_5729


1

CL\_2067


1

CL\_2067


1

CL\_2095


1

CL\_5729


1

CL\_5729


1

CL\_2339


1

CL\_2821


1

CL\_5729


1

CL\_5729


1

CL\_2821


1

CL\_2067


1

CL\_5729


1

CL\_5729


1

CL\_5729


1

CL\_5729


1

CL\_5729


1

CL\_1638


1

CL\_2821

fGI ID


CL\_INS\_368
CL\_INS\_368
CL\_INS\_368
CL\_INS\_368
CL\_INS\_368
CL\_INS\_368
CL\_INS\_368
CL\_INS\_368
CL\_INS\_368
CL\_INS\_368
CL\_INS\_368
CL\_INS\_368
CL\_INS\_368
CL\_INS\_368
CL\_INS\_325
CL\_INS\_368
CL\_INS\_368
CL\_INS\_368
CL\_INS\_368
CL\_INS\_366
CL\_INS\_368
CL\_INS\_368
CL\_INS\_141
CL\_INS\_368
CL\_INS\_141
CL\_INS\_141
CL\_INS\_141
CL\_INS\_141
CL\_INS\_141
CL\_INS\_368
CL\_INS\_141
CL\_INS\_141
CL\_INS\_141
CL\_INS\_141
CL\_INS\_141
CL\_INS\_141
CL\_INS\_141
CL\_INS\_368
CL\_INS\_141
CL\_INS\_368
CL\_INS\_141
CL\_INS\_325
CL\_INS\_325
CL\_INS\_141
CL\_INS\_141
CL\_INS\_141
CL\_INS\_368
CL\_INS\_368
CL\_INS\_141
CL\_INS\_368
CL\_INS\_368
CL\_INS\_141
CL\_INS\_141
CL\_INS\_141
CL\_INS\_141
CL\_INS\_141
CL\_INS\_141
CL\_INS\_325
CL\_INS\_325
CL\_INS\_325
CL\_INS\_325
CL\_INS\_325
CL\_INS\_325
CL\_INS\_325
CL\_INS\_325
CL\_INS\_325
CL\_INS\_325
CL\_INS\_325
CL\_INS\_325
CL\_INS\_325
CL\_INS\_325
CL\_INS\_325
CL\_INS\_325
CL\_INS\_325
CL\_INS\_368
CL\_INS\_325
CL\_INS\_325
CL\_INS\_141
CL\_INS\_325
CL\_INS\_325
CL\_INS\_325
CL\_INS\_325
CL\_INS\_325
CL\_INS\_141
CL\_INS\_141
CL\_INS\_141
CL\_INS\_325
CL\_INS\_325
CL\_INS\_366
CL\_INS\_368
CL\_INS\_368
CL\_INS\_368
CL\_INS\_368
CL\_INS\_204
CL\_INS\_204
CL\_INS\_204
CL\_INS\_204
CL\_INS\_204
CL\_INS\_204
CL\_INS\_204
CL\_INS\_204
CL\_INS\_204
CL\_INS\_368
CL\_INS\_368
CL\_INS\_141
CL\_INS\_141
CL\_INS\_115
CL\_INS\_141
CL\_INS\_204
CL\_INS\_204
CL\_INS\_204
CL\_INS\_204
CL\_INS\_141
CL\_INS\_141
CL\_INS\_141
CL\_INS\_115
CL\_INS\_141
CL\_INS\_141
CL\_INS\_141
CL\_INS\_141
CL\_INS\_141
CL\_INS\_141
CL\_INS\_141
CL\_INS\_141
CL\_INS\_141
CL\_INS\_141
CL\_INS\_141
CL\_INS\_141
CL\_INS\_141
CL\_INS\_141
CL\_INS\_141
CL\_INS\_141
CL\_INS\_141
CL\_INS\_141
CL\_INS\_141
CL\_INS\_141
CL\_INS\_141
CL\_INS\_141
CL\_INS\_141
CL\_INS\_141
CL\_INS\_141
CL\_INS\_141
CL\_INS\_115
CL\_INS\_115
CL\_INS\_141
CL\_INS\_141
CL\_INS\_141
CL\_INS\_141
CL\_INS\_365
CL\_INS\_368
CL\_INS\_365
CL\_INS\_365
CL\_INS\_365
CL\_INS\_141
CL\_INS\_141
CL\_INS\_365
CL\_INS\_365
CL\_INS\_365
CL\_INS\_365
CL\_INS\_365
CL\_INS\_365
CL\_INS\_365
CL\_INS\_365
CL\_INS\_365
CL\_INS\_365
CL\_INS\_365
CL\_INS\_365
CL\_INS\_365
CL\_INS\_365
CL\_INS\_365
CL\_INS\_368
CL\_INS\_368
CL\_INS\_368
CL\_INS\_365
CL\_INS\_365
CL\_INS\_365
CL\_INS\_365
CL\_INS\_368
CL\_INS\_368
CL\_INS\_365
CL\_INS\_365
CL\_INS\_365
CL\_INS\_368
CL\_INS\_141
CL\_INS\_141
CL\_INS\_141
CL\_INS\_141
CL\_INS\_141
CL\_INS\_141
CL\_INS\_141
CL\_INS\_141
CL\_INS\_141
CL\_INS\_141
CL\_INS\_141
CL\_INS\_141
CL\_INS\_141
CL\_INS\_141
CL\_INS\_368
CL\_INS\_141
CL\_INS\_141
CL\_INS\_141
CL\_INS\_132
CL\_INS\_141
CL\_INS\_141
CL\_INS\_204
CL\_INS\_204
CL\_INS\_204
CL\_INS\_141
CL\_INS\_204
CL\_INS\_204
CL\_INS\_204
CL\_INS\_204
CL\_INS\_204
CL\_INS\_204
CL\_INS\_204
CL\_INS\_204
CL\_INS\_204
CL\_INS\_204
CL\_INS\_204
CL\_INS\_204
CL\_INS\_204
CL\_INS\_204
CL\_INS\_204
CL\_INS\_204
CL\_INS\_204
CL\_INS\_204
CL\_INS\_204
CL\_INS\_204
CL\_INS\_368
CL\_INS\_368
CL\_INS\_141
CL\_INS\_141
CL\_INS\_141
CL\_INS\_141
CL\_INS\_141
CL\_INS\_141
CL\_INS\_141
CL\_INS\_141
CL\_INS\_325
CL\_INS\_325
CL\_INS\_325
CL\_INS\_325
CL\_INS\_325
CL\_INS\_325
CL\_INS\_368
CL\_INS\_141
CL\_INS\_141
CL\_INS\_141
CL\_INS\_368
CL\_INS\_325
CL\_INS\_141
CL\_INS\_141
CL\_INS\_141
CL\_INS\_368
CL\_INS\_141
CL\_INS\_141
CL\_INS\_141
CL\_INS\_141
CL\_INS\_368
CL\_INS\_368
CL\_INS\_368
CL\_INS\_368
CL\_INS\_325
CL\_INS\_141
CL\_INS\_141
CL\_INS\_141
CL\_INS\_141
CL\_INS\_141
CL\_INS\_366
CL\_INS\_366
CL\_INS\_366
CL\_INS\_366
CL\_INS\_141
CL\_INS\_325
CL\_INS\_141
CL\_INS\_141
CL\_INS\_141
CL\_INS\_325
CL\_INS\_141
CL\_INS\_141
CL\_INS\_141
CL\_INS\_141
CL\_INS\_115
CL\_INS\_115
CL\_INS\_141
CL\_INS\_325
CL\_INS\_141
CL\_INS\_141
CL\_INS\_325
CL\_INS\_141
CL\_INS\_141
CL\_INS\_115
CL\_INS\_325
CL\_INS\_368
CL\_INS\_141
CL\_INS\_366
CL\_INS\_368
CL\_INS\_368
CL\_INS\_368
CL\_INS\_270
CL\_INS\_368
CL\_INS\_141
CL\_INS\_325
CL\_INS\_141
CL\_INS\_141
CL\_INS\_141
CL\_INS\_325
CL\_INS\_141
CL\_INS\_141
CL\_INS\_141
CL\_INS\_141
CL\_INS\_141
CL\_INS\_141
CL\_INS\_141
CL\_INS\_141
CL\_INS\_141
CL\_INS\_141
CL\_INS\_141
CL\_INS\_141
CL\_INS\_141
CL\_INS\_325
CL\_INS\_366
CL\_INS\_366
CL\_INS\_366
CL\_INS\_141
CL\_INS\_141
CL\_INS\_141
CL\_INS\_141
CL\_INS\_141
CL\_INS\_325
CL\_INS\_141
CL\_INS\_141
CL\_INS\_325
CL\_INS\_325
CL\_INS\_325
CL\_INS\_325
CL\_INS\_366
CL\_INS\_368
CL\_INS\_366
CL\_INS\_366
CL\_INS\_366
CL\_INS\_366
CL\_INS\_366
CL\_INS\_366
CL\_INS\_366
CL\_INS\_366
CL\_INS\_366
CL\_INS\_368
CL\_INS\_366
CL\_INS\_366
CL\_INS\_366
CL\_INS\_366
CL\_INS\_366
CL\_INS\_366
CL\_INS\_368
CL\_INS\_368
CL\_INS\_368
CL\_INS\_368
CL\_INS\_368
CL\_INS\_368
CL\_INS\_368
CL\_INS\_368
CL\_INS\_368
CL\_INS\_270
CL\_INS\_325
CL\_INS\_325
CL\_INS\_141
CL\_INS\_141
CL\_INS\_325
CL\_INS\_141
CL\_INS\_141
CL\_INS\_141
CL\_INS\_141
CL\_INS\_325
CL\_INS\_204
CL\_INS\_204
CL\_INS\_204
CL\_INS\_204
CL\_INS\_368
CL\_INS\_368
CL\_INS\_204
CL\_INS\_204
CL\_INS\_204
CL\_INS\_204
CL\_INS\_204
CL\_INS\_141
CL\_INS\_204
CL\_INS\_204
CL\_INS\_204
CL\_INS\_204
CL\_INS\_204
CL\_INS\_204
CL\_INS\_204
CL\_INS\_204
CL\_INS\_204
CL\_INS\_204
CL\_INS\_368
CL\_INS\_368
CL\_INS\_368
CL\_INS\_368
CL\_INS\_368
CL\_INS\_368
CL\_INS\_368
CL\_INS\_368
CL\_INS\_368
CL\_INS\_368
CL\_INS\_368
CL\_INS\_366
CL\_INS\_366
CL\_INS\_366
CL\_INS\_368
CL\_INS\_368
CL\_INS\_368
CL\_INS\_368
CL\_INS\_366
CL\_INS\_366
CL\_INS\_368
CL\_INS\_366
CL\_INS\_368
CL\_INS\_368
CL\_INS\_368
CL\_INS\_178
CL\_INS\_270
CL\_INS\_368
CL\_INS\_368
CL\_INS\_366
CL\_INS\_366
CL\_INS\_366
CL\_INS\_366
CL\_INS\_366
CL\_INS\_366
CL\_INS\_178
CL\_INS\_368
CL\_INS\_368
CL\_INS\_368
CL\_INS\_368
CL\_INS\_368
CL\_INS\_368
CL\_INS\_368
CL\_INS\_368
CL\_INS\_368
CL\_INS\_368
CL\_INS\_368
CL\_INS\_368
CL\_INS\_368
CL\_INS\_368
CL\_INS\_368
CL\_INS\_368
CL\_INS\_368
CL\_INS\_368
CL\_INS\_368
CL\_INS\_368
CL\_INS\_368
CL\_INS\_368
CL\_INS\_178
CL\_INS\_368
CL\_INS\_368
CL\_INS\_368
CL\_INS\_368
CL\_INS\_178
CL\_INS\_368
CL\_INS\_368
CL\_INS\_368
CL\_INS\_368
CL\_INS\_368
CL\_INS\_368
CL\_INS\_368
CL\_INS\_368
CL\_INS\_368
CL\_INS\_368
CL\_INS\_368
CL\_INS\_368
CL\_INS\_368
CL\_INS\_368
CL\_INS\_368
CL\_INS\_368
CL\_INS\_368
CL\_INS\_368
CL\_INS\_368
CL\_INS\_368
CL\_INS\_368
CL\_INS\_368
CL\_INS\_368
CL\_INS\_368
CL\_INS\_368
CL\_INS\_368
CL\_INS\_368
CL\_INS\_368
CL\_INS\_368
CL\_INS\_368
CL\_INS\_368
CL\_INS\_368
CL\_INS\_368
CL\_INS\_368
CL\_INS\_368
CL\_INS\_368
CL\_INS\_368
CL\_INS\_368
CL\_INS\_368
CL\_INS\_368
CL\_INS\_368
CL\_INS\_368
CL\_INS\_368
CL\_INS\_368
CL\_INS\_368
CL\_INS\_368
CL\_INS\_368
CL\_INS\_368
CL\_INS\_368
CL\_INS\_368
CL\_INS\_368
CL\_INS\_368
CL\_INS\_368
CL\_INS\_368
CL\_INS\_368
CL\_INS\_368
CL\_INS\_368
CL\_INS\_368
CL\_INS\_368
CL\_INS\_368
CL\_INS\_368
CL\_INS\_114
CL\_INS\_114
CL\_INS\_114
CL\_INS\_114
CL\_INS\_114
CL\_INS\_114
CL\_INS\_114
CL\_INS\_114
CL\_INS\_114
CL\_INS\_114
CL\_INS\_114
CL\_INS\_114
CL\_INS\_114
CL\_INS\_114
CL\_INS\_114
CL\_INS\_114
CL\_INS\_114
CL\_INS\_114
CL\_INS\_114
CL\_INS\_114
CL\_INS\_114
CL\_INS\_114
CL\_INS\_114
CL\_INS\_114
CL\_INS\_114
CL\_INS\_114
CL\_INS\_114
CL\_INS\_114
CL\_INS\_114
CL\_INS\_114
CL\_INS\_114
CL\_INS\_114
CL\_INS\_114
CL\_INS\_114
CL\_INS\_114
CL\_INS\_114
CL\_INS\_114
CL\_INS\_114
CL\_INS\_114
CL\_INS\_114
CL\_INS\_114
CL\_INS\_114
CL\_INS\_114
CL\_INS\_114
CL\_INS\_114
CL\_INS\_114
CL\_INS\_114
CL\_INS\_114
CL\_INS\_114
CL\_INS\_114
CL\_INS\_114
CL\_INS\_368
CL\_INS\_325
CL\_INS\_325
CL\_INS\_141
CL\_INS\_368
CL\_INS\_325
CL\_INS\_368
CL\_INS\_368
CL\_INS\_325
CL\_INS\_325
CL\_INS\_368
CL\_INS\_368
CL\_INS\_368
CL\_INS\_368
CL\_INS\_368
CL\_INS\_368
CL\_INS\_368
CL\_INS\_368
CL\_INS\_368
CL\_INS\_368
CL\_INS\_368
CL\_INS\_368
CL\_INS\_368
CL\_INS\_368
CL\_INS\_368
CL\_INS\_368
Cluster ID


CL\_10545
CL\_10546
CL\_10547
CL\_10548
CL\_10549
CL\_10550
CL\_10551
CL\_10552
CL\_12671
CL\_12672
CL\_11970
CL\_12673
CL\_12674
CL\_11972
CL\_10089
CL\_9398
CL\_9399
CL\_12487
CL\_12486
CL\_12485
CL\_12484
CL\_12483
CL\_1280
CL\_1281
CL\_9266
CL\_11973
CL\_1285
CL\_7221
CL\_7220
CL\_1287
CL\_1288
CL\_1289
CL\_5270
CL\_1292
CL\_2023
CL\_2024
CL\_2025
CL\_12482
CL\_1297
CL\_12481
CL\_2029
CL\_11859
CL\_11860
CL\_2030
CL\_7960
CL\_12480
CL\_12479
CL\_12478
CL\_10451
CL\_12477
CL\_12476
CL\_9410
CL\_9411
CL\_9412
CL\_9413
CL\_9414
CL\_5282
CL\_5283
CL\_5284
CL\_5285
CL\_5286
CL\_5287
CL\_5288
CL\_5289
CL\_11861
CL\_5290
CL\_11862
CL\_5292
CL\_5293
CL\_5294
CL\_5295
CL\_5296
CL\_5297
CL\_5299
CL\_11863
CL\_5687
CL\_5688
CL\_5689
CL\_10088
CL\_10087
CL\_10086
CL\_11856
CL\_11857
CL\_5690
CL\_5691
CL\_2007
CL\_5692
CL\_5693
CL\_12035
CL\_2865
CL\_12230
CL\_12229
CL\_12228
CL\_7020
CL\_7021
CL\_5275
CL\_5276
CL\_5277
CL\_5278
CL\_5279
CL\_5280
CL\_7022
CL\_9461
CL\_9462
CL\_2864
CL\_2863
CL\_12782
CL\_10820
CL\_2862
CL\_6124
CL\_6123
CL\_2861
CL\_10821
CL\_10822
CL\_10823
CL\_12783
CL\_10824
CL\_10825
CL\_10826
CL\_10827
CL\_10828
CL\_10829
CL\_10830
CL\_10831
CL\_10832
CL\_10833
CL\_10834
CL\_10835
CL\_10836
CL\_10837
CL\_10838
CL\_10839
CL\_11723
CL\_10840
CL\_10841
CL\_10842
CL\_10843
CL\_10844
CL\_10845
CL\_11722
CL\_10846
CL\_10847
CL\_12784
CL\_12785
CL\_2062
CL\_11721
CL\_10848
CL\_2860
CL\_6122
CL\_12649
CL\_8748
CL\_8784
CL\_6121
CL\_2859
CL\_2858
CL\_2857
CL\_2856
CL\_2855
CL\_6120
CL\_2854
CL\_2853
CL\_2852
CL\_2851
CL\_2850
CL\_2849
CL\_2848
CL\_6119
CL\_2847
CL\_2846
CL\_2845
CL\_13730
CL\_13731
CL\_13732
CL\_2844
CL\_2843
CL\_2842
CL\_6118
CL\_6117
CL\_6116
CL\_8788
CL\_6060
CL\_8787
CL\_12648
CL\_2841
CL\_2840
CL\_2839
CL\_2838
CL\_2837
CL\_2836
CL\_2835
CL\_2834
CL\_2833
CL\_2832
CL\_2831
CL\_2830
CL\_2829
CL\_2828
CL\_2827
CL\_2826
CL\_2825
CL\_2824
CL\_2823
CL\_7941
CL\_7942
CL\_7205
CL\_7204
CL\_9273
CL\_7203
CL\_7202
CL\_7201
CL\_7200
CL\_9274
CL\_9275
CL\_9276
CL\_9277
CL\_9278
CL\_9279
CL\_9280
CL\_9281
CL\_9282
CL\_9283
CL\_9284
CL\_9285
CL\_9286
CL\_11977
CL\_12405
CL\_12406
CL\_12407
CL\_12680
CL\_12681
CL\_7199
CL\_7198
CL\_7197
CL\_7196
CL\_1314
CL\_1315
CL\_8151
CL\_1316
CL\_8164
CL\_8163
CL\_8162
CL\_8161
CL\_8160
CL\_8159
CL\_8158
CL\_1317
CL\_1318
CL\_1319
CL\_12227
CL\_10080
CL\_7943
CL\_9010
CL\_9009
CL\_9008
CL\_7944
CL\_7945
CL\_3143
CL\_3142
CL\_12226
CL\_12225
CL\_12224
CL\_12223
CL\_5727
CL\_2059
CL\_2060
CL\_6140
CL\_2061
CL\_1326
CL\_6137
CL\_7147
CL\_9449
CL\_5694
CL\_5695
CL\_5696
CL\_5697
CL\_7223
CL\_7222
CL\_8185
CL\_5267
CL\_5268
CL\_5269
CL\_5698
CL\_5699
CL\_5700
CL\_5701
CL\_8184
CL\_5702
CL\_5703
CL\_8183
CL\_2022
CL\_7219
CL\_7218
CL\_8182
CL\_12675
CL\_7211
CL\_7210
CL\_12676
CL\_12677
CL\_12678
CL\_5402
CL\_12679
CL\_7209
CL\_8181
CL\_7208
CL\_7207
CL\_8180
CL\_8179
CL\_8178
CL\_8177
CL\_8176
CL\_8175
CL\_8174
CL\_8173
CL\_8172
CL\_8171
CL\_8170
CL\_8169
CL\_8168
CL\_8167
CL\_8166
CL\_8165
CL\_5704
CL\_5705
CL\_5706
CL\_5707
CL\_7217
CL\_10085
CL\_10084
CL\_5708
CL\_11858
CL\_9440
CL\_10083
CL\_5273
CL\_10082
CL\_10081
CL\_5709
CL\_7146
CL\_13058
CL\_9047
CL\_9450
CL\_7144
CL\_9043
CL\_9451
CL\_9040
CL\_9452
CL\_10490
CL\_10491
CL\_10492
CL\_10493
CL\_9038
CL\_9037
CL\_7136
CL\_9036
CL\_9035
CL\_10494
CL\_10495
CL\_10496
CL\_10497
CL\_10498
CL\_10499
CL\_10500
CL\_10501
CL\_10502
CL\_5710
CL\_5711
CL\_5712
CL\_3162
CL\_3161
CL\_5713
CL\_3159
CL\_3158
CL\_3157
CL\_3156
CL\_5714
CL\_5715
CL\_5716
CL\_3154
CL\_5717
CL\_12475
CL\_12474
CL\_5718
CL\_5719
CL\_5720
CL\_7023
CL\_7024
CL\_12045
CL\_5721
CL\_7025
CL\_7026
CL\_7027
CL\_7028
CL\_7029
CL\_7030
CL\_7031
CL\_7032
CL\_7033
CL\_12473
CL\_12472
CL\_12471
CL\_12470
CL\_10503
CL\_10504
CL\_10505
CL\_10506
CL\_10507
CL\_10508
CL\_10509
CL\_9046
CL\_9045
CL\_9044
CL\_13057
CL\_13056
CL\_13055
CL\_13054
CL\_10536
CL\_10537
CL\_12034
CL\_10538
CL\_12033
CL\_12032
CL\_12031
CL\_10882
CL\_10881
CL\_12030
CL\_13053
CL\_9455
CL\_9456
CL\_7130
CL\_7129
CL\_7128
CL\_7127
CL\_10878
CL\_12029
CL\_10479
CL\_9032
CL\_9031
CL\_9030
CL\_9029
CL\_9028
CL\_10481
CL\_12028
CL\_12027
CL\_13052
CL\_13051
CL\_13050
CL\_13049
CL\_13048
CL\_13047
CL\_13046
CL\_13045
CL\_10589
CL\_10588
CL\_10587
CL\_5397
CL\_7115
CL\_10586
CL\_10585
CL\_10584
CL\_10583
CL\_5390
CL\_7108
CL\_13717
CL\_13718
CL\_13719
CL\_13720
CL\_13721
CL\_13722
CL\_13723
CL\_13724
CL\_13725
CL\_13726
CL\_13727
CL\_13728
CL\_13729
CL\_10510
CL\_9027
CL\_9026
CL\_9463
CL\_9025
CL\_9024
CL\_10511
CL\_10512
CL\_10513
CL\_10514
CL\_10515
CL\_10516
CL\_9023
CL\_13360
CL\_9022
CL\_9021
CL\_9464
CL\_9020
CL\_9019
CL\_9018
CL\_10553
CL\_10554
CL\_9017
CL\_9016
CL\_10517
CL\_10518
CL\_10519
CL\_10520
CL\_10521
CL\_9467
CL\_9468
CL\_9469
CL\_9470
CL\_10522
CL\_9015
CL\_9014
CL\_9465
CL\_9013
CL\_10555
CL\_10556
CL\_10557
CL\_10558
CL\_9012
CL\_9466
CL\_9011
CL\_12026
CL\_12025
CL\_9334
CL\_13044
CL\_13043
CL\_9338
CL\_9339
CL\_9340
CL\_9341
CL\_9342
CL\_9343
CL\_9344
CL\_9346
CL\_9347
CL\_9348
CL\_9349
CL\_9350
CL\_9351
CL\_9352
CL\_9353
CL\_9354
CL\_9355
CL\_13042
CL\_9357
CL\_9358
CL\_9359
CL\_9360
CL\_10569
CL\_9362
CL\_9363
CL\_9364
CL\_9365
CL\_9366
CL\_9367
CL\_9368
CL\_9369
CL\_9370
CL\_9371
CL\_9372
CL\_9373
CL\_9374
CL\_13041
CL\_9378
CL\_13040
CL\_9379
CL\_10574
CL\_9380
CL\_9381
CL\_9382
CL\_9383
CL\_9384
CL\_9385
CL\_13039
CL\_13038
CL\_5722
CL\_5723
CL\_3145
CL\_10523
CL\_5724
CL\_10559
CL\_9007
CL\_5725
CL\_5726
CL\_9471
CL\_10560
CL\_10561
CL\_10562
CL\_10563
CL\_10564
CL\_10565
CL\_10566
CL\_9006
CL\_12024
CL\_9472
CL\_9473
CL\_10524
CL\_10525
CL\_13037
CL\_5728
