## Supplementary material for "A novel method for integrating genomic and Tn-Seq data to identify common *in vivo* fitness mechanisms across multiple bacterial species": S4 Dataset: CL_INS_369.html

Legend

 Mobile +extrachromosomalelementfunctions
 Hypothetical
 DNA Metabolism
 AntibioticResistance
 Other
 All VFDB Genes
 Transport +binding proteins

FULL


WINDOWSVGPNG

Trim RowsRemove SingletonsSave Fasta

CL\_5729


CL\_5729


CL\_5729


CL\_5729


CL\_5729


CL\_5729


CL\_5729


CL\_5729


CL\_5729


CL\_5729


CL\_5729

HighlightSelectShow Genomes


6

CL\_2095


4

CL\_4532


2

CL\_2095


1

CL\_2067


1

CL\_2246


1

CL\_1955


1

CL\_2095


1

CL\_2095


1

CL\_6159


1

CL\_2095


1

CL\_2814

fGI ID


CL\_INS\_115
CL\_INS\_369
CL\_INS\_369
CL\_INS\_369
CL\_INS\_369
CL\_INS\_369
CL\_INS\_369
CL\_INS\_369
CL\_INS\_369
CL\_INS\_141
CL\_INS\_141
CL\_INS\_369
CL\_INS\_369
CL\_INS\_369
CL\_INS\_132
CL\_INS\_132
CL\_INS\_132
CL\_INS\_132
CL\_INS\_132
CL\_INS\_132
CL\_INS\_132
CL\_INS\_132
CL\_INS\_132
CL\_INS\_132
CL\_INS\_132
CL\_INS\_132
Cluster ID


CL\_1637
CL\_12587
CL\_6115
CL\_6114
CL\_6113
CL\_6112
CL\_6111
CL\_12682
CL\_5730
CL\_9005
CL\_2064
CL\_9474
CL\_9475
CL\_9476
CL\_10788
CL\_10789
CL\_10790
CL\_2823
CL\_10791
CL\_10792
CL\_10793
CL\_10794
CL\_10795
CL\_10796
CL\_10797
CL\_10798
