## Supplementary material for "A novel method for integrating genomic and Tn-Seq data to identify common *in vivo* fitness mechanisms across multiple bacterial species": S4 Dataset: CL_INS_370.html

Legend

 Mobile +extrachromosomalelementfunctions
 Regulatoryfunctions
 Hypothetical
 DNA Metabolism
 Cell Envelope
 Proteinsynthesis/fate
 Other
 All VFDB Genes
 Transport +binding proteins

FULL


WINDOWSVGPNG

Trim RowsRemove SingletonsSave Fasta

CL\_5661


CL\_5661


CL\_5661


CL\_5661


CL\_5661


CL\_5661


CL\_5661


CL\_5661


CL\_5661


CL\_5661


CL\_5661


CL\_5661


CL\_5661


CL\_5661


CL\_5661


CL\_5661


CL\_5661


CL\_5661


CL\_5661


CL\_5661

HighlightSelectShow Genomes


3

CL\_665


3

CL\_3871


2

CL\_255


1

CL\_912


1

CL\_256


1

CL\_771


1

CL\_3969


1

CL\_256


1

CL\_74


1

CL\_665


1

CL\_1214


1

CL\_4315


1

CL\_4315


1

CL\_912


1

CL\_4455


1

CL\_3969


1

CL\_665


1

CL\_4315


1

CL\_4456


1

CL\_912

fGI ID


CL\_INS\_49
CL\_INS\_49
CL\_INS\_49
CL\_INS\_365
CL\_INS\_6
CL\_INS\_6
CL\_INS\_6
CL\_INS\_6
CL\_INS\_6
CL\_INS\_6
CL\_INS\_6
CL\_INS\_6
CL\_INS\_297
CL\_INS\_297
CL\_INS\_297
CL\_INS\_297
CL\_INS\_297
CL\_INS\_297
CL\_INS\_297
CL\_INS\_297
CL\_INS\_297
CL\_INS\_297
CL\_INS\_365
CL\_INS\_365
CL\_INS\_49
CL\_INS\_365
CL\_INS\_365
CL\_INS\_365
CL\_INS\_365
CL\_INS\_365
CL\_INS\_365
CL\_INS\_365
CL\_INS\_365
CL\_INS\_365
CL\_INS\_365
CL\_INS\_365
CL\_INS\_365
CL\_INS\_365
CL\_INS\_365
CL\_INS\_365
CL\_INS\_365
CL\_INS\_365
CL\_INS\_365
CL\_INS\_365
CL\_INS\_365
CL\_INS\_365
CL\_INS\_365
CL\_INS\_365
CL\_INS\_365
CL\_INS\_49
CL\_INS\_49
CL\_INS\_370
CL\_INS\_297
CL\_INS\_297
CL\_INS\_49
CL\_INS\_49
CL\_INS\_49
CL\_INS\_49
CL\_INS\_49
CL\_INS\_49
CL\_INS\_49
CL\_INS\_49
CL\_INS\_49
CL\_INS\_49
CL\_INS\_49
CL\_INS\_49
CL\_INS\_49
CL\_INS\_49
CL\_INS\_49
CL\_INS\_49
CL\_INS\_49
CL\_INS\_49
CL\_INS\_49
CL\_INS\_49
CL\_INS\_49
CL\_INS\_49
CL\_INS\_49
CL\_INS\_365
CL\_INS\_365
CL\_INS\_365
CL\_INS\_365
CL\_INS\_365
CL\_INS\_365
CL\_INS\_365
CL\_INS\_312
CL\_INS\_365
CL\_INS\_365
CL\_INS\_365
CL\_INS\_312
CL\_INS\_370
CL\_INS\_175
CL\_INS\_175
CL\_INS\_175
CL\_INS\_175
CL\_INS\_175
CL\_INS\_175
CL\_INS\_175
CL\_INS\_365
CL\_INS\_365
CL\_INS\_365
CL\_INS\_365
CL\_INS\_365
CL\_INS\_365
CL\_INS\_365
CL\_INS\_365
CL\_INS\_365
CL\_INS\_365
CL\_INS\_365
CL\_INS\_365
CL\_INS\_365
CL\_INS\_365
CL\_INS\_365
CL\_INS\_365
CL\_INS\_365
CL\_INS\_365
CL\_INS\_312
CL\_INS\_270
CL\_INS\_270
CL\_INS\_270
CL\_INS\_270
CL\_INS\_270
CL\_INS\_270
CL\_INS\_270
CL\_INS\_270
CL\_INS\_270
CL\_INS\_270
CL\_INS\_270
CL\_INS\_270
CL\_INS\_270
CL\_INS\_370
CL\_INS\_365
CL\_INS\_270
CL\_INS\_370
CL\_INS\_365
CL\_INS\_270
CL\_INS\_270
CL\_INS\_270
CL\_INS\_270
CL\_INS\_270
CL\_INS\_270
CL\_INS\_270
CL\_INS\_270
CL\_INS\_370
CL\_INS\_370
CL\_INS\_370
CL\_INS\_370
CL\_INS\_370
CL\_INS\_370
CL\_INS\_370
CL\_INS\_370
CL\_INS\_370
CL\_INS\_370
CL\_INS\_370
CL\_INS\_370
CL\_INS\_370
CL\_INS\_370
CL\_INS\_370
CL\_INS\_370
CL\_INS\_370
CL\_INS\_370
CL\_INS\_370
CL\_INS\_370
CL\_INS\_270
CL\_INS\_270
CL\_INS\_270
CL\_INS\_270
CL\_INS\_270
CL\_INS\_270
CL\_INS\_270
CL\_INS\_270
CL\_INS\_270
CL\_INS\_270
CL\_INS\_270
CL\_INS\_270
CL\_INS\_270
CL\_INS\_270
CL\_INS\_270
CL\_INS\_270
CL\_INS\_270
CL\_INS\_270
CL\_INS\_270
CL\_INS\_270
CL\_INS\_270
CL\_INS\_270
CL\_INS\_270
CL\_INS\_270
CL\_INS\_270
CL\_INS\_270
CL\_INS\_270
CL\_INS\_270
CL\_INS\_270
CL\_INS\_270
CL\_INS\_270
CL\_INS\_270
CL\_INS\_270
CL\_INS\_270
CL\_INS\_270
CL\_INS\_270
CL\_INS\_270
CL\_INS\_270
CL\_INS\_270
CL\_INS\_270
CL\_INS\_270
CL\_INS\_270
CL\_INS\_49
CL\_INS\_49
CL\_INS\_49
CL\_INS\_49
CL\_INS\_370
CL\_INS\_270
CL\_INS\_270
CL\_INS\_270
CL\_INS\_270
CL\_INS\_270
CL\_INS\_270
CL\_INS\_270
CL\_INS\_270
CL\_INS\_270
CL\_INS\_270
CL\_INS\_270
CL\_INS\_270
CL\_INS\_365
CL\_INS\_270
CL\_INS\_270
CL\_INS\_270
CL\_INS\_270
CL\_INS\_297
CL\_INS\_297
CL\_INS\_297
CL\_INS\_270
CL\_INS\_270
CL\_INS\_270
CL\_INS\_270
CL\_INS\_270
CL\_INS\_6
CL\_INS\_6
CL\_INS\_297
CL\_INS\_297
CL\_INS\_297
CL\_INS\_297
CL\_INS\_297
CL\_INS\_297
CL\_INS\_6
CL\_INS\_6
CL\_INS\_6
CL\_INS\_6
CL\_INS\_6
CL\_INS\_6
CL\_INS\_49
CL\_INS\_49
CL\_INS\_49
CL\_INS\_49
CL\_INS\_6
CL\_INS\_6
CL\_INS\_6
CL\_INS\_6
CL\_INS\_6
CL\_INS\_6
CL\_INS\_6
CL\_INS\_6
CL\_INS\_6
CL\_INS\_6
CL\_INS\_370
CL\_INS\_370
CL\_INS\_270
CL\_INS\_370
CL\_INS\_270
CL\_INS\_270
CL\_INS\_270
CL\_INS\_270
CL\_INS\_270
CL\_INS\_270
CL\_INS\_270
CL\_INS\_270
CL\_INS\_270
CL\_INS\_270
CL\_INS\_270
CL\_INS\_270
CL\_INS\_270
CL\_INS\_270
CL\_INS\_270
CL\_INS\_270
CL\_INS\_270
CL\_INS\_270
CL\_INS\_270
CL\_INS\_270
CL\_INS\_270
CL\_INS\_270
CL\_INS\_270
CL\_INS\_270
CL\_INS\_270
CL\_INS\_270
CL\_INS\_270
CL\_INS\_270
CL\_INS\_270
CL\_INS\_270
CL\_INS\_270
CL\_INS\_270
CL\_INS\_270
CL\_INS\_270
CL\_INS\_270
CL\_INS\_270
CL\_INS\_270
CL\_INS\_370
CL\_INS\_370
CL\_INS\_370
CL\_INS\_270
CL\_INS\_270
CL\_INS\_370
CL\_INS\_9
CL\_INS\_370
CL\_INS\_370
CL\_INS\_370
CL\_INS\_370
CL\_INS\_370
CL\_INS\_370
CL\_INS\_370
CL\_INS\_270
CL\_INS\_270
CL\_INS\_270
CL\_INS\_270
CL\_INS\_370
CL\_INS\_9
CL\_INS\_9
CL\_INS\_270
CL\_INS\_270
CL\_INS\_270
CL\_INS\_270
CL\_INS\_370
CL\_INS\_370
CL\_INS\_312
CL\_INS\_312
CL\_INS\_312
CL\_INS\_312
CL\_INS\_297
CL\_INS\_297
CL\_INS\_297
CL\_INS\_312
CL\_INS\_312
CL\_INS\_312
CL\_INS\_312
CL\_INS\_312
CL\_INS\_312
CL\_INS\_312
CL\_INS\_312
CL\_INS\_312
CL\_INS\_312
CL\_INS\_312
CL\_INS\_312
CL\_INS\_312
CL\_INS\_312
CL\_INS\_312
CL\_INS\_312
CL\_INS\_312
CL\_INS\_312
CL\_INS\_312
CL\_INS\_312
CL\_INS\_312
CL\_INS\_312
CL\_INS\_312
CL\_INS\_312
CL\_INS\_312
CL\_INS\_312
CL\_INS\_312
CL\_INS\_312
CL\_INS\_312
CL\_INS\_312
CL\_INS\_312
CL\_INS\_312
CL\_INS\_312
CL\_INS\_312
CL\_INS\_312
CL\_INS\_312
CL\_INS\_312
CL\_INS\_312
CL\_INS\_312
CL\_INS\_312
CL\_INS\_312
CL\_INS\_312
CL\_INS\_312
CL\_INS\_312
CL\_INS\_312
CL\_INS\_312
CL\_INS\_312
CL\_INS\_312
CL\_INS\_312
CL\_INS\_312
CL\_INS\_312
CL\_INS\_312
CL\_INS\_312
CL\_INS\_312
CL\_INS\_312
CL\_INS\_312
CL\_INS\_312
CL\_INS\_312
CL\_INS\_312
CL\_INS\_312
CL\_INS\_365
Cluster ID


CL\_5660
CL\_12121
CL\_12122
CL\_10146
CL\_5659
CL\_5658
CL\_5657
CL\_5656
CL\_5655
CL\_5654
CL\_5653
CL\_5652
CL\_12375
CL\_12374
CL\_12373
CL\_12372
CL\_12371
CL\_12370
CL\_12369
CL\_12368
CL\_12367
CL\_12366
CL\_9131
CL\_10145
CL\_9130
CL\_9129
CL\_9128
CL\_10144
CL\_10143
CL\_10142
CL\_10141
CL\_10140
CL\_10139
CL\_10138
CL\_10401
CL\_10137
CL\_259
CL\_258
CL\_10136
CL\_10135
CL\_10134
CL\_10133
CL\_10132
CL\_10131
CL\_7272
CL\_7273
CL\_7274
CL\_6654
CL\_6655
CL\_9127
CL\_9126
CL\_9125
CL\_12377
CL\_12376
CL\_11838
CL\_11839
CL\_11840
CL\_11841
CL\_11842
CL\_11843
CL\_11844
CL\_11845
CL\_11846
CL\_11847
CL\_11848
CL\_11849
CL\_11850
CL\_11851
CL\_11852
CL\_5213
CL\_5212
CL\_5211
CL\_7981
CL\_12127
CL\_11853
CL\_5208
CL\_5207
CL\_6326
CL\_6455
CL\_6454
CL\_6453
CL\_8832
CL\_8833
CL\_8834
CL\_8835
CL\_11835
CL\_8836
CL\_8837
CL\_8838
CL\_11011
CL\_8840
CL\_8841
CL\_8842
CL\_8843
CL\_8844
CL\_8845
CL\_8846
CL\_13775
CL\_13774
CL\_238
CL\_239
CL\_240
CL\_241
CL\_242
CL\_243
CL\_244
CL\_245
CL\_246
CL\_247
CL\_248
CL\_249
CL\_250
CL\_251
CL\_252
CL\_253
CL\_9787
CL\_6327
CL\_3896
CL\_3895
CL\_3894
CL\_3892
CL\_12853
CL\_12854
CL\_12855
CL\_12856
CL\_12857
CL\_12645
CL\_5751
CL\_5752
CL\_6328
CL\_5753
CL\_5754
CL\_6329
CL\_5755
CL\_12644
CL\_12643
CL\_12642
CL\_12641
CL\_5756
CL\_3891
CL\_5757
CL\_7320
CL\_7321
CL\_7322
CL\_7323
CL\_7324
CL\_7325
CL\_7326
CL\_7327
CL\_7328
CL\_7329
CL\_7330
CL\_7331
CL\_7332
CL\_7333
CL\_7334
CL\_7335
CL\_7336
CL\_7337
CL\_7338
CL\_7339
CL\_7340
CL\_5758
CL\_5759
CL\_5760
CL\_13170
CL\_13169
CL\_13168
CL\_13167
CL\_13166
CL\_13165
CL\_13164
CL\_13163
CL\_13162
CL\_13161
CL\_13160
CL\_13159
CL\_9151
CL\_10694
CL\_3886
CL\_3885
CL\_3884
CL\_3883
CL\_3882
CL\_5998
CL\_3878
CL\_3877
CL\_3876
CL\_3875
CL\_3874
CL\_6000
CL\_3872
CL\_12157
CL\_13158
CL\_7191
CL\_13157
CL\_13156
CL\_7187
CL\_7186
CL\_7185
CL\_7184
CL\_12160
CL\_7183
CL\_12161
CL\_6330
CL\_7978
CL\_7979
CL\_7980
CL\_6331
CL\_5651
CL\_12638
CL\_12637
CL\_12636
CL\_12635
CL\_12634
CL\_12633
CL\_12632
CL\_12631
CL\_12630
CL\_12629
CL\_12628
CL\_7680
CL\_12627
CL\_7678
CL\_10687
CL\_10688
CL\_10689
CL\_10690
CL\_10691
CL\_12626
CL\_12625
CL\_12624
CL\_12623
CL\_12622
CL\_5650
CL\_5649
CL\_13589
CL\_13588
CL\_13587
CL\_13586
CL\_13210
CL\_13209
CL\_5648
CL\_5647
CL\_5646
CL\_5645
CL\_5644
CL\_5643
CL\_12123
CL\_12124
CL\_12125
CL\_12126
CL\_5642
CL\_5641
CL\_5640
CL\_5639
CL\_5638
CL\_5637
CL\_5636
CL\_5635
CL\_5634
CL\_5633
CL\_6332
CL\_6333
CL\_6334
CL\_6335
CL\_6336
CL\_6337
CL\_6338
CL\_6339
CL\_6340
CL\_6341
CL\_12858
CL\_12859
CL\_6342
CL\_12640
CL\_12639
CL\_6343
CL\_6344
CL\_6345
CL\_6346
CL\_6347
CL\_6348
CL\_6349
CL\_6350
CL\_6351
CL\_12860
CL\_12862
CL\_12861
CL\_12863
CL\_12864
CL\_12865
CL\_12866
CL\_12867
CL\_12868
CL\_12869
CL\_12870
CL\_6352
CL\_6353
CL\_6354
CL\_12871
CL\_12872
CL\_12873
CL\_6355
CL\_6356
CL\_6357
CL\_6358
CL\_6359
CL\_6360
CL\_6361
CL\_6362
CL\_6363
CL\_6364
CL\_6365
CL\_6366
CL\_6367
CL\_6368
CL\_6369
CL\_6370
CL\_6371
CL\_6372
CL\_6373
CL\_6374
CL\_6375
CL\_6376
CL\_6377
CL\_6378
CL\_6379
CL\_6380
CL\_6381
CL\_9786
CL\_9785
CL\_9784
CL\_9783
CL\_5604
CL\_5605
CL\_5606
CL\_9782
CL\_9781
CL\_9780
CL\_9779
CL\_9778
CL\_9777
CL\_9776
CL\_9775
CL\_9774
CL\_9773
CL\_9772
CL\_9771
CL\_9141
CL\_9142
CL\_9143
CL\_9770
CL\_9145
CL\_9146
CL\_9147
CL\_9148
CL\_9769
CL\_9768
CL\_9767
CL\_9766
CL\_9765
CL\_9764
CL\_9763
CL\_9762
CL\_9761
CL\_9760
CL\_9759
CL\_9758
CL\_9757
CL\_9756
CL\_9755
CL\_9754
CL\_9753
CL\_9752
CL\_9751
CL\_9750
CL\_9749
CL\_9748
CL\_9747
CL\_9746
CL\_9745
CL\_9744
CL\_9743
CL\_9742
CL\_9741
CL\_9740
CL\_9739
CL\_9738
CL\_9737
CL\_9736
CL\_9735
CL\_9734
CL\_9733
CL\_9732
CL\_9731
CL\_9136
CL\_9135
