## Supplementary material for "A novel method for integrating genomic and Tn-Seq data to identify common *in vivo* fitness mechanisms across multiple bacterial species": S5 Dataset: CL_INS_1.html

Legend

 Hypothetical
 Serritia FitnessGene
 All VFDB Genes
/local/ifs2\_projdata/0800/projects/UMIsepsis/Serra\_Prot\_70/centroids.cluster\_roles.txt

FULL


WINDOWSVGPNG

Trim RowsRemove SingletonsSave Fasta

CL\_1014


CL\_1014


CL\_1014


CL\_1014


CL\_1014


CL\_1014


CL\_1014


CL\_1014


CL\_1014


CL\_1014


CL\_1014


CL\_1014


CL\_1014


CL\_1014


CL\_1014


CL\_1014


CL\_1014

HighlightSelectShow Genomes


14

CL\_1023


6

CL\_1023


4

CL\_1023


4

CL\_1023


3

CL\_1023


3

CL\_1023


3

CL\_1023


2

CL\_1023


1

CL\_1023


1

CL\_1023


1

CL\_1023


1

CL\_1023


1

CL\_1023


1

CL\_1023


1

CL\_1023


1

CL\_1023


1

CL\_1023

fGI ID


CL\_INS\_1
CL\_INS\_1
CL\_INS\_1
CL\_INS\_1
CL\_INS\_1
CL\_INS\_1
CL\_INS\_1
CL\_INS\_1
CL\_INS\_1
CL\_INS\_1
CL\_INS\_1
CL\_INS\_1
CL\_INS\_1
CL\_INS\_1
CL\_INS\_381
CL\_INS\_381
CL\_INS\_1
CL\_INS\_1
CL\_INS\_1
CL\_INS\_1
CL\_INS\_1
CL\_INS\_1
CL\_INS\_1
CL\_INS\_1
CL\_INS\_1
CL\_INS\_1
CL\_INS\_1
CL\_INS\_1
CL\_INS\_1
CL\_INS\_1
CL\_INS\_1
CL\_INS\_1
CL\_INS\_1
CL\_INS\_1
CL\_INS\_1
CL\_INS\_1
CL\_INS\_1
CL\_INS\_1
CL\_INS\_1
CL\_INS\_1
CL\_INS\_1
CL\_INS\_1
CL\_INS\_1
CL\_INS\_1
CL\_INS\_1
CL\_INS\_1
CL\_INS\_1
CL\_INS\_1
CL\_INS\_1
CL\_INS\_1
CL\_INS\_1
CL\_INS\_1
CL\_INS\_1
CL\_INS\_1
CL\_INS\_1
CL\_INS\_1
CL\_INS\_1
CL\_INS\_1
CL\_INS\_1
CL\_INS\_1
CL\_INS\_1
Cluster ID


CL\_5157
CL\_5158
CL\_5159
CL\_5160
CL\_5161
CL\_6516
CL\_6165
CL\_6166
CL\_6167
CL\_6168
CL\_6169
CL\_6170
CL\_6171
CL\_6172
CL\_6437
CL\_6438
CL\_9060
CL\_9061
CL\_5672
CL\_5671
CL\_10878
CL\_7708
CL\_7707
CL\_7025
CL\_5670
CL\_5669
CL\_5668
CL\_5667
CL\_5666
CL\_5665
CL\_10879
CL\_10880
CL\_7024
CL\_7181
CL\_7182
CL\_7183
CL\_7184
CL\_7185
CL\_11025
CL\_11026
CL\_11027
CL\_7706
CL\_7023
CL\_7022
CL\_11028
CL\_5664
CL\_1015
CL\_1016
CL\_1017
CL\_1018
CL\_1019
CL\_1020
CL\_1021
CL\_1022
CL\_9062
CL\_9063
CL\_9064
CL\_9065
CL\_9066
CL\_9067
CL\_9068
