## Supplementary material for "A novel method for integrating genomic and Tn-Seq data to identify common *in vivo* fitness mechanisms across multiple bacterial species": S5 Dataset: CL_INS_2.html

Legend

 Hypothetical
 Serritia FitnessGene
 All VFDB Genes
/local/ifs2\_projdata/0800/projects/UMIsepsis/Serra\_Prot\_70/centroids.cluster\_roles.txt

FULL


WINDOWSVGPNG

Trim RowsRemove SingletonsSave Fasta

CL\_1023


CL\_1023


CL\_1023


CL\_1023


CL\_1023


CL\_1023

HighlightSelectShow Genomes


35

CL\_1026


5

CL\_1026


4

CL\_1026


2

CL\_1026


1

CL\_1026


1

CL\_1026

fGI ID


CL\_INS\_2
CL\_INS\_2
CL\_INS\_2
CL\_INS\_2
CL\_INS\_2
CL\_INS\_2
CL\_INS\_2
CL\_INS\_2
CL\_INS\_2
CL\_INS\_2
CL\_INS\_2
Cluster ID


CL\_9069
CL\_7705
CL\_1024
CL\_1025
CL\_8868
CL\_8869
CL\_8870
CL\_11935
CL\_11936
CL\_11937
CL\_11938
