## Supplementary material for "A novel method for integrating genomic and Tn-Seq data to identify common *in vivo* fitness mechanisms across multiple bacterial species": S5 Dataset: CL_INS_3.html

Legend

 Serritia FitnessGene
 All VFDB Genes
/local/ifs2\_projdata/0800/projects/UMIsepsis/Serra\_Prot\_70/centroids.cluster\_roles.txt

FULL


WINDOWSVGPNG

Trim RowsRemove SingletonsSave Fasta

CL\_1034


CL\_1034

HighlightSelectShow Genomes


27

CL\_1035


21

CL\_1035

fGI ID


CL\_INS\_3
CL\_INS\_3
Cluster ID


CL\_5162
CL\_5163
