## Supplementary material for "A novel method for integrating genomic and Tn-Seq data to identify common *in vivo* fitness mechanisms across multiple bacterial species": S5 Dataset: CL_INS_5.html

Legend

 Hypothetical
 All VFDB Genes
/local/ifs2\_projdata/0800/projects/UMIsepsis/Serra\_Prot\_70/centroids.cluster\_roles.txt

FULL


WINDOWSVGPNG

Trim RowsRemove SingletonsSave Fasta

CL\_1044


CL\_1044


CL\_1044

HighlightSelectShow Genomes


37

CL\_1045


9

CL\_1045


1

CL\_1046

fGI ID

CL\_INS\_5
Cluster ID

CL\_8871
