## Supplementary material for "A novel method for integrating genomic and Tn-Seq data to identify common *in vivo* fitness mechanisms across multiple bacterial species": S5 Dataset: CL_INS_12.html

Legend

 Hypothetical
 All EssentialGenes

FULL


WINDOWSVGPNG

Trim RowsRemove SingletonsSave Fasta

CL\_1130


CL\_1130


CL\_1130

HighlightSelectShow Genomes


38

CL\_1133


6

CL\_1133


3

CL\_1133

fGI ID


CL\_INS\_12
CL\_INS\_12
Cluster ID


CL\_1131
CL\_1132
