## Supplementary material for "A novel method for integrating genomic and Tn-Seq data to identify common *in vivo* fitness mechanisms across multiple bacterial species": S5 Dataset: CL_INS_28.html

Legend

 Hypothetical
 Serritia FitnessGene
 All EssentialGenes
/local/ifs2\_projdata/0800/projects/UMIsepsis/Serra\_Prot\_70/centroids.cluster\_roles.txt

FULL


WINDOWSVGPNG

Trim RowsRemove SingletonsSave Fasta

CL\_1341


CL\_1341


CL\_1341


CL\_1341

HighlightSelectShow Genomes


35

CL\_1342


7

CL\_1342


2

CL\_1342


2

CL\_1342

fGI ID


CL\_INS\_28
CL\_INS\_28
CL\_INS\_28
Cluster ID


CL\_8715
CL\_6523
CL\_10001
