## Supplementary material for "A novel method for integrating genomic and Tn-Seq data to identify common *in vivo* fitness mechanisms across multiple bacterial species": S5 Dataset: CL_INS_29.html

Legend

 Hypothetical
 All VFDB Genes
/local/ifs2\_projdata/0800/projects/UMIsepsis/Serra\_Prot\_70/centroids.cluster\_roles.txt

FULL


WINDOWSVGPNG

Trim RowsRemove SingletonsSave Fasta

CL\_1357


CL\_1357


CL\_1357


CL\_1357


CL\_1357


CL\_1357


CL\_1357


CL\_1357


CL\_1357


CL\_1357


CL\_1357


CL\_1357


CL\_1357


CL\_1357

HighlightSelectShow Genomes


31

CL\_1358


2

CL\_1358


1

CL\_1359


1

CL\_1358


1

CL\_1358


1

CL\_1358


1

CL\_1358


1

CL\_1358


1

CL\_1359


1

CL\_1358


1

CL\_1359


1

CL\_1359


1

CL\_1359


1

CL\_1358

fGI ID


CL\_INS\_29
CL\_INS\_29
CL\_INS\_186
CL\_INS\_259
CL\_INS\_259
CL\_INS\_259
CL\_INS\_259
CL\_INS\_259
CL\_INS\_259
CL\_INS\_259
CL\_INS\_259
CL\_INS\_259
CL\_INS\_259
CL\_INS\_259
CL\_INS\_259
CL\_INS\_259
CL\_INS\_259
CL\_INS\_259
CL\_INS\_259
CL\_INS\_259
CL\_INS\_259
CL\_INS\_259
CL\_INS\_259
CL\_INS\_259
CL\_INS\_259
CL\_INS\_259
CL\_INS\_29
CL\_INS\_29
CL\_INS\_29
CL\_INS\_304
CL\_INS\_29
CL\_INS\_304
CL\_INS\_304
CL\_INS\_304
CL\_INS\_304
CL\_INS\_304
CL\_INS\_186
CL\_INS\_35
CL\_INS\_35
CL\_INS\_186
CL\_INS\_29
CL\_INS\_304
CL\_INS\_304
CL\_INS\_186
CL\_INS\_186
CL\_INS\_29
CL\_INS\_186
CL\_INS\_29
CL\_INS\_29
CL\_INS\_304
CL\_INS\_29
CL\_INS\_304
CL\_INS\_186
CL\_INS\_186
CL\_INS\_304
CL\_INS\_304
CL\_INS\_304
CL\_INS\_304
CL\_INS\_29
CL\_INS\_29
CL\_INS\_186
CL\_INS\_29
CL\_INS\_29
CL\_INS\_29
CL\_INS\_29
CL\_INS\_259
CL\_INS\_29
CL\_INS\_29
CL\_INS\_29
CL\_INS\_29
CL\_INS\_29
CL\_INS\_29
CL\_INS\_29
CL\_INS\_29
CL\_INS\_29
CL\_INS\_29
CL\_INS\_29
CL\_INS\_29
CL\_INS\_29
CL\_INS\_29
CL\_INS\_186
CL\_INS\_186
CL\_INS\_186
CL\_INS\_29
CL\_INS\_29
CL\_INS\_29
CL\_INS\_186
CL\_INS\_29
CL\_INS\_29
CL\_INS\_29
CL\_INS\_29
CL\_INS\_29
CL\_INS\_29
CL\_INS\_29
CL\_INS\_29
CL\_INS\_29
CL\_INS\_29
CL\_INS\_29
CL\_INS\_29
CL\_INS\_29
CL\_INS\_29
CL\_INS\_29
CL\_INS\_29
Cluster ID


CL\_9097
CL\_7688
CL\_4432
CL\_5878
CL\_5877
CL\_5876
CL\_5875
CL\_5874
CL\_5873
CL\_5872
CL\_5871
CL\_5870
CL\_5869
CL\_7890
CL\_5868
CL\_5867
CL\_5866
CL\_5865
CL\_5864
CL\_5863
CL\_5862
CL\_5861
CL\_5860
CL\_5859
CL\_5858
CL\_5857
CL\_9081
CL\_11036
CL\_9082
CL\_4449
CL\_9083
CL\_4450
CL\_4451
CL\_4452
CL\_4453
CL\_4454
CL\_8681
CL\_10408
CL\_10409
CL\_5801
CL\_9084
CL\_4446
CL\_4445
CL\_5799
CL\_5798
CL\_9085
CL\_5797
CL\_10795
CL\_9086
CL\_4439
CL\_9087
CL\_4437
CL\_8688
CL\_4435
CL\_5476
CL\_5477
CL\_5478
CL\_4430
CL\_8876
CL\_8877
CL\_8691
CL\_8878
CL\_6186
CL\_8473
CL\_8474
CL\_5856
CL\_8475
CL\_8476
CL\_8477
CL\_8478
CL\_8479
CL\_8480
CL\_8481
CL\_8482
CL\_8483
CL\_8484
CL\_9088
CL\_9892
CL\_9089
CL\_9090
CL\_8695
CL\_8697
CL\_8698
CL\_10796
CL\_10797
CL\_9091
CL\_8699
CL\_9092
CL\_8485
CL\_8486
CL\_6187
CL\_9480
CL\_6188
CL\_6189
CL\_6190
CL\_8879
CL\_8880
CL\_8487
CL\_10798
CL\_9093
CL\_9094
CL\_9095
CL\_9096
