## Supplementary material for "A novel method for integrating genomic and Tn-Seq data to identify common *in vivo* fitness mechanisms across multiple bacterial species": S5 Dataset: CL_INS_35.html

Legend

 Hypothetical
 All EssentialGenes
 All VFDB Genes
/local/ifs2\_projdata/0800/projects/UMIsepsis/Serra\_Prot\_70/centroids.cluster\_roles.txt

FULL


WINDOWSVGPNG

Trim RowsRemove SingletonsSave Fasta

CL\_1390


CL\_1390


CL\_1390


CL\_1390


CL\_1390


CL\_1390


CL\_1390


CL\_1390


CL\_1390


CL\_1390


CL\_1390


CL\_1390


CL\_1390


CL\_1390


CL\_1390


CL\_1390


CL\_1390


CL\_1390


CL\_1390


CL\_1390

HighlightSelectShow Genomes


24

CL\_1391


2

CL\_1391


2

CL\_1394


2

CL\_1394


2

CL\_1391


2

CL\_1391


1

CL\_1391


1

CL\_1391


1

CL\_1391


1

CL\_1391


1

CL\_1391


1

CL\_1394


1

CL\_1391


1

CL\_1391


1

CL\_1391


1

CL\_1391


1

CL\_1391


1

CL\_1394


1

CL\_1394


1

CL\_1391

fGI ID


CL\_INS\_35
CL\_INS\_35
CL\_INS\_36
CL\_INS\_13
CL\_INS\_35
CL\_INS\_35
CL\_INS\_35
CL\_INS\_36
CL\_INS\_36
CL\_INS\_36
CL\_INS\_36
CL\_INS\_36
CL\_INS\_36
CL\_INS\_36
CL\_INS\_36
CL\_INS\_36
CL\_INS\_36
CL\_INS\_36
CL\_INS\_36
CL\_INS\_36
CL\_INS\_36
CL\_INS\_36
CL\_INS\_36
CL\_INS\_36
CL\_INS\_36
CL\_INS\_36
CL\_INS\_36
CL\_INS\_36
CL\_INS\_13
CL\_INS\_13
CL\_INS\_13
CL\_INS\_13
CL\_INS\_13
CL\_INS\_35
CL\_INS\_35
CL\_INS\_13
CL\_INS\_13
CL\_INS\_13
CL\_INS\_13
CL\_INS\_13
CL\_INS\_13
CL\_INS\_13
CL\_INS\_13
CL\_INS\_13
CL\_INS\_35
CL\_INS\_35
CL\_INS\_13
CL\_INS\_13
CL\_INS\_35
CL\_INS\_13
CL\_INS\_36
CL\_INS\_35
CL\_INS\_36
CL\_INS\_35
CL\_INS\_35
CL\_INS\_35
CL\_INS\_35
CL\_INS\_35
CL\_INS\_36
CL\_INS\_36
CL\_INS\_35
CL\_INS\_36
CL\_INS\_35
CL\_INS\_35
CL\_INS\_35
CL\_INS\_36
CL\_INS\_36
CL\_INS\_36
CL\_INS\_36
CL\_INS\_36
CL\_INS\_36
CL\_INS\_36
CL\_INS\_35
CL\_INS\_35
CL\_INS\_35
CL\_INS\_36
CL\_INS\_36
CL\_INS\_36
CL\_INS\_35
CL\_INS\_35
CL\_INS\_35
CL\_INS\_35
CL\_INS\_35
CL\_INS\_35
CL\_INS\_35
CL\_INS\_35
CL\_INS\_35
CL\_INS\_35
CL\_INS\_35
CL\_INS\_36
CL\_INS\_35
CL\_INS\_35
CL\_INS\_70
CL\_INS\_35
CL\_INS\_35
CL\_INS\_35
CL\_INS\_215
CL\_INS\_35
CL\_INS\_36
CL\_INS\_35
CL\_INS\_35
CL\_INS\_35
CL\_INS\_35
CL\_INS\_35
CL\_INS\_35
CL\_INS\_35
CL\_INS\_35
CL\_INS\_36
CL\_INS\_36
CL\_INS\_35
CL\_INS\_35
CL\_INS\_36
CL\_INS\_35
CL\_INS\_36
CL\_INS\_36
CL\_INS\_36
CL\_INS\_36
CL\_INS\_36
CL\_INS\_36
CL\_INS\_36
CL\_INS\_36
CL\_INS\_36
CL\_INS\_36
CL\_INS\_36
CL\_INS\_36
CL\_INS\_36
CL\_INS\_36
CL\_INS\_36
CL\_INS\_36
CL\_INS\_36
CL\_INS\_36
CL\_INS\_35
CL\_INS\_36
CL\_INS\_36
CL\_INS\_36
CL\_INS\_35
CL\_INS\_35
CL\_INS\_35
CL\_INS\_35
CL\_INS\_35
CL\_INS\_35
CL\_INS\_35
CL\_INS\_35
CL\_INS\_35
CL\_INS\_35
CL\_INS\_35
CL\_INS\_35
CL\_INS\_35
CL\_INS\_36
CL\_INS\_36
CL\_INS\_35
CL\_INS\_36
CL\_INS\_36
CL\_INS\_36
CL\_INS\_36
CL\_INS\_36
CL\_INS\_36
CL\_INS\_36
CL\_INS\_36
CL\_INS\_36
CL\_INS\_36
CL\_INS\_36
CL\_INS\_36
CL\_INS\_36
CL\_INS\_36
CL\_INS\_36
CL\_INS\_35
CL\_INS\_35
CL\_INS\_35
CL\_INS\_35
CL\_INS\_35
CL\_INS\_35
CL\_INS\_35
CL\_INS\_35
CL\_INS\_35
CL\_INS\_35
CL\_INS\_35
CL\_INS\_35
CL\_INS\_35
CL\_INS\_35
CL\_INS\_35
CL\_INS\_35
CL\_INS\_35
CL\_INS\_35
CL\_INS\_35
CL\_INS\_35
CL\_INS\_35
CL\_INS\_35
CL\_INS\_35
CL\_INS\_35
CL\_INS\_35
CL\_INS\_35
CL\_INS\_35
CL\_INS\_35
CL\_INS\_35
CL\_INS\_35
CL\_INS\_35
CL\_INS\_35
CL\_INS\_35
CL\_INS\_70
CL\_INS\_70
CL\_INS\_70
CL\_INS\_35
CL\_INS\_70
CL\_INS\_70
CL\_INS\_70
CL\_INS\_70
CL\_INS\_70
CL\_INS\_70
CL\_INS\_70
CL\_INS\_70
CL\_INS\_70
CL\_INS\_70
CL\_INS\_70
CL\_INS\_70
CL\_INS\_70
CL\_INS\_70
CL\_INS\_35
CL\_INS\_70
CL\_INS\_35
CL\_INS\_35
CL\_INS\_70
CL\_INS\_70
CL\_INS\_70
CL\_INS\_70
CL\_INS\_70
CL\_INS\_35
CL\_INS\_35
CL\_INS\_35
CL\_INS\_35
CL\_INS\_35
CL\_INS\_35
CL\_INS\_35
CL\_INS\_35
CL\_INS\_35
CL\_INS\_35
CL\_INS\_35
CL\_INS\_35
CL\_INS\_35
CL\_INS\_35
CL\_INS\_35
CL\_INS\_35
CL\_INS\_35
CL\_INS\_35
CL\_INS\_35
CL\_INS\_35
CL\_INS\_35
CL\_INS\_35
CL\_INS\_35
CL\_INS\_35
CL\_INS\_35
CL\_INS\_35
CL\_INS\_35
CL\_INS\_35
CL\_INS\_35
CL\_INS\_35
CL\_INS\_35
CL\_INS\_35
CL\_INS\_35
CL\_INS\_35
CL\_INS\_35
CL\_INS\_35
CL\_INS\_186
CL\_INS\_186
CL\_INS\_186
CL\_INS\_35
CL\_INS\_186
CL\_INS\_186
CL\_INS\_186
CL\_INS\_186
CL\_INS\_186
CL\_INS\_35
CL\_INS\_35
CL\_INS\_186
CL\_INS\_35
CL\_INS\_304
CL\_INS\_304
CL\_INS\_186
CL\_INS\_186
CL\_INS\_304
CL\_INS\_304
CL\_INS\_304
CL\_INS\_304
CL\_INS\_35
CL\_INS\_35
CL\_INS\_35
CL\_INS\_304
CL\_INS\_186
CL\_INS\_186
CL\_INS\_304
CL\_INS\_304
CL\_INS\_186
CL\_INS\_304
CL\_INS\_35
CL\_INS\_304
CL\_INS\_304
CL\_INS\_304
CL\_INS\_304
CL\_INS\_35
CL\_INS\_35
CL\_INS\_35
CL\_INS\_35
CL\_INS\_35
CL\_INS\_304
CL\_INS\_35
CL\_INS\_35
CL\_INS\_35
CL\_INS\_35
CL\_INS\_35
CL\_INS\_35
CL\_INS\_35
CL\_INS\_35
CL\_INS\_35
CL\_INS\_304
CL\_INS\_35
CL\_INS\_304
CL\_INS\_35
CL\_INS\_35
CL\_INS\_35
CL\_INS\_35
CL\_INS\_35
Cluster ID


CL\_10003
CL\_7685
CL\_9098
CL\_1193
CL\_6192
CL\_6193
CL\_8881
CL\_12312
CL\_12313
CL\_12314
CL\_12315
CL\_12316
CL\_12317
CL\_12318
CL\_12319
CL\_12320
CL\_12321
CL\_12322
CL\_12323
CL\_12324
CL\_12325
CL\_12326
CL\_12327
CL\_12328
CL\_4690
CL\_4689
CL\_12329
CL\_6762
CL\_1195
CL\_1197
CL\_1198
CL\_1199
CL\_1200
CL\_12367
CL\_12368
CL\_1202
CL\_1203
CL\_1204
CL\_1205
CL\_1206
CL\_1207
CL\_6140
CL\_6139
CL\_1208
CL\_12369
CL\_12370
CL\_1209
CL\_6135
CL\_10302
CL\_1214
CL\_6763
CL\_9627
CL\_6764
CL\_6765
CL\_9647
CL\_9648
CL\_9649
CL\_11943
CL\_9628
CL\_6766
CL\_11944
CL\_6767
CL\_6768
CL\_6769
CL\_9629
CL\_8017
CL\_8018
CL\_6770
CL\_9630
CL\_10626
CL\_6771
CL\_6772
CL\_6773
CL\_9631
CL\_9632
CL\_7163
CL\_6774
CL\_6134
CL\_8019
CL\_8020
CL\_9656
CL\_9657
CL\_9658
CL\_9660
CL\_11945
CL\_11946
CL\_11947
CL\_11948
CL\_11949
CL\_6133
CL\_8882
CL\_8883
CL\_2762
CL\_11347
CL\_11348
CL\_12371
CL\_541
CL\_10303
CL\_6775
CL\_6776
CL\_6777
CL\_6778
CL\_6779
CL\_6780
CL\_6781
CL\_6782
CL\_9633
CL\_10627
CL\_10304
CL\_11349
CL\_11350
CL\_5178
CL\_5179
CL\_542
CL\_544
CL\_5177
CL\_6783
CL\_6784
CL\_6785
CL\_6786
CL\_6787
CL\_6788
CL\_6789
CL\_6790
CL\_6791
CL\_6792
CL\_6793
CL\_6794
CL\_6795
CL\_6796
CL\_6797
CL\_6798
CL\_6799
CL\_6800
CL\_6801
CL\_9634
CL\_9635
CL\_9636
CL\_6806
CL\_9637
CL\_6808
CL\_6809
CL\_6810
CL\_6811
CL\_9638
CL\_9639
CL\_9640
CL\_9641
CL\_6802
CL\_6803
CL\_6804
CL\_10628
CL\_10629
CL\_6805
CL\_10630
CL\_6807
CL\_10631
CL\_10632
CL\_10633
CL\_10634
CL\_6812
CL\_7233
CL\_10635
CL\_10636
CL\_10637
CL\_10638
CL\_11950
CL\_11951
CL\_9669
CL\_9670
CL\_545
CL\_6130
CL\_12372
CL\_6129
CL\_6128
CL\_6127
CL\_6126
CL\_6125
CL\_6124
CL\_6123
CL\_6122
CL\_6121
CL\_6120
CL\_6119
CL\_6118
CL\_6117
CL\_6116
CL\_8030
CL\_6115
CL\_6114
CL\_6113
CL\_6112
CL\_6111
CL\_6110
CL\_6109
CL\_6108
CL\_6107
CL\_11716
CL\_10305
CL\_547
CL\_549
CL\_550
CL\_551
CL\_552
CL\_553
CL\_554
CL\_555
CL\_556
CL\_557
CL\_558
CL\_8499
CL\_559
CL\_560
CL\_561
CL\_562
CL\_8500
CL\_563
CL\_10306
CL\_8503
CL\_10307
CL\_10308
CL\_7230
CL\_568
CL\_569
CL\_570
CL\_571
CL\_10309
CL\_11952
CL\_9673
CL\_11953
CL\_9675
CL\_9676
CL\_9677
CL\_9678
CL\_9679
CL\_9680
CL\_11954
CL\_9681
CL\_9682
CL\_9683
CL\_9684
CL\_9685
CL\_9686
CL\_9687
CL\_9688
CL\_9690
CL\_11955
CL\_11956
CL\_9691
CL\_9692
CL\_9693
CL\_9694
CL\_9695
CL\_9696
CL\_9697
CL\_9698
CL\_11957
CL\_9701
CL\_11958
CL\_11959
CL\_11960
CL\_10406
CL\_8678
CL\_5809
CL\_5808
CL\_10407
CL\_5806
CL\_5805
CL\_5804
CL\_5803
CL\_8681
CL\_10408
CL\_10409
CL\_5801
CL\_12291
CL\_4446
CL\_4445
CL\_5799
CL\_5798
CL\_4442
CL\_4441
CL\_6620
CL\_4440
CL\_10410
CL\_10411
CL\_10412
CL\_4437
CL\_8688
CL\_4435
CL\_5476
CL\_5477
CL\_4432
CL\_4430
CL\_10413
CL\_4433
CL\_5478
CL\_10732
CL\_5479
CL\_10414
CL\_10415
CL\_10416
CL\_10417
CL\_10418
CL\_8693
CL\_10419
CL\_10420
CL\_10421
CL\_12292
CL\_12293
CL\_12294
CL\_10422
CL\_10423
CL\_10424
CL\_5489
CL\_10425
CL\_6625
CL\_10426
CL\_10427
CL\_10428
CL\_10429
CL\_10430
