## Supplementary material for "A novel method for integrating genomic and Tn-Seq data to identify common *in vivo* fitness mechanisms across multiple bacterial species": S5 Dataset: CL_INS_36.html

Legend

 Hypothetical
 All EssentialGenes
 All VFDB Genes
/local/ifs2\_projdata/0800/projects/UMIsepsis/Serra\_Prot\_70/centroids.cluster\_roles.txt

FULL


WINDOWSVGPNG

Trim RowsRemove SingletonsSave Fasta

CL\_1391


CL\_1391


CL\_1391


CL\_1391


CL\_1390


CL\_1390


CL\_1391


CL\_1390


CL\_1390


CL\_1390

HighlightSelectShow Genomes


15

CL\_1394


11

CL\_1394


10

CL\_1394


4

CL\_1394


2

CL\_1394


2

CL\_1394


1

CL\_1394


1

CL\_1394


1

CL\_1394


1

CL\_1394

fGI ID


CL\_INS\_36
CL\_INS\_36
CL\_INS\_36
CL\_INS\_36
CL\_INS\_36
CL\_INS\_36
CL\_INS\_36
CL\_INS\_36
CL\_INS\_36
CL\_INS\_36
CL\_INS\_36
CL\_INS\_36
CL\_INS\_36
CL\_INS\_36
CL\_INS\_36
CL\_INS\_36
CL\_INS\_36
CL\_INS\_36
CL\_INS\_36
CL\_INS\_36
CL\_INS\_36
CL\_INS\_36
CL\_INS\_36
CL\_INS\_36
CL\_INS\_36
CL\_INS\_36
CL\_INS\_36
CL\_INS\_36
CL\_INS\_36
CL\_INS\_36
CL\_INS\_36
CL\_INS\_36
CL\_INS\_36
CL\_INS\_36
CL\_INS\_36
CL\_INS\_36
CL\_INS\_36
CL\_INS\_36
CL\_INS\_36
CL\_INS\_36
CL\_INS\_36
CL\_INS\_36
CL\_INS\_36
CL\_INS\_36
CL\_INS\_36
CL\_INS\_36
CL\_INS\_36
CL\_INS\_36
CL\_INS\_36
CL\_INS\_36
CL\_INS\_36
CL\_INS\_36
CL\_INS\_36
CL\_INS\_36
CL\_INS\_36
CL\_INS\_36
CL\_INS\_36
CL\_INS\_36
CL\_INS\_36
CL\_INS\_36
CL\_INS\_36
CL\_INS\_36
CL\_INS\_36
CL\_INS\_36
CL\_INS\_36
CL\_INS\_36
CL\_INS\_36
CL\_INS\_36
CL\_INS\_36
CL\_INS\_36
CL\_INS\_36
CL\_INS\_36
CL\_INS\_36
CL\_INS\_36
CL\_INS\_36
CL\_INS\_36
CL\_INS\_36
CL\_INS\_36
CL\_INS\_36
CL\_INS\_36
CL\_INS\_36
CL\_INS\_36
CL\_INS\_36
CL\_INS\_36
CL\_INS\_36
Cluster ID


CL\_9098
CL\_5178
CL\_1392
CL\_5180
CL\_1393
CL\_7197
CL\_12312
CL\_12313
CL\_12314
CL\_12315
CL\_12316
CL\_12317
CL\_12318
CL\_12319
CL\_12320
CL\_12321
CL\_12322
CL\_12323
CL\_12324
CL\_12325
CL\_12326
CL\_12327
CL\_12328
CL\_4690
CL\_4689
CL\_12329
CL\_12330
CL\_6762
CL\_6763
CL\_6764
CL\_9628
CL\_6766
CL\_6767
CL\_8017
CL\_8018
CL\_6770
CL\_9630
CL\_10626
CL\_6771
CL\_6772
CL\_7163
CL\_6774
CL\_6134
CL\_6133
CL\_6775
CL\_10627
CL\_10304
CL\_542
CL\_544
CL\_5177
CL\_6783
CL\_6784
CL\_6785
CL\_6786
CL\_6787
CL\_6788
CL\_6789
CL\_6790
CL\_6791
CL\_6792
CL\_6793
CL\_6794
CL\_6795
CL\_6796
CL\_6797
CL\_6799
CL\_6800
CL\_6801
CL\_6802
CL\_6803
CL\_10628
CL\_10629
CL\_6805
CL\_10630
CL\_6807
CL\_10631
CL\_10632
CL\_10633
CL\_10634
CL\_6812
CL\_7233
CL\_10635
CL\_10636
CL\_10637
CL\_10638
