## Supplementary material for "A novel method for integrating genomic and Tn-Seq data to identify common *in vivo* fitness mechanisms across multiple bacterial species": S5 Dataset: CL_INS_39.html

Legend

 Hypothetical
 All VFDB Genes

FULL


WINDOWSVGPNG

Trim RowsRemove SingletonsSave Fasta

CL\_1397


CL\_1397


CL\_1395

HighlightSelectShow Genomes


30

CL\_1398


17

CL\_1398


1

CL\_1398

fGI ID


CL\_INS\_38
CL\_INS\_38
CL\_INS\_38
CL\_INS\_39
Cluster ID


CL\_6196
CL\_6197
CL\_6198
CL\_6199
