## Supplementary material for "A novel method for integrating genomic and Tn-Seq data to identify common *in vivo* fitness mechanisms across multiple bacterial species": S5 Dataset: CL_INS_45.html

Legend

 Hypothetical
 All VFDB Genes
/local/ifs2\_projdata/0800/projects/UMIsepsis/Serra\_Prot\_70/centroids.cluster\_roles.txt

FULL


WINDOWSVGPNG

Trim RowsRemove SingletonsSave Fasta

CL\_1436


CL\_1436


CL\_1436

HighlightSelectShow Genomes


38

CL\_1437


9

CL\_1437


1

CL\_1437

fGI ID


CL\_INS\_45
CL\_INS\_45
Cluster ID


CL\_11226
CL\_10310
