## Supplementary material for "A novel method for integrating genomic and Tn-Seq data to identify common *in vivo* fitness mechanisms across multiple bacterial species": S5 Dataset: CL_INS_47.html

Legend

 Hypothetical
 AntibioticResistance
 All EssentialGenes
 All VFDB Genes

FULL


WINDOWSVGPNG

Trim RowsRemove SingletonsSave Fasta

CL\_1445


CL\_1445


CL\_1445


CL\_1445


CL\_1445


CL\_1445


CL\_1445

HighlightSelectShow Genomes


25

CL\_1447


11

CL\_1447


6

CL\_1447


3

CL\_1447


1

CL\_1447


1

CL\_1447


1

CL\_1447

fGI ID


CL\_INS\_47
CL\_INS\_47
CL\_INS\_47
CL\_INS\_47
CL\_INS\_47
Cluster ID


CL\_6205
CL\_6206
CL\_1446
CL\_7664
CL\_7663
