## Supplementary material for "A novel method for integrating genomic and Tn-Seq data to identify common *in vivo* fitness mechanisms across multiple bacterial species": S5 Dataset: CL_INS_50.html

Legend

 Hypothetical
 All VFDB Genes
/local/ifs2\_projdata/0800/projects/UMIsepsis/Serra\_Prot\_70/centroids.cluster\_roles.txt

FULL


WINDOWSVGPNG

Trim RowsRemove SingletonsSave Fasta

CL\_1458


CL\_1458


CL\_1458

HighlightSelectShow Genomes


37

CL\_1459


9

CL\_1459


2

CL\_1459

fGI ID


CL\_INS\_50
CL\_INS\_50
Cluster ID


CL\_11971
CL\_6525
