## Supplementary material for "A novel method for integrating genomic and Tn-Seq data to identify common *in vivo* fitness mechanisms across multiple bacterial species": S5 Dataset: CL_INS_52.html

Legend

 Hypothetical
 All EssentialGenes
 All VFDB Genes
/local/ifs2\_projdata/0800/projects/UMIsepsis/Serra\_Prot\_70/centroids.cluster\_roles.txt

FULL


WINDOWSVGPNG

Trim RowsRemove SingletonsSave Fasta

CL\_1461


CL\_1461


CL\_1461


CL\_1461


CL\_1461


CL\_1461


CL\_1461


CL\_1461


CL\_1461


CL\_1461

HighlightSelectShow Genomes


35

CL\_1460


3

CL\_1460


2

CL\_1459


2

CL\_1460


1

CL\_1459


1

CL\_1459


1

CL\_1459


1

CL\_1459


1

CL\_1459


1

CL\_1459

fGI ID


CL\_INS\_52
CL\_INS\_52
CL\_INS\_52
CL\_INS\_52
CL\_INS\_52
CL\_INS\_52
CL\_INS\_52
CL\_INS\_52
CL\_INS\_52
CL\_INS\_52
CL\_INS\_52
CL\_INS\_52
CL\_INS\_52
CL\_INS\_52
CL\_INS\_52
CL\_INS\_52
CL\_INS\_52
Cluster ID


CL\_9099
CL\_5956
CL\_5957
CL\_9644
CL\_9643
CL\_6816
CL\_12373
CL\_10012
CL\_10011
CL\_10010
CL\_5648
CL\_5649
CL\_5650
CL\_5651
CL\_5652
CL\_5653
CL\_5654
