## Supplementary material for "A novel method for integrating genomic and Tn-Seq data to identify common *in vivo* fitness mechanisms across multiple bacterial species": S5 Dataset: CL_INS_53.html

Legend

 Hypothetical
/local/ifs2\_projdata/0800/projects/UMIsepsis/Serra\_Prot\_70/centroids.cluster\_roles.txt

FULL


WINDOWSVGPNG

Trim RowsRemove SingletonsSave Fasta

CL\_1465


CL\_1465

HighlightSelectShow Genomes


44

CL\_1466


4

CL\_1466

fGI ID


CL\_INS\_53
CL\_INS\_53
CL\_INS\_53
Cluster ID


CL\_6207
CL\_6208
CL\_6209
