## Supplementary material for "A novel method for integrating genomic and Tn-Seq data to identify common *in vivo* fitness mechanisms across multiple bacterial species": S5 Dataset: CL_INS_58.html

Legend

 Hypothetical
 All VFDB Genes
/local/ifs2\_projdata/0800/projects/UMIsepsis/Serra\_Prot\_70/centroids.cluster\_roles.txt

FULL


WINDOWSVGPNG

Trim RowsRemove SingletonsSave Fasta

CL\_1483


CL\_1483


CL\_1483


CL\_1483


CL\_1483

HighlightSelectShow Genomes


29

CL\_1486


7

CL\_1486


6

CL\_1486


3

CL\_1486


1

CL\_1486

fGI ID


CL\_INS\_58
CL\_INS\_58
CL\_INS\_58
CL\_INS\_58
CL\_INS\_58
CL\_INS\_58
Cluster ID


CL\_1484
CL\_1485
CL\_5198
CL\_6818
CL\_6819
CL\_6820
