## Supplementary material for "A novel method for integrating genomic and Tn-Seq data to identify common *in vivo* fitness mechanisms across multiple bacterial species": S5 Dataset: CL_INS_59.html

Legend

 Hypothetical
 All VFDB Genes
/local/ifs2\_projdata/0800/projects/UMIsepsis/Serra\_Prot\_70/centroids.cluster\_roles.txt

FULL


WINDOWSVGPNG

Trim RowsRemove SingletonsSave Fasta

CL\_1488


CL\_1488


CL\_1488


CL\_1488

HighlightSelectShow Genomes


31

CL\_1489


13

CL\_1489


1

CL\_1489


1

CL\_1489

fGI ID


CL\_INS\_59
CL\_INS\_59
CL\_INS\_59
CL\_INS\_59
Cluster ID


CL\_5199
CL\_12126
CL\_5200
CL\_5201
