## Supplementary material for "A novel method for integrating genomic and Tn-Seq data to identify common *in vivo* fitness mechanisms across multiple bacterial species": S5 Dataset: CL_INS_63.html

Legend

 Hypothetical
 All VFDB Genes
/local/ifs2\_projdata/0800/projects/UMIsepsis/Serra\_Prot\_70/centroids.cluster\_roles.txt

FULL


WINDOWSVGPNG

Trim RowsRemove SingletonsSave Fasta

CL\_1520


CL\_1520


CL\_1520

HighlightSelectShow Genomes


21

CL\_5208


15

CL\_5208


1

CL\_5208

fGI ID


CL\_INS\_63
CL\_INS\_63
Cluster ID


CL\_10017
CL\_5207
