## Supplementary material for "A novel method for integrating genomic and Tn-Seq data to identify common *in vivo* fitness mechanisms across multiple bacterial species": S5 Dataset: CL_INS_69.html

Legend

 Hypothetical
/local/ifs2\_projdata/0800/projects/UMIsepsis/Serra\_Prot\_70/centroids.cluster\_roles.txt

FULL


WINDOWSVGPNG

Trim RowsRemove SingletonsSave Fasta

CL\_1565


CL\_1565


CL\_1565

HighlightSelectShow Genomes


19

CL\_1566


16

CL\_1566


8

CL\_1567

fGI ID

CL\_INS\_69
Cluster ID

CL\_5213
