## Supplementary material for "A novel method for integrating genomic and Tn-Seq data to identify common *in vivo* fitness mechanisms across multiple bacterial species": S5 Dataset: CL_INS_81.html

Legend

 All VFDB Genes
/local/ifs2\_projdata/0800/projects/UMIsepsis/Serra\_Prot\_70/centroids.cluster\_roles.txt

FULL


WINDOWSVGPNG

Trim RowsRemove SingletonsSave Fasta

CL\_1732


CL\_1732


CL\_1731

HighlightSelectShow Genomes


24

CL\_1734


19

CL\_1734


1

CL\_1734

fGI ID

CL\_INS\_81
Cluster ID

CL\_1733
