## Supplementary material for "A novel method for integrating genomic and Tn-Seq data to identify common *in vivo* fitness mechanisms across multiple bacterial species": S5 Dataset: CL_INS_83.html

Legend

 Hypothetical
 All VFDB Genes
/local/ifs2\_projdata/0800/projects/UMIsepsis/Serra\_Prot\_70/centroids.cluster\_roles.txt

FULL


WINDOWSVGPNG

Trim RowsRemove SingletonsSave Fasta

CL\_1740


CL\_1735


CL\_1740


CL\_1735


CL\_1735


CL\_1740

HighlightSelectShow Genomes


30

CL\_1741


11

CL\_1741


2

CL\_2435


2

CL\_1741


1

CL\_1741


1

CL\_2669

fGI ID


CL\_INS\_83
CL\_INS\_82
CL\_INS\_207
CL\_INS\_207
CL\_INS\_182
CL\_INS\_182
CL\_INS\_181
CL\_INS\_182
CL\_INS\_181
CL\_INS\_181
CL\_INS\_182
CL\_INS\_182
Cluster ID


CL\_6218
CL\_5221
CL\_6004
CL\_6005
CL\_9146
CL\_9145
CL\_9144
CL\_9143
CL\_9142
CL\_9141
CL\_5317
CL\_5316
