## Supplementary material for "A novel method for integrating genomic and Tn-Seq data to identify common *in vivo* fitness mechanisms across multiple bacterial species": S5 Dataset: CL_INS_85.html

Legend

 Hypothetical
 All VFDB Genes
/local/ifs2\_projdata/0800/projects/UMIsepsis/Serra\_Prot\_70/centroids.cluster\_roles.txt

FULL


WINDOWSVGPNG

Trim RowsRemove SingletonsSave Fasta

CL\_1745


CL\_1745


CL\_1745


CL\_1745


CL\_1745


Break

HighlightSelectShow Genomes


25

CL\_1746


8

CL\_1746


7

CL\_1746


4

CL\_1746


2

CL\_1746


1

CL\_1746

fGI ID


CL\_INS\_85
CL\_INS\_70
CL\_INS\_85
CL\_INS\_85
CL\_INS\_85
CL\_INS\_85
CL\_INS\_85
CL\_INS\_85
CL\_INS\_85
CL\_INS\_85
CL\_INS\_85
CL\_INS\_85
CL\_INS\_85
CL\_INS\_85
CL\_INS\_85
CL\_INS\_85
CL\_INS\_85
CL\_INS\_85
Cluster ID


CL\_7235
CL\_8503
CL\_10641
CL\_10642
CL\_7236
CL\_7237
CL\_7238
CL\_7239
CL\_7240
CL\_7241
CL\_7242
CL\_7243
CL\_7244
CL\_7245
CL\_7246
CL\_7247
CL\_5646
CL\_5645
