## Supplementary material for "A novel method for integrating genomic and Tn-Seq data to identify common *in vivo* fitness mechanisms across multiple bacterial species": S5 Dataset: CL_INS_94.html

Legend

 Hypothetical
 All VFDB Genes
/local/ifs2\_projdata/0800/projects/UMIsepsis/Serra\_Prot\_70/centroids.cluster\_roles.txt

FULL


WINDOWSVGPNG

Trim RowsRemove SingletonsSave Fasta

CL\_1794


CL\_1794


CL\_1791


CL\_1791

HighlightSelectShow Genomes


36

CL\_1795


6

CL\_1795


5

CL\_1795


1

CL\_1795

fGI ID


CL\_INS\_94
CL\_INS\_94
CL\_INS\_94
Cluster ID


CL\_8075
CL\_6234
CL\_6235
