## Supplementary material for "A novel method for integrating genomic and Tn-Seq data to identify common *in vivo* fitness mechanisms across multiple bacterial species": S5 Dataset: CL_INS_98.html

Legend

 Hypothetical
 All VFDB Genes
/local/ifs2\_projdata/0800/projects/UMIsepsis/Serra\_Prot\_70/centroids.cluster\_roles.txt

FULL


WINDOWSVGPNG

Trim RowsRemove SingletonsSave Fasta

CL\_1820


CL\_1820


CL\_1820


CL\_1820


CL\_1820


CL\_1820

HighlightSelectShow Genomes


28

CL\_1821


14

CL\_1821


2

CL\_1821


1

CL\_1821


1

CL\_1821


1

CL\_1821

fGI ID


CL\_INS\_98
CL\_INS\_98
CL\_INS\_98
CL\_INS\_98
CL\_INS\_98
CL\_INS\_98
CL\_INS\_98
CL\_INS\_98
CL\_INS\_98
CL\_INS\_98
CL\_INS\_98
CL\_INS\_98
CL\_INS\_98
CL\_INS\_98
CL\_INS\_98
CL\_INS\_98
CL\_INS\_98
Cluster ID


CL\_9106
CL\_12489
CL\_5238
CL\_5239
CL\_5240
CL\_11364
CL\_11365
CL\_11366
CL\_11367
CL\_11368
CL\_11369
CL\_11370
CL\_11371
CL\_11372
CL\_11373
CL\_11374
CL\_11375
