## Supplementary material for "A novel method for integrating genomic and Tn-Seq data to identify common *in vivo* fitness mechanisms across multiple bacterial species": S5 Dataset: CL_INS_107.html

Legend

 Hypothetical
 AntibioticResistance
 All VFDB Genes
/local/ifs2\_projdata/0800/projects/UMIsepsis/Serra\_Prot\_70/centroids.cluster\_roles.txt

FULL


WINDOWSVGPNG

Trim RowsRemove SingletonsSave Fasta

CL\_1849


CL\_1847


CL\_1849


CL\_1849


CL\_1847


CL\_1833


CL\_1833


CL\_1847


CL\_1847


Break

HighlightSelectShow Genomes


33

CL\_1850


4

CL\_1850


3

CL\_1850


1

CL\_1850


1

CL\_1850


1

CL\_1850


1

CL\_1850


1

CL\_1850


1

CL\_1850


1

CL\_1850

fGI ID


CL\_INS\_106
CL\_INS\_106
CL\_INS\_106
CL\_INS\_106
CL\_INS\_106
CL\_INS\_106
CL\_INS\_44
CL\_INS\_107
CL\_INS\_107
CL\_INS\_107
CL\_INS\_107
CL\_INS\_107
CL\_INS\_107
CL\_INS\_107
CL\_INS\_107
CL\_INS\_104
CL\_INS\_104
CL\_INS\_104
CL\_INS\_104
CL\_INS\_106
CL\_INS\_107
CL\_INS\_104
CL\_INS\_104
CL\_INS\_104
CL\_INS\_104
CL\_INS\_107
CL\_INS\_107
CL\_INS\_107
CL\_INS\_107
CL\_INS\_107
CL\_INS\_107
CL\_INS\_107
CL\_INS\_107
CL\_INS\_107
CL\_INS\_107
CL\_INS\_107
CL\_INS\_107
CL\_INS\_107
CL\_INS\_107
CL\_INS\_107
CL\_INS\_107
CL\_INS\_107
CL\_INS\_107
CL\_INS\_107
CL\_INS\_107
CL\_INS\_107
CL\_INS\_107
CL\_INS\_107
CL\_INS\_107
CL\_INS\_107
CL\_INS\_107
CL\_INS\_107
CL\_INS\_107
CL\_INS\_107
CL\_INS\_107
CL\_INS\_107
CL\_INS\_107
CL\_INS\_107
CL\_INS\_107
CL\_INS\_107
CL\_INS\_107
CL\_INS\_107
CL\_INS\_107
CL\_INS\_107
CL\_INS\_107
CL\_INS\_107
CL\_INS\_107
CL\_INS\_107
CL\_INS\_107
CL\_INS\_107
CL\_INS\_107
CL\_INS\_104
CL\_INS\_104
CL\_INS\_104
CL\_INS\_104
CL\_INS\_104
CL\_INS\_107
Cluster ID


CL\_8706
CL\_9896
CL\_9108
CL\_9109
CL\_9110
CL\_9111
CL\_5256
CL\_5257
CL\_5258
CL\_5259
CL\_5260
CL\_5261
CL\_5262
CL\_11039
CL\_11040
CL\_8058
CL\_11674
CL\_11673
CL\_11672
CL\_1848
CL\_9112
CL\_8057
CL\_8056
CL\_8055
CL\_8054
CL\_11763
CL\_11762
CL\_11761
CL\_11760
CL\_11759
CL\_11758
CL\_11757
CL\_11756
CL\_11755
CL\_11754
CL\_11753
CL\_11752
CL\_11751
CL\_11750
CL\_11749
CL\_11748
CL\_11747
CL\_11746
CL\_11745
CL\_11744
CL\_11743
CL\_11742
CL\_11741
CL\_11740
CL\_11739
CL\_11738
CL\_11737
CL\_11736
CL\_11735
CL\_11734
CL\_11733
CL\_11732
CL\_11731
CL\_11730
CL\_11729
CL\_11728
CL\_11727
CL\_11726
CL\_11725
CL\_11724
CL\_11723
CL\_11722
CL\_11721
CL\_11720
CL\_11719
CL\_11718
CL\_8053
CL\_8052
CL\_8051
CL\_8050
CL\_8049
CL\_8048
