## Supplementary material for "A novel method for integrating genomic and Tn-Seq data to identify common *in vivo* fitness mechanisms across multiple bacterial species": S5 Dataset: CL_INS_109.html

Legend

 Hypothetical
 All VFDB Genes
/local/ifs2\_projdata/0800/projects/UMIsepsis/Serra\_Prot\_70/centroids.cluster\_roles.txt

FULL


WINDOWSVGPNG

Trim RowsRemove SingletonsSave Fasta

CL\_1855


CL\_1855


CL\_1855


CL\_1855


CL\_1855


CL\_1855


CL\_1855


CL\_1855


CL\_1855


CL\_1855


CL\_1854

HighlightSelectShow Genomes


18

CL\_1857


10

CL\_1857


6

CL\_1857


4

CL\_1857


3

CL\_1857


2

CL\_1857


1

CL\_1858


1

CL\_1857


1

CL\_1858


1

CL\_1857


1

CL\_1857

fGI ID


CL\_INS\_109
CL\_INS\_109
CL\_INS\_109
CL\_INS\_109
CL\_INS\_109
CL\_INS\_109
CL\_INS\_109
Cluster ID


CL\_1856
CL\_6014
CL\_5264
CL\_5265
CL\_5266
CL\_6839
CL\_6840
