## Supplementary material for "A novel method for integrating genomic and Tn-Seq data to identify common *in vivo* fitness mechanisms across multiple bacterial species": S5 Dataset: CL_INS_120.html

Legend

 All VFDB Genes
/local/ifs2\_projdata/0800/projects/UMIsepsis/Serra\_Prot\_70/centroids.cluster\_roles.txt

FULL


WINDOWSVGPNG

Trim RowsRemove SingletonsSave Fasta

CL\_1890


CL\_1890


CL\_1890


CL\_1890


CL\_1890


CL\_1890

HighlightSelectShow Genomes


22

CL\_1895


21

CL\_1895


2

CL\_1896


1

CL\_1895


1

CL\_1895


1

CL\_1896

fGI ID


CL\_INS\_120
CL\_INS\_121
CL\_INS\_120
CL\_INS\_120
Cluster ID


CL\_1891
CL\_1892
CL\_1893
CL\_1894
