## Supplementary material for "A novel method for integrating genomic and Tn-Seq data to identify common *in vivo* fitness mechanisms across multiple bacterial species": S5 Dataset: CL_INS_122.html

Legend

 Hypothetical
 All EssentialGenes
 All VFDB Genes
/local/ifs2\_projdata/0800/projects/UMIsepsis/Serra\_Prot\_70/centroids.cluster\_roles.txt

FULL


WINDOWSVGPNG

Trim RowsRemove SingletonsSave Fasta

CL\_1898


CL\_1898


CL\_1898


CL\_1898


CL\_1898


CL\_1898


CL\_1898


CL\_1897


CL\_1898


CL\_1898


CL\_1898


CL\_1898

HighlightSelectShow Genomes


14

CL\_5278


11

CL\_5278


11

CL\_5278


3

CL\_5278


2

CL\_5278


1

CL\_5278


1

CL\_1907


1

CL\_5278


1

CL\_1907


1

CL\_5278


1

CL\_5278


1

CL\_5278

fGI ID


CL\_INS\_122
CL\_INS\_122
CL\_INS\_122
CL\_INS\_122
CL\_INS\_122
CL\_INS\_122
CL\_INS\_122
CL\_INS\_122
CL\_INS\_122
CL\_INS\_122
CL\_INS\_122
CL\_INS\_122
CL\_INS\_122
Cluster ID


CL\_6243
CL\_5276
CL\_6845
CL\_5277
CL\_6541
CL\_1899
CL\_1900
CL\_1901
CL\_1902
CL\_1903
CL\_1904
CL\_1905
CL\_1906
