## Supplementary material for "A novel method for integrating genomic and Tn-Seq data to identify common *in vivo* fitness mechanisms across multiple bacterial species": S5 Dataset: CL_INS_123.html

Legend

 Hypothetical
 Serritia FitnessGene
 All VFDB Genes
/local/ifs2\_projdata/0800/projects/UMIsepsis/Serra\_Prot\_70/centroids.cluster\_roles.txt

FULL


WINDOWSVGPNG

Trim RowsRemove SingletonsSave Fasta

CL\_1908


CL\_1908


CL\_1908


CL\_1908


CL\_1908


CL\_1908


CL\_1908


CL\_1908

HighlightSelectShow Genomes


17

CL\_1911


15

CL\_1911


8

CL\_1911


2

CL\_1911


1

CL\_1911


1

CL\_1912


1

CL\_1912


1

CL\_1912

fGI ID


CL\_INS\_123
CL\_INS\_123
CL\_INS\_123
CL\_INS\_123
Cluster ID


CL\_10036
CL\_5279
CL\_1909
CL\_1910
