## Supplementary material for "A novel method for integrating genomic and Tn-Seq data to identify common *in vivo* fitness mechanisms across multiple bacterial species": S5 Dataset: CL_INS_136.html

Legend

 Hypothetical
 All VFDB Genes
/local/ifs2\_projdata/0800/projects/UMIsepsis/Serra\_Prot\_70/centroids.cluster\_roles.txt

FULL


WINDOWSVGPNG

Trim RowsRemove SingletonsSave Fasta

CL\_1979


CL\_1979


CL\_1978

HighlightSelectShow Genomes


29

CL\_1982


17

CL\_1982


2

CL\_1982

fGI ID


CL\_INS\_136
CL\_INS\_136
Cluster ID


CL\_1980
CL\_1981
