## Supplementary material for "A novel method for integrating genomic and Tn-Seq data to identify common *in vivo* fitness mechanisms across multiple bacterial species": S5 Dataset: CL_INS_137.html

Legend

 Hypothetical
 All EssentialGenes
 All VFDB Genes
/local/ifs2\_projdata/0800/projects/UMIsepsis/Serra\_Prot\_70/centroids.cluster\_roles.txt

FULL


WINDOWSVGPNG

Trim RowsRemove SingletonsSave Fasta

CL\_1986


CL\_1986


CL\_1986


CL\_1986


CL\_1986


CL\_1986


CL\_1986


CL\_1986

HighlightSelectShow Genomes


27

CL\_1987


11

CL\_1987


4

CL\_1987


2

CL\_1987


1

CL\_1987


1

CL\_1987


1

CL\_1987


1

CL\_1987

fGI ID


CL\_INS\_137
CL\_INS\_137
CL\_INS\_137
CL\_INS\_137
CL\_INS\_35
CL\_INS\_35
CL\_INS\_137
CL\_INS\_13
CL\_INS\_137
CL\_INS\_137
CL\_INS\_210
CL\_INS\_137
CL\_INS\_137
CL\_INS\_35
CL\_INS\_35
CL\_INS\_35
CL\_INS\_35
CL\_INS\_35
CL\_INS\_35
CL\_INS\_35
CL\_INS\_35
CL\_INS\_35
CL\_INS\_35
CL\_INS\_35
CL\_INS\_35
CL\_INS\_35
CL\_INS\_35
CL\_INS\_35
CL\_INS\_35
CL\_INS\_35
CL\_INS\_35
CL\_INS\_35
CL\_INS\_35
CL\_INS\_35
CL\_INS\_35
CL\_INS\_35
CL\_INS\_210
CL\_INS\_210
CL\_INS\_210
CL\_INS\_210
CL\_INS\_210
CL\_INS\_137
CL\_INS\_210
CL\_INS\_217
CL\_INS\_217
CL\_INS\_137
CL\_INS\_261
CL\_INS\_261
CL\_INS\_261
CL\_INS\_137
CL\_INS\_137
CL\_INS\_137
CL\_INS\_137
CL\_INS\_137
CL\_INS\_137
CL\_INS\_137
CL\_INS\_137
CL\_INS\_137
CL\_INS\_35
CL\_INS\_137
CL\_INS\_137
CL\_INS\_434
CL\_INS\_106
CL\_INS\_35
CL\_INS\_36
CL\_INS\_36
CL\_INS\_215
CL\_INS\_137
CL\_INS\_137
CL\_INS\_36
CL\_INS\_137
CL\_INS\_137
CL\_INS\_35
CL\_INS\_137
CL\_INS\_137
CL\_INS\_36
CL\_INS\_137
CL\_INS\_137
CL\_INS\_137
CL\_INS\_137
CL\_INS\_137
CL\_INS\_137
CL\_INS\_137
CL\_INS\_36
CL\_INS\_137
CL\_INS\_137
CL\_INS\_137
CL\_INS\_35
CL\_INS\_35
CL\_INS\_36
CL\_INS\_36
CL\_INS\_137
CL\_INS\_137
CL\_INS\_36
CL\_INS\_137
CL\_INS\_36
CL\_INS\_137
CL\_INS\_137
CL\_INS\_137
CL\_INS\_137
CL\_INS\_137
CL\_INS\_137
CL\_INS\_137
CL\_INS\_137
CL\_INS\_137
CL\_INS\_137
CL\_INS\_137
CL\_INS\_137
Cluster ID


CL\_10323
CL\_10324
CL\_8032
CL\_8031
CL\_11716
CL\_6107
CL\_6245
CL\_6102
CL\_10824
CL\_10825
CL\_8907
CL\_10457
CL\_6106
CL\_6108
CL\_6109
CL\_6110
CL\_6111
CL\_6112
CL\_6113
CL\_6114
CL\_6115
CL\_8030
CL\_6116
CL\_6117
CL\_6118
CL\_6119
CL\_6120
CL\_6121
CL\_6122
CL\_6123
CL\_6124
CL\_6125
CL\_6126
CL\_6127
CL\_6128
CL\_6129
CL\_8519
CL\_2747
CL\_2748
CL\_2749
CL\_2750
CL\_11264
CL\_2752
CL\_2753
CL\_2754
CL\_8793
CL\_10499
CL\_10500
CL\_10502
CL\_10503
CL\_10505
CL\_10506
CL\_10507
CL\_11265
CL\_11266
CL\_10511
CL\_11267
CL\_10513
CL\_6130
CL\_11268
CL\_11269
CL\_8029
CL\_6318
CL\_545
CL\_544
CL\_542
CL\_541
CL\_10826
CL\_6132
CL\_6133
CL\_8028
CL\_8027
CL\_6776
CL\_8026
CL\_8025
CL\_6134
CL\_8024
CL\_10827
CL\_10516
CL\_9661
CL\_8023
CL\_10828
CL\_10829
CL\_6771
CL\_8022
CL\_8021
CL\_10830
CL\_8020
CL\_8019
CL\_8018
CL\_9630
CL\_10519
CL\_10520
CL\_8017
CL\_8016
CL\_6767
CL\_8015
CL\_10831
CL\_11270
CL\_8014
CL\_10523
CL\_10524
CL\_8013
CL\_8012
CL\_8011
CL\_11271
CL\_11272
CL\_11273
