## Supplementary material for "A novel method for integrating genomic and Tn-Seq data to identify common *in vivo* fitness mechanisms across multiple bacterial species": S5 Dataset: CL_INS_140.html

Legend

 Hypothetical
 All VFDB Genes
/local/ifs2\_projdata/0800/projects/UMIsepsis/Serra\_Prot\_70/centroids.cluster\_roles.txt

FULL


WINDOWSVGPNG

Trim RowsRemove SingletonsSave Fasta

CL\_2025


CL\_2025


CL\_2025


CL\_2025


CL\_2025


CL\_2025


CL\_2025


CL\_2025


CL\_2025


CL\_2025


CL\_2025


CL\_2025


CL\_2025


CL\_2025


CL\_2025


CL\_2025


CL\_2025


CL\_2025


CL\_2025


CL\_2025

HighlightSelectShow Genomes


25

CL\_2029


3

CL\_2029


2

CL\_2029


2

CL\_2029


1

CL\_2029


1

CL\_2029


1

CL\_2029


1

CL\_2029


1

CL\_2029


1

CL\_2029


1

CL\_2029


1

CL\_2029


1

CL\_2029


1

CL\_2029


1

CL\_2029


1

CL\_2029


1

CL\_2029


1

CL\_2029


1

CL\_2029


1

CL\_2029

fGI ID


CL\_INS\_140
CL\_INS\_140
CL\_INS\_140
CL\_INS\_140
CL\_INS\_140
CL\_INS\_140
CL\_INS\_140
CL\_INS\_140
CL\_INS\_140
CL\_INS\_140
CL\_INS\_140
CL\_INS\_140
CL\_INS\_140
CL\_INS\_140
CL\_INS\_140
CL\_INS\_140
CL\_INS\_140
CL\_INS\_140
CL\_INS\_140
CL\_INS\_140
CL\_INS\_140
CL\_INS\_140
CL\_INS\_140
CL\_INS\_140
CL\_INS\_140
CL\_INS\_140
CL\_INS\_140
CL\_INS\_140
CL\_INS\_140
CL\_INS\_140
CL\_INS\_140
CL\_INS\_140
CL\_INS\_140
CL\_INS\_140
CL\_INS\_140
CL\_INS\_140
CL\_INS\_140
CL\_INS\_140
CL\_INS\_140
CL\_INS\_140
CL\_INS\_140
Cluster ID


CL\_2026
CL\_6013
CL\_8121
CL\_7254
CL\_7255
CL\_9118
CL\_11715
CL\_8010
CL\_8009
CL\_8008
CL\_9119
CL\_8007
CL\_8006
CL\_8005
CL\_12491
CL\_12492
CL\_12493
CL\_8119
CL\_8118
CL\_9120
CL\_9121
CL\_9125
CL\_9126
CL\_9124
CL\_9123
CL\_6012
CL\_6547
CL\_6992
CL\_6991
CL\_6990
CL\_8120
CL\_6548
CL\_9122
CL\_9897
CL\_9127
CL\_9128
CL\_2027
CL\_10326
CL\_6993
CL\_11714
CL\_2028
