## Supplementary material for "A novel method for integrating genomic and Tn-Seq data to identify common *in vivo* fitness mechanisms across multiple bacterial species": S5 Dataset: CL_INS_144.html

Legend

 All EssentialGenes
 All VFDB Genes

FULL


WINDOWSVGPNG

Trim RowsRemove SingletonsSave Fasta

CL\_2095


CL\_2095


CL\_2095

HighlightSelectShow Genomes


38

CL\_2096


8

CL\_2096


1

CL\_2097

fGI ID

CL\_INS\_144
Cluster ID

CL\_5296
