## Supplementary material for "A novel method for integrating genomic and Tn-Seq data to identify common *in vivo* fitness mechanisms across multiple bacterial species": S5 Dataset: CL_INS_148.html

Legend

 Hypothetical
 All VFDB Genes
/local/ifs2\_projdata/0800/projects/UMIsepsis/Serra\_Prot\_70/centroids.cluster\_roles.txt

FULL


WINDOWSVGPNG

Trim RowsRemove SingletonsSave Fasta

CL\_2108


CL\_2108


CL\_2098


CL\_2098

HighlightSelectShow Genomes


35

CL\_2109


6

CL\_2109


4

CL\_2109


1

CL\_2109

fGI ID

CL\_INS\_148
Cluster ID

CL\_8001
