## Supplementary material for "A novel method for integrating genomic and Tn-Seq data to identify common *in vivo* fitness mechanisms across multiple bacterial species": S5 Dataset: CL_INS_154.html

Legend

 Hypothetical
 All EssentialGenes
 All VFDB Genes

FULL


WINDOWSVGPNG

Trim RowsRemove SingletonsSave Fasta

CL\_2206


CL\_2206


CL\_2206


CL\_2206

HighlightSelectShow Genomes


31

CL\_2207


14

CL\_2207


1

CL\_2207


1

CL\_2207

fGI ID


CL\_INS\_154
CL\_INS\_154
CL\_INS\_154
Cluster ID


CL\_11296
CL\_6248
CL\_5839
