## Supplementary material for "A novel method for integrating genomic and Tn-Seq data to identify common *in vivo* fitness mechanisms across multiple bacterial species": S5 Dataset: CL_INS_164.html

Legend

 Hypothetical
 All VFDB Genes

FULL


WINDOWSVGPNG

Trim RowsRemove SingletonsSave Fasta

CL\_2308


CL\_2308


CL\_2308


CL\_2308

HighlightSelectShow Genomes


26

CL\_2310


15

CL\_2310


5

CL\_2310


2

CL\_2310

fGI ID


CL\_INS\_164
CL\_INS\_164
Cluster ID


CL\_2309
CL\_5624
