## Supplementary material for "A novel method for integrating genomic and Tn-Seq data to identify common *in vivo* fitness mechanisms across multiple bacterial species": S5 Dataset: CL_INS_165.html

Legend

 Hypothetical
 All VFDB Genes
/local/ifs2\_projdata/0800/projects/UMIsepsis/Serra\_Prot\_70/centroids.cluster\_roles.txt

FULL


WINDOWSVGPNG

Trim RowsRemove SingletonsSave Fasta

CL\_2317


CL\_2317


CL\_2317


CL\_2317


CL\_2317


CL\_2317


CL\_2317


CL\_2317

HighlightSelectShow Genomes


38

CL\_2318


4

CL\_2318


1

CL\_2318


1

CL\_2318


1

CL\_2318


1

CL\_2318


1

CL\_2318


1

CL\_2318

fGI ID


CL\_INS\_165
CL\_INS\_165
CL\_INS\_165
CL\_INS\_165
CL\_INS\_165
CL\_INS\_165
CL\_INS\_165
CL\_INS\_165
CL\_INS\_165
CL\_INS\_165
CL\_INS\_165
CL\_INS\_165
CL\_INS\_165
CL\_INS\_165
CL\_INS\_165
CL\_INS\_165
CL\_INS\_165
CL\_INS\_304
CL\_INS\_304
CL\_INS\_304
CL\_INS\_165
CL\_INS\_165
CL\_INS\_165
CL\_INS\_165
CL\_INS\_165
CL\_INS\_165
CL\_INS\_165
CL\_INS\_165
CL\_INS\_165
CL\_INS\_165
CL\_INS\_165
CL\_INS\_165
CL\_INS\_165
CL\_INS\_186
CL\_INS\_186
CL\_INS\_304
CL\_INS\_186
CL\_INS\_304
CL\_INS\_165
CL\_INS\_165
CL\_INS\_165
CL\_INS\_165
CL\_INS\_165
CL\_INS\_165
CL\_INS\_165
CL\_INS\_165
CL\_INS\_304
CL\_INS\_304
CL\_INS\_165
CL\_INS\_165
CL\_INS\_165
CL\_INS\_165
CL\_INS\_165
CL\_INS\_165
Cluster ID


CL\_11520
CL\_12218
CL\_12219
CL\_12220
CL\_7989
CL\_7988
CL\_7987
CL\_7986
CL\_7985
CL\_8758
CL\_8757
CL\_8756
CL\_8755
CL\_8754
CL\_8753
CL\_8752
CL\_8751
CL\_5470
CL\_5472
CL\_4447
CL\_8750
CL\_8749
CL\_8748
CL\_8747
CL\_8746
CL\_8745
CL\_8744
CL\_8743
CL\_8742
CL\_8741
CL\_8740
CL\_8739
CL\_8738
CL\_4435
CL\_4434
CL\_4433
CL\_4432
CL\_4430
CL\_6864
CL\_6865
CL\_6866
CL\_6867
CL\_8737
CL\_8736
CL\_8735
CL\_4427
CL\_4426
CL\_4425
CL\_4424
CL\_4423
CL\_4422
CL\_4420
CL\_4419
CL\_4418
