## Supplementary material for "A novel method for integrating genomic and Tn-Seq data to identify common *in vivo* fitness mechanisms across multiple bacterial species": S5 Dataset: CL_INS_171.html

Legend

 Hypothetical
 All VFDB Genes
/local/ifs2\_projdata/0800/projects/UMIsepsis/Serra\_Prot\_70/centroids.cluster\_roles.txt

FULL


WINDOWSVGPNG

Trim RowsRemove SingletonsSave Fasta

CL\_2374


CL\_2374


CL\_2374


CL\_2374

HighlightSelectShow Genomes


28

CL\_2377


17

CL\_2377


1

CL\_2377


1

CL\_2377

fGI ID


CL\_INS\_171
CL\_INS\_171
CL\_INS\_171
Cluster ID


CL\_6558
CL\_2375
CL\_2376
