## Supplementary material for "A novel method for integrating genomic and Tn-Seq data to identify common *in vivo* fitness mechanisms across multiple bacterial species": S5 Dataset: CL_INS_181.html

Legend

 Hypothetical
 All VFDB Genes
/local/ifs2\_projdata/0800/projects/UMIsepsis/Serra\_Prot\_70/centroids.cluster\_roles.txt

FULL


WINDOWSVGPNG

Trim RowsRemove SingletonsSave Fasta

CL\_2435


CL\_2435


CL\_2434


CL\_2435


CL\_2435


CL\_2435


CL\_2434


CL\_2435

HighlightSelectShow Genomes


18

CL\_2438


16

CL\_2438


2

CL\_2438


2

CL\_1740


1

CL\_2438


1

CL\_2438


1

CL\_2438


1

CL\_2438

fGI ID


CL\_INS\_181
CL\_INS\_181
CL\_INS\_182
CL\_INS\_181
CL\_INS\_182
CL\_INS\_182
CL\_INS\_181
CL\_INS\_181
CL\_INS\_182
CL\_INS\_181
CL\_INS\_182
CL\_INS\_181
CL\_INS\_181
CL\_INS\_181
CL\_INS\_181
CL\_INS\_181
CL\_INS\_181
CL\_INS\_181
CL\_INS\_181
CL\_INS\_181
CL\_INS\_181
CL\_INS\_181
CL\_INS\_181
Cluster ID


CL\_2436
CL\_2437
CL\_12133
CL\_5315
CL\_5316
CL\_5317
CL\_9141
CL\_9142
CL\_9143
CL\_9144
CL\_9145
CL\_2934
CL\_2933
CL\_2932
CL\_5621
CL\_5620
CL\_5619
CL\_5618
CL\_5617
CL\_5616
CL\_5615
CL\_5614
CL\_5613
