## Supplementary material for "A novel method for integrating genomic and Tn-Seq data to identify common *in vivo* fitness mechanisms across multiple bacterial species": S5 Dataset: CL_INS_184.html

Legend

 Hypothetical
 All EssentialGenes
 All VFDB Genes
/local/ifs2\_projdata/0800/projects/UMIsepsis/Serra\_Prot\_70/centroids.cluster\_roles.txt

FULL


WINDOWSVGPNG

Trim RowsRemove SingletonsSave Fasta

CL\_5320


CL\_5320

HighlightSelectShow Genomes


34

CL\_2447


3

CL\_2447

fGI ID


CL\_INS\_184
CL\_INS\_184
CL\_INS\_184
CL\_INS\_184
CL\_INS\_184
CL\_INS\_184
CL\_INS\_184
CL\_INS\_193
CL\_INS\_193
CL\_INS\_193
CL\_INS\_193
CL\_INS\_193
Cluster ID


CL\_6882
CL\_6881
CL\_6880
CL\_6879
CL\_6878
CL\_6877
CL\_6876
CL\_2531
CL\_2529
CL\_2528
CL\_2527
CL\_2526
