## Supplementary material for "A novel method for integrating genomic and Tn-Seq data to identify common *in vivo* fitness mechanisms across multiple bacterial species": S5 Dataset: CL_INS_189.html

Legend

 Hypothetical
 All VFDB Genes
/local/ifs2\_projdata/0800/projects/UMIsepsis/Serra\_Prot\_70/centroids.cluster\_roles.txt

FULL


WINDOWSVGPNG

Trim RowsRemove SingletonsSave Fasta

CL\_2502


CL\_2502


CL\_2497


CL\_2497

HighlightSelectShow Genomes


26

CL\_2504


13

CL\_2504


3

CL\_2504


1

CL\_2504

fGI ID


CL\_INS\_189
CL\_INS\_189
Cluster ID


CL\_2503
CL\_10065
