## Supplementary material for "A novel method for integrating genomic and Tn-Seq data to identify common *in vivo* fitness mechanisms across multiple bacterial species": S5 Dataset: CL_INS_190.html

Legend

 Hypothetical
 AntibioticResistance
 All VFDB Genes

FULL


WINDOWSVGPNG

Trim RowsRemove SingletonsSave Fasta

CL\_2507


CL\_2507


CL\_2507


CL\_2507


CL\_2507


CL\_2507


CL\_2507


CL\_2507


CL\_2507


CL\_2507


CL\_2507


CL\_2507

HighlightSelectShow Genomes


18

CL\_2520


15

CL\_2520


2

CL\_2520


2

CL\_2520


1

CL\_2520


1

CL\_2520


1

CL\_2520


1

CL\_2520


1

CL\_2520


1

CL\_2521


1

CL\_2520


1

CL\_2520

fGI ID


CL\_INS\_190
CL\_INS\_190
CL\_INS\_190
CL\_INS\_190
CL\_INS\_190
CL\_INS\_190
CL\_INS\_190
CL\_INS\_190
CL\_INS\_190
CL\_INS\_190
CL\_INS\_190
CL\_INS\_190
CL\_INS\_190
CL\_INS\_190
CL\_INS\_190
CL\_INS\_190
CL\_INS\_190
CL\_INS\_190
CL\_INS\_190
CL\_INS\_190
Cluster ID


CL\_11839
CL\_11840
CL\_11841
CL\_11842
CL\_11843
CL\_2508
CL\_7940
CL\_2509
CL\_6561
CL\_2510
CL\_2511
CL\_2512
CL\_2513
CL\_2514
CL\_2515
CL\_2516
CL\_2517
CL\_12222
CL\_2518
CL\_2519
