## Supplementary material for "A novel method for integrating genomic and Tn-Seq data to identify common *in vivo* fitness mechanisms across multiple bacterial species": S5 Dataset: CL_INS_196.html

Legend

 Hypothetical
 All VFDB Genes
/local/ifs2\_projdata/0800/projects/UMIsepsis/Serra\_Prot\_70/centroids.cluster\_roles.txt

FULL


WINDOWSVGPNG

Trim RowsRemove SingletonsSave Fasta

CL\_2557


CL\_2557


CL\_2557


CL\_2557


CL\_2557


CL\_2557


CL\_2557


CL\_2557


CL\_2557


CL\_2557


CL\_2557


CL\_2557


CL\_2557

HighlightSelectShow Genomes


20

CL\_2565


7

CL\_2565


4

CL\_2565


4

CL\_2565


3

CL\_2565


3

CL\_2565


1

CL\_2565


1

CL\_2565


1

CL\_2565


1

CL\_2565


1

CL\_2565


1

CL\_2565


1

CL\_2565

fGI ID


CL\_INS\_196
CL\_INS\_196
CL\_INS\_196
CL\_INS\_196
CL\_INS\_196
CL\_INS\_196
CL\_INS\_196
CL\_INS\_196
CL\_INS\_196
CL\_INS\_196
CL\_INS\_196
CL\_INS\_196
CL\_INS\_196
CL\_INS\_196
CL\_INS\_196
CL\_INS\_196
CL\_INS\_196
CL\_INS\_196
CL\_INS\_196
CL\_INS\_196
CL\_INS\_196
CL\_INS\_196
CL\_INS\_196
CL\_INS\_196
CL\_INS\_196
CL\_INS\_196
CL\_INS\_196
CL\_INS\_196
CL\_INS\_196
CL\_INS\_345
CL\_INS\_196
CL\_INS\_196
CL\_INS\_196
CL\_INS\_196
CL\_INS\_196
CL\_INS\_196
CL\_INS\_196
Cluster ID


CL\_5328
CL\_5329
CL\_5330
CL\_5331
CL\_5332
CL\_5333
CL\_5334
CL\_5335
CL\_5336
CL\_2558
CL\_2559
CL\_2560
CL\_2561
CL\_2562
CL\_5337
CL\_10069
CL\_2563
CL\_2564
CL\_6269
CL\_6270
CL\_6271
CL\_6272
CL\_6273
CL\_6274
CL\_6275
CL\_6276
CL\_6277
CL\_5606
CL\_6567
CL\_5528
CL\_5527
CL\_6278
CL\_6279
CL\_6280
CL\_6281
CL\_6282
CL\_6283
