## Supplementary material for "A novel method for integrating genomic and Tn-Seq data to identify common *in vivo* fitness mechanisms across multiple bacterial species": S5 Dataset: CL_INS_199.html

Legend

 Hypothetical
 All EssentialGenes
 All VFDB Genes
/local/ifs2\_projdata/0800/projects/UMIsepsis/Serra\_Prot\_70/centroids.cluster\_roles.txt

FULL


WINDOWSVGPNG

Trim RowsRemove SingletonsSave Fasta

CL\_2616


CL\_2616


CL\_2616


CL\_2616


CL\_2584


CL\_2583


CL\_2616


CL\_2616


CL\_2584


CL\_2616


CL\_2584


CL\_2616


CL\_2616


CL\_2616


CL\_2616


CL\_2616


CL\_2584


CL\_2584


CL\_2616


CL\_2616


CL\_2616


CL\_2616


CL\_2584


CL\_2584


CL\_2616


CL\_2616

HighlightSelectShow Genomes


12

CL\_2617


4

CL\_2617


3

CL\_2617


3

CL\_2617


3

CL\_2617


2

CL\_2617


1

CL\_2617


1

CL\_2617


1

CL\_2617


1

CL\_2617


1

CL\_2617


1

CL\_2617


1

CL\_2617


1

CL\_2617


1

CL\_2617


1

CL\_2617


1

CL\_2617


1

CL\_2617


1

CL\_2617


1

CL\_2617


1

CL\_2618


1

CL\_2617


1

CL\_2617


1

CL\_2617


1

CL\_2617


1

CL\_2617

fGI ID


CL\_INS\_199
CL\_INS\_199
CL\_INS\_199
CL\_INS\_199
CL\_INS\_199
CL\_INS\_199
CL\_INS\_199
CL\_INS\_304
CL\_INS\_304
CL\_INS\_199
CL\_INS\_8
CL\_INS\_199
CL\_INS\_199
CL\_INS\_199
CL\_INS\_199
CL\_INS\_199
CL\_INS\_199
CL\_INS\_199
CL\_INS\_199
CL\_INS\_358
CL\_INS\_199
CL\_INS\_199
CL\_INS\_199
CL\_INS\_199
CL\_INS\_199
CL\_INS\_199
CL\_INS\_199
CL\_INS\_199
CL\_INS\_199
CL\_INS\_199
CL\_INS\_199
CL\_INS\_199
CL\_INS\_199
CL\_INS\_199
CL\_INS\_199
CL\_INS\_199
CL\_INS\_199
CL\_INS\_199
CL\_INS\_199
CL\_INS\_199
CL\_INS\_199
CL\_INS\_199
CL\_INS\_199
CL\_INS\_199
CL\_INS\_199
CL\_INS\_199
CL\_INS\_199
CL\_INS\_199
CL\_INS\_199
CL\_INS\_199
CL\_INS\_199
CL\_INS\_199
CL\_INS\_199
CL\_INS\_199
CL\_INS\_199
CL\_INS\_199
CL\_INS\_8
CL\_INS\_8
CL\_INS\_199
CL\_INS\_199
CL\_INS\_199
CL\_INS\_199
CL\_INS\_199
CL\_INS\_199
CL\_INS\_199
CL\_INS\_208
CL\_INS\_208
CL\_INS\_208
CL\_INS\_208
CL\_INS\_208
CL\_INS\_377
CL\_INS\_199
Cluster ID


CL\_7149
CL\_5338
CL\_5339
CL\_11042
CL\_11043
CL\_11975
CL\_11976
CL\_5480
CL\_8470
CL\_9733
CL\_279
CL\_8512
CL\_8513
CL\_8514
CL\_8515
CL\_8516
CL\_10549
CL\_10550
CL\_10551
CL\_9516
CL\_10552
CL\_7931
CL\_7930
CL\_7929
CL\_9325
CL\_9522
CL\_9521
CL\_12375
CL\_12376
CL\_12377
CL\_12378
CL\_12379
CL\_11154
CL\_11155
CL\_11156
CL\_11157
CL\_11158
CL\_11159
CL\_11160
CL\_11161
CL\_11162
CL\_11163
CL\_11164
CL\_11165
CL\_9174
CL\_9175
CL\_9176
CL\_9177
CL\_9178
CL\_9179
CL\_9180
CL\_9181
CL\_9182
CL\_9183
CL\_9184
CL\_6920
CL\_280
CL\_281
CL\_6284
CL\_6921
CL\_6922
CL\_6923
CL\_6924
CL\_6925
CL\_10662
CL\_7148
CL\_6007
CL\_6008
CL\_7147
CL\_6009
CL\_7055
CL\_6926
