## Supplementary material for "A novel method for integrating genomic and Tn-Seq data to identify common *in vivo* fitness mechanisms across multiple bacterial species": S5 Dataset: CL_INS_204.html

Legend

 Hypothetical
 All VFDB Genes
/local/ifs2\_projdata/0800/projects/UMIsepsis/Serra\_Prot\_70/centroids.cluster\_roles.txt

FULL


WINDOWSVGPNG

Trim RowsRemove SingletonsSave Fasta

CL\_2647


CL\_2647


CL\_2647


CL\_2647


CL\_2647


CL\_2647


CL\_2647

HighlightSelectShow Genomes


35

CL\_2648


5

CL\_2651


2

CL\_2649


1

CL\_2651


1

CL\_2651


1

CL\_2648


1

CL\_2649

fGI ID


CL\_INS\_204
CL\_INS\_204
CL\_INS\_204
CL\_INS\_204
CL\_INS\_204
CL\_INS\_204
CL\_INS\_204
CL\_INS\_204
CL\_INS\_204
CL\_INS\_204
CL\_INS\_204
CL\_INS\_204
CL\_INS\_204
CL\_INS\_204
CL\_INS\_204
CL\_INS\_204
CL\_INS\_204
CL\_INS\_204
CL\_INS\_204
CL\_INS\_204
CL\_INS\_204
CL\_INS\_204
CL\_INS\_204
CL\_INS\_204
CL\_INS\_204
CL\_INS\_204
CL\_INS\_204
CL\_INS\_204
CL\_INS\_204
CL\_INS\_204
CL\_INS\_204
CL\_INS\_204
CL\_INS\_204
CL\_INS\_204
CL\_INS\_204
CL\_INS\_204
CL\_INS\_204
CL\_INS\_204
CL\_INS\_204
CL\_INS\_204
CL\_INS\_204
CL\_INS\_204
CL\_INS\_204
CL\_INS\_204
CL\_INS\_204
CL\_INS\_204
CL\_INS\_204
CL\_INS\_204
CL\_INS\_204
CL\_INS\_204
CL\_INS\_204
CL\_INS\_204
CL\_INS\_204
CL\_INS\_204
CL\_INS\_204
CL\_INS\_204
CL\_INS\_204
Cluster ID


CL\_6927
CL\_6928
CL\_6929
CL\_6930
CL\_6931
CL\_6932
CL\_6933
CL\_6934
CL\_5983
CL\_5984
CL\_5985
CL\_5986
CL\_5987
CL\_5988
CL\_5989
CL\_5990
CL\_5991
CL\_5992
CL\_5993
CL\_5994
CL\_5995
CL\_5996
CL\_5997
CL\_5998
CL\_9902
CL\_9903
CL\_9904
CL\_9905
CL\_9906
CL\_9907
CL\_9908
CL\_9909
CL\_9910
CL\_9911
CL\_9912
CL\_9913
CL\_9914
CL\_2601
CL\_7934
CL\_2604
CL\_2605
CL\_9168
CL\_2607
CL\_9915
CL\_9916
CL\_9917
CL\_9918
CL\_9919
CL\_9920
CL\_9921
CL\_9922
CL\_9923
CL\_7928
CL\_7927
CL\_7926
CL\_9520
CL\_9519
