## Supplementary material for "A novel method for integrating genomic and Tn-Seq data to identify common *in vivo* fitness mechanisms across multiple bacterial species": S5 Dataset: CL_INS_207.html

Legend

 Hypothetical
 All VFDB Genes
/local/ifs2\_projdata/0800/projects/UMIsepsis/Serra\_Prot\_70/centroids.cluster\_roles.txt

FULL


WINDOWSVGPNG

Trim RowsRemove SingletonsSave Fasta

CL\_2667


CL\_2667


CL\_2667


CL\_2667


CL\_2667


CL\_2667


CL\_2667


CL\_2667


CL\_2667


CL\_2667


CL\_2667


CL\_2667


CL\_2653


CL\_2667


CL\_2667


CL\_1740


CL\_2667


CL\_2667


CL\_2667


CL\_2653


CL\_2667


CL\_2653

HighlightSelectShow Genomes


6

CL\_2669


6

CL\_2669


6

CL\_2669


4

CL\_2669


3

CL\_2669


3

CL\_2669


2

CL\_2669


2

CL\_2669


2

CL\_2669


1

CL\_2673


1

CL\_2669


1

CL\_2669


1

CL\_2669


1

CL\_2669


1

CL\_2669


1

CL\_2669


1

CL\_2669


1

CL\_1739


1

CL\_2669


1

CL\_2669


1

CL\_2669


1

CL\_2669

fGI ID


CL\_INS\_207
CL\_INS\_207
CL\_INS\_82
CL\_INS\_207
CL\_INS\_207
CL\_INS\_207
CL\_INS\_207
CL\_INS\_207
CL\_INS\_207
CL\_INS\_207
CL\_INS\_207
CL\_INS\_207
CL\_INS\_207
CL\_INS\_207
CL\_INS\_207
Cluster ID


CL\_2668
CL\_11395
CL\_11396
CL\_5344
CL\_8430
CL\_8429
CL\_6285
CL\_6000
CL\_6001
CL\_6002
CL\_6003
CL\_6004
CL\_6005
CL\_6006
CL\_7923
