## Supplementary material for "A novel method for integrating genomic and Tn-Seq data to identify common *in vivo* fitness mechanisms across multiple bacterial species": S5 Dataset: CL_INS_208.html

Legend

 Hypothetical
 All VFDB Genes
/local/ifs2\_projdata/0800/projects/UMIsepsis/Serra\_Prot\_70/centroids.cluster\_roles.txt

FULL


WINDOWSVGPNG

Trim RowsRemove SingletonsSave Fasta

CL\_2670


CL\_2670


CL\_2670


CL\_2670


CL\_2653


CL\_2670


CL\_2670


CL\_2670


CL\_2670


CL\_2670


CL\_2670


CL\_2670


CL\_2670


CL\_2670


CL\_2670


CL\_2670


CL\_2670


CL\_2670


CL\_2670


CL\_2670


CL\_2670


CL\_2670


CL\_2670


CL\_2670


CL\_2670


CL\_2670


CL\_2670


CL\_2670


CL\_2670


CL\_2670


CL\_2670


CL\_2670


CL\_2670


CL\_2667


CL\_2670


CL\_2670


CL\_2670

HighlightSelectShow Genomes


4

CL\_2673


3

CL\_2673


2

CL\_2673


2

CL\_2673


2

CL\_2673


2

CL\_2673


2

CL\_2673


2

CL\_2673


1

CL\_2673


1

CL\_2673


1

CL\_2673


1

CL\_2673


1

CL\_2673


1

CL\_2673


1

CL\_2673


1

CL\_2673


1

CL\_2673


1

CL\_2673


1

CL\_2673


1

CL\_2673


1

CL\_2673


1

CL\_2673


1

CL\_2673


1

CL\_2673


1

CL\_2673


1

CL\_2673


1

CL\_2673


1

CL\_2673


1

CL\_2673


1

CL\_2673


1

CL\_2673


1

CL\_2673


1

CL\_2673


1

CL\_2673


1

CL\_2673


1

CL\_2673


1

CL\_2673

fGI ID


CL\_INS\_208
CL\_INS\_208
CL\_INS\_208
CL\_INS\_208
CL\_INS\_208
CL\_INS\_208
CL\_INS\_377
CL\_INS\_208
CL\_INS\_208
CL\_INS\_208
CL\_INS\_208
CL\_INS\_208
CL\_INS\_208
CL\_INS\_208
CL\_INS\_208
CL\_INS\_208
CL\_INS\_208
CL\_INS\_208
CL\_INS\_208
CL\_INS\_208
CL\_INS\_208
CL\_INS\_208
CL\_INS\_358
CL\_INS\_208
CL\_INS\_208
CL\_INS\_208
CL\_INS\_208
CL\_INS\_208
CL\_INS\_208
CL\_INS\_208
CL\_INS\_208
CL\_INS\_208
CL\_INS\_208
CL\_INS\_208
CL\_INS\_208
CL\_INS\_208
CL\_INS\_208
CL\_INS\_208
CL\_INS\_208
CL\_INS\_208
CL\_INS\_208
CL\_INS\_208
CL\_INS\_208
CL\_INS\_208
CL\_INS\_208
CL\_INS\_208
CL\_INS\_208
CL\_INS\_208
CL\_INS\_208
CL\_INS\_208
CL\_INS\_207
CL\_INS\_208
CL\_INS\_208
CL\_INS\_208
CL\_INS\_208
CL\_INS\_208
CL\_INS\_208
CL\_INS\_208
CL\_INS\_208
CL\_INS\_208
CL\_INS\_207
CL\_INS\_207
CL\_INS\_208
CL\_INS\_207
CL\_INS\_208
CL\_INS\_208
CL\_INS\_208
CL\_INS\_208
CL\_INS\_208
CL\_INS\_208
CL\_INS\_208
CL\_INS\_208
CL\_INS\_208
CL\_INS\_208
CL\_INS\_208
CL\_INS\_208
CL\_INS\_208
CL\_INS\_208
CL\_INS\_208
CL\_INS\_208
CL\_INS\_208
CL\_INS\_208
CL\_INS\_208
CL\_INS\_208
CL\_INS\_208
CL\_INS\_208
CL\_INS\_208
CL\_INS\_208
CL\_INS\_208
CL\_INS\_208
CL\_INS\_208
CL\_INS\_208
CL\_INS\_208
CL\_INS\_208
CL\_INS\_208
CL\_INS\_208
Cluster ID


CL\_6935
CL\_6286
CL\_11977
CL\_5602
CL\_7922
CL\_10553
CL\_5821
CL\_11044
CL\_8677
CL\_2671
CL\_2672
CL\_11844
CL\_11845
CL\_5345
CL\_5346
CL\_11166
CL\_11167
CL\_12223
CL\_12224
CL\_9734
CL\_9735
CL\_9517
CL\_9516
CL\_8903
CL\_8904
CL\_10075
CL\_10076
CL\_11521
CL\_11522
CL\_10832
CL\_10833
CL\_8734
CL\_8733
CL\_8732
CL\_7271
CL\_7272
CL\_8517
CL\_8428
CL\_8427
CL\_8426
CL\_10663
CL\_11300
CL\_6569
CL\_6570
CL\_11137
CL\_11138
CL\_11139
CL\_12380
CL\_12381
CL\_12382
CL\_6004
CL\_10894
CL\_10895
CL\_10896
CL\_10897
CL\_7148
CL\_6007
CL\_6008
CL\_7147
CL\_6009
CL\_6006
CL\_7923
CL\_10664
CL\_6005
CL\_9185
CL\_9186
CL\_9187
CL\_9188
CL\_10177
CL\_10178
CL\_10179
CL\_10180
CL\_10181
CL\_10182
CL\_10183
CL\_10184
CL\_10185
CL\_10186
CL\_6571
CL\_10187
CL\_10328
CL\_10329
CL\_10330
CL\_10331
CL\_10332
CL\_10333
CL\_10334
CL\_10335
CL\_10336
CL\_10337
CL\_10338
CL\_10339
CL\_10340
CL\_10341
CL\_10342
CL\_10343
