## Supplementary material for "A novel method for integrating genomic and Tn-Seq data to identify common *in vivo* fitness mechanisms across multiple bacterial species": S5 Dataset: CL_INS_213.html

Legend

 Hypothetical
 All VFDB Genes
/local/ifs2\_projdata/0800/projects/UMIsepsis/Serra\_Prot\_70/centroids.cluster\_roles.txt

FULL


WINDOWSVGPNG

Trim RowsRemove SingletonsSave Fasta

CL\_2720


CL\_2720


CL\_2720

HighlightSelectShow Genomes


35

CL\_7145


1

CL\_2721


1

CL\_7145

fGI ID


CL\_INS\_213
CL\_INS\_213
CL\_INS\_213
CL\_INS\_214
CL\_INS\_214
CL\_INS\_214
CL\_INS\_214
CL\_INS\_214
CL\_INS\_214
CL\_INS\_213
CL\_INS\_213
CL\_INS\_213
CL\_INS\_213
CL\_INS\_213
CL\_INS\_213
CL\_INS\_213
CL\_INS\_213
CL\_INS\_213
CL\_INS\_214
CL\_INS\_214
CL\_INS\_214
CL\_INS\_214
CL\_INS\_214
CL\_INS\_214
CL\_INS\_214
CL\_INS\_27
CL\_INS\_214
CL\_INS\_214
CL\_INS\_214
CL\_INS\_214
CL\_INS\_214
CL\_INS\_213
CL\_INS\_213
CL\_INS\_213
CL\_INS\_213
CL\_INS\_213
CL\_INS\_213
CL\_INS\_213
CL\_INS\_213
CL\_INS\_213
CL\_INS\_213
CL\_INS\_213
CL\_INS\_213
CL\_INS\_213
CL\_INS\_213
CL\_INS\_213
CL\_INS\_213
CL\_INS\_213
CL\_INS\_213
CL\_INS\_213
CL\_INS\_214
CL\_INS\_213
CL\_INS\_214
Cluster ID


CL\_12543
CL\_12544
CL\_5349
CL\_5350
CL\_5351
CL\_5352
CL\_5353
CL\_5354
CL\_5355
CL\_12545
CL\_12546
CL\_12547
CL\_12548
CL\_12549
CL\_12550
CL\_12551
CL\_12552
CL\_12553
CL\_5356
CL\_5357
CL\_5358
CL\_5359
CL\_5360
CL\_5361
CL\_5362
CL\_5363
CL\_5364
CL\_5365
CL\_5366
CL\_5367
CL\_5368
CL\_12554
CL\_12555
CL\_12556
CL\_12557
CL\_12558
CL\_12559
CL\_12560
CL\_12561
CL\_12562
CL\_12563
CL\_12564
CL\_12565
CL\_12566
CL\_12567
CL\_12568
CL\_11406
CL\_12569
CL\_12570
CL\_11409
CL\_5369
CL\_12571
CL\_5370
