## Supplementary material for "A novel method for integrating genomic and Tn-Seq data to identify common *in vivo* fitness mechanisms across multiple bacterial species": S5 Dataset: CL_INS_218.html

Legend

 Serritia FitnessGene
 All VFDB Genes

FULL


WINDOWSVGPNG

Trim RowsRemove SingletonsSave Fasta

CL\_5373


CL\_5373


CL\_5373


CL\_5373


CL\_2778


CL\_2778


CL\_5373

HighlightSelectShow Genomes


24

CL\_2780


14

CL\_2780


3

CL\_2780


2

CL\_2780


1

CL\_2780


1

CL\_2780


1

CL\_2780

fGI ID


CL\_INS\_218
CL\_INS\_218
CL\_INS\_218
Cluster ID


CL\_9190
CL\_2779
CL\_6347
