## Supplementary material for "A novel method for integrating genomic and Tn-Seq data to identify common *in vivo* fitness mechanisms across multiple bacterial species": S5 Dataset: CL_INS_220.html

Legend

 Hypothetical
 Serritia FitnessGene
 All VFDB Genes
/local/ifs2\_projdata/0800/projects/UMIsepsis/Serra\_Prot\_70/centroids.cluster\_roles.txt

FULL


WINDOWSVGPNG

Trim RowsRemove SingletonsSave Fasta

CL\_2818


CL\_2818


CL\_2818


CL\_2818


CL\_2818


CL\_2818


CL\_2818


CL\_2818


CL\_2818


CL\_2818


CL\_2818


CL\_2815


CL\_2818


CL\_2818


CL\_2818

HighlightSelectShow Genomes


12

CL\_2821


8

CL\_2821


8

CL\_2821


4

CL\_2821


4

CL\_2821


2

CL\_2821


1

CL\_2821


1

CL\_2821


1

CL\_2821


1

CL\_2821


1

CL\_2821


1

CL\_2821


1

CL\_2821


1

CL\_2821


1

Break

fGI ID


CL\_INS\_220
CL\_INS\_220
CL\_INS\_220
CL\_INS\_220
CL\_INS\_220
CL\_INS\_220
CL\_INS\_220
CL\_INS\_220
Cluster ID


CL\_11398
CL\_6576
CL\_12337
CL\_5601
CL\_7144
CL\_2819
CL\_7902
CL\_2820
