## Supplementary material for "A novel method for integrating genomic and Tn-Seq data to identify common *in vivo* fitness mechanisms across multiple bacterial species": S5 Dataset: CL_INS_222.html

Legend

 Hypothetical
 All VFDB Genes
/local/ifs2\_projdata/0800/projects/UMIsepsis/Serra\_Prot\_70/centroids.cluster\_roles.txt

FULL


WINDOWSVGPNG

Trim RowsRemove SingletonsSave Fasta

CL\_2893


CL\_2893


CL\_2893


CL\_2893


CL\_2893


CL\_2893


CL\_2893


CL\_2893


CL\_2893

HighlightSelectShow Genomes


36

CL\_2894


2

CL\_2894


2

CL\_2894


1

CL\_2894


1

CL\_2894


1

CL\_2894


1

CL\_2894


1

CL\_2895


1

CL\_2894

fGI ID


CL\_INS\_222
CL\_INS\_222
CL\_INS\_222
CL\_INS\_222
CL\_INS\_222
CL\_INS\_222
CL\_INS\_222
CL\_INS\_222
CL\_INS\_222
CL\_INS\_222
CL\_INS\_222
CL\_INS\_222
CL\_INS\_222
CL\_INS\_222
CL\_INS\_222
CL\_INS\_222
CL\_INS\_222
CL\_INS\_222
CL\_INS\_222
CL\_INS\_222
CL\_INS\_222
Cluster ID


CL\_5379
CL\_5380
CL\_11045
CL\_9515
CL\_9514
CL\_10344
CL\_11523
CL\_11524
CL\_11525
CL\_10898
CL\_10899
CL\_11168
CL\_11169
CL\_10900
CL\_10901
CL\_10902
CL\_10903
CL\_10904
CL\_10905
CL\_10906
CL\_10907
