## Supplementary material for "A novel method for integrating genomic and Tn-Seq data to identify common *in vivo* fitness mechanisms across multiple bacterial species": S5 Dataset: CL_INS_223.html

Legend

 Hypothetical
 All VFDB Genes

FULL


WINDOWSVGPNG

Trim RowsRemove SingletonsSave Fasta

CL\_2899


CL\_2899


CL\_2899


CL\_2899

HighlightSelectShow Genomes


34

CL\_2900


9

CL\_2900


2

CL\_2900


2

CL\_2900

fGI ID


CL\_INS\_223
CL\_INS\_223
CL\_INS\_223
CL\_INS\_223
Cluster ID


CL\_10085
CL\_7901
CL\_7900
CL\_7899
