## Supplementary material for "A novel method for integrating genomic and Tn-Seq data to identify common *in vivo* fitness mechanisms across multiple bacterial species": S5 Dataset: CL_INS_229.html

Legend

 Hypothetical
 All VFDB Genes
/local/ifs2\_projdata/0800/projects/UMIsepsis/Serra\_Prot\_70/centroids.cluster\_roles.txt

FULL


WINDOWSVGPNG

Trim RowsRemove SingletonsSave Fasta

CL\_2977


CL\_2977


CL\_2977

HighlightSelectShow Genomes


43

CL\_2978


3

CL\_2978


1

CL\_2978

fGI ID


CL\_INS\_229
CL\_INS\_229
CL\_INS\_229
Cluster ID


CL\_11170
CL\_11171
CL\_11172
