## Supplementary material for "A novel method for integrating genomic and Tn-Seq data to identify common *in vivo* fitness mechanisms across multiple bacterial species": S5 Dataset: CL_INS_230.html

Legend

 Hypothetical
 All VFDB Genes
/local/ifs2\_projdata/0800/projects/UMIsepsis/Serra\_Prot\_70/centroids.cluster\_roles.txt

FULL


WINDOWSVGPNG

Trim RowsRemove SingletonsSave Fasta

CL\_2983


CL\_2983


CL\_2983

HighlightSelectShow Genomes


27

CL\_2984


19

CL\_2984


2

CL\_2984

fGI ID


CL\_INS\_230
CL\_INS\_230
CL\_INS\_230
CL\_INS\_230
Cluster ID


CL\_5383
CL\_7139
CL\_6400
CL\_5384
