## Supplementary material for "A novel method for integrating genomic and Tn-Seq data to identify common *in vivo* fitness mechanisms across multiple bacterial species": S5 Dataset: CL_INS_232.html

Legend

 Hypothetical
 All VFDB Genes
/local/ifs2\_projdata/0800/projects/UMIsepsis/Serra\_Prot\_70/centroids.cluster\_roles.txt

FULL


WINDOWSVGPNG

Trim RowsRemove SingletonsSave Fasta

CL\_2990


CL\_2990


CL\_2990


CL\_2990


CL\_2990


CL\_2990


CL\_2989

HighlightSelectShow Genomes


22

CL\_2992


17

CL\_2992


3

CL\_2992


2

CL\_2992


2

CL\_2992


1

CL\_2992


1

CL\_2992

fGI ID


CL\_INS\_232
CL\_INS\_232
CL\_INS\_232
CL\_INS\_232
CL\_INS\_232
CL\_INS\_232
CL\_INS\_232
CL\_INS\_232
CL\_INS\_232
Cluster ID


CL\_7893
CL\_5385
CL\_5386
CL\_5387
CL\_5388
CL\_5389
CL\_5390
CL\_2991
CL\_6349
