## Supplementary material for "A novel method for integrating genomic and Tn-Seq data to identify common *in vivo* fitness mechanisms across multiple bacterial species": S5 Dataset: CL_INS_237.html

Legend

 All VFDB Genes
/local/ifs2\_projdata/0800/projects/UMIsepsis/Serra\_Prot\_70/centroids.cluster\_roles.txt

FULL


WINDOWSVGPNG

Trim RowsRemove SingletonsSave Fasta

CL\_3014


CL\_3014


CL\_3014


CL\_3014


CL\_3014

HighlightSelectShow Genomes


23

CL\_3017


17

CL\_3017


6

CL\_3017


1

CL\_3017


1

CL\_3017

fGI ID


CL\_INS\_237
CL\_INS\_237
CL\_INS\_237
CL\_INS\_237
Cluster ID


CL\_5395
CL\_5396
CL\_3015
CL\_3016
