## Supplementary material for "A novel method for integrating genomic and Tn-Seq data to identify common *in vivo* fitness mechanisms across multiple bacterial species": S5 Dataset: CL_INS_239.html

Legend

 Hypothetical
 All EssentialGenes
/local/ifs2\_projdata/0800/projects/UMIsepsis/Serra\_Prot\_70/centroids.cluster\_roles.txt

FULL


WINDOWSVGPNG

Trim RowsRemove SingletonsSave Fasta

CL\_3023


CL\_3023


CL\_3023


CL\_3022


CL\_3023

HighlightSelectShow Genomes


28

CL\_5397


15

CL\_5397


3

CL\_5397


1

CL\_5397


1

CL\_3026

fGI ID


CL\_INS\_239
CL\_INS\_239
Cluster ID


CL\_3024
CL\_3025
