## Supplementary material for "A novel method for integrating genomic and Tn-Seq data to identify common *in vivo* fitness mechanisms across multiple bacterial species": S5 Dataset: CL_INS_241.html

Legend

 Hypothetical
 All VFDB Genes
/local/ifs2\_projdata/0800/projects/UMIsepsis/Serra\_Prot\_70/centroids.cluster\_roles.txt

FULL


WINDOWSVGPNG

Trim RowsRemove SingletonsSave Fasta

CL\_3032


CL\_3032


CL\_3032


CL\_3032

HighlightSelectShow Genomes


43

CL\_3033


2

CL\_3033


2

CL\_3033


1

CL\_3033

fGI ID


CL\_INS\_241
CL\_INS\_241
CL\_INS\_241
CL\_INS\_241
CL\_INS\_241
CL\_INS\_241
CL\_INS\_241
CL\_INS\_241
CL\_INS\_241
CL\_INS\_241
CL\_INS\_241
CL\_INS\_241
CL\_INS\_241
CL\_INS\_241
CL\_INS\_241
CL\_INS\_241
CL\_INS\_241
CL\_INS\_241
Cluster ID


CL\_12225
CL\_9513
CL\_9512
CL\_9511
CL\_9510
CL\_9509
CL\_9508
CL\_9507
CL\_9506
CL\_9505
CL\_9504
CL\_9503
CL\_9502
CL\_9501
CL\_10345
CL\_9500
CL\_9499
CL\_9498
