## Supplementary material for "A novel method for integrating genomic and Tn-Seq data to identify common *in vivo* fitness mechanisms across multiple bacterial species": S5 Dataset: CL_INS_242.html

Legend

 Hypothetical
 All VFDB Genes
/local/ifs2\_projdata/0800/projects/UMIsepsis/Serra\_Prot\_70/centroids.cluster\_roles.txt

FULL


WINDOWSVGPNG

Trim RowsRemove SingletonsSave Fasta

CL\_3033


CL\_3033


CL\_3033


CL\_3033


CL\_3033


CL\_3033

HighlightSelectShow Genomes


31

CL\_3035


13

CL\_3035


1

CL\_3043


1

CL\_3043


1

CL\_3043


1

CL\_3035

fGI ID


CL\_INS\_242
CL\_INS\_242
CL\_INS\_243
CL\_INS\_243
CL\_INS\_243
CL\_INS\_243
CL\_INS\_243
CL\_INS\_243
CL\_INS\_243
Cluster ID


CL\_10096
CL\_3034
CL\_3037
CL\_6042
CL\_3039
CL\_10188
CL\_12136
CL\_3040
CL\_3041
