## Supplementary material for "A novel method for integrating genomic and Tn-Seq data to identify common *in vivo* fitness mechanisms across multiple bacterial species": S5 Dataset: CL_INS_248.html

Legend

 Hypothetical
 All VFDB Genes
/local/ifs2\_projdata/0800/projects/UMIsepsis/Serra\_Prot\_70/centroids.cluster\_roles.txt

FULL


WINDOWSVGPNG

Trim RowsRemove SingletonsSave Fasta

CL\_3067


CL\_3067


CL\_3067


CL\_3067

HighlightSelectShow Genomes


23

CL\_3068


21

CL\_3068


2

CL\_3068


2

CL\_3068

fGI ID


CL\_INS\_248
CL\_INS\_248
CL\_INS\_248
CL\_INS\_248
CL\_INS\_248
CL\_INS\_248
Cluster ID


CL\_5401
CL\_5402
CL\_5403
CL\_5404
CL\_5405
CL\_5406
