## Supplementary material for "A novel method for integrating genomic and Tn-Seq data to identify common *in vivo* fitness mechanisms across multiple bacterial species": S5 Dataset: CL_INS_249.html

Legend

 Hypothetical
/local/ifs2\_projdata/0800/projects/UMIsepsis/Serra\_Prot\_70/centroids.cluster\_roles.txt

FULL


WINDOWSVGPNG

Trim RowsRemove SingletonsSave Fasta

CL\_3068


CL\_3068


CL\_3068

HighlightSelectShow Genomes


27

CL\_3070


19

CL\_3070


2

CL\_3070

fGI ID


CL\_INS\_249
CL\_INS\_249
Cluster ID


CL\_3069
CL\_5888
