## Supplementary material for "A novel method for integrating genomic and Tn-Seq data to identify common *in vivo* fitness mechanisms across multiple bacterial species": S5 Dataset: CL_INS_253.html

Legend

 Hypothetical
 All VFDB Genes
/local/ifs2\_projdata/0800/projects/UMIsepsis/Serra\_Prot\_70/centroids.cluster\_roles.txt

FULL


WINDOWSVGPNG

Trim RowsRemove SingletonsSave Fasta

CL\_3085


CL\_3085


CL\_3085


CL\_3085

HighlightSelectShow Genomes


32

CL\_3086


10

CL\_3086


3

CL\_3086


1

CL\_3086

fGI ID


CL\_INS\_253
CL\_INS\_253
CL\_INS\_253
CL\_INS\_253
Cluster ID


CL\_7136
CL\_7135
CL\_11423
CL\_11424
