## Supplementary material for "A novel method for integrating genomic and Tn-Seq data to identify common *in vivo* fitness mechanisms across multiple bacterial species": S5 Dataset: CL_INS_257.html

Legend

 Hypothetical
 All VFDB Genes
/local/ifs2\_projdata/0800/projects/UMIsepsis/Serra\_Prot\_70/centroids.cluster\_roles.txt

FULL


WINDOWSVGPNG

Trim RowsRemove SingletonsSave Fasta

CL\_3139


CL\_3139


CL\_3139

HighlightSelectShow Genomes


34

CL\_3140


13

CL\_3140


1

CL\_3140

fGI ID


CL\_INS\_257
CL\_INS\_257
Cluster ID


CL\_7130
CL\_10346
