## Supplementary material for "A novel method for integrating genomic and Tn-Seq data to identify common *in vivo* fitness mechanisms across multiple bacterial species": S5 Dataset: CL_INS_261.html

Legend

 Hypothetical
 All EssentialGenes
/local/ifs2\_projdata/0800/projects/UMIsepsis/Serra\_Prot\_70/centroids.cluster\_roles.txt

FULL


WINDOWSVGPNG

Trim RowsRemove SingletonsSave Fasta

CL\_3201


CL\_3201

HighlightSelectShow Genomes


43

CL\_3200


4

CL\_3200

fGI ID


CL\_INS\_261
CL\_INS\_261
CL\_INS\_261
CL\_INS\_261
CL\_INS\_261
CL\_INS\_261
CL\_INS\_261
CL\_INS\_217
CL\_INS\_217
CL\_INS\_210
CL\_INS\_210
CL\_INS\_210
CL\_INS\_217
CL\_INS\_261
Cluster ID


CL\_11985
CL\_11984
CL\_11983
CL\_10502
CL\_10500
CL\_10499
CL\_11982
CL\_2754
CL\_2753
CL\_2752
CL\_2750
CL\_2749
CL\_2746
CL\_11981
