## Supplementary material for "A novel method for integrating genomic and Tn-Seq data to identify common *in vivo* fitness mechanisms across multiple bacterial species": S5 Dataset: CL_INS_263.html

Legend

 Hypothetical
/local/ifs2\_projdata/0800/projects/UMIsepsis/Serra\_Prot\_70/centroids.cluster\_roles.txt

FULL


WINDOWSVGPNG

Trim RowsRemove SingletonsSave Fasta

CL\_3208


CL\_3208


CL\_3208

HighlightSelectShow Genomes


40

CL\_3209


5

CL\_3211


3

CL\_3209

fGI ID

CL\_INS\_263
Cluster ID

CL\_5425
