## Supplementary material for "A novel method for integrating genomic and Tn-Seq data to identify common *in vivo* fitness mechanisms across multiple bacterial species": S5 Dataset: CL_INS_264.html

Legend

/local/ifs2\_projdata/0800/projects/UMIsepsis/Serra\_Prot\_70/centroids.cluster\_roles.txt

FULL


WINDOWSVGPNG

Trim RowsRemove SingletonsSave Fasta

CL\_3209


CL\_3209


CL\_3208


CL\_3209

HighlightSelectShow Genomes


36

CL\_3211


6

CL\_3211


5

CL\_3211


1

CL\_3211

fGI ID


CL\_INS\_264
CL\_INS\_264
Cluster ID


CL\_3210
CL\_10726
