## Supplementary material for "A novel method for integrating genomic and Tn-Seq data to identify common *in vivo* fitness mechanisms across multiple bacterial species": S5 Dataset: CL_INS_271.html

Legend

 Hypothetical
 All VFDB Genes
/local/ifs2\_projdata/0800/projects/UMIsepsis/Serra\_Prot\_70/centroids.cluster\_roles.txt

FULL


WINDOWSVGPNG

Trim RowsRemove SingletonsSave Fasta

CL\_3322


CL\_3322


CL\_3322


CL\_3321

HighlightSelectShow Genomes


31

CL\_3323


12

CL\_3323


3

CL\_3323


2

CL\_3323

fGI ID


CL\_INS\_271
CL\_INS\_271
CL\_INS\_270
CL\_INS\_270
CL\_INS\_270
Cluster ID


CL\_8374
CL\_6596
CL\_6358
CL\_6359
CL\_6360
