## Supplementary material for "A novel method for integrating genomic and Tn-Seq data to identify common *in vivo* fitness mechanisms across multiple bacterial species": S5 Dataset: CL_INS_272.html

Legend

 Hypothetical
 All VFDB Genes
/local/ifs2\_projdata/0800/projects/UMIsepsis/Serra\_Prot\_70/centroids.cluster\_roles.txt

FULL


WINDOWSVGPNG

Trim RowsRemove SingletonsSave Fasta

CL\_3323


CL\_3323

HighlightSelectShow Genomes


36

CL\_3324


4

CL\_3326

fGI ID


CL\_INS\_272
CL\_INS\_272
CL\_INS\_272
CL\_INS\_272
CL\_INS\_272
CL\_INS\_272
CL\_INS\_272
CL\_INS\_272
CL\_INS\_272
CL\_INS\_272
CL\_INS\_272
CL\_INS\_272
Cluster ID


CL\_11988
CL\_11989
CL\_11990
CL\_11991
CL\_11992
CL\_11993
CL\_11994
CL\_11995
CL\_11996
CL\_11997
CL\_11998
CL\_11999
