## Supplementary material for "A novel method for integrating genomic and Tn-Seq data to identify common *in vivo* fitness mechanisms across multiple bacterial species": S5 Dataset: CL_INS_274.html

Legend

 All VFDB Genes
/local/ifs2\_projdata/0800/projects/UMIsepsis/Serra\_Prot\_70/centroids.cluster\_roles.txt

FULL


WINDOWSVGPNG

Trim RowsRemove SingletonsSave Fasta

CL\_5433


CL\_3446


CL\_3446


CL\_7082


CL\_3446


CL\_3446

HighlightSelectShow Genomes


30

CL\_5916


3

CL\_5916


1

CL\_5916


1

CL\_5916


1

CL\_5916


1

CL\_5916

fGI ID


CL\_INS\_291
CL\_INS\_291
CL\_INS\_291
CL\_INS\_291
CL\_INS\_291
CL\_INS\_274
CL\_INS\_274
CL\_INS\_274
CL\_INS\_274
CL\_INS\_274
CL\_INS\_274
Cluster ID


CL\_6364
CL\_6365
CL\_6366
CL\_6367
CL\_6368
CL\_5922
CL\_5921
CL\_5920
CL\_5919
CL\_5918
CL\_5917
