## Supplementary material for "A novel method for integrating genomic and Tn-Seq data to identify common *in vivo* fitness mechanisms across multiple bacterial species": S5 Dataset: CL_INS_275.html

Legend

 Hypothetical
 All EssentialGenes
 All VFDB Genes
/local/ifs2\_projdata/0800/projects/UMIsepsis/Serra\_Prot\_70/centroids.cluster\_roles.txt

FULL


WINDOWSVGPNG

Trim RowsRemove SingletonsSave Fasta

CL\_5916


CL\_5916


CL\_5916


CL\_5916


CL\_5916


CL\_5916


CL\_3336

HighlightSelectShow Genomes


28

CL\_3339


4

CL\_3447


2

CL\_3447


1

CL\_3339


1

CL\_3339


1

CL\_3339


1

CL\_3339

fGI ID


CL\_INS\_275
CL\_INS\_275
CL\_INS\_275
CL\_INS\_275
CL\_INS\_275
CL\_INS\_275
CL\_INS\_275
CL\_INS\_275
CL\_INS\_275
CL\_INS\_275
CL\_INS\_275
CL\_INS\_275
CL\_INS\_275
CL\_INS\_275
CL\_INS\_275
CL\_INS\_275
CL\_INS\_275
CL\_INS\_275
CL\_INS\_275
CL\_INS\_275
CL\_INS\_275
CL\_INS\_275
Cluster ID


CL\_7865
CL\_5915
CL\_5914
CL\_5913
CL\_5912
CL\_10844
CL\_10845
CL\_10846
CL\_10847
CL\_10848
CL\_10849
CL\_10850
CL\_10851
CL\_10852
CL\_5911
CL\_5910
CL\_5909
CL\_5908
CL\_5907
CL\_3337
CL\_3338
CL\_6598
