## Supplementary material for "A novel method for integrating genomic and Tn-Seq data to identify common *in vivo* fitness mechanisms across multiple bacterial species": S5 Dataset: CL_INS_280.html

Legend

 Hypothetical
 All VFDB Genes
/local/ifs2\_projdata/0800/projects/UMIsepsis/Serra\_Prot\_70/centroids.cluster\_roles.txt

FULL


WINDOWSVGPNG

Trim RowsRemove SingletonsSave Fasta

CL\_3353


CL\_3353


CL\_3352


CL\_3353


CL\_3352


CL\_3353


CL\_3352


CL\_3353

HighlightSelectShow Genomes


19

CL\_3354


11

CL\_3354


9

CL\_3354


5

CL\_3354


1

CL\_3354


1

CL\_3354


1

CL\_3354


1

CL\_3355

fGI ID


CL\_INS\_280
CL\_INS\_279
CL\_INS\_279
CL\_INS\_280
CL\_INS\_280
Cluster ID


CL\_7297
CL\_10728
CL\_5435
CL\_5436
CL\_5437
