## Supplementary material for "A novel method for integrating genomic and Tn-Seq data to identify common *in vivo* fitness mechanisms across multiple bacterial species": S5 Dataset: CL_INS_282.html

Legend

 Hypothetical
 All EssentialGenes
 All VFDB Genes
/local/ifs2\_projdata/0800/projects/UMIsepsis/Serra\_Prot\_70/centroids.cluster\_roles.txt

FULL


WINDOWSVGPNG

Trim RowsRemove SingletonsSave Fasta

CL\_3357


CL\_3355


CL\_3357


CL\_3357


CL\_3355


CL\_3357


CL\_3357


CL\_3355


CL\_3357


CL\_3357


CL\_3355


CL\_3357


CL\_3357


CL\_3357


CL\_3355


CL\_3357


CL\_3355

HighlightSelectShow Genomes


23

CL\_3358


8

CL\_3358


2

CL\_3358


2

CL\_3358


2

CL\_3358


1

CL\_3358


1

CL\_3358


1

CL\_3358


1

CL\_4235


1

CL\_3358


1

CL\_3358


1

CL\_3358


1

CL\_3358


1

CL\_3358


1

CL\_3358


1

CL\_3358


1

CL\_3358

fGI ID


CL\_INS\_282
CL\_INS\_358
CL\_INS\_358
CL\_INS\_358
CL\_INS\_282
CL\_INS\_282
CL\_INS\_282
CL\_INS\_282
CL\_INS\_282
CL\_INS\_282
CL\_INS\_282
CL\_INS\_282
CL\_INS\_282
CL\_INS\_298
CL\_INS\_298
CL\_INS\_298
CL\_INS\_282
CL\_INS\_282
CL\_INS\_282
CL\_INS\_282
CL\_INS\_282
CL\_INS\_282
CL\_INS\_282
CL\_INS\_282
CL\_INS\_282
CL\_INS\_282
CL\_INS\_282
CL\_INS\_282
CL\_INS\_282
CL\_INS\_282
CL\_INS\_282
CL\_INS\_282
CL\_INS\_282
CL\_INS\_282
CL\_INS\_282
CL\_INS\_282
CL\_INS\_282
CL\_INS\_282
CL\_INS\_282
CL\_INS\_282
CL\_INS\_282
CL\_INS\_282
CL\_INS\_282
CL\_INS\_282
CL\_INS\_282
CL\_INS\_282
CL\_INS\_282
CL\_INS\_282
CL\_INS\_282
CL\_INS\_282
CL\_INS\_259
CL\_INS\_282
CL\_INS\_282
CL\_INS\_304
CL\_INS\_282
CL\_INS\_282
CL\_INS\_282
CL\_INS\_282
CL\_INS\_282
CL\_INS\_282
CL\_INS\_282
CL\_INS\_282
CL\_INS\_282
CL\_INS\_282
CL\_INS\_282
CL\_INS\_282
CL\_INS\_282
CL\_INS\_282
CL\_INS\_282
CL\_INS\_282
CL\_INS\_282
CL\_INS\_282
CL\_INS\_282
CL\_INS\_282
CL\_INS\_282
CL\_INS\_282
CL\_INS\_282
CL\_INS\_282
CL\_INS\_282
CL\_INS\_282
CL\_INS\_282
CL\_INS\_282
CL\_INS\_282
CL\_INS\_282
CL\_INS\_282
CL\_INS\_282
CL\_INS\_282
CL\_INS\_282
CL\_INS\_282
CL\_INS\_282
CL\_INS\_282
CL\_INS\_282
CL\_INS\_282
CL\_INS\_282
CL\_INS\_282
CL\_INS\_282
CL\_INS\_282
CL\_INS\_282
CL\_INS\_282
CL\_INS\_282
CL\_INS\_282
CL\_INS\_282
CL\_INS\_282
CL\_INS\_282
CL\_INS\_282
CL\_INS\_282
CL\_INS\_282
CL\_INS\_282
CL\_INS\_282
CL\_INS\_282
CL\_INS\_282
CL\_INS\_282
CL\_INS\_282
CL\_INS\_282
CL\_INS\_282
CL\_INS\_282
CL\_INS\_282
CL\_INS\_282
CL\_INS\_282
CL\_INS\_282
CL\_INS\_282
CL\_INS\_282
Cluster ID


CL\_11085
CL\_11084
CL\_11083
CL\_11082
CL\_9496
CL\_9495
CL\_12238
CL\_12239
CL\_12240
CL\_12241
CL\_8924
CL\_8925
CL\_12014
CL\_5439
CL\_5440
CL\_5441
CL\_10114
CL\_5442
CL\_5443
CL\_5444
CL\_10115
CL\_11173
CL\_11174
CL\_11175
CL\_11176
CL\_11177
CL\_11178
CL\_11179
CL\_11066
CL\_11067
CL\_11068
CL\_11069
CL\_11070
CL\_11071
CL\_11072
CL\_11073
CL\_11074
CL\_11075
CL\_11076
CL\_11077
CL\_11078
CL\_11079
CL\_11080
CL\_11545
CL\_11546
CL\_11547
CL\_8926
CL\_8927
CL\_8928
CL\_8929
CL\_10431
CL\_6600
CL\_12137
CL\_6601
CL\_8930
CL\_11548
CL\_8931
CL\_8932
CL\_12015
CL\_8933
CL\_6602
CL\_6603
CL\_11549
CL\_11550
CL\_11551
CL\_11552
CL\_11553
CL\_11554
CL\_11555
CL\_11556
CL\_6604
CL\_11557
CL\_8934
CL\_6605
CL\_11558
CL\_12016
CL\_12017
CL\_7884
CL\_7883
CL\_7882
CL\_12138
CL\_8419
CL\_8418
CL\_7881
CL\_7880
CL\_12139
CL\_12140
CL\_7879
CL\_7878
CL\_7877
CL\_12141
CL\_7876
CL\_7875
CL\_7874
CL\_11697
CL\_7873
CL\_7872
CL\_11696
CL\_11695
CL\_6606
CL\_6607
CL\_6608
CL\_8935
CL\_12018
CL\_11559
CL\_7871
CL\_7870
CL\_6609
CL\_12747
CL\_12748
CL\_12749
CL\_12750
CL\_12751
CL\_12752
CL\_12753
CL\_12754
CL\_12755
CL\_12756
CL\_12757
CL\_12758
CL\_12759
CL\_12760
