## Supplementary material for "A novel method for integrating genomic and Tn-Seq data to identify common *in vivo* fitness mechanisms across multiple bacterial species": S5 Dataset: CL_INS_286.html

Legend

 Hypothetical
 All VFDB Genes
/local/ifs2\_projdata/0800/projects/UMIsepsis/Serra\_Prot\_70/centroids.cluster\_roles.txt

FULL


WINDOWSVGPNG

Trim RowsRemove SingletonsSave Fasta

CL\_3380


CL\_3380


CL\_3380

HighlightSelectShow Genomes


28

CL\_3386


16

CL\_3386


1

CL\_3386

fGI ID


CL\_INS\_286
CL\_INS\_286
CL\_INS\_286
CL\_INS\_286
CL\_INS\_286
Cluster ID


CL\_3381
CL\_3382
CL\_3383
CL\_3384
CL\_3385
