## Supplementary material for "A novel method for integrating genomic and Tn-Seq data to identify common *in vivo* fitness mechanisms across multiple bacterial species": S5 Dataset: CL_INS_288.html

Legend

 Hypothetical
 All VFDB Genes
/local/ifs2\_projdata/0800/projects/UMIsepsis/Serra\_Prot\_70/centroids.cluster\_roles.txt

FULL


WINDOWSVGPNG

Trim RowsRemove SingletonsSave Fasta

CL\_3412


CL\_3412


CL\_3412

HighlightSelectShow Genomes


39

CL\_3413


7

CL\_3413


2

CL\_3413

fGI ID


CL\_INS\_288
CL\_INS\_288
Cluster ID


CL\_9755
CL\_9199
