## Supplementary material for "A novel method for integrating genomic and Tn-Seq data to identify common *in vivo* fitness mechanisms across multiple bacterial species": S5 Dataset: CL_INS_289.html

Legend

 All VFDB Genes
/local/ifs2\_projdata/0800/projects/UMIsepsis/Serra\_Prot\_70/centroids.cluster\_roles.txt

FULL


WINDOWSVGPNG

Trim RowsRemove SingletonsSave Fasta

CL\_3417


CL\_3417


CL\_3417


CL\_3417

HighlightSelectShow Genomes


24

CL\_3424


20

CL\_3424


3

CL\_3424


1

CL\_3424

fGI ID


CL\_INS\_289
CL\_INS\_289
CL\_INS\_289
CL\_INS\_289
CL\_INS\_289
CL\_INS\_289
Cluster ID


CL\_3418
CL\_3419
CL\_3420
CL\_3421
CL\_3422
CL\_3423
