## Supplementary material for "A novel method for integrating genomic and Tn-Seq data to identify common *in vivo* fitness mechanisms across multiple bacterial species": S5 Dataset: CL_INS_294.html

Legend

 Hypothetical
 All EssentialGenes
 All VFDB Genes
/local/ifs2\_projdata/0800/projects/UMIsepsis/Serra\_Prot\_70/centroids.cluster\_roles.txt

FULL


WINDOWSVGPNG

Trim RowsRemove SingletonsSave Fasta

CL\_3461


CL\_3461


CL\_3461

HighlightSelectShow Genomes


40

CL\_3464


7

CL\_3464


1

CL\_3464

fGI ID


CL\_INS\_294
CL\_INS\_294
CL\_INS\_294
CL\_INS\_294
CL\_INS\_294
CL\_INS\_294
CL\_INS\_294
Cluster ID


CL\_3462
CL\_3463
CL\_5829
CL\_5830
CL\_5831
CL\_5832
CL\_5833
