## Supplementary material for "A novel method for integrating genomic and Tn-Seq data to identify common *in vivo* fitness mechanisms across multiple bacterial species": S5 Dataset: CL_INS_299.html

Legend

 Hypothetical
 All EssentialGenes
 All VFDB Genes
/local/ifs2\_projdata/0800/projects/UMIsepsis/Serra\_Prot\_70/centroids.cluster\_roles.txt

FULL


WINDOWSVGPNG

Trim RowsRemove SingletonsSave Fasta

CL\_3538


CL\_3538


CL\_3538


CL\_3538

HighlightSelectShow Genomes


25

CL\_3543


12

CL\_3543


2

CL\_3543


1

CL\_3543

fGI ID


CL\_INS\_299
CL\_INS\_299
CL\_INS\_299
CL\_INS\_299
CL\_INS\_299
Cluster ID


CL\_3539
CL\_10118
CL\_3540
CL\_3541
CL\_3542
