## Supplementary material for "A novel method for integrating genomic and Tn-Seq data to identify common *in vivo* fitness mechanisms across multiple bacterial species": S5 Dataset: CL_INS_300.html

Legend

 All VFDB Genes

FULL


WINDOWSVGPNG

Trim RowsRemove SingletonsSave Fasta

CL\_3551


CL\_3551


CL\_3551

HighlightSelectShow Genomes


30

CL\_3552


16

CL\_3552


2

CL\_3552

fGI ID


CL\_INS\_300
CL\_INS\_300
Cluster ID


CL\_5467
CL\_5468
