## Supplementary material for "A novel method for integrating genomic and Tn-Seq data to identify common *in vivo* fitness mechanisms across multiple bacterial species": S5 Dataset: CL_INS_301.html

Legend

 Hypothetical
 All EssentialGenes
 All VFDB Genes
/local/ifs2\_projdata/0800/projects/UMIsepsis/Serra\_Prot\_70/centroids.cluster\_roles.txt

FULL


WINDOWSVGPNG

Trim RowsRemove SingletonsSave Fasta

CL\_3555


CL\_3555


CL\_3555


CL\_3555

HighlightSelectShow Genomes


45

CL\_3556


1

CL\_3556


1

CL\_3556


1

CL\_3556

fGI ID


CL\_INS\_301
CL\_INS\_301
CL\_INS\_301
CL\_INS\_301
CL\_INS\_301
CL\_INS\_301
CL\_INS\_301
CL\_INS\_301
CL\_INS\_301
CL\_INS\_301
CL\_INS\_301
CL\_INS\_301
CL\_INS\_301
CL\_INS\_301
CL\_INS\_301
CL\_INS\_301
CL\_INS\_301
CL\_INS\_301
CL\_INS\_301
CL\_INS\_301
CL\_INS\_301
CL\_INS\_301
CL\_INS\_301
CL\_INS\_301
CL\_INS\_301
CL\_INS\_301
CL\_INS\_301
CL\_INS\_301
CL\_INS\_301
CL\_INS\_301
CL\_INS\_301
CL\_INS\_301
CL\_INS\_301
CL\_INS\_301
CL\_INS\_301
CL\_INS\_301
CL\_INS\_301
CL\_INS\_301
CL\_INS\_301
CL\_INS\_301
CL\_INS\_301
CL\_INS\_301
CL\_INS\_301
CL\_INS\_301
CL\_INS\_301
CL\_INS\_301
CL\_INS\_301
CL\_INS\_301
CL\_INS\_301
CL\_INS\_301
CL\_INS\_301
CL\_INS\_301
CL\_INS\_301
CL\_INS\_301
CL\_INS\_301
CL\_INS\_301
CL\_INS\_369
CL\_INS\_369
CL\_INS\_301
CL\_INS\_301
CL\_INS\_301
CL\_INS\_369
CL\_INS\_301
CL\_INS\_301
CL\_INS\_301
CL\_INS\_301
CL\_INS\_301
CL\_INS\_301
CL\_INS\_301
CL\_INS\_301
CL\_INS\_301
CL\_INS\_301
CL\_INS\_301
CL\_INS\_301
CL\_INS\_301
CL\_INS\_301
CL\_INS\_301
CL\_INS\_301
CL\_INS\_358
CL\_INS\_358
CL\_INS\_358
CL\_INS\_301
CL\_INS\_304
CL\_INS\_301
CL\_INS\_301
CL\_INS\_301
CL\_INS\_301
CL\_INS\_301
CL\_INS\_301
CL\_INS\_301
CL\_INS\_301
CL\_INS\_301
CL\_INS\_301
CL\_INS\_301
CL\_INS\_301
CL\_INS\_301
CL\_INS\_301
CL\_INS\_301
CL\_INS\_301
CL\_INS\_301
CL\_INS\_301
CL\_INS\_301
CL\_INS\_259
CL\_INS\_259
CL\_INS\_301
CL\_INS\_301
CL\_INS\_301
CL\_INS\_301
CL\_INS\_301
CL\_INS\_301
Cluster ID


CL\_8275
CL\_9925
CL\_8274
CL\_8273
CL\_8272
CL\_8271
CL\_8270
CL\_8269
CL\_8268
CL\_8267
CL\_8266
CL\_9205
CL\_7497
CL\_7496
CL\_7495
CL\_7494
CL\_9206
CL\_9207
CL\_9208
CL\_9209
CL\_9210
CL\_9211
CL\_9212
CL\_9213
CL\_9214
CL\_9215
CL\_9216
CL\_9217
CL\_9218
CL\_9219
CL\_9220
CL\_9221
CL\_9222
CL\_9223
CL\_9224
CL\_8265
CL\_7493
CL\_7492
CL\_7491
CL\_7490
CL\_8264
CL\_7652
CL\_7651
CL\_7650
CL\_7649
CL\_7648
CL\_7647
CL\_7646
CL\_7645
CL\_7644
CL\_7643
CL\_8263
CL\_8262
CL\_7579
CL\_8261
CL\_7578
CL\_7577
CL\_7576
CL\_7575
CL\_7573
CL\_7572
CL\_7571
CL\_8260
CL\_8259
CL\_8258
CL\_8257
CL\_8256
CL\_8255
CL\_8254
CL\_8253
CL\_8252
CL\_8251
CL\_8250
CL\_8249
CL\_4704
CL\_8248
CL\_8247
CL\_8246
CL\_8245
CL\_8244
CL\_8243
CL\_8242
CL\_8241
CL\_5563
CL\_8240
CL\_8239
CL\_8238
CL\_8237
CL\_8236
CL\_8235
CL\_8234
CL\_8233
CL\_9225
CL\_8232
CL\_8231
CL\_9226
CL\_8230
CL\_9227
CL\_9228
CL\_8229
CL\_8228
CL\_8227
CL\_7641
CL\_7642
CL\_8226
CL\_8225
CL\_9229
CL\_8224
CL\_8223
CL\_8222
