## Supplementary material for "A novel method for integrating genomic and Tn-Seq data to identify common *in vivo* fitness mechanisms across multiple bacterial species": S5 Dataset: CL_INS_303.html

Legend

 Hypothetical
 All VFDB Genes
/local/ifs2\_projdata/0800/projects/UMIsepsis/Serra\_Prot\_70/centroids.cluster\_roles.txt

FULL


WINDOWSVGPNG

Trim RowsRemove SingletonsSave Fasta

CL\_3598


CL\_3598


CL\_3598

HighlightSelectShow Genomes


21

CL\_3599


21

CL\_3599


6

CL\_3599

fGI ID


CL\_INS\_303
CL\_INS\_303
Cluster ID


CL\_7861
CL\_5469
