## Supplementary material for "A novel method for integrating genomic and Tn-Seq data to identify common *in vivo* fitness mechanisms across multiple bacterial species": S5 Dataset: CL_INS_311.html

Legend

 Hypothetical
 All VFDB Genes
/local/ifs2\_projdata/0800/projects/UMIsepsis/Serra\_Prot\_70/centroids.cluster\_roles.txt

FULL


WINDOWSVGPNG

Trim RowsRemove SingletonsSave Fasta

CL\_3696


CL\_3696


CL\_3696

HighlightSelectShow Genomes


24

CL\_3699


22

CL\_3699


1

CL\_3699

fGI ID


CL\_INS\_311
CL\_INS\_311
Cluster ID


CL\_3697
CL\_3698
