## Supplementary material for "A novel method for integrating genomic and Tn-Seq data to identify common *in vivo* fitness mechanisms across multiple bacterial species": S5 Dataset: CL_INS_313.html

Legend

 Hypothetical

FULL


WINDOWSVGPNG

Trim RowsRemove SingletonsSave Fasta

CL\_3725


CL\_3725


CL\_3725

HighlightSelectShow Genomes


41

CL\_3726


5

CL\_3726


2

CL\_3726

fGI ID


CL\_INS\_313
CL\_INS\_313
Cluster ID


CL\_12019
CL\_10124
