## Supplementary material for "A novel method for integrating genomic and Tn-Seq data to identify common *in vivo* fitness mechanisms across multiple bacterial species": S5 Dataset: CL_INS_318.html

Legend

 Hypothetical
 All EssentialGenes
 All VFDB Genes
/local/ifs2\_projdata/0800/projects/UMIsepsis/Serra\_Prot\_70/centroids.cluster\_roles.txt

FULL


WINDOWSVGPNG

Trim RowsRemove SingletonsSave Fasta

CL\_3762


CL\_3762


CL\_3762


CL\_3762


CL\_3762


CL\_3762


CL\_3762

HighlightSelectShow Genomes


23

CL\_3763


17

CL\_3763


2

CL\_3763


2

CL\_3763


2

CL\_3763


1

CL\_3763


1

CL\_3763

fGI ID


CL\_INS\_318
CL\_INS\_318
CL\_INS\_318
CL\_INS\_318
CL\_INS\_318
CL\_INS\_318
CL\_INS\_318
CL\_INS\_318
CL\_INS\_318
CL\_INS\_318
CL\_INS\_318
CL\_INS\_318
CL\_INS\_318
CL\_INS\_318
CL\_INS\_318
CL\_INS\_318
Cluster ID


CL\_7834
CL\_5503
CL\_5504
CL\_12349
CL\_12350
CL\_11561
CL\_11562
CL\_11563
CL\_9231
CL\_9232
CL\_9233
CL\_9234
CL\_9235
CL\_9236
CL\_9237
CL\_9238
