## Supplementary material for "A novel method for integrating genomic and Tn-Seq data to identify common *in vivo* fitness mechanisms across multiple bacterial species": S5 Dataset: CL_INS_322.html

Legend

 AntibioticResistance
 All VFDB Genes
/local/ifs2\_projdata/0800/projects/UMIsepsis/Serra\_Prot\_70/centroids.cluster\_roles.txt

FULL


WINDOWSVGPNG

Trim RowsRemove SingletonsSave Fasta

CL\_3799


CL\_3799

HighlightSelectShow Genomes


45

CL\_3800


2

CL\_3801

fGI ID


CL\_INS\_322
CL\_INS\_322
CL\_INS\_322
CL\_INS\_322
CL\_INS\_322
CL\_INS\_322
CL\_INS\_322
CL\_INS\_322
CL\_INS\_322
CL\_INS\_322
CL\_INS\_322
CL\_INS\_322
CL\_INS\_322
CL\_INS\_322
CL\_INS\_322
CL\_INS\_322
Cluster ID


CL\_9239
CL\_9240
CL\_9241
CL\_9242
CL\_9243
CL\_9244
CL\_5592
CL\_7350
CL\_4873
CL\_7349
CL\_7311
CL\_9245
CL\_9246
CL\_9247
CL\_9248
CL\_9249
