## Supplementary material for "A novel method for integrating genomic and Tn-Seq data to identify common *in vivo* fitness mechanisms across multiple bacterial species": S5 Dataset: CL_INS_323.html

Legend

 Hypothetical
 All VFDB Genes
/local/ifs2\_projdata/0800/projects/UMIsepsis/Serra\_Prot\_70/centroids.cluster\_roles.txt

FULL


WINDOWSVGPNG

Trim RowsRemove SingletonsSave Fasta

CL\_3898


CL\_3898


CL\_3898


CL\_3898

HighlightSelectShow Genomes


22

CL\_3901


15

CL\_3901


10

CL\_3901


1

CL\_3901

fGI ID


CL\_INS\_323
CL\_INS\_323
CL\_INS\_323
Cluster ID


CL\_3899
CL\_3900
CL\_6636
