## Supplementary material for "A novel method for integrating genomic and Tn-Seq data to identify common *in vivo* fitness mechanisms across multiple bacterial species": S5 Dataset: CL_INS_326.html

Legend

 Hypothetical
/local/ifs2\_projdata/0800/projects/UMIsepsis/Serra\_Prot\_70/centroids.cluster\_roles.txt

FULL


WINDOWSVGPNG

Trim RowsRemove SingletonsSave Fasta

CL\_3910


CL\_3910


CL\_3910

HighlightSelectShow Genomes


26

CL\_3911


18

CL\_3911


3

CL\_3911

fGI ID


CL\_INS\_326
CL\_INS\_326
Cluster ID


CL\_7074
CL\_6638
