## Supplementary material for "A novel method for integrating genomic and Tn-Seq data to identify common *in vivo* fitness mechanisms across multiple bacterial species": S5 Dataset: CL_INS_328.html

Legend

 Hypothetical
 Serritia FitnessGene
 All VFDB Genes
/local/ifs2\_projdata/0800/projects/UMIsepsis/Serra\_Prot\_70/centroids.cluster\_roles.txt

FULL


WINDOWSVGPNG

Trim RowsRemove SingletonsSave Fasta

CL\_3963


CL\_3963


CL\_3963


CL\_3963


CL\_4509

HighlightSelectShow Genomes


41

CL\_3964


4

CL\_3964


2

CL\_3964


1

CL\_4508


1

CL\_3964

fGI ID


CL\_INS\_328
CL\_INS\_328
CL\_INS\_328
CL\_INS\_328
CL\_INS\_328
CL\_INS\_328
CL\_INS\_328
Cluster ID


CL\_10855
CL\_10853
CL\_5906
CL\_5905
CL\_5904
CL\_5903
CL\_9252
