## Supplementary material for "A novel method for integrating genomic and Tn-Seq data to identify common *in vivo* fitness mechanisms across multiple bacterial species": S5 Dataset: CL_INS_330.html

Legend

 Hypothetical
 All VFDB Genes
/local/ifs2\_projdata/0800/projects/UMIsepsis/Serra\_Prot\_70/centroids.cluster\_roles.txt

FULL


WINDOWSVGPNG

Trim RowsRemove SingletonsSave Fasta

CL\_3973


CL\_3973


CL\_3973


CL\_3973

HighlightSelectShow Genomes


37

CL\_5509


9

CL\_5509


1

CL\_5509


1

CL\_3983

fGI ID


CL\_INS\_330
CL\_INS\_330
CL\_INS\_330
CL\_INS\_330
CL\_INS\_330
CL\_INS\_330
CL\_INS\_330
CL\_INS\_330
CL\_INS\_330
CL\_INS\_330
Cluster ID


CL\_3974
CL\_3975
CL\_3976
CL\_3977
CL\_3978
CL\_3979
CL\_3980
CL\_3981
CL\_3982
CL\_6639
