## Supplementary material for "A novel method for integrating genomic and Tn-Seq data to identify common *in vivo* fitness mechanisms across multiple bacterial species": S5 Dataset: CL_INS_331.html

Legend

 Hypothetical
/local/ifs2\_projdata/0800/projects/UMIsepsis/Serra\_Prot\_70/centroids.cluster\_roles.txt

FULL


WINDOWSVGPNG

Trim RowsRemove SingletonsSave Fasta

CL\_3990


CL\_3990

HighlightSelectShow Genomes


32

CL\_3991


16

CL\_3991

fGI ID


CL\_INS\_331
CL\_INS\_331
Cluster ID


CL\_5510
CL\_5511
