## Supplementary material for "A novel method for integrating genomic and Tn-Seq data to identify common *in vivo* fitness mechanisms across multiple bacterial species": S5 Dataset: CL_INS_334.html

Legend

 Hypothetical
 All VFDB Genes
/local/ifs2\_projdata/0800/projects/UMIsepsis/Serra\_Prot\_70/centroids.cluster\_roles.txt

FULL


WINDOWSVGPNG

Trim RowsRemove SingletonsSave Fasta

CL\_4006


CL\_4006


CL\_4006


CL\_4006


CL\_4006

HighlightSelectShow Genomes


32

CL\_4007


11

CL\_4007


3

CL\_4007


1

CL\_4007


1

CL\_4007

fGI ID


CL\_INS\_334
CL\_INS\_334
CL\_INS\_334
CL\_INS\_334
CL\_INS\_334
CL\_INS\_334
CL\_INS\_334
CL\_INS\_334
CL\_INS\_334
CL\_INS\_334
CL\_INS\_334
CL\_INS\_334
CL\_INS\_334
CL\_INS\_334
CL\_INS\_334
Cluster ID


CL\_8948
CL\_941
CL\_940
CL\_939
CL\_8949
CL\_10133
CL\_10383
CL\_6641
CL\_6642
CL\_6643
CL\_6644
CL\_6645
CL\_6646
CL\_6647
CL\_6648
