## Supplementary material for "A novel method for integrating genomic and Tn-Seq data to identify common *in vivo* fitness mechanisms across multiple bacterial species": S5 Dataset: CL_INS_335.html

Legend

 Hypothetical
 All VFDB Genes
/local/ifs2\_projdata/0800/projects/UMIsepsis/Serra\_Prot\_70/centroids.cluster\_roles.txt

FULL


WINDOWSVGPNG

Trim RowsRemove SingletonsSave Fasta

CL\_4010


CL\_4010

HighlightSelectShow Genomes


46

CL\_4009


1

CL\_4008

fGI ID


CL\_INS\_335
CL\_INS\_335
CL\_INS\_221
CL\_INS\_221
CL\_INS\_221
CL\_INS\_221
CL\_INS\_221
CL\_INS\_221
CL\_INS\_221
CL\_INS\_221
CL\_INS\_221
CL\_INS\_221
CL\_INS\_221
CL\_INS\_221
CL\_INS\_221
CL\_INS\_221
CL\_INS\_221
CL\_INS\_221
CL\_INS\_221
CL\_INS\_221
CL\_INS\_221
CL\_INS\_221
CL\_INS\_221
CL\_INS\_221
CL\_INS\_221
CL\_INS\_221
CL\_INS\_221
CL\_INS\_221
CL\_INS\_221
CL\_INS\_221
CL\_INS\_161
CL\_INS\_221
CL\_INS\_221
CL\_INS\_335
CL\_INS\_221
CL\_INS\_221
CL\_INS\_221
CL\_INS\_221
CL\_INS\_221
CL\_INS\_335
CL\_INS\_221
CL\_INS\_221
CL\_INS\_221
CL\_INS\_221
Cluster ID


CL\_8181
CL\_8182
CL\_8183
CL\_8184
CL\_8185
CL\_7948
CL\_7949
CL\_7950
CL\_8186
CL\_8187
CL\_8188
CL\_7954
CL\_7955
CL\_7956
CL\_8189
CL\_8190
CL\_8191
CL\_8192
CL\_8193
CL\_8194
CL\_8195
CL\_8196
CL\_8197
CL\_8198
CL\_8199
CL\_8200
CL\_7965
CL\_8201
CL\_8202
CL\_8203
CL\_8204
CL\_8205
CL\_8206
CL\_8207
CL\_8208
CL\_8209
CL\_8210
CL\_8211
CL\_8212
CL\_8213
CL\_8214
CL\_8215
CL\_8216
CL\_8217
