## Supplementary material for "A novel method for integrating genomic and Tn-Seq data to identify common *in vivo* fitness mechanisms across multiple bacterial species": S5 Dataset: CL_INS_338.html

Legend

 Hypothetical
 All VFDB Genes

FULL


WINDOWSVGPNG

Trim RowsRemove SingletonsSave Fasta

CL\_4016


CL\_4016

HighlightSelectShow Genomes


33

CL\_4020


15

CL\_4020

fGI ID


CL\_INS\_338
CL\_INS\_338
CL\_INS\_338
Cluster ID


CL\_4017
CL\_4018
CL\_4019
