## Supplementary material for "A novel method for integrating genomic and Tn-Seq data to identify common *in vivo* fitness mechanisms across multiple bacterial species": S5 Dataset: CL_INS_346.html

Legend

 Hypothetical
 Serritia FitnessGene

FULL


WINDOWSVGPNG

Trim RowsRemove SingletonsSave Fasta

CL\_4359


CL\_4359


CL\_4359

HighlightSelectShow Genomes


25

CL\_4358


22

CL\_4358


1

Break

fGI ID

CL\_INS\_346
Cluster ID

CL\_7072
