## Supplementary material for "A novel method for integrating genomic and Tn-Seq data to identify common *in vivo* fitness mechanisms across multiple bacterial species": S5 Dataset: CL_INS_348.html

Legend

 Hypothetical
 All EssentialGenes
 All VFDB Genes
/local/ifs2\_projdata/0800/projects/UMIsepsis/Serra\_Prot\_70/centroids.cluster\_roles.txt

FULL


WINDOWSVGPNG

Trim RowsRemove SingletonsSave Fasta

CL\_4301


CL\_4301


CL\_4301


CL\_4301


CL\_4301


CL\_4301


CL\_4301

HighlightSelectShow Genomes


21

CL\_4299


11

CL\_4299


5

Break


5

CL\_4299


3

Break


2

Break


1

CL\_4299

fGI ID


CL\_INS\_348
CL\_INS\_348
CL\_INS\_348
CL\_INS\_348
CL\_INS\_348
CL\_INS\_348
CL\_INS\_348
Cluster ID


CL\_4300
CL\_5530
CL\_5531
CL\_5532
CL\_7298
CL\_7299
CL\_7300
