## Supplementary material for "A novel method for integrating genomic and Tn-Seq data to identify common *in vivo* fitness mechanisms across multiple bacterial species": S5 Dataset: CL_INS_352.html

Legend

 Hypothetical
 All EssentialGenes
 All VFDB Genes

FULL


WINDOWSVGPNG

Trim RowsRemove SingletonsSave Fasta

CL\_4269


CL\_4269


CL\_4269


CL\_4269


CL\_4269


CL\_4269

HighlightSelectShow Genomes


37

CL\_4268


7

CL\_4268


1

CL\_4268


1

CL\_4268


1

CL\_4268


1

CL\_4268

fGI ID


CL\_INS\_352
CL\_INS\_352
CL\_INS\_352
CL\_INS\_352
CL\_INS\_352
Cluster ID


CL\_12022
CL\_9935
CL\_11777
CL\_6446
CL\_8806
