## Supplementary material for "A novel method for integrating genomic and Tn-Seq data to identify common *in vivo* fitness mechanisms across multiple bacterial species": S5 Dataset: CL_INS_353.html

Legend

 Hypothetical
/local/ifs2\_projdata/0800/projects/UMIsepsis/Serra\_Prot\_70/centroids.cluster\_roles.txt

FULL


WINDOWSVGPNG

Trim RowsRemove SingletonsSave Fasta

CL\_4266


CL\_4266

HighlightSelectShow Genomes


43

CL\_4265


5

CL\_4265

fGI ID


CL\_INS\_353
CL\_INS\_353
CL\_INS\_353
Cluster ID


CL\_8657
CL\_8656
CL\_8655
