## Supplementary material for "A novel method for integrating genomic and Tn-Seq data to identify common *in vivo* fitness mechanisms across multiple bacterial species": S5 Dataset: CL_INS_357.html

Legend

 Hypothetical
 All VFDB Genes
/local/ifs2\_projdata/0800/projects/UMIsepsis/Serra\_Prot\_70/centroids.cluster\_roles.txt

FULL


WINDOWSVGPNG

Trim RowsRemove SingletonsSave Fasta

CL\_4240


CL\_4240


CL\_4240


CL\_4240


CL\_4240


CL\_4240

HighlightSelectShow Genomes


28

CL\_4237


13

CL\_4237


3

CL\_4237


2

CL\_4237


1

CL\_4237


1

CL\_4237

fGI ID


CL\_INS\_357
CL\_INS\_357
CL\_INS\_357
CL\_INS\_357
CL\_INS\_357
CL\_INS\_357
CL\_INS\_357
CL\_INS\_357
CL\_INS\_382
CL\_INS\_357
CL\_INS\_357
CL\_INS\_357
CL\_INS\_369
CL\_INS\_357
CL\_INS\_357
CL\_INS\_357
CL\_INS\_357
CL\_INS\_357
CL\_INS\_357
CL\_INS\_357
CL\_INS\_357
CL\_INS\_357
CL\_INS\_357
CL\_INS\_357
CL\_INS\_357
CL\_INS\_357
CL\_INS\_357
Cluster ID


CL\_11314
CL\_11315
CL\_11316
CL\_11317
CL\_11318
CL\_5966
CL\_5054
CL\_11190
CL\_5972
CL\_11191
CL\_11192
CL\_11193
CL\_5060
CL\_11194
CL\_11195
CL\_11196
CL\_11197
CL\_5073
CL\_5074
CL\_5075
CL\_5076
CL\_11198
CL\_11199
CL\_11200
CL\_11201
CL\_4239
CL\_4238
