## Supplementary material for "A novel method for integrating genomic and Tn-Seq data to identify common *in vivo* fitness mechanisms across multiple bacterial species": S5 Dataset: CL_INS_367.html

Legend

 Hypothetical
/local/ifs2\_projdata/0800/projects/UMIsepsis/Serra\_Prot\_70/centroids.cluster\_roles.txt

FULL


WINDOWSVGPNG

Trim RowsRemove SingletonsSave Fasta

CL\_4081


CL\_4081


CL\_4081


CL\_4081

HighlightSelectShow Genomes


34

CL\_4079


11

CL\_4079


2

CL\_4079


1

CL\_4079

fGI ID


CL\_INS\_367
CL\_INS\_367
Cluster ID


CL\_6405
CL\_4080
