## Supplementary material for "A novel method for integrating genomic and Tn-Seq data to identify common *in vivo* fitness mechanisms across multiple bacterial species": S5 Dataset: CL_INS_368.html

Legend

 Hypothetical
/local/ifs2\_projdata/0800/projects/UMIsepsis/Serra\_Prot\_70/centroids.cluster\_roles.txt

FULL


WINDOWSVGPNG

Trim RowsRemove SingletonsSave Fasta

CL\_4065


CL\_4065


Break


CL\_4065


CL\_4065

HighlightSelectShow Genomes


37

CL\_4499


7

CL\_4499


1

CL\_4499


1

Break


1

CL\_4499

fGI ID


CL\_INS\_368
CL\_INS\_368
CL\_INS\_368
CL\_INS\_368
Cluster ID


CL\_9584
CL\_8561
CL\_5933
CL\_5934
