## Supplementary material for "A novel method for integrating genomic and Tn-Seq data to identify common *in vivo* fitness mechanisms across multiple bacterial species": S5 Dataset: CL_INS_374.html

Legend

 Hypothetical
 All EssentialGenes
 All VFDB Genes
/local/ifs2\_projdata/0800/projects/UMIsepsis/Serra\_Prot\_70/centroids.cluster\_roles.txt

FULL


WINDOWSVGPNG

Trim RowsRemove SingletonsSave Fasta

CL\_4625


CL\_4625


CL\_4625


CL\_4625


CL\_4625


CL\_4625


CL\_4625

HighlightSelectShow Genomes


23

CL\_4627


8

CL\_4627


7

CL\_4627


5

CL\_4627


3

CL\_4627


1

CL\_4627


1

CL\_4627

fGI ID


CL\_INS\_374
CL\_INS\_374
CL\_INS\_374
CL\_INS\_374
CL\_INS\_374
CL\_INS\_374
CL\_INS\_374
CL\_INS\_374
CL\_INS\_374
CL\_INS\_374
CL\_INS\_374
CL\_INS\_374
CL\_INS\_298
CL\_INS\_374
CL\_INS\_44
CL\_INS\_44
CL\_INS\_374
CL\_INS\_358
CL\_INS\_358
CL\_INS\_374
CL\_INS\_374
CL\_INS\_374
CL\_INS\_374
CL\_INS\_374
CL\_INS\_374
CL\_INS\_374
CL\_INS\_374
CL\_INS\_374
CL\_INS\_374
CL\_INS\_374
Cluster ID


CL\_4626
CL\_5020
CL\_7801
CL\_7800
CL\_12437
CL\_12438
CL\_12439
CL\_12440
CL\_12441
CL\_12442
CL\_12443
CL\_12444
CL\_6418
CL\_6419
CL\_5187
CL\_5188
CL\_6420
CL\_5891
CL\_6421
CL\_6422
CL\_6423
CL\_6424
CL\_6425
CL\_6426
CL\_6427
CL\_6428
CL\_6429
CL\_6430
CL\_6431
CL\_6432
