## Supplementary material for "A novel method for integrating genomic and Tn-Seq data to identify common *in vivo* fitness mechanisms across multiple bacterial species": S5 Dataset: CL_INS_375.html

Legend

 Hypothetical
 All EssentialGenes
 All VFDB Genes

FULL


WINDOWSVGPNG

Trim RowsRemove SingletonsSave Fasta

CL\_4631


CL\_4631

HighlightSelectShow Genomes


40

CL\_4632


8

CL\_4632

fGI ID

CL\_INS\_375
Cluster ID

CL\_5021
