## Supplementary material for "A novel method for integrating genomic and Tn-Seq data to identify common *in vivo* fitness mechanisms across multiple bacterial species": S5 Dataset: CL_INS_379.html

Legend

 Hypothetical
 All EssentialGenes
 All VFDB Genes
/local/ifs2\_projdata/0800/projects/UMIsepsis/Serra\_Prot\_70/centroids.cluster\_roles.txt

FULL


WINDOWSVGPNG

Trim RowsRemove SingletonsSave Fasta

CL\_4658


CL\_4658


CL\_4658


CL\_4658


CL\_4658

HighlightSelectShow Genomes


33

CL\_4660


7

CL\_4660


6

CL\_4660


1

CL\_4660


1

CL\_4660

fGI ID


CL\_INS\_379
CL\_INS\_379
CL\_INS\_379
CL\_INS\_379
CL\_INS\_379
CL\_INS\_379
CL\_INS\_379
Cluster ID


CL\_4659
CL\_7053
CL\_7052
CL\_6683
CL\_6684
CL\_6685
CL\_6686
