## Supplementary material for "A novel method for integrating genomic and Tn-Seq data to identify common *in vivo* fitness mechanisms across multiple bacterial species": S5 Dataset: CL_INS_380.html

Legend

 Hypothetical
 Serritia FitnessGene
 All VFDB Genes
/local/ifs2\_projdata/0800/projects/UMIsepsis/Serra\_Prot\_70/centroids.cluster\_roles.txt

FULL


WINDOWSVGPNG

Trim RowsRemove SingletonsSave Fasta

CL\_4673


CL\_4673


CL\_4673

HighlightSelectShow Genomes


25

CL\_4675


22

CL\_4675


1

CL\_4698

fGI ID


CL\_INS\_380
CL\_INS\_382
CL\_INS\_382
CL\_INS\_382
CL\_INS\_382
CL\_INS\_382
CL\_INS\_382
CL\_INS\_382
CL\_INS\_381
CL\_INS\_381
CL\_INS\_381
Cluster ID


CL\_4674
CL\_4677
CL\_4678
CL\_4679
CL\_4680
CL\_4681
CL\_4682
CL\_4683
CL\_7051
CL\_12445
CL\_7049
