## Supplementary material for "A novel method for integrating genomic and Tn-Seq data to identify common *in vivo* fitness mechanisms across multiple bacterial species": S5 Dataset: CL_INS_381.html

Legend

 Hypothetical
 Serritia FitnessGene
 All VFDB Genes
/local/ifs2\_projdata/0800/projects/UMIsepsis/Serra\_Prot\_70/centroids.cluster\_roles.txt

FULL


WINDOWSVGPNG

Trim RowsRemove SingletonsSave Fasta

CL\_4675


CL\_4675


CL\_4675


CL\_4675


CL\_4675


CL\_4675


CL\_4675


CL\_4675


CL\_4675


CL\_4675


CL\_4675


CL\_4675


CL\_4675


CL\_4675


CL\_4675


CL\_4675


CL\_4673


CL\_4675


CL\_4675


CL\_4675


CL\_4675


CL\_4675


CL\_4675


CL\_4675


CL\_4675


CL\_4675


CL\_4675


CL\_4675


CL\_4675


CL\_4675


CL\_4675


CL\_4675


CL\_4675


CL\_4675


CL\_4675


CL\_4675


CL\_4675


CL\_4675


CL\_4675


CL\_4675


CL\_4675

HighlightSelectShow Genomes


5

CL\_4698


4

CL\_4698


1

CL\_4717


1

CL\_4699


1

CL\_4698


1

CL\_4699


1

CL\_4699


1

CL\_4698


1

CL\_4698


1

CL\_4698


1

CL\_4698


1

CL\_4698


1

CL\_4698


1

CL\_4698


1

CL\_4698


1

CL\_4699


1

CL\_4698


1

CL\_4698


1

CL\_4698


1

CL\_4699


1

CL\_4717


1

CL\_4698


1

CL\_4698


1

CL\_4717


1

CL\_4698


1

CL\_4698


1

CL\_4698


1

CL\_4698


1

CL\_4698


1

CL\_4698


1

CL\_4698


1

CL\_4698


1

CL\_4699


1

CL\_4717


1

CL\_4698


1

CL\_4698


1

CL\_4699


1

CL\_4698


1

CL\_4699


1

CL\_4698


1

CL\_4698

fGI ID


CL\_INS\_381
CL\_INS\_381
CL\_INS\_381
CL\_INS\_381
CL\_INS\_381
CL\_INS\_381
CL\_INS\_381
CL\_INS\_381
CL\_INS\_381
CL\_INS\_381
CL\_INS\_381
CL\_INS\_381
CL\_INS\_381
CL\_INS\_381
CL\_INS\_381
CL\_INS\_381
CL\_INS\_381
CL\_INS\_381
CL\_INS\_381
CL\_INS\_382
CL\_INS\_382
CL\_INS\_382
CL\_INS\_382
CL\_INS\_381
CL\_INS\_381
CL\_INS\_381
CL\_INS\_381
CL\_INS\_381
CL\_INS\_382
CL\_INS\_382
CL\_INS\_382
CL\_INS\_381
CL\_INS\_381
CL\_INS\_381
CL\_INS\_381
CL\_INS\_381
CL\_INS\_381
CL\_INS\_381
CL\_INS\_381
CL\_INS\_381
CL\_INS\_381
CL\_INS\_381
CL\_INS\_381
CL\_INS\_381
CL\_INS\_381
CL\_INS\_381
CL\_INS\_381
CL\_INS\_381
CL\_INS\_381
CL\_INS\_381
CL\_INS\_381
CL\_INS\_259
CL\_INS\_381
CL\_INS\_381
CL\_INS\_381
CL\_INS\_381
CL\_INS\_381
CL\_INS\_381
CL\_INS\_381
CL\_INS\_381
CL\_INS\_381
CL\_INS\_357
CL\_INS\_357
CL\_INS\_382
CL\_INS\_382
CL\_INS\_382
CL\_INS\_382
CL\_INS\_382
CL\_INS\_382
CL\_INS\_382
CL\_INS\_382
CL\_INS\_381
CL\_INS\_381
CL\_INS\_382
CL\_INS\_382
CL\_INS\_358
CL\_INS\_358
CL\_INS\_358
CL\_INS\_358
CL\_INS\_381
CL\_INS\_381
CL\_INS\_381
CL\_INS\_381
CL\_INS\_381
CL\_INS\_381
CL\_INS\_381
CL\_INS\_44
CL\_INS\_381
CL\_INS\_381
CL\_INS\_381
CL\_INS\_44
CL\_INS\_44
CL\_INS\_381
CL\_INS\_381
CL\_INS\_381
CL\_INS\_44
CL\_INS\_382
CL\_INS\_382
CL\_INS\_382
CL\_INS\_382
CL\_INS\_381
CL\_INS\_381
CL\_INS\_381
CL\_INS\_381
CL\_INS\_382
CL\_INS\_382
CL\_INS\_382
CL\_INS\_382
CL\_INS\_382
CL\_INS\_382
CL\_INS\_382
CL\_INS\_382
CL\_INS\_382
CL\_INS\_382
CL\_INS\_382
CL\_INS\_382
CL\_INS\_382
CL\_INS\_382
CL\_INS\_382
CL\_INS\_382
CL\_INS\_382
CL\_INS\_381
CL\_INS\_381
CL\_INS\_381
CL\_INS\_381
CL\_INS\_382
CL\_INS\_381
CL\_INS\_381
CL\_INS\_381
CL\_INS\_381
CL\_INS\_381
CL\_INS\_381
CL\_INS\_381
CL\_INS\_382
CL\_INS\_382
CL\_INS\_381
CL\_INS\_381
CL\_INS\_381
CL\_INS\_381
CL\_INS\_381
CL\_INS\_381
CL\_INS\_381
CL\_INS\_358
CL\_INS\_358
CL\_INS\_358
CL\_INS\_358
CL\_INS\_381
CL\_INS\_381
CL\_INS\_381
CL\_INS\_381
CL\_INS\_381
CL\_INS\_381
CL\_INS\_381
CL\_INS\_381
CL\_INS\_381
CL\_INS\_381
CL\_INS\_381
CL\_INS\_381
CL\_INS\_381
CL\_INS\_381
CL\_INS\_381
CL\_INS\_381
CL\_INS\_381
CL\_INS\_381
CL\_INS\_381
CL\_INS\_381
CL\_INS\_381
CL\_INS\_381
CL\_INS\_381
CL\_INS\_381
CL\_INS\_381
CL\_INS\_381
CL\_INS\_381
CL\_INS\_382
CL\_INS\_382
CL\_INS\_382
CL\_INS\_382
CL\_INS\_382
CL\_INS\_382
CL\_INS\_381
CL\_INS\_259
CL\_INS\_381
CL\_INS\_381
CL\_INS\_358
CL\_INS\_358
CL\_INS\_382
CL\_INS\_381
CL\_INS\_381
CL\_INS\_381
CL\_INS\_381
CL\_INS\_381
CL\_INS\_381
CL\_INS\_381
CL\_INS\_381
CL\_INS\_381
CL\_INS\_381
CL\_INS\_381
CL\_INS\_381
CL\_INS\_381
CL\_INS\_381
CL\_INS\_259
CL\_INS\_259
CL\_INS\_259
CL\_INS\_259
CL\_INS\_259
CL\_INS\_259
CL\_INS\_304
CL\_INS\_434
CL\_INS\_434
CL\_INS\_434
CL\_INS\_434
CL\_INS\_36
CL\_INS\_36
CL\_INS\_381
CL\_INS\_381
CL\_INS\_381
CL\_INS\_27
CL\_INS\_381
CL\_INS\_381
CL\_INS\_381
CL\_INS\_381
CL\_INS\_381
CL\_INS\_381
CL\_INS\_381
CL\_INS\_381
CL\_INS\_381
CL\_INS\_381
CL\_INS\_381
CL\_INS\_381
CL\_INS\_381
CL\_INS\_381
CL\_INS\_381
CL\_INS\_381
CL\_INS\_382
CL\_INS\_382
CL\_INS\_382
CL\_INS\_381
CL\_INS\_381
CL\_INS\_381
CL\_INS\_381
CL\_INS\_381
CL\_INS\_381
CL\_INS\_381
CL\_INS\_381
CL\_INS\_381
Cluster ID


CL\_6435
CL\_6436
CL\_10921
CL\_10922
CL\_10923
CL\_10924
CL\_11505
CL\_11506
CL\_11507
CL\_11508
CL\_11330
CL\_11331
CL\_11332
CL\_9490
CL\_9489
CL\_10400
CL\_10401
CL\_10402
CL\_4676
CL\_4677
CL\_4678
CL\_4679
CL\_4680
CL\_10206
CL\_10207
CL\_10208
CL\_10209
CL\_6011
CL\_4681
CL\_4682
CL\_4683
CL\_7051
CL\_12445
CL\_7050
CL\_7049
CL\_8178
CL\_8177
CL\_8176
CL\_8175
CL\_8174
CL\_8173
CL\_8172
CL\_4706
CL\_4707
CL\_4708
CL\_9488
CL\_6437
CL\_6438
CL\_6439
CL\_6440
CL\_4684
CL\_7513
CL\_4685
CL\_8980
CL\_8981
CL\_11870
CL\_11871
CL\_11872
CL\_4686
CL\_4687
CL\_4688
CL\_5966
CL\_5054
CL\_5055
CL\_5972
CL\_8650
CL\_7065
CL\_7064
CL\_11787
CL\_11788
CL\_11789
CL\_11790
CL\_11791
CL\_12203
CL\_12204
CL\_5056
CL\_5057
CL\_5033
CL\_5034
CL\_7797
CL\_7796
CL\_7795
CL\_7794
CL\_7793
CL\_11686
CL\_7792
CL\_5035
CL\_5036
CL\_5037
CL\_5038
CL\_5039
CL\_5040
CL\_5041
CL\_7303
CL\_7304
CL\_5042
CL\_5940
CL\_5939
CL\_7305
CL\_9603
CL\_8818
CL\_8819
CL\_8820
CL\_8821
CL\_11307
CL\_11308
CL\_11309
CL\_11310
CL\_11311
CL\_11312
CL\_5938
CL\_5937
CL\_12207
CL\_7791
CL\_8954
CL\_8955
CL\_8956
CL\_8957
CL\_8958
CL\_8959
CL\_8960
CL\_10565
CL\_10566
CL\_10567
CL\_10568
CL\_6693
CL\_7790
CL\_7789
CL\_7788
CL\_11685
CL\_7787
CL\_7786
CL\_7785
CL\_7306
CL\_7307
CL\_11144
CL\_11145
CL\_11146
CL\_11147
CL\_11208
CL\_11209
CL\_11210
CL\_6461
CL\_6462
CL\_6463
CL\_6464
CL\_6465
CL\_6466
CL\_6467
CL\_6468
CL\_6469
CL\_6470
CL\_6471
CL\_6472
CL\_6473
CL\_6474
CL\_6475
CL\_6476
CL\_6477
CL\_5025
CL\_10860
CL\_10861
CL\_10862
CL\_10863
CL\_10864
CL\_11115
CL\_11116
CL\_11117
CL\_11118
CL\_11119
CL\_11120
CL\_11121
CL\_5026
CL\_12355
CL\_12356
CL\_8652
CL\_12202
CL\_6027
CL\_6028
CL\_11207
CL\_11131
CL\_7798
CL\_7302
CL\_5027
CL\_5028
CL\_5029
CL\_5030
CL\_5031
CL\_5032
CL\_6478
CL\_6479
CL\_8456
CL\_6480
CL\_6481
CL\_6482
CL\_8648
CL\_8647
CL\_8646
CL\_8645
CL\_8644
CL\_8643
CL\_8642
CL\_8641
CL\_8640
CL\_8639
CL\_8638
CL\_3609
CL\_3607
CL\_3606
CL\_3604
CL\_3603
CL\_4689
CL\_4690
CL\_8457
CL\_8458
CL\_4691
CL\_4692
CL\_4693
CL\_4694
CL\_6687
CL\_9947
CL\_9948
CL\_6688
CL\_6689
CL\_6690
CL\_6691
CL\_6692
CL\_4695
CL\_6483
CL\_6484
CL\_4696
CL\_4697
CL\_6485
CL\_5820
CL\_5819
CL\_5818
CL\_5817
CL\_5816
CL\_5815
CL\_5814
CL\_7308
CL\_7309
CL\_8637
CL\_8636
CL\_8635
