## Supplementary material for "A novel method for integrating genomic and Tn-Seq data to identify common *in vivo* fitness mechanisms across multiple bacterial species": S5 Dataset: CL_INS_383.html

Legend

 Hypothetical
 All VFDB Genes
/local/ifs2\_projdata/0800/projects/UMIsepsis/Serra\_Prot\_70/centroids.cluster\_roles.txt

FULL


WINDOWSVGPNG

Trim RowsRemove SingletonsSave Fasta

CL\_4699


CL\_4699


CL\_4699


CL\_4699


CL\_4699


CL\_4699


CL\_4699


CL\_4699


CL\_4699


CL\_4699


CL\_4675


CL\_4699


CL\_4699


CL\_4699


CL\_4675


CL\_4699


CL\_4699


CL\_4699


CL\_4699


CL\_4675


CL\_4699


CL\_4699


CL\_4675


CL\_4699


CL\_4699


CL\_4699


CL\_4699


CL\_4699


CL\_4699


CL\_4699


CL\_4699


CL\_4699


CL\_4699

HighlightSelectShow Genomes


6

CL\_4717


4

CL\_4717


3

CL\_4717


2

CL\_4717


2

CL\_4717


2

CL\_4717


2

CL\_4717


2

CL\_4717


1

CL\_4717


1

CL\_4717


1

CL\_4717


1

CL\_4717


1

CL\_4717


1

CL\_4717


1

CL\_4717


1

CL\_4717


1

CL\_4717


1

CL\_4717


1

CL\_4717


1

CL\_4717


1

CL\_4717


1

CL\_4717


1

CL\_4717


1

CL\_4717


1

CL\_4717


1

CL\_4717


1

CL\_4717


1

CL\_4717


1

CL\_4718


1

CL\_4717


1

CL\_4717


1

CL\_4717


1

CL\_4717

fGI ID


CL\_INS\_383
CL\_INS\_381
CL\_INS\_381
CL\_INS\_381
CL\_INS\_383
CL\_INS\_383
CL\_INS\_383
CL\_INS\_383
CL\_INS\_383
CL\_INS\_383
CL\_INS\_383
CL\_INS\_383
CL\_INS\_383
CL\_INS\_383
CL\_INS\_383
CL\_INS\_301
CL\_INS\_298
CL\_INS\_382
CL\_INS\_382
CL\_INS\_382
CL\_INS\_382
CL\_INS\_381
CL\_INS\_381
CL\_INS\_381
CL\_INS\_381
CL\_INS\_382
CL\_INS\_382
CL\_INS\_382
CL\_INS\_381
CL\_INS\_381
CL\_INS\_381
CL\_INS\_381
CL\_INS\_381
CL\_INS\_381
CL\_INS\_381
CL\_INS\_381
CL\_INS\_383
CL\_INS\_383
CL\_INS\_383
CL\_INS\_381
CL\_INS\_381
CL\_INS\_381
CL\_INS\_383
CL\_INS\_383
CL\_INS\_383
CL\_INS\_383
CL\_INS\_383
CL\_INS\_383
CL\_INS\_383
CL\_INS\_383
CL\_INS\_383
CL\_INS\_381
CL\_INS\_381
CL\_INS\_381
CL\_INS\_381
CL\_INS\_383
Cluster ID


CL\_11873
CL\_9489
CL\_10401
CL\_10402
CL\_8962
CL\_8963
CL\_5043
CL\_10925
CL\_6694
CL\_4700
CL\_4701
CL\_4702
CL\_10865
CL\_5044
CL\_4703
CL\_4704
CL\_4705
CL\_4677
CL\_4678
CL\_4679
CL\_4680
CL\_10206
CL\_10207
CL\_10208
CL\_10209
CL\_4681
CL\_4682
CL\_4683
CL\_8178
CL\_8177
CL\_8176
CL\_8175
CL\_8174
CL\_8173
CL\_8172
CL\_4706
CL\_8459
CL\_7784
CL\_7783
CL\_4707
CL\_4708
CL\_9488
CL\_9487
CL\_4709
CL\_4710
CL\_4711
CL\_4712
CL\_4713
CL\_4714
CL\_4715
CL\_4716
CL\_6437
CL\_6438
CL\_6439
CL\_6440
CL\_6441
