## Supplementary material for "A novel method for integrating genomic and Tn-Seq data to identify common *in vivo* fitness mechanisms across multiple bacterial species": S5 Dataset: CL_INS_384.html

Legend

 All VFDB Genes
/local/ifs2\_projdata/0800/projects/UMIsepsis/Serra\_Prot\_70/centroids.cluster\_roles.txt

FULL


WINDOWSVGPNG

Trim RowsRemove SingletonsSave Fasta

CL\_4719


CL\_4719


CL\_4719

HighlightSelectShow Genomes


26

CL\_4720


15

CL\_4720


5

CL\_4726

fGI ID

CL\_INS\_384
Cluster ID

CL\_5045
