## Supplementary material for "A novel method for integrating genomic and Tn-Seq data to identify common *in vivo* fitness mechanisms across multiple bacterial species": S5 Dataset: CL_INS_385.html

Legend

 Hypothetical
 All VFDB Genes

FULL


WINDOWSVGPNG

Trim RowsRemove SingletonsSave Fasta

CL\_4729


CL\_4729


CL\_4729

HighlightSelectShow Genomes


35

CL\_4730


11

CL\_4736


1

CL\_4736

fGI ID


CL\_INS\_385
CL\_INS\_385
Cluster ID


CL\_5046
CL\_5047
