## Supplementary material for "A novel method for integrating genomic and Tn-Seq data to identify common *in vivo* fitness mechanisms across multiple bacterial species": S5 Dataset: CL_INS_388.html

Legend

 Hypothetical
 All VFDB Genes
/local/ifs2\_projdata/0800/projects/UMIsepsis/Serra\_Prot\_70/centroids.cluster\_roles.txt

FULL


WINDOWSVGPNG

Trim RowsRemove SingletonsSave Fasta

CL\_4747


CL\_4747


CL\_4747


CL\_4747


CL\_4747


CL\_4747


CL\_4747


CL\_4747


CL\_4747


CL\_4747


CL\_4747


CL\_4747

HighlightSelectShow Genomes


10

CL\_4750


7

CL\_4750


6

CL\_4750


6

CL\_4750


5

CL\_4750


4

CL\_4750


3

CL\_4750


2

CL\_4750


2

CL\_4750


1

CL\_4750


1

CL\_4750


1

CL\_4750

fGI ID


CL\_INS\_388
CL\_INS\_388
CL\_INS\_388
CL\_INS\_388
CL\_INS\_224
CL\_INS\_388
CL\_INS\_388
CL\_INS\_388
CL\_INS\_388
CL\_INS\_388
CL\_INS\_388
CL\_INS\_388
CL\_INS\_388
CL\_INS\_388
CL\_INS\_388
CL\_INS\_388
Cluster ID


CL\_5813
CL\_5048
CL\_5049
CL\_8964
CL\_2911
CL\_7048
CL\_4748
CL\_8634
CL\_6443
CL\_4749
CL\_6695
CL\_6696
CL\_6697
CL\_6698
CL\_6699
CL\_6700
