## Supplementary material for "A novel method for integrating genomic and Tn-Seq data to identify common *in vivo* fitness mechanisms across multiple bacterial species": S5 Dataset: CL_INS_396.html

Legend

 All VFDB Genes
/local/ifs2\_projdata/0800/projects/UMIsepsis/Serra\_Prot\_70/centroids.cluster\_roles.txt

FULL


WINDOWSVGPNG

Trim RowsRemove SingletonsSave Fasta

CL\_4798


CL\_4798


CL\_4798

HighlightSelectShow Genomes


32

CL\_4800


1

CL\_4800


1

Break

fGI ID

CL\_INS\_396
Cluster ID

CL\_4799
