## Supplementary material for "A novel method for integrating genomic and Tn-Seq data to identify common *in vivo* fitness mechanisms across multiple bacterial species": S5 Dataset: CL_INS_399.html

Legend

/local/ifs2\_projdata/0800/projects/UMIsepsis/Serra\_Prot\_70/centroids.cluster\_roles.txt

FULL


WINDOWSVGPNG

Trim RowsRemove SingletonsSave Fasta

CL\_4819


CL\_4819

HighlightSelectShow Genomes


25

CL\_4820


19

CL\_4820

fGI ID

CL\_INS\_399
Cluster ID

CL\_5812
