## Supplementary material for "A novel method for integrating genomic and Tn-Seq data to identify common *in vivo* fitness mechanisms across multiple bacterial species": S5 Dataset: CL_INS_400.html

Legend

 All VFDB Genes

FULL


WINDOWSVGPNG

Trim RowsRemove SingletonsSave Fasta

CL\_4826


CL\_4826


CL\_4826


CL\_4826


CL\_4826


CL\_4826

HighlightSelectShow Genomes


19

CL\_4830


12

CL\_4830


4

CL\_4830


4

Break


4

CL\_4830


1

Break

fGI ID


CL\_INS\_400
CL\_INS\_400
CL\_INS\_400
Cluster ID


CL\_4827
CL\_4828
CL\_4829
