## Supplementary material for "A novel method for integrating genomic and Tn-Seq data to identify common *in vivo* fitness mechanisms across multiple bacterial species": S5 Dataset: CL_INS_403.html

Legend

 AntibioticResistance
/local/ifs2\_projdata/0800/projects/UMIsepsis/Serra\_Prot\_70/centroids.cluster\_roles.txt

FULL


WINDOWSVGPNG

Trim RowsRemove SingletonsSave Fasta

CL\_58


CL\_58

HighlightSelectShow Genomes


32

CL\_59


16

CL\_59

fGI ID

CL\_INS\_403
Cluster ID

CL\_8719
