## Supplementary material for "A novel method for integrating genomic and Tn-Seq data to identify common *in vivo* fitness mechanisms across multiple bacterial species": S5 Dataset: CL_INS_408.html

Legend

 Hypothetical
/local/ifs2\_projdata/0800/projects/UMIsepsis/Serra\_Prot\_70/centroids.cluster\_roles.txt

FULL


WINDOWSVGPNG

Trim RowsRemove SingletonsSave Fasta

CL\_170


CL\_170


CL\_170


CL\_170


CL\_170


CL\_170


CL\_170


CL\_170

HighlightSelectShow Genomes


38

CL\_171


3

CL\_173


2

CL\_173


1

CL\_173


1

CL\_173


1

CL\_173


1

CL\_173


1

CL\_173

fGI ID


CL\_INS\_186
CL\_INS\_186
CL\_INS\_409
CL\_INS\_409
CL\_INS\_186
CL\_INS\_186
CL\_INS\_186
CL\_INS\_186
CL\_INS\_409
CL\_INS\_409
CL\_INS\_304
CL\_INS\_304
CL\_INS\_186
CL\_INS\_409
CL\_INS\_186
CL\_INS\_186
CL\_INS\_186
CL\_INS\_186
CL\_INS\_409
CL\_INS\_186
CL\_INS\_409
CL\_INS\_409
CL\_INS\_409
CL\_INS\_409
Cluster ID


CL\_5809
CL\_5808
CL\_10569
CL\_5807
CL\_5806
CL\_5805
CL\_5804
CL\_5803
CL\_10866
CL\_5802
CL\_5472
CL\_4447
CL\_5801
CL\_5800
CL\_5799
CL\_5798
CL\_5797
CL\_5796
CL\_5795
CL\_5794
CL\_5793
CL\_5792
CL\_5935
CL\_5791
