## Supplementary material for "A novel method for integrating genomic and Tn-Seq data to identify common *in vivo* fitness mechanisms across multiple bacterial species": S5 Dataset: CL_INS_414.html

Legend

 Hypothetical
 All VFDB Genes
/local/ifs2\_projdata/0800/projects/UMIsepsis/Serra\_Prot\_70/centroids.cluster\_roles.txt

FULL


WINDOWSVGPNG

Trim RowsRemove SingletonsSave Fasta

CL\_259


CL\_259


CL\_259


CL\_259


CL\_259


CL\_259


CL\_259


CL\_259


CL\_259


CL\_259

HighlightSelectShow Genomes


17

CL\_260


13

CL\_260


5

CL\_260


5

CL\_260


2

CL\_260


2

CL\_260


1

CL\_261


1

CL\_260


1

CL\_260


1

CL\_260

fGI ID


CL\_INS\_414
CL\_INS\_414
CL\_INS\_414
CL\_INS\_414
CL\_INS\_414
CL\_INS\_414
Cluster ID


CL\_12447
CL\_7767
CL\_5084
CL\_5082
CL\_5083
CL\_5965
