## Supplementary material for "A novel method for integrating genomic and Tn-Seq data to identify common *in vivo* fitness mechanisms across multiple bacterial species": S5 Dataset: CL_INS_416.html

Legend

 Hypothetical
 All VFDB Genes

FULL


WINDOWSVGPNG

Trim RowsRemove SingletonsSave Fasta

CL\_283


CL\_283

HighlightSelectShow Genomes


42

CL\_284


6

CL\_284

fGI ID


CL\_INS\_416
CL\_INS\_416
Cluster ID


CL\_6729
CL\_6730
