## Supplementary material for "A novel method for integrating genomic and Tn-Seq data to identify common *in vivo* fitness mechanisms across multiple bacterial species": S5 Dataset: CL_INS_421.html

Legend

 All EssentialGenes
/local/ifs2\_projdata/0800/projects/UMIsepsis/Serra\_Prot\_70/centroids.cluster\_roles.txt

FULL


WINDOWSVGPNG

Trim RowsRemove SingletonsSave Fasta

CL\_312


CL\_312

HighlightSelectShow Genomes


29

CL\_313


19

CL\_313

fGI ID

CL\_INS\_421
Cluster ID

CL\_5779
