## Supplementary material for "A novel method for integrating genomic and Tn-Seq data to identify common *in vivo* fitness mechanisms across multiple bacterial species": S5 Dataset: CL_INS_424.html

Legend

 Hypothetical
 All VFDB Genes
/local/ifs2\_projdata/0800/projects/UMIsepsis/Serra\_Prot\_70/centroids.cluster\_roles.txt

FULL


WINDOWSVGPNG

Trim RowsRemove SingletonsSave Fasta

CL\_355


CL\_355


CL\_355

HighlightSelectShow Genomes


30

CL\_356


14

CL\_356


4

CL\_356

fGI ID


CL\_INS\_424
CL\_INS\_424
Cluster ID


CL\_5087
CL\_5088
