## Supplementary material for "A novel method for integrating genomic and Tn-Seq data to identify common *in vivo* fitness mechanisms across multiple bacterial species": S5 Dataset: CL_INS_426.html

Legend

 Hypothetical
 All VFDB Genes
/local/ifs2\_projdata/0800/projects/UMIsepsis/Serra\_Prot\_70/centroids.cluster\_roles.txt

FULL


WINDOWSVGPNG

Trim RowsRemove SingletonsSave Fasta

CL\_412


CL\_412


CL\_412


CL\_412


CL\_412


CL\_412


CL\_412


CL\_412


CL\_411


CL\_412

HighlightSelectShow Genomes


25

CL\_432


12

CL\_432


2

CL\_432


2

CL\_432


2

CL\_432


1

CL\_432


1

CL\_432


1

CL\_432


1

CL\_432


1

CL\_432

fGI ID


CL\_INS\_426
CL\_INS\_426
CL\_INS\_426
CL\_INS\_377
CL\_INS\_377
CL\_INS\_426
CL\_INS\_426
CL\_INS\_426
CL\_INS\_426
CL\_INS\_426
CL\_INS\_426
CL\_INS\_426
CL\_INS\_426
CL\_INS\_426
CL\_INS\_426
CL\_INS\_426
CL\_INS\_426
CL\_INS\_426
CL\_INS\_426
CL\_INS\_426
CL\_INS\_426
CL\_INS\_426
CL\_INS\_426
CL\_INS\_426
CL\_INS\_426
CL\_INS\_426
CL\_INS\_426
CL\_INS\_426
CL\_INS\_426
CL\_INS\_426
CL\_INS\_426
CL\_INS\_426
CL\_INS\_426
Cluster ID


CL\_9024
CL\_9025
CL\_9026
CL\_9027
CL\_9028
CL\_413
CL\_7762
CL\_9967
CL\_7761
CL\_9968
CL\_7760
CL\_7759
CL\_7758
CL\_9969
CL\_9607
CL\_414
CL\_415
CL\_416
CL\_417
CL\_418
CL\_419
CL\_420
CL\_421
CL\_422
CL\_423
CL\_424
CL\_425
CL\_426
CL\_427
CL\_428
CL\_429
CL\_430
CL\_431
