## Supplementary material for "A novel method for integrating genomic and Tn-Seq data to identify common *in vivo* fitness mechanisms across multiple bacterial species": S5 Dataset: CL_INS_428.html

Legend

/local/ifs2\_projdata/0800/projects/UMIsepsis/Serra\_Prot\_70/centroids.cluster\_roles.txt

FULL


WINDOWSVGPNG

Trim RowsRemove SingletonsSave Fasta

CL\_443


CL\_443

HighlightSelectShow Genomes


30

CL\_444


18

CL\_444

fGI ID

CL\_INS\_428
Cluster ID

CL\_5089
