## Supplementary material for "A novel method for integrating genomic and Tn-Seq data to identify common *in vivo* fitness mechanisms across multiple bacterial species": S5 Dataset: CL_INS_431.html

Legend

 Hypothetical
 All VFDB Genes
/local/ifs2\_projdata/0800/projects/UMIsepsis/Serra\_Prot\_70/centroids.cluster\_roles.txt

FULL


WINDOWSVGPNG

Trim RowsRemove SingletonsSave Fasta

CL\_472


CL\_472


CL\_472

HighlightSelectShow Genomes


40

CL\_473


5

CL\_473


2

CL\_473

fGI ID


CL\_INS\_431
CL\_INS\_431
CL\_INS\_431
CL\_INS\_431
CL\_INS\_431
CL\_INS\_431
CL\_INS\_431
CL\_INS\_431
CL\_INS\_431
CL\_INS\_431
CL\_INS\_431
CL\_INS\_431
CL\_INS\_431
CL\_INS\_431
CL\_INS\_431
CL\_INS\_431
Cluster ID


CL\_5090
CL\_5091
CL\_5092
CL\_5093
CL\_5094
CL\_5095
CL\_5096
CL\_5097
CL\_5098
CL\_5099
CL\_5100
CL\_9972
CL\_5101
CL\_5102
CL\_5103
CL\_9973
