## Supplementary material for "A novel method for integrating genomic and Tn-Seq data to identify common *in vivo* fitness mechanisms across multiple bacterial species": S5 Dataset: CL_INS_436.html

Legend

 Hypothetical
 All VFDB Genes
/local/ifs2\_projdata/0800/projects/UMIsepsis/Serra\_Prot\_70/centroids.cluster\_roles.txt

FULL


WINDOWSVGPNG

Trim RowsRemove SingletonsSave Fasta

CL\_583


CL\_583


CL\_583


CL\_583

HighlightSelectShow Genomes


45

CL\_584


1

CL\_584


1

CL\_584


1

CL\_584

fGI ID


CL\_INS\_469
CL\_INS\_469
CL\_INS\_469
CL\_INS\_469
CL\_INS\_469
CL\_INS\_469
CL\_INS\_469
CL\_INS\_469
CL\_INS\_469
CL\_INS\_469
CL\_INS\_469
CL\_INS\_469
CL\_INS\_469
CL\_INS\_469
CL\_INS\_469
CL\_INS\_469
CL\_INS\_436
CL\_INS\_436
CL\_INS\_469
CL\_INS\_469
CL\_INS\_469
CL\_INS\_469
CL\_INS\_469
CL\_INS\_469
CL\_INS\_469
CL\_INS\_469
CL\_INS\_469
CL\_INS\_469
CL\_INS\_469
CL\_INS\_469
CL\_INS\_469
CL\_INS\_469
CL\_INS\_217
CL\_INS\_469
CL\_INS\_469
CL\_INS\_469
CL\_INS\_436
CL\_INS\_469
CL\_INS\_469
CL\_INS\_469
CL\_INS\_469
CL\_INS\_469
CL\_INS\_469
CL\_INS\_469
CL\_INS\_469
CL\_INS\_469
CL\_INS\_469
CL\_INS\_469
CL\_INS\_469
CL\_INS\_469
CL\_INS\_469
CL\_INS\_469
Cluster ID


CL\_5777
CL\_5776
CL\_5775
CL\_5774
CL\_5773
CL\_5772
CL\_5771
CL\_9046
CL\_5770
CL\_5769
CL\_5768
CL\_5767
CL\_5766
CL\_5765
CL\_5764
CL\_9047
CL\_7737
CL\_7736
CL\_5763
CL\_5762
CL\_5761
CL\_9048
CL\_5760
CL\_5759
CL\_5758
CL\_5757
CL\_5756
CL\_5755
CL\_7735
CL\_5754
CL\_5753
CL\_5752
CL\_5751
CL\_5750
CL\_7734
CL\_5749
CL\_5748
CL\_5747
CL\_5746
CL\_5745
CL\_5744
CL\_5743
CL\_5742
CL\_5741
CL\_5740
CL\_5739
CL\_5738
CL\_5737
CL\_5736
CL\_5735
CL\_5734
CL\_5733
