## Supplementary material for "A novel method for integrating genomic and Tn-Seq data to identify common *in vivo* fitness mechanisms across multiple bacterial species": S5 Dataset: CL_INS_437.html

Legend

 Hypothetical
 All VFDB Genes
/local/ifs2\_projdata/0800/projects/UMIsepsis/Serra\_Prot\_70/centroids.cluster\_roles.txt

FULL


WINDOWSVGPNG

Trim RowsRemove SingletonsSave Fasta

CL\_589


CL\_589


CL\_589

HighlightSelectShow Genomes


31

CL\_590


16

CL\_590


1

CL\_590

fGI ID


CL\_INS\_437
CL\_INS\_437
Cluster ID


CL\_10403
CL\_5110
