## Supplementary material for "A novel method for integrating genomic and Tn-Seq data to identify common *in vivo* fitness mechanisms across multiple bacterial species": S5 Dataset: CL_INS_443.html

Legend

 Hypothetical
 All VFDB Genes
/local/ifs2\_projdata/0800/projects/UMIsepsis/Serra\_Prot\_70/centroids.cluster\_roles.txt

FULL


WINDOWSVGPNG

Trim RowsRemove SingletonsSave Fasta

CL\_731


CL\_731


CL\_736


CL\_731

HighlightSelectShow Genomes


43

CL\_730


3

CL\_683


1

CL\_730


1

CL\_1755

fGI ID


CL\_INS\_443
CL\_INS\_443
CL\_INS\_443
Cluster ID


CL\_5951
CL\_5952
CL\_5953
