## Supplementary material for "A novel method for integrating genomic and Tn-Seq data to identify common *in vivo* fitness mechanisms across multiple bacterial species": S5 Dataset: CL_INS_446.html

Legend

 All VFDB Genes
/local/ifs2\_projdata/0800/projects/UMIsepsis/Serra\_Prot\_70/centroids.cluster\_roles.txt

FULL


WINDOWSVGPNG

Trim RowsRemove SingletonsSave Fasta

CL\_752


CL\_752


CL\_751

HighlightSelectShow Genomes


38

CL\_753


8

CL\_753


1

CL\_753

fGI ID

CL\_INS\_446
Cluster ID

CL\_9044
