## Supplementary material for "A novel method for integrating genomic and Tn-Seq data to identify common *in vivo* fitness mechanisms across multiple bacterial species": S5 Dataset: CL_INS_447.html

Legend

/local/ifs2\_projdata/0800/projects/UMIsepsis/Serra\_Prot\_70/centroids.cluster\_roles.txt

FULL


WINDOWSVGPNG

Trim RowsRemove SingletonsSave Fasta

CL\_760


CL\_760


CL\_760

HighlightSelectShow Genomes


28

CL\_761


15

CL\_761


3

CL\_761

fGI ID


CL\_INS\_447
CL\_INS\_447
Cluster ID


CL\_6152
CL\_6153
