## Supplementary material for "A novel method for integrating genomic and Tn-Seq data to identify common *in vivo* fitness mechanisms across multiple bacterial species": S5 Dataset: CL_INS_450.html

Legend

 Hypothetical
 All VFDB Genes
/local/ifs2\_projdata/0800/projects/UMIsepsis/Serra\_Prot\_70/centroids.cluster\_roles.txt

FULL


WINDOWSVGPNG

Trim RowsRemove SingletonsSave Fasta

CL\_782


CL\_782


CL\_782


CL\_782


CL\_782


CL\_782

HighlightSelectShow Genomes


23

CL\_783


17

CL\_783


4

CL\_788


2

CL\_788


1

CL\_788


1

CL\_788

fGI ID


CL\_INS\_450
CL\_INS\_450
CL\_INS\_450
CL\_INS\_450
CL\_INS\_450
Cluster ID


CL\_5679
CL\_5121
CL\_8862
CL\_8863
CL\_9484
