## Supplementary material for "A novel method for integrating genomic and Tn-Seq data to identify common *in vivo* fitness mechanisms across multiple bacterial species": S5 Dataset: CL_INS_452.html

Legend

 All VFDB Genes
/local/ifs2\_projdata/0800/projects/UMIsepsis/Serra\_Prot\_70/centroids.cluster\_roles.txt

FULL


WINDOWSVGPNG

Trim RowsRemove SingletonsSave Fasta

CL\_789


CL\_789


CL\_789


CL\_789

HighlightSelectShow Genomes


24

CL\_794


22

CL\_794


1

CL\_795


1

CL\_794

fGI ID


CL\_INS\_452
CL\_INS\_452
CL\_INS\_452
CL\_INS\_452
Cluster ID


CL\_790
CL\_791
CL\_792
CL\_793
