## Supplementary material for "A novel method for integrating genomic and Tn-Seq data to identify common *in vivo* fitness mechanisms across multiple bacterial species": S5 Dataset: CL_INS_461.html

Legend

 Hypothetical
 All VFDB Genes
/local/ifs2\_projdata/0800/projects/UMIsepsis/Serra\_Prot\_70/centroids.cluster\_roles.txt

FULL


WINDOWSVGPNG

Trim RowsRemove SingletonsSave Fasta

CL\_866


CL\_866


CL\_866


CL\_866

HighlightSelectShow Genomes


28

CL\_867


16

CL\_867


2

CL\_867


2

CL\_867

fGI ID


CL\_INS\_461
CL\_INS\_461
CL\_INS\_461
CL\_INS\_461
Cluster ID


CL\_5950
CL\_6156
CL\_6157
CL\_6158
