## Supplementary material for "A novel method for integrating genomic and Tn-Seq data to identify common *in vivo* fitness mechanisms across multiple bacterial species": S5 Dataset: CL_INS_462.html

Legend

 Hypothetical
 All VFDB Genes
/local/ifs2\_projdata/0800/projects/UMIsepsis/Serra\_Prot\_70/centroids.cluster\_roles.txt

FULL


WINDOWSVGPNG

Trim RowsRemove SingletonsSave Fasta

CL\_868


CL\_868


CL\_868

HighlightSelectShow Genomes


41

CL\_869


6

CL\_869


1

CL\_869

fGI ID


CL\_INS\_462
CL\_INS\_462
CL\_INS\_462
CL\_INS\_462
CL\_INS\_462
CL\_INS\_462
Cluster ID


CL\_7031
CL\_10148
CL\_10149
CL\_10150
CL\_10151
CL\_10152
