## Supplementary material for "A novel method for integrating genomic and Tn-Seq data to identify common *in vivo* fitness mechanisms across multiple bacterial species": S5 Dataset: CL_INS_463.html

Legend

 Hypothetical
 All VFDB Genes

FULL


WINDOWSVGPNG

Trim RowsRemove SingletonsSave Fasta

CL\_889


CL\_889


CL\_889

HighlightSelectShow Genomes


37

CL\_890


6

CL\_890


4

CL\_890

fGI ID


CL\_INS\_463
CL\_INS\_463
Cluster ID


CL\_7717
CL\_7716
