## Supplementary material for "A novel method for integrating genomic and Tn-Seq data to identify common *in vivo* fitness mechanisms across multiple bacterial species": S5 Dataset: CL_INS_466.html

Legend

 Hypothetical
 All VFDB Genes
/local/ifs2\_projdata/0800/projects/UMIsepsis/Serra\_Prot\_70/centroids.cluster\_roles.txt

FULL


WINDOWSVGPNG

Trim RowsRemove SingletonsSave Fasta

CL\_912


CL\_912


CL\_912


CL\_912

HighlightSelectShow Genomes


27

CL\_913


7

CL\_913


4

CL\_926


1

CL\_914

fGI ID


CL\_INS\_466
CL\_INS\_467
CL\_INS\_467
CL\_INS\_467
CL\_INS\_467
CL\_INS\_467
CL\_INS\_467
Cluster ID


CL\_5676
CL\_919
CL\_920
CL\_921
CL\_922
CL\_923
CL\_924
