## Supplementary material for "A novel method for integrating genomic and Tn-Seq data to identify common *in vivo* fitness mechanisms across multiple bacterial species": S5 Dataset: CL_INS_469.html

Legend

 Hypothetical
 All VFDB Genes
/local/ifs2\_projdata/0800/projects/UMIsepsis/Serra\_Prot\_70/centroids.cluster\_roles.txt

FULL


WINDOWSVGPNG

Trim RowsRemove SingletonsSave Fasta

CL\_943


CL\_943


CL\_943


CL\_943


CL\_943

HighlightSelectShow Genomes


39

CL\_5139


5

CL\_938


2

CL\_5139


1

CL\_938


1

CL\_5139

fGI ID


CL\_INS\_469
CL\_INS\_469
CL\_INS\_334
CL\_INS\_334
CL\_INS\_469
CL\_INS\_469
CL\_INS\_469
CL\_INS\_469
CL\_INS\_469
CL\_INS\_469
CL\_INS\_469
CL\_INS\_469
CL\_INS\_469
CL\_INS\_469
CL\_INS\_469
CL\_INS\_469
CL\_INS\_469
CL\_INS\_469
CL\_INS\_469
CL\_INS\_469
CL\_INS\_469
CL\_INS\_469
CL\_INS\_469
CL\_INS\_469
CL\_INS\_469
CL\_INS\_469
CL\_INS\_469
CL\_INS\_469
CL\_INS\_469
CL\_INS\_469
CL\_INS\_469
CL\_INS\_469
CL\_INS\_469
CL\_INS\_469
CL\_INS\_469
CL\_INS\_469
CL\_INS\_469
CL\_INS\_469
CL\_INS\_469
CL\_INS\_469
CL\_INS\_469
CL\_INS\_469
CL\_INS\_469
CL\_INS\_469
CL\_INS\_469
CL\_INS\_469
CL\_INS\_469
CL\_INS\_469
CL\_INS\_469
CL\_INS\_469
CL\_INS\_469
CL\_INS\_469
Cluster ID


CL\_6514
CL\_942
CL\_941
CL\_940
CL\_5733
CL\_5734
CL\_5735
CL\_5736
CL\_5737
CL\_5738
CL\_5739
CL\_5740
CL\_5741
CL\_5742
CL\_5743
CL\_5744
CL\_5745
CL\_5746
CL\_5747
CL\_5749
CL\_7734
CL\_5750
CL\_5752
CL\_5753
CL\_5754
CL\_7735
CL\_5755
CL\_5756
CL\_5757
CL\_5758
CL\_5759
CL\_5760
CL\_9048
CL\_5761
CL\_5762
CL\_5763
CL\_5764
CL\_9047
CL\_5765
CL\_5766
CL\_5767
CL\_5768
CL\_5769
CL\_5770
CL\_9046
CL\_5771
CL\_5772
CL\_5773
CL\_5774
CL\_5775
CL\_5776
CL\_5777
