## Supplementary material for "A novel method for integrating genomic and Tn-Seq data to identify common *in vivo* fitness mechanisms across multiple bacterial species": S5 Dataset: CL_INS_473.html

Legend

 Hypothetical
/local/ifs2\_projdata/0800/projects/UMIsepsis/Serra\_Prot\_70/centroids.cluster\_roles.txt

FULL


WINDOWSVGPNG

Trim RowsRemove SingletonsSave Fasta

CL\_964


CL\_964


CL\_964


CL\_964

HighlightSelectShow Genomes


26

CL\_967


19

CL\_967


2

CL\_967


1

CL\_967

fGI ID


CL\_INS\_473
CL\_INS\_473
Cluster ID


CL\_965
CL\_966
