## Supplementary material for "A novel method for integrating genomic and Tn-Seq data to identify common *in vivo* fitness mechanisms across multiple bacterial species": S5 Dataset: CL_INS_478.html

Legend

 Hypothetical
 Serritia FitnessGene
 All VFDB Genes
/local/ifs2\_projdata/0800/projects/UMIsepsis/Serra\_Prot\_70/centroids.cluster\_roles.txt

FULL


WINDOWSVGPNG

Trim RowsRemove SingletonsSave Fasta

CL\_996


CL\_996


CL\_996


CL\_996


CL\_996


CL\_996


CL\_996


CL\_996


CL\_996


CL\_996


CL\_996


CL\_996


CL\_996


CL\_996


CL\_996


CL\_995


CL\_996


CL\_996


CL\_996


CL\_996

HighlightSelectShow Genomes


10

CL\_5154


5

CL\_1008


5

CL\_5154


4

CL\_5154


3

CL\_5154


2

CL\_5154


2

CL\_5154


2

CL\_1006


2

CL\_5154


2

CL\_5154


1

CL\_5154


1

CL\_1008


1

CL\_1008


1

CL\_1006


1

CL\_5154


1

CL\_5154


1

CL\_1006


1

CL\_5154


1

CL\_5154


1

CL\_1009

fGI ID


CL\_INS\_478
CL\_INS\_478
CL\_INS\_478
CL\_INS\_478
CL\_INS\_478
CL\_INS\_478
CL\_INS\_478
CL\_INS\_478
CL\_INS\_478
CL\_INS\_478
CL\_INS\_478
CL\_INS\_478
CL\_INS\_478
CL\_INS\_478
CL\_INS\_478
CL\_INS\_478
CL\_INS\_478
CL\_INS\_478
CL\_INS\_478
CL\_INS\_478
CL\_INS\_478
CL\_INS\_478
CL\_INS\_483
CL\_INS\_478
CL\_INS\_483
CL\_INS\_479
CL\_INS\_479
CL\_INS\_479
CL\_INS\_479
CL\_INS\_479
CL\_INS\_479
CL\_INS\_483
CL\_INS\_483
CL\_INS\_483
CL\_INS\_483
CL\_INS\_483
CL\_INS\_483
Cluster ID


CL\_9483
CL\_9991
CL\_997
CL\_998
CL\_999
CL\_1000
CL\_1001
CL\_1002
CL\_1003
CL\_1004
CL\_5146
CL\_5147
CL\_5148
CL\_5149
CL\_5675
CL\_5150
CL\_5151
CL\_5152
CL\_6750
CL\_9055
CL\_5153
CL\_11213
CL\_10155
CL\_10273
CL\_10156
CL\_10157
CL\_10158
CL\_10159
CL\_10160
CL\_10161
CL\_10162
CL\_10274
CL\_10275
CL\_7026
CL\_10276
CL\_10277
CL\_10278
