## Supplementary material for "A novel method for integrating genomic and Tn-Seq data to identify common *in vivo* fitness mechanisms across multiple bacterial species": S5 Dataset: CL_INS_482.html

Legend

 Hypothetical
 Serritia FitnessGene
 All VFDB Genes
/local/ifs2\_projdata/0800/projects/UMIsepsis/Serra\_Prot\_70/centroids.cluster\_roles.txt

FULL


WINDOWSVGPNG

Trim RowsRemove SingletonsSave Fasta

CL\_1008


CL\_1008


CL\_1006


CL\_996

HighlightSelectShow Genomes


29

CL\_1009


16

CL\_1006


2

CL\_1009


1

CL\_1009

fGI ID


CL\_INS\_483
CL\_INS\_483
CL\_INS\_483
CL\_INS\_483
CL\_INS\_483
CL\_INS\_483
CL\_INS\_483
CL\_INS\_483
Cluster ID


CL\_9058
CL\_9059
CL\_10156
CL\_7026
CL\_10276
CL\_10277
CL\_10278
CL\_10279
