## Supplementary material for "A novel method for integrating genomic and Tn-Seq data to identify common *in vivo* fitness mechanisms across multiple bacterial species": S5 Dataset: CL_INS_484.html

Legend

 Hypothetical
 All EssentialGenes
 All VFDB Genes
/local/ifs2\_projdata/0800/projects/UMIsepsis/Serra\_Prot\_70/centroids.cluster\_roles.txt

FULL


WINDOWSVGPNG

Trim RowsRemove SingletonsSave Fasta

CL\_3518


CL\_3518


CL\_3518


CL\_3518


CL\_3518


CL\_3518


CL\_3518


CL\_3518


CL\_3518


CL\_3518


CL\_3518


CL\_3518


CL\_3518

HighlightSelectShow Genomes


9

CL\_3512


4

CL\_4520


4

CL\_3516


3

CL\_3512


2

CL\_3512


1

CL\_4520


1

CL\_4520


1

CL\_3512


1

CL\_3512


1

CL\_4520


1

CL\_4520


1

CL\_4520


1

CL\_3512

fGI ID


CL\_INS\_484
CL\_INS\_369
CL\_INS\_369
CL\_INS\_369
CL\_INS\_369
CL\_INS\_369
CL\_INS\_369
CL\_INS\_369
CL\_INS\_369
CL\_INS\_369
CL\_INS\_369
CL\_INS\_369
CL\_INS\_369
CL\_INS\_369
CL\_INS\_369
CL\_INS\_369
CL\_INS\_369
CL\_INS\_369
CL\_INS\_369
CL\_INS\_369
CL\_INS\_369
CL\_INS\_298
CL\_INS\_298
CL\_INS\_298
CL\_INS\_298
CL\_INS\_298
CL\_INS\_298
CL\_INS\_298
CL\_INS\_298
CL\_INS\_298
CL\_INS\_298
CL\_INS\_369
CL\_INS\_369
CL\_INS\_369
CL\_INS\_369
CL\_INS\_369
CL\_INS\_369
CL\_INS\_369
CL\_INS\_369
CL\_INS\_369
CL\_INS\_369
CL\_INS\_369
CL\_INS\_369
CL\_INS\_369
CL\_INS\_369
CL\_INS\_369
CL\_INS\_369
CL\_INS\_369
CL\_INS\_369
CL\_INS\_369
CL\_INS\_369
CL\_INS\_369
CL\_INS\_369
CL\_INS\_369
CL\_INS\_369
CL\_INS\_369
CL\_INS\_369
CL\_INS\_369
CL\_INS\_369
CL\_INS\_369
CL\_INS\_369
CL\_INS\_369
CL\_INS\_369
CL\_INS\_369
CL\_INS\_369
CL\_INS\_369
CL\_INS\_369
CL\_INS\_369
CL\_INS\_369
CL\_INS\_369
CL\_INS\_369
CL\_INS\_369
CL\_INS\_369
CL\_INS\_369
CL\_INS\_369
CL\_INS\_369
CL\_INS\_369
CL\_INS\_369
CL\_INS\_369
CL\_INS\_369
CL\_INS\_369
CL\_INS\_369
CL\_INS\_369
CL\_INS\_369
CL\_INS\_369
CL\_INS\_369
CL\_INS\_369
CL\_INS\_369
CL\_INS\_369
CL\_INS\_369
CL\_INS\_369
CL\_INS\_369
CL\_INS\_369
CL\_INS\_369
CL\_INS\_369
CL\_INS\_369
CL\_INS\_369
CL\_INS\_369
CL\_INS\_369
CL\_INS\_369
CL\_INS\_369
CL\_INS\_369
CL\_INS\_369
CL\_INS\_369
CL\_INS\_369
CL\_INS\_369
CL\_INS\_369
CL\_INS\_369
CL\_INS\_369
CL\_INS\_369
CL\_INS\_369
CL\_INS\_369
CL\_INS\_369
CL\_INS\_369
CL\_INS\_369
CL\_INS\_369
CL\_INS\_369
CL\_INS\_369
CL\_INS\_369
CL\_INS\_369
CL\_INS\_369
CL\_INS\_369
CL\_INS\_369
CL\_INS\_369
CL\_INS\_369
CL\_INS\_369
CL\_INS\_369
CL\_INS\_369
CL\_INS\_369
CL\_INS\_369
CL\_INS\_369
CL\_INS\_369
CL\_INS\_369
CL\_INS\_369
CL\_INS\_369
CL\_INS\_369
CL\_INS\_369
CL\_INS\_369
CL\_INS\_369
CL\_INS\_369
CL\_INS\_369
CL\_INS\_369
CL\_INS\_369
CL\_INS\_369
CL\_INS\_369
Cluster ID


CL\_11859
CL\_5017
CL\_5018
CL\_6672
CL\_5949
CL\_5948
CL\_5947
CL\_5946
CL\_11779
CL\_11780
CL\_9939
CL\_6673
CL\_5019
CL\_6674
CL\_6675
CL\_6676
CL\_6677
CL\_6678
CL\_6679
CL\_6680
CL\_3513
CL\_12769
CL\_12768
CL\_12767
CL\_12766
CL\_12765
CL\_12764
CL\_12763
CL\_12762
CL\_7862
CL\_12761
CL\_8276
CL\_9585
CL\_8277
CL\_8278
CL\_8279
CL\_9586
CL\_8280
CL\_8281
CL\_8282
CL\_8283
CL\_8284
CL\_8285
CL\_8290
CL\_8291
CL\_9587
CL\_8292
CL\_8293
CL\_8294
CL\_8295
CL\_8296
CL\_8297
CL\_8609
CL\_8608
CL\_8298
CL\_8299
CL\_8607
CL\_8301
CL\_8302
CL\_9588
CL\_8304
CL\_8305
CL\_8306
CL\_8307
CL\_8308
CL\_8309
CL\_8310
CL\_8312
CL\_8313
CL\_8314
CL\_9589
CL\_9590
CL\_9591
CL\_8606
CL\_8605
CL\_8604
CL\_8318
CL\_8319
CL\_8320
CL\_8321
CL\_8322
CL\_8603
CL\_9592
CL\_8602
CL\_8601
CL\_8600
CL\_8599
CL\_8598
CL\_8597
CL\_8596
CL\_8595
CL\_8594
CL\_8593
CL\_8592
CL\_8591
CL\_8590
CL\_8589
CL\_8588
CL\_8587
CL\_8586
CL\_8585
CL\_8584
CL\_8583
CL\_8582
CL\_8581
CL\_8580
CL\_8328
CL\_9593
CL\_9594
CL\_9595
CL\_9596
CL\_9597
CL\_8579
CL\_8578
CL\_8577
CL\_8576
CL\_4845
CL\_8575
CL\_7571
CL\_8574
CL\_7576
CL\_7577
CL\_8573
CL\_8572
CL\_8571
CL\_8570
CL\_8363
CL\_8569
CL\_8364
CL\_8366
CL\_8568
CL\_8567
CL\_8367
CL\_9598
CL\_9599
CL\_9600
CL\_9601
CL\_9602
CL\_8368
CL\_8566
CL\_8565
CL\_8564
CL\_8563
CL\_8370
CL\_8371
