## Supplementary material for "A novel method for integrating genomic and Tn-Seq data to identify common *in vivo* fitness mechanisms across multiple bacterial species": S6 Dataset: CL_INS_1.html

Legend

 Hypothetical
 All EssentialGenes
 All Fitness Genes
 Other

FULL


WINDOWSVGPNG

Trim RowsRemove SingletonsSave Fasta

CL\_1917


CL\_1917


CL\_1917

HighlightSelectShow Genomes


3

CL\_1915


1

CL\_1915


1

CL\_1915

fGI ID


CL\_INS\_1
CL\_INS\_1
Cluster ID


CL\_5878
CL\_1916
