## Supplementary material for "A novel method for integrating genomic and Tn-Seq data to identify common *in vivo* fitness mechanisms across multiple bacterial species": S6 Dataset: CL_INS_2.html

Legend

 Hypothetical
 Regulatoryfunctions
 All Fitness Genes
 Other
 Transport +binding proteins

FULL


WINDOWSVGPNG

Trim RowsRemove SingletonsSave Fasta

CL\_1910


CL\_1910


CL\_1910

HighlightSelectShow Genomes


2

CL\_1908


2

CL\_1908


1

CL\_1908

fGI ID


CL\_INS\_2
CL\_INS\_2
CL\_INS\_2
CL\_INS\_2
CL\_INS\_2
CL\_INS\_2
CL\_INS\_2
Cluster ID


CL\_1909
CL\_4445
CL\_4444
CL\_4443
CL\_4442
CL\_4441
CL\_4440
