## Supplementary material for "A novel method for integrating genomic and Tn-Seq data to identify common *in vivo* fitness mechanisms across multiple bacterial species": S6 Dataset: CL_INS_3.html

Legend

 Regulatoryfunctions
 Hypothetical
 All Fitness Genes

FULL


WINDOWSVGPNG

Trim RowsRemove SingletonsSave Fasta

CL\_1877


CL\_1877


CL\_1877

HighlightSelectShow Genomes


2

CL\_1875


1

CL\_1866


1

CL\_1866

fGI ID


CL\_INS\_3
CL\_INS\_3
CL\_INS\_3
CL\_INS\_3
CL\_INS\_3
Cluster ID


CL\_1876
CL\_4454
CL\_4453
CL\_4452
CL\_4450
