## Supplementary material for "A novel method for integrating genomic and Tn-Seq data to identify common *in vivo* fitness mechanisms across multiple bacterial species": S6 Dataset: CL_INS_4.html

Legend

 Regulatoryfunctions
 Hypothetical
 All Fitness Genes
 SignalTransduction +Quorum Sensing
 Other
 Transport +binding proteins

FULL


WINDOWSVGPNG

Trim RowsRemove SingletonsSave Fasta

CL\_1866


CL\_1866


CL\_1866


CL\_4100


CL\_1866

HighlightSelectShow Genomes


2

CL\_1867


1

CL\_1877


1

CL\_1867


1

CL\_1867


1

CL\_1877

fGI ID


CL\_INS\_4
CL\_INS\_150
CL\_INS\_3
CL\_INS\_3
CL\_INS\_3
CL\_INS\_4
CL\_INS\_4
CL\_INS\_4
CL\_INS\_4
CL\_INS\_4
CL\_INS\_4
CL\_INS\_4
CL\_INS\_4
CL\_INS\_4
CL\_INS\_4
CL\_INS\_4
CL\_INS\_4
CL\_INS\_4
CL\_INS\_4
CL\_INS\_4
CL\_INS\_4
Cluster ID


CL\_6707
CL\_4449
CL\_4450
CL\_4452
CL\_4453
CL\_4102
CL\_4103
CL\_4104
CL\_4105
CL\_4106
CL\_4107
CL\_4108
CL\_4109
CL\_4110
CL\_4111
CL\_4112
CL\_4113
CL\_4114
CL\_4115
CL\_4116
CL\_4117
