## Supplementary material for "A novel method for integrating genomic and Tn-Seq data to identify common *in vivo* fitness mechanisms across multiple bacterial species": S6 Dataset: CL_INS_5.html

Legend

 Regulatoryfunctions
 Other
 EnergyMetabolism
 Transport +binding proteins

FULL


WINDOWSVGPNG

Trim RowsRemove SingletonsSave Fasta

CL\_4533


CL\_4533

HighlightSelectShow Genomes


3

CL\_1830


1

CL\_1699

fGI ID


CL\_INS\_5
CL\_INS\_5
CL\_INS\_5
Cluster ID


CL\_5876
CL\_4534
CL\_4535
