## Supplementary material for "A novel method for integrating genomic and Tn-Seq data to identify common *in vivo* fitness mechanisms across multiple bacterial species": S6 Dataset: CL_INS_6.html

Legend

 Hypothetical
 Regulatoryfunctions
 All EssentialGenes
 All Fitness Genes
 SignalTransduction +Quorum Sensing
 Other
 EnergyMetabolism
 Transport +binding proteins

FULL


WINDOWSVGPNG

Trim RowsRemove SingletonsSave Fasta

CL\_1814


CL\_1814


CL\_1829


CL\_1411


CL\_1814

HighlightSelectShow Genomes


2

CL\_1812


1

CL\_1955


1

CL\_1812


1

CL\_1812


1

CL\_273

fGI ID


CL\_INS\_138
CL\_INS\_182
CL\_INS\_182
CL\_INS\_182
CL\_INS\_182
CL\_INS\_182
CL\_INS\_6
CL\_INS\_136
CL\_INS\_136
CL\_INS\_6
CL\_INS\_107
CL\_INS\_107
CL\_INS\_107
CL\_INS\_107
CL\_INS\_107
CL\_INS\_107
CL\_INS\_107
CL\_INS\_6
CL\_INS\_6
CL\_INS\_6
CL\_INS\_6
CL\_INS\_6
CL\_INS\_6
CL\_INS\_6
CL\_INS\_6
CL\_INS\_6
CL\_INS\_6
CL\_INS\_6
CL\_INS\_6
CL\_INS\_6
CL\_INS\_6
CL\_INS\_6
CL\_INS\_6
CL\_INS\_6
CL\_INS\_6
CL\_INS\_6
CL\_INS\_6
CL\_INS\_6
CL\_INS\_6
CL\_INS\_6
CL\_INS\_6
CL\_INS\_6
CL\_INS\_6
CL\_INS\_6
Cluster ID


CL\_1813
CL\_4427
CL\_4426
CL\_4425
CL\_4424
CL\_4423
CL\_5624
CL\_4494
CL\_4495
CL\_5623
CL\_4952
CL\_4951
CL\_4950
CL\_4949
CL\_4948
CL\_4947
CL\_4946
CL\_5702
CL\_6579
CL\_6578
CL\_6577
CL\_6576
CL\_6575
CL\_6574
CL\_6573
CL\_6572
CL\_6571
CL\_6570
CL\_6569
CL\_6568
CL\_6567
CL\_6566
CL\_6565
CL\_6564
CL\_6563
CL\_6562
CL\_6561
CL\_6560
CL\_6559
CL\_6558
CL\_6557
CL\_6556
CL\_5089
CL\_6555
