## Supplementary material for "A novel method for integrating genomic and Tn-Seq data to identify common *in vivo* fitness mechanisms across multiple bacterial species": S6 Dataset: CL_INS_8.html

Legend

 Regulatoryfunctions
 Hypothetical
 SignalTransduction +Quorum Sensing
 Other
 Transport +binding proteins

FULL


WINDOWSVGPNG

Trim RowsRemove SingletonsSave Fasta

CL\_1761


CL\_1761


CL\_1761


CL\_1761


CL\_1182

HighlightSelectShow Genomes


1

CL\_1763


1

CL\_1763


1

CL\_1763


1

CL\_1763


1

CL\_1763

fGI ID


CL\_INS\_8
CL\_INS\_149
CL\_INS\_8
CL\_INS\_8
CL\_INS\_8
CL\_INS\_8
CL\_INS\_8
CL\_INS\_8
CL\_INS\_8
CL\_INS\_8
CL\_INS\_8
CL\_INS\_8
CL\_INS\_8
CL\_INS\_8
CL\_INS\_8
CL\_INS\_8
Cluster ID


CL\_1762
CL\_4572
CL\_3141
CL\_3142
CL\_5116
CL\_4884
CL\_5182
CL\_5181
CL\_5180
CL\_5179
CL\_5178
CL\_5177
CL\_5176
CL\_5175
CL\_5174
CL\_5173
