## Supplementary material for "A novel method for integrating genomic and Tn-Seq data to identify common *in vivo* fitness mechanisms across multiple bacterial species": S6 Dataset: CL_INS_9.html

Legend

 Regulatoryfunctions
 SignalTransduction +Quorum Sensing

FULL


WINDOWSVGPNG

Trim RowsRemove SingletonsSave Fasta

CL\_1750


CL\_1750

HighlightSelectShow Genomes


3

CL\_1748


1

CL\_1748

fGI ID

CL\_INS\_9
Cluster ID

CL\_1749
