## Supplementary material for "A novel method for integrating genomic and Tn-Seq data to identify common *in vivo* fitness mechanisms across multiple bacterial species": S6 Dataset: CL_INS_10.html

Legend

 Regulatoryfunctions
 Hypothetical
 All Fitness Genes
 Other

FULL


WINDOWSVGPNG

Trim RowsRemove SingletonsSave Fasta

CL\_1726


CL\_947


CL\_1726


CL\_1726

HighlightSelectShow Genomes


2

CL\_1722


1

CL\_1722


1

CL\_2230


1

CL\_1722

fGI ID


CL\_INS\_10
CL\_INS\_10
CL\_INS\_10
CL\_INS\_10
CL\_INS\_10
CL\_INS\_10
Cluster ID


CL\_1725
CL\_1724
CL\_1723
CL\_3840
CL\_3839
CL\_3838
