## Supplementary material for "A novel method for integrating genomic and Tn-Seq data to identify common *in vivo* fitness mechanisms across multiple bacterial species": S6 Dataset: CL_INS_11.html

Legend

 Hypothetical
 Other
 Transport +binding proteins

FULL


WINDOWSVGPNG

Trim RowsRemove SingletonsSave Fasta

CL\_1722


CL\_1722

HighlightSelectShow Genomes


2

CL\_1720


1

CL\_1720

fGI ID


CL\_INS\_11
CL\_INS\_11
Cluster ID


CL\_1721
CL\_7071
