## Supplementary material for "A novel method for integrating genomic and Tn-Seq data to identify common *in vivo* fitness mechanisms across multiple bacterial species": S6 Dataset: CL_INS_12.html

Legend

 Hypothetical
 All EssentialGenes
 All Fitness Genes
 Other

FULL


WINDOWSVGPNG

Trim RowsRemove SingletonsSave Fasta

CL\_1710


CL\_1710


CL\_1710

HighlightSelectShow Genomes


2

CL\_1708


2

CL\_1708


1

CL\_1708

fGI ID


CL\_INS\_12
CL\_INS\_12
CL\_INS\_12
Cluster ID


CL\_1709
CL\_4541
CL\_4540
