## Supplementary material for "A novel method for integrating genomic and Tn-Seq data to identify common *in vivo* fitness mechanisms across multiple bacterial species": S6 Dataset: CL_INS_13.html

Legend

 Regulatoryfunctions
 Biosynthesis ofcofactors,prostheticgroups, +carriers
 Other

FULL


WINDOWSVGPNG

Trim RowsRemove SingletonsSave Fasta

CL\_1698


CL\_1736


CL\_1698

HighlightSelectShow Genomes


3

CL\_1696


1

CL\_1696


1

CL\_1696

fGI ID

CL\_INS\_13
Cluster ID

CL\_1697
