## Supplementary material for "A novel method for integrating genomic and Tn-Seq data to identify common *in vivo* fitness mechanisms across multiple bacterial species": S6 Dataset: CL_INS_14.html

Legend

 Hypothetical
 All Fitness Genes
 Cell Envelope
 Other
 All VFDB Genes

FULL


WINDOWSVGPNG

Trim RowsRemove SingletonsSave Fasta

CL\_1690


CL\_604


CL\_1690

HighlightSelectShow Genomes


3

CL\_1679


1

CL\_1679


1

Break

fGI ID


CL\_INS\_14
CL\_INS\_14
CL\_INS\_14
CL\_INS\_14
CL\_INS\_14
CL\_INS\_14
CL\_INS\_14
CL\_INS\_14
CL\_INS\_14
CL\_INS\_14
CL\_INS\_14
CL\_INS\_14
CL\_INS\_14
CL\_INS\_14
Cluster ID


CL\_5325
CL\_3692
CL\_3691
CL\_3690
CL\_1689
CL\_1688
CL\_1687
CL\_1686
CL\_1685
CL\_1684
CL\_1683
CL\_1682
CL\_1681
CL\_1680
