## Supplementary material for "A novel method for integrating genomic and Tn-Seq data to identify common *in vivo* fitness mechanisms across multiple bacterial species": S6 Dataset: CL_INS_15.html

Legend

 Hypothetical
 All Fitness Genes
 Other
 All VFDB Genes

FULL


WINDOWSVGPNG

Trim RowsRemove SingletonsSave Fasta

CL\_1677

HighlightSelectShow Genomes


3

CL\_1672

fGI ID


CL\_INS\_15
CL\_INS\_15
CL\_INS\_15
CL\_INS\_15
Cluster ID


CL\_1676
CL\_1675
CL\_1674
CL\_1673
