## Supplementary material for "A novel method for integrating genomic and Tn-Seq data to identify common *in vivo* fitness mechanisms across multiple bacterial species": S6 Dataset: CL_INS_16.html

Legend

 All Fitness Genes
 Cell Envelope
 Other
 All VFDB Genes

FULL


WINDOWSVGPNG

Trim RowsRemove SingletonsSave Fasta

CL\_4170


CL\_4170


CL\_4170


CL\_1670

HighlightSelectShow Genomes


2

CL\_1668


1

CL\_1025


1

CL\_1668


1

CL\_1668

fGI ID


CL\_INS\_16
CL\_INS\_16
Cluster ID


CL\_6679
CL\_1669
