## Supplementary material for "A novel method for integrating genomic and Tn-Seq data to identify common *in vivo* fitness mechanisms across multiple bacterial species": S6 Dataset: CL_INS_17.html

Legend

 Regulatoryfunctions
 Hypothetical
 All EssentialGenes
 Other
 All VFDB Genes
 Transport +binding proteins

FULL


WINDOWSVGPNG

Trim RowsRemove SingletonsSave Fasta

CL\_1635


CL\_1635


CL\_1635


CL\_1635


CL\_1635

HighlightSelectShow Genomes


1

CL\_1643


1

CL\_3646


1

CL\_1637


1

CL\_1643


1

CL\_5047

fGI ID


CL\_INS\_17
CL\_INS\_17
CL\_INS\_17
CL\_INS\_17
CL\_INS\_17
CL\_INS\_17
CL\_INS\_17
CL\_INS\_17
CL\_INS\_17
CL\_INS\_17
CL\_INS\_17
CL\_INS\_17
CL\_INS\_17
CL\_INS\_17
CL\_INS\_17
CL\_INS\_17
Cluster ID


CL\_6664
CL\_7077
CL\_7076
CL\_7075
CL\_5860
CL\_5861
CL\_5862
CL\_5863
CL\_4338
CL\_4339
CL\_4340
CL\_4341
CL\_4342
CL\_4343
CL\_4344
CL\_4345
