## Supplementary material for "A novel method for integrating genomic and Tn-Seq data to identify common *in vivo* fitness mechanisms across multiple bacterial species": S6 Dataset: CL_INS_18.html

Legend

 Regulatoryfunctions
 Hypothetical
 All EssentialGenes
 SignalTransduction +Quorum Sensing
 Other

FULL


WINDOWSVGPNG

Trim RowsRemove SingletonsSave Fasta

CL\_1635


CL\_1635


CL\_1635


CL\_1632


CL\_3478

HighlightSelectShow Genomes


1

CL\_4390


1

CL\_1627


1

CL\_1627


1

CL\_1627


1

CL\_1627

fGI ID


CL\_INS\_199
CL\_INS\_18
CL\_INS\_207
CL\_INS\_18
CL\_INS\_18
CL\_INS\_18
Cluster ID


CL\_5856
CL\_4135
CL\_4392
CL\_1630
CL\_1629
CL\_1628
