## Supplementary material for "A novel method for integrating genomic and Tn-Seq data to identify common *in vivo* fitness mechanisms across multiple bacterial species": S6 Dataset: CL_INS_19.html

Legend

 Hypothetical
 All Fitness Genes

FULL


WINDOWSVGPNG

Trim RowsRemove SingletonsSave Fasta

CL\_1621


CL\_1621


CL\_1621


CL\_1621

HighlightSelectShow Genomes


2

CL\_1617


1

CL\_3082


1

CL\_1617


1

CL\_1617

fGI ID


CL\_INS\_19
CL\_INS\_19
CL\_INS\_19
Cluster ID


CL\_1620
CL\_1619
CL\_1618
