## Supplementary material for "A novel method for integrating genomic and Tn-Seq data to identify common *in vivo* fitness mechanisms across multiple bacterial species": S6 Dataset: CL_INS_20.html

Legend

 Regulatoryfunctions
 Hypothetical
 All EssentialGenes
 All Fitness Genes
 Cell Envelope
 All VFDB Genes

FULL


WINDOWSVGPNG

Trim RowsRemove SingletonsSave Fasta

CL\_1598


CL\_1615


CL\_1243


CL\_1598


CL\_1601

HighlightSelectShow Genomes


2

CL\_1596


1

CL\_1596


1

CL\_1596


1

CL\_1251


1

CL\_1596

fGI ID


CL\_INS\_20
CL\_INS\_20
CL\_INS\_20
CL\_INS\_20
CL\_INS\_20
Cluster ID


CL\_4203
CL\_1597
CL\_5217
CL\_5218
CL\_5219
