## Supplementary material for "A novel method for integrating genomic and Tn-Seq data to identify common *in vivo* fitness mechanisms across multiple bacterial species": S6 Dataset: CL_INS_21.html

Legend

 Regulatoryfunctions
 Hypothetical
 All EssentialGenes
 Other
 Transport +binding proteins

FULL


WINDOWSVGPNG

Trim RowsRemove SingletonsSave Fasta

CL\_1579


CL\_1579


CL\_1579


CL\_1579

HighlightSelectShow Genomes


1

CL\_1580


1

CL\_1580


1

CL\_1580


1

CL\_1580

fGI ID


CL\_INS\_21
CL\_INS\_21
CL\_INS\_21
CL\_INS\_21
CL\_INS\_21
CL\_INS\_21
CL\_INS\_21
CL\_INS\_90
CL\_INS\_90
CL\_INS\_90
CL\_INS\_90
CL\_INS\_90
CL\_INS\_21
CL\_INS\_21
CL\_INS\_21
Cluster ID


CL\_6661
CL\_7083
CL\_7082
CL\_7081
CL\_7080
CL\_7079
CL\_7078
CL\_4387
CL\_4386
CL\_4385
CL\_4384
CL\_4383
CL\_5854
CL\_4870
CL\_5855
