## Supplementary material for "A novel method for integrating genomic and Tn-Seq data to identify common *in vivo* fitness mechanisms across multiple bacterial species": S6 Dataset: CL_INS_22.html

Legend

 Hypothetical
 Other

FULL


WINDOWSVGPNG

Trim RowsRemove SingletonsSave Fasta

CL\_1576


CL\_1576


CL\_1576


CL\_1576

HighlightSelectShow Genomes


1

CL\_1573


1

CL\_1573


1

CL\_1573


1

CL\_1573

fGI ID


CL\_INS\_22
CL\_INS\_22
CL\_INS\_22
CL\_INS\_22
CL\_INS\_22
Cluster ID


CL\_6660
CL\_1575
CL\_5853
CL\_5852
CL\_1574
