## Supplementary material for "A novel method for integrating genomic and Tn-Seq data to identify common *in vivo* fitness mechanisms across multiple bacterial species": S6 Dataset: CL_INS_24.html

Legend

 All Fitness Genes
 Other

FULL


WINDOWSVGPNG

Trim RowsRemove SingletonsSave Fasta

CL\_1517


CL\_1517


CL\_1517

HighlightSelectShow Genomes


3

CL\_1515


1

CL\_1515


1

CL\_1515

fGI ID


CL\_INS\_24
CL\_INS\_24
CL\_INS\_24
Cluster ID


CL\_1516
CL\_3806
CL\_3807
