## Supplementary material for "A novel method for integrating genomic and Tn-Seq data to identify common *in vivo* fitness mechanisms across multiple bacterial species": S6 Dataset: CL_INS_25.html

Legend

 All Fitness Genes
 Other

FULL


WINDOWSVGPNG

Trim RowsRemove SingletonsSave Fasta

CL\_1495


CL\_1495


CL\_1495

HighlightSelectShow Genomes


3

CL\_1494


1

CL\_1494


1

CL\_1494

fGI ID


CL\_INS\_25
CL\_INS\_25
Cluster ID


CL\_4193
CL\_5843
