## Supplementary material for "A novel method for integrating genomic and Tn-Seq data to identify common *in vivo* fitness mechanisms across multiple bacterial species": S6 Dataset: CL_INS_26.html

Legend

 Hypothetical
 Regulatoryfunctions
 All EssentialGenes
 All Fitness Genes
 Other
 EnergyMetabolism
 Transport +binding proteins

FULL


WINDOWSVGPNG

Trim RowsRemove SingletonsSave Fasta

CL\_1476


CL\_960


CL\_1476


CL\_1476

HighlightSelectShow Genomes


3

CL\_969


1

CL\_969


1

CL\_1471


1

CL\_969

fGI ID


CL\_INS\_32
CL\_INS\_32
CL\_INS\_32
CL\_INS\_26
CL\_INS\_26
CL\_INS\_26
CL\_INS\_26
CL\_INS\_26
CL\_INS\_26
CL\_INS\_26
CL\_INS\_26
Cluster ID


CL\_1475
CL\_1474
CL\_1473
CL\_1472
CL\_962
CL\_963
CL\_964
CL\_965
CL\_966
CL\_967
CL\_968
