## Supplementary material for "A novel method for integrating genomic and Tn-Seq data to identify common *in vivo* fitness mechanisms across multiple bacterial species": S6 Dataset: CL_INS_27.html

Legend

 Hypothetical
 Biosynthesis ofcofactors,prostheticgroups, +carriers
 All Fitness Genes
 Other
 EnergyMetabolism
 Transport +binding proteins

FULL


WINDOWSVGPNG

Trim RowsRemove SingletonsSave Fasta

CL\_987


CL\_987


CL\_987


CL\_987


CL\_987

HighlightSelectShow Genomes


1

CL\_990


1

CL\_990


1

CL\_990


1

CL\_1239


1

CL\_990

fGI ID


CL\_INS\_27
CL\_INS\_27
CL\_INS\_27
CL\_INS\_27
CL\_INS\_27
CL\_INS\_27
CL\_INS\_27
CL\_INS\_27
CL\_INS\_27
CL\_INS\_27
CL\_INS\_27
CL\_INS\_27
CL\_INS\_27
CL\_INS\_27
Cluster ID


CL\_6653
CL\_3901
CL\_3902
CL\_3903
CL\_3904
CL\_988
CL\_5842
CL\_989
CL\_7084
CL\_7085
CL\_7086
CL\_7087
CL\_7088
CL\_7089
