## Supplementary material for "A novel method for integrating genomic and Tn-Seq data to identify common *in vivo* fitness mechanisms across multiple bacterial species": S6 Dataset: CL_INS_28.html

Legend

 Regulatoryfunctions
 Hypothetical
 All EssentialGenes
 Cell Envelope
 Other
 Centralintermediarymetabolism

FULL


WINDOWSVGPNG

Trim RowsRemove SingletonsSave Fasta

CL\_2554


CL\_2554


CL\_2554


CL\_2554


CL\_2554

HighlightSelectShow Genomes


1

CL\_2549


1

CL\_2549


1

CL\_2549


1

CL\_2549


1

CL\_2549

fGI ID


CL\_INS\_28
CL\_INS\_28
CL\_INS\_28
CL\_INS\_28
CL\_INS\_28
CL\_INS\_28
CL\_INS\_28
CL\_INS\_28
CL\_INS\_28
CL\_INS\_28
CL\_INS\_28
CL\_INS\_28
CL\_INS\_28
CL\_INS\_28
CL\_INS\_28
Cluster ID


CL\_6845
CL\_5343
CL\_5342
CL\_5341
CL\_5340
CL\_5339
CL\_5338
CL\_5337
CL\_5336
CL\_6844
CL\_5335
CL\_2553
CL\_2552
CL\_2551
CL\_2550
