## Supplementary material for "A novel method for integrating genomic and Tn-Seq data to identify common *in vivo* fitness mechanisms across multiple bacterial species": S6 Dataset: CL_INS_29.html

Legend

 Regulatoryfunctions
 Hypothetical
 All EssentialGenes
 Proteinsynthesis/fate
 Other
 EnergyMetabolism
 All VFDB Genes
 Transport +binding proteins

FULL


WINDOWSVGPNG

Trim RowsRemove SingletonsSave Fasta

Break


CL\_2504


CL\_1196


CL\_2504


CL\_2504


CL\_2504

HighlightSelectShow Genomes


1

CL\_2502


1

CL\_2502


1

CL\_2502


1

CL\_2490


1

CL\_2502


1

CL\_2453

fGI ID


CL\_INS\_29
CL\_INS\_29
CL\_INS\_29
CL\_INS\_29
CL\_INS\_29
CL\_INS\_29
CL\_INS\_29
CL\_INS\_29
CL\_INS\_29
CL\_INS\_29
CL\_INS\_29
CL\_INS\_29
CL\_INS\_29
CL\_INS\_29
CL\_INS\_29
CL\_INS\_29
CL\_INS\_29
CL\_INS\_29
CL\_INS\_29
CL\_INS\_29
CL\_INS\_29
CL\_INS\_31
CL\_INS\_31
CL\_INS\_155
CL\_INS\_31
CL\_INS\_31
CL\_INS\_31
CL\_INS\_31
CL\_INS\_31
CL\_INS\_31
CL\_INS\_31
CL\_INS\_31
CL\_INS\_31
CL\_INS\_31
CL\_INS\_31
CL\_INS\_31
Cluster ID


CL\_2503
CL\_6843
CL\_6842
CL\_4019
CL\_4018
CL\_4017
CL\_5948
CL\_5947
CL\_5946
CL\_2501
CL\_2500
CL\_5945
CL\_2499
CL\_2498
CL\_2497
CL\_2496
CL\_2495
CL\_2494
CL\_2493
CL\_2492
CL\_4693
CL\_4692
CL\_4691
CL\_4690
CL\_4689
CL\_4688
CL\_4687
CL\_4686
CL\_4685
CL\_4684
CL\_4683
CL\_4682
CL\_4681
CL\_4680
CL\_4679
CL\_4678
