## Supplementary material for "A novel method for integrating genomic and Tn-Seq data to identify common *in vivo* fitness mechanisms across multiple bacterial species": S6 Dataset: CL_INS_30.html

Legend

 Hypothetical
 Regulatoryfunctions
 Biosynthesis ofcofactors,prostheticgroups, +carriers
 All Fitness Genes
 SignalTransduction +Quorum Sensing
 Other
 EnergyMetabolism
 Transport +binding proteins
 All VFDB Genes

FULL


WINDOWSVGPNG

Trim RowsRemove SingletonsSave Fasta

CL\_2454


CL\_2454


CL\_2454


CL\_2454

HighlightSelectShow Genomes


1

CL\_2489


1

CL\_4481


1

CL\_2489


1

CL\_2489

fGI ID


CL\_INS\_30
CL\_INS\_30
CL\_INS\_30
CL\_INS\_30
CL\_INS\_30
CL\_INS\_30
CL\_INS\_30
CL\_INS\_30
CL\_INS\_30
CL\_INS\_30
CL\_INS\_30
CL\_INS\_30
CL\_INS\_30
CL\_INS\_30
CL\_INS\_30
CL\_INS\_30
CL\_INS\_30
CL\_INS\_30
CL\_INS\_30
CL\_INS\_30
CL\_INS\_30
CL\_INS\_30
CL\_INS\_30
CL\_INS\_30
CL\_INS\_30
CL\_INS\_30
CL\_INS\_30
CL\_INS\_30
CL\_INS\_30
CL\_INS\_30
CL\_INS\_30
CL\_INS\_30
CL\_INS\_30
CL\_INS\_30
CL\_INS\_30
CL\_INS\_30
CL\_INS\_30
CL\_INS\_30
CL\_INS\_30
CL\_INS\_30
CL\_INS\_30
CL\_INS\_30
CL\_INS\_30
CL\_INS\_30
CL\_INS\_30
Cluster ID


CL\_4549
CL\_4548
CL\_4547
CL\_4546
CL\_3709
CL\_3708
CL\_5903
CL\_5904
CL\_6839
CL\_6840
CL\_4896
CL\_2455
CL\_2456
CL\_2457
CL\_2458
CL\_2459
CL\_2460
CL\_2461
CL\_2462
CL\_2463
CL\_2464
CL\_2465
CL\_2466
CL\_2467
CL\_2468
CL\_2469
CL\_2470
CL\_2471
CL\_2472
CL\_2473
CL\_2474
CL\_2475
CL\_2476
CL\_2477
CL\_2478
CL\_2479
CL\_2480
CL\_2481
CL\_2482
CL\_2483
CL\_2484
CL\_2485
CL\_2486
CL\_2487
CL\_2488
