## Supplementary material for "A novel method for integrating genomic and Tn-Seq data to identify common *in vivo* fitness mechanisms across multiple bacterial species": S6 Dataset: CL_INS_31.html

Legend

 Regulatoryfunctions
 Hypothetical
 Other
 Proteinsynthesis/fate
 EnergyMetabolism
 Transport +binding proteins
 All VFDB Genes

FULL


WINDOWSVGPNG

Trim RowsRemove SingletonsSave Fasta

CL\_2453


CL\_2453

HighlightSelectShow Genomes


4

CL\_2454


1

CL\_2504

fGI ID


CL\_INS\_31
CL\_INS\_31
CL\_INS\_31
CL\_INS\_31
CL\_INS\_31
CL\_INS\_31
CL\_INS\_31
CL\_INS\_31
CL\_INS\_31
CL\_INS\_31
CL\_INS\_31
CL\_INS\_31
CL\_INS\_31
CL\_INS\_155
CL\_INS\_31
CL\_INS\_31
Cluster ID


CL\_4677
CL\_4678
CL\_4679
CL\_4680
CL\_4681
CL\_4682
CL\_4683
CL\_4684
CL\_4685
CL\_4686
CL\_4687
CL\_4688
CL\_4689
CL\_4690
CL\_4691
CL\_4692
