## Supplementary material for "A novel method for integrating genomic and Tn-Seq data to identify common *in vivo* fitness mechanisms across multiple bacterial species": S6 Dataset: CL_INS_32.html

Legend

 Regulatoryfunctions
 Hypothetical
 All EssentialGenes
 All Fitness Genes
 SignalTransduction +Quorum Sensing
 Cell Envelope
 Other
 EnergyMetabolism
 Transport +binding proteins

FULL


WINDOWSVGPNG

Trim RowsRemove SingletonsSave Fasta

CL\_3425


CL\_827


CL\_827


CL\_4092


CL\_827


CL\_1091

HighlightSelectShow Genomes


1

CL\_804


1

CL\_804


1

CL\_3465


1

CL\_804


1

CL\_804


1

CL\_804

fGI ID


CL\_INS\_32
CL\_INS\_32
CL\_INS\_32
CL\_INS\_32
CL\_INS\_32
CL\_INS\_32
CL\_INS\_32
CL\_INS\_32
CL\_INS\_32
CL\_INS\_32
CL\_INS\_32
CL\_INS\_32
CL\_INS\_32
CL\_INS\_32
CL\_INS\_32
CL\_INS\_32
CL\_INS\_32
CL\_INS\_32
CL\_INS\_32
CL\_INS\_32
CL\_INS\_32
CL\_INS\_32
CL\_INS\_32
CL\_INS\_32
CL\_INS\_32
CL\_INS\_32
CL\_INS\_32
CL\_INS\_32
CL\_INS\_32
CL\_INS\_32
CL\_INS\_32
CL\_INS\_32
CL\_INS\_32
CL\_INS\_32
CL\_INS\_32
CL\_INS\_32
CL\_INS\_32
CL\_INS\_32
CL\_INS\_32
CL\_INS\_32
CL\_INS\_32
CL\_INS\_32
CL\_INS\_32
CL\_INS\_32
CL\_INS\_32
CL\_INS\_32
CL\_INS\_32
CL\_INS\_32
CL\_INS\_32
CL\_INS\_32
CL\_INS\_32
Cluster ID


CL\_5988
CL\_5987
CL\_5986
CL\_3474
CL\_3740
CL\_3739
CL\_3738
CL\_3737
CL\_5670
CL\_1473
CL\_6220
CL\_1474
CL\_6219
CL\_1475
CL\_6218
CL\_6217
CL\_7197
CL\_7198
CL\_7199
CL\_7200
CL\_7201
CL\_5669
CL\_5668
CL\_5667
CL\_7202
CL\_7203
CL\_7204
CL\_7205
CL\_5666
CL\_826
CL\_825
CL\_824
CL\_823
CL\_822
CL\_821
CL\_820
CL\_819
CL\_818
CL\_817
CL\_816
CL\_815
CL\_814
CL\_813
CL\_812
CL\_811
CL\_810
CL\_809
CL\_808
CL\_807
CL\_806
CL\_805
