## Supplementary material for "A novel method for integrating genomic and Tn-Seq data to identify common *in vivo* fitness mechanisms across multiple bacterial species": S6 Dataset: CL_INS_33.html

Legend

 Hypothetical
 Proteinsynthesis/fate
 Other

FULL


WINDOWSVGPNG

Trim RowsRemove SingletonsSave Fasta

CL\_684


CL\_684

HighlightSelectShow Genomes


4

CL\_683


1

CL\_682

fGI ID


CL\_INS\_30
CL\_INS\_30
CL\_INS\_187
CL\_INS\_33
Cluster ID


CL\_3709
CL\_3708
CL\_3707
CL\_3706
