## Supplementary material for "A novel method for integrating genomic and Tn-Seq data to identify common *in vivo* fitness mechanisms across multiple bacterial species": S6 Dataset: CL_INS_34.html

Legend

 Regulatoryfunctions
 Other
 Transport +binding proteins

FULL


WINDOWSVGPNG

Trim RowsRemove SingletonsSave Fasta

CL\_666


CL\_666

HighlightSelectShow Genomes


2

CL\_665


2

CL\_665

fGI ID


CL\_INS\_34
CL\_INS\_34
CL\_INS\_34
Cluster ID


CL\_4552
CL\_4551
CL\_4550
