## Supplementary material for "A novel method for integrating genomic and Tn-Seq data to identify common *in vivo* fitness mechanisms across multiple bacterial species": S6 Dataset: CL_INS_36.html

Legend

 Regulatoryfunctions
 Hypothetical
 Proteinsynthesis/fate
 Other
 EnergyMetabolism
 Transport +binding proteins

FULL


WINDOWSVGPNG

Trim RowsRemove SingletonsSave Fasta

CL\_3869


CL\_3869

HighlightSelectShow Genomes


3

CL\_617


1

CL\_1509

fGI ID


CL\_INS\_36
CL\_INS\_36
CL\_INS\_36
CL\_INS\_36
CL\_INS\_36
CL\_INS\_36
CL\_INS\_36
CL\_INS\_36
CL\_INS\_36
CL\_INS\_36
CL\_INS\_36
CL\_INS\_36
CL\_INS\_36
Cluster ID


CL\_5640
CL\_5639
CL\_3870
CL\_3871
CL\_3872
CL\_3873
CL\_3874
CL\_3875
CL\_3876
CL\_3877
CL\_3878
CL\_3879
CL\_3880
