## Supplementary material for "A novel method for integrating genomic and Tn-Seq data to identify common *in vivo* fitness mechanisms across multiple bacterial species": S6 Dataset: CL_INS_37.html

Legend

 Regulatoryfunctions
 Hypothetical
 Other

FULL


WINDOWSVGPNG

Trim RowsRemove SingletonsSave Fasta

CL\_571


CL\_1230

HighlightSelectShow Genomes


3

CL\_5152


1

CL\_5152

fGI ID


CL\_INS\_37
CL\_INS\_37
CL\_INS\_187
CL\_INS\_187
CL\_INS\_187
CL\_INS\_187
CL\_INS\_187
CL\_INS\_187
CL\_INS\_187
CL\_INS\_37
Cluster ID


CL\_5154
CL\_5153
CL\_3935
CL\_5774
CL\_5775
CL\_5776
CL\_4032
CL\_5777
CL\_5778
CL\_5779
