## Supplementary material for "A novel method for integrating genomic and Tn-Seq data to identify common *in vivo* fitness mechanisms across multiple bacterial species": S6 Dataset: CL_INS_38.html

Legend

 Hypothetical
 All EssentialGenes
 Other

FULL


WINDOWSVGPNG

Trim RowsRemove SingletonsSave Fasta

CL\_567


CL\_567


CL\_567


CL\_567

HighlightSelectShow Genomes


2

CL\_565


1

CL\_565


1

CL\_565


1

CL\_565

fGI ID


CL\_INS\_38
CL\_INS\_38
CL\_INS\_38
Cluster ID


CL\_5151
CL\_566
CL\_7228
