## Supplementary material for "A novel method for integrating genomic and Tn-Seq data to identify common *in vivo* fitness mechanisms across multiple bacterial species": S6 Dataset: CL_INS_39.html

Legend

 Regulatoryfunctions
 Hypothetical
 All Fitness Genes
 Other

FULL


WINDOWSVGPNG

Trim RowsRemove SingletonsSave Fasta

CL\_557


CL\_557


CL\_557


CL\_557


CL\_557

HighlightSelectShow Genomes


1

CL\_554


1

CL\_554


1

CL\_554


1

CL\_554


1

CL\_554

fGI ID


CL\_INS\_39
CL\_INS\_39
CL\_INS\_39
CL\_INS\_39
CL\_INS\_39
CL\_INS\_39
Cluster ID


CL\_6196
CL\_5149
CL\_5148
CL\_4923
CL\_556
CL\_555
