## Supplementary material for "A novel method for integrating genomic and Tn-Seq data to identify common *in vivo* fitness mechanisms across multiple bacterial species": S6 Dataset: CL_INS_40.html

Legend

 Hypothetical
 All EssentialGenes
 Other

FULL


WINDOWSVGPNG

Trim RowsRemove SingletonsSave Fasta

CL\_519


CL\_519


CL\_519

HighlightSelectShow Genomes


2

CL\_517


2

CL\_517


1

CL\_517

fGI ID


CL\_INS\_40
CL\_INS\_40
Cluster ID


CL\_6816
CL\_518
