## Supplementary material for "A novel method for integrating genomic and Tn-Seq data to identify common *in vivo* fitness mechanisms across multiple bacterial species": S6 Dataset: CL_INS_42.html

Legend

 Regulatoryfunctions
 Hypothetical
 Biosynthesis ofcofactors,prostheticgroups, +carriers
 Other

FULL


WINDOWSVGPNG

Trim RowsRemove SingletonsSave Fasta

CL\_500


CL\_278


CL\_500


CL\_500


CL\_500

HighlightSelectShow Genomes


2

CL\_499


1

CL\_499


1

CL\_495


1

CL\_499


1

CL\_2204

fGI ID


CL\_INS\_42
CL\_INS\_42
CL\_INS\_42
CL\_INS\_42
CL\_INS\_42
CL\_INS\_42
CL\_INS\_42
Cluster ID


CL\_4591
CL\_5943
CL\_4590
CL\_4589
CL\_4588
CL\_4587
CL\_4586
