## Supplementary material for "A novel method for integrating genomic and Tn-Seq data to identify common *in vivo* fitness mechanisms across multiple bacterial species": S6 Dataset: CL_INS_45.html

Legend

 Regulatoryfunctions
 Hypothetical
 Other
 Transport +binding proteins

FULL


WINDOWSVGPNG

Trim RowsRemove SingletonsSave Fasta

CL\_483


CL\_483


CL\_483


CL\_483

HighlightSelectShow Genomes


2

CL\_482


1

CL\_209


1

CL\_482


1

CL\_482

fGI ID


CL\_INS\_45
CL\_INS\_45
CL\_INS\_45
CL\_INS\_45
CL\_INS\_45
CL\_INS\_45
CL\_INS\_45
CL\_INS\_45
CL\_INS\_45
CL\_INS\_45
CL\_INS\_45
CL\_INS\_45
CL\_INS\_45
CL\_INS\_45
CL\_INS\_45
CL\_INS\_45
CL\_INS\_45
CL\_INS\_45
CL\_INS\_45
CL\_INS\_45
CL\_INS\_45
CL\_INS\_45
CL\_INS\_45
CL\_INS\_45
Cluster ID


CL\_5053
CL\_5054
CL\_6192
CL\_6191
CL\_6190
CL\_6189
CL\_6188
CL\_6187
CL\_6186
CL\_6185
CL\_5052
CL\_6184
CL\_6183
CL\_6182
CL\_6181
CL\_4089
CL\_6180
CL\_6179
CL\_5055
CL\_5056
CL\_5628
CL\_5322
CL\_4411
CL\_4410
