## Supplementary material for "A novel method for integrating genomic and Tn-Seq data to identify common *in vivo* fitness mechanisms across multiple bacterial species": S6 Dataset: CL_INS_46.html

Legend

 Regulatoryfunctions
 Hypothetical
 Other
 Transport +binding proteins

FULL


WINDOWSVGPNG

Trim RowsRemove SingletonsSave Fasta

CL\_461


CL\_461


CL\_461

HighlightSelectShow Genomes


2

CL\_460


1

CL\_460


1

CL\_460

fGI ID


CL\_INS\_46
CL\_INS\_106
CL\_INS\_46
CL\_INS\_46
CL\_INS\_46
Cluster ID


CL\_5618
CL\_4063
CL\_5617
CL\_5616
CL\_5615
