## Supplementary material for "A novel method for integrating genomic and Tn-Seq data to identify common *in vivo* fitness mechanisms across multiple bacterial species": S6 Dataset: CL_INS_47.html

Legend

 Hypothetical
 Regulatoryfunctions
 All EssentialGenes
 Transport +binding proteins

FULL


WINDOWSVGPNG

Trim RowsRemove SingletonsSave Fasta

CL\_446


CL\_446


CL\_446

HighlightSelectShow Genomes


2

CL\_444


2

CL\_444


1

CL\_444

fGI ID


CL\_INS\_47
CL\_INS\_47
CL\_INS\_47
Cluster ID


CL\_5598
CL\_5597
CL\_445
