## Supplementary material for "A novel method for integrating genomic and Tn-Seq data to identify common *in vivo* fitness mechanisms across multiple bacterial species": S6 Dataset: CL_INS_48.html

Legend

 Regulatoryfunctions
 Hypothetical
 DNA Metabolism
 Proteinsynthesis/fate
 Other
 All VFDB Genes
 Transport +binding proteins

FULL


WINDOWSVGPNG

Trim RowsRemove SingletonsSave Fasta

CL\_235


CL\_380


CL\_235


CL\_235


CL\_235

HighlightSelectShow Genomes


2

CL\_383


1

CL\_383


1

CL\_228


1

CL\_386


1

CL\_383

fGI ID


CL\_INS\_49
CL\_INS\_48
CL\_INS\_48
CL\_INS\_156
CL\_INS\_156
CL\_INS\_156
CL\_INS\_183
CL\_INS\_113
CL\_INS\_113
CL\_INS\_113
CL\_INS\_113
Cluster ID


CL\_382
CL\_5594
CL\_5595
CL\_5125
CL\_5124
CL\_5123
CL\_234
CL\_233
CL\_232
CL\_231
CL\_230
