## Supplementary material for "A novel method for integrating genomic and Tn-Seq data to identify common *in vivo* fitness mechanisms across multiple bacterial species": S6 Dataset: CL_INS_49.html

Legend

 Hypothetical
 Regulatoryfunctions
 All EssentialGenes
 Proteinsynthesis/fate
 Other
 EnergyMetabolism
 Transport +binding proteins
 All VFDB Genes

FULL


WINDOWSVGPNG

Trim RowsRemove SingletonsSave Fasta

CL\_236


CL\_236


CL\_1468


CL\_236


CL\_278


CL\_236

HighlightSelectShow Genomes


1

CL\_278


1

CL\_4297


1

CL\_4297


1

CL\_248


1

CL\_4297


1

CL\_4297

fGI ID


CL\_INS\_49
CL\_INS\_49
CL\_INS\_49
CL\_INS\_49
CL\_INS\_50
CL\_INS\_49
CL\_INS\_192
CL\_INS\_192
CL\_INS\_192
CL\_INS\_192
CL\_INS\_192
CL\_INS\_192
CL\_INS\_192
CL\_INS\_192
CL\_INS\_192
CL\_INS\_49
CL\_INS\_49
CL\_INS\_49
CL\_INS\_49
CL\_INS\_49
CL\_INS\_49
CL\_INS\_49
CL\_INS\_49
CL\_INS\_49
CL\_INS\_49
CL\_INS\_49
CL\_INS\_49
CL\_INS\_49
CL\_INS\_49
Cluster ID


CL\_4300
CL\_4299
CL\_4298
CL\_382
CL\_381
CL\_237
CL\_238
CL\_239
CL\_240
CL\_241
CL\_242
CL\_243
CL\_244
CL\_245
CL\_246
CL\_5593
CL\_6124
CL\_6123
CL\_6122
CL\_6121
CL\_6120
CL\_6119
CL\_6118
CL\_6117
CL\_6116
CL\_1728
CL\_6115
CL\_5592
CL\_5591
