## Supplementary material for "A novel method for integrating genomic and Tn-Seq data to identify common *in vivo* fitness mechanisms across multiple bacterial species": S6 Dataset: CL_INS_50.html

Legend

 Hypothetical
 DNA Metabolism
 All EssentialGenes

FULL


WINDOWSVGPNG

Trim RowsRemove SingletonsSave Fasta

CL\_4297


CL\_383


CL\_4297


CL\_4297

HighlightSelectShow Genomes


2

CL\_380


1

CL\_380


1

CL\_1616


1

CL\_380

fGI ID


CL\_INS\_50
CL\_INS\_50
CL\_INS\_50
CL\_INS\_50
Cluster ID


CL\_4296
CL\_381
CL\_5590
CL\_6114
