## Supplementary material for "A novel method for integrating genomic and Tn-Seq data to identify common *in vivo* fitness mechanisms across multiple bacterial species": S6 Dataset: CL_INS_51.html

Legend

 Hypothetical
 Regulatoryfunctions
 All EssentialGenes
 Other

FULL


WINDOWSVGPNG

Trim RowsRemove SingletonsSave Fasta

CL\_337


CL\_337

HighlightSelectShow Genomes


3

CL\_335


1

CL\_3869

fGI ID


CL\_INS\_51
CL\_INS\_51
CL\_INS\_51
CL\_INS\_51
CL\_INS\_51
CL\_INS\_51
CL\_INS\_51
Cluster ID


CL\_5584
CL\_5583
CL\_3863
CL\_3864
CL\_3865
CL\_3866
CL\_3867
