## Supplementary material for "A novel method for integrating genomic and Tn-Seq data to identify common *in vivo* fitness mechanisms across multiple bacterial species": S6 Dataset: CL_INS_52.html

Legend

 Hypothetical
 Other
 Transport +binding proteins

FULL


WINDOWSVGPNG

Trim RowsRemove SingletonsSave Fasta

CL\_329


CL\_329


CL\_329


CL\_5258

HighlightSelectShow Genomes


2

CL\_5260


1

CL\_5120


1

CL\_5120


1

CL\_5120

fGI ID


CL\_INS\_52
CL\_INS\_52
CL\_INS\_52
CL\_INS\_52
CL\_INS\_52
CL\_INS\_52
CL\_INS\_52
Cluster ID


CL\_5119
CL\_5581
CL\_5582
CL\_6099
CL\_6100
CL\_6101
CL\_6102
