## Supplementary material for "A novel method for integrating genomic and Tn-Seq data to identify common *in vivo* fitness mechanisms across multiple bacterial species": S6 Dataset: CL_INS_53.html

Legend

 Hypothetical
 All EssentialGenes
 All Fitness Genes

FULL


WINDOWSVGPNG

Trim RowsRemove SingletonsSave Fasta

CL\_313


CL\_313

HighlightSelectShow Genomes


3

CL\_311


1

CL\_311

fGI ID

CL\_INS\_53
Cluster ID

CL\_312
