## Supplementary material for "A novel method for integrating genomic and Tn-Seq data to identify common *in vivo* fitness mechanisms across multiple bacterial species": S6 Dataset: CL_INS_54.html

Legend

 Regulatoryfunctions
 Hypothetical
 All EssentialGenes
 Proteinsynthesis/fate
 Other
 All VFDB Genes

FULL


WINDOWSVGPNG

Trim RowsRemove SingletonsSave Fasta

CL\_303


CL\_303


CL\_303

HighlightSelectShow Genomes


3

CL\_300


1

CL\_664


1

CL\_300

fGI ID


CL\_INS\_54
CL\_INS\_54
CL\_INS\_54
CL\_INS\_54
CL\_INS\_54
CL\_INS\_54
CL\_INS\_54
CL\_INS\_54
CL\_INS\_54
CL\_INS\_54
CL\_INS\_54
CL\_INS\_54
CL\_INS\_54
CL\_INS\_54
CL\_INS\_54
CL\_INS\_54
CL\_INS\_54
CL\_INS\_54
CL\_INS\_54
CL\_INS\_54
CL\_INS\_54
CL\_INS\_54
CL\_INS\_54
CL\_INS\_54
CL\_INS\_54
CL\_INS\_54
CL\_INS\_54
CL\_INS\_54
CL\_INS\_54
CL\_INS\_54
CL\_INS\_54
CL\_INS\_54
CL\_INS\_54
CL\_INS\_54
CL\_INS\_54
CL\_INS\_54
CL\_INS\_54
CL\_INS\_54
Cluster ID


CL\_302
CL\_301
CL\_4993
CL\_4992
CL\_4991
CL\_4990
CL\_4989
CL\_4988
CL\_4987
CL\_4986
CL\_4985
CL\_4984
CL\_4983
CL\_4982
CL\_4981
CL\_4980
CL\_4979
CL\_4978
CL\_4977
CL\_4976
CL\_4975
CL\_4974
CL\_4973
CL\_4972
CL\_4971
CL\_4970
CL\_4969
CL\_4968
CL\_4967
CL\_4966
CL\_4965
CL\_4964
CL\_4963
CL\_4962
CL\_4961
CL\_4960
CL\_4959
CL\_4958
