## Supplementary material for "A novel method for integrating genomic and Tn-Seq data to identify common *in vivo* fitness mechanisms across multiple bacterial species": S6 Dataset: CL_INS_55.html

Legend

 Regulatoryfunctions
 Hypothetical
 All EssentialGenes
 Other

FULL


WINDOWSVGPNG

Trim RowsRemove SingletonsSave Fasta

CL\_3465


CL\_3465


CL\_3465


CL\_3465

HighlightSelectShow Genomes


2

CL\_3468


1

CL\_827


1

CL\_3475


1

CL\_3477

fGI ID


CL\_INS\_55
CL\_INS\_55
CL\_INS\_55
CL\_INS\_55
CL\_INS\_55
CL\_INS\_32
CL\_INS\_32
CL\_INS\_32
Cluster ID


CL\_3466
CL\_3774
CL\_3773
CL\_3772
CL\_3473
CL\_3474
CL\_5986
CL\_5987
