## Supplementary material for "A novel method for integrating genomic and Tn-Seq data to identify common *in vivo* fitness mechanisms across multiple bacterial species": S6 Dataset: CL_INS_57.html

Legend

 Hypothetical
 All EssentialGenes
 Other
 Transport +binding proteins

FULL


WINDOWSVGPNG

Trim RowsRemove SingletonsSave Fasta

CL\_3461


CL\_3461

HighlightSelectShow Genomes


3

CL\_3457


1

CL\_3457

fGI ID


CL\_INS\_57
CL\_INS\_57
CL\_INS\_57
Cluster ID


CL\_3460
CL\_3459
CL\_3458
