## Supplementary material for "A novel method for integrating genomic and Tn-Seq data to identify common *in vivo* fitness mechanisms across multiple bacterial species": S6 Dataset: CL_INS_58.html

Legend

 Hypothetical
 Other

FULL


WINDOWSVGPNG

Trim RowsRemove SingletonsSave Fasta

CL\_3449


CL\_3449


CL\_3449

HighlightSelectShow Genomes


3

CL\_3447


1

CL\_3447


1

CL\_3447

fGI ID


CL\_INS\_58
CL\_INS\_58
Cluster ID


CL\_3448
CL\_4924
