## Supplementary material for "A novel method for integrating genomic and Tn-Seq data to identify common *in vivo* fitness mechanisms across multiple bacterial species": S6 Dataset: CL_INS_59.html

Legend

 Regulatoryfunctions
 Hypothetical
 SignalTransduction +Quorum Sensing
 Other
 Transport +binding proteins

FULL


WINDOWSVGPNG

Trim RowsRemove SingletonsSave Fasta

CL\_3446


CL\_3446


CL\_3446


CL\_3446

HighlightSelectShow Genomes


1

CL\_3439


1

CL\_3439


1

CL\_3439


1

CL\_3439

fGI ID


CL\_INS\_59
CL\_INS\_59
CL\_INS\_59
CL\_INS\_59
CL\_INS\_59
CL\_INS\_59
CL\_INS\_59
CL\_INS\_59
CL\_INS\_59
CL\_INS\_59
CL\_INS\_59
CL\_INS\_59
CL\_INS\_59
CL\_INS\_59
CL\_INS\_59
CL\_INS\_59
CL\_INS\_59
CL\_INS\_59
Cluster ID


CL\_3445
CL\_5506
CL\_7315
CL\_5505
CL\_5504
CL\_5503
CL\_5502
CL\_5501
CL\_5500
CL\_5985
CL\_5984
CL\_5499
CL\_5498
CL\_3444
CL\_3443
CL\_3442
CL\_3441
CL\_3440
