## Supplementary material for "A novel method for integrating genomic and Tn-Seq data to identify common *in vivo* fitness mechanisms across multiple bacterial species": S6 Dataset: CL_INS_60.html

Legend

 Regulatoryfunctions
 Hypothetical
 Other

FULL


WINDOWSVGPNG

Trim RowsRemove SingletonsSave Fasta

CL\_3411


CL\_3411


CL\_3411


CL\_3411

HighlightSelectShow Genomes


2

CL\_3410


1

CL\_3410


1

CL\_3410


1

CL\_3410

fGI ID


CL\_INS\_60
CL\_INS\_60
CL\_INS\_60
Cluster ID


CL\_4747
CL\_5496
CL\_5495
