## Supplementary material for "A novel method for integrating genomic and Tn-Seq data to identify common *in vivo* fitness mechanisms across multiple bacterial species": S6 Dataset: CL_INS_61.html

Legend

 Hypothetical
 All EssentialGenes
 All Fitness Genes
 Other

FULL


WINDOWSVGPNG

Trim RowsRemove SingletonsSave Fasta

CL\_3396


CL\_3396


CL\_3396


CL\_3396


CL\_3396

HighlightSelectShow Genomes


1

CL\_3390


1

CL\_3390


1

CL\_3390


1

CL\_3390


1

CL\_3390

fGI ID


CL\_INS\_61
CL\_INS\_61
CL\_INS\_61
CL\_INS\_61
CL\_INS\_61
CL\_INS\_61
CL\_INS\_61
CL\_INS\_61
CL\_INS\_61
CL\_INS\_61
CL\_INS\_61
CL\_INS\_61
Cluster ID


CL\_4752
CL\_3395
CL\_3394
CL\_3393
CL\_3392
CL\_3391
CL\_5963
CL\_5962
CL\_5961
CL\_5960
CL\_5959
CL\_5195
