## Supplementary material for "A novel method for integrating genomic and Tn-Seq data to identify common *in vivo* fitness mechanisms across multiple bacterial species": S6 Dataset: CL_INS_62.html

Legend

 Regulatoryfunctions
 Hypothetical
 Other
 All VFDB Genes

FULL


WINDOWSVGPNG

Trim RowsRemove SingletonsSave Fasta

CL\_3389


CL\_3389


CL\_3389


CL\_3389

HighlightSelectShow Genomes


2

CL\_3388


1

CL\_3388


1

CL\_3385


1

CL\_3388

fGI ID


CL\_INS\_62
CL\_INS\_62
CL\_INS\_62
CL\_INS\_62
CL\_INS\_62
CL\_INS\_62
Cluster ID


CL\_4754
CL\_4755
CL\_7316
CL\_7317
CL\_7318
CL\_7319
