## Supplementary material for "A novel method for integrating genomic and Tn-Seq data to identify common *in vivo* fitness mechanisms across multiple bacterial species": S6 Dataset: CL_INS_64.html

Legend

 Regulatoryfunctions
 All EssentialGenes
 All Fitness Genes
 SignalTransduction +Quorum Sensing

FULL


WINDOWSVGPNG

Trim RowsRemove SingletonsSave Fasta

CL\_3376


CL\_3376

HighlightSelectShow Genomes


3

CL\_3372


2

CL\_3372

fGI ID


CL\_INS\_64
CL\_INS\_64
CL\_INS\_64
Cluster ID


CL\_3375
CL\_3374
CL\_3373
