## Supplementary material for "A novel method for integrating genomic and Tn-Seq data to identify common *in vivo* fitness mechanisms across multiple bacterial species": S6 Dataset: CL_INS_65.html

Legend

 Hypothetical
 All EssentialGenes
 Other
 Transport +binding proteins

FULL


WINDOWSVGPNG

Trim RowsRemove SingletonsSave Fasta

CL\_3360


CL\_3360


CL\_3360

HighlightSelectShow Genomes


2

CL\_4770


1

CL\_3359


1

CL\_4770

fGI ID


CL\_INS\_32
CL\_INS\_65
CL\_INS\_65
CL\_INS\_65
CL\_INS\_65
CL\_INS\_65
Cluster ID


CL\_816
CL\_5494
CL\_5493
CL\_4907
CL\_4906
CL\_4905
