## Supplementary material for "A novel method for integrating genomic and Tn-Seq data to identify common *in vivo* fitness mechanisms across multiple bacterial species": S6 Dataset: CL_INS_67.html

Legend

 Hypothetical
 All EssentialGenes
 Other
 Transport +binding proteins

FULL


WINDOWSVGPNG

Trim RowsRemove SingletonsSave Fasta

CL\_3285


CL\_3285


CL\_3285

HighlightSelectShow Genomes


2

CL\_3284


1

CL\_3284


1

CL\_3284

fGI ID


CL\_INS\_67
CL\_INS\_67
CL\_INS\_67
CL\_INS\_67
CL\_INS\_67
Cluster ID


CL\_5410
CL\_5411
CL\_5412
CL\_5413
CL\_5414
