## Supplementary material for "A novel method for integrating genomic and Tn-Seq data to identify common *in vivo* fitness mechanisms across multiple bacterial species": S6 Dataset: CL_INS_68.html

Legend

 Regulatoryfunctions
 Hypothetical
 Other
 EnergyMetabolism

FULL


WINDOWSVGPNG

Trim RowsRemove SingletonsSave Fasta

CL\_4077


CL\_4077

HighlightSelectShow Genomes


3

CL\_3283


1

CL\_4083

fGI ID


CL\_INS\_68
CL\_INS\_68
CL\_INS\_68
CL\_INS\_68
CL\_INS\_68
CL\_INS\_68
CL\_INS\_68
Cluster ID


CL\_5418
CL\_5419
CL\_5420
CL\_4078
CL\_4079
CL\_4080
CL\_4081
