## Supplementary material for "A novel method for integrating genomic and Tn-Seq data to identify common *in vivo* fitness mechanisms across multiple bacterial species": S6 Dataset: CL_INS_69.html

Legend

 Regulatoryfunctions
 Hypothetical
 Other

FULL


WINDOWSVGPNG

Trim RowsRemove SingletonsSave Fasta

CL\_3280


CL\_3280


CL\_3280


CL\_3280

HighlightSelectShow Genomes


2

CL\_3277


1

CL\_3277


1

CL\_3277


1

CL\_3021

fGI ID


CL\_INS\_69
CL\_INS\_69
CL\_INS\_69
CL\_INS\_69
Cluster ID


CL\_5949
CL\_3279
CL\_3278
CL\_5421
