## Supplementary material for "A novel method for integrating genomic and Tn-Seq data to identify common *in vivo* fitness mechanisms across multiple bacterial species": S6 Dataset: CL_INS_70.html

Legend

 Hypothetical
 Regulatoryfunctions
 All Fitness Genes
 Other
 Transport +binding proteins

FULL


WINDOWSVGPNG

Trim RowsRemove SingletonsSave Fasta

CL\_3274


CL\_3274


CL\_3274


CL\_3274

HighlightSelectShow Genomes


2

CL\_3268


1

CL\_3268


1

CL\_3268


1

CL\_3268

fGI ID


CL\_INS\_70
CL\_INS\_70
CL\_INS\_70
CL\_INS\_70
CL\_INS\_70
CL\_INS\_70
CL\_INS\_70
CL\_INS\_70
CL\_INS\_70
CL\_INS\_70
CL\_INS\_70
Cluster ID


CL\_3273
CL\_3272
CL\_3271
CL\_3270
CL\_3269
CL\_5422
CL\_7334
CL\_5423
CL\_5424
CL\_5425
CL\_5426
