## Supplementary material for "A novel method for integrating genomic and Tn-Seq data to identify common *in vivo* fitness mechanisms across multiple bacterial species": S6 Dataset: CL_INS_71.html

Legend

 Hypothetical
 All EssentialGenes
 All Fitness Genes
 Other
 Transport +binding proteins

FULL


WINDOWSVGPNG

Trim RowsRemove SingletonsSave Fasta

CL\_3143


CL\_4852


CL\_3143


CL\_3143

HighlightSelectShow Genomes


1

CL\_3136


1

CL\_3136


1

CL\_3136


1

CL\_3136

fGI ID


CL\_INS\_71
CL\_INS\_8
CL\_INS\_8
CL\_INS\_71
CL\_INS\_71
CL\_INS\_71
CL\_INS\_71
Cluster ID


CL\_4854
CL\_3142
CL\_3141
CL\_3140
CL\_3139
CL\_3138
CL\_3137
