## Supplementary material for "A novel method for integrating genomic and Tn-Seq data to identify common *in vivo* fitness mechanisms across multiple bacterial species": S6 Dataset: CL_INS_72.html

Legend

 Regulatoryfunctions
 Hypothetical
 Other

FULL


WINDOWSVGPNG

Trim RowsRemove SingletonsSave Fasta

CL\_3116


CL\_3116


CL\_3116


CL\_3116


CL\_3116

HighlightSelectShow Genomes


1

CL\_3108


1

CL\_3108


1

CL\_3108


1

CL\_3108


1

CL\_3108

fGI ID


CL\_INS\_72
CL\_INS\_72
CL\_INS\_72
CL\_INS\_72
CL\_INS\_72
CL\_INS\_72
CL\_INS\_72
CL\_INS\_72
CL\_INS\_72
CL\_INS\_72
CL\_INS\_72
Cluster ID


CL\_4857
CL\_4858
CL\_3115
CL\_6926
CL\_6925
CL\_3114
CL\_3113
CL\_3112
CL\_3111
CL\_3110
CL\_3109
