## Supplementary material for "A novel method for integrating genomic and Tn-Seq data to identify common *in vivo* fitness mechanisms across multiple bacterial species": S6 Dataset: CL_INS_73.html

Legend

 Regulatoryfunctions
 Hypothetical
 All EssentialGenes
 SignalTransduction +Quorum Sensing
 Other

FULL


WINDOWSVGPNG

Trim RowsRemove SingletonsSave Fasta

CL\_3098


CL\_1969


CL\_3098


CL\_3098

HighlightSelectShow Genomes


2

CL\_1957


1

CL\_1957


1

CL\_3096


1

CL\_1957

fGI ID


CL\_INS\_73
CL\_INS\_73
CL\_INS\_73
CL\_INS\_73
CL\_INS\_73
CL\_INS\_73
CL\_INS\_73
CL\_INS\_73
CL\_INS\_149
CL\_INS\_73
CL\_INS\_73
Cluster ID


CL\_3097
CL\_1967
CL\_1966
CL\_1965
CL\_1964
CL\_1963
CL\_1962
CL\_1961
CL\_1960
CL\_1959
CL\_1958
