## Supplementary material for "A novel method for integrating genomic and Tn-Seq data to identify common *in vivo* fitness mechanisms across multiple bacterial species": S6 Dataset: CL_INS_74.html

Legend

 Regulatoryfunctions
 Hypothetical
 All EssentialGenes
 Other
 Transport +binding proteins

FULL


WINDOWSVGPNG

Trim RowsRemove SingletonsSave Fasta

CL\_1957


CL\_1957


CL\_1956

HighlightSelectShow Genomes


3

CL\_4474


1

CL\_4088


1

CL\_4474

fGI ID


CL\_INS\_74
CL\_INS\_74
CL\_INS\_50
CL\_INS\_74
CL\_INS\_74
CL\_INS\_74
CL\_INS\_74
CL\_INS\_74
CL\_INS\_74
CL\_INS\_74
Cluster ID


CL\_5430
CL\_4084
CL\_381
CL\_4085
CL\_4086
CL\_4469
CL\_4470
CL\_4471
CL\_4472
CL\_4473
