## Supplementary material for "A novel method for integrating genomic and Tn-Seq data to identify common *in vivo* fitness mechanisms across multiple bacterial species": S6 Dataset: CL_INS_75.html

Legend

 Other

FULL


WINDOWSVGPNG

Trim RowsRemove SingletonsSave Fasta

CL\_3096


CL\_3096


CL\_3096


CL\_3096

HighlightSelectShow Genomes


2

CL\_3095


1

CL\_3095


1

CL\_3095


1

CL\_3306

fGI ID


CL\_INS\_75
CL\_INS\_75
Cluster ID


CL\_5431
CL\_5432
