## Supplementary material for "A novel method for integrating genomic and Tn-Seq data to identify common *in vivo* fitness mechanisms across multiple bacterial species": S6 Dataset: CL_INS_76.html

Legend

 Regulatoryfunctions
 Hypothetical
 Other
 EnergyMetabolism
 Transport +binding proteins

FULL


WINDOWSVGPNG

Trim RowsRemove SingletonsSave Fasta

CL\_3082


CL\_3082


CL\_3082


CL\_3082


CL\_3081


CL\_3082

HighlightSelectShow Genomes


1

CL\_3084


1

CL\_3084


1

CL\_2013


1

CL\_3081


1

CL\_3084


1

CL\_1758

fGI ID


CL\_INS\_76
CL\_INS\_76
CL\_INS\_76
CL\_INS\_147
CL\_INS\_147
CL\_INS\_147
CL\_INS\_147
CL\_INS\_76
CL\_INS\_76
CL\_INS\_76
CL\_INS\_76
CL\_INS\_76
CL\_INS\_76
CL\_INS\_76
CL\_INS\_76
CL\_INS\_76
CL\_INS\_76
CL\_INS\_147
CL\_INS\_147
CL\_INS\_147
CL\_INS\_76
CL\_INS\_147
CL\_INS\_147
CL\_INS\_147
CL\_INS\_147
CL\_INS\_76
CL\_INS\_76
CL\_INS\_76
CL\_INS\_76
CL\_INS\_76
CL\_INS\_76
CL\_INS\_76
CL\_INS\_76
CL\_INS\_76
CL\_INS\_76
CL\_INS\_76
CL\_INS\_76
CL\_INS\_76
CL\_INS\_76
CL\_INS\_76
CL\_INS\_76
CL\_INS\_76
CL\_INS\_76
CL\_INS\_76
CL\_INS\_76
CL\_INS\_76
Cluster ID


CL\_5193
CL\_5194
CL\_6910
CL\_6911
CL\_4429
CL\_6912
CL\_6913
CL\_6949
CL\_6948
CL\_6947
CL\_6946
CL\_6945
CL\_6944
CL\_6943
CL\_6942
CL\_6941
CL\_3083
CL\_5446
CL\_5445
CL\_5444
CL\_5443
CL\_4071
CL\_4070
CL\_4069
CL\_5442
CL\_5441
CL\_6915
CL\_6916
CL\_6917
CL\_6918
CL\_6919
CL\_6920
CL\_6921
CL\_5440
CL\_5439
CL\_5438
CL\_5042
CL\_5437
CL\_5436
CL\_3888
CL\_3889
CL\_3890
CL\_3891
CL\_3892
CL\_3893
CL\_3894
