## Supplementary material for "A novel method for integrating genomic and Tn-Seq data to identify common *in vivo* fitness mechanisms across multiple bacterial species": S6 Dataset: CL_INS_78.html

Legend

 Hypothetical
 All EssentialGenes
 All Fitness Genes
 Cell Envelope
 Other

FULL


WINDOWSVGPNG

Trim RowsRemove SingletonsSave Fasta

CL\_3055


CL\_3055


CL\_3055


CL\_3055


CL\_3055

HighlightSelectShow Genomes


1

CL\_3048


1

CL\_3048


1

CL\_3048


1

CL\_3048


1

CL\_3048

fGI ID


CL\_INS\_78
CL\_INS\_78
CL\_INS\_78
CL\_INS\_78
CL\_INS\_78
CL\_INS\_78
CL\_INS\_78
CL\_INS\_78
CL\_INS\_78
CL\_INS\_78
CL\_INS\_78
CL\_INS\_78
CL\_INS\_78
CL\_INS\_78
CL\_INS\_78
CL\_INS\_78
CL\_INS\_78
CL\_INS\_78
CL\_INS\_78
CL\_INS\_78
CL\_INS\_78
CL\_INS\_78
Cluster ID


CL\_3054
CL\_3053
CL\_3052
CL\_3051
CL\_3050
CL\_3049
CL\_5486
CL\_4845
CL\_4844
CL\_4843
CL\_4842
CL\_4841
CL\_6893
CL\_6892
CL\_6891
CL\_5487
CL\_5488
CL\_5489
CL\_4840
CL\_4839
CL\_4838
CL\_4837
