## Supplementary material for "A novel method for integrating genomic and Tn-Seq data to identify common *in vivo* fitness mechanisms across multiple bacterial species": S6 Dataset: CL_INS_79.html

Legend

 Regulatoryfunctions
 Hypothetical
 Transport +binding proteins

FULL


WINDOWSVGPNG

Trim RowsRemove SingletonsSave Fasta

CL\_3028


CL\_3028


CL\_3028

HighlightSelectShow Genomes


3

CL\_3026


1

CL\_3026


1

CL\_3026

fGI ID


CL\_INS\_79
CL\_INS\_79
CL\_INS\_79
Cluster ID


CL\_3027
CL\_4866
CL\_4865
