## Supplementary material for "A novel method for integrating genomic and Tn-Seq data to identify common *in vivo* fitness mechanisms across multiple bacterial species": S6 Dataset: CL_INS_80.html

Legend

 Regulatoryfunctions
 Hypothetical
 Other
 Transport +binding proteins

FULL


WINDOWSVGPNG

Trim RowsRemove SingletonsSave Fasta

CL\_3015


CL\_3015


CL\_3015

HighlightSelectShow Genomes


2

CL\_3007


1

CL\_1648


1

CL\_3007

fGI ID


CL\_INS\_80
CL\_INS\_80
CL\_INS\_80
CL\_INS\_80
CL\_INS\_80
CL\_INS\_80
CL\_INS\_80
CL\_INS\_80
CL\_INS\_80
CL\_INS\_80
CL\_INS\_80
CL\_INS\_80
CL\_INS\_80
Cluster ID


CL\_4397
CL\_4398
CL\_4399
CL\_4400
CL\_3014
CL\_3013
CL\_5401
CL\_5400
CL\_3012
CL\_3011
CL\_3010
CL\_3009
CL\_3008
