## Supplementary material for "A novel method for integrating genomic and Tn-Seq data to identify common *in vivo* fitness mechanisms across multiple bacterial species": S6 Dataset: CL_INS_81.html

Legend

 Regulatoryfunctions
 Hypothetical
 Other
 Transport +binding proteins

FULL


WINDOWSVGPNG

Trim RowsRemove SingletonsSave Fasta

CL\_3002


CL\_3002

HighlightSelectShow Genomes


3

CL\_2998


1

CL\_2998

fGI ID


CL\_INS\_81
CL\_INS\_81
CL\_INS\_81
Cluster ID


CL\_3001
CL\_3000
CL\_2999
