## Supplementary material for "A novel method for integrating genomic and Tn-Seq data to identify common *in vivo* fitness mechanisms across multiple bacterial species": S6 Dataset: CL_INS_82.html

Legend

 Hypothetical
 Regulatoryfunctions
 Other

FULL


WINDOWSVGPNG

Trim RowsRemove SingletonsSave Fasta

CL\_2997


CL\_260


CL\_2997


CL\_2997

HighlightSelectShow Genomes


2

CL\_2994


1

CL\_2994


1

CL\_2994


1

CL\_2994

fGI ID


CL\_INS\_82
CL\_INS\_82
CL\_INS\_82
Cluster ID


CL\_4872
CL\_2996
CL\_2995
