## Supplementary material for "A novel method for integrating genomic and Tn-Seq data to identify common *in vivo* fitness mechanisms across multiple bacterial species": S6 Dataset: CL_INS_83.html

Legend

 Hypothetical
 Regulatoryfunctions
 Other

FULL


WINDOWSVGPNG

Trim RowsRemove SingletonsSave Fasta

CL\_2954


CL\_2954


CL\_2954


CL\_2954

HighlightSelectShow Genomes


1

CL\_4484


1

CL\_4484


1

CL\_2952


1

CL\_4484

fGI ID


CL\_INS\_83
CL\_INS\_83
CL\_INS\_83
CL\_INS\_83
CL\_INS\_83
Cluster ID


CL\_5399
CL\_6968
CL\_6969
CL\_2953
CL\_5398
