## Supplementary material for "A novel method for integrating genomic and Tn-Seq data to identify common *in vivo* fitness mechanisms across multiple bacterial species": S6 Dataset: CL_INS_85.html

Legend

 Regulatoryfunctions
 Hypothetical
 SignalTransduction +Quorum Sensing
 Cell Envelope
 Other
 Transport +binding proteins

FULL


WINDOWSVGPNG

Trim RowsRemove SingletonsSave Fasta

CL\_4092


CL\_4092


CL\_2914


CL\_4092


CL\_451


CL\_415

HighlightSelectShow Genomes


2

CL\_2925


1

CL\_804


1

CL\_2925


1

CL\_4088


1

CL\_2925


1

CL\_2925

fGI ID


CL\_INS\_85
CL\_INS\_85
CL\_INS\_85
CL\_INS\_85
CL\_INS\_32
CL\_INS\_32
CL\_INS\_32
CL\_INS\_32
CL\_INS\_86
CL\_INS\_86
CL\_INS\_86
CL\_INS\_86
CL\_INS\_86
CL\_INS\_86
CL\_INS\_86
CL\_INS\_86
CL\_INS\_85
CL\_INS\_85
CL\_INS\_85
CL\_INS\_85
CL\_INS\_85
CL\_INS\_85
CL\_INS\_85
CL\_INS\_85
CL\_INS\_85
CL\_INS\_85
CL\_INS\_85
CL\_INS\_85
CL\_INS\_85
CL\_INS\_85
CL\_INS\_85
Cluster ID


CL\_5395
CL\_4091
CL\_4090
CL\_5671
CL\_5670
CL\_5669
CL\_5668
CL\_5667
CL\_2916
CL\_2917
CL\_2918
CL\_2919
CL\_2920
CL\_2921
CL\_2922
CL\_2923
CL\_2924
CL\_4792
CL\_4793
CL\_4794
CL\_4795
CL\_4796
CL\_4797
CL\_4798
CL\_4799
CL\_4800
CL\_4801
CL\_4802
CL\_4803
CL\_4804
CL\_4805
