## Supplementary material for "A novel method for integrating genomic and Tn-Seq data to identify common *in vivo* fitness mechanisms across multiple bacterial species": S6 Dataset: CL_INS_86.html

Legend

 Regulatoryfunctions
 Hypothetical
 Other
 EnergyMetabolism
 Transport +binding proteins

FULL


WINDOWSVGPNG

Trim RowsRemove SingletonsSave Fasta

CL\_4092


CL\_4092


CL\_4092


CL\_4092

HighlightSelectShow Genomes


1

CL\_2914


1

CL\_829


1

CL\_1869


1

CL\_3560

fGI ID


CL\_INS\_86
CL\_INS\_86
CL\_INS\_86
CL\_INS\_86
CL\_INS\_86
CL\_INS\_86
CL\_INS\_86
CL\_INS\_86
CL\_INS\_86
CL\_INS\_86
CL\_INS\_86
CL\_INS\_86
CL\_INS\_86
CL\_INS\_86
CL\_INS\_86
Cluster ID


CL\_2923
CL\_2922
CL\_2921
CL\_4093
CL\_4094
CL\_4095
CL\_4096
CL\_4097
CL\_4098
CL\_2920
CL\_2919
CL\_2918
CL\_2917
CL\_2916
CL\_5387
