## Supplementary material for "A novel method for integrating genomic and Tn-Seq data to identify common *in vivo* fitness mechanisms across multiple bacterial species": S6 Dataset: CL_INS_87.html

Legend

 Regulatoryfunctions
 Other

FULL


WINDOWSVGPNG

Trim RowsRemove SingletonsSave Fasta

CL\_2560


CL\_2560


CL\_2560

HighlightSelectShow Genomes


2

CL\_2562


1

CL\_2562


1

CL\_2562

fGI ID


CL\_INS\_87
CL\_INS\_87
Cluster ID


CL\_5345
CL\_2561
