## Supplementary material for "A novel method for integrating genomic and Tn-Seq data to identify common *in vivo* fitness mechanisms across multiple bacterial species": S6 Dataset: CL_INS_88.html

Legend

 Hypothetical
 Regulatoryfunctions
 All EssentialGenes
 All Fitness Genes
 Other
 All VFDB Genes
 Transport +binding proteins

FULL


WINDOWSVGPNG

Trim RowsRemove SingletonsSave Fasta

CL\_2562


CL\_1229


CL\_1092


CL\_1229


CL\_1229

HighlightSelectShow Genomes


2

CL\_2570


2

CL\_2570


1

CL\_2570


1

CL\_1208


1

CL\_336

fGI ID


CL\_INS\_88
CL\_INS\_88
CL\_INS\_88
CL\_INS\_88
CL\_INS\_88
CL\_INS\_88
CL\_INS\_88
CL\_INS\_88
CL\_INS\_88
CL\_INS\_88
CL\_INS\_88
CL\_INS\_88
CL\_INS\_88
CL\_INS\_88
CL\_INS\_88
CL\_INS\_88
CL\_INS\_88
CL\_INS\_88
CL\_INS\_88
CL\_INS\_88
CL\_INS\_88
CL\_INS\_88
CL\_INS\_88
CL\_INS\_88
CL\_INS\_88
CL\_INS\_88
CL\_INS\_88
CL\_INS\_88
CL\_INS\_88
CL\_INS\_88
CL\_INS\_88
CL\_INS\_88
Cluster ID


CL\_4707
CL\_2564
CL\_2565
CL\_2566
CL\_2567
CL\_2568
CL\_2569
CL\_1228
CL\_1227
CL\_5348
CL\_5349
CL\_5350
CL\_5351
CL\_5215
CL\_5214
CL\_1226
CL\_1225
CL\_1224
CL\_1223
CL\_1222
CL\_1221
CL\_1220
CL\_1219
CL\_1218
CL\_1217
CL\_1216
CL\_1215
CL\_1214
CL\_1213
CL\_1212
CL\_1211
CL\_1210
