## Supplementary material for "A novel method for integrating genomic and Tn-Seq data to identify common *in vivo* fitness mechanisms across multiple bacterial species": S6 Dataset: CL_INS_89.html

Legend

 Hypothetical
 Other
 Transport +binding proteins

FULL


WINDOWSVGPNG

Trim RowsRemove SingletonsSave Fasta

CL\_2577


CL\_2577

HighlightSelectShow Genomes


3

CL\_2579


2

CL\_2579

fGI ID

CL\_INS\_89
Cluster ID

CL\_2578
