## Supplementary material for "A novel method for integrating genomic and Tn-Seq data to identify common *in vivo* fitness mechanisms across multiple bacterial species": S6 Dataset: CL_INS_90.html

Legend

 Regulatoryfunctions
 Hypothetical
 All EssentialGenes
 Other
 EnergyMetabolism
 Transport +binding proteins

FULL


WINDOWSVGPNG

Trim RowsRemove SingletonsSave Fasta

CL\_2594


CL\_4390


CL\_2594


CL\_2594

HighlightSelectShow Genomes


2

CL\_2600


1

CL\_2600


1

CL\_2607


1

CL\_2600

fGI ID


CL\_INS\_90
CL\_INS\_90
CL\_INS\_90
CL\_INS\_90
CL\_INS\_90
CL\_INS\_90
CL\_INS\_149
CL\_INS\_90
CL\_INS\_90
CL\_INS\_90
CL\_INS\_90
CL\_INS\_90
CL\_INS\_90
CL\_INS\_90
Cluster ID


CL\_6863
CL\_2595
CL\_2596
CL\_2597
CL\_2598
CL\_2599
CL\_4573
CL\_6644
CL\_6645
CL\_4383
CL\_4384
CL\_4385
CL\_4386
CL\_4387
