## Supplementary material for "A novel method for integrating genomic and Tn-Seq data to identify common *in vivo* fitness mechanisms across multiple bacterial species": S6 Dataset: CL_INS_91.html

Legend

 Hypothetical
 Other
 EnergyMetabolism

FULL


WINDOWSVGPNG

Trim RowsRemove SingletonsSave Fasta

CL\_2609


CL\_2609


CL\_2609


CL\_2609

HighlightSelectShow Genomes


1

CL\_2611


1

CL\_2611


1

CL\_2611


1

CL\_2611

fGI ID


CL\_INS\_91
CL\_INS\_91
CL\_INS\_91
Cluster ID


CL\_2610
CL\_6865
CL\_6993
