## Supplementary material for "A novel method for integrating genomic and Tn-Seq data to identify common *in vivo* fitness mechanisms across multiple bacterial species": S6 Dataset: CL_INS_92.html

Legend

 Regulatoryfunctions
 Hypothetical
 Other
 Transport +binding proteins

FULL


WINDOWSVGPNG

Trim RowsRemove SingletonsSave Fasta

CL\_2619


CL\_4119


CL\_2619

HighlightSelectShow Genomes


3

CL\_2622


1

CL\_2622


1

CL\_2350

fGI ID


CL\_INS\_92
CL\_INS\_92
CL\_INS\_92
CL\_INS\_92
CL\_INS\_92
CL\_INS\_92
CL\_INS\_92
CL\_INS\_92
CL\_INS\_92
CL\_INS\_92
CL\_INS\_92
CL\_INS\_92
CL\_INS\_92
Cluster ID


CL\_2620
CL\_2621
CL\_4607
CL\_4606
CL\_4605
CL\_4604
CL\_4121
CL\_4122
CL\_4123
CL\_4124
CL\_4125
CL\_4126
CL\_4127
