## Supplementary material for "A novel method for integrating genomic and Tn-Seq data to identify common *in vivo* fitness mechanisms across multiple bacterial species": S6 Dataset: CL_INS_99.html

Legend

 Regulatoryfunctions
 Hypothetical
 All EssentialGenes
 Other
 EnergyMetabolism

FULL


WINDOWSVGPNG

Trim RowsRemove SingletonsSave Fasta

CL\_1092


CL\_1092


CL\_1092


CL\_1092


CL\_2899

HighlightSelectShow Genomes


1

CL\_4852


1

CL\_2570


1

CL\_2901


1

CL\_2901


1

CL\_2901

fGI ID


CL\_INS\_99
CL\_INS\_99
CL\_INS\_99
CL\_INS\_99
CL\_INS\_99
CL\_INS\_99
CL\_INS\_99
CL\_INS\_99
CL\_INS\_99
Cluster ID


CL\_4894
CL\_4706
CL\_7157
CL\_4560
CL\_7158
CL\_7159
CL\_7160
CL\_7161
CL\_7162
