## Supplementary material for "A novel method for integrating genomic and Tn-Seq data to identify common *in vivo* fitness mechanisms across multiple bacterial species": S6 Dataset: CL_INS_100.html

Legend

 Regulatoryfunctions
 Hypothetical
 Other
 Transport +binding proteins

FULL


WINDOWSVGPNG

Trim RowsRemove SingletonsSave Fasta

CL\_2914


CL\_2914


CL\_2914

HighlightSelectShow Genomes


1

CL\_456


1

CL\_4092


1

CL\_2925

fGI ID


CL\_INS\_100
CL\_INS\_100
CL\_INS\_100
CL\_INS\_100
CL\_INS\_100
CL\_INS\_100
CL\_INS\_100
CL\_INS\_100
CL\_INS\_100
CL\_INS\_100
CL\_INS\_100
CL\_INS\_100
CL\_INS\_100
CL\_INS\_100
CL\_INS\_100
CL\_INS\_100
CL\_INS\_100
CL\_INS\_86
CL\_INS\_86
CL\_INS\_86
CL\_INS\_86
CL\_INS\_86
CL\_INS\_86
CL\_INS\_86
CL\_INS\_86
CL\_INS\_100
CL\_INS\_86
CL\_INS\_149
CL\_INS\_100
CL\_INS\_100
CL\_INS\_100
CL\_INS\_100
Cluster ID


CL\_5370
CL\_5371
CL\_5372
CL\_5373
CL\_5374
CL\_5375
CL\_5376
CL\_5377
CL\_5378
CL\_5379
CL\_5380
CL\_5381
CL\_5382
CL\_5383
CL\_5384
CL\_5385
CL\_2915
CL\_2916
CL\_2917
CL\_2918
CL\_2919
CL\_2920
CL\_2921
CL\_2922
CL\_2923
CL\_5386
CL\_5387
CL\_5388
CL\_5389
CL\_5390
CL\_5391
CL\_5392
