## Supplementary material for "A novel method for integrating genomic and Tn-Seq data to identify common *in vivo* fitness mechanisms across multiple bacterial species": S6 Dataset: CL_INS_101.html

Legend

 Hypothetical
 All EssentialGenes
 Other
 EnergyMetabolism

FULL


WINDOWSVGPNG

Trim RowsRemove SingletonsSave Fasta

CL\_36


CL\_36


CL\_36


CL\_36

HighlightSelectShow Genomes


2

CL\_3846


1

CL\_1468


1

CL\_1139


1

CL\_1139

fGI ID


CL\_INS\_101
CL\_INS\_101
CL\_INS\_101
CL\_INS\_101
CL\_INS\_101
CL\_INS\_101
CL\_INS\_101
CL\_INS\_101
CL\_INS\_101
CL\_INS\_101
CL\_INS\_126
CL\_INS\_126
CL\_INS\_126
CL\_INS\_126
CL\_INS\_126
CL\_INS\_126
CL\_INS\_126
CL\_INS\_126
CL\_INS\_126
CL\_INS\_126
CL\_INS\_126
CL\_INS\_126
CL\_INS\_126
CL\_INS\_126
CL\_INS\_126
CL\_INS\_126
CL\_INS\_126
CL\_INS\_126
CL\_INS\_101
CL\_INS\_101
CL\_INS\_101
CL\_INS\_101
CL\_INS\_101
Cluster ID


CL\_37
CL\_38
CL\_39
CL\_40
CL\_41
CL\_42
CL\_43
CL\_44
CL\_45
CL\_4320
CL\_4319
CL\_4318
CL\_4317
CL\_4316
CL\_4315
CL\_4314
CL\_4313
CL\_4312
CL\_4311
CL\_4310
CL\_4309
CL\_4308
CL\_5731
CL\_5730
CL\_5729
CL\_6311
CL\_5622
CL\_5081
CL\_4307
CL\_4306
CL\_4305
CL\_4304
CL\_4303
