## Supplementary material for "A novel method for integrating genomic and Tn-Seq data to identify common *in vivo* fitness mechanisms across multiple bacterial species": S6 Dataset: CL_INS_102.html

Legend

 Regulatoryfunctions
 Hypothetical
 All EssentialGenes
 Other

FULL


WINDOWSVGPNG

Trim RowsRemove SingletonsSave Fasta

CL\_32


CL\_32


CL\_32


CL\_32

HighlightSelectShow Genomes


2

CL\_30


1

CL\_30


1

CL\_30


1

CL\_30

fGI ID


CL\_INS\_102
CL\_INS\_102
CL\_INS\_102
CL\_INS\_102
CL\_INS\_102
CL\_INS\_102
CL\_INS\_102
CL\_INS\_102
Cluster ID


CL\_5019
CL\_5018
CL\_5017
CL\_5016
CL\_4353
CL\_4354
CL\_6060
CL\_31
