## Supplementary material for "A novel method for integrating genomic and Tn-Seq data to identify common *in vivo* fitness mechanisms across multiple bacterial species": S6 Dataset: CL_INS_103.html

Legend

 Hypothetical
 All EssentialGenes
 Other

FULL


WINDOWSVGPNG

Trim RowsRemove SingletonsSave Fasta

CL\_19


CL\_19

HighlightSelectShow Genomes


3

CL\_16


2

CL\_16

fGI ID


CL\_INS\_103
CL\_INS\_103
Cluster ID


CL\_18
CL\_17
