## Supplementary material for "A novel method for integrating genomic and Tn-Seq data to identify common *in vivo* fitness mechanisms across multiple bacterial species": S6 Dataset: CL_INS_104.html

Legend

 Hypothetical
 All EssentialGenes
 Other

FULL


WINDOWSVGPNG

Trim RowsRemove SingletonsSave Fasta

CL\_3667


CL\_3667


Break

HighlightSelectShow Genomes


2

CL\_3664


2

CL\_3664


1

CL\_3664

fGI ID


CL\_INS\_104
CL\_INS\_104
CL\_INS\_104
Cluster ID


CL\_5010
CL\_3666
CL\_3665
