## Supplementary material for "A novel method for integrating genomic and Tn-Seq data to identify common *in vivo* fitness mechanisms across multiple bacterial species": S6 Dataset: CL_INS_106.html

Legend

 Regulatoryfunctions
 Hypothetical
 Other
 EnergyMetabolism
 Transport +binding proteins

FULL


WINDOWSVGPNG

Trim RowsRemove SingletonsSave Fasta

CL\_3654


CL\_4077

HighlightSelectShow Genomes


3

CL\_4052


1

CL\_4052

fGI ID


CL\_INS\_106
CL\_INS\_106
CL\_INS\_106
CL\_INS\_106
CL\_INS\_106
CL\_INS\_147
CL\_INS\_147
CL\_INS\_147
CL\_INS\_106
CL\_INS\_106
CL\_INS\_106
CL\_INS\_106
CL\_INS\_106
CL\_INS\_106
CL\_INS\_106
CL\_INS\_106
CL\_INS\_106
CL\_INS\_106
CL\_INS\_106
CL\_INS\_106
CL\_INS\_106
CL\_INS\_106
CL\_INS\_106
CL\_INS\_106
Cluster ID


CL\_5559
CL\_4075
CL\_4074
CL\_4073
CL\_4072
CL\_4071
CL\_4070
CL\_4069
CL\_4068
CL\_4067
CL\_4066
CL\_4065
CL\_4064
CL\_4063
CL\_4062
CL\_4061
CL\_4060
CL\_4059
CL\_4058
CL\_4057
CL\_4056
CL\_4055
CL\_4054
CL\_4053
