## Supplementary material for "A novel method for integrating genomic and Tn-Seq data to identify common *in vivo* fitness mechanisms across multiple bacterial species": S6 Dataset: CL_INS_107.html

Legend

 Hypothetical
 Regulatoryfunctions
 Other
 Centralintermediarymetabolism
 Transport +binding proteins
 All VFDB Genes

FULL


WINDOWSVGPNG

Trim RowsRemove SingletonsSave Fasta

CL\_3646


CL\_3646


CL\_3646


CL\_664


CL\_3646

HighlightSelectShow Genomes


1

CL\_3641


1

CL\_3641


1

CL\_3641


1

CL\_3641


1

CL\_3641

fGI ID


CL\_INS\_107
CL\_INS\_107
CL\_INS\_107
CL\_INS\_107
CL\_INS\_107
CL\_INS\_107
CL\_INS\_107
CL\_INS\_107
CL\_INS\_107
CL\_INS\_107
CL\_INS\_107
CL\_INS\_107
CL\_INS\_107
CL\_INS\_107
CL\_INS\_107
CL\_INS\_107
CL\_INS\_107
CL\_INS\_107
CL\_INS\_107
CL\_INS\_107
CL\_INS\_107
CL\_INS\_107
CL\_INS\_107
CL\_INS\_107
CL\_INS\_107
CL\_INS\_107
CL\_INS\_107
CL\_INS\_107
CL\_INS\_107
CL\_INS\_107
CL\_INS\_107
CL\_INS\_107
CL\_INS\_107
CL\_INS\_107
CL\_INS\_107
CL\_INS\_107
CL\_INS\_107
CL\_INS\_107
CL\_INS\_107
CL\_INS\_107
CL\_INS\_107
CL\_INS\_107
CL\_INS\_107
CL\_INS\_107
CL\_INS\_107
CL\_INS\_107
CL\_INS\_107
CL\_INS\_54
CL\_INS\_54
CL\_INS\_54
CL\_INS\_54
CL\_INS\_107
CL\_INS\_54
CL\_INS\_54
CL\_INS\_54
CL\_INS\_54
CL\_INS\_54
CL\_INS\_54
CL\_INS\_54
CL\_INS\_107
CL\_INS\_107
CL\_INS\_107
Cluster ID


CL\_7297
CL\_3645
CL\_3644
CL\_3643
CL\_7298
CL\_5558
CL\_3642
CL\_4955
CL\_4954
CL\_4953
CL\_4952
CL\_4951
CL\_4950
CL\_4949
CL\_4948
CL\_4947
CL\_4946
CL\_4945
CL\_4944
CL\_4943
CL\_4942
CL\_4941
CL\_4940
CL\_4939
CL\_4938
CL\_4937
CL\_4936
CL\_4935
CL\_6055
CL\_6054
CL\_6053
CL\_6052
CL\_6051
CL\_6050
CL\_6049
CL\_6048
CL\_6047
CL\_6046
CL\_6045
CL\_6044
CL\_6043
CL\_6042
CL\_6041
CL\_6040
CL\_6039
CL\_6038
CL\_3760
CL\_4982
CL\_4983
CL\_4984
CL\_4985
CL\_5557
CL\_4987
CL\_4988
CL\_4989
CL\_4990
CL\_4991
CL\_4992
CL\_4993
CL\_5556
CL\_6037
CL\_6036
