## Supplementary material for "A novel method for integrating genomic and Tn-Seq data to identify common *in vivo* fitness mechanisms across multiple bacterial species": S6 Dataset: CL_INS_108.html

Legend

 Regulatoryfunctions
 Hypothetical
 All Fitness Genes
 Other
 Transport +binding proteins

FULL


WINDOWSVGPNG

Trim RowsRemove SingletonsSave Fasta

CL\_3639


CL\_5254


CL\_3639


CL\_3639


CL\_3639


CL\_3639

HighlightSelectShow Genomes


1

CL\_5254


1

CL\_3635


1

CL\_3604


1

CL\_3635


1

CL\_3635


1

CL\_3635

fGI ID


CL\_INS\_108
CL\_INS\_108
CL\_INS\_201
CL\_INS\_108
CL\_INS\_201
CL\_INS\_201
CL\_INS\_201
CL\_INS\_108
CL\_INS\_108
CL\_INS\_108
CL\_INS\_108
CL\_INS\_108
CL\_INS\_108
CL\_INS\_108
CL\_INS\_108
CL\_INS\_108
CL\_INS\_108
CL\_INS\_108
CL\_INS\_108
Cluster ID


CL\_4934
CL\_4933
CL\_3638
CL\_3637
CL\_5552
CL\_5551
CL\_5550
CL\_5098
CL\_5555
CL\_5554
CL\_5553
CL\_6035
CL\_6034
CL\_7299
CL\_6033
CL\_3636
CL\_7300
CL\_7301
CL\_7302
