## Supplementary material for "A novel method for integrating genomic and Tn-Seq data to identify common *in vivo* fitness mechanisms across multiple bacterial species": S6 Dataset: CL_INS_111.html

Legend

 Proteinsynthesis/fate
 Other

FULL


WINDOWSVGPNG

Trim RowsRemove SingletonsSave Fasta

CL\_3550


CL\_3550


CL\_3550

HighlightSelectShow Genomes


3

CL\_3549


1

CL\_3541


1

CL\_3549

fGI ID

CL\_INS\_111
Cluster ID

CL\_4023
