## Supplementary material for "A novel method for integrating genomic and Tn-Seq data to identify common *in vivo* fitness mechanisms across multiple bacterial species": S6 Dataset: CL_INS_112.html

Legend

 Hypothetical
 Regulatoryfunctions
 Other

FULL


WINDOWSVGPNG

Trim RowsRemove SingletonsSave Fasta

CL\_3507


CL\_3507


CL\_3507

HighlightSelectShow Genomes


2

CL\_3505


2

CL\_3505


1

CL\_3505

fGI ID


CL\_INS\_112
CL\_INS\_112
Cluster ID


CL\_5512
CL\_3506
