## Supplementary material for "A novel method for integrating genomic and Tn-Seq data to identify common *in vivo* fitness mechanisms across multiple bacterial species": S6 Dataset: CL_INS_113.html

Legend

 Regulatoryfunctions
 Hypothetical
 All EssentialGenes
 Other
 EnergyMetabolism
 All VFDB Genes

FULL


WINDOWSVGPNG

Trim RowsRemove SingletonsSave Fasta

CL\_226


CL\_226


CL\_226


CL\_226


CL\_226

HighlightSelectShow Genomes


1

CL\_3850


1

CL\_5074


1

CL\_1091


1

CL\_228


1

CL\_2189

fGI ID


CL\_INS\_113
CL\_INS\_113
CL\_INS\_113
CL\_INS\_113
CL\_INS\_113
CL\_INS\_113
CL\_INS\_113
CL\_INS\_113
CL\_INS\_113
CL\_INS\_113
CL\_INS\_113
CL\_INS\_113
CL\_INS\_113
CL\_INS\_113
CL\_INS\_113
CL\_INS\_113
CL\_INS\_149
CL\_INS\_113
CL\_INS\_113
CL\_INS\_113
CL\_INS\_113
CL\_INS\_113
CL\_INS\_113
CL\_INS\_113
CL\_INS\_113
CL\_INS\_113
CL\_INS\_113
CL\_INS\_113
CL\_INS\_113
CL\_INS\_113
CL\_INS\_113
CL\_INS\_113
CL\_INS\_113
CL\_INS\_113
CL\_INS\_32
CL\_INS\_32
CL\_INS\_32
CL\_INS\_32
CL\_INS\_32
CL\_INS\_32
CL\_INS\_32
CL\_INS\_32
Cluster ID


CL\_5067
CL\_5068
CL\_5069
CL\_5070
CL\_5071
CL\_5072
CL\_6085
CL\_6084
CL\_6083
CL\_6082
CL\_6081
CL\_6080
CL\_5575
CL\_5576
CL\_5577
CL\_5578
CL\_2187
CL\_7285
CL\_233
CL\_232
CL\_231
CL\_230
CL\_229
CL\_7284
CL\_7283
CL\_7282
CL\_7281
CL\_7280
CL\_4765
CL\_4766
CL\_4767
CL\_7279
CL\_7278
CL\_7277
CL\_813
CL\_812
CL\_811
CL\_810
CL\_809
CL\_808
CL\_807
CL\_806
