## Supplementary material for "A novel method for integrating genomic and Tn-Seq data to identify common *in vivo* fitness mechanisms across multiple bacterial species": S6 Dataset: CL_INS_114.html

Legend

 Hypothetical
 DNA Metabolism
 Other
 EnergyMetabolism

FULL


WINDOWSVGPNG

Trim RowsRemove SingletonsSave Fasta

CL\_216


CL\_216


CL\_216


CL\_216


CL\_216

HighlightSelectShow Genomes


1

CL\_212


1

CL\_212


1

CL\_212


1

CL\_212


1

CL\_212

fGI ID


CL\_INS\_114
CL\_INS\_114
CL\_INS\_114
CL\_INS\_114
CL\_INS\_114
CL\_INS\_114
CL\_INS\_114
CL\_INS\_114
CL\_INS\_114
CL\_INS\_114
CL\_INS\_114
CL\_INS\_114
CL\_INS\_114
Cluster ID


CL\_7286
CL\_215
CL\_214
CL\_213
CL\_5066
CL\_5065
CL\_5064
CL\_5063
CL\_5062
CL\_5061
CL\_5060
CL\_5059
CL\_5058
