## Supplementary material for "A novel method for integrating genomic and Tn-Seq data to identify common *in vivo* fitness mechanisms across multiple bacterial species": S6 Dataset: CL_INS_116.html

Legend

 Regulatoryfunctions
 All EssentialGenes
 Other

FULL


WINDOWSVGPNG

Trim RowsRemove SingletonsSave Fasta

CL\_183


CL\_183

HighlightSelectShow Genomes


4

CL\_184


1

CL\_184

fGI ID


CL\_INS\_116
CL\_INS\_164
CL\_INS\_164
CL\_INS\_164
Cluster ID


CL\_3488
CL\_3487
CL\_3486
CL\_3485
