## Supplementary material for "A novel method for integrating genomic and Tn-Seq data to identify common *in vivo* fitness mechanisms across multiple bacterial species": S6 Dataset: CL_INS_117.html

Legend

 Hypothetical
 All Fitness Genes
 Other

FULL


WINDOWSVGPNG

Trim RowsRemove SingletonsSave Fasta

CL\_114


CL\_114


CL\_114


CL\_114

HighlightSelectShow Genomes


1

CL\_113


1

CL\_113


1

CL\_113


1

CL\_113

fGI ID


CL\_INS\_117
CL\_INS\_117
CL\_INS\_117
Cluster ID


CL\_5567
CL\_7291
CL\_6067
