## Supplementary material for "A novel method for integrating genomic and Tn-Seq data to identify common *in vivo* fitness mechanisms across multiple bacterial species": S6 Dataset: CL_INS_118.html

Legend

 Regulatoryfunctions
 Hypothetical
 Other
 EnergyMetabolism

FULL


WINDOWSVGPNG

Trim RowsRemove SingletonsSave Fasta

CL\_109


CL\_109


CL\_109

HighlightSelectShow Genomes


3

CL\_108


1

CL\_108


1

CL\_2432

fGI ID


CL\_INS\_118
CL\_INS\_118
CL\_INS\_118
CL\_INS\_118
CL\_INS\_118
CL\_INS\_118
CL\_INS\_118
CL\_INS\_118
Cluster ID


CL\_5566
CL\_5565
CL\_4669
CL\_4670
CL\_4671
CL\_4672
CL\_4673
CL\_4674
