## Supplementary material for "A novel method for integrating genomic and Tn-Seq data to identify common *in vivo* fitness mechanisms across multiple bacterial species": S6 Dataset: CL_INS_119.html

Legend

 Other
 Hypothetical

FULL


WINDOWSVGPNG

Trim RowsRemove SingletonsSave Fasta

CL\_1011


CL\_1011


CL\_1011

HighlightSelectShow Genomes


2

CL\_1012


1

CL\_1012


1

CL\_1012

fGI ID


CL\_INS\_119
CL\_INS\_119
Cluster ID


CL\_5835
CL\_5834
