## Supplementary material for "A novel method for integrating genomic and Tn-Seq data to identify common *in vivo* fitness mechanisms across multiple bacterial species": S6 Dataset: CL_INS_121.html

Legend

 Other
 Proteinsynthesis/fate
 All VFDB Genes

FULL


WINDOWSVGPNG

Trim RowsRemove SingletonsSave Fasta

CL\_228


CL\_228

HighlightSelectShow Genomes


3

CL\_1044


1

CL\_235

fGI ID


CL\_INS\_113
CL\_INS\_113
CL\_INS\_113
CL\_INS\_113
CL\_INS\_113
Cluster ID


CL\_229
CL\_230
CL\_231
CL\_232
CL\_233
