## Supplementary material for "A novel method for integrating genomic and Tn-Seq data to identify common *in vivo* fitness mechanisms across multiple bacterial species": S6 Dataset: CL_INS_122.html

Legend

 Other
 Regulatoryfunctions
 Hypothetical

FULL


WINDOWSVGPNG

Trim RowsRemove SingletonsSave Fasta

CL\_1048


CL\_1048


CL\_1048


CL\_1048

HighlightSelectShow Genomes


1

CL\_1044


1

CL\_1044


1

CL\_1044


1

CL\_1044

fGI ID


CL\_INS\_122
CL\_INS\_122
CL\_INS\_122
CL\_INS\_122
CL\_INS\_122
CL\_INS\_122
CL\_INS\_122
CL\_INS\_122
CL\_INS\_122
CL\_INS\_122
CL\_INS\_122
CL\_INS\_35
CL\_INS\_35
Cluster ID


CL\_4155
CL\_1047
CL\_1046
CL\_1045
CL\_7100
CL\_7099
CL\_7098
CL\_1639
CL\_6257
CL\_6256
CL\_6255
CL\_649
CL\_648
