## Supplementary material for "A novel method for integrating genomic and Tn-Seq data to identify common *in vivo* fitness mechanisms across multiple bacterial species": S6 Dataset: CL_INS_123.html

Legend

 Other
 Regulatoryfunctions
 Hypothetical
 All EssentialGenes
 All Fitness Genes
 Transport +binding proteins

FULL


WINDOWSVGPNG

Trim RowsRemove SingletonsSave Fasta

CL\_1097


CL\_1097


CL\_1097


CL\_1097

HighlightSelectShow Genomes


2

CL\_1111


1

CL\_1111


1

CL\_1111


1

CL\_1111

fGI ID


CL\_INS\_123
CL\_INS\_123
CL\_INS\_123
CL\_INS\_123
CL\_INS\_123
CL\_INS\_123
CL\_INS\_123
CL\_INS\_123
CL\_INS\_123
CL\_INS\_123
CL\_INS\_123
CL\_INS\_123
CL\_INS\_123
Cluster ID


CL\_1098
CL\_1099
CL\_1100
CL\_1101
CL\_1102
CL\_1103
CL\_1104
CL\_1105
CL\_1106
CL\_1107
CL\_1108
CL\_1109
CL\_1110
