## Supplementary material for "A novel method for integrating genomic and Tn-Seq data to identify common *in vivo* fitness mechanisms across multiple bacterial species": S6 Dataset: CL_INS_124.html

Legend

 Proteinsynthesis/fate
 Other
 Regulatoryfunctions
 Hypothetical
 All Fitness Genes
 All VFDB Genes
 Transport +binding proteins

FULL


WINDOWSVGPNG

Trim RowsRemove SingletonsSave Fasta

CL\_1123


CL\_1128


CL\_1128


CL\_261


CL\_1128


CL\_1128


CL\_1128

HighlightSelectShow Genomes


1

CL\_1120


1

CL\_1120


1

CL\_1126


1

CL\_1120


1

CL\_1120


1

CL\_1126


1

CL\_1120

fGI ID


CL\_INS\_204
CL\_INS\_124
CL\_INS\_124
CL\_INS\_124
CL\_INS\_124
CL\_INS\_124
CL\_INS\_124
CL\_INS\_6
CL\_INS\_6
CL\_INS\_124
CL\_INS\_124
CL\_INS\_124
CL\_INS\_127
CL\_INS\_127
CL\_INS\_124
CL\_INS\_124
CL\_INS\_124
CL\_INS\_124
CL\_INS\_124
CL\_INS\_124
CL\_INS\_124
CL\_INS\_124
CL\_INS\_124
CL\_INS\_124
CL\_INS\_124
CL\_INS\_124
CL\_INS\_124
CL\_INS\_124
CL\_INS\_124
CL\_INS\_124
CL\_INS\_124
CL\_INS\_124
CL\_INS\_124
CL\_INS\_124
CL\_INS\_124
CL\_INS\_124
CL\_INS\_124
CL\_INS\_124
CL\_INS\_124
CL\_INS\_124
CL\_INS\_124
CL\_INS\_124
CL\_INS\_124
CL\_INS\_124
CL\_INS\_124
CL\_INS\_124
CL\_INS\_124
CL\_INS\_124
CL\_INS\_124
CL\_INS\_124
CL\_INS\_106
CL\_INS\_124
CL\_INS\_124
CL\_INS\_124
CL\_INS\_32
CL\_INS\_124
CL\_INS\_124
CL\_INS\_124
CL\_INS\_124
CL\_INS\_124
CL\_INS\_124
CL\_INS\_124
Cluster ID


CL\_1121
CL\_4137
CL\_4138
CL\_4139
CL\_5821
CL\_7107
CL\_5822
CL\_5089
CL\_6555
CL\_7105
CL\_7104
CL\_7103
CL\_4009
CL\_4010
CL\_6308
CL\_6307
CL\_6306
CL\_4745
CL\_6305
CL\_6304
CL\_6303
CL\_6302
CL\_6301
CL\_6300
CL\_6299
CL\_6298
CL\_6297
CL\_6296
CL\_6295
CL\_6294
CL\_6293
CL\_6292
CL\_6291
CL\_6290
CL\_6289
CL\_3593
CL\_6288
CL\_6287
CL\_6286
CL\_6285
CL\_6284
CL\_6283
CL\_6282
CL\_6281
CL\_6280
CL\_6279
CL\_6278
CL\_6277
CL\_6276
CL\_5823
CL\_4067
CL\_6275
CL\_6274
CL\_6273
CL\_817
CL\_6272
CL\_6271
CL\_6270
CL\_5719
CL\_5720
CL\_6269
CL\_6268
