## Supplementary material for "A novel method for integrating genomic and Tn-Seq data to identify common *in vivo* fitness mechanisms across multiple bacterial species": S6 Dataset: CL_INS_125.html

Legend

 Other
 Hypothetical
 All Fitness Genes

FULL


WINDOWSVGPNG

Trim RowsRemove SingletonsSave Fasta

CL\_1130


CL\_1130


CL\_1130

HighlightSelectShow Genomes


2

CL\_1132


1

CL\_1132


1

CL\_1132

fGI ID


CL\_INS\_125
CL\_INS\_125
CL\_INS\_125
CL\_INS\_125
Cluster ID


CL\_1131
CL\_6309
CL\_5820
CL\_5819
