## Supplementary material for "A novel method for integrating genomic and Tn-Seq data to identify common *in vivo* fitness mechanisms across multiple bacterial species": S6 Dataset: CL_INS_126.html

Legend

 Other
 Regulatoryfunctions
 Hypothetical
 All EssentialGenes
 EnergyMetabolism
 All Fitness Genes
 Cell Envelope

FULL


WINDOWSVGPNG

Trim RowsRemove SingletonsSave Fasta

CL\_1139


CL\_1139


CL\_1139


CL\_1139


CL\_36


CL\_1139

HighlightSelectShow Genomes


1

CL\_36


1

CL\_1144


1

CL\_1144


1

CL\_36


1

CL\_1144


1

CL\_1144

fGI ID


CL\_INS\_126
CL\_INS\_126
CL\_INS\_126
CL\_INS\_126
CL\_INS\_126
CL\_INS\_126
CL\_INS\_126
CL\_INS\_126
CL\_INS\_126
CL\_INS\_126
CL\_INS\_126
CL\_INS\_126
CL\_INS\_126
CL\_INS\_126
CL\_INS\_126
CL\_INS\_126
CL\_INS\_126
CL\_INS\_126
CL\_INS\_126
CL\_INS\_126
CL\_INS\_126
CL\_INS\_126
CL\_INS\_126
CL\_INS\_126
Cluster ID


CL\_7109
CL\_1140
CL\_1141
CL\_1142
CL\_1143
CL\_5080
CL\_5081
CL\_5622
CL\_6311
CL\_5729
CL\_5730
CL\_5731
CL\_4308
CL\_4309
CL\_4310
CL\_4311
CL\_4312
CL\_4313
CL\_4314
CL\_4315
CL\_4316
CL\_4317
CL\_4318
CL\_4319
