## Supplementary material for "A novel method for integrating genomic and Tn-Seq data to identify common *in vivo* fitness mechanisms across multiple bacterial species": S6 Dataset: CL_INS_127.html

Legend

 Other
 Regulatoryfunctions
 Hypothetical
 DNA Metabolism
 EnergyMetabolism
 All EssentialGenes
 All Fitness Genes
 Transport +binding proteins

FULL


WINDOWSVGPNG

Trim RowsRemove SingletonsSave Fasta

CL\_1149


CL\_1149


CL\_978


CL\_1149


CL\_296

HighlightSelectShow Genomes


1

CL\_290


1

CL\_1153


1

CL\_290


1

CL\_290


1

CL\_290

fGI ID


CL\_INS\_127
CL\_INS\_127
CL\_INS\_127
CL\_INS\_127
CL\_INS\_127
CL\_INS\_127
CL\_INS\_127
CL\_INS\_127
CL\_INS\_127
CL\_INS\_127
CL\_INS\_127
CL\_INS\_127
CL\_INS\_127
CL\_INS\_127
CL\_INS\_127
CL\_INS\_127
CL\_INS\_127
CL\_INS\_127
CL\_INS\_127
CL\_INS\_127
CL\_INS\_127
CL\_INS\_86
CL\_INS\_149
CL\_INS\_127
Cluster ID


CL\_1150
CL\_1151
CL\_294
CL\_293
CL\_292
CL\_291
CL\_7112
CL\_7113
CL\_7114
CL\_7115
CL\_4010
CL\_4009
CL\_5748
CL\_5749
CL\_5750
CL\_5751
CL\_5752
CL\_5753
CL\_5754
CL\_5755
CL\_5756
CL\_4098
CL\_5757
CL\_5758
