## Supplementary material for "A novel method for integrating genomic and Tn-Seq data to identify common *in vivo* fitness mechanisms across multiple bacterial species": S6 Dataset: CL_INS_128.html

Legend

 Other
 Hypothetical
 EnergyMetabolism
 All EssentialGenes
 All Fitness Genes

FULL


WINDOWSVGPNG

Trim RowsRemove SingletonsSave Fasta

CL\_290


CL\_290


CL\_1149


CL\_290

HighlightSelectShow Genomes


2

CL\_1153


1

CL\_1153


1

CL\_1153


1

CL\_288

fGI ID


CL\_INS\_128
CL\_INS\_127
CL\_INS\_128
Cluster ID


CL\_6537
CL\_1151
CL\_1152
