## Supplementary material for "A novel method for integrating genomic and Tn-Seq data to identify common *in vivo* fitness mechanisms across multiple bacterial species": S6 Dataset: CL_INS_129.html

Legend

 Other
 Hypothetical
 All Fitness Genes

FULL


WINDOWSVGPNG

Trim RowsRemove SingletonsSave Fasta

CL\_1172


CL\_1172


CL\_1172


CL\_1172

HighlightSelectShow Genomes


2

CL\_1174


1

CL\_1470


1

CL\_2123


1

CL\_1174

fGI ID


CL\_INS\_99
CL\_INS\_129
CL\_INS\_129
CL\_INS\_129
CL\_INS\_129
CL\_INS\_129
Cluster ID


CL\_4560
CL\_1469
CL\_6531
CL\_6530
CL\_6529
CL\_1173
