## Supplementary material for "A novel method for integrating genomic and Tn-Seq data to identify common *in vivo* fitness mechanisms across multiple bacterial species": S6 Dataset: CL_INS_131.html

Legend

 Other
 Regulatoryfunctions
 Hypothetical
 Transport +binding proteins

FULL


WINDOWSVGPNG

Trim RowsRemove SingletonsSave Fasta

CL\_5137


CL\_4337


CL\_5137


CL\_5137


CL\_5137

HighlightSelectShow Genomes


1

CL\_1183


1

CL\_1183


1

CL\_4337


1

CL\_5135


1

CL\_1183

fGI ID


CL\_INS\_131
CL\_INS\_131
CL\_INS\_131
CL\_INS\_131
CL\_INS\_131
CL\_INS\_131
CL\_INS\_131
CL\_INS\_131
CL\_INS\_131
CL\_INS\_131
CL\_INS\_131
Cluster ID


CL\_4099
CL\_5809
CL\_7119
CL\_7120
CL\_3800
CL\_5813
CL\_5812
CL\_6523
CL\_6522
CL\_6521
CL\_6520
