## Supplementary material for "A novel method for integrating genomic and Tn-Seq data to identify common *in vivo* fitness mechanisms across multiple bacterial species": S6 Dataset: CL_INS_132.html

Legend

 Other
 Hypothetical
 Regulatoryfunctions
 EnergyMetabolism
 All Fitness Genes
 All VFDB Genes

FULL


WINDOWSVGPNG

Trim RowsRemove SingletonsSave Fasta

CL\_1190


CL\_1190


CL\_337


CL\_1190


CL\_1190

HighlightSelectShow Genomes


1

CL\_1192


1

CL\_1192


1

CL\_1192


1

CL\_1192


1

CL\_1192

fGI ID


CL\_INS\_132
CL\_INS\_132
CL\_INS\_132
CL\_INS\_132
CL\_INS\_132
CL\_INS\_132
CL\_INS\_132
CL\_INS\_132
CL\_INS\_132
CL\_INS\_132
CL\_INS\_132
CL\_INS\_132
Cluster ID


CL\_5808
CL\_5807
CL\_7121
CL\_1191
CL\_6516
CL\_6515
CL\_6514
CL\_6513
CL\_6512
CL\_6511
CL\_6510
CL\_6509
