## Supplementary material for "A novel method for integrating genomic and Tn-Seq data to identify common *in vivo* fitness mechanisms across multiple bacterial species": S6 Dataset: CL_INS_133.html

Legend

 Regulatoryfunctions
 Hypothetical
 Other
 EnergyMetabolism

FULL


WINDOWSVGPNG

Trim RowsRemove SingletonsSave Fasta

CL\_1969


CL\_1956


CL\_4481


CL\_1969

HighlightSelectShow Genomes


2

CL\_1971


1

CL\_1971


1

CL\_1971


1

CL\_1971

fGI ID


CL\_INS\_133
CL\_INS\_133
CL\_INS\_133
CL\_INS\_133
CL\_INS\_133
CL\_INS\_133
CL\_INS\_133
Cluster ID


CL\_7055
CL\_4483
CL\_6709
CL\_6710
CL\_6711
CL\_6712
CL\_1970
