## Supplementary material for "A novel method for integrating genomic and Tn-Seq data to identify common *in vivo* fitness mechanisms across multiple bacterial species": S6 Dataset: CL_INS_134.html

Legend

 Regulatoryfunctions
 Hypothetical
 All EssentialGenes
 All Fitness Genes
 Proteinsynthesis/fate
 Other
 EnergyMetabolism
 Transport +binding proteins

FULL


WINDOWSVGPNG

Trim RowsRemove SingletonsSave Fasta

CL\_1973


CL\_1973


CL\_1973


CL\_1973

HighlightSelectShow Genomes


2

CL\_1980


1

CL\_1980


1

CL\_1980


1

CL\_1980

fGI ID


CL\_INS\_134
CL\_INS\_134
CL\_INS\_134
CL\_INS\_134
CL\_INS\_134
CL\_INS\_134
CL\_INS\_134
CL\_INS\_134
CL\_INS\_134
CL\_INS\_134
Cluster ID


CL\_4486
CL\_4487
CL\_1974
CL\_7054
CL\_7053
CL\_1975
CL\_1976
CL\_1977
CL\_1978
CL\_1979
