## Supplementary material for "A novel method for integrating genomic and Tn-Seq data to identify common *in vivo* fitness mechanisms across multiple bacterial species": S6 Dataset: CL_INS_135.html

Legend

 Hypothetical
 All EssentialGenes

FULL


WINDOWSVGPNG

Trim RowsRemove SingletonsSave Fasta

CL\_1984


CL\_1984


CL\_1984

HighlightSelectShow Genomes


3

CL\_1986


1

CL\_1988


1

CL\_1986

fGI ID

CL\_INS\_135
Cluster ID

CL\_1985
