## Supplementary material for "A novel method for integrating genomic and Tn-Seq data to identify common *in vivo* fitness mechanisms across multiple bacterial species": S6 Dataset: CL_INS_136.html

Legend

 Hypothetical
 Regulatoryfunctions
 All EssentialGenes
 Other
 EnergyMetabolism
 All VFDB Genes
 Transport +binding proteins

FULL


WINDOWSVGPNG

Trim RowsRemove SingletonsSave Fasta

CL\_1988


CL\_1988


CL\_1988


CL\_1988


CL\_470

HighlightSelectShow Genomes


1

CL\_1994


1

CL\_1994


1

CL\_1994


1

CL\_1994


1

CL\_1994

fGI ID


CL\_INS\_136
CL\_INS\_136
CL\_INS\_136
CL\_INS\_136
CL\_INS\_136
CL\_INS\_136
CL\_INS\_136
CL\_INS\_136
CL\_INS\_136
CL\_INS\_136
CL\_INS\_136
CL\_INS\_136
CL\_INS\_136
Cluster ID


CL\_1989
CL\_1990
CL\_1991
CL\_1992
CL\_5879
CL\_1993
CL\_4494
CL\_4495
CL\_4496
CL\_4497
CL\_4498
CL\_4499
CL\_4500
