## Supplementary material for "A novel method for integrating genomic and Tn-Seq data to identify common *in vivo* fitness mechanisms across multiple bacterial species": S6 Dataset: CL_INS_137.html

Legend

 Hypothetical
 Regulatoryfunctions
 All EssentialGenes
 All Fitness Genes
 Other
 EnergyMetabolism

FULL


WINDOWSVGPNG

Trim RowsRemove SingletonsSave Fasta

CL\_2020


CL\_2020

HighlightSelectShow Genomes


3

CL\_2038


1

CL\_2038

fGI ID


CL\_INS\_137
CL\_INS\_137
CL\_INS\_137
CL\_INS\_137
CL\_INS\_137
CL\_INS\_137
CL\_INS\_137
CL\_INS\_137
CL\_INS\_137
CL\_INS\_137
CL\_INS\_137
CL\_INS\_137
CL\_INS\_137
CL\_INS\_137
CL\_INS\_137
CL\_INS\_137
CL\_INS\_137
Cluster ID


CL\_2021
CL\_2022
CL\_2023
CL\_2024
CL\_2025
CL\_2026
CL\_2027
CL\_2028
CL\_2029
CL\_2030
CL\_2031
CL\_2032
CL\_2033
CL\_2034
CL\_2035
CL\_2036
CL\_2037
