## Supplementary material for "A novel method for integrating genomic and Tn-Seq data to identify common *in vivo* fitness mechanisms across multiple bacterial species": S6 Dataset: CL_INS_138.html

Legend

 Hypothetical
 Regulatoryfunctions
 SignalTransduction +Quorum Sensing
 Other

FULL


WINDOWSVGPNG

Trim RowsRemove SingletonsSave Fasta

CL\_2043


CL\_2042


CL\_2043

HighlightSelectShow Genomes


3

CL\_2045


1

CL\_2045


1

CL\_2045

fGI ID


CL\_INS\_138
CL\_INS\_138
CL\_INS\_138
Cluster ID


CL\_2044
CL\_1813
CL\_4516
