## Supplementary material for "A novel method for integrating genomic and Tn-Seq data to identify common *in vivo* fitness mechanisms across multiple bacterial species": S6 Dataset: CL_INS_139.html

Legend

 Regulatoryfunctions
 Hypothetical
 All EssentialGenes
 Other

FULL


WINDOWSVGPNG

Trim RowsRemove SingletonsSave Fasta

CL\_2045


CL\_2045


CL\_2045

HighlightSelectShow Genomes


3

CL\_2047


1

CL\_2047


1

CL\_2047

fGI ID


CL\_INS\_139
CL\_INS\_139
CL\_INS\_139
Cluster ID


CL\_2046
CL\_4517
CL\_4518
