## Supplementary material for "A novel method for integrating genomic and Tn-Seq data to identify common *in vivo* fitness mechanisms across multiple bacterial species": S6 Dataset: CL_INS_140.html

Legend

 Hypothetical
 Other

FULL


WINDOWSVGPNG

Trim RowsRemove SingletonsSave Fasta

CL\_2057


CL\_2057


CL\_2057

HighlightSelectShow Genomes


3

CL\_2064


1

CL\_2064


1

CL\_2064

fGI ID


CL\_INS\_140
CL\_INS\_140
CL\_INS\_140
CL\_INS\_140
CL\_INS\_140
CL\_INS\_140
CL\_INS\_140
CL\_INS\_140
CL\_INS\_140
Cluster ID


CL\_3626
CL\_3627
CL\_3628
CL\_2058
CL\_2059
CL\_2060
CL\_2061
CL\_2062
CL\_2063
