## Supplementary material for "A novel method for integrating genomic and Tn-Seq data to identify common *in vivo* fitness mechanisms across multiple bacterial species": S6 Dataset: CL_INS_141.html

Legend

 Hypothetical
 All Fitness Genes
 Other

FULL


WINDOWSVGPNG

Trim RowsRemove SingletonsSave Fasta

CL\_2069


CL\_2069


CL\_2069

HighlightSelectShow Genomes


2

CL\_2070


2

CL\_2070


1

CL\_2070

fGI ID


CL\_INS\_141
CL\_INS\_141
Cluster ID


CL\_5884
CL\_6718
