## Supplementary material for "A novel method for integrating genomic and Tn-Seq data to identify common *in vivo* fitness mechanisms across multiple bacterial species": S6 Dataset: CL_INS_142.html

Legend

 Regulatoryfunctions
 Hypothetical
 Transcription
 All Fitness Genes
 SignalTransduction +Quorum Sensing
 Other

FULL


WINDOWSVGPNG

Trim RowsRemove SingletonsSave Fasta

CL\_2078


CL\_2078


CL\_2078


CL\_2078


CL\_2078

HighlightSelectShow Genomes


1

CL\_2075


1

CL\_2075


1

CL\_2075


1

CL\_3387


1

CL\_2075

fGI ID


CL\_INS\_142
CL\_INS\_142
CL\_INS\_142
CL\_INS\_30
CL\_INS\_30
CL\_INS\_30
CL\_INS\_142
CL\_INS\_142
CL\_INS\_142
CL\_INS\_142
CL\_INS\_170
CL\_INS\_170
CL\_INS\_170
CL\_INS\_170
CL\_INS\_142
CL\_INS\_142
CL\_INS\_142
CL\_INS\_142
CL\_INS\_142
Cluster ID


CL\_7051
CL\_2077
CL\_2076
CL\_4549
CL\_4548
CL\_4547
CL\_6722
CL\_6721
CL\_6720
CL\_6719
CL\_3804
CL\_3803
CL\_3802
CL\_3801
CL\_5889
CL\_5888
CL\_5887
CL\_5886
CL\_5885
