## Supplementary material for "A novel method for integrating genomic and Tn-Seq data to identify common *in vivo* fitness mechanisms across multiple bacterial species": S6 Dataset: CL_INS_143.html

Legend

 Regulatoryfunctions
 Hypothetical
 All EssentialGenes
 Other
 Transport +binding proteins

FULL


WINDOWSVGPNG

Trim RowsRemove SingletonsSave Fasta

CL\_2106


CL\_2106


CL\_2106

HighlightSelectShow Genomes


3

CL\_4558


1

CL\_2104


1

CL\_4558

fGI ID


CL\_INS\_84
CL\_INS\_174
CL\_INS\_174
CL\_INS\_174
CL\_INS\_174
CL\_INS\_174
Cluster ID


CL\_4813
CL\_4812
CL\_4811
CL\_4810
CL\_4809
CL\_4808
