## Supplementary material for "A novel method for integrating genomic and Tn-Seq data to identify common *in vivo* fitness mechanisms across multiple bacterial species": S6 Dataset: CL_INS_144.html

Legend

 Regulatoryfunctions
 Hypothetical
 All EssentialGenes
 Other
 Transport +binding proteins

FULL


WINDOWSVGPNG

Trim RowsRemove SingletonsSave Fasta

CL\_2107


CL\_2107


CL\_2107


CL\_2107

HighlightSelectShow Genomes


1

CL\_2114


1

CL\_2122


1

CL\_2114


1

CL\_2114

fGI ID


CL\_INS\_144
CL\_INS\_144
CL\_INS\_144
CL\_INS\_144
CL\_INS\_144
CL\_INS\_144
CL\_INS\_144
CL\_INS\_144
CL\_INS\_144
CL\_INS\_144
Cluster ID


CL\_6726
CL\_6727
CL\_2120
CL\_2121
CL\_2108
CL\_2109
CL\_2110
CL\_2111
CL\_2112
CL\_2113
