## Supplementary material for "A novel method for integrating genomic and Tn-Seq data to identify common *in vivo* fitness mechanisms across multiple bacterial species": S6 Dataset: CL_INS_147.html

Legend

 Hypothetical
 Regulatoryfunctions
 Other

FULL


WINDOWSVGPNG

Trim RowsRemove SingletonsSave Fasta

CL\_2013


CL\_2013


CL\_2013

HighlightSelectShow Genomes


3

CL\_2132


1

CL\_2010


1

CL\_3082

fGI ID


CL\_INS\_147
CL\_INS\_147
CL\_INS\_147
CL\_INS\_147
CL\_INS\_147
CL\_INS\_147
CL\_INS\_147
CL\_INS\_147
CL\_INS\_147
CL\_INS\_147
CL\_INS\_147
CL\_INS\_147
Cluster ID


CL\_2012
CL\_5442
CL\_4069
CL\_4070
CL\_4071
CL\_5444
CL\_5445
CL\_5446
CL\_6913
CL\_6912
CL\_4429
CL\_6911
