## Supplementary material for "A novel method for integrating genomic and Tn-Seq data to identify common *in vivo* fitness mechanisms across multiple bacterial species": S6 Dataset: CL_INS_149.html

Legend

 Mobile +extrachromosomalelementfunctions
 Regulatoryfunctions
 Hypothetical
 All EssentialGenes
 All Fitness Genes
 SignalTransduction +Quorum Sensing
 Cell Envelope
 Other
 EnergyMetabolism
 All VFDB Genes
 Transport +binding proteins

FULL


WINDOWSVGPNG

Trim RowsRemove SingletonsSave Fasta

CL\_2186


CL\_2186


CL\_2186


CL\_2186


CL\_2186

HighlightSelectShow Genomes


1

CL\_2189


1

CL\_260


1

CL\_273


1

CL\_4576


1

CL\_3017

fGI ID


CL\_INS\_149
CL\_INS\_149
CL\_INS\_149
CL\_INS\_149
CL\_INS\_149
CL\_INS\_149
CL\_INS\_149
CL\_INS\_149
CL\_INS\_149
CL\_INS\_149
CL\_INS\_170
CL\_INS\_170
CL\_INS\_170
CL\_INS\_170
CL\_INS\_149
CL\_INS\_149
CL\_INS\_149
CL\_INS\_149
CL\_INS\_149
CL\_INS\_149
CL\_INS\_149
CL\_INS\_149
CL\_INS\_149
CL\_INS\_149
CL\_INS\_149
CL\_INS\_149
CL\_INS\_149
CL\_INS\_149
CL\_INS\_149
CL\_INS\_149
CL\_INS\_149
CL\_INS\_149
CL\_INS\_149
CL\_INS\_149
CL\_INS\_149
CL\_INS\_30
CL\_INS\_30
CL\_INS\_30
CL\_INS\_149
CL\_INS\_149
CL\_INS\_149
CL\_INS\_149
CL\_INS\_149
CL\_INS\_149
CL\_INS\_149
CL\_INS\_149
CL\_INS\_149
CL\_INS\_149
CL\_INS\_149
CL\_INS\_149
CL\_INS\_149
CL\_INS\_149
CL\_INS\_149
CL\_INS\_149
CL\_INS\_149
CL\_INS\_149
CL\_INS\_149
CL\_INS\_149
CL\_INS\_149
CL\_INS\_149
CL\_INS\_149
CL\_INS\_149
CL\_INS\_149
Cluster ID


CL\_2187
CL\_4569
CL\_4570
CL\_4571
CL\_4572
CL\_4573
CL\_4574
CL\_6729
CL\_6730
CL\_6731
CL\_3804
CL\_3803
CL\_3802
CL\_3801
CL\_6732
CL\_3700
CL\_6733
CL\_6734
CL\_6735
CL\_6736
CL\_6737
CL\_6738
CL\_5388
CL\_5757
CL\_6739
CL\_6740
CL\_6741
CL\_6742
CL\_6743
CL\_6744
CL\_7048
CL\_7047
CL\_5901
CL\_5902
CL\_408
CL\_4896
CL\_5903
CL\_5904
CL\_7046
CL\_4787
CL\_7045
CL\_4785
CL\_5257
CL\_7044
CL\_7043
CL\_7042
CL\_7041
CL\_4784
CL\_4788
CL\_4786
CL\_2938
CL\_2946
CL\_2947
CL\_6851
CL\_6850
CL\_6849
CL\_6848
CL\_1960
CL\_7040
CL\_7039
CL\_7038
CL\_7037
CL\_7036
