## Supplementary material for "A novel method for integrating genomic and Tn-Seq data to identify common *in vivo* fitness mechanisms across multiple bacterial species": S6 Dataset: CL_INS_150.html

Legend

 Regulatoryfunctions
 Hypothetical
 All EssentialGenes
 All Fitness Genes
 Cell Envelope
 SignalTransduction +Quorum Sensing
 Other
 Transport +binding proteins
 All VFDB Genes

FULL


WINDOWSVGPNG

Trim RowsRemove SingletonsSave Fasta

CL\_2189


CL\_2189


CL\_2189


CL\_2189

HighlightSelectShow Genomes


1

CL\_2186


1

CL\_273


1

CL\_36


1

CL\_226

fGI ID


CL\_INS\_150
CL\_INS\_150
CL\_INS\_150
CL\_INS\_150
CL\_INS\_150
CL\_INS\_150
CL\_INS\_150
CL\_INS\_150
CL\_INS\_149
CL\_INS\_113
CL\_INS\_113
CL\_INS\_113
CL\_INS\_113
CL\_INS\_113
CL\_INS\_113
CL\_INS\_113
CL\_INS\_150
CL\_INS\_150
CL\_INS\_150
CL\_INS\_150
CL\_INS\_30
CL\_INS\_30
CL\_INS\_3
CL\_INS\_3
CL\_INS\_3
CL\_INS\_3
CL\_INS\_150
CL\_INS\_150
CL\_INS\_150
CL\_INS\_150
CL\_INS\_150
CL\_INS\_150
CL\_INS\_150
CL\_INS\_150
CL\_INS\_150
CL\_INS\_150
CL\_INS\_150
CL\_INS\_150
CL\_INS\_150
CL\_INS\_150
CL\_INS\_150
CL\_INS\_150
CL\_INS\_150
Cluster ID


CL\_4328
CL\_4327
CL\_4326
CL\_4325
CL\_4324
CL\_4323
CL\_4322
CL\_2188
CL\_2187
CL\_5576
CL\_5575
CL\_6080
CL\_6081
CL\_6082
CL\_6083
CL\_6084
CL\_7014
CL\_7015
CL\_7016
CL\_7017
CL\_5904
CL\_5903
CL\_4454
CL\_4453
CL\_4452
CL\_4450
CL\_4449
CL\_7018
CL\_7019
CL\_7020
CL\_7021
CL\_7022
CL\_7023
CL\_7024
CL\_7025
CL\_7026
CL\_7027
CL\_7028
CL\_7029
CL\_7030
CL\_7031
CL\_7032
CL\_7033
