## Supplementary material for "A novel method for integrating genomic and Tn-Seq data to identify common *in vivo* fitness mechanisms across multiple bacterial species": S6 Dataset: CL_INS_151.html

Legend

 Regulatoryfunctions
 Hypothetical
 All EssentialGenes
 SignalTransduction +Quorum Sensing
 Cell Envelope
 Other
 All VFDB Genes
 Transport +binding proteins

FULL


WINDOWSVGPNG

Trim RowsRemove SingletonsSave Fasta

CL\_2189


CL\_2524


CL\_2189


CL\_830


CL\_1208


CL\_2189

HighlightSelectShow Genomes


2

CL\_2193


1

CL\_2193


1

CL\_223


1

CL\_2193


1

CL\_2193


1

CL\_1119

fGI ID


CL\_INS\_151
CL\_INS\_20
CL\_INS\_151
CL\_INS\_151
CL\_INS\_151
CL\_INS\_151
CL\_INS\_151
CL\_INS\_151
CL\_INS\_151
CL\_INS\_151
CL\_INS\_151
CL\_INS\_151
CL\_INS\_151
CL\_INS\_151
CL\_INS\_151
CL\_INS\_151
CL\_INS\_151
CL\_INS\_151
CL\_INS\_151
CL\_INS\_151
CL\_INS\_151
CL\_INS\_151
CL\_INS\_151
CL\_INS\_151
CL\_INS\_151
CL\_INS\_151
CL\_INS\_151
CL\_INS\_151
CL\_INS\_151
CL\_INS\_151
CL\_INS\_151
CL\_INS\_151
CL\_INS\_151
CL\_INS\_151
CL\_INS\_151
CL\_INS\_151
CL\_INS\_151
CL\_INS\_151
CL\_INS\_151
CL\_INS\_151
CL\_INS\_151
CL\_INS\_151
CL\_INS\_151
CL\_INS\_151
CL\_INS\_151
CL\_INS\_151
CL\_INS\_151
CL\_INS\_151
CL\_INS\_151
CL\_INS\_151
CL\_INS\_151
CL\_INS\_151
CL\_INS\_151
CL\_INS\_151
Cluster ID


CL\_4329
CL\_4203
CL\_4204
CL\_5330
CL\_5329
CL\_5328
CL\_2190
CL\_2191
CL\_2192
CL\_6389
CL\_6390
CL\_6391
CL\_6392
CL\_5202
CL\_5203
CL\_6393
CL\_6394
CL\_6395
CL\_6396
CL\_6397
CL\_6398
CL\_6399
CL\_6400
CL\_6401
CL\_6402
CL\_6403
CL\_6404
CL\_6405
CL\_6406
CL\_6407
CL\_6408
CL\_6409
CL\_6410
CL\_6411
CL\_6412
CL\_6413
CL\_6414
CL\_6415
CL\_4626
CL\_4628
CL\_6416
CL\_6417
CL\_6418
CL\_6419
CL\_6420
CL\_6421
CL\_6422
CL\_6423
CL\_6424
CL\_6425
CL\_6426
CL\_6427
CL\_6428
CL\_6429
