## Supplementary material for "A novel method for integrating genomic and Tn-Seq data to identify common *in vivo* fitness mechanisms across multiple bacterial species": S6 Dataset: CL_INS_152.html

Legend

 Hypothetical
 Regulatoryfunctions
 All EssentialGenes
 All Fitness Genes
 Other

FULL


WINDOWSVGPNG

Trim RowsRemove SingletonsSave Fasta

CL\_2194


CL\_2194


CL\_2194


CL\_2186


CL\_3017


CL\_2194

HighlightSelectShow Genomes


1

CL\_1147


1

CL\_2197


1

CL\_4576


1

CL\_4576


1

CL\_4576


1

CL\_4576

fGI ID


CL\_INS\_152
CL\_INS\_152
CL\_INS\_152
CL\_INS\_153
CL\_INS\_152
CL\_INS\_149
CL\_INS\_149
CL\_INS\_149
CL\_INS\_149
CL\_INS\_149
CL\_INS\_152
Cluster ID


CL\_6747
CL\_4242
CL\_2195
CL\_2196
CL\_4239
CL\_4570
CL\_4571
CL\_4572
CL\_4573
CL\_4574
CL\_4575
