## Supplementary material for "A novel method for integrating genomic and Tn-Seq data to identify common *in vivo* fitness mechanisms across multiple bacterial species": S6 Dataset: CL_INS_153.html

Legend

 Regulatoryfunctions
 All EssentialGenes
 Other

FULL


WINDOWSVGPNG

Trim RowsRemove SingletonsSave Fasta

CL\_2197


CL\_2197

HighlightSelectShow Genomes


3

CL\_4576


1

CL\_2194

fGI ID

CL\_INS\_153
Cluster ID

CL\_2196
