## Supplementary material for "A novel method for integrating genomic and Tn-Seq data to identify common *in vivo* fitness mechanisms across multiple bacterial species": S6 Dataset: CL_INS_154.html

Legend

 Regulatoryfunctions
 All EssentialGenes
 Other

FULL


WINDOWSVGPNG

Trim RowsRemove SingletonsSave Fasta

CL\_2200


CL\_2200


CL\_2199

HighlightSelectShow Genomes


2

CL\_2201


2

CL\_2201


1

CL\_2201

fGI ID


CL\_INS\_154
CL\_INS\_154
Cluster ID


CL\_4583
CL\_4584
