## Supplementary material for "A novel method for integrating genomic and Tn-Seq data to identify common *in vivo* fitness mechanisms across multiple bacterial species": S6 Dataset: CL_INS_155.html

Legend

 Regulatoryfunctions
 Hypothetical
 Other
 All VFDB Genes

FULL


WINDOWSVGPNG

Trim RowsRemove SingletonsSave Fasta

CL\_2213


CL\_2213


CL\_2213

HighlightSelectShow Genomes


2

CL\_2214


1

CL\_2214


1

CL\_2214

fGI ID


CL\_INS\_155
CL\_INS\_155
CL\_INS\_155
Cluster ID


CL\_4690
CL\_3782
CL\_3783
