## Supplementary material for "A novel method for integrating genomic and Tn-Seq data to identify common *in vivo* fitness mechanisms across multiple bacterial species": S6 Dataset: CL_INS_156.html

Legend

 Other
 Transport +binding proteins

FULL


WINDOWSVGPNG

Trim RowsRemove SingletonsSave Fasta

CL\_2222


CL\_2222


CL\_2222

HighlightSelectShow Genomes


2

CL\_2223


2

CL\_2223


1

CL\_1405

fGI ID


CL\_INS\_156
CL\_INS\_156
CL\_INS\_156
Cluster ID


CL\_5125
CL\_5124
CL\_5123
