## Supplementary material for "A novel method for integrating genomic and Tn-Seq data to identify common *in vivo* fitness mechanisms across multiple bacterial species": S6 Dataset: CL_INS_157.html

Legend

 Hypothetical
 Regulatoryfunctions
 All EssentialGenes
 Other
 All VFDB Genes
 Transport +binding proteins

FULL


WINDOWSVGPNG

Trim RowsRemove SingletonsSave Fasta

CL\_2255


CL\_2255


CL\_2255


CL\_2255


CL\_2255

HighlightSelectShow Genomes


1

CL\_2263


1

CL\_2263


1

CL\_2263


1

CL\_2263


1

CL\_2263

fGI ID


CL\_INS\_157
CL\_INS\_157
CL\_INS\_157
CL\_INS\_157
CL\_INS\_157
CL\_INS\_157
CL\_INS\_157
CL\_INS\_157
CL\_INS\_157
CL\_INS\_157
CL\_INS\_157
CL\_INS\_157
CL\_INS\_157
CL\_INS\_157
CL\_INS\_157
Cluster ID


CL\_2256
CL\_2257
CL\_2258
CL\_2259
CL\_2260
CL\_2261
CL\_2262
CL\_2276
CL\_5908
CL\_5909
CL\_5910
CL\_5911
CL\_4579
CL\_4580
CL\_4581
