## Supplementary material for "A novel method for integrating genomic and Tn-Seq data to identify common *in vivo* fitness mechanisms across multiple bacterial species": S6 Dataset: CL_INS_158.html

Legend

 Biosynthesis ofcofactors,prostheticgroups, +carriers
 Other

FULL


WINDOWSVGPNG

Trim RowsRemove SingletonsSave Fasta

CL\_2274


CL\_2274

HighlightSelectShow Genomes


3

CL\_2275


2

CL\_2275

fGI ID

CL\_INS\_158
Cluster ID

CL\_5034
