## Supplementary material for "A novel method for integrating genomic and Tn-Seq data to identify common *in vivo* fitness mechanisms across multiple bacterial species": S6 Dataset: CL_INS_159.html

Legend

 Regulatoryfunctions
 Hypothetical
 All EssentialGenes
 Other
 EnergyMetabolism
 Transport +binding proteins

FULL


WINDOWSVGPNG

Trim RowsRemove SingletonsSave Fasta

CL\_2291


CL\_2291


CL\_2291


CL\_2291


CL\_5031

HighlightSelectShow Genomes


2

CL\_2293


1

CL\_5032


1

CL\_2293


1

CL\_2293


1

CL\_2293

fGI ID


CL\_INS\_159
CL\_INS\_159
CL\_INS\_159
CL\_INS\_159
CL\_INS\_159
CL\_INS\_159
CL\_INS\_159
CL\_INS\_159
Cluster ID


CL\_4404
CL\_2292
CL\_5029
CL\_5028
CL\_5027
CL\_3730
CL\_3729
CL\_3728
