## Supplementary material for "A novel method for integrating genomic and Tn-Seq data to identify common *in vivo* fitness mechanisms across multiple bacterial species": S6 Dataset: CL_INS_161.html

Legend

 Regulatoryfunctions
 Hypothetical
 All Fitness Genes
 Other
 Transport +binding proteins

FULL


WINDOWSVGPNG

Trim RowsRemove SingletonsSave Fasta

CL\_2348


CL\_2348


CL\_2348


CL\_2348

HighlightSelectShow Genomes


2

CL\_2349


1

CL\_2349


1

CL\_2349


1

CL\_2349

fGI ID


CL\_INS\_161
CL\_INS\_161
CL\_INS\_161
CL\_INS\_161
CL\_INS\_161
CL\_INS\_161
CL\_INS\_161
Cluster ID


CL\_4601
CL\_4602
CL\_5914
CL\_5915
CL\_5916
CL\_5917
CL\_5918
