## Supplementary material for "A novel method for integrating genomic and Tn-Seq data to identify common *in vivo* fitness mechanisms across multiple bacterial species": S6 Dataset: CL_INS_163.html

Legend

 Hypothetical
 Other

FULL


WINDOWSVGPNG

Trim RowsRemove SingletonsSave Fasta

CL\_2353


CL\_2353


CL\_2353

HighlightSelectShow Genomes


2

CL\_2355


1

CL\_2355


1

CL\_2355

fGI ID


CL\_INS\_163
CL\_INS\_163
CL\_INS\_163
Cluster ID


CL\_2354
CL\_5925
CL\_5926
