## Supplementary material for "A novel method for integrating genomic and Tn-Seq data to identify common *in vivo* fitness mechanisms across multiple bacterial species": S6 Dataset: CL_INS_164.html

Legend

 Hypothetical
 Regulatoryfunctions
 All EssentialGenes
 All Fitness Genes
 Proteinsynthesis/fate
 Other
 Transport +binding proteins
 All VFDB Genes

FULL


WINDOWSVGPNG

Trim RowsRemove SingletonsSave Fasta

CL\_3481


CL\_3481


CL\_3481


CL\_260


CL\_3481

HighlightSelectShow Genomes


1

CL\_3489


1

CL\_2356


1

CL\_2356


1

CL\_2356


1

CL\_260

fGI ID


CL\_INS\_49
CL\_INS\_202
CL\_INS\_202
CL\_INS\_202
CL\_INS\_202
CL\_INS\_202
CL\_INS\_202
CL\_INS\_164
CL\_INS\_164
CL\_INS\_164
CL\_INS\_164
CL\_INS\_164
CL\_INS\_164
CL\_INS\_164
CL\_INS\_164
CL\_INS\_164
CL\_INS\_164
CL\_INS\_164
CL\_INS\_164
CL\_INS\_164
CL\_INS\_164
CL\_INS\_164
CL\_INS\_164
CL\_INS\_164
CL\_INS\_164
CL\_INS\_164
CL\_INS\_164
CL\_INS\_164
CL\_INS\_164
CL\_INS\_164
CL\_INS\_164
CL\_INS\_164
CL\_INS\_164
CL\_INS\_164
CL\_INS\_164
CL\_INS\_126
Cluster ID


CL\_1728
CL\_253
CL\_255
CL\_256
CL\_257
CL\_258
CL\_5905
CL\_3482
CL\_3483
CL\_3484
CL\_3485
CL\_3486
CL\_3487
CL\_6829
CL\_3618
CL\_3617
CL\_3616
CL\_3615
CL\_3614
CL\_3613
CL\_3612
CL\_3611
CL\_3610
CL\_5508
CL\_6828
CL\_6827
CL\_6826
CL\_6825
CL\_6824
CL\_6823
CL\_6822
CL\_6821
CL\_6820
CL\_6819
CL\_6818
CL\_1140
