## Supplementary material for "A novel method for integrating genomic and Tn-Seq data to identify common *in vivo* fitness mechanisms across multiple bacterial species": S6 Dataset: CL_INS_165.html

Legend

 Hypothetical
 All EssentialGenes
 Other

FULL


WINDOWSVGPNG

Trim RowsRemove SingletonsSave Fasta

CL\_3747


CL\_832


CL\_3747

HighlightSelectShow Genomes


2

CL\_840


1

CL\_840


1

CL\_840

fGI ID


CL\_INS\_165
CL\_INS\_165
CL\_INS\_165
CL\_INS\_165
CL\_INS\_165
CL\_INS\_165
CL\_INS\_165
Cluster ID


CL\_833
CL\_834
CL\_835
CL\_836
CL\_837
CL\_838
CL\_839
