## Supplementary material for "A novel method for integrating genomic and Tn-Seq data to identify common *in vivo* fitness mechanisms across multiple bacterial species": S6 Dataset: CL_INS_167.html

Legend

 Hypothetical
 Other
 Transport +binding proteins

FULL


WINDOWSVGPNG

Trim RowsRemove SingletonsSave Fasta

CL\_843


CL\_843

HighlightSelectShow Genomes


3

CL\_844


2

CL\_844

fGI ID


CL\_INS\_167
CL\_INS\_167
Cluster ID


CL\_4849
CL\_4848
