## Supplementary material for "A novel method for integrating genomic and Tn-Seq data to identify common *in vivo* fitness mechanisms across multiple bacterial species": S6 Dataset: CL_INS_169.html

Legend

 Regulatoryfunctions
 All Fitness Genes
 Other
 All VFDB Genes

FULL


WINDOWSVGPNG

Trim RowsRemove SingletonsSave Fasta

CL\_900


CL\_879


CL\_886


CL\_900

HighlightSelectShow Genomes


2

CL\_901


1

CL\_901


1

CL\_901


1

CL\_901

fGI ID


CL\_INS\_169
CL\_INS\_169
Cluster ID


CL\_5674
CL\_5675
