## Supplementary material for "A novel method for integrating genomic and Tn-Seq data to identify common *in vivo* fitness mechanisms across multiple bacterial species": S6 Dataset: CL_INS_170.html

Legend

 Regulatoryfunctions
 All EssentialGenes
 All Fitness Genes
 Other

FULL


WINDOWSVGPNG

Trim RowsRemove SingletonsSave Fasta

CL\_926


CL\_926

HighlightSelectShow Genomes


2

CL\_927


1

CL\_851

fGI ID


CL\_INS\_170
CL\_INS\_170
CL\_INS\_170
CL\_INS\_170
Cluster ID


CL\_3801
CL\_3802
CL\_3803
CL\_3804
