## Supplementary material for "A novel method for integrating genomic and Tn-Seq data to identify common *in vivo* fitness mechanisms across multiple bacterial species": S6 Dataset: CL_INS_172.html

Legend

 Regulatoryfunctions
 Hypothetical
 All EssentialGenes
 Other
 Transport +binding proteins

FULL


WINDOWSVGPNG

Trim RowsRemove SingletonsSave Fasta

CL\_1468


CL\_1468


CL\_1471


CL\_1468


CL\_1468

HighlightSelectShow Genomes


2

CL\_1466


1

CL\_4297


1

CL\_1466


1

CL\_1466


1

CL\_1359

fGI ID


CL\_INS\_172
CL\_INS\_172
CL\_INS\_172
CL\_INS\_49
CL\_INS\_49
CL\_INS\_172
CL\_INS\_194
CL\_INS\_194
CL\_INS\_181
CL\_INS\_32
CL\_INS\_32
CL\_INS\_32
CL\_INS\_181
CL\_INS\_181
CL\_INS\_181
CL\_INS\_181
CL\_INS\_181
CL\_INS\_181
CL\_INS\_181
CL\_INS\_181
Cluster ID


CL\_6246
CL\_1467
CL\_4301
CL\_4300
CL\_4299
CL\_6621
CL\_4643
CL\_4644
CL\_6620
CL\_5667
CL\_5668
CL\_5669
CL\_6619
CL\_6618
CL\_6617
CL\_6616
CL\_6615
CL\_6614
CL\_6613
CL\_6612
