## Supplementary material for "A novel method for integrating genomic and Tn-Seq data to identify common *in vivo* fitness mechanisms across multiple bacterial species": S6 Dataset: CL_INS_174.html

Legend

 Hypothetical
 All Fitness Genes
 Other
 EnergyMetabolism
 Transport +binding proteins

FULL


WINDOWSVGPNG

Trim RowsRemove SingletonsSave Fasta

CL\_1433


CL\_1433


CL\_1433

HighlightSelectShow Genomes


3

CL\_1434


1

CL\_2930


1

CL\_1434

fGI ID


CL\_INS\_174
CL\_INS\_174
CL\_INS\_174
CL\_INS\_174
CL\_INS\_174
CL\_INS\_174
Cluster ID


CL\_6254
CL\_4808
CL\_4809
CL\_4810
CL\_4811
CL\_4812
