## Supplementary material for "A novel method for integrating genomic and Tn-Seq data to identify common *in vivo* fitness mechanisms across multiple bacterial species": S6 Dataset: CL_INS_175.html

Legend

 Hypothetical
 All EssentialGenes

FULL


WINDOWSVGPNG

Trim RowsRemove SingletonsSave Fasta

CL\_1426


CL\_1426


CL\_1426


CL\_1426

HighlightSelectShow Genomes


1

CL\_1421


1

CL\_1421


1

CL\_1421


1

CL\_1421

fGI ID


CL\_INS\_175
CL\_INS\_175
CL\_INS\_175
CL\_INS\_175
CL\_INS\_175
Cluster ID


CL\_5683
CL\_1425
CL\_1424
CL\_1423
CL\_1422
