## Supplementary material for "A novel method for integrating genomic and Tn-Seq data to identify common *in vivo* fitness mechanisms across multiple bacterial species": S6 Dataset: CL_INS_176.html

Legend

 All EssentialGenes
 All Fitness Genes
 Other
 Transport +binding proteins

FULL


WINDOWSVGPNG

Trim RowsRemove SingletonsSave Fasta

CL\_1415


CL\_985


CL\_1415

HighlightSelectShow Genomes


3

CL\_1413


1

CL\_1413


1

CL\_1413

fGI ID


CL\_INS\_176
CL\_INS\_176
Cluster ID


CL\_1414
CL\_3778
