## Supplementary material for "A novel method for integrating genomic and Tn-Seq data to identify common *in vivo* fitness mechanisms across multiple bacterial species": S6 Dataset: CL_INS_179.html

Legend

 Hypothetical
 All EssentialGenes
 All Fitness Genes

FULL


WINDOWSVGPNG

Trim RowsRemove SingletonsSave Fasta

CL\_1383


CL\_3671


CL\_1383

HighlightSelectShow Genomes


3

CL\_1381


1

CL\_1381


1

CL\_1381

fGI ID

CL\_INS\_179
Cluster ID

CL\_1382
