## Supplementary material for "A novel method for integrating genomic and Tn-Seq data to identify common *in vivo* fitness mechanisms across multiple bacterial species": S6 Dataset: CL_INS_180.html

Legend

 Hypothetical
 All EssentialGenes
 Other
 All VFDB Genes

FULL


WINDOWSVGPNG

Trim RowsRemove SingletonsSave Fasta

CL\_3913


CL\_3913

HighlightSelectShow Genomes


3

CL\_1380


1

CL\_2254

fGI ID


CL\_INS\_180
CL\_INS\_180
CL\_INS\_29
CL\_INS\_180
CL\_INS\_180
Cluster ID


CL\_3914
CL\_6763
CL\_4017
CL\_6762
CL\_6761
