## Supplementary material for "A novel method for integrating genomic and Tn-Seq data to identify common *in vivo* fitness mechanisms across multiple bacterial species": S6 Dataset: CL_INS_181.html

Legend

 Hypothetical
 Regulatoryfunctions
 All EssentialGenes
 Cell Envelope
 Other
 Transport +binding proteins

FULL


WINDOWSVGPNG

Trim RowsRemove SingletonsSave Fasta

CL\_1359


CL\_1359


CL\_960


CL\_1359


CL\_1359

HighlightSelectShow Genomes


1

CL\_1361


1

CL\_1360


1

CL\_1360


1

CL\_1468


1

CL\_1360

fGI ID


CL\_INS\_185
CL\_INS\_185
CL\_INS\_181
CL\_INS\_181
CL\_INS\_181
CL\_INS\_181
CL\_INS\_181
CL\_INS\_181
CL\_INS\_181
CL\_INS\_181
CL\_INS\_125
CL\_INS\_181
CL\_INS\_181
CL\_INS\_181
CL\_INS\_181
CL\_INS\_181
CL\_INS\_181
CL\_INS\_181
CL\_INS\_181
CL\_INS\_32
CL\_INS\_32
CL\_INS\_32
CL\_INS\_181
CL\_INS\_194
CL\_INS\_194
Cluster ID


CL\_5801
CL\_5802
CL\_5696
CL\_5695
CL\_5694
CL\_5693
CL\_5692
CL\_5691
CL\_5690
CL\_5689
CL\_5820
CL\_6612
CL\_6613
CL\_6614
CL\_6615
CL\_6616
CL\_6617
CL\_6618
CL\_6619
CL\_5669
CL\_5668
CL\_5667
CL\_6620
CL\_4644
CL\_4643
