## Supplementary material for "A novel method for integrating genomic and Tn-Seq data to identify common *in vivo* fitness mechanisms across multiple bacterial species": S6 Dataset: CL_INS_182.html

Legend

 Regulatoryfunctions
 Hypothetical
 SignalTransduction +Quorum Sensing
 Other
 EnergyMetabolism
 Transport +binding proteins

FULL


WINDOWSVGPNG

Trim RowsRemove SingletonsSave Fasta

CL\_1351


CL\_1351


CL\_1351


CL\_1351


CL\_1351

HighlightSelectShow Genomes


1

CL\_1350


1

CL\_1340


1

CL\_1350


1

CL\_1350


1

CL\_1350

fGI ID


CL\_INS\_182
CL\_INS\_30
CL\_INS\_30
CL\_INS\_182
CL\_INS\_182
CL\_INS\_182
CL\_INS\_182
CL\_INS\_182
CL\_INS\_182
CL\_INS\_182
CL\_INS\_182
CL\_INS\_182
CL\_INS\_182
CL\_INS\_182
Cluster ID


CL\_6611
CL\_3708
CL\_3709
CL\_7165
CL\_7164
CL\_7163
CL\_5227
CL\_5226
CL\_5225
CL\_4423
CL\_4424
CL\_4425
CL\_4426
CL\_4427
