## Supplementary material for "A novel method for integrating genomic and Tn-Seq data to identify common *in vivo* fitness mechanisms across multiple bacterial species": S6 Dataset: CL_INS_183.html

Legend

 Regulatoryfunctions
 Hypothetical
 All EssentialGenes
 All Fitness Genes
 Other
 All VFDB Genes
 Transport +binding proteins

FULL


WINDOWSVGPNG

Trim RowsRemove SingletonsSave Fasta

CL\_3136


CL\_1350


CL\_1350


CL\_224


CL\_1350


CL\_1350

HighlightSelectShow Genomes


1

CL\_4852


1

CL\_1343


1

CL\_4852


1

CL\_4852


1

CL\_645


1

CL\_4852

fGI ID


CL\_INS\_183
CL\_INS\_183
CL\_INS\_183
CL\_INS\_183
CL\_INS\_183
CL\_INS\_183
CL\_INS\_183
CL\_INS\_183
CL\_INS\_183
CL\_INS\_183
CL\_INS\_183
CL\_INS\_183
CL\_INS\_183
CL\_INS\_183
CL\_INS\_113
CL\_INS\_113
CL\_INS\_113
CL\_INS\_113
CL\_INS\_113
CL\_INS\_183
CL\_INS\_183
CL\_INS\_183
CL\_INS\_183
CL\_INS\_183
CL\_INS\_183
CL\_INS\_183
CL\_INS\_183
CL\_INS\_183
CL\_INS\_183
CL\_INS\_183
CL\_INS\_194
CL\_INS\_194
CL\_INS\_194
CL\_INS\_194
CL\_INS\_194
CL\_INS\_194
CL\_INS\_194
CL\_INS\_194
CL\_INS\_183
CL\_INS\_183
CL\_INS\_183
CL\_INS\_183
CL\_INS\_183
CL\_INS\_183
CL\_INS\_183
CL\_INS\_183
CL\_INS\_183
CL\_INS\_183
CL\_INS\_183
CL\_INS\_183
CL\_INS\_183
CL\_INS\_183
CL\_INS\_183
CL\_INS\_183
CL\_INS\_183
CL\_INS\_183
CL\_INS\_183
CL\_INS\_183
CL\_INS\_183
CL\_INS\_183
CL\_INS\_183
CL\_INS\_183
CL\_INS\_183
Cluster ID


CL\_4853
CL\_1349
CL\_1348
CL\_1347
CL\_1346
CL\_1345
CL\_1344
CL\_6087
CL\_6088
CL\_5237
CL\_5236
CL\_5235
CL\_5234
CL\_234
CL\_233
CL\_232
CL\_231
CL\_230
CL\_229
CL\_5698
CL\_5699
CL\_5700
CL\_6610
CL\_6609
CL\_6608
CL\_6607
CL\_6606
CL\_6605
CL\_6604
CL\_6603
CL\_280
CL\_281
CL\_282
CL\_283
CL\_284
CL\_285
CL\_286
CL\_287
CL\_6602
CL\_6601
CL\_6600
CL\_6599
CL\_6598
CL\_6597
CL\_6596
CL\_6595
CL\_6594
CL\_6593
CL\_6592
CL\_6591
CL\_6590
CL\_6589
CL\_6588
CL\_6587
CL\_5037
CL\_5036
CL\_5035
CL\_6586
CL\_6585
CL\_6584
CL\_6583
CL\_6582
CL\_6581
