## Supplementary material for "A novel method for integrating genomic and Tn-Seq data to identify common *in vivo* fitness mechanisms across multiple bacterial species": S6 Dataset: CL_INS_184.html

Legend

 Regulatoryfunctions
 Hypothetical
 Other
 EnergyMetabolism

FULL


WINDOWSVGPNG

Trim RowsRemove SingletonsSave Fasta

CL\_1298


CL\_1298


CL\_1251


CL\_1298

HighlightSelectShow Genomes


3

CL\_1297


1

CL\_1297


1

CL\_1297


1

CL\_1149

fGI ID


CL\_INS\_184
CL\_INS\_184
CL\_INS\_184
CL\_INS\_187
CL\_INS\_187
CL\_INS\_184
Cluster ID


CL\_6541
CL\_6540
CL\_6539
CL\_3935
CL\_3936
CL\_3937
