## Supplementary material for "A novel method for integrating genomic and Tn-Seq data to identify common *in vivo* fitness mechanisms across multiple bacterial species": S6 Dataset: CL_INS_185.html

Legend

 Regulatoryfunctions
 Hypothetical
 All Fitness Genes
 Proteinsynthesis/fate
 Other

FULL


WINDOWSVGPNG

Trim RowsRemove SingletonsSave Fasta

CL\_1198


CL\_1084


CL\_1198


CL\_1198


CL\_1198

HighlightSelectShow Genomes


1

CL\_1200


1

CL\_1200


1

CL\_1200


1

CL\_1084


1

CL\_1200

fGI ID


CL\_INS\_185
CL\_INS\_185
CL\_INS\_185
CL\_INS\_185
CL\_INS\_185
CL\_INS\_185
CL\_INS\_185
CL\_INS\_185
CL\_INS\_185
CL\_INS\_185
CL\_INS\_185
CL\_INS\_185
CL\_INS\_185
Cluster ID


CL\_1199
CL\_5803
CL\_5802
CL\_5801
CL\_6496
CL\_6495
CL\_6494
CL\_6493
CL\_5930
CL\_5931
CL\_5933
CL\_6492
CL\_6491
