## Supplementary material for "A novel method for integrating genomic and Tn-Seq data to identify common *in vivo* fitness mechanisms across multiple bacterial species": S6 Dataset: CL_INS_186.html

Legend

 Regulatoryfunctions
 Hypothetical
 All EssentialGenes
 All Fitness Genes
 Proteinsynthesis/fate
 Other
 EnergyMetabolism
 Transport +binding proteins

FULL


WINDOWSVGPNG

Trim RowsRemove SingletonsSave Fasta

CL\_1200


CL\_1200


CL\_1200


CL\_1200


CL\_5171

HighlightSelectShow Genomes


1

CL\_1208


1

CL\_1231


1

CL\_1208


1

CL\_1208


1

CL\_1208

fGI ID


CL\_INS\_186
CL\_INS\_186
CL\_INS\_186
CL\_INS\_186
CL\_INS\_186
CL\_INS\_186
CL\_INS\_186
CL\_INS\_186
CL\_INS\_186
CL\_INS\_186
CL\_INS\_186
CL\_INS\_186
CL\_INS\_186
CL\_INS\_186
CL\_INS\_88
CL\_INS\_107
CL\_INS\_107
CL\_INS\_107
CL\_INS\_107
CL\_INS\_107
CL\_INS\_107
CL\_INS\_107
CL\_INS\_186
CL\_INS\_186
CL\_INS\_186
CL\_INS\_186
CL\_INS\_186
CL\_INS\_186
CL\_INS\_186
CL\_INS\_186
CL\_INS\_186
CL\_INS\_186
CL\_INS\_186
CL\_INS\_186
CL\_INS\_186
CL\_INS\_186
CL\_INS\_186
CL\_INS\_186
CL\_INS\_186
CL\_INS\_186
CL\_INS\_31
CL\_INS\_186
CL\_INS\_31
CL\_INS\_31
CL\_INS\_29
CL\_INS\_186
CL\_INS\_186
CL\_INS\_186
CL\_INS\_186
Cluster ID


CL\_5229
CL\_6490
CL\_6489
CL\_6377
CL\_6378
CL\_6379
CL\_6380
CL\_6381
CL\_6382
CL\_6383
CL\_6384
CL\_6385
CL\_6386
CL\_6387
CL\_1225
CL\_4952
CL\_4951
CL\_4950
CL\_4949
CL\_4948
CL\_4947
CL\_4946
CL\_5800
CL\_5799
CL\_1201
CL\_1202
CL\_1203
CL\_1204
CL\_5798
CL\_1205
CL\_1206
CL\_1207
CL\_5797
CL\_7122
CL\_7123
CL\_7124
CL\_7125
CL\_7126
CL\_5796
CL\_5795
CL\_4689
CL\_5794
CL\_4691
CL\_4692
CL\_4693
CL\_5793
CL\_5792
CL\_5791
CL\_5790
