## Supplementary material for "A novel method for integrating genomic and Tn-Seq data to identify common *in vivo* fitness mechanisms across multiple bacterial species": S6 Dataset: CL_INS_187.html

Legend

 Hypothetical
 Regulatoryfunctions
 Cell Envelope
 Other
 Transport +binding proteins

FULL


WINDOWSVGPNG

Trim RowsRemove SingletonsSave Fasta

CL\_1230


CL\_1230


CL\_5152


CL\_1230

HighlightSelectShow Genomes


1

CL\_5152


1

CL\_5171


1

CL\_5171


1

CL\_5171

fGI ID


CL\_INS\_187
CL\_INS\_187
CL\_INS\_187
CL\_INS\_187
CL\_INS\_187
CL\_INS\_187
CL\_INS\_187
CL\_INS\_187
CL\_INS\_187
CL\_INS\_187
CL\_INS\_187
CL\_INS\_187
CL\_INS\_187
CL\_INS\_187
CL\_INS\_187
CL\_INS\_187
CL\_INS\_187
CL\_INS\_187
CL\_INS\_187
CL\_INS\_187
CL\_INS\_187
CL\_INS\_187
CL\_INS\_187
CL\_INS\_187
CL\_INS\_187
CL\_INS\_187
CL\_INS\_187
CL\_INS\_187
CL\_INS\_187
CL\_INS\_187
CL\_INS\_187
CL\_INS\_187
CL\_INS\_187
CL\_INS\_187
CL\_INS\_187
CL\_INS\_187
CL\_INS\_187
CL\_INS\_187
Cluster ID


CL\_3936
CL\_3935
CL\_5774
CL\_5775
CL\_5776
CL\_4032
CL\_5777
CL\_5778
CL\_5781
CL\_5782
CL\_5783
CL\_5784
CL\_5785
CL\_5786
CL\_5787
CL\_5788
CL\_5789
CL\_6367
CL\_6368
CL\_6369
CL\_6370
CL\_6371
CL\_6372
CL\_6373
CL\_6374
CL\_6375
CL\_3707
CL\_7134
CL\_7133
CL\_7132
CL\_7131
CL\_7130
CL\_3830
CL\_3831
CL\_3832
CL\_7129
CL\_7128
CL\_7127
