## Supplementary material for "A novel method for integrating genomic and Tn-Seq data to identify common *in vivo* fitness mechanisms across multiple bacterial species": S6 Dataset: CL_INS_188.html

Legend

 Hypothetical
 All EssentialGenes
 All Fitness Genes
 Other

FULL


WINDOWSVGPNG

Trim RowsRemove SingletonsSave Fasta

CL\_1232


CL\_1232


CL\_1232


CL\_1836


CL\_1232

HighlightSelectShow Genomes


1

CL\_1234


1

CL\_1234


1

CL\_1234


1

CL\_1234


1

CL\_1234

fGI ID


CL\_INS\_188
CL\_INS\_188
CL\_INS\_188
Cluster ID


CL\_5768
CL\_5767
CL\_1233
