## Supplementary material for "A novel method for integrating genomic and Tn-Seq data to identify common *in vivo* fitness mechanisms across multiple bacterial species": S6 Dataset: CL_INS_189.html

Legend

 Regulatoryfunctions
 Hypothetical
 Other
 Transport +binding proteins

FULL


WINDOWSVGPNG

Trim RowsRemove SingletonsSave Fasta

CL\_1238


CL\_1238


CL\_1238


CL\_1238

HighlightSelectShow Genomes


2

CL\_1237


1

CL\_1237


1

CL\_4390


1

CL\_1237

fGI ID


CL\_INS\_189
CL\_INS\_189
CL\_INS\_189
CL\_INS\_189
CL\_INS\_189
CL\_INS\_189
CL\_INS\_189
CL\_INS\_189
CL\_INS\_189
CL\_INS\_189
CL\_INS\_189
CL\_INS\_189
CL\_INS\_90
CL\_INS\_90
CL\_INS\_90
CL\_INS\_90
CL\_INS\_90
CL\_INS\_189
Cluster ID


CL\_5764
CL\_5765
CL\_6357
CL\_6358
CL\_6359
CL\_6360
CL\_6361
CL\_6362
CL\_6363
CL\_4380
CL\_4381
CL\_4382
CL\_4383
CL\_4384
CL\_4385
CL\_4386
CL\_4387
CL\_4388
