## Supplementary material for "A novel method for integrating genomic and Tn-Seq data to identify common *in vivo* fitness mechanisms across multiple bacterial species": S6 Dataset: CL_INS_190.html

Legend

 Regulatoryfunctions
 Hypothetical
 All Fitness Genes
 Proteinsynthesis/fate
 Other
 Transport +binding proteins

FULL


WINDOWSVGPNG

Trim RowsRemove SingletonsSave Fasta

CL\_1244


CL\_5254


CL\_1244


CL\_1244

HighlightSelectShow Genomes


1

CL\_5245


1

CL\_5245


1

CL\_5245


1

CL\_1248

fGI ID


CL\_INS\_190
CL\_INS\_190
CL\_INS\_190
CL\_INS\_190
CL\_INS\_190
CL\_INS\_190
CL\_INS\_190
CL\_INS\_190
CL\_INS\_190
CL\_INS\_190
Cluster ID


CL\_6356
CL\_1245
CL\_1246
CL\_5252
CL\_5251
CL\_5250
CL\_5249
CL\_5248
CL\_5247
CL\_5246
