## Supplementary material for "A novel method for integrating genomic and Tn-Seq data to identify common *in vivo* fitness mechanisms across multiple bacterial species": S6 Dataset: CL_INS_191.html

Legend

 All EssentialGenes
 All Fitness Genes
 Other

FULL


WINDOWSVGPNG

Trim RowsRemove SingletonsSave Fasta

CL\_1252


CL\_1252


CL\_1252

HighlightSelectShow Genomes


2

CL\_1256


1

CL\_1256


1

CL\_1256

fGI ID


CL\_INS\_191
CL\_INS\_191
CL\_INS\_191
CL\_INS\_191
Cluster ID


CL\_6339
CL\_1253
CL\_1254
CL\_1255
