## Supplementary material for "A novel method for integrating genomic and Tn-Seq data to identify common *in vivo* fitness mechanisms across multiple bacterial species": S6 Dataset: CL_INS_192.html

Legend

 Hypothetical
 Regulatoryfunctions
 Cell Envelope
 Proteinsynthesis/fate
 Other
 Transport +binding proteins
 All VFDB Genes

FULL


WINDOWSVGPNG

Trim RowsRemove SingletonsSave Fasta

CL\_248


CL\_248


CL\_248


CL\_248

HighlightSelectShow Genomes


1

CL\_3424


1

CL\_5135


1

CL\_236


1

CL\_462

fGI ID


CL\_INS\_192
CL\_INS\_192
CL\_INS\_192
CL\_INS\_192
CL\_INS\_192
CL\_INS\_192
CL\_INS\_192
CL\_INS\_192
CL\_INS\_192
CL\_INS\_192
CL\_INS\_192
CL\_INS\_192
CL\_INS\_192
CL\_INS\_192
CL\_INS\_192
CL\_INS\_192
CL\_INS\_192
CL\_INS\_192
CL\_INS\_192
CL\_INS\_192
CL\_INS\_192
CL\_INS\_192
CL\_INS\_192
CL\_INS\_192
CL\_INS\_192
CL\_INS\_192
CL\_INS\_192
CL\_INS\_192
CL\_INS\_192
CL\_INS\_192
Cluster ID


CL\_5306
CL\_5307
CL\_5308
CL\_5309
CL\_5310
CL\_5311
CL\_247
CL\_246
CL\_245
CL\_244
CL\_243
CL\_242
CL\_241
CL\_240
CL\_239
CL\_238
CL\_5625
CL\_4237
CL\_4236
CL\_4235
CL\_4234
CL\_6166
CL\_6165
CL\_6164
CL\_6163
CL\_6162
CL\_6161
CL\_6160
CL\_6159
CL\_6158
