## Supplementary material for "A novel method for integrating genomic and Tn-Seq data to identify common *in vivo* fitness mechanisms across multiple bacterial species": S6 Dataset: CL_INS_193.html

Legend

 Regulatoryfunctions
 Hypothetical
 DNA Metabolism
 All Fitness Genes
 Proteinsynthesis/fate
 Other
 EnergyMetabolism
 Cellularprocesses
 All VFDB Genes
 Transport +binding proteins

FULL


WINDOWSVGPNG

Trim RowsRemove SingletonsSave Fasta

CL\_978


CL\_978


CL\_978


CL\_978

HighlightSelectShow Genomes


1

CL\_1455


1

CL\_1178


1

CL\_290


1

CL\_977

fGI ID


CL\_INS\_193
CL\_INS\_127
CL\_INS\_127
CL\_INS\_127
CL\_INS\_127
CL\_INS\_186
CL\_INS\_186
CL\_INS\_193
CL\_INS\_193
CL\_INS\_193
CL\_INS\_193
CL\_INS\_193
CL\_INS\_193
CL\_INS\_193
CL\_INS\_193
CL\_INS\_193
CL\_INS\_193
CL\_INS\_193
CL\_INS\_193
CL\_INS\_193
CL\_INS\_193
CL\_INS\_193
CL\_INS\_193
CL\_INS\_193
CL\_INS\_193
CL\_INS\_193
CL\_INS\_193
CL\_INS\_193
CL\_INS\_193
CL\_INS\_193
CL\_INS\_193
CL\_INS\_193
CL\_INS\_193
CL\_INS\_193
CL\_INS\_193
CL\_INS\_193
CL\_INS\_193
CL\_INS\_193
CL\_INS\_193
CL\_INS\_193
CL\_INS\_193
CL\_INS\_193
CL\_INS\_193
CL\_INS\_193
CL\_INS\_193
CL\_INS\_193
CL\_INS\_193
CL\_INS\_193
CL\_INS\_193
CL\_INS\_193
CL\_INS\_193
CL\_INS\_193
CL\_INS\_193
CL\_INS\_193
CL\_INS\_193
Cluster ID


CL\_7111
CL\_7112
CL\_7113
CL\_7114
CL\_7115
CL\_5800
CL\_5799
CL\_6249
CL\_6250
CL\_6251
CL\_3915
CL\_4031
CL\_4030
CL\_4029
CL\_4028
CL\_4027
CL\_4026
CL\_4025
CL\_4282
CL\_4281
CL\_4280
CL\_4279
CL\_4278
CL\_4277
CL\_4276
CL\_4275
CL\_4274
CL\_4273
CL\_4272
CL\_4271
CL\_4270
CL\_4269
CL\_4268
CL\_4267
CL\_4266
CL\_4265
CL\_4264
CL\_4263
CL\_4262
CL\_4261
CL\_4260
CL\_4259
CL\_4258
CL\_4257
CL\_4256
CL\_4255
CL\_4254
CL\_4253
CL\_4252
CL\_4251
CL\_4250
CL\_4249
CL\_4248
CL\_4247
CL\_4246
