## Supplementary material for "A novel method for integrating genomic and Tn-Seq data to identify common *in vivo* fitness mechanisms across multiple bacterial species": S6 Dataset: CL_INS_194.html

Legend

 Other
 Hypothetical
 Regulatoryfunctions
 All EssentialGenes
 All Fitness Genes
 All VFDB Genes

FULL


WINDOWSVGPNG

Trim RowsRemove SingletonsSave Fasta

CL\_288


CL\_288


CL\_288


CL\_288


CL\_288

HighlightSelectShow Genomes


1

CL\_36


1

CL\_278


1

CL\_995


1

CL\_2415


1

CL\_4337

fGI ID


CL\_INS\_194
CL\_INS\_194
CL\_INS\_194
CL\_INS\_194
CL\_INS\_194
CL\_INS\_194
CL\_INS\_194
CL\_INS\_194
CL\_INS\_194
CL\_INS\_194
CL\_INS\_194
CL\_INS\_194
CL\_INS\_194
CL\_INS\_194
CL\_INS\_194
CL\_INS\_194
CL\_INS\_194
CL\_INS\_194
CL\_INS\_194
CL\_INS\_194
CL\_INS\_194
CL\_INS\_194
CL\_INS\_194
CL\_INS\_194
CL\_INS\_194
CL\_INS\_194
CL\_INS\_194
CL\_INS\_194
CL\_INS\_194
CL\_INS\_194
CL\_INS\_194
CL\_INS\_194
CL\_INS\_194
CL\_INS\_194
CL\_INS\_194
Cluster ID


CL\_287
CL\_286
CL\_285
CL\_284
CL\_283
CL\_282
CL\_281
CL\_280
CL\_6642
CL\_6641
CL\_6640
CL\_6639
CL\_6638
CL\_6637
CL\_6636
CL\_6635
CL\_6634
CL\_6633
CL\_6632
CL\_6631
CL\_3819
CL\_4643
CL\_4644
CL\_5744
CL\_5743
CL\_5742
CL\_5741
CL\_5740
CL\_5739
CL\_5738
CL\_5737
CL\_5736
CL\_5735
CL\_5734
CL\_5733
