## Supplementary material for "A novel method for integrating genomic and Tn-Seq data to identify common *in vivo* fitness mechanisms across multiple bacterial species": S6 Dataset: CL_INS_195.html

Legend

 Regulatoryfunctions
 All EssentialGenes
 SignalTransduction +Quorum Sensing
 Other

FULL


WINDOWSVGPNG

Trim RowsRemove SingletonsSave Fasta

CL\_3468


CL\_3468


CL\_3468

HighlightSelectShow Genomes


2

CL\_3465


2

CL\_2663


1

CL\_624

fGI ID

CL\_INS\_195
Cluster ID

CL\_3467
