## Supplementary material for "A novel method for integrating genomic and Tn-Seq data to identify common *in vivo* fitness mechanisms across multiple bacterial species": S6 Dataset: CL_INS_196.html

Legend

 Hypothetical
 Regulatoryfunctions
 All EssentialGenes
 Other

FULL


WINDOWSVGPNG

Trim RowsRemove SingletonsSave Fasta

CL\_3469


CL\_3469


CL\_3469

HighlightSelectShow Genomes


1

CL\_3475


1

CL\_1324


1

CL\_3475

fGI ID


CL\_INS\_196
CL\_INS\_196
CL\_INS\_196
CL\_INS\_196
CL\_INS\_55
CL\_INS\_32
CL\_INS\_32
Cluster ID


CL\_3919
CL\_3470
CL\_3471
CL\_3472
CL\_3473
CL\_3474
CL\_5986
