## Supplementary material for "A novel method for integrating genomic and Tn-Seq data to identify common *in vivo* fitness mechanisms across multiple bacterial species": S6 Dataset: CL_INS_197.html

Legend

 Regulatoryfunctions
 Hypothetical
 SignalTransduction +Quorum Sensing
 Other

FULL


WINDOWSVGPNG

Trim RowsRemove SingletonsSave Fasta

CL\_5258


CL\_5254

HighlightSelectShow Genomes


3

CL\_5260


1

CL\_5260

fGI ID


CL\_INS\_197
CL\_INS\_197
CL\_INS\_197
CL\_INS\_197
CL\_INS\_197
CL\_INS\_197
CL\_INS\_197
CL\_INS\_197
CL\_INS\_197
CL\_INS\_197
CL\_INS\_197
Cluster ID


CL\_5259
CL\_5544
CL\_5543
CL\_5542
CL\_5541
CL\_5540
CL\_5539
CL\_5538
CL\_5537
CL\_5536
CL\_5535
