## Supplementary material for "A novel method for integrating genomic and Tn-Seq data to identify common *in vivo* fitness mechanisms across multiple bacterial species": S6 Dataset: CL_INS_199.html

Legend

 Hypothetical
 Regulatoryfunctions
 All Fitness Genes
 Cell Envelope
 Other
 EnergyMetabolism
 All VFDB Genes
 Transport +binding proteins

FULL


WINDOWSVGPNG

Trim RowsRemove SingletonsSave Fasta

CL\_3478


CL\_3478


CL\_3478


CL\_3478


CL\_3478

HighlightSelectShow Genomes


1

CL\_1276


1

CL\_1627


1

CL\_3480


1

CL\_995


1

CL\_3489

fGI ID


CL\_INS\_199
CL\_INS\_199
CL\_INS\_199
CL\_INS\_199
CL\_INS\_199
CL\_INS\_199
CL\_INS\_199
CL\_INS\_199
CL\_INS\_199
CL\_INS\_199
CL\_INS\_199
CL\_INS\_199
CL\_INS\_199
CL\_INS\_199
CL\_INS\_199
CL\_INS\_199
CL\_INS\_199
CL\_INS\_199
CL\_INS\_194
CL\_INS\_199
CL\_INS\_199
CL\_INS\_199
CL\_INS\_199
CL\_INS\_199
CL\_INS\_199
CL\_INS\_199
CL\_INS\_199
CL\_INS\_199
CL\_INS\_199
CL\_INS\_187
CL\_INS\_187
CL\_INS\_187
CL\_INS\_199
CL\_INS\_199
Cluster ID


CL\_3479
CL\_4156
CL\_3814
CL\_6335
CL\_5856
CL\_6336
CL\_5358
CL\_5359
CL\_5360
CL\_5361
CL\_5362
CL\_5363
CL\_5364
CL\_5365
CL\_3815
CL\_3816
CL\_3817
CL\_3818
CL\_3819
CL\_3820
CL\_3821
CL\_3822
CL\_3823
CL\_3824
CL\_3825
CL\_3826
CL\_3827
CL\_3828
CL\_3829
CL\_3830
CL\_3831
CL\_3832
CL\_3833
CL\_3834
