## Supplementary material for "A novel method for integrating genomic and Tn-Seq data to identify common *in vivo* fitness mechanisms across multiple bacterial species": S6 Dataset: CL_INS_200.html

Legend

 Regulatoryfunctions
 Hypothetical
 Other
 Transport +binding proteins

FULL


WINDOWSVGPNG

Trim RowsRemove SingletonsSave Fasta

CL\_271


CL\_271

HighlightSelectShow Genomes


1

CL\_261


1

CL\_2693

fGI ID


CL\_INS\_200
CL\_INS\_204
CL\_INS\_204
CL\_INS\_204
CL\_INS\_204
CL\_INS\_204
CL\_INS\_204
CL\_INS\_204
Cluster ID


CL\_270
CL\_269
CL\_268
CL\_267
CL\_266
CL\_265
CL\_264
CL\_263
