## Supplementary material for "A novel method for integrating genomic and Tn-Seq data to identify common *in vivo* fitness mechanisms across multiple bacterial species": S6 Dataset: CL_INS_201.html

Legend

 Regulatoryfunctions
 Hypothetical
 Other

FULL


WINDOWSVGPNG

Trim RowsRemove SingletonsSave Fasta

CL\_5254


CL\_5254


CL\_5254


CL\_5254

HighlightSelectShow Genomes


1

CL\_3639


1

CL\_5258


1

CL\_3635


1

CL\_3635

fGI ID


CL\_INS\_201
CL\_INS\_201
CL\_INS\_201
CL\_INS\_201
CL\_INS\_201
CL\_INS\_201
CL\_INS\_201
CL\_INS\_201
CL\_INS\_201
CL\_INS\_201
CL\_INS\_201
CL\_INS\_201
CL\_INS\_201
CL\_INS\_201
CL\_INS\_201
Cluster ID


CL\_5255
CL\_5256
CL\_5546
CL\_5547
CL\_5548
CL\_5315
CL\_5549
CL\_3638
CL\_5550
CL\_5551
CL\_5552
CL\_3632
CL\_3633
CL\_3634
CL\_6032
