## Supplementary material for "A novel method for integrating genomic and Tn-Seq data to identify common *in vivo* fitness mechanisms across multiple bacterial species": S6 Dataset: CL_INS_202.html

Legend

 Regulatoryfunctions
 Hypothetical
 All EssentialGenes
 Other

FULL


WINDOWSVGPNG

Trim RowsRemove SingletonsSave Fasta

CL\_260


CL\_260


CL\_260


CL\_260

HighlightSelectShow Genomes


1

CL\_3481


1

CL\_2194


1

CL\_251


1

CL\_1333

fGI ID


CL\_INS\_202
CL\_INS\_202
CL\_INS\_202
CL\_INS\_202
CL\_INS\_202
CL\_INS\_202
CL\_INS\_202
CL\_INS\_202
CL\_INS\_202
CL\_INS\_202
CL\_INS\_202
CL\_INS\_202
CL\_INS\_202
CL\_INS\_202
CL\_INS\_202
CL\_INS\_202
Cluster ID


CL\_259
CL\_5905
CL\_258
CL\_257
CL\_256
CL\_255
CL\_254
CL\_253
CL\_4874
CL\_4875
CL\_4876
CL\_4877
CL\_4878
CL\_4879
CL\_4880
CL\_4881
