## Supplementary material for "A novel method for integrating genomic and Tn-Seq data to identify common *in vivo* fitness mechanisms across multiple bacterial species": S6 Dataset: CL_INS_204.html

Legend

 Hypothetical
 Regulatoryfunctions
 All Fitness Genes
 Other
 Transport +binding proteins

FULL


WINDOWSVGPNG

Trim RowsRemove SingletonsSave Fasta

CL\_261


CL\_261


CL\_261

HighlightSelectShow Genomes


1

CL\_1123


1

CL\_3082


1

CL\_271

fGI ID


CL\_INS\_204
CL\_INS\_204
CL\_INS\_204
CL\_INS\_204
CL\_INS\_204
CL\_INS\_204
CL\_INS\_204
CL\_INS\_204
CL\_INS\_204
CL\_INS\_204
CL\_INS\_204
CL\_INS\_204
Cluster ID


CL\_1121
CL\_5463
CL\_5462
CL\_6908
CL\_262
CL\_263
CL\_264
CL\_265
CL\_266
CL\_267
CL\_268
CL\_269
