## Supplementary material for "A novel method for integrating genomic and Tn-Seq data to identify common *in vivo* fitness mechanisms across multiple bacterial species": S6 Dataset: CL_INS_205.html

Legend

 Regulatoryfunctions
 Hypothetical
 All Fitness Genes
 Other

FULL


WINDOWSVGPNG

Trim RowsRemove SingletonsSave Fasta

CL\_1084


CL\_1084

HighlightSelectShow Genomes


1

CL\_278


1

CL\_1200

fGI ID


CL\_INS\_204
CL\_INS\_204
CL\_INS\_205
CL\_INS\_205
CL\_INS\_185
CL\_INS\_185
CL\_INS\_185
CL\_INS\_185
CL\_INS\_185
CL\_INS\_185
CL\_INS\_185
CL\_INS\_185
Cluster ID


CL\_5462
CL\_5463
CL\_7266
CL\_6497
CL\_6496
CL\_6495
CL\_6494
CL\_6493
CL\_5930
CL\_5931
CL\_5933
CL\_6492
