## Supplementary material for "A novel method for integrating genomic and Tn-Seq data to identify common *in vivo* fitness mechanisms across multiple bacterial species": S6 Dataset: CL_INS_206.html

Legend

 Regulatoryfunctions
 Hypothetical
 All Fitness Genes
 Other
 EnergyMetabolism
 Transport +binding proteins

FULL


WINDOWSVGPNG

Trim RowsRemove SingletonsSave Fasta

CL\_4390


CL\_4390


CL\_4390


CL\_4390

HighlightSelectShow Genomes


1

CL\_493


1

CL\_2600


1

CL\_1238


1

CL\_548

fGI ID


CL\_INS\_206
CL\_INS\_206
CL\_INS\_207
CL\_INS\_207
CL\_INS\_206
CL\_INS\_149
CL\_INS\_90
CL\_INS\_90
CL\_INS\_206
CL\_INS\_189
CL\_INS\_90
CL\_INS\_90
CL\_INS\_90
CL\_INS\_90
CL\_INS\_90
CL\_INS\_189
CL\_INS\_189
Cluster ID


CL\_7231
CL\_7230
CL\_4394
CL\_4395
CL\_6643
CL\_4573
CL\_6644
CL\_6645
CL\_4389
CL\_4388
CL\_4387
CL\_4386
CL\_4385
CL\_4384
CL\_4383
CL\_4382
CL\_4381
