## Supplementary material for "A novel method for integrating genomic and Tn-Seq data to identify common *in vivo* fitness mechanisms across multiple bacterial species": S6 Dataset: CL_INS_207.html

Legend

 Regulatoryfunctions
 Hypothetical
 All EssentialGenes
 All Fitness Genes
 Biosynthesis ofcofactors,prostheticgroups, +carriers
 Cell Envelope
 Other
 EnergyMetabolism

FULL


WINDOWSVGPNG

Trim RowsRemove SingletonsSave Fasta

CL\_4390


CL\_4390

HighlightSelectShow Genomes


2

CL\_1635


1

CL\_3016

fGI ID


CL\_INS\_207
CL\_INS\_207
CL\_INS\_207
CL\_INS\_30
CL\_INS\_30
CL\_INS\_30
CL\_INS\_30
CL\_INS\_30
CL\_INS\_30
CL\_INS\_30
CL\_INS\_30
CL\_INS\_30
CL\_INS\_30
CL\_INS\_30
CL\_INS\_30
CL\_INS\_30
CL\_INS\_207
CL\_INS\_207
Cluster ID


CL\_4391
CL\_4392
CL\_4393
CL\_2475
CL\_2476
CL\_2477
CL\_2478
CL\_2479
CL\_2487
CL\_2486
CL\_2485
CL\_2484
CL\_2483
CL\_2482
CL\_2481
CL\_2480
CL\_4394
CL\_4395
