## Supplementary material for "A novel method for integrating genomic and Tn-Seq data to identify common *in vivo* fitness mechanisms across multiple bacterial species": S6 Dataset: CL_INS_208.html

Legend

 Hypothetical
 Regulatoryfunctions
 All EssentialGenes
 Other
 Transport +binding proteins

FULL


WINDOWSVGPNG

Trim RowsRemove SingletonsSave Fasta

CL\_5135


CL\_5135


CL\_5135


CL\_5135

HighlightSelectShow Genomes


1

CL\_4507


1

CL\_248


1

CL\_4390


1

CL\_447

fGI ID


CL\_INS\_207
CL\_INS\_208
CL\_INS\_208
CL\_INS\_208
CL\_INS\_192
CL\_INS\_192
CL\_INS\_192
CL\_INS\_192
CL\_INS\_194
CL\_INS\_194
CL\_INS\_194
CL\_INS\_194
CL\_INS\_194
CL\_INS\_194
CL\_INS\_194
Cluster ID


CL\_4392
CL\_5134
CL\_5133
CL\_5132
CL\_4234
CL\_4235
CL\_4236
CL\_4237
CL\_280
CL\_281
CL\_282
CL\_283
CL\_284
CL\_285
CL\_286
